## Supplementary material for "Nerve-associated macrophages control adipose homeostasis across lifespan and restrain age-related inflammation": Table S4

| p_val | avg_log2FC | pct.1 | pct.2 | p_val_adj | cluster | gene | pct.diff | condition |
|---|---|---|---|---|---|---|---|---|
| 1.83E-39 | 3.178281 | 0.129 | 0.016 | 5.67E-35 | 0 | Xist | 0.113 | Old |
| 7.92E-29 | 3.263704 | 0.083 | 0.008 | 2.46E-24 | 0 | Cd55 | 0.075 | Old |
| 1.97E-27 | 1.222965 | 0.412 | 0.217 | 6.13E-23 | 0 | Vcam1 | 0.195 | Old |
| 4.53E-27 | 2.291799 | 0.149 | 0.038 | 1.41E-22 | 0 | Cxcl13 | 0.111 | Old |
| 2.90E-26 | 0.372334 | 0.986 | 0.968 | 9.01E-22 | 0 | H2-Aa | 0.018 | Old |
| 1.79E-25 | 0.390788 | 0.966 | 0.933 | 5.55E-21 | 0 | B2m | 0.033 | Old |
| 2.30E-25 | 0.842193 | 0.657 | 0.483 | 7.15E-21 | 0 | Gda | 0.174 | Old |
| 2.19E-21 | 0.630919 | 0.938 | 0.899 | 6.79E-17 | 0 | H3f3b | 0.039 | Old |
| 2.42E-21 | 1.125774 | 0.514 | 0.338 | 7.50E-17 | 0 | Ccl8 | 0.176 | Old |
| 2.10E-19 | 0.361535 | 0.947 | 0.895 | 6.52E-15 | 0 | H2-K1 | 0.052 | Old |
| 8.93E-19 | 0.379952 | 0.963 | 0.926 | 2.77E-14 | 0 | H2-Eb1 | 0.037 | Old |
| 4.02E-16 | 0.307074 | 0.974 | 0.964 | 1.25E-11 | 0 | H2-Ab1 | 0.01 | Old |
| 4.16E-16 | 0.214073 | 1 | 1 | 1.29E-11 | 0 | Actb | 0 | Old |
| 1.75E-14 | 0.778832 | 0.438 | 0.292 | 5.45E-10 | 0 | Plbd1 | 0.146 | Old |
| 2.42E-13 | 0.314468 | 0.914 | 0.821 | 7.53E-09 | 0 | Hsp90b1 | 0.093 | Old |
| 3.71E-13 | 1.835671 | 0.1 | 0.034 | 1.15E-08 | 0 | C3 | 0.066 | Old |
| 1.65E-12 | 0.138424 | 1 | 1 | 5.13E-08 | 0 | mt-Co1 | 0 | Old |
| 2.70E-12 | 0.303641 | 0.963 | 0.955 | 8.39E-08 | 0 | Actg1 | 0.008 | Old |
| 3.88E-12 | 0.168551 | 1 | 1 | 1.21E-07 | 0 | mt-Co3 | 0 | Old |
| 9.13E-11 | 0.669897 | 0.343 | 0.225 | 2.84E-06 | 0 | Lfng | 0.118 | Old |
| 1.04E-10 | 1.522447 | 0.094 | 0.036 | 3.23E-06 | 0 | Mylk | 0.058 | Old |
| 1.11E-10 | 0.281877 | 0.912 | 0.878 | 3.43E-06 | 0 | H2-D1 | 0.034 | Old |
| 1.81E-10 | 0.342411 | 0.926 | 0.894 | 5.62E-06 | 0 | Ccl6 | 0.032 | Old |
| 1.88E-10 | 0.520201 | 0.523 | 0.399 | 5.82E-06 | 0 | Mpeg1 | 0.124 | Old |
| 2.46E-10 | 0.492087 | 0.598 | 0.485 | 7.63E-06 | 0 | Vmp1 | 0.113 | Old |
| 3.52E-10 | 0.357885 | 0.865 | 0.847 | 1.09E-05 | 0 | Ccl9 | 0.018 | Old |
| 4.30E-10 | 0.431581 | 0.715 | 0.628 | 1.34E-05 | 0 | Tmem176a | 0.087 | Old |
| 4.30E-10 | 1.087128 | 0.171 | 0.09 | 1.34E-05 | 0 | Il2rg | 0.081 | Old |
| 1.08E-09 | 0.174842 | 0.996 | 0.999 | 3.37E-05 | 0 | mt-Atp6 | -0.003 | Old |
| 3.16E-09 | 0.27153 | 0.947 | 0.922 | 9.80E-05 | 0 | Fos | 0.025 | Old |
| 3.89E-09 | 0.572321 | 0.505 | 0.398 | 0.000121 | 0 | Apobec3 | 0.107 | Old |
| 7.61E-09 | 1.256801 | 0.161 | 0.088 | 0.000236 | 0 | Rgs1 | 0.073 | Old |
| 9.04E-09 | 0.249975 | 0.923 | 0.88 | 0.000281 | 0 | Ptma | 0.043 | Old |
| 1.12E-08 | 0.876546 | 0.327 | 0.227 | 0.000348 | 0 | Anxa1 | 0.1 | Old |
| 1.44E-08 | 0.284604 | 0.891 | 0.83 | 0.000449 | 0 | mt-Nd3 | 0.061 | Old |
| 2.12E-08 | 1.756379 | 0.041 | 0.01 | 0.000657 | 0 | Gdf3 | 0.031 | Old |
| 2.16E-08 | 0.19428 | 0.99 | 0.986 | 0.000672 | 0 | mt-Cytb | 0.004 | Old |
| 2.44E-08 | 0.406501 | 0.577 | 0.464 | 0.000758 | 0 | Xbp1 | 0.113 | Old |
| 3.75E-08 | 0.807721 | 0.202 | 0.124 | 0.001166 | 0 | Rab20 | 0.078 | Old |
| 4.09E-08 | 0.590395 | 0.382 | 0.278 | 0.001271 | 0 | Lgals3 | 0.104 | Old |
| 5.36E-08 | 0.352942 | 0.796 | 0.744 | 0.001664 | 0 | Vim | 0.052 | Old |
| 7.35E-08 | 0.180051 | 0.992 | 0.984 | 0.002282 | 0 | Psap | 0.008 | Old |
| 2.43E-07 | 0.290321 | 0.918 | 0.885 | 0.007533 | 0 | Neat1 | 0.033 | Old |
| 2.61E-07 | 0.31634 | 0.759 | 0.679 | 0.008101 | 0 | Hspa5 | 0.08 | Old |

| | | | | | | | | |
|---|---|---|---|---|---|---|---|---|
| 3.15E-07 | 0.31894 | 0.751 | 0.66 | 0.009793 | 0 | C5ar1 | 0.091 | Old |
| 3.60E-07 | 0.488245 | 0.382 | 0.279 | 0.011175 | 0 | Olfm1 | 0.103 | Old |
| 3.60E-07 | 0.289634 | 0.778 | 0.687 | 0.011188 | 0 | Cfp | 0.091 | Old |
| 6.68E-07 | 0.355883 | 0.837 | 0.799 | 0.020749 | 0 | Egr1 | 0.038 | Old |
| 7.65E-07 | 0.348084 | 0.736 | 0.649 | 0.023743 | 0 | Sdc3 | 0.087 | Old |
| 9.18E-07 | 0.431089 | 0.342 | 0.246 | 0.028515 | 0 | Surf4 | 0.096 | Old |
| 9.20E-07 | 0.491606 | 0.51 | 0.417 | 0.028581 | 0 | Myadm | 0.093 | Old |
| 1.09E-06 | 0.336009 | 0.673 | 0.58 | 0.033758 | 0 | Cd47 | 0.093 | Old |
| 1.09E-06 | 0.709237 | 0.219 | 0.144 | 0.033867 | 0 | Lgals3bp | 0.075 | Old |
| 1.17E-06 | 2.077546 | 0.016 | 0.002 | 0.036393 | 0 | Gm10800 | 0.014 | Old |
| 1.34E-06 | 0.138295 | 0.997 | 0.996 | 0.041686 | 0 | C1qa | 0.001 | Old |
| 1.35E-06 | 0.509994 | 0.276 | 0.19 | 0.041863 | 0 | Tuba1c | 0.086 | Old |
| 1.67E-06 | 0.279136 | 0.919 | 0.872 | 0.051791 | 0 | Klf6 | 0.047 | Old |
| 1.85E-06 | 1.36423 | 0.048 | 0.017 | 0.05737 | 0 | Gm26870 | 0.031 | Old |
| 1.88E-06 | 0.385991 | 0.585 | 0.481 | 0.058304 | 0 | Ms4a6c | 0.104 | Old |
| 2.10E-06 | 0.747278 | 0.179 | 0.114 | 0.065104 | 0 | Gfpt1 | 0.065 | Old |
| 2.50E-06 | 0.280009 | 0.845 | 0.795 | 0.077775 | 0 | Ctss | 0.05 | Old |
| 2.64E-06 | 0.379015 | 0.713 | 0.656 | 0.081852 | 0 | Ly6e | 0.057 | Old |
| 2.93E-06 | 0.540382 | 0.297 | 0.214 | 0.091092 | 0 | Acer3 | 0.083 | Old |
| 3.14E-06 | 0.272474 | 0.631 | 0.529 | 0.097368 | 0 | Ssr3 | 0.102 | Old |
| 3.17E-06 | 0.683226 | 0.155 | 0.093 | 0.098364 | 0 | Dhrs1 | 0.062 | Old |
| 3.22E-06 | 0.328747 | 0.549 | 0.451 | 0.099836 | 0 | Prkcd | 0.098 | Old |
| 3.67E-06 | 0.952242 | 0.286 | 0.213 | 0.11394 | 0 | Ccl12 | 0.073 | Old |
| 4.03E-06 | 0.40814 | 0.687 | 0.62 | 0.125033 | 0 | Fosb | 0.067 | Old |
| 4.78E-06 | 0.510097 | 0.22 | 0.147 | 0.148303 | 0 | Csnk2b | 0.073 | Old |
| 4.97E-06 | 0.331005 | 0.517 | 0.423 | 0.154199 | 0 | Itgam | 0.094 | Old |
| 5.15E-06 | 0.453139 | 0.442 | 0.354 | 0.159894 | 0 | Pla2g7 | 0.088 | Old |
| 6.60E-06 | 0.752012 | 0.218 | 0.15 | 0.204988 | 0 | Fgl2 | 0.068 | Old |
| 6.69E-06 | 0.598267 | 0.174 | 0.11 | 0.207686 | 0 | Ckap4 | 0.064 | Old |
| 7.43E-06 | 0.429587 | 0.423 | 0.329 | 0.230875 | 0 | Wsb1 | 0.094 | Old |
| 7.74E-06 | 0.267003 | 0.655 | 0.552 | 0.240294 | 0 | Efhd2 | 0.103 | Old |
| 8.98E-06 | 0.430009 | 0.432 | 0.35 | 0.278947 | 0 | Gm26532 | 0.082 | Old |
| 1.23E-05 | 0.186318 | 0.949 | 0.93 | 0.381703 | 0 | Junb | 0.019 | Old |
| 1.28E-05 | 0.367316 | 0.449 | 0.362 | 0.396219 | 0 | Cstb | 0.087 | Old |
| 1.29E-05 | 0.269504 | 0.761 | 0.697 | 0.401714 | 0 | Sirpa | 0.064 | Old |
| 1.30E-05 | 0.691948 | 0.131 | 0.078 | 0.404352 | 0 | Tmem51 | 0.053 | Old |
| 1.37E-05 | 0.675256 | 0.189 | 0.126 | 0.426781 | 0 | Grk3 | 0.063 | Old |
| 1.45E-05 | 0.296789 | 0.576 | 0.485 | 0.451387 | 0 | Pdia3 | 0.091 | Old |
| 1.46E-05 | 0.190973 | 0.833 | 0.765 | 0.45368 | 0 | BC005537 | 0.068 | Old |
| 1.56E-05 | 0.710711 | 0.152 | 0.096 | 0.485821 | 0 | Noct | 0.056 | Old |
| 1.87E-05 | 0.350756 | 0.479 | 0.394 | 0.58113 | 0 | Atp6ap2 | 0.085 | Old |
| 1.95E-05 | 0.857238 | 0.077 | 0.038 | 0.604223 | 0 | Fcgr4 | 0.039 | Old |
| 1.96E-05 | 1.966819 | 0.01 | 0 | 0.609296 | 0 | Gm15675 | 0.01 | Old |
| 2.12E-05 | 0.284961 | 0.579 | 0.485 | 0.657512 | 0 | H2-DMb1 | 0.094 | Old |
| 2.16E-05 | 0.563163 | 0.101 | 0.056 | 0.671275 | 0 | Pqlc3 | 0.045 | Old |

| | | | | | | | | |
|---|---|---|---|---|---|---|---|---|
| 2.16E-05 | 1.261345 | 0.04 | 0.014 | 0.672004 | 0 | Rbm12b2 | 0.026 | Old |
| 2.31E-05 | 0.269648 | 0.653 | 0.568 | 0.718331 | 0 | Pim1 | 0.085 | Old |
| 2.43E-05 | 0.809751 | 0.111 | 0.064 | 0.753297 | 0 | Yipf6 | 0.047 | Old |
| 2.67E-05 | 0.232171 | 0.491 | 0.386 | 0.829702 | 0 | Rpn2 | 0.105 | Old |
| 2.73E-05 | 0.492802 | 0.424 | 0.349 | 0.846359 | 0 | Lmna | 0.075 | Old |
| 2.98E-05 | 0.398015 | 0.226 | 0.156 | 0.924526 | 0 | Slc12a7 | 0.07 | Old |
| 3.04E-05 | 0.364572 | 0.509 | 0.424 | 0.94548 | 0 | Naaa | 0.085 | Old |
| 3.11E-05 | 0.207255 | 0.724 | 0.642 | 0.965959 | 0 | Actr2 | 0.082 | Old |
| 3.11E-05 | 0.242403 | 0.705 | 0.618 | 0.96728 | 0 | Hnrnpf | 0.087 | Old |
| 3.12E-05 | 0.319921 | 0.595 | 0.503 | 0.967436 | 0 | Gm26917 | 0.092 | Old |
| 3.16E-05 | 0.22976 | 0.813 | 0.761 | 0.982774 | 0 | Pfn1 | 0.052 | Old |
| 3.61E-05 | 0.216824 | 0.938 | 0.93 | 1 | 0 | Dusp1 | 0.008 | Old |
| 3.69E-05 | 0.241559 | 0.8 | 0.75 | 1 | 0 | Sdcbp | 0.05 | Old |
| 3.77E-05 | 0.314093 | 0.778 | 0.743 | 1 | 0 | Mgl2 | 0.035 | Old |
| 4.34E-05 | 0.707153 | 0.155 | 0.101 | 1 | 0 | AW112010 | 0.054 | Old |
| 4.43E-05 | 0.420984 | 0.487 | 0.406 | 1 | 0 | Mmp9 | 0.081 | Old |
| 4.50E-05 | 0.288832 | 0.43 | 0.334 | 1 | 0 | Arf4 | 0.096 | Old |
| 4.59E-05 | 0.444103 | 0.304 | 0.23 | 1 | 0 | Pld3 | 0.074 | Old |
| 4.80E-05 | 0.506299 | 0.238 | 0.171 | 1 | 0 | Basp1 | 0.067 | Old |
| 5.45E-05 | 0.282352 | 0.643 | 0.565 | 1 | 0 | Ctsz | 0.078 | Old |
| 5.69E-05 | 1.268331 | 0.019 | 0.004 | 1 | 0 | Csprs | 0.015 | Old |
| 6.17E-05 | 0.529895 | 0.179 | 0.12 | 1 | 0 | Plekhm2 | 0.059 | Old |
| 6.35E-05 | 0.258832 | 0.769 | 0.714 | 1 | 0 | Btg2 | 0.055 | Old |
| 6.48E-05 | 0.81474 | 0.067 | 0.033 | 1 | 0 | Hax1 | 0.034 | Old |
| 6.50E-05 | 0.268334 | 0.535 | 0.444 | 1 | 0 | Rbm3 | 0.091 | Old |
| 7.07E-05 | 0.135521 | 0.979 | 0.956 | 1 | 0 | Lgmn | 0.023 | Old |
| 7.29E-05 | 0.492042 | 0.235 | 0.169 | 1 | 0 | Gga1 | 0.066 | Old |
| 7.52E-05 | 0.417765 | 0.451 | 0.378 | 1 | 0 | Metrnl | 0.073 | Old |
| 7.80E-05 | 0.3359 | 0.294 | 0.219 | 1 | 0 | Tmsb10 | 0.075 | Old |
| 8.51E-05 | 0.97438 | 0.088 | 0.049 | 1 | 0 | Fabp5 | 0.039 | Old |
| 8.78E-05 | 0.350988 | 0.401 | 0.32 | 1 | 0 | Clec4a2 | 0.081 | Old |
| 8.79E-05 | 0.541703 | 0.17 | 0.114 | 1 | 0 | H2-Q7 | 0.056 | Old |
| 0.000104 | 0.617853 | 0.167 | 0.112 | 1 | 0 | Il11ra1 | 0.055 | Old |
| 0.000106 | 0.259192 | 0.587 | 0.491 | 1 | 0 | Cox6a1 | 0.096 | Old |
| 0.000112 | 1.370063 | 0.04 | 0.016 | 1 | 0 | Ttyh2 | 0.024 | Old |
| 0.000118 | 0.512813 | 0.449 | 0.394 | 1 | 0 | Ccr2 | 0.055 | Old |
| 0.000118 | 1.664054 | 0.011 | 0.001 | 1 | 0 | Anxa9 | 0.01 | Old |
| 0.00012 | 0.339102 | 0.306 | 0.23 | 1 | 0 | Emb | 0.076 | Old |
| 0.000122 | 0.782393 | 0.067 | 0.034 | 1 | 0 | Aspa | 0.033 | Old |
| 0.000123 | 0.402297 | 0.283 | 0.213 | 1 | 0 | Plec | 0.07 | Old |
| 0.000127 | 0.540114 | 0.167 | 0.113 | 1 | 0 | Slc16a10 | 0.054 | Old |
| 0.000129 | 1.166805 | 0.057 | 0.028 | 1 | 0 | Tspyl2 | 0.029 | Old |
| 0.000132 | 0.803176 | 0.041 | 0.017 | 1 | 0 | Cd274 | 0.024 | Old |
| 0.00014 | 0.269561 | 0.479 | 0.386 | 1 | 0 | Srsf2 | 0.093 | Old |
| 0.000147 | 0.382081 | 0.283 | 0.214 | 1 | 0 | Anxa4 | 0.069 | Old |

| | | | | | | | | |
|---|---|---|---|---|---|---|---|---|
| 0.000151 | 0.405767 | 0.341 | 0.266 | 1 | 0 | Ddx39b | 0.075 | Old |
| 0.000153 | 0.459716 | 0.23 | 0.165 | 1 | 0 | C5ar2 | 0.065 | Old |
| 0.00016 | 0.223976 | 0.636 | 0.546 | 1 | 0 | Plin2 | 0.09 | Old |
| 0.000165 | 0.372466 | 0.209 | 0.148 | 1 | 0 | Slc39a7 | 0.061 | Old |
| 0.000168 | 0.283639 | 0.529 | 0.447 | 1 | 0 | Pdia6 | 0.082 | Old |
| 0.000169 | 0.340299 | 0.347 | 0.272 | 1 | 0 | Man2a1 | 0.075 | Old |
| 0.000169 | 0.245474 | 0.387 | 0.305 | 1 | 0 | Plxnd1 | 0.082 | Old |
| 0.000184 | 0.231237 | 0.636 | 0.566 | 1 | 0 | Calr | 0.07 | Old |
| 0.000196 | 0.309005 | 0.315 | 0.239 | 1 | 0 | Nme1 | 0.076 | Old |
| 0.0002 | 0.778412 | 0.182 | 0.13 | 1 | 0 | Rnd3 | 0.052 | Old |
| 0.000203 | 0.759996 | 0.107 | 0.066 | 1 | 0 | Vta1 | 0.041 | Old |
| 0.000222 | 0.730472 | 0.156 | 0.107 | 1 | 0 | Slc20a1 | 0.049 | Old |
| 0.000222 | 0.321125 | 0.269 | 0.2 | 1 | 0 | Ms4a6d | 0.069 | Old |
| 0.000229 | 0.447497 | 0.274 | 0.209 | 1 | 0 | Sdf2 | 0.065 | Old |
| 0.000232 | 1.394141 | 0.04 | 0.017 | 1 | 0 | Cep78 | 0.023 | Old |
| 0.000255 | 0.3947 | 0.562 | 0.5 | 1 | 0 | Rhob | 0.062 | Old |
| 0.000264 | 1.726302 | 0.016 | 0.004 | 1 | 0 | Igf2r | 0.012 | Old |
| 0.000265 | 0.365696 | 0.342 | 0.273 | 1 | 0 | Susd6 | 0.069 | Old |
| 0.00027 | 0.990372 | 0.049 | 0.023 | 1 | 0 | Dnaaf3 | 0.026 | Old |
| 0.00027 | 0.371613 | 0.499 | 0.42 | 1 | 0 | AC149090. | 0.079 | Old |
| 0.000277 | 0.527959 | 0.131 | 0.085 | 1 | 0 | Etv6 | 0.046 | Old |
| 0.00028 | 1.25151 | 0.016 | 0.004 | 1 | 0 | Pde8b | 0.012 | Old |
| 0.000282 | 0.126655 | 0.992 | 0.985 | 1 | 0 | Cd74 | 0.007 | Old |
| 0.000285 | 0.347995 | 0.342 | 0.27 | 1 | 0 | Psme1 | 0.072 | Old |
| 0.000295 | 0.31717 | 0.341 | 0.267 | 1 | 0 | Gsn | 0.074 | Old |
| 0.000299 | 0.356676 | 0.494 | 0.422 | 1 | 0 | Trps1 | 0.072 | Old |
| 0.0003 | 0.265021 | 0.41 | 0.327 | 1 | 0 | Arpc4 | 0.083 | Old |
| 0.000316 | 0.211051 | 0.702 | 0.629 | 1 | 0 | Mob1a | 0.073 | Old |
| 0.000323 | 0.648862 | 0.109 | 0.069 | 1 | 0 | Banp | 0.04 | Old |
| 0.000339 | 0.577254 | 0.183 | 0.13 | 1 | 0 | A4galt | 0.053 | Old |
| 0.000351 | 1.000666 | 0.09 | 0.054 | 1 | 0 | Mta3 | 0.036 | Old |
| 0.000351 | 0.233261 | 0.605 | 0.532 | 1 | 0 | Tuba1b | 0.073 | Old |
| 0.000388 | 1.100692 | 0.03 | 0.011 | 1 | 0 | Tnni3 | 0.019 | Old |
| 0.000398 | 1.086499 | 0.034 | 0.014 | 1 | 0 | Il27ra | 0.02 | Old |
| 0.0004 | 1.545917 | 0.015 | 0.003 | 1 | 0 | C130026I2 | 0.012 | Old |
| 0.000402 | 0.467896 | 0.134 | 0.088 | 1 | 0 | Dpagt1 | 0.046 | Old |
| 0.000405 | 0.41966 | 0.249 | 0.19 | 1 | 0 | Hnrnpa1 | 0.059 | Old |
| 0.000407 | 0.386224 | 0.309 | 0.244 | 1 | 0 | Arcn1 | 0.065 | Old |
| 0.00041 | 0.260081 | 0.328 | 0.254 | 1 | 0 | Psmd2 | 0.074 | Old |
| 0.000415 | 0.796393 | 0.068 | 0.037 | 1 | 0 | Pam | 0.031 | Old |
| 0.000425 | 0.972496 | 0.1 | 0.062 | 1 | 0 | Gpx3 | 0.038 | Old |
| 0.000427 | 0.326883 | 0.279 | 0.215 | 1 | 0 | Tmem123 | 0.064 | Old |
| 0.000428 | 0.584311 | 0.118 | 0.076 | 1 | 0 | Map3k14 | 0.042 | Old |
| 0.000439 | 0.685405 | 0.075 | 0.043 | 1 | 0 | Nup93 | 0.032 | Old |
| 0.00044 | 0.160727 | 0.679 | 0.586 | 1 | 0 | Nrros | 0.093 | Old |

| | | | | | | | | |
|---|---|---|---|---|---|---|---|---|
| 0.000458 | 0.303314 | 0.339 | 0.267 | 1 | 0 | Elf1 | 0.072 | Old |
| 0.000469 | 0.316272 | 0.383 | 0.312 | 1 | 0 | P4hb | 0.071 | Old |
| 0.000475 | 0.248816 | 0.512 | 0.434 | 1 | 0 | Pirb | 0.078 | Old |
| 0.000491 | 0.289318 | 0.312 | 0.243 | 1 | 0 | Ddi2 | 0.069 | Old |
| 0.0005 | 0.234178 | 0.421 | 0.342 | 1 | 0 | Lyn | 0.079 | Old |
| 0.000509 | 0.439188 | 0.171 | 0.12 | 1 | 0 | Ufm1 | 0.051 | Old |
| 0.000527 | 1.759558 | 0.01 | 0.001 | 1 | 0 | Cttn | 0.009 | Old |
| 0.000528 | 0.220193 | 0.603 | 0.527 | 1 | 0 | Clec4a1 | 0.076 | Old |
| 0.000532 | 1.178474 | 0.026 | 0.009 | 1 | 0 | Zfp111 | 0.017 | Old |
| 0.000533 | 0.961931 | 0.01 | 0.001 | 1 | 0 | Zfp955a | 0.009 | Old |
| 0.000539 | 0.239509 | 0.134 | 0.088 | 1 | 0 | Gm36161 | 0.046 | Old |
| 0.000543 | 0.20509 | 0.605 | 0.519 | 1 | 0 | Pcbp1 | 0.086 | Old |
| 0.000548 | 0.605784 | 0.101 | 0.063 | 1 | 0 | Atp8b4 | 0.038 | Old |
| 0.000551 | 0.203045 | 0.564 | 0.478 | 1 | 0 | Rap1a | 0.086 | Old |
| 0.000569 | 0.405632 | 0.254 | 0.196 | 1 | 0 | Tle3 | 0.058 | Old |
| 0.000582 | 0.272801 | 0.361 | 0.291 | 1 | 0 | Degs1 | 0.07 | Old |
| 0.000583 | 0.180556 | 0.758 | 0.694 | 1 | 0 | Capzb | 0.064 | Old |
| 0.000587 | 0.552993 | 0.115 | 0.074 | 1 | 0 | Stk25 | 0.041 | Old |
| 0.000596 | 1.022676 | 0.057 | 0.03 | 1 | 0 | Kcnn4 | 0.027 | Old |
| 0.000596 | 0.683105 | 0.163 | 0.116 | 1 | 0 | Fbxw2 | 0.047 | Old |
| 0.000602 | 0.632616 | 0.088 | 0.053 | 1 | 0 | Cdk2ap1 | 0.035 | Old |
| 0.000608 | 0.382154 | 0.226 | 0.168 | 1 | 0 | Prelid1 | 0.058 | Old |
| 0.000609 | 0.985662 | 0.031 | 0.013 | 1 | 0 | Psph | 0.018 | Old |
| 0.000609 | 1.050952 | 0.044 | 0.021 | 1 | 0 | Zfp715 | 0.023 | Old |
| 0.000618 | 0.591637 | 0.17 | 0.122 | 1 | 0 | Dck | 0.048 | Old |
| 0.00064 | 0.322679 | 0.443 | 0.374 | 1 | 0 | Atp5g3 | 0.069 | Old |
| 0.000642 | 0.329515 | 0.464 | 0.402 | 1 | 0 | Anxa2 | 0.062 | Old |
| 0.000663 | 0.364887 | 0.248 | 0.187 | 1 | 0 | Gdi1 | 0.061 | Old |
| 0.000673 | 0.341339 | 0.282 | 0.221 | 1 | 0 | Casp8 | 0.061 | Old |
| 0.000675 | 0.320964 | 0.309 | 0.244 | 1 | 0 | Pdia4 | 0.065 | Old |
| 0.000699 | 0.799584 | 0.053 | 0.027 | 1 | 0 | Plp2 | 0.026 | Old |
| 0.000703 | 0.507688 | 0.096 | 0.059 | 1 | 0 | Igip | 0.037 | Old |
| 0.000704 | 0.331945 | 0.249 | 0.188 | 1 | 0 | Rc3h2 | 0.061 | Old |
| 0.000711 | 0.365916 | 0.319 | 0.252 | 1 | 0 | Cd52 | 0.067 | Old |
| 0.000722 | 0.30843 | 0.332 | 0.265 | 1 | 0 | Sp100 | 0.067 | Old |
| 0.000727 | 0.778714 | 0.052 | 0.027 | 1 | 0 | Cstf1 | 0.025 | Old |
| 0.000727 | 0.608144 | 0.082 | 0.049 | 1 | 0 | N4bp2 | 0.033 | Old |
| 0.000729 | 0.434583 | 0.088 | 0.053 | 1 | 0 | Farsb | 0.035 | Old |
| 0.000733 | 0.385049 | 0.323 | 0.259 | 1 | 0 | Jarid2 | 0.064 | Old |
| 0.000741 | 0.376464 | 0.172 | 0.123 | 1 | 0 | Wbp1l | 0.049 | Old |
| 0.000742 | 0.330845 | 0.43 | 0.37 | 1 | 0 | Cd44 | 0.06 | Old |
| 0.000749 | 0.214245 | 0.398 | 0.319 | 1 | 0 | Tmed5 | 0.079 | Old |
| 0.000751 | 1.61009 | 0.018 | 0.005 | 1 | 0 | Oas1g | 0.013 | Old |
| 0.000757 | 0.327585 | 0.238 | 0.179 | 1 | 0 | Tspan32 | 0.059 | Old |
| 0.000759 | 1.147419 | 0.023 | 0.008 | 1 | 0 | Acsf2 | 0.015 | Old |

| | | | | | | | | |
|---|---|---|---|---|---|---|---|---|
| 0.000812 | 0.345283 | 0.327 | 0.262 | 1 | 0 | Leprot | 0.065 | Old |
| 0.000828 | 0.818273 | 0.073 | 0.042 | 1 | 0 | Fkbp5 | 0.031 | Old |
| 0.000843 | 0.602976 | 0.059 | 0.032 | 1 | 0 | Usp10 | 0.027 | Old |
| 0.000855 | 0.392964 | 0.146 | 0.101 | 1 | 0 | Ttc39b | 0.045 | Old |
| 0.000863 | 0.188941 | 0.647 | 0.583 | 1 | 0 | H2-DMa | 0.064 | Old |
| 0.000868 | 0.515198 | 0.098 | 0.062 | 1 | 0 | Rnf168 | 0.036 | Old |
| 0.000879 | 0.310366 | 0.332 | 0.265 | 1 | 0 | Hspa9 | 0.067 | Old |
| 0.000883 | 0.467596 | 0.19 | 0.139 | 1 | 0 | Mgat2 | 0.051 | Old |
| 0.000885 | 0.498018 | 0.14 | 0.096 | 1 | 0 | Senp2 | 0.044 | Old |
| 0.000886 | 0.315075 | 0.398 | 0.33 | 1 | 0 | Prpf38b | 0.068 | Old |
| 0.000891 | 0.246853 | 0.538 | 0.461 | 1 | 0 | Rpn1 | 0.077 | Old |
| 0.000896 | 0.317368 | 0.375 | 0.31 | 1 | 0 | Slc39a1 | 0.065 | Old |
| 0.000937 | 0.595554 | 0.075 | 0.044 | 1 | 0 | Zfp738 | 0.031 | Old |
| 0.000955 | 0.231671 | 0.278 | 0.213 | 1 | 0 | Tspan31 | 0.065 | Old |
| 0.001019 | 0.413281 | 0.168 | 0.12 | 1 | 0 | Tmem65 | 0.048 | Old |
| 0.00103 | 0.243779 | 0.302 | 0.238 | 1 | 0 | N4bp1 | 0.064 | Old |
| 0.001037 | 0.245791 | 0.439 | 0.361 | 1 | 0 | Plekhb2 | 0.078 | Old |
| 0.001046 | 0.288991 | 0.424 | 0.354 | 1 | 0 | Cd84 | 0.07 | Old |
| 0.001069 | 0.284396 | 0.271 | 0.21 | 1 | 0 | Il4ra | 0.061 | Old |
| 0.001077 | 0.461778 | 0.135 | 0.093 | 1 | 0 | 2210016F1 | 0.042 | Old |
| 0.001078 | 0.198435 | 0.471 | 0.391 | 1 | 0 | Colgalt1 | 0.08 | Old |
| 0.001083 | 1.255201 | 0.021 | 0.007 | 1 | 0 | Timp3 | 0.014 | Old |
| 0.001089 | 1.029225 | 0.021 | 0.007 | 1 | 0 | Bmpr1a | 0.014 | Old |
| 0.001105 | 0.6974 | 0.085 | 0.052 | 1 | 0 | Traf3 | 0.033 | Old |
| 0.001112 | 0.269542 | 0.347 | 0.281 | 1 | 0 | Hmgb2 | 0.066 | Old |
| 0.001153 | 0.329417 | 0.324 | 0.262 | 1 | 0 | Ppp1r12c | 0.062 | Old |
| 0.001192 | 0.422743 | 0.131 | 0.09 | 1 | 0 | Cstf3 | 0.041 | Old |
| 0.001202 | 2.073718 | 0.011 | 0.002 | 1 | 0 | Gm44777 | 0.009 | Old |
| 0.001206 | 0.677349 | 0.083 | 0.051 | 1 | 0 | 1700025G( | 0.032 | Old |
| 0.001224 | 0.356568 | 0.152 | 0.107 | 1 | 0 | Atad2b | 0.045 | Old |
| 0.001244 | 0.739131 | 0.075 | 0.045 | 1 | 0 | Pira2 | 0.03 | Old |
| 0.001263 | 1.139198 | 0.031 | 0.013 | 1 | 0 | Odf2l | 0.018 | Old |
| 0.001272 | 0.373595 | 0.724 | 0.683 | 1 | 0 | Ier2 | 0.041 | Old |
| 0.001277 | 0.311503 | 0.256 | 0.197 | 1 | 0 | Birc6 | 0.059 | Old |
| 0.001283 | 0.274289 | 0.319 | 0.252 | 1 | 0 | Nono | 0.067 | Old |
| 0.001294 | 0.204242 | 0.472 | 0.396 | 1 | 0 | Arhgap30 | 0.076 | Old |
| 0.001297 | 0.257336 | 0.454 | 0.382 | 1 | 0 | Stxbp3 | 0.072 | Old |
| 0.001311 | 0.703856 | 0.059 | 0.032 | 1 | 0 | Klhl2 | 0.027 | Old |
| 0.001345 | 1.149109 | 0.014 | 0.003 | 1 | 0 | Gm4070 | 0.011 | Old |
| 0.00135 | 1.027491 | 0.026 | 0.01 | 1 | 0 | Rpl9-ps6 | 0.016 | Old |
| 0.001388 | 0.401247 | 0.259 | 0.202 | 1 | 0 | Sp3 | 0.057 | Old |
| 0.001389 | 0.265389 | 0.393 | 0.325 | 1 | 0 | Ndufc2 | 0.068 | Old |
| 0.001405 | 0.24253 | 0.253 | 0.193 | 1 | 0 | Hadhb | 0.06 | Old |
| 0.001406 | 0.658841 | 0.057 | 0.032 | 1 | 0 | Zfp865 | 0.025 | Old |
| 0.001483 | 0.168411 | 0.763 | 0.706 | 1 | 0 | Tmem176b | 0.057 | Old |

| | | | | | | | | |
|---|---|---|---|---|---|---|---|---|
| 0.001499 | 0.34944 | 0.334 | 0.274 | 1 | 0 | Shtn1 | 0.06 | Old |
| 0.00151 | 0.519814 | 0.097 | 0.062 | 1 | 0 | E2f1 | 0.035 | Old |
| 0.001521 | 0.344763 | 0.22 | 0.166 | 1 | 0 | Pgam1 | 0.054 | Old |
| 0.001527 | 0.314519 | 0.308 | 0.245 | 1 | 0 | Cd300a | 0.063 | Old |
| 0.00153 | 0.190648 | 0.527 | 0.444 | 1 | 0 | Pdcd6ip | 0.083 | Old |
| 0.001549 | 1.974028 | 0.019 | 0.006 | 1 | 0 | Paqr9 | 0.013 | Old |
| 0.001576 | 0.656161 | 0.064 | 0.037 | 1 | 0 | Vopp1 | 0.027 | Old |
| 0.001581 | 0.383099 | 0.098 | 0.063 | 1 | 0 | Slc35b1 | 0.035 | Old |
| 0.001608 | 0.606027 | 0.06 | 0.034 | 1 | 0 | Mterf3 | 0.026 | Old |
| 0.001668 | 0.623863 | 0.055 | 0.03 | 1 | 0 | Pde8a | 0.025 | Old |
| 0.001681 | 0.406078 | 0.15 | 0.107 | 1 | 0 | Tuba1a | 0.043 | Old |
| 0.001689 | 1.543092 | 0.01 | 0.002 | 1 | 0 | Lrp4 | 0.008 | Old |
| 0.001692 | 1.124329 | 0.01 | 0.002 | 1 | 0 | F7 | 0.008 | Old |
| 0.001702 | 0.538175 | 0.137 | 0.095 | 1 | 0 | Oasl2 | 0.042 | Old |
| 0.001711 | 0.788879 | 0.01 | 0.002 | 1 | 0 | Gm15448 | 0.008 | Old |
| 0.001718 | 0.608751 | 0.082 | 0.051 | 1 | 0 | Casp6 | 0.031 | Old |
| 0.001724 | 0.160612 | 0.585 | 0.508 | 1 | 0 | Arpc3 | 0.077 | Old |
| 0.001745 | 0.236053 | 0.363 | 0.295 | 1 | 0 | Ran | 0.068 | Old |
| 0.001755 | 0.57822 | 0.146 | 0.105 | 1 | 0 | U2af1 | 0.041 | Old |
| 0.001787 | 0.373614 | 0.122 | 0.083 | 1 | 0 | Adrm1 | 0.039 | Old |
| 0.001793 | 1.592518 | 0.022 | 0.008 | 1 | 0 | Aurkb | 0.014 | Old |
| 0.001796 | 0.604681 | 0.055 | 0.03 | 1 | 0 | Agpat2 | 0.025 | Old |
| 0.001797 | 0.79508 | 0.15 | 0.111 | 1 | 0 | Clec4n | 0.039 | Old |
| 0.001798 | 1.16946 | 0.015 | 0.004 | 1 | 0 | Slc22a21 | 0.011 | Old |
| 0.001872 | 0.368258 | 0.215 | 0.163 | 1 | 0 | Hsd17b12 | 0.052 | Old |
| 0.001903 | 0.149694 | 0.844 | 0.786 | 1 | 0 | Ap2a2 | 0.058 | Old |
| 0.001905 | 0.35202 | 0.202 | 0.152 | 1 | 0 | Arf6 | 0.05 | Old |
| 0.001926 | 0.280767 | 0.259 | 0.201 | 1 | 0 | Dock7 | 0.058 | Old |
| 0.001946 | 0.429168 | 0.115 | 0.078 | 1 | 0 | Cul4a | 0.037 | Old |
| 0.001951 | 0.211471 | 0.249 | 0.19 | 1 | 0 | Ripk1 | 0.059 | Old |
| 0.00196 | 0.300502 | 0.298 | 0.238 | 1 | 0 | Trim8 | 0.06 | Old |
| 0.001963 | 0.228757 | 0.527 | 0.451 | 1 | 0 | Man1a | 0.076 | Old |
| 0.001997 | 1.004299 | 0.026 | 0.011 | 1 | 0 | Ankzf1 | 0.015 | Old |
| 0.002044 | 0.86033 | 0.025 | 0.01 | 1 | 0 | Pigb | 0.015 | Old |
| 0.002088 | 0.771272 | 0.068 | 0.041 | 1 | 0 | Armc10 | 0.027 | Old |
| 0.002146 | 0.365285 | 0.114 | 0.076 | 1 | 0 | Hyou1 | 0.038 | Old |
| 0.002156 | 0.322039 | 0.595 | 0.537 | 1 | 0 | Klf4 | 0.058 | Old |
| 0.002158 | 0.403181 | 0.153 | 0.11 | 1 | 0 | Plod3 | 0.043 | Old |
| 0.002189 | 0.306719 | 0.171 | 0.125 | 1 | 0 | Hipk2 | 0.046 | Old |
| 0.00219 | 0.35238 | 0.253 | 0.199 | 1 | 0 | Tomm22 | 0.054 | Old |
| 0.002238 | 0.174613 | 0.673 | 0.619 | 1 | 0 | Slc25a3 | 0.054 | Old |
| 0.002281 | 0.339973 | 0.207 | 0.158 | 1 | 0 | Ldlr | 0.049 | Old |
| 0.002308 | 0.541198 | 0.146 | 0.105 | 1 | 0 | Ly9 | 0.041 | Old |
| 0.002312 | 0.493109 | 0.218 | 0.169 | 1 | 0 | Eif3d | 0.049 | Old |
| 0.002371 | 0.280114 | 0.172 | 0.126 | 1 | 0 | Tmem33 | 0.046 | Old |

| | | | | | | | | |
|---|---|---|---|---|---|---|---|---|
| 0.002381 | 0.914895 | 0.048 | 0.026 | 1 | 0 | Exoc6 | 0.022 | Old |
| 0.002389 | 0.283407 | 0.194 | 0.145 | 1 | 0 | Cdk6 | 0.049 | Old |
| 0.002399 | 0.290278 | 0.291 | 0.233 | 1 | 0 | Larp1 | 0.058 | Old |
| 0.002455 | 0.836338 | 0.048 | 0.026 | 1 | 0 | Hotairm1 | 0.022 | Old |
| 0.002489 | 0.229388 | 0.486 | 0.426 | 1 | 0 | Ywhah | 0.06 | Old |
| 0.002496 | 0.394974 | 0.186 | 0.14 | 1 | 0 | Dpp8 | 0.046 | Old |
| 0.00251 | 0.960215 | 0.042 | 0.022 | 1 | 0 | Fdft1 | 0.02 | Old |
| 0.002541 | 0.377389 | 0.134 | 0.094 | 1 | 0 | Manba | 0.04 | Old |
| 0.002622 | 0.713694 | 0.075 | 0.047 | 1 | 0 | Helb | 0.028 | Old |
| 0.002634 | 0.299676 | 0.198 | 0.15 | 1 | 0 | Api5 | 0.048 | Old |
| 0.002649 | 1.212567 | 0.037 | 0.018 | 1 | 0 | Ralgps1 | 0.019 | Old |
| 0.002678 | 0.853877 | 0.047 | 0.025 | 1 | 0 | Pomk | 0.022 | Old |
| 0.002697 | 0.122689 | 0.899 | 0.862 | 1 | 0 | Cfl1 | 0.037 | Old |
| 0.002718 | 0.286789 | 0.357 | 0.299 | 1 | 0 | Lbh | 0.058 | Old |
| 0.002736 | 0.410754 | 0.098 | 0.065 | 1 | 0 | Josd1 | 0.033 | Old |
| 0.00278 | 0.908621 | 0.019 | 0.007 | 1 | 0 | Zfp169 | 0.012 | Old |
| 0.002788 | 0.293314 | 0.271 | 0.215 | 1 | 0 | Psma3 | 0.056 | Old |
| 0.002809 | 0.223438 | 0.451 | 0.382 | 1 | 0 | Cd300ld | 0.069 | Old |
| 0.002815 | 0.351084 | 0.239 | 0.19 | 1 | 0 | Ggh | 0.049 | Old |
| 0.002831 | 0.92532 | 0.029 | 0.013 | 1 | 0 | Glb1l | 0.016 | Old |
| 0.002897 | 0.658032 | 0.212 | 0.169 | 1 | 0 | Phlda1 | 0.043 | Old |
| 0.002897 | 1.278059 | 0.022 | 0.008 | 1 | 0 | Cadm1 | 0.014 | Old |
| 0.002905 | 0.252509 | 0.265 | 0.207 | 1 | 0 | Psma4 | 0.058 | Old |
| 0.003001 | 1.107807 | 0.011 | 0.003 | 1 | 0 | Tspan7 | 0.008 | Old |
| 0.003031 | 0.106548 | 0.985 | 0.977 | 1 | 0 | C1qc | 0.008 | Old |
| 0.003033 | 0.982307 | 0.011 | 0.003 | 1 | 0 | Cdh5 | 0.008 | Old |
| 0.003038 | 0.337471 | 0.222 | 0.172 | 1 | 0 | Tcirg1 | 0.05 | Old |
| 0.00306 | 0.475716 | 0.105 | 0.071 | 1 | 0 | Plau | 0.034 | Old |
| 0.003074 | 0.900719 | 0.034 | 0.016 | 1 | 0 | Spon1 | 0.018 | Old |
| 0.003083 | 0.41159 | 0.256 | 0.207 | 1 | 0 | Setd1b | 0.049 | Old |
| 0.003108 | 0.35062 | 0.293 | 0.242 | 1 | 0 | Nufip2 | 0.051 | Old |
| 0.003122 | 0.225135 | 0.328 | 0.265 | 1 | 0 | Fermt3 | 0.063 | Old |
| 0.003134 | 0.278078 | 0.219 | 0.168 | 1 | 0 | Igsf6 | 0.051 | Old |
| 0.003249 | 0.998725 | 0.038 | 0.019 | 1 | 0 | Polr3d | 0.019 | Old |
| 0.00326 | 0.838129 | 0.053 | 0.03 | 1 | 0 | Ankrd54 | 0.023 | Old |
| 0.003266 | 0.432603 | 0.114 | 0.078 | 1 | 0 | Clcn7 | 0.036 | Old |
| 0.003319 | 0.556595 | 0.071 | 0.044 | 1 | 0 | Med22 | 0.027 | Old |
| 0.003348 | 0.676833 | 0.088 | 0.057 | 1 | 0 | Vars | 0.031 | Old |
| 0.003376 | 0.229969 | 0.694 | 0.634 | 1 | 0 | Picalm | 0.06 | Old |
| 0.003421 | 0.539153 | 0.04 | 0.02 | 1 | 0 | Ice2 | 0.02 | Old |
| 0.003429 | 0.951518 | 0.049 | 0.027 | 1 | 0 | Cd200r4 | 0.022 | Old |
| 0.003444 | 0.217426 | 0.369 | 0.305 | 1 | 0 | Bst2 | 0.064 | Old |
| 0.003463 | 0.262007 | 0.285 | 0.227 | 1 | 0 | Bag1 | 0.058 | Old |
| 0.003476 | 0.574147 | 0.093 | 0.061 | 1 | 0 | Afg3l2 | 0.032 | Old |
| 0.003485 | 0.536766 | 0.133 | 0.096 | 1 | 0 | Snd1 | 0.037 | Old |

| | | | | | | | | |
|---|---|---|---|---|---|---|---|---|
| 0.003497 | 0.589056 | 0.082 | 0.053 | 1 | 0 | Brix1 | 0.029 | Old |
| 0.003498 | 0.450488 | 0.148 | 0.108 | 1 | 0 | Arhgap25 | 0.04 | Old |
| 0.003523 | 0.600194 | 0.079 | 0.051 | 1 | 0 | Mitd1 | 0.028 | Old |
| 0.003536 | 0.493325 | 0.107 | 0.073 | 1 | 0 | Zfp655 | 0.034 | Old |
| 0.00362 | 0.387604 | 0.044 | 0.023 | 1 | 0 | Ranbp6 | 0.021 | Old |
| 0.003627 | 0.297452 | 0.253 | 0.202 | 1 | 0 | Fem1c | 0.051 | Old |
| 0.003655 | 0.891079 | 0.059 | 0.035 | 1 | 0 | Fbxl18 | 0.024 | Old |
| 0.003727 | 0.315922 | 0.231 | 0.181 | 1 | 0 | Fcgr1 | 0.05 | Old |
| 0.0038 | 0.485923 | 0.134 | 0.096 | 1 | 0 | Zmpste24 | 0.038 | Old |
| 0.003804 | 0.545585 | 0.062 | 0.037 | 1 | 0 | Gnpat | 0.025 | Old |
| 0.003853 | 0.152385 | 0.714 | 0.623 | 1 | 0 | Ywhaz | 0.091 | Old |
| 0.003875 | 0.39336 | 0.124 | 0.088 | 1 | 0 | Gm34084 | 0.036 | Old |
| 0.003877 | 0.492233 | 0.07 | 0.043 | 1 | 0 | Zdhhc18 | 0.027 | Old |
| 0.003946 | 0.184333 | 0.487 | 0.42 | 1 | 0 | Ifngr1 | 0.067 | Old |
| 0.003961 | 0.150443 | 0.672 | 0.592 | 1 | 0 | Prcp | 0.08 | Old |
| 0.003962 | 0.844021 | 0.059 | 0.035 | 1 | 0 | Uckl1 | 0.024 | Old |
| 0.003972 | 0.244838 | 0.293 | 0.235 | 1 | 0 | Wdfy2 | 0.058 | Old |
| 0.003983 | 0.451312 | 0.138 | 0.1 | 1 | 0 | Txndc5 | 0.038 | Old |
| 0.004064 | 0.178751 | 0.672 | 0.608 | 1 | 0 | Akr1a1 | 0.064 | Old |
| 0.004072 | 0.284677 | 0.334 | 0.275 | 1 | 0 | Arl8a | 0.059 | Old |
| 0.004076 | 0.754597 | 0.071 | 0.045 | 1 | 0 | Wdr7 | 0.026 | Old |
| 0.004085 | 0.510628 | 0.131 | 0.095 | 1 | 0 | Chmp1b | 0.036 | Old |
| 0.004089 | 0.292896 | 0.662 | 0.617 | 1 | 0 | Ier5 | 0.045 | Old |
| 0.004151 | 0.175013 | 0.219 | 0.167 | 1 | 0 | Eif3l | 0.052 | Old |
| 0.004166 | 0.206948 | 0.412 | 0.352 | 1 | 0 | Fosl2 | 0.06 | Old |
| 0.004181 | 1.136166 | 0.026 | 0.011 | 1 | 0 | Tmem129 | 0.015 | Old |
| 0.004201 | 0.272716 | 0.332 | 0.274 | 1 | 0 | Morf4l2 | 0.058 | Old |
| 0.004213 | 0.871045 | 0.034 | 0.017 | 1 | 0 | Ndnf | 0.017 | Old |
| 0.004219 | 0.356137 | 0.079 | 0.051 | 1 | 0 | Cyth2 | 0.028 | Old |
| 0.004232 | 0.485339 | 0.144 | 0.105 | 1 | 0 | Tpd52l2 | 0.039 | Old |
| 0.004239 | 1.004064 | 0.012 | 0.003 | 1 | 0 | Ankrd9 | 0.009 | Old |
| 0.004265 | 0.385301 | 0.163 | 0.121 | 1 | 0 | Slain2 | 0.042 | Old |
| 0.004271 | 0.287883 | 0.235 | 0.184 | 1 | 0 | 9-Sep | 0.051 | Old |
| 0.004355 | 0.601654 | 0.053 | 0.031 | 1 | 0 | Gng11 | 0.022 | Old |
| 0.004368 | 0.166913 | 0.316 | 0.257 | 1 | 0 | Prdx2 | 0.059 | Old |
| 0.004381 | 0.401609 | 0.164 | 0.123 | 1 | 0 | Esyt1 | 0.041 | Old |
| 0.004453 | 0.28151 | 0.159 | 0.117 | 1 | 0 | Kif1c | 0.042 | Old |
| 0.004471 | 0.984684 | 0.023 | 0.01 | 1 | 0 | Zfp976 | 0.013 | Old |
| 0.004485 | 0.891525 | 0.023 | 0.01 | 1 | 0 | Trmt12 | 0.013 | Old |
| 0.004572 | 0.335809 | 0.287 | 0.237 | 1 | 0 | Pcmtd1 | 0.05 | Old |
| 0.004602 | 0.726847 | 0.1 | 0.069 | 1 | 0 | Cops3 | 0.031 | Old |
| 0.004727 | 0.476223 | 0.108 | 0.075 | 1 | 0 | Siglec1 | 0.033 | Old |
| 0.004749 | 0.215188 | 0.512 | 0.45 | 1 | 0 | B4galt6 | 0.062 | Old |
| 0.004753 | 0.445557 | 0.118 | 0.083 | 1 | 0 | Slc38a1 | 0.035 | Old |
| 0.004788 | 0.237072 | 0.126 | 0.088 | 1 | 0 | Msmo1 | 0.038 | Old |

| | | | | | | | | |
|---|---|---|---|---|---|---|---|---|
| 0.004836 | 0.435817 | 0.166 | 0.124 | 1 | 0 | Brox | 0.042 | Old |
| 0.004851 | 0.494526 | 0.088 | 0.058 | 1 | 0 | Cfap20 | 0.03 | Old |
| 0.00492 | 0.461971 | 0.062 | 0.037 | 1 | 0 | Fbxo6 | 0.025 | Old |
| 0.004935 | 0.727095 | 0.041 | 0.022 | 1 | 0 | Rfx3 | 0.019 | Old |
| 0.005001 | 0.252705 | 0.219 | 0.17 | 1 | 0 | Psma1 | 0.049 | Old |
| 0.005013 | 0.447081 | 0.104 | 0.071 | 1 | 0 | Polr3c | 0.033 | Old |
| 0.005047 | 1.523825 | 0.014 | 0.004 | 1 | 0 | Smoc2 | 0.01 | Old |
| 0.005059 | 0.698614 | 0.051 | 0.029 | 1 | 0 | Elk1 | 0.022 | Old |
| 0.005119 | 0.253807 | 0.178 | 0.134 | 1 | 0 | Rps6ka1 | 0.044 | Old |
| 0.005143 | 0.438659 | 0.16 | 0.12 | 1 | 0 | Jdp2 | 0.04 | Old |
| 0.005181 | 0.487361 | 0.124 | 0.088 | 1 | 0 | Trim30d | 0.036 | Old |
| 0.005224 | 0.221007 | 0.246 | 0.195 | 1 | 0 | Ube2q1 | 0.051 | Old |
| 0.005295 | 0.207672 | 0.49 | 0.419 | 1 | 0 | Cops9 | 0.071 | Old |
| 0.005305 | 0.379982 | 0.248 | 0.2 | 1 | 0 | Rnf19b | 0.048 | Old |
| 0.005313 | 0.518501 | 0.13 | 0.095 | 1 | 0 | Bak1 | 0.035 | Old |
| 0.005351 | 0.322788 | 0.241 | 0.193 | 1 | 0 | Dnajb9 | 0.048 | Old |
| 0.005358 | 0.299394 | 0.204 | 0.158 | 1 | 0 | Cux1 | 0.046 | Old |
| 0.005446 | 0.239489 | 0.265 | 0.213 | 1 | 0 | Hnrnpdl | 0.052 | Old |
| 0.005482 | 0.344398 | 0.176 | 0.134 | 1 | 0 | Gtpbp2 | 0.042 | Old |
| 0.00552 | 0.203139 | 0.328 | 0.271 | 1 | 0 | Cct5 | 0.057 | Old |
| 0.005523 | 0.652372 | 0.036 | 0.018 | 1 | 0 | Ppp3cc | 0.018 | Old |
| 0.005624 | 0.534242 | 0.093 | 0.063 | 1 | 0 | Rabggtb | 0.03 | Old |
| 0.005657 | 0.28207 | 0.304 | 0.252 | 1 | 0 | Anp32b | 0.052 | Old |
| 0.005666 | 0.537918 | 0.112 | 0.08 | 1 | 0 | Aph1c | 0.032 | Old |
| 0.005676 | 0.166394 | 0.46 | 0.39 | 1 | 0 | Serinc1 | 0.07 | Old |
| 0.005684 | 0.56916 | 0.075 | 0.048 | 1 | 0 | Aig1 | 0.027 | Old |
| 0.005691 | 0.272007 | 0.248 | 0.198 | 1 | 0 | Klc1 | 0.05 | Old |
| 0.0057 | 0.55913 | 0.056 | 0.033 | 1 | 0 | Spred2 | 0.023 | Old |
| 0.005838 | 0.487622 | 0.104 | 0.072 | 1 | 0 | Oas2 | 0.032 | Old |
| 0.005871 | 0.237765 | 0.337 | 0.283 | 1 | 0 | Gnpda1 | 0.054 | Old |
| 0.005919 | 0.185343 | 0.525 | 0.463 | 1 | 0 | Clec4a3 | 0.062 | Old |
| 0.005921 | 0.228944 | 0.211 | 0.164 | 1 | 0 | Uap1l1 | 0.047 | Old |
| 0.005921 | 0.64368 | 0.082 | 0.054 | 1 | 0 | Cavin1 | 0.028 | Old |
| 0.005923 | 0.532404 | 0.089 | 0.06 | 1 | 0 | Ccnd2 | 0.029 | Old |
| 0.00595 | 0.968892 | 0.026 | 0.012 | 1 | 0 | Slc44a2 | 0.014 | Old |
| 0.005972 | 0.389477 | 0.112 | 0.079 | 1 | 0 | Rnmt | 0.033 | Old |
| 0.006219 | 0.363107 | 0.06 | 0.037 | 1 | 0 | Lrrc42 | 0.023 | Old |
| 0.006272 | 0.30377 | 0.155 | 0.115 | 1 | 0 | Tor1aip2 | 0.04 | Old |
| 0.006319 | 1.317768 | 0.016 | 0.006 | 1 | 0 | Tle6 | 0.01 | Old |
| 0.00637 | 0.170549 | 0.554 | 0.487 | 1 | 0 | Tpd52 | 0.067 | Old |
| 0.006467 | 0.561996 | 0.089 | 0.06 | 1 | 0 | Zbtb22 | 0.029 | Old |
| 0.006549 | 0.306977 | 0.163 | 0.124 | 1 | 0 | Ndufa10 | 0.039 | Old |
| 0.006555 | 0.393117 | 0.082 | 0.054 | 1 | 0 | Nqo2 | 0.028 | Old |
| 0.006612 | 0.462953 | 0.016 | 0.006 | 1 | 0 | Dmpk | 0.01 | Old |
| 0.006629 | 1.159329 | 0.038 | 0.021 | 1 | 0 | Zfp709 | 0.017 | Old |

| | | | | | | | | |
|---|---|---|---|---|---|---|---|---|
| 0.006681 | 0.23056 | 0.728 | 0.678 | 1 | 0 | Srsf5 | 0.05 | Old |
| 0.006692 | 0.151897 | 0.353 | 0.288 | 1 | 0 | Camk1d | 0.065 | Old |
| 0.006698 | 0.198876 | 0.189 | 0.144 | 1 | 0 | Rbbp4 | 0.045 | Old |
| 0.00675 | 0.231029 | 0.368 | 0.312 | 1 | 0 | Ptpn1 | 0.056 | Old |
| 0.006792 | 0.456148 | 0.092 | 0.062 | 1 | 0 | Sympk | 0.03 | Old |
| 0.006839 | 0.175097 | 0.41 | 0.35 | 1 | 0 | Notch2 | 0.06 | Old |
| 0.006849 | 0.122918 | 0.491 | 0.417 | 1 | 0 | Tnfaip8 | 0.074 | Old |
| 0.006851 | 0.138438 | 0.531 | 0.46 | 1 | 0 | Ctsl | 0.071 | Old |
| 0.006891 | 0.529696 | 0.063 | 0.039 | 1 | 0 | Cars2 | 0.024 | Old |
| 0.0069 | 0.156802 | 0.647 | 0.576 | 1 | 0 | Ifi207 | 0.071 | Old |
| 0.006975 | 0.275862 | 0.13 | 0.094 | 1 | 0 | Pik3r5 | 0.036 | Old |
| 0.007041 | 0.323837 | 0.194 | 0.151 | 1 | 0 | Gpr146 | 0.043 | Old |
| 0.007058 | 0.32175 | 0.19 | 0.148 | 1 | 0 | Ap3d1 | 0.042 | Old |
| 0.007072 | 0.868922 | 0.034 | 0.018 | 1 | 0 | Asah2 | 0.016 | Old |
| 0.007077 | 0.438914 | 0.156 | 0.119 | 1 | 0 | Hadh | 0.037 | Old |
| 0.007089 | 0.267373 | 0.176 | 0.135 | 1 | 0 | Alcam | 0.041 | Old |
| 0.007124 | 0.297129 | 0.213 | 0.168 | 1 | 0 | Map4 | 0.045 | Old |
| 0.007169 | 0.392214 | 0.104 | 0.072 | 1 | 0 | Papss1 | 0.032 | Old |
| 0.007186 | 0.618721 | 0.111 | 0.079 | 1 | 0 | Stard7 | 0.032 | Old |
| 0.007212 | 0.880056 | 0.038 | 0.021 | 1 | 0 | Ints4 | 0.017 | Old |
| 0.007278 | 0.211796 | 0.531 | 0.468 | 1 | 0 | Srsf3 | 0.063 | Old |
| 0.007286 | 0.49044 | 0.156 | 0.119 | 1 | 0 | Zmiz2 | 0.037 | Old |
| 0.007292 | 0.879043 | 0.073 | 0.048 | 1 | 0 | Fgr | 0.025 | Old |
| 0.007371 | 0.206737 | 0.473 | 0.415 | 1 | 0 | Rassf2 | 0.058 | Old |
| 0.007585 | 0.847092 | 0.044 | 0.025 | 1 | 0 | Mrpl22 | 0.019 | Old |
| 0.007589 | 0.947492 | 0.037 | 0.02 | 1 | 0 | Nr1d1 | 0.017 | Old |
| 0.007591 | 0.760116 | 0.049 | 0.029 | 1 | 0 | Mthfd2 | 0.02 | Old |
| 0.007602 | 0.461904 | 0.075 | 0.049 | 1 | 0 | Sc5d | 0.026 | Old |
| 0.007626 | 0.137704 | 0.658 | 0.595 | 1 | 0 | Celf2 | 0.063 | Old |
| 0.007832 | 0.397743 | 0.089 | 0.06 | 1 | 0 | Tada1 | 0.029 | Old |
| 0.007915 | 0.258451 | 0.138 | 0.102 | 1 | 0 | Bicd2 | 0.036 | Old |
| 0.007985 | 0.68794 | 0.033 | 0.017 | 1 | 0 | Fdps | 0.016 | Old |
| 0.007995 | 0.34017 | 0.208 | 0.166 | 1 | 0 | Brd7 | 0.042 | Old |
| 0.008012 | 1.078291 | 0.012 | 0.004 | 1 | 0 | Pomc | 0.008 | Old |
| 0.008038 | 1.249651 | 0.012 | 0.004 | 1 | 0 | Zfp493 | 0.008 | Old |
| 0.008046 | 0.308519 | 0.308 | 0.262 | 1 | 0 | AC160336. | 0.046 | Old |
| 0.008073 | 0.489729 | 0.161 | 0.123 | 1 | 0 | Chka | 0.038 | Old |
| 0.008089 | 0.821444 | 0.012 | 0.004 | 1 | 0 | Arv1 | 0.008 | Old |
| 0.008096 | 0.311705 | 0.103 | 0.072 | 1 | 0 | Prpf38a | 0.031 | Old |
| 0.008147 | 0.301931 | 0.298 | 0.248 | 1 | 0 | Axl | 0.05 | Old |
| 0.00817 | 0.239713 | 0.304 | 0.253 | 1 | 0 | Eif1a | 0.051 | Old |
| 0.008226 | 0.240091 | 0.227 | 0.179 | 1 | 0 | Psmb10 | 0.048 | Old |
| 0.008241 | 1.083314 | 0.031 | 0.016 | 1 | 0 | Rrp12 | 0.015 | Old |
| 0.008279 | 0.682111 | 0.052 | 0.031 | 1 | 0 | Adprhl2 | 0.021 | Old |
| 0.008312 | 0.176991 | 0.399 | 0.339 | 1 | 0 | Tm9sf2 | 0.06 | Old |

| | | | | | | | | |
|---|---|---|---|---|---|---|---|---|
| 0.008342 | 0.547587 | 0.071 | 0.046 | 1 | 0 | Usp21 | 0.025 | Old |
| 0.008396 | 0.433766 | 0.062 | 0.038 | 1 | 0 | Snapc3 | 0.024 | Old |
| 0.008435 | 0.549669 | 0.071 | 0.046 | 1 | 0 | Zfyve26 | 0.025 | Old |
| 0.00846 | 0.30415 | 0.234 | 0.187 | 1 | 0 | Rap2b | 0.047 | Old |
| 0.008509 | 0.227625 | 0.276 | 0.227 | 1 | 0 | Apobec1 | 0.049 | Old |
| 0.008525 | 0.755667 | 0.026 | 0.012 | 1 | 0 | Tbc1d19 | 0.014 | Old |
| 0.008595 | 0.300797 | 0.207 | 0.163 | 1 | 0 | Csnk1g3 | 0.044 | Old |
| 0.008607 | 0.493486 | 0.116 | 0.084 | 1 | 0 | Naa20 | 0.032 | Old |
| 0.008685 | 0.17292 | 0.518 | 0.456 | 1 | 0 | Clec2d | 0.062 | Old |
| 0.008688 | 0.399727 | 0.15 | 0.114 | 1 | 0 | Fbxw11 | 0.036 | Old |
| 0.008714 | 0.310716 | 0.17 | 0.131 | 1 | 0 | Elmo2 | 0.039 | Old |
| 0.008826 | 0.292241 | 0.254 | 0.21 | 1 | 0 | Atxn2l | 0.044 | Old |
| 0.008834 | 0.236714 | 0.363 | 0.31 | 1 | 0 | Trip12 | 0.053 | Old |
| 0.008836 | 0.168566 | 0.546 | 0.484 | 1 | 0 | Btg1 | 0.062 | Old |
| 0.008849 | 0.613278 | 0.059 | 0.037 | 1 | 0 | Smad3 | 0.022 | Old |
| 0.008886 | 1.821792 | 0.014 | 0.005 | 1 | 0 | Ptrh1 | 0.009 | Old |
| 0.008932 | 0.857773 | 0.025 | 0.011 | 1 | 0 | Nlrp1b | 0.014 | Old |
| 0.008945 | 0.280162 | 0.237 | 0.192 | 1 | 0 | Sec31a | 0.045 | Old |
| 0.008999 | 1.333304 | 0.014 | 0.005 | 1 | 0 | Plppr4 | 0.009 | Old |
| 0.009007 | 0.410102 | 0.101 | 0.071 | 1 | 0 | Rbm14 | 0.03 | Old |
| 0.009048 | 0.238934 | 0.491 | 0.432 | 1 | 0 | Ptms | 0.059 | Old |
| 0.00906 | 1.268451 | 0.014 | 0.005 | 1 | 0 | Zfp39 | 0.009 | Old |
| 0.009071 | 0.449431 | 0.105 | 0.075 | 1 | 0 | Plcg2 | 0.03 | Old |
| 0.009104 | 0.204088 | 0.142 | 0.105 | 1 | 0 | Znfx1 | 0.037 | Old |
| 0.009106 | 0.220143 | 0.15 | 0.113 | 1 | 0 | Dennd1c | 0.037 | Old |
| 0.009146 | 0.343721 | 0.096 | 0.066 | 1 | 0 | Wdr5 | 0.03 | Old |
| 0.009191 | 0.894271 | 0.034 | 0.018 | 1 | 0 | Tbrg4 | 0.016 | Old |
| 0.009269 | 0.188139 | 0.427 | 0.365 | 1 | 0 | Dync1h1 | 0.062 | Old |
| 0.009269 | 0.206804 | 0.29 | 0.238 | 1 | 0 | Drap1 | 0.052 | Old |
| 0.009284 | 0.243835 | 0.111 | 0.078 | 1 | 0 | Slc35c2 | 0.033 | Old |
| 0.009293 | 0.599379 | 0.109 | 0.08 | 1 | 0 | Cdkn2aipnl | 0.029 | Old |
| 0.009363 | 0.204407 | 0.521 | 0.463 | 1 | 0 | Tmem106a | 0.058 | Old |
| 0.009429 | 0.411699 | 0.067 | 0.043 | 1 | 0 | Ankrd49 | 0.024 | Old |
| 0.009449 | 0.177758 | 0.335 | 0.276 | 1 | 0 | Stx7 | 0.059 | Old |
| 0.009455 | 0.759786 | 0.089 | 0.062 | 1 | 0 | Il21r | 0.027 | Old |
| 0.0097 | 0.072349 | 0.449 | 0.375 | 1 | 0 | Mapre1 | 0.074 | Old |
| 0.009752 | 1.222596 | 0.021 | 0.009 | 1 | 0 | Efnb1 | 0.012 | Old |
| 0.009817 | 0.764105 | 0.015 | 0.005 | 1 | 0 | Gm17435 | 0.01 | Old |
| 0.009817 | 0.231956 | 0.301 | 0.25 | 1 | 0 | Cmtm3 | 0.051 | Old |
| 0.009875 | 1.032729 | 0.021 | 0.009 | 1 | 0 | Akap17b | 0.012 | Old |
| 0.009876 | 0.224945 | 0.368 | 0.312 | 1 | 0 | Tkt | 0.056 | Old |
| 0.009881 | 0.302717 | 0.175 | 0.136 | 1 | 0 | Btbd1 | 0.039 | Old |
| 0.009887 | 0.446903 | 0.092 | 0.064 | 1 | 0 | Parp11 | 0.028 | Old |
| 0.009909 | 0.739289 | 0.029 | 0.014 | 1 | 0 | Nt5c3b | 0.015 | Old |
| 0.009956 | 0.125129 | 0.476 | 0.41 | 1 | 0 | Cflar | 0.066 | Old |

| | | | | | | | | |
|---|---|---|---|---|---|---|---|---|
| 0.010059 | 0.984954 | 0.021 | 0.009 | 1 | 0 | Padi4 | 0.012 | Old |
| 0.010103 | 0.21584 | 0.261 | 0.211 | 1 | 0 | Tmed3 | 0.05 | Old |
| 0.010146 | 1.064186 | 0.027 | 0.013 | 1 | 0 | Cep19 | 0.014 | Old |
| 0.010172 | 0.718785 | 0.073 | 0.048 | 1 | 0 | St7l | 0.025 | Old |
| 0.010173 | 0.137434 | 0.674 | 0.611 | 1 | 0 | Ly86 | 0.063 | Old |
| 0.010269 | 0.137657 | 0.528 | 0.47 | 1 | 0 | Reep5 | 0.058 | Old |
| 0.010336 | 0.565626 | 0.066 | 0.043 | 1 | 0 | Med29 | 0.023 | Old |
| 0.010349 | 0.202509 | 0.427 | 0.362 | 1 | 0 | Sbno2 | 0.065 | Old |
| 0.010423 | 1.005075 | 0.027 | 0.013 | 1 | 0 | Plcg1 | 0.014 | Old |
| 0.010424 | 0.749781 | 0.019 | 0.008 | 1 | 0 | Hgh1 | 0.011 | Old |
| 0.010473 | 0.320289 | 0.118 | 0.086 | 1 | 0 | Dapk3 | 0.032 | Old |
| 0.0105 | 0.147498 | 0.297 | 0.243 | 1 | 0 | Vapa | 0.054 | Old |
| 0.010553 | 0.533034 | 0.098 | 0.07 | 1 | 0 | Adpgk | 0.028 | Old |
| 0.010673 | 0.149274 | 0.216 | 0.171 | 1 | 0 | Emg1 | 0.045 | Old |
| 0.010711 | 0.261979 | 0.219 | 0.175 | 1 | 0 | Ino80d | 0.044 | Old |
| 0.010754 | 0.490612 | 0.105 | 0.075 | 1 | 0 | Lypla1 | 0.03 | Old |
| 0.010772 | 0.10639 | 0.254 | 0.204 | 1 | 0 | Card19 | 0.05 | Old |
| 0.010954 | 0.25397 | 0.253 | 0.207 | 1 | 0 | Mfsd14b | 0.046 | Old |
| 0.010966 | 0.381036 | 0.06 | 0.038 | 1 | 0 | Zfp84 | 0.022 | Old |
| 0.011147 | 0.204189 | 0.464 | 0.406 | 1 | 0 | Prkar1a | 0.058 | Old |
| 0.011162 | 0.100022 | 0.711 | 0.643 | 1 | 0 | Cd53 | 0.068 | Old |
| 0.011173 | 0.510167 | 0.068 | 0.045 | 1 | 0 | Zfp398 | 0.023 | Old |
| 0.011248 | 0.189951 | 0.524 | 0.46 | 1 | 0 | Tapbp | 0.064 | Old |
| 0.011309 | 0.644864 | 0.036 | 0.019 | 1 | 0 | Rrs1 | 0.017 | Old |
| 0.011341 | 0.504221 | 0.122 | 0.091 | 1 | 0 | Ccdc22 | 0.031 | Old |
| 0.011392 | 0.11334 | 0.955 | 0.938 | 1 | 0 | Zfp36 | 0.017 | Old |
| 0.0114 | 0.126468 | 0.364 | 0.298 | 1 | 0 | Tmem256 | 0.066 | Old |
| 0.011456 | 0.299017 | 0.052 | 0.032 | 1 | 0 | Maip1 | 0.02 | Old |
| 0.011498 | 0.213856 | 0.218 | 0.174 | 1 | 0 | Dbnl | 0.044 | Old |
| 0.01151 | 0.140469 | 0.543 | 0.47 | 1 | 0 | Ap1b1 | 0.073 | Old |
| 0.011547 | 1.221017 | 0.025 | 0.012 | 1 | 0 | Lrmda | 0.013 | Old |
| 0.011566 | 0.307381 | 0.075 | 0.05 | 1 | 0 | Rgs14 | 0.025 | Old |
| 0.011641 | 0.220745 | 0.129 | 0.094 | 1 | 0 | Cep120 | 0.035 | Old |
| 0.011666 | 0.197753 | 0.275 | 0.226 | 1 | 0 | Eif4g1 | 0.049 | Old |
| 0.01167 | 0.433342 | 0.104 | 0.075 | 1 | 0 | Gskip | 0.029 | Old |
| 0.011693 | 0.234821 | 0.133 | 0.098 | 1 | 0 | Gatd1 | 0.035 | Old |
| 0.011712 | 0.567765 | 0.026 | 0.013 | 1 | 0 | Ccdc130 | 0.013 | Old |
| 0.011761 | 0.301416 | 0.231 | 0.188 | 1 | 0 | 2310035C2 | 0.043 | Old |
| 0.011794 | 0.661608 | 0.057 | 0.036 | 1 | 0 | H2-Q6 | 0.021 | Old |
| 0.011817 | 0.562393 | 0.062 | 0.04 | 1 | 0 | Tmem199 | 0.022 | Old |
| 0.011946 | 0.469643 | 0.063 | 0.04 | 1 | 0 | Vhl | 0.023 | Old |
| 0.011953 | 1.114769 | 0.025 | 0.012 | 1 | 0 | Clec1a | 0.013 | Old |
| 0.011982 | 0.382215 | 0.104 | 0.075 | 1 | 0 | Hk1 | 0.029 | Old |
| 0.01205 | 0.210367 | 0.331 | 0.278 | 1 | 0 | Atp5b | 0.053 | Old |
| 0.012275 | 0.468484 | 0.057 | 0.036 | 1 | 0 | Fkrp | 0.021 | Old |

| | | | | | | | | |
|---|---|---|---|---|---|---|---|---|
| 0.012291 | 1.453168 | 0.011 | 0.003 | 1 | 0 | Hspa1l | 0.008 | Old |
| 0.012298 | 0.351844 | 0.115 | 0.084 | 1 | 0 | Tnfrsf21 | 0.031 | Old |
| 0.012342 | 0.491558 | 0.097 | 0.069 | 1 | 0 | Agap1 | 0.028 | Old |
| 0.012385 | 0.40598 | 0.144 | 0.11 | 1 | 0 | Tmem230 | 0.034 | Old |
| 0.012419 | 0.442178 | 0.073 | 0.048 | 1 | 0 | Gtf2f1 | 0.025 | Old |
| 0.01245 | 1.027718 | 0.011 | 0.003 | 1 | 0 | BC048403 | 0.008 | Old |
| 0.012483 | 0.474657 | 0.097 | 0.07 | 1 | 0 | Ecd | 0.027 | Old |
| 0.012528 | 0.128896 | 0.773 | 0.714 | 1 | 0 | Hnrnpa2b1 | 0.059 | Old |
| 0.012543 | 1.121341 | 0.029 | 0.015 | 1 | 0 | Inf2 | 0.014 | Old |
| 0.012583 | 0.439796 | 0.083 | 0.057 | 1 | 0 | Dcaf10 | 0.026 | Old |
| 0.012669 | 0.24959 | 0.215 | 0.171 | 1 | 0 | Nceh1 | 0.044 | Old |
| 0.012697 | 0.144176 | 0.286 | 0.235 | 1 | 0 | Ccnd1 | 0.051 | Old |
| 0.012786 | 0.294824 | 0.093 | 0.065 | 1 | 0 | Fuca2 | 0.028 | Old |
| 0.01289 | 1.3146 | 0.022 | 0.01 | 1 | 0 | Sqle | 0.012 | Old |
| 0.012912 | 0.298352 | 0.123 | 0.091 | 1 | 0 | Atxn3 | 0.032 | Old |
| 0.012935 | 0.208194 | 0.213 | 0.17 | 1 | 0 | Borcs6 | 0.043 | Old |
| 0.012955 | 0.788275 | 0.075 | 0.051 | 1 | 0 | Dcn | 0.024 | Old |
| 0.013139 | 0.137575 | 0.557 | 0.491 | 1 | 0 | Sfpq | 0.066 | Old |
| 0.013386 | 0.223511 | 0.3 | 0.249 | 1 | 0 | Zfp703 | 0.051 | Old |
| 0.013449 | 0.3891 | 0.108 | 0.078 | 1 | 0 | Nop58 | 0.03 | Old |
| 0.013452 | 0.236533 | 0.412 | 0.364 | 1 | 0 | Flna | 0.048 | Old |
| 0.013569 | 0.552586 | 0.06 | 0.039 | 1 | 0 | Slc13a3 | 0.021 | Old |
| 0.01358 | 0.581064 | 0.033 | 0.018 | 1 | 0 | Sbk1 | 0.015 | Old |
| 0.013583 | 0.185471 | 0.223 | 0.179 | 1 | 0 | Mgat4b | 0.044 | Old |
| 0.013586 | 0.61874 | 0.098 | 0.071 | 1 | 0 | Gm28875 | 0.027 | Old |
| 0.013587 | 0.392559 | 0.093 | 0.066 | 1 | 0 | Trappc3 | 0.027 | Old |
| 0.0136 | 0.254144 | 0.157 | 0.121 | 1 | 0 | Tomm34 | 0.036 | Old |
| 0.013636 | 0.219969 | 0.16 | 0.122 | 1 | 0 | B4galt5 | 0.038 | Old |
| 0.013661 | 0.183686 | 0.585 | 0.532 | 1 | 0 | Gnb1 | 0.053 | Old |
| 0.013699 | 0.708367 | 0.022 | 0.01 | 1 | 0 | Loxl3 | 0.012 | Old |
| 0.013748 | 0.207457 | 0.246 | 0.199 | 1 | 0 | Acly | 0.047 | Old |
| 0.013794 | 0.228373 | 0.323 | 0.272 | 1 | 0 | Ranbp2 | 0.051 | Old |
| 0.013814 | 0.332274 | 0.168 | 0.131 | 1 | 0 | Pdlim4 | 0.037 | Old |
| 0.013817 | 0.381627 | 0.189 | 0.151 | 1 | 0 | Dpy19l1 | 0.038 | Old |
| 0.013824 | 0.416732 | 0.112 | 0.083 | 1 | 0 | Prkag2 | 0.029 | Old |
| 0.013929 | 0.42281 | 0.148 | 0.115 | 1 | 0 | Wdr37 | 0.033 | Old |
| 0.013953 | 0.686605 | 0.09 | 0.064 | 1 | 0 | Avpi1 | 0.026 | Old |
| 0.01396 | 1.533334 | 0.012 | 0.004 | 1 | 0 | Rnls | 0.008 | Old |
| 0.014016 | 0.250142 | 0.282 | 0.236 | 1 | 0 | Tmod3 | 0.046 | Old |
| 0.01405 | 0.106424 | 0.925 | 0.902 | 1 | 0 | Rpsa | 0.023 | Old |
| 0.014096 | 0.725988 | 0.038 | 0.022 | 1 | 0 | Xylt1 | 0.016 | Old |
| 0.014282 | 0.654127 | 0.031 | 0.017 | 1 | 0 | Tlr8 | 0.014 | Old |
| 0.01429 | 0.812155 | 0.031 | 0.017 | 1 | 0 | Bcar3 | 0.014 | Old |
| 0.014415 | 0.457871 | 0.012 | 0.004 | 1 | 0 | Gm48529 | 0.008 | Old |
| 0.014427 | 0.250207 | 0.186 | 0.147 | 1 | 0 | Arnt | 0.039 | Old |

| | | | | | | | | |
|---|---|---|---|---|---|---|---|---|
| 0.014449 | 0.278481 | 0.213 | 0.173 | 1 | 0 | Mdh2 | 0.04 | Old |
| 0.014462 | 0.150688 | 0.205 | 0.162 | 1 | 0 | Bscl2 | 0.043 | Old |
| 0.01448 | 0.417219 | 0.141 | 0.108 | 1 | 0 | Arl6ip5 | 0.033 | Old |
| 0.014564 | 0.561003 | 0.021 | 0.009 | 1 | 0 | Acyp2 | 0.012 | Old |
| 0.014586 | 0.459042 | 0.09 | 0.064 | 1 | 0 | Acads | 0.026 | Old |
| 0.014705 | 0.461911 | 0.064 | 0.042 | 1 | 0 | Gm26782 | 0.022 | Old |
| 0.014745 | 0.334237 | 0.153 | 0.119 | 1 | 0 | Strap | 0.034 | Old |
| 0.014773 | 0.95023 | 0.019 | 0.008 | 1 | 0 | Vcpkmt | 0.011 | Old |
| 0.014845 | 0.41025 | 0.088 | 0.061 | 1 | 0 | Gan | 0.027 | Old |
| 0.014871 | 0.26419 | 0.109 | 0.079 | 1 | 0 | Rpe | 0.03 | Old |
| 0.014935 | 0.569256 | 0.056 | 0.036 | 1 | 0 | Enoph1 | 0.02 | Old |
| 0.014941 | 1.442701 | 0.018 | 0.008 | 1 | 0 | Dennd2d | 0.01 | Old |
| 0.014943 | 0.152052 | 0.642 | 0.586 | 1 | 0 | Ube2d3 | 0.056 | Old |
| 0.015017 | 0.557168 | 0.063 | 0.041 | 1 | 0 | Socs5 | 0.022 | Old |
| 0.015024 | 1.125611 | 0.014 | 0.005 | 1 | 0 | Camk2n1 | 0.009 | Old |
| 0.015065 | 1.064824 | 0.014 | 0.005 | 1 | 0 | Med18 | 0.009 | Old |
| 0.015072 | 0.12946 | 0.603 | 0.534 | 1 | 0 | Vamp8 | 0.069 | Old |
| 0.015093 | 1.119786 | 0.033 | 0.018 | 1 | 0 | AC127341. | 0.015 | Old |
| 0.015153 | 0.550155 | 0.067 | 0.045 | 1 | 0 | Wdr77 | 0.022 | Old |
| 0.015176 | 0.360353 | 0.114 | 0.084 | 1 | 0 | Cfap43 | 0.03 | Old |
| 0.015188 | 0.338218 | 0.101 | 0.073 | 1 | 0 | Ergic1 | 0.028 | Old |
| 0.015197 | 1.286785 | 0.016 | 0.007 | 1 | 0 | Xlr4a | 0.009 | Old |
| 0.015216 | 0.201075 | 0.178 | 0.138 | 1 | 0 | Ogfr | 0.04 | Old |
| 0.015276 | 0.17419 | 0.269 | 0.223 | 1 | 0 | Rasa1 | 0.046 | Old |
| 0.015341 | 0.16936 | 0.135 | 0.102 | 1 | 0 | 1810030O( | 0.033 | Old |
| 0.015365 | 1.024679 | 0.042 | 0.025 | 1 | 0 | Glipr1 | 0.017 | Old |
| 0.015394 | 0.712598 | 0.034 | 0.019 | 1 | 0 | Pwp2 | 0.015 | Old |
| 0.015397 | 1.04957 | 0.016 | 0.007 | 1 | 0 | Chpf | 0.009 | Old |
| 0.015447 | 0.692998 | 0.014 | 0.005 | 1 | 0 | Pdss1 | 0.009 | Old |
| 0.015491 | 0.453088 | 0.116 | 0.086 | 1 | 0 | Naa35 | 0.03 | Old |
| 0.015503 | 0.258911 | 0.239 | 0.196 | 1 | 0 | Pdxk | 0.043 | Old |
| 0.0156 | 0.894396 | 0.016 | 0.007 | 1 | 0 | Simc1 | 0.009 | Old |
| 0.015616 | 0.281824 | 0.178 | 0.139 | 1 | 0 | Spag7 | 0.039 | Old |
| 0.015641 | 0.115811 | 0.932 | 0.929 | 1 | 0 | Rps26 | 0.003 | Old |
| 0.015662 | 0.830927 | 0.016 | 0.007 | 1 | 0 | Ddb2 | 0.009 | Old |
| 0.015699 | 0.619656 | 0.016 | 0.007 | 1 | 0 | Serpine1 | 0.009 | Old |
| 0.015729 | 0.726077 | 0.048 | 0.029 | 1 | 0 | Gm12840 | 0.019 | Old |
| 0.015798 | 0.406359 | 0.141 | 0.108 | 1 | 0 | Zfp263 | 0.033 | Old |
| 0.015847 | 0.326997 | 0.108 | 0.079 | 1 | 0 | Trak1 | 0.029 | Old |
| 0.015933 | 0.176975 | 0.338 | 0.283 | 1 | 0 | Nras | 0.055 | Old |
| 0.015937 | 0.917507 | 0.015 | 0.006 | 1 | 0 | Cdc7 | 0.009 | Old |
| 0.016006 | 0.859071 | 0.033 | 0.018 | 1 | 0 | Snupn | 0.015 | Old |
| 0.016024 | 0.456229 | 0.085 | 0.059 | 1 | 0 | Med12 | 0.026 | Old |
| 0.01607 | 0.165993 | 0.241 | 0.195 | 1 | 0 | D17Wsu92 | 0.046 | Old |
| 0.016075 | 0.255046 | 0.111 | 0.081 | 1 | 0 | Pilrb2 | 0.03 | Old |

| | | | | | | | | |
|---|---|---|---|---|---|---|---|---|
| 0.01611 | 0.461075 | 0.081 | 0.056 | 1 | 0 | Dap3 | 0.025 | Old |
| 0.016126 | 0.346811 | 0.124 | 0.093 | 1 | 0 | Bcat2 | 0.031 | Old |
| 0.016219 | 0.118529 | 0.772 | 0.707 | 1 | 0 | Capza2 | 0.065 | Old |
| 0.016235 | 0.417676 | 0.211 | 0.173 | 1 | 0 | Pmaip1 | 0.038 | Old |
| 0.016238 | 0.40377 | 0.089 | 0.063 | 1 | 0 | Strn | 0.026 | Old |
| 0.016316 | 0.333622 | 0.176 | 0.139 | 1 | 0 | Gpat4 | 0.037 | Old |
| 0.016345 | 0.327509 | 0.145 | 0.112 | 1 | 0 | Pim3 | 0.033 | Old |
| 0.016499 | 0.101348 | 0.958 | 0.929 | 1 | 0 | C1qb | 0.029 | Old |
| 0.016652 | 0.257751 | 0.108 | 0.079 | 1 | 0 | Rps6ka4 | 0.029 | Old |
| 0.016665 | 0.195325 | 0.38 | 0.323 | 1 | 0 | Wipf1 | 0.057 | Old |
| 0.016669 | 0.18352 | 0.391 | 0.337 | 1 | 0 | Rnf141 | 0.054 | Old |
| 0.016767 | 0.271092 | 0.215 | 0.175 | 1 | 0 | Nsf | 0.04 | Old |
| 0.016793 | 0.455063 | 0.037 | 0.021 | 1 | 0 | Tnfsf13os | 0.016 | Old |
| 0.016811 | 1.171635 | 0.023 | 0.011 | 1 | 0 | Dppa3 | 0.012 | Old |
| 0.016851 | 0.568378 | 0.094 | 0.068 | 1 | 0 | Odc1 | 0.026 | Old |
| 0.017019 | 0.22319 | 0.278 | 0.232 | 1 | 0 | Actn4 | 0.046 | Old |
| 0.017037 | 0.23499 | 0.373 | 0.328 | 1 | 0 | Cmpk1 | 0.045 | Old |
| 0.017066 | 0.532375 | 0.04 | 0.023 | 1 | 0 | Tsr1 | 0.017 | Old |
| 0.017116 | 0.302944 | 0.246 | 0.204 | 1 | 0 | Pgk1 | 0.042 | Old |
| 0.017168 | 0.146726 | 0.338 | 0.281 | 1 | 0 | Ube2r2 | 0.057 | Old |
| 0.017173 | 0.404495 | 0.083 | 0.059 | 1 | 0 | Atic | 0.024 | Old |
| 0.017279 | 0.130206 | 0.473 | 0.417 | 1 | 0 | Atp6v1b2 | 0.056 | Old |
| 0.017412 | 0.505367 | 0.082 | 0.057 | 1 | 0 | Herc6 | 0.025 | Old |
| 0.017527 | 0.717615 | 0.044 | 0.027 | 1 | 0 | Rtca | 0.017 | Old |
| 0.017563 | 0.635466 | 0.036 | 0.02 | 1 | 0 | Mdm1 | 0.016 | Old |
| 0.017676 | 0.258975 | 0.167 | 0.132 | 1 | 0 | Sdhb | 0.035 | Old |
| 0.017791 | 0.2736 | 0.397 | 0.35 | 1 | 0 | Atox1 | 0.047 | Old |
| 0.017808 | 0.081089 | 0.244 | 0.196 | 1 | 0 | Atp13a3 | 0.048 | Old |
| 0.017826 | 0.447032 | 0.122 | 0.092 | 1 | 0 | Atg2a | 0.03 | Old |
| 0.017849 | 0.405503 | 0.114 | 0.085 | 1 | 0 | Ltn1 | 0.029 | Old |
| 0.01786 | 0.217622 | 0.317 | 0.27 | 1 | 0 | Tbc1d14 | 0.047 | Old |
| 0.017904 | 0.170949 | 0.35 | 0.298 | 1 | 0 | Snx9 | 0.052 | Old |
| 0.017953 | 0.330021 | 0.146 | 0.114 | 1 | 0 | Fam192a | 0.032 | Old |
| 0.018089 | 0.344595 | 0.079 | 0.055 | 1 | 0 | Tmx2 | 0.024 | Old |
| 0.018104 | 0.375974 | 0.078 | 0.054 | 1 | 0 | Taf2 | 0.024 | Old |
| 0.018135 | 0.195342 | 0.1 | 0.072 | 1 | 0 | Stard5 | 0.028 | Old |
| 0.018184 | 0.502121 | 0.036 | 0.02 | 1 | 0 | Nr1h3 | 0.016 | Old |
| 0.018186 | 0.202628 | 0.326 | 0.276 | 1 | 0 | Spcs1 | 0.05 | Old |
| 0.018337 | 0.197858 | 0.275 | 0.225 | 1 | 0 | Ctdnep1 | 0.05 | Old |
| 0.018352 | 0.069947 | 0.424 | 0.36 | 1 | 0 | Prrc2c | 0.064 | Old |
| 0.01842 | 0.222741 | 0.182 | 0.144 | 1 | 0 | Fam96a | 0.038 | Old |
| 0.018445 | 0.44274 | 0.155 | 0.121 | 1 | 0 | Insig1 | 0.034 | Old |
| 0.01854 | 0.180328 | 0.282 | 0.233 | 1 | 0 | Rtraf | 0.049 | Old |
| 0.018602 | 0.654793 | 0.027 | 0.014 | 1 | 0 | Gk5 | 0.013 | Old |
| 0.018625 | 0.519473 | 0.074 | 0.051 | 1 | 0 | Sfxn1 | 0.023 | Old |

| | | | | | | | | |
|---|---|---|---|---|---|---|---|---|
| 0.018663 | 0.095355 | 0.343 | 0.289 | 1 | 0 | 9530068E0 | 0.054 | Old |
| 0.018784 | 0.498111 | 0.09 | 0.065 | 1 | 0 | Slc36a4 | 0.025 | Old |
| 0.018856 | 0.299125 | 0.55 | 0.511 | 1 | 0 | Sdc4 | 0.039 | Old |
| 0.018981 | 0.252587 | 0.269 | 0.226 | 1 | 0 | 1110008F1 | 0.043 | Old |
| 0.019098 | 0.222801 | 0.226 | 0.184 | 1 | 0 | Copb1 | 0.042 | Old |
| 0.019231 | 0.320756 | 0.157 | 0.123 | 1 | 0 | Por | 0.034 | Old |
| 0.019271 | 0.422786 | 0.104 | 0.077 | 1 | 0 | Kdm5c | 0.027 | Old |
| 0.019298 | 1.255898 | 0.021 | 0.01 | 1 | 0 | Atp6v0c | 0.011 | Old |
| 0.019304 | 0.247404 | 0.168 | 0.132 | 1 | 0 | Dido1 | 0.036 | Old |
| 0.019319 | 0.201069 | 0.309 | 0.259 | 1 | 0 | Mcrip1 | 0.05 | Old |
| 0.019398 | 0.860883 | 0.01 | 0.003 | 1 | 0 | Ascl2 | 0.007 | Old |
| 0.019416 | 0.266957 | 0.215 | 0.174 | 1 | 0 | Ubash3b | 0.041 | Old |
| 0.019419 | 0.198783 | 0.205 | 0.165 | 1 | 0 | Snx4 | 0.04 | Old |
| 0.01944 | 0.651721 | 0.034 | 0.019 | 1 | 0 | Drosha | 0.015 | Old |
| 0.019592 | 0.187544 | 0.13 | 0.098 | 1 | 0 | Gmps | 0.032 | Old |
| 0.019605 | 0.178056 | 0.215 | 0.174 | 1 | 0 | Szrd1 | 0.041 | Old |
| 0.019631 | 0.765994 | 0.01 | 0.003 | 1 | 0 | A830052D1 | 0.007 | Old |
| 0.019685 | 0.133714 | 0.156 | 0.12 | 1 | 0 | Sqor | 0.036 | Old |
| 0.019717 | 0.487123 | 0.01 | 0.003 | 1 | 0 | Gnal | 0.007 | Old |
| 0.019733 | 0.20254 | 0.317 | 0.268 | 1 | 0 | Idh1 | 0.049 | Old |
| 0.01974 | 0.83797 | 0.044 | 0.027 | 1 | 0 | St8sia6 | 0.017 | Old |
| 0.019767 | 0.346806 | 0.108 | 0.08 | 1 | 0 | Ube3b | 0.028 | Old |
| 0.019981 | 0.592991 | 0.045 | 0.028 | 1 | 0 | Alad | 0.017 | Old |
| 0.019985 | 0.315214 | 0.12 | 0.091 | 1 | 0 | Acvrl1 | 0.029 | Old |
| 0.020035 | 0.365858 | 0.118 | 0.088 | 1 | 0 | Preb | 0.03 | Old |
| 0.02016 | 1.292108 | 0.019 | 0.009 | 1 | 0 | Fam78b | 0.01 | Old |
| 0.020175 | 0.416672 | 0.096 | 0.07 | 1 | 0 | Mpdu1 | 0.026 | Old |
| 0.020299 | 0.31735 | 0.2 | 0.161 | 1 | 0 | Mff | 0.039 | Old |
| 0.020376 | 0.217342 | 0.271 | 0.226 | 1 | 0 | Dynll2 | 0.045 | Old |
| 0.020427 | 0.511418 | 0.021 | 0.01 | 1 | 0 | Dcaf4 | 0.011 | Old |
| 0.020581 | 0.396525 | 0.105 | 0.078 | 1 | 0 | Taf11 | 0.027 | Old |
| 0.020632 | 0.149054 | 0.26 | 0.212 | 1 | 0 | Antxr2 | 0.048 | Old |
| 0.020683 | 0.220658 | 0.306 | 0.262 | 1 | 0 | Twf1 | 0.044 | Old |
| 0.020705 | 0.439918 | 0.115 | 0.086 | 1 | 0 | Tbc1d22a | 0.029 | Old |
| 0.020825 | 0.28665 | 0.073 | 0.05 | 1 | 0 | Nolc1 | 0.023 | Old |
| 0.020939 | 0.133282 | 0.793 | 0.76 | 1 | 0 | Pabpc1 | 0.033 | Old |
| 0.021003 | 0.125955 | 0.501 | 0.438 | 1 | 0 | Atp1a1 | 0.063 | Old |
| 0.02102 | 1.075278 | 0.025 | 0.013 | 1 | 0 | Arap3 | 0.012 | Old |
| 0.02105 | 0.931485 | 0.019 | 0.009 | 1 | 0 | Nck2 | 0.01 | Old |
| 0.021155 | 0.2851 | 0.198 | 0.162 | 1 | 0 | Ppp6c | 0.036 | Old |
| 0.021226 | 0.175456 | 0.163 | 0.127 | 1 | 0 | Htt | 0.036 | Old |
| 0.021282 | 0.235557 | 0.202 | 0.164 | 1 | 0 | Ssfa2 | 0.038 | Old |
| 0.021332 | 0.048378 | 0.047 | 0.029 | 1 | 0 | Drg2 | 0.018 | Old |
| 0.02136 | 0.254714 | 0.161 | 0.127 | 1 | 0 | Rnf213 | 0.034 | Old |
| 0.02137 | 0.303722 | 0.179 | 0.144 | 1 | 0 | Dr1 | 0.035 | Old |

| | | | | | | | | |
|---|---|---|---|---|---|---|---|---|
| 0.021542 | 0.289053 | 0.101 | 0.074 | 1 | 0 | Pigm | 0.027 | Old |
| 0.021604 | 0.348985 | 0.208 | 0.172 | 1 | 0 | Skp1a | 0.036 | Old |
| 0.021612 | 0.08998 | 0.332 | 0.282 | 1 | 0 | Ldha | 0.05 | Old |
| 0.021705 | 0.396763 | 0.134 | 0.104 | 1 | 0 | Cdk4 | 0.03 | Old |
| 0.021774 | 0.153785 | 0.491 | 0.43 | 1 | 0 | Rgl1 | 0.061 | Old |
| 0.0219 | 0.352564 | 0.103 | 0.076 | 1 | 0 | Slc35f5 | 0.027 | Old |
| 0.021967 | 0.205799 | 0.15 | 0.117 | 1 | 0 | Gsk3a | 0.033 | Old |
| 0.021975 | 0.53351 | 0.104 | 0.078 | 1 | 0 | Zfp512 | 0.026 | Old |
| 0.021994 | 0.194746 | 0.172 | 0.136 | 1 | 0 | Marveld1 | 0.036 | Old |
| 0.022081 | 0.274623 | 0.16 | 0.125 | 1 | 0 | Gabpb2 | 0.035 | Old |
| 0.022208 | 0.688546 | 0.041 | 0.025 | 1 | 0 | Dennd2a | 0.016 | Old |
| 0.022234 | 0.113325 | 0.075 | 0.052 | 1 | 0 | Rasa2 | 0.023 | Old |
| 0.022553 | 0.212683 | 0.227 | 0.185 | 1 | 0 | Samd4b | 0.042 | Old |
| 0.022772 | 1.372666 | 0.012 | 0.005 | 1 | 0 | Tm7sf2 | 0.007 | Old |
| 0.022876 | 0.380848 | 0.13 | 0.1 | 1 | 0 | Mrpl13 | 0.03 | Old |
| 0.022896 | 0.192494 | 0.409 | 0.358 | 1 | 0 | Cd68 | 0.051 | Old |
| 0.02291 | 0.337028 | 0.178 | 0.144 | 1 | 0 | Hist1h1c | 0.034 | Old |
| 0.022917 | 0.197069 | 0.328 | 0.281 | 1 | 0 | Lamtor1 | 0.047 | Old |
| 0.022937 | 0.238846 | 0.131 | 0.1 | 1 | 0 | Babam2 | 0.031 | Old |
| 0.023125 | 0.385572 | 0.085 | 0.061 | 1 | 0 | Lpcat3 | 0.024 | Old |
| 0.023156 | 0.215147 | 0.334 | 0.288 | 1 | 0 | Mcub | 0.046 | Old |
| 0.02317 | 0.200426 | 0.267 | 0.223 | 1 | 0 | Cyp4f18 | 0.044 | Old |
| 0.023193 | 0.217724 | 0.256 | 0.213 | 1 | 0 | Cox17 | 0.043 | Old |
| 0.023289 | 0.163239 | 0.192 | 0.152 | 1 | 0 | Srsf10 | 0.04 | Old |
| 0.023315 | 0.159993 | 0.12 | 0.09 | 1 | 0 | Hlcs | 0.03 | Old |
| 0.023402 | 0.07762 | 0.96 | 0.942 | 1 | 0 | Laptm5 | 0.018 | Old |
| 0.023438 | 0.228647 | 0.167 | 0.131 | 1 | 0 | Cd300lb | 0.036 | Old |
| 0.023534 | 0.785427 | 0.012 | 0.005 | 1 | 0 | C230037L1 | 0.007 | Old |
| 0.023698 | 0.227812 | 0.074 | 0.051 | 1 | 0 | Dlat | 0.023 | Old |
| 0.023712 | 0.470889 | 0.073 | 0.051 | 1 | 0 | Nln | 0.022 | Old |
| 0.023767 | 0.074193 | 0.919 | 0.915 | 1 | 0 | Grn | 0.004 | Old |
| 0.023817 | 1.098787 | 0.014 | 0.005 | 1 | 0 | Gins1 | 0.009 | Old |
| 0.023876 | 0.9265 | 0.014 | 0.005 | 1 | 0 | Slc22a5 | 0.009 | Old |
| 0.023892 | 0.404491 | 0.012 | 0.005 | 1 | 0 | Xlr3b | 0.007 | Old |
| 0.023939 | 0.714831 | 0.015 | 0.006 | 1 | 0 | Plag1 | 0.009 | Old |
| 0.024062 | 0.372205 | 0.012 | 0.005 | 1 | 0 | Agbl5 | 0.007 | Old |
| 0.024087 | 0.568037 | 0.034 | 0.02 | 1 | 0 | Mtfr1 | 0.014 | Old |
| 0.024094 | 0.626982 | 0.014 | 0.005 | 1 | 0 | Acat2 | 0.009 | Old |
| 0.024094 | 0.578256 | 0.014 | 0.005 | 1 | 0 | Kif9 | 0.009 | Old |
| 0.024155 | 0.069214 | 0.499 | 0.428 | 1 | 0 | Eef1g | 0.071 | Old |
| 0.024176 | 0.440175 | 0.045 | 0.028 | 1 | 0 | Acvr2b | 0.017 | Old |
| 0.024219 | 0.410592 | 0.063 | 0.043 | 1 | 0 | Tmem161a | 0.02 | Old |
| 0.024257 | 0.349058 | 0.257 | 0.22 | 1 | 0 | Msl2 | 0.037 | Old |
| 0.024296 | 0.244296 | 0.129 | 0.098 | 1 | 0 | Trnau1ap | 0.031 | Old |
| 0.024296 | 0.307256 | 0.223 | 0.184 | 1 | 0 | Cct3 | 0.039 | Old |

| | | | | | | | | |
|---|---|---|---|---|---|---|---|---|
| 0.024307 | 0.736162 | 0.057 | 0.038 | 1 | 0 | Hps1 | 0.019 | Old |
| 0.024349 | 0.160472 | 0.282 | 0.237 | 1 | 0 | Hivep2 | 0.045 | Old |
| 0.024617 | 0.117737 | 0.739 | 0.693 | 1 | 0 | Tagln2 | 0.046 | Old |
| 0.024628 | 0.870278 | 0.03 | 0.017 | 1 | 0 | Tdp1 | 0.013 | Old |
| 0.024859 | 0.519751 | 0.06 | 0.04 | 1 | 0 | Rnaset2b | 0.02 | Old |
| 0.024886 | 0.249018 | 0.078 | 0.055 | 1 | 0 | Dolpp1 | 0.023 | Old |
| 0.024887 | 0.364949 | 0.085 | 0.061 | 1 | 0 | Cav2 | 0.024 | Old |
| 0.024904 | 0.49359 | 0.027 | 0.015 | 1 | 0 | Hpn | 0.012 | Old |
| 0.024929 | 0.1221 | 0.2 | 0.161 | 1 | 0 | Lgals9 | 0.039 | Old |
| 0.024996 | 0.295514 | 0.079 | 0.056 | 1 | 0 | Xab2 | 0.023 | Old |
| 0.025079 | 0.420842 | 0.052 | 0.034 | 1 | 0 | Vps8 | 0.018 | Old |
| 0.025169 | 0.425955 | 0.103 | 0.077 | 1 | 0 | Ndufs8 | 0.026 | Old |
| 0.02521 | 0.209724 | 0.417 | 0.372 | 1 | 0 | Nipa2 | 0.045 | Old |
| 0.025358 | 0.338268 | 0.137 | 0.105 | 1 | 0 | Snx12 | 0.032 | Old |
| 0.025426 | 0.283424 | 0.137 | 0.106 | 1 | 0 | Smim13 | 0.031 | Old |
| 0.025442 | 0.153857 | 0.027 | 0.015 | 1 | 0 | Rps6kb2 | 0.012 | Old |
| 0.025475 | 0.361786 | 0.129 | 0.099 | 1 | 0 | Tmem170b | 0.03 | Old |
| 0.025638 | 0.243592 | 0.111 | 0.083 | 1 | 0 | Nf1 | 0.028 | Old |
| 0.025664 | 0.361116 | 0.075 | 0.053 | 1 | 0 | Sun1 | 0.022 | Old |
| 0.025665 | 0.333426 | 0.183 | 0.148 | 1 | 0 | Usp16 | 0.035 | Old |
| 0.025669 | 0.148285 | 0.338 | 0.291 | 1 | 0 | G3bp1 | 0.047 | Old |
| 0.025694 | 0.126889 | 0.272 | 0.224 | 1 | 0 | Pkig | 0.048 | Old |
| 0.02577 | 0.401631 | 0.063 | 0.043 | 1 | 0 | Nfrkb | 0.02 | Old |
| 0.025804 | 0.483521 | 0.049 | 0.032 | 1 | 0 | Hars2 | 0.017 | Old |
| 0.025842 | 0.285931 | 0.192 | 0.156 | 1 | 0 | Eif2a | 0.036 | Old |
| 0.025863 | 0.113825 | 0.826 | 0.789 | 1 | 0 | Arpc2 | 0.037 | Old |
| 0.025912 | 0.540065 | 0.03 | 0.017 | 1 | 0 | Arhgap22 | 0.013 | Old |
| 0.02594 | 0.121006 | 0.39 | 0.338 | 1 | 0 | Cep170 | 0.052 | Old |
| 0.026088 | 0.127225 | 0.354 | 0.308 | 1 | 0 | Irf2 | 0.046 | Old |
| 0.026262 | 0.34427 | 0.088 | 0.063 | 1 | 0 | Rai1 | 0.025 | Old |
| 0.026343 | 0.092813 | 0.108 | 0.08 | 1 | 0 | Ptgr2 | 0.028 | Old |
| 0.02675 | 0.139021 | 0.347 | 0.297 | 1 | 0 | Lims1 | 0.05 | Old |
| 0.026827 | 0.114915 | 0.424 | 0.37 | 1 | 0 | Khdrbs1 | 0.054 | Old |
| 0.02683 | 0.775782 | 0.077 | 0.055 | 1 | 0 | Med20 | 0.022 | Old |
| 0.026975 | 0.21378 | 0.175 | 0.141 | 1 | 0 | AB124611 | 0.034 | Old |
| 0.026998 | 0.209967 | 0.386 | 0.337 | 1 | 0 | Prr13 | 0.049 | Old |
| 0.027022 | 0.494924 | 0.122 | 0.095 | 1 | 0 | Chsy1 | 0.027 | Old |
| 0.027023 | 0.279379 | 0.17 | 0.136 | 1 | 0 | Bcl3 | 0.034 | Old |
| 0.027065 | 0.152501 | 0.435 | 0.378 | 1 | 0 | Rbm25 | 0.057 | Old |
| 0.027079 | 0.317994 | 0.1 | 0.074 | 1 | 0 | Mfsd12 | 0.026 | Old |
| 0.027189 | 0.271373 | 0.212 | 0.175 | 1 | 0 | Rnf44 | 0.037 | Old |
| 0.027398 | 0.207608 | 0.312 | 0.267 | 1 | 0 | Ccdc88a | 0.045 | Old |
| 0.027551 | 0.605407 | 0.081 | 0.059 | 1 | 0 | Tlr1 | 0.022 | Old |
| 0.02759 | 0.055274 | 0.848 | 0.823 | 1 | 0 | Cdc42 | 0.025 | Old |
| 0.027645 | 0.640915 | 0.021 | 0.01 | 1 | 0 | Sag | 0.011 | Old |

| | | | | | | | | |
|---|---|---|---|---|---|---|---|---|
| 0.027669 | 0.115652 | 0.644 | 0.591 | 1 | 0 | Fcgr3 | 0.053 | Old |
| 0.02767 | 0.120838 | 0.81 | 0.754 | 1 | 0 | H3f3a | 0.056 | Old |
| 0.027784 | 0.727156 | 0.042 | 0.027 | 1 | 0 | Mblac2 | 0.015 | Old |
| 0.02786 | 0.582386 | 0.056 | 0.037 | 1 | 0 | Ltbp3 | 0.019 | Old |
| 0.028043 | 1.175656 | 0.019 | 0.009 | 1 | 0 | Tmem132a | 0.01 | Old |
| 0.028105 | 0.178498 | 0.516 | 0.46 | 1 | 0 | Eif4a1 | 0.056 | Old |
| 0.028246 | 0.212826 | 0.198 | 0.16 | 1 | 0 | Tbc1d5 | 0.038 | Old |
| 0.028301 | 0.315512 | 0.101 | 0.076 | 1 | 0 | Tmem41b | 0.025 | Old |
| 0.028388 | 0.120131 | 0.316 | 0.266 | 1 | 0 | Itch | 0.05 | Old |
| 0.028475 | 0.117215 | 0.807 | 0.767 | 1 | 0 | Wnk1 | 0.04 | Old |
| 0.028561 | 0.185886 | 0.148 | 0.115 | 1 | 0 | Eif4ebp1 | 0.033 | Old |
| 0.028649 | 0.256619 | 0.446 | 0.403 | 1 | 0 | Dnaja1 | 0.043 | Old |
| 0.028683 | 0.702022 | 0.037 | 0.022 | 1 | 0 | Ppat | 0.015 | Old |
| 0.028869 | 0.236166 | 0.086 | 0.062 | 1 | 0 | Cd151 | 0.024 | Old |
| 0.028883 | 0.181022 | 0.222 | 0.184 | 1 | 0 | Setd2 | 0.038 | Old |
| 0.02892 | 0.40142 | 0.133 | 0.103 | 1 | 0 | Tiam1 | 0.03 | Old |
| 0.029037 | 0.392987 | 0.088 | 0.064 | 1 | 0 | Erg28 | 0.024 | Old |
| 0.029089 | 0.613093 | 0.04 | 0.024 | 1 | 0 | Borcs5 | 0.016 | Old |
| 0.029104 | 0.314616 | 0.134 | 0.104 | 1 | 0 | Wwc2 | 0.03 | Old |
| 0.029122 | 0.393771 | 0.071 | 0.05 | 1 | 0 | Ppid | 0.021 | Old |
| 0.029391 | 0.258867 | 0.066 | 0.045 | 1 | 0 | Mtmr2 | 0.021 | Old |
| 0.029451 | 0.387506 | 0.049 | 0.032 | 1 | 0 | Scyl3 | 0.017 | Old |
| 0.029457 | 0.362574 | 0.14 | 0.11 | 1 | 0 | Dicer1 | 0.03 | Old |
| 0.029542 | 0.284297 | 0.159 | 0.127 | 1 | 0 | Srrt | 0.032 | Old |
| 0.029634 | 0.201937 | 0.14 | 0.108 | 1 | 0 | Vps13d | 0.032 | Old |
| 0.029645 | 0.551833 | 0.019 | 0.009 | 1 | 0 | Epas1 | 0.01 | Old |
| 0.029686 | 0.423008 | 0.031 | 0.018 | 1 | 0 | Ift27 | 0.013 | Old |
| 0.029761 | 0.150686 | 0.523 | 0.463 | 1 | 0 | Grk2 | 0.06 | Old |
| 0.029771 | 0.190414 | 0.335 | 0.288 | 1 | 0 | Atp2a2 | 0.047 | Old |
| 0.029874 | 0.411614 | 0.019 | 0.009 | 1 | 0 | Car5b | 0.01 | Old |
| 0.029901 | 0.985844 | 0.038 | 0.024 | 1 | 0 | Syne1 | 0.014 | Old |
| 0.030004 | 0.226977 | 0.327 | 0.282 | 1 | 0 | Dnmt3a | 0.045 | Old |
| 0.030096 | 0.127802 | 0.068 | 0.047 | 1 | 0 | Zfp362 | 0.021 | Old |
| 0.03011 | 0.269657 | 0.096 | 0.07 | 1 | 0 | Cog7 | 0.026 | Old |
| 0.03012 | 0.625955 | 0.044 | 0.028 | 1 | 0 | Isg15 | 0.016 | Old |
| 0.030121 | 0.319864 | 0.141 | 0.111 | 1 | 0 | Xdh | 0.03 | Old |
| 0.030151 | 0.151508 | 0.246 | 0.203 | 1 | 0 | mt-Atp8 | 0.043 | Old |
| 0.03032 | 0.960954 | 0.036 | 0.021 | 1 | 0 | Etaa1 | 0.015 | Old |
| 0.030351 | 0.186646 | 0.142 | 0.111 | 1 | 0 | Lrrc8c | 0.031 | Old |
| 0.030378 | 0.23449 | 0.067 | 0.046 | 1 | 0 | Hpse | 0.021 | Old |
| 0.030433 | 1.05511 | 0.018 | 0.008 | 1 | 0 | Gba2 | 0.01 | Old |
| 0.030436 | 0.357209 | 0.074 | 0.052 | 1 | 0 | Prpf19 | 0.022 | Old |
| 0.030448 | 0.164458 | 0.075 | 0.053 | 1 | 0 | Fbxl4 | 0.022 | Old |
| 0.030616 | 0.674021 | 0.027 | 0.015 | 1 | 0 | Kdm4b | 0.012 | Old |
| 0.030916 | 0.409994 | 0.067 | 0.047 | 1 | 0 | Prkch | 0.02 | Old |

| | | | | | | | | |
|---|---|---|---|---|---|---|---|---|
| 0.030956 | 0.211662 | 0.146 | 0.115 | 1 | 0 | Cstf2t | 0.031 | Old |
| 0.03101 | 0.355522 | 0.122 | 0.094 | 1 | 0 | Rangap1 | 0.028 | Old |
| 0.031052 | 0.449139 | 0.057 | 0.039 | 1 | 0 | Heatr1 | 0.018 | Old |
| 0.031132 | 0.715427 | 0.018 | 0.008 | 1 | 0 | Gm10521 | 0.01 | Old |
| 0.031139 | 0.240009 | 0.089 | 0.065 | 1 | 0 | A430005L1 | 0.024 | Old |
| 0.031188 | 0.114264 | 0.304 | 0.254 | 1 | 0 | Stag2 | 0.05 | Old |
| 0.031188 | 0.445138 | 0.027 | 0.015 | 1 | 0 | Vars2 | 0.012 | Old |
| 0.031264 | 0.655168 | 0.098 | 0.075 | 1 | 0 | Apmap | 0.023 | Old |
| 0.031269 | 0.507674 | 0.027 | 0.015 | 1 | 0 | Smug1 | 0.012 | Old |
| 0.031299 | 0.619663 | 0.018 | 0.008 | 1 | 0 | Ss18l1 | 0.01 | Old |
| 0.031304 | 0.335286 | 0.07 | 0.049 | 1 | 0 | Scmh1 | 0.021 | Old |
| 0.03131 | 0.329306 | 0.053 | 0.035 | 1 | 0 | Stk16 | 0.018 | Old |
| 0.031333 | 0.14393 | 0.14 | 0.108 | 1 | 0 | 2310011J0 | 0.032 | Old |
| 0.031457 | 0.837146 | 0.038 | 0.024 | 1 | 0 | Setdb2 | 0.014 | Old |
| 0.031475 | 0.149033 | 0.093 | 0.068 | 1 | 0 | Hdac4 | 0.025 | Old |
| 0.03148 | 0.268999 | 0.138 | 0.108 | 1 | 0 | Piezo1 | 0.03 | Old |
| 0.031508 | 0.358246 | 0.13 | 0.102 | 1 | 0 | Slc4a2 | 0.028 | Old |
| 0.031519 | 0.568415 | 0.066 | 0.046 | 1 | 0 | Rnf121 | 0.02 | Old |
| 0.031653 | 0.527072 | 0.081 | 0.059 | 1 | 0 | Slamf8 | 0.022 | Old |
| 0.03171 | 0.231547 | 0.126 | 0.097 | 1 | 0 | Kcnab2 | 0.029 | Old |
| 0.031711 | 0.828554 | 0.042 | 0.027 | 1 | 0 | Gm26881 | 0.015 | Old |
| 0.03172 | 0.091387 | 0.283 | 0.237 | 1 | 0 | Hdlbp | 0.046 | Old |
| 0.031741 | 0.286912 | 0.144 | 0.112 | 1 | 0 | Aco1 | 0.032 | Old |
| 0.031819 | 0.254 | 0.118 | 0.09 | 1 | 0 | Katna1 | 0.028 | Old |
| 0.031845 | 0.226708 | 0.171 | 0.138 | 1 | 0 | Zcchc7 | 0.033 | Old |
| 0.032021 | 0.170575 | 0.138 | 0.108 | 1 | 0 | Txlna | 0.03 | Old |
| 0.032032 | 0.128271 | 0.301 | 0.259 | 1 | 0 | Mapre2 | 0.042 | Old |
| 0.032089 | 0.505714 | 0.023 | 0.012 | 1 | 0 | Akr1c12 | 0.011 | Old |
| 0.032093 | 0.176754 | 0.235 | 0.195 | 1 | 0 | Dcaf12 | 0.04 | Old |
| 0.032093 | 0.220316 | 0.123 | 0.094 | 1 | 0 | Svil | 0.029 | Old |
| 0.032106 | 0.193187 | 0.145 | 0.113 | 1 | 0 | Mtmr10 | 0.032 | Old |
| 0.032136 | 0.118711 | 0.256 | 0.213 | 1 | 0 | Mfsd14a | 0.043 | Old |
| 0.032178 | 0.377097 | 0.114 | 0.087 | 1 | 0 | Supt6 | 0.027 | Old |
| 0.032296 | 0.467028 | 0.083 | 0.061 | 1 | 0 | Sipa1l3 | 0.022 | Old |
| 0.032398 | 1.038894 | 0.016 | 0.008 | 1 | 0 | Fsd2 | 0.008 | Old |
| 0.032491 | 0.109733 | 0.692 | 0.655 | 1 | 0 | Arpc5 | 0.037 | Old |
| 0.032506 | 0.1411 | 0.272 | 0.23 | 1 | 0 | Aldoa | 0.042 | Old |
| 0.032532 | 0.355842 | 0.053 | 0.035 | 1 | 0 | Ipo13 | 0.018 | Old |
| 0.032572 | 0.205482 | 0.219 | 0.182 | 1 | 0 | Ergic3 | 0.037 | Old |
| 0.032613 | 0.247773 | 0.379 | 0.341 | 1 | 0 | Cdkn1a | 0.038 | Old |
| 0.032623 | 0.314456 | 0.189 | 0.155 | 1 | 0 | Ndufa11 | 0.034 | Old |
| 0.032655 | 0.355441 | 0.074 | 0.053 | 1 | 0 | Ikbke | 0.021 | Old |
| 0.032972 | 0.18494 | 0.419 | 0.366 | 1 | 0 | P2rx4 | 0.053 | Old |
| 0.033142 | 0.263903 | 0.328 | 0.284 | 1 | 0 | Atp6v1e1 | 0.044 | Old |
| 0.033294 | 0.338379 | 0.163 | 0.132 | 1 | 0 | Mdm2 | 0.031 | Old |

| | | | | | | | | |
|---|---|---|---|---|---|---|---|---|
| 0.033327 | 0.443231 | 0.105 | 0.08 | 1 | 0 | A630001G2 | 0.025 | Old |
| 0.033428 | 0.388068 | 0.038 | 0.024 | 1 | 0 | Ift22 | 0.014 | Old |
| 0.033566 | 0.360575 | 0.175 | 0.142 | 1 | 0 | Ndufa8 | 0.033 | Old |
| 0.033568 | 0.459042 | 0.038 | 0.024 | 1 | 0 | Tram2 | 0.014 | Old |
| 0.033645 | 0.289472 | 0.174 | 0.142 | 1 | 0 | Lat2 | 0.032 | Old |
| 0.033702 | 0.857049 | 0.034 | 0.021 | 1 | 0 | Gpr68 | 0.013 | Old |
| 0.033755 | 0.545735 | 0.057 | 0.039 | 1 | 0 | Slc33a1 | 0.018 | Old |
| 0.033807 | 0.241504 | 0.097 | 0.072 | 1 | 0 | Tax1bp3 | 0.025 | Old |
| 0.033872 | 0.096729 | 0.408 | 0.349 | 1 | 0 | Hnrnpab | 0.059 | Old |
| 0.033882 | 0.886878 | 0.031 | 0.019 | 1 | 0 | Tmem115 | 0.012 | Old |
| 0.033904 | 0.138844 | 0.077 | 0.054 | 1 | 0 | C1galt1c1 | 0.023 | Old |
| 0.034028 | 0.234692 | 0.211 | 0.174 | 1 | 0 | Polr2k | 0.037 | Old |
| 0.034031 | 0.155307 | 0.224 | 0.186 | 1 | 0 | Rcc2 | 0.038 | Old |
| 0.03406 | 0.865958 | 0.029 | 0.016 | 1 | 0 | Mus81 | 0.013 | Old |
| 0.034154 | 1.035595 | 0.015 | 0.007 | 1 | 0 | Rabl2 | 0.008 | Old |
| 0.034396 | 0.080883 | 0.417 | 0.36 | 1 | 0 | Kif5b | 0.057 | Old |
| 0.03446 | 0.346619 | 0.083 | 0.061 | 1 | 0 | Mrps6 | 0.022 | Old |
| 0.034507 | 0.174191 | 0.193 | 0.157 | 1 | 0 | Fam49a | 0.036 | Old |
| 0.03453 | 0.059223 | 0.915 | 0.869 | 1 | 0 | Lamp1 | 0.046 | Old |
| 0.034628 | 0.160805 | 0.352 | 0.31 | 1 | 0 | Tram1 | 0.042 | Old |
| 0.034681 | 0.173082 | 0.371 | 0.324 | 1 | 0 | Capns1 | 0.047 | Old |
| 0.034758 | 0.59279 | 0.01 | 0.003 | 1 | 0 | 2010013B2 | 0.007 | Old |
| 0.034774 | 0.158254 | 0.295 | 0.253 | 1 | 0 | Strn3 | 0.042 | Old |
| 0.03483 | 0.535015 | 0.015 | 0.007 | 1 | 0 | Cpeb1 | 0.008 | Old |
| 0.034846 | 0.783444 | 0.034 | 0.021 | 1 | 0 | Asb6 | 0.013 | Old |
| 0.034866 | 0.502587 | 0.063 | 0.044 | 1 | 0 | Zfp938 | 0.019 | Old |
| 0.034952 | 0.106491 | 0.343 | 0.291 | 1 | 0 | Ywhaq | 0.052 | Old |
| 0.035339 | 0.143613 | 0.16 | 0.127 | 1 | 0 | Srp19 | 0.033 | Old |
| 0.035364 | 0.386613 | 0.119 | 0.093 | 1 | 0 | Spg20 | 0.026 | Old |
| 0.035391 | 0.930392 | 0.025 | 0.013 | 1 | 0 | Pigf | 0.012 | Old |
| 0.035396 | 0.632816 | 0.034 | 0.021 | 1 | 0 | Sh3bp5l | 0.013 | Old |
| 0.035427 | 0.437425 | 0.049 | 0.032 | 1 | 0 | Nlrc5 | 0.017 | Old |
| 0.035428 | 1.030426 | 0.021 | 0.011 | 1 | 0 | Prepl | 0.01 | Old |
| 0.035503 | 1.068304 | 0.014 | 0.006 | 1 | 0 | Tmem144 | 0.008 | Old |
| 0.035503 | 0.96011 | 0.014 | 0.006 | 1 | 0 | Rsph3b | 0.008 | Old |
| 0.035541 | 0.643494 | 0.029 | 0.016 | 1 | 0 | Nol10 | 0.013 | Old |
| 0.035587 | 0.471775 | 0.075 | 0.054 | 1 | 0 | Irf2bp1 | 0.021 | Old |
| 0.035829 | 0.21621 | 0.268 | 0.227 | 1 | 0 | Smg1 | 0.041 | Old |
| 0.035862 | 0.499171 | 0.029 | 0.016 | 1 | 0 | Ercc2 | 0.013 | Old |
| 0.035916 | 0.775729 | 0.011 | 0.004 | 1 | 0 | Nat8l | 0.007 | Old |
| 0.036135 | 0.109056 | 0.888 | 0.848 | 1 | 0 | Mcl1 | 0.04 | Old |
| 0.036235 | 0.638635 | 0.048 | 0.032 | 1 | 0 | Cdc42ep4 | 0.016 | Old |
| 0.036394 | 0.324852 | 0.083 | 0.061 | 1 | 0 | Fbxo28 | 0.022 | Old |
| 0.036442 | 0.401161 | 0.12 | 0.094 | 1 | 0 | Traf1 | 0.026 | Old |
| 0.036633 | 0.183752 | 0.361 | 0.318 | 1 | 0 | Pias1 | 0.043 | Old |

| | | | | | | | | |
|---|---|---|---|---|---|---|---|---|
| 0.036641 | 0.09611 | 0.807 | 0.75 | 1 | 0 | Rtn4 | 0.057 | Old |
| 0.036676 | 0.289313 | 0.155 | 0.125 | 1 | 0 | Rcor1 | 0.03 | Old |
| 0.036771 | 0.185294 | 0.295 | 0.254 | 1 | 0 | Nedd8 | 0.041 | Old |
| 0.036851 | 0.169345 | 0.416 | 0.362 | 1 | 0 | Ap2s1 | 0.054 | Old |
| 0.036858 | 0.135026 | 0.795 | 0.763 | 1 | 0 | Irf2bp2 | 0.032 | Old |
| 0.037028 | 0.46799 | 0.012 | 0.005 | 1 | 0 | Cyb5rl | 0.007 | Old |
| 0.037028 | 0.38857 | 0.012 | 0.005 | 1 | 0 | Ctps | 0.007 | Old |
| 0.037098 | 0.172064 | 0.175 | 0.142 | 1 | 0 | Dcun1d5 | 0.033 | Old |
| 0.037164 | 0.149777 | 0.297 | 0.252 | 1 | 0 | Tmem55b | 0.045 | Old |
| 0.037173 | 0.140448 | 0.398 | 0.35 | 1 | 0 | Irf8 | 0.048 | Old |
| 0.037175 | 0.437965 | 0.059 | 0.04 | 1 | 0 | Gtdc1 | 0.019 | Old |
| 0.037267 | 0.388015 | 0.09 | 0.067 | 1 | 0 | Ubac2 | 0.023 | Old |
| 0.037417 | 0.163738 | 0.643 | 0.589 | 1 | 0 | Zfand5 | 0.054 | Old |
| 0.037473 | 0.233772 | 0.152 | 0.122 | 1 | 0 | Usp22 | 0.03 | Old |
| 0.037489 | 0.658723 | 0.038 | 0.024 | 1 | 0 | Dnajc12 | 0.014 | Old |
| 0.037558 | 0.440415 | 0.107 | 0.082 | 1 | 0 | Imp4 | 0.025 | Old |
| 0.037744 | 0.178735 | 0.107 | 0.081 | 1 | 0 | Elp2 | 0.026 | Old |
| 0.037858 | 0.258319 | 0.103 | 0.078 | 1 | 0 | Dpp3 | 0.025 | Old |
| 0.037903 | 0.187035 | 0.181 | 0.146 | 1 | 0 | Gba | 0.035 | Old |
| 0.037916 | 0.134569 | 0.741 | 0.698 | 1 | 0 | Rhoa | 0.043 | Old |
| 0.038114 | 0.766667 | 0.036 | 0.022 | 1 | 0 | Emilin1 | 0.014 | Old |
| 0.03816 | 0.275211 | 0.123 | 0.096 | 1 | 0 | Syvn1 | 0.027 | Old |
| 0.038269 | 0.276784 | 0.131 | 0.103 | 1 | 0 | Plgrkt | 0.028 | Old |
| 0.038279 | 0.166454 | 0.141 | 0.11 | 1 | 0 | Eif4a3 | 0.031 | Old |
| 0.038301 | 0.287453 | 0.115 | 0.089 | 1 | 0 | Mrps9 | 0.026 | Old |
| 0.038319 | 0.45141 | 0.073 | 0.052 | 1 | 0 | Smarcal1 | 0.021 | Old |
| 0.038588 | 0.224936 | 0.327 | 0.279 | 1 | 0 | Tpm4 | 0.048 | Old |
| 0.038604 | 0.577294 | 0.063 | 0.044 | 1 | 0 | Vps45 | 0.019 | Old |
| 0.038606 | 0.923783 | 0.019 | 0.01 | 1 | 0 | Pvt1 | 0.009 | Old |
| 0.038631 | 0.420123 | 0.066 | 0.046 | 1 | 0 | Lonp2 | 0.02 | Old |
| 0.038788 | 0.343388 | 0.048 | 0.032 | 1 | 0 | Magohb | 0.016 | Old |
| 0.038848 | 0.244221 | 0.368 | 0.328 | 1 | 0 | Csf2ra | 0.04 | Old |
| 0.039033 | 0.277331 | 0.213 | 0.178 | 1 | 0 | Galc | 0.035 | Old |
| 0.039218 | 0.484685 | 0.045 | 0.029 | 1 | 0 | Ptger2 | 0.016 | Old |
| 0.039221 | 0.133571 | 0.256 | 0.215 | 1 | 0 | R3hdm4 | 0.041 | Old |
| 0.039223 | 0.129226 | 0.144 | 0.113 | 1 | 0 | Fam20b | 0.031 | Old |
| 0.039241 | 0.124663 | 0.305 | 0.258 | 1 | 0 | Pik3cd | 0.047 | Old |
| 0.0393 | 0.262058 | 0.15 | 0.12 | 1 | 0 | Tor1b | 0.03 | Old |
| 0.039306 | 0.217973 | 0.29 | 0.249 | 1 | 0 | Lrrc25 | 0.041 | Old |
| 0.03941 | 0.110548 | 0.209 | 0.171 | 1 | 0 | Ppp4c | 0.038 | Old |
| 0.039507 | 0.12559 | 0.501 | 0.449 | 1 | 0 | Lgals1 | 0.052 | Old |
| 0.039537 | 0.109138 | 0.989 | 0.991 | 1 | 0 | Lyz2 | -0.002 | Old |
| 0.0396 | 0.266481 | 0.14 | 0.111 | 1 | 0 | Dyrk1a | 0.029 | Old |
| 0.039643 | 0.476291 | 0.118 | 0.092 | 1 | 0 | Usp3 | 0.026 | Old |
| 0.039879 | 0.53244 | 0.085 | 0.063 | 1 | 0 | Stard4 | 0.022 | Old |

| | | | | | | | | |
|---|---|---|---|---|---|---|---|---|
| 0.039981 | 0.419768 | 0.027 | 0.016 | 1 | 0 | Zfp934 | 0.011 | Old |
| 0.040079 | 0.241541 | 0.144 | 0.115 | 1 | 0 | Adss | 0.029 | Old |
| 0.040284 | 0.375823 | 0.056 | 0.038 | 1 | 0 | Pcnx3 | 0.018 | Old |
| 0.040359 | 0.792813 | 0.052 | 0.035 | 1 | 0 | Nudt7 | 0.017 | Old |
| 0.040384 | 0.133815 | 0.148 | 0.117 | 1 | 0 | Ubr3 | 0.031 | Old |
| 0.040458 | 0.387046 | 0.057 | 0.04 | 1 | 0 | Gmcl1 | 0.017 | Old |
| 0.040508 | 0.302386 | 0.056 | 0.038 | 1 | 0 | Abhd5 | 0.018 | Old |
| 0.040578 | 0.595531 | 0.124 | 0.099 | 1 | 0 | Ipmk | 0.025 | Old |
| 0.040605 | 0.671252 | 0.023 | 0.013 | 1 | 0 | Cep41 | 0.01 | Old |
| 0.040633 | 0.502495 | 0.03 | 0.018 | 1 | 0 | Adgrl2 | 0.012 | Old |
| 0.040659 | 0.351561 | 0.06 | 0.042 | 1 | 0 | Papd7 | 0.018 | Old |
| 0.040781 | 0.24555 | 0.135 | 0.107 | 1 | 0 | Emc2 | 0.028 | Old |
| 0.040898 | 0.617247 | 0.042 | 0.027 | 1 | 0 | D930016D( | 0.015 | Old |
| 0.041007 | 0.5048 | 0.042 | 0.027 | 1 | 0 | Ints8 | 0.015 | Old |
| 0.041026 | 0.685162 | 0.037 | 0.023 | 1 | 0 | Ndufaf3 | 0.014 | Old |
| 0.041151 | 0.417642 | 0.103 | 0.079 | 1 | 0 | Polg | 0.024 | Old |
| 0.041191 | 0.213573 | 0.118 | 0.091 | 1 | 0 | Hacd3 | 0.027 | Old |
| 0.041197 | 0.183949 | 0.167 | 0.135 | 1 | 0 | Otub1 | 0.032 | Old |
| 0.041254 | 0.346743 | 0.126 | 0.099 | 1 | 0 | Akna | 0.027 | Old |
| 0.041482 | 0.664103 | 0.037 | 0.023 | 1 | 0 | Cdt1 | 0.014 | Old |
| 0.041679 | 0.477854 | 0.051 | 0.034 | 1 | 0 | Ammecr1 | 0.017 | Old |
| 0.041859 | 0.263021 | 0.112 | 0.087 | 1 | 0 | Pdgfb | 0.025 | Old |
| 0.041888 | 0.340487 | 0.094 | 0.071 | 1 | 0 | Oxsr1 | 0.023 | Old |
| 0.041905 | 0.172691 | 0.105 | 0.08 | 1 | 0 | Afg3l1 | 0.025 | Old |
| 0.041956 | 0.243599 | 0.134 | 0.106 | 1 | 0 | Lysmd3 | 0.028 | Old |
| 0.042121 | 0.677232 | 0.044 | 0.029 | 1 | 0 | Ppip5k1 | 0.015 | Old |
| 0.042192 | 0.420917 | 0.118 | 0.093 | 1 | 0 | St6galnac6 | 0.025 | Old |
| 0.042321 | 1.044485 | 0.022 | 0.012 | 1 | 0 | Pradc1 | 0.01 | Old |
| 0.042327 | 0.429006 | 0.083 | 0.062 | 1 | 0 | Ap1ar | 0.021 | Old |
| 0.042356 | 0.642473 | 0.042 | 0.027 | 1 | 0 | Utp4 | 0.015 | Old |
| 0.04245 | 0.469954 | 0.044 | 0.029 | 1 | 0 | Clcn6 | 0.015 | Old |
| 0.042457 | 0.267349 | 0.174 | 0.142 | 1 | 0 | Alyref | 0.032 | Old |
| 0.042498 | 0.176077 | 0.06 | 0.042 | 1 | 0 | Rufy2 | 0.018 | Old |
| 0.042525 | 0.061811 | 0.657 | 0.598 | 1 | 0 | Chchd2 | 0.059 | Old |
| 0.042555 | 0.455363 | 0.074 | 0.054 | 1 | 0 | Grpel1 | 0.02 | Old |
| 0.042571 | 0.141684 | 0.35 | 0.304 | 1 | 0 | Acp2 | 0.046 | Old |
| 0.042575 | 0.215759 | 0.155 | 0.125 | 1 | 0 | Cand1 | 0.03 | Old |
| 0.042615 | 0.716539 | 0.018 | 0.009 | 1 | 0 | Tcea3 | 0.009 | Old |
| 0.042661 | 0.134718 | 0.521 | 0.461 | 1 | 0 | Csnk1a1 | 0.06 | Old |
| 0.042726 | 0.664732 | 0.034 | 0.021 | 1 | 0 | I830077J02 | 0.013 | Old |
| 0.042839 | 0.216574 | 0.186 | 0.154 | 1 | 0 | Yme1l1 | 0.032 | Old |
| 0.043138 | 0.085184 | 0.486 | 0.423 | 1 | 0 | Atf4 | 0.063 | Old |
| 0.043139 | 0.241833 | 0.079 | 0.058 | 1 | 0 | Sipa1l2 | 0.021 | Old |
| 0.043182 | 0.19574 | 0.168 | 0.136 | 1 | 0 | Zkscan3 | 0.032 | Old |
| 0.043208 | 0.325403 | 0.049 | 0.033 | 1 | 0 | Paxx | 0.016 | Old |

| | | | | | | | | |
|---|---|---|---|---|---|---|---|---|
| 0.043406 | 0.102635 | 0.729 | 0.677 | 1 | 0 | Ptprc | 0.052 | Old |
| 0.04342 | 0.377961 | 0.097 | 0.074 | 1 | 0 | Mmadhc | 0.023 | Old |
| 0.043434 | 0.226578 | 0.335 | 0.297 | 1 | 0 | Fam111a | 0.038 | Old |
| 0.043506 | 0.17967 | 0.168 | 0.136 | 1 | 0 | Trio | 0.032 | Old |
| 0.043594 | 0.485339 | 0.048 | 0.032 | 1 | 0 | Zbtb6 | 0.016 | Old |
| 0.0436 | 0.235593 | 0.208 | 0.172 | 1 | 0 | Smarcc2 | 0.036 | Old |
| 0.043602 | 0.769394 | 0.038 | 0.024 | 1 | 0 | Gm16104 | 0.014 | Old |
| 0.04361 | 0.520449 | 0.044 | 0.029 | 1 | 0 | Ppp2r3a | 0.015 | Old |
| 0.043693 | 0.234594 | 0.042 | 0.027 | 1 | 0 | Atpaf2 | 0.015 | Old |
| 0.043715 | 0.371351 | 0.055 | 0.037 | 1 | 0 | Slc39a13 | 0.018 | Old |
| 0.043876 | 0.274604 | 0.063 | 0.044 | 1 | 0 | Lsm10 | 0.019 | Old |
| 0.044051 | 0.404141 | 0.034 | 0.021 | 1 | 0 | 2010111I0 | 0.013 | Old |
| 0.044056 | 0.130209 | 0.698 | 0.64 | 1 | 0 | Ifitm2 | 0.058 | Old |
| 0.044116 | 0.093571 | 0.294 | 0.249 | 1 | 0 | Ap3s1 | 0.045 | Old |
| 0.044215 | 0.271531 | 0.083 | 0.061 | 1 | 0 | Creb3 | 0.022 | Old |
| 0.044264 | 0.199493 | 0.443 | 0.398 | 1 | 0 | Ppp2ca | 0.045 | Old |
| 0.044345 | 0.252118 | 0.071 | 0.051 | 1 | 0 | Ppdpf | 0.02 | Old |
| 0.044515 | 0.128679 | 0.077 | 0.056 | 1 | 0 | Timm10 | 0.021 | Old |
| 0.044588 | 0.064226 | 0.49 | 0.42 | 1 | 0 | Ostf1 | 0.07 | Old |
| 0.044795 | 0.138069 | 0.389 | 0.34 | 1 | 0 | Necap2 | 0.049 | Old |
| 0.044827 | 0.226043 | 0.201 | 0.167 | 1 | 0 | Kansl1 | 0.034 | Old |
| 0.045087 | 0.887884 | 0.025 | 0.014 | 1 | 0 | Ttc39c | 0.011 | Old |
| 0.045097 | 0.2972 | 0.16 | 0.13 | 1 | 0 | Slc27a1 | 0.03 | Old |
| 0.045154 | 1.061475 | 0.016 | 0.008 | 1 | 0 | Ifit3 | 0.008 | Old |
| 0.045183 | 0.989216 | 0.016 | 0.008 | 1 | 0 | Rab11fip3 | 0.008 | Old |
| 0.045187 | 0.03714 | 0.253 | 0.21 | 1 | 0 | Fxr1 | 0.043 | Old |
| 0.045198 | 0.222362 | 0.263 | 0.225 | 1 | 0 | Ifnar1 | 0.038 | Old |
| 0.045372 | 0.1825 | 0.133 | 0.104 | 1 | 0 | Zc3h4 | 0.029 | Old |
| 0.045405 | 0.51372 | 0.063 | 0.045 | 1 | 0 | Tmem104 | 0.018 | Old |
| 0.045433 | 0.26081 | 0.308 | 0.268 | 1 | 0 | Cerk | 0.04 | Old |
| 0.04547 | 0.302785 | 0.155 | 0.126 | 1 | 0 | Tmem5 | 0.029 | Old |
| 0.045578 | 0.109862 | 0.81 | 0.765 | 1 | 0 | Cebpb | 0.045 | Old |
| 0.045665 | 0.545002 | 0.038 | 0.024 | 1 | 0 | Amigo1 | 0.014 | Old |
| 0.045775 | 0.219234 | 0.083 | 0.062 | 1 | 0 | Rnf181 | 0.021 | Old |
| 0.04578 | 0.838779 | 0.025 | 0.014 | 1 | 0 | Gnb5 | 0.011 | Old |
| 0.045837 | 0.274562 | 0.126 | 0.099 | 1 | 0 | Mrps17 | 0.027 | Old |
| 0.045901 | 0.262388 | 0.037 | 0.023 | 1 | 0 | Dmac2 | 0.014 | Old |
| 0.045929 | 0.17494 | 0.204 | 0.169 | 1 | 0 | Btk | 0.035 | Old |
| 0.04599 | 0.411658 | 0.041 | 0.027 | 1 | 0 | Atm | 0.014 | Old |
| 0.046103 | 0.601986 | 0.016 | 0.008 | 1 | 0 | Cdca8 | 0.008 | Old |
| 0.046153 | 0.486325 | 0.074 | 0.054 | 1 | 0 | Aars | 0.02 | Old |
| 0.046282 | 0.133502 | 0.28 | 0.238 | 1 | 0 | Ddost | 0.042 | Old |
| 0.046335 | 0.389816 | 0.112 | 0.088 | 1 | 0 | Snx24 | 0.024 | Old |
| 0.046358 | 0.08678 | 0.399 | 0.344 | 1 | 0 | Eif3e | 0.055 | Old |
| 0.046382 | 0.314491 | 0.097 | 0.074 | 1 | 0 | Ddx58 | 0.023 | Old |

| | | | | | | | | |
|---|---|---|---|---|---|---|---|---|
| 0.04645 | 0.206996 | 0.155 | 0.125 | 1 | 0 | Rheb | 0.03 | Old |
| 0.046513 | 0.149331 | 0.216 | 0.182 | 1 | 0 | Prkx | 0.034 | Old |
| 0.046629 | 0.190186 | 0.26 | 0.222 | 1 | 0 | Bola2 | 0.038 | Old |
| 0.046636 | 0.438477 | 0.509 | 0.492 | 1 | 0 | Ccl7 | 0.017 | Old |
| 0.04669 | 0.092293 | 0.528 | 0.467 | 1 | 0 | Serbp1 | 0.061 | Old |
| 0.046739 | 0.28042 | 0.103 | 0.079 | 1 | 0 | Tango2 | 0.024 | Old |
| 0.046757 | 0.159116 | 0.034 | 0.021 | 1 | 0 | Cog2 | 0.013 | Old |
| 0.046832 | 0.53901 | 0.092 | 0.07 | 1 | 0 | Pmpca | 0.022 | Old |
| 0.046951 | 0.429716 | 0.07 | 0.051 | 1 | 0 | Vps26b | 0.019 | Old |
| 0.047042 | 0.671458 | 0.03 | 0.018 | 1 | 0 | Prdm10 | 0.012 | Old |
| 0.047049 | 0.291886 | 0.074 | 0.054 | 1 | 0 | Tufm | 0.02 | Old |
| 0.047176 | 0.178614 | 0.06 | 0.042 | 1 | 0 | Taf12 | 0.018 | Old |
| 0.047208 | 0.312661 | 0.116 | 0.091 | 1 | 0 | Eif2s1 | 0.025 | Old |
| 0.047279 | 0.189999 | 0.23 | 0.195 | 1 | 0 | Ostc | 0.035 | Old |
| 0.047318 | 0.059659 | 0.211 | 0.173 | 1 | 0 | Ppm1a | 0.038 | Old |
| 0.047334 | 0.351394 | 0.053 | 0.037 | 1 | 0 | Kantr | 0.016 | Old |
| 0.047494 | 0.445601 | 0.06 | 0.043 | 1 | 0 | Fyn | 0.017 | Old |
| 0.047511 | 0.147294 | 0.194 | 0.16 | 1 | 0 | Rhoc | 0.034 | Old |
| 0.047724 | 0.116073 | 0.317 | 0.272 | 1 | 0 | Dnajc13 | 0.045 | Old |
| 0.047842 | 0.165062 | 0.197 | 0.163 | 1 | 0 | Egln2 | 0.034 | Old |
| 0.048129 | 0.53783 | 0.027 | 0.016 | 1 | 0 | Unc13d | 0.011 | Old |
| 0.048148 | 0.142925 | 0.086 | 0.064 | 1 | 0 | 2310001H1 | 0.022 | Old |
| 0.048175 | 0.607079 | 0.021 | 0.011 | 1 | 0 | Coq8a | 0.01 | Old |
| 0.048314 | 0.408787 | 0.112 | 0.088 | 1 | 0 | Mbd6 | 0.024 | Old |
| 0.048324 | 0.44677 | 0.021 | 0.011 | 1 | 0 | Efcab2 | 0.01 | Old |
| 0.048368 | 0.150166 | 0.071 | 0.051 | 1 | 0 | 3110082I1 | 0.02 | Old |
| 0.048471 | 1.228514 | 0.015 | 0.007 | 1 | 0 | E430018J2 | 0.008 | Old |
| 0.048666 | 0.583175 | 0.057 | 0.04 | 1 | 0 | Cdca4 | 0.017 | Old |
| 0.048783 | 0.247246 | 0.073 | 0.053 | 1 | 0 | Dph5 | 0.02 | Old |
| 0.048863 | 0.149956 | 0.13 | 0.102 | 1 | 0 | Mrps18c | 0.028 | Old |
| 0.048945 | 0.144788 | 0.293 | 0.25 | 1 | 0 | Cct8 | 0.043 | Old |
| 0.048947 | 1.23377 | 0.026 | 0.015 | 1 | 0 | Sco1 | 0.011 | Old |
| 0.049016 | 0.252176 | 0.083 | 0.062 | 1 | 0 | Nup214 | 0.021 | Old |
| 0.049031 | 0.750605 | 0.015 | 0.007 | 1 | 0 | Ift88 | 0.008 | Old |
| 0.049213 | 0.042494 | 0.44 | 0.382 | 1 | 0 | Psmb4 | 0.058 | Old |
| 0.049253 | 0.311952 | 0.116 | 0.091 | 1 | 0 | Ube2j1 | 0.025 | Old |
| 0.049288 | 0.511586 | 0.094 | 0.073 | 1 | 0 | Cyp51 | 0.021 | Old |
| 0.049394 | 0.238147 | 0.122 | 0.095 | 1 | 0 | Ppp4r1 | 0.027 | Old |
| 0.049506 | 0.126579 | 0.471 | 0.426 | 1 | 0 | Tmem30a | 0.045 | Old |
| 0.049516 | 0.374451 | 0.07 | 0.051 | 1 | 0 | Mrps5 | 0.019 | Old |
| 0.049614 | 0.304855 | 0.116 | 0.091 | 1 | 0 | Cops8 | 0.025 | Old |
| 0.049701 | 0.291904 | 0.055 | 0.038 | 1 | 0 | Ubac1 | 0.017 | Old |
| 0.049913 | 0.286072 | 0.03 | 0.018 | 1 | 0 | Sdhaf3 | 0.012 | Old |
| 0.050036 | 0.431732 | 0.052 | 0.036 | 1 | 0 | Cox19 | 0.016 | Old |
| 0.050103 | 0.268833 | 0.075 | 0.055 | 1 | 0 | Uspl1 | 0.02 | Old |

| | | | | | | | | |
|---|---|---|---|---|---|---|---|---|
| 0.050399 | 0.781602 | 0.034 | 0.021 | 1 | 0 | Mrps18b | 0.013 | Old |
| 0.050415 | 0.138129 | 0.364 | 0.32 | 1 | 0 | Ms4a6b | 0.044 | Old |
| 0.050425 | 0.372887 | 0.055 | 0.038 | 1 | 0 | Scly | 0.017 | Old |
| 0.050476 | 0.257616 | 0.092 | 0.07 | 1 | 0 | Smarcb1 | 0.022 | Old |
| 0.050482 | 0.0884 | 0.988 | 0.988 | 1 | 0 | mt-Nd4 | 0 | Old |
| 0.050499 | 0.238108 | 0.295 | 0.258 | 1 | 0 | Slfn5 | 0.037 | Old |
| 0.050567 | 0.110937 | 0.079 | 0.058 | 1 | 0 | Dhps | 0.021 | Old |
| 0.050613 | 0.155385 | 0.405 | 0.368 | 1 | 0 | Tmed2 | 0.037 | Old |
| 0.050823 | 1.034185 | 0.014 | 0.006 | 1 | 0 | Zscan22 | 0.008 | Old |
| 0.05083 | 0.345785 | 0.145 | 0.119 | 1 | 0 | Cd300c2 | 0.026 | Old |
| 0.050862 | 0.322744 | 0.15 | 0.123 | 1 | 0 | Cyb5b | 0.027 | Old |
| 0.050925 | 0.093082 | 0.114 | 0.088 | 1 | 0 | Aldh3a2 | 0.026 | Old |
| 0.050934 | 0.403442 | 0.085 | 0.064 | 1 | 0 | Prpf6 | 0.021 | Old |
| 0.051109 | 0.380722 | 0.048 | 0.032 | 1 | 0 | Fpr2 | 0.016 | Old |
| 0.051117 | 0.276063 | 0.056 | 0.039 | 1 | 0 | Ttc4 | 0.017 | Old |
| 0.051136 | 0.978669 | 0.029 | 0.017 | 1 | 0 | Cox10 | 0.012 | Old |
| 0.05119 | 0.216098 | 0.107 | 0.083 | 1 | 0 | Aldh16a1 | 0.024 | Old |
| 0.051231 | 0.207227 | 0.301 | 0.265 | 1 | 0 | Tmed9 | 0.036 | Old |
| 0.051254 | 0.354364 | 0.085 | 0.064 | 1 | 0 | Scaf4 | 0.021 | Old |
| 0.051325 | 0.166125 | 0.269 | 0.23 | 1 | 0 | Dram2 | 0.039 | Old |
| 0.051423 | 0.239465 | 0.137 | 0.11 | 1 | 0 | Fundc1 | 0.027 | Old |
| 0.051502 | 0.914065 | 0.026 | 0.015 | 1 | 0 | Zfp46 | 0.011 | Old |
| 0.051611 | 0.077899 | 0.443 | 0.39 | 1 | 0 | Ppp1r18 | 0.053 | Old |
| 0.051865 | 0.110311 | 0.186 | 0.151 | 1 | 0 | Yipf5 | 0.035 | Old |
| 0.051878 | 0.455308 | 0.07 | 0.051 | 1 | 0 | Fem1a | 0.019 | Old |
| 0.052101 | 0.322095 | 0.109 | 0.086 | 1 | 0 | Unc119 | 0.023 | Old |
| 0.052102 | 0.23572 | 0.112 | 0.088 | 1 | 0 | Echs1 | 0.024 | Old |
| 0.052134 | 0.14432 | 0.516 | 0.468 | 1 | 0 | Rab1a | 0.048 | Old |
| 0.052311 | 0.299323 | 0.06 | 0.043 | 1 | 0 | Pde4dip | 0.017 | Old |
| 0.052373 | 0.581316 | 0.059 | 0.042 | 1 | 0 | Coq2 | 0.017 | Old |
| 0.052404 | 0.054136 | 0.343 | 0.296 | 1 | 0 | Ier3ip1 | 0.047 | Old |
| 0.052452 | 0.327801 | 0.052 | 0.036 | 1 | 0 | Tmem9 | 0.016 | Old |
| 0.05259 | 0.349887 | 0.04 | 0.026 | 1 | 0 | Nectin4 | 0.014 | Old |
| 0.052613 | 0.653783 | 0.026 | 0.015 | 1 | 0 | Rps6ka2 | 0.011 | Old |
| 0.052697 | 0.152007 | 0.39 | 0.348 | 1 | 0 | Csrnp1 | 0.042 | Old |
| 0.052707 | 0.349373 | 0.052 | 0.036 | 1 | 0 | Mrto4 | 0.016 | Old |
| 0.052768 | 0.335665 | 0.014 | 0.006 | 1 | 0 | Arl16 | 0.008 | Old |
| 0.052792 | 0.489575 | 0.019 | 0.01 | 1 | 0 | Trim30b | 0.009 | Old |
| 0.052794 | 0.531239 | 0.041 | 0.027 | 1 | 0 | Ctu2 | 0.014 | Old |
| 0.052898 | 0.000205 | 0.295 | 0.246 | 1 | 0 | Usp7 | 0.049 | Old |
| 0.053126 | 0.516518 | 0.019 | 0.01 | 1 | 0 | Myo10 | 0.009 | Old |
| 0.053794 | 0.297745 | 0.123 | 0.098 | 1 | 0 | Luc7l | 0.025 | Old |
| 0.053801 | 0.615437 | 0.042 | 0.028 | 1 | 0 | Arel1 | 0.014 | Old |
| 0.053875 | 0.174748 | 0.164 | 0.134 | 1 | 0 | Abcf1 | 0.03 | Old |
| 0.053888 | 0.993644 | 0.012 | 0.005 | 1 | 0 | Nr6a1 | 0.007 | Old |

| | | | | | | | | |
|---|---|---|---|---|---|---|---|---|
| 0.053985 | 0.248078 | 0.019 | 0.01 | 1 | 0 | Tyw5 | 0.009 | Old |
| 0.054021 | 0.256548 | 0.141 | 0.114 | 1 | 0 | Tpst2 | 0.027 | Old |
| 0.054046 | 0.146414 | 0.405 | 0.364 | 1 | 0 | Spred1 | 0.041 | Old |
| 0.054083 | 0.119157 | 0.282 | 0.241 | 1 | 0 | Cers2 | 0.041 | Old |
| 0.054097 | 0.408974 | 0.067 | 0.049 | 1 | 0 | Heatr3 | 0.018 | Old |
| 0.05413 | 0.183388 | 0.193 | 0.16 | 1 | 0 | Trim12c | 0.033 | Old |
| 0.054131 | 0.723213 | 0.012 | 0.005 | 1 | 0 | Tlr11 | 0.007 | Old |
| 0.054204 | 0.308271 | 0.064 | 0.046 | 1 | 0 | Mfge8 | 0.018 | Old |
| 0.054212 | 0.820446 | 0.012 | 0.005 | 1 | 0 | Nlrp10 | 0.007 | Old |
| 0.054294 | 0.796325 | 0.012 | 0.005 | 1 | 0 | C1qtnf6 | 0.007 | Old |
| 0.054416 | 0.191978 | 0.423 | 0.378 | 1 | 0 | Socs3 | 0.045 | Old |
| 0.054499 | 0.07759 | 0.995 | 0.982 | 1 | 0 | Jund | 0.013 | Old |
| 0.054504 | 0.313797 | 0.107 | 0.083 | 1 | 0 | Tmem222 | 0.024 | Old |
| 0.054542 | 0.092026 | 0.368 | 0.318 | 1 | 0 | Eif4b | 0.05 | Old |
| 0.054622 | 0.384228 | 0.026 | 0.015 | 1 | 0 | Atg4a | 0.011 | Old |
| 0.054724 | 0.290421 | 0.145 | 0.118 | 1 | 0 | Mbp | 0.027 | Old |
| 0.054824 | 0.576381 | 0.012 | 0.005 | 1 | 0 | Hoxb3 | 0.007 | Old |
| 0.054854 | 0.242945 | 0.083 | 0.063 | 1 | 0 | Gsap | 0.02 | Old |
| 0.055096 | 0.264957 | 0.179 | 0.149 | 1 | 0 | Plod1 | 0.03 | Old |
| 0.055116 | 0.096117 | 0.624 | 0.565 | 1 | 0 | Ccnl1 | 0.059 | Old |
| 0.055247 | 0.872584 | 0.022 | 0.012 | 1 | 0 | Magee1 | 0.01 | Old |
| 0.0553 | 0.135102 | 0.267 | 0.227 | 1 | 0 | Supt16 | 0.04 | Old |
| 0.055315 | 0.323212 | 0.196 | 0.165 | 1 | 0 | Mospd2 | 0.031 | Old |
| 0.055434 | 0.203408 | 0.168 | 0.139 | 1 | 0 | Commd7 | 0.029 | Old |
| 0.055459 | 0.157071 | 0.244 | 0.207 | 1 | 0 | Ccnt2 | 0.037 | Old |
| 0.055605 | 0.457629 | 0.094 | 0.073 | 1 | 0 | Cog5 | 0.021 | Old |
| 0.055696 | 0.149308 | 0.213 | 0.179 | 1 | 0 | Vasp | 0.034 | Old |
| 0.055769 | 0.191122 | 0.506 | 0.458 | 1 | 0 | Atp5g2 | 0.048 | Old |
| 0.0558 | 0.506493 | 0.022 | 0.012 | 1 | 0 | Il18r1 | 0.01 | Old |
| 0.055878 | 0.140065 | 0.145 | 0.117 | 1 | 0 | Lmbrd2 | 0.028 | Old |
| 0.055884 | 0.240565 | 0.031 | 0.019 | 1 | 0 | Sort1 | 0.012 | Old |
| 0.055903 | 0.828185 | 0.011 | 0.005 | 1 | 0 | Cmbl | 0.006 | Old |
| 0.056032 | 0.267316 | 0.059 | 0.042 | 1 | 0 | Ppfibp1 | 0.017 | Old |
| 0.056065 | 1.005059 | 0.01 | 0.004 | 1 | 0 | A530032D1 | 0.006 | Old |
| 0.056232 | 0.190294 | 0.253 | 0.217 | 1 | 0 | Zmynd11 | 0.036 | Old |
| 0.056263 | 0.780988 | 0.011 | 0.005 | 1 | 0 | C330027C0 | 0.006 | Old |
| 0.056264 | 0.838222 | 0.037 | 0.024 | 1 | 0 | Exosc1 | 0.013 | Old |
| 0.056276 | 0.329085 | 0.181 | 0.151 | 1 | 0 | Baz1a | 0.03 | Old |
| 0.056307 | 0.07097 | 0.227 | 0.19 | 1 | 0 | Usp19 | 0.037 | Old |
| 0.056353 | 0.514432 | 0.011 | 0.005 | 1 | 0 | Pou5f1 | 0.006 | Old |
| 0.056375 | 0.227436 | 0.103 | 0.08 | 1 | 0 | Hck | 0.023 | Old |
| 0.056457 | 0.637736 | 0.01 | 0.004 | 1 | 0 | Tmc4 | 0.006 | Old |
| 0.056562 | 0.065677 | 0.512 | 0.451 | 1 | 0 | Eid1 | 0.061 | Old |
| 0.056605 | 0.427069 | 0.01 | 0.004 | 1 | 0 | Mettl18 | 0.006 | Old |
| 0.056755 | 0.165653 | 0.535 | 0.495 | 1 | 0 | Vsir | 0.04 | Old |

| | | | | | | | | |
|---|---|---|---|---|---|---|---|---|
| 0.05685 | 0.154312 | 0.282 | 0.241 | 1 | 0 | Dad1 | 0.041 | Old |
| 0.056862 | 0.305647 | 0.242 | 0.211 | 1 | 0 | Furin | 0.031 | Old |
| 0.0569 | 0.304009 | 0.155 | 0.126 | 1 | 0 | Derl2 | 0.029 | Old |
| 0.057049 | 0.306029 | 0.071 | 0.052 | 1 | 0 | Anapc10 | 0.019 | Old |
| 0.057382 | 0.144327 | 0.155 | 0.126 | 1 | 0 | Stk24 | 0.029 | Old |
| 0.057442 | 0.393095 | 0.068 | 0.05 | 1 | 0 | Sepsecs | 0.018 | Old |
| 0.057734 | 0.213771 | 0.19 | 0.16 | 1 | 0 | Hs6st1 | 0.03 | Old |
| 0.057768 | 0.643589 | 0.025 | 0.014 | 1 | 0 | Oas3 | 0.011 | Old |
| 0.057923 | 0.181316 | 0.063 | 0.045 | 1 | 0 | Tbc1d10a | 0.018 | Old |
| 0.058354 | 0.127189 | 0.406 | 0.362 | 1 | 0 | Stat3 | 0.044 | Old |
| 0.058538 | 0.102321 | 0.053 | 0.037 | 1 | 0 | Rce1 | 0.016 | Old |
| 0.058758 | 0.406651 | 0.101 | 0.079 | 1 | 0 | Mbd1 | 0.022 | Old |
| 0.058764 | 0.116497 | 0.205 | 0.17 | 1 | 0 | Uqcrc2 | 0.035 | Old |
| 0.058792 | 0.289272 | 0.052 | 0.036 | 1 | 0 | Hilpda | 0.016 | Old |
| 0.059086 | 0.328389 | 0.056 | 0.04 | 1 | 0 | Gramd1b | 0.016 | Old |
| 0.059088 | 0.090602 | 0.192 | 0.159 | 1 | 0 | Kmt5b | 0.033 | Old |
| 0.059247 | 0.257887 | 0.185 | 0.156 | 1 | 0 | Atg3 | 0.029 | Old |
| 0.059337 | 0.300054 | 0.085 | 0.064 | 1 | 0 | Slc41a2 | 0.021 | Old |
| 0.059364 | 0.20169 | 0.116 | 0.092 | 1 | 0 | Nfu1 | 0.024 | Old |
| 0.059518 | 0.184507 | 0.104 | 0.08 | 1 | 0 | Mpnd | 0.024 | Old |
| 0.059569 | 0.148091 | 0.111 | 0.087 | 1 | 0 | Fam118b | 0.024 | Old |
| 0.05957 | 0.23092 | 0.066 | 0.048 | 1 | 0 | H2-Ke6 | 0.018 | Old |
| 0.059723 | 0.714791 | 0.034 | 0.022 | 1 | 0 | Cipc | 0.012 | Old |
| 0.059752 | 0.199053 | 0.208 | 0.176 | 1 | 0 | Snrpg | 0.032 | Old |
| 0.059765 | 0.375834 | 0.04 | 0.026 | 1 | 0 | Wdfy1 | 0.014 | Old |
| 0.060076 | 0.204925 | 0.176 | 0.147 | 1 | 0 | Smarca2 | 0.029 | Old |
| 0.060158 | 0.117206 | 0.358 | 0.313 | 1 | 0 | H2afy | 0.045 | Old |
| 0.060162 | 0.405266 | 0.071 | 0.053 | 1 | 0 | Utp6 | 0.018 | Old |
| 0.060455 | 0.17731 | 0.033 | 0.021 | 1 | 0 | Akr1b8 | 0.012 | Old |
| 0.060486 | 0.328733 | 0.115 | 0.091 | 1 | 0 | Ormdl2 | 0.024 | Old |
| 0.060528 | 0.140354 | 0.313 | 0.273 | 1 | 0 | B4galt1 | 0.04 | Old |
| 0.060531 | 0.301623 | 0.093 | 0.072 | 1 | 0 | Rfc2 | 0.021 | Old |
| 0.060678 | 0.063589 | 0.382 | 0.332 | 1 | 0 | Dpysl2 | 0.05 | Old |
| 0.060696 | 0.102713 | 0.384 | 0.337 | 1 | 0 | Selenot | 0.047 | Old |
| 0.060795 | 0.16865 | 0.267 | 0.232 | 1 | 0 | Ppp1r15b | 0.035 | Old |
| 0.060832 | 0.285113 | 0.115 | 0.091 | 1 | 0 | Prdx3 | 0.024 | Old |
| 0.060843 | 0.102802 | 0.254 | 0.217 | 1 | 0 | Gm16286 | 0.037 | Old |
| 0.06087 | 0.56832 | 0.062 | 0.045 | 1 | 0 | Arhgef11 | 0.017 | Old |
| 0.06107 | 0.245382 | 0.328 | 0.288 | 1 | 0 | Trir | 0.04 | Old |
| 0.061118 | 0.010255 | 0.133 | 0.105 | 1 | 0 | Txndc15 | 0.028 | Old |
| 0.061657 | 0.588473 | 0.021 | 0.011 | 1 | 0 | Ercc8 | 0.01 | Old |
| 0.061726 | 0.173434 | 0.166 | 0.136 | 1 | 0 | Tiprl | 0.03 | Old |
| 0.06192 | 0.248158 | 0.13 | 0.104 | 1 | 0 | Pptc7 | 0.026 | Old |
| 0.061946 | 0.225666 | 0.131 | 0.106 | 1 | 0 | Rprd2 | 0.025 | Old |
| 0.061985 | 0.140903 | 0.364 | 0.321 | 1 | 0 | Bmp2k | 0.043 | Old |

| | | | | | | | | |
|---|---|---|---|---|---|---|---|---|
| 0.062 | 0.146531 | 0.218 | 0.184 | 1 | 0 | Stat6 | 0.034 | Old |
| 0.06205 | 0.322784 | 0.164 | 0.136 | 1 | 0 | Naa15 | 0.028 | Old |
| 0.062175 | 0.03816 | 0.157 | 0.127 | 1 | 0 | Erlin1 | 0.03 | Old |
| 0.062279 | 0.103142 | 0.213 | 0.18 | 1 | 0 | Atxn2 | 0.033 | Old |
| 0.062365 | 0.415331 | 0.016 | 0.008 | 1 | 0 | Zfp949 | 0.008 | Old |
| 0.062398 | 0.065024 | 0.245 | 0.207 | 1 | 0 | Glyr1 | 0.038 | Old |
| 0.062435 | 0.173236 | 0.161 | 0.132 | 1 | 0 | Bmt2 | 0.029 | Old |
| 0.062487 | 0.058466 | 0.079 | 0.059 | 1 | 0 | Ctbs | 0.02 | Old |
| 0.062532 | 0.125903 | 0.891 | 0.855 | 1 | 0 | Clec10a | 0.036 | Old |
| 0.062627 | 0.201405 | 0.063 | 0.045 | 1 | 0 | Dgcr8 | 0.018 | Old |
| 0.062675 | 0.348739 | 0.066 | 0.048 | 1 | 0 | Naglu | 0.018 | Old |
| 0.062706 | 0.203687 | 0.172 | 0.144 | 1 | 0 | Rsbn1l | 0.028 | Old |
| 0.062742 | 0.473821 | 0.062 | 0.045 | 1 | 0 | D430042O( | 0.017 | Old |
| 0.06299 | 0.333422 | 0.064 | 0.047 | 1 | 0 | 1190002N1 | 0.017 | Old |
| 0.063196 | 0.452769 | 0.057 | 0.041 | 1 | 0 | Haus8 | 0.016 | Old |
| 0.063334 | 0.092129 | 0.208 | 0.173 | 1 | 0 | Smarcd2 | 0.035 | Old |
| 0.063336 | 0.206291 | 0.105 | 0.083 | 1 | 0 | Senp1 | 0.022 | Old |
| 0.063361 | 0.383801 | 0.067 | 0.049 | 1 | 0 | Lyst | 0.018 | Old |
| 0.063467 | 0.296777 | 0.346 | 0.313 | 1 | 0 | Tfrc | 0.033 | Old |
| 0.063564 | 0.407798 | 0.075 | 0.056 | 1 | 0 | Gm13212 | 0.019 | Old |
| 0.06357 | 0.488846 | 0.071 | 0.053 | 1 | 0 | Daxx | 0.018 | Old |
| 0.063983 | 0.091092 | 0.547 | 0.492 | 1 | 0 | Rab8b | 0.055 | Old |
| 0.06416 | 0.378158 | 0.075 | 0.056 | 1 | 0 | Pias2 | 0.019 | Old |
| 0.064193 | 0.14634 | 0.105 | 0.082 | 1 | 0 | Dld | 0.023 | Old |
| 0.064254 | 0.102215 | 0.416 | 0.369 | 1 | 0 | Rab11b | 0.047 | Old |
| 0.064331 | 0.194161 | 0.119 | 0.096 | 1 | 0 | Utp3 | 0.023 | Old |
| 0.064347 | 0.258066 | 0.094 | 0.073 | 1 | 0 | Snapc1 | 0.021 | Old |
| 0.064753 | 0.123416 | 0.324 | 0.284 | 1 | 0 | Tmbim1 | 0.04 | Old |
| 0.064822 | 0.035232 | 0.118 | 0.092 | 1 | 0 | Mcrs1 | 0.026 | Old |
| 0.064892 | 0.118986 | 0.726 | 0.667 | 1 | 0 | Rbms1 | 0.059 | Old |
| 0.064931 | 0.144435 | 0.44 | 0.392 | 1 | 0 | Nktr | 0.048 | Old |
| 0.064937 | 0.208614 | 0.256 | 0.222 | 1 | 0 | Mib1 | 0.034 | Old |
| 0.064977 | 0.142075 | 0.345 | 0.301 | 1 | 0 | Cdc42se2 | 0.044 | Old |
| 0.065111 | 0.206935 | 0.369 | 0.335 | 1 | 0 | S100a4 | 0.034 | Old |
| 0.065124 | 0.778919 | 0.041 | 0.028 | 1 | 0 | Mllt11 | 0.013 | Old |
| 0.065124 | 0.462793 | 0.031 | 0.02 | 1 | 0 | Dnajc18 | 0.011 | Old |
| 0.065465 | 0.115534 | 0.421 | 0.372 | 1 | 0 | Tmed7 | 0.049 | Old |
| 0.065471 | 0.164897 | 0.045 | 0.031 | 1 | 0 | Gmpr2 | 0.014 | Old |
| 0.065502 | 0.658681 | 0.033 | 0.021 | 1 | 0 | Osgin1 | 0.012 | Old |
| 0.065563 | 0.127425 | 0.123 | 0.098 | 1 | 0 | Cmas | 0.025 | Old |
| 0.065612 | 0.155139 | 0.053 | 0.037 | 1 | 0 | Nt5c3 | 0.016 | Old |
| 0.065711 | 0.114779 | 0.754 | 0.728 | 1 | 0 | Lamp2 | 0.026 | Old |
| 0.065787 | 0.516478 | 0.048 | 0.033 | 1 | 0 | Cblb | 0.015 | Old |
| 0.065883 | 0.137822 | 0.442 | 0.396 | 1 | 0 | Gng2 | 0.046 | Old |
| 0.065885 | 0.227661 | 0.068 | 0.05 | 1 | 0 | Ppp1r8 | 0.018 | Old |

| | | | | | | | | |
|---|---|---|---|---|---|---|---|---|
| 0.065963 | 0.800321 | 0.015 | 0.008 | 1 | 0 | Hps6 | 0.007 | Old |
| 0.06612 | 0.390948 | 0.086 | 0.067 | 1 | 0 | B4galnt1 | 0.019 | Old |
| 0.066131 | 0.730902 | 0.033 | 0.021 | 1 | 0 | Trpm2 | 0.012 | Old |
| 0.066223 | 0.632798 | 0.04 | 0.027 | 1 | 0 | Smg8 | 0.013 | Old |
| 0.066397 | 0.111375 | 0.557 | 0.498 | 1 | 0 | Tubb5 | 0.059 | Old |
| 0.066504 | 0.820559 | 0.019 | 0.011 | 1 | 0 | Cpne8 | 0.008 | Old |
| 0.066747 | 0.059409 | 0.395 | 0.344 | 1 | 0 | Usf2 | 0.051 | Old |
| 0.067321 | 0.183203 | 0.345 | 0.305 | 1 | 0 | Hsd17b11 | 0.04 | Old |
| 0.067334 | 0.11196 | 0.261 | 0.224 | 1 | 0 | Kpnb1 | 0.037 | Old |
| 0.067363 | 0.352214 | 0.097 | 0.076 | 1 | 0 | Pdcd11 | 0.021 | Old |
| 0.067448 | 0.048715 | 0.479 | 0.428 | 1 | 0 | Laptm4a | 0.051 | Old |
| 0.067474 | 0.24966 | 0.168 | 0.141 | 1 | 0 | Tsg101 | 0.027 | Old |
| 0.067563 | 0.118132 | 0.141 | 0.113 | 1 | 0 | Cnot8 | 0.028 | Old |
| 0.067605 | 0.583086 | 0.015 | 0.008 | 1 | 0 | Kcnj2 | 0.007 | Old |
| 0.067667 | 0.403922 | 0.088 | 0.067 | 1 | 0 | Lipo3 | 0.021 | Old |
| 0.067669 | 0.397105 | 0.066 | 0.048 | 1 | 0 | Gm44751 | 0.018 | Old |
| 0.067702 | 0.089579 | 0.051 | 0.07 | 1 | 0 | Ptgs2 | -0.019 | Old |
| 0.067814 | 0.080751 | 0.239 | 0.203 | 1 | 0 | Nbeal1 | 0.036 | Old |
| 0.067861 | 0.317657 | 0.186 | 0.16 | 1 | 0 | Uggt1 | 0.026 | Old |
| 0.068077 | 0.488265 | 0.033 | 0.021 | 1 | 0 | Fam117a | 0.012 | Old |
| 0.068201 | 0.320475 | 0.105 | 0.083 | 1 | 0 | Zdhhc21 | 0.022 | Old |
| 0.068462 | 0.897054 | 0.027 | 0.017 | 1 | 0 | Lrrc20 | 0.01 | Old |
| 0.06849 | 0.118203 | 0.234 | 0.199 | 1 | 0 | Csnk1e | 0.035 | Old |
| 0.068629 | 0.306885 | 0.079 | 0.06 | 1 | 0 | Taf5l | 0.019 | Old |
| 0.068738 | 0.0854 | 0.271 | 0.233 | 1 | 0 | Lman2 | 0.038 | Old |
| 0.068745 | 0.41119 | 0.163 | 0.136 | 1 | 0 | Ide | 0.027 | Old |
| 0.068862 | 0.69919 | 0.037 | 0.024 | 1 | 0 | Klhl15 | 0.013 | Old |
| 0.068902 | 0.439577 | 0.089 | 0.069 | 1 | 0 | Lrpprc | 0.02 | Old |
| 0.069138 | 0.231626 | 0.17 | 0.142 | 1 | 0 | Zfp516 | 0.028 | Old |
| 0.069205 | 0.27586 | 0.114 | 0.091 | 1 | 0 | Prkag1 | 0.023 | Old |
| 0.069476 | 0.098831 | 0.829 | 0.8 | 1 | 0 | Rap1b | 0.029 | Old |
| 0.069666 | 0.274469 | 0.152 | 0.125 | 1 | 0 | Frg1 | 0.027 | Old |
| 0.069689 | 0.596763 | 0.051 | 0.036 | 1 | 0 | Dvl2 | 0.015 | Old |
| 0.070133 | 0.226255 | 0.092 | 0.071 | 1 | 0 | Rtp4 | 0.021 | Old |
| 0.070196 | 0.109621 | 0.237 | 0.202 | 1 | 0 | Exoc5 | 0.035 | Old |
| 0.070233 | 0.339764 | 0.04 | 0.027 | 1 | 0 | B230307C2 | 0.013 | Old |
| 0.070416 | 0.398881 | 0.033 | 0.021 | 1 | 0 | Lrrc61 | 0.012 | Old |
| 0.070441 | 0.445526 | 0.092 | 0.072 | 1 | 0 | Anapc4 | 0.02 | Old |
| 0.070656 | 0.198385 | 0.215 | 0.182 | 1 | 0 | Dynlt3 | 0.033 | Old |
| 0.070733 | 0.424466 | 0.03 | 0.019 | 1 | 0 | Irf7 | 0.011 | Old |
| 0.070807 | 0.898589 | 0.06 | 0.044 | 1 | 0 | Mmp19 | 0.016 | Old |
| 0.070876 | 0.288177 | 0.114 | 0.091 | 1 | 0 | Cyp20a1 | 0.023 | Old |
| 0.071026 | 0.136936 | 0.192 | 0.16 | 1 | 0 | Sav1 | 0.032 | Old |
| 0.071073 | 0.860453 | 0.025 | 0.015 | 1 | 0 | Gm20300 | 0.01 | Old |
| 0.071121 | 0.37945 | 0.103 | 0.081 | 1 | 0 | Selenon | 0.022 | Old |

| | | | | | | | | |
|---|---|---|---|---|---|---|---|---|
| 0.071157 | 0.116139 | 0.107 | 0.083 | 1 | 0 | Rbm12 | 0.024 | Old |
| 0.071167 | 0.113139 | 0.17 | 0.14 | 1 | 0 | Ociad1 | 0.03 | Old |
| 0.071252 | 0.133449 | 0.265 | 0.23 | 1 | 0 | Cct4 | 0.035 | Old |
| 0.071258 | 0.467297 | 0.398 | 0.361 | 1 | 0 | Lpl | 0.037 | Old |
| 0.071285 | 0.113593 | 0.436 | 0.388 | 1 | 0 | Sppl2a | 0.048 | Old |
| 0.071403 | 0.233309 | 0.161 | 0.135 | 1 | 0 | Nabp1 | 0.026 | Old |
| 0.071437 | 0.282497 | 0.078 | 0.059 | 1 | 0 | Wsb2 | 0.019 | Old |
| 0.071461 | 0.273586 | 0.118 | 0.094 | 1 | 0 | Dnmt1 | 0.024 | Old |
| 0.071544 | 0.458826 | 0.057 | 0.042 | 1 | 0 | Evi2 | 0.015 | Old |
| 0.071553 | 0.228169 | 0.155 | 0.127 | 1 | 0 | Cs | 0.028 | Old |
| 0.07159 | 0.197526 | 0.111 | 0.088 | 1 | 0 | Kif13b | 0.023 | Old |
| 0.071632 | 0.067735 | 0.213 | 0.18 | 1 | 0 | Dhx9 | 0.033 | Old |
| 0.071723 | 0.199724 | 0.209 | 0.179 | 1 | 0 | Cnih4 | 0.03 | Old |
| 0.071939 | 0.19677 | 0.12 | 0.096 | 1 | 0 | Aip | 0.024 | Old |
| 0.071978 | 0.181658 | 0.068 | 0.051 | 1 | 0 | Actr8 | 0.017 | Old |
| 0.072052 | 0.270854 | 0.063 | 0.046 | 1 | 0 | Faap100 | 0.017 | Old |
| 0.07214 | 0.179468 | 0.208 | 0.178 | 1 | 0 | Ubxn4 | 0.03 | Old |
| 0.072191 | 0.09371 | 0.383 | 0.338 | 1 | 0 | Ubap2l | 0.045 | Old |
| 0.07221 | 0.43659 | 0.096 | 0.075 | 1 | 0 | Zfp322a | 0.021 | Old |
| 0.072274 | 0.102881 | 0.32 | 0.279 | 1 | 0 | Spcs2 | 0.041 | Old |
| 0.072451 | 0.134651 | 0.16 | 0.132 | 1 | 0 | Ncf4 | 0.028 | Old |
| 0.072649 | 0.310915 | 0.104 | 0.083 | 1 | 0 | Zcchc17 | 0.021 | Old |
| 0.072853 | 0.107552 | 0.32 | 0.28 | 1 | 0 | Ndufa3 | 0.04 | Old |
| 0.072918 | 0.101876 | 0.421 | 0.37 | 1 | 0 | Abi1 | 0.051 | Old |
| 0.073222 | 0.461681 | 0.014 | 0.007 | 1 | 0 | Ripply3 | 0.007 | Old |
| 0.073257 | 0.390601 | 0.073 | 0.055 | 1 | 0 | Hat1 | 0.018 | Old |
| 0.073461 | 0.057446 | 0.208 | 0.174 | 1 | 0 | 4931406C0 | 0.034 | Old |
| 0.073541 | 0.124251 | 0.116 | 0.092 | 1 | 0 | Ppm1h | 0.024 | Old |
| 0.073674 | 0.229862 | 0.142 | 0.117 | 1 | 0 | Setdb1 | 0.025 | Old |
| 0.073675 | 0.297561 | 0.185 | 0.157 | 1 | 0 | Atf7ip | 0.028 | Old |
| 0.073815 | 0.893131 | 0.018 | 0.01 | 1 | 0 | Stox2 | 0.008 | Old |
| 0.073961 | 0.154625 | 0.254 | 0.219 | 1 | 0 | Arhgef2 | 0.035 | Old |
| 0.074062 | 0.088836 | 0.399 | 0.352 | 1 | 0 | Fam174a | 0.047 | Old |
| 0.074204 | 0.452545 | 0.037 | 0.024 | 1 | 0 | Zfp113 | 0.013 | Old |
| 0.074217 | 0.256781 | 0.064 | 0.047 | 1 | 0 | Slc10a7 | 0.017 | Old |
| 0.074267 | 0.196593 | 0.668 | 0.636 | 1 | 0 | Ddx3x | 0.032 | Old |
| 0.074349 | 0.69894 | 0.056 | 0.041 | 1 | 0 | Abhd2 | 0.015 | Old |
| 0.07448 | 0.413658 | 0.052 | 0.037 | 1 | 0 | Clk2 | 0.015 | Old |
| 0.074608 | 0.110007 | 0.265 | 0.229 | 1 | 0 | Myo9b | 0.036 | Old |
| 0.074631 | 0.164898 | 0.32 | 0.278 | 1 | 0 | Tsc22d1 | 0.042 | Old |
| 0.07464 | 0.512589 | 0.018 | 0.01 | 1 | 0 | Gm15859 | 0.008 | Old |
| 0.074686 | 0.187334 | 0.098 | 0.077 | 1 | 0 | Pogz | 0.021 | Old |
| 0.074752 | 0.158271 | 0.118 | 0.094 | 1 | 0 | Psme3 | 0.024 | Old |
| 0.074891 | 0.030069 | 0.052 | 0.037 | 1 | 0 | Naf1 | 0.015 | Old |
| 0.075013 | 0.820683 | 0.029 | 0.018 | 1 | 0 | Ddx20 | 0.011 | Old |

| | | | | | | | | |
|---|---|---|---|---|---|---|---|---|
| 0.075181 | 0.558632 | 0.092 | 0.072 | 1 | 0 | Crebzf | 0.02 | Old |
| 0.075435 | 0.115086 | 0.393 | 0.349 | 1 | 0 | Synj1 | 0.044 | Old |
| 0.075563 | 0.246611 | 0.097 | 0.076 | 1 | 0 | Suclg1 | 0.021 | Old |
| 0.075652 | 0.590754 | 0.04 | 0.027 | 1 | 0 | Med27 | 0.013 | Old |
| 0.075751 | 0.148913 | 0.135 | 0.11 | 1 | 0 | Chfr | 0.025 | Old |
| 0.075935 | 0.139983 | 0.491 | 0.443 | 1 | 0 | Ctdsp2 | 0.048 | Old |
| 0.075936 | 0.430021 | 0.092 | 0.072 | 1 | 0 | Entpd7 | 0.02 | Old |
| 0.076201 | 0.047679 | 0.037 | 0.024 | 1 | 0 | F8a | 0.013 | Old |
| 0.076382 | 0.310348 | 0.137 | 0.113 | 1 | 0 | Ube2v2 | 0.024 | Old |
| 0.076564 | 0.808149 | 0.012 | 0.006 | 1 | 0 | Phldb3 | 0.006 | Old |
| 0.076595 | 0.200901 | 0.146 | 0.121 | 1 | 0 | Shoc2 | 0.025 | Old |
| 0.076612 | 0.179594 | 0.09 | 0.07 | 1 | 0 | Slc46a3 | 0.02 | Old |
| 0.076628 | 0.244967 | 0.034 | 0.022 | 1 | 0 | Bloc1s3 | 0.012 | Old |
| 0.076782 | 0.157842 | 0.287 | 0.251 | 1 | 0 | Sgk3 | 0.036 | Old |
| 0.076936 | 0.289697 | 0.047 | 0.032 | 1 | 0 | Zw10 | 0.015 | Old |
| 0.077101 | 0.087095 | 0.347 | 0.303 | 1 | 0 | Txn2 | 0.044 | Old |
| 0.0772 | 0.758945 | 0.012 | 0.006 | 1 | 0 | 0610040B1 | 0.006 | Old |
| 0.077227 | 0.193396 | 0.302 | 0.269 | 1 | 0 | Klf7 | 0.033 | Old |
| 0.077533 | 0.046376 | 0.555 | 0.503 | 1 | 0 | G3bp2 | 0.052 | Old |
| 0.077589 | 0.064497 | 0.172 | 0.142 | 1 | 0 | Aup1 | 0.03 | Old |
| 0.077592 | 0.177157 | 0.387 | 0.344 | 1 | 0 | Nfkb1 | 0.043 | Old |
| 0.077783 | 0.174594 | 0.137 | 0.111 | 1 | 0 | Pdxdc1 | 0.026 | Old |
| 0.077841 | 0.493665 | 0.012 | 0.006 | 1 | 0 | Siah1b | 0.006 | Old |
| 0.077879 | 0.269877 | 0.038 | 0.026 | 1 | 0 | Tmem184c | 0.012 | Old |
| 0.078043 | 0.174596 | 0.131 | 0.107 | 1 | 0 | Fundc2 | 0.024 | Old |
| 0.07817 | 0.562502 | 0.029 | 0.018 | 1 | 0 | Ttc21b | 0.011 | Old |
| 0.078259 | 0.327985 | 0.14 | 0.116 | 1 | 0 | Psmd14 | 0.024 | Old |
| 0.078267 | 0.016121 | 0.114 | 0.09 | 1 | 0 | Trnt1 | 0.024 | Old |
| 0.078314 | 0.250628 | 0.131 | 0.107 | 1 | 0 | Osbp | 0.024 | Old |
| 0.078358 | 0.085192 | 0.655 | 0.598 | 1 | 0 | Rac1 | 0.057 | Old |
| 0.078451 | 0.094888 | 0.098 | 0.077 | 1 | 0 | Zbtb37 | 0.021 | Old |
| 0.078507 | 0.562642 | 0.036 | 0.024 | 1 | 0 | D17H6S53I | 0.012 | Old |
| 0.078858 | 0.174725 | 0.101 | 0.08 | 1 | 0 | Mvb12a | 0.021 | Old |
| 0.078874 | 0.317759 | 0.109 | 0.088 | 1 | 0 | Fam98b | 0.021 | Old |
| 0.078885 | 0.293429 | 0.064 | 0.048 | 1 | 0 | Slc9a8 | 0.016 | Old |
| 0.078887 | 0.15941 | 0.161 | 0.134 | 1 | 0 | Chd3 | 0.027 | Old |
| 0.078938 | 0.153181 | 0.265 | 0.231 | 1 | 0 | Fndc3b | 0.034 | Old |
| 0.078972 | 0.324692 | 0.119 | 0.096 | 1 | 0 | Nipsnap3b | 0.023 | Old |
| 0.079186 | 0.641712 | 0.023 | 0.014 | 1 | 0 | Apba3 | 0.009 | Old |
| 0.079229 | 1.101146 | 0.016 | 0.009 | 1 | 0 | Cacna1a | 0.007 | Old |
| 0.079231 | 0.0932 | 0.194 | 0.162 | 1 | 0 | Rufy3 | 0.032 | Old |
| 0.079297 | 0.161405 | 0.092 | 0.071 | 1 | 0 | Tnip2 | 0.021 | Old |
| 0.0793 | 0.29569 | 0.14 | 0.115 | 1 | 0 | Arrdc4 | 0.025 | Old |
| 0.079328 | 0.321318 | 0.04 | 0.027 | 1 | 0 | Dbf4 | 0.013 | Old |
| 0.07937 | 0.222436 | 0.063 | 0.046 | 1 | 0 | Decr2 | 0.017 | Old |

| | | | | | | | | |
|---|---|---|---|---|---|---|---|---|
| 0.079392 | 0.319148 | 0.044 | 0.03 | 1 | 0 | Tmem18 | 0.014 | Old |
| 0.079471 | 0.266122 | 0.115 | 0.092 | 1 | 0 | Mrpl17 | 0.023 | Old |
| 0.079543 | 0.137134 | 0.424 | 0.382 | 1 | 0 | Gls | 0.042 | Old |
| 0.079674 | 0.414417 | 0.045 | 0.032 | 1 | 0 | Fam35a | 0.013 | Old |
| 0.079714 | 0.051343 | 0.175 | 0.145 | 1 | 0 | C330007P( | 0.03 | Old |
| 0.07978 | 0.622138 | 0.023 | 0.014 | 1 | 0 | Htr1b | 0.009 | Old |
| 0.080003 | 0.51284 | 0.023 | 0.014 | 1 | 0 | Fmnl3 | 0.009 | Old |
| 0.080045 | 0.435228 | 0.051 | 0.036 | 1 | 0 | Rrn3 | 0.015 | Old |
| 0.080066 | 0.254335 | 0.129 | 0.105 | 1 | 0 | Tmem131l | 0.024 | Old |
| 0.080248 | 0.187511 | 0.275 | 0.241 | 1 | 0 | Adgre5 | 0.034 | Old |
| 0.080456 | 0.122756 | 0.207 | 0.175 | 1 | 0 | Cept1 | 0.032 | Old |
| 0.080464 | 0.93135 | 0.016 | 0.009 | 1 | 0 | Grhpr | 0.007 | Old |
| 0.080699 | 0.361235 | 0.029 | 0.018 | 1 | 0 | Nup107 | 0.011 | Old |
| 0.08074 | 1.040548 | 0.016 | 0.009 | 1 | 0 | Parp16 | 0.007 | Old |
| 0.080753 | 0.433197 | 0.056 | 0.041 | 1 | 0 | Pabpc4 | 0.015 | Old |
| 0.080864 | 0.281635 | 0.048 | 0.034 | 1 | 0 | Ppm1d | 0.014 | Old |
| 0.080885 | 0.366869 | 0.157 | 0.132 | 1 | 0 | Prpf39 | 0.025 | Old |
| 0.081109 | 0.096829 | 0.259 | 0.22 | 1 | 0 | H2-M3 | 0.039 | Old |
| 0.081221 | 0.623023 | 0.011 | 0.005 | 1 | 0 | Hist1h2ap | 0.006 | Old |
| 0.081253 | 0.636523 | 0.115 | 0.094 | 1 | 0 | Osm | 0.021 | Old |
| 0.081269 | 0.696654 | 0.042 | 0.029 | 1 | 0 | Adcy3 | 0.013 | Old |
| 0.081329 | 0.204071 | 0.036 | 0.024 | 1 | 0 | Pcyox1l | 0.012 | Old |
| 0.08134 | 0.84762 | 0.011 | 0.005 | 1 | 0 | Igfbp7 | 0.006 | Old |
| 0.081346 | 0.099582 | 0.07 | 0.052 | 1 | 0 | Ambra1 | 0.018 | Old |
| 0.081369 | 0.084918 | 0.389 | 0.343 | 1 | 0 | Ets2 | 0.046 | Old |
| 0.081403 | 0.193906 | 0.036 | 0.024 | 1 | 0 | Nt5m | 0.012 | Old |
| 0.081516 | 0.334349 | 0.026 | 0.016 | 1 | 0 | 1110004E( | 0.01 | Old |
| 0.081637 | 0.886733 | 0.011 | 0.005 | 1 | 0 | Cyp2r1 | 0.006 | Old |
| 0.081876 | 0.74979 | 0.011 | 0.005 | 1 | 0 | Ankrd66 | 0.006 | Old |
| 0.081904 | 0.104656 | 0.419 | 0.375 | 1 | 0 | Ube2d2a | 0.044 | Old |
| 0.08201 | 0.281094 | 0.134 | 0.111 | 1 | 0 | Xpot | 0.023 | Old |
| 0.08202 | 0.090963 | 0.601 | 0.552 | 1 | 0 | Emp3 | 0.049 | Old |
| 0.082083 | 0.279303 | 0.127 | 0.105 | 1 | 0 | Abhd17a | 0.022 | Old |
| 0.082227 | 0.438835 | 0.016 | 0.009 | 1 | 0 | Msh5 | 0.007 | Old |
| 0.082358 | 0.181991 | 0.159 | 0.133 | 1 | 0 | Rnf114 | 0.026 | Old |
| 0.082383 | 0.154277 | 0.222 | 0.191 | 1 | 0 | Nus1 | 0.031 | Old |
| 0.082416 | 0.417703 | 0.011 | 0.005 | 1 | 0 | Gm12689 | 0.006 | Old |
| 0.08265 | 0.215016 | 0.23 | 0.201 | 1 | 0 | Smim7 | 0.029 | Old |
| 0.08266 | 0.210726 | 0.044 | 0.03 | 1 | 0 | Nat6 | 0.014 | Old |
| 0.082864 | 0.623547 | 0.03 | 0.019 | 1 | 0 | Hdac10 | 0.011 | Old |
| 0.083114 | 0.106382 | 0.077 | 0.058 | 1 | 0 | Stx2 | 0.019 | Old |
| 0.083148 | 0.317561 | 0.14 | 0.117 | 1 | 0 | Gripap1 | 0.023 | Old |
| 0.08329 | 0.323548 | 0.081 | 0.062 | 1 | 0 | Mrpl9 | 0.019 | Old |
| 0.083493 | 0.544982 | 0.03 | 0.019 | 1 | 0 | Emc9 | 0.011 | Old |
| 0.083576 | 0.171314 | 0.207 | 0.177 | 1 | 0 | Cdk11b | 0.03 | Old |

| | | | | | | | | |
|---|---|---|---|---|---|---|---|---|
| 0.083887 | 0.700881 | 0.049 | 0.035 | 1 | 0 | Dnajc30 | 0.014 | Old |
| 0.083974 | 0.063928 | 0.174 | 0.144 | 1 | 0 | Mrpl14 | 0.03 | Old |
| 0.084143 | 0.810198 | 0.025 | 0.015 | 1 | 0 | Sirpb1c | 0.01 | Old |
| 0.084173 | 0.255451 | 0.092 | 0.072 | 1 | 0 | Terf2 | 0.02 | Old |
| 0.084367 | 0.588795 | 0.027 | 0.017 | 1 | 0 | Ccdc167 | 0.01 | Old |
| 0.084383 | 0.296917 | 0.104 | 0.083 | 1 | 0 | Trmt1l | 0.021 | Old |
| 0.084539 | 0.638661 | 0.038 | 0.026 | 1 | 0 | Zfp511 | 0.012 | Old |
| 0.084551 | 0.569879 | 0.034 | 0.023 | 1 | 0 | Vwa8 | 0.011 | Old |
| 0.084642 | 0.089206 | 0.328 | 0.29 | 1 | 0 | Osbpl9 | 0.038 | Old |
| 0.084683 | 0.085613 | 0.172 | 0.143 | 1 | 0 | Tmem9b | 0.029 | Old |
| 0.084691 | 0.346716 | 0.041 | 0.028 | 1 | 0 | Atp6v0d2 | 0.013 | Old |
| 0.08482 | 0.062246 | 0.081 | 0.062 | 1 | 0 | Nde1 | 0.019 | Old |
| 0.084844 | 0.114129 | 0.248 | 0.215 | 1 | 0 | Sel1l | 0.033 | Old |
| 0.084906 | 0.203004 | 0.033 | 0.021 | 1 | 0 | Pparg | 0.012 | Old |
| 0.085086 | 0.315621 | 0.089 | 0.07 | 1 | 0 | Cops6 | 0.019 | Old |
| 0.085185 | 0.159777 | 0.04 | 0.027 | 1 | 0 | Mccc1 | 0.013 | Old |
| 0.085196 | 0.171601 | 0.163 | 0.136 | 1 | 0 | Fam76b | 0.027 | Old |
| 0.085413 | 0.11902 | 0.339 | 0.299 | 1 | 0 | Tubb6 | 0.04 | Old |
| 0.085486 | 0.209422 | 0.189 | 0.162 | 1 | 0 | Trpv4 | 0.027 | Old |
| 0.085607 | 0.930512 | 0.01 | 0.004 | 1 | 0 | Gpd1 | 0.006 | Old |
| 0.085674 | 0.59018 | 0.01 | 0.004 | 1 | 0 | Ddr2 | 0.006 | Old |
| 0.085809 | 0.986742 | 0.01 | 0.004 | 1 | 0 | Il4 | 0.006 | Old |
| 0.085864 | 0.470347 | 0.038 | 0.026 | 1 | 0 | Bnip1 | 0.012 | Old |
| 0.085878 | 0.452768 | 0.03 | 0.019 | 1 | 0 | Tfpt | 0.011 | Old |
| 0.086006 | 0.164387 | 0.239 | 0.206 | 1 | 0 | 2610001J0 | 0.033 | Old |
| 0.086014 | 0.234555 | 0.213 | 0.185 | 1 | 0 | Sh3bp1 | 0.028 | Old |
| 0.086077 | 0.319775 | 0.064 | 0.048 | 1 | 0 | Csf2rb2 | 0.016 | Old |
| 0.086083 | 0.206776 | 0.067 | 0.05 | 1 | 0 | Thoc3 | 0.017 | Old |
| 0.086161 | 0.449338 | 0.038 | 0.026 | 1 | 0 | Rbm19 | 0.012 | Old |
| 0.086268 | 0.206693 | 0.045 | 0.032 | 1 | 0 | Homer3 | 0.013 | Old |
| 0.086289 | 0.25182 | 0.144 | 0.12 | 1 | 0 | Ubqln1 | 0.024 | Old |
| 0.086294 | 0.262375 | 0.056 | 0.041 | 1 | 0 | Ubl4a | 0.015 | Old |
| 0.086511 | 0.167775 | 0.23 | 0.2 | 1 | 0 | Vma21 | 0.03 | Old |
| 0.08654 | 1.115335 | 0.015 | 0.008 | 1 | 0 | Ccdc114 | 0.007 | Old |
| 0.086552 | 0.505614 | 0.01 | 0.004 | 1 | 0 | Cryab | 0.006 | Old |
| 0.086722 | 0.1678 | 0.197 | 0.168 | 1 | 0 | Azin1 | 0.029 | Old |
| 0.086928 | 0.132542 | 0.22 | 0.191 | 1 | 0 | Arid5a | 0.029 | Old |
| 0.086974 | 0.293605 | 0.037 | 0.025 | 1 | 0 | Zbtb46 | 0.012 | Old |
| 0.087076 | 0.285425 | 0.1 | 0.08 | 1 | 0 | Copg1 | 0.02 | Old |
| 0.087225 | 0.242077 | 0.047 | 0.033 | 1 | 0 | Pex16 | 0.014 | Old |
| 0.087294 | 0.103589 | 0.049 | 0.035 | 1 | 0 | Usp53 | 0.014 | Old |
| 0.08734 | 0.473157 | 0.027 | 0.017 | 1 | 0 | 1700066M | 0.01 | Old |
| 0.087392 | 0.420786 | 0.062 | 0.046 | 1 | 0 | Calcrl | 0.016 | Old |
| 0.087413 | 0.454265 | 0.027 | 0.017 | 1 | 0 | Aste1 | 0.01 | Old |
| 0.087465 | 0.951513 | 0.015 | 0.008 | 1 | 0 | Gm16973 | 0.007 | Old |

| | | | | | | | | |
|---|---|---|---|---|---|---|---|---|
| 0.087618 | 0.191736 | 0.071 | 0.054 | 1 | 0 | Fam96b | 0.017 | Old |
| 0.087658 | 0.15378 | 0.105 | 0.084 | 1 | 0 | Prmt1 | 0.021 | Old |
| 0.087816 | 0.172197 | 0.212 | 0.181 | 1 | 0 | Uvrag | 0.031 | Old |
| 0.087886 | 0.341602 | 0.027 | 0.017 | 1 | 0 | Ercc1 | 0.01 | Old |
| 0.088048 | 0.012752 | 0.196 | 0.164 | 1 | 0 | Fes | 0.032 | Old |
| 0.08818 | 0.145586 | 0.358 | 0.317 | 1 | 0 | Usp9x | 0.041 | Old |
| 0.088279 | 0.130701 | 0.208 | 0.177 | 1 | 0 | Sf3b3 | 0.031 | Old |
| 0.088467 | 0.052175 | 0.354 | 0.31 | 1 | 0 | Eif4ebp2 | 0.044 | Old |
| 0.088475 | 0.564044 | 0.059 | 0.044 | 1 | 0 | Nol4l | 0.015 | Old |
| 0.088535 | 0.255333 | 0.161 | 0.135 | 1 | 0 | Cpeb2 | 0.026 | Old |
| 0.088651 | 0.154176 | 0.09 | 0.071 | 1 | 0 | Sec16a | 0.019 | Old |
| 0.088658 | 0.647437 | 0.015 | 0.008 | 1 | 0 | Rpgr | 0.007 | Old |
| 0.088661 | 0.270712 | 0.09 | 0.071 | 1 | 0 | Dcaf17 | 0.019 | Old |
| 0.088671 | 0.15185 | 0.245 | 0.214 | 1 | 0 | Ubald2 | 0.031 | Old |
| 0.088705 | 0.075978 | 0.135 | 0.111 | 1 | 0 | Uqcc2 | 0.024 | Old |
| 0.088711 | 0.241347 | 0.286 | 0.257 | 1 | 0 | Tmem189 | 0.029 | Old |
| 0.088738 | 0.072637 | 0.198 | 0.168 | 1 | 0 | Zfx | 0.03 | Old |
| 0.088742 | 0.329726 | 0.078 | 0.06 | 1 | 0 | Tpra1 | 0.018 | Old |
| 0.088776 | 0.409282 | 0.257 | 0.23 | 1 | 0 | Mt2 | 0.027 | Old |
| 0.088911 | 0.35332 | 0.048 | 0.034 | 1 | 0 | Etfdh | 0.014 | Old |
| 0.088961 | 0.087845 | 0.269 | 0.235 | 1 | 0 | Dock10 | 0.034 | Old |
| 0.089074 | 0.256994 | 0.042 | 0.029 | 1 | 0 | Borcs8 | 0.013 | Old |
| 0.089096 | 0.172291 | 0.321 | 0.288 | 1 | 0 | Cdk2ap2 | 0.033 | Old |
| 0.089172 | 0.341827 | 0.098 | 0.079 | 1 | 0 | Vezt | 0.019 | Old |
| 0.089254 | 0.316098 | 0.164 | 0.139 | 1 | 0 | Nudt16l1 | 0.025 | Old |
| 0.089692 | 0.127621 | 0.16 | 0.133 | 1 | 0 | Tbc1d23 | 0.027 | Old |
| 0.089956 | 0.484986 | 0.057 | 0.043 | 1 | 0 | Snhg20 | 0.014 | Old |
| 0.090264 | 0.373739 | 0.025 | 0.015 | 1 | 0 | Mageh1 | 0.01 | Old |
| 0.090587 | 0.25166 | 0.103 | 0.082 | 1 | 0 | Sgk1 | 0.021 | Old |
| 0.090613 | 0.333119 | 0.081 | 0.063 | 1 | 0 | Cyp4f16 | 0.018 | Old |
| 0.09064 | 0.296726 | 0.094 | 0.075 | 1 | 0 | Med16 | 0.019 | Old |
| 0.090666 | 0.350711 | 0.049 | 0.035 | 1 | 0 | Tipin | 0.014 | Old |
| 0.090698 | 0.124415 | 0.166 | 0.138 | 1 | 0 | Ss18 | 0.028 | Old |
| 0.090827 | 0.131492 | 0.239 | 0.207 | 1 | 0 | Aldh9a1 | 0.032 | Old |
| 0.090855 | 0.065941 | 0.175 | 0.147 | 1 | 0 | Cct7 | 0.028 | Old |
| 0.090923 | 0.104768 | 0.409 | 0.364 | 1 | 0 | Runx1 | 0.045 | Old |
| 0.090977 | 0.431562 | 0.015 | 0.008 | 1 | 0 | Syn1 | 0.007 | Old |
| 0.091077 | 0.151837 | 0.226 | 0.194 | 1 | 0 | Il3ra | 0.032 | Old |
| 0.091281 | 0.27499 | 0.048 | 0.034 | 1 | 0 | Acap3 | 0.014 | Old |
| 0.091319 | 0.105383 | 0.28 | 0.243 | 1 | 0 | Daglb | 0.037 | Old |
| 0.091333 | 0.211643 | 0.038 | 0.026 | 1 | 0 | Zfp81 | 0.012 | Old |
| 0.091555 | 0.196789 | 0.176 | 0.149 | 1 | 0 | Gpatch8 | 0.027 | Old |
| 0.091787 | 0.070694 | 0.207 | 0.175 | 1 | 0 | Usp4 | 0.032 | Old |
| 0.091958 | 0.126508 | 0.14 | 0.115 | 1 | 0 | Yipf1 | 0.025 | Old |
| 0.092028 | 0.346404 | 0.092 | 0.073 | 1 | 0 | Tmem88 | 0.019 | Old |

| | | | | | | | | |
|---|---|---|---|---|---|---|---|---|
| 0.092067 | 0.310783 | 0.09 | 0.072 | 1 | 0 | Prelid3b | 0.018 | Old |
| 0.092092 | 0.155349 | 0.226 | 0.196 | 1 | 0 | Mrpl28 | 0.03 | Old |
| 0.092145 | 0.315994 | 0.105 | 0.085 | 1 | 0 | Riok1 | 0.02 | Old |
| 0.092329 | 0.135544 | 0.435 | 0.4 | 1 | 0 | Slc3a2 | 0.035 | Old |
| 0.092514 | 0.152879 | 0.308 | 0.275 | 1 | 0 | Rbx1 | 0.033 | Old |
| 0.092521 | 0.230203 | 0.168 | 0.143 | 1 | 0 | Ap1s1 | 0.025 | Old |
| 0.092696 | 0.044777 | 0.26 | 0.225 | 1 | 0 | Arglu1 | 0.035 | Old |
| 0.092712 | 0.149789 | 0.067 | 0.051 | 1 | 0 | Apeh | 0.016 | Old |
| 0.092721 | 0.126492 | 0.218 | 0.186 | 1 | 0 | Map3k8 | 0.032 | Old |
| 0.092906 | 0.462227 | 0.041 | 0.029 | 1 | 0 | Ino80b | 0.012 | Old |
| 0.093017 | 0.458509 | 0.045 | 0.032 | 1 | 0 | Cpsf1 | 0.013 | Old |
| 0.093044 | 0.170593 | 0.144 | 0.119 | 1 | 0 | Rps12-ps3 | 0.025 | Old |
| 0.093122 | 0.239299 | 0.135 | 0.112 | 1 | 0 | Zscan26 | 0.023 | Old |
| 0.093128 | 0.402341 | 0.077 | 0.059 | 1 | 0 | Bcl2l2 | 0.018 | Old |
| 0.093234 | 0.142722 | 0.692 | 0.672 | 1 | 0 | Cd63 | 0.02 | Old |
| 0.093238 | 0.185094 | 0.044 | 0.031 | 1 | 0 | Zkscan8 | 0.013 | Old |
| 0.093367 | 0.198638 | 0.074 | 0.056 | 1 | 0 | Rae1 | 0.018 | Old |
| 0.093449 | 0.149228 | 0.048 | 0.034 | 1 | 0 | Kif19a | 0.014 | Old |
| 0.093556 | 0.098424 | 0.156 | 0.13 | 1 | 0 | Mif | 0.026 | Old |
| 0.09381 | 0.102038 | 0.17 | 0.142 | 1 | 0 | Spopl | 0.028 | Old |
| 0.093915 | 0.233211 | 0.055 | 0.04 | 1 | 0 | Prr14l | 0.015 | Old |
| 0.093981 | 0.154368 | 0.436 | 0.398 | 1 | 0 | 2900097C1 | 0.038 | Old |
| 0.094019 | 0.502825 | 0.068 | 0.052 | 1 | 0 | Med25 | 0.016 | Old |
| 0.094097 | 0.318609 | 0.138 | 0.116 | 1 | 0 | Tnfaip2 | 0.022 | Old |
| 0.094236 | 0.31091 | 0.06 | 0.045 | 1 | 0 | Tmem161b | 0.015 | Old |
| 0.094503 | 0.098528 | 0.17 | 0.142 | 1 | 0 | Igbp1 | 0.028 | Old |
| 0.094755 | 0.341744 | 0.073 | 0.056 | 1 | 0 | Eya3 | 0.017 | Old |
| 0.094942 | 0.113244 | 0.36 | 0.321 | 1 | 0 | Nipbl | 0.039 | Old |
| 0.095007 | 0.118988 | 0.249 | 0.214 | 1 | 0 | Atp5o.1 | 0.035 | Old |
| 0.095092 | 0.124215 | 0.357 | 0.315 | 1 | 0 | Ppp1r9b | 0.042 | Old |
| 0.09516 | 0.159564 | 0.301 | 0.267 | 1 | 0 | Smim15 | 0.034 | Old |
| 0.095473 | 0.141419 | 0.371 | 0.329 | 1 | 0 | Trim25 | 0.042 | Old |
| 0.095903 | 0.733577 | 0.021 | 0.012 | 1 | 0 | Zfp764 | 0.009 | Old |
| 0.096001 | 0.269698 | 0.018 | 0.01 | 1 | 0 | 3110009E1 | 0.008 | Old |
| 0.096437 | 0.325256 | 0.073 | 0.056 | 1 | 0 | Umad1 | 0.017 | Old |
| 0.09678 | 0.222752 | 0.119 | 0.097 | 1 | 0 | Lrrc8d | 0.022 | Old |
| 0.096879 | 0.654002 | 0.051 | 0.037 | 1 | 0 | 9130401M | 0.014 | Old |
| 0.097362 | 0.352897 | 0.018 | 0.01 | 1 | 0 | Adal | 0.008 | Old |
| 0.097674 | 0.164502 | 0.138 | 0.114 | 1 | 0 | Ptprs | 0.024 | Old |
| 0.097723 | 0.234691 | 0.093 | 0.074 | 1 | 0 | 1600012H0 | 0.019 | Old |
| 0.097829 | 0.069188 | 0.313 | 0.273 | 1 | 0 | Hspe1 | 0.04 | Old |
| 0.097861 | 0.341569 | 0.033 | 0.022 | 1 | 0 | Rnf135 | 0.011 | Old |
| 0.098094 | 0.129511 | 0.078 | 0.06 | 1 | 0 | Abcd3 | 0.018 | Old |
| 0.098243 | 0.081465 | 0.209 | 0.179 | 1 | 0 | Tank | 0.03 | Old |
| 0.098305 | 0.612734 | 0.034 | 0.023 | 1 | 0 | Eid3 | 0.011 | Old |

| | | | | | | | | |
|---|---|---|---|---|---|---|---|---|
| 0.098434 | 0.266377 | 0.135 | 0.113 | 1 | 0 | C1d | 0.022 | Old |
| 0.098688 | 0.137505 | 0.38 | 0.341 | 1 | 0 | Fxyd5 | 0.039 | Old |
| 0.098753 | 0.110193 | 0.198 | 0.17 | 1 | 0 | Pank3 | 0.028 | Old |
| 0.09903 | 0.2736 | 0.041 | 0.029 | 1 | 0 | Gm45902 | 0.012 | Old |
| 0.099051 | 0.434834 | 0.056 | 0.042 | 1 | 0 | AL590144.1 | 0.014 | Old |
| 0.099067 | 0.178404 | 0.213 | 0.183 | 1 | 0 | Mcur1 | 0.03 | Old |
| 0.099146 | 0.117362 | 0.153 | 0.127 | 1 | 0 | Lpgat1 | 0.026 | Old |
| 0.099155 | 0.141822 | 0.15 | 0.126 | 1 | 0 | Ccdc115 | 0.024 | Old |
| 0.099192 | 0.13161 | 0.135 | 0.112 | 1 | 0 | Dnajb11 | 0.023 | Old |
| 0.099218 | 0.078498 | 0.631 | 0.602 | 1 | 0 | Pitpna | 0.029 | Old |
| 0.099219 | 0.328949 | 0.073 | 0.056 | 1 | 0 | Dhdds | 0.017 | Old |
| 0.099585 | 0.086917 | 0.105 | 0.084 | 1 | 0 | Mcoln1 | 0.021 | Old |
| 0.099589 | 0.26959 | 0.104 | 0.084 | 1 | 0 | Lrrc41 | 0.02 | Old |
| 0.099591 | 0.285585 | 0.141 | 0.118 | 1 | 0 | Wdr89 | 0.023 | Old |
| 0.099601 | 0.169116 | 0.216 | 0.186 | 1 | 0 | Ago3 | 0.03 | Old |
| 0.099658 | 0.074475 | 0.156 | 0.13 | 1 | 0 | Tia1 | 0.026 | Old |
| 0.099744 | 0.074255 | 0.14 | 0.115 | 1 | 0 | Lamtor5 | 0.025 | Old |
| 0.099773 | 0.380013 | 0.088 | 0.07 | 1 | 0 | Rbm15 | 0.018 | Old |
| 0.099841 | 0.263537 | 0.037 | 0.025 | 1 | 0 | Ms4a14 | 0.012 | Old |
| 0.099858 | 0.302222 | 0.101 | 0.082 | 1 | 0 | Usp32 | 0.019 | Old |
| 0.099878 | 0.375534 | 0.055 | 0.04 | 1 | 0 | Alg9 | 0.015 | Old |
| 0.099971 | 0.127283 | 0.204 | 0.174 | 1 | 0 | Stt3a | 0.03 | Old |
| 0.100064 | 0.20773 | 0.014 | 0.007 | 1 | 0 | Cep97 | 0.007 | Old |
| 0.10019 | 0.274411 | 0.149 | 0.126 | 1 | 0 | Pole4 | 0.023 | Old |
| 0.100309 | 0.626476 | 0.027 | 0.018 | 1 | 0 | Kbtbd4 | 0.009 | Old |
| 0.100565 | 0.357673 | 0.077 | 0.06 | 1 | 0 | Kif13a | 0.017 | Old |
| 0.100829 | 0.178875 | 0.148 | 0.123 | 1 | 0 | Nmt1 | 0.025 | Old |
| 0.100877 | 0.28381 | 0.063 | 0.048 | 1 | 0 | Pogk | 0.015 | Old |
| 0.100934 | 0.318043 | 0.037 | 0.025 | 1 | 0 | Ccdc61 | 0.012 | Old |
| 0.100946 | 0.378986 | 0.056 | 0.042 | 1 | 0 | Slc39a6 | 0.014 | Old |
| 0.101103 | 0.532751 | 0.052 | 0.038 | 1 | 0 | Metrn | 0.014 | Old |
| 0.101274 | 0.243563 | 0.145 | 0.121 | 1 | 0 | Mitf | 0.024 | Old |
| 0.101325 | 0.194099 | 0.192 | 0.165 | 1 | 0 | Rbm6 | 0.027 | Old |
| 0.101439 | 0.208109 | 0.052 | 0.038 | 1 | 0 | Dnaja3 | 0.014 | Old |
| 0.10161 | 0.212784 | 0.048 | 0.035 | 1 | 0 | 2510009E0 | 0.013 | Old |
| 0.101658 | 0.174343 | 0.078 | 0.06 | 1 | 0 | Mrpl10 | 0.018 | Old |
| 0.101664 | 0.02543 | 0.096 | 0.075 | 1 | 0 | Mboat7 | 0.021 | Old |
| 0.10178 | 0.151994 | 0.123 | 0.101 | 1 | 0 | Adgrg6 | 0.022 | Old |
| 0.101883 | 0.565404 | 0.051 | 0.037 | 1 | 0 | Lilra6 | 0.014 | Old |
| 0.102082 | 0.175621 | 0.03 | 0.02 | 1 | 0 | Mvd | 0.01 | Old |
| 0.102256 | 0.581543 | 0.027 | 0.018 | 1 | 0 | Prelid3a | 0.009 | Old |
| 0.102273 | 0.114012 | 0.074 | 0.057 | 1 | 0 | Rpa3 | 0.017 | Old |
| 0.102291 | 0.003751 | 0.146 | 0.12 | 1 | 0 | Rbm22 | 0.026 | Old |
| 0.102541 | 0.188796 | 0.133 | 0.109 | 1 | 0 | Ndufa12 | 0.024 | Old |
| 0.102721 | 0.055076 | 0.231 | 0.198 | 1 | 0 | Stard9 | 0.033 | Old |

| | | | | | | | | |
|---|---|---|---|---|---|---|---|---|
| 0.102741 | 0.017987 | 0.421 | 0.373 | 1 | 0 | Itpripl2 | 0.048 | Old |
| 0.102768 | 0.165087 | 0.092 | 0.072 | 1 | 0 | Cep250 | 0.02 | Old |
| 0.102817 | 0.649926 | 0.04 | 0.028 | 1 | 0 | Mid1 | 0.012 | Old |
| 0.103024 | 0.178317 | 0.098 | 0.079 | 1 | 0 | Asb8 | 0.019 | Old |
| 0.103306 | 0.110218 | 0.357 | 0.32 | 1 | 0 | Ewsr1 | 0.037 | Old |
| 0.103352 | 0.171322 | 0.204 | 0.174 | 1 | 0 | Sdhd | 0.03 | Old |
| 0.103391 | 0.065265 | 0.33 | 0.289 | 1 | 0 | Capza1 | 0.041 | Old |
| 0.103445 | 0.340667 | 0.03 | 0.02 | 1 | 0 | Spindoc | 0.01 | Old |
| 0.103469 | 0.215415 | 0.122 | 0.099 | 1 | 0 | Crtap | 0.023 | Old |
| 0.103485 | 0.418809 | 0.036 | 0.024 | 1 | 0 | Pigh | 0.012 | Old |
| 0.10385 | 0.196546 | 0.094 | 0.075 | 1 | 0 | Fubp3 | 0.019 | Old |
| 0.103932 | 0.250253 | 0.048 | 0.035 | 1 | 0 | Usp50 | 0.013 | Old |
| 0.104221 | 0.170838 | 0.06 | 0.045 | 1 | 0 | Gpn1 | 0.015 | Old |
| 0.104221 | 0.476125 | 0.052 | 0.038 | 1 | 0 | Nt5dc2 | 0.014 | Old |
| 0.104252 | 0.590963 | 0.036 | 0.024 | 1 | 0 | Prmt9 | 0.012 | Old |
| 0.10431 | 0.098228 | 0.074 | 0.057 | 1 | 0 | Ttc1 | 0.017 | Old |
| 0.104343 | 0.248589 | 0.055 | 0.04 | 1 | 0 | Narf | 0.015 | Old |
| 0.104591 | 0.257344 | 0.092 | 0.073 | 1 | 0 | Mxra7 | 0.019 | Old |
| 0.104694 | 0.476424 | 0.063 | 0.048 | 1 | 0 | Cacna1d | 0.015 | Old |
| 0.104695 | 0.42486 | 0.097 | 0.079 | 1 | 0 | Dst | 0.018 | Old |
| 0.104826 | 0.101702 | 0.152 | 0.126 | 1 | 0 | Ei24 | 0.026 | Old |
| 0.104864 | 0.182326 | 0.119 | 0.098 | 1 | 0 | Fam50a | 0.021 | Old |
| 0.104866 | 0.712933 | 0.025 | 0.016 | 1 | 0 | Lzts2 | 0.009 | Old |
| 0.105141 | 0.607365 | 0.016 | 0.009 | 1 | 0 | Thg1l | 0.007 | Old |
| 0.105272 | 0.864085 | 0.012 | 0.006 | 1 | 0 | Cdkl3 | 0.006 | Old |
| 0.105339 | 0.380011 | 0.012 | 0.006 | 1 | 0 | Car9 | 0.006 | Old |
| 0.105745 | 0.522431 | 0.055 | 0.041 | 1 | 0 | Pno1 | 0.014 | Old |
| 0.105752 | 0.027929 | 0.393 | 0.344 | 1 | 0 | Spi1 | 0.049 | Old |
| 0.105769 | 0.074196 | 0.311 | 0.275 | 1 | 0 | U2af2 | 0.036 | Old |
| 0.105842 | 0.54829 | 0.025 | 0.016 | 1 | 0 | Phf7 | 0.009 | Old |
| 0.106269 | 0.450378 | 0.016 | 0.009 | 1 | 0 | Gm26542 | 0.007 | Old |
| 0.106954 | 0.06169 | 0.012 | 0.006 | 1 | 0 | Btbd17 | 0.006 | Old |
| 0.107035 | 0.185873 | 0.1 | 0.08 | 1 | 0 | Hcfc2 | 0.02 | Old |
| 0.107124 | 0.237093 | 0.142 | 0.12 | 1 | 0 | Ube2f | 0.022 | Old |
| 0.107187 | 0.091129 | 0.289 | 0.253 | 1 | 0 | Derl1 | 0.036 | Old |
| 0.107203 | 0.247811 | 0.089 | 0.071 | 1 | 0 | Tbck | 0.018 | Old |
| 0.107321 | 0.41964 | 0.025 | 0.016 | 1 | 0 | Tspyl3 | 0.009 | Old |
| 0.10741 | 0.099876 | 0.077 | 0.059 | 1 | 0 | Nop56 | 0.018 | Old |
| 0.107564 | 0.33979 | 0.031 | 0.021 | 1 | 0 | Ssh3 | 0.01 | Old |
| 0.107703 | 0.270884 | 0.126 | 0.104 | 1 | 0 | Nup62 | 0.022 | Old |
| 0.107738 | 0.599898 | 0.022 | 0.013 | 1 | 0 | Crtc1 | 0.009 | Old |
| 0.107739 | 0.144467 | 0.164 | 0.139 | 1 | 0 | S100a11 | 0.025 | Old |
| 0.107808 | 0.095638 | 0.594 | 0.54 | 1 | 0 | Btf3 | 0.054 | Old |
| 0.107876 | 0.163521 | 0.092 | 0.073 | 1 | 0 | Psmc2 | 0.019 | Old |
| 0.108202 | 0.246065 | 0.079 | 0.062 | 1 | 0 | 1700037H( | 0.017 | Old |

| | | | | | | | | |
|---|---|---|---|---|---|---|---|---|
| 0.108418 | 0.63763 | 0.068 | 0.053 | 1 | 0 | Acot2 | 0.015 | Old |
| 0.108508 | 0.337185 | 0.052 | 0.038 | 1 | 0 | Dnttip1 | 0.014 | Old |
| 0.108531 | 0.16411 | 0.089 | 0.07 | 1 | 0 | Pdrg1 | 0.019 | Old |
| 0.108556 | 0.079374 | 0.492 | 0.447 | 1 | 0 | Plekho2 | 0.045 | Old |
| 0.108582 | 0.127579 | 0.375 | 0.334 | 1 | 0 | Rhog | 0.041 | Old |
| 0.108767 | 0.173842 | 0.22 | 0.19 | 1 | 0 | Rer1 | 0.03 | Old |
| 0.109044 | 0.086249 | 0.12 | 0.098 | 1 | 0 | Hibadh | 0.022 | Old |
| 0.10913 | 0.227818 | 0.197 | 0.171 | 1 | 0 | Cisd2 | 0.026 | Old |
| 0.109215 | 0.100592 | 0.256 | 0.225 | 1 | 0 | Rap1gds1 | 0.031 | Old |
| 0.109218 | 0.023067 | 0.202 | 0.172 | 1 | 0 | Thoc2 | 0.03 | Old |
| 0.109294 | 0.480157 | 0.056 | 0.042 | 1 | 0 | Tbc1d8 | 0.014 | Old |
| 0.109358 | 0.070669 | 0.201 | 0.172 | 1 | 0 | Ppp2r2a | 0.029 | Old |
| 0.109533 | 0.163798 | 0.088 | 0.07 | 1 | 0 | Ptgr1 | 0.018 | Old |
| 0.109705 | 0.281135 | 0.353 | 0.333 | 1 | 0 | Ccr5 | 0.02 | Old |
| 0.109819 | 0.714787 | 0.026 | 0.017 | 1 | 0 | Pank4 | 0.009 | Old |
| 0.109855 | 0.15407 | 0.189 | 0.163 | 1 | 0 | Mga | 0.026 | Old |
| 0.109902 | 0.16983 | 0.031 | 0.021 | 1 | 0 | Xrcc6 | 0.01 | Old |
| 0.109947 | 0.536637 | 0.089 | 0.072 | 1 | 0 | Rnpc3 | 0.017 | Old |
| 0.110324 | 0.259067 | 0.022 | 0.013 | 1 | 0 | Eif2d | 0.009 | Old |
| 0.110523 | 0.576927 | 0.044 | 0.032 | 1 | 0 | Rrp15 | 0.012 | Old |
| 0.11062 | 0.022079 | 0.529 | 0.47 | 1 | 0 | Reep3 | 0.059 | Old |
| 0.110629 | 0.458566 | 0.048 | 0.035 | 1 | 0 | Stmn1 | 0.013 | Old |
| 0.110638 | 0.081985 | 0.449 | 0.405 | 1 | 0 | Bsg | 0.044 | Old |
| 0.110707 | 0.093132 | 0.059 | 0.044 | 1 | 0 | 9330160F1 | 0.015 | Old |
| 0.110969 | 0.245608 | 0.234 | 0.204 | 1 | 0 | Rela | 0.03 | Old |
| 0.110995 | 0.308026 | 0.033 | 0.022 | 1 | 0 | Psmg1 | 0.011 | Old |
| 0.111115 | 0.888528 | 0.03 | 0.02 | 1 | 0 | Haus6 | 0.01 | Old |
| 0.111232 | 0.117226 | 0.019 | 0.011 | 1 | 0 | Chac2 | 0.008 | Old |
| 0.111274 | 0.37929 | 0.029 | 0.019 | 1 | 0 | Phykpl | 0.01 | Old |
| 0.11133 | 0.313319 | 0.062 | 0.047 | 1 | 0 | Srprb | 0.015 | Old |
| 0.111559 | 0.260318 | 0.059 | 0.044 | 1 | 0 | Zswim4 | 0.015 | Old |
| 0.111617 | 0.324026 | 0.077 | 0.06 | 1 | 0 | Bin3 | 0.017 | Old |
| 0.111618 | 0.175706 | 0.196 | 0.169 | 1 | 0 | Mbd2 | 0.027 | Old |
| 0.111621 | 0.179502 | 0.176 | 0.15 | 1 | 0 | Mgrn1 | 0.026 | Old |
| 0.111775 | 0.012605 | 0.168 | 0.141 | 1 | 0 | Stt3b | 0.027 | Old |
| 0.111832 | 0.128552 | 0.302 | 0.27 | 1 | 0 | Syncrip | 0.032 | Old |
| 0.111876 | 0.193812 | 0.096 | 0.077 | 1 | 0 | Ddx52 | 0.019 | Old |
| 0.111959 | 0.104646 | 0.07 | 0.053 | 1 | 0 | Ptpn9 | 0.017 | Old |
| 0.112307 | 0.35176 | 0.077 | 0.06 | 1 | 0 | Lsm2 | 0.017 | Old |
| 0.112557 | 0.157899 | 0.057 | 0.043 | 1 | 0 | Atp11a | 0.014 | Old |
| 0.112743 | 0.266763 | 0.111 | 0.091 | 1 | 0 | Acadsb | 0.02 | Old |
| 0.112787 | 0.397964 | 0.088 | 0.07 | 1 | 0 | Gopc | 0.018 | Old |
| 0.11301 | 0.076574 | 0.088 | 0.069 | 1 | 0 | Tbc1d1 | 0.019 | Old |
| 0.113313 | 0.654497 | 0.03 | 0.02 | 1 | 0 | Hoxb4 | 0.01 | Old |
| 0.113323 | 0.135884 | 0.168 | 0.143 | 1 | 0 | Stat5b | 0.025 | Old |

| | | | | | | | | |
|---|---|---|---|---|---|---|---|---|
| 0.113411 | 0.181554 | 0.041 | 0.029 | 1 | 0 | Wdr75 | 0.012 | Old |
| 0.113537 | 1.068359 | 0.015 | 0.008 | 1 | 0 | Ntmt1 | 0.007 | Old |
| 0.113541 | 0.284144 | 0.066 | 0.051 | 1 | 0 | Nomo1 | 0.015 | Old |
| 0.1138 | 0.250011 | 0.233 | 0.206 | 1 | 0 | Npc1 | 0.027 | Old |
| 0.113942 | 0.078982 | 0.237 | 0.204 | 1 | 0 | Ilk | 0.033 | Old |
| 0.114058 | 0.535212 | 0.047 | 0.034 | 1 | 0 | Map4k1 | 0.013 | Old |
| 0.114142 | 0.818434 | 0.034 | 0.024 | 1 | 0 | Tma16 | 0.01 | Old |
| 0.114166 | 0.178988 | 0.083 | 0.066 | 1 | 0 | Lnpk | 0.017 | Old |
| 0.114244 | 0.039609 | 0.368 | 0.321 | 1 | 0 | Gsk3b | 0.047 | Old |
| 0.114281 | 0.232533 | 0.152 | 0.128 | 1 | 0 | Adprh | 0.024 | Old |
| 0.114506 | 0.475504 | 0.026 | 0.017 | 1 | 0 | Ttc27 | 0.009 | Old |
| 0.114895 | 0.198869 | 0.189 | 0.163 | 1 | 0 | Pip5k1c | 0.026 | Old |
| 0.115006 | 0.136401 | 0.066 | 0.05 | 1 | 0 | Isoc1 | 0.016 | Old |
| 0.115036 | 0.704167 | 0.023 | 0.015 | 1 | 0 | Fastkd2 | 0.008 | Old |
| 0.11508 | 0.736725 | 0.011 | 0.005 | 1 | 0 | Cenpk | 0.006 | Old |
| 0.11508 | 0.634376 | 0.011 | 0.005 | 1 | 0 | Mks1 | 0.006 | Old |
| 0.115156 | 0.718018 | 0.011 | 0.005 | 1 | 0 | Afap1 | 0.006 | Old |
| 0.11534 | 0.122735 | 0.287 | 0.254 | 1 | 0 | Lasp1 | 0.033 | Old |
| 0.115366 | 0.471896 | 0.045 | 0.033 | 1 | 0 | Zbp1 | 0.012 | Old |
| 0.115462 | 0.143403 | 0.156 | 0.133 | 1 | 0 | Setd5 | 0.023 | Old |
| 0.115535 | 0.135656 | 0.279 | 0.247 | 1 | 0 | Adam17 | 0.032 | Old |
| 0.115667 | 0.178483 | 0.13 | 0.108 | 1 | 0 | Ptdss1 | 0.022 | Old |
| 0.115704 | 0.199077 | 0.067 | 0.051 | 1 | 0 | Plekha3 | 0.016 | Old |
| 0.115742 | 0.61011 | 0.015 | 0.008 | 1 | 0 | Nectin2 | 0.007 | Old |
| 0.116057 | 0.055876 | 0.181 | 0.152 | 1 | 0 | Coro1b | 0.029 | Old |
| 0.116165 | 0.242296 | 0.026 | 0.017 | 1 | 0 | Stk11ip | 0.009 | Old |
| 0.116247 | 0.114659 | 0.531 | 0.488 | 1 | 0 | Srgn | 0.043 | Old |
| 0.116255 | 0.631002 | 0.036 | 0.025 | 1 | 0 | Lncpint | 0.011 | Old |
| 0.116272 | 0.535045 | 0.04 | 0.028 | 1 | 0 | Sephs1 | 0.012 | Old |
| 0.11642 | 0.012847 | 0.249 | 0.214 | 1 | 0 | Tmem258 | 0.035 | Old |
| 0.116454 | 0.166306 | 0.098 | 0.08 | 1 | 0 | Flvcr1 | 0.018 | Old |
| 0.116467 | 0.139922 | 0.15 | 0.127 | 1 | 0 | Nsfl1c | 0.023 | Old |
| 0.116577 | 0.128662 | 0.094 | 0.075 | 1 | 0 | Kmt2b | 0.019 | Old |
| 0.116596 | 0.037828 | 0.249 | 0.217 | 1 | 0 | Psme2 | 0.032 | Old |
| 0.11662 | 0.157992 | 0.011 | 0.005 | 1 | 0 | Lysmd4 | 0.006 | Old |
| 0.116629 | 0.312853 | 0.026 | 0.017 | 1 | 0 | Zfp868 | 0.009 | Old |
| 0.116803 | 0.120444 | 0.146 | 0.123 | 1 | 0 | Psmb2 | 0.023 | Old |
| 0.116841 | 0.332299 | 0.059 | 0.045 | 1 | 0 | Pgp | 0.014 | Old |
| 0.11693 | 0.18716 | 0.011 | 0.005 | 1 | 0 | 2210408I2 | 0.006 | Old |
| 0.116998 | 0.595914 | 0.041 | 0.029 | 1 | 0 | Mrpl1 | 0.012 | Old |
| 0.11702 | 0.366034 | 0.044 | 0.032 | 1 | 0 | Ints2 | 0.012 | Old |
| 0.11706 | 0.07692 | 0.141 | 0.118 | 1 | 0 | Ak2 | 0.023 | Old |
| 0.117346 | 0.494677 | 0.053 | 0.04 | 1 | 0 | Atg4c | 0.013 | Old |
| 0.11735 | 0.138348 | 0.079 | 0.062 | 1 | 0 | Sms | 0.017 | Old |
| 0.117912 | 0.118308 | 0.044 | 0.032 | 1 | 0 | Cd226 | 0.012 | Old |

| | | | | | | | | |
|---|---|---|---|---|---|---|---|---|
| 0.117982 | 0.198301 | 0.07 | 0.054 | 1 | 0 | Tsga10 | 0.016 | Old |
| 0.118105 | 0.256177 | 0.063 | 0.048 | 1 | 0 | Crls1 | 0.015 | Old |
| 0.118139 | 0.194708 | 0.083 | 0.066 | 1 | 0 | Nabp2 | 0.017 | Old |
| 0.118158 | 0.16237 | 0.074 | 0.058 | 1 | 0 | Nip7 | 0.016 | Old |
| 0.118236 | 0.209219 | 0.172 | 0.149 | 1 | 0 | Mllt10 | 0.023 | Old |
| 0.118244 | 0.188014 | 0.212 | 0.184 | 1 | 0 | Tmem167 | 0.028 | Old |
| 0.118361 | 0.618581 | 0.021 | 0.013 | 1 | 0 | Gm26853 | 0.008 | Old |
| 0.118422 | 0.278078 | 0.07 | 0.054 | 1 | 0 | Ubl7 | 0.016 | Old |
| 0.11852 | 0.167671 | 0.142 | 0.119 | 1 | 0 | Zcrb1 | 0.023 | Old |
| 0.118589 | 0.249399 | 0.068 | 0.053 | 1 | 0 | Mbtps2 | 0.015 | Old |
| 0.11863 | 0.591346 | 0.021 | 0.013 | 1 | 0 | Eps8l1 | 0.008 | Old |
| 0.118656 | 0.338053 | 0.098 | 0.08 | 1 | 0 | Ngdn | 0.018 | Old |
| 0.118756 | 0.267061 | 0.09 | 0.072 | 1 | 0 | Med14 | 0.018 | Old |
| 0.118804 | 0.124578 | 0.228 | 0.201 | 1 | 0 | Vdac3 | 0.027 | Old |
| 0.118811 | 0.092041 | 0.059 | 0.044 | 1 | 0 | Ptpn23 | 0.015 | Old |
| 0.119106 | 0.514588 | 0.018 | 0.011 | 1 | 0 | Traf3ip3 | 0.007 | Old |
| 0.119194 | 0.235107 | 0.365 | 0.333 | 1 | 0 | Ptger4 | 0.032 | Old |
| 0.119326 | 0.084949 | 0.564 | 0.512 | 1 | 0 | Ubl3 | 0.052 | Old |
| 0.119331 | 0.0763 | 0.321 | 0.283 | 1 | 0 | Fgfr1op2 | 0.038 | Old |
| 0.119493 | 0.609432 | 0.021 | 0.013 | 1 | 0 | Fam26f | 0.008 | Old |
| 0.119515 | 0.166988 | 0.097 | 0.078 | 1 | 0 | Hmg20a | 0.019 | Old |
| 0.119572 | 0.317244 | 0.2 | 0.177 | 1 | 0 | Peak1 | 0.023 | Old |
| 0.119585 | 0.123411 | 0.224 | 0.193 | 1 | 0 | Ttyh3 | 0.031 | Old |
| 0.119657 | 0.420764 | 0.218 | 0.196 | 1 | 0 | Bmp2 | 0.022 | Old |
| 0.119947 | 0.076204 | 0.264 | 0.232 | 1 | 0 | Parvb | 0.032 | Old |
| 0.120014 | 0.268347 | 0.064 | 0.049 | 1 | 0 | Rab10os | 0.015 | Old |
| 0.120022 | 0.119425 | 0.179 | 0.153 | 1 | 0 | Gigyf1 | 0.026 | Old |
| 0.120144 | 0.499548 | 0.021 | 0.013 | 1 | 0 | Ppif | 0.008 | Old |
| 0.120152 | 0.220948 | 0.045 | 0.033 | 1 | 0 | Dpep2 | 0.012 | Old |
| 0.120277 | 0.276146 | 0.142 | 0.12 | 1 | 0 | Kcnq1ot1 | 0.022 | Old |
| 0.120374 | 0.118812 | 0.204 | 0.176 | 1 | 0 | Pik3ca | 0.028 | Old |
| 0.120459 | 0.038492 | 0.112 | 0.091 | 1 | 0 | Smurf2 | 0.021 | Old |
| 0.120562 | 0.168093 | 0.062 | 0.047 | 1 | 0 | Calcoco1 | 0.015 | Old |
| 0.120634 | 0.085334 | 0.275 | 0.241 | 1 | 0 | Wdr83os | 0.034 | Old |
| 0.120644 | 0.277216 | 0.023 | 0.015 | 1 | 0 | Zfp946 | 0.008 | Old |
| 0.120765 | 0.097276 | 0.282 | 0.251 | 1 | 0 | Mef2d | 0.031 | Old |
| 0.120805 | 0.334448 | 0.053 | 0.04 | 1 | 0 | Rtl5 | 0.013 | Old |
| 0.120809 | 0.224415 | 0.122 | 0.101 | 1 | 0 | Hnrnpll | 0.021 | Old |
| 0.120936 | 0.36722 | 0.018 | 0.011 | 1 | 0 | Zfp790 | 0.007 | Old |
| 0.121096 | 0.190236 | 0.049 | 0.036 | 1 | 0 | Cnot11 | 0.013 | Old |
| 0.121196 | 0.124625 | 0.053 | 0.04 | 1 | 0 | Ksr1 | 0.013 | Old |
| 0.121328 | 0.181128 | 0.051 | 0.037 | 1 | 0 | Nectin1 | 0.014 | Old |
| 0.121446 | 0.090225 | 0.267 | 0.233 | 1 | 0 | Aff4 | 0.034 | Old |
| 0.121471 | 0.321934 | 0.018 | 0.011 | 1 | 0 | Ydjc | 0.007 | Old |
| 0.12165 | 0.437081 | 0.018 | 0.011 | 1 | 0 | Gm14296 | 0.007 | Old |

| | | | | | | | | |
|---|---|---|---|---|---|---|---|---|
| 0.121679 | 0.111576 | 0.105 | 0.086 | 1 | 0 | Ube4a | 0.019 | Old |
| 0.121823 | 0.059007 | 0.728 | 0.678 | 1 | 0 | Eif4g2 | 0.05 | Old |
| 0.122057 | 0.538282 | 0.027 | 0.018 | 1 | 0 | Gm15445 | 0.009 | Old |
| 0.122145 | 0.174594 | 0.075 | 0.059 | 1 | 0 | Atp6v0a2 | 0.016 | Old |
| 0.122167 | 0.175195 | 0.09 | 0.072 | 1 | 0 | Slc38a7 | 0.018 | Old |
| 0.122221 | 0.182517 | 0.272 | 0.243 | 1 | 0 | Anxa7 | 0.029 | Old |
| 0.122438 | 0.26513 | 0.135 | 0.114 | 1 | 0 | Bcl2l1 | 0.021 | Old |
| 0.122841 | 0.232893 | 0.053 | 0.04 | 1 | 0 | Lsm1 | 0.013 | Old |
| 0.122848 | 0.080176 | 0.302 | 0.268 | 1 | 0 | Dnm2 | 0.034 | Old |
| 0.122877 | 0.112274 | 0.335 | 0.302 | 1 | 0 | Cd302 | 0.033 | Old |
| 0.122987 | 0.112344 | 0.149 | 0.126 | 1 | 0 | Slc30a5 | 0.023 | Old |
| 0.123321 | 1.098645 | 0.01 | 0.005 | 1 | 0 | Gm48420 | 0.005 | Old |
| 0.123407 | 0.202046 | 0.159 | 0.135 | 1 | 0 | Arl1 | 0.024 | Old |
| 0.123409 | 0.74455 | 0.01 | 0.005 | 1 | 0 | Usp54 | 0.005 | Old |
| 0.123496 | 1.225783 | 0.01 | 0.005 | 1 | 0 | Tiam2 | 0.005 | Old |
| 0.123671 | 0.886995 | 0.01 | 0.005 | 1 | 0 | E430024I0 | 0.005 | Old |
| 0.124241 | 0.134865 | 0.315 | 0.286 | 1 | 0 | Tspan3 | 0.029 | Old |
| 0.124285 | 0.14372 | 0.01 | 0.005 | 1 | 0 | 1700056E2 | 0.005 | Old |
| 0.124425 | 0.155199 | 0.212 | 0.184 | 1 | 0 | Adam9 | 0.028 | Old |
| 0.124755 | 0.085334 | 0.421 | 0.382 | 1 | 0 | Fyb | 0.039 | Old |
| 0.124813 | 0.383461 | 0.01 | 0.005 | 1 | 0 | Nptxr | 0.005 | Old |
| 0.124989 | 0.304925 | 0.01 | 0.005 | 1 | 0 | Hddc3 | 0.005 | Old |
| 0.125431 | 0.275606 | 0.01 | 0.005 | 1 | 0 | Serpinh1 | 0.005 | Old |
| 0.125482 | 0.224907 | 0.096 | 0.078 | 1 | 0 | Snrnp48 | 0.018 | Old |
| 0.125776 | 0.013234 | 0.186 | 0.158 | 1 | 0 | Mcts1 | 0.028 | Old |
| 0.125843 | 0.021367 | 0.226 | 0.196 | 1 | 0 | Dynlrb1 | 0.03 | Old |
| 0.125848 | 0.132192 | 0.144 | 0.121 | 1 | 0 | Snrnp200 | 0.023 | Old |
| 0.126071 | 0.458299 | 0.037 | 0.026 | 1 | 0 | Mbip | 0.011 | Old |
| 0.126395 | 0.445922 | 0.037 | 0.026 | 1 | 0 | Prim2 | 0.011 | Old |
| 0.126439 | 0.407543 | 0.042 | 0.031 | 1 | 0 | Prkdc | 0.011 | Old |
| 0.126447 | 0.128136 | 0.259 | 0.23 | 1 | 0 | Tbl1xr1 | 0.029 | Old |
| 0.126749 | 0.206118 | 0.271 | 0.244 | 1 | 0 | Gpr132 | 0.027 | Old |
| 0.126834 | 0.206589 | 0.153 | 0.131 | 1 | 0 | Glb1 | 0.022 | Old |
| 0.126837 | 0.079448 | 0.313 | 0.277 | 1 | 0 | Lbr | 0.036 | Old |
| 0.127086 | 0.625351 | 0.014 | 0.008 | 1 | 0 | Nr3c2 | 0.006 | Old |
| 0.12722 | 0.383424 | 0.079 | 0.063 | 1 | 0 | Snrpa1 | 0.016 | Old |
| 0.127291 | 0.041211 | 0.706 | 0.656 | 1 | 0 | Ptbp3 | 0.05 | Old |
| 0.127596 | 0.07832 | 0.308 | 0.277 | 1 | 0 | Tomm20 | 0.031 | Old |
| 0.127713 | 0.144647 | 0.227 | 0.2 | 1 | 0 | Tbcb | 0.027 | Old |
| 0.127714 | 0.253698 | 0.193 | 0.169 | 1 | 0 | Eif3b | 0.024 | Old |
| 0.12805 | 0.068793 | 0.118 | 0.096 | 1 | 0 | Rpl7l1 | 0.022 | Old |
| 0.128145 | 0.120749 | 0.107 | 0.087 | 1 | 0 | Fbxw7 | 0.02 | Old |
| 0.128208 | 0.450879 | 0.029 | 0.019 | 1 | 0 | Echdc1 | 0.01 | Old |
| 0.128213 | 0.124396 | 0.115 | 0.094 | 1 | 0 | Slc35a5 | 0.021 | Old |
| 0.128285 | 0.238149 | 0.157 | 0.134 | 1 | 0 | Gatad1 | 0.023 | Old |

| | | | | | | | | |
|---|---|---|---|---|---|---|---|---|
| 0.128349 | 0.614813 | 0.029 | 0.019 | 1 | 0 | Zmym6 | 0.01 | Old |
| 0.128546 | 0.067853 | 0.293 | 0.257 | 1 | 0 | Mcfd2 | 0.036 | Old |
| 0.12863 | 0.108205 | 0.098 | 0.08 | 1 | 0 | Polr2b | 0.018 | Old |
| 0.128651 | 0.080052 | 0.497 | 0.451 | 1 | 0 | 2010107E0 | 0.046 | Old |
| 0.128858 | 0.326676 | 0.056 | 0.043 | 1 | 0 | Agpat4 | 0.013 | Old |
| 0.128918 | 0.124983 | 0.249 | 0.218 | 1 | 0 | Ifngr2 | 0.031 | Old |
| 0.129059 | 0.039197 | 0.085 | 0.067 | 1 | 0 | Ddx1 | 0.018 | Old |
| 0.129486 | 0.866577 | 0.026 | 0.017 | 1 | 0 | Eif5a2 | 0.009 | Old |
| 0.129662 | 0.500725 | 0.062 | 0.048 | 1 | 0 | 9430038I0 | 0.014 | Old |
| 0.129668 | 0.068344 | 0.144 | 0.12 | 1 | 0 | Mast3 | 0.024 | Old |
| 0.129802 | 0.157793 | 0.252 | 0.225 | 1 | 0 | Hexb | 0.027 | Old |
| 0.129806 | 0.201694 | 0.209 | 0.182 | 1 | 0 | Usp15 | 0.027 | Old |
| 0.129813 | 0.085211 | 0.242 | 0.209 | 1 | 0 | Cat | 0.033 | Old |
| 0.129973 | 0.181427 | 0.073 | 0.057 | 1 | 0 | E2f4 | 0.016 | Old |
| 0.130084 | 0.185398 | 0.161 | 0.137 | 1 | 0 | Asxl2 | 0.024 | Old |
| 0.130293 | 0.478053 | 0.074 | 0.059 | 1 | 0 | Aga | 0.015 | Old |
| 0.130504 | 0.253299 | 0.14 | 0.118 | 1 | 0 | Rapgef2 | 0.022 | Old |
| 0.130544 | 0.031915 | 0.152 | 0.128 | 1 | 0 | Zfp217 | 0.024 | Old |
| 0.13061 | 0.14039 | 0.127 | 0.106 | 1 | 0 | Rps27rt | 0.021 | Old |
| 0.130679 | 0.224175 | 0.111 | 0.091 | 1 | 0 | Snrpa | 0.02 | Old |
| 0.130791 | 0.299064 | 0.033 | 0.023 | 1 | 0 | Ttc38 | 0.01 | Old |
| 0.130863 | 0.246025 | 0.063 | 0.048 | 1 | 0 | Eci2 | 0.015 | Old |
| 0.130914 | 0.158824 | 0.209 | 0.182 | 1 | 0 | Pabpn1 | 0.027 | Old |
| 0.130944 | 0.547344 | 0.034 | 0.024 | 1 | 0 | Rrnad1 | 0.01 | Old |
| 0.131246 | 0.051986 | 0.323 | 0.286 | 1 | 0 | Fuca1 | 0.037 | Old |
| 0.131433 | 0.038602 | 0.291 | 0.257 | 1 | 0 | Rsu1 | 0.034 | Old |
| 0.131453 | 0.347609 | 0.037 | 0.026 | 1 | 0 | Mier2 | 0.011 | Old |
| 0.131489 | 0.258294 | 0.198 | 0.176 | 1 | 0 | Yeats4 | 0.022 | Old |
| 0.13199 | 0.565828 | 0.019 | 0.012 | 1 | 0 | Tada2a | 0.007 | Old |
| 0.132004 | 0.386751 | 0.068 | 0.053 | 1 | 0 | Sirt1 | 0.015 | Old |
| 0.132028 | 0.506422 | 0.016 | 0.01 | 1 | 0 | Fitm2 | 0.006 | Old |
| 0.132475 | 0.149223 | 0.167 | 0.142 | 1 | 0 | Clptm1 | 0.025 | Old |
| 0.132624 | 0.101517 | 0.458 | 0.418 | 1 | 0 | Arf1 | 0.04 | Old |
| 0.132642 | 0.604384 | 0.026 | 0.017 | 1 | 0 | Cluap1 | 0.009 | Old |
| 0.132695 | 0.546152 | 0.047 | 0.035 | 1 | 0 | Gfm1 | 0.012 | Old |
| 0.132718 | 0.448221 | 0.019 | 0.012 | 1 | 0 | Crocc | 0.007 | Old |
| 0.132817 | 0.163432 | 0.168 | 0.145 | 1 | 0 | Rbfa | 0.023 | Old |
| 0.133163 | 0.092189 | 0.063 | 0.048 | 1 | 0 | Atg4d | 0.015 | Old |
| 0.133244 | 0.053017 | 0.135 | 0.113 | 1 | 0 | Dync1li1 | 0.022 | Old |
| 0.133291 | 0.518397 | 0.016 | 0.01 | 1 | 0 | Fam135a | 0.006 | Old |
| 0.133293 | 0.468712 | 0.081 | 0.065 | 1 | 0 | Pygo2 | 0.016 | Old |
| 0.133525 | 0.489481 | 0.016 | 0.01 | 1 | 0 | Tmem41a | 0.006 | Old |
| 0.133603 | 0.164654 | 0.14 | 0.117 | 1 | 0 | Sh3gl1 | 0.023 | Old |
| 0.133751 | 0.063606 | 0.053 | 0.04 | 1 | 0 | Aasdhppt | 0.013 | Old |
| 0.133762 | 0.28868 | 0.111 | 0.092 | 1 | 0 | Zfp281 | 0.019 | Old |

| | | | | | | | | |
|---|---|---|---|---|---|---|---|---|
| 0.133892 | 0.376844 | 0.016 | 0.01 | 1 | 0 | Pecr | 0.006 | Old |
| 0.133943 | 0.125686 | 0.071 | 0.056 | 1 | 0 | Soga1 | 0.015 | Old |
| 0.134005 | 0.396074 | 0.053 | 0.04 | 1 | 0 | Slc35b3 | 0.013 | Old |
| 0.134015 | 0.116497 | 0.209 | 0.182 | 1 | 0 | Mtch1 | 0.027 | Old |
| 0.134029 | 0.200568 | 0.044 | 0.032 | 1 | 0 | Tcp11l1 | 0.012 | Old |
| 0.134204 | 0.118311 | 0.205 | 0.179 | 1 | 0 | Ppp3cb | 0.026 | Old |
| 0.13428 | 0.12297 | 0.428 | 0.389 | 1 | 0 | Tra2a | 0.039 | Old |
| 0.134295 | 0.379865 | 0.016 | 0.01 | 1 | 0 | Palm | 0.006 | Old |
| 0.134295 | 0.287233 | 0.016 | 0.01 | 1 | 0 | Zfp605 | 0.006 | Old |
| 0.134429 | 0.285114 | 0.016 | 0.01 | 1 | 0 | Chchd6 | 0.006 | Old |
| 0.134438 | 0.266926 | 0.089 | 0.072 | 1 | 0 | Ptpn22 | 0.017 | Old |
| 0.134486 | 0.015641 | 0.352 | 0.311 | 1 | 0 | Aamp | 0.041 | Old |
| 0.134612 | 0.080289 | 0.332 | 0.298 | 1 | 0 | Cap1 | 0.034 | Old |
| 0.13498 | 0.063533 | 0.365 | 0.328 | 1 | 0 | Ssb | 0.037 | Old |
| 0.135002 | 0.472456 | 0.026 | 0.017 | 1 | 0 | Fbxw9 | 0.009 | Old |
| 0.135351 | 0.362869 | 0.019 | 0.012 | 1 | 0 | Mfsd4b4 | 0.007 | Old |
| 0.135403 | 0.058016 | 0.19 | 0.165 | 1 | 0 | Ube2i | 0.025 | Old |
| 0.135468 | 0.117387 | 0.141 | 0.118 | 1 | 0 | Iws1 | 0.023 | Old |
| 0.135526 | 0.152103 | 0.146 | 0.125 | 1 | 0 | Herc2 | 0.021 | Old |
| 0.135847 | 0.197797 | 0.1 | 0.082 | 1 | 0 | Iah1 | 0.018 | Old |
| 0.135852 | 0.091659 | 0.253 | 0.221 | 1 | 0 | Raly | 0.032 | Old |
| 0.135887 | 0.090494 | 0.109 | 0.09 | 1 | 0 | Ssr2 | 0.019 | Old |
| 0.136203 | 0.215063 | 0.14 | 0.118 | 1 | 0 | Hspa14 | 0.022 | Old |
| 0.136217 | 0.135235 | 0.019 | 0.012 | 1 | 0 | Elovl6 | 0.007 | Old |
| 0.136497 | 0.082582 | 0.257 | 0.225 | 1 | 0 | Ikzf1 | 0.032 | Old |
| 0.136624 | 0.254394 | 0.137 | 0.117 | 1 | 0 | Oxr1 | 0.02 | Old |
| 0.136853 | 0.084014 | 0.073 | 0.057 | 1 | 0 | Mb21d1 | 0.016 | Old |
| 0.137005 | 0.144875 | 0.109 | 0.091 | 1 | 0 | Nhp2 | 0.018 | Old |
| 0.137332 | 0.298621 | 0.04 | 0.029 | 1 | 0 | Nek1 | 0.011 | Old |
| 0.137343 | 0.29246 | 0.034 | 0.024 | 1 | 0 | Wdr47 | 0.01 | Old |
| 0.137472 | 0.115635 | 0.182 | 0.157 | 1 | 0 | Lpxn | 0.025 | Old |
| 0.137744 | 0.241136 | 0.126 | 0.107 | 1 | 0 | Pqbp1 | 0.019 | Old |
| 0.138016 | 0.110561 | 0.133 | 0.111 | 1 | 0 | Plpbp | 0.022 | Old |
| 0.138038 | 0.31531 | 0.053 | 0.04 | 1 | 0 | Letmd1 | 0.013 | Old |
| 0.138219 | 0.588816 | 0.027 | 0.019 | 1 | 0 | Laptm4b | 0.008 | Old |
| 0.138332 | 0.089888 | 0.176 | 0.151 | 1 | 0 | Nab1 | 0.025 | Old |
| 0.138411 | 0.143279 | 0.218 | 0.19 | 1 | 0 | Huwe1 | 0.028 | Old |
| 0.138591 | 0.345148 | 0.041 | 0.03 | 1 | 0 | Pramef8 | 0.011 | Old |
| 0.13861 | 0.150217 | 0.104 | 0.086 | 1 | 0 | Htatsf1 | 0.018 | Old |
| 0.138767 | 0.043331 | 0.148 | 0.125 | 1 | 0 | Timm10b | 0.023 | Old |
| 0.138825 | 0.262529 | 0.167 | 0.145 | 1 | 0 | Lmo4 | 0.022 | Old |
| 0.138985 | 0.623164 | 0.027 | 0.019 | 1 | 0 | Ulk3 | 0.008 | Old |
| 0.139081 | 0.286208 | 0.096 | 0.078 | 1 | 0 | Cnrip1 | 0.018 | Old |
| 0.139245 | 0.116381 | 0.189 | 0.163 | 1 | 0 | Eif2s3x | 0.026 | Old |
| 0.139437 | 0.479506 | 0.07 | 0.056 | 1 | 0 | Fam222b | 0.014 | Old |

| | | | | | | | | |
|---|---|---|---|---|---|---|---|---|
| 0.139638 | 0.280842 | 0.041 | 0.03 | 1 | 0 | Megf8 | 0.011 | Old |
| 0.139753 | 0.369104 | 0.048 | 0.036 | 1 | 0 | Fmn1 | 0.012 | Old |
| 0.13978 | 0.65869 | 0.027 | 0.019 | 1 | 0 | Mon1a | 0.008 | Old |
| 0.139951 | 0.230469 | 0.062 | 0.048 | 1 | 0 | Slc25a12 | 0.014 | Old |
| 0.140003 | 0.165075 | 0.115 | 0.096 | 1 | 0 | Gid8 | 0.019 | Old |
| 0.140003 | 0.909334 | 0.012 | 0.007 | 1 | 0 | Rflnb | 0.005 | Old |
| 0.140037 | 0.076893 | 0.197 | 0.171 | 1 | 0 | Ube2s | 0.026 | Old |
| 0.140534 | 0.037953 | 0.138 | 0.116 | 1 | 0 | Myo1g | 0.022 | Old |
| 0.140985 | 0.222736 | 0.062 | 0.048 | 1 | 0 | Sfxn2 | 0.014 | Old |
| 0.14137 | 0.073879 | 0.321 | 0.288 | 1 | 0 | Vps4b | 0.033 | Old |
| 0.141918 | 0.580977 | 0.038 | 0.028 | 1 | 0 | Rhno1 | 0.01 | Old |
| 0.141918 | 0.482626 | 0.081 | 0.065 | 1 | 0 | Pnkd | 0.016 | Old |
| 0.141924 | 0.26884 | 0.077 | 0.061 | 1 | 0 | Ddx56 | 0.016 | Old |
| 0.141956 | 0.197999 | 0.041 | 0.03 | 1 | 0 | Scimp | 0.011 | Old |
| 0.142021 | 0.097183 | 0.244 | 0.214 | 1 | 0 | Med13l | 0.03 | Old |
| 0.142057 | 0.062427 | 0.386 | 0.345 | 1 | 0 | Atp6v0e | 0.041 | Old |
| 0.142059 | 0.209132 | 0.133 | 0.112 | 1 | 0 | Nans | 0.021 | Old |
| 0.142318 | 0.073188 | 0.436 | 0.392 | 1 | 0 | Tspo | 0.044 | Old |
| 0.142395 | 0.32059 | 0.097 | 0.08 | 1 | 0 | Hbs1l | 0.017 | Old |
| 0.14248 | 0.218194 | 0.062 | 0.048 | 1 | 0 | Dubr | 0.014 | Old |
| 0.142906 | 0.114209 | 0.304 | 0.27 | 1 | 0 | Snx8 | 0.034 | Old |
| 0.14306 | 0.816102 | 0.021 | 0.013 | 1 | 0 | Ptch1 | 0.008 | Old |
| 0.143097 | 0.417633 | 0.049 | 0.037 | 1 | 0 | Hectd4 | 0.012 | Old |
| 0.143293 | 0.091771 | 0.09 | 0.073 | 1 | 0 | Yrdc | 0.017 | Old |
| 0.143393 | 0.163997 | 0.023 | 0.015 | 1 | 0 | Trit1 | 0.008 | Old |
| 0.143419 | 0.100141 | 0.071 | 0.056 | 1 | 0 | Matk | 0.015 | Old |
| 0.143584 | 0.605785 | 0.021 | 0.013 | 1 | 0 | Slc6a8 | 0.008 | Old |
| 0.143878 | 0.144309 | 0.053 | 0.04 | 1 | 0 | Mrps28 | 0.013 | Old |
| 0.143893 | 0.703085 | 0.021 | 0.013 | 1 | 0 | Disc1 | 0.008 | Old |
| 0.144252 | 0.059406 | 0.311 | 0.276 | 1 | 0 | Myl12b | 0.035 | Old |
| 0.144276 | 0.495209 | 0.137 | 0.117 | 1 | 0 | Tnip3 | 0.02 | Old |
| 0.144327 | 0.644232 | 0.021 | 0.013 | 1 | 0 | Miga2 | 0.008 | Old |
| 0.144464 | 0.864388 | 0.015 | 0.009 | 1 | 0 | Sac3d1 | 0.006 | Old |
| 0.14464 | 0.152089 | 0.104 | 0.086 | 1 | 0 | Itfg1 | 0.018 | Old |
| 0.144761 | 0.966935 | 0.015 | 0.009 | 1 | 0 | Gpr85 | 0.006 | Old |
| 0.144796 | 0.213748 | 0.144 | 0.122 | 1 | 0 | Keap1 | 0.022 | Old |
| 0.145062 | 0.060015 | 0.599 | 0.543 | 1 | 0 | Aif1 | 0.056 | Old |
| 0.14517 | 0.745438 | 0.015 | 0.009 | 1 | 0 | Gcat | 0.006 | Old |
| 0.145321 | 0.519069 | 0.021 | 0.013 | 1 | 0 | Noa1 | 0.008 | Old |
| 0.145452 | 0.060791 | 0.417 | 0.38 | 1 | 0 | Pou2f2 | 0.037 | Old |
| 0.145513 | 0.109433 | 0.097 | 0.079 | 1 | 0 | Zfp646 | 0.018 | Old |
| 0.145546 | 0.353096 | 0.031 | 0.022 | 1 | 0 | Zbtb42 | 0.009 | Old |
| 0.145739 | 0.121191 | 0.178 | 0.154 | 1 | 0 | Rbm27 | 0.024 | Old |
| 0.145792 | 0.261424 | 0.031 | 0.022 | 1 | 0 | Tab1 | 0.009 | Old |
| 0.145804 | 0.784817 | 0.015 | 0.009 | 1 | 0 | Zc3h11a | 0.006 | Old |

| | | | | | | | | |
|---|---|---|---|---|---|---|---|---|
| 0.145863 | 0.080249 | 0.415 | 0.373 | 1 | 0 | Set | 0.042 | Old |
| 0.146151 | 0.198919 | 0.156 | 0.134 | 1 | 0 | Rab5b | 0.022 | Old |
| 0.146158 | 0.055128 | 0.067 | 0.052 | 1 | 0 | Fhl3 | 0.015 | Old |
| 0.146338 | 0.098965 | 0.271 | 0.242 | 1 | 0 | Gna13 | 0.029 | Old |
| 0.146513 | 0.134109 | 0.194 | 0.168 | 1 | 0 | Trim47 | 0.026 | Old |
| 0.146536 | 0.16823 | 0.167 | 0.144 | 1 | 0 | Dock1 | 0.023 | Old |
| 0.146537 | 0.183943 | 0.456 | 0.425 | 1 | 0 | Birc3 | 0.031 | Old |
| 0.146696 | 0.068074 | 0.471 | 0.428 | 1 | 0 | Cox6b1 | 0.043 | Old |
| 0.146737 | 0.029555 | 0.997 | 1 | 1 | 0 | mt-Co2 | -0.003 | Old |
| 0.146828 | 0.281478 | 0.119 | 0.1 | 1 | 0 | Lrrfip1 | 0.019 | Old |
| 0.146846 | 0.1893 | 0.096 | 0.079 | 1 | 0 | Prr14 | 0.017 | Old |
| 0.146971 | 0.247078 | 0.037 | 0.027 | 1 | 0 | Iqce | 0.01 | Old |
| 0.147078 | 0.541708 | 0.015 | 0.009 | 1 | 0 | Ficd | 0.006 | Old |
| 0.147432 | 0.521462 | 0.018 | 0.011 | 1 | 0 | Yars2 | 0.007 | Old |
| 0.147432 | 0.063659 | 0.114 | 0.094 | 1 | 0 | Slc39a9 | 0.02 | Old |
| 0.147569 | 0.469591 | 0.018 | 0.011 | 1 | 0 | Tyms | 0.007 | Old |
| 0.147823 | 0.205961 | 0.042 | 0.031 | 1 | 0 | 2410004B1 | 0.011 | Old |
| 0.147911 | 0.471095 | 0.018 | 0.011 | 1 | 0 | Acp5 | 0.007 | Old |
| 0.147992 | 0.263201 | 0.037 | 0.027 | 1 | 0 | Cry1 | 0.01 | Old |
| 0.14808 | 0.29009 | 0.079 | 0.064 | 1 | 0 | Camk2g | 0.015 | Old |
| 0.148143 | 0.196751 | 0.056 | 0.043 | 1 | 0 | Casp9 | 0.013 | Old |
| 0.148254 | 0.288507 | 0.092 | 0.075 | 1 | 0 | Rab43 | 0.017 | Old |
| 0.148261 | 0.074183 | 0.384 | 0.347 | 1 | 0 | Klhl9 | 0.037 | Old |
| 0.148478 | 0.170378 | 0.135 | 0.115 | 1 | 0 | Unc50 | 0.02 | Old |
| 0.148507 | 0.106696 | 0.187 | 0.162 | 1 | 0 | Csnk2a1 | 0.025 | Old |
| 0.148597 | 0.493886 | 0.018 | 0.011 | 1 | 0 | Pop1 | 0.007 | Old |
| 0.148641 | 0.235702 | 0.13 | 0.111 | 1 | 0 | Gnl1 | 0.019 | Old |
| 0.14879 | 0.237255 | 0.052 | 0.04 | 1 | 0 | Nr1d2 | 0.012 | Old |
| 0.148843 | 0.124541 | 0.082 | 0.066 | 1 | 0 | Zfp24 | 0.016 | Old |
| 0.148942 | 0.078969 | 0.234 | 0.206 | 1 | 0 | 2810474O1 | 0.028 | Old |
| 0.148976 | 0.118577 | 0.311 | 0.279 | 1 | 0 | Snrpb2 | 0.032 | Old |
| 0.149082 | 0.245371 | 0.015 | 0.009 | 1 | 0 | Haus5 | 0.006 | Old |
| 0.149286 | 0.091884 | 0.906 | 0.89 | 1 | 0 | Gpx1 | 0.016 | Old |
| 0.14935 | 0.227095 | 0.097 | 0.08 | 1 | 0 | Virma | 0.017 | Old |
| 0.14952 | 0.367748 | 0.093 | 0.077 | 1 | 0 | Tmem128 | 0.016 | Old |
| 0.149573 | 0.069514 | 0.093 | 0.075 | 1 | 0 | Faim | 0.018 | Old |
| 0.149701 | 0.241341 | 0.018 | 0.011 | 1 | 0 | Acsf3 | 0.007 | Old |
| 0.149787 | 0.150841 | 0.105 | 0.087 | 1 | 0 | 2310022A1 | 0.018 | Old |
| 0.150233 | 0.120941 | 0.213 | 0.188 | 1 | 0 | Sri | 0.025 | Old |
| 0.150322 | 0.340073 | 0.053 | 0.041 | 1 | 0 | Ddx41 | 0.012 | Old |
| 0.15043 | 0.101391 | 0.127 | 0.107 | 1 | 0 | Klhl28 | 0.02 | Old |
| 0.150744 | 0.189022 | 0.189 | 0.167 | 1 | 0 | Eif4e | 0.022 | Old |
| 0.150761 | 0.280963 | 0.075 | 0.06 | 1 | 0 | Phactr4 | 0.015 | Old |
| 0.150837 | 0.252306 | 0.088 | 0.072 | 1 | 0 | Rbmxl1 | 0.016 | Old |
| 0.150926 | 0.171322 | 0.192 | 0.167 | 1 | 0 | Eif3i | 0.025 | Old |

| | | | | | | | | |
|---|---|---|---|---|---|---|---|---|
| 0.150979 | 0.107548 | 0.43 | 0.393 | 1 | 0 | Litaf | 0.037 | Old |
| 0.151019 | 0.165784 | 0.018 | 0.011 | 1 | 0 | Zfp316 | 0.007 | Old |
| 0.151035 | 0.819411 | 0.026 | 0.018 | 1 | 0 | Poll | 0.008 | Old |
| 0.151203 | 0.10033 | 0.116 | 0.097 | 1 | 0 | Nudt4 | 0.019 | Old |
| 0.15124 | 0.456233 | 0.064 | 0.051 | 1 | 0 | Sass6 | 0.013 | Old |
| 0.151442 | 0.076398 | 0.393 | 0.358 | 1 | 0 | Gm2a | 0.035 | Old |
| 0.15148 | 0.107441 | 0.464 | 0.422 | 1 | 0 | Brk1 | 0.042 | Old |
| 0.151499 | 0.247729 | 0.06 | 0.047 | 1 | 0 | Rasgrp1 | 0.013 | Old |
| 0.151538 | 0.162278 | 0.311 | 0.282 | 1 | 0 | Ogdh | 0.029 | Old |
| 0.15168 | 0.756413 | 0.022 | 0.014 | 1 | 0 | Map2k3os | 0.008 | Old |
| 0.151862 | 0.289449 | 0.029 | 0.02 | 1 | 0 | Rnaseh1 | 0.009 | Old |
| 0.152135 | 0.145717 | 0.168 | 0.145 | 1 | 0 | Rab22a | 0.023 | Old |
| 0.152512 | 0.116602 | 0.104 | 0.085 | 1 | 0 | Abcd2 | 0.019 | Old |
| 0.152542 | 0.402218 | 0.081 | 0.066 | 1 | 0 | 4930455G( | 0.015 | Old |
| 0.152748 | 0.076709 | 0.092 | 0.075 | 1 | 0 | Usp6nl | 0.017 | Old |
| 0.152761 | 1.087059 | 0.011 | 0.006 | 1 | 0 | Ube2c | 0.005 | Old |
| 0.152777 | 0.202291 | 0.094 | 0.078 | 1 | 0 | Anpep | 0.016 | Old |
| 0.152834 | 0.52093 | 0.126 | 0.109 | 1 | 0 | Errfi1 | 0.017 | Old |
| 0.152958 | 0.023964 | 0.083 | 0.067 | 1 | 0 | Trpv2 | 0.016 | Old |
| 0.152999 | 0.221925 | 0.071 | 0.056 | 1 | 0 | Fbxl17 | 0.015 | Old |
| 0.153094 | 0.066698 | 0.291 | 0.256 | 1 | 0 | Mdh1 | 0.035 | Old |
| 0.153256 | 0.056772 | 0.391 | 0.352 | 1 | 0 | Taok1 | 0.039 | Old |
| 0.153358 | 0.064709 | 0.319 | 0.289 | 1 | 0 | Camk1 | 0.03 | Old |
| 0.153461 | 0.359816 | 0.041 | 0.03 | 1 | 0 | Snhg9 | 0.011 | Old |
| 0.153508 | 1.181977 | 0.011 | 0.006 | 1 | 0 | BC052040 | 0.005 | Old |
| 0.153614 | 0.070781 | 0.261 | 0.23 | 1 | 0 | Swap70 | 0.031 | Old |
| 0.153824 | 0.176059 | 0.075 | 0.06 | 1 | 0 | Fam220a | 0.015 | Old |
| 0.153883 | 0.747644 | 0.011 | 0.006 | 1 | 0 | Dus4l | 0.005 | Old |
| 0.154111 | 0.350719 | 0.098 | 0.082 | 1 | 0 | Ccdc88b | 0.016 | Old |
| 0.15441 | 0.113245 | 0.575 | 0.526 | 1 | 0 | Fam49b | 0.049 | Old |
| 0.154446 | 0.815454 | 0.011 | 0.006 | 1 | 0 | Pcbp3 | 0.005 | Old |
| 0.154453 | 0.113978 | 0.111 | 0.093 | 1 | 0 | Dusp16 | 0.018 | Old |
| 0.154457 | 0.093682 | 0.198 | 0.173 | 1 | 0 | Ctbp1 | 0.025 | Old |
| 0.154668 | 0.54997 | 0.022 | 0.014 | 1 | 0 | Qtrt2 | 0.008 | Old |
| 0.154816 | 0.28168 | 0.111 | 0.093 | 1 | 0 | Bst1 | 0.018 | Old |
| 0.155427 | 0.411391 | 0.049 | 0.037 | 1 | 0 | Pprc1 | 0.012 | Old |
| 0.155604 | 0.148187 | 0.238 | 0.211 | 1 | 0 | Pon2 | 0.027 | Old |
| 0.155789 | 0.05896 | 0.352 | 0.315 | 1 | 0 | Rab3il1 | 0.037 | Old |
| 0.156126 | 0.596139 | 0.026 | 0.018 | 1 | 0 | Gin1 | 0.008 | Old |
| 0.156174 | 0.337492 | 0.062 | 0.048 | 1 | 0 | Fam129b | 0.014 | Old |
| 0.156187 | 0.043233 | 0.089 | 0.072 | 1 | 0 | Prrc1 | 0.017 | Old |
| 0.156218 | 0.157481 | 0.197 | 0.172 | 1 | 0 | Dennd6a | 0.025 | Old |
| 0.156292 | 0.193065 | 0.126 | 0.107 | 1 | 0 | Ndufs4 | 0.019 | Old |
| 0.156296 | 0.026313 | 0.094 | 0.077 | 1 | 0 | Bccip | 0.017 | Old |
| 0.156345 | 0.416868 | 0.109 | 0.092 | 1 | 0 | Itpr2 | 0.017 | Old |

| | | | | | | | | |
|---|---|---|---|---|---|---|---|---|
| 0.156665 | 0.23985 | 0.134 | 0.115 | 1 | 0 | Ipo7 | 0.019 | Old |
| 0.156709 | 0.234005 | 0.034 | 0.024 | 1 | 0 | Zbtb5 | 0.01 | Old |
| 0.156835 | 0.215077 | 0.149 | 0.128 | 1 | 0 | Xpo1 | 0.021 | Old |
| 0.156925 | 0.366938 | 0.137 | 0.118 | 1 | 0 | Rbbp7 | 0.019 | Old |
| 0.157096 | 0.236458 | 0.011 | 0.006 | 1 | 0 | Sncg | 0.005 | Old |
| 0.157391 | 0.357324 | 0.097 | 0.081 | 1 | 0 | Zfp397 | 0.016 | Old |
| 0.157785 | 0.228208 | 0.089 | 0.073 | 1 | 0 | Gars | 0.016 | Old |
| 0.157999 | 0.078589 | 0.621 | 0.587 | 1 | 0 | Rnf130 | 0.034 | Old |
| 0.158003 | 0.061768 | 0.479 | 0.434 | 1 | 0 | Clic1 | 0.045 | Old |
| 0.158059 | 0.215573 | 0.104 | 0.086 | 1 | 0 | Nsmce1 | 0.018 | Old |
| 0.158444 | 0.297698 | 0.157 | 0.137 | 1 | 0 | Aoah | 0.02 | Old |
| 0.158544 | 0.409466 | 0.03 | 0.021 | 1 | 0 | Zfp959 | 0.009 | Old |
| 0.159273 | 0.107675 | 0.053 | 0.041 | 1 | 0 | Nop9 | 0.012 | Old |
| 0.159491 | 0.157068 | 0.119 | 0.099 | 1 | 0 | Nub1 | 0.02 | Old |
| 0.159622 | 0.001708 | 0.26 | 0.227 | 1 | 0 | Csk | 0.033 | Old |
| 0.160099 | 0.096649 | 0.811 | 0.783 | 1 | 0 | Lcp1 | 0.028 | Old |
| 0.160369 | 0.075772 | 0.286 | 0.254 | 1 | 0 | Cpt1a | 0.032 | Old |
| 0.160418 | 0.040117 | 0.079 | 0.064 | 1 | 0 | Cdk2 | 0.015 | Old |
| 0.160556 | 0.117123 | 0.181 | 0.156 | 1 | 0 | Stap1 | 0.025 | Old |
| 0.160947 | 0.715922 | 0.027 | 0.019 | 1 | 0 | Hsd17b7 | 0.008 | Old |
| 0.161072 | 0.443113 | 0.031 | 0.022 | 1 | 0 | Smyd5 | 0.009 | Old |
| 0.161081 | 0.724151 | 0.023 | 0.016 | 1 | 0 | Zfp931 | 0.007 | Old |
| 0.16115 | 0.127521 | 0.406 | 0.373 | 1 | 0 | Rpl22l1 | 0.033 | Old |
| 0.161204 | 0.166357 | 0.13 | 0.11 | 1 | 0 | Samd8 | 0.02 | Old |
| 0.161294 | 0.23678 | 0.056 | 0.043 | 1 | 0 | Ebag9 | 0.013 | Old |
| 0.161574 | 0.093012 | 0.331 | 0.299 | 1 | 0 | Hnrnph2 | 0.032 | Old |
| 0.161744 | 0.089049 | 0.256 | 0.227 | 1 | 0 | Kmt2a | 0.029 | Old |
| 0.161748 | 0.210204 | 0.03 | 0.021 | 1 | 0 | Gm26601 | 0.009 | Old |
| 0.161825 | 0.179619 | 0.036 | 0.026 | 1 | 0 | Lrch4 | 0.01 | Old |
| 0.161968 | 0.003013 | 0.152 | 0.128 | 1 | 0 | Stip1 | 0.024 | Old |
| 0.161985 | 0.792456 | 0.016 | 0.01 | 1 | 0 | Dtnb | 0.006 | Old |
| 0.162004 | 0.462998 | 0.019 | 0.012 | 1 | 0 | Wdr73 | 0.007 | Old |
| 0.162651 | 0.464853 | 0.037 | 0.027 | 1 | 0 | Comtd1 | 0.01 | Old |
| 0.162974 | 0.041193 | 0.096 | 0.078 | 1 | 0 | Fblim1 | 0.018 | Old |
| 0.163065 | 0.334137 | 0.037 | 0.027 | 1 | 0 | Fbxw17 | 0.01 | Old |
| 0.163134 | 0.750738 | 0.016 | 0.01 | 1 | 0 | Gm12166 | 0.006 | Old |
| 0.163266 | 0.145053 | 0.098 | 0.081 | 1 | 0 | Glrx | 0.017 | Old |
| 0.163565 | 0.270125 | 0.036 | 0.026 | 1 | 0 | Acad9 | 0.01 | Old |
| 0.163906 | 0.129235 | 0.04 | 0.029 | 1 | 0 | Pdp2 | 0.011 | Old |
| 0.164198 | 0.076448 | 0.097 | 0.08 | 1 | 0 | Trip4 | 0.017 | Old |
| 0.164288 | 0.491718 | 0.014 | 0.008 | 1 | 0 | Stat4 | 0.006 | Old |
| 0.164623 | 0.439059 | 0.038 | 0.028 | 1 | 0 | AI839979 | 0.01 | Old |
| 0.164717 | 0.332432 | 0.014 | 0.008 | 1 | 0 | C030006K1 | 0.006 | Old |
| 0.164737 | 0.10379 | 0.23 | 0.203 | 1 | 0 | Mat2b | 0.027 | Old |
| 0.164769 | 0.443866 | 0.042 | 0.032 | 1 | 0 | Gabpb1 | 0.01 | Old |

| | | | | | | | | |
|---|---|---|---|---|---|---|---|---|
| 0.165117 | 0.055035 | 0.036 | 0.026 | 1 | 0 | Gm46430 | 0.01 | Old |
| 0.165139 | 0.509362 | 0.016 | 0.01 | 1 | 0 | Jag1 | 0.006 | Old |
| 0.165242 | 0.136393 | 0.056 | 0.043 | 1 | 0 | Tfdp1 | 0.013 | Old |
| 0.16534 | 0.317757 | 0.063 | 0.05 | 1 | 0 | Nr2c1 | 0.013 | Old |
| 0.165414 | 0.116139 | 0.326 | 0.294 | 1 | 0 | Bcl10 | 0.032 | Old |
| 0.165605 | 0.383808 | 0.016 | 0.01 | 1 | 0 | Gpatch3 | 0.006 | Old |
| 0.165999 | 0.087976 | 0.108 | 0.09 | 1 | 0 | Naga | 0.018 | Old |
| 0.166032 | 0.150028 | 0.09 | 0.074 | 1 | 0 | Wdr91 | 0.016 | Old |
| 0.166826 | 0.002241 | 0.03 | 0.021 | 1 | 0 | Zfp810 | 0.009 | Old |
| 0.166841 | 0.384306 | 0.059 | 0.046 | 1 | 0 | Mrpl37 | 0.013 | Old |
| 0.167075 | 0.056189 | 0.711 | 0.667 | 1 | 0 | Hnrnpk | 0.044 | Old |
| 0.167114 | 0.421437 | 0.027 | 0.019 | 1 | 0 | Lhpp | 0.008 | Old |
| 0.167496 | 0.167866 | 0.133 | 0.113 | 1 | 0 | BC003331 | 0.02 | Old |
| 0.167571 | 0.187145 | 0.088 | 0.072 | 1 | 0 | Ebna1bp2 | 0.016 | Old |
| 0.167623 | 0.248796 | 0.205 | 0.182 | 1 | 0 | Cxcr4 | 0.023 | Old |
| 0.16772 | 0.232379 | 0.023 | 0.016 | 1 | 0 | Klhl42 | 0.007 | Old |
| 0.167834 | 0.328301 | 0.066 | 0.052 | 1 | 0 | Rcan3 | 0.014 | Old |
| 0.16786 | 0.127956 | 0.1 | 0.083 | 1 | 0 | Pi4ka | 0.017 | Old |
| 0.168568 | 0.099536 | 0.144 | 0.123 | 1 | 0 | Ktn1 | 0.021 | Old |
| 0.168632 | 0.422385 | 0.063 | 0.05 | 1 | 0 | Rgs3 | 0.013 | Old |
| 0.16875 | 0.296459 | 0.023 | 0.016 | 1 | 0 | Rtn4ip1 | 0.007 | Old |
| 0.16875 | 0.089353 | 0.097 | 0.08 | 1 | 0 | Amn1 | 0.017 | Old |
| 0.168901 | 0.067867 | 0.279 | 0.248 | 1 | 0 | Ssr1 | 0.031 | Old |
| 0.168951 | 0.068775 | 0.219 | 0.194 | 1 | 0 | Atp13a2 | 0.025 | Old |
| 0.169009 | 0.115545 | 0.1 | 0.083 | 1 | 0 | Mrps16 | 0.017 | Old |
| 0.169144 | 0.078449 | 0.185 | 0.162 | 1 | 0 | Vamp2 | 0.023 | Old |
| 0.169151 | 0.089951 | 0.308 | 0.28 | 1 | 0 | Caprin1 | 0.028 | Old |
| 0.169169 | 0.727845 | 0.021 | 0.013 | 1 | 0 | Papss2 | 0.008 | Old |
| 0.169317 | 0.315311 | 0.027 | 0.019 | 1 | 0 | Slc25a40 | 0.008 | Old |
| 0.169549 | 0.276614 | 0.073 | 0.059 | 1 | 0 | Gart | 0.014 | Old |
| 0.170007 | 0.073405 | 0.112 | 0.093 | 1 | 0 | Kcnk13 | 0.019 | Old |
| 0.170031 | 0.292761 | 0.057 | 0.045 | 1 | 0 | Mlycd | 0.012 | Old |
| 0.170193 | 0.145995 | 0.164 | 0.142 | 1 | 0 | Appbp2 | 0.022 | Old |
| 0.170195 | 0.558361 | 0.01 | 0.005 | 1 | 0 | Gm33973 | 0.005 | Old |
| 0.170228 | 0.176277 | 0.234 | 0.209 | 1 | 0 | Capg | 0.025 | Old |
| 0.170378 | 0.073293 | 0.104 | 0.086 | 1 | 0 | Tob1 | 0.018 | Old |
| 0.170433 | 0.008336 | 0.023 | 0.016 | 1 | 0 | Haus1 | 0.007 | Old |
| 0.170835 | 0.27083 | 0.038 | 0.028 | 1 | 0 | Mlh1 | 0.01 | Old |
| 0.170849 | 0.54492 | 0.01 | 0.005 | 1 | 0 | Ctif | 0.005 | Old |
| 0.170971 | 0.128994 | 0.375 | 0.347 | 1 | 0 | 7-Sep | 0.028 | Old |
| 0.171184 | 0.643353 | 0.021 | 0.013 | 1 | 0 | Ccnyl1 | 0.008 | Old |
| 0.171558 | 0.160028 | 0.112 | 0.094 | 1 | 0 | Sfswap | 0.018 | Old |
| 0.171743 | 0.557117 | 0.021 | 0.013 | 1 | 0 | Tepsin | 0.008 | Old |
| 0.171867 | 0.087024 | 0.023 | 0.016 | 1 | 0 | 2700081O1 | 0.007 | Old |
| 0.172162 | 0.497733 | 0.01 | 0.005 | 1 | 0 | Top1mt | 0.005 | Old |

| | | | | | | | | |
|---|---|---|---|---|---|---|---|---|
| 0.172376 | 0.058165 | 0.152 | 0.13 | 1 | 0 | Slc38a9 | 0.022 | Old |
| 0.17239 | 0.152571 | 0.159 | 0.138 | 1 | 0 | Map2k1 | 0.021 | Old |
| 0.172614 | 0.175257 | 0.07 | 0.056 | 1 | 0 | Fbxo30 | 0.014 | Old |
| 0.172634 | 0.196303 | 0.133 | 0.114 | 1 | 0 | Mrpl42 | 0.019 | Old |
| 0.172822 | 0.153585 | 0.01 | 0.005 | 1 | 0 | Kdr | 0.005 | Old |
| 0.172859 | 0.09873 | 0.15 | 0.128 | 1 | 0 | Samm50 | 0.022 | Old |
| 0.17291 | 0.166803 | 0.044 | 0.033 | 1 | 0 | E2f3 | 0.011 | Old |
| 0.173154 | 0.1424 | 0.134 | 0.114 | 1 | 0 | Nudt21 | 0.02 | Old |
| 0.173175 | 0.104254 | 0.22 | 0.194 | 1 | 0 | Psma2 | 0.026 | Old |
| 0.173182 | 0.471071 | 0.021 | 0.013 | 1 | 0 | Zfp518b | 0.008 | Old |
| 0.173324 | 0.56899 | 0.044 | 0.033 | 1 | 0 | Cstf2 | 0.011 | Old |
| 0.173584 | 0.32437 | 0.047 | 0.035 | 1 | 0 | Dvl1 | 0.012 | Old |
| 0.173689 | 0.481507 | 0.057 | 0.045 | 1 | 0 | Sertad3 | 0.012 | Old |
| 0.173723 | 0.040045 | 0.17 | 0.146 | 1 | 0 | Sec24b | 0.024 | Old |
| 0.173818 | 0.023971 | 0.027 | 0.019 | 1 | 0 | Btg3 | 0.008 | Old |
| 0.173993 | 0.325807 | 0.021 | 0.013 | 1 | 0 | Gadd45a | 0.008 | Old |
| 0.174465 | 0.188095 | 0.149 | 0.129 | 1 | 0 | Ehmt2 | 0.02 | Old |
| 0.174923 | 0.430773 | 0.025 | 0.017 | 1 | 0 | Cdpf1 | 0.008 | Old |
| 0.175166 | 0.103592 | 0.077 | 0.062 | 1 | 0 | Timm44 | 0.015 | Old |
| 0.175174 | 0.041944 | 0.291 | 0.26 | 1 | 0 | Ppp1cc | 0.031 | Old |
| 0.175306 | 0.120632 | 0.144 | 0.123 | 1 | 0 | Ubr4 | 0.021 | Old |
| 0.17537 | 0.298834 | 0.041 | 0.031 | 1 | 0 | Mov10 | 0.01 | Old |
| 0.175411 | 0.35598 | 0.021 | 0.013 | 1 | 0 | Tmco4 | 0.008 | Old |
| 0.175567 | 0.44849 | 0.036 | 0.026 | 1 | 0 | Cd209e | 0.01 | Old |
| 0.175673 | 0.30683 | 0.062 | 0.049 | 1 | 0 | Timm29 | 0.013 | Old |
| 0.175978 | 0.034991 | 0.197 | 0.171 | 1 | 0 | Araf | 0.026 | Old |
| 0.176047 | 0.112852 | 0.093 | 0.077 | 1 | 0 | Map2k4 | 0.016 | Old |
| 0.176071 | 0.148924 | 0.108 | 0.091 | 1 | 0 | Arl6ip6 | 0.017 | Old |
| 0.176173 | 0.003326 | 0.189 | 0.164 | 1 | 0 | Becn1 | 0.025 | Old |
| 0.176211 | 0.147857 | 0.114 | 0.096 | 1 | 0 | Ppp2r5e | 0.018 | Old |
| 0.17633 | 0.099041 | 0.152 | 0.13 | 1 | 0 | Cers5 | 0.022 | Old |
| 0.176769 | 0.213157 | 0.152 | 0.132 | 1 | 0 | Cdc73 | 0.02 | Old |
| 0.17692 | 0.329015 | 0.078 | 0.064 | 1 | 0 | Mcm6 | 0.014 | Old |
| 0.177448 | 0.153281 | 0.108 | 0.091 | 1 | 0 | Nfkb2 | 0.017 | Old |
| 0.177511 | 0.18271 | 0.041 | 0.031 | 1 | 0 | Dus1l | 0.01 | Old |
| 0.177563 | 0.579125 | 0.018 | 0.011 | 1 | 0 | Vill | 0.007 | Old |
| 0.177745 | 0.204286 | 0.034 | 0.025 | 1 | 0 | Foxred1 | 0.009 | Old |
| 0.178066 | 0.497554 | 0.051 | 0.04 | 1 | 0 | Slc7a1 | 0.011 | Old |
| 0.178094 | 0.088544 | 0.053 | 0.041 | 1 | 0 | Adi1 | 0.012 | Old |
| 0.178301 | 0.17887 | 0.17 | 0.149 | 1 | 0 | Cyld | 0.021 | Old |
| 0.178395 | 0.903259 | 0.012 | 0.007 | 1 | 0 | Mast4 | 0.005 | Old |
| 0.178428 | 0.13225 | 0.233 | 0.207 | 1 | 0 | Supt5 | 0.026 | Old |
| 0.178436 | 0.10683 | 0.317 | 0.286 | 1 | 0 | Atp6v0d1 | 0.031 | Old |
| 0.178502 | 0.09274 | 0.469 | 0.433 | 1 | 0 | Ms4a7 | 0.036 | Old |
| 0.178506 | 0.15802 | 0.029 | 0.02 | 1 | 0 | Ifi213 | 0.009 | Old |

| | | | | | | | | |
|---|---|---|---|---|---|---|---|---|
| 0.178831 | 0.145214 | 0.124 | 0.105 | 1 | 0 | Pip4k2b | 0.019 | Old |
| 0.178927 | 0.15692 | 0.127 | 0.108 | 1 | 0 | Ahsa1 | 0.019 | Old |
| 0.178965 | 0.526287 | 0.018 | 0.011 | 1 | 0 | 9130221H1 | 0.007 | Old |
| 0.178989 | 0.151644 | 0.06 | 0.047 | 1 | 0 | Memo1 | 0.013 | Old |
| 0.179043 | 0.519728 | 0.018 | 0.011 | 1 | 0 | Zfp873 | 0.007 | Old |
| 0.179091 | 0.333487 | 0.044 | 0.033 | 1 | 0 | Bcl2l13 | 0.011 | Old |
| 0.179179 | 0.154786 | 0.085 | 0.07 | 1 | 0 | 0610010F0 | 0.015 | Old |
| 0.179312 | 0.196354 | 0.122 | 0.104 | 1 | 0 | Psen2 | 0.018 | Old |
| 0.179467 | 0.1341 | 0.098 | 0.082 | 1 | 0 | Gaa | 0.016 | Old |
| 0.179487 | 0.101681 | 0.297 | 0.267 | 1 | 0 | Papola | 0.03 | Old |
| 0.179508 | 0.076572 | 0.244 | 0.217 | 1 | 0 | Eif4g3 | 0.027 | Old |
| 0.179626 | 0.151536 | 0.15 | 0.131 | 1 | 0 | Fbxo11 | 0.019 | Old |
| 0.179702 | 0.275407 | 0.064 | 0.051 | 1 | 0 | Cd79b | 0.013 | Old |
| 0.17977 | 0.104547 | 0.174 | 0.151 | 1 | 0 | Emsy | 0.023 | Old |
| 0.17978 | 0.026368 | 0.313 | 0.279 | 1 | 0 | Map7d1 | 0.034 | Old |
| 0.179877 | 0.094561 | 0.413 | 0.376 | 1 | 0 | Nptn | 0.037 | Old |
| 0.180012 | 0.092455 | 0.227 | 0.202 | 1 | 0 | Ap2b1 | 0.025 | Old |
| 0.180061 | 0.255004 | 0.018 | 0.011 | 1 | 0 | Pcdhga12 | 0.007 | Old |
| 0.180215 | 0.108953 | 0.312 | 0.285 | 1 | 0 | Lcp2 | 0.027 | Old |
| 0.180526 | 0.6534 | 0.012 | 0.007 | 1 | 0 | B130006D( | 0.005 | Old |
| 0.180687 | 0.149212 | 0.059 | 0.046 | 1 | 0 | Dis3l2 | 0.013 | Old |
| 0.18073 | 0.185754 | 0.157 | 0.137 | 1 | 0 | Tspan4 | 0.02 | Old |
| 0.180929 | 0.062735 | 0.175 | 0.152 | 1 | 0 | Taz | 0.023 | Old |
| 0.180977 | 0.658429 | 0.022 | 0.015 | 1 | 0 | Ccdc137 | 0.007 | Old |
| 0.181063 | 0.087152 | 0.737 | 0.682 | 1 | 0 | Ehd4 | 0.055 | Old |
| 0.181081 | 0.456614 | 0.015 | 0.009 | 1 | 0 | Adora2a | 0.006 | Old |
| 0.181208 | 0.334468 | 0.068 | 0.055 | 1 | 0 | AI606181 | 0.013 | Old |
| 0.18123 | 0.249272 | 0.079 | 0.065 | 1 | 0 | Pld2 | 0.014 | Old |
| 0.181281 | 0.102118 | 0.235 | 0.211 | 1 | 0 | Ik | 0.024 | Old |
| 0.181339 | 0.09361 | 0.163 | 0.14 | 1 | 0 | Pigk | 0.023 | Old |
| 0.18153 | 0.015476 | 0.134 | 0.113 | 1 | 0 | D5Ertd579 | 0.021 | Old |
| 0.181758 | 0.13787 | 0.083 | 0.068 | 1 | 0 | Ddx47 | 0.015 | Old |
| 0.181808 | 0.231479 | 0.06 | 0.048 | 1 | 0 | Hace1 | 0.012 | Old |
| 0.181956 | 0.239525 | 0.123 | 0.105 | 1 | 0 | Sarnp | 0.018 | Old |
| 0.181996 | 0.182647 | 0.231 | 0.207 | 1 | 0 | Gxylt1 | 0.024 | Old |
| 0.182287 | 0.142747 | 0.092 | 0.076 | 1 | 0 | Dok1 | 0.016 | Old |
| 0.182682 | 0.108024 | 0.059 | 0.046 | 1 | 0 | Eef1e1 | 0.013 | Old |
| 0.182829 | 0.064248 | 0.841 | 0.805 | 1 | 0 | Rpl8 | 0.036 | Old |
| 0.182868 | 0.707128 | 0.033 | 0.024 | 1 | 0 | Mad2l1bp | 0.009 | Old |
| 0.183069 | 0.187302 | 0.012 | 0.007 | 1 | 0 | Gm14548 | 0.005 | Old |
| 0.183098 | 0.072595 | 0.089 | 0.073 | 1 | 0 | Gtf3c6 | 0.016 | Old |
| 0.183365 | 0.211421 | 0.012 | 0.007 | 1 | 0 | Plac8 | 0.005 | Old |
| 0.183508 | 0.084627 | 0.538 | 0.498 | 1 | 0 | Serp1 | 0.04 | Old |
| 0.183562 | 0.305275 | 0.012 | 0.007 | 1 | 0 | Mocos | 0.005 | Old |
| 0.183691 | 0.245965 | 0.052 | 0.04 | 1 | 0 | Myef2 | 0.012 | Old |

| | | | | | | | | |
|---|---|---|---|---|---|---|---|---|
| 0.183753 | 0.405838 | 0.022 | 0.015 | 1 | 0 | Gm42715 | 0.007 | Old |
| 0.183834 | 0.319021 | 0.056 | 0.044 | 1 | 0 | Tpcn2 | 0.012 | Old |
| 0.183932 | 0.208506 | 0.122 | 0.104 | 1 | 0 | Sh3bp2 | 0.018 | Old |
| 0.184027 | 0.190228 | 0.067 | 0.054 | 1 | 0 | Bloc1s4 | 0.013 | Old |
| 0.184142 | 0.356139 | 0.022 | 0.015 | 1 | 0 | Ripk3 | 0.007 | Old |
| 0.184299 | 0.328097 | 0.063 | 0.05 | 1 | 0 | Riok2 | 0.013 | Old |
| 0.184683 | 0.207001 | 0.085 | 0.07 | 1 | 0 | Arsa | 0.015 | Old |
| 0.184764 | 0.151162 | 0.098 | 0.082 | 1 | 0 | Arhgap39 | 0.016 | Old |
| 0.18478 | 0.39609 | 0.022 | 0.015 | 1 | 0 | Nat2 | 0.007 | Old |
| 0.184944 | 0.054208 | 0.375 | 0.34 | 1 | 0 | Eif5a | 0.035 | Old |
| 0.185029 | 0.105145 | 0.079 | 0.064 | 1 | 0 | Fam3a | 0.015 | Old |
| 0.18521 | 0.138056 | 0.32 | 0.289 | 1 | 0 | Syngr2 | 0.031 | Old |
| 0.18528 | 0.443272 | 0.038 | 0.029 | 1 | 0 | Cacfd1 | 0.009 | Old |
| 0.185369 | 0.272451 | 0.026 | 0.018 | 1 | 0 | Nlrx1 | 0.008 | Old |
| 0.185718 | 0.183545 | 0.015 | 0.009 | 1 | 0 | 3-Mar | 0.006 | Old |
| 0.18608 | 0.081957 | 0.126 | 0.107 | 1 | 0 | Parp8 | 0.019 | Old |
| 0.186172 | 0.073162 | 0.285 | 0.255 | 1 | 0 | Myl12a | 0.03 | Old |
| 0.1862 | 0.080966 | 0.093 | 0.077 | 1 | 0 | Hsph1 | 0.016 | Old |
| 0.186248 | 0.032462 | 0.015 | 0.009 | 1 | 0 | Neil1 | 0.006 | Old |
| 0.186346 | 0.109626 | 0.011 | 0.018 | 1 | 0 | Atp5s | -0.007 | Old |
| 0.186426 | 0.03425 | 0.093 | 0.077 | 1 | 0 | Cyhr1 | 0.016 | Old |
| 0.186766 | 0.098738 | 0.116 | 0.099 | 1 | 0 | Rnh1 | 0.017 | Old |
| 0.186809 | 0.129207 | 0.144 | 0.123 | 1 | 0 | Tap1 | 0.021 | Old |
| 0.186836 | 0.109465 | 0.145 | 0.125 | 1 | 0 | Ireb2 | 0.02 | Old |
| 0.186846 | 0.064286 | 0.28 | 0.251 | 1 | 0 | Emc10 | 0.029 | Old |
| 0.187179 | 0.085897 | 0.066 | 0.052 | 1 | 0 | Cdadc1 | 0.014 | Old |
| 0.187214 | 0.021874 | 0.237 | 0.208 | 1 | 0 | Mapk1ip1l | 0.029 | Old |
| 0.18742 | 0.077308 | 0.347 | 0.315 | 1 | 0 | Srsf6 | 0.032 | Old |
| 0.18778 | 0.407727 | 0.064 | 0.052 | 1 | 0 | Kctd6 | 0.012 | Old |
| 0.187813 | 0.116607 | 0.282 | 0.257 | 1 | 0 | 11-Sep | 0.025 | Old |
| 0.187856 | 0.312452 | 0.057 | 0.045 | 1 | 0 | Rnf40 | 0.012 | Old |
| 0.188002 | 0.146938 | 0.149 | 0.129 | 1 | 0 | Rnft1 | 0.02 | Old |
| 0.188468 | 0.240354 | 0.073 | 0.059 | 1 | 0 | Pla2g4a | 0.014 | Old |
| 0.18854 | 0.347204 | 0.047 | 0.036 | 1 | 0 | Scarf1 | 0.011 | Old |
| 0.188925 | 0.278457 | 0.064 | 0.052 | 1 | 0 | Snta1 | 0.012 | Old |
| 0.188989 | 0.059489 | 0.446 | 0.407 | 1 | 0 | Tgoln1 | 0.039 | Old |
| 0.189141 | 0.144705 | 0.042 | 0.032 | 1 | 0 | Rdh11 | 0.01 | Old |
| 0.189146 | 0.143693 | 0.026 | 0.018 | 1 | 0 | Dgcr14 | 0.008 | Old |
| 0.189448 | 0.606138 | 0.047 | 0.036 | 1 | 0 | Fastk | 0.011 | Old |
| 0.189577 | 0.213565 | 0.202 | 0.181 | 1 | 0 | Immt | 0.021 | Old |
| 0.189668 | 0.139416 | 0.163 | 0.142 | 1 | 0 | Dguok | 0.021 | Old |
| 0.189688 | 0.1341 | 0.294 | 0.268 | 1 | 0 | Wapl | 0.026 | Old |
| 0.189989 | 0.104353 | 0.118 | 0.1 | 1 | 0 | Emc6 | 0.018 | Old |
| 0.190014 | 0.330124 | 0.027 | 0.019 | 1 | 0 | Dusp10 | 0.008 | Old |
| 0.190031 | 0.107159 | 0.207 | 0.182 | 1 | 0 | Lsm4 | 0.025 | Old |

| | | | | | | | | |
|---|---|---|---|---|---|---|---|---|
| 0.190397 | 0.226012 | 0.07 | 0.056 | 1 | 0 | Ints1 | 0.014 | Old |
| 0.190546 | 0.163754 | 0.067 | 0.054 | 1 | 0 | Tmem63b | 0.013 | Old |
| 0.190759 | 0.047517 | 0.278 | 0.249 | 1 | 0 | Tnpo3 | 0.029 | Old |
| 0.190776 | 0.404869 | 0.042 | 0.032 | 1 | 0 | Styx | 0.01 | Old |
| 0.19084 | 0.135609 | 0.055 | 0.043 | 1 | 0 | Mrpl40 | 0.012 | Old |
| 0.1912 | 0.448881 | 0.051 | 0.04 | 1 | 0 | Zdhhc8 | 0.011 | Old |
| 0.191292 | 0.255455 | 0.126 | 0.11 | 1 | 0 | Sik1 | 0.016 | Old |
| 0.191475 | 0.330385 | 0.059 | 0.047 | 1 | 0 | Cog8 | 0.012 | Old |
| 0.191812 | 0.078986 | 0.482 | 0.44 | 1 | 0 | Atp5f1 | 0.042 | Old |
| 0.192117 | 0.353503 | 0.047 | 0.036 | 1 | 0 | Hebp1 | 0.011 | Old |
| 0.19213 | 0.408083 | 0.027 | 0.019 | 1 | 0 | Gtf3c5 | 0.008 | Old |
| 0.19274 | 0.520491 | 0.019 | 0.013 | 1 | 0 | Nipal3 | 0.006 | Old |
| 0.192811 | 0.241133 | 0.661 | 0.642 | 1 | 0 | Ccl2 | 0.019 | Old |
| 0.193098 | 0.092873 | 0.709 | 0.682 | 1 | 0 | Actr3 | 0.027 | Old |
| 0.193647 | 0.420204 | 0.019 | 0.013 | 1 | 0 | Gm45871 | 0.006 | Old |
| 0.193777 | 0.086707 | 0.153 | 0.133 | 1 | 0 | Plin3 | 0.02 | Old |
| 0.19384 | 0.006549 | 0.137 | 0.117 | 1 | 0 | Atp9b | 0.02 | Old |
| 0.193887 | 0.184917 | 0.186 | 0.165 | 1 | 0 | Sh3pxd2b | 0.021 | Old |
| 0.19402 | 0.070965 | 0.234 | 0.21 | 1 | 0 | Tsen34 | 0.024 | Old |
| 0.195376 | 0.481769 | 0.029 | 0.021 | 1 | 0 | Chst14 | 0.008 | Old |
| 0.195402 | 0.222473 | 0.124 | 0.107 | 1 | 0 | Cdk13 | 0.017 | Old |
| 0.195578 | 0.143749 | 0.193 | 0.171 | 1 | 0 | Ugp2 | 0.022 | Old |
| 0.19578 | 0.05793 | 0.261 | 0.234 | 1 | 0 | Sf3b5 | 0.027 | Old |
| 0.195922 | 0.154503 | 0.142 | 0.123 | 1 | 0 | Taf1d | 0.019 | Old |
| 0.196025 | 0.655432 | 0.037 | 0.028 | 1 | 0 | Cyb561d1 | 0.009 | Old |
| 0.19631 | 0.01906 | 0.242 | 0.214 | 1 | 0 | Mlf2 | 0.028 | Old |
| 0.196327 | 0.050799 | 0.107 | 0.09 | 1 | 0 | Drg1 | 0.017 | Old |
| 0.196389 | 0.382861 | 0.057 | 0.046 | 1 | 0 | Ak6 | 0.011 | Old |
| 0.196496 | 0.041372 | 0.175 | 0.153 | 1 | 0 | Azi2 | 0.022 | Old |
| 0.196505 | 0.228696 | 0.133 | 0.115 | 1 | 0 | Acadl | 0.018 | Old |
| 0.196616 | 0.196669 | 0.112 | 0.096 | 1 | 0 | Tyk2 | 0.016 | Old |
| 0.196786 | 0.128626 | 0.468 | 0.433 | 1 | 0 | Ehd1 | 0.035 | Old |
| 0.196836 | 0.219799 | 0.123 | 0.106 | 1 | 0 | Epc1 | 0.017 | Old |
| 0.19717 | 0.063573 | 0.157 | 0.136 | 1 | 0 | Zhx1 | 0.021 | Old |
| 0.197182 | 0.737752 | 0.016 | 0.011 | 1 | 0 | Iqcc | 0.005 | Old |
| 0.197362 | 0.142625 | 0.201 | 0.179 | 1 | 0 | Rilpl2 | 0.022 | Old |
| 0.198306 | 0.041212 | 0.098 | 0.082 | 1 | 0 | Golga2 | 0.016 | Old |
| 0.198349 | 0.007166 | 0.295 | 0.261 | 1 | 0 | Tnks2 | 0.034 | Old |
| 0.198358 | 0.150452 | 0.2 | 0.178 | 1 | 0 | Syf2 | 0.022 | Old |
| 0.198544 | 0.509159 | 0.03 | 0.022 | 1 | 0 | Cd3eap | 0.008 | Old |
| 0.198567 | 0.1556 | 0.062 | 0.049 | 1 | 0 | Gpn3 | 0.013 | Old |
| 0.198616 | 0.018598 | 0.064 | 0.051 | 1 | 0 | Lyrm2 | 0.013 | Old |
| 0.198837 | 0.676874 | 0.025 | 0.017 | 1 | 0 | Nbas | 0.008 | Old |
| 0.19901 | 0.356711 | 0.029 | 0.021 | 1 | 0 | Gbp2b | 0.008 | Old |
| 0.199027 | 0.157901 | 0.04 | 0.03 | 1 | 0 | Slc25a13 | 0.01 | Old |

| | | | | | | | | |
|---|---|---|---|---|---|---|---|---|
| 0.199203 | 0.169708 | 0.034 | 0.025 | 1 | 0 | Alg8 | 0.009 | Old |
| 0.19957 | 0.314457 | 0.112 | 0.097 | 1 | 0 | Mrps15 | 0.015 | Old |
| 0.199574 | 0.166915 | 0.263 | 0.24 | 1 | 0 | Fn1 | 0.023 | Old |
| 0.199764 | 0.04275 | 0.123 | 0.104 | 1 | 0 | Arhgap10 | 0.019 | Old |
| 0.199841 | 0.229279 | 0.07 | 0.057 | 1 | 0 | Srebf1 | 0.013 | Old |
| 0.199867 | 0.228088 | 0.098 | 0.083 | 1 | 0 | Mras | 0.015 | Old |
| 0.19991 | 0.070425 | 0.718 | 0.688 | 1 | 0 | Samhd1 | 0.03 | Old |
| 0.200025 | 0.58823 | 0.025 | 0.017 | 1 | 0 | Zfp788 | 0.008 | Old |
| 0.200242 | 0.10937 | 0.13 | 0.111 | 1 | 0 | Slc35a3 | 0.019 | Old |
| 0.200427 | 0.881307 | 0.014 | 0.008 | 1 | 0 | Gm31243 | 0.006 | Old |
| 0.200475 | 0.034761 | 0.223 | 0.198 | 1 | 0 | Spcs3 | 0.025 | Old |
| 0.200607 | 0.20091 | 0.055 | 0.043 | 1 | 0 | Srxn1 | 0.012 | Old |
| 0.200674 | 0.036851 | 0.096 | 0.08 | 1 | 0 | Cul2 | 0.016 | Old |
| 0.200958 | 0.279934 | 0.016 | 0.011 | 1 | 0 | Fastkd5 | 0.005 | Old |
| 0.200998 | 0.75967 | 0.011 | 0.006 | 1 | 0 | Zfp446 | 0.005 | Old |
| 0.201037 | 0.222139 | 0.062 | 0.049 | 1 | 0 | Snrnp25 | 0.013 | Old |
| 0.20115 | 0.063791 | 0.089 | 0.074 | 1 | 0 | Hmgcr | 0.015 | Old |
| 0.201238 | 0.189714 | 0.078 | 0.064 | 1 | 0 | Zdhhc5 | 0.014 | Old |
| 0.201253 | 0.144869 | 0.198 | 0.176 | 1 | 0 | Ppp4r2 | 0.022 | Old |
| 0.201269 | 0.084655 | 0.265 | 0.236 | 1 | 0 | Psmd13 | 0.029 | Old |
| 0.201374 | 0.088465 | 0.049 | 0.038 | 1 | 0 | Zbtb16 | 0.011 | Old |
| 0.201446 | 0.830952 | 0.011 | 0.006 | 1 | 0 | Gm10384 | 0.005 | Old |
| 0.201908 | 0.085629 | 0.094 | 0.079 | 1 | 0 | Mrpl4 | 0.015 | Old |
| 0.202134 | 0.489245 | 0.025 | 0.017 | 1 | 0 | Suv39h1 | 0.008 | Old |
| 0.202266 | 0.275842 | 0.096 | 0.081 | 1 | 0 | Hmgn2 | 0.015 | Old |
| 0.202692 | 0.183639 | 0.029 | 0.021 | 1 | 0 | Creld1 | 0.008 | Old |
| 0.203245 | 0.307573 | 0.011 | 0.006 | 1 | 0 | Acot11 | 0.005 | Old |
| 0.203281 | 0.735594 | 0.014 | 0.008 | 1 | 0 | Mrm1 | 0.006 | Old |
| 0.203305 | 0.513555 | 0.066 | 0.053 | 1 | 0 | Gbp2 | 0.013 | Old |
| 0.203534 | 0.346117 | 0.016 | 0.011 | 1 | 0 | Etl4 | 0.005 | Old |
| 0.203663 | 0.158207 | 0.144 | 0.125 | 1 | 0 | Yipf4 | 0.019 | Old |
| 0.203776 | 0.853578 | 0.014 | 0.008 | 1 | 0 | Asf1b | 0.006 | Old |
| 0.20381 | 0.368945 | 0.011 | 0.006 | 1 | 0 | Mttp | 0.005 | Old |
| 0.203959 | 0.434882 | 0.03 | 0.022 | 1 | 0 | 5730480H( | 0.008 | Old |
| 0.204086 | 0.370357 | 0.03 | 0.022 | 1 | 0 | Acaca | 0.008 | Old |
| 0.20418 | 0.058704 | 0.312 | 0.283 | 1 | 0 | Cln8 | 0.029 | Old |
| 0.204233 | 0.435462 | 0.038 | 0.029 | 1 | 0 | Piga | 0.009 | Old |
| 0.204259 | 0.431007 | 0.021 | 0.014 | 1 | 0 | Ptgfrn | 0.007 | Old |
| 0.204277 | 0.220995 | 0.135 | 0.118 | 1 | 0 | Ythdf2 | 0.017 | Old |
| 0.204496 | 0.546374 | 0.021 | 0.014 | 1 | 0 | Rab38 | 0.007 | Old |
| 0.204741 | 0.144375 | 0.049 | 0.038 | 1 | 0 | Kctd3 | 0.011 | Old |
| 0.205145 | 0.261584 | 0.045 | 0.035 | 1 | 0 | Pigv | 0.01 | Old |
| 0.205352 | 0.049353 | 0.285 | 0.257 | 1 | 0 | Usp25 | 0.028 | Old |
| 0.20539 | 0.114859 | 0.529 | 0.5 | 1 | 0 | Sulf2 | 0.029 | Old |
| 0.20542 | 0.248954 | 0.114 | 0.097 | 1 | 0 | Gm49336 | 0.017 | Old |

| | | | | | | | | |
|---|---|---|---|---|---|---|---|---|
| 0.205512 | 0.174969 | 0.052 | 0.041 | 1 | 0 | Trap1 | 0.011 | Old |
| 0.205665 | 0.205854 | 0.014 | 0.008 | 1 | 0 | Neurl1b | 0.006 | Old |
| 0.205701 | 0.348566 | 0.037 | 0.028 | 1 | 0 | Tmem39b | 0.009 | Old |
| 0.205922 | 0.30655 | 0.092 | 0.078 | 1 | 0 | Adgre4 | 0.014 | Old |
| 0.206164 | 0.348621 | 0.014 | 0.008 | 1 | 0 | Nupl2 | 0.006 | Old |
| 0.206207 | 0.126383 | 0.137 | 0.118 | 1 | 0 | Kctd20 | 0.019 | Old |
| 0.206322 | 0.51371 | 0.064 | 0.053 | 1 | 0 | Rpf1 | 0.011 | Old |
| 0.206386 | 0.137868 | 0.174 | 0.153 | 1 | 0 | Elk4 | 0.021 | Old |
| 0.206544 | 0.309887 | 0.025 | 0.017 | 1 | 0 | Osbpl3 | 0.008 | Old |
| 0.207136 | 0.32728 | 0.037 | 0.028 | 1 | 0 | Fbxl19 | 0.009 | Old |
| 0.207491 | 0.05447 | 0.249 | 0.222 | 1 | 0 | Osbpl11 | 0.027 | Old |
| 0.207945 | 0.212356 | 0.025 | 0.017 | 1 | 0 | Smyd2 | 0.008 | Old |
| 0.208021 | 0.069106 | 0.302 | 0.271 | 1 | 0 | Ywhag | 0.031 | Old |
| 0.208036 | 0.011894 | 0.156 | 0.135 | 1 | 0 | Rnf169 | 0.021 | Old |
| 0.208126 | 0.077134 | 0.025 | 0.017 | 1 | 0 | Bphl | 0.008 | Old |
| 0.208294 | 0.459163 | 0.092 | 0.078 | 1 | 0 | Zbtb1 | 0.014 | Old |
| 0.208364 | 0.110475 | 0.397 | 0.367 | 1 | 0 | Slfn2 | 0.03 | Old |
| 0.208387 | 1.0253 | 0.029 | 0.021 | 1 | 0 | 2610044O1 | 0.008 | Old |
| 0.20854 | 0.13241 | 0.052 | 0.041 | 1 | 0 | Ccdc152 | 0.011 | Old |
| 0.208548 | 0.095532 | 0.376 | 0.349 | 1 | 0 | Ppp2r5c | 0.027 | Old |
| 0.20887 | 0.047129 | 0.836 | 0.788 | 1 | 0 | Rack1 | 0.048 | Old |
| 0.208885 | 0.093709 | 0.346 | 0.321 | 1 | 0 | Dynll1 | 0.025 | Old |
| 0.208885 | 0.122442 | 0.204 | 0.182 | 1 | 0 | Dnajc8 | 0.022 | Old |
| 0.208989 | 0.063902 | 0.404 | 0.372 | 1 | 0 | Epsti1 | 0.032 | Old |
| 0.209009 | 0.497007 | 0.026 | 0.019 | 1 | 0 | Dcun1d2 | 0.007 | Old |
| 0.209569 | 0.12836 | 0.049 | 0.038 | 1 | 0 | Pigz | 0.011 | Old |
| 0.209614 | 0.368965 | 0.052 | 0.041 | 1 | 0 | Brms1 | 0.011 | Old |
| 0.209638 | 0.189354 | 0.057 | 0.046 | 1 | 0 | Nf2 | 0.011 | Old |
| 0.210308 | 0.084432 | 0.212 | 0.188 | 1 | 0 | Ftl1-ps1 | 0.024 | Old |
| 0.210419 | 0.025847 | 0.209 | 0.185 | 1 | 0 | Zfp445 | 0.024 | Old |
| 0.21057 | 0.049105 | 0.301 | 0.27 | 1 | 0 | Pebp1 | 0.031 | Old |
| 0.210728 | 0.264169 | 0.073 | 0.06 | 1 | 0 | Smg9 | 0.013 | Old |
| 0.210982 | 0.213669 | 0.12 | 0.104 | 1 | 0 | Ccdc71l | 0.016 | Old |
| 0.211315 | 0.244318 | 0.13 | 0.113 | 1 | 0 | Cln6 | 0.017 | Old |
| 0.21133 | 0.058549 | 0.108 | 0.091 | 1 | 0 | Ppie | 0.017 | Old |
| 0.211795 | 0.238902 | 0.059 | 0.047 | 1 | 0 | Mtpap | 0.012 | Old |
| 0.211912 | 0.101591 | 0.131 | 0.113 | 1 | 0 | Raf1 | 0.018 | Old |
| 0.212308 | 0.040096 | 0.501 | 0.455 | 1 | 0 | Msn | 0.046 | Old |
| 0.212395 | 0.219761 | 0.185 | 0.166 | 1 | 0 | Dcaf7 | 0.019 | Old |
| 0.212613 | 0.275217 | 0.038 | 0.029 | 1 | 0 | Ubxn2b | 0.009 | Old |
| 0.212674 | 0.132096 | 0.146 | 0.127 | 1 | 0 | Fnbp1l | 0.019 | Old |
| 0.212814 | 0.03011 | 0.488 | 0.443 | 1 | 0 | Ncf2 | 0.045 | Old |
| 0.212889 | 0.176479 | 0.049 | 0.039 | 1 | 0 | Camta1 | 0.01 | Old |
| 0.213077 | 0.038697 | 0.176 | 0.155 | 1 | 0 | Inpp5k | 0.021 | Old |
| 0.213238 | 0.019341 | 0.279 | 0.25 | 1 | 0 | Gm26740 | 0.029 | Old |

| | | | | | | | | |
|---|---|---|---|---|---|---|---|---|
| 0.213395 | 0.195315 | 0.178 | 0.159 | 1 | 0 | Qdpr | 0.019 | Old |
| 0.213665 | 0.067831 | 0.239 | 0.215 | 1 | 0 | Rlim | 0.024 | Old |
| 0.213791 | 0.536171 | 0.022 | 0.015 | 1 | 0 | Gm49359 | 0.007 | Old |
| 0.213909 | 0.110468 | 0.415 | 0.385 | 1 | 0 | Mapk3 | 0.03 | Old |
| 0.214187 | 0.090439 | 0.064 | 0.052 | 1 | 0 | Per3 | 0.012 | Old |
| 0.214293 | 0.08661 | 0.3 | 0.274 | 1 | 0 | Metap2 | 0.026 | Old |
| 0.214428 | 0.124933 | 0.083 | 0.07 | 1 | 0 | Slc5a3 | 0.013 | Old |
| 0.214677 | 0.488179 | 0.018 | 0.012 | 1 | 0 | Nmrk1 | 0.006 | Old |
| 0.215116 | 0.140946 | 0.098 | 0.083 | 1 | 0 | Hist1h2bc | 0.015 | Old |
| 0.215517 | 0.193009 | 0.109 | 0.094 | 1 | 0 | Wdr33 | 0.015 | Old |
| 0.215656 | 0.22656 | 0.129 | 0.112 | 1 | 0 | Wdfy4 | 0.017 | Old |
| 0.215726 | 0.311107 | 0.085 | 0.072 | 1 | 0 | Slc30a1 | 0.013 | Old |
| 0.21579 | 0.006982 | 0.096 | 0.08 | 1 | 0 | Gcn1l1 | 0.016 | Old |
| 0.217014 | 0.394819 | 0.022 | 0.015 | 1 | 0 | Galm | 0.007 | Old |
| 0.217633 | 0.058918 | 0.142 | 0.123 | 1 | 0 | Vrk2 | 0.019 | Old |
| 0.217665 | 0.269252 | 0.068 | 0.056 | 1 | 0 | Jmy | 0.012 | Old |
| 0.21786 | 0.028632 | 0.305 | 0.273 | 1 | 0 | Manf | 0.032 | Old |
| 0.218084 | 0.464508 | 0.048 | 0.038 | 1 | 0 | Tmem175 | 0.01 | Old |
| 0.218185 | 0.026465 | 0.425 | 0.383 | 1 | 0 | Rab5c | 0.042 | Old |
| 0.21819 | 0.083285 | 0.674 | 0.639 | 1 | 0 | Arf5 | 0.035 | Old |
| 0.218274 | 0.088594 | 0.1 | 0.084 | 1 | 0 | Fbrs | 0.016 | Old |
| 0.218469 | 0.043518 | 0.068 | 0.056 | 1 | 0 | Manbal | 0.012 | Old |
| 0.218736 | 0.101987 | 0.166 | 0.147 | 1 | 0 | Atp6v1d | 0.019 | Old |
| 0.218741 | 0.184155 | 0.145 | 0.127 | 1 | 0 | Rras | 0.018 | Old |
| 0.219067 | 0.122147 | 0.286 | 0.257 | 1 | 0 | Copz1 | 0.029 | Old |
| 0.219142 | 0.200971 | 0.131 | 0.115 | 1 | 0 | Cd200r1 | 0.016 | Old |
| 0.219149 | 0.303263 | 0.078 | 0.065 | 1 | 0 | Mtf1 | 0.013 | Old |
| 0.219434 | 0.794006 | 0.023 | 0.016 | 1 | 0 | Il18bp | 0.007 | Old |
| 0.219551 | 0.182724 | 0.175 | 0.156 | 1 | 0 | Wtap | 0.019 | Old |
| 0.219691 | 0.126251 | 0.124 | 0.107 | 1 | 0 | Zwint | 0.017 | Old |
| 0.219868 | 0.130848 | 0.085 | 0.071 | 1 | 0 | Nelfa | 0.014 | Old |
| 0.219925 | 0.163398 | 0.018 | 0.012 | 1 | 0 | Tubb2b | 0.006 | Old |
| 0.220074 | 0.001663 | 0.066 | 0.053 | 1 | 0 | Ap1g2 | 0.013 | Old |
| 0.220088 | 0.049661 | 0.215 | 0.19 | 1 | 0 | Dnajc7 | 0.025 | Old |
| 0.220163 | 0.359477 | 0.051 | 0.04 | 1 | 0 | Mief1 | 0.011 | Old |
| 0.220168 | 0.195837 | 0.127 | 0.11 | 1 | 0 | Rbm42 | 0.017 | Old |
| 0.220381 | 0.140793 | 0.09 | 0.076 | 1 | 0 | Brip1os | 0.014 | Old |
| 0.220529 | 0.220292 | 0.114 | 0.098 | 1 | 0 | Sos2 | 0.016 | Old |
| 0.220536 | 0.057379 | 0.181 | 0.158 | 1 | 0 | Ifnar2 | 0.023 | Old |
| 0.220571 | 0.430425 | 0.034 | 0.026 | 1 | 0 | Pqlc2 | 0.008 | Old |
| 0.220892 | 0.236699 | 0.033 | 0.024 | 1 | 0 | Syne3 | 0.009 | Old |
| 0.220993 | 0.135983 | 0.022 | 0.015 | 1 | 0 | Gm21781 | 0.007 | Old |
| 0.221598 | 0.274951 | 0.098 | 0.084 | 1 | 0 | Rab29 | 0.014 | Old |
| 0.221598 | 0.685182 | 0.015 | 0.01 | 1 | 0 | Mmaa | 0.005 | Old |
| 0.221836 | 0.309239 | 0.168 | 0.152 | 1 | 0 | Got1 | 0.016 | Old |

| | | | | | | | | |
|---|---|---|---|---|---|---|---|---|
| 0.222072 | 0.243785 | 0.092 | 0.078 | 1 | 0 | Alkbh1 | 0.014 | Old |
| 0.22213 | 0.151755 | 0.35 | 0.324 | 1 | 0 | Clic4 | 0.026 | Old |
| 0.222144 | 0.188753 | 0.053 | 0.043 | 1 | 0 | Gtf2f2 | 0.01 | Old |
| 0.22226 | 0.033276 | 0.112 | 0.095 | 1 | 0 | Elf4 | 0.017 | Old |
| 0.222478 | 0.189279 | 0.134 | 0.116 | 1 | 0 | Dctn5 | 0.018 | Old |
| 0.222763 | 0.108276 | 0.041 | 0.032 | 1 | 0 | Dedd | 0.009 | Old |
| 0.222925 | 0.188727 | 0.149 | 0.131 | 1 | 0 | Hipk3 | 0.018 | Old |
| 0.223043 | 0.094429 | 0.073 | 0.059 | 1 | 0 | Nvl | 0.014 | Old |
| 0.223193 | 0.005414 | 0.109 | 0.093 | 1 | 0 | Ahctf1 | 0.016 | Old |
| 0.223306 | 1.111703 | 0.012 | 0.008 | 1 | 0 | Jaml | 0.004 | Old |
| 0.223452 | 0.13283 | 0.06 | 0.048 | 1 | 0 | Nhlrc2 | 0.012 | Old |
| 0.22355 | 0.229588 | 0.131 | 0.115 | 1 | 0 | Bckdk | 0.016 | Old |
| 0.223783 | 0.042613 | 0.007 | 0.012 | 1 | 0 | Trim16 | -0.005 | Old |
| 0.224182 | 0.442424 | 0.015 | 0.01 | 1 | 0 | Sprypd4 | 0.005 | Old |
| 0.224194 | 0.156522 | 0.071 | 0.059 | 1 | 0 | Gmppa | 0.012 | Old |
| 0.224381 | 0.393653 | 0.015 | 0.01 | 1 | 0 | Ddx59 | 0.005 | Old |
| 0.22441 | 0.202291 | 0.034 | 0.026 | 1 | 0 | Dhx35 | 0.008 | Old |
| 0.224752 | 0.529065 | 0.038 | 0.029 | 1 | 0 | Klhl7 | 0.009 | Old |
| 0.224918 | 0.133638 | 0.202 | 0.181 | 1 | 0 | Srpr | 0.021 | Old |
| 0.224931 | 0.561469 | 0.019 | 0.013 | 1 | 0 | Tmem154 | 0.006 | Old |
| 0.224949 | 0.075285 | 0.315 | 0.284 | 1 | 0 | Dnase2a | 0.031 | Old |
| 0.225044 | 0.287288 | 0.034 | 0.026 | 1 | 0 | Zdhhc13 | 0.008 | Old |
| 0.225288 | 0.186018 | 0.055 | 0.044 | 1 | 0 | Eipr1 | 0.011 | Old |
| 0.225486 | 0.041888 | 0.26 | 0.233 | 1 | 0 | Ppp6r3 | 0.027 | Old |
| 0.225775 | 0.061839 | 0.161 | 0.141 | 1 | 0 | Fto | 0.02 | Old |
| 0.225782 | 0.277027 | 0.015 | 0.01 | 1 | 0 | 2700062C0 | 0.005 | Old |
| 0.226165 | 0.037554 | 0.074 | 0.061 | 1 | 0 | Fam98c | 0.013 | Old |
| 0.226219 | 0.751008 | 0.012 | 0.008 | 1 | 0 | Nthl1 | 0.004 | Old |
| 0.226229 | 0.039592 | 0.074 | 0.061 | 1 | 0 | Wipi2 | 0.013 | Old |
| 0.226261 | 0.317893 | 0.01 | 0.005 | 1 | 0 | Gm13205 | 0.005 | Old |
| 0.226391 | 0.369533 | 0.01 | 0.005 | 1 | 0 | Kif15 | 0.005 | Old |
| 0.226398 | 0.037555 | 0.137 | 0.118 | 1 | 0 | Tbc1d2b | 0.019 | Old |
| 0.226469 | 0.087723 | 0.15 | 0.132 | 1 | 0 | Eny2 | 0.018 | Old |
| 0.226653 | 0.430475 | 0.01 | 0.005 | 1 | 0 | Zfp189 | 0.005 | Old |
| 0.226915 | 0.378101 | 0.01 | 0.005 | 1 | 0 | Gm36839 | 0.005 | Old |
| 0.227235 | 0.315039 | 0.036 | 0.027 | 1 | 0 | Ccdc86 | 0.009 | Old |
| 0.227246 | 0.166436 | 0.19 | 0.17 | 1 | 0 | Tial1 | 0.02 | Old |
| 0.227445 | 0.096049 | 0.238 | 0.214 | 1 | 0 | Ccar1 | 0.024 | Old |
| 0.227599 | 0.018331 | 0.442 | 0.397 | 1 | 0 | M6pr | 0.045 | Old |
| 0.227717 | 0.324226 | 0.029 | 0.021 | 1 | 0 | Fggy | 0.008 | Old |
| 0.227742 | 0.197266 | 0.098 | 0.084 | 1 | 0 | Sec13 | 0.014 | Old |
| 0.227828 | 0.102542 | 0.263 | 0.237 | 1 | 0 | Sdhc | 0.026 | Old |
| 0.227861 | 0.288952 | 0.036 | 0.027 | 1 | 0 | Enho | 0.009 | Old |
| 0.228079 | 0.073946 | 0.293 | 0.265 | 1 | 0 | Smchd1 | 0.028 | Old |
| 0.228147 | 0.139642 | 0.097 | 0.082 | 1 | 0 | Casp1 | 0.015 | Old |

| | | | | | | | | |
|---|---|---|---|---|---|---|---|---|
| 0.228179 | 0.083368 | 0.397 | 0.364 | 1 | 0 | Larp4b | 0.033 | Old |
| 0.228249 | 0.405438 | 0.037 | 0.028 | 1 | 0 | 3110002H1 | 0.009 | Old |
| 0.228317 | 0.085923 | 0.119 | 0.103 | 1 | 0 | Ptpn2 | 0.016 | Old |
| 0.228389 | 0.458082 | 0.019 | 0.013 | 1 | 0 | Zfp580 | 0.006 | Old |
| 0.228448 | 0.076061 | 0.624 | 0.599 | 1 | 0 | Arhgdia | 0.025 | Old |
| 0.228658 | 0.512946 | 0.062 | 0.051 | 1 | 0 | Hvcn1 | 0.011 | Old |
| 0.228743 | 0.416118 | 0.081 | 0.068 | 1 | 0 | Rnf144b | 0.013 | Old |
| 0.228932 | 0.526645 | 0.012 | 0.008 | 1 | 0 | Pim2 | 0.004 | Old |
| 0.229016 | 0.124264 | 0.01 | 0.005 | 1 | 0 | Tmlhe | 0.005 | Old |
| 0.229029 | 0.105826 | 0.137 | 0.119 | 1 | 0 | Ddx21 | 0.018 | Old |
| 0.229222 | 0.118224 | 0.138 | 0.12 | 1 | 0 | Sp110 | 0.018 | Old |
| 0.229241 | 0.268976 | 0.036 | 0.027 | 1 | 0 | Zbtb45 | 0.009 | Old |
| 0.229251 | 0.218343 | 0.114 | 0.099 | 1 | 0 | Idh3b | 0.015 | Old |
| 0.229273 | 0.478578 | 0.012 | 0.008 | 1 | 0 | Tmem14a | 0.004 | Old |
| 0.229497 | 0.176474 | 0.116 | 0.1 | 1 | 0 | Htra2 | 0.016 | Old |
| 0.230045 | 0.346898 | 0.085 | 0.072 | 1 | 0 | Ccdc9 | 0.013 | Old |
| 0.230183 | 0.312448 | 0.012 | 0.008 | 1 | 0 | Trmo | 0.004 | Old |
| 0.230718 | 0.007196 | 0.324 | 0.288 | 1 | 0 | Sdf4 | 0.036 | Old |
| 0.230731 | 0.333279 | 0.019 | 0.013 | 1 | 0 | Zfp60 | 0.006 | Old |
| 0.230833 | 0.326413 | 0.137 | 0.121 | 1 | 0 | Nxpe5 | 0.016 | Old |
| 0.230875 | 0.137255 | 0.077 | 0.064 | 1 | 0 | Atg7 | 0.013 | Old |
| 0.231062 | 0.523696 | 0.031 | 0.024 | 1 | 0 | Fancg | 0.007 | Old |
| 0.231134 | 0.030541 | 0.049 | 0.039 | 1 | 0 | Prkrip1 | 0.01 | Old |
| 0.231175 | 0.29177 | 0.019 | 0.013 | 1 | 0 | Homer2 | 0.006 | Old |
| 0.231554 | 0.25568 | 0.012 | 0.008 | 1 | 0 | Rexo5 | 0.004 | Old |
| 0.231878 | 0.274298 | 0.089 | 0.075 | 1 | 0 | Edem2 | 0.014 | Old |
| 0.231878 | 0.045986 | 0.156 | 0.137 | 1 | 0 | Srd5a3 | 0.019 | Old |
| 0.232236 | 0.024045 | 0.189 | 0.167 | 1 | 0 | Edf1 | 0.022 | Old |
| 0.232254 | 0.142835 | 0.094 | 0.08 | 1 | 0 | Aak1 | 0.014 | Old |
| 0.232304 | 0.136892 | 0.045 | 0.035 | 1 | 0 | Ints11 | 0.01 | Old |
| 0.232496 | 0.073668 | 0.036 | 0.027 | 1 | 0 | Pym1 | 0.009 | Old |
| 0.232684 | 0.101786 | 0.156 | 0.137 | 1 | 0 | Birc2 | 0.019 | Old |
| 0.232853 | 0.261519 | 0.067 | 0.055 | 1 | 0 | Rpl10-ps3 | 0.012 | Old |
| 0.233243 | 0.180717 | 0.029 | 0.021 | 1 | 0 | D3Ertd254 | 0.008 | Old |
| 0.233332 | 0.336236 | 0.051 | 0.04 | 1 | 0 | Ikzf2 | 0.011 | Old |
| 0.233353 | 0.122809 | 0.222 | 0.199 | 1 | 0 | Rbm8a | 0.023 | Old |
| 0.233505 | 0.001729 | 0.012 | 0.008 | 1 | 0 | 4930481A1 | 0.004 | Old |
| 0.23366 | 0.05154 | 0.034 | 0.026 | 1 | 0 | Prmt7 | 0.008 | Old |
| 0.233877 | 0.100809 | 0.118 | 0.102 | 1 | 0 | Snx1 | 0.016 | Old |
| 0.233939 | 0.009667 | 0.019 | 0.013 | 1 | 0 | Slc17a9 | 0.006 | Old |
| 0.234258 | 0.280805 | 0.037 | 0.028 | 1 | 0 | Zfp143 | 0.009 | Old |
| 0.234618 | 0.318583 | 0.025 | 0.018 | 1 | 0 | Slc22a15 | 0.007 | Old |
| 0.234657 | 0.051286 | 0.019 | 0.013 | 1 | 0 | Tgm1 | 0.006 | Old |
| 0.234739 | 0.068772 | 0.042 | 0.033 | 1 | 0 | Isca1 | 0.009 | Old |
| 0.234748 | 0.095194 | 0.189 | 0.168 | 1 | 0 | Fam91a1 | 0.021 | Old |

| | | | | | | | | |
|---|---|---|---|---|---|---|---|---|
| 0.23482 | 0.096491 | 0.228 | 0.205 | 1 | 0 | Cnn2 | 0.023 | Old |
| 0.234827 | 0.14123 | 0.238 | 0.217 | 1 | 0 | Pitpnc1 | 0.021 | Old |
| 0.234894 | 0.264494 | 0.1 | 0.086 | 1 | 0 | Tmem71 | 0.014 | Old |
| 0.23508 | 0.134712 | 0.092 | 0.077 | 1 | 0 | Zfp984 | 0.015 | Old |
| 0.235255 | 0.201355 | 0.111 | 0.096 | 1 | 0 | Cask | 0.015 | Old |
| 0.235324 | 0.21637 | 0.178 | 0.16 | 1 | 0 | Ranbp1 | 0.018 | Old |
| 0.235462 | 0.038224 | 0.361 | 0.324 | 1 | 0 | Arid1a | 0.037 | Old |
| 0.235541 | 0.053919 | 0.248 | 0.221 | 1 | 0 | Ppp3r1 | 0.027 | Old |
| 0.235556 | 0.077431 | 0.323 | 0.292 | 1 | 0 | Kdelr1 | 0.031 | Old |
| 0.235753 | 0.191074 | 0.029 | 0.021 | 1 | 0 | Tcim | 0.008 | Old |
| 0.235858 | 0.296757 | 0.155 | 0.138 | 1 | 0 | Creb5 | 0.017 | Old |
| 0.235919 | 0.235035 | 0.14 | 0.123 | 1 | 0 | Rara | 0.017 | Old |
| 0.23593 | 0.372336 | 0.06 | 0.049 | 1 | 0 | Poldip2 | 0.011 | Old |
| 0.236246 | 0.281403 | 0.051 | 0.04 | 1 | 0 | Spice1 | 0.011 | Old |
| 0.236401 | 0.453501 | 0.033 | 0.025 | 1 | 0 | Asf1a | 0.008 | Old |
| 0.236449 | 0.080983 | 0.078 | 0.065 | 1 | 0 | Tfeb | 0.013 | Old |
| 0.236526 | 0.093871 | 0.297 | 0.27 | 1 | 0 | Psmb1 | 0.027 | Old |
| 0.236541 | 0.724209 | 0.016 | 0.011 | 1 | 0 | A930007I1 | 0.005 | Old |
| 0.237072 | 0.061385 | 0.512 | 0.472 | 1 | 0 | Slc25a5 | 0.04 | Old |
| 0.237621 | 0.097773 | 0.443 | 0.416 | 1 | 0 | Top1 | 0.027 | Old |
| 0.237774 | 0.445361 | 0.033 | 0.025 | 1 | 0 | Trim37 | 0.008 | Old |
| 0.237964 | 0.254727 | 0.049 | 0.039 | 1 | 0 | Slc39a14 | 0.01 | Old |
| 0.238029 | 0.353664 | 0.014 | 0.021 | 1 | 0 | Serf1 | -0.007 | Old |
| 0.238062 | 0.085016 | 0.261 | 0.238 | 1 | 0 | Ubtf | 0.023 | Old |
| 0.238069 | 0.173761 | 0.085 | 0.071 | 1 | 0 | Pon3 | 0.014 | Old |
| 0.238073 | 0.244914 | 0.071 | 0.059 | 1 | 0 | Ccdc25 | 0.012 | Old |
| 0.238268 | 0.158204 | 0.168 | 0.149 | 1 | 0 | Napa | 0.019 | Old |
| 0.238436 | 0.096703 | 0.446 | 0.414 | 1 | 0 | Atp1b3 | 0.032 | Old |
| 0.238494 | 0.377629 | 0.104 | 0.09 | 1 | 0 | Agpat5 | 0.014 | Old |
| 0.238733 | 0.306495 | 0.025 | 0.018 | 1 | 0 | Tbl2 | 0.007 | Old |
| 0.238911 | 0.3529 | 0.042 | 0.033 | 1 | 0 | Sema4d | 0.009 | Old |
| 0.238924 | 0.078977 | 0.192 | 0.17 | 1 | 0 | Stx16 | 0.022 | Old |
| 0.239192 | 0.116833 | 0.183 | 0.163 | 1 | 0 | Ube2e3 | 0.02 | Old |
| 0.239201 | 0.640421 | 0.016 | 0.011 | 1 | 0 | Amotl2 | 0.005 | Old |
| 0.239285 | 0.101049 | 0.133 | 0.116 | 1 | 0 | Hpf1 | 0.017 | Old |
| 0.239618 | 0.088977 | 0.124 | 0.107 | 1 | 0 | Oaz2 | 0.017 | Old |
| 0.23969 | 0.080641 | 0.109 | 0.094 | 1 | 0 | Stk40 | 0.015 | Old |
| 0.239874 | 0.186785 | 0.068 | 0.056 | 1 | 0 | Mkl2 | 0.012 | Old |
| 0.240307 | 0.096805 | 0.057 | 0.046 | 1 | 0 | Xaf1 | 0.011 | Old |
| 0.240449 | 0.143283 | 0.048 | 0.038 | 1 | 0 | Hectd3 | 0.01 | Old |
| 0.24062 | 0.126255 | 0.075 | 0.063 | 1 | 0 | Ncbp1 | 0.012 | Old |
| 0.240641 | 0.367908 | 0.14 | 0.124 | 1 | 0 | Ahr | 0.016 | Old |
| 0.240701 | 0.028573 | 0.081 | 0.067 | 1 | 0 | Zpr1 | 0.014 | Old |
| 0.240751 | 0.146849 | 0.025 | 0.018 | 1 | 0 | Sgsm2 | 0.007 | Old |
| 0.240927 | 0.025459 | 0.167 | 0.146 | 1 | 0 | Nampt | 0.021 | Old |

| | | | | | | | | |
|---|---|---|---|---|---|---|---|---|
| 0.241102 | 0.57458 | 0.056 | 0.046 | 1 | 0 | Coro2a | 0.01 | Old |
| 0.241169 | 0.10128 | 0.233 | 0.21 | 1 | 0 | Pgls | 0.023 | Old |
| 0.241331 | 0.05116 | 0.272 | 0.245 | 1 | 0 | Ocrl | 0.027 | Old |
| 0.241435 | 0.099662 | 0.378 | 0.345 | 1 | 0 | Neurl3 | 0.033 | Old |
| 0.241972 | 0.001586 | 0.263 | 0.236 | 1 | 0 | Smim10l1 | 0.027 | Old |
| 0.242032 | 0.149862 | 0.1 | 0.086 | 1 | 0 | Lta4h | 0.014 | Old |
| 0.242034 | 0.355906 | 0.021 | 0.014 | 1 | 0 | AW209491 | 0.007 | Old |
| 0.242103 | 0.10052 | 0.133 | 0.115 | 1 | 0 | Crat | 0.018 | Old |
| 0.24233 | 0.033027 | 0.03 | 0.022 | 1 | 0 | Mettl8 | 0.008 | Old |
| 0.24269 | 0.135335 | 0.135 | 0.118 | 1 | 0 | Psen1 | 0.017 | Old |
| 0.2428 | 0.053173 | 0.274 | 0.248 | 1 | 0 | Amfr | 0.026 | Old |
| 0.242858 | 0.293503 | 0.059 | 0.048 | 1 | 0 | Ndufaf4 | 0.011 | Old |
| 0.243168 | 0.436808 | 0.042 | 0.033 | 1 | 0 | Bcam | 0.009 | Old |
| 0.243212 | 0.045307 | 0.208 | 0.186 | 1 | 0 | Nfat5 | 0.022 | Old |
| 0.243696 | 0.144232 | 0.031 | 0.024 | 1 | 0 | Repin1 | 0.007 | Old |
| 0.243791 | 0.087831 | 0.562 | 0.531 | 1 | 0 | Coro1a | 0.031 | Old |
| 0.243796 | 0.075367 | 0.683 | 0.634 | 1 | 0 | Sh3bgrl3 | 0.049 | Old |
| 0.243991 | 0.135196 | 0.038 | 0.029 | 1 | 0 | Ogfod2 | 0.009 | Old |
| 0.244114 | 0.052459 | 0.088 | 0.074 | 1 | 0 | Impa1 | 0.014 | Old |
| 0.244778 | 0.312992 | 0.042 | 0.033 | 1 | 0 | Copz2 | 0.009 | Old |
| 0.245172 | 0.122834 | 0.361 | 0.328 | 1 | 0 | Dusp6 | 0.033 | Old |
| 0.245372 | 0.369935 | 0.049 | 0.04 | 1 | 0 | Tusc1 | 0.009 | Old |
| 0.245956 | 0.38053 | 0.044 | 0.035 | 1 | 0 | Pex14 | 0.009 | Old |
| 0.246126 | 0.175469 | 0.071 | 0.059 | 1 | 0 | Sntb2 | 0.012 | Old |
| 0.246231 | 0.037388 | 0.518 | 0.475 | 1 | 0 | Atp6ap1 | 0.043 | Old |
| 0.246245 | 0.186377 | 0.042 | 0.033 | 1 | 0 | Pdpr | 0.009 | Old |
| 0.246288 | 0.254867 | 0.026 | 0.019 | 1 | 0 | Prickle3 | 0.007 | Old |
| 0.246379 | 0.04217 | 0.843 | 0.818 | 1 | 0 | Myl6 | 0.025 | Old |
| 0.246462 | 0.477545 | 0.036 | 0.027 | 1 | 0 | Pi4kb | 0.009 | Old |
| 0.247058 | 0.100921 | 0.153 | 0.135 | 1 | 0 | Cacybp | 0.018 | Old |
| 0.247077 | 0.111629 | 0.152 | 0.134 | 1 | 0 | Rab24 | 0.018 | Old |
| 0.247305 | 0.074969 | 0.319 | 0.289 | 1 | 0 | Trim30a | 0.03 | Old |
| 0.247809 | 0.218225 | 0.042 | 0.033 | 1 | 0 | Slc35e2 | 0.009 | Old |
| 0.247959 | 0.327187 | 0.059 | 0.048 | 1 | 0 | Slc7a13 | 0.011 | Old |
| 0.24805 | 0.134933 | 0.137 | 0.12 | 1 | 0 | Amdhd2 | 0.017 | Old |
| 0.248388 | 0.1555 | 0.111 | 0.096 | 1 | 0 | Washc1 | 0.015 | Old |
| 0.248544 | 0.102766 | 0.082 | 0.069 | 1 | 0 | Ppm1k | 0.013 | Old |
| 0.24871 | 0.485936 | 0.037 | 0.029 | 1 | 0 | Lrrc14 | 0.008 | Old |
| 0.248844 | 0.077508 | 0.211 | 0.188 | 1 | 0 | Calu | 0.023 | Old |
| 0.249245 | 0.129636 | 0.068 | 0.057 | 1 | 0 | Bloc1s2 | 0.011 | Old |
| 0.2496 | 0.065446 | 0.119 | 0.103 | 1 | 0 | Pofut1 | 0.016 | Old |
| 0.249608 | 0.217837 | 0.155 | 0.138 | 1 | 0 | Myd88 | 0.017 | Old |
| 0.249629 | 0.269538 | 0.045 | 0.036 | 1 | 0 | Sik2 | 0.009 | Old |
| 0.24986 | 0.243187 | 0.057 | 0.047 | 1 | 0 | Itpa | 0.01 | Old |
| 0.249897 | 0.122333 | 0.059 | 0.048 | 1 | 0 | Lrrc57 | 0.011 | Old |

| | | | | | | | | |
|---|---|---|---|---|---|---|---|---|
| 0.250074 | 0.189568 | 0.124 | 0.109 | 1 | 0 | Krt80 | 0.015 | Old |
| 0.25013 | 0.066121 | 0.202 | 0.181 | 1 | 0 | Prkcsh | 0.021 | Old |
| 0.250152 | 0.410285 | 0.014 | 0.009 | 1 | 0 | Oas1c | 0.005 | Old |
| 0.250423 | 0.429769 | 0.034 | 0.026 | 1 | 0 | Cdc25a | 0.008 | Old |
| 0.250524 | 0.19481 | 0.044 | 0.035 | 1 | 0 | Tpgs1 | 0.009 | Old |
| 0.250604 | 0.111134 | 0.185 | 0.166 | 1 | 0 | Mdm4 | 0.019 | Old |
| 0.250747 | 0.148884 | 0.179 | 0.161 | 1 | 0 | Snrpc | 0.018 | Old |
| 0.251094 | 0.019951 | 0.647 | 0.599 | 1 | 0 | Ppp3ca | 0.048 | Old |
| 0.251313 | 0.124675 | 0.122 | 0.106 | 1 | 0 | Tars | 0.016 | Old |
| 0.251419 | 0.620658 | 0.018 | 0.012 | 1 | 0 | Kyat1 | 0.006 | Old |
| 0.251533 | 0.418904 | 0.036 | 0.027 | 1 | 0 | Rmnd1 | 0.009 | Old |
| 0.251735 | 0.449977 | 0.014 | 0.009 | 1 | 0 | Cpt1c | 0.005 | Old |
| 0.251756 | 0.330868 | 0.049 | 0.04 | 1 | 0 | Mettl21a | 0.009 | Old |
| 0.251795 | 0.060429 | 0.133 | 0.115 | 1 | 0 | Plxnc1 | 0.018 | Old |
| 0.251809 | 0.056243 | 0.145 | 0.126 | 1 | 0 | Ubald1 | 0.019 | Old |
| 0.251863 | 0.134196 | 0.271 | 0.25 | 1 | 0 | Egr2 | 0.021 | Old |
| 0.252154 | 0.052448 | 0.222 | 0.199 | 1 | 0 | Uhmk1 | 0.023 | Old |
| 0.252493 | 0.622628 | 0.018 | 0.012 | 1 | 0 | Oasl1 | 0.006 | Old |
| 0.252649 | 0.038494 | 0.051 | 0.04 | 1 | 0 | Borcs7 | 0.011 | Old |
| 0.252672 | 0.077909 | 0.175 | 0.156 | 1 | 0 | Tox4 | 0.019 | Old |
| 0.252884 | 0.638512 | 0.018 | 0.012 | 1 | 0 | Nubpl | 0.006 | Old |
| 0.252991 | 0.030159 | 0.114 | 0.097 | 1 | 0 | Ints6 | 0.017 | Old |
| 0.253069 | 0.192277 | 0.027 | 0.02 | 1 | 0 | Ercc4 | 0.007 | Old |
| 0.253089 | 0.167211 | 0.09 | 0.077 | 1 | 0 | Crybg1 | 0.013 | Old |
| 0.253186 | 0.11549 | 0.413 | 0.387 | 1 | 0 | Herpud1 | 0.026 | Old |
| 0.253235 | 0.625033 | 0.03 | 0.023 | 1 | 0 | Tmem38a | 0.007 | Old |
| 0.253408 | 0.155111 | 0.092 | 0.078 | 1 | 0 | Rars | 0.014 | Old |
| 0.25357 | 0.178836 | 0.022 | 0.016 | 1 | 0 | Ssx2ip | 0.006 | Old |
| 0.253573 | 0.115681 | 0.104 | 0.089 | 1 | 0 | S1pr2 | 0.015 | Old |
| 0.25398 | 0.224789 | 0.166 | 0.149 | 1 | 0 | Rap2c | 0.017 | Old |
| 0.254075 | 0.121376 | 0.027 | 0.02 | 1 | 0 | Etfbkmt | 0.007 | Old |
| 0.254147 | 0.001203 | 0.516 | 0.537 | 1 | 0 | Hsp90aa1 | -0.021 | Old |
| 0.254774 | 0.04415 | 0.081 | 0.068 | 1 | 0 | Sf3a2 | 0.013 | Old |
| 0.254833 | 0.026228 | 0.186 | 0.164 | 1 | 0 | Ythdf1 | 0.022 | Old |
| 0.255048 | 0.023787 | 0.088 | 0.074 | 1 | 0 | Pop5 | 0.014 | Old |
| 0.255176 | 0.590807 | 0.023 | 0.017 | 1 | 0 | Txnrd3 | 0.006 | Old |
| 0.255217 | 0.120714 | 0.034 | 0.026 | 1 | 0 | A530017D: | 0.008 | Old |
| 0.255439 | 0.532565 | 0.03 | 0.023 | 1 | 0 | Orai2 | 0.007 | Old |
| 0.255634 | 0.211576 | 0.14 | 0.124 | 1 | 0 | Limd2 | 0.016 | Old |
| 0.25568 | 0.156769 | 0.079 | 0.067 | 1 | 0 | Usp45 | 0.012 | Old |
| 0.255813 | 0.075864 | 0.631 | 0.615 | 1 | 0 | Ctsa | 0.016 | Old |
| 0.255817 | 0.339104 | 0.029 | 0.021 | 1 | 0 | Oraov1 | 0.008 | Old |
| 0.25611 | 0.090662 | 0.168 | 0.15 | 1 | 0 | Ppp6r1 | 0.018 | Old |
| 0.256459 | 0.511285 | 0.09 | 0.079 | 1 | 0 | Creb3l2 | 0.011 | Old |
| 0.256495 | 0.116711 | 0.254 | 0.231 | 1 | 0 | Edem1 | 0.023 | Old |

| | | | | | | | | |
|---|---|---|---|---|---|---|---|---|
| 0.256556 | 0.611204 | 0.011 | 0.007 | 1 | 0 | Tmem216 | 0.004 | Old |
| 0.25736 | 0.078072 | 0.034 | 0.026 | 1 | 0 | Yipf2 | 0.008 | Old |
| 0.257365 | 0.062515 | 0.194 | 0.174 | 1 | 0 | Sptlc2 | 0.02 | Old |
| 0.257862 | 0.453985 | 0.011 | 0.007 | 1 | 0 | Rccd1 | 0.004 | Old |
| 0.257993 | 0.270435 | 0.011 | 0.007 | 1 | 0 | Cad | 0.004 | Old |
| 0.258043 | 0.26473 | 0.037 | 0.029 | 1 | 0 | Fam151b | 0.008 | Old |
| 0.25811 | 0.080118 | 0.438 | 0.41 | 1 | 0 | Ifi204 | 0.028 | Old |
| 0.25838 | 0.001782 | 0.178 | 0.158 | 1 | 0 | St6galnac4 | 0.02 | Old |
| 0.258485 | 0.223486 | 0.114 | 0.099 | 1 | 0 | Atad1 | 0.015 | Old |
| 0.258485 | 0.126731 | 0.067 | 0.056 | 1 | 0 | Eef1akmt1 | 0.011 | Old |
| 0.25891 | 0.133644 | 0.011 | 0.007 | 1 | 0 | Oxld1 | 0.004 | Old |
| 0.258963 | 0.072403 | 0.358 | 0.334 | 1 | 0 | Fubp1 | 0.024 | Old |
| 0.25899 | 0.025881 | 0.027 | 0.02 | 1 | 0 | Lancl2 | 0.007 | Old |
| 0.259041 | 0.12535 | 0.011 | 0.007 | 1 | 0 | Ehhadh | 0.004 | Old |
| 0.259082 | 0.011592 | 0.2 | 0.178 | 1 | 0 | Rpl36al | 0.022 | Old |
| 0.259091 | 0.103103 | 0.129 | 0.112 | 1 | 0 | Ift20 | 0.017 | Old |
| 0.259256 | 0.02908 | 0.142 | 0.124 | 1 | 0 | Fkbp4 | 0.018 | Old |
| 0.259444 | 0.216434 | 0.097 | 0.084 | 1 | 0 | Kifc3 | 0.013 | Old |
| 0.259544 | 0.252781 | 0.056 | 0.046 | 1 | 0 | Kin | 0.01 | Old |
| 0.260258 | 0.405685 | 0.03 | 0.023 | 1 | 0 | Ppp1r16a | 0.007 | Old |
| 0.260339 | 0.119217 | 0.09 | 0.077 | 1 | 0 | Nsrp1 | 0.013 | Old |
| 0.260339 | 0.133972 | 0.071 | 0.059 | 1 | 0 | Plxna4os1 | 0.012 | Old |
| 0.260494 | 0.203964 | 0.09 | 0.078 | 1 | 0 | Abcb7 | 0.012 | Old |
| 0.260556 | 0.643563 | 0.025 | 0.018 | 1 | 0 | Gemin6 | 0.007 | Old |
| 0.260754 | 0.259728 | 0.066 | 0.055 | 1 | 0 | Mrvi1 | 0.011 | Old |
| 0.260844 | 0.130155 | 0.034 | 0.026 | 1 | 0 | Akip1 | 0.008 | Old |
| 0.26097 | 0.0937 | 0.089 | 0.076 | 1 | 0 | Erap1 | 0.013 | Old |
| 0.261231 | 0.042052 | 0.17 | 0.15 | 1 | 0 | Senp6 | 0.02 | Old |
| 0.261245 | 0.185936 | 0.122 | 0.107 | 1 | 0 | 1810013L2 | 0.015 | Old |
| 0.261409 | 0.013525 | 0.133 | 0.116 | 1 | 0 | Atg16l1 | 0.017 | Old |
| 0.26142 | 0.118667 | 0.083 | 0.071 | 1 | 0 | Faap20 | 0.012 | Old |
| 0.261618 | 0.183691 | 0.077 | 0.064 | 1 | 0 | Lman2l | 0.013 | Old |
| 0.262301 | 0.115548 | 0.052 | 0.042 | 1 | 0 | Wdyhv1 | 0.01 | Old |
| 0.262374 | 0.106716 | 0.062 | 0.051 | 1 | 0 | Rngtt | 0.011 | Old |
| 0.262377 | 0.165355 | 0.119 | 0.104 | 1 | 0 | B4galt3 | 0.015 | Old |
| 0.262623 | 0.173842 | 0.023 | 0.017 | 1 | 0 | Gmppb | 0.006 | Old |
| 0.262897 | 0.048334 | 0.408 | 0.369 | 1 | 0 | Hdgf | 0.039 | Old |
| 0.2629 | 0.30153 | 0.059 | 0.049 | 1 | 0 | Rab11fip1 | 0.01 | Old |
| 0.26306 | 0.168483 | 0.092 | 0.078 | 1 | 0 | Cfdp1 | 0.014 | Old |
| 0.263106 | 0.166421 | 0.048 | 0.038 | 1 | 0 | Hmces | 0.01 | Old |
| 0.263134 | 0.244501 | 0.094 | 0.082 | 1 | 0 | Ssbp1 | 0.012 | Old |
| 0.263143 | 0.752016 | 0.025 | 0.018 | 1 | 0 | Slc25a38 | 0.007 | Old |
| 0.263712 | 0.157728 | 0.067 | 0.056 | 1 | 0 | Tmem167b | 0.011 | Old |
| 0.264004 | 0.173466 | 0.201 | 0.182 | 1 | 0 | Diaph2 | 0.019 | Old |
| 0.265183 | 0.143037 | 0.073 | 0.061 | 1 | 0 | Usp38 | 0.012 | Old |

| | | | | | | | | |
|---|---|---|---|---|---|---|---|---|
| 0.26538 | 0.015103 | 0.048 | 0.038 | 1 | 0 | Ndc1 | 0.01 | Old |
| 0.265597 | 0.762063 | 0.026 | 0.019 | 1 | 0 | Gga2 | 0.007 | Old |
| 0.265699 | 0.094874 | 0.234 | 0.211 | 1 | 0 | Tmx3 | 0.023 | Old |
| 0.265715 | 0.071382 | 0.029 | 0.021 | 1 | 0 | Cox15 | 0.008 | Old |
| 0.266082 | 0.072095 | 0.148 | 0.13 | 1 | 0 | Chtf8 | 0.018 | Old |
| 0.266443 | 0.017966 | 0.354 | 0.321 | 1 | 0 | Macf1 | 0.033 | Old |
| 0.266446 | 0.153836 | 0.023 | 0.017 | 1 | 0 | Dph6 | 0.006 | Old |
| 0.266691 | 0.078164 | 0.186 | 0.166 | 1 | 0 | Plbd2 | 0.02 | Old |
| 0.266761 | 0.04103 | 0.101 | 0.087 | 1 | 0 | Ccz1 | 0.014 | Old |
| 0.26691 | 0.084721 | 0.068 | 0.057 | 1 | 0 | 6720427I0 | 0.011 | Old |
| 0.267688 | 0.03147 | 0.311 | 0.28 | 1 | 0 | Pacs2 | 0.031 | Old |
| 0.26781 | 0.431894 | 0.019 | 0.013 | 1 | 0 | Pms2 | 0.006 | Old |
| 0.268717 | 0.082525 | 0.213 | 0.194 | 1 | 0 | Asap1 | 0.019 | Old |
| 0.268819 | 0.200707 | 0.067 | 0.056 | 1 | 0 | Isca2 | 0.011 | Old |
| 0.269108 | 0.176016 | 0.056 | 0.046 | 1 | 0 | Ifi211 | 0.01 | Old |
| 0.269213 | 0.128107 | 0.053 | 0.043 | 1 | 0 | Hira | 0.01 | Old |
| 0.26936 | 0.126441 | 0.142 | 0.125 | 1 | 0 | Pin4 | 0.017 | Old |
| 0.269615 | 0.316259 | 0.044 | 0.035 | 1 | 0 | Ptrh2 | 0.009 | Old |
| 0.269634 | 0.157143 | 0.04 | 0.031 | 1 | 0 | Timm21 | 0.009 | Old |
| 0.270063 | 0.383089 | 0.057 | 0.048 | 1 | 0 | Bop1 | 0.009 | Old |
| 0.270333 | 0.288233 | 0.041 | 0.032 | 1 | 0 | Gm47754 | 0.009 | Old |
| 0.270339 | 0.047502 | 0.503 | 0.482 | 1 | 0 | C4b | 0.021 | Old |
| 0.270548 | 0.108241 | 0.17 | 0.152 | 1 | 0 | 2510039O1 | 0.018 | Old |
| 0.270709 | 0.362529 | 0.127 | 0.113 | 1 | 0 | Slf2 | 0.014 | Old |
| 0.270835 | 0.064566 | 0.047 | 0.037 | 1 | 0 | Atg10 | 0.01 | Old |
| 0.270891 | 0.130497 | 0.482 | 0.455 | 1 | 0 | Gnas | 0.027 | Old |
| 0.271012 | 0.132549 | 0.142 | 0.127 | 1 | 0 | Atg5 | 0.015 | Old |
| 0.27121 | 0.033981 | 0.298 | 0.269 | 1 | 0 | Slc12a6 | 0.029 | Old |
| 0.271257 | 0.254764 | 0.049 | 0.04 | 1 | 0 | Alpk1 | 0.009 | Old |
| 0.271361 | 0.148992 | 0.059 | 0.048 | 1 | 0 | Tmem80 | 0.011 | Old |
| 0.271406 | 0.069441 | 0.516 | 0.476 | 1 | 0 | Ddx6 | 0.04 | Old |
| 0.27142 | 0.017366 | 0.068 | 0.057 | 1 | 0 | Hs1bp3 | 0.011 | Old |
| 0.271495 | 0.194563 | 0.052 | 0.042 | 1 | 0 | Traf5 | 0.01 | Old |
| 0.271804 | 0.268592 | 0.033 | 0.025 | 1 | 0 | Zfp821 | 0.008 | Old |
| 0.2719 | 0.365993 | 0.066 | 0.055 | 1 | 0 | Usp36 | 0.011 | Old |
| 0.272006 | 0.004064 | 0.187 | 0.167 | 1 | 0 | Man2a2 | 0.02 | Old |
| 0.272049 | 0.073534 | 0.038 | 0.03 | 1 | 0 | Cul4b | 0.008 | Old |
| 0.272124 | 0.08395 | 0.141 | 0.124 | 1 | 0 | Chd6 | 0.017 | Old |
| 0.272174 | 0.033096 | 0.015 | 0.01 | 1 | 0 | Adat2 | 0.005 | Old |
| 0.272183 | 0.17044 | 0.078 | 0.066 | 1 | 0 | Iars | 0.012 | Old |
| 0.272478 | 0.193513 | 0.09 | 0.078 | 1 | 0 | Pak1ip1 | 0.012 | Old |
| 0.272637 | 0.140449 | 0.025 | 0.018 | 1 | 0 | Oxnad1 | 0.007 | Old |
| 0.272716 | 0.238987 | 0.081 | 0.069 | 1 | 0 | Dcaf15 | 0.012 | Old |
| 0.272894 | 0.183504 | 0.096 | 0.083 | 1 | 0 | Ak3 | 0.013 | Old |
| 0.272905 | 0.011382 | 0.2 | 0.178 | 1 | 0 | Tmem87b | 0.022 | Old |

| | | | | | | | | |
|---|---|---|---|---|---|---|---|---|
| 0.273348 | 0.148414 | 0.752 | 0.74 | 1 | 0 | Klf2 | 0.012 | Old |
| 0.273449 | 0.577352 | 0.021 | 0.015 | 1 | 0 | B3galt6 | 0.006 | Old |
| 0.273779 | 0.180177 | 0.042 | 0.034 | 1 | 0 | Mpv17l2 | 0.008 | Old |
| 0.274779 | 0.182494 | 0.06 | 0.05 | 1 | 0 | Golga5 | 0.01 | Old |
| 0.274797 | 0.152615 | 0.103 | 0.089 | 1 | 0 | Casd1 | 0.014 | Old |
| 0.275157 | 0.23415 | 0.074 | 0.062 | 1 | 0 | Gsr | 0.012 | Old |
| 0.275222 | 0.201279 | 0.066 | 0.055 | 1 | 0 | Acot9 | 0.011 | Old |
| 0.275398 | 0.034816 | 0.276 | 0.25 | 1 | 0 | Vdac2 | 0.026 | Old |
| 0.275519 | 0.321681 | 0.048 | 0.039 | 1 | 0 | Rhbdd1 | 0.009 | Old |
| 0.275908 | 0.003115 | 0.131 | 0.114 | 1 | 0 | Utrn | 0.017 | Old |
| 0.276308 | 0.083293 | 0.166 | 0.147 | 1 | 0 | Dap | 0.019 | Old |
| 0.276633 | 0.034586 | 0.17 | 0.151 | 1 | 0 | Slc48a1 | 0.019 | Old |
| 0.276716 | 0.320419 | 0.034 | 0.027 | 1 | 0 | Zfp758 | 0.007 | Old |
| 0.277163 | 0.622605 | 0.021 | 0.015 | 1 | 0 | Nedd4 | 0.006 | Old |
| 0.277275 | 0.104199 | 0.382 | 0.352 | 1 | 0 | Tet3 | 0.03 | Old |
| 0.277518 | 0.291973 | 0.073 | 0.061 | 1 | 0 | Gt(ROSA)2( | 0.012 | Old |
| 0.277701 | 0.167027 | 0.07 | 0.059 | 1 | 0 | Fam8a1 | 0.011 | Old |
| 0.277712 | 0.360244 | 0.036 | 0.028 | 1 | 0 | Ubxn8 | 0.008 | Old |
| 0.277855 | 0.141216 | 0.033 | 0.025 | 1 | 0 | Def6 | 0.008 | Old |
| 0.278265 | 0.760034 | 0.012 | 0.008 | 1 | 0 | Phf19 | 0.004 | Old |
| 0.279035 | 0.207284 | 0.074 | 0.062 | 1 | 0 | Rev1 | 0.012 | Old |
| 0.279087 | 0.074664 | 0.042 | 0.034 | 1 | 0 | Gtpbp1 | 0.008 | Old |
| 0.279425 | 0.247486 | 0.118 | 0.104 | 1 | 0 | Wdr41 | 0.014 | Old |
| 0.279757 | 0.401566 | 0.067 | 0.057 | 1 | 0 | Ubqln4 | 0.01 | Old |
| 0.279929 | 0.649467 | 0.012 | 0.008 | 1 | 0 | Nsl1 | 0.004 | Old |
| 0.280048 | 0.041285 | 0.454 | 0.413 | 1 | 0 | Fli1 | 0.041 | Old |
| 0.280205 | 0.031846 | 0.424 | 0.387 | 1 | 0 | 2610507B1 | 0.037 | Old |
| 0.280817 | 0.109026 | 0.086 | 0.074 | 1 | 0 | Ddx19a | 0.012 | Old |
| 0.280877 | 0.036492 | 0.033 | 0.025 | 1 | 0 | Gcdh | 0.008 | Old |
| 0.280984 | 0.11213 | 0.073 | 0.061 | 1 | 0 | Tom1 | 0.012 | Old |
| 0.281017 | 0.203604 | 0.118 | 0.103 | 1 | 0 | Prdx6 | 0.015 | Old |
| 0.28111 | 0.606795 | 0.029 | 0.022 | 1 | 0 | Smpd2 | 0.007 | Old |
| 0.2813 | 0.009286 | 0.122 | 0.106 | 1 | 0 | Zdhhc3 | 0.016 | Old |
| 0.281515 | 0.067077 | 0.725 | 0.682 | 1 | 0 | Fcgr2b | 0.043 | Old |
| 0.281878 | 0.167928 | 0.021 | 0.015 | 1 | 0 | Hdac6 | 0.006 | Old |
| 0.281965 | 0.155883 | 0.085 | 0.072 | 1 | 0 | Ssh1 | 0.013 | Old |
| 0.28251 | 0.093401 | 0.155 | 0.138 | 1 | 0 | Paf1 | 0.017 | Old |
| 0.282962 | 0.209893 | 0.127 | 0.112 | 1 | 0 | Slc25a28 | 0.015 | Old |
| 0.282984 | 0.016437 | 0.265 | 0.239 | 1 | 0 | Ciita | 0.026 | Old |
| 0.283018 | 0.29249 | 0.012 | 0.008 | 1 | 0 | Gm38251 | 0.004 | Old |
| 0.283121 | 0.564634 | 0.029 | 0.022 | 1 | 0 | Tmem38b | 0.007 | Old |
| 0.283434 | 0.087484 | 0.026 | 0.019 | 1 | 0 | Mrpl39 | 0.007 | Old |
| 0.284178 | 0.147746 | 0.187 | 0.17 | 1 | 0 | Bcl2l11 | 0.017 | Old |
| 0.284275 | 0.026099 | 0.166 | 0.147 | 1 | 0 | Arid1b | 0.019 | Old |
| 0.28474 | 1.072862 | 0.01 | 0.006 | 1 | 0 | Gm31597 | 0.004 | Old |

| | | | | | | | | |
|---|---|---|---|---|---|---|---|---|
| 0.285039 | 0.477543 | 0.052 | 0.043 | 1 | 0 | Tlr13 | 0.009 | Old |
| 0.285263 | 0.326409 | 0.029 | 0.022 | 1 | 0 | Serpinb9 | 0.007 | Old |
| 0.285322 | 0.238063 | 0.086 | 0.074 | 1 | 0 | Tmem19 | 0.012 | Old |
| 0.285445 | 0.050594 | 0.461 | 0.428 | 1 | 0 | Bach1 | 0.033 | Old |
| 0.285689 | 0.239867 | 0.067 | 0.056 | 1 | 0 | Krr1 | 0.011 | Old |
| 0.285694 | 0.046008 | 0.966 | 0.969 | 1 | 0 | Rpl41 | -0.003 | Old |
| 0.285807 | 0.072925 | 0.269 | 0.247 | 1 | 0 | Srsf1 | 0.022 | Old |
| 0.285837 | 0.128564 | 0.051 | 0.041 | 1 | 0 | Prcc | 0.01 | Old |
| 0.286182 | 0.119772 | 0.047 | 0.037 | 1 | 0 | Dhrs4 | 0.01 | Old |
| 0.286929 | 0.240041 | 0.064 | 0.054 | 1 | 0 | Mrps18a | 0.01 | Old |
| 0.287344 | 0.201024 | 0.053 | 0.044 | 1 | 0 | Zfp414 | 0.009 | Old |
| 0.287449 | 0.085375 | 0.802 | 0.762 | 1 | 0 | Dazap2 | 0.04 | Old |
| 0.287466 | 0.133873 | 0.114 | 0.099 | 1 | 0 | Cetn3 | 0.015 | Old |
| 0.287524 | 0.259053 | 0.07 | 0.059 | 1 | 0 | Brwd3 | 0.011 | Old |
| 0.28796 | 0.622324 | 0.023 | 0.017 | 1 | 0 | Adam22 | 0.006 | Old |
| 0.288055 | 0.15887 | 0.077 | 0.065 | 1 | 0 | Nfkbib | 0.012 | Old |
| 0.288128 | 0.117905 | 0.357 | 0.332 | 1 | 0 | Il10rb | 0.025 | Old |
| 0.288176 | 0.005213 | 0.086 | 0.073 | 1 | 0 | Arfrp1 | 0.013 | Old |
| 0.288249 | 0.080554 | 0.045 | 0.036 | 1 | 0 | Brcc3 | 0.009 | Old |
| 0.288374 | 0.067127 | 0.134 | 0.119 | 1 | 0 | Ube2m | 0.015 | Old |
| 0.288498 | 0.513169 | 0.01 | 0.006 | 1 | 0 | Gm43614 | 0.004 | Old |
| 0.288583 | 0.15716 | 0.17 | 0.152 | 1 | 0 | Ly96 | 0.018 | Old |
| 0.288951 | 0.572023 | 0.01 | 0.006 | 1 | 0 | Dnph1 | 0.004 | Old |
| 0.289724 | 0.49711 | 0.031 | 0.024 | 1 | 0 | Adck1 | 0.007 | Old |
| 0.289989 | 0.108807 | 0.093 | 0.08 | 1 | 0 | Vps54 | 0.013 | Old |
| 0.290041 | 0.352985 | 0.187 | 0.174 | 1 | 0 | Cyth1 | 0.013 | Old |
| 0.290097 | 0.203716 | 0.016 | 0.011 | 1 | 0 | Lins1 | 0.005 | Old |
| 0.290504 | 0.151804 | 0.051 | 0.041 | 1 | 0 | Tdp2 | 0.01 | Old |
| 0.290839 | 0.085291 | 0.051 | 0.041 | 1 | 0 | Tomm6 | 0.01 | Old |
| 0.290996 | 0.1629 | 0.01 | 0.006 | 1 | 0 | Gm27003 | 0.004 | Old |
| 0.291259 | 0.125395 | 0.037 | 0.029 | 1 | 0 | Tor4a | 0.008 | Old |
| 0.291627 | 0.18618 | 0.15 | 0.136 | 1 | 0 | Fam104a | 0.014 | Old |
| 0.291652 | 0.051306 | 0.079 | 0.067 | 1 | 0 | Jak2 | 0.012 | Old |
| 0.29168 | 0.053446 | 0.01 | 0.006 | 1 | 0 | Mmp12 | 0.004 | Old |
| 0.291832 | 0.013795 | 0.01 | 0.006 | 1 | 0 | Pus7l | 0.004 | Old |
| 0.291839 | 0.055783 | 0.075 | 0.064 | 1 | 0 | Ftsj3 | 0.011 | Old |
| 0.292035 | 0.039466 | 0.603 | 0.564 | 1 | 0 | Pld4 | 0.039 | Old |
| 0.293194 | 0.012217 | 0.323 | 0.295 | 1 | 0 | Rnpep | 0.028 | Old |
| 0.293384 | 0.150814 | 0.097 | 0.084 | 1 | 0 | Crkl | 0.013 | Old |
| 0.293827 | 0.74639 | 0.018 | 0.013 | 1 | 0 | Wdr4 | 0.005 | Old |
| 0.293833 | 0.032074 | 0.09 | 0.077 | 1 | 0 | Fech | 0.013 | Old |
| 0.293871 | 0.064871 | 0.212 | 0.191 | 1 | 0 | N4bp2l2 | 0.021 | Old |
| 0.294003 | 0.073362 | 0.007 | 0.011 | 1 | 0 | Prpf40b | -0.004 | Old |
| 0.294136 | 0.080688 | 0.107 | 0.093 | 1 | 0 | Lrrc59 | 0.014 | Old |
| 0.294334 | 0.036701 | 0.022 | 0.016 | 1 | 0 | 2810006K2 | 0.006 | Old |

| | | | | | | | | |
|---|---|---|---|---|---|---|---|---|
| 0.294639 | 0.025934 | 0.248 | 0.224 | 1 | 0 | Nfix | 0.024 | Old |
| 0.294653 | 0.056834 | 0.037 | 0.029 | 1 | 0 | Phospho2 | 0.008 | Old |
| 0.294793 | 0.680713 | 0.018 | 0.013 | 1 | 0 | Cep131 | 0.005 | Old |
| 0.29501 | 0.163461 | 0.041 | 0.033 | 1 | 0 | Rhoj | 0.008 | Old |
| 0.295215 | 0.043194 | 0.174 | 0.154 | 1 | 0 | Ssbp3 | 0.02 | Old |
| 0.295236 | 0.034839 | 0.193 | 0.173 | 1 | 0 | Lpin1 | 0.02 | Old |
| 0.295276 | 0.040595 | 0.029 | 0.037 | 1 | 0 | Sema6a | -0.008 | Old |
| 0.29539 | 0.092188 | 0.068 | 0.058 | 1 | 0 | Psmd9 | 0.01 | Old |
| 0.295524 | 0.016794 | 0.073 | 0.061 | 1 | 0 | Usf3 | 0.012 | Old |
| 0.295565 | 0.010565 | 0.705 | 0.666 | 1 | 0 | Cd81 | 0.039 | Old |
| 0.296029 | 0.734026 | 0.044 | 0.036 | 1 | 0 | Rpf2 | 0.008 | Old |
| 0.296353 | 0.048589 | 0.022 | 0.016 | 1 | 0 | Rbm4 | 0.006 | Old |
| 0.296508 | 0.05763 | 0.244 | 0.221 | 1 | 0 | Myo1f | 0.023 | Old |
| 0.296642 | 0.229372 | 0.101 | 0.088 | 1 | 0 | Pttg1 | 0.013 | Old |
| 0.296907 | 0.024961 | 0.145 | 0.129 | 1 | 0 | Abcd1 | 0.016 | Old |
| 0.297163 | 0.053895 | 0.274 | 0.251 | 1 | 0 | Ptp4a1 | 0.023 | Old |
| 0.297253 | 0.040499 | 0.008 | 0.013 | 1 | 0 | Ndufaf1 | -0.005 | Old |
| 0.297304 | 0.033086 | 0.248 | 0.224 | 1 | 0 | Chmp5 | 0.024 | Old |
| 0.297345 | 0.042833 | 0.997 | 0.997 | 1 | 0 | Itm2b | 0 | Old |
| 0.297379 | 0.618043 | 0.018 | 0.013 | 1 | 0 | Tmem107 | 0.005 | Old |
| 0.297487 | 0.494026 | 0.018 | 0.013 | 1 | 0 | Surf6 | 0.005 | Old |
| 0.297658 | 0.30449 | 0.031 | 0.024 | 1 | 0 | Fance | 0.007 | Old |
| 0.297977 | 0.385514 | 0.033 | 0.026 | 1 | 0 | Rhbdd2 | 0.007 | Old |
| 0.297986 | 0.036477 | 0.127 | 0.112 | 1 | 0 | Chml | 0.015 | Old |
| 0.298071 | 0.135638 | 0.041 | 0.033 | 1 | 0 | Nos1ap | 0.008 | Old |
| 0.298256 | 0.163426 | 0.096 | 0.083 | 1 | 0 | Trappc1 | 0.013 | Old |
| 0.298347 | 0.203967 | 0.041 | 0.033 | 1 | 0 | Polr3f | 0.008 | Old |
| 0.29909 | 0.159833 | 0.077 | 0.065 | 1 | 0 | Pfdn4 | 0.012 | Old |
| 0.299286 | 0.189121 | 0.044 | 0.035 | 1 | 0 | Accs | 0.009 | Old |
| 0.299312 | 0.063191 | 0.321 | 0.291 | 1 | 0 | Snap23 | 0.03 | Old |
| 0.299341 | 0.060271 | 0.064 | 0.054 | 1 | 0 | Nagpa | 0.01 | Old |
| 0.299436 | 0.188819 | 0.075 | 0.064 | 1 | 0 | Naxe | 0.011 | Old |
| 0.299546 | 0.239387 | 0.018 | 0.013 | 1 | 0 | Mad2l1 | 0.005 | Old |
| 0.299718 | 0.058291 | 0.382 | 0.351 | 1 | 0 | Shisa5 | 0.031 | Old |
| 0.300198 | 0.309215 | 0.018 | 0.013 | 1 | 0 | Zfp688 | 0.005 | Old |
| 0.300537 | 0.044077 | 0.295 | 0.271 | 1 | 0 | Ube2n | 0.024 | Old |
| 0.300575 | 0.160433 | 0.042 | 0.034 | 1 | 0 | Fam160a2 | 0.008 | Old |
| 0.300836 | 0.045807 | 0.023 | 0.017 | 1 | 0 | Zc3h12d | 0.006 | Old |
| 0.300851 | 0.269661 | 0.018 | 0.013 | 1 | 0 | 1810044D( | 0.005 | Old |
| 0.300928 | 0.033429 | 0.135 | 0.153 | 1 | 0 | Smad1 | -0.018 | Old |
| 0.301062 | 0.168186 | 0.048 | 0.039 | 1 | 0 | Ctnnbl1 | 0.009 | Old |
| 0.301534 | 0.187958 | 0.142 | 0.128 | 1 | 0 | Evl | 0.014 | Old |
| 0.301735 | 0.097928 | 0.062 | 0.051 | 1 | 0 | Cd2bp2 | 0.011 | Old |
| 0.301825 | 0.160673 | 0.101 | 0.088 | 1 | 0 | Nuak1 | 0.013 | Old |
| 0.301966 | 0.09814 | 0.073 | 0.061 | 1 | 0 | Nup50 | 0.012 | Old |

| | | | | | | | | |
|---|---|---|---|---|---|---|---|---|
| 0.302143 | 0.260096 | 0.014 | 0.009 | 1 | 0 | Camk2a | 0.005 | Old |
| 0.302248 | 0.126072 | 0.023 | 0.017 | 1 | 0 | Zfp953 | 0.006 | Old |
| 0.302615 | 0.02901 | 0.041 | 0.033 | 1 | 0 | Zfp617 | 0.008 | Old |
| 0.303101 | 0.022719 | 0.449 | 0.411 | 1 | 0 | Pomp | 0.038 | Old |
| 0.303115 | 0.112198 | 0.765 | 0.747 | 1 | 0 | Nfkbia | 0.018 | Old |
| 0.303203 | 0.03899 | 0.286 | 0.264 | 1 | 0 | Prex1 | 0.022 | Old |
| 0.303219 | 0.177723 | 0.103 | 0.089 | 1 | 0 | Sdf2l1 | 0.014 | Old |
| 0.303245 | 0.061544 | 0.466 | 0.432 | 1 | 0 | Asah1 | 0.034 | Old |
| 0.303408 | 0.029241 | 0.186 | 0.167 | 1 | 0 | Efcab14 | 0.019 | Old |
| 0.303505 | 0.034546 | 0.514 | 0.478 | 1 | 0 | Ndufa4 | 0.036 | Old |
| 0.303549 | 0.066165 | 0.096 | 0.083 | 1 | 0 | Ppt2 | 0.013 | Old |
| 0.303709 | 0.268465 | 0.033 | 0.026 | 1 | 0 | Eef1akmt2 | 0.007 | Old |
| 0.303758 | 0.137228 | 0.112 | 0.099 | 1 | 0 | Sgta | 0.013 | Old |
| 0.304225 | 0.009514 | 0.192 | 0.171 | 1 | 0 | Cnppd1 | 0.021 | Old |
| 0.304464 | 0.17739 | 0.044 | 0.035 | 1 | 0 | Stx11 | 0.009 | Old |
| 0.304498 | 0.065874 | 0.15 | 0.134 | 1 | 0 | Vamp4 | 0.016 | Old |
| 0.304584 | 0.063195 | 0.349 | 0.321 | 1 | 0 | Eif4a2 | 0.028 | Old |
| 0.304793 | 0.089404 | 0.116 | 0.102 | 1 | 0 | Atf6b | 0.014 | Old |
| 0.304859 | 0.052551 | 0.395 | 0.364 | 1 | 0 | Skap2 | 0.031 | Old |
| 0.304892 | 0.027706 | 0.042 | 0.034 | 1 | 0 | Epg5 | 0.008 | Old |
| 0.304964 | 0.015417 | 0.186 | 0.167 | 1 | 0 | Fam173a | 0.019 | Old |
| 0.305012 | 0.117808 | 0.079 | 0.068 | 1 | 0 | Lemd3 | 0.011 | Old |
| 0.305407 | 0.237795 | 0.052 | 0.043 | 1 | 0 | Mrpl58 | 0.009 | Old |
| 0.305547 | 0.053079 | 0.044 | 0.035 | 1 | 0 | Gnl3l | 0.009 | Old |
| 0.305851 | 0.015297 | 0.201 | 0.182 | 1 | 0 | R3hdm2 | 0.019 | Old |
| 0.305924 | 0.106544 | 0.131 | 0.117 | 1 | 0 | Galnt7 | 0.014 | Old |
| 0.306385 | 0.380496 | 0.038 | 0.031 | 1 | 0 | Galns | 0.007 | Old |
| 0.306473 | 0.122246 | 0.078 | 0.067 | 1 | 0 | Pspc1 | 0.011 | Old |
| 0.307238 | 0.571118 | 0.019 | 0.014 | 1 | 0 | Zbed3 | 0.005 | Old |
| 0.307245 | 0.072085 | 0.044 | 0.035 | 1 | 0 | Rab13 | 0.009 | Old |
| 0.307312 | 0.684662 | 0.027 | 0.021 | 1 | 0 | Nudt18 | 0.006 | Old |
| 0.307364 | 0.177002 | 0.176 | 0.161 | 1 | 0 | Sertad1 | 0.015 | Old |
| 0.307421 | 0.061332 | 0.144 | 0.128 | 1 | 0 | Dennd4b | 0.016 | Old |
| 0.307653 | 0.025679 | 0.286 | 0.261 | 1 | 0 | Itgb2 | 0.025 | Old |
| 0.307978 | 0.531225 | 0.019 | 0.014 | 1 | 0 | Zfp456 | 0.005 | Old |
| 0.308879 | 0.522593 | 0.019 | 0.014 | 1 | 0 | Exo5 | 0.005 | Old |
| 0.308967 | 0.552256 | 0.027 | 0.021 | 1 | 0 | Svip | 0.006 | Old |
| 0.309098 | 0.487554 | 0.027 | 0.021 | 1 | 0 | Zfp637 | 0.006 | Old |
| 0.309597 | 0.000314 | 0.163 | 0.147 | 1 | 0 | Il1rl1 | 0.016 | Old |
| 0.310456 | 0.065315 | 0.263 | 0.24 | 1 | 0 | 7-Mar | 0.023 | Old |
| 0.310909 | 0.046116 | 0.15 | 0.135 | 1 | 0 | Rnf216 | 0.015 | Old |
| 0.311032 | 0.126895 | 0.557 | 0.536 | 1 | 0 | Hnrnph1 | 0.021 | Old |
| 0.311112 | 0.22119 | 0.019 | 0.014 | 1 | 0 | Fam216a | 0.005 | Old |
| 0.311199 | 0.392988 | 0.04 | 0.032 | 1 | 0 | Fbxo45 | 0.008 | Old |
| 0.311718 | 0.111461 | 0.192 | 0.174 | 1 | 0 | Taf15 | 0.018 | Old |

| | | | | | | | | |
|---|---|---|---|---|---|---|---|---|
| 0.312172 | 0.15921 | 0.209 | 0.192 | 1 | 0 | Sp1 | 0.017 | Old |
| 0.312195 | 0.025245 | 0.29 | 0.266 | 1 | 0 | Mndal | 0.024 | Old |
| 0.312361 | 0.000724 | 0.278 | 0.252 | 1 | 0 | Stk38 | 0.026 | Old |
| 0.312401 | 0.149337 | 0.053 | 0.044 | 1 | 0 | Ccnc | 0.009 | Old |
| 0.312949 | 0.115664 | 0.034 | 0.027 | 1 | 0 | Pnpt1 | 0.007 | Old |
| 0.313038 | 0.23643 | 0.082 | 0.071 | 1 | 0 | Cetn2 | 0.011 | Old |
| 0.313698 | 0.163578 | 0.098 | 0.086 | 1 | 0 | Wdr11 | 0.012 | Old |
| 0.313828 | 0.136242 | 0.093 | 0.081 | 1 | 0 | Dpp9 | 0.012 | Old |
| 0.313997 | 0.45903 | 0.038 | 0.031 | 1 | 0 | Zmat3 | 0.007 | Old |
| 0.314504 | 0.181944 | 0.192 | 0.176 | 1 | 0 | Nrbp1 | 0.016 | Old |
| 0.314643 | 0.852472 | 0.015 | 0.011 | 1 | 0 | Trdmt1 | 0.004 | Old |
| 0.314826 | 0.095734 | 0.053 | 0.044 | 1 | 0 | Ciapin1 | 0.009 | Old |
| 0.314914 | 0.158736 | 0.098 | 0.086 | 1 | 0 | Akap10 | 0.012 | Old |
| 0.314965 | 0.051175 | 0.051 | 0.042 | 1 | 0 | Rrad | 0.009 | Old |
| 0.315505 | 0.268569 | 0.037 | 0.029 | 1 | 0 | Mon1b | 0.008 | Old |
| 0.315595 | 0.014328 | 0.189 | 0.168 | 1 | 0 | Gspt1 | 0.021 | Old |
| 0.31604 | 0.404608 | 0.045 | 0.037 | 1 | 0 | Nostrin | 0.008 | Old |
| 0.316196 | 0.166989 | 0.063 | 0.053 | 1 | 0 | Ttc13 | 0.01 | Old |
| 0.316262 | 0.109144 | 0.804 | 0.788 | 1 | 0 | Glul | 0.016 | Old |
| 0.316714 | 0.017965 | 0.108 | 0.094 | 1 | 0 | Nup153 | 0.014 | Old |
| 0.316733 | 0.298977 | 0.053 | 0.045 | 1 | 0 | Polr1a | 0.008 | Old |
| 0.317206 | 0.182674 | 0.082 | 0.071 | 1 | 0 | Ampd3 | 0.011 | Old |
| 0.317613 | 0.047624 | 0.086 | 0.074 | 1 | 0 | Mrps23 | 0.012 | Old |
| 0.31774 | 0.668574 | 0.033 | 0.026 | 1 | 0 | Ankrd26 | 0.007 | Old |
| 0.318291 | 0.311481 | 0.011 | 0.007 | 1 | 0 | Cyp2d22 | 0.004 | Old |
| 0.318438 | 0.133348 | 0.011 | 0.007 | 1 | 0 | Sirpb1b | 0.004 | Old |
| 0.318854 | 0.135299 | 0.182 | 0.166 | 1 | 0 | Sike1 | 0.016 | Old |
| 0.318945 | 0.031769 | 0.049 | 0.04 | 1 | 0 | 6030458C1 | 0.009 | Old |
| 0.318965 | 0.098108 | 0.119 | 0.106 | 1 | 0 | Nucb2 | 0.013 | Old |
| 0.319027 | 0.083644 | 0.181 | 0.163 | 1 | 0 | Bub3 | 0.018 | Old |
| 0.319071 | 0.024704 | 0.123 | 0.109 | 1 | 0 | Anapc1 | 0.014 | Old |
| 0.319489 | 0.094463 | 0.179 | 0.163 | 1 | 0 | Mydgf | 0.016 | Old |
| 0.319757 | 0.363577 | 0.021 | 0.015 | 1 | 0 | Optn | 0.006 | Old |
| 0.320109 | 0.03157 | 0.112 | 0.099 | 1 | 0 | Slfn8 | 0.013 | Old |
| 0.320326 | 0.238289 | 0.041 | 0.033 | 1 | 0 | Psmf1 | 0.008 | Old |
| 0.320383 | 0.18219 | 0.021 | 0.015 | 1 | 0 | Mpst | 0.006 | Old |
| 0.320513 | 0.052149 | 0.011 | 0.007 | 1 | 0 | Rpp38 | 0.004 | Old |
| 0.320572 | 0.126466 | 0.038 | 0.031 | 1 | 0 | Wwox | 0.007 | Old |
| 0.320591 | 0.031937 | 0.118 | 0.104 | 1 | 0 | Rnps1 | 0.014 | Old |
| 0.320728 | 0.093249 | 0.103 | 0.09 | 1 | 0 | Tmem184b | 0.013 | Old |
| 0.321383 | 0.04804 | 0.051 | 0.042 | 1 | 0 | Mir17hg | 0.009 | Old |
| 0.32206 | 0.153267 | 0.027 | 0.021 | 1 | 0 | 2310057M | 0.006 | Old |
| 0.322448 | 0.336106 | 0.029 | 0.022 | 1 | 0 | Prpf3 | 0.007 | Old |
| 0.322672 | 0.189089 | 0.071 | 0.061 | 1 | 0 | Prmt2 | 0.01 | Old |
| 0.322703 | 0.202636 | 0.108 | 0.096 | 1 | 0 | D10Jhu81e | 0.012 | Old |

| | | | | | | | | |
|---|---|---|---|---|---|---|---|---|
| 0.322921 | 0.252114 | 0.074 | 0.064 | 1 | 0 | E2f5 | 0.01 | Old |
| 0.323292 | 0.011108 | 0.015 | 0.011 | 1 | 0 | Mrps22 | 0.004 | Old |
| 0.324189 | 0.064979 | 0.616 | 0.572 | 1 | 0 | Cotl1 | 0.044 | Old |
| 0.324749 | 0.267646 | 0.052 | 0.043 | 1 | 0 | Gsto1 | 0.009 | Old |
| 0.324932 | 0.228372 | 0.056 | 0.047 | 1 | 0 | Kctd5 | 0.009 | Old |
| 0.324966 | 0.211576 | 0.111 | 0.098 | 1 | 0 | Cycs | 0.013 | Old |
| 0.325198 | 0.121527 | 0.13 | 0.116 | 1 | 0 | Paip1 | 0.014 | Old |
| 0.325798 | 0.317917 | 0.049 | 0.041 | 1 | 0 | Aldh6a1 | 0.008 | Old |
| 0.326125 | 0.103198 | 0.041 | 0.033 | 1 | 0 | Ppa1 | 0.008 | Old |
| 0.326737 | 0.298178 | 0.022 | 0.016 | 1 | 0 | Plcb2 | 0.006 | Old |
| 0.326818 | 0.094157 | 0.041 | 0.033 | 1 | 0 | Gm11808 | 0.008 | Old |
| 0.327383 | 0.042047 | 0.097 | 0.085 | 1 | 0 | B3galnt1 | 0.012 | Old |
| 0.327524 | 0.102105 | 0.168 | 0.152 | 1 | 0 | Inppl1 | 0.016 | Old |
| 0.327687 | 0.025087 | 0.261 | 0.24 | 1 | 0 | Ugcg | 0.021 | Old |
| 0.327792 | 0.182991 | 0.066 | 0.056 | 1 | 0 | Sdhaf2 | 0.01 | Old |
| 0.327801 | 0.000519 | 0.1 | 0.087 | 1 | 0 | Cnep1r1 | 0.013 | Old |
| 0.327903 | 0.400879 | 0.114 | 0.102 | 1 | 0 | Sowahc | 0.012 | Old |
| 0.328091 | 0.183354 | 0.063 | 0.053 | 1 | 0 | Sil1 | 0.01 | Old |
| 0.328329 | 0.249082 | 0.057 | 0.048 | 1 | 0 | Rnf138 | 0.009 | Old |
| 0.32836 | 0.026894 | 0.055 | 0.045 | 1 | 0 | Klhdc4 | 0.01 | Old |
| 0.328472 | 0.2593 | 0.022 | 0.016 | 1 | 0 | Top2a | 0.006 | Old |
| 0.328686 | 0.108892 | 0.123 | 0.11 | 1 | 0 | Ykt6 | 0.013 | Old |
| 0.32888 | 0.162249 | 0.03 | 0.024 | 1 | 0 | Lonrf1 | 0.006 | Old |
| 0.328943 | 0.197961 | 0.052 | 0.043 | 1 | 0 | Ubtd1 | 0.009 | Old |
| 0.32929 | 0.287858 | 0.022 | 0.016 | 1 | 0 | Syp | 0.006 | Old |
| 0.329304 | 0.040371 | 0.276 | 0.252 | 1 | 0 | Atp5c1 | 0.024 | Old |
| 0.329428 | 0.024883 | 0.248 | 0.227 | 1 | 0 | Ndufs6 | 0.021 | Old |
| 0.329509 | 0.04608 | 0.412 | 0.383 | 1 | 0 | Plxnb2 | 0.029 | Old |
| 0.329591 | 0.312852 | 0.056 | 0.047 | 1 | 0 | Msra | 0.009 | Old |
| 0.329881 | 0.640066 | 0.016 | 0.012 | 1 | 0 | Mrm2 | 0.004 | Old |
| 0.329906 | 0.050479 | 0.074 | 0.063 | 1 | 0 | Thoc6 | 0.011 | Old |
| 0.330191 | 0.615848 | 0.023 | 0.018 | 1 | 0 | Urb1 | 0.005 | Old |
| 0.330552 | 0.382779 | 0.063 | 0.054 | 1 | 0 | Grk5 | 0.009 | Old |
| 0.330742 | 0.042221 | 0.153 | 0.138 | 1 | 0 | Gpbp1l1 | 0.015 | Old |
| 0.330923 | 0.300484 | 0.049 | 0.041 | 1 | 0 | Dcps | 0.008 | Old |
| 0.331322 | 0.435204 | 0.016 | 0.012 | 1 | 0 | AI480526 | 0.004 | Old |
| 0.331707 | 0.097073 | 0.045 | 0.037 | 1 | 0 | Pphln1 | 0.008 | Old |
| 0.332393 | 0.034837 | 0.241 | 0.221 | 1 | 0 | Srsf9 | 0.02 | Old |
| 0.332566 | 0.155848 | 0.09 | 0.079 | 1 | 0 | Cass4 | 0.011 | Old |
| 0.332682 | 0.031394 | 0.163 | 0.146 | 1 | 0 | R3hdm1 | 0.017 | Old |
| 0.332966 | 0.177893 | 0.06 | 0.051 | 1 | 0 | Chid1 | 0.009 | Old |
| 0.333012 | 0.128657 | 0.122 | 0.109 | 1 | 0 | Lrch3 | 0.013 | Old |
| 0.333349 | 0.124485 | 0.075 | 0.065 | 1 | 0 | Scfd1 | 0.01 | Old |
| 0.333793 | 0.10821 | 0.016 | 0.012 | 1 | 0 | Ly6a | 0.004 | Old |
| 0.334059 | 0.072137 | 0.451 | 0.424 | 1 | 0 | Arap1 | 0.027 | Old |

| | | | | | | | | |
|---|---|---|---|---|---|---|---|---|
| 0.334319 | 0.014555 | 0.094 | 0.082 | 1 | 0 | Rnf103 | 0.012 | Old |
| 0.334453 | 0.337674 | 0.038 | 0.031 | 1 | 0 | Ptcd2 | 0.007 | Old |
| 0.334458 | 0.197909 | 0.016 | 0.012 | 1 | 0 | Ncapd2 | 0.004 | Old |
| 0.334751 | 0.135434 | 0.068 | 0.059 | 1 | 0 | Clec4e | 0.009 | Old |
| 0.334901 | 0.226523 | 0.101 | 0.09 | 1 | 0 | Bms1 | 0.011 | Old |
| 0.334917 | 0.207279 | 0.033 | 0.026 | 1 | 0 | Fen1 | 0.007 | Old |
| 0.335731 | 0.205698 | 0.06 | 0.051 | 1 | 0 | Fiz1 | 0.009 | Old |
| 0.33587 | 0.283293 | 0.023 | 0.018 | 1 | 0 | AW146154 | 0.005 | Old |
| 0.336183 | 0.000306 | 0.278 | 0.254 | 1 | 0 | Tppp3 | 0.024 | Old |
| 0.33635 | 0.131447 | 0.094 | 0.083 | 1 | 0 | Plekhf2 | 0.011 | Old |
| 0.336391 | 0.150787 | 0.064 | 0.055 | 1 | 0 | Gnpda2 | 0.009 | Old |
| 0.336714 | 0.052067 | 0.042 | 0.035 | 1 | 0 | Chchd4 | 0.007 | Old |
| 0.33678 | 0.14048 | 0.174 | 0.158 | 1 | 0 | Ppp4r3a | 0.016 | Old |
| 0.336802 | 0.115381 | 0.149 | 0.134 | 1 | 0 | Yipf3 | 0.015 | Old |
| 0.336856 | 0.015782 | 0.115 | 0.102 | 1 | 0 | Ppm1b | 0.013 | Old |
| 0.337484 | 0.152064 | 0.064 | 0.055 | 1 | 0 | Cd82 | 0.009 | Old |
| 0.337734 | 0.014278 | 0.016 | 0.012 | 1 | 0 | Rhobtb2 | 0.004 | Old |
| 0.337846 | 0.128443 | 0.101 | 0.089 | 1 | 0 | Nr1h2 | 0.012 | Old |
| 0.337886 | 0.049244 | 0.042 | 0.035 | 1 | 0 | Tdrkh | 0.007 | Old |
| 0.338101 | 0.551793 | 0.012 | 0.008 | 1 | 0 | Map7d3 | 0.004 | Old |
| 0.338187 | 0.177258 | 0.092 | 0.08 | 1 | 0 | Kdm3b | 0.012 | Old |
| 0.338359 | 0.016979 | 0.534 | 0.488 | 1 | 0 | Cbl | 0.046 | Old |
| 0.338387 | 0.40753 | 0.012 | 0.008 | 1 | 0 | Tedc2 | 0.004 | Old |
| 0.338387 | 0.295202 | 0.012 | 0.008 | 1 | 0 | Sardh | 0.004 | Old |
| 0.338559 | 0.095139 | 0.066 | 0.056 | 1 | 0 | Plaur | 0.01 | Old |
| 0.338959 | 0.329437 | 0.012 | 0.008 | 1 | 0 | Enpp4 | 0.004 | Old |
| 0.339014 | 0.728183 | 0.018 | 0.013 | 1 | 0 | Acss2 | 0.005 | Old |
| 0.339042 | 0.178026 | 0.112 | 0.1 | 1 | 0 | Asxl1 | 0.012 | Old |
| 0.33915 | 0.313441 | 0.037 | 0.03 | 1 | 0 | Ccdc71 | 0.007 | Old |
| 0.339572 | 0.179927 | 0.108 | 0.096 | 1 | 0 | Ostm1 | 0.012 | Old |
| 0.33984 | 0.269236 | 0.034 | 0.027 | 1 | 0 | Pkdcc | 0.007 | Old |
| 0.339903 | 0.107883 | 0.055 | 0.046 | 1 | 0 | Mrpl38 | 0.009 | Old |
| 0.340374 | 0.204777 | 0.034 | 0.027 | 1 | 0 | Ust | 0.007 | Old |
| 0.34057 | 0.03161 | 0.375 | 0.348 | 1 | 0 | Galnt1 | 0.027 | Old |
| 0.340673 | 0.406395 | 0.044 | 0.036 | 1 | 0 | Zbtb33 | 0.008 | Old |
| 0.340818 | 0.131208 | 0.189 | 0.172 | 1 | 0 | Vps26a | 0.017 | Old |
| 0.34088 | 0.605516 | 0.018 | 0.013 | 1 | 0 | Polrmt | 0.005 | Old |
| 0.340909 | 0.077981 | 0.031 | 0.025 | 1 | 0 | Rmdn3 | 0.006 | Old |
| 0.341113 | 0.245048 | 0.036 | 0.029 | 1 | 0 | Chkb | 0.007 | Old |
| 0.34117 | 0.060599 | 0.575 | 0.532 | 1 | 0 | Hnrnpu | 0.043 | Old |
| 0.341489 | 0.224481 | 0.033 | 0.026 | 1 | 0 | Carf | 0.007 | Old |
| 0.341586 | 0.039948 | 0.104 | 0.091 | 1 | 0 | Rnf139 | 0.013 | Old |
| 0.341618 | 0.119771 | 0.082 | 0.071 | 1 | 0 | Mis12 | 0.011 | Old |
| 0.341713 | 0.201287 | 0.023 | 0.018 | 1 | 0 | Polm | 0.005 | Old |
| 0.341762 | 0.089209 | 0.15 | 0.135 | 1 | 0 | Ddhd2 | 0.015 | Old |

| | | | | | | | | |
|---|---|---|---|---|---|---|---|---|
| 0.342031 | 0.304253 | 0.07 | 0.06 | 1 | 0 | Asb7 | 0.01 | Old |
| 0.342179 | 0.088662 | 0.07 | 0.06 | 1 | 0 | Ifih1 | 0.01 | Old |
| 0.342402 | 0.045211 | 0.146 | 0.131 | 1 | 0 | Mark3 | 0.015 | Old |
| 0.342504 | 0.052762 | 0.218 | 0.198 | 1 | 0 | Gpsm3 | 0.02 | Old |
| 0.3428 | 0.090374 | 0.141 | 0.126 | 1 | 0 | Fbxo33 | 0.015 | Old |
| 0.343322 | 0.204712 | 0.135 | 0.123 | 1 | 0 | Pde3b | 0.012 | Old |
| 0.34343 | 0.058734 | 0.057 | 0.048 | 1 | 0 | Eif2ak3 | 0.009 | Old |
| 0.343673 | 0.148587 | 0.034 | 0.027 | 1 | 0 | Isyna1 | 0.007 | Old |
| 0.344044 | 0.362922 | 0.018 | 0.013 | 1 | 0 | Bpgm | 0.005 | Old |
| 0.344064 | 0.115985 | 0.14 | 0.126 | 1 | 0 | Cops4 | 0.014 | Old |
| 0.344131 | 0.133382 | 0.063 | 0.053 | 1 | 0 | Patz1 | 0.01 | Old |
| 0.344657 | 0.013729 | 0.033 | 0.026 | 1 | 0 | Klhl22 | 0.007 | Old |
| 0.344764 | 0.062968 | 0.053 | 0.045 | 1 | 0 | Hmbs | 0.008 | Old |
| 0.344769 | 0.211629 | 0.037 | 0.03 | 1 | 0 | 4930503L1 | 0.007 | Old |
| 0.344844 | 0.040165 | 0.1 | 0.088 | 1 | 0 | Slirp | 0.012 | Old |
| 0.344886 | 0.161795 | 0.038 | 0.031 | 1 | 0 | Snx19 | 0.007 | Old |
| 0.344929 | 0.193337 | 0.064 | 0.055 | 1 | 0 | Caml | 0.009 | Old |
| 0.346143 | 0.033493 | 0.249 | 0.227 | 1 | 0 | Adam19 | 0.022 | Old |
| 0.346199 | 0.821206 | 0.085 | 0.076 | 1 | 0 | Gdf15 | 0.009 | Old |
| 0.346282 | 0.270503 | 0.018 | 0.013 | 1 | 0 | 5830416I1! | 0.005 | Old |
| 0.346606 | 0.254405 | 0.038 | 0.031 | 1 | 0 | Pex1 | 0.007 | Old |
| 0.347094 | 0.118937 | 0.149 | 0.135 | 1 | 0 | Nrd1 | 0.014 | Old |
| 0.347105 | 0.280939 | 0.045 | 0.037 | 1 | 0 | Gon7 | 0.008 | Old |
| 0.347255 | 0.078612 | 0.025 | 0.019 | 1 | 0 | Ppil1 | 0.006 | Old |
| 0.347304 | 0.048082 | 0.025 | 0.019 | 1 | 0 | Rcc1 | 0.006 | Old |
| 0.34743 | 0.076569 | 0.067 | 0.057 | 1 | 0 | Zfp639 | 0.01 | Old |
| 0.347582 | 0.197411 | 0.018 | 0.013 | 1 | 0 | Zfp12 | 0.005 | Old |
| 0.34768 | 0.256605 | 0.045 | 0.037 | 1 | 0 | Orc5 | 0.008 | Old |
| 0.3477 | 0.151884 | 0.018 | 0.013 | 1 | 0 | Alg3 | 0.005 | Old |
| 0.348407 | 0.046169 | 0.313 | 0.291 | 1 | 0 | Crlf3 | 0.022 | Old |
| 0.348765 | 0.031452 | 0.23 | 0.211 | 1 | 0 | Mvp | 0.019 | Old |
| 0.349062 | 0.128204 | 0.018 | 0.013 | 1 | 0 | Dync2h1 | 0.005 | Old |
| 0.349212 | 0.182803 | 0.026 | 0.02 | 1 | 0 | Fdxr | 0.006 | Old |
| 0.349278 | 0.190962 | 0.142 | 0.128 | 1 | 0 | Nbr1 | 0.014 | Old |
| 0.349355 | 0.085751 | 0.105 | 0.092 | 1 | 0 | Actr10 | 0.013 | Old |
| 0.349398 | 0.213063 | 0.042 | 0.035 | 1 | 0 | Parn | 0.007 | Old |
| 0.349398 | 0.212347 | 0.042 | 0.035 | 1 | 0 | Slc9a1 | 0.007 | Old |
| 0.349478 | 0.094028 | 0.018 | 0.013 | 1 | 0 | Tbc1d8b | 0.005 | Old |
| 0.349629 | 0.017508 | 0.198 | 0.18 | 1 | 0 | Baiap2 | 0.018 | Old |
| 0.349937 | 0.153015 | 0.183 | 0.168 | 1 | 0 | Srebf2 | 0.015 | Old |
| 0.349999 | 0.013564 | 0.038 | 0.047 | 1 | 0 | Wipi1 | -0.009 | Old |
| 0.350911 | 0.087169 | 0.253 | 0.236 | 1 | 0 | Bcap31 | 0.017 | Old |
| 0.351304 | 0.030804 | 0.787 | 0.734 | 1 | 0 | Rbpj | 0.053 | Old |
| 0.352081 | 0.181555 | 0.059 | 0.05 | 1 | 0 | Adap1 | 0.009 | Old |
| 0.352902 | 0.282542 | 0.055 | 0.046 | 1 | 0 | Idh3a | 0.009 | Old |

| | | | | | | | | |
|---|---|---|---|---|---|---|---|---|
| 0.353186 | 0.160576 | 0.037 | 0.03 | 1 | 0 | Cyp2j6 | 0.007 | Old |
| 0.353461 | 0.211536 | 0.078 | 0.068 | 1 | 0 | Cfl2 | 0.01 | Old |
| 0.353757 | 0.22119 | 0.107 | 0.095 | 1 | 0 | Tsnax | 0.012 | Old |
| 0.353882 | 0.136881 | 0.124 | 0.112 | 1 | 0 | BC031181 | 0.012 | Old |
| 0.35397 | 0.242233 | 0.051 | 0.043 | 1 | 0 | Atr | 0.008 | Old |
| 0.354302 | 0.080135 | 0.269 | 0.249 | 1 | 0 | Fam129a | 0.02 | Old |
| 0.354325 | 0.16634 | 0.079 | 0.069 | 1 | 0 | Nasp | 0.01 | Old |
| 0.354477 | 0.271368 | 0.019 | 0.014 | 1 | 0 | Rnf123 | 0.005 | Old |
| 0.355008 | 0.036927 | 0.241 | 0.221 | 1 | 0 | Dennd1a | 0.02 | Old |
| 0.355052 | 0.303827 | 0.019 | 0.014 | 1 | 0 | Zgpat | 0.005 | Old |
| 0.355361 | 0.013304 | 0.157 | 0.141 | 1 | 0 | Hltf | 0.016 | Old |
| 0.355763 | 0.080248 | 0.038 | 0.031 | 1 | 0 | Fpgt | 0.007 | Old |
| 0.355775 | 0.043837 | 0.239 | 0.219 | 1 | 0 | Phf21a | 0.02 | Old |
| 0.356391 | 0.007729 | 0.109 | 0.097 | 1 | 0 | Spast | 0.012 | Old |
| 0.356465 | 0.174025 | 0.144 | 0.131 | 1 | 0 | Ccny | 0.013 | Old |
| 0.356554 | 0.430818 | 0.014 | 0.01 | 1 | 0 | Pex11g | 0.004 | Old |
| 0.356671 | 0.027283 | 0.257 | 0.237 | 1 | 0 | Eif1b | 0.02 | Old |
| 0.356864 | 0.184771 | 0.044 | 0.036 | 1 | 0 | Hsf1 | 0.008 | Old |
| 0.357013 | 0.203998 | 0.049 | 0.041 | 1 | 0 | Rffl | 0.008 | Old |
| 0.35735 | 0.359898 | 0.03 | 0.024 | 1 | 0 | Ttll4 | 0.006 | Old |
| 0.357509 | 0.510598 | 0.021 | 0.016 | 1 | 0 | Ifi206 | 0.005 | Old |
| 0.357576 | 0.005863 | 0.122 | 0.108 | 1 | 0 | Rnf20 | 0.014 | Old |
| 0.357594 | 0.331114 | 0.014 | 0.01 | 1 | 0 | Ctla2a | 0.004 | Old |
| 0.358113 | 0.461493 | 0.01 | 0.006 | 1 | 0 | Zfp7 | 0.004 | Old |
| 0.35827 | 0.150629 | 0.052 | 0.044 | 1 | 0 | Sbf1 | 0.008 | Old |
| 0.358596 | 0.017703 | 0.038 | 0.031 | 1 | 0 | Cdk5rap2 | 0.007 | Old |
| 0.358622 | 0.122277 | 0.211 | 0.198 | 1 | 0 | Tnfsf9 | 0.013 | Old |
| 0.358648 | 0.008131 | 0.361 | 0.333 | 1 | 0 | Ppp1ca | 0.028 | Old |
| 0.359069 | 0.419493 | 0.03 | 0.024 | 1 | 0 | B4galt7 | 0.006 | Old |
| 0.359117 | 0.413487 | 0.031 | 0.025 | 1 | 0 | Hmgxb4 | 0.006 | Old |
| 0.359549 | 0.033076 | 0.019 | 0.014 | 1 | 0 | Mfsd4a | 0.005 | Old |
| 0.359644 | 0.43248 | 0.01 | 0.006 | 1 | 0 | Mfsd9 | 0.004 | Old |
| 0.35968 | 0.272029 | 0.014 | 0.01 | 1 | 0 | Gdf9 | 0.004 | Old |
| 0.359707 | 0.246523 | 0.079 | 0.07 | 1 | 0 | Upf3a | 0.009 | Old |
| 0.359814 | 0.318013 | 0.01 | 0.006 | 1 | 0 | Ccdc94 | 0.004 | Old |
| 0.359814 | 0.208347 | 0.01 | 0.006 | 1 | 0 | Icam2 | 0.004 | Old |
| 0.359985 | 0.305222 | 0.01 | 0.006 | 1 | 0 | Gm11423 | 0.004 | Old |
| 0.360476 | 0.097675 | 0.019 | 0.014 | 1 | 0 | Rab23 | 0.005 | Old |
| 0.360496 | 0.316113 | 0.01 | 0.006 | 1 | 0 | Slc19a2 | 0.004 | Old |
| 0.360507 | 0.113495 | 0.097 | 0.086 | 1 | 0 | Atg12 | 0.011 | Old |
| 0.360507 | 0.238376 | 0.059 | 0.05 | 1 | 0 | Akr7a5 | 0.009 | Old |
| 0.360623 | 0.109878 | 0.042 | 0.035 | 1 | 0 | Pop4 | 0.007 | Old |
| 0.360667 | 0.29182 | 0.01 | 0.006 | 1 | 0 | Cmtr1 | 0.004 | Old |
| 0.360667 | 0.277348 | 0.01 | 0.006 | 1 | 0 | Aplnr | 0.004 | Old |
| 0.360936 | 0.166958 | 0.014 | 0.01 | 1 | 0 | Spef1 | 0.004 | Old |

| | | | | | | | | |
|---|---|---|---|---|---|---|---|---|
| 0.361008 | 0.109376 | 0.01 | 0.006 | 1 | 0 | Nipsnap1 | 0.004 | Old |
| 0.361049 | 0.40382 | 0.077 | 0.067 | 1 | 0 | Tshz1 | 0.01 | Old |
| 0.361179 | 0.177024 | 0.01 | 0.006 | 1 | 0 | Fzd1 | 0.004 | Old |
| 0.361179 | 0.136442 | 0.01 | 0.006 | 1 | 0 | Frg2f1 | 0.004 | Old |
| 0.36164 | 0.120015 | 0.074 | 0.064 | 1 | 0 | Acbd6 | 0.01 | Old |
| 0.363047 | 0.039946 | 0.192 | 0.174 | 1 | 0 | Apc | 0.018 | Old |
| 0.363262 | 0.27527 | 0.034 | 0.028 | 1 | 0 | Selenoo | 0.006 | Old |
| 0.363402 | 0.708989 | 0.042 | 0.035 | 1 | 0 | Yae1d1 | 0.007 | Old |
| 0.363666 | 0.177678 | 0.056 | 0.048 | 1 | 0 | Hook2 | 0.008 | Old |
| 0.363745 | 0.455927 | 0.059 | 0.051 | 1 | 0 | 4930453N2 | 0.008 | Old |
| 0.36385 | 0.011751 | 0.019 | 0.014 | 1 | 0 | Slc16a13 | 0.005 | Old |
| 0.363902 | 0.381031 | 0.036 | 0.029 | 1 | 0 | Klc4 | 0.007 | Old |
| 0.363931 | 0.577123 | 0.022 | 0.017 | 1 | 0 | 4930430F0 | 0.005 | Old |
| 0.364096 | 0.470467 | 0.041 | 0.034 | 1 | 0 | Zfp629 | 0.007 | Old |
| 0.364151 | 0.055307 | 0.321 | 0.297 | 1 | 0 | Zfand6 | 0.024 | Old |
| 0.364638 | 0.144938 | 0.059 | 0.05 | 1 | 0 | Pctp | 0.009 | Old |
| 0.364681 | 0.039567 | 0.845 | 0.818 | 1 | 0 | Ppia | 0.027 | Old |
| 0.365437 | 0.041958 | 0.079 | 0.069 | 1 | 0 | Kansl2 | 0.01 | Old |
| 0.366091 | 0.062933 | 0.267 | 0.246 | 1 | 0 | Ensa | 0.021 | Old |
| 0.3662 | 0.165568 | 0.104 | 0.093 | 1 | 0 | Fam69a | 0.011 | Old |
| 0.366773 | 0.11321 | 0.163 | 0.149 | 1 | 0 | Gas2l1 | 0.014 | Old |
| 0.3675 | 0.13177 | 0.082 | 0.072 | 1 | 0 | Nudcd2 | 0.01 | Old |
| 0.36762 | 0.052812 | 0.048 | 0.04 | 1 | 0 | Carm1 | 0.008 | Old |
| 0.367675 | 0.265118 | 0.022 | 0.017 | 1 | 0 | Spata5 | 0.005 | Old |
| 0.367765 | 0.032484 | 0.048 | 0.04 | 1 | 0 | Hic2 | 0.008 | Old |
| 0.367924 | 0.001159 | 0.082 | 0.071 | 1 | 0 | Abraxas2 | 0.011 | Old |
| 0.368122 | 0.043656 | 0.116 | 0.104 | 1 | 0 | Nsun2 | 0.012 | Old |
| 0.368829 | 0.184801 | 0.096 | 0.085 | 1 | 0 | BC035044 | 0.011 | Old |
| 0.369047 | 0.020199 | 0.12 | 0.107 | 1 | 0 | Tbc1d9b | 0.013 | Old |
| 0.369157 | 0.103962 | 0.059 | 0.05 | 1 | 0 | Ccdc93 | 0.009 | Old |
| 0.369744 | 0.249349 | 0.031 | 0.025 | 1 | 0 | Pigg | 0.006 | Old |
| 0.370065 | 0.065335 | 0.163 | 0.148 | 1 | 0 | Hmbox1 | 0.015 | Old |
| 0.370186 | 0.06748 | 0.163 | 0.148 | 1 | 0 | Trappc6b | 0.015 | Old |
| 0.370195 | 0.459828 | 0.015 | 0.011 | 1 | 0 | Trim7 | 0.004 | Old |
| 0.370422 | 0.328177 | 0.034 | 0.028 | 1 | 0 | Poc5 | 0.006 | Old |
| 0.370456 | 0.216249 | 0.074 | 0.064 | 1 | 0 | Rpp21 | 0.01 | Old |
| 0.370498 | 0.218262 | 0.068 | 0.059 | 1 | 0 | Rab9 | 0.009 | Old |
| 0.370968 | 0.039224 | 0.462 | 0.425 | 1 | 0 | Atp6v1g1 | 0.037 | Old |
| 0.371066 | 0.076203 | 0.064 | 0.055 | 1 | 0 | Naa40 | 0.009 | Old |
| 0.371095 | 0.261331 | 0.056 | 0.048 | 1 | 0 | B3galnt2 | 0.008 | Old |
| 0.371371 | 0.16167 | 0.033 | 0.027 | 1 | 0 | Fam122b | 0.006 | Old |
| 0.371419 | 0.267945 | 0.045 | 0.038 | 1 | 0 | Nbn | 0.007 | Old |
| 0.371771 | 0.217964 | 0.022 | 0.017 | 1 | 0 | Slc25a53 | 0.005 | Old |
| 0.371966 | 0.051479 | 0.114 | 0.102 | 1 | 0 | Mrpl34 | 0.012 | Old |
| 0.37203 | 0.047393 | 0.029 | 0.023 | 1 | 0 | B3gnt8 | 0.006 | Old |

| | | | | | | | | |
|---|---|---|---|---|---|---|---|---|
| 0.373067 | 0.414588 | 0.023 | 0.018 | 1 | 0 | Sumf2 | 0.005 | Old |
| 0.373197 | 0.256905 | 0.022 | 0.017 | 1 | 0 | Ets1 | 0.005 | Old |
| 0.373203 | 0.088535 | 0.105 | 0.093 | 1 | 0 | Casp8ap2 | 0.012 | Old |
| 0.373617 | 0.034707 | 0.051 | 0.043 | 1 | 0 | Pou5f2 | 0.008 | Old |
| 0.373941 | 0.356758 | 0.057 | 0.049 | 1 | 0 | Rnf19a | 0.008 | Old |
| 0.374623 | 0.34379 | 0.042 | 0.035 | 1 | 0 | Gm15987 | 0.007 | Old |
| 0.374781 | 0.063009 | 0.337 | 0.315 | 1 | 0 | Spop | 0.022 | Old |
| 0.375003 | 0.006774 | 0.062 | 0.072 | 1 | 0 | Xpa | -0.01 | Old |
| 0.375762 | 0.084628 | 0.241 | 0.223 | 1 | 0 | Brd1 | 0.018 | Old |
| 0.37724 | 0.143424 | 0.083 | 0.073 | 1 | 0 | Wdr61 | 0.01 | Old |
| 0.377674 | 0.029402 | 0.601 | 0.562 | 1 | 0 | Snx3 | 0.039 | Old |
| 0.377716 | 0.126117 | 0.073 | 0.063 | 1 | 0 | Frmd8 | 0.01 | Old |
| 0.377778 | 0.408972 | 0.025 | 0.019 | 1 | 0 | Sorl1 | 0.006 | Old |
| 0.37785 | 0.254948 | 0.055 | 0.047 | 1 | 0 | Mfn1 | 0.008 | Old |
| 0.378095 | 0.285971 | 0.073 | 0.064 | 1 | 0 | Slc35e1 | 0.009 | Old |
| 0.378933 | 0.214313 | 0.025 | 0.031 | 1 | 0 | Bambi | -0.006 | Old |
| 0.37921 | 0.126639 | 0.038 | 0.032 | 1 | 0 | Stx18 | 0.006 | Old |
| 0.379533 | 1.157055 | 0.011 | 0.008 | 1 | 0 | Rnase2a | 0.003 | Old |
| 0.379533 | 1.127406 | 0.011 | 0.008 | 1 | 0 | Smad7 | 0.003 | Old |
| 0.37958 | 0.003614 | 0.06 | 0.051 | 1 | 0 | Traf7 | 0.009 | Old |
| 0.379643 | 0.325783 | 0.047 | 0.039 | 1 | 0 | Zfp597 | 0.008 | Old |
| 0.379858 | 0.613045 | 0.011 | 0.008 | 1 | 0 | Abcb9 | 0.003 | Old |
| 0.379939 | 0.783756 | 0.011 | 0.008 | 1 | 0 | Polr3g | 0.003 | Old |
| 0.38025 | 0.40541 | 0.026 | 0.021 | 1 | 0 | Nbeal2 | 0.005 | Old |
| 0.380427 | 0.222953 | 0.047 | 0.039 | 1 | 0 | Mir142hg | 0.008 | Old |
| 0.38068 | 0.181406 | 0.045 | 0.038 | 1 | 0 | Dip2a | 0.007 | Old |
| 0.380777 | 0.099071 | 0.161 | 0.147 | 1 | 0 | Slc4a7 | 0.014 | Old |
| 0.380864 | 0.199874 | 0.034 | 0.028 | 1 | 0 | Saal1 | 0.006 | Old |
| 0.381043 | 0.018328 | 0.228 | 0.211 | 1 | 0 | Sf3b6 | 0.017 | Old |
| 0.381243 | 0.167174 | 0.041 | 0.034 | 1 | 0 | Ccdc122 | 0.007 | Old |
| 0.381323 | 0.176738 | 0.041 | 0.034 | 1 | 0 | Jpt2 | 0.007 | Old |
| 0.381896 | 0.402661 | 0.064 | 0.056 | 1 | 0 | Selenoi | 0.008 | Old |
| 0.381913 | 0.107609 | 0.112 | 0.1 | 1 | 0 | Lats1 | 0.012 | Old |
| 0.38212 | 0.079879 | 0.238 | 0.219 | 1 | 0 | Ist1 | 0.019 | Old |
| 0.382174 | 0.070134 | 0.386 | 0.362 | 1 | 0 | Rab7 | 0.024 | Old |
| 0.382296 | 0.456503 | 0.011 | 0.008 | 1 | 0 | Cep76 | 0.003 | Old |
| 0.382621 | 0.491867 | 0.011 | 0.008 | 1 | 0 | Pms1 | 0.003 | Old |
| 0.382784 | 0.471369 | 0.011 | 0.008 | 1 | 0 | Srgap3 | 0.003 | Old |
| 0.38311 | 0.288703 | 0.011 | 0.008 | 1 | 0 | Gm17259 | 0.003 | Old |
| 0.383436 | 0.388112 | 0.011 | 0.008 | 1 | 0 | Tyw3 | 0.003 | Old |
| 0.383501 | 0.071519 | 0.248 | 0.228 | 1 | 0 | Lypla2 | 0.02 | Old |
| 0.384126 | 0.45038 | 0.016 | 0.012 | 1 | 0 | Nmb | 0.004 | Old |
| 0.384201 | 0.076686 | 0.041 | 0.034 | 1 | 0 | Mzt2 | 0.007 | Old |
| 0.384319 | 0.008832 | 0.363 | 0.332 | 1 | 0 | Slc38a2 | 0.031 | Old |
| 0.38434 | 0.268103 | 0.037 | 0.03 | 1 | 0 | Asna1 | 0.007 | Old |

| | | | | | | | | |
|---|---|---|---|---|---|---|---|---|
| 0.384401 | 0.009135 | 0.54 | 0.513 | 1 | 0 | Son | 0.027 | Old |
| 0.385041 | 0.125906 | 0.167 | 0.152 | 1 | 0 | Psmb7 | 0.015 | Old |
| 0.385058 | 0.061653 | 0.324 | 0.301 | 1 | 0 | Mrpl33 | 0.023 | Old |
| 0.38517 | 0.355741 | 0.016 | 0.012 | 1 | 0 | Ccdc73 | 0.004 | Old |
| 0.385234 | 0.094097 | 0.056 | 0.048 | 1 | 0 | Rap1gap2 | 0.008 | Old |
| 0.385274 | 0.153184 | 0.051 | 0.043 | 1 | 0 | Mfsd8 | 0.008 | Old |
| 0.385318 | 0.102825 | 0.185 | 0.17 | 1 | 0 | Chtop | 0.015 | Old |
| 0.385451 | 0.185332 | 0.047 | 0.039 | 1 | 0 | Fam213a | 0.008 | Old |
| 0.385865 | 0.182679 | 0.052 | 0.044 | 1 | 0 | Ikbkap | 0.008 | Old |
| 0.386085 | 0.275424 | 0.016 | 0.012 | 1 | 0 | Sgms2 | 0.004 | Old |
| 0.38638 | 0.063825 | 0.011 | 0.008 | 1 | 0 | Zfp251 | 0.003 | Old |
| 0.38638 | 0.058626 | 0.011 | 0.008 | 1 | 0 | Fkbpl | 0.003 | Old |
| 0.386708 | 0.146353 | 0.011 | 0.008 | 1 | 0 | Zfp449 | 0.003 | Old |
| 0.386751 | 0.309517 | 0.026 | 0.021 | 1 | 0 | Gmnn | 0.005 | Old |
| 0.38683 | 0.056348 | 0.107 | 0.095 | 1 | 0 | Tmem43 | 0.012 | Old |
| 0.387132 | 0.174232 | 0.016 | 0.012 | 1 | 0 | Fbxl8 | 0.004 | Old |
| 0.387158 | 0.070098 | 0.141 | 0.127 | 1 | 0 | Etfa | 0.014 | Old |
| 0.387394 | 0.124062 | 0.016 | 0.012 | 1 | 0 | Zscan12 | 0.004 | Old |
| 0.387517 | 0.241577 | 0.026 | 0.021 | 1 | 0 | Cdc42ep2 | 0.005 | Old |
| 0.387602 | 0.233364 | 0.074 | 0.064 | 1 | 0 | Arl6ip4 | 0.01 | Old |
| 0.387727 | 0.230924 | 0.129 | 0.118 | 1 | 0 | Tfg | 0.011 | Old |
| 0.387787 | 0.322885 | 0.016 | 0.012 | 1 | 0 | 1810011H1 | 0.004 | Old |
| 0.387857 | 0.078898 | 0.011 | 0.008 | 1 | 0 | Gm15564 | 0.003 | Old |
| 0.388014 | 0.13345 | 0.044 | 0.037 | 1 | 0 | Dcp1a | 0.007 | Old |
| 0.388379 | 0.021342 | 0.104 | 0.092 | 1 | 0 | Mms19 | 0.012 | Old |
| 0.388706 | 0.103873 | 0.041 | 0.034 | 1 | 0 | Tlcd2 | 0.007 | Old |
| 0.388789 | 0.021225 | 0.042 | 0.035 | 1 | 0 | Tpgs2 | 0.007 | Old |
| 0.3891 | 0.044295 | 0.016 | 0.012 | 1 | 0 | Synj2 | 0.004 | Old |
| 0.389317 | 0.067435 | 0.599 | 0.583 | 1 | 0 | S100a6 | 0.016 | Old |
| 0.389391 | 0.143028 | 0.094 | 0.084 | 1 | 0 | Tor1a | 0.01 | Old |
| 0.389528 | 0.08407 | 0.163 | 0.148 | 1 | 0 | Trim28 | 0.015 | Old |
| 0.389902 | 0.06228 | 0.159 | 0.144 | 1 | 0 | Bcl7b | 0.015 | Old |
| 0.390381 | 0.030424 | 0.614 | 0.575 | 1 | 0 | Itm2c | 0.039 | Old |
| 0.390557 | 0.066616 | 0.025 | 0.019 | 1 | 0 | Ccdc84 | 0.006 | Old |
| 0.390923 | 0.615141 | 0.018 | 0.013 | 1 | 0 | Mlkl | 0.005 | Old |
| 0.391193 | 0.090493 | 0.088 | 0.077 | 1 | 0 | Hhex | 0.011 | Old |
| 0.391363 | 0.159244 | 0.026 | 0.021 | 1 | 0 | Rbm38 | 0.005 | Old |
| 0.391379 | 0.095703 | 0.175 | 0.161 | 1 | 0 | Rasa3 | 0.014 | Old |
| 0.391394 | 0.028533 | 0.156 | 0.142 | 1 | 0 | Psma5 | 0.014 | Old |
| 0.391457 | 0.074794 | 0.041 | 0.034 | 1 | 0 | Spsb2 | 0.007 | Old |
| 0.391501 | 0.080248 | 0.294 | 0.277 | 1 | 0 | Anxa3 | 0.017 | Old |
| 0.392079 | 0.226972 | 0.114 | 0.102 | 1 | 0 | Galnt11 | 0.012 | Old |
| 0.392188 | 0.032804 | 0.369 | 0.345 | 1 | 0 | Atp6v1a | 0.024 | Old |
| 0.392347 | 0.447669 | 0.04 | 0.033 | 1 | 0 | AI597479 | 0.007 | Old |
| 0.392436 | 0.099446 | 0.044 | 0.037 | 1 | 0 | Tbp | 0.007 | Old |

| | | | | | | | | |
|---|---|---|---|---|---|---|---|---|
| 0.392598 | 0.169021 | 0.026 | 0.021 | 1 | 0 | Per2 | 0.005 | Old |
| 0.392757 | 0.20578 | 0.118 | 0.107 | 1 | 0 | Erlec1 | 0.011 | Old |
| 0.392909 | 0.480318 | 0.027 | 0.022 | 1 | 0 | Zfp606 | 0.005 | Old |
| 0.393209 | 0.092715 | 0.042 | 0.035 | 1 | 0 | Mospd1 | 0.007 | Old |
| 0.393346 | 0.063989 | 0.059 | 0.051 | 1 | 0 | Ppp2cb | 0.008 | Old |
| 0.393488 | 0.18165 | 0.06 | 0.052 | 1 | 0 | Lin54 | 0.008 | Old |
| 0.393952 | 0.083733 | 0.025 | 0.019 | 1 | 0 | Kbtbd7 | 0.006 | Old |
| 0.394014 | 0.29707 | 0.018 | 0.013 | 1 | 0 | Purg | 0.005 | Old |
| 0.394095 | 0.049648 | 0.174 | 0.159 | 1 | 0 | Tmbim4 | 0.015 | Old |
| 0.394183 | 0.015023 | 0.161 | 0.146 | 1 | 0 | Dscr3 | 0.015 | Old |
| 0.394415 | 0.171256 | 0.027 | 0.022 | 1 | 0 | Kremen1 | 0.005 | Old |
| 0.394786 | 0.081963 | 0.183 | 0.168 | 1 | 0 | Srp72 | 0.015 | Old |
| 0.394809 | 0.010002 | 0.042 | 0.035 | 1 | 0 | Med31 | 0.007 | Old |
| 0.394992 | 0.000166 | 0.014 | 0.019 | 1 | 0 | Zfp763 | -0.005 | Old |
| 0.395249 | 0.01215 | 0.208 | 0.191 | 1 | 0 | Akirin2 | 0.017 | Old |
| 0.395899 | 0.173775 | 0.153 | 0.14 | 1 | 0 | Stat1 | 0.013 | Old |
| 0.396025 | 0.034331 | 0.253 | 0.236 | 1 | 0 | Ankrd12 | 0.017 | Old |
| 0.396173 | 0.013023 | 0.19 | 0.173 | 1 | 0 | Vps13a | 0.017 | Old |
| 0.396209 | 0.055314 | 0.241 | 0.224 | 1 | 0 | Rab21 | 0.017 | Old |
| 0.396463 | 0.023905 | 0.311 | 0.291 | 1 | 0 | Hif1a | 0.02 | Old |
| 0.396702 | 0.028392 | 0.23 | 0.212 | 1 | 0 | Etnk1 | 0.018 | Old |
| 0.396915 | 0.012025 | 0.174 | 0.158 | 1 | 0 | Rad23a | 0.016 | Old |
| 0.396986 | 0.117288 | 0.078 | 0.069 | 1 | 0 | Crnkl1 | 0.009 | Old |
| 0.397148 | 0.079296 | 0.026 | 0.021 | 1 | 0 | D2hgdh | 0.005 | Old |
| 0.397294 | 0.019507 | 0.752 | 0.709 | 1 | 0 | Gabarap | 0.043 | Old |
| 0.397397 | 0.130788 | 0.066 | 0.057 | 1 | 0 | Heatr6 | 0.009 | Old |
| 0.397538 | 0.118763 | 0.116 | 0.105 | 1 | 0 | Ncbp3 | 0.011 | Old |
| 0.397552 | 0.59638 | 0.089 | 0.102 | 1 | 0 | Egr3 | -0.013 | Old |
| 0.397633 | 0.043659 | 0.029 | 0.023 | 1 | 0 | Slc12a4 | 0.006 | Old |
| 0.397744 | 0.173069 | 0.055 | 0.047 | 1 | 0 | Josd2 | 0.008 | Old |
| 0.397845 | 0.04131 | 0.119 | 0.107 | 1 | 0 | Clcc1 | 0.012 | Old |
| 0.397992 | 0.168925 | 0.027 | 0.022 | 1 | 0 | Lmln | 0.005 | Old |
| 0.398435 | 0.033979 | 0.062 | 0.053 | 1 | 0 | Immp1l | 0.009 | Old |
| 0.398661 | 0.05845 | 0.052 | 0.044 | 1 | 0 | Zbtb14 | 0.008 | Old |
| 0.399223 | 0.000715 | 0.047 | 0.039 | 1 | 0 | Cers4 | 0.008 | Old |
| 0.399348 | 0.11424 | 0.018 | 0.013 | 1 | 0 | Fadd | 0.005 | Old |
| 0.399454 | 0.221215 | 0.047 | 0.039 | 1 | 0 | Osgin2 | 0.008 | Old |
| 0.4001 | 0.45366 | 0.012 | 0.009 | 1 | 0 | Glcci1 | 0.003 | Old |
| 0.400135 | 0.082772 | 0.16 | 0.146 | 1 | 0 | Nek9 | 0.014 | Old |
| 0.400184 | 0.330567 | 0.019 | 0.015 | 1 | 0 | Trub1 | 0.004 | Old |
| 0.400558 | 0.035464 | 0.914 | 0.922 | 1 | 0 | Rps28 | -0.008 | Old |
| 0.400689 | 0.190684 | 0.029 | 0.023 | 1 | 0 | Prdm15 | 0.006 | Old |
| 0.400922 | 0.02494 | 0.163 | 0.147 | 1 | 0 | Slc25a24 | 0.016 | Old |
| 0.400927 | 0.267057 | 0.063 | 0.055 | 1 | 0 | Isg20l2 | 0.008 | Old |
| 0.401354 | 0.441711 | 0.042 | 0.036 | 1 | 0 | Bag4 | 0.006 | Old |

| | | | | | | | | |
|---|---|---|---|---|---|---|---|---|
| 0.401355 | 0.739065 | 0.021 | 0.016 | 1 | 0 | P2ry10b | 0.005 | Old |
| 0.401425 | 0.207401 | 0.052 | 0.045 | 1 | 0 | Pgm1 | 0.007 | Old |
| 0.401505 | 0.453119 | 0.012 | 0.009 | 1 | 0 | Cnbd2 | 0.003 | Old |
| 0.401983 | 0.051782 | 0.181 | 0.166 | 1 | 0 | Vav1 | 0.015 | Old |
| 0.402063 | 0.017505 | 0.726 | 0.7 | 1 | 0 | Adgre1 | 0.026 | Old |
| 0.402092 | 0.024223 | 0.193 | 0.175 | 1 | 0 | Cnot6 | 0.018 | Old |
| 0.402287 | 0.394704 | 0.012 | 0.009 | 1 | 0 | Cxcl3 | 0.003 | Old |
| 0.402297 | 0.237495 | 0.052 | 0.045 | 1 | 0 | Tmc6 | 0.007 | Old |
| 0.402324 | 0.060969 | 0.513 | 0.485 | 1 | 0 | Sub1 | 0.028 | Old |
| 0.403383 | 0.227376 | 0.012 | 0.009 | 1 | 0 | Ppp1r3e | 0.003 | Old |
| 0.40363 | 0.031611 | 0.19 | 0.174 | 1 | 0 | Ssu72 | 0.016 | Old |
| 0.403718 | 0.108389 | 0.134 | 0.121 | 1 | 0 | Tm2d1 | 0.013 | Old |
| 0.403744 | 0.645703 | 0.019 | 0.015 | 1 | 0 | Pex11a | 0.004 | Old |
| 0.403854 | 0.168059 | 0.012 | 0.009 | 1 | 0 | Impa2 | 0.003 | Old |
| 0.403929 | 0.252002 | 0.019 | 0.015 | 1 | 0 | Tmem186 | 0.004 | Old |
| 0.40399 | 0.131362 | 0.019 | 0.015 | 1 | 0 | Cnr2 | 0.004 | Old |
| 0.40436 | 0.028151 | 0.019 | 0.015 | 1 | 0 | AC163018. | 0.004 | Old |
| 0.404602 | 0.027534 | 0.776 | 0.76 | 1 | 0 | Aplp2 | 0.016 | Old |
| 0.404814 | 0.130783 | 0.03 | 0.024 | 1 | 0 | Dus3l | 0.006 | Old |
| 0.404846 | 0.227166 | 0.04 | 0.033 | 1 | 0 | Bpnt1 | 0.007 | Old |
| 0.404857 | 0.218917 | 0.045 | 0.038 | 1 | 0 | Apex1 | 0.007 | Old |
| 0.404985 | 0.051696 | 0.446 | 0.416 | 1 | 0 | Jpt1 | 0.03 | Old |
| 0.405039 | 0.063687 | 0.311 | 0.292 | 1 | 0 | Rab2a | 0.019 | Old |
| 0.405141 | 0.577204 | 0.021 | 0.016 | 1 | 0 | Zfp748 | 0.005 | Old |
| 0.405238 | 0.082371 | 0.119 | 0.107 | 1 | 0 | Kdm6a | 0.012 | Old |
| 0.405414 | 0.139369 | 0.164 | 0.152 | 1 | 0 | Psmb3 | 0.012 | Old |
| 0.40569 | 0.059508 | 0.172 | 0.158 | 1 | 0 | Sppl3 | 0.014 | Old |
| 0.405774 | 0.126911 | 0.104 | 0.094 | 1 | 0 | Mrps25 | 0.01 | Old |
| 0.406443 | 0.03325 | 0.238 | 0.222 | 1 | 0 | Ccnt1 | 0.016 | Old |
| 0.406814 | 0.020263 | 0.078 | 0.069 | 1 | 0 | Auh | 0.009 | Old |
| 0.408212 | 0.300111 | 0.038 | 0.032 | 1 | 0 | Wdr70 | 0.006 | Old |
| 0.408349 | 0.060668 | 0.28 | 0.262 | 1 | 0 | Prkcb | 0.018 | Old |
| 0.408988 | 0.534285 | 0.022 | 0.017 | 1 | 0 | 2700038G2 | 0.005 | Old |
| 0.410061 | 0.121358 | 0.048 | 0.041 | 1 | 0 | Man2c1 | 0.007 | Old |
| 0.410159 | 0.083526 | 0.064 | 0.056 | 1 | 0 | Taf4 | 0.008 | Old |
| 0.410412 | 0.083253 | 0.074 | 0.065 | 1 | 0 | Sh3glb2 | 0.009 | Old |
| 0.41049 | 0.030936 | 0.847 | 0.808 | 1 | 0 | Mbnl1 | 0.039 | Old |
| 0.410693 | 0.075833 | 0.175 | 0.161 | 1 | 0 | Trpm7 | 0.014 | Old |
| 0.411635 | 0.071096 | 0.03 | 0.024 | 1 | 0 | Eogt | 0.006 | Old |
| 0.412172 | 0.00582 | 0.419 | 0.39 | 1 | 0 | Itsn1 | 0.029 | Old |
| 0.412682 | 0.350092 | 0.022 | 0.017 | 1 | 0 | Rabif | 0.005 | Old |
| 0.412856 | 0.061708 | 0.126 | 0.113 | 1 | 0 | Gch1 | 0.013 | Old |
| 0.413004 | 0.062478 | 0.39 | 0.366 | 1 | 0 | Nfe2l2 | 0.024 | Old |
| 0.41314 | 0.214701 | 0.042 | 0.036 | 1 | 0 | Skp2 | 0.006 | Old |
| 0.413247 | 0.011244 | 0.078 | 0.068 | 1 | 0 | Gpaa1 | 0.01 | Old |

| | | | | | | | | |
|---|---|---|---|---|---|---|---|---|
| 0.413409 | 0.141965 | 0.068 | 0.06 | 1 | 0 | Uri1 | 0.008 | Old |
| 0.413419 | 0.131675 | 0.063 | 0.055 | 1 | 0 | Elof1 | 0.008 | Old |
| 0.413438 | 0.091229 | 0.034 | 0.028 | 1 | 0 | 1190007I0' | 0.006 | Old |
| 0.413482 | 0.138935 | 0.104 | 0.094 | 1 | 0 | Fbxo22 | 0.01 | Old |
| 0.413576 | 0.043927 | 0.227 | 0.21 | 1 | 0 | U2surp | 0.017 | Old |
| 0.413618 | 0.184344 | 0.021 | 0.016 | 1 | 0 | Qtrt1 | 0.005 | Old |
| 0.414218 | 0.172412 | 0.109 | 0.099 | 1 | 0 | Isy1 | 0.01 | Old |
| 0.414306 | 0.020166 | 0.187 | 0.171 | 1 | 0 | Sh2b3 | 0.016 | Old |
| 0.415078 | 0.463532 | 0.014 | 0.01 | 1 | 0 | Mfsd3 | 0.004 | Old |
| 0.415125 | 0.009743 | 0.105 | 0.094 | 1 | 0 | As3mt | 0.011 | Old |
| 0.4152 | 0.068872 | 0.134 | 0.122 | 1 | 0 | Nup88 | 0.012 | Old |
| 0.41535 | 0.354371 | 0.022 | 0.017 | 1 | 0 | Slc19a1 | 0.005 | Old |
| 0.415697 | 0.042798 | 0.811 | 0.801 | 1 | 0 | Anxa5 | 0.01 | Old |
| 0.416127 | 0.426697 | 0.014 | 0.01 | 1 | 0 | Pinx1 | 0.004 | Old |
| 0.416342 | 0.19853 | 0.048 | 0.041 | 1 | 0 | Uqcc1 | 0.007 | Old |
| 0.416458 | 0.177917 | 0.109 | 0.099 | 1 | 0 | Ccng1 | 0.01 | Old |
| 0.416923 | 0.126683 | 0.09 | 0.081 | 1 | 0 | Sin3a | 0.009 | Old |
| 0.417235 | 0.01868 | 0.109 | 0.098 | 1 | 0 | Kmt5a | 0.011 | Old |
| 0.417291 | 0.212878 | 0.118 | 0.108 | 1 | 0 | Mif4gd | 0.01 | Old |
| 0.417357 | 0.145771 | 0.044 | 0.037 | 1 | 0 | Rpain | 0.007 | Old |
| 0.417449 | 0.071483 | 0.057 | 0.05 | 1 | 0 | Grcc10 | 0.007 | Old |
| 0.417663 | 0.107223 | 0.042 | 0.036 | 1 | 0 | Hinfp | 0.006 | Old |
| 0.418296 | 0.070123 | 0.097 | 0.087 | 1 | 0 | Kif2a | 0.01 | Old |
| 0.418421 | 0.374397 | 0.023 | 0.019 | 1 | 0 | Tmem191c | 0.004 | Old |
| 0.418753 | 0.021124 | 0.22 | 0.205 | 1 | 0 | Nucb1 | 0.015 | Old |
| 0.419311 | 0.174344 | 0.022 | 0.017 | 1 | 0 | Zfp653 | 0.005 | Old |
| 0.419435 | 0.079673 | 0.119 | 0.107 | 1 | 0 | Ctso | 0.012 | Old |
| 0.420437 | 0.038078 | 0.326 | 0.311 | 1 | 0 | Selenos | 0.015 | Old |
| 0.420587 | 0.036774 | 0.227 | 0.21 | 1 | 0 | Dhx15 | 0.017 | Old |
| 0.420687 | 0.097697 | 0.06 | 0.052 | 1 | 0 | Frs2 | 0.008 | Old |
| 0.420852 | 0.084029 | 0.038 | 0.032 | 1 | 0 | Nol6 | 0.006 | Old |
| 0.421097 | 0.097895 | 0.014 | 0.01 | 1 | 0 | Zfp58 | 0.004 | Old |
| 0.421215 | 0.024329 | 0.242 | 0.224 | 1 | 0 | Arf2 | 0.018 | Old |
| 0.421621 | 0.803195 | 0.015 | 0.011 | 1 | 0 | Zfp458 | 0.004 | Old |
| 0.421776 | 0.028781 | 0.172 | 0.159 | 1 | 0 | Fam172a | 0.013 | Old |
| 0.421795 | 0.350231 | 0.025 | 0.02 | 1 | 0 | Gm26778 | 0.005 | Old |
| 0.422156 | 0.304408 | 0.023 | 0.019 | 1 | 0 | Cyb5d2 | 0.004 | Old |
| 0.422337 | 0.675646 | 0.015 | 0.011 | 1 | 0 | Nfib | 0.004 | Old |
| 0.42249 | 0.010632 | 0.1 | 0.089 | 1 | 0 | Cbx4 | 0.011 | Old |
| 0.422621 | 0.006046 | 0.187 | 0.171 | 1 | 0 | Phf14 | 0.016 | Old |
| 0.423031 | 0.015828 | 0.781 | 0.73 | 1 | 0 | Rpl3 | 0.051 | Old |
| 0.423081 | 0.058535 | 0.137 | 0.124 | 1 | 0 | Dnajc15 | 0.013 | Old |
| 0.423178 | 0.184999 | 0.023 | 0.019 | 1 | 0 | Gstm4 | 0.004 | Old |
| 0.42321 | 0.272779 | 0.074 | 0.066 | 1 | 0 | Togaram1 | 0.008 | Old |
| 0.423519 | 0.22658 | 0.023 | 0.019 | 1 | 0 | Tysnd1 | 0.004 | Old |

| | | | | | | | | |
|---|---|---|---|---|---|---|---|---|
| 0.423951 | 0.041498 | 0.088 | 0.078 | 1 | 0 | Fnbp4 | 0.01 | Old |
| 0.424003 | 0.517613 | 0.029 | 0.024 | 1 | 0 | Gm7694 | 0.005 | Old |
| 0.4242 | 0.01253 | 0.146 | 0.133 | 1 | 0 | Uba6 | 0.013 | Old |
| 0.424656 | 0.127454 | 0.023 | 0.019 | 1 | 0 | Rttn | 0.004 | Old |
| 0.424833 | 0.021228 | 0.562 | 0.529 | 1 | 0 | Arl6ip1 | 0.033 | Old |
| 0.42486 | 0.384391 | 0.026 | 0.021 | 1 | 0 | Racgap1 | 0.005 | Old |
| 0.425501 | 0.246649 | 0.06 | 0.053 | 1 | 0 | Dram1 | 0.007 | Old |
| 0.426458 | 0.124353 | 0.098 | 0.088 | 1 | 0 | Spg7 | 0.01 | Old |
| 0.426538 | 0.018644 | 0.29 | 0.268 | 1 | 0 | Peli1 | 0.022 | Old |
| 0.426633 | 0.035348 | 0.057 | 0.05 | 1 | 0 | Decr1 | 0.007 | Old |
| 0.426649 | 0.477629 | 0.015 | 0.011 | 1 | 0 | Cmtr2 | 0.004 | Old |
| 0.426842 | 0.000156 | 0.287 | 0.309 | 1 | 0 | Ccr1 | -0.022 | Old |
| 0.427351 | 0.039519 | 0.079 | 0.07 | 1 | 0 | Cdk8 | 0.009 | Old |
| 0.427461 | 0.55142 | 0.185 | 0.176 | 1 | 0 | Ccl4 | 0.009 | Old |
| 0.427514 | 0.401733 | 0.015 | 0.011 | 1 | 0 | Ipp | 0.004 | Old |
| 0.427593 | 0.025458 | 0.085 | 0.075 | 1 | 0 | Prorsd1 | 0.01 | Old |
| 0.427764 | 0.091667 | 0.088 | 0.078 | 1 | 0 | Tmem55a | 0.01 | Old |
| 0.427771 | 0.051793 | 0.085 | 0.075 | 1 | 0 | Poc1b | 0.01 | Old |
| 0.427884 | 0.26168 | 0.007 | 0.01 | 1 | 0 | Agbl3 | -0.003 | Old |
| 0.428724 | 0.090419 | 0.512 | 0.479 | 1 | 0 | Tm9sf3 | 0.033 | Old |
| 0.428814 | 0.346304 | 0.015 | 0.011 | 1 | 0 | Smim1 | 0.004 | Old |
| 0.429313 | 0.002609 | 0.022 | 0.017 | 1 | 0 | Akap1 | 0.005 | Old |
| 0.429378 | 0.250349 | 0.026 | 0.021 | 1 | 0 | Clip2 | 0.005 | Old |
| 0.429643 | 0.096125 | 0.122 | 0.111 | 1 | 0 | Sf3a1 | 0.011 | Old |
| 0.430007 | 0.065592 | 0.167 | 0.153 | 1 | 0 | Pdcl3 | 0.014 | Old |
| 0.430142 | 0.04787 | 0.157 | 0.144 | 1 | 0 | Cop1 | 0.013 | Old |
| 0.430878 | 0.001119 | 0.172 | 0.158 | 1 | 0 | Pi4k2a | 0.014 | Old |
| 0.431309 | 0.080936 | 0.123 | 0.112 | 1 | 0 | Smarcad1 | 0.011 | Old |
| 0.431457 | 0.044433 | 0.059 | 0.051 | 1 | 0 | Zfp994 | 0.008 | Old |
| 0.431659 | 0.009473 | 0.071 | 0.063 | 1 | 0 | Bad | 0.008 | Old |
| 0.43185 | 0.062795 | 0.083 | 0.074 | 1 | 0 | Abcb1b | 0.009 | Old |
| 0.431851 | 0.148562 | 0.064 | 0.056 | 1 | 0 | Akr1e1 | 0.008 | Old |
| 0.43204 | 0.015658 | 0.205 | 0.19 | 1 | 0 | Cenpb | 0.015 | Old |
| 0.432423 | 0.337191 | 0.03 | 0.025 | 1 | 0 | Slc20a2 | 0.005 | Old |
| 0.432504 | 0.000813 | 0.111 | 0.1 | 1 | 0 | Sde2 | 0.011 | Old |
| 0.432729 | 0.077116 | 0.081 | 0.072 | 1 | 0 | Nsmce3 | 0.009 | Old |
| 0.433024 | 0.223646 | 0.03 | 0.025 | 1 | 0 | Park2 | 0.005 | Old |
| 0.433088 | 0.07656 | 0.053 | 0.046 | 1 | 0 | Slc2a8 | 0.007 | Old |
| 0.433406 | 0.338413 | 0.033 | 0.027 | 1 | 0 | Atad3a | 0.006 | Old |
| 0.433521 | 0.045726 | 0.204 | 0.187 | 1 | 0 | Sidt2 | 0.017 | Old |
| 0.433841 | 0.501884 | 0.01 | 0.007 | 1 | 0 | Gemin8 | 0.003 | Old |
| 0.433861 | 0.214891 | 0.064 | 0.056 | 1 | 0 | Uvssa | 0.008 | Old |
| 0.434195 | 0.176033 | 0.026 | 0.021 | 1 | 0 | Trim14 | 0.005 | Old |
| 0.434214 | 0.289618 | 0.01 | 0.007 | 1 | 0 | Gm11696 | 0.003 | Old |
| 0.434774 | 0.313226 | 0.01 | 0.007 | 1 | 0 | Eya4 | 0.003 | Old |

| | | | | | | | | |
|---|---|---|---|---|---|---|---|---|
| 0.435581 | 0.28278 | 0.047 | 0.04 | 1 | 0 | Zxdc | 0.007 | Old |
| 0.435896 | 0.286774 | 0.01 | 0.007 | 1 | 0 | Dlg3 | 0.003 | Old |
| 0.436158 | 0.012104 | 0.114 | 0.102 | 1 | 0 | Fam193a | 0.012 | Old |
| 0.436323 | 0.092943 | 0.027 | 0.022 | 1 | 0 | Cbr3 | 0.005 | Old |
| 0.43637 | 0.059298 | 0.026 | 0.021 | 1 | 0 | Snx7 | 0.005 | Old |
| 0.436564 | 0.153559 | 0.19 | 0.179 | 1 | 0 | Mapk6 | 0.011 | Old |
| 0.437021 | 0.068273 | 0.412 | 0.391 | 1 | 0 | Mfsd1 | 0.021 | Old |
| 0.437157 | 0.153748 | 0.077 | 0.068 | 1 | 0 | Rhbdf2 | 0.009 | Old |
| 0.437303 | 0.06461 | 0.083 | 0.074 | 1 | 0 | Tm7sf3 | 0.009 | Old |
| 0.43755 | 0.145338 | 0.03 | 0.025 | 1 | 0 | Tmub1 | 0.005 | Old |
| 0.437984 | 0.260944 | 0.086 | 0.078 | 1 | 0 | Igsf9 | 0.008 | Old |
| 0.438322 | 0.206558 | 0.041 | 0.035 | 1 | 0 | Ppp2r1b | 0.006 | Old |
| 0.438351 | 0.13831 | 0.092 | 0.083 | 1 | 0 | Elovl1 | 0.009 | Old |
| 0.438745 | 0.284852 | 0.016 | 0.013 | 1 | 0 | Ppcs | 0.003 | Old |
| 0.438794 | 0.354733 | 0.06 | 0.053 | 1 | 0 | Tstd2 | 0.007 | Old |
| 0.438832 | 0.008666 | 0.171 | 0.158 | 1 | 0 | Erp44 | 0.013 | Old |
| 0.439043 | 0.117218 | 0.026 | 0.021 | 1 | 0 | Olfr920 | 0.005 | Old |
| 0.439262 | 0.270277 | 0.04 | 0.034 | 1 | 0 | Pwp1 | 0.006 | Old |
| 0.439539 | 0.113522 | 0.089 | 0.08 | 1 | 0 | Gpatch2 | 0.009 | Old |
| 0.439575 | 0.11788 | 0.06 | 0.053 | 1 | 0 | Gid4 | 0.007 | Old |
| 0.439582 | 0.311833 | 0.016 | 0.013 | 1 | 0 | Tmem138 | 0.003 | Old |
| 0.440036 | 0.067902 | 0.142 | 0.131 | 1 | 0 | Chordc1 | 0.011 | Old |
| 0.440404 | 0.167306 | 0.092 | 0.083 | 1 | 0 | Tmem245 | 0.009 | Old |
| 0.440985 | 0.028467 | 0.115 | 0.104 | 1 | 0 | Arid2 | 0.011 | Old |
| 0.441402 | 0.196387 | 0.033 | 0.027 | 1 | 0 | C1ra | 0.006 | Old |
| 0.441875 | 0.149881 | 0.047 | 0.04 | 1 | 0 | Bloc1s6 | 0.007 | Old |
| 0.441936 | 0.217306 | 0.063 | 0.056 | 1 | 0 | Ccdc134 | 0.007 | Old |
| 0.442058 | 0.104955 | 0.059 | 0.051 | 1 | 0 | Chmp6 | 0.008 | Old |
| 0.442083 | 0.02016 | 0.673 | 0.65 | 1 | 0 | mt-Nd4l | 0.023 | Old |
| 0.442958 | 0.124778 | 0.081 | 0.072 | 1 | 0 | Spns1 | 0.009 | Old |
| 0.442972 | 0.093374 | 0.066 | 0.058 | 1 | 0 | Stoml2 | 0.008 | Old |
| 0.443025 | 0.101717 | 0.185 | 0.172 | 1 | 0 | Dlst | 0.013 | Old |
| 0.44307 | 0.196679 | 0.048 | 0.041 | 1 | 0 | Tarbp2 | 0.007 | Old |
| 0.443212 | 0.311139 | 0.04 | 0.034 | 1 | 0 | Xlr4b | 0.006 | Old |
| 0.443257 | 0.077246 | 0.275 | 0.261 | 1 | 0 | Midn | 0.014 | Old |
| 0.443508 | 0.089344 | 0.13 | 0.119 | 1 | 0 | Ap3m1 | 0.011 | Old |
| 0.443602 | 0.148363 | 0.034 | 0.029 | 1 | 0 | Hist2h2be | 0.005 | Old |
| 0.443931 | 0.030895 | 0.189 | 0.176 | 1 | 0 | Ssr4 | 0.013 | Old |
| 0.444623 | 0.004977 | 0.016 | 0.013 | 1 | 0 | Ndufaf5 | 0.003 | Old |
| 0.444979 | 0.222768 | 0.018 | 0.014 | 1 | 0 | Tsr3 | 0.004 | Old |
| 0.445105 | 0.006081 | 0.031 | 0.026 | 1 | 0 | Mettl1 | 0.005 | Old |
| 0.445919 | 0.11128 | 0.052 | 0.045 | 1 | 0 | Pop7 | 0.007 | Old |
| 0.446055 | 0.267598 | 0.018 | 0.014 | 1 | 0 | Lekr1 | 0.004 | Old |
| 0.446162 | 0.192475 | 0.033 | 0.027 | 1 | 0 | Slc35g1 | 0.006 | Old |
| 0.446357 | 0.471106 | 0.047 | 0.04 | 1 | 0 | Ttc17 | 0.007 | Old |

| | | | | | | | | |
|---|---|---|---|---|---|---|---|---|
| 0.446593 | 0.057491 | 0.063 | 0.055 | 1 | 0 | Nkap | 0.008 | Old |
| 0.446599 | 0.074098 | 0.031 | 0.026 | 1 | 0 | Pcca | 0.005 | Old |
| 0.446776 | 0.003249 | 0.146 | 0.134 | 1 | 0 | Cers6 | 0.012 | Old |
| 0.446887 | 0.06228 | 0.554 | 0.535 | 1 | 0 | Ifi27l2a | 0.019 | Old |
| 0.446955 | 0.027482 | 0.078 | 0.069 | 1 | 0 | Ap4s1 | 0.009 | Old |
| 0.447264 | 0.016583 | 0.103 | 0.092 | 1 | 0 | H2-T22 | 0.011 | Old |
| 0.447289 | 0.078534 | 0.253 | 0.237 | 1 | 0 | Rab6a | 0.016 | Old |
| 0.447445 | 0.008372 | 0.918 | 0.917 | 1 | 0 | Dab2 | 0.001 | Old |
| 0.447494 | 0.150057 | 0.045 | 0.039 | 1 | 0 | Zfyve1 | 0.006 | Old |
| 0.447502 | 0.084586 | 0.198 | 0.184 | 1 | 0 | Dennd1b | 0.014 | Old |
| 0.447603 | 0.022255 | 0.075 | 0.067 | 1 | 0 | Rbpms | 0.008 | Old |
| 0.447875 | 0.096323 | 0.14 | 0.128 | 1 | 0 | Gtpbp4 | 0.012 | Old |
| 0.448072 | 0.452526 | 0.019 | 0.015 | 1 | 0 | Dcp1b | 0.004 | Old |
| 0.448422 | 0.067565 | 0.279 | 0.262 | 1 | 0 | 1-Mar | 0.017 | Old |
| 0.448594 | 0.060777 | 0.038 | 0.032 | 1 | 0 | Ppox | 0.006 | Old |
| 0.448717 | 0.110736 | 0.059 | 0.051 | 1 | 0 | Setd1a | 0.008 | Old |
| 0.448886 | 0.243431 | 0.018 | 0.014 | 1 | 0 | L3mbtl2 | 0.004 | Old |
| 0.449182 | 0.069116 | 0.104 | 0.094 | 1 | 0 | CAAA0111{ | 0.01 | Old |
| 0.449313 | 0.076191 | 0.067 | 0.059 | 1 | 0 | Tceanc2 | 0.008 | Old |
| 0.449741 | 0.13973 | 0.189 | 0.178 | 1 | 0 | Tmem164 | 0.011 | Old |
| 0.450176 | 0.051506 | 0.175 | 0.163 | 1 | 0 | Tmem168 | 0.012 | Old |
| 0.450363 | 0.036806 | 0.033 | 0.027 | 1 | 0 | Eif2b3 | 0.006 | Old |
| 0.450703 | 0.062695 | 0.019 | 0.024 | 1 | 0 | Pik3ap1 | -0.005 | Old |
| 0.451148 | 0.110683 | 0.208 | 0.194 | 1 | 0 | Ifitm6 | 0.014 | Old |
| 0.451712 | 0.065629 | 0.094 | 0.085 | 1 | 0 | Cbx5 | 0.009 | Old |
| 0.451999 | 0.115226 | 0.018 | 0.014 | 1 | 0 | Ecsit | 0.004 | Old |
| 0.452134 | 0.11341 | 0.018 | 0.014 | 1 | 0 | Gm13431 | 0.004 | Old |
| 0.45235 | 0.022269 | 0.075 | 0.067 | 1 | 0 | Ankrd27 | 0.008 | Old |
| 0.452617 | 0.181247 | 0.019 | 0.015 | 1 | 0 | Ndrg1 | 0.004 | Old |
| 0.452634 | 0.036946 | 0.083 | 0.074 | 1 | 0 | Vamp7 | 0.009 | Old |
| 0.452833 | 0.058082 | 0.133 | 0.122 | 1 | 0 | Vps28 | 0.011 | Old |
| 0.452935 | 0.031382 | 0.375 | 0.35 | 1 | 0 | Sf3b2 | 0.025 | Old |
| 0.453308 | 0.146491 | 0.155 | 0.142 | 1 | 0 | Slc25a39 | 0.013 | Old |
| 0.45333 | 0.599976 | 0.023 | 0.019 | 1 | 0 | Ube2e2 | 0.004 | Old |
| 0.453709 | 0.138825 | 0.078 | 0.07 | 1 | 0 | Slc1a5 | 0.008 | Old |
| 0.453818 | 0.012983 | 0.048 | 0.041 | 1 | 0 | Senp3 | 0.007 | Old |
| 0.454347 | 0.257794 | 0.011 | 0.008 | 1 | 0 | Dnaaf5 | 0.003 | Old |
| 0.454454 | 0.004139 | 0.352 | 0.326 | 1 | 0 | Cast | 0.026 | Old |
| 0.454662 | 0.070618 | 0.038 | 0.032 | 1 | 0 | Dhx58 | 0.006 | Old |
| 0.45494 | 0.048354 | 0.036 | 0.03 | 1 | 0 | Nob1 | 0.006 | Old |
| 0.455072 | 0.082733 | 0.064 | 0.056 | 1 | 0 | Mafk | 0.008 | Old |
| 0.455449 | 0.151688 | 0.052 | 0.045 | 1 | 0 | Rptor | 0.007 | Old |
| 0.455533 | 0.050018 | 0.09 | 0.081 | 1 | 0 | Stx6 | 0.009 | Old |
| 0.455704 | 0.190679 | 0.074 | 0.066 | 1 | 0 | Hikeshi | 0.008 | Old |
| 0.45666 | 0.089703 | 0.064 | 0.057 | 1 | 0 | Shc1 | 0.007 | Old |

| | | | | | | | | |
|---|---|---|---|---|---|---|---|---|
| 0.456864 | 0.062352 | 0.037 | 0.031 | 1 | 0 | Alkbh3 | 0.006 | Old |
| 0.456916 | 0.145481 | 0.227 | 0.213 | 1 | 0 | Rere | 0.014 | Old |
| 0.457006 | 0.127266 | 0.011 | 0.008 | 1 | 0 | Tut1 | 0.003 | Old |
| 0.457484 | 0.218077 | 0.082 | 0.074 | 1 | 0 | Ppfia4 | 0.008 | Old |
| 0.458106 | 0.129553 | 0.019 | 0.015 | 1 | 0 | Zbtb12 | 0.004 | Old |
| 0.45843 | 0.075164 | 0.086 | 0.077 | 1 | 0 | Appl2 | 0.009 | Old |
| 0.458523 | 0.016665 | 0.141 | 0.128 | 1 | 0 | Ncoa2 | 0.013 | Old |
| 0.458622 | 0.196347 | 0.081 | 0.073 | 1 | 0 | Hps5 | 0.008 | Old |
| 0.459118 | 0.042307 | 0.134 | 0.123 | 1 | 0 | Nap1l4 | 0.011 | Old |
| 0.459534 | 0.062036 | 0.166 | 0.152 | 1 | 0 | 5031425E2 | 0.014 | Old |
| 0.459684 | 0.190776 | 0.057 | 0.051 | 1 | 0 | Zbtb21 | 0.006 | Old |
| 0.460831 | 0.072118 | 0.149 | 0.137 | 1 | 0 | Itgb5 | 0.012 | Old |
| 0.461061 | 0.006954 | 0.055 | 0.048 | 1 | 0 | Clmp | 0.007 | Old |
| 0.461292 | 0.07234 | 0.148 | 0.136 | 1 | 0 | Ralgds | 0.012 | Old |
| 0.461363 | 0.211443 | 0.063 | 0.056 | 1 | 0 | Ralgapb | 0.007 | Old |
| 0.461759 | 0.075255 | 0.075 | 0.067 | 1 | 0 | Alas1 | 0.008 | Old |
| 0.462266 | 0.045413 | 0.71 | 0.699 | 1 | 0 | Ifitm3 | 0.011 | Old |
| 0.46251 | 0.063906 | 0.09 | 0.081 | 1 | 0 | Tm2d3 | 0.009 | Old |
| 0.462562 | 0.039275 | 0.15 | 0.139 | 1 | 0 | Dcp2 | 0.011 | Old |
| 0.462978 | 0.021345 | 0.093 | 0.084 | 1 | 0 | Cnot10 | 0.009 | Old |
| 0.463193 | 0.4227 | 0.033 | 0.028 | 1 | 0 | Sufu | 0.005 | Old |
| 0.463398 | 0.047217 | 0.287 | 0.269 | 1 | 0 | Comt | 0.018 | Old |
| 0.464679 | 0.286373 | 0.057 | 0.051 | 1 | 0 | Med23 | 0.006 | Old |
| 0.464832 | 0.105806 | 0.092 | 0.083 | 1 | 0 | Atxn10 | 0.009 | Old |
| 0.464868 | 0.007982 | 0.14 | 0.128 | 1 | 0 | Gtf3c2 | 0.012 | Old |
| 0.46496 | 0.289896 | 0.047 | 0.04 | 1 | 0 | Psd4 | 0.007 | Old |
| 0.46523 | 0.067811 | 0.108 | 0.099 | 1 | 0 | Mrps12 | 0.009 | Old |
| 0.465354 | 0.070426 | 0.379 | 0.359 | 1 | 0 | Hcls1 | 0.02 | Old |
| 0.465862 | 0.402956 | 0.022 | 0.018 | 1 | 0 | Limk2 | 0.004 | Old |
| 0.46622 | 0.021013 | 0.163 | 0.15 | 1 | 0 | Asph | 0.013 | Old |
| 0.46631 | 0.283542 | 0.025 | 0.02 | 1 | 0 | Leng9 | 0.005 | Old |
| 0.466467 | 0.161315 | 0.078 | 0.07 | 1 | 0 | Fry | 0.008 | Old |
| 0.466883 | 0.310313 | 0.026 | 0.021 | 1 | 0 | Fasn | 0.005 | Old |
| 0.466996 | 0.082125 | 0.264 | 0.247 | 1 | 0 | Rgs18 | 0.017 | Old |
| 0.467054 | 0.13554 | 0.083 | 0.075 | 1 | 0 | Commd1 | 0.008 | Old |
| 0.467196 | 0.126638 | 0.164 | 0.153 | 1 | 0 | Polr2e | 0.011 | Old |
| 0.467524 | 0.552852 | 0.012 | 0.009 | 1 | 0 | Smco4 | 0.003 | Old |
| 0.467588 | 0.381598 | 0.031 | 0.027 | 1 | 0 | Cd9 | 0.004 | Old |
| 0.467656 | 0.032294 | 0.008 | 0.011 | 1 | 0 | Zbtb49 | -0.003 | Old |
| 0.468076 | 0.173634 | 0.051 | 0.044 | 1 | 0 | AI837181 | 0.007 | Old |
| 0.468894 | 0.019918 | 0.171 | 0.158 | 1 | 0 | 9030624J0 | 0.013 | Old |
| 0.469205 | 0.17963 | 0.012 | 0.009 | 1 | 0 | Gmpr | 0.003 | Old |
| 0.469499 | 0.398167 | 0.029 | 0.024 | 1 | 0 | Ppp2r5b | 0.005 | Old |
| 0.469524 | 0.287333 | 0.064 | 0.057 | 1 | 0 | Dnpep | 0.007 | Old |
| 0.469678 | 0.061707 | 0.124 | 0.114 | 1 | 0 | Uap1 | 0.01 | Old |

| | | | | | | | | |
|---|---|---|---|---|---|---|---|---|
| 0.469932 | 0.071427 | 0.022 | 0.018 | 1 | 0 | Kptn | 0.004 | Old |
| 0.470047 | 0.265462 | 0.012 | 0.009 | 1 | 0 | Zfp870 | 0.003 | Old |
| 0.470215 | 0.093388 | 0.012 | 0.009 | 1 | 0 | Coq4 | 0.003 | Old |
| 0.470318 | 0.053835 | 0.424 | 0.406 | 1 | 0 | Selenof | 0.018 | Old |
| 0.470733 | 0.011123 | 0.073 | 0.064 | 1 | 0 | Dusp23 | 0.009 | Old |
| 0.470933 | 0.054678 | 0.598 | 0.575 | 1 | 0 | C3ar1 | 0.023 | Old |
| 0.471485 | 0.030631 | 0.178 | 0.165 | 1 | 0 | Rhoq | 0.013 | Old |
| 0.472226 | 0.001533 | 0.142 | 0.131 | 1 | 0 | Sertad2 | 0.011 | Old |
| 0.472341 | 0.222112 | 0.025 | 0.02 | 1 | 0 | Gm20186 | 0.005 | Old |
| 0.472366 | 0.005885 | 0.171 | 0.158 | 1 | 0 | Slc36a1 | 0.013 | Old |
| 0.472713 | 0.278611 | 0.029 | 0.024 | 1 | 0 | Atpaf1 | 0.005 | Old |
| 0.473094 | 0.112995 | 0.023 | 0.019 | 1 | 0 | Clybl | 0.004 | Old |
| 0.473324 | 0.690758 | 0.014 | 0.011 | 1 | 0 | Brca2 | 0.003 | Old |
| 0.473477 | 0.13233 | 0.089 | 0.08 | 1 | 0 | Gpr137b | 0.009 | Old |
| 0.474441 | 0.229846 | 0.13 | 0.122 | 1 | 0 | Tomm70a | 0.008 | Old |
| 0.474538 | 0.029687 | 0.503 | 0.479 | 1 | 0 | Creg1 | 0.024 | Old |
| 0.475115 | 0.20754 | 0.041 | 0.035 | 1 | 0 | Ddt | 0.006 | Old |
| 0.475432 | 0.27768 | 0.037 | 0.032 | 1 | 0 | Nudt2 | 0.005 | Old |
| 0.475705 | 0.041535 | 0.332 | 0.309 | 1 | 0 | Klf13 | 0.023 | Old |
| 0.475923 | 0.177888 | 0.04 | 0.034 | 1 | 0 | Naip6 | 0.006 | Old |
| 0.477035 | 0.214445 | 0.145 | 0.135 | 1 | 0 | Nr4a2 | 0.01 | Old |
| 0.477221 | 0.224809 | 0.041 | 0.035 | 1 | 0 | Tbc1d32 | 0.006 | Old |
| 0.477232 | 0.161692 | 0.094 | 0.086 | 1 | 0 | Topors | 0.008 | Old |
| 0.477584 | 0.160547 | 0.063 | 0.056 | 1 | 0 | Psmd10 | 0.007 | Old |
| 0.477789 | 0.242485 | 0.034 | 0.029 | 1 | 0 | Mlh3 | 0.005 | Old |
| 0.47849 | 0.083746 | 0.051 | 0.044 | 1 | 0 | Lrch1 | 0.007 | Old |
| 0.4795 | 0.197634 | 0.029 | 0.024 | 1 | 0 | Zfand2a | 0.005 | Old |
| 0.480198 | 0.473381 | 0.014 | 0.011 | 1 | 0 | Ggnbp1 | 0.003 | Old |
| 0.480248 | 0.048781 | 0.131 | 0.12 | 1 | 0 | Pex6 | 0.011 | Old |
| 0.480373 | 0.080748 | 0.134 | 0.123 | 1 | 0 | Ythdc2 | 0.011 | Old |
| 0.480949 | 0.114091 | 0.326 | 0.31 | 1 | 0 | Ptpro | 0.016 | Old |
| 0.481098 | 0.012859 | 0.483 | 0.453 | 1 | 0 | Tpp1 | 0.03 | Old |
| 0.481377 | 0.745575 | 0.015 | 0.012 | 1 | 0 | Mvk | 0.003 | Old |
| 0.481475 | 0.119272 | 0.122 | 0.112 | 1 | 0 | Sf3a3 | 0.01 | Old |
| 0.48152 | 0.066623 | 0.304 | 0.289 | 1 | 0 | Ehbp1l1 | 0.015 | Old |
| 0.481613 | 0.05919 | 0.144 | 0.133 | 1 | 0 | Tlnrd1 | 0.011 | Old |
| 0.481991 | 0.167314 | 0.041 | 0.035 | 1 | 0 | Nup160 | 0.006 | Old |
| 0.482043 | 0.049462 | 0.073 | 0.065 | 1 | 0 | Trim34a | 0.008 | Old |
| 0.482053 | 0.141802 | 0.156 | 0.144 | 1 | 0 | Fam213b | 0.012 | Old |
| 0.482769 | 0.228156 | 0.014 | 0.011 | 1 | 0 | Homez | 0.003 | Old |
| 0.482941 | 0.202726 | 0.038 | 0.033 | 1 | 0 | Dclre1a | 0.005 | Old |
| 0.482954 | 0.056047 | 0.048 | 0.042 | 1 | 0 | Mgat5 | 0.006 | Old |
| 0.484123 | 0.72169 | 0.015 | 0.012 | 1 | 0 | Mettl2 | 0.003 | Old |
| 0.484258 | 0.034877 | 0.338 | 0.321 | 1 | 0 | Rp2 | 0.017 | Old |
| 0.484728 | 0.013783 | 0.331 | 0.308 | 1 | 0 | Tnrc18 | 0.023 | Old |

| | | | | | | | | |
|---|---|---|---|---|---|---|---|---|
| 0.484867 | 0.083735 | 0.034 | 0.029 | 1 | 0 | Slc10a3 | 0.005 | Old |
| 0.485603 | 0.061554 | 0.067 | 0.06 | 1 | 0 | Vps9d1 | 0.007 | Old |
| 0.48576 | 0.030034 | 0.189 | 0.176 | 1 | 0 | Tmem165 | 0.013 | Old |
| 0.486174 | 0.021361 | 0.317 | 0.302 | 1 | 0 | Zfp207 | 0.015 | Old |
| 0.486381 | 0.040396 | 0.171 | 0.158 | 1 | 0 | Txnl1 | 0.013 | Old |
| 0.48647 | 0.051247 | 0.041 | 0.035 | 1 | 0 | Noc4l | 0.006 | Old |
| 0.486594 | 0.075092 | 0.161 | 0.15 | 1 | 0 | Acbd3 | 0.011 | Old |
| 0.487353 | 0.056103 | 0.042 | 0.037 | 1 | 0 | Mrs2 | 0.005 | Old |
| 0.487576 | 0.653354 | 0.016 | 0.013 | 1 | 0 | Trp53rkb | 0.003 | Old |
| 0.487822 | 0.155803 | 0.108 | 0.099 | 1 | 0 | Fem1b | 0.009 | Old |
| 0.487831 | 0.094058 | 0.041 | 0.035 | 1 | 0 | Numb | 0.006 | Old |
| 0.487893 | 0.285155 | 0.092 | 0.085 | 1 | 0 | Ppm1g | 0.007 | Old |
| 0.487955 | 0.129047 | 0.086 | 0.078 | 1 | 0 | Dcaf11 | 0.008 | Old |
| 0.488311 | 0.004291 | 0.212 | 0.198 | 1 | 0 | Vdac1 | 0.014 | Old |
| 0.488387 | 0.086362 | 0.092 | 0.083 | 1 | 0 | Atp11b | 0.009 | Old |
| 0.488695 | 0.091186 | 0.03 | 0.025 | 1 | 0 | Cdk7 | 0.005 | Old |
| 0.48871 | 0.052103 | 0.1 | 0.091 | 1 | 0 | Palld | 0.009 | Old |
| 0.489141 | 0.042332 | 0.13 | 0.119 | 1 | 0 | Socs6 | 0.011 | Old |
| 0.489794 | 0.280405 | 0.015 | 0.012 | 1 | 0 | Zdhhc1 | 0.003 | Old |
| 0.490382 | 0.042306 | 0.029 | 0.024 | 1 | 0 | Mtor | 0.005 | Old |
| 0.491825 | 0.36743 | 0.016 | 0.013 | 1 | 0 | Eml6 | 0.003 | Old |
| 0.491836 | 0.024808 | 0.204 | 0.191 | 1 | 0 | Zfand3 | 0.013 | Old |
| 0.492253 | 0.007513 | 0.022 | 0.027 | 1 | 0 | Cnnm4 | -0.005 | Old |
| 0.492574 | 0.513573 | 0.036 | 0.031 | 1 | 0 | Cactin | 0.005 | Old |
| 0.49376 | 0.029483 | 0.279 | 0.26 | 1 | 0 | Acin1 | 0.019 | Old |
| 0.494109 | 0.110843 | 0.015 | 0.012 | 1 | 0 | Gm28151 | 0.003 | Old |
| 0.494662 | 0.183305 | 0.052 | 0.046 | 1 | 0 | Rwdd4a | 0.006 | Old |
| 0.494868 | 0.71556 | 0.018 | 0.014 | 1 | 0 | Tjap1 | 0.004 | Old |
| 0.495356 | 0.477899 | 0.016 | 0.013 | 1 | 0 | BC030499 | 0.003 | Old |
| 0.4957 | 0.024598 | 0.324 | 0.302 | 1 | 0 | Ctnna1 | 0.022 | Old |
| 0.495821 | 0.068558 | 0.313 | 0.296 | 1 | 0 | Znrf2 | 0.017 | Old |
| 0.496415 | 0.326642 | 0.063 | 0.056 | 1 | 0 | Rinl | 0.007 | Old |
| 0.496584 | 0.162879 | 0.19 | 0.18 | 1 | 0 | Atf7 | 0.01 | Old |
| 0.496703 | 0.424821 | 0.018 | 0.014 | 1 | 0 | Cks1b | 0.004 | Old |
| 0.496757 | 0.293297 | 0.016 | 0.013 | 1 | 0 | Traf3ip1 | 0.003 | Old |
| 0.497131 | 0.025689 | 0.527 | 0.499 | 1 | 0 | Hnrnpa0 | 0.028 | Old |
| 0.497396 | 0.325167 | 0.042 | 0.037 | 1 | 0 | Ric8b | 0.005 | Old |
| 0.497849 | 0.699485 | 0.019 | 0.016 | 1 | 0 | Tmem218 | 0.003 | Old |
| 0.49819 | 0.01636 | 0.201 | 0.186 | 1 | 0 | Tlr7 | 0.015 | Old |
| 0.4984 | 0.333851 | 0.018 | 0.014 | 1 | 0 | Wars2 | 0.004 | Old |
| 0.498522 | 0.070031 | 0.115 | 0.105 | 1 | 0 | Eif4enif1 | 0.01 | Old |
| 0.49858 | 0.002163 | 0.263 | 0.243 | 1 | 0 | Vamp3 | 0.02 | Old |
| 0.498648 | 0.088066 | 0.079 | 0.072 | 1 | 0 | Larp7 | 0.007 | Old |
| 0.499031 | 0.035033 | 0.098 | 0.09 | 1 | 0 | Siah2 | 0.008 | Old |
| 0.499196 | 0.172717 | 0.016 | 0.013 | 1 | 0 | Ms4a4c | 0.003 | Old |

| | | | | | | | | |
|---|---|---|---|---|---|---|---|---|
| 0.49924 | 0.118662 | 0.103 | 0.094 | 1 | 0 | Mocs2 | 0.009 | Old |
| 0.499635 | 0.003161 | 0.29 | 0.271 | 1 | 0 | Hnrnpl | 0.019 | Old |
| 0.500149 | 0.078902 | 0.115 | 0.106 | 1 | 0 | Dnajc2 | 0.009 | Old |
| 0.500453 | 0.090401 | 0.074 | 0.067 | 1 | 0 | Ugdh | 0.007 | Old |
| 0.500543 | 0.240747 | 0.066 | 0.059 | 1 | 0 | Vcl | 0.007 | Old |
| 0.500888 | 0.014042 | 0.068 | 0.061 | 1 | 0 | Polr2j | 0.007 | Old |
| 0.501123 | 0.294239 | 0.016 | 0.013 | 1 | 0 | Rbak | 0.003 | Old |
| 0.501185 | 0.256223 | 0.063 | 0.056 | 1 | 0 | Tdg | 0.007 | Old |
| 0.502227 | 0.604896 | 0.029 | 0.024 | 1 | 0 | Tmem268 | 0.005 | Old |
| 0.502285 | 0.279927 | 0.04 | 0.035 | 1 | 0 | Cinp | 0.005 | Old |
| 0.502301 | 0.283781 | 0.018 | 0.014 | 1 | 0 | Pex12 | 0.004 | Old |
| 0.503016 | 0.036593 | 0.133 | 0.123 | 1 | 0 | Nr2c2 | 0.01 | Old |
| 0.503035 | 0.171857 | 0.115 | 0.107 | 1 | 0 | Nfil3 | 0.008 | Old |
| 0.503225 | 0.191295 | 0.018 | 0.014 | 1 | 0 | Fads6 | 0.004 | Old |
| 0.50387 | 0.293925 | 0.053 | 0.048 | 1 | 0 | Sdccag8 | 0.005 | Old |
| 0.504079 | 0.200533 | 0.018 | 0.014 | 1 | 0 | Ttll1 | 0.004 | Old |
| 0.504122 | 0.115937 | 0.073 | 0.065 | 1 | 0 | Ttc33 | 0.008 | Old |
| 0.504474 | 0.018564 | 0.15 | 0.163 | 1 | 0 | Ep400 | -0.013 | Old |
| 0.50456 | 0.442964 | 0.021 | 0.017 | 1 | 0 | Msl3l2 | 0.004 | Old |
| 0.50503 | 0.059745 | 0.287 | 0.271 | 1 | 0 | Fam120a | 0.016 | Old |
| 0.505121 | 0.021665 | 0.098 | 0.09 | 1 | 0 | Patl1 | 0.008 | Old |
| 0.505232 | 0.209708 | 0.111 | 0.102 | 1 | 0 | Cspp1 | 0.009 | Old |
| 0.50549 | 0.422753 | 0.019 | 0.016 | 1 | 0 | Fam20a | 0.003 | Old |
| 0.506012 | 0.334534 | 0.021 | 0.017 | 1 | 0 | Ift57 | 0.004 | Old |
| 0.506098 | 0.039091 | 0.238 | 0.224 | 1 | 0 | Srrm1 | 0.014 | Old |
| 0.506606 | 0.143304 | 0.174 | 0.163 | 1 | 0 | Ddx50 | 0.011 | Old |
| 0.506735 | 0.028223 | 0.053 | 0.047 | 1 | 0 | Rpusd4 | 0.006 | Old |
| 0.506887 | 0.308426 | 0.038 | 0.033 | 1 | 0 | Dglucy | 0.005 | Old |
| 0.507517 | 0.280399 | 0.037 | 0.032 | 1 | 0 | Taf1b | 0.005 | Old |
| 0.50768 | 0.180901 | 0.12 | 0.112 | 1 | 0 | Frmd4a | 0.008 | Old |
| 0.507715 | 0.014432 | 0.272 | 0.256 | 1 | 0 | Fam208a | 0.016 | Old |
| 0.508106 | 0.365231 | 0.022 | 0.018 | 1 | 0 | Gstt2 | 0.004 | Old |
| 0.508218 | 0.113792 | 0.018 | 0.014 | 1 | 0 | Rnf152 | 0.004 | Old |
| 0.50826 | 0.388047 | 0.021 | 0.017 | 1 | 0 | Zfp609 | 0.004 | Old |
| 0.50879 | 0.067896 | 0.018 | 0.014 | 1 | 0 | Prmt6 | 0.004 | Old |
| 0.508874 | 0.325462 | 0.022 | 0.018 | 1 | 0 | Ankrd39 | 0.004 | Old |
| 0.509329 | 0.041518 | 0.222 | 0.209 | 1 | 0 | Rab18 | 0.013 | Old |
| 0.509599 | 0.004191 | 0.071 | 0.079 | 1 | 0 | Dda1 | -0.008 | Old |
| 0.509917 | 0.418163 | 0.029 | 0.024 | 1 | 0 | Flrt2 | 0.005 | Old |
| 0.510138 | 0.075686 | 0.045 | 0.04 | 1 | 0 | Cpsf4 | 0.005 | Old |
| 0.510316 | 0.742794 | 0.01 | 0.007 | 1 | 0 | Tkfc | 0.003 | Old |
| 0.510848 | 0.188245 | 0.019 | 0.016 | 1 | 0 | Tbc1d22b | 0.003 | Old |
| 0.51099 | 0.027073 | 0.148 | 0.137 | 1 | 0 | Cacul1 | 0.011 | Old |
| 0.511025 | 0.419321 | 0.03 | 0.026 | 1 | 0 | Tbccd1 | 0.004 | Old |
| 0.511066 | 0.248519 | 0.041 | 0.036 | 1 | 0 | Eri3 | 0.005 | Old |

| | | | | | | | | |
|---|---|---|---|---|---|---|---|---|
| 0.511096 | 0.078733 | 0.118 | 0.109 | 1 | 0 | Pfkfb3 | 0.009 | Old |
| 0.511315 | 0.599145 | 0.01 | 0.007 | 1 | 0 | Zfp30 | 0.003 | Old |
| 0.511515 | 0.51609 | 0.01 | 0.007 | 1 | 0 | 4930579G2 | 0.003 | Old |
| 0.511692 | 0.19405 | 0.141 | 0.133 | 1 | 0 | Zc3h12a | 0.008 | Old |
| 0.511751 | 0.395476 | 0.026 | 0.022 | 1 | 0 | Pianp | 0.004 | Old |
| 0.511916 | 0.203322 | 0.01 | 0.013 | 1 | 0 | 6-Sep | -0.003 | Old |
| 0.511951 | 0.041179 | 0.019 | 0.016 | 1 | 0 | Sec22c | 0.003 | Old |
| 0.512474 | 0.358105 | 0.034 | 0.029 | 1 | 0 | Znhit6 | 0.005 | Old |
| 0.512772 | 0.11296 | 0.021 | 0.017 | 1 | 0 | Mccc2 | 0.004 | Old |
| 0.513107 | 0.250386 | 0.022 | 0.018 | 1 | 0 | Rcl1 | 0.004 | Old |
| 0.513318 | 0.490029 | 0.01 | 0.007 | 1 | 0 | Chaf1a | 0.003 | Old |
| 0.513463 | 0.006917 | 0.576 | 0.539 | 1 | 0 | Tm6sf1 | 0.037 | Old |
| 0.513469 | 0.179881 | 0.019 | 0.016 | 1 | 0 | Klhl12 | 0.003 | Old |
| 0.513518 | 0.140636 | 0.01 | 0.007 | 1 | 0 | Dock5 | 0.003 | Old |
| 0.513703 | 0.139327 | 0.021 | 0.017 | 1 | 0 | Zbtb8a | 0.004 | Old |
| 0.513745 | 0.04164 | 0.019 | 0.016 | 1 | 0 | BC024978 | 0.003 | Old |
| 0.514926 | 0.148182 | 0.023 | 0.019 | 1 | 0 | Itpr1 | 0.004 | Old |
| 0.51522 | 0.211742 | 0.077 | 0.07 | 1 | 0 | Cmc2 | 0.007 | Old |
| 0.515324 | 0.454274 | 0.01 | 0.007 | 1 | 0 | A630072M | 0.003 | Old |
| 0.515328 | 0.048895 | 0.398 | 0.377 | 1 | 0 | Ifrd1 | 0.021 | Old |
| 0.515779 | 0.259964 | 0.042 | 0.037 | 1 | 0 | Rad51d | 0.005 | Old |
| 0.515839 | 0.039347 | 0.082 | 0.075 | 1 | 0 | Pex2 | 0.007 | Old |
| 0.515927 | 0.173146 | 0.01 | 0.007 | 1 | 0 | Adgra3 | 0.003 | Old |
| 0.516035 | 0.364572 | 0.031 | 0.027 | 1 | 0 | Slc2a9 | 0.004 | Old |
| 0.516351 | 0.1889 | 0.034 | 0.029 | 1 | 0 | Gnl2 | 0.005 | Old |
| 0.516405 | 0.073218 | 0.142 | 0.133 | 1 | 0 | Mnt | 0.009 | Old |
| 0.51653 | 0.244734 | 0.01 | 0.007 | 1 | 0 | AC129328. | 0.003 | Old |
| 0.516562 | 0.159349 | 0.074 | 0.067 | 1 | 0 | Fam208b | 0.007 | Old |
| 0.516712 | 0.015434 | 0.192 | 0.178 | 1 | 0 | Ssrp1 | 0.014 | Old |
| 0.51678 | 0.140306 | 0.036 | 0.031 | 1 | 0 | Ddx19b | 0.005 | Old |
| 0.516828 | 0.172586 | 0.029 | 0.024 | 1 | 0 | Bclaf3 | 0.005 | Old |
| 0.517017 | 0.01213 | 0.114 | 0.104 | 1 | 0 | Chmp2b | 0.01 | Old |
| 0.517288 | 0.141467 | 0.03 | 0.026 | 1 | 0 | Kctd11 | 0.004 | Old |
| 0.517335 | 0.094916 | 0.01 | 0.007 | 1 | 0 | 9530082P2 | 0.003 | Old |
| 0.51809 | 0.026203 | 0.116 | 0.107 | 1 | 0 | Rab3gap2 | 0.009 | Old |
| 0.518514 | 0.114303 | 0.025 | 0.021 | 1 | 0 | Bloc1s1 | 0.004 | Old |
| 0.518783 | 0.143787 | 0.098 | 0.09 | 1 | 0 | Map3k5 | 0.008 | Old |
| 0.518984 | 0.182831 | 0.027 | 0.023 | 1 | 0 | Nat10 | 0.004 | Old |
| 0.519425 | 0.187084 | 0.023 | 0.019 | 1 | 0 | Phka1 | 0.004 | Old |
| 0.51987 | 0.001894 | 0.086 | 0.078 | 1 | 0 | Zeb2os | 0.008 | Old |
| 0.519884 | 0.341668 | 0.034 | 0.029 | 1 | 0 | Alkbh6 | 0.005 | Old |
| 0.52007 | 0.070135 | 0.436 | 0.412 | 1 | 0 | Dock2 | 0.024 | Old |
| 0.521529 | 0.002995 | 0.16 | 0.148 | 1 | 0 | Arhgap12 | 0.012 | Old |
| 0.521636 | 0.199991 | 0.111 | 0.103 | 1 | 0 | Taok2 | 0.008 | Old |
| 0.521707 | 0.017091 | 0.111 | 0.101 | 1 | 0 | Cdk17 | 0.01 | Old |

| | | | | | | | | |
|---|---|---|---|---|---|---|---|---|
| 0.522315 | 0.106002 | 0.071 | 0.064 | 1 | 0 | Nom1 | 0.007 | Old |
| 0.522446 | 0.049791 | 0.042 | 0.037 | 1 | 0 | Il18 | 0.005 | Old |
| 0.523131 | 0.151401 | 0.027 | 0.023 | 1 | 0 | Kctd13 | 0.004 | Old |
| 0.523269 | 0.019769 | 0.008 | 0.011 | 1 | 0 | Gm28557 | -0.003 | Old |
| 0.523567 | 0.633312 | 0.011 | 0.008 | 1 | 0 | Dzip1 | 0.003 | Old |
| 0.523791 | 0.072554 | 0.145 | 0.134 | 1 | 0 | Vbp1 | 0.011 | Old |
| 0.524478 | 0.118052 | 0.036 | 0.031 | 1 | 0 | Ap5s1 | 0.005 | Old |
| 0.524506 | 0.453556 | 0.011 | 0.008 | 1 | 0 | 4933406I1 | 0.003 | Old |
| 0.524634 | 0.061626 | 0.063 | 0.056 | 1 | 0 | Med7 | 0.007 | Old |
| 0.524913 | 0.000547 | 0.074 | 0.067 | 1 | 0 | Itgav | 0.007 | Old |
| 0.52498 | 0.144423 | 0.031 | 0.027 | 1 | 0 | Haus7 | 0.004 | Old |
| 0.525069 | 0.658573 | 0.011 | 0.008 | 1 | 0 | Map3k6 | 0.003 | Old |
| 0.525087 | 0.101212 | 0.031 | 0.027 | 1 | 0 | Vps37c | 0.004 | Old |
| 0.525481 | 0.022245 | 0.821 | 0.797 | 1 | 0 | Npc2 | 0.024 | Old |
| 0.52596 | 0.01101 | 0.023 | 0.019 | 1 | 0 | Zfp995 | 0.004 | Old |
| 0.526342 | 0.045764 | 0.03 | 0.026 | 1 | 0 | Zcchc4 | 0.004 | Old |
| 0.526497 | 0.182946 | 0.138 | 0.128 | 1 | 0 | Acaa2 | 0.01 | Old |
| 0.526573 | 0.450793 | 0.011 | 0.008 | 1 | 0 | Slc7a4 | 0.003 | Old |
| 0.52674 | 0.018354 | 0.817 | 0.811 | 1 | 0 | Calm1 | 0.006 | Old |
| 0.526997 | 0.084097 | 0.231 | 0.221 | 1 | 0 | Pan3 | 0.01 | Old |
| 0.527064 | 0.057059 | 0.027 | 0.023 | 1 | 0 | Lig3 | 0.004 | Old |
| 0.527586 | 0.148067 | 0.026 | 0.022 | 1 | 0 | Mgmt | 0.004 | Old |
| 0.527752 | 0.012371 | 0.038 | 0.033 | 1 | 0 | G6pc3 | 0.005 | Old |
| 0.527909 | 0.093429 | 0.116 | 0.108 | 1 | 0 | Dpf2 | 0.008 | Old |
| 0.528101 | 0.178699 | 0.086 | 0.079 | 1 | 0 | Maff | 0.007 | Old |
| 0.528786 | 0.030781 | 0.105 | 0.097 | 1 | 0 | Tnpo2 | 0.008 | Old |
| 0.528884 | 0.130638 | 0.181 | 0.171 | 1 | 0 | F630028O1 | 0.01 | Old |
| 0.529069 | 0.060746 | 0.082 | 0.075 | 1 | 0 | Creld2 | 0.007 | Old |
| 0.529211 | 0.389539 | 0.011 | 0.008 | 1 | 0 | Tnfrsf23 | 0.003 | Old |
| 0.529211 | 0.260903 | 0.011 | 0.008 | 1 | 0 | B3galt4 | 0.003 | Old |
| 0.529234 | 0.05859 | 0.07 | 0.063 | 1 | 0 | Iqsec1 | 0.007 | Old |
| 0.529291 | 0.021805 | 0.182 | 0.171 | 1 | 0 | Ctdsp1 | 0.011 | Old |
| 0.529377 | 0.185216 | 0.092 | 0.084 | 1 | 0 | Adh5 | 0.008 | Old |
| 0.529388 | 0.140173 | 0.064 | 0.058 | 1 | 0 | Pdlim2 | 0.006 | Old |
| 0.529468 | 0.33012 | 0.042 | 0.037 | 1 | 0 | Hsf2 | 0.005 | Old |
| 0.529563 | 0.057917 | 0.031 | 0.027 | 1 | 0 | Kbtbd3 | 0.004 | Old |
| 0.529777 | 0.263673 | 0.011 | 0.008 | 1 | 0 | Snx32 | 0.003 | Old |
| 0.529836 | 0.056376 | 0.094 | 0.086 | 1 | 0 | Flot1 | 0.008 | Old |
| 0.53059 | 0.00114 | 0.031 | 0.027 | 1 | 0 | B230354K1 | 0.004 | Old |
| 0.530896 | 0.124453 | 0.093 | 0.086 | 1 | 0 | Senp7 | 0.007 | Old |
| 0.530946 | 0.111061 | 0.1 | 0.092 | 1 | 0 | Retreg1 | 0.008 | Old |
| 0.531393 | 0.02306 | 0.316 | 0.301 | 1 | 0 | Vps35 | 0.015 | Old |
| 0.532579 | 0.009424 | 0.093 | 0.085 | 1 | 0 | Eaf1 | 0.008 | Old |
| 0.533079 | 0.068973 | 0.228 | 0.215 | 1 | 0 | Wdr45b | 0.013 | Old |
| 0.533638 | 0.013652 | 0.074 | 0.067 | 1 | 0 | Manea | 0.007 | Old |

| | | | | | | | | |
|---|---|---|---|---|---|---|---|---|
| 0.53376 | 0.686193 | 0.029 | 0.025 | 1 | 0 | Zfp846 | 0.004 | Old |
| 0.534185 | 0.027884 | 0.083 | 0.076 | 1 | 0 | Fibp | 0.007 | Old |
| 0.535153 | 0.093703 | 0.071 | 0.064 | 1 | 0 | Tmem11 | 0.007 | Old |
| 0.535651 | 0.020233 | 0.049 | 0.055 | 1 | 0 | Sfi1 | -0.006 | Old |
| 0.536183 | 0.09185 | 0.015 | 0.019 | 1 | 0 | A230050P2 | -0.004 | Old |
| 0.536877 | 0.187817 | 0.049 | 0.044 | 1 | 0 | Tex10 | 0.005 | Old |
| 0.537053 | 0.565597 | 0.012 | 0.01 | 1 | 0 | Gtf2h3 | 0.002 | Old |
| 0.537115 | 0.005623 | 0.271 | 0.255 | 1 | 0 | Ppig | 0.016 | Old |
| 0.537572 | 0.062634 | 0.052 | 0.046 | 1 | 0 | Gadd45gip | 0.006 | Old |
| 0.537585 | 0.343396 | 0.048 | 0.043 | 1 | 0 | Nsun6 | 0.005 | Old |
| 0.537862 | 0.133205 | 0.059 | 0.053 | 1 | 0 | Yif1b | 0.006 | Old |
| 0.538943 | 0.000399 | 0.365 | 0.346 | 1 | 0 | Dnajb6 | 0.019 | Old |
| 0.53933 | 0.009395 | 0.105 | 0.097 | 1 | 0 | Rnf220 | 0.008 | Old |
| 0.539515 | 0.06954 | 0.089 | 0.082 | 1 | 0 | Cabin1 | 0.007 | Old |
| 0.540515 | 0.060915 | 0.059 | 0.066 | 1 | 0 | Sik3 | -0.007 | Old |
| 0.540721 | 0.087136 | 0.022 | 0.026 | 1 | 0 | Mamld1 | -0.004 | Old |
| 0.540875 | 0.014423 | 0.471 | 0.436 | 1 | 0 | Selenok | 0.035 | Old |
| 0.542986 | 0.463068 | 0.014 | 0.011 | 1 | 0 | Dhfr | 0.003 | Old |
| 0.543415 | 0.575514 | 0.015 | 0.012 | 1 | 0 | Ch25h | 0.003 | Old |
| 0.543438 | 0.111528 | 0.112 | 0.104 | 1 | 0 | Ylpm1 | 0.008 | Old |
| 0.543955 | 0.019297 | 0.118 | 0.109 | 1 | 0 | Lrrk1 | 0.009 | Old |
| 0.544592 | 0.173554 | 0.037 | 0.032 | 1 | 0 | Zfp160 | 0.005 | Old |
| 0.544821 | 0.004596 | 0.127 | 0.118 | 1 | 0 | Ganab | 0.009 | Old |
| 0.545173 | 0.024816 | 0.049 | 0.044 | 1 | 0 | Ppm1m | 0.005 | Old |
| 0.545719 | 0.147713 | 0.057 | 0.052 | 1 | 0 | Gps2 | 0.005 | Old |
| 0.546262 | 0.033299 | 0.2 | 0.188 | 1 | 0 | Bod1l | 0.012 | Old |
| 0.546366 | 0.187064 | 0.014 | 0.011 | 1 | 0 | Usp27x | 0.003 | Old |
| 0.547063 | 0.097268 | 0.152 | 0.144 | 1 | 0 | Aida | 0.008 | Old |
| 0.547607 | 0.095883 | 0.119 | 0.11 | 1 | 0 | Atf1 | 0.009 | Old |
| 0.548355 | 0.307552 | 0.071 | 0.065 | 1 | 0 | Il1rn | 0.006 | Old |
| 0.548716 | 0.011448 | 0.085 | 0.078 | 1 | 0 | Dcun1d1 | 0.007 | Old |
| 0.548738 | 0.04738 | 0.014 | 0.011 | 1 | 0 | Focad | 0.003 | Old |
| 0.549234 | 0.015584 | 0.047 | 0.041 | 1 | 0 | Nup205 | 0.006 | Old |
| 0.549247 | 0.092238 | 0.014 | 0.011 | 1 | 0 | Map2k6 | 0.003 | Old |
| 0.54936 | 0.546788 | 0.015 | 0.012 | 1 | 0 | Lrrc45 | 0.003 | Old |
| 0.549725 | 0.182887 | 0.078 | 0.072 | 1 | 0 | Qrich1 | 0.006 | Old |
| 0.549926 | 0.090529 | 0.014 | 0.011 | 1 | 0 | D830025C( | 0.003 | Old |
| 0.550569 | 0.324222 | 0.015 | 0.012 | 1 | 0 | Cobll1 | 0.003 | Old |
| 0.551429 | 0.029775 | 0.43 | 0.406 | 1 | 0 | Fcho2 | 0.024 | Old |
| 0.551559 | 0.555824 | 0.016 | 0.013 | 1 | 0 | Rsad1 | 0.003 | Old |
| 0.551657 | 0.158197 | 0.097 | 0.091 | 1 | 0 | Sh3pxd2a | 0.006 | Old |
| 0.551773 | 0.426332 | 0.027 | 0.024 | 1 | 0 | Entpd6 | 0.003 | Old |
| 0.551775 | 0.078089 | 0.166 | 0.156 | 1 | 0 | Gorasp2 | 0.01 | Old |
| 0.551981 | 0.113099 | 0.109 | 0.102 | 1 | 0 | Mtfr1l | 0.007 | Old |
| 0.552175 | 0.437525 | 0.016 | 0.013 | 1 | 0 | Arhgef18 | 0.003 | Old |

| | | | | | | | | |
|---|---|---|---|---|---|---|---|---|
| 0.55223 | 0.337254 | 0.067 | 0.061 | 1 | 0 | C6 | 0.006 | Old |
| 0.55304 | 0.060615 | 0.068 | 0.062 | 1 | 0 | Actr1b | 0.006 | Old |
| 0.553661 | 0.3474 | 0.031 | 0.027 | 1 | 0 | Wdr46 | 0.004 | Old |
| 0.553714 | 0.017126 | 0.253 | 0.237 | 1 | 0 | Uba1 | 0.016 | Old |
| 0.553769 | 0.42193 | 0.025 | 0.021 | 1 | 0 | Gde1 | 0.004 | Old |
| 0.553772 | 0.219156 | 0.034 | 0.03 | 1 | 0 | 1700021F0 | 0.004 | Old |
| 0.553844 | 0.075522 | 0.135 | 0.126 | 1 | 0 | Ndufs3 | 0.009 | Old |
| 0.553872 | 0.521797 | 0.016 | 0.013 | 1 | 0 | Rgcc | 0.003 | Old |
| 0.553949 | 0.20534 | 0.096 | 0.089 | 1 | 0 | Lrrc47 | 0.007 | Old |
| 0.554238 | 0.431876 | 0.018 | 0.015 | 1 | 0 | Herc3 | 0.003 | Old |
| 0.55466 | 0.140516 | 0.036 | 0.031 | 1 | 0 | Slc16a3 | 0.005 | Old |
| 0.554689 | 0.000167 | 0.015 | 0.012 | 1 | 0 | Dmwd | 0.003 | Old |
| 0.55476 | 0.251827 | 0.031 | 0.027 | 1 | 0 | Atp8b2 | 0.004 | Old |
| 0.555219 | 0.076948 | 0.118 | 0.11 | 1 | 0 | Brd8 | 0.008 | Old |
| 0.555363 | 0.073751 | 0.038 | 0.034 | 1 | 0 | Recql | 0.004 | Old |
| 0.555715 | 0.033961 | 0.239 | 0.25 | 1 | 0 | Rgs2 | -0.011 | Old |
| 0.55614 | 0.102171 | 0.051 | 0.045 | 1 | 0 | Pcnx | 0.006 | Old |
| 0.556484 | 0.092285 | 0.417 | 0.405 | 1 | 0 | Etv3 | 0.012 | Old |
| 0.556634 | 0.060352 | 0.015 | 0.012 | 1 | 0 | Fntb | 0.003 | Old |
| 0.556682 | 0.037864 | 0.15 | 0.141 | 1 | 0 | Cript | 0.009 | Old |
| 0.556738 | 0.464185 | 0.019 | 0.016 | 1 | 0 | Zmynd15 | 0.003 | Old |
| 0.556859 | 0.030121 | 0.399 | 0.378 | 1 | 0 | Inpp5d | 0.021 | Old |
| 0.556972 | 0.034178 | 0.085 | 0.078 | 1 | 0 | Pes1 | 0.007 | Old |
| 0.557036 | 0.008297 | 0.923 | 0.92 | 1 | 0 | Zfp36l1 | 0.003 | Old |
| 0.557266 | 0.371094 | 0.027 | 0.024 | 1 | 0 | Cdk20 | 0.003 | Old |
| 0.557282 | 0.148897 | 0.342 | 0.339 | 1 | 0 | Cd38 | 0.003 | Old |
| 0.557492 | 0.031841 | 0.501 | 0.485 | 1 | 0 | Tnfaip3 | 0.016 | Old |
| 0.557499 | 0.386432 | 0.023 | 0.02 | 1 | 0 | Gm29340 | 0.003 | Old |
| 0.557622 | 0.017181 | 0.56 | 0.533 | 1 | 0 | Eif3a | 0.027 | Old |
| 0.558389 | 0.385871 | 0.025 | 0.021 | 1 | 0 | Cutc | 0.004 | Old |
| 0.558427 | 0.029281 | 0.014 | 0.017 | 1 | 0 | Zkscan6 | -0.003 | Old |
| 0.558648 | 0.057059 | 0.157 | 0.148 | 1 | 0 | Guk1 | 0.009 | Old |
| 0.559054 | 0.060674 | 0.104 | 0.096 | 1 | 0 | Acox1 | 0.008 | Old |
| 0.559779 | 0.138406 | 0.078 | 0.072 | 1 | 0 | Scyl2 | 0.006 | Old |
| 0.560167 | 0.042539 | 0.137 | 0.128 | 1 | 0 | Atp11c | 0.009 | Old |
| 0.560695 | 0.080825 | 0.033 | 0.029 | 1 | 0 | 1600010M | 0.004 | Old |
| 0.561062 | 0.195965 | 0.018 | 0.015 | 1 | 0 | Xndc1 | 0.003 | Old |
| 0.561787 | 0.092439 | 0.202 | 0.194 | 1 | 0 | Pde4b | 0.008 | Old |
| 0.562091 | 0.03015 | 0.446 | 0.425 | 1 | 0 | Dusp3 | 0.021 | Old |
| 0.562356 | 0.078351 | 0.122 | 0.114 | 1 | 0 | Pih1d1 | 0.008 | Old |
| 0.562586 | 0.135249 | 0.129 | 0.121 | 1 | 0 | Gnl3 | 0.008 | Old |
| 0.562723 | 0.068727 | 0.081 | 0.074 | 1 | 0 | Eif2ak1 | 0.007 | Old |
| 0.562751 | 5.69E-05 | 0.289 | 0.27 | 1 | 0 | Kras | 0.019 | Old |
| 0.563592 | 0.005963 | 0.01 | 0.012 | 1 | 0 | Tsen54 | -0.002 | Old |
| 0.564536 | 0.228056 | 0.025 | 0.021 | 1 | 0 | Rasgrf2 | 0.004 | Old |

| | | | | | | | | |
|---|---|---|---|---|---|---|---|---|
| 0.565361 | 0.22509 | 0.045 | 0.04 | 1 | 0 | Fdx1 | 0.005 | Old |
| 0.565606 | 0.103474 | 0.022 | 0.019 | 1 | 0 | Spout1 | 0.003 | Old |
| 0.565739 | 0.202368 | 0.022 | 0.019 | 1 | 0 | Mtmr11 | 0.003 | Old |
| 0.566302 | 0.149459 | 0.048 | 0.043 | 1 | 0 | Gtf3c4 | 0.005 | Old |
| 0.566552 | 0.028331 | 0.726 | 0.691 | 1 | 0 | Tmbim6 | 0.035 | Old |
| 0.566805 | 0.117949 | 0.029 | 0.025 | 1 | 0 | Lmbr1l | 0.004 | Old |
| 0.56748 | 0.077029 | 0.031 | 0.027 | 1 | 0 | Tmem94 | 0.004 | Old |
| 0.567562 | 0.232918 | 0.062 | 0.056 | 1 | 0 | 2610002M | 0.006 | Old |
| 0.567711 | 0.007945 | 0.034 | 0.03 | 1 | 0 | Med19 | 0.004 | Old |
| 0.567808 | 0.023545 | 0.059 | 0.053 | 1 | 0 | Ankrd50 | 0.006 | Old |
| 0.568013 | 0.044141 | 0.021 | 0.017 | 1 | 0 | Zfp472 | 0.004 | Old |
| 0.568085 | 0.188423 | 0.022 | 0.019 | 1 | 0 | Maged2 | 0.003 | Old |
| 0.568156 | 0.212375 | 0.078 | 0.072 | 1 | 0 | Fbxl14 | 0.006 | Old |
| 0.568927 | 0.194546 | 0.031 | 0.027 | 1 | 0 | Mrgbp | 0.004 | Old |
| 0.569326 | 0.021307 | 0.025 | 0.021 | 1 | 0 | Hibch | 0.004 | Old |
| 0.569378 | 0.013236 | 0.207 | 0.196 | 1 | 0 | Zc3hav1 | 0.011 | Old |
| 0.569718 | 0.107043 | 0.048 | 0.043 | 1 | 0 | Zfp369 | 0.005 | Old |
| 0.570084 | 0.048969 | 0.025 | 0.021 | 1 | 0 | Bbs7 | 0.004 | Old |
| 0.571376 | 0.062001 | 0.022 | 0.019 | 1 | 0 | Dffa | 0.003 | Old |
| 0.571393 | 0.199417 | 0.092 | 0.086 | 1 | 0 | Plk3 | 0.006 | Old |
| 0.57182 | 0.075461 | 0.131 | 0.123 | 1 | 0 | Smim27 | 0.008 | Old |
| 0.572465 | 0.088469 | 0.063 | 0.057 | 1 | 0 | Vamp5 | 0.006 | Old |
| 0.574429 | 0.025604 | 0.037 | 0.042 | 1 | 0 | Ctdp1 | -0.005 | Old |
| 0.575137 | 0.241632 | 0.041 | 0.037 | 1 | 0 | Urgcp | 0.004 | Old |
| 0.575315 | 0.139085 | 0.085 | 0.079 | 1 | 0 | Ear2 | 0.006 | Old |
| 0.575745 | 0.038092 | 0.051 | 0.045 | 1 | 0 | Cdc27 | 0.006 | Old |
| 0.576113 | 0.021606 | 0.242 | 0.23 | 1 | 0 | Ubxn1 | 0.012 | Old |
| 0.576779 | 0.074802 | 0.254 | 0.243 | 1 | 0 | Ids | 0.011 | Old |
| 0.576916 | 0.109948 | 0.077 | 0.071 | 1 | 0 | Prkra | 0.006 | Old |
| 0.577751 | 0.028825 | 0.311 | 0.293 | 1 | 0 | Sypl | 0.018 | Old |
| 0.579025 | 0.128771 | 0.048 | 0.043 | 1 | 0 | Tirap | 0.005 | Old |
| 0.57933 | 0.09965 | 0.166 | 0.156 | 1 | 0 | Tecr | 0.01 | Old |
| 0.579371 | 0.184102 | 0.063 | 0.058 | 1 | 0 | Dtymk | 0.005 | Old |
| 0.579489 | 0.127145 | 0.107 | 0.1 | 1 | 0 | St3gal5 | 0.007 | Old |
| 0.579584 | 0.153893 | 0.044 | 0.039 | 1 | 0 | Tmem39a | 0.005 | Old |
| 0.580233 | 0.044849 | 0.223 | 0.211 | 1 | 0 | Eif3j1 | 0.012 | Old |
| 0.580675 | 0.147714 | 0.067 | 0.061 | 1 | 0 | 0610037L1 | 0.006 | Old |
| 0.58092 | 0.083769 | 0.613 | 0.591 | 1 | 0 | Cd14 | 0.022 | Old |
| 0.582678 | 0.142203 | 0.073 | 0.067 | 1 | 0 | Gcc1 | 0.006 | Old |
| 0.583025 | 0.039329 | 0.015 | 0.018 | 1 | 0 | Rad9a | -0.003 | Old |
| 0.583041 | 0.25677 | 0.04 | 0.035 | 1 | 0 | Zc3h8 | 0.005 | Old |
| 0.583154 | 0.03464 | 0.587 | 0.562 | 1 | 0 | Clk1 | 0.025 | Old |
| 0.58406 | 0.086832 | 0.089 | 0.083 | 1 | 0 | Eed | 0.006 | Old |
| 0.585301 | 0.125039 | 0.109 | 0.102 | 1 | 0 | Ptbp2 | 0.007 | Old |
| 0.585574 | 0.051959 | 0.008 | 0.011 | 1 | 0 | Tasp1 | -0.003 | Old |

| | | | | | | | | |
|---|---|---|---|---|---|---|---|---|
| 0.586749 | 0.108042 | 0.041 | 0.037 | 1 | 0 | Mib2 | 0.004 | Old |
| 0.587417 | 0.002942 | 0.149 | 0.14 | 1 | 0 | Rabl6 | 0.009 | Old |
| 0.587574 | 0.165738 | 0.036 | 0.032 | 1 | 0 | Nubp1 | 0.004 | Old |
| 0.589246 | 0.019149 | 0.044 | 0.039 | 1 | 0 | Extl2 | 0.005 | Old |
| 0.589619 | 0.002791 | 0.047 | 0.042 | 1 | 0 | Sec22a | 0.005 | Old |
| 0.589639 | 0.157361 | 0.053 | 0.048 | 1 | 0 | Mak16 | 0.005 | Old |
| 0.590543 | 0.003465 | 0.301 | 0.286 | 1 | 0 | Ube2l3 | 0.015 | Old |
| 0.59157 | 0.024572 | 0.083 | 0.077 | 1 | 0 | Mdp1 | 0.006 | Old |
| 0.592056 | 0.123279 | 0.04 | 0.035 | 1 | 0 | Naa25 | 0.005 | Old |
| 0.592765 | 0.132507 | 0.037 | 0.033 | 1 | 0 | Bysl | 0.004 | Old |
| 0.592938 | 0.246328 | 0.056 | 0.051 | 1 | 0 | Utp14b | 0.005 | Old |
| 0.593267 | 0.039837 | 0.447 | 0.427 | 1 | 0 | Scp2 | 0.02 | Old |
| 0.593278 | 0.133254 | 0.085 | 0.079 | 1 | 0 | Ipo5 | 0.006 | Old |
| 0.593987 | 0.118836 | 0.036 | 0.032 | 1 | 0 | Tnfrsf12a | 0.004 | Old |
| 0.594714 | 0.051891 | 0.2 | 0.19 | 1 | 0 | Rnf4 | 0.01 | Old |
| 0.594779 | 0.228246 | 0.07 | 0.064 | 1 | 0 | Taf3 | 0.006 | Old |
| 0.594895 | 0.424498 | 0.01 | 0.008 | 1 | 0 | H2-Q10 | 0.002 | Old |
| 0.595104 | 0.041787 | 0.14 | 0.131 | 1 | 0 | Sptlc1 | 0.009 | Old |
| 0.595214 | 0.009749 | 0.462 | 0.433 | 1 | 0 | Gnaq | 0.029 | Old |
| 0.595317 | 0.483937 | 0.01 | 0.008 | 1 | 0 | Tnfrsf22 | 0.002 | Old |
| 0.595622 | 0.092798 | 0.108 | 0.101 | 1 | 0 | Nagk | 0.007 | Old |
| 0.595951 | 0.357991 | 0.01 | 0.008 | 1 | 0 | Cenpq | 0.002 | Old |
| 0.596091 | 0.05093 | 0.265 | 0.254 | 1 | 0 | Gnai3 | 0.011 | Old |
| 0.596349 | 0.032975 | 0.108 | 0.101 | 1 | 0 | Cdip1 | 0.007 | Old |
| 0.596585 | 0.254029 | 0.01 | 0.008 | 1 | 0 | Pfkfb2 | 0.002 | Old |
| 0.596602 | 0.059424 | 0.101 | 0.095 | 1 | 0 | Ube4b | 0.006 | Old |
| 0.596621 | 0.104762 | 0.038 | 0.034 | 1 | 0 | Myc | 0.004 | Old |
| 0.598148 | 0.070461 | 0.627 | 0.61 | 1 | 0 | Atf3 | 0.017 | Old |
| 0.598605 | 0.177383 | 0.096 | 0.09 | 1 | 0 | Clasp1 | 0.006 | Old |
| 0.599147 | 0.023227 | 0.131 | 0.123 | 1 | 0 | Mbtps1 | 0.008 | Old |
| 0.59923 | 0.050225 | 0.01 | 0.008 | 1 | 0 | Rwdd2b | 0.002 | Old |
| 0.599405 | 0.087578 | 0.156 | 0.148 | 1 | 0 | Txnrd1 | 0.008 | Old |
| 0.599593 | 0.123451 | 0.066 | 0.06 | 1 | 0 | Snx14 | 0.006 | Old |
| 0.599831 | 0.281688 | 0.062 | 0.056 | 1 | 0 | Nbdy | 0.006 | Old |
| 0.60082 | 0.052639 | 0.01 | 0.008 | 1 | 0 | Hist1h4h | 0.002 | Old |
| 0.600973 | 0.107017 | 0.031 | 0.036 | 1 | 0 | Zfp386 | -0.005 | Old |
| 0.601802 | 0.482784 | 0.023 | 0.02 | 1 | 0 | Abcb10 | 0.003 | Old |
| 0.602567 | 0.24944 | 0.026 | 0.023 | 1 | 0 | Clec4d | 0.003 | Old |
| 0.602574 | 0.009588 | 0.97 | 0.968 | 1 | 0 | Ctsc | 0.002 | Old |
| 0.602715 | 0.063667 | 0.034 | 0.03 | 1 | 0 | Kctd18 | 0.004 | Old |
| 0.603687 | 0.5542 | 0.011 | 0.009 | 1 | 0 | Gm15446 | 0.002 | Old |
| 0.604229 | 0.001341 | 0.294 | 0.275 | 1 | 0 | Rpl27 | 0.019 | Old |
| 0.604633 | 0.150929 | 0.064 | 0.059 | 1 | 0 | Gdap2 | 0.005 | Old |
| 0.604675 | 0.182163 | 0.011 | 0.009 | 1 | 0 | Thop1 | 0.002 | Old |
| 0.604713 | 0.072516 | 0.064 | 0.059 | 1 | 0 | Mea1 | 0.005 | Old |

| | | | | | | | | |
|---|---|---|---|---|---|---|---|---|
| 0.604993 | 0.37592 | 0.027 | 0.024 | 1 | 0 | Amd2 | 0.003 | Old |
| 0.605173 | 0.05409 | 0.187 | 0.177 | 1 | 0 | Stx12 | 0.01 | Old |
| 0.605226 | 0.112129 | 0.089 | 0.083 | 1 | 0 | Lrif1 | 0.006 | Old |
| 0.605641 | 0.29628 | 0.025 | 0.021 | 1 | 0 | Armt1 | 0.004 | Old |
| 0.605774 | 0.071207 | 0.264 | 0.254 | 1 | 0 | Tceal9 | 0.01 | Old |
| 0.605949 | 0.084266 | 0.138 | 0.13 | 1 | 0 | Taf10 | 0.008 | Old |
| 0.60611 | 0.505659 | 0.012 | 0.01 | 1 | 0 | Il15ra | 0.002 | Old |
| 0.60611 | 0.102012 | 0.03 | 0.027 | 1 | 0 | Ggta1 | 0.003 | Old |
| 0.606245 | 0.124722 | 0.081 | 0.075 | 1 | 0 | Akap7 | 0.006 | Old |
| 0.606266 | 0.095817 | 0.034 | 0.03 | 1 | 0 | Habp4 | 0.004 | Old |
| 0.606344 | 0.140257 | 0.03 | 0.027 | 1 | 0 | Sppl2b | 0.003 | Old |
| 0.606803 | 0.263402 | 0.025 | 0.021 | 1 | 0 | Pus7 | 0.004 | Old |
| 0.607018 | 0.041691 | 0.04 | 0.035 | 1 | 0 | 2310061I0[illegible] | 0.005 | Old |
| 0.607077 | 0.02526 | 0.731 | 0.707 | 1 | 0 | Pid1 | 0.024 | Old |
| 0.607314 | 0.52028 | 0.015 | 0.013 | 1 | 0 | Lig4 | 0.002 | Old |
| 0.607432 | 0.062776 | 0.242 | 0.23 | 1 | 0 | Cd37 | 0.012 | Old |
| 0.607851 | 0.092875 | 0.144 | 0.136 | 1 | 0 | Stk3 | 0.008 | Old |
| 0.608674 | 0.017007 | 0.088 | 0.081 | 1 | 0 | Seh1l | 0.007 | Old |
| 0.609149 | 0.004295 | 0.115 | 0.108 | 1 | 0 | Samsn1 | 0.007 | Old |
| 0.611099 | 0.048782 | 0.111 | 0.104 | 1 | 0 | Gadd45g | 0.007 | Old |
| 0.611202 | 0.15211 | 0.025 | 0.021 | 1 | 0 | Mcm7 | 0.004 | Old |
| 0.611248 | 0.51888 | 0.016 | 0.014 | 1 | 0 | Knstrn | 0.002 | Old |
| 0.611411 | 0.132635 | 0.011 | 0.009 | 1 | 0 | Nudt15 | 0.002 | Old |
| 0.611968 | 0.245853 | 0.012 | 0.01 | 1 | 0 | Klhdc1 | 0.002 | Old |
| 0.61284 | 0.200148 | 0.027 | 0.024 | 1 | 0 | Mrpl44 | 0.003 | Old |
| 0.613148 | 0.157258 | 0.027 | 0.024 | 1 | 0 | Zbtb24 | 0.003 | Old |
| 0.613394 | 0.310251 | 0.018 | 0.015 | 1 | 0 | Plscr1 | 0.003 | Old |
| 0.61349 | 0.142871 | 0.016 | 0.019 | 1 | 0 | Tmem238 | -0.003 | Old |
| 0.613556 | 0.050148 | 0.945 | 0.932 | 1 | 0 | Crip1 | 0.013 | Old |
| 0.613806 | 0.016274 | 0.337 | 0.316 | 1 | 0 | Itsn2 | 0.021 | Old |
| 0.61444 | 0.192741 | 0.027 | 0.024 | 1 | 0 | Micall1 | 0.003 | Old |
| 0.614504 | 0.121032 | 0.022 | 0.019 | 1 | 0 | Heatr5b | 0.003 | Old |
| 0.614611 | 0.142557 | 0.044 | 0.04 | 1 | 0 | Farsa | 0.004 | Old |
| 0.614641 | 0.490149 | 0.019 | 0.016 | 1 | 0 | Ubiad1 | 0.003 | Old |
| 0.614721 | 0.346593 | 0.014 | 0.011 | 1 | 0 | Nicn1 | 0.003 | Old |
| 0.614834 | 0.11505 | 0.025 | 0.021 | 1 | 0 | Ogfod3 | 0.004 | Old |
| 0.614897 | 0.183476 | 0.014 | 0.011 | 1 | 0 | Nup43 | 0.003 | Old |
| 0.615347 | 0.393091 | 0.015 | 0.013 | 1 | 0 | Fam102a | 0.002 | Old |
| 0.615972 | 0.293354 | 0.019 | 0.016 | 1 | 0 | Ipo4 | 0.003 | Old |
| 0.616186 | 0.237673 | 0.015 | 0.013 | 1 | 0 | Pard6a | 0.002 | Old |
| 0.616368 | 0.200261 | 0.022 | 0.019 | 1 | 0 | Hus1 | 0.003 | Old |
| 0.616468 | 0.279946 | 0.018 | 0.015 | 1 | 0 | Shq1 | 0.003 | Old |
| 0.616515 | 0.06186 | 0.126 | 0.118 | 1 | 0 | Polr2g | 0.008 | Old |
| 0.616905 | 0.187593 | 0.052 | 0.048 | 1 | 0 | Fgd3 | 0.004 | Old |
| 0.61722 | 0.020827 | 0.335 | 0.321 | 1 | 0 | Git2 | 0.014 | Old |

| | | | | | | | | |
|---|---|---|---|---|---|---|---|---|
| 0.61737 | 0.326494 | 0.014 | 0.011 | 1 | 0 | Armc6 | 0.003 | Old |
| 0.617438 | 0.204751 | 0.021 | 0.018 | 1 | 0 | Kctd17 | 0.003 | Old |
| 0.617581 | 0.144742 | 0.021 | 0.018 | 1 | 0 | Gcfc2 | 0.003 | Old |
| 0.617588 | 0.075988 | 0.056 | 0.051 | 1 | 0 | Arhgef7 | 0.005 | Old |
| 0.617693 | 0.024865 | 0.022 | 0.025 | 1 | 0 | Dnajc4 | -0.003 | Old |
| 0.617735 | 0.209136 | 0.168 | 0.16 | 1 | 0 | Id3 | 0.008 | Old |
| 0.618539 | 0.186485 | 0.015 | 0.013 | 1 | 0 | Cep152 | 0.002 | Old |
| 0.618607 | 0.175196 | 0.025 | 0.021 | 1 | 0 | Gm5086 | 0.004 | Old |
| 0.619044 | 0.14644 | 0.015 | 0.013 | 1 | 0 | Slc26a6 | 0.002 | Old |
| 0.619626 | 0.032587 | 0.07 | 0.064 | 1 | 0 | Copg2 | 0.006 | Old |
| 0.619702 | 0.098795 | 0.018 | 0.015 | 1 | 0 | Tctn3 | 0.003 | Old |
| 0.620106 | 0.077605 | 0.17 | 0.161 | 1 | 0 | Elmsan1 | 0.009 | Old |
| 0.620791 | 0.020069 | 0.112 | 0.105 | 1 | 0 | Tm9sf4 | 0.007 | Old |
| 0.621233 | 0.000198 | 0.015 | 0.013 | 1 | 0 | Ogg1 | 0.002 | Old |
| 0.621514 | 0.059856 | 0.016 | 0.014 | 1 | 0 | Mtfmt | 0.002 | Old |
| 0.621907 | 0.014807 | 0.581 | 0.544 | 1 | 0 | Atp5a1 | 0.037 | Old |
| 0.622611 | 0.043841 | 0.042 | 0.038 | 1 | 0 | Nup155 | 0.004 | Old |
| 0.623132 | 0.116391 | 0.06 | 0.055 | 1 | 0 | Stam | 0.005 | Old |
| 0.623303 | 0.007275 | 0.352 | 0.338 | 1 | 0 | Rnf149 | 0.014 | Old |
| 0.623448 | 0.050483 | 0.016 | 0.014 | 1 | 0 | Ada | 0.002 | Old |
| 0.623945 | 0.092842 | 0.074 | 0.069 | 1 | 0 | Exosc3 | 0.005 | Old |
| 0.624042 | 0.028322 | 0.438 | 0.421 | 1 | 0 | Ap2m1 | 0.017 | Old |
| 0.624512 | 0.109439 | 0.116 | 0.11 | 1 | 0 | Aph1b | 0.006 | Old |
| 0.625126 | 0.011932 | 0.041 | 0.045 | 1 | 0 | Mrps34 | -0.004 | Old |
| 0.625289 | 0.014298 | 0.042 | 0.038 | 1 | 0 | Edc3 | 0.004 | Old |
| 0.625705 | 0.148008 | 0.116 | 0.11 | 1 | 0 | Snf8 | 0.006 | Old |
| 0.626015 | 0.006532 | 0.06 | 0.055 | 1 | 0 | Tomm40 | 0.005 | Old |
| 0.627328 | 0.202766 | 0.044 | 0.04 | 1 | 0 | Ece1 | 0.004 | Old |
| 0.6276 | 0.006314 | 0.055 | 0.05 | 1 | 0 | 2310033P0 | 0.005 | Old |
| 0.627982 | 0.045858 | 0.245 | 0.234 | 1 | 0 | Hpcal1 | 0.011 | Old |
| 0.628333 | 0.097268 | 0.041 | 0.037 | 1 | 0 | Zfp943 | 0.004 | Old |
| 0.628443 | 0.182057 | 0.133 | 0.126 | 1 | 0 | Brd3 | 0.007 | Old |
| 0.629762 | 0.083206 | 0.114 | 0.107 | 1 | 0 | Zfp280d | 0.007 | Old |
| 0.62998 | 0.173101 | 0.031 | 0.035 | 1 | 0 | Rrp8 | -0.004 | Old |
| 0.630077 | 0.01584 | 0.183 | 0.175 | 1 | 0 | Atp6v1h | 0.008 | Old |
| 0.630274 | 0.108389 | 0.012 | 0.015 | 1 | 0 | Rrp1b | -0.003 | Old |
| 0.630611 | 0.054127 | 0.111 | 0.104 | 1 | 0 | Cramp1l | 0.007 | Old |
| 0.632115 | 0.017406 | 0.105 | 0.099 | 1 | 0 | Prdm1 | 0.006 | Old |
| 0.63239 | 0.003587 | 0.204 | 0.193 | 1 | 0 | Retreg2 | 0.011 | Old |
| 0.632417 | 0.020146 | 0.015 | 0.018 | 1 | 0 | Zfp35 | -0.003 | Old |
| 0.633037 | 0.132988 | 0.074 | 0.069 | 1 | 0 | Relt | 0.005 | Old |
| 0.633349 | 0.015737 | 0.166 | 0.157 | 1 | 0 | Bfar | 0.009 | Old |
| 0.634326 | 0.009712 | 0.056 | 0.051 | 1 | 0 | Mrps7 | 0.005 | Old |
| 0.634379 | 0.063596 | 0.149 | 0.142 | 1 | 0 | Itga9 | 0.007 | Old |
| 0.635393 | 0.124557 | 0.036 | 0.032 | 1 | 0 | Npat | 0.004 | Old |

| | | | | | | | | |
|---|---|---|---|---|---|---|---|---|
| 0.636418 | 0.077464 | 0.192 | 0.183 | 1 | 0 | Dhrs7 | 0.009 | Old |
| 0.637895 | 0.089361 | 0.052 | 0.057 | 1 | 0 | Cd72 | -0.005 | Old |
| 0.638285 | 0.310281 | 0.031 | 0.028 | 1 | 0 | Clp1 | 0.003 | Old |
| 0.63832 | 0.290217 | 0.012 | 0.015 | 1 | 0 | Coq6 | -0.003 | Old |
| 0.638591 | 0.284662 | 0.212 | 0.209 | 1 | 0 | Fam43a | 0.003 | Old |
| 0.6388 | 0.010117 | 0.105 | 0.112 | 1 | 0 | Eri1 | -0.007 | Old |
| 0.639639 | 0.036814 | 0.051 | 0.046 | 1 | 0 | Xpnpep1 | 0.005 | Old |
| 0.641066 | 0.062504 | 0.092 | 0.086 | 1 | 0 | Pdcd7 | 0.006 | Old |
| 0.641199 | 0.228522 | 0.077 | 0.072 | 1 | 0 | Opa3 | 0.005 | Old |
| 0.641246 | 0.040744 | 0.086 | 0.08 | 1 | 0 | Magoh | 0.006 | Old |
| 0.641251 | 0.020113 | 0.817 | 0.798 | 1 | 0 | Nfkbiz | 0.019 | Old |
| 0.641287 | 0.129298 | 0.089 | 0.084 | 1 | 0 | Slc30a9 | 0.005 | Old |
| 0.64139 | 0.10521 | 0.129 | 0.122 | 1 | 0 | Spin1 | 0.007 | Old |
| 0.642077 | 0.166378 | 0.034 | 0.031 | 1 | 0 | Gatc | 0.003 | Old |
| 0.642078 | 0.153052 | 0.131 | 0.125 | 1 | 0 | Tmem208 | 0.006 | Old |
| 0.642999 | 0.130351 | 0.047 | 0.043 | 1 | 0 | Thoc1 | 0.004 | Old |
| 0.643603 | 0.189571 | 0.03 | 0.027 | 1 | 0 | Alg6 | 0.003 | Old |
| 0.64479 | 0.121437 | 0.023 | 0.027 | 1 | 0 | Pcgf6 | -0.004 | Old |
| 0.644812 | 0.001904 | 0.105 | 0.099 | 1 | 0 | Meaf6 | 0.006 | Old |
| 0.644874 | 0.094593 | 0.051 | 0.046 | 1 | 0 | Pdgfa | 0.005 | Old |
| 0.645173 | 0.257249 | 0.068 | 0.064 | 1 | 0 | Kpna6 | 0.004 | Old |
| 0.646275 | 0.005947 | 0.116 | 0.109 | 1 | 0 | Ash2l | 0.007 | Old |
| 0.64796 | 0.001511 | 0.077 | 0.071 | 1 | 0 | Tmub2 | 0.006 | Old |
| 0.648291 | 0.052393 | 0.1 | 0.094 | 1 | 0 | Phf8 | 0.006 | Old |
| 0.648777 | 0.14199 | 0.008 | 0.01 | 1 | 0 | Ftx | -0.002 | Old |
| 0.648887 | 0.111591 | 0.085 | 0.079 | 1 | 0 | Kansl3 | 0.006 | Old |
| 0.650163 | 0.370814 | 0.037 | 0.034 | 1 | 0 | Supt7l | 0.003 | Old |
| 0.650982 | 0.177034 | 0.008 | 0.01 | 1 | 0 | Mefv | -0.002 | Old |
| 0.65124 | 0.14545 | 0.042 | 0.039 | 1 | 0 | Acvr1b | 0.003 | Old |
| 0.652443 | 0.010164 | 0.031 | 0.028 | 1 | 0 | Tsr2 | 0.003 | Old |
| 0.652669 | 0.110546 | 0.078 | 0.073 | 1 | 0 | Zfp120 | 0.005 | Old |
| 0.652753 | 0.266984 | 0.026 | 0.023 | 1 | 0 | Pdcd2 | 0.003 | Old |
| 0.65319 | 0.49605 | 0.008 | 0.01 | 1 | 0 | Mterf1a | -0.002 | Old |
| 0.653293 | 0.166218 | 0.049 | 0.045 | 1 | 0 | Cdyl | 0.004 | Old |
| 0.653729 | 0.027516 | 0.334 | 0.321 | 1 | 0 | Casp4 | 0.013 | Old |
| 0.654341 | 0.014119 | 0.048 | 0.052 | 1 | 0 | Surf1 | -0.004 | Old |
| 0.654567 | 0.177479 | 0.096 | 0.091 | 1 | 0 | 4931406P1 | 0.005 | Old |
| 0.655622 | 0.185791 | 0.096 | 0.091 | 1 | 0 | Pdcl | 0.005 | Old |
| 0.655858 | 0.02403 | 0.176 | 0.168 | 1 | 0 | Ppp1r2 | 0.008 | Old |
| 0.65659 | 0.23656 | 0.038 | 0.035 | 1 | 0 | Abt1 | 0.003 | Old |
| 0.657008 | 0.241202 | 0.026 | 0.023 | 1 | 0 | Tmem101 | 0.003 | Old |
| 0.657081 | 0.147169 | 0.055 | 0.051 | 1 | 0 | Dgat1 | 0.004 | Old |
| 0.658947 | 0.003553 | 0.107 | 0.101 | 1 | 0 | Slc39a10 | 0.006 | Old |
| 0.660731 | 0.032534 | 0.204 | 0.195 | 1 | 0 | Eif3m | 0.009 | Old |
| 0.661305 | 0.159584 | 0.073 | 0.068 | 1 | 0 | Sec24d | 0.005 | Old |

| | | | | | | | | |
|---|---|---|---|---|---|---|---|---|
| 0.661313 | 0.026777 | 0.016 | 0.019 | 1 | 0 | Ifit1 | -0.003 | Old |
| 0.662248 | 0.187445 | 0.025 | 0.022 | 1 | 0 | Mipep | 0.003 | Old |
| 0.662595 | 0.022259 | 0.052 | 0.048 | 1 | 0 | Fbxl6 | 0.004 | Old |
| 0.662924 | 0.035747 | 0.126 | 0.12 | 1 | 0 | Nudt16 | 0.006 | Old |
| 0.663685 | 0.001229 | 0.073 | 0.078 | 1 | 0 | Acsl3 | -0.005 | Old |
| 0.664057 | 0.009652 | 0.914 | 0.915 | 1 | 0 | Hspa8 | -0.001 | Old |
| 0.664136 | 0.03806 | 0.014 | 0.016 | 1 | 0 | Phlpp1 | -0.002 | Old |
| 0.66419 | 0.14127 | 0.026 | 0.023 | 1 | 0 | Usp46 | 0.003 | Old |
| 0.664258 | 0.128008 | 0.049 | 0.045 | 1 | 0 | 2510002D2 | 0.004 | Old |
| 0.66503 | 0.10023 | 0.053 | 0.049 | 1 | 0 | Pip4k2c | 0.004 | Old |
| 0.665555 | 0.148108 | 0.081 | 0.076 | 1 | 0 | Ptk2b | 0.005 | Old |
| 0.665669 | 0.09185 | 0.049 | 0.045 | 1 | 0 | Galk1 | 0.004 | Old |
| 0.66574 | 0.272944 | 0.014 | 0.016 | 1 | 0 | Nup210 | -0.002 | Old |
| 0.666266 | 0.028628 | 0.026 | 0.023 | 1 | 0 | Pdss2 | 0.003 | Old |
| 0.668591 | 0.042906 | 0.06 | 0.056 | 1 | 0 | Csf3r | 0.004 | Old |
| 0.668795 | 0.112008 | 0.126 | 0.12 | 1 | 0 | Arsk | 0.006 | Old |
| 0.669051 | 0.025784 | 0.174 | 0.166 | 1 | 0 | Cbfb | 0.008 | Old |
| 0.669627 | 0.193084 | 0.037 | 0.034 | 1 | 0 | Polr2d | 0.003 | Old |
| 0.669715 | 0.218355 | 0.022 | 0.019 | 1 | 0 | 3110001I2 | 0.003 | Old |
| 0.670041 | 0.094076 | 0.029 | 0.032 | 1 | 0 | Lrwd1 | -0.003 | Old |
| 0.670539 | 0.078815 | 0.038 | 0.035 | 1 | 0 | Alg11 | 0.003 | Old |
| 0.670624 | 0.015692 | 0.047 | 0.051 | 1 | 0 | Mfsd7a | -0.004 | Old |
| 0.670631 | 0.007914 | 0.086 | 0.081 | 1 | 0 | Dhrs7b | 0.005 | Old |
| 0.670787 | 0.068287 | 0.025 | 0.022 | 1 | 0 | Slc25a23 | 0.003 | Old |
| 0.671134 | 0.231811 | 0.022 | 0.019 | 1 | 0 | Acpp | 0.003 | Old |
| 0.671407 | 0.246371 | 0.021 | 0.018 | 1 | 0 | Plpp5 | 0.003 | Old |
| 0.671623 | 0.061889 | 0.14 | 0.133 | 1 | 0 | Mta2 | 0.007 | Old |
| 0.672206 | 0.073033 | 0.023 | 0.021 | 1 | 0 | Fpgs | 0.002 | Old |
| 0.672842 | 0.473273 | 0.041 | 0.038 | 1 | 0 | Rdh14 | 0.003 | Old |
| 0.673315 | 0.096839 | 0.021 | 0.018 | 1 | 0 | Slc25a15 | 0.003 | Old |
| 0.673322 | 0.040171 | 0.271 | 0.261 | 1 | 0 | Map4k4 | 0.01 | Old |
| 0.67337 | 0.551602 | 0.015 | 0.013 | 1 | 0 | Cenpo | 0.002 | Old |
| 0.675074 | 0.107002 | 0.037 | 0.034 | 1 | 0 | Tomm40l | 0.003 | Old |
| 0.675082 | 0.487787 | 0.033 | 0.03 | 1 | 0 | Gpr35 | 0.003 | Old |
| 0.675092 | 0.015734 | 0.308 | 0.295 | 1 | 0 | Sft2d2 | 0.013 | Old |
| 0.675666 | 0.215239 | 0.036 | 0.032 | 1 | 0 | Zc3hc1 | 0.004 | Old |
| 0.675683 | 0.124915 | 0.022 | 0.019 | 1 | 0 | Riox1 | 0.003 | Old |
| 0.675995 | 0.116366 | 0.097 | 0.091 | 1 | 0 | 4833420G1 | 0.006 | Old |
| 0.676258 | 0.05382 | 0.079 | 0.075 | 1 | 0 | Nelfcd | 0.004 | Old |
| 0.676766 | 0.190185 | 0.071 | 0.067 | 1 | 0 | Rab28 | 0.004 | Old |
| 0.678101 | 0.006485 | 0.581 | 0.559 | 1 | 0 | Rab14 | 0.022 | Old |
| 0.67885 | 0.180047 | 0.044 | 0.04 | 1 | 0 | Tjp2 | 0.004 | Old |
| 0.678952 | 0.451522 | 0.01 | 0.008 | 1 | 0 | Pkd1l2 | 0.002 | Old |
| 0.679244 | 0.003647 | 0.981 | 0.98 | 1 | 0 | Eef1a1 | 0.001 | Old |
| 0.679482 | 0.016132 | 0.577 | 0.595 | 1 | 0 | Nr4a1 | -0.018 | Old |

| | | | | | | | | |
|---|---|---|---|---|---|---|---|---|
| 0.679505 | 0.1738 | 0.036 | 0.032 | 1 | 0 | Pik3cb | 0.004 | Old |
| 0.679648 | 0.149814 | 0.114 | 0.109 | 1 | 0 | Lcorl | 0.005 | Old |
| 0.679843 | 0.024947 | 0.049 | 0.045 | 1 | 0 | Mrps26 | 0.004 | Old |
| 0.679924 | 0.23785 | 0.015 | 0.013 | 1 | 0 | Nme6 | 0.002 | Old |
| 0.68001 | 0.283017 | 0.015 | 0.013 | 1 | 0 | Cdyl2 | 0.002 | Old |
| 0.680593 | 0.633774 | 0.014 | 0.012 | 1 | 0 | Cox18 | 0.002 | Old |
| 0.680623 | 0.198828 | 0.016 | 0.014 | 1 | 0 | Mtrr | 0.002 | Old |
| 0.680811 | 0.217629 | 0.012 | 0.011 | 1 | 0 | Otof | 0.001 | Old |
| 0.681362 | 0.337764 | 0.01 | 0.008 | 1 | 0 | Nim1k | 0.002 | Old |
| 0.682112 | 0.257797 | 0.016 | 0.014 | 1 | 0 | Wdhd1 | 0.002 | Old |
| 0.683308 | 0.343934 | 0.01 | 0.011 | 1 | 0 | Nckap5l | -0.001 | Old |
| 0.683683 | 0.086998 | 0.014 | 0.012 | 1 | 0 | Gm10762 | 0.002 | Old |
| 0.683857 | 0.003634 | 0.378 | 0.358 | 1 | 0 | Rab11a | 0.02 | Old |
| 0.684215 | 0.129034 | 0.01 | 0.008 | 1 | 0 | Them6 | 0.002 | Old |
| 0.684363 | 0.263273 | 0.03 | 0.027 | 1 | 0 | Traf2 | 0.003 | Old |
| 0.684423 | 0.251639 | 0.015 | 0.013 | 1 | 0 | Gm30329 | 0.002 | Old |
| 0.684435 | 0.074809 | 0.088 | 0.083 | 1 | 0 | Erlin2 | 0.005 | Old |
| 0.684677 | 0.104826 | 0.034 | 0.031 | 1 | 0 | H2-T24 | 0.003 | Old |
| 0.684725 | 0.145411 | 0.019 | 0.017 | 1 | 0 | Trmt10a | 0.002 | Old |
| 0.684735 | 0.058434 | 0.114 | 0.108 | 1 | 0 | Ncbp2 | 0.006 | Old |
| 0.684874 | 0.092791 | 0.01 | 0.008 | 1 | 0 | Ccdc14 | 0.002 | Old |
| 0.68522 | 0.009141 | 0.103 | 0.097 | 1 | 0 | Brpf1 | 0.006 | Old |
| 0.685231 | 0.194183 | 0.012 | 0.011 | 1 | 0 | Hdhd5 | 0.001 | Old |
| 0.685246 | 0.058037 | 0.034 | 0.031 | 1 | 0 | C2cd2l | 0.003 | Old |
| 0.686255 | 0.028714 | 0.011 | 0.009 | 1 | 0 | Hells | 0.002 | Old |
| 0.68646 | 0.192842 | 0.011 | 0.009 | 1 | 0 | Gm20632 | 0.002 | Old |
| 0.686578 | 0.079098 | 0.012 | 0.011 | 1 | 0 | Tmem69 | 0.001 | Old |
| 0.686671 | 0.068004 | 0.034 | 0.031 | 1 | 0 | Actr5 | 0.003 | Old |
| 0.687059 | 0.031318 | 0.012 | 0.011 | 1 | 0 | Gins2 | 0.001 | Old |
| 0.687074 | 0.055031 | 0.011 | 0.009 | 1 | 0 | Armc2 | 0.002 | Old |
| 0.687993 | 0.165173 | 0.021 | 0.023 | 1 | 0 | 3222401L1 | -0.002 | Old |
| 0.688125 | 0.005264 | 0.357 | 0.344 | 1 | 0 | Il10ra | 0.013 | Old |
| 0.688886 | 0.16815 | 0.101 | 0.097 | 1 | 0 | Ccdc34 | 0.004 | Old |
| 0.688954 | 0.141911 | 0.101 | 0.097 | 1 | 0 | Chrac1 | 0.004 | Old |
| 0.689275 | 0.071097 | 0.012 | 0.011 | 1 | 0 | Zmynd19 | 0.001 | Old |
| 0.689739 | 0.066832 | 0.758 | 0.729 | 1 | 0 | Ucp2 | 0.029 | Old |
| 0.69015 | 0.049187 | 0.011 | 0.009 | 1 | 0 | Gm10642 | 0.002 | Old |
| 0.69113 | 0.009054 | 0.044 | 0.04 | 1 | 0 | Armc7 | 0.004 | Old |
| 0.691157 | 0.096194 | 0.145 | 0.14 | 1 | 0 | Irf2bpl | 0.005 | Old |
| 0.691203 | 0.163186 | 0.085 | 0.091 | 1 | 0 | Tbcel | -0.006 | Old |
| 0.692715 | 0.33456 | 0.027 | 0.025 | 1 | 0 | Snapc2 | 0.002 | Old |
| 0.692787 | 0.243849 | 0.031 | 0.029 | 1 | 0 | Sco2 | 0.002 | Old |
| 0.693659 | 0.097474 | 0.063 | 0.059 | 1 | 0 | Wdr48 | 0.004 | Old |
| 0.695479 | 0.026516 | 0.044 | 0.04 | 1 | 0 | Eif2b2 | 0.004 | Old |
| 0.696227 | 0.131172 | 0.051 | 0.047 | 1 | 0 | Smim26 | 0.004 | Old |

| | | | | | | | | |
|---|---|---|---|---|---|---|---|---|
| 0.699205 | 0.072686 | 0.124 | 0.119 | 1 | 0 | Srp54a | 0.005 | Old |
| 0.699388 | 0.00183 | 0.795 | 0.797 | 1 | 0 | Iqgap1 | -0.002 | Old |
| 0.699481 | 0.075463 | 0.064 | 0.061 | 1 | 0 | Rin3 | 0.003 | Old |
| 0.699627 | 0.083498 | 0.038 | 0.042 | 1 | 0 | Snx25 | -0.004 | Old |
| 0.700741 | 0.461036 | 0.025 | 0.022 | 1 | 0 | Fam136a | 0.003 | Old |
| 0.701031 | 0.164205 | 0.027 | 0.025 | 1 | 0 | Ifitm5 | 0.002 | Old |
| 0.701199 | 0.032527 | 0.031 | 0.035 | 1 | 0 | Pls3 | -0.004 | Old |
| 0.702852 | 0.150718 | 0.094 | 0.09 | 1 | 0 | Exoc2 | 0.004 | Old |
| 0.703236 | 0.004549 | 0.081 | 0.076 | 1 | 0 | Zfp992 | 0.005 | Old |
| 0.703977 | 0.034661 | 0.107 | 0.102 | 1 | 0 | Nupr1 | 0.005 | Old |
| 0.704831 | 0.044622 | 0.038 | 0.035 | 1 | 0 | Cxcl10 | 0.003 | Old |
| 0.704854 | 0.012878 | 0.033 | 0.03 | 1 | 0 | Erich1 | 0.003 | Old |
| 0.705541 | 0.053192 | 0.126 | 0.131 | 1 | 0 | Tnf | -0.005 | Old |
| 0.706132 | 0.042968 | 0.141 | 0.135 | 1 | 0 | Cdk12 | 0.006 | Old |
| 0.706133 | 0.120017 | 0.038 | 0.035 | 1 | 0 | Ermard | 0.003 | Old |
| 0.709653 | 0.033404 | 0.077 | 0.072 | 1 | 0 | Pfdn6 | 0.005 | Old |
| 0.709827 | 0.218874 | 0.038 | 0.035 | 1 | 0 | Alg1 | 0.003 | Old |
| 0.710606 | 0.03777 | 0.027 | 0.025 | 1 | 0 | Gstt3 | 0.002 | Old |
| 0.710865 | 0.10704 | 0.105 | 0.101 | 1 | 0 | Rabgef1 | 0.004 | Old |
| 0.71196 | 0.144737 | 0.055 | 0.051 | 1 | 0 | Batf | 0.004 | Old |
| 0.71218 | 0.015278 | 0.235 | 0.226 | 1 | 0 | Tnpo1 | 0.009 | Old |
| 0.712331 | 0.166964 | 0.04 | 0.037 | 1 | 0 | Fam13b | 0.003 | Old |
| 0.712768 | 0.029447 | 0.219 | 0.212 | 1 | 0 | Dusp11 | 0.007 | Old |
| 0.713053 | 0.001658 | 0.175 | 0.168 | 1 | 0 | Sec63 | 0.007 | Old |
| 0.713208 | 0.029525 | 0.201 | 0.195 | 1 | 0 | Copb2 | 0.006 | Old |
| 0.713792 | 0.023472 | 0.239 | 0.231 | 1 | 0 | Tcea1 | 0.008 | Old |
| 0.713981 | 0.060516 | 0.231 | 0.225 | 1 | 0 | Neu1 | 0.006 | Old |
| 0.714123 | 0.01921 | 0.449 | 0.427 | 1 | 0 | Lnpep | 0.022 | Old |
| 0.714667 | 0.061604 | 0.161 | 0.156 | 1 | 0 | Rbm7 | 0.005 | Old |
| 0.714709 | 0.043459 | 0.06 | 0.064 | 1 | 0 | Znrf1 | -0.004 | Old |
| 0.715324 | 0.001765 | 0.446 | 0.425 | 1 | 0 | Jmjd1c | 0.021 | Old |
| 0.715575 | 0.349738 | 0.022 | 0.02 | 1 | 0 | Cdan1 | 0.002 | Old |
| 0.716503 | 0.13247 | 0.038 | 0.042 | 1 | 0 | Zdhhc23 | -0.004 | Old |
| 0.718029 | 0.269791 | 0.022 | 0.02 | 1 | 0 | Cfap45 | 0.002 | Old |
| 0.718741 | 0.000704 | 0.242 | 0.233 | 1 | 0 | Mrps33 | 0.009 | Old |
| 0.719563 | 0.003774 | 0.711 | 0.708 | 1 | 0 | mt-Nd5 | 0.003 | Old |
| 0.720741 | 0.071394 | 0.064 | 0.061 | 1 | 0 | Selenoh | 0.003 | Old |
| 0.720943 | 0.01878 | 0.055 | 0.059 | 1 | 0 | Txnrd2 | -0.004 | Old |
| 0.720963 | 0.140519 | 0.036 | 0.033 | 1 | 0 | Ccdc32 | 0.003 | Old |
| 0.721292 | 0.084618 | 0.068 | 0.064 | 1 | 0 | Pja1 | 0.004 | Old |
| 0.721349 | 0.092845 | 0.023 | 0.021 | 1 | 0 | Znhit2 | 0.002 | Old |
| 0.721724 | 0.164813 | 0.011 | 0.013 | 1 | 0 | Mars2 | -0.002 | Old |
| 0.722051 | 0.014697 | 0.023 | 0.021 | 1 | 0 | Nphp3 | 0.002 | Old |
| 0.72256 | 0.012878 | 0.048 | 0.045 | 1 | 0 | Pithd1 | 0.003 | Old |
| 0.72378 | 0.011631 | 0.219 | 0.21 | 1 | 0 | Mfsd5 | 0.009 | Old |

| | | | | | | | | |
|---|---|---|---|---|---|---|---|---|
| 0.724688 | 0.077029 | 0.166 | 0.16 | 1 | 0 | Btbd7 | 0.006 | Old |
| 0.724931 | 0.064026 | 0.023 | 0.021 | 1 | 0 | Dhx37 | 0.002 | Old |
| 0.725462 | 0.327462 | 0.03 | 0.028 | 1 | 0 | Cnnm2 | 0.002 | Old |
| 0.725879 | 0.000325 | 0.057 | 0.054 | 1 | 0 | Cln5 | 0.003 | Old |
| 0.726195 | 0.119667 | 0.051 | 0.048 | 1 | 0 | Pde12 | 0.003 | Old |
| 0.728176 | 0.440239 | 0.019 | 0.017 | 1 | 0 | Arl15 | 0.002 | Old |
| 0.728453 | 0.149595 | 0.022 | 0.02 | 1 | 0 | Hemk1 | 0.002 | Old |
| 0.729528 | 0.039279 | 0.048 | 0.045 | 1 | 0 | Acot13 | 0.003 | Old |
| 0.730035 | 0.016501 | 0.233 | 0.227 | 1 | 0 | Zranb2 | 0.006 | Old |
| 0.73005 | 0.094064 | 0.042 | 0.04 | 1 | 0 | Ubap2 | 0.002 | Old |
| 0.732435 | 0.03324 | 0.062 | 0.058 | 1 | 0 | Gtf3c1 | 0.004 | Old |
| 0.733195 | 0.034255 | 0.156 | 0.15 | 1 | 0 | Lsm6 | 0.006 | Old |
| 0.733233 | 0.244087 | 0.03 | 0.028 | 1 | 0 | Slc35d2 | 0.002 | Old |
| 0.734891 | 0.04483 | 0.152 | 0.145 | 1 | 0 | 8-Mar | 0.007 | Old |
| 0.735047 | 0.03577 | 0.272 | 0.266 | 1 | 0 | Notch1 | 0.006 | Old |
| 0.735529 | 0.429379 | 0.016 | 0.015 | 1 | 0 | Ears2 | 0.001 | Old |
| 0.736413 | 0.034152 | 0.042 | 0.04 | 1 | 0 | Ercc5 | 0.002 | Old |
| 0.737065 | 0.028867 | 0.086 | 0.091 | 1 | 0 | Kmo | -0.005 | Old |
| 0.737184 | 0.029865 | 0.339 | 0.327 | 1 | 0 | Bclaf1 | 0.012 | Old |
| 0.739063 | 0.393958 | 0.016 | 0.015 | 1 | 0 | Tbc1d7 | 0.001 | Old |
| 0.739634 | 0.005963 | 0.186 | 0.179 | 1 | 0 | Vps13c | 0.007 | Old |
| 0.741007 | 0.1353 | 0.07 | 0.067 | 1 | 0 | Cdkn2d | 0.003 | Old |
| 0.741105 | 0.009294 | 0.681 | 0.665 | 1 | 0 | Sat1 | 0.016 | Old |
| 0.741128 | 0.020238 | 0.108 | 0.104 | 1 | 0 | Cyth3 | 0.004 | Old |
| 0.741662 | 0.004319 | 0.044 | 0.041 | 1 | 0 | Acad8 | 0.003 | Old |
| 0.742532 | 0.044076 | 0.042 | 0.04 | 1 | 0 | Mthfs | 0.002 | Old |
| 0.742587 | 0.01546 | 0.07 | 0.066 | 1 | 0 | Faf2 | 0.004 | Old |
| 0.743188 | 0.117945 | 0.085 | 0.09 | 1 | 0 | Mybbp1a | -0.005 | Old |
| 0.74379 | 0.031951 | 0.213 | 0.219 | 1 | 0 | Cp | -0.006 | Old |
| 0.745303 | 0.160016 | 0.016 | 0.015 | 1 | 0 | Pdk2 | 0.001 | Old |
| 0.745387 | 0.110943 | 0.093 | 0.089 | 1 | 0 | Rsl24d1 | 0.004 | Old |
| 0.745667 | 0.472201 | 0.015 | 0.013 | 1 | 0 | Med26 | 0.002 | Old |
| 0.745954 | 0.161896 | 0.049 | 0.046 | 1 | 0 | Cul9 | 0.003 | Old |
| 0.746071 | 0.123636 | 0.249 | 0.243 | 1 | 0 | Mertk | 0.006 | Old |
| 0.746403 | 0.053268 | 0.103 | 0.098 | 1 | 0 | Pou2f1 | 0.005 | Old |
| 0.746599 | 0.011677 | 0.09 | 0.086 | 1 | 0 | Pfkfb4 | 0.004 | Old |
| 0.746702 | 0.035686 | 0.26 | 0.251 | 1 | 0 | Srsf11 | 0.009 | Old |
| 0.74675 | 0.082634 | 0.033 | 0.035 | 1 | 0 | Rbmx2 | -0.002 | Old |
| 0.746823 | 0.088366 | 0.016 | 0.015 | 1 | 0 | Acp6 | 0.001 | Old |
| 0.746876 | 0.251976 | 0.044 | 0.041 | 1 | 0 | Dnajc24 | 0.003 | Old |
| 0.747118 | 0.066579 | 0.04 | 0.037 | 1 | 0 | Leo1 | 0.003 | Old |
| 0.747571 | 0.092518 | 0.048 | 0.045 | 1 | 0 | Xrcc1 | 0.003 | Old |
| 0.747603 | 0.054071 | 0.068 | 0.065 | 1 | 0 | Zcchc9 | 0.003 | Old |
| 0.748319 | 0.059891 | 0.083 | 0.08 | 1 | 0 | Vgll4 | 0.003 | Old |
| 0.748705 | 0.027289 | 0.051 | 0.048 | 1 | 0 | Flt3l | 0.003 | Old |

| | | | | | | | | |
|---|---|---|---|---|---|---|---|---|
| 0.748873 | 0.055522 | 0.012 | 0.014 | 1 | 0 | 4632404H1 | -0.002 | Old |
| 0.74888 | 0.227495 | 0.036 | 0.033 | 1 | 0 | Sft2d3 | 0.003 | Old |
| 0.749056 | 0.13469 | 0.651 | 0.654 | 1 | 0 | Stk17b | -0.003 | Old |
| 0.749562 | 0.08596 | 0.098 | 0.094 | 1 | 0 | S100pbp | 0.004 | Old |
| 0.749729 | 0.153683 | 0.015 | 0.013 | 1 | 0 | Mcrip2 | 0.002 | Old |
| 0.750712 | 0.13655 | 0.016 | 0.015 | 1 | 0 | Slc12a5 | 0.001 | Old |
| 0.75086 | 0.029029 | 0.079 | 0.076 | 1 | 0 | 4930523C0 | 0.003 | Old |
| 0.751462 | 0.182913 | 0.06 | 0.057 | 1 | 0 | Slc9a6 | 0.003 | Old |
| 0.751982 | 0.418381 | 0.014 | 0.012 | 1 | 0 | Gm26982 | 0.002 | Old |
| 0.752119 | 0.00093 | 0.122 | 0.116 | 1 | 0 | Cse1l | 0.006 | Old |
| 0.752316 | 0.363044 | 0.034 | 0.032 | 1 | 0 | Tbcd | 0.002 | Old |
| 0.753436 | 0.12605 | 0.112 | 0.108 | 1 | 0 | Snrk | 0.004 | Old |
| 0.753526 | 0.219925 | 0.012 | 0.014 | 1 | 0 | Ppp1r3d | -0.002 | Old |
| 0.754508 | 0.000998 | 0.041 | 0.038 | 1 | 0 | Rtl8b | 0.003 | Old |
| 0.754546 | 0.129792 | 0.047 | 0.044 | 1 | 0 | AI987944 | 0.003 | Old |
| 0.754577 | 0.620037 | 0.012 | 0.011 | 1 | 0 | A430106G1 | 0.001 | Old |
| 0.754597 | 0.023793 | 0.057 | 0.054 | 1 | 0 | Las1l | 0.003 | Old |
| 0.755906 | 0.154052 | 0.026 | 0.024 | 1 | 0 | Tatdn1 | 0.002 | Old |
| 0.75637 | 0.019774 | 0.051 | 0.048 | 1 | 0 | Terf2ip | 0.003 | Old |
| 0.756433 | 0.488146 | 0.011 | 0.01 | 1 | 0 | Trip13 | 0.001 | Old |
| 0.756442 | 0.135183 | 0.014 | 0.012 | 1 | 0 | Szt2 | 0.002 | Old |
| 0.756607 | 0.332669 | 0.053 | 0.051 | 1 | 0 | Scai | 0.002 | Old |
| 0.756648 | 0.047729 | 0.075 | 0.072 | 1 | 0 | Rps6kc1 | 0.003 | Old |
| 0.756776 | 0.000991 | 0.06 | 0.057 | 1 | 0 | Snrnp40 | 0.003 | Old |
| 0.757321 | 0.042893 | 0.16 | 0.156 | 1 | 0 | Pdpk1 | 0.004 | Old |
| 0.757523 | 0.083662 | 0.108 | 0.104 | 1 | 0 | Hmgcs1 | 0.004 | Old |
| 0.7576 | 0.143072 | 0.055 | 0.052 | 1 | 0 | Rbm4b | 0.003 | Old |
| 0.758116 | 0.02589 | 0.014 | 0.012 | 1 | 0 | Phactr1 | 0.002 | Old |
| 0.758305 | 0.188643 | 0.019 | 0.021 | 1 | 0 | Zfp770 | -0.002 | Old |
| 0.758608 | 0.023702 | 0.092 | 0.088 | 1 | 0 | Rhot1 | 0.004 | Old |
| 0.759074 | 0.027563 | 0.037 | 0.035 | 1 | 0 | Wdr45 | 0.002 | Old |
| 0.759112 | 0.136932 | 0.026 | 0.028 | 1 | 0 | Cbr1 | -0.002 | Old |
| 0.759589 | 0.124651 | 0.03 | 0.032 | 1 | 0 | Acrbp | -0.002 | Old |
| 0.759877 | 0.159684 | 0.012 | 0.011 | 1 | 0 | Rnf217 | 0.001 | Old |
| 0.760348 | 0.161258 | 0.085 | 0.082 | 1 | 0 | Mcm3 | 0.003 | Old |
| 0.760367 | 0.051446 | 0.053 | 0.051 | 1 | 0 | Ino80c | 0.002 | Old |
| 0.760844 | 0.21982 | 0.068 | 0.066 | 1 | 0 | Mettl16 | 0.002 | Old |
| 0.761204 | 0.072591 | 0.083 | 0.08 | 1 | 0 | Zfp518a | 0.003 | Old |
| 0.761892 | 0.043773 | 0.037 | 0.035 | 1 | 0 | Lrrc40 | 0.002 | Old |
| 0.762044 | 0.026486 | 0.038 | 0.036 | 1 | 0 | Smpd3 | 0.002 | Old |
| 0.763288 | 0.101962 | 0.026 | 0.024 | 1 | 0 | Rbm48 | 0.002 | Old |
| 0.763525 | 0.028377 | 0.078 | 0.082 | 1 | 0 | Abhd4 | -0.004 | Old |
| 0.765281 | 0.077113 | 0.078 | 0.075 | 1 | 0 | Cep192 | 0.003 | Old |
| 0.76692 | 0.061009 | 0.083 | 0.08 | 1 | 0 | Psmc4 | 0.003 | Old |
| 0.767834 | 0.05193 | 0.134 | 0.129 | 1 | 0 | B3gat3 | 0.005 | Old |

| | | | | | | | | |
|---|---|---|---|---|---|---|---|---|
| 0.767978 | 0.284636 | 0.01 | 0.008 | 1 | 0 | Cenpe | 0.002 | Old |
| 0.76798 | 0.060486 | 0.112 | 0.108 | 1 | 0 | Abl2 | 0.004 | Old |
| 0.76852 | 0.004413 | 0.259 | 0.252 | 1 | 0 | Gbp7 | 0.007 | Old |
| 0.768747 | 0.062316 | 0.025 | 0.023 | 1 | 0 | Ptgir | 0.002 | Old |
| 0.769451 | 0.005105 | 0.09 | 0.087 | 1 | 0 | Trim65 | 0.003 | Old |
| 0.769548 | 0.048813 | 0.01 | 0.008 | 1 | 0 | Slc2a3 | 0.002 | Old |
| 0.769772 | 0.108929 | 0.01 | 0.008 | 1 | 0 | Zfp623 | 0.002 | Old |
| 0.770319 | 0.203437 | 0.098 | 0.096 | 1 | 0 | Rsrc1 | 0.002 | Old |
| 0.770445 | 0.047256 | 0.01 | 0.008 | 1 | 0 | 1700037C1 | 0.002 | Old |
| 0.771857 | 0.038717 | 0.034 | 0.032 | 1 | 0 | E130309D( | 0.002 | Old |
| 0.772444 | 0.002434 | 0.034 | 0.032 | 1 | 0 | Zfand2b | 0.002 | Old |
| 0.772786 | 0.057378 | 0.044 | 0.041 | 1 | 0 | Tnip1 | 0.003 | Old |
| 0.773633 | 0.085791 | 0.116 | 0.112 | 1 | 0 | Gdap10 | 0.004 | Old |
| 0.774165 | 0.006076 | 0.111 | 0.107 | 1 | 0 | Cope | 0.004 | Old |
| 0.775554 | 0.018163 | 0.127 | 0.123 | 1 | 0 | Rnf146 | 0.004 | Old |
| 0.777184 | 0.003547 | 0.09 | 0.087 | 1 | 0 | Ttc5 | 0.003 | Old |
| 0.777808 | 0.155733 | 0.103 | 0.099 | 1 | 0 | Mtmr12 | 0.004 | Old |
| 0.778228 | 0.282311 | 0.037 | 0.035 | 1 | 0 | Fam58b | 0.002 | Old |
| 0.77864 | 0.004795 | 0.144 | 0.139 | 1 | 0 | Ahcyl2 | 0.005 | Old |
| 0.778888 | 0.046437 | 0.197 | 0.203 | 1 | 0 | Mid1ip1 | -0.006 | Old |
| 0.7789 | 0.038156 | 0.021 | 0.022 | 1 | 0 | Coq3 | -0.001 | Old |
| 0.779305 | 0.004751 | 0.059 | 0.056 | 1 | 0 | Tecpr1 | 0.003 | Old |
| 0.77986 | 0.334595 | 0.021 | 0.019 | 1 | 0 | Adora2b | 0.002 | Old |
| 0.780063 | 0.144964 | 0.022 | 0.02 | 1 | 0 | Armcx5 | 0.002 | Old |
| 0.781289 | 0.218628 | 0.031 | 0.029 | 1 | 0 | Aimp2 | 0.002 | Old |
| 0.781297 | 0.036763 | 0.234 | 0.227 | 1 | 0 | Rreb1 | 0.007 | Old |
| 0.782125 | 0.143921 | 0.014 | 0.015 | 1 | 0 | Cnn3 | -0.001 | Old |
| 0.782276 | 0.026199 | 0.041 | 0.039 | 1 | 0 | Morc2a | 0.002 | Old |
| 0.782332 | 0.007651 | 0.226 | 0.219 | 1 | 0 | Arih1 | 0.007 | Old |
| 0.782374 | 0.082309 | 0.074 | 0.071 | 1 | 0 | Siae | 0.003 | Old |
| 0.782747 | 0.158712 | 0.021 | 0.019 | 1 | 0 | Slco3a1 | 0.002 | Old |
| 0.782924 | 0.028872 | 0.052 | 0.049 | 1 | 0 | Sprydu7 | 0.003 | Old |
| 0.783103 | 0.077261 | 0.071 | 0.068 | 1 | 0 | Rbsn | 0.003 | Old |
| 0.78352 | 0.251423 | 0.051 | 0.048 | 1 | 0 | Edem3 | 0.003 | Old |
| 0.783596 | 0.305513 | 0.022 | 0.02 | 1 | 0 | Gstk1 | 0.002 | Old |
| 0.783738 | 0.023876 | 0.07 | 0.073 | 1 | 0 | Med21 | -0.003 | Old |
| 0.784794 | 0.024323 | 0.025 | 0.027 | 1 | 0 | Dynlt1f | -0.002 | Old |
| 0.784889 | 0.375991 | 0.019 | 0.018 | 1 | 0 | 4933427D1 | 0.001 | Old |
| 0.784996 | 0.142082 | 0.022 | 0.02 | 1 | 0 | Slc46a1 | 0.002 | Old |
| 0.785028 | 0.22495 | 0.021 | 0.019 | 1 | 0 | Dhcr24 | 0.002 | Old |
| 0.785051 | 0.00187 | 0.171 | 0.175 | 1 | 0 | Atp6v1c1 | -0.004 | Old |
| 0.785133 | 0.018144 | 0.011 | 0.012 | 1 | 0 | A830010M | -0.001 | Old |
| 0.78521 | 0.078533 | 0.146 | 0.142 | 1 | 0 | Cuedc2 | 0.004 | Old |
| 0.785484 | 0.195998 | 0.021 | 0.019 | 1 | 0 | Inpp5e | 0.002 | Old |
| 0.785659 | 0.035274 | 0.022 | 0.02 | 1 | 0 | Aasdh | 0.002 | Old |

| | | | | | | | | |
|---|---|---|---|---|---|---|---|---|
| 0.787149 | 0.014284 | 0.056 | 0.053 | 1 | 0 | Gphn | 0.003 | Old |
| 0.788212 | 0.518028 | 0.027 | 0.026 | 1 | 0 | Lpcat2 | 0.001 | Old |
| 0.788823 | 0.182502 | 0.036 | 0.038 | 1 | 0 | Prune1 | -0.002 | Old |
| 0.789771 | 0.239415 | 0.019 | 0.018 | 1 | 0 | Rfesd | 0.001 | Old |
| 0.790181 | 0.000187 | 0.077 | 0.08 | 1 | 0 | Bcap29 | -0.003 | Old |
| 0.790796 | 0.017077 | 0.211 | 0.206 | 1 | 0 | Arfgef1 | 0.005 | Old |
| 0.790869 | 0.11051 | 0.036 | 0.034 | 1 | 0 | Dhx29 | 0.002 | Old |
| 0.791125 | 0.060236 | 0.725 | 0.727 | 1 | 0 | Marcksl1 | -0.002 | Old |
| 0.791671 | 0.034844 | 0.047 | 0.049 | 1 | 0 | Epor | -0.002 | Old |
| 0.794241 | 0.114713 | 0.059 | 0.056 | 1 | 0 | Gtf3c3 | 0.003 | Old |
| 0.794721 | 0.232746 | 0.053 | 0.051 | 1 | 0 | Prpf31 | 0.002 | Old |
| 0.794789 | 0.045641 | 0.038 | 0.036 | 1 | 0 | Plpp6 | 0.002 | Old |
| 0.795159 | 0.027166 | 0.166 | 0.171 | 1 | 0 | Map2k3 | -0.005 | Old |
| 0.795241 | 0.11034 | 0.018 | 0.016 | 1 | 0 | Lrrc8b | 0.002 | Old |
| 0.795399 | 0.093517 | 0.066 | 0.063 | 1 | 0 | Mtmr14 | 0.003 | Old |
| 0.795428 | 0.18808 | 0.018 | 0.019 | 1 | 0 | Mad2l2 | -0.001 | Old |
| 0.797047 | 0.000666 | 0.275 | 0.263 | 1 | 0 | Slc7a8 | 0.012 | Old |
| 0.797207 | 0.271489 | 0.018 | 0.016 | 1 | 0 | AU041133 | 0.002 | Old |
| 0.797246 | 0.026406 | 0.022 | 0.024 | 1 | 0 | Usp35 | -0.002 | Old |
| 0.79758 | 0.212938 | 0.034 | 0.032 | 1 | 0 | 10-Sep | 0.002 | Old |
| 0.797583 | 0.058694 | 0.056 | 0.054 | 1 | 0 | Plscr4 | 0.002 | Old |
| 0.798994 | 0.015327 | 0.049 | 0.047 | 1 | 0 | Cd1d1 | 0.002 | Old |
| 0.799256 | 0.044241 | 0.079 | 0.076 | 1 | 0 | Acadvl | 0.003 | Old |
| 0.800233 | 0.104706 | 0.037 | 0.035 | 1 | 0 | Srm | 0.002 | Old |
| 0.800387 | 0.178899 | 0.03 | 0.032 | 1 | 0 | Timm8a1 | -0.002 | Old |
| 0.800834 | 0.551592 | 0.016 | 0.015 | 1 | 0 | Hspb1 | 0.001 | Old |
| 0.8013 | 0.011818 | 0.019 | 0.018 | 1 | 0 | 2900076A( | 0.001 | Old |
| 0.801535 | 0.070529 | 0.105 | 0.102 | 1 | 0 | Hdac2 | 0.003 | Old |
| 0.803984 | 0.1221 | 0.026 | 0.024 | 1 | 0 | Zkscan17 | 0.002 | Old |
| 0.805027 | 0.056598 | 0.038 | 0.036 | 1 | 0 | Sema4c | 0.002 | Old |
| 0.805485 | 0.059179 | 0.083 | 0.081 | 1 | 0 | Abcg1 | 0.002 | Old |
| 0.805727 | 0.058049 | 0.07 | 0.067 | 1 | 0 | Steap3 | 0.003 | Old |
| 0.8063 | 0.298251 | 0.016 | 0.015 | 1 | 0 | Vamp1 | 0.001 | Old |
| 0.806946 | 0.086432 | 0.027 | 0.026 | 1 | 0 | BC002059 | 0.001 | Old |
| 0.807653 | 0.20489 | 0.053 | 0.051 | 1 | 0 | Tmem87a | 0.002 | Old |
| 0.809341 | 0.029286 | 0.034 | 0.032 | 1 | 0 | Ryr1 | 0.002 | Old |
| 0.809795 | 0.017884 | 0.152 | 0.148 | 1 | 0 | Tmpo | 0.004 | Old |
| 0.810552 | 0.0205 | 0.012 | 0.013 | 1 | 0 | Gm7160 | -0.001 | Old |
| 0.810908 | 0.039878 | 0.534 | 0.511 | 1 | 0 | Tmed10 | 0.023 | Old |
| 0.811149 | 0.023973 | 0.12 | 0.116 | 1 | 0 | Dctn2 | 0.004 | Old |
| 0.812446 | 0.455682 | 0.015 | 0.014 | 1 | 0 | Kcng2 | 0.001 | Old |
| 0.812485 | 0.002316 | 0.263 | 0.254 | 1 | 0 | Itpkb | 0.009 | Old |
| 0.813242 | 0.039061 | 0.067 | 0.07 | 1 | 0 | Rexo1 | -0.003 | Old |
| 0.813354 | 0.066449 | 0.034 | 0.036 | 1 | 0 | Tmem126a | -0.002 | Old |
| 0.814002 | 0.015625 | 0.016 | 0.015 | 1 | 0 | Tceal1 | 0.001 | Old |

| | | | | | | | | |
|---|---|---|---|---|---|---|---|---|
| 0.814974 | 0.210032 | 0.041 | 0.039 | 1 | 0 | Pdk3 | 0.002 | Old |
| 0.815224 | 0.04865 | 0.101 | 0.099 | 1 | 0 | Khnyn | 0.002 | Old |
| 0.815894 | 0.03038 | 0.057 | 0.055 | 1 | 0 | Sart3 | 0.002 | Old |
| 0.816135 | 0.165792 | 0.026 | 0.024 | 1 | 0 | Qpctl | 0.002 | Old |
| 0.816203 | 0.207595 | 0.015 | 0.014 | 1 | 0 | C1rl | 0.001 | Old |
| 0.816293 | 0.154749 | 0.012 | 0.013 | 1 | 0 | Mir155hg | -0.001 | Old |
| 0.816382 | 0.169935 | 0.015 | 0.014 | 1 | 0 | Glmn | 0.001 | Old |
| 0.816585 | 0.084837 | 0.088 | 0.085 | 1 | 0 | Chpt1 | 0.003 | Old |
| 0.816708 | 0.010029 | 0.111 | 0.114 | 1 | 0 | Aurkaip1 | -0.003 | Old |
| 0.817034 | 0.222307 | 0.012 | 0.013 | 1 | 0 | Msrb2 | -0.001 | Old |
| 0.817261 | 0.316774 | 0.019 | 0.021 | 1 | 0 | Rrp9 | -0.002 | Old |
| 0.817515 | 0.161075 | 0.066 | 0.064 | 1 | 0 | Pum3 | 0.002 | Old |
| 0.818474 | 0.002192 | 0.848 | 0.833 | 1 | 0 | Ubc | 0.015 | Old |
| 0.818582 | 0.020117 | 0.252 | 0.245 | 1 | 0 | Slc25a37 | 0.007 | Old |
| 0.818889 | 0.043507 | 0.015 | 0.014 | 1 | 0 | Tmem53 | 0.001 | Old |
| 0.818993 | 0.032244 | 0.026 | 0.024 | 1 | 0 | Zc3h6 | 0.002 | Old |
| 0.8197 | 0.055642 | 0.052 | 0.054 | 1 | 0 | Vps16 | -0.002 | Old |
| 0.820888 | 0.073152 | 0.016 | 0.018 | 1 | 0 | Intu | -0.002 | Old |
| 0.822637 | 0.374265 | 0.029 | 0.027 | 1 | 0 | Rfx2 | 0.002 | Old |
| 0.822967 | 0.012818 | 0.049 | 0.047 | 1 | 0 | 2310022B0 | 0.002 | Old |
| 0.823005 | 0.010915 | 0.041 | 0.039 | 1 | 0 | Iqsec2 | 0.002 | Old |
| 0.82389 | 0.135878 | 0.033 | 0.031 | 1 | 0 | Tecpr2 | 0.002 | Old |
| 0.824265 | 0.104197 | 0.074 | 0.077 | 1 | 0 | Cbx6 | -0.003 | Old |
| 0.824825 | 0.010601 | 0.971 | 0.962 | 1 | 0 | Marcks | 0.009 | Old |
| 0.825138 | 0.063379 | 0.083 | 0.081 | 1 | 0 | Mplkip | 0.002 | Old |
| 0.825161 | 0.282678 | 0.014 | 0.013 | 1 | 0 | Acot7 | 0.001 | Old |
| 0.825324 | 0.137253 | 0.062 | 0.064 | 1 | 0 | Clns1a | -0.002 | Old |
| 0.825348 | 0.034185 | 0.196 | 0.193 | 1 | 0 | Stag1 | 0.003 | Old |
| 0.826078 | 0.092344 | 0.016 | 0.018 | 1 | 0 | Tmem150b | -0.002 | Old |
| 0.826958 | 0.058251 | 0.025 | 0.023 | 1 | 0 | Hmgn5 | 0.002 | Old |
| 0.827008 | 0.049205 | 0.289 | 0.283 | 1 | 0 | Lair1 | 0.006 | Old |
| 0.827331 | 0.034559 | 0.313 | 0.307 | 1 | 0 | Lrrc58 | 0.006 | Old |
| 0.827419 | 0.102163 | 0.014 | 0.013 | 1 | 0 | Urb2 | 0.001 | Old |
| 0.828033 | 0.058118 | 0.074 | 0.072 | 1 | 0 | Mrpl15 | 0.002 | Old |
| 0.828345 | 0.19705 | 0.03 | 0.029 | 1 | 0 | Hscb | 0.001 | Old |
| 0.82852 | 0.022467 | 0.063 | 0.061 | 1 | 0 | Zmat5 | 0.002 | Old |
| 0.829129 | 0.074647 | 0.16 | 0.156 | 1 | 0 | Ube2g2 | 0.004 | Old |
| 0.829679 | 0.014389 | 0.014 | 0.013 | 1 | 0 | Trim68 | 0.001 | Old |
| 0.830127 | 0.185711 | 0.037 | 0.035 | 1 | 0 | Gpr180 | 0.002 | Old |
| 0.830172 | 0.028907 | 0.056 | 0.059 | 1 | 0 | Mill2 | -0.003 | Old |
| 0.830244 | 0.541493 | 0.012 | 0.011 | 1 | 0 | Znrf3 | 0.001 | Old |
| 0.830405 | 0.243403 | 0.021 | 0.022 | 1 | 0 | L3mbtl3 | -0.001 | Old |
| 0.830413 | 0.135373 | 0.037 | 0.035 | 1 | 0 | Adck5 | 0.002 | Old |
| 0.830828 | 0.118551 | 0.045 | 0.043 | 1 | 0 | Rrp36 | 0.002 | Old |
| 0.831055 | 0.179851 | 0.063 | 0.061 | 1 | 0 | Nfyc | 0.002 | Old |

| | | | | | | | | |
|---|---|---|---|---|---|---|---|---|
| 0.832088 | 0.356629 | 0.016 | 0.018 | 1 | 0 | Spn | -0.002 | Old |
| 0.832229 | 0.258845 | 0.012 | 0.011 | 1 | 0 | L2hgdh | 0.001 | Old |
| 0.832334 | 0.23755 | 0.031 | 0.033 | 1 | 0 | Plpp2 | -0.002 | Old |
| 0.832427 | 0.15518 | 0.012 | 0.011 | 1 | 0 | BC147527 | 0.001 | Old |
| 0.833193 | 0.016552 | 0.346 | 0.349 | 1 | 0 | Pura | -0.003 | Old |
| 0.833236 | 0.332323 | 0.027 | 0.026 | 1 | 0 | Scrn3 | 0.001 | Old |
| 0.834367 | 0.090547 | 0.022 | 0.021 | 1 | 0 | Zc3h12c | 0.001 | Old |
| 0.835959 | 0.199772 | 0.062 | 0.06 | 1 | 0 | Mrpl16 | 0.002 | Old |
| 0.836207 | 0.544164 | 0.011 | 0.01 | 1 | 0 | Fam185a | 0.001 | Old |
| 0.836572 | 0.333334 | 0.014 | 0.015 | 1 | 0 | Gngt2 | -0.001 | Old |
| 0.837338 | 0.113993 | 0.022 | 0.021 | 1 | 0 | Twsg1 | 0.001 | Old |
| 0.841102 | 0.394247 | 0.019 | 0.018 | 1 | 0 | Nup188 | 0.001 | Old |
| 0.841477 | 0.314435 | 0.011 | 0.01 | 1 | 0 | Zfp677 | 0.001 | Old |
| 0.842677 | 0.053858 | 0.045 | 0.043 | 1 | 0 | N6amt1 | 0.002 | Old |
| 0.842882 | 0.114819 | 0.051 | 0.049 | 1 | 0 | Zfp26 | 0.002 | Old |
| 0.843567 | 0.060184 | 0.3 | 0.297 | 1 | 0 | Copa | 0.003 | Old |
| 0.843587 | 0.124708 | 0.011 | 0.01 | 1 | 0 | Rbks | 0.001 | Old |
| 0.844854 | 0.323741 | 0.011 | 0.01 | 1 | 0 | 1810032O( | 0.001 | Old |
| 0.845313 | 0.053438 | 0.021 | 0.019 | 1 | 0 | Mtg2 | 0.002 | Old |
| 0.845738 | 0.13394 | 0.051 | 0.049 | 1 | 0 | Ppard | 0.002 | Old |
| 0.846701 | 0.096238 | 0.06 | 0.062 | 1 | 0 | Trappc12 | -0.002 | Old |
| 0.847377 | 0.025929 | 0.083 | 0.086 | 1 | 0 | B3glct | -0.003 | Old |
| 0.847417 | 0.006099 | 0.042 | 0.044 | 1 | 0 | Cep63 | -0.002 | Old |
| 0.847554 | 0.006825 | 0.216 | 0.222 | 1 | 0 | Snx10 | -0.006 | Old |
| 0.847567 | 0.020449 | 0.156 | 0.159 | 1 | 0 | Ogfrl1 | -0.003 | Old |
| 0.847614 | 0.22978 | 0.019 | 0.018 | 1 | 0 | Nek4 | 0.001 | Old |
| 0.847875 | 0.073943 | 0.057 | 0.056 | 1 | 0 | Slc30a4 | 0.001 | Old |
| 0.848312 | 0.026604 | 0.027 | 0.026 | 1 | 0 | Hmgxb3 | 0.001 | Old |
| 0.848825 | 0.142719 | 0.066 | 0.064 | 1 | 0 | Ascc1 | 0.002 | Old |
| 0.849044 | 0.175327 | 0.019 | 0.018 | 1 | 0 | Aacs | 0.001 | Old |
| 0.8499 | 0.059671 | 0.045 | 0.047 | 1 | 0 | Tyw1 | -0.002 | Old |
| 0.850109 | 0.129305 | 0.027 | 0.026 | 1 | 0 | Mios | 0.001 | Old |
| 0.850686 | 0.244683 | 0.026 | 0.025 | 1 | 0 | Zcchc10 | 0.001 | Old |
| 0.851112 | 0.14322 | 0.019 | 0.018 | 1 | 0 | Cald1 | 0.001 | Old |
| 0.851758 | 0.495814 | 0.01 | 0.009 | 1 | 0 | Tnfaip8l1 | 0.001 | Old |
| 0.852133 | 0.095238 | 0.011 | 0.012 | 1 | 0 | Ankrd33b | -0.001 | Old |
| 0.852662 | 0.132845 | 0.01 | 0.009 | 1 | 0 | Sh2d1b1 | 0.001 | Old |
| 0.853131 | 0.008367 | 0.011 | 0.012 | 1 | 0 | Tprn | -0.001 | Old |
| 0.853675 | 0.150296 | 0.077 | 0.075 | 1 | 0 | Spata2 | 0.002 | Old |
| 0.854389 | 0.18473 | 0.055 | 0.057 | 1 | 0 | Ifi35 | -0.002 | Old |
| 0.854788 | 0.378735 | 0.037 | 0.036 | 1 | 0 | Pcyt2 | 0.001 | Old |
| 0.855149 | 0.025333 | 0.01 | 0.009 | 1 | 0 | Col3a1 | 0.001 | Old |
| 0.855455 | 0.009584 | 0.119 | 0.121 | 1 | 0 | Twistnb | -0.002 | Old |
| 0.85549 | 0.114152 | 0.04 | 0.041 | 1 | 0 | Xrcc4 | -0.001 | Old |
| 0.855505 | 0.267857 | 0.027 | 0.026 | 1 | 0 | Vsig4 | 0.001 | Old |

| | | | | | | | | |
|---|---|---|---|---|---|---|---|---|
| 0.855943 | 0.126108 | 0.088 | 0.086 | 1 | 0 | Mrpl32 | 0.002 | Old |
| 0.856796 | 0.169358 | 0.114 | 0.118 | 1 | 0 | Mylip | -0.004 | Old |
| 0.856845 | 0.128299 | 0.01 | 0.009 | 1 | 0 | Zbtb25 | 0.001 | Old |
| 0.857111 | 0.010142 | 0.071 | 0.069 | 1 | 0 | Mcee | 0.002 | Old |
| 0.857972 | 0.055132 | 0.027 | 0.026 | 1 | 0 | Hist1h4i | 0.001 | Old |
| 0.859889 | 0.068665 | 0.127 | 0.131 | 1 | 0 | Otulin | -0.004 | Old |
| 0.860245 | 0.111682 | 0.126 | 0.125 | 1 | 0 | Cxcl1 | 0.001 | Old |
| 0.860357 | 0.008923 | 0.078 | 0.076 | 1 | 0 | Nt5c2 | 0.002 | Old |
| 0.860834 | 0.227776 | 0.031 | 0.03 | 1 | 0 | Tmem242 | 0.001 | Old |
| 0.862753 | 0.076373 | 0.04 | 0.038 | 1 | 0 | Map2k5 | 0.002 | Old |
| 0.865044 | 0.097961 | 0.129 | 0.126 | 1 | 0 | Zzz3 | 0.003 | Old |
| 0.865577 | 0.025517 | 0.134 | 0.131 | 1 | 0 | Maea | 0.003 | Old |
| 0.867033 | 0.218296 | 0.016 | 0.016 | 1 | 0 | Tmco6 | 0 | Old |
| 0.867179 | 0.203661 | 0.025 | 0.024 | 1 | 0 | Zeb1 | 0.001 | Old |
| 0.86739 | 0.140178 | 0.019 | 0.02 | 1 | 0 | Slc10a6 | -0.001 | Old |
| 0.867672 | 0.034639 | 0.948 | 0.949 | 1 | 0 | Mafb | -0.001 | Old |
| 0.86817 | 0.141163 | 0.131 | 0.13 | 1 | 0 | Fads1 | 0.001 | Old |
| 0.868273 | 0.019198 | 0.029 | 0.03 | 1 | 0 | Ublcp1 | -0.001 | Old |
| 0.868646 | 0.06565 | 0.144 | 0.142 | 1 | 0 | Luzp1 | 0.002 | Old |
| 0.869125 | 0.006379 | 0.018 | 0.017 | 1 | 0 | Strbp | 0.001 | Old |
| 0.869442 | 0.146324 | 0.016 | 0.016 | 1 | 0 | Gm1673 | 0 | Old |
| 0.872272 | 0.011777 | 0.033 | 0.034 | 1 | 0 | Slc18b1 | -0.001 | Old |
| 0.872351 | 0.023047 | 0.105 | 0.107 | 1 | 0 | Cbx7 | -0.002 | Old |
| 0.872662 | 0.080497 | 0.025 | 0.024 | 1 | 0 | Rad1 | 0.001 | Old |
| 0.873241 | 0.015439 | 0.308 | 0.303 | 1 | 0 | Nfic | 0.005 | Old |
| 0.873573 | 0.017178 | 0.016 | 0.016 | 1 | 0 | Tango6 | 0 | Old |
| 0.87446 | 0.07252 | 0.071 | 0.07 | 1 | 0 | Fkbp1b | 0.001 | Old |
| 0.874755 | 0.02603 | 0.012 | 0.013 | 1 | 0 | Lysmd1 | -0.001 | Old |
| 0.874833 | 0.045704 | 0.105 | 0.103 | 1 | 0 | Il17ra | 0.002 | Old |
| 0.875133 | 0.201518 | 0.047 | 0.045 | 1 | 0 | B9d2 | 0.002 | Old |
| 0.875747 | 0.493347 | 0.015 | 0.014 | 1 | 0 | Fsd1l | 0.001 | Old |
| 0.876254 | 0.119729 | 0.077 | 0.075 | 1 | 0 | Ubap1 | 0.002 | Old |
| 0.876735 | 0.416546 | 0.015 | 0.014 | 1 | 0 | Pex26 | 0.001 | Old |
| 0.876911 | 0.005458 | 0.12 | 0.123 | 1 | 0 | Tbc1d10b | -0.003 | Old |
| 0.877527 | 0.061251 | 0.074 | 0.072 | 1 | 0 | Hivep3 | 0.002 | Old |
| 0.879457 | 0.106315 | 0.033 | 0.034 | 1 | 0 | Map4k5 | -0.001 | Old |
| 0.879699 | 0.243898 | 0.029 | 0.028 | 1 | 0 | Rundc1 | 0.001 | Old |
| 0.88055 | 0.028633 | 0.029 | 0.03 | 1 | 0 | Ccar2 | -0.001 | Old |
| 0.881148 | 0.029875 | 0.059 | 0.057 | 1 | 0 | Lrba | 0.002 | Old |
| 0.881261 | 0.120561 | 0.042 | 0.041 | 1 | 0 | Hagh | 0.001 | Old |
| 0.881609 | 0.028482 | 0.114 | 0.111 | 1 | 0 | Otud4 | 0.003 | Old |
| 0.881643 | 0.175606 | 0.082 | 0.081 | 1 | 0 | Pex3 | 0.001 | Old |
| 0.881748 | 0.003764 | 0.082 | 0.08 | 1 | 0 | Gpd1l | 0.002 | Old |
| 0.882306 | 0.092825 | 0.064 | 0.066 | 1 | 0 | Ppp1r13b | -0.002 | Old |
| 0.882468 | 0.112851 | 0.016 | 0.017 | 1 | 0 | Ctla2b | -0.001 | Old |

| | | | | | | | | |
|---|---|---|---|---|---|---|---|---|
| 0.882766 | 0.096696 | 0.082 | 0.081 | 1 | 0 | Agtpbp1 | 0.001 | Old |
| 0.882848 | 0.078327 | 0.015 | 0.014 | 1 | 0 | Klf8 | 0.001 | Old |
| 0.883962 | 0.128934 | 0.016 | 0.017 | 1 | 0 | Fndc7 | -0.001 | Old |
| 0.884323 | 0.082861 | 0.056 | 0.055 | 1 | 0 | Slc35e4 | 0.001 | Old |
| 0.884631 | 0.030261 | 0.077 | 0.075 | 1 | 0 | Cdr2 | 0.002 | Old |
| 0.884648 | 0.038375 | 0.015 | 0.014 | 1 | 0 | Rnf122 | 0.001 | Old |
| 0.884804 | 0.026615 | 0.083 | 0.082 | 1 | 0 | Ikzf5 | 0.001 | Old |
| 0.885383 | 0.012699 | 0.057 | 0.056 | 1 | 0 | Pitpnm1 | 0.001 | Old |
| 0.88552 | 0.020993 | 0.419 | 0.42 | 1 | 0 | Trib1 | -0.001 | Old |
| 0.886522 | 0.516534 | 0.014 | 0.013 | 1 | 0 | Ggct | 0.001 | Old |
| 0.887032 | 0.066638 | 0.094 | 0.097 | 1 | 0 | Gpr108 | -0.003 | Old |
| 0.888993 | 0.052918 | 0.094 | 0.096 | 1 | 0 | 3110043O2 | -0.002 | Old |
| 0.889381 | 0.064595 | 0.105 | 0.107 | 1 | 0 | Map3k7 | -0.002 | Old |
| 0.889869 | 0.082718 | 0.015 | 0.014 | 1 | 0 | Hectd2 | 0.001 | Old |
| 0.890611 | 0.078747 | 0.055 | 0.053 | 1 | 0 | Cdc42ep3 | 0.002 | Old |
| 0.890853 | 0.0044 | 0.093 | 0.095 | 1 | 0 | Mrpl43 | -0.002 | Old |
| 0.891425 | 0.370353 | 0.014 | 0.013 | 1 | 0 | Fchsd1 | 0.001 | Old |
| 0.892888 | 0.275403 | 0.021 | 0.02 | 1 | 0 | Hspbap1 | 0.001 | Old |
| 0.89342 | 0.09239 | 0.059 | 0.058 | 1 | 0 | Fam160b2 | 0.001 | Old |
| 0.895609 | 0.183039 | 0.033 | 0.032 | 1 | 0 | Mthfsl | 0.001 | Old |
| 0.895681 | 0.004383 | 0.347 | 0.344 | 1 | 0 | Cd164 | 0.003 | Old |
| 0.896037 | 0.020301 | 0.862 | 0.837 | 1 | 0 | Ddx5 | 0.025 | Old |
| 0.896301 | 0.060475 | 0.034 | 0.033 | 1 | 0 | Mcm4 | 0.001 | Old |
| 0.896329 | 0.043762 | 0.048 | 0.049 | 1 | 0 | Gla | -0.001 | Old |
| 0.898887 | 0.05665 | 0.103 | 0.102 | 1 | 0 | Kpna3 | 0.001 | Old |
| 0.899327 | 0.012685 | 0.042 | 0.041 | 1 | 0 | Rfng | 0.001 | Old |
| 0.899862 | 0.026238 | 0.153 | 0.151 | 1 | 0 | Vps13b | 0.002 | Old |
| 0.899936 | 0.015508 | 0.04 | 0.039 | 1 | 0 | Stambpl1 | 0.001 | Old |
| 0.901074 | 0.143353 | 0.157 | 0.156 | 1 | 0 | Pcmtd2 | 0.001 | Old |
| 0.901362 | 0.035861 | 0.021 | 0.02 | 1 | 0 | Tbc1d16 | 0.001 | Old |
| 0.90173 | 0.079829 | 0.018 | 0.019 | 1 | 0 | D3Ertd751 | -0.001 | Old |
| 0.902324 | 0.11275 | 0.014 | 0.014 | 1 | 0 | Patj | 0 | Old |
| 0.902846 | 0.177936 | 0.01 | 0.01 | 1 | 0 | Stxbp4 | 0 | Old |
| 0.902992 | 0.454466 | 0.012 | 0.012 | 1 | 0 | Rgs5 | 0 | Old |
| 0.90426 | 0.1712 | 0.022 | 0.023 | 1 | 0 | Pycrl | -0.001 | Old |
| 0.904413 | 0.016982 | 0.185 | 0.181 | 1 | 0 | Trip11 | 0.004 | Old |
| 0.904514 | 0.101165 | 0.014 | 0.014 | 1 | 0 | Fam167b | 0 | Old |
| 0.904582 | 0.231712 | 0.01 | 0.01 | 1 | 0 | Hr | 0 | Old |
| 0.905105 | 0.175156 | 0.018 | 0.019 | 1 | 0 | Slc2a6 | -0.001 | Old |
| 0.90634 | 0.26067 | 0.014 | 0.014 | 1 | 0 | Als2 | 0 | Old |
| 0.906788 | 0.181521 | 0.064 | 0.064 | 1 | 0 | Tvp23b | 0 | Old |
| 0.90701 | 0.169982 | 0.146 | 0.145 | 1 | 0 | Klf11 | 0.001 | Old |
| 0.907336 | 0.015431 | 0.033 | 0.032 | 1 | 0 | Zfp954 | 0.001 | Old |
| 0.90801 | 0.09696 | 0.036 | 0.037 | 1 | 0 | Abcg2 | -0.001 | Old |
| 0.908299 | 0.147609 | 0.019 | 0.019 | 1 | 0 | H2-Ob | 0 | Old |

| | | | | | | | | |
|---|---|---|---|---|---|---|---|---|
| 0.910692 | 0.111488 | 0.019 | 0.019 | 1 | 0 | Mettl13 | 0 | Old |
| 0.910931 | 0.020218 | 0.067 | 0.066 | 1 | 0 | Csrp1 | 0.001 | Old |
| 0.9113 | 0.064532 | 0.055 | 0.056 | 1 | 0 | Srpk1 | -0.001 | Old |
| 0.911341 | 0.01641 | 0.023 | 0.024 | 1 | 0 | Sergef | -0.001 | Old |
| 0.911746 | 0.017948 | 0.012 | 0.012 | 1 | 0 | Ugt1a6a | 0 | Old |
| 0.911809 | 0.201131 | 0.019 | 0.019 | 1 | 0 | Gm41307 | 0 | Old |
| 0.91333 | 0.025057 | 0.034 | 0.033 | 1 | 0 | Papolg | 0.001 | Old |
| 0.913538 | 0.018209 | 0.012 | 0.012 | 1 | 0 | Tigar | 0 | Old |
| 0.913559 | 0.04093 | 0.101 | 0.1 | 1 | 0 | Sgsh | 0.001 | Old |
| 0.914032 | 0.047949 | 0.023 | 0.024 | 1 | 0 | Clasrp | -0.001 | Old |
| 0.915 | 0.074022 | 0.019 | 0.019 | 1 | 0 | Fdxacb1 | 0 | Old |
| 0.916616 | 0.021386 | 0.075 | 0.073 | 1 | 0 | Stx17 | 0.002 | Old |
| 0.917062 | 0.189094 | 0.057 | 0.059 | 1 | 0 | Dusp5 | -0.002 | Old |
| 0.917102 | 0.301984 | 0.018 | 0.017 | 1 | 0 | Sall2 | 0.001 | Old |
| 0.917505 | 0.066416 | 0.038 | 0.039 | 1 | 0 | Bach2 | -0.001 | Old |
| 0.918525 | 0.084857 | 0.103 | 0.102 | 1 | 0 | Esf1 | 0.001 | Old |
| 0.918715 | 0.182915 | 0.031 | 0.032 | 1 | 0 | Zfp280c | -0.001 | Old |
| 0.918854 | 0.072905 | 0.037 | 0.036 | 1 | 0 | Ttl | 0.001 | Old |
| 0.919011 | 0.048406 | 0.09 | 0.089 | 1 | 0 | Ppfia1 | 0.001 | Old |
| 0.91948 | 0.101625 | 0.051 | 0.05 | 1 | 0 | Hbegf | 0.001 | Old |
| 0.919649 | 0.037639 | 0.155 | 0.156 | 1 | 0 | Lats2 | -0.001 | Old |
| 0.919801 | 0.000939 | 0.025 | 0.024 | 1 | 0 | Pcyt1b | 0.001 | Old |
| 0.919831 | 0.213461 | 0.018 | 0.017 | 1 | 0 | Map3k15 | 0.001 | Old |
| 0.922178 | 0.026096 | 0.666 | 0.63 | 1 | 0 | Atp5l | 0.036 | Old |
| 0.922396 | 0.099409 | 0.063 | 0.062 | 1 | 0 | Ino80 | 0.001 | Old |
| 0.922544 | 0.240974 | 0.011 | 0.011 | 1 | 0 | Tubg1 | 0 | Old |
| 0.922759 | 0.00052 | 0.025 | 0.024 | 1 | 0 | 4632415L0 | 0.001 | Old |
| 0.922922 | 0.084229 | 0.103 | 0.104 | 1 | 0 | 4833439L1 | -0.001 | Old |
| 0.923569 | 0.086212 | 0.015 | 0.016 | 1 | 0 | 2810454H( | -0.001 | Old |
| 0.92381 | 0.089853 | 0.011 | 0.011 | 1 | 0 | Prr7 | 0 | Old |
| 0.926439 | 0.024691 | 0.502 | 0.5 | 1 | 0 | Fcrls | 0.002 | Old |
| 0.927514 | 0.001641 | 0.156 | 0.155 | 1 | 0 | Suz12 | 0.001 | Old |
| 0.927823 | 0.018839 | 0.044 | 0.045 | 1 | 0 | Abr | -0.001 | Old |
| 0.927978 | 0.059373 | 0.049 | 0.048 | 1 | 0 | Fam98a | 0.001 | Old |
| 0.92813 | 0.280723 | 0.015 | 0.016 | 1 | 0 | Numbl | -0.001 | Old |
| 0.928701 | 0.12537 | 0.096 | 0.098 | 1 | 0 | Slc15a2 | -0.002 | Old |
| 0.929214 | 0.029765 | 0.729 | 0.739 | 1 | 0 | Mt1 | -0.01 | Old |
| 0.929725 | 0.062181 | 0.088 | 0.087 | 1 | 0 | Impdh1 | 0.001 | Old |
| 0.93132 | 0.362283 | 0.011 | 0.011 | 1 | 0 | Dnajc28 | 0 | Old |
| 0.9326 | 0.141141 | 0.021 | 0.021 | 1 | 0 | Rfc5 | 0 | Old |
| 0.932824 | 0.146111 | 0.016 | 0.016 | 1 | 0 | Gm31718 | 0 | Old |
| 0.933558 | 0.010413 | 0.023 | 0.023 | 1 | 0 | Actn1 | 0 | Old |
| 0.933878 | 0.017948 | 0.03 | 0.029 | 1 | 0 | Fam122a | 0.001 | Old |
| 0.934658 | 0.386949 | 0.01 | 0.009 | 1 | 0 | Pdia5 | 0.001 | Old |
| 0.935405 | 0.171422 | 0.016 | 0.016 | 1 | 0 | Ccdc69 | 0 | Old |

| | | | | | | | | |
|---|---|---|---|---|---|---|---|---|
| 0.935749 | 0.031962 | 0.056 | 0.057 | 1 | 0 | Mdc1 | -0.001 | Old |
| 0.936505 | 0.219112 | 0.027 | 0.027 | 1 | 0 | Hdac7 | 0 | Old |
| 0.938663 | 0.123332 | 0.016 | 0.017 | 1 | 0 | Nmnat1 | -0.001 | Old |
| 0.938712 | 0.199759 | 0.01 | 0.009 | 1 | 0 | Mansc1 | 0.001 | Old |
| 0.93894 | 0.108259 | 0.041 | 0.04 | 1 | 0 | Nop16 | 0.001 | Old |
| 0.939163 | 0.215488 | 0.01 | 0.009 | 1 | 0 | Wrap53 | 0.001 | Old |
| 0.94046 | 0.100857 | 0.048 | 0.049 | 1 | 0 | Tfam | -0.001 | Old |
| 0.940469 | 0.207583 | 0.021 | 0.021 | 1 | 0 | Pfkm | 0 | Old |
| 0.940481 | 0.007009 | 0.083 | 0.084 | 1 | 0 | Mex3c | -0.001 | Old |
| 0.941223 | 0.085546 | 0.575 | 0.589 | 1 | 0 | Cxcl2 | -0.014 | Old |
| 0.943158 | 0.208949 | 0.012 | 0.013 | 1 | 0 | Plpp3 | -0.001 | Old |
| 0.943553 | 0.04055 | 0.066 | 0.065 | 1 | 0 | Ppme1 | 0.001 | Old |
| 0.943633 | 0.02364 | 0.986 | 0.986 | 1 | 0 | Ctsb | 0 | Old |
| 0.944139 | 0.013785 | 0.867 | 0.821 | 1 | 0 | Oaz1 | 0.046 | Old |
| 0.944165 | 0.148751 | 0.036 | 0.036 | 1 | 0 | Reep4 | 0 | Old |
| 0.946445 | 0.035338 | 0.164 | 0.163 | 1 | 0 | 1500011K1 | 0.001 | Old |
| 0.948034 | 0.019668 | 0.097 | 0.096 | 1 | 0 | Rbm18 | 0.001 | Old |
| 0.948463 | 0.177983 | 0.033 | 0.032 | 1 | 0 | Wdr12 | 0.001 | Old |
| 0.949062 | 0.014715 | 0.181 | 0.177 | 1 | 0 | Rraga | 0.004 | Old |
| 0.949307 | 0.376975 | 0.016 | 0.017 | 1 | 0 | Inpp1 | -0.001 | Old |
| 0.949563 | 0.040198 | 0.033 | 0.032 | 1 | 0 | Zfp367 | 0.001 | Old |
| 0.950019 | 0.100267 | 0.015 | 0.015 | 1 | 0 | Nmnat3 | 0 | Old |
| 0.951684 | 0.0856 | 0.051 | 0.051 | 1 | 0 | Bid | 0 | Old |
| 0.951762 | 0.032033 | 0.052 | 0.053 | 1 | 0 | Mpg | -0.001 | Old |
| 0.952622 | 0.141532 | 0.015 | 0.015 | 1 | 0 | Gja1 | 0 | Old |
| 0.952824 | 0.016625 | 0.068 | 0.068 | 1 | 0 | Ddit3 | 0 | Old |
| 0.9532 | 0.149327 | 0.074 | 0.073 | 1 | 0 | Ift52 | 0.001 | Old |
| 0.95348 | 0.309628 | 0.012 | 0.013 | 1 | 0 | Dus2 | -0.001 | Old |
| 0.953529 | 0.010281 | 0.442 | 0.436 | 1 | 0 | Sgpp1 | 0.006 | Old |
| 0.953744 | 0.004952 | 0.049 | 0.05 | 1 | 0 | 4921524J1 | -0.001 | Old |
| 0.954052 | 0.051999 | 0.063 | 0.063 | 1 | 0 | Nudt13 | 0 | Old |
| 0.954204 | 0.050747 | 0.027 | 0.028 | 1 | 0 | Zfp579 | -0.001 | Old |
| 0.954224 | 0.049809 | 0.09 | 0.09 | 1 | 0 | Akap8 | 0 | Old |
| 0.954747 | 0.005742 | 0.027 | 0.027 | 1 | 0 | 2900089D1 | 0 | Old |
| 0.957427 | 0.038415 | 0.152 | 0.151 | 1 | 0 | Rasa4 | 0.001 | Old |
| 0.957479 | 0.139991 | 0.066 | 0.067 | 1 | 0 | Uty | -0.001 | Old |
| 0.957515 | 0.021428 | 0.038 | 0.038 | 1 | 0 | Mtmr9 | 0 | Old |
| 0.959495 | 0.058761 | 0.044 | 0.044 | 1 | 0 | Tmem181a | 0 | Old |
| 0.959596 | 0.203092 | 0.018 | 0.018 | 1 | 0 | Nudt22 | 0 | Old |
| 0.959845 | 0.531467 | 0.014 | 0.013 | 1 | 0 | Crip2 | 0.001 | Old |
| 0.961064 | 0.231785 | 0.018 | 0.018 | 1 | 0 | Rbm41 | 0 | Old |
| 0.961161 | 0.573281 | 0.014 | 0.013 | 1 | 0 | Fam212a | 0.001 | Old |
| 0.96139 | 0.246398 | 0.018 | 0.018 | 1 | 0 | Lrrc28 | 0 | Old |
| 0.96235 | 0.199503 | 0.031 | 0.032 | 1 | 0 | Prr3 | -0.001 | Old |
| 0.962742 | 0.016936 | 0.16 | 0.161 | 1 | 0 | Cdc37l1 | -0.001 | Old |

| | | | | | | | | |
|---|---|---|---|---|---|---|---|---|
| 0.962931 | 0.01606 | 0.083 | 0.083 | 1 | 0 | Hdac3 | 0 | Old |
| 0.964203 | 0.08877 | 0.07 | 0.07 | 1 | 0 | Metap1 | 0 | Old |
| 0.966904 | 0.015708 | 0.014 | 0.014 | 1 | 0 | Rasal3 | 0 | Old |
| 0.967752 | 0.127597 | 0.03 | 0.03 | 1 | 0 | Smpd4 | 0 | Old |
| 0.968985 | 0.00367 | 0.157 | 0.157 | 1 | 0 | Taf1 | 0 | Old |
| 0.969192 | 0.039167 | 0.053 | 0.054 | 1 | 0 | Exosc9 | -0.001 | Old |
| 0.969698 | 0.049545 | 0.025 | 0.024 | 1 | 0 | Cd2 | 0.001 | Old |
| 0.970944 | 0.210739 | 0.033 | 0.033 | 1 | 0 | Tada3 | 0 | Old |
| 0.971319 | 0.222833 | 0.023 | 0.024 | 1 | 0 | Gpn2 | -0.001 | Old |
| 0.971474 | 0.051684 | 0.034 | 0.035 | 1 | 0 | Usp31 | -0.001 | Old |
| 0.972262 | 0.079202 | 0.036 | 0.035 | 1 | 0 | Eef2kmt | 0.001 | Old |
| 0.972649 | 0.084129 | 0.025 | 0.024 | 1 | 0 | Smo | 0.001 | Old |
| 0.972909 | 0.009844 | 0.037 | 0.037 | 1 | 0 | Cnpy4 | 0 | Old |
| 0.973841 | 0.010639 | 0.447 | 0.447 | 1 | 0 | Eif3f | 0 | Old |
| 0.973967 | 0.334875 | 0.014 | 0.014 | 1 | 0 | Gstm5 | 0 | Old |
| 0.974429 | 0.183044 | 0.025 | 0.025 | 1 | 0 | Zhx3 | 0 | Old |
| 0.974516 | 0.199188 | 0.023 | 0.023 | 1 | 0 | Sh3d19 | 0 | Old |
| 0.976053 | 0.043899 | 0.045 | 0.045 | 1 | 0 | Bckdha | 0 | Old |
| 0.976615 | 0.024083 | 0.62 | 0.62 | 1 | 0 | Sh3bgrl | 0 | Old |
| 0.976875 | 0.020773 | 0.238 | 0.238 | 1 | 0 | Timm13 | 0 | Old |
| 0.97708 | 0.164465 | 0.019 | 0.019 | 1 | 0 | Csrp2 | 0 | Old |
| 0.977228 | 0.045993 | 0.01 | 0.01 | 1 | 0 | Nod2 | 0 | Old |
| 0.978291 | 0.040206 | 0.059 | 0.059 | 1 | 0 | Rnf170 | 0 | Old |
| 0.979449 | 0.08839 | 0.01 | 0.01 | 1 | 0 | Src | 0 | Old |
| 0.980332 | 0.138317 | 0.012 | 0.012 | 1 | 0 | Cox11 | 0 | Old |
| 0.980547 | 0.075024 | 0.651 | 0.642 | 1 | 0 | Ctsd | 0.009 | Old |
| 0.980925 | 0.241137 | 0.012 | 0.012 | 1 | 0 | Zfp280b | 0 | Old |
| 0.981077 | 0.00986 | 0.263 | 0.261 | 1 | 0 | Ankfy1 | 0.002 | Old |
| 0.981466 | 0.009919 | 0.047 | 0.047 | 1 | 0 | Angptl4 | 0 | Old |
| 0.98167 | 0.17633 | 0.01 | 0.01 | 1 | 0 | Cd320 | 0 | Old |
| 0.981892 | 0.178729 | 0.01 | 0.01 | 1 | 0 | Slc2a1 | 0 | Old |
| 0.982712 | 0.112316 | 0.134 | 0.136 | 1 | 0 | Eif2s3y | -0.002 | Old |
| 0.983635 | 0.009625 | 0.097 | 0.097 | 1 | 0 | Opa1 | 0 | Old |
| 0.983703 | 0.004517 | 0.066 | 0.065 | 1 | 0 | Prps2 | 0.001 | Old |
| 0.983867 | 0.012291 | 0.164 | 0.164 | 1 | 0 | Gak | 0 | Old |
| 0.983892 | 0.345202 | 0.01 | 0.01 | 1 | 0 | Ercc6 | 0 | Old |
| 0.984246 | 0.008285 | 0.047 | 0.047 | 1 | 0 | E4f1 | 0 | Old |
| 0.984535 | 0.124191 | 0.071 | 0.072 | 1 | 0 | Idi1 | -0.001 | Old |
| 0.984595 | 0.234668 | 0.018 | 0.018 | 1 | 0 | Mgp | 0 | Old |
| 0.984686 | 0.121358 | 0.12 | 0.121 | 1 | 0 | Arhgap19 | -0.001 | Old |
| 0.985188 | 0.035197 | 0.085 | 0.085 | 1 | 0 | Dph3 | 0 | Old |
| 0.985224 | 0.275203 | 0.03 | 0.03 | 1 | 0 | Ripk2 | 0 | Old |
| 0.986 | 0.158667 | 0.042 | 0.043 | 1 | 0 | Ttc19 | -0.001 | Old |
| 0.986271 | 0.030558 | 0.09 | 0.091 | 1 | 0 | Stim1 | -0.001 | Old |
| 0.987349 | 0.13673 | 0.064 | 0.064 | 1 | 0 | Psmd5 | 0 | Old |

| | | | | | | | | |
|---|---|---|---|---|---|---|---|---|
| 0.987629 | 0.06845 | 0.015 | 0.015 | 1 | 0 | Nme7 | 0 | Old |
| 0.988097 | 0.082622 | 0.055 | 0.055 | 1 | 0 | Brpf3 | 0 | Old |
| 0.989592 | 0.184272 | 0.096 | 0.096 | 1 | 0 | Slc16a6 | 0 | Old |
| 0.98989 | 0.096562 | 0.041 | 0.041 | 1 | 0 | Tpst1 | 0 | Old |
| 0.990278 | 0.002188 | 0.027 | 0.027 | 1 | 0 | Dalrd3 | 0 | Old |
| 0.993246 | 0.246643 | 0.016 | 0.016 | 1 | 0 | Srr | 0 | Old |
| 0.994313 | 0.236813 | 0.045 | 0.045 | 1 | 0 | Elp3 | 0 | Old |
| 0.997264 | 0.125366 | 0.016 | 0.016 | 1 | 0 | Fam222a | 0 | Old |
| 0.997326 | 0.008528 | 0.055 | 0.055 | 1 | 0 | Spic | 0 | Old |
| 0.997995 | 0.073333 | 0.022 | 0.022 | 1 | 0 | Pde6d | 0 | Old |
| 0.998016 | 0.29051 | 0.011 | 0.011 | 1 | 0 | Zfp119b | 0 | Old |
| 0.99829 | 0.049954 | 0.016 | 0.016 | 1 | 0 | Gm17586 | 0 | Old |
| 0.998412 | 0.073055 | 0.077 | 0.077 | 1 | 0 | Fnip2 | 0 | Old |
| 0.999029 | 0.025013 | 0.093 | 0.093 | 1 | 0 | Eftud2 | 0 | Old |
| 0.999581 | 0.000893 | 0.034 | 0.034 | 1 | 0 | Mkrn2 | 0 | Old |
| 0.999687 | 0.095147 | 0.011 | 0.011 | 1 | 0 | Rwdd2a | 0 | Old |
| 0.999896 | 0.073869 | 0.011 | 0.011 | 1 | 0 | Ska2 | 0 | Old |
| 2.02E-31 | 0.911136 | 0.725 | 0.524 | 6.27E-27 | 0 | Ccl24 | 0.201 | Young |
| 3.70E-24 | 0.336783 | 0.989 | 0.986 | 1.15E-19 | 0 | Cst3 | 0.003 | Young |
| 5.66E-24 | 0.4313 | 0.524 | 0.291 | 1.76E-19 | 0 | Hspa1b | 0.233 | Young |
| 1.29E-23 | 0.38421 | 0.955 | 0.907 | 4.00E-19 | 0 | Rps14 | 0.048 | Young |
| 8.09E-21 | 0.214119 | 0.999 | 1 | 2.51E-16 | 0 | Rps29 | -0.001 | Young |
| 1.02E-18 | 0.170781 | 1 | 1 | 3.18E-14 | 0 | Gm42418 | 0 | Young |
| 1.74E-16 | 0.231488 | 0.998 | 0.989 | 5.40E-12 | 0 | Selenop | 0.009 | Young |
| 1.09E-15 | 0.696837 | 0.558 | 0.408 | 3.40E-11 | 0 | Ednrb | 0.15 | Young |
| 1.81E-15 | 0.237286 | 0.986 | 0.978 | 5.62E-11 | 0 | Rplp1 | 0.008 | Young |
| 2.17E-15 | 0.394786 | 0.843 | 0.756 | 6.73E-11 | 0 | Igfbp4 | 0.087 | Young |
| 2.20E-15 | 0.405472 | 0.89 | 0.803 | 6.83E-11 | 0 | Retnla | 0.087 | Young |
| 1.79E-14 | 0.382121 | 0.541 | 0.371 | 5.57E-10 | 0 | Hspa1a | 0.17 | Young |
| 3.46E-14 | 0.617721 | 0.546 | 0.397 | 1.08E-09 | 0 | Ltc4s | 0.149 | Young |
| 7.83E-14 | 0.282852 | 0.973 | 0.956 | 2.43E-09 | 0 | F13a1 | 0.017 | Young |
| 1.15E-13 | 0.277011 | 0.984 | 0.975 | 3.58E-09 | 0 | Pf4 | 0.009 | Young |
| 3.51E-13 | 0.266033 | 0.991 | 0.971 | 1.09E-08 | 0 | Apoe | 0.02 | Young |
| 4.82E-12 | 0.308667 | 0.885 | 0.839 | 1.50E-07 | 0 | Csf1r | 0.046 | Young |
| 5.31E-12 | 0.766897 | 0.426 | 0.291 | 1.65E-07 | 0 | Hpgd | 0.135 | Young |
| 1.07E-11 | 0.266255 | 0.927 | 0.911 | 3.34E-07 | 0 | Rps20 | 0.016 | Young |
| 1.54E-11 | 0.283147 | 0.913 | 0.9 | 4.79E-07 | 0 | Rpl30 | 0.013 | Young |
| 1.57E-11 | 0.829677 | 0.669 | 0.58 | 4.87E-07 | 0 | Ighm | 0.089 | Young |
| 2.13E-11 | 0.321716 | 0.856 | 0.843 | 6.61E-07 | 0 | Tgfbi | 0.013 | Young |
| 2.20E-11 | 0.370474 | 0.797 | 0.763 | 6.84E-07 | 0 | Rpl34 | 0.034 | Young |
| 2.83E-11 | 2.350712 | 0.092 | 0.018 | 8.79E-07 | 0 | Colec12 | 0.074 | Young |
| 2.09E-10 | 0.231785 | 0.934 | 0.938 | 6.50E-06 | 0 | Rps23 | -0.004 | Young |
| 2.89E-10 | 0.288276 | 0.901 | 0.844 | 8.98E-06 | 0 | Cd36 | 0.057 | Young |
| 5.28E-10 | 0.379396 | 0.796 | 0.714 | 1.64E-05 | 0 | Folr2 | 0.082 | Young |
| 6.10E-10 | 0.278343 | 0.881 | 0.858 | 1.89E-05 | 0 | Rps4x | 0.023 | Young |

| | | | | | | | | |
|---|---|---|---|---|---|---|---|---|
| 6.27E-10 | 1.983472 | 0.016 | 0.057 | 1.95E-05 | 0 | Igkc | -0.041 | Young |
| 8.73E-10 | 0.484355 | 0.578 | 0.491 | 2.71E-05 | 0 | Gmfg | 0.087 | Young |
| 9.76E-10 | 0.347167 | 0.797 | 0.761 | 3.03E-05 | 0 | Rpl26 | 0.036 | Young |
| 1.49E-09 | 0.207238 | 0.973 | 0.966 | 4.63E-05 | 0 | Rps24 | 0.007 | Young |
| 2.74E-09 | 0.171826 | 0.985 | 0.978 | 8.50E-05 | 0 | Rpl35a | 0.007 | Young |
| 5.69E-09 | 0.793591 | 0.273 | 0.166 | 0.000177 | 0 | Sh3bp5 | 0.107 | Young |
| 7.21E-09 | 0.405301 | 0.647 | 0.588 | 0.000224 | 0 | Rpl35 | 0.059 | Young |
| 1.15E-08 | 0.203494 | 0.967 | 0.952 | 0.000356 | 0 | Fau | 0.015 | Young |
| 1.31E-08 | 0.195794 | 0.955 | 0.959 | 0.000408 | 0 | Rps8 | -0.004 | Young |
| 1.40E-08 | 0.317057 | 0.816 | 0.785 | 0.000436 | 0 | Rpl18a | 0.031 | Young |
| 2.98E-08 | 0.568146 | 0.472 | 0.371 | 0.000926 | 0 | Fgfr1 | 0.101 | Young |
| 5.80E-08 | 0.184576 | 0.961 | 0.958 | 0.0018 | 0 | Rps12 | 0.003 | Young |
| 6.98E-08 | 0.273958 | 0.717 | 0.591 | 0.002166 | 0 | Cd163 | 0.126 | Young |
| 7.13E-08 | 0.354717 | 0.628 | 0.536 | 0.002215 | 0 | Tsc22d3 | 0.092 | Young |
| 8.69E-08 | 0.257424 | 0.84 | 0.81 | 0.002699 | 0 | Rps13 | 0.03 | Young |
| 8.70E-08 | 0.15511 | 1 | 0.997 | 0.002702 | 0 | Malat1 | 0.003 | Young |
| 8.74E-08 | 0.156318 | 0.984 | 0.984 | 0.002713 | 0 | Ftl1 | 0 | Young |
| 9.70E-08 | 0.318152 | 0.735 | 0.679 | 0.003013 | 0 | Rpl32 | 0.056 | Young |
| 1.01E-07 | 0.370437 | 0.455 | 0.338 | 0.003131 | 0 | Cd209g | 0.117 | Young |
| 1.12E-07 | 0.222319 | 0.925 | 0.902 | 0.003472 | 0 | Rpl23 | 0.023 | Young |
| 1.44E-07 | 0.230662 | 0.892 | 0.829 | 0.004483 | 0 | Txnip | 0.063 | Young |
| 1.48E-07 | 0.214271 | 0.904 | 0.902 | 0.004605 | 0 | Rpl13 | 0.002 | Young |
| 1.65E-07 | 0.393215 | 0.594 | 0.472 | 0.005129 | 0 | Lyve1 | 0.122 | Young |
| 2.01E-07 | 0.292132 | 0.635 | 0.529 | 0.006232 | 0 | Rnase4 | 0.106 | Young |
| 3.27E-07 | 0.600531 | 0.436 | 0.343 | 0.010149 | 0 | Fcna | 0.093 | Young |
| 4.10E-07 | 0.364089 | 0.685 | 0.64 | 0.012728 | 0 | Rpl11 | 0.045 | Young |
| 5.40E-07 | 0.407313 | 0.43 | 0.316 | 0.016769 | 0 | Cd209d | 0.114 | Young |
| 5.62E-07 | 0.289181 | 0.781 | 0.732 | 0.017456 | 0 | Rpl18 | 0.049 | Young |
| 6.67E-07 | 0.166629 | 0.992 | 0.988 | 0.020709 | 0 | Fth1 | 0.004 | Young |
| 6.76E-07 | 0.272563 | 0.655 | 0.583 | 0.020998 | 0 | Ier3 | 0.072 | Young |
| 8.56E-07 | 1.135839 | 0.14 | 0.074 | 0.026584 | 0 | Gypc | 0.066 | Young |
| 1.83E-06 | 0.654007 | 0.297 | 0.213 | 0.056901 | 0 | Gpr34 | 0.084 | Young |
| 1.92E-06 | 2.881545 | 0.044 | 0.007 | 0.059726 | 0 | Tln2 | 0.037 | Young |
| 2.49E-06 | 0.687479 | 0.198 | 0.12 | 0.077436 | 0 | Slc40a1 | 0.078 | Young |
| 2.80E-06 | 0.406858 | 0.468 | 0.379 | 0.087093 | 0 | Gatm | 0.089 | Young |
| 2.87E-06 | 0.240994 | 0.785 | 0.724 | 0.088985 | 0 | Rsrp1 | 0.061 | Young |
| 3.81E-06 | 0.296305 | 0.697 | 0.665 | 0.118332 | 0 | Rpl21 | 0.032 | Young |
| 3.81E-06 | 0.274257 | 0.82 | 0.802 | 0.118333 | 0 | Fcgrt | 0.018 | Young |
| 3.91E-06 | 1.59411 | 0.086 | 0.036 | 0.121531 | 0 | Timd4 | 0.05 | Young |
| 4.68E-06 | 0.223006 | 0.587 | 0.472 | 0.145423 | 0 | Cd209f | 0.115 | Young |
| 4.94E-06 | 0.978862 | 0.21 | 0.14 | 0.153376 | 0 | Rasgrp3 | 0.07 | Young |
| 6.85E-06 | 0.489802 | 0.456 | 0.391 | 0.212664 | 0 | Pfdn5 | 0.065 | Young |
| 7.03E-06 | 0.295409 | 0.703 | 0.644 | 0.218402 | 0 | Mtss1 | 0.059 | Young |
| 7.21E-06 | 0.207298 | 0.867 | 0.869 | 0.223834 | 0 | Rps16 | -0.002 | Young |
| 9.32E-06 | 0.269469 | 0.743 | 0.679 | 0.289349 | 0 | Cfh | 0.064 | Young |

| | | | | | | | | |
|---|---|---|---|---|---|---|---|---|
| 1.91E-05 | 0.589261 | 0.288 | 0.213 | 0.593061 | 0 | Tanc2 | 0.075 | Young |
| 1.98E-05 | 0.306112 | 0.584 | 0.516 | 0.615966 | 0 | Atp5e | 0.068 | Young |
| 2.12E-05 | 0.595767 | 0.258 | 0.186 | 0.659246 | 0 | Sptbn1 | 0.072 | Young |
| 2.75E-05 | 0.409182 | 0.508 | 0.431 | 0.853825 | 0 | Tns1 | 0.077 | Young |
| 3.16E-05 | 1.243996 | 0.094 | 0.047 | 0.981705 | 0 | Tppp | 0.047 | Young |
| 3.34E-05 | 0.907604 | 0.145 | 0.088 | 1 | 0 | Car3 | 0.057 | Young |
| 3.52E-05 | 0.305521 | 0.639 | 0.609 | 1 | 0 | Rps3 | 0.03 | Young |
| 3.69E-05 | 0.765362 | 0.167 | 0.107 | 1 | 0 | Arsb | 0.06 | Young |
| 5.16E-05 | 1.70582 | 0.059 | 0.022 | 1 | 0 | Cfd | 0.037 | Young |
| 5.73E-05 | 1.277467 | 0.053 | 0.018 | 1 | 0 | P2ry13 | 0.035 | Young |
| 6.14E-05 | 0.492071 | 0.404 | 0.341 | 1 | 0 | Lipa | 0.063 | Young |
| 6.53E-05 | 0.310344 | 0.627 | 0.595 | 1 | 0 | Cox7c | 0.032 | Young |
| 6.94E-05 | 0.398659 | 0.715 | 0.683 | 1 | 0 | Wfdc17 | 0.032 | Young |
| 9.68E-05 | 0.403943 | 0.462 | 0.408 | 1 | 0 | Rgs10 | 0.054 | Young |
| 0.000126 | 0.220712 | 0.773 | 0.752 | 1 | 0 | Rps15a | 0.021 | Young |
| 0.00014 | 0.546642 | 0.299 | 0.235 | 1 | 0 | Jup | 0.064 | Young |
| 0.000146 | 1.059122 | 0.095 | 0.051 | 1 | 0 | Plxna4 | 0.044 | Young |
| 0.000163 | 0.540148 | 0.311 | 0.246 | 1 | 0 | Cmah | 0.065 | Young |
| 0.000176 | 3.866165 | 0.022 | 0.001 | 1 | 0 | Cyr61 | 0.021 | Young |
| 0.000205 | 0.585721 | 0.288 | 0.228 | 1 | 0 | Nfxl1 | 0.06 | Young |
| 0.00022 | 0.163309 | 0.88 | 0.881 | 1 | 0 | Rpl37 | -0.001 | Young |
| 0.00023 | 0.234274 | 0.687 | 0.647 | 1 | 0 | Rpl36 | 0.04 | Young |
| 0.00025 | 0.392123 | 0.421 | 0.364 | 1 | 0 | Blvrb | 0.057 | Young |
| 0.000256 | 0.192053 | 0.789 | 0.77 | 1 | 0 | Wwp1 | 0.019 | Young |
| 0.000263 | 0.253588 | 0.586 | 0.557 | 1 | 0 | Gng5 | 0.029 | Young |
| 0.000288 | 0.181631 | 0.864 | 0.825 | 1 | 0 | Gas6 | 0.039 | Young |
| 0.000292 | 0.297575 | 0.495 | 0.421 | 1 | 0 | Fxyd2 | 0.074 | Young |
| 0.000311 | 0.632827 | 0.181 | 0.126 | 1 | 0 | Ralgps2 | 0.055 | Young |
| 0.000312 | 0.257611 | 0.59 | 0.524 | 1 | 0 | Serpinb6a | 0.066 | Young |
| 0.000325 | 0.98632 | 0.097 | 0.055 | 1 | 0 | Nr4a3 | 0.042 | Young |
| 0.000335 | 0.192048 | 0.815 | 0.81 | 1 | 0 | Rpl19 | 0.005 | Young |
| 0.00035 | 0.248968 | 0.622 | 0.575 | 1 | 0 | Pepd | 0.047 | Young |
| 0.000462 | 0.482313 | 0.334 | 0.275 | 1 | 0 | Bin1 | 0.059 | Young |
| 0.000481 | 0.18569 | 0.818 | 0.807 | 1 | 0 | Sem1 | 0.011 | Young |
| 0.000506 | 0.222698 | 0.662 | 0.632 | 1 | 0 | Rpl10a | 0.03 | Young |
| 0.00052 | 0.754821 | 0.138 | 0.09 | 1 | 0 | Ctsf | 0.048 | Young |
| 0.000529 | 0.22036 | 0.651 | 0.601 | 1 | 0 | Zcchc6 | 0.05 | Young |
| 0.000531 | 1.202698 | 0.076 | 0.04 | 1 | 0 | Mtus1 | 0.036 | Young |
| 0.000548 | 0.388132 | 0.348 | 0.287 | 1 | 0 | Dek | 0.061 | Young |
| 0.000558 | 0.14532 | 0.687 | 0.627 | 1 | 0 | Stab1 | 0.06 | Young |
| 0.000581 | 0.170499 | 0.846 | 0.834 | 1 | 0 | Rps19 | 0.012 | Young |
| 0.000589 | 0.200611 | 0.801 | 0.776 | 1 | 0 | Tyrobp | 0.025 | Young |
| 0.000594 | 0.511808 | 0.179 | 0.124 | 1 | 0 | Gprc5c | 0.055 | Young |
| 0.000595 | 0.652582 | 0.237 | 0.185 | 1 | 0 | Tmem234 | 0.052 | Young |
| 0.000599 | 0.313705 | 0.547 | 0.506 | 1 | 0 | Mrfap1 | 0.041 | Young |

| | | | | | | | | |
|---|---|---|---|---|---|---|---|---|
| 0.00061 | 0.256379 | 0.639 | 0.596 | 1 | 0 | Eps15 | 0.043 | Young |
| 0.000646 | 0.169912 | 0.866 | 0.878 | 1 | 0 | Rpl17 | -0.012 | Young |
| 0.000664 | 0.350618 | 0.442 | 0.386 | 1 | 0 | Aldh2 | 0.056 | Young |
| 0.000678 | 0.46763 | 0.154 | 0.103 | 1 | 0 | Frmd6 | 0.051 | Young |
| 0.000731 | 0.257625 | 0.574 | 0.546 | 1 | 0 | Tln1 | 0.028 | Young |
| 0.000784 | 1.219905 | 0.066 | 0.033 | 1 | 0 | Tcf7l2 | 0.033 | Young |
| 0.000834 | 0.599058 | 0.163 | 0.112 | 1 | 0 | Tslp | 0.051 | Young |
| 0.000841 | 0.133956 | 0.929 | 0.907 | 1 | 0 | Maf | 0.022 | Young |
| 0.000861 | 0.25885 | 0.62 | 0.581 | 1 | 0 | Foxp1 | 0.039 | Young |
| 0.000877 | 0.235985 | 0.676 | 0.643 | 1 | 0 | Fam46a | 0.033 | Young |
| 0.000893 | 0.726958 | 0.141 | 0.094 | 1 | 0 | Cx3cr1 | 0.047 | Young |
| 0.00091 | 0.577606 | 0.256 | 0.202 | 1 | 0 | Rcn3 | 0.054 | Young |
| 0.001008 | 0.557454 | 0.199 | 0.146 | 1 | 0 | Ang | 0.053 | Young |
| 0.001096 | 0.277636 | 0.58 | 0.535 | 1 | 0 | Nrp1 | 0.045 | Young |
| 0.001119 | 0.765889 | 0.119 | 0.077 | 1 | 0 | Ripor2 | 0.042 | Young |
| 0.0012 | 0.195956 | 0.781 | 0.765 | 1 | 0 | Cybb | 0.016 | Young |
| 0.001217 | 0.890683 | 0.111 | 0.071 | 1 | 0 | Tmem229b | 0.04 | Young |
| 0.001262 | 0.323515 | 0.396 | 0.338 | 1 | 0 | Slc9a9 | 0.058 | Young |
| 0.001296 | 0.218367 | 0.715 | 0.694 | 1 | 0 | Rpl28 | 0.021 | Young |
| 0.001393 | 0.104791 | 0.959 | 0.953 | 1 | 0 | Rps21 | 0.006 | Young |
| 0.00153 | 0.848527 | 0.109 | 0.07 | 1 | 0 | Fam126b | 0.039 | Young |
| 0.001586 | 0.417193 | 0.287 | 0.231 | 1 | 0 | S1pr1 | 0.056 | Young |
| 0.001734 | 0.280166 | 0.434 | 0.378 | 1 | 0 | Cd33 | 0.056 | Young |
| 0.001742 | 0.273673 | 0.541 | 0.51 | 1 | 0 | Atp5h | 0.031 | Young |
| 0.001785 | 0.182071 | 0.749 | 0.732 | 1 | 0 | Calm2 | 0.017 | Young |
| 0.001815 | 0.140831 | 0.832 | 0.836 | 1 | 0 | Rpl23a | -0.004 | Young |
| 0.001834 | 0.387858 | 0.397 | 0.35 | 1 | 0 | Rnf150 | 0.047 | Young |
| 0.001868 | 0.150406 | 0.872 | 0.865 | 1 | 0 | Rpl39 | 0.007 | Young |
| 0.001925 | 0.227572 | 0.501 | 0.451 | 1 | 0 | Sema4a | 0.05 | Young |
| 0.001951 | 0.569334 | 0.203 | 0.157 | 1 | 0 | Rnf145 | 0.046 | Young |
| 0.002252 | 0.669783 | 0.111 | 0.073 | 1 | 0 | Rasgrp2 | 0.038 | Young |
| 0.002294 | 0.270573 | 0.507 | 0.462 | 1 | 0 | Rpl31 | 0.045 | Young |
| 0.002311 | 0.288795 | 0.573 | 0.555 | 1 | 0 | Abca9 | 0.018 | Young |
| 0.002368 | 1.597192 | 0.032 | 0.011 | 1 | 0 | Ccdc106 | 0.021 | Young |
| 0.002385 | 0.528789 | 0.314 | 0.269 | 1 | 0 | Dnajb14 | 0.045 | Young |
| 0.002392 | 0.239977 | 0.534 | 0.494 | 1 | 0 | Ccdc50 | 0.04 | Young |
| 0.002419 | 0.532429 | 0.154 | 0.109 | 1 | 0 | Fam118a | 0.045 | Young |
| 0.002467 | 0.179258 | 0.737 | 0.725 | 1 | 0 | Tcf4 | 0.012 | Young |
| 0.002489 | 0.409356 | 0.215 | 0.164 | 1 | 0 | Smim11 | 0.051 | Young |
| 0.002515 | 0.375887 | 0.178 | 0.129 | 1 | 0 | Cd209b | 0.049 | Young |
| 0.002753 | 0.148679 | 0.818 | 0.802 | 1 | 0 | Cd83 | 0.016 | Young |
| 0.002781 | 0.555501 | 0.253 | 0.205 | 1 | 0 | Dennd4a | 0.048 | Young |
| 0.002856 | 0.102371 | 0.95 | 0.959 | 1 | 0 | Rps3a1 | -0.009 | Young |
| 0.002925 | 0.170526 | 0.722 | 0.709 | 1 | 0 | Mef2c | 0.013 | Young |
| 0.002946 | 0.412343 | 0.347 | 0.3 | 1 | 0 | Ankrd44 | 0.047 | Young |

| | | | | | | | | |
|---|---|---|---|---|---|---|---|---|
| 0.002963 | 0.285617 | 0.524 | 0.484 | 1 | 0 | Nr3c1 | 0.04 | Young |
| 0.003086 | 0.742168 | 0.112 | 0.075 | 1 | 0 | Pcbd2 | 0.037 | Young |
| 0.003171 | 1.99908 | 0.02 | 0.004 | 1 | 0 | Itgal | 0.016 | Young |
| 0.003279 | 0.235772 | 0.561 | 0.52 | 1 | 0 | Pnrc1 | 0.041 | Young |
| 0.003313 | 1.449114 | 0.017 | 0.003 | 1 | 0 | Cox6b2 | 0.014 | Young |
| 0.003387 | 0.206954 | 0.695 | 0.687 | 1 | 0 | Rps6 | 0.008 | Young |
| 0.003427 | 1.867712 | 0.032 | 0.012 | 1 | 0 | Btnl2 | 0.02 | Young |
| 0.003451 | 0.345255 | 0.402 | 0.357 | 1 | 0 | Gadd45b | 0.045 | Young |
| 0.003492 | 0.51463 | 0.214 | 0.17 | 1 | 0 | Rfk | 0.044 | Young |
| 0.003882 | 0.154898 | 0.744 | 0.717 | 1 | 0 | Serf2 | 0.027 | Young |
| 0.003912 | 0.433154 | 0.244 | 0.196 | 1 | 0 | Arhgef3 | 0.048 | Young |
| 0.003986 | 0.257163 | 0.519 | 0.488 | 1 | 0 | Rpl15 | 0.031 | Young |
| 0.003997 | 0.62839 | 0.178 | 0.135 | 1 | 0 | Aaed1 | 0.043 | Young |
| 0.004156 | 0.265302 | 0.458 | 0.41 | 1 | 0 | Lrp6 | 0.048 | Young |
| 0.004171 | 0.446767 | 0.286 | 0.239 | 1 | 0 | Sesn1 | 0.047 | Young |
| 0.004224 | 0.162564 | 0.761 | 0.769 | 1 | 0 | Rps5 | -0.008 | Young |
| 0.004319 | 0.191037 | 0.678 | 0.663 | 1 | 0 | Atp6v0b | 0.015 | Young |
| 0.004341 | 0.68938 | 0.095 | 0.062 | 1 | 0 | Il6 | 0.033 | Young |
| 0.004393 | 0.140944 | 0.846 | 0.847 | 1 | 0 | Rps10 | -0.001 | Young |
| 0.004428 | 0.13747 | 0.779 | 0.756 | 1 | 0 | Rpl5 | 0.023 | Young |
| 0.004448 | 0.821409 | 0.056 | 0.03 | 1 | 0 | Cks2 | 0.026 | Young |
| 0.004749 | 0.632588 | 0.139 | 0.101 | 1 | 0 | Jade1 | 0.038 | Young |
| 0.004858 | 0.525836 | 0.216 | 0.172 | 1 | 0 | Mctp1 | 0.044 | Young |
| 0.005219 | 0.118071 | 0.326 | 0.264 | 1 | 0 | Cd209a | 0.062 | Young |
| 0.005263 | 0.335162 | 0.486 | 0.453 | 1 | 0 | Slc43a2 | 0.033 | Young |
| 0.005284 | 0.091575 | 0.39 | 0.323 | 1 | 0 | Il1b | 0.067 | Young |
| 0.005474 | 0.185957 | 0.393 | 0.339 | 1 | 0 | Cited2 | 0.054 | Young |
| 0.005723 | 0.155122 | 0.698 | 0.68 | 1 | 0 | Uqcrh | 0.018 | Young |
| 0.005869 | 0.478851 | 0.227 | 0.183 | 1 | 0 | Dnajc19 | 0.044 | Young |
| 0.005932 | 0.318729 | 0.281 | 0.231 | 1 | 0 | Use1 | 0.05 | Young |
| 0.005957 | 0.884984 | 0.045 | 0.022 | 1 | 0 | Enpep | 0.023 | Young |
| 0.006074 | 1.194028 | 0.073 | 0.045 | 1 | 0 | Tnfrsf25 | 0.028 | Young |
| 0.006196 | 1.266742 | 0.033 | 0.014 | 1 | 0 | Cpt2 | 0.019 | Young |
| 0.006478 | 0.749926 | 0.099 | 0.067 | 1 | 0 | Prpsap2 | 0.032 | Young |
| 0.00651 | 1.51376 | 0.045 | 0.023 | 1 | 0 | Aff3 | 0.022 | Young |
| 0.006773 | 0.736504 | 0.069 | 0.041 | 1 | 0 | Pank1 | 0.028 | Young |
| 0.006837 | 0.218593 | 0.661 | 0.662 | 1 | 0 | Rps7 | -0.001 | Young |
| 0.00707 | 0.438689 | 0.115 | 0.079 | 1 | 0 | Zmym2 | 0.036 | Young |
| 0.007101 | 1.277518 | 0.078 | 0.051 | 1 | 0 | Mamdc2 | 0.027 | Young |
| 0.007323 | 2.093919 | 0.013 | 0.001 | 1 | 0 | Il1rl2 | 0.012 | Young |
| 0.007516 | 0.475328 | 0.242 | 0.198 | 1 | 0 | Nrip1 | 0.044 | Young |
| 0.007573 | 2.265527 | 0.01 | 0 | 1 | 0 | Hp | 0.01 | Young |
| 0.007653 | 0.496838 | 0.165 | 0.124 | 1 | 0 | Lilrb4a | 0.041 | Young |
| 0.007801 | 1.145695 | 0.028 | 0.011 | 1 | 0 | Lrrc49 | 0.017 | Young |
| 0.00806 | 0.084165 | 0.946 | 0.952 | 1 | 0 | Rps27a | -0.006 | Young |

| | | | | | | | | |
|---|---|---|---|---|---|---|---|---|
| 0.008325 | 1.505577 | 0.03 | 0.012 | 1 | 0 | Xlr | 0.018 | Young |
| 0.008371 | 0.703152 | 0.124 | 0.09 | 1 | 0 | Kbtbd11 | 0.034 | Young |
| 0.008512 | 0.125086 | 0.912 | 0.896 | 1 | 0 | Atp2b1 | 0.016 | Young |
| 0.008684 | 0.417461 | 0.217 | 0.174 | 1 | 0 | Gpcpd1 | 0.043 | Young |
| 0.008887 | 0.228466 | 0.417 | 0.373 | 1 | 0 | Kitl | 0.044 | Young |
| 0.00897 | 0.163405 | 0.754 | 0.743 | 1 | 0 | Hexa | 0.011 | Young |
| 0.008992 | 0.763157 | 0.043 | 0.022 | 1 | 0 | Rbm12b1 | 0.021 | Young |
| 0.009156 | 0.477678 | 0.278 | 0.242 | 1 | 0 | B230219D2 | 0.036 | Young |
| 0.009529 | 0.747504 | 0.064 | 0.038 | 1 | 0 | Stxbp1 | 0.026 | Young |
| 0.009592 | 0.070874 | 0.985 | 0.981 | 1 | 0 | Rps9 | 0.004 | Young |
| 0.009645 | 0.712307 | 0.104 | 0.073 | 1 | 0 | Zfp592 | 0.031 | Young |
| 0.009806 | 0.439965 | 0.158 | 0.12 | 1 | 0 | Selenbp1 | 0.038 | Young |
| 0.010127 | 0.979726 | 0.037 | 0.018 | 1 | 0 | 5430427O1 | 0.019 | Young |
| 0.010145 | 0.272516 | 0.334 | 0.287 | 1 | 0 | Reps2 | 0.047 | Young |
| 0.010277 | 0.283934 | 0.468 | 0.436 | 1 | 0 | 1810037I1 | 0.032 | Young |
| 0.010379 | 1.273709 | 0.04 | 0.021 | 1 | 0 | Zfhx2 | 0.019 | Young |
| 0.010403 | 1.733545 | 0.019 | 0.005 | 1 | 0 | Slc22a18 | 0.014 | Young |
| 0.010874 | 1.299123 | 0.023 | 0.008 | 1 | 0 | Cmc4 | 0.015 | Young |
| 0.010989 | 1.248096 | 0.021 | 0.007 | 1 | 0 | Kiz | 0.014 | Young |
| 0.011021 | 0.20015 | 0.603 | 0.587 | 1 | 0 | Rpl12 | 0.016 | Young |
| 0.011174 | 0.150438 | 0.78 | 0.754 | 1 | 0 | Cbr2 | 0.026 | Young |
| 0.012115 | 0.295327 | 0.469 | 0.447 | 1 | 0 | Sf3b1 | 0.022 | Young |
| 0.012404 | 0.701612 | 0.102 | 0.073 | 1 | 0 | Tmod1 | 0.029 | Young |
| 0.012439 | 1.022823 | 0.04 | 0.021 | 1 | 0 | Mark4 | 0.019 | Young |
| 0.012504 | 0.676735 | 0.031 | 0.014 | 1 | 0 | Tbkbp1 | 0.017 | Young |
| 0.012975 | 0.405596 | 0.147 | 0.111 | 1 | 0 | Il16 | 0.036 | Young |
| 0.013244 | 0.638988 | 0.134 | 0.101 | 1 | 0 | Chchd7 | 0.033 | Young |
| 0.013288 | 1.032133 | 0.047 | 0.026 | 1 | 0 | E130307A1 | 0.021 | Young |
| 0.013301 | 0.177964 | 0.574 | 0.534 | 1 | 0 | Stard8 | 0.04 | Young |
| 0.013574 | 0.580003 | 0.122 | 0.09 | 1 | 0 | Trim26 | 0.032 | Young |
| 0.013649 | 0.402607 | 0.158 | 0.123 | 1 | 0 | Tbc1d4 | 0.035 | Young |
| 0.013652 | 1.075041 | 0.027 | 0.011 | 1 | 0 | Aven | 0.016 | Young |
| 0.014059 | 0.313219 | 0.303 | 0.261 | 1 | 0 | Per1 | 0.042 | Young |
| 0.014164 | 0.100548 | 0.904 | 0.915 | 1 | 0 | Rpl7 | -0.011 | Young |
| 0.014415 | 0.512776 | 0.024 | 0.01 | 1 | 0 | Tuba4a | 0.014 | Young |
| 0.014492 | 0.350722 | 0.149 | 0.112 | 1 | 0 | Pgm2l1 | 0.037 | Young |
| 0.014535 | 0.232896 | 0.468 | 0.438 | 1 | 0 | Fkbp1a | 0.03 | Young |
| 0.014666 | 1.699101 | 0.013 | 0.003 | 1 | 0 | Lrfn1 | 0.01 | Young |
| 0.014693 | 0.724385 | 0.101 | 0.073 | 1 | 0 | Adam33 | 0.028 | Young |
| 0.014855 | 0.354657 | 0.259 | 0.219 | 1 | 0 | Chp2 | 0.04 | Young |
| 0.015315 | 0.617277 | 0.108 | 0.078 | 1 | 0 | Wdr36 | 0.03 | Young |
| 0.015605 | 1.110447 | 0.026 | 0.011 | 1 | 0 | Tfap4 | 0.015 | Young |
| 0.015753 | 0.769813 | 0.048 | 0.027 | 1 | 0 | Cxxc5 | 0.021 | Young |
| 0.015981 | 0.787772 | 0.048 | 0.027 | 1 | 0 | Polr3b | 0.021 | Young |
| 0.015984 | 0.653501 | 0.053 | 0.031 | 1 | 0 | Pxdc1 | 0.022 | Young |

| | | | | | | | | |
|---|---|---|---|---|---|---|---|---|
| 0.016028 | 1.915598 | 0.016 | 0.004 | 1 | 0 | Spp1 | 0.012 | Young |
| 0.016599 | 0.909249 | 0.064 | 0.041 | 1 | 0 | Ramp1 | 0.023 | Young |
| 0.01674 | 0.795084 | 0.108 | 0.079 | 1 | 0 | Prkar2b | 0.029 | Young |
| 0.017112 | 0.644467 | 0.106 | 0.077 | 1 | 0 | Tagap | 0.029 | Young |
| 0.017384 | 0.822552 | 0.02 | 0.007 | 1 | 0 | 2010320M | 0.013 | Young |
| 0.017408 | 1.686995 | 0.011 | 0.001 | 1 | 0 | Nhej1 | 0.01 | Young |
| 0.017711 | 0.131554 | 0.746 | 0.722 | 1 | 0 | Cox4i1 | 0.024 | Young |
| 0.01784 | 0.674943 | 0.124 | 0.094 | 1 | 0 | Rnf2 | 0.03 | Young |
| 0.018 | 0.333288 | 0.28 | 0.244 | 1 | 0 | Ncoa4 | 0.036 | Young |
| 0.018054 | 0.896499 | 0.051 | 0.03 | 1 | 0 | Cela1 | 0.021 | Young |
| 0.018073 | 0.664803 | 0.213 | 0.182 | 1 | 0 | Smc3 | 0.031 | Young |
| 0.018452 | 0.715894 | 0.061 | 0.038 | 1 | 0 | Mrpl27 | 0.023 | Young |
| 0.01854 | 1.412353 | 0.024 | 0.01 | 1 | 0 | Zfp74 | 0.014 | Young |
| 0.018674 | 1.778021 | 0.015 | 0.004 | 1 | 0 | Caprin2 | 0.011 | Young |
| 0.018747 | 0.269008 | 0.33 | 0.29 | 1 | 0 | Smdt1 | 0.04 | Young |
| 0.018932 | 0.090001 | 0.831 | 0.832 | 1 | 0 | Hsp90ab1 | -0.001 | Young |
| 0.019031 | 1.47527 | 0.021 | 0.008 | 1 | 0 | Retn | 0.013 | Young |
| 0.019186 | 0.780038 | 0.024 | 0.01 | 1 | 0 | Dars2 | 0.014 | Young |
| 0.019423 | 0.37788 | 0.193 | 0.156 | 1 | 0 | Arpc1a | 0.037 | Young |
| 0.019424 | 0.42922 | 0.125 | 0.093 | 1 | 0 | Gtf2i | 0.032 | Young |
| 0.019653 | 0.422012 | 0.205 | 0.168 | 1 | 0 | Tbl1x | 0.037 | Young |
| 0.019797 | 0.063593 | 0.35 | 0.293 | 1 | 0 | Pla2g2d | 0.057 | Young |
| 0.019974 | 0.342341 | 0.257 | 0.219 | 1 | 0 | Adap2 | 0.038 | Young |
| 0.020298 | 0.504992 | 0.123 | 0.093 | 1 | 0 | Ammecr1l | 0.03 | Young |
| 0.020349 | 0.114672 | 0.861 | 0.854 | 1 | 0 | Rps27 | 0.007 | Young |
| 0.02063 | 1.578547 | 0.013 | 0.003 | 1 | 0 | C2 | 0.01 | Young |
| 0.020725 | 0.060035 | 0.914 | 0.908 | 1 | 0 | Kctd12 | 0.006 | Young |
| 0.020878 | 0.123619 | 0.89 | 0.895 | 1 | 0 | Timp2 | -0.005 | Young |
| 0.020949 | 1.109784 | 0.063 | 0.041 | 1 | 0 | Plekhg2 | 0.022 | Young |
| 0.021055 | 0.541436 | 0.119 | 0.089 | 1 | 0 | Mrpl54 | 0.03 | Young |
| 0.021232 | 1.403187 | 0.043 | 0.025 | 1 | 0 | Zfp687 | 0.018 | Young |
| 0.021334 | 0.094822 | 0.821 | 0.818 | 1 | 0 | Rpl22 | 0.003 | Young |
| 0.021632 | 0.396402 | 0.293 | 0.26 | 1 | 0 | Ttc3 | 0.033 | Young |
| 0.02221 | 0.441052 | 0.111 | 0.082 | 1 | 0 | Cntrl | 0.029 | Young |
| 0.022434 | 0.587965 | 0.127 | 0.098 | 1 | 0 | Ccrl2 | 0.029 | Young |
| 0.023141 | 0.314721 | 0.317 | 0.28 | 1 | 0 | Srp14 | 0.037 | Young |
| 0.023433 | 0.998188 | 0.025 | 0.011 | 1 | 0 | Kat8 | 0.014 | Young |
| 0.023647 | 0.991793 | 0.046 | 0.027 | 1 | 0 | Gbp5 | 0.019 | Young |
| 0.023824 | 0.376042 | 0.256 | 0.219 | 1 | 0 | Etv1 | 0.037 | Young |
| 0.024279 | 0.389477 | 0.283 | 0.252 | 1 | 0 | Mrps21 | 0.031 | Young |
| 0.024519 | 0.66554 | 0.093 | 0.067 | 1 | 0 | Cdc14b | 0.026 | Young |
| 0.024625 | 0.348619 | 0.206 | 0.171 | 1 | 0 | Igf1r | 0.035 | Young |
| 0.025299 | 0.509943 | 0.159 | 0.127 | 1 | 0 | Mgst1 | 0.032 | Young |
| 0.025336 | 0.253333 | 0.261 | 0.226 | 1 | 0 | Fabp4 | 0.035 | Young |
| 0.025603 | 0.462768 | 0.246 | 0.216 | 1 | 0 | Elmo1 | 0.03 | Young |

| | | | | | | | | |
|---|---|---|---|---|---|---|---|---|
| 0.025773 | 0.268888 | 0.34 | 0.298 | 1 | 0 | Ddx3y | 0.042 | Young |
| 0.025853 | 1.446997 | 0.014 | 0.004 | 1 | 0 | Rnf26 | 0.01 | Young |
| 0.025863 | 0.368987 | 0.171 | 0.137 | 1 | 0 | Dstyk | 0.034 | Young |
| 0.02587 | 1.543102 | 0.014 | 0.004 | 1 | 0 | 5430403G1 | 0.01 | Young |
| 0.025997 | 0.427051 | 0.156 | 0.123 | 1 | 0 | Orai3 | 0.033 | Young |
| 0.026046 | 0.644979 | 0.056 | 0.036 | 1 | 0 | Lix1l | 0.02 | Young |
| 0.026451 | 0.277137 | 0.288 | 0.249 | 1 | 0 | Agtrap | 0.039 | Young |
| 0.026671 | 0.49061 | 0.061 | 0.04 | 1 | 0 | Epn2 | 0.021 | Young |
| 0.026708 | 0.243017 | 0.491 | 0.465 | 1 | 0 | Raph1 | 0.026 | Young |
| 0.026841 | 0.186233 | 0.559 | 0.538 | 1 | 0 | Rps18 | 0.021 | Young |
| 0.026907 | 0.114142 | 0.529 | 0.49 | 1 | 0 | Nfkbid | 0.039 | Young |
| 0.027406 | 0.17473 | 0.631 | 0.629 | 1 | 0 | Rpl10 | 0.002 | Young |
| 0.027427 | 0.677026 | 0.049 | 0.03 | 1 | 0 | Fra10ac1 | 0.019 | Young |
| 0.028289 | 0.056886 | 0.994 | 0.99 | 1 | 0 | Tpt1 | 0.004 | Young |
| 0.028336 | 0.344731 | 0.31 | 0.278 | 1 | 0 | Mier1 | 0.032 | Young |
| 0.028937 | 0.385303 | 0.175 | 0.142 | 1 | 0 | Pink1 | 0.033 | Young |
| 0.029125 | 1.029379 | 0.042 | 0.025 | 1 | 0 | Fam207a | 0.017 | Young |
| 0.029305 | 0.193333 | 0.454 | 0.428 | 1 | 0 | Tomm7 | 0.026 | Young |
| 0.029481 | 0.888508 | 0.026 | 0.012 | 1 | 0 | Zfp952 | 0.014 | Young |
| 0.029498 | 0.44417 | 0.126 | 0.097 | 1 | 0 | Cyp27a1 | 0.029 | Young |
| 0.029963 | 0.34584 | 0.073 | 0.049 | 1 | 0 | Ptp4a3 | 0.024 | Young |
| 0.02997 | 0.297527 | 0.217 | 0.181 | 1 | 0 | Cbfa2t3 | 0.036 | Young |
| 0.030039 | 1.020967 | 0.033 | 0.018 | 1 | 0 | Mettl7a1 | 0.015 | Young |
| 0.030077 | 0.221914 | 0.248 | 0.207 | 1 | 0 | Ecm1 | 0.041 | Young |
| 0.030246 | 1.001571 | 0.033 | 0.018 | 1 | 0 | Bcl2a1a | 0.015 | Young |
| 0.030328 | 0.844523 | 0.045 | 0.027 | 1 | 0 | Cebpzos | 0.018 | Young |
| 0.030412 | 0.344297 | 0.353 | 0.328 | 1 | 0 | Mapk14 | 0.025 | Young |
| 0.030416 | 0.072086 | 0.886 | 0.888 | 1 | 0 | Rps15 | -0.002 | Young |
| 0.030447 | 0.52545 | 0.077 | 0.053 | 1 | 0 | Sp3os | 0.024 | Young |
| 0.030454 | 0.393908 | 0.28 | 0.25 | 1 | 0 | Rb1cc1 | 0.03 | Young |
| 0.030579 | 0.879333 | 0.033 | 0.018 | 1 | 0 | Uaca | 0.015 | Young |
| 0.030879 | 0.897005 | 0.052 | 0.033 | 1 | 0 | Stoml1 | 0.019 | Young |
| 0.030978 | 1.025591 | 0.018 | 0.007 | 1 | 0 | Dnmt3b | 0.011 | Young |
| 0.031002 | 0.240878 | 0.373 | 0.339 | 1 | 0 | Rin2 | 0.034 | Young |
| 0.031165 | 1.254907 | 0.029 | 0.015 | 1 | 0 | Tspan13 | 0.014 | Young |
| 0.031399 | 0.925303 | 0.016 | 0.005 | 1 | 0 | Fgf9 | 0.011 | Young |
| 0.031542 | 0.267849 | 0.454 | 0.435 | 1 | 0 | Hint1 | 0.019 | Young |
| 0.031555 | 0.451929 | 0.175 | 0.144 | 1 | 0 | Mpc2 | 0.031 | Young |
| 0.032058 | 0.544469 | 0.059 | 0.038 | 1 | 0 | Ap3s2 | 0.021 | Young |
| 0.032429 | 0.738667 | 0.075 | 0.052 | 1 | 0 | Zcchc14 | 0.023 | Young |
| 0.032469 | 0.357209 | 0.302 | 0.275 | 1 | 0 | Entpd1 | 0.027 | Young |
| 0.033148 | 0.31777 | 0.174 | 0.141 | 1 | 0 | Pdgfc | 0.033 | Young |
| 0.033293 | 2.014542 | 0.011 | 0.003 | 1 | 0 | Fam109b | 0.008 | Young |
| 0.033314 | 0.897671 | 0.026 | 0.012 | 1 | 0 | Ankrd52 | 0.014 | Young |
| 0.033411 | 1.949115 | 0.011 | 0.003 | 1 | 0 | Napb | 0.008 | Young |

| | | | | | | | | |
|---|---|---|---|---|---|---|---|---|
| 0.033615 | 1.469405 | 0.02 | 0.008 | 1 | 0 | Ap4b1 | 0.012 | Young |
| 0.033677 | 0.640446 | 0.043 | 0.026 | 1 | 0 | Bet1l | 0.017 | Young |
| 0.033802 | 0.300471 | 0.413 | 0.393 | 1 | 0 | Rell1 | 0.02 | Young |
| 0.033857 | 0.297465 | 0.348 | 0.317 | 1 | 0 | 9930111J2 | 0.031 | Young |
| 0.033875 | 0.667524 | 0.099 | 0.074 | 1 | 0 | Adcy9 | 0.025 | Young |
| 0.034099 | 0.851522 | 0.011 | 0.003 | 1 | 0 | Mcemp1 | 0.008 | Young |
| 0.034167 | 0.79123 | 0.058 | 0.038 | 1 | 0 | Faf1 | 0.02 | Young |
| 0.034697 | 0.990391 | 0.051 | 0.033 | 1 | 0 | Abtb1 | 0.018 | Young |
| 0.034756 | 0.503604 | 0.154 | 0.124 | 1 | 0 | Pdcd4 | 0.03 | Young |
| 0.034893 | 0.434911 | 0.174 | 0.144 | 1 | 0 | Gm10073 | 0.03 | Young |
| 0.035241 | 0.876738 | 0.051 | 0.033 | 1 | 0 | Dhx8 | 0.018 | Young |
| 0.035992 | 0.292895 | 0.238 | 0.204 | 1 | 0 | Adrb2 | 0.034 | Young |
| 0.036114 | 0.252572 | 0.128 | 0.1 | 1 | 0 | Arhgap5 | 0.028 | Young |
| 0.036297 | 0.674162 | 0.077 | 0.055 | 1 | 0 | Zfp945 | 0.022 | Young |
| 0.036511 | 0.855871 | 0.029 | 0.015 | 1 | 0 | Mturn | 0.014 | Young |
| 0.036668 | 0.13467 | 0.644 | 0.625 | 1 | 0 | Eef1b2 | 0.019 | Young |
| 0.036669 | 0.428325 | 0.152 | 0.123 | 1 | 0 | Rab1b | 0.029 | Young |
| 0.036685 | 0.541073 | 0.1 | 0.075 | 1 | 0 | Cmc1 | 0.025 | Young |
| 0.036876 | 0.448342 | 0.116 | 0.089 | 1 | 0 | Eef2k | 0.027 | Young |
| 0.037908 | 0.682209 | 0.048 | 0.03 | 1 | 0 | Tfpi | 0.018 | Young |
| 0.038615 | 0.442864 | 0.202 | 0.174 | 1 | 0 | Vapb | 0.028 | Young |
| 0.0387 | 0.430178 | 0.107 | 0.081 | 1 | 0 | Tceal8 | 0.026 | Young |
| 0.03881 | 0.400698 | 0.274 | 0.246 | 1 | 0 | Hmox2 | 0.028 | Young |
| 0.039024 | 0.766867 | 0.032 | 0.018 | 1 | 0 | Tsen15 | 0.014 | Young |
| 0.039049 | 1.255162 | 0.019 | 0.008 | 1 | 0 | Zkscan14 | 0.011 | Young |
| 0.039965 | 0.118114 | 0.77 | 0.774 | 1 | 0 | Rps25 | -0.004 | Young |
| 0.040426 | 0.940416 | 0.043 | 0.026 | 1 | 0 | Tle1 | 0.017 | Young |
| 0.040524 | 1.055547 | 0.051 | 0.033 | 1 | 0 | Gm38843 | 0.018 | Young |
| 0.040767 | 1.425232 | 0.017 | 0.007 | 1 | 0 | Hexdc | 0.01 | Young |
| 0.041006 | 0.185994 | 0.63 | 0.627 | 1 | 0 | Tmcc1 | 0.003 | Young |
| 0.04128 | 0.370246 | 0.269 | 0.241 | 1 | 0 | Cxcl12 | 0.028 | Young |
| 0.041421 | 0.352394 | 0.181 | 0.15 | 1 | 0 | Rcsd1 | 0.031 | Young |
| 0.041777 | 1.066729 | 0.03 | 0.016 | 1 | 0 | Cdkn1c | 0.014 | Young |
| 0.04199 | 1.266593 | 0.015 | 0.005 | 1 | 0 | Slc8a1 | 0.01 | Young |
| 0.042274 | 1.283805 | 0.042 | 0.026 | 1 | 0 | Pam16 | 0.016 | Young |
| 0.042346 | 0.246012 | 0.39 | 0.369 | 1 | 0 | Minos1 | 0.021 | Young |
| 0.044035 | 0.218868 | 0.417 | 0.39 | 1 | 0 | Sec14l1 | 0.027 | Young |
| 0.044867 | 0.732447 | 0.086 | 0.064 | 1 | 0 | Srp68 | 0.022 | Young |
| 0.04506 | 0.773257 | 0.035 | 0.021 | 1 | 0 | Aifm1 | 0.014 | Young |
| 0.045677 | 0.707659 | 0.049 | 0.031 | 1 | 0 | Peli2 | 0.018 | Young |
| 0.045989 | 0.692101 | 0.039 | 0.023 | 1 | 0 | Cd24a | 0.016 | Young |
| 0.04599 | 0.960486 | 0.019 | 0.008 | 1 | 0 | Tmem206 | 0.011 | Young |
| 0.046099 | 0.387584 | 0.105 | 0.079 | 1 | 0 | Ckap5 | 0.026 | Young |
| 0.046355 | 0.143223 | 0.633 | 0.627 | 1 | 0 | Snx5 | 0.006 | Young |
| 0.047403 | 1.239359 | 0.017 | 0.007 | 1 | 0 | Ccl22 | 0.01 | Young |

| | | | | | | | | |
|---|---|---|---|---|---|---|---|---|
| 0.047587 | 0.1065 | 0.864 | 0.866 | 1 | 0 | Rpl27a | -0.002 | Young |
| 0.047671 | 0.296133 | 0.356 | 0.327 | 1 | 0 | Ifi203 | 0.029 | Young |
| 0.047881 | 0.468032 | 0.165 | 0.138 | 1 | 0 | Gabarapl1 | 0.027 | Young |
| 0.047961 | 0.905696 | 0.055 | 0.037 | 1 | 0 | Acvr1 | 0.018 | Young |
| 0.047993 | 0.212929 | 0.155 | 0.124 | 1 | 0 | Eapp | 0.031 | Young |
| 0.04803 | 0.782744 | 0.017 | 0.007 | 1 | 0 | Rgs7bp | 0.01 | Young |
| 0.048474 | 0.896548 | 0.024 | 0.012 | 1 | 0 | Rabep2 | 0.012 | Young |
| 0.048527 | 1.564568 | 0.015 | 0.005 | 1 | 0 | Bcr | 0.01 | Young |
| 0.048557 | 0.293323 | 0.321 | 0.29 | 1 | 0 | Rel | 0.031 | Young |
| 0.048722 | 1.453212 | 0.013 | 0.004 | 1 | 0 | Dhrs11 | 0.009 | Young |
| 0.048741 | 1.151036 | 0.021 | 0.01 | 1 | 0 | Dcbld2 | 0.011 | Young |
| 0.048752 | 1.572693 | 0.013 | 0.004 | 1 | 0 | Dtd1 | 0.009 | Young |
| 0.048779 | 1.204533 | 0.015 | 0.005 | 1 | 0 | Gm44067 | 0.01 | Young |
| 0.049057 | 1.361158 | 0.013 | 0.004 | 1 | 0 | A430033K( | 0.009 | Young |
| 0.049517 | 1.049124 | 0.013 | 0.004 | 1 | 0 | Tgif2 | 0.009 | Young |
| 0.049601 | 0.228354 | 0.37 | 0.341 | 1 | 0 | Rab31 | 0.029 | Young |
| 0.049617 | 0.24787 | 0.34 | 0.312 | 1 | 0 | Lamtor4 | 0.028 | Young |
| 0.049738 | 0.362097 | 0.193 | 0.164 | 1 | 0 | Zfp644 | 0.029 | Young |
| 0.049978 | 0.504053 | 0.111 | 0.086 | 1 | 0 | Ccdc186 | 0.025 | Young |
| 0.050453 | 0.408618 | 0.134 | 0.107 | 1 | 0 | Slc38a6 | 0.027 | Young |
| 0.050463 | 0.67481 | 0.033 | 0.019 | 1 | 0 | Ptpn21 | 0.014 | Young |
| 0.050815 | 0.674101 | 0.101 | 0.078 | 1 | 0 | Cox16 | 0.023 | Young |
| 0.051474 | 0.328322 | 0.163 | 0.134 | 1 | 0 | Acss1 | 0.029 | Young |
| 0.05167 | 1.2374 | 0.019 | 0.008 | 1 | 0 | Immp2l | 0.011 | Young |
| 0.051847 | 1.098593 | 0.029 | 0.016 | 1 | 0 | Wfdc18 | 0.013 | Young |
| 0.052006 | 0.748123 | 0.028 | 0.015 | 1 | 0 | Barx2 | 0.013 | Young |
| 0.052051 | 0.497426 | 0.08 | 0.059 | 1 | 0 | Lap3 | 0.021 | Young |
| 0.05238 | 0.191457 | 0.401 | 0.375 | 1 | 0 | Sult1a1 | 0.026 | Young |
| 0.052529 | 0.492844 | 0.077 | 0.056 | 1 | 0 | Mpp5 | 0.021 | Young |
| 0.052601 | 0.138293 | 0.65 | 0.655 | 1 | 0 | Rpl14 | -0.005 | Young |
| 0.053375 | 0.370046 | 0.326 | 0.305 | 1 | 0 | Mxd4 | 0.021 | Young |
| 0.054167 | 0.274624 | 0.187 | 0.156 | 1 | 0 | Smagp | 0.031 | Young |
| 0.054277 | 0.465571 | 0.126 | 0.101 | 1 | 0 | Dnajc1 | 0.025 | Young |
| 0.055302 | 0.403378 | 0.173 | 0.145 | 1 | 0 | Pik3c2a | 0.028 | Young |
| 0.055598 | 0.416767 | 0.064 | 0.045 | 1 | 0 | Exosc8 | 0.019 | Young |
| 0.055695 | 1.356449 | 0.01 | 0.003 | 1 | 0 | Gm26756 | 0.007 | Young |
| 0.055974 | 0.312938 | 0.312 | 0.289 | 1 | 0 | H2afj | 0.023 | Young |
| 0.056037 | 0.265206 | 0.099 | 0.075 | 1 | 0 | Smim4 | 0.024 | Young |
| 0.056175 | 0.058974 | 0.875 | 0.855 | 1 | 0 | Pltp | 0.02 | Young |
| 0.057607 | 0.166562 | 0.587 | 0.579 | 1 | 0 | Ptpn18 | 0.008 | Young |
| 0.057718 | 0.675487 | 0.038 | 0.023 | 1 | 0 | Otud1 | 0.015 | Young |
| 0.058441 | 0.663018 | 0.075 | 0.055 | 1 | 0 | Rfx7 | 0.02 | Young |
| 0.058484 | 0.679992 | 0.078 | 0.057 | 1 | 0 | Ints3 | 0.021 | Young |
| 0.058676 | 1.236666 | 0.022 | 0.011 | 1 | 0 | Ftsj1 | 0.011 | Young |
| 0.058771 | 0.190199 | 0.031 | 0.018 | 1 | 0 | Napsa | 0.013 | Young |

| | | | | | | | | |
|---|---|---|---|---|---|---|---|---|
| 0.059034 | 0.313695 | 0.056 | 0.038 | 1 | 0 | Cgrrf1 | 0.018 | Young |
| 0.059178 | 0.440119 | 0.134 | 0.108 | 1 | 0 | Foxred2 | 0.026 | Young |
| 0.05939 | 0.506114 | 0.062 | 0.044 | 1 | 0 | AI504432 | 0.018 | Young |
| 0.059691 | 0.424983 | 0.163 | 0.137 | 1 | 0 | Phactr2 | 0.026 | Young |
| 0.059739 | 0.417262 | 0.117 | 0.093 | 1 | 0 | Ipo11 | 0.024 | Young |
| 0.059816 | 0.894779 | 0.029 | 0.016 | 1 | 0 | Slx1b | 0.013 | Young |
| 0.059846 | 1.166828 | 0.018 | 0.008 | 1 | 0 | Zfp974 | 0.01 | Young |
| 0.059939 | 0.770068 | 0.048 | 0.031 | 1 | 0 | 2010315B( | 0.017 | Young |
| 0.06065 | 0.779385 | 0.085 | 0.064 | 1 | 0 | Rabgap1l | 0.021 | Young |
| 0.060802 | 0.269103 | 0.394 | 0.369 | 1 | 0 | Cmtm6 | 0.025 | Young |
| 0.061445 | 0.500909 | 0.161 | 0.137 | 1 | 0 | Casc3 | 0.024 | Young |
| 0.061467 | 0.650428 | 0.018 | 0.008 | 1 | 0 | Mvb12b | 0.01 | Young |
| 0.061821 | 0.711598 | 0.036 | 0.022 | 1 | 0 | Cry2 | 0.014 | Young |
| 0.062027 | 0.672656 | 0.031 | 0.018 | 1 | 0 | Rfc3 | 0.013 | Young |
| 0.062051 | 0.720103 | 0.072 | 0.053 | 1 | 0 | Kif23 | 0.019 | Young |
| 0.062452 | 0.285966 | 0.138 | 0.112 | 1 | 0 | Npl | 0.026 | Young |
| 0.062784 | 1.42038 | 0.016 | 0.007 | 1 | 0 | Zbtb26 | 0.009 | Young |
| 0.063126 | 0.208723 | 0.164 | 0.135 | 1 | 0 | C1qtnf1 | 0.029 | Young |
| 0.063331 | 0.586924 | 0.068 | 0.049 | 1 | 0 | Vps37b | 0.019 | Young |
| 0.063382 | 0.246317 | 0.375 | 0.358 | 1 | 0 | Atxn7l3b | 0.017 | Young |
| 0.063614 | 0.235965 | 0.361 | 0.335 | 1 | 0 | Pnisr | 0.026 | Young |
| 0.063721 | 0.473409 | 0.118 | 0.094 | 1 | 0 | Mrpl30 | 0.024 | Young |
| 0.063849 | 0.607577 | 0.123 | 0.1 | 1 | 0 | Srek1ip1 | 0.023 | Young |
| 0.064154 | 0.257155 | 0.22 | 0.189 | 1 | 0 | Prune2 | 0.031 | Young |
| 0.064355 | 0.265858 | 0.264 | 0.233 | 1 | 0 | Idh2 | 0.031 | Young |
| 0.065077 | 0.584943 | 0.089 | 0.068 | 1 | 0 | Irak3 | 0.021 | Young |
| 0.065136 | 0.485667 | 0.046 | 0.03 | 1 | 0 | Pmepa1 | 0.016 | Young |
| 0.065586 | 0.458397 | 0.193 | 0.168 | 1 | 0 | Cdc37 | 0.025 | Young |
| 0.065627 | 0.025191 | 0.128 | 0.159 | 1 | 0 | Dtx3l | -0.031 | Young |
| 0.06573 | 0.880473 | 0.027 | 0.015 | 1 | 0 | Zfp874a | 0.012 | Young |
| 0.065919 | 1.266567 | 0.01 | 0.003 | 1 | 0 | Cdkn2a | 0.007 | Young |
| 0.066083 | 0.496298 | 0.048 | 0.031 | 1 | 0 | Tlr5 | 0.017 | Young |
| 0.066113 | 1.070951 | 0.014 | 0.005 | 1 | 0 | Tmem143 | 0.009 | Young |
| 0.066226 | 0.966039 | 0.027 | 0.015 | 1 | 0 | Hepacam2 | 0.012 | Young |
| 0.066373 | 1.002851 | 0.014 | 0.005 | 1 | 0 | Ccl27a | 0.009 | Young |
| 0.066521 | 1.076082 | 0.014 | 0.005 | 1 | 0 | Mblac1 | 0.009 | Young |
| 0.066841 | 1.050867 | 0.021 | 0.011 | 1 | 0 | E130102H2 | 0.01 | Young |
| 0.066958 | 0.407796 | 0.183 | 0.157 | 1 | 0 | Ndufs5 | 0.026 | Young |
| 0.067324 | 0.412251 | 0.079 | 0.059 | 1 | 0 | Ryk | 0.02 | Young |
| 0.067348 | 1.009976 | 0.021 | 0.011 | 1 | 0 | Pald1 | 0.01 | Young |
| 0.067735 | 0.109649 | 0.797 | 0.814 | 1 | 0 | Rps11 | -0.017 | Young |
| 0.067947 | 0.136914 | 0.501 | 0.475 | 1 | 0 | Man2b1 | 0.026 | Young |
| 0.068146 | 0.624337 | 0.101 | 0.079 | 1 | 0 | Sord | 0.022 | Young |
| 0.068763 | 0.863907 | 0.018 | 0.008 | 1 | 0 | Twnk | 0.01 | Young |
| 0.068877 | 0.319215 | 0.012 | 0.004 | 1 | 0 | Hist1h1e | 0.008 | Young |

| | | | | | | | | |
|---|---|---|---|---|---|---|---|---|
| 0.069048 | 0.813103 | 0.056 | 0.04 | 1 | 0 | Rora | 0.016 | Young |
| 0.069124 | 0.189333 | 0.508 | 0.501 | 1 | 0 | Tpr | 0.007 | Young |
| 0.069525 | 0.082262 | 0.811 | 0.811 | 1 | 0 | Rpl9 | 0 | Young |
| 0.06975 | 0.386183 | 0.154 | 0.129 | 1 | 0 | Thra | 0.025 | Young |
| 0.070013 | 0.314804 | 0.251 | 0.223 | 1 | 0 | Zfp395 | 0.028 | Young |
| 0.070025 | 0.649294 | 0.058 | 0.041 | 1 | 0 | Ddah2 | 0.017 | Young |
| 0.070029 | 0.641094 | 0.053 | 0.037 | 1 | 0 | Bcl2a1d | 0.016 | Young |
| 0.070265 | 0.104149 | 0.635 | 0.616 | 1 | 0 | Igf1 | 0.019 | Young |
| 0.070599 | 0.500733 | 0.061 | 0.044 | 1 | 0 | Slc9a3r2 | 0.017 | Young |
| 0.070923 | 0.665049 | 0.03 | 0.018 | 1 | 0 | Lix1 | 0.012 | Young |
| 0.07155 | 0.368982 | 0.123 | 0.1 | 1 | 0 | Samd4 | 0.023 | Young |
| 0.071961 | 0.666119 | 0.023 | 0.012 | 1 | 0 | 1600020E0 | 0.011 | Young |
| 0.072073 | 0.426256 | 0.044 | 0.029 | 1 | 0 | Rbms2 | 0.015 | Young |
| 0.072483 | 0.965168 | 0.019 | 0.01 | 1 | 0 | Ildr2 | 0.009 | Young |
| 0.072708 | 0.40844 | 0.055 | 0.038 | 1 | 0 | Afdn | 0.017 | Young |
| 0.072996 | 0.585585 | 0.088 | 0.068 | 1 | 0 | Slc12a9 | 0.02 | Young |
| 0.073357 | 0.271903 | 0.136 | 0.111 | 1 | 0 | Crim1 | 0.025 | Young |
| 0.07359 | 0.740492 | 0.067 | 0.049 | 1 | 0 | Nudt5 | 0.018 | Young |
| 0.073666 | 0.842766 | 0.038 | 0.025 | 1 | 0 | Zfp760 | 0.013 | Young |
| 0.073774 | 0.593671 | 0.045 | 0.03 | 1 | 0 | Cd28 | 0.015 | Young |
| 0.074086 | 0.367043 | 0.066 | 0.048 | 1 | 0 | Osbpl2 | 0.018 | Young |
| 0.074136 | 0.459342 | 0.052 | 0.036 | 1 | 0 | Fmc1 | 0.016 | Young |
| 0.074478 | 0.787152 | 0.019 | 0.01 | 1 | 0 | 1110065P2 | 0.009 | Young |
| 0.074847 | 0.207733 | 0.177 | 0.148 | 1 | 0 | 1700017B0 | 0.029 | Young |
| 0.074885 | 0.962977 | 0.032 | 0.019 | 1 | 0 | Map4k3 | 0.013 | Young |
| 0.074952 | 0.228824 | 0.288 | 0.261 | 1 | 0 | Lacc1 | 0.027 | Young |
| 0.075014 | 1.908354 | 0.013 | 0.005 | 1 | 0 | Mocs3 | 0.008 | Young |
| 0.075114 | 0.519325 | 0.11 | 0.088 | 1 | 0 | Bmyc | 0.022 | Young |
| 0.075379 | 0.46125 | 0.069 | 0.051 | 1 | 0 | Uchl3 | 0.018 | Young |
| 0.075413 | 0.427568 | 0.175 | 0.15 | 1 | 0 | Trem2 | 0.025 | Young |
| 0.075526 | 0.200873 | 0.425 | 0.399 | 1 | 0 | Kdm7a | 0.026 | Young |
| 0.07585 | 1.574837 | 0.013 | 0.005 | 1 | 0 | Ighmbp2 | 0.008 | Young |
| 0.075975 | 0.095862 | 0.918 | 0.908 | 1 | 0 | Cltc | 0.01 | Young |
| 0.07627 | 0.274787 | 0.207 | 0.181 | 1 | 0 | Ulk2 | 0.026 | Young |
| 0.077008 | 0.528663 | 0.055 | 0.038 | 1 | 0 | Gm46224 | 0.017 | Young |
| 0.077118 | 1.08191 | 0.013 | 0.005 | 1 | 0 | Exog | 0.008 | Young |
| 0.077215 | 0.29288 | 0.256 | 0.231 | 1 | 0 | Nars | 0.025 | Young |
| 0.077545 | 0.784576 | 0.013 | 0.005 | 1 | 0 | Ugt1a7c | 0.008 | Young |
| 0.077673 | 0.829864 | 0.013 | 0.005 | 1 | 0 | Gm36266 | 0.008 | Young |
| 0.077783 | 0.262492 | 0.438 | 0.427 | 1 | 0 | Anp32a | 0.011 | Young |
| 0.078224 | 0.772703 | 0.081 | 0.063 | 1 | 0 | Pot1a | 0.018 | Young |
| 0.078333 | 1.510112 | 0.011 | 0.004 | 1 | 0 | Olr1 | 0.007 | Young |
| 0.078346 | 0.42224 | 0.08 | 0.06 | 1 | 0 | Rlf | 0.02 | Young |
| 0.078664 | 1.224917 | 0.011 | 0.004 | 1 | 0 | Ctnnd2 | 0.007 | Young |
| 0.078995 | 0.43128 | 0.148 | 0.124 | 1 | 0 | Angel2 | 0.024 | Young |

| | | | | | | | | |
|---|---|---|---|---|---|---|---|---|
| 0.079615 | 0.735462 | 0.011 | 0.004 | 1 | 0 | Cwf19l1 | 0.007 | Young |
| 0.079751 | 0.315868 | 0.399 | 0.387 | 1 | 0 | Ralbp1 | 0.012 | Young |
| 0.080058 | 0.532537 | 0.054 | 0.038 | 1 | 0 | Nup85 | 0.016 | Young |
| 0.080114 | 0.666001 | 0.017 | 0.008 | 1 | 0 | Col14a1 | 0.009 | Young |
| 0.080323 | 0.244366 | 0.313 | 0.287 | 1 | 0 | Iqgap2 | 0.026 | Young |
| 0.080448 | 0.811217 | 0.023 | 0.012 | 1 | 0 | Trim39 | 0.011 | Young |
| 0.080487 | 0.047596 | 0.985 | 0.995 | 1 | 0 | Mrc1 | -0.01 | Young |
| 0.080856 | 0.977267 | 0.038 | 0.025 | 1 | 0 | Phlpp2 | 0.013 | Young |
| 0.081115 | 0.626153 | 0.07 | 0.052 | 1 | 0 | Wtip | 0.018 | Young |
| 0.081684 | 0.561844 | 0.045 | 0.03 | 1 | 0 | Bmf | 0.015 | Young |
| 0.0818 | 0.031624 | 0.056 | 0.075 | 1 | 0 | Dnajc16 | -0.019 | Young |
| 0.08252 | 0.943435 | 0.026 | 0.015 | 1 | 0 | Prkd2 | 0.011 | Young |
| 0.082618 | 1.010069 | 0.028 | 0.016 | 1 | 0 | Gstp1 | 0.012 | Young |
| 0.08277 | 0.9935 | 0.055 | 0.04 | 1 | 0 | Gpr137 | 0.015 | Young |
| 0.08287 | 0.464225 | 0.056 | 0.04 | 1 | 0 | Nacc1 | 0.016 | Young |
| 0.083072 | 0.151003 | 0.084 | 0.063 | 1 | 0 | Kank2 | 0.021 | Young |
| 0.083371 | 1.348619 | 0.015 | 0.007 | 1 | 0 | Fam46c | 0.008 | Young |
| 0.083527 | 0.379081 | 0.136 | 0.112 | 1 | 0 | Dot1l | 0.024 | Young |
| 0.083618 | 0.553962 | 0.068 | 0.051 | 1 | 0 | Rps6ka5 | 0.017 | Young |
| 0.083627 | 1.372208 | 0.015 | 0.007 | 1 | 0 | Dtwd1 | 0.008 | Young |
| 0.084354 | 0.861346 | 0.044 | 0.03 | 1 | 0 | Rfx5 | 0.014 | Young |
| 0.084552 | 0.418986 | 0.224 | 0.201 | 1 | 0 | Pcnp | 0.023 | Young |
| 0.08484 | 0.229355 | 0.486 | 0.48 | 1 | 0 | Klf3 | 0.006 | Young |
| 0.085739 | 1.138737 | 0.021 | 0.011 | 1 | 0 | Tmed1 | 0.01 | Young |
| 0.085828 | 0.824644 | 0.029 | 0.018 | 1 | 0 | Tcta | 0.011 | Young |
| 0.086147 | 1.397741 | 0.021 | 0.011 | 1 | 0 | Mmp13 | 0.01 | Young |
| 0.086545 | 0.859303 | 0.043 | 0.029 | 1 | 0 | Gimap6 | 0.014 | Young |
| 0.086564 | 0.128761 | 0.326 | 0.297 | 1 | 0 | Cebpd | 0.029 | Young |
| 0.086807 | 0.370511 | 0.102 | 0.081 | 1 | 0 | Apopt1 | 0.021 | Young |
| 0.086938 | 0.59266 | 0.083 | 0.064 | 1 | 0 | Crot | 0.019 | Young |
| 0.087741 | 0.460332 | 0.132 | 0.109 | 1 | 0 | Rasgef1b | 0.023 | Young |
| 0.087922 | 0.828677 | 0.041 | 0.027 | 1 | 0 | Sssca1 | 0.014 | Young |
| 0.088182 | 0.226866 | 0.382 | 0.364 | 1 | 0 | Gpi1 | 0.018 | Young |
| 0.088259 | 0.492389 | 0.102 | 0.082 | 1 | 0 | mt-Nd6 | 0.02 | Young |
| 0.088293 | 0.764385 | 0.063 | 0.047 | 1 | 0 | Nxt2 | 0.016 | Young |
| 0.088409 | 0.61473 | 0.078 | 0.06 | 1 | 0 | Zdhhc17 | 0.018 | Young |
| 0.089142 | 0.586988 | 0.046 | 0.031 | 1 | 0 | Depdc5 | 0.015 | Young |
| 0.089279 | 0.276737 | 0.18 | 0.155 | 1 | 0 | Mfsd6 | 0.025 | Young |
| 0.089581 | 0.264122 | 0.352 | 0.33 | 1 | 0 | Kmt2e | 0.022 | Young |
| 0.089658 | 0.475836 | 0.049 | 0.034 | 1 | 0 | Atg2b | 0.015 | Young |
| 0.089922 | 0.801429 | 0.013 | 0.005 | 1 | 0 | Thumpd2 | 0.008 | Young |
| 0.090453 | 1.05796 | 0.017 | 0.008 | 1 | 0 | Lmbr1 | 0.009 | Young |
| 0.090856 | 0.550505 | 0.046 | 0.031 | 1 | 0 | Zfp68 | 0.015 | Young |
| 0.091188 | 0.353728 | 0.171 | 0.146 | 1 | 0 | Lyz1 | 0.025 | Young |
| 0.0912 | 0.534392 | 0.143 | 0.122 | 1 | 0 | Aagab | 0.021 | Young |

| | | | | | | | | |
|---|---|---|---|---|---|---|---|---|
| 0.091531 | 0.868094 | 0.037 | 0.025 | 1 | 0 | Acot8 | 0.012 | Young |
| 0.091944 | 0.383789 | 0.189 | 0.166 | 1 | 0 | Plekha2 | 0.023 | Young |
| 0.091954 | 1.284251 | 0.011 | 0.004 | 1 | 0 | Nlrc3 | 0.007 | Young |
| 0.092337 | 0.859732 | 0.011 | 0.004 | 1 | 0 | L3hypdh | 0.007 | Young |
| 0.09246 | 0.565361 | 0.068 | 0.051 | 1 | 0 | Irgm1 | 0.017 | Young |
| 0.092525 | 0.503262 | 0.129 | 0.108 | 1 | 0 | Grap | 0.021 | Young |
| 0.092692 | 0.24673 | 0.473 | 0.475 | 1 | 0 | Myo5a | -0.002 | Young |
| 0.093054 | 0.441403 | 0.111 | 0.09 | 1 | 0 | Vrk1 | 0.021 | Young |
| 0.093056 | 0.764195 | 0.056 | 0.041 | 1 | 0 | Snx11 | 0.015 | Young |
| 0.093539 | 0.299089 | 0.188 | 0.163 | 1 | 0 | Nfia | 0.025 | Young |
| 0.093554 | 0.809597 | 0.026 | 0.015 | 1 | 0 | B230118H( | 0.011 | Young |
| 0.093896 | 1.103715 | 0.019 | 0.01 | 1 | 0 | Rida | 0.009 | Young |
| 0.093939 | 0.990822 | 0.019 | 0.01 | 1 | 0 | Zfp628 | 0.009 | Young |
| 0.094712 | 1.012017 | 0.039 | 0.026 | 1 | 0 | Mical2 | 0.013 | Young |
| 0.094727 | 0.778389 | 0.026 | 0.015 | 1 | 0 | Spats2 | 0.011 | Young |
| 0.094788 | 0.15998 | 0.609 | 0.61 | 1 | 0 | Ogt | -0.001 | Young |
| 0.094805 | 0.665264 | 0.045 | 0.031 | 1 | 0 | Gcsh | 0.014 | Young |
| 0.094995 | 1.029817 | 0.019 | 0.01 | 1 | 0 | Thada | 0.009 | Young |
| 0.095065 | 0.637684 | 0.047 | 0.033 | 1 | 0 | Crbn | 0.014 | Young |
| 0.095127 | 0.653787 | 0.045 | 0.031 | 1 | 0 | Foxo4 | 0.014 | Young |
| 0.095276 | 0.786176 | 0.029 | 0.018 | 1 | 0 | Mtrf1l | 0.011 | Young |
| 0.095304 | 0.397222 | 0.105 | 0.085 | 1 | 0 | Wars | 0.02 | Young |
| 0.095442 | 0.104979 | 0.691 | 0.694 | 1 | 0 | Clta | -0.003 | Young |
| 0.095803 | 0.784408 | 0.019 | 0.01 | 1 | 0 | Zfp667 | 0.009 | Young |
| 0.096421 | 0.377578 | 0.128 | 0.105 | 1 | 0 | Zcchc2 | 0.023 | Young |
| 0.096495 | 1.122926 | 0.015 | 0.007 | 1 | 0 | Tmem81 | 0.008 | Young |
| 0.096546 | 0.28401 | 0.087 | 0.067 | 1 | 0 | Kmt5c | 0.02 | Young |
| 0.096738 | 0.261537 | 0.345 | 0.32 | 1 | 0 | Cmklr1 | 0.025 | Young |
| 0.096834 | 1.106633 | 0.015 | 0.007 | 1 | 0 | E330009J0 | 0.008 | Young |
| 0.096936 | 0.231231 | 0.19 | 0.164 | 1 | 0 | Golga4 | 0.026 | Young |
| 0.097265 | 0.594321 | 0.019 | 0.01 | 1 | 0 | Cox7a1 | 0.009 | Young |
| 0.097533 | 0.388926 | 0.043 | 0.029 | 1 | 0 | Bcl2l12 | 0.014 | Young |
| 0.098088 | 1.042749 | 0.02 | 0.011 | 1 | 0 | Rab39 | 0.009 | Young |
| 0.098297 | 0.316692 | 0.029 | 0.018 | 1 | 0 | Mlst8 | 0.011 | Young |
| 0.09843 | 0.594315 | 0.072 | 0.055 | 1 | 0 | Kif3a | 0.017 | Young |
| 0.098455 | 0.459952 | 0.031 | 0.019 | 1 | 0 | Zxdb | 0.012 | Young |
| 0.098504 | 0.931511 | 0.02 | 0.011 | 1 | 0 | BC049352 | 0.009 | Young |
| 0.099262 | 0.513908 | 0.084 | 0.066 | 1 | 0 | AY036118 | 0.018 | Young |
| 0.099317 | 0.293529 | 0.167 | 0.142 | 1 | 0 | Phc2 | 0.025 | Young |
| 0.099882 | 0.082668 | 0.007 | 0.014 | 1 | 0 | Phf11a | -0.007 | Young |
| 0.099992 | 0.236031 | 0.399 | 0.383 | 1 | 0 | Ssh2 | 0.016 | Young |
| 0.10001 | 1.167752 | 0.024 | 0.014 | 1 | 0 | Cmpk2 | 0.01 | Young |
| 0.100391 | 1.03705 | 0.022 | 0.012 | 1 | 0 | Hyal1 | 0.01 | Young |
| 0.101075 | 0.610643 | 0.02 | 0.011 | 1 | 0 | Zdhhc24 | 0.009 | Young |
| 0.101301 | 0.084696 | 0.712 | 0.71 | 1 | 0 | Rps17 | 0.002 | Young |

| | | | | | | | | |
|---|---|---|---|---|---|---|---|---|
| 0.101441 | 0.673846 | 0.079 | 0.062 | 1 | 0 | Plxdc2 | 0.017 | Young |
| 0.101934 | 0.507922 | 0.056 | 0.041 | 1 | 0 | Ttc28 | 0.015 | Young |
| 0.102199 | 0.0559 | 0.158 | 0.13 | 1 | 0 | Siglech | 0.028 | Young |
| 0.102471 | 1.087692 | 0.013 | 0.005 | 1 | 0 | Tmem246 | 0.008 | Young |
| 0.10258 | 0.197247 | 0.482 | 0.472 | 1 | 0 | Cox6c | 0.01 | Young |
| 0.10278 | 0.168979 | 0.513 | 0.503 | 1 | 0 | Eef2 | 0.01 | Young |
| 0.102973 | 0.861093 | 0.032 | 0.021 | 1 | 0 | Lin9 | 0.011 | Young |
| 0.103406 | 0.515622 | 0.05 | 0.036 | 1 | 0 | Trub2 | 0.014 | Young |
| 0.103515 | 0.0797 | 0.022 | 0.012 | 1 | 0 | Runx3 | 0.01 | Young |
| 0.103558 | 0.699697 | 0.037 | 0.025 | 1 | 0 | Afap1l1 | 0.012 | Young |
| 0.103594 | 0.273432 | 0.291 | 0.265 | 1 | 0 | Ypel3 | 0.026 | Young |
| 0.103674 | 0.093137 | 0.64 | 0.627 | 1 | 0 | Abca1 | 0.013 | Young |
| 0.104085 | 0.336226 | 0.158 | 0.134 | 1 | 0 | Cnot4 | 0.024 | Young |
| 0.104117 | 0.769764 | 0.034 | 0.022 | 1 | 0 | Stradb | 0.012 | Young |
| 0.104468 | 0.69498 | 0.013 | 0.005 | 1 | 0 | Slc30a6 | 0.008 | Young |
| 0.105001 | 0.549901 | 0.035 | 0.023 | 1 | 0 | Hint3 | 0.012 | Young |
| 0.105031 | 0.065955 | 0.03 | 0.042 | 1 | 0 | Glipr2 | -0.012 | Young |
| 0.105084 | 0.505417 | 0.013 | 0.005 | 1 | 0 | Tamm41 | 0.008 | Young |
| 0.106099 | 0.718938 | 0.043 | 0.03 | 1 | 0 | Pafah1b3 | 0.013 | Young |
| 0.106245 | 0.788933 | 0.029 | 0.018 | 1 | 0 | Zdhhc12 | 0.011 | Young |
| 0.106867 | 1.401012 | 0.011 | 0.004 | 1 | 0 | Zfyve9 | 0.007 | Young |
| 0.106924 | 0.612966 | 0.035 | 0.023 | 1 | 0 | Zc3h7b | 0.012 | Young |
| 0.107685 | 1.105445 | 0.011 | 0.004 | 1 | 0 | Mctp2 | 0.007 | Young |
| 0.107695 | 0.655125 | 0.029 | 0.018 | 1 | 0 | Ccl25 | 0.011 | Young |
| 0.107778 | 0.305222 | 0.078 | 0.06 | 1 | 0 | Pilra | 0.018 | Young |
| 0.107786 | 0.210098 | 0.34 | 0.317 | 1 | 0 | Irf1 | 0.023 | Young |
| 0.107871 | 0.955608 | 0.018 | 0.01 | 1 | 0 | Slc37a3 | 0.008 | Young |
| 0.108225 | 0.308878 | 0.259 | 0.233 | 1 | 0 | Arl4c | 0.026 | Young |
| 0.108633 | 0.521484 | 0.027 | 0.016 | 1 | 0 | Wdr83 | 0.011 | Young |
| 0.10867 | 0.498477 | 0.1 | 0.081 | 1 | 0 | Rtn1 | 0.019 | Young |
| 0.109131 | 0.243468 | 0.374 | 0.36 | 1 | 0 | Abhd12 | 0.014 | Young |
| 0.109332 | 0.153243 | 0.11 | 0.088 | 1 | 0 | Polr3gl | 0.022 | Young |
| 0.109375 | 0.257668 | 0.218 | 0.192 | 1 | 0 | Il6st | 0.026 | Young |
| 0.110746 | 0.530975 | 0.018 | 0.01 | 1 | 0 | Det1 | 0.008 | Young |
| 0.111345 | 0.617194 | 0.109 | 0.09 | 1 | 0 | Setd7 | 0.019 | Young |
| 0.111514 | 0.589303 | 0.102 | 0.083 | 1 | 0 | Hk2 | 0.019 | Young |
| 0.111812 | 0.991751 | 0.055 | 0.041 | 1 | 0 | Agfg2 | 0.014 | Young |
| 0.112054 | 0.506167 | 0.043 | 0.03 | 1 | 0 | Kat14 | 0.013 | Young |
| 0.112178 | 0.251956 | 0.067 | 0.051 | 1 | 0 | Pbxip1 | 0.016 | Young |
| 0.112389 | 0.166374 | 0.54 | 0.535 | 1 | 0 | Csde1 | 0.005 | Young |
| 0.112754 | 1.003787 | 0.027 | 0.016 | 1 | 0 | Zc2hc1a | 0.011 | Young |
| 0.11328 | 0.721288 | 0.035 | 0.023 | 1 | 0 | Rtn4rl1 | 0.012 | Young |
| 0.11344 | 0.402198 | 0.137 | 0.116 | 1 | 0 | Pnrc2 | 0.021 | Young |
| 0.113581 | 0.476355 | 0.048 | 0.034 | 1 | 0 | Sorbs3 | 0.014 | Young |
| 0.114362 | 0.082807 | 0.035 | 0.048 | 1 | 0 | Narfl | -0.013 | Young |

| | | | | | | | | |
|---|---|---|---|---|---|---|---|---|
| 0.114404 | 1.034051 | 0.025 | 0.015 | 1 | 0 | Prr5 | 0.01 | Young |
| 0.115311 | 0.311882 | 0.294 | 0.276 | 1 | 0 | Cox14 | 0.018 | Young |
| 0.115336 | 0.399159 | 0.062 | 0.047 | 1 | 0 | Orc4 | 0.015 | Young |
| 0.115368 | 0.04575 | 0.104 | 0.127 | 1 | 0 | Dgkz | -0.023 | Young |
| 0.115983 | 0.286346 | 0.274 | 0.254 | 1 | 0 | Ankrd17 | 0.02 | Young |
| 0.116456 | 0.479591 | 0.083 | 0.066 | 1 | 0 | Tmem183a | 0.017 | Young |
| 0.116459 | 0.617639 | 0.035 | 0.023 | 1 | 0 | Kdm1b | 0.012 | Young |
| 0.116597 | 0.056197 | 0.026 | 0.037 | 1 | 0 | Capn15 | -0.011 | Young |
| 0.116659 | 2.009062 | 0.012 | 0.005 | 1 | 0 | C730034F0 | 0.007 | Young |
| 0.117146 | 1.080985 | 0.016 | 0.008 | 1 | 0 | Suox | 0.008 | Young |
| 0.117415 | 0.595305 | 0.023 | 0.014 | 1 | 0 | Cish | 0.009 | Young |
| 0.117812 | 0.681525 | 0.025 | 0.015 | 1 | 0 | Sccpdh | 0.01 | Young |
| 0.11782 | 0.896949 | 0.04 | 0.027 | 1 | 0 | Sclt1 | 0.013 | Young |
| 0.11835 | 0.467433 | 0.092 | 0.074 | 1 | 0 | Pkd2 | 0.018 | Young |
| 0.118739 | 0.661352 | 0.03 | 0.019 | 1 | 0 | Ezh2 | 0.011 | Young |
| 0.119535 | 0.699486 | 0.016 | 0.008 | 1 | 0 | Ccdc91 | 0.008 | Young |
| 0.120075 | 0.208012 | 0.095 | 0.075 | 1 | 0 | Mxi1 | 0.02 | Young |
| 0.120113 | 0.325963 | 0.038 | 0.026 | 1 | 0 | 2310039H0 | 0.012 | Young |
| 0.120311 | 0.59157 | 0.027 | 0.016 | 1 | 0 | Zc3h3 | 0.011 | Young |
| 0.120359 | 0.656177 | 0.016 | 0.008 | 1 | 0 | Fscn1 | 0.008 | Young |
| 0.12102 | 0.192735 | 0.086 | 0.067 | 1 | 0 | Csnk1g1 | 0.019 | Young |
| 0.121161 | 0.276659 | 0.192 | 0.167 | 1 | 0 | Xpr1 | 0.025 | Young |
| 0.121337 | 0.844857 | 0.036 | 0.025 | 1 | 0 | Casp2 | 0.011 | Young |
| 0.122275 | 0.251056 | 0.106 | 0.086 | 1 | 0 | Ccm2 | 0.02 | Young |
| 0.122659 | 0.397317 | 0.027 | 0.016 | 1 | 0 | Ppm1f | 0.011 | Young |
| 0.123408 | 0.009264 | 0.168 | 0.197 | 1 | 0 | Riok3 | -0.029 | Young |
| 0.123753 | 0.3213 | 0.051 | 0.037 | 1 | 0 | Dnlz | 0.014 | Young |
| 0.124132 | 0.270588 | 0.099 | 0.079 | 1 | 0 | Fam133b | 0.02 | Young |
| 0.124658 | 0.32079 | 0.062 | 0.047 | 1 | 0 | Pold4 | 0.015 | Young |
| 0.124876 | 0.325508 | 0.142 | 0.12 | 1 | 0 | Mlec | 0.022 | Young |
| 0.125179 | 0.324315 | 0.099 | 0.081 | 1 | 0 | Cldnd1 | 0.018 | Young |
| 0.125227 | 1.270913 | 0.01 | 0.004 | 1 | 0 | Wdr34 | 0.006 | Young |
| 0.125734 | 1.135074 | 0.01 | 0.004 | 1 | 0 | Phldb1 | 0.006 | Young |
| 0.126024 | 0.949054 | 0.01 | 0.004 | 1 | 0 | D930030I0 | 0.006 | Young |
| 0.126083 | 0.325667 | 0.2 | 0.178 | 1 | 0 | Mfsd11 | 0.022 | Young |
| 0.126121 | 0.231656 | 0.214 | 0.192 | 1 | 0 | P2rx7 | 0.022 | Young |
| 0.126125 | 0.495367 | 0.038 | 0.026 | 1 | 0 | Tnfrsf17 | 0.012 | Young |
| 0.126317 | 0.127143 | 0.778 | 0.798 | 1 | 0 | Ptafr | -0.02 | Young |
| 0.126482 | 0.585891 | 0.132 | 0.114 | 1 | 0 | Cd180 | 0.018 | Young |
| 0.12692 | 1.188303 | 0.023 | 0.014 | 1 | 0 | Ncapd3 | 0.009 | Young |
| 0.126944 | 0.827783 | 0.019 | 0.011 | 1 | 0 | Itgax | 0.008 | Young |
| 0.126945 | 1.094177 | 0.026 | 0.016 | 1 | 0 | F830016B0 | 0.01 | Young |
| 0.127412 | 0.833425 | 0.019 | 0.011 | 1 | 0 | Plekhh3 | 0.008 | Young |
| 0.127425 | 0.257183 | 0.048 | 0.034 | 1 | 0 | Bola1 | 0.014 | Young |
| 0.127516 | 0.71738 | 0.019 | 0.011 | 1 | 0 | Inpp5b | 0.008 | Young |

| | | | | | | | | |
|---|---|---|---|---|---|---|---|---|
| 0.127529 | 0.138351 | 0.509 | 0.49 | 1 | 0 | Ptprj | 0.019 | Young |
| 0.12791 | 0.980992 | 0.024 | 0.015 | 1 | 0 | Armc3 | 0.009 | Young |
| 0.127992 | 0.300059 | 0.267 | 0.25 | 1 | 0 | Prdx5 | 0.017 | Young |
| 0.128611 | 0.792097 | 0.021 | 0.012 | 1 | 0 | Mrpl47 | 0.009 | Young |
| 0.128666 | 0.382588 | 0.04 | 0.027 | 1 | 0 | Dctpp1 | 0.013 | Young |
| 0.128815 | 0.022068 | 0.136 | 0.163 | 1 | 0 | Zdhhc9 | -0.027 | Young |
| 0.128851 | 0.821608 | 0.014 | 0.007 | 1 | 0 | Mapkapk5 | 0.007 | Young |
| 0.129038 | 0.860327 | 0.014 | 0.007 | 1 | 0 | Zfp882 | 0.007 | Young |
| 0.129063 | 0.595152 | 0.021 | 0.012 | 1 | 0 | Natd1 | 0.009 | Young |
| 0.129766 | 0.409204 | 0.088 | 0.071 | 1 | 0 | Mfap1a | 0.017 | Young |
| 0.130405 | 0.374497 | 0.16 | 0.14 | 1 | 0 | Ndufaf8 | 0.02 | Young |
| 0.130457 | 0.317987 | 0.019 | 0.011 | 1 | 0 | Lyrm4 | 0.008 | Young |
| 0.130613 | 0.303489 | 0.074 | 0.057 | 1 | 0 | Ttll12 | 0.017 | Young |
| 0.130999 | 0.562984 | 0.094 | 0.077 | 1 | 0 | Zbtb11 | 0.017 | Young |
| 0.131072 | 0.299227 | 0.167 | 0.145 | 1 | 0 | Pcm1 | 0.022 | Young |
| 0.131209 | 0.570064 | 0.116 | 0.098 | 1 | 0 | Pdzd8 | 0.018 | Young |
| 0.131935 | 0.597753 | 0.045 | 0.033 | 1 | 0 | Xpc | 0.012 | Young |
| 0.132312 | 0.401325 | 0.229 | 0.211 | 1 | 0 | Eif3h | 0.018 | Young |
| 0.132483 | 0.418576 | 0.046 | 0.033 | 1 | 0 | AU020206 | 0.013 | Young |
| 0.132489 | 0.196167 | 0.274 | 0.248 | 1 | 0 | Dclre1c | 0.026 | Young |
| 0.132799 | 0.38522 | 0.204 | 0.183 | 1 | 0 | Phc3 | 0.021 | Young |
| 0.133094 | 0.451834 | 0.023 | 0.014 | 1 | 0 | Zfp275 | 0.009 | Young |
| 0.133434 | 0.462243 | 0.078 | 0.062 | 1 | 0 | Myh10 | 0.016 | Young |
| 0.133469 | 0.418095 | 0.031 | 0.021 | 1 | 0 | Enpp1 | 0.01 | Young |
| 0.133858 | 1.095665 | 0.016 | 0.008 | 1 | 0 | Bco2 | 0.008 | Young |
| 0.133918 | 0.446516 | 0.029 | 0.019 | 1 | 0 | Unc119b | 0.01 | Young |
| 0.134127 | 0.484811 | 0.09 | 0.073 | 1 | 0 | Pex5 | 0.017 | Young |
| 0.134226 | 0.184485 | 0.229 | 0.204 | 1 | 0 | Mob3c | 0.025 | Young |
| 0.134268 | 0.660848 | 0.068 | 0.053 | 1 | 0 | Icosl | 0.015 | Young |
| 0.134318 | 0.07963 | 0.058 | 0.074 | 1 | 0 | Amz2 | -0.016 | Young |
| 0.134825 | 0.24995 | 0.29 | 0.271 | 1 | 0 | Tiparp | 0.019 | Young |
| 0.134947 | 0.293279 | 0.075 | 0.059 | 1 | 0 | Ttf1 | 0.016 | Young |
| 0.135271 | 0.305664 | 0.127 | 0.107 | 1 | 0 | Irf4 | 0.02 | Young |
| 0.135473 | 0.538971 | 0.064 | 0.049 | 1 | 0 | Abcc3 | 0.015 | Young |
| 0.135479 | 0.25264 | 0.262 | 0.239 | 1 | 0 | Hp1bp3 | 0.023 | Young |
| 0.135553 | 0.337166 | 0.028 | 0.018 | 1 | 0 | Stau2 | 0.01 | Young |
| 0.135866 | 0.282671 | 0.195 | 0.174 | 1 | 0 | 4932438A1 | 0.021 | Young |
| 0.136171 | 0.748863 | 0.016 | 0.008 | 1 | 0 | Tal1 | 0.008 | Young |
| 0.136476 | 0.365034 | 0.024 | 0.015 | 1 | 0 | Osbpl1a | 0.009 | Young |
| 0.137323 | 0.196907 | 0.226 | 0.204 | 1 | 0 | Acadm | 0.022 | Young |
| 0.137388 | 0.607369 | 0.068 | 0.053 | 1 | 0 | Upf2 | 0.015 | Young |
| 0.13791 | 0.54451 | 0.044 | 0.031 | 1 | 0 | 1500011B0 | 0.013 | Young |
| 0.138145 | 0.393694 | 0.131 | 0.112 | 1 | 0 | Tnks | 0.019 | Young |
| 0.13823 | 0.179829 | 0.01 | 0.016 | 1 | 0 | Sap130 | -0.006 | Young |
| 0.138241 | 0.365512 | 0.231 | 0.213 | 1 | 0 | Map2k2 | 0.018 | Young |

| | | | | | | | | |
|---|---|---|---|---|---|---|---|---|
| 0.138991 | 0.45513 | 0.155 | 0.135 | 1 | 0 | Anapc13 | 0.02 | Young |
| 0.13924 | 0.45954 | 0.134 | 0.115 | 1 | 0 | Dlgap4 | 0.019 | Young |
| 0.139764 | 0.044957 | 0.1 | 0.122 | 1 | 0 | Wdr82 | -0.022 | Young |
| 0.140722 | 0.406908 | 0.158 | 0.138 | 1 | 0 | Rnpepl1 | 0.02 | Young |
| 0.140936 | 0.46174 | 0.012 | 0.005 | 1 | 0 | Spata7 | 0.007 | Young |
| 0.141418 | 0.372853 | 0.122 | 0.103 | 1 | 0 | Mrpl57 | 0.019 | Young |
| 0.141522 | 0.61075 | 0.045 | 0.033 | 1 | 0 | Nop14 | 0.012 | Young |
| 0.141987 | 0.663769 | 0.059 | 0.045 | 1 | 0 | Cntln | 0.014 | Young |
| 0.141991 | 1.074711 | 0.021 | 0.012 | 1 | 0 | Izumo4 | 0.009 | Young |
| 0.142066 | 1.108664 | 0.022 | 0.014 | 1 | 0 | Lppos | 0.008 | Young |
| 0.142119 | 0.35642 | 0.07 | 0.055 | 1 | 0 | Ikbip | 0.015 | Young |
| 0.142383 | 0.252012 | 0.179 | 0.157 | 1 | 0 | Ndufs2 | 0.022 | Young |
| 0.142392 | 2.141242 | 0.007 | 0.012 | 1 | 0 | Jchain | -0.005 | Young |
| 0.142645 | 0.531578 | 0.058 | 0.044 | 1 | 0 | Serhl | 0.014 | Young |
| 0.142671 | 0.25208 | 0.35 | 0.334 | 1 | 0 | Txn1 | 0.016 | Young |
| 0.142812 | 1.043205 | 0.021 | 0.012 | 1 | 0 | 1810024B0 | 0.009 | Young |
| 0.142882 | 0.358804 | 0.017 | 0.01 | 1 | 0 | Zfp933 | 0.007 | Young |
| 0.143181 | 0.28922 | 0.189 | 0.167 | 1 | 0 | Adipor2 | 0.022 | Young |
| 0.143373 | 0.300195 | 0.203 | 0.182 | 1 | 0 | Tpcn1 | 0.021 | Young |
| 0.14354 | 0.166036 | 0.418 | 0.398 | 1 | 0 | Ndufa6 | 0.02 | Young |
| 0.143641 | 0.342679 | 0.131 | 0.111 | 1 | 0 | Epb41 | 0.02 | Young |
| 0.143779 | 0.23728 | 0.459 | 0.451 | 1 | 0 | Mtdh | 0.008 | Young |
| 0.144093 | 0.26414 | 0.188 | 0.166 | 1 | 0 | Cyb5a | 0.022 | Young |
| 0.145341 | 0.174027 | 0.032 | 0.022 | 1 | 0 | Clpp | 0.01 | Young |
| 0.145402 | 0.205405 | 0.203 | 0.181 | 1 | 0 | Lyl1 | 0.022 | Young |
| 0.145534 | 0.701876 | 0.029 | 0.019 | 1 | 0 | H2-DMb2 | 0.01 | Young |
| 0.145991 | 1.490776 | 0.03 | 0.021 | 1 | 0 | Mthfd1 | 0.009 | Young |
| 0.146095 | 0.643338 | 0.029 | 0.019 | 1 | 0 | C330018D2 | 0.01 | Young |
| 0.146879 | 1.009342 | 0.01 | 0.004 | 1 | 0 | Kdelc1 | 0.006 | Young |
| 0.14706 | 0.002602 | 0.144 | 0.17 | 1 | 0 | Poldip3 | -0.026 | Young |
| 0.147061 | 0.415759 | 0.187 | 0.17 | 1 | 0 | D1Ertd622 | 0.017 | Young |
| 0.147062 | 0.077167 | 0.758 | 0.74 | 1 | 0 | Ctsh | 0.018 | Young |
| 0.147213 | 0.832716 | 0.01 | 0.004 | 1 | 0 | Mmab | 0.006 | Young |
| 0.147222 | 0.944216 | 0.013 | 0.007 | 1 | 0 | Pafah2 | 0.006 | Young |
| 0.147289 | 0.624762 | 0.06 | 0.047 | 1 | 0 | Lrrcc1 | 0.013 | Young |
| 0.147547 | 0.806546 | 0.01 | 0.004 | 1 | 0 | Cldn10 | 0.006 | Young |
| 0.147552 | 0.172411 | 0.015 | 0.023 | 1 | 0 | BC017158 | -0.008 | Young |
| 0.147717 | 0.411087 | 0.095 | 0.078 | 1 | 0 | Naa60 | 0.017 | Young |
| 0.14782 | 0.626711 | 0.022 | 0.014 | 1 | 0 | Ccdc58 | 0.008 | Young |
| 0.148216 | 0.57776 | 0.01 | 0.004 | 1 | 0 | Alkbh4 | 0.006 | Young |
| 0.148569 | 0.525232 | 0.065 | 0.051 | 1 | 0 | Ubr1 | 0.014 | Young |
| 0.148593 | 0.522212 | 0.05 | 0.037 | 1 | 0 | Klf16 | 0.013 | Young |
| 0.148891 | 0.66864 | 0.035 | 0.025 | 1 | 0 | Noc3l | 0.01 | Young |
| 0.149123 | 0.700132 | 0.013 | 0.007 | 1 | 0 | Tnks1bp1 | 0.006 | Young |
| 0.149183 | 0.198469 | 0.358 | 0.342 | 1 | 0 | Rock1 | 0.016 | Young |

| | | | | | | | | |
|---|---|---|---|---|---|---|---|---|
| 0.14997 | 0.7153 | 0.035 | 0.025 | 1 | 0 | Adprm | 0.01 | Young |
| 0.150782 | 0.241442 | 0.308 | 0.287 | 1 | 0 | Abracl | 0.021 | Young |
| 0.150929 | 0.102189 | 0.53 | 0.513 | 1 | 0 | S100a10 | 0.017 | Young |
| 0.150994 | 0.259168 | 0.289 | 0.268 | 1 | 0 | Arhgap15 | 0.021 | Young |
| 0.151045 | 0.330589 | 0.085 | 0.068 | 1 | 0 | Adamtsl5 | 0.017 | Young |
| 0.151249 | 0.336681 | 0.242 | 0.224 | 1 | 0 | Dusp22 | 0.018 | Young |
| 0.151735 | 1.111872 | 0.015 | 0.008 | 1 | 0 | Cd209c | 0.007 | Young |
| 0.151792 | 0.139809 | 0.646 | 0.657 | 1 | 0 | Gdi2 | -0.011 | Young |
| 0.151904 | 0.200132 | 0.027 | 0.018 | 1 | 0 | Dgkq | 0.009 | Young |
| 0.152281 | 0.198359 | 0.209 | 0.186 | 1 | 0 | Arid4a | 0.023 | Young |
| 0.152743 | 0.247847 | 0.243 | 0.22 | 1 | 0 | Akt1 | 0.023 | Young |
| 0.152912 | 0.58163 | 0.046 | 0.034 | 1 | 0 | Alg2 | 0.012 | Young |
| 0.153209 | 0.612975 | 0.051 | 0.038 | 1 | 0 | Zkscan1 | 0.013 | Young |
| 0.153326 | 0.38075 | 0.144 | 0.126 | 1 | 0 | Vezf1 | 0.018 | Young |
| 0.153608 | 0.405571 | 0.037 | 0.026 | 1 | 0 | Ripor1 | 0.011 | Young |
| 0.153699 | 0.205635 | 0.112 | 0.093 | 1 | 0 | Gpr183 | 0.019 | Young |
| 0.153704 | 0.23937 | 0.256 | 0.234 | 1 | 0 | Snx29 | 0.022 | Young |
| 0.153707 | 0.525461 | 0.115 | 0.098 | 1 | 0 | Ankib1 | 0.017 | Young |
| 0.153834 | 0.823799 | 0.015 | 0.008 | 1 | 0 | Tspyl4 | 0.007 | Young |
| 0.153872 | 0.027495 | 0.083 | 0.101 | 1 | 0 | Pygb | -0.018 | Young |
| 0.154008 | 0.016626 | 0.062 | 0.078 | 1 | 0 | Ola1 | -0.016 | Young |
| 0.154106 | 0.892252 | 0.015 | 0.008 | 1 | 0 | Cavin2 | 0.007 | Young |
| 0.15431 | 0.206829 | 0.175 | 0.153 | 1 | 0 | Rabep1 | 0.022 | Young |
| 0.154311 | 0.776939 | 0.015 | 0.008 | 1 | 0 | Trmt61a | 0.007 | Young |
| 0.154372 | 0.415159 | 0.056 | 0.042 | 1 | 0 | Ppwd1 | 0.014 | Young |
| 0.154926 | 0.960412 | 0.015 | 0.008 | 1 | 0 | Hcar2 | 0.007 | Young |
| 0.15498 | 0.592194 | 0.032 | 0.022 | 1 | 0 | Mad1l1 | 0.01 | Young |
| 0.155037 | 0.276022 | 0.129 | 0.109 | 1 | 0 | Slc16a7 | 0.02 | Young |
| 0.155108 | 0.012583 | 0.051 | 0.066 | 1 | 0 | Chmp7 | -0.015 | Young |
| 0.155318 | 0.391063 | 0.108 | 0.09 | 1 | 0 | Smc5 | 0.018 | Young |
| 0.155611 | 0.62179 | 0.015 | 0.008 | 1 | 0 | Mettl25 | 0.007 | Young |
| 0.155729 | 0.424699 | 0.075 | 0.06 | 1 | 0 | Usp1 | 0.015 | Young |
| 0.155917 | 0.298694 | 0.045 | 0.033 | 1 | 0 | Rnf214 | 0.012 | Young |
| 0.156102 | 0.526328 | 0.046 | 0.034 | 1 | 0 | Rnf31 | 0.012 | Young |
| 0.156298 | 0.463377 | 0.035 | 0.025 | 1 | 0 | Slf1 | 0.01 | Young |
| 0.158097 | 0.159768 | 0.294 | 0.275 | 1 | 0 | Msl1 | 0.019 | Young |
| 0.158127 | 0.345443 | 0.098 | 0.081 | 1 | 0 | Ppil4 | 0.017 | Young |
| 0.158218 | 0.539608 | 0.051 | 0.038 | 1 | 0 | Tom1l2 | 0.013 | Young |
| 0.158408 | 0.187022 | 0.243 | 0.222 | 1 | 0 | Cdkn1b | 0.021 | Young |
| 0.15845 | 0.622828 | 0.038 | 0.027 | 1 | 0 | Gprasp1 | 0.011 | Young |
| 0.158739 | 0.147521 | 0.174 | 0.152 | 1 | 0 | Smap1 | 0.022 | Young |
| 0.158942 | 1.1891 | 0.011 | 0.005 | 1 | 0 | Brf2 | 0.006 | Young |
| 0.159023 | 1.158412 | 0.011 | 0.005 | 1 | 0 | Zbtb7c | 0.006 | Young |
| 0.159276 | 0.602434 | 0.063 | 0.049 | 1 | 0 | Ranbp3 | 0.014 | Young |
| 0.159785 | 0.109772 | 0.651 | 0.647 | 1 | 0 | Ybx1 | 0.004 | Young |

| | | | | | | | | |
|---|---|---|---|---|---|---|---|---|
| 0.159825 | 0.027636 | 0.222 | 0.256 | 1 | 0 | Dctn1 | -0.034 | Young |
| 0.16018 | 0.220926 | 0.383 | 0.367 | 1 | 0 | Bzw1 | 0.016 | Young |
| 0.160243 | 0.780284 | 0.011 | 0.005 | 1 | 0 | Socs1 | 0.006 | Young |
| 0.16057 | 0.831467 | 0.011 | 0.005 | 1 | 0 | Gm42722 | 0.006 | Young |
| 0.160623 | 0.230109 | 0.149 | 0.129 | 1 | 0 | Uhrf1bp1l | 0.02 | Young |
| 0.160623 | 0.118263 | 0.016 | 0.025 | 1 | 0 | Ncaph | -0.009 | Young |
| 0.161265 | 0.364424 | 0.232 | 0.215 | 1 | 0 | Rab5a | 0.017 | Young |
| 0.161267 | 0.269489 | 0.267 | 0.248 | 1 | 0 | Ash1l | 0.019 | Young |
| 0.161424 | 0.407581 | 0.066 | 0.052 | 1 | 0 | Hrh1 | 0.014 | Young |
| 0.161568 | 0.006668 | 0.147 | 0.172 | 1 | 0 | Slc15a4 | -0.025 | Young |
| 0.161586 | 0.316213 | 0.198 | 0.179 | 1 | 0 | Tifa | 0.019 | Young |
| 0.161616 | 0.509423 | 0.019 | 0.011 | 1 | 0 | Gem | 0.008 | Young |
| 0.162055 | 0.150265 | 0.367 | 0.345 | 1 | 0 | Evi2a | 0.022 | Young |
| 0.162102 | 0.333526 | 0.078 | 0.063 | 1 | 0 | Knop1 | 0.015 | Young |
| 0.162129 | 0.431349 | 0.084 | 0.068 | 1 | 0 | Paxbp1 | 0.016 | Young |
| 0.162155 | 0.405247 | 0.06 | 0.047 | 1 | 0 | Foxk2 | 0.013 | Young |
| 0.162185 | 0.504537 | 0.043 | 0.031 | 1 | 0 | Ccdc88c | 0.012 | Young |
| 0.162191 | 0.348093 | 0.032 | 0.022 | 1 | 0 | Wrap73 | 0.01 | Young |
| 0.162246 | 0.400864 | 0.03 | 0.021 | 1 | 0 | Smim3 | 0.009 | Young |
| 0.162318 | 0.182913 | 0.135 | 0.116 | 1 | 0 | Slc14a1 | 0.019 | Young |
| 0.162344 | 0.409394 | 0.045 | 0.033 | 1 | 0 | Gm5617 | 0.012 | Young |
| 0.16239 | 0.020023 | 0.163 | 0.19 | 1 | 0 | Plekhg3 | -0.027 | Young |
| 0.163021 | 0.013228 | 0.416 | 0.464 | 1 | 0 | 5031439G( | -0.048 | Young |
| 0.163276 | 0.47413 | 0.019 | 0.011 | 1 | 0 | Gm16124 | 0.008 | Young |
| 0.164041 | 0.013609 | 0.071 | 0.056 | 1 | 0 | Hes1 | 0.015 | Young |
| 0.164186 | 0.018363 | 0.042 | 0.055 | 1 | 0 | Ppil3 | -0.013 | Young |
| 0.164223 | 0.512497 | 0.117 | 0.1 | 1 | 0 | Tcerg1 | 0.017 | Young |
| 0.16435 | 0.46823 | 0.069 | 0.055 | 1 | 0 | Usp40 | 0.014 | Young |
| 0.164428 | 0.109053 | 0.611 | 0.595 | 1 | 0 | Ninj1 | 0.016 | Young |
| 0.164627 | 0.082564 | 0.567 | 0.553 | 1 | 0 | Cyfip1 | 0.014 | Young |
| 0.16487 | 0.546654 | 0.02 | 0.012 | 1 | 0 | Eefsec | 0.008 | Young |
| 0.164872 | 0.669877 | 0.033 | 0.023 | 1 | 0 | Zfp839 | 0.01 | Young |
| 0.165163 | 0.008962 | 0.507 | 0.557 | 1 | 0 | Dse | -0.05 | Young |
| 0.165228 | 0.67777 | 0.064 | 0.051 | 1 | 0 | Fmo5 | 0.013 | Young |
| 0.165233 | 0.178946 | 0.329 | 0.312 | 1 | 0 | Scaf11 | 0.017 | Young |
| 0.165257 | 0.110247 | 0.552 | 0.546 | 1 | 0 | Rpl7a | 0.006 | Young |
| 0.165371 | 0.538804 | 0.043 | 0.031 | 1 | 0 | Mthfd2l | 0.012 | Young |
| 0.165391 | 0.18527 | 0.334 | 0.316 | 1 | 0 | Tlr2 | 0.018 | Young |
| 0.165576 | 0.460912 | 0.082 | 0.067 | 1 | 0 | Armc1 | 0.015 | Young |
| 0.16586 | 0.464971 | 0.057 | 0.044 | 1 | 0 | Med11 | 0.013 | Young |
| 0.166105 | 0.375933 | 0.025 | 0.016 | 1 | 0 | Pdk4 | 0.009 | Young |
| 0.166214 | 0.057983 | 0.511 | 0.498 | 1 | 0 | Ndufa2 | 0.013 | Young |
| 0.166551 | 0.02448 | 0.027 | 0.037 | 1 | 0 | Tenm4 | -0.01 | Young |
| 0.166631 | 0.229718 | 0.419 | 0.416 | 1 | 0 | Hmgb1 | 0.003 | Young |
| 0.16742 | 0.569958 | 0.063 | 0.049 | 1 | 0 | Zfp467 | 0.014 | Young |

| | | | | | | | | |
|---|---|---|---|---|---|---|---|---|
| 0.167633 | 1.185448 | 0.013 | 0.007 | 1 | 0 | R3hcc1 | 0.006 | Young |
| 0.167841 | 0.433568 | 0.079 | 0.064 | 1 | 0 | Ice1 | 0.015 | Young |
| 0.167853 | 0.246386 | 0.02 | 0.012 | 1 | 0 | Gemin5 | 0.008 | Young |
| 0.168499 | 0.067947 | 0.045 | 0.059 | 1 | 0 | Phf13 | -0.014 | Young |
| 0.168617 | 0.109803 | 0.693 | 0.705 | 1 | 0 | Rpl24 | -0.012 | Young |
| 0.169285 | 0.966414 | 0.013 | 0.007 | 1 | 0 | Snapc4 | 0.006 | Young |
| 0.169285 | 0.861513 | 0.013 | 0.007 | 1 | 0 | Atp1a2 | 0.006 | Young |
| 0.169332 | 0.216823 | 0.027 | 0.018 | 1 | 0 | Lclat1 | 0.009 | Young |
| 0.17006 | 0.303083 | 0.136 | 0.118 | 1 | 0 | Serpinf1 | 0.018 | Young |
| 0.170312 | 0.124847 | 0.132 | 0.112 | 1 | 0 | Cir1 | 0.02 | Young |
| 0.170551 | 0.646702 | 0.013 | 0.007 | 1 | 0 | Fastkd1 | 0.006 | Young |
| 0.170584 | 0.310019 | 0.134 | 0.115 | 1 | 0 | P3h2 | 0.019 | Young |
| 0.171659 | 0.27958 | 0.35 | 0.337 | 1 | 0 | Arhgap17 | 0.013 | Young |
| 0.172416 | 0.561821 | 0.033 | 0.023 | 1 | 0 | Cpeb3 | 0.01 | Young |
| 0.172433 | 0.274679 | 0.247 | 0.23 | 1 | 0 | Cdv3 | 0.017 | Young |
| 0.172815 | 0.399949 | 0.208 | 0.192 | 1 | 0 | Gpr160 | 0.016 | Young |
| 0.172932 | 0.367662 | 0.005 | 0.01 | 1 | 0 | Pi16 | -0.005 | Young |
| 0.173001 | 0.52404 | 0.059 | 0.047 | 1 | 0 | Exoc7 | 0.012 | Young |
| 0.173237 | 0.503165 | 0.044 | 0.033 | 1 | 0 | Btbd3 | 0.011 | Young |
| 0.173484 | 0.009536 | 0.075 | 0.092 | 1 | 0 | Zbtb7b | -0.017 | Young |
| 0.173883 | 0.186303 | 0.337 | 0.32 | 1 | 0 | Ski | 0.017 | Young |
| 0.174007 | 0.254037 | 0.057 | 0.044 | 1 | 0 | Prpf4 | 0.013 | Young |
| 0.174456 | 0.373729 | 0.041 | 0.03 | 1 | 0 | Rasal2 | 0.011 | Young |
| 0.175617 | 0.066524 | 0.043 | 0.056 | 1 | 0 | Kat2a | -0.013 | Young |
| 0.175643 | 0.094557 | 0.175 | 0.152 | 1 | 0 | Fbxw4 | 0.023 | Young |
| 0.175692 | 0.874562 | 0.025 | 0.016 | 1 | 0 | Spryd3 | 0.009 | Young |
| 0.175693 | 0.5552 | 0.047 | 0.036 | 1 | 0 | Fuom | 0.011 | Young |
| 0.176115 | 0.337707 | 0.03 | 0.021 | 1 | 0 | 1500015A( | 0.009 | Young |
| 0.176145 | 0.457437 | 0.058 | 0.045 | 1 | 0 | Gab3 | 0.013 | Young |
| 0.176197 | 0.278218 | 0.061 | 0.048 | 1 | 0 | Ccdc117 | 0.013 | Young |
| 0.176441 | 0.388745 | 0.156 | 0.138 | 1 | 0 | Ankrd40 | 0.018 | Young |
| 0.176766 | 0.213758 | 0.271 | 0.252 | 1 | 0 | Numa1 | 0.019 | Young |
| 0.176768 | 0.440097 | 0.167 | 0.15 | 1 | 0 | N4bp2l1 | 0.017 | Young |
| 0.177014 | 0.49575 | 0.07 | 0.056 | 1 | 0 | Ormdl3 | 0.014 | Young |
| 0.177354 | 0.619781 | 0.036 | 0.026 | 1 | 0 | Rnf8 | 0.01 | Young |
| 0.177443 | 0.22526 | 0.352 | 0.338 | 1 | 0 | Akr1b3 | 0.014 | Young |
| 0.17768 | 0.47749 | 0.056 | 0.044 | 1 | 0 | Incenp | 0.012 | Young |
| 0.177765 | 0.121377 | 0.465 | 0.447 | 1 | 0 | Rnasel | 0.018 | Young |
| 0.177784 | 0.124774 | 0.052 | 0.066 | 1 | 0 | Gm43848 | -0.014 | Young |
| 0.177838 | 0.51777 | 0.028 | 0.019 | 1 | 0 | Ptcd3 | 0.009 | Young |
| 0.178315 | 0.718221 | 0.035 | 0.025 | 1 | 0 | Batf2 | 0.01 | Young |
| 0.178636 | 0.544718 | 0.052 | 0.04 | 1 | 0 | Scyl1 | 0.012 | Young |
| 0.178806 | 0.474996 | 0.065 | 0.052 | 1 | 0 | Sp140 | 0.013 | Young |
| 0.178853 | 0.007155 | 0.034 | 0.045 | 1 | 0 | Polr3e | -0.011 | Young |
| 0.178977 | 0.339957 | 0.187 | 0.17 | 1 | 0 | Krtcap2 | 0.017 | Young |

| | | | | | | | | |
|---|---|---|---|---|---|---|---|---|
| 0.179218 | 0.013813 | 0.095 | 0.114 | 1 | 0 | Fam219a | -0.019 | Young |
| 0.179482 | 0.193125 | 0.494 | 0.492 | 1 | 0 | Atpif1 | 0.002 | Young |
| 0.180082 | 0.017014 | 0.209 | 0.238 | 1 | 0 | Epb41l2 | -0.029 | Young |
| 0.181139 | 0.793585 | 0.021 | 0.014 | 1 | 0 | Dedd2 | 0.007 | Young |
| 0.18123 | 0.729241 | 0.023 | 0.015 | 1 | 0 | Nipa1 | 0.008 | Young |
| 0.181321 | 0.495176 | 0.055 | 0.042 | 1 | 0 | Slc12a2 | 0.013 | Young |
| 0.181457 | 0.732775 | 0.025 | 0.016 | 1 | 0 | Ighd | 0.009 | Young |
| 0.181514 | 0.805922 | 0.02 | 0.012 | 1 | 0 | 9030025P2 | 0.008 | Young |
| 0.181941 | 0.695068 | 0.021 | 0.014 | 1 | 0 | Cd4 | 0.007 | Young |
| 0.18299 | 0.711575 | 0.02 | 0.012 | 1 | 0 | Btf3l4 | 0.008 | Young |
| 0.18328 | 0.417183 | 0.091 | 0.075 | 1 | 0 | Cytip | 0.016 | Young |
| 0.183335 | 0.541407 | 0.023 | 0.015 | 1 | 0 | 1110012L1 | 0.008 | Young |
| 0.183362 | 0.321901 | 0.148 | 0.13 | 1 | 0 | Gabarapl2 | 0.018 | Young |
| 0.183419 | 0.492701 | 0.088 | 0.074 | 1 | 0 | Hdac5 | 0.014 | Young |
| 0.18355 | 1.206669 | 0.011 | 0.005 | 1 | 0 | Cyp2e1 | 0.006 | Young |
| 0.183596 | 0.658007 | 0.02 | 0.012 | 1 | 0 | Plxna3 | 0.008 | Young |
| 0.184039 | 0.261623 | 0.307 | 0.293 | 1 | 0 | Pten | 0.014 | Young |
| 0.184077 | 0.01321 | 0.175 | 0.202 | 1 | 0 | AI314180 | -0.027 | Young |
| 0.184098 | 0.174612 | 0.421 | 0.408 | 1 | 0 | Rabac1 | 0.013 | Young |
| 0.184469 | 0.941729 | 0.011 | 0.005 | 1 | 0 | Armc9 | 0.006 | Young |
| 0.184519 | 0.073486 | 0.07 | 0.055 | 1 | 0 | Cables1 | 0.015 | Young |
| 0.184561 | 1.02602 | 0.011 | 0.005 | 1 | 0 | Fancm | 0.006 | Young |
| 0.184676 | 0.551376 | 0.087 | 0.073 | 1 | 0 | Mgat4a | 0.014 | Young |
| 0.184851 | 0.327113 | 0.018 | 0.011 | 1 | 0 | Mpv17l | 0.007 | Young |
| 0.184962 | 0.433506 | 0.126 | 0.109 | 1 | 0 | Flot2 | 0.017 | Young |
| 0.185239 | 0.171985 | 0.027 | 0.018 | 1 | 0 | Cep70 | 0.009 | Young |
| 0.185796 | 0.059768 | 0.214 | 0.246 | 1 | 0 | Slc11a1 | -0.032 | Young |
| 0.186275 | 0.429341 | 0.023 | 0.015 | 1 | 0 | Zfp317 | 0.008 | Young |
| 0.186501 | 0.844386 | 0.05 | 0.038 | 1 | 0 | Arl3 | 0.012 | Young |
| 0.187887 | 0.278838 | 0.097 | 0.081 | 1 | 0 | Mrpl12 | 0.016 | Young |
| 0.187991 | 0.18022 | 0.217 | 0.194 | 1 | 0 | Coq10b | 0.023 | Young |
| 0.188098 | 0.51832 | 0.062 | 0.049 | 1 | 0 | Ufd1 | 0.013 | Young |
| 0.188404 | 0.122749 | 0.029 | 0.04 | 1 | 0 | Mrps2 | -0.011 | Young |
| 0.189169 | 0.494131 | 0.031 | 0.022 | 1 | 0 | Tmem237 | 0.009 | Young |
| 0.189603 | 0.315688 | 0.117 | 0.1 | 1 | 0 | Pqlc1 | 0.017 | Young |
| 0.189634 | 0.518507 | 0.031 | 0.022 | 1 | 0 | Dkc1 | 0.009 | Young |
| 0.189649 | 0.740469 | 0.029 | 0.021 | 1 | 0 | Taf8 | 0.008 | Young |
| 0.189717 | 0.260447 | 0.228 | 0.208 | 1 | 0 | Alox5 | 0.02 | Young |
| 0.189721 | 0.260413 | 0.035 | 0.025 | 1 | 0 | Rnf113a2 | 0.01 | Young |
| 0.189893 | 0.628502 | 0.088 | 0.074 | 1 | 0 | Tsc1 | 0.014 | Young |
| 0.190248 | 0.51668 | 0.059 | 0.047 | 1 | 0 | Hsd17b10 | 0.012 | Young |
| 0.190526 | 0.305746 | 0.097 | 0.082 | 1 | 0 | Tmem37 | 0.015 | Young |
| 0.190867 | 0.199205 | 0.414 | 0.416 | 1 | 0 | Lrp1 | -0.002 | Young |
| 0.191138 | 0.755021 | 0.04 | 0.03 | 1 | 0 | Myg1 | 0.01 | Young |
| 0.192514 | 0.129937 | 0.018 | 0.026 | 1 | 0 | Trmt10c | -0.008 | Young |

| | | | | | | | | |
|---|---|---|---|---|---|---|---|---|
| 0.192592 | 1.044652 | 0.013 | 0.007 | 1 | 0 | Tmem177 | 0.006 | Young |
| 0.192955 | 0.003602 | 0.124 | 0.144 | 1 | 0 | Uqcrc1 | -0.02 | Young |
| 0.193298 | 0.748284 | 0.013 | 0.007 | 1 | 0 | Ybey | 0.006 | Young |
| 0.1933 | 0.011472 | 0.233 | 0.264 | 1 | 0 | Atf6 | -0.031 | Young |
| 0.19332 | 0.199452 | 0.132 | 0.114 | 1 | 0 | Thbd | 0.018 | Young |
| 0.193722 | 0.216049 | 0.119 | 0.103 | 1 | 0 | Srpk2 | 0.016 | Young |
| 0.193828 | 0.855734 | 0.013 | 0.007 | 1 | 0 | Setd4 | 0.006 | Young |
| 0.193851 | 0.76225 | 0.034 | 0.025 | 1 | 0 | Ccdc28b | 0.009 | Young |
| 0.193967 | 0.471658 | 0.029 | 0.021 | 1 | 0 | Ppargc1b | 0.008 | Young |
| 0.19425 | 0.398873 | 0.172 | 0.156 | 1 | 0 | Mtmr6 | 0.016 | Young |
| 0.194549 | 0.438307 | 0.059 | 0.047 | 1 | 0 | Dcaf1 | 0.012 | Young |
| 0.194899 | 0.235506 | 0.288 | 0.272 | 1 | 0 | Psenen | 0.016 | Young |
| 0.195426 | 0.280659 | 0.039 | 0.029 | 1 | 0 | Tmem126b | 0.01 | Young |
| 0.1956 | 0.403173 | 0.126 | 0.109 | 1 | 0 | Fam114a2 | 0.017 | Young |
| 0.195644 | 0.463495 | 0.101 | 0.086 | 1 | 0 | Fmr1 | 0.015 | Young |
| 0.195646 | 0.224919 | 0.091 | 0.075 | 1 | 0 | Aggf1 | 0.016 | Young |
| 0.196515 | 0.602641 | 0.039 | 0.029 | 1 | 0 | Brms1l | 0.01 | Young |
| 0.197022 | 0.344313 | 0.242 | 0.227 | 1 | 0 | Rapgef6 | 0.015 | Young |
| 0.197085 | 0.188567 | 0.378 | 0.363 | 1 | 0 | Slco2b1 | 0.015 | Young |
| 0.197302 | 0.264973 | 0.05 | 0.038 | 1 | 0 | Slc25a25 | 0.012 | Young |
| 0.197361 | 0.273496 | 0.28 | 0.265 | 1 | 0 | Fam102b | 0.015 | Young |
| 0.197933 | 0.118632 | 0.037 | 0.027 | 1 | 0 | Rcc1l | 0.01 | Young |
| 0.198194 | 0.633244 | 0.034 | 0.025 | 1 | 0 | Sparc | 0.009 | Young |
| 0.199827 | 0.278294 | 0.254 | 0.237 | 1 | 0 | Stub1 | 0.017 | Young |
| 0.199889 | 0.479271 | 0.04 | 0.03 | 1 | 0 | Nmi | 0.01 | Young |
| 0.200283 | 0.310194 | 0.126 | 0.109 | 1 | 0 | Pbx3 | 0.017 | Young |
| 0.200342 | 0.439196 | 0.051 | 0.04 | 1 | 0 | Rassf8 | 0.011 | Young |
| 0.200395 | 0.154979 | 0.073 | 0.059 | 1 | 0 | Zscan29 | 0.014 | Young |
| 0.200436 | 0.295085 | 0.162 | 0.144 | 1 | 0 | Helz | 0.018 | Young |
| 0.20053 | 0.841084 | 0.04 | 0.03 | 1 | 0 | Gna11 | 0.01 | Young |
| 0.20091 | 0.452243 | 0.034 | 0.025 | 1 | 0 | 2300009A0 | 0.009 | Young |
| 0.201039 | 0.859628 | 0.016 | 0.01 | 1 | 0 | Ncs1 | 0.006 | Young |
| 0.201164 | 0.610576 | 0.014 | 0.008 | 1 | 0 | Bend6 | 0.006 | Young |
| 0.201184 | 0.743443 | 0.021 | 0.014 | 1 | 0 | Zbtb11os1 | 0.007 | Young |
| 0.201347 | 0.558715 | 0.075 | 0.062 | 1 | 0 | Scaf1 | 0.013 | Young |
| 0.201682 | 0.739955 | 0.016 | 0.01 | 1 | 0 | Gm3448 | 0.006 | Young |
| 0.201752 | 0.501965 | 0.032 | 0.023 | 1 | 0 | Abhd10 | 0.009 | Young |
| 0.201791 | 0.32269 | 0.132 | 0.115 | 1 | 0 | 2410015M | 0.017 | Young |
| 0.202 | 0.954923 | 0.029 | 0.021 | 1 | 0 | Ints5.1 | 0.008 | Young |
| 0.202248 | 0.04767 | 0.076 | 0.092 | 1 | 0 | Fam199x | -0.016 | Young |
| 0.202365 | 0.201283 | 0.1 | 0.083 | 1 | 0 | Plcl2 | 0.017 | Young |
| 0.202462 | 0.002819 | 0.599 | 0.648 | 1 | 0 | Ywhae | -0.049 | Young |
| 0.202771 | 0.822145 | 0.018 | 0.011 | 1 | 0 | Cbx8 | 0.007 | Young |
| 0.202976 | 0.227621 | 0.047 | 0.036 | 1 | 0 | Gm48065 | 0.011 | Young |
| 0.203305 | 0.361487 | 0.146 | 0.13 | 1 | 0 | Plk2 | 0.016 | Young |

| | | | | | | | | |
|---|---|---|---|---|---|---|---|---|
| 0.203378 | 0.798063 | 0.019 | 0.012 | 1 | 0 | Zfp276 | 0.007 | Young |
| 0.203451 | 0.878568 | 0.019 | 0.012 | 1 | 0 | Vmac | 0.007 | Young |
| 0.203571 | 0.384981 | 0.034 | 0.025 | 1 | 0 | Mfng | 0.009 | Young |
| 0.203942 | 0.378049 | 0.04 | 0.03 | 1 | 0 | Zfp830 | 0.01 | Young |
| 0.204456 | 0.482391 | 0.059 | 0.047 | 1 | 0 | Dgcr6 | 0.012 | Young |
| 0.204928 | 0.652661 | 0.018 | 0.011 | 1 | 0 | Mtif3 | 0.007 | Young |
| 0.204943 | 0.027192 | 0.006 | 0.011 | 1 | 0 | Nlrp1a | -0.005 | Young |
| 0.205203 | 0.191494 | 0.171 | 0.152 | 1 | 0 | Themis2 | 0.019 | Young |
| 0.205224 | 0.882101 | 0.029 | 0.021 | 1 | 0 | Il1rap | 0.008 | Young |
| 0.205469 | 0.788598 | 0.048 | 0.037 | 1 | 0 | Mlx | 0.011 | Young |
| 0.205668 | 0.471196 | 0.023 | 0.015 | 1 | 0 | 2610035D1 | 0.008 | Young |
| 0.206025 | 0.268495 | 0.232 | 0.215 | 1 | 0 | Cpne3 | 0.017 | Young |
| 0.206396 | 0.395104 | 0.078 | 0.064 | 1 | 0 | Atl2 | 0.014 | Young |
| 0.206734 | 0.179487 | 0.372 | 0.36 | 1 | 0 | Mafg | 0.012 | Young |
| 0.206845 | 0.077413 | 0.054 | 0.067 | 1 | 0 | Minpp1 | -0.013 | Young |
| 0.206857 | 0.226867 | 0.222 | 0.204 | 1 | 0 | Mrpl23 | 0.018 | Young |
| 0.207393 | 0.248816 | 0.12 | 0.103 | 1 | 0 | Emd | 0.017 | Young |
| 0.207677 | 0.095361 | 0.419 | 0.405 | 1 | 0 | Ptpra | 0.014 | Young |
| 0.207936 | 0.22274 | 0.261 | 0.245 | 1 | 0 | Top2b | 0.016 | Young |
| 0.208213 | 0.098247 | 0.73 | 0.739 | 1 | 0 | Ptp4a2 | -0.009 | Young |
| 0.208273 | 0.021419 | 0.076 | 0.092 | 1 | 0 | Tbc1d15 | -0.016 | Young |
| 0.20873 | 0.199799 | 0.289 | 0.272 | 1 | 0 | Ndufa13 | 0.017 | Young |
| 0.208789 | 0.377672 | 0.029 | 0.021 | 1 | 0 | Flywch1 | 0.008 | Young |
| 0.208814 | 0.219654 | 0.196 | 0.176 | 1 | 0 | Usp34 | 0.02 | Young |
| 0.209035 | 0.487499 | 0.046 | 0.036 | 1 | 0 | Pin1 | 0.01 | Young |
| 0.20904 | 0.320331 | 0.07 | 0.056 | 1 | 0 | Sap30l | 0.014 | Young |
| 0.20908 | 0.249728 | 0.077 | 0.063 | 1 | 0 | Pkp4 | 0.014 | Young |
| 0.209088 | 0.376253 | 0.066 | 0.053 | 1 | 0 | Lztfl1 | 0.013 | Young |
| 0.209667 | 0.03113 | 0.085 | 0.101 | 1 | 0 | Ppfibp2 | -0.016 | Young |
| 0.209674 | 0.346722 | 0.056 | 0.044 | 1 | 0 | Fbxo4 | 0.012 | Young |
| 0.210528 | 0.31116 | 0.102 | 0.086 | 1 | 0 | Ube2a | 0.016 | Young |
| 0.210828 | 0.302826 | 0.088 | 0.074 | 1 | 0 | Otud6b | 0.014 | Young |
| 0.211041 | 0.203742 | 0.121 | 0.104 | 1 | 0 | Txndc16 | 0.017 | Young |
| 0.211498 | 0.414477 | 0.145 | 0.13 | 1 | 0 | Skiv2l | 0.015 | Young |
| 0.211531 | 0.30592 | 0.22 | 0.202 | 1 | 0 | Arpp19 | 0.018 | Young |
| 0.211778 | 0.356338 | 0.197 | 0.181 | 1 | 0 | Ggt5 | 0.016 | Young |
| 0.212267 | 0.269988 | 0.051 | 0.04 | 1 | 0 | Slc26a11 | 0.011 | Young |
| 0.212545 | 0.337145 | 0.067 | 0.055 | 1 | 0 | Ints12 | 0.012 | Young |
| 0.212774 | 0.661784 | 0.027 | 0.019 | 1 | 0 | Nrg2 | 0.008 | Young |
| 0.212835 | 0.873656 | 0.041 | 0.031 | 1 | 0 | Golga1 | 0.01 | Young |
| 0.21337 | 0.185533 | 0.065 | 0.052 | 1 | 0 | Erf | 0.013 | Young |
| 0.213985 | 0.707843 | 0.011 | 0.005 | 1 | 0 | Klhl3 | 0.006 | Young |
| 0.214091 | 0.238526 | 0.26 | 0.242 | 1 | 0 | Nfe2l1 | 0.018 | Young |
| 0.214194 | 0.652469 | 0.011 | 0.005 | 1 | 0 | Dpp4 | 0.006 | Young |
| 0.21436 | 0.473617 | 0.069 | 0.056 | 1 | 0 | Cpped1 | 0.013 | Young |

| | | | | | | | | |
|---|---|---|---|---|---|---|---|---|
| 0.21519 | 1.216704 | 0.029 | 0.021 | 1 | 0 | Dis3l | 0.008 | Young |
| 0.215398 | 0.486973 | 0.011 | 0.005 | 1 | 0 | Slc25a42 | 0.006 | Young |
| 0.215485 | 0.559364 | 0.056 | 0.045 | 1 | 0 | Med4 | 0.011 | Young |
| 0.215978 | 0.225553 | 0.18 | 0.16 | 1 | 0 | Gpr65 | 0.02 | Young |
| 0.216376 | 0.026384 | 0.212 | 0.241 | 1 | 0 | Stk4 | -0.029 | Young |
| 0.216473 | 0.056998 | 0.176 | 0.202 | 1 | 0 | Ankrd13a | -0.026 | Young |
| 0.217049 | 0.013058 | 0.27 | 0.3 | 1 | 0 | Stx4a | -0.03 | Young |
| 0.217122 | 0.170482 | 0.324 | 0.305 | 1 | 0 | Itga4 | 0.019 | Young |
| 0.217556 | 0.454343 | 0.128 | 0.114 | 1 | 0 | Rftn1 | 0.014 | Young |
| 0.217724 | 0.56841 | 0.038 | 0.029 | 1 | 0 | Snip1 | 0.009 | Young |
| 0.217995 | 0.220071 | 0.313 | 0.297 | 1 | 0 | Sptssa | 0.016 | Young |
| 0.219257 | 0.26705 | 0.201 | 0.182 | 1 | 0 | Psme4 | 0.019 | Young |
| 0.220019 | 0.902153 | 0.012 | 0.007 | 1 | 0 | Lcmt2 | 0.005 | Young |
| 0.220099 | 0.225209 | 0.316 | 0.304 | 1 | 0 | Ttc14 | 0.012 | Young |
| 0.220218 | 0.02042 | 0.995 | 0.989 | 1 | 0 | Rpl37a | 0.006 | Young |
| 0.220851 | 0.131256 | 0.565 | 0.576 | 1 | 0 | Akap13 | -0.011 | Young |
| 0.220953 | 0.255547 | 0.104 | 0.089 | 1 | 0 | Pdcd2l | 0.015 | Young |
| 0.221052 | 0.55079 | 0.083 | 0.07 | 1 | 0 | Mnat1 | 0.013 | Young |
| 0.221192 | 0.234087 | 0.169 | 0.152 | 1 | 0 | Ccdc12 | 0.017 | Young |
| 0.221773 | 0.473667 | 0.085 | 0.071 | 1 | 0 | Rad50 | 0.014 | Young |
| 0.221933 | 0.43351 | 0.09 | 0.077 | 1 | 0 | Prkab2 | 0.013 | Young |
| 0.222753 | 0.231849 | 0.247 | 0.227 | 1 | 0 | Hpgds | 0.02 | Young |
| 0.223086 | 0.57028 | 0.012 | 0.007 | 1 | 0 | Tnnt1 | 0.005 | Young |
| 0.223111 | 0.289131 | 0.072 | 0.059 | 1 | 0 | Exosc5 | 0.013 | Young |
| 0.223184 | 0.437407 | 0.102 | 0.088 | 1 | 0 | Cep68 | 0.014 | Young |
| 0.223432 | 0.388483 | 0.053 | 0.042 | 1 | 0 | Map3k3 | 0.011 | Young |
| 0.223688 | 1.32272 | 0.016 | 0.01 | 1 | 0 | Wdr24 | 0.006 | Young |
| 0.224171 | 0.320314 | 0.012 | 0.018 | 1 | 0 | Fn3krp | -0.006 | Young |
| 0.224392 | 0.152821 | 0.167 | 0.148 | 1 | 0 | P2ry12 | 0.019 | Young |
| 0.224446 | 0.354972 | 0.15 | 0.134 | 1 | 0 | Map3k2 | 0.016 | Young |
| 0.224747 | 0.187936 | 0.233 | 0.213 | 1 | 0 | Specc1l | 0.02 | Young |
| 0.224943 | 0.976713 | 0.019 | 0.012 | 1 | 0 | Phtf1 | 0.007 | Young |
| 0.225011 | 0.671505 | 0.047 | 0.037 | 1 | 0 | Slc39a3 | 0.01 | Young |
| 0.225041 | 1.030131 | 0.014 | 0.008 | 1 | 0 | Ak8 | 0.006 | Young |
| 0.225269 | 0.449806 | 0.113 | 0.098 | 1 | 0 | Otud5 | 0.015 | Young |
| 0.22564 | 0.03094 | 0.854 | 0.87 | 1 | 0 | Zfp36l2 | -0.016 | Young |
| 0.225897 | 0.531122 | 0.037 | 0.027 | 1 | 0 | Arhgef5 | 0.01 | Young |
| 0.226165 | 0.75745 | 0.014 | 0.008 | 1 | 0 | Lmnb2 | 0.006 | Young |
| 0.226198 | 0.373013 | 0.064 | 0.052 | 1 | 0 | Mroh1 | 0.012 | Young |
| 0.226272 | 0.327665 | 0.15 | 0.134 | 1 | 0 | Zranb1 | 0.016 | Young |
| 0.226451 | 0.502479 | 0.115 | 0.101 | 1 | 0 | Clpx | 0.014 | Young |
| 0.226688 | 0.841589 | 0.017 | 0.011 | 1 | 0 | Hirip3 | 0.006 | Young |
| 0.227293 | 0.653301 | 0.014 | 0.008 | 1 | 0 | Zfp41 | 0.006 | Young |
| 0.227392 | 0.0133 | 0.02 | 0.027 | 1 | 0 | Alg13 | -0.007 | Young |
| 0.227817 | 0.314098 | 0.022 | 0.015 | 1 | 0 | Zfp40 | 0.007 | Young |

| | | | | | | | | |
|---|---|---|---|---|---|---|---|---|
| 0.227996 | 0.537524 | 0.093 | 0.079 | 1 | 0 | Tex264 | 0.014 | Young |
| 0.228763 | 0.502973 | 0.04 | 0.03 | 1 | 0 | Dnajb2 | 0.01 | Young |
| 0.228854 | 0.340559 | 0.022 | 0.015 | 1 | 0 | Wee1 | 0.007 | Young |
| 0.228882 | 0.280448 | 0.038 | 0.029 | 1 | 0 | Stx3 | 0.009 | Young |
| 0.229019 | 0.037203 | 0.184 | 0.166 | 1 | 0 | Dnajb1 | 0.018 | Young |
| 0.22943 | 0.52451 | 0.04 | 0.03 | 1 | 0 | Dym | 0.01 | Young |
| 0.229442 | 0.603083 | 0.016 | 0.01 | 1 | 0 | Sec16b | 0.006 | Young |
| 0.229638 | 0.111477 | 0.02 | 0.027 | 1 | 0 | Galt | -0.007 | Young |
| 0.230157 | 0.496366 | 0.016 | 0.01 | 1 | 0 | Fxn | 0.006 | Young |
| 0.230395 | 0.027427 | 0.304 | 0.338 | 1 | 0 | Tax1bp1 | -0.034 | Young |
| 0.230432 | 0.225319 | 0.31 | 0.298 | 1 | 0 | Ndufb11 | 0.012 | Young |
| 0.230518 | 0.219221 | 0.128 | 0.111 | 1 | 0 | Fam103a1 | 0.017 | Young |
| 0.230931 | 0.245812 | 0.275 | 0.26 | 1 | 0 | Cpsf6 | 0.015 | Young |
| 0.231053 | 0.571874 | 0.019 | 0.012 | 1 | 0 | Il1a | 0.007 | Young |
| 0.231284 | 0.224839 | 0.184 | 0.167 | 1 | 0 | Crlf2 | 0.017 | Young |
| 0.23134 | 0.294634 | 0.177 | 0.16 | 1 | 0 | Trappc4 | 0.017 | Young |
| 0.231592 | 0.409463 | 0.016 | 0.01 | 1 | 0 | Phc1 | 0.006 | Young |
| 0.231857 | 0.339074 | 0.165 | 0.149 | 1 | 0 | Mrpl24 | 0.016 | Young |
| 0.23205 | 0.343554 | 0.07 | 0.057 | 1 | 0 | Rad54l2 | 0.013 | Young |
| 0.232296 | 0.99339 | 0.027 | 0.019 | 1 | 0 | Sdc1 | 0.008 | Young |
| 0.232521 | 0.202454 | 0.384 | 0.373 | 1 | 0 | Psd3 | 0.011 | Young |
| 0.232815 | 0.088942 | 0.008 | 0.012 | 1 | 0 | Eno3 | -0.004 | Young |
| 0.233246 | 0.257996 | 0.207 | 0.189 | 1 | 0 | Evi5 | 0.018 | Young |
| 0.233328 | 0.307302 | 0.097 | 0.083 | 1 | 0 | Phf1 | 0.014 | Young |
| 0.233493 | 0.396196 | 0.118 | 0.103 | 1 | 0 | Slc38a10 | 0.015 | Young |
| 0.233615 | 0.166407 | 0.524 | 0.52 | 1 | 0 | Lifr | 0.004 | Young |
| 0.233764 | 0.134721 | 0.059 | 0.048 | 1 | 0 | Tmx4 | 0.011 | Young |
| 0.233996 | 0.384422 | 0.107 | 0.092 | 1 | 0 | Nuak2 | 0.015 | Young |
| 0.234014 | 0.38443 | 0.123 | 0.108 | 1 | 0 | Mrps24 | 0.015 | Young |
| 0.23405 | 0.1752 | 0.476 | 0.479 | 1 | 0 | Rps27l | -0.003 | Young |
| 0.234413 | 0.32751 | 0.16 | 0.145 | 1 | 0 | Mepce | 0.015 | Young |
| 0.234464 | 0.091924 | 0.016 | 0.023 | 1 | 0 | Rmdn1 | -0.007 | Young |
| 0.234545 | 0.043566 | 0.016 | 0.023 | 1 | 0 | Cenpl | -0.007 | Young |
| 0.234635 | 0.027884 | 0.034 | 0.044 | 1 | 0 | Pstk | -0.01 | Young |
| 0.234888 | 0.138983 | 0.434 | 0.415 | 1 | 0 | Clec12a | 0.019 | Young |
| 0.234889 | 0.166719 | 0.019 | 0.012 | 1 | 0 | Wbp1 | 0.007 | Young |
| 0.235383 | 0.010138 | 0.375 | 0.413 | 1 | 0 | Rtn3 | -0.038 | Young |
| 0.236027 | 0.779182 | 0.025 | 0.018 | 1 | 0 | Ccdc28a | 0.007 | Young |
| 0.236041 | 0.083649 | 0.019 | 0.012 | 1 | 0 | AC163032. | 0.007 | Young |
| 0.236093 | 0.399562 | 0.114 | 0.1 | 1 | 0 | Commd3 | 0.014 | Young |
| 0.236145 | 0.330561 | 0.065 | 0.053 | 1 | 0 | Fcf1 | 0.012 | Young |
| 0.236214 | 0.236042 | 0.199 | 0.182 | 1 | 0 | Tcf12 | 0.017 | Young |
| 0.236473 | 0.560854 | 0.027 | 0.019 | 1 | 0 | Fkbp7 | 0.008 | Young |
| 0.236802 | 0.464801 | 0.029 | 0.021 | 1 | 0 | Zfp961 | 0.008 | Young |
| 0.238028 | 0.141012 | 0.016 | 0.023 | 1 | 0 | Mkks | -0.007 | Young |

| | | | | | | | | |
|---|---|---|---|---|---|---|---|---|
| 0.23841 | 0.145603 | 0.366 | 0.353 | 1 | 0 | Atp6v1f | 0.013 | Young |
| 0.238505 | 0.067173 | 0.165 | 0.187 | 1 | 0 | Leprotl1 | -0.022 | Young |
| 0.238524 | 0.348511 | 0.09 | 0.077 | 1 | 0 | Slu7 | 0.013 | Young |
| 0.238979 | 0.218442 | 0.112 | 0.097 | 1 | 0 | Trove2 | 0.015 | Young |
| 0.239334 | 0.028998 | 0.076 | 0.09 | 1 | 0 | Tatdn2 | -0.014 | Young |
| 0.239465 | 0.227805 | 0.241 | 0.224 | 1 | 0 | Tnrc6b | 0.017 | Young |
| 0.239617 | 0.335134 | 0.137 | 0.122 | 1 | 0 | Smarca4 | 0.015 | Young |
| 0.240329 | 0.223197 | 0.159 | 0.142 | 1 | 0 | Psmc6 | 0.017 | Young |
| 0.240381 | 0.347332 | 0.044 | 0.034 | 1 | 0 | Gins4 | 0.01 | Young |
| 0.241598 | 0.443835 | 0.116 | 0.103 | 1 | 0 | Trmt1 | 0.013 | Young |
| 0.242778 | 0.294022 | 0.083 | 0.07 | 1 | 0 | Tef | 0.013 | Young |
| 0.243481 | 0.016313 | 0.165 | 0.186 | 1 | 0 | Sec11a | -0.021 | Young |
| 0.24364 | 0.144816 | 0.411 | 0.402 | 1 | 0 | Pum2 | 0.009 | Young |
| 0.243975 | 0.453905 | 0.025 | 0.018 | 1 | 0 | Trp53rka | 0.007 | Young |
| 0.244684 | 0.024436 | 0.011 | 0.016 | 1 | 0 | Fbxo10 | -0.005 | Young |
| 0.244762 | 0.600035 | 0.035 | 0.026 | 1 | 0 | Clstn1 | 0.009 | Young |
| 0.244956 | 0.477049 | 0.054 | 0.044 | 1 | 0 | Rabggta | 0.01 | Young |
| 0.244989 | 0.736727 | 0.01 | 0.005 | 1 | 0 | Slc22a23 | 0.005 | Young |
| 0.245106 | 0.895421 | 0.01 | 0.005 | 1 | 0 | Adap2os | 0.005 | Young |
| 0.245106 | 0.727587 | 0.01 | 0.005 | 1 | 0 | Chst1 | 0.005 | Young |
| 0.245638 | 0.302677 | 0.093 | 0.079 | 1 | 0 | Tubgcp4 | 0.014 | Young |
| 0.246027 | 0.194491 | 0.018 | 0.025 | 1 | 0 | D030056L2 | -0.007 | Young |
| 0.246043 | 0.629935 | 0.01 | 0.005 | 1 | 0 | Uhrf1bp1 | 0.005 | Young |
| 0.246383 | 0.186242 | 0.197 | 0.181 | 1 | 0 | Agfg1 | 0.016 | Young |
| 0.246427 | 3.99E-05 | 0.117 | 0.135 | 1 | 0 | Sec23b | -0.018 | Young |
| 0.246549 | 0.107673 | 0.109 | 0.127 | 1 | 0 | Dazap1 | -0.018 | Young |
| 0.246598 | 0.486018 | 0.069 | 0.057 | 1 | 0 | Tusc2 | 0.012 | Young |
| 0.246866 | 0.533748 | 0.01 | 0.005 | 1 | 0 | Aplf | 0.005 | Young |
| 0.246878 | 0.428001 | 0.056 | 0.045 | 1 | 0 | Rab11fip2 | 0.011 | Young |
| 0.246948 | 0.029793 | 0.056 | 0.068 | 1 | 0 | Fbxo8 | -0.012 | Young |
| 0.24699 | 0.397838 | 0.108 | 0.094 | 1 | 0 | Sptan1 | 0.014 | Young |
| 0.247111 | 0.224098 | 0.192 | 0.175 | 1 | 0 | St3gal6 | 0.017 | Young |
| 0.247205 | 0.450247 | 0.024 | 0.016 | 1 | 0 | Gmds | 0.008 | Young |
| 0.247244 | 1.024416 | 0.02 | 0.014 | 1 | 0 | Cidec | 0.006 | Young |
| 0.247556 | 0.275806 | 0.038 | 0.029 | 1 | 0 | Zc3h10 | 0.009 | Young |
| 0.247572 | 0.380925 | 0.01 | 0.005 | 1 | 0 | Rad18 | 0.005 | Young |
| 0.247777 | 0.295648 | 0.124 | 0.109 | 1 | 0 | Lgals8 | 0.015 | Young |
| 0.247969 | 0.278879 | 0.101 | 0.086 | 1 | 0 | Phf5a | 0.015 | Young |
| 0.247985 | 0.0073 | 0.278 | 0.311 | 1 | 0 | Fgd2 | -0.033 | Young |
| 0.248009 | 0.250774 | 0.134 | 0.119 | 1 | 0 | Trappc5 | 0.015 | Young |
| 0.248295 | 0.263351 | 0.218 | 0.204 | 1 | 0 | Zfp652 | 0.014 | Young |
| 0.248392 | 0.422913 | 0.141 | 0.127 | 1 | 0 | Cc2d1b | 0.014 | Young |
| 0.248421 | 0.271337 | 0.011 | 0.016 | 1 | 0 | Slfn1 | -0.005 | Young |
| 0.248713 | 0.1954 | 0.076 | 0.063 | 1 | 0 | Dnase1l3 | 0.013 | Young |
| 0.249373 | 1.24172 | 0.012 | 0.007 | 1 | 0 | Frat2 | 0.005 | Young |

| | | | | | | | | |
|---|---|---|---|---|---|---|---|---|
| 0.249499 | 0.410714 | 0.044 | 0.034 | 1 | 0 | Sugp2 | 0.01 | Young |
| 0.249611 | 0.486195 | 0.057 | 0.047 | 1 | 0 | BC005561 | 0.01 | Young |
| 0.249755 | 0.958862 | 0.012 | 0.007 | 1 | 0 | Mr1 | 0.005 | Young |
| 0.250411 | 1.085084 | 0.012 | 0.007 | 1 | 0 | Fhit | 0.005 | Young |
| 0.250562 | 0.002022 | 0.088 | 0.104 | 1 | 0 | Pex7 | -0.016 | Young |
| 0.251537 | 0.387485 | 0.099 | 0.086 | 1 | 0 | Zc3h14 | 0.013 | Young |
| 0.251597 | 0.392996 | 0.054 | 0.044 | 1 | 0 | Dpm2 | 0.01 | Young |
| 0.251659 | 0.312773 | 0.051 | 0.041 | 1 | 0 | Gtf3a | 0.01 | Young |
| 0.251725 | 0.046962 | 0.173 | 0.197 | 1 | 0 | Bcas2 | -0.024 | Young |
| 0.25222 | 0.553034 | 0.035 | 0.026 | 1 | 0 | Supt3 | 0.009 | Young |
| 0.252248 | 0.390238 | 0.077 | 0.064 | 1 | 0 | Zfp384 | 0.013 | Young |
| 0.252317 | 0.328651 | 0.068 | 0.056 | 1 | 0 | Gm4951 | 0.012 | Young |
| 0.252703 | 0.077804 | 0.04 | 0.03 | 1 | 0 | Skint3 | 0.01 | Young |
| 0.252793 | 0.449905 | 0.035 | 0.026 | 1 | 0 | Phf11d | 0.009 | Young |
| 0.252967 | 0.348399 | 0.107 | 0.093 | 1 | 0 | Dirc2 | 0.014 | Young |
| 0.253066 | 0.218442 | 0.265 | 0.25 | 1 | 0 | Arhgap45 | 0.015 | Young |
| 0.253407 | 0.372329 | 0.022 | 0.015 | 1 | 0 | Zfp768 | 0.007 | Young |
| 0.25349 | 0.552856 | 0.012 | 0.007 | 1 | 0 | Capn5 | 0.005 | Young |
| 0.253706 | 0.999196 | 0.034 | 0.026 | 1 | 0 | Apool | 0.008 | Young |
| 0.253743 | 0.118817 | 0.073 | 0.06 | 1 | 0 | Qser1 | 0.013 | Young |
| 0.254145 | 0.822113 | 0.017 | 0.011 | 1 | 0 | Gm6563 | 0.006 | Young |
| 0.25415 | 0.455837 | 0.047 | 0.037 | 1 | 0 | Bola3 | 0.01 | Young |
| 0.254154 | 0.527598 | 0.012 | 0.007 | 1 | 0 | Acot1 | 0.005 | Young |
| 0.254599 | 0.35087 | 0.115 | 0.101 | 1 | 0 | Naa38 | 0.014 | Young |
| 0.254719 | 0.355111 | 0.147 | 0.133 | 1 | 0 | Foxn2 | 0.014 | Young |
| 0.254818 | 0.451227 | 0.012 | 0.007 | 1 | 0 | Aaas | 0.005 | Young |
| 0.25487 | 0.011943 | 0.134 | 0.153 | 1 | 0 | Tlr4 | -0.019 | Young |
| 0.254877 | 0.95422 | 0.013 | 0.008 | 1 | 0 | Coa7 | 0.005 | Young |
| 0.254929 | 0.432832 | 0.012 | 0.007 | 1 | 0 | Tmem67 | 0.005 | Young |
| 0.255291 | 0.796692 | 0.013 | 0.008 | 1 | 0 | Bcl7a | 0.005 | Young |
| 0.255409 | 0.027001 | 0.157 | 0.178 | 1 | 0 | Aprt | -0.021 | Young |
| 0.255617 | 0.193882 | 0.092 | 0.078 | 1 | 0 | Pknox1 | 0.014 | Young |
| 0.255711 | 0.729992 | 0.017 | 0.011 | 1 | 0 | Colca2 | 0.006 | Young |
| 0.255788 | 0.092656 | 0.032 | 0.041 | 1 | 0 | 0610009B2 | -0.009 | Young |
| 0.255916 | 0.087638 | 0.052 | 0.041 | 1 | 0 | Cdkn2aip | 0.011 | Young |
| 0.256223 | 0.748455 | 0.013 | 0.008 | 1 | 0 | A430046D1 | 0.005 | Young |
| 0.256847 | 0.630685 | 0.013 | 0.008 | 1 | 0 | Cep112 | 0.005 | Young |
| 0.25693 | 0.719816 | 0.015 | 0.01 | 1 | 0 | Gtf2ird2 | 0.005 | Young |
| 0.257125 | 0.68763 | 0.015 | 0.01 | 1 | 0 | Pigo | 0.005 | Young |
| 0.257642 | 0.447993 | 0.142 | 0.129 | 1 | 0 | Zfp451 | 0.013 | Young |
| 0.257701 | 0.290067 | 0.196 | 0.182 | 1 | 0 | Snx20 | 0.014 | Young |
| 0.257914 | 0.048005 | 0.238 | 0.265 | 1 | 0 | Rpl13a | -0.027 | Young |
| 0.258168 | 0.405837 | 0.028 | 0.021 | 1 | 0 | Lsm11 | 0.007 | Young |
| 0.258452 | 0.177586 | 0.052 | 0.041 | 1 | 0 | Larp1b | 0.011 | Young |
| 0.258541 | 0.006599 | 0.026 | 0.034 | 1 | 0 | Rpap3 | -0.008 | Young |

| | | | | | | | | |
|---|---|---|---|---|---|---|---|---|
| 0.258722 | 0.369779 | 0.013 | 0.008 | 1 | 0 | Map3k12 | 0.005 | Young |
| 0.258906 | 0.284933 | 0.088 | 0.075 | 1 | 0 | Ing3 | 0.013 | Young |
| 0.259611 | 0.413629 | 0.017 | 0.011 | 1 | 0 | Katnb1 | 0.006 | Young |
| 0.259644 | 0.509273 | 0.048 | 0.038 | 1 | 0 | Ero1lb | 0.01 | Young |
| 0.260422 | 0.220771 | 0.145 | 0.129 | 1 | 0 | Arid4b | 0.016 | Young |
| 0.260442 | 0.179311 | 0.147 | 0.131 | 1 | 0 | Rgs19 | 0.016 | Young |
| 0.260448 | 0.155225 | 0.012 | 0.018 | 1 | 0 | Zfp668 | -0.006 | Young |
| 0.260858 | 0.12034 | 0.092 | 0.078 | 1 | 0 | Zbtb18 | 0.014 | Young |
| 0.26115 | 0.459652 | 0.053 | 0.042 | 1 | 0 | Rprd1a | 0.011 | Young |
| 0.261288 | 0.24523 | 0.222 | 0.208 | 1 | 0 | Fkbp8 | 0.014 | Young |
| 0.261437 | 0.293845 | 0.227 | 0.215 | 1 | 0 | Usp24 | 0.012 | Young |
| 0.261707 | 0.157122 | 0.118 | 0.103 | 1 | 0 | Fryl | 0.015 | Young |
| 0.261817 | 0.046906 | 0.904 | 0.903 | 1 | 0 | Eif1 | 0.001 | Young |
| 0.262093 | 0.216387 | 0.159 | 0.142 | 1 | 0 | S100a1 | 0.017 | Young |
| 0.262825 | 0.321497 | 0.11 | 0.097 | 1 | 0 | Mtr | 0.013 | Young |
| 0.263345 | 0.114465 | 0.4 | 0.389 | 1 | 0 | Tsc22d4 | 0.011 | Young |
| 0.263716 | 0.656866 | 0.039 | 0.03 | 1 | 0 | Rubcnl | 0.009 | Young |
| 0.263892 | 0.343468 | 0.085 | 0.073 | 1 | 0 | Ccng2 | 0.012 | Young |
| 0.263931 | 0.432387 | 0.025 | 0.018 | 1 | 0 | Msh3 | 0.007 | Young |
| 0.26422 | 0.210518 | 0.075 | 0.063 | 1 | 0 | Msi2 | 0.012 | Young |
| 0.26523 | 0.466671 | 0.063 | 0.052 | 1 | 0 | Hspb11 | 0.011 | Young |
| 0.265332 | 0.136162 | 0.059 | 0.048 | 1 | 0 | Arl10 | 0.011 | Young |
| 0.265651 | 0.81108 | 0.046 | 0.037 | 1 | 0 | Rcan1 | 0.009 | Young |
| 0.267208 | 0.326969 | 0.187 | 0.174 | 1 | 0 | Micu1 | 0.013 | Young |
| 0.267291 | 0.639398 | 0.029 | 0.022 | 1 | 0 | Pan2 | 0.007 | Young |
| 0.267725 | 0.248175 | 0.119 | 0.104 | 1 | 0 | Rbm28 | 0.015 | Young |
| 0.267859 | 0.355202 | 0.075 | 0.063 | 1 | 0 | Nup54 | 0.012 | Young |
| 0.268051 | 0.210461 | 0.126 | 0.112 | 1 | 0 | Phf10 | 0.014 | Young |
| 0.268084 | 0.420352 | 0.036 | 0.027 | 1 | 0 | Vipas39 | 0.009 | Young |
| 0.268245 | 0.280247 | 0.154 | 0.14 | 1 | 0 | Xpo7 | 0.014 | Young |
| 0.268335 | 0.363851 | 0.12 | 0.107 | 1 | 0 | Hspa4l | 0.013 | Young |
| 0.268784 | 0.493963 | 0.126 | 0.114 | 1 | 0 | Tnfaip1 | 0.012 | Young |
| 0.269143 | 0.753204 | 0.028 | 0.021 | 1 | 0 | Zfp319 | 0.007 | Young |
| 0.269259 | 0.143185 | 0.258 | 0.241 | 1 | 0 | Ppp1r21 | 0.017 | Young |
| 0.269573 | 0.146897 | 0.037 | 0.029 | 1 | 0 | Thoc5 | 0.008 | Young |
| 0.269739 | 0.263794 | 0.297 | 0.291 | 1 | 0 | Ikbkb | 0.006 | Young |
| 0.270156 | 0.196448 | 0.057 | 0.047 | 1 | 0 | Chst7 | 0.01 | Young |
| 0.270554 | 0.0469 | 0.248 | 0.228 | 1 | 0 | Kat2b | 0.02 | Young |
| 0.270859 | 0.483183 | 0.048 | 0.038 | 1 | 0 | Usp39 | 0.01 | Young |
| 0.271253 | 0.180552 | 0.171 | 0.155 | 1 | 0 | Ccdc47 | 0.016 | Young |
| 0.272056 | 0.450379 | 0.094 | 0.082 | 1 | 0 | Phf2 | 0.012 | Young |
| 0.272357 | 0.378724 | 0.043 | 0.034 | 1 | 0 | Dock4 | 0.009 | Young |
| 0.273236 | 0.363696 | 0.089 | 0.077 | 1 | 0 | Bag6 | 0.012 | Young |
| 0.273579 | 0.294023 | 0.182 | 0.167 | 1 | 0 | Hcfc1r1 | 0.015 | Young |
| 0.273609 | 0.053803 | 0.128 | 0.146 | 1 | 0 | Ubqln2 | -0.018 | Young |

| | | | | | | | | |
|---|---|---|---|---|---|---|---|---|
| 0.273652 | 0.542214 | 0.045 | 0.036 | 1 | 0 | Cryzl2 | 0.009 | Young |
| 0.27388 | 0.345165 | 0.046 | 0.037 | 1 | 0 | Gamt | 0.009 | Young |
| 0.273922 | 0.342644 | 0.036 | 0.027 | 1 | 0 | Pola1 | 0.009 | Young |
| 0.27406 | 0.419516 | 0.069 | 0.057 | 1 | 0 | Coasy | 0.012 | Young |
| 0.274447 | 0.569681 | 0.021 | 0.015 | 1 | 0 | Aldh4a1 | 0.006 | Young |
| 0.275879 | 0.237333 | 0.08 | 0.068 | 1 | 0 | Gm38399 | 0.012 | Young |
| 0.276466 | 0.56184 | 0.051 | 0.041 | 1 | 0 | Cep57 | 0.01 | Young |
| 0.276636 | 0.030692 | 0.054 | 0.066 | 1 | 0 | Pycr2 | -0.012 | Young |
| 0.276689 | 0.005567 | 0.058 | 0.07 | 1 | 0 | Tmem2 | -0.012 | Young |
| 0.277063 | 0.008416 | 0.126 | 0.144 | 1 | 0 | Pcna | -0.018 | Young |
| 0.277692 | 0.269554 | 0.144 | 0.13 | 1 | 0 | Nhlrc3 | 0.014 | Young |
| 0.277698 | 0.386598 | 0.061 | 0.051 | 1 | 0 | Fktn | 0.01 | Young |
| 0.278368 | 0.552909 | 0.02 | 0.014 | 1 | 0 | Cdc6 | 0.006 | Young |
| 0.278628 | 0.224367 | 0.304 | 0.291 | 1 | 0 | Spag9 | 0.013 | Young |
| 0.278659 | 0.212107 | 0.021 | 0.015 | 1 | 0 | Padi2 | 0.006 | Young |
| 0.278931 | 0.12507 | 0.105 | 0.122 | 1 | 0 | Rnf38 | -0.017 | Young |
| 0.278999 | 0.773642 | 0.02 | 0.014 | 1 | 0 | Btd | 0.006 | Young |
| 0.279761 | 0.550412 | 0.045 | 0.036 | 1 | 0 | Tns4 | 0.009 | Young |
| 0.279998 | 1.235691 | 0.01 | 0.005 | 1 | 0 | Tnni2 | 0.005 | Young |
| 0.279998 | 1.037173 | 0.01 | 0.005 | 1 | 0 | Lag3 | 0.005 | Young |
| 0.280126 | 0.322449 | 0.052 | 0.042 | 1 | 0 | Bhlhe40 | 0.01 | Young |
| 0.280262 | 0.531943 | 0.02 | 0.014 | 1 | 0 | Tnfsfm13 | 0.006 | Young |
| 0.280269 | 0.747586 | 0.044 | 0.036 | 1 | 0 | Rnf185 | 0.008 | Young |
| 0.280321 | 0.111553 | 0.533 | 0.534 | 1 | 0 | Hmox1 | -0.001 | Young |
| 0.280372 | 0.281916 | 0.231 | 0.22 | 1 | 0 | Sirt2 | 0.011 | Young |
| 0.281139 | 0.00116 | 0.025 | 0.033 | 1 | 0 | Maml3 | -0.008 | Young |
| 0.281149 | 0.276189 | 0.061 | 0.051 | 1 | 0 | Fkbp9 | 0.01 | Young |
| 0.281226 | 0.082383 | 0.57 | 0.558 | 1 | 0 | Bri3 | 0.012 | Young |
| 0.281314 | 0.571436 | 0.074 | 0.063 | 1 | 0 | Bmi1 | 0.011 | Young |
| 0.281705 | 0.233785 | 0.241 | 0.228 | 1 | 0 | Gfra2 | 0.013 | Young |
| 0.281969 | 0.34218 | 0.081 | 0.07 | 1 | 0 | Lpar6 | 0.011 | Young |
| 0.282542 | 0.416288 | 0.083 | 0.071 | 1 | 0 | Fam120b | 0.012 | Young |
| 0.283317 | 0.010464 | 0.103 | 0.119 | 1 | 0 | Atp7a | -0.016 | Young |
| 0.283803 | 0.472035 | 0.01 | 0.005 | 1 | 0 | Gm26624 | 0.005 | Young |
| 0.283821 | 0.027097 | 1 | 1 | 1 | 0 | Tmsb4x | 0 | Young |
| 0.28386 | 0.244664 | 0.018 | 0.012 | 1 | 0 | Pde1b | 0.006 | Young |
| 0.283935 | 0.26819 | 0.01 | 0.005 | 1 | 0 | Gatsl3 | 0.005 | Young |
| 0.283983 | 0.080598 | 0.042 | 0.052 | 1 | 0 | Zfp101 | -0.01 | Young |
| 0.284175 | 0.352753 | 0.156 | 0.142 | 1 | 0 | Nfatc1 | 0.014 | Young |
| 0.284337 | 0.708442 | 0.018 | 0.012 | 1 | 0 | Pfas | 0.006 | Young |
| 0.284641 | 0.384326 | 0.114 | 0.101 | 1 | 0 | Pigp | 0.013 | Young |
| 0.284981 | 0.489858 | 0.026 | 0.019 | 1 | 0 | Gm26669 | 0.007 | Young |
| 0.285489 | 0.249632 | 0.285 | 0.275 | 1 | 0 | Ppp2r5a | 0.01 | Young |
| 0.285688 | 0.035936 | 0.1 | 0.086 | 1 | 0 | Mfap3 | 0.014 | Young |
| 0.285856 | 0.404442 | 0.026 | 0.019 | 1 | 0 | Dis3 | 0.007 | Young |

| | | | | | | | | |
|---|---|---|---|---|---|---|---|---|
| 0.286133 | 0.870099 | 0.011 | 0.007 | 1 | 0 | Ispd | 0.004 | Young |
| 0.286377 | 0.797403 | 0.016 | 0.011 | 1 | 0 | Cby1 | 0.005 | Young |
| 0.286564 | 0.091418 | 0.127 | 0.145 | 1 | 0 | 4933434E2 | -0.018 | Young |
| 0.286843 | 0.115159 | 0.015 | 0.021 | 1 | 0 | Mettl17 | -0.006 | Young |
| 0.286891 | 0.439331 | 0.026 | 0.019 | 1 | 0 | Fam217b | 0.007 | Young |
| 0.28723 | 0.009067 | 0.063 | 0.075 | 1 | 0 | Mex3d | -0.012 | Young |
| 0.287353 | 0.597197 | 0.011 | 0.007 | 1 | 0 | 2510046G1 | 0.004 | Young |
| 0.287454 | 0.484599 | 0.058 | 0.048 | 1 | 0 | Camsap1 | 0.01 | Young |
| 0.287847 | 0.019533 | 0.134 | 0.152 | 1 | 0 | Srek1 | -0.018 | Young |
| 0.288339 | 0.447697 | 0.043 | 0.034 | 1 | 0 | Mpi | 0.009 | Young |
| 0.289088 | 0.26242 | 0.066 | 0.055 | 1 | 0 | Maml2 | 0.011 | Young |
| 0.289156 | 0.190868 | 0.115 | 0.101 | 1 | 0 | Csad | 0.014 | Young |
| 0.289283 | 0.060241 | 0.665 | 0.666 | 1 | 0 | Rpl36a | -0.001 | Young |
| 0.289429 | 0.055048 | 0.011 | 0.016 | 1 | 0 | Eral1 | -0.005 | Young |
| 0.289629 | 0.329714 | 0.056 | 0.047 | 1 | 0 | Anks3 | 0.009 | Young |
| 0.289681 | 0.359374 | 0.011 | 0.007 | 1 | 0 | Rai14 | 0.004 | Young |
| 0.289734 | 0.267274 | 0.179 | 0.164 | 1 | 0 | Dhx40 | 0.015 | Young |
| 0.289793 | 0.421042 | 0.056 | 0.047 | 1 | 0 | Klhl21 | 0.009 | Young |
| 0.28988 | 0.04798 | 0.046 | 0.056 | 1 | 0 | Trim36 | -0.01 | Young |
| 0.289935 | 0.118527 | 0.119 | 0.104 | 1 | 0 | Actr1a | 0.015 | Young |
| 0.290334 | 0.17651 | 0.042 | 0.033 | 1 | 0 | Pomgnt1 | 0.009 | Young |
| 0.290914 | 0.404618 | 0.015 | 0.01 | 1 | 0 | Sdr42e1 | 0.005 | Young |
| 0.290998 | 0.476091 | 0.08 | 0.068 | 1 | 0 | Atg4b | 0.012 | Young |
| 0.291051 | 0.254528 | 0.138 | 0.124 | 1 | 0 | Arhgap26 | 0.014 | Young |
| 0.291158 | 0.413452 | 0.011 | 0.007 | 1 | 0 | Zfp652os | 0.004 | Young |
| 0.29121 | 0.595815 | 0.029 | 0.022 | 1 | 0 | Kti12 | 0.007 | Young |
| 0.291391 | 0.331325 | 0.176 | 0.161 | 1 | 0 | Cnpy2 | 0.015 | Young |
| 0.291683 | 0.251819 | 0.128 | 0.115 | 1 | 0 | Kctd2 | 0.013 | Young |
| 0.291833 | 0.425726 | 0.049 | 0.04 | 1 | 0 | Mre11a | 0.009 | Young |
| 0.291842 | 0.401758 | 0.087 | 0.075 | 1 | 0 | Lsm7 | 0.012 | Young |
| 0.291957 | 0.316358 | 0.137 | 0.123 | 1 | 0 | Ssbp4 | 0.014 | Young |
| 0.291971 | 0.002079 | 0.273 | 0.298 | 1 | 0 | Hprt | -0.025 | Young |
| 0.292038 | 0.463121 | 0.072 | 0.062 | 1 | 0 | Tigd2 | 0.01 | Young |
| 0.292491 | 0.023469 | 0.157 | 0.176 | 1 | 0 | Pcf11 | -0.019 | Young |
| 0.292508 | 0.422186 | 0.013 | 0.008 | 1 | 0 | Gpsm1 | 0.005 | Young |
| 0.292735 | 0.372199 | 0.035 | 0.027 | 1 | 0 | Thyn1 | 0.008 | Young |
| 0.292836 | 0.028205 | 0.155 | 0.175 | 1 | 0 | Ccnl2 | -0.02 | Young |
| 0.292975 | 0.493866 | 0.034 | 0.026 | 1 | 0 | Fam149b | 0.008 | Young |
| 0.29328 | 0.016303 | 0.155 | 0.175 | 1 | 0 | Trim35 | -0.02 | Young |
| 0.293535 | 0.211028 | 0.183 | 0.167 | 1 | 0 | Slamf7 | 0.016 | Young |
| 0.293801 | 0.082862 | 0.476 | 0.469 | 1 | 0 | Hacd4 | 0.007 | Young |
| 0.293926 | 0.141185 | 0.322 | 0.305 | 1 | 0 | Leng8 | 0.017 | Young |
| 0.293927 | 0.283552 | 0.196 | 0.183 | 1 | 0 | Col4a3bp | 0.013 | Young |
| 0.293991 | 0.459896 | 0.112 | 0.1 | 1 | 0 | Bcl7c | 0.012 | Young |
| 0.294011 | 0.477978 | 0.024 | 0.018 | 1 | 0 | Msto1 | 0.006 | Young |

| | | | | | | | | |
|---|---|---|---|---|---|---|---|---|
| 0.294081 | 0.194726 | 0.075 | 0.089 | 1 | 0 | Swt1 | -0.014 | Young |
| 0.294561 | 0.210226 | 0.085 | 0.073 | 1 | 0 | Usp12 | 0.012 | Young |
| 0.294944 | 0.006463 | 0.122 | 0.138 | 1 | 0 | Npepl1 | -0.016 | Young |
| 0.29598 | 0.170983 | 0.201 | 0.186 | 1 | 0 | Ppt1 | 0.015 | Young |
| 0.296994 | 0.537463 | 0.05 | 0.041 | 1 | 0 | Kif21b | 0.009 | Young |
| 0.297154 | 0.486436 | 0.023 | 0.016 | 1 | 0 | Gm9493 | 0.007 | Young |
| 0.297181 | 0.170929 | 0.203 | 0.185 | 1 | 0 | Trafd1 | 0.018 | Young |
| 0.297222 | 0.106337 | 0.016 | 0.022 | 1 | 0 | Usp20 | -0.006 | Young |
| 0.297408 | 0.7433 | 0.057 | 0.048 | 1 | 0 | Atn1 | 0.009 | Young |
| 0.297942 | 0.600491 | 0.023 | 0.016 | 1 | 0 | Tnk2 | 0.007 | Young |
| 0.298013 | 0.498661 | 0.08 | 0.07 | 1 | 0 | Tgfbr1 | 0.01 | Young |
| 0.29811 | 0.099294 | 0.037 | 0.047 | 1 | 0 | Slc25a19 | -0.01 | Young |
| 0.298792 | 0.319781 | 0.062 | 0.052 | 1 | 0 | Zfp329 | 0.01 | Young |
| 0.299218 | 0.177041 | 0.455 | 0.464 | 1 | 0 | Arhgap18 | -0.009 | Young |
| 0.299703 | 0.333485 | 0.086 | 0.074 | 1 | 0 | Gkap1 | 0.012 | Young |
| 0.30008 | 0.103746 | 0.561 | 0.566 | 1 | 0 | H2afz | -0.005 | Young |
| 0.300238 | 0.645014 | 0.027 | 0.021 | 1 | 0 | Gorasp1 | 0.006 | Young |
| 0.300893 | 0.197198 | 0.259 | 0.245 | 1 | 0 | Adcy7 | 0.014 | Young |
| 0.301277 | 0.370613 | 0.078 | 0.067 | 1 | 0 | Upf3b | 0.011 | Young |
| 0.301565 | 0.281653 | 0.068 | 0.057 | 1 | 0 | Dok3 | 0.011 | Young |
| 0.301777 | 0.006237 | 0.105 | 0.12 | 1 | 0 | Sdccag3 | -0.015 | Young |
| 0.301884 | 0.007237 | 0.251 | 0.278 | 1 | 0 | Myof | -0.027 | Young |
| 0.301925 | 0.599041 | 0.027 | 0.021 | 1 | 0 | Adrb1 | 0.006 | Young |
| 0.302119 | 0.029361 | 0.095 | 0.109 | 1 | 0 | Tbc1d17 | -0.014 | Young |
| 0.302746 | 0.46039 | 0.079 | 0.068 | 1 | 0 | Thap12 | 0.011 | Young |
| 0.302892 | 1.127412 | 0.029 | 0.022 | 1 | 0 | Usp28 | 0.007 | Young |
| 0.302958 | 0.719464 | 0.03 | 0.023 | 1 | 0 | Mul1 | 0.007 | Young |
| 0.303427 | 0.560916 | 0.04 | 0.031 | 1 | 0 | Slc37a4 | 0.009 | Young |
| 0.304021 | 0.077993 | 0.12 | 0.137 | 1 | 0 | Abca3 | -0.017 | Young |
| 0.304451 | 0.292279 | 0.235 | 0.224 | 1 | 0 | Fkbp15 | 0.011 | Young |
| 0.304697 | 0.052808 | 0.198 | 0.22 | 1 | 0 | Tle4 | -0.022 | Young |
| 0.304702 | 0.166994 | 0.086 | 0.1 | 1 | 0 | Gm8186 | -0.014 | Young |
| 0.304882 | 0.334082 | 0.072 | 0.062 | 1 | 0 | Cog3 | 0.01 | Young |
| 0.305076 | 0.513732 | 0.04 | 0.031 | 1 | 0 | Nfkbil1 | 0.009 | Young |
| 0.305124 | 0.234381 | 0.203 | 0.192 | 1 | 0 | Oxct1 | 0.011 | Young |
| 0.305865 | 0.294012 | 0.064 | 0.053 | 1 | 0 | Zfp524 | 0.011 | Young |
| 0.306521 | 0.491397 | 0.032 | 0.025 | 1 | 0 | Zfp65 | 0.007 | Young |
| 0.306629 | 0.789442 | 0.019 | 0.014 | 1 | 0 | Erc1 | 0.005 | Young |
| 0.306715 | 0.407668 | 0.044 | 0.036 | 1 | 0 | Arfgap3 | 0.008 | Young |
| 0.306822 | 0.269161 | 0.086 | 0.074 | 1 | 0 | Scoc | 0.012 | Young |
| 0.307486 | 0.009936 | 0.033 | 0.041 | 1 | 0 | Mrps10 | -0.008 | Young |
| 0.307538 | 0.232235 | 0.231 | 0.218 | 1 | 0 | Cbx3 | 0.013 | Young |
| 0.307677 | 0.117883 | 0.043 | 0.034 | 1 | 0 | Pskh1 | 0.009 | Young |
| 0.307978 | 0.009386 | 0.237 | 0.263 | 1 | 0 | Etf1 | -0.026 | Young |
| 0.307987 | 0.766405 | 0.019 | 0.014 | 1 | 0 | Apbb1 | 0.005 | Young |

| | | | | | | | | |
|---|---|---|---|---|---|---|---|---|
| 0.308404 | 0.126497 | 0.172 | 0.156 | 1 | 0 | Zfc3h1 | 0.016 | Young |
| 0.308567 | 0.011286 | 0.117 | 0.134 | 1 | 0 | Retreg3 | -0.017 | Young |
| 0.308864 | 0.409146 | 0.092 | 0.081 | 1 | 0 | Grsf1 | 0.011 | Young |
| 0.309511 | 0.45469 | 0.026 | 0.019 | 1 | 0 | Rpap1 | 0.007 | Young |
| 0.309722 | 0.435334 | 0.075 | 0.064 | 1 | 0 | G6pdx | 0.011 | Young |
| 0.309969 | 0.099405 | 0.57 | 0.573 | 1 | 0 | Glud1 | -0.003 | Young |
| 0.310422 | 0.524487 | 0.019 | 0.014 | 1 | 0 | Platr25 | 0.005 | Young |
| 0.310453 | 0.062429 | 0.112 | 0.129 | 1 | 0 | Ncaph2 | -0.017 | Young |
| 0.31054 | 0.081482 | 0.617 | 0.622 | 1 | 0 | Sh3glb1 | -0.005 | Young |
| 0.310778 | 0.203216 | 0.218 | 0.204 | 1 | 0 | Zfp148 | 0.014 | Young |
| 0.31094 | 0.001434 | 0.318 | 0.346 | 1 | 0 | Hook3 | -0.028 | Young |
| 0.311758 | 0.02129 | 0.241 | 0.264 | 1 | 0 | Cuta | -0.023 | Young |
| 0.312029 | 0.054193 | 0.719 | 0.728 | 1 | 0 | Canx | -0.009 | Young |
| 0.312477 | 0.312108 | 0.092 | 0.081 | 1 | 0 | Fnta | 0.011 | Young |
| 0.312923 | 0.382815 | 0.131 | 0.119 | 1 | 0 | Synrg | 0.012 | Young |
| 0.312984 | 0.402 | 0.066 | 0.056 | 1 | 0 | Prr12 | 0.01 | Young |
| 0.313754 | 0.346234 | 0.026 | 0.019 | 1 | 0 | Dxo | 0.007 | Young |
| 0.313804 | 0.265033 | 0.181 | 0.168 | 1 | 0 | Cr1l | 0.013 | Young |
| 0.313983 | 0.324941 | 0.15 | 0.138 | 1 | 0 | Crybg3 | 0.012 | Young |
| 0.314048 | 0.350329 | 0.019 | 0.014 | 1 | 0 | Ifi208 | 0.005 | Young |
| 0.31517 | 0.242183 | 0.071 | 0.06 | 1 | 0 | Tmem60 | 0.011 | Young |
| 0.315288 | 0.338581 | 0.051 | 0.042 | 1 | 0 | Sgcb | 0.009 | Young |
| 0.315628 | 0.027539 | 0.105 | 0.12 | 1 | 0 | Cd40 | -0.015 | Young |
| 0.315685 | 0.150643 | 0.055 | 0.045 | 1 | 0 | Gprc5b | 0.01 | Young |
| 0.31611 | 0.039796 | 0.088 | 0.103 | 1 | 0 | Nae1 | -0.015 | Young |
| 0.316212 | 0.021008 | 0.059 | 0.07 | 1 | 0 | Tbce | -0.011 | Young |
| 0.316435 | 0.208068 | 0.222 | 0.208 | 1 | 0 | Pacsin2 | 0.014 | Young |
| 0.316459 | 0.225416 | 0.159 | 0.145 | 1 | 0 | Chd8 | 0.014 | Young |
| 0.316521 | 0.377295 | 0.069 | 0.059 | 1 | 0 | Zfp318 | 0.01 | Young |
| 0.316597 | 0.498876 | 0.067 | 0.057 | 1 | 0 | Nelfe | 0.01 | Young |
| 0.317067 | 0.355094 | 0.053 | 0.044 | 1 | 0 | Zfyve21 | 0.009 | Young |
| 0.317385 | 0.651661 | 0.042 | 0.034 | 1 | 0 | Ddx55 | 0.008 | Young |
| 0.317434 | 0.033328 | 0.24 | 0.265 | 1 | 0 | Ptpa | -0.025 | Young |
| 0.318079 | 0.055132 | 0.186 | 0.207 | 1 | 0 | Rb1 | -0.021 | Young |
| 0.318152 | 0.381983 | 0.07 | 0.06 | 1 | 0 | Rgmb | 0.01 | Young |
| 0.318242 | 0.106928 | 0.349 | 0.331 | 1 | 0 | Snx6 | 0.018 | Young |
| 0.318505 | 0.336922 | 0.118 | 0.105 | 1 | 0 | Trim44 | 0.013 | Young |
| 0.318681 | 0.270742 | 0.019 | 0.025 | 1 | 0 | Ulbp1 | -0.006 | Young |
| 0.318716 | 0.540148 | 0.018 | 0.012 | 1 | 0 | Tatdn3 | 0.006 | Young |
| 0.319064 | 0.333867 | 0.014 | 0.019 | 1 | 0 | Dusp28 | -0.005 | Young |
| 0.319241 | 0.109314 | 0.085 | 0.098 | 1 | 0 | Fbrsl1 | -0.013 | Young |
| 0.319512 | 0.011133 | 0.039 | 0.048 | 1 | 0 | Tmem70 | -0.009 | Young |
| 0.319636 | 0.09721 | 0.13 | 0.148 | 1 | 0 | Chmp1a | -0.018 | Young |
| 0.319648 | 0.19313 | 0.195 | 0.181 | 1 | 0 | Uba2 | 0.014 | Young |
| 0.3197 | 0.227453 | 0.29 | 0.282 | 1 | 0 | Wdfy3 | 0.008 | Young |

| | | | | | | | | |
|---|---|---|---|---|---|---|---|---|
| 0.3199 | 0.194531 | 0.188 | 0.174 | 1 | 0 | Chd9 | 0.014 | Young |
| 0.320239 | 0.066777 | 0.049 | 0.04 | 1 | 0 | Rbmx | 0.009 | Young |
| 0.32043 | 0.344558 | 0.016 | 0.011 | 1 | 0 | Upb1 | 0.005 | Young |
| 0.320513 | 0.042812 | 0.007 | 0.011 | 1 | 0 | Magi3 | -0.004 | Young |
| 0.320592 | 0.121184 | 0.018 | 0.012 | 1 | 0 | Gm20604 | 0.006 | Young |
| 0.320627 | 0.187845 | 0.189 | 0.174 | 1 | 0 | Pcyt1a | 0.015 | Young |
| 0.320645 | 0.114616 | 0.489 | 0.495 | 1 | 0 | Wdr26 | -0.006 | Young |
| 0.321482 | 0.151639 | 0.393 | 0.383 | 1 | 0 | Psma7 | 0.01 | Young |
| 0.321551 | 0.437336 | 0.061 | 0.052 | 1 | 0 | Cwc25 | 0.009 | Young |
| 0.321553 | 0.123961 | 0.007 | 0.011 | 1 | 0 | Tceanc | -0.004 | Young |
| 0.321593 | 0.948639 | 0.014 | 0.01 | 1 | 0 | 1700008J0 | 0.004 | Young |
| 0.321611 | 0.418443 | 0.024 | 0.018 | 1 | 0 | 2310040G2 | 0.006 | Young |
| 0.32197 | 0.519388 | 0.016 | 0.011 | 1 | 0 | Bahcc1 | 0.005 | Young |
| 0.322176 | 0.897207 | 0.014 | 0.01 | 1 | 0 | Fut10 | 0.004 | Young |
| 0.322734 | 0.444285 | 0.053 | 0.044 | 1 | 0 | Tmem140 | 0.009 | Young |
| 0.323136 | 0.319034 | 0.064 | 0.055 | 1 | 0 | Serinc5 | 0.009 | Young |
| 0.323331 | 1.063187 | 0.011 | 0.007 | 1 | 0 | Zfp235 | 0.004 | Young |
| 0.323613 | 0.240249 | 0.053 | 0.044 | 1 | 0 | Scd1 | 0.009 | Young |
| 0.323949 | 0.227813 | 0.281 | 0.271 | 1 | 0 | Polr2a | 0.01 | Young |
| 0.324127 | 0.369675 | 0.029 | 0.022 | 1 | 0 | Pik3r2 | 0.007 | Young |
| 0.324242 | 0.671833 | 0.022 | 0.016 | 1 | 0 | Iba57 | 0.006 | Young |
| 0.324396 | 0.285916 | 0.024 | 0.018 | 1 | 0 | Taf9b | 0.006 | Young |
| 0.32445 | 0.105509 | 0.054 | 0.045 | 1 | 0 | Kdm4c | 0.009 | Young |
| 0.324488 | 0.293692 | 0.132 | 0.119 | 1 | 0 | Arntl | 0.013 | Young |
| 0.324665 | 1.03405 | 0.013 | 0.008 | 1 | 0 | Zfp747 | 0.005 | Young |
| 0.325209 | 0.165821 | 0.481 | 0.492 | 1 | 0 | Uqcr11 | -0.011 | Young |
| 0.325285 | 0.24816 | 0.14 | 0.127 | 1 | 0 | Mrpl51 | 0.013 | Young |
| 0.325328 | 0.215788 | 0.173 | 0.16 | 1 | 0 | Nck1 | 0.013 | Young |
| 0.325475 | 0.354336 | 0.044 | 0.036 | 1 | 0 | Ss18l2 | 0.008 | Young |
| 0.325569 | 0.327006 | 0.024 | 0.018 | 1 | 0 | Whamm | 0.006 | Young |
| 0.325674 | 0.31051 | 0.096 | 0.085 | 1 | 0 | Atg101 | 0.011 | Young |
| 0.326059 | 0.059371 | 0.028 | 0.036 | 1 | 0 | Lrp5 | -0.008 | Young |
| 0.326063 | 0.027144 | 0.13 | 0.146 | 1 | 0 | Nemf | -0.016 | Young |
| 0.326133 | 0.044221 | 0.135 | 0.152 | 1 | 0 | Iscu | -0.017 | Young |
| 0.326571 | 0.030858 | 0.104 | 0.119 | 1 | 0 | Eml4 | -0.015 | Young |
| 0.326764 | 0.048619 | 0.719 | 0.725 | 1 | 0 | Arpc1b | -0.006 | Young |
| 0.32693 | 0.056452 | 0.244 | 0.269 | 1 | 0 | Erbin | -0.025 | Young |
| 0.327235 | 0.283037 | 0.069 | 0.059 | 1 | 0 | Stx8 | 0.01 | Young |
| 0.327241 | 0.436402 | 0.032 | 0.025 | 1 | 0 | Bloc1s5 | 0.007 | Young |
| 0.327253 | 0.121967 | 0.395 | 0.389 | 1 | 0 | Ctnnb1 | 0.006 | Young |
| 0.327431 | 0.019401 | 0.255 | 0.28 | 1 | 0 | Slc29a3 | -0.025 | Young |
| 0.32757 | 0.15888 | 0.014 | 0.01 | 1 | 0 | Jade3 | 0.004 | Young |
| 0.327958 | 0.150751 | 0.238 | 0.224 | 1 | 0 | Sumo1 | 0.014 | Young |
| 0.327993 | 0.288442 | 0.171 | 0.159 | 1 | 0 | Cpsf2 | 0.012 | Young |
| 0.328187 | 0.645992 | 0.011 | 0.007 | 1 | 0 | Gm49368 | 0.004 | Young |

| | | | | | | | | |
|---|---|---|---|---|---|---|---|---|
| 0.328904 | 0.219376 | 0.202 | 0.19 | 1 | 0 | Rybp | 0.012 | Young |
| 0.328988 | 0.072618 | 0.167 | 0.187 | 1 | 0 | Fam32a | -0.02 | Young |
| 0.329043 | 0.267756 | 0.079 | 0.068 | 1 | 0 | Kdm3a | 0.011 | Young |
| 0.329066 | 0.266496 | 0.061 | 0.052 | 1 | 0 | Brwd1 | 0.009 | Young |
| 0.329324 | 0.398292 | 0.067 | 0.057 | 1 | 0 | Egfr | 0.01 | Young |
| 0.329772 | 0.441366 | 0.032 | 0.025 | 1 | 0 | 2700097O( | 0.007 | Young |
| 0.330025 | 0.703451 | 0.03 | 0.023 | 1 | 0 | Zscan21 | 0.007 | Young |
| 0.3302 | 0.465411 | 0.093 | 0.082 | 1 | 0 | Engase | 0.011 | Young |
| 0.330675 | 0.099565 | 0.045 | 0.037 | 1 | 0 | Oxsm | 0.008 | Young |
| 0.330867 | 0.192057 | 0.121 | 0.108 | 1 | 0 | Dnttip2 | 0.013 | Young |
| 0.331429 | 0.024351 | 0.131 | 0.148 | 1 | 0 | Tmem219 | -0.017 | Young |
| 0.331709 | 0.390515 | 0.027 | 0.021 | 1 | 0 | Fzd5 | 0.006 | Young |
| 0.33219 | 0.14996 | 0.014 | 0.01 | 1 | 0 | Cxcr3 | 0.004 | Young |
| 0.332289 | 0.02457 | 0.053 | 0.063 | 1 | 0 | Lztr1 | -0.01 | Young |
| 0.332338 | 0.318153 | 0.092 | 0.081 | 1 | 0 | Wdr3 | 0.011 | Young |
| 0.332389 | 0.039687 | 0.032 | 0.04 | 1 | 0 | Aifm2 | -0.008 | Young |
| 0.332454 | 0.261673 | 0.06 | 0.051 | 1 | 0 | Dera | 0.009 | Young |
| 0.333002 | 0.231503 | 0.285 | 0.278 | 1 | 0 | Gapdh | 0.007 | Young |
| 0.33309 | 0.008194 | 0.878 | 0.878 | 1 | 0 | Jun | 0 | Young |
| 0.333193 | 0.314615 | 0.032 | 0.025 | 1 | 0 | Gabbr1 | 0.007 | Young |
| 0.333279 | 0.281784 | 0.057 | 0.048 | 1 | 0 | Kif1bp | 0.009 | Young |
| 0.333461 | 0.23041 | 0.211 | 0.197 | 1 | 0 | Rc3h1 | 0.014 | Young |
| 0.333738 | 0.259026 | 0.122 | 0.109 | 1 | 0 | Psmg4 | 0.013 | Young |
| 0.334223 | 0.351515 | 0.035 | 0.027 | 1 | 0 | Mrpl46 | 0.008 | Young |
| 0.334534 | 0.092614 | 0.553 | 0.561 | 1 | 0 | Dbi | -0.008 | Young |
| 0.334662 | 0.046591 | 0.199 | 0.223 | 1 | 0 | Psmd8 | -0.024 | Young |
| 0.334995 | 0.068183 | 0.024 | 0.03 | 1 | 0 | Cog1 | -0.006 | Young |
| 0.335521 | 0.126145 | 0.193 | 0.176 | 1 | 0 | Arrdc3 | 0.017 | Young |
| 0.335677 | 0.874841 | 0.019 | 0.014 | 1 | 0 | P2ry14 | 0.005 | Young |
| 0.335882 | 0.280246 | 0.096 | 0.085 | 1 | 0 | Scn1b | 0.011 | Young |
| 0.336138 | 0.231822 | 0.079 | 0.068 | 1 | 0 | Tspan17 | 0.011 | Young |
| 0.336337 | 0.384423 | 0.08 | 0.07 | 1 | 0 | Kifap3 | 0.01 | Young |
| 0.336975 | 0.045016 | 0.374 | 0.417 | 1 | 0 | Arrb2 | -0.043 | Young |
| 0.336988 | 0.083452 | 0.066 | 0.077 | 1 | 0 | Tubb4b | -0.011 | Young |
| 0.337003 | 0.217546 | 0.05 | 0.041 | 1 | 0 | Cep164 | 0.009 | Young |
| 0.337125 | 0.107541 | 0.102 | 0.089 | 1 | 0 | Glrx2 | 0.013 | Young |
| 0.337248 | 0.48759 | 0.025 | 0.019 | 1 | 0 | B3gnt3 | 0.006 | Young |
| 0.337553 | 0.585563 | 0.052 | 0.044 | 1 | 0 | Ciao1 | 0.008 | Young |
| 0.337749 | 0.209829 | 0.334 | 0.331 | 1 | 0 | Washc4 | 0.003 | Young |
| 0.338043 | 0.328759 | 0.051 | 0.042 | 1 | 0 | Rexo4 | 0.009 | Young |
| 0.338364 | 0.458224 | 0.03 | 0.023 | 1 | 0 | Ndst1 | 0.007 | Young |
| 0.338415 | 0.132304 | 0.534 | 0.539 | 1 | 0 | Naca | -0.005 | Young |
| 0.338421 | 0.18548 | 0.053 | 0.044 | 1 | 0 | D16Ertd47: | 0.009 | Young |
| 0.338672 | 0.193538 | 0.157 | 0.144 | 1 | 0 | Denr | 0.013 | Young |
| 0.339034 | 0.218485 | 0.076 | 0.066 | 1 | 0 | Mapk8 | 0.01 | Young |

| | | | | | | | | |
|---|---|---|---|---|---|---|---|---|
| 0.33933 | 0.369595 | 0.081 | 0.071 | 1 | 0 | Itgb1bp1 | 0.01 | Young |
| 0.339513 | 0.210165 | 0.25 | 0.239 | 1 | 0 | Atp5d | 0.011 | Young |
| 0.339619 | 0.191144 | 0.368 | 0.364 | 1 | 0 | Nfam1 | 0.004 | Young |
| 0.339864 | 0.194998 | 0.256 | 0.246 | 1 | 0 | Lamtor2 | 0.01 | Young |
| 0.33994 | 0.066731 | 0.612 | 0.614 | 1 | 0 | Srrm2 | -0.002 | Young |
| 0.340169 | 0.071184 | 0.701 | 0.699 | 1 | 0 | Morf4l1 | 0.002 | Young |
| 0.341284 | 0.332887 | 0.042 | 0.034 | 1 | 0 | Lsg1 | 0.008 | Young |
| 0.341305 | 0.55588 | 0.019 | 0.014 | 1 | 0 | Itgb7 | 0.005 | Young |
| 0.341452 | 0.287579 | 0.146 | 0.134 | 1 | 0 | Polr2f | 0.012 | Young |
| 0.341551 | 0.445063 | 0.04 | 0.033 | 1 | 0 | Nt5dc1 | 0.007 | Young |
| 0.341829 | 0.553311 | 0.019 | 0.014 | 1 | 0 | Zbtb10 | 0.005 | Young |
| 0.343304 | 0.021367 | 0.351 | 0.383 | 1 | 0 | Calm3 | -0.032 | Young |
| 0.343417 | 0.159205 | 0.021 | 0.015 | 1 | 0 | Gm42031 | 0.006 | Young |
| 0.343537 | 0.025295 | 0.185 | 0.208 | 1 | 0 | Ndufb4 | -0.023 | Young |
| 0.34358 | 0.149875 | 0.485 | 0.488 | 1 | 0 | Nisch | -0.003 | Young |
| 0.343825 | 0.395209 | 0.019 | 0.014 | 1 | 0 | Gm16638 | 0.005 | Young |
| 0.344141 | 0.558239 | 0.019 | 0.014 | 1 | 0 | Pus3 | 0.005 | Young |
| 0.344255 | 0.002482 | 0.14 | 0.156 | 1 | 0 | Eprs | -0.016 | Young |
| 0.344316 | 0.646677 | 0.028 | 0.022 | 1 | 0 | Nmral1 | 0.006 | Young |
| 0.34444 | 0.190582 | 0.301 | 0.29 | 1 | 0 | H13 | 0.011 | Young |
| 0.344587 | 0.284845 | 0.037 | 0.03 | 1 | 0 | Zfp729b | 0.007 | Young |
| 0.345145 | 0.072276 | 0.008 | 0.012 | 1 | 0 | 1700006J1 | -0.004 | Young |
| 0.345586 | 0.429403 | 0.058 | 0.049 | 1 | 0 | Otud7b | 0.009 | Young |
| 0.346014 | 0.235844 | 0.008 | 0.012 | 1 | 0 | Rtl6 | -0.004 | Young |
| 0.346232 | 0.178418 | 0.211 | 0.198 | 1 | 0 | Snx27 | 0.013 | Young |
| 0.346236 | 0.237661 | 0.193 | 0.181 | 1 | 0 | Plekhg5 | 0.012 | Young |
| 0.346293 | 0.094329 | 0.056 | 0.066 | 1 | 0 | Arhgap4 | -0.01 | Young |
| 0.346304 | 0.013801 | 0.361 | 0.384 | 1 | 0 | Lst1 | -0.023 | Young |
| 0.346458 | 0.203455 | 0.235 | 0.223 | 1 | 0 | Prpf4b | 0.012 | Young |
| 0.346647 | 0.278383 | 0.127 | 0.115 | 1 | 0 | Ano6 | 0.012 | Young |
| 0.346752 | 0.27372 | 0.03 | 0.023 | 1 | 0 | Bri3bp | 0.007 | Young |
| 0.347138 | 0.212287 | 0.059 | 0.049 | 1 | 0 | Cds1 | 0.01 | Young |
| 0.347184 | 0.076869 | 0.192 | 0.175 | 1 | 0 | Ddx24 | 0.017 | Young |
| 0.347193 | 0.17316 | 0.279 | 0.271 | 1 | 0 | Mbnl2 | 0.008 | Young |
| 0.347555 | 0.102066 | 0.158 | 0.178 | 1 | 0 | Pcmt1 | -0.02 | Young |
| 0.347801 | 0.028059 | 0.325 | 0.354 | 1 | 0 | Sec61g | -0.029 | Young |
| 0.348158 | 0.281645 | 0.18 | 0.17 | 1 | 0 | Impact | 0.01 | Young |
| 0.348618 | 0.025233 | 0.027 | 0.034 | 1 | 0 | Endog | -0.007 | Young |
| 0.348789 | 0.278893 | 0.077 | 0.067 | 1 | 0 | Akr1b10 | 0.01 | Young |
| 0.348979 | 0.005774 | 0.102 | 0.116 | 1 | 0 | Clk3 | -0.014 | Young |
| 0.349008 | 0.066354 | 0.55 | 0.539 | 1 | 0 | Adam15 | 0.011 | Young |
| 0.349022 | 0.192842 | 0.151 | 0.138 | 1 | 0 | Rxra | 0.013 | Young |
| 0.349614 | 0.469746 | 0.045 | 0.037 | 1 | 0 | Mtmr4 | 0.008 | Young |
| 0.350711 | 0.104036 | 0.053 | 0.063 | 1 | 0 | Zfp626 | -0.01 | Young |
| 0.350721 | 0.354923 | 0.136 | 0.124 | 1 | 0 | Ank | 0.012 | Young |

| | | | | | | | | |
|---|---|---|---|---|---|---|---|---|
| 0.350934 | 0.035868 | 0.019 | 0.025 | 1 | 0 | Abhd11 | -0.006 | Young |
| 0.351355 | 0.539796 | 0.048 | 0.04 | 1 | 0 | Mphosph6 | 0.008 | Young |
| 0.353364 | 0.308646 | 0.017 | 0.012 | 1 | 0 | Cnst | 0.005 | Young |
| 0.353532 | 0.487726 | 0.017 | 0.012 | 1 | 0 | Fignl2 | 0.005 | Young |
| 0.353581 | 0.282239 | 0.131 | 0.119 | 1 | 0 | Hgsnat | 0.012 | Young |
| 0.353812 | 0.399102 | 0.14 | 0.13 | 1 | 0 | Slc25a11 | 0.01 | Young |
| 0.353824 | 0.272396 | 0.037 | 0.045 | 1 | 0 | Myo1e | -0.008 | Young |
| 0.354178 | 0.186221 | 0.163 | 0.15 | 1 | 0 | Cpeb4 | 0.013 | Young |
| 0.354466 | 0.126169 | 0.123 | 0.111 | 1 | 0 | Rnaseh2c | 0.012 | Young |
| 0.354514 | 0.811273 | 0.016 | 0.011 | 1 | 0 | Slfn9 | 0.005 | Young |
| 0.354886 | 0.444128 | 0.024 | 0.018 | 1 | 0 | Ltb4r1 | 0.006 | Young |
| 0.354934 | 0.240017 | 0.282 | 0.275 | 1 | 0 | Adam10 | 0.007 | Young |
| 0.355285 | 0.36714 | 0.098 | 0.088 | 1 | 0 | Cenpa | 0.01 | Young |
| 0.355434 | 0.038782 | 0.046 | 0.055 | 1 | 0 | Cdc16 | -0.009 | Young |
| 0.355529 | 0.101609 | 0.07 | 0.081 | 1 | 0 | Uba3 | -0.011 | Young |
| 0.356294 | 0.523966 | 0.016 | 0.011 | 1 | 0 | Plcl1 | 0.005 | Young |
| 0.356311 | 0.015139 | 0.283 | 0.309 | 1 | 0 | Fgd4 | -0.026 | Young |
| 0.357226 | 0.546421 | 0.031 | 0.025 | 1 | 0 | Mbd4 | 0.006 | Young |
| 0.357376 | 0.069238 | 0.364 | 0.349 | 1 | 0 | S100a13 | 0.015 | Young |
| 0.357391 | 0.074546 | 0.693 | 0.703 | 1 | 0 | Tmem50a | -0.01 | Young |
| 0.357395 | 0.363366 | 0.094 | 0.083 | 1 | 0 | Relb | 0.011 | Young |
| 0.357495 | 0.264281 | 0.055 | 0.047 | 1 | 0 | Zfp991 | 0.008 | Young |
| 0.357535 | 0.020068 | 0.021 | 0.027 | 1 | 0 | Thap1 | -0.006 | Young |
| 0.357669 | 0.547592 | 0.034 | 0.027 | 1 | 0 | Haus4 | 0.007 | Young |
| 0.357781 | 0.00106 | 0.05 | 0.059 | 1 | 0 | Prkab1 | -0.009 | Young |
| 0.358183 | 0.078 | 0.308 | 0.339 | 1 | 0 | Taldo1 | -0.031 | Young |
| 0.35819 | 0.411039 | 0.034 | 0.027 | 1 | 0 | Cep135 | 0.007 | Young |
| 0.359509 | 0.508553 | 0.058 | 0.049 | 1 | 0 | Fam214a | 0.009 | Young |
| 0.359916 | 0.348041 | 0.042 | 0.034 | 1 | 0 | 1110034G2 | 0.008 | Young |
| 0.360016 | 0.205621 | 0.194 | 0.182 | 1 | 0 | Phf20l1 | 0.012 | Young |
| 0.360029 | 0.104665 | 0.02 | 0.026 | 1 | 0 | Nradd | -0.006 | Young |
| 0.360088 | 0.124504 | 0.351 | 0.342 | 1 | 0 | Cd2ap | 0.009 | Young |
| 0.360107 | 0.039256 | 0.048 | 0.057 | 1 | 0 | Mrps30 | -0.009 | Young |
| 0.360532 | 0.348251 | 0.091 | 0.081 | 1 | 0 | Fbxo38 | 0.01 | Young |
| 0.360685 | 0.287817 | 0.027 | 0.021 | 1 | 0 | Zfp932 | 0.006 | Young |
| 0.360825 | 0.274631 | 0.166 | 0.155 | 1 | 0 | Renbp | 0.011 | Young |
| 0.361331 | 0.123813 | 0.075 | 0.086 | 1 | 0 | Cyp4f13 | -0.011 | Young |
| 0.361365 | 0.356309 | 0.022 | 0.016 | 1 | 0 | Nynrin | 0.006 | Young |
| 0.361375 | 0.275855 | 0.102 | 0.092 | 1 | 0 | Ccdc6 | 0.01 | Young |
| 0.361479 | 0.3194 | 0.056 | 0.048 | 1 | 0 | Poglut1 | 0.008 | Young |
| 0.361521 | 0.104888 | 0.006 | 0.01 | 1 | 0 | Asap2 | -0.004 | Young |
| 0.361604 | 0.102851 | 0.088 | 0.1 | 1 | 0 | Ncor2 | -0.012 | Young |
| 0.362024 | 0.118239 | 0.404 | 0.398 | 1 | 0 | Ptpre | 0.006 | Young |
| 0.362027 | 0.139159 | 0.085 | 0.074 | 1 | 0 | Mzt1 | 0.011 | Young |
| 0.362269 | 0.225433 | 0.059 | 0.051 | 1 | 0 | Cd80 | 0.008 | Young |

| | | | | | | | | |
|---|---|---|---|---|---|---|---|---|
| 0.362384 | 0.409631 | 0.162 | 0.153 | 1 | 0 | Rgl2 | 0.009 | Young |
| 0.362665 | 0.015587 | 0.073 | 0.083 | 1 | 0 | Stk19 | -0.01 | Young |
| 0.362976 | 0.45831 | 0.027 | 0.021 | 1 | 0 | Alkbh7 | 0.006 | Young |
| 0.363092 | 0.043377 | 0.069 | 0.079 | 1 | 0 | Parp2 | -0.01 | Young |
| 0.363855 | 0.016814 | 0.021 | 0.027 | 1 | 0 | Atg13 | -0.006 | Young |
| 0.364004 | 0.003512 | 0.067 | 0.077 | 1 | 0 | Ech1 | -0.01 | Young |
| 0.364503 | 0.005135 | 0.208 | 0.228 | 1 | 0 | Saraf | -0.02 | Young |
| 0.365003 | 0.1905 | 0.039 | 0.031 | 1 | 0 | Trmt13 | 0.008 | Young |
| 0.365069 | 0.409346 | 0.014 | 0.01 | 1 | 0 | Osgepl1 | 0.004 | Young |
| 0.365464 | 0.15899 | 0.022 | 0.016 | 1 | 0 | Rbm10 | 0.006 | Young |
| 0.365953 | 0.133434 | 0.014 | 0.019 | 1 | 0 | Snx33 | -0.005 | Young |
| 0.366015 | 0.004503 | 0.34 | 0.368 | 1 | 0 | Map1lc3b | -0.028 | Young |
| 0.36626 | 0.410649 | 0.049 | 0.041 | 1 | 0 | Rpa1 | 0.008 | Young |
| 0.366606 | 0.071421 | 0.193 | 0.179 | 1 | 0 | Tmem8 | 0.014 | Young |
| 0.366663 | 0.214252 | 0.155 | 0.144 | 1 | 0 | Rtcb | 0.011 | Young |
| 0.366753 | 0.395664 | 0.029 | 0.023 | 1 | 0 | Pdk1 | 0.006 | Young |
| 0.367297 | 0.195398 | 0.039 | 0.031 | 1 | 0 | Adnp2 | 0.008 | Young |
| 0.367501 | 0.287363 | 0.177 | 0.166 | 1 | 0 | Cebpz | 0.011 | Young |
| 0.367557 | 0.812436 | 0.012 | 0.008 | 1 | 0 | Lrg1 | 0.004 | Young |
| 0.367602 | 0.004456 | 0.957 | 0.97 | 1 | 0 | Rplp2 | -0.013 | Young |
| 0.367694 | 0.70348 | 0.012 | 0.008 | 1 | 0 | Cdo1 | 0.004 | Young |
| 0.367694 | 0.648514 | 0.012 | 0.008 | 1 | 0 | Rmdn2 | 0.004 | Young |
| 0.36777 | 0.21342 | 0.027 | 0.021 | 1 | 0 | Ano8 | 0.006 | Young |
| 0.368089 | 0.071157 | 0.167 | 0.186 | 1 | 0 | Ube2k | -0.019 | Young |
| 0.368789 | 0.228153 | 0.014 | 0.01 | 1 | 0 | 4930522L1 | 0.004 | Young |
| 0.369044 | 0.236824 | 0.144 | 0.133 | 1 | 0 | Sec22b | 0.011 | Young |
| 0.369153 | 0.056992 | 0.684 | 0.68 | 1 | 0 | Rrbp1 | 0.004 | Young |
| 0.369618 | 0.467875 | 0.012 | 0.008 | 1 | 0 | Slc22a17 | 0.004 | Young |
| 0.369645 | 0.881487 | 0.011 | 0.007 | 1 | 0 | Cd276 | 0.004 | Young |
| 0.369645 | 0.698471 | 0.011 | 0.007 | 1 | 0 | Kif21a | 0.004 | Young |
| 0.36987 | 0.419859 | 0.02 | 0.015 | 1 | 0 | Ttf2 | 0.005 | Young |
| 0.370078 | 0.078129 | 0.118 | 0.133 | 1 | 0 | Acp1 | -0.015 | Young |
| 0.37009 | 0.723246 | 0.011 | 0.007 | 1 | 0 | Thrsp | 0.004 | Young |
| 0.370227 | 0.07054 | 0.531 | 0.536 | 1 | 0 | Mef2a | -0.005 | Young |
| 0.370286 | 0.387233 | 0.037 | 0.03 | 1 | 0 | Arl2 | 0.007 | Young |
| 0.370407 | 0.149115 | 0.202 | 0.19 | 1 | 0 | Prrc2b | 0.012 | Young |
| 0.37048 | 0.547416 | 0.081 | 0.073 | 1 | 0 | Itga5 | 0.008 | Young |
| 0.370972 | 0.024875 | 0.105 | 0.118 | 1 | 0 | Pfkp | -0.013 | Young |
| 0.371076 | 0.009781 | 0.054 | 0.063 | 1 | 0 | Arhgap21 | -0.009 | Young |
| 0.371223 | 0.239327 | 0.212 | 0.201 | 1 | 0 | Psmb6 | 0.011 | Young |
| 0.371577 | 0.83071 | 0.019 | 0.014 | 1 | 0 | Inpp4b | 0.005 | Young |
| 0.371728 | 0.576214 | 0.011 | 0.007 | 1 | 0 | Ptpdc1 | 0.004 | Young |
| 0.371775 | 0.261897 | 0.212 | 0.202 | 1 | 0 | Tomm5 | 0.01 | Young |
| 0.37179 | 0.155668 | 0.133 | 0.122 | 1 | 0 | Ddx54 | 0.011 | Young |
| 0.371819 | 0.270159 | 0.084 | 0.074 | 1 | 0 | Rxrb | 0.01 | Young |

| | | | | | | | | |
|---|---|---|---|---|---|---|---|---|
| 0.371963 | 0.449774 | 0.059 | 0.051 | 1 | 0 | Cib2 | 0.008 | Young |
| 0.372027 | 0.570522 | 0.011 | 0.007 | 1 | 0 | Dnajb13 | 0.004 | Young |
| 0.372147 | 0.05088 | 0.348 | 0.384 | 1 | 0 | Tor1aip1 | -0.036 | Young |
| 0.372305 | 0.347086 | 0.04 | 0.033 | 1 | 0 | Sharpin | 0.007 | Young |
| 0.372549 | 0.426587 | 0.092 | 0.082 | 1 | 0 | Washc5 | 0.01 | Young |
| 0.372859 | 0.287217 | 0.02 | 0.015 | 1 | 0 | Trmt11 | 0.005 | Young |
| 0.372921 | 0.121118 | 0.448 | 0.442 | 1 | 0 | Cox8a | 0.006 | Young |
| 0.373214 | 0.024049 | 0.07 | 0.081 | 1 | 0 | Pik3c3 | -0.011 | Young |
| 0.373221 | 0.214343 | 0.011 | 0.007 | 1 | 0 | Zfp229 | 0.004 | Young |
| 0.373819 | 0.324931 | 0.011 | 0.007 | 1 | 0 | Isg20 | 0.004 | Young |
| 0.374682 | 0.239699 | 0.065 | 0.056 | 1 | 0 | Prps1 | 0.009 | Young |
| 0.374702 | 0.023558 | 0.221 | 0.242 | 1 | 0 | Mgea5 | -0.021 | Young |
| 0.37473 | 0.172736 | 0.056 | 0.048 | 1 | 0 | Cep83 | 0.008 | Young |
| 0.374984 | 0.365822 | 0.043 | 0.036 | 1 | 0 | Nat9 | 0.007 | Young |
| 0.37507 | 0.108861 | 0.327 | 0.313 | 1 | 0 | Add3 | 0.014 | Young |
| 0.375161 | 0.233157 | 0.053 | 0.045 | 1 | 0 | Xpo4 | 0.008 | Young |
| 0.375209 | 0.250276 | 0.131 | 0.119 | 1 | 0 | Stxbp2 | 0.012 | Young |
| 0.375444 | 0.140646 | 0.07 | 0.081 | 1 | 0 | Cpsf3 | -0.011 | Young |
| 0.375671 | 0.268027 | 0.124 | 0.114 | 1 | 0 | Ethe1 | 0.01 | Young |
| 0.375691 | 0.036801 | 0.027 | 0.021 | 1 | 0 | Umps | 0.006 | Young |
| 0.375953 | 0.146438 | 0.222 | 0.209 | 1 | 0 | Ppp1r10 | 0.013 | Young |
| 0.376333 | 0.474513 | 0.041 | 0.034 | 1 | 0 | Ap2a1 | 0.007 | Young |
| 0.376406 | 0.0782 | 0.169 | 0.187 | 1 | 0 | Polr2m | -0.018 | Young |
| 0.376419 | 0.023687 | 0.148 | 0.164 | 1 | 0 | Smarce1 | -0.016 | Young |
| 0.376435 | 0.28872 | 0.07 | 0.062 | 1 | 0 | Il15 | 0.008 | Young |
| 0.376523 | 0.698747 | 0.023 | 0.018 | 1 | 0 | Ppp1r12b | 0.005 | Young |
| 0.376576 | 0.248268 | 0.302 | 0.298 | 1 | 0 | Eif5 | 0.004 | Young |
| 0.37677 | 0.08842 | 0.044 | 0.052 | 1 | 0 | Fam173b | -0.008 | Young |
| 0.376817 | 0.37322 | 0.096 | 0.086 | 1 | 0 | Casp3 | 0.01 | Young |
| 0.376984 | 0.452405 | 0.094 | 0.085 | 1 | 0 | Aldh7a1 | 0.009 | Young |
| 0.377006 | 0.267639 | 0.149 | 0.138 | 1 | 0 | Glrx5 | 0.011 | Young |
| 0.377495 | 0.663647 | 0.031 | 0.025 | 1 | 0 | Gdpd3 | 0.006 | Young |
| 0.37759 | 0.088386 | 0.025 | 0.031 | 1 | 0 | Ttbk2 | -0.006 | Young |
| 0.377823 | 0.382133 | 0.108 | 0.098 | 1 | 0 | Mink1 | 0.01 | Young |
| 0.377995 | 0.046361 | 0.314 | 0.341 | 1 | 0 | Coro1c | -0.027 | Young |
| 0.378065 | 0.281416 | 0.24 | 0.231 | 1 | 0 | Cdipt | 0.009 | Young |
| 0.378138 | 0.214623 | 0.024 | 0.03 | 1 | 0 | Ints10 | -0.006 | Young |
| 0.378363 | 0.496431 | 0.019 | 0.014 | 1 | 0 | Ift140 | 0.005 | Young |
| 0.378568 | 0.145329 | 0.305 | 0.295 | 1 | 0 | Ncor1 | 0.01 | Young |
| 0.378958 | 0.329355 | 0.068 | 0.059 | 1 | 0 | Esrra | 0.009 | Young |
| 0.378971 | 0.04603 | 0.262 | 0.287 | 1 | 0 | Tubb2a | -0.025 | Young |
| 0.379057 | 0.148988 | 0.09 | 0.103 | 1 | 0 | Gemin7 | -0.013 | Young |
| 0.379415 | 0.684597 | 0.034 | 0.027 | 1 | 0 | Gtpbp8 | 0.007 | Young |
| 0.379722 | 0.438579 | 0.038 | 0.031 | 1 | 0 | Gna15 | 0.007 | Young |
| 0.379807 | 0.352215 | 0.115 | 0.105 | 1 | 0 | Coa3 | 0.01 | Young |

| | | | | | | | | |
|---|---|---|---|---|---|---|---|---|
| 0.379993 | 0.172187 | 0.122 | 0.109 | 1 | 0 | Anapc16 | 0.013 | Young |
| 0.380035 | 0.096558 | 0.249 | 0.234 | 1 | 0 | Washc2 | 0.015 | Young |
| 0.380103 | 0.15631 | 0.315 | 0.309 | 1 | 0 | Cox7a2 | 0.006 | Young |
| 0.380547 | 0.024294 | 0.131 | 0.146 | 1 | 0 | Ndfip2 | -0.015 | Young |
| 0.380744 | 0.03812 | 0.118 | 0.131 | 1 | 0 | Ube2j2 | -0.013 | Young |
| 0.381686 | 0.46566 | 0.084 | 0.075 | 1 | 0 | Hmg20b | 0.009 | Young |
| 0.381879 | 0.237734 | 0.111 | 0.1 | 1 | 0 | Clip1 | 0.011 | Young |
| 0.382258 | 0.306851 | 0.157 | 0.146 | 1 | 0 | Emc3 | 0.011 | Young |
| 0.382377 | 0.172136 | 0.389 | 0.39 | 1 | 0 | Dennd5a | -0.001 | Young |
| 0.38255 | 0.254564 | 0.019 | 0.014 | 1 | 0 | Gys1 | 0.005 | Young |
| 0.382994 | 0.133475 | 0.38 | 0.372 | 1 | 0 | Rpl29 | 0.008 | Young |
| 0.383475 | 0.532843 | 0.112 | 0.104 | 1 | 0 | Coro7 | 0.008 | Young |
| 0.383496 | 0.378525 | 0.024 | 0.03 | 1 | 0 | Mapk7 | -0.006 | Young |
| 0.383629 | 0.056775 | 0.165 | 0.183 | 1 | 0 | Smad4 | -0.018 | Young |
| 0.383723 | 0.525631 | 0.031 | 0.025 | 1 | 0 | Gnptab | 0.006 | Young |
| 0.383901 | 0.321997 | 0.066 | 0.057 | 1 | 0 | Scrib | 0.009 | Young |
| 0.384382 | 0.545071 | 0.037 | 0.03 | 1 | 0 | Arid3b | 0.007 | Young |
| 0.384819 | 0.279413 | 0.024 | 0.03 | 1 | 0 | Acad12 | -0.006 | Young |
| 0.38507 | 0.559218 | 0.037 | 0.03 | 1 | 0 | Scfd2 | 0.007 | Young |
| 0.385095 | 0.092979 | 0.08 | 0.092 | 1 | 0 | Commd2 | -0.012 | Young |
| 0.385156 | 0.193841 | 0.221 | 0.209 | 1 | 0 | Med13 | 0.012 | Young |
| 0.38565 | 0.289732 | 0.055 | 0.047 | 1 | 0 | Klhdc2 | 0.008 | Young |
| 0.385841 | 0.25536 | 0.038 | 0.031 | 1 | 0 | Nufip1 | 0.007 | Young |
| 0.385884 | 0.175648 | 0.055 | 0.047 | 1 | 0 | Svbp | 0.008 | Young |
| 0.387083 | 0.171836 | 0.028 | 0.034 | 1 | 0 | Wrb | -0.006 | Young |
| 0.387155 | 0.426966 | 0.017 | 0.012 | 1 | 0 | Letm2 | 0.005 | Young |
| 0.387244 | 0.24838 | 0.182 | 0.171 | 1 | 0 | Uqcrfs1 | 0.011 | Young |
| 0.387366 | 0.183191 | 0.157 | 0.145 | 1 | 0 | Ddx42 | 0.012 | Young |
| 0.387416 | 0.28908 | 0.096 | 0.086 | 1 | 0 | Qpct | 0.01 | Young |
| 0.387657 | 0.270079 | 0.191 | 0.181 | 1 | 0 | Spg21 | 0.01 | Young |
| 0.38772 | 0.215418 | 0.067 | 0.057 | 1 | 0 | Zcchc8 | 0.01 | Young |
| 0.388129 | 0.156676 | 0.324 | 0.316 | 1 | 0 | Scand1 | 0.008 | Young |
| 0.388181 | 0.031317 | 0.065 | 0.075 | 1 | 0 | Cdc34 | -0.01 | Young |
| 0.388186 | 0.070653 | 0.008 | 0.011 | 1 | 0 | Tcaf1 | -0.003 | Young |
| 0.388215 | 0.261602 | 0.035 | 0.029 | 1 | 0 | Unc45a | 0.006 | Young |
| 0.388266 | 0.085045 | 0.33 | 0.317 | 1 | 0 | Emp1 | 0.013 | Young |
| 0.38865 | 0.138406 | 0.401 | 0.399 | 1 | 0 | Mpp1 | 0.002 | Young |
| 0.388683 | 0.544199 | 0.037 | 0.03 | 1 | 0 | Zfp866 | 0.007 | Young |
| 0.388954 | 0.120779 | 0.317 | 0.308 | 1 | 0 | Zfp91 | 0.009 | Young |
| 0.389253 | 0.451814 | 0.066 | 0.057 | 1 | 0 | Ctc1 | 0.009 | Young |
| 0.389309 | 0.228678 | 0.023 | 0.018 | 1 | 0 | Pde4d | 0.005 | Young |
| 0.389448 | 0.265186 | 0.11 | 0.1 | 1 | 0 | Cirbp | 0.01 | Young |
| 0.389667 | 0.33951 | 0.06 | 0.052 | 1 | 0 | Tes | 0.008 | Young |
| 0.390169 | 0.233436 | 0.123 | 0.112 | 1 | 0 | Rab32 | 0.011 | Young |
| 0.390332 | 0.212471 | 0.31 | 0.305 | 1 | 0 | Fam168b | 0.005 | Young |

| | | | | | | | | |
|---|---|---|---|---|---|---|---|---|
| 0.390437 | 0.133933 | 0.359 | 0.36 | 1 | 0 | Selenow | -0.001 | Young |
| 0.39073 | 0.044174 | 0.105 | 0.118 | 1 | 0 | Eif6 | -0.013 | Young |
| 0.390999 | 0.332273 | 0.048 | 0.041 | 1 | 0 | Mcc | 0.007 | Young |
| 0.391084 | 0.131896 | 0.031 | 0.025 | 1 | 0 | Pla2g12a | 0.006 | Young |
| 0.391217 | 0.425961 | 0.078 | 0.07 | 1 | 0 | Ubxn2a | 0.008 | Young |
| 0.392075 | 0.930197 | 0.015 | 0.011 | 1 | 0 | Acy1 | 0.004 | Young |
| 0.392128 | 0.695515 | 0.041 | 0.034 | 1 | 0 | Scnm1 | 0.007 | Young |
| 0.392164 | 0.241925 | 0.05 | 0.042 | 1 | 0 | Gbe1 | 0.008 | Young |
| 0.392261 | 0.381765 | 0.034 | 0.027 | 1 | 0 | Anln | 0.007 | Young |
| 0.392293 | 0.103627 | 0.566 | 0.566 | 1 | 0 | Frmd4b | 0 | Young |
| 0.392698 | 0.320213 | 0.163 | 0.155 | 1 | 0 | Srsf7 | 0.008 | Young |
| 0.392731 | 0.196722 | 0.222 | 0.209 | 1 | 0 | Lpp | 0.013 | Young |
| 0.393018 | 0.305589 | 0.15 | 0.141 | 1 | 0 | Mettl9 | 0.009 | Young |
| 0.393024 | 0.405162 | 0.021 | 0.016 | 1 | 0 | Ptpn4 | 0.005 | Young |
| 0.393122 | 0.402594 | 0.026 | 0.021 | 1 | 0 | Tubgcp6 | 0.005 | Young |
| 0.393183 | 0.270674 | 0.017 | 0.012 | 1 | 0 | Asb10 | 0.005 | Young |
| 0.393583 | 0.214421 | 0.012 | 0.016 | 1 | 0 | Sh3rf1 | -0.004 | Young |
| 0.393842 | 0.145642 | 0.073 | 0.083 | 1 | 0 | Dtnbp1 | -0.01 | Young |
| 0.394418 | 0.265878 | 0.178 | 0.168 | 1 | 0 | Maf1 | 0.01 | Young |
| 0.394427 | 0.156577 | 0.111 | 0.1 | 1 | 0 | Gramd1a | 0.011 | Young |
| 0.394516 | 0.265746 | 0.164 | 0.153 | 1 | 0 | Vcpip1 | 0.011 | Young |
| 0.394651 | 0.182865 | 0.27 | 0.26 | 1 | 0 | Klf10 | 0.01 | Young |
| 0.394817 | 0.110558 | 0.032 | 0.038 | 1 | 0 | Mapkapk3 | -0.006 | Young |
| 0.395108 | 0.044339 | 0.118 | 0.131 | 1 | 0 | Uba5 | -0.013 | Young |
| 0.395498 | 0.012905 | 0.105 | 0.093 | 1 | 0 | Ppp1r11 | 0.012 | Young |
| 0.396117 | 0.035302 | 0.073 | 0.083 | 1 | 0 | Katnbl1 | -0.01 | Young |
| 0.396358 | 0.139136 | 0.05 | 0.042 | 1 | 0 | Dhx32 | 0.008 | Young |
| 0.396361 | 0.566152 | 0.015 | 0.011 | 1 | 0 | Fam13a | 0.004 | Young |
| 0.396731 | 0.224499 | 0.056 | 0.048 | 1 | 0 | 2310009B1 | 0.008 | Young |
| 0.396877 | 0.002652 | 0.109 | 0.122 | 1 | 0 | Dhx38 | -0.013 | Young |
| 0.397399 | 0.0285 | 0.229 | 0.248 | 1 | 0 | BC017643 | -0.019 | Young |
| 0.397438 | 0.165505 | 0.041 | 0.034 | 1 | 0 | Ndufaf2 | 0.007 | Young |
| 0.397838 | 0.549225 | 0.015 | 0.011 | 1 | 0 | Zfp408 | 0.004 | Young |
| 0.397907 | 0.231644 | 0.175 | 0.164 | 1 | 0 | Tspyl1 | 0.011 | Young |
| 0.398042 | 0.044229 | 0.022 | 0.027 | 1 | 0 | Oas1a | -0.005 | Young |
| 0.398224 | 0.437645 | 0.015 | 0.011 | 1 | 0 | Polr3h | 0.004 | Young |
| 0.398306 | 0.55393 | 0.047 | 0.04 | 1 | 0 | Kat5 | 0.007 | Young |
| 0.398515 | 0.320258 | 0.164 | 0.155 | 1 | 0 | Csnk2a2 | 0.009 | Young |
| 0.398691 | 0.226877 | 0.07 | 0.062 | 1 | 0 | Dpm1 | 0.008 | Young |
| 0.398739 | 0.346619 | 0.015 | 0.011 | 1 | 0 | Tacc3 | 0.004 | Young |
| 0.398846 | 0.273482 | 0.118 | 0.108 | 1 | 0 | Vps50 | 0.01 | Young |
| 0.399513 | 0.248973 | 0.015 | 0.011 | 1 | 0 | Ptar1 | 0.004 | Young |
| 0.399514 | 0.68899 | 0.045 | 0.038 | 1 | 0 | Tmem97 | 0.007 | Young |
| 0.400545 | 0.327618 | 0.015 | 0.011 | 1 | 0 | Mical1 | 0.004 | Young |
| 0.400569 | 0.008274 | 0.061 | 0.07 | 1 | 0 | Wdr43 | -0.009 | Young |

| | | | | | | | | |
|---|---|---|---|---|---|---|---|---|
| 0.400581 | 0.27426 | 0.021 | 0.026 | 1 | 0 | Ccp110 | -0.005 | Young |
| 0.400683 | 0.545087 | 0.038 | 0.031 | 1 | 0 | Kank3 | 0.007 | Young |
| 0.40073 | 0.228315 | 0.172 | 0.161 | 1 | 0 | Lrpap1 | 0.011 | Young |
| 0.400759 | 0.014597 | 0.414 | 0.445 | 1 | 0 | Tra2b | -0.031 | Young |
| 0.400972 | 0.816295 | 0.02 | 0.015 | 1 | 0 | Areg | 0.005 | Young |
| 0.401726 | 0.312544 | 0.154 | 0.144 | 1 | 0 | Fez2 | 0.01 | Young |
| 0.401807 | 0.005383 | 0.231 | 0.25 | 1 | 0 | Aph1a | -0.019 | Young |
| 0.402264 | 0.190951 | 0.153 | 0.142 | 1 | 0 | Tbc1d9 | 0.011 | Young |
| 0.402513 | 0.272198 | 0.115 | 0.105 | 1 | 0 | Psmd12 | 0.01 | Young |
| 0.402921 | 0.142923 | 0.076 | 0.067 | 1 | 0 | Mphosph8 | 0.009 | Young |
| 0.404616 | 0.00306 | 0.215 | 0.234 | 1 | 0 | Cct2 | -0.019 | Young |
| 0.405142 | 0.655962 | 0.03 | 0.025 | 1 | 0 | Neurl1a | 0.005 | Young |
| 0.405859 | 0.239969 | 0.103 | 0.093 | 1 | 0 | Trrap | 0.01 | Young |
| 0.40606 | 0.243484 | 0.033 | 0.04 | 1 | 0 | Orc2 | -0.007 | Young |
| 0.406505 | 0.504155 | 0.013 | 0.01 | 1 | 0 | 1110051M | 0.003 | Young |
| 0.406789 | 0.316138 | 0.024 | 0.019 | 1 | 0 | Smyd3 | 0.005 | Young |
| 0.406898 | 0.117419 | 0.458 | 0.454 | 1 | 0 | Pip4k2a | 0.004 | Young |
| 0.406929 | 0.346954 | 0.132 | 0.123 | 1 | 0 | Man1a2 | 0.009 | Young |
| 0.407058 | 0.179464 | 0.013 | 0.01 | 1 | 0 | Amacr | 0.003 | Young |
| 0.407131 | 0.041994 | 0.087 | 0.098 | 1 | 0 | Med8 | -0.011 | Young |
| 0.407436 | 0.405584 | 0.044 | 0.037 | 1 | 0 | Zfp180 | 0.007 | Young |
| 0.407612 | 0.381101 | 0.013 | 0.01 | 1 | 0 | Gm48066 | 0.003 | Young |
| 0.407783 | 0.329868 | 0.099 | 0.09 | 1 | 0 | Pnpla7 | 0.009 | Young |
| 0.408232 | 0.325687 | 0.02 | 0.015 | 1 | 0 | Mllt3 | 0.005 | Young |
| 0.40834 | 0.048731 | 0.023 | 0.029 | 1 | 0 | Zmym1 | -0.006 | Young |
| 0.40837 | 0.144571 | 0.073 | 0.064 | 1 | 0 | Zfp598 | 0.009 | Young |
| 0.408702 | 0.376309 | 0.044 | 0.037 | 1 | 0 | Ccs | 0.007 | Young |
| 0.408809 | 0.006304 | 0.015 | 0.019 | 1 | 0 | Meis3 | -0.004 | Young |
| 0.408815 | 0.067733 | 0.086 | 0.097 | 1 | 0 | Cdk16 | -0.011 | Young |
| 0.409037 | 0.126366 | 0.116 | 0.13 | 1 | 0 | Aqr | -0.014 | Young |
| 0.409381 | 0.032754 | 0.206 | 0.226 | 1 | 0 | Sbno1 | -0.02 | Young |
| 0.409523 | 0.153676 | 0.009 | 0.012 | 1 | 0 | Zfp820 | -0.003 | Young |
| 0.409543 | 0.29575 | 0.027 | 0.022 | 1 | 0 | Dbp | 0.005 | Young |
| 0.409617 | 0.145623 | 0.278 | 0.265 | 1 | 0 | Pikfyve | 0.013 | Young |
| 0.409818 | 0.400462 | 0.123 | 0.115 | 1 | 0 | Slc50a1 | 0.008 | Young |
| 0.409937 | 0.021718 | 0.107 | 0.119 | 1 | 0 | Gcc2 | -0.012 | Young |
| 0.410151 | 0.937081 | 0.012 | 0.008 | 1 | 0 | Zfp607a | 0.004 | Young |
| 0.410758 | 0.150642 | 0.286 | 0.274 | 1 | 0 | Dip2b | 0.012 | Young |
| 0.410824 | 0.056453 | 0.096 | 0.108 | 1 | 0 | Hlx | -0.012 | Young |
| 0.410923 | 0.080605 | 0.056 | 0.064 | 1 | 0 | Slc25a30 | -0.008 | Young |
| 0.410956 | 0.002246 | 0.39 | 0.421 | 1 | 0 | Nap1l1 | -0.031 | Young |
| 0.410975 | 0.169458 | 0.029 | 0.023 | 1 | 0 | Zfp944 | 0.006 | Young |
| 0.411169 | 0.007723 | 0.015 | 0.019 | 1 | 0 | Abca2 | -0.004 | Young |
| 0.41135 | 0.061287 | 0.092 | 0.104 | 1 | 0 | Ubfd1 | -0.012 | Young |
| 0.411399 | 0.724378 | 0.018 | 0.014 | 1 | 0 | Myom1 | 0.004 | Young |

| | | | | | | | | |
|---|---|---|---|---|---|---|---|---|
| 0.41153 | 0.00898 | 0.067 | 0.077 | 1 | 0 | Golga3 | -0.01 | Young |
| 0.411531 | 0.672777 | 0.023 | 0.018 | 1 | 0 | Nudt1 | 0.005 | Young |
| 0.411789 | 0.706589 | 0.012 | 0.008 | 1 | 0 | Zswim7 | 0.004 | Young |
| 0.411846 | 0.391246 | 0.052 | 0.045 | 1 | 0 | Fgfr1op | 0.007 | Young |
| 0.411938 | 0.048804 | 0.218 | 0.205 | 1 | 0 | Ptpn12 | 0.013 | Young |
| 0.412065 | 0.527805 | 0.023 | 0.018 | 1 | 0 | Mapkbp1 | 0.005 | Young |
| 0.412294 | 0.313225 | 0.066 | 0.057 | 1 | 0 | Pmpcb | 0.009 | Young |
| 0.41255 | 0.099907 | 0.288 | 0.276 | 1 | 0 | Scarb2 | 0.012 | Young |
| 0.41277 | 0.075489 | 0.057 | 0.066 | 1 | 0 | Ap4m1 | -0.009 | Young |
| 0.412896 | 0.014101 | 0.034 | 0.027 | 1 | 0 | Sprtn | 0.007 | Young |
| 0.412973 | 0.198449 | 0.06 | 0.07 | 1 | 0 | Sos1 | -0.01 | Young |
| 0.413602 | 0.218817 | 0.032 | 0.026 | 1 | 0 | Trim32 | 0.006 | Young |
| 0.413622 | 0.362568 | 0.03 | 0.025 | 1 | 0 | Cep128 | 0.005 | Young |
| 0.413678 | 0.577532 | 0.018 | 0.014 | 1 | 0 | Isoc2b | 0.004 | Young |
| 0.413731 | 0.55642 | 0.012 | 0.008 | 1 | 0 | Gm340 | 0.004 | Young |
| 0.413757 | 0.168874 | 0.023 | 0.029 | 1 | 0 | Ift74 | -0.006 | Young |
| 0.414015 | 0.145312 | 0.02 | 0.015 | 1 | 0 | Mars | 0.005 | Young |
| 0.414285 | 0.222072 | 0.069 | 0.079 | 1 | 0 | Abcg3 | -0.01 | Young |
| 0.414329 | 0.553606 | 0.012 | 0.008 | 1 | 0 | Alkbh2 | 0.004 | Young |
| 0.415267 | 0.459496 | 0.033 | 0.027 | 1 | 0 | Ctdspl | 0.006 | Young |
| 0.4154 | 0.195347 | 0.153 | 0.142 | 1 | 0 | Tgs1 | 0.011 | Young |
| 0.415667 | 0.236006 | 0.056 | 0.048 | 1 | 0 | Lrp12 | 0.008 | Young |
| 0.415827 | 0.407959 | 0.012 | 0.008 | 1 | 0 | Gm14326 | 0.004 | Young |
| 0.415977 | 0.359259 | 0.012 | 0.008 | 1 | 0 | Apex2 | 0.004 | Young |
| 0.416456 | 0.239512 | 0.326 | 0.327 | 1 | 0 | Atp5j | -0.001 | Young |
| 0.416785 | 0.240907 | 0.16 | 0.149 | 1 | 0 | Clcn4 | 0.011 | Young |
| 0.416835 | 0.325181 | 0.05 | 0.042 | 1 | 0 | Mgst3 | 0.008 | Young |
| 0.41699 | 0.100622 | 0.016 | 0.021 | 1 | 0 | Gm35154 | -0.005 | Young |
| 0.417236 | 0.414619 | 0.01 | 0.007 | 1 | 0 | Npr1 | 0.003 | Young |
| 0.417645 | 0.179626 | 0.197 | 0.185 | 1 | 0 | Heca | 0.012 | Young |
| 0.418024 | 0.014521 | 0.044 | 0.037 | 1 | 0 | Cdc14a | 0.007 | Young |
| 0.418375 | 0.809117 | 0.01 | 0.007 | 1 | 0 | Wdr35 | 0.003 | Young |
| 0.418401 | 0.009496 | 0.329 | 0.353 | 1 | 0 | Tep1 | -0.024 | Young |
| 0.418409 | 0.084688 | 0.043 | 0.036 | 1 | 0 | Gm5547 | 0.007 | Young |
| 0.418549 | 0.350109 | 0.09 | 0.081 | 1 | 0 | Adssl1 | 0.009 | Young |
| 0.418961 | 0.139079 | 0.25 | 0.242 | 1 | 0 | Nol7 | 0.008 | Young |
| 0.419889 | 0.202502 | 0.23 | 0.223 | 1 | 0 | Ddb1 | 0.007 | Young |
| 0.420005 | 0.774829 | 0.01 | 0.007 | 1 | 0 | Gm15853 | 0.003 | Young |
| 0.420031 | 0.084504 | 0.152 | 0.14 | 1 | 0 | Prkd3 | 0.012 | Young |
| 0.420185 | 0.451811 | 0.045 | 0.038 | 1 | 0 | Rfxap | 0.007 | Young |
| 0.420304 | 0.047386 | 0.074 | 0.083 | 1 | 0 | Uba52 | -0.009 | Young |
| 0.420416 | 0.320289 | 0.045 | 0.038 | 1 | 0 | Yif1a | 0.007 | Young |
| 0.420459 | 0.173122 | 0.142 | 0.13 | 1 | 0 | Zc3h15 | 0.012 | Young |
| 0.420717 | 0.236368 | 0.146 | 0.135 | 1 | 0 | Enox2 | 0.011 | Young |
| 0.422064 | 0.216365 | 0.083 | 0.074 | 1 | 0 | Atl3 | 0.009 | Young |

| | | | | | | | | |
|---|---|---|---|---|---|---|---|---|
| 0.422139 | 0.170237 | 0.294 | 0.286 | 1 | 0 | Ndufb10 | 0.008 | Young |
| 0.422912 | 0.755305 | 0.021 | 0.016 | 1 | 0 | Paqr4 | 0.005 | Young |
| 0.422912 | 0.055259 | 0.01 | 0.014 | 1 | 0 | Nars2 | -0.004 | Young |
| 0.42323 | 0.071102 | 0.636 | 0.642 | 1 | 0 | Mat2a | -0.006 | Young |
| 0.423294 | 0.395429 | 0.059 | 0.052 | 1 | 0 | Cog6 | 0.007 | Young |
| 0.423428 | 0.275931 | 0.077 | 0.068 | 1 | 0 | Sfmbt1 | 0.009 | Young |
| 0.423908 | 0.145918 | 0.102 | 0.092 | 1 | 0 | Supt4a | 0.01 | Young |
| 0.423963 | 0.025633 | 0.193 | 0.211 | 1 | 0 | Pet100 | -0.018 | Young |
| 0.424176 | 0.459969 | 0.016 | 0.012 | 1 | 0 | Snn | 0.004 | Young |
| 0.424271 | 0.08922 | 0.08 | 0.09 | 1 | 0 | Ulk1 | -0.01 | Young |
| 0.424358 | 0.274145 | 0.195 | 0.186 | 1 | 0 | Snx17 | 0.009 | Young |
| 0.424516 | 0.118951 | 0.069 | 0.06 | 1 | 0 | Rfwd3 | 0.009 | Young |
| 0.424686 | 0.158368 | 0.384 | 0.386 | 1 | 0 | Mycbp2 | -0.002 | Young |
| 0.424717 | 0.5632 | 0.021 | 0.016 | 1 | 0 | Dnajb5 | 0.005 | Young |
| 0.424765 | 0.433688 | 0.092 | 0.083 | 1 | 0 | Eif3g | 0.009 | Young |
| 0.425412 | 0.105149 | 0.01 | 0.007 | 1 | 0 | 0610040J0 | 0.003 | Young |
| 0.425749 | 0.147588 | 0.323 | 0.319 | 1 | 0 | Eif3c | 0.004 | Young |
| 0.42581 | 0.307953 | 0.026 | 0.031 | 1 | 0 | Vps33b | -0.005 | Young |
| 0.425961 | 0.514037 | 0.021 | 0.016 | 1 | 0 | Zfp512b | 0.005 | Young |
| 0.426003 | 0.22649 | 0.017 | 0.022 | 1 | 0 | Jpx | -0.005 | Young |
| 0.427316 | 0.281728 | 0.054 | 0.047 | 1 | 0 | Rnf34 | 0.007 | Young |
| 0.427383 | 0.361996 | 0.026 | 0.021 | 1 | 0 | Med6 | 0.005 | Young |
| 0.427685 | 0.209539 | 0.145 | 0.135 | 1 | 0 | Slc28a2 | 0.01 | Young |
| 0.427805 | 0.002218 | 0.254 | 0.274 | 1 | 0 | Arhgap9 | -0.02 | Young |
| 0.428072 | 0.170426 | 0.05 | 0.042 | 1 | 0 | Rad51b | 0.008 | Young |
| 0.428242 | 0.322816 | 0.142 | 0.133 | 1 | 0 | Rest | 0.009 | Young |
| 0.428804 | 0.065644 | 0.095 | 0.085 | 1 | 0 | Rbbp8 | 0.01 | Young |
| 0.429045 | 0.311629 | 0.114 | 0.105 | 1 | 0 | Mmd | 0.009 | Young |
| 0.429129 | 0.208144 | 0.127 | 0.116 | 1 | 0 | Banf1 | 0.011 | Young |
| 0.429454 | 0.529625 | 0.04 | 0.034 | 1 | 0 | Ubr7 | 0.006 | Young |
| 0.429748 | 1.17794 | 0.015 | 0.011 | 1 | 0 | Zfp930 | 0.004 | Young |
| 0.429995 | 0.008726 | 0.06 | 0.068 | 1 | 0 | Ormdl1 | -0.008 | Young |
| 0.430003 | 0.430734 | 0.032 | 0.026 | 1 | 0 | Mllt1 | 0.006 | Young |
| 0.430024 | 0.121769 | 0.299 | 0.289 | 1 | 0 | Grina | 0.01 | Young |
| 0.430245 | 0.28605 | 0.048 | 0.056 | 1 | 0 | Ermp1 | -0.008 | Young |
| 0.430304 | 0.206212 | 0.198 | 0.189 | 1 | 0 | Ivns1abp | 0.009 | Young |
| 0.430636 | 0.218138 | 0.115 | 0.105 | 1 | 0 | Mien1 | 0.01 | Young |
| 0.430823 | 0.169563 | 0.158 | 0.148 | 1 | 0 | Lpar1 | 0.01 | Young |
| 0.430882 | 0.107888 | 0.102 | 0.114 | 1 | 0 | Nsd2 | -0.012 | Young |
| 0.43144 | 0.036215 | 0.315 | 0.343 | 1 | 0 | Prrc2a | -0.028 | Young |
| 0.431897 | 0.190922 | 0.016 | 0.012 | 1 | 0 | Wdr76 | 0.004 | Young |
| 0.432009 | 0.029273 | 0.146 | 0.16 | 1 | 0 | Cyb5r4 | -0.014 | Young |
| 0.43211 | 0.638262 | 0.024 | 0.019 | 1 | 0 | Fig4 | 0.005 | Young |
| 0.432169 | 0.075847 | 0.031 | 0.037 | 1 | 0 | Vps72 | -0.006 | Young |
| 0.432554 | 0.201875 | 0.021 | 0.016 | 1 | 0 | Gm17745 | 0.005 | Young |

| | | | | | | | | |
|---|---|---|---|---|---|---|---|---|
| 0.432797 | 0.009187 | 0.059 | 0.067 | 1 | 0 | Ipo9 | -0.008 | Young |
| 0.432798 | 0.064075 | 0.104 | 0.093 | 1 | 0 | Idnk | 0.011 | Young |
| 0.433011 | 0.356281 | 0.021 | 0.016 | 1 | 0 | Dmap1 | 0.005 | Young |
| 0.433807 | 0.369248 | 0.048 | 0.041 | 1 | 0 | Zfp358 | 0.007 | Young |
| 0.433969 | 0.041609 | 0.016 | 0.012 | 1 | 0 | Ccdc171 | 0.004 | Young |
| 0.434509 | 0.154214 | 0.138 | 0.127 | 1 | 0 | Hdac1 | 0.011 | Young |
| 0.434522 | 0.645913 | 0.027 | 0.022 | 1 | 0 | Edrf1 | 0.005 | Young |
| 0.434621 | 0.092896 | 0.079 | 0.07 | 1 | 0 | Elp5 | 0.009 | Young |
| 0.434748 | 0.14775 | 0.016 | 0.012 | 1 | 0 | Grwd1 | 0.004 | Young |
| 0.43492 | 0.052449 | 0.078 | 0.088 | 1 | 0 | Itpk1 | -0.01 | Young |
| 0.435546 | 0.324361 | 0.171 | 0.163 | 1 | 0 | Exoc4 | 0.008 | Young |
| 0.435857 | 0.504362 | 0.024 | 0.019 | 1 | 0 | Gm20703 | 0.005 | Young |
| 0.436149 | 0.028878 | 0.157 | 0.171 | 1 | 0 | Mrps14 | -0.014 | Young |
| 0.436475 | 0.070593 | 0.463 | 0.464 | 1 | 0 | Tma7 | -0.001 | Young |
| 0.436587 | 0.75761 | 0.015 | 0.011 | 1 | 0 | Zfp553 | 0.004 | Young |
| 0.436645 | 0.253694 | 0.007 | 0.01 | 1 | 0 | Cacna1f | -0.003 | Young |
| 0.436734 | 0.050638 | 0.578 | 0.565 | 1 | 0 | Pmp22 | 0.013 | Young |
| 0.437435 | 0.016579 | 0.163 | 0.178 | 1 | 0 | Rassf5 | -0.015 | Young |
| 0.437721 | 0.049547 | 0.178 | 0.194 | 1 | 0 | Ms4a8a | -0.016 | Young |
| 0.437869 | 0.254531 | 0.019 | 0.023 | 1 | 0 | Bcorl1 | -0.004 | Young |
| 0.437923 | 0.393887 | 0.069 | 0.062 | 1 | 0 | Bag5 | 0.007 | Young |
| 0.438069 | 0.324478 | 0.067 | 0.059 | 1 | 0 | Letm1 | 0.008 | Young |
| 0.438101 | 0.686486 | 0.015 | 0.011 | 1 | 0 | Sirt5 | 0.004 | Young |
| 0.438711 | 0.248034 | 0.064 | 0.056 | 1 | 0 | Sesn3 | 0.008 | Young |
| 0.438881 | 0.388981 | 0.032 | 0.026 | 1 | 0 | Steap4 | 0.006 | Young |
| 0.439054 | 0.142861 | 0.055 | 0.048 | 1 | 0 | Trp53inp1 | 0.007 | Young |
| 0.439314 | 0.022336 | 0.59 | 0.631 | 1 | 0 | Prdx1 | -0.041 | Young |
| 0.439375 | 0.310356 | 0.12 | 0.111 | 1 | 0 | Ing1 | 0.009 | Young |
| 0.439447 | 0.104126 | 0.066 | 0.057 | 1 | 0 | Hjurp | 0.009 | Young |
| 0.439484 | 0.095762 | 0.229 | 0.216 | 1 | 0 | Crebrf | 0.013 | Young |
| 0.43954 | 0.416268 | 0.035 | 0.029 | 1 | 0 | Spsb3 | 0.006 | Young |
| 0.439975 | 0.230818 | 0.312 | 0.311 | 1 | 0 | Ndufb3 | 0.001 | Young |
| 0.4401 | 0.485786 | 0.015 | 0.011 | 1 | 0 | Nnt | 0.004 | Young |
| 0.440131 | 0.331881 | 0.043 | 0.037 | 1 | 0 | Lancl1 | 0.006 | Young |
| 0.44065 | 0.416339 | 0.065 | 0.057 | 1 | 0 | Pmvk | 0.008 | Young |
| 0.441048 | 0.360716 | 0.04 | 0.034 | 1 | 0 | Rnaseh2b | 0.006 | Young |
| 0.441071 | 0.204841 | 0.14 | 0.13 | 1 | 0 | Tdrd7 | 0.01 | Young |
| 0.441354 | 0.068648 | 0.782 | 0.788 | 1 | 0 | Gnai2 | -0.006 | Young |
| 0.441456 | 0.281321 | 0.037 | 0.031 | 1 | 0 | Acvr2a | 0.006 | Young |
| 0.441537 | 0.134542 | 0.158 | 0.146 | 1 | 0 | Ddx46 | 0.012 | Young |
| 0.441675 | 0.358761 | 0.024 | 0.019 | 1 | 0 | Cd300ld3 | 0.005 | Young |
| 0.441693 | 0.131168 | 0.19 | 0.181 | 1 | 0 | Slc25a36 | 0.009 | Young |
| 0.441758 | 0.005655 | 0.289 | 0.312 | 1 | 0 | Mia2 | -0.023 | Young |
| 0.44189 | 0.166368 | 0.021 | 0.026 | 1 | 0 | Stn1 | -0.005 | Young |
| 0.442035 | 0.301009 | 0.015 | 0.011 | 1 | 0 | Klra2 | 0.004 | Young |

| | | | | | | | | |
|---|---|---|---|---|---|---|---|---|
| 0.442116 | 0.115967 | 0.445 | 0.447 | 1 | 0 | Fnbp1 | -0.002 | Young |
| 0.442121 | 0.28551 | 0.236 | 0.23 | 1 | 0 | Dock8 | 0.006 | Young |
| 0.442325 | 0.129053 | 0.25 | 0.275 | 1 | 0 | Pkn1 | -0.025 | Young |
| 0.442402 | 0.030337 | 0.024 | 0.029 | 1 | 0 | Ppan | -0.005 | Young |
| 0.442491 | 0.14175 | 0.05 | 0.057 | 1 | 0 | Cep295 | -0.007 | Young |
| 0.442834 | 0.277857 | 0.043 | 0.037 | 1 | 0 | Slc41a1 | 0.006 | Young |
| 0.442859 | 0.002397 | 0.307 | 0.332 | 1 | 0 | Tmco1 | -0.025 | Young |
| 0.44342 | 0.143828 | 0.265 | 0.254 | 1 | 0 | Fchsd2 | 0.011 | Young |
| 0.443468 | 0.123504 | 0.077 | 0.068 | 1 | 0 | Ncoa5 | 0.009 | Young |
| 0.44347 | 0.189598 | 0.068 | 0.06 | 1 | 0 | Stard3 | 0.008 | Young |
| 0.443728 | 0.299181 | 0.145 | 0.135 | 1 | 0 | Blvra | 0.01 | Young |
| 0.444003 | 0.340555 | 0.024 | 0.019 | 1 | 0 | Stk38l | 0.005 | Young |
| 0.444123 | 0.005747 | 0.18 | 0.196 | 1 | 0 | Safb2 | -0.016 | Young |
| 0.444253 | 0.160646 | 0.015 | 0.011 | 1 | 0 | Lipt2 | 0.004 | Young |
| 0.444306 | 0.05116 | 0.61 | 0.61 | 1 | 0 | Snx2 | 0 | Young |
| 0.444596 | 0.14736 | 0.307 | 0.3 | 1 | 0 | Cysltr1 | 0.007 | Young |
| 0.444769 | 0.042431 | 0.112 | 0.124 | 1 | 0 | Ccndbp1 | -0.012 | Young |
| 0.444978 | 0.018876 | 0.087 | 0.097 | 1 | 0 | Nfx1 | -0.01 | Young |
| 0.44519 | 0.308885 | 0.086 | 0.078 | 1 | 0 | Plcb3 | 0.008 | Young |
| 0.445705 | 0.131258 | 0.368 | 0.368 | 1 | 0 | Ubl5 | 0 | Young |
| 0.446251 | 0.202102 | 0.049 | 0.042 | 1 | 0 | Stambp | 0.007 | Young |
| 0.446477 | 0.184952 | 0.015 | 0.011 | 1 | 0 | Slc9a4 | 0.004 | Young |
| 0.446838 | 0.278556 | 0.046 | 0.04 | 1 | 0 | Ppp2r3c | 0.006 | Young |
| 0.446877 | 0.209014 | 0.072 | 0.064 | 1 | 0 | Topbp1 | 0.008 | Young |
| 0.446894 | 0.283582 | 0.015 | 0.011 | 1 | 0 | Tns2 | 0.004 | Young |
| 0.447296 | 0.230923 | 0.062 | 0.055 | 1 | 0 | Rcor3 | 0.007 | Young |
| 0.447646 | 0.169955 | 0.024 | 0.019 | 1 | 0 | Fbxo21 | 0.005 | Young |
| 0.447808 | 0.258675 | 0.207 | 0.198 | 1 | 0 | Nhsl2 | 0.009 | Young |
| 0.44782 | 0.217162 | 0.043 | 0.037 | 1 | 0 | Guf1 | 0.006 | Young |
| 0.448189 | 0.021771 | 0.252 | 0.272 | 1 | 0 | Ndufa1 | -0.02 | Young |
| 0.448193 | 0.315427 | 0.036 | 0.03 | 1 | 0 | Plekhn1 | 0.006 | Young |
| 0.448449 | 0.133429 | 0.019 | 0.015 | 1 | 0 | A930033H1 | 0.004 | Young |
| 0.448566 | 0.085487 | 0.024 | 0.029 | 1 | 0 | Jak3 | -0.005 | Young |
| 0.44906 | 0.278522 | 0.067 | 0.059 | 1 | 0 | Rac3 | 0.008 | Young |
| 0.449567 | 0.141615 | 0.125 | 0.115 | 1 | 0 | Slc44a1 | 0.01 | Young |
| 0.44963 | 0.420355 | 0.027 | 0.022 | 1 | 0 | Pyurf | 0.005 | Young |
| 0.449703 | 0.322563 | 0.174 | 0.167 | 1 | 0 | Pgrmc1 | 0.007 | Young |
| 0.449922 | 0.760182 | 0.013 | 0.01 | 1 | 0 | Sesn2 | 0.003 | Young |
| 0.45019 | 0.18042 | 0.058 | 0.051 | 1 | 0 | Ascc2 | 0.007 | Young |
| 0.4503 | 0.087929 | 0.107 | 0.119 | 1 | 0 | Txndc11 | -0.012 | Young |
| 0.450613 | 0.091569 | 0.547 | 0.547 | 1 | 0 | Ywhab | 0 | Young |
| 0.450755 | 0.300907 | 0.03 | 0.025 | 1 | 0 | Rdh10 | 0.005 | Young |
| 0.450861 | 0.340607 | 0.066 | 0.059 | 1 | 0 | Siah1a | 0.007 | Young |
| 0.450968 | 0.466539 | 0.022 | 0.018 | 1 | 0 | Usp49 | 0.004 | Young |
| 0.451431 | 0.063917 | 0.019 | 0.015 | 1 | 0 | Map3k10 | 0.004 | Young |

| | | | | | | | | |
|---|---|---|---|---|---|---|---|---|
| 0.451457 | 0.006302 | 0.107 | 0.118 | 1 | 0 | Srf | -0.011 | Young |
| 0.451509 | 0.041449 | 0.241 | 0.261 | 1 | 0 | Cwc15 | -0.02 | Young |
| 0.451525 | 0.073769 | 0.45 | 0.45 | 1 | 0 | Ncl | 0 | Young |
| 0.451563 | 0.293739 | 0.104 | 0.096 | 1 | 0 | Wasl | 0.008 | Young |
| 0.451834 | 0.269242 | 0.081 | 0.073 | 1 | 0 | Naa30 | 0.008 | Young |
| 0.451974 | 0.222757 | 0.059 | 0.052 | 1 | 0 | Ccdc97 | 0.007 | Young |
| 0.452023 | 0.166576 | 0.105 | 0.096 | 1 | 0 | Zfp62 | 0.009 | Young |
| 0.452253 | 0.188399 | 0.157 | 0.175 | 1 | 0 | Dennd4c | -0.018 | Young |
| 0.452304 | 0.566436 | 0.037 | 0.031 | 1 | 0 | Atg16l2 | 0.006 | Young |
| 0.452418 | 0.019414 | 0.034 | 0.04 | 1 | 0 | Rusc1 | -0.006 | Young |
| 0.45242 | 0.721886 | 0.028 | 0.023 | 1 | 0 | Uxt | 0.005 | Young |
| 0.452601 | 0.399459 | 0.013 | 0.01 | 1 | 0 | Ift43 | 0.003 | Young |
| 0.452819 | 0.664918 | 0.018 | 0.014 | 1 | 0 | Tgds | 0.004 | Young |
| 0.452879 | 0.324592 | 0.092 | 0.083 | 1 | 0 | Kctd10 | 0.009 | Young |
| 0.453055 | 0.030262 | 0.026 | 0.031 | 1 | 0 | Zfp142 | -0.005 | Young |
| 0.453143 | 0.10118 | 0.17 | 0.187 | 1 | 0 | Sema6d | -0.017 | Young |
| 0.453379 | 0.213127 | 0.046 | 0.053 | 1 | 0 | Ube2l6 | -0.007 | Young |
| 0.453762 | 0.049223 | 0.097 | 0.108 | 1 | 0 | Eps15l1 | -0.011 | Young |
| 0.45386 | 0.136441 | 0.025 | 0.03 | 1 | 0 | Pyroxd1 | -0.005 | Young |
| 0.453938 | 0.039607 | 0.029 | 0.034 | 1 | 0 | Plekhg1 | -0.005 | Young |
| 0.453972 | 0.031194 | 0.032 | 0.026 | 1 | 0 | Mettl26 | 0.006 | Young |
| 0.454037 | 0.589939 | 0.028 | 0.023 | 1 | 0 | Btrc | 0.005 | Young |
| 0.454093 | 0.569293 | 0.013 | 0.01 | 1 | 0 | Vegfa | 0.003 | Young |
| 0.454126 | 0.042528 | 0.056 | 0.064 | 1 | 0 | Tspan5 | -0.008 | Young |
| 0.454186 | 0.29923 | 0.061 | 0.053 | 1 | 0 | Timm22 | 0.008 | Young |
| 0.454195 | 0.198886 | 0.127 | 0.118 | 1 | 0 | Irak4 | 0.009 | Young |
| 0.454409 | 0.011629 | 0.129 | 0.141 | 1 | 0 | Xpo6 | -0.012 | Young |
| 0.45497 | 0.375618 | 0.138 | 0.13 | 1 | 0 | Nfatc3 | 0.008 | Young |
| 0.454991 | 0.196477 | 0.118 | 0.108 | 1 | 0 | Bcl6 | 0.01 | Young |
| 0.455094 | 0.002671 | 0.034 | 0.04 | 1 | 0 | Ap5z1 | -0.006 | Young |
| 0.455353 | 0.289276 | 0.102 | 0.093 | 1 | 0 | Hmgcl | 0.009 | Young |
| 0.455362 | 0.185896 | 0.098 | 0.089 | 1 | 0 | Dopey1 | 0.009 | Young |
| 0.455374 | 0.173001 | 0.03 | 0.025 | 1 | 0 | Tdrd3 | 0.005 | Young |
| 0.455892 | 0.585856 | 0.018 | 0.014 | 1 | 0 | Rpap2 | 0.004 | Young |
| 0.456195 | 0.332381 | 0.062 | 0.055 | 1 | 0 | Ralgapa1 | 0.007 | Young |
| 0.456287 | 0.162675 | 0.065 | 0.057 | 1 | 0 | Chchd5 | 0.008 | Young |
| 0.456406 | 0.09416 | 0.137 | 0.152 | 1 | 0 | Ankrd13c | -0.015 | Young |
| 0.456693 | 0.31098 | 0.082 | 0.074 | 1 | 0 | Rcbtb2 | 0.008 | Young |
| 0.45672 | 0.076825 | 0.147 | 0.161 | 1 | 0 | Fam76a | -0.014 | Young |
| 0.45689 | 0.103692 | 0.05 | 0.057 | 1 | 0 | Esr1 | -0.007 | Young |
| 0.456987 | 0.063642 | 0.06 | 0.068 | 1 | 0 | Exosc10 | -0.008 | Young |
| 0.457014 | 0.037773 | 0.13 | 0.142 | 1 | 0 | Lsm5 | -0.012 | Young |
| 0.457236 | 0.133822 | 0.013 | 0.01 | 1 | 0 | Nemp2 | 0.003 | Young |
| 0.457334 | 0.264632 | 0.14 | 0.131 | 1 | 0 | Impad1 | 0.009 | Young |
| 0.457587 | 0.201914 | 0.119 | 0.109 | 1 | 0 | Rabgap1 | 0.01 | Young |

| | | | | | | | | |
|---|---|---|---|---|---|---|---|---|
| 0.457646 | 0.356886 | 0.049 | 0.042 | 1 | 0 | Nosip | 0.007 | Young |
| 0.45767 | 0.033421 | 0.098 | 0.108 | 1 | 0 | Maml1 | -0.01 | Young |
| 0.457743 | 0.096789 | 0.026 | 0.031 | 1 | 0 | Pnpla6 | -0.005 | Young |
| 0.457805 | 0.137747 | 0.343 | 0.339 | 1 | 0 | Matr3 | 0.004 | Young |
| 0.457938 | 0.110766 | 0.138 | 0.127 | 1 | 0 | Stk11 | 0.011 | Young |
| 0.458272 | 0.460196 | 0.064 | 0.057 | 1 | 0 | Wrn | 0.007 | Young |
| 0.45845 | 0.322836 | 0.147 | 0.14 | 1 | 0 | Stxbp6 | 0.007 | Young |
| 0.458605 | 0.490104 | 0.04 | 0.034 | 1 | 0 | Fas | 0.006 | Young |
| 0.458783 | 0.068475 | 0.008 | 0.011 | 1 | 0 | Tagap1 | -0.003 | Young |
| 0.459061 | 0.036947 | 0.269 | 0.286 | 1 | 0 | Camkk2 | -0.017 | Young |
| 0.459286 | 0.464722 | 0.081 | 0.074 | 1 | 0 | Fam219b | 0.007 | Young |
| 0.459323 | 0.242992 | 0.043 | 0.037 | 1 | 0 | Sephs2 | 0.006 | Young |
| 0.459472 | 0.443529 | 0.144 | 0.138 | 1 | 0 | Anxa11 | 0.006 | Young |
| 0.459594 | 0.201198 | 0.148 | 0.138 | 1 | 0 | Ufsp2 | 0.01 | Young |
| 0.460156 | 0.398483 | 0.025 | 0.021 | 1 | 0 | Ank2 | 0.004 | Young |
| 0.460485 | 0.016953 | 0.071 | 0.079 | 1 | 0 | Oxa1l | -0.008 | Young |
| 0.460684 | 0.470555 | 0.011 | 0.008 | 1 | 0 | Mtcp1 | 0.003 | Young |
| 0.460745 | 0.154215 | 0.145 | 0.16 | 1 | 0 | Psme2b | -0.015 | Young |
| 0.46129 | 0.226811 | 0.218 | 0.209 | 1 | 0 | Fam234a | 0.009 | Young |
| 0.4613 | 0.070379 | 0.054 | 0.062 | 1 | 0 | Ston2 | -0.008 | Young |
| 0.461873 | 0.310454 | 0.056 | 0.049 | 1 | 0 | Gramd3 | 0.007 | Young |
| 0.461977 | 0.76292 | 0.011 | 0.008 | 1 | 0 | Zfp958 | 0.003 | Young |
| 0.462221 | 0.469912 | 0.035 | 0.041 | 1 | 0 | Tor2a | -0.006 | Young |
| 0.462529 | 0.148811 | 0.288 | 0.283 | 1 | 0 | Add1 | 0.005 | Young |
| 0.462948 | 0.554276 | 0.011 | 0.008 | 1 | 0 | AC151836. | 0.003 | Young |
| 0.463118 | 0.197257 | 0.089 | 0.081 | 1 | 0 | Nkiras2 | 0.008 | Young |
| 0.463432 | 0.078203 | 0.04 | 0.047 | 1 | 0 | Sugp1 | -0.007 | Young |
| 0.463725 | 0.116858 | 0.029 | 0.034 | 1 | 0 | Mettl3 | -0.005 | Young |
| 0.46392 | 0.568019 | 0.011 | 0.008 | 1 | 0 | 5830432E0 | 0.003 | Young |
| 0.464094 | 0.0945 | 0.031 | 0.037 | 1 | 0 | Hdhd2 | -0.006 | Young |
| 0.464158 | 0.2568 | 0.097 | 0.089 | 1 | 0 | Spr | 0.008 | Young |
| 0.464309 | 0.051933 | 0.036 | 0.042 | 1 | 0 | Cnot9 | -0.006 | Young |
| 0.464326 | 0.367704 | 0.122 | 0.114 | 1 | 0 | Kdm4a | 0.008 | Young |
| 0.464424 | 0.081494 | 0.147 | 0.137 | 1 | 0 | Anks1 | 0.01 | Young |
| 0.464569 | 0.527057 | 0.011 | 0.008 | 1 | 0 | Nphp1 | 0.003 | Young |
| 0.464731 | 0.449634 | 0.011 | 0.008 | 1 | 0 | Mag | 0.003 | Young |
| 0.464784 | 0.127503 | 0.407 | 0.41 | 1 | 0 | Atrx | -0.003 | Young |
| 0.465395 | 0.106961 | 0.319 | 0.312 | 1 | 0 | Tcn2 | 0.007 | Young |
| 0.465597 | 0.726739 | 0.016 | 0.012 | 1 | 0 | Ap5b1 | 0.004 | Young |
| 0.465651 | 0.550825 | 0.021 | 0.016 | 1 | 0 | Fndc4 | 0.005 | Young |
| 0.466233 | 0.373382 | 0.037 | 0.031 | 1 | 0 | Fam114a1 | 0.006 | Young |
| 0.466254 | 0.335438 | 0.073 | 0.066 | 1 | 0 | Tmed4 | 0.007 | Young |
| 0.466425 | 0.004453 | 0.78 | 0.782 | 1 | 0 | Trf | -0.002 | Young |
| 0.466534 | 0.290797 | 0.031 | 0.037 | 1 | 0 | Bod1 | -0.006 | Young |
| 0.466667 | 0.063448 | 0.379 | 0.376 | 1 | 0 | Mapkapk2 | 0.003 | Young |

| | | | | | | | | |
|---|---|---|---|---|---|---|---|---|
| 0.466701 | 0.286878 | 0.188 | 0.181 | 1 | 0 | Tnfrsf13b | 0.007 | Young |
| 0.466877 | 0.487236 | 0.047 | 0.041 | 1 | 0 | Nemp1 | 0.006 | Young |
| 0.467204 | 0.157312 | 0.152 | 0.142 | 1 | 0 | Msrb1 | 0.01 | Young |
| 0.467236 | 0.610951 | 0.016 | 0.012 | 1 | 0 | Spred3 | 0.004 | Young |
| 0.467399 | 0.045287 | 0.016 | 0.021 | 1 | 0 | Pold2 | -0.005 | Young |
| 0.467478 | 0.245504 | 0.046 | 0.04 | 1 | 0 | Rhbdf1 | 0.006 | Young |
| 0.46778 | 0.0617 | 0.168 | 0.157 | 1 | 0 | Maoa | 0.011 | Young |
| 0.468287 | 0.055064 | 0.076 | 0.085 | 1 | 0 | Sae1 | -0.009 | Young |
| 0.468399 | 0.268609 | 0.043 | 0.037 | 1 | 0 | Utp15 | 0.006 | Young |
| 0.468449 | 0.054769 | 0.147 | 0.135 | 1 | 0 | Ubxn6 | 0.012 | Young |
| 0.46856 | 0.373249 | 0.034 | 0.029 | 1 | 0 | Haus3 | 0.005 | Young |
| 0.468658 | 0.062735 | 0.053 | 0.06 | 1 | 0 | Epm2aip1 | -0.007 | Young |
| 0.468688 | 0.354735 | 0.163 | 0.157 | 1 | 0 | Hbp1 | 0.006 | Young |
| 0.468752 | 0.135176 | 0.117 | 0.108 | 1 | 0 | Epc2 | 0.009 | Young |
| 0.468986 | 0.209651 | 0.025 | 0.021 | 1 | 0 | Mapk1ip1 | 0.004 | Young |
| 0.469036 | 0.309481 | 0.04 | 0.034 | 1 | 0 | Zdhhc16 | 0.006 | Young |
| 0.46908 | 0.014221 | 0.117 | 0.129 | 1 | 0 | Gpalpp1 | -0.012 | Young |
| 0.469087 | 0.13982 | 0.25 | 0.242 | 1 | 0 | Nacc2 | 0.008 | Young |
| 0.469437 | 0.075517 | 0.16 | 0.176 | 1 | 0 | Golga7 | -0.016 | Young |
| 0.469449 | 0.165121 | 0.011 | 0.008 | 1 | 0 | Ehbp1 | 0.003 | Young |
| 0.469603 | 0.058 | 0.034 | 0.04 | 1 | 0 | Pot1b | -0.006 | Young |
| 0.46971 | 0.129952 | 0.084 | 0.094 | 1 | 0 | Slc17a5 | -0.01 | Young |
| 0.470173 | 0.639926 | 0.029 | 0.025 | 1 | 0 | Elac2 | 0.004 | Young |
| 0.470194 | 0.212526 | 0.233 | 0.227 | 1 | 0 | Hps3 | 0.006 | Young |
| 0.47027 | 0.003697 | 0.178 | 0.193 | 1 | 0 | Lman1 | -0.015 | Young |
| 0.471053 | 0.271158 | 0.109 | 0.101 | 1 | 0 | Suds3 | 0.008 | Young |
| 0.471395 | 0.00478 | 0.009 | 0.012 | 1 | 0 | C130013H( | -0.003 | Young |
| 0.471545 | 0.238655 | 0.191 | 0.183 | 1 | 0 | Chic2 | 0.008 | Young |
| 0.471576 | 0.010519 | 0.061 | 0.068 | 1 | 0 | Snapc5 | -0.007 | Young |
| 0.471758 | 0.469505 | 0.06 | 0.053 | 1 | 0 | Lrig2 | 0.007 | Young |
| 0.472292 | 0.266068 | 0.069 | 0.062 | 1 | 0 | Slc35a2 | 0.007 | Young |
| 0.472689 | 0.140691 | 0.217 | 0.207 | 1 | 0 | Kcnk6 | 0.01 | Young |
| 0.472992 | 0.051003 | 0.059 | 0.067 | 1 | 0 | Hras | -0.008 | Young |
| 0.473053 | 0.569898 | 0.027 | 0.022 | 1 | 0 | Mcoln2 | 0.005 | Young |
| 0.473519 | 0.188286 | 0.073 | 0.066 | 1 | 0 | Got2 | 0.007 | Young |
| 0.473649 | 0.122429 | 0.422 | 0.427 | 1 | 0 | Rnf13 | -0.005 | Young |
| 0.473667 | 0.7793 | 0.01 | 0.007 | 1 | 0 | Hist2h3c1 | 0.003 | Young |
| 0.473684 | 0.391004 | 0.016 | 0.012 | 1 | 0 | Wdr59 | 0.004 | Young |
| 0.473798 | 0.138999 | 0.388 | 0.383 | 1 | 0 | Sec62 | 0.005 | Young |
| 0.473815 | 0.006643 | 0.019 | 0.023 | 1 | 0 | Mcm2 | -0.004 | Young |
| 0.474052 | 0.18171 | 0.276 | 0.274 | 1 | 0 | Ndufa7 | 0.002 | Young |
| 0.474111 | 0.030213 | 0.084 | 0.093 | 1 | 0 | Cpne2 | -0.009 | Young |
| 0.475033 | 0.273543 | 0.063 | 0.056 | 1 | 0 | Btbd10 | 0.007 | Young |
| 0.475045 | 0.097367 | 0.061 | 0.053 | 1 | 0 | Sigmar1 | 0.008 | Young |
| 0.475283 | 0.205158 | 0.009 | 0.012 | 1 | 0 | Catip | -0.003 | Young |

| | | | | | | | | |
|---|---|---|---|---|---|---|---|---|
| 0.475301 | 0.301382 | 0.037 | 0.031 | 1 | 0 | Tinf2 | 0.006 | Young |
| 0.475982 | 0.485193 | 0.01 | 0.007 | 1 | 0 | 1110008L1 | 0.003 | Young |
| 0.475982 | 0.4164 | 0.01 | 0.007 | 1 | 0 | Dand5 | 0.003 | Young |
| 0.47607 | 0.194765 | 0.198 | 0.187 | 1 | 0 | Mknk2 | 0.011 | Young |
| 0.476075 | 0.195097 | 0.044 | 0.051 | 1 | 0 | Srcap | -0.007 | Young |
| 0.476138 | 0.46276 | 0.053 | 0.047 | 1 | 0 | Zfp746 | 0.006 | Young |
| 0.476161 | 0.454498 | 0.01 | 0.007 | 1 | 0 | Arl4d | 0.003 | Young |
| 0.476235 | 0.015646 | 0.17 | 0.183 | 1 | 0 | Exoc3 | -0.013 | Young |
| 0.476339 | 0.400346 | 0.01 | 0.007 | 1 | 0 | Trmu | 0.003 | Young |
| 0.476446 | 0.159577 | 0.217 | 0.205 | 1 | 0 | Zfp638 | 0.012 | Young |
| 0.47658 | 0.284276 | 0.016 | 0.012 | 1 | 0 | Mms22l | 0.004 | Young |
| 0.476678 | 0.136871 | 0.046 | 0.04 | 1 | 0 | Terf1 | 0.006 | Young |
| 0.47679 | 0.069846 | 0.149 | 0.163 | 1 | 0 | Bsdc1 | -0.014 | Young |
| 0.477053 | 0.032133 | 0.01 | 0.007 | 1 | 0 | Peg10 | 0.003 | Young |
| 0.477352 | 0.369771 | 0.096 | 0.089 | 1 | 0 | Pak1 | 0.007 | Young |
| 0.477371 | 0.298196 | 0.118 | 0.109 | 1 | 0 | Ccdc59 | 0.009 | Young |
| 0.477799 | 0.07074 | 0.147 | 0.16 | 1 | 0 | Vps29 | -0.013 | Young |
| 0.477894 | 0.018932 | 0.093 | 0.103 | 1 | 0 | 9930021J0 | -0.01 | Young |
| 0.478536 | 0.160468 | 0.096 | 0.088 | 1 | 0 | Usp37 | 0.008 | Young |
| 0.478983 | 0.025026 | 0.152 | 0.164 | 1 | 0 | Kdm2b | -0.012 | Young |
| 0.47902 | 0.243977 | 0.01 | 0.007 | 1 | 0 | Eaf2 | 0.003 | Young |
| 0.4796 | 0.043758 | 0.364 | 0.389 | 1 | 0 | Diaph1 | -0.025 | Young |
| 0.480131 | 0.041153 | 0.021 | 0.026 | 1 | 0 | Rnf157 | -0.005 | Young |
| 0.480808 | 0.154993 | 0.116 | 0.107 | 1 | 0 | Aimp1 | 0.009 | Young |
| 0.480821 | 0.994662 | 0.014 | 0.011 | 1 | 0 | Gbp9 | 0.003 | Young |
| 0.481058 | 0.211104 | 0.202 | 0.196 | 1 | 0 | Kcmf1 | 0.006 | Young |
| 0.481235 | 0.161308 | 0.053 | 0.047 | 1 | 0 | Trim3 | 0.006 | Young |
| 0.48133 | 0.296166 | 0.024 | 0.019 | 1 | 0 | Polr3a | 0.005 | Young |
| 0.481415 | 0.388083 | 0.019 | 0.015 | 1 | 0 | Cmss1 | 0.004 | Young |
| 0.481526 | 0.271649 | 0.098 | 0.09 | 1 | 0 | Nckipsd | 0.008 | Young |
| 0.481851 | 0.874871 | 0.031 | 0.026 | 1 | 0 | Tradd | 0.005 | Young |
| 0.482539 | 0.215109 | 0.05 | 0.044 | 1 | 0 | Cep290 | 0.006 | Young |
| 0.482578 | 0.093412 | 0.053 | 0.047 | 1 | 0 | Susd1 | 0.006 | Young |
| 0.482731 | 0.728808 | 0.014 | 0.011 | 1 | 0 | Arhgap35 | 0.003 | Young |
| 0.483359 | 0.194143 | 0.075 | 0.067 | 1 | 0 | Ankrd46 | 0.008 | Young |
| 0.483457 | 0.465948 | 0.019 | 0.015 | 1 | 0 | Oplah | 0.004 | Young |
| 0.483506 | 0.556989 | 0.073 | 0.067 | 1 | 0 | Mum1 | 0.006 | Young |
| 0.484106 | 0.035328 | 0.365 | 0.391 | 1 | 0 | Zmiz1 | -0.026 | Young |
| 0.484236 | 0.05449 | 0.085 | 0.094 | 1 | 0 | Hsdl1 | -0.009 | Young |
| 0.484241 | 0.279838 | 0.051 | 0.045 | 1 | 0 | Cog4 | 0.006 | Young |
| 0.484242 | 0.294568 | 0.057 | 0.051 | 1 | 0 | Snhg12 | 0.006 | Young |
| 0.484467 | 0.344233 | 0.029 | 0.025 | 1 | 0 | Mmgt2 | 0.004 | Young |
| 0.484479 | 0.244438 | 0.019 | 0.015 | 1 | 0 | Tanc1 | 0.004 | Young |
| 0.484675 | 0.002034 | 0.12 | 0.131 | 1 | 0 | Ric8a | -0.011 | Young |
| 0.485619 | 0.05588 | 0.205 | 0.223 | 1 | 0 | Wbp2 | -0.018 | Young |

| | | | | | | | | |
|---|---|---|---|---|---|---|---|---|
| 0.486415 | 0.538787 | 0.014 | 0.011 | 1 | 0 | Slamf6 | 0.003 | Young |
| 0.486472 | 0.164212 | 0.092 | 0.083 | 1 | 0 | Cmip | 0.009 | Young |
| 0.487597 | 0.350967 | 0.014 | 0.011 | 1 | 0 | Bivm | 0.003 | Young |
| 0.488041 | 0.388473 | 0.014 | 0.011 | 1 | 0 | Ppp1r18os | 0.003 | Young |
| 0.488068 | 0.309919 | 0.019 | 0.015 | 1 | 0 | Itgb3bp | 0.004 | Young |
| 0.48816 | 0.05837 | 0.266 | 0.283 | 1 | 0 | Wac | -0.017 | Young |
| 0.488189 | 0.384311 | 0.014 | 0.011 | 1 | 0 | Cldn34c1 | 0.003 | Young |
| 0.488302 | 0.330597 | 0.06 | 0.053 | 1 | 0 | Akt1s1 | 0.007 | Young |
| 0.488682 | 0.281223 | 0.029 | 0.025 | 1 | 0 | Clpb | 0.004 | Young |
| 0.489077 | 0.438281 | 0.014 | 0.011 | 1 | 0 | Adat1 | 0.003 | Young |
| 0.489581 | 0.448437 | 0.034 | 0.029 | 1 | 0 | Parp6 | 0.005 | Young |
| 0.489762 | 0.333504 | 0.06 | 0.053 | 1 | 0 | Bcl2 | 0.007 | Young |
| 0.490124 | 0.0479 | 0.063 | 0.071 | 1 | 0 | Ctps2 | -0.008 | Young |
| 0.490369 | 0.228753 | 0.094 | 0.086 | 1 | 0 | Axin1 | 0.008 | Young |
| 0.490486 | 0.172145 | 0.085 | 0.077 | 1 | 0 | Akr1c13 | 0.008 | Young |
| 0.490486 | 0.041527 | 0.187 | 0.204 | 1 | 0 | Clptm1l | -0.017 | Young |
| 0.490665 | 0.05024 | 0.246 | 0.234 | 1 | 0 | Smap2 | 0.012 | Young |
| 0.491114 | 0.137907 | 0.195 | 0.186 | 1 | 0 | Map3k11 | 0.009 | Young |
| 0.491341 | 0.081194 | 0.238 | 0.259 | 1 | 0 | Nop10 | -0.021 | Young |
| 0.491526 | 0.025747 | 0.121 | 0.133 | 1 | 0 | Fbxo3 | -0.012 | Young |
| 0.491537 | 0.124403 | 0.29 | 0.319 | 1 | 0 | Adipor1 | -0.029 | Young |
| 0.491539 | 0.32376 | 0.109 | 0.101 | 1 | 0 | Mbd3 | 0.008 | Young |
| 0.491702 | 0.539253 | 0.051 | 0.045 | 1 | 0 | Ccdc107 | 0.006 | Young |
| 0.491734 | 0.028882 | 0.042 | 0.048 | 1 | 0 | Ganc | -0.006 | Young |
| 0.491918 | 0.037786 | 0.186 | 0.201 | 1 | 0 | Was | -0.015 | Young |
| 0.491967 | 0.34455 | 0.081 | 0.074 | 1 | 0 | Ilvbl | 0.007 | Young |
| 0.492188 | 0.043637 | 0.185 | 0.2 | 1 | 0 | Fmnl1 | -0.015 | Young |
| 0.4922 | 0.193512 | 0.08 | 0.073 | 1 | 0 | Bnip3 | 0.007 | Young |
| 0.492204 | 0.058182 | 0.048 | 0.042 | 1 | 0 | 1110059E2 | 0.006 | Young |
| 0.492299 | 0.134471 | 0.027 | 0.031 | 1 | 0 | Rhbdd3 | -0.004 | Young |
| 0.492422 | 0.551934 | 0.031 | 0.026 | 1 | 0 | Tex30 | 0.005 | Young |
| 0.492589 | 0.418742 | 0.058 | 0.052 | 1 | 0 | Ppp6r2 | 0.006 | Young |
| 0.492593 | 0.261938 | 0.136 | 0.129 | 1 | 0 | Hadha | 0.007 | Young |
| 0.492613 | 0.021594 | 0.035 | 0.041 | 1 | 0 | Aar2 | -0.006 | Young |
| 0.492628 | 0.225915 | 0.102 | 0.094 | 1 | 0 | Gps1 | 0.008 | Young |
| 0.492942 | 0.150166 | 0.288 | 0.282 | 1 | 0 | Arhgap31 | 0.006 | Young |
| 0.493156 | 0.192407 | 0.086 | 0.078 | 1 | 0 | Nploc4 | 0.008 | Young |
| 0.493184 | 0.112013 | 0.448 | 0.451 | 1 | 0 | Tgfb1 | -0.003 | Young |
| 0.493234 | 0.29127 | 0.014 | 0.011 | 1 | 0 | Ccdc181 | 0.003 | Young |
| 0.493842 | 0.022014 | 0.375 | 0.398 | 1 | 0 | Ncf1 | -0.023 | Young |
| 0.494583 | 0.407925 | 0.093 | 0.086 | 1 | 0 | Kctd12b | 0.007 | Young |
| 0.494769 | 0.018919 | 0.106 | 0.116 | 1 | 0 | Ibtk | -0.01 | Young |
| 0.495015 | 0.047008 | 0.129 | 0.119 | 1 | 0 | Zfp704 | 0.01 | Young |
| 0.495238 | 0.149696 | 0.085 | 0.094 | 1 | 0 | Cdc42bpb | -0.009 | Young |
| 0.495347 | 0.295508 | 0.063 | 0.056 | 1 | 0 | Ginm1 | 0.007 | Young |

| | | | | | | | | |
|---|---|---|---|---|---|---|---|---|
| 0.495466 | 0.405889 | 0.037 | 0.031 | 1 | 0 | Zfp444 | 0.006 | Young |
| 0.496516 | 0.191078 | 0.033 | 0.038 | 1 | 0 | Itpkc | -0.005 | Young |
| 0.4966 | 0.013028 | 0.153 | 0.166 | 1 | 0 | Epb41l1 | -0.013 | Young |
| 0.497008 | 0.911733 | 0.013 | 0.01 | 1 | 0 | Gm26802 | 0.003 | Young |
| 0.497129 | 0.220617 | 0.148 | 0.14 | 1 | 0 | Uqcc3 | 0.008 | Young |
| 0.497173 | 0.078388 | 0.135 | 0.148 | 1 | 0 | Polb | -0.013 | Young |
| 0.497662 | 0.07199 | 0.297 | 0.321 | 1 | 0 | Map3k1 | -0.024 | Young |
| 0.497675 | 0.007264 | 0.071 | 0.079 | 1 | 0 | Cryzl1 | -0.008 | Young |
| 0.497975 | 0.130773 | 0.102 | 0.112 | 1 | 0 | Hnrnph3 | -0.01 | Young |
| 0.498067 | 0.113291 | 0.206 | 0.197 | 1 | 0 | Pik3cg | 0.009 | Young |
| 0.498121 | 0.060656 | 0.198 | 0.187 | 1 | 0 | Zfp710 | 0.011 | Young |
| 0.498406 | 0.378791 | 0.028 | 0.023 | 1 | 0 | Atf5 | 0.005 | Young |
| 0.4986 | 0.756869 | 0.013 | 0.01 | 1 | 0 | 4931414P1 | 0.003 | Young |
| 0.498751 | 0.172612 | 0.035 | 0.041 | 1 | 0 | Bbc3 | -0.006 | Young |
| 0.498899 | 0.057313 | 0.037 | 0.042 | 1 | 0 | Psmg2 | -0.005 | Young |
| 0.49891 | 0.290156 | 0.025 | 0.021 | 1 | 0 | Pou6f1 | 0.004 | Young |
| 0.499127 | 0.024558 | 0.16 | 0.174 | 1 | 0 | Atp2c1 | -0.014 | Young |
| 0.49925 | 0.257129 | 0.025 | 0.021 | 1 | 0 | Upp2 | 0.004 | Young |
| 0.500044 | 0.285388 | 0.025 | 0.021 | 1 | 0 | Ppih | 0.004 | Young |
| 0.50074 | 0.446296 | 0.059 | 0.053 | 1 | 0 | Dnajb12 | 0.006 | Young |
| 0.500744 | 0.338614 | 0.034 | 0.029 | 1 | 0 | Abhd6 | 0.005 | Young |
| 0.500986 | 0.12829 | 0.271 | 0.264 | 1 | 0 | Mapk1 | 0.007 | Young |
| 0.501116 | 0.365782 | 0.017 | 0.014 | 1 | 0 | Gtf2h2 | 0.003 | Young |
| 0.501211 | 0.152591 | 0.111 | 0.103 | 1 | 0 | 0610030E2 | 0.008 | Young |
| 0.501246 | 0.209006 | 0.089 | 0.1 | 1 | 0 | Exoc1 | -0.011 | Young |
| 0.50157 | 0.407747 | 0.041 | 0.036 | 1 | 0 | Haus2 | 0.005 | Young |
| 0.501624 | 0.268297 | 0.134 | 0.126 | 1 | 0 | Zbtb44 | 0.008 | Young |
| 0.502376 | 0.024883 | 0.096 | 0.105 | 1 | 0 | Dgcr2 | -0.009 | Young |
| 0.502983 | 0.187532 | 0.146 | 0.138 | 1 | 0 | Ncoa7 | 0.008 | Young |
| 0.503287 | 0.373537 | 0.034 | 0.029 | 1 | 0 | Ifitm1 | 0.005 | Young |
| 0.503684 | 0.252453 | 0.025 | 0.021 | 1 | 0 | Fars2 | 0.004 | Young |
| 0.504031 | 0.009841 | 0.244 | 0.263 | 1 | 0 | Elavl1 | -0.019 | Young |
| 0.504038 | 0.342383 | 0.031 | 0.026 | 1 | 0 | Phf6 | 0.005 | Young |
| 0.50421 | 0.253888 | 0.242 | 0.239 | 1 | 0 | Rnf7 | 0.003 | Young |
| 0.504442 | 0.101667 | 0.057 | 0.064 | 1 | 0 | Wdr20 | -0.007 | Young |
| 0.504478 | 0.735436 | 0.029 | 0.025 | 1 | 0 | 0610009O2 | 0.004 | Young |
| 0.504834 | 0.400136 | 0.013 | 0.01 | 1 | 0 | Vsig10l | 0.003 | Young |
| 0.504846 | 0.05612 | 0.05 | 0.056 | 1 | 0 | Cd300lg | -0.006 | Young |
| 0.504996 | 0.09649 | 0.123 | 0.114 | 1 | 0 | Zfp740 | 0.009 | Young |
| 0.505109 | 0.465881 | 0.048 | 0.042 | 1 | 0 | Eml3 | 0.006 | Young |
| 0.505327 | 0.038098 | 0.069 | 0.077 | 1 | 0 | Smox | -0.008 | Young |
| 0.505752 | 0.184437 | 0.043 | 0.037 | 1 | 0 | Stk35 | 0.006 | Young |
| 0.505781 | 0.089172 | 0.13 | 0.142 | 1 | 0 | Smad5 | -0.012 | Young |
| 0.50649 | 0.310236 | 0.217 | 0.212 | 1 | 0 | Baz2b | 0.005 | Young |
| 0.506861 | 0.189925 | 0.017 | 0.014 | 1 | 0 | Pcgf1 | 0.003 | Young |

| | | | | | | | | |
|---|---|---|---|---|---|---|---|---|
| 0.50739 | 0.290996 | 0.057 | 0.051 | 1 | 0 | Abcf3 | 0.006 | Young |
| 0.507971 | 0.339572 | 0.028 | 0.033 | 1 | 0 | Gnpnat1 | -0.005 | Young |
| 0.508231 | 0.323606 | 0.201 | 0.194 | 1 | 0 | Zc3h7a | 0.007 | Young |
| 0.50829 | 0.000285 | 0.114 | 0.124 | 1 | 0 | Tmem131 | -0.01 | Young |
| 0.508344 | 0.082863 | 0.113 | 0.124 | 1 | 0 | Fut11 | -0.011 | Young |
| 0.508665 | 0.296276 | 0.131 | 0.123 | 1 | 0 | Thoc7 | 0.008 | Young |
| 0.50871 | 0.183685 | 0.161 | 0.152 | 1 | 0 | Slc25a4 | 0.009 | Young |
| 0.508854 | 0.025256 | 0.068 | 0.077 | 1 | 0 | Thap11 | -0.009 | Young |
| 0.509523 | 0.093802 | 0.077 | 0.07 | 1 | 0 | Vps53 | 0.007 | Young |
| 0.509688 | 0.05443 | 0.237 | 0.257 | 1 | 0 | Snu13 | -0.02 | Young |
| 0.50985 | 0.513216 | 0.032 | 0.027 | 1 | 0 | Ddx31 | 0.005 | Young |
| 0.509901 | 0.074552 | 0.289 | 0.279 | 1 | 0 | Bptf | 0.01 | Young |
| 0.510066 | 0.025335 | 0.064 | 0.071 | 1 | 0 | Thap2 | -0.007 | Young |
| 0.510086 | 0.067204 | 0.025 | 0.021 | 1 | 0 | Usp11 | 0.004 | Young |
| 0.510578 | 0.16388 | 0.204 | 0.196 | 1 | 0 | Nlrp3 | 0.008 | Young |
| 0.510682 | 0.028014 | 0.069 | 0.077 | 1 | 0 | Zfp236 | -0.008 | Young |
| 0.511246 | 0.082134 | 0.057 | 0.064 | 1 | 0 | Ssna1 | -0.007 | Young |
| 0.511298 | 0.35035 | 0.086 | 0.079 | 1 | 0 | Gle1 | 0.007 | Young |
| 0.51155 | 0.338425 | 0.115 | 0.108 | 1 | 0 | Gbp3 | 0.007 | Young |
| 0.511634 | 0.0324 | 0.247 | 0.265 | 1 | 0 | Sh3kbp1 | -0.018 | Young |
| 0.511742 | 0.021104 | 0.159 | 0.171 | 1 | 0 | Pts | -0.012 | Young |
| 0.511905 | 0.06456 | 0.057 | 0.051 | 1 | 0 | Oma1 | 0.006 | Young |
| 0.512158 | 0.004824 | 0.087 | 0.096 | 1 | 0 | Ctdspl2 | -0.009 | Young |
| 0.512884 | 0.424324 | 0.061 | 0.055 | 1 | 0 | Mier3 | 0.006 | Young |
| 0.514161 | 0.472597 | 0.053 | 0.048 | 1 | 0 | Rfx1 | 0.005 | Young |
| 0.51439 | 0.427519 | 0.026 | 0.022 | 1 | 0 | Iqcb1 | 0.004 | Young |
| 0.514471 | 0.088499 | 0.084 | 0.077 | 1 | 0 | Pcnt | 0.007 | Young |
| 0.514538 | 0.069801 | 0.286 | 0.309 | 1 | 0 | Magt1 | -0.023 | Young |
| 0.514671 | 0.503549 | 0.026 | 0.022 | 1 | 0 | Dcun1d4 | 0.004 | Young |
| 0.515348 | 0.450993 | 0.083 | 0.077 | 1 | 0 | Nsmf | 0.006 | Young |
| 0.515702 | 0.009394 | 0.017 | 0.021 | 1 | 0 | Rsl1 | -0.004 | Young |
| 0.51573 | 0.013266 | 0.214 | 0.23 | 1 | 0 | Morc3 | -0.016 | Young |
| 0.51574 | 0.141022 | 0.02 | 0.016 | 1 | 0 | Itga6 | 0.004 | Young |
| 0.516295 | 0.153455 | 0.219 | 0.211 | 1 | 0 | Dnaja2 | 0.008 | Young |
| 0.516329 | 0.02416 | 0.007 | 0.01 | 1 | 0 | Gm19951 | -0.003 | Young |
| 0.516332 | 0.068425 | 0.307 | 0.295 | 1 | 0 | Pik3r1 | 0.012 | Young |
| 0.516468 | 0.232144 | 0.023 | 0.019 | 1 | 0 | Dnajc27 | 0.004 | Young |
| 0.517346 | 0.17526 | 0.101 | 0.093 | 1 | 0 | Snap29 | 0.008 | Young |
| 0.517805 | 0.07285 | 0.069 | 0.062 | 1 | 0 | Chpf2 | 0.007 | Young |
| 0.517841 | 0.234891 | 0.19 | 0.183 | 1 | 0 | Xrn2 | 0.007 | Young |
| 0.518129 | 0.145033 | 0.067 | 0.075 | 1 | 0 | Ccser2 | -0.008 | Young |
| 0.518238 | 0.165107 | 0.314 | 0.312 | 1 | 0 | 1700020I1 | 0.002 | Young |
| 0.518518 | 0.189177 | 0.104 | 0.096 | 1 | 0 | Pbx1 | 0.008 | Young |
| 0.51852 | 0.475707 | 0.046 | 0.041 | 1 | 0 | Aatf | 0.005 | Young |
| 0.518658 | 0.13845 | 0.566 | 0.587 | 1 | 0 | Erp29 | -0.021 | Young |

| | | | | | | | | |
|---|---|---|---|---|---|---|---|---|
| 0.518715 | 0.466574 | 0.011 | 0.008 | 1 | 0 | Zfp707 | 0.003 | Young |
| 0.518948 | 0.427172 | 0.026 | 0.022 | 1 | 0 | Msh6 | 0.004 | Young |
| 0.518978 | 0.449658 | 0.011 | 0.008 | 1 | 0 | Dock6 | 0.003 | Young |
| 0.519065 | 0.489013 | 0.011 | 0.008 | 1 | 0 | Fam177a | 0.003 | Young |
| 0.51937 | 0.025049 | 0.235 | 0.253 | 1 | 0 | Higd2a | -0.018 | Young |
| 0.51969 | 0.277913 | 0.083 | 0.093 | 1 | 0 | Zfp110 | -0.01 | Young |
| 0.519753 | 0.099579 | 0.007 | 0.01 | 1 | 0 | Car13 | -0.003 | Young |
| 0.519939 | 0.350759 | 0.011 | 0.008 | 1 | 0 | E2f6 | 0.003 | Young |
| 0.520063 | 0.288164 | 0.045 | 0.04 | 1 | 0 | Apip | 0.005 | Young |
| 0.520438 | 0.101189 | 0.061 | 0.068 | 1 | 0 | Tmem159 | -0.007 | Young |
| 0.520524 | 0.297734 | 0.029 | 0.025 | 1 | 0 | Zfp658 | 0.004 | Young |
| 0.520783 | 0.130065 | 0.11 | 0.12 | 1 | 0 | Med1 | -0.01 | Young |
| 0.520785 | 0.098845 | 0.123 | 0.134 | 1 | 0 | Arid3a | -0.011 | Young |
| 0.52084 | 0.376552 | 0.045 | 0.04 | 1 | 0 | Anapc7 | 0.005 | Young |
| 0.521233 | 0.046449 | 0.073 | 0.081 | 1 | 0 | Ino80e | -0.008 | Young |
| 0.52129 | 0.045824 | 0.191 | 0.205 | 1 | 0 | 6-Mar | -0.014 | Young |
| 0.521293 | 0.099373 | 0.104 | 0.096 | 1 | 0 | Rogdi | 0.008 | Young |
| 0.521313 | 0.140046 | 0.134 | 0.126 | 1 | 0 | Cyb5r3 | 0.008 | Young |
| 0.521431 | 0.216386 | 0.16 | 0.153 | 1 | 0 | 0610012G( | 0.007 | Young |
| 0.521571 | 0.324297 | 0.016 | 0.012 | 1 | 0 | Nif3l1 | 0.004 | Young |
| 0.521575 | 0.256298 | 0.045 | 0.04 | 1 | 0 | Ankmy2 | 0.005 | Young |
| 0.521691 | 0.337085 | 0.011 | 0.008 | 1 | 0 | Mmp14 | 0.003 | Young |
| 0.521792 | 0.062278 | 0.535 | 0.543 | 1 | 0 | Pea15a | -0.008 | Young |
| 0.522003 | 0.193235 | 0.099 | 0.092 | 1 | 0 | Abi2 | 0.007 | Young |
| 0.52217 | 0.007721 | 0.086 | 0.094 | 1 | 0 | Il10 | -0.008 | Young |
| 0.522185 | 0.105041 | 0.096 | 0.105 | 1 | 0 | Slc35a4 | -0.009 | Young |
| 0.522193 | 0.267577 | 0.223 | 0.222 | 1 | 0 | Cox5b | 0.001 | Young |
| 0.52226 | 0.631717 | 0.019 | 0.015 | 1 | 0 | Pced1a | 0.004 | Young |
| 0.522297 | 0.209448 | 0.061 | 0.055 | 1 | 0 | Prkaa1 | 0.006 | Young |
| 0.522326 | 0.325293 | 0.052 | 0.047 | 1 | 0 | Diras2 | 0.005 | Young |
| 0.522409 | 0.0575 | 0.02 | 0.016 | 1 | 0 | Rnf180 | 0.004 | Young |
| 0.522459 | 0.294731 | 0.023 | 0.019 | 1 | 0 | Mki67 | 0.004 | Young |
| 0.522543 | 0.012908 | 0.103 | 0.112 | 1 | 0 | Rhoh | -0.009 | Young |
| 0.522674 | 0.254921 | 0.067 | 0.06 | 1 | 0 | Ttc9c | 0.007 | Young |
| 0.522815 | 0.414492 | 0.077 | 0.086 | 1 | 0 | Gon4l | -0.009 | Young |
| 0.522891 | 0.285263 | 0.016 | 0.012 | 1 | 0 | Zfp1 | 0.004 | Young |
| 0.523028 | 0.260971 | 0.042 | 0.037 | 1 | 0 | Pdik1l | 0.005 | Young |
| 0.52305 | 0.178357 | 0.101 | 0.093 | 1 | 0 | Sars | 0.008 | Young |
| 0.523066 | 0.57356 | 0.019 | 0.015 | 1 | 0 | Il17rd | 0.004 | Young |
| 0.52307 | 0.002439 | 0.086 | 0.094 | 1 | 0 | Rab3d | -0.008 | Young |
| 0.523438 | 0.043062 | 0.064 | 0.071 | 1 | 0 | Rnf215 | -0.007 | Young |
| 0.523798 | 0.048608 | 0.161 | 0.174 | 1 | 0 | Psmc5 | -0.013 | Young |
| 0.523913 | 0.112421 | 0.094 | 0.103 | 1 | 0 | Ddx18 | -0.009 | Young |
| 0.523963 | 0.073918 | 0.018 | 0.022 | 1 | 0 | Zfand1 | -0.004 | Young |
| 0.524055 | 0.021345 | 0.096 | 0.088 | 1 | 0 | Rap2a | 0.008 | Young |

| | | | | | | | | |
|---|---|---|---|---|---|---|---|---|
| 0.52406 | 0.270315 | 0.076 | 0.07 | 1 | 0 | Mpv17 | 0.006 | Young |
| 0.524275 | 0.095472 | 0.474 | 0.472 | 1 | 0 | Plek | 0.002 | Young |
| 0.524373 | 0.106391 | 0.081 | 0.09 | 1 | 0 | Nsmaf | -0.009 | Young |
| 0.524652 | 0.215382 | 0.016 | 0.012 | 1 | 0 | E130308A1 | 0.004 | Young |
| 0.524657 | 0.398091 | 0.036 | 0.031 | 1 | 0 | Ccdc126 | 0.005 | Young |
| 0.525302 | 0.122716 | 0.053 | 0.06 | 1 | 0 | Malsu1 | -0.007 | Young |
| 0.525485 | 0.267261 | 0.127 | 0.12 | 1 | 0 | Rfc1 | 0.007 | Young |
| 0.525686 | 0.136126 | 0.162 | 0.153 | 1 | 0 | Tmem106b | 0.009 | Young |
| 0.525944 | 0.056593 | 0.05 | 0.056 | 1 | 0 | Tctex1d2 | -0.006 | Young |
| 0.526022 | 0.065725 | 0.023 | 0.027 | 1 | 0 | Dsel | -0.004 | Young |
| 0.526323 | 0.181862 | 0.147 | 0.14 | 1 | 0 | Sugt1 | 0.007 | Young |
| 0.526794 | 0.179736 | 0.061 | 0.055 | 1 | 0 | Arl11 | 0.006 | Young |
| 0.526817 | 0.442588 | 0.072 | 0.066 | 1 | 0 | Smg6 | 0.006 | Young |
| 0.526925 | 0.486511 | 0.036 | 0.031 | 1 | 0 | Yars | 0.005 | Young |
| 0.527691 | 0.111801 | 0.11 | 0.103 | 1 | 0 | Pgrmc2 | 0.007 | Young |
| 0.528073 | 0.031122 | 0.029 | 0.034 | 1 | 0 | Lig1 | -0.005 | Young |
| 0.528134 | 0.220786 | 0.065 | 0.059 | 1 | 0 | Map3k4 | 0.006 | Young |
| 0.528148 | 0.04218 | 0.117 | 0.127 | 1 | 0 | Cpd | -0.01 | Young |
| 0.528303 | 0.016908 | 0.024 | 0.029 | 1 | 0 | Rilp | -0.005 | Young |
| 0.528476 | 0.103523 | 0.031 | 0.036 | 1 | 0 | Pygl | -0.005 | Young |
| 0.528499 | 0.636636 | 0.027 | 0.023 | 1 | 0 | Gmeb2 | 0.004 | Young |
| 0.528522 | 0.35274 | 0.019 | 0.015 | 1 | 0 | Zfp619 | 0.004 | Young |
| 0.528674 | 0.139784 | 0.078 | 0.086 | 1 | 0 | Ppip5k2 | -0.008 | Young |
| 0.528819 | 0.191472 | 0.071 | 0.064 | 1 | 0 | Arfip1 | 0.007 | Young |
| 0.529037 | 0.070658 | 0.067 | 0.074 | 1 | 0 | Arl2bp | -0.007 | Young |
| 0.5292 | 0.130699 | 0.195 | 0.186 | 1 | 0 | Snw1 | 0.009 | Young |
| 0.529673 | 0.083254 | 0.177 | 0.193 | 1 | 0 | Tmem259 | -0.016 | Young |
| 0.529692 | 0.002276 | 0.162 | 0.174 | 1 | 0 | Dnase1l1 | -0.012 | Young |
| 0.529732 | 0.114678 | 0.022 | 0.026 | 1 | 0 | Slx4 | -0.004 | Young |
| 0.529795 | 0.115842 | 0.031 | 0.036 | 1 | 0 | Tmem110 | -0.005 | Young |
| 0.529961 | 0.022349 | 0.155 | 0.167 | 1 | 0 | Agps | -0.012 | Young |
| 0.530124 | 0.401202 | 0.033 | 0.029 | 1 | 0 | Acyp1 | 0.004 | Young |
| 0.530374 | 0.397869 | 0.104 | 0.098 | 1 | 0 | Trim33 | 0.006 | Young |
| 0.530709 | 0.091959 | 0.031 | 0.036 | 1 | 0 | Mrps11 | -0.005 | Young |
| 0.530817 | 0.170366 | 0.273 | 0.268 | 1 | 0 | Pdlim5 | 0.005 | Young |
| 0.530835 | 0.447618 | 0.027 | 0.023 | 1 | 0 | Mettl5 | 0.004 | Young |
| 0.530947 | 0.029757 | 0.055 | 0.062 | 1 | 0 | Sp2 | -0.007 | Young |
| 0.53138 | 0.179696 | 0.084 | 0.093 | 1 | 0 | Edc4 | -0.009 | Young |
| 0.531477 | 0.016195 | 0.008 | 0.011 | 1 | 0 | 4833418N( | -0.003 | Young |
| 0.531762 | 0.072089 | 0.176 | 0.19 | 1 | 0 | Ndufab1 | -0.014 | Young |
| 0.531908 | 0.467753 | 0.021 | 0.018 | 1 | 0 | Coq9 | 0.003 | Young |
| 0.53193 | 0.182753 | 0.024 | 0.029 | 1 | 0 | Gstp3 | -0.005 | Young |
| 0.532279 | 0.105493 | 0.054 | 0.048 | 1 | 0 | Haghl | 0.006 | Young |
| 0.532405 | 0.337697 | 0.08 | 0.074 | 1 | 0 | Phf11b | 0.006 | Young |
| 0.532523 | 0.007232 | 0.114 | 0.123 | 1 | 0 | Myo7a | -0.009 | Young |

| | | | | | | | | |
|---|---|---|---|---|---|---|---|---|
| 0.532702 | 0.108694 | 0.029 | 0.025 | 1 | 0 | Zfp426 | 0.004 | Young |
| 0.532801 | 0.104315 | 0.008 | 0.011 | 1 | 0 | 6330562C2 | -0.003 | Young |
| 0.532812 | 0.167244 | 0.052 | 0.047 | 1 | 0 | Erh | 0.005 | Young |
| 0.53314 | 0.119655 | 0.379 | 0.378 | 1 | 0 | Pros1 | 0.001 | Young |
| 0.533161 | 0.055241 | 0.1 | 0.109 | 1 | 0 | Dhx30 | -0.009 | Young |
| 0.533193 | 0.186398 | 0.019 | 0.015 | 1 | 0 | Naif1 | 0.004 | Young |
| 0.533447 | 0.03481 | 0.088 | 0.096 | 1 | 0 | Tmem57 | -0.008 | Young |
| 0.533516 | 0.043897 | 0.25 | 0.268 | 1 | 0 | Pxn | -0.018 | Young |
| 0.533857 | 0.063347 | 0.082 | 0.09 | 1 | 0 | Ago1 | -0.008 | Young |
| 0.534247 | 0.399655 | 0.059 | 0.053 | 1 | 0 | Prpsap1 | 0.006 | Young |
| 0.534488 | 0.171602 | 0.136 | 0.129 | 1 | 0 | Tcp11l2 | 0.007 | Young |
| 0.534554 | 0.478653 | 0.021 | 0.018 | 1 | 0 | Kctd9 | 0.003 | Young |
| 0.534589 | 0.157459 | 0.132 | 0.124 | 1 | 0 | Dapp1 | 0.008 | Young |
| 0.534871 | 0.136898 | 0.179 | 0.17 | 1 | 0 | Rhobtb1 | 0.009 | Young |
| 0.534953 | 0.76047 | 0.014 | 0.011 | 1 | 0 | Gm3336 | 0.003 | Young |
| 0.535075 | 0.296271 | 0.008 | 0.011 | 1 | 0 | Gm26510 | -0.003 | Young |
| 0.535227 | 0.097388 | 0.024 | 0.029 | 1 | 0 | Trpm4 | -0.005 | Young |
| 0.535277 | 0.155155 | 0.036 | 0.041 | 1 | 0 | Thrb | -0.005 | Young |
| 0.535559 | 0.203633 | 0.023 | 0.027 | 1 | 0 | Mtg1 | -0.004 | Young |
| 0.535606 | 0.095028 | 0.148 | 0.14 | 1 | 0 | Slc15a3 | 0.008 | Young |
| 0.535947 | 0.331521 | 0.067 | 0.062 | 1 | 0 | Ahsa2 | 0.005 | Young |
| 0.536201 | 0.039219 | 0.27 | 0.289 | 1 | 0 | Pigt | -0.019 | Young |
| 0.536735 | 0.063908 | 0.187 | 0.202 | 1 | 0 | Mob4 | -0.015 | Young |
| 0.53714 | 0.317542 | 0.051 | 0.045 | 1 | 0 | Fam110b | 0.006 | Young |
| 0.537237 | 0.008313 | 0.099 | 0.092 | 1 | 0 | Ctr9 | 0.007 | Young |
| 0.537839 | 0.288583 | 0.021 | 0.018 | 1 | 0 | Zbtb40 | 0.003 | Young |
| 0.537918 | 0.127076 | 0.027 | 0.031 | 1 | 0 | Ttc7b | -0.004 | Young |
| 0.538379 | 0.245926 | 0.066 | 0.06 | 1 | 0 | BC003965 | 0.006 | Young |
| 0.538436 | 0.368036 | 0.027 | 0.023 | 1 | 0 | Mcm5 | 0.004 | Young |
| 0.538517 | 0.127371 | 0.027 | 0.031 | 1 | 0 | Coq7 | -0.004 | Young |
| 0.53879 | 0.220406 | 0.049 | 0.044 | 1 | 0 | Fbxo34 | 0.005 | Young |
| 0.539087 | 0.2466 | 0.024 | 0.021 | 1 | 0 | Ncapg2 | 0.003 | Young |
| 0.539562 | 0.031941 | 0.051 | 0.057 | 1 | 0 | Triap1 | -0.006 | Young |
| 0.539755 | 0.143627 | 0.087 | 0.096 | 1 | 0 | Wdr13 | -0.009 | Young |
| 0.539974 | 0.126619 | 0.126 | 0.118 | 1 | 0 | Bcor | 0.008 | Young |
| 0.540086 | 0.011122 | 0.084 | 0.092 | 1 | 0 | Ifi47 | -0.008 | Young |
| 0.540149 | 0.254261 | 0.014 | 0.011 | 1 | 0 | Fahd2a | 0.003 | Young |
| 0.540157 | 0.107505 | 0.177 | 0.17 | 1 | 0 | Clasp2 | 0.007 | Young |
| 0.540699 | 0.299776 | 0.06 | 0.055 | 1 | 0 | Hnmt | 0.005 | Young |
| 0.540737 | 0.394353 | 0.048 | 0.042 | 1 | 0 | Sh2b1 | 0.006 | Young |
| 0.5409 | 0.060233 | 0.343 | 0.368 | 1 | 0 | Selplg | -0.025 | Young |
| 0.541142 | 0.001514 | 0.489 | 0.523 | 1 | 0 | Tab2 | -0.034 | Young |
| 0.541247 | 0.035995 | 0.285 | 0.305 | 1 | 0 | Cebpg | -0.02 | Young |
| 0.541545 | 0.251229 | 0.078 | 0.071 | 1 | 0 | Pex13 | 0.007 | Young |
| 0.541785 | 0.362414 | 0.145 | 0.14 | 1 | 0 | Sod2 | 0.005 | Young |

| | | | | | | | | |
|---|---|---|---|---|---|---|---|---|
| 0.542031 | 0.016085 | 0.109 | 0.118 | 1 | 0 | Dlg1 | -0.009 | Young |
| 0.542043 | 0.320992 | 0.055 | 0.049 | 1 | 0 | Dopey2 | 0.006 | Young |
| 0.542068 | 0.285765 | 0.04 | 0.036 | 1 | 0 | Ticam1 | 0.004 | Young |
| 0.542178 | 0.120404 | 0.339 | 0.338 | 1 | 0 | Ep300 | 0.001 | Young |
| 0.542408 | 0.152966 | 0.032 | 0.037 | 1 | 0 | Dhx33 | -0.005 | Young |
| 0.542418 | 0.242869 | 0.024 | 0.021 | 1 | 0 | Ap4e1 | 0.003 | Young |
| 0.542664 | 0.107063 | 0.052 | 0.047 | 1 | 0 | Abhd13 | 0.005 | Young |
| 0.542848 | 0.223746 | 0.037 | 0.042 | 1 | 0 | 2410002F2 | -0.005 | Young |
| 0.542898 | 0.170638 | 0.036 | 0.031 | 1 | 0 | Fancl | 0.005 | Young |
| 0.54318 | 0.748902 | 0.017 | 0.014 | 1 | 0 | Znhit3 | 0.003 | Young |
| 0.543346 | 0.173853 | 0.033 | 0.029 | 1 | 0 | Fbxl12 | 0.004 | Young |
| 0.543485 | 0.120754 | 0.033 | 0.038 | 1 | 0 | Retsat | -0.005 | Young |
| 0.543506 | 0.378841 | 0.086 | 0.079 | 1 | 0 | Zfyve16 | 0.007 | Young |
| 0.5443 | 0.522076 | 0.05 | 0.045 | 1 | 0 | Pacs1 | 0.005 | Young |
| 0.544438 | 0.12153 | 0.021 | 0.018 | 1 | 0 | Slc52a2 | 0.003 | Young |
| 0.544568 | 0.00217 | 0.147 | 0.157 | 1 | 0 | Mpc1 | -0.01 | Young |
| 0.544975 | 0.02044 | 0.931 | 0.947 | 1 | 0 | Rpl38 | -0.016 | Young |
| 0.54503 | 0.460022 | 0.042 | 0.037 | 1 | 0 | Miga1 | 0.005 | Young |
| 0.545828 | 0.162627 | 0.414 | 0.42 | 1 | 0 | Sod1 | -0.006 | Young |
| 0.545872 | 0.190372 | 0.258 | 0.256 | 1 | 0 | Nek7 | 0.002 | Young |
| 0.546201 | 0.006205 | 0.079 | 0.086 | 1 | 0 | Pias4 | -0.007 | Young |
| 0.546315 | 0.056636 | 0.826 | 0.822 | 1 | 0 | Unc93b1 | 0.004 | Young |
| 0.546447 | 0.064322 | 0.583 | 0.588 | 1 | 0 | Cox7a2l | -0.005 | Young |
| 0.546474 | 0.090745 | 0.406 | 0.413 | 1 | 0 | Sec61b | -0.007 | Young |
| 0.546554 | 0.626322 | 0.02 | 0.016 | 1 | 0 | Cyp2ab1 | 0.004 | Young |
| 0.546677 | 0.232343 | 0.164 | 0.157 | 1 | 0 | Cdc40 | 0.007 | Young |
| 0.546686 | 0.06083 | 0.21 | 0.226 | 1 | 0 | Rdx | -0.016 | Young |
| 0.547153 | 0.2061 | 0.072 | 0.066 | 1 | 0 | Med17 | 0.006 | Young |
| 0.547597 | 0.064036 | 0.058 | 0.052 | 1 | 0 | Ddhd1 | 0.006 | Young |
| 0.547603 | 0.110893 | 0.149 | 0.161 | 1 | 0 | Fam168a | -0.012 | Young |
| 0.54766 | 0.260717 | 0.147 | 0.141 | 1 | 0 | Ezh1 | 0.006 | Young |
| 0.547886 | 0.307979 | 0.053 | 0.048 | 1 | 0 | Tapbpl | 0.005 | Young |
| 0.547983 | 0.513399 | 0.043 | 0.038 | 1 | 0 | Nadk2 | 0.005 | Young |
| 0.548116 | 0.407135 | 0.045 | 0.04 | 1 | 0 | Mcm9 | 0.005 | Young |
| 0.548274 | 0.237904 | 0.108 | 0.101 | 1 | 0 | Ntpcr | 0.007 | Young |
| 0.54851 | 0.063643 | 0.334 | 0.358 | 1 | 0 | Tnfrsf1b | -0.024 | Young |
| 0.548557 | 0.347538 | 0.037 | 0.033 | 1 | 0 | Cnksr3 | 0.004 | Young |
| 0.548756 | 0.020167 | 0.102 | 0.111 | 1 | 0 | Vps4a | -0.009 | Young |
| 0.548805 | 0.173269 | 0.027 | 0.023 | 1 | 0 | Ampd2 | 0.004 | Young |
| 0.548923 | 0.334216 | 0.035 | 0.03 | 1 | 0 | Unc5a | 0.005 | Young |
| 0.549758 | 0.03912 | 0.07 | 0.077 | 1 | 0 | Dcaf13 | -0.007 | Young |
| 0.550444 | 0.129808 | 0.03 | 0.026 | 1 | 0 | Zfp608 | 0.004 | Young |
| 0.55092 | 0.355612 | 0.06 | 0.055 | 1 | 0 | Gm15417 | 0.005 | Young |
| 0.550988 | 0.449795 | 0.029 | 0.025 | 1 | 0 | 1700096K1 | 0.004 | Young |
| 0.551431 | 0.410198 | 0.029 | 0.025 | 1 | 0 | Dolk | 0.004 | Young |

| | | | | | | | | |
|---|---|---|---|---|---|---|---|---|
| 0.551887 | 0.315751 | 0.047 | 0.053 | 1 | 0 | Atg14 | -0.006 | Young |
| 0.551901 | 0.535882 | 0.023 | 0.019 | 1 | 0 | Eif2ak4 | 0.004 | Young |
| 0.552523 | 0.479112 | 0.023 | 0.019 | 1 | 0 | Thap7 | 0.004 | Young |
| 0.55264 | 0.656843 | 0.01 | 0.012 | 1 | 0 | Gm38832 | -0.002 | Young |
| 0.552726 | 0.155821 | 0.094 | 0.103 | 1 | 0 | Pigx | -0.009 | Young |
| 0.552772 | 0.371648 | 0.023 | 0.019 | 1 | 0 | Sacs | 0.004 | Young |
| 0.553049 | 0.20281 | 0.086 | 0.079 | 1 | 0 | Ube2w | 0.007 | Young |
| 0.553349 | 0.183044 | 0.069 | 0.063 | 1 | 0 | Nifk | 0.006 | Young |
| 0.553614 | 0.098784 | 0.128 | 0.138 | 1 | 0 | Phb2 | -0.01 | Young |
| 0.553846 | 0.02317 | 0.056 | 0.063 | 1 | 0 | Mrpl19 | -0.007 | Young |
| 0.554093 | 0.139876 | 0.069 | 0.077 | 1 | 0 | Yeats2 | -0.008 | Young |
| 0.554129 | 0.202073 | 0.148 | 0.141 | 1 | 0 | Baz2a | 0.007 | Young |
| 0.5544 | 0.076168 | 0.071 | 0.078 | 1 | 0 | Reps1 | -0.007 | Young |
| 0.555879 | 0.056525 | 0.05 | 0.056 | 1 | 0 | Dpysl3 | -0.006 | Young |
| 0.556014 | 0.380598 | 0.023 | 0.019 | 1 | 0 | Lnx2 | 0.004 | Young |
| 0.556055 | 0.006068 | 0.011 | 0.014 | 1 | 0 | Dact3 | -0.003 | Young |
| 0.556077 | 0.570757 | 0.023 | 0.019 | 1 | 0 | Pola2 | 0.004 | Young |
| 0.556174 | 0.085893 | 0.017 | 0.014 | 1 | 0 | Bckdhb | 0.003 | Young |
| 0.556284 | 0.255717 | 0.066 | 0.06 | 1 | 0 | Llgl1 | 0.006 | Young |
| 0.556397 | 0.570867 | 0.012 | 0.01 | 1 | 0 | Cbwd1 | 0.002 | Young |
| 0.556666 | 0.191971 | 0.149 | 0.142 | 1 | 0 | Stk10 | 0.007 | Young |
| 0.557121 | 0.135559 | 0.298 | 0.298 | 1 | 0 | Bnip3l | 0 | Young |
| 0.557447 | 0.421028 | 0.061 | 0.056 | 1 | 0 | Slc4a1ap | 0.005 | Young |
| 0.557723 | 0.169607 | 0.304 | 0.302 | 1 | 0 | Ankrd11 | 0.002 | Young |
| 0.557818 | 0.019802 | 0.041 | 0.047 | 1 | 0 | Ints13 | -0.006 | Young |
| 0.558278 | 0.179955 | 0.29 | 0.29 | 1 | 0 | Ndufb9 | 0 | Young |
| 0.558908 | 0.029368 | 0.032 | 0.037 | 1 | 0 | Bin2 | -0.005 | Young |
| 0.558974 | 0.071984 | 0.043 | 0.048 | 1 | 0 | Gapt | -0.005 | Young |
| 0.559603 | 0.16238 | 0.144 | 0.137 | 1 | 0 | Mknk1 | 0.007 | Young |
| 0.559646 | 0.075455 | 0.092 | 0.085 | 1 | 0 | Gpr155 | 0.007 | Young |
| 0.559646 | 0.292952 | 0.012 | 0.01 | 1 | 0 | Gpx7 | 0.002 | Young |
| 0.559973 | 0.181385 | 0.02 | 0.016 | 1 | 0 | Sh3bp4 | 0.004 | Young |
| 0.560015 | 0.499222 | 0.033 | 0.029 | 1 | 0 | Gpank1 | 0.004 | Young |
| 0.560332 | 0.250398 | 0.012 | 0.01 | 1 | 0 | Afg1l | 0.002 | Young |
| 0.560802 | 0.057414 | 0.29 | 0.311 | 1 | 0 | Rassf1 | -0.021 | Young |
| 0.561069 | 0.041867 | 0.34 | 0.363 | 1 | 0 | Snrpb | -0.023 | Young |
| 0.561128 | 0.111221 | 0.213 | 0.205 | 1 | 0 | Gtf2h5 | 0.008 | Young |
| 0.561129 | 0.327948 | 0.026 | 0.022 | 1 | 0 | Ankrd13d | 0.004 | Young |
| 0.561425 | 0.286824 | 0.042 | 0.037 | 1 | 0 | Efl1 | 0.005 | Young |
| 0.561647 | 0.216588 | 0.023 | 0.019 | 1 | 0 | Mcph1 | 0.004 | Young |
| 0.562082 | 0.179563 | 0.269 | 0.265 | 1 | 0 | H2afv | 0.004 | Young |
| 0.562322 | 0.326709 | 0.072 | 0.066 | 1 | 0 | Pik3r4 | 0.006 | Young |
| 0.56238 | 0.022678 | 0.043 | 0.048 | 1 | 0 | Msantd2 | -0.005 | Young |
| 0.562999 | 0.311593 | 0.084 | 0.078 | 1 | 0 | Vrk3 | 0.006 | Young |
| 0.563077 | 0.098883 | 0.012 | 0.01 | 1 | 0 | Harbi1 | 0.002 | Young |

| | | | | | | | | |
|---|---|---|---|---|---|---|---|---|
| 0.563569 | 0.018686 | 0.086 | 0.078 | 1 | 0 | Parp4 | 0.008 | Young |
| 0.564013 | 0.053643 | 0.095 | 0.104 | 1 | 0 | Zufsp | -0.009 | Young |
| 0.564285 | 0.197164 | 0.023 | 0.019 | 1 | 0 | Bzw2 | 0.004 | Young |
| 0.564556 | 0.371971 | 0.054 | 0.049 | 1 | 0 | Ercc3 | 0.005 | Young |
| 0.564853 | 0.461004 | 0.015 | 0.012 | 1 | 0 | Jmjd8 | 0.003 | Young |
| 0.564911 | 0.172193 | 0.073 | 0.081 | 1 | 0 | Guca1a | -0.008 | Young |
| 0.565079 | 0.039923 | 0.045 | 0.051 | 1 | 0 | Asgr2 | -0.006 | Young |
| 0.565108 | 0.025476 | 0.224 | 0.238 | 1 | 0 | Usp8 | -0.014 | Young |
| 0.565318 | 0.222867 | 0.111 | 0.104 | 1 | 0 | Lsm8 | 0.007 | Young |
| 0.565922 | 0.207036 | 0.123 | 0.116 | 1 | 0 | Vps11 | 0.007 | Young |
| 0.566213 | 0.40201 | 0.078 | 0.073 | 1 | 0 | Suclg2 | 0.005 | Young |
| 0.566359 | 0.190058 | 0.073 | 0.067 | 1 | 0 | Commd10 | 0.006 | Young |
| 0.566381 | 0.020905 | 0.267 | 0.283 | 1 | 0 | Sgms1 | -0.016 | Young |
| 0.566541 | 0.304526 | 0.074 | 0.068 | 1 | 0 | Vac14 | 0.006 | Young |
| 0.566899 | 0.002418 | 0.321 | 0.345 | 1 | 0 | Gna12 | -0.024 | Young |
| 0.567185 | 0.30415 | 0.054 | 0.049 | 1 | 0 | Tuft1 | 0.005 | Young |
| 0.567298 | 0.014706 | 0.23 | 0.244 | 1 | 0 | Parp14 | -0.014 | Young |
| 0.567618 | 0.212053 | 0.091 | 0.085 | 1 | 0 | Cd99l2 | 0.006 | Young |
| 0.568042 | 0.105073 | 0.075 | 0.068 | 1 | 0 | Zfp330 | 0.007 | Young |
| 0.568327 | 0.406499 | 0.04 | 0.036 | 1 | 0 | Pus10 | 0.004 | Young |
| 0.568335 | 0.461655 | 0.027 | 0.023 | 1 | 0 | Pwwp2b | 0.004 | Young |
| 0.569352 | 0.104806 | 0.132 | 0.144 | 1 | 0 | Taf6l | -0.012 | Young |
| 0.569686 | 0.195514 | 0.03 | 0.034 | 1 | 0 | Rnf126 | -0.004 | Young |
| 0.569825 | 0.149036 | 0.227 | 0.222 | 1 | 0 | Tspan14 | 0.005 | Young |
| 0.570085 | 0.053477 | 0.163 | 0.175 | 1 | 0 | Noc2l | -0.012 | Young |
| 0.570346 | 0.126736 | 0.049 | 0.055 | 1 | 0 | Pole3 | -0.006 | Young |
| 0.57045 | 0.011036 | 0.121 | 0.114 | 1 | 0 | Ccl3 | 0.007 | Young |
| 0.570454 | 0.032697 | 0.992 | 0.993 | 1 | 0 | Serinc3 | -0.001 | Young |
| 0.570569 | 0.04437 | 0.019 | 0.022 | 1 | 0 | Ddx28 | -0.003 | Young |
| 0.570663 | 0.159075 | 0.036 | 0.031 | 1 | 0 | Ralgapa2 | 0.005 | Young |
| 0.570787 | 0.134802 | 0.157 | 0.149 | 1 | 0 | Rchy1 | 0.008 | Young |
| 0.570853 | 0.145572 | 0.187 | 0.179 | 1 | 0 | Nfatc2 | 0.008 | Young |
| 0.570982 | 0.256558 | 0.222 | 0.222 | 1 | 0 | Lima1 | 0 | Young |
| 0.571561 | 0.225785 | 0.07 | 0.064 | 1 | 0 | Dcxr | 0.006 | Young |
| 0.571765 | 0.251655 | 0.134 | 0.127 | 1 | 0 | Ap1g1 | 0.007 | Young |
| 0.571865 | 0.142935 | 0.137 | 0.13 | 1 | 0 | Gosr2 | 0.007 | Young |
| 0.571895 | 0.133262 | 0.036 | 0.041 | 1 | 0 | Spidr | -0.005 | Young |
| 0.571969 | 0.093415 | 0.089 | 0.097 | 1 | 0 | Atp13a1 | -0.008 | Young |
| 0.57229 | 0.051184 | 0.025 | 0.029 | 1 | 0 | Zfp346 | -0.004 | Young |
| 0.572389 | 0.235387 | 0.069 | 0.077 | 1 | 0 | Mavs | -0.008 | Young |
| 0.572397 | 0.075459 | 0.026 | 0.022 | 1 | 0 | Shmt2 | 0.004 | Young |
| 0.572659 | 0.483084 | 0.061 | 0.056 | 1 | 0 | Ivd | 0.005 | Young |
| 0.572675 | 0.173218 | 0.076 | 0.07 | 1 | 0 | Eng | 0.006 | Young |
| 0.572791 | 0.258304 | 0.104 | 0.097 | 1 | 0 | Washc3 | 0.007 | Young |
| 0.572795 | 0.098067 | 0.093 | 0.101 | 1 | 0 | Braf | -0.008 | Young |

| | | | | | | | | |
|---|---|---|---|---|---|---|---|---|
| 0.572927 | 0.12037 | 0.088 | 0.081 | 1 | 0 | Insig2 | 0.007 | Young |
| 0.572991 | 0.090241 | 0.019 | 0.022 | 1 | 0 | Zfp335 | -0.003 | Young |
| 0.573001 | 0.013634 | 0.107 | 0.115 | 1 | 0 | Ranbp9 | -0.008 | Young |
| 0.573342 | 0.079394 | 0.214 | 0.23 | 1 | 0 | Psmd11 | -0.016 | Young |
| 0.573415 | 0.243416 | 0.118 | 0.112 | 1 | 0 | Dpy19l4 | 0.006 | Young |
| 0.573586 | 0.332307 | 0.04 | 0.036 | 1 | 0 | Tfb2m | 0.004 | Young |
| 0.574301 | 0.008789 | 0.209 | 0.222 | 1 | 0 | Tcp1 | -0.013 | Young |
| 0.574552 | 0.177409 | 0.017 | 0.021 | 1 | 0 | Tmem220 | -0.004 | Young |
| 0.575038 | 0.237276 | 0.066 | 0.06 | 1 | 0 | Fam45a | 0.006 | Young |
| 0.575277 | 0.035391 | 0.021 | 0.025 | 1 | 0 | Plekha8 | -0.004 | Young |
| 0.57535 | 0.497901 | 0.021 | 0.018 | 1 | 0 | Elmod3 | 0.003 | Young |
| 0.575378 | 0.133711 | 0.015 | 0.012 | 1 | 0 | Vsig10 | 0.003 | Young |
| 0.575622 | 0.132025 | 0.034 | 0.038 | 1 | 0 | Slx4ip | -0.004 | Young |
| 0.575628 | 0.130293 | 0.086 | 0.094 | 1 | 0 | Nr2f6 | -0.008 | Young |
| 0.575736 | 0.037041 | 0.02 | 0.023 | 1 | 0 | Ctu1 | -0.003 | Young |
| 0.575836 | 0.123812 | 0.109 | 0.118 | 1 | 0 | Fxr2 | -0.009 | Young |
| 0.576087 | 0.193072 | 0.25 | 0.25 | 1 | 0 | St6gal1 | 0 | Young |
| 0.576188 | 0.875489 | 0.011 | 0.008 | 1 | 0 | 1700030KC | 0.003 | Young |
| 0.576228 | 0.173624 | 0.019 | 0.022 | 1 | 0 | Rpgrip1 | -0.003 | Young |
| 0.576363 | 0.302883 | 0.077 | 0.071 | 1 | 0 | Sacm1l | 0.006 | Young |
| 0.576524 | 0.141473 | 0.131 | 0.124 | 1 | 0 | Tubgcp5 | 0.007 | Young |
| 0.576637 | 0.046379 | 0.054 | 0.06 | 1 | 0 | Trim11 | -0.006 | Young |
| 0.576721 | 0.022073 | 0.249 | 0.267 | 1 | 0 | 8-Sep | -0.018 | Young |
| 0.576731 | 0.064032 | 0.209 | 0.224 | 1 | 0 | Hfe | -0.015 | Young |
| 0.577188 | 0.027618 | 0.061 | 0.067 | 1 | 0 | Rrm2b | -0.006 | Young |
| 0.577419 | 0.193779 | 0.027 | 0.031 | 1 | 0 | Rabepk | -0.004 | Young |
| 0.57746 | 0.307085 | 0.018 | 0.015 | 1 | 0 | Qrsl1 | 0.003 | Young |
| 0.577691 | 0.079048 | 0.054 | 0.06 | 1 | 0 | Tprkb | -0.006 | Young |
| 0.577807 | 0.500731 | 0.086 | 0.081 | 1 | 0 | Ccdc90b | 0.005 | Young |
| 0.578009 | 0.300022 | 0.129 | 0.123 | 1 | 0 | Clcn3 | 0.006 | Young |
| 0.578024 | 0.01893 | 0.015 | 0.012 | 1 | 0 | Dgke | 0.003 | Young |
| 0.578056 | 0.192096 | 0.208 | 0.202 | 1 | 0 | Mfhas1 | 0.006 | Young |
| 0.578655 | 0.148136 | 0.469 | 0.503 | 1 | 0 | Rpl4 | -0.034 | Young |
| 0.579008 | 0.291987 | 0.05 | 0.045 | 1 | 0 | Dvl3 | 0.005 | Young |
| 0.579371 | 0.270466 | 0.011 | 0.008 | 1 | 0 | Fam109a | 0.003 | Young |
| 0.579808 | 0.459752 | 0.015 | 0.018 | 1 | 0 | Fads3 | -0.003 | Young |
| 0.579839 | 0.3289 | 0.021 | 0.018 | 1 | 0 | Atp6v1g2 | 0.003 | Young |
| 0.579873 | 0.254736 | 0.201 | 0.198 | 1 | 0 | Akt2 | 0.003 | Young |
| 0.580268 | 0.111066 | 0.136 | 0.129 | 1 | 0 | Ufc1 | 0.007 | Young |
| 0.580286 | 0.245032 | 0.019 | 0.022 | 1 | 0 | C030034I2 | -0.003 | Young |
| 0.580341 | 0.017833 | 0.111 | 0.119 | 1 | 0 | Klhdc3 | -0.008 | Young |
| 0.580409 | 0.22226 | 0.043 | 0.038 | 1 | 0 | Adam8 | 0.005 | Young |
| 0.580648 | 0.124934 | 0.026 | 0.03 | 1 | 0 | Zfp874b | -0.004 | Young |
| 0.580684 | 0.315405 | 0.011 | 0.008 | 1 | 0 | Slc7a6 | 0.003 | Young |
| 0.580871 | 0.296193 | 0.078 | 0.073 | 1 | 0 | Hspa13 | 0.005 | Young |

| | | | | | | | | |
|---|---|---|---|---|---|---|---|---|
| 0.580936 | 0.231418 | 0.072 | 0.079 | 1 | 0 | Emc8 | -0.007 | Young |
| 0.581565 | 0.251945 | 0.025 | 0.029 | 1 | 0 | Pex11b | -0.004 | Young |
| 0.581615 | 0.128535 | 0.23 | 0.224 | 1 | 0 | Trpc4ap | 0.006 | Young |
| 0.581674 | 0.290695 | 0.064 | 0.059 | 1 | 0 | Tmem214 | 0.005 | Young |
| 0.581735 | 0.311886 | 0.027 | 0.023 | 1 | 0 | Chd1l | 0.004 | Young |
| 0.582017 | 0.006184 | 0.096 | 0.104 | 1 | 0 | Polr2i | -0.008 | Young |
| 0.582592 | 0.439518 | 0.019 | 0.022 | 1 | 0 | 6330418K0 | -0.003 | Young |
| 0.582677 | 0.16716 | 0.233 | 0.227 | 1 | 0 | Pdcd5 | 0.006 | Young |
| 0.582882 | 0.064934 | 0.018 | 0.015 | 1 | 0 | Tmem42 | 0.003 | Young |
| 0.582937 | 0.364357 | 0.093 | 0.088 | 1 | 0 | 1110038F1 | 0.005 | Young |
| 0.582981 | 0.201859 | 0.231 | 0.226 | 1 | 0 | Rbbp6 | 0.005 | Young |
| 0.583127 | 0.116238 | 0.011 | 0.008 | 1 | 0 | Timeless | 0.003 | Young |
| 0.583168 | 0.003094 | 0.018 | 0.015 | 1 | 0 | Atp2a3 | 0.003 | Young |
| 0.58331 | 0.0374 | 0.204 | 0.218 | 1 | 0 | Sar1a | -0.014 | Young |
| 0.583327 | 0.501661 | 0.038 | 0.034 | 1 | 0 | Tnfsf13b | 0.004 | Young |
| 0.58353 | 0.272181 | 0.065 | 0.06 | 1 | 0 | Rcbtb1 | 0.005 | Young |
| 0.583907 | 0.24996 | 0.075 | 0.07 | 1 | 0 | Chst12 | 0.005 | Young |
| 0.583993 | 0.203648 | 0.063 | 0.07 | 1 | 0 | Golph3l | -0.007 | Young |
| 0.584063 | 0.013377 | 0.497 | 0.523 | 1 | 0 | Itgb1 | -0.026 | Young |
| 0.584071 | 0.069913 | 0.143 | 0.153 | 1 | 0 | Abhd17b | -0.01 | Young |
| 0.584211 | 0.06234 | 0.325 | 0.35 | 1 | 0 | Ctnnd1 | -0.025 | Young |
| 0.584579 | 0.354104 | 0.075 | 0.07 | 1 | 0 | Pofut2 | 0.005 | Young |
| 0.584617 | 0.158622 | 0.027 | 0.031 | 1 | 0 | Nprl2 | -0.004 | Young |
| 0.584676 | 0.07428 | 0.332 | 0.326 | 1 | 0 | Sumo3 | 0.006 | Young |
| 0.58492 | 0.064791 | 0.206 | 0.22 | 1 | 0 | Pum1 | -0.014 | Young |
| 0.584921 | 0.112133 | 0.102 | 0.094 | 1 | 0 | Sirt7 | 0.008 | Young |
| 0.584985 | 0.198367 | 0.088 | 0.082 | 1 | 0 | Ipo8 | 0.006 | Young |
| 0.585009 | 0.148794 | 0.011 | 0.008 | 1 | 0 | Dffb | 0.003 | Young |
| 0.585045 | 0.165056 | 0.04 | 0.036 | 1 | 0 | Gtpbp6 | 0.004 | Young |
| 0.585321 | 0.168179 | 0.078 | 0.073 | 1 | 0 | Mosmo | 0.005 | Young |
| 0.585406 | 0.163513 | 0.021 | 0.018 | 1 | 0 | Ccdc191 | 0.003 | Young |
| 0.585469 | 0.065295 | 0.13 | 0.141 | 1 | 0 | Ubp1 | -0.011 | Young |
| 0.585569 | 0.011384 | 0.059 | 0.064 | 1 | 0 | Urm1 | -0.005 | Young |
| 0.586179 | 0.037121 | 0.045 | 0.051 | 1 | 0 | Zdhhc6 | -0.006 | Young |
| 0.587045 | 0.035643 | 0.257 | 0.274 | 1 | 0 | Cmtm7 | -0.017 | Young |
| 0.58717 | 0.302186 | 0.034 | 0.03 | 1 | 0 | Limk1 | 0.004 | Young |
| 0.587277 | 0.411033 | 0.038 | 0.034 | 1 | 0 | Zfp672 | 0.004 | Young |
| 0.587746 | 0.188007 | 0.14 | 0.152 | 1 | 0 | Ap3b1 | -0.012 | Young |
| 0.587867 | 0.158006 | 0.093 | 0.101 | 1 | 0 | Nrf1 | -0.008 | Young |
| 0.588158 | 0.050011 | 0.19 | 0.182 | 1 | 0 | Gm6377 | 0.008 | Young |
| 0.588595 | 0.041657 | 0.147 | 0.157 | 1 | 0 | BC005624 | -0.01 | Young |
| 0.588622 | 0.056181 | 0.069 | 0.075 | 1 | 0 | BC029722 | -0.006 | Young |
| 0.588683 | 0.001552 | 0.157 | 0.168 | 1 | 0 | Tmcc3 | -0.011 | Young |
| 0.589401 | 0.051443 | 0.269 | 0.287 | 1 | 0 | Abcc1 | -0.018 | Young |
| 0.58946 | 0.049196 | 0.524 | 0.539 | 1 | 0 | Zfhx3 | -0.015 | Young |

| | | | | | | | | |
|---|---|---|---|---|---|---|---|---|
| 0.589732 | 0.241111 | 0.11 | 0.104 | 1 | 0 | Pde7a | 0.006 | Young |
| 0.59023 | 0.406794 | 0.028 | 0.025 | 1 | 0 | Lin52 | 0.003 | Young |
| 0.590765 | 0.128787 | 0.04 | 0.036 | 1 | 0 | Smim8 | 0.004 | Young |
| 0.590887 | 0.447729 | 0.025 | 0.022 | 1 | 0 | Ccdc66 | 0.003 | Young |
| 0.590978 | 0.181339 | 0.068 | 0.063 | 1 | 0 | Phrf1 | 0.005 | Young |
| 0.591169 | 0.186287 | 0.084 | 0.078 | 1 | 0 | Tmem63a | 0.006 | Young |
| 0.591397 | 0.160918 | 0.099 | 0.093 | 1 | 0 | Smim20 | 0.006 | Young |
| 0.591417 | 0.038369 | 0.183 | 0.196 | 1 | 0 | Sec61a1 | -0.013 | Young |
| 0.592098 | 0.134221 | 0.045 | 0.051 | 1 | 0 | Ift46 | -0.006 | Young |
| 0.592315 | 0.214166 | 0.125 | 0.119 | 1 | 0 | Hcfc1 | 0.006 | Young |
| 0.592753 | 0.676448 | 0.016 | 0.014 | 1 | 0 | B4galt4 | 0.002 | Young |
| 0.593348 | 0.291964 | 0.13 | 0.124 | 1 | 0 | Cracr2b | 0.006 | Young |
| 0.593693 | 0.306653 | 0.064 | 0.059 | 1 | 0 | Txndc12 | 0.005 | Young |
| 0.594038 | 0.504326 | 0.028 | 0.025 | 1 | 0 | Thbs1 | 0.003 | Young |
| 0.594096 | 0.044625 | 0.083 | 0.09 | 1 | 0 | Trappc9 | -0.007 | Young |
| 0.594479 | 0.192095 | 0.111 | 0.105 | 1 | 0 | Ctbp2 | 0.006 | Young |
| 0.595458 | 0.216513 | 0.061 | 0.067 | 1 | 0 | Pmf1 | -0.006 | Young |
| 0.595624 | 0.104032 | 0.116 | 0.109 | 1 | 0 | Carhsp1 | 0.007 | Young |
| 0.595888 | 0.332261 | 0.025 | 0.022 | 1 | 0 | Slc16a1 | 0.003 | Young |
| 0.596054 | 0.413001 | 0.047 | 0.042 | 1 | 0 | Cops7b | 0.005 | Young |
| 0.59608 | 0.449037 | 0.019 | 0.016 | 1 | 0 | Tsen2 | 0.003 | Young |
| 0.596122 | 0.135315 | 0.061 | 0.067 | 1 | 0 | Qsox1 | -0.006 | Young |
| 0.596158 | 0.346112 | 0.036 | 0.041 | 1 | 0 | Ddx49 | -0.005 | Young |
| 0.596661 | 0.074785 | 0.068 | 0.074 | 1 | 0 | Cdk10 | -0.006 | Young |
| 0.596689 | 0.338862 | 0.016 | 0.014 | 1 | 0 | Cav1 | 0.002 | Young |
| 0.596937 | 0.034067 | 0.497 | 0.509 | 1 | 0 | Ubb | -0.012 | Young |
| 0.597102 | 0.113084 | 0.151 | 0.144 | 1 | 0 | Rsf1 | 0.007 | Young |
| 0.598165 | 0.087595 | 0.2 | 0.215 | 1 | 0 | Tnfrsf11a | -0.015 | Young |
| 0.599186 | 0.063023 | 0.065 | 0.071 | 1 | 0 | Pgpep1 | -0.006 | Young |
| 0.599515 | 0.233721 | 0.082 | 0.077 | 1 | 0 | Tmem260 | 0.005 | Young |
| 0.599725 | 0.388352 | 0.016 | 0.014 | 1 | 0 | Foxp4 | 0.002 | Young |
| 0.600921 | 0.123647 | 0.129 | 0.123 | 1 | 0 | Foxo3 | 0.006 | Young |
| 0.600966 | 0.234088 | 0.059 | 0.055 | 1 | 0 | Irak2 | 0.004 | Young |
| 0.601091 | 0.292452 | 0.025 | 0.022 | 1 | 0 | Gss | 0.003 | Young |
| 0.601216 | 0.181497 | 0.085 | 0.093 | 1 | 0 | Nudt9 | -0.008 | Young |
| 0.601251 | 0.341655 | 0.019 | 0.016 | 1 | 0 | Negr1 | 0.003 | Young |
| 0.601342 | 0.23023 | 0.022 | 0.019 | 1 | 0 | Klhl13 | 0.003 | Young |
| 0.601386 | 0.162522 | 0.103 | 0.097 | 1 | 0 | Ufl1 | 0.006 | Young |
| 0.601603 | 0.156588 | 0.063 | 0.057 | 1 | 0 | Cyp4v3 | 0.006 | Young |
| 0.601617 | 0.17261 | 0.066 | 0.073 | 1 | 0 | Sphk2 | -0.007 | Young |
| 0.601742 | 0.085074 | 0.047 | 0.052 | 1 | 0 | Prdm4 | -0.005 | Young |
| 0.602125 | 0.327797 | 0.022 | 0.019 | 1 | 0 | Acd | 0.003 | Young |
| 0.602158 | 0.208876 | 0.016 | 0.014 | 1 | 0 | Slc25a33 | 0.002 | Young |
| 0.602698 | 0.123222 | 0.084 | 0.078 | 1 | 0 | Slc25a1 | 0.006 | Young |
| 0.603164 | 0.155632 | 0.252 | 0.249 | 1 | 0 | Rassf3 | 0.003 | Young |

| | | | | | | | | |
|---|---|---|---|---|---|---|---|---|
| 0.603213 | 0.277235 | 0.019 | 0.016 | 1 | 0 | Cep57l1 | 0.003 | Young |
| 0.603282 | 0.025778 | 0.181 | 0.171 | 1 | 0 | Smc1a | 0.01 | Young |
| 0.603455 | 0.043546 | 0.156 | 0.167 | 1 | 0 | Kdm1a | -0.011 | Young |
| 0.60354 | 0.020628 | 0.037 | 0.033 | 1 | 0 | Trim23 | 0.004 | Young |
| 0.603727 | 0.112365 | 0.091 | 0.085 | 1 | 0 | Abca7 | 0.006 | Young |
| 0.604005 | 0.363008 | 0.008 | 0.01 | 1 | 0 | Sars2 | -0.002 | Young |
| 0.604225 | 0.298965 | 0.08 | 0.075 | 1 | 0 | Arpc5l | 0.005 | Young |
| 0.604346 | 0.21947 | 0.025 | 0.022 | 1 | 0 | St7 | 0.003 | Young |
| 0.604734 | 0.251563 | 0.099 | 0.093 | 1 | 0 | Shprh | 0.006 | Young |
| 0.605563 | 0.078002 | 0.251 | 0.267 | 1 | 0 | Eloc | -0.016 | Young |
| 0.605842 | 0.05148 | 0.327 | 0.354 | 1 | 0 | Atp5j2 | -0.027 | Young |
| 0.606056 | 0.027632 | 0.055 | 0.06 | 1 | 0 | D6Wsu163 | -0.005 | Young |
| 0.606149 | 0.013976 | 0.081 | 0.088 | 1 | 0 | Rnf14 | -0.007 | Young |
| 0.606162 | 0.544024 | 0.045 | 0.041 | 1 | 0 | Ints14 | 0.004 | Young |
| 0.606165 | 0.218214 | 0.048 | 0.053 | 1 | 0 | Fhod1 | -0.005 | Young |
| 0.606211 | 0.021172 | 0.03 | 0.034 | 1 | 0 | Ncdn | -0.004 | Young |
| 0.606707 | 0.19142 | 0.043 | 0.038 | 1 | 0 | Fbxo42 | 0.005 | Young |
| 0.606961 | 0.078582 | 0.087 | 0.081 | 1 | 0 | Hccs | 0.006 | Young |
| 0.607152 | 0.266781 | 0.1 | 0.094 | 1 | 0 | Pom121 | 0.006 | Young |
| 0.607197 | 0.065839 | 0.127 | 0.135 | 1 | 0 | Gpr107 | -0.008 | Young |
| 0.607337 | 0.330131 | 0.04 | 0.036 | 1 | 0 | St3gal2 | 0.004 | Young |
| 0.607399 | 0.049078 | 0.353 | 0.373 | 1 | 0 | Rab11fip5 | -0.02 | Young |
| 0.607532 | 0.243059 | 0.188 | 0.183 | 1 | 0 | Hnrnpr | 0.005 | Young |
| 0.607828 | 0.122318 | 0.261 | 0.259 | 1 | 0 | Blnk | 0.002 | Young |
| 0.607879 | 0.958321 | 0.012 | 0.01 | 1 | 0 | Adipoq | 0.002 | Young |
| 0.608171 | 0.048838 | 0.064 | 0.07 | 1 | 0 | Gtf2b | -0.006 | Young |
| 0.608234 | 0.085695 | 0.077 | 0.083 | 1 | 0 | Mettl23 | -0.006 | Young |
| 0.608428 | 0.146614 | 0.045 | 0.041 | 1 | 0 | Hps4 | 0.004 | Young |
| 0.608602 | 0.260482 | 0.147 | 0.142 | 1 | 0 | Cnp | 0.005 | Young |
| 0.608697 | 0.052755 | 0.241 | 0.254 | 1 | 0 | Irf5 | -0.013 | Young |
| 0.608735 | 0.02764 | 0.131 | 0.141 | 1 | 0 | Scamp1 | -0.01 | Young |
| 0.608814 | 0.01492 | 0.048 | 0.053 | 1 | 0 | H6pd | -0.005 | Young |
| 0.60903 | 0.218213 | 0.009 | 0.011 | 1 | 0 | Ipcef1 | -0.002 | Young |
| 0.609619 | 0.012138 | 0.309 | 0.326 | 1 | 0 | Gatad2b | -0.017 | Young |
| 0.609741 | 0.092289 | 0.123 | 0.131 | 1 | 0 | Ankle2 | -0.008 | Young |
| 0.609776 | 0.078304 | 0.059 | 0.064 | 1 | 0 | Nup133 | -0.005 | Young |
| 0.609858 | 0.279454 | 0.04 | 0.045 | 1 | 0 | Aamdc | -0.005 | Young |
| 0.609881 | 0.107076 | 0.083 | 0.077 | 1 | 0 | Pggt1b | 0.006 | Young |
| 0.610142 | 0.142211 | 0.143 | 0.137 | 1 | 0 | Glrx3 | 0.006 | Young |
| 0.610258 | 0.237945 | 0.24 | 0.239 | 1 | 0 | Inpp4a | 0.001 | Young |
| 0.610391 | 0.072647 | 0.073 | 0.079 | 1 | 0 | Ndst2 | -0.006 | Young |
| 0.610419 | 0.202007 | 0.047 | 0.052 | 1 | 0 | Tlr6 | -0.005 | Young |
| 0.611113 | 0.041319 | 0.028 | 0.031 | 1 | 0 | Ndufaf7 | -0.003 | Young |
| 0.611149 | 0.239036 | 0.043 | 0.038 | 1 | 0 | C030034L1 | 0.005 | Young |
| 0.611227 | 0.817953 | 0.021 | 0.018 | 1 | 0 | Orc6 | 0.003 | Young |

| | | | | | | | | |
|---|---|---|---|---|---|---|---|---|
| 0.611368 | 0.274489 | 0.117 | 0.112 | 1 | 0 | Ldah | 0.005 | Young |
| 0.611577 | 0.019488 | 0.109 | 0.118 | 1 | 0 | Plagl2 | -0.009 | Young |
| 0.612102 | 0.005472 | 0.045 | 0.049 | 1 | 0 | Nol8 | -0.004 | Young |
| 0.612303 | 0.008853 | 0.222 | 0.235 | 1 | 0 | Kat6a | -0.013 | Young |
| 0.61272 | 0.208381 | 0.08 | 0.075 | 1 | 0 | Pla2g16 | 0.005 | Young |
| 0.612791 | 0.070359 | 0.107 | 0.115 | 1 | 0 | Tcof1 | -0.008 | Young |
| 0.613 | 0.031174 | 0.009 | 0.011 | 1 | 0 | Slc16a9 | -0.002 | Young |
| 0.613258 | 0.283844 | 0.044 | 0.04 | 1 | 0 | Fam110a | 0.004 | Young |
| 0.613316 | 0.06572 | 0.116 | 0.124 | 1 | 0 | Slc37a2 | -0.008 | Young |
| 0.613792 | 0.194185 | 0.039 | 0.044 | 1 | 0 | AC165953. | -0.005 | Young |
| 0.613986 | 0.119402 | 0.099 | 0.107 | 1 | 0 | Rnf166 | -0.008 | Young |
| 0.613988 | 0.07738 | 0.202 | 0.216 | 1 | 0 | Arrdc1 | -0.014 | Young |
| 0.614208 | 0.168338 | 0.175 | 0.17 | 1 | 0 | Slc23a2 | 0.005 | Young |
| 0.614425 | 0.162852 | 0.039 | 0.044 | 1 | 0 | Slc7a6os | -0.005 | Young |
| 0.614474 | 0.808089 | 0.015 | 0.012 | 1 | 0 | Usp18 | 0.003 | Young |
| 0.614592 | 0.180323 | 0.009 | 0.011 | 1 | 0 | Tnfaip8l3 | -0.002 | Young |
| 0.614783 | 0.013254 | 0.276 | 0.293 | 1 | 0 | Ap1s2 | -0.017 | Young |
| 0.615118 | 0.58922 | 0.012 | 0.01 | 1 | 0 | Cenpj | 0.002 | Young |
| 0.615168 | 0.012947 | 0.103 | 0.111 | 1 | 0 | Ranbp10 | -0.008 | Young |
| 0.615701 | 0.013339 | 0.01 | 0.012 | 1 | 0 | Ifrd2 | -0.002 | Young |
| 0.616219 | 0.138158 | 0.131 | 0.124 | 1 | 0 | Zbtb38 | 0.007 | Young |
| 0.616515 | 0.089139 | 0.418 | 0.425 | 1 | 0 | Syk | -0.007 | Young |
| 0.616822 | 0.126551 | 0.01 | 0.012 | 1 | 0 | Zc4h2 | -0.002 | Young |
| 0.616903 | 0.167421 | 0.131 | 0.124 | 1 | 0 | Slc35f6 | 0.007 | Young |
| 0.617083 | 0.821392 | 0.018 | 0.015 | 1 | 0 | Ptcd1 | 0.003 | Young |
| 0.617201 | 0.268032 | 0.053 | 0.049 | 1 | 0 | Syap1 | 0.004 | Young |
| 0.61723 | 0.213251 | 0.044 | 0.04 | 1 | 0 | Kif16b | 0.004 | Young |
| 0.617248 | 0.550018 | 0.021 | 0.018 | 1 | 0 | Fut4 | 0.003 | Young |
| 0.617301 | 0.019405 | 0.023 | 0.026 | 1 | 0 | Bud13 | -0.003 | Young |
| 0.617311 | 0.236321 | 0.056 | 0.052 | 1 | 0 | Zfp780b | 0.004 | Young |
| 0.617384 | 0.007424 | 0.224 | 0.235 | 1 | 0 | Nsd3 | -0.011 | Young |
| 0.617659 | 0.082591 | 0.112 | 0.12 | 1 | 0 | BC004004 | -0.008 | Young |
| 0.618208 | 0.110419 | 0.029 | 0.033 | 1 | 0 | Rpp30 | -0.004 | Young |
| 0.618249 | 0.260117 | 0.079 | 0.074 | 1 | 0 | Tusc3 | 0.005 | Young |
| 0.618329 | 0.216319 | 0.128 | 0.122 | 1 | 0 | Fh1 | 0.006 | Young |
| 0.61839 | 0.143097 | 0.092 | 0.086 | 1 | 0 | Dmtf1 | 0.006 | Young |
| 0.618398 | 0.05331 | 0.057 | 0.052 | 1 | 0 | Coq8b | 0.005 | Young |
| 0.618445 | 0.132614 | 0.143 | 0.137 | 1 | 0 | Ncoa6 | 0.006 | Young |
| 0.618529 | 0.476534 | 0.015 | 0.012 | 1 | 0 | Klhl8 | 0.003 | Young |
| 0.618642 | 0.120726 | 0.242 | 0.237 | 1 | 0 | Vat1 | 0.005 | Young |
| 0.618667 | 0.565614 | 0.043 | 0.04 | 1 | 0 | Mmgt1 | 0.003 | Young |
| 0.618868 | 0.038818 | 0.02 | 0.023 | 1 | 0 | Atp1a3 | -0.003 | Young |
| 0.61908 | 0.088765 | 0.053 | 0.048 | 1 | 0 | Zfp407 | 0.005 | Young |
| 0.619297 | 0.258761 | 0.012 | 0.01 | 1 | 0 | Adamtsl4 | 0.002 | Young |
| 0.619362 | 0.373388 | 0.024 | 0.021 | 1 | 0 | Grpel2 | 0.003 | Young |

| | | | | | | | | |
|---|---|---|---|---|---|---|---|---|
| 0.619631 | 0.388896 | 0.01 | 0.012 | 1 | 0 | Gm26890 | -0.002 | Young |
| 0.619679 | 0.236531 | 0.087 | 0.082 | 1 | 0 | C2cd5 | 0.005 | Young |
| 0.619896 | 0.107653 | 0.069 | 0.075 | 1 | 0 | Pgap1 | -0.006 | Young |
| 0.619941 | 0.001647 | 0.02 | 0.023 | 1 | 0 | Zfp809 | -0.003 | Young |
| 0.620025 | 0.235693 | 0.012 | 0.01 | 1 | 0 | Senp8 | 0.002 | Young |
| 0.620119 | 0.10573 | 0.098 | 0.105 | 1 | 0 | Eif2ak2 | -0.007 | Young |
| 0.620229 | 0.121127 | 0.048 | 0.053 | 1 | 0 | Tmco3 | -0.005 | Young |
| 0.620268 | 0.531008 | 0.021 | 0.018 | 1 | 0 | Cgnl1 | 0.003 | Young |
| 0.62027 | 0.145727 | 0.055 | 0.06 | 1 | 0 | Gosr1 | -0.005 | Young |
| 0.620378 | 0.047569 | 0.144 | 0.153 | 1 | 0 | Xrn1 | -0.009 | Young |
| 0.620755 | 0.31307 | 0.01 | 0.012 | 1 | 0 | Gm26522 | -0.002 | Young |
| 0.620962 | 0.113312 | 0.065 | 0.071 | 1 | 0 | Tubgcp3 | -0.006 | Young |
| 0.621044 | 0.074949 | 0.073 | 0.079 | 1 | 0 | Qars | -0.006 | Young |
| 0.621193 | 0.347576 | 0.088 | 0.083 | 1 | 0 | Celf4 | 0.005 | Young |
| 0.621446 | 0.057482 | 0.018 | 0.021 | 1 | 0 | Panx1 | -0.003 | Young |
| 0.621505 | 0.610902 | 0.021 | 0.018 | 1 | 0 | Zbtb17 | 0.003 | Young |
| 0.621554 | 0.037174 | 0.047 | 0.042 | 1 | 0 | 5730455P1 | 0.005 | Young |
| 0.621863 | 0.459522 | 0.015 | 0.012 | 1 | 0 | Nsun5 | 0.003 | Young |
| 0.621903 | 0.01184 | 0.016 | 0.019 | 1 | 0 | Casz1 | -0.003 | Young |
| 0.622086 | 0.21086 | 0.08 | 0.075 | 1 | 0 | Akap8l | 0.005 | Young |
| 0.622394 | 0.068953 | 0.012 | 0.01 | 1 | 0 | Apoo | 0.002 | Young |
| 0.622751 | 0.020054 | 0.013 | 0.015 | 1 | 0 | Ippk | -0.002 | Young |
| 0.622782 | 0.055609 | 0.051 | 0.056 | 1 | 0 | Fyco1 | -0.005 | Young |
| 0.62329 | 0.20739 | 0.069 | 0.075 | 1 | 0 | Dzip3 | -0.006 | Young |
| 0.623306 | 0.088966 | 0.043 | 0.038 | 1 | 0 | Zfp593 | 0.005 | Young |
| 0.62339 | 0.192492 | 0.153 | 0.149 | 1 | 0 | Hcst | 0.004 | Young |
| 0.623473 | 0.481224 | 0.046 | 0.042 | 1 | 0 | Exosc4 | 0.004 | Young |
| 0.623932 | 0.225326 | 0.127 | 0.137 | 1 | 0 | Cnot3 | -0.01 | Young |
| 0.623977 | 0.248657 | 0.181 | 0.178 | 1 | 0 | Clk4 | 0.003 | Young |
| 0.623982 | 0.30829 | 0.044 | 0.049 | 1 | 0 | Akt3 | -0.005 | Young |
| 0.624096 | 0.023916 | 0.046 | 0.051 | 1 | 0 | Slc35a1 | -0.005 | Young |
| 0.624187 | 0.145285 | 0.114 | 0.108 | 1 | 0 | Cops2 | 0.006 | Young |
| 0.624462 | 0.130845 | 0.052 | 0.057 | 1 | 0 | Neurl4 | -0.005 | Young |
| 0.624589 | 0.248255 | 0.044 | 0.04 | 1 | 0 | Pus1 | 0.004 | Young |
| 0.624611 | 0.100218 | 0.019 | 0.022 | 1 | 0 | Gm26518 | -0.003 | Young |
| 0.624738 | 0.017075 | 0.014 | 0.016 | 1 | 0 | Atp23 | -0.002 | Young |
| 0.62476 | 0.198649 | 0.096 | 0.104 | 1 | 0 | Tpm1 | -0.008 | Young |
| 0.6248 | 0.057438 | 0.57 | 0.59 | 1 | 0 | Hnrnpa3 | -0.02 | Young |
| 0.625152 | 0.215928 | 0.217 | 0.213 | 1 | 0 | Hspa4 | 0.004 | Young |
| 0.625332 | 0.2545 | 0.065 | 0.06 | 1 | 0 | Sap30 | 0.005 | Young |
| 0.625357 | 0.203759 | 0.203 | 0.201 | 1 | 0 | Ophn1 | 0.002 | Young |
| 0.625527 | 0.231811 | 0.011 | 0.014 | 1 | 0 | 2010007H( | -0.003 | Young |
| 0.625665 | 0.109489 | 0.17 | 0.182 | 1 | 0 | Tap2 | -0.012 | Young |
| 0.62572 | 0.029789 | 0.31 | 0.327 | 1 | 0 | Brd2 | -0.017 | Young |
| 0.625725 | 0.078103 | 0.02 | 0.023 | 1 | 0 | Inafm2 | -0.003 | Young |

| | | | | | | | | |
|---|---|---|---|---|---|---|---|---|
| 0.625796 | 0.152628 | 0.093 | 0.088 | 1 | 0 | Klhl18 | 0.005 | Young |
| 0.625837 | 0.199118 | 0.194 | 0.19 | 1 | 0 | Smc6 | 0.004 | Young |
| 0.626029 | 0.306882 | 0.126 | 0.122 | 1 | 0 | Sun2 | 0.004 | Young |
| 0.626042 | 0.113916 | 0.113 | 0.122 | 1 | 0 | Bbip1 | -0.009 | Young |
| 0.626323 | 0.309836 | 0.072 | 0.067 | 1 | 0 | Naip5 | 0.005 | Young |
| 0.626432 | 0.219097 | 0.027 | 0.03 | 1 | 0 | Dcakd | -0.003 | Young |
| 0.626479 | 0.060004 | 0.279 | 0.274 | 1 | 0 | Hnrnpul2 | 0.005 | Young |
| 0.626603 | 0.122142 | 0.021 | 0.018 | 1 | 0 | Hddc2 | 0.003 | Young |
| 0.626822 | 0.088164 | 0.072 | 0.078 | 1 | 0 | Phax | -0.006 | Young |
| 0.626837 | 0.13585 | 0.014 | 0.016 | 1 | 0 | Gsdme | -0.002 | Young |
| 0.627046 | 0.029244 | 0.018 | 0.021 | 1 | 0 | Eid2b | -0.003 | Young |
| 0.62723 | 0.079832 | 0.021 | 0.025 | 1 | 0 | Tti2 | -0.004 | Young |
| 0.627434 | 0.101348 | 0.067 | 0.073 | 1 | 0 | G2e3 | -0.006 | Young |
| 0.627513 | 0.091574 | 0.052 | 0.048 | 1 | 0 | Tfec | 0.004 | Young |
| 0.627611 | 0.161193 | 0.024 | 0.027 | 1 | 0 | Prrg1 | -0.003 | Young |
| 0.62769 | 0.094857 | 0.047 | 0.042 | 1 | 0 | Lsr | 0.005 | Young |
| 0.627989 | 0.019395 | 0.066 | 0.071 | 1 | 0 | Lipe | -0.005 | Young |
| 0.628136 | 0.042398 | 0.227 | 0.238 | 1 | 0 | Smim14 | -0.011 | Young |
| 0.628399 | 0.230192 | 0.021 | 0.018 | 1 | 0 | Fmnl2 | 0.003 | Young |
| 0.628472 | 0.048593 | 0.444 | 0.48 | 1 | 0 | Vcp | -0.036 | Young |
| 0.628567 | 0.001928 | 0.087 | 0.093 | 1 | 0 | Sestd1 | -0.006 | Young |
| 0.628997 | 0.238116 | 0.204 | 0.202 | 1 | 0 | Rab35 | 0.002 | Young |
| 0.629097 | 0.066864 | 0.271 | 0.29 | 1 | 0 | Slc29a1 | -0.019 | Young |
| 0.629262 | 0.148465 | 0.044 | 0.04 | 1 | 0 | Timm17b | 0.004 | Young |
| 0.629291 | 0.123607 | 0.015 | 0.012 | 1 | 0 | Tmem221 | 0.003 | Young |
| 0.629433 | 0.03954 | 0.468 | 0.464 | 1 | 0 | Sgpl1 | 0.004 | Young |
| 0.629595 | 0.272584 | 0.155 | 0.152 | 1 | 0 | Scamp5 | 0.003 | Young |
| 0.629777 | 0.207567 | 0.185 | 0.182 | 1 | 0 | 1110004F1 | 0.003 | Young |
| 0.629782 | 0.102573 | 0.015 | 0.012 | 1 | 0 | 6330403L0 | 0.003 | Young |
| 0.630398 | 0.13098 | 0.016 | 0.019 | 1 | 0 | Gstz1 | -0.003 | Young |
| 0.630404 | 0.260909 | 0.021 | 0.018 | 1 | 0 | Arfip2 | 0.003 | Young |
| 0.630472 | 0.173799 | 0.069 | 0.075 | 1 | 0 | Mrpl2 | -0.006 | Young |
| 0.630771 | 0.275755 | 0.018 | 0.015 | 1 | 0 | Abhd14a | 0.003 | Young |
| 0.630894 | 0.231516 | 0.041 | 0.037 | 1 | 0 | Scaper | 0.004 | Young |
| 0.631116 | 0.350841 | 0.038 | 0.034 | 1 | 0 | Fam210a | 0.004 | Young |
| 0.631189 | 0.122297 | 0.096 | 0.09 | 1 | 0 | Tlk2 | 0.006 | Young |
| 0.631784 | 0.261169 | 0.031 | 0.027 | 1 | 0 | Ube2o | 0.004 | Young |
| 0.63183 | 0.0079 | 0.11 | 0.116 | 1 | 0 | Pgap2 | -0.006 | Young |
| 0.632297 | 0.138807 | 0.125 | 0.134 | 1 | 0 | 2810403A0 | -0.009 | Young |
| 0.6325 | 0.253492 | 0.053 | 0.049 | 1 | 0 | Atrn | 0.004 | Young |
| 0.632552 | 0.214242 | 0.019 | 0.022 | 1 | 0 | Glis2 | -0.003 | Young |
| 0.632622 | 0.009853 | 0.091 | 0.097 | 1 | 0 | Gabpa | -0.006 | Young |
| 0.632727 | 0.113471 | 0.189 | 0.182 | 1 | 0 | Gstm1 | 0.007 | Young |
| 0.632751 | 0.255701 | 0.031 | 0.027 | 1 | 0 | Hacd2 | 0.004 | Young |
| 0.633009 | 0.033649 | 0.062 | 0.057 | 1 | 0 | Glce | 0.005 | Young |

| | | | | | | | | |
|---|---|---|---|---|---|---|---|---|
| 0.633247 | 0.039105 | 0.388 | 0.409 | 1 | 0 | Nav1 | -0.021 | Young |
| 0.633249 | 0.294991 | 0.142 | 0.138 | 1 | 0 | Lars2 | 0.004 | Young |
| 0.633468 | 0.191201 | 0.051 | 0.056 | 1 | 0 | Rmi1 | -0.005 | Young |
| 0.633902 | 0.092607 | 0.288 | 0.309 | 1 | 0 | Mgat1 | -0.021 | Young |
| 0.634197 | 0.141854 | 0.041 | 0.045 | 1 | 0 | Rps19bp1 | -0.004 | Young |
| 0.634404 | 0.095172 | 0.064 | 0.059 | 1 | 0 | Ttc32 | 0.005 | Young |
| 0.63448 | 0.285461 | 0.076 | 0.071 | 1 | 0 | Glod4 | 0.005 | Young |
| 0.634547 | 0.252985 | 0.118 | 0.114 | 1 | 0 | Pxk | 0.004 | Young |
| 0.634816 | 0.133552 | 0.138 | 0.148 | 1 | 0 | Kmt2d | -0.01 | Young |
| 0.635064 | 0.319817 | 0.122 | 0.118 | 1 | 0 | Ezr | 0.004 | Young |
| 0.635328 | 0.56944 | 0.04 | 0.037 | 1 | 0 | Trmt2b | 0.003 | Young |
| 0.635519 | 0.073653 | 0.048 | 0.044 | 1 | 0 | Cluh | 0.004 | Young |
| 0.635794 | 0.141015 | 0.364 | 0.367 | 1 | 0 | Aftph | -0.003 | Young |
| 0.635989 | 0.010498 | 0.047 | 0.042 | 1 | 0 | Dclre1b | 0.005 | Young |
| 0.636033 | 0.139265 | 0.164 | 0.176 | 1 | 0 | Blmh | -0.012 | Young |
| 0.636217 | 0.402203 | 0.025 | 0.022 | 1 | 0 | Katnal1 | 0.003 | Young |
| 0.636308 | 0.011453 | 0.221 | 0.213 | 1 | 0 | Tsc22d2 | 0.008 | Young |
| 0.636976 | 0.00737 | 0.541 | 0.572 | 1 | 0 | Eps8 | -0.031 | Young |
| 0.637072 | 0.432006 | 0.088 | 0.083 | 1 | 0 | Fam204a | 0.005 | Young |
| 0.637173 | 0.14422 | 0.099 | 0.107 | 1 | 0 | Mob3a | -0.008 | Young |
| 0.637186 | 0.145918 | 0.115 | 0.123 | 1 | 0 | Plscr3 | -0.008 | Young |
| 0.63721 | 0.34864 | 0.088 | 0.083 | 1 | 0 | Btaf1 | 0.005 | Young |
| 0.637249 | 0.00576 | 0.066 | 0.071 | 1 | 0 | Ggps1 | -0.005 | Young |
| 0.637393 | 0.037208 | 0.055 | 0.06 | 1 | 0 | Smcr8 | -0.005 | Young |
| 0.637594 | 0.212275 | 0.069 | 0.064 | 1 | 0 | Usp5 | 0.005 | Young |
| 0.637827 | 0.003448 | 0.036 | 0.04 | 1 | 0 | Eci1 | -0.004 | Young |
| 0.637855 | 0.321573 | 0.015 | 0.018 | 1 | 0 | Asb1 | -0.003 | Young |
| 0.638072 | 0.364714 | 0.063 | 0.059 | 1 | 0 | Samd1 | 0.004 | Young |
| 0.638196 | 0.841281 | 0.01 | 0.008 | 1 | 0 | Wdpcp | 0.002 | Young |
| 0.638198 | 0.119756 | 0.122 | 0.13 | 1 | 0 | Slc31a1 | -0.008 | Young |
| 0.638216 | 0.101764 | 0.155 | 0.166 | 1 | 0 | Tug1 | -0.011 | Young |
| 0.638395 | 0.211832 | 0.045 | 0.041 | 1 | 0 | Gzf1 | 0.004 | Young |
| 0.638793 | 0.904823 | 0.01 | 0.008 | 1 | 0 | Ahi1 | 0.002 | Young |
| 0.639319 | 0.118193 | 0.073 | 0.079 | 1 | 0 | Irf3 | -0.006 | Young |
| 0.639505 | 0.19605 | 0.06 | 0.066 | 1 | 0 | Rnaset2a | -0.006 | Young |
| 0.63976 | 0.175905 | 0.203 | 0.219 | 1 | 0 | Chmp3 | -0.016 | Young |
| 0.639996 | 0.650512 | 0.029 | 0.026 | 1 | 0 | Pgm3 | 0.003 | Young |
| 0.640187 | 0.755831 | 0.01 | 0.008 | 1 | 0 | Naprt | 0.002 | Young |
| 0.640188 | 0.218791 | 0.087 | 0.082 | 1 | 0 | Ing5 | 0.005 | Young |
| 0.640255 | 0.14541 | 0.082 | 0.089 | 1 | 0 | Esco1 | -0.007 | Young |
| 0.64045 | 0.052873 | 0.037 | 0.033 | 1 | 0 | Dnm1 | 0.004 | Young |
| 0.640533 | 0.031481 | 0.982 | 0.974 | 1 | 0 | mt-Nd2 | 0.008 | Young |
| 0.640606 | 0.323692 | 0.064 | 0.06 | 1 | 0 | Rpp25l | 0.004 | Young |
| 0.640913 | 0.008182 | 0.059 | 0.055 | 1 | 0 | Ocel1 | 0.004 | Young |
| 0.641045 | 0.043206 | 0.035 | 0.038 | 1 | 0 | Gm45606 | -0.003 | Young |

| | | | | | | | | |
|---|---|---|---|---|---|---|---|---|
| 0.641071 | 0.131535 | 0.148 | 0.142 | 1 | 0 | Fbl | 0.006 | Young |
| 0.641141 | 0.041765 | 0.036 | 0.04 | 1 | 0 | Nubp2 | -0.004 | Young |
| 0.64132 | 0.176451 | 0.061 | 0.056 | 1 | 0 | Dmxl2 | 0.005 | Young |
| 0.641786 | 0.434055 | 0.049 | 0.045 | 1 | 0 | Dennd2c | 0.004 | Young |
| 0.642364 | 0.430069 | 0.036 | 0.033 | 1 | 0 | Txnl4a | 0.003 | Young |
| 0.64238 | 0.467921 | 0.01 | 0.008 | 1 | 0 | Gm16867 | 0.002 | Young |
| 0.642492 | 0.044843 | 0.094 | 0.101 | 1 | 0 | Ebpl | -0.007 | Young |
| 0.642649 | 0.100734 | 0.235 | 0.233 | 1 | 0 | Emilin2 | 0.002 | Young |
| 0.642761 | 0.208834 | 0.107 | 0.115 | 1 | 0 | Hdac9 | -0.008 | Young |
| 0.643068 | 0.474978 | 0.085 | 0.081 | 1 | 0 | Man1b1 | 0.004 | Young |
| 0.643216 | 0.119749 | 0.125 | 0.119 | 1 | 0 | Klhl6 | 0.006 | Young |
| 0.643406 | 0.05214 | 0.462 | 0.464 | 1 | 0 | Serpinb8 | -0.002 | Young |
| 0.643437 | 0.303313 | 0.12 | 0.116 | 1 | 0 | Ergic2 | 0.004 | Young |
| 0.643521 | 0.440636 | 0.022 | 0.019 | 1 | 0 | Nsdhl | 0.003 | Young |
| 0.643829 | 0.077895 | 0.109 | 0.116 | 1 | 0 | Sf3b4 | -0.007 | Young |
| 0.644048 | 0.415488 | 0.019 | 0.016 | 1 | 0 | Ttc8 | 0.003 | Young |
| 0.644104 | 0.247987 | 0.066 | 0.062 | 1 | 0 | Lilra5 | 0.004 | Young |
| 0.64419 | 0.088035 | 0.084 | 0.079 | 1 | 0 | Zswim8 | 0.005 | Young |
| 0.644233 | 0.318043 | 0.032 | 0.029 | 1 | 0 | Lmf1 | 0.003 | Young |
| 0.644233 | 0.010177 | 0.08 | 0.086 | 1 | 0 | Trappc6a | -0.006 | Young |
| 0.644585 | 0.243261 | 0.15 | 0.146 | 1 | 0 | Klhl5 | 0.004 | Young |
| 0.64474 | 0.361791 | 0.022 | 0.019 | 1 | 0 | Taf1c | 0.003 | Young |
| 0.644855 | 0.127783 | 0.231 | 0.248 | 1 | 0 | Ppib | -0.017 | Young |
| 0.64527 | 0.73382 | 0.013 | 0.011 | 1 | 0 | Gm13091 | 0.002 | Young |
| 0.645395 | 0.338867 | 0.052 | 0.048 | 1 | 0 | Uba7 | 0.004 | Young |
| 0.645583 | 0.272888 | 0.051 | 0.047 | 1 | 0 | Man2b2 | 0.004 | Young |
| 0.645998 | 0.085465 | 0.071 | 0.077 | 1 | 0 | Lyar | -0.006 | Young |
| 0.646002 | 0.106428 | 0.108 | 0.103 | 1 | 0 | Tmem248 | 0.005 | Young |
| 0.646063 | 0.093059 | 0.165 | 0.175 | 1 | 0 | Elf2 | -0.01 | Young |
| 0.64611 | 0.012298 | 0.103 | 0.109 | 1 | 0 | Frrs1 | -0.006 | Young |
| 0.646122 | 0.192388 | 0.025 | 0.022 | 1 | 0 | Arrdc2 | 0.003 | Young |
| 0.646475 | 0.289669 | 0.01 | 0.008 | 1 | 0 | Gm35363 | 0.002 | Young |
| 0.646608 | 0.006596 | 0.093 | 0.098 | 1 | 0 | Cyc1 | -0.005 | Young |
| 0.646657 | 0.118944 | 0.178 | 0.19 | 1 | 0 | Ptov1 | -0.012 | Young |
| 0.647079 | 0.498204 | 0.054 | 0.051 | 1 | 0 | Paxip1 | 0.003 | Young |
| 0.647452 | 0.038294 | 0.299 | 0.29 | 1 | 0 | Cggbp1 | 0.009 | Young |
| 0.647609 | 0.100439 | 0.031 | 0.034 | 1 | 0 | Espn | -0.003 | Young |
| 0.647675 | 0.073162 | 0.028 | 0.025 | 1 | 0 | Zfp595 | 0.003 | Young |
| 0.647699 | 0.051606 | 0.061 | 0.066 | 1 | 0 | Ruvbl1 | -0.005 | Young |
| 0.647809 | 0.394879 | 0.068 | 0.064 | 1 | 0 | Clec5a | 0.004 | Young |
| 0.648274 | 0.413531 | 0.019 | 0.016 | 1 | 0 | Abcb8 | 0.003 | Young |
| 0.648408 | 0.584279 | 0.026 | 0.023 | 1 | 0 | Gm5431 | 0.003 | Young |
| 0.648609 | 0.710805 | 0.013 | 0.011 | 1 | 0 | Dyrk1b | 0.002 | Young |
| 0.648658 | 0.470499 | 0.016 | 0.014 | 1 | 0 | Btbd6 | 0.002 | Young |
| 0.648786 | 0.152046 | 0.054 | 0.059 | 1 | 0 | Coa6 | -0.005 | Young |

| | | | | | | | | |
|---|---|---|---|---|---|---|---|---|
| 0.649156 | 0.153788 | 0.123 | 0.118 | 1 | 0 | Agpat3 | 0.005 | Young |
| 0.649239 | 0.278709 | 0.06 | 0.056 | 1 | 0 | Sipa1l1 | 0.004 | Young |
| 0.649352 | 0.13897 | 0.065 | 0.06 | 1 | 0 | Fer | 0.005 | Young |
| 0.64944 | 0.16878 | 0.058 | 0.063 | 1 | 0 | Irgm2 | -0.005 | Young |
| 0.649452 | 0.222592 | 0.074 | 0.07 | 1 | 0 | Senp5 | 0.004 | Young |
| 0.649597 | 0.115902 | 0.179 | 0.174 | 1 | 0 | Ip6k1 | 0.005 | Young |
| 0.649637 | 0.374225 | 0.059 | 0.055 | 1 | 0 | Parp12 | 0.004 | Young |
| 0.64969 | 0.101619 | 0.061 | 0.066 | 1 | 0 | Ell2 | -0.005 | Young |
| 0.649718 | 0.113336 | 0.118 | 0.112 | 1 | 0 | Cwf19l2 | 0.006 | Young |
| 0.649841 | 0.564967 | 0.013 | 0.011 | 1 | 0 | Gm8369 | 0.002 | Young |
| 0.649851 | 0.068169 | 0.092 | 0.098 | 1 | 0 | 3830406C1 | -0.006 | Young |
| 0.650033 | 0.181629 | 0.022 | 0.019 | 1 | 0 | Srrd | 0.003 | Young |
| 0.650296 | 0.056463 | 0.282 | 0.297 | 1 | 0 | Ralb | -0.015 | Young |
| 0.650604 | 0.450683 | 0.033 | 0.03 | 1 | 0 | Nudt14 | 0.003 | Young |
| 0.651152 | 0.15735 | 0.107 | 0.101 | 1 | 0 | Susd3 | 0.006 | Young |
| 0.651314 | 0.052656 | 0.648 | 0.673 | 1 | 0 | Rps2 | -0.025 | Young |
| 0.651359 | 0.096116 | 0.497 | 0.514 | 1 | 0 | Tacc1 | -0.017 | Young |
| 0.65139 | 0.072162 | 0.174 | 0.185 | 1 | 0 | Appl1 | -0.011 | Young |
| 0.651801 | 0.085757 | 0.022 | 0.019 | 1 | 0 | Gpr157 | 0.003 | Young |
| 0.652005 | 0.203524 | 0.022 | 0.019 | 1 | 0 | 2810001G2 | 0.003 | Young |
| 0.652528 | 0.010861 | 0.284 | 0.295 | 1 | 0 | Tex261 | -0.011 | Young |
| 0.65266 | 0.292242 | 0.013 | 0.011 | 1 | 0 | Slc25a26 | 0.002 | Young |
| 0.652811 | 0.304123 | 0.111 | 0.107 | 1 | 0 | Ppp2r2d | 0.004 | Young |
| 0.652826 | 0.08317 | 0.294 | 0.291 | 1 | 0 | Hsbp1 | 0.003 | Young |
| 0.652879 | 0.119801 | 0.098 | 0.093 | 1 | 0 | Hs2st1 | 0.005 | Young |
| 0.652962 | 0.22838 | 0.073 | 0.068 | 1 | 0 | Hopx | 0.005 | Young |
| 0.652987 | 0.289597 | 0.069 | 0.064 | 1 | 0 | Timm17a | 0.005 | Young |
| 0.653152 | 0.384145 | 0.057 | 0.053 | 1 | 0 | Mapk8ip3 | 0.004 | Young |
| 0.653907 | 0.28814 | 0.016 | 0.014 | 1 | 0 | Slc5a6 | 0.002 | Young |
| 0.654066 | 0.317112 | 0.016 | 0.014 | 1 | 0 | AI427809 | 0.002 | Young |
| 0.6541 | 0.050522 | 0.046 | 0.051 | 1 | 0 | Fbxo25 | -0.005 | Young |
| 0.654577 | 0.334365 | 0.026 | 0.023 | 1 | 0 | Wdr92 | 0.003 | Young |
| 0.654805 | 0.225917 | 0.127 | 0.123 | 1 | 0 | Rictor | 0.004 | Young |
| 0.655278 | 0.034404 | 0.05 | 0.055 | 1 | 0 | Kri1 | -0.005 | Young |
| 0.655403 | 0.184258 | 0.112 | 0.107 | 1 | 0 | Cdc123 | 0.005 | Young |
| 0.655537 | 0.169485 | 0.254 | 0.254 | 1 | 0 | Eea1 | 0 | Young |
| 0.655576 | 0.162957 | 0.13 | 0.124 | 1 | 0 | Ccdc82 | 0.006 | Young |
| 0.655657 | 0.015842 | 0.156 | 0.164 | 1 | 0 | Esd | -0.008 | Young |
| 0.65579 | 0.196329 | 0.093 | 0.088 | 1 | 0 | Polr2l | 0.005 | Young |
| 0.65593 | 0.17699 | 0.042 | 0.038 | 1 | 0 | Triobp | 0.004 | Young |
| 0.65629 | 0.196849 | 0.201 | 0.197 | 1 | 0 | Madd | 0.004 | Young |
| 0.656668 | 0.593678 | 0.023 | 0.021 | 1 | 0 | Alyref2 | 0.002 | Young |
| 0.656752 | 0.240005 | 0.084 | 0.079 | 1 | 0 | Zfp622 | 0.005 | Young |
| 0.656955 | 0.001159 | 0.028 | 0.031 | 1 | 0 | Pef1 | -0.003 | Young |
| 0.657184 | 0.1625 | 0.195 | 0.19 | 1 | 0 | Ndufa5 | 0.005 | Young |

| | | | | | | | | |
|---|---|---|---|---|---|---|---|---|
| 0.657199 | 0.607868 | 0.023 | 0.021 | 1 | 0 | Nudcd1 | 0.002 | Young |
| 0.65723 | 0.154124 | 0.029 | 0.026 | 1 | 0 | Gm20707 | 0.003 | Young |
| 0.657605 | 0.168174 | 0.22 | 0.218 | 1 | 0 | Pnpla8 | 0.002 | Young |
| 0.65777 | 0.189636 | 0.073 | 0.068 | 1 | 0 | Rnase6 | 0.005 | Young |
| 0.657958 | 0.240806 | 0.184 | 0.181 | 1 | 0 | Anapc11 | 0.003 | Young |
| 0.658141 | 0.223745 | 0.077 | 0.073 | 1 | 0 | Gbf1 | 0.004 | Young |
| 0.658214 | 0.163797 | 0.179 | 0.176 | 1 | 0 | Ubr2 | 0.003 | Young |
| 0.658486 | 0.144217 | 0.123 | 0.118 | 1 | 0 | Ndufv2 | 0.005 | Young |
| 0.658754 | 0.152973 | 0.036 | 0.033 | 1 | 0 | Nr2c2ap | 0.003 | Young |
| 0.658831 | 0.017247 | 0.029 | 0.033 | 1 | 0 | Rbm43 | -0.004 | Young |
| 0.658984 | 0.00659 | 0.135 | 0.144 | 1 | 0 | Mrpl3 | -0.009 | Young |
| 0.659057 | 0.114869 | 0.086 | 0.092 | 1 | 0 | Galnt10 | -0.006 | Young |
| 0.659317 | 0.030717 | 0.094 | 0.1 | 1 | 0 | Zc3h18 | -0.006 | Young |
| 0.659748 | 0.048625 | 0.083 | 0.089 | 1 | 0 | Bex3 | -0.006 | Young |
| 0.659867 | 0.014001 | 0.091 | 0.097 | 1 | 0 | Slc30a7 | -0.006 | Young |
| 0.660426 | 0.2039 | 0.048 | 0.044 | 1 | 0 | Igtp | 0.004 | Young |
| 0.660652 | 0.379663 | 0.023 | 0.021 | 1 | 0 | Mrpl49 | 0.002 | Young |
| 0.6609 | 0.436662 | 0.065 | 0.062 | 1 | 0 | Kdsr | 0.003 | Young |
| 0.661488 | 0.072682 | 0.108 | 0.103 | 1 | 0 | Ilf3 | 0.005 | Young |
| 0.661585 | 0.301247 | 0.026 | 0.023 | 1 | 0 | Brat1 | 0.003 | Young |
| 0.661846 | 0.082974 | 0.075 | 0.081 | 1 | 0 | Commd4 | -0.006 | Young |
| 0.662576 | 0.402062 | 0.05 | 0.047 | 1 | 0 | Rad17 | 0.003 | Young |
| 0.662945 | 0.03587 | 0.204 | 0.216 | 1 | 0 | Smpdl3a | -0.012 | Young |
| 0.662999 | 0.061506 | 0.057 | 0.062 | 1 | 0 | Slc25a20 | -0.005 | Young |
| 0.663079 | 0.014773 | 0.019 | 0.016 | 1 | 0 | Rfc4 | 0.003 | Young |
| 0.663599 | 0.075775 | 0.026 | 0.029 | 1 | 0 | Actr6 | -0.003 | Young |
| 0.66365 | 0.155795 | 0.348 | 0.35 | 1 | 0 | Foxn3 | -0.002 | Young |
| 0.663745 | 0.020342 | 0.178 | 0.186 | 1 | 0 | Clec4b1 | -0.008 | Young |
| 0.66432 | 0.022269 | 0.455 | 0.486 | 1 | 0 | Purb | -0.031 | Young |
| 0.664544 | 0.09757 | 0.107 | 0.115 | 1 | 0 | Sft2d1 | -0.008 | Young |
| 0.664591 | 0.048996 | 0.056 | 0.06 | 1 | 0 | Dip2c | -0.004 | Young |
| 0.664891 | 0.093784 | 0.123 | 0.118 | 1 | 0 | Gigyf2 | 0.005 | Young |
| 0.665047 | 0.157575 | 0.08 | 0.075 | 1 | 0 | Tmem250- | 0.005 | Young |
| 0.665216 | 0.159616 | 0.357 | 0.367 | 1 | 0 | Cnot6l | -0.01 | Young |
| 0.665321 | 0.273275 | 0.045 | 0.049 | 1 | 0 | Lonp1 | -0.004 | Young |
| 0.665382 | 0.05576 | 0.065 | 0.07 | 1 | 0 | Pcsk7 | -0.005 | Young |
| 0.665523 | 0.203867 | 0.16 | 0.157 | 1 | 0 | Rtf2 | 0.003 | Young |
| 0.66572 | 0.042781 | 0.115 | 0.122 | 1 | 0 | Hist3h2a | -0.007 | Young |
| 0.665776 | 0.059243 | 0.23 | 0.224 | 1 | 0 | Fkbp2 | 0.006 | Young |
| 0.665863 | 0.117559 | 0.112 | 0.107 | 1 | 0 | Oser1 | 0.005 | Young |
| 0.665922 | 0.464479 | 0.043 | 0.04 | 1 | 0 | Emc4 | 0.003 | Young |
| 0.666091 | 0.056832 | 0.131 | 0.14 | 1 | 0 | Parp1 | -0.009 | Young |
| 0.666152 | 0.002965 | 0.471 | 0.498 | 1 | 0 | Cox5a | -0.027 | Young |
| 0.666174 | 0.09948 | 0.056 | 0.06 | 1 | 0 | Mtmr1 | -0.004 | Young |
| 0.666377 | 0.561821 | 0.027 | 0.025 | 1 | 0 | Aptx | 0.002 | Young |

| | | | | | | | | |
|---|---|---|---|---|---|---|---|---|
| 0.66638 | 0.040258 | 0.033 | 0.03 | 1 | 0 | Xpnpep3 | 0.003 | Young |
| 0.666538 | 0.194048 | 0.125 | 0.12 | 1 | 0 | Pisd | 0.005 | Young |
| 0.666664 | 0.17065 | 0.061 | 0.057 | 1 | 0 | Ppp1r9a | 0.004 | Young |
| 0.66673 | 0.089924 | 0.265 | 0.282 | 1 | 0 | Mia3 | -0.017 | Young |
| 0.666822 | 0.190447 | 0.123 | 0.119 | 1 | 0 | Smpdl3b | 0.004 | Young |
| 0.666959 | 0.325033 | 0.071 | 0.067 | 1 | 0 | Coro2b | 0.004 | Young |
| 0.667115 | 0.130339 | 0.113 | 0.108 | 1 | 0 | Prpf8 | 0.005 | Young |
| 0.667355 | 0.164305 | 0.169 | 0.166 | 1 | 0 | Setx | 0.003 | Young |
| 0.667432 | 0.239298 | 0.045 | 0.041 | 1 | 0 | Phkg2 | 0.004 | Young |
| 0.667526 | 0.300538 | 0.134 | 0.131 | 1 | 0 | Safb | 0.003 | Young |
| 0.667668 | 0.016749 | 0.345 | 0.363 | 1 | 0 | Man1c1 | -0.018 | Young |
| 0.667686 | 0.051889 | 0.105 | 0.112 | 1 | 0 | Fbxo18 | -0.007 | Young |
| 0.667923 | 0.144962 | 0.225 | 0.241 | 1 | 0 | Glmp | -0.016 | Young |
| 0.667948 | 0.21058 | 0.066 | 0.062 | 1 | 0 | Dctn6 | 0.004 | Young |
| 0.668105 | 0.055586 | 0.069 | 0.064 | 1 | 0 | 4930402H2 | 0.005 | Young |
| 0.668118 | 0.233008 | 0.138 | 0.134 | 1 | 0 | Eif1ax | 0.004 | Young |
| 0.668291 | 0.050478 | 0.026 | 0.029 | 1 | 0 | C2cd2 | -0.003 | Young |
| 0.668308 | 0.631338 | 0.032 | 0.029 | 1 | 0 | Diexf | 0.003 | Young |
| 0.668485 | 0.167448 | 0.07 | 0.066 | 1 | 0 | Arfgef2 | 0.004 | Young |
| 0.668531 | 0.150752 | 0.209 | 0.204 | 1 | 0 | Rasgrp4 | 0.005 | Young |
| 0.668672 | 0.408001 | 0.065 | 0.062 | 1 | 0 | Lmf2 | 0.003 | Young |
| 0.668751 | 0.174387 | 0.049 | 0.053 | 1 | 0 | Ankrd16 | -0.004 | Young |
| 0.668979 | 0.151534 | 0.06 | 0.056 | 1 | 0 | Lsm14b | 0.004 | Young |
| 0.669075 | 0.246653 | 0.05 | 0.055 | 1 | 0 | Yod1 | -0.005 | Young |
| 0.669467 | 0.321193 | 0.03 | 0.027 | 1 | 0 | Klhl33 | 0.003 | Young |
| 0.669552 | 0.136975 | 0.054 | 0.059 | 1 | 0 | Ddx27 | -0.005 | Young |
| 0.669582 | 0.084332 | 0.325 | 0.346 | 1 | 0 | Pttg1ip | -0.021 | Young |
| 0.669648 | 0.222669 | 0.245 | 0.246 | 1 | 0 | Acat1 | -0.001 | Young |
| 0.67019 | 0.058385 | 0.239 | 0.253 | 1 | 0 | Kpna4 | -0.014 | Young |
| 0.670777 | 0.026813 | 0.197 | 0.208 | 1 | 0 | Pnn | -0.011 | Young |
| 0.670977 | 0.119799 | 0.374 | 0.383 | 1 | 0 | Arhgef6 | -0.009 | Young |
| 0.671052 | 0.150256 | 0.083 | 0.089 | 1 | 0 | Fam214b | -0.006 | Young |
| 0.671253 | 0.072339 | 0.053 | 0.057 | 1 | 0 | Sdad1 | -0.004 | Young |
| 0.671345 | 0.008104 | 0.051 | 0.055 | 1 | 0 | Ttc37 | -0.004 | Young |
| 0.671661 | 0.01432 | 0.17 | 0.178 | 1 | 0 | Anapc5 | -0.008 | Young |
| 0.672543 | 0.189451 | 0.078 | 0.083 | 1 | 0 | Trak2 | -0.005 | Young |
| 0.672603 | 0.401463 | 0.031 | 0.034 | 1 | 0 | Mtap | -0.003 | Young |
| 0.672643 | 0.097316 | 0.033 | 0.03 | 1 | 0 | Agk | 0.003 | Young |
| 0.673151 | 0.143271 | 0.024 | 0.027 | 1 | 0 | Zfp429 | -0.003 | Young |
| 0.673171 | 0.18751 | 0.049 | 0.053 | 1 | 0 | Mrpl45 | -0.004 | Young |
| 0.673684 | 0.070409 | 0.11 | 0.104 | 1 | 0 | Fam160b1 | 0.006 | Young |
| 0.673905 | 0.202754 | 0.088 | 0.094 | 1 | 0 | Tmem86a | -0.006 | Young |
| 0.673907 | 0.017786 | 0.19 | 0.2 | 1 | 0 | Zmat2 | -0.01 | Young |
| 0.674229 | 0.216236 | 0.182 | 0.179 | 1 | 0 | Tgif1 | 0.003 | Young |
| 0.674288 | 0.027624 | 0.135 | 0.142 | 1 | 0 | Atp8a1 | -0.007 | Young |

| | | | | | | | | |
|---|---|---|---|---|---|---|---|---|
| 0.67434 | 0.191342 | 0.042 | 0.038 | 1 | 0 | Tbc1d12 | 0.004 | Young |
| 0.674645 | 0.006362 | 0.062 | 0.067 | 1 | 0 | Bicra | -0.005 | Young |
| 0.67467 | 0.031083 | 0.184 | 0.194 | 1 | 0 | Aebp2 | -0.01 | Young |
| 0.674872 | 0.689775 | 0.011 | 0.01 | 1 | 0 | Gm15706 | 0.001 | Young |
| 0.67517 | 0.068065 | 0.29 | 0.286 | 1 | 0 | Ptgs1 | 0.004 | Young |
| 0.675541 | 0.056354 | 0.019 | 0.022 | 1 | 0 | Shb | -0.003 | Young |
| 0.675863 | 0.274463 | 0.056 | 0.062 | 1 | 0 | Vkorc1l1 | -0.006 | Young |
| 0.676073 | 0.149731 | 0.16 | 0.171 | 1 | 0 | Akirin1 | -0.011 | Young |
| 0.6763 | 0.236186 | 0.044 | 0.048 | 1 | 0 | Tmcc2 | -0.004 | Young |
| 0.676765 | 0.013935 | 0.066 | 0.062 | 1 | 0 | Zfp560 | 0.004 | Young |
| 0.676818 | 0.400641 | 0.017 | 0.015 | 1 | 0 | Dut | 0.002 | Young |
| 0.676912 | 0.152458 | 0.08 | 0.075 | 1 | 0 | Zadh2 | 0.005 | Young |
| 0.676923 | 0.527998 | 0.014 | 0.012 | 1 | 0 | Endov | 0.002 | Young |
| 0.677413 | 0.242871 | 0.104 | 0.1 | 1 | 0 | Napg | 0.004 | Young |
| 0.677755 | 0.378858 | 0.027 | 0.03 | 1 | 0 | Sgf29 | -0.003 | Young |
| 0.67777 | 0.155565 | 0.053 | 0.049 | 1 | 0 | Rpia | 0.004 | Young |
| 0.677809 | 0.102609 | 0.295 | 0.293 | 1 | 0 | Wls | 0.002 | Young |
| 0.677935 | 0.510264 | 0.011 | 0.01 | 1 | 0 | Fam189b | 0.001 | Young |
| 0.677973 | 0.116512 | 0.097 | 0.104 | 1 | 0 | Mllt6 | -0.007 | Young |
| 0.677993 | 0.038997 | 0.326 | 0.346 | 1 | 0 | Eif5b | -0.02 | Young |
| 0.678001 | 0.00996 | 0.162 | 0.17 | 1 | 0 | Rab8a | -0.008 | Young |
| 0.678019 | 0.058222 | 0.091 | 0.097 | 1 | 0 | Maged1 | -0.006 | Young |
| 0.678061 | 0.370191 | 0.017 | 0.015 | 1 | 0 | Amer1 | 0.002 | Young |
| 0.678318 | 0.493317 | 0.011 | 0.01 | 1 | 0 | Ak1 | 0.001 | Young |
| 0.678319 | 0.039692 | 0.051 | 0.055 | 1 | 0 | Ccdc43 | -0.004 | Young |
| 0.678499 | 0.411171 | 0.03 | 0.027 | 1 | 0 | Dna2 | 0.003 | Young |
| 0.679378 | 0.494118 | 0.054 | 0.051 | 1 | 0 | Mdn1 | 0.003 | Young |
| 0.679457 | 0.136509 | 0.147 | 0.141 | 1 | 0 | Wwp2 | 0.006 | Young |
| 0.67949 | 0.032522 | 0.157 | 0.164 | 1 | 0 | Nudcd3 | -0.007 | Young |
| 0.679654 | 0.429096 | 0.014 | 0.012 | 1 | 0 | Ccdc138 | 0.002 | Young |
| 0.679943 | 0.102247 | 0.1 | 0.107 | 1 | 0 | Parl | -0.007 | Young |
| 0.68065 | 0.086121 | 0.166 | 0.175 | 1 | 0 | Ube2h | -0.009 | Young |
| 0.680679 | 0.323739 | 0.014 | 0.012 | 1 | 0 | Chek2 | 0.002 | Young |
| 0.680697 | 0.091193 | 0.036 | 0.04 | 1 | 0 | Atg9a | -0.004 | Young |
| 0.680927 | 0.078684 | 0.027 | 0.025 | 1 | 0 | Ift80 | 0.002 | Young |
| 0.680951 | 0.051297 | 0.049 | 0.053 | 1 | 0 | Inip | -0.004 | Young |
| 0.681322 | 0.386905 | 0.032 | 0.036 | 1 | 0 | Pelp1 | -0.004 | Young |
| 0.681352 | 0.041988 | 0.153 | 0.148 | 1 | 0 | Phf23 | 0.005 | Young |
| 0.681355 | 0.125788 | 0.152 | 0.146 | 1 | 0 | Ptpn11 | 0.006 | Young |
| 0.681387 | 0.294589 | 0.011 | 0.01 | 1 | 0 | Mtrf1 | 0.001 | Young |
| 0.682008 | 0.03845 | 0.299 | 0.316 | 1 | 0 | Herc1 | -0.017 | Young |
| 0.68206 | 0.289063 | 0.022 | 0.025 | 1 | 0 | Mto1 | -0.003 | Young |
| 0.682129 | 0.062892 | 0.233 | 0.23 | 1 | 0 | Nup98 | 0.003 | Young |
| 0.682155 | 0.313622 | 0.011 | 0.01 | 1 | 0 | Wdr25 | 0.001 | Young |
| 0.682263 | 0.263027 | 0.056 | 0.052 | 1 | 0 | Pitrm1 | 0.004 | Young |

| | | | | | | | | |
|---|---|---|---|---|---|---|---|---|
| 0.682347 | 0.234961 | 0.011 | 0.01 | 1 | 0 | Faap24 | 0.001 | Young |
| 0.682417 | 0.17138 | 0.017 | 0.015 | 1 | 0 | Trp53i13 | 0.002 | Young |
| 0.682499 | 0.009993 | 0.227 | 0.237 | 1 | 0 | Aco2 | -0.01 | Young |
| 0.682736 | 0.068171 | 0.017 | 0.019 | 1 | 0 | Plk4 | -0.002 | Young |
| 0.683038 | 0.061727 | 0.037 | 0.034 | 1 | 0 | Trp53bp1 | 0.003 | Young |
| 0.683116 | 0.253898 | 0.011 | 0.01 | 1 | 0 | Mpp7 | 0.001 | Young |
| 0.683134 | 0.341006 | 0.165 | 0.164 | 1 | 0 | Stom | 0.001 | Young |
| 0.683219 | 0.092106 | 0.027 | 0.025 | 1 | 0 | Fahd1 | 0.002 | Young |
| 0.683501 | 0.360122 | 0.014 | 0.012 | 1 | 0 | Tfb1m | 0.002 | Young |
| 0.683753 | 0.256885 | 0.101 | 0.097 | 1 | 0 | Nedd9 | 0.004 | Young |
| 0.683808 | 0.09955 | 0.216 | 0.228 | 1 | 0 | Acbd5 | -0.012 | Young |
| 0.683823 | 0.027948 | 0.191 | 0.201 | 1 | 0 | Pgd | -0.01 | Young |
| 0.684077 | 0.188035 | 0.011 | 0.01 | 1 | 0 | Srfbp1 | 0.001 | Young |
| 0.684125 | 0.064079 | 0.066 | 0.062 | 1 | 0 | Ero1l | 0.004 | Young |
| 0.684215 | 0.037611 | 0.008 | 0.01 | 1 | 0 | Ldlrad4 | -0.002 | Young |
| 0.684265 | 0.085194 | 0.102 | 0.108 | 1 | 0 | Zfp90 | -0.006 | Young |
| 0.684474 | 0.042165 | 0.155 | 0.163 | 1 | 0 | Tet2 | -0.008 | Young |
| 0.684503 | 0.239172 | 0.056 | 0.06 | 1 | 0 | Iffo2 | -0.004 | Young |
| 0.684691 | 0.316557 | 0.067 | 0.063 | 1 | 0 | Kars | 0.004 | Young |
| 0.684725 | 0.144553 | 0.017 | 0.019 | 1 | 0 | Depdc7 | -0.002 | Young |
| 0.684961 | 0.194996 | 0.297 | 0.3 | 1 | 0 | Ndufc1 | -0.003 | Young |
| 0.685067 | 0.10582 | 0.094 | 0.089 | 1 | 0 | Gtf2h1 | 0.005 | Young |
| 0.685241 | 0.094714 | 0.201 | 0.194 | 1 | 0 | Rbm26 | 0.007 | Young |
| 0.68547 | 0.14529 | 0.014 | 0.012 | 1 | 0 | Arvcf | 0.002 | Young |
| 0.685584 | 0.145605 | 0.232 | 0.228 | 1 | 0 | Fam126a | 0.004 | Young |
| 0.685663 | 0.286084 | 0.061 | 0.057 | 1 | 0 | Phb | 0.004 | Young |
| 0.685783 | 0.140413 | 0.16 | 0.156 | 1 | 0 | Chd2 | 0.004 | Young |
| 0.685813 | 0.065186 | 0.014 | 0.012 | 1 | 0 | Net1 | 0.002 | Young |
| 0.685999 | 0.117559 | 0.094 | 0.1 | 1 | 0 | Micu2 | -0.006 | Young |
| 0.68605 | 0.232254 | 0.009 | 0.011 | 1 | 0 | Platr17 | -0.002 | Young |
| 0.686058 | 0.061424 | 0.096 | 0.103 | 1 | 0 | Nipsnap2 | -0.007 | Young |
| 0.686079 | 0.252401 | 0.041 | 0.045 | 1 | 0 | Gm20696 | -0.004 | Young |
| 0.686288 | 0.25725 | 0.018 | 0.021 | 1 | 0 | Wdr55 | -0.003 | Young |
| 0.686705 | 0.232355 | 0.036 | 0.033 | 1 | 0 | Abhd18 | 0.003 | Young |
| 0.686705 | 0.081774 | 0.085 | 0.081 | 1 | 0 | Smg5 | 0.004 | Young |
| 0.68678 | 0.15405 | 0.072 | 0.077 | 1 | 0 | Me2 | -0.005 | Young |
| 0.686851 | 0.025254 | 0.287 | 0.298 | 1 | 0 | Ndel1 | -0.011 | Young |
| 0.687101 | 0.02916 | 0.172 | 0.181 | 1 | 0 | Bud31 | -0.009 | Young |
| 0.687156 | 0.03187 | 0.083 | 0.078 | 1 | 0 | Mtf2 | 0.005 | Young |
| 0.687202 | 0.470726 | 0.033 | 0.03 | 1 | 0 | Nek6 | 0.003 | Young |
| 0.687326 | 0.010275 | 0.012 | 0.014 | 1 | 0 | Zhx2 | -0.002 | Young |
| 0.687482 | 0.351172 | 0.053 | 0.057 | 1 | 0 | Rab40c | -0.004 | Young |
| 0.68753 | 0.042684 | 0.086 | 0.092 | 1 | 0 | Ybx3 | -0.006 | Young |
| 0.687926 | 0.064134 | 0.011 | 0.01 | 1 | 0 | Asb4 | 0.001 | Young |
| 0.688311 | 0.068798 | 0.011 | 0.01 | 1 | 0 | Proser3 | 0.001 | Young |

| | | | | | | | | |
|---|---|---|---|---|---|---|---|---|
| 0.688608 | 0.041204 | 0.048 | 0.052 | 1 | 0 | Prmt3 | -0.004 | Young |
| 0.688834 | 0.002681 | 0.235 | 0.227 | 1 | 0 | Bank1 | 0.008 | Young |
| 0.688904 | 0.077099 | 0.044 | 0.048 | 1 | 0 | Micu3 | -0.004 | Young |
| 0.688933 | 0.069964 | 0.013 | 0.015 | 1 | 0 | Mb21d2 | -0.002 | Young |
| 0.689203 | 0.240256 | 0.024 | 0.022 | 1 | 0 | Dexi | 0.002 | Young |
| 0.689246 | 0.179659 | 0.077 | 0.073 | 1 | 0 | Phf12 | 0.004 | Young |
| 0.689275 | 0.125408 | 0.011 | 0.012 | 1 | 0 | C130050O1 | -0.001 | Young |
| 0.68941 | 0.85931 | 0.019 | 0.016 | 1 | 0 | Taf5 | 0.003 | Young |
| 0.689415 | 0.024031 | 0.014 | 0.012 | 1 | 0 | 2410131K1 | 0.002 | Young |
| 0.689493 | 0.238679 | 0.008 | 0.01 | 1 | 0 | C920006O1 | -0.002 | Young |
| 0.69018 | 0.106074 | 0.053 | 0.049 | 1 | 0 | Prep | 0.004 | Young |
| 0.690236 | 0.155559 | 0.014 | 0.016 | 1 | 0 | Zfp141 | -0.002 | Young |
| 0.690262 | 0.290885 | 0.019 | 0.022 | 1 | 0 | Fgf11 | -0.003 | Young |
| 0.690406 | 0.267124 | 0.019 | 0.022 | 1 | 0 | Golm1 | -0.003 | Young |
| 0.690741 | 0.0863 | 0.054 | 0.051 | 1 | 0 | Rbbp5 | 0.003 | Young |
| 0.690838 | 0.03754 | 0.031 | 0.034 | 1 | 0 | Txlng | -0.003 | Young |
| 0.690846 | 0.025488 | 0.104 | 0.109 | 1 | 0 | Rnf5 | -0.005 | Young |
| 0.690934 | 0.149699 | 0.065 | 0.062 | 1 | 0 | Mtif2 | 0.003 | Young |
| 0.690961 | 0.040925 | 0.014 | 0.012 | 1 | 0 | Tti1 | 0.002 | Young |
| 0.690971 | 0.207258 | 0.009 | 0.011 | 1 | 0 | Plppr3 | -0.002 | Young |
| 0.691025 | 0.439125 | 0.05 | 0.047 | 1 | 0 | Mcm3ap | 0.003 | Young |
| 0.691263 | 0.15761 | 0.049 | 0.053 | 1 | 0 | Nlrc4 | -0.004 | Young |
| 0.691265 | 0.001285 | 0.118 | 0.124 | 1 | 0 | Sla | -0.006 | Young |
| 0.691501 | 0.154305 | 0.118 | 0.114 | 1 | 0 | Slc45a4 | 0.004 | Young |
| 0.691526 | 0.001578 | 0.063 | 0.067 | 1 | 0 | Smim12 | -0.004 | Young |
| 0.691695 | 0.200425 | 0.047 | 0.044 | 1 | 0 | Extl3 | 0.003 | Young |
| 0.691733 | 0.04074 | 0.227 | 0.237 | 1 | 0 | Sipa1 | -0.01 | Young |
| 0.691939 | 0.236635 | 0.072 | 0.068 | 1 | 0 | Atad2 | 0.004 | Young |
| 0.692356 | 0.124447 | 0.082 | 0.078 | 1 | 0 | Ctns | 0.004 | Young |
| 0.692745 | 0.365645 | 0.07 | 0.075 | 1 | 0 | Sdhaf4 | -0.005 | Young |
| 0.692781 | 0.040652 | 0.098 | 0.104 | 1 | 0 | Stat2 | -0.006 | Young |
| 0.692879 | 0.01454 | 0.114 | 0.108 | 1 | 0 | Syne2 | 0.006 | Young |
| 0.693018 | 0.178472 | 0.226 | 0.227 | 1 | 0 | Cox7b | -0.001 | Young |
| 0.693115 | 0.125157 | 0.121 | 0.129 | 1 | 0 | Znhit1 | -0.008 | Young |
| 0.693229 | 0.203194 | 0.052 | 0.056 | 1 | 0 | Sap30bp | -0.004 | Young |
| 0.693353 | 0.284657 | 0.05 | 0.047 | 1 | 0 | Rpa2 | 0.003 | Young |
| 0.694055 | 0.033798 | 0.179 | 0.189 | 1 | 0 | Cul1 | -0.01 | Young |
| 0.694264 | 0.022335 | 0.081 | 0.086 | 1 | 0 | Gga3 | -0.005 | Young |
| 0.694506 | 0.175311 | 0.071 | 0.067 | 1 | 0 | Cmtm4 | 0.004 | Young |
| 0.694879 | 0.136108 | 0.104 | 0.1 | 1 | 0 | Atmin | 0.004 | Young |
| 0.695003 | 0.080982 | 0.044 | 0.048 | 1 | 0 | Dnajc9 | -0.004 | Young |
| 0.695017 | 0.020215 | 0.074 | 0.07 | 1 | 0 | Ate1 | 0.004 | Young |
| 0.695063 | 0.115984 | 0.21 | 0.205 | 1 | 0 | Klf9 | 0.005 | Young |
| 0.695124 | 0.458007 | 0.052 | 0.049 | 1 | 0 | Gpkow | 0.003 | Young |
| 0.695519 | 0.151846 | 0.035 | 0.038 | 1 | 0 | Mycbp | -0.003 | Young |

| | | | | | | | | |
|---|---|---|---|---|---|---|---|---|
| 0.695912 | 0.1149 | 0.185 | 0.196 | 1 | 0 | Npepps | -0.011 | Young |
| 0.696451 | 0.085294 | 0.298 | 0.32 | 1 | 0 | D8Ertd738 | -0.022 | Young |
| 0.696641 | 0.20928 | 0.107 | 0.103 | 1 | 0 | Zfp800 | 0.004 | Young |
| 0.697081 | 0.15144 | 0.142 | 0.15 | 1 | 0 | Chp1 | -0.008 | Young |
| 0.697189 | 0.032776 | 0.29 | 0.305 | 1 | 0 | Polr1d | -0.015 | Young |
| 0.697192 | 0.141363 | 0.105 | 0.111 | 1 | 0 | Arfgap2 | -0.006 | Young |
| 0.697449 | 0.094994 | 0.344 | 0.367 | 1 | 0 | Epn1 | -0.023 | Young |
| 0.697503 | 0.15174 | 0.173 | 0.17 | 1 | 0 | Snx13 | 0.003 | Young |
| 0.698409 | 0.069308 | 0.118 | 0.124 | 1 | 0 | Atxn1 | -0.006 | Young |
| 0.698424 | 0.219848 | 0.146 | 0.144 | 1 | 0 | Abcc5 | 0.002 | Young |
| 0.698808 | 0.105499 | 0.032 | 0.029 | 1 | 0 | Rbbp9 | 0.003 | Young |
| 0.698981 | 0.114397 | 0.183 | 0.194 | 1 | 0 | Rex1bd | -0.011 | Young |
| 0.698996 | 0.436503 | 0.052 | 0.049 | 1 | 0 | Elac1 | 0.003 | Young |
| 0.699085 | 0.426266 | 0.019 | 0.016 | 1 | 0 | Fbxo32 | 0.003 | Young |
| 0.699158 | 0.328741 | 0.044 | 0.041 | 1 | 0 | Gramd4 | 0.003 | Young |
| 0.699166 | 0.220699 | 0.032 | 0.036 | 1 | 0 | Ttll3 | -0.004 | Young |
| 0.699365 | 0.001587 | 0.292 | 0.306 | 1 | 0 | Rps6ka3 | -0.014 | Young |
| 0.699479 | 0.18953 | 0.296 | 0.301 | 1 | 0 | Smarca5 | -0.005 | Young |
| 0.699499 | 0.266358 | 0.115 | 0.112 | 1 | 0 | Vps37a | 0.003 | Young |
| 0.699523 | 0.146015 | 0.067 | 0.063 | 1 | 0 | Pign | 0.004 | Young |
| 0.699573 | 0.168686 | 0.034 | 0.037 | 1 | 0 | Rdm1 | -0.003 | Young |
| 0.699829 | 0.019496 | 0.131 | 0.138 | 1 | 0 | Mecp2 | -0.007 | Young |
| 0.700419 | 0.108089 | 0.191 | 0.187 | 1 | 0 | Il13ra1 | 0.004 | Young |
| 0.700586 | 0.112989 | 0.044 | 0.048 | 1 | 0 | Ypel2 | -0.004 | Young |
| 0.700678 | 0.390619 | 0.026 | 0.023 | 1 | 0 | Nfatc2ip | 0.003 | Young |
| 0.700758 | 0.216118 | 0.087 | 0.083 | 1 | 0 | Impdh2 | 0.004 | Young |
| 0.700905 | 0.385576 | 0.019 | 0.016 | 1 | 0 | Telo2 | 0.003 | Young |
| 0.701497 | 0.034724 | 0.047 | 0.051 | 1 | 0 | Gdpd1 | -0.004 | Young |
| 0.701554 | 0.108326 | 0.079 | 0.075 | 1 | 0 | Pcgf3 | 0.004 | Young |
| 0.70161 | 0.216713 | 0.159 | 0.156 | 1 | 0 | Ebi3 | 0.003 | Young |
| 0.701683 | 0.191119 | 0.056 | 0.06 | 1 | 0 | Nfyb | -0.004 | Young |
| 0.701713 | 0.096097 | 0.072 | 0.077 | 1 | 0 | Secisbp2 | -0.005 | Young |
| 0.70187 | 0.28041 | 0.075 | 0.073 | 1 | 0 | Zfpl1 | 0.002 | Young |
| 0.702485 | 0.396726 | 0.026 | 0.023 | 1 | 0 | Snx21 | 0.003 | Young |
| 0.70252 | 0.003822 | 0.158 | 0.166 | 1 | 0 | Slc35b2 | -0.008 | Young |
| 0.702697 | 0.37597 | 0.089 | 0.086 | 1 | 0 | 1810022K0 | 0.003 | Young |
| 0.703252 | 0.278328 | 0.055 | 0.052 | 1 | 0 | Aspscr1 | 0.003 | Young |
| 0.703556 | 0.069996 | 0.334 | 0.354 | 1 | 0 | Eif2s2 | -0.02 | Young |
| 0.703566 | 0.211481 | 0.074 | 0.079 | 1 | 0 | Ppp1r7 | -0.005 | Young |
| 0.703647 | 0.215587 | 0.054 | 0.059 | 1 | 0 | Mettl14 | -0.005 | Young |
| 0.703739 | 0.009101 | 0.112 | 0.119 | 1 | 0 | Cln3 | -0.007 | Young |
| 0.703967 | 0.015716 | 0.07 | 0.074 | 1 | 0 | Fam53c | -0.004 | Young |
| 0.704238 | 0.156518 | 0.069 | 0.074 | 1 | 0 | Iffo1 | -0.005 | Young |
| 0.704412 | 0.242748 | 0.067 | 0.063 | 1 | 0 | Pigq | 0.004 | Young |
| 0.704462 | 0.390451 | 0.081 | 0.078 | 1 | 0 | Mindy3 | 0.003 | Young |

| | | | | | | | | |
|---|---|---|---|---|---|---|---|---|
| 0.704546 | 0.11608 | 0.123 | 0.118 | 1 | 0 | Mtch2 | 0.005 | Young |
| 0.704843 | 0.132262 | 0.059 | 0.063 | 1 | 0 | Tbpl1 | -0.004 | Young |
| 0.705238 | 0.130899 | 0.17 | 0.181 | 1 | 0 | Osgep | -0.011 | Young |
| 0.705274 | 0.141951 | 0.081 | 0.086 | 1 | 0 | Igsf8 | -0.005 | Young |
| 0.705444 | 0.062523 | 0.429 | 0.439 | 1 | 0 | Rab7b | -0.01 | Young |
| 0.705594 | 0.088459 | 0.045 | 0.049 | 1 | 0 | Rbm15b | -0.004 | Young |
| 0.705613 | 0.289235 | 0.019 | 0.016 | 1 | 0 | Arsg | 0.003 | Young |
| 0.705727 | 0.201577 | 0.08 | 0.086 | 1 | 0 | Phyh | -0.006 | Young |
| 0.705924 | 0.149091 | 0.083 | 0.079 | 1 | 0 | Sart1 | 0.004 | Young |
| 0.705964 | 0.012687 | 0.126 | 0.131 | 1 | 0 | Mob1b | -0.005 | Young |
| 0.706696 | 0.053906 | 0.3 | 0.319 | 1 | 0 | Kif1b | -0.019 | Young |
| 0.706856 | 0.028225 | 0.211 | 0.207 | 1 | 0 | H1f0 | 0.004 | Young |
| 0.707436 | 0.088969 | 0.127 | 0.122 | 1 | 0 | Cdk19 | 0.005 | Young |
| 0.707458 | 0.166538 | 0.251 | 0.252 | 1 | 0 | Llph | -0.001 | Young |
| 0.707682 | 0.327137 | 0.03 | 0.027 | 1 | 0 | Cdkn2c | 0.003 | Young |
| 0.707784 | 0.100575 | 0.061 | 0.066 | 1 | 0 | Brap | -0.005 | Young |
| 0.707899 | 0.101513 | 0.063 | 0.059 | 1 | 0 | Ints9 | 0.004 | Young |
| 0.708 | 0.026373 | 0.047 | 0.044 | 1 | 0 | Cfap36 | 0.003 | Young |
| 0.708391 | 0.052119 | 0.134 | 0.131 | 1 | 0 | Ephx1 | 0.003 | Young |
| 0.708399 | 0.162483 | 0.126 | 0.122 | 1 | 0 | Pip5k1a | 0.004 | Young |
| 0.708468 | 0.18753 | 0.226 | 0.227 | 1 | 0 | Ndufb8 | -0.001 | Young |
| 0.708498 | 0.07977 | 0.137 | 0.133 | 1 | 0 | Mrpl48 | 0.004 | Young |
| 0.708625 | 0.125871 | 0.111 | 0.107 | 1 | 0 | Eif4e2 | 0.004 | Young |
| 0.708761 | 0.344493 | 0.034 | 0.031 | 1 | 0 | Exosc7 | 0.003 | Young |
| 0.709076 | 0.123916 | 0.115 | 0.122 | 1 | 0 | Cnpy3 | -0.007 | Young |
| 0.70914 | 0.10156 | 0.027 | 0.03 | 1 | 0 | Ier5l | -0.003 | Young |
| 0.709443 | 0.038933 | 0.309 | 0.305 | 1 | 0 | Mkln1 | 0.004 | Young |
| 0.709689 | 0.180312 | 0.192 | 0.189 | 1 | 0 | Pdlim1 | 0.003 | Young |
| 0.709707 | 0.114269 | 0.233 | 0.231 | 1 | 0 | Secisbp2l | 0.002 | Young |
| 0.709791 | 0.576908 | 0.013 | 0.011 | 1 | 0 | Synpo | 0.002 | Young |
| 0.709817 | 0.063688 | 0.052 | 0.056 | 1 | 0 | Slc27a4 | -0.004 | Young |
| 0.709865 | 0.229296 | 0.041 | 0.038 | 1 | 0 | Cep85l | 0.003 | Young |
| 0.709974 | 0.212734 | 0.314 | 0.327 | 1 | 0 | Aes | -0.013 | Young |
| 0.710198 | 0.357526 | 0.053 | 0.051 | 1 | 0 | Tmtc3 | 0.002 | Young |
| 0.710415 | 0.074558 | 0.08 | 0.085 | 1 | 0 | Ppp2r3d | -0.005 | Young |
| 0.710504 | 0.318928 | 0.069 | 0.066 | 1 | 0 | Tmem203 | 0.003 | Young |
| 0.710734 | 0.081115 | 0.228 | 0.224 | 1 | 0 | Herc4 | 0.004 | Young |
| 0.710893 | 0.097548 | 0.205 | 0.201 | 1 | 0 | Icam1 | 0.004 | Young |
| 0.711048 | 0.332149 | 0.03 | 0.027 | 1 | 0 | Trip10 | 0.003 | Young |
| 0.711276 | 0.590911 | 0.01 | 0.008 | 1 | 0 | Abhd8 | 0.002 | Young |
| 0.711716 | 0.184245 | 0.054 | 0.051 | 1 | 0 | Hsdl2 | 0.003 | Young |
| 0.711987 | 0.102333 | 0.153 | 0.161 | 1 | 0 | Usp48 | -0.008 | Young |
| 0.712185 | 0.106274 | 0.039 | 0.042 | 1 | 0 | Exoc8 | -0.003 | Young |
| 0.712189 | 0.092892 | 0.124 | 0.131 | 1 | 0 | Fbxl20 | -0.007 | Young |
| 0.712543 | 0.456132 | 0.01 | 0.008 | 1 | 0 | Jmjd4 | 0.002 | Young |

| | | | | | | | | |
|---|---|---|---|---|---|---|---|---|
| 0.712543 | 0.413236 | 0.01 | 0.008 | 1 | 0 | Rab26os | 0.002 | Young |
| 0.712612 | 0.17909 | 0.128 | 0.124 | 1 | 0 | Fkbp3 | 0.004 | Young |
| 0.7127 | 0.160131 | 0.102 | 0.097 | 1 | 0 | 6430548M | 0.005 | Young |
| 0.712794 | 0.346428 | 0.116 | 0.114 | 1 | 0 | Mesd | 0.002 | Young |
| 0.713032 | 0.143824 | 0.016 | 0.014 | 1 | 0 | Olfr1167 | 0.002 | Young |
| 0.713096 | 0.012357 | 0.595 | 0.61 | 1 | 0 | Dhrs3 | -0.015 | Young |
| 0.713176 | 0.289657 | 0.01 | 0.008 | 1 | 0 | Grk4 | 0.002 | Young |
| 0.713229 | 0.062689 | 0.192 | 0.185 | 1 | 0 | Krit1 | 0.007 | Young |
| 0.71326 | 0.152823 | 0.275 | 0.275 | 1 | 0 | Elk3 | 0 | Young |
| 0.713938 | 0.039115 | 0.112 | 0.118 | 1 | 0 | Mospd3 | -0.006 | Young |
| 0.713971 | 0.147205 | 0.272 | 0.275 | 1 | 0 | Pla2g15 | -0.003 | Young |
| 0.714004 | 0.200658 | 0.044 | 0.048 | 1 | 0 | Rusc2 | -0.004 | Young |
| 0.714049 | 0.116367 | 0.136 | 0.144 | 1 | 0 | Cant1 | -0.008 | Young |
| 0.714546 | 0.017886 | 0.452 | 0.451 | 1 | 0 | Tob2 | 0.001 | Young |
| 0.714605 | 0.016226 | 0.278 | 0.289 | 1 | 0 | Clint1 | -0.011 | Young |
| 0.714876 | 0.174064 | 0.094 | 0.09 | 1 | 0 | Zfp131 | 0.004 | Young |
| 0.715076 | 0.233593 | 0.187 | 0.186 | 1 | 0 | Gtf2a1 | 0.001 | Young |
| 0.715078 | 0.308901 | 0.01 | 0.008 | 1 | 0 | Rab3ip | 0.002 | Young |
| 0.715197 | 0.082518 | 0.038 | 0.041 | 1 | 0 | Polr2h | -0.003 | Young |
| 0.715244 | 0.183549 | 0.163 | 0.16 | 1 | 0 | Samd9l | 0.003 | Young |
| 0.715324 | 0.129839 | 0.05 | 0.047 | 1 | 0 | Nfs1 | 0.003 | Young |
| 0.715501 | 0.28971 | 0.01 | 0.008 | 1 | 0 | Mpzl1 | 0.002 | Young |
| 0.715811 | 0.143131 | 0.131 | 0.127 | 1 | 0 | Spty2d1 | 0.004 | Young |
| 0.716 | 0.042029 | 0.278 | 0.291 | 1 | 0 | Sec11c | -0.013 | Young |
| 0.716028 | 0.027797 | 0.355 | 0.371 | 1 | 0 | Hnrnpc | -0.016 | Young |
| 0.716299 | 0.122722 | 0.09 | 0.096 | 1 | 0 | Tmem205 | -0.006 | Young |
| 0.716471 | 0.482598 | 0.037 | 0.034 | 1 | 0 | Nol9 | 0.003 | Young |
| 0.716724 | 0.113834 | 0.126 | 0.122 | 1 | 0 | Tbc1d20 | 0.004 | Young |
| 0.7168 | 0.143524 | 0.068 | 0.073 | 1 | 0 | Fam107b | -0.005 | Young |
| 0.716832 | 0.207264 | 0.071 | 0.067 | 1 | 0 | Ado | 0.004 | Young |
| 0.716981 | 0.1016 | 0.01 | 0.008 | 1 | 0 | Ajuba | 0.002 | Young |
| 0.717209 | 0.032228 | 0.05 | 0.047 | 1 | 0 | Slc35b4 | 0.003 | Young |
| 0.717239 | 0.266306 | 0.064 | 0.06 | 1 | 0 | Diablo | 0.004 | Young |
| 0.71724 | 0.226536 | 0.074 | 0.071 | 1 | 0 | Iars2 | 0.003 | Young |
| 0.71725 | 0.11207 | 0.037 | 0.034 | 1 | 0 | Pgghg | 0.003 | Young |
| 0.717339 | 0.088096 | 0.372 | 0.373 | 1 | 0 | St8sia4 | -0.001 | Young |
| 0.717649 | 0.10874 | 0.058 | 0.062 | 1 | 0 | Pbdc1 | -0.004 | Young |
| 0.717694 | 0.194267 | 0.023 | 0.021 | 1 | 0 | Fam210b | 0.002 | Young |
| 0.718018 | 0.086295 | 0.105 | 0.111 | 1 | 0 | Bag3 | -0.006 | Young |
| 0.718185 | 0.132224 | 0.134 | 0.13 | 1 | 0 | Arih2 | 0.004 | Young |
| 0.718358 | 0.200095 | 0.034 | 0.031 | 1 | 0 | 2810402E2 | 0.003 | Young |
| 0.718387 | 0.072195 | 0.013 | 0.011 | 1 | 0 | Slc37a1 | 0.002 | Young |
| 0.718446 | 0.012665 | 0.134 | 0.127 | 1 | 0 | Siglece | 0.007 | Young |
| 0.719082 | 0.118074 | 0.099 | 0.105 | 1 | 0 | Inpp5f | -0.006 | Young |
| 0.719322 | 0.079289 | 0.496 | 0.492 | 1 | 0 | Jak1 | 0.004 | Young |

| | | | | | | | | |
|---|---|---|---|---|---|---|---|---|
| 0.719799 | 0.048551 | 0.056 | 0.06 | 1 | 0 | Traf6 | -0.004 | Young |
| 0.719805 | 0.023971 | 0.051 | 0.048 | 1 | 0 | Ngrn | 0.003 | Young |
| 0.720137 | 0.026452 | 0.16 | 0.168 | 1 | 0 | Eno1 | -0.008 | Young |
| 0.720184 | 0.180112 | 0.039 | 0.042 | 1 | 0 | Exd2 | -0.003 | Young |
| 0.720682 | 0.34806 | 0.044 | 0.041 | 1 | 0 | Utp18 | 0.003 | Young |
| 0.720752 | 0.197893 | 0.121 | 0.129 | 1 | 0 | Nudc | -0.008 | Young |
| 0.720787 | 0.057193 | 0.323 | 0.342 | 1 | 0 | Chmp4b | -0.019 | Young |
| 0.720836 | 0.331291 | 0.069 | 0.066 | 1 | 0 | Trp53inp2 | 0.003 | Young |
| 0.721232 | 0.163422 | 0.044 | 0.041 | 1 | 0 | Cenpc1 | 0.003 | Young |
| 0.721235 | 0.180041 | 0.09 | 0.086 | 1 | 0 | Glo1 | 0.004 | Young |
| 0.721351 | 0.175784 | 0.319 | 0.326 | 1 | 0 | Sdha | -0.007 | Young |
| 0.721426 | 0.097261 | 0.038 | 0.036 | 1 | 0 | Tsta3 | 0.002 | Young |
| 0.721508 | 0.142854 | 0.061 | 0.066 | 1 | 0 | C2cd3 | -0.005 | Young |
| 0.721617 | 0.739879 | 0.017 | 0.015 | 1 | 0 | Cenpv | 0.002 | Young |
| 0.721806 | 0.115433 | 0.09 | 0.086 | 1 | 0 | Brd9 | 0.004 | Young |
| 0.722309 | 0.088813 | 0.112 | 0.108 | 1 | 0 | Zyg11b | 0.004 | Young |
| 0.722502 | 0.114518 | 0.089 | 0.094 | 1 | 0 | Smim19 | -0.005 | Young |
| 0.72269 | 0.324411 | 0.029 | 0.031 | 1 | 0 | Trim21 | -0.002 | Young |
| 0.722977 | 0.139021 | 0.174 | 0.171 | 1 | 0 | Max | 0.003 | Young |
| 0.723068 | 0.262518 | 0.027 | 0.03 | 1 | 0 | 1300002E1 | -0.003 | Young |
| 0.723997 | 0.035497 | 0.202 | 0.209 | 1 | 0 | Ubn1 | -0.007 | Young |
| 0.72427 | 0.125613 | 0.116 | 0.112 | 1 | 0 | Mrpl36 | 0.004 | Young |
| 0.724459 | 0.229638 | 0.091 | 0.088 | 1 | 0 | Malt1 | 0.003 | Young |
| 0.724477 | 0.168951 | 0.096 | 0.093 | 1 | 0 | Fam53b | 0.003 | Young |
| 0.724519 | 0.152698 | 0.122 | 0.129 | 1 | 0 | P4ha1 | -0.007 | Young |
| 0.724524 | 0.155055 | 0.055 | 0.059 | 1 | 0 | Aktip | -0.004 | Young |
| 0.724663 | 0.037979 | 0.231 | 0.226 | 1 | 0 | Arhgef1 | 0.005 | Young |
| 0.724744 | 0.1161 | 0.091 | 0.088 | 1 | 0 | Mta1 | 0.003 | Young |
| 0.724771 | 0.27568 | 0.065 | 0.07 | 1 | 0 | Btbd9 | -0.005 | Young |
| 0.724798 | 0.231076 | 0.027 | 0.025 | 1 | 0 | Dcaf6 | 0.002 | Young |
| 0.725198 | 0.186794 | 0.067 | 0.071 | 1 | 0 | Pmm2 | -0.004 | Young |
| 0.725302 | 0.053659 | 0.249 | 0.245 | 1 | 0 | Klhl24 | 0.004 | Young |
| 0.72544 | 0.071097 | 0.366 | 0.386 | 1 | 0 | Bnip2 | -0.02 | Young |
| 0.7259 | 0.225063 | 0.047 | 0.044 | 1 | 0 | Msh2 | 0.003 | Young |
| 0.726528 | 0.071269 | 0.333 | 0.328 | 1 | 0 | Zcchc24 | 0.005 | Young |
| 0.726612 | 0.037203 | 0.078 | 0.082 | 1 | 0 | Selenom | -0.004 | Young |
| 0.727022 | 0.053156 | 0.087 | 0.092 | 1 | 0 | D730003I1 | -0.005 | Young |
| 0.727128 | 0.078098 | 0.056 | 0.06 | 1 | 0 | Klhl20 | -0.004 | Young |
| 0.727163 | 0.123345 | 0.051 | 0.048 | 1 | 0 | Rnaseh2a | 0.003 | Young |
| 0.727532 | 0.139763 | 0.088 | 0.093 | 1 | 0 | Myo18a | -0.005 | Young |
| 0.727675 | 0.180751 | 0.203 | 0.201 | 1 | 0 | Mpp6 | 0.002 | Young |
| 0.727688 | 0.146022 | 0.111 | 0.118 | 1 | 0 | Uxs1 | -0.007 | Young |
| 0.727951 | 0.38629 | 0.052 | 0.056 | 1 | 0 | Abcf2 | -0.004 | Young |
| 0.728306 | 0.065378 | 0.034 | 0.037 | 1 | 0 | Scap | -0.003 | Young |
| 0.728619 | 0.078623 | 0.402 | 0.415 | 1 | 0 | Hnrnpm | -0.013 | Young |

| | | | | | | | | |
|---|---|---|---|---|---|---|---|---|
| 0.728667 | 0.051348 | 0.238 | 0.233 | 1 | 0 | Snx18 | 0.005 | Young |
| 0.72869 | 0.395681 | 0.017 | 0.015 | 1 | 0 | Zrsr1 | 0.002 | Young |
| 0.728781 | 0.010173 | 0.169 | 0.176 | 1 | 0 | Heatr5a | -0.007 | Young |
| 0.728851 | 0.491535 | 0.017 | 0.015 | 1 | 0 | Flad1 | 0.002 | Young |
| 0.728999 | 0.233729 | 0.048 | 0.045 | 1 | 0 | Etfrf1 | 0.003 | Young |
| 0.729055 | 0.239719 | 0.02 | 0.018 | 1 | 0 | Gm20045 | 0.002 | Young |
| 0.729217 | 0.031471 | 0.106 | 0.101 | 1 | 0 | Ccnh | 0.005 | Young |
| 0.729703 | 0.371596 | 0.024 | 0.026 | 1 | 0 | Nit2 | -0.002 | Young |
| 0.730102 | 0.096726 | 0.066 | 0.07 | 1 | 0 | 2210016L2 | -0.004 | Young |
| 0.730259 | 0.059025 | 0.199 | 0.209 | 1 | 0 | Ncstn | -0.01 | Young |
| 0.730367 | 0.365594 | 0.094 | 0.092 | 1 | 0 | Prdx4 | 0.002 | Young |
| 0.730462 | 0.540648 | 0.017 | 0.015 | 1 | 0 | Zfp825 | 0.002 | Young |
| 0.730481 | 0.272187 | 0.085 | 0.082 | 1 | 0 | Rab6b | 0.003 | Young |
| 0.730539 | 0.161722 | 0.027 | 0.025 | 1 | 0 | Ahdc1 | 0.002 | Young |
| 0.730543 | 0.114271 | 0.259 | 0.259 | 1 | 0 | Ost4 | 0 | Young |
| 0.730843 | 0.290808 | 0.056 | 0.06 | 1 | 0 | Tbc1d31 | -0.004 | Young |
| 0.730889 | 0.904834 | 0.014 | 0.012 | 1 | 0 | Dpy19l3 | 0.002 | Young |
| 0.731345 | 0.054085 | 0.069 | 0.066 | 1 | 0 | Ppp1r37 | 0.003 | Young |
| 0.731358 | 0.030064 | 0.03 | 0.027 | 1 | 0 | Nsun3 | 0.003 | Young |
| 0.731543 | 0.056664 | 0.397 | 0.399 | 1 | 0 | Dnajc3 | -0.002 | Young |
| 0.731747 | 0.014548 | 0.169 | 0.175 | 1 | 0 | Dctn4 | -0.006 | Young |
| 0.732137 | 0.048656 | 0.249 | 0.245 | 1 | 0 | Orai1 | 0.004 | Young |
| 0.732151 | 0.067146 | 0.146 | 0.141 | 1 | 0 | Ube2q2 | 0.005 | Young |
| 0.732571 | 0.070878 | 0.298 | 0.311 | 1 | 0 | Ythdc1 | -0.013 | Young |
| 0.732965 | 0.446998 | 0.063 | 0.06 | 1 | 0 | Ppp5c | 0.003 | Young |
| 0.733134 | 0.003019 | 0.469 | 0.49 | 1 | 0 | Wdr1 | -0.021 | Young |
| 0.733163 | 0.028689 | 0.083 | 0.088 | 1 | 0 | Nmd3 | -0.005 | Young |
| 0.733315 | 0.228346 | 0.097 | 0.094 | 1 | 0 | Chm | 0.003 | Young |
| 0.73335 | 0.090875 | 0.033 | 0.036 | 1 | 0 | Zmym3 | -0.003 | Young |
| 0.733382 | 0.029817 | 0.055 | 0.059 | 1 | 0 | Rnf41 | -0.004 | Young |
| 0.733718 | 0.058764 | 0.116 | 0.122 | 1 | 0 | Plxna1 | -0.006 | Young |
| 0.733831 | 0.078289 | 0.038 | 0.036 | 1 | 0 | Zbtb41 | 0.002 | Young |
| 0.733903 | 0.111889 | 0.152 | 0.16 | 1 | 0 | Ube2e1 | -0.008 | Young |
| 0.733977 | 0.099889 | 0.033 | 0.036 | 1 | 0 | Gpatch11 | -0.003 | Young |
| 0.734 | 0.114531 | 0.035 | 0.038 | 1 | 0 | Rsad2 | -0.003 | Young |
| 0.734223 | 0.322984 | 0.092 | 0.089 | 1 | 0 | Hk3 | 0.003 | Young |
| 0.734308 | 0.110828 | 0.297 | 0.316 | 1 | 0 | Paip2 | -0.019 | Young |
| 0.734849 | 0.155347 | 0.02 | 0.022 | 1 | 0 | Gstcd | -0.002 | Young |
| 0.734937 | 0.108592 | 0.361 | 0.368 | 1 | 0 | Myh9 | -0.007 | Young |
| 0.734975 | 0.276252 | 0.028 | 0.026 | 1 | 0 | Trim2 | 0.002 | Young |
| 0.73529 | 0.230596 | 0.021 | 0.023 | 1 | 0 | Mocs1 | -0.002 | Young |
| 0.735408 | 0.133409 | 0.034 | 0.037 | 1 | 0 | Asb3 | -0.003 | Young |
| 0.735447 | 0.000601 | 0.038 | 0.036 | 1 | 0 | Mphosph1 | 0.002 | Young |
| 0.735589 | 0.099186 | 0.134 | 0.13 | 1 | 0 | Smu1 | 0.004 | Young |
| 0.735612 | 0.213559 | 0.022 | 0.025 | 1 | 0 | Dusp18 | -0.003 | Young |

| | | | | | | | | |
|---|---|---|---|---|---|---|---|---|
| 0.735679 | 0.289779 | 0.118 | 0.115 | 1 | 0 | Rab3gap1 | 0.003 | Young |
| 0.735847 | 0.012357 | 0.061 | 0.064 | 1 | 0 | Arl4a | -0.003 | Young |
| 0.736101 | 0.004655 | 0.017 | 0.019 | 1 | 0 | Rabl3 | -0.002 | Young |
| 0.736109 | 0.306769 | 0.017 | 0.015 | 1 | 0 | Prr5l | 0.002 | Young |
| 0.736297 | 0.275201 | 0.095 | 0.101 | 1 | 0 | Cyb5r1 | -0.006 | Young |
| 0.736601 | 0.061516 | 0.047 | 0.044 | 1 | 0 | Mkl1 | 0.003 | Young |
| 0.736874 | 0.132587 | 0.031 | 0.029 | 1 | 0 | Utp23 | 0.002 | Young |
| 0.737049 | 0.093187 | 0.042 | 0.045 | 1 | 0 | Metap1d | -0.003 | Young |
| 0.737216 | 0.046265 | 0.39 | 0.413 | 1 | 0 | Kdm6b | -0.023 | Young |
| 0.737333 | 0.13008 | 0.33 | 0.331 | 1 | 0 | Zdhhc20 | -0.001 | Young |
| 0.737561 | 0.111097 | 0.019 | 0.021 | 1 | 0 | Paip2b | -0.002 | Young |
| 0.737693 | 0.13642 | 0.355 | 0.36 | 1 | 0 | Gm21188 | -0.005 | Young |
| 0.737862 | 0.05473 | 0.045 | 0.042 | 1 | 0 | Yaf2 | 0.003 | Young |
| 0.737935 | 0.213021 | 0.085 | 0.09 | 1 | 0 | Stx5a | -0.005 | Young |
| 0.737975 | 0.590917 | 0.059 | 0.056 | 1 | 0 | Chn2 | 0.003 | Young |
| 0.738072 | 0.13849 | 0.085 | 0.09 | 1 | 0 | Crtc2 | -0.005 | Young |
| 0.738706 | 0.596873 | 0.014 | 0.012 | 1 | 0 | Coil | 0.002 | Young |
| 0.739044 | 0.01282 | 0.134 | 0.129 | 1 | 0 | Arhgap1 | 0.005 | Young |
| 0.739237 | 0.034624 | 0.139 | 0.145 | 1 | 0 | Plekha1 | -0.006 | Young |
| 0.739366 | 0.309191 | 0.049 | 0.047 | 1 | 0 | Bap1 | 0.002 | Young |
| 0.739522 | 0.034063 | 0.03 | 0.033 | 1 | 0 | Slc25a22 | -0.003 | Young |
| 0.739774 | 0.403433 | 0.014 | 0.012 | 1 | 0 | Gm43936 | 0.002 | Young |
| 0.740058 | 0.107072 | 0.079 | 0.075 | 1 | 0 | Ccdc174 | 0.004 | Young |
| 0.740169 | 0.086852 | 0.287 | 0.286 | 1 | 0 | Cab39 | 0.001 | Young |
| 0.740897 | 0.091328 | 0.109 | 0.105 | 1 | 0 | Pank2 | 0.004 | Young |
| 0.740988 | 0.189241 | 0.076 | 0.073 | 1 | 0 | Rif1 | 0.003 | Young |
| 0.74108 | 0.150898 | 0.024 | 0.022 | 1 | 0 | Top3a | 0.002 | Young |
| 0.741115 | 0.113364 | 0.032 | 0.034 | 1 | 0 | Qsox2 | -0.002 | Young |
| 0.741165 | 0.107644 | 0.152 | 0.148 | 1 | 0 | Ptbp1 | 0.004 | Young |
| 0.741208 | 0.150737 | 0.045 | 0.048 | 1 | 0 | Inpp5a | -0.003 | Young |
| 0.741513 | 0.385065 | 0.021 | 0.019 | 1 | 0 | Dnal4 | 0.002 | Young |
| 0.741661 | 0.267488 | 0.049 | 0.047 | 1 | 0 | Prrg2 | 0.002 | Young |
| 0.742082 | 0.085695 | 0.198 | 0.208 | 1 | 0 | Cnih1 | -0.01 | Young |
| 0.742222 | 0.201509 | 0.191 | 0.19 | 1 | 0 | Bmpr2 | 0.001 | Young |
| 0.742278 | 0.065229 | 0.042 | 0.045 | 1 | 0 | Arl14ep | -0.003 | Young |
| 0.742312 | 0.429005 | 0.057 | 0.055 | 1 | 0 | Tmem198b | 0.002 | Young |
| 0.742366 | 0.088078 | 0.028 | 0.026 | 1 | 0 | Pck2 | 0.002 | Young |
| 0.742418 | 0.133976 | 0.128 | 0.124 | 1 | 0 | Strn4 | 0.004 | Young |
| 0.742603 | 0.366578 | 0.052 | 0.049 | 1 | 0 | Carnmt1 | 0.003 | Young |
| 0.74267 | 0.156497 | 0.055 | 0.059 | 1 | 0 | Tbc1d24 | -0.004 | Young |
| 0.742721 | 0.167396 | 0.04 | 0.037 | 1 | 0 | Git1 | 0.003 | Young |
| 0.74321 | 0.203674 | 0.094 | 0.092 | 1 | 0 | Sec23ip | 0.002 | Young |
| 0.743241 | 0.335472 | 0.134 | 0.133 | 1 | 0 | Cdk14 | 0.001 | Young |
| 0.74327 | 0.139685 | 0.131 | 0.127 | 1 | 0 | Vps41 | 0.004 | Young |
| 0.743574 | 0.059929 | 0.163 | 0.17 | 1 | 0 | Mtm1 | -0.007 | Young |

| | | | | | | | | |
|---|---|---|---|---|---|---|---|---|
| 0.743875 | 0.106085 | 0.029 | 0.031 | 1 | 0 | Cables2 | -0.002 | Young |
| 0.744054 | 0.689949 | 0.018 | 0.016 | 1 | 0 | Abcc4 | 0.002 | Young |
| 0.744203 | 0.135815 | 0.123 | 0.13 | 1 | 0 | Camta2 | -0.007 | Young |
| 0.744238 | 0.098131 | 0.323 | 0.327 | 1 | 0 | Sfr1 | -0.004 | Young |
| 0.744614 | 0.089235 | 0.032 | 0.034 | 1 | 0 | Rmnd5b | -0.002 | Young |
| 0.744636 | 0.041529 | 0.599 | 0.61 | 1 | 0 | Ddx17 | -0.011 | Young |
| 0.744758 | 0.50154 | 0.011 | 0.01 | 1 | 0 | Adgrg1 | 0.001 | Young |
| 0.744977 | 0.151054 | 0.15 | 0.146 | 1 | 0 | Psmd6 | 0.004 | Young |
| 0.74505 | 0.061217 | 0.108 | 0.104 | 1 | 0 | Flcn | 0.004 | Young |
| 0.745122 | 0.214448 | 0.014 | 0.012 | 1 | 0 | Snap47 | 0.002 | Young |
| 0.745396 | 0.029492 | 0.154 | 0.16 | 1 | 0 | Mrpl20 | -0.006 | Young |
| 0.745472 | 0.152542 | 0.3 | 0.306 | 1 | 0 | Ndufb7 | -0.006 | Young |
| 0.74548 | 0.047971 | 0.074 | 0.07 | 1 | 0 | Snx15 | 0.004 | Young |
| 0.745562 | 0.268764 | 0.011 | 0.01 | 1 | 0 | Trim59 | 0.001 | Young |
| 0.745764 | 0.417421 | 0.011 | 0.01 | 1 | 0 | Tcaim | 0.001 | Young |
| 0.745787 | 0.060246 | 0.214 | 0.209 | 1 | 0 | Snx30 | 0.005 | Young |
| 0.745938 | 0.618043 | 0.02 | 0.022 | 1 | 0 | Nedd4l | -0.002 | Young |
| 0.745986 | 0.065722 | 0.016 | 0.018 | 1 | 0 | Ms4a4a | -0.002 | Young |
| 0.746028 | 0.216645 | 0.102 | 0.108 | 1 | 0 | Npm3 | -0.006 | Young |
| 0.746083 | 0.102847 | 0.119 | 0.115 | 1 | 0 | Dnajc10 | 0.004 | Young |
| 0.746205 | 0.121618 | 0.184 | 0.194 | 1 | 0 | Ythdf3 | -0.01 | Young |
| 0.746246 | 0.052479 | 0.028 | 0.03 | 1 | 0 | Med9 | -0.002 | Young |
| 0.746367 | 0.388252 | 0.011 | 0.01 | 1 | 0 | 1110046J0 | 0.001 | Young |
| 0.746568 | 0.666754 | 0.011 | 0.01 | 1 | 0 | Wdr19 | 0.001 | Young |
| 0.746589 | 0.329655 | 0.021 | 0.019 | 1 | 0 | E130311K1 | 0.002 | Young |
| 0.746693 | 0.061868 | 0.229 | 0.239 | 1 | 0 | Psma6 | -0.01 | Young |
| 0.746693 | 0.119659 | 0.113 | 0.109 | 1 | 0 | Pdap1 | 0.004 | Young |
| 0.747184 | 0.388222 | 0.029 | 0.027 | 1 | 0 | Ghdc | 0.002 | Young |
| 0.74735 | 0.213537 | 0.037 | 0.034 | 1 | 0 | Snx16 | 0.003 | Young |
| 0.747476 | 0.160245 | 0.063 | 0.06 | 1 | 0 | Jade2 | 0.003 | Young |
| 0.747823 | 0.270762 | 0.021 | 0.019 | 1 | 0 | Pomt2 | 0.002 | Young |
| 0.747854 | 0.143649 | 0.118 | 0.124 | 1 | 0 | Chchd1 | -0.006 | Young |
| 0.747979 | 0.05137 | 0.372 | 0.398 | 1 | 0 | Nucks1 | -0.026 | Young |
| 0.748029 | 0.104118 | 0.085 | 0.082 | 1 | 0 | Ap5m1 | 0.003 | Young |
| 0.74844 | 0.180815 | 0.062 | 0.059 | 1 | 0 | Cops7a | 0.003 | Young |
| 0.748497 | 0.205082 | 0.084 | 0.081 | 1 | 0 | Dusp2 | 0.003 | Young |
| 0.748583 | 0.167662 | 0.016 | 0.018 | 1 | 0 | Casp7 | -0.002 | Young |
| 0.748652 | 0.113757 | 0.126 | 0.133 | 1 | 0 | Cul5 | -0.007 | Young |
| 0.748995 | 0.022921 | 0.13 | 0.135 | 1 | 0 | Tnfaip8l2 | -0.005 | Young |
| 0.749141 | 0.031627 | 0.069 | 0.073 | 1 | 0 | Tmem120a | -0.004 | Young |
| 0.749338 | 0.053391 | 0.064 | 0.06 | 1 | 0 | Brf1 | 0.004 | Young |
| 0.749599 | 0.041676 | 0.551 | 0.59 | 1 | 0 | Npm1 | -0.039 | Young |
| 0.749825 | 0.134283 | 0.025 | 0.023 | 1 | 0 | Krt10 | 0.002 | Young |
| 0.749991 | 0.311302 | 0.011 | 0.01 | 1 | 0 | Cdiptos | 0.001 | Young |
| 0.75011 | 0.057085 | 0.407 | 0.434 | 1 | 0 | Scpep1 | -0.027 | Young |

| | | | | | | | | |
|---|---|---|---|---|---|---|---|---|
| 0.750137 | 0.411857 | 0.029 | 0.027 | 1 | 0 | Rft1 | 0.002 | Young |
| 0.750154 | 0.311974 | 0.018 | 0.016 | 1 | 0 | Clec16a | 0.002 | Young |
| 0.750205 | 0.074856 | 0.015 | 0.016 | 1 | 0 | Fancc | -0.001 | Young |
| 0.750259 | 0.196788 | 0.054 | 0.057 | 1 | 0 | Cpox | -0.003 | Young |
| 0.750355 | 0.169368 | 0.237 | 0.237 | 1 | 0 | Clcn5 | 0 | Young |
| 0.750374 | 0.055233 | 0.015 | 0.016 | 1 | 0 | Fam131a | -0.001 | Young |
| 0.750389 | 0.060121 | 0.114 | 0.109 | 1 | 0 | Zfp422 | 0.005 | Young |
| 0.750399 | 0.142294 | 0.032 | 0.03 | 1 | 0 | Tars2 | 0.002 | Young |
| 0.750563 | 0.00147 | 0.12 | 0.126 | 1 | 0 | Grk6 | -0.006 | Young |
| 0.750681 | 0.143785 | 0.34 | 0.347 | 1 | 0 | Prkacb | -0.007 | Young |
| 0.750712 | 0.253817 | 0.107 | 0.104 | 1 | 0 | Arhgap11a | 0.003 | Young |
| 0.750842 | 0.080392 | 0.09 | 0.094 | 1 | 0 | Med10 | -0.004 | Young |
| 0.751007 | 0.137019 | 0.128 | 0.124 | 1 | 0 | Msl3 | 0.004 | Young |
| 0.751038 | 0.444002 | 0.039 | 0.037 | 1 | 0 | Ccdc112 | 0.002 | Young |
| 0.751123 | 0.388537 | 0.029 | 0.027 | 1 | 0 | Cdc25b | 0.002 | Young |
| 0.75121 | 0.10278 | 0.391 | 0.399 | 1 | 0 | Mindy2 | -0.008 | Young |
| 0.751314 | 0.159902 | 0.063 | 0.06 | 1 | 0 | Kdm5b | 0.003 | Young |
| 0.751487 | 0.06043 | 0.135 | 0.141 | 1 | 0 | Clec2i | -0.006 | Young |
| 0.751506 | 0.088579 | 0.224 | 0.22 | 1 | 0 | Sbds | 0.004 | Young |
| 0.751746 | 0.101588 | 0.162 | 0.17 | 1 | 0 | Ankhd1 | -0.008 | Young |
| 0.751803 | 0.153471 | 0.053 | 0.051 | 1 | 0 | Zfp942 | 0.002 | Young |
| 0.751961 | 0.038921 | 0.085 | 0.089 | 1 | 0 | Ddx10 | -0.004 | Young |
| 0.752267 | 0.270254 | 0.046 | 0.044 | 1 | 0 | Khk | 0.002 | Young |
| 0.75235 | 0.257196 | 0.025 | 0.023 | 1 | 0 | Dusp12 | 0.002 | Young |
| 0.752368 | 0.048416 | 0.12 | 0.116 | 1 | 0 | Nfkbie | 0.004 | Young |
| 0.752505 | 0.226114 | 0.018 | 0.016 | 1 | 0 | A130010J1 | 0.002 | Young |
| 0.752662 | 0.438818 | 0.018 | 0.016 | 1 | 0 | Klhl36 | 0.002 | Young |
| 0.752738 | 0.021081 | 0.215 | 0.223 | 1 | 0 | Thrap3 | -0.008 | Young |
| 0.753083 | 0.130858 | 0.015 | 0.016 | 1 | 0 | Tmtc4 | -0.001 | Young |
| 0.753083 | 0.001295 | 0.015 | 0.016 | 1 | 0 | Tldc1 | -0.001 | Young |
| 0.753153 | 0.018863 | 0.028 | 0.03 | 1 | 0 | Osbpl7 | -0.002 | Young |
| 0.753183 | 0.376509 | 0.052 | 0.056 | 1 | 0 | Pigu | -0.004 | Young |
| 0.753299 | 0.119215 | 0.127 | 0.123 | 1 | 0 | Bbx | 0.004 | Young |
| 0.753329 | 0.102264 | 0.069 | 0.066 | 1 | 0 | Dbt | 0.003 | Young |
| 0.753341 | 0.237547 | 0.029 | 0.027 | 1 | 0 | Toe1 | 0.002 | Young |
| 0.753591 | 0.244592 | 0.015 | 0.016 | 1 | 0 | Pold1 | -0.001 | Young |
| 0.753839 | 0.019634 | 0.195 | 0.19 | 1 | 0 | Etfb | 0.005 | Young |
| 0.753887 | 0.135213 | 0.156 | 0.166 | 1 | 0 | Mcmbp | -0.01 | Young |
| 0.754154 | 0.110252 | 0.196 | 0.193 | 1 | 0 | Ndufs7 | 0.003 | Young |
| 0.754171 | 0.066225 | 0.062 | 0.059 | 1 | 0 | 3110062M | 0.003 | Young |
| 0.754338 | 0.09689 | 0.028 | 0.03 | 1 | 0 | Cc2d1a | -0.002 | Young |
| 0.754345 | 0.144768 | 0.053 | 0.051 | 1 | 0 | Asnsd1 | 0.002 | Young |
| 0.754364 | 0.201242 | 0.169 | 0.167 | 1 | 0 | Rala | 0.002 | Young |
| 0.754439 | 0.14404 | 0.015 | 0.016 | 1 | 0 | Lrsam1 | -0.001 | Young |
| 0.755038 | 0.037401 | 0.013 | 0.015 | 1 | 0 | A130077B1 | -0.002 | Young |

| | | | | | | | | |
|---|---|---|---|---|---|---|---|---|
| 0.755235 | 0.014662 | 0.011 | 0.01 | 1 | 0 | Enc1 | 0.001 | Young |
| 0.755522 | 0.043334 | 0.304 | 0.319 | 1 | 0 | Cd86 | -0.015 | Young |
| 0.75566 | 0.488839 | 0.027 | 0.025 | 1 | 0 | Nsun4 | 0.002 | Young |
| 0.756071 | 0.177307 | 0.121 | 0.118 | 1 | 0 | Rbck1 | 0.003 | Young |
| 0.756187 | 0.231176 | 0.071 | 0.075 | 1 | 0 | Slc35e3 | -0.004 | Young |
| 0.756353 | 0.027577 | 0.353 | 0.364 | 1 | 0 | Dync1i2 | -0.011 | Young |
| 0.756442 | 0.021141 | 0.012 | 0.014 | 1 | 0 | Gk | -0.002 | Young |
| 0.756473 | 0.002828 | 0.323 | 0.335 | 1 | 0 | Uqcrb | -0.012 | Young |
| 0.75658 | 0.137885 | 0.382 | 0.399 | 1 | 0 | Tmem59 | -0.017 | Young |
| 0.756601 | 0.177251 | 0.146 | 0.145 | 1 | 0 | Kidins220 | 0.001 | Young |
| 0.757218 | 0.257439 | 0.159 | 0.159 | 1 | 0 | Jtb | 0 | Young |
| 0.757264 | 0.328057 | 0.103 | 0.101 | 1 | 0 | Tlk1 | 0.002 | Young |
| 0.757302 | 0.400641 | 0.072 | 0.07 | 1 | 0 | Ccdc127 | 0.002 | Young |
| 0.757578 | 0.040308 | 0.253 | 0.265 | 1 | 0 | Dgkd | -0.012 | Young |
| 0.757593 | 0.428165 | 0.083 | 0.081 | 1 | 0 | Adk | 0.002 | Young |
| 0.757637 | 0.129193 | 0.113 | 0.119 | 1 | 0 | Cryl1 | -0.006 | Young |
| 0.757805 | 0.020499 | 0.317 | 0.335 | 1 | 0 | Crebbp | -0.018 | Young |
| 0.758036 | 0.064183 | 0.664 | 0.679 | 1 | 0 | Tpm3 | -0.015 | Young |
| 0.758101 | 0.076268 | 0.106 | 0.111 | 1 | 0 | Tmem50b | -0.005 | Young |
| 0.758127 | 0.225165 | 0.118 | 0.115 | 1 | 0 | Gab2 | 0.003 | Young |
| 0.758221 | 0.127241 | 0.185 | 0.196 | 1 | 0 | Mbtd1 | -0.011 | Young |
| 0.758221 | 0.202438 | 0.041 | 0.044 | 1 | 0 | Rgp1 | -0.003 | Young |
| 0.758437 | 0.170328 | 0.251 | 0.25 | 1 | 0 | Stard3nl | 0.001 | Young |
| 0.75871 | 0.079321 | 0.12 | 0.116 | 1 | 0 | Ago4 | 0.004 | Young |
| 0.758854 | 0.039859 | 0.277 | 0.272 | 1 | 0 | Efr3a | 0.005 | Young |
| 0.758859 | 0.129708 | 0.066 | 0.07 | 1 | 0 | Necap1 | -0.004 | Young |
| 0.75904 | 0.379448 | 0.027 | 0.025 | 1 | 0 | Tfip11 | 0.002 | Young |
| 0.759062 | 0.003326 | 0.101 | 0.105 | 1 | 0 | Spire1 | -0.004 | Young |
| 0.759174 | 0.465613 | 0.031 | 0.029 | 1 | 0 | Gpatch1 | 0.002 | Young |
| 0.759374 | 0.127626 | 0.118 | 0.115 | 1 | 0 | Rnf115 | 0.003 | Young |
| 0.75953 | 0.204408 | 0.015 | 0.016 | 1 | 0 | Zswim9 | -0.001 | Young |
| 0.759551 | 0.26551 | 0.034 | 0.031 | 1 | 0 | Ofd1 | 0.003 | Young |
| 0.759606 | 0.045208 | 0.012 | 0.014 | 1 | 0 | Gnb4 | -0.002 | Young |
| 0.759885 | 0.494875 | 0.017 | 0.019 | 1 | 0 | Cryz | -0.002 | Young |
| 0.760005 | 0.108491 | 0.027 | 0.029 | 1 | 0 | Dhcr7 | -0.002 | Young |
| 0.760292 | 0.074673 | 0.092 | 0.089 | 1 | 0 | Ilf2 | 0.003 | Young |
| 0.760351 | 0.081991 | 0.012 | 0.014 | 1 | 0 | Them4 | -0.002 | Young |
| 0.760462 | 0.057811 | 0.138 | 0.134 | 1 | 0 | Gcnt1 | 0.004 | Young |
| 0.760478 | 0.137684 | 0.086 | 0.082 | 1 | 0 | Tmem251 | 0.004 | Young |
| 0.760921 | 0.271943 | 0.039 | 0.037 | 1 | 0 | Tacc2 | 0.002 | Young |
| 0.761296 | 0.047882 | 0.139 | 0.145 | 1 | 0 | Nsmce4a | -0.006 | Young |
| 0.761359 | 0.046412 | 0.15 | 0.155 | 1 | 0 | Flii | -0.005 | Young |
| 0.761419 | 0.263263 | 0.139 | 0.137 | 1 | 0 | Ifi27 | 0.002 | Young |
| 0.76159 | 0.079121 | 0.152 | 0.159 | 1 | 0 | Snrpd1 | -0.007 | Young |
| 0.76174 | 0.227004 | 0.029 | 0.027 | 1 | 0 | Dtd2 | 0.002 | Young |

| | | | | | | | | |
|---|---|---|---|---|---|---|---|---|
| 0.761767 | 0.141016 | 0.216 | 0.216 | 1 | 0 | Pycard | 0 | Young |
| 0.762585 | 0.060638 | 0.09 | 0.094 | 1 | 0 | Ube3c | -0.004 | Young |
| 0.762638 | 0.136838 | 0.038 | 0.036 | 1 | 0 | Timm9 | 0.002 | Young |
| 0.76296 | 0.170242 | 0.012 | 0.014 | 1 | 0 | Klhl25 | -0.002 | Young |
| 0.76332 | 0.126861 | 0.025 | 0.027 | 1 | 0 | Gtf2h4 | -0.002 | Young |
| 0.763557 | 0.229647 | 0.092 | 0.089 | 1 | 0 | Dnm1l | 0.003 | Young |
| 0.763762 | 0.438983 | 0.045 | 0.042 | 1 | 0 | Mecr | 0.003 | Young |
| 0.763893 | 0.256414 | 0.012 | 0.014 | 1 | 0 | Smim10l2a | -0.002 | Young |
| 0.763898 | 0.456027 | 0.031 | 0.029 | 1 | 0 | Miip | 0.002 | Young |
| 0.76393 | 0.129777 | 0.099 | 0.096 | 1 | 0 | Lactb | 0.003 | Young |
| 0.764044 | 0.013753 | 0.123 | 0.119 | 1 | 0 | Cklf | 0.004 | Young |
| 0.764236 | 0.154863 | 0.022 | 0.021 | 1 | 0 | Nup35 | 0.001 | Young |
| 0.764898 | 0.099706 | 0.18 | 0.179 | 1 | 0 | Slbp | 0.001 | Young |
| 0.765156 | 0.597983 | 0.013 | 0.015 | 1 | 0 | Gorab | -0.002 | Young |
| 0.765195 | 0.222052 | 0.107 | 0.104 | 1 | 0 | Ndufs1 | 0.003 | Young |
| 0.765403 | 0.218383 | 0.074 | 0.078 | 1 | 0 | Pkd1 | -0.004 | Young |
| 0.765573 | 0.502514 | 0.012 | 0.014 | 1 | 0 | Pigw | -0.002 | Young |
| 0.765612 | 0.121951 | 0.046 | 0.044 | 1 | 0 | Lias | 0.002 | Young |
| 0.765708 | 0.038676 | 0.024 | 0.026 | 1 | 0 | Zfyve19 | -0.002 | Young |
| 0.765743 | 0.11935 | 0.069 | 0.073 | 1 | 0 | Smurf1 | -0.004 | Young |
| 0.765745 | 0.055034 | 0.358 | 0.376 | 1 | 0 | Gapvd1 | -0.018 | Young |
| 0.765975 | 0.045596 | 0.379 | 0.404 | 1 | 0 | Zfp706 | -0.025 | Young |
| 0.766014 | 0.255844 | 0.093 | 0.09 | 1 | 0 | Ticam2 | 0.003 | Young |
| 0.766023 | 0.211556 | 0.126 | 0.123 | 1 | 0 | Atf2 | 0.003 | Young |
| 0.766192 | 0.168543 | 0.094 | 0.09 | 1 | 0 | Pml | 0.004 | Young |
| 0.766266 | 0.18299 | 0.072 | 0.07 | 1 | 0 | Akap11 | 0.002 | Young |
| 0.766369 | 0.118529 | 0.011 | 0.012 | 1 | 0 | Capn10 | -0.001 | Young |
| 0.766763 | 0.211444 | 0.011 | 0.012 | 1 | 0 | Kctd21 | -0.001 | Young |
| 0.766812 | 0.031498 | 0.082 | 0.086 | 1 | 0 | Tbrg1 | -0.004 | Young |
| 0.766827 | 0.135138 | 0.147 | 0.144 | 1 | 0 | Trim56 | 0.003 | Young |
| 0.766861 | 0.212838 | 0.011 | 0.012 | 1 | 0 | 1600014C1 | -0.001 | Young |
| 0.76696 | 0.197155 | 0.011 | 0.012 | 1 | 0 | Bbs9 | -0.001 | Young |
| 0.76705 | 0.07347 | 0.064 | 0.067 | 1 | 0 | Tmem185b | -0.003 | Young |
| 0.767265 | 0.123635 | 0.091 | 0.088 | 1 | 0 | Suco | 0.003 | Young |
| 0.767477 | 0.224959 | 0.031 | 0.029 | 1 | 0 | Smim24 | 0.002 | Young |
| 0.767587 | 0.164399 | 0.038 | 0.041 | 1 | 0 | Hes6 | -0.003 | Young |
| 0.767608 | 0.307243 | 0.043 | 0.047 | 1 | 0 | Tsc2 | -0.004 | Young |
| 0.767695 | 0.219171 | 0.053 | 0.056 | 1 | 0 | Lxn | -0.003 | Young |
| 0.767796 | 0.184331 | 0.23 | 0.234 | 1 | 0 | Arhgef12 | -0.004 | Young |
| 0.767859 | 0.255683 | 0.022 | 0.021 | 1 | 0 | 3110040N1 | 0.001 | Young |
| 0.767882 | 0.217251 | 0.043 | 0.041 | 1 | 0 | Fbxw5 | 0.002 | Young |
| 0.768337 | 0.051644 | 0.01 | 0.011 | 1 | 0 | Zfp799 | -0.001 | Young |
| 0.768715 | 0.123797 | 0.034 | 0.031 | 1 | 0 | Nt5dc3 | 0.003 | Young |
| 0.76883 | 0.069242 | 0.063 | 0.06 | 1 | 0 | Trappc11 | 0.003 | Young |
| 0.768878 | 0.168245 | 0.038 | 0.041 | 1 | 0 | Ercc6l2 | -0.003 | Young |

| | | | | | | | | |
|---|---|---|---|---|---|---|---|---|
| 0.768965 | 0.309437 | 0.01 | 0.011 | 1 | 0 | 8030462N1 | -0.001 | Young |
| 0.769162 | 0.327198 | 0.057 | 0.055 | 1 | 0 | Urod | 0.002 | Young |
| 0.769342 | 0.215683 | 0.015 | 0.014 | 1 | 0 | B930036N1 | 0.001 | Young |
| 0.769374 | 0.123994 | 0.147 | 0.144 | 1 | 0 | Cpsf7 | 0.003 | Young |
| 0.769523 | 0.442062 | 0.011 | 0.012 | 1 | 0 | Zfp146 | -0.001 | Young |
| 0.769584 | 0.164576 | 0.056 | 0.053 | 1 | 0 | Cisd1 | 0.003 | Young |
| 0.769683 | 0.651092 | 0.012 | 0.014 | 1 | 0 | Bcs1l | -0.002 | Young |
| 0.769785 | 0.030331 | 0.12 | 0.116 | 1 | 0 | Abce1 | 0.004 | Young |
| 0.770052 | 0.043763 | 0.034 | 0.031 | 1 | 0 | Gfer | 0.003 | Young |
| 0.770066 | 0.047215 | 0.25 | 0.259 | 1 | 0 | Hspd1 | -0.009 | Young |
| 0.770278 | 0.212343 | 0.025 | 0.027 | 1 | 0 | 1810043G0 | -0.002 | Young |
| 0.770339 | 0.211907 | 0.078 | 0.075 | 1 | 0 | Sucla2 | 0.003 | Young |
| 0.77036 | 0.228934 | 0.052 | 0.049 | 1 | 0 | Crebl2 | 0.003 | Young |
| 0.770676 | 0.45634 | 0.07 | 0.068 | 1 | 0 | Prkci | 0.002 | Young |
| 0.770763 | 0.030571 | 0.112 | 0.116 | 1 | 0 | Crem | -0.004 | Young |
| 0.770807 | 0.097221 | 0.038 | 0.036 | 1 | 0 | Gm16536 | 0.002 | Young |
| 0.771009 | 0.11862 | 0.409 | 0.421 | 1 | 0 | Cnbp | -0.012 | Young |
| 0.771096 | 0.182515 | 0.097 | 0.103 | 1 | 0 | Dnajc14 | -0.006 | Young |
| 0.771118 | 0.03969 | 0.008 | 0.01 | 1 | 0 | Zfp202 | -0.002 | Young |
| 0.771244 | 0.404342 | 0.042 | 0.04 | 1 | 0 | Gipc1 | 0.002 | Young |
| 0.771342 | 0.087067 | 0.008 | 0.01 | 1 | 0 | Zfp51 | -0.002 | Young |
| 0.772318 | 0.288728 | 0.01 | 0.011 | 1 | 0 | Gpr84 | -0.001 | Young |
| 0.772483 | 0.134028 | 0.044 | 0.047 | 1 | 0 | Desi1 | -0.003 | Young |
| 0.772586 | 0.120021 | 0.046 | 0.044 | 1 | 0 | Mbd5 | 0.002 | Young |
| 0.772967 | 0.260469 | 0.038 | 0.041 | 1 | 0 | Rapgef1 | -0.003 | Young |
| 0.773215 | 0.04914 | 0.265 | 0.261 | 1 | 0 | Zyx | 0.004 | Young |
| 0.773251 | 0.149681 | 0.2 | 0.201 | 1 | 0 | Psmd4 | -0.001 | Young |
| 0.773383 | 0.103623 | 0.107 | 0.112 | 1 | 0 | Cpq | -0.005 | Young |
| 0.773812 | 0.145531 | 0.008 | 0.01 | 1 | 0 | Zfp420 | -0.002 | Young |
| 0.774037 | 0.115213 | 0.008 | 0.01 | 1 | 0 | Mx1 | -0.002 | Young |
| 0.774314 | 0.131418 | 0.132 | 0.138 | 1 | 0 | Golgb1 | -0.006 | Young |
| 0.774394 | 0.080563 | 0.176 | 0.183 | 1 | 0 | Clock | -0.007 | Young |
| 0.774424 | 0.132685 | 0.074 | 0.078 | 1 | 0 | Sra1 | -0.004 | Young |
| 0.774517 | 0.038697 | 0.015 | 0.014 | 1 | 0 | Rbfox2 | 0.001 | Young |
| 0.775554 | 0.126304 | 0.089 | 0.086 | 1 | 0 | Rnf167 | 0.003 | Young |
| 0.775577 | 0.111602 | 0.078 | 0.082 | 1 | 0 | Tcf3 | -0.004 | Young |
| 0.775756 | 0.110879 | 0.22 | 0.218 | 1 | 0 | Psmb8 | 0.002 | Young |
| 0.775757 | 0.013122 | 0.752 | 0.763 | 1 | 0 | Rpl6 | -0.011 | Young |
| 0.775952 | 0.142428 | 0.089 | 0.086 | 1 | 0 | Proser1 | 0.003 | Young |
| 0.775995 | 0.048316 | 0.12 | 0.126 | 1 | 0 | Itpripl1 | -0.006 | Young |
| 0.776285 | 0.120362 | 0.349 | 0.358 | 1 | 0 | Tbca | -0.009 | Young |
| 0.77645 | 0.069905 | 0.078 | 0.075 | 1 | 0 | Parp9 | 0.003 | Young |
| 0.776477 | 0.075679 | 0.031 | 0.029 | 1 | 0 | Pgm2 | 0.002 | Young |
| 0.776538 | 0.222158 | 0.064 | 0.068 | 1 | 0 | 2810004N2 | -0.004 | Young |
| 0.776558 | 0.002121 | 0.061 | 0.064 | 1 | 0 | Idua | -0.003 | Young |

| | | | | | | | | |
|---|---|---|---|---|---|---|---|---|
| 0.776654 | 0.243475 | 0.012 | 0.011 | 1 | 0 | Pmm1 | 0.001 | Young |
| 0.776757 | 0.081033 | 0.331 | 0.337 | 1 | 0 | Sash1 | -0.006 | Young |
| 0.776902 | 0.026787 | 0.021 | 0.023 | 1 | 0 | Zmat1 | -0.002 | Young |
| 0.776982 | 0.242583 | 0.061 | 0.059 | 1 | 0 | Polk | 0.002 | Young |
| 0.777092 | 0.056149 | 0.069 | 0.073 | 1 | 0 | Vps52 | -0.004 | Young |
| 0.777358 | 0.129487 | 0.075 | 0.073 | 1 | 0 | Gse1 | 0.002 | Young |
| 0.777739 | 0.125784 | 0.096 | 0.093 | 1 | 0 | Plaa | 0.003 | Young |
| 0.777828 | 0.051792 | 0.153 | 0.149 | 1 | 0 | Zmym5 | 0.004 | Young |
| 0.77798 | 0.255621 | 0.024 | 0.026 | 1 | 0 | Card9 | -0.002 | Young |
| 0.777996 | 0.08974 | 0.226 | 0.226 | 1 | 0 | Uhrf2 | 0 | Young |
| 0.778162 | 0.19483 | 0.097 | 0.094 | 1 | 0 | Tmem243 | 0.003 | Young |
| 0.778378 | 0.326168 | 0.032 | 0.03 | 1 | 0 | Cep44 | 0.002 | Young |
| 0.778538 | 0.110267 | 0.11 | 0.115 | 1 | 0 | Dctn3 | -0.005 | Young |
| 0.778725 | 0.041608 | 0.152 | 0.159 | 1 | 0 | Pdcd6 | -0.007 | Young |
| 0.779304 | 0.1326 | 0.06 | 0.057 | 1 | 0 | Agl | 0.003 | Young |
| 0.779499 | 0.077181 | 0.125 | 0.13 | 1 | 0 | Lin7c | -0.005 | Young |
| 0.779752 | 0.401844 | 0.07 | 0.068 | 1 | 0 | Phka2 | 0.002 | Young |
| 0.779783 | 0.255337 | 0.061 | 0.059 | 1 | 0 | Mtx1 | 0.002 | Young |
| 0.779973 | 0.368222 | 0.024 | 0.022 | 1 | 0 | Large1 | 0.002 | Young |
| 0.780007 | 0.088307 | 0.122 | 0.127 | 1 | 0 | Csnk1g2 | -0.005 | Young |
| 0.780917 | 0.067744 | 0.199 | 0.196 | 1 | 0 | Papd4 | 0.003 | Young |
| 0.781153 | 0.386169 | 0.024 | 0.022 | 1 | 0 | Gm27010 | 0.002 | Young |
| 0.781179 | 0.003507 | 0.034 | 0.031 | 1 | 0 | C87436 | 0.003 | Young |
| 0.781217 | 0.459444 | 0.048 | 0.047 | 1 | 0 | Fbxo7 | 0.001 | Young |
| 0.781716 | 0.265633 | 0.131 | 0.13 | 1 | 0 | Plekhm3 | 0.001 | Young |
| 0.781984 | 0.249311 | 0.05 | 0.048 | 1 | 0 | Nelfb | 0.002 | Young |
| 0.782128 | 0.247286 | 0.05 | 0.048 | 1 | 0 | Champ1 | 0.002 | Young |
| 0.782464 | 0.290231 | 0.075 | 0.074 | 1 | 0 | Dusp8 | 0.001 | Young |
| 0.782719 | 0.133291 | 0.329 | 0.341 | 1 | 0 | Ube2b | -0.012 | Young |
| 0.782729 | 0.230007 | 0.058 | 0.062 | 1 | 0 | Mob2 | -0.004 | Young |
| 0.78305 | 0.156897 | 0.324 | 0.332 | 1 | 0 | Kdm2a | -0.008 | Young |
| 0.783077 | 0.047709 | 0.179 | 0.185 | 1 | 0 | Cdc5l | -0.006 | Young |
| 0.78331 | 0.03461 | 0.04 | 0.042 | 1 | 0 | Cep162 | -0.002 | Young |
| 0.783321 | 0.032323 | 0.337 | 0.352 | 1 | 0 | Gusb | -0.015 | Young |
| 0.783509 | 0.211519 | 0.118 | 0.115 | 1 | 0 | Pwwp2a | 0.003 | Young |
| 0.783633 | 0.310234 | 0.087 | 0.085 | 1 | 0 | Fam162a | 0.002 | Young |
| 0.783893 | 0.296331 | 0.057 | 0.055 | 1 | 0 | Mtx2 | 0.002 | Young |
| 0.783976 | 0.058969 | 0.012 | 0.011 | 1 | 0 | Gemin2 | 0.001 | Young |
| 0.784018 | 0.502942 | 0.016 | 0.015 | 1 | 0 | Adnp | 0.001 | Young |
| 0.78436 | 0.103847 | 0.075 | 0.079 | 1 | 0 | Milr1 | -0.004 | Young |
| 0.784575 | 0.254124 | 0.036 | 0.034 | 1 | 0 | Pag1 | 0.002 | Young |
| 0.784605 | 0.157165 | 0.049 | 0.052 | 1 | 0 | Zfyve27 | -0.003 | Young |
| 0.784759 | 0.182173 | 0.021 | 0.023 | 1 | 0 | Dbr1 | -0.002 | Young |
| 0.784979 | 0.29165 | 0.019 | 0.018 | 1 | 0 | Lmo1 | 0.001 | Young |
| 0.78504 | 0.081525 | 0.086 | 0.083 | 1 | 0 | Ssbp2 | 0.003 | Young |

| | | | | | | | | |
|---|---|---|---|---|---|---|---|---|
| 0.785114 | 0.232369 | 0.035 | 0.037 | 1 | 0 | Ppp1r3b | -0.002 | Young |
| 0.785181 | 0.229211 | 0.205 | 0.207 | 1 | 0 | Psmc3 | -0.002 | Young |
| 0.785203 | 0.021375 | 0.059 | 0.056 | 1 | 0 | Apobr | 0.003 | Young |
| 0.785268 | 0.028327 | 0.38 | 0.393 | 1 | 0 | Pbrm1 | -0.013 | Young |
| 0.785512 | 0.013314 | 0.02 | 0.022 | 1 | 0 | 4933433G1 | -0.002 | Young |
| 0.785683 | 0.311326 | 0.107 | 0.105 | 1 | 0 | Smad2 | 0.002 | Young |
| 0.785829 | 0.243719 | 0.087 | 0.085 | 1 | 0 | Lactb2 | 0.002 | Young |
| 0.786134 | 0.080434 | 0.19 | 0.186 | 1 | 0 | Dapk1 | 0.004 | Young |
| 0.786225 | 0.059023 | 0.098 | 0.103 | 1 | 0 | Pitpnb | -0.005 | Young |
| 0.786713 | 0.087446 | 0.305 | 0.323 | 1 | 0 | Atp6v0a1 | -0.018 | Young |
| 0.786963 | 0.005091 | 0.027 | 0.025 | 1 | 0 | Gm10138 | 0.002 | Young |
| 0.78706 | 0.127902 | 0.122 | 0.127 | 1 | 0 | Pigs | -0.005 | Young |
| 0.787442 | 0.069431 | 0.072 | 0.075 | 1 | 0 | Rprd1b | -0.003 | Young |
| 0.7878 | 0.031744 | 0.401 | 0.419 | 1 | 0 | Cd93 | -0.018 | Young |
| 0.787876 | 0.111022 | 0.354 | 0.363 | 1 | 0 | Wasf2 | -0.009 | Young |
| 0.78803 | 0.052575 | 0.268 | 0.264 | 1 | 0 | 2-Mar | 0.004 | Young |
| 0.788088 | 0.057976 | 0.652 | 0.67 | 1 | 0 | Arhgdib | -0.018 | Young |
| 0.788102 | 0.180169 | 0.057 | 0.06 | 1 | 0 | Sec23a | -0.003 | Young |
| 0.78827 | 0.100499 | 0.106 | 0.111 | 1 | 0 | Dars | -0.005 | Young |
| 0.788495 | 0.055711 | 0.674 | 0.689 | 1 | 0 | Gnb2 | -0.015 | Young |
| 0.78859 | 0.138269 | 0.112 | 0.109 | 1 | 0 | Map1lc3a | 0.003 | Young |
| 0.789012 | 0.20565 | 0.12 | 0.118 | 1 | 0 | Mysm1 | 0.002 | Young |
| 0.789512 | 0.437899 | 0.016 | 0.015 | 1 | 0 | Rab3a | 0.001 | Young |
| 0.789615 | 0.183821 | 0.111 | 0.116 | 1 | 0 | Tmem106c | -0.005 | Young |
| 0.789899 | 0.082852 | 0.261 | 0.272 | 1 | 0 | Rsrc2 | -0.011 | Young |
| 0.789921 | 0.162469 | 0.028 | 0.026 | 1 | 0 | Cep104 | 0.002 | Young |
| 0.79035 | 0.134662 | 0.149 | 0.146 | 1 | 0 | Slc25a51 | 0.003 | Young |
| 0.791098 | 0.389538 | 0.033 | 0.031 | 1 | 0 | Cdc23 | 0.002 | Young |
| 0.791518 | 0.119583 | 0.036 | 0.034 | 1 | 0 | Cep85 | 0.002 | Young |
| 0.791728 | 0.182507 | 0.05 | 0.048 | 1 | 0 | Adamts10 | 0.002 | Young |
| 0.791797 | 0.095441 | 0.382 | 0.393 | 1 | 0 | Mtpn | -0.011 | Young |
| 0.791883 | 0.230923 | 0.047 | 0.045 | 1 | 0 | Nid2 | 0.002 | Young |
| 0.791965 | 0.113378 | 0.186 | 0.185 | 1 | 0 | Cox20 | 0.001 | Young |
| 0.791982 | 0.036168 | 0.039 | 0.041 | 1 | 0 | Fam241a | -0.002 | Young |
| 0.792504 | 0.022951 | 0.051 | 0.053 | 1 | 0 | Pxmp4 | -0.002 | Young |
| 0.792763 | 0.0356 | 0.016 | 0.015 | 1 | 0 | Wdr60 | 0.001 | Young |
| 0.793711 | 0.034665 | 0.019 | 0.021 | 1 | 0 | Pcnx4 | -0.002 | Young |
| 0.793957 | 0.056289 | 0.343 | 0.357 | 1 | 0 | Phip | -0.014 | Young |
| 0.794076 | 0.060625 | 0.093 | 0.097 | 1 | 0 | Parg | -0.004 | Young |
| 0.794473 | 0.042239 | 0.15 | 0.146 | 1 | 0 | Vav3 | 0.004 | Young |
| 0.794476 | 0.102818 | 0.137 | 0.142 | 1 | 0 | Tmem127 | -0.005 | Young |
| 0.794503 | 0.103531 | 0.151 | 0.157 | 1 | 0 | Csgalnact2 | -0.006 | Young |
| 0.794506 | 0.061667 | 0.261 | 0.275 | 1 | 0 | Elob | -0.014 | Young |
| 0.794585 | 0.024034 | 0.099 | 0.103 | 1 | 0 | Stau1 | -0.004 | Young |
| 0.795199 | 0.263827 | 0.025 | 0.023 | 1 | 0 | Ptdss2 | 0.002 | Young |

| | | | | | | | | |
|---|---|---|---|---|---|---|---|---|
| 0.795339 | 0.038453 | 0.127 | 0.131 | 1 | 0 | Vps36 | -0.004 | Young |
| 0.795396 | 0.210437 | 0.042 | 0.04 | 1 | 0 | Mettl6 | 0.002 | Young |
| 0.795976 | 0.124685 | 0.072 | 0.075 | 1 | 0 | Camsap2 | -0.003 | Young |
| 0.796073 | 0.303829 | 0.048 | 0.047 | 1 | 0 | Tmem209 | 0.001 | Young |
| 0.797114 | 0.105907 | 0.249 | 0.248 | 1 | 0 | Acap2 | 0.001 | Young |
| 0.797374 | 0.122067 | 0.149 | 0.146 | 1 | 0 | Myo9a | 0.003 | Young |
| 0.797402 | 0.000109 | 0.149 | 0.155 | 1 | 0 | Ndufb6 | -0.006 | Young |
| 0.797416 | 0.051162 | 0.626 | 0.648 | 1 | 0 | Pcbp2 | -0.022 | Young |
| 0.797815 | 0.162506 | 0.175 | 0.174 | 1 | 0 | Marf1 | 0.001 | Young |
| 0.797833 | 0.100726 | 0.103 | 0.1 | 1 | 0 | Jkamp | 0.003 | Young |
| 0.79791 | 0.141754 | 0.064 | 0.067 | 1 | 0 | Cnot7 | -0.003 | Young |
| 0.798072 | 0.232147 | 0.053 | 0.051 | 1 | 0 | Phtf2 | 0.002 | Young |
| 0.798102 | 0.156469 | 0.047 | 0.049 | 1 | 0 | Wiz | -0.002 | Young |
| 0.798398 | 0.314546 | 0.039 | 0.037 | 1 | 0 | Wfs1 | 0.002 | Young |
| 0.798528 | 0.092401 | 0.18 | 0.189 | 1 | 0 | Slc8b1 | -0.009 | Young |
| 0.798836 | 0.374722 | 0.039 | 0.037 | 1 | 0 | Tubgcp2 | 0.002 | Young |
| 0.799056 | 0.34998 | 0.019 | 0.021 | 1 | 0 | Nop2 | -0.002 | Young |
| 0.799113 | 0.051841 | 0.516 | 0.534 | 1 | 0 | H2-T23 | -0.018 | Young |
| 0.799137 | 0.305044 | 0.037 | 0.036 | 1 | 0 | Tada2b | 0.001 | Young |
| 0.799275 | 0.043893 | 0.116 | 0.12 | 1 | 0 | Ltbr | -0.004 | Young |
| 0.799453 | 0.101112 | 0.045 | 0.048 | 1 | 0 | Zdhhc4 | -0.003 | Young |
| 0.79958 | 0.062508 | 0.324 | 0.338 | 1 | 0 | Zcchc11 | -0.014 | Young |
| 0.799821 | 0.241617 | 0.019 | 0.021 | 1 | 0 | Dynlt1a | -0.002 | Young |
| 0.800044 | 0.181179 | 0.043 | 0.041 | 1 | 0 | Zfp513 | 0.002 | Young |
| 0.800287 | 0.06919 | 0.12 | 0.124 | 1 | 0 | Kdelr2 | -0.004 | Young |
| 0.800339 | 0.087876 | 0.056 | 0.059 | 1 | 0 | Kat6b | -0.003 | Young |
| 0.800693 | 0.021489 | 0.113 | 0.118 | 1 | 0 | Slc25a45 | -0.005 | Young |
| 0.800791 | 0.090992 | 0.16 | 0.156 | 1 | 0 | Bcl2a1b | 0.004 | Young |
| 0.800866 | 0.629477 | 0.032 | 0.03 | 1 | 0 | Usb1 | 0.002 | Young |
| 0.801446 | 0.172969 | 0.056 | 0.053 | 1 | 0 | Trp53bp2 | 0.003 | Young |
| 0.801555 | 0.158248 | 0.232 | 0.233 | 1 | 0 | Tnfsf12 | -0.001 | Young |
| 0.801616 | 0.052061 | 0.04 | 0.038 | 1 | 0 | Abhd17c | 0.002 | Young |
| 0.801674 | 0.037437 | 0.145 | 0.142 | 1 | 0 | Fam3c | 0.003 | Young |
| 0.80173 | 0.013987 | 0.246 | 0.254 | 1 | 0 | Ubn2 | -0.008 | Young |
| 0.801934 | 0.008152 | 0.398 | 0.412 | 1 | 0 | Sumo2 | -0.014 | Young |
| 0.802143 | 0.260271 | 0.025 | 0.023 | 1 | 0 | Blzf1 | 0.002 | Young |
| 0.802597 | 0.065571 | 0.267 | 0.278 | 1 | 0 | Crk | -0.011 | Young |
| 0.80288 | 0.132701 | 0.037 | 0.04 | 1 | 0 | Srbd1 | -0.003 | Young |
| 0.803204 | 0.328331 | 0.026 | 0.025 | 1 | 0 | Zfp87 | 0.001 | Young |
| 0.803318 | 0.255282 | 0.037 | 0.036 | 1 | 0 | Rpl27-ps3 | 0.001 | Young |
| 0.803415 | 0.032388 | 0.129 | 0.134 | 1 | 0 | Ccnd3 | -0.005 | Young |
| 0.803854 | 0.099464 | 0.382 | 0.387 | 1 | 0 | Tbxas1 | -0.005 | Young |
| 0.804083 | 0.368627 | 0.013 | 0.012 | 1 | 0 | Fuz | 0.001 | Young |
| 0.804135 | 0.511438 | 0.026 | 0.025 | 1 | 0 | Leng1 | 0.001 | Young |
| 0.804501 | 0.193132 | 0.062 | 0.06 | 1 | 0 | Rtl8a | 0.002 | Young |

| | | | | | | | | |
|---|---|---|---|---|---|---|---|---|
| 0.80459 | 0.083221 | 0.016 | 0.018 | 1 | 0 | Deaf1 | -0.002 | Young |
| 0.80486 | 0.04322 | 0.355 | 0.369 | 1 | 0 | Ube3a | -0.014 | Young |
| 0.805209 | 0.062885 | 0.206 | 0.212 | 1 | 0 | Sltm | -0.006 | Young |
| 0.805386 | 0.053096 | 0.027 | 0.029 | 1 | 0 | Slc35d1 | -0.002 | Young |
| 0.805557 | 0.0499 | 0.068 | 0.066 | 1 | 0 | Rufy1 | 0.002 | Young |
| 0.805583 | 0.341902 | 0.039 | 0.037 | 1 | 0 | 2610524H( | 0.002 | Young |
| 0.805661 | 0.085213 | 0.157 | 0.163 | 1 | 0 | Gng12 | -0.006 | Young |
| 0.805686 | 0.378657 | 0.069 | 0.067 | 1 | 0 | Eif2b5 | 0.002 | Young |
| 0.805833 | 0.053789 | 0.16 | 0.157 | 1 | 0 | Fip1l1 | 0.003 | Young |
| 0.806115 | 0.357982 | 0.013 | 0.012 | 1 | 0 | Blm | 0.001 | Young |
| 0.806244 | 0.007947 | 0.09 | 0.093 | 1 | 0 | AU040320 | -0.003 | Young |
| 0.806481 | 0.059356 | 0.206 | 0.216 | 1 | 0 | Pja2 | -0.01 | Young |
| 0.806541 | 0.126649 | 0.156 | 0.155 | 1 | 0 | Psmc1 | 0.001 | Young |
| 0.807442 | 0.09884 | 0.128 | 0.134 | 1 | 0 | Lrrc8a | -0.006 | Young |
| 0.807539 | 0.376678 | 0.091 | 0.09 | 1 | 0 | Trim5 | 0.001 | Young |
| 0.80784 | 0.008194 | 0.542 | 0.565 | 1 | 0 | Gas7 | -0.023 | Young |
| 0.808157 | 0.350904 | 0.03 | 0.029 | 1 | 0 | Cyb561d2 | 0.001 | Young |
| 0.808289 | 0.064754 | 0.118 | 0.122 | 1 | 0 | Rcn2 | -0.004 | Young |
| 0.808296 | 0.115384 | 0.097 | 0.101 | 1 | 0 | Commd6 | -0.004 | Young |
| 0.808338 | 0.042258 | 0.029 | 0.027 | 1 | 0 | Fbxo31 | 0.002 | Young |
| 0.808417 | 0.030508 | 0.021 | 0.019 | 1 | 0 | Hgf | 0.002 | Young |
| 0.808533 | 0.321358 | 0.085 | 0.083 | 1 | 0 | AI413582 | 0.002 | Young |
| 0.808535 | 0.272013 | 0.016 | 0.018 | 1 | 0 | Gtpbp3 | -0.002 | Young |
| 0.808591 | 0.134236 | 0.029 | 0.027 | 1 | 0 | Usp30 | 0.002 | Young |
| 0.808669 | 0.076934 | 0.035 | 0.037 | 1 | 0 | Dtx3 | -0.002 | Young |
| 0.808835 | 0.121205 | 0.177 | 0.175 | 1 | 0 | Psmd1 | 0.002 | Young |
| 0.80949 | 0.066823 | 0.182 | 0.178 | 1 | 0 | Eloa | 0.004 | Young |
| 0.809887 | 0.430241 | 0.018 | 0.016 | 1 | 0 | Fam84b | 0.002 | Young |
| 0.809911 | 0.089583 | 0.228 | 0.227 | 1 | 0 | Anxa6 | 0.001 | Young |
| 0.810054 | 0.150938 | 0.152 | 0.15 | 1 | 0 | Acsl5 | 0.002 | Young |
| 0.810055 | 0.11971 | 0.135 | 0.133 | 1 | 0 | Stim2 | 0.002 | Young |
| 0.810243 | 0.067471 | 0.276 | 0.275 | 1 | 0 | Cdk9 | 0.001 | Young |
| 0.810737 | 0.143836 | 0.013 | 0.012 | 1 | 0 | Taf1a | 0.001 | Young |
| 0.810853 | 0.120654 | 0.093 | 0.097 | 1 | 0 | Ncln | -0.004 | Young |
| 0.810904 | 0.273251 | 0.04 | 0.042 | 1 | 0 | 1600002K( | -0.002 | Young |
| 0.811091 | 0.138537 | 0.094 | 0.098 | 1 | 0 | BC037034 | -0.004 | Young |
| 0.811236 | 0.091594 | 0.348 | 0.368 | 1 | 0 | Nsa2 | -0.02 | Young |
| 0.81124 | 0.010724 | 0.353 | 0.354 | 1 | 0 | Sqstm1 | -0.001 | Young |
| 0.811277 | 0.060225 | 0.071 | 0.068 | 1 | 0 | Zbtb8os | 0.003 | Young |
| 0.812325 | 0.134061 | 0.313 | 0.317 | 1 | 0 | Cd48 | -0.004 | Young |
| 0.812641 | 0.202889 | 0.028 | 0.03 | 1 | 0 | Zfp212 | -0.002 | Young |
| 0.813134 | 0.370199 | 0.041 | 0.044 | 1 | 0 | Gtf2e1 | -0.003 | Young |
| 0.813231 | 0.205009 | 0.08 | 0.078 | 1 | 0 | Zdhhc7 | 0.002 | Young |
| 0.813333 | 0.120517 | 0.226 | 0.226 | 1 | 0 | Hnrnpd | 0 | Young |
| 0.813542 | 0.16484 | 0.094 | 0.098 | 1 | 0 | Pld1 | -0.004 | Young |

| | | | | | | | | |
|---|---|---|---|---|---|---|---|---|
| 0.813584 | 0.310596 | 0.083 | 0.081 | 1 | 0 | Sec24a | 0.002 | Young |
| 0.813926 | 0.284089 | 0.072 | 0.07 | 1 | 0 | Trappc13 | 0.002 | Young |
| 0.814562 | 0.143606 | 0.151 | 0.159 | 1 | 0 | Rab12 | -0.008 | Young |
| 0.814574 | 0.644955 | 0.011 | 0.01 | 1 | 0 | Thnsl1 | 0.001 | Young |
| 0.815099 | 0.223849 | 0.087 | 0.085 | 1 | 0 | Papd5 | 0.002 | Young |
| 0.81513 | 0.029333 | 0.523 | 0.518 | 1 | 0 | Zbtb20 | 0.005 | Young |
| 0.815727 | 0.018287 | 0.024 | 0.026 | 1 | 0 | Mrps31 | -0.002 | Young |
| 0.816276 | 0.045469 | 0.112 | 0.116 | 1 | 0 | Gab1 | -0.004 | Young |
| 0.816354 | 0.190445 | 0.018 | 0.016 | 1 | 0 | Smyd4 | 0.002 | Young |
| 0.816652 | 0.056669 | 0.084 | 0.088 | 1 | 0 | Chd7 | -0.004 | Young |
| 0.816798 | 0.107149 | 0.251 | 0.252 | 1 | 0 | Hipk1 | -0.001 | Young |
| 0.816964 | 0.061531 | 0.201 | 0.208 | 1 | 0 | Tpp2 | -0.007 | Young |
| 0.817275 | 0.322881 | 0.052 | 0.051 | 1 | 0 | Pgs1 | 0.001 | Young |
| 0.817716 | 0.517365 | 0.011 | 0.01 | 1 | 0 | Cenpw | 0.001 | Young |
| 0.817909 | 0.11562 | 0.179 | 0.187 | 1 | 0 | Nenf | -0.008 | Young |
| 0.817946 | 0.219266 | 0.015 | 0.016 | 1 | 0 | Ccdc77 | -0.001 | Young |
| 0.818019 | 0.061368 | 0.167 | 0.166 | 1 | 0 | Arhgef10l | 0.001 | Young |
| 0.818436 | 0.077925 | 0.102 | 0.105 | 1 | 0 | Pdha1 | -0.003 | Young |
| 0.818596 | 0.136648 | 0.071 | 0.068 | 1 | 0 | Apba1 | 0.003 | Young |
| 0.818654 | 0.299505 | 0.054 | 0.057 | 1 | 0 | Daam1 | -0.003 | Young |
| 0.818717 | 0.173233 | 0.08 | 0.078 | 1 | 0 | Ndrg3 | 0.002 | Young |
| 0.818878 | 0.011181 | 0.048 | 0.051 | 1 | 0 | Ddx39 | -0.003 | Young |
| 0.81891 | 0.049105 | 0.647 | 0.684 | 1 | 0 | Gm10076 | -0.037 | Young |
| 0.818919 | 0.147579 | 0.107 | 0.104 | 1 | 0 | C1qbp | 0.003 | Young |
| 0.819418 | 0.093525 | 0.035 | 0.037 | 1 | 0 | Ldlrap1 | -0.002 | Young |
| 0.819469 | 0.165053 | 0.026 | 0.025 | 1 | 0 | Fbxl15 | 0.001 | Young |
| 0.819683 | 0.12268 | 0.06 | 0.063 | 1 | 0 | Zbtb34 | -0.003 | Young |
| 0.819823 | 0.093802 | 0.023 | 0.025 | 1 | 0 | Lzic | -0.002 | Young |
| 0.820277 | 0.085677 | 0.035 | 0.037 | 1 | 0 | Mgme1 | -0.002 | Young |
| 0.820444 | 0.130174 | 0.086 | 0.083 | 1 | 0 | Sumf1 | 0.003 | Young |
| 0.820616 | 0.014221 | 0.056 | 0.059 | 1 | 0 | Slc41a3 | -0.003 | Young |
| 0.820746 | 0.058577 | 0.03 | 0.029 | 1 | 0 | Coq10a | 0.001 | Young |
| 0.820761 | 0.098065 | 0.29 | 0.304 | 1 | 0 | Mrpl52 | -0.014 | Young |
| 0.820764 | 0.250749 | 0.08 | 0.085 | 1 | 0 | Adar | -0.005 | Young |
| 0.82085 | 0.3312 | 0.039 | 0.041 | 1 | 0 | Plrg1 | -0.002 | Young |
| 0.820882 | 0.135743 | 0.034 | 0.036 | 1 | 0 | Pnkp | -0.002 | Young |
| 0.820984 | 0.322139 | 0.047 | 0.045 | 1 | 0 | Gm2000 | 0.002 | Young |
| 0.821105 | 0.092999 | 0.305 | 0.306 | 1 | 0 | Cxcl16 | -0.001 | Young |
| 0.82151 | 0.114351 | 0.111 | 0.116 | 1 | 0 | Pdhb | -0.005 | Young |
| 0.821553 | 0.018545 | 0.227 | 0.235 | 1 | 0 | Tnfrsf1a | -0.008 | Young |
| 0.821606 | 0.087177 | 0.126 | 0.13 | 1 | 0 | Snapin | -0.004 | Young |
| 0.821678 | 0.20943 | 0.03 | 0.029 | 1 | 0 | Med24 | 0.001 | Young |
| 0.821792 | 0.175543 | 0.076 | 0.079 | 1 | 0 | Actl6a | -0.003 | Young |
| 0.82182 | 0.051441 | 0.039 | 0.037 | 1 | 0 | Parp10 | 0.002 | Young |
| 0.822028 | 0.176488 | 0.04 | 0.038 | 1 | 0 | Pgam5 | 0.002 | Young |

| | | | | | | | | |
|---|---|---|---|---|---|---|---|---|
| 0.822041 | 0.008347 | 0.039 | 0.037 | 1 | 0 | Pdhx | 0.002 | Young |
| 0.822405 | 0.02788 | 0.102 | 0.105 | 1 | 0 | Rsbn1 | -0.003 | Young |
| 0.822473 | 0.128018 | 0.014 | 0.015 | 1 | 0 | Gm4631 | -0.001 | Young |
| 0.822753 | 0.324486 | 0.015 | 0.016 | 1 | 0 | Rdh13 | -0.001 | Young |
| 0.822799 | 0.175661 | 0.094 | 0.092 | 1 | 0 | Filip1l | 0.002 | Young |
| 0.822959 | 0.099208 | 0.011 | 0.01 | 1 | 0 | Rab34 | 0.001 | Young |
| 0.823097 | 0.21913 | 0.015 | 0.016 | 1 | 0 | Mterf4 | -0.001 | Young |
| 0.823255 | 0.028622 | 0.118 | 0.115 | 1 | 0 | Hivep1 | 0.003 | Young |
| 0.82337 | 0.106533 | 0.014 | 0.015 | 1 | 0 | Klhl26 | -0.001 | Young |
| 0.823549 | 0.18189 | 0.014 | 0.015 | 1 | 0 | Tnfrsf26 | -0.001 | Young |
| 0.823799 | 0.142084 | 0.011 | 0.01 | 1 | 0 | Megf9 | 0.001 | Young |
| 0.823799 | 0.101098 | 0.011 | 0.01 | 1 | 0 | Alg12 | 0.001 | Young |
| 0.823967 | 0.393087 | 0.043 | 0.041 | 1 | 0 | Fbxw8 | 0.002 | Young |
| 0.823973 | 0.409513 | 0.103 | 0.103 | 1 | 0 | Ifi209 | 0 | Young |
| 0.824418 | 0.088711 | 0.313 | 0.319 | 1 | 0 | Sbf2 | -0.006 | Young |
| 0.824647 | 0.017958 | 0.194 | 0.189 | 1 | 0 | Zbtb4 | 0.005 | Young |
| 0.824765 | 0.055246 | 0.108 | 0.112 | 1 | 0 | Trappc8 | -0.004 | Young |
| 0.824988 | 0.39459 | 0.037 | 0.04 | 1 | 0 | Commd9 | -0.003 | Young |
| 0.825012 | 0.154052 | 0.13 | 0.129 | 1 | 0 | Egln1 | 0.001 | Young |
| 0.825189 | 0.714458 | 0.016 | 0.018 | 1 | 0 | Amz1 | -0.002 | Young |
| 0.825199 | 0.173951 | 0.072 | 0.07 | 1 | 0 | Gmeb1 | 0.002 | Young |
| 0.825293 | 0.096458 | 0.022 | 0.021 | 1 | 0 | Cbr4 | 0.001 | Young |
| 0.825567 | 0.163061 | 0.214 | 0.226 | 1 | 0 | Os9 | -0.012 | Young |
| 0.826026 | 0.102627 | 0.275 | 0.287 | 1 | 0 | Snrpf | -0.012 | Young |
| 0.826241 | 0.079047 | 0.014 | 0.015 | 1 | 0 | Vwf | -0.001 | Young |
| 0.82625 | 0.04473 | 0.322 | 0.332 | 1 | 0 | Sf1 | -0.01 | Young |
| 0.826584 | 0.027691 | 0.353 | 0.361 | 1 | 0 | Nrp2 | -0.008 | Young |
| 0.827513 | 0.117326 | 0.05 | 0.052 | 1 | 0 | Golt1b | -0.002 | Young |
| 0.827907 | 0.063997 | 0.04 | 0.042 | 1 | 0 | Dtx2 | -0.002 | Young |
| 0.827968 | 0.093303 | 0.2 | 0.208 | 1 | 0 | Pdcd10 | -0.008 | Young |
| 0.828253 | 0.413443 | 0.033 | 0.031 | 1 | 0 | Zbed4 | 0.002 | Young |
| 0.828306 | 0.104513 | 0.249 | 0.26 | 1 | 0 | AI467606 | -0.011 | Young |
| 0.8284 | 0.254003 | 0.024 | 0.026 | 1 | 0 | Tmem29 | -0.002 | Young |
| 0.828576 | 0.234358 | 0.014 | 0.015 | 1 | 0 | Hspa2 | -0.001 | Young |
| 0.828623 | 0.161654 | 0.131 | 0.129 | 1 | 0 | Pbx2 | 0.002 | Young |
| 0.828649 | 0.066804 | 0.273 | 0.274 | 1 | 0 | Lsp1 | -0.001 | Young |
| 0.828748 | 0.290368 | 0.127 | 0.127 | 1 | 0 | Exoc6b | 0 | Young |
| 0.828841 | 0.231881 | 0.077 | 0.075 | 1 | 0 | Pold3 | 0.002 | Young |
| 0.828913 | 0.10194 | 0.053 | 0.056 | 1 | 0 | 2410089E0 | -0.003 | Young |
| 0.828962 | 0.10176 | 0.055 | 0.057 | 1 | 0 | Mrpl35 | -0.002 | Young |
| 0.82928 | 0.290007 | 0.015 | 0.014 | 1 | 0 | Tmem86b | 0.001 | Young |
| 0.829567 | 0.11838 | 0.109 | 0.112 | 1 | 0 | Utp11 | -0.003 | Young |
| 0.829569 | 0.192896 | 0.134 | 0.141 | 1 | 0 | Ing4 | -0.007 | Young |
| 0.829635 | 0.175189 | 0.168 | 0.176 | 1 | 0 | Psmb5 | -0.008 | Young |
| 0.829741 | 0.221021 | 0.035 | 0.037 | 1 | 0 | Slc35c1 | -0.002 | Young |

| | | | | | | | | |
|---|---|---|---|---|---|---|---|---|
| 0.830008 | 0.123283 | 0.197 | 0.205 | 1 | 0 | Lamtor3 | -0.008 | Young |
| 0.830014 | 0.081506 | 0.03 | 0.031 | 1 | 0 | Cwc27 | -0.001 | Young |
| 0.830055 | 0.050773 | 0.305 | 0.32 | 1 | 0 | Tcf25 | -0.015 | Young |
| 0.830071 | 0.086804 | 0.358 | 0.376 | 1 | 0 | Rab10 | -0.018 | Young |
| 0.830109 | 0.288782 | 0.027 | 0.026 | 1 | 0 | 1110032A( | 0.001 | Young |
| 0.830172 | 0.053739 | 0.069 | 0.071 | 1 | 0 | Thumpd1 | -0.002 | Young |
| 0.830199 | 0.065047 | 0.069 | 0.067 | 1 | 0 | Med30 | 0.002 | Young |
| 0.830299 | 0.297064 | 0.043 | 0.041 | 1 | 0 | Mrpl50 | 0.002 | Young |
| 0.830512 | 0.218326 | 0.037 | 0.036 | 1 | 0 | 2610008E1 | 0.001 | Young |
| 0.830602 | 0.243138 | 0.023 | 0.025 | 1 | 0 | Tbl3 | -0.002 | Young |
| 0.830763 | 0.212942 | 0.123 | 0.122 | 1 | 0 | Rnf11 | 0.001 | Young |
| 0.830801 | 0.099553 | 0.04 | 0.042 | 1 | 0 | Dgkh | -0.002 | Young |
| 0.830884 | 0.248908 | 0.019 | 0.018 | 1 | 0 | Abca6 | 0.001 | Young |
| 0.831022 | 0.058903 | 0.023 | 0.025 | 1 | 0 | Nprl3 | -0.002 | Young |
| 0.831137 | 0.176283 | 0.259 | 0.264 | 1 | 0 | Fndc3a | -0.005 | Young |
| 0.831262 | 0.067758 | 0.158 | 0.161 | 1 | 0 | Arl5b | -0.003 | Young |
| 0.831384 | 0.035325 | 0.291 | 0.305 | 1 | 0 | P2ry6 | -0.014 | Young |
| 0.831502 | 0.201713 | 0.26 | 0.264 | 1 | 0 | Rbm5 | -0.004 | Young |
| 0.831678 | 0.04503 | 0.054 | 0.056 | 1 | 0 | Galnt4 | -0.002 | Young |
| 0.831751 | 0.152321 | 0.013 | 0.014 | 1 | 0 | Ift81 | -0.001 | Young |
| 0.831806 | 0.07462 | 0.272 | 0.282 | 1 | 0 | Gmfb | -0.01 | Young |
| 0.832045 | 0.158315 | 0.118 | 0.123 | 1 | 0 | Tbc1d13 | -0.005 | Young |
| 0.832425 | 0.181005 | 0.086 | 0.085 | 1 | 0 | Trabd | 0.001 | Young |
| 0.832468 | 0.072998 | 0.206 | 0.207 | 1 | 0 | Fis1 | -0.001 | Young |
| 0.83276 | 0.107138 | 0.045 | 0.044 | 1 | 0 | Gm45716 | 0.001 | Young |
| 0.832882 | 0.040585 | 0.013 | 0.014 | 1 | 0 | Arpin | -0.001 | Young |
| 0.832971 | 0.182223 | 0.102 | 0.1 | 1 | 0 | Cnot2 | 0.002 | Young |
| 0.833265 | 0.238697 | 0.096 | 0.1 | 1 | 0 | Psmd3 | -0.004 | Young |
| 0.833298 | 0.20052 | 0.023 | 0.022 | 1 | 0 | Cul7 | 0.001 | Young |
| 0.833429 | 0.292359 | 0.014 | 0.015 | 1 | 0 | Pstpip2 | -0.001 | Young |
| 0.834073 | 0.189735 | 0.04 | 0.042 | 1 | 0 | Anapc15 | -0.002 | Young |
| 0.834159 | 0.096694 | 0.081 | 0.079 | 1 | 0 | Naip2 | 0.002 | Young |
| 0.834175 | 0.213564 | 0.056 | 0.055 | 1 | 0 | Eif2b4 | 0.001 | Young |
| 0.83422 | 0.413358 | 0.034 | 0.033 | 1 | 0 | Tex2 | 0.001 | Young |
| 0.834323 | 0.097425 | 0.039 | 0.041 | 1 | 0 | Zer1 | -0.002 | Young |
| 0.834378 | 0.101297 | 0.022 | 0.023 | 1 | 0 | 9930104L0 | -0.001 | Young |
| 0.834657 | 0.078306 | 0.048 | 0.051 | 1 | 0 | Vti1a | -0.003 | Young |
| 0.834739 | 0.002745 | 0.663 | 0.7 | 1 | 0 | Slc6a6 | -0.037 | Young |
| 0.834767 | 0.29116 | 0.013 | 0.014 | 1 | 0 | Zfp948 | -0.001 | Young |
| 0.835095 | 0.266996 | 0.024 | 0.026 | 1 | 0 | Cdc42bpg | -0.002 | Young |
| 0.835152 | 0.035916 | 0.658 | 0.688 | 1 | 0 | Qk | -0.03 | Young |
| 0.835225 | 0.001727 | 0.073 | 0.071 | 1 | 0 | Cbll1 | 0.002 | Young |
| 0.835716 | 0.294009 | 0.023 | 0.022 | 1 | 0 | Def8 | 0.001 | Young |
| 0.835898 | 0.187033 | 0.155 | 0.163 | 1 | 0 | Rnf6 | -0.008 | Young |
| 0.83618 | 0.05642 | 0.273 | 0.287 | 1 | 0 | Eef1d | -0.014 | Young |

| | | | | | | | | |
|---|---|---|---|---|---|---|---|---|
| 0.83632 | 0.473504 | 0.024 | 0.023 | 1 | 0 | Tmed8 | 0.001 | Young |
| 0.836324 | 0.075166 | 0.046 | 0.048 | 1 | 0 | Rpp14 | -0.002 | Young |
| 0.836425 | 0.039976 | 0.126 | 0.13 | 1 | 0 | Pik3r6 | -0.004 | Young |
| 0.836643 | 0.283064 | 0.055 | 0.053 | 1 | 0 | Wdtc1 | 0.002 | Young |
| 0.836749 | 0.073693 | 0.086 | 0.083 | 1 | 0 | Pilrb1 | 0.003 | Young |
| 0.836752 | 0.116334 | 0.058 | 0.056 | 1 | 0 | Ceacam1 | 0.002 | Young |
| 0.837011 | 0.031228 | 0.181 | 0.178 | 1 | 0 | Cebpa | 0.003 | Young |
| 0.837239 | 0.104568 | 0.05 | 0.052 | 1 | 0 | Alg14 | -0.002 | Young |
| 0.837338 | 0.129368 | 0.171 | 0.178 | 1 | 0 | Tollip | -0.007 | Young |
| 0.837597 | 0.364755 | 0.013 | 0.014 | 1 | 0 | Tle2 | -0.001 | Young |
| 0.837598 | 0.013922 | 0.058 | 0.06 | 1 | 0 | Zfp277 | -0.002 | Young |
| 0.838064 | 0.222144 | 0.051 | 0.049 | 1 | 0 | Smn1 | 0.002 | Young |
| 0.838299 | 0.141118 | 0.027 | 0.026 | 1 | 0 | Tab3 | 0.001 | Young |
| 0.838369 | 0.20576 | 0.037 | 0.036 | 1 | 0 | D230025D: | 0.001 | Young |
| 0.838501 | 0.039131 | 0.096 | 0.094 | 1 | 0 | Stam2 | 0.002 | Young |
| 0.838634 | 0.085292 | 0.038 | 0.04 | 1 | 0 | Zfp157 | -0.002 | Young |
| 0.838741 | 0.126088 | 0.232 | 0.234 | 1 | 0 | Trim41 | -0.002 | Young |
| 0.838891 | 0.270767 | 0.034 | 0.036 | 1 | 0 | AU022252 | -0.002 | Young |
| 0.839023 | 0.320033 | 0.023 | 0.025 | 1 | 0 | Taf6 | -0.002 | Young |
| 0.839198 | 0.044365 | 0.054 | 0.052 | 1 | 0 | Gpd2 | 0.002 | Young |
| 0.839886 | 0.360252 | 0.067 | 0.066 | 1 | 0 | Mrpl21 | 0.001 | Young |
| 0.839959 | 0.162059 | 0.091 | 0.089 | 1 | 0 | Utp14a | 0.002 | Young |
| 0.840238 | 0.00428 | 0.681 | 0.711 | 1 | 0 | Alox5ap | -0.03 | Young |
| 0.840745 | 0.089981 | 0.203 | 0.213 | 1 | 0 | Rragc | -0.01 | Young |
| 0.840805 | 0.094709 | 0.248 | 0.259 | 1 | 0 | Chmp2a | -0.011 | Young |
| 0.84111 | 0.110424 | 0.067 | 0.07 | 1 | 0 | Ap1m1 | -0.003 | Young |
| 0.841674 | 0.129594 | 0.086 | 0.089 | 1 | 0 | Acsl1 | -0.003 | Young |
| 0.841725 | 0.405686 | 0.02 | 0.019 | 1 | 0 | Zfp777 | 0.001 | Young |
| 0.841825 | 0.187669 | 0.086 | 0.085 | 1 | 0 | Mapkap1 | 0.001 | Young |
| 0.841969 | 0.554713 | 0.012 | 0.011 | 1 | 0 | Ttc30b | 0.001 | Young |
| 0.842014 | 0.264624 | 0.146 | 0.146 | 1 | 0 | Aldh3b1 | 0 | Young |
| 0.84218 | 0.088744 | 0.166 | 0.166 | 1 | 0 | Paics | 0 | Young |
| 0.842384 | 0.233305 | 0.123 | 0.122 | 1 | 0 | Tcf20 | 0.001 | Young |
| 0.842537 | 0.201978 | 0.059 | 0.057 | 1 | 0 | Zfp219 | 0.002 | Young |
| 0.842606 | 0.186256 | 0.068 | 0.071 | 1 | 0 | Hypk | -0.003 | Young |
| 0.842613 | 0.113953 | 0.225 | 0.226 | 1 | 0 | Ddrgk1 | -0.001 | Young |
| 0.842765 | 0.343385 | 0.011 | 0.012 | 1 | 0 | Praf2 | -0.001 | Young |
| 0.843119 | 0.088312 | 0.204 | 0.201 | 1 | 0 | Fam105a | 0.003 | Young |
| 0.843123 | 0.063769 | 0.023 | 0.022 | 1 | 0 | Dgka | 0.001 | Young |
| 0.843355 | 0.03718 | 0.113 | 0.111 | 1 | 0 | Rbl2 | 0.002 | Young |
| 0.843557 | 0.257707 | 0.096 | 0.094 | 1 | 0 | Timm8b | 0.002 | Young |
| 0.843761 | 0.307979 | 0.011 | 0.012 | 1 | 0 | Pter | -0.001 | Young |
| 0.843824 | 0.219209 | 0.067 | 0.07 | 1 | 0 | Gucd1 | -0.003 | Young |
| 0.844059 | 0.192487 | 0.081 | 0.079 | 1 | 0 | Nmt2 | 0.002 | Young |
| 0.844177 | 0.207451 | 0.247 | 0.261 | 1 | 0 | Rad21 | -0.014 | Young |

| | | | | | | | | |
|---|---|---|---|---|---|---|---|---|
| 0.844557 | 0.070586 | 0.333 | 0.337 | 1 | 0 | Eif3k | -0.004 | Young |
| 0.84457 | 0.174973 | 0.23 | 0.233 | 1 | 0 | Ppp2r1a | -0.003 | Young |
| 0.844582 | 0.103935 | 0.401 | 0.415 | 1 | 0 | Tgfbr2 | -0.014 | Young |
| 0.844717 | 0.058689 | 0.098 | 0.101 | 1 | 0 | Spen | -0.003 | Young |
| 0.844965 | 0.092858 | 0.158 | 0.163 | 1 | 0 | Tmx1 | -0.005 | Young |
| 0.84504 | 0.110266 | 0.184 | 0.19 | 1 | 0 | Krcc1 | -0.006 | Young |
| 0.84505 | 0.214109 | 0.031 | 0.033 | 1 | 0 | Atat1 | -0.002 | Young |
| 0.845357 | 0.153294 | 0.076 | 0.079 | 1 | 0 | Ppp1r35 | -0.003 | Young |
| 0.845391 | 0.243799 | 0.061 | 0.064 | 1 | 0 | Icmt | -0.003 | Young |
| 0.845461 | 0.048157 | 0.574 | 0.614 | 1 | 0 | Dstn | -0.04 | Young |
| 0.845605 | 0.34799 | 0.1 | 0.1 | 1 | 0 | Smg7 | 0 | Young |
| 0.845687 | 0.102271 | 0.133 | 0.138 | 1 | 0 | Nfya | -0.005 | Young |
| 0.845691 | 0.128182 | 0.106 | 0.104 | 1 | 0 | Capn1 | 0.002 | Young |
| 0.845846 | 0.05261 | 0.436 | 0.439 | 1 | 0 | Chd4 | -0.003 | Young |
| 0.845998 | 0.002909 | 0.136 | 0.14 | 1 | 0 | Usp47 | -0.004 | Young |
| 0.846113 | 0.020783 | 0.027 | 0.026 | 1 | 0 | Trim30c | 0.001 | Young |
| 0.846147 | 0.195426 | 0.074 | 0.073 | 1 | 0 | Tmem192 | 0.001 | Young |
| 0.84647 | 0.037605 | 0.265 | 0.276 | 1 | 0 | Anp32e | -0.011 | Young |
| 0.846944 | 0.115561 | 0.077 | 0.075 | 1 | 0 | Avl9 | 0.002 | Young |
| 0.846966 | 0.064467 | 0.01 | 0.011 | 1 | 0 | Sgsm3 | -0.001 | Young |
| 0.847003 | 0.325545 | 0.073 | 0.073 | 1 | 0 | Glt8d1 | 0 | Young |
| 0.847513 | 0.037756 | 0.112 | 0.109 | 1 | 0 | Zdhhc14 | 0.003 | Young |
| 0.847608 | 0.0568 | 0.021 | 0.022 | 1 | 0 | Eif2b1 | -0.001 | Young |
| 0.847851 | 0.093065 | 0.23 | 0.241 | 1 | 0 | Mkrn1 | -0.011 | Young |
| 0.847895 | 0.20769 | 0.094 | 0.093 | 1 | 0 | Aim2 | 0.001 | Young |
| 0.847928 | 0.109035 | 0.147 | 0.146 | 1 | 0 | Galk2 | 0.001 | Young |
| 0.848008 | 0.186235 | 0.058 | 0.056 | 1 | 0 | Alg10b | 0.002 | Young |
| 0.848543 | 0.186583 | 0.012 | 0.011 | 1 | 0 | 9430015G1 | 0.001 | Young |
| 0.848661 | 0.12052 | 0.054 | 0.056 | 1 | 0 | Pigyl | -0.002 | Young |
| 0.848742 | 0.196115 | 0.012 | 0.011 | 1 | 0 | Bhlhe41 | 0.001 | Young |
| 0.848756 | 0.434943 | 0.016 | 0.015 | 1 | 0 | Sirt3 | 0.001 | Young |
| 0.84883 | 0.120667 | 0.033 | 0.031 | 1 | 0 | Wrnip1 | 0.002 | Young |
| 0.848925 | 0.059814 | 0.019 | 0.021 | 1 | 0 | Cdc42bpa | -0.002 | Young |
| 0.849047 | 0.070022 | 0.1 | 0.103 | 1 | 0 | Ccdc124 | -0.003 | Young |
| 0.84906 | 0.563694 | 0.031 | 0.03 | 1 | 0 | Paqr7 | 0.001 | Young |
| 0.849088 | 0.044782 | 0.287 | 0.295 | 1 | 0 | Tprgl | -0.008 | Young |
| 0.849126 | 0.27516 | 0.029 | 0.027 | 1 | 0 | Tmem35b | 0.002 | Young |
| 0.849451 | 0.027858 | 0.115 | 0.112 | 1 | 0 | Map1s | 0.003 | Young |
| 0.849498 | 0.066598 | 0.125 | 0.129 | 1 | 0 | Higd1a | -0.004 | Young |
| 0.849757 | 0.159953 | 0.157 | 0.157 | 1 | 0 | Myo1c | 0 | Young |
| 0.849911 | 0.048147 | 0.059 | 0.062 | 1 | 0 | Blcap | -0.003 | Young |
| 0.850136 | 0.137178 | 0.01 | 0.011 | 1 | 0 | Zfp971 | -0.001 | Young |
| 0.850565 | 0.212249 | 0.079 | 0.082 | 1 | 0 | Atrnl1 | -0.003 | Young |
| 0.85057 | 0.24153 | 0.038 | 0.04 | 1 | 0 | Ogfod1 | -0.002 | Young |
| 0.850727 | 0.482153 | 0.021 | 0.021 | 1 | 0 | Prnp | 0 | Young |

| | | | | | | | | |
|---|---|---|---|---|---|---|---|---|
| 0.850855 | 0.051822 | 0.042 | 0.044 | 1 | 0 | Slc25a44 | -0.002 | Young |
| 0.850917 | 0.149489 | 0.052 | 0.051 | 1 | 0 | Ptrhd1 | 0.001 | Young |
| 0.851179 | 0.259389 | 0.02 | 0.019 | 1 | 0 | Evi5l | 0.001 | Young |
| 0.851186 | 0.0912 | 0.045 | 0.047 | 1 | 0 | Dnajc25 | -0.002 | Young |
| 0.851413 | 0.049424 | 0.251 | 0.25 | 1 | 0 | Batf3 | 0.001 | Young |
| 0.851434 | 0.156225 | 0.069 | 0.067 | 1 | 0 | Tmem68 | 0.002 | Young |
| 0.852032 | 0.088395 | 0.175 | 0.174 | 1 | 0 | Phf20 | 0.001 | Young |
| 0.85223 | 0.120501 | 0.128 | 0.126 | 1 | 0 | Kbtbd2 | 0.002 | Young |
| 0.852456 | 0.041939 | 0.082 | 0.085 | 1 | 0 | Mrpl18 | -0.003 | Young |
| 0.852528 | 0.144804 | 0.221 | 0.222 | 1 | 0 | Dmxl1 | -0.001 | Young |
| 0.852533 | 0.120809 | 0.082 | 0.085 | 1 | 0 | Agap3 | -0.003 | Young |
| 0.85257 | 0.144088 | 0.066 | 0.064 | 1 | 0 | Tor3a | 0.002 | Young |
| 0.852623 | 0.110696 | 0.072 | 0.07 | 1 | 0 | Oard1 | 0.002 | Young |
| 0.852706 | 0.113685 | 0.024 | 0.023 | 1 | 0 | Mapre3 | 0.001 | Young |
| 0.852736 | 0.208447 | 0.059 | 0.062 | 1 | 0 | Spaca6 | -0.003 | Young |
| 0.852885 | 0.425944 | 0.01 | 0.011 | 1 | 0 | Gm4673 | -0.001 | Young |
| 0.852904 | 0.04339 | 0.629 | 0.657 | 1 | 0 | Cyba | -0.028 | Young |
| 0.853056 | 0.226595 | 0.103 | 0.101 | 1 | 0 | Epb41l4aos | 0.002 | Young |
| 0.853114 | 0.003254 | 0.248 | 0.252 | 1 | 0 | Soat1 | -0.004 | Young |
| 0.853161 | 0.005648 | 0.274 | 0.282 | 1 | 0 | Tsn | -0.008 | Young |
| 0.85334 | 0.16988 | 0.009 | 0.01 | 1 | 0 | Spink2 | -0.001 | Young |
| 0.853436 | 0.118944 | 0.037 | 0.038 | 1 | 0 | Lyrm9 | -0.001 | Young |
| 0.853574 | 0.032497 | 0.115 | 0.112 | 1 | 0 | Irs2 | 0.003 | Young |
| 0.853759 | 0.184386 | 0.029 | 0.027 | 1 | 0 | Invs | 0.002 | Young |
| 0.85379 | 0.208846 | 0.048 | 0.049 | 1 | 0 | Iigp1 | -0.001 | Young |
| 0.853873 | 0.094241 | 0.16 | 0.159 | 1 | 0 | Zfp292 | 0.001 | Young |
| 0.854507 | 0.001531 | 0.1 | 0.103 | 1 | 0 | Dhdh | -0.003 | Young |
| 0.854845 | 0.035871 | 0.168 | 0.172 | 1 | 0 | Alkbh5 | -0.004 | Young |
| 0.855045 | 0.109582 | 0.08 | 0.083 | 1 | 0 | Mis18a | -0.003 | Young |
| 0.85513 | 0.064518 | 0.061 | 0.059 | 1 | 0 | Pigc | 0.002 | Young |
| 0.855243 | 0.109382 | 0.069 | 0.067 | 1 | 0 | Taf7 | 0.002 | Young |
| 0.855375 | 0.132242 | 0.009 | 0.01 | 1 | 0 | Kif20b | -0.001 | Young |
| 0.855454 | 0.173936 | 0.19 | 0.198 | 1 | 0 | Nudt3 | -0.008 | Young |
| 0.855553 | 0.128992 | 0.08 | 0.083 | 1 | 0 | Tssc4 | -0.003 | Young |
| 0.855728 | 0.044402 | 0.018 | 0.019 | 1 | 0 | Aqp9 | -0.001 | Young |
| 0.855833 | 0.230808 | 0.051 | 0.049 | 1 | 0 | Slc25a16 | 0.002 | Young |
| 0.855849 | 0.188313 | 0.056 | 0.059 | 1 | 0 | Htatip2 | -0.003 | Young |
| 0.856125 | 0.007887 | 0.671 | 0.698 | 1 | 0 | Rplp0 | -0.027 | Young |
| 0.856263 | 0.10555 | 0.052 | 0.051 | 1 | 0 | Bcl9 | 0.001 | Young |
| 0.856437 | 0.302669 | 0.046 | 0.045 | 1 | 0 | Spg11 | 0.001 | Young |
| 0.856518 | 0.113217 | 0.037 | 0.036 | 1 | 0 | Irgq | 0.001 | Young |
| 0.856581 | 0.160234 | 0.088 | 0.09 | 1 | 0 | Mxd1 | -0.002 | Young |
| 0.856712 | 0.05913 | 0.184 | 0.19 | 1 | 0 | Gpbp1 | -0.006 | Young |
| 0.856904 | 0.049219 | 0.128 | 0.131 | 1 | 0 | Rit1 | -0.003 | Young |
| 0.85695 | 0.049272 | 0.169 | 0.167 | 1 | 0 | Khsrp | 0.002 | Young |

| | | | | | | | | |
|---|---|---|---|---|---|---|---|---|
| 0.857241 | 0.009522 | 0.061 | 0.059 | 1 | 0 | Dtx4 | 0.002 | Young |
| 0.857849 | 0.085707 | 0.095 | 0.098 | 1 | 0 | Naxd | -0.003 | Young |
| 0.857881 | 0.010366 | 0.029 | 0.027 | 1 | 0 | Rnf25 | 0.002 | Young |
| 0.858125 | 0.219916 | 0.049 | 0.048 | 1 | 0 | Map3k20 | 0.001 | Young |
| 0.85843 | 0.148627 | 0.215 | 0.218 | 1 | 0 | Rac2 | -0.003 | Young |
| 0.858452 | 0.068035 | 0.224 | 0.234 | 1 | 0 | Ndufv3 | -0.01 | Young |
| 0.858585 | 0.043212 | 0.506 | 0.516 | 1 | 0 | Scamp2 | -0.01 | Young |
| 0.858615 | 0.197753 | 0.079 | 0.082 | 1 | 0 | Uchl5 | -0.003 | Young |
| 0.858633 | 0.041779 | 0.119 | 0.122 | 1 | 0 | Atxn7l1 | -0.003 | Young |
| 0.858823 | 0.130773 | 0.086 | 0.085 | 1 | 0 | Ngly1 | 0.001 | Young |
| 0.85895 | 0.200315 | 0.247 | 0.253 | 1 | 0 | Zmynd8 | -0.006 | Young |
| 0.859359 | 0.140956 | 0.192 | 0.194 | 1 | 0 | Med28 | -0.002 | Young |
| 0.859377 | 0.092322 | 0.235 | 0.244 | 1 | 0 | Cul3 | -0.009 | Young |
| 0.859623 | 0.184974 | 0.214 | 0.223 | 1 | 0 | Tns3 | -0.009 | Young |
| 0.859797 | 0.185364 | 0.051 | 0.049 | 1 | 0 | Tpk1 | 0.002 | Young |
| 0.859939 | 0.066596 | 0.1 | 0.103 | 1 | 0 | Fut8 | -0.003 | Young |
| 0.860026 | 0.332785 | 0.067 | 0.066 | 1 | 0 | Asl | 0.001 | Young |
| 0.860035 | 0.120882 | 0.396 | 0.41 | 1 | 0 | Vwa5a | -0.014 | Young |
| 0.860126 | 0.215893 | 0.009 | 0.01 | 1 | 0 | Dusp19 | -0.001 | Young |
| 0.860179 | 0.038682 | 0.086 | 0.083 | 1 | 0 | Gclc | 0.003 | Young |
| 0.860327 | 0.12108 | 0.069 | 0.067 | 1 | 0 | Ccnk | 0.002 | Young |
| 0.860575 | 0.338029 | 0.088 | 0.092 | 1 | 0 | Kdelc2 | -0.004 | Young |
| 0.860827 | 0.095534 | 0.388 | 0.404 | 1 | 0 | Glg1 | -0.016 | Young |
| 0.86105 | 0.07308 | 0.251 | 0.26 | 1 | 0 | Ghitm | -0.009 | Young |
| 0.861092 | 0.34559 | 0.019 | 0.021 | 1 | 0 | Ip6k2 | -0.002 | Young |
| 0.861104 | 0.053654 | 0.016 | 0.015 | 1 | 0 | Wnt11 | 0.001 | Young |
| 0.861169 | 0.008581 | 0.029 | 0.027 | 1 | 0 | Zfp719 | 0.002 | Young |
| 0.861172 | 0.062726 | 0.153 | 0.152 | 1 | 0 | Ctcf | 0.001 | Young |
| 0.861938 | 0.263715 | 0.009 | 0.01 | 1 | 0 | Zfr2 | -0.001 | Young |
| 0.861964 | 0.107304 | 0.187 | 0.193 | 1 | 0 | Csf2rb | -0.006 | Young |
| 0.862115 | 0.529589 | 0.027 | 0.026 | 1 | 0 | Adsl | 0.001 | Young |
| 0.862164 | 0.315611 | 0.009 | 0.01 | 1 | 0 | Carmil1 | -0.001 | Young |
| 0.862545 | 0.086483 | 0.515 | 0.531 | 1 | 0 | Rassf4 | -0.016 | Young |
| 0.862776 | 0.109871 | 0.026 | 0.027 | 1 | 0 | Ift172 | -0.001 | Young |
| 0.862786 | 0.205647 | 0.121 | 0.12 | 1 | 0 | Prpf18 | 0.001 | Young |
| 0.862903 | 0.227481 | 0.165 | 0.166 | 1 | 0 | Rtf1 | -0.001 | Young |
| 0.863135 | 0.161567 | 0.117 | 0.116 | 1 | 0 | Mlxip | 0.001 | Young |
| 0.863523 | 0.845386 | 0.009 | 0.01 | 1 | 0 | Clcf1 | -0.001 | Young |
| 0.863871 | 0.032357 | 0.066 | 0.064 | 1 | 0 | Ube2g1 | 0.002 | Young |
| 0.863961 | 0.299837 | 0.076 | 0.075 | 1 | 0 | Chchd3 | 0.001 | Young |
| 0.864251 | 0.093574 | 0.188 | 0.193 | 1 | 0 | Oat | -0.005 | Young |
| 0.86438 | 0.204019 | 0.115 | 0.115 | 1 | 0 | Supt20 | 0 | Young |
| 0.864631 | 0.136003 | 0.13 | 0.134 | 1 | 0 | Scamp3 | -0.004 | Young |
| 0.864844 | 0.240822 | 0.046 | 0.048 | 1 | 0 | Smarcd1 | -0.002 | Young |
| 0.865018 | 0.059822 | 0.056 | 0.055 | 1 | 0 | Hdgfl2 | 0.001 | Young |

| | | | | | | | | |
|---|---|---|---|---|---|---|---|---|
| 0.865095 | 0.073001 | 0.04 | 0.038 | 1 | 0 | Zfp771 | 0.002 | Young |
| 0.865261 | 0.056384 | 0.346 | 0.347 | 1 | 0 | Lmo2 | -0.001 | Young |
| 0.865358 | 0.175863 | 0.059 | 0.062 | 1 | 0 | 5031425F1 | -0.003 | Young |
| 0.865453 | 0.262548 | 0.018 | 0.019 | 1 | 0 | Trip6 | -0.001 | Young |
| 0.865798 | 0.093169 | 0.279 | 0.291 | 1 | 0 | Limd1 | -0.012 | Young |
| 0.86625 | 0.035548 | 0.048 | 0.049 | 1 | 0 | Fbxo9 | -0.001 | Young |
| 0.866467 | 0.042346 | 0.072 | 0.07 | 1 | 0 | Arl5c | 0.002 | Young |
| 0.86664 | 0.179699 | 0.209 | 0.218 | 1 | 0 | Osbpl8 | -0.009 | Young |
| 0.867042 | 0.011601 | 0.146 | 0.149 | 1 | 0 | Slc31a2 | -0.003 | Young |
| 0.867288 | 0.041971 | 0.219 | 0.226 | 1 | 0 | Ppp1r14b | -0.007 | Young |
| 0.867541 | 0.016001 | 0.077 | 0.075 | 1 | 0 | Nlk | 0.002 | Young |
| 0.86781 | 0.039329 | 0.281 | 0.293 | 1 | 0 | Ptpn6 | -0.012 | Young |
| 0.867888 | 0.117431 | 0.05 | 0.052 | 1 | 0 | Nit1 | -0.002 | Young |
| 0.867893 | 0.128464 | 0.112 | 0.116 | 1 | 0 | Mut | -0.004 | Young |
| 0.868009 | 0.136599 | 0.078 | 0.081 | 1 | 0 | Lage3 | -0.003 | Young |
| 0.868325 | 0.034202 | 0.052 | 0.053 | 1 | 0 | Eif1ad | -0.001 | Young |
| 0.868408 | 0.128733 | 0.063 | 0.062 | 1 | 0 | Inafm1 | 0.001 | Young |
| 0.868486 | 0.368265 | 0.018 | 0.019 | 1 | 0 | Ggcx | -0.001 | Young |
| 0.868781 | 0.073437 | 0.149 | 0.153 | 1 | 0 | Zfp950 | -0.004 | Young |
| 0.868816 | 0.122734 | 0.091 | 0.094 | 1 | 0 | Scaf8 | -0.003 | Young |
| 0.868839 | 0.094905 | 0.041 | 0.042 | 1 | 0 | Nrde2 | -0.001 | Young |
| 0.868918 | 0.141284 | 0.045 | 0.047 | 1 | 0 | Cherp | -0.002 | Young |
| 0.869238 | 0.523052 | 0.013 | 0.012 | 1 | 0 | Zbtb39 | 0.001 | Young |
| 0.869375 | 0.283738 | 0.041 | 0.04 | 1 | 0 | Ltv1 | 0.001 | Young |
| 0.869847 | 0.070833 | 0.08 | 0.082 | 1 | 0 | Foxo1 | -0.002 | Young |
| 0.869907 | 0.266981 | 0.042 | 0.041 | 1 | 0 | Hyal2 | 0.001 | Young |
| 0.870111 | 0.02711 | 0.263 | 0.269 | 1 | 0 | Dmac1 | -0.006 | Young |
| 0.870375 | 0.023209 | 0.124 | 0.122 | 1 | 0 | Rspry1 | 0.002 | Young |
| 0.87058 | 0.215793 | 0.087 | 0.086 | 1 | 0 | Uimc1 | 0.001 | Young |
| 0.870907 | 0.101376 | 0.021 | 0.021 | 1 | 0 | Tchp | 0 | Young |
| 0.870939 | 0.174279 | 0.031 | 0.03 | 1 | 0 | Btbd2 | 0.001 | Young |
| 0.871264 | 0.335908 | 0.046 | 0.048 | 1 | 0 | Mynn | -0.002 | Young |
| 0.87145 | 0.011955 | 0.275 | 0.285 | 1 | 0 | Slk | -0.01 | Young |
| 0.871804 | 0.264474 | 0.091 | 0.09 | 1 | 0 | Sar1b | 0.001 | Young |
| 0.871868 | 0.020221 | 0.092 | 0.094 | 1 | 0 | Capn7 | -0.002 | Young |
| 0.871983 | 0.227183 | 0.041 | 0.04 | 1 | 0 | Eepd1 | 0.001 | Young |
| 0.872007 | 0.121869 | 0.095 | 0.098 | 1 | 0 | Tifab | -0.003 | Young |
| 0.872231 | 0.014183 | 0.99 | 0.992 | 1 | 0 | mt-Nd1 | -0.002 | Young |
| 0.872484 | 0.052799 | 0.04 | 0.041 | 1 | 0 | Zbtb2 | -0.001 | Young |
| 0.873069 | 0.035207 | 0.026 | 0.027 | 1 | 0 | 2900093K2 | -0.001 | Young |
| 0.87367 | 0.296344 | 0.097 | 0.097 | 1 | 0 | Cib1 | 0 | Young |
| 0.873702 | 0.203863 | 0.152 | 0.152 | 1 | 0 | Psmb9 | 0 | Young |
| 0.874508 | 0.156164 | 0.017 | 0.016 | 1 | 0 | Fbxo46 | 0.001 | Young |
| 0.874784 | 0.112174 | 0.139 | 0.138 | 1 | 0 | Esyt2 | 0.001 | Young |
| 0.8748 | 0.004201 | 0.938 | 0.94 | 1 | 0 | Ahnak | -0.002 | Young |

| | | | | | | | | |
|---|---|---|---|---|---|---|---|---|
| 0.874845 | 0.019629 | 0.643 | 0.674 | 1 | 0 | Rbm39 | -0.031 | Young |
| 0.874922 | 0.25128 | 0.017 | 0.018 | 1 | 0 | Nkiras1 | -0.001 | Young |
| 0.87504 | 0.079163 | 0.062 | 0.064 | 1 | 0 | Rras2 | -0.002 | Young |
| 0.875076 | 0.149502 | 0.233 | 0.234 | 1 | 0 | Nadk | -0.001 | Young |
| 0.875327 | 0.02872 | 0.272 | 0.275 | 1 | 0 | Ppp1cb | -0.003 | Young |
| 0.875493 | 0.178979 | 0.161 | 0.167 | 1 | 0 | Ube2z | -0.006 | Young |
| 0.875829 | 0.110019 | 0.053 | 0.055 | 1 | 0 | Rrp7a | -0.002 | Young |
| 0.87606 | 0.080096 | 0.273 | 0.28 | 1 | 0 | Uqcr10 | -0.007 | Young |
| 0.876093 | 0.416121 | 0.043 | 0.042 | 1 | 0 | Pak4 | 0.001 | Young |
| 0.876093 | 0.225668 | 0.075 | 0.074 | 1 | 0 | Abhd16a | 0.001 | Young |
| 0.876247 | 0.103619 | 0.225 | 0.227 | 1 | 0 | Irak1 | -0.002 | Young |
| 0.876347 | 0.077272 | 0.537 | 0.565 | 1 | 0 | Snrnp70 | -0.028 | Young |
| 0.876895 | 0.070094 | 0.304 | 0.316 | 1 | 0 | Cdc42se1 | -0.012 | Young |
| 0.877864 | 0.390217 | 0.023 | 0.022 | 1 | 0 | Tapt1 | 0.001 | Young |
| 0.878166 | 0.209216 | 0.09 | 0.089 | 1 | 0 | Sap18 | 0.001 | Young |
| 0.878273 | 0.080198 | 0.045 | 0.047 | 1 | 0 | Caap1 | -0.002 | Young |
| 0.878404 | 0.104464 | 0.159 | 0.159 | 1 | 0 | Camk2d | 0 | Young |
| 0.878677 | 0.238424 | 0.079 | 0.078 | 1 | 0 | Lcmt1 | 0.001 | Young |
| 0.878751 | 0.116447 | 0.067 | 0.066 | 1 | 0 | Gpatch2l | 0.001 | Young |
| 0.878918 | 0.109387 | 0.224 | 0.23 | 1 | 0 | Twf2 | -0.006 | Young |
| 0.879026 | 0.086916 | 0.152 | 0.15 | 1 | 0 | Atp5g1 | 0.002 | Young |
| 0.879158 | 0.162962 | 0.032 | 0.031 | 1 | 0 | Dnmbp | 0.001 | Young |
| 0.879415 | 0.039493 | 0.111 | 0.109 | 1 | 0 | Upf1 | 0.002 | Young |
| 0.879455 | 0.149748 | 0.201 | 0.209 | 1 | 0 | Tmem14c | -0.008 | Young |
| 0.879466 | 0.145671 | 0.082 | 0.081 | 1 | 0 | Ubxn7 | 0.001 | Young |
| 0.87953 | 0.070838 | 0.103 | 0.107 | 1 | 0 | Lars | -0.004 | Young |
| 0.879667 | 0.157621 | 0.027 | 0.026 | 1 | 0 | Crcp | 0.001 | Young |
| 0.879706 | 0.313029 | 0.025 | 0.026 | 1 | 0 | Usp42 | -0.001 | Young |
| 0.879805 | 0.56189 | 0.047 | 0.047 | 1 | 0 | Serpinb1a | 0 | Young |
| 0.879897 | 0.095458 | 0.361 | 0.373 | 1 | 0 | Atp5k | -0.012 | Young |
| 0.880331 | 0.243772 | 0.123 | 0.123 | 1 | 0 | Ptpn7 | 0 | Young |
| 0.880352 | 0.003813 | 0.321 | 0.332 | 1 | 0 | Eif4h | -0.011 | Young |
| 0.880511 | 0.074706 | 0.185 | 0.19 | 1 | 0 | Dyrk2 | -0.005 | Young |
| 0.880614 | 0.289544 | 0.023 | 0.022 | 1 | 0 | Zfp574 | 0.001 | Young |
| 0.881291 | 0.087377 | 0.061 | 0.063 | 1 | 0 | Skiv2l2 | -0.002 | Young |
| 0.881312 | 0.091915 | 0.238 | 0.239 | 1 | 0 | Dpm3 | -0.001 | Young |
| 0.881956 | 0.23758 | 0.041 | 0.042 | 1 | 0 | Tpbgl | -0.001 | Young |
| 0.882306 | 0.02058 | 0.258 | 0.265 | 1 | 0 | Ptges3 | -0.007 | Young |
| 0.882443 | 0.085607 | 0.03 | 0.031 | 1 | 0 | Zfp64 | -0.001 | Young |
| 0.882473 | 0.235879 | 0.095 | 0.094 | 1 | 0 | Srsf4 | 0.001 | Young |
| 0.88252 | 0.243455 | 0.115 | 0.119 | 1 | 0 | Smpd1 | -0.004 | Young |
| 0.88254 | 0.210808 | 0.033 | 0.034 | 1 | 0 | Lin37 | -0.001 | Young |
| 0.882543 | 0.096435 | 0.255 | 0.259 | 1 | 0 | Apbb1ip | -0.004 | Young |
| 0.883017 | 0.033162 | 0.091 | 0.093 | 1 | 0 | Hint2 | -0.002 | Young |
| 0.883066 | 0.211421 | 0.056 | 0.057 | 1 | 0 | Alg5 | -0.001 | Young |

| | | | | | | | | |
|---|---|---|---|---|---|---|---|---|
| 0.88351 | 0.030757 | 0.022 | 0.023 | 1 | 0 | Thap4 | -0.001 | Young |
| 0.883744 | 0.092704 | 0.068 | 0.067 | 1 | 0 | Sh3tc1 | 0.001 | Young |
| 0.883947 | 0.012404 | 0.927 | 0.926 | 1 | 0 | Zeb2 | 0.001 | Young |
| 0.884274 | 0.087852 | 0.393 | 0.401 | 1 | 0 | Ckb | -0.008 | Young |
| 0.884481 | 0.110986 | 0.037 | 0.036 | 1 | 0 | Nrm | 0.001 | Young |
| 0.884852 | 0.13497 | 0.219 | 0.226 | 1 | 0 | Snrpd3 | -0.007 | Young |
| 0.884878 | 0.099612 | 0.03 | 0.031 | 1 | 0 | Nxt1 | -0.001 | Young |
| 0.885043 | 0.34054 | 0.019 | 0.018 | 1 | 0 | Timmdc1 | 0.001 | Young |
| 0.885278 | 0.214767 | 0.092 | 0.092 | 1 | 0 | Zmym4 | 0 | Young |
| 0.885337 | 0.006026 | 0.029 | 0.03 | 1 | 0 | Rars2 | -0.001 | Young |
| 0.885668 | 0.049761 | 0.051 | 0.052 | 1 | 0 | Trappc2 | -0.001 | Young |
| 0.885684 | 0.053662 | 0.152 | 0.15 | 1 | 0 | Herpud2 | 0.002 | Young |
| 0.885809 | 0.238303 | 0.095 | 0.094 | 1 | 0 | Hars | 0.001 | Young |
| 0.885813 | 0.117465 | 0.067 | 0.066 | 1 | 0 | 1700123O2 | 0.001 | Young |
| 0.885827 | 0.085316 | 0.14 | 0.144 | 1 | 0 | Mtmr3 | -0.004 | Young |
| 0.886012 | 0.072131 | 0.049 | 0.051 | 1 | 0 | Coq5 | -0.002 | Young |
| 0.886496 | 0.088691 | 0.22 | 0.222 | 1 | 0 | Tmem160 | -0.002 | Young |
| 0.886497 | 0.012515 | 0.641 | 0.674 | 1 | 0 | Gltp | -0.033 | Young |
| 0.886535 | 0.017342 | 0.134 | 0.133 | 1 | 0 | Trappc2l | 0.001 | Young |
| 0.886604 | 0.330396 | 0.028 | 0.027 | 1 | 0 | Pdp1 | 0.001 | Young |
| 0.886748 | 0.023547 | 0.135 | 0.138 | 1 | 0 | Foxj2 | -0.003 | Young |
| 0.887132 | 0.055685 | 0.022 | 0.023 | 1 | 0 | Scd2 | -0.001 | Young |
| 0.887188 | 0.130915 | 0.03 | 0.031 | 1 | 0 | Exosc2 | -0.001 | Young |
| 0.887314 | 0.077122 | 0.029 | 0.03 | 1 | 0 | Aen | -0.001 | Young |
| 0.887375 | 0.204222 | 0.114 | 0.118 | 1 | 0 | Paox | -0.004 | Young |
| 0.887376 | 0.056813 | 0.03 | 0.031 | 1 | 0 | Strada | -0.001 | Young |
| 0.887433 | 0.007042 | 0.174 | 0.178 | 1 | 0 | Rnf111 | -0.004 | Young |
| 0.887469 | 0.082134 | 0.06 | 0.059 | 1 | 0 | Pde4a | 0.001 | Young |
| 0.887884 | 0.077337 | 0.11 | 0.112 | 1 | 0 | Lemd2 | -0.002 | Young |
| 0.887952 | 0.436531 | 0.029 | 0.029 | 1 | 0 | Mypopos | 0 | Young |
| 0.888147 | 0.179642 | 0.022 | 0.023 | 1 | 0 | Surf2 | -0.001 | Young |
| 0.888486 | 0.262748 | 0.078 | 0.078 | 1 | 0 | Thap3 | 0 | Young |
| 0.888733 | 0.134667 | 0.188 | 0.189 | 1 | 0 | Cic | -0.001 | Young |
| 0.888844 | 0.075797 | 0.148 | 0.152 | 1 | 0 | Fyttd1 | -0.004 | Young |
| 0.888969 | 0.30949 | 0.045 | 0.044 | 1 | 0 | Zbtb43 | 0.001 | Young |
| 0.889509 | 0.155733 | 0.014 | 0.015 | 1 | 0 | Zfat | -0.001 | Young |
| 0.889561 | 0.018417 | 0.06 | 0.062 | 1 | 0 | Ankrd10 | -0.002 | Young |
| 0.889763 | 0.110445 | 0.167 | 0.174 | 1 | 0 | Insr | -0.007 | Young |
| 0.890151 | 0.29942 | 0.058 | 0.057 | 1 | 0 | Arhgap23 | 0.001 | Young |
| 0.890706 | 0.800491 | 0.01 | 0.01 | 1 | 0 | 9130008F2 | 0 | Young |
| 0.890706 | 0.427669 | 0.01 | 0.01 | 1 | 0 | Cep89 | 0 | Young |
| 0.890836 | 0.636991 | 0.014 | 0.014 | 1 | 0 | Elp6 | 0 | Young |
| 0.89095 | 0.075931 | 0.014 | 0.015 | 1 | 0 | C1qtnf12 | -0.001 | Young |
| 0.890953 | 0.081868 | 0.12 | 0.123 | 1 | 0 | Ahcyl1 | -0.003 | Young |
| 0.891068 | 0.031129 | 0.08 | 0.082 | 1 | 0 | Trim27 | -0.002 | Young |

| | | | | | | | | |
|---|---|---|---|---|---|---|---|---|
| 0.891236 | 0.053203 | 0.13 | 0.133 | 1 | 0 | Lsm12 | -0.003 | Young |
| 0.891394 | 0.118135 | 0.107 | 0.105 | 1 | 0 | Mprip | 0.002 | Young |
| 0.891576 | 0.070816 | 0.042 | 0.041 | 1 | 0 | Saysd1 | 0.001 | Young |
| 0.891614 | 0.14274 | 0.109 | 0.112 | 1 | 0 | Pfdn1 | -0.003 | Young |
| 0.891634 | 0.186989 | 0.061 | 0.06 | 1 | 0 | Strip1 | 0.001 | Young |
| 0.891911 | 0.103654 | 0.166 | 0.172 | 1 | 0 | Coa5 | -0.006 | Young |
| 0.891969 | 0.378697 | 0.024 | 0.023 | 1 | 0 | Ddx51 | 0.001 | Young |
| 0.892097 | 0.140993 | 0.019 | 0.018 | 1 | 0 | Dimt1 | 0.001 | Young |
| 0.89212 | 0.094029 | 0.175 | 0.179 | 1 | 0 | Tpi1 | -0.004 | Young |
| 0.892184 | 0.166884 | 0.059 | 0.06 | 1 | 0 | Gclm | -0.001 | Young |
| 0.892185 | 0.067526 | 0.125 | 0.127 | 1 | 0 | Mogs | -0.002 | Young |
| 0.892349 | 0.057278 | 0.061 | 0.06 | 1 | 0 | Stat5a | 0.001 | Young |
| 0.892355 | 0.150646 | 0.238 | 0.248 | 1 | 0 | Emc7 | -0.01 | Young |
| 0.892476 | 0.283725 | 0.014 | 0.014 | 1 | 0 | Psat1 | 0 | Young |
| 0.892626 | 0.121752 | 0.107 | 0.109 | 1 | 0 | Ehmt1 | -0.002 | Young |
| 0.892739 | 0.12754 | 0.019 | 0.018 | 1 | 0 | Zfp568 | 0.001 | Young |
| 0.892866 | 0.217805 | 0.039 | 0.038 | 1 | 0 | Rab2b | 0.001 | Young |
| 0.892922 | 0.271684 | 0.039 | 0.038 | 1 | 0 | Cisd3 | 0.001 | Young |
| 0.892941 | 0.168756 | 0.054 | 0.056 | 1 | 0 | Atxn1l | -0.002 | Young |
| 0.892955 | 0.027002 | 0.234 | 0.233 | 1 | 0 | Arrb1 | 0.001 | Young |
| 0.893088 | 0.505297 | 0.01 | 0.01 | 1 | 0 | Mgll | 0 | Young |
| 0.893117 | 0.083867 | 0.092 | 0.09 | 1 | 0 | Cltb | 0.002 | Young |
| 0.893282 | 0.090644 | 0.059 | 0.057 | 1 | 0 | Cdk5rap3 | 0.002 | Young |
| 0.893473 | 0.191375 | 0.014 | 0.015 | 1 | 0 | Setd6 | -0.001 | Young |
| 0.893712 | 0.409185 | 0.054 | 0.053 | 1 | 0 | Trmt2a | 0.001 | Young |
| 0.893812 | 0.233047 | 0.046 | 0.045 | 1 | 0 | Utp20 | 0.001 | Young |
| 0.893857 | 0.131513 | 0.036 | 0.037 | 1 | 0 | Sema4b | -0.001 | Young |
| 0.893862 | 0.137454 | 0.13 | 0.129 | 1 | 0 | Vti1b | 0.001 | Young |
| 0.893955 | 0.499948 | 0.01 | 0.01 | 1 | 0 | Dhodh | 0 | Young |
| 0.893988 | 0.046161 | 0.225 | 0.23 | 1 | 0 | Tnrc6a | -0.005 | Young |
| 0.894532 | 0.257264 | 0.076 | 0.075 | 1 | 0 | Cnnm3 | 0.001 | Young |
| 0.894604 | 0.102091 | 0.288 | 0.297 | 1 | 0 | Srgap2 | -0.009 | Young |
| 0.894976 | 0.090953 | 0.139 | 0.144 | 1 | 0 | Arl5a | -0.005 | Young |
| 0.895146 | 0.161061 | 0.019 | 0.018 | 1 | 0 | 4930555A( | 0.001 | Young |
| 0.895312 | 0.03099 | 0.161 | 0.164 | 1 | 0 | Irf9 | -0.003 | Young |
| 0.895327 | 0.2487 | 0.133 | 0.138 | 1 | 0 | Idh3g | -0.005 | Young |
| 0.895392 | 0.208467 | 0.014 | 0.014 | 1 | 0 | Gpatch4 | 0 | Young |
| 0.89567 | 0.143192 | 0.032 | 0.031 | 1 | 0 | Slc39a11 | 0.001 | Young |
| 0.895689 | 0.294495 | 0.01 | 0.01 | 1 | 0 | Ccdc51 | 0 | Young |
| 0.896057 | 0.151342 | 0.086 | 0.085 | 1 | 0 | Gatsl2 | 0.001 | Young |
| 0.896248 | 0.079752 | 0.021 | 0.022 | 1 | 0 | Bcdin3d | -0.001 | Young |
| 0.896338 | 0.054762 | 0.161 | 0.16 | 1 | 0 | Etv5 | 0.001 | Young |
| 0.896547 | 0.057113 | 0.021 | 0.022 | 1 | 0 | Rbm45 | -0.001 | Young |
| 0.896627 | 0.165997 | 0.077 | 0.079 | 1 | 0 | Rarg | -0.002 | Young |
| 0.896753 | 0.252353 | 0.03 | 0.031 | 1 | 0 | Dcun1d3 | -0.001 | Young |

| | | | | | | | | |
|---|---|---|---|---|---|---|---|---|
| 0.897089 | 0.085136 | 0.112 | 0.115 | 1 | 0 | Wbp4 | -0.003 | Young |
| 0.897092 | 0.039495 | 0.036 | 0.037 | 1 | 0 | Riox2 | -0.001 | Young |
| 0.897154 | 0.034156 | 0.467 | 0.488 | 1 | 0 | Gns | -0.021 | Young |
| 0.897238 | 0.118306 | 0.028 | 0.029 | 1 | 0 | Cdkal1 | -0.001 | Young |
| 0.897454 | 0.183704 | 0.121 | 0.122 | 1 | 0 | Stxbp5 | -0.001 | Young |
| 0.897471 | 0.456764 | 0.034 | 0.036 | 1 | 0 | Tlr3 | -0.002 | Young |
| 0.897485 | 0.264779 | 0.045 | 0.044 | 1 | 0 | Sdhaf1 | 0.001 | Young |
| 0.897653 | 0.00406 | 0.013 | 0.014 | 1 | 0 | Vps51 | -0.001 | Young |
| 0.897881 | 0.240816 | 0.057 | 0.056 | 1 | 0 | R3hcc1l | 0.001 | Young |
| 0.897944 | 0.075566 | 0.325 | 0.332 | 1 | 0 | Plekho1 | -0.007 | Young |
| 0.897945 | 0.106028 | 0.014 | 0.014 | 1 | 0 | Adck2 | 0 | Young |
| 0.897945 | 0.097454 | 0.014 | 0.014 | 1 | 0 | Zfp52 | 0 | Young |
| 0.898222 | 0.00802 | 0.035 | 0.036 | 1 | 0 | Tpm3-rs7 | -0.001 | Young |
| 0.898607 | 0.333246 | 0.046 | 0.045 | 1 | 0 | Agpat1 | 0.001 | Young |
| 0.898803 | 0.009358 | 0.064 | 0.063 | 1 | 0 | Odf2 | 0.001 | Young |
| 0.898941 | 0.151824 | 0.01 | 0.01 | 1 | 0 | 1810034E1 | 0 | Young |
| 0.898968 | 0.041905 | 0.037 | 0.036 | 1 | 0 | Map4k2 | 0.001 | Young |
| 0.899015 | 0.096039 | 0.3 | 0.313 | 1 | 0 | Nckap1l | -0.013 | Young |
| 0.899039 | 0.048737 | 0.014 | 0.014 | 1 | 0 | Fgfbp3 | 0 | Young |
| 0.899061 | 0.429158 | 0.053 | 0.052 | 1 | 0 | Scarb1 | 0.001 | Young |
| 0.899111 | 0.380752 | 0.042 | 0.041 | 1 | 0 | Slc25a32 | 0.001 | Young |
| 0.899164 | 0.097261 | 0.013 | 0.014 | 1 | 0 | Phpt1 | -0.001 | Young |
| 0.89921 | 0.18422 | 0.049 | 0.048 | 1 | 0 | Prmt5 | 0.001 | Young |
| 0.899825 | 0.111422 | 0.013 | 0.014 | 1 | 0 | BC051226 | -0.001 | Young |
| 0.899956 | 0.187942 | 0.029 | 0.03 | 1 | 0 | Macrod1 | -0.001 | Young |
| 0.900123 | 0.033088 | 0.141 | 0.144 | 1 | 0 | Rbm33 | -0.003 | Young |
| 0.900413 | 0.041429 | 0.508 | 0.534 | 1 | 0 | Rock2 | -0.026 | Young |
| 0.900555 | 0.21326 | 0.096 | 0.096 | 1 | 0 | Xylt2 | 0 | Young |
| 0.900676 | 0.005292 | 0.01 | 0.01 | 1 | 0 | Ahrr | 0 | Young |
| 0.900749 | 0.052092 | 0.1 | 0.098 | 1 | 0 | Desi2 | 0.002 | Young |
| 0.900864 | 0.127218 | 0.013 | 0.014 | 1 | 0 | Abcb6 | -0.001 | Young |
| 0.901075 | 0.163876 | 0.167 | 0.168 | 1 | 0 | Hexim1 | -0.001 | Young |
| 0.901331 | 0.102927 | 0.113 | 0.116 | 1 | 0 | Gtf2a2 | -0.003 | Young |
| 0.901358 | 0.120844 | 0.124 | 0.127 | 1 | 0 | Eif4e3 | -0.003 | Young |
| 0.90146 | 0.378161 | 0.101 | 0.101 | 1 | 0 | Acox3 | 0 | Young |
| 0.9015 | 0.074188 | 0.037 | 0.036 | 1 | 0 | 1110008P1 | 0.001 | Young |
| 0.90168 | 0.256471 | 0.1 | 0.1 | 1 | 0 | Dusp7 | 0 | Young |
| 0.901737 | 0.14468 | 0.105 | 0.108 | 1 | 0 | Znrd1 | -0.003 | Young |
| 0.90193 | 0.113326 | 0.227 | 0.23 | 1 | 0 | Sin3b | -0.003 | Young |
| 0.901933 | 0.124709 | 0.034 | 0.033 | 1 | 0 | Recql5 | 0.001 | Young |
| 0.901959 | 0.1128 | 0.286 | 0.298 | 1 | 0 | Mdfic | -0.012 | Young |
| 0.901982 | 0.20831 | 0.057 | 0.056 | 1 | 0 | Flnb | 0.001 | Young |
| 0.90214 | 0.103461 | 0.107 | 0.109 | 1 | 0 | Chuk | -0.002 | Young |
| 0.902473 | 0.149252 | 0.051 | 0.052 | 1 | 0 | Cwc22 | -0.001 | Young |
| 0.902588 | 0.511211 | 0.032 | 0.033 | 1 | 0 | Ints7 | -0.001 | Young |

| | | | | | | | | |
|---|---|---|---|---|---|---|---|---|
| 0.902648 | 0.126435 | 0.126 | 0.126 | 1 | 0 | Dag1 | 0 | Young |
| 0.902745 | 0.165002 | 0.145 | 0.146 | 1 | 0 | Ncoa1 | -0.001 | Young |
| 0.902885 | 0.116619 | 0.093 | 0.096 | 1 | 0 | Rubcn | -0.003 | Young |
| 0.902943 | 0.010548 | 0.067 | 0.066 | 1 | 0 | Hif1an | 0.001 | Young |
| 0.903074 | 0.064396 | 0.08 | 0.082 | 1 | 0 | Tmem135 | -0.002 | Young |
| 0.903151 | 0.134266 | 0.231 | 0.235 | 1 | 0 | Lmbrd1 | -0.004 | Young |
| 0.903321 | 0.136841 | 0.013 | 0.014 | 1 | 0 | Mthfd1l | -0.001 | Young |
| 0.903548 | 0.096537 | 0.067 | 0.066 | 1 | 0 | Nol11 | 0.001 | Young |
| 0.904652 | 0.398473 | 0.058 | 0.057 | 1 | 0 | Rab33b | 0.001 | Young |
| 0.905326 | 0.154629 | 0.024 | 0.023 | 1 | 0 | Abhd14b | 0.001 | Young |
| 0.905549 | 0.059402 | 0.14 | 0.144 | 1 | 0 | Bdp1 | -0.004 | Young |
| 0.905783 | 0.039233 | 0.205 | 0.208 | 1 | 0 | Rmnd5a | -0.003 | Young |
| 0.906563 | 0.230111 | 0.02 | 0.019 | 1 | 0 | Chst15 | 0.001 | Young |
| 0.906729 | 0.077932 | 0.034 | 0.033 | 1 | 0 | Gtpbp10 | 0.001 | Young |
| 0.907224 | 0.084748 | 0.14 | 0.14 | 1 | 0 | Lpcat1 | 0 | Young |
| 0.907765 | 0.047302 | 0.012 | 0.012 | 1 | 0 | Plcb1 | 0 | Young |
| 0.907964 | 0.007301 | 0.012 | 0.012 | 1 | 0 | Fam206a | 0 | Young |
| 0.908271 | 0.110434 | 0.067 | 0.066 | 1 | 0 | Psip1 | 0.001 | Young |
| 0.908357 | 0.214495 | 0.067 | 0.066 | 1 | 0 | Zfp787 | 0.001 | Young |
| 0.908403 | 0.029654 | 0.217 | 0.218 | 1 | 0 | Pnp | -0.001 | Young |
| 0.908687 | 0.183988 | 0.049 | 0.051 | 1 | 0 | Rhot2 | -0.002 | Young |
| 0.908725 | 0.186107 | 0.133 | 0.137 | 1 | 0 | Mark2 | -0.004 | Young |
| 0.908896 | 0.058878 | 0.136 | 0.14 | 1 | 0 | Ascc3 | -0.004 | Young |
| 0.908984 | 0.17406 | 0.046 | 0.045 | 1 | 0 | Ppcdc | 0.001 | Young |
| 0.909409 | 0.047538 | 0.052 | 0.053 | 1 | 0 | Pfkl | -0.001 | Young |
| 0.909458 | 0.113575 | 0.098 | 0.097 | 1 | 0 | Mob3b | 0.001 | Young |
| 0.909608 | 0.248181 | 0.07 | 0.07 | 1 | 0 | Dcaf5 | 0 | Young |
| 0.909862 | 0.067593 | 0.088 | 0.088 | 1 | 0 | Get4 | 0 | Young |
| 0.909868 | 0.031117 | 0.192 | 0.193 | 1 | 0 | 1810058I2 | -0.001 | Young |
| 0.910197 | 0.161708 | 0.163 | 0.168 | 1 | 0 | Rexo2 | -0.005 | Young |
| 0.910644 | 0.040092 | 0.095 | 0.094 | 1 | 0 | Fam193b | 0.001 | Young |
| 0.910705 | 0.0792 | 0.162 | 0.166 | 1 | 0 | Setd3 | -0.004 | Young |
| 0.91095 | 0.224338 | 0.012 | 0.012 | 1 | 0 | Sh2b2 | 0 | Young |
| 0.910987 | 0.081118 | 0.109 | 0.108 | 1 | 0 | Nt5c | 0.001 | Young |
| 0.911081 | 0.151905 | 0.195 | 0.197 | 1 | 0 | Tnrc6c | -0.002 | Young |
| 0.911314 | 0.132591 | 0.173 | 0.175 | 1 | 0 | Prkar2a | -0.002 | Young |
| 0.911547 | 0.11896 | 0.012 | 0.012 | 1 | 0 | Wdr90 | 0 | Young |
| 0.911703 | 0.26514 | 0.097 | 0.097 | 1 | 0 | Gsdmd | 0 | Young |
| 0.911922 | 0.153358 | 0.148 | 0.149 | 1 | 0 | Smarcc1 | -0.001 | Young |
| 0.912008 | 0.106407 | 0.02 | 0.019 | 1 | 0 | Wdr53 | 0.001 | Young |
| 0.91225 | 0.134353 | 0.092 | 0.094 | 1 | 0 | AW549877 | -0.002 | Young |
| 0.912283 | 0.122318 | 0.4 | 0.423 | 1 | 0 | Srp9 | -0.023 | Young |
| 0.912546 | 0.065203 | 0.088 | 0.09 | 1 | 0 | Homer1 | -0.002 | Young |
| 0.912651 | 0.103309 | 0.049 | 0.048 | 1 | 0 | Dohh | 0.001 | Young |
| 0.91329 | 0.194773 | 0.13 | 0.134 | 1 | 0 | Ndufb2 | -0.004 | Young |

| | | | | | | | | |
|---|---|---|---|---|---|---|---|---|
| 0.913435 | 0.020016 | 0.152 | 0.155 | 1 | 0 | Smndc1 | -0.003 | Young |
| 0.913728 | 0.040008 | 0.22 | 0.224 | 1 | 0 | Rbm47 | -0.004 | Young |
| 0.91373 | 0.033777 | 0.088 | 0.086 | 1 | 0 | Lsm3 | 0.002 | Young |
| 0.913936 | 0.053662 | 0.012 | 0.012 | 1 | 0 | Nup37 | 0 | Young |
| 0.914277 | 0.48592 | 0.021 | 0.022 | 1 | 0 | Ptges2 | -0.001 | Young |
| 0.914381 | 0.017706 | 0.048 | 0.047 | 1 | 0 | Wdr18 | 0.001 | Young |
| 0.914466 | 0.097242 | 0.196 | 0.196 | 1 | 0 | Hsd17b4 | 0 | Young |
| 0.914524 | 0.090608 | 0.025 | 0.026 | 1 | 0 | Tstd3 | -0.001 | Young |
| 0.914814 | 0.088563 | 0.102 | 0.101 | 1 | 0 | Ilkap | 0.001 | Young |
| 0.915065 | 0.181491 | 0.035 | 0.036 | 1 | 0 | Prkca | -0.001 | Young |
| 0.915449 | 0.153292 | 0.168 | 0.171 | 1 | 0 | Atraid | -0.003 | Young |
| 0.915701 | 0.168686 | 0.06 | 0.062 | 1 | 0 | Shkbp1 | -0.002 | Young |
| 0.915903 | 0.248837 | 0.065 | 0.064 | 1 | 0 | Thumpd3 | 0.001 | Young |
| 0.915928 | 0.339948 | 0.012 | 0.012 | 1 | 0 | Gm29291 | 0 | Young |
| 0.916326 | 0.143761 | 0.166 | 0.171 | 1 | 0 | Zzef1 | -0.005 | Young |
| 0.91634 | 0.148198 | 0.108 | 0.108 | 1 | 0 | Tbk1 | 0 | Young |
| 0.916442 | 0.308433 | 0.097 | 0.097 | 1 | 0 | Cds2 | 0 | Young |
| 0.916725 | 0.362902 | 0.012 | 0.012 | 1 | 0 | Gstp2 | 0 | Young |
| 0.916789 | 0.159587 | 0.209 | 0.213 | 1 | 0 | Cnot1 | -0.004 | Young |
| 0.916896 | 0.057936 | 0.533 | 0.549 | 1 | 0 | Cyth4 | -0.016 | Young |
| 0.917258 | 0.058027 | 0.016 | 0.015 | 1 | 0 | Supv3l1 | 0.001 | Young |
| 0.917471 | 0.126619 | 0.144 | 0.148 | 1 | 0 | Tmem173 | -0.004 | Young |
| 0.917629 | 0.013165 | 0.57 | 0.579 | 1 | 0 | Luc7l2 | -0.009 | Young |
| 0.918034 | 0.134454 | 0.019 | 0.019 | 1 | 0 | 2-Sep | 0 | Young |
| 0.918119 | 0.358212 | 0.012 | 0.012 | 1 | 0 | Pnpla1 | 0 | Young |
| 0.918163 | 0.171438 | 0.131 | 0.135 | 1 | 0 | Dhx36 | -0.004 | Young |
| 0.918624 | 0.061869 | 0.14 | 0.144 | 1 | 0 | Puf60 | -0.004 | Young |
| 0.91866 | 0.002614 | 0.016 | 0.015 | 1 | 0 | Sirt4 | 0.001 | Young |
| 0.918667 | 0.088823 | 0.134 | 0.138 | 1 | 0 | Rps6kb1 | -0.004 | Young |
| 0.919083 | 0.365632 | 0.039 | 0.038 | 1 | 0 | Top3b | 0.001 | Young |
| 0.91927 | 0.198337 | 0.02 | 0.021 | 1 | 0 | 2700049A( | -0.001 | Young |
| 0.919358 | 0.052749 | 0.093 | 0.094 | 1 | 0 | Ndufa9 | -0.001 | Young |
| 0.91945 | 0.087189 | 0.032 | 0.033 | 1 | 0 | Prelid2 | -0.001 | Young |
| 0.919636 | 0.629399 | 0.011 | 0.011 | 1 | 0 | Apold1 | 0 | Young |
| 0.919637 | 0.092299 | 0.181 | 0.186 | 1 | 0 | Apaf1 | -0.005 | Young |
| 0.919887 | 0.075967 | 0.016 | 0.015 | 1 | 0 | Gm37305 | 0.001 | Young |
| 0.91993 | 0.211212 | 0.049 | 0.051 | 1 | 0 | Zfp266 | -0.002 | Young |
| 0.920298 | 0.085556 | 0.039 | 0.04 | 1 | 0 | Cbfa2t2 | -0.001 | Young |
| 0.920311 | 0.476127 | 0.012 | 0.012 | 1 | 0 | AC162302. | 0 | Young |
| 0.920503 | 0.009413 | 0.061 | 0.06 | 1 | 0 | Pvr | 0.001 | Young |
| 0.920682 | 0.021065 | 0.125 | 0.127 | 1 | 0 | Wipf2 | -0.002 | Young |
| 0.920826 | 0.235118 | 0.113 | 0.116 | 1 | 0 | Usp14 | -0.003 | Young |
| 0.921114 | 0.123462 | 0.084 | 0.083 | 1 | 0 | Kansl1l | 0.001 | Young |
| 0.921305 | 0.064949 | 0.217 | 0.22 | 1 | 0 | Ypel5 | -0.003 | Young |
| 0.921441 | 0.170765 | 0.076 | 0.075 | 1 | 0 | Foxk1 | 0.001 | Young |

| | | | | | | | | |
|---|---|---|---|---|---|---|---|---|
| 0.921516 | 0.193028 | 0.115 | 0.118 | 1 | 0 | Tk2 | -0.003 | Young |
| 0.92169 | 0.12123 | 0.054 | 0.053 | 1 | 0 | Tsfm | 0.001 | Young |
| 0.921773 | 0.284687 | 0.074 | 0.074 | 1 | 0 | Tmem64 | 0 | Young |
| 0.921803 | 0.1243 | 0.06 | 0.059 | 1 | 0 | Aldh1a2 | 0.001 | Young |
| 0.921838 | 0.163149 | 0.155 | 0.156 | 1 | 0 | Wbp11 | -0.001 | Young |
| 0.921851 | 0.179456 | 0.144 | 0.145 | 1 | 0 | Kxd1 | -0.001 | Young |
| 0.921942 | 0.091441 | 0.021 | 0.021 | 1 | 0 | Wdr6 | 0 | Young |
| 0.921977 | 0.085454 | 0.351 | 0.36 | 1 | 0 | Uqcrq | -0.009 | Young |
| 0.922096 | 0.38863 | 0.044 | 0.045 | 1 | 0 | Cdk5 | -0.001 | Young |
| 0.922321 | 0.320606 | 0.041 | 0.042 | 1 | 0 | Parp3 | -0.001 | Young |
| 0.92268 | 0.058974 | 0.096 | 0.098 | 1 | 0 | Cab39l | -0.002 | Young |
| 0.922994 | 0.055099 | 0.072 | 0.071 | 1 | 0 | Ttpal | 0.001 | Young |
| 0.923264 | 0.085644 | 0.077 | 0.078 | 1 | 0 | Mrpl41 | -0.001 | Young |
| 0.923311 | 0.03178 | 0.502 | 0.521 | 1 | 0 | Gpx4 | -0.019 | Young |
| 0.923465 | 0.26009 | 0.019 | 0.019 | 1 | 0 | Trmt10b | 0 | Young |
| 0.923734 | 0.099587 | 0.011 | 0.011 | 1 | 0 | Spaca9 | 0 | Young |
| 0.923885 | 0.141655 | 0.226 | 0.23 | 1 | 0 | Ppp4r3b | -0.004 | Young |
| 0.92398 | 0.135634 | 0.206 | 0.212 | 1 | 0 | Commd8 | -0.006 | Young |
| 0.924015 | 0.174305 | 0.036 | 0.036 | 1 | 0 | Gpr89 | 0 | Young |
| 0.924099 | 0.524178 | 0.012 | 0.012 | 1 | 0 | Hist3h2ba | 0 | Young |
| 0.924318 | 0.055224 | 0.051 | 0.051 | 1 | 0 | Calml4 | 0 | Young |
| 0.9246 | 0.12588 | 0.039 | 0.04 | 1 | 0 | Pcolce | -0.001 | Young |
| 0.924938 | 0.117052 | 0.329 | 0.342 | 1 | 0 | Taok3 | -0.013 | Young |
| 0.924971 | 0.038711 | 0.119 | 0.12 | 1 | 0 | Siva1 | -0.001 | Young |
| 0.925063 | 0.317473 | 0.019 | 0.019 | 1 | 0 | Itfg2 | 0 | Young |
| 0.925407 | 0.264724 | 0.05 | 0.049 | 1 | 0 | Elmod2 | 0.001 | Young |
| 0.925606 | 0.07308 | 0.133 | 0.131 | 1 | 0 | Ppil2 | 0.002 | Young |
| 0.925641 | 0.010201 | 0.266 | 0.271 | 1 | 0 | Mon2 | -0.005 | Young |
| 0.925819 | 0.071328 | 0.124 | 0.124 | 1 | 0 | Sec24c | 0 | Young |
| 0.925932 | 0.169992 | 0.131 | 0.131 | 1 | 0 | Churc1 | 0 | Young |
| 0.92615 | 0.218367 | 0.04 | 0.041 | 1 | 0 | Cxxc1 | -0.001 | Young |
| 0.926182 | 0.286793 | 0.019 | 0.019 | 1 | 0 | Coa4 | 0 | Young |
| 0.926413 | 0.028013 | 0.172 | 0.172 | 1 | 0 | C1galt1 | 0 | Young |
| 0.926458 | 0.099724 | 0.126 | 0.124 | 1 | 0 | Pa2g4 | 0.002 | Young |
| 0.926645 | 0.01541 | 0.04 | 0.04 | 1 | 0 | Coprs | 0 | Young |
| 0.927189 | 0.144502 | 0.053 | 0.052 | 1 | 0 | Gtf2e2 | 0.001 | Young |
| 0.92722 | 0.042366 | 0.031 | 0.031 | 1 | 0 | Sgce | 0 | Young |
| 0.927441 | 0.157351 | 0.12 | 0.12 | 1 | 0 | Chd1 | 0 | Young |
| 0.927573 | 0.241926 | 0.096 | 0.098 | 1 | 0 | Tmem147 | -0.002 | Young |
| 0.928092 | 0.297119 | 0.095 | 0.096 | 1 | 0 | Ctnnbip1 | -0.001 | Young |
| 0.928256 | 0.18312 | 0.085 | 0.085 | 1 | 0 | Pccb | 0 | Young |
| 0.929549 | 0.252453 | 0.078 | 0.079 | 1 | 0 | Wdr81 | -0.001 | Young |
| 0.929585 | 0.030427 | 0.036 | 0.036 | 1 | 0 | Fam20c | 0 | Young |
| 0.929607 | 0.092268 | 0.087 | 0.086 | 1 | 0 | Cops5 | 0.001 | Young |
| 0.929615 | 0.125206 | 0.053 | 0.052 | 1 | 0 | Zfp410 | 0.001 | Young |

| | | | | | | | | |
|---|---|---|---|---|---|---|---|---|
| 0.929639 | 0.009322 | 0.896 | 0.917 | 1 | 0 | Fcer1g | -0.021 | Young |
| 0.929661 | 0.173463 | 0.168 | 0.171 | 1 | 0 | Lrmp | -0.003 | Young |
| 0.929686 | 0.1447 | 0.08 | 0.079 | 1 | 0 | Foxj3 | 0.001 | Young |
| 0.929689 | 0.312702 | 0.039 | 0.038 | 1 | 0 | St3gal3 | 0.001 | Young |
| 0.929819 | 0.076533 | 0.257 | 0.263 | 1 | 0 | Msr1 | -0.006 | Young |
| 0.929931 | 0.265166 | 0.011 | 0.011 | 1 | 0 | Gas8 | 0 | Young |
| 0.930114 | 0.098783 | 0.223 | 0.226 | 1 | 0 | Ndufb5 | -0.003 | Young |
| 0.930425 | 0.192918 | 0.066 | 0.066 | 1 | 0 | Ric1 | 0 | Young |
| 0.930458 | 0.383563 | 0.033 | 0.034 | 1 | 0 | Bend4 | -0.001 | Young |
| 0.930494 | 0.017406 | 0.029 | 0.03 | 1 | 0 | Fam60a | -0.001 | Young |
| 0.9307 | 0.013289 | 0.28 | 0.28 | 1 | 0 | Hmgn1 | 0 | Young |
| 0.930889 | 0.030814 | 0.214 | 0.218 | 1 | 0 | Csnk1d | -0.004 | Young |
| 0.931368 | 0.122465 | 0.027 | 0.026 | 1 | 0 | Rnf219 | 0.001 | Young |
| 0.931419 | 0.176527 | 0.044 | 0.044 | 1 | 0 | Zfp729a | 0 | Young |
| 0.931562 | 0.096515 | 0.036 | 0.037 | 1 | 0 | Rbl1 | -0.001 | Young |
| 0.931682 | 0.193482 | 0.028 | 0.027 | 1 | 0 | Lamc1 | 0.001 | Young |
| 0.93179 | 0.191494 | 0.103 | 0.105 | 1 | 0 | Pstpip1 | -0.002 | Young |
| 0.931833 | 0.13514 | 0.081 | 0.081 | 1 | 0 | Zfp260 | 0 | Young |
| 0.931847 | 0.05808 | 0.062 | 0.062 | 1 | 0 | Nsmce2 | 0 | Young |
| 0.931948 | 0.144553 | 0.194 | 0.196 | 1 | 0 | Map2k7 | -0.002 | Young |
| 0.932162 | 0.164312 | 0.243 | 0.25 | 1 | 0 | Ggnbp2 | -0.007 | Young |
| 0.932348 | 0.147854 | 0.031 | 0.031 | 1 | 0 | Cradd | 0 | Young |
| 0.93248 | 0.351946 | 0.041 | 0.041 | 1 | 0 | Dhx57 | 0 | Young |
| 0.932658 | 0.155789 | 0.101 | 0.101 | 1 | 0 | Pds5b | 0 | Young |
| 0.932746 | 0.137157 | 0.138 | 0.14 | 1 | 0 | Slc26a2 | -0.002 | Young |
| 0.932774 | 0.487072 | 0.035 | 0.036 | 1 | 0 | Syt11 | -0.001 | Young |
| 0.93289 | 0.052391 | 0.131 | 0.134 | 1 | 0 | Tgfbrap1 | -0.003 | Young |
| 0.932906 | 0.110669 | 0.226 | 0.23 | 1 | 0 | Fam117b | -0.004 | Young |
| 0.932911 | 0.114424 | 0.086 | 0.088 | 1 | 0 | Dpy30 | -0.002 | Young |
| 0.93294 | 0.05641 | 0.063 | 0.064 | 1 | 0 | Mapk9 | -0.001 | Young |
| 0.933284 | 0.036128 | 0.228 | 0.227 | 1 | 0 | Rad23b | 0.001 | Young |
| 0.933369 | 0.103522 | 0.092 | 0.092 | 1 | 0 | 2610037D( | 0 | Young |
| 0.933411 | 0.28088 | 0.064 | 0.066 | 1 | 0 | Phkb | -0.002 | Young |
| 0.93344 | 0.095355 | 0.109 | 0.111 | 1 | 0 | St3gal4 | -0.002 | Young |
| 0.933738 | 0.050313 | 0.117 | 0.116 | 1 | 0 | Bet1 | 0.001 | Young |
| 0.933786 | 0.220329 | 0.115 | 0.118 | 1 | 0 | Crtc3 | -0.003 | Young |
| 0.933956 | 0.03071 | 0.035 | 0.034 | 1 | 0 | Ccdc163 | 0.001 | Young |
| 0.934581 | 0.019644 | 0.35 | 0.354 | 1 | 0 | Xiap | -0.004 | Young |
| 0.934793 | 0.129789 | 0.228 | 0.231 | 1 | 0 | Prpf40a | -0.003 | Young |
| 0.934903 | 0.322777 | 0.022 | 0.022 | 1 | 0 | Mrrf | 0 | Young |
| 0.934976 | 0.094183 | 0.096 | 0.096 | 1 | 0 | Nab2 | 0 | Young |
| 0.935082 | 0.061474 | 0.027 | 0.026 | 1 | 0 | AW554918 | 0.001 | Young |
| 0.935084 | 0.273614 | 0.035 | 0.034 | 1 | 0 | Mtx3 | 0.001 | Young |
| 0.935236 | 0.073077 | 0.104 | 0.105 | 1 | 0 | H2-Q4 | -0.001 | Young |
| 0.935484 | 0.091428 | 0.181 | 0.183 | 1 | 0 | Ccni | -0.002 | Young |

| | | | | | | | | |
|---|---|---|---|---|---|---|---|---|
| 0.936 | 0.057703 | 0.227 | 0.231 | 1 | 0 | Baz1b | -0.004 | Young |
| 0.936117 | 0.004565 | 0.472 | 0.49 | 1 | 0 | Celf1 | -0.018 | Young |
| 0.936119 | 0.278703 | 0.112 | 0.114 | 1 | 0 | Slc7a7 | -0.002 | Young |
| 0.936467 | 0.042544 | 0.017 | 0.016 | 1 | 0 | Gm13373 | 0.001 | Young |
| 0.93677 | 0.131814 | 0.059 | 0.059 | 1 | 0 | Ndufv1 | 0 | Young |
| 0.936918 | 0.086782 | 0.143 | 0.146 | 1 | 0 | Txndc9 | -0.003 | Young |
| 0.937011 | 0.041183 | 0.12 | 0.119 | 1 | 0 | Pcgf5 | 0.001 | Young |
| 0.937181 | 0.127775 | 0.023 | 0.023 | 1 | 0 | Pick1 | 0 | Young |
| 0.937327 | 0.008059 | 0.029 | 0.03 | 1 | 0 | Tmem185a | -0.001 | Young |
| 0.93738 | 0.080522 | 0.143 | 0.146 | 1 | 0 | Med15 | -0.003 | Young |
| 0.937649 | 0.011333 | 0.017 | 0.016 | 1 | 0 | Tesk2 | 0.001 | Young |
| 0.937769 | 0.024226 | 0.276 | 0.278 | 1 | 0 | Luc7l3 | -0.002 | Young |
| 0.938095 | 0.07969 | 0.067 | 0.068 | 1 | 0 | Kat7 | -0.001 | Young |
| 0.938248 | 0.150378 | 0.089 | 0.089 | 1 | 0 | Fgd6 | 0 | Young |
| 0.938284 | 0.138128 | 0.059 | 0.059 | 1 | 0 | Fam78a | 0 | Young |
| 0.938592 | 0.776445 | 0.011 | 0.011 | 1 | 0 | Cdkl2 | 0 | Young |
| 0.938876 | 0.145797 | 0.028 | 0.027 | 1 | 0 | Gm47675 | 0.001 | Young |
| 0.938899 | 0.17826 | 0.109 | 0.109 | 1 | 0 | Pafah1b2 | 0 | Young |
| 0.939116 | 0.048911 | 0.029 | 0.03 | 1 | 0 | Sema6b | -0.001 | Young |
| 0.939191 | 0.174103 | 0.167 | 0.172 | 1 | 0 | Deptor | -0.005 | Young |
| 0.939816 | 0.017417 | 0.093 | 0.092 | 1 | 0 | Socs7 | 0.001 | Young |
| 0.939966 | 0.14998 | 0.053 | 0.055 | 1 | 0 | Xxylt1 | -0.002 | Young |
| 0.940077 | 0.174779 | 0.11 | 0.111 | 1 | 0 | Trp53 | -0.001 | Young |
| 0.940246 | 0.012326 | 0.277 | 0.282 | 1 | 0 | Usmg5 | -0.005 | Young |
| 0.940285 | 0.542753 | 0.017 | 0.018 | 1 | 0 | Kif5c | -0.001 | Young |
| 0.940558 | 0.229033 | 0.068 | 0.07 | 1 | 0 | Bahd1 | -0.002 | Young |
| 0.94057 | 0.049287 | 0.016 | 0.016 | 1 | 0 | Atad5 | 0 | Young |
| 0.941016 | 0.150544 | 0.013 | 0.012 | 1 | 0 | Mphosph9 | 0.001 | Young |
| 0.941049 | 0.06348 | 0.09 | 0.092 | 1 | 0 | Trim24 | -0.002 | Young |
| 0.941133 | 0.092894 | 0.086 | 0.088 | 1 | 0 | Slc25a46 | -0.002 | Young |
| 0.941191 | 0.069682 | 0.009 | 0.01 | 1 | 0 | E230032D2 | -0.001 | Young |
| 0.941211 | 0.17357 | 0.013 | 0.012 | 1 | 0 | Magi2 | 0.001 | Young |
| 0.941705 | 0.017912 | 0.079 | 0.078 | 1 | 0 | Ern1 | 0.001 | Young |
| 0.941793 | 0.069131 | 0.042 | 0.042 | 1 | 0 | Ccdc85b | 0 | Young |
| 0.942259 | 0.337095 | 0.032 | 0.033 | 1 | 0 | Pkib | -0.001 | Young |
| 0.942362 | 0.030459 | 0.028 | 0.029 | 1 | 0 | Tfdp2 | -0.001 | Young |
| 0.942393 | 0.460457 | 0.041 | 0.042 | 1 | 0 | Nol12 | -0.001 | Young |
| 0.942737 | 0.170585 | 0.101 | 0.103 | 1 | 0 | Acaa1a | -0.002 | Young |
| 0.943041 | 0.1323 | 0.085 | 0.086 | 1 | 0 | Usp33 | -0.001 | Young |
| 0.943044 | 0.242793 | 0.052 | 0.052 | 1 | 0 | Bcl9l | 0 | Young |
| 0.94316 | 0.126344 | 0.213 | 0.216 | 1 | 0 | Dcaf8 | -0.003 | Young |
| 0.943185 | 0.029561 | 0.251 | 0.252 | 1 | 0 | Swi5 | -0.001 | Young |
| 0.943309 | 0.315484 | 0.029 | 0.029 | 1 | 0 | Entpd5 | 0 | Young |
| 0.943362 | 0.207184 | 0.031 | 0.031 | 1 | 0 | Naa16 | 0 | Young |
| 0.943459 | 0.208773 | 0.163 | 0.166 | 1 | 0 | Gmip | -0.003 | Young |

| | | | | | | | | |
|---|---|---|---|---|---|---|---|---|
| 0.943558 | 0.038216 | 0.626 | 0.651 | 1 | 0 | Fus | -0.025 | Young |
| 0.943621 | 0.191665 | 0.128 | 0.13 | 1 | 0 | Rsl1d1 | -0.002 | Young |
| 0.943936 | 0.025738 | 0.013 | 0.012 | 1 | 0 | Zswim1 | 0.001 | Young |
| 0.944054 | 0.050352 | 0.108 | 0.109 | 1 | 0 | Lpin2 | -0.001 | Young |
| 0.944201 | 0.026948 | 0.15 | 0.149 | 1 | 0 | Ing2 | 0.001 | Young |
| 0.944904 | 0.006434 | 0.419 | 0.423 | 1 | 0 | Dok2 | -0.004 | Young |
| 0.945073 | 0.390754 | 0.048 | 0.049 | 1 | 0 | Smc2 | -0.001 | Young |
| 0.945073 | 0.117559 | 0.142 | 0.145 | 1 | 0 | Bax | -0.003 | Young |
| 0.94511 | 0.017432 | 0.051 | 0.052 | 1 | 0 | Mthfr | -0.001 | Young |
| 0.945156 | 0.145862 | 0.023 | 0.023 | 1 | 0 | Syce2 | 0 | Young |
| 0.9457 | 0.265798 | 0.029 | 0.029 | 1 | 0 | Asb13 | 0 | Young |
| 0.946345 | 0.260394 | 0.079 | 0.079 | 1 | 0 | Fzr1 | 0 | Young |
| 0.946848 | 0.207381 | 0.258 | 0.267 | 1 | 0 | Hip1 | -0.009 | Young |
| 0.947152 | 0.126833 | 0.136 | 0.137 | 1 | 0 | Imp3 | -0.001 | Young |
| 0.947284 | 0.04179 | 0.168 | 0.17 | 1 | 0 | Snrpd2 | -0.002 | Young |
| 0.947803 | 0.115702 | 0.088 | 0.089 | 1 | 0 | Pex19 | -0.001 | Young |
| 0.947862 | 0.112703 | 0.05 | 0.051 | 1 | 0 | D10Wsu10 | -0.001 | Young |
| 0.947865 | 0.213967 | 0.091 | 0.093 | 1 | 0 | Dnajb4 | -0.002 | Young |
| 0.948136 | 0.326134 | 0.056 | 0.056 | 1 | 0 | Pi4k2b | 0 | Young |
| 0.948149 | 0.258518 | 0.016 | 0.016 | 1 | 0 | Lbx2 | 0 | Young |
| 0.948207 | 0.112992 | 0.046 | 0.047 | 1 | 0 | Arfgap1 | -0.001 | Young |
| 0.94826 | 0.121454 | 0.061 | 0.06 | 1 | 0 | Slc25a10 | 0.001 | Young |
| 0.948321 | 0.287993 | 0.016 | 0.016 | 1 | 0 | Polr1b | 0 | Young |
| 0.948478 | 0.050938 | 0.072 | 0.073 | 1 | 0 | Atxn7l3 | -0.001 | Young |
| 0.948585 | 0.069625 | 0.033 | 0.033 | 1 | 0 | Pcid2 | 0 | Young |
| 0.948586 | 0.201996 | 0.155 | 0.159 | 1 | 0 | Lrp10 | -0.004 | Young |
| 0.94871 | 0.077307 | 0.04 | 0.04 | 1 | 0 | Ppa2 | 0 | Young |
| 0.948843 | 0.136447 | 0.145 | 0.145 | 1 | 0 | Ldb1 | 0 | Young |
| 0.94888 | 0.076348 | 0.374 | 0.386 | 1 | 0 | Cndp2 | -0.012 | Young |
| 0.948948 | 0.08164 | 0.096 | 0.097 | 1 | 0 | Zfp385a | -0.001 | Young |
| 0.949354 | 0.159806 | 0.07 | 0.07 | 1 | 0 | Mfap1b | 0 | Young |
| 0.949614 | 0.174269 | 0.033 | 0.033 | 1 | 0 | Wdcp | 0 | Young |
| 0.949657 | 0.269519 | 0.078 | 0.078 | 1 | 0 | Mrpl11 | 0 | Young |
| 0.949807 | 0.089228 | 0.139 | 0.14 | 1 | 0 | Rbm17 | -0.001 | Young |
| 0.950401 | 0.006796 | 0.043 | 0.042 | 1 | 0 | Pdzd11 | 0.001 | Young |
| 0.950537 | 0.168292 | 0.04 | 0.04 | 1 | 0 | Neurl2 | 0 | Young |
| 0.950923 | 0.04848 | 0.22 | 0.22 | 1 | 0 | Creb1 | 0 | Young |
| 0.950998 | 0.204507 | 0.05 | 0.051 | 1 | 0 | Ano10 | -0.001 | Young |
| 0.951 | 0.096586 | 0.326 | 0.339 | 1 | 0 | Txndc17 | -0.013 | Young |
| 0.951206 | 0.218899 | 0.193 | 0.198 | 1 | 0 | Dock11 | -0.005 | Young |
| 0.95149 | 0.153645 | 0.067 | 0.068 | 1 | 0 | Rbm34 | -0.001 | Young |
| 0.951551 | 0.032134 | 0.316 | 0.332 | 1 | 0 | Snrpe | -0.016 | Young |
| 0.951769 | 0.144509 | 0.018 | 0.018 | 1 | 0 | Stk39 | 0 | Young |
| 0.952033 | 0.231358 | 0.075 | 0.077 | 1 | 0 | Sash3 | -0.002 | Young |
| 0.952274 | 0.099926 | 0.405 | 0.424 | 1 | 0 | Pafah1b1 | -0.019 | Young |

| | | | | | | | | |
|---|---|---|---|---|---|---|---|---|
| 0.952298 | 0.069493 | 0.35 | 0.36 | 1 | 0 | Far1 | -0.01 | Young |
| 0.952313 | 0.010151 | 0.021 | 0.022 | 1 | 0 | Gatb | -0.001 | Young |
| 0.952323 | 0.109803 | 0.177 | 0.179 | 1 | 0 | Kdm5a | -0.002 | Young |
| 0.952629 | 0.200341 | 0.106 | 0.108 | 1 | 0 | Abl1 | -0.002 | Young |
| 0.952634 | 0.08036 | 0.174 | 0.175 | 1 | 0 | Tmem141 | -0.001 | Young |
| 0.952997 | 0.199974 | 0.042 | 0.042 | 1 | 0 | Nudt19 | 0 | Young |
| 0.953051 | 0.157329 | 0.024 | 0.023 | 1 | 0 | Fzd7 | 0.001 | Young |
| 0.953446 | 0.042101 | 0.051 | 0.052 | 1 | 0 | Jrkl | -0.001 | Young |
| 0.95367 | 0.157703 | 0.084 | 0.083 | 1 | 0 | Arid5b | 0.001 | Young |
| 0.953673 | 0.12443 | 0.283 | 0.289 | 1 | 0 | Zfp106 | -0.006 | Young |
| 0.954252 | 0.159431 | 0.091 | 0.093 | 1 | 0 | Lcor | -0.002 | Young |
| 0.954262 | 0.178155 | 0.174 | 0.176 | 1 | 0 | St3gal1 | -0.002 | Young |
| 0.954364 | 0.019248 | 0.098 | 0.097 | 1 | 0 | Zfp871 | 0.001 | Young |
| 0.954418 | 8.52E-06 | 0.015 | 0.015 | 1 | 0 | Dcstamp | 0 | Young |
| 0.954814 | 0.006824 | 0.027 | 0.027 | 1 | 0 | Primpol | 0 | Young |
| 0.954829 | 0.474635 | 0.014 | 0.014 | 1 | 0 | Elp4 | 0 | Young |
| 0.954901 | 0.30636 | 0.043 | 0.044 | 1 | 0 | Orc3 | -0.001 | Young |
| 0.955054 | 0.16623 | 0.118 | 0.12 | 1 | 0 | Vkorc1 | -0.002 | Young |
| 0.955195 | 0.041402 | 0.028 | 0.027 | 1 | 0 | Sec61a2 | 0.001 | Young |
| 0.955203 | 0.109066 | 0.093 | 0.094 | 1 | 0 | Anapc2 | -0.001 | Young |
| 0.95538 | 0.252107 | 0.07 | 0.07 | 1 | 0 | Taf13 | 0 | Young |
| 0.955476 | 0.043376 | 0.137 | 0.137 | 1 | 0 | Pcyox1 | 0 | Young |
| 0.955829 | 0.148393 | 0.053 | 0.053 | 1 | 0 | Ell | 0 | Young |
| 0.955939 | 0.015298 | 0.255 | 0.26 | 1 | 0 | Nsd1 | -0.005 | Young |
| 0.956098 | 0.167946 | 0.135 | 0.138 | 1 | 0 | Ebp | -0.003 | Young |
| 0.956617 | 0.212747 | 0.143 | 0.145 | 1 | 0 | Gm37494 | -0.002 | Young |
| 0.956711 | 0.047342 | 0.155 | 0.156 | 1 | 0 | Rwdd1 | -0.001 | Young |
| 0.956751 | 0.150502 | 0.167 | 0.17 | 1 | 0 | Rrp1 | -0.003 | Young |
| 0.956754 | 0.050846 | 0.015 | 0.015 | 1 | 0 | Cenpt | 0 | Young |
| 0.956872 | 0.251992 | 0.014 | 0.014 | 1 | 0 | Hdac8 | 0 | Young |
| 0.956897 | 0.284525 | 0.042 | 0.042 | 1 | 0 | Ppp2r5d | 0 | Young |
| 0.957113 | 0.315799 | 0.015 | 0.015 | 1 | 0 | Gm30978 | 0 | Young |
| 0.957207 | 0.093955 | 0.075 | 0.075 | 1 | 0 | Arl13b | 0 | Young |
| 0.957333 | 0.172599 | 0.12 | 0.12 | 1 | 0 | Trappc10 | 0 | Young |
| 0.957362 | 0.097718 | 0.169 | 0.171 | 1 | 0 | Fbxl5 | -0.002 | Young |
| 0.957582 | 0.040706 | 0.267 | 0.269 | 1 | 0 | Dnajc5 | -0.002 | Young |
| 0.958007 | 0.053045 | 0.04 | 0.041 | 1 | 0 | Dlg4 | -0.001 | Young |
| 0.958011 | 0.399998 | 0.015 | 0.015 | 1 | 0 | Pomt1 | 0 | Young |
| 0.958227 | 0.120457 | 0.229 | 0.23 | 1 | 0 | Ppp1r12a | -0.001 | Young |
| 0.958308 | 0.344224 | 0.056 | 0.057 | 1 | 0 | Psmg3 | -0.001 | Young |
| 0.958586 | 0.49745 | 0.041 | 0.042 | 1 | 0 | Klhdc10 | -0.001 | Young |
| 0.958609 | 0.164747 | 0.094 | 0.094 | 1 | 0 | Snrnp27 | 0 | Young |
| 0.958723 | 0.004574 | 0.112 | 0.112 | 1 | 0 | Tec | 0 | Young |
| 0.958822 | 0.192635 | 0.014 | 0.014 | 1 | 0 | Gm13479 | 0 | Young |
| 0.959083 | 0.049877 | 0.115 | 0.116 | 1 | 0 | Smc4 | -0.001 | Young |

| | | | | | | | | |
|---|---|---|---|---|---|---|---|---|
| 0.959089 | 0.17548 | 0.015 | 0.015 | 1 | 0 | Pcbp4 | 0 | Young |
| 0.959118 | 0.359546 | 0.043 | 0.044 | 1 | 0 | Jagn1 | -0.001 | Young |
| 0.959249 | 0.226992 | 0.047 | 0.048 | 1 | 0 | Nedd1 | -0.001 | Young |
| 0.959456 | 0.10075 | 0.201 | 0.204 | 1 | 0 | Kmt2c | -0.003 | Young |
| 0.959934 | 0.275161 | 0.09 | 0.092 | 1 | 0 | Babam1 | -0.002 | Young |
| 0.959988 | 0.328357 | 0.015 | 0.015 | 1 | 0 | Mrps27 | 0 | Young |
| 0.960099 | 0.245526 | 0.063 | 0.064 | 1 | 0 | Slc25a17 | -0.001 | Young |
| 0.960167 | 0.271914 | 0.015 | 0.015 | 1 | 0 | Mex3b | 0 | Young |
| 0.960338 | 0.04547 | 0.036 | 0.036 | 1 | 0 | Uck1 | 0 | Young |
| 0.960443 | 0.070618 | 0.197 | 0.201 | 1 | 0 | Ints6l | -0.004 | Young |
| 0.960481 | 0.041741 | 0.184 | 0.186 | 1 | 0 | Rp9 | -0.002 | Young |
| 0.960527 | 0.064753 | 0.015 | 0.015 | 1 | 0 | Alms1 | 0 | Young |
| 0.960549 | 0.170578 | 0.161 | 0.164 | 1 | 0 | Ubr5 | -0.003 | Young |
| 0.960703 | 0.173947 | 0.18 | 0.183 | 1 | 0 | Tulp4 | -0.003 | Young |
| 0.960821 | 0.115805 | 0.201 | 0.205 | 1 | 0 | Emc1 | -0.004 | Young |
| 0.960839 | 0.129464 | 0.229 | 0.234 | 1 | 0 | Nop53 | -0.005 | Young |
| 0.961108 | 0.053806 | 0.124 | 0.123 | 1 | 0 | BC028528 | 0.001 | Young |
| 0.961339 | 0.10438 | 0.113 | 0.115 | 1 | 0 | Akap9 | -0.002 | Young |
| 0.961378 | 0.217219 | 0.102 | 0.103 | 1 | 0 | Rnf10 | -0.001 | Young |
| 0.961477 | 0.031887 | 0.029 | 0.029 | 1 | 0 | Card6 | 0 | Young |
| 0.961484 | 0.048208 | 0.278 | 0.28 | 1 | 0 | Slamf9 | -0.002 | Young |
| 0.961569 | 0.225057 | 0.027 | 0.027 | 1 | 0 | Tfcp2 | 0 | Young |
| 0.961573 | 0.291001 | 0.021 | 0.022 | 1 | 0 | Txnl4b | -0.001 | Young |
| 0.961656 | 0.127683 | 0.152 | 0.153 | 1 | 0 | Tm2d2 | -0.001 | Young |
| 0.96174 | 0.274677 | 0.051 | 0.051 | 1 | 0 | Vps39 | 0 | Young |
| 0.962143 | 0.216889 | 0.063 | 0.063 | 1 | 0 | Pibf1 | 0 | Young |
| 0.962555 | 0.119412 | 0.02 | 0.021 | 1 | 0 | Ldlrad3 | -0.001 | Young |
| 0.962932 | 0.061853 | 0.164 | 0.167 | 1 | 0 | Rnf187 | -0.003 | Young |
| 0.963187 | 0.067096 | 0.242 | 0.244 | 1 | 0 | Hectd1 | -0.002 | Young |
| 0.96371 | 0.020008 | 0.045 | 0.045 | 1 | 0 | Wdsub1 | 0 | Young |
| 0.963898 | 0.120816 | 0.058 | 0.057 | 1 | 0 | 0610010K1 | 0.001 | Young |
| 0.964265 | 0.126742 | 0.077 | 0.077 | 1 | 0 | Mfsd10 | 0 | Young |
| 0.964634 | 0.117672 | 0.02 | 0.021 | 1 | 0 | Zfp53 | -0.001 | Young |
| 0.964872 | 0.133133 | 0.041 | 0.041 | 1 | 0 | Snhg6 | 0 | Young |
| 0.965019 | 0.402042 | 0.015 | 0.015 | 1 | 0 | Zfp523 | 0 | Young |
| 0.965046 | 0.106006 | 0.014 | 0.014 | 1 | 0 | Nepro | 0 | Young |
| 0.965107 | 0.159173 | 0.143 | 0.146 | 1 | 0 | Synj2bp | -0.003 | Young |
| 0.965517 | 0.076826 | 0.367 | 0.379 | 1 | 0 | Tardbp | -0.012 | Young |
| 0.965671 | 0.180012 | 0.019 | 0.019 | 1 | 0 | H2afx | 0 | Young |
| 0.966144 | 0.221253 | 0.019 | 0.019 | 1 | 0 | Acad11 | 0 | Young |
| 0.966219 | 0.085373 | 0.108 | 0.108 | 1 | 0 | Mau2 | 0 | Young |
| 0.966226 | 0.106179 | 0.155 | 0.159 | 1 | 0 | Slc9a3r1 | -0.004 | Young |
| 0.966236 | 0.013166 | 0.035 | 0.034 | 1 | 0 | Rint1 | 0.001 | Young |
| 0.966281 | 0.045557 | 0.134 | 0.135 | 1 | 0 | Aff1 | -0.001 | Young |
| 0.966329 | 0.179237 | 0.025 | 0.025 | 1 | 0 | Amd1 | 0 | Young |

| | | | | | | | | |
|---|---|---|---|---|---|---|---|---|
| 0.966743 | 0.03587 | 0.099 | 0.098 | 1 | 0 | Pcif1 | 0.001 | Young |
| 0.966884 | 0.130129 | 0.04 | 0.04 | 1 | 0 | Alkbh8 | 0 | Young |
| 0.966929 | 0.026846 | 0.391 | 0.397 | 1 | 0 | St13 | -0.006 | Young |
| 0.966988 | 0.659325 | 0.015 | 0.015 | 1 | 0 | Btbd19 | 0 | Young |
| 0.967486 | 0.117588 | 0.166 | 0.168 | 1 | 0 | Uso1 | -0.002 | Young |
| 0.967493 | 0.195927 | 0.041 | 0.041 | 1 | 0 | Bcas3 | 0 | Young |
| 0.967862 | 0.162917 | 0.074 | 0.075 | 1 | 0 | Dnajc21 | -0.001 | Young |
| 0.968002 | 0.202106 | 0.063 | 0.063 | 1 | 0 | Arhgap6 | 0 | Young |
| 0.968013 | 0.167055 | 0.109 | 0.109 | 1 | 0 | Cenpx | 0 | Young |
| 0.9682 | 0.06949 | 0.035 | 0.034 | 1 | 0 | Endod1 | 0.001 | Young |
| 0.968209 | 0.160783 | 0.034 | 0.034 | 1 | 0 | Mcu | 0 | Young |
| 0.96821 | 0.223882 | 0.044 | 0.044 | 1 | 0 | Gm17018 | 0 | Young |
| 0.968258 | 0.059873 | 0.262 | 0.269 | 1 | 0 | Pkm | -0.007 | Young |
| 0.968339 | 0.150295 | 0.117 | 0.119 | 1 | 0 | Prdm2 | -0.002 | Young |
| 0.968435 | 0.642597 | 0.015 | 0.015 | 1 | 0 | Tefm | 0 | Young |
| 0.968462 | 0.088962 | 0.075 | 0.075 | 1 | 0 | Pnpla2 | 0 | Young |
| 0.968467 | 0.090178 | 0.268 | 0.274 | 1 | 0 | Arf3 | -0.006 | Young |
| 0.968567 | 0.699565 | 0.01 | 0.01 | 1 | 0 | D130040H | 0 | Young |
| 0.968844 | 0.036902 | 0.081 | 0.082 | 1 | 0 | Armc5 | -0.001 | Young |
| 0.968919 | 0.069095 | 0.305 | 0.311 | 1 | 0 | Ppp1r15a | -0.006 | Young |
| 0.968968 | 0.201196 | 0.096 | 0.097 | 1 | 0 | Kpna1 | -0.001 | Young |
| 0.969322 | 0.154241 | 0.086 | 0.088 | 1 | 0 | Usf1 | -0.002 | Young |
| 0.96957 | 0.032265 | 0.093 | 0.093 | 1 | 0 | Mrps36 | 0 | Young |
| 0.969712 | 0.182301 | 0.149 | 0.153 | 1 | 0 | Ndfip1 | -0.004 | Young |
| 0.970215 | 0.186085 | 0.026 | 0.026 | 1 | 0 | Bud23 | 0 | Young |
| 0.970338 | 0.242504 | 0.057 | 0.057 | 1 | 0 | Gyg | 0 | Young |
| 0.970373 | 0.085148 | 0.106 | 0.107 | 1 | 0 | Socs4 | -0.001 | Young |
| 0.970916 | 0.072386 | 0.061 | 0.062 | 1 | 0 | Trmt6 | -0.001 | Young |
| 0.971256 | 0.07982 | 0.079 | 0.079 | 1 | 0 | Ptk2 | 0 | Young |
| 0.971312 | 0.141059 | 0.013 | 0.014 | 1 | 0 | Polg2 | -0.001 | Young |
| 0.971357 | 0.019956 | 0.025 | 0.025 | 1 | 0 | Snrnp35 | 0 | Young |
| 0.971436 | 0.184202 | 0.091 | 0.092 | 1 | 0 | Elovl5 | -0.001 | Young |
| 0.971446 | 0.086273 | 0.063 | 0.063 | 1 | 0 | Dpp7 | 0 | Young |
| 0.971468 | 0.068833 | 0.03 | 0.03 | 1 | 0 | Pnpo | 0 | Young |
| 0.971584 | 0.162821 | 0.09 | 0.09 | 1 | 0 | Espl1 | 0 | Young |
| 0.971684 | 0.082253 | 0.203 | 0.208 | 1 | 0 | Acsl4 | -0.005 | Young |
| 0.971717 | 0.021809 | 0.351 | 0.361 | 1 | 0 | Romo1 | -0.01 | Young |
| 0.971723 | 0.071152 | 0.052 | 0.052 | 1 | 0 | Pias3 | 0 | Young |
| 0.972252 | 0.29736 | 0.013 | 0.014 | 1 | 0 | Dpt | -0.001 | Young |
| 0.972352 | 0.051097 | 0.158 | 0.16 | 1 | 0 | Ddx60 | -0.002 | Young |
| 0.972472 | 0.047913 | 0.071 | 0.071 | 1 | 0 | Tesk1 | 0 | Young |
| 0.972629 | 0.27491 | 0.013 | 0.014 | 1 | 0 | Cep95 | -0.001 | Young |
| 0.97294 | 0.058791 | 0.241 | 0.244 | 1 | 0 | Ncoa3 | -0.003 | Young |
| 0.973275 | 0.135546 | 0.08 | 0.081 | 1 | 0 | 1110059G | -0.001 | Young |
| 0.973539 | 0.081517 | 0.233 | 0.238 | 1 | 0 | Golph3 | -0.005 | Young |

| | | | | | | | | |
|---|---|---|---|---|---|---|---|---|
| 0.973698 | 0.325917 | 0.067 | 0.068 | 1 | 0 | Fopnl | -0.001 | Young |
| 0.973736 | 0.145729 | 0.026 | 0.026 | 1 | 0 | Tbcc | 0 | Young |
| 0.973782 | 0.076857 | 0.23 | 0.233 | 1 | 0 | Naa50 | -0.003 | Young |
| 0.974201 | 0.026573 | 0.035 | 0.036 | 1 | 0 | Gfm2 | -0.001 | Young |
| 0.974377 | 0.198213 | 0.08 | 0.081 | 1 | 0 | Zfp664 | -0.001 | Young |
| 0.974411 | 0.227523 | 0.053 | 0.053 | 1 | 0 | Mrpl55 | 0 | Young |
| 0.974568 | 0.087805 | 0.259 | 0.263 | 1 | 0 | Hnrnpul1 | -0.004 | Young |
| 0.9747 | 0.138145 | 0.193 | 0.197 | 1 | 0 | Larp4 | -0.004 | Young |
| 0.974811 | 0.131708 | 0.147 | 0.148 | 1 | 0 | Ext1 | -0.001 | Young |
| 0.97507 | 0.181165 | 0.063 | 0.063 | 1 | 0 | Dhx16 | 0 | Young |
| 0.975391 | 0.048789 | 0.043 | 0.044 | 1 | 0 | Rev3l | -0.001 | Young |
| 0.975489 | 0.075506 | 0.019 | 0.019 | 1 | 0 | 2610021A( | 0 | Young |
| 0.975564 | 0.297032 | 0.027 | 0.027 | 1 | 0 | Donson | 0 | Young |
| 0.975772 | 0.148848 | 0.109 | 0.109 | 1 | 0 | Ccpg1 | 0 | Young |
| 0.976168 | 0.197084 | 0.044 | 0.044 | 1 | 0 | Prkce | 0 | Young |
| 0.976462 | 0.140686 | 0.097 | 0.098 | 1 | 0 | Hspbp1 | -0.001 | Young |
| 0.976744 | 0.18657 | 0.133 | 0.134 | 1 | 0 | Zrsr2 | -0.001 | Young |
| 0.976747 | 0.102513 | 0.179 | 0.182 | 1 | 0 | Fbxl3 | -0.003 | Young |
| 0.977039 | 0.193915 | 0.092 | 0.093 | 1 | 0 | Ext2 | -0.001 | Young |
| 0.977278 | 0.171995 | 0.085 | 0.086 | 1 | 0 | Bicral | -0.001 | Young |
| 0.977407 | 0.076514 | 0.109 | 0.109 | 1 | 0 | Trim12a | 0 | Young |
| 0.977452 | 0.04393 | 0.155 | 0.156 | 1 | 0 | Zfr | -0.001 | Young |
| 0.977744 | 0.183478 | 0.111 | 0.112 | 1 | 0 | Psmd7 | -0.001 | Young |
| 0.977796 | 0.142727 | 0.239 | 0.245 | 1 | 0 | Yy1 | -0.006 | Young |
| 0.977975 | 0.030432 | 0.067 | 0.067 | 1 | 0 | Polr3k | 0 | Young |
| 0.978338 | 0.313615 | 0.01 | 0.01 | 1 | 0 | Taco1os | 0 | Young |
| 0.978388 | 0.113201 | 0.098 | 0.098 | 1 | 0 | Gm47283 | 0 | Young |
| 0.97841 | 0.078391 | 0.059 | 0.059 | 1 | 0 | Gpt2 | 0 | Young |
| 0.978594 | 0.017699 | 0.242 | 0.246 | 1 | 0 | Phf3 | -0.004 | Young |
| 0.978624 | 0.167105 | 0.043 | 0.042 | 1 | 0 | Cars | 0.001 | Young |
| 0.978704 | 0.213566 | 0.031 | 0.031 | 1 | 0 | Mri1 | 0 | Young |
| 0.978811 | 0.163939 | 0.072 | 0.073 | 1 | 0 | AU019823 | -0.001 | Young |
| 0.978915 | 0.196823 | 0.192 | 0.197 | 1 | 0 | Sfxn3 | -0.005 | Young |
| 0.97899 | 0.177643 | 0.068 | 0.068 | 1 | 0 | Naa10 | 0 | Young |
| 0.979009 | 0.215965 | 0.107 | 0.109 | 1 | 0 | Gatad2a | -0.002 | Young |
| 0.979539 | 0.111902 | 0.043 | 0.042 | 1 | 0 | Gar1 | 0.001 | Young |
| 0.979798 | 0.200994 | 0.015 | 0.015 | 1 | 0 | Syt3 | 0 | Young |
| 0.979824 | 0.232706 | 0.033 | 0.033 | 1 | 0 | Ick | 0 | Young |
| 0.979925 | 0.055209 | 0.27 | 0.274 | 1 | 0 | Nxf1 | -0.004 | Young |
| 0.98029 | 0.326109 | 0.027 | 0.027 | 1 | 0 | Nod1 | 0 | Young |
| 0.980383 | 0.058622 | 0.042 | 0.042 | 1 | 0 | B4gat1 | 0 | Young |
| 0.980733 | 0.268079 | 0.046 | 0.047 | 1 | 0 | Ap3m2 | -0.001 | Young |
| 0.981087 | 0.046861 | 0.083 | 0.083 | 1 | 0 | Ankra2 | 0 | Young |
| 0.981147 | 0.055577 | 0.142 | 0.144 | 1 | 0 | Dync1li2 | -0.002 | Young |
| 0.981294 | 0.123958 | 0.135 | 0.138 | 1 | 0 | Sys1 | -0.003 | Young |

| | | | | | | | | |
|---|---|---|---|---|---|---|---|---|
| 0.981382 | 0.193836 | 0.102 | 0.103 | 1 | 0 | Plekhm1 | -0.001 | Young |
| 0.981392 | 0.050175 | 0.11 | 0.111 | 1 | 0 | Lrrfip2 | -0.001 | Young |
| 0.981608 | 0.072466 | 0.168 | 0.168 | 1 | 0 | Gas2l3 | 0 | Young |
| 0.981992 | 0.109203 | 0.122 | 0.123 | 1 | 0 | Zfp326 | -0.001 | Young |
| 0.982234 | 0.182598 | 0.073 | 0.074 | 1 | 0 | Msantd4 | -0.001 | Young |
| 0.982337 | 0.196079 | 0.171 | 0.175 | 1 | 0 | Maz | -0.004 | Young |
| 0.982557 | 0.155368 | 0.195 | 0.2 | 1 | 0 | Cbx1 | -0.005 | Young |
| 0.982873 | 0.245938 | 0.074 | 0.075 | 1 | 0 | Vps33a | -0.001 | Young |
| 0.983582 | 0.126106 | 0.265 | 0.276 | 1 | 0 | Tmem109 | -0.011 | Young |
| 0.983604 | 0.221326 | 0.102 | 0.104 | 1 | 0 | Specc1 | -0.002 | Young |
| 0.983713 | 0.179445 | 0.015 | 0.015 | 1 | 0 | Cptp | 0 | Young |
| 0.983998 | 0.121598 | 0.324 | 0.338 | 1 | 0 | Tfe3 | -0.014 | Young |
| 0.984016 | 0.060927 | 0.021 | 0.021 | 1 | 0 | 1110019D1 | 0 | Young |
| 0.984157 | 0.0106 | 0.197 | 0.198 | 1 | 0 | Cttnbp2nl | -0.001 | Young |
| 0.984215 | 0.018002 | 0.032 | 0.031 | 1 | 0 | 5730409E0 | 0.001 | Young |
| 0.984438 | 0.236652 | 0.136 | 0.141 | 1 | 0 | Ube2d1 | -0.005 | Young |
| 0.984454 | 0.07001 | 0.224 | 0.226 | 1 | 0 | Armc8 | -0.002 | Young |
| 0.984705 | 0.164576 | 0.08 | 0.081 | 1 | 0 | Ddx23 | -0.001 | Young |
| 0.984878 | 0.265471 | 0.019 | 0.019 | 1 | 0 | Pgap3 | 0 | Young |
| 0.984977 | 0.059971 | 0.608 | 0.637 | 1 | 0 | Il6ra | -0.029 | Young |
| 0.985073 | 0.236702 | 0.031 | 0.031 | 1 | 0 | Mrps35 | 0 | Young |
| 0.985181 | 0.067534 | 0.496 | 0.52 | 1 | 0 | Grb2 | -0.024 | Young |
| 0.985237 | 0.284247 | 0.053 | 0.053 | 1 | 0 | Zfp282 | 0 | Young |
| 0.985356 | 0.050434 | 0.244 | 0.248 | 1 | 0 | Skil | -0.004 | Young |
| 0.985398 | 0.125036 | 0.209 | 0.212 | 1 | 0 | Ago2 | -0.003 | Young |
| 0.985469 | 0.008814 | 0.182 | 0.182 | 1 | 0 | Id2 | 0 | Young |
| 0.985493 | 0.168047 | 0.169 | 0.172 | 1 | 0 | Lsm14a | -0.003 | Young |
| 0.985546 | 0.163514 | 0.061 | 0.062 | 1 | 0 | Timm50 | -0.001 | Young |
| 0.985551 | 0.107499 | 0.123 | 0.124 | 1 | 0 | Nin | -0.001 | Young |
| 0.985652 | 0.094454 | 0.163 | 0.166 | 1 | 0 | Zc3h13 | -0.003 | Young |
| 0.985784 | 0.090826 | 0.449 | 0.468 | 1 | 0 | Pak2 | -0.019 | Young |
| 0.985857 | 0.071142 | 0.149 | 0.15 | 1 | 0 | 2-Mar | -0.001 | Young |
| 0.985903 | 0.696615 | 0.011 | 0.011 | 1 | 0 | Ttc12 | 0 | Young |
| 0.985946 | 0.068514 | 0.043 | 0.042 | 1 | 0 | Vegfb | 0.001 | Young |
| 0.986041 | 0.080526 | 0.033 | 0.033 | 1 | 0 | Rem1 | 0 | Young |
| 0.986279 | 0.058852 | 0.19 | 0.193 | 1 | 0 | Atxn7 | -0.003 | Young |
| 0.986569 | 0.102471 | 0.024 | 0.025 | 1 | 0 | Unk | -0.001 | Young |
| 0.9867 | 0.168462 | 0.107 | 0.108 | 1 | 0 | Wdr44 | -0.001 | Young |
| 0.986884 | 0.207517 | 0.031 | 0.031 | 1 | 0 | Mthfsd | 0 | Young |
| 0.986948 | 0.207127 | 0.027 | 0.027 | 1 | 0 | Men1 | 0 | Young |
| 0.986962 | 0.298837 | 0.096 | 0.097 | 1 | 0 | Jmjd6 | -0.001 | Young |
| 0.986999 | 0.038671 | 0.058 | 0.057 | 1 | 0 | Cfap97 | 0.001 | Young |
| 0.987077 | 0.170585 | 0.265 | 0.275 | 1 | 0 | Brd4 | -0.01 | Young |
| 0.987113 | 0.055465 | 0.092 | 0.093 | 1 | 0 | Agmo | -0.001 | Young |
| 0.987153 | 0.337991 | 0.059 | 0.06 | 1 | 0 | Dnajc11 | -0.001 | Young |

| | | | | | | | | |
|---|---|---|---|---|---|---|---|---|
| 0.987383 | 0.118917 | 0.045 | 0.045 | 1 | 0 | Sp4 | 0 | Young |
| 0.987396 | 0.190779 | 0.018 | 0.018 | 1 | 0 | Lmnb1 | 0 | Young |
| 0.987701 | 0.205525 | 0.106 | 0.108 | 1 | 0 | Ten1 | -0.002 | Young |
| 0.987901 | 0.012086 | 0.187 | 0.187 | 1 | 0 | Pkn2 | 0 | Young |
| 0.988356 | 0.054734 | 0.049 | 0.049 | 1 | 0 | Arhgef10 | 0 | Young |
| 0.988357 | 0.157608 | 0.192 | 0.196 | 1 | 0 | Cep350 | -0.004 | Young |
| 0.98905 | 0.14495 | 0.034 | 0.034 | 1 | 0 | Zfp182 | 0 | Young |
| 0.989174 | 0.125723 | 0.113 | 0.115 | 1 | 0 | Cdc26 | -0.002 | Young |
| 0.989178 | 0.004001 | 0.144 | 0.144 | 1 | 0 | Armcx3 | 0 | Young |
| 0.989443 | 0.028602 | 0.071 | 0.071 | 1 | 0 | Mfn2 | 0 | Young |
| 0.989757 | 0.120241 | 0.254 | 0.26 | 1 | 0 | Capn2 | -0.006 | Young |
| 0.989774 | 0.051965 | 0.06 | 0.06 | 1 | 0 | Ciz1 | 0 | Young |
| 0.98982 | 0.347981 | 0.012 | 0.012 | 1 | 0 | Tctn1 | 0 | Young |
| 0.98982 | 0.304517 | 0.012 | 0.012 | 1 | 0 | Mcat | 0 | Young |
| 0.989836 | 0.245146 | 0.133 | 0.135 | 1 | 0 | Slmap | -0.002 | Young |
| 0.989949 | 0.142048 | 0.018 | 0.018 | 1 | 0 | Prdm11 | 0 | Young |
| 0.990004 | 0.102757 | 0.089 | 0.09 | 1 | 0 | 5-Mar | -0.001 | Young |
| 0.990234 | 0.129579 | 0.185 | 0.189 | 1 | 0 | Mindy1 | -0.004 | Young |
| 0.990296 | 0.125369 | 0.224 | 0.23 | 1 | 0 | Arl8b | -0.006 | Young |
| 0.990332 | 0.184313 | 0.15 | 0.153 | 1 | 0 | Spata6 | -0.003 | Young |
| 0.990366 | 0.776651 | 0.02 | 0.021 | 1 | 0 | Kcnq1 | -0.001 | Young |
| 0.990537 | 0.373922 | 0.035 | 0.036 | 1 | 0 | Fam53a | -0.001 | Young |
| 0.990719 | 0.247041 | 0.022 | 0.022 | 1 | 0 | Eva1b | 0 | Young |
| 0.991458 | 0.546376 | 0.047 | 0.048 | 1 | 0 | Golim4 | -0.001 | Young |
| 0.991462 | 0.136059 | 0.022 | 0.022 | 1 | 0 | Chd3os | 0 | Young |
| 0.992377 | 0.351547 | 0.011 | 0.011 | 1 | 0 | Zfp867 | 0 | Young |
| 0.992591 | 0.184831 | 0.17 | 0.174 | 1 | 0 | Park7 | -0.004 | Young |
| 0.992647 | 0.264416 | 0.016 | 0.016 | 1 | 0 | Rpl36a-ps1 | 0 | Young |
| 0.992897 | 0.155111 | 0.024 | 0.025 | 1 | 0 | Rgl3 | -0.001 | Young |
| 0.992964 | 0.044055 | 0.176 | 0.176 | 1 | 0 | Plekhj1 | 0 | Young |
| 0.993003 | 0.357187 | 0.011 | 0.011 | 1 | 0 | Opn3 | 0 | Young |
| 0.993091 | 0.05771 | 0.228 | 0.231 | 1 | 0 | Fnip1 | -0.003 | Young |
| 0.993106 | 0.154547 | 0.152 | 0.155 | 1 | 0 | Ntan1 | -0.003 | Young |
| 0.99322 | 0.064347 | 0.168 | 0.17 | 1 | 0 | Pfdn2 | -0.002 | Young |
| 0.993234 | 0.069053 | 0.033 | 0.033 | 1 | 0 | Fbf1 | 0 | Young |
| 0.993575 | 0.622138 | 0.012 | 0.012 | 1 | 0 | Zfp951 | 0 | Young |
| 0.993688 | 0.094847 | 0.067 | 0.067 | 1 | 0 | Unkl | 0 | Young |
| 0.993718 | 0.198815 | 0.023 | 0.023 | 1 | 0 | Uevld | 0 | Young |
| 0.993888 | 0.04925 | 0.244 | 0.246 | 1 | 0 | Pds5a | -0.002 | Young |
| 0.993891 | 0.087483 | 0.09 | 0.09 | 1 | 0 | Vav2 | 0 | Young |
| 0.994116 | 0.229661 | 0.045 | 0.045 | 1 | 0 | Nme4 | 0 | Young |
| 0.994283 | 0.093708 | 0.022 | 0.022 | 1 | 0 | Rfxank | 0 | Young |
| 0.994465 | 0.354885 | 0.011 | 0.011 | 1 | 0 | Kpna2 | 0 | Young |
| 0.994559 | 0.019407 | 0.064 | 0.064 | 1 | 0 | Polr1c | 0 | Young |
| 0.994697 | 0.026938 | 0.122 | 0.122 | 1 | 0 | Vps18 | 0 | Young |

| | | | | | | | | |
|---|---|---|---|---|---|---|---|---|
| 0.994773 | 0.329413 | 0.03 | 0.03 | 1 | 0 | Ruvbl2 | 0 | Young |
| 0.994824 | 0.018153 | 0.348 | 0.353 | 1 | 0 | Zbtb7a | -0.005 | Young |
| 0.995004 | 0.129655 | 0.035 | 0.036 | 1 | 0 | Gm37233 | -0.001 | Young |
| 0.995108 | 0.203314 | 0.087 | 0.088 | 1 | 0 | Prkaca | -0.001 | Young |
| 0.995139 | 0.322746 | 0.018 | 0.018 | 1 | 0 | Atrip | 0 | Young |
| 0.995392 | 0.111219 | 0.041 | 0.041 | 1 | 0 | Gm20342 | 0 | Young |
| 0.995488 | 0.039852 | 0.377 | 0.383 | 1 | 0 | Lilr4b | -0.006 | Young |
| 0.995684 | 0.038797 | 0.049 | 0.049 | 1 | 0 | Gnptg | 0 | Young |
| 0.995738 | 0.078338 | 0.027 | 0.027 | 1 | 0 | Tmem241 | 0 | Young |
| 0.995884 | 0.362543 | 0.023 | 0.023 | 1 | 0 | Efemp2 | 0 | Young |
| 0.995927 | 0.293142 | 0.011 | 0.011 | 1 | 0 | Gm9796 | 0 | Young |
| 0.995927 | 0.18941 | 0.011 | 0.011 | 1 | 0 | Lss | 0 | Young |
| 0.996943 | 0.055559 | 0.068 | 0.068 | 1 | 0 | Abcd4 | 0 | Young |
| 0.996972 | 0.085349 | 0.011 | 0.011 | 1 | 0 | Pyroxd2 | 0 | Young |
| 0.997027 | 0.199803 | 0.042 | 0.042 | 1 | 0 | Ring1 | 0 | Young |
| 0.99712 | 0.10476 | 0.093 | 0.093 | 1 | 0 | Calhm2 | 0 | Young |
| 0.997171 | 0.179678 | 0.111 | 0.112 | 1 | 0 | Polr2c | -0.001 | Young |
| 0.997502 | 0.07799 | 0.048 | 0.048 | 1 | 0 | Ikbkg | 0 | Young |
| 0.997578 | 0.261345 | 0.03 | 0.03 | 1 | 0 | Mast2 | 0 | Young |
| 0.997644 | 0.026898 | 0.627 | 0.647 | 1 | 0 | App | -0.02 | Young |
| 0.997715 | 0.233782 | 0.083 | 0.083 | 1 | 0 | Nrbf2 | 0 | Young |
| 0.997785 | 0.04153 | 0.061 | 0.062 | 1 | 0 | Slc11a2 | -0.001 | Young |
| 0.998158 | 0.085844 | 0.104 | 0.105 | 1 | 0 | Kif3b | -0.001 | Young |
| 0.998225 | 0.097395 | 0.011 | 0.011 | 1 | 0 | Tarsl2 | 0 | Young |
| 0.998225 | 0.047252 | 0.011 | 0.011 | 1 | 0 | P2ry1 | 0 | Young |
| 0.998738 | 0.138751 | 0.212 | 0.218 | 1 | 0 | Trmt112 | -0.006 | Young |
| 0.998766 | 0.339429 | 0.035 | 0.036 | 1 | 0 | Dhx34 | -0.001 | Young |
| 0.99883 | 0.005333 | 0.206 | 0.209 | 1 | 0 | Ttc7 | -0.003 | Young |
| 0.998857 | 0.146147 | 0.071 | 0.071 | 1 | 0 | Rrm1 | 0 | Young |
| 0.99886 | 0.011182 | 0.174 | 0.175 | 1 | 0 | Tmf1 | -0.001 | Young |
| 0.998947 | 0.182458 | 0.054 | 0.055 | 1 | 0 | Zfp654 | -0.001 | Young |
| 0.999044 | 0.05108 | 0.067 | 0.067 | 1 | 0 | Kdm5d | 0 | Young |
| 0.999269 | 0.339688 | 0.011 | 0.011 | 1 | 0 | Heg1 | 0 | Young |
| 1.40E-48 | 3.457939 | 0.237 | 0.034 | 4.35E-44 | 1 | Cxcl13 | 0.203 | Old |
| 9.68E-35 | 2.479375 | 0.364 | 0.146 | 3.01E-30 | 1 | Ccl8 | 0.218 | Old |
| 4.29E-30 | 4.954236 | 0.113 | 0.007 | 1.33E-25 | 1 | Cd55 | 0.106 | Old |
| 1.48E-22 | 0.504442 | 0.892 | 0.857 | 4.59E-18 | 1 | B2m | 0.035 | Old |
| 2.15E-22 | 3.023182 | 0.14 | 0.03 | 6.68E-18 | 1 | C3 | 0.11 | Old |
| 4.51E-16 | 0.502499 | 0.886 | 0.824 | 1.40E-11 | 1 | H2-K1 | 0.062 | Old |
| 1.98E-14 | 0.662656 | 0.605 | 0.466 | 6.14E-10 | 1 | Clec10a | 0.139 | Old |
| 6.53E-14 | 0.604195 | 0.703 | 0.616 | 2.03E-09 | 1 | Ccl6 | 0.087 | Old |
| 3.84E-13 | 0.458551 | 0.855 | 0.784 | 1.19E-08 | 1 | Crip1 | 0.071 | Old |
| 4.50E-13 | 1.202052 | 0.267 | 0.152 | 1.40E-08 | 1 | Mmp9 | 0.115 | Old |
| 4.84E-13 | 0.377878 | 0.861 | 0.816 | 1.50E-08 | 1 | Ptma | 0.045 | Old |
| 1.30E-12 | 0.895408 | 0.415 | 0.294 | 4.03E-08 | 1 | Gda | 0.121 | Old |

| | | | | | | | | |
|---|---|---|---|---|---|---|---|---|
| 1.83E-12 | 0.565346 | 0.704 | 0.653 | 5.67E-08 | 1 | Pfn1 | 0.051 | Old |
| 1.99E-11 | 0.356578 | 0.956 | 0.95 | 6.18E-07 | 1 | Psap | 0.006 | Old |
| 1.20E-10 | 0.21325 | 0.995 | 0.996 | 3.72E-06 | 1 | Actb | -0.001 | Old |
| 4.30E-10 | 0.847934 | 0.483 | 0.394 | 1.33E-05 | 1 | Ly6e | 0.089 | Old |
| 1.12E-09 | 3.731456 | 0.032 | 0.001 | 3.49E-05 | 1 | Dmwd | 0.031 | Old |
| 1.13E-09 | 1.091725 | 0.24 | 0.142 | 3.52E-05 | 1 | Cd209f | 0.098 | Old |
| 2.02E-09 | 0.476771 | 0.742 | 0.702 | 6.26E-05 | 1 | Hsp90b1 | 0.04 | Old |
| 2.66E-09 | 0.486086 | 0.683 | 0.619 | 8.25E-05 | 1 | Ccl9 | 0.064 | Old |
| 4.80E-09 | 1.28197 | 0.184 | 0.098 | 0.000149 | 1 | Anxa1 | 0.086 | Old |
| 7.04E-09 | 0.33504 | 0.925 | 0.914 | 0.000218 | 1 | Actg1 | 0.011 | Old |
| 7.72E-09 | 0.655506 | 0.598 | 0.524 | 0.00024 | 1 | Chchd2 | 0.074 | Old |
| 2.73E-08 | 0.892368 | 0.504 | 0.413 | 0.000847 | 1 | Lpl | 0.091 | Old |
| 9.06E-08 | 2.366171 | 0.058 | 0.016 | 0.002813 | 1 | Gm26870 | 0.042 | Old |
| 9.10E-08 | 1.093441 | 0.184 | 0.106 | 0.002825 | 1 | Cd209g | 0.078 | Old |
| 3.12E-07 | 0.700709 | 0.342 | 0.257 | 0.009676 | 1 | Cstb | 0.085 | Old |
| 3.65E-07 | 0.719841 | 0.409 | 0.323 | 0.011348 | 1 | Naaa | 0.086 | Old |
| 4.74E-07 | 1.681839 | 0.062 | 0.02 | 0.014734 | 1 | Xist | 0.042 | Old |
| 7.00E-07 | 0.336287 | 0.744 | 0.73 | 0.021725 | 1 | H2-D1 | 0.014 | Old |
| 7.58E-07 | 0.336179 | 0.865 | 0.859 | 0.023528 | 1 | Ctsb | 0.006 | Old |
| 1.71E-06 | 0.515117 | 0.553 | 0.5 | 0.053024 | 1 | Hnrnpf | 0.053 | Old |
| 1.71E-06 | 2.456562 | 0.026 | 0.003 | 0.053035 | 1 | Adgrg1 | 0.023 | Old |
| 2.36E-06 | 1.87834 | 0.04 | 0.01 | 0.073432 | 1 | C6 | 0.03 | Old |
| 3.76E-06 | 0.261744 | 0.883 | 0.942 | 0.116864 | 1 | Junb | -0.059 | Old |
| 4.87E-06 | 0.129891 | 0.991 | 0.999 | 0.151271 | 1 | mt-Co3 | -0.008 | Old |
| 6.09E-06 | 0.129471 | 0.995 | 0.996 | 0.189127 | 1 | mt-Co2 | -0.001 | Old |
| 7.56E-06 | 0.264889 | 0.937 | 0.938 | 0.234751 | 1 | Jund | -0.001 | Old |
| 8.99E-06 | 0.648904 | 0.378 | 0.301 | 0.279151 | 1 | Anxa2 | 0.077 | Old |
| 1.12E-05 | 0.246003 | 0.798 | 0.764 | 0.348883 | 1 | Ahnak | 0.034 | Old |
| 1.39E-05 | 0.320202 | 0.754 | 0.747 | 0.432022 | 1 | Grn | 0.007 | Old |
| 1.64E-05 | 0.472884 | 0.617 | 0.593 | 0.508305 | 1 | Sdc3 | 0.024 | Old |
| 2.74E-05 | 0.837227 | 0.263 | 0.196 | 0.85078 | 1 | Nme1 | 0.067 | Old |
| 2.79E-05 | 0.676022 | 0.406 | 0.352 | 0.867058 | 1 | Pdia6 | 0.054 | Old |
| 2.85E-05 | 0.863933 | 0.269 | 0.203 | 0.884156 | 1 | Lgals3 | 0.066 | Old |
| 3.04E-05 | 1.056862 | 0.18 | 0.121 | 0.943625 | 1 | Pla2g2d | 0.059 | Old |
| 3.24E-05 | 1.31619 | 0.09 | 0.046 | 1 | 1 | Akr1c13 | 0.044 | Old |
| 3.27E-05 | 3.703575 | 0.019 | 0.002 | 1 | 1 | Colq | 0.017 | Old |
| 4.14E-05 | 0.703735 | 0.351 | 0.288 | 1 | 1 | Atp1a1 | 0.063 | Old |
| 5.13E-05 | 0.541506 | 0.415 | 0.357 | 1 | 1 | Plbd1 | 0.058 | Old |
| 6.83E-05 | 0.555927 | 0.431 | 0.369 | 1 | 1 | Tuba1b | 0.062 | Old |
| 6.98E-05 | 0.491798 | 0.512 | 0.453 | 1 | 1 | Mgl2 | 0.059 | Old |
| 7.53E-05 | 1.066005 | 0.132 | 0.081 | 1 | 1 | Smpdl3a | 0.051 | Old |
| 7.77E-05 | 0.639913 | 0.231 | 0.166 | 1 | 1 | Cdk2ap2 | 0.065 | Old |
| 8.35E-05 | 0.395295 | 0.593 | 0.553 | 1 | 1 | Vim | 0.04 | Old |
| 8.64E-05 | 0.234916 | 0.924 | 0.905 | 1 | 1 | H2-Eb1 | 0.019 | Old |
| 9.65E-05 | 0.734203 | 0.202 | 0.142 | 1 | 1 | Nrp2 | 0.06 | Old |

| | | | | | | | | |
|---|---|---|---|---|---|---|---|---|
| 0.000101 | 0.884851 | 0.221 | 0.162 | 1 | 1 | Man2a1 | 0.059 | Old |
| 0.000109 | 0.784935 | 0.129 | 0.078 | 1 | 1 | Fcna | 0.051 | Old |
| 0.000121 | 0.574631 | 0.402 | 0.341 | 1 | 1 | Klf4 | 0.061 | Old |
| 0.000135 | 0.304727 | 0.639 | 0.603 | 1 | 1 | Timp2 | 0.036 | Old |
| 0.000144 | 0.842375 | 0.267 | 0.21 | 1 | 1 | Sulf2 | 0.057 | Old |
| 0.000236 | 0.429772 | 0.374 | 0.308 | 1 | 1 | Plin2 | 0.066 | Old |
| 0.000244 | 0.384961 | 0.478 | 0.431 | 1 | 1 | Cd63 | 0.047 | Old |
| 0.000374 | 0.080732 | 0.044 | 0.018 | 1 | 1 | Igkc | 0.026 | Old |
| 0.000424 | 0.496596 | 0.433 | 0.397 | 1 | 1 | Cfp | 0.036 | Old |
| 0.000435 | 0.634742 | 0.303 | 0.245 | 1 | 1 | Ctsl | 0.058 | Old |
| 0.000435 | 3.204349 | 0.015 | 0.002 | 1 | 1 | Dmpk | 0.013 | Old |
| 0.000534 | 1.837516 | 0.026 | 0.007 | 1 | 1 | Ly6a | 0.019 | Old |
| 0.000636 | 0.88686 | 0.138 | 0.092 | 1 | 1 | Ckap4 | 0.046 | Old |
| 0.000687 | 1.231752 | 0.126 | 0.084 | 1 | 1 | Fam43a | 0.042 | Old |
| 0.000734 | 0.159629 | 0.965 | 0.958 | 1 | 1 | H2-Aa | 0.007 | Old |
| 0.000747 | 0.749891 | 0.203 | 0.152 | 1 | 1 | Sf3b6 | 0.051 | Old |
| 0.0008 | 0.616717 | 0.424 | 0.381 | 1 | 1 | Ccl7 | 0.043 | Old |
| 0.001064 | 2.788064 | 0.01 | 0.001 | 1 | 1 | Cdh6 | 0.009 | Old |
| 0.001307 | 0.951874 | 0.125 | 0.084 | 1 | 1 | Cd209d | 0.041 | Old |
| 0.001313 | 2.206762 | 0.015 | 0.003 | 1 | 1 | A530032D1 | 0.012 | Old |
| 0.001387 | 0.249374 | 0.744 | 0.762 | 1 | 1 | Gpx1 | -0.018 | Old |
| 0.001463 | 0.959724 | 0.06 | 0.032 | 1 | 1 | A4galt | 0.028 | Old |
| 0.001589 | 0.474956 | 0.51 | 0.478 | 1 | 1 | Klf2 | 0.032 | Old |
| 0.00164 | 0.406392 | 0.508 | 0.493 | 1 | 1 | Tmem176a | 0.015 | Old |
| 0.001691 | 0.837577 | 0.107 | 0.069 | 1 | 1 | Il2rg | 0.038 | Old |
| 0.001709 | 2.429108 | 0.018 | 0.004 | 1 | 1 | Mcoln2 | 0.014 | Old |
| 0.001711 | 0.62003 | 0.335 | 0.289 | 1 | 1 | Apobec3 | 0.046 | Old |
| 0.001753 | 1.92854 | 0.012 | 0.001 | 1 | 1 | Rtl6 | 0.011 | Old |
| 0.001759 | 0.801954 | 0.166 | 0.122 | 1 | 1 | Pdia4 | 0.044 | Old |
| 0.001767 | 0.808655 | 0.1 | 0.064 | 1 | 1 | Ms4a8a | 0.036 | Old |
| 0.001791 | 0.364227 | 0.436 | 0.38 | 1 | 1 | Clec4a1 | 0.056 | Old |
| 0.001847 | 1.507914 | 0.028 | 0.01 | 1 | 1 | Mmp12 | 0.018 | Old |
| 0.001984 | 1.413114 | 0.028 | 0.01 | 1 | 1 | Hpse | 0.018 | Old |
| 0.002029 | 1.539034 | 0.054 | 0.028 | 1 | 1 | Kcnab2 | 0.026 | Old |
| 0.002269 | 0.722703 | 0.064 | 0.035 | 1 | 1 | Gm36161 | 0.029 | Old |
| 0.002342 | 0.465648 | 0.438 | 0.399 | 1 | 1 | Prdx1 | 0.039 | Old |
| 0.00236 | 1.053759 | 0.109 | 0.073 | 1 | 1 | Dusp5 | 0.036 | Old |
| 0.002399 | 0.66925 | 0.308 | 0.268 | 1 | 1 | Rhob | 0.04 | Old |
| 0.002506 | 0.182247 | 0.889 | 0.918 | 1 | 1 | C1qa | -0.029 | Old |
| 0.002782 | 0.711313 | 0.114 | 0.076 | 1 | 1 | Il1rn | 0.038 | Old |
| 0.003084 | 0.416399 | 0.545 | 0.521 | 1 | 1 | Arf5 | 0.024 | Old |
| 0.003118 | 1.114216 | 0.045 | 0.022 | 1 | 1 | Steap3 | 0.023 | Old |
| 0.003262 | 2.472063 | 0.015 | 0.004 | 1 | 1 | Gm10800 | 0.011 | Old |
| 0.003278 | 0.233798 | 0.704 | 0.716 | 1 | 1 | Myl6 | -0.012 | Old |
| 0.003846 | 1.383211 | 0.035 | 0.016 | 1 | 1 | Nr1h3 | 0.019 | Old |

| | | | | | | | | |
|---|---|---|---|---|---|---|---|---|
| 0.003903 | 2.063886 | 0.022 | 0.007 | 1 | 1 | Igf2r | 0.015 | Old |
| 0.003951 | 1.845118 | 0.019 | 0.006 | 1 | 1 | Sema6a | 0.013 | Old |
| 0.003988 | 1.675856 | 0.022 | 0.007 | 1 | 1 | Prmt6 | 0.015 | Old |
| 0.003994 | 1.117737 | 0.035 | 0.016 | 1 | 1 | Rrp15 | 0.019 | Old |
| 0.004016 | 0.128477 | 0.964 | 0.947 | 1 | 1 | H2-Ab1 | 0.017 | Old |
| 0.004062 | 0.715984 | 0.161 | 0.12 | 1 | 1 | Tfrc | 0.041 | Old |
| 0.004122 | 2.984277 | 0.01 | 0.001 | 1 | 1 | Gm15675 | 0.009 | Old |
| 0.004171 | 2.546847 | 0.01 | 0.001 | 1 | 1 | Hsd17b14 | 0.009 | Old |
| 0.00497 | 0.712781 | 0.058 | 0.032 | 1 | 1 | Anpep | 0.026 | Old |
| 0.004974 | 0.540058 | 0.357 | 0.323 | 1 | 1 | Pdia3 | 0.034 | Old |
| 0.005425 | 0.206018 | 0.842 | 0.886 | 1 | 1 | Dusp1 | -0.044 | Old |
| 0.005438 | 2.210636 | 0.012 | 0.002 | 1 | 1 | Adcy3 | 0.01 | Old |
| 0.005457 | 0.714342 | 0.118 | 0.083 | 1 | 1 | Etfa | 0.035 | Old |
| 0.005617 | 0.599308 | 0.263 | 0.222 | 1 | 1 | Ninj1 | 0.041 | Old |
| 0.006101 | 1.779544 | 0.026 | 0.01 | 1 | 1 | Panx1 | 0.016 | Old |
| 0.006767 | 0.240647 | 0.591 | 0.592 | 1 | 1 | Ptp4a2 | -0.001 | Old |
| 0.006813 | 0.479705 | 0.348 | 0.311 | 1 | 1 | Arpc4 | 0.037 | Old |
| 0.006933 | 0.751585 | 0.144 | 0.108 | 1 | 1 | Klc1 | 0.036 | Old |
| 0.007118 | 1.682535 | 0.014 | 0.004 | 1 | 1 | Mgll | 0.01 | Old |
| 0.007223 | 0.801874 | 0.147 | 0.11 | 1 | 1 | Tspan3 | 0.037 | Old |
| 0.007767 | 0.533901 | 0.999 | 1 | 1 | 1 | Gm42418 | -0.001 | Old |
| 0.007799 | 0.982398 | 0.037 | 0.018 | 1 | 1 | Cdkn2c | 0.019 | Old |
| 0.007953 | 1.02856 | 0.017 | 0.005 | 1 | 1 | Cald1 | 0.012 | Old |
| 0.008592 | 1.078086 | 0.067 | 0.041 | 1 | 1 | Pqlc3 | 0.026 | Old |
| 0.008666 | 1.379673 | 0.059 | 0.035 | 1 | 1 | Slc35a2 | 0.024 | Old |
| 0.008734 | 0.357763 | 0.383 | 0.347 | 1 | 1 | Xbp1 | 0.036 | Old |
| 0.008841 | 0.335264 | 0.53 | 0.521 | 1 | 1 | Hspa5 | 0.009 | Old |
| 0.008857 | 1.426687 | 0.048 | 0.027 | 1 | 1 | Pira2 | 0.021 | Old |
| 0.009234 | 0.17849 | 0.817 | 0.857 | 1 | 1 | C1qc | -0.04 | Old |
| 0.009493 | 1.659096 | 0.033 | 0.016 | 1 | 1 | Gng11 | 0.017 | Old |
| 0.009558 | 1.340367 | 0.027 | 0.012 | 1 | 1 | Cd200r4 | 0.015 | Old |
| 0.010336 | 0.202906 | 0.758 | 0.783 | 1 | 1 | mt-Nd3 | -0.025 | Old |
| 0.010415 | 0.260704 | 0.513 | 0.479 | 1 | 1 | Ywhaz | 0.034 | Old |
| 0.01079 | 0.649428 | 0.132 | 0.098 | 1 | 1 | Ms4a6d | 0.034 | Old |
| 0.01093 | 0.650274 | 0.132 | 0.098 | 1 | 1 | Uap1l1 | 0.034 | Old |
| 0.011275 | 0.911843 | 0.089 | 0.061 | 1 | 1 | Slc27a1 | 0.028 | Old |
| 0.011377 | 0.327846 | 0.555 | 0.544 | 1 | 1 | Arpc5 | 0.011 | Old |
| 0.011438 | 0.649953 | 0.111 | 0.079 | 1 | 1 | Kxd1 | 0.032 | Old |
| 0.011693 | 0.954111 | 0.068 | 0.044 | 1 | 1 | Gpx3 | 0.024 | Old |
| 0.01229 | 0.439985 | 0.279 | 0.241 | 1 | 1 | Lbh | 0.038 | Old |
| 0.01251 | 1.575223 | 0.022 | 0.009 | 1 | 1 | Gbp2b | 0.013 | Old |
| 0.012682 | 1.369007 | 0.054 | 0.032 | 1 | 1 | Ddx56 | 0.022 | Old |
| 0.012969 | 1.714396 | 0.021 | 0.008 | 1 | 1 | Mov10 | 0.013 | Old |
| 0.01317 | 1.697771 | 0.021 | 0.008 | 1 | 1 | Atpaf1 | 0.013 | Old |
| 0.013209 | 1.049248 | 0.035 | 0.018 | 1 | 1 | Fkbp9 | 0.017 | Old |

| | | | | | | | | |
|---|---|---|---|---|---|---|---|---|
| 0.013452 | 0.389596 | 0.497 | 0.485 | 1 | 1 | Mob1a | 0.012 | Old |
| 0.013767 | 1.302683 | 0.042 | 0.024 | 1 | 1 | Ptpn23 | 0.018 | Old |
| 0.013798 | 2.097375 | 0.012 | 0.003 | 1 | 1 | Arhgef5 | 0.009 | Old |
| 0.013819 | 2.311241 | 0.012 | 0.003 | 1 | 1 | Cit | 0.009 | Old |
| 0.013819 | 1.501521 | 0.033 | 0.017 | 1 | 1 | Rars2 | 0.016 | Old |
| 0.013945 | 1.667729 | 0.012 | 0.003 | 1 | 1 | Timeless | 0.009 | Old |
| 0.01449 | 0.808063 | 0.089 | 0.061 | 1 | 1 | Lrpprc | 0.028 | Old |
| 0.014665 | 2.123919 | 0.017 | 0.006 | 1 | 1 | Ptgfrn | 0.011 | Old |
| 0.014702 | 0.672985 | 0.058 | 0.035 | 1 | 1 | Bst1 | 0.023 | Old |
| 0.014751 | 1.232763 | 0.027 | 0.013 | 1 | 1 | E2f3 | 0.014 | Old |
| 0.014765 | 0.979448 | 0.018 | 0.007 | 1 | 1 | Tcim | 0.011 | Old |
| 0.014897 | 1.629723 | 0.013 | 0.004 | 1 | 1 | Kif2c | 0.009 | Old |
| 0.01496 | 1.368826 | 0.013 | 0.004 | 1 | 1 | Rab23 | 0.009 | Old |
| 0.014989 | 0.541892 | 0.193 | 0.157 | 1 | 1 | Ppp3r1 | 0.036 | Old |
| 0.015019 | 1.10649 | 0.055 | 0.034 | 1 | 1 | Cd209b | 0.021 | Old |
| 0.015294 | 1.183735 | 0.014 | 0.004 | 1 | 1 | Ggct | 0.01 | Old |
| 0.015448 | 0.828308 | 0.05 | 0.03 | 1 | 1 | Zfpl1 | 0.02 | Old |
| 0.015579 | 1.269967 | 0.064 | 0.041 | 1 | 1 | Fabp5 | 0.023 | Old |
| 0.015874 | 0.904075 | 0.199 | 0.163 | 1 | 1 | Fabp4 | 0.036 | Old |
| 0.016224 | 0.184615 | 0.679 | 0.676 | 1 | 1 | Lcp1 | 0.003 | Old |
| 0.016229 | 0.004391 | 0.211 | 0.267 | 1 | 1 | Tet3 | -0.056 | Old |
| 0.017336 | 0.46586 | 0.323 | 0.294 | 1 | 1 | Atp5g3 | 0.029 | Old |
| 0.018115 | 1.320386 | 0.046 | 0.027 | 1 | 1 | Bcl2l13 | 0.019 | Old |
| 0.018377 | 0.220258 | 0.629 | 0.636 | 1 | 1 | Calm1 | -0.007 | Old |
| 0.018691 | 0.539481 | 0.17 | 0.134 | 1 | 1 | Chmp5 | 0.036 | Old |
| 0.018738 | 1.561959 | 0.023 | 0.01 | 1 | 1 | 2-Sep | 0.013 | Old |
| 0.018746 | 0.4169 | 0.344 | 0.309 | 1 | 1 | Tmem30a | 0.035 | Old |
| 0.019632 | 0.221858 | 0.548 | 0.545 | 1 | 1 | Pltp | 0.003 | Old |
| 0.019937 | 0.791737 | 0.125 | 0.095 | 1 | 1 | Ftl1-ps1 | 0.03 | Old |
| 0.020214 | 1.079003 | 0.033 | 0.018 | 1 | 1 | Gdf3 | 0.015 | Old |
| 0.020224 | 0.469838 | 0.317 | 0.286 | 1 | 1 | Ap1b1 | 0.031 | Old |
| 0.020454 | 0.611528 | 0.103 | 0.075 | 1 | 1 | Rnd3 | 0.028 | Old |
| 0.020665 | 0.574094 | 0.199 | 0.165 | 1 | 1 | Copa | 0.034 | Old |
| 0.020875 | 1.164908 | 0.037 | 0.021 | 1 | 1 | Rrp12 | 0.016 | Old |
| 0.021429 | 0.801397 | 0.105 | 0.078 | 1 | 1 | Pcna | 0.027 | Old |
| 0.021514 | 0.764748 | 0.055 | 0.035 | 1 | 1 | Ifi211 | 0.02 | Old |
| 0.021841 | 0.960698 | 0.041 | 0.024 | 1 | 1 | Traf6 | 0.017 | Old |
| 0.022035 | 1.432088 | 0.021 | 0.009 | 1 | 1 | Gtf3c5 | 0.012 | Old |
| 0.022146 | 0.022594 | 0.059 | 0.088 | 1 | 1 | Polr2f | -0.029 | Old |
| 0.022207 | 1.437294 | 0.021 | 0.009 | 1 | 1 | Ttc27 | 0.012 | Old |
| 0.022221 | 0.62419 | 0.217 | 0.185 | 1 | 1 | Actn4 | 0.032 | Old |
| 0.022274 | 0.597969 | 0.087 | 0.061 | 1 | 1 | Slc25a24 | 0.026 | Old |
| 0.022294 | 0.965801 | 0.021 | 0.009 | 1 | 1 | Dusp18 | 0.012 | Old |
| 0.022933 | 0.507466 | 0.256 | 0.22 | 1 | 1 | Selenos | 0.036 | Old |
| 0.023378 | 0.903609 | 0.063 | 0.041 | 1 | 1 | Celf4 | 0.022 | Old |

| | | | | | | | | |
|---|---|---|---|---|---|---|---|---|
| 0.0234 | 0.931819 | 0.048 | 0.029 | 1 | 1 | Mmp19 | 0.019 | Old |
| 0.023714 | 0.602302 | 0.143 | 0.112 | 1 | 1 | Fbl | 0.031 | Old |
| 0.023872 | 0.991693 | 0.075 | 0.052 | 1 | 1 | Rab29 | 0.023 | Old |
| 0.024672 | 0.725227 | 0.066 | 0.044 | 1 | 1 | Cul4a | 0.022 | Old |
| 0.024935 | 0.77639 | 0.066 | 0.044 | 1 | 1 | Thoc6 | 0.022 | Old |
| 0.025079 | 0.434461 | 0.08 | 0.055 | 1 | 1 | Polr2l | 0.025 | Old |
| 0.025081 | 1.570408 | 0.018 | 0.007 | 1 | 1 | Kctd17 | 0.011 | Old |
| 0.025373 | 0.580774 | 0.208 | 0.176 | 1 | 1 | Kdelr1 | 0.032 | Old |
| 0.025506 | 0.509372 | 0.064 | 0.042 | 1 | 1 | Nabp2 | 0.022 | Old |
| 0.025736 | 1.197881 | 0.035 | 0.019 | 1 | 1 | Txnl4a | 0.016 | Old |
| 0.026105 | 0.449356 | 0.332 | 0.301 | 1 | 1 | Creg1 | 0.031 | Old |
| 0.02681 | 0.475151 | 0.235 | 0.202 | 1 | 1 | Psmd2 | 0.033 | Old |
| 0.026913 | 1.334939 | 0.017 | 0.007 | 1 | 1 | Meis3 | 0.01 | Old |
| 0.026999 | 1.26836 | 0.017 | 0.007 | 1 | 1 | Ulbp1 | 0.01 | Old |
| 0.027524 | 0.675775 | 0.017 | 0.007 | 1 | 1 | Kti12 | 0.01 | Old |
| 0.027808 | 1.76384 | 0.015 | 0.006 | 1 | 1 | Akr1c12 | 0.009 | Old |
| 0.027818 | 1.010833 | 0.084 | 0.06 | 1 | 1 | Emc2 | 0.024 | Old |
| 0.027839 | 1.756478 | 0.015 | 0.006 | 1 | 1 | Nectin2 | 0.009 | Old |
| 0.028541 | 0.60658 | 0.175 | 0.143 | 1 | 1 | 11-Sep | 0.032 | Old |
| 0.02917 | 2.493913 | 0.01 | 0.003 | 1 | 1 | C130026I2 | 0.007 | Old |
| 0.029171 | 1.362091 | 0.014 | 0.005 | 1 | 1 | Ighmbp2 | 0.009 | Old |
| 0.029253 | 2.05555 | 0.01 | 0.003 | 1 | 1 | Thrsp | 0.007 | Old |
| 0.029569 | 0.496458 | 0.303 | 0.276 | 1 | 1 | Folr2 | 0.027 | Old |
| 0.029607 | 0.817821 | 0.045 | 0.027 | 1 | 1 | Slc39a6 | 0.018 | Old |
| 0.029928 | 1.84204 | 0.012 | 0.004 | 1 | 1 | Sh3bgrl2 | 0.008 | Old |
| 0.029929 | 0.12771 | 0.01 | 0.003 | 1 | 1 | Nat8f1 | 0.007 | Old |
| 0.030015 | 1.460817 | 0.013 | 0.004 | 1 | 1 | Gpnmb | 0.009 | Old |
| 0.03059 | 1.105402 | 0.028 | 0.015 | 1 | 1 | Nectin1 | 0.013 | Old |
| 0.030725 | 0.206606 | 0.87 | 0.888 | 1 | 1 | Lgmn | -0.018 | Old |
| 0.031033 | 0.014688 | 0.113 | 0.151 | 1 | 1 | Cox7b | -0.038 | Old |
| 0.031866 | 0.421224 | 0.285 | 0.253 | 1 | 1 | Dync1h1 | 0.032 | Old |
| 0.03241 | 0.559101 | 0.154 | 0.125 | 1 | 1 | Dck | 0.029 | Old |
| 0.033402 | 0.45408 | 0.239 | 0.207 | 1 | 1 | Vcam1 | 0.032 | Old |
| 0.033855 | 0.559291 | 0.068 | 0.047 | 1 | 1 | Eif2s1 | 0.021 | Old |
| 0.033951 | 1.486196 | 0.031 | 0.017 | 1 | 1 | Ubl4a | 0.014 | Old |
| 0.033997 | 0.622367 | 0.064 | 0.044 | 1 | 1 | Vta1 | 0.02 | Old |
| 0.034249 | 0.535636 | 0.224 | 0.192 | 1 | 1 | Atp6v0a1 | 0.032 | Old |
| 0.034932 | 1.368583 | 0.021 | 0.01 | 1 | 1 | Gadd45a | 0.011 | Old |
| 0.034934 | 0.653321 | 0.084 | 0.06 | 1 | 1 | Htra2 | 0.024 | Old |
| 0.035569 | 0.557119 | 0.158 | 0.128 | 1 | 1 | Sdf2 | 0.03 | Old |
| 0.03567 | 0.722699 | 0.064 | 0.044 | 1 | 1 | Mfsd12 | 0.02 | Old |
| 0.036226 | 0.533477 | 0.152 | 0.122 | 1 | 1 | Sec31a | 0.03 | Old |
| 0.036251 | 0.615621 | 0.126 | 0.098 | 1 | 1 | Vat1 | 0.028 | Old |
| 0.036712 | 0.014491 | 0.176 | 0.221 | 1 | 1 | Usp9x | -0.045 | Old |
| 0.036892 | 0.132983 | 0.153 | 0.196 | 1 | 1 | Wapl | -0.043 | Old |

| | | | | | | | | |
|---|---|---|---|---|---|---|---|---|
| 0.03759 | 0.277073 | 0.613 | 0.629 | 1 | 1 | Anxa5 | -0.016 | Old |
| 0.037808 | 1.308429 | 0.019 | 0.009 | 1 | 1 | Nol10 | 0.01 | Old |
| 0.037829 | 0.627905 | 0.161 | 0.131 | 1 | 1 | Rab20 | 0.03 | Old |
| 0.037862 | 0.673465 | 0.163 | 0.136 | 1 | 1 | 9-Sep | 0.027 | Old |
| 0.039056 | 0.078834 | 0.138 | 0.177 | 1 | 1 | Rbm5 | -0.039 | Old |
| 0.039583 | 1.21031 | 0.024 | 0.013 | 1 | 1 | Fam160a2 | 0.011 | Old |
| 0.040068 | 0.615521 | 0.166 | 0.138 | 1 | 1 | Neu1 | 0.028 | Old |
| 0.040167 | 0.602361 | 0.117 | 0.091 | 1 | 1 | Arhgef10l | 0.026 | Old |
| 0.040557 | 0.839731 | 0.059 | 0.04 | 1 | 1 | Taf4 | 0.019 | Old |
| 0.041308 | 0.000986 | 0.109 | 0.143 | 1 | 1 | Tmem160 | -0.034 | Old |
| 0.041864 | 0.356564 | 0.123 | 0.095 | 1 | 1 | Rap2c | 0.028 | Old |
| 0.04255 | 0.773607 | 0.075 | 0.053 | 1 | 1 | Bcat2 | 0.022 | Old |
| 0.042868 | 0.730794 | 0.046 | 0.03 | 1 | 1 | Utp20 | 0.016 | Old |
| 0.043695 | 1.6584 | 0.017 | 0.007 | 1 | 1 | Atg4a | 0.01 | Old |
| 0.043915 | 0.436344 | 0.195 | 0.165 | 1 | 1 | Arf3 | 0.03 | Old |
| 0.044432 | 0.460275 | 0.208 | 0.177 | 1 | 1 | U2af2 | 0.031 | Old |
| 0.04467 | 0.905618 | 0.032 | 0.018 | 1 | 1 | Lxn | 0.014 | Old |
| 0.044764 | 0.345854 | 0.262 | 0.229 | 1 | 1 | Serpinb6a | 0.033 | Old |
| 0.045307 | 0.497404 | 0.098 | 0.074 | 1 | 1 | St6galnac4 | 0.024 | Old |
| 0.046142 | 0.854563 | 0.032 | 0.018 | 1 | 1 | Atp8b2 | 0.014 | Old |
| 0.046354 | 0.42874 | 0.085 | 0.062 | 1 | 1 | Ear2 | 0.023 | Old |
| 0.046674 | 0.183137 | 0.672 | 0.68 | 1 | 1 | Mbnl1 | -0.008 | Old |
| 0.046776 | 1.118 | 0.031 | 0.018 | 1 | 1 | Ciapin1 | 0.013 | Old |
| 0.046901 | 0.402217 | 0.087 | 0.064 | 1 | 1 | Msmo1 | 0.023 | Old |
| 0.047427 | 0.240038 | 0.512 | 0.49 | 1 | 1 | Cyba | 0.022 | Old |
| 0.047498 | 0.776644 | 0.066 | 0.046 | 1 | 1 | Rpl10-ps3 | 0.02 | Old |
| 0.047739 | 0.254247 | 0.539 | 0.528 | 1 | 1 | Canx | 0.011 | Old |
| 0.047896 | 1.583422 | 0.015 | 0.007 | 1 | 1 | Tmco6 | 0.008 | Old |
| 0.048337 | 1.104255 | 0.022 | 0.011 | 1 | 1 | Tmem18 | 0.011 | Old |
| 0.04838 | 1.000469 | 0.015 | 0.007 | 1 | 1 | Efcab2 | 0.008 | Old |
| 0.048637 | 0.353208 | 0.134 | 0.106 | 1 | 1 | Tppp3 | 0.028 | Old |
| 0.048693 | 1.035892 | 0.022 | 0.011 | 1 | 1 | Ulk3 | 0.011 | Old |
| 0.049375 | 0.709946 | 0.027 | 0.015 | 1 | 1 | Yars | 0.012 | Old |
| 0.049784 | 0.016272 | 0.113 | 0.148 | 1 | 1 | Osbpl8 | -0.035 | Old |
| 0.049877 | 0.675112 | 0.174 | 0.147 | 1 | 1 | Azin1 | 0.027 | Old |
| 0.049925 | 0.855977 | 0.134 | 0.109 | 1 | 1 | Hist1h1c | 0.025 | Old |
| 0.050029 | 0.197644 | 0.654 | 0.675 | 1 | 1 | BC005537 | -0.021 | Old |
| 0.050238 | 0.432491 | 0.239 | 0.211 | 1 | 1 | Atp5k | 0.028 | Old |
| 0.050434 | 1.896937 | 0.014 | 0.006 | 1 | 1 | Gatb | 0.008 | Old |
| 0.051716 | 1.176635 | 0.014 | 0.006 | 1 | 1 | Gm26982 | 0.008 | Old |
| 0.05177 | 0.907811 | 0.014 | 0.006 | 1 | 1 | Intu | 0.008 | Old |
| 0.051875 | 1.266479 | 0.033 | 0.02 | 1 | 1 | Pop7 | 0.013 | Old |
| 0.051891 | 0.030315 | 0.258 | 0.309 | 1 | 1 | Cox8a | -0.051 | Old |
| 0.052258 | 0.392911 | 0.337 | 0.314 | 1 | 1 | Itgam | 0.023 | Old |
| 0.052969 | 1.011292 | 0.049 | 0.032 | 1 | 1 | Mfn2 | 0.017 | Old |

| | | | | | | | | |
|---|---|---|---|---|---|---|---|---|
| 0.053239 | 0.259282 | 0.378 | 0.348 | 1 | 1 | Ctsd | 0.03 | Old |
| 0.053895 | 0.833924 | 0.021 | 0.01 | 1 | 1 | Phlpp1 | 0.011 | Old |
| 0.054011 | 1.57299 | 0.013 | 0.005 | 1 | 1 | Gprc5b | 0.008 | Old |
| 0.05407 | 1.65417 | 0.013 | 0.005 | 1 | 1 | Sgce | 0.008 | Old |
| 0.054098 | 0.117552 | 0.118 | 0.153 | 1 | 1 | Nop10 | -0.035 | Old |
| 0.054709 | 0.362483 | 0.365 | 0.346 | 1 | 1 | Tubb5 | 0.019 | Old |
| 0.054709 | 0.627276 | 0.1 | 0.078 | 1 | 1 | BC031181 | 0.022 | Old |
| 0.05549 | 0.385968 | 0.208 | 0.179 | 1 | 1 | Tmem123 | 0.029 | Old |
| 0.056139 | 0.550867 | 0.21 | 0.184 | 1 | 1 | Cdc42se2 | 0.026 | Old |
| 0.056605 | 1.910538 | 0.012 | 0.004 | 1 | 1 | Fitm2 | 0.008 | Old |
| 0.056626 | 0.860708 | 0.077 | 0.057 | 1 | 1 | 2310011J0 | 0.02 | Old |
| 0.056736 | 1.895114 | 0.012 | 0.004 | 1 | 1 | Lum | 0.008 | Old |
| 0.057113 | 0.971765 | 0.024 | 0.013 | 1 | 1 | Bcas3 | 0.011 | Old |
| 0.057727 | 1.347675 | 0.012 | 0.004 | 1 | 1 | Gemin5 | 0.008 | Old |
| 0.057833 | 0.58087 | 0.201 | 0.174 | 1 | 1 | Pros1 | 0.027 | Old |
| 0.058156 | 1.037014 | 0.019 | 0.01 | 1 | 1 | Wdhd1 | 0.009 | Old |
| 0.058194 | 0.8218 | 0.012 | 0.004 | 1 | 1 | Gpr162 | 0.008 | Old |
| 0.058219 | 1.104606 | 0.028 | 0.016 | 1 | 1 | Larp1b | 0.012 | Old |
| 0.058501 | 0.858278 | 0.019 | 0.01 | 1 | 1 | Inpp5e | 0.009 | Old |
| 0.058595 | 0.115677 | 0.147 | 0.185 | 1 | 1 | Hnrnpdl | -0.038 | Old |
| 0.058955 | 2.081568 | 0.01 | 0.004 | 1 | 1 | Sparcl1 | 0.006 | Old |
| 0.059491 | 0.813854 | 0.028 | 0.016 | 1 | 1 | Smpd3 | 0.012 | Old |
| 0.059759 | 1.39482 | 0.01 | 0.004 | 1 | 1 | Tmlhe | 0.006 | Old |
| 0.060099 | 0.371153 | 0.019 | 0.01 | 1 | 1 | Tacc2 | 0.009 | Old |
| 0.06012 | 0.416845 | 0.105 | 0.081 | 1 | 1 | Il11ra1 | 0.024 | Old |
| 0.060264 | 1.539719 | 0.031 | 0.018 | 1 | 1 | Pola1 | 0.013 | Old |
| 0.060702 | 0.078333 | 0.905 | 0.948 | 1 | 1 | mt-Nd1 | -0.043 | Old |
| 0.06119 | 1.431814 | 0.023 | 0.013 | 1 | 1 | Dnaaf3 | 0.01 | Old |
| 0.061308 | 0.891846 | 0.048 | 0.032 | 1 | 1 | Nup93 | 0.016 | Old |
| 0.062056 | 0.753836 | 0.06 | 0.043 | 1 | 1 | Cavin1 | 0.017 | Old |
| 0.062815 | 1.155497 | 0.023 | 0.013 | 1 | 1 | Trip10 | 0.01 | Old |
| 0.064033 | 0.517471 | 0.039 | 0.024 | 1 | 1 | Kpna6 | 0.015 | Old |
| 0.064252 | 0.204085 | 0.567 | 0.575 | 1 | 1 | Actr3 | -0.008 | Old |
| 0.064284 | 1.121592 | 0.023 | 0.013 | 1 | 1 | Narfl | 0.01 | Old |
| 0.064332 | 1.033141 | 0.023 | 0.013 | 1 | 1 | Rrnad1 | 0.01 | Old |
| 0.064521 | 0.592266 | 0.195 | 0.171 | 1 | 1 | C4b | 0.024 | Old |
| 0.064656 | 0.523512 | 0.048 | 0.032 | 1 | 1 | Pum3 | 0.016 | Old |
| 0.064795 | 0.386384 | 0.093 | 0.07 | 1 | 1 | Gstm1 | 0.023 | Old |
| 0.064839 | 0.479512 | 0.172 | 0.146 | 1 | 1 | Snx6 | 0.026 | Old |
| 0.065168 | 0.938908 | 0.044 | 0.029 | 1 | 1 | Cfap36 | 0.015 | Old |
| 0.065458 | 0.041929 | 0.275 | 0.325 | 1 | 1 | Slc25a5 | -0.05 | Old |
| 0.065972 | 0.739403 | 0.063 | 0.045 | 1 | 1 | Insig2 | 0.018 | Old |
| 0.067329 | 0.721758 | 0.027 | 0.016 | 1 | 1 | 6030458C1 | 0.011 | Old |
| 0.068676 | 0.155487 | 0.883 | 0.908 | 1 | 1 | Itm2b | -0.025 | Old |
| 0.069646 | 0.806619 | 0.063 | 0.045 | 1 | 1 | Bms1 | 0.018 | Old |

| | | | | | | | | |
|---|---|---|---|---|---|---|---|---|
| 0.070069 | 0.491377 | 0.23 | 0.204 | 1 | 1 | Edem1 | 0.026 | Old |
| 0.070073 | 0.006557 | 0.265 | 0.311 | 1 | 1 | Gnas | -0.046 | Old |
| 0.07011 | 1.129296 | 0.03 | 0.018 | 1 | 1 | Lsm1 | 0.012 | Old |
| 0.070849 | 1.23241 | 0.026 | 0.015 | 1 | 1 | Dnttip1 | 0.011 | Old |
| 0.071061 | 0.713637 | 0.075 | 0.055 | 1 | 1 | Odc1 | 0.02 | Old |
| 0.071489 | 0.471639 | 0.022 | 0.012 | 1 | 1 | Actn1 | 0.01 | Old |
| 0.07159 | 0.919128 | 0.03 | 0.018 | 1 | 1 | Mfsd7a | 0.012 | Old |
| 0.071711 | 0.556174 | 0.125 | 0.101 | 1 | 1 | Sri | 0.024 | Old |
| 0.07196 | 0.672251 | 0.054 | 0.038 | 1 | 1 | Klhdc4 | 0.016 | Old |
| 0.072091 | 0.484845 | 0.049 | 0.033 | 1 | 1 | Tada1 | 0.016 | Old |
| 0.072362 | 0.723265 | 0.03 | 0.018 | 1 | 1 | Rnf31 | 0.012 | Old |
| 0.073117 | 0.361368 | 0.306 | 0.281 | 1 | 1 | Pirb | 0.025 | Old |
| 0.073327 | 0.105006 | 0.013 | 0.024 | 1 | 1 | Cep164 | -0.011 | Old |
| 0.073387 | 0.668908 | 0.093 | 0.072 | 1 | 1 | B4galt5 | 0.021 | Old |
| 0.073911 | 0.737097 | 0.026 | 0.015 | 1 | 1 | Rdm1 | 0.011 | Old |
| 0.074012 | 0.414872 | 0.523 | 0.515 | 1 | 1 | Ccl2 | 0.008 | Old |
| 0.074174 | 0.568059 | 0.132 | 0.109 | 1 | 1 | Lamtor3 | 0.023 | Old |
| 0.074514 | 0.909672 | 0.068 | 0.05 | 1 | 1 | Nup153 | 0.018 | Old |
| 0.074626 | 1.417298 | 0.021 | 0.011 | 1 | 1 | Mill2 | 0.01 | Old |
| 0.075121 | 0.914006 | 0.055 | 0.039 | 1 | 1 | Parp11 | 0.016 | Old |
| 0.075179 | 0.562638 | 0.126 | 0.103 | 1 | 1 | Fam49a | 0.023 | Old |
| 0.075474 | 0.9581 | 0.045 | 0.03 | 1 | 1 | Trappc13 | 0.015 | Old |
| 0.075598 | 0.416863 | 0.303 | 0.282 | 1 | 1 | Rpn1 | 0.021 | Old |
| 0.076459 | 0.651179 | 0.066 | 0.048 | 1 | 1 | Pygo2 | 0.018 | Old |
| 0.076934 | 0.545444 | 0.015 | 0.007 | 1 | 1 | F10 | 0.008 | Old |
| 0.077571 | 1.006749 | 0.021 | 0.011 | 1 | 1 | Klhl15 | 0.01 | Old |
| 0.078673 | 0.190588 | 0.656 | 0.666 | 1 | 1 | Pabpc1 | -0.01 | Old |
| 0.078864 | 1.128368 | 0.024 | 0.014 | 1 | 1 | Alg8 | 0.01 | Old |
| 0.079206 | 0.29508 | 0.409 | 0.39 | 1 | 1 | Sh3bgrl | 0.019 | Old |
| 0.079518 | 1.016181 | 0.024 | 0.014 | 1 | 1 | Ceacam1 | 0.01 | Old |
| 0.079683 | 0.464671 | 0.224 | 0.2 | 1 | 1 | S100a4 | 0.024 | Old |
| 0.079745 | 0.017818 | 0.054 | 0.075 | 1 | 1 | Chtf8 | -0.021 | Old |
| 0.079747 | 0.528546 | 0.103 | 0.081 | 1 | 1 | Cpeb4 | 0.022 | Old |
| 0.08052 | 0.001585 | 0.06 | 0.082 | 1 | 1 | Ahcyl1 | -0.022 | Old |
| 0.080626 | 0.288269 | 0.321 | 0.298 | 1 | 1 | Flna | 0.023 | Old |
| 0.080956 | 0.517078 | 0.118 | 0.096 | 1 | 1 | Csnk1g2 | 0.022 | Old |
| 0.081616 | 0.481004 | 0.053 | 0.037 | 1 | 1 | Vps53 | 0.016 | Old |
| 0.081687 | 0.134598 | 0.048 | 0.067 | 1 | 1 | Zbtb11 | -0.019 | Old |
| 0.082083 | 0.081396 | 0.145 | 0.178 | 1 | 1 | Ndufb10 | -0.033 | Old |
| 0.082749 | 1.472955 | 0.014 | 0.007 | 1 | 1 | Papss2 | 0.007 | Old |
| 0.083056 | 0.521301 | 0.19 | 0.165 | 1 | 1 | Tsc22d1 | 0.025 | Old |
| 0.083137 | 1.602348 | 0.014 | 0.007 | 1 | 1 | Cdc6 | 0.007 | Old |
| 0.083274 | 0.549801 | 0.148 | 0.124 | 1 | 1 | Rcc2 | 0.024 | Old |
| 0.083341 | 0.881395 | 0.031 | 0.019 | 1 | 1 | Tsc2 | 0.012 | Old |
| 0.083684 | 0.475282 | 0.147 | 0.123 | 1 | 1 | Aco2 | 0.024 | Old |

| | | | | | | | | |
|---|---|---|---|---|---|---|---|---|
| 0.083958 | 0.844125 | 0.067 | 0.049 | 1 | 1 | Sptan1 | 0.018 | Old |
| 0.084627 | 0.714645 | 0.014 | 0.007 | 1 | 1 | 2010320M | 0.007 | Old |
| 0.084848 | 0.518423 | 0.091 | 0.071 | 1 | 1 | Ralgds | 0.02 | Old |
| 0.085022 | 0.622805 | 0.014 | 0.007 | 1 | 1 | Gstm4 | 0.007 | Old |
| 0.085348 | 0.570177 | 0.093 | 0.072 | 1 | 1 | Chuk | 0.021 | Old |
| 0.08545 | 0.208589 | 0.621 | 0.66 | 1 | 1 | Neat1 | -0.039 | Old |
| 0.085488 | 1.070092 | 0.042 | 0.029 | 1 | 1 | Neurl4 | 0.013 | Old |
| 0.086274 | 0.903312 | 0.055 | 0.04 | 1 | 1 | Tmem185b | 0.015 | Old |
| 0.087098 | 0.028912 | 0.064 | 0.086 | 1 | 1 | Dhrs1 | -0.022 | Old |
| 0.087342 | 0.009824 | 0.117 | 0.145 | 1 | 1 | Tnrc6a | -0.028 | Old |
| 0.088111 | 1.106091 | 0.042 | 0.029 | 1 | 1 | Iars2 | 0.013 | Old |
| 0.088561 | 0.005551 | 0.017 | 0.029 | 1 | 1 | Arl11 | -0.012 | Old |
| 0.088673 | 0.652759 | 0.095 | 0.075 | 1 | 1 | Vbp1 | 0.02 | Old |
| 0.088824 | 1.002782 | 0.045 | 0.031 | 1 | 1 | Tarbp2 | 0.014 | Old |
| 0.089662 | 0.09593 | 0.326 | 0.378 | 1 | 1 | Mat2a | -0.052 | Old |
| 0.08974 | 0.749068 | 0.066 | 0.049 | 1 | 1 | Mrpl17 | 0.017 | Old |
| 0.090175 | 0.748821 | 0.042 | 0.029 | 1 | 1 | Pam | 0.013 | Old |
| 0.090532 | 1.34307 | 0.013 | 0.006 | 1 | 1 | Gins2 | 0.007 | Old |
| 0.090978 | 0.302825 | 0.521 | 0.534 | 1 | 1 | Sat1 | -0.013 | Old |
| 0.091009 | 0.05551 | 0.09 | 0.115 | 1 | 1 | Ssr4 | -0.025 | Old |
| 0.091126 | 0.51444 | 0.213 | 0.189 | 1 | 1 | Hspa9 | 0.024 | Old |
| 0.091864 | 0.768414 | 0.06 | 0.044 | 1 | 1 | C1galt1c1 | 0.016 | Old |
| 0.092035 | 0.86682 | 0.013 | 0.006 | 1 | 1 | Cela1 | 0.007 | Old |
| 0.093106 | 1.40212 | 0.018 | 0.01 | 1 | 1 | Ppox | 0.008 | Old |
| 0.09371 | 0.290424 | 0.251 | 0.223 | 1 | 1 | Rgl1 | 0.028 | Old |
| 0.094288 | 0.998187 | 0.018 | 0.01 | 1 | 1 | 1810024B0 | 0.008 | Old |
| 0.094618 | 0.66206 | 0.094 | 0.074 | 1 | 1 | Rnf114 | 0.02 | Old |
| 0.094667 | 0.054967 | 0.162 | 0.194 | 1 | 1 | Vps4b | -0.032 | Old |
| 0.094763 | 0.022132 | 0.219 | 0.259 | 1 | 1 | Ptpra | -0.04 | Old |
| 0.094823 | 0.542737 | 0.138 | 0.116 | 1 | 1 | Smarcd2 | 0.022 | Old |
| 0.095316 | 1.54693 | 0.035 | 0.023 | 1 | 1 | Mrpl38 | 0.012 | Old |
| 0.095601 | 0.191139 | 0.026 | 0.04 | 1 | 1 | Appl2 | -0.014 | Old |
| 0.095637 | 0.016268 | 0.495 | 0.564 | 1 | 1 | Hnrnpk | -0.069 | Old |
| 0.095933 | 0.524395 | 0.045 | 0.031 | 1 | 1 | Tspan17 | 0.014 | Old |
| 0.09625 | 0.065969 | 0.184 | 0.222 | 1 | 1 | Ptpn6 | -0.038 | Old |
| 0.097231 | 0.010832 | 0.132 | 0.163 | 1 | 1 | Ccr1 | -0.031 | Old |
| 0.097247 | 0.801995 | 0.039 | 0.026 | 1 | 1 | Tbc1d10a | 0.013 | Old |
| 0.098162 | 0.905353 | 0.026 | 0.016 | 1 | 1 | Adora2b | 0.01 | Old |
| 0.09829 | 0.258928 | 0.018 | 0.01 | 1 | 1 | Qtrt1 | 0.008 | Old |
| 0.098439 | 0.885798 | 0.022 | 0.013 | 1 | 1 | Gpn2 | 0.009 | Old |
| 0.098908 | 0.023113 | 0.203 | 0.242 | 1 | 1 | Cmtm6 | -0.039 | Old |
| 0.098996 | 0.793322 | 0.072 | 0.055 | 1 | 1 | Casd1 | 0.017 | Old |
| 0.099488 | 0.755127 | 0.089 | 0.07 | 1 | 1 | Prmt1 | 0.019 | Old |
| 0.099902 | 0.010164 | 0.149 | 0.182 | 1 | 1 | Uqcr10 | -0.033 | Old |
| 0.100003 | 0.340512 | 0.27 | 0.247 | 1 | 1 | H2afy | 0.023 | Old |

| | | | | | | | | |
|---|---|---|---|---|---|---|---|---|
| 0.100011 | 0.038394 | 0.03 | 0.044 | 1 | 1 | Commd10 | -0.014 | Old |
| 0.100117 | 0.515783 | 0.126 | 0.104 | 1 | 1 | Hsd17b12 | 0.022 | Old |
| 0.100303 | 0.551913 | 0.026 | 0.016 | 1 | 1 | Nup160 | 0.01 | Old |
| 0.100409 | 0.079241 | 0.068 | 0.089 | 1 | 1 | Dguok | -0.021 | Old |
| 0.101401 | 0.486482 | 0.233 | 0.21 | 1 | 1 | Trip12 | 0.023 | Old |
| 0.101452 | 0.492509 | 0.026 | 0.016 | 1 | 1 | Akt3 | 0.01 | Old |
| 0.101539 | 0.012925 | 0.337 | 0.384 | 1 | 1 | Zfand5 | -0.047 | Old |
| 0.102968 | 0.335954 | 0.103 | 0.082 | 1 | 1 | Atp2c1 | 0.021 | Old |
| 0.103128 | 0.948357 | 0.035 | 0.023 | 1 | 1 | Reep4 | 0.012 | Old |
| 0.103206 | 0.378749 | 0.217 | 0.196 | 1 | 1 | Morf4l2 | 0.021 | Old |
| 0.103794 | 0.780497 | 0.032 | 0.021 | 1 | 1 | Pparg | 0.011 | Old |
| 0.1038 | 0.084707 | 0.022 | 0.013 | 1 | 1 | Ptger2 | 0.009 | Old |
| 0.103848 | 0.35687 | 0.144 | 0.121 | 1 | 1 | Lmbrd1 | 0.023 | Old |
| 0.10455 | 1.045934 | 0.017 | 0.009 | 1 | 1 | Zbtb42 | 0.008 | Old |
| 0.104887 | 0.639575 | 0.099 | 0.08 | 1 | 1 | Elmo2 | 0.019 | Old |
| 0.104948 | 0.618831 | 0.148 | 0.126 | 1 | 1 | Polr2k | 0.022 | Old |
| 0.105339 | 0.752541 | 0.035 | 0.023 | 1 | 1 | Akr1e1 | 0.012 | Old |
| 0.10544 | 1.125441 | 0.024 | 0.015 | 1 | 1 | Coprs | 0.009 | Old |
| 0.106336 | 0.018985 | 0.234 | 0.275 | 1 | 1 | Ostf1 | -0.041 | Old |
| 0.106433 | 1.652746 | 0.01 | 0.004 | 1 | 1 | Cdan1 | 0.006 | Old |
| 0.106512 | 0.806898 | 0.055 | 0.041 | 1 | 1 | Hebp1 | 0.014 | Old |
| 0.106537 | 0.102508 | 0.05 | 0.069 | 1 | 1 | Flot2 | -0.019 | Old |
| 0.107001 | 0.355601 | 0.017 | 0.009 | 1 | 1 | Mon1b | 0.008 | Old |
| 0.107228 | 1.154092 | 0.01 | 0.004 | 1 | 1 | Bmpr1a | 0.006 | Old |
| 0.10757 | 0.06865 | 0.976 | 0.995 | 1 | 1 | mt-Atp6 | -0.019 | Old |
| 0.107597 | 0.75106 | 0.028 | 0.018 | 1 | 1 | Espn | 0.01 | Old |
| 0.109253 | 0.878289 | 0.024 | 0.015 | 1 | 1 | Mccc1 | 0.009 | Old |
| 0.110972 | 0.909574 | 0.033 | 0.022 | 1 | 1 | Cdc27 | 0.011 | Old |
| 0.111338 | 0.671879 | 0.073 | 0.057 | 1 | 1 | Bhlhe40 | 0.016 | Old |
| 0.111384 | 0.377292 | 0.067 | 0.05 | 1 | 1 | Snx1 | 0.017 | Old |
| 0.111713 | 0.298383 | 0.465 | 0.459 | 1 | 1 | Nrros | 0.006 | Old |
| 0.111969 | 0.174954 | 0.021 | 0.012 | 1 | 1 | Zcchc10 | 0.009 | Old |
| 0.112618 | 0.011107 | 0.044 | 0.061 | 1 | 1 | Gabpa | -0.017 | Old |
| 0.112792 | 0.770217 | 0.062 | 0.047 | 1 | 1 | Dcn | 0.015 | Old |
| 0.113539 | 0.228852 | 0.054 | 0.039 | 1 | 1 | Mras | 0.015 | Old |
| 0.1137 | 0.649513 | 0.048 | 0.034 | 1 | 1 | Mzt1 | 0.014 | Old |
| 0.114163 | 0.786062 | 0.053 | 0.038 | 1 | 1 | Casp1 | 0.015 | Old |
| 0.115345 | 0.542417 | 0.093 | 0.074 | 1 | 1 | Cyb5b | 0.019 | Old |
| 0.115424 | 0.975855 | 0.015 | 0.008 | 1 | 1 | Asb6 | 0.007 | Old |
| 0.115456 | 1.010096 | 0.049 | 0.035 | 1 | 1 | Ccdc25 | 0.014 | Old |
| 0.115716 | 0.153397 | 0.021 | 0.032 | 1 | 1 | Arid3b | -0.011 | Old |
| 0.116087 | 0.86373 | 0.015 | 0.008 | 1 | 1 | Mars | 0.007 | Old |
| 0.116178 | 0.607061 | 0.033 | 0.022 | 1 | 1 | Usp50 | 0.011 | Old |
| 0.11625 | 0.613129 | 0.069 | 0.053 | 1 | 1 | Tnfaip1 | 0.016 | Old |
| 0.116663 | 1.641653 | 0.019 | 0.011 | 1 | 1 | Ogfod2 | 0.008 | Old |

| | | | | | | | | |
|---|---|---|---|---|---|---|---|---|
| 0.117261 | 0.017433 | 0.14 | 0.171 | 1 | 1 | Pdlim5 | -0.031 | Old |
| 0.117551 | 0.05093 | 0.03 | 0.044 | 1 | 1 | Ap1ar | -0.014 | Old |
| 0.117623 | 1.205972 | 0.023 | 0.014 | 1 | 1 | Sgcb | 0.009 | Old |
| 0.117714 | 0.125411 | 0.085 | 0.108 | 1 | 1 | Dhx9 | -0.023 | Old |
| 0.11792 | 1.301316 | 0.019 | 0.011 | 1 | 1 | Gm47754 | 0.008 | Old |
| 0.119145 | 0.030508 | 0.095 | 0.12 | 1 | 1 | Tle4 | -0.025 | Old |
| 0.119443 | 1.153073 | 0.019 | 0.011 | 1 | 1 | Retsat | 0.008 | Old |
| 0.119649 | 0.787571 | 0.062 | 0.047 | 1 | 1 | Mtmr10 | 0.015 | Old |
| 0.119848 | 0.124818 | 0.021 | 0.032 | 1 | 1 | Trp53inp1 | -0.011 | Old |
| 0.119929 | 0.333817 | 0.384 | 0.363 | 1 | 1 | Cbr2 | 0.021 | Old |
| 0.120391 | 0.139103 | 0.022 | 0.034 | 1 | 1 | Coa6 | -0.012 | Old |
| 0.120755 | 0.480624 | 0.131 | 0.11 | 1 | 1 | Pdxk | 0.021 | Old |
| 0.120757 | 0.084263 | 0.125 | 0.153 | 1 | 1 | Fndc3b | -0.028 | Old |
| 0.121 | 0.112522 | 0.073 | 0.095 | 1 | 1 | Timm10b | -0.022 | Old |
| 0.123024 | 0.510494 | 0.243 | 0.227 | 1 | 1 | AC149090. | 0.016 | Old |
| 0.123107 | 0.465115 | 0.276 | 0.26 | 1 | 1 | Trim25 | 0.016 | Old |
| 0.123231 | 0.501409 | 0.019 | 0.011 | 1 | 1 | Foxred1 | 0.008 | Old |
| 0.123883 | 0.032856 | 0.071 | 0.092 | 1 | 1 | Rogdi | -0.021 | Old |
| 0.124126 | 0.26808 | 0.375 | 0.364 | 1 | 1 | Gnb1 | 0.011 | Old |
| 0.124421 | 0.062334 | 0.104 | 0.13 | 1 | 1 | Ulk2 | -0.026 | Old |
| 0.124435 | 0.45695 | 0.078 | 0.061 | 1 | 1 | Dnajc14 | 0.017 | Old |
| 0.124751 | 1.082209 | 0.026 | 0.016 | 1 | 1 | Mylk | 0.01 | Old |
| 0.125329 | 0.05136 | 0.134 | 0.16 | 1 | 1 | Clint1 | -0.026 | Old |
| 0.125367 | 0.450724 | 0.039 | 0.027 | 1 | 1 | Dubr | 0.012 | Old |
| 0.125475 | 0.032277 | 0.045 | 0.061 | 1 | 1 | Fam208b | -0.016 | Old |
| 0.127315 | 0.496593 | 0.145 | 0.126 | 1 | 1 | Snx8 | 0.019 | Old |
| 0.12758 | 0.934008 | 0.014 | 0.007 | 1 | 1 | Cmpk2 | 0.007 | Old |
| 0.127861 | 0.001401 | 0.341 | 0.389 | 1 | 1 | Ckb | -0.048 | Old |
| 0.128006 | 0.07649 | 0.073 | 0.094 | 1 | 1 | Eif4e3 | -0.021 | Old |
| 0.128007 | 1.279534 | 0.014 | 0.007 | 1 | 1 | Hectd2 | 0.007 | Old |
| 0.128055 | 0.475108 | 0.054 | 0.04 | 1 | 1 | Tmem250- | 0.014 | Old |
| 0.128064 | 0.083081 | 0.086 | 0.109 | 1 | 1 | Tifa | -0.023 | Old |
| 0.128333 | 1.401139 | 0.028 | 0.018 | 1 | 1 | Dcstamp | 0.01 | Old |
| 0.128849 | 0.098556 | 0.027 | 0.04 | 1 | 1 | Upf2 | -0.013 | Old |
| 0.129621 | 0.532758 | 0.014 | 0.007 | 1 | 1 | Ldlrad3 | 0.007 | Old |
| 0.130056 | 0.671535 | 0.026 | 0.016 | 1 | 1 | Timm8a1 | 0.01 | Old |
| 0.130066 | 0.450611 | 0.032 | 0.021 | 1 | 1 | Magohb | 0.011 | Old |
| 0.130312 | 0.767442 | 0.035 | 0.024 | 1 | 1 | Ascc2 | 0.011 | Old |
| 0.130386 | 0.221769 | 0.041 | 0.029 | 1 | 1 | Ltv1 | 0.012 | Old |
| 0.131186 | 0.244452 | 0.285 | 0.266 | 1 | 1 | Clec4a2 | 0.019 | Old |
| 0.131332 | 0.030809 | 0.013 | 0.022 | 1 | 1 | Gga2 | -0.009 | Old |
| 0.13149 | 0.229152 | 0.143 | 0.12 | 1 | 1 | Bub3 | 0.023 | Old |
| 0.131567 | 0.375866 | 0.226 | 0.203 | 1 | 1 | Atp6v1e1 | 0.023 | Old |
| 0.131759 | 0.483066 | 0.026 | 0.016 | 1 | 1 | Pex16 | 0.01 | Old |
| 0.131965 | 0.533929 | 0.147 | 0.127 | 1 | 1 | Snx9 | 0.02 | Old |

| | | | | | | | | |
|---|---|---|---|---|---|---|---|---|
| 0.131987 | 1.161464 | 0.018 | 0.01 | 1 | 1 | Dis3 | 0.008 | Old |
| 0.131995 | 0.78259 | 0.042 | 0.03 | 1 | 1 | Sec24d | 0.012 | Old |
| 0.132416 | 0.604792 | 0.035 | 0.024 | 1 | 1 | Tyw1 | 0.011 | Old |
| 0.132427 | 1.026962 | 0.051 | 0.038 | 1 | 1 | Cd24a | 0.013 | Old |
| 0.132464 | 0.611199 | 0.162 | 0.143 | 1 | 1 | Stub1 | 0.019 | Old |
| 0.13249 | 0.071768 | 0.051 | 0.069 | 1 | 1 | Gtf2i | -0.018 | Old |
| 0.132757 | 0.326533 | 0.059 | 0.044 | 1 | 1 | Dph3 | 0.015 | Old |
| 0.133659 | 0.726694 | 0.042 | 0.03 | 1 | 1 | Hvcn1 | 0.012 | Old |
| 0.13382 | 0.011451 | 0.096 | 0.119 | 1 | 1 | Fbxl3 | -0.023 | Old |
| 0.133899 | 0.916944 | 0.018 | 0.01 | 1 | 1 | Ppm1f | 0.008 | Old |
| 0.134961 | 0.48697 | 0.131 | 0.112 | 1 | 1 | Eif2s3x | 0.019 | Old |
| 0.135191 | 0.611159 | 0.022 | 0.013 | 1 | 1 | Slc35d2 | 0.009 | Old |
| 0.135513 | 0.063996 | 0.167 | 0.198 | 1 | 1 | Zdhhc20 | -0.031 | Old |
| 0.135599 | 0.598457 | 0.05 | 0.037 | 1 | 1 | Llgl1 | 0.013 | Old |
| 0.13561 | 0.295661 | 0.343 | 0.326 | 1 | 1 | Cd44 | 0.017 | Old |
| 0.135742 | 0.585075 | 0.055 | 0.041 | 1 | 1 | Sucla2 | 0.014 | Old |
| 0.135838 | 0.340435 | 0.044 | 0.031 | 1 | 1 | Zbtb21 | 0.013 | Old |
| 0.136028 | 0.812412 | 0.018 | 0.01 | 1 | 1 | Gm26778 | 0.008 | Old |
| 0.136465 | 0.510694 | 0.069 | 0.054 | 1 | 1 | Rangap1 | 0.015 | Old |
| 0.137887 | 0.333769 | 0.018 | 0.01 | 1 | 1 | Mpst | 0.008 | Old |
| 0.138692 | 0.700388 | 0.064 | 0.049 | 1 | 1 | Siah2 | 0.015 | Old |
| 0.140125 | 0.623242 | 0.096 | 0.079 | 1 | 1 | Sik1 | 0.017 | Old |
| 0.141797 | 0.34652 | 0.028 | 0.018 | 1 | 1 | Cd1d1 | 0.01 | Old |
| 0.141971 | 0.144659 | 0.006 | 0.013 | 1 | 1 | Dhx37 | -0.007 | Old |
| 0.14214 | 0.289801 | 0.014 | 0.024 | 1 | 1 | Zfp715 | -0.01 | Old |
| 0.142276 | 0.87524 | 0.013 | 0.007 | 1 | 1 | Acss2 | 0.006 | Old |
| 0.142452 | 0.583478 | 0.102 | 0.084 | 1 | 1 | Trappc2l | 0.018 | Old |
| 0.142521 | 0.736583 | 0.013 | 0.007 | 1 | 1 | Birc5 | 0.006 | Old |
| 0.142643 | 0.798407 | 0.013 | 0.007 | 1 | 1 | Shb | 0.006 | Old |
| 0.143127 | 0.576378 | 0.1 | 0.083 | 1 | 1 | Stt3b | 0.017 | Old |
| 0.143378 | 0.653835 | 0.013 | 0.007 | 1 | 1 | 9430015G1 | 0.006 | Old |
| 0.143559 | 0.785181 | 0.069 | 0.055 | 1 | 1 | Snrpa | 0.014 | Old |
| 0.143581 | 0.021585 | 0.438 | 0.505 | 1 | 1 | Igfbp4 | -0.067 | Old |
| 0.143593 | 0.390177 | 0.05 | 0.037 | 1 | 1 | Fyn | 0.013 | Old |
| 0.143709 | 1.102597 | 0.027 | 0.018 | 1 | 1 | Snupn | 0.009 | Old |
| 0.14387 | 0.252296 | 0.013 | 0.007 | 1 | 1 | Kif3c | 0.006 | Old |
| 0.143874 | 0.315829 | 0.289 | 0.27 | 1 | 1 | Bnip2 | 0.019 | Old |
| 0.144236 | 0.239032 | 0.42 | 0.417 | 1 | 1 | Cfh | 0.003 | Old |
| 0.14449 | 0.605264 | 0.024 | 0.016 | 1 | 1 | Slco3a1 | 0.008 | Old |
| 0.145677 | 0.437299 | 0.135 | 0.115 | 1 | 1 | Emilin2 | 0.02 | Old |
| 0.145931 | 0.056799 | 0.171 | 0.202 | 1 | 1 | Prkcb | -0.031 | Old |
| 0.146261 | 0.182968 | 0.015 | 0.025 | 1 | 1 | Pi4kb | -0.01 | Old |
| 0.146699 | 0.018246 | 0.091 | 0.112 | 1 | 1 | Cbfa2t3 | -0.021 | Old |
| 0.146738 | 1.259466 | 0.017 | 0.01 | 1 | 1 | Tbc1d22b | 0.007 | Old |
| 0.146815 | 0.530083 | 0.166 | 0.148 | 1 | 1 | Sec11c | 0.018 | Old |

| | | | | | | | | |
|---|---|---|---|---|---|---|---|---|
| 0.147116 | 0.484105 | 0.059 | 0.045 | 1 | 1 | Gpr108 | 0.014 | Old |
| 0.14786 | 0.750185 | 0.085 | 0.069 | 1 | 1 | Tiam1 | 0.016 | Old |
| 0.148114 | 0.973704 | 0.021 | 0.013 | 1 | 1 | Inf2 | 0.008 | Old |
| 0.148336 | 0.275097 | 0.077 | 0.061 | 1 | 1 | Actr1a | 0.016 | Old |
| 0.148781 | 0.320444 | 0.294 | 0.277 | 1 | 1 | Tkt | 0.017 | Old |
| 0.148905 | 0.680618 | 0.039 | 0.027 | 1 | 1 | Selenoi | 0.012 | Old |
| 0.149272 | 0.104069 | 0.127 | 0.153 | 1 | 1 | Dram2 | -0.026 | Old |
| 0.150382 | 0.587662 | 0.139 | 0.12 | 1 | 1 | Tmem14c | 0.019 | Old |
| 0.151091 | 0.499055 | 0.036 | 0.025 | 1 | 1 | Nt5c3 | 0.011 | Old |
| 0.151244 | 0.335354 | 0.033 | 0.023 | 1 | 1 | Rtp4 | 0.01 | Old |
| 0.151517 | 0.460299 | 0.066 | 0.051 | 1 | 1 | Sms | 0.015 | Old |
| 0.15179 | 0.359427 | 0.021 | 0.013 | 1 | 1 | Dym | 0.008 | Old |
| 0.152085 | 0.433441 | 0.089 | 0.072 | 1 | 1 | Rab12 | 0.017 | Old |
| 0.152291 | 0.127738 | 0.102 | 0.123 | 1 | 1 | Rex1bd | -0.021 | Old |
| 0.153239 | 0.103634 | 0.09 | 0.111 | 1 | 1 | Snw1 | -0.021 | Old |
| 0.153381 | 0.015129 | 0.181 | 0.213 | 1 | 1 | Snrpb2 | -0.032 | Old |
| 0.15352 | 0.43792 | 0.039 | 0.027 | 1 | 1 | Gbp2 | 0.012 | Old |
| 0.153907 | 0.610447 | 0.055 | 0.042 | 1 | 1 | Slc41a2 | 0.013 | Old |
| 0.155028 | 1.574241 | 0.012 | 0.006 | 1 | 1 | Mdm1 | 0.006 | Old |
| 0.15526 | 0.61545 | 0.089 | 0.072 | 1 | 1 | Scamp1 | 0.017 | Old |
| 0.156006 | 0.135713 | 0.171 | 0.204 | 1 | 1 | Elf1 | -0.033 | Old |
| 0.156071 | 0.056915 | 0.221 | 0.258 | 1 | 1 | Plxnb2 | -0.037 | Old |
| 0.156294 | 0.356758 | 0.185 | 0.165 | 1 | 1 | Rbx1 | 0.02 | Old |
| 0.15641 | 1.10146 | 0.012 | 0.006 | 1 | 1 | Cyb5d2 | 0.006 | Old |
| 0.156688 | 1.095059 | 0.012 | 0.006 | 1 | 1 | Rbfox2 | 0.006 | Old |
| 0.156748 | 0.208679 | 0.45 | 0.441 | 1 | 1 | Gltp | 0.009 | Old |
| 0.156826 | 0.967934 | 0.012 | 0.006 | 1 | 1 | Pi16 | 0.006 | Old |
| 0.157175 | 0.00249 | 0.116 | 0.14 | 1 | 1 | St6gal1 | -0.024 | Old |
| 0.157814 | 0.118093 | 0.076 | 0.095 | 1 | 1 | Ssu72 | -0.019 | Old |
| 0.158297 | 0.707929 | 0.032 | 0.022 | 1 | 1 | Fbxw8 | 0.01 | Old |
| 0.159016 | 0.480113 | 0.067 | 0.052 | 1 | 1 | Rab11fip1 | 0.015 | Old |
| 0.159704 | 0.000782 | 0.04 | 0.054 | 1 | 1 | Ttc9c | -0.014 | Old |
| 0.159773 | 0.698075 | 0.062 | 0.048 | 1 | 1 | Nfx1 | 0.014 | Old |
| 0.159923 | 0.533277 | 0.103 | 0.086 | 1 | 1 | Slc16a10 | 0.017 | Old |
| 0.160195 | 0.009496 | 0.13 | 0.153 | 1 | 1 | Ttc3 | -0.023 | Old |
| 0.160315 | 0.779964 | 0.035 | 0.024 | 1 | 1 | Ttc13 | 0.011 | Old |
| 0.160507 | 0.962025 | 0.042 | 0.031 | 1 | 1 | Tmem2 | 0.011 | Old |
| 0.16119 | 0.763195 | 0.035 | 0.024 | 1 | 1 | Galk1 | 0.011 | Old |
| 0.161556 | 0.490469 | 0.023 | 0.015 | 1 | 1 | Map4k5 | 0.008 | Old |
| 0.162047 | 0.378376 | 0.145 | 0.126 | 1 | 1 | Anxa4 | 0.019 | Old |
| 0.162223 | 0.633526 | 0.078 | 0.064 | 1 | 1 | Trim30d | 0.014 | Old |
| 0.162314 | 0.059208 | 0.005 | 0.011 | 1 | 1 | Bphl | -0.006 | Old |
| 0.162363 | 0.104978 | 0.138 | 0.164 | 1 | 1 | Atp5c1 | -0.026 | Old |
| 0.162577 | 0.199442 | 0.022 | 0.032 | 1 | 1 | Pkib | -0.01 | Old |
| 0.163622 | 0.223351 | 0.036 | 0.049 | 1 | 1 | Nt5c2 | -0.013 | Old |

| | | | | | | | | |
|---|---|---|---|---|---|---|---|---|
| 0.163792 | 0.720969 | 0.04 | 0.029 | 1 | 1 | Tmem60 | 0.011 | Old |
| 0.164311 | 1.125961 | 0.015 | 0.009 | 1 | 1 | Unc5a | 0.006 | Old |
| 0.164334 | 0.701288 | 0.019 | 0.012 | 1 | 1 | Acp5 | 0.007 | Old |
| 0.164458 | 0.021416 | 0.064 | 0.082 | 1 | 1 | Ndufa10 | -0.018 | Old |
| 0.164643 | 0.142776 | 0.022 | 0.032 | 1 | 1 | Ppp2r3c | -0.01 | Old |
| 0.164671 | 0.493553 | 0.032 | 0.022 | 1 | 1 | Casp9 | 0.01 | Old |
| 0.164731 | 0.05885 | 0.093 | 0.114 | 1 | 1 | Tmem259 | -0.021 | Old |
| 0.165406 | 0.561049 | 0.019 | 0.012 | 1 | 1 | Loxl3 | 0.007 | Old |
| 0.166016 | 1.006031 | 0.015 | 0.009 | 1 | 1 | Pank4 | 0.006 | Old |
| 0.166598 | 0.511503 | 0.075 | 0.06 | 1 | 1 | Dscr3 | 0.015 | Old |
| 0.166878 | 0.00603 | 0.136 | 0.161 | 1 | 1 | Ndufa13 | -0.025 | Old |
| 0.167294 | 0.300882 | 0.177 | 0.156 | 1 | 1 | Eif1a | 0.021 | Old |
| 0.167338 | 1.050123 | 0.055 | 0.043 | 1 | 1 | Yipf6 | 0.012 | Old |
| 0.167746 | 0.563663 | 0.026 | 0.017 | 1 | 1 | Rbm45 | 0.009 | Old |
| 0.167892 | 0.616492 | 0.019 | 0.012 | 1 | 1 | Gss | 0.007 | Old |
| 0.168179 | 0.085468 | 0.107 | 0.131 | 1 | 1 | Npepps | -0.024 | Old |
| 0.168238 | 0.101403 | 0.096 | 0.118 | 1 | 1 | Dock11 | -0.022 | Old |
| 0.168394 | 0.255442 | 0.009 | 0.016 | 1 | 1 | Def6 | -0.007 | Old |
| 0.168672 | 0.495255 | 0.026 | 0.017 | 1 | 1 | Cdt1 | 0.009 | Old |
| 0.168732 | 0.24972 | 0.009 | 0.016 | 1 | 1 | Cyb561d2 | -0.007 | Old |
| 0.168872 | 0.313131 | 0.019 | 0.012 | 1 | 1 | Wrb | 0.007 | Old |
| 0.169008 | 0.011851 | 0.064 | 0.082 | 1 | 1 | Chmp1a | -0.018 | Old |
| 0.169706 | 0.618612 | 0.051 | 0.039 | 1 | 1 | Secisbp2 | 0.012 | Old |
| 0.169843 | 0.064206 | 0.023 | 0.034 | 1 | 1 | Exoc6 | -0.011 | Old |
| 0.169963 | 0.865285 | 0.046 | 0.035 | 1 | 1 | Ruvbl1 | 0.011 | Old |
| 0.170202 | 0.258806 | 0.009 | 0.016 | 1 | 1 | Unk | -0.007 | Old |
| 0.170237 | 0.331079 | 0.171 | 0.15 | 1 | 1 | Tspan31 | 0.021 | Old |
| 0.170764 | 1.827875 | 0.01 | 0.005 | 1 | 1 | Camk2a | 0.005 | Old |
| 0.171189 | 0.246816 | 0.362 | 0.351 | 1 | 1 | Rab8b | 0.011 | Old |
| 0.171197 | 0.316086 | 0.048 | 0.035 | 1 | 1 | Bad | 0.013 | Old |
| 0.172079 | 0.175415 | 0.015 | 0.024 | 1 | 1 | Mrpl1 | -0.009 | Old |
| 0.172086 | 0.54341 | 0.057 | 0.044 | 1 | 1 | Elovl1 | 0.013 | Old |
| 0.172158 | 0.148283 | 0.645 | 0.67 | 1 | 1 | Arpc2 | -0.025 | Old |
| 0.172504 | 1.311522 | 0.01 | 0.005 | 1 | 1 | Gm14286 | 0.005 | Old |
| 0.172663 | 1.263774 | 0.01 | 0.005 | 1 | 1 | Cep78 | 0.005 | Old |
| 0.172761 | 0.03733 | 0.059 | 0.075 | 1 | 1 | Phf10 | -0.016 | Old |
| 0.172822 | 1.077931 | 0.01 | 0.005 | 1 | 1 | Atp5s | 0.005 | Old |
| 0.172836 | 0.196051 | 0.095 | 0.117 | 1 | 1 | Il6st | -0.022 | Old |
| 0.173619 | 1.137918 | 0.01 | 0.005 | 1 | 1 | Mtg2 | 0.005 | Old |
| 0.173938 | 0.770981 | 0.01 | 0.005 | 1 | 1 | Ifi206 | 0.005 | Old |
| 0.174098 | 0.906702 | 0.01 | 0.005 | 1 | 1 | Tcaim | 0.005 | Old |
| 0.174223 | 0.086989 | 0.099 | 0.121 | 1 | 1 | Fchsd2 | -0.022 | Old |
| 0.175334 | 0.02117 | 0.06 | 0.077 | 1 | 1 | Vps36 | -0.017 | Old |
| 0.175703 | 0.358475 | 0.01 | 0.005 | 1 | 1 | Rad54l | 0.005 | Old |
| 0.17572 | 0.067231 | 0.028 | 0.04 | 1 | 1 | Dcxr | -0.012 | Old |

| | | | | | | | | |
|---|---|---|---|---|---|---|---|---|
| 0.176045 | 0.007908 | 0.121 | 0.145 | 1 | 1 | Creb1 | -0.024 | Old |
| 0.176878 | 0.576679 | 0.028 | 0.019 | 1 | 1 | Gm26782 | 0.009 | Old |
| 0.176937 | 0.817721 | 0.044 | 0.032 | 1 | 1 | Hikeshi | 0.012 | Old |
| 0.176967 | 0.027741 | 0.095 | 0.116 | 1 | 1 | Ep400 | -0.021 | Old |
| 0.177261 | 0.605874 | 0.06 | 0.047 | 1 | 1 | Nlk | 0.013 | Old |
| 0.17881 | 0.097807 | 0.01 | 0.018 | 1 | 1 | Dut | -0.008 | Old |
| 0.179255 | 0.544598 | 0.163 | 0.148 | 1 | 1 | Supt5 | 0.015 | Old |
| 0.179594 | 0.025088 | 0.708 | 0.71 | 1 | 1 | Ctss | -0.002 | Old |
| 0.179726 | 0.747215 | 0.058 | 0.045 | 1 | 1 | Usp6nl | 0.013 | Old |
| 0.179754 | 0.629993 | 0.077 | 0.063 | 1 | 1 | Crybg1 | 0.014 | Old |
| 0.180862 | 1.613278 | 0.014 | 0.008 | 1 | 1 | Wfdc18 | 0.006 | Old |
| 0.180968 | 0.611324 | 0.089 | 0.074 | 1 | 1 | Cyb5r3 | 0.015 | Old |
| 0.18112 | 0.944924 | 0.018 | 0.011 | 1 | 1 | Ccar2 | 0.007 | Old |
| 0.181444 | 0.041104 | 0.064 | 0.081 | 1 | 1 | Akirin1 | -0.017 | Old |
| 0.181715 | 1.023746 | 0.018 | 0.011 | 1 | 1 | Tradd | 0.007 | Old |
| 0.181827 | 0.545674 | 0.033 | 0.024 | 1 | 1 | Aig1 | 0.009 | Old |
| 0.182145 | 0.272504 | 0.084 | 0.068 | 1 | 1 | Gfra2 | 0.016 | Old |
| 0.182457 | 0.575916 | 0.039 | 0.028 | 1 | 1 | Selenom | 0.011 | Old |
| 0.182898 | 0.220394 | 0.51 | 0.514 | 1 | 1 | Samhd1 | -0.004 | Old |
| 0.182977 | 0.052747 | 0.055 | 0.071 | 1 | 1 | Aimp1 | -0.016 | Old |
| 0.183303 | 0.855132 | 0.024 | 0.016 | 1 | 1 | Elac2 | 0.008 | Old |
| 0.18346 | 0.116863 | 0.042 | 0.056 | 1 | 1 | Cbx5 | -0.014 | Old |
| 0.183562 | 0.62198 | 0.039 | 0.028 | 1 | 1 | Zfp787 | 0.011 | Old |
| 0.183628 | 0.833563 | 0.018 | 0.011 | 1 | 1 | Lrrc28 | 0.007 | Old |
| 0.183669 | 0.249053 | 0.474 | 0.479 | 1 | 1 | Srsf5 | -0.005 | Old |
| 0.183868 | 0.783757 | 0.018 | 0.011 | 1 | 1 | Phf7 | 0.007 | Old |
| 0.184169 | 0.513707 | 0.018 | 0.011 | 1 | 1 | Rtl8b | 0.007 | Old |
| 0.184326 | 0.405417 | 0.044 | 0.032 | 1 | 1 | Minpp1 | 0.012 | Old |
| 0.184484 | 0.224585 | 0.391 | 0.388 | 1 | 1 | Pcbp1 | 0.003 | Old |
| 0.184597 | 0.414934 | 0.141 | 0.123 | 1 | 1 | Snrpd2 | 0.018 | Old |
| 0.18465 | 0.016923 | 0.067 | 0.083 | 1 | 1 | Inppl1 | -0.016 | Old |
| 0.185067 | 0.356861 | 0.244 | 0.227 | 1 | 1 | Olfm1 | 0.017 | Old |
| 0.185265 | 0.008759 | 0.046 | 0.061 | 1 | 1 | Zfyve16 | -0.015 | Old |
| 0.185284 | 0.2715 | 0.068 | 0.054 | 1 | 1 | Adgrg6 | 0.014 | Old |
| 0.185394 | 0.887675 | 0.014 | 0.008 | 1 | 1 | Ssh3 | 0.006 | Old |
| 0.185898 | 0.021122 | 0.035 | 0.047 | 1 | 1 | Zfp518a | -0.012 | Old |
| 0.185982 | 0.090642 | 0.147 | 0.172 | 1 | 1 | Ndufc1 | -0.025 | Old |
| 0.186561 | 0.51064 | 0.024 | 0.016 | 1 | 1 | Scarf1 | 0.008 | Old |
| 0.186803 | 0.05178 | 0.04 | 0.053 | 1 | 1 | Zbtb37 | -0.013 | Old |
| 0.187408 | 0.474697 | 0.118 | 0.102 | 1 | 1 | Apaf1 | 0.016 | Old |
| 0.187503 | 0.523202 | 0.104 | 0.088 | 1 | 1 | Chchd1 | 0.016 | Old |
| 0.187686 | 0.160712 | 0.049 | 0.064 | 1 | 1 | Slc31a1 | -0.015 | Old |
| 0.188682 | 0.185739 | 0.04 | 0.053 | 1 | 1 | Ssbp2 | -0.013 | Old |
| 0.188743 | 0.298039 | 0.014 | 0.008 | 1 | 1 | Tab1 | 0.006 | Old |
| 0.1897 | 0.517803 | 0.068 | 0.055 | 1 | 1 | Gse1 | 0.013 | Old |

| | | | | | | | | |
|---|---|---|---|---|---|---|---|---|
| 0.189959 | 0.476632 | 0.082 | 0.067 | 1 | 1 | Fgr | 0.015 | Old |
| 0.191033 | 0.170968 | 0.526 | 0.523 | 1 | 1 | Ap2a2 | 0.003 | Old |
| 0.191484 | 0.316767 | 0.13 | 0.112 | 1 | 1 | Ankrd13a | 0.018 | Old |
| 0.191635 | 0.011436 | 0.023 | 0.033 | 1 | 1 | Ksr1 | -0.01 | Old |
| 0.192047 | 0.17783 | 0.499 | 0.501 | 1 | 1 | Lamp2 | -0.002 | Old |
| 0.192221 | 0.403507 | 0.12 | 0.103 | 1 | 1 | Psma5 | 0.017 | Old |
| 0.193437 | 0.240198 | 0.105 | 0.088 | 1 | 1 | Fes | 0.017 | Old |
| 0.193732 | 0.457597 | 0.067 | 0.053 | 1 | 1 | Rbm12 | 0.014 | Old |
| 0.194443 | 0.532527 | 0.111 | 0.095 | 1 | 1 | Alyref | 0.016 | Old |
| 0.194443 | 0.469572 | 0.12 | 0.103 | 1 | 1 | 9030624J0 | 0.017 | Old |
| 0.194462 | 0.489525 | 0.021 | 0.013 | 1 | 1 | Dnm1 | 0.008 | Old |
| 0.194641 | 0.482401 | 0.077 | 0.063 | 1 | 1 | Aco1 | 0.014 | Old |
| 0.194945 | 0.35123 | 0.085 | 0.069 | 1 | 1 | Pqbp1 | 0.016 | Old |
| 0.195086 | 0.601722 | 0.051 | 0.04 | 1 | 1 | Creld2 | 0.011 | Old |
| 0.195312 | 0.438423 | 0.113 | 0.097 | 1 | 1 | Rasa4 | 0.016 | Old |
| 0.195606 | 0.424823 | 0.202 | 0.185 | 1 | 1 | Cct8 | 0.017 | Old |
| 0.195762 | 0.566376 | 0.035 | 0.025 | 1 | 1 | Mpv17l2 | 0.01 | Old |
| 0.19601 | 0.099778 | 0.028 | 0.039 | 1 | 1 | Pgs1 | -0.011 | Old |
| 0.196342 | 0.154948 | 0.598 | 0.623 | 1 | 1 | Rap1b | -0.025 | Old |
| 0.19641 | 0.286292 | 0.085 | 0.069 | 1 | 1 | Abcc5 | 0.016 | Old |
| 0.196647 | 0.556542 | 0.093 | 0.078 | 1 | 1 | Dync1li1 | 0.015 | Old |
| 0.196766 | 0.03127 | 0.076 | 0.061 | 1 | 1 | Cd300lb | 0.015 | Old |
| 0.197097 | 0.509275 | 0.093 | 0.078 | 1 | 1 | Ccdc71l | 0.015 | Old |
| 0.197218 | 0.019937 | 0.026 | 0.036 | 1 | 1 | Cenpc1 | -0.01 | Old |
| 0.197762 | 0.646694 | 0.127 | 0.112 | 1 | 1 | Pfdn2 | 0.015 | Old |
| 0.198361 | 0.530918 | 0.045 | 0.034 | 1 | 1 | Abhd2 | 0.011 | Old |
| 0.198429 | 0.491461 | 0.021 | 0.013 | 1 | 1 | Uxt | 0.008 | Old |
| 0.199666 | 0.217529 | 0.067 | 0.053 | 1 | 1 | Ahctf1 | 0.014 | Old |
| 0.199761 | 0.031887 | 0.242 | 0.277 | 1 | 1 | Laptm4a | -0.035 | Old |
| 0.200198 | 0.475667 | 0.021 | 0.013 | 1 | 1 | Zscan21 | 0.008 | Old |
| 0.20079 | 0.320469 | 0.021 | 0.013 | 1 | 1 | Gna11 | 0.008 | Old |
| 0.200824 | 0.472068 | 0.082 | 0.068 | 1 | 1 | Ecm1 | 0.014 | Old |
| 0.201152 | 1.140607 | 0.026 | 0.018 | 1 | 1 | Rbm38 | 0.008 | Old |
| 0.201596 | 0.075703 | 0.09 | 0.109 | 1 | 1 | Ubr2 | -0.019 | Old |
| 0.20192 | 0.001915 | 0.032 | 0.044 | 1 | 1 | Pex13 | -0.012 | Old |
| 0.202797 | 1.230082 | 0.017 | 0.01 | 1 | 1 | 1810011H1 | 0.007 | Old |
| 0.203013 | 0.50018 | 0.09 | 0.075 | 1 | 1 | Plaa | 0.015 | Old |
| 0.203522 | 0.464851 | 0.261 | 0.248 | 1 | 1 | Stxbp3 | 0.013 | Old |
| 0.203567 | 0.780516 | 0.023 | 0.016 | 1 | 1 | Recql | 0.007 | Old |
| 0.203567 | 0.650185 | 0.023 | 0.016 | 1 | 1 | Usp53 | 0.007 | Old |
| 0.203584 | 0.589787 | 0.06 | 0.048 | 1 | 1 | H2-T22 | 0.012 | Old |
| 0.20371 | 0.152796 | 0.067 | 0.083 | 1 | 1 | Nemf | -0.016 | Old |
| 0.203993 | 0.002135 | 0.102 | 0.122 | 1 | 1 | Fkbp8 | -0.02 | Old |
| 0.204091 | 0.039273 | 0.048 | 0.061 | 1 | 1 | Hmgcr | -0.013 | Old |
| 0.204202 | 0.210497 | 0.411 | 0.402 | 1 | 1 | S100a6 | 0.009 | Old |

| | | | | | | | | |
|---|---|---|---|---|---|---|---|---|
| 0.204677 | 0.014755 | 0.125 | 0.147 | 1 | 1 | Nus1 | -0.022 | Old |
| 0.204737 | 1.2086 | 0.013 | 0.007 | 1 | 1 | Ppif | 0.006 | Old |
| 0.204893 | 1.318329 | 0.013 | 0.007 | 1 | 1 | Gcat | 0.006 | Old |
| 0.205106 | 0.034277 | 0.058 | 0.073 | 1 | 1 | Vrk1 | -0.015 | Old |
| 0.205174 | 0.043754 | 0.039 | 0.051 | 1 | 1 | Brwd3 | -0.012 | Old |
| 0.2055 | 0.483613 | 0.08 | 0.066 | 1 | 1 | Nhsl2 | 0.014 | Old |
| 0.206147 | 1.032562 | 0.013 | 0.007 | 1 | 1 | Neil1 | 0.006 | Old |
| 0.206228 | 0.518713 | 0.118 | 0.103 | 1 | 1 | Smc6 | 0.015 | Old |
| 0.206671 | 0.106909 | 0.067 | 0.083 | 1 | 1 | Chfr | -0.016 | Old |
| 0.206776 | 1.022857 | 0.013 | 0.007 | 1 | 1 | Ube2c | 0.006 | Old |
| 0.206971 | 0.085087 | 0.281 | 0.319 | 1 | 1 | Atf4 | -0.038 | Old |
| 0.207091 | 0.867159 | 0.013 | 0.007 | 1 | 1 | Dpt | 0.006 | Old |
| 0.207406 | 0.58579 | 0.013 | 0.007 | 1 | 1 | Gstp1 | 0.006 | Old |
| 0.207652 | 0.297364 | 0.009 | 0.016 | 1 | 1 | Tbcc | -0.007 | Old |
| 0.208039 | 0.024447 | 0.198 | 0.226 | 1 | 1 | Tram1 | -0.028 | Old |
| 0.208104 | 0.363926 | 0.152 | 0.134 | 1 | 1 | Cnn2 | 0.018 | Old |
| 0.208512 | 0.553145 | 0.013 | 0.007 | 1 | 1 | Fam216a | 0.006 | Old |
| 0.208687 | 0.072794 | 0.215 | 0.25 | 1 | 1 | Fkbp1a | -0.035 | Old |
| 0.209119 | 0.148972 | 0.03 | 0.041 | 1 | 1 | Smim3 | -0.011 | Old |
| 0.209557 | 0.932395 | 0.036 | 0.027 | 1 | 1 | Rfx1 | 0.009 | Old |
| 0.209729 | 0.406045 | 0.109 | 0.093 | 1 | 1 | Mdm2 | 0.016 | Old |
| 0.209813 | 0.002408 | 0.212 | 0.239 | 1 | 1 | Nipbl | -0.027 | Old |
| 0.210316 | 0.017278 | 0.143 | 0.165 | 1 | 1 | Nadk | -0.022 | Old |
| 0.210412 | 0.865963 | 0.028 | 0.02 | 1 | 1 | Acvr2a | 0.008 | Old |
| 0.210473 | 0.130245 | 0.009 | 0.016 | 1 | 1 | Aasdh | -0.007 | Old |
| 0.210511 | 0.773443 | 0.026 | 0.018 | 1 | 1 | Zfp367 | 0.008 | Old |
| 0.210536 | 0.625291 | 0.039 | 0.029 | 1 | 1 | Hax1 | 0.01 | Old |
| 0.210736 | 0.526036 | 0.049 | 0.038 | 1 | 1 | Abhd16a | 0.011 | Old |
| 0.210743 | 0.591069 | 0.009 | 0.016 | 1 | 1 | 2700062C0 | -0.007 | Old |
| 0.210796 | 0.151578 | 0.62 | 0.643 | 1 | 1 | Dazap2 | -0.023 | Old |
| 0.210813 | 0.015013 | 0.072 | 0.089 | 1 | 1 | Ndufa8 | -0.017 | Old |
| 0.210819 | 0.887766 | 0.033 | 0.024 | 1 | 1 | Tprkb | 0.009 | Old |
| 0.210996 | 0.088336 | 0.005 | 0.01 | 1 | 1 | Il15ra | -0.005 | Old |
| 0.211472 | 0.367922 | 0.317 | 0.308 | 1 | 1 | Ifi27l2a | 0.009 | Old |
| 0.2116 | 0.300619 | 0.017 | 0.025 | 1 | 1 | Slc35a1 | -0.008 | Old |
| 0.212121 | 0.53368 | 0.044 | 0.033 | 1 | 1 | Commd1 | 0.011 | Old |
| 0.213415 | 0.166014 | 0.051 | 0.066 | 1 | 1 | Zbtb38 | -0.015 | Old |
| 0.213955 | 0.689853 | 0.067 | 0.055 | 1 | 1 | Traf3 | 0.012 | Old |
| 0.214277 | 0.983857 | 0.019 | 0.013 | 1 | 1 | Tmem175 | 0.006 | Old |
| 0.214683 | 0.461538 | 0.041 | 0.031 | 1 | 1 | Irf2bp1 | 0.01 | Old |
| 0.214769 | 0.025852 | 0.017 | 0.025 | 1 | 1 | Mtmr4 | -0.008 | Old |
| 0.215499 | 0.583738 | 0.033 | 0.024 | 1 | 1 | Eri3 | 0.009 | Old |
| 0.21607 | 0.387693 | 0.064 | 0.052 | 1 | 1 | Phf2 | 0.012 | Old |
| 0.216133 | 0.653091 | 0.031 | 0.022 | 1 | 1 | Leo1 | 0.009 | Old |
| 0.216586 | 0.019385 | 0.066 | 0.081 | 1 | 1 | Lpin1 | -0.015 | Old |

| | | | | | | | | |
|---|---|---|---|---|---|---|---|---|
| 0.216591 | 0.939687 | 0.019 | 0.013 | 1 | 1 | Wdr47 | 0.006 | Old |
| 0.217517 | 0.144128 | 0.064 | 0.08 | 1 | 1 | Mif4gd | -0.016 | Old |
| 0.217712 | 0.222285 | 0.238 | 0.218 | 1 | 1 | Pla2g7 | 0.02 | Old |
| 0.217863 | 0.025714 | 0.188 | 0.216 | 1 | 1 | Rab11a | -0.028 | Old |
| 0.217884 | 0.780954 | 0.019 | 0.013 | 1 | 1 | Wdr46 | 0.006 | Old |
| 0.218503 | 0.416965 | 0.098 | 0.083 | 1 | 1 | Yipf5 | 0.015 | Old |
| 0.218792 | 0.182517 | 0.06 | 0.075 | 1 | 1 | Uqcrc1 | -0.015 | Old |
| 0.21883 | 0.597116 | 0.053 | 0.041 | 1 | 1 | St8sia6 | 0.012 | Old |
| 0.219002 | 0.51061 | 0.098 | 0.083 | 1 | 1 | Tmem9b | 0.015 | Old |
| 0.219505 | 0.103409 | 0.278 | 0.318 | 1 | 1 | Pak2 | -0.04 | Old |
| 0.219756 | 0.012457 | 0.386 | 0.438 | 1 | 1 | Ccnl1 | -0.052 | Old |
| 0.219971 | 0.01907 | 0.284 | 0.322 | 1 | 1 | Fxyd5 | -0.038 | Old |
| 0.220049 | 0.079044 | 0.125 | 0.147 | 1 | 1 | Mib1 | -0.022 | Old |
| 0.220055 | 0.383616 | 0.147 | 0.129 | 1 | 1 | Smim7 | 0.018 | Old |
| 0.220524 | 0.349688 | 0.293 | 0.283 | 1 | 1 | Ndufa4 | 0.01 | Old |
| 0.220835 | 0.28863 | 0.415 | 0.418 | 1 | 1 | Cd47 | -0.003 | Old |
| 0.221629 | 0.04852 | 0.05 | 0.064 | 1 | 1 | Cenpa | -0.014 | Old |
| 0.221719 | 0.04554 | 0.076 | 0.093 | 1 | 1 | Zhx1 | -0.017 | Old |
| 0.221728 | 0.046936 | 0.215 | 0.244 | 1 | 1 | Eif4b | -0.029 | Old |
| 0.221855 | 0.021186 | 0.064 | 0.079 | 1 | 1 | Gm49336 | -0.015 | Old |
| 0.221985 | 0.006127 | 0.099 | 0.118 | 1 | 1 | Psma1 | -0.019 | Old |
| 0.222944 | 0.436706 | 0.055 | 0.044 | 1 | 1 | Cyc1 | 0.011 | Old |
| 0.223432 | 0.224501 | 0.036 | 0.027 | 1 | 1 | Dnajc9 | 0.009 | Old |
| 0.224111 | 0.470675 | 0.059 | 0.047 | 1 | 1 | Vcl | 0.012 | Old |
| 0.224717 | 0.998863 | 0.024 | 0.017 | 1 | 1 | Pdp1 | 0.007 | Old |
| 0.225044 | 1.420567 | 0.015 | 0.01 | 1 | 1 | Tenm4 | 0.005 | Old |
| 0.225989 | 0.003857 | 0.125 | 0.145 | 1 | 1 | 1110008F1 | -0.02 | Old |
| 0.226621 | 1.09358 | 0.015 | 0.01 | 1 | 1 | Heatr5b | 0.005 | Old |
| 0.226995 | 0.591365 | 0.054 | 0.043 | 1 | 1 | Hk1 | 0.011 | Old |
| 0.226998 | 0.745463 | 0.015 | 0.01 | 1 | 1 | Macrod1 | 0.005 | Old |
| 0.22705 | 0.060662 | 0.006 | 0.012 | 1 | 1 | Jaml | -0.006 | Old |
| 0.227299 | 0.85016 | 0.015 | 0.01 | 1 | 1 | Gmppb | 0.005 | Old |
| 0.22761 | 0.202963 | 0.059 | 0.074 | 1 | 1 | Arhgap11a | -0.015 | Old |
| 0.228084 | 0.020682 | 0.134 | 0.155 | 1 | 1 | Ago2 | -0.021 | Old |
| 0.228102 | 0.059987 | 0.053 | 0.066 | 1 | 1 | Atp11c | -0.013 | Old |
| 0.228224 | 1.293854 | 0.012 | 0.007 | 1 | 1 | Coq3 | 0.005 | Old |
| 0.228439 | 0.080651 | 0.085 | 0.103 | 1 | 1 | Rbm33 | -0.018 | Old |
| 0.228859 | 0.597791 | 0.022 | 0.015 | 1 | 1 | Kctd11 | 0.007 | Old |
| 0.228984 | 0.515912 | 0.022 | 0.015 | 1 | 1 | Fastk | 0.007 | Old |
| 0.2294 | 0.129946 | 0.14 | 0.163 | 1 | 1 | Rtraf | -0.023 | Old |
| 0.22969 | 0.058802 | 0.055 | 0.069 | 1 | 1 | Ctnnbip1 | -0.014 | Old |
| 0.229989 | 0.126483 | 0.154 | 0.179 | 1 | 1 | Usp7 | -0.025 | Old |
| 0.23006 | 0.00789 | 0.195 | 0.219 | 1 | 1 | Lims1 | -0.024 | Old |
| 0.230088 | 0.599941 | 0.058 | 0.047 | 1 | 1 | Arhgap10 | 0.011 | Old |
| 0.230633 | 0.388151 | 0.085 | 0.071 | 1 | 1 | Ppil2 | 0.014 | Old |

| | | | | | | | | |
|---|---|---|---|---|---|---|---|---|
| 0.230761 | 0.528761 | 0.082 | 0.069 | 1 | 1 | Sf3a1 | 0.013 | Old |
| 0.230895 | 0.253749 | 0.398 | 0.387 | 1 | 1 | Ssr3 | 0.011 | Old |
| 0.231085 | 1.142536 | 0.012 | 0.007 | 1 | 1 | Tmem115 | 0.005 | Old |
| 0.231138 | 0.160093 | 0.125 | 0.146 | 1 | 1 | Med13l | -0.021 | Old |
| 0.231245 | 0.770056 | 0.032 | 0.024 | 1 | 1 | Cops7b | 0.008 | Old |
| 0.232191 | 0.03514 | 0.093 | 0.11 | 1 | 1 | Zc3h13 | -0.017 | Old |
| 0.232241 | 0.112187 | 0.037 | 0.049 | 1 | 1 | Ubxn7 | -0.012 | Old |
| 0.232525 | 0.907707 | 0.012 | 0.007 | 1 | 1 | Adat2 | 0.005 | Old |
| 0.232525 | 0.324275 | 0.012 | 0.007 | 1 | 1 | Asf1b | 0.005 | Old |
| 0.232774 | 0.272171 | 0.715 | 0.758 | 1 | 1 | Retnla | -0.043 | Old |
| 0.233067 | 0.704056 | 0.012 | 0.007 | 1 | 1 | Pcgf1 | 0.005 | Old |
| 0.233487 | 0.673978 | 0.027 | 0.019 | 1 | 1 | Fbxl6 | 0.008 | Old |
| 0.233826 | 0.621414 | 0.027 | 0.019 | 1 | 1 | Dhcr24 | 0.008 | Old |
| 0.234319 | 0.18234 | 0.039 | 0.05 | 1 | 1 | Mis18a | -0.011 | Old |
| 0.234786 | 0.751436 | 0.027 | 0.019 | 1 | 1 | Rpl27-ps3 | 0.008 | Old |
| 0.234882 | 0.039336 | 0.136 | 0.157 | 1 | 1 | Arih1 | -0.021 | Old |
| 0.235024 | 0.80394 | 0.035 | 0.026 | 1 | 1 | Rgs3 | 0.009 | Old |
| 0.23506 | 0.219364 | 0.012 | 0.007 | 1 | 1 | Slc37a1 | 0.005 | Old |
| 0.235066 | 0.073074 | 0.058 | 0.072 | 1 | 1 | Oxr1 | -0.014 | Old |
| 0.236539 | 0.758105 | 0.05 | 0.04 | 1 | 1 | Timm50 | 0.01 | Old |
| 0.236881 | 0.023355 | 0.338 | 0.38 | 1 | 1 | Bri3 | -0.042 | Old |
| 0.236911 | 0.571915 | 0.046 | 0.036 | 1 | 1 | Cpsf3 | 0.01 | Old |
| 0.237953 | 0.705825 | 0.03 | 0.021 | 1 | 1 | Tmtc3 | 0.009 | Old |
| 0.238083 | 0.333987 | 0.066 | 0.053 | 1 | 1 | Ctso | 0.013 | Old |
| 0.238239 | 0.291002 | 0.134 | 0.117 | 1 | 1 | Antxr2 | 0.017 | Old |
| 0.238754 | 0.444828 | 0.075 | 0.062 | 1 | 1 | Capn1 | 0.013 | Old |
| 0.238937 | 0.095634 | 0.069 | 0.085 | 1 | 1 | Paip1 | -0.016 | Old |
| 0.239719 | 0.186676 | 0.014 | 0.021 | 1 | 1 | Man2c1 | -0.007 | Old |
| 0.240009 | 0.467284 | 0.06 | 0.049 | 1 | 1 | Cyp20a1 | 0.011 | Old |
| 0.240249 | 0.048098 | 0.08 | 0.096 | 1 | 1 | Zfc3h1 | -0.016 | Old |
| 0.240748 | 0.345541 | 0.333 | 0.325 | 1 | 1 | Pmp22 | 0.008 | Old |
| 0.241093 | 0.024875 | 0.222 | 0.252 | 1 | 1 | Arf4 | -0.03 | Old |
| 0.241384 | 0.632744 | 0.041 | 0.032 | 1 | 1 | Mdc1 | 0.009 | Old |
| 0.241418 | 0.379031 | 0.035 | 0.026 | 1 | 1 | Fam160b2 | 0.009 | Old |
| 0.241696 | 0.254651 | 0.393 | 0.388 | 1 | 1 | Calr | 0.005 | Old |
| 0.241964 | 0.200226 | 0.014 | 0.021 | 1 | 1 | Cd274 | -0.007 | Old |
| 0.241964 | 0.006851 | 0.014 | 0.021 | 1 | 1 | Orai2 | -0.007 | Old |
| 0.241993 | 0.643921 | 0.055 | 0.044 | 1 | 1 | Faap20 | 0.011 | Old |
| 0.242079 | 0.001838 | 0.203 | 0.229 | 1 | 1 | Tmed5 | -0.026 | Old |
| 0.242732 | 0.243535 | 0.035 | 0.026 | 1 | 1 | Sdhaf1 | 0.009 | Old |
| 0.242841 | 0.144858 | 0.036 | 0.047 | 1 | 1 | Upf3b | -0.011 | Old |
| 0.243029 | 0.355719 | 0.202 | 0.188 | 1 | 1 | Surf4 | 0.014 | Old |
| 0.243965 | 0.115216 | 0.098 | 0.116 | 1 | 1 | Ndufb4 | -0.018 | Old |
| 0.2446 | 0.078678 | 0.008 | 0.013 | 1 | 1 | Tcea3 | -0.005 | Old |
| 0.24462 | 0.244143 | 0.111 | 0.095 | 1 | 1 | Max | 0.016 | Old |

| | | | | | | | | |
|---|---|---|---|---|---|---|---|---|
| 0.244691 | 0.687269 | 0.05 | 0.04 | 1 | 1 | Bloc1s2 | 0.01 | Old |
| 0.244897 | 1.086587 | 0.041 | 0.032 | 1 | 1 | Stx11 | 0.009 | Old |
| 0.244938 | 0.314179 | 0.04 | 0.03 | 1 | 1 | Gpd2 | 0.01 | Old |
| 0.245055 | 0.24353 | 0.111 | 0.095 | 1 | 1 | Cul1 | 0.016 | Old |
| 0.24585 | 0.222225 | 0.181 | 0.164 | 1 | 1 | Ids | 0.017 | Old |
| 0.245958 | 0.575434 | 0.062 | 0.05 | 1 | 1 | Iah1 | 0.012 | Old |
| 0.247864 | 0.05819 | 0.125 | 0.146 | 1 | 1 | Huwe1 | -0.021 | Old |
| 0.248868 | 0.556747 | 0.084 | 0.071 | 1 | 1 | Klhl28 | 0.013 | Old |
| 0.249451 | 0.01456 | 0.093 | 0.109 | 1 | 1 | Immt | -0.016 | Old |
| 0.249907 | 0.037059 | 0.077 | 0.092 | 1 | 1 | Polr2e | -0.015 | Old |
| 0.250159 | 0.151216 | 0.65 | 0.685 | 1 | 1 | Irf2bp2 | -0.035 | Old |
| 0.250268 | 0.021116 | 0.039 | 0.049 | 1 | 1 | Fblim1 | -0.01 | Old |
| 0.251705 | 0.387259 | 0.033 | 0.044 | 1 | 1 | Aga | -0.011 | Old |
| 0.251803 | 0.775598 | 0.021 | 0.014 | 1 | 1 | Cks1b | 0.007 | Old |
| 0.252137 | 0.337631 | 0.229 | 0.218 | 1 | 1 | Gm26532 | 0.011 | Old |
| 0.252202 | 0.884705 | 0.014 | 0.009 | 1 | 1 | Twnk | 0.005 | Old |
| 0.252849 | 0.305588 | 0.12 | 0.105 | 1 | 1 | Snx27 | 0.015 | Old |
| 0.253661 | 0.623453 | 0.048 | 0.038 | 1 | 1 | Gpr137b | 0.01 | Old |
| 0.253836 | 0.042565 | 0.046 | 0.058 | 1 | 1 | Fzr1 | -0.012 | Old |
| 0.25415 | 0.236212 | 0.019 | 0.027 | 1 | 1 | Ccdc122 | -0.008 | Old |
| 0.255039 | 0.118894 | 0.028 | 0.038 | 1 | 1 | Med23 | -0.01 | Old |
| 0.255168 | 0.291732 | 0.049 | 0.038 | 1 | 1 | Hypk | 0.011 | Old |
| 0.255223 | 0.325584 | 0.207 | 0.191 | 1 | 1 | Jarid2 | 0.016 | Old |
| 0.255523 | 0.05328 | 0.009 | 0.015 | 1 | 1 | Coq10a | -0.006 | Old |
| 0.255709 | 0.151979 | 0.054 | 0.067 | 1 | 1 | Akap9 | -0.013 | Old |
| 0.256093 | 0.035173 | 0.158 | 0.182 | 1 | 1 | Adipor1 | -0.024 | Old |
| 0.256519 | 0.676211 | 0.093 | 0.081 | 1 | 1 | Ubr4 | 0.012 | Old |
| 0.256611 | 0.024194 | 0.176 | 0.202 | 1 | 1 | Ptpro | -0.026 | Old |
| 0.256627 | 0.847833 | 0.014 | 0.009 | 1 | 1 | Gm41307 | 0.005 | Old |
| 0.256783 | 0.16092 | 0.021 | 0.014 | 1 | 1 | Otud7b | 0.007 | Old |
| 0.256894 | 0.551539 | 0.048 | 0.038 | 1 | 1 | Hmg20b | 0.01 | Old |
| 0.257201 | 0.312819 | 0.021 | 0.014 | 1 | 1 | Bysl | 0.007 | Old |
| 0.257225 | 1.149828 | 0.01 | 0.006 | 1 | 1 | Ftsj1 | 0.004 | Old |
| 0.257701 | 0.00554 | 0.046 | 0.058 | 1 | 1 | Nxpe5 | -0.012 | Old |
| 0.257897 | 0.046265 | 0.009 | 0.015 | 1 | 1 | Col1a2 | -0.006 | Old |
| 0.258588 | 0.590313 | 0.048 | 0.038 | 1 | 1 | Scn1b | 0.01 | Old |
| 0.258812 | 0.612678 | 0.023 | 0.016 | 1 | 1 | Dhx35 | 0.007 | Old |
| 0.259077 | 0.633503 | 0.01 | 0.006 | 1 | 1 | Ccdc91 | 0.004 | Old |
| 0.259205 | 0.153408 | 0.014 | 0.009 | 1 | 1 | Ccdc77 | 0.005 | Old |
| 0.26011 | 0.890384 | 0.01 | 0.006 | 1 | 1 | Zfp12 | 0.004 | Old |
| 0.260141 | 0.324868 | 0.104 | 0.09 | 1 | 1 | Ap1s1 | 0.014 | Old |
| 0.260481 | 0.066803 | 0.33 | 0.371 | 1 | 1 | Ifi207 | -0.041 | Old |
| 0.260724 | 0.041463 | 0.023 | 0.032 | 1 | 1 | Hectd3 | -0.009 | Old |
| 0.260939 | 0.30643 | 0.01 | 0.006 | 1 | 1 | Gm16973 | 0.004 | Old |
| 0.261015 | 0.60586 | 0.026 | 0.018 | 1 | 1 | Arl2 | 0.008 | Old |

| | | | | | | | | |
|---|---|---|---|---|---|---|---|---|
| 0.261065 | 0.024164 | 0.454 | 0.504 | 1 | 1 | Gabarap | -0.05 | Old |
| 0.261146 | 0.456324 | 0.01 | 0.006 | 1 | 1 | Mfsd3 | 0.004 | Old |
| 0.261354 | 0.495215 | 0.01 | 0.006 | 1 | 1 | Aatk | 0.004 | Old |
| 0.261354 | 0.290973 | 0.01 | 0.006 | 1 | 1 | Rsl1 | 0.004 | Old |
| 0.261402 | 0.069794 | 0.039 | 0.049 | 1 | 1 | Camsap2 | -0.01 | Old |
| 0.261888 | 0.627025 | 0.026 | 0.018 | 1 | 1 | Ofd1 | 0.008 | Old |
| 0.262385 | 0.243287 | 0.019 | 0.027 | 1 | 1 | Zfp865 | -0.008 | Old |
| 0.26334 | 0.504331 | 0.036 | 0.027 | 1 | 1 | Hsdl1 | 0.009 | Old |
| 0.263467 | 0.154808 | 0.03 | 0.039 | 1 | 1 | Ccdc97 | -0.009 | Old |
| 0.263651 | 0.316457 | 0.189 | 0.174 | 1 | 1 | Tuba1c | 0.015 | Old |
| 0.264672 | 0.45189 | 0.112 | 0.098 | 1 | 1 | Sppl3 | 0.014 | Old |
| 0.264888 | 0.179754 | 0.01 | 0.016 | 1 | 1 | Foxp4 | -0.006 | Old |
| 0.265215 | 0.166971 | 0.036 | 0.047 | 1 | 1 | Rnf170 | -0.011 | Old |
| 0.265492 | 0.122585 | 0.032 | 0.042 | 1 | 1 | Mfap1a | -0.01 | Old |
| 0.265777 | 0.368864 | 0.026 | 0.018 | 1 | 1 | Dvl3 | 0.008 | Old |
| 0.265873 | 0.181514 | 0.491 | 0.498 | 1 | 1 | Ptafr | -0.007 | Old |
| 0.266335 | 0.464172 | 0.123 | 0.109 | 1 | 1 | Stat1 | 0.014 | Old |
| 0.266575 | 0.135085 | 0.352 | 0.394 | 1 | 1 | Ywhab | -0.042 | Old |
| 0.266658 | 0.054713 | 0.005 | 0.01 | 1 | 1 | Cep89 | -0.005 | Old |
| 0.266664 | 0.003957 | 0.167 | 0.187 | 1 | 1 | Ppp2r5a | -0.02 | Old |
| 0.267016 | 0.167088 | 0.388 | 0.388 | 1 | 1 | Ptms | 0 | Old |
| 0.26709 | 0.010653 | 0.353 | 0.39 | 1 | 1 | Snx5 | -0.037 | Old |
| 0.267124 | 0.057822 | 0.175 | 0.199 | 1 | 1 | Psmb1 | -0.024 | Old |
| 0.267179 | 1.250224 | 0.017 | 0.011 | 1 | 1 | Mvd | 0.006 | Old |
| 0.267228 | 0.597526 | 0.166 | 0.154 | 1 | 1 | Sdhc | 0.012 | Old |
| 0.267277 | 0.492194 | 0.064 | 0.053 | 1 | 1 | Hbs1l | 0.011 | Old |
| 0.26783 | 0.406389 | 0.069 | 0.058 | 1 | 1 | Parl | 0.011 | Old |
| 0.267904 | 0.270758 | 0.085 | 0.101 | 1 | 1 | Esd | -0.016 | Old |
| 0.267985 | 0.461067 | 0.031 | 0.023 | 1 | 1 | Akt1s1 | 0.008 | Old |
| 0.268058 | 1.040605 | 0.017 | 0.011 | 1 | 1 | Gm26669 | 0.006 | Old |
| 0.268263 | 1.136681 | 0.032 | 0.024 | 1 | 1 | Arhgap23 | 0.008 | Old |
| 0.268501 | 0.002141 | 0.005 | 0.01 | 1 | 1 | Mrm1 | -0.005 | Old |
| 0.268549 | 0.177369 | 0.035 | 0.045 | 1 | 1 | Auh | -0.01 | Old |
| 0.268862 | 0.539773 | 0.06 | 0.049 | 1 | 1 | Tmem43 | 0.011 | Old |
| 0.268948 | 0.562904 | 0.049 | 0.039 | 1 | 1 | Dcaf15 | 0.01 | Old |
| 0.269139 | 0.051833 | 0.339 | 0.38 | 1 | 1 | Rnf130 | -0.041 | Old |
| 0.269502 | 0.594764 | 0.017 | 0.011 | 1 | 1 | Atp6v0d2 | 0.006 | Old |
| 0.270284 | 0.072727 | 0.24 | 0.273 | 1 | 1 | Sppl2a | -0.033 | Old |
| 0.270614 | 0.039138 | 0.049 | 0.061 | 1 | 1 | Eif2ak2 | -0.012 | Old |
| 0.27071 | 0.712153 | 0.017 | 0.011 | 1 | 1 | D17H6S53I | 0.006 | Old |
| 0.270816 | 0.695395 | 0.059 | 0.049 | 1 | 1 | Nhp2 | 0.01 | Old |
| 0.270937 | 0.334324 | 0.057 | 0.046 | 1 | 1 | Rbbp8 | 0.011 | Old |
| 0.270937 | 0.15731 | 0.156 | 0.178 | 1 | 1 | Srsf9 | -0.022 | Old |
| 0.270982 | 0.197137 | 0.068 | 0.056 | 1 | 1 | Ubfd1 | 0.012 | Old |
| 0.272022 | 0.103675 | 0.068 | 0.083 | 1 | 1 | Acss1 | -0.015 | Old |

| | | | | | | | | |
|---|---|---|---|---|---|---|---|---|
| 0.272224 | 1.449131 | 0.019 | 0.013 | 1 | 1 | Rnf113a2 | 0.006 | Old |
| 0.272341 | 0.004673 | 0.105 | 0.122 | 1 | 1 | Ccnl2 | -0.017 | Old |
| 0.272973 | 0.369985 | 0.017 | 0.011 | 1 | 1 | Nipal3 | 0.006 | Old |
| 0.273519 | 0.096979 | 0.094 | 0.11 | 1 | 1 | Smim11 | -0.016 | Old |
| 0.273614 | 0.06411 | 0.123 | 0.143 | 1 | 1 | Tbcb | -0.02 | Old |
| 0.273625 | 0.935359 | 0.035 | 0.027 | 1 | 1 | Heatr1 | 0.008 | Old |
| 0.273801 | 0.033332 | 0.135 | 0.155 | 1 | 1 | Tceal9 | -0.02 | Old |
| 0.2744 | 0.153396 | 0.172 | 0.198 | 1 | 1 | Rpl27 | -0.026 | Old |
| 0.274594 | 0.324556 | 0.103 | 0.089 | 1 | 1 | Nup88 | 0.014 | Old |
| 0.274645 | 0.682846 | 0.077 | 0.066 | 1 | 1 | Raf1 | 0.011 | Old |
| 0.274769 | 0.053892 | 0.012 | 0.018 | 1 | 1 | Slc35d1 | -0.006 | Old |
| 0.275106 | 0.075945 | 0.131 | 0.151 | 1 | 1 | Acap2 | -0.02 | Old |
| 0.275813 | 0.009574 | 0.162 | 0.184 | 1 | 1 | Drap1 | -0.022 | Old |
| 0.276803 | 0.599599 | 0.035 | 0.027 | 1 | 1 | Slc35b3 | 0.008 | Old |
| 0.276929 | 0.787164 | 0.03 | 0.022 | 1 | 1 | Cars2 | 0.008 | Old |
| 0.277903 | 0.279973 | 0.252 | 0.242 | 1 | 1 | Hnrnpc | 0.01 | Old |
| 0.278248 | 0.798115 | 0.03 | 0.022 | 1 | 1 | Actr8 | 0.008 | Old |
| 0.278997 | 0.322521 | 0.012 | 0.018 | 1 | 1 | Nos1ap | -0.006 | Old |
| 0.279092 | 0.799515 | 0.044 | 0.035 | 1 | 1 | Samd1 | 0.009 | Old |
| 0.279421 | 0.826772 | 0.022 | 0.016 | 1 | 1 | 1110034G2 | 0.006 | Old |
| 0.279867 | 0.573092 | 0.032 | 0.024 | 1 | 1 | Camta1 | 0.008 | Old |
| 0.280387 | 0.065772 | 0.111 | 0.129 | 1 | 1 | Usp34 | -0.018 | Old |
| 0.280597 | 0.999388 | 0.027 | 0.02 | 1 | 1 | Zbtb6 | 0.007 | Old |
| 0.280891 | 0.023947 | 0.046 | 0.058 | 1 | 1 | 4833439L1 | -0.012 | Old |
| 0.281312 | 0.39482 | 0.198 | 0.187 | 1 | 1 | Emc10 | 0.011 | Old |
| 0.281502 | 0.019721 | 0.188 | 0.211 | 1 | 1 | Ap2m1 | -0.023 | Old |
| 0.282301 | 0.983242 | 0.024 | 0.018 | 1 | 1 | Pde12 | 0.006 | Old |
| 0.282319 | 1.494231 | 0.013 | 0.008 | 1 | 1 | Sec22c | 0.005 | Old |
| 0.282376 | 0.731818 | 0.027 | 0.02 | 1 | 1 | Ctu2 | 0.007 | Old |
| 0.282646 | 0.424107 | 0.032 | 0.024 | 1 | 1 | Adamtsl5 | 0.008 | Old |
| 0.283529 | 0.417734 | 0.086 | 0.074 | 1 | 1 | Tmem141 | 0.012 | Old |
| 0.283684 | 0.204345 | 0.045 | 0.056 | 1 | 1 | Wdr61 | -0.011 | Old |
| 0.283729 | 0.27308 | 0.019 | 0.013 | 1 | 1 | Synj2 | 0.006 | Old |
| 0.283791 | 0.014013 | 0.147 | 0.167 | 1 | 1 | Ypel3 | -0.02 | Old |
| 0.2838 | 0.009066 | 0.379 | 0.429 | 1 | 1 | Ctsa | -0.05 | Old |
| 0.284007 | 0.311106 | 0.156 | 0.142 | 1 | 1 | Tnpo3 | 0.014 | Old |
| 0.284176 | 0.267934 | 0.352 | 0.346 | 1 | 1 | Tapbp | 0.006 | Old |
| 0.284646 | 0.35861 | 0.075 | 0.063 | 1 | 1 | Impad1 | 0.012 | Old |
| 0.285005 | 0.876518 | 0.013 | 0.008 | 1 | 1 | Crocc | 0.005 | Old |
| 0.285632 | 0.007386 | 0.251 | 0.278 | 1 | 1 | Asah1 | -0.027 | Old |
| 0.285956 | 0.664848 | 0.027 | 0.02 | 1 | 1 | Tjp2 | 0.007 | Old |
| 0.286281 | 0.73267 | 0.022 | 0.016 | 1 | 1 | Ano10 | 0.006 | Old |
| 0.286355 | 0.8335 | 0.013 | 0.008 | 1 | 1 | Kdelc1 | 0.005 | Old |
| 0.286355 | 0.771383 | 0.013 | 0.008 | 1 | 1 | Zfp141 | 0.005 | Old |
| 0.286356 | 0.238861 | 0.037 | 0.029 | 1 | 1 | Akr7a5 | 0.008 | Old |

| | | | | | | | | |
|---|---|---|---|---|---|---|---|---|
| 0.286418 | 0.369321 | 0.145 | 0.131 | 1 | 1 | Orai1 | 0.014 | Old |
| 0.286741 | 0.406168 | 0.013 | 0.008 | 1 | 1 | Gm42715 | 0.005 | Old |
| 0.286799 | 0.01291 | 0.586 | 0.666 | 1 | 1 | Ddx5 | -0.08 | Old |
| 0.286844 | 0.600429 | 0.067 | 0.056 | 1 | 1 | Ibtk | 0.011 | Old |
| 0.286935 | 0.004861 | 0.143 | 0.164 | 1 | 1 | Tex261 | -0.021 | Old |
| 0.287307 | 0.086128 | 0.036 | 0.046 | 1 | 1 | Plcl2 | -0.01 | Old |
| 0.287355 | 0.055965 | 0.036 | 0.046 | 1 | 1 | Slc25a25 | -0.01 | Old |
| 0.287412 | 0.240382 | 0.154 | 0.139 | 1 | 1 | Mlf2 | 0.015 | Old |
| 0.28754 | 0.621269 | 0.03 | 0.022 | 1 | 1 | Rnf40 | 0.008 | Old |
| 0.288452 | 0.075216 | 0.015 | 0.022 | 1 | 1 | Noc3l | -0.007 | Old |
| 0.288508 | 0.492726 | 0.044 | 0.035 | 1 | 1 | Trmt6 | 0.009 | Old |
| 0.288571 | 0.226554 | 0.085 | 0.101 | 1 | 1 | Fam3c | -0.016 | Old |
| 0.288812 | 0.420298 | 0.049 | 0.039 | 1 | 1 | Zbtb7b | 0.01 | Old |
| 0.289038 | 0.049301 | 0.247 | 0.276 | 1 | 1 | Lsp1 | -0.029 | Old |
| 0.289048 | 0.438364 | 0.121 | 0.108 | 1 | 1 | Tap2 | 0.013 | Old |
| 0.289196 | 0.297494 | 0.014 | 0.021 | 1 | 1 | Abhd10 | -0.007 | Old |
| 0.289261 | 0.252719 | 0.013 | 0.008 | 1 | 1 | Plac8 | 0.005 | Old |
| 0.289514 | 0.067432 | 0.037 | 0.029 | 1 | 1 | Atg7 | 0.008 | Old |
| 0.290161 | 0.092485 | 0.105 | 0.123 | 1 | 1 | Srp72 | -0.018 | Old |
| 0.290398 | 0.040235 | 0.234 | 0.261 | 1 | 1 | Sf3b2 | -0.027 | Old |
| 0.290451 | 0.366506 | 0.143 | 0.129 | 1 | 1 | Parvb | 0.014 | Old |
| 0.291014 | 0.164596 | 0.118 | 0.137 | 1 | 1 | Aprt | -0.019 | Old |
| 0.291025 | 0.009773 | 0.201 | 0.223 | 1 | 1 | Wipf1 | -0.022 | Old |
| 0.29108 | 0.045022 | 0.015 | 0.022 | 1 | 1 | Plpp6 | -0.007 | Old |
| 0.291353 | 0.051508 | 0.089 | 0.104 | 1 | 1 | Pbx2 | -0.015 | Old |
| 0.291405 | 0.116687 | 0.013 | 0.008 | 1 | 1 | Flad1 | 0.005 | Old |
| 0.292516 | 0.005608 | 0.114 | 0.132 | 1 | 1 | Golph3 | -0.018 | Old |
| 0.2936 | 0.091154 | 0.373 | 0.422 | 1 | 1 | Atp5e | -0.049 | Old |
| 0.293848 | 0.010594 | 0.067 | 0.08 | 1 | 1 | Senp2 | -0.013 | Old |
| 0.293871 | 0.085606 | 0.269 | 0.3 | 1 | 1 | Atox1 | -0.031 | Old |
| 0.294058 | 0.229369 | 0.018 | 0.025 | 1 | 1 | Ripk2 | -0.007 | Old |
| 0.294067 | 0.129837 | 0.032 | 0.024 | 1 | 1 | Pde4a | 0.008 | Old |
| 0.294602 | 0.034607 | 0.55 | 0.606 | 1 | 1 | Hnrnpa2b1 | -0.056 | Old |
| 0.29493 | 0.026725 | 0.019 | 0.027 | 1 | 1 | Setd1a | -0.008 | Old |
| 0.295453 | 0.209634 | 0.145 | 0.167 | 1 | 1 | Usmg5 | -0.022 | Old |
| 0.295497 | 0.214221 | 0.486 | 0.497 | 1 | 1 | Cd53 | -0.011 | Old |
| 0.295534 | 0.823317 | 0.053 | 0.044 | 1 | 1 | Sos1 | 0.009 | Old |
| 0.295811 | 0.001649 | 0.066 | 0.078 | 1 | 1 | Cdk13 | -0.012 | Old |
| 0.295829 | 0.329152 | 0.027 | 0.02 | 1 | 1 | AC127341. | 0.007 | Old |
| 0.296279 | 0.688734 | 0.048 | 0.038 | 1 | 1 | Mta3 | 0.01 | Old |
| 0.296593 | 0.05616 | 0.18 | 0.205 | 1 | 1 | Lrp6 | -0.025 | Old |
| 0.296716 | 0.332409 | 0.019 | 0.027 | 1 | 1 | Cep85 | -0.008 | Old |
| 0.296735 | 0.414441 | 0.095 | 0.083 | 1 | 1 | Stk40 | 0.012 | Old |
| 0.29694 | 0.268863 | 0.042 | 0.053 | 1 | 1 | Fam204a | -0.011 | Old |
| 0.297115 | 0.023937 | 0.1 | 0.116 | 1 | 1 | Mospd2 | -0.016 | Old |

| | | | | | | | | |
|---|---|---|---|---|---|---|---|---|
| 0.297586 | 0.050149 | 0.467 | 0.528 | 1 | 1 | Tmem50a | -0.061 | Old |
| 0.297793 | 1.242284 | 0.015 | 0.01 | 1 | 1 | Ccdc69 | 0.005 | Old |
| 0.297867 | 0.273666 | 0.019 | 0.027 | 1 | 1 | Naglu | -0.008 | Old |
| 0.298064 | 0.578254 | 0.054 | 0.044 | 1 | 1 | Gaa | 0.01 | Old |
| 0.298081 | 0.007897 | 0.045 | 0.055 | 1 | 1 | Adrm1 | -0.01 | Old |
| 0.298228 | 0.393298 | 0.105 | 0.092 | 1 | 1 | Txnrd1 | 0.013 | Old |
| 0.298733 | 0.091079 | 0.039 | 0.049 | 1 | 1 | Tmem63a | -0.01 | Old |
| 0.298943 | 0.060284 | 0.09 | 0.106 | 1 | 1 | Jup | -0.016 | Old |
| 0.298969 | 0.075547 | 0.108 | 0.126 | 1 | 1 | Retreg2 | -0.018 | Old |
| 0.299402 | 0.833678 | 0.015 | 0.01 | 1 | 1 | Dynlt1f | 0.005 | Old |
| 0.299793 | 0.033733 | 0.563 | 0.64 | 1 | 1 | Iqgap1 | -0.077 | Old |
| 0.299929 | 0.132082 | 0.1 | 0.086 | 1 | 1 | Gpat4 | 0.014 | Old |
| 0.30006 | 0.265137 | 0.165 | 0.189 | 1 | 1 | Pgk1 | -0.024 | Old |
| 0.300073 | 0.302432 | 0.125 | 0.111 | 1 | 1 | Plec | 0.014 | Old |
| 0.300134 | 0.436892 | 0.099 | 0.086 | 1 | 1 | Glrx3 | 0.013 | Old |
| 0.300369 | 0.165996 | 0.027 | 0.035 | 1 | 1 | Usf3 | -0.008 | Old |
| 0.300376 | 0.123694 | 0.841 | 0.891 | 1 | 1 | Ftl1 | -0.05 | Old |
| 0.300407 | 0.009778 | 0.008 | 0.013 | 1 | 1 | Il18r1 | -0.005 | Old |
| 0.300656 | 0.975547 | 0.015 | 0.01 | 1 | 1 | Trim39 | 0.005 | Old |
| 0.300688 | 0.120993 | 0.021 | 0.028 | 1 | 1 | Ercc6l2 | -0.007 | Old |
| 0.300781 | 0.939704 | 0.04 | 0.032 | 1 | 1 | Snap29 | 0.008 | Old |
| 0.301456 | 0.045777 | 0.096 | 0.112 | 1 | 1 | Psmc6 | -0.016 | Old |
| 0.301541 | 0.280954 | 0.059 | 0.049 | 1 | 1 | Rras2 | 0.01 | Old |
| 0.301568 | 0.705558 | 0.033 | 0.026 | 1 | 1 | Shkbp1 | 0.007 | Old |
| 0.301915 | 0.507002 | 0.015 | 0.01 | 1 | 1 | Tnni3 | 0.005 | Old |
| 0.301935 | 0.093348 | 0.008 | 0.013 | 1 | 1 | Prkca | -0.005 | Old |
| 0.30224 | 0.17291 | 0.031 | 0.04 | 1 | 1 | Cops7a | -0.009 | Old |
| 0.302291 | 0.025042 | 0.087 | 0.102 | 1 | 1 | Rtf1 | -0.015 | Old |
| 0.302537 | 0.435399 | 0.104 | 0.092 | 1 | 1 | Psmc1 | 0.012 | Old |
| 0.302572 | 0.368382 | 0.15 | 0.136 | 1 | 1 | Ldlr | 0.014 | Old |
| 0.302611 | 0.304064 | 0.222 | 0.21 | 1 | 1 | Capns1 | 0.012 | Old |
| 0.302777 | 0.068323 | 0.087 | 0.103 | 1 | 1 | Aldh9a1 | -0.016 | Old |
| 0.302942 | 0.260876 | 0.057 | 0.047 | 1 | 1 | Drg1 | 0.01 | Old |
| 0.303795 | 0.028034 | 0.026 | 0.034 | 1 | 1 | Rngtt | -0.008 | Old |
| 0.303804 | 1.22108 | 0.018 | 0.013 | 1 | 1 | Atat1 | 0.005 | Old |
| 0.303816 | 0.316896 | 0.05 | 0.041 | 1 | 1 | Dph5 | 0.009 | Old |
| 0.30509 | 0.035783 | 0.094 | 0.109 | 1 | 1 | Ube2f | -0.015 | Old |
| 0.305632 | 0.094647 | 0.192 | 0.217 | 1 | 1 | Gapvd1 | -0.025 | Old |
| 0.306074 | 0.481902 | 0.015 | 0.01 | 1 | 1 | Cav1 | 0.005 | Old |
| 0.306198 | 0.005559 | 0.086 | 0.1 | 1 | 1 | Ppp6r1 | -0.014 | Old |
| 0.306464 | 0.287158 | 0.156 | 0.14 | 1 | 1 | Tubb2a | 0.016 | Old |
| 0.306797 | 0.822536 | 0.028 | 0.021 | 1 | 1 | Trim23 | 0.007 | Old |
| 0.307963 | 0.013286 | 0.324 | 0.357 | 1 | 1 | Msn | -0.033 | Old |
| 0.308224 | 0.057632 | 0.163 | 0.185 | 1 | 1 | Ski | -0.022 | Old |
| 0.308274 | 0.416225 | 0.075 | 0.064 | 1 | 1 | Ipo5 | 0.011 | Old |

| | | | | | | | | |
|---|---|---|---|---|---|---|---|---|
| 0.308569 | 0.894806 | 0.021 | 0.015 | 1 | 1 | Tbcd | 0.006 | Old |
| 0.308969 | 0.619083 | 0.033 | 0.026 | 1 | 1 | Rasgrp2 | 0.007 | Old |
| 0.309474 | 0.985334 | 0.018 | 0.013 | 1 | 1 | Gm37233 | 0.005 | Old |
| 0.309696 | 0.791583 | 0.026 | 0.019 | 1 | 1 | Prmt5 | 0.007 | Old |
| 0.310299 | 0.511896 | 0.066 | 0.055 | 1 | 1 | Cmas | 0.011 | Old |
| 0.310624 | 0.399123 | 0.108 | 0.095 | 1 | 1 | Gas2l3 | 0.013 | Old |
| 0.311323 | 0.581724 | 0.018 | 0.013 | 1 | 1 | Prkd2 | 0.005 | Old |
| 0.311464 | 0.12088 | 0.009 | 0.014 | 1 | 1 | Rbm41 | -0.005 | Old |
| 0.311663 | 0.103352 | 0.095 | 0.111 | 1 | 1 | Hcfc1r1 | -0.016 | Old |
| 0.312053 | 0.496558 | 0.028 | 0.021 | 1 | 1 | Ndc1 | 0.007 | Old |
| 0.312087 | 0.801807 | 0.023 | 0.017 | 1 | 1 | Plekhn1 | 0.006 | Old |
| 0.312166 | 0.52945 | 0.018 | 0.013 | 1 | 1 | Dus1l | 0.005 | Old |
| 0.312222 | 0.072773 | 0.084 | 0.097 | 1 | 1 | Ddx21 | -0.013 | Old |
| 0.312251 | 0.695482 | 0.018 | 0.013 | 1 | 1 | Entpd5 | 0.005 | Old |
| 0.312256 | 0.44411 | 0.099 | 0.087 | 1 | 1 | Zmiz2 | 0.012 | Old |
| 0.312335 | 0.48486 | 0.018 | 0.013 | 1 | 1 | Zmym6 | 0.005 | Old |
| 0.312887 | 0.935255 | 0.026 | 0.019 | 1 | 1 | Srbd1 | 0.007 | Old |
| 0.313675 | 0.476158 | 0.009 | 0.014 | 1 | 1 | Bpgm | -0.005 | Old |
| 0.313675 | 0.069351 | 0.009 | 0.014 | 1 | 1 | Fars2 | -0.005 | Old |
| 0.314262 | 0.44017 | 0.117 | 0.105 | 1 | 1 | Nenf | 0.012 | Old |
| 0.314275 | 0.176547 | 0.059 | 0.049 | 1 | 1 | Ccm2 | 0.01 | Old |
| 0.314445 | 0.574044 | 0.023 | 0.017 | 1 | 1 | Polr2d | 0.006 | Old |
| 0.314621 | 0.074693 | 0.263 | 0.297 | 1 | 1 | Ube2d2a | -0.034 | Old |
| 0.31472 | 0.913533 | 0.037 | 0.03 | 1 | 1 | Scaper | 0.007 | Old |
| 0.315567 | 0.218656 | 0.105 | 0.092 | 1 | 1 | Zc3h12a | 0.013 | Old |
| 0.316304 | 0.396973 | 0.135 | 0.123 | 1 | 1 | 8-Sep | 0.012 | Old |
| 0.316707 | 0.029553 | 0.132 | 0.151 | 1 | 1 | Timm13 | -0.019 | Old |
| 0.316984 | 0.14751 | 0.149 | 0.171 | 1 | 1 | Papola | -0.022 | Old |
| 0.317291 | 0.150694 | 0.031 | 0.024 | 1 | 1 | Farsa | 0.007 | Old |
| 0.317597 | 0.724198 | 0.037 | 0.03 | 1 | 1 | Ube2l6 | 0.007 | Old |
| 0.317679 | 0.417593 | 0.08 | 0.069 | 1 | 1 | Slc48a1 | 0.011 | Old |
| 0.317782 | 0.634583 | 0.023 | 0.017 | 1 | 1 | Top2a | 0.006 | Old |
| 0.31784 | 0.03643 | 0.086 | 0.1 | 1 | 1 | Gtpbp2 | -0.014 | Old |
| 0.318208 | 0.388012 | 0.026 | 0.019 | 1 | 1 | Fmn1 | 0.007 | Old |
| 0.318239 | 0.29842 | 0.089 | 0.077 | 1 | 1 | Tmem208 | 0.012 | Old |
| 0.318305 | 0.565823 | 0.028 | 0.021 | 1 | 1 | Nqo2 | 0.007 | Old |
| 0.31836 | 0.237113 | 0.163 | 0.148 | 1 | 1 | Hpcal1 | 0.015 | Old |
| 0.318792 | 0.480855 | 0.021 | 0.015 | 1 | 1 | Rem1 | 0.006 | Old |
| 0.318897 | 0.557232 | 0.067 | 0.057 | 1 | 1 | Snapc1 | 0.01 | Old |
| 0.318949 | 0.000143 | 0.418 | 0.458 | 1 | 1 | Ctsz | -0.04 | Old |
| 0.319587 | 0.473736 | 0.021 | 0.015 | 1 | 1 | Ap2a1 | 0.006 | Old |
| 0.319778 | 0.308703 | 0.18 | 0.165 | 1 | 1 | Ppp1r12c | 0.015 | Old |
| 0.319913 | 0.011248 | 0.071 | 0.083 | 1 | 1 | Strap | -0.012 | Old |
| 0.320353 | 1.131033 | 0.012 | 0.007 | 1 | 1 | Jade3 | 0.005 | Old |
| 0.320803 | 0.075197 | 0.057 | 0.068 | 1 | 1 | Mrpl36 | -0.011 | Old |

| | | | | | | | | |
|---|---|---|---|---|---|---|---|---|
| 0.321022 | 0.344614 | 0.021 | 0.015 | 1 | 1 | Etfdh | 0.006 | Old |
| 0.321142 | 0.383892 | 0.023 | 0.017 | 1 | 1 | Dalrd3 | 0.006 | Old |
| 0.322324 | 0.78101 | 0.012 | 0.007 | 1 | 1 | Rfx2 | 0.005 | Old |
| 0.322353 | 0.701801 | 0.08 | 0.069 | 1 | 1 | Fam104a | 0.011 | Old |
| 0.322476 | 0.826949 | 0.081 | 0.071 | 1 | 1 | BC005624 | 0.01 | Old |
| 0.322544 | 0.534826 | 0.012 | 0.007 | 1 | 1 | Tmem132a | 0.005 | Old |
| 0.322656 | 0.65907 | 0.078 | 0.068 | 1 | 1 | Sde2 | 0.01 | Old |
| 0.322971 | 0.123874 | 0.01 | 0.016 | 1 | 1 | Osbpl7 | -0.006 | Old |
| 0.323141 | 0.028899 | 0.051 | 0.062 | 1 | 1 | Atp11b | -0.011 | Old |
| 0.323643 | 0.491716 | 0.012 | 0.007 | 1 | 1 | Dgke | 0.005 | Old |
| 0.32372 | 0.418064 | 0.288 | 0.284 | 1 | 1 | Pomp | 0.004 | Old |
| 0.323863 | 0.513703 | 0.012 | 0.007 | 1 | 1 | Slc35g1 | 0.005 | Old |
| 0.323933 | 0.716513 | 0.054 | 0.045 | 1 | 1 | Rbm34 | 0.009 | Old |
| 0.324221 | 0.211521 | 0.01 | 0.016 | 1 | 1 | Vps33b | -0.006 | Old |
| 0.324451 | 0.358149 | 0.107 | 0.095 | 1 | 1 | Exoc3 | 0.012 | Old |
| 0.324745 | 0.530418 | 0.123 | 0.112 | 1 | 1 | Nfix | 0.011 | Old |
| 0.324985 | 0.040171 | 0.199 | 0.221 | 1 | 1 | Calm3 | -0.022 | Old |
| 0.325825 | 0.022347 | 0.163 | 0.183 | 1 | 1 | Magt1 | -0.02 | Old |
| 0.326523 | 0.184889 | 0.123 | 0.141 | 1 | 1 | Lamtor2 | -0.018 | Old |
| 0.327227 | 0.466531 | 0.143 | 0.132 | 1 | 1 | Tnfaip2 | 0.011 | Old |
| 0.327242 | 0.240485 | 0.229 | 0.216 | 1 | 1 | Zfp706 | 0.013 | Old |
| 0.327403 | 0.509609 | 0.071 | 0.061 | 1 | 1 | Vps37a | 0.01 | Old |
| 0.327473 | 0.28729 | 0.012 | 0.017 | 1 | 1 | Plpp2 | -0.005 | Old |
| 0.328596 | 0.056466 | 0.037 | 0.047 | 1 | 1 | Mcoln1 | -0.01 | Old |
| 0.328698 | 0.55397 | 0.072 | 0.062 | 1 | 1 | Idi1 | 0.01 | Old |
| 0.328796 | 0.47176 | 0.098 | 0.086 | 1 | 1 | Plbd2 | 0.012 | Old |
| 0.329458 | 0.399678 | 0.093 | 0.081 | 1 | 1 | Ybx3 | 0.012 | Old |
| 0.329541 | 0.132496 | 0.071 | 0.06 | 1 | 1 | Tars | 0.011 | Old |
| 0.329852 | 0.043045 | 0.228 | 0.252 | 1 | 1 | 2610507B1 | -0.024 | Old |
| 0.330067 | 0.467378 | 0.089 | 0.078 | 1 | 1 | Cers6 | 0.011 | Old |
| 0.330832 | 0.4856 | 0.179 | 0.168 | 1 | 1 | Ptges3 | 0.011 | Old |
| 0.331008 | 0.210105 | 0.19 | 0.176 | 1 | 1 | Shtn1 | 0.014 | Old |
| 0.33117 | 0.633331 | 0.032 | 0.025 | 1 | 1 | Ormdl3 | 0.007 | Old |
| 0.331418 | 0.049594 | 0.095 | 0.109 | 1 | 1 | Vps26a | -0.014 | Old |
| 0.332933 | 0.162048 | 0.263 | 0.294 | 1 | 1 | Srp9 | -0.031 | Old |
| 0.333501 | 0.129348 | 0.024 | 0.032 | 1 | 1 | Ppp1r3b | -0.008 | Old |
| 0.3337 | 0.042006 | 0.013 | 0.018 | 1 | 1 | Ints10 | -0.005 | Old |
| 0.334128 | 0.019563 | 0.028 | 0.036 | 1 | 1 | Caap1 | -0.008 | Old |
| 0.334261 | 0.525973 | 0.06 | 0.051 | 1 | 1 | Rras | 0.009 | Old |
| 0.334356 | 0.062102 | 0.057 | 0.068 | 1 | 1 | Cbx7 | -0.011 | Old |
| 0.335162 | 0.015233 | 0.032 | 0.041 | 1 | 1 | Ambra1 | -0.009 | Old |
| 0.335431 | 0.069473 | 0.438 | 0.487 | 1 | 1 | Ddx17 | -0.049 | Old |
| 0.335503 | 0.014689 | 0.03 | 0.038 | 1 | 1 | Gramd4 | -0.008 | Old |
| 0.335524 | 0.318639 | 0.243 | 0.233 | 1 | 1 | Taok1 | 0.01 | Old |
| 0.336788 | 0.036981 | 0.347 | 0.385 | 1 | 1 | Ndufa2 | -0.038 | Old |

| | | | | | | | | |
|---|---|---|---|---|---|---|---|---|
| 0.336798 | 0.029553 | 0.026 | 0.033 | 1 | 1 | Ppp2r5d | -0.007 | Old |
| 0.336951 | 1.019869 | 0.014 | 0.01 | 1 | 1 | Ddx28 | 0.004 | Old |
| 0.337555 | 0.558251 | 0.014 | 0.01 | 1 | 1 | Elovl6 | 0.004 | Old |
| 0.337573 | 0.571556 | 0.057 | 0.047 | 1 | 1 | Phf11b | 0.01 | Old |
| 0.337728 | 0.030713 | 0.032 | 0.041 | 1 | 1 | Phyh | -0.009 | Old |
| 0.337843 | 0.490752 | 0.03 | 0.023 | 1 | 1 | Rnf144b | 0.007 | Old |
| 0.337924 | 0.640749 | 0.058 | 0.049 | 1 | 1 | Cds2 | 0.009 | Old |
| 0.338362 | 0.800008 | 0.014 | 0.01 | 1 | 1 | Dmap1 | 0.004 | Old |
| 0.338728 | 0.110037 | 0.031 | 0.039 | 1 | 1 | H2-Ke6 | -0.008 | Old |
| 0.338823 | 0.336647 | 0.067 | 0.057 | 1 | 1 | Tmem131 | 0.01 | Old |
| 0.339271 | 0.492892 | 0.014 | 0.01 | 1 | 1 | Tubgcp6 | 0.004 | Old |
| 0.339361 | 0.436323 | 0.054 | 0.045 | 1 | 1 | Smim19 | 0.009 | Old |
| 0.339589 | 0.745907 | 0.049 | 0.041 | 1 | 1 | Dpp9 | 0.008 | Old |
| 0.33964 | 0.029645 | 0.089 | 0.103 | 1 | 1 | D1Ertd622 | -0.014 | Old |
| 0.339695 | 0.576759 | 0.051 | 0.043 | 1 | 1 | Ipo11 | 0.008 | Old |
| 0.33973 | 0.334094 | 0.044 | 0.035 | 1 | 1 | Cd151 | 0.009 | Old |
| 0.339894 | 0.106229 | 0.018 | 0.024 | 1 | 1 | Plcb2 | -0.006 | Old |
| 0.340181 | 0.488761 | 0.014 | 0.01 | 1 | 1 | Ptcd1 | 0.004 | Old |
| 0.34035 | 0.962379 | 0.022 | 0.016 | 1 | 1 | Lias | 0.006 | Old |
| 0.34143 | 0.004349 | 0.045 | 0.055 | 1 | 1 | Fam133b | -0.01 | Old |
| 0.341658 | 0.022748 | 0.334 | 0.372 | 1 | 1 | Sub1 | -0.038 | Old |
| 0.342006 | 0.203295 | 0.014 | 0.01 | 1 | 1 | Zdhhc24 | 0.004 | Old |
| 0.342306 | 0.791757 | 0.024 | 0.018 | 1 | 1 | Ino80b | 0.006 | Old |
| 0.342411 | 0.233755 | 0.015 | 0.021 | 1 | 1 | Asb13 | -0.006 | Old |
| 0.342575 | 0.0888 | 0.098 | 0.113 | 1 | 1 | Ndufa5 | -0.015 | Old |
| 0.342907 | 0.973832 | 0.022 | 0.016 | 1 | 1 | Tomm40l | 0.006 | Old |
| 0.342949 | 0.151818 | 0.217 | 0.245 | 1 | 1 | Aamp | -0.028 | Old |
| 0.342964 | 0.124206 | 0.118 | 0.136 | 1 | 1 | Pum1 | -0.018 | Old |
| 0.342974 | 1.136899 | 0.017 | 0.012 | 1 | 1 | Dtd2 | 0.005 | Old |
| 0.343706 | 0.04916 | 0.104 | 0.119 | 1 | 1 | Bod1l | -0.015 | Old |
| 0.343865 | 0.45192 | 0.032 | 0.025 | 1 | 1 | Pphln1 | 0.007 | Old |
| 0.344028 | 0.243441 | 0.279 | 0.269 | 1 | 1 | Cd300ld | 0.01 | Old |
| 0.344062 | 0.187587 | 0.041 | 0.033 | 1 | 1 | Mtx1 | 0.008 | Old |
| 0.344831 | 0.797619 | 0.017 | 0.012 | 1 | 1 | 6330418K0 | 0.005 | Old |
| 0.345299 | 0.388796 | 0.095 | 0.084 | 1 | 1 | Ociad1 | 0.011 | Old |
| 0.34539 | 0.984406 | 0.017 | 0.012 | 1 | 1 | Psmg1 | 0.005 | Old |
| 0.34545 | 0.334223 | 0.482 | 0.505 | 1 | 1 | Ier2 | -0.023 | Old |
| 0.345762 | 0.907994 | 0.017 | 0.012 | 1 | 1 | Mrpl22 | 0.005 | Old |
| 0.345874 | 0.108518 | 0.051 | 0.062 | 1 | 1 | Uimc1 | -0.011 | Old |
| 0.345889 | 0.190273 | 0.018 | 0.024 | 1 | 1 | Thoc1 | -0.006 | Old |
| 0.346229 | 0.127866 | 0.17 | 0.191 | 1 | 1 | Caprin1 | -0.021 | Old |
| 0.346632 | 0.157564 | 0.054 | 0.065 | 1 | 1 | Zscan26 | -0.011 | Old |
| 0.346695 | 0.802621 | 0.017 | 0.012 | 1 | 1 | Cavin2 | 0.005 | Old |
| 0.346897 | 0.284182 | 0.206 | 0.195 | 1 | 1 | Ralb | 0.011 | Old |
| 0.347135 | 0.400641 | 0.147 | 0.134 | 1 | 1 | Ptpa | 0.013 | Old |

| | | | | | | | | |
|---|---|---|---|---|---|---|---|---|
| 0.347208 | 0.126202 | 0.053 | 0.064 | 1 | 1 | Samd8 | -0.011 | Old |
| 0.347835 | 0.370773 | 0.027 | 0.021 | 1 | 1 | Galns | 0.006 | Old |
| 0.348224 | 0.495807 | 0.125 | 0.114 | 1 | 1 | Lsm4 | 0.011 | Old |
| 0.348344 | 0.542787 | 0.198 | 0.189 | 1 | 1 | Tmsb10 | 0.009 | Old |
| 0.348466 | 0.825216 | 0.036 | 0.029 | 1 | 1 | Trappc9 | 0.007 | Old |
| 0.34856 | 0.127275 | 0.019 | 0.026 | 1 | 1 | Mtmr9 | -0.007 | Old |
| 0.349127 | 0.340528 | 0.017 | 0.012 | 1 | 1 | Mtfr1 | 0.005 | Old |
| 0.349231 | 0.285536 | 0.315 | 0.308 | 1 | 1 | Herpud1 | 0.007 | Old |
| 0.349641 | 0.505658 | 0.208 | 0.2 | 1 | 1 | Degs1 | 0.008 | Old |
| 0.350309 | 0.420571 | 0.054 | 0.045 | 1 | 1 | Fam118b | 0.009 | Old |
| 0.350309 | 0.079666 | 0.042 | 0.052 | 1 | 1 | St6galnac6 | -0.01 | Old |
| 0.350734 | 0.318344 | 0.027 | 0.021 | 1 | 1 | Atp11a | 0.006 | Old |
| 0.350782 | 0.148113 | 0.078 | 0.091 | 1 | 1 | Mecp2 | -0.013 | Old |
| 0.350836 | 0.446277 | 0.062 | 0.052 | 1 | 1 | Tnks | 0.01 | Old |
| 0.351333 | 0.514931 | 0.036 | 0.029 | 1 | 1 | Cdk5rap3 | 0.007 | Old |
| 0.351444 | 0.056223 | 0.476 | 0.532 | 1 | 1 | Gas6 | -0.056 | Old |
| 0.351926 | 0.146675 | 0.068 | 0.081 | 1 | 1 | Smim27 | -0.013 | Old |
| 0.352115 | 0.06022 | 0.077 | 0.09 | 1 | 1 | Vrk2 | -0.013 | Old |
| 0.352416 | 0.062926 | 0.167 | 0.19 | 1 | 1 | Aldoa | -0.023 | Old |
| 0.35424 | 0.278072 | 0.307 | 0.3 | 1 | 1 | Clec4a3 | 0.007 | Old |
| 0.354463 | 0.419339 | 0.051 | 0.043 | 1 | 1 | Zfp384 | 0.008 | Old |
| 0.354628 | 0.287382 | 0.022 | 0.029 | 1 | 1 | Kank2 | -0.007 | Old |
| 0.355955 | 0.015274 | 0.08 | 0.092 | 1 | 1 | Ank | -0.012 | Old |
| 0.355969 | 0.724595 | 0.031 | 0.024 | 1 | 1 | Klra2 | 0.007 | Old |
| 0.35597 | 0.403971 | 0.208 | 0.199 | 1 | 1 | P4hb | 0.009 | Old |
| 0.356051 | 0.473656 | 0.066 | 0.056 | 1 | 1 | Coa3 | 0.01 | Old |
| 0.356125 | 0.632293 | 0.033 | 0.027 | 1 | 1 | Dpp7 | 0.006 | Old |
| 0.356256 | 0.636445 | 0.033 | 0.027 | 1 | 1 | Alg14 | 0.006 | Old |
| 0.356282 | 0.229739 | 0.019 | 0.014 | 1 | 1 | Slc12a4 | 0.005 | Old |
| 0.356357 | 0.083791 | 0.098 | 0.112 | 1 | 1 | Rp9 | -0.014 | Old |
| 0.356387 | 0.653214 | 0.033 | 0.027 | 1 | 1 | Sirpb1c | 0.006 | Old |
| 0.357093 | 0.015572 | 0.033 | 0.041 | 1 | 1 | Virma | -0.008 | Old |
| 0.357225 | 0.286383 | 0.077 | 0.066 | 1 | 1 | Chordc1 | 0.011 | Old |
| 0.357376 | 0.091982 | 0.247 | 0.274 | 1 | 1 | Sqstm1 | -0.027 | Old |
| 0.357763 | 0.564828 | 0.063 | 0.054 | 1 | 1 | Spg7 | 0.009 | Old |
| 0.358134 | 0.059656 | 0.154 | 0.174 | 1 | 1 | Ndufa7 | -0.02 | Old |
| 0.358233 | 0.126711 | 0.054 | 0.064 | 1 | 1 | Acaa2 | -0.01 | Old |
| 0.358299 | 0.491758 | 0.059 | 0.05 | 1 | 1 | Riok1 | 0.009 | Old |
| 0.358449 | 0.213019 | 0.117 | 0.135 | 1 | 1 | Rragc | -0.018 | Old |
| 0.358456 | 0.450551 | 0.006 | 0.01 | 1 | 1 | Parp3 | -0.004 | Old |
| 0.358503 | 0.331748 | 0.058 | 0.049 | 1 | 1 | Rab3gap2 | 0.009 | Old |
| 0.358769 | 1.408523 | 0.01 | 0.007 | 1 | 1 | Plppr4 | 0.003 | Old |
| 0.358799 | 0.043333 | 0.166 | 0.187 | 1 | 1 | Naa50 | -0.021 | Old |
| 0.359165 | 0.258177 | 0.077 | 0.066 | 1 | 1 | Ly96 | 0.011 | Old |
| 0.359386 | 0.815699 | 0.031 | 0.024 | 1 | 1 | Epg5 | 0.007 | Old |

| | | | | | | | | |
|---|---|---|---|---|---|---|---|---|
| 0.360048 | 0.331295 | 0.068 | 0.058 | 1 | 1 | Sec24c | 0.01 | Old |
| 0.360057 | 0.31173 | 0.066 | 0.056 | 1 | 1 | Atp13a1 | 0.01 | Old |
| 0.360339 | 0.082008 | 0.006 | 0.01 | 1 | 1 | Mfsd4a | -0.004 | Old |
| 0.360415 | 0.610574 | 0.078 | 0.069 | 1 | 1 | Puf60 | 0.009 | Old |
| 0.36076 | 0.902888 | 0.01 | 0.007 | 1 | 1 | Pex26 | 0.003 | Old |
| 0.360899 | 0.546233 | 0.067 | 0.058 | 1 | 1 | Hars | 0.009 | Old |
| 0.361009 | 0.964196 | 0.01 | 0.007 | 1 | 1 | Thop1 | 0.003 | Old |
| 0.361073 | 0.113678 | 0.058 | 0.069 | 1 | 1 | Smarcad1 | -0.011 | Old |
| 0.36167 | 0.440635 | 0.12 | 0.109 | 1 | 1 | Calu | 0.011 | Old |
| 0.362418 | 0.02866 | 0.039 | 0.047 | 1 | 1 | Kif13b | -0.008 | Old |
| 0.362508 | 0.678731 | 0.01 | 0.007 | 1 | 1 | Tmem218 | 0.003 | Old |
| 0.362556 | 0.588939 | 0.04 | 0.032 | 1 | 1 | Haus8 | 0.008 | Old |
| 0.362669 | 0.539882 | 0.123 | 0.113 | 1 | 1 | Tmem168 | 0.01 | Old |
| 0.362949 | 0.040033 | 0.04 | 0.049 | 1 | 1 | Alas1 | -0.009 | Old |
| 0.363045 | 0.198854 | 0.429 | 0.435 | 1 | 1 | Ifitm2 | -0.006 | Old |
| 0.363911 | 0.054873 | 0.162 | 0.181 | 1 | 1 | Dnm2 | -0.019 | Old |
| 0.36401 | 0.450903 | 0.01 | 0.007 | 1 | 1 | Zfp280b | 0.003 | Old |
| 0.364205 | 0.692876 | 0.031 | 0.024 | 1 | 1 | Phtf2 | 0.007 | Old |
| 0.36426 | 0.723181 | 0.01 | 0.007 | 1 | 1 | Gm26853 | 0.003 | Old |
| 0.36426 | 0.661224 | 0.01 | 0.007 | 1 | 1 | Aldh18a1 | 0.003 | Old |
| 0.364409 | 0.160259 | 0.127 | 0.114 | 1 | 1 | Map4 | 0.013 | Old |
| 0.364511 | 0.718718 | 0.01 | 0.007 | 1 | 1 | Itfg2 | 0.003 | Old |
| 0.36462 | 0.385615 | 0.031 | 0.024 | 1 | 1 | Cep57 | 0.007 | Old |
| 0.364695 | 0.409908 | 0.033 | 0.027 | 1 | 1 | Gtf2f2 | 0.006 | Old |
| 0.365053 | 0.511048 | 0.08 | 0.07 | 1 | 1 | U2af1 | 0.01 | Old |
| 0.366752 | 0.420144 | 0.122 | 0.112 | 1 | 1 | Slc20a1 | 0.01 | Old |
| 0.367122 | 0.215861 | 0.141 | 0.162 | 1 | 1 | Pcmtd1 | -0.021 | Old |
| 0.367786 | 0.083685 | 0.461 | 0.518 | 1 | 1 | Pcbp2 | -0.057 | Old |
| 0.367958 | 0.380862 | 0.084 | 0.073 | 1 | 1 | Gatad1 | 0.011 | Old |
| 0.367962 | 0.363639 | 0.076 | 0.066 | 1 | 1 | Ifi27 | 0.01 | Old |
| 0.368012 | 0.426845 | 0.04 | 0.032 | 1 | 1 | Slc25a10 | 0.008 | Old |
| 0.368026 | 0.165594 | 0.189 | 0.213 | 1 | 1 | Tnrc18 | -0.024 | Old |
| 0.368762 | 0.459071 | 0.068 | 0.059 | 1 | 1 | Tpd52l2 | 0.009 | Old |
| 0.368828 | 0.472292 | 0.044 | 0.036 | 1 | 1 | Pnpla2 | 0.008 | Old |
| 0.369817 | 0.129648 | 0.055 | 0.066 | 1 | 1 | Cetn3 | -0.011 | Old |
| 0.370256 | 0.567874 | 0.048 | 0.04 | 1 | 1 | Rbbp5 | 0.008 | Old |
| 0.370288 | 0.035692 | 0.13 | 0.146 | 1 | 1 | Ubald2 | -0.016 | Old |
| 0.371006 | 0.182914 | 0.433 | 0.432 | 1 | 1 | Itm2c | 0.001 | Old |
| 0.371498 | 0.108158 | 0.008 | 0.012 | 1 | 1 | 2310057M | -0.004 | Old |
| 0.371723 | 0.077317 | 0.05 | 0.06 | 1 | 1 | Slc1a5 | -0.01 | Old |
| 0.372211 | 0.324327 | 0.108 | 0.097 | 1 | 1 | Chp1 | 0.011 | Old |
| 0.372671 | 0.255915 | 0.094 | 0.083 | 1 | 1 | Isy1 | 0.011 | Old |
| 0.373292 | 0.467999 | 0.039 | 0.047 | 1 | 1 | Gtf3c6 | -0.008 | Old |
| 0.373822 | 0.35548 | 0.314 | 0.32 | 1 | 1 | Cox5a | -0.006 | Old |
| 0.373939 | 0.051903 | 0.276 | 0.303 | 1 | 1 | Ccr5 | -0.027 | Old |

| | | | | | | | | |
|---|---|---|---|---|---|---|---|---|
| 0.373963 | 0.480802 | 0.06 | 0.052 | 1 | 1 | A430005L1 | 0.008 | Old |
| 0.374286 | 0.553443 | 0.111 | 0.1 | 1 | 1 | Atg3 | 0.011 | Old |
| 0.374315 | 0.446533 | 0.031 | 0.024 | 1 | 1 | Tmem167b | 0.007 | Old |
| 0.374449 | 0.009466 | 0.121 | 0.136 | 1 | 1 | Cct7 | -0.015 | Old |
| 0.374649 | 0.01782 | 0.033 | 0.041 | 1 | 1 | Mrpl10 | -0.008 | Old |
| 0.374919 | 0.42377 | 0.1 | 0.089 | 1 | 1 | Lgals3bp | 0.011 | Old |
| 0.375187 | 0.39238 | 0.028 | 0.022 | 1 | 1 | Pdhx | 0.006 | Old |
| 0.375391 | 0.758639 | 0.026 | 0.02 | 1 | 1 | Pcnx3 | 0.006 | Old |
| 0.375769 | 0.998429 | 0.021 | 0.016 | 1 | 1 | Zfp770 | 0.005 | Old |
| 0.375984 | 0.576594 | 0.008 | 0.012 | 1 | 1 | Pigg | -0.004 | Old |
| 0.376132 | 0.137279 | 0.551 | 0.566 | 1 | 1 | Capza2 | -0.015 | Old |
| 0.376196 | 0.168491 | 0.105 | 0.12 | 1 | 1 | Krit1 | -0.015 | Old |
| 0.376434 | 0.909935 | 0.013 | 0.009 | 1 | 1 | 2700049A0 | 0.004 | Old |
| 0.376434 | 0.898774 | 0.013 | 0.009 | 1 | 1 | Epas1 | 0.004 | Old |
| 0.376603 | 0.528189 | 0.09 | 0.081 | 1 | 1 | Srp54a | 0.009 | Old |
| 0.376759 | 0.261942 | 0.069 | 0.06 | 1 | 1 | Eif4a3 | 0.009 | Old |
| 0.377118 | 0.335167 | 0.086 | 0.076 | 1 | 1 | Cdk14 | 0.01 | Old |
| 0.377336 | 0.718156 | 0.013 | 0.009 | 1 | 1 | Pdss2 | 0.004 | Old |
| 0.377555 | 0.070351 | 0.135 | 0.151 | 1 | 1 | Il13ra1 | -0.016 | Old |
| 0.377622 | 0.140664 | 0.067 | 0.078 | 1 | 1 | Ankle2 | -0.011 | Old |
| 0.377676 | 0.055553 | 0.031 | 0.038 | 1 | 1 | Atrnl1 | -0.007 | Old |
| 0.37874 | 0.074338 | 0.062 | 0.072 | 1 | 1 | Tbc1d20 | -0.01 | Old |
| 0.379095 | 0.040176 | 0.163 | 0.18 | 1 | 1 | Tnks2 | -0.017 | Old |
| 0.379349 | 1.075456 | 0.021 | 0.016 | 1 | 1 | Rmdn3 | 0.005 | Old |
| 0.379678 | 0.041306 | 0.121 | 0.136 | 1 | 1 | Chd9 | -0.015 | Old |
| 0.379901 | 0.022151 | 0.171 | 0.19 | 1 | 1 | Zfp106 | -0.019 | Old |
| 0.380008 | 0.129824 | 0.046 | 0.055 | 1 | 1 | Bag6 | -0.009 | Old |
| 0.380314 | 0.962609 | 0.021 | 0.016 | 1 | 1 | Rab2b | 0.005 | Old |
| 0.380388 | 0.029749 | 0.031 | 0.038 | 1 | 1 | BC003965 | -0.007 | Old |
| 0.380515 | 0.019868 | 0.009 | 0.013 | 1 | 1 | Pde4d | -0.004 | Old |
| 0.380645 | 0.11004 | 0.21 | 0.233 | 1 | 1 | Rp2 | -0.023 | Old |
| 0.380665 | 0.822264 | 0.021 | 0.016 | 1 | 1 | Igtp | 0.005 | Old |
| 0.38073 | 0.493312 | 0.013 | 0.009 | 1 | 1 | Tnk2 | 0.004 | Old |
| 0.381529 | 0.314276 | 0.24 | 0.233 | 1 | 1 | Ddx39b | 0.007 | Old |
| 0.381638 | 0.165873 | 0.069 | 0.06 | 1 | 1 | Piezo1 | 0.009 | Old |
| 0.381681 | 1.299021 | 0.015 | 0.011 | 1 | 1 | Pyurf | 0.004 | Old |
| 0.381721 | 0.472568 | 0.058 | 0.049 | 1 | 1 | Lrrc8a | 0.009 | Old |
| 0.381908 | 0.287779 | 0.035 | 0.028 | 1 | 1 | Dhx8 | 0.007 | Old |
| 0.38236 | 1.041276 | 0.018 | 0.013 | 1 | 1 | Znhit6 | 0.005 | Old |
| 0.382377 | 0.296455 | 0.037 | 0.03 | 1 | 1 | Ptgr1 | 0.007 | Old |
| 0.382915 | 1.267393 | 0.015 | 0.011 | 1 | 1 | Gm37305 | 0.004 | Old |
| 0.382933 | 0.184837 | 0.667 | 0.712 | 1 | 1 | Egr1 | -0.045 | Old |
| 0.383125 | 0.643144 | 0.032 | 0.026 | 1 | 1 | Wdr7 | 0.006 | Old |
| 0.383145 | 0.485794 | 0.041 | 0.034 | 1 | 1 | Lrig2 | 0.007 | Old |
| 0.384104 | 0.041446 | 0.01 | 0.015 | 1 | 1 | Zfp275 | -0.005 | Old |

| | | | | | | | | |
|---|---|---|---|---|---|---|---|---|
| 0.384288 | 0.408565 | 0.053 | 0.063 | 1 | 1 | Rxra | -0.01 | Old |
| 0.384682 | 0.337901 | 0.082 | 0.072 | 1 | 1 | Sptlc1 | 0.01 | Old |
| 0.384769 | 0.940014 | 0.015 | 0.011 | 1 | 1 | 2700097O( | 0.004 | Old |
| 0.384771 | 0.515973 | 0.082 | 0.073 | 1 | 1 | Ubr3 | 0.009 | Old |
| 0.384823 | 0.345446 | 0.023 | 0.018 | 1 | 1 | Ammecr1 | 0.005 | Old |
| 0.384867 | 0.070734 | 0.06 | 0.071 | 1 | 1 | S100a1 | -0.011 | Old |
| 0.385174 | 0.266954 | 0.05 | 0.042 | 1 | 1 | Rpa3 | 0.008 | Old |
| 0.385751 | 0.029506 | 0.117 | 0.131 | 1 | 1 | Gbp7 | -0.014 | Old |
| 0.385933 | 0.623672 | 0.055 | 0.047 | 1 | 1 | Ddx19a | 0.008 | Old |
| 0.386008 | 0.590613 | 0.015 | 0.011 | 1 | 1 | Src | 0.004 | Old |
| 0.386064 | 0.097443 | 0.028 | 0.035 | 1 | 1 | Skint3 | -0.007 | Old |
| 0.386367 | 0.111683 | 0.185 | 0.205 | 1 | 1 | Kdm2a | -0.02 | Old |
| 0.386428 | 0.024113 | 0.072 | 0.083 | 1 | 1 | Chp2 | -0.011 | Old |
| 0.386618 | 0.109145 | 0.023 | 0.03 | 1 | 1 | Wdtc1 | -0.007 | Old |
| 0.386836 | 0.455312 | 0.015 | 0.011 | 1 | 1 | Ccdc130 | 0.004 | Old |
| 0.386892 | 0.12778 | 0.044 | 0.052 | 1 | 1 | Pdcd11 | -0.008 | Old |
| 0.386933 | 0.046249 | 0.028 | 0.022 | 1 | 1 | Usp10 | 0.006 | Old |
| 0.387326 | 0.091687 | 0.082 | 0.094 | 1 | 1 | Tap1 | -0.012 | Old |
| 0.387617 | 0.051874 | 0.145 | 0.162 | 1 | 1 | Ubxn1 | -0.017 | Old |
| 0.387726 | 0.664185 | 0.021 | 0.016 | 1 | 1 | Men1 | 0.005 | Old |
| 0.387736 | 0.334172 | 0.023 | 0.018 | 1 | 1 | Tsr1 | 0.005 | Old |
| 0.388377 | 0.036421 | 0.026 | 0.032 | 1 | 1 | Tipin | -0.006 | Old |
| 0.388445 | 0.004098 | 0.026 | 0.032 | 1 | 1 | Lrrc42 | -0.006 | Old |
| 0.388574 | 0.220383 | 0.01 | 0.015 | 1 | 1 | Enpp4 | -0.005 | Old |
| 0.388716 | 0.019824 | 0.012 | 0.016 | 1 | 1 | Glmn | -0.004 | Old |
| 0.38887 | 0.42567 | 0.041 | 0.034 | 1 | 1 | Rarg | 0.007 | Old |
| 0.389024 | 0.026088 | 0.342 | 0.374 | 1 | 1 | Eif3a | -0.032 | Old |
| 0.389068 | 0.515462 | 0.058 | 0.049 | 1 | 1 | Trak1 | 0.009 | Old |
| 0.389203 | 0.338021 | 0.018 | 0.013 | 1 | 1 | Rmnd1 | 0.005 | Old |
| 0.389311 | 0.15631 | 0.03 | 0.037 | 1 | 1 | Traf7 | -0.007 | Old |
| 0.389372 | 0.824469 | 0.03 | 0.024 | 1 | 1 | Gpn3 | 0.006 | Old |
| 0.389394 | 0.451006 | 0.018 | 0.013 | 1 | 1 | Zfand1 | 0.005 | Old |
| 0.389533 | 0.273306 | 0.015 | 0.011 | 1 | 1 | Plekha8 | 0.004 | Old |
| 0.389594 | 0.118898 | 0.01 | 0.015 | 1 | 1 | Strbp | -0.005 | Old |
| 0.389667 | 0.047406 | 0.018 | 0.024 | 1 | 1 | 5031425F1 | -0.006 | Old |
| 0.38992 | 0.574391 | 0.104 | 0.095 | 1 | 1 | Mydgf | 0.009 | Old |
| 0.390032 | 0.118228 | 0.103 | 0.117 | 1 | 1 | Krcc1 | -0.014 | Old |
| 0.390076 | 0.106484 | 0.012 | 0.016 | 1 | 1 | Nmral1 | -0.004 | Old |
| 0.390144 | 0.361719 | 0.06 | 0.052 | 1 | 1 | Map3k14 | 0.008 | Old |
| 0.390157 | 0.067097 | 0.015 | 0.011 | 1 | 1 | Mocs1 | 0.004 | Old |
| 0.390303 | 0.07351 | 0.231 | 0.254 | 1 | 1 | Cd84 | -0.023 | Old |
| 0.390566 | 0.063354 | 0.24 | 0.267 | 1 | 1 | 5031439G( | -0.027 | Old |
| 0.390709 | 0.249094 | 0.152 | 0.14 | 1 | 1 | Ocrl | 0.012 | Old |
| 0.390757 | 0.048308 | 0.023 | 0.03 | 1 | 1 | Crnkl1 | -0.007 | Old |
| 0.391033 | 0.071161 | 0.015 | 0.021 | 1 | 1 | Gypc | -0.006 | Old |

| | | | | | | | | |
|---|---|---|---|---|---|---|---|---|
| 0.39126 | 0.160063 | 0.024 | 0.031 | 1 | 1 | Dusp23 | -0.007 | Old |
| 0.391574 | 0.772577 | 0.03 | 0.024 | 1 | 1 | Pno1 | 0.006 | Old |
| 0.391745 | 0.208497 | 0.041 | 0.049 | 1 | 1 | Ccdc90b | -0.008 | Old |
| 0.39231 | 0.013213 | 0.024 | 0.031 | 1 | 1 | Bmi1 | -0.007 | Old |
| 0.392415 | 0.231018 | 0.035 | 0.028 | 1 | 1 | Rnf181 | 0.007 | Old |
| 0.392688 | 0.144249 | 0.041 | 0.049 | 1 | 1 | Med30 | -0.008 | Old |
| 0.393012 | 0.031502 | 0.023 | 0.018 | 1 | 1 | Bet1l | 0.005 | Old |
| 0.393298 | 0.339258 | 0.058 | 0.049 | 1 | 1 | BC035044 | 0.009 | Old |
| 0.393339 | 0.019628 | 0.023 | 0.03 | 1 | 1 | Mak16 | -0.007 | Old |
| 0.393486 | 0.535893 | 0.01 | 0.015 | 1 | 1 | Wwox | -0.005 | Old |
| 0.39359 | 0.303284 | 0.103 | 0.118 | 1 | 1 | Dr1 | -0.015 | Old |
| 0.393779 | 0.051523 | 0.012 | 0.016 | 1 | 1 | Scfd2 | -0.004 | Old |
| 0.393944 | 0.496842 | 0.037 | 0.046 | 1 | 1 | Dpy30 | -0.009 | Old |
| 0.394361 | 0.02499 | 0.051 | 0.061 | 1 | 1 | Ano6 | -0.01 | Old |
| 0.394995 | 0.121201 | 0.015 | 0.021 | 1 | 1 | Pnkp | -0.006 | Old |
| 0.395039 | 0.161155 | 0.143 | 0.16 | 1 | 1 | Ppp1r21 | -0.017 | Old |
| 0.395218 | 0.332547 | 0.233 | 0.227 | 1 | 1 | P2rx4 | 0.006 | Old |
| 0.395462 | 0.226892 | 0.107 | 0.123 | 1 | 1 | Rapgef6 | -0.016 | Old |
| 0.395614 | 0.167388 | 0.035 | 0.028 | 1 | 1 | D10Wsu10 | 0.007 | Old |
| 0.395631 | 0.31 | 0.032 | 0.026 | 1 | 1 | Gps2 | 0.006 | Old |
| 0.3961 | 0.136089 | 0.018 | 0.024 | 1 | 1 | Trappc2 | -0.006 | Old |
| 0.396146 | 0.366948 | 0.041 | 0.034 | 1 | 1 | Nup214 | 0.007 | Old |
| 0.396444 | 0.070209 | 0.075 | 0.086 | 1 | 1 | Ldb1 | -0.011 | Old |
| 0.396453 | 0.317069 | 0.31 | 0.309 | 1 | 1 | Atp6v1b2 | 0.001 | Old |
| 0.396508 | 0.093266 | 0.013 | 0.018 | 1 | 1 | Sephs1 | -0.005 | Old |
| 0.396818 | 0.114697 | 0.022 | 0.028 | 1 | 1 | Mrps5 | -0.006 | Old |
| 0.396892 | 0.690365 | 0.03 | 0.024 | 1 | 1 | Thyn1 | 0.006 | Old |
| 0.397035 | 0.453082 | 0.055 | 0.047 | 1 | 1 | Lipo3 | 0.008 | Old |
| 0.397211 | 0.181029 | 0.073 | 0.085 | 1 | 1 | Abce1 | -0.012 | Old |
| 0.397253 | 0.475698 | 0.148 | 0.14 | 1 | 1 | Tbc1d14 | 0.008 | Old |
| 0.397279 | 0.290115 | 0.153 | 0.142 | 1 | 1 | Nudt3 | 0.011 | Old |
| 0.397301 | 0.062959 | 0.019 | 0.025 | 1 | 1 | Trap1 | -0.006 | Old |
| 0.397411 | 0.025011 | 0.022 | 0.028 | 1 | 1 | Gm43848 | -0.006 | Old |
| 0.397731 | 0.199839 | 0.044 | 0.052 | 1 | 1 | Aak1 | -0.008 | Old |
| 0.397863 | 0.739403 | 0.04 | 0.033 | 1 | 1 | Snhg9 | 0.007 | Old |
| 0.39795 | 0.291153 | 0.184 | 0.174 | 1 | 1 | Hspd1 | 0.01 | Old |
| 0.398011 | 0.146522 | 0.013 | 0.018 | 1 | 1 | Trmt10a | -0.005 | Old |
| 0.398275 | 0.347831 | 0.202 | 0.194 | 1 | 1 | Fam120a | 0.008 | Old |
| 0.398391 | 0.221222 | 0.482 | 0.485 | 1 | 1 | Tagln2 | -0.003 | Old |
| 0.398543 | 0.386441 | 0.073 | 0.064 | 1 | 1 | Ptdss1 | 0.009 | Old |
| 0.398572 | 0.019417 | 0.042 | 0.051 | 1 | 1 | Maml1 | -0.009 | Old |
| 0.398697 | 0.033802 | 0.26 | 0.285 | 1 | 1 | Arap1 | -0.025 | Old |
| 0.398766 | 0.096532 | 0.044 | 0.036 | 1 | 1 | Ssh1 | 0.008 | Old |
| 0.399188 | 0.241841 | 0.019 | 0.025 | 1 | 1 | Chchd4 | -0.006 | Old |
| 0.399721 | 0.15961 | 0.035 | 0.028 | 1 | 1 | Eipr1 | 0.007 | Old |

| | | | | | | | | |
|---|---|---|---|---|---|---|---|---|
| 0.399973 | 0.073982 | 0.238 | 0.264 | 1 | 1 | Syk | -0.026 | Old |
| 0.400044 | 0.00582 | 0.134 | 0.149 | 1 | 1 | Wdfy2 | -0.015 | Old |
| 0.400222 | 0.017627 | 0.067 | 0.078 | 1 | 1 | Epc2 | -0.011 | Old |
| 0.400501 | 0.145117 | 0.533 | 0.547 | 1 | 1 | Capzb | -0.014 | Old |
| 0.400778 | 0.707224 | 0.027 | 0.021 | 1 | 1 | Noc4l | 0.006 | Old |
| 0.401307 | 0.007478 | 0.33 | 0.363 | 1 | 1 | Csnk1a1 | -0.033 | Old |
| 0.402025 | 0.177003 | 0.071 | 0.061 | 1 | 1 | Sgta | 0.01 | Old |
| 0.402256 | 0.207465 | 0.06 | 0.052 | 1 | 1 | Dhx38 | 0.008 | Old |
| 0.402403 | 0.254992 | 0.058 | 0.068 | 1 | 1 | Bbx | -0.01 | Old |
| 0.402676 | 0.430167 | 0.026 | 0.032 | 1 | 1 | Ap1m1 | -0.006 | Old |
| 0.403616 | 0.01233 | 0.103 | 0.116 | 1 | 1 | Dcaf8 | -0.013 | Old |
| 0.403654 | 0.086906 | 0.126 | 0.143 | 1 | 1 | Kat2b | -0.017 | Old |
| 0.404799 | 0.495695 | 0.022 | 0.028 | 1 | 1 | Aldh7a1 | -0.006 | Old |
| 0.40501 | 0.143951 | 0.04 | 0.048 | 1 | 1 | Zfp330 | -0.008 | Old |
| 0.405578 | 0.196715 | 0.402 | 0.456 | 1 | 1 | Ddx3x | -0.054 | Old |
| 0.406943 | 0.154978 | 0.166 | 0.186 | 1 | 1 | Uqcrb | -0.02 | Old |
| 0.407658 | 0.056589 | 0.706 | 0.781 | 1 | 1 | Klf6 | -0.075 | Old |
| 0.408646 | 0.187397 | 0.13 | 0.148 | 1 | 1 | Vamp3 | -0.018 | Old |
| 0.408699 | 0.918607 | 0.022 | 0.017 | 1 | 1 | Nudcd1 | 0.005 | Old |
| 0.408763 | 0.547812 | 0.068 | 0.06 | 1 | 1 | Hyou1 | 0.008 | Old |
| 0.409249 | 0.117126 | 0.032 | 0.026 | 1 | 1 | Sirt1 | 0.006 | Old |
| 0.409257 | 0.112981 | 0.122 | 0.137 | 1 | 1 | Myo1f | -0.015 | Old |
| 0.409406 | 0.342557 | 0.059 | 0.051 | 1 | 1 | Erap1 | 0.008 | Old |
| 0.409515 | 0.097583 | 0.063 | 0.073 | 1 | 1 | Utp11 | -0.01 | Old |
| 0.410101 | 0.580948 | 0.04 | 0.033 | 1 | 1 | Serinc5 | 0.007 | Old |
| 0.410115 | 0.240031 | 0.114 | 0.104 | 1 | 1 | Mrpl28 | 0.01 | Old |
| 0.410126 | 0.154425 | 0.213 | 0.24 | 1 | 1 | Hmgb2 | -0.027 | Old |
| 0.410428 | 0.404499 | 0.082 | 0.073 | 1 | 1 | Psme3 | 0.009 | Old |
| 0.410829 | 0.548367 | 0.072 | 0.064 | 1 | 1 | Nfu1 | 0.008 | Old |
| 0.411009 | 0.414687 | 0.078 | 0.069 | 1 | 1 | Sdccag3 | 0.009 | Old |
| 0.411041 | 0.082495 | 0.068 | 0.079 | 1 | 1 | Synrg | -0.011 | Old |
| 0.412601 | 0.384082 | 0.027 | 0.021 | 1 | 1 | Polr3f | 0.006 | Old |
| 0.412751 | 0.179918 | 0.098 | 0.111 | 1 | 1 | Dctn4 | -0.013 | Old |
| 0.412754 | 0.049283 | 0.105 | 0.119 | 1 | 1 | Zfand3 | -0.014 | Old |
| 0.413295 | 0.007923 | 0.072 | 0.083 | 1 | 1 | Lmbrd2 | -0.011 | Old |
| 0.413502 | 0.497953 | 0.13 | 0.12 | 1 | 1 | Dock7 | 0.01 | Old |
| 0.41401 | 1.082268 | 0.022 | 0.017 | 1 | 1 | Ahdc1 | 0.005 | Old |
| 0.414167 | 0.11363 | 0.03 | 0.024 | 1 | 1 | Surf1 | 0.006 | Old |
| 0.414273 | 0.574228 | 0.024 | 0.019 | 1 | 1 | Rps19bp1 | 0.005 | Old |
| 0.414535 | 0.256219 | 0.204 | 0.194 | 1 | 1 | Notch2 | 0.01 | Old |
| 0.414796 | 0.327389 | 0.299 | 0.3 | 1 | 1 | Colgalt1 | -0.001 | Old |
| 0.415256 | 0.766302 | 0.031 | 0.025 | 1 | 1 | Mief1 | 0.006 | Old |
| 0.415459 | 0.144634 | 0.036 | 0.044 | 1 | 1 | Bcl2l2 | -0.008 | Old |
| 0.415571 | 0.345955 | 0.217 | 0.208 | 1 | 1 | Atp5b | 0.009 | Old |
| 0.415841 | 0.20703 | 0.09 | 0.103 | 1 | 1 | Ubr5 | -0.013 | Old |

| | | | | | | | | |
|---|---|---|---|---|---|---|---|---|
| 0.416206 | 0.137035 | 0.049 | 0.058 | 1 | 1 | Fubp3 | -0.009 | Old |
| 0.416627 | 0.775411 | 0.024 | 0.019 | 1 | 1 | Zfyve1 | 0.005 | Old |
| 0.41686 | 0.745349 | 0.022 | 0.017 | 1 | 1 | Elk1 | 0.005 | Old |
| 0.417499 | 0.014195 | 0.044 | 0.052 | 1 | 1 | Mtf2 | -0.008 | Old |
| 0.417897 | 0.122018 | 0.139 | 0.156 | 1 | 1 | Cmtm3 | -0.017 | Old |
| 0.418418 | 0.300514 | 0.104 | 0.094 | 1 | 1 | Tollip | 0.01 | Old |
| 0.419154 | 0.365136 | 0.087 | 0.078 | 1 | 1 | Usp14 | 0.009 | Old |
| 0.419561 | 0.039704 | 0.059 | 0.069 | 1 | 1 | Zwint | -0.01 | Old |
| 0.419691 | 0.108017 | 0.032 | 0.039 | 1 | 1 | Hif1an | -0.007 | Old |
| 0.419705 | 0.020049 | 0.064 | 0.074 | 1 | 1 | Rbm42 | -0.01 | Old |
| 0.420457 | 0.450383 | 0.05 | 0.043 | 1 | 1 | Sec23ip | 0.007 | Old |
| 0.420851 | 0.434019 | 0.04 | 0.033 | 1 | 1 | Sash3 | 0.007 | Old |
| 0.420896 | 0.565203 | 0.048 | 0.041 | 1 | 1 | Vars | 0.007 | Old |
| 0.420919 | 0.108483 | 0.027 | 0.021 | 1 | 1 | Cars | 0.006 | Old |
| 0.42119 | 0.40298 | 0.121 | 0.112 | 1 | 1 | Rab18 | 0.009 | Old |
| 0.421495 | 0.972822 | 0.019 | 0.015 | 1 | 1 | Gnpnat1 | 0.004 | Old |
| 0.421638 | 0.124155 | 0.046 | 0.055 | 1 | 1 | Zcchc17 | -0.009 | Old |
| 0.422165 | 0.3557 | 0.04 | 0.033 | 1 | 1 | Ankrd27 | 0.007 | Old |
| 0.422261 | 0.251877 | 0.1 | 0.09 | 1 | 1 | Tmem165 | 0.01 | Old |
| 0.422654 | 1.023292 | 0.012 | 0.008 | 1 | 1 | Etfbkmt | 0.004 | Old |
| 0.422678 | 0.272644 | 0.162 | 0.151 | 1 | 1 | Gnpda1 | 0.011 | Old |
| 0.422949 | 0.137143 | 0.136 | 0.151 | 1 | 1 | Kpna4 | -0.015 | Old |
| 0.423019 | 0.041307 | 0.118 | 0.131 | 1 | 1 | Pgd | -0.013 | Old |
| 0.423086 | 0.494903 | 0.04 | 0.033 | 1 | 1 | Tbrg1 | 0.007 | Old |
| 0.423161 | 1.343695 | 0.012 | 0.008 | 1 | 1 | 4930555A( | 0.004 | Old |
| 0.423237 | 0.506239 | 0.057 | 0.049 | 1 | 1 | Pex2 | 0.008 | Old |
| 0.423692 | 0.073954 | 0.036 | 0.03 | 1 | 1 | Abhd17c | 0.006 | Old |
| 0.42372 | 0.342573 | 0.109 | 0.099 | 1 | 1 | Xpr1 | 0.01 | Old |
| 0.423734 | 0.254104 | 0.035 | 0.042 | 1 | 1 | Fbxo30 | -0.007 | Old |
| 0.423943 | 0.266482 | 0.27 | 0.267 | 1 | 1 | Usf2 | 0.003 | Old |
| 0.423955 | 0.030032 | 0.058 | 0.067 | 1 | 1 | Dnmt1 | -0.009 | Old |
| 0.424551 | 0.148477 | 0.035 | 0.042 | 1 | 1 | Map3k4 | -0.007 | Old |
| 0.424683 | 0.919175 | 0.012 | 0.008 | 1 | 1 | Tctn3 | 0.004 | Old |
| 0.425006 | 0.349349 | 0.057 | 0.049 | 1 | 1 | Cask | 0.008 | Old |
| 0.425052 | 0.24968 | 0.019 | 0.015 | 1 | 1 | Aptx | 0.004 | Old |
| 0.425154 | 0.130707 | 0.042 | 0.035 | 1 | 1 | Nr1h2 | 0.007 | Old |
| 0.425445 | 0.54506 | 0.012 | 0.008 | 1 | 1 | Coa4 | 0.004 | Old |
| 0.425726 | 0.075353 | 0.104 | 0.117 | 1 | 1 | Kansl1 | -0.013 | Old |
| 0.425909 | 0.009022 | 0.027 | 0.033 | 1 | 1 | Rabggta | -0.006 | Old |
| 0.426019 | 0.375484 | 0.022 | 0.017 | 1 | 1 | Abt1 | 0.005 | Old |
| 0.426279 | 0.361875 | 0.078 | 0.069 | 1 | 1 | Myo1c | 0.009 | Old |
| 0.426462 | 0.927501 | 0.012 | 0.008 | 1 | 1 | Bub1b | 0.004 | Old |
| 0.426792 | 0.256158 | 0.03 | 0.036 | 1 | 1 | Gtf2f1 | -0.006 | Old |
| 0.426944 | 0.674903 | 0.041 | 0.035 | 1 | 1 | Ufd1 | 0.006 | Old |
| 0.427204 | 0.046709 | 0.373 | 0.412 | 1 | 1 | App | -0.039 | Old |

| | | | | | | | | |
|---|---|---|---|---|---|---|---|---|
| 0.427698 | 0.081445 | 0.228 | 0.247 | 1 | 1 | Tpp1 | -0.019 | Old |
| 0.427736 | 0.664404 | 0.012 | 0.008 | 1 | 1 | Thap7 | 0.004 | Old |
| 0.427736 | 0.206014 | 0.012 | 0.008 | 1 | 1 | Snx33 | 0.004 | Old |
| 0.428016 | 0.122207 | 0.058 | 0.067 | 1 | 1 | Otud4 | -0.009 | Old |
| 0.428501 | 0.35332 | 0.012 | 0.008 | 1 | 1 | Gpsm1 | 0.004 | Old |
| 0.428529 | 0.547532 | 0.019 | 0.015 | 1 | 1 | Rhbdf1 | 0.004 | Old |
| 0.428742 | 0.056951 | 0.121 | 0.134 | 1 | 1 | Tmem258 | -0.013 | Old |
| 0.428784 | 0.442271 | 0.017 | 0.013 | 1 | 1 | Tuft1 | 0.004 | Old |
| 0.428878 | 0.725182 | 0.014 | 0.01 | 1 | 1 | Ranbp6 | 0.004 | Old |
| 0.428878 | 0.681732 | 0.014 | 0.01 | 1 | 1 | 4930430F0 | 0.004 | Old |
| 0.42909 | 0.113905 | 0.066 | 0.057 | 1 | 1 | Clcc1 | 0.009 | Old |
| 0.429268 | 0.026967 | 0.012 | 0.008 | 1 | 1 | Pex12 | 0.004 | Old |
| 0.429523 | 0.237577 | 0.012 | 0.008 | 1 | 1 | Arrdc2 | 0.004 | Old |
| 0.429564 | 0.46703 | 0.014 | 0.01 | 1 | 1 | Rsad1 | 0.004 | Old |
| 0.429597 | 0.236894 | 0.073 | 0.064 | 1 | 1 | Dpy19l4 | 0.009 | Old |
| 0.429636 | 0.74785 | 0.053 | 0.046 | 1 | 1 | Gps1 | 0.007 | Old |
| 0.429642 | 0.280036 | 0.022 | 0.017 | 1 | 1 | Gamt | 0.005 | Old |
| 0.429912 | 0.097857 | 0.122 | 0.136 | 1 | 1 | Ttyh3 | -0.014 | Old |
| 0.429953 | 0.27184 | 0.036 | 0.044 | 1 | 1 | Phactr4 | -0.008 | Old |
| 0.430022 | 0.91203 | 0.014 | 0.01 | 1 | 1 | Dimt1 | 0.004 | Old |
| 0.430177 | 0.380103 | 0.019 | 0.015 | 1 | 1 | Ubxn8 | 0.004 | Old |
| 0.430249 | 0.603492 | 0.017 | 0.013 | 1 | 1 | Slc25a13 | 0.004 | Old |
| 0.43048 | 0.655992 | 0.028 | 0.023 | 1 | 1 | Slc39a13 | 0.005 | Old |
| 0.430553 | 0.180074 | 0.071 | 0.081 | 1 | 1 | Rnf38 | -0.01 | Old |
| 0.430671 | 0.176178 | 0.046 | 0.055 | 1 | 1 | Scaf8 | -0.009 | Old |
| 0.430856 | 0.508408 | 0.019 | 0.015 | 1 | 1 | Suv39h1 | 0.004 | Old |
| 0.431266 | 0.21797 | 0.068 | 0.079 | 1 | 1 | Cln6 | -0.011 | Old |
| 0.431277 | 0.463444 | 0.05 | 0.043 | 1 | 1 | Tmem205 | 0.007 | Old |
| 0.431459 | 0.196944 | 0.057 | 0.066 | 1 | 1 | Irak3 | -0.009 | Old |
| 0.431631 | 0.271852 | 0.031 | 0.038 | 1 | 1 | Dnajc11 | -0.007 | Old |
| 0.431926 | 0.6035 | 0.017 | 0.013 | 1 | 1 | Ap5s1 | 0.004 | Old |
| 0.431931 | 0.356535 | 0.261 | 0.258 | 1 | 1 | Wsb1 | 0.003 | Old |
| 0.432321 | 0.196282 | 0.104 | 0.118 | 1 | 1 | Dcaf7 | -0.014 | Old |
| 0.432321 | 0.167495 | 0.434 | 0.446 | 1 | 1 | H2-DMb1 | -0.012 | Old |
| 0.432499 | 0.198211 | 0.045 | 0.053 | 1 | 1 | Fam162a | -0.008 | Old |
| 0.432546 | 0.520037 | 0.014 | 0.01 | 1 | 1 | Irgq | 0.004 | Old |
| 0.433005 | 0.011129 | 0.014 | 0.01 | 1 | 1 | Tmem101 | 0.004 | Old |
| 0.433165 | 0.531424 | 0.086 | 0.078 | 1 | 1 | Pigk | 0.008 | Old |
| 0.433296 | 0.152582 | 0.046 | 0.055 | 1 | 1 | Trim27 | -0.009 | Old |
| 0.433857 | 0.380296 | 0.127 | 0.118 | 1 | 1 | Mob4 | 0.009 | Old |
| 0.433925 | 0.2624 | 0.19 | 0.181 | 1 | 1 | Lamtor1 | 0.009 | Old |
| 0.434659 | 0.395753 | 0.017 | 0.013 | 1 | 1 | Mad1l1 | 0.004 | Old |
| 0.43467 | 0.3042 | 0.031 | 0.025 | 1 | 1 | Faap100 | 0.006 | Old |
| 0.435078 | 0.21424 | 0.014 | 0.01 | 1 | 1 | Atp1a2 | 0.004 | Old |
| 0.435355 | 0.150038 | 0.174 | 0.161 | 1 | 1 | Limd1 | 0.013 | Old |

| | | | | | | | | |
|---|---|---|---|---|---|---|---|---|
| 0.435533 | 0.211211 | 0.019 | 0.015 | 1 | 1 | Fbf1 | 0.004 | Old |
| 0.436496 | 0.034554 | 0.026 | 0.032 | 1 | 1 | Fbxo4 | -0.006 | Old |
| 0.436756 | 0.45811 | 0.041 | 0.035 | 1 | 1 | Stoml2 | 0.006 | Old |
| 0.436861 | 0.346862 | 0.058 | 0.05 | 1 | 1 | Nucb2 | 0.008 | Old |
| 0.437163 | 0.337957 | 0.044 | 0.052 | 1 | 1 | AI413582 | -0.008 | Old |
| 0.437191 | 0.247144 | 0.017 | 0.013 | 1 | 1 | Rabep2 | 0.004 | Old |
| 0.43736 | 0.454358 | 0.071 | 0.063 | 1 | 1 | Mtr | 0.008 | Old |
| 0.43797 | 0.072657 | 0.911 | 0.922 | 1 | 1 | Lyz2 | -0.011 | Old |
| 0.438438 | 0.244303 | 0.054 | 0.047 | 1 | 1 | Tnfrsf21 | 0.007 | Old |
| 0.439041 | 0.05149 | 0.129 | 0.143 | 1 | 1 | Cdipt | -0.014 | Old |
| 0.439605 | 0.397366 | 0.069 | 0.061 | 1 | 1 | Fkbp4 | 0.008 | Old |
| 0.439797 | 0.800323 | 0.045 | 0.038 | 1 | 1 | Ext2 | 0.007 | Old |
| 0.439812 | 0.071086 | 0.15 | 0.166 | 1 | 1 | Itch | -0.016 | Old |
| 0.440042 | 0.118253 | 0.086 | 0.097 | 1 | 1 | Csnk2a1 | -0.011 | Old |
| 0.440326 | 0.085733 | 0.838 | 0.873 | 1 | 1 | Selenop | -0.035 | Old |
| 0.440638 | 0.744936 | 0.039 | 0.032 | 1 | 1 | E2f1 | 0.007 | Old |
| 0.440827 | 0.354785 | 0.069 | 0.061 | 1 | 1 | Cdk17 | 0.008 | Old |
| 0.440851 | 0.325199 | 0.176 | 0.167 | 1 | 1 | Ssr1 | 0.009 | Old |
| 0.441476 | 0.56932 | 0.026 | 0.021 | 1 | 1 | Acot2 | 0.005 | Old |
| 0.441501 | 0.077502 | 0.871 | 0.874 | 1 | 1 | Mrc1 | -0.003 | Old |
| 0.441756 | 0.094648 | 0.037 | 0.044 | 1 | 1 | Asl | -0.007 | Old |
| 0.441891 | 0.188561 | 0.06 | 0.07 | 1 | 1 | Srek1ip1 | -0.01 | Old |
| 0.441981 | 0.346396 | 0.075 | 0.066 | 1 | 1 | Klf11 | 0.009 | Old |
| 0.442155 | 0.174342 | 0.087 | 0.099 | 1 | 1 | Rps27rt | -0.012 | Old |
| 0.44233 | 0.293065 | 0.138 | 0.128 | 1 | 1 | Ptpn12 | 0.01 | Old |
| 0.442571 | 0.381527 | 0.081 | 0.072 | 1 | 1 | Arl5a | 0.009 | Old |
| 0.442579 | 0.559674 | 0.045 | 0.038 | 1 | 1 | Ppm1k | 0.007 | Old |
| 0.442584 | 0.000949 | 0.071 | 0.081 | 1 | 1 | Stip1 | -0.01 | Old |
| 0.442945 | 0.061595 | 0.171 | 0.187 | 1 | 1 | Klf13 | -0.016 | Old |
| 0.442972 | 0.117635 | 0.072 | 0.082 | 1 | 1 | Rheb | -0.01 | Old |
| 0.443411 | 0.128069 | 0.144 | 0.16 | 1 | 1 | Derl1 | -0.016 | Old |
| 0.443708 | 0.313578 | 0.006 | 0.01 | 1 | 1 | Lag3 | -0.004 | Old |
| 0.444185 | 1.102714 | 0.023 | 0.018 | 1 | 1 | Pgm2 | 0.005 | Old |
| 0.444269 | 0.01102 | 0.176 | 0.194 | 1 | 1 | Pik3cd | -0.018 | Old |
| 0.444729 | 0.364526 | 0.098 | 0.089 | 1 | 1 | Cops4 | 0.009 | Old |
| 0.444858 | 0.694385 | 0.026 | 0.021 | 1 | 1 | Ermard | 0.005 | Old |
| 0.444892 | 0.059744 | 0.276 | 0.302 | 1 | 1 | Clic1 | -0.026 | Old |
| 0.444936 | 0.28402 | 0.172 | 0.163 | 1 | 1 | Aes | 0.009 | Old |
| 0.445045 | 0.007119 | 0.312 | 0.333 | 1 | 1 | Wdr1 | -0.021 | Old |
| 0.445081 | 0.535685 | 0.006 | 0.01 | 1 | 1 | Mcrip2 | -0.004 | Old |
| 0.445116 | 0.510537 | 0.045 | 0.038 | 1 | 1 | Oas2 | 0.007 | Old |
| 0.445206 | 0.142807 | 0.054 | 0.047 | 1 | 1 | Npl | 0.007 | Old |
| 0.445242 | 0.039023 | 0.219 | 0.236 | 1 | 1 | Rab11b | -0.017 | Old |
| 0.445242 | 0.023405 | 0.051 | 0.06 | 1 | 1 | Ppfia1 | -0.009 | Old |
| 0.445289 | 0.18062 | 0.028 | 0.035 | 1 | 1 | Ubac1 | -0.007 | Old |

| | | | | | | | | |
|---|---|---|---|---|---|---|---|---|
| 0.445356 | 0.176293 | 0.006 | 0.01 | 1 | 1 | Sumf2 | -0.004 | Old |
| 0.445376 | 0.017324 | 0.023 | 0.029 | 1 | 1 | Metap1d | -0.006 | Old |
| 0.445631 | 0.229479 | 0.006 | 0.01 | 1 | 1 | Sardh | -0.004 | Old |
| 0.445907 | 0.45014 | 0.006 | 0.01 | 1 | 1 | Fam78b | -0.004 | Old |
| 0.445992 | 0.304415 | 0.027 | 0.033 | 1 | 1 | Prorsd1 | -0.006 | Old |
| 0.446024 | 0.718296 | 0.032 | 0.027 | 1 | 1 | Ptrhd1 | 0.005 | Old |
| 0.446182 | 0.088222 | 0.006 | 0.01 | 1 | 1 | Ctu1 | -0.004 | Old |
| 0.446474 | 0.875726 | 0.03 | 0.024 | 1 | 1 | Lrrc57 | 0.006 | Old |
| 0.446475 | 0.670617 | 0.055 | 0.049 | 1 | 1 | Strn | 0.006 | Old |
| 0.446484 | 0.399534 | 0.123 | 0.115 | 1 | 1 | Smarcc2 | 0.008 | Old |
| 0.446528 | 0.753343 | 0.06 | 0.053 | 1 | 1 | Tatdn2 | 0.007 | Old |
| 0.446598 | 0.045369 | 0.158 | 0.174 | 1 | 1 | Irf5 | -0.016 | Old |
| 0.446894 | 0.420957 | 0.026 | 0.021 | 1 | 1 | Mrpl40 | 0.005 | Old |
| 0.447292 | 0.246612 | 0.026 | 0.032 | 1 | 1 | Fbxl4 | -0.006 | Old |
| 0.447328 | 0.189813 | 0.023 | 0.029 | 1 | 1 | Bcl9 | -0.006 | Old |
| 0.447389 | 0.066446 | 0.134 | 0.148 | 1 | 1 | Arhgef1 | -0.014 | Old |
| 0.44778 | 0.917051 | 0.036 | 0.03 | 1 | 1 | Eif1ad | 0.006 | Old |
| 0.447943 | 0.140501 | 0.174 | 0.19 | 1 | 1 | Slc29a3 | -0.016 | Old |
| 0.447953 | 0.133345 | 0.023 | 0.029 | 1 | 1 | Usp31 | -0.006 | Old |
| 0.448126 | 0.772545 | 0.023 | 0.018 | 1 | 1 | Xrcc4 | 0.005 | Old |
| 0.44865 | 0.638681 | 0.049 | 0.042 | 1 | 1 | Kcnn4 | 0.007 | Old |
| 0.44875 | 0.077595 | 0.032 | 0.038 | 1 | 1 | Gkap1 | -0.006 | Old |
| 0.448831 | 0.080659 | 0.294 | 0.323 | 1 | 1 | Ppp1r18 | -0.029 | Old |
| 0.448847 | 0.298543 | 0.077 | 0.069 | 1 | 1 | Tm2d1 | 0.008 | Old |
| 0.448873 | 0.415762 | 0.109 | 0.101 | 1 | 1 | Tomm70a | 0.008 | Old |
| 0.448971 | 0.131743 | 0.023 | 0.029 | 1 | 1 | Metrn | -0.006 | Old |
| 0.449492 | 0.8164 | 0.006 | 0.01 | 1 | 1 | Rrp1b | -0.004 | Old |
| 0.449598 | 0.267385 | 0.024 | 0.03 | 1 | 1 | Rrp7a | -0.006 | Old |
| 0.44961 | 0.295684 | 0.078 | 0.07 | 1 | 1 | Ube2j2 | 0.008 | Old |
| 0.449625 | 0.053051 | 0.077 | 0.087 | 1 | 1 | Rnpepl1 | -0.01 | Old |
| 0.450025 | 0.60968 | 0.046 | 0.04 | 1 | 1 | Dbt | 0.006 | Old |
| 0.450547 | 0.264306 | 0.022 | 0.027 | 1 | 1 | Smim24 | -0.005 | Old |
| 0.450618 | 0.139805 | 0.023 | 0.029 | 1 | 1 | Lmnb1 | -0.006 | Old |
| 0.450694 | 0.07491 | 0.037 | 0.044 | 1 | 1 | Cmc2 | -0.007 | Old |
| 0.450841 | 0.293578 | 0.175 | 0.165 | 1 | 1 | Cd48 | 0.01 | Old |
| 0.451177 | 0.087825 | 0.141 | 0.157 | 1 | 1 | Kif1b | -0.016 | Old |
| 0.451365 | 0.895737 | 0.023 | 0.018 | 1 | 1 | Dusp10 | 0.005 | Old |
| 0.451524 | 0.578079 | 0.057 | 0.049 | 1 | 1 | Mcrs1 | 0.008 | Old |
| 0.451688 | 0.006293 | 0.036 | 0.043 | 1 | 1 | Zfp110 | -0.007 | Old |
| 0.452215 | 0.026881 | 0.037 | 0.044 | 1 | 1 | Zbtb22 | -0.007 | Old |
| 0.452225 | 0.022621 | 0.008 | 0.011 | 1 | 1 | Pdlim7 | -0.003 | Old |
| 0.452336 | 0.540058 | 0.039 | 0.032 | 1 | 1 | Tmem97 | 0.007 | Old |
| 0.452443 | 0.019712 | 0.055 | 0.064 | 1 | 1 | Gtf2b | -0.009 | Old |
| 0.452455 | 0.23934 | 0.105 | 0.096 | 1 | 1 | Snx4 | 0.009 | Old |
| 0.452628 | 0.56761 | 0.023 | 0.018 | 1 | 1 | Ublcp1 | 0.005 | Old |

| | | | | | | | | |
|---|---|---|---|---|---|---|---|---|
| 0.452971 | 0.335631 | 0.123 | 0.115 | 1 | 1 | Bcas2 | 0.008 | Old |
| 0.453123 | 0.163169 | 0.024 | 0.03 | 1 | 1 | Traf2 | -0.006 | Old |
| 0.453623 | 0.213812 | 0.057 | 0.066 | 1 | 1 | Lysmd3 | -0.009 | Old |
| 0.453959 | 0.047178 | 0.165 | 0.181 | 1 | 1 | Pten | -0.016 | Old |
| 0.454231 | 0.421155 | 0.077 | 0.069 | 1 | 1 | Sec23b | 0.008 | Old |
| 0.454541 | 0.068124 | 0.008 | 0.011 | 1 | 1 | Uevld | -0.003 | Old |
| 0.454681 | 0.394342 | 0.066 | 0.058 | 1 | 1 | Chml | 0.008 | Old |
| 0.454794 | 0.464451 | 0.049 | 0.042 | 1 | 1 | Ngly1 | 0.007 | Old |
| 0.454894 | 1.112061 | 0.021 | 0.016 | 1 | 1 | Cul4b | 0.005 | Old |
| 0.454935 | 0.522849 | 0.086 | 0.078 | 1 | 1 | Ccdc115 | 0.008 | Old |
| 0.455426 | 0.121249 | 0.072 | 0.082 | 1 | 1 | Aida | -0.01 | Old |
| 0.455445 | 0.330411 | 0.059 | 0.052 | 1 | 1 | Naa20 | 0.007 | Old |
| 0.455561 | 0.282599 | 0.055 | 0.048 | 1 | 1 | Zbtb8os | 0.007 | Old |
| 0.455763 | 0.20525 | 0.073 | 0.083 | 1 | 1 | Mllt6 | -0.01 | Old |
| 0.455812 | 0.037742 | 0.01 | 0.014 | 1 | 1 | Mical1 | -0.004 | Old |
| 0.455859 | 0.124989 | 0.015 | 0.02 | 1 | 1 | Rtca | -0.005 | Old |
| 0.456257 | 0.128863 | 0.017 | 0.021 | 1 | 1 | Isg15 | -0.004 | Old |
| 0.456536 | 0.198047 | 0.18 | 0.2 | 1 | 1 | Ubl5 | -0.02 | Old |
| 0.456881 | 0.548159 | 0.112 | 0.104 | 1 | 1 | Setx | 0.008 | Old |
| 0.456906 | 0.192416 | 0.051 | 0.044 | 1 | 1 | Cfdp1 | 0.007 | Old |
| 0.457151 | 0.064606 | 0.091 | 0.102 | 1 | 1 | 4931406C0 | -0.011 | Old |
| 0.457401 | 0.104405 | 0.111 | 0.124 | 1 | 1 | Hexim1 | -0.013 | Old |
| 0.457507 | 0.097312 | 0.254 | 0.278 | 1 | 1 | Mapre1 | -0.024 | Old |
| 0.457585 | 0.17964 | 0.019 | 0.024 | 1 | 1 | 1500011B0 | -0.005 | Old |
| 0.458198 | 0.405932 | 0.032 | 0.027 | 1 | 1 | Pcsk7 | 0.005 | Old |
| 0.4587 | 0.387044 | 0.125 | 0.116 | 1 | 1 | Ssrp1 | 0.009 | Old |
| 0.45887 | 0.179589 | 0.069 | 0.061 | 1 | 1 | Fuca2 | 0.008 | Old |
| 0.458938 | 0.46476 | 0.073 | 0.066 | 1 | 1 | Rara | 0.007 | Old |
| 0.459016 | 0.622025 | 0.03 | 0.024 | 1 | 1 | Aar2 | 0.006 | Old |
| 0.4592 | 0.647733 | 0.03 | 0.036 | 1 | 1 | Sugp2 | -0.006 | Old |
| 0.459287 | 0.30255 | 0.406 | 0.423 | 1 | 1 | Atp5a1 | -0.017 | Old |
| 0.459319 | 0.511142 | 0.05 | 0.044 | 1 | 1 | Vps45 | 0.006 | Old |
| 0.459548 | 0.671786 | 0.044 | 0.038 | 1 | 1 | Nelfe | 0.006 | Old |
| 0.460014 | 0.078057 | 0.352 | 0.39 | 1 | 1 | Fosl2 | -0.038 | Old |
| 0.460119 | 0.281948 | 0.009 | 0.013 | 1 | 1 | Ttc39c | -0.004 | Old |
| 0.460339 | 0.252767 | 0.219 | 0.21 | 1 | 1 | Chmp4b | 0.009 | Old |
| 0.460437 | 0.193914 | 0.201 | 0.224 | 1 | 1 | Rab7 | -0.023 | Old |
| 0.460771 | 0.100811 | 0.108 | 0.12 | 1 | 1 | Ccnt2 | -0.012 | Old |
| 0.461072 | 0.115018 | 0.01 | 0.014 | 1 | 1 | Olfr920 | -0.004 | Old |
| 0.461117 | 0.170329 | 0.032 | 0.027 | 1 | 1 | Kdelc2 | 0.005 | Old |
| 0.461287 | 0.37854 | 0.161 | 0.153 | 1 | 1 | Lypla2 | 0.008 | Old |
| 0.461296 | 0.321759 | 0.019 | 0.024 | 1 | 1 | Fam114a1 | -0.005 | Old |
| 0.461658 | 0.81893 | 0.104 | 0.098 | 1 | 1 | Gfpt1 | 0.006 | Old |
| 0.461968 | 0.002109 | 0.087 | 0.097 | 1 | 1 | Polr2m | -0.01 | Old |
| 0.46199 | 0.095976 | 0.01 | 0.014 | 1 | 1 | Eif5a2 | -0.004 | Old |

| | | | | | | | | |
|---|---|---|---|---|---|---|---|---|
| 0.462223 | 0.431112 | 0.03 | 0.024 | 1 | 1 | Tinf2 | 0.006 | Old |
| 0.462258 | 0.314998 | 0.023 | 0.018 | 1 | 1 | Hmgxb4 | 0.005 | Old |
| 0.462392 | 0.442114 | 0.024 | 0.03 | 1 | 1 | Pgm1 | -0.006 | Old |
| 0.462973 | 0.017379 | 0.113 | 0.125 | 1 | 1 | Mdh2 | -0.012 | Old |
| 0.463178 | 0.390168 | 0.212 | 0.208 | 1 | 1 | Ap2s1 | 0.004 | Old |
| 0.463312 | 0.335265 | 0.015 | 0.02 | 1 | 1 | Zfp142 | -0.005 | Old |
| 0.463413 | 0.15505 | 0.181 | 0.2 | 1 | 1 | Stx7 | -0.019 | Old |
| 0.463419 | 0.324863 | 0.057 | 0.049 | 1 | 1 | Commd2 | 0.008 | Old |
| 0.463572 | 0.369849 | 0.184 | 0.177 | 1 | 1 | Klhl9 | 0.007 | Old |
| 0.463989 | 0.648844 | 0.021 | 0.016 | 1 | 1 | Clpb | 0.005 | Old |
| 0.464045 | 0.103651 | 0.179 | 0.196 | 1 | 1 | Hsd17b11 | -0.017 | Old |
| 0.4646 | 0.198976 | 0.051 | 0.06 | 1 | 1 | Aph1c | -0.009 | Old |
| 0.464768 | 0.545023 | 0.021 | 0.016 | 1 | 1 | Rgp1 | 0.005 | Old |
| 0.464825 | 0.12859 | 0.171 | 0.188 | 1 | 1 | Ppp1r15b | -0.017 | Old |
| 0.464847 | 0.043612 | 0.013 | 0.017 | 1 | 1 | Serpine1 | -0.004 | Old |
| 0.464847 | 0.197711 | 0.013 | 0.017 | 1 | 1 | Plk4 | -0.004 | Old |
| 0.464912 | 0.078657 | 0.194 | 0.213 | 1 | 1 | Hspe1 | -0.019 | Old |
| 0.465128 | 0.191168 | 0.032 | 0.027 | 1 | 1 | Mterf3 | 0.005 | Old |
| 0.46565 | 0.334109 | 0.014 | 0.018 | 1 | 1 | Tsr2 | -0.004 | Old |
| 0.466064 | 0.271227 | 0.084 | 0.095 | 1 | 1 | Ing2 | -0.011 | Old |
| 0.466254 | 0.330039 | 0.012 | 0.016 | 1 | 1 | Cnnm2 | -0.004 | Old |
| 0.466456 | 0.249497 | 0.08 | 0.091 | 1 | 1 | Rhoq | -0.011 | Old |
| 0.466547 | 0.171256 | 0.032 | 0.038 | 1 | 1 | Immp1l | -0.006 | Old |
| 0.466828 | 0.708656 | 0.027 | 0.022 | 1 | 1 | Asgr2 | 0.005 | Old |
| 0.466907 | 0.152818 | 0.036 | 0.043 | 1 | 1 | Wdr20 | -0.007 | Old |
| 0.466962 | 0.689194 | 0.04 | 0.034 | 1 | 1 | Sfxn1 | 0.006 | Old |
| 0.467984 | 0.537073 | 0.01 | 0.014 | 1 | 1 | Mus81 | -0.004 | Old |
| 0.468009 | 0.346274 | 0.012 | 0.016 | 1 | 1 | Tram2 | -0.004 | Old |
| 0.468628 | 0.224043 | 0.053 | 0.061 | 1 | 1 | Tlk2 | -0.008 | Old |
| 0.468897 | 0.470384 | 0.015 | 0.02 | 1 | 1 | Srcap | -0.005 | Old |
| 0.468928 | 0.213008 | 0.091 | 0.103 | 1 | 1 | Sptbn1 | -0.012 | Old |
| 0.468971 | 0.374949 | 0.149 | 0.141 | 1 | 1 | Mrps33 | 0.008 | Old |
| 0.469145 | 0.087247 | 0.046 | 0.054 | 1 | 1 | Syvn1 | -0.008 | Old |
| 0.469517 | 0.769157 | 0.027 | 0.033 | 1 | 1 | Snhg20 | -0.006 | Old |
| 0.469689 | 0.121301 | 0.184 | 0.203 | 1 | 1 | Vdac2 | -0.019 | Old |
| 0.469743 | 0.093257 | 0.159 | 0.175 | 1 | 1 | Dnase2a | -0.016 | Old |
| 0.469792 | 0.121954 | 0.31 | 0.298 | 1 | 1 | Prkar1a | 0.012 | Old |
| 0.469807 | 0.385651 | 0.03 | 0.024 | 1 | 1 | Fbxo25 | 0.006 | Old |
| 0.470011 | 0.407932 | 0.104 | 0.096 | 1 | 1 | Mindy1 | 0.008 | Old |
| 0.470041 | 0.286808 | 0.042 | 0.036 | 1 | 1 | Polr3k | 0.006 | Old |
| 0.470059 | 0.098911 | 0.139 | 0.154 | 1 | 1 | Hps3 | -0.015 | Old |
| 0.470195 | 0.393553 | 0.036 | 0.043 | 1 | 1 | Usp40 | -0.007 | Old |
| 0.470364 | 0.100327 | 0.183 | 0.2 | 1 | 1 | Ubap2l | -0.017 | Old |
| 0.470762 | 0.191331 | 0.113 | 0.126 | 1 | 1 | Eif3d | -0.013 | Old |
| 0.471026 | 0.318041 | 0.015 | 0.02 | 1 | 1 | Fpgt | -0.005 | Old |

| | | | | | | | | |
|---|---|---|---|---|---|---|---|---|
| 0.471108 | 0.002689 | 0.152 | 0.166 | 1 | 1 | Tcea1 | -0.014 | Old |
| 0.472373 | 0.578309 | 0.084 | 0.076 | 1 | 1 | Pik3r5 | 0.008 | Old |
| 0.472595 | 0.209611 | 0.021 | 0.016 | 1 | 1 | Gm20342 | 0.005 | Old |
| 0.472608 | 0.451198 | 0.018 | 0.014 | 1 | 1 | Ppat | 0.004 | Old |
| 0.472737 | 0.760645 | 0.044 | 0.038 | 1 | 1 | Fbxo22 | 0.006 | Old |
| 0.472766 | 0.464518 | 0.027 | 0.022 | 1 | 1 | Piga | 0.005 | Old |
| 0.47342 | 0.083227 | 0.121 | 0.134 | 1 | 1 | Spcs3 | -0.013 | Old |
| 0.473973 | 0.421015 | 0.111 | 0.102 | 1 | 1 | Noct | 0.009 | Old |
| 0.473989 | 0.849416 | 0.015 | 0.012 | 1 | 1 | Rmnd5b | 0.003 | Old |
| 0.474224 | 0.433096 | 0.046 | 0.04 | 1 | 1 | Rgmb | 0.006 | Old |
| 0.474423 | 0.29577 | 0.036 | 0.043 | 1 | 1 | Nfyc | -0.007 | Old |
| 0.474489 | 0.451802 | 0.05 | 0.044 | 1 | 1 | Dpagt1 | 0.006 | Old |
| 0.474537 | 0.147681 | 0.076 | 0.086 | 1 | 1 | Vps28 | -0.01 | Old |
| 0.47461 | 0.011732 | 0.031 | 0.037 | 1 | 1 | Wdr91 | -0.006 | Old |
| 0.475318 | 0.145357 | 0.481 | 0.499 | 1 | 1 | Slc25a3 | -0.018 | Old |
| 0.475323 | 0.489114 | 0.027 | 0.022 | 1 | 1 | Cep63 | 0.005 | Old |
| 0.476199 | 0.295518 | 0.05 | 0.058 | 1 | 1 | Arsk | -0.008 | Old |
| 0.476315 | 0.295387 | 0.03 | 0.024 | 1 | 1 | Dpep2 | 0.006 | Old |
| 0.476395 | 0.581945 | 0.015 | 0.012 | 1 | 1 | Npat | 0.003 | Old |
| 0.476612 | 0.802061 | 0.01 | 0.007 | 1 | 1 | Zfp619 | 0.003 | Old |
| 0.476663 | 1.122353 | 0.013 | 0.01 | 1 | 1 | Prrg1 | 0.003 | Old |
| 0.477027 | 0.467558 | 0.098 | 0.09 | 1 | 1 | Chd3 | 0.008 | Old |
| 0.477084 | 0.760279 | 0.015 | 0.012 | 1 | 1 | Nubpl | 0.003 | Old |
| 0.477148 | 0.137244 | 0.064 | 0.073 | 1 | 1 | Echs1 | -0.009 | Old |
| 0.477668 | 0.393182 | 0.06 | 0.053 | 1 | 1 | Kctd10 | 0.007 | Old |
| 0.477717 | 0.433697 | 0.027 | 0.022 | 1 | 1 | Epn2 | 0.005 | Old |
| 0.477929 | 0.830732 | 0.013 | 0.01 | 1 | 1 | Cpt2 | 0.003 | Old |
| 0.478177 | 0.08626 | 0.031 | 0.037 | 1 | 1 | Unkl | -0.006 | Old |
| 0.478257 | 0.052294 | 0.062 | 0.07 | 1 | 1 | Idh3b | -0.008 | Old |
| 0.47834 | 0.100722 | 0.03 | 0.035 | 1 | 1 | Itgax | -0.005 | Old |
| 0.478405 | 0.348955 | 0.168 | 0.16 | 1 | 1 | Zyx | 0.008 | Old |
| 0.478459 | 0.278546 | 0.086 | 0.078 | 1 | 1 | Nudt16l1 | 0.008 | Old |
| 0.478706 | 0.206623 | 0.091 | 0.103 | 1 | 1 | Stag1 | -0.012 | Old |
| 0.479153 | 0.55965 | 0.015 | 0.012 | 1 | 1 | Tab3 | 0.003 | Old |
| 0.479186 | 0.737532 | 0.01 | 0.007 | 1 | 1 | Nat10 | 0.003 | Old |
| 0.479212 | 0.127946 | 0.054 | 0.062 | 1 | 1 | Gsdmd | -0.008 | Old |
| 0.479704 | 1.216366 | 0.013 | 0.01 | 1 | 1 | Eefsec | 0.003 | Old |
| 0.479844 | 0.513141 | 0.015 | 0.012 | 1 | 1 | Zfp426 | 0.003 | Old |
| 0.479908 | 0.096694 | 0.085 | 0.095 | 1 | 1 | Edf1 | -0.01 | Old |
| 0.479966 | 0.367693 | 0.138 | 0.129 | 1 | 1 | Comt | 0.009 | Old |
| 0.480014 | 0.207126 | 0.018 | 0.014 | 1 | 1 | Tstd3 | 0.004 | Old |
| 0.480046 | 0.227647 | 0.102 | 0.092 | 1 | 1 | Mfsd5 | 0.01 | Old |
| 0.480075 | 0.382708 | 0.015 | 0.012 | 1 | 1 | Tnfsf13os | 0.003 | Old |
| 0.480212 | 0.647666 | 0.013 | 0.01 | 1 | 1 | Thg1l | 0.003 | Old |
| 0.48047 | 0.083079 | 0.059 | 0.067 | 1 | 1 | Tor1aip2 | -0.008 | Old |

| | | | | | | | | |
|---|---|---|---|---|---|---|---|---|
| 0.480619 | 0.592537 | 0.01 | 0.007 | 1 | 1 | Fn3krp | 0.003 | Old |
| 0.480766 | 0.432221 | 0.015 | 0.012 | 1 | 1 | Trp53rka | 0.003 | Old |
| 0.480809 | 0.264537 | 0.195 | 0.186 | 1 | 1 | Hnrnph2 | 0.009 | Old |
| 0.481498 | 0.28117 | 0.126 | 0.141 | 1 | 1 | Dnajc7 | -0.015 | Old |
| 0.48161 | 0.826981 | 0.013 | 0.01 | 1 | 1 | Ehf | 0.003 | Old |
| 0.481626 | 0.447894 | 0.107 | 0.099 | 1 | 1 | Oat | 0.008 | Old |
| 0.481736 | 0.023903 | 0.424 | 0.46 | 1 | 1 | Wwp1 | -0.036 | Old |
| 0.482055 | 0.167278 | 0.01 | 0.007 | 1 | 1 | Zfp974 | 0.003 | Old |
| 0.482144 | 0.476233 | 0.156 | 0.149 | 1 | 1 | Emc1 | 0.007 | Old |
| 0.482342 | 0.62384 | 0.01 | 0.007 | 1 | 1 | Ripk3 | 0.003 | Old |
| 0.482342 | 0.266543 | 0.01 | 0.007 | 1 | 1 | Cebpe | 0.003 | Old |
| 0.482501 | 0.677963 | 0.013 | 0.01 | 1 | 1 | Tcta | 0.003 | Old |
| 0.482501 | 0.672261 | 0.013 | 0.01 | 1 | 1 | Mrrf | 0.003 | Old |
| 0.482533 | 0.599127 | 0.031 | 0.026 | 1 | 1 | Ccdc88c | 0.005 | Old |
| 0.482607 | 0.250486 | 0.033 | 0.028 | 1 | 1 | Cdadc1 | 0.005 | Old |
| 0.482755 | 0.607354 | 0.013 | 0.01 | 1 | 1 | Dennd2a | 0.003 | Old |
| 0.482982 | 0.076974 | 0.04 | 0.047 | 1 | 1 | Pofut2 | -0.007 | Old |
| 0.483075 | 0.24116 | 0.015 | 0.012 | 1 | 1 | Srrd | 0.003 | Old |
| 0.483326 | 0.595389 | 0.041 | 0.035 | 1 | 1 | Ndufaf4 | 0.006 | Old |
| 0.483425 | 0.045394 | 0.407 | 0.441 | 1 | 1 | Arhgdia | -0.034 | Old |
| 0.483664 | 0.123378 | 0.228 | 0.252 | 1 | 1 | Minos1 | -0.024 | Old |
| 0.483775 | 0.438554 | 0.013 | 0.01 | 1 | 1 | Phc1 | 0.003 | Old |
| 0.483786 | 0.05603 | 0.432 | 0.469 | 1 | 1 | Aif1 | -0.037 | Old |
| 0.483841 | 0.087225 | 0.095 | 0.106 | 1 | 1 | Ctbp1 | -0.011 | Old |
| 0.484069 | 0.227252 | 0.01 | 0.007 | 1 | 1 | Efnb1 | 0.003 | Old |
| 0.484088 | 0.03395 | 0.311 | 0.337 | 1 | 1 | Tmed10 | -0.026 | Old |
| 0.484178 | 0.104316 | 0.152 | 0.166 | 1 | 1 | Eif1b | -0.014 | Old |
| 0.484258 | 0.197352 | 0.046 | 0.04 | 1 | 1 | Daxx | 0.006 | Old |
| 0.484551 | 0.047773 | 0.05 | 0.058 | 1 | 1 | Polg | -0.008 | Old |
| 0.484575 | 0.016559 | 0.028 | 0.034 | 1 | 1 | Tom1 | -0.006 | Old |
| 0.484614 | 0.345742 | 0.033 | 0.028 | 1 | 1 | Ndufv1 | 0.005 | Old |
| 0.485424 | 0.204471 | 0.271 | 0.262 | 1 | 1 | Eif5a | 0.009 | Old |
| 0.485626 | 0.126291 | 0.028 | 0.034 | 1 | 1 | Apba1 | -0.006 | Old |
| 0.485854 | 0.048593 | 0.015 | 0.012 | 1 | 1 | Nudt18 | 0.003 | Old |
| 0.486063 | 1.189106 | 0.022 | 0.018 | 1 | 1 | Prkdc | 0.004 | Old |
| 0.486073 | 0.195852 | 0.013 | 0.01 | 1 | 1 | Gramd3 | 0.003 | Old |
| 0.486077 | 0.267808 | 0.028 | 0.034 | 1 | 1 | Urm1 | -0.006 | Old |
| 0.486292 | 0.085834 | 0.058 | 0.066 | 1 | 1 | mt-Nd6 | -0.008 | Old |
| 0.486329 | 0.12281 | 0.013 | 0.01 | 1 | 1 | Gpr85 | 0.003 | Old |
| 0.486329 | 0.075213 | 0.013 | 0.01 | 1 | 1 | Dcun1d4 | 0.003 | Old |
| 0.48652 | 0.284279 | 0.01 | 0.007 | 1 | 1 | Xlr4a | 0.003 | Old |
| 0.48666 | 0.05292 | 0.122 | 0.134 | 1 | 1 | Csnk1e | -0.012 | Old |
| 0.487181 | 0.305288 | 0.206 | 0.199 | 1 | 1 | Zfp207 | 0.007 | Old |
| 0.487665 | 0.060896 | 0.063 | 0.072 | 1 | 1 | Lrch3 | -0.009 | Old |
| 0.488121 | 0.014852 | 0.027 | 0.032 | 1 | 1 | Fam222b | -0.005 | Old |

| | | | | | | | | |
|---|---|---|---|---|---|---|---|---|
| 0.488223 | 0.049142 | 0.027 | 0.022 | 1 | 1 | Dtx3 | 0.005 | Old |
| 0.488736 | 0.025072 | 0.048 | 0.055 | 1 | 1 | Tm9sf4 | -0.007 | Old |
| 0.488814 | 0.543242 | 0.045 | 0.039 | 1 | 1 | Zfp664 | 0.006 | Old |
| 0.490537 | 0.240612 | 0.041 | 0.048 | 1 | 1 | Med7 | -0.007 | Old |
| 0.491186 | 0.069428 | 0.108 | 0.119 | 1 | 1 | Ggh | -0.011 | Old |
| 0.491508 | 0.218188 | 0.064 | 0.072 | 1 | 1 | Snf8 | -0.008 | Old |
| 0.49176 | 0.909393 | 0.028 | 0.024 | 1 | 1 | Rnaset2b | 0.004 | Old |
| 0.492256 | 0.378304 | 0.024 | 0.02 | 1 | 1 | Tpm3-rs7 | 0.004 | Old |
| 0.492361 | 0.510995 | 0.107 | 0.099 | 1 | 1 | Gspt1 | 0.008 | Old |
| 0.492368 | 0.135777 | 0.24 | 0.263 | 1 | 1 | Lmna | -0.023 | Old |
| 0.492437 | 0.364007 | 0.105 | 0.097 | 1 | 1 | AI467606 | 0.008 | Old |
| 0.492551 | 0.166945 | 0.131 | 0.122 | 1 | 1 | Fgfr1 | 0.009 | Old |
| 0.492606 | 0.062168 | 0.049 | 0.056 | 1 | 1 | Akap8 | -0.007 | Old |
| 0.492651 | 0.039392 | 0.371 | 0.41 | 1 | 1 | Hnrnpa3 | -0.039 | Old |
| 0.492768 | 0.203625 | 0.041 | 0.048 | 1 | 1 | Mrpl16 | -0.007 | Old |
| 0.493017 | 0.528431 | 0.028 | 0.034 | 1 | 1 | Gm11808 | -0.006 | Old |
| 0.493398 | 0.019015 | 0.086 | 0.095 | 1 | 1 | Cyb5a | -0.009 | Old |
| 0.493427 | 0.096647 | 0.114 | 0.106 | 1 | 1 | Sod2 | 0.008 | Old |
| 0.493665 | 0.020816 | 0.163 | 0.178 | 1 | 1 | Dgkd | -0.015 | Old |
| 0.493986 | 0.121997 | 0.157 | 0.146 | 1 | 1 | Csk | 0.011 | Old |
| 0.494104 | 0.086338 | 0.246 | 0.267 | 1 | 1 | Rock1 | -0.021 | Old |
| 0.495171 | 0.012965 | 0.315 | 0.34 | 1 | 1 | Cox6a1 | -0.025 | Old |
| 0.495922 | 0.073061 | 0.049 | 0.056 | 1 | 1 | Mrps36 | -0.007 | Old |
| 0.496604 | 0.123959 | 0.059 | 0.067 | 1 | 1 | Ppm1h | -0.008 | Old |
| 0.496623 | 0.343243 | 0.126 | 0.117 | 1 | 1 | Pycard | 0.009 | Old |
| 0.49731 | 0.164901 | 0.505 | 0.533 | 1 | 1 | Rhoa | -0.028 | Old |
| 0.498641 | 0.005753 | 0.023 | 0.028 | 1 | 1 | Anks3 | -0.005 | Old |
| 0.499239 | 0.075029 | 0.406 | 0.448 | 1 | 1 | Ywhae | -0.042 | Old |
| 0.499264 | 0.188531 | 0.1 | 0.112 | 1 | 1 | Rtf2 | -0.012 | Old |
| 0.499672 | 0.173892 | 0.107 | 0.097 | 1 | 1 | Pdlim1 | 0.01 | Old |
| 0.500251 | 0.026811 | 0.024 | 0.02 | 1 | 1 | Rhot2 | 0.004 | Old |
| 0.50054 | 0.144439 | 0.077 | 0.069 | 1 | 1 | Dennd4c | 0.008 | Old |
| 0.500579 | 0.019249 | 0.105 | 0.116 | 1 | 1 | Nucb1 | -0.011 | Old |
| 0.500905 | 0.094655 | 0.033 | 0.028 | 1 | 1 | Arfgef2 | 0.005 | Old |
| 0.501018 | 0.439072 | 0.049 | 0.057 | 1 | 1 | Stau1 | -0.008 | Old |
| 0.50113 | 0.447008 | 0.063 | 0.056 | 1 | 1 | H2-Q4 | 0.007 | Old |
| 0.501416 | 0.010091 | 0.1 | 0.111 | 1 | 1 | Mrpl24 | -0.011 | Old |
| 0.502361 | 0.254448 | 0.022 | 0.018 | 1 | 1 | Acot8 | 0.004 | Old |
| 0.502419 | 0.024035 | 0.208 | 0.225 | 1 | 1 | Eif2s2 | -0.017 | Old |
| 0.502467 | 0.281456 | 0.089 | 0.081 | 1 | 1 | Gorasp2 | 0.008 | Old |
| 0.502524 | 0.028557 | 0.422 | 0.459 | 1 | 1 | Atp6v0b | -0.037 | Old |
| 0.503566 | 0.39826 | 0.041 | 0.035 | 1 | 1 | Tomm40 | 0.006 | Old |
| 0.503634 | 0.036772 | 0.035 | 0.041 | 1 | 1 | Sap30l | -0.006 | Old |
| 0.504107 | 0.028777 | 0.442 | 0.482 | 1 | 1 | Hexa | -0.04 | Old |
| 0.504129 | 0.161586 | 0.369 | 0.369 | 1 | 1 | Grk2 | 0 | Old |

| | | | | | | | | |
|---|---|---|---|---|---|---|---|---|
| 0.504613 | 0.051989 | 0.022 | 0.027 | 1 | 1 | Trp53bp1 | -0.005 | Old |
| 0.504618 | 0.062524 | 0.084 | 0.093 | 1 | 1 | Lpgat1 | -0.009 | Old |
| 0.504705 | 0.38172 | 0.028 | 0.024 | 1 | 1 | Plrg1 | 0.004 | Old |
| 0.504831 | 0.163556 | 0.055 | 0.049 | 1 | 1 | Slc36a4 | 0.006 | Old |
| 0.50494 | 0.204099 | 0.063 | 0.072 | 1 | 1 | Polr2g | -0.009 | Old |
| 0.504974 | 0.018443 | 0.166 | 0.179 | 1 | 1 | Etf1 | -0.013 | Old |
| 0.504996 | 0.058038 | 0.071 | 0.079 | 1 | 1 | Dst | -0.008 | Old |
| 0.505049 | 0.766833 | 0.032 | 0.027 | 1 | 1 | Sigmar1 | 0.005 | Old |
| 0.50519 | 0.508005 | 0.159 | 0.154 | 1 | 1 | Rnpep | 0.005 | Old |
| 0.506083 | 0.683837 | 0.045 | 0.039 | 1 | 1 | Cfap20 | 0.006 | Old |
| 0.506099 | 0.13803 | 0.046 | 0.053 | 1 | 1 | Jmjd6 | -0.007 | Old |
| 0.506345 | 0.100227 | 0.022 | 0.027 | 1 | 1 | Ipcef1 | -0.005 | Old |
| 0.506463 | 0.153326 | 0.174 | 0.188 | 1 | 1 | Sypl | -0.014 | Old |
| 0.506672 | 0.164885 | 0.091 | 0.101 | 1 | 1 | Lsm6 | -0.01 | Old |
| 0.506714 | 0.38201 | 0.156 | 0.149 | 1 | 1 | Bag1 | 0.007 | Old |
| 0.506718 | 0.036639 | 0.382 | 0.422 | 1 | 1 | Btf3 | -0.04 | Old |
| 0.506837 | 0.197681 | 0.046 | 0.053 | 1 | 1 | Glrx2 | -0.007 | Old |
| 0.507101 | 0.243487 | 0.045 | 0.039 | 1 | 1 | Sh3glb2 | 0.006 | Old |
| 0.507109 | 0.621904 | 0.028 | 0.024 | 1 | 1 | Dvl1 | 0.004 | Old |
| 0.507133 | 0.276943 | 0.121 | 0.134 | 1 | 1 | Gtf2h5 | -0.013 | Old |
| 0.507283 | 0.717782 | 0.036 | 0.031 | 1 | 1 | Manea | 0.005 | Old |
| 0.507476 | 0.035612 | 0.072 | 0.081 | 1 | 1 | Srd5a3 | -0.009 | Old |
| 0.507809 | 0.006636 | 0.023 | 0.028 | 1 | 1 | Timm17b | -0.005 | Old |
| 0.508051 | 0.059391 | 0.081 | 0.09 | 1 | 1 | Rnf187 | -0.009 | Old |
| 0.508204 | 0.385788 | 0.121 | 0.113 | 1 | 1 | Blmh | 0.008 | Old |
| 0.50841 | 0.275228 | 0.035 | 0.03 | 1 | 1 | Klhl20 | 0.005 | Old |
| 0.508694 | 0.183965 | 0.062 | 0.069 | 1 | 1 | Pdhb | -0.007 | Old |
| 0.508733 | 0.129723 | 0.073 | 0.066 | 1 | 1 | Ly9 | 0.007 | Old |
| 0.508954 | 0.374324 | 0.064 | 0.058 | 1 | 1 | Ngdn | 0.006 | Old |
| 0.510209 | 0.066398 | 0.497 | 0.556 | 1 | 1 | Efhd2 | -0.059 | Old |
| 0.510458 | 0.298358 | 0.165 | 0.157 | 1 | 1 | Atp2a2 | 0.008 | Old |
| 0.510746 | 0.204842 | 0.28 | 0.276 | 1 | 1 | Dusp3 | 0.004 | Old |
| 0.511016 | 0.75197 | 0.026 | 0.021 | 1 | 1 | Mlycd | 0.005 | Old |
| 0.511042 | 0.004287 | 0.021 | 0.025 | 1 | 1 | Card9 | -0.004 | Old |
| 0.511401 | 0.010588 | 0.021 | 0.025 | 1 | 1 | Ciao1 | -0.004 | Old |
| 0.511622 | 0.166107 | 0.063 | 0.071 | 1 | 1 | Elovl5 | -0.008 | Old |
| 0.511981 | 1.093181 | 0.017 | 0.013 | 1 | 1 | Zmym1 | 0.004 | Old |
| 0.512013 | 0.187587 | 0.339 | 0.345 | 1 | 1 | Pdcd6ip | -0.006 | Old |
| 0.512473 | 0.058331 | 0.141 | 0.155 | 1 | 1 | Tmem109 | -0.014 | Old |
| 0.512571 | 0.26949 | 0.293 | 0.295 | 1 | 1 | Tmem106a | -0.002 | Old |
| 0.512699 | 0.398151 | 0.026 | 0.031 | 1 | 1 | Pkp4 | -0.005 | Old |
| 0.512734 | 0.551854 | 0.019 | 0.016 | 1 | 1 | Kctd3 | 0.003 | Old |
| 0.512808 | 0.191337 | 0.131 | 0.145 | 1 | 1 | Dctn1 | -0.014 | Old |
| 0.513506 | 0.164075 | 0.058 | 0.066 | 1 | 1 | Seh1l | -0.008 | Old |
| 0.513826 | 0.400097 | 0.076 | 0.069 | 1 | 1 | 8-Mar | 0.007 | Old |

| | | | | | | | | |
|---|---|---|---|---|---|---|---|---|
| 0.5144 | 0.307975 | 0.035 | 0.041 | 1 | 1 | Pmvk | -0.006 | Old |
| 0.514422 | 0.592641 | 0.019 | 0.016 | 1 | 1 | Lzic | 0.003 | Old |
| 0.514703 | 0.186109 | 0.022 | 0.027 | 1 | 1 | Gab3 | -0.005 | Old |
| 0.51491 | 0.450176 | 0.05 | 0.044 | 1 | 1 | Acox1 | 0.006 | Old |
| 0.515592 | 0.238047 | 0.246 | 0.238 | 1 | 1 | Prr13 | 0.008 | Old |
| 0.515616 | 0.91334 | 0.017 | 0.013 | 1 | 1 | Zfp874b | 0.004 | Old |
| 0.515715 | 0.30396 | 0.076 | 0.069 | 1 | 1 | Rnf11 | 0.007 | Old |
| 0.515994 | 0.051452 | 0.388 | 0.424 | 1 | 1 | Akr1a1 | -0.036 | Old |
| 0.516154 | 0.08651 | 0.199 | 0.216 | 1 | 1 | Trir | -0.017 | Old |
| 0.516537 | 0.349744 | 0.019 | 0.016 | 1 | 1 | Aspa | 0.003 | Old |
| 0.516537 | 0.329712 | 0.185 | 0.179 | 1 | 1 | Copz1 | 0.006 | Old |
| 0.516733 | 0.267851 | 0.08 | 0.089 | 1 | 1 | Ide | -0.009 | Old |
| 0.516749 | 0.267242 | 0.019 | 0.016 | 1 | 1 | Myom1 | 0.003 | Old |
| 0.517155 | 0.109679 | 0.021 | 0.025 | 1 | 1 | Ubtd1 | -0.004 | Old |
| 0.517885 | 0.135781 | 0.114 | 0.126 | 1 | 1 | Fis1 | -0.012 | Old |
| 0.518593 | 0.054063 | 0.107 | 0.117 | 1 | 1 | Lrmp | -0.01 | Old |
| 0.519012 | 0.340107 | 0.026 | 0.021 | 1 | 1 | Scly | 0.005 | Old |
| 0.519413 | 0.265649 | 0.12 | 0.112 | 1 | 1 | Tcn2 | 0.008 | Old |
| 0.52072 | 0.26444 | 0.068 | 0.077 | 1 | 1 | Sft2d1 | -0.009 | Old |
| 0.521224 | 0.368342 | 0.032 | 0.027 | 1 | 1 | Parp12 | 0.005 | Old |
| 0.521334 | 0.104285 | 0.042 | 0.049 | 1 | 1 | Tvp23b | -0.007 | Old |
| 0.521418 | 0.321102 | 0.084 | 0.077 | 1 | 1 | Cyp51 | 0.007 | Old |
| 0.52155 | 0.339982 | 0.032 | 0.027 | 1 | 1 | Mettl14 | 0.005 | Old |
| 0.521591 | 0.540747 | 0.019 | 0.024 | 1 | 1 | Lncpint | -0.005 | Old |
| 0.522006 | 0.119242 | 0.051 | 0.058 | 1 | 1 | Pak1 | -0.007 | Old |
| 0.522327 | 0.18699 | 0.257 | 0.251 | 1 | 1 | Tardbp | 0.006 | Old |
| 0.522506 | 0.051429 | 0.042 | 0.049 | 1 | 1 | Ugdh | -0.007 | Old |
| 0.52253 | 0.063237 | 0.91 | 0.94 | 1 | 1 | mt-Cytb | -0.03 | Old |
| 0.523007 | 0.049035 | 0.039 | 0.033 | 1 | 1 | Ado | 0.006 | Old |
| 0.523641 | 0.040028 | 0.068 | 0.076 | 1 | 1 | Stk3 | -0.008 | Old |
| 0.523733 | 0.099834 | 0.645 | 0.681 | 1 | 1 | Ppia | -0.036 | Old |
| 0.523878 | 0.128033 | 0.072 | 0.081 | 1 | 1 | Fbrsl1 | -0.009 | Old |
| 0.524225 | 0.059837 | 0.452 | 0.51 | 1 | 1 | Tmbim6 | -0.058 | Old |
| 0.524534 | 0.016531 | 0.017 | 0.021 | 1 | 1 | Snip1 | -0.004 | Old |
| 0.525101 | 0.004064 | 0.044 | 0.05 | 1 | 1 | Rnf5 | -0.006 | Old |
| 0.525224 | 0.274879 | 0.017 | 0.013 | 1 | 1 | Taf6 | 0.004 | Old |
| 0.525545 | 0.270256 | 0.116 | 0.108 | 1 | 1 | Akt2 | 0.008 | Old |
| 0.525648 | 0.783151 | 0.014 | 0.011 | 1 | 1 | Gorasp1 | 0.003 | Old |
| 0.525785 | 0.3579 | 0.084 | 0.077 | 1 | 1 | Pts | 0.007 | Old |
| 0.526081 | 0.250088 | 0.019 | 0.024 | 1 | 1 | Tars2 | -0.005 | Old |
| 0.526135 | 0.263429 | 0.161 | 0.154 | 1 | 1 | R3hdm4 | 0.007 | Old |
| 0.526401 | 0.8799 | 0.014 | 0.011 | 1 | 1 | Slc7a13 | 0.003 | Old |
| 0.527134 | 0.754725 | 0.023 | 0.019 | 1 | 1 | Ccs | 0.004 | Old |
| 0.527489 | 0.440163 | 0.019 | 0.024 | 1 | 1 | Mrps11 | -0.005 | Old |
| 0.527673 | 0.273838 | 0.111 | 0.103 | 1 | 1 | Tmem167 | 0.008 | Old |

| | | | | | | | | |
|---|---|---|---|---|---|---|---|---|
| 0.527793 | 0.04029 | 0.05 | 0.057 | 1 | 1 | Flcn | -0.007 | Old |
| 0.527948 | 0.57687 | 0.04 | 0.035 | 1 | 1 | Psmd9 | 0.005 | Old |
| 0.528746 | 0.028146 | 0.017 | 0.021 | 1 | 1 | Fam213a | -0.004 | Old |
| 0.529172 | 0.69325 | 0.023 | 0.019 | 1 | 1 | Cep104 | 0.004 | Old |
| 0.529583 | 0.181174 | 0.039 | 0.033 | 1 | 1 | Prkci | 0.006 | Old |
| 0.529669 | 0.850533 | 0.014 | 0.011 | 1 | 1 | E130311K1 | 0.003 | Old |
| 0.529756 | 0.172504 | 0.09 | 0.1 | 1 | 1 | Fam96a | -0.01 | Old |
| 0.529989 | 1.078516 | 0.035 | 0.03 | 1 | 1 | Pja1 | 0.005 | Old |
| 0.530138 | 0.200566 | 0.044 | 0.05 | 1 | 1 | Fnip2 | -0.006 | Old |
| 0.530172 | 0.587958 | 0.014 | 0.011 | 1 | 1 | Coro2b | 0.003 | Old |
| 0.530234 | 0.35568 | 0.062 | 0.055 | 1 | 1 | Dctn5 | 0.007 | Old |
| 0.530415 | 0.263745 | 0.064 | 0.072 | 1 | 1 | Sf3a3 | -0.008 | Old |
| 0.530743 | 0.308287 | 0.135 | 0.127 | 1 | 1 | Psma4 | 0.008 | Old |
| 0.530934 | 0.069845 | 0.039 | 0.033 | 1 | 1 | Lrp12 | 0.006 | Old |
| 0.531144 | 0.130038 | 0.08 | 0.089 | 1 | 1 | 4933434E2 | -0.009 | Old |
| 0.531684 | 1.015977 | 0.014 | 0.011 | 1 | 1 | Ncs1 | 0.003 | Old |
| 0.531776 | 0.225526 | 0.14 | 0.132 | 1 | 1 | Ezr | 0.008 | Old |
| 0.531882 | 0.101233 | 0.048 | 0.054 | 1 | 1 | Lsr | -0.006 | Old |
| 0.531906 | 0.09503 | 0.112 | 0.123 | 1 | 1 | Chmp3 | -0.011 | Old |
| 0.531991 | 0.641269 | 0.041 | 0.036 | 1 | 1 | Banp | 0.005 | Old |
| 0.532189 | 0.841461 | 0.014 | 0.011 | 1 | 1 | Stn1 | 0.003 | Old |
| 0.532296 | 0.112018 | 0.239 | 0.261 | 1 | 1 | Selenow | -0.022 | Old |
| 0.532509 | 0.350078 | 0.125 | 0.118 | 1 | 1 | Api5 | 0.007 | Old |
| 0.533082 | 0.303299 | 0.064 | 0.072 | 1 | 1 | Mrpl13 | -0.008 | Old |
| 0.533127 | 0.021466 | 0.015 | 0.019 | 1 | 1 | Mcm7 | -0.004 | Old |
| 0.533239 | 0.008357 | 0.063 | 0.07 | 1 | 1 | Dyrk1a | -0.007 | Old |
| 0.533247 | 0.201012 | 0.154 | 0.146 | 1 | 1 | Med28 | 0.008 | Old |
| 0.533546 | 0.19318 | 0.015 | 0.019 | 1 | 1 | Eid3 | -0.004 | Old |
| 0.533692 | 0.104905 | 0.054 | 0.061 | 1 | 1 | Homer1 | -0.007 | Old |
| 0.533783 | 0.340411 | 0.017 | 0.021 | 1 | 1 | Zfp954 | -0.004 | Old |
| 0.534193 | 0.20752 | 0.188 | 0.181 | 1 | 1 | Itsn1 | 0.007 | Old |
| 0.534588 | 0.269568 | 0.091 | 0.101 | 1 | 1 | Mta2 | -0.01 | Old |
| 0.534737 | 0.033904 | 0.057 | 0.064 | 1 | 1 | Mob3a | -0.007 | Old |
| 0.534914 | 0.149224 | 0.042 | 0.037 | 1 | 1 | Gsto1 | 0.005 | Old |
| 0.535036 | 0.136874 | 0.19 | 0.208 | 1 | 1 | Atp6v1a | -0.018 | Old |
| 0.535078 | 0.440872 | 0.055 | 0.049 | 1 | 1 | Nsmce1 | 0.006 | Old |
| 0.535211 | 0.580169 | 0.023 | 0.019 | 1 | 1 | 2010111I0 | 0.004 | Old |
| 0.535225 | 0.159446 | 0.015 | 0.019 | 1 | 1 | Bzw2 | -0.004 | Old |
| 0.5355 | 0.083639 | 0.059 | 0.066 | 1 | 1 | Atxn3 | -0.007 | Old |
| 0.535924 | 0.020026 | 0.055 | 0.062 | 1 | 1 | Rnf220 | -0.007 | Old |
| 0.536188 | 0.325837 | 0.023 | 0.019 | 1 | 1 | Apip | 0.004 | Old |
| 0.536579 | 0.320079 | 0.018 | 0.022 | 1 | 1 | Zer1 | -0.004 | Old |
| 0.537228 | 0.177376 | 0.212 | 0.23 | 1 | 1 | Anp32b | -0.018 | Old |
| 0.537405 | 0.058216 | 0.179 | 0.192 | 1 | 1 | Nedd8 | -0.013 | Old |
| 0.537538 | 0.1543 | 0.053 | 0.06 | 1 | 1 | Arl6ip6 | -0.007 | Old |

| | | | | | | | | |
|---|---|---|---|---|---|---|---|---|
| 0.53771 | 0.647367 | 0.012 | 0.009 | 1 | 1 | Arhgef18 | 0.003 | Old |
| 0.537777 | 0.773519 | 0.027 | 0.023 | 1 | 1 | Itpa | 0.004 | Old |
| 0.537789 | 0.044709 | 0.042 | 0.037 | 1 | 1 | Ints3 | 0.005 | Old |
| 0.537909 | 0.012735 | 0.028 | 0.033 | 1 | 1 | Lmf2 | -0.005 | Old |
| 0.537911 | 0.277632 | 0.04 | 0.035 | 1 | 1 | Cdc42ep3 | 0.005 | Old |
| 0.538389 | 0.404025 | 0.03 | 0.025 | 1 | 1 | Mrpl19 | 0.005 | Old |
| 0.538487 | 0.07874 | 0.076 | 0.084 | 1 | 1 | Med1 | -0.008 | Old |
| 0.538556 | 0.989369 | 0.012 | 0.009 | 1 | 1 | Gstm5 | 0.003 | Old |
| 0.538589 | 0.117606 | 0.015 | 0.019 | 1 | 1 | Clp1 | -0.004 | Old |
| 0.538617 | 0.151248 | 0.069 | 0.078 | 1 | 1 | Ifitm6 | -0.009 | Old |
| 0.538745 | 0.896517 | 0.021 | 0.017 | 1 | 1 | Hdhd2 | 0.004 | Old |
| 0.538953 | 1.002261 | 0.021 | 0.017 | 1 | 1 | Sufu | 0.004 | Old |
| 0.539023 | 0.443481 | 0.033 | 0.029 | 1 | 1 | 4930455G( | 0.004 | Old |
| 0.539113 | 0.134128 | 0.45 | 0.453 | 1 | 1 | Morf4l1 | -0.003 | Old |
| 0.539212 | 0.319318 | 0.013 | 0.016 | 1 | 1 | Mag | -0.003 | Old |
| 0.539262 | 0.273776 | 0.049 | 0.055 | 1 | 1 | Gpd1l | -0.006 | Old |
| 0.539278 | 0.282855 | 0.014 | 0.011 | 1 | 1 | Kbtbd4 | 0.003 | Old |
| 0.539366 | 0.541616 | 0.033 | 0.039 | 1 | 1 | Zfp560 | -0.006 | Old |
| 0.539403 | 0.720179 | 0.012 | 0.009 | 1 | 1 | Deaf1 | 0.003 | Old |
| 0.539517 | 0.251 | 0.023 | 0.019 | 1 | 1 | Wdr45 | 0.004 | Old |
| 0.53957 | 0.288068 | 0.031 | 0.036 | 1 | 1 | Timm22 | -0.005 | Old |
| 0.539822 | 0.292779 | 0.014 | 0.018 | 1 | 1 | Fancl | -0.004 | Old |
| 0.540005 | 0.122771 | 0.13 | 0.141 | 1 | 1 | 2610001J0 | -0.011 | Old |
| 0.540064 | 0.391866 | 0.015 | 0.019 | 1 | 1 | Abhd18 | -0.004 | Old |
| 0.540267 | 0.140408 | 0.077 | 0.085 | 1 | 1 | Lats2 | -0.008 | Old |
| 0.540278 | 0.592552 | 0.041 | 0.036 | 1 | 1 | Slc25a20 | 0.005 | Old |
| 0.540589 | 0.072773 | 0.013 | 0.016 | 1 | 1 | Kbtbd3 | -0.003 | Old |
| 0.540798 | 0.206962 | 0.103 | 0.095 | 1 | 1 | S100a11 | 0.008 | Old |
| 0.540982 | 0.162806 | 0.03 | 0.025 | 1 | 1 | Fam20c | 0.005 | Old |
| 0.541523 | 0.374616 | 0.012 | 0.009 | 1 | 1 | Gm20513 | 0.003 | Old |
| 0.541577 | 0.001431 | 0.063 | 0.056 | 1 | 1 | Mrpl57 | 0.007 | Old |
| 0.541874 | 0.12271 | 0.023 | 0.019 | 1 | 1 | Lyrm2 | 0.004 | Old |
| 0.541927 | 0.622995 | 0.045 | 0.04 | 1 | 1 | Cul2 | 0.005 | Old |
| 0.542089 | 0.280392 | 0.012 | 0.009 | 1 | 1 | Ehbp1 | 0.003 | Old |
| 0.542514 | 0.820463 | 0.012 | 0.009 | 1 | 1 | Inpp1 | 0.003 | Old |
| 0.5427 | 0.422198 | 0.08 | 0.073 | 1 | 1 | Lrrk1 | 0.007 | Old |
| 0.542797 | 0.198072 | 0.012 | 0.009 | 1 | 1 | Med26 | 0.003 | Old |
| 0.543357 | 0.094773 | 0.145 | 0.158 | 1 | 1 | Wdr45b | -0.013 | Old |
| 0.543563 | 0.325913 | 0.063 | 0.057 | 1 | 1 | Gtf2h1 | 0.006 | Old |
| 0.54362 | 0.033509 | 0.073 | 0.081 | 1 | 1 | Vps13d | -0.008 | Old |
| 0.543691 | 0.01006 | 0.062 | 0.069 | 1 | 1 | Ptbp2 | -0.007 | Old |
| 0.543785 | 0.18994 | 0.04 | 0.035 | 1 | 1 | Ero1lb | 0.005 | Old |
| 0.544048 | 0.941558 | 0.024 | 0.021 | 1 | 1 | Mettl21a | 0.003 | Old |
| 0.544214 | 0.568332 | 0.012 | 0.009 | 1 | 1 | Mrps31 | 0.003 | Old |
| 0.544328 | 0.091458 | 0.012 | 0.015 | 1 | 1 | Spsb3 | -0.003 | Old |

| | | | | | | | | |
|---|---|---|---|---|---|---|---|---|
| 0.544562 | 0.105007 | 0.026 | 0.03 | 1 | 1 | Foxk2 | -0.004 | Old |
| 0.544623 | 0.084381 | 0.03 | 0.025 | 1 | 1 | Lilra6 | 0.005 | Old |
| 0.544632 | 0.216794 | 0.028 | 0.033 | 1 | 1 | Pabpc4 | -0.005 | Old |
| 0.54487 | 0.28702 | 0.048 | 0.042 | 1 | 1 | Gbf1 | 0.006 | Old |
| 0.545004 | 0.04225 | 0.04 | 0.046 | 1 | 1 | Pmpcb | -0.006 | Old |
| 0.545054 | 0.241011 | 0.012 | 0.015 | 1 | 1 | Tdp1 | -0.003 | Old |
| 0.545162 | 0.030546 | 0.08 | 0.087 | 1 | 1 | Yipf3 | -0.007 | Old |
| 0.545212 | 0.150251 | 0.058 | 0.065 | 1 | 1 | Mrpl51 | -0.007 | Old |
| 0.545599 | 0.147915 | 0.072 | 0.08 | 1 | 1 | Hpf1 | -0.008 | Old |
| 0.545819 | 1.202657 | 0.018 | 0.015 | 1 | 1 | Cd4 | 0.003 | Old |
| 0.545917 | 0.587146 | 0.012 | 0.009 | 1 | 1 | Gemin6 | 0.003 | Old |
| 0.546156 | 0.093779 | 0.066 | 0.073 | 1 | 1 | Dgkz | -0.007 | Old |
| 0.546575 | 0.170958 | 0.008 | 0.01 | 1 | 1 | Gm43936 | -0.002 | Old |
| 0.546731 | 0.106842 | 0.041 | 0.047 | 1 | 1 | Adam33 | -0.006 | Old |
| 0.54684 | 0.327646 | 0.051 | 0.046 | 1 | 1 | Prkaca | 0.005 | Old |
| 0.546925 | 0.316602 | 0.096 | 0.108 | 1 | 1 | Stx12 | -0.012 | Old |
| 0.547298 | 0.216611 | 0.009 | 0.012 | 1 | 1 | Cnst | -0.003 | Old |
| 0.547449 | 0.189982 | 0.008 | 0.01 | 1 | 1 | Wars2 | -0.002 | Old |
| 0.547469 | 0.166609 | 0.247 | 0.242 | 1 | 1 | Sumo2 | 0.005 | Old |
| 0.54776 | 0.242925 | 0.026 | 0.03 | 1 | 1 | Miip | -0.004 | Old |
| 0.547927 | 0.185775 | 0.139 | 0.153 | 1 | 1 | Smim10l1 | -0.014 | Old |
| 0.547933 | 0.603758 | 0.021 | 0.017 | 1 | 1 | Gpt2 | 0.004 | Old |
| 0.548032 | 0.296229 | 0.008 | 0.01 | 1 | 1 | BC017158 | -0.002 | Old |
| 0.54812 | 0.425833 | 0.069 | 0.064 | 1 | 1 | Mrps15 | 0.005 | Old |
| 0.548323 | 4.87E-05 | 0.008 | 0.01 | 1 | 1 | Herc3 | -0.002 | Old |
| 0.548332 | 0.27428 | 0.028 | 0.033 | 1 | 1 | Slc9a8 | -0.005 | Old |
| 0.548354 | 0.340375 | 0.044 | 0.05 | 1 | 1 | Fibp | -0.006 | Old |
| 0.548557 | 0.364833 | 0.063 | 0.057 | 1 | 1 | Lrrfip2 | 0.006 | Old |
| 0.548591 | 0.30209 | 0.01 | 0.013 | 1 | 1 | Gys1 | -0.003 | Old |
| 0.548591 | 0.113546 | 0.01 | 0.013 | 1 | 1 | Ift27 | -0.003 | Old |
| 0.548615 | 0.033915 | 0.008 | 0.01 | 1 | 1 | Snapc2 | -0.002 | Old |
| 0.54891 | 0.291602 | 0.03 | 0.035 | 1 | 1 | Yod1 | -0.005 | Old |
| 0.549192 | 0.245754 | 0.021 | 0.017 | 1 | 1 | Rcc1l | 0.004 | Old |
| 0.549203 | 0.324977 | 0.033 | 0.029 | 1 | 1 | Rcan1 | 0.004 | Old |
| 0.549533 | 0.19008 | 0.225 | 0.221 | 1 | 1 | Tnfrsf1b | 0.004 | Old |
| 0.549693 | 0.102403 | 0.158 | 0.171 | 1 | 1 | Cebpg | -0.013 | Old |
| 0.550043 | 0.336189 | 0.042 | 0.049 | 1 | 1 | Hacd3 | -0.007 | Old |
| 0.550074 | 0.152596 | 0.008 | 0.01 | 1 | 1 | Gmpr | -0.002 | Old |
| 0.5501 | 0.323763 | 0.203 | 0.199 | 1 | 1 | Mcrip1 | 0.004 | Old |
| 0.55024 | 0.45773 | 0.12 | 0.114 | 1 | 1 | Etfb | 0.006 | Old |
| 0.550246 | 0.029989 | 0.17 | 0.183 | 1 | 1 | Cd164 | -0.013 | Old |
| 0.55039 | 0.468019 | 0.012 | 0.015 | 1 | 1 | Dnajc27 | -0.003 | Old |
| 0.550553 | 0.494171 | 0.08 | 0.073 | 1 | 1 | Pfkp | 0.007 | Old |
| 0.550969 | 0.584848 | 0.013 | 0.016 | 1 | 1 | Ankrd54 | -0.003 | Old |
| 0.551032 | 0.018319 | 0.062 | 0.069 | 1 | 1 | Nae1 | -0.007 | Old |

| | | | | | | | | |
|---|---|---|---|---|---|---|---|---|
| 0.551172 | 0.102784 | 0.037 | 0.043 | 1 | 1 | Tmub2 | -0.006 | Old |
| 0.551354 | 0.578048 | 0.035 | 0.03 | 1 | 1 | Slc25a16 | 0.005 | Old |
| 0.551457 | 0.124046 | 0.077 | 0.086 | 1 | 1 | Msrb1 | -0.009 | Old |
| 0.551536 | 0.615796 | 0.008 | 0.01 | 1 | 1 | Ccdc138 | -0.002 | Old |
| 0.551581 | 0.199067 | 0.027 | 0.023 | 1 | 1 | Zbed4 | 0.004 | Old |
| 0.551674 | 0.402342 | 0.077 | 0.071 | 1 | 1 | Psen2 | 0.006 | Old |
| 0.552028 | 0.014191 | 0.085 | 0.093 | 1 | 1 | Eno1 | -0.008 | Old |
| 0.552723 | 0.065474 | 0.188 | 0.202 | 1 | 1 | Ikbkb | -0.014 | Old |
| 0.552796 | 0.596315 | 0.054 | 0.049 | 1 | 1 | Naa35 | 0.005 | Old |
| 0.552941 | 0.039636 | 0.162 | 0.174 | 1 | 1 | Pacs2 | -0.012 | Old |
| 0.55325 | 0.149692 | 0.036 | 0.041 | 1 | 1 | Slc5a3 | -0.005 | Old |
| 0.553294 | 0.048969 | 0.263 | 0.284 | 1 | 1 | Rab1a | -0.021 | Old |
| 0.553504 | 0.065469 | 0.026 | 0.03 | 1 | 1 | Eya3 | -0.004 | Old |
| 0.553618 | 0.325298 | 0.027 | 0.032 | 1 | 1 | Msra | -0.005 | Old |
| 0.553895 | 0.204096 | 0.15 | 0.142 | 1 | 1 | Eloc | 0.008 | Old |
| 0.554267 | 0.108086 | 0.026 | 0.03 | 1 | 1 | Trim3 | -0.004 | Old |
| 0.554361 | 0.778454 | 0.018 | 0.015 | 1 | 1 | Vegfa | 0.003 | Old |
| 0.554687 | 0.037131 | 0.206 | 0.225 | 1 | 1 | Tnip3 | -0.019 | Old |
| 0.555076 | 0.18197 | 0.073 | 0.081 | 1 | 1 | Maff | -0.008 | Old |
| 0.555117 | 0.060268 | 0.111 | 0.12 | 1 | 1 | Ppp4r2 | -0.009 | Old |
| 0.555211 | 0.376487 | 0.009 | 0.012 | 1 | 1 | Mcc | -0.003 | Old |
| 0.555928 | 0.188301 | 0.18 | 0.172 | 1 | 1 | Gna13 | 0.008 | Old |
| 0.556014 | 0.326396 | 0.033 | 0.029 | 1 | 1 | Golim4 | 0.004 | Old |
| 0.556124 | 0.30075 | 0.039 | 0.044 | 1 | 1 | Ddx39 | -0.005 | Old |
| 0.556413 | 0.199734 | 0.116 | 0.128 | 1 | 1 | Gdi1 | -0.012 | Old |
| 0.55702 | 0.442406 | 0.042 | 0.049 | 1 | 1 | Bex3 | -0.007 | Old |
| 0.55705 | 0.13015 | 0.111 | 0.103 | 1 | 1 | Riok3 | 0.008 | Old |
| 0.557229 | 0.241221 | 0.028 | 0.033 | 1 | 1 | Uba7 | -0.005 | Old |
| 0.557466 | 0.095311 | 0.036 | 0.041 | 1 | 1 | Nol11 | -0.005 | Old |
| 0.557495 | 0.369337 | 0.068 | 0.062 | 1 | 1 | Zc3h4 | 0.006 | Old |
| 0.557892 | 0.20464 | 0.229 | 0.222 | 1 | 1 | Rnf141 | 0.007 | Old |
| 0.558513 | 0.173802 | 0.075 | 0.083 | 1 | 1 | Esyt2 | -0.008 | Old |
| 0.5588 | 0.168414 | 0.175 | 0.191 | 1 | 1 | Strn3 | -0.016 | Old |
| 0.559012 | 0.182016 | 0.086 | 0.079 | 1 | 1 | Smg7 | 0.007 | Old |
| 0.559119 | 0.301083 | 0.041 | 0.036 | 1 | 1 | Plcb3 | 0.005 | Old |
| 0.559565 | 0.083501 | 0.024 | 0.029 | 1 | 1 | Plekha3 | -0.005 | Old |
| 0.559565 | 0.100605 | 0.024 | 0.029 | 1 | 1 | Ccdc117 | -0.005 | Old |
| 0.559787 | 0.274245 | 0.041 | 0.036 | 1 | 1 | Isca2 | 0.005 | Old |
| 0.559851 | 0.841342 | 0.032 | 0.028 | 1 | 1 | Gm5547 | 0.004 | Old |
| 0.560361 | 0.308574 | 0.057 | 0.051 | 1 | 1 | Ccz1 | 0.006 | Old |
| 0.560469 | 0.743773 | 0.018 | 0.015 | 1 | 1 | Mllt1 | 0.003 | Old |
| 0.56048 | 0.041005 | 0.027 | 0.023 | 1 | 1 | Parp10 | 0.004 | Old |
| 0.560928 | 0.172564 | 0.078 | 0.086 | 1 | 1 | Gm47283 | -0.008 | Old |
| 0.561153 | 0.36811 | 0.026 | 0.03 | 1 | 1 | Nom1 | -0.004 | Old |
| 0.561232 | 0.072455 | 0.093 | 0.101 | 1 | 1 | Chtop | -0.008 | Old |

| | | | | | | | | |
|---|---|---|---|---|---|---|---|---|
| 0.561356 | 0.304477 | 0.123 | 0.117 | 1 | 1 | Uba2 | 0.006 | Old |
| 0.561566 | 0.111613 | 0.073 | 0.066 | 1 | 1 | Slc37a2 | 0.007 | Old |
| 0.561681 | 0.290348 | 0.254 | 0.254 | 1 | 1 | Hdgf | 0 | Old |
| 0.561704 | 0.199655 | 0.138 | 0.131 | 1 | 1 | Mcfd2 | 0.007 | Old |
| 0.56209 | 0.530079 | 0.027 | 0.032 | 1 | 1 | Narf | -0.005 | Old |
| 0.562092 | 0.66618 | 0.028 | 0.024 | 1 | 1 | Iigp1 | 0.004 | Old |
| 0.562122 | 0.534122 | 0.024 | 0.021 | 1 | 1 | Tex2 | 0.003 | Old |
| 0.562265 | 0.33311 | 0.051 | 0.046 | 1 | 1 | Copg1 | 0.005 | Old |
| 0.562285 | 0.730314 | 0.018 | 0.015 | 1 | 1 | Atp1a3 | 0.003 | Old |
| 0.562428 | 0.008939 | 0.033 | 0.038 | 1 | 1 | Agpat1 | -0.005 | Old |
| 0.563529 | 0.204274 | 0.023 | 0.027 | 1 | 1 | 2310009B1 | -0.004 | Old |
| 0.563673 | 0.428054 | 0.024 | 0.021 | 1 | 1 | Ppil3 | 0.003 | Old |
| 0.56368 | 0.07793 | 0.105 | 0.115 | 1 | 1 | Tulp4 | -0.01 | Old |
| 0.563779 | 0.270976 | 0.127 | 0.12 | 1 | 1 | Gpbp1 | 0.007 | Old |
| 0.563888 | 0.136471 | 0.023 | 0.027 | 1 | 1 | Zscan29 | -0.004 | Old |
| 0.564017 | 0.279598 | 0.031 | 0.027 | 1 | 1 | Kif1bp | 0.004 | Old |
| 0.5646 | 0.346871 | 0.039 | 0.044 | 1 | 1 | Mrpl9 | -0.005 | Old |
| 0.564644 | 0.396751 | 0.024 | 0.021 | 1 | 1 | Alg11 | 0.003 | Old |
| 0.565301 | 0.008603 | 0.067 | 0.075 | 1 | 1 | Ankib1 | -0.008 | Old |
| 0.565474 | 0.293245 | 0.117 | 0.111 | 1 | 1 | Plekhg3 | 0.006 | Old |
| 0.565514 | 0.13633 | 0.024 | 0.029 | 1 | 1 | Hook2 | -0.005 | Old |
| 0.565657 | 0.12728 | 0.221 | 0.239 | 1 | 1 | Bzw1 | -0.018 | Old |
| 0.565977 | 0.173371 | 0.098 | 0.107 | 1 | 1 | Wdr89 | -0.009 | Old |
| 0.566005 | 0.233131 | 0.045 | 0.04 | 1 | 1 | Faf2 | 0.005 | Old |
| 0.566587 | 0.478025 | 0.024 | 0.021 | 1 | 1 | B4gat1 | 0.003 | Old |
| 0.567123 | 0.255111 | 0.031 | 0.027 | 1 | 1 | Gnpda2 | 0.004 | Old |
| 0.567222 | 0.093491 | 0.154 | 0.165 | 1 | 1 | Batf3 | -0.011 | Old |
| 0.56727 | 0.408427 | 0.024 | 0.029 | 1 | 1 | Yae1d1 | -0.005 | Old |
| 0.567293 | 0.94687 | 0.015 | 0.013 | 1 | 1 | Oxnad1 | 0.002 | Old |
| 0.567297 | 0.200885 | 0.068 | 0.062 | 1 | 1 | Atg5 | 0.006 | Old |
| 0.567536 | 0.21103 | 0.123 | 0.116 | 1 | 1 | Agtrap | 0.007 | Old |
| 0.568145 | 0.329097 | 0.024 | 0.021 | 1 | 1 | Arl14ep | 0.003 | Old |
| 0.568306 | 0.067202 | 0.037 | 0.032 | 1 | 1 | Gyg | 0.005 | Old |
| 0.568359 | 0.213298 | 0.035 | 0.04 | 1 | 1 | Rab28 | -0.005 | Old |
| 0.568812 | 0.099778 | 0.069 | 0.063 | 1 | 1 | Slc28a2 | 0.006 | Old |
| 0.569195 | 0.20414 | 0.073 | 0.067 | 1 | 1 | Rhobtb1 | 0.006 | Old |
| 0.5701 | 0.443123 | 0.125 | 0.119 | 1 | 1 | Dbnl | 0.006 | Old |
| 0.570185 | 0.017707 | 0.033 | 0.038 | 1 | 1 | Phb | -0.005 | Old |
| 0.570521 | 0.405739 | 0.035 | 0.03 | 1 | 1 | Mrps18a | 0.005 | Old |
| 0.570768 | 0.352876 | 0.105 | 0.1 | 1 | 1 | Skp1a | 0.005 | Old |
| 0.570798 | 0.22833 | 0.221 | 0.218 | 1 | 1 | Cndp2 | 0.003 | Old |
| 0.571701 | 0.537041 | 0.064 | 0.059 | 1 | 1 | Arrdc4 | 0.005 | Old |
| 0.571894 | 0.067752 | 0.107 | 0.116 | 1 | 1 | Usp4 | -0.009 | Old |
| 0.571963 | 0.081386 | 0.194 | 0.209 | 1 | 1 | Slk | -0.015 | Old |
| 0.571984 | 0.571387 | 0.055 | 0.05 | 1 | 1 | Cab39l | 0.005 | Old |

| | | | | | | | | |
|---|---|---|---|---|---|---|---|---|
| 0.572169 | 0.096372 | 0.694 | 0.73 | 1 | 1 | Cebpb | -0.036 | Old |
| 0.572933 | 0.16184 | 0.042 | 0.048 | 1 | 1 | Hcfc2 | -0.006 | Old |
| 0.573167 | 0.108775 | 0.022 | 0.026 | 1 | 1 | Flt3l | -0.004 | Old |
| 0.57323 | 1.191566 | 0.015 | 0.013 | 1 | 1 | Ptgir | 0.002 | Old |
| 0.574098 | 0.428572 | 0.015 | 0.013 | 1 | 1 | Zfp944 | 0.002 | Old |
| 0.574362 | 0.459564 | 0.032 | 0.028 | 1 | 1 | Anapc15 | 0.004 | Old |
| 0.574605 | 0.214207 | 0.035 | 0.03 | 1 | 1 | Ppp1r13b | 0.005 | Old |
| 0.574651 | 0.145817 | 0.117 | 0.128 | 1 | 1 | Phf20l1 | -0.011 | Old |
| 0.575096 | 0.342241 | 0.035 | 0.03 | 1 | 1 | Crls1 | 0.005 | Old |
| 0.575283 | 0.11205 | 0.077 | 0.085 | 1 | 1 | Ap3m1 | -0.008 | Old |
| 0.575905 | 0.153977 | 0.522 | 0.558 | 1 | 1 | Tmem176b | -0.036 | Old |
| 0.576019 | 0.466264 | 0.036 | 0.032 | 1 | 1 | Gopc | 0.004 | Old |
| 0.576459 | 0.251124 | 0.102 | 0.095 | 1 | 1 | Zzef1 | 0.007 | Old |
| 0.576629 | 0.300326 | 0.077 | 0.072 | 1 | 1 | Plekhg5 | 0.005 | Old |
| 0.577042 | 0.358523 | 0.102 | 0.095 | 1 | 1 | Mfhas1 | 0.007 | Old |
| 0.577311 | 0.102475 | 0.058 | 0.064 | 1 | 1 | Abca7 | -0.006 | Old |
| 0.577472 | 0.359325 | 0.049 | 0.055 | 1 | 1 | Ctns | -0.006 | Old |
| 0.577629 | 0.424007 | 0.028 | 0.024 | 1 | 1 | Zfp839 | 0.004 | Old |
| 0.577789 | 0.391893 | 0.163 | 0.16 | 1 | 1 | Cers2 | 0.003 | Old |
| 0.578065 | 0.100179 | 0.05 | 0.056 | 1 | 1 | Ilkap | -0.006 | Old |
| 0.578111 | 0.072744 | 0.022 | 0.026 | 1 | 1 | Zfp994 | -0.004 | Old |
| 0.578441 | 0.013931 | 0.057 | 0.063 | 1 | 1 | Osbp | -0.006 | Old |
| 0.578449 | 0.508789 | 0.015 | 0.013 | 1 | 1 | Tmem185a | 0.002 | Old |
| 0.578702 | 0.159435 | 0.035 | 0.03 | 1 | 1 | Ppa1 | 0.005 | Old |
| 0.578709 | 0.239929 | 0.073 | 0.081 | 1 | 1 | Cdk12 | -0.008 | Old |
| 0.578919 | 0.055832 | 0.039 | 0.044 | 1 | 1 | C2cd5 | -0.005 | Old |
| 0.5797 | 0.156879 | 0.293 | 0.291 | 1 | 1 | M6pr | 0.002 | Old |
| 0.580589 | 0.006859 | 0.077 | 0.084 | 1 | 1 | Mark3 | -0.007 | Old |
| 0.580693 | 0.401823 | 0.015 | 0.013 | 1 | 1 | Rtn4rl1 | 0.002 | Old |
| 0.581169 | 0.074057 | 0.021 | 0.024 | 1 | 1 | B3gnt8 | -0.003 | Old |
| 0.581441 | 0.899295 | 0.026 | 0.022 | 1 | 1 | Cep295 | 0.004 | Old |
| 0.581789 | 0.043229 | 0.031 | 0.035 | 1 | 1 | Abcd4 | -0.004 | Old |
| 0.58255 | 0.149993 | 0.17 | 0.185 | 1 | 1 | Dusp6 | -0.015 | Old |
| 0.582555 | 0.191707 | 0.076 | 0.083 | 1 | 1 | Trio | -0.007 | Old |
| 0.582836 | 0.219077 | 0.13 | 0.123 | 1 | 1 | H2-Q7 | 0.007 | Old |
| 0.582944 | 0.007244 | 0.019 | 0.023 | 1 | 1 | Zfp830 | -0.004 | Old |
| 0.582994 | 0.365607 | 0.058 | 0.052 | 1 | 1 | Kdm3a | 0.006 | Old |
| 0.583041 | 0.279994 | 0.053 | 0.059 | 1 | 1 | Itfg1 | -0.006 | Old |
| 0.583874 | 0.198092 | 0.095 | 0.088 | 1 | 1 | Safb | 0.007 | Old |
| 0.583905 | 1.210463 | 0.013 | 0.01 | 1 | 1 | Rbm10 | 0.003 | Old |
| 0.583969 | 0.085006 | 0.021 | 0.024 | 1 | 1 | Bcl2l12 | -0.003 | Old |
| 0.584633 | 0.171123 | 0.109 | 0.119 | 1 | 1 | Mtch1 | -0.01 | Old |
| 0.585268 | 0.78678 | 0.026 | 0.022 | 1 | 1 | Rufy2 | 0.004 | Old |
| 0.585442 | 0.346899 | 0.015 | 0.013 | 1 | 1 | 0610010K1 | 0.002 | Old |
| 0.585578 | 0.160347 | 0.163 | 0.177 | 1 | 1 | Slfn5 | -0.014 | Old |

| | | | | | | | | |
|---|---|---|---|---|---|---|---|---|
| 0.585663 | 0.194959 | 0.153 | 0.167 | 1 | 1 | Man1c1 | -0.014 | Old |
| 0.585914 | 0.441418 | 0.096 | 0.091 | 1 | 1 | Dcp2 | 0.005 | Old |
| 0.586132 | 0.448118 | 0.032 | 0.028 | 1 | 1 | Rtl5 | 0.004 | Old |
| 0.586141 | 0.058857 | 0.233 | 0.248 | 1 | 1 | Abi1 | -0.015 | Old |
| 0.586181 | 0.021992 | 0.076 | 0.083 | 1 | 1 | Paics | -0.007 | Old |
| 0.586407 | 0.305945 | 0.057 | 0.064 | 1 | 1 | Soga1 | -0.007 | Old |
| 0.586452 | 0.428327 | 0.044 | 0.039 | 1 | 1 | Tmem11 | 0.005 | Old |
| 0.586527 | 0.466211 | 0.04 | 0.035 | 1 | 1 | BC029722 | 0.005 | Old |
| 0.587056 | 0.476323 | 0.103 | 0.097 | 1 | 1 | 2510039O1 | 0.006 | Old |
| 0.587238 | 0.27294 | 0.319 | 0.326 | 1 | 1 | Ywhah | -0.007 | Old |
| 0.587467 | 0.152702 | 0.171 | 0.182 | 1 | 1 | Ranbp2 | -0.011 | Old |
| 0.587478 | 0.039119 | 0.04 | 0.045 | 1 | 1 | Nelfa | -0.005 | Old |
| 0.587633 | 0.010811 | 0.096 | 0.105 | 1 | 1 | Kdm2b | -0.009 | Old |
| 0.587847 | 0.365805 | 0.13 | 0.124 | 1 | 1 | Rer1 | 0.006 | Old |
| 0.588135 | 0.161396 | 0.019 | 0.023 | 1 | 1 | Ndufaf2 | -0.004 | Old |
| 0.588183 | 0.068923 | 0.073 | 0.081 | 1 | 1 | Pin4 | -0.008 | Old |
| 0.588328 | 0.193368 | 0.104 | 0.097 | 1 | 1 | Itga9 | 0.007 | Old |
| 0.588586 | 0.298351 | 0.093 | 0.086 | 1 | 1 | Tspan32 | 0.007 | Old |
| 0.588779 | 0.30936 | 0.051 | 0.047 | 1 | 1 | Get4 | 0.004 | Old |
| 0.58891 | 0.570995 | 0.021 | 0.024 | 1 | 1 | Gfm1 | -0.003 | Old |
| 0.588978 | 0.265999 | 0.09 | 0.098 | 1 | 1 | Ddx24 | -0.008 | Old |
| 0.589167 | 0.360558 | 0.062 | 0.056 | 1 | 1 | Lrrc59 | 0.006 | Old |
| 0.589418 | 0.145684 | 0.13 | 0.14 | 1 | 1 | Atp5o.1 | -0.01 | Old |
| 0.589536 | 0.01384 | 0.019 | 0.023 | 1 | 1 | Ikzf2 | -0.004 | Old |
| 0.589736 | 0.232362 | 0.032 | 0.028 | 1 | 1 | 2310033P0 | 0.004 | Old |
| 0.589846 | 0.123393 | 0.075 | 0.082 | 1 | 1 | Psmb2 | -0.007 | Old |
| 0.590372 | 0.001127 | 0.116 | 0.124 | 1 | 1 | Sar1a | -0.008 | Old |
| 0.590475 | 0.310426 | 0.051 | 0.047 | 1 | 1 | Epb41l1 | 0.004 | Old |
| 0.590514 | 0.695523 | 0.013 | 0.01 | 1 | 1 | Them4 | 0.003 | Old |
| 0.590658 | 0.297349 | 0.021 | 0.024 | 1 | 1 | Flnb | -0.003 | Old |
| 0.590956 | 0.158981 | 0.066 | 0.072 | 1 | 1 | Lrrc8d | -0.006 | Old |
| 0.590964 | 0.016446 | 0.015 | 0.013 | 1 | 1 | Dolk | 0.002 | Old |
| 0.59114 | 0.333871 | 0.019 | 0.023 | 1 | 1 | Cluh | -0.004 | Old |
| 0.591435 | 0.347457 | 0.018 | 0.021 | 1 | 1 | Ubap2 | -0.003 | Old |
| 0.591481 | 0.557542 | 0.013 | 0.01 | 1 | 1 | Ercc8 | 0.003 | Old |
| 0.591716 | 0.029872 | 0.096 | 0.105 | 1 | 1 | Atp8a1 | -0.009 | Old |
| 0.591895 | 0.702895 | 0.013 | 0.01 | 1 | 1 | Sccpdh | 0.003 | Old |
| 0.591909 | 0.696392 | 0.042 | 0.038 | 1 | 1 | Zdhhc7 | 0.004 | Old |
| 0.59215 | 0.60906 | 0.085 | 0.081 | 1 | 1 | Slf2 | 0.004 | Old |
| 0.592171 | 0.778875 | 0.013 | 0.01 | 1 | 1 | Mapkbp1 | 0.003 | Old |
| 0.592171 | 0.486469 | 0.013 | 0.01 | 1 | 1 | Zfyve21 | 0.003 | Old |
| 0.59268 | 0.101441 | 0.018 | 0.021 | 1 | 1 | Wfs1 | -0.003 | Old |
| 0.592765 | 0.521871 | 0.026 | 0.022 | 1 | 1 | Akap7 | 0.004 | Old |
| 0.593001 | 0.21275 | 0.013 | 0.01 | 1 | 1 | Urb1 | 0.003 | Old |
| 0.593054 | 0.475036 | 0.026 | 0.022 | 1 | 1 | Rnaseh2a | 0.004 | Old |

| | | | | | | | | |
|---|---|---|---|---|---|---|---|---|
| 0.593178 | 0.093192 | 0.058 | 0.052 | 1 | 1 | Rsl24d1 | 0.006 | Old |
| 0.593706 | 0.285887 | 0.216 | 0.213 | 1 | 1 | Dnajc3 | 0.003 | Old |
| 0.59375 | 0.40429 | 0.019 | 0.023 | 1 | 1 | Dnajb2 | -0.004 | Old |
| 0.59385 | 0.240096 | 0.112 | 0.121 | 1 | 1 | Evl | -0.009 | Old |
| 0.594358 | 0.136261 | 0.342 | 0.377 | 1 | 1 | Serbp1 | -0.035 | Old |
| 0.594368 | 0.134168 | 0.054 | 0.049 | 1 | 1 | Lpar1 | 0.005 | Old |
| 0.594401 | 0.160498 | 0.058 | 0.052 | 1 | 1 | Cops8 | 0.006 | Old |
| 0.595216 | 0.468785 | 0.013 | 0.01 | 1 | 1 | Sco1 | 0.003 | Old |
| 0.595216 | 0.357194 | 0.013 | 0.01 | 1 | 1 | Btg3 | 0.003 | Old |
| 0.595447 | 0.395011 | 0.057 | 0.052 | 1 | 1 | Oasl2 | 0.005 | Old |
| 0.595707 | 0.578966 | 0.03 | 0.026 | 1 | 1 | Ccdc43 | 0.004 | Old |
| 0.595795 | 0.190622 | 0.031 | 0.035 | 1 | 1 | Sec23a | -0.004 | Old |
| 0.595988 | 0.327721 | 0.037 | 0.042 | 1 | 1 | Cfl2 | -0.005 | Old |
| 0.596555 | 0.37584 | 0.126 | 0.12 | 1 | 1 | Ubash3b | 0.006 | Old |
| 0.596712 | 0.035044 | 0.109 | 0.118 | 1 | 1 | Suz12 | -0.009 | Old |
| 0.596818 | 0.074134 | 0.159 | 0.17 | 1 | 1 | Arhgap9 | -0.011 | Old |
| 0.596969 | 0.250454 | 0.019 | 0.023 | 1 | 1 | Atg2b | -0.004 | Old |
| 0.597005 | 0.39654 | 0.165 | 0.161 | 1 | 1 | Smim15 | 0.004 | Old |
| 0.597018 | 0.152503 | 0.013 | 0.01 | 1 | 1 | Serpinb1a | 0.003 | Old |
| 0.597077 | 0.24485 | 0.017 | 0.02 | 1 | 1 | Mettl3 | -0.003 | Old |
| 0.597226 | 0.779874 | 0.019 | 0.016 | 1 | 1 | Timm21 | 0.003 | Old |
| 0.597226 | 0.59189 | 0.019 | 0.016 | 1 | 1 | Bola1 | 0.003 | Old |
| 0.597482 | 0.165252 | 0.256 | 0.278 | 1 | 1 | Nucks1 | -0.022 | Old |
| 0.598031 | 0.373785 | 0.036 | 0.032 | 1 | 1 | Rpf1 | 0.004 | Old |
| 0.598039 | 0.556491 | 0.064 | 0.059 | 1 | 1 | Htatsf1 | 0.005 | Old |
| 0.598251 | 0.277914 | 0.049 | 0.055 | 1 | 1 | Babam1 | -0.006 | Old |
| 0.598309 | 1.403169 | 0.01 | 0.008 | 1 | 1 | Timp3 | 0.002 | Old |
| 0.598504 | 0.24482 | 0.018 | 0.021 | 1 | 1 | Tmem184c | -0.003 | Old |
| 0.598585 | 0.263295 | 0.094 | 0.088 | 1 | 1 | Nck1 | 0.006 | Old |
| 0.599099 | 0.056945 | 0.121 | 0.131 | 1 | 1 | Rnf7 | -0.01 | Old |
| 0.599657 | 0.338513 | 0.013 | 0.01 | 1 | 1 | Poc5 | 0.003 | Old |
| 0.599678 | 0.037265 | 0.075 | 0.082 | 1 | 1 | Atf1 | -0.007 | Old |
| 0.599868 | 0.057315 | 0.122 | 0.131 | 1 | 1 | Thrap3 | -0.009 | Old |
| 0.599949 | 0.04049 | 0.262 | 0.279 | 1 | 1 | Set | -0.017 | Old |
| 0.600157 | 0.274977 | 0.068 | 0.063 | 1 | 1 | Suds3 | 0.005 | Old |
| 0.600543 | 0.490686 | 0.071 | 0.066 | 1 | 1 | Qpct | 0.005 | Old |
| 0.600593 | 0.546475 | 0.019 | 0.016 | 1 | 1 | Tmed8 | 0.003 | Old |
| 0.601148 | 0.332633 | 0.031 | 0.035 | 1 | 1 | Rpp25l | -0.004 | Old |
| 0.60171 | 0.316884 | 0.19 | 0.188 | 1 | 1 | Tmem256 | 0.002 | Old |
| 0.601947 | 0.494846 | 0.041 | 0.037 | 1 | 1 | Gsr | 0.004 | Old |
| 0.602261 | 0.456368 | 0.018 | 0.021 | 1 | 1 | Clpp | -0.003 | Old |
| 0.602635 | 0.171631 | 0.026 | 0.022 | 1 | 1 | Gprasp1 | 0.004 | Old |
| 0.602955 | 0.22421 | 0.243 | 0.243 | 1 | 1 | Csf2ra | 0 | Old |
| 0.603681 | 0.152284 | 0.135 | 0.128 | 1 | 1 | Pld3 | 0.007 | Old |
| 0.60456 | 1.2632 | 0.01 | 0.008 | 1 | 1 | Fdxacb1 | 0.002 | Old |

| | | | | | | | | |
|---|---|---|---|---|---|---|---|---|
| 0.604755 | 0.156723 | 0.028 | 0.032 | 1 | 1 | Hmces | -0.004 | Old |
| 0.604968 | 0.242811 | 0.026 | 0.022 | 1 | 1 | Tpcn2 | 0.004 | Old |
| 0.605096 | 0.30997 | 0.019 | 0.016 | 1 | 1 | Tbl3 | 0.003 | Old |
| 0.605268 | 0.329246 | 0.03 | 0.026 | 1 | 1 | Prmt3 | 0.004 | Old |
| 0.605268 | 0.265998 | 0.03 | 0.026 | 1 | 1 | Mrpl55 | 0.004 | Old |
| 0.6055 | 0.699396 | 0.01 | 0.008 | 1 | 1 | Slc12a5 | 0.002 | Old |
| 0.605518 | 0.309765 | 0.104 | 0.099 | 1 | 1 | Ogfr | 0.005 | Old |
| 0.605551 | 0.063083 | 0.026 | 0.022 | 1 | 1 | Exoc8 | 0.004 | Old |
| 0.605678 | 0.206883 | 0.08 | 0.074 | 1 | 1 | Znrd1 | 0.006 | Old |
| 0.60569 | 0.102384 | 0.028 | 0.032 | 1 | 1 | Arhgef7 | -0.004 | Old |
| 0.605735 | 0.374045 | 0.017 | 0.02 | 1 | 1 | Drg2 | -0.003 | Old |
| 0.605772 | 0.311065 | 0.019 | 0.016 | 1 | 1 | Tmem80 | 0.003 | Old |
| 0.606211 | 0.337967 | 0.107 | 0.101 | 1 | 1 | Vapb | 0.006 | Old |
| 0.606308 | 0.586046 | 0.041 | 0.037 | 1 | 1 | Tbpl1 | 0.004 | Old |
| 0.606441 | 0.721488 | 0.01 | 0.008 | 1 | 1 | Elp4 | 0.002 | Old |
| 0.606675 | 0.094027 | 0.015 | 0.018 | 1 | 1 | Fdxr | -0.003 | Old |
| 0.606754 | 0.857795 | 0.01 | 0.008 | 1 | 1 | Lancl2 | 0.002 | Old |
| 0.607382 | 0.978258 | 0.01 | 0.008 | 1 | 1 | Pcbp4 | 0.002 | Old |
| 0.607696 | 0.668472 | 0.01 | 0.008 | 1 | 1 | Slc6a8 | 0.002 | Old |
| 0.607876 | 0.216197 | 0.082 | 0.09 | 1 | 1 | Pgrmc1 | -0.008 | Old |
| 0.60801 | 0.531091 | 0.01 | 0.008 | 1 | 1 | Fkbp7 | 0.002 | Old |
| 0.608206 | 0.115534 | 0.409 | 0.464 | 1 | 1 | Cotl1 | -0.055 | Old |
| 0.608372 | 0.233761 | 0.03 | 0.034 | 1 | 1 | Ikbip | -0.004 | Old |
| 0.608451 | 0.000286 | 0.166 | 0.177 | 1 | 1 | Mrpl33 | -0.011 | Old |
| 0.608638 | 0.369818 | 0.01 | 0.008 | 1 | 1 | C1qtnf12 | 0.002 | Old |
| 0.608751 | 0.032569 | 0.186 | 0.198 | 1 | 1 | Ptp4a1 | -0.012 | Old |
| 0.609022 | 0.30969 | 0.094 | 0.089 | 1 | 1 | Slc23a2 | 0.005 | Old |
| 0.609267 | 0.270618 | 0.01 | 0.008 | 1 | 1 | Megf9 | 0.002 | Old |
| 0.609386 | 0.211235 | 0.087 | 0.096 | 1 | 1 | Rab24 | -0.009 | Old |
| 0.609478 | 0.280112 | 0.027 | 0.031 | 1 | 1 | Slc41a1 | -0.004 | Old |
| 0.609491 | 0.369888 | 0.131 | 0.126 | 1 | 1 | Dapk1 | 0.005 | Old |
| 0.609658 | 0.145325 | 0.037 | 0.042 | 1 | 1 | Fopnl | -0.005 | Old |
| 0.609896 | 0.540277 | 0.01 | 0.008 | 1 | 1 | Ndufaf1 | 0.002 | Old |
| 0.610223 | 0.042462 | 0.13 | 0.14 | 1 | 1 | U2surp | -0.01 | Old |
| 0.610269 | 0.269561 | 0.064 | 0.059 | 1 | 1 | Ctbp2 | 0.005 | Old |
| 0.610461 | 0.335378 | 0.033 | 0.03 | 1 | 1 | Gbe1 | 0.003 | Old |
| 0.610621 | 0.327418 | 0.111 | 0.105 | 1 | 1 | Pcyt1a | 0.006 | Old |
| 0.610641 | 0.031062 | 0.026 | 0.03 | 1 | 1 | Cog8 | -0.004 | Old |
| 0.610718 | 0.465226 | 0.105 | 0.1 | 1 | 1 | Rbbp4 | 0.005 | Old |
| 0.611071 | 0.155301 | 0.045 | 0.05 | 1 | 1 | Lipe | -0.005 | Old |
| 0.611425 | 0.220104 | 0.015 | 0.018 | 1 | 1 | Zmym3 | -0.003 | Old |
| 0.611588 | 0.261986 | 0.057 | 0.052 | 1 | 1 | Hdac3 | 0.005 | Old |
| 0.612099 | 0.151415 | 0.01 | 0.008 | 1 | 1 | Slx4 | 0.002 | Old |
| 0.613056 | 0.111994 | 0.026 | 0.03 | 1 | 1 | Utp15 | -0.004 | Old |
| 0.61314 | 0.083869 | 0.686 | 0.736 | 1 | 1 | Cfl1 | -0.05 | Old |

| | | | | | | | | |
|---|---|---|---|---|---|---|---|---|
| 0.613497 | 0.110124 | 0.224 | 0.241 | 1 | 1 | Dnajb6 | -0.017 | Old |
| 0.613695 | 0.199661 | 0.045 | 0.041 | 1 | 1 | 6430548M | 0.004 | Old |
| 0.613714 | 0.20755 | 0.08 | 0.088 | 1 | 1 | Fyttd1 | -0.008 | Old |
| 0.613999 | 0.148668 | 0.046 | 0.052 | 1 | 1 | Hspa13 | -0.006 | Old |
| 0.614379 | 0.01862 | 0.095 | 0.103 | 1 | 1 | Btbd1 | -0.008 | Old |
| 0.614583 | 0.086816 | 0.263 | 0.282 | 1 | 1 | Nktr | -0.019 | Old |
| 0.614772 | 0.338138 | 0.037 | 0.042 | 1 | 1 | Adh5 | -0.005 | Old |
| 0.615247 | 0.185384 | 0.027 | 0.031 | 1 | 1 | Mphosph6 | -0.004 | Old |
| 0.615534 | 0.321921 | 0.041 | 0.037 | 1 | 1 | Tubgcp3 | 0.004 | Old |
| 0.615964 | 0.330308 | 0.015 | 0.018 | 1 | 1 | Bcar3 | -0.003 | Old |
| 0.616474 | 0.218753 | 0.027 | 0.031 | 1 | 1 | Spic | -0.004 | Old |
| 0.616528 | 0.163013 | 0.121 | 0.131 | 1 | 1 | 6-Mar | -0.01 | Old |
| 0.616952 | 0.267288 | 0.064 | 0.059 | 1 | 1 | Eif4enif1 | 0.005 | Old |
| 0.617101 | 0.1621 | 0.015 | 0.018 | 1 | 1 | Tmem268 | -0.003 | Old |
| 0.617199 | 0.067709 | 0.413 | 0.448 | 1 | 1 | C5ar1 | -0.035 | Old |
| 0.617306 | 0.491037 | 0.028 | 0.032 | 1 | 1 | Dolpp1 | -0.004 | Old |
| 0.617551 | 0.445495 | 0.017 | 0.014 | 1 | 1 | Arhgap22 | 0.003 | Old |
| 0.617721 | 0.065135 | 0.026 | 0.03 | 1 | 1 | Foxo4 | -0.004 | Old |
| 0.617761 | 0.420903 | 0.09 | 0.085 | 1 | 1 | Bbip1 | 0.005 | Old |
| 0.617992 | 0.441717 | 0.028 | 0.032 | 1 | 1 | Hes1 | -0.004 | Old |
| 0.618062 | 0.005074 | 0.147 | 0.155 | 1 | 1 | Rbm47 | -0.008 | Old |
| 0.618086 | 0.43593 | 0.076 | 0.071 | 1 | 1 | Mettl9 | 0.005 | Old |
| 0.619092 | 0.352923 | 0.045 | 0.05 | 1 | 1 | Atg12 | -0.005 | Old |
| 0.619234 | 0.177637 | 0.014 | 0.017 | 1 | 1 | Nek1 | -0.003 | Old |
| 0.619287 | 0.414093 | 0.055 | 0.051 | 1 | 1 | Ranbp9 | 0.004 | Old |
| 0.619785 | 0.143873 | 0.355 | 0.386 | 1 | 1 | Hnrnpu | -0.031 | Old |
| 0.620015 | 0.200777 | 0.087 | 0.095 | 1 | 1 | Insig1 | -0.008 | Old |
| 0.620242 | 0.503001 | 0.017 | 0.014 | 1 | 1 | Zfp719 | 0.003 | Old |
| 0.620489 | 0.252968 | 0.045 | 0.041 | 1 | 1 | Thoc3 | 0.004 | Old |
| 0.620787 | 0.123131 | 0.127 | 0.137 | 1 | 1 | Mbtd1 | -0.01 | Old |
| 0.620805 | 0.206932 | 0.023 | 0.02 | 1 | 1 | Nfs1 | 0.003 | Old |
| 0.620915 | 0.26843 | 0.148 | 0.143 | 1 | 1 | Psmb10 | 0.005 | Old |
| 0.620982 | 0.010933 | 0.132 | 0.141 | 1 | 1 | Trim41 | -0.009 | Old |
| 0.621086 | 0.051888 | 0.082 | 0.089 | 1 | 1 | Abcf1 | -0.007 | Old |
| 0.621234 | 0.126128 | 0.276 | 0.275 | 1 | 1 | Khdrbs1 | 0.001 | Old |
| 0.621323 | 0.045004 | 0.013 | 0.016 | 1 | 1 | D230025D: | -0.003 | Old |
| 0.621492 | 0.311943 | 0.014 | 0.017 | 1 | 1 | Dxo | -0.003 | Old |
| 0.62243 | 0.467264 | 0.046 | 0.042 | 1 | 1 | Fut8 | 0.004 | Old |
| 0.622623 | 0.482908 | 0.035 | 0.031 | 1 | 1 | Ppp1r8 | 0.004 | Old |
| 0.622682 | 0.37322 | 0.014 | 0.017 | 1 | 1 | Mettl5 | -0.003 | Old |
| 0.622925 | 0.222281 | 0.051 | 0.057 | 1 | 1 | Tcf3 | -0.006 | Old |
| 0.622947 | 0.179863 | 0.024 | 0.028 | 1 | 1 | Skiv2l2 | -0.004 | Old |
| 0.623417 | 0.089258 | 0.044 | 0.049 | 1 | 1 | Med16 | -0.005 | Old |
| 0.623639 | 0.286609 | 0.075 | 0.082 | 1 | 1 | Dnajb11 | -0.007 | Old |
| 0.623675 | 0.469192 | 0.017 | 0.014 | 1 | 1 | Zfp617 | 0.003 | Old |

| | | | | | | | | |
|---|---|---|---|---|---|---|---|---|
| 0.624313 | 0.239347 | 0.013 | 0.016 | 1 | 1 | Atg13 | -0.003 | Old |
| 0.624628 | 0.598704 | 0.015 | 0.018 | 1 | 1 | Pop4 | -0.003 | Old |
| 0.624794 | 0.073852 | 0.163 | 0.175 | 1 | 1 | Manf | -0.012 | Old |
| 0.624821 | 0.239581 | 0.12 | 0.114 | 1 | 1 | Trim12c | 0.006 | Old |
| 0.625189 | 0.14704 | 0.105 | 0.114 | 1 | 1 | Prelid1 | -0.009 | Old |
| 0.625303 | 0.176812 | 0.014 | 0.017 | 1 | 1 | 0610009B2 | -0.003 | Old |
| 0.62613 | 0.34344 | 0.085 | 0.08 | 1 | 1 | Gabarapl1 | 0.005 | Old |
| 0.62631 | 0.417353 | 0.013 | 0.016 | 1 | 1 | Lig3 | -0.003 | Old |
| 0.626623 | 0.029735 | 0.017 | 0.014 | 1 | 1 | Dgcr14 | 0.003 | Old |
| 0.626717 | 0.215211 | 0.189 | 0.185 | 1 | 1 | Rnf19b | 0.004 | Old |
| 0.627059 | 0.172403 | 0.013 | 0.016 | 1 | 1 | Gcfc2 | -0.003 | Old |
| 0.627639 | 0.188484 | 0.026 | 0.03 | 1 | 1 | Nxt1 | -0.004 | Old |
| 0.627655 | 0.33592 | 0.084 | 0.078 | 1 | 1 | Pdpk1 | 0.006 | Old |
| 0.627659 | 0.064084 | 0.023 | 0.02 | 1 | 1 | Rpp14 | 0.003 | Old |
| 0.627794 | 0.078012 | 0.159 | 0.17 | 1 | 1 | Smchd1 | -0.011 | Old |
| 0.628 | 0.23171 | 0.067 | 0.062 | 1 | 1 | Ufl1 | 0.005 | Old |
| 0.628864 | 0.209128 | 0.041 | 0.037 | 1 | 1 | Agmo | 0.004 | Old |
| 0.62905 | 0.044595 | 0.05 | 0.055 | 1 | 1 | Lypla1 | -0.005 | Old |
| 0.629232 | 0.217358 | 0.18 | 0.195 | 1 | 1 | Spcs1 | -0.015 | Old |
| 0.629405 | 0.090775 | 0.299 | 0.319 | 1 | 1 | Gas7 | -0.02 | Old |
| 0.629715 | 0.163127 | 0.033 | 0.038 | 1 | 1 | Haghl | -0.005 | Old |
| 0.629909 | 0.637041 | 0.044 | 0.04 | 1 | 1 | Phrf1 | 0.004 | Old |
| 0.630063 | 0.295323 | 0.027 | 0.024 | 1 | 1 | Mrps26 | 0.003 | Old |
| 0.630112 | 0.100636 | 0.1 | 0.109 | 1 | 1 | Cyth1 | -0.009 | Old |
| 0.630466 | 0.337607 | 0.112 | 0.107 | 1 | 1 | Card19 | 0.005 | Old |
| 0.630734 | 0.277931 | 0.041 | 0.037 | 1 | 1 | Usp1 | 0.004 | Old |
| 0.631292 | 0.456737 | 0.031 | 0.027 | 1 | 1 | Stat5a | 0.004 | Old |
| 0.631487 | 0.625209 | 0.021 | 0.018 | 1 | 1 | Ift46 | 0.003 | Old |
| 0.631768 | 0.01178 | 0.956 | 0.956 | 1 | 1 | Rps9 | 0 | Old |
| 0.632077 | 0.171804 | 0.041 | 0.046 | 1 | 1 | Qrich1 | -0.005 | Old |
| 0.632168 | 0.277261 | 0.017 | 0.014 | 1 | 1 | Mul1 | 0.003 | Old |
| 0.632696 | 0.097346 | 0.096 | 0.104 | 1 | 1 | 1110004F1 | -0.008 | Old |
| 0.632744 | 0.953383 | 0.014 | 0.012 | 1 | 1 | Zbtb26 | 0.002 | Old |
| 0.632944 | 0.616574 | 0.013 | 0.016 | 1 | 1 | Batf2 | -0.003 | Old |
| 0.633113 | 0.163366 | 0.044 | 0.049 | 1 | 1 | Mocs2 | -0.005 | Old |
| 0.633255 | 0.199981 | 0.012 | 0.014 | 1 | 1 | Akap1 | -0.002 | Old |
| 0.633387 | 0.083075 | 0.041 | 0.037 | 1 | 1 | 1700123O2 | 0.004 | Old |
| 0.633521 | 0.279599 | 0.15 | 0.145 | 1 | 1 | Aph1a | 0.005 | Old |
| 0.633728 | 0.250704 | 0.054 | 0.049 | 1 | 1 | Tbc1d13 | 0.005 | Old |
| 0.633951 | 0.030893 | 0.045 | 0.041 | 1 | 1 | Mtx2 | 0.004 | Old |
| 0.634576 | 0.498926 | 0.013 | 0.016 | 1 | 1 | Rbmx2 | -0.003 | Old |
| 0.634704 | 0.12436 | 0.37 | 0.374 | 1 | 1 | Dbi | -0.004 | Old |
| 0.634807 | 0.362515 | 0.035 | 0.031 | 1 | 1 | Mpp5 | 0.004 | Old |
| 0.635133 | 0.187309 | 0.032 | 0.036 | 1 | 1 | Nup133 | -0.004 | Old |
| 0.635526 | 0.269422 | 0.031 | 0.027 | 1 | 1 | Elac1 | 0.004 | Old |

| | | | | | | | | |
|---|---|---|---|---|---|---|---|---|
| 0.635543 | 0.16902 | 0.072 | 0.078 | 1 | 1 | Slc36a1 | -0.006 | Old |
| 0.635694 | 0.546762 | 0.021 | 0.018 | 1 | 1 | P2ry10b | 0.003 | Old |
| 0.636507 | 0.293317 | 0.035 | 0.031 | 1 | 1 | Gcsh | 0.004 | Old |
| 0.636822 | 0.27537 | 0.012 | 0.014 | 1 | 1 | Bnip1 | -0.002 | Old |
| 0.636886 | 0.370954 | 0.053 | 0.058 | 1 | 1 | Zbtb1 | -0.005 | Old |
| 0.637086 | 0.230321 | 0.012 | 0.014 | 1 | 1 | Adam22 | -0.002 | Old |
| 0.637148 | 0.080471 | 0.01 | 0.013 | 1 | 1 | Abhd11 | -0.003 | Old |
| 0.637586 | 0.704251 | 0.014 | 0.012 | 1 | 1 | Ttc21b | 0.002 | Old |
| 0.637826 | 0.098471 | 0.04 | 0.044 | 1 | 1 | Asb8 | -0.004 | Old |
| 0.637914 | 0.482348 | 0.021 | 0.018 | 1 | 1 | Mtap | 0.003 | Old |
| 0.637993 | 0.265554 | 0.051 | 0.057 | 1 | 1 | Hccs | -0.006 | Old |
| 0.638198 | 0.692513 | 0.032 | 0.029 | 1 | 1 | Neurl2 | 0.003 | Old |
| 0.638209 | 0.252688 | 0.035 | 0.031 | 1 | 1 | Heatr3 | 0.004 | Old |
| 0.638369 | 0.567444 | 0.044 | 0.04 | 1 | 1 | Psmc4 | 0.004 | Old |
| 0.638939 | 0.322338 | 0.012 | 0.014 | 1 | 1 | Trmt61a | -0.002 | Old |
| 0.638975 | 0.016368 | 0.039 | 0.043 | 1 | 1 | Thap3 | -0.004 | Old |
| 0.639025 | 0.532783 | 0.021 | 0.018 | 1 | 1 | Fam122b | 0.003 | Old |
| 0.639126 | 0.000922 | 0.039 | 0.043 | 1 | 1 | Slc25a17 | -0.004 | Old |
| 0.639204 | 0.404486 | 0.012 | 0.014 | 1 | 1 | Mcph1 | -0.002 | Old |
| 0.63941 | 0.125469 | 0.032 | 0.036 | 1 | 1 | B3galnt1 | -0.004 | Old |
| 0.639469 | 0.622829 | 0.012 | 0.014 | 1 | 1 | 1600020E0 | -0.002 | Old |
| 0.639625 | 0.210548 | 0.033 | 0.038 | 1 | 1 | Rhbdf2 | -0.005 | Old |
| 0.639905 | 0.127646 | 0.032 | 0.036 | 1 | 1 | Golt1b | -0.004 | Old |
| 0.640173 | 0.435399 | 0.04 | 0.036 | 1 | 1 | Tmx2 | 0.004 | Old |
| 0.640229 | 0.173243 | 0.01 | 0.013 | 1 | 1 | Snx7 | -0.003 | Old |
| 0.640299 | 0.048413 | 0.031 | 0.035 | 1 | 1 | Kri1 | -0.004 | Old |
| 0.640765 | 0.221579 | 0.283 | 0.284 | 1 | 1 | 1810037I1 | -0.001 | Old |
| 0.640901 | 0.088294 | 0.059 | 0.064 | 1 | 1 | Adgre4 | -0.005 | Old |
| 0.64102 | 0.175083 | 0.094 | 0.101 | 1 | 1 | Tmpo | -0.007 | Old |
| 0.641317 | 0.00712 | 0.113 | 0.121 | 1 | 1 | Clptm1l | -0.008 | Old |
| 0.641351 | 0.172258 | 0.01 | 0.013 | 1 | 1 | Klhl36 | -0.003 | Old |
| 0.641455 | 0.169917 | 0.324 | 0.328 | 1 | 1 | Emp3 | -0.004 | Old |
| 0.64152 | 0.390141 | 0.136 | 0.132 | 1 | 1 | Snrpg | 0.004 | Old |
| 0.641801 | 0.225709 | 0.032 | 0.036 | 1 | 1 | Prpf19 | -0.004 | Old |
| 0.642016 | 0.359586 | 0.044 | 0.049 | 1 | 1 | Aggf1 | -0.005 | Old |
| 0.642121 | 0.986739 | 0.012 | 0.014 | 1 | 1 | Cnn3 | -0.002 | Old |
| 0.642205 | 0.100479 | 0.212 | 0.208 | 1 | 1 | Tm9sf2 | 0.004 | Old |
| 0.642619 | 0.170268 | 0.054 | 0.049 | 1 | 1 | 3830406C1 | 0.005 | Old |
| 0.642714 | 0.199859 | 0.014 | 0.012 | 1 | 1 | Eid2b | 0.002 | Old |
| 0.643054 | 0.0594 | 0.112 | 0.119 | 1 | 1 | Zmat2 | -0.007 | Old |
| 0.643622 | 0.278463 | 0.108 | 0.117 | 1 | 1 | Ppp4c | -0.009 | Old |
| 0.643706 | 0.004879 | 0.132 | 0.14 | 1 | 1 | Uba1 | -0.008 | Old |
| 0.643714 | 0.506151 | 0.012 | 0.014 | 1 | 1 | Rida | -0.002 | Old |
| 0.643835 | 1.063462 | 0.018 | 0.016 | 1 | 1 | Emilin1 | 0.002 | Old |
| 0.643868 | 0.415641 | 0.028 | 0.025 | 1 | 1 | Rpusd4 | 0.003 | Old |

| | | | | | | | | |
|---|---|---|---|---|---|---|---|---|
| 0.644399 | 0.118682 | 0.068 | 0.074 | 1 | 1 | Sdhb | -0.006 | Old |
| 0.645484 | 0.169211 | 0.021 | 0.018 | 1 | 1 | Jak3 | 0.003 | Old |
| 0.646217 | 1.012284 | 0.018 | 0.016 | 1 | 1 | Ccdc71 | 0.002 | Old |
| 0.646353 | 0.040549 | 0.08 | 0.086 | 1 | 1 | Smad5 | -0.006 | Old |
| 0.646638 | 0.394727 | 0.024 | 0.021 | 1 | 1 | Zfp160 | 0.003 | Old |
| 0.646836 | 0.084912 | 0.126 | 0.135 | 1 | 1 | Ddost | -0.009 | Old |
| 0.647493 | 0.35718 | 0.113 | 0.109 | 1 | 1 | R3hdm2 | 0.004 | Old |
| 0.647815 | 0.375879 | 0.022 | 0.025 | 1 | 1 | Srprb | -0.003 | Old |
| 0.647815 | 0.155187 | 0.022 | 0.025 | 1 | 1 | Tpgs2 | -0.003 | Old |
| 0.647894 | 0.46845 | 0.049 | 0.045 | 1 | 1 | Gid4 | 0.004 | Old |
| 0.648068 | 0.357943 | 0.024 | 0.021 | 1 | 1 | Atg4c | 0.003 | Old |
| 0.648289 | 0.147239 | 0.035 | 0.031 | 1 | 1 | Vps39 | 0.004 | Old |
| 0.648671 | 0.195238 | 0.087 | 0.082 | 1 | 1 | Sec22b | 0.005 | Old |
| 0.648842 | 0.967373 | 0.018 | 0.016 | 1 | 1 | Fam149b | 0.002 | Old |
| 0.648871 | 0.122293 | 0.091 | 0.086 | 1 | 1 | Spag7 | 0.005 | Old |
| 0.649023 | 0.306818 | 0.266 | 0.273 | 1 | 1 | Gns | -0.007 | Old |
| 0.649091 | 0.375509 | 0.024 | 0.021 | 1 | 1 | Smim8 | 0.003 | Old |
| 0.649097 | 0.385421 | 0.073 | 0.069 | 1 | 1 | Pdap1 | 0.004 | Old |
| 0.649119 | 0.20036 | 0.04 | 0.044 | 1 | 1 | Pold3 | -0.004 | Old |
| 0.649702 | 0.269851 | 0.022 | 0.025 | 1 | 1 | Zbp1 | -0.003 | Old |
| 0.64993 | 0.348167 | 0.048 | 0.052 | 1 | 1 | Timm8b | -0.004 | Old |
| 0.650114 | 0.324422 | 0.024 | 0.021 | 1 | 1 | Cog4 | 0.003 | Old |
| 0.650655 | 0.027976 | 0.037 | 0.041 | 1 | 1 | Atl2 | -0.004 | Old |
| 0.650854 | 0.108839 | 0.009 | 0.011 | 1 | 1 | Chd3os | -0.002 | Old |
| 0.651088 | 0.279978 | 0.082 | 0.089 | 1 | 1 | Fam173a | -0.007 | Old |
| 0.651191 | 0.15768 | 0.067 | 0.062 | 1 | 1 | Prdx3 | 0.005 | Old |
| 0.651589 | 0.642905 | 0.033 | 0.03 | 1 | 1 | Fbxo8 | 0.003 | Old |
| 0.652057 | 0.072841 | 0.009 | 0.011 | 1 | 1 | Laptm4b | -0.002 | Old |
| 0.652358 | 0.192478 | 0.009 | 0.011 | 1 | 1 | Rasgrf2 | -0.002 | Old |
| 0.65247 | 0.311669 | 0.129 | 0.124 | 1 | 1 | Mfsd14b | 0.005 | Old |
| 0.652655 | 0.360289 | 0.055 | 0.051 | 1 | 1 | Ppp2r3d | 0.004 | Old |
| 0.653488 | 0.27845 | 0.081 | 0.077 | 1 | 1 | Frrs1 | 0.004 | Old |
| 0.653864 | 0.162761 | 0.009 | 0.011 | 1 | 1 | Zfp961 | -0.002 | Old |
| 0.653925 | 0.448804 | 0.028 | 0.025 | 1 | 1 | Gzf1 | 0.003 | Old |
| 0.654213 | 0.120889 | 0.073 | 0.068 | 1 | 1 | Kif3b | 0.005 | Old |
| 0.654548 | 0.072749 | 0.086 | 0.093 | 1 | 1 | Fam32a | -0.007 | Old |
| 0.654717 | 0.103234 | 0.497 | 0.521 | 1 | 1 | Actr2 | -0.024 | Old |
| 0.654984 | 0.591202 | 0.045 | 0.041 | 1 | 1 | Nfyb | 0.004 | Old |
| 0.655069 | 0.336379 | 0.009 | 0.011 | 1 | 1 | Tnfrsf17 | -0.002 | Old |
| 0.655091 | 0.065419 | 0.054 | 0.059 | 1 | 1 | Naga | -0.005 | Old |
| 0.655097 | 0.043811 | 0.067 | 0.072 | 1 | 1 | Clec4n | -0.005 | Old |
| 0.655371 | 0.391759 | 0.009 | 0.011 | 1 | 1 | Mccc2 | -0.002 | Old |
| 0.65558 | 0.065486 | 0.008 | 0.01 | 1 | 1 | Tnni2 | -0.002 | Old |
| 0.655735 | 0.144059 | 0.03 | 0.033 | 1 | 1 | Uchl3 | -0.003 | Old |
| 0.655904 | 0.221882 | 0.008 | 0.01 | 1 | 1 | Tmem238 | -0.002 | Old |

| | | | | | | | | |
|---|---|---|---|---|---|---|---|---|
| 0.65593 | 0.40316 | 0.055 | 0.051 | 1 | 1 | Timm17a | 0.004 | Old |
| 0.656015 | 0.129509 | 0.154 | 0.164 | 1 | 1 | Tsn | -0.01 | Old |
| 0.656171 | 0.059481 | 0.069 | 0.075 | 1 | 1 | Gigyf2 | -0.006 | Old |
| 0.656418 | 0.192561 | 0.044 | 0.048 | 1 | 1 | Jade1 | -0.004 | Old |
| 0.65649 | 0.148481 | 0.206 | 0.22 | 1 | 1 | Sptssa | -0.014 | Old |
| 0.656553 | 0.140625 | 0.008 | 0.01 | 1 | 1 | Bahcc1 | -0.002 | Old |
| 0.656574 | 0.242722 | 0.332 | 0.36 | 1 | 1 | Wfdc17 | -0.028 | Old |
| 0.656626 | 0.03352 | 0.406 | 0.431 | 1 | 1 | Snrnp70 | -0.025 | Old |
| 0.65679 | 0.061237 | 0.053 | 0.058 | 1 | 1 | Ubald1 | -0.005 | Old |
| 0.656879 | 0.450999 | 0.009 | 0.011 | 1 | 1 | Gk5 | -0.002 | Old |
| 0.657052 | 0.128932 | 0.046 | 0.051 | 1 | 1 | Zfp236 | -0.005 | Old |
| 0.657062 | 0.239828 | 0.067 | 0.072 | 1 | 1 | Pilra | -0.005 | Old |
| 0.657202 | 0.146382 | 0.008 | 0.01 | 1 | 1 | Olr1 | -0.002 | Old |
| 0.657785 | 0.594457 | 0.009 | 0.011 | 1 | 1 | Tysnd1 | -0.002 | Old |
| 0.658176 | 0.23826 | 0.008 | 0.01 | 1 | 1 | Wdr90 | -0.002 | Old |
| 0.658753 | 0.121235 | 0.095 | 0.102 | 1 | 1 | Nfkbie | -0.007 | Old |
| 0.658904 | 0.066157 | 0.037 | 0.041 | 1 | 1 | Cdk8 | -0.004 | Old |
| 0.659034 | 0.130087 | 0.059 | 0.064 | 1 | 1 | Itpr2 | -0.005 | Old |
| 0.6594 | 0.019271 | 0.816 | 0.863 | 1 | 1 | Fos | -0.047 | Old |
| 0.659598 | 0.405016 | 0.012 | 0.01 | 1 | 1 | Bcdin3d | 0.002 | Old |
| 0.65962 | 0.440807 | 0.154 | 0.153 | 1 | 1 | Rad21 | 0.001 | Old |
| 0.659801 | 0.252686 | 0.008 | 0.01 | 1 | 1 | Snap47 | -0.002 | Old |
| 0.660125 | 0.491457 | 0.037 | 0.034 | 1 | 1 | Sertad3 | 0.003 | Old |
| 0.660126 | 0.398646 | 0.008 | 0.01 | 1 | 1 | B3gnt3 | -0.002 | Old |
| 0.660266 | 0.26568 | 0.04 | 0.044 | 1 | 1 | Rexo1 | -0.004 | Old |
| 0.660307 | 0.390694 | 0.04 | 0.036 | 1 | 1 | Gm13212 | 0.004 | Old |
| 0.660505 | 0.716344 | 0.012 | 0.01 | 1 | 1 | Ints4 | 0.002 | Old |
| 0.660677 | 0.197406 | 0.175 | 0.187 | 1 | 1 | Sp100 | -0.012 | Old |
| 0.660693 | 0.412183 | 0.021 | 0.024 | 1 | 1 | Zfp943 | -0.003 | Old |
| 0.660802 | 0.240324 | 0.228 | 0.249 | 1 | 1 | Tspo | -0.021 | Old |
| 0.660808 | 0.333786 | 0.012 | 0.01 | 1 | 1 | Grasp | 0.002 | Old |
| 0.660878 | 0.09927 | 0.064 | 0.069 | 1 | 1 | Oser1 | -0.005 | Old |
| 0.66154 | 0.398443 | 0.087 | 0.095 | 1 | 1 | Rps12-ps3 | -0.008 | Old |
| 0.661731 | 0.045244 | 0.147 | 0.155 | 1 | 1 | Ifi203 | -0.008 | Old |
| 0.662018 | 0.584503 | 0.012 | 0.01 | 1 | 1 | Tyw5 | 0.002 | Old |
| 0.662018 | 0.674896 | 0.012 | 0.01 | 1 | 1 | Zdhhc1 | 0.002 | Old |
| 0.66227 | 0.124045 | 0.32 | 0.321 | 1 | 1 | Rap1a | -0.001 | Old |
| 0.662321 | 0.775954 | 0.012 | 0.01 | 1 | 1 | Mx1 | 0.002 | Old |
| 0.662321 | 0.307146 | 0.012 | 0.01 | 1 | 1 | Eps8l1 | 0.002 | Old |
| 0.662895 | 0.199288 | 0.045 | 0.049 | 1 | 1 | Med14 | -0.004 | Old |
| 0.662995 | 0.405971 | 0.018 | 0.016 | 1 | 1 | Eci1 | 0.002 | Old |
| 0.663044 | 0.260428 | 0.019 | 0.022 | 1 | 1 | Pdlim2 | -0.003 | Old |
| 0.663056 | 0.338379 | 0.008 | 0.01 | 1 | 1 | Ctla2a | -0.002 | Old |
| 0.663088 | 0.821313 | 0.048 | 0.044 | 1 | 1 | Ldlrap1 | 0.004 | Old |
| 0.663274 | 0.191733 | 0.063 | 0.069 | 1 | 1 | Commd3 | -0.006 | Old |

| | | | | | | | | |
|---|---|---|---|---|---|---|---|---|
| 0.663286 | 0.148557 | 0.29 | 0.289 | 1 | 1 | Ehd1 | 0.001 | Old |
| 0.663561 | 0.128517 | 0.359 | 0.388 | 1 | 1 | Csde1 | -0.029 | Old |
| 0.663707 | 0.457275 | 0.008 | 0.01 | 1 | 1 | Sirt5 | -0.002 | Old |
| 0.663836 | 0.396372 | 0.012 | 0.01 | 1 | 1 | Nek4 | 0.002 | Old |
| 0.663843 | 0.182589 | 0.481 | 0.532 | 1 | 1 | Btg1 | -0.051 | Old |
| 0.664229 | 0.051934 | 0.064 | 0.069 | 1 | 1 | Ppm1b | -0.005 | Old |
| 0.664233 | 0.039203 | 0.852 | 0.888 | 1 | 1 | Laptm5 | -0.036 | Old |
| 0.664304 | 0.021257 | 0.054 | 0.049 | 1 | 1 | Atp8b4 | 0.005 | Old |
| 0.66477 | 0.018488 | 0.035 | 0.038 | 1 | 1 | Dnajb12 | -0.003 | Old |
| 0.664921 | 0.289649 | 0.084 | 0.091 | 1 | 1 | Mrps14 | -0.007 | Old |
| 0.665165 | 0.197817 | 0.018 | 0.016 | 1 | 1 | Pomgnt1 | 0.002 | Old |
| 0.665352 | 0.48992 | 0.012 | 0.01 | 1 | 1 | Trim30c | 0.002 | Old |
| 0.665418 | 0.117002 | 0.051 | 0.056 | 1 | 1 | Ccdc6 | -0.005 | Old |
| 0.665601 | 0.212438 | 0.019 | 0.022 | 1 | 1 | Rrp36 | -0.003 | Old |
| 0.666097 | 0.510708 | 0.057 | 0.052 | 1 | 1 | Sars | 0.005 | Old |
| 0.666098 | 0.077808 | 0.037 | 0.041 | 1 | 1 | Rabgap1l | -0.004 | Old |
| 0.666235 | 0.082682 | 0.132 | 0.14 | 1 | 1 | Yy1 | -0.008 | Old |
| 0.666359 | 0.53093 | 0.054 | 0.05 | 1 | 1 | Csrp1 | 0.004 | Old |
| 0.666613 | 0.023564 | 0.018 | 0.016 | 1 | 1 | Prdm15 | 0.002 | Old |
| 0.666627 | 0.54954 | 0.04 | 0.044 | 1 | 1 | Sympk | -0.004 | Old |
| 0.667272 | 0.253218 | 0.1 | 0.095 | 1 | 1 | Kdm1a | 0.005 | Old |
| 0.667522 | 0.237203 | 0.019 | 0.022 | 1 | 1 | Triobp | -0.003 | Old |
| 0.66778 | 0.298516 | 0.134 | 0.131 | 1 | 1 | Dcaf12 | 0.003 | Old |
| 0.667813 | 0.057404 | 0.261 | 0.276 | 1 | 1 | Rnf149 | -0.015 | Old |
| 0.6682 | 0.086479 | 0.144 | 0.153 | 1 | 1 | Sf3b5 | -0.009 | Old |
| 0.668717 | 0.172013 | 0.053 | 0.058 | 1 | 1 | Ing5 | -0.005 | Old |
| 0.668953 | 0.741492 | 0.044 | 0.041 | 1 | 1 | Papss1 | 0.003 | Old |
| 0.668986 | 0.029303 | 0.042 | 0.047 | 1 | 1 | Phf12 | -0.005 | Old |
| 0.669021 | 0.032634 | 0.134 | 0.143 | 1 | 1 | Rbm8a | -0.009 | Old |
| 0.669025 | 0.100205 | 0.203 | 0.214 | 1 | 1 | Ifi204 | -0.011 | Old |
| 0.669156 | 0.469868 | 0.03 | 0.027 | 1 | 1 | Bambi | 0.003 | Old |
| 0.669202 | 0.091572 | 0.067 | 0.072 | 1 | 1 | Gpr146 | -0.005 | Old |
| 0.669333 | 0.188099 | 0.121 | 0.13 | 1 | 1 | Akt1 | -0.009 | Old |
| 0.670126 | 0.047794 | 0.098 | 0.104 | 1 | 1 | Nacc2 | -0.006 | Old |
| 0.670229 | 0.153282 | 0.147 | 0.157 | 1 | 1 | Klf7 | -0.01 | Old |
| 0.67048 | 0.048518 | 0.018 | 0.016 | 1 | 1 | 1110008P1 | 0.002 | Old |
| 0.670584 | 0.065151 | 0.087 | 0.094 | 1 | 1 | Zcchc7 | -0.007 | Old |
| 0.670621 | 0.199817 | 0.059 | 0.064 | 1 | 1 | Krt80 | -0.005 | Old |
| 0.670655 | 0.522529 | 0.03 | 0.027 | 1 | 1 | Zfp780b | 0.003 | Old |
| 0.670686 | 0.039418 | 0.722 | 0.764 | 1 | 1 | Eif1 | -0.042 | Old |
| 0.670972 | 0.083048 | 0.488 | 0.506 | 1 | 1 | Gnb2 | -0.018 | Old |
| 0.671652 | 0.167299 | 0.028 | 0.032 | 1 | 1 | Slc10a7 | -0.004 | Old |
| 0.671772 | 0.13855 | 0.05 | 0.055 | 1 | 1 | Slu7 | -0.005 | Old |
| 0.67185 | 0.108019 | 0.134 | 0.142 | 1 | 1 | Tpi1 | -0.008 | Old |
| 0.672032 | 0.048268 | 0.112 | 0.118 | 1 | 1 | Capn2 | -0.006 | Old |

| | | | | | | | | |
|---|---|---|---|---|---|---|---|---|
| 0.672343 | 0.437274 | 0.03 | 0.027 | 1 | 1 | Prkce | 0.003 | Old |
| 0.672812 | 0.012704 | 0.072 | 0.066 | 1 | 1 | Glb1 | 0.006 | Old |
| 0.673267 | 0.141259 | 0.355 | 0.36 | 1 | 1 | Snx3 | -0.005 | Old |
| 0.673401 | 0.21481 | 0.082 | 0.089 | 1 | 1 | Hnrnpll | -0.007 | Old |
| 0.673537 | 0.189141 | 0.1 | 0.107 | 1 | 1 | N4bp2l1 | -0.007 | Old |
| 0.673802 | 0.194829 | 0.018 | 0.021 | 1 | 1 | Tbccd1 | -0.003 | Old |
| 0.674463 | 0.055409 | 0.086 | 0.092 | 1 | 1 | Trpm7 | -0.006 | Old |
| 0.674617 | 0.387954 | 0.048 | 0.052 | 1 | 1 | Bcap29 | -0.004 | Old |
| 0.675018 | 0.144505 | 0.027 | 0.03 | 1 | 1 | Rnf121 | -0.003 | Old |
| 0.675366 | 0.077918 | 0.048 | 0.044 | 1 | 1 | Fam3a | 0.004 | Old |
| 0.675391 | 0.410993 | 0.104 | 0.1 | 1 | 1 | Tecr | 0.004 | Old |
| 0.675415 | 0.402666 | 0.028 | 0.032 | 1 | 1 | Pigyl | -0.004 | Old |
| 0.675457 | 0.215327 | 0.148 | 0.159 | 1 | 1 | Cyp4f18 | -0.011 | Old |
| 0.675629 | 0.362507 | 0.03 | 0.027 | 1 | 1 | Dcaf1 | 0.003 | Old |
| 0.676218 | 0.370749 | 0.112 | 0.109 | 1 | 1 | Ppp6c | 0.003 | Old |
| 0.676322 | 0.757354 | 0.015 | 0.013 | 1 | 1 | Gdpd1 | 0.002 | Old |
| 0.676512 | 0.009675 | 0.246 | 0.259 | 1 | 1 | Lilr4b | -0.013 | Old |
| 0.676691 | 0.51941 | 0.042 | 0.039 | 1 | 1 | Gpaa1 | 0.003 | Old |
| 0.676745 | 0.194688 | 0.113 | 0.109 | 1 | 1 | Cept1 | 0.004 | Old |
| 0.676846 | 0.929303 | 0.015 | 0.013 | 1 | 1 | Usp35 | 0.002 | Old |
| 0.677208 | 0.011827 | 0.06 | 0.065 | 1 | 1 | Ahsa1 | -0.005 | Old |
| 0.67765 | 0.19216 | 0.116 | 0.124 | 1 | 1 | Nabp1 | -0.008 | Old |
| 0.677686 | 0.195284 | 0.018 | 0.021 | 1 | 1 | Alkbh8 | -0.003 | Old |
| 0.678062 | 0.102517 | 0.026 | 0.023 | 1 | 1 | Rsad2 | 0.003 | Old |
| 0.678461 | 0.23044 | 0.086 | 0.093 | 1 | 1 | Hmbox1 | -0.007 | Old |
| 0.678691 | 0.043107 | 0.027 | 0.03 | 1 | 1 | Pex3 | -0.003 | Old |
| 0.678715 | 0.219833 | 0.24 | 0.241 | 1 | 1 | Psma7 | -0.001 | Old |
| 0.680549 | 0.438944 | 0.118 | 0.115 | 1 | 1 | Tmem234 | 0.003 | Old |
| 0.680626 | 0.098495 | 0.325 | 0.349 | 1 | 1 | Plekho2 | -0.024 | Old |
| 0.680894 | 0.140861 | 0.113 | 0.121 | 1 | 1 | Arid5a | -0.008 | Old |
| 0.680999 | 0.185095 | 0.154 | 0.166 | 1 | 1 | Mia3 | -0.012 | Old |
| 0.681035 | 0.272992 | 0.132 | 0.128 | 1 | 1 | Psmd8 | 0.004 | Old |
| 0.68147 | 0.222572 | 0.134 | 0.129 | 1 | 1 | Oxct1 | 0.005 | Old |
| 0.681719 | 0.335733 | 0.063 | 0.069 | 1 | 1 | Mrpl12 | -0.006 | Old |
| 0.682357 | 0.386665 | 0.015 | 0.013 | 1 | 1 | Riox2 | 0.002 | Old |
| 0.682505 | 0.089151 | 0.04 | 0.044 | 1 | 1 | Sfxn2 | -0.004 | Old |
| 0.682751 | 0.297006 | 0.073 | 0.079 | 1 | 1 | Ufm1 | -0.006 | Old |
| 0.683256 | 0.252822 | 0.017 | 0.019 | 1 | 1 | Cables1 | -0.002 | Old |
| 0.683463 | 0.27724 | 0.303 | 0.315 | 1 | 1 | Vcp | -0.012 | Old |
| 0.683506 | 0.201009 | 0.041 | 0.038 | 1 | 1 | Rbm4b | 0.003 | Old |
| 0.683947 | 0.20712 | 0.049 | 0.045 | 1 | 1 | Plxna1 | 0.004 | Old |
| 0.68406 | 0.287631 | 0.094 | 0.089 | 1 | 1 | Stxbp2 | 0.005 | Old |
| 0.684461 | 0.463592 | 0.015 | 0.013 | 1 | 1 | Stk11ip | 0.002 | Old |
| 0.684705 | 0.255859 | 0.123 | 0.132 | 1 | 1 | Uhmk1 | -0.009 | Old |
| 0.685002 | 0.271098 | 0.05 | 0.055 | 1 | 1 | Jak2 | -0.005 | Old |

| | | | | | | | | |
|---|---|---|---|---|---|---|---|---|
| 0.685045 | 0.556594 | 0.031 | 0.028 | 1 | 1 | Sap30bp | 0.003 | Old |
| 0.685203 | 0.277288 | 0.033 | 0.03 | 1 | 1 | Isoc1 | 0.003 | Old |
| 0.685382 | 0.447081 | 0.015 | 0.013 | 1 | 1 | Adck1 | 0.002 | Old |
| 0.685413 | 0.178201 | 0.058 | 0.054 | 1 | 1 | Mrpl32 | 0.004 | Old |
| 0.685564 | 0.466854 | 0.039 | 0.035 | 1 | 1 | Fam8a1 | 0.004 | Old |
| 0.685761 | 0.261709 | 0.332 | 0.339 | 1 | 1 | Dstn | -0.007 | Old |
| 0.685981 | 0.19612 | 0.121 | 0.129 | 1 | 1 | Fkbp15 | -0.008 | Old |
| 0.686041 | 0.653694 | 0.015 | 0.013 | 1 | 1 | Bloc1s1 | 0.002 | Old |
| 0.686165 | 0.096264 | 0.026 | 0.023 | 1 | 1 | 2810402E2 | 0.003 | Old |
| 0.686567 | 0.066387 | 0.03 | 0.027 | 1 | 1 | 2610002M | 0.003 | Old |
| 0.686575 | 0.055436 | 0.049 | 0.053 | 1 | 1 | Pdzd8 | -0.004 | Old |
| 0.686607 | 0.308783 | 0.234 | 0.237 | 1 | 1 | Plxnd1 | -0.003 | Old |
| 0.686774 | 0.165106 | 0.026 | 0.023 | 1 | 1 | Champ1 | 0.003 | Old |
| 0.686851 | 0.162619 | 0.026 | 0.029 | 1 | 1 | Yif1a | -0.003 | Old |
| 0.686851 | 0.015401 | 0.026 | 0.029 | 1 | 1 | Dera | -0.003 | Old |
| 0.687334 | 0.225589 | 0.098 | 0.092 | 1 | 1 | Adss | 0.006 | Old |
| 0.687807 | 0.248587 | 0.063 | 0.069 | 1 | 1 | Cdip1 | -0.006 | Old |
| 0.687963 | 0.147649 | 0.224 | 0.239 | 1 | 1 | Il10rb | -0.015 | Old |
| 0.688009 | 0.036389 | 0.199 | 0.21 | 1 | 1 | Lcp2 | -0.011 | Old |
| 0.688234 | 0.534763 | 0.035 | 0.032 | 1 | 1 | Med25 | 0.003 | Old |
| 0.6883 | 0.268946 | 0.129 | 0.124 | 1 | 1 | Zfp148 | 0.005 | Old |
| 0.688327 | 0.05285 | 0.185 | 0.195 | 1 | 1 | Abracl | -0.01 | Old |
| 0.688413 | 0.139255 | 0.015 | 0.013 | 1 | 1 | Ppp1r16a | 0.002 | Old |
| 0.688933 | 0.375037 | 0.026 | 0.029 | 1 | 1 | E4f1 | -0.003 | Old |
| 0.688939 | 0.273864 | 0.033 | 0.03 | 1 | 1 | Fgd6 | 0.003 | Old |
| 0.689489 | 0.283293 | 0.019 | 0.017 | 1 | 1 | Zfp597 | 0.002 | Old |
| 0.689597 | 0.245621 | 0.033 | 0.037 | 1 | 1 | Nosip | -0.004 | Old |
| 0.690042 | 0.726721 | 0.044 | 0.041 | 1 | 1 | Sdad1 | 0.003 | Old |
| 0.690077 | 0.211367 | 0.069 | 0.075 | 1 | 1 | Rgs19 | -0.006 | Old |
| 0.690173 | 0.221485 | 0.063 | 0.059 | 1 | 1 | Pigp | 0.004 | Old |
| 0.690194 | 0.021984 | 0.024 | 0.027 | 1 | 1 | Siae | -0.003 | Old |
| 0.690423 | 0.17561 | 0.127 | 0.137 | 1 | 1 | Nampt | -0.01 | Old |
| 0.690496 | 0.398053 | 0.027 | 0.03 | 1 | 1 | Ap3m2 | -0.003 | Old |
| 0.690621 | 0.052793 | 0.094 | 0.1 | 1 | 1 | Rabep1 | -0.006 | Old |
| 0.691274 | 0.146083 | 0.033 | 0.037 | 1 | 1 | Fpr2 | -0.004 | Old |
| 0.691276 | 0.195502 | 0.135 | 0.13 | 1 | 1 | Hdlbp | 0.005 | Old |
| 0.69161 | 0.196209 | 0.033 | 0.037 | 1 | 1 | Dcaf10 | -0.004 | Old |
| 0.691944 | 0.325751 | 0.017 | 0.019 | 1 | 1 | Tmcc2 | -0.002 | Old |
| 0.692074 | 0.525351 | 0.019 | 0.017 | 1 | 1 | Ifitm5 | 0.002 | Old |
| 0.692307 | 0.172015 | 0.051 | 0.056 | 1 | 1 | Plau | -0.005 | Old |
| 0.692545 | 0.431188 | 0.019 | 0.017 | 1 | 1 | 1300002E1 | 0.002 | Old |
| 0.692597 | 0.233234 | 0.039 | 0.035 | 1 | 1 | Ifi47 | 0.004 | Old |
| 0.693076 | 0.043096 | 0.045 | 0.041 | 1 | 1 | Ptpn9 | 0.004 | Old |
| 0.693166 | 0.209984 | 0.015 | 0.013 | 1 | 1 | Prmt7 | 0.002 | Old |
| 0.693245 | 0.213883 | 0.098 | 0.093 | 1 | 1 | Nsfl1c | 0.005 | Old |

| | | | | | | | | |
|---|---|---|---|---|---|---|---|---|
| 0.693487 | 0.291185 | 0.102 | 0.097 | 1 | 1 | Mgat2 | 0.005 | Old |
| 0.693574 | 0.1504 | 0.114 | 0.123 | 1 | 1 | Pnpla8 | -0.009 | Old |
| 0.693644 | 0.50372 | 0.035 | 0.032 | 1 | 1 | Fbxo28 | 0.003 | Old |
| 0.693675 | 0.438529 | 0.039 | 0.035 | 1 | 1 | Ppdpf | 0.004 | Old |
| 0.693695 | 0.053295 | 0.015 | 0.013 | 1 | 1 | Tacc3 | 0.002 | Old |
| 0.693746 | 0.179336 | 0.111 | 0.106 | 1 | 1 | Bcl3 | 0.005 | Old |
| 0.694093 | 0.633457 | 0.023 | 0.021 | 1 | 1 | Tmem70 | 0.002 | Old |
| 0.695135 | 0.32859 | 0.019 | 0.017 | 1 | 1 | Unc45a | 0.002 | Old |
| 0.695389 | 0.271847 | 0.093 | 0.089 | 1 | 1 | Ak2 | 0.004 | Old |
| 0.695755 | 0.460989 | 0.109 | 0.106 | 1 | 1 | Sfxn3 | 0.003 | Old |
| 0.69583 | 0.281257 | 0.035 | 0.032 | 1 | 1 | Anapc10 | 0.003 | Old |
| 0.696116 | 0.137076 | 0.147 | 0.156 | 1 | 1 | Efr3a | -0.009 | Old |
| 0.696485 | 0.106182 | 0.032 | 0.035 | 1 | 1 | Eef1e1 | -0.003 | Old |
| 0.696595 | 0.527491 | 0.017 | 0.019 | 1 | 1 | Fbxw5 | -0.002 | Old |
| 0.696676 | 0.483674 | 0.048 | 0.044 | 1 | 1 | Hhex | 0.004 | Old |
| 0.697179 | 0.312437 | 0.154 | 0.152 | 1 | 1 | Gm16286 | 0.002 | Old |
| 0.69732 | 0.512783 | 0.057 | 0.053 | 1 | 1 | Asxl1 | 0.004 | Old |
| 0.697401 | 0.073635 | 0.024 | 0.027 | 1 | 1 | Rfx5 | -0.003 | Old |
| 0.69772 | 0.300944 | 0.104 | 0.112 | 1 | 1 | Mcts1 | -0.008 | Old |
| 0.697988 | 0.238273 | 0.177 | 0.174 | 1 | 1 | Rab31 | 0.003 | Old |
| 0.698043 | 0.202397 | 0.033 | 0.03 | 1 | 1 | Trappc12 | 0.003 | Old |
| 0.698595 | 0.209352 | 0.015 | 0.018 | 1 | 1 | Bloc1s6 | -0.003 | Old |
| 0.699044 | 0.288498 | 0.252 | 0.256 | 1 | 1 | Lgals1 | -0.004 | Old |
| 0.699164 | 0.291434 | 0.069 | 0.066 | 1 | 1 | Amdhd2 | 0.003 | Old |
| 0.699193 | 0.26801 | 0.122 | 0.119 | 1 | 1 | Cnih1 | 0.003 | Old |
| 0.699218 | 0.410803 | 0.036 | 0.04 | 1 | 1 | Cdc14b | -0.004 | Old |
| 0.699354 | 0.03479 | 0.024 | 0.027 | 1 | 1 | Sun1 | -0.003 | Old |
| 0.700048 | 0.05811 | 0.039 | 0.042 | 1 | 1 | Evi2 | -0.003 | Old |
| 0.700229 | 0.172132 | 0.27 | 0.293 | 1 | 1 | Atp6v1g1 | -0.023 | Old |
| 0.700263 | 0.200462 | 0.032 | 0.035 | 1 | 1 | Tbc1d8 | -0.003 | Old |
| 0.700368 | 0.302564 | 0.049 | 0.053 | 1 | 1 | Tbc1d4 | -0.004 | Old |
| 0.700869 | 0.08079 | 0.402 | 0.435 | 1 | 1 | Fcgrt | -0.033 | Old |
| 0.700981 | 0.488853 | 0.031 | 0.028 | 1 | 1 | Lztfl1 | 0.003 | Old |
| 0.701005 | 0.316209 | 0.027 | 0.024 | 1 | 1 | Mfsd8 | 0.003 | Old |
| 0.701323 | 0.210594 | 0.076 | 0.072 | 1 | 1 | Lrrc8c | 0.004 | Old |
| 0.701353 | 0.40016 | 0.031 | 0.028 | 1 | 1 | Ipo13 | 0.003 | Old |
| 0.701995 | 0.064204 | 0.068 | 0.064 | 1 | 1 | Bet1 | 0.004 | Old |
| 0.702217 | 1.101632 | 0.013 | 0.011 | 1 | 1 | Zmat1 | 0.002 | Old |
| 0.702234 | 0.199703 | 0.015 | 0.018 | 1 | 1 | Dph6 | -0.003 | Old |
| 0.702259 | 0.31934 | 0.023 | 0.021 | 1 | 1 | Med27 | 0.002 | Old |
| 0.702994 | 0.248742 | 0.068 | 0.074 | 1 | 1 | Derl2 | -0.006 | Old |
| 0.703113 | 0.124828 | 0.197 | 0.208 | 1 | 1 | Zbtb7a | -0.011 | Old |
| 0.70312 | 0.263853 | 0.046 | 0.043 | 1 | 1 | Slc25a1 | 0.003 | Old |
| 0.703282 | 0.425574 | 0.035 | 0.032 | 1 | 1 | Kin | 0.003 | Old |
| 0.70356 | 0.062633 | 0.315 | 0.337 | 1 | 1 | Eif4a1 | -0.022 | Old |

| | | | | | | | | |
|---|---|---|---|---|---|---|---|---|
| 0.703851 | 0.419717 | 0.183 | 0.182 | 1 | 1 | Lfng | 0.001 | Old |
| 0.703899 | 0.510229 | 0.033 | 0.037 | 1 | 1 | Nop56 | -0.004 | Old |
| 0.703935 | 0.701766 | 0.015 | 0.018 | 1 | 1 | Naa16 | -0.003 | Old |
| 0.703935 | 0.355694 | 0.015 | 0.018 | 1 | 1 | Cep70 | -0.003 | Old |
| 0.704736 | 0.765879 | 0.036 | 0.033 | 1 | 1 | Mrps7 | 0.003 | Old |
| 0.704989 | 0.255962 | 0.027 | 0.024 | 1 | 1 | Prkar2b | 0.003 | Old |
| 0.705168 | 0.578456 | 0.066 | 0.063 | 1 | 1 | Ell2 | 0.003 | Old |
| 0.7053 | 0.040438 | 0.014 | 0.016 | 1 | 1 | Umps | -0.002 | Old |
| 0.706123 | 0.338369 | 0.078 | 0.075 | 1 | 1 | Abhd17b | 0.003 | Old |
| 0.706141 | 0.317012 | 0.082 | 0.078 | 1 | 1 | Marveld1 | 0.004 | Old |
| 0.70657 | 0.188239 | 0.014 | 0.016 | 1 | 1 | Ogfod3 | -0.002 | Old |
| 0.707019 | 0.198854 | 0.049 | 0.053 | 1 | 1 | Fam199x | -0.004 | Old |
| 0.707332 | 1.273963 | 0.017 | 0.015 | 1 | 1 | Fahd1 | 0.002 | Old |
| 0.707426 | 0.13661 | 0.019 | 0.017 | 1 | 1 | Aasdhppt | 0.002 | Old |
| 0.707587 | 0.136916 | 0.014 | 0.016 | 1 | 1 | Dbf4 | -0.002 | Old |
| 0.707744 | 0.227896 | 0.039 | 0.035 | 1 | 1 | Pml | 0.004 | Old |
| 0.707795 | 0.152854 | 0.071 | 0.075 | 1 | 1 | Ythdc2 | -0.004 | Old |
| 0.707841 | 0.183928 | 0.014 | 0.016 | 1 | 1 | Fam53a | -0.002 | Old |
| 0.708029 | 0.26278 | 0.051 | 0.048 | 1 | 1 | Ergic1 | 0.003 | Old |
| 0.708062 | 0.237335 | 0.174 | 0.171 | 1 | 1 | H2-M3 | 0.003 | Old |
| 0.708078 | 0.047882 | 0.062 | 0.058 | 1 | 1 | 2210016F1 | 0.004 | Old |
| 0.708157 | 0.076255 | 0.073 | 0.069 | 1 | 1 | Slc35a3 | 0.004 | Old |
| 0.708243 | 0.261002 | 0.094 | 0.1 | 1 | 1 | Smarce1 | -0.006 | Old |
| 0.708375 | 0.088193 | 0.019 | 0.017 | 1 | 1 | Dusp12 | 0.002 | Old |
| 0.708622 | 0.938012 | 0.013 | 0.011 | 1 | 1 | Fdps | 0.002 | Old |
| 0.708919 | 0.157548 | 0.076 | 0.081 | 1 | 1 | Srp19 | -0.005 | Old |
| 0.708992 | 0.05724 | 0.039 | 0.042 | 1 | 1 | Rfwd3 | -0.003 | Old |
| 0.709146 | 0.163459 | 0.022 | 0.024 | 1 | 1 | Ogfod1 | -0.002 | Old |
| 0.709206 | 0.470455 | 0.013 | 0.011 | 1 | 1 | Gm20300 | 0.002 | Old |
| 0.709327 | 0.121773 | 0.095 | 0.101 | 1 | 1 | Dpy19l1 | -0.006 | Old |
| 0.709354 | 0.053466 | 0.022 | 0.024 | 1 | 1 | Zdhhc6 | -0.002 | Old |
| 0.710233 | 0.412688 | 0.035 | 0.032 | 1 | 1 | Mum1 | 0.003 | Old |
| 0.710428 | 0.356544 | 0.069 | 0.066 | 1 | 1 | Dirc2 | 0.003 | Old |
| 0.710571 | 0.438413 | 0.091 | 0.088 | 1 | 1 | Ptprs | 0.003 | Old |
| 0.710665 | 0.46208 | 0.013 | 0.011 | 1 | 1 | Tnfrsf25 | 0.002 | Old |
| 0.710841 | 0.269926 | 0.067 | 0.064 | 1 | 1 | Exoc6b | 0.003 | Old |
| 0.710943 | 0.106622 | 0.058 | 0.062 | 1 | 1 | Mitf | -0.004 | Old |
| 0.711006 | 0.360526 | 0.058 | 0.055 | 1 | 1 | Cdk4 | 0.003 | Old |
| 0.711214 | 0.174101 | 0.224 | 0.222 | 1 | 1 | Ran | 0.002 | Old |
| 0.711548 | 0.212645 | 0.103 | 0.11 | 1 | 1 | Mertk | -0.007 | Old |
| 0.712167 | 0.088792 | 0.014 | 0.016 | 1 | 1 | Tfcp2 | -0.002 | Old |
| 0.712231 | 0.021635 | 0.03 | 0.032 | 1 | 1 | Slc38a7 | -0.002 | Old |
| 0.712421 | 0.045097 | 0.123 | 0.129 | 1 | 1 | Ap3s1 | -0.006 | Old |
| 0.712677 | 1.049648 | 0.017 | 0.015 | 1 | 1 | Tmem38b | 0.002 | Old |
| 0.712789 | 0.03016 | 0.058 | 0.062 | 1 | 1 | Itgb1bp1 | -0.004 | Old |

| | | | | | | | | |
|---|---|---|---|---|---|---|---|---|
| 0.712875 | 0.227101 | 0.1 | 0.096 | 1 | 1 | Yme1l1 | 0.004 | Old |
| 0.713002 | 0.330346 | 0.013 | 0.011 | 1 | 1 | Zfp952 | 0.002 | Old |
| 0.713361 | 0.21096 | 0.039 | 0.042 | 1 | 1 | Xdh | -0.003 | Old |
| 0.713505 | 0.192584 | 0.05 | 0.047 | 1 | 1 | Suclg1 | 0.003 | Old |
| 0.713883 | 0.079498 | 0.027 | 0.024 | 1 | 1 | Fktn | 0.003 | Old |
| 0.714199 | 0.028941 | 0.093 | 0.088 | 1 | 1 | Myd88 | 0.005 | Old |
| 0.714564 | 0.124731 | 0.127 | 0.135 | 1 | 1 | 2310035C2 | -0.008 | Old |
| 0.714723 | 0.223444 | 0.051 | 0.055 | 1 | 1 | Suco | -0.004 | Old |
| 0.714748 | 0.043662 | 0.05 | 0.054 | 1 | 1 | Pfdn1 | -0.004 | Old |
| 0.714799 | 0.819184 | 0.021 | 0.018 | 1 | 1 | Dgcr6 | 0.003 | Old |
| 0.714972 | 0.397352 | 0.049 | 0.053 | 1 | 1 | Akap11 | -0.004 | Old |
| 0.714992 | 0.470948 | 0.073 | 0.07 | 1 | 1 | Phf20 | 0.003 | Old |
| 0.715242 | 0.094418 | 0.05 | 0.054 | 1 | 1 | Med10 | -0.004 | Old |
| 0.715276 | 0.084869 | 0.046 | 0.043 | 1 | 1 | Fbxw7 | 0.003 | Old |
| 0.715359 | 0.422487 | 0.036 | 0.033 | 1 | 1 | Mthfd2 | 0.003 | Old |
| 0.71699 | 0.036954 | 0.027 | 0.024 | 1 | 1 | Rusc1 | 0.003 | Old |
| 0.717178 | 0.408833 | 0.036 | 0.033 | 1 | 1 | Cd99l2 | 0.003 | Old |
| 0.717214 | 0.074227 | 0.036 | 0.039 | 1 | 1 | Lnpk | -0.003 | Old |
| 0.717916 | 0.119288 | 0.031 | 0.034 | 1 | 1 | Prpf31 | -0.003 | Old |
| 0.717974 | 0.231752 | 0.113 | 0.12 | 1 | 1 | Rdx | -0.007 | Old |
| 0.718011 | 0.842293 | 0.021 | 0.018 | 1 | 1 | Hscb | 0.003 | Old |
| 0.718025 | 0.286993 | 0.086 | 0.083 | 1 | 1 | Rbfa | 0.003 | Old |
| 0.718346 | 0.277978 | 0.046 | 0.043 | 1 | 1 | Pla2g4a | 0.003 | Old |
| 0.718349 | 0.061084 | 0.05 | 0.054 | 1 | 1 | Sra1 | -0.004 | Old |
| 0.718664 | 0.44548 | 0.075 | 0.072 | 1 | 1 | Ncoa1 | 0.003 | Old |
| 0.718713 | 0.136205 | 0.053 | 0.057 | 1 | 1 | Setdb1 | -0.004 | Old |
| 0.718934 | 0.063251 | 0.039 | 0.042 | 1 | 1 | Entpd7 | -0.003 | Old |
| 0.719057 | 0.510887 | 0.014 | 0.016 | 1 | 1 | Zbtb33 | -0.002 | Old |
| 0.720335 | 0.369052 | 0.017 | 0.015 | 1 | 1 | Lin52 | 0.002 | Old |
| 0.7207 | 0.015944 | 0.042 | 0.046 | 1 | 1 | Prpf6 | -0.004 | Old |
| 0.720828 | 0.068975 | 0.139 | 0.134 | 1 | 1 | Dennd1a | 0.005 | Old |
| 0.721382 | 0.416673 | 0.045 | 0.042 | 1 | 1 | Zpr1 | 0.003 | Old |
| 0.721408 | 0.128601 | 0.105 | 0.1 | 1 | 1 | Kcnk6 | 0.005 | Old |
| 0.721481 | 0.095541 | 0.195 | 0.206 | 1 | 1 | Rps6ka3 | -0.011 | Old |
| 0.721933 | 0.130524 | 0.013 | 0.015 | 1 | 1 | D030056L2 | -0.002 | Old |
| 0.721961 | 0.265655 | 0.089 | 0.085 | 1 | 1 | Pcmt1 | 0.004 | Old |
| 0.722013 | 0.196061 | 0.325 | 0.334 | 1 | 1 | Reep5 | -0.009 | Old |
| 0.722292 | 0.146044 | 0.203 | 0.201 | 1 | 1 | Spop | 0.002 | Old |
| 0.722761 | 0.32908 | 0.049 | 0.046 | 1 | 1 | Tbc1d15 | 0.003 | Old |
| 0.722763 | 0.055486 | 0.057 | 0.061 | 1 | 1 | Cstf3 | -0.004 | Old |
| 0.723135 | 0.049341 | 0.028 | 0.031 | 1 | 1 | Tapbpl | -0.003 | Old |
| 0.723275 | 0.131591 | 0.013 | 0.015 | 1 | 1 | Mospd1 | -0.002 | Old |
| 0.7233 | 0.515643 | 0.021 | 0.018 | 1 | 1 | Alg2 | 0.003 | Old |
| 0.723352 | 0.494404 | 0.028 | 0.026 | 1 | 1 | Usp39 | 0.002 | Old |
| 0.723379 | 0.502395 | 0.046 | 0.044 | 1 | 1 | Togaram1 | 0.002 | Old |

| | | | | | | | | |
|---|---|---|---|---|---|---|---|---|
| 0.723522 | 0.630219 | 0.024 | 0.022 | 1 | 1 | Fas | 0.002 | Old |
| 0.723897 | 0.282505 | 0.197 | 0.199 | 1 | 1 | Sumo3 | -0.002 | Old |
| 0.724081 | 0.087762 | 0.013 | 0.015 | 1 | 1 | Alyref2 | -0.002 | Old |
| 0.724136 | 0.422423 | 0.028 | 0.026 | 1 | 1 | Crebl2 | 0.002 | Old |
| 0.724344 | 0.321999 | 0.023 | 0.026 | 1 | 1 | Maip1 | -0.003 | Old |
| 0.724711 | 0.113008 | 0.028 | 0.031 | 1 | 1 | Osbpl2 | -0.003 | Old |
| 0.72475 | 0.64876 | 0.023 | 0.026 | 1 | 1 | Mrto4 | -0.003 | Old |
| 0.725062 | 0.502841 | 0.06 | 0.058 | 1 | 1 | Mex3c | 0.002 | Old |
| 0.725105 | 0.108808 | 0.149 | 0.157 | 1 | 1 | Ist1 | -0.008 | Old |
| 0.725201 | 0.296265 | 0.017 | 0.015 | 1 | 1 | Mthfsd | 0.002 | Old |
| 0.725297 | 0.571968 | 0.059 | 0.056 | 1 | 1 | Txndc5 | 0.003 | Old |
| 0.725776 | 0.292081 | 0.036 | 0.033 | 1 | 1 | Cpox | 0.003 | Old |
| 0.725965 | 0.201949 | 0.053 | 0.057 | 1 | 1 | Hdac9 | -0.004 | Old |
| 0.726195 | 0.083315 | 0.83 | 0.858 | 1 | 1 | Ctsc | -0.028 | Old |
| 0.72628 | 0.113219 | 0.276 | 0.292 | 1 | 1 | Arf1 | -0.016 | Old |
| 0.726379 | 0.092769 | 0.051 | 0.048 | 1 | 1 | C1qtnf1 | 0.003 | Old |
| 0.726681 | 0.164905 | 0.076 | 0.072 | 1 | 1 | Nop58 | 0.004 | Old |
| 0.727017 | 0.330619 | 0.087 | 0.084 | 1 | 1 | Psmd6 | 0.003 | Old |
| 0.727213 | 0.095632 | 0.071 | 0.075 | 1 | 1 | Katna1 | -0.004 | Old |
| 0.72722 | 0.527312 | 0.042 | 0.047 | 1 | 1 | Gmppa | -0.005 | Old |
| 0.727253 | 0.837597 | 0.017 | 0.015 | 1 | 1 | Sdc1 | 0.002 | Old |
| 0.727305 | 0.421115 | 0.013 | 0.015 | 1 | 1 | Cipc | -0.002 | Old |
| 0.727305 | 0.334425 | 0.013 | 0.015 | 1 | 1 | Zfp574 | -0.002 | Old |
| 0.727605 | 0.323973 | 0.036 | 0.033 | 1 | 1 | Nrm | 0.003 | Old |
| 0.727637 | 0.67059 | 0.017 | 0.015 | 1 | 1 | Inafm2 | 0.002 | Old |
| 0.727651 | 0.214728 | 0.089 | 0.095 | 1 | 1 | Rasa3 | -0.006 | Old |
| 0.727679 | 0.181948 | 0.028 | 0.031 | 1 | 1 | Gm46224 | -0.003 | Old |
| 0.727703 | 0.285086 | 0.071 | 0.067 | 1 | 1 | Slc50a1 | 0.004 | Old |
| 0.727774 | 0.027622 | 0.062 | 0.066 | 1 | 1 | Ykt6 | -0.004 | Old |
| 0.728113 | 0.139107 | 0.224 | 0.22 | 1 | 1 | Atp6v0e | 0.004 | Old |
| 0.729058 | 0.029671 | 0.021 | 0.023 | 1 | 1 | Nudt14 | -0.002 | Old |
| 0.729078 | 0.136688 | 0.035 | 0.038 | 1 | 1 | Grpel1 | -0.003 | Old |
| 0.729196 | 0.361188 | 0.081 | 0.087 | 1 | 1 | Maz | -0.006 | Old |
| 0.729792 | 0.15245 | 0.144 | 0.152 | 1 | 1 | Tomm22 | -0.008 | Old |
| 0.72982 | 0.620425 | 0.017 | 0.015 | 1 | 1 | Bud23 | 0.002 | Old |
| 0.730138 | 0.370219 | 0.021 | 0.023 | 1 | 1 | Kctd6 | -0.002 | Old |
| 0.730161 | 0.042606 | 0.127 | 0.134 | 1 | 1 | Pikfyve | -0.007 | Old |
| 0.730218 | 0.342241 | 0.021 | 0.018 | 1 | 1 | Gpr89 | 0.003 | Old |
| 0.730591 | 0.056891 | 0.017 | 0.015 | 1 | 1 | 5730480H( | 0.002 | Old |
| 0.730911 | 0.447397 | 0.095 | 0.092 | 1 | 1 | Otulin | 0.003 | Old |
| 0.730915 | 0.131172 | 0.298 | 0.316 | 1 | 1 | Gpx4 | -0.018 | Old |
| 0.731167 | 0.174721 | 0.099 | 0.105 | 1 | 1 | Cyb5r4 | -0.006 | Old |
| 0.731175 | 0.29901 | 0.073 | 0.078 | 1 | 1 | Elf4 | -0.005 | Old |
| 0.731184 | 0.061303 | 0.09 | 0.095 | 1 | 1 | Cdc26 | -0.005 | Old |
| 0.731343 | 0.239557 | 0.013 | 0.015 | 1 | 1 | Pced1a | -0.002 | Old |

| | | | | | | | | |
|---|---|---|---|---|---|---|---|---|
| 0.731908 | 0.456989 | 0.099 | 0.097 | 1 | 1 | Slc15a4 | 0.002 | Old |
| 0.73233 | 0.232474 | 0.075 | 0.08 | 1 | 1 | Camta2 | -0.005 | Old |
| 0.732608 | 0.02345 | 0.012 | 0.013 | 1 | 1 | Tmem191c | -0.001 | Old |
| 0.732804 | 0.048178 | 0.172 | 0.168 | 1 | 1 | Stx4a | 0.004 | Old |
| 0.732905 | 0.012477 | 0.017 | 0.015 | 1 | 1 | Acrbp | 0.002 | Old |
| 0.732956 | 0.008468 | 0.048 | 0.051 | 1 | 1 | Parg | -0.003 | Old |
| 0.733104 | 0.34458 | 0.077 | 0.074 | 1 | 1 | Rab5b | 0.003 | Old |
| 0.733442 | 0.040118 | 0.028 | 0.031 | 1 | 1 | Tfam | -0.003 | Old |
| 0.733489 | 0.150381 | 0.166 | 0.176 | 1 | 1 | Id3 | -0.01 | Old |
| 0.733635 | 1.14796 | 0.01 | 0.009 | 1 | 1 | 2900076A0 | 0.001 | Old |
| 0.73377 | 0.422547 | 0.013 | 0.015 | 1 | 1 | Ccl22 | -0.002 | Old |
| 0.733791 | 0.112074 | 0.066 | 0.069 | 1 | 1 | Plpbp | -0.003 | Old |
| 0.734131 | 0.207867 | 0.172 | 0.171 | 1 | 1 | Pkig | 0.001 | Old |
| 0.734354 | 0.341482 | 0.021 | 0.023 | 1 | 1 | Ifi213 | -0.002 | Old |
| 0.734403 | 0.036542 | 0.035 | 0.038 | 1 | 1 | Cdk10 | -0.003 | Old |
| 0.734932 | 0.261323 | 0.202 | 0.204 | 1 | 1 | H13 | -0.002 | Old |
| 0.735049 | 0.193513 | 0.285 | 0.291 | 1 | 1 | Mapk3 | -0.006 | Old |
| 0.735982 | 0.164037 | 0.036 | 0.033 | 1 | 1 | Gm2000 | 0.003 | Old |
| 0.736489 | 0.238921 | 0.037 | 0.041 | 1 | 1 | Man1b1 | -0.004 | Old |
| 0.736659 | 0.15884 | 0.05 | 0.054 | 1 | 1 | Flvcr1 | -0.004 | Old |
| 0.736663 | 0.088524 | 0.258 | 0.273 | 1 | 1 | Psmb4 | -0.015 | Old |
| 0.736859 | 0.132646 | 0.033 | 0.036 | 1 | 1 | Atrn | -0.003 | Old |
| 0.736872 | 0.118986 | 0.012 | 0.013 | 1 | 1 | Zfp932 | -0.001 | Old |
| 0.737098 | 0.695867 | 0.03 | 0.027 | 1 | 1 | Mfge8 | 0.003 | Old |
| 0.737157 | 0.282235 | 0.012 | 0.013 | 1 | 1 | Lppos | -0.001 | Old |
| 0.73723 | 0.102948 | 0.558 | 0.572 | 1 | 1 | Cd36 | -0.014 | Old |
| 0.737923 | 0.705719 | 0.01 | 0.009 | 1 | 1 | Dsel | 0.001 | Old |
| 0.737923 | 0.587255 | 0.01 | 0.009 | 1 | 1 | Eif2d | 0.001 | Old |
| 0.738004 | 0.162676 | 0.094 | 0.1 | 1 | 1 | 1500011K1 | -0.006 | Old |
| 0.738186 | 0.187578 | 0.095 | 0.101 | 1 | 1 | Vcpip1 | -0.006 | Old |
| 0.738264 | 0.037281 | 0.027 | 0.03 | 1 | 1 | Foxo1 | -0.003 | Old |
| 0.738436 | 0.117155 | 0.197 | 0.209 | 1 | 1 | Hif1a | -0.012 | Old |
| 0.738746 | 0.307745 | 0.085 | 0.091 | 1 | 1 | Agfg1 | -0.006 | Old |
| 0.738798 | 0.121859 | 0.064 | 0.069 | 1 | 1 | Mcm3 | -0.005 | Old |
| 0.739245 | 0.637055 | 0.01 | 0.009 | 1 | 1 | Trim7 | 0.001 | Old |
| 0.739435 | 0.270643 | 0.012 | 0.013 | 1 | 1 | Thrb | -0.001 | Old |
| 0.739575 | 0.805864 | 0.01 | 0.009 | 1 | 1 | Itgb3bp | 0.001 | Old |
| 0.739575 | 0.734828 | 0.01 | 0.009 | 1 | 1 | Dnal4 | 0.001 | Old |
| 0.739593 | 0.239933 | 0.028 | 0.031 | 1 | 1 | Afdn | -0.003 | Old |
| 0.739721 | 0.28152 | 0.18 | 0.179 | 1 | 1 | Metap2 | 0.001 | Old |
| 0.740589 | 0.163222 | 0.036 | 0.039 | 1 | 1 | Pycr2 | -0.003 | Old |
| 0.740774 | 0.063324 | 0.089 | 0.094 | 1 | 1 | Acbd3 | -0.005 | Old |
| 0.740972 | 0.29375 | 0.154 | 0.151 | 1 | 1 | Cwc15 | 0.003 | Old |
| 0.741228 | 0.397839 | 0.01 | 0.009 | 1 | 1 | Fgfbp3 | 0.001 | Old |
| 0.741291 | 0.22277 | 0.283 | 0.305 | 1 | 1 | Jpt1 | -0.022 | Old |

| | | | | | | | | |
|---|---|---|---|---|---|---|---|---|
| 0.74161 | 0.212419 | 0.118 | 0.126 | 1 | 1 | Hnrnpr | -0.008 | Old |
| 0.741889 | 0.396845 | 0.01 | 0.009 | 1 | 1 | Atad5 | 0.001 | Old |
| 0.7419 | 0.339559 | 0.063 | 0.06 | 1 | 1 | Lactb | 0.003 | Old |
| 0.741925 | 0.242762 | 0.143 | 0.153 | 1 | 1 | Pgls | -0.01 | Old |
| 0.742006 | 0.344284 | 0.112 | 0.109 | 1 | 1 | Arf6 | 0.003 | Old |
| 0.742053 | 0.445532 | 0.033 | 0.031 | 1 | 1 | Slc13a3 | 0.002 | Old |
| 0.742078 | 0.014779 | 0.019 | 0.021 | 1 | 1 | Fdx1 | -0.002 | Old |
| 0.742173 | 0.201189 | 0.154 | 0.165 | 1 | 1 | Ubtf | -0.011 | Old |
| 0.742478 | 0.312749 | 0.078 | 0.075 | 1 | 1 | Ube2m | 0.003 | Old |
| 0.742637 | 0.272862 | 0.036 | 0.033 | 1 | 1 | Cwc22 | 0.003 | Old |
| 0.742871 | 0.664766 | 0.026 | 0.024 | 1 | 1 | Nop16 | 0.002 | Old |
| 0.743649 | 0.228292 | 0.019 | 0.021 | 1 | 1 | Mecr | -0.002 | Old |
| 0.743779 | 0.183239 | 0.095 | 0.101 | 1 | 1 | Eif1ax | -0.006 | Old |
| 0.744071 | 0.21157 | 0.035 | 0.038 | 1 | 1 | Mgst1 | -0.003 | Old |
| 0.744406 | 0.170134 | 0.08 | 0.076 | 1 | 1 | Plekhm2 | 0.004 | Old |
| 0.744571 | 0.025236 | 0.105 | 0.111 | 1 | 1 | Hs6st1 | -0.006 | Old |
| 0.74459 | 0.175726 | 0.15 | 0.158 | 1 | 1 | Fxr1 | -0.008 | Old |
| 0.745099 | 0.142628 | 0.216 | 0.228 | 1 | 1 | Sbno2 | -0.012 | Old |
| 0.745456 | 0.325486 | 0.051 | 0.055 | 1 | 1 | Espl1 | -0.004 | Old |
| 0.745496 | 0.622544 | 0.039 | 0.036 | 1 | 1 | Utp6 | 0.003 | Old |
| 0.745821 | 0.354244 | 0.089 | 0.086 | 1 | 1 | Lrrfip1 | 0.003 | Old |
| 0.746281 | 0.831215 | 0.012 | 0.013 | 1 | 1 | Urgcp | -0.001 | Old |
| 0.746298 | 0.162026 | 0.027 | 0.03 | 1 | 1 | Stx3 | -0.003 | Old |
| 0.746745 | 0.504801 | 0.01 | 0.012 | 1 | 1 | Pomk | -0.002 | Old |
| 0.746934 | 0.056669 | 0.067 | 0.071 | 1 | 1 | Stk11 | -0.004 | Old |
| 0.747132 | 0.215846 | 0.104 | 0.1 | 1 | 1 | Atp6v1h | 0.004 | Old |
| 0.747141 | 0.094125 | 0.281 | 0.302 | 1 | 1 | Bach1 | -0.021 | Old |
| 0.747177 | 0.149824 | 0.141 | 0.138 | 1 | 1 | Lyve1 | 0.003 | Old |
| 0.74721 | 0.143381 | 0.314 | 0.321 | 1 | 1 | Snx2 | -0.007 | Old |
| 0.747257 | 0.376553 | 0.082 | 0.08 | 1 | 1 | Ddx42 | 0.002 | Old |
| 0.747296 | 0.31332 | 0.107 | 0.113 | 1 | 1 | Cbfb | -0.006 | Old |
| 0.74732 | 0.088339 | 0.125 | 0.12 | 1 | 1 | Mkrn1 | 0.005 | Old |
| 0.747479 | 0.460611 | 0.049 | 0.052 | 1 | 1 | Ppm1g | -0.003 | Old |
| 0.747486 | 0.436584 | 0.021 | 0.023 | 1 | 1 | Bpnt1 | -0.002 | Old |
| 0.747819 | 0.434154 | 0.072 | 0.078 | 1 | 1 | C1qbp | -0.006 | Old |
| 0.748279 | 0.228285 | 0.158 | 0.168 | 1 | 1 | Hp1bp3 | -0.01 | Old |
| 0.748422 | 0.413205 | 0.107 | 0.103 | 1 | 1 | Ahcyl2 | 0.004 | Old |
| 0.74938 | 0.264149 | 0.231 | 0.233 | 1 | 1 | Sema4a | -0.002 | Old |
| 0.74994 | 0.733679 | 0.014 | 0.013 | 1 | 1 | Btbd3 | 0.001 | Old |
| 0.750079 | 0.424947 | 0.01 | 0.012 | 1 | 1 | Pkdcc | -0.002 | Old |
| 0.750079 | 0.105302 | 0.01 | 0.012 | 1 | 1 | Gm26601 | -0.002 | Old |
| 0.75039 | 0.776101 | 0.018 | 0.016 | 1 | 1 | Limk1 | 0.002 | Old |
| 0.750392 | 0.581843 | 0.022 | 0.02 | 1 | 1 | Ctdspl | 0.002 | Old |
| 0.750481 | 0.151065 | 0.154 | 0.151 | 1 | 1 | Tmem189 | 0.003 | Old |
| 0.750486 | 0.363892 | 0.058 | 0.055 | 1 | 1 | AU040320 | 0.003 | Old |

| | | | | | | | | |
|---|---|---|---|---|---|---|---|---|
| 0.75051 | 0.011517 | 0.045 | 0.048 | 1 | 1 | Pcnt | -0.003 | Old |
| 0.750685 | 0.415975 | 0.01 | 0.012 | 1 | 1 | Tlr8 | -0.002 | Old |
| 0.750714 | 0.144166 | 0.033 | 0.036 | 1 | 1 | Snrnp25 | -0.003 | Old |
| 0.750756 | 0.239189 | 0.172 | 0.171 | 1 | 1 | Mpp1 | 0.001 | Old |
| 0.750798 | 0.398259 | 0.03 | 0.027 | 1 | 1 | Ak6 | 0.003 | Old |
| 0.750851 | 0.487421 | 0.048 | 0.045 | 1 | 1 | Rars | 0.003 | Old |
| 0.751065 | 0.559036 | 0.014 | 0.013 | 1 | 1 | Gstcd | 0.001 | Old |
| 0.751251 | 0.347134 | 0.039 | 0.036 | 1 | 1 | Poglut1 | 0.003 | Old |
| 0.751293 | 0.813398 | 0.022 | 0.02 | 1 | 1 | Mgp | 0.002 | Old |
| 0.751346 | 0.539485 | 0.014 | 0.013 | 1 | 1 | Thada | 0.001 | Old |
| 0.751423 | 0.085612 | 0.055 | 0.059 | 1 | 1 | Ube2a | -0.004 | Old |
| 0.751742 | 0.272632 | 0.041 | 0.044 | 1 | 1 | Pot1a | -0.003 | Old |
| 0.751909 | 0.358891 | 0.014 | 0.013 | 1 | 1 | 5730409E0 | 0.001 | Old |
| 0.752098 | 0.372687 | 0.039 | 0.036 | 1 | 1 | Atxn1l | 0.003 | Old |
| 0.752146 | 0.322561 | 0.094 | 0.091 | 1 | 1 | Pptc7 | 0.003 | Old |
| 0.752395 | 0.34917 | 0.055 | 0.052 | 1 | 1 | Necap1 | 0.003 | Old |
| 0.752471 | 0.512929 | 0.014 | 0.013 | 1 | 1 | Gstt2 | 0.001 | Old |
| 0.752644 | 0.49221 | 0.019 | 0.021 | 1 | 1 | Fbxo31 | -0.002 | Old |
| 0.753114 | 0.73037 | 0.01 | 0.012 | 1 | 1 | Casz1 | -0.002 | Old |
| 0.75332 | 0.71878 | 0.022 | 0.02 | 1 | 1 | Gtf3c4 | 0.002 | Old |
| 0.753454 | 0.287845 | 0.057 | 0.061 | 1 | 1 | Gmps | -0.004 | Old |
| 0.753523 | 0.22571 | 0.058 | 0.062 | 1 | 1 | Sqor | -0.004 | Old |
| 0.753698 | 0.234454 | 0.111 | 0.107 | 1 | 1 | Cd37 | 0.004 | Old |
| 0.753721 | 0.38772 | 0.01 | 0.012 | 1 | 1 | Tmtc4 | -0.002 | Old |
| 0.754016 | 0.23346 | 0.033 | 0.036 | 1 | 1 | Polr1c | -0.003 | Old |
| 0.754221 | 0.703691 | 0.036 | 0.034 | 1 | 1 | Lpcat3 | 0.002 | Old |
| 0.754302 | 0.031723 | 0.663 | 0.713 | 1 | 1 | Oaz1 | -0.05 | Old |
| 0.754895 | 0.444623 | 0.026 | 0.024 | 1 | 1 | Khk | 0.002 | Old |
| 0.755005 | 0.351606 | 0.014 | 0.013 | 1 | 1 | Krt10 | 0.001 | Old |
| 0.75524 | 0.314065 | 0.01 | 0.012 | 1 | 1 | Zfyve19 | -0.002 | Old |
| 0.755527 | 0.253237 | 0.121 | 0.118 | 1 | 1 | Tns3 | 0.003 | Old |
| 0.75611 | 0.422664 | 0.018 | 0.016 | 1 | 1 | Fbxo6 | 0.002 | Old |
| 0.756132 | 0.40527 | 0.014 | 0.013 | 1 | 1 | Amd2 | 0.001 | Old |
| 0.756389 | 0.016206 | 0.107 | 0.112 | 1 | 1 | Gm6377 | -0.005 | Old |
| 0.756413 | 0.367101 | 0.014 | 0.013 | 1 | 1 | Sirpb1b | 0.001 | Old |
| 0.756477 | 0.011123 | 0.039 | 0.041 | 1 | 1 | Arl10 | -0.002 | Old |
| 0.756534 | 0.206294 | 0.112 | 0.117 | 1 | 1 | Slc25a37 | -0.005 | Old |
| 0.756547 | 0.387863 | 0.075 | 0.08 | 1 | 1 | Pbx3 | -0.005 | Old |
| 0.756745 | 0.335643 | 0.109 | 0.107 | 1 | 1 | Usp16 | 0.002 | Old |
| 0.756932 | 0.467295 | 0.044 | 0.041 | 1 | 1 | Cdc34 | 0.003 | Old |
| 0.756969 | 0.117798 | 0.075 | 0.071 | 1 | 1 | Stat5b | 0.004 | Old |
| 0.756977 | 0.37236 | 0.014 | 0.013 | 1 | 1 | Echdc1 | 0.001 | Old |
| 0.757197 | 0.276755 | 0.091 | 0.097 | 1 | 1 | Tug1 | -0.006 | Old |
| 0.757259 | 0.347722 | 0.014 | 0.013 | 1 | 1 | Gm26881 | 0.001 | Old |
| 0.757346 | 0.137831 | 0.113 | 0.109 | 1 | 1 | Elk4 | 0.004 | Old |

| | | | | | | | | |
|---|---|---|---|---|---|---|---|---|
| 0.758102 | 0.416943 | 0.018 | 0.016 | 1 | 1 | Mlh3 | 0.002 | Old |
| 0.758309 | 0.05509 | 0.059 | 0.063 | 1 | 1 | Fbxo3 | -0.004 | Old |
| 0.75832 | 0.099672 | 0.095 | 0.092 | 1 | 1 | Pole4 | 0.003 | Old |
| 0.758352 | 0.286061 | 0.018 | 0.016 | 1 | 1 | Mynn | 0.002 | Old |
| 0.758387 | 0.206508 | 0.014 | 0.013 | 1 | 1 | Rusc2 | 0.001 | Old |
| 0.758727 | 0.013511 | 0.037 | 0.04 | 1 | 1 | Rab10os | -0.003 | Old |
| 0.759414 | 0.272633 | 0.033 | 0.036 | 1 | 1 | Snapc5 | -0.003 | Old |
| 0.759612 | 0.333962 | 0.099 | 0.096 | 1 | 1 | Qdpr | 0.003 | Old |
| 0.759847 | 0.232262 | 0.149 | 0.16 | 1 | 1 | Psenen | -0.011 | Old |
| 0.759911 | 0.209268 | 0.03 | 0.027 | 1 | 1 | Mtmr1 | 0.003 | Old |
| 0.76007 | 0.299813 | 0.107 | 0.104 | 1 | 1 | Eif3b | 0.003 | Old |
| 0.76009 | 0.642999 | 0.019 | 0.021 | 1 | 1 | Orc5 | -0.002 | Old |
| 0.76009 | 0.507464 | 0.019 | 0.021 | 1 | 1 | Pan2 | -0.002 | Old |
| 0.760097 | 0.348469 | 0.018 | 0.016 | 1 | 1 | Naa25 | 0.002 | Old |
| 0.760432 | 0.285283 | 0.051 | 0.049 | 1 | 1 | Mrpl18 | 0.002 | Old |
| 0.761377 | 0.190379 | 0.026 | 0.028 | 1 | 1 | Rpf2 | -0.002 | Old |
| 0.76145 | 0.002335 | 0.031 | 0.033 | 1 | 1 | Vti1a | -0.002 | Old |
| 0.761514 | 0.159629 | 0.156 | 0.152 | 1 | 1 | Tsen34 | 0.004 | Old |
| 0.761556 | 0.293775 | 0.144 | 0.141 | 1 | 1 | Ppp2r1a | 0.003 | Old |
| 0.762049 | 0.170475 | 0.03 | 0.027 | 1 | 1 | Gon7 | 0.003 | Old |
| 0.762243 | 0.181386 | 0.03 | 0.027 | 1 | 1 | Clk2 | 0.003 | Old |
| 0.762419 | 0.718527 | 0.031 | 0.029 | 1 | 1 | Gpr35 | 0.002 | Old |
| 0.762474 | 0.245392 | 0.369 | 0.388 | 1 | 1 | Vamp8 | -0.019 | Old |
| 0.762518 | 0.134259 | 0.024 | 0.027 | 1 | 1 | Camsap1 | -0.003 | Old |
| 0.762805 | 0.073148 | 0.026 | 0.024 | 1 | 1 | Fam13b | 0.002 | Old |
| 0.763163 | 0.42957 | 0.064 | 0.062 | 1 | 1 | Mrps9 | 0.002 | Old |
| 0.763255 | 0.183547 | 0.026 | 0.028 | 1 | 1 | Mycbp | -0.002 | Old |
| 0.763327 | 0.229376 | 0.026 | 0.024 | 1 | 1 | Slc30a4 | 0.002 | Old |
| 0.763348 | 0.186304 | 0.17 | 0.168 | 1 | 1 | Prdx2 | 0.002 | Old |
| 0.763427 | 0.310544 | 0.009 | 0.01 | 1 | 1 | Aqp9 | -0.001 | Old |
| 0.763434 | 0.130439 | 0.18 | 0.19 | 1 | 1 | Rassf3 | -0.01 | Old |
| 0.763848 | 0.162575 | 0.026 | 0.024 | 1 | 1 | Utp18 | 0.002 | Old |
| 0.763927 | 0.347933 | 0.063 | 0.061 | 1 | 1 | Hmgcl | 0.002 | Old |
| 0.764383 | 0.148202 | 0.03 | 0.027 | 1 | 1 | Terf1 | 0.003 | Old |
| 0.764405 | 0.043459 | 0.009 | 0.01 | 1 | 1 | Evi5l | -0.001 | Old |
| 0.764744 | 0.191072 | 0.081 | 0.086 | 1 | 1 | Slc25a45 | -0.005 | Old |
| 0.765082 | 0.233395 | 0.064 | 0.061 | 1 | 1 | Rabgap1 | 0.003 | Old |
| 0.765369 | 0.301085 | 0.108 | 0.105 | 1 | 1 | Psmd1 | 0.003 | Old |
| 0.765478 | 0.016641 | 0.084 | 0.088 | 1 | 1 | Fto | -0.004 | Old |
| 0.7656 | 0.115886 | 0.509 | 0.538 | 1 | 1 | Eif4g2 | -0.029 | Old |
| 0.765612 | 0.165847 | 0.104 | 0.1 | 1 | 1 | Rad23a | 0.004 | Old |
| 0.765718 | 0.158417 | 0.032 | 0.035 | 1 | 1 | Bloc1s4 | -0.003 | Old |
| 0.765831 | 0.241296 | 0.026 | 0.024 | 1 | 1 | Nostrin | 0.002 | Old |
| 0.766108 | 0.042967 | 0.053 | 0.056 | 1 | 1 | Ppie | -0.003 | Old |
| 0.766539 | 0.035853 | 0.022 | 0.02 | 1 | 1 | Prcc | 0.002 | Old |

| | | | | | | | | |
|---|---|---|---|---|---|---|---|---|
| 0.766791 | 0.582599 | 0.027 | 0.03 | 1 | 1 | Tnip1 | -0.003 | Old |
| 0.766826 | 0.024949 | 0.096 | 0.1 | 1 | 1 | Mcmbp | -0.004 | Old |
| 0.766832 | 0.270942 | 0.085 | 0.082 | 1 | 1 | Ccndbp1 | 0.003 | Old |
| 0.767051 | 0.046153 | 0.017 | 0.018 | 1 | 1 | Cdc25a | -0.001 | Old |
| 0.767339 | 0.162366 | 0.009 | 0.01 | 1 | 1 | Fadd | -0.001 | Old |
| 0.767616 | 0.260547 | 0.154 | 0.162 | 1 | 1 | Larp1 | -0.008 | Old |
| 0.767858 | 0.323565 | 0.032 | 0.035 | 1 | 1 | Asnsd1 | -0.003 | Old |
| 0.767992 | 0.220559 | 0.009 | 0.01 | 1 | 1 | Acad11 | -0.001 | Old |
| 0.768108 | 0.278754 | 0.114 | 0.112 | 1 | 1 | Pdlim4 | 0.002 | Old |
| 0.768259 | 0.248269 | 0.067 | 0.071 | 1 | 1 | Sarnp | -0.004 | Old |
| 0.768484 | 0.124955 | 0.059 | 0.063 | 1 | 1 | 4930523C0 | -0.004 | Old |
| 0.768661 | 0.080595 | 0.032 | 0.035 | 1 | 1 | Armc10 | -0.003 | Old |
| 0.768668 | 0.044919 | 0.114 | 0.12 | 1 | 1 | Zfp516 | -0.006 | Old |
| 0.76875 | 0.043403 | 0.032 | 0.035 | 1 | 1 | Stx17 | -0.003 | Old |
| 0.768758 | 0.283934 | 0.033 | 0.036 | 1 | 1 | Dna2 | -0.003 | Old |
| 0.769464 | 0.189056 | 0.389 | 0.411 | 1 | 1 | Arl6ip1 | -0.022 | Old |
| 0.769999 | 0.144703 | 0.09 | 0.095 | 1 | 1 | Eny2 | -0.005 | Old |
| 0.770064 | 0.153507 | 0.054 | 0.058 | 1 | 1 | Ticam2 | -0.004 | Old |
| 0.770104 | 0.256256 | 0.055 | 0.052 | 1 | 1 | Ino80e | 0.003 | Old |
| 0.770205 | 0.307875 | 0.098 | 0.103 | 1 | 1 | Glrx5 | -0.005 | Old |
| 0.770274 | 0.125462 | 0.099 | 0.104 | 1 | 1 | Psmb9 | -0.005 | Old |
| 0.770278 | 0.445736 | 0.009 | 0.01 | 1 | 1 | Fam234b | -0.001 | Old |
| 0.770466 | 0.350359 | 0.054 | 0.052 | 1 | 1 | Parp9 | 0.002 | Old |
| 0.770565 | 0.041785 | 0.045 | 0.048 | 1 | 1 | Mmadhc | -0.003 | Old |
| 0.770665 | 0.019865 | 0.09 | 0.095 | 1 | 1 | Psmc5 | -0.005 | Old |
| 0.77083 | 0.575751 | 0.04 | 0.038 | 1 | 1 | Abhd4 | 0.002 | Old |
| 0.77087 | 0.228581 | 0.045 | 0.048 | 1 | 1 | Glo1 | -0.003 | Old |
| 0.771014 | 0.381593 | 0.166 | 0.168 | 1 | 1 | Spcs2 | -0.002 | Old |
| 0.771047 | 0.292796 | 0.046 | 0.049 | 1 | 1 | Dap3 | -0.003 | Old |
| 0.771368 | 0.074815 | 0.023 | 0.025 | 1 | 1 | Aamdc | -0.002 | Old |
| 0.771587 | 0.182772 | 0.249 | 0.252 | 1 | 1 | Bsg | -0.003 | Old |
| 0.772273 | 0.05091 | 0.024 | 0.027 | 1 | 1 | Slc7a6os | -0.003 | Old |
| 0.772503 | 0.355395 | 0.031 | 0.029 | 1 | 1 | Psmd5 | 0.002 | Old |
| 0.772529 | 0.262672 | 0.035 | 0.032 | 1 | 1 | Psmd10 | 0.003 | Old |
| 0.772735 | 0.227924 | 0.098 | 0.103 | 1 | 1 | Trim28 | -0.005 | Old |
| 0.772767 | 0.146628 | 0.373 | 0.383 | 1 | 1 | Pea15a | -0.01 | Old |
| 0.772785 | 0.411828 | 0.033 | 0.036 | 1 | 1 | Wdr77 | -0.003 | Old |
| 0.772847 | 0.173292 | 0.096 | 0.101 | 1 | 1 | Myo9a | -0.005 | Old |
| 0.77319 | 0.357372 | 0.024 | 0.027 | 1 | 1 | Golph3l | -0.003 | Old |
| 0.773459 | 0.048634 | 0.022 | 0.02 | 1 | 1 | Sephs2 | 0.002 | Old |
| 0.774646 | 0.138005 | 0.059 | 0.056 | 1 | 1 | Zfp385a | 0.003 | Old |
| 0.774735 | 0.376948 | 0.032 | 0.035 | 1 | 1 | Ncbp1 | -0.003 | Old |
| 0.775099 | 0.244663 | 0.017 | 0.018 | 1 | 1 | Cbfa2t2 | -0.001 | Old |
| 0.775808 | 0.166676 | 0.053 | 0.056 | 1 | 1 | Il6 | -0.003 | Old |
| 0.775853 | 0.761948 | 0.032 | 0.03 | 1 | 1 | Fmc1 | 0.002 | Old |

| | | | | | | | | |
|---|---|---|---|---|---|---|---|---|
| 0.776573 | 0.056421 | 0.059 | 0.062 | 1 | 1 | Tor1a | -0.003 | Old |
| 0.777023 | 0.139703 | 0.023 | 0.025 | 1 | 1 | Mib2 | -0.002 | Old |
| 0.778259 | 0.36829 | 0.095 | 0.093 | 1 | 1 | Pip5k1c | 0.002 | Old |
| 0.778586 | 0.009567 | 0.049 | 0.052 | 1 | 1 | Ipo8 | -0.003 | Old |
| 0.778919 | 0.331915 | 0.049 | 0.047 | 1 | 1 | Ppip5k2 | 0.002 | Old |
| 0.779075 | 0.220703 | 0.039 | 0.041 | 1 | 1 | Slc7a1 | -0.002 | Old |
| 0.779312 | 0.335596 | 0.031 | 0.033 | 1 | 1 | Med29 | -0.002 | Old |
| 0.779663 | 0.345161 | 0.054 | 0.058 | 1 | 1 | Pttg1 | -0.004 | Old |
| 0.779726 | 0.147381 | 0.037 | 0.04 | 1 | 1 | Ccdc134 | -0.003 | Old |
| 0.78019 | 0.172567 | 0.073 | 0.07 | 1 | 1 | Map2k1 | 0.003 | Old |
| 0.780302 | 0.114702 | 0.184 | 0.182 | 1 | 1 | Sgms1 | 0.002 | Old |
| 0.780589 | 0.001714 | 0.023 | 0.025 | 1 | 1 | Rfxap | -0.002 | Old |
| 0.78089 | 0.109354 | 0.114 | 0.12 | 1 | 1 | Bcl6 | -0.006 | Old |
| 0.780976 | 0.317364 | 0.049 | 0.047 | 1 | 1 | Srsf4 | 0.002 | Old |
| 0.781073 | 0.206035 | 0.08 | 0.084 | 1 | 1 | Phf23 | -0.004 | Old |
| 0.781875 | 0.443698 | 0.048 | 0.051 | 1 | 1 | Adssl1 | -0.003 | Old |
| 0.782005 | 1.00712 | 0.012 | 0.01 | 1 | 1 | Usp18 | 0.002 | Old |
| 0.782394 | 0.398707 | 0.086 | 0.084 | 1 | 1 | Slc8b1 | 0.002 | Old |
| 0.782437 | 0.295913 | 0.017 | 0.018 | 1 | 1 | Hacd2 | -0.001 | Old |
| 0.782674 | 0.227899 | 0.051 | 0.055 | 1 | 1 | Slc39a9 | -0.004 | Old |
| 0.782815 | 0.199774 | 0.058 | 0.061 | 1 | 1 | Fam213b | -0.003 | Old |
| 0.782934 | 0.531581 | 0.04 | 0.038 | 1 | 1 | Slc35b1 | 0.002 | Old |
| 0.783306 | 0.034009 | 0.036 | 0.038 | 1 | 1 | Lsm2 | -0.002 | Old |
| 0.783515 | 0.070645 | 0.03 | 0.032 | 1 | 1 | Relt | -0.002 | Old |
| 0.78366 | 0.258657 | 0.08 | 0.084 | 1 | 1 | C1d | -0.004 | Old |
| 0.783795 | 0.778874 | 0.019 | 0.018 | 1 | 1 | Alkbh7 | 0.001 | Old |
| 0.78414 | 0.384691 | 0.045 | 0.043 | 1 | 1 | Stk19 | 0.002 | Old |
| 0.784515 | 0.778153 | 0.012 | 0.01 | 1 | 1 | Rnf219 | 0.002 | Old |
| 0.784784 | 0.144309 | 0.015 | 0.017 | 1 | 1 | Abhd5 | -0.002 | Old |
| 0.784814 | 0.287261 | 0.054 | 0.052 | 1 | 1 | Pfdn4 | 0.002 | Old |
| 0.785305 | 0.154418 | 0.031 | 0.029 | 1 | 1 | Zfp593 | 0.002 | Old |
| 0.785337 | 0.183563 | 0.094 | 0.098 | 1 | 1 | Ctdsp1 | -0.004 | Old |
| 0.785631 | 0.247991 | 0.023 | 0.025 | 1 | 1 | Cd226 | -0.002 | Old |
| 0.785732 | 0.118973 | 0.158 | 0.165 | 1 | 1 | Ube2r2 | -0.007 | Old |
| 0.786057 | 0.177989 | 0.035 | 0.032 | 1 | 1 | Zdhhc18 | 0.003 | Old |
| 0.786061 | 0.031486 | 0.015 | 0.017 | 1 | 1 | Nedd4l | -0.002 | Old |
| 0.786232 | 0.260861 | 0.266 | 0.274 | 1 | 1 | Atp6ap2 | -0.008 | Old |
| 0.786272 | 0.05384 | 0.078 | 0.075 | 1 | 1 | Srf | 0.003 | Old |
| 0.786493 | 0.571616 | 0.015 | 0.014 | 1 | 1 | Angptl4 | 0.001 | Old |
| 0.786571 | 0.003916 | 0.015 | 0.017 | 1 | 1 | Commd9 | -0.002 | Old |
| 0.786868 | 0.455808 | 0.023 | 0.021 | 1 | 1 | Rnf214 | 0.002 | Old |
| 0.786963 | 0.035337 | 0.141 | 0.146 | 1 | 1 | Slc29a1 | -0.005 | Old |
| 0.787046 | 0.662993 | 0.053 | 0.051 | 1 | 1 | Naxd | 0.002 | Old |
| 0.787735 | 0.083128 | 0.023 | 0.025 | 1 | 1 | Slc20a2 | -0.002 | Old |
| 0.787785 | 0.210246 | 0.105 | 0.112 | 1 | 1 | AW112010 | -0.007 | Old |

| | | | | | | | | |
|---|---|---|---|---|---|---|---|---|
| 0.787849 | 0.220103 | 0.015 | 0.017 | 1 | 1 | Zfp444 | -0.002 | Old |
| 0.78797 | 0.247879 | 0.012 | 0.01 | 1 | 1 | Etaa1 | 0.002 | Old |
| 0.788274 | 0.196403 | 0.062 | 0.059 | 1 | 1 | Babam2 | 0.003 | Old |
| 0.788571 | 0.173849 | 0.022 | 0.024 | 1 | 1 | Flywch1 | -0.002 | Old |
| 0.788688 | 0.762201 | 0.028 | 0.027 | 1 | 1 | Bid | 0.001 | Old |
| 0.788913 | 0.300312 | 0.012 | 0.01 | 1 | 1 | Rad9a | 0.002 | Old |
| 0.788934 | 0.749321 | 0.015 | 0.014 | 1 | 1 | Polr3a | 0.001 | Old |
| 0.788998 | 0.379076 | 0.023 | 0.025 | 1 | 1 | Ganc | -0.002 | Old |
| 0.789081 | 0.225561 | 0.199 | 0.201 | 1 | 1 | Nckap1l | -0.002 | Old |
| 0.789411 | 0.396945 | 0.023 | 0.021 | 1 | 1 | Edc3 | 0.002 | Old |
| 0.789638 | 0.027163 | 0.015 | 0.017 | 1 | 1 | Atg16l2 | -0.002 | Old |
| 0.789767 | 0.47861 | 0.06 | 0.058 | 1 | 1 | Hopx | 0.002 | Old |
| 0.790543 | 0.12503 | 0.062 | 0.065 | 1 | 1 | Cd200r1 | -0.003 | Old |
| 0.790984 | 0.190739 | 0.22 | 0.232 | 1 | 1 | Plekhb2 | -0.012 | Old |
| 0.791115 | 0.235441 | 0.012 | 0.01 | 1 | 1 | Hirip3 | 0.002 | Old |
| 0.791151 | 0.230755 | 0.049 | 0.052 | 1 | 1 | Impa1 | -0.003 | Old |
| 0.791214 | 0.247969 | 0.054 | 0.052 | 1 | 1 | Calhm2 | 0.002 | Old |
| 0.791466 | 0.131354 | 0.045 | 0.048 | 1 | 1 | Cog3 | -0.003 | Old |
| 0.791567 | 0.25729 | 0.194 | 0.208 | 1 | 1 | Epsti1 | -0.014 | Old |
| 0.791734 | 0.401945 | 0.055 | 0.053 | 1 | 1 | Mvb12a | 0.002 | Old |
| 0.792098 | 0.226954 | 0.122 | 0.129 | 1 | 1 | Osbpl9 | -0.007 | Old |
| 0.792202 | 0.080166 | 0.575 | 0.645 | 1 | 1 | H3f3a | -0.07 | Old |
| 0.79267 | 0.262861 | 0.064 | 0.068 | 1 | 1 | Ganab | -0.004 | Old |
| 0.793007 | 0.41282 | 0.015 | 0.014 | 1 | 1 | Zkscan6 | 0.001 | Old |
| 0.793057 | 0.355137 | 0.072 | 0.07 | 1 | 1 | Cstf2t | 0.002 | Old |
| 0.793483 | 0.167411 | 0.044 | 0.041 | 1 | 1 | Nagk | 0.003 | Old |
| 0.793731 | 0.055035 | 0.015 | 0.017 | 1 | 1 | Ms4a4c | -0.002 | Old |
| 0.794094 | 0.320669 | 0.015 | 0.014 | 1 | 1 | Zbtb17 | 0.001 | Old |
| 0.794218 | 0.122705 | 0.093 | 0.089 | 1 | 1 | Rgl2 | 0.004 | Old |
| 0.794365 | 0.235909 | 0.015 | 0.014 | 1 | 1 | Adsl | 0.001 | Old |
| 0.794741 | 0.16371 | 0.055 | 0.058 | 1 | 1 | Mbd1 | -0.003 | Old |
| 0.794845 | 0.046736 | 0.06 | 0.064 | 1 | 1 | Smad2 | -0.004 | Old |
| 0.795292 | 0.27927 | 0.072 | 0.069 | 1 | 1 | Fundc2 | 0.003 | Old |
| 0.795296 | 0.033568 | 0.022 | 0.024 | 1 | 1 | Pex14 | -0.002 | Old |
| 0.795379 | 0.128144 | 0.027 | 0.025 | 1 | 1 | Urod | 0.002 | Old |
| 0.795593 | 0.336653 | 0.117 | 0.114 | 1 | 1 | 1810058I2 | 0.003 | Old |
| 0.79565 | 0.014344 | 0.035 | 0.037 | 1 | 1 | Klhl21 | -0.002 | Old |
| 0.795735 | 0.348478 | 0.068 | 0.066 | 1 | 1 | Lamtor5 | 0.002 | Old |
| 0.795795 | 0.173004 | 0.127 | 0.125 | 1 | 1 | Prkar2a | 0.002 | Old |
| 0.795974 | 0.130035 | 0.244 | 0.256 | 1 | 1 | Selenot | -0.012 | Old |
| 0.795987 | 0.161976 | 0.03 | 0.032 | 1 | 1 | Mrs2 | -0.002 | Old |
| 0.79641 | 0.305656 | 0.072 | 0.076 | 1 | 1 | Fbxw11 | -0.004 | Old |
| 0.796555 | 0.639232 | 0.033 | 0.032 | 1 | 1 | Jrkl | 0.001 | Old |
| 0.796596 | 0.271217 | 0.171 | 0.182 | 1 | 1 | Lbr | -0.011 | Old |
| 0.796813 | 0.279856 | 0.015 | 0.014 | 1 | 1 | Atg9a | 0.001 | Old |

| | | | | | | | | |
|---|---|---|---|---|---|---|---|---|
| 0.797204 | 0.102459 | 0.041 | 0.044 | 1 | 1 | Zfp984 | -0.003 | Old |
| 0.797386 | 0.111268 | 0.023 | 0.021 | 1 | 1 | Zfp608 | 0.002 | Old |
| 0.797959 | 0.22623 | 0.08 | 0.078 | 1 | 1 | Tomm34 | 0.002 | Old |
| 0.798127 | 0.028062 | 0.062 | 0.065 | 1 | 1 | Ccdc59 | -0.003 | Old |
| 0.798578 | 0.138673 | 0.054 | 0.052 | 1 | 1 | Spg20 | 0.002 | Old |
| 0.798609 | 0.337168 | 0.036 | 0.038 | 1 | 1 | Aspscr1 | -0.002 | Old |
| 0.798649 | 0.408006 | 0.067 | 0.071 | 1 | 1 | Mrps17 | -0.004 | Old |
| 0.798822 | 0.093783 | 0.054 | 0.057 | 1 | 1 | Hps5 | -0.003 | Old |
| 0.798984 | 0.062791 | 0.053 | 0.05 | 1 | 1 | Pdcd7 | 0.003 | Old |
| 0.799235 | 0.300876 | 0.078 | 0.076 | 1 | 1 | Mfsd11 | 0.002 | Old |
| 0.799262 | 0.157309 | 0.015 | 0.014 | 1 | 1 | Gtpbp8 | 0.001 | Old |
| 0.799686 | 0.079 | 0.847 | 0.875 | 1 | 1 | Rpsa | -0.028 | Old |
| 0.799787 | 0.090864 | 0.67 | 0.723 | 1 | 1 | Hsp90ab1 | -0.053 | Old |
| 0.799957 | 0.299673 | 0.145 | 0.144 | 1 | 1 | Cdk11b | 0.001 | Old |
| 0.800079 | 0.124446 | 0.015 | 0.014 | 1 | 1 | Dnajc18 | 0.001 | Old |
| 0.800182 | 0.097107 | 0.062 | 0.059 | 1 | 1 | Pigx | 0.003 | Old |
| 0.801053 | 0.033016 | 0.028 | 0.03 | 1 | 1 | Dcp1a | -0.002 | Old |
| 0.801384 | 0.196922 | 0.063 | 0.066 | 1 | 1 | D5Ertd579 | -0.003 | Old |
| 0.801557 | 0.052318 | 0.656 | 0.66 | 1 | 1 | F13a1 | -0.004 | Old |
| 0.802383 | 0.276555 | 0.036 | 0.038 | 1 | 1 | Ints9 | -0.002 | Old |
| 0.802526 | 0.023786 | 0.206 | 0.212 | 1 | 1 | Srsf6 | -0.006 | Old |
| 0.802972 | 0.081244 | 0.053 | 0.055 | 1 | 1 | A630001G | -0.002 | Old |
| 0.803312 | 0.126625 | 0.082 | 0.08 | 1 | 1 | Pa2g4 | 0.002 | Old |
| 0.804083 | 0.187115 | 0.05 | 0.048 | 1 | 1 | Umad1 | 0.002 | Old |
| 0.804197 | 0.208279 | 0.242 | 0.246 | 1 | 1 | Serinc1 | -0.004 | Old |
| 0.8042 | 0.211384 | 0.048 | 0.05 | 1 | 1 | Cops3 | -0.002 | Old |
| 0.804557 | 0.033423 | 0.069 | 0.072 | 1 | 1 | Npm3 | -0.003 | Old |
| 0.804594 | 0.018568 | 0.014 | 0.016 | 1 | 1 | Clstn1 | -0.002 | Old |
| 0.804715 | 0.122758 | 0.021 | 0.022 | 1 | 1 | Cgrrf1 | -0.001 | Old |
| 0.805127 | 0.336496 | 0.024 | 0.023 | 1 | 1 | Rprd1a | 0.001 | Old |
| 0.805213 | 0.284278 | 0.064 | 0.062 | 1 | 1 | B3gat3 | 0.002 | Old |
| 0.805259 | 0.322007 | 0.035 | 0.037 | 1 | 1 | Vps9d1 | -0.002 | Old |
| 0.805678 | 0.093063 | 0.028 | 0.03 | 1 | 1 | Arfgap3 | -0.002 | Old |
| 0.806039 | 0.449793 | 0.042 | 0.041 | 1 | 1 | Creb3l2 | 0.001 | Old |
| 0.806105 | 0.170904 | 0.14 | 0.147 | 1 | 1 | Ndufa1 | -0.007 | Old |
| 0.806212 | 0.173136 | 0.085 | 0.089 | 1 | 1 | Guk1 | -0.004 | Old |
| 0.806257 | 0.171737 | 0.028 | 0.03 | 1 | 1 | Sik3 | -0.002 | Old |
| 0.806359 | 0.337901 | 0.051 | 0.049 | 1 | 1 | Glod4 | 0.002 | Old |
| 0.806395 | 0.851609 | 0.03 | 0.028 | 1 | 1 | Hdac7 | 0.002 | Old |
| 0.806432 | 0.22543 | 0.081 | 0.085 | 1 | 1 | Slc12a7 | -0.004 | Old |
| 0.80645 | 0.1651 | 0.028 | 0.03 | 1 | 1 | Mthfr | -0.002 | Old |
| 0.806458 | 0.043252 | 0.06 | 0.058 | 1 | 1 | Slc39a10 | 0.002 | Old |
| 0.806554 | 0.231485 | 0.059 | 0.057 | 1 | 1 | Cnpy3 | 0.002 | Old |
| 0.807011 | 0.2611 | 0.014 | 0.016 | 1 | 1 | Nt5m | -0.002 | Old |
| 0.807513 | 0.215096 | 0.104 | 0.102 | 1 | 1 | Becn1 | 0.002 | Old |

| | | | | | | | | |
|---|---|---|---|---|---|---|---|---|
| 0.80761 | 0.174856 | 0.099 | 0.097 | 1 | 1 | Traf1 | 0.002 | Old |
| 0.807781 | 0.092232 | 0.136 | 0.142 | 1 | 1 | Psmd4 | -0.006 | Old |
| 0.808793 | 0.414548 | 0.022 | 0.024 | 1 | 1 | Cwc27 | -0.002 | Old |
| 0.809615 | 0.053648 | 0.098 | 0.095 | 1 | 1 | Jdp2 | 0.003 | Old |
| 0.809774 | 0.03256 | 0.022 | 0.024 | 1 | 1 | Pdp2 | -0.002 | Old |
| 0.809951 | 0.085446 | 0.338 | 0.357 | 1 | 1 | Itgb1 | -0.019 | Old |
| 0.810047 | 0.117645 | 0.094 | 0.098 | 1 | 1 | Ugp2 | -0.004 | Old |
| 0.81031 | 0.285109 | 0.028 | 0.03 | 1 | 1 | Dip2c | -0.002 | Old |
| 0.810577 | 0.09797 | 0.129 | 0.126 | 1 | 1 | Clec4b1 | 0.003 | Old |
| 0.810966 | 0.346232 | 0.032 | 0.03 | 1 | 1 | Mex3d | 0.002 | Old |
| 0.811045 | 0.215332 | 0.014 | 0.016 | 1 | 1 | Map4k3 | -0.002 | Old |
| 0.811083 | 0.078994 | 0.064 | 0.067 | 1 | 1 | Cep120 | -0.003 | Old |
| 0.811238 | 0.097555 | 0.021 | 0.022 | 1 | 1 | Supt3 | -0.001 | Old |
| 0.811566 | 0.175628 | 0.111 | 0.108 | 1 | 1 | Osgep | 0.003 | Old |
| 0.811953 | 0.342052 | 0.041 | 0.039 | 1 | 1 | Mosmo | 0.002 | Old |
| 0.812273 | 0.543874 | 0.051 | 0.049 | 1 | 1 | Gm8186 | 0.002 | Old |
| 0.812446 | 0.232426 | 0.158 | 0.157 | 1 | 1 | Psma2 | 0.001 | Old |
| 0.812617 | 0.313291 | 0.037 | 0.035 | 1 | 1 | Hras | 0.002 | Old |
| 0.812622 | 0.291907 | 0.041 | 0.039 | 1 | 1 | Josd1 | 0.002 | Old |
| 0.812721 | 0.111806 | 0.112 | 0.116 | 1 | 1 | Ddb1 | -0.004 | Old |
| 0.812815 | 0.192531 | 0.033 | 0.035 | 1 | 1 | Pgp | -0.002 | Old |
| 0.813001 | 0.16158 | 0.112 | 0.117 | 1 | 1 | Sertad1 | -0.005 | Old |
| 0.81344 | 0.242851 | 0.075 | 0.078 | 1 | 1 | Synj2bp | -0.003 | Old |
| 0.813531 | 0.204265 | 0.153 | 0.151 | 1 | 1 | Vapa | 0.002 | Old |
| 0.813766 | 0.121616 | 0.398 | 0.425 | 1 | 1 | Ifitm3 | -0.027 | Old |
| 0.813873 | 0.032226 | 0.549 | 0.572 | 1 | 1 | Sdcbp | -0.023 | Old |
| 0.814257 | 0.155504 | 0.027 | 0.029 | 1 | 1 | Nol4l | -0.002 | Old |
| 0.81428 | 0.093538 | 0.053 | 0.055 | 1 | 1 | Gcnt1 | -0.002 | Old |
| 0.814337 | 0.055701 | 0.076 | 0.079 | 1 | 1 | Tmem50b | -0.003 | Old |
| 0.814457 | 0.647068 | 0.021 | 0.019 | 1 | 1 | Jpt2 | 0.002 | Old |
| 0.814833 | 0.096705 | 0.129 | 0.133 | 1 | 1 | Vdac3 | -0.004 | Old |
| 0.815192 | 0.871106 | 0.017 | 0.016 | 1 | 1 | Nfatc2ip | 0.001 | Old |
| 0.815284 | 0.323832 | 0.024 | 0.023 | 1 | 1 | Sclt1 | 0.001 | Old |
| 0.815294 | 0.303398 | 0.369 | 0.391 | 1 | 1 | Gm26917 | -0.022 | Old |
| 0.815453 | 0.951717 | 0.017 | 0.016 | 1 | 1 | Prim2 | 0.001 | Old |
| 0.816108 | 0.044284 | 0.076 | 0.079 | 1 | 1 | Ccny | -0.003 | Old |
| 0.816205 | 0.124625 | 0.153 | 0.158 | 1 | 1 | Tcirg1 | -0.005 | Old |
| 0.816505 | 0.385217 | 0.082 | 0.081 | 1 | 1 | Cs | 0.001 | Old |
| 0.816532 | 0.034851 | 0.045 | 0.043 | 1 | 1 | Tufm | 0.002 | Old |
| 0.8166 | 0.656246 | 0.03 | 0.028 | 1 | 1 | Fam45a | 0.002 | Old |
| 0.816661 | 0.571883 | 0.039 | 0.037 | 1 | 1 | Creb3 | 0.002 | Old |
| 0.816761 | 0.950238 | 0.017 | 0.016 | 1 | 1 | Mbip | 0.001 | Old |
| 0.816815 | 0.358658 | 0.021 | 0.019 | 1 | 1 | Pramef8 | 0.002 | Old |
| 0.816843 | 1.051817 | 0.013 | 0.012 | 1 | 1 | Copz2 | 0.001 | Old |
| 0.817003 | 0.051119 | 0.045 | 0.047 | 1 | 1 | Nupr1 | -0.002 | Old |

| | | | | | | | | |
|---|---|---|---|---|---|---|---|---|
| 0.817168 | 0.499644 | 0.021 | 0.019 | 1 | 1 | Zfp746 | 0.002 | Old |
| 0.81733 | 0.091252 | 0.08 | 0.077 | 1 | 1 | Ndfip2 | 0.003 | Old |
| 0.817519 | 0.046754 | 0.068 | 0.071 | 1 | 1 | Arl1 | -0.003 | Old |
| 0.817719 | 0.2482 | 0.132 | 0.131 | 1 | 1 | Dnaja2 | 0.001 | Old |
| 0.817754 | 0.132133 | 0.072 | 0.069 | 1 | 1 | Gas2l1 | 0.003 | Old |
| 0.817778 | 0.654817 | 0.014 | 0.016 | 1 | 1 | Mad2l1bp | -0.002 | Old |
| 0.817778 | 0.333121 | 0.014 | 0.016 | 1 | 1 | Spice1 | -0.002 | Old |
| 0.817796 | 0.145846 | 0.099 | 0.097 | 1 | 1 | Nceh1 | 0.002 | Old |
| 0.817811 | 0.061554 | 0.041 | 0.039 | 1 | 1 | Naip5 | 0.002 | Old |
| 0.817884 | 0.31135 | 0.05 | 0.053 | 1 | 1 | Dram1 | -0.003 | Old |
| 0.818159 | 0.118021 | 0.09 | 0.094 | 1 | 1 | Tsg101 | -0.004 | Old |
| 0.819209 | 0.540338 | 0.013 | 0.014 | 1 | 1 | Pclaf | -0.001 | Old |
| 0.819414 | 0.333274 | 0.09 | 0.095 | 1 | 1 | Emd | -0.005 | Old |
| 0.819417 | 0.194033 | 0.06 | 0.064 | 1 | 1 | Plaur | -0.004 | Old |
| 0.819619 | 0.308275 | 0.048 | 0.05 | 1 | 1 | Igsf8 | -0.002 | Old |
| 0.819961 | 0.128453 | 0.28 | 0.293 | 1 | 1 | Myadm | -0.013 | Old |
| 0.820518 | 0.012559 | 0.062 | 0.064 | 1 | 1 | Stom | -0.002 | Old |
| 0.820653 | 0.228325 | 0.296 | 0.303 | 1 | 1 | Igf1 | -0.007 | Old |
| 0.820708 | 0.372177 | 0.021 | 0.019 | 1 | 1 | Mrpl46 | 0.002 | Old |
| 0.82118 | 0.477446 | 0.021 | 0.019 | 1 | 1 | Map4k2 | 0.002 | Old |
| 0.821222 | 0.136444 | 0.279 | 0.295 | 1 | 1 | Rpn2 | -0.016 | Old |
| 0.821229 | 0.191091 | 0.033 | 0.035 | 1 | 1 | G2e3 | -0.002 | Old |
| 0.82131 | 0.372397 | 0.035 | 0.037 | 1 | 1 | Hdgfl2 | -0.002 | Old |
| 0.821386 | 0.300173 | 0.021 | 0.022 | 1 | 1 | Ttc17 | -0.001 | Old |
| 0.821421 | 0.224175 | 0.211 | 0.222 | 1 | 1 | Rhog | -0.011 | Old |
| 0.821838 | 0.284292 | 0.021 | 0.022 | 1 | 1 | Cnpy4 | -0.001 | Old |
| 0.8219 | 0.256476 | 0.013 | 0.014 | 1 | 1 | Pola2 | -0.001 | Old |
| 0.822138 | 0.01444 | 0.129 | 0.133 | 1 | 1 | Cln8 | -0.004 | Old |
| 0.822271 | 0.222076 | 0.048 | 0.05 | 1 | 1 | Grsf1 | -0.002 | Old |
| 0.822318 | 0.124386 | 0.051 | 0.054 | 1 | 1 | Rfc2 | -0.003 | Old |
| 0.822318 | 0.112188 | 0.122 | 0.126 | 1 | 1 | Tmem164 | -0.004 | Old |
| 0.822419 | 0.339881 | 0.033 | 0.032 | 1 | 1 | Stam | 0.001 | Old |
| 0.822521 | 0.339235 | 0.017 | 0.016 | 1 | 1 | Tbrg4 | 0.001 | Old |
| 0.822566 | 0.206469 | 0.073 | 0.071 | 1 | 1 | Plin3 | 0.002 | Old |
| 0.82263 | 0.276497 | 0.035 | 0.037 | 1 | 1 | Hilpda | -0.002 | Old |
| 0.822826 | 0.243025 | 0.042 | 0.041 | 1 | 1 | Ccnk | 0.001 | Old |
| 0.823092 | 0.509376 | 0.03 | 0.028 | 1 | 1 | Sh3tc1 | 0.002 | Old |
| 0.823168 | 0.148886 | 0.032 | 0.034 | 1 | 1 | Mrps6 | -0.002 | Old |
| 0.823643 | 0.443436 | 0.089 | 0.094 | 1 | 1 | Cenpb | -0.005 | Old |
| 0.823645 | 0.861048 | 0.021 | 0.022 | 1 | 1 | Med22 | -0.001 | Old |
| 0.823701 | 0.490385 | 0.017 | 0.016 | 1 | 1 | Sorbs3 | 0.001 | Old |
| 0.823736 | 0.475921 | 0.024 | 0.023 | 1 | 1 | Spred2 | 0.001 | Old |
| 0.823843 | 0.123089 | 0.252 | 0.26 | 1 | 1 | S100a10 | -0.008 | Old |
| 0.824181 | 0.105201 | 0.026 | 0.027 | 1 | 1 | Klhl2 | -0.001 | Old |
| 0.824356 | 0.607292 | 0.017 | 0.016 | 1 | 1 | Wdr83 | 0.001 | Old |

| | | | | | | | | |
|---|---|---|---|---|---|---|---|---|
| 0.824504 | 0.305784 | 0.063 | 0.066 | 1 | 1 | Trpv4 | -0.003 | Old |
| 0.82519 | 0.055991 | 0.041 | 0.039 | 1 | 1 | Mta1 | 0.002 | Old |
| 0.825363 | 0.431446 | 0.027 | 0.029 | 1 | 1 | Senp3 | -0.002 | Old |
| 0.8256 | 0.054668 | 0.093 | 0.096 | 1 | 1 | Dynlt3 | -0.003 | Old |
| 0.825713 | 0.058223 | 0.033 | 0.035 | 1 | 1 | Zfp524 | -0.002 | Old |
| 0.825727 | 0.067354 | 0.013 | 0.014 | 1 | 1 | Egfr | -0.001 | Old |
| 0.825733 | 0.270573 | 0.046 | 0.044 | 1 | 1 | Nasp | 0.002 | Old |
| 0.825844 | 0.244982 | 0.046 | 0.049 | 1 | 1 | Nudcd2 | -0.003 | Old |
| 0.825906 | 0.145377 | 0.021 | 0.019 | 1 | 1 | Lamc1 | 0.002 | Old |
| 0.825936 | 0.115771 | 0.051 | 0.049 | 1 | 1 | Tmem183a | 0.002 | Old |
| 0.826075 | 0.504328 | 0.049 | 0.047 | 1 | 1 | Snrpa1 | 0.002 | Old |
| 0.826112 | 0.500628 | 0.013 | 0.012 | 1 | 1 | H2-T24 | 0.001 | Old |
| 0.826168 | 0.252314 | 0.078 | 0.083 | 1 | 1 | Xpot | -0.005 | Old |
| 0.826195 | 0.288235 | 0.082 | 0.086 | 1 | 1 | Lsm5 | -0.004 | Old |
| 0.826199 | 0.024068 | 0.102 | 0.105 | 1 | 1 | Ppp1r2 | -0.003 | Old |
| 0.826559 | 0.079718 | 0.08 | 0.083 | 1 | 1 | Ireb2 | -0.003 | Old |
| 0.826688 | 0.311084 | 0.051 | 0.049 | 1 | 1 | Trmt1l | 0.002 | Old |
| 0.826922 | 0.120981 | 0.171 | 0.177 | 1 | 1 | Gmfb | -0.006 | Old |
| 0.826989 | 0.111793 | 0.051 | 0.049 | 1 | 1 | Tbc1d22a | 0.002 | Old |
| 0.827196 | 0.109771 | 0.032 | 0.034 | 1 | 1 | Mroh1 | -0.002 | Old |
| 0.82731 | 0.425314 | 0.013 | 0.012 | 1 | 1 | Ttf2 | 0.001 | Old |
| 0.827705 | 0.031464 | 0.211 | 0.218 | 1 | 1 | Eif5b | -0.007 | Old |
| 0.827766 | 0.366368 | 0.017 | 0.016 | 1 | 1 | Rrp8 | 0.001 | Old |
| 0.828142 | 0.163132 | 0.154 | 0.154 | 1 | 1 | Camkk2 | 0 | Old |
| 0.828168 | 0.46574 | 0.021 | 0.022 | 1 | 1 | Atp2a3 | -0.001 | Old |
| 0.828341 | 0.502141 | 0.053 | 0.056 | 1 | 1 | Gon4l | -0.003 | Old |
| 0.828513 | 0.247518 | 0.024 | 0.023 | 1 | 1 | Srm | 0.001 | Old |
| 0.828675 | 0.191942 | 0.063 | 0.061 | 1 | 1 | Kdm5c | 0.002 | Old |
| 0.828855 | 0.164548 | 0.332 | 0.34 | 1 | 1 | Adam15 | -0.008 | Old |
| 0.828867 | 0.102143 | 0.026 | 0.027 | 1 | 1 | Eml3 | -0.001 | Old |
| 0.829162 | 0.028249 | 0.054 | 0.052 | 1 | 1 | Exosc3 | 0.002 | Old |
| 0.829341 | 0.098952 | 0.017 | 0.016 | 1 | 1 | Tmem9 | 0.001 | Old |
| 0.830006 | 0.378088 | 0.013 | 0.012 | 1 | 1 | Rasal2 | 0.001 | Old |
| 0.830129 | 0.131277 | 0.017 | 0.016 | 1 | 1 | Nod1 | 0.001 | Old |
| 0.830497 | 0.201315 | 0.048 | 0.046 | 1 | 1 | Wdr11 | 0.002 | Old |
| 0.830929 | 0.363648 | 0.027 | 0.029 | 1 | 1 | Memo1 | -0.002 | Old |
| 0.831179 | 0.194485 | 0.017 | 0.016 | 1 | 1 | 3110040N1 | 0.001 | Old |
| 0.831321 | 0.354521 | 0.063 | 0.061 | 1 | 1 | Rprd2 | 0.002 | Old |
| 0.831334 | 0.171954 | 0.042 | 0.041 | 1 | 1 | Nr2f6 | 0.001 | Old |
| 0.831505 | 0.263269 | 0.013 | 0.012 | 1 | 1 | Rnf135 | 0.001 | Old |
| 0.831505 | 0.099902 | 0.013 | 0.012 | 1 | 1 | Spryd3 | 0.001 | Old |
| 0.831675 | 0.280133 | 0.072 | 0.07 | 1 | 1 | Cope | 0.002 | Old |
| 0.831705 | 0.22177 | 0.017 | 0.016 | 1 | 1 | Pex11b | 0.001 | Old |
| 0.831871 | 0.185654 | 0.071 | 0.069 | 1 | 1 | Cln3 | 0.002 | Old |
| 0.832105 | 0.45707 | 0.013 | 0.012 | 1 | 1 | Coq9 | 0.001 | Old |

| | | | | | | | | |
|---|---|---|---|---|---|---|---|---|
| 0.832359 | 0.162333 | 0.03 | 0.028 | 1 | 1 | Rpa2 | 0.002 | Old |
| 0.832756 | 0.146889 | 0.017 | 0.016 | 1 | 1 | Ruvbl2 | 0.001 | Old |
| 0.832913 | 0.100191 | 0.033 | 0.032 | 1 | 1 | Cdkn2d | 0.001 | Old |
| 0.833005 | 0.035647 | 0.013 | 0.012 | 1 | 1 | Trdmt1 | 0.001 | Old |
| 0.833206 | 0.578229 | 0.05 | 0.049 | 1 | 1 | Ube3b | 0.001 | Old |
| 0.833281 | 0.088591 | 0.017 | 0.016 | 1 | 1 | Amz1 | 0.001 | Old |
| 0.833285 | 0.187024 | 0.221 | 0.233 | 1 | 1 | Synj1 | -0.012 | Old |
| 0.833395 | 0.493946 | 0.013 | 0.014 | 1 | 1 | Cdk7 | -0.001 | Old |
| 0.834146 | 0.082261 | 0.042 | 0.041 | 1 | 1 | Polr2b | 0.001 | Old |
| 0.834247 | 0.550033 | 0.013 | 0.014 | 1 | 1 | Fam117a | -0.001 | Old |
| 0.834505 | 0.096761 | 0.013 | 0.012 | 1 | 1 | Plekhh3 | 0.001 | Old |
| 0.834641 | 0.332535 | 0.045 | 0.047 | 1 | 1 | Ftsj3 | -0.002 | Old |
| 0.834893 | 0.304382 | 0.019 | 0.021 | 1 | 1 | Ncdn | -0.002 | Old |
| 0.834947 | 0.129811 | 0.024 | 0.026 | 1 | 1 | Mettl1 | -0.002 | Old |
| 0.835457 | 0.132291 | 0.333 | 0.355 | 1 | 1 | Clec2d | -0.022 | Old |
| 0.835566 | 0.155772 | 0.239 | 0.242 | 1 | 1 | Sgpp1 | -0.003 | Old |
| 0.835822 | 0.247121 | 0.19 | 0.19 | 1 | 1 | Scand1 | 0 | Old |
| 0.836069 | 0.626973 | 0.022 | 0.021 | 1 | 1 | Uck1 | 0.001 | Old |
| 0.83617 | 0.129777 | 0.033 | 0.032 | 1 | 1 | Polr2h | 0.001 | Old |
| 0.836198 | 0.122043 | 0.176 | 0.182 | 1 | 1 | Myl12a | -0.006 | Old |
| 0.836413 | 0.711406 | 0.022 | 0.021 | 1 | 1 | Rab13 | 0.001 | Old |
| 0.836533 | 0.141382 | 0.019 | 0.021 | 1 | 1 | Eif2b2 | -0.002 | Old |
| 0.836731 | 0.152762 | 0.037 | 0.035 | 1 | 1 | Decr1 | 0.002 | Old |
| 0.836807 | 1.062427 | 0.013 | 0.014 | 1 | 1 | Ttll4 | -0.001 | Old |
| 0.836841 | 0.096897 | 0.077 | 0.08 | 1 | 1 | Cxcl12 | -0.003 | Old |
| 0.837129 | 0.246632 | 0.039 | 0.041 | 1 | 1 | Naa60 | -0.002 | Old |
| 0.837219 | 0.576128 | 0.045 | 0.044 | 1 | 1 | Esrra | 0.001 | Old |
| 0.837242 | 0.481623 | 0.04 | 0.038 | 1 | 1 | Metap1 | 0.002 | Old |
| 0.838051 | 0.242466 | 0.103 | 0.101 | 1 | 1 | Pdcd6 | 0.002 | Old |
| 0.838202 | 0.260457 | 0.102 | 0.1 | 1 | 1 | Ddx46 | 0.002 | Old |
| 0.838645 | 0.168009 | 0.129 | 0.134 | 1 | 1 | Exoc5 | -0.005 | Old |
| 0.838766 | 0.016947 | 0.06 | 0.063 | 1 | 1 | Slc30a7 | -0.003 | Old |
| 0.838895 | 0.468415 | 0.027 | 0.029 | 1 | 1 | Slc39a3 | -0.002 | Old |
| 0.839007 | 0.223478 | 0.1 | 0.098 | 1 | 1 | Rchy1 | 0.002 | Old |
| 0.839041 | 0.06347 | 0.263 | 0.27 | 1 | 1 | Rnf13 | -0.007 | Old |
| 0.839067 | 0.38176 | 0.026 | 0.024 | 1 | 1 | Rrn3 | 0.002 | Old |
| 0.839213 | 0.249935 | 0.032 | 0.034 | 1 | 1 | Desi1 | -0.002 | Old |
| 0.839364 | 0.370979 | 0.04 | 0.042 | 1 | 1 | Stk25 | -0.002 | Old |
| 0.840122 | 0.171906 | 0.048 | 0.046 | 1 | 1 | 1110038F1 | 0.002 | Old |
| 0.840638 | 0.431231 | 0.019 | 0.021 | 1 | 1 | 10-Sep | -0.002 | Old |
| 0.840662 | 0.074466 | 0.012 | 0.013 | 1 | 1 | Smyd5 | -0.001 | Old |
| 0.840865 | 0.164587 | 0.14 | 0.139 | 1 | 1 | Ctdnep1 | 0.001 | Old |
| 0.840954 | 0.040478 | 0.06 | 0.063 | 1 | 1 | Crat | -0.003 | Old |
| 0.841263 | 0.195802 | 0.012 | 0.013 | 1 | 1 | Rasal3 | -0.001 | Old |
| 0.841296 | 0.272903 | 0.078 | 0.077 | 1 | 1 | Cnot4 | 0.001 | Old |

| | | | | | | | | |
|---|---|---|---|---|---|---|---|---|
| 0.841425 | 0.186323 | 0.086 | 0.089 | 1 | 1 | Cse1l | -0.003 | Old |
| 0.841488 | 0.359361 | 0.053 | 0.051 | 1 | 1 | Chm | 0.002 | Old |
| 0.841864 | 0.022291 | 0.012 | 0.013 | 1 | 1 | Ttc7b | -0.001 | Old |
| 0.842442 | 0.41044 | 0.03 | 0.028 | 1 | 1 | Scmh1 | 0.002 | Old |
| 0.842466 | 0.036079 | 0.012 | 0.013 | 1 | 1 | Sdhaf3 | -0.001 | Old |
| 0.842838 | 0.20915 | 0.035 | 0.033 | 1 | 1 | Gclm | 0.002 | Old |
| 0.843107 | 0.145112 | 0.272 | 0.275 | 1 | 1 | Hmox1 | -0.003 | Old |
| 0.843173 | 0.240314 | 0.084 | 0.082 | 1 | 1 | Shoc2 | 0.002 | Old |
| 0.843493 | 0.054082 | 0.048 | 0.046 | 1 | 1 | Med12 | 0.002 | Old |
| 0.84397 | 0.332278 | 0.012 | 0.013 | 1 | 1 | E130308A1 | -0.001 | Old |
| 0.844001 | 0.250249 | 0.283 | 0.295 | 1 | 1 | Sec61b | -0.012 | Old |
| 0.844164 | 0.092349 | 0.775 | 0.819 | 1 | 1 | Rpl7 | -0.044 | Old |
| 0.844872 | 0.172181 | 0.012 | 0.013 | 1 | 1 | Tfec | -0.001 | Old |
| 0.84494 | 0.101304 | 0.049 | 0.051 | 1 | 1 | Arfrp1 | -0.002 | Old |
| 0.845001 | 0.202433 | 0.026 | 0.024 | 1 | 1 | Ptp4a3 | 0.002 | Old |
| 0.845345 | 0.138382 | 0.24 | 0.252 | 1 | 1 | Mtpn | -0.012 | Old |
| 0.845537 | 0.159769 | 0.031 | 0.032 | 1 | 1 | Sbf1 | -0.001 | Old |
| 0.845635 | 0.134556 | 0.207 | 0.216 | 1 | 1 | Sec61g | -0.009 | Old |
| 0.84569 | 0.422233 | 0.054 | 0.057 | 1 | 1 | Dag1 | -0.003 | Old |
| 0.845775 | 0.273799 | 0.012 | 0.013 | 1 | 1 | Acaca | -0.001 | Old |
| 0.84603 | 0.157179 | 0.018 | 0.019 | 1 | 1 | Ccdc85b | -0.001 | Old |
| 0.846122 | 0.136187 | 0.13 | 0.134 | 1 | 1 | Irak1 | -0.004 | Old |
| 0.846274 | 0.003776 | 0.018 | 0.019 | 1 | 1 | Areg | -0.001 | Old |
| 0.846311 | 0.172378 | 0.059 | 0.061 | 1 | 1 | Cops5 | -0.002 | Old |
| 0.846315 | 0.242643 | 0.145 | 0.145 | 1 | 1 | Ndufs6 | 0 | Old |
| 0.846761 | 0.105521 | 0.018 | 0.019 | 1 | 1 | Fasn | -0.001 | Old |
| 0.846844 | 0.183103 | 0.121 | 0.119 | 1 | 1 | Dynlrb1 | 0.002 | Old |
| 0.847335 | 0.284414 | 0.026 | 0.024 | 1 | 1 | Scyl1 | 0.002 | Old |
| 0.847335 | 0.143753 | 0.024 | 0.026 | 1 | 1 | Fam98a | -0.002 | Old |
| 0.84742 | 0.170911 | 0.114 | 0.119 | 1 | 1 | Parp14 | -0.005 | Old |
| 0.847581 | 0.476012 | 0.012 | 0.013 | 1 | 1 | Mthfd1l | -0.001 | Old |
| 0.847892 | 0.215232 | 0.174 | 0.174 | 1 | 1 | Elavl1 | 0 | Old |
| 0.847977 | 0.22296 | 0.129 | 0.127 | 1 | 1 | Rad23b | 0.002 | Old |
| 0.848782 | 0.155532 | 0.03 | 0.028 | 1 | 1 | Arfgap1 | 0.002 | Old |
| 0.848821 | 0.334432 | 0.026 | 0.024 | 1 | 1 | Pigq | 0.002 | Old |
| 0.848891 | 0.209049 | 0.058 | 0.061 | 1 | 1 | Ccpg1 | -0.003 | Old |
| 0.848961 | 0.25419 | 0.058 | 0.061 | 1 | 1 | Taf13 | -0.003 | Old |
| 0.849194 | 0.133012 | 0.081 | 0.084 | 1 | 1 | Tmem173 | -0.003 | Old |
| 0.849374 | 0.272057 | 0.19 | 0.192 | 1 | 1 | Cct5 | -0.002 | Old |
| 0.849388 | 0.5163 | 0.012 | 0.013 | 1 | 1 | Abcc4 | -0.001 | Old |
| 0.849468 | 0.205058 | 0.114 | 0.114 | 1 | 1 | Lman1 | 0 | Old |
| 0.849643 | 0.241952 | 0.126 | 0.126 | 1 | 1 | Sltm | 0 | Old |
| 0.849689 | 0.438788 | 0.012 | 0.013 | 1 | 1 | Asah2 | -0.001 | Old |
| 0.849985 | 0.124353 | 0.096 | 0.094 | 1 | 1 | Mff | 0.002 | Old |
| 0.850021 | 0.500416 | 0.033 | 0.035 | 1 | 1 | Chchd5 | -0.002 | Old |

| | | | | | | | | |
|---|---|---|---|---|---|---|---|---|
| 0.850107 | 0.531118 | 0.041 | 0.04 | 1 | 1 | Avl9 | 0.001 | Old |
| 0.850261 | 0.204158 | 0.249 | 0.258 | 1 | 1 | Syngr2 | -0.009 | Old |
| 0.850281 | 0.105375 | 0.024 | 0.026 | 1 | 1 | Map2k5 | -0.002 | Old |
| 0.850404 | 0.215074 | 0.135 | 0.134 | 1 | 1 | Snrpd3 | 0.001 | Old |
| 0.850466 | 0.198122 | 0.089 | 0.092 | 1 | 1 | Ddhd2 | -0.003 | Old |
| 0.850696 | 0.121769 | 0.103 | 0.101 | 1 | 1 | Arnt | 0.002 | Old |
| 0.851146 | 0.148687 | 0.018 | 0.019 | 1 | 1 | Lrrc14 | -0.001 | Old |
| 0.851158 | 0.144539 | 0.107 | 0.104 | 1 | 1 | Alox5 | 0.003 | Old |
| 0.851315 | 0.59137 | 0.04 | 0.042 | 1 | 1 | Sp140 | -0.002 | Old |
| 0.851381 | 0.18566 | 0.314 | 0.323 | 1 | 1 | Srsf2 | -0.009 | Old |
| 0.851664 | 0.257227 | 0.04 | 0.038 | 1 | 1 | Taf2 | 0.002 | Old |
| 0.852083 | 0.263856 | 0.117 | 0.117 | 1 | 1 | Adam9 | 0 | Old |
| 0.852693 | 0.06855 | 0.036 | 0.038 | 1 | 1 | Akap10 | -0.002 | Old |
| 0.852871 | 0.228274 | 0.058 | 0.061 | 1 | 1 | Mink1 | -0.003 | Old |
| 0.852878 | 0.285558 | 0.113 | 0.112 | 1 | 1 | Arhgef12 | 0.001 | Old |
| 0.852904 | 0.23582 | 0.22 | 0.225 | 1 | 1 | Tomm20 | -0.005 | Old |
| 0.852947 | 0.011998 | 0.03 | 0.028 | 1 | 1 | Cln5 | 0.002 | Old |
| 0.853005 | 0.68117 | 0.018 | 0.017 | 1 | 1 | Qsox2 | 0.001 | Old |
| 0.853151 | 0.3481 | 0.06 | 0.059 | 1 | 1 | Snrnp27 | 0.001 | Old |
| 0.853171 | 0.0907 | 0.107 | 0.11 | 1 | 1 | Emp1 | -0.003 | Old |
| 0.853484 | 0.023942 | 0.591 | 0.636 | 1 | 1 | Sirpa | -0.045 | Old |
| 0.853873 | 0.452898 | 0.046 | 0.045 | 1 | 1 | Pi4ka | 0.001 | Old |
| 0.854055 | 0.311094 | 0.066 | 0.069 | 1 | 1 | Crim1 | -0.003 | Old |
| 0.854652 | 0.124087 | 0.461 | 0.487 | 1 | 1 | Fam49b | -0.026 | Old |
| 0.854927 | 0.197326 | 0.249 | 0.26 | 1 | 1 | Snrpb | -0.011 | Old |
| 0.855145 | 0.343379 | 0.104 | 0.109 | 1 | 1 | Mogs | -0.005 | Old |
| 0.855645 | 0.245109 | 0.162 | 0.163 | 1 | 1 | Klf10 | -0.001 | Old |
| 0.855714 | 0.327908 | 0.031 | 0.03 | 1 | 1 | Dgcr8 | 0.001 | Old |
| 0.855793 | 0.269969 | 0.018 | 0.017 | 1 | 1 | Prelid2 | 0.001 | Old |
| 0.855809 | 0.314863 | 0.082 | 0.081 | 1 | 1 | Renbp | 0.001 | Old |
| 0.856286 | 0.034324 | 0.054 | 0.056 | 1 | 1 | Trabd | -0.002 | Old |
| 0.856296 | 0.272746 | 0.049 | 0.051 | 1 | 1 | Hdac5 | -0.002 | Old |
| 0.856349 | 0.256881 | 0.044 | 0.046 | 1 | 1 | Micu2 | -0.002 | Old |
| 0.856868 | 0.252998 | 0.078 | 0.077 | 1 | 1 | Npepl1 | 0.001 | Old |
| 0.857177 | 0.443311 | 0.053 | 0.055 | 1 | 1 | Tmem57 | -0.002 | Old |
| 0.857279 | 0.24815 | 0.219 | 0.225 | 1 | 1 | Ndufc2 | -0.006 | Old |
| 0.857653 | 0.143958 | 0.149 | 0.154 | 1 | 1 | Myof | -0.005 | Old |
| 0.857657 | 0.235482 | 0.17 | 0.178 | 1 | 1 | Acer3 | -0.008 | Old |
| 0.857734 | 0.190117 | 0.179 | 0.185 | 1 | 1 | Psmd13 | -0.006 | Old |
| 0.858068 | 0.263814 | 0.199 | 0.202 | 1 | 1 | Pmaip1 | -0.003 | Old |
| 0.858221 | 0.250385 | 0.018 | 0.019 | 1 | 1 | Gpatch11 | -0.001 | Old |
| 0.858409 | 0.334197 | 0.134 | 0.134 | 1 | 1 | Rgs18 | 0 | Old |
| 0.858521 | 0.057397 | 0.085 | 0.087 | 1 | 1 | Slc25a28 | -0.002 | Old |
| 0.858857 | 0.287294 | 0.207 | 0.211 | 1 | 1 | Anp32e | -0.004 | Old |
| 0.85888 | 0.266 | 0.042 | 0.044 | 1 | 1 | Nudt9 | -0.002 | Old |

| | | | | | | | | |
|---|---|---|---|---|---|---|---|---|
| 0.858881 | 0.185875 | 0.118 | 0.123 | 1 | 1 | Uqcrfs1 | -0.005 | Old |
| 0.859512 | 0.363327 | 0.036 | 0.035 | 1 | 1 | Pign | 0.001 | Old |
| 0.859644 | 0.159017 | 0.145 | 0.151 | 1 | 1 | Camk1 | -0.006 | Old |
| 0.859947 | 0.253181 | 0.239 | 0.247 | 1 | 1 | Gsn | -0.008 | Old |
| 0.860764 | 0.160381 | 0.082 | 0.081 | 1 | 1 | Vps29 | 0.001 | Old |
| 0.861121 | 0.299802 | 0.018 | 0.017 | 1 | 1 | Pym1 | 0.001 | Old |
| 0.861121 | 0.287292 | 0.018 | 0.017 | 1 | 1 | B230307C2 | 0.001 | Old |
| 0.861172 | 0.161241 | 0.071 | 0.073 | 1 | 1 | Pik3r6 | -0.002 | Old |
| 0.861229 | 0.258747 | 0.181 | 0.183 | 1 | 1 | Pebp1 | -0.002 | Old |
| 0.861351 | 0.467347 | 0.014 | 0.013 | 1 | 1 | Hist1h4i | 0.001 | Old |
| 0.861849 | 0.058072 | 0.04 | 0.038 | 1 | 1 | Krr1 | 0.002 | Old |
| 0.862193 | 0.338363 | 0.109 | 0.109 | 1 | 1 | Ube2z | 0 | Old |
| 0.862259 | 0.200179 | 0.081 | 0.084 | 1 | 1 | Nap1l4 | -0.003 | Old |
| 0.862533 | 0.203266 | 0.102 | 0.1 | 1 | 1 | Hadhb | 0.002 | Old |
| 0.863432 | 0.150768 | 0.224 | 0.232 | 1 | 1 | Slc39a1 | -0.008 | Old |
| 0.863535 | 0.190046 | 0.072 | 0.075 | 1 | 1 | Slain2 | -0.003 | Old |
| 0.863661 | 0.348773 | 0.017 | 0.018 | 1 | 1 | Mark4 | -0.001 | Old |
| 0.863791 | 0.005277 | 0.067 | 0.069 | 1 | 1 | Tbc1d2b | -0.002 | Old |
| 0.864065 | 0.303282 | 0.027 | 0.026 | 1 | 1 | Lonp2 | 0.001 | Old |
| 0.864085 | 0.689231 | 0.018 | 0.019 | 1 | 1 | Sass6 | -0.001 | Old |
| 0.86444 | 0.03261 | 0.057 | 0.058 | 1 | 1 | Crtc2 | -0.001 | Old |
| 0.864687 | 0.274851 | 0.027 | 0.026 | 1 | 1 | Pias3 | 0.001 | Old |
| 0.864716 | 0.297339 | 0.057 | 0.055 | 1 | 1 | Crtap | 0.002 | Old |
| 0.864911 | 0.790613 | 0.028 | 0.027 | 1 | 1 | Zfp729a | 0.001 | Old |
| 0.864931 | 0.176094 | 0.017 | 0.018 | 1 | 1 | 6-Sep | -0.001 | Old |
| 0.865567 | 0.025632 | 0.01 | 0.011 | 1 | 1 | Sgf29 | -0.001 | Old |
| 0.865574 | 0.208397 | 0.301 | 0.312 | 1 | 1 | Nap1l1 | -0.011 | Old |
| 0.865639 | 0.24206 | 0.099 | 0.097 | 1 | 1 | Sp110 | 0.002 | Old |
| 0.86591 | 0.097548 | 0.104 | 0.108 | 1 | 1 | Cyld | -0.004 | Old |
| 0.866209 | 0.043437 | 0.01 | 0.011 | 1 | 1 | Tmem237 | -0.001 | Old |
| 0.866248 | 0.29011 | 0.081 | 0.08 | 1 | 1 | Adprh | 0.001 | Old |
| 0.866456 | 0.03912 | 0.018 | 0.017 | 1 | 1 | Zfp64 | 0.001 | Old |
| 0.866482 | 0.097997 | 0.085 | 0.087 | 1 | 1 | Ncoa2 | -0.002 | Old |
| 0.86651 | 0.16098 | 0.127 | 0.126 | 1 | 1 | Tle3 | 0.001 | Old |
| 0.866829 | 0.98363 | 0.01 | 0.01 | 1 | 1 | Tmem143 | 0 | Old |
| 0.866906 | 0.367918 | 0.023 | 0.024 | 1 | 1 | Wdsub1 | -0.001 | Old |
| 0.867043 | 0.229189 | 0.127 | 0.129 | 1 | 1 | Impact | -0.002 | Old |
| 0.867151 | 0.071737 | 0.104 | 0.102 | 1 | 1 | Gga1 | 0.002 | Old |
| 0.867272 | 0.335872 | 0.165 | 0.168 | 1 | 1 | Tns1 | -0.003 | Old |
| 0.867444 | 0.085605 | 0.081 | 0.083 | 1 | 1 | 2-Mar | -0.002 | Old |
| 0.867469 | 0.327132 | 0.118 | 0.117 | 1 | 1 | Cpne3 | 0.001 | Old |
| 0.86783 | 0.120548 | 0.071 | 0.073 | 1 | 1 | Nsun2 | -0.002 | Old |
| 0.867837 | 1.007906 | 0.01 | 0.01 | 1 | 1 | Sema4b | 0 | Old |
| 0.868175 | 0.02237 | 0.04 | 0.041 | 1 | 1 | Aars | -0.001 | Old |
| 0.86823 | 0.093057 | 0.075 | 0.077 | 1 | 1 | Map3k5 | -0.002 | Old |

| | | | | | | | | |
|---|---|---|---|---|---|---|---|---|
| 0.868426 | 0.896885 | 0.023 | 0.022 | 1 | 1 | Bri3bp | 0.001 | Old |
| 0.868554 | 0.033185 | 0.049 | 0.047 | 1 | 1 | Mrpl41 | 0.002 | Old |
| 0.868689 | 0.04244 | 0.028 | 0.03 | 1 | 1 | Atg14 | -0.002 | Old |
| 0.868716 | 0.091424 | 0.035 | 0.036 | 1 | 1 | Snhg12 | -0.001 | Old |
| 0.86878 | 0.255072 | 0.01 | 0.011 | 1 | 1 | Nipa1 | -0.001 | Old |
| 0.869061 | 0.248631 | 0.057 | 0.059 | 1 | 1 | Cib1 | -0.002 | Old |
| 0.869102 | 0.337413 | 0.01 | 0.011 | 1 | 1 | Slc19a1 | -0.001 | Old |
| 0.869164 | 0.149887 | 0.035 | 0.036 | 1 | 1 | Mapk9 | -0.001 | Old |
| 0.869243 | 0.026131 | 0.842 | 0.914 | 1 | 1 | Zfp36 | -0.072 | Old |
| 0.869371 | 0.249383 | 0.041 | 0.043 | 1 | 1 | Mpnd | -0.002 | Old |
| 0.869373 | 0.086676 | 0.051 | 0.053 | 1 | 1 | Nploc4 | -0.002 | Old |
| 0.86944 | 0.135533 | 0.19 | 0.196 | 1 | 1 | Mcub | -0.006 | Old |
| 0.869674 | 0.191583 | 0.081 | 0.083 | 1 | 1 | Etv6 | -0.002 | Old |
| 0.87 | 0.259216 | 0.153 | 0.154 | 1 | 1 | Supt16 | -0.001 | Old |
| 0.870069 | 0.522092 | 0.037 | 0.036 | 1 | 1 | Tpra1 | 0.001 | Old |
| 0.870416 | 0.019065 | 0.148 | 0.151 | 1 | 1 | D17Wsu92 | -0.003 | Old |
| 0.87052 | 0.034949 | 0.014 | 0.013 | 1 | 1 | Trmt10c | 0.001 | Old |
| 0.870524 | 0.469179 | 0.017 | 0.018 | 1 | 1 | Rps6kb2 | -0.001 | Old |
| 0.87071 | 0.737525 | 0.01 | 0.011 | 1 | 1 | Unc13d | -0.001 | Old |
| 0.870799 | 0.183578 | 0.09 | 0.093 | 1 | 1 | Cytip | -0.003 | Old |
| 0.870864 | 0.628767 | 0.01 | 0.01 | 1 | 1 | Lrrc45 | 0 | Old |
| 0.870932 | 0.018605 | 0.113 | 0.111 | 1 | 1 | Tmed3 | 0.002 | Old |
| 0.871031 | 0.401551 | 0.01 | 0.011 | 1 | 1 | Hspa2 | -0.001 | Old |
| 0.871223 | 0.167434 | 0.069 | 0.072 | 1 | 1 | Nfya | -0.003 | Old |
| 0.871252 | 0.505095 | 0.03 | 0.031 | 1 | 1 | Syap1 | -0.001 | Old |
| 0.8713 | 0.002311 | 0.063 | 0.061 | 1 | 1 | Nipsnap3b | 0.002 | Old |
| 0.87163 | 0.160089 | 0.077 | 0.08 | 1 | 1 | Cpeb2 | -0.003 | Old |
| 0.871667 | 0.230628 | 0.014 | 0.013 | 1 | 1 | Agk | 0.001 | Old |
| 0.87177 | 0.186916 | 0.114 | 0.119 | 1 | 1 | Prkcsh | -0.005 | Old |
| 0.871781 | 0.395369 | 0.054 | 0.053 | 1 | 1 | Arntl | 0.001 | Old |
| 0.872305 | 0.165654 | 0.017 | 0.018 | 1 | 1 | Supt7l | -0.001 | Old |
| 0.872318 | 0.642659 | 0.01 | 0.011 | 1 | 1 | 1600014C1 | -0.001 | Old |
| 0.872318 | 0.375946 | 0.01 | 0.011 | 1 | 1 | A130010J1 | -0.001 | Old |
| 0.872559 | 0.01124 | 0.018 | 0.017 | 1 | 1 | Zkscan1 | 0.001 | Old |
| 0.872993 | 0.129536 | 0.076 | 0.078 | 1 | 1 | Tmem33 | -0.002 | Old |
| 0.873038 | 0.000699 | 0.085 | 0.087 | 1 | 1 | Ezh1 | -0.002 | Old |
| 0.873332 | 0.11449 | 0.108 | 0.107 | 1 | 1 | Osbpl11 | 0.001 | Old |
| 0.873605 | 0.596781 | 0.01 | 0.011 | 1 | 1 | Zfp568 | -0.001 | Old |
| 0.873927 | 0.456882 | 0.01 | 0.011 | 1 | 1 | Cyp2j6 | -0.001 | Old |
| 0.874566 | 0.512184 | 0.01 | 0.01 | 1 | 1 | Mrpl47 | 0 | Old |
| 0.874566 | 0.388967 | 0.01 | 0.01 | 1 | 1 | Taf5 | 0 | Old |
| 0.874848 | 0.044338 | 0.203 | 0.207 | 1 | 1 | Shisa5 | -0.004 | Old |
| 0.874973 | 0.394435 | 0.121 | 0.128 | 1 | 1 | Lmo4 | -0.007 | Old |
| 0.875077 | 0.263925 | 0.035 | 0.036 | 1 | 1 | Ell | -0.001 | Old |
| 0.875553 | 0.202953 | 0.041 | 0.04 | 1 | 1 | Sc5d | 0.001 | Old |

| | | | | | | | | |
|---|---|---|---|---|---|---|---|---|
| 0.875576 | 0.150285 | 0.01 | 0.01 | 1 | 1 | Slfn9 | 0 | Old |
| 0.875646 | 0.001927 | 0.068 | 0.07 | 1 | 1 | Chmp2b | -0.002 | Old |
| 0.875869 | 0.517576 | 0.017 | 0.018 | 1 | 1 | Cbr1 | -0.001 | Old |
| 0.876581 | 0.429563 | 0.066 | 0.068 | 1 | 1 | Cops2 | -0.002 | Old |
| 0.87726 | 0.537063 | 0.01 | 0.01 | 1 | 1 | Zfp688 | 0 | Old |
| 0.87726 | 0.408075 | 0.01 | 0.01 | 1 | 1 | Oas1a | 0 | Old |
| 0.877412 | 0.22022 | 0.041 | 0.04 | 1 | 1 | Eif2b5 | 0.001 | Old |
| 0.877844 | 0.129544 | 0.066 | 0.068 | 1 | 1 | Fam192a | -0.002 | Old |
| 0.878271 | 0.170529 | 0.01 | 0.01 | 1 | 1 | Dhrs11 | 0 | Old |
| 0.878944 | 0.652221 | 0.01 | 0.01 | 1 | 1 | Plcb1 | 0 | Old |
| 0.879396 | 0.279098 | 0.066 | 0.064 | 1 | 1 | B4galnt1 | 0.002 | Old |
| 0.879507 | 0.220575 | 0.183 | 0.183 | 1 | 1 | Ppig | 0 | Old |
| 0.879742 | 0.043001 | 0.022 | 0.023 | 1 | 1 | Gfer | -0.001 | Old |
| 0.879865 | 0.603438 | 0.039 | 0.038 | 1 | 1 | Mtmr14 | 0.001 | Old |
| 0.879955 | 0.123823 | 0.01 | 0.01 | 1 | 1 | D3Ertd254 | 0 | Old |
| 0.88011 | 0.102428 | 0.183 | 0.182 | 1 | 1 | Zcchc24 | 0.001 | Old |
| 0.880165 | 0.319107 | 0.032 | 0.031 | 1 | 1 | Fkbp5 | 0.001 | Old |
| 0.8802 | 0.123997 | 0.077 | 0.075 | 1 | 1 | Fem1b | 0.002 | Old |
| 0.88109 | 0.336804 | 0.087 | 0.091 | 1 | 1 | Ubxn6 | -0.004 | Old |
| 0.881322 | 0.123208 | 0.027 | 0.028 | 1 | 1 | Stard5 | -0.001 | Old |
| 0.881537 | 0.316091 | 0.022 | 0.023 | 1 | 1 | Polr3b | -0.001 | Old |
| 0.882045 | 0.240745 | 0.028 | 0.03 | 1 | 1 | Snx11 | -0.002 | Old |
| 0.882923 | 0.252923 | 0.075 | 0.073 | 1 | 1 | Ube2v2 | 0.002 | Old |
| 0.883108 | 0.146944 | 0.022 | 0.023 | 1 | 1 | Gramd1b | -0.001 | Old |
| 0.883162 | 0.135328 | 0.041 | 0.04 | 1 | 1 | Ipo9 | 0.001 | Old |
| 0.884235 | 0.072132 | 0.183 | 0.188 | 1 | 1 | Stag2 | -0.005 | Old |
| 0.8843 | 0.728804 | 0.051 | 0.051 | 1 | 1 | Arpc5l | 0 | Old |
| 0.884744 | 0.406649 | 0.032 | 0.031 | 1 | 1 | Rpa1 | 0.001 | Old |
| 0.884936 | 0.093635 | 0.14 | 0.144 | 1 | 1 | Mgea5 | -0.004 | Old |
| 0.885186 | 0.318146 | 0.087 | 0.09 | 1 | 1 | Chka | -0.003 | Old |
| 0.885443 | 0.205691 | 0.289 | 0.298 | 1 | 1 | Gls | -0.009 | Old |
| 0.885455 | 0.236812 | 0.033 | 0.035 | 1 | 1 | Shc1 | -0.002 | Old |
| 0.885631 | 0.117302 | 0.287 | 0.291 | 1 | 1 | Dock2 | -0.004 | Old |
| 0.88594 | 0.456075 | 0.08 | 0.08 | 1 | 1 | Mbtps1 | 0 | Old |
| 0.885969 | 0.329529 | 0.042 | 0.041 | 1 | 1 | Thumpd1 | 0.001 | Old |
| 0.886017 | 0.499246 | 0.019 | 0.018 | 1 | 1 | Morc2a | 0.001 | Old |
| 0.886108 | 0.368859 | 0.028 | 0.03 | 1 | 1 | Pik3cb | -0.002 | Old |
| 0.886171 | 0.184497 | 0.077 | 0.076 | 1 | 1 | Ubp1 | 0.001 | Old |
| 0.886497 | 0.18102 | 0.04 | 0.041 | 1 | 1 | Atn1 | -0.001 | Old |
| 0.886541 | 0.067929 | 0.054 | 0.052 | 1 | 1 | Ddx10 | 0.002 | Old |
| 0.886589 | 0.035179 | 0.023 | 0.022 | 1 | 1 | Cdc42ep2 | 0.001 | Old |
| 0.886617 | 0.260752 | 0.442 | 0.486 | 1 | 1 | Trf | -0.044 | Old |
| 0.887066 | 0.181554 | 0.184 | 0.192 | 1 | 1 | S100a13 | -0.008 | Old |
| 0.887221 | 0.200464 | 0.287 | 0.301 | 1 | 1 | Cops9 | -0.014 | Old |
| 0.887344 | 0.544432 | 0.044 | 0.046 | 1 | 1 | Cyp4f16 | -0.002 | Old |

| | | | | | | | | |
|---|---|---|---|---|---|---|---|---|
| 0.887824 | 0.258705 | 0.022 | 0.023 | 1 | 1 | Mthfd1 | -0.001 | Old |
| 0.888115 | 0.441767 | 0.063 | 0.062 | 1 | 1 | Wdr82 | 0.001 | Old |
| 0.888363 | 0.272633 | 0.035 | 0.036 | 1 | 1 | Myo1e | -0.001 | Old |
| 0.888654 | 0.193012 | 0.219 | 0.229 | 1 | 1 | Ssb | -0.01 | Old |
| 0.888845 | 0.358779 | 0.033 | 0.035 | 1 | 1 | C2cd3 | -0.002 | Old |
| 0.888884 | 0.154224 | 0.042 | 0.041 | 1 | 1 | Commd4 | 0.001 | Old |
| 0.889207 | 0.111062 | 0.039 | 0.04 | 1 | 1 | Gpatch2 | -0.001 | Old |
| 0.889733 | 0.124575 | 0.1 | 0.103 | 1 | 1 | Copb1 | -0.003 | Old |
| 0.890236 | 0.23567 | 0.067 | 0.066 | 1 | 1 | Gba | 0.001 | Old |
| 0.890445 | 0.001212 | 0.073 | 0.075 | 1 | 1 | Slc25a51 | -0.002 | Old |
| 0.89069 | 0.026571 | 0.009 | 0.01 | 1 | 1 | Fam136a | -0.001 | Old |
| 0.890716 | 0.191926 | 0.203 | 0.212 | 1 | 1 | Taldo1 | -0.009 | Old |
| 0.890791 | 0.271078 | 0.08 | 0.083 | 1 | 1 | Tcp11l2 | -0.003 | Old |
| 0.890836 | 0.170356 | 0.105 | 0.105 | 1 | 1 | Psmb3 | 0 | Old |
| 0.891103 | 0.027829 | 0.037 | 0.038 | 1 | 1 | Tspan5 | -0.001 | Old |
| 0.891302 | 0.114471 | 0.071 | 0.069 | 1 | 1 | Fam20b | 0.002 | Old |
| 0.891728 | 0.259602 | 0.009 | 0.01 | 1 | 1 | Alg3 | -0.001 | Old |
| 0.892002 | 0.36362 | 0.122 | 0.123 | 1 | 1 | Armc8 | -0.001 | Old |
| 0.892074 | 0.076714 | 0.009 | 0.01 | 1 | 1 | Dnajc4 | -0.001 | Old |
| 0.892115 | 0.170663 | 0.051 | 0.053 | 1 | 1 | Slc35c2 | -0.002 | Old |
| 0.892383 | 0.040706 | 0.111 | 0.113 | 1 | 1 | Mat2b | -0.002 | Old |
| 0.892757 | 0.422852 | 0.033 | 0.032 | 1 | 1 | Alg5 | 0.001 | Old |
| 0.892803 | 0.377712 | 0.175 | 0.181 | 1 | 1 | Ndufa3 | -0.006 | Old |
| 0.893112 | 0.063705 | 0.009 | 0.01 | 1 | 1 | Blzf1 | -0.001 | Old |
| 0.893149 | 0.299859 | 0.024 | 0.024 | 1 | 1 | Mri1 | 0 | Old |
| 0.893324 | 0.04282 | 0.041 | 0.04 | 1 | 1 | Cyp4f13 | 0.001 | Old |
| 0.893458 | 0.744997 | 0.009 | 0.01 | 1 | 1 | Izumo4 | -0.001 | Old |
| 0.893586 | 0.391058 | 0.024 | 0.024 | 1 | 1 | Aktip | 0 | Old |
| 0.893836 | 0.30336 | 0.053 | 0.052 | 1 | 1 | Senp1 | 0.001 | Old |
| 0.894151 | 0.420765 | 0.009 | 0.01 | 1 | 1 | Grwd1 | -0.001 | Old |
| 0.894456 | 0.307088 | 0.049 | 0.048 | 1 | 1 | Lactb2 | 0.001 | Old |
| 0.894781 | 0.112225 | 0.166 | 0.171 | 1 | 1 | Ddi2 | -0.005 | Old |
| 0.894939 | 0.765458 | 0.03 | 0.029 | 1 | 1 | Pskh1 | 0.001 | Old |
| 0.895043 | 0.250493 | 0.046 | 0.048 | 1 | 1 | Foxred2 | -0.002 | Old |
| 0.895616 | 0.208508 | 0.089 | 0.088 | 1 | 1 | Anxa6 | 0.001 | Old |
| 0.895712 | 0.216319 | 0.053 | 0.052 | 1 | 1 | Nup62 | 0.001 | Old |
| 0.895882 | 0.776462 | 0.009 | 0.01 | 1 | 1 | Ptpn4 | -0.001 | Old |
| 0.895952 | 0.189047 | 0.027 | 0.028 | 1 | 1 | Iqsec1 | -0.001 | Old |
| 0.896058 | 0.439034 | 0.015 | 0.015 | 1 | 1 | Rgcc | 0 | Old |
| 0.896089 | 0.162438 | 0.033 | 0.035 | 1 | 1 | Slc35e3 | -0.002 | Old |
| 0.896155 | 0.143322 | 0.028 | 0.027 | 1 | 1 | Phkg2 | 0.001 | Old |
| 0.896228 | 0.757372 | 0.009 | 0.01 | 1 | 1 | Flrt3 | -0.001 | Old |
| 0.896257 | 0.056593 | 0.087 | 0.086 | 1 | 1 | Rnf213 | 0.001 | Old |
| 0.896776 | 0.323492 | 0.288 | 0.308 | 1 | 1 | Fn1 | -0.02 | Old |
| 0.89692 | 0.462192 | 0.009 | 0.01 | 1 | 1 | Map2k3os | -0.001 | Old |

| | | | | | | | | |
|---|---|---|---|---|---|---|---|---|
| 0.897412 | 0.068435 | 0.049 | 0.05 | 1 | 1 | Fut11 | -0.001 | Old |
| 0.898139 | 0.103072 | 0.084 | 0.082 | 1 | 1 | Pim3 | 0.002 | Old |
| 0.898257 | 0.207637 | 0.062 | 0.064 | 1 | 1 | D10Jhu81e | -0.002 | Old |
| 0.898702 | 0.207191 | 0.087 | 0.086 | 1 | 1 | Smu1 | 0.001 | Old |
| 0.898719 | 0.011615 | 0.026 | 0.027 | 1 | 1 | Csf2rb2 | -0.001 | Old |
| 0.898794 | 0.146193 | 0.049 | 0.05 | 1 | 1 | Adar | -0.001 | Old |
| 0.898956 | 0.365604 | 0.064 | 0.064 | 1 | 1 | Acaa1a | 0 | Old |
| 0.899131 | 0.165158 | 0.108 | 0.112 | 1 | 1 | Rnf44 | -0.004 | Old |
| 0.899141 | 0.248065 | 0.057 | 0.055 | 1 | 1 | Cltb | 0.002 | Old |
| 0.899226 | 0.154842 | 0.093 | 0.095 | 1 | 1 | Frg1 | -0.002 | Old |
| 0.899284 | 0.201424 | 0.046 | 0.048 | 1 | 1 | Pdgfa | -0.002 | Old |
| 0.899484 | 0.330774 | 0.024 | 0.024 | 1 | 1 | Gar1 | 0 | Old |
| 0.899604 | 0.294377 | 0.096 | 0.096 | 1 | 1 | Pdcl3 | 0 | Old |
| 0.8997 | 0.328208 | 0.081 | 0.081 | 1 | 1 | Galnt7 | 0 | Old |
| 0.90035 | 0.141647 | 0.12 | 0.119 | 1 | 1 | Pnn | 0.001 | Old |
| 0.900405 | 0.194548 | 0.057 | 0.058 | 1 | 1 | Zfp592 | -0.001 | Old |
| 0.901373 | 0.26391 | 0.033 | 0.032 | 1 | 1 | Xpo4 | 0.001 | Old |
| 0.901453 | 0.204934 | 0.064 | 0.066 | 1 | 1 | Ctsf | -0.002 | Old |
| 0.901542 | 0.101897 | 0.021 | 0.021 | 1 | 1 | Slc39a11 | 0 | Old |
| 0.901774 | 0.065814 | 0.021 | 0.021 | 1 | 1 | Il18 | 0 | Old |
| 0.902113 | 0.546592 | 0.048 | 0.047 | 1 | 1 | Terf2 | 0.001 | Old |
| 0.902283 | 0.251793 | 0.048 | 0.047 | 1 | 1 | Fam214b | 0.001 | Old |
| 0.902659 | 0.219029 | 0.175 | 0.177 | 1 | 1 | Ube2l3 | -0.002 | Old |
| 0.902664 | 0.162579 | 0.152 | 0.156 | 1 | 1 | Sh3kbp1 | -0.004 | Old |
| 0.903061 | 0.054127 | 0.075 | 0.076 | 1 | 1 | Rcn2 | -0.001 | Old |
| 0.903419 | 0.255636 | 0.024 | 0.024 | 1 | 1 | Fyco1 | 0 | Old |
| 0.903611 | 0.29706 | 0.015 | 0.015 | 1 | 1 | Ccdc28b | 0 | Old |
| 0.903637 | 0.164223 | 0.024 | 0.024 | 1 | 1 | Papolg | 0 | Old |
| 0.90377 | 0.24234 | 0.057 | 0.058 | 1 | 1 | Preb | -0.001 | Old |
| 0.903817 | 0.205736 | 0.042 | 0.041 | 1 | 1 | Fam120b | 0.001 | Old |
| 0.90396 | 0.278997 | 0.039 | 0.04 | 1 | 1 | Mafk | -0.001 | Old |
| 0.904298 | 0.281945 | 0.015 | 0.015 | 1 | 1 | Fam26f | 0 | Old |
| 0.904298 | 0.240907 | 0.015 | 0.015 | 1 | 1 | Plcg1 | 0 | Old |
| 0.904475 | 0.306689 | 0.039 | 0.04 | 1 | 1 | Brd9 | -0.001 | Old |
| 0.904671 | 0.229503 | 0.189 | 0.193 | 1 | 1 | Nras | -0.004 | Old |
| 0.905541 | 0.190088 | 0.185 | 0.194 | 1 | 1 | Polr1d | -0.009 | Old |
| 0.90575 | 0.151517 | 0.046 | 0.045 | 1 | 1 | Tmem88 | 0.001 | Old |
| 0.906016 | 0.17156 | 0.049 | 0.05 | 1 | 1 | Usp32 | -0.001 | Old |
| 0.906155 | 0.15809 | 0.032 | 0.033 | 1 | 1 | Xrcc1 | -0.001 | Old |
| 0.906175 | 0.193417 | 0.117 | 0.117 | 1 | 1 | Atp6v1c1 | 0 | Old |
| 0.906193 | 0.339391 | 0.039 | 0.04 | 1 | 1 | Tmem120a | -0.001 | Old |
| 0.906223 | 0.114302 | 0.015 | 0.015 | 1 | 1 | Exosc1 | 0 | Old |
| 0.906295 | 0.460257 | 0.039 | 0.038 | 1 | 1 | Tmem55a | 0.001 | Old |
| 0.90647 | 0.183516 | 0.037 | 0.038 | 1 | 1 | Abcd3 | -0.001 | Old |
| 0.906494 | 0.114132 | 0.342 | 0.357 | 1 | 1 | Atp5g2 | -0.015 | Old |

| | | | | | | | | |
|---|---|---|---|---|---|---|---|---|
| 0.906498 | 0.140609 | 0.015 | 0.015 | 1 | 1 | Hrh1 | 0 | Old |
| 0.907017 | 0.043369 | 0.068 | 0.069 | 1 | 1 | Bak1 | -0.001 | Old |
| 0.907058 | 0.189876 | 0.326 | 0.343 | 1 | 1 | Birc3 | -0.017 | Old |
| 0.908145 | 0.136137 | 0.09 | 0.092 | 1 | 1 | Iscu | -0.002 | Old |
| 0.908284 | 0.178881 | 0.042 | 0.044 | 1 | 1 | Rab9 | -0.002 | Old |
| 0.908451 | 0.262187 | 0.09 | 0.089 | 1 | 1 | Gm34084 | 0.001 | Old |
| 0.908532 | 0.080258 | 0.686 | 0.737 | 1 | 1 | Cltc | -0.051 | Old |
| 0.908662 | 0.417187 | 0.014 | 0.015 | 1 | 1 | Slc9a3r2 | -0.001 | Old |
| 0.909534 | 0.148812 | 0.066 | 0.067 | 1 | 1 | Aaed1 | -0.001 | Old |
| 0.909573 | 0.752778 | 0.012 | 0.011 | 1 | 1 | 1810043G( | 0.001 | Old |
| 0.90989 | 0.587266 | 0.012 | 0.011 | 1 | 1 | Sbk1 | 0.001 | Old |
| 0.909968 | 0.40712 | 0.039 | 0.038 | 1 | 1 | Mdn1 | 0.001 | Old |
| 0.91029 | 0.159036 | 0.135 | 0.135 | 1 | 1 | Fndc3a | 0 | Old |
| 0.910323 | 0.119132 | 0.127 | 0.126 | 1 | 1 | Slc11a1 | 0.001 | Old |
| 0.910618 | 0.175661 | 0.014 | 0.015 | 1 | 1 | Myg1 | -0.001 | Old |
| 0.910712 | 0.174413 | 0.184 | 0.188 | 1 | 1 | Srp14 | -0.004 | Old |
| 0.910821 | 0.458963 | 0.027 | 0.028 | 1 | 1 | E2f5 | -0.001 | Old |
| 0.911688 | 0.00823 | 0.499 | 0.516 | 1 | 1 | Glul | -0.017 | Old |
| 0.911701 | 0.406614 | 0.021 | 0.02 | 1 | 1 | Rnf25 | 0.001 | Old |
| 0.912027 | 0.35007 | 0.071 | 0.073 | 1 | 1 | Prpf18 | -0.002 | Old |
| 0.912437 | 0.221062 | 0.03 | 0.029 | 1 | 1 | Fam118a | 0.001 | Old |
| 0.912603 | 0.287603 | 0.059 | 0.058 | 1 | 1 | Fbrs | 0.001 | Old |
| 0.912692 | 0.272826 | 0.275 | 0.292 | 1 | 1 | Pepd | -0.017 | Old |
| 0.913486 | 0.443575 | 0.021 | 0.02 | 1 | 1 | Tlr3 | 0.001 | Old |
| 0.913707 | 0.327966 | 0.136 | 0.138 | 1 | 1 | Ndufb5 | -0.002 | Old |
| 0.91371 | 0.262693 | 0.168 | 0.172 | 1 | 1 | Ogdh | -0.004 | Old |
| 0.913972 | 0.451076 | 0.014 | 0.015 | 1 | 1 | Ddx51 | -0.001 | Old |
| 0.914094 | 0.056434 | 0.071 | 0.072 | 1 | 1 | Cant1 | -0.001 | Old |
| 0.914217 | 0.158862 | 0.031 | 0.032 | 1 | 1 | Malsu1 | -0.001 | Old |
| 0.914472 | 0.130331 | 0.036 | 0.037 | 1 | 1 | Lin54 | -0.001 | Old |
| 0.914645 | 0.41453 | 0.104 | 0.105 | 1 | 1 | Etv5 | -0.001 | Old |
| 0.915034 | 0.380054 | 0.021 | 0.02 | 1 | 1 | Dhx58 | 0.001 | Old |
| 0.91509 | 0.557598 | 0.014 | 0.015 | 1 | 1 | Pdcd2 | -0.001 | Old |
| 0.915282 | 0.59453 | 0.012 | 0.011 | 1 | 1 | Casp7 | 0.001 | Old |
| 0.915444 | 0.016414 | 0.044 | 0.043 | 1 | 1 | Pop5 | 0.001 | Old |
| 0.915504 | 0.253607 | 0.145 | 0.147 | 1 | 1 | Add3 | -0.002 | Old |
| 0.915628 | 0.271824 | 0.089 | 0.088 | 1 | 1 | Tmx1 | 0.001 | Old |
| 0.915883 | 0.249801 | 0.134 | 0.138 | 1 | 1 | Ythdf3 | -0.004 | Old |
| 0.915917 | 0.078855 | 0.012 | 0.011 | 1 | 1 | C1ra | 0.001 | Old |
| 0.916209 | 0.492622 | 0.014 | 0.015 | 1 | 1 | Pycrl | -0.001 | Old |
| 0.916711 | 0.15631 | 0.031 | 0.032 | 1 | 1 | Diablo | -0.001 | Old |
| 0.917239 | 0.309908 | 0.026 | 0.025 | 1 | 1 | Pgam5 | 0.001 | Old |
| 0.917743 | 0.201582 | 0.102 | 0.101 | 1 | 1 | Araf | 0.001 | Old |
| 0.917821 | 0.103836 | 0.012 | 0.011 | 1 | 1 | Ccdc66 | 0.001 | Old |
| 0.918217 | 0.047559 | 0.156 | 0.155 | 1 | 1 | Gnai3 | 0.001 | Old |

| | | | | | | | | |
|---|---|---|---|---|---|---|---|---|
| 0.918368 | 0.209009 | 0.021 | 0.02 | 1 | 1 | Wdr75 | 0.001 | Old |
| 0.918493 | 0.100429 | 0.048 | 0.047 | 1 | 1 | Dtx4 | 0.001 | Old |
| 0.918598 | 0.216081 | 0.183 | 0.19 | 1 | 1 | Fgl2 | -0.007 | Old |
| 0.919559 | 0.216931 | 0.021 | 0.02 | 1 | 1 | Tex30 | 0.001 | Old |
| 0.919903 | 0.383391 | 0.026 | 0.025 | 1 | 1 | Mmgt1 | 0.001 | Old |
| 0.919986 | 0.706154 | 0.014 | 0.015 | 1 | 1 | Ttl | -0.001 | Old |
| 0.920488 | 0.222226 | 0.054 | 0.053 | 1 | 1 | Washc3 | 0.001 | Old |
| 0.920678 | 0.063435 | 0.012 | 0.011 | 1 | 1 | Nup188 | 0.001 | Old |
| 0.920877 | 0.165479 | 0.048 | 0.049 | 1 | 1 | Hist1h2bc | -0.001 | Old |
| 0.921076 | 0.187635 | 0.141 | 0.142 | 1 | 1 | Idh1 | -0.001 | Old |
| 0.923115 | 0.047496 | 0.416 | 0.439 | 1 | 1 | Prcp | -0.023 | Old |
| 0.923129 | 0.237106 | 0.042 | 0.042 | 1 | 1 | Nkap | 0 | Old |
| 0.923433 | 0.23111 | 0.301 | 0.321 | 1 | 1 | Ppp2ca | -0.02 | Old |
| 0.923571 | 0.41973 | 0.019 | 0.02 | 1 | 1 | Net1 | -0.001 | Old |
| 0.923689 | 0.552831 | 0.032 | 0.032 | 1 | 1 | Slc33a1 | 0 | Old |
| 0.92385 | 0.007692 | 0.057 | 0.058 | 1 | 1 | Zdhhc3 | -0.001 | Old |
| 0.92386 | 0.369981 | 0.031 | 0.03 | 1 | 1 | Mre11a | 0.001 | Old |
| 0.924717 | 0.123044 | 0.067 | 0.068 | 1 | 1 | Maoa | -0.001 | Old |
| 0.924774 | 0.387346 | 0.031 | 0.032 | 1 | 1 | Mcee | -0.001 | Old |
| 0.924906 | 0.247997 | 0.036 | 0.037 | 1 | 1 | Coq8b | -0.001 | Old |
| 0.925447 | 0.127759 | 0.026 | 0.025 | 1 | 1 | Ndnf | 0.001 | Old |
| 0.925782 | 0.239045 | 0.084 | 0.083 | 1 | 1 | Taf1 | 0.001 | Old |
| 0.926035 | 0.116693 | 0.127 | 0.131 | 1 | 1 | Emc7 | -0.004 | Old |
| 0.92622 | 0.186027 | 0.121 | 0.122 | 1 | 1 | Ube2q1 | -0.001 | Old |
| 0.926566 | 0.113921 | 0.058 | 0.059 | 1 | 1 | Ddx60 | -0.001 | Old |
| 0.926706 | 0.246326 | 0.019 | 0.02 | 1 | 1 | Dpysl3 | -0.001 | Old |
| 0.926979 | 0.409373 | 0.031 | 0.03 | 1 | 1 | 2810004N2 | 0.001 | Old |
| 0.927624 | 0.047689 | 0.051 | 0.052 | 1 | 1 | Abi2 | -0.001 | Old |
| 0.927877 | 0.247174 | 0.105 | 0.106 | 1 | 1 | Zfx | -0.001 | Old |
| 0.928627 | 0.043449 | 0.066 | 0.066 | 1 | 1 | Eif4e2 | 0 | Old |
| 0.928676 | 0.270057 | 0.144 | 0.149 | 1 | 1 | Atp5d | -0.005 | Old |
| 0.928879 | 0.022827 | 0.045 | 0.046 | 1 | 1 | Uspl1 | -0.001 | Old |
| 0.929305 | 0.289295 | 0.046 | 0.046 | 1 | 1 | Stmn1 | 0 | Old |
| 0.929333 | 0.025171 | 0.021 | 0.02 | 1 | 1 | Mkrn2 | 0.001 | Old |
| 0.929776 | 0.418472 | 0.099 | 0.1 | 1 | 1 | Lima1 | -0.001 | Old |
| 0.929832 | 0.096029 | 0.075 | 0.076 | 1 | 1 | Mtch2 | -0.001 | Old |
| 0.929885 | 0.355964 | 0.068 | 0.07 | 1 | 1 | Pip4k2b | -0.002 | Old |
| 0.930198 | 0.331504 | 0.042 | 0.044 | 1 | 1 | Hsph1 | -0.002 | Old |
| 0.930265 | 0.334574 | 0.046 | 0.046 | 1 | 1 | Topbp1 | 0 | Old |
| 0.930451 | 0.264556 | 0.013 | 0.013 | 1 | 1 | Psat1 | 0 | Old |
| 0.930746 | 0.17864 | 0.013 | 0.013 | 1 | 1 | Il27ra | 0 | Old |
| 0.930811 | 0.373142 | 0.017 | 0.016 | 1 | 1 | Gmds | 0.001 | Old |
| 0.931146 | 0.124108 | 0.185 | 0.186 | 1 | 1 | Dnmt3a | -0.001 | Old |
| 0.931156 | 0.302586 | 0.099 | 0.103 | 1 | 1 | Glmp | -0.004 | Old |
| 0.931276 | 0.528911 | 0.022 | 0.021 | 1 | 1 | Mfn1 | 0.001 | Old |

| | | | | | | | | |
|---|---|---|---|---|---|---|---|---|
| 0.93131 | 0.423459 | 0.053 | 0.052 | 1 | 1 | As3mt | 0.001 | Old |
| 0.931339 | 0.367475 | 0.017 | 0.016 | 1 | 1 | Zfand2a | 0.001 | Old |
| 0.932342 | 0.208449 | 0.162 | 0.168 | 1 | 1 | Fuca1 | -0.006 | Old |
| 0.932354 | 0.09701 | 0.086 | 0.088 | 1 | 1 | Mpc1 | -0.002 | Old |
| 0.932678 | 0.449322 | 0.048 | 0.047 | 1 | 1 | Tm7sf3 | 0.001 | Old |
| 0.932923 | 0.463376 | 0.017 | 0.016 | 1 | 1 | Sco2 | 0.001 | Old |
| 0.932991 | 0.566528 | 0.027 | 0.027 | 1 | 1 | Ccdc93 | 0 | Old |
| 0.933173 | 0.593034 | 0.037 | 0.037 | 1 | 1 | Scaf4 | 0 | Old |
| 0.933696 | 0.305785 | 0.113 | 0.114 | 1 | 1 | Prrc2b | -0.001 | Old |
| 0.933971 | 0.245319 | 0.05 | 0.049 | 1 | 1 | Aldh3a2 | 0.001 | Old |
| 0.933987 | 0.103436 | 0.013 | 0.013 | 1 | 1 | Mettl17 | 0 | Old |
| 0.934131 | 0.476635 | 0.042 | 0.042 | 1 | 1 | Dnm1l | 0 | Old |
| 0.934576 | 0.234367 | 0.013 | 0.013 | 1 | 1 | Dppa3 | 0 | Old |
| 0.934831 | 0.013821 | 0.067 | 0.066 | 1 | 1 | Rnmt | 0.001 | Old |
| 0.934871 | 0.361557 | 0.013 | 0.013 | 1 | 1 | Mlst8 | 0 | Old |
| 0.93506 | 0.095651 | 0.123 | 0.125 | 1 | 1 | Ppp1r14b | -0.002 | Old |
| 0.935104 | 0.247999 | 0.024 | 0.025 | 1 | 1 | Nr2c2ap | -0.001 | Old |
| 0.935392 | 0.246904 | 0.163 | 0.165 | 1 | 1 | Ywhag | -0.002 | Old |
| 0.935395 | 0.106575 | 0.174 | 0.177 | 1 | 1 | Tmbim1 | -0.003 | Old |
| 0.935461 | 0.173239 | 0.013 | 0.013 | 1 | 1 | Acad12 | 0 | Old |
| 0.935621 | 0.50966 | 0.036 | 0.037 | 1 | 1 | Wdr43 | -0.001 | Old |
| 0.935893 | 0.343767 | 0.022 | 0.021 | 1 | 1 | Esr1 | 0.001 | Old |
| 0.935921 | 0.091773 | 0.064 | 0.064 | 1 | 1 | Bcl7c | 0 | Old |
| 0.936106 | 0.139394 | 0.353 | 0.369 | 1 | 1 | Ms4a6c | -0.016 | Old |
| 0.937008 | 0.033303 | 0.075 | 0.075 | 1 | 1 | Spata6 | 0 | Old |
| 0.937575 | 0.125724 | 0.256 | 0.266 | 1 | 1 | Vwa5a | -0.01 | Old |
| 0.937584 | 0.024517 | 0.072 | 0.071 | 1 | 1 | Lrpap1 | 0.001 | Old |
| 0.937748 | 0.166234 | 0.136 | 0.14 | 1 | 1 | Taf15 | -0.004 | Old |
| 0.938113 | 0.734265 | 0.013 | 0.013 | 1 | 1 | Hr | 0 | Old |
| 0.938381 | 0.060778 | 0.198 | 0.2 | 1 | 1 | Dynll1 | -0.002 | Old |
| 0.938629 | 0.246116 | 0.094 | 0.096 | 1 | 1 | Mrpl14 | -0.002 | Old |
| 0.938934 | 0.176948 | 0.127 | 0.131 | 1 | 1 | Tank | -0.004 | Old |
| 0.939007 | 0.30055 | 0.072 | 0.072 | 1 | 1 | Socs4 | 0 | Old |
| 0.939566 | 0.268087 | 0.041 | 0.042 | 1 | 1 | Cks2 | -0.001 | Old |
| 0.939735 | 0.167809 | 0.013 | 0.013 | 1 | 1 | Nbeal2 | 0 | Old |
| 0.939779 | 0.060617 | 0.139 | 0.142 | 1 | 1 | Dnajc13 | -0.003 | Old |
| 0.939841 | 0.100956 | 0.436 | 0.462 | 1 | 1 | Arhgdib | -0.026 | Old |
| 0.939882 | 0.636558 | 0.013 | 0.013 | 1 | 1 | AU022252 | 0 | Old |
| 0.94027 | 0.082605 | 0.018 | 0.018 | 1 | 1 | Dcaf6 | 0 | Old |
| 0.940297 | 0.308974 | 0.166 | 0.168 | 1 | 1 | Ywhaq | -0.002 | Old |
| 0.940745 | 0.305435 | 0.072 | 0.072 | 1 | 1 | Por | 0 | Old |
| 0.940986 | 0.287936 | 0.165 | 0.167 | 1 | 1 | Fxyd2 | -0.002 | Old |
| 0.941121 | 0.144672 | 0.066 | 0.065 | 1 | 1 | Usf1 | 0.001 | Old |
| 0.941291 | 0.152101 | 0.099 | 0.101 | 1 | 1 | Atxn2 | -0.002 | Old |
| 0.94136 | 0.160149 | 0.051 | 0.052 | 1 | 1 | Wars | -0.001 | Old |

| | | | | | | | | |
|---|---|---|---|---|---|---|---|---|
| 0.941728 | 0.282466 | 0.027 | 0.027 | 1 | 1 | Atg10 | 0 | Old |
| 0.94247 | 0.066614 | 0.057 | 0.058 | 1 | 1 | Plk2 | -0.001 | Old |
| 0.942598 | 0.234339 | 0.102 | 0.102 | 1 | 1 | Cdc40 | 0 | Old |
| 0.942901 | 0.323877 | 0.018 | 0.018 | 1 | 1 | Rabepk | 0 | Old |
| 0.942975 | 0.254893 | 0.06 | 0.06 | 1 | 1 | Eif3g | 0 | Old |
| 0.943121 | 0.354308 | 0.037 | 0.037 | 1 | 1 | Tm2d3 | 0 | Old |
| 0.943387 | 0.22036 | 0.046 | 0.047 | 1 | 1 | Itpk1 | -0.001 | Old |
| 0.943484 | 0.179391 | 0.067 | 0.068 | 1 | 1 | Hspa14 | -0.001 | Old |
| 0.943497 | 0.074771 | 0.033 | 0.034 | 1 | 1 | Arsa | -0.001 | Old |
| 0.943688 | 0.221506 | 0.045 | 0.046 | 1 | 1 | S1pr2 | -0.001 | Old |
| 0.94421 | 0.343012 | 0.022 | 0.021 | 1 | 1 | Pdzd11 | 0.001 | Old |
| 0.944493 | 0.133075 | 0.122 | 0.122 | 1 | 1 | Sp3 | 0 | Old |
| 0.944714 | 0.199933 | 0.03 | 0.03 | 1 | 1 | Inpp5a | 0 | Old |
| 0.944716 | 0.084189 | 0.13 | 0.129 | 1 | 1 | Psmd11 | 0.001 | Old |
| 0.94479 | 0.308145 | 0.135 | 0.137 | 1 | 1 | Chmp2a | -0.002 | Old |
| 0.944905 | 0.264276 | 0.018 | 0.018 | 1 | 1 | Lsm10 | 0 | Old |
| 0.945216 | 0.171756 | 0.057 | 0.058 | 1 | 1 | Adap1 | -0.001 | Old |
| 0.945468 | 0.680289 | 0.018 | 0.018 | 1 | 1 | Zcchc4 | 0 | Old |
| 0.945794 | 0.238956 | 0.057 | 0.058 | 1 | 1 | Slirp | -0.001 | Old |
| 0.945848 | 0.232775 | 0.046 | 0.047 | 1 | 1 | Otud6b | -0.001 | Old |
| 0.945879 | 0.043332 | 0.023 | 0.024 | 1 | 1 | Nudt19 | -0.001 | Old |
| 0.94629 | 0.166006 | 0.022 | 0.021 | 1 | 1 | Hmbs | 0.001 | Old |
| 0.946553 | 0.491307 | 0.049 | 0.05 | 1 | 1 | Prkra | -0.001 | Old |
| 0.946951 | 0.287139 | 0.041 | 0.042 | 1 | 1 | Plekhm1 | -0.001 | Old |
| 0.94724 | 0.25683 | 0.028 | 0.029 | 1 | 1 | Pigc | -0.001 | Old |
| 0.947412 | 0.296351 | 0.018 | 0.018 | 1 | 1 | Ccdc152 | 0 | Old |
| 0.947663 | 0.042826 | 0.055 | 0.055 | 1 | 1 | Crkl | 0 | Old |
| 0.947805 | 0.606908 | 0.013 | 0.013 | 1 | 1 | Rps6ka2 | 0 | Old |
| 0.947973 | 0.147055 | 0.027 | 0.027 | 1 | 1 | Dnajc30 | 0 | Old |
| 0.948009 | 0.207522 | 0.051 | 0.052 | 1 | 1 | Sae1 | -0.001 | Old |
| 0.948186 | 0.080554 | 0.039 | 0.039 | 1 | 1 | Vps33a | 0 | Old |
| 0.948333 | 0.455773 | 0.062 | 0.064 | 1 | 1 | Erlin2 | -0.002 | Old |
| 0.948832 | 0.287633 | 0.06 | 0.062 | 1 | 1 | Plgrkt | -0.002 | Old |
| 0.949307 | 0.015753 | 0.013 | 0.013 | 1 | 1 | 2610044O1 | 0 | Old |
| 0.949339 | 0.394737 | 0.03 | 0.03 | 1 | 1 | Cnot9 | 0 | Old |
| 0.949465 | 0.224651 | 0.032 | 0.032 | 1 | 1 | Uri1 | 0 | Old |
| 0.949739 | 0.43577 | 0.048 | 0.047 | 1 | 1 | Ddit3 | 0.001 | Old |
| 0.949784 | 0.624702 | 0.018 | 0.018 | 1 | 1 | Wnt11 | 0 | Old |
| 0.950208 | 0.318178 | 0.013 | 0.013 | 1 | 1 | Tango6 | 0 | Old |
| 0.950964 | 0.078467 | 0.627 | 0.665 | 1 | 1 | Ubc | -0.038 | Old |
| 0.951054 | 0.598011 | 0.018 | 0.018 | 1 | 1 | Cryzl2 | 0 | Old |
| 0.951478 | 0.176321 | 0.084 | 0.085 | 1 | 1 | Atp6v1d | -0.001 | Old |
| 0.951973 | 0.149134 | 0.041 | 0.041 | 1 | 1 | Smg6 | 0 | Old |
| 0.952175 | 0.622433 | 0.018 | 0.018 | 1 | 1 | Alkbh6 | 0 | Old |
| 0.952177 | 0.13614 | 0.028 | 0.029 | 1 | 1 | Nrde2 | -0.001 | Old |

| | | | | | | | | |
|---|---|---|---|---|---|---|---|---|
| 0.952761 | 0.344166 | 0.023 | 0.024 | 1 | 1 | Gna15 | -0.001 | Old |
| 0.952872 | 0.098938 | 0.266 | 0.271 | 1 | 1 | Eps8 | -0.005 | Old |
| 0.953084 | 0.231895 | 0.028 | 0.029 | 1 | 1 | Dkc1 | -0.001 | Old |
| 0.953275 | 0.406763 | 0.057 | 0.057 | 1 | 1 | Mut | 0 | Old |
| 0.953514 | 0.037001 | 0.013 | 0.013 | 1 | 1 | Gm46430 | 0 | Old |
| 0.953649 | 0.343547 | 0.023 | 0.024 | 1 | 1 | Dhrs4 | -0.001 | Old |
| 0.953655 | 0.343604 | 0.102 | 0.103 | 1 | 1 | Igbp1 | -0.001 | Old |
| 0.953825 | 0.032058 | 0.046 | 0.046 | 1 | 1 | Crebzf | 0 | Old |
| 0.954683 | 0.638069 | 0.018 | 0.018 | 1 | 1 | Zfp414 | 0 | Old |
| 0.954759 | 0.457444 | 0.023 | 0.024 | 1 | 1 | Dnlz | -0.001 | Old |
| 0.954921 | 0.30576 | 0.067 | 0.068 | 1 | 1 | Ufc1 | -0.001 | Old |
| 0.954945 | 0.316418 | 0.132 | 0.137 | 1 | 1 | Cd38 | -0.005 | Old |
| 0.955775 | 0.271149 | 0.032 | 0.032 | 1 | 1 | Nop14 | 0 | Old |
| 0.955825 | 0.319569 | 0.105 | 0.106 | 1 | 1 | Stt3a | -0.001 | Old |
| 0.955907 | 0.1145 | 0.071 | 0.072 | 1 | 1 | Cldnd1 | -0.001 | Old |
| 0.956158 | 0.401749 | 0.032 | 0.032 | 1 | 1 | Klf16 | 0 | Old |
| 0.956519 | 0.037577 | 0.013 | 0.013 | 1 | 1 | Trim32 | 0 | Old |
| 0.957785 | 0.138351 | 0.018 | 0.018 | 1 | 1 | Bcl9l | 0 | Old |
| 0.957912 | 0.320384 | 0.018 | 0.018 | 1 | 1 | Rhbdd2 | 0 | Old |
| 0.957918 | 0.232278 | 0.216 | 0.226 | 1 | 1 | Tmed7 | -0.01 | Old |
| 0.958093 | 0.300743 | 0.033 | 0.033 | 1 | 1 | Dctn6 | 0 | Old |
| 0.959018 | 0.543491 | 0.059 | 0.061 | 1 | 1 | Snrnp48 | -0.002 | Old |
| 0.959181 | 0.059347 | 0.058 | 0.058 | 1 | 1 | Fam98b | 0 | Old |
| 0.959601 | 0.252098 | 0.032 | 0.032 | 1 | 1 | Slc4a1ap | 0 | Old |
| 0.959684 | 0.120698 | 0.093 | 0.092 | 1 | 1 | Zfp263 | 0.001 | Old |
| 0.959699 | 0.746463 | 0.018 | 0.018 | 1 | 1 | Tbc1d32 | 0 | Old |
| 0.959826 | 0.128855 | 0.114 | 0.114 | 1 | 1 | Noc2l | 0 | Old |
| 0.959833 | 0.060671 | 0.012 | 0.012 | 1 | 1 | Pus7 | 0 | Old |
| 0.960147 | 0.197633 | 0.096 | 0.097 | 1 | 1 | Ap3d1 | -0.001 | Old |
| 0.960258 | 0.050675 | 0.109 | 0.11 | 1 | 1 | Sec11a | -0.001 | Old |
| 0.960564 | 0.02727 | 0.017 | 0.017 | 1 | 1 | Psmg2 | 0 | Old |
| 0.960656 | 0.192428 | 0.107 | 0.109 | 1 | 1 | Fam126a | -0.002 | Old |
| 0.961032 | 0.303276 | 0.069 | 0.07 | 1 | 1 | Ubqln2 | -0.001 | Old |
| 0.961193 | 0.544794 | 0.035 | 0.035 | 1 | 1 | Slc35e1 | 0 | Old |
| 0.961707 | 0.108082 | 0.012 | 0.012 | 1 | 1 | Ints5.1 | 0 | Old |
| 0.962283 | 0.336698 | 0.033 | 0.034 | 1 | 1 | Actr1b | -0.001 | Old |
| 0.962318 | 0.229744 | 0.093 | 0.095 | 1 | 1 | Nek9 | -0.002 | Old |
| 0.962332 | 0.254166 | 0.012 | 0.012 | 1 | 1 | Ctnnd2 | 0 | Old |
| 0.962387 | 0.546027 | 0.023 | 0.023 | 1 | 1 | Zxdc | 0 | Old |
| 0.962441 | 0.428449 | 0.066 | 0.067 | 1 | 1 | Vti1b | -0.001 | Old |
| 0.962838 | 0.457004 | 0.059 | 0.061 | 1 | 1 | Snd1 | -0.002 | Old |
| 0.963199 | 0.183724 | 0.284 | 0.298 | 1 | 1 | Cox6b1 | -0.014 | Old |
| 0.963247 | 0.104777 | 0.109 | 0.109 | 1 | 1 | Sdhd | 0 | Old |
| 0.963256 | 0.014352 | 0.078 | 0.079 | 1 | 1 | Sema6d | -0.001 | Old |
| 0.963659 | 0.066879 | 0.062 | 0.061 | 1 | 1 | Lsm8 | 0.001 | Old |

| | | | | | | | | |
|---|---|---|---|---|---|---|---|---|
| 0.964232 | 0.372675 | 0.168 | 0.176 | 1 | 1 | Cct4 | -0.008 | Old |
| 0.964479 | 0.110652 | 0.017 | 0.017 | 1 | 1 | Mamld1 | 0 | Old |
| 0.964653 | 0.059365 | 0.116 | 0.116 | 1 | 1 | Pgam1 | 0 | Old |
| 0.964831 | 0.361618 | 0.012 | 0.012 | 1 | 1 | Pvt1 | 0 | Old |
| 0.964981 | 0.191981 | 0.274 | 0.287 | 1 | 1 | Scamp2 | -0.013 | Old |
| 0.965456 | 0.365035 | 0.012 | 0.012 | 1 | 1 | Lrch4 | 0 | Old |
| 0.965586 | 0.219059 | 0.028 | 0.029 | 1 | 1 | Zfp398 | -0.001 | Old |
| 0.965653 | 0.178007 | 0.037 | 0.038 | 1 | 1 | Pex5 | -0.001 | Old |
| 0.96598 | 0.231335 | 0.157 | 0.16 | 1 | 1 | Swi5 | -0.003 | Old |
| 0.966254 | 0.242335 | 0.138 | 0.14 | 1 | 1 | Psme2 | -0.002 | Old |
| 0.966548 | 0.514597 | 0.085 | 0.087 | 1 | 1 | Otub1 | -0.002 | Old |
| 0.966705 | 0.155308 | 0.012 | 0.012 | 1 | 1 | Gm10138 | 0 | Old |
| 0.966759 | 0.584547 | 0.035 | 0.035 | 1 | 1 | Sgsh | 0 | Old |
| 0.967087 | 0.224399 | 0.033 | 0.033 | 1 | 1 | Tusc1 | 0 | Old |
| 0.967316 | 0.067878 | 0.018 | 0.018 | 1 | 1 | Slc9a1 | 0 | Old |
| 0.967498 | 0.058025 | 0.107 | 0.106 | 1 | 1 | Anapc11 | 0.001 | Old |
| 0.967544 | 0.173485 | 0.039 | 0.038 | 1 | 1 | Mis12 | 0.001 | Old |
| 0.967549 | 0.266804 | 0.023 | 0.023 | 1 | 1 | Ryr1 | 0 | Old |
| 0.968338 | 0.175489 | 0.152 | 0.155 | 1 | 1 | Dhx15 | -0.003 | Old |
| 0.96858 | 0.456069 | 0.012 | 0.012 | 1 | 1 | Klhl42 | 0 | Old |
| 0.968928 | 0.077635 | 0.063 | 0.064 | 1 | 1 | Atg16l1 | -0.001 | Old |
| 0.969047 | 0.124268 | 0.114 | 0.114 | 1 | 1 | Uqcrc2 | 0 | Old |
| 0.969317 | 0.146317 | 0.076 | 0.077 | 1 | 1 | Tcerg1 | -0.001 | Old |
| 0.969416 | 0.141551 | 0.058 | 0.058 | 1 | 1 | Esf1 | 0 | Old |
| 0.969511 | 0.031981 | 0.081 | 0.081 | 1 | 1 | Acvrl1 | 0 | Old |
| 0.969692 | 0.381306 | 0.121 | 0.123 | 1 | 1 | Cct3 | -0.002 | Old |
| 0.9697 | 0.36688 | 0.017 | 0.017 | 1 | 1 | Fen1 | 0 | Old |
| 0.969826 | 0.341017 | 0.064 | 0.064 | 1 | 1 | Tuba1a | 0 | Old |
| 0.970013 | 0.219421 | 0.049 | 0.049 | 1 | 1 | Clcn7 | 0 | Old |
| 0.97104 | 0.042731 | 0.066 | 0.066 | 1 | 1 | Sec13 | 0 | Old |
| 0.97108 | 0.841942 | 0.012 | 0.012 | 1 | 1 | Sec61a2 | 0 | Old |
| 0.971083 | 0.048619 | 0.032 | 0.032 | 1 | 1 | Mpv17 | 0 | Old |
| 0.971205 | 0.688488 | 0.019 | 0.019 | 1 | 1 | Gpr157 | 0 | Old |
| 0.971393 | 0.914226 | 0.012 | 0.012 | 1 | 1 | Ankrd52 | 0 | Old |
| 0.97185 | 0.333353 | 0.125 | 0.128 | 1 | 1 | Papd4 | -0.003 | Old |
| 0.971878 | 0.339958 | 0.072 | 0.072 | 1 | 1 | Fcgr4 | 0 | Old |
| 0.972323 | 0.204495 | 0.087 | 0.089 | 1 | 1 | Tor1b | -0.002 | Old |
| 0.972982 | 0.187622 | 0.225 | 0.23 | 1 | 1 | Rab10 | -0.005 | Old |
| 0.973546 | 0.336749 | 0.033 | 0.034 | 1 | 1 | Rab33b | -0.001 | Old |
| 0.974013 | 0.169599 | 0.054 | 0.054 | 1 | 1 | Prelid3b | 0 | Old |
| 0.974292 | 0.233338 | 0.09 | 0.092 | 1 | 1 | Socs6 | -0.002 | Old |
| 0.974568 | 0.962845 | 0.014 | 0.014 | 1 | 1 | Ncapd3 | 0 | Old |
| 0.974568 | 0.148711 | 0.014 | 0.014 | 1 | 1 | Zfp81 | 0 | Old |
| 0.974661 | 0.375711 | 0.017 | 0.017 | 1 | 1 | Mllt11 | 0 | Old |
| 0.974811 | 0.303005 | 0.08 | 0.081 | 1 | 1 | Ncaph2 | -0.001 | Old |

| | | | | | | | | |
|---|---|---|---|---|---|---|---|---|
| 0.97513 | 0.270836 | 0.022 | 0.022 | 1 | 1 | Nop9 | 0 | Old |
| 0.975406 | 0.064313 | 0.023 | 0.023 | 1 | 1 | AI987944 | 0 | Old |
| 0.975967 | 0.386311 | 0.077 | 0.078 | 1 | 1 | Vps41 | -0.001 | Old |
| 0.976095 | 0.087102 | 0.037 | 0.038 | 1 | 1 | Yeats2 | -0.001 | Old |
| 0.976279 | 0.241816 | 0.067 | 0.068 | 1 | 1 | Atf2 | -0.001 | Old |
| 0.976392 | 0.036964 | 0.031 | 0.031 | 1 | 1 | Igip | 0 | Old |
| 0.976743 | 0.226341 | 0.279 | 0.29 | 1 | 1 | Dse | -0.011 | Old |
| 0.976812 | 0.318817 | 0.072 | 0.072 | 1 | 1 | Fam114a2 | 0 | Old |
| 0.976936 | 0.169282 | 0.044 | 0.044 | 1 | 1 | Ptgr2 | 0 | Old |
| 0.977098 | 0.286087 | 0.129 | 0.131 | 1 | 1 | Arrdc1 | -0.002 | Old |
| 0.977339 | 0.11776 | 0.039 | 0.038 | 1 | 1 | Thap2 | 0.001 | Old |
| 0.977954 | 0.499778 | 0.067 | 0.068 | 1 | 1 | Ei24 | -0.001 | Old |
| 0.97805 | 0.316877 | 0.201 | 0.21 | 1 | 1 | Atxn7l3b | -0.009 | Old |
| 0.978063 | 0.078797 | 0.198 | 0.202 | 1 | 1 | Paip2 | -0.004 | Old |
| 0.978225 | 0.221544 | 0.032 | 0.032 | 1 | 1 | Tbce | 0 | Old |
| 0.978334 | 0.044712 | 0.031 | 0.031 | 1 | 1 | Rpia | 0 | Old |
| 0.978406 | 0.049926 | 0.713 | 0.75 | 1 | 1 | Mafb | -0.037 | Old |
| 0.978689 | 0.251708 | 0.116 | 0.117 | 1 | 1 | Tomm5 | -0.001 | Old |
| 0.978711 | 0.181868 | 0.033 | 0.033 | 1 | 1 | Cryzl1 | 0 | Old |
| 0.979281 | 0.236506 | 0.014 | 0.014 | 1 | 1 | Ppip5k1 | 0 | Old |
| 0.979926 | 0.546382 | 0.024 | 0.024 | 1 | 1 | Pus10 | 0 | Old |
| 0.980206 | 0.166785 | 0.156 | 0.157 | 1 | 1 | Coro1c | -0.001 | Old |
| 0.980471 | 0.121494 | 0.031 | 0.031 | 1 | 1 | Ppp5c | 0 | Old |
| 0.980567 | 0.26882 | 0.014 | 0.014 | 1 | 1 | 1700021F0 | 0 | Old |
| 0.98068 | 0.000908 | 0.033 | 0.033 | 1 | 1 | Slc25a12 | 0 | Old |
| 0.980853 | 0.250761 | 0.014 | 0.014 | 1 | 1 | Chd1l | 0 | Old |
| 0.980926 | 0.107337 | 0.039 | 0.038 | 1 | 1 | Ttc33 | 0.001 | Old |
| 0.982281 | 0.093761 | 0.014 | 0.014 | 1 | 1 | Lin37 | 0 | Old |
| 0.982496 | 0.931211 | 0.017 | 0.017 | 1 | 1 | Sergef | 0 | Old |
| 0.982828 | 0.111225 | 0.171 | 0.172 | 1 | 1 | Arl8a | -0.001 | Old |
| 0.982853 | 0.190117 | 0.014 | 0.014 | 1 | 1 | Def8 | 0 | Old |
| 0.98351 | 0.15201 | 0.05 | 0.05 | 1 | 1 | Edem2 | 0 | Old |
| 0.984069 | 0.462337 | 0.027 | 0.027 | 1 | 1 | Tmem181a | 0 | Old |
| 0.984503 | 0.360724 | 0.05 | 0.051 | 1 | 1 | Gab1 | -0.001 | Old |
| 0.984657 | 0.574899 | 0.05 | 0.051 | 1 | 1 | Fh1 | -0.001 | Old |
| 0.984969 | 0.192258 | 0.234 | 0.243 | 1 | 1 | Mfsd1 | -0.009 | Old |
| 0.985266 | 0.155166 | 0.373 | 0.398 | 1 | 1 | Prkcd | -0.025 | Old |
| 0.98531 | 0.380987 | 0.027 | 0.027 | 1 | 1 | Sil1 | 0 | Old |
| 0.985322 | 0.324812 | 0.051 | 0.052 | 1 | 1 | Psmd12 | -0.001 | Old |
| 0.985655 | 0.027889 | 0.114 | 0.114 | 1 | 1 | Slc35b2 | 0 | Old |
| 0.985684 | 0.149542 | 0.015 | 0.016 | 1 | 1 | Tfip11 | -0.001 | Old |
| 0.985747 | 0.453563 | 0.033 | 0.034 | 1 | 1 | Rps6kc1 | -0.001 | Old |
| 0.985897 | 0.515758 | 0.107 | 0.109 | 1 | 1 | Ndufab1 | -0.002 | Old |
| 0.986037 | 0.223501 | 0.156 | 0.157 | 1 | 1 | Itgb2 | -0.001 | Old |
| 0.986187 | 0.184455 | 0.063 | 0.064 | 1 | 1 | Cycs | -0.001 | Old |

| | | | | | | | | |
|---|---|---|---|---|---|---|---|---|
| 0.986335 | 0.068275 | 0.163 | 0.164 | 1 | 1 | Cmklr1 | -0.001 | Old |
| 0.986623 | 0.067036 | 0.021 | 0.021 | 1 | 1 | Mrps18b | 0 | Old |
| 0.986642 | 0.240957 | 0.019 | 0.019 | 1 | 1 | Acvr1b | 0 | Old |
| 0.986869 | 0.276971 | 0.078 | 0.08 | 1 | 1 | 1810030O( | -0.002 | Old |
| 0.987132 | 0.09409 | 0.019 | 0.019 | 1 | 1 | Kif19a | 0 | Old |
| 0.987139 | 0.207487 | 0.014 | 0.014 | 1 | 1 | Unc119b | 0 | Old |
| 0.987212 | 0.625755 | 0.039 | 0.039 | 1 | 1 | Atad2 | 0 | Old |
| 0.988411 | 0.291012 | 0.015 | 0.016 | 1 | 1 | Ppp3cc | -0.001 | Old |
| 0.988481 | 0.355219 | 0.068 | 0.069 | 1 | 1 | Ndufs4 | -0.001 | Old |
| 0.988624 | 0.359049 | 0.062 | 0.062 | 1 | 1 | Utp3 | 0 | Old |
| 0.989037 | 0.467597 | 0.049 | 0.049 | 1 | 1 | Proser1 | 0 | Old |
| 0.989107 | 0.158179 | 0.348 | 0.367 | 1 | 1 | Rbm3 | -0.019 | Old |
| 0.989507 | 0.060456 | 0.01 | 0.01 | 1 | 1 | Platr25 | 0 | Old |
| 0.989579 | 0.065664 | 0.078 | 0.079 | 1 | 1 | Trim44 | -0.001 | Old |
| 0.990255 | 0.208865 | 0.035 | 0.035 | 1 | 1 | Phax | 0 | Old |
| 0.990529 | 0.312619 | 0.021 | 0.021 | 1 | 1 | Fmnl2 | 0 | Old |
| 0.990617 | 0.312518 | 0.024 | 0.024 | 1 | 1 | Hs1bp3 | 0 | Old |
| 0.990868 | 0.246706 | 0.091 | 0.092 | 1 | 1 | Ndufa11 | -0.001 | Old |
| 0.991281 | 0.135154 | 0.071 | 0.071 | 1 | 1 | Dctn2 | 0 | Old |
| 0.991354 | 0.409802 | 0.06 | 0.061 | 1 | 1 | Rfc1 | -0.001 | Old |
| 0.99141 | 0.137562 | 0.015 | 0.016 | 1 | 1 | Oraov1 | -0.001 | Old |
| 0.99143 | 0.280635 | 0.181 | 0.187 | 1 | 1 | Ifngr2 | -0.006 | Old |
| 0.992153 | 0.334605 | 0.036 | 0.036 | 1 | 1 | Pogk | 0 | Old |
| 0.992615 | 0.398678 | 0.071 | 0.072 | 1 | 1 | Dnajc2 | -0.001 | Old |
| 0.992712 | 0.330658 | 0.031 | 0.031 | 1 | 1 | Nup85 | 0 | Old |
| 0.992712 | 0.192657 | 0.031 | 0.031 | 1 | 1 | Acot9 | 0 | Old |
| 0.992838 | 0.661882 | 0.01 | 0.01 | 1 | 1 | Uhrf1 | 0 | Old |
| 0.992865 | 0.115421 | 0.292 | 0.302 | 1 | 1 | Jak1 | -0.01 | Old |
| 0.993015 | 0.670069 | 0.021 | 0.021 | 1 | 1 | Tmem29 | 0 | Old |
| 0.99348 | 0.26056 | 0.087 | 0.089 | 1 | 1 | Rnft1 | -0.002 | Old |
| 0.993725 | 0.169374 | 0.021 | 0.021 | 1 | 1 | Fdft1 | 0 | Old |
| 0.993725 | 0.237633 | 0.021 | 0.021 | 1 | 1 | Kctd13 | 0 | Old |
| 0.993981 | 0.034921 | 0.103 | 0.103 | 1 | 1 | Arpc1a | 0 | Old |
| 0.993986 | 0.023253 | 0.156 | 0.155 | 1 | 1 | Gm21188 | 0.001 | Old |
| 0.994244 | 0.271139 | 0.135 | 0.138 | 1 | 1 | Bcap31 | -0.003 | Old |
| 0.994401 | 0.031871 | 0.045 | 0.045 | 1 | 1 | Lyar | 0 | Old |
| 0.994682 | 0.865074 | 0.015 | 0.016 | 1 | 1 | Sqle | -0.001 | Old |
| 0.994682 | 0.448566 | 0.015 | 0.016 | 1 | 1 | Slc16a1 | -0.001 | Old |
| 0.994941 | 0.38307 | 0.064 | 0.065 | 1 | 1 | Ccnd3 | -0.001 | Old |
| 0.995199 | 0.024322 | 0.024 | 0.024 | 1 | 1 | Bach2 | 0 | Old |
| 0.9954 | 0.238064 | 0.058 | 0.058 | 1 | 1 | Tmem184b | 0 | Old |
| 0.995458 | 0.151019 | 0.122 | 0.124 | 1 | 1 | Rnf4 | -0.002 | Old |
| 0.995838 | 0.408769 | 0.042 | 0.043 | 1 | 1 | Glipr1 | -0.001 | Old |
| 0.996169 | 0.388046 | 0.01 | 0.01 | 1 | 1 | Clasrp | 0 | Old |
| 0.996642 | 0.112361 | 0.046 | 0.047 | 1 | 1 | Sumf1 | -0.001 | Old |

| | | | | | | | | |
|---|---|---|---|---|---|---|---|---|
| 0.996835 | 0.84492 | 0.01 | 0.01 | 1 | 1 | Cmss1 | 0 | Old |
| 0.997375 | 0.234324 | 0.063 | 0.064 | 1 | 1 | Mtmr12 | -0.001 | Old |
| 0.997835 | 0.930206 | 0.01 | 0.01 | 1 | 1 | Cd209e | 0 | Old |
| 0.997875 | 0.032383 | 0.12 | 0.119 | 1 | 1 | Rb1 | 0.001 | Old |
| 0.998117 | 0.316263 | 0.089 | 0.09 | 1 | 1 | Lpcat1 | -0.001 | Old |
| 0.998121 | 0.210825 | 0.084 | 0.084 | 1 | 1 | Gosr2 | 0 | Old |
| 0.9985 | 0.462093 | 0.015 | 0.016 | 1 | 1 | Ppih | -0.001 | Old |
| 0.998501 | 0.476916 | 0.01 | 0.01 | 1 | 1 | Nup35 | 0 | Old |
| 0.998698 | 0.445694 | 0.021 | 0.021 | 1 | 1 | D6Wsu163 | 0 | Old |
| 0.998912 | 0.31202 | 0.13 | 0.134 | 1 | 1 | Capg | -0.004 | Old |
| 0.998917 | 0.274044 | 0.066 | 0.066 | 1 | 1 | Maea | 0 | Old |
| 0.99936 | 0.388782 | 0.059 | 0.06 | 1 | 1 | Polr2i | -0.001 | Old |
| 0.999369 | 0.327598 | 0.048 | 0.048 | 1 | 1 | Naa30 | 0 | Old |
| 0.999519 | 0.284459 | 0.175 | 0.18 | 1 | 1 | Mndal | -0.005 | Old |
| 1.09E-27 | 1.438213 | 0.371 | 0.15 | 3.38E-23 | 1 | Cx3cr1 | 0.221 | Young |
| 3.49E-24 | 0.347605 | 0.976 | 0.934 | 1.08E-19 | 1 | Rps29 | 0.042 | Young |
| 3.58E-23 | 1.125594 | 0.435 | 0.217 | 1.11E-18 | 1 | Gatm | 0.218 | Young |
| 3.88E-22 | 0.492862 | 0.927 | 0.82 | 1.20E-17 | 1 | Apoe | 0.107 | Young |
| 4.35E-20 | 0.786135 | 0.606 | 0.41 | 1.35E-15 | 1 | Il1b | 0.196 | Young |
| 3.67E-19 | 0.383947 | 0.925 | 0.869 | 1.14E-14 | 1 | Rpl35a | 0.056 | Young |
| 9.76E-19 | 0.339504 | 0.965 | 0.851 | 3.03E-14 | 1 | Malat1 | 0.114 | Young |
| 1.69E-18 | 0.944359 | 0.343 | 0.162 | 5.24E-14 | 1 | Hspa1b | 0.181 | Young |
| 3.27E-18 | 0.438611 | 0.838 | 0.695 | 1.01E-13 | 1 | Rps12 | 0.143 | Young |
| 3.94E-18 | 0.565394 | 0.699 | 0.515 | 1.22E-13 | 1 | Rpl18a | 0.184 | Young |
| 8.23E-18 | 0.446708 | 0.791 | 0.634 | 2.56E-13 | 1 | Rpl17 | 0.157 | Young |
| 1.65E-17 | 0.446931 | 0.86 | 0.749 | 5.11E-13 | 1 | Rps14 | 0.111 | Young |
| 2.27E-17 | 0.500876 | 0.754 | 0.602 | 7.06E-13 | 1 | Rpl30 | 0.152 | Young |
| 1.46E-16 | 0.448547 | 0.81 | 0.686 | 4.53E-12 | 1 | Rpl23 | 0.124 | Young |
| 1.53E-15 | 0.442053 | 0.775 | 0.622 | 4.76E-11 | 1 | Nfkbiz | 0.153 | Young |
| 3.17E-15 | 0.491457 | 0.722 | 0.567 | 9.86E-11 | 1 | Rps27 | 0.155 | Young |
| 3.26E-15 | 0.452776 | 0.824 | 0.686 | 1.01E-10 | 1 | Cd83 | 0.138 | Young |
| 3.93E-15 | 0.345536 | 0.905 | 0.833 | 1.22E-10 | 1 | Fau | 0.072 | Young |
| 2.17E-14 | 0.676745 | 0.534 | 0.366 | 6.75E-10 | 1 | Mef2c | 0.168 | Young |
| 2.27E-14 | 0.319331 | 0.911 | 0.842 | 7.06E-10 | 1 | Rps24 | 0.069 | Young |
| 4.52E-14 | 0.400508 | 0.792 | 0.683 | 1.40E-09 | 1 | Rps20 | 0.109 | Young |
| 5.13E-14 | 0.692509 | 0.341 | 0.185 | 1.59E-09 | 1 | Hspa1a | 0.156 | Young |
| 5.63E-14 | 0.630043 | 0.557 | 0.386 | 1.75E-09 | 1 | Cd14 | 0.171 | Young |
| 7.47E-14 | 0.721322 | 0.501 | 0.333 | 2.32E-09 | 1 | Ier3 | 0.168 | Young |
| 1.76E-13 | 0.566655 | 0.518 | 0.351 | 5.45E-09 | 1 | Rpl11 | 0.167 | Young |
| 2.49E-13 | 1.969915 | 0.129 | 0.033 | 7.73E-09 | 1 | Itga6 | 0.096 | Young |
| 1.25E-12 | 1.08181 | 0.264 | 0.136 | 3.90E-08 | 1 | Gpr65 | 0.128 | Young |
| 3.09E-12 | 0.46985 | 0.492 | 0.33 | 9.60E-08 | 1 | Rpl35 | 0.162 | Young |
| 3.23E-12 | 0.585472 | 0.317 | 0.172 | 1.00E-07 | 1 | Cited2 | 0.145 | Young |
| 2.37E-11 | 0.223296 | 0.979 | 0.954 | 7.36E-07 | 1 | Rpl37a | 0.025 | Young |
| 4.33E-11 | 0.366772 | 0.737 | 0.602 | 1.35E-06 | 1 | Rpl39 | 0.135 | Young |

| | | | | | | | | |
|---|---|---|---|---|---|---|---|---|
| 4.95E-11 | 0.48493 | 0.648 | 0.512 | 1.54E-06 | 1 | Rpl18 | 0.136 | Young |
| 7.26E-11 | 0.332768 | 0.782 | 0.657 | 2.26E-06 | 1 | Rps4x | 0.125 | Young |
| 7.32E-11 | 0.370202 | 0.753 | 0.644 | 2.27E-06 | 1 | Rps16 | 0.109 | Young |
| 1.14E-10 | 0.36999 | 0.721 | 0.607 | 3.54E-06 | 1 | Rps10 | 0.114 | Young |
| 1.40E-10 | 1.790059 | 0.093 | 0.021 | 4.35E-06 | 1 | Pmepa1 | 0.072 | Young |
| 1.42E-10 | 0.319381 | 0.771 | 0.643 | 4.40E-06 | 1 | Rps15 | 0.128 | Young |
| 1.66E-10 | 0.491611 | 0.52 | 0.368 | 5.15E-06 | 1 | Rpl12 | 0.152 | Young |
| 1.90E-10 | 1.0025 | 0.262 | 0.144 | 5.91E-06 | 1 | Dennd4a | 0.118 | Young |
| 2.00E-10 | 0.770003 | 0.344 | 0.213 | 6.22E-06 | 1 | Ptpre | 0.131 | Young |
| 2.21E-10 | 0.70888 | 0.346 | 0.212 | 6.87E-06 | 1 | Pou2f2 | 0.134 | Young |
| 3.13E-10 | 0.982503 | 0.25 | 0.135 | 9.73E-06 | 1 | Hpgd | 0.115 | Young |
| 3.22E-10 | 0.37096 | 0.497 | 0.337 | 9.99E-06 | 1 | Atf3 | 0.16 | Young |
| 3.27E-10 | 0.367798 | 0.661 | 0.508 | 1.02E-05 | 1 | Rpl9 | 0.153 | Young |
| 3.30E-10 | 0.320117 | 0.835 | 0.71 | 1.02E-05 | 1 | Kctd12 | 0.125 | Young |
| 3.31E-10 | 0.565845 | 0.656 | 0.532 | 1.03E-05 | 1 | Cxcl2 | 0.124 | Young |
| 4.05E-10 | 0.354082 | 0.63 | 0.49 | 1.26E-05 | 1 | Csf1r | 0.14 | Young |
| 4.63E-10 | 0.674818 | 0.347 | 0.22 | 1.44E-05 | 1 | Skil | 0.127 | Young |
| 4.63E-10 | 1.310413 | 0.121 | 0.04 | 1.44E-05 | 1 | Tgfbr1 | 0.081 | Young |
| 4.66E-10 | 1.070702 | 0.174 | 0.076 | 1.45E-05 | 1 | Ccl3 | 0.098 | Young |
| 4.90E-10 | 0.32245 | 0.815 | 0.693 | 1.52E-05 | 1 | Rpl13 | 0.122 | Young |
| 6.98E-10 | 0.801297 | 0.304 | 0.179 | 2.17E-05 | 1 | Cd33 | 0.125 | Young |
| 8.19E-10 | 2.635621 | 0.076 | 0.014 | 2.54E-05 | 1 | Col14a1 | 0.062 | Young |
| 8.51E-10 | 0.35139 | 0.629 | 0.485 | 2.64E-05 | 1 | Tyrobp | 0.144 | Young |
| 1.03E-09 | 0.358322 | 0.801 | 0.681 | 3.21E-05 | 1 | Pf4 | 0.12 | Young |
| 1.09E-09 | 0.390541 | 0.636 | 0.494 | 3.38E-05 | 1 | Rpl26 | 0.142 | Young |
| 1.16E-09 | 0.910455 | 0.213 | 0.107 | 3.60E-05 | 1 | Hpgds | 0.106 | Young |
| 1.20E-09 | 3.282598 | 0.068 | 0.01 | 3.74E-05 | 1 | F11r | 0.058 | Young |
| 2.25E-09 | 0.426216 | 0.648 | 0.527 | 6.98E-05 | 1 | Rpl34 | 0.121 | Young |
| 2.27E-09 | 0.303612 | 0.769 | 0.641 | 7.06E-05 | 1 | Rpl37 | 0.128 | Young |
| 2.57E-09 | 0.3163 | 0.729 | 0.596 | 7.97E-05 | 1 | Rps19 | 0.133 | Young |
| 3.08E-09 | 1.037383 | 0.335 | 0.221 | 9.56E-05 | 1 | Hexb | 0.114 | Young |
| 4.67E-09 | 0.926548 | 0.189 | 0.093 | 0.000145 | 1 | Dyrk2 | 0.096 | Young |
| 5.17E-09 | 0.377949 | 0.629 | 0.483 | 0.00016 | 1 | Rps15a | 0.146 | Young |
| 5.80E-09 | 0.425886 | 0.567 | 0.433 | 0.00018 | 1 | Rpl10a | 0.134 | Young |
| 7.93E-09 | 0.683879 | 0.289 | 0.177 | 0.000246 | 1 | Arl4c | 0.112 | Young |
| 9.64E-09 | 0.652129 | 0.315 | 0.199 | 0.000299 | 1 | Tlr2 | 0.116 | Young |
| 1.00E-08 | 0.400803 | 0.617 | 0.485 | 0.000311 | 1 | Rpl32 | 0.132 | Young |
| 1.28E-08 | 0.382077 | 0.681 | 0.563 | 0.000396 | 1 | Rps13 | 0.118 | Young |
| 1.67E-08 | 0.549993 | 0.405 | 0.28 | 0.000517 | 1 | Mef2a | 0.125 | Young |
| 1.91E-08 | 0.443331 | 0.453 | 0.319 | 0.000592 | 1 | Rps18 | 0.134 | Young |
| 2.36E-08 | 0.230796 | 0.886 | 0.817 | 0.000731 | 1 | Cst3 | 0.069 | Young |
| 4.17E-08 | 0.579975 | 0.315 | 0.202 | 0.001294 | 1 | Runx1 | 0.113 | Young |
| 5.42E-08 | 0.371969 | 0.636 | 0.521 | 0.001683 | 1 | Rps25 | 0.115 | Young |
| 6.60E-08 | 0.742284 | 0.253 | 0.154 | 0.002051 | 1 | Itga4 | 0.099 | Young |
| 6.70E-08 | 1.039032 | 0.106 | 0.039 | 0.002082 | 1 | AU020206 | 0.067 | Young |

| | | | | | | | | |
|---|---|---|---|---|---|---|---|---|
| 6.92E-08 | 0.460706 | 0.437 | 0.311 | 0.002148 | 1 | Stab1 | 0.126 | Young |
| 7.76E-08 | 0.905086 | 0.2 | 0.111 | 0.00241 | 1 | Aff4 | 0.089 | Young |
| 8.41E-08 | 0.82372 | 0.299 | 0.193 | 0.002612 | 1 | Ccl4 | 0.106 | Young |
| 9.62E-08 | 0.313062 | 0.581 | 0.445 | 0.002987 | 1 | Rps6 | 0.136 | Young |
| 1.13E-07 | 0.436485 | 0.348 | 0.228 | 0.003498 | 1 | Tsc22d3 | 0.12 | Young |
| 1.85E-07 | 1.804684 | 0.099 | 0.037 | 0.00575 | 1 | Lpcat2 | 0.062 | Young |
| 1.86E-07 | 0.688129 | 0.293 | 0.192 | 0.005773 | 1 | Arrb2 | 0.101 | Young |
| 2.15E-07 | 0.286256 | 0.673 | 0.553 | 0.006681 | 1 | Rpl22 | 0.12 | Young |
| 2.23E-07 | 0.608189 | 0.324 | 0.217 | 0.00694 | 1 | Ubb | 0.107 | Young |
| 2.31E-07 | 1.03724 | 0.151 | 0.075 | 0.007182 | 1 | Cd72 | 0.076 | Young |
| 2.43E-07 | 0.469174 | 0.444 | 0.323 | 0.007559 | 1 | Nfkbid | 0.121 | Young |
| 2.47E-07 | 0.52426 | 0.388 | 0.274 | 0.007663 | 1 | Gadd45b | 0.114 | Young |
| 2.73E-07 | 0.405504 | 0.667 | 0.548 | 0.008475 | 1 | Cybb | 0.119 | Young |
| 3.12E-07 | 0.553275 | 0.392 | 0.276 | 0.009684 | 1 | Ighm | 0.116 | Young |
| 3.12E-07 | 1.491819 | 0.089 | 0.031 | 0.009687 | 1 | Tagap | 0.058 | Young |
| 3.49E-07 | 0.229658 | 0.821 | 0.738 | 0.010842 | 1 | Rps23 | 0.083 | Young |
| 3.51E-07 | 0.590902 | 0.238 | 0.147 | 0.010909 | 1 | Stap1 | 0.091 | Young |
| 3.56E-07 | 0.521017 | 0.273 | 0.171 | 0.011051 | 1 | Ssh2 | 0.102 | Young |
| 3.99E-07 | 0.632845 | 0.194 | 0.108 | 0.012398 | 1 | Ptgs1 | 0.086 | Young |
| 4.54E-07 | 0.435181 | 0.398 | 0.284 | 0.014087 | 1 | Rpl31 | 0.114 | Young |
| 5.46E-07 | 0.479874 | 0.4 | 0.29 | 0.016959 | 1 | Zfhx3 | 0.11 | Young |
| 6.58E-07 | 0.382938 | 0.415 | 0.299 | 0.020439 | 1 | Pnrc1 | 0.116 | Young |
| 7.39E-07 | 0.2593 | 0.688 | 0.557 | 0.022963 | 1 | Rps11 | 0.131 | Young |
| 7.71E-07 | 0.421991 | 0.49 | 0.365 | 0.023945 | 1 | Coro1a | 0.125 | Young |
| 7.71E-07 | 1.238421 | 0.097 | 0.039 | 0.023956 | 1 | Lilra5 | 0.058 | Young |
| 9.00E-07 | 0.880343 | 0.174 | 0.096 | 0.02794 | 1 | Arhgap45 | 0.078 | Young |
| 9.21E-07 | 0.458277 | 0.282 | 0.183 | 0.028609 | 1 | Nlrp3 | 0.099 | Young |
| 1.09E-06 | 1.077707 | 0.155 | 0.082 | 0.033827 | 1 | Adap2 | 0.073 | Young |
| 1.16E-06 | 1.303951 | 0.114 | 0.051 | 0.036099 | 1 | Nr4a3 | 0.063 | Young |
| 1.41E-06 | 0.59533 | 0.199 | 0.116 | 0.04382 | 1 | Prdx5 | 0.083 | Young |
| 1.46E-06 | 0.64158 | 0.186 | 0.105 | 0.045197 | 1 | Gpr34 | 0.081 | Young |
| 1.56E-06 | 0.413234 | 0.402 | 0.287 | 0.048579 | 1 | Cxcl16 | 0.115 | Young |
| 1.57E-06 | 0.210188 | 0.874 | 0.824 | 0.048772 | 1 | Rps21 | 0.05 | Young |
| 1.66E-06 | 0.3386 | 0.441 | 0.325 | 0.051437 | 1 | Abca1 | 0.116 | Young |
| 1.81E-06 | 0.886923 | 0.219 | 0.138 | 0.056164 | 1 | Bcl2a1b | 0.081 | Young |
| 1.87E-06 | 0.369887 | 0.338 | 0.233 | 0.05793 | 1 | Mrfap1 | 0.105 | Young |
| 1.87E-06 | 0.808259 | 0.188 | 0.111 | 0.058138 | 1 | Elmo1 | 0.077 | Young |
| 2.03E-06 | 0.333152 | 0.388 | 0.278 | 0.062903 | 1 | Atp5h | 0.11 | Young |
| 2.03E-06 | 0.624667 | 0.233 | 0.148 | 0.063059 | 1 | Itpkb | 0.085 | Young |
| 2.36E-06 | 0.617869 | 0.279 | 0.185 | 0.073298 | 1 | Lair1 | 0.094 | Young |
| 2.48E-06 | 0.297679 | 0.504 | 0.383 | 0.076857 | 1 | Rps2 | 0.121 | Young |
| 2.62E-06 | 0.856241 | 0.16 | 0.087 | 0.081384 | 1 | Cttnbp2nl | 0.073 | Young |
| 2.93E-06 | 0.428377 | 0.356 | 0.253 | 0.090849 | 1 | Rel | 0.103 | Young |
| 2.96E-06 | 0.649646 | 0.241 | 0.154 | 0.092011 | 1 | Pnisr | 0.087 | Young |
| 3.03E-06 | 0.908832 | 0.171 | 0.098 | 0.093982 | 1 | Ccrl2 | 0.073 | Young |

| | | | | | | | | |
|---|---|---|---|---|---|---|---|---|
| 3.66E-06 | 0.321264 | 0.29 | 0.192 | 0.113673 | 1 | Nsa2 | 0.098 | Young |
| 3.96E-06 | 0.573704 | 0.201 | 0.121 | 0.122865 | 1 | Nrip1 | 0.08 | Young |
| 4.02E-06 | 0.886177 | 0.163 | 0.093 | 0.124979 | 1 | Slc15a3 | 0.07 | Young |
| 4.51E-06 | 0.551659 | 0.226 | 0.141 | 0.14019 | 1 | Dek | 0.085 | Young |
| 4.62E-06 | 0.269192 | 0.471 | 0.348 | 0.143512 | 1 | Rbm39 | 0.123 | Young |
| 4.79E-06 | 0.38746 | 0.439 | 0.333 | 0.148846 | 1 | Srrm2 | 0.106 | Young |
| 4.84E-06 | 0.774312 | 0.151 | 0.084 | 0.150378 | 1 | Tnf | 0.067 | Young |
| 5.40E-06 | 0.213734 | 0.892 | 0.857 | 0.167564 | 1 | Rps27a | 0.035 | Young |
| 5.73E-06 | 0.259523 | 0.502 | 0.384 | 0.178022 | 1 | Rps3 | 0.118 | Young |
| 6.12E-06 | 0.243758 | 0.73 | 0.609 | 0.190122 | 1 | Fcer1g | 0.121 | Young |
| 6.17E-06 | 0.914731 | 0.118 | 0.058 | 0.191495 | 1 | Cpd | 0.06 | Young |
| 6.33E-06 | 2.915112 | 0.042 | 0.008 | 0.196702 | 1 | Spp1 | 0.034 | Young |
| 6.46E-06 | 0.445508 | 0.49 | 0.384 | 0.200715 | 1 | Mpeg1 | 0.106 | Young |
| 6.64E-06 | 1.007761 | 0.081 | 0.031 | 0.206315 | 1 | Icosl | 0.05 | Young |
| 6.65E-06 | 1.198104 | 0.097 | 0.044 | 0.206465 | 1 | Nuak2 | 0.053 | Young |
| 6.95E-06 | 0.524529 | 0.207 | 0.125 | 0.215966 | 1 | Cdkn1b | 0.082 | Young |
| 7.03E-06 | 1.644532 | 0.066 | 0.022 | 0.218193 | 1 | Tmem86a | 0.044 | Young |
| 7.03E-06 | 0.770089 | 0.186 | 0.112 | 0.218408 | 1 | Coq10b | 0.074 | Young |
| 7.63E-06 | 0.285189 | 0.628 | 0.521 | 0.236908 | 1 | Rpl3 | 0.107 | Young |
| 8.11E-06 | 0.242432 | 0.655 | 0.546 | 0.251778 | 1 | Rps5 | 0.109 | Young |
| 8.28E-06 | 0.448314 | 0.357 | 0.26 | 0.257039 | 1 | Clec12a | 0.097 | Young |
| 8.57E-06 | 0.259362 | 0.547 | 0.428 | 0.266199 | 1 | Rpl36a | 0.119 | Young |
| 8.61E-06 | 0.243242 | 0.717 | 0.602 | 0.267417 | 1 | Dab2 | 0.115 | Young |
| 8.72E-06 | 1.516792 | 0.061 | 0.019 | 0.270835 | 1 | Mtus1 | 0.042 | Young |
| 9.19E-06 | 0.185738 | 0.778 | 0.703 | 0.285517 | 1 | Rpl27a | 0.075 | Young |
| 9.50E-06 | 0.399698 | 0.318 | 0.221 | 0.294909 | 1 | Rnase4 | 0.097 | Young |
| 9.78E-06 | 0.388922 | 0.364 | 0.263 | 0.303851 | 1 | Tmcc1 | 0.101 | Young |
| 1.08E-05 | 0.21101 | 0.459 | 0.338 | 0.334136 | 1 | Rpl10 | 0.121 | Young |
| 1.11E-05 | 0.736973 | 0.235 | 0.156 | 0.34612 | 1 | Tnfsf9 | 0.079 | Young |
| 1.13E-05 | 0.451677 | 0.303 | 0.211 | 0.350672 | 1 | Slc3a2 | 0.092 | Young |
| 1.16E-05 | 3.292615 | 0.03 | 0.003 | 0.359932 | 1 | Tmem119 | 0.027 | Young |
| 1.23E-05 | 2.359064 | 0.03 | 0.003 | 0.382411 | 1 | Cxcl14 | 0.027 | Young |
| 1.28E-05 | 0.448758 | 0.343 | 0.249 | 0.397104 | 1 | Ms4a7 | 0.094 | Young |
| 1.30E-05 | 0.537921 | 0.239 | 0.158 | 0.403945 | 1 | Phip | 0.081 | Young |
| 1.61E-05 | 0.892843 | 0.173 | 0.105 | 0.500479 | 1 | Itgb5 | 0.068 | Young |
| 1.65E-05 | 0.156584 | 0.886 | 0.815 | 0.51174 | 1 | Rpl41 | 0.071 | Young |
| 1.73E-05 | 0.443618 | 0.351 | 0.26 | 0.537238 | 1 | Rassf4 | 0.091 | Young |
| 2.10E-05 | 0.550611 | 0.177 | 0.107 | 0.652422 | 1 | Tanc2 | 0.07 | Young |
| 2.20E-05 | 0.693494 | 0.126 | 0.067 | 0.683366 | 1 | Arsb | 0.059 | Young |
| 2.22E-05 | 0.264708 | 0.699 | 0.599 | 0.690738 | 1 | Rpl19 | 0.1 | Young |
| 2.36E-05 | 0.362161 | 0.282 | 0.193 | 0.733707 | 1 | Tbca | 0.089 | Young |
| 2.43E-05 | 0.287476 | 0.601 | 0.499 | 0.755526 | 1 | Rplp0 | 0.102 | Young |
| 2.48E-05 | 0.931024 | 0.116 | 0.06 | 0.771098 | 1 | AI504432 | 0.056 | Young |
| 2.50E-05 | 0.513936 | 0.273 | 0.189 | 0.777296 | 1 | Axl | 0.084 | Young |
| 2.59E-05 | 0.248791 | 0.724 | 0.622 | 0.803365 | 1 | Rpl23a | 0.102 | Young |

| | | | | | | | | |
|---|---|---|---|---|---|---|---|---|
| 2.70E-05 | 0.480734 | 0.242 | 0.159 | 0.837066 | 1 | Spag9 | 0.083 | Young |
| 2.74E-05 | 1.201072 | 0.058 | 0.019 | 0.852307 | 1 | Mmp14 | 0.039 | Young |
| 2.83E-05 | 0.461987 | 0.253 | 0.171 | 0.879809 | 1 | Zmiz1 | 0.082 | Young |
| 2.95E-05 | 0.28253 | 0.624 | 0.519 | 0.914851 | 1 | Marcksl1 | 0.105 | Young |
| 3.00E-05 | 0.327528 | 0.527 | 0.425 | 0.932118 | 1 | Adgre1 | 0.102 | Young |
| 3.02E-05 | 0.363295 | 0.343 | 0.251 | 0.937787 | 1 | Rpl4 | 0.092 | Young |
| 3.48E-05 | 0.745489 | 0.165 | 0.1 | 1 | 1 | Arrb1 | 0.065 | Young |
| 3.48E-05 | 0.233349 | 0.793 | 0.681 | 1 | 1 | Zfp36l1 | 0.112 | Young |
| 3.85E-05 | 0.569028 | 0.142 | 0.081 | 1 | 1 | Arpp19 | 0.061 | Young |
| 3.90E-05 | 1.00472 | 0.097 | 0.048 | 1 | 1 | Sema4d | 0.049 | Young |
| 4.00E-05 | 0.858383 | 0.16 | 0.096 | 1 | 1 | Fcgr1 | 0.064 | Young |
| 4.10E-05 | 0.360568 | 0.399 | 0.306 | 1 | 1 | Rpl15 | 0.093 | Young |
| 4.17E-05 | 0.969288 | 0.094 | 0.045 | 1 | 1 | Rap2a | 0.049 | Young |
| 4.34E-05 | 0.546693 | 0.281 | 0.197 | 1 | 1 | Ncf1 | 0.084 | Young |
| 4.38E-05 | 0.439193 | 0.193 | 0.121 | 1 | 1 | B4galt1 | 0.072 | Young |
| 4.40E-05 | 1.106428 | 0.072 | 0.03 | 1 | 1 | Slc14a1 | 0.042 | Young |
| 4.83E-05 | 0.283488 | 0.609 | 0.501 | 1 | 1 | Rpl24 | 0.108 | Young |
| 5.02E-05 | 0.527307 | 0.153 | 0.09 | 1 | 1 | Sirt2 | 0.063 | Young |
| 5.15E-05 | 0.555243 | 0.234 | 0.162 | 1 | 1 | Ankrd11 | 0.072 | Young |
| 5.35E-05 | 0.265097 | 0.634 | 0.513 | 1 | 1 | Tgfbi | 0.121 | Young |
| 5.40E-05 | 0.644099 | 0.204 | 0.135 | 1 | 1 | Zcchc11 | 0.069 | Young |
| 5.42E-05 | 0.999376 | 0.106 | 0.055 | 1 | 1 | Dnajb1 | 0.051 | Young |
| 5.56E-05 | 0.364464 | 0.264 | 0.18 | 1 | 1 | Ddx3y | 0.084 | Young |
| 5.66E-05 | 0.857115 | 0.13 | 0.073 | 1 | 1 | Prdm1 | 0.057 | Young |
| 5.90E-05 | 0.145287 | 0.891 | 0.832 | 1 | 1 | Rps8 | 0.059 | Young |
| 6.29E-05 | 0.335871 | 0.445 | 0.347 | 1 | 1 | Ubl3 | 0.098 | Young |
| 6.30E-05 | 0.68918 | 0.177 | 0.112 | 1 | 1 | St3gal6 | 0.065 | Young |
| 6.35E-05 | 0.269633 | 0.533 | 0.429 | 1 | 1 | Rps7 | 0.104 | Young |
| 6.37E-05 | 0.532753 | 0.278 | 0.199 | 1 | 1 | Rbm25 | 0.079 | Young |
| 6.43E-05 | 0.153122 | 0.967 | 0.954 | 1 | 1 | Cd74 | 0.013 | Young |
| 6.65E-05 | 0.604996 | 0.19 | 0.122 | 1 | 1 | Ifnar1 | 0.068 | Young |
| 6.86E-05 | 0.414754 | 0.222 | 0.148 | 1 | 1 | Gapdh | 0.074 | Young |
| 7.30E-05 | 1.115529 | 0.105 | 0.055 | 1 | 1 | Retreg1 | 0.05 | Young |
| 7.36E-05 | 0.207459 | 0.389 | 0.292 | 1 | 1 | Cd52 | 0.097 | Young |
| 7.39E-05 | 0.296363 | 0.441 | 0.342 | 1 | 1 | Rpl7a | 0.099 | Young |
| 7.50E-05 | 1.214979 | 0.069 | 0.028 | 1 | 1 | Cxcl10 | 0.041 | Young |
| 7.94E-05 | 0.08292 | 0.995 | 0.999 | 1 | 1 | Tmsb4x | -0.004 | Young |
| 8.25E-05 | 0.226189 | 0.273 | 0.19 | 1 | 1 | Id2 | 0.083 | Young |
| 8.70E-05 | 0.353219 | 0.305 | 0.224 | 1 | 1 | Neurl3 | 0.081 | Young |
| 8.70E-05 | 0.5619 | 0.153 | 0.093 | 1 | 1 | Cd300c2 | 0.06 | Young |
| 8.87E-05 | 0.30277 | 0.292 | 0.206 | 1 | 1 | Irf8 | 0.086 | Young |
| 8.88E-05 | 1.419199 | 0.047 | 0.014 | 1 | 1 | Sema4c | 0.033 | Young |
| 9.34E-05 | 0.904412 | 0.117 | 0.066 | 1 | 1 | Ppp2r2a | 0.051 | Young |
| 9.60E-05 | 0.434328 | 0.304 | 0.222 | 1 | 1 | Slamf9 | 0.082 | Young |
| 9.67E-05 | 0.135815 | 0.696 | 0.576 | 1 | 1 | Btg2 | 0.12 | Young |

| | | | | | | | | |
|---|---|---|---|---|---|---|---|---|
| 9.69E-05 | 1.002289 | 0.066 | 0.027 | 1 | 1 | Zfp871 | 0.039 | Young |
| 9.83E-05 | 0.085188 | 0.939 | 0.915 | 1 | 1 | Rplp1 | 0.024 | Young |
| 9.84E-05 | 0.566047 | 0.196 | 0.129 | 1 | 1 | Use1 | 0.067 | Young |
| 0.000101 | 1.483166 | 0.076 | 0.035 | 1 | 1 | Zfp740 | 0.041 | Young |
| 0.000103 | 0.153394 | 0.806 | 0.699 | 1 | 1 | Hspa8 | 0.107 | Young |
| 0.000104 | 0.289864 | 0.38 | 0.288 | 1 | 1 | Tob2 | 0.092 | Young |
| 0.000105 | 1.116549 | 0.083 | 0.04 | 1 | 1 | Pstpip1 | 0.043 | Young |
| 0.000107 | 0.423606 | 0.183 | 0.117 | 1 | 1 | Numa1 | 0.066 | Young |
| 0.000109 | 0.622375 | 0.171 | 0.108 | 1 | 1 | Arhgap31 | 0.063 | Young |
| 0.00011 | 0.181819 | 0.564 | 0.445 | 1 | 1 | Rpl36 | 0.119 | Young |
| 0.000113 | 0.753823 | 0.105 | 0.055 | 1 | 1 | Gpr183 | 0.05 | Young |
| 0.000115 | 0.181997 | 0.674 | 0.575 | 1 | 1 | Rpl5 | 0.099 | Young |
| 0.00012 | 0.277605 | 0.201 | 0.131 | 1 | 1 | Phlda1 | 0.07 | Young |
| 0.00012 | 1.826161 | 0.035 | 0.008 | 1 | 1 | Tln2 | 0.027 | Young |
| 0.000123 | 0.206424 | 0.609 | 0.508 | 1 | 1 | Rpl28 | 0.101 | Young |
| 0.000127 | 1.488711 | 0.027 | 0.004 | 1 | 1 | Inpp4b | 0.023 | Young |
| 0.000129 | 0.93855 | 0.093 | 0.048 | 1 | 1 | Chd7 | 0.045 | Young |
| 0.00013 | 0.313257 | 0.304 | 0.221 | 1 | 1 | Tra2b | 0.083 | Young |
| 0.000138 | 0.459945 | 0.191 | 0.126 | 1 | 1 | Sash1 | 0.065 | Young |
| 0.000141 | 0.516822 | 0.205 | 0.138 | 1 | 1 | Crebbp | 0.067 | Young |
| 0.000141 | 0.428276 | 0.255 | 0.183 | 1 | 1 | Glg1 | 0.072 | Young |
| 0.000146 | 3.41952 | 0.021 | 0.001 | 1 | 1 | Cd69 | 0.02 | Young |
| 0.000146 | 0.738937 | 0.126 | 0.073 | 1 | 1 | Ptbp1 | 0.053 | Young |
| 0.000147 | 0.542078 | 0.16 | 0.099 | 1 | 1 | Tsc22d2 | 0.061 | Young |
| 0.000151 | 0.124022 | 0.927 | 0.904 | 1 | 1 | Eef1a1 | 0.023 | Young |
| 0.000155 | 0.42998 | 0.236 | 0.163 | 1 | 1 | Hacd4 | 0.073 | Young |
| 0.000161 | 0.2753 | 0.456 | 0.359 | 1 | 1 | Srgn | 0.097 | Young |
| 0.000162 | 0.176044 | 0.321 | 0.23 | 1 | 1 | Uqcr11 | 0.091 | Young |
| 0.000163 | 0.517148 | 0.152 | 0.094 | 1 | 1 | Trem2 | 0.058 | Young |
| 0.000163 | 0.429393 | 0.205 | 0.136 | 1 | 1 | Hook3 | 0.069 | Young |
| 0.000166 | 0.357915 | 0.292 | 0.212 | 1 | 1 | Basp1 | 0.08 | Young |
| 0.000172 | 0.373494 | 0.27 | 0.193 | 1 | 1 | St8sia4 | 0.077 | Young |
| 0.000173 | 0.221736 | 0.711 | 0.625 | 1 | 1 | Maf | 0.086 | Young |
| 0.000184 | 0.694296 | 0.119 | 0.068 | 1 | 1 | Atf7 | 0.051 | Young |
| 0.000185 | 0.696683 | 0.078 | 0.036 | 1 | 1 | Atxn1 | 0.042 | Young |
| 0.000199 | 0.158719 | 0.848 | 0.771 | 1 | 1 | Rpl38 | 0.077 | Young |
| 0.0002 | 1.844318 | 0.04 | 0.012 | 1 | 1 | Spryd7 | 0.028 | Young |
| 0.000206 | 0.389908 | 0.301 | 0.22 | 1 | 1 | Nisch | 0.081 | Young |
| 0.000209 | 0.608207 | 0.113 | 0.063 | 1 | 1 | 1700017B0 | 0.05 | Young |
| 0.000212 | 0.904001 | 0.131 | 0.08 | 1 | 1 | Pdgfb | 0.051 | Young |
| 0.000222 | 0.271209 | 0.388 | 0.293 | 1 | 1 | Luc7l2 | 0.095 | Young |
| 0.000222 | 0.215987 | 0.383 | 0.288 | 1 | 1 | Kdm6b | 0.095 | Young |
| 0.000232 | 0.781571 | 0.142 | 0.089 | 1 | 1 | Ubxn4 | 0.053 | Young |
| 0.000238 | 0.255064 | 0.357 | 0.265 | 1 | 1 | Fnbp1 | 0.092 | Young |
| 0.000249 | 0.541084 | 0.123 | 0.072 | 1 | 1 | Gm10073 | 0.051 | Young |

| | | | | | | | | |
|---|---|---|---|---|---|---|---|---|
| 0.000252 | 0.829435 | 0.117 | 0.068 | 1 | 1 | Eif2a | 0.049 | Young |
| 0.000257 | 0.327213 | 0.278 | 0.202 | 1 | 1 | Gpi1 | 0.076 | Young |
| 0.000259 | 0.274087 | 0.328 | 0.242 | 1 | 1 | Clic4 | 0.086 | Young |
| 0.000261 | 0.231158 | 0.582 | 0.476 | 1 | 1 | Gnai2 | 0.106 | Young |
| 0.000272 | 0.280945 | 0.283 | 0.203 | 1 | 1 | Mycbp2 | 0.08 | Young |
| 0.000273 | 0.164094 | 0.349 | 0.261 | 1 | 1 | Ncl | 0.088 | Young |
| 0.000275 | 0.952854 | 0.104 | 0.058 | 1 | 1 | Pdcd4 | 0.046 | Young |
| 0.000284 | 0.53971 | 0.117 | 0.067 | 1 | 1 | Serpinf1 | 0.05 | Young |
| 0.000289 | 0.453712 | 0.148 | 0.093 | 1 | 1 | Gpsm3 | 0.055 | Young |
| 0.000291 | 0.617414 | 0.177 | 0.118 | 1 | 1 | Tiparp | 0.059 | Young |
| 0.000291 | 0.203179 | 0.529 | 0.422 | 1 | 1 | Rpl14 | 0.107 | Young |
| 0.000295 | 0.349191 | 0.222 | 0.154 | 1 | 1 | Mrpl52 | 0.068 | Young |
| 0.000299 | 1.086765 | 0.035 | 0.009 | 1 | 1 | Pxdc1 | 0.026 | Young |
| 0.000304 | 0.558319 | 0.106 | 0.059 | 1 | 1 | Arrdc3 | 0.047 | Young |
| 0.000306 | 0.521569 | 0.173 | 0.113 | 1 | 1 | Atf6 | 0.06 | Young |
| 0.000308 | 0.526407 | 0.184 | 0.123 | 1 | 1 | Cox14 | 0.061 | Young |
| 0.000317 | 0.594505 | 0.106 | 0.059 | 1 | 1 | Mylip | 0.047 | Young |
| 0.000321 | 0.317232 | 0.513 | 0.424 | 1 | 1 | Sdc4 | 0.089 | Young |
| 0.000322 | 0.250155 | 0.346 | 0.263 | 1 | 1 | Mtdh | 0.083 | Young |
| 0.000327 | 0.414941 | 0.231 | 0.162 | 1 | 1 | Mafg | 0.069 | Young |
| 0.000328 | 0.299017 | 0.36 | 0.275 | 1 | 1 | Nfe2l2 | 0.085 | Young |
| 0.000334 | 0.642314 | 0.21 | 0.149 | 1 | 1 | Map1lc3b | 0.061 | Young |
| 0.000343 | 0.398478 | 0.188 | 0.125 | 1 | 1 | Swap70 | 0.063 | Young |
| 0.000344 | 0.245612 | 0.479 | 0.384 | 1 | 1 | Qk | 0.095 | Young |
| 0.000349 | 0.500754 | 0.202 | 0.14 | 1 | 1 | Brd2 | 0.062 | Young |
| 0.000366 | 1.591954 | 0.034 | 0.009 | 1 | 1 | Git1 | 0.025 | Young |
| 0.000367 | 1.395397 | 0.054 | 0.022 | 1 | 1 | Ago1 | 0.032 | Young |
| 0.000367 | 0.484592 | 0.205 | 0.144 | 1 | 1 | Ttc14 | 0.061 | Young |
| 0.000401 | 0.183009 | 0.564 | 0.461 | 1 | 1 | Eef1b2 | 0.103 | Young |
| 0.000412 | 0.541903 | 0.08 | 0.04 | 1 | 1 | Bmyc | 0.04 | Young |
| 0.000415 | 0.643165 | 0.131 | 0.081 | 1 | 1 | Ankrd12 | 0.05 | Young |
| 0.000417 | 0.286917 | 0.301 | 0.224 | 1 | 1 | Inpp5d | 0.077 | Young |
| 0.000423 | 0.47474 | 0.199 | 0.139 | 1 | 1 | Vasp | 0.06 | Young |
| 0.000434 | 0.412942 | 0.189 | 0.129 | 1 | 1 | Zfp395 | 0.06 | Young |
| 0.000435 | 1.610037 | 0.046 | 0.017 | 1 | 1 | Zcchc9 | 0.029 | Young |
| 0.000444 | 0.33375 | 0.279 | 0.21 | 1 | 1 | Il10ra | 0.069 | Young |
| 0.000451 | 0.686472 | 0.091 | 0.049 | 1 | 1 | Dnajc1 | 0.042 | Young |
| 0.000452 | 0.578668 | 0.109 | 0.063 | 1 | 1 | Gigyf1 | 0.046 | Young |
| 0.000456 | 0.3293 | 0.404 | 0.325 | 1 | 1 | Ogt | 0.079 | Young |
| 0.000482 | 0.357944 | 0.309 | 0.238 | 1 | 1 | Top1 | 0.071 | Young |
| 0.000483 | 0.504199 | 0.148 | 0.095 | 1 | 1 | Zfp638 | 0.053 | Young |
| 0.000484 | 0.439939 | 0.182 | 0.122 | 1 | 1 | Nop53 | 0.06 | Young |
| 0.000484 | 0.877424 | 0.082 | 0.042 | 1 | 1 | Pcyox1 | 0.04 | Young |
| 0.000495 | 0.297652 | 0.137 | 0.085 | 1 | 1 | Kdm5a | 0.052 | Young |
| 0.000496 | 0.457151 | 0.177 | 0.118 | 1 | 1 | Top2b | 0.059 | Young |

| | | | | | | | | |
|---|---|---|---|---|---|---|---|---|
| 0.000497 | 0.550651 | 0.123 | 0.073 | 1 | 1 | Ophn1 | 0.05 | Young |
| 0.000498 | 0.470197 | 0.216 | 0.152 | 1 | 1 | Gatad2b | 0.064 | Young |
| 0.000499 | 0.376413 | 0.167 | 0.111 | 1 | 1 | Smg1 | 0.056 | Young |
| 0.000499 | 0.2962 | 0.311 | 0.238 | 1 | 1 | Rgs10 | 0.073 | Young |
| 0.000517 | 0.517414 | 0.163 | 0.107 | 1 | 1 | Msr1 | 0.056 | Young |
| 0.000522 | 1.105476 | 0.074 | 0.037 | 1 | 1 | Gm48065 | 0.037 | Young |
| 0.000533 | 1.117308 | 0.069 | 0.033 | 1 | 1 | Tspan13 | 0.036 | Young |
| 0.000536 | 0.859671 | 0.084 | 0.045 | 1 | 1 | Frmd6 | 0.039 | Young |
| 0.000541 | 0.673342 | 0.057 | 0.024 | 1 | 1 | Cacna1d | 0.033 | Young |
| 0.000552 | 0.194375 | 0.637 | 0.542 | 1 | 1 | Rpl6 | 0.095 | Young |
| 0.00056 | 0.294557 | 0.194 | 0.131 | 1 | 1 | Ndufb11 | 0.063 | Young |
| 0.00056 | 0.316574 | 0.289 | 0.216 | 1 | 1 | Cd93 | 0.073 | Young |
| 0.000564 | 0.51648 | 0.194 | 0.135 | 1 | 1 | Arhgap15 | 0.059 | Young |
| 0.000565 | 0.421155 | 0.225 | 0.161 | 1 | 1 | Prex1 | 0.064 | Young |
| 0.000573 | 1.386529 | 0.045 | 0.017 | 1 | 1 | Wipi2 | 0.028 | Young |
| 0.000573 | 0.335116 | 0.238 | 0.171 | 1 | 1 | Cnot6l | 0.067 | Young |
| 0.000574 | 0.28816 | 0.403 | 0.317 | 1 | 1 | C3ar1 | 0.086 | Young |
| 0.000582 | 0.478051 | 0.156 | 0.102 | 1 | 1 | Med13 | 0.054 | Young |
| 0.000584 | 0.312859 | 0.301 | 0.229 | 1 | 1 | Nr3c1 | 0.072 | Young |
| 0.000584 | 0.798701 | 0.106 | 0.062 | 1 | 1 | AI314180 | 0.044 | Young |
| 0.000585 | 0.495593 | 0.202 | 0.143 | 1 | 1 | Hivep2 | 0.059 | Young |
| 0.000593 | 0.510364 | 0.19 | 0.132 | 1 | 1 | Cd86 | 0.058 | Young |
| 0.000596 | 0.300633 | 0.222 | 0.156 | 1 | 1 | Snx18 | 0.066 | Young |
| 0.000602 | 0.521586 | 0.207 | 0.147 | 1 | 1 | Mxd4 | 0.06 | Young |
| 0.00061 | 0.129148 | 0.911 | 0.862 | 1 | 1 | Marcks | 0.049 | Young |
| 0.000619 | 0.291679 | 0.284 | 0.213 | 1 | 1 | Chd4 | 0.071 | Young |
| 0.000621 | 0.75204 | 0.114 | 0.069 | 1 | 1 | Vezf1 | 0.045 | Young |
| 0.00063 | 0.579489 | 0.125 | 0.076 | 1 | 1 | Sh3bp5 | 0.049 | Young |
| 0.000632 | 0.593768 | 0.143 | 0.091 | 1 | 1 | Ube2h | 0.052 | Young |
| 0.000643 | 0.268111 | 0.391 | 0.308 | 1 | 1 | Cyth4 | 0.083 | Young |
| 0.000653 | 0.834555 | 0.069 | 0.033 | 1 | 1 | Nmt2 | 0.036 | Young |
| 0.000657 | 0.796809 | 0.084 | 0.045 | 1 | 1 | Smc4 | 0.039 | Young |
| 0.000676 | 0.435285 | 0.166 | 0.111 | 1 | 1 | Rac2 | 0.055 | Young |
| 0.000682 | 0.108654 | 0.933 | 0.902 | 1 | 1 | mt-Nd4 | 0.031 | Young |
| 0.000692 | 2.89377 | 0.018 | 0.001 | 1 | 1 | Lhfpl2 | 0.017 | Young |
| 0.000692 | 0.415622 | 0.202 | 0.141 | 1 | 1 | Icam1 | 0.061 | Young |
| 0.0007 | 0.356342 | 0.336 | 0.261 | 1 | 1 | Pfdn5 | 0.075 | Young |
| 0.000705 | 0.249375 | 0.309 | 0.231 | 1 | 1 | Ets2 | 0.078 | Young |
| 0.000707 | 0.814528 | 0.047 | 0.018 | 1 | 1 | Dnase1l3 | 0.029 | Young |
| 0.000721 | 0.704469 | 0.089 | 0.049 | 1 | 1 | Dido1 | 0.04 | Young |
| 0.000729 | 0.396616 | 0.167 | 0.112 | 1 | 1 | Nars | 0.055 | Young |
| 0.000734 | 0.251979 | 0.315 | 0.235 | 1 | 1 | Tpr | 0.08 | Young |
| 0.000739 | 0.370947 | 0.188 | 0.129 | 1 | 1 | Kras | 0.059 | Young |
| 0.000744 | 0.125549 | 0.578 | 0.472 | 1 | 1 | Rpl21 | 0.106 | Young |
| 0.000794 | 0.364531 | 0.298 | 0.226 | 1 | 1 | Ccl24 | 0.072 | Young |

| | | | | | | | | |
|---|---|---|---|---|---|---|---|---|
| 0.000806 | 0.42653 | 0.164 | 0.109 | 1 | 1 | Snx20 | 0.055 | Young |
| 0.000808 | 0.442249 | 0.149 | 0.096 | 1 | 1 | Ifnar2 | 0.053 | Young |
| 0.000809 | 2.185292 | 0.03 | 0.008 | 1 | 1 | Lrp5 | 0.022 | Young |
| 0.000819 | 0.478103 | 0.185 | 0.129 | 1 | 1 | Slc9a9 | 0.056 | Young |
| 0.000821 | 1.09188 | 0.052 | 0.022 | 1 | 1 | Pfkl | 0.03 | Young |
| 0.000849 | 1.573787 | 0.058 | 0.027 | 1 | 1 | Plxdc2 | 0.031 | Young |
| 0.000864 | 0.2176 | 0.493 | 0.404 | 1 | 1 | mt-Nd4l | 0.089 | Young |
| 0.000876 | 0.227926 | 0.312 | 0.237 | 1 | 1 | Tma7 | 0.075 | Young |
| 0.000888 | 0.530181 | 0.078 | 0.04 | 1 | 1 | Arhgap19 | 0.038 | Young |
| 0.000902 | 1.717283 | 0.038 | 0.013 | 1 | 1 | Mir155hg | 0.025 | Young |
| 0.000906 | 0.578201 | 0.107 | 0.063 | 1 | 1 | Fbxw4 | 0.044 | Young |
| 0.000907 | 3.22632 | 0.014 | 0 | 1 | 1 | Olfr1167 | 0.014 | Young |
| 0.000913 | 0.359479 | 0.369 | 0.299 | 1 | 1 | Fcgr3 | 0.07 | Young |
| 0.000921 | 0.283887 | 0.401 | 0.32 | 1 | 1 | Tln1 | 0.081 | Young |
| 0.000943 | 1.154689 | 0.058 | 0.027 | 1 | 1 | Smox | 0.031 | Young |
| 0.000947 | 0.62986 | 0.103 | 0.06 | 1 | 1 | Fam172a | 0.043 | Young |
| 0.000948 | 0.184716 | 0.384 | 0.302 | 1 | 1 | Gmfg | 0.082 | Young |
| 0.000957 | 0.350981 | 0.229 | 0.167 | 1 | 1 | Lacc1 | 0.062 | Young |
| 0.00096 | 0.206526 | 0.353 | 0.275 | 1 | 1 | Atpif1 | 0.078 | Young |
| 0.000967 | 0.238711 | 0.36 | 0.279 | 1 | 1 | Wdr26 | 0.081 | Young |
| 0.000974 | 0.340196 | 0.218 | 0.157 | 1 | 1 | Mef2d | 0.061 | Young |
| 0.000993 | 0.846601 | 0.049 | 0.021 | 1 | 1 | Arap3 | 0.028 | Young |
| 0.001011 | 0.552197 | 0.129 | 0.081 | 1 | 1 | Tnfrsf13b | 0.048 | Young |
| 0.001018 | 0.257359 | 0.342 | 0.27 | 1 | 1 | Tnfaip8 | 0.072 | Young |
| 0.001022 | 0.213444 | 0.411 | 0.329 | 1 | 1 | Clk1 | 0.082 | Young |
| 0.001023 | 0.717876 | 0.1 | 0.059 | 1 | 1 | Bmp2 | 0.041 | Young |
| 0.001023 | 0.321805 | 0.36 | 0.285 | 1 | 1 | Eef1g | 0.075 | Young |
| 0.001053 | 0.862113 | 0.082 | 0.045 | 1 | 1 | 1700025G( | 0.037 | Young |
| 0.001055 | 0.806588 | 0.083 | 0.045 | 1 | 1 | Mbp | 0.038 | Young |
| 0.001059 | 0.763095 | 0.066 | 0.032 | 1 | 1 | Hs2st1 | 0.034 | Young |
| 0.001061 | 0.305766 | 0.273 | 0.203 | 1 | 1 | Tgfbr2 | 0.07 | Young |
| 0.001065 | 0.29361 | 0.186 | 0.127 | 1 | 1 | Crlf3 | 0.059 | Young |
| 0.001081 | 0.361417 | 0.253 | 0.19 | 1 | 1 | Spi1 | 0.063 | Young |
| 0.001086 | 0.115676 | 0.558 | 0.455 | 1 | 1 | Rps17 | 0.103 | Young |
| 0.001095 | 0.331164 | 0.304 | 0.234 | 1 | 1 | Tra2a | 0.07 | Young |
| 0.001163 | 1.026813 | 0.077 | 0.041 | 1 | 1 | Tubgcp5 | 0.036 | Young |
| 0.001172 | 0.145015 | 0.518 | 0.42 | 1 | 1 | Gm10076 | 0.098 | Young |
| 0.001174 | 0.562828 | 0.145 | 0.096 | 1 | 1 | Ivns1abp | 0.049 | Young |
| 0.001184 | 0.263247 | 0.416 | 0.341 | 1 | 1 | Erp29 | 0.075 | Young |
| 0.001198 | 0.257225 | 0.147 | 0.095 | 1 | 1 | Sh3bp1 | 0.052 | Young |
| 0.001222 | 0.789129 | 0.107 | 0.066 | 1 | 1 | Tifab | 0.041 | Young |
| 0.001226 | 0.965065 | 0.083 | 0.046 | 1 | 1 | Tmem127 | 0.037 | Young |
| 0.001231 | 0.441182 | 0.165 | 0.112 | 1 | 1 | Ndufb9 | 0.053 | Young |
| 0.001243 | 1.428307 | 0.055 | 0.026 | 1 | 1 | Poc1b | 0.029 | Young |
| 0.001245 | 0.246332 | 0.386 | 0.307 | 1 | 1 | Il6ra | 0.079 | Young |

| | | | | | | | | |
|---|---|---|---|---|---|---|---|---|
| 0.001247 | 0.803418 | 0.078 | 0.042 | 1 | 1 | Slc45a4 | 0.036 | Young |
| 0.001273 | 0.411532 | 0.235 | 0.175 | 1 | 1 | Cd2ap | 0.06 | Young |
| 0.001291 | 0.208202 | 0.516 | 0.423 | 1 | 1 | Aplp2 | 0.093 | Young |
| 0.001329 | 0.925234 | 0.047 | 0.019 | 1 | 1 | Abr | 0.028 | Young |
| 0.001336 | 0.356166 | 0.206 | 0.148 | 1 | 1 | Rab3il1 | 0.058 | Young |
| 0.001337 | 1.913253 | 0.021 | 0.004 | 1 | 1 | Utp23 | 0.017 | Young |
| 0.00134 | 0.770036 | 0.107 | 0.066 | 1 | 1 | Pde3b | 0.041 | Young |
| 0.001392 | 0.366081 | 0.22 | 0.161 | 1 | 1 | Ankrd44 | 0.059 | Young |
| 0.001401 | 0.98719 | 0.069 | 0.036 | 1 | 1 | D730003I1 | 0.033 | Young |
| 0.001411 | 0.565734 | 0.095 | 0.055 | 1 | 1 | Dhx36 | 0.04 | Young |
| 0.001419 | 0.429087 | 0.144 | 0.095 | 1 | 1 | Ubn1 | 0.049 | Young |
| 0.00143 | 0.282606 | 0.308 | 0.234 | 1 | 1 | Nfkb1 | 0.074 | Young |
| 0.001433 | 0.425783 | 0.21 | 0.153 | 1 | 1 | Cysltr1 | 0.057 | Young |
| 0.001443 | 0.802679 | 0.044 | 0.018 | 1 | 1 | Selenoh | 0.026 | Young |
| 0.001445 | 0.162743 | 0.473 | 0.383 | 1 | 1 | Rrbp1 | 0.09 | Young |
| 0.001453 | 1.30903 | 0.038 | 0.014 | 1 | 1 | Il1a | 0.024 | Young |
| 0.001455 | 0.69274 | 0.128 | 0.084 | 1 | 1 | Atxn7 | 0.044 | Young |
| 0.001482 | 0.239179 | 0.327 | 0.252 | 1 | 1 | Pafah1b1 | 0.075 | Young |
| 0.001485 | 1.592187 | 0.028 | 0.008 | 1 | 1 | Pald1 | 0.02 | Young |
| 0.001502 | 1.301465 | 0.046 | 0.019 | 1 | 1 | Sorl1 | 0.027 | Young |
| 0.001503 | 0.574499 | 0.078 | 0.042 | 1 | 1 | Idnk | 0.036 | Young |
| 0.001513 | 0.726928 | 0.055 | 0.026 | 1 | 1 | Sfmbt1 | 0.029 | Young |
| 0.001541 | 0.511991 | 0.104 | 0.063 | 1 | 1 | Tox4 | 0.041 | Young |
| 0.001542 | 0.359154 | 0.208 | 0.15 | 1 | 1 | Rnasel | 0.058 | Young |
| 0.001542 | 0.438498 | 0.148 | 0.099 | 1 | 1 | Rtcb | 0.049 | Young |
| 0.001574 | 0.693581 | 0.117 | 0.075 | 1 | 1 | Tnfrsf1a | 0.042 | Young |
| 0.001609 | 0.156558 | 0.448 | 0.36 | 1 | 1 | Tcf4 | 0.088 | Young |
| 0.001637 | 0.448582 | 0.161 | 0.111 | 1 | 1 | Fam208a | 0.05 | Young |
| 0.001638 | 0.720516 | 0.078 | 0.042 | 1 | 1 | Milr1 | 0.036 | Young |
| 0.001654 | 0.302448 | 0.188 | 0.132 | 1 | 1 | Vps35 | 0.056 | Young |
| 0.00167 | 0.327804 | 0.13 | 0.084 | 1 | 1 | Snx29 | 0.046 | Young |
| 0.001676 | 0.63781 | 0.105 | 0.064 | 1 | 1 | Tlr7 | 0.041 | Young |
| 0.001694 | 0.503104 | 0.117 | 0.073 | 1 | 1 | Ssfa2 | 0.044 | Young |
| 0.001695 | 0.241505 | 0.352 | 0.276 | 1 | 1 | Tacc1 | 0.076 | Young |
| 0.001713 | 0.344007 | 0.144 | 0.095 | 1 | 1 | Tmcc3 | 0.049 | Young |
| 0.001722 | 0.339427 | 0.17 | 0.117 | 1 | 1 | Etnk1 | 0.053 | Young |
| 0.00173 | 0.255995 | 0.277 | 0.207 | 1 | 1 | Slc38a2 | 0.07 | Young |
| 0.001747 | 0.414 | 0.156 | 0.105 | 1 | 1 | Acbd5 | 0.051 | Young |
| 0.001777 | 2.811533 | 0.021 | 0.004 | 1 | 1 | AA467197 | 0.017 | Young |
| 0.001798 | 0.566903 | 0.131 | 0.085 | 1 | 1 | Cmah | 0.046 | Young |
| 0.001799 | 0.186134 | 0.339 | 0.262 | 1 | 1 | Cox6c | 0.077 | Young |
| 0.001802 | 0.551037 | 0.142 | 0.095 | 1 | 1 | Eif4g3 | 0.047 | Young |
| 0.00182 | 0.712716 | 0.114 | 0.073 | 1 | 1 | Lpar6 | 0.041 | Young |
| 0.00183 | 0.270973 | 0.205 | 0.148 | 1 | 1 | Srsf11 | 0.057 | Young |
| 0.001838 | 0.12351 | 0.629 | 0.517 | 1 | 1 | Pim1 | 0.112 | Young |

| | | | | | | | | |
|---|---|---|---|---|---|---|---|---|
| 0.00184 | 0.440942 | 0.295 | 0.234 | 1 | 1 | Ptger4 | 0.061 | Young |
| 0.001848 | 0.415222 | 0.163 | 0.113 | 1 | 1 | Rb1cc1 | 0.05 | Young |
| 0.001849 | 0.893558 | 0.055 | 0.026 | 1 | 1 | Bnip3 | 0.029 | Young |
| 0.001858 | 0.233265 | 0.31 | 0.244 | 1 | 1 | Ndufa6 | 0.066 | Young |
| 0.001858 | 2.411437 | 0.025 | 0.006 | 1 | 1 | Nes | 0.019 | Young |
| 0.001868 | 0.791822 | 0.078 | 0.044 | 1 | 1 | P2ry12 | 0.034 | Young |
| 0.001881 | 0.258022 | 0.261 | 0.195 | 1 | 1 | Tgoln1 | 0.066 | Young |
| 0.001884 | 0.542278 | 0.117 | 0.075 | 1 | 1 | Tbl1x | 0.042 | Young |
| 0.001898 | 0.733742 | 0.077 | 0.042 | 1 | 1 | Ccng2 | 0.035 | Young |
| 0.001924 | 0.153099 | 0.242 | 0.179 | 1 | 1 | Fcho2 | 0.063 | Young |
| 0.001928 | 0.209143 | 0.451 | 0.365 | 1 | 1 | Cox7a2l | 0.086 | Young |
| 0.001944 | 0.570054 | 0.07 | 0.037 | 1 | 1 | Il17ra | 0.033 | Young |
| 0.001949 | 1.254789 | 0.041 | 0.017 | 1 | 1 | Fry | 0.024 | Young |
| 0.001967 | 0.400373 | 0.167 | 0.116 | 1 | 1 | Fam105a | 0.051 | Young |
| 0.002004 | 0.526854 | 0.137 | 0.091 | 1 | 1 | Dclre1c | 0.046 | Young |
| 0.002033 | 0.473042 | 0.124 | 0.08 | 1 | 1 | Foxj2 | 0.044 | Young |
| 0.00204 | 0.713622 | 0.137 | 0.093 | 1 | 1 | 2810474O1 | 0.044 | Young |
| 0.00205 | 0.217452 | 0.367 | 0.29 | 1 | 1 | Akap13 | 0.077 | Young |
| 0.002061 | 0.405747 | 0.16 | 0.111 | 1 | 1 | Phf3 | 0.049 | Young |
| 0.002134 | 0.432754 | 0.147 | 0.1 | 1 | 1 | Mrpl23 | 0.047 | Young |
| 0.002142 | 0.410902 | 0.202 | 0.147 | 1 | 1 | Eef1d | 0.055 | Young |
| 0.002154 | 0.297344 | 0.182 | 0.129 | 1 | 1 | Erbin | 0.053 | Young |
| 0.002168 | 0.230245 | 0.294 | 0.228 | 1 | 1 | Hint1 | 0.066 | Young |
| 0.002174 | 0.509133 | 0.156 | 0.108 | 1 | 1 | Dnajc5 | 0.048 | Young |
| 0.002211 | 0.36116 | 0.129 | 0.084 | 1 | 1 | Chic2 | 0.045 | Young |
| 0.002241 | 0.710604 | 0.095 | 0.058 | 1 | 1 | Smagp | 0.037 | Young |
| 0.002253 | 0.576406 | 0.127 | 0.084 | 1 | 1 | Baz1a | 0.043 | Young |
| 0.002277 | 0.542538 | 0.106 | 0.066 | 1 | 1 | Ripk1 | 0.04 | Young |
| 0.002281 | 0.129332 | 0.245 | 0.181 | 1 | 1 | Ube3a | 0.064 | Young |
| 0.002304 | 0.327118 | 0.19 | 0.136 | 1 | 1 | Scaf11 | 0.054 | Young |
| 0.002318 | 0.415951 | 0.143 | 0.096 | 1 | 1 | Fam117b | 0.047 | Young |
| 0.002338 | 0.775028 | 0.066 | 0.035 | 1 | 1 | Ang | 0.031 | Young |
| 0.002346 | 3.174031 | 0.012 | 0 | 1 | 1 | Hcar2 | 0.012 | Young |
| 0.002346 | 2.209672 | 0.012 | 0 | 1 | 1 | Grap2 | 0.012 | Young |
| 0.002357 | 0.869066 | 0.09 | 0.054 | 1 | 1 | Brox | 0.036 | Young |
| 0.00236 | 0.141965 | 0.406 | 0.324 | 1 | 1 | Eef2 | 0.082 | Young |
| 0.002373 | 0.320396 | 0.169 | 0.12 | 1 | 1 | Tnrc6b | 0.049 | Young |
| 0.002376 | 3.152641 | 0.015 | 0.001 | 1 | 1 | Prr5 | 0.014 | Young |
| 0.002417 | 2.624717 | 0.015 | 0.001 | 1 | 1 | Rapgef5 | 0.014 | Young |
| 0.002434 | 1.704312 | 0.029 | 0.009 | 1 | 1 | St3gal2 | 0.02 | Young |
| 0.00244 | 0.371639 | 0.196 | 0.144 | 1 | 1 | Dock8 | 0.052 | Young |
| 0.002455 | 0.329331 | 0.288 | 0.225 | 1 | 1 | Slc43a2 | 0.063 | Young |
| 0.002464 | 0.662197 | 0.107 | 0.068 | 1 | 1 | Ccdc12 | 0.039 | Young |
| 0.00248 | 1.060029 | 0.094 | 0.058 | 1 | 1 | Asxl2 | 0.036 | Young |
| 0.002496 | 0.992213 | 0.069 | 0.037 | 1 | 1 | Slc35a4 | 0.032 | Young |

| | | | | | | | | |
|---|---|---|---|---|---|---|---|---|
| 0.002503 | 0.655231 | 0.127 | 0.085 | 1 | 1 | Map3k8 | 0.042 | Young |
| 0.002507 | 1.59236 | 0.022 | 0.005 | 1 | 1 | Sugp1 | 0.017 | Young |
| 0.002509 | 0.974685 | 0.092 | 0.057 | 1 | 1 | Heatr5a | 0.035 | Young |
| 0.00251 | 0.347835 | 0.168 | 0.117 | 1 | 1 | Eif3h | 0.051 | Young |
| 0.002522 | 0.231051 | 0.382 | 0.31 | 1 | 1 | Son | 0.072 | Young |
| 0.002536 | 0.255721 | 0.256 | 0.194 | 1 | 1 | Macf1 | 0.062 | Young |
| 0.002553 | 0.599415 | 0.12 | 0.078 | 1 | 1 | Clcn5 | 0.042 | Young |
| 0.00256 | 0.57235 | 0.067 | 0.036 | 1 | 1 | Dock4 | 0.031 | Young |
| 0.002564 | 0.111476 | 0.477 | 0.378 | 1 | 1 | Stk17b | 0.099 | Young |
| 0.002592 | 1.448605 | 0.022 | 0.005 | 1 | 1 | Pik3r2 | 0.017 | Young |
| 0.002592 | 0.535892 | 0.143 | 0.098 | 1 | 1 | Tnrc6c | 0.045 | Young |
| 0.002628 | 0.139728 | 0.248 | 0.185 | 1 | 1 | Pik3r1 | 0.063 | Young |
| 0.002649 | 0.525357 | 0.15 | 0.104 | 1 | 1 | Adgre5 | 0.046 | Young |
| 0.002666 | 0.74066 | 0.079 | 0.045 | 1 | 1 | Slc12a9 | 0.034 | Young |
| 0.002682 | 0.302248 | 0.19 | 0.135 | 1 | 1 | Lyz1 | 0.055 | Young |
| 0.002701 | 0.146802 | 0.357 | 0.284 | 1 | 1 | Gng5 | 0.073 | Young |
| 0.002738 | 0.170794 | 0.561 | 0.465 | 1 | 1 | Pid1 | 0.096 | Young |
| 0.002753 | 0.531884 | 0.108 | 0.068 | 1 | 1 | Smad4 | 0.04 | Young |
| 0.00277 | 0.449717 | 0.092 | 0.055 | 1 | 1 | Itgav | 0.037 | Young |
| 0.002778 | 0.400369 | 0.189 | 0.138 | 1 | 1 | Pde4b | 0.051 | Young |
| 0.002781 | 0.460798 | 0.089 | 0.053 | 1 | 1 | P4ha1 | 0.036 | Young |
| 0.002886 | 0.452647 | 0.142 | 0.096 | 1 | 1 | Inpp4a | 0.046 | Young |
| 0.002892 | 0.326219 | 0.214 | 0.158 | 1 | 1 | Gsk3b | 0.056 | Young |
| 0.002897 | 0.337182 | 0.224 | 0.166 | 1 | 1 | Map4k4 | 0.058 | Young |
| 0.002899 | 0.523754 | 0.079 | 0.045 | 1 | 1 | Ndufs3 | 0.034 | Young |
| 0.002919 | 2.193846 | 0.017 | 0.003 | 1 | 1 | Zbtb5 | 0.014 | Young |
| 0.002932 | 0.163871 | 0.542 | 0.459 | 1 | 1 | mt-Nd5 | 0.083 | Young |
| 0.002951 | 0.658338 | 0.064 | 0.033 | 1 | 1 | Abl1 | 0.031 | Young |
| 0.002966 | 0.480826 | 0.129 | 0.086 | 1 | 1 | Ino80d | 0.043 | Young |
| 0.002967 | 0.442367 | 0.117 | 0.076 | 1 | 1 | Zfp292 | 0.041 | Young |
| 0.002976 | 0.31806 | 0.131 | 0.087 | 1 | 1 | Madd | 0.044 | Young |
| 0.003003 | 0.878804 | 0.055 | 0.027 | 1 | 1 | Wasl | 0.028 | Young |
| 0.003036 | 0.480743 | 0.23 | 0.177 | 1 | 1 | Ccdc88a | 0.053 | Young |
| 0.003048 | 1.359199 | 0.036 | 0.014 | 1 | 1 | Patz1 | 0.022 | Young |
| 0.003052 | 0.16896 | 0.595 | 0.492 | 1 | 1 | Ptprc | 0.103 | Young |
| 0.003067 | 0.458978 | 0.167 | 0.12 | 1 | 1 | Srsf1 | 0.047 | Young |
| 0.003083 | 0.745698 | 0.06 | 0.031 | 1 | 1 | Cog5 | 0.029 | Young |
| 0.003108 | 0.732845 | 0.078 | 0.045 | 1 | 1 | Lsm7 | 0.033 | Young |
| 0.003149 | 0.430913 | 0.111 | 0.071 | 1 | 1 | Map3k11 | 0.04 | Young |
| 0.003152 | 0.786547 | 0.06 | 0.031 | 1 | 1 | Tor3a | 0.029 | Young |
| 0.003166 | 0.391339 | 0.119 | 0.077 | 1 | 1 | Taz | 0.042 | Young |
| 0.003176 | 1.731983 | 0.028 | 0.009 | 1 | 1 | Bmf | 0.019 | Young |
| 0.00318 | 0.758966 | 0.055 | 0.027 | 1 | 1 | Afg3l1 | 0.028 | Young |
| 0.003196 | 0.487359 | 0.06 | 0.031 | 1 | 1 | Wdr3 | 0.029 | Young |
| 0.003201 | 0.71318 | 0.083 | 0.049 | 1 | 1 | Herpud2 | 0.034 | Young |

| | | | | | | | | |
|---|---|---|---|---|---|---|---|---|
| 0.003221 | 1.188943 | 0.049 | 0.023 | 1 | 1 | Dcaf17 | 0.026 | Young |
| 0.003227 | 2.91796 | 0.011 | 0 | 1 | 1 | Fkbpl | 0.011 | Young |
| 0.003298 | 0.314592 | 0.232 | 0.176 | 1 | 1 | Atp5j2 | 0.056 | Young |
| 0.0033 | 0.264654 | 0.157 | 0.107 | 1 | 1 | Ltc4s | 0.05 | Young |
| 0.003314 | 1.417334 | 0.045 | 0.021 | 1 | 1 | Ap1g2 | 0.024 | Young |
| 0.003343 | 1.643041 | 0.026 | 0.008 | 1 | 1 | Ets1 | 0.018 | Young |
| 0.003372 | 1.743547 | 0.021 | 0.005 | 1 | 1 | Cox10 | 0.016 | Young |
| 0.003449 | 0.648993 | 0.1 | 0.063 | 1 | 1 | Man1a2 | 0.037 | Young |
| 0.003454 | 0.478526 | 0.12 | 0.08 | 1 | 1 | Cux1 | 0.04 | Young |
| 0.003477 | 0.287874 | 0.259 | 0.201 | 1 | 1 | Myh9 | 0.058 | Young |
| 0.003486 | 0.397783 | 0.194 | 0.141 | 1 | 1 | Ikzf1 | 0.053 | Young |
| 0.003518 | 1.401389 | 0.032 | 0.012 | 1 | 1 | Itgal | 0.02 | Young |
| 0.00352 | 0.177418 | 0.454 | 0.371 | 1 | 1 | Plek | 0.083 | Young |
| 0.00358 | 0.718363 | 0.081 | 0.048 | 1 | 1 | Ylpm1 | 0.033 | Young |
| 0.003665 | 0.160605 | 0.157 | 0.108 | 1 | 1 | Bin1 | 0.049 | Young |
| 0.003677 | 1.023511 | 0.038 | 0.015 | 1 | 1 | Epm2aip1 | 0.023 | Young |
| 0.003679 | 0.235016 | 0.328 | 0.261 | 1 | 1 | Trib1 | 0.067 | Young |
| 0.003695 | 0.72535 | 0.049 | 0.023 | 1 | 1 | Abcc3 | 0.026 | Young |
| 0.003699 | 0.177467 | 0.16 | 0.112 | 1 | 1 | Cnot1 | 0.048 | Young |
| 0.003717 | 0.227116 | 0.232 | 0.176 | 1 | 1 | Dync1i2 | 0.056 | Young |
| 0.003721 | 0.17448 | 0.577 | 0.485 | 1 | 1 | Rsrp1 | 0.092 | Young |
| 0.00373 | 0.412427 | 0.177 | 0.13 | 1 | 1 | Asph | 0.047 | Young |
| 0.003732 | 0.754704 | 0.081 | 0.048 | 1 | 1 | Mrps24 | 0.033 | Young |
| 0.003738 | 0.729319 | 0.069 | 0.039 | 1 | 1 | Tbk1 | 0.03 | Young |
| 0.00382 | 0.617706 | 0.129 | 0.087 | 1 | 1 | Pik3c2a | 0.042 | Young |
| 0.003849 | 0.455251 | 0.12 | 0.08 | 1 | 1 | Lilrb4a | 0.04 | Young |
| 0.003908 | 0.337915 | 0.162 | 0.116 | 1 | 1 | Nek7 | 0.046 | Young |
| 0.003915 | 0.427528 | 0.061 | 0.032 | 1 | 1 | Cd79b | 0.029 | Young |
| 0.003979 | 0.837529 | 0.092 | 0.058 | 1 | 1 | Arhgef3 | 0.034 | Young |
| 0.003985 | 0.411625 | 0.157 | 0.111 | 1 | 1 | Fem1c | 0.046 | Young |
| 0.004002 | 2.488122 | 0.016 | 0.003 | 1 | 1 | Cyr61 | 0.013 | Young |
| 0.004012 | 0.345762 | 0.165 | 0.117 | 1 | 1 | Morc3 | 0.048 | Young |
| 0.004013 | 1.060724 | 0.064 | 0.035 | 1 | 1 | Spen | 0.029 | Young |
| 0.004038 | 0.062172 | 0.391 | 0.307 | 1 | 1 | Serp1 | 0.084 | Young |
| 0.004047 | 0.384502 | 0.227 | 0.174 | 1 | 1 | Slco2b1 | 0.053 | Young |
| 0.004054 | 0.810179 | 0.062 | 0.033 | 1 | 1 | Ubac2 | 0.029 | Young |
| 0.004056 | 1.862222 | 0.027 | 0.009 | 1 | 1 | Ankrd26 | 0.018 | Young |
| 0.00408 | 0.907608 | 0.057 | 0.03 | 1 | 1 | Ctc1 | 0.027 | Young |
| 0.004132 | 0.047746 | 0.484 | 0.4 | 1 | 1 | H2afz | 0.084 | Young |
| 0.004135 | 0.431479 | 0.153 | 0.108 | 1 | 1 | Sipa1 | 0.045 | Young |
| 0.004245 | 0.578701 | 0.095 | 0.059 | 1 | 1 | Ing4 | 0.036 | Young |
| 0.004262 | 1.456948 | 0.041 | 0.018 | 1 | 1 | D16Ertd47 | 0.023 | Young |
| 0.00429 | 0.397086 | 0.134 | 0.091 | 1 | 1 | Rasgef1b | 0.043 | Young |
| 0.004329 | 0.799042 | 0.052 | 0.026 | 1 | 1 | Ttc28 | 0.026 | Young |
| 0.004337 | 0.554936 | 0.061 | 0.032 | 1 | 1 | Tbc1d17 | 0.029 | Young |

| | | | | | | | | |
|---|---|---|---|---|---|---|---|---|
| 0.004347 | 2.005568 | 0.023 | 0.006 | 1 | 1 | Mblac2 | 0.017 | Young |
| 0.004378 | 0.27936 | 0.239 | 0.184 | 1 | 1 | Git2 | 0.055 | Young |
| 0.004386 | 1.409788 | 0.041 | 0.018 | 1 | 1 | 5430427O1 | 0.023 | Young |
| 0.004395 | 2.299744 | 0.021 | 0.005 | 1 | 1 | Pick1 | 0.016 | Young |
| 0.004402 | 0.526505 | 0.092 | 0.057 | 1 | 1 | Clock | 0.035 | Young |
| 0.004404 | 0.413789 | 0.138 | 0.095 | 1 | 1 | Slc25a36 | 0.043 | Young |
| 0.004423 | 0.798286 | 0.065 | 0.036 | 1 | 1 | Vav2 | 0.029 | Young |
| 0.004448 | 0.314971 | 0.15 | 0.105 | 1 | 1 | Trafd1 | 0.045 | Young |
| 0.004458 | 2.090894 | 0.021 | 0.005 | 1 | 1 | Fbxo21 | 0.016 | Young |
| 0.004462 | 0.101552 | 0.678 | 0.576 | 1 | 1 | Nr4a1 | 0.102 | Young |
| 0.004464 | 2.94459 | 0.013 | 0.001 | 1 | 1 | Fgf11 | 0.012 | Young |
| 0.004476 | 0.225617 | 0.22 | 0.165 | 1 | 1 | Acp2 | 0.055 | Young |
| 0.004483 | 1.778942 | 0.023 | 0.006 | 1 | 1 | Ccl25 | 0.017 | Young |
| 0.004518 | 0.347831 | 0.117 | 0.077 | 1 | 1 | Ncoa4 | 0.04 | Young |
| 0.004551 | 0.472592 | 0.064 | 0.035 | 1 | 1 | Svbp | 0.029 | Young |
| 0.004566 | 0.285259 | 0.172 | 0.123 | 1 | 1 | Pkn1 | 0.049 | Young |
| 0.004611 | 0.857032 | 0.069 | 0.04 | 1 | 1 | Kansl3 | 0.029 | Young |
| 0.004632 | 0.149011 | 0.49 | 0.407 | 1 | 1 | Npm1 | 0.083 | Young |
| 0.004679 | 0.780587 | 0.037 | 0.015 | 1 | 1 | Tmod1 | 0.022 | Young |
| 0.004708 | 0.184517 | 0.417 | 0.342 | 1 | 1 | Fam46a | 0.075 | Young |
| 0.004763 | 0.42665 | 0.12 | 0.081 | 1 | 1 | Elf2 | 0.039 | Young |
| 0.004764 | 1.097823 | 0.046 | 0.022 | 1 | 1 | Bicra | 0.024 | Young |
| 0.004799 | 0.809226 | 0.049 | 0.024 | 1 | 1 | 0610010F0 | 0.025 | Young |
| 0.004803 | 0.577613 | 0.14 | 0.098 | 1 | 1 | Peak1 | 0.042 | Young |
| 0.004819 | 0.197952 | 0.452 | 0.369 | 1 | 1 | Slc6a6 | 0.083 | Young |
| 0.00486 | 0.242764 | 0.25 | 0.19 | 1 | 1 | Dock10 | 0.06 | Young |
| 0.004871 | 0.561214 | 0.158 | 0.114 | 1 | 1 | Creb5 | 0.044 | Young |
| 0.004885 | 0.307893 | 0.081 | 0.048 | 1 | 1 | Rnf216 | 0.033 | Young |
| 0.004906 | 0.674246 | 0.069 | 0.04 | 1 | 1 | Aim2 | 0.029 | Young |
| 0.00498 | 0.70648 | 0.031 | 0.012 | 1 | 1 | P2ry13 | 0.019 | Young |
| 0.005086 | 0.230296 | 0.34 | 0.279 | 1 | 1 | Cflar | 0.061 | Young |
| 0.005095 | 0.109536 | 0.302 | 0.234 | 1 | 1 | Rgs2 | 0.068 | Young |
| 0.005128 | 0.25237 | 0.184 | 0.135 | 1 | 1 | Rilpl2 | 0.049 | Young |
| 0.00514 | 0.698749 | 0.042 | 0.019 | 1 | 1 | Qser1 | 0.023 | Young |
| 0.005173 | 0.654184 | 0.112 | 0.075 | 1 | 1 | Ube2e1 | 0.037 | Young |
| 0.005282 | 0.152903 | 0.077 | 0.045 | 1 | 1 | Trim33 | 0.032 | Young |
| 0.005315 | 0.449707 | 0.075 | 0.044 | 1 | 1 | Sfswap | 0.031 | Young |
| 0.005437 | 0.280139 | 0.101 | 0.064 | 1 | 1 | Hbp1 | 0.037 | Young |
| 0.005486 | 0.765351 | 0.051 | 0.026 | 1 | 1 | Sfi1 | 0.025 | Young |
| 0.005493 | 0.943066 | 0.044 | 0.021 | 1 | 1 | Brpf3 | 0.023 | Young |
| 0.00555 | 0.390222 | 0.143 | 0.102 | 1 | 1 | Hspa4 | 0.041 | Young |
| 0.005579 | 0.220702 | 0.103 | 0.067 | 1 | 1 | Osm | 0.036 | Young |
| 0.005588 | 0.6035 | 0.128 | 0.089 | 1 | 1 | Leprotl1 | 0.039 | Young |
| 0.005629 | 0.521298 | 0.089 | 0.055 | 1 | 1 | Aagab | 0.034 | Young |
| 0.00564 | 0.614741 | 0.074 | 0.044 | 1 | 1 | Wipf2 | 0.03 | Young |

| | | | | | | | | |
|---|---|---|---|---|---|---|---|---|
| 0.005675 | 0.155983 | 0.302 | 0.238 | 1 | 1 | Sod1 | 0.064 | Young |
| 0.005707 | 0.246615 | 0.258 | 0.203 | 1 | 1 | St13 | 0.055 | Young |
| 0.00576 | 1.173037 | 0.045 | 0.022 | 1 | 1 | Extl3 | 0.023 | Young |
| 0.005801 | 0.29132 | 0.051 | 0.026 | 1 | 1 | Ulk1 | 0.025 | Young |
| 0.005811 | 0.521413 | 0.069 | 0.04 | 1 | 1 | Slc4a2 | 0.029 | Young |
| 0.005815 | 1.006089 | 0.072 | 0.042 | 1 | 1 | Wdr44 | 0.03 | Young |
| 0.005841 | 0.304204 | 0.21 | 0.161 | 1 | 1 | Wasf2 | 0.049 | Young |
| 0.005897 | 0.86004 | 0.04 | 0.018 | 1 | 1 | Nit1 | 0.022 | Young |
| 0.005901 | 0.230158 | 0.231 | 0.177 | 1 | 1 | Bmp2k | 0.054 | Young |
| 0.005901 | 0.255975 | 0.209 | 0.157 | 1 | 1 | Bcl10 | 0.052 | Young |
| 0.005907 | 0.190208 | 0.575 | 0.51 | 1 | 1 | Cox4i1 | 0.065 | Young |
| 0.005913 | 0.968223 | 0.072 | 0.042 | 1 | 1 | Slc16a6 | 0.03 | Young |
| 0.005924 | 0.154843 | 0.311 | 0.247 | 1 | 1 | Nrp1 | 0.064 | Young |
| 0.005932 | 0.679085 | 0.032 | 0.013 | 1 | 1 | Hps4 | 0.019 | Young |
| 0.005952 | 0.416673 | 0.194 | 0.145 | 1 | 1 | Wac | 0.049 | Young |
| 0.005959 | 0.726391 | 0.054 | 0.028 | 1 | 1 | Apmap | 0.026 | Young |
| 0.005978 | 0.49357 | 0.126 | 0.086 | 1 | 1 | Ube2k | 0.04 | Young |
| 0.005999 | 0.552283 | 0.066 | 0.037 | 1 | 1 | AW549877 | 0.029 | Young |
| 0.006007 | 1.059143 | 0.041 | 0.019 | 1 | 1 | Nifk | 0.022 | Young |
| 0.006019 | 0.419662 | 0.138 | 0.098 | 1 | 1 | Arl5b | 0.04 | Young |
| 0.00604 | 0.61076 | 0.104 | 0.068 | 1 | 1 | Arl5c | 0.036 | Young |
| 0.00606 | 0.245739 | 0.089 | 0.055 | 1 | 1 | Ufsp2 | 0.034 | Young |
| 0.006081 | 0.863816 | 0.062 | 0.035 | 1 | 1 | 5-Mar | 0.027 | Young |
| 0.006127 | 0.271146 | 0.249 | 0.195 | 1 | 1 | Nipa2 | 0.054 | Young |
| 0.006134 | 2.446528 | 0.01 | 0 | 1 | 1 | Kcnip3 | 0.01 | Young |
| 0.006136 | 0.657344 | 0.044 | 0.021 | 1 | 1 | Nup50 | 0.023 | Young |
| 0.006143 | 2.829297 | 0.013 | 0.001 | 1 | 1 | E2f6 | 0.012 | Young |
| 0.006145 | 0.266717 | 0.101 | 0.066 | 1 | 1 | Krtcap2 | 0.035 | Young |
| 0.006158 | 0.238946 | 0.206 | 0.154 | 1 | 1 | Anxa3 | 0.052 | Young |
| 0.006161 | 0.543485 | 0.135 | 0.095 | 1 | 1 | Higd2a | 0.04 | Young |
| 0.006181 | 0.540486 | 0.095 | 0.06 | 1 | 1 | Csnk2a2 | 0.035 | Young |
| 0.0062 | 0.728809 | 0.078 | 0.048 | 1 | 1 | Csad | 0.03 | Young |
| 0.006205 | 1.113706 | 0.03 | 0.012 | 1 | 1 | Per3 | 0.018 | Young |
| 0.006221 | 2.368261 | 0.013 | 0.001 | 1 | 1 | Vamp1 | 0.012 | Young |
| 0.006225 | 0.814632 | 0.072 | 0.042 | 1 | 1 | Setd7 | 0.03 | Young |
| 0.00626 | 0.540628 | 0.117 | 0.08 | 1 | 1 | Map2k7 | 0.037 | Young |
| 0.006283 | 0.274386 | 0.178 | 0.131 | 1 | 1 | Pds5a | 0.047 | Young |
| 0.006294 | 0.971048 | 0.038 | 0.017 | 1 | 1 | Fgd3 | 0.021 | Young |
| 0.006324 | 0.563403 | 0.059 | 0.032 | 1 | 1 | Zfp90 | 0.027 | Young |
| 0.006335 | 1.53801 | 0.028 | 0.01 | 1 | 1 | Peli2 | 0.018 | Young |
| 0.006417 | 0.181797 | 0.185 | 0.138 | 1 | 1 | Gm26740 | 0.047 | Young |
| 0.00645 | 0.149345 | 0.248 | 0.19 | 1 | 1 | Irf1 | 0.058 | Young |
| 0.006457 | 0.556126 | 0.105 | 0.069 | 1 | 1 | Aup1 | 0.036 | Young |
| 0.006459 | 0.333138 | 0.145 | 0.103 | 1 | 1 | Ubn2 | 0.042 | Young |
| 0.006483 | 0.45705 | 0.096 | 0.062 | 1 | 1 | Sh2b3 | 0.034 | Young |

| | | | | | | | | |
|---|---|---|---|---|---|---|---|---|
| 0.006487 | 0.894891 | 0.058 | 0.032 | 1 | 1 | Tubgcp4 | 0.026 | Young |
| 0.006489 | 0.117167 | 0.747 | 0.658 | 1 | 1 | Atp2b1 | 0.089 | Young |
| 0.006554 | 0.098218 | 0.935 | 0.91 | 1 | 1 | Fth1 | 0.025 | Young |
| 0.006605 | 1.121663 | 0.048 | 0.024 | 1 | 1 | Cep68 | 0.024 | Young |
| 0.006711 | 1.621385 | 0.032 | 0.013 | 1 | 1 | Ttc4 | 0.019 | Young |
| 0.00675 | 0.687711 | 0.103 | 0.068 | 1 | 1 | Mark2 | 0.035 | Young |
| 0.006774 | 1.819806 | 0.026 | 0.009 | 1 | 1 | Iqce | 0.017 | Young |
| 0.006778 | 0.730567 | 0.064 | 0.036 | 1 | 1 | Tssc4 | 0.028 | Young |
| 0.006784 | 0.36106 | 0.142 | 0.1 | 1 | 1 | Kmt2a | 0.042 | Young |
| 0.006786 | 0.419936 | 0.159 | 0.116 | 1 | 1 | Trim8 | 0.043 | Young |
| 0.006797 | 0.293151 | 0.187 | 0.14 | 1 | 1 | P2ry6 | 0.047 | Young |
| 0.006799 | 0.18377 | 0.213 | 0.162 | 1 | 1 | Eif4ebp2 | 0.051 | Young |
| 0.006919 | 0.652774 | 0.064 | 0.036 | 1 | 1 | Cpne2 | 0.028 | Young |
| 0.00693 | 0.464471 | 0.103 | 0.068 | 1 | 1 | Arid4b | 0.035 | Young |
| 0.00695 | 1.069895 | 0.032 | 0.013 | 1 | 1 | Pdk1 | 0.019 | Young |
| 0.006956 | 0.458905 | 0.092 | 0.058 | 1 | 1 | Emc6 | 0.034 | Young |
| 0.006967 | 0.424729 | 0.157 | 0.114 | 1 | 1 | Crk | 0.043 | Young |
| 0.006992 | 0.23144 | 0.038 | 0.017 | 1 | 1 | Pfdn6 | 0.021 | Young |
| 0.007011 | 0.518652 | 0.11 | 0.075 | 1 | 1 | Maf1 | 0.035 | Young |
| 0.007047 | 0.304566 | 0.273 | 0.217 | 1 | 1 | Ccnd1 | 0.056 | Young |
| 0.007056 | 0.582782 | 0.028 | 0.01 | 1 | 1 | Msantd2 | 0.018 | Young |
| 0.007077 | 0.607384 | 0.081 | 0.05 | 1 | 1 | Epb41l4ao | 0.031 | Young |
| 0.007198 | 0.138179 | 0.313 | 0.249 | 1 | 1 | Cnbp | 0.064 | Young |
| 0.007249 | 2.225637 | 0.015 | 0.003 | 1 | 1 | Tmem206 | 0.012 | Young |
| 0.007285 | 0.852316 | 0.035 | 0.015 | 1 | 1 | Ch25h | 0.02 | Young |
| 0.007317 | 0.734404 | 0.078 | 0.048 | 1 | 1 | Ago4 | 0.03 | Young |
| 0.007344 | 1.086456 | 0.044 | 0.022 | 1 | 1 | 1190002N1 | 0.022 | Young |
| 0.007378 | 0.729132 | 0.106 | 0.072 | 1 | 1 | Rwdd1 | 0.034 | Young |
| 0.007441 | 1.653031 | 0.015 | 0.003 | 1 | 1 | Ildr2 | 0.012 | Young |
| 0.007445 | 0.435266 | 0.058 | 0.032 | 1 | 1 | Sin3a | 0.026 | Young |
| 0.007445 | 0.813396 | 0.055 | 0.03 | 1 | 1 | Rif1 | 0.025 | Young |
| 0.007445 | 0.614834 | 0.055 | 0.03 | 1 | 1 | Bcl2 | 0.025 | Young |
| 0.007461 | 0.965762 | 0.074 | 0.045 | 1 | 1 | Gsk3a | 0.029 | Young |
| 0.007469 | 1.04341 | 0.015 | 0.003 | 1 | 1 | Fam102a | 0.012 | Young |
| 0.007518 | 1.049685 | 0.046 | 0.023 | 1 | 1 | Mgat5 | 0.023 | Young |
| 0.007526 | 0.604437 | 0.043 | 0.021 | 1 | 1 | Samd4 | 0.022 | Young |
| 0.007567 | 0.276311 | 0.331 | 0.272 | 1 | 1 | Ptprj | 0.059 | Young |
| 0.007583 | 0.645616 | 0.099 | 0.066 | 1 | 1 | Mllt10 | 0.033 | Young |
| 0.007626 | 1.087807 | 0.024 | 0.008 | 1 | 1 | Capn15 | 0.016 | Young |
| 0.007634 | 1.232175 | 0.024 | 0.008 | 1 | 1 | Miga1 | 0.016 | Young |
| 0.007663 | 0.215561 | 0.174 | 0.127 | 1 | 1 | Aebp2 | 0.047 | Young |
| 0.007669 | 0.371029 | 0.203 | 0.156 | 1 | 1 | Fgd2 | 0.047 | Young |
| 0.007698 | 0.290491 | 0.202 | 0.153 | 1 | 1 | Msl1 | 0.049 | Young |
| 0.007734 | 1.284954 | 0.037 | 0.017 | 1 | 1 | Syt11 | 0.02 | Young |
| 0.007744 | 0.623741 | 0.039 | 0.018 | 1 | 1 | Ggps1 | 0.021 | Young |

| | | | | | | | | |
|---|---|---|---|---|---|---|---|---|
| 0.007749 | 0.057079 | 0.345 | 0.274 | 1 | 1 | Eps15 | 0.071 | Young |
| 0.007765 | 0.836786 | 0.058 | 0.032 | 1 | 1 | Wdr13 | 0.026 | Young |
| 0.007821 | 1.180756 | 0.037 | 0.017 | 1 | 1 | BC005561 | 0.02 | Young |
| 0.007823 | 0.261132 | 0.306 | 0.248 | 1 | 1 | Abca9 | 0.058 | Young |
| 0.007847 | 0.760656 | 0.061 | 0.035 | 1 | 1 | Kansl1l | 0.026 | Young |
| 0.007885 | 0.227094 | 0.14 | 0.099 | 1 | 1 | Psmb6 | 0.041 | Young |
| 0.007913 | 1.00292 | 0.052 | 0.028 | 1 | 1 | Klhl18 | 0.024 | Young |
| 0.007965 | 0.595245 | 0.068 | 0.04 | 1 | 1 | Tmem8 | 0.028 | Young |
| 0.007974 | 0.141491 | 0.353 | 0.283 | 1 | 1 | Pip4k2a | 0.07 | Young |
| 0.00798 | 0.423321 | 0.098 | 0.064 | 1 | 1 | Rala | 0.034 | Young |
| 0.00798 | 0.533717 | 0.144 | 0.105 | 1 | 1 | Bmpr2 | 0.039 | Young |
| 0.008015 | 0.327568 | 0.227 | 0.177 | 1 | 1 | Diaph1 | 0.05 | Young |
| 0.008046 | 0.150828 | 0.129 | 0.089 | 1 | 1 | Lpxn | 0.04 | Young |
| 0.008073 | 0.617546 | 0.099 | 0.066 | 1 | 1 | Gtf3c2 | 0.033 | Young |
| 0.008144 | 0.28403 | 0.134 | 0.094 | 1 | 1 | Brd1 | 0.04 | Young |
| 0.008163 | 0.55394 | 0.081 | 0.05 | 1 | 1 | Chd6 | 0.031 | Young |
| 0.008172 | 0.960297 | 0.042 | 0.021 | 1 | 1 | Slc16a3 | 0.021 | Young |
| 0.008198 | 1.589625 | 0.027 | 0.01 | 1 | 1 | Hepacam2 | 0.017 | Young |
| 0.008227 | 0.736815 | 0.072 | 0.044 | 1 | 1 | Ccng1 | 0.028 | Young |
| 0.008264 | 0.728207 | 0.056 | 0.031 | 1 | 1 | Herc6 | 0.025 | Young |
| 0.008394 | 0.177363 | 0.156 | 0.111 | 1 | 1 | Mt2 | 0.045 | Young |
| 0.00846 | 2.508298 | 0.012 | 0.001 | 1 | 1 | Nckap5l | 0.011 | Young |
| 0.008473 | 0.230587 | 0.13 | 0.09 | 1 | 1 | Dnajc19 | 0.04 | Young |
| 0.00848 | 1.227406 | 0.027 | 0.01 | 1 | 1 | Slc25a33 | 0.017 | Young |
| 0.008545 | 0.481528 | 0.157 | 0.116 | 1 | 1 | B230219D2 | 0.041 | Young |
| 0.008558 | 1.438568 | 0.031 | 0.013 | 1 | 1 | Tmem161a | 0.018 | Young |
| 0.008579 | 0.809128 | 0.054 | 0.03 | 1 | 1 | Ap5m1 | 0.024 | Young |
| 0.008601 | 0.84809 | 0.086 | 0.055 | 1 | 1 | Armcx3 | 0.031 | Young |
| 0.008606 | 1.456067 | 0.031 | 0.013 | 1 | 1 | Efl1 | 0.018 | Young |
| 0.008611 | 0.250418 | 0.159 | 0.116 | 1 | 1 | Sin3b | 0.043 | Young |
| 0.008629 | 0.11739 | 0.546 | 0.463 | 1 | 1 | Uqcrh | 0.083 | Young |
| 0.008639 | 0.756299 | 0.064 | 0.037 | 1 | 1 | Tle1 | 0.027 | Young |
| 0.00866 | 2.266043 | 0.025 | 0.009 | 1 | 1 | Zfp182 | 0.016 | Young |
| 0.008679 | 0.672162 | 0.069 | 0.041 | 1 | 1 | Kbtbd2 | 0.028 | Young |
| 0.008683 | 0.682495 | 0.097 | 0.064 | 1 | 1 | Epb41 | 0.033 | Young |
| 0.008715 | 0.229906 | 0.225 | 0.175 | 1 | 1 | Cap1 | 0.05 | Young |
| 0.008728 | 0.247234 | 0.117 | 0.08 | 1 | 1 | Clk4 | 0.037 | Young |
| 0.008762 | 0.438543 | 0.203 | 0.159 | 1 | 1 | Lipa | 0.044 | Young |
| 0.008832 | 0.880166 | 0.047 | 0.024 | 1 | 1 | Hjurp | 0.023 | Young |
| 0.008835 | 0.594253 | 0.052 | 0.028 | 1 | 1 | Mrpl15 | 0.024 | Young |
| 0.008916 | 1.166791 | 0.031 | 0.013 | 1 | 1 | Dcakd | 0.018 | Young |
| 0.008922 | 0.280185 | 0.211 | 0.163 | 1 | 1 | Ralbp1 | 0.048 | Young |
| 0.008938 | 0.173366 | 0.292 | 0.237 | 1 | 1 | Ctdsp2 | 0.055 | Young |
| 0.008993 | 0.365333 | 0.094 | 0.06 | 1 | 1 | Kcnq1ot1 | 0.034 | Young |
| 0.009017 | 0.735956 | 0.069 | 0.041 | 1 | 1 | Stx5a | 0.028 | Young |

| | | | | | | | | |
|---|---|---|---|---|---|---|---|---|
| 0.00902 | 0.292048 | 0.163 | 0.12 | 1 | 1 | Nfe2l1 | 0.043 | Young |
| 0.009048 | 0.814081 | 0.072 | 0.044 | 1 | 1 | Casp3 | 0.028 | Young |
| 0.009049 | 0.197387 | 0.167 | 0.122 | 1 | 1 | Msl2 | 0.045 | Young |
| 0.009065 | 0.343716 | 0.117 | 0.081 | 1 | 1 | Sec63 | 0.036 | Young |
| 0.009072 | 0.325249 | 0.158 | 0.117 | 1 | 1 | Stk4 | 0.041 | Young |
| 0.009197 | 0.127167 | 0.209 | 0.158 | 1 | 1 | Rab2a | 0.051 | Young |
| 0.009214 | 0.504588 | 0.086 | 0.055 | 1 | 1 | Ndufa12 | 0.031 | Young |
| 0.009222 | 1.184628 | 0.025 | 0.009 | 1 | 1 | Zfp629 | 0.016 | Young |
| 0.009249 | 0.244036 | 0.136 | 0.096 | 1 | 1 | Trmt112 | 0.04 | Young |
| 0.009279 | 0.329721 | 0.179 | 0.134 | 1 | 1 | Cab39 | 0.045 | Young |
| 0.009306 | 0.666527 | 0.067 | 0.04 | 1 | 1 | Acox3 | 0.027 | Young |
| 0.009367 | 0.284161 | 0.204 | 0.157 | 1 | 1 | Syncrip | 0.047 | Young |
| 0.009393 | 0.337783 | 0.1 | 0.066 | 1 | 1 | Tgs1 | 0.034 | Young |
| 0.009417 | 0.255929 | 0.155 | 0.113 | 1 | 1 | Glyr1 | 0.042 | Young |
| 0.009495 | 0.626111 | 0.1 | 0.067 | 1 | 1 | Adipor2 | 0.033 | Young |
| 0.009496 | 0.377448 | 0.134 | 0.095 | 1 | 1 | Igsf6 | 0.039 | Young |
| 0.009507 | 0.381547 | 0.072 | 0.044 | 1 | 1 | Gpr155 | 0.028 | Young |
| 0.009579 | 0.302803 | 0.155 | 0.113 | 1 | 1 | Vav1 | 0.042 | Young |
| 0.009661 | 0.775671 | 0.04 | 0.019 | 1 | 1 | Rcbtb1 | 0.021 | Young |
| 0.009662 | 1.531334 | 0.023 | 0.008 | 1 | 1 | E130309D( | 0.015 | Young |
| 0.009772 | 0.184951 | 0.131 | 0.091 | 1 | 1 | Smc3 | 0.04 | Young |
| 0.009775 | 0.398831 | 0.097 | 0.064 | 1 | 1 | Rbm17 | 0.033 | Young |
| 0.009797 | 0.501847 | 0.098 | 0.066 | 1 | 1 | Snx13 | 0.032 | Young |
| 0.009836 | 2.337125 | 0.014 | 0.003 | 1 | 1 | Prmt9 | 0.011 | Young |
| 0.009853 | 0.196046 | 0.106 | 0.071 | 1 | 1 | Wdfy4 | 0.035 | Young |
| 0.009964 | 0.392078 | 0.14 | 0.1 | 1 | 1 | Rbm26 | 0.04 | Young |
| 0.010065 | 0.35014 | 0.194 | 0.148 | 1 | 1 | Adam10 | 0.046 | Young |
| 0.010111 | 0.344464 | 0.084 | 0.054 | 1 | 1 | Nudc | 0.03 | Young |
| 0.010159 | 0.604965 | 0.057 | 0.032 | 1 | 1 | Slc7a7 | 0.025 | Young |
| 0.010177 | 0.435452 | 0.062 | 0.036 | 1 | 1 | Trnt1 | 0.026 | Young |
| 0.010241 | 0.913632 | 0.06 | 0.035 | 1 | 1 | Hist3h2a | 0.025 | Young |
| 0.01028 | 2.265742 | 0.018 | 0.005 | 1 | 1 | Alms1 | 0.013 | Young |
| 0.010297 | 0.141819 | 0.703 | 0.634 | 1 | 1 | Rpl8 | 0.069 | Young |
| 0.010335 | 0.401021 | 0.072 | 0.044 | 1 | 1 | Mxd1 | 0.028 | Young |
| 0.010359 | 0.325775 | 0.173 | 0.129 | 1 | 1 | Ppp1cc | 0.044 | Young |
| 0.010416 | 0.662029 | 0.058 | 0.033 | 1 | 1 | Cep250 | 0.025 | Young |
| 0.010444 | 0.689804 | 0.099 | 0.067 | 1 | 1 | Stx16 | 0.032 | Young |
| 0.010487 | 1.862991 | 0.018 | 0.005 | 1 | 1 | Rbm43 | 0.013 | Young |
| 0.010492 | 0.483285 | 0.148 | 0.109 | 1 | 1 | Stard9 | 0.039 | Young |
| 0.010526 | 0.791081 | 0.06 | 0.035 | 1 | 1 | Retreg3 | 0.025 | Young |
| 0.010529 | 1.940447 | 0.016 | 0.004 | 1 | 1 | Odf2l | 0.012 | Young |
| 0.010547 | 0.67076 | 0.068 | 0.041 | 1 | 1 | Jkamp | 0.027 | Young |
| 0.010551 | 0.408106 | 0.089 | 0.058 | 1 | 1 | Mast3 | 0.031 | Young |
| 0.010675 | 0.226736 | 0.134 | 0.095 | 1 | 1 | Prune2 | 0.039 | Young |
| 0.010685 | 1.485875 | 0.027 | 0.01 | 1 | 1 | Nemp1 | 0.017 | Young |

| | | | | | | | | |
|---|---|---|---|---|---|---|---|---|
| 0.010698 | 0.777801 | 0.151 | 0.112 | 1 | 1 | Ptgs2 | 0.039 | Young |
| 0.010793 | 1.503424 | 0.03 | 0.013 | 1 | 1 | Fem1a | 0.017 | Young |
| 0.010839 | 0.632099 | 0.076 | 0.048 | 1 | 1 | Tmem170b | 0.028 | Young |
| 0.01084 | 1.282507 | 0.027 | 0.01 | 1 | 1 | Cdk5 | 0.017 | Young |
| 0.01085 | 0.519657 | 0.05 | 0.027 | 1 | 1 | Rfx7 | 0.023 | Young |
| 0.010867 | 0.14426 | 0.392 | 0.325 | 1 | 1 | Mtss1 | 0.067 | Young |
| 0.010897 | 0.412897 | 0.14 | 0.102 | 1 | 1 | Zmynd11 | 0.038 | Young |
| 0.010898 | 0.233821 | 0.156 | 0.116 | 1 | 1 | Secisbp2l | 0.04 | Young |
| 0.010905 | 0.157862 | 0.349 | 0.287 | 1 | 1 | Purb | 0.062 | Young |
| 0.01094 | 0.455155 | 0.087 | 0.057 | 1 | 1 | Smap1 | 0.03 | Young |
| 0.01098 | 1.368323 | 0.027 | 0.01 | 1 | 1 | Kat5 | 0.017 | Young |
| 0.011032 | 0.548237 | 0.068 | 0.041 | 1 | 1 | Tmem251 | 0.027 | Young |
| 0.01106 | 0.326608 | 0.199 | 0.152 | 1 | 1 | Tep1 | 0.047 | Young |
| 0.011185 | 0.544005 | 0.12 | 0.085 | 1 | 1 | Rhoc | 0.035 | Young |
| 0.011188 | 0.94467 | 0.048 | 0.026 | 1 | 1 | Mnat1 | 0.022 | Young |
| 0.011199 | 0.860043 | 0.043 | 0.022 | 1 | 1 | Tpbgl | 0.021 | Young |
| 0.011299 | 0.533669 | 0.032 | 0.014 | 1 | 1 | Dennd2c | 0.018 | Young |
| 0.011413 | 0.324625 | 0.041 | 0.021 | 1 | 1 | Tsga10 | 0.02 | Young |
| 0.011531 | 0.245136 | 0.171 | 0.127 | 1 | 1 | 7-Mar | 0.044 | Young |
| 0.011566 | 0.155523 | 0.154 | 0.112 | 1 | 1 | Mon2 | 0.042 | Young |
| 0.011568 | 2.85099 | 0.011 | 0.001 | 1 | 1 | Zfp599 | 0.01 | Young |
| 0.011578 | 0.60195 | 0.043 | 0.022 | 1 | 1 | Unc119 | 0.021 | Young |
| 0.011584 | 2.605486 | 0.011 | 0.001 | 1 | 1 | 4632415L0 | 0.01 | Young |
| 0.011634 | 0.353221 | 0.119 | 0.084 | 1 | 1 | Ago3 | 0.035 | Young |
| 0.011664 | 0.494563 | 0.089 | 0.058 | 1 | 1 | Specc1 | 0.031 | Young |
| 0.011704 | 0.243905 | 0.192 | 0.145 | 1 | 1 | Pura | 0.047 | Young |
| 0.011705 | 0.89202 | 0.049 | 0.027 | 1 | 1 | Ero1l | 0.022 | Young |
| 0.011714 | 0.997591 | 0.044 | 0.023 | 1 | 1 | Cdr2 | 0.021 | Young |
| 0.011767 | 0.160863 | 0.194 | 0.147 | 1 | 1 | Apbb1ip | 0.047 | Young |
| 0.011773 | 0.601561 | 0.038 | 0.018 | 1 | 1 | Dusp7 | 0.02 | Young |
| 0.011787 | 0.849776 | 0.067 | 0.041 | 1 | 1 | Tmem128 | 0.026 | Young |
| 0.011797 | 0.478258 | 0.122 | 0.086 | 1 | 1 | Tspyl1 | 0.036 | Young |
| 0.011809 | 0.726757 | 0.064 | 0.039 | 1 | 1 | AY036118 | 0.025 | Young |
| 0.011812 | 1.152668 | 0.011 | 0.001 | 1 | 1 | Wdr73 | 0.01 | Young |
| 0.011812 | 1.124989 | 0.011 | 0.001 | 1 | 1 | Natd1 | 0.01 | Young |
| 0.011889 | 0.705126 | 0.072 | 0.045 | 1 | 1 | Ipmk | 0.027 | Young |
| 0.011997 | 0.211079 | 0.068 | 0.041 | 1 | 1 | Tlk1 | 0.027 | Young |
| 0.012115 | 0.330834 | 0.15 | 0.111 | 1 | 1 | Iqgap2 | 0.039 | Young |
| 0.012129 | 0.232668 | 0.078 | 0.049 | 1 | 1 | Ramp1 | 0.029 | Young |
| 0.01215 | 0.467098 | 0.081 | 0.051 | 1 | 1 | Irs2 | 0.03 | Young |
| 0.012188 | 0.37573 | 0.243 | 0.195 | 1 | 1 | Ndel1 | 0.048 | Young |
| 0.012214 | 0.369891 | 0.061 | 0.036 | 1 | 1 | Mxi1 | 0.025 | Young |
| 0.012289 | 0.299018 | 0.12 | 0.085 | 1 | 1 | Smc1a | 0.035 | Young |
| 0.0123 | 0.43798 | 0.123 | 0.087 | 1 | 1 | Stk10 | 0.036 | Young |
| 0.012317 | 2.054156 | 0.022 | 0.008 | 1 | 1 | Upb1 | 0.014 | Young |

| | | | | | | | | |
|---|---|---|---|---|---|---|---|---|
| 0.012318 | 0.37385 | 0.123 | 0.086 | 1 | 1 | Galc | 0.037 | Young |
| 0.012432 | 0.719921 | 0.069 | 0.042 | 1 | 1 | Lrif1 | 0.027 | Young |
| 0.012441 | 0.379845 | 0.136 | 0.099 | 1 | 1 | Bcl2l11 | 0.037 | Young |
| 0.012455 | 1.275478 | 0.032 | 0.014 | 1 | 1 | 4930453N2 | 0.018 | Young |
| 0.012463 | 0.339563 | 0.086 | 0.057 | 1 | 1 | Eml4 | 0.029 | Young |
| 0.012555 | 0.414256 | 0.123 | 0.087 | 1 | 1 | Bank1 | 0.036 | Young |
| 0.012578 | 1.144155 | 0.032 | 0.014 | 1 | 1 | Acvr1 | 0.018 | Young |
| 0.01262 | 0.911113 | 0.035 | 0.017 | 1 | 1 | Scd1 | 0.018 | Young |
| 0.012646 | 0.244675 | 0.217 | 0.17 | 1 | 1 | Txn1 | 0.047 | Young |
| 0.012698 | 0.122483 | 0.422 | 0.352 | 1 | 1 | Cox7c | 0.07 | Young |
| 0.012766 | 0.795888 | 0.039 | 0.019 | 1 | 1 | Mvb12b | 0.02 | Young |
| 0.012858 | 0.440507 | 0.074 | 0.046 | 1 | 1 | Btaf1 | 0.028 | Young |
| 0.012915 | 0.92514 | 0.032 | 0.014 | 1 | 1 | Pcyt2 | 0.018 | Young |
| 0.012964 | 0.186211 | 0.352 | 0.29 | 1 | 1 | Cbl | 0.062 | Young |
| 0.01304 | 0.563058 | 0.039 | 0.019 | 1 | 1 | Nf2 | 0.02 | Young |
| 0.013046 | 0.563203 | 0.131 | 0.095 | 1 | 1 | Appl1 | 0.036 | Young |
| 0.013169 | 0.522196 | 0.094 | 0.063 | 1 | 1 | Htt | 0.031 | Young |
| 0.013174 | 0.156902 | 0.278 | 0.222 | 1 | 1 | Pum2 | 0.056 | Young |
| 0.013174 | 1.263535 | 0.037 | 0.018 | 1 | 1 | Chkb | 0.019 | Young |
| 0.013188 | 0.213843 | 0.319 | 0.261 | 1 | 1 | Kdm7a | 0.058 | Young |
| 0.013221 | 0.163373 | 0.233 | 0.181 | 1 | 1 | Prrc2a | 0.052 | Young |
| 0.013225 | 0.438812 | 0.175 | 0.134 | 1 | 1 | Psmb8 | 0.041 | Young |
| 0.013276 | 0.585647 | 0.107 | 0.075 | 1 | 1 | Ext1 | 0.032 | Young |
| 0.013404 | 0.179234 | 0.298 | 0.243 | 1 | 1 | Hcls1 | 0.055 | Young |
| 0.013411 | 2.123858 | 0.013 | 0.003 | 1 | 1 | Tmco4 | 0.01 | Young |
| 0.013502 | 0.870972 | 0.047 | 0.026 | 1 | 1 | Med17 | 0.021 | Young |
| 0.013515 | 0.189129 | 0.38 | 0.319 | 1 | 1 | Sfpq | 0.061 | Young |
| 0.01354 | 0.332771 | 0.117 | 0.082 | 1 | 1 | Btk | 0.035 | Young |
| 0.013553 | 0.194928 | 0.255 | 0.202 | 1 | 1 | Fam174a | 0.053 | Young |
| 0.013599 | 0.223347 | 0.345 | 0.288 | 1 | 1 | Eif3f | 0.057 | Young |
| 0.013631 | 0.103746 | 0.176 | 0.132 | 1 | 1 | Sesn1 | 0.044 | Young |
| 0.013644 | 1.073051 | 0.026 | 0.01 | 1 | 1 | Tmem94 | 0.016 | Young |
| 0.013666 | 0.081167 | 0.261 | 0.204 | 1 | 1 | Tor1aip1 | 0.057 | Young |
| 0.013724 | 0.921371 | 0.042 | 0.022 | 1 | 1 | Kat7 | 0.02 | Young |
| 0.013743 | 2.362507 | 0.016 | 0.004 | 1 | 1 | Neurl1b | 0.012 | Young |
| 0.013759 | 1.214691 | 0.026 | 0.01 | 1 | 1 | Carf | 0.016 | Young |
| 0.013776 | 1.271005 | 0.026 | 0.01 | 1 | 1 | Lmf1 | 0.016 | Young |
| 0.013883 | 0.57245 | 0.052 | 0.03 | 1 | 1 | Dgkh | 0.022 | Young |
| 0.013948 | 0.354185 | 0.135 | 0.098 | 1 | 1 | Cat | 0.037 | Young |
| 0.013961 | 1.131735 | 0.033 | 0.015 | 1 | 1 | Cstf2 | 0.018 | Young |
| 0.014014 | 1.168204 | 0.026 | 0.01 | 1 | 1 | Rbm12b1 | 0.016 | Young |
| 0.014087 | 1.27685 | 0.03 | 0.013 | 1 | 1 | Stambp | 0.017 | Young |
| 0.014098 | 0.794727 | 0.065 | 0.04 | 1 | 1 | Dhx30 | 0.025 | Young |
| 0.014099 | 0.641947 | 0.099 | 0.068 | 1 | 1 | Ss18 | 0.031 | Young |
| 0.014104 | 0.630265 | 0.072 | 0.045 | 1 | 1 | Khnyn | 0.027 | Young |

| | | | | | | | | |
|---|---|---|---|---|---|---|---|---|
| 0.014185 | 0.169567 | 0.351 | 0.287 | 1 | 1 | Trps1 | 0.064 | Young |
| 0.014188 | 1.127429 | 0.018 | 0.005 | 1 | 1 | Msto1 | 0.013 | Young |
| 0.014235 | 0.846514 | 0.047 | 0.026 | 1 | 1 | AI606181 | 0.021 | Young |
| 0.01425 | 0.063208 | 0.48 | 0.398 | 1 | 1 | Ehd4 | 0.082 | Young |
| 0.014252 | 0.71468 | 0.037 | 0.018 | 1 | 1 | Anapc4 | 0.019 | Young |
| 0.014263 | 0.115692 | 0.11 | 0.076 | 1 | 1 | Acsl5 | 0.034 | Young |
| 0.014281 | 0.304151 | 0.09 | 0.059 | 1 | 1 | Anapc13 | 0.031 | Young |
| 0.014286 | 0.307701 | 0.051 | 0.028 | 1 | 1 | Ern1 | 0.023 | Young |
| 0.014292 | 0.32522 | 0.1 | 0.068 | 1 | 1 | Rbm7 | 0.032 | Young |
| 0.01439 | 0.253638 | 0.234 | 0.188 | 1 | 1 | Eif3k | 0.046 | Young |
| 0.014457 | 0.204577 | 0.154 | 0.112 | 1 | 1 | Uhrf2 | 0.042 | Young |
| 0.014496 | 0.473772 | 0.077 | 0.049 | 1 | 1 | Brd8 | 0.028 | Young |
| 0.014628 | 0.66179 | 0.037 | 0.018 | 1 | 1 | Polk | 0.019 | Young |
| 0.01464 | 0.584333 | 0.054 | 0.031 | 1 | 1 | Eaf1 | 0.023 | Young |
| 0.014672 | 0.231949 | 0.399 | 0.342 | 1 | 1 | Cdkn1a | 0.057 | Young |
| 0.014676 | 0.499803 | 0.105 | 0.073 | 1 | 1 | Gadd45g | 0.032 | Young |
| 0.014692 | 0.095439 | 0.798 | 0.716 | 1 | 1 | Mcl1 | 0.082 | Young |
| 0.014695 | 0.42524 | 0.055 | 0.032 | 1 | 1 | Ltbr | 0.023 | Young |
| 0.014741 | 0.626033 | 0.061 | 0.036 | 1 | 1 | Fhod1 | 0.025 | Young |
| 0.014818 | 0.30791 | 0.133 | 0.096 | 1 | 1 | Setd3 | 0.037 | Young |
| 0.01489 | 0.384173 | 0.137 | 0.1 | 1 | 1 | Ppp3cb | 0.037 | Young |
| 0.014894 | 0.257341 | 0.097 | 0.066 | 1 | 1 | Gpr160 | 0.031 | Young |
| 0.015058 | 1.343911 | 0.024 | 0.009 | 1 | 1 | Mtx3 | 0.015 | Young |
| 0.015068 | 0.148263 | 0.357 | 0.299 | 1 | 1 | Dhrs3 | 0.058 | Young |
| 0.015072 | 0.532961 | 0.054 | 0.031 | 1 | 1 | Pkd2 | 0.023 | Young |
| 0.015097 | 0.978383 | 0.05 | 0.028 | 1 | 1 | Tshz1 | 0.022 | Young |
| 0.01517 | 0.269049 | 0.181 | 0.139 | 1 | 1 | Acin1 | 0.042 | Young |
| 0.015179 | 0.454542 | 0.083 | 0.054 | 1 | 1 | Dtx3l | 0.029 | Young |
| 0.015205 | 0.36151 | 0.109 | 0.076 | 1 | 1 | Crem | 0.033 | Young |
| 0.015239 | 0.160117 | 0.198 | 0.152 | 1 | 1 | Smim14 | 0.046 | Young |
| 0.015241 | 0.13705 | 0.487 | 0.409 | 1 | 1 | H2-DMa | 0.078 | Young |
| 0.015286 | 0.098429 | 0.255 | 0.203 | 1 | 1 | Eif3e | 0.052 | Young |
| 0.015311 | 0.393263 | 0.129 | 0.094 | 1 | 1 | Pik3cg | 0.035 | Young |
| 0.015325 | 0.104327 | 0.089 | 0.058 | 1 | 1 | Ktn1 | 0.031 | Young |
| 0.01536 | 0.301011 | 0.067 | 0.041 | 1 | 1 | Stam2 | 0.026 | Young |
| 0.015612 | 0.480085 | 0.099 | 0.068 | 1 | 1 | Sun2 | 0.031 | Young |
| 0.015625 | 1.32094 | 0.041 | 0.022 | 1 | 1 | Gtf3c1 | 0.019 | Young |
| 0.015677 | 0.545226 | 0.167 | 0.13 | 1 | 1 | Arhgef2 | 0.037 | Young |
| 0.015798 | 1.214784 | 0.027 | 0.012 | 1 | 1 | Lmo1 | 0.015 | Young |
| 0.015841 | 0.550546 | 0.078 | 0.05 | 1 | 1 | Zfp280d | 0.028 | Young |
| 0.015852 | 0.236097 | 0.227 | 0.179 | 1 | 1 | Camk1d | 0.048 | Young |
| 0.015853 | 0.608059 | 0.055 | 0.032 | 1 | 1 | Wsb2 | 0.023 | Young |
| 0.015876 | 0.353572 | 0.069 | 0.042 | 1 | 1 | Tlr1 | 0.027 | Young |
| 0.015886 | 0.108566 | 0.133 | 0.095 | 1 | 1 | Errfi1 | 0.038 | Young |
| 0.015947 | 2.246819 | 0.01 | 0.001 | 1 | 1 | Gm8369 | 0.009 | Young |

| | | | | | | | | |
|---|---|---|---|---|---|---|---|---|
| 0.015954 | 0.4831 | 0.09 | 0.06 | 1 | 1 | Hck | 0.03 | Young |
| 0.01603 | 0.729188 | 0.061 | 0.037 | 1 | 1 | Tlr13 | 0.024 | Young |
| 0.016035 | 1.165985 | 0.058 | 0.035 | 1 | 1 | Ncor2 | 0.023 | Young |
| 0.016061 | 0.732687 | 0.035 | 0.017 | 1 | 1 | Spaca6 | 0.018 | Young |
| 0.016073 | 0.136038 | 0.261 | 0.21 | 1 | 1 | Arhgef6 | 0.051 | Young |
| 0.016119 | 0.801086 | 0.04 | 0.021 | 1 | 1 | Pik3c3 | 0.019 | Young |
| 0.016146 | 1.747231 | 0.01 | 0.001 | 1 | 1 | Gdpgp1 | 0.009 | Young |
| 0.016191 | 1.853855 | 0.01 | 0.001 | 1 | 1 | Hes7 | 0.009 | Young |
| 0.016233 | 0.197666 | 0.168 | 0.126 | 1 | 1 | Twf2 | 0.042 | Young |
| 0.016251 | 0.151974 | 0.196 | 0.152 | 1 | 1 | Snrpf | 0.044 | Young |
| 0.01629 | 1.663983 | 0.021 | 0.008 | 1 | 1 | Cnot11 | 0.013 | Young |
| 0.016359 | 0.039133 | 0.083 | 0.054 | 1 | 1 | Sgk1 | 0.029 | Young |
| 0.01644 | 0.741151 | 0.068 | 0.042 | 1 | 1 | Pigm | 0.026 | Young |
| 0.016492 | 0.545927 | 0.053 | 0.031 | 1 | 1 | Pias2 | 0.022 | Young |
| 0.016517 | 0.142721 | 0.523 | 0.452 | 1 | 1 | Tnfaip3 | 0.071 | Young |
| 0.016618 | 1.301667 | 0.021 | 0.008 | 1 | 1 | Stambpl1 | 0.013 | Young |
| 0.016627 | 0.28088 | 0.123 | 0.089 | 1 | 1 | P2rx7 | 0.034 | Young |
| 0.01663 | 1.048952 | 0.059 | 0.036 | 1 | 1 | Cmc1 | 0.023 | Young |
| 0.016641 | 0.492044 | 0.094 | 0.064 | 1 | 1 | Eri1 | 0.03 | Young |
| 0.016665 | 0.136473 | 0.346 | 0.287 | 1 | 1 | Ddx6 | 0.059 | Young |
| 0.01671 | 0.63543 | 0.031 | 0.014 | 1 | 1 | Donson | 0.017 | Young |
| 0.016745 | 0.292173 | 0.188 | 0.147 | 1 | 1 | Fgfr1op2 | 0.041 | Young |
| 0.016752 | 0.363488 | 0.068 | 0.042 | 1 | 1 | Ammecr1l | 0.026 | Young |
| 0.016825 | 0.577248 | 0.071 | 0.045 | 1 | 1 | Arhgap39 | 0.026 | Young |
| 0.016831 | 0.460569 | 0.1 | 0.069 | 1 | 1 | R3hdm1 | 0.031 | Young |
| 0.016869 | 0.247782 | 0.055 | 0.032 | 1 | 1 | Mb21d1 | 0.023 | Young |
| 0.01688 | 0.752218 | 0.096 | 0.067 | 1 | 1 | Zrsr2 | 0.029 | Young |
| 0.016938 | 0.482885 | 0.103 | 0.072 | 1 | 1 | Cnppd1 | 0.031 | Young |
| 0.01706 | 0.504306 | 0.071 | 0.045 | 1 | 1 | Impdh2 | 0.026 | Young |
| 0.017094 | 0.108828 | 0.19 | 0.145 | 1 | 1 | Mier1 | 0.045 | Young |
| 0.017153 | 0.553933 | 0.135 | 0.1 | 1 | 1 | Ccnd2 | 0.035 | Young |
| 0.017201 | 0.651688 | 0.053 | 0.031 | 1 | 1 | Hint2 | 0.022 | Young |
| 0.01723 | 1.982156 | 0.019 | 0.006 | 1 | 1 | Zfp113 | 0.013 | Young |
| 0.017242 | 0.98034 | 0.049 | 0.028 | 1 | 1 | Gm45716 | 0.021 | Young |
| 0.017252 | 1.729574 | 0.039 | 0.021 | 1 | 1 | Ino80 | 0.018 | Young |
| 0.017261 | 0.289959 | 0.218 | 0.176 | 1 | 1 | Smdt1 | 0.042 | Young |
| 0.017356 | 0.438815 | 0.164 | 0.127 | 1 | 1 | Slamf7 | 0.037 | Young |
| 0.017499 | 0.192781 | 0.211 | 0.165 | 1 | 1 | Atp5j | 0.046 | Young |
| 0.017528 | 0.05105 | 0.272 | 0.219 | 1 | 1 | Ccl12 | 0.053 | Young |
| 0.017698 | 0.5941 | 0.113 | 0.081 | 1 | 1 | Clasp2 | 0.032 | Young |
| 0.017756 | 0.370677 | 0.131 | 0.096 | 1 | 1 | Mcur1 | 0.035 | Young |
| 0.017764 | 0.583653 | 0.036 | 0.018 | 1 | 1 | Cdc23 | 0.018 | Young |
| 0.017792 | 0.560125 | 0.078 | 0.051 | 1 | 1 | Psmg4 | 0.027 | Young |
| 0.017985 | 1.144354 | 0.029 | 0.013 | 1 | 1 | Ttbk2 | 0.016 | Young |
| 0.018034 | 1.258806 | 0.054 | 0.032 | 1 | 1 | Bcl2a1a | 0.022 | Young |

| | | | | | | | | |
|---|---|---|---|---|---|---|---|---|
| 0.018043 | 0.392624 | 0.123 | 0.09 | 1 | 1 | Chd2 | 0.033 | Young |
| 0.018057 | 2.430162 | 0.017 | 0.005 | 1 | 1 | Fancc | 0.012 | Young |
| 0.018067 | 0.519539 | 0.094 | 0.064 | 1 | 1 | Nhlrc3 | 0.03 | Young |
| 0.018075 | 0.453331 | 0.071 | 0.045 | 1 | 1 | Zzz3 | 0.026 | Young |
| 0.018094 | 0.407087 | 0.114 | 0.082 | 1 | 1 | Apc | 0.032 | Young |
| 0.018095 | 0.241448 | 0.168 | 0.127 | 1 | 1 | Emb | 0.041 | Young |
| 0.0181 | 0.245923 | 0.141 | 0.104 | 1 | 1 | Zfp710 | 0.037 | Young |
| 0.018104 | 0.263014 | 0.066 | 0.041 | 1 | 1 | Ppp1r11 | 0.025 | Young |
| 0.018119 | 0.49245 | 0.092 | 0.063 | 1 | 1 | Cc2d1b | 0.029 | Young |
| 0.018133 | 0.206043 | 0.195 | 0.15 | 1 | 1 | Snap23 | 0.045 | Young |
| 0.018183 | 0.334223 | 0.069 | 0.044 | 1 | 1 | Itpripl1 | 0.025 | Young |
| 0.018227 | 0.105842 | 0.468 | 0.392 | 1 | 1 | Clta | 0.076 | Young |
| 0.018305 | 0.791851 | 0.059 | 0.036 | 1 | 1 | Arhgap1 | 0.023 | Young |
| 0.018305 | 0.087691 | 0.935 | 0.891 | 1 | 1 | Rplp2 | 0.044 | Young |
| 0.018348 | 0.828143 | 0.013 | 0.003 | 1 | 1 | Brat1 | 0.01 | Young |
| 0.018415 | 1.293662 | 0.013 | 0.003 | 1 | 1 | Itpr3 | 0.01 | Young |
| 0.01842 | 0.352887 | 0.115 | 0.082 | 1 | 1 | Gak | 0.033 | Young |
| 0.018459 | 1.411469 | 0.013 | 0.003 | 1 | 1 | Cep57l1 | 0.01 | Young |
| 0.018494 | 0.459837 | 0.126 | 0.093 | 1 | 1 | Zfp644 | 0.033 | Young |
| 0.018502 | 0.304284 | 0.158 | 0.12 | 1 | 1 | Psma3 | 0.038 | Young |
| 0.01851 | 0.263417 | 0.15 | 0.112 | 1 | 1 | Cuta | 0.038 | Young |
| 0.018584 | 0.244673 | 0.14 | 0.104 | 1 | 1 | Cpt1a | 0.036 | Young |
| 0.018639 | 0.072835 | 0.485 | 0.404 | 1 | 1 | Cd81 | 0.081 | Young |
| 0.018659 | 0.804757 | 0.083 | 0.057 | 1 | 1 | Gpr107 | 0.026 | Young |
| 0.018666 | 0.489402 | 0.067 | 0.042 | 1 | 1 | Kpna3 | 0.025 | Young |
| 0.018768 | 1.221133 | 0.017 | 0.005 | 1 | 1 | Zfp429 | 0.012 | Young |
| 0.018801 | 1.666366 | 0.015 | 0.004 | 1 | 1 | Cep152 | 0.011 | Young |
| 0.018842 | 0.135281 | 0.132 | 0.096 | 1 | 1 | Pcnp | 0.036 | Young |
| 0.018904 | 1.623283 | 0.015 | 0.004 | 1 | 1 | Col3a1 | 0.011 | Young |
| 0.018917 | 1.006985 | 0.034 | 0.017 | 1 | 1 | Snapc3 | 0.017 | Young |
| 0.01892 | 0.095524 | 0.586 | 0.503 | 1 | 1 | Rbpj | 0.083 | Young |
| 0.018945 | 0.445327 | 0.066 | 0.041 | 1 | 1 | Hivep3 | 0.025 | Young |
| 0.018945 | 1.478871 | 0.015 | 0.004 | 1 | 1 | Nradd | 0.011 | Young |
| 0.018952 | 0.585535 | 0.139 | 0.105 | 1 | 1 | Nfat5 | 0.034 | Young |
| 0.018963 | 0.404401 | 0.091 | 0.062 | 1 | 1 | Cirbp | 0.029 | Young |
| 0.018989 | 0.674401 | 0.058 | 0.035 | 1 | 1 | Kcnk13 | 0.023 | Young |
| 0.019061 | 0.579252 | 0.09 | 0.062 | 1 | 1 | Zfp704 | 0.028 | Young |
| 0.0191 | 1.195011 | 0.034 | 0.017 | 1 | 1 | Ppwd1 | 0.017 | Young |
| 0.019101 | 0.167214 | 0.284 | 0.231 | 1 | 1 | Stat3 | 0.053 | Young |
| 0.019172 | 0.624409 | 0.017 | 0.005 | 1 | 1 | Tsen15 | 0.012 | Young |
| 0.019215 | 0.500682 | 0.067 | 0.042 | 1 | 1 | Mrps18c | 0.025 | Young |
| 0.019255 | 0.889799 | 0.052 | 0.031 | 1 | 1 | Atxn7l3 | 0.021 | Young |
| 0.019263 | 0.109812 | 0.871 | 0.798 | 1 | 1 | Serinc3 | 0.073 | Young |
| 0.019312 | 0.198627 | 0.186 | 0.143 | 1 | 1 | Blvrb | 0.043 | Young |
| 0.019384 | 0.412323 | 0.137 | 0.103 | 1 | 1 | Siglece | 0.034 | Young |

| | | | | | | | | |
|---|---|---|---|---|---|---|---|---|
| 0.019418 | 0.397106 | 0.09 | 0.062 | 1 | 1 | Klhdc3 | 0.028 | Young |
| 0.019427 | 1.235507 | 0.046 | 0.026 | 1 | 1 | Trappc6a | 0.02 | Young |
| 0.019428 | 0.918338 | 0.015 | 0.004 | 1 | 1 | Tal1 | 0.011 | Young |
| 0.019444 | 0.263953 | 0.231 | 0.186 | 1 | 1 | Dennd5a | 0.045 | Young |
| 0.019461 | 0.375011 | 0.101 | 0.071 | 1 | 1 | Nfatc1 | 0.03 | Young |
| 0.019587 | 0.117821 | 0.16 | 0.12 | 1 | 1 | Nfxl1 | 0.04 | Young |
| 0.019687 | 0.448018 | 0.081 | 0.054 | 1 | 1 | Kidins220 | 0.027 | Young |
| 0.019789 | 0.146245 | 0.203 | 0.158 | 1 | 1 | Slc12a6 | 0.045 | Young |
| 0.019891 | 0.820887 | 0.034 | 0.017 | 1 | 1 | Ppp1r9a | 0.017 | Young |
| 0.019959 | 0.688312 | 0.078 | 0.051 | 1 | 1 | Filip1l | 0.027 | Young |
| 0.019988 | 0.234863 | 0.188 | 0.145 | 1 | 1 | Luc7l3 | 0.043 | Young |
| 0.020007 | 0.644856 | 0.069 | 0.044 | 1 | 1 | Atf6b | 0.025 | Young |
| 0.02007 | 0.170431 | 0.239 | 0.19 | 1 | 1 | Scpep1 | 0.049 | Young |
| 0.020099 | 0.671713 | 0.062 | 0.039 | 1 | 1 | Cmip | 0.023 | Young |
| 0.020108 | 0.062409 | 0.628 | 0.542 | 1 | 1 | Rtn4 | 0.086 | Young |
| 0.020141 | 0.046943 | 0.19 | 0.147 | 1 | 1 | Ppp1cb | 0.043 | Young |
| 0.020203 | 0.484658 | 0.058 | 0.035 | 1 | 1 | Tceal8 | 0.023 | Young |
| 0.020222 | 0.479648 | 0.085 | 0.058 | 1 | 1 | Bcl2a1d | 0.027 | Young |
| 0.020256 | 0.356031 | 0.098 | 0.068 | 1 | 1 | Gng12 | 0.03 | Young |
| 0.020264 | 0.827382 | 0.035 | 0.018 | 1 | 1 | Adamts10 | 0.017 | Young |
| 0.020354 | 0.795554 | 0.03 | 0.014 | 1 | 1 | Plscr1 | 0.016 | Young |
| 0.020359 | 0.39098 | 0.123 | 0.09 | 1 | 1 | Rab5a | 0.033 | Young |
| 0.02038 | 0.317058 | 0.066 | 0.041 | 1 | 1 | Utrn | 0.025 | Young |
| 0.020383 | 0.775173 | 0.03 | 0.014 | 1 | 1 | Rffl | 0.016 | Young |
| 0.02042 | 0.1135 | 0.476 | 0.409 | 1 | 1 | Fus | 0.067 | Young |
| 0.02042 | 0.231673 | 0.266 | 0.217 | 1 | 1 | Ppp1r15a | 0.049 | Young |
| 0.020548 | 0.462491 | 0.102 | 0.072 | 1 | 1 | Mnt | 0.03 | Young |
| 0.020591 | 0.180043 | 0.074 | 0.048 | 1 | 1 | Lats1 | 0.026 | Young |
| 0.020644 | 0.447361 | 0.07 | 0.045 | 1 | 1 | Dnajb4 | 0.025 | Young |
| 0.020653 | 0.49024 | 0.087 | 0.059 | 1 | 1 | Nuak1 | 0.028 | Young |
| 0.020809 | 0.302199 | 0.08 | 0.053 | 1 | 1 | Dpf2 | 0.027 | Young |
| 0.020871 | 0.62858 | 0.064 | 0.04 | 1 | 1 | Slc35a5 | 0.024 | Young |
| 0.020896 | 0.582885 | 0.094 | 0.066 | 1 | 1 | Emsy | 0.028 | Young |
| 0.021172 | 0.583682 | 0.076 | 0.05 | 1 | 1 | Pfkfb4 | 0.026 | Young |
| 0.021237 | 0.095812 | 0.497 | 0.419 | 1 | 1 | Alox5ap | 0.078 | Young |
| 0.021371 | 0.685811 | 0.046 | 0.026 | 1 | 1 | Trim34a | 0.02 | Young |
| 0.021388 | 0.283999 | 0.097 | 0.067 | 1 | 1 | Inpp5k | 0.03 | Young |
| 0.021415 | 1.086246 | 0.042 | 0.023 | 1 | 1 | Axin1 | 0.019 | Young |
| 0.021422 | 0.154843 | 0.179 | 0.139 | 1 | 1 | Bptf | 0.04 | Young |
| 0.02143 | 0.28502 | 0.174 | 0.135 | 1 | 1 | Nol7 | 0.039 | Young |
| 0.021444 | 0.917016 | 0.049 | 0.028 | 1 | 1 | Vrk3 | 0.021 | Young |
| 0.021445 | 0.573183 | 0.054 | 0.032 | 1 | 1 | Rnase6 | 0.022 | Young |
| 0.021449 | 0.449722 | 0.078 | 0.051 | 1 | 1 | Wbp1l | 0.027 | Young |
| 0.021501 | 0.834715 | 0.041 | 0.022 | 1 | 1 | Kdm5d | 0.019 | Young |
| 0.021556 | 1.227541 | 0.021 | 0.008 | 1 | 1 | Utp4 | 0.013 | Young |

| | | | | | | | | |
|---|---|---|---|---|---|---|---|---|
| 0.02156 | 0.375348 | 0.099 | 0.069 | 1 | 1 | Hsd17b4 | 0.03 | Young |
| 0.02156 | 0.412884 | 0.157 | 0.121 | 1 | 1 | Ncoa3 | 0.036 | Young |
| 0.021661 | 0.039709 | 0.497 | 0.414 | 1 | 1 | Picalm | 0.083 | Young |
| 0.021801 | 1.064216 | 0.044 | 0.024 | 1 | 1 | Tes | 0.02 | Young |
| 0.02186 | 0.442562 | 0.078 | 0.051 | 1 | 1 | Psmd3 | 0.027 | Young |
| 0.021886 | 0.102948 | 0.541 | 0.47 | 1 | 1 | Ccr2 | 0.071 | Young |
| 0.021926 | 0.824655 | 0.071 | 0.046 | 1 | 1 | Ift20 | 0.025 | Young |
| 0.021983 | 2.351075 | 0.01 | 0.001 | 1 | 1 | Zfp691 | 0.009 | Young |
| 0.022013 | 2.325284 | 0.01 | 0.001 | 1 | 1 | AU041133 | 0.009 | Young |
| 0.022044 | 0.654218 | 0.075 | 0.05 | 1 | 1 | Pdcd2l | 0.025 | Young |
| 0.022058 | 0.11333 | 0.459 | 0.387 | 1 | 1 | Calm2 | 0.072 | Young |
| 0.022074 | 2.261315 | 0.01 | 0.001 | 1 | 1 | Hdac8 | 0.009 | Young |
| 0.022074 | 2.141522 | 0.01 | 0.001 | 1 | 1 | Pigb | 0.009 | Young |
| 0.022104 | 1.99684 | 0.01 | 0.001 | 1 | 1 | Ydjc | 0.009 | Young |
| 0.022105 | 0.374976 | 0.151 | 0.117 | 1 | 1 | Hsbp1 | 0.034 | Young |
| 0.022145 | 0.286683 | 0.197 | 0.156 | 1 | 1 | Lmo2 | 0.041 | Young |
| 0.022165 | 2.014114 | 0.01 | 0.001 | 1 | 1 | Stat4 | 0.009 | Young |
| 0.022165 | 1.95052 | 0.01 | 0.001 | 1 | 1 | Gm38394 | 0.009 | Young |
| 0.022195 | 2.071477 | 0.01 | 0.001 | 1 | 1 | Gm26624 | 0.009 | Young |
| 0.022216 | 0.526493 | 0.021 | 0.008 | 1 | 1 | Spata13 | 0.013 | Young |
| 0.022218 | 0.928566 | 0.045 | 0.026 | 1 | 1 | Opa3 | 0.019 | Young |
| 0.022226 | 1.867102 | 0.01 | 0.001 | 1 | 1 | Abhd8 | 0.009 | Young |
| 0.022266 | 0.073472 | 0.399 | 0.33 | 1 | 1 | Pld4 | 0.069 | Young |
| 0.022287 | 1.410194 | 0.024 | 0.01 | 1 | 1 | Anln | 0.014 | Young |
| 0.022322 | 0.100488 | 0.134 | 0.098 | 1 | 1 | Mif | 0.036 | Young |
| 0.022326 | 0.443537 | 0.076 | 0.05 | 1 | 1 | Ergic2 | 0.026 | Young |
| 0.022413 | 0.302319 | 0.096 | 0.067 | 1 | 1 | Tia1 | 0.029 | Young |
| 0.022484 | 0.641357 | 0.032 | 0.015 | 1 | 1 | Endod1 | 0.017 | Young |
| 0.022504 | 1.322238 | 0.024 | 0.01 | 1 | 1 | Strada | 0.014 | Young |
| 0.022567 | 0.239682 | 0.189 | 0.148 | 1 | 1 | Pkm | 0.041 | Young |
| 0.022609 | 1.756264 | 0.018 | 0.006 | 1 | 1 | Stau2 | 0.012 | Young |
| 0.022635 | 1.245532 | 0.028 | 0.013 | 1 | 1 | Zbtb16 | 0.015 | Young |
| 0.02267 | 0.232979 | 0.16 | 0.122 | 1 | 1 | Hnrnpul1 | 0.038 | Young |
| 0.022822 | 0.045862 | 0.557 | 0.478 | 1 | 1 | Serf2 | 0.079 | Young |
| 0.022952 | 1.37842 | 0.018 | 0.006 | 1 | 1 | Trim21 | 0.012 | Young |
| 0.022978 | 0.304587 | 0.121 | 0.089 | 1 | 1 | Il1rl1 | 0.032 | Young |
| 0.022998 | 0.80553 | 0.024 | 0.01 | 1 | 1 | Mfng | 0.014 | Young |
| 0.023002 | 0.223552 | 0.119 | 0.086 | 1 | 1 | Kmt5b | 0.033 | Young |
| 0.023012 | 0.34856 | 0.035 | 0.018 | 1 | 1 | Elmod2 | 0.017 | Young |
| 0.023146 | 0.442971 | 0.112 | 0.081 | 1 | 1 | Ralgps2 | 0.031 | Young |
| 0.023216 | 1.031398 | 0.045 | 0.026 | 1 | 1 | Reps1 | 0.019 | Young |
| 0.023229 | 0.267788 | 0.054 | 0.032 | 1 | 1 | Mrpl43 | 0.022 | Young |
| 0.023283 | 0.047289 | 0.155 | 0.117 | 1 | 1 | Il4ra | 0.038 | Young |
| 0.023296 | 0.920923 | 0.024 | 0.01 | 1 | 1 | Adi1 | 0.014 | Young |
| 0.023305 | 0.538258 | 0.092 | 0.064 | 1 | 1 | Taf1d | 0.028 | Young |

| | | | | | | | | |
|---|---|---|---|---|---|---|---|---|
| 0.02332 | 1.118008 | 0.018 | 0.006 | 1 | 1 | Spats2 | 0.012 | Young |
| 0.023351 | 0.403226 | 0.064 | 0.04 | 1 | 1 | Zdhhc14 | 0.024 | Young |
| 0.023405 | 0.52791 | 0.089 | 0.062 | 1 | 1 | St3gal1 | 0.027 | Young |
| 0.023458 | 0.667791 | 0.028 | 0.013 | 1 | 1 | Fance | 0.015 | Young |
| 0.023531 | 0.624756 | 0.06 | 0.037 | 1 | 1 | Rbl2 | 0.023 | Young |
| 0.023599 | 0.653158 | 0.072 | 0.048 | 1 | 1 | Pitpnb | 0.024 | Young |
| 0.023636 | 2.28504 | 0.022 | 0.009 | 1 | 1 | Haus2 | 0.013 | Young |
| 0.023667 | 0.263836 | 0.118 | 0.086 | 1 | 1 | Copb2 | 0.032 | Young |
| 0.02376 | 0.056334 | 0.12 | 0.086 | 1 | 1 | Eloa | 0.034 | Young |
| 0.023828 | 0.162005 | 0.329 | 0.274 | 1 | 1 | Fcrls | 0.055 | Young |
| 0.023849 | 0.690005 | 0.018 | 0.006 | 1 | 1 | Mbd4 | 0.012 | Young |
| 0.02386 | 0.259792 | 0.065 | 0.041 | 1 | 1 | Mesd | 0.024 | Young |
| 0.023932 | 0.915851 | 0.038 | 0.021 | 1 | 1 | Rev1 | 0.017 | Young |
| 0.023965 | 0.200809 | 0.138 | 0.104 | 1 | 1 | Adrb2 | 0.034 | Young |
| 0.024013 | 0.436181 | 0.095 | 0.067 | 1 | 1 | Tiprl | 0.028 | Young |
| 0.024035 | 0.257068 | 0.134 | 0.1 | 1 | 1 | Dlst | 0.034 | Young |
| 0.02415 | 0.437824 | 0.037 | 0.019 | 1 | 1 | Fbxo34 | 0.018 | Young |
| 0.024163 | 0.264899 | 0.141 | 0.107 | 1 | 1 | Ppp1r10 | 0.034 | Young |
| 0.024267 | 2.624661 | 0.012 | 0.003 | 1 | 1 | Cd300ld3 | 0.009 | Young |
| 0.024273 | 0.882468 | 0.033 | 0.017 | 1 | 1 | Pou5f2 | 0.016 | Young |
| 0.0243 | 0.342429 | 0.114 | 0.084 | 1 | 1 | Mknk2 | 0.03 | Young |
| 0.024366 | 1.568085 | 0.016 | 0.005 | 1 | 1 | Kbtbd7 | 0.011 | Young |
| 0.024383 | 1.174804 | 0.047 | 0.027 | 1 | 1 | Qsox1 | 0.02 | Young |
| 0.024383 | 2.790402 | 0.012 | 0.003 | 1 | 1 | Nsdhl | 0.009 | Young |
| 0.02439 | 0.519239 | 0.075 | 0.05 | 1 | 1 | Smurf2 | 0.025 | Young |
| 0.024463 | 0.709466 | 0.045 | 0.026 | 1 | 1 | Uvssa | 0.019 | Young |
| 0.024618 | 2.194821 | 0.012 | 0.003 | 1 | 1 | Olfml3 | 0.009 | Young |
| 0.024726 | 1.003518 | 0.04 | 0.022 | 1 | 1 | Kmo | 0.018 | Young |
| 0.024736 | 0.426563 | 0.094 | 0.066 | 1 | 1 | Atad2b | 0.028 | Young |
| 0.024743 | 0.859366 | 0.047 | 0.027 | 1 | 1 | Cisd1 | 0.02 | Young |
| 0.024856 | 0.547582 | 0.052 | 0.031 | 1 | 1 | Parp4 | 0.021 | Young |
| 0.024857 | 0.072122 | 0.171 | 0.13 | 1 | 1 | Hipk1 | 0.041 | Young |
| 0.024988 | 0.547836 | 0.066 | 0.042 | 1 | 1 | Usp12 | 0.024 | Young |
| 0.025006 | 0.832828 | 0.016 | 0.005 | 1 | 1 | Klhl25 | 0.011 | Young |
| 0.025015 | 0.271073 | 0.085 | 0.058 | 1 | 1 | Atxn7l1 | 0.027 | Young |
| 0.025057 | 0.576683 | 0.093 | 0.066 | 1 | 1 | Foxn2 | 0.027 | Young |
| 0.025108 | 0.669454 | 0.038 | 0.021 | 1 | 1 | Tnfsf13b | 0.017 | Young |
| 0.025171 | 0.605713 | 0.038 | 0.021 | 1 | 1 | Taf5l | 0.017 | Young |
| 0.025217 | 0.6467 | 0.075 | 0.05 | 1 | 1 | Chsy1 | 0.025 | Young |
| 0.025341 | 1.366588 | 0.014 | 0.004 | 1 | 1 | Ppp2r5b | 0.01 | Young |
| 0.025417 | 0.220134 | 0.194 | 0.153 | 1 | 1 | Mdh1 | 0.041 | Young |
| 0.025425 | 0.182505 | 0.136 | 0.1 | 1 | 1 | Elmsan1 | 0.036 | Young |
| 0.025451 | 0.935859 | 0.014 | 0.004 | 1 | 1 | Wbp1 | 0.01 | Young |
| 0.025533 | 1.070735 | 0.014 | 0.004 | 1 | 1 | Slc25a23 | 0.01 | Young |
| 0.025974 | 0.377475 | 0.117 | 0.086 | 1 | 1 | Snrpd1 | 0.031 | Young |

| | | | | | | | | |
|---|---|---|---|---|---|---|---|---|
| 0.025999 | 0.315416 | 0.078 | 0.053 | 1 | 1 | Rcor1 | 0.025 | Young |
| 0.026051 | 0.8902 | 0.035 | 0.018 | 1 | 1 | Nxt2 | 0.017 | Young |
| 0.026109 | 0.151345 | 0.25 | 0.203 | 1 | 1 | Dpysl2 | 0.047 | Young |
| 0.026111 | 0.248345 | 0.165 | 0.129 | 1 | 1 | Cdk6 | 0.036 | Young |
| 0.026188 | 0.736863 | 0.083 | 0.058 | 1 | 1 | Rab32 | 0.025 | Young |
| 0.026338 | 0.154023 | 0.192 | 0.149 | 1 | 1 | Peli1 | 0.043 | Young |
| 0.026488 | 0.503516 | 0.069 | 0.045 | 1 | 1 | Pcif1 | 0.024 | Young |
| 0.026523 | 0.095276 | 0.422 | 0.36 | 1 | 1 | Tm6sf1 | 0.062 | Young |
| 0.02655 | 0.155074 | 0.185 | 0.144 | 1 | 1 | Mia2 | 0.041 | Young |
| 0.026572 | 0.365888 | 0.052 | 0.031 | 1 | 1 | Aph1b | 0.021 | Young |
| 0.026664 | 0.483163 | 0.081 | 0.055 | 1 | 1 | Skiv2l | 0.026 | Young |
| 0.026828 | 0.127292 | 0.499 | 0.432 | 1 | 1 | Tpm3 | 0.067 | Young |
| 0.02686 | 0.391363 | 0.081 | 0.055 | 1 | 1 | Ndfip1 | 0.026 | Young |
| 0.026886 | 1.695422 | 0.02 | 0.008 | 1 | 1 | Paqr4 | 0.012 | Young |
| 0.026909 | 0.174202 | 0.223 | 0.181 | 1 | 1 | Irf2 | 0.042 | Young |
| 0.026932 | 0.614036 | 0.048 | 0.028 | 1 | 1 | Pkd1 | 0.02 | Young |
| 0.026976 | 0.585354 | 0.092 | 0.064 | 1 | 1 | Cd180 | 0.028 | Young |
| 0.026977 | 0.284499 | 0.166 | 0.13 | 1 | 1 | Elob | 0.036 | Young |
| 0.026978 | 0.160002 | 0.107 | 0.076 | 1 | 1 | Cpsf2 | 0.031 | Young |
| 0.027129 | 0.214489 | 0.319 | 0.272 | 1 | 1 | Lnpep | 0.047 | Young |
| 0.027198 | 0.066488 | 0.189 | 0.147 | 1 | 1 | Lat2 | 0.042 | Young |
| 0.027257 | 0.119243 | 0.18 | 0.14 | 1 | 1 | Hnrnpd | 0.04 | Young |
| 0.027281 | 0.117302 | 0.359 | 0.306 | 1 | 1 | Grb2 | 0.053 | Young |
| 0.027283 | 0.705608 | 0.041 | 0.023 | 1 | 1 | Rai1 | 0.018 | Young |
| 0.027296 | 0.470861 | 0.081 | 0.055 | 1 | 1 | Smpdl3b | 0.026 | Young |
| 0.027371 | 0.909811 | 0.031 | 0.015 | 1 | 1 | Mrpl58 | 0.016 | Young |
| 0.027566 | 0.271965 | 0.106 | 0.076 | 1 | 1 | Col4a3bp | 0.03 | Young |
| 0.027588 | 0.751789 | 0.043 | 0.024 | 1 | 1 | Ddx1 | 0.019 | Young |
| 0.027632 | 0.313184 | 0.112 | 0.081 | 1 | 1 | Dpp8 | 0.031 | Young |
| 0.027712 | 1.7458 | 0.024 | 0.01 | 1 | 1 | Cxxc5 | 0.014 | Young |
| 0.027794 | 1.049725 | 0.02 | 0.008 | 1 | 1 | Mical2 | 0.012 | Young |
| 0.02785 | 1.272061 | 0.027 | 0.013 | 1 | 1 | Dctpp1 | 0.014 | Young |
| 0.027875 | 0.265373 | 0.063 | 0.04 | 1 | 1 | Gskip | 0.023 | Young |
| 0.027895 | 0.268683 | 0.035 | 0.018 | 1 | 1 | Wtip | 0.017 | Young |
| 0.027908 | 0.588937 | 0.049 | 0.03 | 1 | 1 | Abraxas2 | 0.019 | Young |
| 0.027965 | 1.229321 | 0.044 | 0.026 | 1 | 1 | Stx8 | 0.018 | Young |
| 0.028074 | 0.423018 | 0.123 | 0.093 | 1 | 1 | Rpl36al | 0.03 | Young |
| 0.028146 | 0.156129 | 0.34 | 0.285 | 1 | 1 | Tm9sf3 | 0.055 | Young |
| 0.028192 | 0.257028 | 0.217 | 0.177 | 1 | 1 | Foxn3 | 0.04 | Young |
| 0.028241 | 1.257558 | 0.027 | 0.013 | 1 | 1 | Sparc | 0.014 | Young |
| 0.02826 | 0.860781 | 0.077 | 0.053 | 1 | 1 | Brd3 | 0.024 | Young |
| 0.028296 | 0.364505 | 0.041 | 0.023 | 1 | 1 | Calcrl | 0.018 | Young |
| 0.02841 | 0.046535 | 0.205 | 0.159 | 1 | 1 | Smarca5 | 0.046 | Young |
| 0.02849 | 0.555407 | 0.079 | 0.054 | 1 | 1 | Nsmce4a | 0.025 | Young |
| 0.028493 | 0.34804 | 0.064 | 0.041 | 1 | 1 | Trove2 | 0.023 | Young |

| | | | | | | | | |
|---|---|---|---|---|---|---|---|---|
| 0.028519 | 0.715065 | 0.051 | 0.031 | 1 | 1 | Tcf7l2 | 0.02 | Young |
| 0.028531 | 0.376605 | 0.067 | 0.044 | 1 | 1 | Cbx4 | 0.023 | Young |
| 0.028617 | 0.376861 | 0.048 | 0.028 | 1 | 1 | Kdm3b | 0.02 | Young |
| 0.028624 | 0.217422 | 0.137 | 0.103 | 1 | 1 | Ttc7 | 0.034 | Young |
| 0.028652 | 0.024876 | 0.203 | 0.159 | 1 | 1 | Usp25 | 0.044 | Young |
| 0.028748 | 0.40082 | 0.151 | 0.117 | 1 | 1 | Nr4a2 | 0.034 | Young |
| 0.028837 | 1.294604 | 0.039 | 0.022 | 1 | 1 | Rad54l2 | 0.017 | Young |
| 0.028907 | 0.186692 | 0.107 | 0.077 | 1 | 1 | Atraid | 0.03 | Young |
| 0.02901 | 0.790035 | 0.044 | 0.026 | 1 | 1 | Nvl | 0.018 | Young |
| 0.02903 | 0.085753 | 0.186 | 0.145 | 1 | 1 | Notch1 | 0.041 | Young |
| 0.029082 | 0.180946 | 0.239 | 0.195 | 1 | 1 | Galnt1 | 0.044 | Young |
| 0.029148 | 0.442647 | 0.031 | 0.015 | 1 | 1 | Neurl1a | 0.016 | Young |
| 0.029571 | 0.546427 | 0.125 | 0.095 | 1 | 1 | Kmt2c | 0.03 | Young |
| 0.029589 | 0.293309 | 0.099 | 0.071 | 1 | 1 | Btbd7 | 0.028 | Young |
| 0.029605 | 0.28232 | 0.058 | 0.036 | 1 | 1 | Rrm2b | 0.022 | Young |
| 0.029697 | 0.416366 | 0.059 | 0.037 | 1 | 1 | Desi2 | 0.022 | Young |
| 0.029741 | 0.295949 | 0.218 | 0.176 | 1 | 1 | Apobec1 | 0.042 | Young |
| 0.029762 | 0.596333 | 0.062 | 0.04 | 1 | 1 | Pvr | 0.022 | Young |
| 0.029849 | 0.455856 | 0.027 | 0.013 | 1 | 1 | Mcm9 | 0.014 | Young |
| 0.029877 | 0.110137 | 0.239 | 0.193 | 1 | 1 | Epb41l2 | 0.046 | Young |
| 0.029898 | 0.431873 | 0.043 | 0.024 | 1 | 1 | Kif21b | 0.019 | Young |
| 0.029899 | 1.708045 | 0.018 | 0.006 | 1 | 1 | Traf3ip3 | 0.012 | Young |
| 0.02995 | 0.337721 | 0.151 | 0.117 | 1 | 1 | Zranb2 | 0.034 | Young |
| 0.030022 | 0.230263 | 0.128 | 0.095 | 1 | 1 | Csf2rb | 0.033 | Young |
| 0.030064 | 1.207407 | 0.018 | 0.006 | 1 | 1 | Aifm1 | 0.012 | Young |
| 0.030091 | 1.108919 | 0.039 | 0.022 | 1 | 1 | Plxna4os1 | 0.017 | Young |
| 0.030177 | 0.862153 | 0.058 | 0.037 | 1 | 1 | Faim | 0.021 | Young |
| 0.030187 | 0.415271 | 0.064 | 0.041 | 1 | 1 | Trp53 | 0.023 | Young |
| 0.030197 | 0.406684 | 0.061 | 0.039 | 1 | 1 | Gsap | 0.022 | Young |
| 0.030245 | 1.25309 | 0.034 | 0.018 | 1 | 1 | Depdc5 | 0.016 | Young |
| 0.030286 | 0.704317 | 0.068 | 0.045 | 1 | 1 | P3h2 | 0.023 | Young |
| 0.03034 | 0.163792 | 0.165 | 0.129 | 1 | 1 | Cmtm7 | 0.036 | Young |
| 0.030368 | 1.316281 | 0.029 | 0.014 | 1 | 1 | Ddt | 0.015 | Young |
| 0.030397 | 0.990281 | 0.018 | 0.006 | 1 | 1 | Slx1b | 0.012 | Young |
| 0.030401 | 0.662966 | 0.032 | 0.017 | 1 | 1 | Smad7 | 0.015 | Young |
| 0.030428 | 0.18187 | 0.325 | 0.272 | 1 | 1 | Vsir | 0.053 | Young |
| 0.030515 | 0.106246 | 0.14 | 0.105 | 1 | 1 | Arfgef1 | 0.035 | Young |
| 0.030578 | 0.096763 | 0.304 | 0.249 | 1 | 1 | Selenok | 0.055 | Young |
| 0.030591 | 0.110904 | 0.356 | 0.297 | 1 | 1 | Srsf3 | 0.059 | Young |
| 0.030632 | 0.281642 | 0.061 | 0.039 | 1 | 1 | Usp3 | 0.022 | Young |
| 0.030737 | 0.421235 | 0.11 | 0.081 | 1 | 1 | Tbc1d5 | 0.029 | Young |
| 0.030803 | 0.476051 | 0.062 | 0.04 | 1 | 1 | Zfp62 | 0.022 | Young |
| 0.030816 | 0.59879 | 0.055 | 0.035 | 1 | 1 | Cd82 | 0.02 | Young |
| 0.031035 | 0.313462 | 0.095 | 0.067 | 1 | 1 | Zranb1 | 0.028 | Young |
| 0.031037 | 0.301155 | 0.143 | 0.109 | 1 | 1 | Commd8 | 0.034 | Young |

| | | | | | | | | |
|---|---|---|---|---|---|---|---|---|
| 0.031048 | 0.396844 | 0.157 | 0.123 | 1 | 1 | Ankrd17 | 0.034 | Young |
| 0.031143 | 0.532274 | 0.065 | 0.042 | 1 | 1 | Myo7a | 0.023 | Young |
| 0.03118 | 0.286933 | 0.153 | 0.118 | 1 | 1 | Rassf5 | 0.035 | Young |
| 0.031208 | 0.256555 | 0.198 | 0.158 | 1 | 1 | Mbnl2 | 0.04 | Young |
| 0.031228 | 0.188255 | 0.216 | 0.174 | 1 | 1 | Prkacb | 0.042 | Young |
| 0.031265 | 1.014232 | 0.029 | 0.014 | 1 | 1 | Fam46c | 0.015 | Young |
| 0.031443 | 0.298039 | 0.158 | 0.123 | 1 | 1 | Tmod3 | 0.035 | Young |
| 0.031453 | 1.000084 | 0.034 | 0.018 | 1 | 1 | Txnrd2 | 0.016 | Young |
| 0.031471 | 0.330886 | 0.081 | 0.055 | 1 | 1 | Sh3bp2 | 0.026 | Young |
| 0.031493 | 0.646898 | 0.034 | 0.018 | 1 | 1 | Zmynd15 | 0.016 | Young |
| 0.031566 | 0.52629 | 0.092 | 0.066 | 1 | 1 | S1pr1 | 0.026 | Young |
| 0.031695 | 0.708474 | 0.034 | 0.018 | 1 | 1 | Zbtb34 | 0.016 | Young |
| 0.031716 | 0.746529 | 0.029 | 0.014 | 1 | 1 | Crbn | 0.015 | Young |
| 0.03177 | 0.629726 | 0.06 | 0.039 | 1 | 1 | Zfp131 | 0.021 | Young |
| 0.031805 | 0.135125 | 0.135 | 0.102 | 1 | 1 | Phf21a | 0.033 | Young |
| 0.031869 | 0.432984 | 0.101 | 0.073 | 1 | 1 | Nfatc3 | 0.028 | Young |
| 0.031947 | 1.156078 | 0.025 | 0.012 | 1 | 1 | Pck2 | 0.013 | Young |
| 0.032043 | 0.842821 | 0.025 | 0.012 | 1 | 1 | Negr1 | 0.013 | Young |
| 0.032122 | 0.498703 | 0.082 | 0.057 | 1 | 1 | Orai3 | 0.025 | Young |
| 0.032225 | 0.041814 | 0.147 | 0.111 | 1 | 1 | Slbp | 0.036 | Young |
| 0.032317 | 0.342681 | 0.083 | 0.058 | 1 | 1 | Pcgf5 | 0.025 | Young |
| 0.032343 | 1.5006 | 0.016 | 0.005 | 1 | 1 | Colec12 | 0.011 | Young |
| 0.032384 | 0.142803 | 0.323 | 0.267 | 1 | 1 | Jmjd1c | 0.056 | Young |
| 0.03247 | 1.260119 | 0.016 | 0.005 | 1 | 1 | Slc25a38 | 0.011 | Young |
| 0.032566 | 1.23092 | 0.016 | 0.005 | 1 | 1 | Diexf | 0.011 | Young |
| 0.032957 | 0.267722 | 0.129 | 0.098 | 1 | 1 | Plekhj1 | 0.031 | Young |
| 0.033029 | 0.055486 | 0.146 | 0.111 | 1 | 1 | Cox5b | 0.035 | Young |
| 0.033037 | 0.091651 | 0.456 | 0.389 | 1 | 1 | Ube2d3 | 0.067 | Young |
| 0.033062 | 0.192978 | 0.14 | 0.105 | 1 | 1 | Cox20 | 0.035 | Young |
| 0.033082 | 0.177915 | 0.021 | 0.009 | 1 | 1 | Disc1 | 0.012 | Young |
| 0.033095 | 0.438382 | 0.071 | 0.048 | 1 | 1 | Ephx1 | 0.023 | Young |
| 0.03316 | 0.119693 | 0.25 | 0.203 | 1 | 1 | Gusb | 0.047 | Young |
| 0.03319 | 0.315968 | 0.021 | 0.009 | 1 | 1 | AC163018. | 0.012 | Young |
| 0.033368 | 0.227848 | 0.029 | 0.014 | 1 | 1 | Lin9 | 0.015 | Young |
| 0.033385 | 0.682422 | 0.05 | 0.031 | 1 | 1 | Zfp512 | 0.019 | Young |
| 0.033386 | 0.080905 | 0.273 | 0.225 | 1 | 1 | Arhgap30 | 0.048 | Young |
| 0.033442 | 0.197822 | 0.148 | 0.114 | 1 | 1 | Ap2b1 | 0.034 | Young |
| 0.033453 | 0.358854 | 0.068 | 0.045 | 1 | 1 | Cep192 | 0.023 | Young |
| 0.033464 | 1.877521 | 0.011 | 0.003 | 1 | 1 | Endov | 0.008 | Young |
| 0.033481 | 0.401381 | 0.11 | 0.081 | 1 | 1 | Slc26a2 | 0.029 | Young |
| 0.033493 | 0.91019 | 0.03 | 0.015 | 1 | 1 | Ercc5 | 0.015 | Young |
| 0.033504 | 1.709609 | 0.011 | 0.003 | 1 | 1 | Iqcb1 | 0.008 | Young |
| 0.033516 | 0.959655 | 0.03 | 0.015 | 1 | 1 | Fmnl3 | 0.015 | Young |
| 0.033543 | 1.803844 | 0.011 | 0.003 | 1 | 1 | Prag1 | 0.008 | Young |
| 0.033552 | 0.012451 | 0.123 | 0.09 | 1 | 1 | Cd209a | 0.033 | Young |

| | | | | | | | | |
|---|---|---|---|---|---|---|---|---|
| 0.033565 | 1.493358 | 0.013 | 0.004 | 1 | 1 | Mad2l2 | 0.009 | Young |
| 0.033641 | 0.103961 | 0.117 | 0.086 | 1 | 1 | Rere | 0.031 | Young |
| 0.033707 | 0.179158 | 0.276 | 0.23 | 1 | 1 | Atrx | 0.046 | Young |
| 0.033856 | 0.586982 | 0.041 | 0.023 | 1 | 1 | Tmem209 | 0.018 | Young |
| 0.033861 | 0.681396 | 0.011 | 0.003 | 1 | 1 | Polr1e | 0.008 | Young |
| 0.033941 | 1.231938 | 0.011 | 0.003 | 1 | 1 | Smo | 0.008 | Young |
| 0.033941 | 0.516688 | 0.011 | 0.003 | 1 | 1 | Cdpf1 | 0.008 | Young |
| 0.033955 | 0.154149 | 0.193 | 0.156 | 1 | 1 | Fam111a | 0.037 | Young |
| 0.034063 | 0.149897 | 0.085 | 0.059 | 1 | 1 | Ap1g1 | 0.026 | Young |
| 0.03414 | 1.229168 | 0.013 | 0.004 | 1 | 1 | Zfp60 | 0.009 | Young |
| 0.034193 | 0.249904 | 0.123 | 0.091 | 1 | 1 | Mbd2 | 0.032 | Young |
| 0.034244 | 0.404925 | 0.086 | 0.06 | 1 | 1 | Dusp16 | 0.026 | Young |
| 0.034284 | 0.148884 | 0.013 | 0.004 | 1 | 1 | Rab3ip | 0.009 | Young |
| 0.034333 | 0.089773 | 0.649 | 0.571 | 1 | 1 | Jun | 0.078 | Young |
| 0.034333 | 0.223658 | 0.21 | 0.171 | 1 | 1 | Selplg | 0.039 | Young |
| 0.034338 | 0.203579 | 0.165 | 0.13 | 1 | 1 | Lrrc25 | 0.035 | Young |
| 0.034354 | 0.047632 | 0.886 | 0.85 | 1 | 1 | Rps3a1 | 0.036 | Young |
| 0.034378 | 0.218683 | 0.151 | 0.116 | 1 | 1 | Nfic | 0.035 | Young |
| 0.034529 | 0.120018 | 0.108 | 0.078 | 1 | 1 | Limd2 | 0.03 | Young |
| 0.034603 | 0.335894 | 0.163 | 0.129 | 1 | 1 | Pxn | 0.034 | Young |
| 0.034669 | 0.218096 | 0.24 | 0.199 | 1 | 1 | Epn1 | 0.041 | Young |
| 0.034716 | 0.052102 | 0.612 | 0.519 | 1 | 1 | Zfp36l2 | 0.093 | Young |
| 0.034987 | 1.590371 | 0.019 | 0.008 | 1 | 1 | Plpp5 | 0.011 | Young |
| 0.035167 | 0.488459 | 0.035 | 0.019 | 1 | 1 | Stxbp1 | 0.016 | Young |
| 0.035436 | 0.182554 | 0.069 | 0.046 | 1 | 1 | Commd6 | 0.023 | Young |
| 0.035443 | 0.346288 | 0.132 | 0.102 | 1 | 1 | Srsf7 | 0.03 | Young |
| 0.035527 | 0.437099 | 0.086 | 0.06 | 1 | 1 | Txlna | 0.026 | Young |
| 0.035714 | 0.131998 | 0.229 | 0.185 | 1 | 1 | Sec62 | 0.044 | Young |
| 0.035718 | 0.425923 | 0.06 | 0.039 | 1 | 1 | St3gal5 | 0.021 | Young |
| 0.03572 | 0.245449 | 0.092 | 0.066 | 1 | 1 | Pcmtd2 | 0.026 | Young |
| 0.035848 | 0.122785 | 0.21 | 0.168 | 1 | 1 | Rell1 | 0.042 | Young |
| 0.035862 | 0.423781 | 0.135 | 0.104 | 1 | 1 | Rc3h2 | 0.031 | Young |
| 0.036184 | 0.787908 | 0.056 | 0.036 | 1 | 1 | Lrrc41 | 0.02 | Young |
| 0.036187 | 0.460853 | 0.067 | 0.045 | 1 | 1 | Prkab2 | 0.022 | Young |
| 0.036236 | 0.267272 | 0.095 | 0.068 | 1 | 1 | Rnf2 | 0.027 | Young |
| 0.036282 | 0.256326 | 0.148 | 0.116 | 1 | 1 | Samd4b | 0.032 | Young |
| 0.036298 | 1.237299 | 0.04 | 0.023 | 1 | 1 | Xab2 | 0.017 | Young |
| 0.036328 | 0.316459 | 0.058 | 0.037 | 1 | 1 | Utp14b | 0.021 | Young |
| 0.036343 | 0.904574 | 0.032 | 0.017 | 1 | 1 | Nat9 | 0.015 | Young |
| 0.036363 | 0.510015 | 0.027 | 0.013 | 1 | 1 | Gabpb1 | 0.014 | Young |
| 0.036478 | 0.703653 | 0.045 | 0.027 | 1 | 1 | Amz2 | 0.018 | Young |
| 0.036483 | 0.908162 | 0.019 | 0.008 | 1 | 1 | Galnt12 | 0.011 | Young |
| 0.036519 | 0.500662 | 0.027 | 0.013 | 1 | 1 | Exosc4 | 0.014 | Young |
| 0.03652 | 0.416622 | 0.042 | 0.024 | 1 | 1 | Asb7 | 0.018 | Young |
| 0.036609 | 0.589829 | 0.069 | 0.046 | 1 | 1 | Trrap | 0.023 | Young |

| | | | | | | | | |
|---|---|---|---|---|---|---|---|---|
| 0.036647 | 0.591799 | 0.073 | 0.05 | 1 | 1 | Phf1 | 0.023 | Young |
| 0.036682 | 0.123938 | 0.25 | 0.207 | 1 | 1 | Xiap | 0.043 | Young |
| 0.036696 | 0.435256 | 0.064 | 0.042 | 1 | 1 | 4833420G1 | 0.022 | Young |
| 0.036728 | 0.545587 | 0.027 | 0.013 | 1 | 1 | Slc25a44 | 0.014 | Young |
| 0.036742 | 0.146678 | 0.153 | 0.118 | 1 | 1 | Ccnt1 | 0.035 | Young |
| 0.036822 | 0.216947 | 0.171 | 0.136 | 1 | 1 | Tnpo1 | 0.035 | Young |
| 0.036825 | 0.541009 | 0.072 | 0.049 | 1 | 1 | Arid2 | 0.023 | Young |
| 0.036843 | 0.88025 | 0.023 | 0.01 | 1 | 1 | Klhl22 | 0.013 | Young |
| 0.036877 | 0.418245 | 0.083 | 0.058 | 1 | 1 | Samm50 | 0.025 | Young |
| 0.036924 | 0.091644 | 0.127 | 0.095 | 1 | 1 | Dpm3 | 0.032 | Young |
| 0.037016 | 0.211205 | 0.264 | 0.22 | 1 | 1 | Klf3 | 0.044 | Young |
| 0.037043 | 0.107824 | 0.361 | 0.305 | 1 | 1 | Rock2 | 0.056 | Young |
| 0.037064 | 0.655872 | 0.078 | 0.054 | 1 | 1 | 1810013L2 | 0.024 | Young |
| 0.037083 | 0.49916 | 0.07 | 0.048 | 1 | 1 | Tnfaip8l2 | 0.022 | Young |
| 0.037202 | 0.838665 | 0.028 | 0.014 | 1 | 1 | Gdpd3 | 0.014 | Young |
| 0.037226 | 0.507971 | 0.044 | 0.026 | 1 | 1 | Larp7 | 0.018 | Young |
| 0.037294 | 0.423746 | 0.089 | 0.063 | 1 | 1 | C1galt1 | 0.026 | Young |
| 0.037389 | 0.196236 | 0.134 | 0.103 | 1 | 1 | Ppp6r3 | 0.031 | Young |
| 0.037404 | 0.141661 | 0.208 | 0.168 | 1 | 1 | Cpsf6 | 0.04 | Young |
| 0.037556 | 0.068154 | 0.213 | 0.171 | 1 | 1 | Dnajb14 | 0.042 | Young |
| 0.037735 | 0.210164 | 0.152 | 0.118 | 1 | 1 | Acat1 | 0.034 | Young |
| 0.037763 | 0.63721 | 0.045 | 0.027 | 1 | 1 | Uty | 0.018 | Young |
| 0.037877 | 0.182432 | 0.071 | 0.048 | 1 | 1 | Hlx | 0.023 | Young |
| 0.037883 | 2.003068 | 0.017 | 0.006 | 1 | 1 | Zc3h10 | 0.011 | Young |
| 0.038075 | 0.319412 | 0.063 | 0.041 | 1 | 1 | Rnf169 | 0.022 | Young |
| 0.038206 | 0.337713 | 0.108 | 0.08 | 1 | 1 | Bsdc1 | 0.028 | Young |
| 0.038207 | 0.385119 | 0.032 | 0.017 | 1 | 1 | Rad17 | 0.015 | Young |
| 0.038231 | 0.701187 | 0.068 | 0.046 | 1 | 1 | Atad1 | 0.022 | Young |
| 0.038372 | 0.319634 | 0.05 | 0.031 | 1 | 1 | Gprc5c | 0.019 | Young |
| 0.038387 | 0.49058 | 0.079 | 0.055 | 1 | 1 | Tbc1d10b | 0.024 | Young |
| 0.038435 | 1.910052 | 0.017 | 0.006 | 1 | 1 | C330018D2 | 0.011 | Young |
| 0.038469 | 0.031631 | 0.335 | 0.281 | 1 | 1 | Atp6ap1 | 0.054 | Young |
| 0.038507 | 0.766459 | 0.033 | 0.018 | 1 | 1 | Elof1 | 0.015 | Young |
| 0.038523 | 0.031313 | 0.188 | 0.149 | 1 | 1 | Pttg1ip | 0.039 | Young |
| 0.038556 | 0.604077 | 0.061 | 0.04 | 1 | 1 | Ola1 | 0.021 | Young |
| 0.038559 | 0.028914 | 0.222 | 0.179 | 1 | 1 | Uqcrq | 0.043 | Young |
| 0.038643 | 0.037458 | 0.312 | 0.261 | 1 | 1 | Brk1 | 0.051 | Young |
| 0.038644 | 1.677586 | 0.017 | 0.006 | 1 | 1 | 3110001I2 | 0.011 | Young |
| 0.038677 | 0.384426 | 0.1 | 0.073 | 1 | 1 | Map3k2 | 0.027 | Young |
| 0.038713 | 1.651671 | 0.017 | 0.006 | 1 | 1 | Invs | 0.011 | Young |
| 0.038748 | 1.578474 | 0.017 | 0.006 | 1 | 1 | Plscr4 | 0.011 | Young |
| 0.038859 | 0.164573 | 0.097 | 0.071 | 1 | 1 | Ahr | 0.026 | Young |
| 0.038923 | 0.434536 | 0.094 | 0.068 | 1 | 1 | Nfia | 0.026 | Young |
| 0.03899 | 0.132091 | 0.052 | 0.032 | 1 | 1 | Engase | 0.02 | Young |
| 0.039011 | 0.448482 | 0.055 | 0.035 | 1 | 1 | Acbd6 | 0.02 | Young |

| | | | | | | | | |
|---|---|---|---|---|---|---|---|---|
| 0.039013 | 0.098612 | 0.359 | 0.303 | 1 | 1 | Zcchc6 | 0.056 | Young |
| 0.039146 | 0.007047 | 0.072 | 0.049 | 1 | 1 | Lcorl | 0.023 | Young |
| 0.039334 | 0.678758 | 0.072 | 0.05 | 1 | 1 | Slc35f6 | 0.022 | Young |
| 0.039393 | 0.153756 | 0.177 | 0.14 | 1 | 1 | Hnrnpul2 | 0.037 | Young |
| 0.039416 | 0.334679 | 0.066 | 0.044 | 1 | 1 | Pbx1 | 0.022 | Young |
| 0.039515 | 0.104272 | 0.174 | 0.136 | 1 | 1 | Cct2 | 0.038 | Young |
| 0.039703 | 0.339192 | 0.106 | 0.078 | 1 | 1 | Dapp1 | 0.028 | Young |
| 0.039716 | 0.595391 | 0.078 | 0.054 | 1 | 1 | B4galt3 | 0.024 | Young |
| 0.039767 | 0.379926 | 0.12 | 0.09 | 1 | 1 | Idh2 | 0.03 | Young |
| 0.039858 | 0.687107 | 0.081 | 0.058 | 1 | 1 | Crtc3 | 0.023 | Young |
| 0.039916 | 0.762149 | 0.017 | 0.006 | 1 | 1 | 9930104L0 | 0.011 | Young |
| 0.039923 | 0.716324 | 0.077 | 0.054 | 1 | 1 | Chd1 | 0.023 | Young |
| 0.040022 | 1.054845 | 0.024 | 0.012 | 1 | 1 | Bend4 | 0.012 | Young |
| 0.040052 | 0.167228 | 0.25 | 0.208 | 1 | 1 | Itpripl2 | 0.042 | Young |
| 0.040052 | 0.649541 | 0.033 | 0.018 | 1 | 1 | Klhdc10 | 0.015 | Young |
| 0.040094 | 0.267836 | 0.092 | 0.066 | 1 | 1 | Arid1b | 0.026 | Young |
| 0.040095 | 0.685272 | 0.017 | 0.006 | 1 | 1 | Slc2a1 | 0.011 | Young |
| 0.040201 | 0.337375 | 0.075 | 0.051 | 1 | 1 | Pom121 | 0.024 | Young |
| 0.040222 | 0.751219 | 0.049 | 0.031 | 1 | 1 | 2510009E0 | 0.018 | Young |
| 0.04034 | 0.598564 | 0.04 | 0.023 | 1 | 1 | Pnkd | 0.017 | Young |
| 0.040362 | 0.111634 | 0.265 | 0.219 | 1 | 1 | Skap2 | 0.046 | Young |
| 0.040391 | 1.282612 | 0.024 | 0.012 | 1 | 1 | Brcc3 | 0.012 | Young |
| 0.040575 | 0.439393 | 0.059 | 0.039 | 1 | 1 | Stx6 | 0.02 | Young |
| 0.040595 | 0.22223 | 0.217 | 0.177 | 1 | 1 | Pias1 | 0.04 | Young |
| 0.040616 | 1.240761 | 0.021 | 0.009 | 1 | 1 | Cish | 0.012 | Young |
| 0.040649 | 1.231503 | 0.021 | 0.009 | 1 | 1 | Lsg1 | 0.012 | Young |
| 0.040714 | 1.366169 | 0.021 | 0.009 | 1 | 1 | Zkscan8 | 0.012 | Young |
| 0.040716 | 1.083463 | 0.03 | 0.015 | 1 | 1 | Fam110a | 0.015 | Young |
| 0.040748 | 0.043032 | 0.252 | 0.206 | 1 | 1 | Ppp2r5c | 0.046 | Young |
| 0.04109 | 0.063896 | 0.055 | 0.035 | 1 | 1 | Rad50 | 0.02 | Young |
| 0.041406 | 0.256746 | 0.07 | 0.048 | 1 | 1 | Sos2 | 0.022 | Young |
| 0.041455 | 0.782109 | 0.024 | 0.012 | 1 | 1 | Ndrg1 | 0.012 | Young |
| 0.04153 | 0.245712 | 0.185 | 0.149 | 1 | 1 | Pigt | 0.036 | Young |
| 0.041614 | 0.112057 | 0.467 | 0.405 | 1 | 1 | Gdi2 | 0.062 | Young |
| 0.041681 | 0.321269 | 0.059 | 0.039 | 1 | 1 | Fmr1 | 0.02 | Young |
| 0.041728 | 0.67896 | 0.024 | 0.012 | 1 | 1 | Parp6 | 0.012 | Young |
| 0.041779 | 0.125733 | 0.312 | 0.263 | 1 | 1 | Lrp1 | 0.049 | Young |
| 0.04178 | 0.132661 | 0.19 | 0.153 | 1 | 1 | Ndufb3 | 0.037 | Young |
| 0.041781 | 0.313555 | 0.098 | 0.072 | 1 | 1 | Srrt | 0.026 | Young |
| 0.041908 | 0.159681 | 0.29 | 0.247 | 1 | 1 | Rpl29 | 0.043 | Young |
| 0.042003 | 0.568122 | 0.024 | 0.012 | 1 | 1 | Usp42 | 0.012 | Young |
| 0.042062 | 0.368854 | 0.048 | 0.03 | 1 | 1 | Chst12 | 0.018 | Young |
| 0.042104 | 0.547706 | 0.066 | 0.045 | 1 | 1 | Phf5a | 0.021 | Young |
| 0.042147 | 0.465905 | 0.069 | 0.048 | 1 | 1 | Kctd2 | 0.021 | Young |
| 0.042158 | 0.349477 | 0.083 | 0.059 | 1 | 1 | Luzp1 | 0.024 | Young |

| | | | | | | | | |
|---|---|---|---|---|---|---|---|---|
| 0.042273 | 0.400751 | 0.021 | 0.009 | 1 | 1 | Hmgn5 | 0.012 | Young |
| 0.042281 | 0.352485 | 0.138 | 0.108 | 1 | 1 | Tpp2 | 0.03 | Young |
| 0.042419 | 1.344538 | 0.015 | 0.005 | 1 | 1 | Bend6 | 0.01 | Young |
| 0.042425 | 0.239624 | 0.138 | 0.107 | 1 | 1 | Washc2 | 0.031 | Young |
| 0.042441 | 0.621104 | 0.021 | 0.009 | 1 | 1 | Tfdp2 | 0.012 | Young |
| 0.042459 | 1.460357 | 0.015 | 0.005 | 1 | 1 | Poll | 0.01 | Young |
| 0.042502 | 0.452997 | 0.035 | 0.019 | 1 | 1 | Zfp266 | 0.016 | Young |
| 0.042543 | 0.619125 | 0.021 | 0.009 | 1 | 1 | Ccdc58 | 0.012 | Young |
| 0.042582 | 0.926896 | 0.015 | 0.005 | 1 | 1 | Gpr84 | 0.01 | Young |
| 0.042613 | 0.554902 | 0.052 | 0.033 | 1 | 1 | Tmx4 | 0.019 | Young |
| 0.042758 | 0.404008 | 0.061 | 0.04 | 1 | 1 | Arl13b | 0.021 | Young |
| 0.042918 | 0.376724 | 0.066 | 0.045 | 1 | 1 | Zyg11b | 0.021 | Young |
| 0.04305 | 0.286438 | 0.107 | 0.08 | 1 | 1 | Dhx40 | 0.027 | Young |
| 0.043054 | 0.064961 | 0.504 | 0.429 | 1 | 1 | Atp5l | 0.075 | Young |
| 0.043088 | 0.463007 | 0.021 | 0.009 | 1 | 1 | Bhlhe41 | 0.012 | Young |
| 0.043098 | 0.480865 | 0.072 | 0.05 | 1 | 1 | Acp1 | 0.022 | Young |
| 0.043155 | 0.616054 | 0.054 | 0.035 | 1 | 1 | Pccb | 0.019 | Young |
| 0.043291 | 0.419989 | 0.12 | 0.093 | 1 | 1 | Csnk2b | 0.027 | Young |
| 0.043331 | 0.250447 | 0.202 | 0.165 | 1 | 1 | Txndc17 | 0.037 | Young |
| 0.043333 | 0.61737 | 0.043 | 0.026 | 1 | 1 | Zfp397 | 0.017 | Young |
| 0.04339 | 0.287822 | 0.123 | 0.094 | 1 | 1 | Trip11 | 0.029 | Young |
| 0.043406 | 1.137435 | 0.026 | 0.013 | 1 | 1 | Zfp729b | 0.013 | Young |
| 0.04344 | 0.442917 | 0.091 | 0.066 | 1 | 1 | Rictor | 0.025 | Young |
| 0.043533 | 0.668408 | 0.036 | 0.021 | 1 | 1 | Napsa | 0.015 | Young |
| 0.043552 | 0.483959 | 0.072 | 0.05 | 1 | 1 | Trim12a | 0.022 | Young |
| 0.043613 | 0.12094 | 0.015 | 0.005 | 1 | 1 | Mfsd4b4 | 0.01 | Young |
| 0.043615 | 0.096148 | 0.169 | 0.132 | 1 | 1 | Llph | 0.037 | Young |
| 0.043632 | 0.328393 | 0.091 | 0.066 | 1 | 1 | Hipk3 | 0.025 | Young |
| 0.04365 | 0.90071 | 0.026 | 0.013 | 1 | 1 | Slc10a3 | 0.013 | Young |
| 0.043682 | 0.323957 | 0.079 | 0.055 | 1 | 1 | Pds5b | 0.024 | Young |
| 0.043751 | 0.088114 | 0.172 | 0.136 | 1 | 1 | Amfr | 0.036 | Young |
| 0.043988 | 1.07498 | 0.026 | 0.013 | 1 | 1 | Tom1l2 | 0.013 | Young |
| 0.044121 | 0.121559 | 0.194 | 0.156 | 1 | 1 | Trim30a | 0.038 | Young |
| 0.044142 | 0.314123 | 0.075 | 0.051 | 1 | 1 | Ube2q2 | 0.024 | Young |
| 0.044199 | 0.835121 | 0.049 | 0.031 | 1 | 1 | Zfp646 | 0.018 | Young |
| 0.044254 | 2.068662 | 0.018 | 0.008 | 1 | 1 | Ercc4 | 0.01 | Young |
| 0.044266 | 2.153869 | 0.013 | 0.004 | 1 | 1 | Ccdc102a | 0.009 | Young |
| 0.044361 | 0.402732 | 0.074 | 0.051 | 1 | 1 | Churc1 | 0.023 | Young |
| 0.044586 | 0.306348 | 0.054 | 0.035 | 1 | 1 | Runx3 | 0.019 | Young |
| 0.044607 | 0.886146 | 0.026 | 0.013 | 1 | 1 | Accs | 0.013 | Young |
| 0.04463 | 0.240778 | 0.109 | 0.081 | 1 | 1 | Phactr2 | 0.028 | Young |
| 0.044631 | 0.229772 | 0.03 | 0.015 | 1 | 1 | Zfp687 | 0.015 | Young |
| 0.044659 | 0.260491 | 0.162 | 0.13 | 1 | 1 | Klhl24 | 0.032 | Young |
| 0.044683 | 1.688186 | 0.013 | 0.004 | 1 | 1 | Mapre3 | 0.009 | Young |
| 0.044702 | 1.860031 | 0.018 | 0.008 | 1 | 1 | Tmed1 | 0.01 | Young |

| | | | | | | | | |
|---|---|---|---|---|---|---|---|---|
| 0.044776 | 1.378907 | 0.013 | 0.004 | 1 | 1 | Ctla2b | 0.009 | Young |
| 0.044777 | 1.616954 | 0.018 | 0.008 | 1 | 1 | Fndc7 | 0.01 | Young |
| 0.044823 | 1.59155 | 0.013 | 0.004 | 1 | 1 | B430306N( | 0.009 | Young |
| 0.044851 | 0.45186 | 0.036 | 0.021 | 1 | 1 | Ttc1 | 0.015 | Young |
| 0.044889 | 0.785058 | 0.049 | 0.031 | 1 | 1 | Mrpl21 | 0.018 | Young |
| 0.04507 | 0.197039 | 0.15 | 0.118 | 1 | 1 | Grk3 | 0.032 | Young |
| 0.045104 | 1.140066 | 0.013 | 0.004 | 1 | 1 | Cep41 | 0.009 | Young |
| 0.045143 | 0.062107 | 0.083 | 0.059 | 1 | 1 | Dnttip2 | 0.024 | Young |
| 0.045343 | 0.674432 | 0.038 | 0.022 | 1 | 1 | Camk2g | 0.016 | Young |
| 0.045351 | 0.513779 | 0.057 | 0.037 | 1 | 1 | Fam160b1 | 0.02 | Young |
| 0.045373 | 0.572326 | 0.097 | 0.072 | 1 | 1 | Ccdc47 | 0.025 | Young |
| 0.045392 | 0.272347 | 0.122 | 0.093 | 1 | 1 | Brd7 | 0.029 | Young |
| 0.045449 | 0.244634 | 0.082 | 0.058 | 1 | 1 | Snrnp200 | 0.024 | Young |
| 0.0455 | 1.397173 | 0.01 | 0.003 | 1 | 1 | Nup43 | 0.007 | Young |
| 0.045504 | 0.796087 | 0.039 | 0.023 | 1 | 1 | Psd4 | 0.016 | Young |
| 0.045586 | 0.677966 | 0.031 | 0.017 | 1 | 1 | 3110062M | 0.014 | Young |
| 0.045598 | 0.200231 | 0.217 | 0.179 | 1 | 1 | Ppp1ca | 0.038 | Young |
| 0.045643 | 0.487314 | 0.031 | 0.017 | 1 | 1 | Zfp410 | 0.014 | Young |
| 0.045669 | 0.258148 | 0.123 | 0.094 | 1 | 1 | Tcf12 | 0.029 | Young |
| 0.045685 | 1.022415 | 0.01 | 0.003 | 1 | 1 | Klhl13 | 0.007 | Young |
| 0.045752 | 0.063503 | 0.552 | 0.468 | 1 | 1 | Ucp2 | 0.084 | Young |
| 0.045831 | 1.152705 | 0.022 | 0.01 | 1 | 1 | Nt5dc1 | 0.012 | Young |
| 0.045835 | 0.138637 | 0.091 | 0.066 | 1 | 1 | Slc38a9 | 0.025 | Young |
| 0.045847 | 0.333561 | 0.072 | 0.05 | 1 | 1 | Atxn10 | 0.022 | Young |
| 0.045865 | 0.196481 | 0.132 | 0.102 | 1 | 1 | Setd1b | 0.03 | Young |
| 0.0459 | 0.096312 | 0.275 | 0.225 | 1 | 1 | Litaf | 0.05 | Young |
| 0.045951 | 0.697712 | 0.01 | 0.003 | 1 | 1 | 2610035D1 | 0.007 | Young |
| 0.045981 | 0.11544 | 0.239 | 0.197 | 1 | 1 | Snrpe | 0.042 | Young |
| 0.046063 | 0.5845 | 0.031 | 0.017 | 1 | 1 | Cpsf1 | 0.014 | Young |
| 0.046184 | 1.013501 | 0.018 | 0.008 | 1 | 1 | Haus6 | 0.01 | Young |
| 0.0463 | 0.91414 | 0.018 | 0.008 | 1 | 1 | Zdhhc4 | 0.01 | Young |
| 0.046327 | 0.318222 | 0.08 | 0.057 | 1 | 1 | Banf1 | 0.023 | Young |
| 0.046401 | 0.828075 | 0.05 | 0.032 | 1 | 1 | Srp68 | 0.018 | Young |
| 0.046443 | 0.012362 | 0.537 | 0.461 | 1 | 1 | Fosb | 0.076 | Young |
| 0.046557 | 0.160545 | 0.239 | 0.199 | 1 | 1 | Gng2 | 0.04 | Young |
| 0.046564 | 0.213912 | 0.179 | 0.144 | 1 | 1 | H2afv | 0.035 | Young |
| 0.046609 | 1.012961 | 0.052 | 0.033 | 1 | 1 | Med21 | 0.019 | Young |
| 0.04661 | 0.081909 | 0.665 | 0.568 | 1 | 1 | Wnk1 | 0.097 | Young |
| 0.046739 | 0.832239 | 0.022 | 0.01 | 1 | 1 | Zfp760 | 0.012 | Young |
| 0.046923 | 0.044179 | 0.054 | 0.035 | 1 | 1 | Thap12 | 0.019 | Young |
| 0.047018 | 0.028966 | 0.136 | 0.104 | 1 | 1 | Cbx1 | 0.032 | Young |
| 0.047067 | 0.404243 | 0.055 | 0.036 | 1 | 1 | Phka2 | 0.019 | Young |
| 0.047113 | 0.223588 | 0.133 | 0.103 | 1 | 1 | Zc3h7a | 0.03 | Young |
| 0.047129 | 0.470608 | 0.063 | 0.042 | 1 | 1 | Lars | 0.021 | Young |
| 0.047202 | 0.947598 | 0.041 | 0.024 | 1 | 1 | Rptor | 0.017 | Young |

| | | | | | | | | |
|---|---|---|---|---|---|---|---|---|
| 0.047394 | 0.378419 | 0.057 | 0.037 | 1 | 1 | Clasp1 | 0.02 | Young |
| 0.047543 | 0.101159 | 0.077 | 0.054 | 1 | 1 | Casp8ap2 | 0.023 | Young |
| 0.047947 | 0.223668 | 0.177 | 0.144 | 1 | 1 | Cdc42se1 | 0.033 | Young |
| 0.04813 | 0.23313 | 0.022 | 0.01 | 1 | 1 | Fzd7 | 0.012 | Young |
| 0.048133 | 0.066284 | 0.346 | 0.287 | 1 | 1 | Ifngr1 | 0.059 | Young |
| 0.048448 | 0.325149 | 0.1 | 0.075 | 1 | 1 | Hipk2 | 0.025 | Young |
| 0.048497 | 0.106394 | 0.376 | 0.319 | 1 | 1 | Arpc3 | 0.057 | Young |
| 0.048603 | 0.651576 | 0.032 | 0.018 | 1 | 1 | Vamp5 | 0.014 | Young |
| 0.048678 | 0.501458 | 0.039 | 0.023 | 1 | 1 | Cep83 | 0.016 | Young |
| 0.048859 | 0.424031 | 0.153 | 0.122 | 1 | 1 | Pan3 | 0.031 | Young |
| 0.049042 | 0.740075 | 0.034 | 0.019 | 1 | 1 | Rbmx | 0.015 | Young |
| 0.049113 | 0.314013 | 0.074 | 0.051 | 1 | 1 | CAAA0111[illegible] | 0.023 | Young |
| 0.049177 | 0.500796 | 0.05 | 0.032 | 1 | 1 | Irak2 | 0.018 | Young |
| 0.049258 | 0.276724 | 0.092 | 0.067 | 1 | 1 | Fam76b | 0.025 | Young |
| 0.049293 | 0.71455 | 0.041 | 0.024 | 1 | 1 | Rbpms | 0.017 | Young |
| 0.049324 | 0.259436 | 0.151 | 0.12 | 1 | 1 | Tcp1 | 0.031 | Young |
| 0.049462 | 0.295133 | 0.095 | 0.069 | 1 | 1 | Rybp | 0.026 | Young |
| 0.049533 | 0.341096 | 0.102 | 0.076 | 1 | 1 | Dap | 0.026 | Young |
| 0.049558 | 0.320649 | 0.032 | 0.018 | 1 | 1 | Tmem203 | 0.014 | Young |
| 0.049614 | 0.860842 | 0.044 | 0.027 | 1 | 1 | Zswim4 | 0.017 | Young |
| 0.049627 | 0.19018 | 0.148 | 0.116 | 1 | 1 | Kpnb1 | 0.032 | Young |
| 0.049655 | 0.467998 | 0.073 | 0.051 | 1 | 1 | Rnf146 | 0.022 | Young |
| 0.049671 | 0.764229 | 0.034 | 0.019 | 1 | 1 | Ttc19 | 0.015 | Young |
| 0.049732 | 1.175079 | 0.024 | 0.012 | 1 | 1 | Saal1 | 0.012 | Young |
| 0.049957 | 1.695254 | 0.016 | 0.006 | 1 | 1 | Fig4 | 0.01 | Young |
| 0.050057 | 0.42449 | 0.066 | 0.045 | 1 | 1 | Vps11 | 0.021 | Young |
| 0.050162 | 0.631512 | 0.024 | 0.012 | 1 | 1 | Xpc | 0.012 | Young |
| 0.05023 | 0.083005 | 0.23 | 0.189 | 1 | 1 | Plekho1 | 0.041 | Young |
| 0.050252 | 0.347697 | 0.1 | 0.075 | 1 | 1 | Tmem219 | 0.025 | Young |
| 0.05031 | 1.319129 | 0.016 | 0.006 | 1 | 1 | Zfp319 | 0.01 | Young |
| 0.050369 | 0.428831 | 0.098 | 0.073 | 1 | 1 | Rabgef1 | 0.025 | Young |
| 0.050407 | 0.293325 | 0.16 | 0.129 | 1 | 1 | Cox17 | 0.031 | Young |
| 0.050466 | 0.073049 | 0.202 | 0.163 | 1 | 1 | Eif4a2 | 0.039 | Young |
| 0.050487 | 1.0878 | 0.024 | 0.012 | 1 | 1 | 2610524H[illegible] | 0.012 | Young |
| 0.050523 | 0.354142 | 0.1 | 0.075 | 1 | 1 | Nin | 0.025 | Young |
| 0.05056 | 0.620048 | 0.065 | 0.045 | 1 | 1 | Dlgap4 | 0.02 | Young |
| 0.050577 | 1.508428 | 0.016 | 0.006 | 1 | 1 | Rbm4 | 0.01 | Young |
| 0.050625 | 0.220339 | 0.165 | 0.132 | 1 | 1 | Nsd3 | 0.033 | Young |
| 0.050686 | 0.099046 | 0.314 | 0.267 | 1 | 1 | Rpl22l1 | 0.047 | Young |
| 0.050703 | 0.453565 | 0.088 | 0.064 | 1 | 1 | Trim56 | 0.024 | Young |
| 0.050709 | 0.944029 | 0.035 | 0.021 | 1 | 1 | Fam207a | 0.014 | Young |
| 0.050711 | 1.061742 | 0.016 | 0.006 | 1 | 1 | BC024978 | 0.01 | Young |
| 0.050746 | 0.517102 | 0.075 | 0.053 | 1 | 1 | Rcbtb2 | 0.022 | Young |
| 0.05083 | 0.090734 | 0.068 | 0.046 | 1 | 1 | Cramp1l | 0.022 | Young |
| 0.050887 | 0.989776 | 0.024 | 0.012 | 1 | 1 | Edrf1 | 0.012 | Young |

| | | | | | | | | |
|---|---|---|---|---|---|---|---|---|
| 0.050923 | 0.412063 | 0.063 | 0.042 | 1 | 1 | Fam53c | 0.021 | Young |
| 0.050967 | 0.160541 | 0.047 | 0.03 | 1 | 1 | Mrpl2 | 0.017 | Young |
| 0.051013 | 1.362845 | 0.02 | 0.009 | 1 | 1 | Lmln | 0.011 | Young |
| 0.051024 | 1.066325 | 0.016 | 0.006 | 1 | 1 | Fscn1 | 0.01 | Young |
| 0.051024 | 0.984452 | 0.016 | 0.006 | 1 | 1 | 1190007I0 | 0.01 | Young |
| 0.051053 | 1.348334 | 0.02 | 0.009 | 1 | 1 | Iqsec2 | 0.011 | Young |
| 0.051083 | 0.112443 | 0.129 | 0.099 | 1 | 1 | Diaph2 | 0.03 | Young |
| 0.051142 | 0.720759 | 0.024 | 0.012 | 1 | 1 | Tns4 | 0.012 | Young |
| 0.051179 | 0.185541 | 0.179 | 0.145 | 1 | 1 | 9930111J2 | 0.034 | Young |
| 0.051271 | 0.219079 | 0.196 | 0.162 | 1 | 1 | Rsrc2 | 0.034 | Young |
| 0.051305 | 0.166062 | 0.332 | 0.284 | 1 | 1 | Eid1 | 0.048 | Young |
| 0.05134 | 0.708942 | 0.068 | 0.048 | 1 | 1 | Cd9 | 0.02 | Young |
| 0.051412 | 0.322788 | 0.279 | 0.24 | 1 | 1 | Socs3 | 0.039 | Young |
| 0.051534 | 0.438806 | 0.094 | 0.069 | 1 | 1 | Pex6 | 0.025 | Young |
| 0.051582 | 0.893363 | 0.029 | 0.015 | 1 | 1 | Chmp6 | 0.014 | Young |
| 0.051616 | 1.04581 | 0.02 | 0.009 | 1 | 1 | Ttll3 | 0.011 | Young |
| 0.051738 | 1.132558 | 0.02 | 0.009 | 1 | 1 | Vill | 0.011 | Young |
| 0.051746 | 0.717774 | 0.016 | 0.006 | 1 | 1 | 0610009O2 | 0.01 | Young |
| 0.051772 | 0.055411 | 0.206 | 0.167 | 1 | 1 | Map3k1 | 0.039 | Young |
| 0.051855 | 0.029663 | 0.205 | 0.165 | 1 | 1 | Cox7a2 | 0.04 | Young |
| 0.051957 | 0.284522 | 0.096 | 0.071 | 1 | 1 | Mctp1 | 0.025 | Young |
| 0.051969 | 0.218599 | 0.163 | 0.131 | 1 | 1 | N4bp1 | 0.032 | Young |
| 0.052012 | 0.452271 | 0.151 | 0.123 | 1 | 1 | Dusp11 | 0.028 | Young |
| 0.052185 | 1.017444 | 0.02 | 0.009 | 1 | 1 | Sharpin | 0.011 | Young |
| 0.052277 | 0.600439 | 0.059 | 0.04 | 1 | 1 | Rlf | 0.019 | Young |
| 0.052594 | 0.798139 | 0.02 | 0.009 | 1 | 1 | Ms4a14 | 0.011 | Young |
| 0.052609 | 1.600857 | 0.025 | 0.013 | 1 | 1 | Taf1b | 0.012 | Young |
| 0.052669 | 0.08515 | 0.28 | 0.237 | 1 | 1 | Gnaq | 0.043 | Young |
| 0.052718 | 0.549595 | 0.029 | 0.015 | 1 | 1 | Slc35e2 | 0.014 | Young |
| 0.05276 | 0.354923 | 0.075 | 0.053 | 1 | 1 | Cklf | 0.022 | Young |
| 0.052857 | 0.650508 | 0.044 | 0.027 | 1 | 1 | Tfeb | 0.017 | Young |
| 0.052962 | 0.277402 | 0.119 | 0.091 | 1 | 1 | Gpcpd1 | 0.028 | Young |
| 0.05298 | 0.09897 | 0.143 | 0.112 | 1 | 1 | Rlim | 0.031 | Young |
| 0.053038 | 0.131469 | 0.077 | 0.054 | 1 | 1 | Gdf15 | 0.023 | Young |
| 0.053255 | 0.645547 | 0.02 | 0.009 | 1 | 1 | Hars2 | 0.011 | Young |
| 0.053301 | 0.073268 | 0.227 | 0.185 | 1 | 1 | Ep300 | 0.042 | Young |
| 0.053371 | 1.163493 | 0.03 | 0.017 | 1 | 1 | Nol12 | 0.013 | Young |
| 0.053437 | 0.862046 | 0.03 | 0.017 | 1 | 1 | Mcm4 | 0.013 | Young |
| 0.05344 | 0.648395 | 0.045 | 0.028 | 1 | 1 | Tmem71 | 0.017 | Young |
| 0.05365 | 0.067721 | 0.202 | 0.163 | 1 | 1 | Fam168b | 0.039 | Young |
| 0.053827 | 0.630091 | 0.061 | 0.041 | 1 | 1 | Nsrp1 | 0.02 | Young |
| 0.053828 | 0.337318 | 0.035 | 0.021 | 1 | 1 | Tigd2 | 0.014 | Young |
| 0.053871 | 0.077984 | 0.143 | 0.112 | 1 | 1 | Pdcd5 | 0.031 | Young |
| 0.054004 | 0.842985 | 0.03 | 0.017 | 1 | 1 | Ercc3 | 0.013 | Young |
| 0.054043 | 0.551008 | 0.086 | 0.063 | 1 | 1 | Parp8 | 0.023 | Young |

| | | | | | | | | |
|---|---|---|---|---|---|---|---|---|
| 0.054074 | 0.566529 | 0.077 | 0.055 | 1 | 1 | S100pbp | 0.022 | Young |
| 0.054124 | 0.205034 | 0.123 | 0.095 | 1 | 1 | Marf1 | 0.028 | Young |
| 0.054188 | 0.265208 | 0.064 | 0.044 | 1 | 1 | Ckap5 | 0.02 | Young |
| 0.054462 | 0.054808 | 0.415 | 0.355 | 1 | 1 | Rac1 | 0.06 | Young |
| 0.054729 | 0.348317 | 0.092 | 0.068 | 1 | 1 | Kif1c | 0.024 | Young |
| 0.054838 | 0.935612 | 0.025 | 0.013 | 1 | 1 | Mmp13 | 0.012 | Young |
| 0.054857 | 0.573017 | 0.045 | 0.028 | 1 | 1 | Gan | 0.017 | Young |
| 0.054907 | 0.208099 | 0.131 | 0.103 | 1 | 1 | Ppib | 0.028 | Young |
| 0.055006 | 0.165262 | 0.141 | 0.111 | 1 | 1 | Fmnl1 | 0.03 | Young |
| 0.055142 | 1.723949 | 0.014 | 0.005 | 1 | 1 | Zfp930 | 0.009 | Young |
| 0.055142 | 1.362374 | 0.014 | 0.005 | 1 | 1 | Numb | 0.009 | Young |
| 0.055209 | 0.266506 | 0.103 | 0.077 | 1 | 1 | Slc25a11 | 0.026 | Young |
| 0.055251 | 0.794686 | 0.025 | 0.013 | 1 | 1 | Dnajc25 | 0.012 | Young |
| 0.05527 | 0.680708 | 0.025 | 0.013 | 1 | 1 | Itpkc | 0.012 | Young |
| 0.055372 | 0.111052 | 0.393 | 0.344 | 1 | 1 | Tab2 | 0.049 | Young |
| 0.055413 | 0.085254 | 0.289 | 0.243 | 1 | 1 | Myo5a | 0.046 | Young |
| 0.055465 | 0.048167 | 0.45 | 0.386 | 1 | 1 | Ppp3ca | 0.064 | Young |
| 0.055596 | 0.089742 | 0.103 | 0.077 | 1 | 1 | Setd5 | 0.026 | Young |
| 0.055626 | 0.119126 | 0.063 | 0.042 | 1 | 1 | Slc38a6 | 0.021 | Young |
| 0.055815 | 0.25458 | 0.07 | 0.049 | 1 | 1 | Ilf2 | 0.021 | Young |
| 0.055838 | 0.333842 | 0.135 | 0.107 | 1 | 1 | Arid4a | 0.028 | Young |
| 0.055895 | 0.795592 | 0.025 | 0.013 | 1 | 1 | Gm5086 | 0.012 | Young |
| 0.055917 | 0.080655 | 0.518 | 0.451 | 1 | 1 | Ctsh | 0.067 | Young |
| 0.05592 | 1.071708 | 0.032 | 0.018 | 1 | 1 | Micu3 | 0.014 | Young |
| 0.055924 | 0.663363 | 0.014 | 0.005 | 1 | 1 | Med9 | 0.009 | Young |
| 0.056072 | 0.668969 | 0.048 | 0.031 | 1 | 1 | Fhl3 | 0.017 | Young |
| 0.056078 | 0.541958 | 0.03 | 0.017 | 1 | 1 | Ppa2 | 0.013 | Young |
| 0.056105 | 0.961286 | 0.04 | 0.024 | 1 | 1 | H2-Q6 | 0.016 | Young |
| 0.056123 | 0.567207 | 0.071 | 0.05 | 1 | 1 | Arid5b | 0.021 | Young |
| 0.056167 | 0.91088 | 0.052 | 0.035 | 1 | 1 | Rasa2 | 0.017 | Young |
| 0.056235 | 0.184016 | 0.066 | 0.045 | 1 | 1 | Senp7 | 0.021 | Young |
| 0.056248 | 0.713366 | 0.048 | 0.031 | 1 | 1 | Ppp2cb | 0.017 | Young |
| 0.056356 | 0.078875 | 0.45 | 0.386 | 1 | 1 | Naca | 0.064 | Young |
| 0.056358 | 1.721546 | 0.021 | 0.01 | 1 | 1 | Tmub1 | 0.011 | Young |
| 0.05642 | 0.47575 | 0.037 | 0.022 | 1 | 1 | Eepd1 | 0.015 | Young |
| 0.056449 | 0.138536 | 0.147 | 0.117 | 1 | 1 | Raly | 0.03 | Young |
| 0.056512 | 0.139485 | 0.131 | 0.102 | 1 | 1 | Ppp4r3b | 0.029 | Young |
| 0.056561 | 0.893334 | 0.032 | 0.018 | 1 | 1 | Cebpzos | 0.014 | Young |
| 0.056594 | 0.514499 | 0.047 | 0.03 | 1 | 1 | Pgap1 | 0.017 | Young |
| 0.056601 | 0.421237 | 0.062 | 0.042 | 1 | 1 | Ehmt1 | 0.02 | Young |
| 0.056606 | 0.475718 | 0.066 | 0.046 | 1 | 1 | Deptor | 0.02 | Young |
| 0.056678 | 0.34443 | 0.044 | 0.027 | 1 | 1 | Pibf1 | 0.017 | Young |
| 0.056796 | 0.326075 | 0.056 | 0.037 | 1 | 1 | Cnot7 | 0.019 | Young |
| 0.056818 | 0.206161 | 0.167 | 0.135 | 1 | 1 | Rpl13a | 0.032 | Young |
| 0.056832 | 0.850127 | 0.032 | 0.018 | 1 | 1 | Slc25a19 | 0.014 | Young |

| | | | | | | | | |
|---|---|---|---|---|---|---|---|---|
| 0.056847 | 0.166991 | 0.228 | 0.188 | 1 | 1 | 7-Sep | 0.04 | Young |
| 0.056986 | 1.328252 | 0.021 | 0.01 | 1 | 1 | Rubcnl | 0.011 | Young |
| 0.057087 | 0.536024 | 0.014 | 0.005 | 1 | 1 | Zbed3 | 0.009 | Young |
| 0.057139 | 1.844451 | 0.018 | 0.008 | 1 | 1 | Rundc1 | 0.01 | Young |
| 0.057209 | 1.385549 | 0.027 | 0.014 | 1 | 1 | Wdr18 | 0.013 | Young |
| 0.057379 | 0.098991 | 0.159 | 0.127 | 1 | 1 | Prpf4b | 0.032 | Young |
| 0.057397 | 0.895065 | 0.027 | 0.014 | 1 | 1 | Nup210 | 0.013 | Young |
| 0.057443 | 0.649512 | 0.049 | 0.032 | 1 | 1 | Cetn2 | 0.017 | Young |
| 0.05745 | 0.138398 | 0.175 | 0.141 | 1 | 1 | Brd4 | 0.034 | Young |
| 0.057509 | 0.455779 | 0.093 | 0.069 | 1 | 1 | Akna | 0.024 | Young |
| 0.057536 | 1.002832 | 0.021 | 0.01 | 1 | 1 | Zbtb45 | 0.011 | Young |
| 0.05757 | 0.410832 | 0.102 | 0.077 | 1 | 1 | Slc30a5 | 0.025 | Young |
| 0.057629 | 0.411956 | 0.062 | 0.042 | 1 | 1 | Arhgap4 | 0.02 | Young |
| 0.057692 | 0.41667 | 0.078 | 0.057 | 1 | 1 | Napg | 0.021 | Young |
| 0.057737 | 0.006976 | 0.752 | 0.671 | 1 | 1 | C1qb | 0.081 | Young |
| 0.057743 | 0.263595 | 0.15 | 0.12 | 1 | 1 | Mrps21 | 0.03 | Young |
| 0.057761 | 0.480923 | 0.048 | 0.031 | 1 | 1 | Ddx47 | 0.017 | Young |
| 0.057934 | 0.390132 | 0.062 | 0.042 | 1 | 1 | Siglech | 0.02 | Young |
| 0.058008 | 0.076353 | 0.177 | 0.143 | 1 | 1 | Dmac1 | 0.034 | Young |
| 0.058033 | 0.069691 | 0.186 | 0.152 | 1 | 1 | Nav1 | 0.034 | Young |
| 0.058367 | 1.547161 | 0.018 | 0.008 | 1 | 1 | Zc3h6 | 0.01 | Young |
| 0.058384 | 0.89122 | 0.027 | 0.014 | 1 | 1 | Sh2b1 | 0.013 | Young |
| 0.058477 | 0.667039 | 0.021 | 0.01 | 1 | 1 | Eva1b | 0.011 | Young |
| 0.058505 | 0.809594 | 0.041 | 0.026 | 1 | 1 | Cadm1 | 0.015 | Young |
| 0.058596 | 0.110604 | 0.287 | 0.244 | 1 | 1 | Man2b1 | 0.043 | Young |
| 0.058606 | 0.435889 | 0.021 | 0.01 | 1 | 1 | Fbxl15 | 0.011 | Young |
| 0.058608 | 0.359932 | 0.157 | 0.127 | 1 | 1 | Dusp2 | 0.03 | Young |
| 0.058644 | 0.89728 | 0.033 | 0.019 | 1 | 1 | Ranbp3 | 0.014 | Young |
| 0.058654 | 0.368799 | 0.092 | 0.068 | 1 | 1 | Ccdc82 | 0.024 | Young |
| 0.058721 | 0.774513 | 0.063 | 0.044 | 1 | 1 | Txndc15 | 0.019 | Young |
| 0.058815 | 0.234497 | 0.073 | 0.051 | 1 | 1 | Uhrf1bp1l | 0.022 | Young |
| 0.058914 | 0.057253 | 0.268 | 0.225 | 1 | 1 | 2900097C1 | 0.043 | Young |
| 0.058984 | 0.264829 | 0.134 | 0.105 | 1 | 1 | Srebf2 | 0.029 | Young |
| 0.059049 | 0.647159 | 0.04 | 0.024 | 1 | 1 | Tceanc2 | 0.016 | Young |
| 0.059204 | 0.528838 | 0.054 | 0.036 | 1 | 1 | Sap30 | 0.018 | Young |
| 0.05921 | 0.090665 | 0.436 | 0.382 | 1 | 1 | Sh3glb1 | 0.054 | Young |
| 0.059214 | 0.125018 | 0.047 | 0.03 | 1 | 1 | Ndrg3 | 0.017 | Young |
| 0.059256 | 0.307557 | 0.032 | 0.018 | 1 | 1 | Mcm3ap | 0.014 | Young |
| 0.059317 | 0.355963 | 0.065 | 0.045 | 1 | 1 | Lsm3 | 0.02 | Young |
| 0.059385 | 0.991398 | 0.027 | 0.014 | 1 | 1 | Parvg | 0.013 | Young |
| 0.059423 | 0.978075 | 0.018 | 0.008 | 1 | 1 | Sesn2 | 0.01 | Young |
| 0.059429 | 0.171955 | 0.17 | 0.138 | 1 | 1 | Sbf2 | 0.032 | Young |
| 0.059782 | 0.332344 | 0.077 | 0.055 | 1 | 1 | Fam107b | 0.022 | Young |
| 0.05981 | 1.179725 | 0.012 | 0.004 | 1 | 1 | Cfap45 | 0.008 | Young |
| 0.059878 | 0.361787 | 0.048 | 0.031 | 1 | 1 | Foxj3 | 0.017 | Young |

| | | | | | | | | |
|---|---|---|---|---|---|---|---|---|
| 0.059933 | 0.459041 | 0.033 | 0.019 | 1 | 1 | Map3k3 | 0.014 | Young |
| 0.060072 | 0.568149 | 0.058 | 0.04 | 1 | 1 | Ccdc124 | 0.018 | Young |
| 0.060117 | 0.238707 | 0.097 | 0.072 | 1 | 1 | C330007P0 | 0.025 | Young |
| 0.060315 | 1.050319 | 0.035 | 0.021 | 1 | 1 | Sdhaf2 | 0.014 | Young |
| 0.060363 | 0.32037 | 0.074 | 0.053 | 1 | 1 | Csnk1g1 | 0.021 | Young |
| 0.060384 | 0.152353 | 0.167 | 0.134 | 1 | 1 | H1f0 | 0.033 | Young |
| 0.06049 | 0.833878 | 0.033 | 0.019 | 1 | 1 | Mrpl45 | 0.014 | Young |
| 0.060519 | 0.166107 | 0.079 | 0.057 | 1 | 1 | Srek1 | 0.022 | Young |
| 0.060609 | 0.069522 | 0.864 | 0.803 | 1 | 1 | Rps28 | 0.061 | Young |
| 0.060654 | 0.402967 | 0.062 | 0.042 | 1 | 1 | Spast | 0.02 | Young |
| 0.060659 | 0.423637 | 0.041 | 0.026 | 1 | 1 | Zfp626 | 0.015 | Young |
| 0.060668 | 0.16293 | 0.012 | 0.004 | 1 | 1 | Chst11 | 0.008 | Young |
| 0.060769 | 0.068109 | 0.349 | 0.301 | 1 | 1 | Hnrnpa0 | 0.048 | Young |
| 0.061044 | 0.0027 | 0.554 | 0.482 | 1 | 1 | Ier5 | 0.072 | Young |
| 0.061119 | 0.492644 | 0.091 | 0.068 | 1 | 1 | Rsf1 | 0.023 | Young |
| 0.061158 | 0.281684 | 0.097 | 0.073 | 1 | 1 | Med15 | 0.024 | Young |
| 0.061183 | 0.381354 | 0.04 | 0.024 | 1 | 1 | Arhgap21 | 0.016 | Young |
| 0.06124 | 0.238419 | 0.151 | 0.121 | 1 | 1 | Stard3nl | 0.03 | Young |
| 0.061273 | 1.768815 | 0.01 | 0.003 | 1 | 1 | Zfp764 | 0.007 | Young |
| 0.06135 | 1.818938 | 0.023 | 0.012 | 1 | 1 | Comtd1 | 0.011 | Young |
| 0.061399 | 0.431931 | 0.051 | 0.033 | 1 | 1 | Brip1os | 0.018 | Young |
| 0.061413 | 0.324004 | 0.051 | 0.033 | 1 | 1 | Tmem243 | 0.018 | Young |
| 0.061433 | 0.229504 | 0.285 | 0.246 | 1 | 1 | Tgfb1 | 0.039 | Young |
| 0.061451 | 0.624569 | 0.043 | 0.027 | 1 | 1 | Pias4 | 0.016 | Young |
| 0.061483 | 1.593007 | 0.01 | 0.003 | 1 | 1 | Col27a1 | 0.007 | Young |
| 0.061496 | 0.153212 | 0.069 | 0.049 | 1 | 1 | Spin1 | 0.02 | Young |
| 0.061624 | 1.378651 | 0.01 | 0.003 | 1 | 1 | Kif20b | 0.007 | Young |
| 0.061694 | 1.177325 | 0.01 | 0.003 | 1 | 1 | 1700037C1 | 0.007 | Young |
| 0.061707 | 0.11453 | 0.216 | 0.179 | 1 | 1 | Leng8 | 0.037 | Young |
| 0.061996 | 0.660255 | 0.043 | 0.027 | 1 | 1 | Erf | 0.016 | Young |
| 0.062008 | 0.339593 | 0.136 | 0.108 | 1 | 1 | Baz2b | 0.028 | Young |
| 0.062024 | 0.146773 | 0.22 | 0.184 | 1 | 1 | Abhd12 | 0.036 | Young |
| 0.062126 | 0.301893 | 0.051 | 0.033 | 1 | 1 | Dda1 | 0.018 | Young |
| 0.062178 | 0.145361 | 0.137 | 0.108 | 1 | 1 | Ndufb8 | 0.029 | Young |
| 0.062194 | 0.360246 | 0.081 | 0.059 | 1 | 1 | Svil | 0.022 | Young |
| 0.062238 | 0.146652 | 0.117 | 0.09 | 1 | 1 | Smarcc1 | 0.027 | Young |
| 0.062329 | 0.602385 | 0.01 | 0.003 | 1 | 1 | Mmaa | 0.007 | Young |
| 0.062348 | 1.130348 | 0.023 | 0.012 | 1 | 1 | Mier2 | 0.011 | Young |
| 0.0624 | 0.589064 | 0.01 | 0.003 | 1 | 1 | Arhgap35 | 0.007 | Young |
| 0.062474 | 0.083965 | 0.136 | 0.107 | 1 | 1 | Snrpc | 0.029 | Young |
| 0.062636 | 0.398841 | 0.055 | 0.037 | 1 | 1 | Pggt1b | 0.018 | Young |
| 0.062687 | 0.184033 | 0.109 | 0.084 | 1 | 1 | Pkn2 | 0.025 | Young |
| 0.062856 | 0.031117 | 0.347 | 0.293 | 1 | 1 | Sf3b1 | 0.054 | Young |
| 0.062973 | 0.786808 | 0.046 | 0.03 | 1 | 1 | Abcd2 | 0.016 | Young |
| 0.06302 | 0.563759 | 0.035 | 0.021 | 1 | 1 | Cbll1 | 0.014 | Young |

| | | | | | | | | |
|---|---|---|---|---|---|---|---|---|
| 0.063187 | 0.245539 | 0.146 | 0.116 | 1 | 1 | Entpd1 | 0.03 | Young |
| 0.063192 | 0.220775 | 0.051 | 0.033 | 1 | 1 | Adam8 | 0.018 | Young |
| 0.06342 | 0.635912 | 0.028 | 0.015 | 1 | 1 | Alpk1 | 0.013 | Young |
| 0.063534 | 0.077635 | 0.224 | 0.184 | 1 | 1 | Hmgn1 | 0.04 | Young |
| 0.063546 | 0.120332 | 0.214 | 0.177 | 1 | 1 | Tfe3 | 0.037 | Young |
| 0.063595 | 0.091819 | 0.18 | 0.145 | 1 | 1 | Slc7a8 | 0.035 | Young |
| 0.063618 | 0.085394 | 0.33 | 0.281 | 1 | 1 | Fyb | 0.049 | Young |
| 0.063625 | 0.858344 | 0.023 | 0.012 | 1 | 1 | Fuom | 0.011 | Young |
| 0.063703 | 0.243259 | 0.151 | 0.121 | 1 | 1 | Pla2g15 | 0.03 | Young |
| 0.063711 | 0.300874 | 0.087 | 0.064 | 1 | 1 | Pih1d1 | 0.023 | Young |
| 0.063845 | 1.370814 | 0.03 | 0.017 | 1 | 1 | 2410089E0 | 0.013 | Young |
| 0.06385 | 0.150538 | 0.228 | 0.192 | 1 | 1 | Ube2b | 0.036 | Young |
| 0.063885 | 0.245916 | 0.129 | 0.102 | 1 | 1 | Akirin2 | 0.027 | Young |
| 0.063981 | 0.789938 | 0.023 | 0.012 | 1 | 1 | Ankmy2 | 0.011 | Young |
| 0.064049 | 0.241691 | 0.1 | 0.076 | 1 | 1 | Lpin2 | 0.024 | Young |
| 0.064122 | 0.609147 | 0.063 | 0.044 | 1 | 1 | Ppp2r5e | 0.019 | Young |
| 0.064226 | 0.149587 | 0.17 | 0.139 | 1 | 1 | Fgd4 | 0.031 | Young |
| 0.064235 | 0.177101 | 0.244 | 0.206 | 1 | 1 | Stard8 | 0.038 | Young |
| 0.064263 | 0.290881 | 0.141 | 0.114 | 1 | 1 | Rela | 0.027 | Young |
| 0.064317 | 0.057386 | 0.199 | 0.163 | 1 | 1 | Eif5 | 0.036 | Young |
| 0.064447 | 0.231537 | 0.097 | 0.073 | 1 | 1 | Rsbn1l | 0.024 | Young |
| 0.064512 | 0.015352 | 0.168 | 0.134 | 1 | 1 | Trpc4ap | 0.034 | Young |
| 0.064607 | 0.436033 | 0.097 | 0.073 | 1 | 1 | Wdr37 | 0.024 | Young |
| 0.064661 | 0.645062 | 0.035 | 0.021 | 1 | 1 | Pld2 | 0.014 | Young |
| 0.064782 | 0.242642 | 0.068 | 0.048 | 1 | 1 | Msi2 | 0.02 | Young |
| 0.065046 | 0.639837 | 0.028 | 0.015 | 1 | 1 | Maml3 | 0.013 | Young |
| 0.065054 | 0.456463 | 0.043 | 0.027 | 1 | 1 | Smim20 | 0.016 | Young |
| 0.065068 | 1.277491 | 0.016 | 0.006 | 1 | 1 | Abi3 | 0.01 | Young |
| 0.065124 | 1.232151 | 0.016 | 0.006 | 1 | 1 | Gpr141 | 0.01 | Young |
| 0.065393 | 0.500323 | 0.046 | 0.03 | 1 | 1 | Cnot10 | 0.016 | Young |
| 0.065478 | 0.038727 | 0.215 | 0.176 | 1 | 1 | 1700020I1 | 0.039 | Young |
| 0.065525 | 1.179702 | 0.019 | 0.009 | 1 | 1 | Tbc1d16 | 0.01 | Young |
| 0.065638 | 0.26123 | 0.115 | 0.09 | 1 | 1 | Csnk1g3 | 0.025 | Young |
| 0.065659 | 0.413566 | 0.092 | 0.069 | 1 | 1 | Cand1 | 0.023 | Young |
| 0.065825 | 0.365418 | 0.095 | 0.072 | 1 | 1 | Ttc39b | 0.023 | Young |
| 0.065897 | 0.292377 | 0.095 | 0.071 | 1 | 1 | Sptlc2 | 0.024 | Young |
| 0.065912 | 0.93099 | 0.016 | 0.006 | 1 | 1 | Zfp580 | 0.01 | Young |
| 0.065968 | 1.020898 | 0.016 | 0.006 | 1 | 1 | Bckdhb | 0.01 | Young |
| 0.065986 | 0.359115 | 0.067 | 0.048 | 1 | 1 | Rps6ka4 | 0.019 | Young |
| 0.066025 | 0.800881 | 0.016 | 0.006 | 1 | 1 | Gtf2h2 | 0.01 | Young |
| 0.066178 | 0.685148 | 0.019 | 0.009 | 1 | 1 | H2afx | 0.01 | Young |
| 0.066268 | 0.321372 | 0.057 | 0.039 | 1 | 1 | Suclg2 | 0.018 | Young |
| 0.066421 | 0.492259 | 0.061 | 0.042 | 1 | 1 | Snx12 | 0.019 | Young |
| 0.066428 | 0.693028 | 0.055 | 0.037 | 1 | 1 | Smim4 | 0.018 | Young |
| 0.066476 | 0.299342 | 0.095 | 0.072 | 1 | 1 | Rbck1 | 0.023 | Young |

| | | | | | | | | |
|---|---|---|---|---|---|---|---|---|
| 0.066496 | 0.25244 | 0.043 | 0.027 | 1 | 1 | Vhl | 0.016 | Young |
| 0.066533 | 0.089741 | 0.142 | 0.112 | 1 | 1 | 1-Mar | 0.03 | Young |
| 0.066661 | 0.301491 | 0.159 | 0.13 | 1 | 1 | Specc1l | 0.029 | Young |
| 0.066692 | 0.10836 | 0.09 | 0.067 | 1 | 1 | Hk2 | 0.023 | Young |
| 0.066703 | 0.20912 | 0.078 | 0.057 | 1 | 1 | Atg2a | 0.021 | Young |
| 0.066709 | 0.17762 | 0.153 | 0.123 | 1 | 1 | Hectd1 | 0.03 | Young |
| 0.066836 | 0.014179 | 0.541 | 0.468 | 1 | 1 | Arpc1b | 0.073 | Young |
| 0.066892 | 0.187242 | 0.208 | 0.174 | 1 | 1 | Per1 | 0.034 | Young |
| 0.066925 | 0.521022 | 0.078 | 0.057 | 1 | 1 | Idh3g | 0.021 | Young |
| 0.06728 | 0.528387 | 0.016 | 0.006 | 1 | 1 | Diras2 | 0.01 | Young |
| 0.0673 | 0.317001 | 0.157 | 0.129 | 1 | 1 | Rab11fip5 | 0.028 | Young |
| 0.067311 | 0.360195 | 0.072 | 0.051 | 1 | 1 | Fam50a | 0.021 | Young |
| 0.067522 | 0.498711 | 0.086 | 0.064 | 1 | 1 | Cnot3 | 0.022 | Young |
| 0.0677 | 0.775087 | 0.039 | 0.024 | 1 | 1 | G6pdx | 0.015 | Young |
| 0.067752 | 0.908869 | 0.024 | 0.013 | 1 | 1 | Pdpr | 0.011 | Young |
| 0.067797 | 0.980704 | 0.024 | 0.013 | 1 | 1 | Nfrkb | 0.011 | Young |
| 0.06781 | 0.119486 | 0.057 | 0.039 | 1 | 1 | Oard1 | 0.018 | Young |
| 0.067825 | 0.02864 | 0.035 | 0.021 | 1 | 1 | Fgfr1op | 0.014 | Young |
| 0.067826 | 0.321928 | 0.03 | 0.017 | 1 | 1 | Lrwd1 | 0.013 | Young |
| 0.067842 | 1.065564 | 0.024 | 0.013 | 1 | 1 | Zfp810 | 0.011 | Young |
| 0.067962 | 0.427866 | 0.019 | 0.009 | 1 | 1 | Tma16 | 0.01 | Young |
| 0.068069 | 0.007146 | 0.071 | 0.05 | 1 | 1 | Mprip | 0.021 | Young |
| 0.068102 | 0.171822 | 0.182 | 0.15 | 1 | 1 | Nsd1 | 0.032 | Young |
| 0.06842 | 0.213664 | 0.134 | 0.105 | 1 | 1 | Eea1 | 0.029 | Young |
| 0.068597 | 0.255677 | 0.136 | 0.108 | 1 | 1 | Snx17 | 0.028 | Young |
| 0.068844 | 0.306397 | 0.078 | 0.057 | 1 | 1 | Cfap43 | 0.021 | Young |
| 0.069039 | 0.136723 | 0.137 | 0.108 | 1 | 1 | Sh3pxd2b | 0.029 | Young |
| 0.069051 | 0.303562 | 0.088 | 0.066 | 1 | 1 | Ptpn11 | 0.022 | Young |
| 0.06907 | 0.567365 | 0.058 | 0.04 | 1 | 1 | Rnf103 | 0.018 | Young |
| 0.06917 | 0.222689 | 0.123 | 0.096 | 1 | 1 | Man2a2 | 0.027 | Young |
| 0.069202 | 0.325299 | 0.086 | 0.064 | 1 | 1 | 2410015M | 0.022 | Young |
| 0.069277 | 0.826369 | 0.039 | 0.024 | 1 | 1 | Vkorc1l1 | 0.015 | Young |
| 0.069392 | 0.533644 | 0.038 | 0.023 | 1 | 1 | Pde4dip | 0.015 | Young |
| 0.069436 | 0.551191 | 0.052 | 0.035 | 1 | 1 | Txndc12 | 0.017 | Young |
| 0.069454 | 1.070954 | 0.042 | 0.027 | 1 | 1 | Dzip3 | 0.015 | Young |
| 0.069493 | 0.183792 | 0.171 | 0.141 | 1 | 1 | Kat6a | 0.03 | Young |
| 0.0695 | 0.068503 | 0.069 | 0.049 | 1 | 1 | Fam126b | 0.02 | Young |
| 0.069664 | 0.327285 | 0.165 | 0.135 | 1 | 1 | Wls | 0.03 | Young |
| 0.069716 | 0.048876 | 0.185 | 0.152 | 1 | 1 | Tgif1 | 0.033 | Young |
| 0.069782 | 0.147894 | 0.081 | 0.059 | 1 | 1 | Nsd2 | 0.022 | Young |
| 0.069834 | 0.415317 | 0.066 | 0.046 | 1 | 1 | Rasgrp3 | 0.02 | Young |
| 0.069855 | 0.50065 | 0.049 | 0.032 | 1 | 1 | Slc35f5 | 0.017 | Young |
| 0.06988 | 0.185663 | 0.091 | 0.068 | 1 | 1 | Igsf9 | 0.023 | Young |
| 0.069928 | 0.078579 | 0.11 | 0.084 | 1 | 1 | Pi4k2a | 0.026 | Young |
| 0.07004 | 0.042005 | 0.138 | 0.108 | 1 | 1 | Mob3c | 0.03 | Young |

| | | | | | | | | |
|---|---|---|---|---|---|---|---|---|
| 0.070283 | 0.26427 | 0.097 | 0.073 | 1 | 1 | Ptov1 | 0.024 | Young |
| 0.070444 | 0.250901 | 0.064 | 0.045 | 1 | 1 | Dnajc15 | 0.019 | Young |
| 0.070529 | 0.245158 | 0.046 | 0.03 | 1 | 1 | Pde7a | 0.016 | Young |
| 0.070691 | 0.141214 | 0.089 | 0.066 | 1 | 1 | Uqcc3 | 0.023 | Young |
| 0.070758 | 0.024488 | 0.261 | 0.219 | 1 | 1 | Far1 | 0.042 | Young |
| 0.070894 | 0.595062 | 0.065 | 0.046 | 1 | 1 | Mlxip | 0.019 | Young |
| 0.071146 | 0.305379 | 0.073 | 0.053 | 1 | 1 | Angel2 | 0.02 | Young |
| 0.071302 | 0.247239 | 0.024 | 0.013 | 1 | 1 | Rnf8 | 0.011 | Young |
| 0.071418 | 0.704199 | 0.041 | 0.026 | 1 | 1 | Pik3r4 | 0.015 | Young |
| 0.071446 | 0.065566 | 0.272 | 0.231 | 1 | 1 | Hmgb1 | 0.041 | Young |
| 0.071534 | 0.907059 | 0.032 | 0.019 | 1 | 1 | Fbxl17 | 0.013 | Young |
| 0.071554 | 0.747916 | 0.032 | 0.019 | 1 | 1 | Rora | 0.013 | Young |
| 0.071572 | 1.342561 | 0.021 | 0.01 | 1 | 1 | Rad51b | 0.011 | Young |
| 0.071594 | 0.012765 | 0.18 | 0.145 | 1 | 1 | Lasp1 | 0.035 | Young |
| 0.071718 | 0.123697 | 0.157 | 0.126 | 1 | 1 | Birc6 | 0.031 | Young |
| 0.071908 | 0.292981 | 0.108 | 0.084 | 1 | 1 | Gmip | 0.024 | Young |
| 0.071915 | 1.820707 | 0.013 | 0.005 | 1 | 1 | A630072M | 0.008 | Young |
| 0.071933 | 1.234954 | 0.021 | 0.01 | 1 | 1 | Pnpo | 0.011 | Young |
| 0.072113 | 1.870584 | 0.013 | 0.005 | 1 | 1 | Zfp276 | 0.008 | Young |
| 0.072133 | 0.225641 | 0.092 | 0.069 | 1 | 1 | Trappc6b | 0.023 | Young |
| 0.072378 | 1.605828 | 0.013 | 0.005 | 1 | 1 | Mypopos | 0.008 | Young |
| 0.072391 | 0.158198 | 0.129 | 0.102 | 1 | 1 | Setd2 | 0.027 | Young |
| 0.07247 | 0.089112 | 0.061 | 0.042 | 1 | 1 | Snx24 | 0.019 | Young |
| 0.072568 | 0.081972 | 0.909 | 0.861 | 1 | 1 | mt-Nd2 | 0.048 | Young |
| 0.072577 | 1.248835 | 0.013 | 0.005 | 1 | 1 | Trp53i13 | 0.008 | Young |
| 0.07266 | 1.104046 | 0.021 | 0.01 | 1 | 1 | Zfp157 | 0.011 | Young |
| 0.072686 | 0.087268 | 0.053 | 0.036 | 1 | 1 | Zfp120 | 0.017 | Young |
| 0.072699 | 0.408191 | 0.114 | 0.09 | 1 | 1 | Fam91a1 | 0.024 | Young |
| 0.07328 | 0.018877 | 0.086 | 0.063 | 1 | 1 | Bdp1 | 0.023 | Young |
| 0.073333 | 0.772374 | 0.032 | 0.019 | 1 | 1 | Vps16 | 0.013 | Young |
| 0.073376 | 0.291767 | 0.091 | 0.068 | 1 | 1 | Pwwp2a | 0.023 | Young |
| 0.073448 | 0.195954 | 0.123 | 0.096 | 1 | 1 | Ints6l | 0.027 | Young |
| 0.073512 | 1.087657 | 0.013 | 0.005 | 1 | 1 | Pqlc2 | 0.008 | Young |
| 0.073513 | 0.376059 | 0.089 | 0.067 | 1 | 1 | 2810403A( | 0.022 | Young |
| 0.073773 | 0.834098 | 0.047 | 0.031 | 1 | 1 | Mrps16 | 0.016 | Young |
| 0.073781 | 1.050515 | 0.013 | 0.005 | 1 | 1 | Ada | 0.008 | Young |
| 0.073984 | 0.702228 | 0.013 | 0.005 | 1 | 1 | Pyroxd1 | 0.008 | Young |
| 0.074123 | 0.043293 | 0.345 | 0.296 | 1 | 1 | Ncf2 | 0.049 | Young |
| 0.074133 | 0.154854 | 0.111 | 0.086 | 1 | 1 | Nmt1 | 0.025 | Young |
| 0.074134 | 0.244906 | 0.097 | 0.073 | 1 | 1 | Csgalnact2 | 0.024 | Young |
| 0.074193 | 0.282824 | 0.128 | 0.102 | 1 | 1 | Pabpn1 | 0.026 | Young |
| 0.074371 | 0.261777 | 0.053 | 0.036 | 1 | 1 | Stim1 | 0.017 | Young |
| 0.074385 | 1.286946 | 0.017 | 0.008 | 1 | 1 | Ubxn2b | 0.009 | Young |
| 0.07439 | 0.245032 | 0.052 | 0.035 | 1 | 1 | Trappc5 | 0.017 | Young |
| 0.074399 | 0.41175 | 0.021 | 0.01 | 1 | 1 | Rpp30 | 0.011 | Young |

| | | | | | | | | |
|---|---|---|---|---|---|---|---|---|
| 0.074402 | 0.165991 | 0.168 | 0.136 | 1 | 1 | Atp13a3 | 0.032 | Young |
| 0.074457 | 0.60278 | 0.013 | 0.005 | 1 | 1 | Selenoo | 0.008 | Young |
| 0.074607 | 0.52643 | 0.026 | 0.014 | 1 | 1 | Rnaseh2b | 0.012 | Young |
| 0.074698 | 0.29362 | 0.151 | 0.123 | 1 | 1 | Polr2a | 0.028 | Young |
| 0.07474 | 0.930897 | 0.017 | 0.008 | 1 | 1 | Clybl | 0.009 | Young |
| 0.074973 | 0.470983 | 0.066 | 0.048 | 1 | 1 | Scoc | 0.018 | Young |
| 0.075038 | 1.167892 | 0.017 | 0.008 | 1 | 1 | Timd4 | 0.009 | Young |
| 0.075107 | 0.305685 | 0.056 | 0.039 | 1 | 1 | Nsmaf | 0.017 | Young |
| 0.075244 | 0.019524 | 0.198 | 0.162 | 1 | 1 | Ndufb7 | 0.036 | Young |
| 0.075396 | 0.764267 | 0.017 | 0.008 | 1 | 1 | Fam210b | 0.009 | Young |
| 0.075523 | 0.324983 | 0.021 | 0.01 | 1 | 1 | Wdfy1 | 0.011 | Young |
| 0.075614 | 0.614975 | 0.044 | 0.028 | 1 | 1 | Trim36 | 0.016 | Young |
| 0.075635 | 0.841057 | 0.017 | 0.008 | 1 | 1 | Ccdc126 | 0.009 | Young |
| 0.075649 | 0.203876 | 0.281 | 0.242 | 1 | 1 | Lifr | 0.039 | Young |
| 0.07582 | 0.094203 | 0.066 | 0.046 | 1 | 1 | Cnp | 0.02 | Young |
| 0.075915 | 0.405375 | 0.062 | 0.044 | 1 | 1 | Pld1 | 0.018 | Young |
| 0.075915 | 0.599403 | 0.045 | 0.03 | 1 | 1 | Osgin2 | 0.015 | Young |
| 0.076072 | 0.174313 | 0.149 | 0.121 | 1 | 1 | Atxn2l | 0.028 | Young |
| 0.076151 | 0.139693 | 0.16 | 0.13 | 1 | 1 | Nup98 | 0.03 | Young |
| 0.076194 | 0.553019 | 0.068 | 0.049 | 1 | 1 | Zbtb18 | 0.019 | Young |
| 0.076206 | 0.330385 | 0.032 | 0.019 | 1 | 1 | Nelfb | 0.013 | Young |
| 0.076252 | 0.025497 | 0.095 | 0.072 | 1 | 1 | Mlec | 0.023 | Young |
| 0.076265 | 0.282305 | 0.109 | 0.085 | 1 | 1 | Ndufs5 | 0.024 | Young |
| 0.076361 | 0.40087 | 0.071 | 0.051 | 1 | 1 | Clip1 | 0.02 | Young |
| 0.076376 | 0.109438 | 0.202 | 0.168 | 1 | 1 | Hnrnpl | 0.034 | Young |
| 0.076379 | 0.022739 | 0.182 | 0.149 | 1 | 1 | Atp6v0d1 | 0.033 | Young |
| 0.07639 | 0.172337 | 0.193 | 0.161 | 1 | 1 | D8Ertd738 | 0.032 | Young |
| 0.076401 | 0.752006 | 0.035 | 0.022 | 1 | 1 | Zfp771 | 0.013 | Young |
| 0.076464 | 0.05653 | 0.443 | 0.383 | 1 | 1 | Vmp1 | 0.06 | Young |
| 0.076516 | 0.361758 | 0.084 | 0.063 | 1 | 1 | Hmgcs1 | 0.021 | Young |
| 0.076552 | 0.570189 | 0.092 | 0.071 | 1 | 1 | Txndc9 | 0.021 | Young |
| 0.076644 | 0.372651 | 0.1 | 0.077 | 1 | 1 | Ankrd13c | 0.023 | Young |
| 0.076779 | 0.310668 | 0.105 | 0.081 | 1 | 1 | Psmb7 | 0.024 | Young |
| 0.076822 | 0.975374 | 0.037 | 0.023 | 1 | 1 | Ginm1 | 0.014 | Young |
| 0.076873 | 0.403453 | 0.087 | 0.066 | 1 | 1 | Crybg3 | 0.021 | Young |
| 0.076886 | 0.320344 | 0.105 | 0.081 | 1 | 1 | Smarca2 | 0.024 | Young |
| 0.07693 | 0.312418 | 0.138 | 0.112 | 1 | 1 | Ncstn | 0.026 | Young |
| 0.076966 | 0.247471 | 0.053 | 0.036 | 1 | 1 | Stard4 | 0.017 | Young |
| 0.076997 | 0.053115 | 0.239 | 0.201 | 1 | 1 | G3bp1 | 0.038 | Young |
| 0.077002 | 0.423898 | 0.083 | 0.062 | 1 | 1 | Abcg1 | 0.021 | Young |
| 0.077294 | 0.13268 | 0.225 | 0.19 | 1 | 1 | Tsc22d4 | 0.035 | Young |
| 0.077298 | 0.996778 | 0.04 | 0.026 | 1 | 1 | Trpm2 | 0.014 | Young |
| 0.07734 | 0.655202 | 0.049 | 0.033 | 1 | 1 | Mfap1b | 0.016 | Young |
| 0.077595 | 0.46571 | 0.069 | 0.05 | 1 | 1 | Nbr1 | 0.019 | Young |
| 0.077636 | 1.077458 | 0.029 | 0.017 | 1 | 1 | Ubr7 | 0.012 | Young |

| | | | | | | | | |
|---|---|---|---|---|---|---|---|---|
| 0.077671 | 0.098123 | 0.211 | 0.176 | 1 | 1 | Capza1 | 0.035 | Young |
| 0.077764 | 0.302773 | 0.131 | 0.105 | 1 | 1 | Wbp2 | 0.026 | Young |
| 0.077838 | 0.913375 | 0.035 | 0.022 | 1 | 1 | Fbxo9 | 0.013 | Young |
| 0.078064 | 0.598754 | 0.075 | 0.055 | 1 | 1 | Trnau1ap | 0.02 | Young |
| 0.078083 | 0.432827 | 0.159 | 0.132 | 1 | 1 | Rab6a | 0.027 | Young |
| 0.078227 | 0.145732 | 0.088 | 0.066 | 1 | 1 | Hcst | 0.022 | Young |
| 0.07826 | 0.335358 | 0.077 | 0.057 | 1 | 1 | F630028O1 | 0.02 | Young |
| 0.078304 | 0.235064 | 0.027 | 0.015 | 1 | 1 | Med4 | 0.012 | Young |
| 0.078341 | 1.061632 | 0.022 | 0.012 | 1 | 1 | Mcm2 | 0.01 | Young |
| 0.078362 | 0.070713 | 0.15 | 0.12 | 1 | 1 | Fnip1 | 0.03 | Young |
| 0.078447 | 0.07182 | 0.027 | 0.015 | 1 | 1 | Zfp513 | 0.012 | Young |
| 0.078541 | 0.511682 | 0.048 | 0.032 | 1 | 1 | Zfp260 | 0.016 | Young |
| 0.078553 | 0.498674 | 0.085 | 0.064 | 1 | 1 | Zcrb1 | 0.021 | Young |
| 0.078616 | 1.828113 | 0.011 | 0.004 | 1 | 1 | Txnrd3 | 0.007 | Young |
| 0.078681 | 0.144428 | 0.092 | 0.069 | 1 | 1 | Ube2g2 | 0.023 | Young |
| 0.078696 | 0.509786 | 0.034 | 0.021 | 1 | 1 | Rnf157 | 0.013 | Young |
| 0.078869 | 0.297483 | 0.017 | 0.008 | 1 | 1 | Hpn | 0.009 | Young |
| 0.078876 | 0.930409 | 0.022 | 0.012 | 1 | 1 | Dus3l | 0.01 | Young |
| 0.078929 | 0.075332 | 0.234 | 0.195 | 1 | 1 | Prpf38b | 0.039 | Young |
| 0.078993 | 0.105601 | 0.123 | 0.096 | 1 | 1 | Agps | 0.027 | Young |
| 0.079095 | 0.304621 | 0.074 | 0.054 | 1 | 1 | Foxo3 | 0.02 | Young |
| 0.079268 | 0.469335 | 0.045 | 0.03 | 1 | 1 | Kdm4c | 0.015 | Young |
| 0.079481 | 0.105457 | 0.096 | 0.073 | 1 | 1 | Slc38a1 | 0.023 | Young |
| 0.079577 | 0.230901 | 0.1 | 0.077 | 1 | 1 | Erp44 | 0.023 | Young |
| 0.07971 | 0.620178 | 0.054 | 0.037 | 1 | 1 | Aurkaip1 | 0.017 | Young |
| 0.079914 | 0.740557 | 0.037 | 0.023 | 1 | 1 | Hira | 0.014 | Young |
| 0.079958 | 0.87723 | 0.011 | 0.004 | 1 | 1 | Arhgap27 | 0.007 | Young |
| 0.080117 | 1.163024 | 0.011 | 0.004 | 1 | 1 | Zfp995 | 0.007 | Young |
| 0.080239 | 0.111132 | 0.075 | 0.054 | 1 | 1 | Tob1 | 0.021 | Young |
| 0.080277 | 0.03959 | 0.17 | 0.138 | 1 | 1 | Tmed9 | 0.032 | Young |
| 0.080394 | 0.629155 | 0.035 | 0.022 | 1 | 1 | Exosc5 | 0.013 | Young |
| 0.080588 | 0.187177 | 0.059 | 0.041 | 1 | 1 | Sipa1l1 | 0.018 | Young |
| 0.080595 | 0.467245 | 0.011 | 0.004 | 1 | 1 | Znrf3 | 0.007 | Young |
| 0.080801 | 0.032263 | 0.28 | 0.239 | 1 | 1 | Lyn | 0.041 | Young |
| 0.081102 | 0.552838 | 0.083 | 0.063 | 1 | 1 | Slfn8 | 0.02 | Young |
| 0.081162 | 0.473874 | 0.052 | 0.036 | 1 | 1 | Atmin | 0.016 | Young |
| 0.081262 | 0.433022 | 0.049 | 0.033 | 1 | 1 | Kmt2b | 0.016 | Young |
| 0.08132 | 0.45609 | 0.022 | 0.012 | 1 | 1 | Wrnip1 | 0.01 | Young |
| 0.081403 | 1.688493 | 0.018 | 0.009 | 1 | 1 | Gtf2h4 | 0.009 | Young |
| 0.081411 | 0.711442 | 0.029 | 0.017 | 1 | 1 | Ankrd49 | 0.012 | Young |
| 0.081539 | 0.638985 | 0.037 | 0.023 | 1 | 1 | Zfyve27 | 0.014 | Young |
| 0.081685 | 0.268855 | 0.066 | 0.048 | 1 | 1 | Cwf19l2 | 0.018 | Young |
| 0.081769 | 0.364591 | 0.092 | 0.071 | 1 | 1 | Dync1li2 | 0.021 | Young |
| 0.082347 | 0.231983 | 0.162 | 0.134 | 1 | 1 | Ciita | 0.028 | Young |
| 0.082529 | 0.043183 | 0.134 | 0.105 | 1 | 1 | Pet100 | 0.029 | Young |

| | | | | | | | | |
|---|---|---|---|---|---|---|---|---|
| 0.082676 | 0.269755 | 0.072 | 0.053 | 1 | 1 | Map1lc3a | 0.019 | Young |
| 0.082758 | 0.302677 | 0.08 | 0.059 | 1 | 1 | Dlg1 | 0.021 | Young |
| 0.082759 | 0.931112 | 0.024 | 0.013 | 1 | 1 | Pafah1b3 | 0.011 | Young |
| 0.082796 | 0.829304 | 0.03 | 0.018 | 1 | 1 | Kif23 | 0.012 | Young |
| 0.082859 | 0.262346 | 0.077 | 0.057 | 1 | 1 | Iws1 | 0.02 | Young |
| 0.082886 | 0.185543 | 0.275 | 0.239 | 1 | 1 | Dnaja1 | 0.036 | Young |
| 0.082891 | 0.117539 | 0.229 | 0.194 | 1 | 1 | Lamtor4 | 0.035 | Young |
| 0.083174 | 0.78604 | 0.03 | 0.018 | 1 | 1 | Nmi | 0.012 | Young |
| 0.083239 | 1.130452 | 0.018 | 0.009 | 1 | 1 | Mms22l | 0.009 | Young |
| 0.083291 | 0.048142 | 0.151 | 0.122 | 1 | 1 | Ost4 | 0.029 | Young |
| 0.083319 | 0.098625 | 0.099 | 0.076 | 1 | 1 | Lgals9 | 0.023 | Young |
| 0.083388 | 1.775756 | 0.015 | 0.006 | 1 | 1 | Trit1 | 0.009 | Young |
| 0.083415 | 0.572998 | 0.04 | 0.026 | 1 | 1 | Tmem214 | 0.014 | Young |
| 0.083528 | 1.041908 | 0.015 | 0.006 | 1 | 1 | Rpap1 | 0.009 | Young |
| 0.08355 | 0.103114 | 0.238 | 0.201 | 1 | 1 | Ctnnb1 | 0.037 | Young |
| 0.08356 | 0.259442 | 0.065 | 0.046 | 1 | 1 | Trim5 | 0.019 | Young |
| 0.083704 | 0.552131 | 0.038 | 0.024 | 1 | 1 | Ggta1 | 0.014 | Young |
| 0.083839 | 0.633258 | 0.03 | 0.018 | 1 | 1 | Snta1 | 0.012 | Young |
| 0.083976 | 0.294797 | 0.048 | 0.032 | 1 | 1 | Gmeb1 | 0.016 | Young |
| 0.084239 | 0.910218 | 0.032 | 0.019 | 1 | 1 | Cntln | 0.013 | Young |
| 0.0843 | 1.296908 | 0.015 | 0.006 | 1 | 1 | Rpgrip1 | 0.009 | Young |
| 0.084346 | 0.431727 | 0.029 | 0.017 | 1 | 1 | Hsdl2 | 0.012 | Young |
| 0.084353 | 0.467795 | 0.067 | 0.049 | 1 | 1 | Ppfia4 | 0.018 | Young |
| 0.084385 | 0.832842 | 0.024 | 0.013 | 1 | 1 | Fkrp | 0.011 | Young |
| 0.084563 | 0.122404 | 0.355 | 0.307 | 1 | 1 | Rps27l | 0.048 | Young |
| 0.084604 | 0.840854 | 0.024 | 0.013 | 1 | 1 | Mthfsl | 0.011 | Young |
| 0.084658 | 0.741775 | 0.024 | 0.013 | 1 | 1 | Zfand2b | 0.011 | Young |
| 0.084799 | 0.967497 | 0.032 | 0.019 | 1 | 1 | Rhbdd1 | 0.013 | Young |
| 0.085004 | 0.484578 | 0.057 | 0.04 | 1 | 1 | Elp5 | 0.017 | Young |
| 0.085071 | 0.16962 | 0.045 | 0.03 | 1 | 1 | Mir142hg | 0.015 | Young |
| 0.085078 | 0.399206 | 0.061 | 0.044 | 1 | 1 | Erlec1 | 0.017 | Young |
| 0.085172 | 0.637961 | 0.018 | 0.009 | 1 | 1 | Ppp1r12b | 0.009 | Young |
| 0.085339 | 0.126408 | 0.048 | 0.032 | 1 | 1 | Brf1 | 0.016 | Young |
| 0.085434 | 0.87598 | 0.015 | 0.006 | 1 | 1 | Sacs | 0.009 | Young |
| 0.085621 | 0.104851 | 0.186 | 0.156 | 1 | 1 | Bola2 | 0.03 | Young |
| 0.08576 | 0.643024 | 0.024 | 0.013 | 1 | 1 | Lyrm9 | 0.011 | Young |
| 0.08576 | 0.615967 | 0.024 | 0.013 | 1 | 1 | Cblb | 0.011 | Young |
| 0.085902 | 0.210929 | 0.157 | 0.129 | 1 | 1 | Rab21 | 0.028 | Young |
| 0.085907 | 0.073608 | 0.189 | 0.156 | 1 | 1 | Adam17 | 0.033 | Young |
| 0.085934 | 0.210393 | 0.196 | 0.165 | 1 | 1 | Zfp703 | 0.031 | Young |
| 0.086083 | 0.118665 | 0.062 | 0.044 | 1 | 1 | Fbxo38 | 0.018 | Young |
| 0.086093 | 0.092866 | 0.037 | 0.023 | 1 | 1 | Gla | 0.014 | Young |
| 0.086138 | 0.209105 | 0.16 | 0.131 | 1 | 1 | Nufip2 | 0.029 | Young |
| 0.086221 | 0.111335 | 0.068 | 0.049 | 1 | 1 | Ten1 | 0.019 | Young |
| 0.086245 | 0.247334 | 0.188 | 0.158 | 1 | 1 | Cggbp1 | 0.03 | Young |

| | | | | | | | | |
|---|---|---|---|---|---|---|---|---|
| 0.086298 | 0.385623 | 0.113 | 0.09 | 1 | 1 | Phc3 | 0.023 | Young |
| 0.086571 | 0.129054 | 0.178 | 0.148 | 1 | 1 | Myo9b | 0.03 | Young |
| 0.08658 | 0.686354 | 0.015 | 0.006 | 1 | 1 | Tada2b | 0.009 | Young |
| 0.086629 | 0.267641 | 0.018 | 0.009 | 1 | 1 | Mrgbp | 0.009 | Young |
| 0.086652 | 0.485046 | 0.015 | 0.006 | 1 | 1 | Cmc4 | 0.009 | Young |
| 0.086684 | 0.146116 | 0.049 | 0.033 | 1 | 1 | Spns1 | 0.016 | Young |
| 0.087199 | 0.343794 | 0.112 | 0.089 | 1 | 1 | Pacsin2 | 0.023 | Young |
| 0.087373 | 0.070918 | 0.15 | 0.121 | 1 | 1 | Tial1 | 0.029 | Young |
| 0.087376 | 0.677581 | 0.071 | 0.053 | 1 | 1 | Chchd7 | 0.018 | Young |
| 0.087432 | 0.037185 | 0.371 | 0.321 | 1 | 1 | Rab7b | 0.05 | Young |
| 0.087452 | 0.065646 | 0.064 | 0.045 | 1 | 1 | Plod1 | 0.019 | Young |
| 0.087552 | 0.101317 | 0.173 | 0.141 | 1 | 1 | Necap2 | 0.032 | Young |
| 0.087705 | 0.375145 | 0.106 | 0.084 | 1 | 1 | Sidt2 | 0.022 | Young |
| 0.087823 | 0.361849 | 0.078 | 0.058 | 1 | 1 | Fbxw2 | 0.02 | Young |
| 0.087879 | 0.302557 | 0.058 | 0.041 | 1 | 1 | Lcor | 0.017 | Young |
| 0.087937 | 0.611649 | 0.041 | 0.027 | 1 | 1 | Ppme1 | 0.014 | Young |
| 0.088058 | 1.198738 | 0.025 | 0.014 | 1 | 1 | Tirap | 0.011 | Young |
| 0.088113 | 0.393083 | 0.064 | 0.046 | 1 | 1 | Klhl6 | 0.018 | Young |
| 0.088387 | 1.035543 | 0.025 | 0.014 | 1 | 1 | Stk16 | 0.011 | Young |
| 0.088393 | 0.341006 | 0.085 | 0.064 | 1 | 1 | Rnps1 | 0.021 | Young |
| 0.088442 | 0.845874 | 0.025 | 0.014 | 1 | 1 | Sprtn | 0.011 | Young |
| 0.088467 | 0.389933 | 0.055 | 0.039 | 1 | 1 | Lap3 | 0.016 | Young |
| 0.088503 | 0.027083 | 0.049 | 0.033 | 1 | 1 | Guca1a | 0.016 | Young |
| 0.088682 | 0.864036 | 0.052 | 0.036 | 1 | 1 | Fam69a | 0.016 | Young |
| 0.088689 | 0.565349 | 0.043 | 0.028 | 1 | 1 | Slc41a3 | 0.015 | Young |
| 0.088695 | 0.083973 | 0.105 | 0.081 | 1 | 1 | Cp | 0.024 | Young |
| 0.088962 | 0.987497 | 0.035 | 0.022 | 1 | 1 | Spn | 0.013 | Young |
| 0.089297 | 0.30618 | 0.032 | 0.019 | 1 | 1 | Timm9 | 0.013 | Young |
| 0.089314 | 0.114889 | 0.03 | 0.018 | 1 | 1 | Ttc37 | 0.012 | Young |
| 0.089378 | 1.01283 | 0.025 | 0.014 | 1 | 1 | Rad51d | 0.011 | Young |
| 0.089464 | 0.612449 | 0.044 | 0.03 | 1 | 1 | Sipa1l3 | 0.014 | Young |
| 0.089525 | 0.033067 | 0.04 | 0.026 | 1 | 1 | Pag1 | 0.014 | Young |
| 0.089759 | 0.094545 | 0.134 | 0.107 | 1 | 1 | Nbeal1 | 0.027 | Young |
| 0.089764 | 0.168595 | 0.247 | 0.213 | 1 | 1 | Evi2a | 0.034 | Young |
| 0.089865 | 0.323179 | 0.16 | 0.134 | 1 | 1 | Adcy7 | 0.026 | Young |
| 0.090085 | 0.113326 | 0.126 | 0.102 | 1 | 1 | Uvrag | 0.024 | Young |
| 0.090294 | 0.017786 | 0.216 | 0.18 | 1 | 1 | Rin2 | 0.036 | Young |
| 0.090399 | 0.120371 | 0.116 | 0.091 | 1 | 1 | C5ar2 | 0.025 | Young |
| 0.090421 | 0.650199 | 0.049 | 0.033 | 1 | 1 | Prpsap2 | 0.016 | Young |
| 0.09051 | 0.338583 | 0.083 | 0.063 | 1 | 1 | Epc1 | 0.02 | Young |
| 0.090532 | 0.319696 | 0.092 | 0.071 | 1 | 1 | Zfp451 | 0.021 | Young |
| 0.090572 | 0.110316 | 0.095 | 0.072 | 1 | 1 | Rbm22 | 0.023 | Young |
| 0.091076 | 0.081814 | 0.135 | 0.108 | 1 | 1 | Pank3 | 0.027 | Young |
| 0.091188 | 0.121302 | 0.218 | 0.184 | 1 | 1 | Eif3c | 0.034 | Young |
| 0.091239 | 0.012463 | 0.213 | 0.179 | 1 | 1 | Tbxas1 | 0.034 | Young |

| | | | | | | | | |
|---|---|---|---|---|---|---|---|---|
| 0.091375 | 0.02787 | 0.145 | 0.117 | 1 | 1 | Gtf2a1 | 0.028 | Young |
| 0.091445 | 0.311779 | 0.025 | 0.014 | 1 | 1 | Sec22a | 0.011 | Young |
| 0.091499 | 0.210002 | 0.06 | 0.042 | 1 | 1 | Dopey1 | 0.018 | Young |
| 0.091505 | 0.83033 | 0.02 | 0.01 | 1 | 1 | Med19 | 0.01 | Young |
| 0.091514 | 0.547234 | 0.033 | 0.021 | 1 | 1 | Ivd | 0.012 | Young |
| 0.091542 | 0.340916 | 0.032 | 0.019 | 1 | 1 | Zfp24 | 0.013 | Young |
| 0.091565 | 0.243613 | 0.109 | 0.086 | 1 | 1 | Ranbp1 | 0.023 | Young |
| 0.091758 | 0.250738 | 0.091 | 0.069 | 1 | 1 | Rps6kb1 | 0.022 | Young |
| 0.091817 | 0.366609 | 0.032 | 0.019 | 1 | 1 | Eif2ak3 | 0.013 | Young |
| 0.091896 | 0.511872 | 0.025 | 0.014 | 1 | 1 | Ndufaf7 | 0.011 | Young |
| 0.091973 | 0.198625 | 0.56 | 0.503 | 1 | 1 | Mt1 | 0.057 | Young |
| 0.091981 | 0.130099 | 0.176 | 0.145 | 1 | 1 | Sft2d2 | 0.031 | Young |
| 0.092117 | 0.066658 | 0.032 | 0.019 | 1 | 1 | 2410002F2 | 0.013 | Young |
| 0.092215 | 0.470842 | 0.075 | 0.057 | 1 | 1 | Hadh | 0.018 | Young |
| 0.092339 | 0.561976 | 0.068 | 0.05 | 1 | 1 | Polr2c | 0.018 | Young |
| 0.09251 | 0.785799 | 0.036 | 0.023 | 1 | 1 | Riok2 | 0.013 | Young |
| 0.092575 | 0.305279 | 0.041 | 0.027 | 1 | 1 | Taf7 | 0.014 | Young |
| 0.092604 | 0.324329 | 0.076 | 0.057 | 1 | 1 | Ccdc88b | 0.019 | Young |
| 0.092609 | 2.500781 | 0.013 | 0.005 | 1 | 1 | Mrpl44 | 0.008 | Young |
| 0.09274 | 0.06696 | 0.511 | 0.445 | 1 | 1 | Ybx1 | 0.066 | Young |
| 0.092803 | 0.384562 | 0.086 | 0.066 | 1 | 1 | Arhgap5 | 0.02 | Young |
| 0.092894 | 1.062916 | 0.027 | 0.015 | 1 | 1 | Eogt | 0.012 | Young |
| 0.092896 | 0.259904 | 0.091 | 0.069 | 1 | 1 | Phb2 | 0.022 | Young |
| 0.092933 | 0.233157 | 0.095 | 0.073 | 1 | 1 | Tmem106b | 0.022 | Young |
| 0.093004 | 0.489922 | 0.047 | 0.032 | 1 | 1 | Ctbs | 0.015 | Young |
| 0.093305 | 0.608096 | 0.02 | 0.01 | 1 | 1 | Kremen1 | 0.01 | Young |
| 0.093413 | 0.819914 | 0.035 | 0.022 | 1 | 1 | Ifitm1 | 0.013 | Young |
| 0.09344 | 0.233782 | 0.151 | 0.125 | 1 | 1 | Cdv3 | 0.026 | Young |
| 0.093696 | 0.017388 | 0.14 | 0.112 | 1 | 1 | Dynll2 | 0.028 | Young |
| 0.094071 | 0.109815 | 0.216 | 0.181 | 1 | 1 | Ewsr1 | 0.035 | Young |
| 0.094196 | 0.379046 | 0.049 | 0.033 | 1 | 1 | Pold4 | 0.016 | Young |
| 0.094674 | 0.273969 | 0.02 | 0.01 | 1 | 1 | Nudt7 | 0.01 | Young |
| 0.094829 | 0.818126 | 0.036 | 0.023 | 1 | 1 | Yif1b | 0.013 | Young |
| 0.094869 | 1.025118 | 0.013 | 0.005 | 1 | 1 | Mrpl49 | 0.008 | Young |
| 0.094927 | 0.053796 | 0.29 | 0.248 | 1 | 1 | Sgpl1 | 0.042 | Young |
| 0.094969 | 0.934428 | 0.016 | 0.008 | 1 | 1 | Nit2 | 0.008 | Young |
| 0.094985 | 0.169901 | 0.082 | 0.062 | 1 | 1 | Ctcf | 0.02 | Young |
| 0.095206 | 0.130002 | 0.253 | 0.221 | 1 | 1 | Anp32a | 0.032 | Young |
| 0.095232 | 0.175813 | 0.253 | 0.219 | 1 | 1 | Kif5b | 0.034 | Young |
| 0.095246 | 0.284967 | 0.136 | 0.111 | 1 | 1 | Saraf | 0.025 | Young |
| 0.09539 | 0.238503 | 0.1 | 0.077 | 1 | 1 | Ndufaf8 | 0.023 | Young |
| 0.095446 | 0.284807 | 0.082 | 0.062 | 1 | 1 | Enox2 | 0.02 | Young |
| 0.095484 | 0.728819 | 0.016 | 0.008 | 1 | 1 | Nav2 | 0.008 | Young |
| 0.095631 | 1.070145 | 0.016 | 0.008 | 1 | 1 | Zfp595 | 0.008 | Young |
| 0.095649 | 0.388306 | 0.084 | 0.064 | 1 | 1 | Ntpcr | 0.02 | Young |

| | | | | | | | | |
|---|---|---|---|---|---|---|---|---|
| 0.095918 | 0.262831 | 0.044 | 0.03 | 1 | 1 | Emc8 | 0.014 | Young |
| 0.095991 | 0.21842 | 0.103 | 0.08 | 1 | 1 | Bmt2 | 0.023 | Young |
| 0.096041 | 0.434513 | 0.058 | 0.041 | 1 | 1 | Ranbp10 | 0.017 | Young |
| 0.096058 | 0.793361 | 0.013 | 0.005 | 1 | 1 | Per2 | 0.008 | Young |
| 0.096143 | 0.826421 | 0.013 | 0.005 | 1 | 1 | Mrps27 | 0.008 | Young |
| 0.096147 | 0.455813 | 0.027 | 0.015 | 1 | 1 | Bbc3 | 0.012 | Young |
| 0.096399 | 0.74452 | 0.013 | 0.005 | 1 | 1 | Zfp607a | 0.008 | Young |
| 0.096431 | 0.21604 | 0.078 | 0.058 | 1 | 1 | Tyk2 | 0.02 | Young |
| 0.096442 | 0.081075 | 0.076 | 0.057 | 1 | 1 | Oaz2 | 0.019 | Young |
| 0.096547 | 0.09825 | 0.134 | 0.108 | 1 | 1 | Psmc3 | 0.026 | Young |
| 0.096594 | 0.049032 | 0.139 | 0.112 | 1 | 1 | Hmox2 | 0.027 | Young |
| 0.096619 | 0.265588 | 0.07 | 0.051 | 1 | 1 | Ash2l | 0.019 | Young |
| 0.096645 | 0.62278 | 0.035 | 0.022 | 1 | 1 | Bap1 | 0.013 | Young |
| 0.096734 | 0.525362 | 0.052 | 0.036 | 1 | 1 | Pak1ip1 | 0.016 | Young |
| 0.096893 | 0.400461 | 0.035 | 0.022 | 1 | 1 | Wdr81 | 0.013 | Young |
| 0.097124 | 0.00425 | 0.479 | 0.423 | 1 | 1 | Celf2 | 0.056 | Young |
| 0.097176 | 1.070296 | 0.03 | 0.018 | 1 | 1 | Rbsn | 0.012 | Young |
| 0.097258 | 0.313394 | 0.013 | 0.005 | 1 | 1 | Gimap6 | 0.008 | Young |
| 0.097258 | 0.254927 | 0.013 | 0.005 | 1 | 1 | Gm1673 | 0.008 | Young |
| 0.097266 | 0.349369 | 0.084 | 0.064 | 1 | 1 | Irak4 | 0.02 | Young |
| 0.09737 | 0.210679 | 0.101 | 0.078 | 1 | 1 | Ankrd40 | 0.023 | Young |
| 0.097425 | 0.39427 | 0.053 | 0.037 | 1 | 1 | Ice1 | 0.016 | Young |
| 0.097584 | 0.358883 | 0.055 | 0.039 | 1 | 1 | Tsc1 | 0.016 | Young |
| 0.097705 | 0.074082 | 0.88 | 0.852 | 1 | 1 | Rps26 | 0.028 | Young |
| 0.097775 | 0.028927 | 0.066 | 0.048 | 1 | 1 | Pxk | 0.018 | Young |
| 0.097795 | 0.122014 | 0.097 | 0.076 | 1 | 1 | Helz | 0.021 | Young |
| 0.097895 | 0.158434 | 0.092 | 0.071 | 1 | 1 | Baiap2 | 0.021 | Young |
| 0.09797 | 0.574838 | 0.028 | 0.017 | 1 | 1 | 4921524J1 | 0.011 | Young |
| 0.097973 | 0.218458 | 0.137 | 0.112 | 1 | 1 | Sf3b3 | 0.025 | Young |
| 0.098192 | 0.305335 | 0.021 | 0.012 | 1 | 1 | Sp4 | 0.009 | Young |
| 0.098271 | 0.112165 | 0.047 | 0.032 | 1 | 1 | Ssbp1 | 0.015 | Young |
| 0.098395 | 0.050544 | 0.089 | 0.067 | 1 | 1 | Eif2s3y | 0.022 | Young |
| 0.09852 | 0.305381 | 0.126 | 0.102 | 1 | 1 | Ccar1 | 0.024 | Young |
| 0.098531 | 0.207864 | 0.11 | 0.087 | 1 | 1 | Zmym5 | 0.023 | Young |
| 0.098648 | 0.573773 | 0.072 | 0.054 | 1 | 1 | Tpst2 | 0.018 | Young |
| 0.098694 | 0.421548 | 0.111 | 0.089 | 1 | 1 | Mid1ip1 | 0.022 | Young |
| 0.098761 | 0.021918 | 0.23 | 0.192 | 1 | 1 | Tmem59 | 0.038 | Young |
| 0.098861 | 0.668388 | 0.039 | 0.026 | 1 | 1 | Smad3 | 0.013 | Young |
| 0.099083 | 0.417756 | 0.072 | 0.054 | 1 | 1 | Rnaseh2c | 0.018 | Young |
| 0.099448 | 0.04709 | 0.357 | 0.308 | 1 | 1 | 2010107E0 | 0.049 | Young |
| 0.099518 | 0.272448 | 0.069 | 0.051 | 1 | 1 | Wbp4 | 0.018 | Young |
| 0.099679 | 0.088953 | 0.091 | 0.069 | 1 | 1 | Chd8 | 0.022 | Young |
| 0.099802 | 0.382327 | 0.055 | 0.039 | 1 | 1 | Clns1a | 0.016 | Young |
| 0.099816 | 0.398293 | 0.061 | 0.044 | 1 | 1 | Tsnax | 0.017 | Young |
| 0.100309 | 0.03767 | 0.182 | 0.15 | 1 | 1 | Zfand6 | 0.032 | Young |

| | | | | | | | | |
|---|---|---|---|---|---|---|---|---|
| 0.100441 | 0.262981 | 0.058 | 0.041 | 1 | 1 | Vps54 | 0.017 | Young |
| 0.100596 | 0.502248 | 0.041 | 0.027 | 1 | 1 | Chpf2 | 0.014 | Young |
| 0.101045 | 0.019621 | 0.055 | 0.039 | 1 | 1 | Mfsd6 | 0.016 | Young |
| 0.10131 | 0.158895 | 0.021 | 0.012 | 1 | 1 | Mpi | 0.009 | Young |
| 0.101399 | 0.104126 | 0.123 | 0.098 | 1 | 1 | Rap2b | 0.025 | Young |
| 0.101496 | 0.554326 | 0.075 | 0.057 | 1 | 1 | Golgb1 | 0.018 | Young |
| 0.101535 | 0.848328 | 0.058 | 0.042 | 1 | 1 | Glrx | 0.016 | Young |
| 0.101594 | 0.240812 | 0.075 | 0.057 | 1 | 1 | Psen1 | 0.018 | Young |
| 0.101915 | 0.259353 | 0.069 | 0.051 | 1 | 1 | Keap1 | 0.018 | Young |
| 0.102311 | 0.593885 | 0.049 | 0.035 | 1 | 1 | Abcb7 | 0.014 | Young |
| 0.102484 | 0.903038 | 0.049 | 0.035 | 1 | 1 | Trpv2 | 0.014 | Young |
| 0.102532 | 0.421548 | 0.111 | 0.089 | 1 | 1 | Bcl2l1 | 0.022 | Young |
| 0.102785 | 0.753593 | 0.045 | 0.031 | 1 | 1 | Snx14 | 0.014 | Young |
| 0.103139 | 0.108105 | 0.194 | 0.163 | 1 | 1 | H2afj | 0.031 | Young |
| 0.103308 | 0.276874 | 0.116 | 0.093 | 1 | 1 | Pitpnc1 | 0.023 | Young |
| 0.103324 | 0.761084 | 0.023 | 0.013 | 1 | 1 | Jagn1 | 0.01 | Young |
| 0.10344 | 0.249038 | 0.083 | 0.063 | 1 | 1 | Tbc1d23 | 0.02 | Young |
| 0.103444 | 0.377727 | 0.072 | 0.054 | 1 | 1 | Brpf1 | 0.018 | Young |
| 0.103588 | 0.159067 | 0.213 | 0.184 | 1 | 1 | Taok3 | 0.029 | Young |
| 0.103632 | 0.287294 | 0.064 | 0.046 | 1 | 1 | Med8 | 0.018 | Young |
| 0.103646 | 0.027325 | 0.256 | 0.217 | 1 | 1 | Larp4b | 0.039 | Young |
| 0.103895 | 0.288632 | 0.052 | 0.036 | 1 | 1 | Nmd3 | 0.016 | Young |
| 0.104041 | 0.231088 | 0.08 | 0.06 | 1 | 1 | Zfp800 | 0.02 | Young |
| 0.104054 | 1.82293 | 0.01 | 0.004 | 1 | 1 | Zfp606 | 0.006 | Young |
| 0.104057 | 0.254371 | 0.09 | 0.069 | 1 | 1 | Spopl | 0.021 | Young |
| 0.104121 | 0.475883 | 0.076 | 0.058 | 1 | 1 | Gatad2a | 0.018 | Young |
| 0.104155 | 1.810252 | 0.01 | 0.004 | 1 | 1 | Cd101 | 0.006 | Young |
| 0.104224 | 0.273989 | 0.114 | 0.091 | 1 | 1 | Eif3l | 0.023 | Young |
| 0.104257 | 1.675784 | 0.01 | 0.004 | 1 | 1 | Tbc1d8b | 0.006 | Young |
| 0.104289 | 0.328225 | 0.143 | 0.118 | 1 | 1 | Map2k3 | 0.025 | Young |
| 0.104344 | 0.020579 | 0.162 | 0.132 | 1 | 1 | Grina | 0.03 | Young |
| 0.104395 | 0.092387 | 0.275 | 0.237 | 1 | 1 | Arhgap18 | 0.038 | Young |
| 0.10444 | 0.506013 | 0.042 | 0.028 | 1 | 1 | Trak2 | 0.014 | Young |
| 0.104529 | 0.593838 | 0.048 | 0.033 | 1 | 1 | Tmed4 | 0.015 | Young |
| 0.104662 | 1.68075 | 0.01 | 0.004 | 1 | 1 | Zfp628 | 0.006 | Young |
| 0.104662 | 1.581826 | 0.01 | 0.004 | 1 | 1 | Apold1 | 0.006 | Young |
| 0.104774 | 0.107793 | 0.168 | 0.14 | 1 | 1 | Ghitm | 0.028 | Young |
| 0.104785 | 0.141996 | 0.182 | 0.153 | 1 | 1 | Mapk14 | 0.029 | Young |
| 0.104833 | 1.023115 | 0.032 | 0.021 | 1 | 1 | Prr3 | 0.011 | Young |
| 0.104839 | 0.464893 | 0.044 | 0.03 | 1 | 1 | Sipa1l2 | 0.014 | Young |
| 0.10496 | 0.290509 | 0.041 | 0.027 | 1 | 1 | Kdsr | 0.014 | Young |
| 0.105058 | 0.437681 | 0.041 | 0.027 | 1 | 1 | Atg4b | 0.014 | Young |
| 0.105228 | 0.018644 | 0.092 | 0.071 | 1 | 1 | Ip6k1 | 0.021 | Young |
| 0.105288 | 0.960471 | 0.018 | 0.009 | 1 | 1 | Dbp | 0.009 | Young |
| 0.105473 | 0.401663 | 0.096 | 0.076 | 1 | 1 | Mpc2 | 0.02 | Young |

| | | | | | | | | |
|---|---|---|---|---|---|---|---|---|
| 0.105476 | 1.307199 | 0.01 | 0.004 | 1 | 1 | Bcs1l | 0.006 | Young |
| 0.105492 | 0.086557 | 0.153 | 0.125 | 1 | 1 | Stk38 | 0.028 | Young |
| 0.105578 | 0.782987 | 0.01 | 0.004 | 1 | 1 | Syp | 0.006 | Young |
| 0.105591 | 0.391449 | 0.018 | 0.009 | 1 | 1 | Mcemp1 | 0.009 | Young |
| 0.105639 | 0.301122 | 0.125 | 0.102 | 1 | 1 | Dennd6a | 0.023 | Young |
| 0.105655 | 0.311973 | 0.086 | 0.067 | 1 | 1 | Zdhhc9 | 0.019 | Young |
| 0.10568 | 0.799152 | 0.01 | 0.004 | 1 | 1 | Ankzf1 | 0.006 | Young |
| 0.105745 | 0.115936 | 0.03 | 0.018 | 1 | 1 | Gm17018 | 0.012 | Young |
| 0.105778 | 0.150362 | 0.092 | 0.071 | 1 | 1 | Fbxo33 | 0.021 | Young |
| 0.105815 | 0.749776 | 0.057 | 0.041 | 1 | 1 | Tango2 | 0.016 | Young |
| 0.105846 | 0.379279 | 0.089 | 0.069 | 1 | 1 | Ncoa7 | 0.02 | Young |
| 0.105985 | 0.047969 | 0.127 | 0.102 | 1 | 1 | Npc1 | 0.025 | Young |
| 0.106196 | 0.6176 | 0.06 | 0.044 | 1 | 1 | Ppt2 | 0.016 | Young |
| 0.106261 | 0.169832 | 0.14 | 0.114 | 1 | 1 | Ik | 0.026 | Young |
| 0.106295 | 0.686377 | 0.01 | 0.004 | 1 | 1 | 0610040J0 | 0.006 | Young |
| 0.106338 | 0.074307 | 0.119 | 0.095 | 1 | 1 | Ogfrl1 | 0.024 | Young |
| 0.10675 | 0.451707 | 0.023 | 0.013 | 1 | 1 | Eef1akmt2 | 0.01 | Young |
| 0.107035 | 0.041533 | 0.179 | 0.149 | 1 | 1 | Cerk | 0.03 | Young |
| 0.107093 | 0.001163 | 0.352 | 0.306 | 1 | 1 | Rab14 | 0.046 | Young |
| 0.107304 | 0.891154 | 0.034 | 0.022 | 1 | 1 | Bola3 | 0.012 | Young |
| 0.107484 | 0.888405 | 0.024 | 0.014 | 1 | 1 | Haus3 | 0.01 | Young |
| 0.107787 | 0.235074 | 0.03 | 0.018 | 1 | 1 | Tmem87a | 0.012 | Young |
| 0.107819 | 1.838039 | 0.014 | 0.006 | 1 | 1 | Dcp1b | 0.008 | Young |
| 0.107957 | 0.224997 | 0.023 | 0.013 | 1 | 1 | Nubp1 | 0.01 | Young |
| 0.107977 | 0.426674 | 0.045 | 0.031 | 1 | 1 | Hk3 | 0.014 | Young |
| 0.108004 | 0.462244 | 0.024 | 0.014 | 1 | 1 | AI837181 | 0.01 | Young |
| 0.108395 | 0.83765 | 0.034 | 0.022 | 1 | 1 | Manbal | 0.012 | Young |
| 0.108425 | 0.075708 | 0.093 | 0.072 | 1 | 1 | Smndc1 | 0.021 | Young |
| 0.108522 | 0.087848 | 0.148 | 0.121 | 1 | 1 | Elk3 | 0.027 | Young |
| 0.108598 | 0.102154 | 0.023 | 0.013 | 1 | 1 | Ankrd39 | 0.01 | Young |
| 0.108653 | 0.192934 | 0.207 | 0.176 | 1 | 1 | Sdha | 0.031 | Young |
| 0.108656 | 0.87397 | 0.024 | 0.014 | 1 | 1 | Klhl7 | 0.01 | Young |
| 0.108662 | 0.227403 | 0.018 | 0.009 | 1 | 1 | Cutc | 0.009 | Young |
| 0.108772 | 0.506208 | 0.075 | 0.057 | 1 | 1 | Map3k7 | 0.018 | Young |
| 0.108852 | 0.247991 | 0.123 | 0.1 | 1 | 1 | Cdc5l | 0.023 | Young |
| 0.10886 | 0.067418 | 0.133 | 0.107 | 1 | 1 | Psme4 | 0.026 | Young |
| 0.108874 | 1.091085 | 0.014 | 0.006 | 1 | 1 | D3Ertd751 | 0.008 | Young |
| 0.108926 | 0.174406 | 0.125 | 0.1 | 1 | 1 | Fbxl5 | 0.025 | Young |
| 0.108971 | 0.163905 | 0.134 | 0.108 | 1 | 1 | Dnajc8 | 0.026 | Young |
| 0.10905 | 0.996737 | 0.014 | 0.006 | 1 | 1 | Pls3 | 0.008 | Young |
| 0.109086 | 0.400103 | 0.049 | 0.035 | 1 | 1 | Cdk16 | 0.014 | Young |
| 0.109111 | 0.453295 | 0.052 | 0.037 | 1 | 1 | Oxsr1 | 0.015 | Young |
| 0.109224 | 0.477937 | 0.055 | 0.04 | 1 | 1 | Relb | 0.015 | Young |
| 0.109283 | 0.323259 | 0.138 | 0.114 | 1 | 1 | Ccni | 0.024 | Young |
| 0.109316 | 1.157164 | 0.014 | 0.006 | 1 | 1 | Mtmr11 | 0.008 | Young |

| | | | | | | | | |
|---|---|---|---|---|---|---|---|---|
| 0.109316 | 1.012368 | 0.014 | 0.006 | 1 | 1 | Armc7 | 0.008 | Young |
| 0.109404 | 1.084009 | 0.014 | 0.006 | 1 | 1 | Havcr2 | 0.008 | Young |
| 0.109404 | 0.793296 | 0.014 | 0.006 | 1 | 1 | Ppp1r3d | 0.008 | Young |
| 0.109474 | 0.250815 | 0.047 | 0.032 | 1 | 1 | Patl1 | 0.015 | Young |
| 0.109495 | 0.526106 | 0.084 | 0.066 | 1 | 1 | Dennd1c | 0.018 | Young |
| 0.109626 | 0.561924 | 0.014 | 0.006 | 1 | 1 | Osbpl1a | 0.008 | Young |
| 0.10964 | 0.486768 | 0.024 | 0.014 | 1 | 1 | Mrps35 | 0.01 | Young |
| 0.109757 | 0.467068 | 0.032 | 0.021 | 1 | 1 | Map4k1 | 0.011 | Young |
| 0.110043 | 0.200018 | 0.072 | 0.054 | 1 | 1 | Uqcc2 | 0.018 | Young |
| 0.110063 | 1.414228 | 0.019 | 0.01 | 1 | 1 | I830077J02 | 0.009 | Young |
| 0.110213 | 0.004496 | 0.23 | 0.194 | 1 | 1 | Arhgap17 | 0.036 | Young |
| 0.110215 | 0.549169 | 0.055 | 0.04 | 1 | 1 | Ptpn22 | 0.015 | Young |
| 0.110367 | 0.259481 | 0.024 | 0.014 | 1 | 1 | Gnptab | 0.01 | Young |
| 0.110408 | 0.236105 | 0.146 | 0.121 | 1 | 1 | Emg1 | 0.025 | Young |
| 0.110462 | 0.281903 | 0.042 | 0.028 | 1 | 1 | Mtmr2 | 0.014 | Young |
| 0.110861 | 0.28923 | 0.083 | 0.064 | 1 | 1 | Nudt21 | 0.019 | Young |
| 0.110863 | 0.022622 | 0.081 | 0.062 | 1 | 1 | Mgat4a | 0.019 | Young |
| 0.111098 | 0.349855 | 0.014 | 0.006 | 1 | 1 | Dnmbp | 0.008 | Young |
| 0.111187 | 0.304757 | 0.014 | 0.006 | 1 | 1 | Pcgf6 | 0.008 | Young |
| 0.111204 | 0.153316 | 0.191 | 0.163 | 1 | 1 | Mkln1 | 0.028 | Young |
| 0.11124 | 0.312865 | 0.088 | 0.068 | 1 | 1 | Mrpl48 | 0.02 | Young |
| 0.111282 | 0.506481 | 0.049 | 0.035 | 1 | 1 | Casp6 | 0.014 | Young |
| 0.111397 | 0.043787 | 0.203 | 0.171 | 1 | 1 | Zfp91 | 0.032 | Young |
| 0.111424 | 0.348728 | 0.072 | 0.054 | 1 | 1 | Rabl6 | 0.018 | Young |
| 0.111506 | 0.241027 | 0.115 | 0.093 | 1 | 1 | Sav1 | 0.022 | Young |
| 0.111525 | 0.376932 | 0.032 | 0.021 | 1 | 1 | Mavs | 0.011 | Young |
| 0.111542 | 0.465736 | 0.061 | 0.045 | 1 | 1 | Plcg2 | 0.016 | Young |
| 0.111563 | 0.494233 | 0.024 | 0.014 | 1 | 1 | Slc12a2 | 0.01 | Young |
| 0.111606 | 0.386988 | 0.049 | 0.035 | 1 | 1 | Plxna4 | 0.014 | Young |
| 0.112003 | 0.207531 | 0.082 | 0.063 | 1 | 1 | Samsn1 | 0.019 | Young |
| 0.112085 | 0.856134 | 0.035 | 0.023 | 1 | 1 | Pctp | 0.012 | Young |
| 0.112253 | 0.095325 | 0.031 | 0.019 | 1 | 1 | Helb | 0.012 | Young |
| 0.112373 | 0.369922 | 0.057 | 0.041 | 1 | 1 | Papd5 | 0.016 | Young |
| 0.112406 | 1.024556 | 0.019 | 0.01 | 1 | 1 | Usp30 | 0.009 | Young |
| 0.112483 | 0.671992 | 0.019 | 0.01 | 1 | 1 | Zfp658 | 0.009 | Young |
| 0.112581 | 1.074223 | 0.038 | 0.026 | 1 | 1 | Cnnm3 | 0.012 | Young |
| 0.112788 | 0.909958 | 0.019 | 0.01 | 1 | 1 | 2300009A0 | 0.009 | Young |
| 0.112921 | 0.299917 | 0.091 | 0.071 | 1 | 1 | Ube2e3 | 0.02 | Young |
| 0.112943 | 0.305697 | 0.1 | 0.08 | 1 | 1 | Arih2 | 0.02 | Young |
| 0.112947 | 0.012295 | 0.504 | 0.44 | 1 | 1 | Sh3bgrl3 | 0.064 | Young |
| 0.113106 | 0.253286 | 0.085 | 0.066 | 1 | 1 | Fkbp3 | 0.019 | Young |
| 0.113242 | 0.427462 | 0.024 | 0.014 | 1 | 1 | Zbtb46 | 0.01 | Young |
| 0.113324 | 0.729812 | 0.019 | 0.01 | 1 | 1 | Ube2o | 0.009 | Young |
| 0.113372 | 0.374161 | 0.032 | 0.021 | 1 | 1 | Gtf2e2 | 0.011 | Young |
| 0.113552 | 0.21952 | 0.088 | 0.068 | 1 | 1 | Exoc4 | 0.02 | Young |

| | | | | | | | | |
|---|---|---|---|---|---|---|---|---|
| 0.113599 | 0.106507 | 0.069 | 0.051 | 1 | 1 | 0610030E2 | 0.018 | Young |
| 0.113644 | 0.074638 | 0.357 | 0.315 | 1 | 1 | Foxp1 | 0.042 | Young |
| 0.113673 | 0.045073 | 0.08 | 0.06 | 1 | 1 | Rtn1 | 0.02 | Young |
| 0.113784 | 0.300029 | 0.024 | 0.014 | 1 | 1 | Syce2 | 0.01 | Young |
| 0.113867 | 0.509849 | 0.026 | 0.015 | 1 | 1 | G6pc3 | 0.011 | Young |
| 0.113894 | 0.298929 | 0.055 | 0.04 | 1 | 1 | Ddx52 | 0.015 | Young |
| 0.114031 | 0.199149 | 0.049 | 0.035 | 1 | 1 | Cabin1 | 0.014 | Young |
| 0.114034 | 0.000501 | 0.272 | 0.233 | 1 | 1 | Mapkapk2 | 0.039 | Young |
| 0.114201 | 0.261003 | 0.061 | 0.045 | 1 | 1 | Cryl1 | 0.016 | Young |
| 0.114249 | 0.67047 | 0.065 | 0.049 | 1 | 1 | Fryl | 0.016 | Young |
| 0.114317 | 0.121634 | 0.099 | 0.077 | 1 | 1 | Pcm1 | 0.022 | Young |
| 0.114776 | 0.10806 | 0.115 | 0.093 | 1 | 1 | Tnfsf12 | 0.022 | Young |
| 0.11478 | 0.134815 | 0.121 | 0.098 | 1 | 1 | Yeats4 | 0.023 | Young |
| 0.114862 | 0.262771 | 0.072 | 0.054 | 1 | 1 | Car3 | 0.018 | Young |
| 0.114922 | 0.080457 | 0.026 | 0.015 | 1 | 1 | Cnnm4 | 0.011 | Young |
| 0.115121 | 0.628315 | 0.026 | 0.015 | 1 | 1 | Cfd | 0.011 | Young |
| 0.115585 | 0.512437 | 0.026 | 0.015 | 1 | 1 | Uckl1 | 0.011 | Young |
| 0.115609 | 0.334779 | 0.058 | 0.042 | 1 | 1 | Selenon | 0.016 | Young |
| 0.115737 | 0.701859 | 0.038 | 0.026 | 1 | 1 | Wrn | 0.012 | Young |
| 0.115785 | 0.532961 | 0.026 | 0.015 | 1 | 1 | Tspyl2 | 0.011 | Young |
| 0.115832 | 0.015908 | 0.137 | 0.112 | 1 | 1 | Slc4a7 | 0.025 | Young |
| 0.116109 | 0.29391 | 0.034 | 0.022 | 1 | 1 | Thap11 | 0.012 | Young |
| 0.116167 | 0.445859 | 0.092 | 0.072 | 1 | 1 | Rfk | 0.02 | Young |
| 0.116216 | 0.334912 | 0.066 | 0.049 | 1 | 1 | Tmem248 | 0.017 | Young |
| 0.116265 | 0.101481 | 0.272 | 0.238 | 1 | 1 | Zbtb20 | 0.034 | Young |
| 0.116803 | 0.196686 | 0.064 | 0.048 | 1 | 1 | Rpl7l1 | 0.016 | Young |
| 0.116867 | 0.016601 | 0.024 | 0.014 | 1 | 1 | 2610008E1 | 0.01 | Young |
| 0.117172 | 0.616442 | 0.037 | 0.024 | 1 | 1 | Kat6b | 0.013 | Young |
| 0.117248 | 0.158418 | 0.051 | 0.036 | 1 | 1 | Pilrb1 | 0.015 | Young |
| 0.117391 | 0.437107 | 0.026 | 0.015 | 1 | 1 | H6pd | 0.011 | Young |
| 0.117518 | 0.132647 | 0.116 | 0.094 | 1 | 1 | Sec61a1 | 0.022 | Young |
| 0.117549 | 0.227441 | 0.108 | 0.086 | 1 | 1 | Ppp4r3a | 0.022 | Young |
| 0.117583 | 0.151229 | 0.247 | 0.212 | 1 | 1 | Raph1 | 0.035 | Young |
| 0.117684 | 0.207835 | 0.019 | 0.01 | 1 | 1 | Pcyox1l | 0.009 | Young |
| 0.117857 | 0.307011 | 0.058 | 0.042 | 1 | 1 | Zc3h14 | 0.016 | Young |
| 0.117899 | 0.497676 | 0.037 | 0.024 | 1 | 1 | Zfp945 | 0.013 | Young |
| 0.117915 | 0.178552 | 0.057 | 0.041 | 1 | 1 | Fam193a | 0.016 | Young |
| 0.118764 | 0.19359 | 0.116 | 0.094 | 1 | 1 | Srsf10 | 0.022 | Young |
| 0.118962 | 0.468895 | 0.038 | 0.026 | 1 | 1 | Jmy | 0.012 | Young |
| 0.119116 | 1.2558 | 0.016 | 0.008 | 1 | 1 | Fut4 | 0.008 | Young |
| 0.119376 | 0.732077 | 0.021 | 0.012 | 1 | 1 | Crcp | 0.009 | Young |
| 0.119478 | 0.351256 | 0.027 | 0.017 | 1 | 1 | Ctnnbl1 | 0.01 | Young |
| 0.11958 | 0.322884 | 0.061 | 0.045 | 1 | 1 | Wdr33 | 0.016 | Young |
| 0.119698 | 0.574092 | 0.061 | 0.045 | 1 | 1 | Uxs1 | 0.016 | Young |
| 0.119833 | 1.015197 | 0.016 | 0.008 | 1 | 1 | Dgkq | 0.008 | Young |

| | | | | | | | | |
|---|---|---|---|---|---|---|---|---|
| 0.119883 | 0.225057 | 0.066 | 0.049 | 1 | 1 | Txndc16 | 0.017 | Young |
| 0.119989 | 0.359442 | 0.113 | 0.093 | 1 | 1 | Xrn2 | 0.02 | Young |
| 0.120012 | 1.25862 | 0.016 | 0.008 | 1 | 1 | Hexdc | 0.008 | Young |
| 0.120051 | 0.129822 | 0.196 | 0.165 | 1 | 1 | Lrrc58 | 0.031 | Young |
| 0.120103 | 0.118023 | 0.117 | 0.094 | 1 | 1 | Pnp | 0.023 | Young |
| 0.120253 | 0.126462 | 0.085 | 0.066 | 1 | 1 | Ntan1 | 0.019 | Young |
| 0.120545 | 0.130325 | 0.058 | 0.042 | 1 | 1 | Nudt4 | 0.016 | Young |
| 0.1206 | 0.273718 | 0.051 | 0.036 | 1 | 1 | Rnf167 | 0.015 | Young |
| 0.120633 | 0.075185 | 0.334 | 0.29 | 1 | 1 | B4galt6 | 0.044 | Young |
| 0.120874 | 0.190861 | 0.106 | 0.085 | 1 | 1 | Eprs | 0.021 | Young |
| 0.120901 | 0.14941 | 0.091 | 0.071 | 1 | 1 | Hltf | 0.02 | Young |
| 0.120906 | 0.068018 | 0.048 | 0.033 | 1 | 1 | Zfp322a | 0.015 | Young |
| 0.120928 | 0.138872 | 0.076 | 0.058 | 1 | 1 | Smim13 | 0.018 | Young |
| 0.120999 | 0.262656 | 0.089 | 0.069 | 1 | 1 | Tpcn1 | 0.02 | Young |
| 0.121065 | 0.855689 | 0.029 | 0.018 | 1 | 1 | Mthfs | 0.011 | Young |
| 0.121259 | 0.092344 | 0.057 | 0.041 | 1 | 1 | Dok3 | 0.016 | Young |
| 0.121382 | 0.707447 | 0.021 | 0.012 | 1 | 1 | Rint1 | 0.009 | Young |
| 0.12141 | 0.317077 | 0.027 | 0.017 | 1 | 1 | Entpd6 | 0.01 | Young |
| 0.121765 | 1.817393 | 0.012 | 0.005 | 1 | 1 | Acpp | 0.007 | Young |
| 0.122081 | 0.1363 | 0.123 | 0.1 | 1 | 1 | Dmxl1 | 0.023 | Young |
| 0.122293 | 1.41007 | 0.012 | 0.005 | 1 | 1 | Zfp825 | 0.007 | Young |
| 0.122499 | 0.118343 | 0.083 | 0.064 | 1 | 1 | Cir1 | 0.019 | Young |
| 0.122523 | 0.1282 | 0.168 | 0.141 | 1 | 1 | Ensa | 0.027 | Young |
| 0.122552 | 0.066513 | 0.129 | 0.105 | 1 | 1 | Ilk | 0.024 | Young |
| 0.122669 | 0.264344 | 0.021 | 0.012 | 1 | 1 | Nprl3 | 0.009 | Young |
| 0.122691 | 0.309636 | 0.095 | 0.075 | 1 | 1 | Gpalpp1 | 0.02 | Young |
| 0.122806 | 0.450287 | 0.049 | 0.035 | 1 | 1 | Tmem192 | 0.014 | Young |
| 0.122823 | 1.59641 | 0.012 | 0.005 | 1 | 1 | AW209491 | 0.007 | Young |
| 0.122865 | 0.428029 | 0.021 | 0.012 | 1 | 1 | Fam58b | 0.009 | Young |
| 0.123266 | 0.406495 | 0.04 | 0.027 | 1 | 1 | Cpped1 | 0.013 | Young |
| 0.123277 | 0.226038 | 0.038 | 0.026 | 1 | 1 | Lrrcc1 | 0.012 | Young |
| 0.123415 | 0.219599 | 0.021 | 0.012 | 1 | 1 | Orc2 | 0.009 | Young |
| 0.123568 | 1.120111 | 0.012 | 0.005 | 1 | 1 | Gnb5 | 0.007 | Young |
| 0.123627 | 0.322575 | 0.075 | 0.058 | 1 | 1 | Ubqln1 | 0.017 | Young |
| 0.123782 | 1.200642 | 0.012 | 0.005 | 1 | 1 | Mturn | 0.007 | Young |
| 0.123782 | 0.805397 | 0.012 | 0.005 | 1 | 1 | Tefm | 0.007 | Young |
| 0.123797 | 1.4531 | 0.022 | 0.013 | 1 | 1 | Gm20703 | 0.009 | Young |
| 0.123955 | 0.201025 | 0.119 | 0.096 | 1 | 1 | Kcmf1 | 0.023 | Young |
| 0.124125 | 0.054708 | 0.108 | 0.086 | 1 | 1 | Atp5g1 | 0.022 | Young |
| 0.124204 | 0.225621 | 0.016 | 0.008 | 1 | 1 | Pfkm | 0.008 | Young |
| 0.124221 | 0.415736 | 0.059 | 0.044 | 1 | 1 | Nrf1 | 0.015 | Young |
| 0.12423 | 0.47341 | 0.056 | 0.041 | 1 | 1 | Pes1 | 0.015 | Young |
| 0.124317 | 0.659795 | 0.012 | 0.005 | 1 | 1 | Kcng2 | 0.007 | Young |
| 0.124502 | 0.077317 | 0.12 | 0.098 | 1 | 1 | Golga4 | 0.022 | Young |
| 0.124639 | 0.54149 | 0.012 | 0.005 | 1 | 1 | Fam131a | 0.007 | Young |

| | | | | | | | | |
|---|---|---|---|---|---|---|---|---|
| 0.124917 | 0.334779 | 0.021 | 0.012 | 1 | 1 | Cyb561d1 | 0.009 | Young |
| 0.124961 | 0.585067 | 0.012 | 0.005 | 1 | 1 | Wdr6 | 0.007 | Young |
| 0.124976 | 0.184714 | 0.128 | 0.105 | 1 | 1 | Ebi3 | 0.023 | Young |
| 0.125321 | 0.12461 | 0.303 | 0.263 | 1 | 1 | Gpr132 | 0.04 | Young |
| 0.125336 | 0.187322 | 0.204 | 0.175 | 1 | 1 | Tax1bp1 | 0.029 | Young |
| 0.125392 | 0.227502 | 0.012 | 0.005 | 1 | 1 | Pex11a | 0.007 | Young |
| 0.125439 | 0.104892 | 0.079 | 0.06 | 1 | 1 | Gramd1a | 0.019 | Young |
| 0.125594 | 0.015869 | 0.747 | 0.658 | 1 | 1 | Nfkbia | 0.089 | Young |
| 0.125683 | 0.459427 | 0.053 | 0.039 | 1 | 1 | Bahd1 | 0.014 | Young |
| 0.125874 | 0.350408 | 0.075 | 0.058 | 1 | 1 | Unc50 | 0.017 | Young |
| 0.126017 | 0.49824 | 0.029 | 0.018 | 1 | 1 | Socs5 | 0.011 | Young |
| 0.126082 | 0.559808 | 0.03 | 0.019 | 1 | 1 | Pigu | 0.011 | Young |
| 0.126209 | 0.080901 | 0.055 | 0.04 | 1 | 1 | Zswim8 | 0.015 | Young |
| 0.126239 | 0.28385 | 0.047 | 0.033 | 1 | 1 | Myh10 | 0.014 | Young |
| 0.126271 | 0.067605 | 0.118 | 0.095 | 1 | 1 | Acly | 0.023 | Young |
| 0.12628 | 0.162914 | 0.164 | 0.138 | 1 | 1 | Smap2 | 0.026 | Young |
| 0.126501 | 0.580142 | 0.032 | 0.021 | 1 | 1 | Gtdc1 | 0.011 | Young |
| 0.12654 | 0.41653 | 0.03 | 0.019 | 1 | 1 | Polr2j | 0.011 | Young |
| 0.126704 | 0.134073 | 0.118 | 0.096 | 1 | 1 | Ddx50 | 0.022 | Young |
| 0.126928 | 0.262488 | 0.1 | 0.08 | 1 | 1 | Vamp2 | 0.02 | Young |
| 0.127026 | 0.208318 | 0.029 | 0.018 | 1 | 1 | Gtf3c3 | 0.011 | Young |
| 0.127097 | 0.497 | 0.049 | 0.035 | 1 | 1 | Nckipsd | 0.014 | Young |
| 0.127234 | 0.059574 | 0.063 | 0.046 | 1 | 1 | Yrdc | 0.017 | Young |
| 0.127274 | 0.71245 | 0.033 | 0.022 | 1 | 1 | Rnf19a | 0.011 | Young |
| 0.127293 | 0.422582 | 0.123 | 0.103 | 1 | 1 | Was | 0.02 | Young |
| 0.127405 | 0.540855 | 0.022 | 0.013 | 1 | 1 | Rrs1 | 0.009 | Young |
| 0.127439 | 0.008048 | 0.209 | 0.176 | 1 | 1 | Kmt2e | 0.033 | Young |
| 0.12752 | 0.035599 | 0.97 | 0.964 | 1 | 1 | Tpt1 | 0.006 | Young |
| 0.127716 | 0.562796 | 0.022 | 0.013 | 1 | 1 | Alg9 | 0.009 | Young |
| 0.12779 | 0.073661 | 0.078 | 0.059 | 1 | 1 | Hdac2 | 0.019 | Young |
| 0.127792 | 0.224102 | 0.097 | 0.077 | 1 | 1 | Cdc73 | 0.02 | Young |
| 0.127831 | 0.098164 | 0.118 | 0.095 | 1 | 1 | N4bp2l2 | 0.023 | Young |
| 0.128174 | 0.33822 | 0.047 | 0.033 | 1 | 1 | Irf3 | 0.014 | Young |
| 0.128417 | 0.778652 | 0.022 | 0.013 | 1 | 1 | Rfc3 | 0.009 | Young |
| 0.128548 | 0.001484 | 0.081 | 0.062 | 1 | 1 | Stard7 | 0.019 | Young |
| 0.128568 | 0.081927 | 0.173 | 0.145 | 1 | 1 | Washc4 | 0.028 | Young |
| 0.128638 | 0.118191 | 0.073 | 0.055 | 1 | 1 | Agpat5 | 0.018 | Young |
| 0.128765 | 0.436576 | 0.056 | 0.041 | 1 | 1 | Smg5 | 0.015 | Young |
| 0.128792 | 0.159049 | 0.164 | 0.138 | 1 | 1 | Ash1l | 0.026 | Young |
| 0.129472 | 0.310577 | 0.052 | 0.037 | 1 | 1 | Ssr2 | 0.015 | Young |
| 0.129483 | 0.198703 | 0.043 | 0.03 | 1 | 1 | Scd2 | 0.013 | Young |
| 0.129656 | 0.162538 | 0.059 | 0.044 | 1 | 1 | Zfp622 | 0.015 | Young |
| 0.129763 | 0.095831 | 0.044 | 0.031 | 1 | 1 | Rin3 | 0.013 | Young |
| 0.129769 | 0.309513 | 0.079 | 0.062 | 1 | 1 | Myo1g | 0.017 | Young |
| 0.129875 | 0.38976 | 0.05 | 0.036 | 1 | 1 | Rnpc3 | 0.014 | Young |

| | | | | | | | | |
|---|---|---|---|---|---|---|---|---|
| 0.13002 | 0.080007 | 0.118 | 0.095 | 1 | 1 | Tmf1 | 0.023 | Young |
| 0.130132 | 0.302594 | 0.047 | 0.033 | 1 | 1 | Stard3 | 0.014 | Young |
| 0.130206 | 0.114257 | 0.061 | 0.045 | 1 | 1 | Sirt7 | 0.016 | Young |
| 0.130516 | 0.181195 | 0.098 | 0.078 | 1 | 1 | Lyl1 | 0.02 | Young |
| 0.131026 | 0.052855 | 0.162 | 0.135 | 1 | 1 | Rsu1 | 0.027 | Young |
| 0.13148 | 0.194503 | 0.05 | 0.036 | 1 | 1 | Upf3a | 0.014 | Young |
| 0.131618 | 0.132733 | 0.041 | 0.028 | 1 | 1 | Ciz1 | 0.013 | Young |
| 0.131636 | 0.898964 | 0.017 | 0.009 | 1 | 1 | Mtrf1l | 0.008 | Young |
| 0.131794 | 0.159909 | 0.049 | 0.035 | 1 | 1 | Grap | 0.014 | Young |
| 0.131819 | 0.678401 | 0.017 | 0.009 | 1 | 1 | Prr7 | 0.008 | Young |
| 0.131921 | 0.330053 | 0.083 | 0.066 | 1 | 1 | Plagl2 | 0.017 | Young |
| 0.132115 | 0.273743 | 0.055 | 0.04 | 1 | 1 | Dhrs7b | 0.015 | Young |
| 0.132187 | 0.531622 | 0.017 | 0.009 | 1 | 1 | Fbxo32 | 0.008 | Young |
| 0.132413 | 0.130401 | 0.092 | 0.073 | 1 | 1 | Slc31a2 | 0.019 | Young |
| 0.132472 | 0.218532 | 0.05 | 0.036 | 1 | 1 | Ric1 | 0.014 | Young |
| 0.132588 | 0.975553 | 0.039 | 0.027 | 1 | 1 | Aldh1a2 | 0.012 | Young |
| 0.132648 | 0.670959 | 0.017 | 0.009 | 1 | 1 | Ccdc28a | 0.008 | Young |
| 0.132714 | 0.460714 | 0.032 | 0.021 | 1 | 1 | Cnrip1 | 0.011 | Young |
| 0.132832 | 0.644262 | 0.017 | 0.009 | 1 | 1 | Rbm19 | 0.008 | Young |
| 0.133033 | 0.20403 | 0.158 | 0.134 | 1 | 1 | Wdfy3 | 0.024 | Young |
| 0.133493 | 0.016411 | 0.235 | 0.202 | 1 | 1 | Eif4h | 0.033 | Young |
| 0.133787 | 0.523736 | 0.033 | 0.022 | 1 | 1 | Hps1 | 0.011 | Young |
| 0.133835 | 0.048501 | 0.179 | 0.152 | 1 | 1 | Cdk9 | 0.027 | Young |
| 0.133858 | 0.170283 | 0.072 | 0.055 | 1 | 1 | Otud5 | 0.017 | Young |
| 0.134332 | 0.415681 | 0.079 | 0.062 | 1 | 1 | Ing1 | 0.017 | Young |
| 0.134417 | 0.283972 | 0.068 | 0.051 | 1 | 1 | Cnot2 | 0.017 | Young |
| 0.134458 | 0.241172 | 0.055 | 0.04 | 1 | 1 | 9930021J0 | 0.015 | Young |
| 0.134503 | 0.06226 | 0.017 | 0.009 | 1 | 1 | Phykpl | 0.008 | Young |
| 0.13455 | 0.018251 | 0.281 | 0.246 | 1 | 1 | Gm2a | 0.035 | Young |
| 0.134562 | 0.383495 | 0.093 | 0.075 | 1 | 1 | Sla | 0.018 | Young |
| 0.134897 | 0.444246 | 0.033 | 0.022 | 1 | 1 | Smim12 | 0.011 | Young |
| 0.135259 | 0.174903 | 0.122 | 0.1 | 1 | 1 | Alkbh5 | 0.022 | Young |
| 0.135824 | 0.187171 | 0.032 | 0.021 | 1 | 1 | AI839979 | 0.011 | Young |
| 0.135888 | 0.125999 | 0.12 | 0.098 | 1 | 1 | Nab1 | 0.022 | Young |
| 0.135979 | 0.236896 | 0.052 | 0.037 | 1 | 1 | Lemd3 | 0.015 | Young |
| 0.136362 | 0.555384 | 0.024 | 0.014 | 1 | 1 | Il1rap | 0.01 | Young |
| 0.136363 | 0.622 | 0.054 | 0.04 | 1 | 1 | Carhsp1 | 0.014 | Young |
| 0.136531 | 0.505722 | 0.038 | 0.026 | 1 | 1 | Tmem159 | 0.012 | Young |
| 0.136551 | 0.138079 | 0.055 | 0.04 | 1 | 1 | Ilvbl | 0.015 | Young |
| 0.136637 | 0.547627 | 0.036 | 0.024 | 1 | 1 | Abcf2 | 0.012 | Young |
| 0.137061 | 0.002601 | 0.227 | 0.195 | 1 | 1 | Cd68 | 0.032 | Young |
| 0.137244 | 0.565649 | 0.052 | 0.039 | 1 | 1 | Tax1bp3 | 0.013 | Young |
| 0.137265 | 1.713104 | 0.013 | 0.006 | 1 | 1 | Acap3 | 0.007 | Young |
| 0.137294 | 0.12167 | 0.336 | 0.299 | 1 | 1 | Csrnp1 | 0.037 | Young |
| 0.137338 | 0.598879 | 0.038 | 0.026 | 1 | 1 | Gpr137 | 0.012 | Young |

| | | | | | | | | |
|---|---|---|---|---|---|---|---|---|
| 0.137523 | 0.657866 | 0.039 | 0.027 | 1 | 1 | Adcy9 | 0.012 | Young |
| 0.137731 | 0.224095 | 0.061 | 0.045 | 1 | 1 | Rsbn1 | 0.016 | Young |
| 0.137772 | 0.365402 | 0.041 | 0.028 | 1 | 1 | Adk | 0.013 | Young |
| 0.137773 | 0.407627 | 0.035 | 0.023 | 1 | 1 | Lig1 | 0.012 | Young |
| 0.138135 | 0.189171 | 0.168 | 0.143 | 1 | 1 | Snu13 | 0.025 | Young |
| 0.138241 | 0.138427 | 0.065 | 0.049 | 1 | 1 | Naa38 | 0.016 | Young |
| 0.138266 | 1.452475 | 0.01 | 0.004 | 1 | 1 | Ddb2 | 0.006 | Young |
| 0.13832 | 0.129629 | 0.074 | 0.057 | 1 | 1 | Ube2j1 | 0.017 | Young |
| 0.138396 | 1.400561 | 0.01 | 0.004 | 1 | 1 | Mapk1ip1 | 0.006 | Young |
| 0.138503 | 0.877223 | 0.042 | 0.03 | 1 | 1 | Nkiras2 | 0.012 | Young |
| 0.138526 | 1.150064 | 0.01 | 0.004 | 1 | 1 | Plcl1 | 0.006 | Young |
| 0.138722 | 1.228983 | 0.01 | 0.004 | 1 | 1 | Mpv17l | 0.006 | Young |
| 0.138787 | 0.649644 | 0.01 | 0.004 | 1 | 1 | Nsun5 | 0.006 | Young |
| 0.138787 | 0.070012 | 0.024 | 0.014 | 1 | 1 | Kdm1b | 0.01 | Young |
| 0.138917 | 1.190499 | 0.01 | 0.004 | 1 | 1 | Glb1l | 0.006 | Young |
| 0.139048 | 0.335581 | 0.01 | 0.004 | 1 | 1 | Itgb8 | 0.006 | Young |
| 0.139125 | 0.525042 | 0.058 | 0.044 | 1 | 1 | Tmem147 | 0.014 | Young |
| 0.139179 | 0.983605 | 0.01 | 0.004 | 1 | 1 | Nt5c3b | 0.006 | Young |
| 0.139179 | 0.646452 | 0.01 | 0.004 | 1 | 1 | Nudt22 | 0.006 | Young |
| 0.13922 | 1.09792 | 0.013 | 0.006 | 1 | 1 | Zfp346 | 0.007 | Young |
| 0.139309 | 1.005195 | 0.01 | 0.004 | 1 | 1 | Lekr1 | 0.006 | Young |
| 0.139331 | 0.045116 | 0.064 | 0.048 | 1 | 1 | Nf1 | 0.016 | Young |
| 0.139367 | 0.176511 | 0.032 | 0.021 | 1 | 1 | Vopp1 | 0.011 | Young |
| 0.139375 | 1.072719 | 0.01 | 0.004 | 1 | 1 | Lyrm4 | 0.006 | Young |
| 0.139571 | 0.860736 | 0.01 | 0.004 | 1 | 1 | Chst14 | 0.006 | Young |
| 0.139571 | 0.53543 | 0.01 | 0.004 | 1 | 1 | Zfp523 | 0.006 | Young |
| 0.139595 | 0.052601 | 0.169 | 0.143 | 1 | 1 | Hnrnpa1 | 0.026 | Young |
| 0.139658 | 0.620432 | 0.013 | 0.006 | 1 | 1 | Atrip | 0.007 | Young |
| 0.139717 | 0.446216 | 0.038 | 0.026 | 1 | 1 | Vps26b | 0.012 | Young |
| 0.139964 | 0.844138 | 0.01 | 0.004 | 1 | 1 | Pdia5 | 0.006 | Young |
| 0.140035 | 0.544167 | 0.025 | 0.015 | 1 | 1 | Snx16 | 0.01 | Young |
| 0.140098 | 0.65537 | 0.057 | 0.042 | 1 | 1 | Mboat7 | 0.015 | Young |
| 0.140186 | 0.189198 | 0.123 | 0.102 | 1 | 1 | Dusp22 | 0.021 | Young |
| 0.140227 | 0.563801 | 0.01 | 0.004 | 1 | 1 | Gm10791 | 0.006 | Young |
| 0.14023 | 0.700748 | 0.018 | 0.01 | 1 | 1 | Saysd1 | 0.008 | Young |
| 0.140316 | 0.735993 | 0.013 | 0.006 | 1 | 1 | Ttyh2 | 0.007 | Young |
| 0.140359 | 0.679453 | 0.01 | 0.004 | 1 | 1 | Cdo1 | 0.006 | Young |
| 0.14049 | 0.104043 | 0.01 | 0.004 | 1 | 1 | Tmem107 | 0.006 | Young |
| 0.14069 | 0.656421 | 0.018 | 0.01 | 1 | 1 | Med6 | 0.008 | Young |
| 0.140911 | 0.684283 | 0.045 | 0.032 | 1 | 1 | Klhdc2 | 0.013 | Young |
| 0.140952 | 0.014748 | 0.056 | 0.041 | 1 | 1 | Acadsb | 0.015 | Young |
| 0.141044 | 0.005658 | 0.105 | 0.084 | 1 | 1 | Usp48 | 0.021 | Young |
| 0.141307 | 0.546369 | 0.013 | 0.006 | 1 | 1 | Msh6 | 0.007 | Young |
| 0.141424 | 0.171754 | 0.212 | 0.185 | 1 | 1 | Sec14l1 | 0.027 | Young |
| 0.14151 | 0.462663 | 0.084 | 0.067 | 1 | 1 | Arhgap25 | 0.017 | Young |

| | | | | | | | | |
|---|---|---|---|---|---|---|---|---|
| 0.141528 | 0.398964 | 0.013 | 0.006 | 1 | 1 | Sirt3 | 0.007 | Young |
| 0.141735 | 0.586888 | 0.051 | 0.037 | 1 | 1 | Tbc1d31 | 0.014 | Young |
| 0.14186 | 0.217508 | 0.013 | 0.006 | 1 | 1 | Bcl11a | 0.007 | Young |
| 0.141984 | 0.42114 | 0.018 | 0.01 | 1 | 1 | Cog1 | 0.008 | Young |
| 0.142293 | 0.312961 | 0.036 | 0.024 | 1 | 1 | Snhg6 | 0.012 | Young |
| 0.142358 | 0.219169 | 0.036 | 0.024 | 1 | 1 | Eef1akmt1 | 0.012 | Young |
| 0.14263 | 0.048585 | 0.271 | 0.234 | 1 | 1 | Rtn3 | 0.037 | Young |
| 0.142681 | 0.314728 | 0.078 | 0.06 | 1 | 1 | Rnf115 | 0.018 | Young |
| 0.143019 | 0.037416 | 0.3 | 0.261 | 1 | 1 | Scp2 | 0.039 | Young |
| 0.143062 | 0.089322 | 0.025 | 0.015 | 1 | 1 | Tcp11l1 | 0.01 | Young |
| 0.143065 | 0.266935 | 0.055 | 0.041 | 1 | 1 | Slc30a9 | 0.014 | Young |
| 0.143083 | 0.12412 | 0.013 | 0.006 | 1 | 1 | Zfp768 | 0.007 | Young |
| 0.143254 | 0.524062 | 0.086 | 0.069 | 1 | 1 | Ankhd1 | 0.017 | Young |
| 0.143405 | 0.234704 | 0.121 | 0.1 | 1 | 1 | Thoc2 | 0.021 | Young |
| 0.143536 | 0.173475 | 0.086 | 0.068 | 1 | 1 | Fam76a | 0.018 | Young |
| 0.143638 | 0.240551 | 0.058 | 0.042 | 1 | 1 | Clpx | 0.016 | Young |
| 0.143902 | 0.562856 | 0.042 | 0.03 | 1 | 1 | Tusc3 | 0.012 | Young |
| 0.143908 | 0.35812 | 0.093 | 0.075 | 1 | 1 | Rps6ka1 | 0.018 | Young |
| 0.144176 | 0.341801 | 0.055 | 0.041 | 1 | 1 | Inpp5f | 0.014 | Young |
| 0.144266 | 0.044684 | 0.049 | 0.035 | 1 | 1 | Abcb1b | 0.014 | Young |
| 0.144394 | 0.132813 | 0.088 | 0.069 | 1 | 1 | Fbxo11 | 0.019 | Young |
| 0.144777 | 0.0489 | 0.128 | 0.105 | 1 | 1 | Dot1l | 0.023 | Young |
| 0.14496 | 0.335228 | 0.038 | 0.026 | 1 | 1 | Slc39a14 | 0.012 | Young |
| 0.145079 | 0.385608 | 0.069 | 0.053 | 1 | 1 | Fam103a1 | 0.016 | Young |
| 0.14512 | 0.126176 | 0.072 | 0.055 | 1 | 1 | Vps50 | 0.017 | Young |
| 0.145386 | 0.04579 | 0.311 | 0.271 | 1 | 1 | H2-T23 | 0.04 | Young |
| 0.145715 | 0.283763 | 0.057 | 0.042 | 1 | 1 | Cnep1r1 | 0.015 | Young |
| 0.146045 | 0.357338 | 0.042 | 0.03 | 1 | 1 | Hspb11 | 0.012 | Young |
| 0.146108 | 0.161307 | 0.122 | 0.1 | 1 | 1 | Nrbp1 | 0.022 | Young |
| 0.146217 | 0.413313 | 0.057 | 0.042 | 1 | 1 | Itga5 | 0.015 | Young |
| 0.146221 | 0.391937 | 0.044 | 0.031 | 1 | 1 | Cdkn2aipnl | 0.013 | Young |
| 0.146243 | 0.109552 | 0.074 | 0.057 | 1 | 1 | Cul5 | 0.017 | Young |
| 0.146312 | 0.394198 | 0.049 | 0.036 | 1 | 1 | Elp2 | 0.013 | Young |
| 0.146462 | 0.135031 | 0.119 | 0.098 | 1 | 1 | Zbtb4 | 0.021 | Young |
| 0.146789 | 0.166568 | 0.164 | 0.14 | 1 | 1 | Acsl4 | 0.024 | Young |
| 0.146868 | 0.121106 | 0.16 | 0.135 | 1 | 1 | Got1 | 0.025 | Young |
| 0.147013 | 0.405202 | 0.06 | 0.045 | 1 | 1 | Agtpbp1 | 0.015 | Young |
| 0.147092 | 0.436425 | 0.058 | 0.044 | 1 | 1 | Mplkip | 0.014 | Young |
| 0.147144 | 0.077032 | 0.072 | 0.055 | 1 | 1 | Imp3 | 0.017 | Young |
| 0.147539 | 0.490526 | 0.028 | 0.018 | 1 | 1 | Tmem161b | 0.01 | Young |
| 0.147612 | 0.794425 | 0.02 | 0.012 | 1 | 1 | Zbtb41 | 0.008 | Young |
| 0.147849 | 0.351694 | 0.074 | 0.058 | 1 | 1 | Cpq | 0.016 | Young |
| 0.147862 | 0.481469 | 0.072 | 0.057 | 1 | 1 | Nab2 | 0.015 | Young |
| 0.147957 | 0.046792 | 0.177 | 0.149 | 1 | 1 | Nono | 0.028 | Young |
| 0.148016 | 0.30988 | 0.044 | 0.031 | 1 | 1 | Prpsap1 | 0.013 | Young |

| | | | | | | | | |
|---|---|---|---|---|---|---|---|---|
| 0.148219 | 0.016271 | 0.027 | 0.017 | 1 | 1 | Prkrip1 | 0.01 | Young |
| 0.148444 | 0.321663 | 0.125 | 0.104 | 1 | 1 | Crebrf | 0.021 | Young |
| 0.14848 | 0.410255 | 0.048 | 0.035 | 1 | 1 | Usp38 | 0.013 | Young |
| 0.148505 | 0.332265 | 0.044 | 0.031 | 1 | 1 | Rab6b | 0.013 | Young |
| 0.148565 | 0.209032 | 0.036 | 0.024 | 1 | 1 | Rbm15b | 0.012 | Young |
| 0.148602 | 0.484519 | 0.051 | 0.037 | 1 | 1 | Il15 | 0.014 | Young |
| 0.148613 | 0.217233 | 0.028 | 0.018 | 1 | 1 | Pithd1 | 0.01 | Young |
| 0.148622 | 0.93693 | 0.02 | 0.012 | 1 | 1 | Med24 | 0.008 | Young |
| 0.148829 | 0.213258 | 0.045 | 0.032 | 1 | 1 | Rufy1 | 0.013 | Young |
| 0.148899 | 0.748935 | 0.02 | 0.012 | 1 | 1 | Twsg1 | 0.008 | Young |
| 0.149355 | 0.004975 | 0.038 | 0.026 | 1 | 1 | Ormdl1 | 0.012 | Young |
| 0.14936 | 0.681218 | 0.02 | 0.012 | 1 | 1 | Timm29 | 0.008 | Young |
| 0.14937 | 0.03496 | 0.082 | 0.064 | 1 | 1 | Scamp3 | 0.018 | Young |
| 0.149457 | 0.256322 | 0.047 | 0.033 | 1 | 1 | Clec5a | 0.014 | Young |
| 0.149579 | 0.050281 | 0.127 | 0.104 | 1 | 1 | Acadm | 0.023 | Young |
| 0.149769 | 0.22461 | 0.057 | 0.042 | 1 | 1 | Ndufa9 | 0.015 | Young |
| 0.149916 | 0.197881 | 0.02 | 0.012 | 1 | 1 | Map3k15 | 0.008 | Young |
| 0.149922 | 0.233117 | 0.137 | 0.116 | 1 | 1 | Snx10 | 0.021 | Young |
| 0.150107 | 0.2857 | 0.069 | 0.053 | 1 | 1 | Slamf8 | 0.016 | Young |
| 0.150299 | 0.202372 | 0.049 | 0.036 | 1 | 1 | Polr3c | 0.013 | Young |
| 0.150522 | 0.112964 | 0.052 | 0.039 | 1 | 1 | Atp6v0a2 | 0.013 | Young |
| 0.150682 | 0.290403 | 0.145 | 0.123 | 1 | 1 | Usp8 | 0.022 | Young |
| 0.151014 | 0.208855 | 0.072 | 0.055 | 1 | 1 | Mospd3 | 0.017 | Young |
| 0.1512 | 0.615131 | 0.03 | 0.019 | 1 | 1 | Apex1 | 0.011 | Young |
| 0.151264 | 0.459221 | 0.02 | 0.012 | 1 | 1 | Fancg | 0.008 | Young |
| 0.151438 | 0.61044 | 0.031 | 0.021 | 1 | 1 | Rac3 | 0.01 | Young |
| 0.151592 | 0.156575 | 0.104 | 0.085 | 1 | 1 | Slc9a3r1 | 0.019 | Young |
| 0.151639 | 0.564017 | 0.072 | 0.057 | 1 | 1 | Pfkfb3 | 0.015 | Young |
| 0.15166 | 0.537021 | 0.031 | 0.021 | 1 | 1 | Mlx | 0.01 | Young |
| 0.151817 | 0.101964 | 0.117 | 0.095 | 1 | 1 | Ndufs2 | 0.022 | Young |
| 0.151951 | 0.269185 | 0.048 | 0.035 | 1 | 1 | Chpt1 | 0.013 | Young |
| 0.1521 | 0.002867 | 0.304 | 0.263 | 1 | 1 | Slfn2 | 0.041 | Young |
| 0.152283 | 0.655989 | 0.062 | 0.048 | 1 | 1 | Tpm1 | 0.014 | Young |
| 0.152601 | 0.014949 | 0.061 | 0.046 | 1 | 1 | Slc38a10 | 0.015 | Young |
| 0.152659 | 0.893104 | 0.015 | 0.008 | 1 | 1 | Ssx2ip | 0.007 | Young |
| 0.152667 | 0.491422 | 0.068 | 0.053 | 1 | 1 | Tmem41b | 0.015 | Young |
| 0.152699 | 0.630506 | 0.031 | 0.021 | 1 | 1 | 2010315B0 | 0.01 | Young |
| 0.15289 | 0.329965 | 0.075 | 0.059 | 1 | 1 | Pip5k1a | 0.016 | Young |
| 0.15297 | 0.173685 | 0.066 | 0.05 | 1 | 1 | Rab3gap1 | 0.016 | Young |
| 0.153334 | 0.737 | 0.041 | 0.03 | 1 | 1 | Syne1 | 0.011 | Young |
| 0.153416 | 0.16635 | 0.051 | 0.037 | 1 | 1 | Kbtbd11 | 0.014 | Young |
| 0.153766 | 1.012774 | 0.021 | 0.013 | 1 | 1 | Itgb7 | 0.008 | Young |
| 0.153865 | 0.685174 | 0.056 | 0.042 | 1 | 1 | Fam193b | 0.014 | Young |
| 0.153876 | 0.514288 | 0.015 | 0.008 | 1 | 1 | Zfp65 | 0.007 | Young |
| 0.154131 | 0.108549 | 0.15 | 0.126 | 1 | 1 | Ppp1r12a | 0.024 | Young |

| | | | | | | | | |
|---|---|---|---|---|---|---|---|---|
| 0.154178 | 0.314536 | 0.03 | 0.019 | 1 | 1 | Zc3h7b | 0.011 | Young |
| 0.15448 | 0.759918 | 0.034 | 0.023 | 1 | 1 | Prmt2 | 0.011 | Young |
| 0.154553 | 0.176563 | 0.047 | 0.033 | 1 | 1 | Washc5 | 0.014 | Young |
| 0.155014 | 0.421023 | 0.064 | 0.049 | 1 | 1 | Hibadh | 0.015 | Young |
| 0.155212 | 0.118058 | 0.015 | 0.008 | 1 | 1 | Znhit2 | 0.007 | Young |
| 0.155417 | 0.043215 | 0.028 | 0.018 | 1 | 1 | Calcoco1 | 0.01 | Young |
| 0.155445 | 0.727039 | 0.035 | 0.024 | 1 | 1 | Scimp | 0.011 | Young |
| 0.1555 | 0.711802 | 0.021 | 0.013 | 1 | 1 | Paip2b | 0.008 | Young |
| 0.155874 | 0.277874 | 0.054 | 0.04 | 1 | 1 | Avpi1 | 0.014 | Young |
| 0.155956 | 0.112061 | 0.264 | 0.231 | 1 | 1 | Fli1 | 0.033 | Young |
| 0.156043 | 0.03116 | 0.051 | 0.037 | 1 | 1 | Rrad | 0.014 | Young |
| 0.156078 | 0.213519 | 0.111 | 0.091 | 1 | 1 | Mgrn1 | 0.02 | Young |
| 0.156104 | 0.090192 | 0.064 | 0.049 | 1 | 1 | Fxr2 | 0.015 | Young |
| 0.156132 | 0.068311 | 0.409 | 0.364 | 1 | 1 | Hnrnph1 | 0.045 | Young |
| 0.156234 | 0.53993 | 0.021 | 0.013 | 1 | 1 | Paxx | 0.008 | Young |
| 0.15626 | 0.231923 | 0.171 | 0.147 | 1 | 1 | Furin | 0.024 | Young |
| 0.156467 | 0.161907 | 0.106 | 0.086 | 1 | 1 | Mtmr6 | 0.02 | Young |
| 0.156497 | 0.031321 | 0.2 | 0.171 | 1 | 1 | Ythdc1 | 0.029 | Young |
| 0.156711 | 0.11809 | 0.15 | 0.126 | 1 | 1 | Tmem55b | 0.024 | Young |
| 0.15675 | 0.116101 | 0.138 | 0.117 | 1 | 1 | Lpp | 0.021 | Young |
| 0.157245 | 0.839977 | 0.044 | 0.032 | 1 | 1 | Lrch1 | 0.012 | Young |
| 0.157299 | 0.004846 | 0.4 | 0.346 | 1 | 1 | Ptpn18 | 0.054 | Young |
| 0.157622 | 0.249145 | 0.081 | 0.064 | 1 | 1 | Dctn3 | 0.017 | Young |
| 0.157642 | 1.565715 | 0.011 | 0.005 | 1 | 1 | Cyb5rl | 0.006 | Young |
| 0.15774 | 0.275864 | 0.11 | 0.09 | 1 | 1 | Cnih4 | 0.02 | Young |
| 0.157853 | 1.143061 | 0.023 | 0.014 | 1 | 1 | Zc3hc1 | 0.009 | Young |
| 0.158092 | 0.111744 | 0.079 | 0.062 | 1 | 1 | Tdrd7 | 0.017 | Young |
| 0.15819 | 0.064853 | 0.052 | 0.039 | 1 | 1 | Pmpca | 0.013 | Young |
| 0.158198 | 0.616163 | 0.04 | 0.028 | 1 | 1 | Mapkapk3 | 0.012 | Young |
| 0.158242 | 0.534423 | 0.035 | 0.024 | 1 | 1 | Ndst2 | 0.011 | Young |
| 0.158254 | 0.266128 | 0.103 | 0.085 | 1 | 1 | Sike1 | 0.018 | Young |
| 0.158436 | 1.634727 | 0.011 | 0.005 | 1 | 1 | Gpt | 0.006 | Young |
| 0.158481 | 0.458576 | 0.061 | 0.046 | 1 | 1 | Zmym2 | 0.015 | Young |
| 0.158545 | 0.15307 | 0.021 | 0.013 | 1 | 1 | Gm38843 | 0.008 | Young |
| 0.158701 | 1.684899 | 0.011 | 0.005 | 1 | 1 | Usp2 | 0.006 | Young |
| 0.158917 | 0.2718 | 0.021 | 0.013 | 1 | 1 | Primpol | 0.008 | Young |
| 0.159099 | 1.627707 | 0.011 | 0.005 | 1 | 1 | Card6 | 0.006 | Young |
| 0.159149 | 0.029443 | 0.199 | 0.17 | 1 | 1 | Akr1b3 | 0.029 | Young |
| 0.15917 | 0.68127 | 0.041 | 0.03 | 1 | 1 | Brwd1 | 0.011 | Young |
| 0.159531 | 0.295614 | 0.074 | 0.058 | 1 | 1 | Rab43 | 0.016 | Young |
| 0.159535 | 0.25894 | 0.087 | 0.069 | 1 | 1 | Fads1 | 0.018 | Young |
| 0.159552 | 0.347572 | 0.034 | 0.023 | 1 | 1 | Afg3l2 | 0.011 | Young |
| 0.159628 | 0.22473 | 0.149 | 0.127 | 1 | 1 | Tspan14 | 0.022 | Young |
| 0.159755 | 0.09392 | 0.147 | 0.125 | 1 | 1 | Arl8b | 0.022 | Young |
| 0.159899 | 1.273573 | 0.011 | 0.005 | 1 | 1 | Rmdn1 | 0.006 | Young |

| | | | | | | | | |
|---|---|---|---|---|---|---|---|---|
| 0.159914 | 0.220222 | 0.072 | 0.057 | 1 | 1 | Hivep1 | 0.015 | Young |
| 0.159918 | 0.604137 | 0.023 | 0.014 | 1 | 1 | Mphosph1 | 0.009 | Young |
| 0.159967 | 0.175486 | 0.134 | 0.112 | 1 | 1 | Cebpd | 0.022 | Young |
| 0.160032 | 1.277108 | 0.011 | 0.005 | 1 | 1 | Slc25a53 | 0.006 | Young |
| 0.160057 | 0.076292 | 0.11 | 0.09 | 1 | 1 | Cdc37 | 0.02 | Young |
| 0.160166 | 1.079488 | 0.011 | 0.005 | 1 | 1 | Cd320 | 0.006 | Young |
| 0.160238 | 0.129489 | 0.104 | 0.085 | 1 | 1 | Bax | 0.019 | Young |
| 0.160265 | 0.446345 | 0.056 | 0.042 | 1 | 1 | Usp33 | 0.014 | Young |
| 0.160299 | 0.800807 | 0.011 | 0.005 | 1 | 1 | Jag1 | 0.006 | Young |
| 0.160355 | 0.188158 | 0.097 | 0.078 | 1 | 1 | Nfatc2 | 0.019 | Young |
| 0.160433 | 0.810127 | 0.011 | 0.005 | 1 | 1 | Gm15445 | 0.006 | Young |
| 0.160434 | 0.028072 | 0.031 | 0.021 | 1 | 1 | Stx2 | 0.01 | Young |
| 0.16046 | 0.574006 | 0.023 | 0.014 | 1 | 1 | Ric8b | 0.009 | Young |
| 0.160527 | 0.136853 | 0.086 | 0.068 | 1 | 1 | Frmd4a | 0.018 | Young |
| 0.160758 | 0.150227 | 0.193 | 0.167 | 1 | 1 | Itsn2 | 0.026 | Young |
| 0.160835 | 0.731129 | 0.011 | 0.005 | 1 | 1 | Wdr24 | 0.006 | Young |
| 0.161093 | 0.299587 | 0.066 | 0.051 | 1 | 1 | Abca3 | 0.015 | Young |
| 0.161438 | 0.293394 | 0.085 | 0.068 | 1 | 1 | Tm2d2 | 0.017 | Young |
| 0.161506 | 0.75426 | 0.011 | 0.005 | 1 | 1 | 1110004E0 | 0.006 | Young |
| 0.16164 | 0.795268 | 0.011 | 0.005 | 1 | 1 | Smyd4 | 0.006 | Young |
| 0.16164 | 0.420926 | 0.011 | 0.005 | 1 | 1 | Thap4 | 0.006 | Young |
| 0.1622 | 0.371387 | 0.035 | 0.024 | 1 | 1 | Myc | 0.011 | Young |
| 0.162277 | 0.500274 | 0.023 | 0.014 | 1 | 1 | Cry1 | 0.009 | Young |
| 0.162369 | 0.716997 | 0.023 | 0.014 | 1 | 1 | Vegfb | 0.009 | Young |
| 0.162385 | 0.404678 | 0.047 | 0.035 | 1 | 1 | Fnbp4 | 0.012 | Young |
| 0.162492 | 0.36974 | 0.052 | 0.039 | 1 | 1 | Rxrb | 0.013 | Young |
| 0.162584 | 0.265625 | 0.011 | 0.005 | 1 | 1 | Abcb8 | 0.006 | Young |
| 0.162668 | 0.485255 | 0.038 | 0.027 | 1 | 1 | 0610037L1 | 0.011 | Young |
| 0.162687 | 0.336092 | 0.1 | 0.082 | 1 | 1 | Nudcd3 | 0.018 | Young |
| 0.162714 | 0.041135 | 0.188 | 0.163 | 1 | 1 | Ehbp1l1 | 0.025 | Young |
| 0.162917 | 0.427122 | 0.023 | 0.014 | 1 | 1 | Cd2 | 0.009 | Young |
| 0.162983 | 0.3544 | 0.038 | 0.027 | 1 | 1 | Nlrc4 | 0.011 | Young |
| 0.163114 | 0.467582 | 0.046 | 0.033 | 1 | 1 | Topors | 0.013 | Young |
| 0.163125 | 0.404456 | 0.011 | 0.005 | 1 | 1 | Zfp386 | 0.006 | Young |
| 0.163286 | 0.223161 | 0.087 | 0.069 | 1 | 1 | Fbxl20 | 0.018 | Young |
| 0.163827 | 0.816702 | 0.016 | 0.009 | 1 | 1 | Dnajc12 | 0.007 | Young |
| 0.163854 | 0.442271 | 0.091 | 0.073 | 1 | 1 | Malt1 | 0.018 | Young |
| 0.163951 | 0.3371 | 0.035 | 0.024 | 1 | 1 | Gtf3a | 0.011 | Young |
| 0.164046 | 0.466551 | 0.04 | 0.028 | 1 | 1 | Dtnbp1 | 0.012 | Young |
| 0.164047 | 0.601223 | 0.016 | 0.009 | 1 | 1 | Spsb2 | 0.007 | Young |
| 0.16423 | 0.041833 | 0.113 | 0.093 | 1 | 1 | Efcab14 | 0.02 | Young |
| 0.164291 | 0.337446 | 0.064 | 0.049 | 1 | 1 | Pex7 | 0.015 | Young |
| 0.164493 | 0.10394 | 0.063 | 0.048 | 1 | 1 | Zmym4 | 0.015 | Young |
| 0.164603 | 0.144184 | 0.109 | 0.09 | 1 | 1 | Rab8a | 0.019 | Young |
| 0.164622 | 0.050003 | 0.109 | 0.089 | 1 | 1 | Cebpz | 0.02 | Young |

| | | | | | | | | |
|---|---|---|---|---|---|---|---|---|
| 0.164755 | 0.026616 | 0.061 | 0.046 | 1 | 1 | Fez2 | 0.015 | Young |
| 0.164784 | 0.368072 | 0.049 | 0.036 | 1 | 1 | Prdx4 | 0.013 | Young |
| 0.164892 | 0.162369 | 0.066 | 0.051 | 1 | 1 | Hdac1 | 0.015 | Young |
| 0.165043 | 0.70154 | 0.016 | 0.009 | 1 | 1 | Dmac2 | 0.007 | Young |
| 0.165056 | 0.271057 | 0.038 | 0.027 | 1 | 1 | Ppcdc | 0.011 | Young |
| 0.165096 | 0.441955 | 0.049 | 0.036 | 1 | 1 | Arl6ip4 | 0.013 | Young |
| 0.165123 | 0.377318 | 0.062 | 0.048 | 1 | 1 | Dnajc10 | 0.014 | Young |
| 0.165189 | 0.277094 | 0.071 | 0.055 | 1 | 1 | Kdm4a | 0.016 | Young |
| 0.165265 | 0.644813 | 0.016 | 0.009 | 1 | 1 | Serf1 | 0.007 | Young |
| 0.165294 | 0.381131 | 0.092 | 0.075 | 1 | 1 | Trim35 | 0.017 | Young |
| 0.165377 | 0.34226 | 0.041 | 0.03 | 1 | 1 | Arhgap6 | 0.011 | Young |
| 0.165598 | 0.079526 | 0.016 | 0.009 | 1 | 1 | Fam122a | 0.007 | Young |
| 0.165657 | 0.528398 | 0.038 | 0.027 | 1 | 1 | Srpk1 | 0.011 | Young |
| 0.16582 | 0.622941 | 0.016 | 0.009 | 1 | 1 | Ccdc84 | 0.007 | Young |
| 0.16582 | 0.418645 | 0.016 | 0.009 | 1 | 1 | Zfp212 | 0.007 | Young |
| 0.165831 | 0.436308 | 0.044 | 0.032 | 1 | 1 | Vps37b | 0.012 | Young |
| 0.16595 | 0.075751 | 0.36 | 0.325 | 1 | 1 | Glud1 | 0.035 | Young |
| 0.166016 | 0.315891 | 0.093 | 0.076 | 1 | 1 | Dnase1l1 | 0.017 | Young |
| 0.166031 | 0.378611 | 0.052 | 0.039 | 1 | 1 | Agap1 | 0.013 | Young |
| 0.166233 | 0.220205 | 0.035 | 0.024 | 1 | 1 | Ctps2 | 0.011 | Young |
| 0.166423 | 0.225601 | 0.037 | 0.026 | 1 | 1 | Pitpnm1 | 0.011 | Young |
| 0.166705 | 0.708784 | 0.024 | 0.015 | 1 | 1 | Rdh10 | 0.009 | Young |
| 0.166828 | 0.282919 | 0.038 | 0.027 | 1 | 1 | Rcor3 | 0.011 | Young |
| 0.166936 | 0.369713 | 0.016 | 0.009 | 1 | 1 | Yipf2 | 0.007 | Young |
| 0.167119 | 0.07402 | 0.015 | 0.023 | 1 | 1 | Rdh14 | -0.008 | Young |
| 0.167217 | 0.025296 | 0.294 | 0.256 | 1 | 1 | Rassf2 | 0.038 | Young |
| 0.167336 | 0.369739 | 0.024 | 0.015 | 1 | 1 | Pak4 | 0.009 | Young |
| 0.167445 | 0.017776 | 0.269 | 0.234 | 1 | 1 | Prrc2c | 0.035 | Young |
| 0.167944 | 0.309413 | 0.016 | 0.009 | 1 | 1 | 1700096K1 | 0.007 | Young |
| 0.168059 | 0.172017 | 0.129 | 0.108 | 1 | 1 | Ostc | 0.021 | Young |
| 0.168503 | 0.240841 | 0.04 | 0.028 | 1 | 1 | Sp3os | 0.012 | Young |
| 0.16858 | 0.15979 | 0.151 | 0.13 | 1 | 1 | Eif4g1 | 0.021 | Young |
| 0.168603 | 0.448102 | 0.024 | 0.015 | 1 | 1 | Polr3e | 0.009 | Young |
| 0.168722 | 0.267305 | 0.056 | 0.042 | 1 | 1 | Kmt5c | 0.014 | Young |
| 0.168967 | 0.00436 | 0.222 | 0.19 | 1 | 1 | Arid1a | 0.032 | Young |
| 0.169176 | 0.017066 | 0.095 | 0.076 | 1 | 1 | Mrpl20 | 0.019 | Young |
| 0.169204 | 0.184179 | 0.16 | 0.136 | 1 | 1 | Add1 | 0.024 | Young |
| 0.169302 | 0.321201 | 0.081 | 0.064 | 1 | 1 | Kctd20 | 0.017 | Young |
| 0.169483 | 0.233275 | 0.061 | 0.046 | 1 | 1 | Nfkbib | 0.015 | Young |
| 0.169772 | 0.279535 | 0.04 | 0.028 | 1 | 1 | Fam220a | 0.012 | Young |
| 0.169796 | 0.298954 | 0.092 | 0.075 | 1 | 1 | Nr2c2 | 0.017 | Young |
| 0.169807 | 0.084979 | 0.168 | 0.145 | 1 | 1 | Herc1 | 0.023 | Young |
| 0.17006 | 0.181291 | 0.024 | 0.015 | 1 | 1 | Oxsm | 0.009 | Young |
| 0.170222 | 0.195963 | 0.131 | 0.111 | 1 | 1 | Rreb1 | 0.02 | Young |
| 0.170243 | 0.160702 | 0.024 | 0.015 | 1 | 1 | Pigv | 0.009 | Young |

| | | | | | | | | |
|---|---|---|---|---|---|---|---|---|
| 0.170286 | 0.209406 | 0.055 | 0.041 | 1 | 1 | Hmgn2 | 0.014 | Young |
| 0.170356 | 0.112635 | 0.097 | 0.078 | 1 | 1 | Aoah | 0.019 | Young |
| 0.170658 | 0.352808 | 0.044 | 0.032 | 1 | 1 | Trp53bp2 | 0.012 | Young |
| 0.170701 | 0.254861 | 0.024 | 0.015 | 1 | 1 | Usp28 | 0.009 | Young |
| 0.170778 | 0.027586 | 0.129 | 0.107 | 1 | 1 | Mgat1 | 0.022 | Young |
| 0.1709 | 0.034877 | 0.361 | 0.321 | 1 | 1 | Ccdc50 | 0.04 | Young |
| 0.170977 | 0.165751 | 0.053 | 0.04 | 1 | 1 | Rnf166 | 0.013 | Young |
| 0.171159 | 0.184495 | 0.024 | 0.015 | 1 | 1 | Lrrc40 | 0.009 | Young |
| 0.171205 | 0.186697 | 0.024 | 0.015 | 1 | 1 | Arel1 | 0.009 | Young |
| 0.171441 | 0.102196 | 0.177 | 0.152 | 1 | 1 | Map7d1 | 0.025 | Young |
| 0.171578 | 0.194025 | 0.059 | 0.045 | 1 | 1 | Hspa4l | 0.014 | Young |
| 0.171685 | 0.664756 | 0.027 | 0.018 | 1 | 1 | Zc3h8 | 0.009 | Young |
| 0.171693 | 0.192166 | 0.075 | 0.059 | 1 | 1 | Sh3pxd2a | 0.016 | Young |
| 0.171794 | 0.202378 | 0.05 | 0.037 | 1 | 1 | Pygb | 0.013 | Young |
| 0.172303 | 0.050962 | 0.08 | 0.063 | 1 | 1 | Ncbp3 | 0.017 | Young |
| 0.172465 | 0.742441 | 0.027 | 0.018 | 1 | 1 | Wiz | 0.009 | Young |
| 0.172534 | 0.194724 | 0.055 | 0.041 | 1 | 1 | Capn7 | 0.014 | Young |
| 0.172642 | 0.240061 | 0.041 | 0.03 | 1 | 1 | Nudt5 | 0.011 | Young |
| 0.172707 | 0.318779 | 0.058 | 0.044 | 1 | 1 | Lta4h | 0.014 | Young |
| 0.172776 | 0.784776 | 0.018 | 0.01 | 1 | 1 | Slamf6 | 0.008 | Young |
| 0.172857 | 0.191148 | 0.046 | 0.033 | 1 | 1 | Tmem64 | 0.013 | Young |
| 0.17294 | 0.743606 | 0.018 | 0.01 | 1 | 1 | Pik3ap1 | 0.008 | Young |
| 0.172941 | 0.030072 | 0.066 | 0.051 | 1 | 1 | Dars | 0.015 | Young |
| 0.173125 | 0.043726 | 0.026 | 0.017 | 1 | 1 | Fra10ac1 | 0.009 | Young |
| 0.173378 | 0.750594 | 0.018 | 0.01 | 1 | 1 | Gm27010 | 0.008 | Young |
| 0.173409 | 0.121051 | 0.04 | 0.028 | 1 | 1 | Scyl2 | 0.012 | Young |
| 0.173574 | 0.342434 | 0.026 | 0.017 | 1 | 1 | Acot13 | 0.009 | Young |
| 0.173977 | 0.33941 | 0.066 | 0.051 | 1 | 1 | Abhd17a | 0.015 | Young |
| 0.174232 | 0.224922 | 0.043 | 0.031 | 1 | 1 | Wdr36 | 0.012 | Young |
| 0.17443 | 0.696725 | 0.029 | 0.019 | 1 | 1 | Mkl1 | 0.01 | Young |
| 0.174477 | 0.187072 | 0.018 | 0.01 | 1 | 1 | Cog2 | 0.008 | Young |
| 0.174515 | 0.411197 | 0.029 | 0.019 | 1 | 1 | Papd7 | 0.01 | Young |
| 0.174639 | 0.171528 | 0.047 | 0.035 | 1 | 1 | Fam98c | 0.012 | Young |
| 0.174973 | 0.148717 | 0.076 | 0.06 | 1 | 1 | Zfp326 | 0.016 | Young |
| 0.175121 | 0.180159 | 0.079 | 0.063 | 1 | 1 | Phf8 | 0.016 | Young |
| 0.175185 | 0.657404 | 0.047 | 0.035 | 1 | 1 | Pcgf3 | 0.012 | Young |
| 0.175186 | 0.433882 | 0.106 | 0.089 | 1 | 1 | Ncf4 | 0.017 | Young |
| 0.175197 | 0.240276 | 0.026 | 0.017 | 1 | 1 | Tppp | 0.009 | Young |
| 0.175696 | 0.420596 | 0.078 | 0.063 | 1 | 1 | Rapgef2 | 0.015 | Young |
| 0.175857 | 0.304362 | 0.067 | 0.053 | 1 | 1 | Eed | 0.014 | Young |
| 0.175879 | 0.379306 | 0.013 | 0.006 | 1 | 1 | Vwf | 0.007 | Young |
| 0.17593 | 0.215668 | 0.078 | 0.062 | 1 | 1 | Zufsp | 0.016 | Young |
| 0.176134 | 0.412829 | 0.018 | 0.01 | 1 | 1 | Zfp709 | 0.008 | Young |
| 0.177134 | 0.060379 | 0.018 | 0.01 | 1 | 1 | Sort1 | 0.008 | Young |
| 0.177418 | 0.281473 | 0.066 | 0.051 | 1 | 1 | Braf | 0.015 | Young |

| | | | | | | | | |
|---|---|---|---|---|---|---|---|---|
| 0.177887 | 0.858173 | 0.013 | 0.006 | 1 | 1 | Xlr | 0.007 | Young |
| 0.178156 | 0.766117 | 0.013 | 0.006 | 1 | 1 | 1810032O( | 0.007 | Young |
| 0.178177 | 0.302537 | 0.075 | 0.059 | 1 | 1 | Palld | 0.016 | Young |
| 0.178315 | 0.005273 | 0.095 | 0.076 | 1 | 1 | Slc39a7 | 0.019 | Young |
| 0.178637 | 0.551428 | 0.057 | 0.044 | 1 | 1 | Esco1 | 0.013 | Young |
| 0.178709 | 0.315595 | 0.086 | 0.069 | 1 | 1 | Pgap2 | 0.017 | Young |
| 0.178812 | 0.120155 | 0.099 | 0.081 | 1 | 1 | Anapc5 | 0.018 | Young |
| 0.178882 | 0.106621 | 0.121 | 0.1 | 1 | 1 | Pja2 | 0.021 | Young |
| 0.178988 | 0.43307 | 0.03 | 0.021 | 1 | 1 | Smcr8 | 0.009 | Young |
| 0.178991 | 0.784288 | 0.036 | 0.026 | 1 | 1 | Dhx57 | 0.01 | Young |
| 0.179032 | 0.787596 | 0.013 | 0.006 | 1 | 1 | Enc1 | 0.007 | Young |
| 0.179169 | 0.490731 | 0.084 | 0.068 | 1 | 1 | Slmap | 0.016 | Young |
| 0.179174 | 0.697864 | 0.032 | 0.022 | 1 | 1 | Ubqln4 | 0.01 | Young |
| 0.179224 | 0.13055 | 0.101 | 0.082 | 1 | 1 | Bscl2 | 0.019 | Young |
| 0.17937 | 0.648869 | 0.013 | 0.006 | 1 | 1 | Nkiras1 | 0.007 | Young |
| 0.179399 | 0.175763 | 0.026 | 0.017 | 1 | 1 | Fmo5 | 0.009 | Young |
| 0.1794 | 0.114403 | 0.195 | 0.168 | 1 | 1 | Rnf150 | 0.027 | Young |
| 0.179409 | 0.601404 | 0.069 | 0.055 | 1 | 1 | Luc7l | 0.014 | Young |
| 0.17964 | 0.466262 | 0.013 | 0.006 | 1 | 1 | Lig4 | 0.007 | Young |
| 0.180103 | 0.098265 | 0.081 | 0.064 | 1 | 1 | Baz2a | 0.017 | Young |
| 0.180375 | 0.154665 | 0.122 | 0.102 | 1 | 1 | Eif4e | 0.02 | Young |
| 0.180855 | 0.119147 | 0.089 | 0.072 | 1 | 1 | Mepce | 0.017 | Young |
| 0.181353 | 0.022409 | 0.089 | 0.072 | 1 | 1 | Xrn1 | 0.017 | Young |
| 0.18147 | 0.265319 | 0.078 | 0.062 | 1 | 1 | Pank2 | 0.016 | Young |
| 0.181553 | 0.296484 | 0.075 | 0.059 | 1 | 1 | Atg101 | 0.016 | Young |
| 0.181625 | 0.360535 | 0.03 | 0.021 | 1 | 1 | Gm5617 | 0.009 | Young |
| 0.181664 | 0.468252 | 0.032 | 0.022 | 1 | 1 | Ppard | 0.01 | Young |
| 0.181908 | 0.598901 | 0.019 | 0.012 | 1 | 1 | Lmbr1l | 0.007 | Young |
| 0.182126 | 0.82456 | 0.019 | 0.012 | 1 | 1 | Dclre1b | 0.007 | Young |
| 0.182379 | 0.406398 | 0.069 | 0.055 | 1 | 1 | Ube3c | 0.014 | Young |
| 0.182426 | 0.082068 | 0.158 | 0.135 | 1 | 1 | Daglb | 0.023 | Young |
| 0.182493 | 0.52533 | 0.055 | 0.042 | 1 | 1 | Rrm1 | 0.013 | Young |
| 0.182516 | 0.006442 | 0.23 | 0.201 | 1 | 1 | Mindy2 | 0.029 | Young |
| 0.182747 | 0.118752 | 0.082 | 0.066 | 1 | 1 | Egln1 | 0.016 | Young |
| 0.182998 | 0.082784 | 0.117 | 0.098 | 1 | 1 | Herc4 | 0.019 | Young |
| 0.183108 | 0.105566 | 0.163 | 0.14 | 1 | 1 | Ankfy1 | 0.023 | Young |
| 0.18335 | 0.357041 | 0.06 | 0.046 | 1 | 1 | Cyp27a1 | 0.014 | Young |
| 0.183431 | 0.188836 | 0.067 | 0.053 | 1 | 1 | Rbm14 | 0.014 | Young |
| 0.18346 | 0.524406 | 0.061 | 0.048 | 1 | 1 | Cntrl | 0.013 | Young |
| 0.183611 | 0.256852 | 0.054 | 0.041 | 1 | 1 | Armc5 | 0.013 | Young |
| 0.183683 | 0.251978 | 0.101 | 0.084 | 1 | 1 | Irf9 | 0.017 | Young |
| 0.183872 | 0.335258 | 0.019 | 0.012 | 1 | 1 | Tfb2m | 0.007 | Young |
| 0.18437 | 0.745346 | 0.036 | 0.026 | 1 | 1 | Zfp318 | 0.01 | Young |
| 0.184802 | 0.399548 | 0.054 | 0.041 | 1 | 1 | Akap8l | 0.013 | Young |
| 0.185016 | 0.042323 | 0.194 | 0.167 | 1 | 1 | Tcf25 | 0.027 | Young |

| | | | | | | | | |
|---|---|---|---|---|---|---|---|---|
| 0.185093 | 0.087695 | 0.225 | 0.197 | 1 | 1 | Cep170 | 0.028 | Young |
| 0.185136 | 0.293357 | 0.067 | 0.053 | 1 | 1 | Nans | 0.014 | Young |
| 0.18523 | 0.087321 | 0.115 | 0.096 | 1 | 1 | Cic | 0.019 | Young |
| 0.18587 | 0.421911 | 0.032 | 0.022 | 1 | 1 | Sdccag8 | 0.01 | Young |
| 0.186064 | 0.527639 | 0.035 | 0.024 | 1 | 1 | Tbc1d24 | 0.011 | Young |
| 0.186417 | 0.747041 | 0.021 | 0.013 | 1 | 1 | Pdk3 | 0.008 | Young |
| 0.186737 | 0.077933 | 0.019 | 0.012 | 1 | 1 | Cep128 | 0.007 | Young |
| 0.186871 | 0.025649 | 0.058 | 0.044 | 1 | 1 | Pdxdc1 | 0.014 | Young |
| 0.186891 | 0.509413 | 0.033 | 0.023 | 1 | 1 | Klc4 | 0.01 | Young |
| 0.186949 | 0.793289 | 0.021 | 0.013 | 1 | 1 | Ccdc163 | 0.008 | Young |
| 0.187002 | 0.836085 | 0.021 | 0.013 | 1 | 1 | Fbxo45 | 0.008 | Young |
| 0.187004 | 0.220239 | 0.06 | 0.046 | 1 | 1 | Cox16 | 0.014 | Young |
| 0.187095 | 0.601527 | 0.039 | 0.028 | 1 | 1 | 2310022B0 | 0.011 | Young |
| 0.187383 | 0.401417 | 0.036 | 0.026 | 1 | 1 | Pi4k2b | 0.01 | Young |
| 0.187965 | 0.110666 | 0.114 | 0.095 | 1 | 1 | Asap1 | 0.019 | Young |
| 0.187991 | 0.177577 | 0.061 | 0.048 | 1 | 1 | Nub1 | 0.013 | Young |
| 0.188016 | 0.619781 | 0.021 | 0.013 | 1 | 1 | C030034L1 | 0.008 | Young |
| 0.188117 | 0.421647 | 0.072 | 0.058 | 1 | 1 | Gm37494 | 0.014 | Young |
| 0.188181 | 0.376707 | 0.035 | 0.024 | 1 | 1 | Tlr6 | 0.011 | Young |
| 0.188283 | 0.753516 | 0.021 | 0.013 | 1 | 1 | Nsun3 | 0.008 | Young |
| 0.188346 | 0.166452 | 0.138 | 0.117 | 1 | 1 | Twf1 | 0.021 | Young |
| 0.188551 | 0.622565 | 0.021 | 0.013 | 1 | 1 | Styx | 0.008 | Young |
| 0.188572 | 0.297195 | 0.045 | 0.033 | 1 | 1 | Hsd17b10 | 0.012 | Young |
| 0.1886 | 0.157561 | 0.032 | 0.022 | 1 | 1 | Gmcl1 | 0.01 | Young |
| 0.188659 | 0.50517 | 0.021 | 0.013 | 1 | 1 | Gm29340 | 0.008 | Young |
| 0.188984 | 0.195168 | 0.038 | 0.027 | 1 | 1 | Ikbkap | 0.011 | Young |
| 0.189454 | 0.07913 | 0.165 | 0.143 | 1 | 1 | Sult1a1 | 0.022 | Young |
| 0.189518 | 0.652234 | 0.021 | 0.013 | 1 | 1 | Nsun4 | 0.008 | Young |
| 0.189577 | 0.355979 | 0.045 | 0.033 | 1 | 1 | Tef | 0.012 | Young |
| 0.189843 | 0.254176 | 0.058 | 0.045 | 1 | 1 | Sdf2l1 | 0.013 | Young |
| 0.190062 | 0.485734 | 0.042 | 0.031 | 1 | 1 | Pcnx | 0.011 | Young |
| 0.190185 | 1.259449 | 0.014 | 0.008 | 1 | 1 | Taf9b | 0.006 | Young |
| 0.190437 | 0.066593 | 0.174 | 0.149 | 1 | 1 | Ugcg | 0.025 | Young |
| 0.190488 | 0.501958 | 0.021 | 0.013 | 1 | 1 | Psmf1 | 0.008 | Young |
| 0.190958 | 0.155947 | 0.033 | 0.023 | 1 | 1 | Rbl1 | 0.01 | Young |
| 0.191115 | 0.788941 | 0.014 | 0.008 | 1 | 1 | Polm | 0.006 | Young |
| 0.191283 | 0.269756 | 0.008 | 0.014 | 1 | 1 | Acp6 | -0.006 | Young |
| 0.191407 | 0.259606 | 0.054 | 0.041 | 1 | 1 | Exoc1 | 0.013 | Young |
| 0.191539 | 0.212227 | 0.097 | 0.08 | 1 | 1 | Denr | 0.017 | Young |
| 0.191718 | 0.278666 | 0.06 | 0.046 | 1 | 1 | Gbp3 | 0.014 | Young |
| 0.191871 | 0.168525 | 0.066 | 0.051 | 1 | 1 | Kdelr2 | 0.015 | Young |
| 0.19188 | 0.051006 | 0.07 | 0.055 | 1 | 1 | Mbd3 | 0.015 | Young |
| 0.192029 | 0.445786 | 0.038 | 0.027 | 1 | 1 | Gtpbp6 | 0.011 | Young |
| 0.192037 | 0.029656 | 0.162 | 0.138 | 1 | 1 | Cbx3 | 0.024 | Young |
| 0.192049 | 0.851615 | 0.022 | 0.014 | 1 | 1 | Ring1 | 0.008 | Young |

| | | | | | | | | |
|---|---|---|---|---|---|---|---|---|
| 0.192151 | 1.177709 | 0.024 | 0.015 | 1 | 1 | Coq7 | 0.009 | Young |
| 0.192182 | 1.047168 | 0.014 | 0.008 | 1 | 1 | Usp49 | 0.006 | Young |
| 0.192182 | 0.804462 | 0.014 | 0.008 | 1 | 1 | Fam212a | 0.006 | Young |
| 0.192288 | 0.486035 | 0.075 | 0.06 | 1 | 1 | Ssbp4 | 0.015 | Young |
| 0.192373 | 0.053663 | 0.289 | 0.254 | 1 | 1 | Nptn | 0.035 | Young |
| 0.192504 | 0.225659 | 0.066 | 0.051 | 1 | 1 | Nudt16 | 0.015 | Young |
| 0.192692 | 0.208742 | 0.081 | 0.066 | 1 | 1 | Samd9l | 0.015 | Young |
| 0.192707 | 0.048549 | 0.208 | 0.181 | 1 | 1 | Sfr1 | 0.027 | Young |
| 0.192717 | 0.429829 | 0.014 | 0.008 | 1 | 1 | Psph | 0.006 | Young |
| 0.192794 | 0.248537 | 0.142 | 0.122 | 1 | 1 | Tet2 | 0.02 | Young |
| 0.192813 | 0.110672 | 0.184 | 0.159 | 1 | 1 | Psd3 | 0.025 | Young |
| 0.192984 | 0.107992 | 0.177 | 0.154 | 1 | 1 | Aldh2 | 0.023 | Young |
| 0.193253 | 0.503016 | 0.014 | 0.008 | 1 | 1 | Btf3l4 | 0.006 | Young |
| 0.193425 | 0.2516 | 0.038 | 0.027 | 1 | 1 | Htatip2 | 0.011 | Young |
| 0.193604 | 0.313949 | 0.101 | 0.085 | 1 | 1 | Szrd1 | 0.016 | Young |
| 0.193668 | 0.240062 | 0.042 | 0.031 | 1 | 1 | Sec16a | 0.011 | Young |
| 0.193826 | 0.34225 | 0.038 | 0.027 | 1 | 1 | St7l | 0.011 | Young |
| 0.193948 | 0.187614 | 0.047 | 0.035 | 1 | 1 | Cdc42bpb | 0.012 | Young |
| 0.194046 | 0.630913 | 0.022 | 0.014 | 1 | 1 | Fbxl18 | 0.008 | Young |
| 0.194147 | 0.443762 | 0.076 | 0.062 | 1 | 1 | Ptpn7 | 0.014 | Young |
| 0.194362 | 0.127717 | 0.041 | 0.03 | 1 | 1 | Kars | 0.011 | Young |
| 0.194383 | 0.089109 | 0.075 | 0.059 | 1 | 1 | Cnpy2 | 0.016 | Young |
| 0.194463 | 0.305038 | 0.014 | 0.008 | 1 | 1 | Upp2 | 0.006 | Young |
| 0.194625 | 0.163591 | 0.066 | 0.051 | 1 | 1 | Snrk | 0.015 | Young |
| 0.194873 | 0.277018 | 0.052 | 0.04 | 1 | 1 | Tmem222 | 0.012 | Young |
| 0.194967 | 0.011376 | 0.142 | 0.12 | 1 | 1 | Cul3 | 0.022 | Young |
| 0.195238 | 0.05249 | 0.036 | 0.026 | 1 | 1 | Paxip1 | 0.01 | Young |
| 0.195287 | 0.207925 | 0.094 | 0.077 | 1 | 1 | Rnf6 | 0.017 | Young |
| 0.195337 | 0.224746 | 0.039 | 0.028 | 1 | 1 | Exosc8 | 0.011 | Young |
| 0.195373 | 0.3363 | 0.055 | 0.042 | 1 | 1 | Ddx18 | 0.013 | Young |
| 0.195398 | 0.158644 | 0.045 | 0.033 | 1 | 1 | Dcaf11 | 0.012 | Young |
| 0.19542 | 0.610198 | 0.022 | 0.014 | 1 | 1 | Zfp280c | 0.008 | Young |
| 0.195518 | 0.09615 | 0.558 | 0.514 | 1 | 1 | Ly86 | 0.044 | Young |
| 0.195756 | 0.026345 | 0.041 | 0.03 | 1 | 1 | Brix1 | 0.011 | Young |
| 0.196468 | 0.333253 | 0.112 | 0.095 | 1 | 1 | Heca | 0.017 | Young |
| 0.196913 | 0.126668 | 0.114 | 0.096 | 1 | 1 | Trappc4 | 0.018 | Young |
| 0.197031 | 0.00206 | 0.131 | 0.109 | 1 | 1 | Irf2bpl | 0.022 | Young |
| 0.197392 | 0.223277 | 0.106 | 0.089 | 1 | 1 | Casc3 | 0.017 | Young |
| 0.197463 | 0.247499 | 0.047 | 0.035 | 1 | 1 | Prps1 | 0.012 | Young |
| 0.197494 | 0.134181 | 0.145 | 0.125 | 1 | 1 | Cd300a | 0.02 | Young |
| 0.197604 | 0.200894 | 0.1 | 0.082 | 1 | 1 | Parp1 | 0.018 | Young |
| 0.197656 | 0.278643 | 0.022 | 0.014 | 1 | 1 | Edem3 | 0.008 | Young |
| 0.197801 | 1.53696 | 0.016 | 0.009 | 1 | 1 | Zxdb | 0.007 | Young |
| 0.197908 | 0.582305 | 0.055 | 0.042 | 1 | 1 | Mindy3 | 0.013 | Young |
| 0.197928 | 0.784752 | 0.024 | 0.015 | 1 | 1 | Gphn | 0.009 | Young |

| | | | | | | | | |
|---|---|---|---|---|---|---|---|---|
| 0.197972 | 0.043039 | 0.723 | 0.652 | 1 | 1 | Cdc42 | 0.071 | Young |
| 0.19829 | 0.754245 | 0.053 | 0.041 | 1 | 1 | Scaf1 | 0.012 | Young |
| 0.198351 | 0.220848 | 0.061 | 0.048 | 1 | 1 | Pcbd2 | 0.013 | Young |
| 0.1984 | 0.070488 | 0.131 | 0.111 | 1 | 1 | Sbds | 0.02 | Young |
| 0.198455 | 0.111674 | 0.069 | 0.054 | 1 | 1 | Arid3a | 0.015 | Young |
| 0.199211 | 0.994594 | 0.025 | 0.017 | 1 | 1 | Tubgcp2 | 0.008 | Young |
| 0.199372 | 0.09323 | 0.022 | 0.014 | 1 | 1 | Nbn | 0.008 | Young |
| 0.199625 | 0.111338 | 0.045 | 0.033 | 1 | 1 | Cbx6 | 0.012 | Young |
| 0.200249 | 0.160393 | 0.048 | 0.036 | 1 | 1 | Tmem19 | 0.012 | Young |
| 0.200446 | 0.775983 | 0.027 | 0.018 | 1 | 1 | Katnal1 | 0.009 | Young |
| 0.200485 | 0.131181 | 0.124 | 0.105 | 1 | 1 | Fnbp1l | 0.019 | Young |
| 0.200576 | 0.2852 | 0.114 | 0.096 | 1 | 1 | Coro1b | 0.018 | Young |
| 0.200586 | 0.019144 | 0.676 | 0.617 | 1 | 1 | Lamp1 | 0.059 | Young |
| 0.200838 | 0.42258 | 0.024 | 0.015 | 1 | 1 | Gm20707 | 0.009 | Young |
| 0.200839 | 0.128822 | 0.049 | 0.037 | 1 | 1 | Plekhf2 | 0.012 | Young |
| 0.20096 | 0.109569 | 0.12 | 0.1 | 1 | 1 | Rasa1 | 0.02 | Young |
| 0.201438 | 1.036983 | 0.016 | 0.009 | 1 | 1 | Kctd18 | 0.007 | Young |
| 0.201478 | 0.376967 | 0.083 | 0.068 | 1 | 1 | Atp7a | 0.015 | Young |
| 0.201532 | 0.176176 | 0.022 | 0.014 | 1 | 1 | Tdrkh | 0.008 | Young |
| 0.201614 | 0.063556 | 0.042 | 0.031 | 1 | 1 | Cd2bp2 | 0.011 | Young |
| 0.20163 | 0.291425 | 0.123 | 0.105 | 1 | 1 | Safb2 | 0.018 | Young |
| 0.201637 | 0.035309 | 0.159 | 0.136 | 1 | 1 | Dad1 | 0.023 | Young |
| 0.201699 | 0.969146 | 0.016 | 0.009 | 1 | 1 | Pex1 | 0.007 | Young |
| 0.201771 | 0.088115 | 0.089 | 0.072 | 1 | 1 | Aff1 | 0.017 | Young |
| 0.201906 | 0.098228 | 0.052 | 0.04 | 1 | 1 | Ube2g1 | 0.012 | Young |
| 0.201908 | 0.072341 | 0.254 | 0.224 | 1 | 1 | Cmpk1 | 0.03 | Young |
| 0.202946 | 0.032976 | 0.06 | 0.046 | 1 | 1 | Erg28 | 0.014 | Young |
| 0.202986 | 0.085368 | 0.196 | 0.174 | 1 | 1 | Mdfic | 0.022 | Young |
| 0.203143 | 0.776081 | 0.016 | 0.009 | 1 | 1 | Dedd | 0.007 | Young |
| 0.203199 | 0.400731 | 0.024 | 0.015 | 1 | 1 | Aldh6a1 | 0.009 | Young |
| 0.203208 | 0.230312 | 0.064 | 0.05 | 1 | 1 | Pigs | 0.014 | Young |
| 0.203213 | 0.510299 | 0.025 | 0.017 | 1 | 1 | C2cd2l | 0.008 | Young |
| 0.203499 | 0.090973 | 0.194 | 0.168 | 1 | 1 | Cxcr4 | 0.026 | Young |
| 0.20372 | 0.001365 | 0.239 | 0.21 | 1 | 1 | Bst2 | 0.029 | Young |
| 0.203888 | 0.104703 | 0.089 | 0.072 | 1 | 1 | Yipf4 | 0.017 | Young |
| 0.204065 | 0.995296 | 0.016 | 0.009 | 1 | 1 | Zkscan17 | 0.007 | Young |
| 0.204225 | 0.120207 | 0.08 | 0.064 | 1 | 1 | Galnt11 | 0.016 | Young |
| 0.20433 | 0.514245 | 0.016 | 0.009 | 1 | 1 | Mapk7 | 0.007 | Young |
| 0.204594 | 0.74962 | 0.016 | 0.009 | 1 | 1 | 1110032A0 | 0.007 | Young |
| 0.204698 | 0.019399 | 0.081 | 0.066 | 1 | 1 | Twistnb | 0.015 | Young |
| 0.204858 | 0.953376 | 0.037 | 0.027 | 1 | 1 | Gm15417 | 0.01 | Young |
| 0.204892 | 0.564084 | 0.027 | 0.018 | 1 | 1 | Pwp1 | 0.009 | Young |
| 0.205314 | 0.215016 | 0.068 | 0.054 | 1 | 1 | Anapc1 | 0.014 | Young |
| 0.205364 | 0.429552 | 0.025 | 0.017 | 1 | 1 | Borcs7 | 0.008 | Young |
| 0.205511 | 0.459007 | 0.083 | 0.068 | 1 | 1 | 5031425E2 | 0.015 | Young |

| | | | | | | | | |
|---|---|---|---|---|---|---|---|---|
| 0.205521 | 0.44526 | 0.016 | 0.009 | 1 | 1 | Slc52a2 | 0.007 | Young |
| 0.205826 | 0.846656 | 0.01 | 0.005 | 1 | 1 | Ybey | 0.005 | Young |
| 0.205981 | 0.329884 | 0.025 | 0.017 | 1 | 1 | Bloc1s5 | 0.008 | Young |
| 0.206156 | 0.074634 | 0.147 | 0.126 | 1 | 1 | Zmynd8 | 0.021 | Young |
| 0.206199 | 0.01045 | 0.052 | 0.04 | 1 | 1 | Mcm6 | 0.012 | Young |
| 0.206488 | 0.84488 | 0.01 | 0.005 | 1 | 1 | Lysmd1 | 0.005 | Young |
| 0.206654 | 0.820471 | 0.01 | 0.005 | 1 | 1 | Phactr1 | 0.005 | Young |
| 0.206673 | 0.161622 | 0.08 | 0.064 | 1 | 1 | Usp37 | 0.016 | Young |
| 0.206731 | 0.091253 | 0.19 | 0.166 | 1 | 1 | Ctnnd1 | 0.024 | Young |
| 0.20682 | 0.742225 | 0.01 | 0.005 | 1 | 1 | Gstp3 | 0.005 | Young |
| 0.20682 | 0.109782 | 0.01 | 0.005 | 1 | 1 | Il1rl2 | 0.005 | Young |
| 0.206986 | 0.839295 | 0.01 | 0.005 | 1 | 1 | Zfp53 | 0.005 | Young |
| 0.207251 | 0.145528 | 0.016 | 0.009 | 1 | 1 | Gnl2 | 0.007 | Young |
| 0.207318 | 0.7442 | 0.01 | 0.005 | 1 | 1 | Cenpo | 0.005 | Young |
| 0.207484 | 0.61932 | 0.01 | 0.005 | 1 | 1 | Sars2 | 0.005 | Young |
| 0.207556 | 0.012161 | 0.09 | 0.073 | 1 | 1 | Prpf39 | 0.017 | Young |
| 0.207666 | 0.813397 | 0.03 | 0.021 | 1 | 1 | Clcn6 | 0.009 | Young |
| 0.207898 | 0.105333 | 0.055 | 0.042 | 1 | 1 | Shprh | 0.013 | Young |
| 0.208085 | 0.11936 | 0.091 | 0.075 | 1 | 1 | 0610012G( | 0.016 | Young |
| 0.208116 | 0.452042 | 0.028 | 0.019 | 1 | 1 | Xaf1 | 0.009 | Young |
| 0.208305 | 0.115139 | 0.102 | 0.085 | 1 | 1 | Cop1 | 0.017 | Young |
| 0.208673 | 0.096346 | 0.025 | 0.017 | 1 | 1 | Dip2a | 0.008 | Young |
| 0.208858 | 0.002943 | 0.016 | 0.009 | 1 | 1 | Inpp5b | 0.007 | Young |
| 0.209136 | 0.119017 | 0.074 | 0.059 | 1 | 1 | Msl3 | 0.015 | Young |
| 0.209329 | 0.335884 | 0.059 | 0.046 | 1 | 1 | Pgm2l1 | 0.013 | Young |
| 0.209486 | 0.383044 | 0.01 | 0.005 | 1 | 1 | Cox15 | 0.005 | Young |
| 0.209486 | 0.060768 | 0.01 | 0.005 | 1 | 1 | Ubiad1 | 0.005 | Young |
| 0.209528 | 0.215541 | 0.062 | 0.049 | 1 | 1 | Ccdc22 | 0.013 | Young |
| 0.209558 | 0.232609 | 0.025 | 0.017 | 1 | 1 | Spidr | 0.008 | Young |
| 0.209604 | 0.023291 | 0.176 | 0.153 | 1 | 1 | Fam102b | 0.023 | Young |
| 0.209689 | 0.166126 | 0.1 | 0.084 | 1 | 1 | Ndufs7 | 0.016 | Young |
| 0.209797 | 0.123891 | 0.134 | 0.114 | 1 | 1 | Tnfrsf11a | 0.02 | Young |
| 0.209821 | 0.090115 | 0.01 | 0.005 | 1 | 1 | Gm11423 | 0.005 | Young |
| 0.209924 | 0.053653 | 0.025 | 0.017 | 1 | 1 | Slc25a22 | 0.008 | Young |
| 0.210012 | 1.091869 | 0.017 | 0.01 | 1 | 1 | Ripor1 | 0.007 | Young |
| 0.210171 | 0.249022 | 0.097 | 0.081 | 1 | 1 | Erlin1 | 0.016 | Young |
| 0.21018 | 0.04661 | 0.164 | 0.141 | 1 | 1 | Prpf40a | 0.023 | Young |
| 0.210257 | 0.08141 | 0.031 | 0.022 | 1 | 1 | H2-DMb2 | 0.009 | Young |
| 0.210278 | 0.035021 | 0.228 | 0.201 | 1 | 1 | Ppp1r9b | 0.027 | Young |
| 0.21043 | 0.481667 | 0.097 | 0.082 | 1 | 1 | Anks1 | 0.015 | Young |
| 0.210654 | 0.094684 | 0.078 | 0.063 | 1 | 1 | Eif6 | 0.015 | Young |
| 0.210656 | 0.40124 | 0.061 | 0.048 | 1 | 1 | Clec2i | 0.013 | Young |
| 0.210782 | 0.969683 | 0.017 | 0.01 | 1 | 1 | Eif2ak4 | 0.007 | Young |
| 0.210884 | 0.504557 | 0.037 | 0.027 | 1 | 1 | Ubr1 | 0.01 | Young |
| 0.210931 | 0.227201 | 0.027 | 0.018 | 1 | 1 | Mrps34 | 0.009 | Young |

| | | | | | | | | |
|---|---|---|---|---|---|---|---|---|
| 0.210968 | 0.50658 | 0.032 | 0.023 | 1 | 1 | N4bp2 | 0.009 | Young |
| 0.211365 | 0.441698 | 0.028 | 0.019 | 1 | 1 | Yaf2 | 0.009 | Young |
| 0.211614 | 0.312911 | 0.095 | 0.08 | 1 | 1 | Tmem87b | 0.015 | Young |
| 0.211812 | 0.665559 | 0.017 | 0.01 | 1 | 1 | Lclat1 | 0.007 | Young |
| 0.212008 | 0.030006 | 0.145 | 0.125 | 1 | 1 | Vma21 | 0.02 | Young |
| 0.212071 | 0.329949 | 0.052 | 0.04 | 1 | 1 | Ttpal | 0.012 | Young |
| 0.212437 | 0.554149 | 0.032 | 0.023 | 1 | 1 | Trim11 | 0.009 | Young |
| 0.21287 | 0.473961 | 0.075 | 0.062 | 1 | 1 | St3gal4 | 0.013 | Young |
| 0.213127 | 0.002831 | 0.489 | 0.437 | 1 | 1 | Ptbp3 | 0.052 | Young |
| 0.213319 | 0.579454 | 0.04 | 0.03 | 1 | 1 | Pmm2 | 0.01 | Young |
| 0.213454 | 0.300614 | 0.095 | 0.08 | 1 | 1 | Gab2 | 0.015 | Young |
| 0.213519 | 0.163418 | 0.085 | 0.069 | 1 | 1 | Tcf20 | 0.016 | Young |
| 0.213634 | 0.129197 | 0.094 | 0.077 | 1 | 1 | Arhgap12 | 0.017 | Young |
| 0.213866 | 0.097419 | 0.219 | 0.194 | 1 | 1 | Gna12 | 0.025 | Young |
| 0.214501 | 0.277111 | 0.059 | 0.046 | 1 | 1 | Upf1 | 0.013 | Young |
| 0.214512 | 0.033566 | 0.027 | 0.018 | 1 | 1 | Rev3l | 0.009 | Young |
| 0.214533 | 0.196398 | 0.017 | 0.01 | 1 | 1 | Slc25a40 | 0.007 | Young |
| 0.214621 | 0.020147 | 0.201 | 0.175 | 1 | 1 | Ms4a6b | 0.026 | Young |
| 0.214682 | 0.419727 | 0.037 | 0.027 | 1 | 1 | Ankrd50 | 0.01 | Young |
| 0.214862 | 0.335265 | 0.055 | 0.042 | 1 | 1 | Lrrc47 | 0.013 | Young |
| 0.214924 | 0.529836 | 0.017 | 0.01 | 1 | 1 | Cul7 | 0.007 | Young |
| 0.215191 | 0.428084 | 0.046 | 0.035 | 1 | 1 | Golga5 | 0.011 | Young |
| 0.21523 | 0.572109 | 0.046 | 0.035 | 1 | 1 | Dcaf5 | 0.011 | Young |
| 0.215315 | 0.007104 | 0.017 | 0.01 | 1 | 1 | Mettl7a1 | 0.007 | Young |
| 0.215381 | 0.224451 | 0.03 | 0.021 | 1 | 1 | Kif13a | 0.009 | Young |
| 0.216099 | 0.313753 | 0.017 | 0.01 | 1 | 1 | Tmem35b | 0.007 | Young |
| 0.216115 | 0.238791 | 0.034 | 0.024 | 1 | 1 | Rasgrp1 | 0.01 | Young |
| 0.216363 | 0.239607 | 0.062 | 0.049 | 1 | 1 | Slc15a2 | 0.013 | Young |
| 0.216885 | 0.239578 | 0.017 | 0.01 | 1 | 1 | Zfp758 | 0.007 | Young |
| 0.216922 | 0.172811 | 0.047 | 0.036 | 1 | 1 | Pdcl | 0.011 | Young |
| 0.217008 | 0.445443 | 0.05 | 0.039 | 1 | 1 | Kansl2 | 0.011 | Young |
| 0.21712 | 0.292193 | 0.038 | 0.028 | 1 | 1 | Zfp219 | 0.01 | Young |
| 0.217675 | 0.450861 | 0.04 | 0.03 | 1 | 1 | Trim65 | 0.01 | Young |
| 0.21768 | 0.014183 | 0.146 | 0.126 | 1 | 1 | Sbno1 | 0.02 | Young |
| 0.217685 | 0.527193 | 0.032 | 0.023 | 1 | 1 | Dglucy | 0.009 | Young |
| 0.21801 | 0.025682 | 0.375 | 0.335 | 1 | 1 | Tpd52 | 0.04 | Young |
| 0.218054 | 0.329724 | 0.046 | 0.035 | 1 | 1 | Prps2 | 0.011 | Young |
| 0.218091 | 0.041993 | 0.086 | 0.071 | 1 | 1 | Cpsf7 | 0.015 | Young |
| 0.218235 | 0.119133 | 0.115 | 0.098 | 1 | 1 | Os9 | 0.017 | Young |
| 0.218316 | 0.299722 | 0.047 | 0.036 | 1 | 1 | Slc11a2 | 0.011 | Young |
| 0.218447 | 0.147191 | 0.034 | 0.024 | 1 | 1 | Tex10 | 0.01 | Young |
| 0.218524 | 0.078884 | 0.03 | 0.021 | 1 | 1 | Dopey2 | 0.009 | Young |
| 0.218576 | 0.32078 | 0.032 | 0.023 | 1 | 1 | Timm10 | 0.009 | Young |
| 0.218578 | 0.14273 | 0.244 | 0.219 | 1 | 1 | Rab5c | 0.025 | Young |
| 0.218612 | 0.150178 | 0.031 | 0.022 | 1 | 1 | Caml | 0.009 | Young |

| | | | | | | | | |
|---|---|---|---|---|---|---|---|---|
| 0.218912 | 0.282597 | 0.052 | 0.04 | 1 | 1 | Ikzf5 | 0.012 | Young |
| 0.218972 | 0.472416 | 0.154 | 0.136 | 1 | 1 | Rgs1 | 0.018 | Young |
| 0.219238 | 0.266258 | 0.103 | 0.086 | 1 | 1 | Abcc1 | 0.017 | Young |
| 0.21995 | 0.365667 | 0.061 | 0.049 | 1 | 1 | Trim26 | 0.012 | Young |
| 0.220534 | 0.219 | 0.089 | 0.073 | 1 | 1 | Taf6l | 0.016 | Young |
| 0.220554 | 0.190746 | 0.075 | 0.06 | 1 | 1 | Clk3 | 0.015 | Young |
| 0.220803 | 0.065222 | 0.074 | 0.059 | 1 | 1 | Cdk19 | 0.015 | Young |
| 0.220834 | 0.324985 | 0.038 | 0.028 | 1 | 1 | Ddx58 | 0.01 | Young |
| 0.220954 | 0.152021 | 0.035 | 0.026 | 1 | 1 | Nsmce2 | 0.009 | Young |
| 0.221217 | 0.271764 | 0.053 | 0.041 | 1 | 1 | Alkbh1 | 0.012 | Young |
| 0.221243 | 0.651795 | 0.018 | 0.012 | 1 | 1 | Zfp959 | 0.006 | Young |
| 0.221498 | 0.488583 | 0.018 | 0.012 | 1 | 1 | Ccp110 | 0.006 | Young |
| 0.222007 | 0.519331 | 0.018 | 0.012 | 1 | 1 | Galt | 0.006 | Young |
| 0.22232 | 1.15367 | 0.012 | 0.006 | 1 | 1 | Tlr12 | 0.006 | Young |
| 0.22238 | 0.252752 | 0.047 | 0.036 | 1 | 1 | Cds1 | 0.011 | Young |
| 0.22256 | 0.170466 | 0.143 | 0.123 | 1 | 1 | Sumo1 | 0.02 | Young |
| 0.222768 | 0.197857 | 0.052 | 0.04 | 1 | 1 | Prkag1 | 0.012 | Young |
| 0.222773 | 0.387385 | 0.018 | 0.012 | 1 | 1 | Smpd4 | 0.006 | Young |
| 0.223139 | 0.232314 | 0.037 | 0.027 | 1 | 1 | Zfp467 | 0.01 | Young |
| 0.223293 | 1.344053 | 0.012 | 0.006 | 1 | 1 | Tbl2 | 0.006 | Young |
| 0.223318 | 0.185647 | 0.035 | 0.026 | 1 | 1 | Kdm5b | 0.009 | Young |
| 0.223436 | 0.042561 | 0.298 | 0.263 | 1 | 1 | Cd163 | 0.035 | Young |
| 0.223497 | 0.372846 | 0.037 | 0.027 | 1 | 1 | Zfp654 | 0.01 | Young |
| 0.223669 | 0.440721 | 0.018 | 0.012 | 1 | 1 | Nol9 | 0.006 | Young |
| 0.223915 | 0.028739 | 0.156 | 0.134 | 1 | 1 | Reps2 | 0.022 | Young |
| 0.22401 | 0.151509 | 0.075 | 0.06 | 1 | 1 | Sys1 | 0.015 | Young |
| 0.224072 | 0.270018 | 0.07 | 0.057 | 1 | 1 | Pbxip1 | 0.013 | Young |
| 0.224154 | 0.022102 | 0.085 | 0.069 | 1 | 1 | Clptm1 | 0.016 | Young |
| 0.224238 | 0.038499 | 0.338 | 0.301 | 1 | 1 | Celf1 | 0.037 | Young |
| 0.224439 | 0.21898 | 0.018 | 0.012 | 1 | 1 | Akip1 | 0.006 | Young |
| 0.22448 | 0.016294 | 0.227 | 0.199 | 1 | 1 | Nfam1 | 0.028 | Young |
| 0.224651 | 0.246593 | 0.032 | 0.023 | 1 | 1 | Daam1 | 0.009 | Young |
| 0.224705 | 0.114846 | 0.213 | 0.19 | 1 | 1 | Sf1 | 0.023 | Young |
| 0.224756 | 0.418251 | 0.085 | 0.071 | 1 | 1 | Dennd4b | 0.014 | Young |
| 0.224758 | 0.847343 | 0.012 | 0.006 | 1 | 1 | Srgap3 | 0.006 | Young |
| 0.225082 | 0.416701 | 0.018 | 0.012 | 1 | 1 | Dcun1d2 | 0.006 | Young |
| 0.225288 | 0.038469 | 0.086 | 0.071 | 1 | 1 | Galk2 | 0.015 | Young |
| 0.22547 | 0.021845 | 0.062 | 0.049 | 1 | 1 | Mms19 | 0.013 | Young |
| 0.225539 | 0.030837 | 0.284 | 0.253 | 1 | 1 | Lst1 | 0.031 | Young |
| 0.225574 | 0.668135 | 0.012 | 0.006 | 1 | 1 | Rnf123 | 0.006 | Young |
| 0.225574 | 0.45887 | 0.012 | 0.006 | 1 | 1 | Maged2 | 0.006 | Young |
| 0.225827 | 0.196686 | 0.038 | 0.028 | 1 | 1 | Rae1 | 0.01 | Young |
| 0.225883 | 0.041816 | 0.035 | 0.026 | 1 | 1 | Tusc2 | 0.009 | Young |
| 0.225936 | 0.124459 | 0.252 | 0.225 | 1 | 1 | Pbrm1 | 0.027 | Young |
| 0.226074 | 0.355865 | 0.044 | 0.033 | 1 | 1 | Eef2k | 0.011 | Young |

| | | | | | | | | |
|---|---|---|---|---|---|---|---|---|
| 0.226399 | 0.124443 | 0.034 | 0.024 | 1 | 1 | Ppfibp1 | 0.01 | Young |
| 0.226693 | 0.066759 | 0.1 | 0.084 | 1 | 1 | Gpbp1l1 | 0.016 | Young |
| 0.226722 | 0.535464 | 0.012 | 0.006 | 1 | 1 | AC163032. | 0.006 | Young |
| 0.226886 | 0.405571 | 0.012 | 0.006 | 1 | 1 | Abhd14b | 0.006 | Young |
| 0.22705 | 0.383226 | 0.012 | 0.006 | 1 | 1 | Amer1 | 0.006 | Young |
| 0.227214 | 0.339782 | 0.012 | 0.006 | 1 | 1 | Atp6v1g2 | 0.006 | Young |
| 0.227296 | 0.375003 | 0.058 | 0.046 | 1 | 1 | Ccdc186 | 0.012 | Young |
| 0.227774 | 0.186037 | 0.109 | 0.093 | 1 | 1 | Pik3ca | 0.016 | Young |
| 0.227912 | 0.00738 | 0.103 | 0.086 | 1 | 1 | Rbm6 | 0.017 | Young |
| 0.228015 | 0.400513 | 0.06 | 0.048 | 1 | 1 | Dmtf1 | 0.012 | Young |
| 0.228234 | 0.666269 | 0.02 | 0.013 | 1 | 1 | Fam210a | 0.007 | Young |
| 0.228367 | 0.358269 | 0.012 | 0.006 | 1 | 1 | Abcb6 | 0.006 | Young |
| 0.228999 | 0.043253 | 0.062 | 0.049 | 1 | 1 | Nt5c | 0.013 | Young |
| 0.229602 | 0.105386 | 0.173 | 0.15 | 1 | 1 | Hprt | 0.023 | Young |
| 0.22969 | 0.240242 | 0.012 | 0.006 | 1 | 1 | Park2 | 0.006 | Young |
| 0.229799 | 0.783679 | 0.021 | 0.014 | 1 | 1 | Trub2 | 0.007 | Young |
| 0.230136 | 0.14366 | 0.115 | 0.098 | 1 | 1 | Syf2 | 0.017 | Young |
| 0.230214 | 0.176326 | 0.107 | 0.09 | 1 | 1 | Borcs6 | 0.017 | Young |
| 0.230367 | 0.127224 | 0.037 | 0.027 | 1 | 1 | B3glct | 0.01 | Young |
| 0.230951 | 1.03782 | 0.023 | 0.015 | 1 | 1 | Micall1 | 0.008 | Young |
| 0.23107 | 0.659819 | 0.073 | 0.06 | 1 | 1 | Stxbp6 | 0.013 | Young |
| 0.231129 | 0.599774 | 0.021 | 0.014 | 1 | 1 | Fiz1 | 0.007 | Young |
| 0.231322 | 0.113554 | 0.177 | 0.156 | 1 | 1 | Bnip3l | 0.021 | Young |
| 0.231328 | 0.293466 | 0.086 | 0.072 | 1 | 1 | Rexo2 | 0.014 | Young |
| 0.231335 | 0.437193 | 0.057 | 0.045 | 1 | 1 | Susd3 | 0.012 | Young |
| 0.231646 | 0.058232 | 0.244 | 0.216 | 1 | 1 | Tpm4 | 0.028 | Young |
| 0.231984 | 0.333866 | 0.007 | 0.012 | 1 | 1 | Jchain | -0.005 | Young |
| 0.232417 | 0.147815 | 0.137 | 0.118 | 1 | 1 | Eif3m | 0.019 | Young |
| 0.232448 | 0.107745 | 0.21 | 0.186 | 1 | 1 | Bclaf1 | 0.024 | Young |
| 0.232708 | 0.523632 | 0.021 | 0.014 | 1 | 1 | Tmem126a | 0.007 | Young |
| 0.232932 | 0.864036 | 0.024 | 0.017 | 1 | 1 | Prdm4 | 0.007 | Young |
| 0.232951 | 0.591525 | 0.021 | 0.014 | 1 | 1 | Trmt2b | 0.007 | Young |
| 0.233034 | 0.003808 | 0.307 | 0.272 | 1 | 1 | Tomm7 | 0.035 | Young |
| 0.233136 | 0.086183 | 0.02 | 0.013 | 1 | 1 | Sepsecs | 0.007 | Young |
| 0.233324 | 0.03239 | 0.168 | 0.145 | 1 | 1 | Tprgl | 0.023 | Young |
| 0.233775 | 0.317067 | 0.057 | 0.045 | 1 | 1 | Hmg20a | 0.012 | Young |
| 0.233822 | 0.032328 | 0.117 | 0.099 | 1 | 1 | Srpr | 0.018 | Young |
| 0.233843 | 1.02602 | 0.024 | 0.017 | 1 | 1 | Mki67 | 0.007 | Young |
| 0.233957 | 1.061997 | 0.024 | 0.017 | 1 | 1 | Tfdp1 | 0.007 | Young |
| 0.234315 | 0.057436 | 0.01 | 0.015 | 1 | 1 | Dcbld2 | -0.005 | Young |
| 0.234956 | 0.279389 | 0.06 | 0.048 | 1 | 1 | Agpat3 | 0.012 | Young |
| 0.235203 | 0.132724 | 0.067 | 0.054 | 1 | 1 | Cdc123 | 0.013 | Young |
| 0.235221 | 0.231931 | 0.063 | 0.05 | 1 | 1 | Dnajc21 | 0.013 | Young |
| 0.235361 | 0.369916 | 0.09 | 0.076 | 1 | 1 | Psmd14 | 0.014 | Young |
| 0.235512 | 0.458802 | 0.066 | 0.054 | 1 | 1 | Rbm28 | 0.012 | Young |

| | | | | | | | | |
|---|---|---|---|---|---|---|---|---|
| 0.235599 | 0.492722 | 0.023 | 0.015 | 1 | 1 | Tbp | 0.008 | Young |
| 0.235617 | 0.005625 | 0.134 | 0.114 | 1 | 1 | Rnf111 | 0.02 | Young |
| 0.236267 | 0.283956 | 0.055 | 0.044 | 1 | 1 | Aldh16a1 | 0.011 | Young |
| 0.23627 | 1.343017 | 0.013 | 0.008 | 1 | 1 | Rpap2 | 0.005 | Young |
| 0.236635 | 0.554377 | 0.041 | 0.031 | 1 | 1 | 1700037H( | 0.01 | Young |
| 0.236904 | 1.33866 | 0.013 | 0.008 | 1 | 1 | Adrb1 | 0.005 | Young |
| 0.236904 | 0.976167 | 0.013 | 0.008 | 1 | 1 | Gpatch4 | 0.005 | Young |
| 0.237736 | 0.244439 | 0.021 | 0.014 | 1 | 1 | Cers4 | 0.007 | Young |
| 0.238174 | 0.828741 | 0.013 | 0.008 | 1 | 1 | Nudt1 | 0.005 | Young |
| 0.238334 | 1.017956 | 0.013 | 0.008 | 1 | 1 | Abhd6 | 0.005 | Young |
| 0.238684 | 0.158901 | 0.084 | 0.069 | 1 | 1 | Cers5 | 0.015 | Young |
| 0.238941 | 0.260678 | 0.075 | 0.062 | 1 | 1 | Rspry1 | 0.013 | Young |
| 0.238962 | 0.068934 | 0.634 | 0.566 | 1 | 1 | Unc93b1 | 0.068 | Young |
| 0.239131 | 0.875809 | 0.013 | 0.008 | 1 | 1 | Enho | 0.005 | Young |
| 0.239527 | 0.196118 | 0.084 | 0.069 | 1 | 1 | Cuedc2 | 0.015 | Young |
| 0.23965 | 0.064707 | 0.069 | 0.055 | 1 | 1 | Rpe | 0.014 | Young |
| 0.23977 | 0.650305 | 0.013 | 0.008 | 1 | 1 | 2900093K2 | 0.005 | Young |
| 0.240073 | 0.449048 | 0.023 | 0.015 | 1 | 1 | Erich1 | 0.008 | Young |
| 0.24009 | 0.611043 | 0.013 | 0.008 | 1 | 1 | Gstt3 | 0.005 | Young |
| 0.240554 | 0.348185 | 0.023 | 0.015 | 1 | 1 | 2510002D2 | 0.008 | Young |
| 0.240573 | 0.63638 | 0.027 | 0.019 | 1 | 1 | Osbpl3 | 0.008 | Young |
| 0.240629 | 0.009494 | 0.101 | 0.085 | 1 | 1 | Rrp1 | 0.016 | Young |
| 0.240705 | 0.360198 | 0.051 | 0.04 | 1 | 1 | Zfp422 | 0.011 | Young |
| 0.240724 | 0.609694 | 0.029 | 0.021 | 1 | 1 | Slc25a32 | 0.008 | Young |
| 0.24073 | 0.529491 | 0.013 | 0.008 | 1 | 1 | Phtf1 | 0.005 | Young |
| 0.240815 | 0.079366 | 0.092 | 0.077 | 1 | 1 | Cr1l | 0.015 | Young |
| 0.240827 | 0.098661 | 0.057 | 0.045 | 1 | 1 | Imp4 | 0.012 | Young |
| 0.240858 | 0.226059 | 0.06 | 0.048 | 1 | 1 | Cyb5r1 | 0.012 | Young |
| 0.241063 | 0.669893 | 0.026 | 0.018 | 1 | 1 | Guf1 | 0.008 | Young |
| 0.241156 | 0.342842 | 0.023 | 0.015 | 1 | 1 | Mrpl27 | 0.008 | Young |
| 0.241517 | 0.147364 | 0.023 | 0.015 | 1 | 1 | Fbxl12 | 0.008 | Young |
| 0.241533 | 0.578126 | 0.013 | 0.008 | 1 | 1 | Hdac6 | 0.005 | Young |
| 0.241879 | 0.343925 | 0.023 | 0.015 | 1 | 1 | Nt5dc2 | 0.008 | Young |
| 0.241964 | 0.505319 | 0.038 | 0.028 | 1 | 1 | Dgat1 | 0.01 | Young |
| 0.242134 | 0.654667 | 0.036 | 0.027 | 1 | 1 | Rapgef1 | 0.009 | Young |
| 0.242157 | 0.280948 | 0.024 | 0.017 | 1 | 1 | Sh2d1b1 | 0.007 | Young |
| 0.242333 | 0.341246 | 0.024 | 0.017 | 1 | 1 | Rhno1 | 0.007 | Young |
| 0.242622 | 0.057391 | 0.063 | 0.05 | 1 | 1 | Sh3gl1 | 0.013 | Young |
| 0.242757 | 0.671924 | 0.029 | 0.021 | 1 | 1 | Syne3 | 0.008 | Young |
| 0.242821 | 0.012099 | 0.013 | 0.008 | 1 | 1 | Snrnp35 | 0.005 | Young |
| 0.243071 | 0.196983 | 0.057 | 0.045 | 1 | 1 | Mrps12 | 0.012 | Young |
| 0.243088 | 0.571018 | 0.036 | 0.027 | 1 | 1 | Ermp1 | 0.009 | Young |
| 0.243102 | 0.108364 | 0.072 | 0.059 | 1 | 1 | Pisd | 0.013 | Young |
| 0.243211 | 0.113014 | 0.094 | 0.078 | 1 | 1 | Micu1 | 0.016 | Young |
| 0.243325 | 0.742224 | 0.03 | 0.022 | 1 | 1 | Megf8 | 0.008 | Young |

| | | | | | | | | |
|---|---|---|---|---|---|---|---|---|
| 0.243394 | 0.048683 | 0.069 | 0.055 | 1 | 1 | Ldah | 0.014 | Young |
| 0.243513 | 0.079428 | 0.023 | 0.015 | 1 | 1 | Nudt2 | 0.008 | Young |
| 0.24379 | 0.080672 | 0.013 | 0.008 | 1 | 1 | Mterf4 | 0.005 | Young |
| 0.244015 | 0.028791 | 0.089 | 0.073 | 1 | 1 | Ap3b1 | 0.016 | Young |
| 0.244682 | 0.211579 | 0.088 | 0.073 | 1 | 1 | Scamp5 | 0.015 | Young |
| 0.244694 | 0.641833 | 0.029 | 0.021 | 1 | 1 | Cnksr3 | 0.008 | Young |
| 0.244755 | 0.449757 | 0.026 | 0.018 | 1 | 1 | Tmem199 | 0.008 | Young |
| 0.244869 | 0.4804 | 0.026 | 0.018 | 1 | 1 | Zfp672 | 0.008 | Young |
| 0.244907 | 0.121189 | 0.057 | 0.045 | 1 | 1 | Acads | 0.012 | Young |
| 0.24521 | 0.240543 | 0.024 | 0.017 | 1 | 1 | Fam217b | 0.007 | Young |
| 0.245548 | 0.625774 | 0.035 | 0.026 | 1 | 1 | Eif2b4 | 0.009 | Young |
| 0.245567 | 1.584513 | 0.015 | 0.009 | 1 | 1 | Leng1 | 0.006 | Young |
| 0.245729 | 0.008305 | 0.814 | 0.751 | 1 | 1 | H3f3b | 0.063 | Young |
| 0.245826 | 0.124927 | 0.023 | 0.015 | 1 | 1 | Ppan | 0.008 | Young |
| 0.24584 | 0.331721 | 0.097 | 0.082 | 1 | 1 | Senp6 | 0.015 | Young |
| 0.245862 | 0.127001 | 0.085 | 0.071 | 1 | 1 | Tlnrd1 | 0.014 | Young |
| 0.246028 | 1.455112 | 0.015 | 0.009 | 1 | 1 | Smyd3 | 0.006 | Young |
| 0.246123 | 0.287662 | 0.049 | 0.039 | 1 | 1 | Ebpl | 0.01 | Young |
| 0.24616 | 0.612846 | 0.033 | 0.024 | 1 | 1 | Bckdha | 0.009 | Young |
| 0.246743 | 0.024678 | 0.023 | 0.015 | 1 | 1 | Cul9 | 0.008 | Young |
| 0.247255 | 0.247835 | 0.057 | 0.045 | 1 | 1 | Ccnh | 0.012 | Young |
| 0.247322 | 0.687821 | 0.035 | 0.026 | 1 | 1 | Batf | 0.009 | Young |
| 0.247403 | 0.478718 | 0.029 | 0.021 | 1 | 1 | Ints7 | 0.008 | Young |
| 0.247413 | 1.151686 | 0.015 | 0.009 | 1 | 1 | Mettl13 | 0.006 | Young |
| 0.247453 | 0.246469 | 0.045 | 0.035 | 1 | 1 | Ston2 | 0.01 | Young |
| 0.247495 | 0.315541 | 0.039 | 0.03 | 1 | 1 | Inafm1 | 0.009 | Young |
| 0.247705 | 0.006558 | 0.066 | 0.053 | 1 | 1 | Prdx6 | 0.013 | Young |
| 0.248073 | 0.128488 | 0.054 | 0.042 | 1 | 1 | Spire1 | 0.012 | Young |
| 0.248106 | 0.140229 | 0.049 | 0.039 | 1 | 1 | 2610037D( | 0.01 | Young |
| 0.248511 | 0.407708 | 0.042 | 0.032 | 1 | 1 | Wdr48 | 0.01 | Young |
| 0.248693 | 0.348723 | 0.032 | 0.023 | 1 | 1 | Hinfp | 0.009 | Young |
| 0.248849 | 0.036563 | 0.612 | 0.541 | 1 | 1 | Sem1 | 0.071 | Young |
| 0.248893 | 0.280635 | 0.038 | 0.028 | 1 | 1 | Ece1 | 0.01 | Young |
| 0.248901 | 0.383586 | 0.032 | 0.023 | 1 | 1 | Naa40 | 0.009 | Young |
| 0.249005 | 0.3674 | 0.035 | 0.026 | 1 | 1 | Nhlrc2 | 0.009 | Young |
| 0.24916 | 0.309944 | 0.062 | 0.05 | 1 | 1 | Ctdspl2 | 0.012 | Young |
| 0.249241 | 0.112804 | 0.114 | 0.098 | 1 | 1 | Uggt1 | 0.016 | Young |
| 0.249268 | 0.989512 | 0.015 | 0.009 | 1 | 1 | Cbx8 | 0.006 | Young |
| 0.249457 | 0.28076 | 0.042 | 0.032 | 1 | 1 | Got2 | 0.01 | Young |
| 0.250022 | 0.302386 | 0.044 | 0.033 | 1 | 1 | Mir17hg | 0.011 | Young |
| 0.250266 | 0.308165 | 0.065 | 0.053 | 1 | 1 | Cspp1 | 0.012 | Young |
| 0.250557 | 0.092754 | 0.21 | 0.185 | 1 | 1 | Sowahc | 0.025 | Young |
| 0.25095 | 0.333281 | 0.03 | 0.022 | 1 | 1 | Asb3 | 0.008 | Young |
| 0.251057 | 0.184997 | 0.03 | 0.022 | 1 | 1 | 1600002K( | 0.008 | Young |
| 0.251197 | 0.130204 | 0.1 | 0.084 | 1 | 1 | Wbp11 | 0.016 | Young |

| | | | | | | | | |
|---|---|---|---|---|---|---|---|---|
| 0.251289 | 0.427685 | 0.015 | 0.009 | 1 | 1 | Gm13373 | 0.006 | Young |
| 0.251445 | 0.251057 | 0.015 | 0.009 | 1 | 1 | Asna1 | 0.006 | Young |
| 0.251593 | 0.282074 | 0.03 | 0.022 | 1 | 1 | Orc3 | 0.008 | Young |
| 0.251601 | 0.299504 | 0.015 | 0.009 | 1 | 1 | Ift80 | 0.006 | Young |
| 0.251601 | 0.293226 | 0.015 | 0.009 | 1 | 1 | Svip | 0.006 | Young |
| 0.251653 | 0.01592 | 0.405 | 0.364 | 1 | 1 | Cyfip1 | 0.041 | Young |
| 0.251819 | 0.150761 | 0.166 | 0.147 | 1 | 1 | Egr2 | 0.019 | Young |
| 0.251846 | 0.219862 | 0.044 | 0.033 | 1 | 1 | Washc1 | 0.011 | Young |
| 0.25207 | 0.641937 | 0.015 | 0.009 | 1 | 1 | Aldh4a1 | 0.006 | Young |
| 0.252089 | 0.060262 | 0.105 | 0.089 | 1 | 1 | Phc2 | 0.016 | Young |
| 0.252358 | 0.230056 | 0.092 | 0.077 | 1 | 1 | Rbm27 | 0.015 | Young |
| 0.252365 | 0.223911 | 0.076 | 0.063 | 1 | 1 | Mtm1 | 0.013 | Young |
| 0.253313 | 0.403968 | 0.03 | 0.022 | 1 | 1 | Exosc9 | 0.008 | Young |
| 0.253436 | 0.068387 | 0.029 | 0.021 | 1 | 1 | Nln | 0.008 | Young |
| 0.253479 | 0.361186 | 0.015 | 0.009 | 1 | 1 | Bod1 | 0.006 | Young |
| 0.253563 | 0.244039 | 0.103 | 0.087 | 1 | 1 | Pcf11 | 0.016 | Young |
| 0.253621 | 0.104573 | 0.152 | 0.134 | 1 | 1 | Dip2b | 0.018 | Young |
| 0.253711 | 0.268101 | 0.032 | 0.023 | 1 | 1 | Cep290 | 0.009 | Young |
| 0.253827 | 0.041662 | 0.171 | 0.15 | 1 | 1 | AC160336. | 0.021 | Young |
| 0.254252 | 0.013052 | 0.445 | 0.406 | 1 | 1 | Rbms1 | 0.039 | Young |
| 0.254322 | 0.225073 | 0.029 | 0.021 | 1 | 1 | Fer | 0.008 | Young |
| 0.254433 | 0.257326 | 0.029 | 0.021 | 1 | 1 | Nadk2 | 0.008 | Young |
| 0.254448 | 0.173104 | 0.032 | 0.023 | 1 | 1 | Prkch | 0.009 | Young |
| 0.254617 | 0.178862 | 0.033 | 0.024 | 1 | 1 | Rexo4 | 0.009 | Young |
| 0.254702 | 0.002996 | 0.06 | 0.048 | 1 | 1 | Dgcr2 | 0.012 | Young |
| 0.254736 | 0.215541 | 0.015 | 0.009 | 1 | 1 | Afap1l1 | 0.006 | Young |
| 0.254853 | 0.38982 | 0.075 | 0.062 | 1 | 1 | Ilf3 | 0.013 | Young |
| 0.255158 | 0.000389 | 0.09 | 0.075 | 1 | 1 | Plekha1 | 0.015 | Young |
| 0.255211 | 0.239327 | 0.106 | 0.09 | 1 | 1 | Mga | 0.016 | Young |
| 0.255723 | 0.220712 | 0.068 | 0.055 | 1 | 1 | Tmem230 | 0.013 | Young |
| 0.25582 | 0.100291 | 0.035 | 0.026 | 1 | 1 | Pmf1 | 0.009 | Young |
| 0.256087 | 1.152543 | 0.016 | 0.01 | 1 | 1 | Pwwp2b | 0.006 | Young |
| 0.256114 | 0.294136 | 0.147 | 0.13 | 1 | 1 | Srrm1 | 0.017 | Young |
| 0.256114 | 0.028425 | 0.148 | 0.127 | 1 | 1 | Cxcl1 | 0.021 | Young |
| 0.256428 | 0.124583 | 0.035 | 0.026 | 1 | 1 | Josd2 | 0.009 | Young |
| 0.256544 | 0.104058 | 0.047 | 0.036 | 1 | 1 | Rps6ka5 | 0.011 | Young |
| 0.256801 | 0.101479 | 0.071 | 0.058 | 1 | 1 | Ptk2b | 0.013 | Young |
| 0.256996 | 0.148725 | 0.033 | 0.024 | 1 | 1 | Pus1 | 0.009 | Young |
| 0.256999 | 0.18031 | 0.069 | 0.057 | 1 | 1 | Taok2 | 0.012 | Young |
| 0.257063 | 0.749576 | 0.016 | 0.01 | 1 | 1 | Pgm3 | 0.006 | Young |
| 0.257255 | 0.068408 | 0.033 | 0.024 | 1 | 1 | Siglec1 | 0.009 | Young |
| 0.257273 | 0.315117 | 0.038 | 0.028 | 1 | 1 | Parp2 | 0.01 | Young |
| 0.257293 | 0.043583 | 0.236 | 0.21 | 1 | 1 | Susd6 | 0.026 | Young |
| 0.257428 | 0.130503 | 0.045 | 0.035 | 1 | 1 | Ap4s1 | 0.01 | Young |
| 0.257489 | 0.227141 | 0.048 | 0.037 | 1 | 1 | Mrps23 | 0.011 | Young |

| | | | | | | | | |
|---|---|---|---|---|---|---|---|---|
| 0.257619 | 0.161985 | 0.033 | 0.024 | 1 | 1 | 9430038I01 | 0.009 | Young |
| 0.258141 | 0.158111 | 0.045 | 0.035 | 1 | 1 | Dcaf13 | 0.01 | Young |
| 0.258518 | 0.405007 | 0.036 | 0.027 | 1 | 1 | Cfap97 | 0.009 | Young |
| 0.258858 | 0.230103 | 0.136 | 0.12 | 1 | 1 | Anxa7 | 0.016 | Young |
| 0.259601 | 0.147262 | 0.039 | 0.03 | 1 | 1 | Sphk2 | 0.009 | Young |
| 0.25964 | 0.11946 | 0.065 | 0.053 | 1 | 1 | Snapin | 0.012 | Young |
| 0.259704 | 0.767029 | 0.016 | 0.01 | 1 | 1 | Ap4e1 | 0.006 | Young |
| 0.260007 | 0.723998 | 0.016 | 0.01 | 1 | 1 | Xpnpep3 | 0.006 | Young |
| 0.260017 | 0.029978 | 0.033 | 0.024 | 1 | 1 | Gtpbp10 | 0.009 | Young |
| 0.260066 | 0.25571 | 0.123 | 0.107 | 1 | 1 | Hfe | 0.016 | Young |
| 0.260089 | 0.260619 | 0.059 | 0.048 | 1 | 1 | Fnta | 0.011 | Young |
| 0.260948 | 0.296246 | 0.058 | 0.046 | 1 | 1 | Atic | 0.012 | Young |
| 0.261106 | 0.058347 | 0.088 | 0.073 | 1 | 1 | Ptpn2 | 0.015 | Young |
| 0.261194 | 0.081128 | 0.069 | 0.057 | 1 | 1 | Tfg | 0.012 | Young |
| 0.261384 | 1.989377 | 0.01 | 0.005 | 1 | 1 | Nod2 | 0.005 | Young |
| 0.26182 | 0.120145 | 0.076 | 0.063 | 1 | 1 | Herc2 | 0.013 | Young |
| 0.261989 | 1.550923 | 0.01 | 0.005 | 1 | 1 | Trim68 | 0.005 | Young |
| 0.262549 | 0.631097 | 0.047 | 0.037 | 1 | 1 | Golga2 | 0.01 | Young |
| 0.262574 | 0.425987 | 0.052 | 0.041 | 1 | 1 | Naa10 | 0.011 | Young |
| 0.262911 | 0.411906 | 0.052 | 0.041 | 1 | 1 | Exoc2 | 0.011 | Young |
| 0.262918 | 0.124482 | 0.035 | 0.026 | 1 | 1 | Impdh1 | 0.009 | Young |
| 0.262961 | 0.391082 | 0.053 | 0.042 | 1 | 1 | Dusp8 | 0.011 | Young |
| 0.263203 | 0.132651 | 0.016 | 0.01 | 1 | 1 | Mast2 | 0.006 | Young |
| 0.263235 | 0.026952 | 0.083 | 0.069 | 1 | 1 | Aldh3b1 | 0.014 | Young |
| 0.26328 | 0.250628 | 0.016 | 0.01 | 1 | 1 | Arfip2 | 0.006 | Young |
| 0.263404 | 1.218889 | 0.01 | 0.005 | 1 | 1 | B3galt6 | 0.005 | Young |
| 0.263404 | 0.905224 | 0.01 | 0.005 | 1 | 1 | Vmac | 0.005 | Young |
| 0.263649 | 0.118938 | 0.106 | 0.09 | 1 | 1 | Phf14 | 0.016 | Young |
| 0.264012 | 1.025925 | 0.01 | 0.005 | 1 | 1 | Zfp653 | 0.005 | Young |
| 0.264012 | 1.005826 | 0.01 | 0.005 | 1 | 1 | Alg12 | 0.005 | Young |
| 0.264054 | 0.407026 | 0.055 | 0.044 | 1 | 1 | Smc5 | 0.011 | Young |
| 0.264129 | 0.062946 | 0.042 | 0.032 | 1 | 1 | Nsmce3 | 0.01 | Young |
| 0.264215 | 0.993286 | 0.01 | 0.005 | 1 | 1 | Cdk1 | 0.005 | Young |
| 0.264246 | 1.06475 | 0.018 | 0.012 | 1 | 1 | Pde1b | 0.006 | Young |
| 0.264277 | 0.01359 | 0.033 | 0.024 | 1 | 1 | Gnptg | 0.009 | Young |
| 0.264392 | 0.748876 | 0.018 | 0.012 | 1 | 1 | Homer3 | 0.006 | Young |
| 0.264418 | 0.886004 | 0.01 | 0.005 | 1 | 1 | 4933427D1 | 0.005 | Young |
| 0.264428 | 0.046116 | 0.016 | 0.01 | 1 | 1 | Ick | 0.006 | Young |
| 0.26446 | 0.408118 | 0.053 | 0.042 | 1 | 1 | Xylt2 | 0.011 | Young |
| 0.264621 | 0.640755 | 0.01 | 0.005 | 1 | 1 | Tnfaip8l1 | 0.005 | Young |
| 0.265028 | 0.841517 | 0.01 | 0.005 | 1 | 1 | Ift172 | 0.005 | Young |
| 0.265122 | 0.001167 | 0.039 | 0.03 | 1 | 1 | Dnpep | 0.009 | Young |
| 0.265196 | 0.06525 | 0.172 | 0.152 | 1 | 1 | 2-Mar | 0.02 | Young |
| 0.265231 | 0.975885 | 0.01 | 0.005 | 1 | 1 | Dock5 | 0.005 | Young |
| 0.265271 | 0.635592 | 0.018 | 0.012 | 1 | 1 | Gatc | 0.006 | Young |

| | | | | | | | | |
|---|---|---|---|---|---|---|---|---|
| 0.265302 | 0.211896 | 0.09 | 0.076 | 1 | 1 | Ehmt2 | 0.014 | Young |
| 0.265509 | 0.396494 | 0.056 | 0.045 | 1 | 1 | Anapc16 | 0.011 | Young |
| 0.265638 | 0.657171 | 0.01 | 0.005 | 1 | 1 | Plekhg1 | 0.005 | Young |
| 0.265816 | 0.083404 | 0.095 | 0.08 | 1 | 1 | Ncoa6 | 0.015 | Young |
| 0.265842 | 0.222693 | 0.01 | 0.005 | 1 | 1 | Pms1 | 0.005 | Young |
| 0.266152 | 0.57141 | 0.018 | 0.012 | 1 | 1 | Sgsm2 | 0.006 | Young |
| 0.26625 | 0.032154 | 0.01 | 0.005 | 1 | 1 | Dynlt1a | 0.005 | Young |
| 0.26641 | 0.069827 | 0.23 | 0.208 | 1 | 1 | Rabac1 | 0.022 | Young |
| 0.266446 | 0.476436 | 0.018 | 0.012 | 1 | 1 | Shmt2 | 0.006 | Young |
| 0.266862 | 0.230782 | 0.01 | 0.005 | 1 | 1 | Dlg3 | 0.005 | Young |
| 0.267067 | 0.228302 | 0.01 | 0.005 | 1 | 1 | Gm14548 | 0.005 | Young |
| 0.26732 | 0.040567 | 0.162 | 0.141 | 1 | 1 | Mapk1 | 0.021 | Young |
| 0.267459 | 1.046358 | 0.019 | 0.013 | 1 | 1 | Nek6 | 0.006 | Young |
| 0.267919 | 0.63995 | 0.018 | 0.012 | 1 | 1 | Ercc2 | 0.006 | Young |
| 0.268362 | 0.483345 | 0.018 | 0.012 | 1 | 1 | Mgst3 | 0.006 | Young |
| 0.26851 | 0.166229 | 0.018 | 0.012 | 1 | 1 | Vps37c | 0.006 | Young |
| 0.26866 | 0.097815 | 0.16 | 0.14 | 1 | 1 | Rbbp6 | 0.02 | Young |
| 0.268954 | 0.034499 | 0.145 | 0.126 | 1 | 1 | Alcam | 0.019 | Young |
| 0.269398 | 0.345455 | 0.018 | 0.012 | 1 | 1 | 2310061I0 | 0.006 | Young |
| 0.269725 | 0.994956 | 0.019 | 0.013 | 1 | 1 | Mcu | 0.006 | Young |
| 0.269886 | 0.018261 | 0.1 | 0.085 | 1 | 1 | Mgat4b | 0.015 | Young |
| 0.270202 | 0.044413 | 0.256 | 0.229 | 1 | 1 | Atp6v1f | 0.027 | Young |
| 0.270294 | 0.628867 | 0.019 | 0.013 | 1 | 1 | Zfp511 | 0.006 | Young |
| 0.271024 | 0.125943 | 0.055 | 0.044 | 1 | 1 | Cd80 | 0.011 | Young |
| 0.271338 | 0.15487 | 0.056 | 0.045 | 1 | 1 | Supt6 | 0.011 | Young |
| 0.271434 | 0.348739 | 0.019 | 0.013 | 1 | 1 | Fam35a | 0.006 | Young |
| 0.271577 | 0.659892 | 0.019 | 0.013 | 1 | 1 | Ccdc86 | 0.006 | Young |
| 0.271744 | 0.578664 | 0.044 | 0.035 | 1 | 1 | Pilrb2 | 0.009 | Young |
| 0.271776 | 0.373673 | 0.05 | 0.04 | 1 | 1 | Syne2 | 0.01 | Young |
| 0.272006 | 0.652686 | 0.019 | 0.013 | 1 | 1 | Mefv | 0.006 | Young |
| 0.272255 | 0.273659 | 0.053 | 0.042 | 1 | 1 | Tmem229b | 0.011 | Young |
| 0.272539 | 0.077996 | 0.097 | 0.082 | 1 | 1 | Zfr | 0.015 | Young |
| 0.272951 | 0.22824 | 0.084 | 0.071 | 1 | 1 | Naa15 | 0.013 | Young |
| 0.273151 | 0.525562 | 0.019 | 0.013 | 1 | 1 | Gm35154 | 0.006 | Young |
| 0.273478 | 0.100838 | 0.148 | 0.13 | 1 | 1 | Mapk6 | 0.018 | Young |
| 0.273569 | 0.012435 | 0.018 | 0.012 | 1 | 1 | Ccdc32 | 0.006 | Young |
| 0.273706 | 1.008096 | 0.028 | 0.021 | 1 | 1 | Acad8 | 0.007 | Young |
| 0.273752 | 0.377555 | 0.046 | 0.036 | 1 | 1 | Zdhhc5 | 0.01 | Young |
| 0.275059 | 0.291258 | 0.066 | 0.054 | 1 | 1 | Mybbp1a | 0.012 | Young |
| 0.275128 | 0.101885 | 0.087 | 0.073 | 1 | 1 | 3110043O2 | 0.014 | Young |
| 0.275482 | 0.381611 | 0.041 | 0.032 | 1 | 1 | Ankrd10 | 0.009 | Young |
| 0.275569 | 0.053902 | 0.112 | 0.096 | 1 | 1 | Coa5 | 0.016 | Young |
| 0.275588 | 0.503782 | 0.04 | 0.031 | 1 | 1 | Ahsa2 | 0.009 | Young |
| 0.275595 | 0.276294 | 0.019 | 0.013 | 1 | 1 | Dhx34 | 0.006 | Young |
| 0.275595 | 0.171064 | 0.019 | 0.013 | 1 | 1 | Lsm11 | 0.006 | Young |

| | | | | | | | | |
|---|---|---|---|---|---|---|---|---|
| 0.276005 | 0.423062 | 0.071 | 0.059 | 1 | 1 | Acsl1 | 0.012 | Young |
| 0.276172 | 0.164572 | 0.019 | 0.013 | 1 | 1 | Toe1 | 0.006 | Young |
| 0.276316 | 0.466132 | 0.019 | 0.013 | 1 | 1 | Cdkn1c | 0.006 | Young |
| 0.276432 | 0.063809 | 0.149 | 0.131 | 1 | 1 | Midn | 0.018 | Young |
| 0.276533 | 0.199313 | 0.067 | 0.055 | 1 | 1 | Thra | 0.012 | Young |
| 0.276592 | 0.100142 | 0.143 | 0.125 | 1 | 1 | Gxylt1 | 0.018 | Young |
| 0.276832 | 0.000552 | 0.229 | 0.206 | 1 | 1 | Matr3 | 0.023 | Young |
| 0.277184 | 0.158543 | 0.019 | 0.013 | 1 | 1 | Tmem126b | 0.006 | Young |
| 0.278308 | 1.653157 | 0.011 | 0.006 | 1 | 1 | Ccdc106 | 0.005 | Young |
| 0.278347 | 0.090543 | 0.056 | 0.045 | 1 | 1 | Spata2 | 0.011 | Young |
| 0.278465 | 0.006118 | 0.094 | 0.08 | 1 | 1 | Evi5 | 0.014 | Young |
| 0.278519 | 0.874959 | 0.03 | 0.022 | 1 | 1 | Hnmt | 0.008 | Young |
| 0.278587 | 0.406931 | 0.021 | 0.014 | 1 | 1 | B4galt7 | 0.007 | Young |
| 0.278591 | 0.789785 | 0.024 | 0.017 | 1 | 1 | Dohh | 0.007 | Young |
| 0.278925 | 0.21563 | 0.019 | 0.013 | 1 | 1 | Nubp2 | 0.006 | Young |
| 0.27967 | 0.532339 | 0.034 | 0.026 | 1 | 1 | Fbxo7 | 0.008 | Young |
| 0.279718 | 0.942397 | 0.022 | 0.015 | 1 | 1 | Gm20186 | 0.007 | Young |
| 0.279915 | 0.036839 | 0.052 | 0.041 | 1 | 1 | Cyhr1 | 0.011 | Young |
| 0.279944 | 0.07181 | 0.059 | 0.048 | 1 | 1 | Ethe1 | 0.011 | Young |
| 0.280056 | 0.018843 | 0.058 | 0.046 | 1 | 1 | Ube4a | 0.012 | Young |
| 0.280201 | 0.073958 | 0.145 | 0.129 | 1 | 1 | Rc3h1 | 0.016 | Young |
| 0.280253 | 1.235599 | 0.011 | 0.006 | 1 | 1 | Hspbap1 | 0.005 | Young |
| 0.280253 | 0.874855 | 0.011 | 0.006 | 1 | 1 | Mettl25 | 0.005 | Young |
| 0.280266 | 0.306171 | 0.021 | 0.014 | 1 | 1 | Zeb1 | 0.007 | Young |
| 0.280799 | 0.489121 | 0.022 | 0.015 | 1 | 1 | Cdc42ep4 | 0.007 | Young |
| 0.280839 | 1.330489 | 0.011 | 0.006 | 1 | 1 | Trim30b | 0.005 | Young |
| 0.280928 | 0.153586 | 0.05 | 0.04 | 1 | 1 | Stat2 | 0.01 | Young |
| 0.280941 | 0.575632 | 0.024 | 0.017 | 1 | 1 | Mrvi1 | 0.007 | Young |
| 0.28107 | 0.260423 | 0.022 | 0.015 | 1 | 1 | Lonrf1 | 0.007 | Young |
| 0.28109 | 0.13441 | 0.08 | 0.067 | 1 | 1 | Mtfr1l | 0.013 | Young |
| 0.281109 | 0.130187 | 0.083 | 0.069 | 1 | 1 | Hcfc1 | 0.014 | Young |
| 0.281123 | 0.118936 | 0.049 | 0.039 | 1 | 1 | Snx15 | 0.01 | Young |
| 0.281269 | 0.570221 | 0.024 | 0.017 | 1 | 1 | Ints8 | 0.007 | Young |
| 0.28134 | 0.065931 | 0.134 | 0.117 | 1 | 1 | Nsf | 0.017 | Young |
| 0.281531 | 0.137871 | 0.062 | 0.05 | 1 | 1 | Ppfibp2 | 0.012 | Young |
| 0.281728 | 0.531595 | 0.024 | 0.017 | 1 | 1 | Fam60a | 0.007 | Young |
| 0.281729 | 0.485381 | 0.037 | 0.028 | 1 | 1 | R3hcc1l | 0.009 | Young |
| 0.281746 | 0.493243 | 0.035 | 0.027 | 1 | 1 | Dmxl2 | 0.008 | Young |
| 0.281884 | 0.405828 | 0.022 | 0.015 | 1 | 1 | Sssca1 | 0.007 | Young |
| 0.282093 | 0.303685 | 0.021 | 0.014 | 1 | 1 | Cry2 | 0.007 | Young |
| 0.282207 | 1.007628 | 0.011 | 0.006 | 1 | 1 | Zbtb10 | 0.005 | Young |
| 0.282292 | 0.346864 | 0.022 | 0.015 | 1 | 1 | Aimp2 | 0.007 | Young |
| 0.282386 | 0.549548 | 0.035 | 0.027 | 1 | 1 | Bin2 | 0.008 | Young |
| 0.282599 | 0.656353 | 0.011 | 0.006 | 1 | 1 | Pomt1 | 0.005 | Young |
| 0.282599 | 0.553785 | 0.011 | 0.006 | 1 | 1 | Tmem129 | 0.005 | Young |

| | | | | | | | | |
|---|---|---|---|---|---|---|---|---|
| 0.282732 | 0.125209 | 0.053 | 0.042 | 1 | 1 | Prkag2 | 0.011 | Young |
| 0.283006 | 0.031519 | 0.055 | 0.044 | 1 | 1 | Ndufs8 | 0.011 | Young |
| 0.283478 | 0.15842 | 0.097 | 0.084 | 1 | 1 | Sec24b | 0.013 | Young |
| 0.283504 | 0.267876 | 0.021 | 0.014 | 1 | 1 | Gm16536 | 0.007 | Young |
| 0.283533 | 0.00508 | 0.303 | 0.27 | 1 | 1 | Etv3 | 0.033 | Young |
| 0.283693 | 0.031249 | 0.087 | 0.073 | 1 | 1 | Smad1 | 0.014 | Young |
| 0.283777 | 0.394521 | 0.011 | 0.006 | 1 | 1 | B930036N1 | 0.005 | Young |
| 0.283851 | 0.16848 | 0.063 | 0.051 | 1 | 1 | Tctex1d2 | 0.012 | Young |
| 0.284008 | 0.225705 | 0.077 | 0.064 | 1 | 1 | Bckdk | 0.013 | Young |
| 0.284153 | 0.142338 | 0.137 | 0.121 | 1 | 1 | Baz1b | 0.016 | Young |
| 0.284313 | 0.152779 | 0.049 | 0.039 | 1 | 1 | Fech | 0.01 | Young |
| 0.284368 | 0.733506 | 0.011 | 0.006 | 1 | 1 | Letm2 | 0.005 | Young |
| 0.284368 | 0.497154 | 0.011 | 0.006 | 1 | 1 | Prelid3a | 0.005 | Young |
| 0.284758 | 0.034257 | 0.403 | 0.369 | 1 | 1 | Fcgr2b | 0.034 | Young |
| 0.285062 | 0.057115 | 0.021 | 0.014 | 1 | 1 | Pnpt1 | 0.007 | Young |
| 0.285087 | 0.317101 | 0.022 | 0.015 | 1 | 1 | Otud1 | 0.007 | Young |
| 0.285418 | 0.51729 | 0.024 | 0.017 | 1 | 1 | Rnf126 | 0.007 | Young |
| 0.285571 | 0.34478 | 0.025 | 0.018 | 1 | 1 | Gmpr2 | 0.007 | Young |
| 0.285669 | 0.051082 | 0.276 | 0.252 | 1 | 1 | Metrnl | 0.024 | Young |
| 0.285748 | 0.350557 | 0.011 | 0.006 | 1 | 1 | Nphp3 | 0.005 | Young |
| 0.285775 | 0.038063 | 0.329 | 0.297 | 1 | 1 | Atp5f1 | 0.032 | Young |
| 0.285836 | 0.154081 | 0.043 | 0.033 | 1 | 1 | Fam214a | 0.01 | Young |
| 0.285946 | 0.298603 | 0.011 | 0.006 | 1 | 1 | Gm17745 | 0.005 | Young |
| 0.28602 | 0.132667 | 0.086 | 0.072 | 1 | 1 | Bfar | 0.014 | Young |
| 0.28611 | 0.246194 | 0.041 | 0.032 | 1 | 1 | Znrf1 | 0.009 | Young |
| 0.286235 | 0.316911 | 0.027 | 0.019 | 1 | 1 | Enoph1 | 0.008 | Young |
| 0.286242 | 0.415483 | 0.031 | 0.023 | 1 | 1 | E2f4 | 0.008 | Young |
| 0.286359 | 0.387852 | 0.027 | 0.019 | 1 | 1 | Vps8 | 0.008 | Young |
| 0.286383 | 0.024368 | 0.055 | 0.044 | 1 | 1 | Gemin7 | 0.011 | Young |
| 0.286526 | 0.045412 | 0.022 | 0.015 | 1 | 1 | Adnp2 | 0.007 | Young |
| 0.286539 | 0.151984 | 0.011 | 0.006 | 1 | 1 | Dis3l | 0.005 | Young |
| 0.286594 | 0.231637 | 0.022 | 0.015 | 1 | 1 | Cd300lg | 0.007 | Young |
| 0.286893 | 0.183198 | 0.035 | 0.027 | 1 | 1 | Oxa1l | 0.008 | Young |
| 0.287095 | 0.11813 | 0.088 | 0.075 | 1 | 1 | Egr3 | 0.013 | Young |
| 0.287307 | 0.089756 | 0.043 | 0.033 | 1 | 1 | Trp53inp2 | 0.01 | Young |
| 0.28747 | 0.242114 | 0.038 | 0.03 | 1 | 1 | Ech1 | 0.008 | Young |
| 0.287589 | 0.919572 | 0.042 | 0.033 | 1 | 1 | Gt(ROSA)2( | 0.009 | Young |
| 0.287609 | 0.132057 | 0.077 | 0.064 | 1 | 1 | Rab22a | 0.013 | Young |
| 0.288145 | 0.242575 | 0.04 | 0.031 | 1 | 1 | Blcap | 0.009 | Young |
| 0.288199 | 0.388494 | 0.037 | 0.028 | 1 | 1 | Sesn3 | 0.009 | Young |
| 0.288437 | 0.414555 | 0.031 | 0.023 | 1 | 1 | Alg10b | 0.008 | Young |
| 0.288452 | 0.14837 | 0.022 | 0.015 | 1 | 1 | Anapc7 | 0.007 | Young |
| 0.28886 | 0.011023 | 0.04 | 0.031 | 1 | 1 | Gngt2 | 0.009 | Young |
| 0.288861 | 0.38651 | 0.032 | 0.024 | 1 | 1 | Rgs14 | 0.008 | Young |
| 0.289101 | 0.167032 | 0.037 | 0.028 | 1 | 1 | Dlat | 0.009 | Young |

| | | | | | | | | |
|---|---|---|---|---|---|---|---|---|
| 0.289114 | 0.527462 | 0.027 | 0.019 | 1 | 1 | Fam241a | 0.008 | Young |
| 0.289313 | 0.476856 | 0.037 | 0.028 | 1 | 1 | Las1l | 0.009 | Young |
| 0.289568 | 0.330935 | 0.03 | 0.022 | 1 | 1 | Abtb1 | 0.008 | Young |
| 0.289768 | 0.004411 | 0.235 | 0.211 | 1 | 1 | Serpinb8 | 0.024 | Young |
| 0.289997 | 0.085861 | 0.194 | 0.174 | 1 | 1 | Nxf1 | 0.02 | Young |
| 0.290563 | 0.301071 | 0.032 | 0.024 | 1 | 1 | Trappc11 | 0.008 | Young |
| 0.290825 | 0.066867 | 0.117 | 0.1 | 1 | 1 | Ndufb6 | 0.017 | Young |
| 0.290969 | 0.314871 | 0.057 | 0.046 | 1 | 1 | Rbm18 | 0.011 | Young |
| 0.291017 | 0.144199 | 0.037 | 0.028 | 1 | 1 | Btbd9 | 0.009 | Young |
| 0.291192 | 0.089493 | 0.072 | 0.06 | 1 | 1 | Paxbp1 | 0.012 | Young |
| 0.291574 | 0.271923 | 0.071 | 0.059 | 1 | 1 | Plekhm3 | 0.012 | Young |
| 0.291732 | 0.17877 | 0.099 | 0.085 | 1 | 1 | Spty2d1 | 0.014 | Young |
| 0.291884 | 0.223972 | 0.035 | 0.027 | 1 | 1 | Tmem39a | 0.008 | Young |
| 0.292265 | 0.098404 | 0.027 | 0.019 | 1 | 1 | Cep85l | 0.008 | Young |
| 0.292966 | 0.028607 | 0.216 | 0.193 | 1 | 1 | Aftph | 0.023 | Young |
| 0.293221 | 0.260871 | 0.06 | 0.049 | 1 | 1 | Tlr4 | 0.011 | Young |
| 0.293317 | 0.214087 | 0.037 | 0.028 | 1 | 1 | Vps52 | 0.009 | Young |
| 0.293409 | 1.051772 | 0.013 | 0.008 | 1 | 1 | Ptpn21 | 0.005 | Young |
| 0.293416 | 0.151335 | 0.213 | 0.194 | 1 | 1 | Ncor1 | 0.019 | Young |
| 0.293766 | 0.160501 | 0.064 | 0.053 | 1 | 1 | Rhoh | 0.011 | Young |
| 0.294542 | 0.072347 | 0.091 | 0.077 | 1 | 1 | Paf1 | 0.014 | Young |
| 0.294611 | 0.128104 | 0.1 | 0.086 | 1 | 1 | Vps13c | 0.014 | Young |
| 0.295075 | 0.110688 | 0.072 | 0.06 | 1 | 1 | Zfp950 | 0.012 | Young |
| 0.295387 | 0.435412 | 0.051 | 0.041 | 1 | 1 | Armc1 | 0.01 | Young |
| 0.295563 | 0.01242 | 0.747 | 0.688 | 1 | 1 | Rack1 | 0.059 | Young |
| 0.295792 | 0.478588 | 0.061 | 0.05 | 1 | 1 | Plk3 | 0.011 | Young |
| 0.296023 | 0.210599 | 0.068 | 0.057 | 1 | 1 | Ndufs1 | 0.011 | Young |
| 0.296048 | 0.061736 | 0.03 | 0.022 | 1 | 1 | Prr12 | 0.008 | Young |
| 0.296444 | 0.010735 | 0.028 | 0.021 | 1 | 1 | Pin1 | 0.007 | Young |
| 0.29655 | 0.052361 | 0.064 | 0.053 | 1 | 1 | Mien1 | 0.011 | Young |
| 0.296551 | 0.007135 | 0.041 | 0.032 | 1 | 1 | Ppp1r37 | 0.009 | Young |
| 0.2968 | 0.590761 | 0.013 | 0.008 | 1 | 1 | Acat2 | 0.005 | Young |
| 0.297033 | 0.064373 | 0.061 | 0.05 | 1 | 1 | Acadl | 0.011 | Young |
| 0.297367 | 0.604774 | 0.013 | 0.008 | 1 | 1 | Plxna3 | 0.005 | Young |
| 0.298125 | 0.821946 | 0.013 | 0.008 | 1 | 1 | 1110065P2 | 0.005 | Young |
| 0.298132 | 0.149728 | 0.038 | 0.03 | 1 | 1 | Mettl16 | 0.008 | Young |
| 0.298315 | 0.391985 | 0.013 | 0.008 | 1 | 1 | Susd1 | 0.005 | Young |
| 0.298364 | 0.186754 | 0.032 | 0.024 | 1 | 1 | Tmem198b | 0.008 | Young |
| 0.298505 | 0.584171 | 0.013 | 0.008 | 1 | 1 | Srr | 0.005 | Young |
| 0.298505 | 0.463918 | 0.013 | 0.008 | 1 | 1 | Tmem86b | 0.005 | Young |
| 0.298728 | 0.044246 | 0.321 | 0.293 | 1 | 1 | Man1a | 0.028 | Young |
| 0.298885 | 0.38798 | 0.013 | 0.008 | 1 | 1 | Repin1 | 0.005 | Young |
| 0.29909 | 0.340508 | 0.081 | 0.069 | 1 | 1 | Gabarapl2 | 0.012 | Young |
| 0.29933 | 0.058935 | 0.075 | 0.063 | 1 | 1 | Bicd2 | 0.012 | Young |
| 0.29935 | 0.086379 | 0.034 | 0.026 | 1 | 1 | Farsb | 0.008 | Young |

| | | | | | | | | |
|---|---|---|---|---|---|---|---|---|
| 0.299455 | 0.628939 | 0.013 | 0.008 | 1 | 1 | Tmem150b | 0.005 | Young |
| 0.299997 | 0.079008 | 0.038 | 0.03 | 1 | 1 | Trappc3 | 0.008 | Young |
| 0.300026 | 0.272883 | 0.013 | 0.008 | 1 | 1 | Dars2 | 0.005 | Young |
| 0.300407 | 0.012646 | 0.013 | 0.008 | 1 | 1 | Chpf | 0.005 | Young |
| 0.300598 | 0.316359 | 0.013 | 0.008 | 1 | 1 | Dnajb5 | 0.005 | Young |
| 0.300682 | 0.062765 | 0.091 | 0.077 | 1 | 1 | Tmbim4 | 0.014 | Young |
| 0.301185 | 0.338742 | 0.084 | 0.072 | 1 | 1 | Ndufv2 | 0.012 | Young |
| 0.301285 | 1.549891 | 0.014 | 0.009 | 1 | 1 | Fbxo42 | 0.005 | Young |
| 0.30206 | 0.261076 | 0.081 | 0.068 | 1 | 1 | Znhit1 | 0.013 | Young |
| 0.302402 | 0.107651 | 0.088 | 0.075 | 1 | 1 | Nrd1 | 0.013 | Young |
| 0.302968 | 0.129678 | 0.205 | 0.185 | 1 | 1 | Znrf2 | 0.02 | Young |
| 0.302971 | 0.058848 | 0.091 | 0.077 | 1 | 1 | Cript | 0.014 | Young |
| 0.303105 | 0.200445 | 0.115 | 0.1 | 1 | 1 | Zfp652 | 0.015 | Young |
| 0.303334 | 0.08417 | 0.055 | 0.045 | 1 | 1 | Dcun1d1 | 0.01 | Young |
| 0.303494 | 0.572473 | 0.042 | 0.033 | 1 | 1 | Smg9 | 0.009 | Young |
| 0.30366 | 0.104507 | 0.013 | 0.008 | 1 | 1 | Procr | 0.005 | Young |
| 0.303987 | 0.100915 | 0.093 | 0.08 | 1 | 1 | Ascc3 | 0.013 | Young |
| 0.304186 | 0.514949 | 0.089 | 0.077 | 1 | 1 | Klf9 | 0.012 | Young |
| 0.304217 | 0.07443 | 0.148 | 0.13 | 1 | 1 | Adam19 | 0.018 | Young |
| 0.304714 | 0.858392 | 0.014 | 0.009 | 1 | 1 | Ncapd2 | 0.005 | Young |
| 0.304944 | 0.848484 | 0.038 | 0.03 | 1 | 1 | Csf3r | 0.008 | Young |
| 0.305849 | 0.394401 | 0.08 | 0.068 | 1 | 1 | Tmem51 | 0.012 | Young |
| 0.306165 | 0.542432 | 0.014 | 0.009 | 1 | 1 | Prdm11 | 0.005 | Young |
| 0.306414 | 0.016744 | 0.057 | 0.046 | 1 | 1 | Chchd3 | 0.011 | Young |
| 0.306532 | 0.228437 | 0.062 | 0.051 | 1 | 1 | Kdm6a | 0.011 | Young |
| 0.306733 | 0.03218 | 0.141 | 0.123 | 1 | 1 | Rap1gds1 | 0.018 | Young |
| 0.306895 | 0.27301 | 0.051 | 0.041 | 1 | 1 | Snrnp40 | 0.01 | Young |
| 0.307198 | 0.12522 | 0.11 | 0.095 | 1 | 1 | Rnf145 | 0.015 | Young |
| 0.30748 | 0.014386 | 0.078 | 0.066 | 1 | 1 | Slc44a1 | 0.012 | Young |
| 0.307937 | 0.132404 | 0.117 | 0.103 | 1 | 1 | Dnajb9 | 0.014 | Young |
| 0.308168 | 0.487714 | 0.014 | 0.009 | 1 | 1 | Btbd2 | 0.005 | Young |
| 0.308391 | 1.41719 | 0.016 | 0.01 | 1 | 1 | Nlrp1b | 0.006 | Young |
| 0.308534 | 0.226552 | 0.086 | 0.073 | 1 | 1 | Mtmr3 | 0.013 | Young |
| 0.309264 | 0.405335 | 0.014 | 0.009 | 1 | 1 | L3mbtl3 | 0.005 | Young |
| 0.309264 | 0.369654 | 0.014 | 0.009 | 1 | 1 | Large1 | 0.005 | Young |
| 0.309689 | 0.085618 | 0.069 | 0.058 | 1 | 1 | Mrpl54 | 0.011 | Young |
| 0.311585 | 0.33792 | 0.078 | 0.067 | 1 | 1 | Rest | 0.011 | Young |
| 0.311754 | 0.243766 | 0.045 | 0.036 | 1 | 1 | Mphosph8 | 0.009 | Young |
| 0.311885 | 0.45969 | 0.055 | 0.045 | 1 | 1 | Mau2 | 0.01 | Young |
| 0.313435 | 0.748541 | 0.016 | 0.01 | 1 | 1 | Ndufaf3 | 0.006 | Young |
| 0.313863 | 0.064604 | 0.047 | 0.037 | 1 | 1 | Senp5 | 0.01 | Young |
| 0.31396 | 0.621966 | 0.016 | 0.01 | 1 | 1 | Gm9493 | 0.006 | Young |
| 0.314113 | 0.317543 | 0.047 | 0.037 | 1 | 1 | Wdr5 | 0.01 | Young |
| 0.314412 | 0.120908 | 0.143 | 0.126 | 1 | 1 | Tmx3 | 0.017 | Young |
| 0.314766 | 0.007378 | 0.208 | 0.185 | 1 | 1 | Romo1 | 0.023 | Young |

| | | | | | | | | |
|---|---|---|---|---|---|---|---|---|
| 0.315208 | 0.147791 | 0.138 | 0.122 | 1 | 1 | Zc3hav1 | 0.016 | Young |
| 0.316338 | 0.119194 | 0.126 | 0.111 | 1 | 1 | Igf1r | 0.015 | Young |
| 0.31665 | 0.051294 | 0.188 | 0.168 | 1 | 1 | Blnk | 0.02 | Young |
| 0.316714 | 0.048035 | 0.047 | 0.037 | 1 | 1 | 2310001H1 | 0.01 | Young |
| 0.316906 | 0.293385 | 0.041 | 0.032 | 1 | 1 | Bicral | 0.009 | Young |
| 0.317116 | 0.378669 | 0.042 | 0.033 | 1 | 1 | Cav2 | 0.009 | Young |
| 0.317121 | 0.553905 | 0.016 | 0.01 | 1 | 1 | Qpctl | 0.006 | Young |
| 0.317422 | 0.021403 | 0.247 | 0.224 | 1 | 1 | Selenof | 0.023 | Young |
| 0.318242 | 0.200883 | 0.19 | 0.174 | 1 | 1 | Mapre2 | 0.016 | Young |
| 0.318685 | 0.088031 | 0.049 | 0.04 | 1 | 1 | Sf3a2 | 0.009 | Young |
| 0.318897 | 0.487973 | 0.064 | 0.054 | 1 | 1 | Supt4a | 0.01 | Young |
| 0.319515 | 0.043098 | 0.099 | 0.085 | 1 | 1 | Wtap | 0.014 | Young |
| 0.319517 | 0.020171 | 0.086 | 0.073 | 1 | 1 | Dazap1 | 0.013 | Young |
| 0.319842 | 0.02996 | 0.137 | 0.121 | 1 | 1 | Mapk1ip1l | 0.016 | Young |
| 0.320248 | 0.69382 | 0.032 | 0.024 | 1 | 1 | Ap3s2 | 0.008 | Young |
| 0.320335 | 0.150459 | 0.013 | 0.018 | 1 | 1 | Cxcl9 | -0.005 | Young |
| 0.320481 | 0.098959 | 0.016 | 0.01 | 1 | 1 | Spindoc | 0.006 | Young |
| 0.320835 | 0.014001 | 0.016 | 0.01 | 1 | 1 | Enpp1 | 0.006 | Young |
| 0.320941 | 0.215655 | 0.076 | 0.064 | 1 | 1 | Supt20 | 0.012 | Young |
| 0.3215 | 1.040643 | 0.02 | 0.014 | 1 | 1 | Cox19 | 0.006 | Young |
| 0.321546 | 0.064207 | 0.016 | 0.01 | 1 | 1 | Pgap3 | 0.006 | Young |
| 0.321724 | 0.224342 | 0.016 | 0.01 | 1 | 1 | Wdr76 | 0.006 | Young |
| 0.321902 | 0.125125 | 0.016 | 0.01 | 1 | 1 | Rbbp9 | 0.006 | Young |
| 0.322423 | 0.017757 | 0.091 | 0.078 | 1 | 1 | Gpatch8 | 0.013 | Young |
| 0.322526 | 0.12421 | 0.056 | 0.046 | 1 | 1 | Rcn3 | 0.01 | Young |
| 0.322526 | 0.027992 | 0.042 | 0.033 | 1 | 1 | Fam96b | 0.009 | Young |
| 0.32285 | 0.235925 | 0.056 | 0.046 | 1 | 1 | Pnpla7 | 0.01 | Young |
| 0.323062 | 0.324168 | 0.041 | 0.032 | 1 | 1 | Vamp7 | 0.009 | Young |
| 0.323301 | 0.370474 | 0.036 | 0.028 | 1 | 1 | Knop1 | 0.008 | Young |
| 0.323362 | 0.30352 | 0.093 | 0.081 | 1 | 1 | Xpo6 | 0.012 | Young |
| 0.324447 | 0.809942 | 0.027 | 0.021 | 1 | 1 | Carnmt1 | 0.006 | Young |
| 0.32478 | 0.565513 | 0.02 | 0.014 | 1 | 1 | Golga1 | 0.006 | Young |
| 0.324812 | 0.432824 | 0.018 | 0.013 | 1 | 1 | Zfp317 | 0.005 | Young |
| 0.32525 | 0.050367 | 0.017 | 0.012 | 1 | 1 | Acyp1 | 0.005 | Young |
| 0.32557 | 0.631463 | 0.03 | 0.023 | 1 | 1 | Tmem63b | 0.007 | Young |
| 0.325573 | 0.017808 | 0.047 | 0.037 | 1 | 1 | Nelfcd | 0.01 | Young |
| 0.32607 | 0.106304 | 0.007 | 0.012 | 1 | 1 | Rad1 | -0.005 | Young |
| 0.32634 | 0.022833 | 0.183 | 0.163 | 1 | 1 | Srgap2 | 0.02 | Young |
| 0.326938 | 0.480214 | 0.018 | 0.013 | 1 | 1 | Mtor | 0.005 | Young |
| 0.326945 | 0.557451 | 0.021 | 0.015 | 1 | 1 | Fam78a | 0.006 | Young |
| 0.326981 | 0.65917 | 0.027 | 0.021 | 1 | 1 | D430042O( | 0.006 | Young |
| 0.327136 | 0.708551 | 0.02 | 0.014 | 1 | 1 | Hgf | 0.006 | Young |
| 0.327593 | 0.433385 | 0.018 | 0.013 | 1 | 1 | Adck5 | 0.005 | Young |
| 0.327822 | 0.757228 | 0.024 | 0.018 | 1 | 1 | Dvl2 | 0.006 | Young |
| 0.328168 | 0.746451 | 0.023 | 0.017 | 1 | 1 | 9330160F1 | 0.006 | Young |

| | | | | | | | | |
|---|---|---|---|---|---|---|---|---|
| 0.328691 | 0.744567 | 0.021 | 0.015 | 1 | 1 | Eef2kmt | 0.006 | Young |
| 0.328744 | 0.196829 | 0.036 | 0.028 | 1 | 1 | Rnf215 | 0.008 | Young |
| 0.328784 | 0.13235 | 0.084 | 0.072 | 1 | 1 | Gnl1 | 0.012 | Young |
| 0.328814 | 0.18099 | 0.1 | 0.087 | 1 | 1 | Jtb | 0.013 | Young |
| 0.328897 | 0.356016 | 0.033 | 0.026 | 1 | 1 | Slc35e4 | 0.007 | Young |
| 0.329049 | 0.213979 | 0.086 | 0.075 | 1 | 1 | Trim47 | 0.011 | Young |
| 0.329255 | 0.102597 | 0.078 | 0.067 | 1 | 1 | Smarca4 | 0.011 | Young |
| 0.329757 | 0.507814 | 0.021 | 0.015 | 1 | 1 | Med31 | 0.006 | Young |
| 0.330053 | 0.237107 | 0.033 | 0.026 | 1 | 1 | Ccnc | 0.007 | Young |
| 0.330124 | 0.550851 | 0.032 | 0.024 | 1 | 1 | Traf5 | 0.008 | Young |
| 0.330212 | 0.025512 | 0.084 | 0.072 | 1 | 1 | Pqlc1 | 0.012 | Young |
| 0.330294 | 0.165223 | 0.02 | 0.014 | 1 | 1 | Cables2 | 0.006 | Young |
| 0.330373 | 0.425805 | 0.032 | 0.024 | 1 | 1 | 5730455P1 | 0.008 | Young |
| 0.330373 | 0.226255 | 0.032 | 0.024 | 1 | 1 | Rinl | 0.008 | Young |
| 0.330375 | 0.645564 | 0.023 | 0.017 | 1 | 1 | Zfyve26 | 0.006 | Young |
| 0.330452 | 0.228513 | 0.02 | 0.014 | 1 | 1 | 1600010M | 0.006 | Young |
| 0.330589 | 0.009098 | 0.076 | 0.064 | 1 | 1 | Tmem135 | 0.012 | Young |
| 0.331426 | 0.104584 | 0.162 | 0.145 | 1 | 1 | Eif3j1 | 0.017 | Young |
| 0.331554 | 0.301946 | 0.03 | 0.023 | 1 | 1 | Ss18l2 | 0.007 | Young |
| 0.331555 | 0.571486 | 0.023 | 0.017 | 1 | 1 | Ypel2 | 0.006 | Young |
| 0.331684 | 0.474869 | 0.027 | 0.021 | 1 | 1 | Gcc1 | 0.006 | Young |
| 0.331784 | 0.142268 | 0.042 | 0.033 | 1 | 1 | Odf2 | 0.009 | Young |
| 0.331896 | 0.370757 | 0.021 | 0.015 | 1 | 1 | Slc2a8 | 0.006 | Young |
| 0.331975 | 0.531263 | 0.026 | 0.019 | 1 | 1 | Gnl3l | 0.007 | Young |
| 0.331987 | 0.136732 | 0.129 | 0.114 | 1 | 1 | Map2k2 | 0.015 | Young |
| 0.332125 | 0.529716 | 0.021 | 0.015 | 1 | 1 | Bag4 | 0.006 | Young |
| 0.3323 | 0.342027 | 0.052 | 0.042 | 1 | 1 | Tbc1d9b | 0.01 | Young |
| 0.332568 | 0.287536 | 0.047 | 0.037 | 1 | 1 | Ltn1 | 0.01 | Young |
| 0.332662 | 0.439724 | 0.021 | 0.015 | 1 | 1 | Gm44751 | 0.006 | Young |
| 0.33287 | 0.031342 | 0.106 | 0.093 | 1 | 1 | Fam168a | 0.013 | Young |
| 0.332949 | 0.402673 | 0.026 | 0.019 | 1 | 1 | Kat14 | 0.007 | Young |
| 0.332985 | 0.161677 | 0.033 | 0.026 | 1 | 1 | Ap4m1 | 0.007 | Young |
| 0.333221 | 0.139445 | 0.03 | 0.023 | 1 | 1 | Smc2 | 0.007 | Young |
| 0.333301 | 0.107738 | 0.123 | 0.108 | 1 | 1 | Fkbp2 | 0.015 | Young |
| 0.333478 | 0.416202 | 0.03 | 0.023 | 1 | 1 | Ttf1 | 0.007 | Young |
| 0.333679 | 0.008011 | 0.103 | 0.089 | 1 | 1 | Etv1 | 0.014 | Young |
| 0.333777 | 0.553963 | 0.023 | 0.017 | 1 | 1 | Tpgs1 | 0.006 | Young |
| 0.333962 | 0.028625 | 0.078 | 0.066 | 1 | 1 | Xpo7 | 0.012 | Young |
| 0.334069 | 0.408709 | 0.036 | 0.028 | 1 | 1 | Slc40a1 | 0.008 | Young |
| 0.334418 | 0.083331 | 0.09 | 0.077 | 1 | 1 | Lrp10 | 0.013 | Young |
| 0.334519 | 0.443359 | 0.023 | 0.017 | 1 | 1 | Irf7 | 0.006 | Young |
| 0.334526 | 0.443906 | 0.027 | 0.021 | 1 | 1 | Kat2a | 0.006 | Young |
| 0.334572 | 0.388855 | 0.03 | 0.023 | 1 | 1 | Ikbkg | 0.007 | Young |
| 0.335135 | 0.159134 | 0.032 | 0.024 | 1 | 1 | Tada3 | 0.008 | Young |
| 0.33546 | 0.208503 | 0.026 | 0.019 | 1 | 1 | Gins4 | 0.007 | Young |

| | | | | | | | | |
|---|---|---|---|---|---|---|---|---|
| 0.335583 | 0.20769 | 0.021 | 0.015 | 1 | 1 | Slf1 | 0.006 | Young |
| 0.336008 | 0.417667 | 0.023 | 0.017 | 1 | 1 | Etfrf1 | 0.006 | Young |
| 0.336441 | 0.262111 | 0.026 | 0.019 | 1 | 1 | Nr2c1 | 0.007 | Young |
| 0.336949 | 0.258994 | 0.111 | 0.098 | 1 | 1 | Atp13a2 | 0.013 | Young |
| 0.336998 | 0.108119 | 0.061 | 0.05 | 1 | 1 | Rhot1 | 0.011 | Young |
| 0.337247 | 0.315148 | 0.027 | 0.021 | 1 | 1 | Nup155 | 0.006 | Young |
| 0.337281 | 0.152386 | 0.021 | 0.015 | 1 | 1 | Pam16 | 0.006 | Young |
| 0.337297 | 0.360443 | 0.063 | 0.053 | 1 | 1 | Gtf2a2 | 0.01 | Young |
| 0.337501 | 0.288007 | 0.023 | 0.017 | 1 | 1 | Rnf185 | 0.006 | Young |
| 0.337573 | 0.201154 | 0.024 | 0.018 | 1 | 1 | AC165953. | 0.006 | Young |
| 0.337718 | 0.155647 | 0.024 | 0.018 | 1 | 1 | Smim26 | 0.006 | Young |
| 0.337738 | 0.219324 | 0.03 | 0.023 | 1 | 1 | Ddx55 | 0.007 | Young |
| 0.337793 | 0.195452 | 0.047 | 0.039 | 1 | 1 | Golga3 | 0.008 | Young |
| 0.337998 | 0.132938 | 0.027 | 0.021 | 1 | 1 | Eci2 | 0.006 | Young |
| 0.338099 | 0.172088 | 0.023 | 0.017 | 1 | 1 | Ralgps1 | 0.006 | Young |
| 0.338515 | 0.152947 | 0.024 | 0.018 | 1 | 1 | Gpank1 | 0.006 | Young |
| 0.33888 | 0.169821 | 0.086 | 0.075 | 1 | 1 | Arl6ip5 | 0.011 | Young |
| 0.339944 | 0.189673 | 0.069 | 0.058 | 1 | 1 | Map1s | 0.011 | Young |
| 0.340499 | 0.216053 | 0.023 | 0.017 | 1 | 1 | Zfp942 | 0.006 | Young |
| 0.340849 | 0.329186 | 0.043 | 0.035 | 1 | 1 | Plekhg2 | 0.008 | Young |
| 0.341015 | 0.24466 | 0.049 | 0.04 | 1 | 1 | Actl6a | 0.009 | Young |
| 0.341661 | 0.062993 | 0.112 | 0.098 | 1 | 1 | Rraga | 0.014 | Young |
| 0.341832 | 0.039429 | 0.171 | 0.153 | 1 | 1 | Ube2n | 0.018 | Young |
| 0.342214 | 0.103545 | 0.1 | 0.087 | 1 | 1 | Bcl7b | 0.013 | Young |
| 0.342786 | 0.003135 | 0.041 | 0.032 | 1 | 1 | Rubcn | 0.009 | Young |
| 0.343361 | 0.135822 | 0.029 | 0.022 | 1 | 1 | Zfp26 | 0.007 | Young |
| 0.3442 | 0.016014 | 0.024 | 0.018 | 1 | 1 | Slc26a11 | 0.006 | Young |
| 0.344579 | 0.422233 | 0.044 | 0.036 | 1 | 1 | Cog7 | 0.008 | Young |
| 0.344687 | 0.480233 | 0.044 | 0.036 | 1 | 1 | Tmem260 | 0.008 | Young |
| 0.345691 | 0.310437 | 0.057 | 0.048 | 1 | 1 | Cyth3 | 0.009 | Young |
| 0.345703 | 0.05511 | 0.03 | 0.023 | 1 | 1 | Gosr1 | 0.007 | Young |
| 0.346607 | 0.08928 | 0.063 | 0.053 | 1 | 1 | Uba52 | 0.01 | Young |
| 0.346829 | 0.407516 | 0.068 | 0.058 | 1 | 1 | Pou2f1 | 0.01 | Young |
| 0.347228 | 0.049172 | 0.27 | 0.246 | 1 | 1 | Atp1b3 | 0.024 | Young |
| 0.347427 | 0.070907 | 0.141 | 0.126 | 1 | 1 | Psma6 | 0.015 | Young |
| 0.347499 | 0.533483 | 0.054 | 0.045 | 1 | 1 | Il16 | 0.009 | Young |
| 0.34899 | 0.177848 | 0.047 | 0.039 | 1 | 1 | Pogz | 0.008 | Young |
| 0.349044 | 0.067533 | 0.09 | 0.078 | 1 | 1 | Prkd3 | 0.012 | Young |
| 0.35006 | 0.065085 | 0.095 | 0.082 | 1 | 1 | Commd7 | 0.013 | Young |
| 0.350127 | 0.287962 | 0.044 | 0.036 | 1 | 1 | Ccdc127 | 0.008 | Young |
| 0.350292 | 1.394015 | 0.01 | 0.006 | 1 | 1 | Socs1 | 0.004 | Young |
| 0.350717 | 0.113641 | 0.061 | 0.051 | 1 | 1 | Arfgap2 | 0.01 | Young |
| 0.351108 | 0.250226 | 0.047 | 0.039 | 1 | 1 | Slc25a46 | 0.008 | Young |
| 0.351219 | 1.126585 | 0.01 | 0.006 | 1 | 1 | Fads6 | 0.004 | Young |
| 0.351219 | 1.325774 | 0.01 | 0.006 | 1 | 1 | Ift140 | 0.004 | Young |

| | | | | | | | | |
|---|---|---|---|---|---|---|---|---|
| 0.351219 | 0.603197 | 0.01 | 0.006 | 1 | 1 | Sox4 | 0.004 | Young |
| 0.352147 | 0.879127 | 0.01 | 0.006 | 1 | 1 | Emc9 | 0.004 | Young |
| 0.352371 | 0.073689 | 0.049 | 0.04 | 1 | 1 | Mrpl4 | 0.009 | Young |
| 0.35331 | 0.80698 | 0.01 | 0.006 | 1 | 1 | Tmem154 | 0.004 | Young |
| 0.353543 | 0.35777 | 0.01 | 0.006 | 1 | 1 | BC049352 | 0.004 | Young |
| 0.354195 | 0.136555 | 0.047 | 0.039 | 1 | 1 | Vgll4 | 0.008 | Young |
| 0.354293 | 0.319463 | 0.043 | 0.035 | 1 | 1 | Zcchc8 | 0.008 | Young |
| 0.354664 | 0.013004 | 0.35 | 0.314 | 1 | 1 | Hsp90aa1 | 0.036 | Young |
| 0.354988 | 0.219231 | 0.04 | 0.032 | 1 | 1 | Akr1b10 | 0.008 | Young |
| 0.355252 | 0.049546 | 0.085 | 0.073 | 1 | 1 | Hadha | 0.012 | Young |
| 0.355602 | 0.279654 | 0.1 | 0.089 | 1 | 1 | Ppm1a | 0.011 | Young |
| 0.356345 | 0.397364 | 0.01 | 0.006 | 1 | 1 | Rai14 | 0.004 | Young |
| 0.356345 | 0.050353 | 0.01 | 0.006 | 1 | 1 | Kcnj2 | 0.004 | Young |
| 0.356571 | 0.027175 | 0.094 | 0.081 | 1 | 1 | Ssbp3 | 0.013 | Young |
| 0.356865 | 0.018171 | 0.099 | 0.086 | 1 | 1 | Dstyk | 0.013 | Young |
| 0.357048 | 0.335451 | 0.01 | 0.006 | 1 | 1 | Hibch | 0.004 | Young |
| 0.357229 | 0.007813 | 0.053 | 0.044 | 1 | 1 | Tgfbrap1 | 0.009 | Young |
| 0.357608 | 0.320892 | 0.04 | 0.032 | 1 | 1 | Gga3 | 0.008 | Young |
| 0.357902 | 0.066315 | 0.075 | 0.064 | 1 | 1 | Vps13b | 0.011 | Young |
| 0.358362 | 0.475613 | 0.049 | 0.041 | 1 | 1 | 4930402H2 | 0.008 | Young |
| 0.358926 | 0.022367 | 0.01 | 0.006 | 1 | 1 | Rabif | 0.004 | Young |
| 0.359338 | 0.090525 | 0.121 | 0.108 | 1 | 1 | Dennd1b | 0.013 | Young |
| 0.359595 | 0.127282 | 0.049 | 0.04 | 1 | 1 | Dpp3 | 0.009 | Young |
| 0.35978 | 0.172238 | 0.085 | 0.073 | 1 | 1 | Lars2 | 0.012 | Young |
| 0.360533 | 0.329313 | 0.04 | 0.032 | 1 | 1 | Dnajc16 | 0.008 | Young |
| 0.36092 | 0.220092 | 0.095 | 0.084 | 1 | 1 | Spg21 | 0.011 | Young |
| 0.361049 | 0.841856 | 0.032 | 0.026 | 1 | 1 | Smn1 | 0.006 | Young |
| 0.361308 | 1.220843 | 0.012 | 0.008 | 1 | 1 | Rnf180 | 0.004 | Young |
| 0.362006 | 0.190798 | 0.131 | 0.118 | 1 | 1 | Ndufv3 | 0.013 | Young |
| 0.362131 | 0.107102 | 0.023 | 0.03 | 1 | 1 | Incenp | -0.007 | Young |
| 0.362884 | 0.367653 | 0.04 | 0.032 | 1 | 1 | Arl4a | 0.008 | Young |
| 0.363054 | 0.063748 | 0.083 | 0.072 | 1 | 1 | Appbp2 | 0.011 | Young |
| 0.363067 | 1.36614 | 0.012 | 0.008 | 1 | 1 | Asap2 | 0.004 | Young |
| 0.363729 | 0.837683 | 0.012 | 0.008 | 1 | 1 | Dlc1 | 0.004 | Young |
| 0.36448 | 0.590615 | 0.047 | 0.039 | 1 | 1 | Pdrg1 | 0.008 | Young |
| 0.364611 | 0.48033 | 0.012 | 0.008 | 1 | 1 | A230050P2 | 0.004 | Young |
| 0.364832 | 0.855805 | 0.012 | 0.008 | 1 | 1 | Dusp19 | 0.004 | Young |
| 0.365053 | 1.069521 | 0.012 | 0.008 | 1 | 1 | Map3k10 | 0.004 | Young |
| 0.365274 | 0.776983 | 0.012 | 0.008 | 1 | 1 | Alkbh2 | 0.004 | Young |
| 0.365535 | 0.172048 | 0.044 | 0.036 | 1 | 1 | Myef2 | 0.008 | Young |
| 0.365717 | 0.529935 | 0.012 | 0.008 | 1 | 1 | Gm20604 | 0.004 | Young |
| 0.365938 | 0.731703 | 0.012 | 0.008 | 1 | 1 | Snapc4 | 0.004 | Young |
| 0.366234 | 0.134527 | 0.043 | 0.035 | 1 | 1 | Fam219a | 0.008 | Young |
| 0.3669 | 0.327927 | 0.072 | 0.062 | 1 | 1 | Hnrnph3 | 0.01 | Young |
| 0.367214 | 0.09634 | 0.074 | 0.063 | 1 | 1 | Ggt5 | 0.011 | Young |

| | | | | | | | | |
|---|---|---|---|---|---|---|---|---|
| 0.36749 | 0.23492 | 0.012 | 0.008 | 1 | 1 | Limk2 | 0.004 | Young |
| 0.367921 | 0.077898 | 0.079 | 0.068 | 1 | 1 | Zbtb44 | 0.011 | Young |
| 0.367934 | 0.144073 | 0.012 | 0.008 | 1 | 1 | Urb2 | 0.004 | Young |
| 0.367944 | 0.51247 | 0.034 | 0.027 | 1 | 1 | Tecpr1 | 0.007 | Young |
| 0.368378 | 0.379993 | 0.012 | 0.008 | 1 | 1 | 2700038G2 | 0.004 | Young |
| 0.368448 | 0.113428 | 0.071 | 0.06 | 1 | 1 | Prdm2 | 0.011 | Young |
| 0.368471 | 0.545782 | 0.031 | 0.024 | 1 | 1 | Nol8 | 0.007 | Young |
| 0.3685 | 1.300236 | 0.013 | 0.009 | 1 | 1 | Ank2 | 0.004 | Young |
| 0.369121 | 0.32501 | 0.051 | 0.042 | 1 | 1 | Stxbp5 | 0.009 | Young |
| 0.369133 | 0.168659 | 0.037 | 0.03 | 1 | 1 | Cyth2 | 0.007 | Young |
| 0.369268 | 0.116351 | 0.012 | 0.008 | 1 | 1 | Zbtb40 | 0.004 | Young |
| 0.369375 | 0.271801 | 0.073 | 0.063 | 1 | 1 | Coro7 | 0.01 | Young |
| 0.369753 | 1.101232 | 0.013 | 0.009 | 1 | 1 | Smug1 | 0.004 | Young |
| 0.369902 | 0.166283 | 0.105 | 0.093 | 1 | 1 | Azi2 | 0.012 | Young |
| 0.370153 | 0.528116 | 0.032 | 0.026 | 1 | 1 | Tmem104 | 0.006 | Young |
| 0.370223 | 0.162546 | 0.137 | 0.123 | 1 | 1 | Ppt1 | 0.014 | Young |
| 0.37027 | 0.262365 | 0.012 | 0.008 | 1 | 1 | Zhx3 | 0.004 | Young |
| 0.370381 | 0.815712 | 0.013 | 0.009 | 1 | 1 | 2810001G2 | 0.004 | Young |
| 0.370393 | 0.420382 | 0.034 | 0.027 | 1 | 1 | Tpst1 | 0.007 | Young |
| 0.370419 | 0.076817 | 0.082 | 0.071 | 1 | 1 | Fam234a | 0.011 | Young |
| 0.371498 | 0.039125 | 0.012 | 0.008 | 1 | 1 | Zfyve9 | 0.004 | Young |
| 0.371611 | 0.214431 | 0.037 | 0.03 | 1 | 1 | Oma1 | 0.007 | Young |
| 0.371639 | 0.926249 | 0.013 | 0.009 | 1 | 1 | Actr6 | 0.004 | Young |
| 0.371849 | 1.046388 | 0.013 | 0.009 | 1 | 1 | Arl15 | 0.004 | Young |
| 0.371876 | 1.502307 | 0.016 | 0.012 | 1 | 1 | Zc3h12d | 0.004 | Young |
| 0.372058 | 0.649699 | 0.013 | 0.009 | 1 | 1 | Fggy | 0.004 | Young |
| 0.372176 | 0.448979 | 0.048 | 0.04 | 1 | 1 | Psmc2 | 0.008 | Young |
| 0.372209 | 0.277284 | 0.035 | 0.028 | 1 | 1 | Msantd4 | 0.007 | Young |
| 0.372531 | 0.118187 | 0.038 | 0.031 | 1 | 1 | Pla2g16 | 0.007 | Young |
| 0.373307 | 0.546158 | 0.028 | 0.022 | 1 | 1 | Ttll12 | 0.006 | Young |
| 0.37335 | 0.293782 | 0.052 | 0.044 | 1 | 1 | Hspbp1 | 0.008 | Young |
| 0.37353 | 0.625434 | 0.013 | 0.009 | 1 | 1 | Gm3448 | 0.004 | Young |
| 0.37374 | 0.600823 | 0.013 | 0.009 | 1 | 1 | Slc25a15 | 0.004 | Young |
| 0.373795 | 0.265044 | 0.032 | 0.026 | 1 | 1 | Letmd1 | 0.006 | Young |
| 0.373928 | 0.388835 | 0.032 | 0.026 | 1 | 1 | Agfg2 | 0.006 | Young |
| 0.37481 | 0.085484 | 0.185 | 0.168 | 1 | 1 | Sdf4 | 0.017 | Young |
| 0.374925 | 0.336133 | 0.032 | 0.026 | 1 | 1 | Lonp1 | 0.006 | Young |
| 0.374927 | 0.005685 | 0.104 | 0.091 | 1 | 1 | Slc25a4 | 0.013 | Young |
| 0.374975 | 0.061264 | 0.038 | 0.031 | 1 | 1 | Dhps | 0.007 | Young |
| 0.375005 | 0.408091 | 0.013 | 0.009 | 1 | 1 | Zfp87 | 0.004 | Young |
| 0.375005 | 0.072858 | 0.013 | 0.009 | 1 | 1 | Zfp74 | 0.004 | Young |
| 0.375125 | 0.310278 | 0.032 | 0.026 | 1 | 1 | Cdca4 | 0.006 | Young |
| 0.375207 | 0.206421 | 0.065 | 0.055 | 1 | 1 | Tmem106c | 0.01 | Young |
| 0.375315 | 0.399647 | 0.031 | 0.024 | 1 | 1 | Usp36 | 0.007 | Young |
| 0.375771 | 0.301698 | 0.035 | 0.028 | 1 | 1 | Hat1 | 0.007 | Young |

| | | | | | | | | |
|---|---|---|---|---|---|---|---|---|
| 0.376061 | 0.483156 | 0.013 | 0.009 | 1 | 1 | Cd276 | 0.004 | Young |
| 0.376192 | 0.222107 | 0.078 | 0.068 | 1 | 1 | Xpo1 | 0.01 | Young |
| 0.377392 | 0.313385 | 0.032 | 0.026 | 1 | 1 | Nudt13 | 0.006 | Young |
| 0.377759 | 0.351946 | 0.055 | 0.046 | 1 | 1 | Wwc2 | 0.009 | Young |
| 0.37806 | 0.23093 | 0.032 | 0.026 | 1 | 1 | Glipr2 | 0.006 | Young |
| 0.378198 | 0.476835 | 0.025 | 0.019 | 1 | 1 | Tnfrsf12a | 0.006 | Young |
| 0.378328 | 0.114611 | 0.032 | 0.026 | 1 | 1 | Msh2 | 0.006 | Young |
| 0.378515 | 0.371538 | 0.047 | 0.039 | 1 | 1 | 4931406P1 | 0.008 | Young |
| 0.378892 | 0.065578 | 0.051 | 0.042 | 1 | 1 | Mapk8 | 0.009 | Young |
| 0.379027 | 0.238497 | 0.013 | 0.009 | 1 | 1 | Tlr5 | 0.004 | Young |
| 0.379092 | 0.915738 | 0.016 | 0.012 | 1 | 1 | B230118H( | 0.004 | Young |
| 0.379465 | 0.891555 | 0.015 | 0.01 | 1 | 1 | Rfc5 | 0.005 | Young |
| 0.379465 | 0.766958 | 0.015 | 0.01 | 1 | 1 | Dffa | 0.005 | Young |
| 0.37995 | 0.735855 | 0.022 | 0.017 | 1 | 1 | Dnaja3 | 0.005 | Young |
| 0.37997 | 0.492359 | 0.047 | 0.039 | 1 | 1 | Ing3 | 0.008 | Young |
| 0.380648 | 0.054357 | 0.114 | 0.1 | 1 | 1 | Rasgrp4 | 0.014 | Young |
| 0.380895 | 0.025772 | 0.247 | 0.226 | 1 | 1 | Tmed2 | 0.021 | Young |
| 0.381133 | 0.034876 | 0.113 | 0.1 | 1 | 1 | Ube2i | 0.013 | Young |
| 0.381241 | 0.210072 | 0.07 | 0.06 | 1 | 1 | Trappc10 | 0.01 | Young |
| 0.38215 | 0.047726 | 0.154 | 0.14 | 1 | 1 | Lman2 | 0.014 | Young |
| 0.38254 | 0.694193 | 0.016 | 0.012 | 1 | 1 | Pou6f1 | 0.004 | Young |
| 0.38254 | 0.640901 | 0.016 | 0.012 | 1 | 1 | Wrap73 | 0.004 | Young |
| 0.382587 | 0.258574 | 0.015 | 0.01 | 1 | 1 | Tor4a | 0.005 | Young |
| 0.382744 | 0.428953 | 0.027 | 0.021 | 1 | 1 | 1110059E2 | 0.006 | Young |
| 0.38289 | 0.296902 | 0.015 | 0.01 | 1 | 1 | Drosha | 0.005 | Young |
| 0.383061 | 0.60319 | 0.042 | 0.035 | 1 | 1 | Cass4 | 0.007 | Young |
| 0.383133 | 0.258721 | 0.031 | 0.024 | 1 | 1 | Ccdc107 | 0.007 | Young |
| 0.383295 | 0.131775 | 0.015 | 0.01 | 1 | 1 | Surf6 | 0.005 | Young |
| 0.383308 | 0.405926 | 0.016 | 0.012 | 1 | 1 | Zfp101 | 0.004 | Young |
| 0.383485 | 0.071154 | 0.062 | 0.053 | 1 | 1 | Mmd | 0.009 | Young |
| 0.383754 | 0.159447 | 0.107 | 0.095 | 1 | 1 | Lsm14a | 0.012 | Young |
| 0.383902 | 0.227313 | 0.015 | 0.01 | 1 | 1 | Xrcc6 | 0.005 | Young |
| 0.383962 | 0.049007 | 0.031 | 0.024 | 1 | 1 | Zdhhc13 | 0.007 | Young |
| 0.384008 | 0.045071 | 0.185 | 0.166 | 1 | 1 | Ctnna1 | 0.019 | Young |
| 0.384061 | 0.700664 | 0.018 | 0.013 | 1 | 1 | Gpr180 | 0.005 | Young |
| 0.38409 | 0.707094 | 0.024 | 0.018 | 1 | 1 | Cep135 | 0.006 | Young |
| 0.384826 | 0.029918 | 0.009 | 0.013 | 1 | 1 | Aven | -0.004 | Young |
| 0.385133 | 0.45149 | 0.044 | 0.036 | 1 | 1 | Med20 | 0.008 | Young |
| 0.385134 | 0.295243 | 0.028 | 0.022 | 1 | 1 | Prep | 0.006 | Young |
| 0.385829 | 0.126065 | 0.034 | 0.027 | 1 | 1 | Dpm1 | 0.007 | Young |
| 0.386052 | 0.072689 | 0.051 | 0.042 | 1 | 1 | Sart1 | 0.009 | Young |
| 0.386093 | 0.132446 | 0.066 | 0.057 | 1 | 1 | Abl2 | 0.009 | Young |
| 0.386456 | 0.060111 | 0.203 | 0.186 | 1 | 1 | Psme1 | 0.017 | Young |
| 0.386663 | 0.044323 | 0.129 | 0.116 | 1 | 1 | Usp19 | 0.013 | Young |
| 0.386722 | 0.108203 | 0.049 | 0.041 | 1 | 1 | Gcc2 | 0.008 | Young |

| | | | | | | | | |
|---|---|---|---|---|---|---|---|---|
| 0.386884 | 0.074948 | 0.028 | 0.022 | 1 | 1 | Stk38l | 0.006 | Young |
| 0.387288 | 0.080455 | 0.031 | 0.024 | 1 | 1 | AL590144.: | 0.007 | Young |
| 0.38733 | 0.126584 | 0.025 | 0.019 | 1 | 1 | 2310039H( | 0.006 | Young |
| 0.387358 | 0.003153 | 0.032 | 0.026 | 1 | 1 | Brap | 0.006 | Young |
| 0.38736 | 0.132362 | 0.016 | 0.012 | 1 | 1 | Zfp512b | 0.004 | Young |
| 0.387607 | 0.574999 | 0.021 | 0.015 | 1 | 1 | Ints2 | 0.006 | Young |
| 0.387625 | 0.051665 | 0.105 | 0.093 | 1 | 1 | Vav3 | 0.012 | Young |
| 0.387834 | 0.093283 | 0.168 | 0.154 | 1 | 1 | Tbl1xr1 | 0.014 | Young |
| 0.388081 | 0.219524 | 0.019 | 0.014 | 1 | 1 | Slc2a6 | 0.005 | Young |
| 0.388147 | 0.320292 | 0.022 | 0.017 | 1 | 1 | Pelp1 | 0.005 | Young |
| 0.38858 | 0.175308 | 0.058 | 0.049 | 1 | 1 | Magoh | 0.009 | Young |
| 0.388807 | 0.461729 | 0.022 | 0.017 | 1 | 1 | Mrps30 | 0.005 | Young |
| 0.388887 | 0.291521 | 0.027 | 0.021 | 1 | 1 | Zfp282 | 0.006 | Young |
| 0.38891 | 0.165508 | 0.106 | 0.095 | 1 | 1 | Stat6 | 0.011 | Young |
| 0.389236 | 0.484477 | 0.019 | 0.014 | 1 | 1 | Thoc5 | 0.005 | Young |
| 0.389376 | 0.19263 | 0.031 | 0.024 | 1 | 1 | Cherp | 0.007 | Young |
| 0.389435 | 0.062259 | 0.081 | 0.071 | 1 | 1 | Atp9b | 0.01 | Young |
| 0.389881 | 0.071366 | 0.016 | 0.012 | 1 | 1 | Gcdh | 0.004 | Young |
| 0.389976 | 0.305707 | 0.018 | 0.013 | 1 | 1 | Zfp809 | 0.005 | Young |
| 0.390003 | 0.302393 | 0.021 | 0.015 | 1 | 1 | Nt5dc3 | 0.006 | Young |
| 0.390003 | 0.276503 | 0.021 | 0.015 | 1 | 1 | D930016D( | 0.006 | Young |
| 0.390216 | 0.499713 | 0.019 | 0.014 | 1 | 1 | Gde1 | 0.005 | Young |
| 0.390295 | 0.129525 | 0.022 | 0.017 | 1 | 1 | Ltb4r1 | 0.005 | Young |
| 0.390297 | 0.217455 | 0.104 | 0.093 | 1 | 1 | Mpp6 | 0.011 | Young |
| 0.390544 | 0.053683 | 0.022 | 0.017 | 1 | 1 | Snx25 | 0.005 | Young |
| 0.390651 | 0.173084 | 0.066 | 0.057 | 1 | 1 | Socs7 | 0.009 | Young |
| 0.390751 | 0.045276 | 0.019 | 0.014 | 1 | 1 | Pcid2 | 0.005 | Young |
| 0.3909 | 0.101074 | 0.03 | 0.023 | 1 | 1 | Slc35b4 | 0.007 | Young |
| 0.39092 | 0.155039 | 0.045 | 0.037 | 1 | 1 | Apobr | 0.008 | Young |
| 0.391124 | 0.380367 | 0.022 | 0.017 | 1 | 1 | Stk35 | 0.005 | Young |
| 0.391892 | 0.251925 | 0.021 | 0.015 | 1 | 1 | 2810006K2 | 0.006 | Young |
| 0.392393 | 0.312644 | 0.024 | 0.018 | 1 | 1 | Decr2 | 0.006 | Young |
| 0.39272 | 0.098229 | 0.088 | 0.077 | 1 | 1 | Rsl1d1 | 0.011 | Young |
| 0.393073 | 0.025635 | 0.019 | 0.014 | 1 | 1 | Gm45902 | 0.005 | Young |
| 0.39368 | 0.265476 | 0.024 | 0.018 | 1 | 1 | Rfx3 | 0.006 | Young |
| 0.3937 | 0.052032 | 0.025 | 0.019 | 1 | 1 | Tor2a | 0.006 | Young |
| 0.394477 | 0.191233 | 0.021 | 0.015 | 1 | 1 | Ppargc1b | 0.006 | Young |
| 0.394605 | 0.436018 | 0.052 | 0.044 | 1 | 1 | Mfap3 | 0.008 | Young |
| 0.394614 | 0.066416 | 0.022 | 0.017 | 1 | 1 | Phf6 | 0.005 | Young |
| 0.394633 | 0.495445 | 0.049 | 0.041 | 1 | 1 | Jade2 | 0.008 | Young |
| 0.394961 | 0.107222 | 0.011 | 0.015 | 1 | 1 | Ice2 | -0.004 | Young |
| 0.395584 | 0.033815 | 0.019 | 0.014 | 1 | 1 | Slc37a4 | 0.005 | Young |
| 0.396124 | 0.055425 | 0.019 | 0.014 | 1 | 1 | Trmt13 | 0.005 | Young |
| 0.396381 | 0.10287 | 0.059 | 0.05 | 1 | 1 | Vps4a | 0.009 | Young |
| 0.396491 | 0.019273 | 0.623 | 0.571 | 1 | 1 | Txnip | 0.052 | Young |

| | | | | | | | | |
|---|---|---|---|---|---|---|---|---|
| 0.396573 | 0.087556 | 0.086 | 0.075 | 1 | 1 | Polb | 0.011 | Young |
| 0.39794 | 0.083698 | 0.091 | 0.08 | 1 | 1 | Dcun1d5 | 0.011 | Young |
| 0.399125 | 0.037541 | 0.158 | 0.143 | 1 | 1 | Scarb2 | 0.015 | Young |
| 0.400298 | 0.124684 | 0.066 | 0.057 | 1 | 1 | Kif2a | 0.009 | Young |
| 0.401303 | 0.555421 | 0.036 | 0.03 | 1 | 1 | Vac14 | 0.006 | Young |
| 0.40178 | 0.091852 | 0.042 | 0.035 | 1 | 1 | Rab40c | 0.007 | Young |
| 0.40201 | 0.353658 | 0.061 | 0.053 | 1 | 1 | Opa1 | 0.008 | Young |
| 0.402389 | 0.109244 | 0.067 | 0.058 | 1 | 1 | Vps18 | 0.009 | Young |
| 0.402548 | 0.033983 | 0.045 | 0.037 | 1 | 1 | Lage3 | 0.008 | Young |
| 0.402848 | 0.01446 | 0.084 | 0.073 | 1 | 1 | Esyt1 | 0.011 | Young |
| 0.403588 | 0.228986 | 0.055 | 0.046 | 1 | 1 | Ctr9 | 0.009 | Young |
| 0.405943 | 0.020473 | 0.099 | 0.087 | 1 | 1 | Egln2 | 0.012 | Young |
| 0.406276 | 0.021674 | 0.028 | 0.035 | 1 | 1 | Ltbp3 | -0.007 | Young |
| 0.407631 | 0.147178 | 0.041 | 0.033 | 1 | 1 | Slc46a3 | 0.008 | Young |
| 0.407744 | 0.058597 | 0.093 | 0.082 | 1 | 1 | Cdc37l1 | 0.011 | Young |
| 0.407964 | 0.045338 | 0.116 | 0.103 | 1 | 1 | Hip1 | 0.013 | Young |
| 0.408193 | 0.035266 | 0.041 | 0.033 | 1 | 1 | Scfd1 | 0.008 | Young |
| 0.408351 | 0.05943 | 0.077 | 0.067 | 1 | 1 | Nfkb2 | 0.01 | Young |
| 0.408504 | 0.299213 | 0.052 | 0.044 | 1 | 1 | Swt1 | 0.008 | Young |
| 0.408903 | 0.022795 | 0.073 | 0.063 | 1 | 1 | Plxnc1 | 0.01 | Young |
| 0.408949 | 0.162346 | 0.109 | 0.096 | 1 | 1 | Ednrb | 0.013 | Young |
| 0.409238 | 0.114773 | 0.223 | 0.207 | 1 | 1 | Dok2 | 0.016 | Young |
| 0.40968 | 0.024881 | 0.232 | 0.212 | 1 | 1 | Ptpn1 | 0.02 | Young |
| 0.409835 | 0.377608 | 0.038 | 0.031 | 1 | 1 | Nsmf | 0.007 | Young |
| 0.409965 | 0.262736 | 0.038 | 0.031 | 1 | 1 | Idua | 0.007 | Young |
| 0.410161 | 0.11982 | 0.108 | 0.096 | 1 | 1 | Rab35 | 0.012 | Young |
| 0.411022 | 0.102861 | 0.111 | 0.099 | 1 | 1 | Prkx | 0.012 | Young |
| 0.411174 | 0.067398 | 0.052 | 0.044 | 1 | 1 | Ccdc34 | 0.008 | Young |
| 0.411593 | 0.230643 | 0.038 | 0.031 | 1 | 1 | Me2 | 0.007 | Young |
| 0.411614 | 0.506818 | 0.033 | 0.027 | 1 | 1 | Glce | 0.006 | Young |
| 0.412094 | 0.076222 | 0.039 | 0.032 | 1 | 1 | Frmd8 | 0.007 | Young |
| 0.412231 | 0.191989 | 0.075 | 0.066 | 1 | 1 | Plod3 | 0.009 | Young |
| 0.412532 | 0.018438 | 0.059 | 0.05 | 1 | 1 | Gclc | 0.009 | Young |
| 0.412943 | 0.051885 | 0.195 | 0.179 | 1 | 1 | Casp4 | 0.016 | Young |
| 0.41395 | 0.02499 | 0.125 | 0.112 | 1 | 1 | Ypel5 | 0.013 | Young |
| 0.414534 | 0.388973 | 0.033 | 0.027 | 1 | 1 | Ppm1m | 0.006 | Young |
| 0.414683 | 0.022065 | 0.042 | 0.035 | 1 | 1 | Zfp277 | 0.007 | Young |
| 0.415432 | 0.114853 | 0.039 | 0.032 | 1 | 1 | Pdgfc | 0.007 | Young |
| 0.416715 | 0.28563 | 0.046 | 0.039 | 1 | 1 | Scrib | 0.007 | Young |
| 0.417293 | 0.005919 | 0.042 | 0.035 | 1 | 1 | Rab3d | 0.007 | Young |
| 0.417477 | 0.234834 | 0.036 | 0.03 | 1 | 1 | Thumpd3 | 0.006 | Young |
| 0.417958 | 0.284118 | 0.133 | 0.122 | 1 | 1 | Sp1 | 0.011 | Young |
| 0.418551 | 0.027864 | 0.036 | 0.03 | 1 | 1 | Ppp1r35 | 0.006 | Young |
| 0.418822 | 0.282221 | 0.035 | 0.028 | 1 | 1 | Ampd3 | 0.007 | Young |
| 0.419659 | 0.067143 | 0.113 | 0.102 | 1 | 1 | Ythdf1 | 0.011 | Young |

| | | | | | | | | |
|---|---|---|---|---|---|---|---|---|
| 0.419988 | 0.398885 | 0.033 | 0.027 | 1 | 1 | Zmat5 | 0.006 | Young |
| 0.421433 | 0.262137 | 0.035 | 0.028 | 1 | 1 | 6720427I0' | 0.007 | Young |
| 0.421713 | 0.010862 | 0.032 | 0.039 | 1 | 1 | Tubb4b | -0.007 | Young |
| 0.422552 | 0.380666 | 0.03 | 0.024 | 1 | 1 | Orc4 | 0.006 | Young |
| 0.423168 | 0.022574 | 0.338 | 0.314 | 1 | 1 | G3bp2 | 0.024 | Young |
| 0.423501 | 0.053158 | 0.035 | 0.028 | 1 | 1 | Taf12 | 0.007 | Young |
| 0.423995 | 0.049545 | 0.052 | 0.044 | 1 | 1 | Mpdu1 | 0.008 | Young |
| 0.424177 | 0.37715 | 0.052 | 0.045 | 1 | 1 | Ostm1 | 0.007 | Young |
| 0.424448 | 0.119038 | 0.06 | 0.051 | 1 | 1 | Bag3 | 0.009 | Young |
| 0.425164 | 0.058932 | 0.036 | 0.03 | 1 | 1 | Nomo1 | 0.006 | Young |
| 0.425287 | 0.314512 | 0.118 | 0.108 | 1 | 1 | Zfp445 | 0.01 | Young |
| 0.425418 | 0.325954 | 0.032 | 0.026 | 1 | 1 | Grcc10 | 0.006 | Young |
| 0.425626 | 0.163087 | 0.033 | 0.027 | 1 | 1 | Fbxl14 | 0.006 | Young |
| 0.425838 | 0.22991 | 0.033 | 0.027 | 1 | 1 | Nacc1 | 0.006 | Young |
| 0.426964 | 0.167972 | 0.082 | 0.072 | 1 | 1 | Gnl3 | 0.01 | Young |
| 0.42698 | 0.04754 | 0.29 | 0.271 | 1 | 1 | Frmd4b | 0.019 | Young |
| 0.427992 | 0.00096 | 0.206 | 0.189 | 1 | 1 | Tmco1 | 0.017 | Young |
| 0.429752 | 0.024304 | 0.147 | 0.132 | 1 | 1 | BC017643 | 0.015 | Young |
| 0.430333 | 0.387844 | 0.043 | 0.036 | 1 | 1 | Ecd | 0.007 | Young |
| 0.430437 | 0.205277 | 0.057 | 0.049 | 1 | 1 | Ube4b | 0.008 | Young |
| 0.430712 | 0.522553 | 0.04 | 0.033 | 1 | 1 | Tesk1 | 0.007 | Young |
| 0.431021 | 0.038218 | 0.071 | 0.062 | 1 | 1 | Ndufb2 | 0.009 | Young |
| 0.431562 | 0.12422 | 0.071 | 0.062 | 1 | 1 | Pink1 | 0.009 | Young |
| 0.431666 | 0.211356 | 0.032 | 0.026 | 1 | 1 | Smarcd1 | 0.006 | Young |
| 0.43204 | 0.056148 | 0.052 | 0.044 | 1 | 1 | Apopt1 | 0.008 | Young |
| 0.432156 | 0.681093 | 0.029 | 0.023 | 1 | 1 | Gapt | 0.006 | Young |
| 0.432597 | 0.119122 | 0.008 | 0.012 | 1 | 1 | B230354K1 | -0.004 | Young |
| 0.432702 | 0.41705 | 0.027 | 0.022 | 1 | 1 | Hsf1 | 0.005 | Young |
| 0.433616 | 0.244751 | 0.085 | 0.075 | 1 | 1 | Il10 | 0.01 | Young |
| 0.43412 | 0.391462 | 0.027 | 0.022 | 1 | 1 | Slc35c1 | 0.005 | Young |
| 0.434353 | 0.525564 | 0.026 | 0.021 | 1 | 1 | Tomm6 | 0.005 | Young |
| 0.434519 | 0.170926 | 0.066 | 0.058 | 1 | 1 | Tspan4 | 0.008 | Young |
| 0.434597 | 0.491754 | 0.026 | 0.021 | 1 | 1 | Ints11 | 0.005 | Young |
| 0.434655 | 0.194185 | 0.073 | 0.064 | 1 | 1 | Psme2b | 0.009 | Young |
| 0.434817 | 0.092973 | 0.1 | 0.089 | 1 | 1 | Rufy3 | 0.011 | Young |
| 0.435875 | 0.10402 | 0.106 | 0.095 | 1 | 1 | AB124611 | 0.011 | Young |
| 0.435995 | 0.329067 | 0.043 | 0.036 | 1 | 1 | Mrps25 | 0.007 | Young |
| 0.436316 | 0.309752 | 0.081 | 0.072 | 1 | 1 | Zc3h15 | 0.009 | Young |
| 0.436768 | 0.27265 | 0.029 | 0.023 | 1 | 1 | Tmc6 | 0.006 | Young |
| 0.437151 | 1.02114 | 0.01 | 0.006 | 1 | 1 | Fam84b | 0.004 | Young |
| 0.437236 | 0.139275 | 0.073 | 0.064 | 1 | 1 | Aqr | 0.009 | Young |
| 0.437321 | 0.274679 | 0.064 | 0.055 | 1 | 1 | Spr | 0.009 | Young |
| 0.437458 | 0.075015 | 0.03 | 0.024 | 1 | 1 | Ngrn | 0.006 | Young |
| 0.437663 | 0.384423 | 0.062 | 0.054 | 1 | 1 | Irf4 | 0.008 | Young |
| 0.437669 | 0.017351 | 0.131 | 0.118 | 1 | 1 | Csnk1d | 0.013 | Young |

| | | | | | | | | |
|---|---|---|---|---|---|---|---|---|
| 0.437831 | 0.98117 | 0.01 | 0.006 | 1 | 1 | Kctd9 | 0.004 | Young |
| 0.438075 | 0.381001 | 0.027 | 0.022 | 1 | 1 | Chid1 | 0.005 | Young |
| 0.438512 | 0.970151 | 0.01 | 0.006 | 1 | 1 | Steap4 | 0.004 | Young |
| 0.438758 | 0.092047 | 0.044 | 0.037 | 1 | 1 | Pon3 | 0.007 | Young |
| 0.439035 | 0.093466 | 0.065 | 0.057 | 1 | 1 | Blvra | 0.008 | Young |
| 0.439401 | 0.38161 | 0.026 | 0.021 | 1 | 1 | Arl3 | 0.005 | Young |
| 0.439706 | 0.571806 | 0.024 | 0.019 | 1 | 1 | Spg11 | 0.005 | Young |
| 0.440149 | 0.392024 | 0.01 | 0.006 | 1 | 1 | Cgnl1 | 0.004 | Young |
| 0.440423 | 0.364282 | 0.01 | 0.006 | 1 | 1 | Tasp1 | 0.004 | Young |
| 0.440709 | 0.293915 | 0.026 | 0.021 | 1 | 1 | Kif16b | 0.005 | Young |
| 0.440969 | 0.434046 | 0.01 | 0.006 | 1 | 1 | Spout1 | 0.004 | Young |
| 0.441404 | 0.094766 | 0.104 | 0.094 | 1 | 1 | Bud31 | 0.01 | Young |
| 0.441461 | 0.085538 | 0.047 | 0.04 | 1 | 1 | Clec4e | 0.007 | Young |
| 0.44179 | 0.426484 | 0.01 | 0.006 | 1 | 1 | Brca2 | 0.004 | Young |
| 0.44179 | 0.182684 | 0.01 | 0.006 | 1 | 1 | Pigf | 0.004 | Young |
| 0.44179 | 0.169968 | 0.01 | 0.006 | 1 | 1 | Otof | 0.004 | Young |
| 0.441856 | 0.306944 | 0.026 | 0.021 | 1 | 1 | Rassf8 | 0.005 | Young |
| 0.441898 | 0.661262 | 0.024 | 0.019 | 1 | 1 | Zbtb14 | 0.005 | Young |
| 0.441986 | 0.01591 | 0.211 | 0.194 | 1 | 1 | Ier3ip1 | 0.017 | Young |
| 0.442235 | 0.447911 | 0.024 | 0.019 | 1 | 1 | Gfm2 | 0.005 | Young |
| 0.442358 | 0.057853 | 0.058 | 0.05 | 1 | 1 | Ncln | 0.008 | Young |
| 0.442681 | 0.269001 | 0.023 | 0.018 | 1 | 1 | Parn | 0.005 | Young |
| 0.442706 | 0.241413 | 0.013 | 0.017 | 1 | 1 | Pigh | -0.004 | Young |
| 0.443434 | 0.014734 | 0.01 | 0.006 | 1 | 1 | Mphosph9 | 0.004 | Young |
| 0.443664 | 0.377035 | 0.069 | 0.06 | 1 | 1 | Ebp | 0.009 | Young |
| 0.443873 | 0.527247 | 0.021 | 0.017 | 1 | 1 | Emc4 | 0.004 | Young |
| 0.443926 | 0.436911 | 0.024 | 0.019 | 1 | 1 | Borcs8 | 0.005 | Young |
| 0.443983 | 0.050971 | 0.01 | 0.006 | 1 | 1 | Sh2b2 | 0.004 | Young |
| 0.44447 | 0.264506 | 0.04 | 0.033 | 1 | 1 | Ralgapa1 | 0.007 | Young |
| 0.444887 | 0.04709 | 0.043 | 0.036 | 1 | 1 | Cdc14a | 0.007 | Young |
| 0.444943 | 1.654368 | 0.014 | 0.01 | 1 | 1 | Mamdc2 | 0.004 | Young |
| 0.445062 | 0.822306 | 0.011 | 0.008 | 1 | 1 | Acd | 0.003 | Young |
| 0.445062 | 0.612094 | 0.011 | 0.008 | 1 | 1 | B4galt4 | 0.003 | Young |
| 0.445799 | 0.17924 | 0.026 | 0.021 | 1 | 1 | Tfpi | 0.005 | Young |
| 0.445826 | 0.9416 | 0.011 | 0.008 | 1 | 1 | Pold1 | 0.003 | Young |
| 0.445967 | 0.064852 | 0.027 | 0.022 | 1 | 1 | Hic2 | 0.005 | Young |
| 0.446207 | 0.196612 | 0.027 | 0.022 | 1 | 1 | Cep44 | 0.005 | Young |
| 0.446317 | 0.286027 | 0.061 | 0.053 | 1 | 1 | Rit1 | 0.008 | Young |
| 0.446591 | 0.794764 | 0.011 | 0.008 | 1 | 1 | Mipep | 0.003 | Young |
| 0.446591 | 0.731486 | 0.011 | 0.008 | 1 | 1 | Pfas | 0.003 | Young |
| 0.446953 | 0.101566 | 0.026 | 0.021 | 1 | 1 | Slx4ip | 0.005 | Young |
| 0.447102 | 0.414358 | 0.011 | 0.008 | 1 | 1 | Traf3ip1 | 0.003 | Young |
| 0.447462 | 0.109855 | 0.065 | 0.057 | 1 | 1 | Ddx54 | 0.008 | Young |
| 0.447613 | 0.526848 | 0.011 | 0.008 | 1 | 1 | Tada2a | 0.003 | Young |
| 0.447613 | 0.191096 | 0.026 | 0.021 | 1 | 1 | Gdap2 | 0.005 | Young |

| | | | | | | | | |
|---|---|---|---|---|---|---|---|---|
| 0.447716 | 0.095845 | 0.04 | 0.033 | 1 | 1 | Coasy | 0.007 | Young |
| 0.448124 | 0.819268 | 0.011 | 0.008 | 1 | 1 | Prkar1b | 0.003 | Young |
| 0.448124 | 0.368128 | 0.011 | 0.008 | 1 | 1 | Mvk | 0.003 | Young |
| 0.448458 | 0.257919 | 0.023 | 0.018 | 1 | 1 | Usb1 | 0.005 | Young |
| 0.448635 | 0.230024 | 0.011 | 0.008 | 1 | 1 | Lrrc20 | 0.003 | Young |
| 0.44866 | 0.569078 | 0.045 | 0.039 | 1 | 1 | Timm44 | 0.006 | Young |
| 0.448792 | 0.171464 | 0.078 | 0.069 | 1 | 1 | BC028528 | 0.009 | Young |
| 0.44904 | 0.709175 | 0.02 | 0.015 | 1 | 1 | Borcs5 | 0.005 | Young |
| 0.449147 | 0.127698 | 0.011 | 0.008 | 1 | 1 | Zfp637 | 0.003 | Young |
| 0.449331 | 0.073807 | 0.073 | 0.064 | 1 | 1 | Pnrc2 | 0.009 | Young |
| 0.449337 | 0.283533 | 0.023 | 0.018 | 1 | 1 | Mettl6 | 0.005 | Young |
| 0.449501 | 0.739508 | 0.013 | 0.009 | 1 | 1 | Cenpl | 0.004 | Young |
| 0.449659 | 0.31324 | 0.011 | 0.008 | 1 | 1 | Cenpv | 0.003 | Young |
| 0.449703 | 0.177942 | 0.024 | 0.019 | 1 | 1 | Zfp362 | 0.005 | Young |
| 0.449983 | 0.618927 | 0.02 | 0.015 | 1 | 1 | Stx18 | 0.005 | Young |
| 0.450428 | 0.410119 | 0.011 | 0.008 | 1 | 1 | Sppl2b | 0.003 | Young |
| 0.450544 | 0.011452 | 0.125 | 0.113 | 1 | 1 | Vdac1 | 0.012 | Young |
| 0.450556 | 0.123759 | 0.024 | 0.019 | 1 | 1 | Wipi1 | 0.005 | Young |
| 0.450585 | 0.936036 | 0.014 | 0.01 | 1 | 1 | Spink2 | 0.004 | Young |
| 0.451081 | 0.17511 | 0.048 | 0.041 | 1 | 1 | Thbd | 0.007 | Young |
| 0.451237 | 0.052225 | 0.043 | 0.036 | 1 | 1 | Sestd1 | 0.007 | Young |
| 0.451454 | 0.253276 | 0.011 | 0.008 | 1 | 1 | Sema6b | 0.003 | Young |
| 0.451562 | 0.223475 | 0.092 | 0.082 | 1 | 1 | Klhl5 | 0.01 | Young |
| 0.45187 | 0.189425 | 0.038 | 0.032 | 1 | 1 | Agpat4 | 0.006 | Young |
| 0.452115 | 0.012492 | 0.043 | 0.05 | 1 | 1 | Mob3b | -0.007 | Young |
| 0.452343 | 0.53096 | 0.018 | 0.014 | 1 | 1 | Trpm4 | 0.004 | Young |
| 0.452623 | 0.479665 | 0.013 | 0.009 | 1 | 1 | Ccdc167 | 0.004 | Young |
| 0.453081 | 0.535305 | 0.014 | 0.01 | 1 | 1 | Ano8 | 0.004 | Young |
| 0.453105 | 0.39515 | 0.013 | 0.009 | 1 | 1 | Gtpbp3 | 0.004 | Young |
| 0.453171 | 0.116702 | 0.043 | 0.036 | 1 | 1 | Thbs1 | 0.007 | Young |
| 0.453308 | 0.669199 | 0.014 | 0.01 | 1 | 1 | Cacfd1 | 0.004 | Young |
| 0.453308 | 0.39051 | 0.014 | 0.01 | 1 | 1 | Whamm | 0.004 | Young |
| 0.453308 | 0.345149 | 0.014 | 0.01 | 1 | 1 | Ahi1 | 0.004 | Young |
| 0.453442 | 0.163218 | 0.051 | 0.044 | 1 | 1 | Polr3gl | 0.007 | Young |
| 0.453587 | 0.467517 | 0.013 | 0.009 | 1 | 1 | Blm | 0.004 | Young |
| 0.453854 | 0.586216 | 0.016 | 0.012 | 1 | 1 | Ipo4 | 0.004 | Young |
| 0.453994 | 0.478959 | 0.053 | 0.046 | 1 | 1 | Sec24a | 0.007 | Young |
| 0.453995 | 0.004413 | 0.295 | 0.272 | 1 | 1 | Hnrnpm | 0.023 | Young |
| 0.454371 | 0.395236 | 0.017 | 0.013 | 1 | 1 | 2410004B1 | 0.004 | Young |
| 0.454445 | 0.592287 | 0.014 | 0.01 | 1 | 1 | Mgmt | 0.004 | Young |
| 0.454749 | 0.008932 | 0.024 | 0.019 | 1 | 1 | Nol6 | 0.005 | Young |
| 0.454807 | 0.028701 | 0.177 | 0.161 | 1 | 1 | Leprot | 0.016 | Young |
| 0.4549 | 0.363246 | 0.014 | 0.01 | 1 | 1 | Slc2a9 | 0.004 | Young |
| 0.454962 | 0.080893 | 0.16 | 0.147 | 1 | 1 | Kitl | 0.013 | Young |
| 0.455003 | 0.160987 | 0.038 | 0.032 | 1 | 1 | Ankrd16 | 0.006 | Young |

| | | | | | | | | |
|---|---|---|---|---|---|---|---|---|
| 0.455034 | 0.058254 | 0.013 | 0.009 | 1 | 1 | Pigz | 0.004 | Young |
| 0.455073 | 0.269979 | 0.067 | 0.059 | 1 | 1 | Hgsnat | 0.008 | Young |
| 0.45515 | 0.627074 | 0.016 | 0.012 | 1 | 1 | Slc44a2 | 0.004 | Young |
| 0.455183 | 0.215246 | 0.037 | 0.031 | 1 | 1 | Kctd12b | 0.006 | Young |
| 0.455288 | 0.303617 | 0.02 | 0.015 | 1 | 1 | St3gal3 | 0.005 | Young |
| 0.455699 | 0.57599 | 0.018 | 0.014 | 1 | 1 | Golm1 | 0.004 | Young |
| 0.455799 | 0.110979 | 0.016 | 0.012 | 1 | 1 | A530017D2 | 0.004 | Young |
| 0.456125 | 0.331663 | 0.017 | 0.013 | 1 | 1 | Cradd | 0.004 | Young |
| 0.456967 | 0.29944 | 0.013 | 0.009 | 1 | 1 | Osgin1 | 0.004 | Young |
| 0.457215 | 0.006478 | 0.427 | 0.392 | 1 | 1 | Pitpna | 0.035 | Young |
| 0.457316 | 0.392586 | 0.016 | 0.012 | 1 | 1 | Tspan33 | 0.004 | Young |
| 0.457365 | 0.038625 | 0.017 | 0.013 | 1 | 1 | Clip2 | 0.004 | Young |
| 0.458183 | 0.114794 | 0.016 | 0.012 | 1 | 1 | Acad9 | 0.004 | Young |
| 0.458193 | 0.02034 | 0.017 | 0.013 | 1 | 1 | Sft2d3 | 0.004 | Young |
| 0.458421 | 0.072422 | 0.038 | 0.032 | 1 | 1 | Ankra2 | 0.006 | Young |
| 0.458473 | 0.284012 | 0.018 | 0.014 | 1 | 1 | Zfhx2 | 0.004 | Young |
| 0.458663 | 0.022417 | 0.013 | 0.009 | 1 | 1 | Nat6 | 0.004 | Young |
| 0.458768 | 0.121166 | 0.07 | 0.062 | 1 | 1 | Bcor | 0.008 | Young |
| 0.459068 | 0.025434 | 0.064 | 0.055 | 1 | 1 | Gatd1 | 0.009 | Young |
| 0.459267 | 0.242449 | 0.018 | 0.014 | 1 | 1 | 2900089D1 | 0.004 | Young |
| 0.459294 | 0.348502 | 0.035 | 0.03 | 1 | 1 | Cib2 | 0.005 | Young |
| 0.459467 | 0.085945 | 0.014 | 0.01 | 1 | 1 | Zfp931 | 0.004 | Young |
| 0.459806 | 0.080366 | 0.16 | 0.148 | 1 | 1 | Wdr83os | 0.012 | Young |
| 0.459852 | 0.230333 | 0.017 | 0.013 | 1 | 1 | Pgghg | 0.004 | Young |
| 0.459863 | 0.168751 | 0.02 | 0.015 | 1 | 1 | Nfkbil1 | 0.005 | Young |
| 0.459876 | 0.020581 | 0.013 | 0.009 | 1 | 1 | Pnpla1 | 0.004 | Young |
| 0.460055 | 0.094318 | 0.02 | 0.015 | 1 | 1 | Scyl3 | 0.005 | Young |
| 0.460069 | 0.093528 | 0.059 | 0.051 | 1 | 1 | Meaf6 | 0.008 | Young |
| 0.460292 | 0.097064 | 0.049 | 0.042 | 1 | 1 | Actr10 | 0.007 | Young |
| 0.460548 | 0.009924 | 0.054 | 0.046 | 1 | 1 | Uchl5 | 0.008 | Young |
| 0.460579 | 0.423575 | 0.035 | 0.03 | 1 | 1 | Psmg3 | 0.005 | Young |
| 0.460613 | 0.376206 | 0.014 | 0.01 | 1 | 1 | Dclre1a | 0.004 | Young |
| 0.460779 | 0.058578 | 0.088 | 0.078 | 1 | 1 | Rbbp7 | 0.01 | Young |
| 0.460781 | 0.032886 | 0.054 | 0.046 | 1 | 1 | Ddx23 | 0.008 | Young |
| 0.46089 | 0.219678 | 0.017 | 0.013 | 1 | 1 | Zfp84 | 0.004 | Young |
| 0.461193 | 0.184427 | 0.088 | 0.078 | 1 | 1 | Wwp2 | 0.01 | Young |
| 0.461803 | 0.056215 | 0.115 | 0.104 | 1 | 1 | Cep350 | 0.011 | Young |
| 0.462318 | 0.195175 | 0.016 | 0.012 | 1 | 1 | Pwp2 | 0.004 | Young |
| 0.462734 | 0.260599 | 0.072 | 0.064 | 1 | 1 | Abcd1 | 0.008 | Young |
| 0.462856 | 0.004923 | 0.081 | 0.072 | 1 | 1 | Insr | 0.009 | Young |
| 0.463078 | 0.596693 | 0.032 | 0.027 | 1 | 1 | Mitd1 | 0.005 | Young |
| 0.4631 | 0.178874 | 0.1 | 0.091 | 1 | 1 | Vps13a | 0.009 | Young |
| 0.463261 | 0.098294 | 0.021 | 0.017 | 1 | 1 | Alg6 | 0.004 | Young |
| 0.46337 | 0.064659 | 0.014 | 0.01 | 1 | 1 | Dexi | 0.004 | Young |
| 0.463874 | 0.110456 | 0.04 | 0.033 | 1 | 1 | Ttc5 | 0.007 | Young |

| | | | | | | | | |
|---|---|---|---|---|---|---|---|---|
| 0.464734 | 0.097926 | 0.104 | 0.094 | 1 | 1 | Cebpa | 0.01 | Young |
| 0.465154 | 0.008613 | 0.049 | 0.042 | 1 | 1 | Taf11 | 0.007 | Young |
| 0.465603 | 0.099235 | 0.108 | 0.098 | 1 | 1 | Nfil3 | 0.01 | Young |
| 0.465685 | 0.122535 | 0.037 | 0.031 | 1 | 1 | Arhgef11 | 0.006 | Young |
| 0.465785 | 0.092717 | 0.081 | 0.072 | 1 | 1 | Mrpl3 | 0.009 | Young |
| 0.46654 | 0.249872 | 0.047 | 0.04 | 1 | 1 | Rprd1b | 0.007 | Young |
| 0.46655 | 0.280213 | 0.055 | 0.048 | 1 | 1 | Tnip2 | 0.007 | Young |
| 0.467559 | 0.367042 | 0.045 | 0.039 | 1 | 1 | Zfp655 | 0.006 | Young |
| 0.468157 | 0.143127 | 0.058 | 0.05 | 1 | 1 | Mdp1 | 0.008 | Young |
| 0.468303 | 0.126702 | 0.131 | 0.12 | 1 | 1 | Cisd2 | 0.011 | Young |
| 0.469925 | 0.043464 | 0.103 | 0.093 | 1 | 1 | Atf7ip | 0.01 | Young |
| 0.470352 | 0.256537 | 0.035 | 0.03 | 1 | 1 | Ino80c | 0.005 | Young |
| 0.471464 | 0.041484 | 0.066 | 0.058 | 1 | 1 | Kpna1 | 0.008 | Young |
| 0.472192 | 0.068435 | 0.049 | 0.042 | 1 | 1 | Cdk2 | 0.007 | Young |
| 0.47226 | 0.391037 | 0.044 | 0.037 | 1 | 1 | Sacm1l | 0.007 | Young |
| 0.47306 | 0.012792 | 0.089 | 0.08 | 1 | 1 | Emc3 | 0.009 | Young |
| 0.473667 | 0.07447 | 0.395 | 0.37 | 1 | 1 | Ifrd1 | 0.025 | Young |
| 0.475282 | 0.108006 | 0.032 | 0.027 | 1 | 1 | Extl2 | 0.005 | Young |
| 0.475629 | 0.101978 | 0.056 | 0.049 | 1 | 1 | Smpd1 | 0.007 | Young |
| 0.475805 | 0.016566 | 0.104 | 0.094 | 1 | 1 | Park7 | 0.01 | Young |
| 0.475855 | 0.010552 | 0.07 | 0.062 | 1 | 1 | Gid8 | 0.008 | Young |
| 0.475886 | 0.022512 | 0.086 | 0.077 | 1 | 1 | Kmt2d | 0.009 | Young |
| 0.475963 | 0.271645 | 0.053 | 0.046 | 1 | 1 | Nipsnap2 | 0.007 | Young |
| 0.475977 | 0.187331 | 0.045 | 0.039 | 1 | 1 | Ubxn2a | 0.006 | Young |
| 0.476452 | 0.247373 | 0.044 | 0.037 | 1 | 1 | Eif2ak1 | 0.007 | Young |
| 0.47716 | 0.294679 | 0.031 | 0.026 | 1 | 1 | Cmtm4 | 0.005 | Young |
| 0.478114 | 0.087144 | 0.064 | 0.057 | 1 | 1 | Mrpl42 | 0.007 | Young |
| 0.479236 | 0.11957 | 0.032 | 0.027 | 1 | 1 | Zfp407 | 0.005 | Young |
| 0.481036 | 0.088667 | 0.066 | 0.058 | 1 | 1 | Trappc8 | 0.008 | Young |
| 0.482443 | 0.0821 | 0.032 | 0.027 | 1 | 1 | Chmp7 | 0.005 | Young |
| 0.482743 | 0.24508 | 0.05 | 0.044 | 1 | 1 | Adpgk | 0.006 | Young |
| 0.483997 | 0.050875 | 0.126 | 0.116 | 1 | 1 | 4932438A1 | 0.01 | Young |
| 0.484437 | 0.008431 | 0.028 | 0.023 | 1 | 1 | Mbd5 | 0.005 | Young |
| 0.485421 | 0.011119 | 0.047 | 0.04 | 1 | 1 | Sp2 | 0.007 | Young |
| 0.487037 | 0.0925 | 0.081 | 0.072 | 1 | 1 | Rnh1 | 0.009 | Young |
| 0.487428 | 0.341453 | 0.049 | 0.042 | 1 | 1 | Nup54 | 0.007 | Young |
| 0.487655 | 0.047038 | 0.056 | 0.049 | 1 | 1 | Lcmt1 | 0.007 | Young |
| 0.487679 | 0.162945 | 0.03 | 0.024 | 1 | 1 | Srxn1 | 0.006 | Young |
| 0.487982 | 0.027872 | 0.237 | 0.222 | 1 | 1 | Fubp1 | 0.015 | Young |
| 0.488276 | 0.125267 | 0.031 | 0.026 | 1 | 1 | Fcf1 | 0.005 | Young |
| 0.488319 | 0.043614 | 0.071 | 0.063 | 1 | 1 | Lemd2 | 0.008 | Young |
| 0.488402 | 0.281456 | 0.028 | 0.023 | 1 | 1 | Nsun6 | 0.005 | Young |
| 0.48891 | 0.540276 | 0.039 | 0.033 | 1 | 1 | Cwc25 | 0.006 | Young |
| 0.488947 | 0.044834 | 0.112 | 0.102 | 1 | 1 | Mdm4 | 0.01 | Young |
| 0.489039 | 0.254986 | 0.042 | 0.036 | 1 | 1 | Psip1 | 0.006 | Young |

| | | | | | | | | |
|---|---|---|---|---|---|---|---|---|
| 0.48934 | 0.098655 | 0.012 | 0.015 | 1 | 1 | Rcl1 | -0.003 | Young |
| 0.489698 | 0.140658 | 0.03 | 0.024 | 1 | 1 | Lrba | 0.006 | Young |
| 0.489806 | 0.271717 | 0.012 | 0.015 | 1 | 1 | Zfp35 | -0.003 | Young |
| 0.491201 | 0.229341 | 0.01 | 0.013 | 1 | 1 | Nup107 | -0.003 | Young |
| 0.491261 | 0.356032 | 0.041 | 0.035 | 1 | 1 | 1110059G1 | 0.006 | Young |
| 0.491716 | 0.188521 | 0.01 | 0.013 | 1 | 1 | Bco2 | -0.003 | Young |
| 0.493089 | 0.492323 | 0.027 | 0.022 | 1 | 1 | Ints13 | 0.005 | Young |
| 0.494458 | 0.562984 | 0.027 | 0.022 | 1 | 1 | Gbp5 | 0.005 | Young |
| 0.494882 | 0.134345 | 0.028 | 0.023 | 1 | 1 | Itpr1 | 0.005 | Young |
| 0.495616 | 0.00203 | 0.202 | 0.185 | 1 | 1 | 9530068E0 | 0.017 | Young |
| 0.495828 | 0.293667 | 0.027 | 0.022 | 1 | 1 | N6amt1 | 0.005 | Young |
| 0.49643 | 0.064863 | 0.03 | 0.024 | 1 | 1 | Lix1l | 0.006 | Young |
| 0.496479 | 0.269735 | 0.038 | 0.032 | 1 | 1 | Exosc10 | 0.006 | Young |
| 0.496858 | 0.370498 | 0.027 | 0.022 | 1 | 1 | Kank3 | 0.005 | Young |
| 0.496986 | 0.599103 | 0.025 | 0.021 | 1 | 1 | Gm20696 | 0.004 | Young |
| 0.497781 | 0.393368 | 0.025 | 0.021 | 1 | 1 | Naip6 | 0.004 | Young |
| 0.497964 | 0.047711 | 0.069 | 0.062 | 1 | 1 | Prpf8 | 0.007 | Young |
| 0.498157 | 0.089958 | 0.042 | 0.036 | 1 | 1 | Uba3 | 0.006 | Young |
| 0.498235 | 0.291102 | 0.055 | 0.049 | 1 | 1 | Zc3h18 | 0.006 | Young |
| 0.498748 | 0.420541 | 0.027 | 0.022 | 1 | 1 | Pank1 | 0.005 | Young |
| 0.498749 | 0.158999 | 0.041 | 0.035 | 1 | 1 | Rap1gap2 | 0.006 | Young |
| 0.498817 | 0.307571 | 0.085 | 0.077 | 1 | 1 | Tmem5 | 0.008 | Young |
| 0.498978 | 0.053793 | 0.042 | 0.036 | 1 | 1 | Ccdc9 | 0.006 | Young |
| 0.499124 | 0.044683 | 0.044 | 0.037 | 1 | 1 | Zadh2 | 0.007 | Young |
| 0.499264 | 0.288534 | 0.027 | 0.022 | 1 | 1 | Naf1 | 0.005 | Young |
| 0.499772 | 0.293991 | 0.039 | 0.033 | 1 | 1 | Rpp21 | 0.006 | Young |
| 0.50013 | 0.100257 | 0.171 | 0.159 | 1 | 1 | Tubb6 | 0.012 | Young |
| 0.501491 | 0.016822 | 0.125 | 0.114 | 1 | 1 | Usp15 | 0.011 | Young |
| 0.502178 | 0.011845 | 0.151 | 0.139 | 1 | 1 | Arf2 | 0.012 | Young |
| 0.503301 | 0.292623 | 0.057 | 0.05 | 1 | 1 | Dicer1 | 0.007 | Young |
| 0.504304 | 0.009253 | 0.197 | 0.183 | 1 | 1 | Txn2 | 0.014 | Young |
| 0.504414 | 0.209571 | 0.02 | 0.024 | 1 | 1 | Ust | -0.004 | Young |
| 0.504713 | 0.421973 | 0.052 | 0.046 | 1 | 1 | Pknox1 | 0.006 | Young |
| 0.504967 | 0.211455 | 0.025 | 0.021 | 1 | 1 | Ddx41 | 0.004 | Young |
| 0.505017 | 0.079748 | 0.038 | 0.032 | 1 | 1 | Triap1 | 0.006 | Young |
| 0.505133 | 0.215451 | 0.027 | 0.022 | 1 | 1 | Tlcd2 | 0.005 | Young |
| 0.506696 | 0.834119 | 0.022 | 0.018 | 1 | 1 | Tpk1 | 0.004 | Young |
| 0.506866 | 0.083578 | 0.027 | 0.022 | 1 | 1 | Dnajc24 | 0.005 | Young |
| 0.506965 | 0.027082 | 0.039 | 0.033 | 1 | 1 | Prrc1 | 0.006 | Young |
| 0.507272 | 0.062764 | 0.038 | 0.032 | 1 | 1 | Sntb2 | 0.006 | Young |
| 0.507823 | 0.09764 | 0.039 | 0.033 | 1 | 1 | Smarcb1 | 0.006 | Young |
| 0.508212 | 0.139093 | 0.091 | 0.082 | 1 | 1 | Poldip3 | 0.009 | Young |
| 0.508682 | 0.133117 | 0.039 | 0.033 | 1 | 1 | Agap3 | 0.006 | Young |
| 0.509105 | 0.072847 | 0.073 | 0.066 | 1 | 1 | Strn4 | 0.007 | Young |
| 0.509314 | 0.051991 | 0.05 | 0.044 | 1 | 1 | Gle1 | 0.006 | Young |

| | | | | | | | | |
|---|---|---|---|---|---|---|---|---|
| 0.509367 | 0.014396 | 0.057 | 0.05 | 1 | 1 | Fbxo18 | 0.007 | Young |
| 0.509735 | 0.003202 | 0.061 | 0.054 | 1 | 1 | Sf3b4 | 0.007 | Young |
| 0.51065 | 0.109769 | 0.049 | 0.042 | 1 | 1 | Rnf14 | 0.007 | Young |
| 0.51119 | 0.177915 | 0.024 | 0.019 | 1 | 1 | Mzt2 | 0.005 | Young |
| 0.51179 | 0.038835 | 0.05 | 0.044 | 1 | 1 | Txndc11 | 0.006 | Young |
| 0.512017 | 0.194133 | 0.073 | 0.066 | 1 | 1 | Tmem131l | 0.007 | Young |
| 0.512093 | 0.220863 | 0.038 | 0.032 | 1 | 1 | Nlrc5 | 0.006 | Young |
| 0.512594 | 0.01568 | 0.018 | 0.022 | 1 | 1 | Gtf2e1 | -0.004 | Young |
| 0.513073 | 0.301052 | 0.036 | 0.031 | 1 | 1 | Serhl | 0.005 | Young |
| 0.513672 | 0.300021 | 0.035 | 0.03 | 1 | 1 | Smarcal1 | 0.005 | Young |
| 0.514184 | 0.081327 | 0.123 | 0.113 | 1 | 1 | Sel1l | 0.01 | Young |
| 0.514198 | 0.102033 | 0.109 | 0.099 | 1 | 1 | Khsrp | 0.01 | Young |
| 0.514336 | 0.108336 | 0.024 | 0.019 | 1 | 1 | Ddah2 | 0.005 | Young |
| 0.515293 | 0.033539 | 0.05 | 0.044 | 1 | 1 | Eftud2 | 0.006 | Young |
| 0.515692 | 0.157412 | 0.036 | 0.031 | 1 | 1 | Prkaa1 | 0.005 | Young |
| 0.516191 | 0.015507 | 0.024 | 0.019 | 1 | 1 | 3110082I1 | 0.005 | Young |
| 0.516623 | 0.032791 | 0.022 | 0.018 | 1 | 1 | Mlh1 | 0.004 | Young |
| 0.51674 | 0.423534 | 0.033 | 0.028 | 1 | 1 | Mtif2 | 0.005 | Young |
| 0.517353 | 0.317071 | 0.021 | 0.017 | 1 | 1 | Zfp358 | 0.004 | Young |
| 0.517582 | 0.31926 | 0.022 | 0.018 | 1 | 1 | Cstf1 | 0.004 | Young |
| 0.517829 | 0.396875 | 0.051 | 0.045 | 1 | 1 | Nde1 | 0.006 | Young |
| 0.518546 | 0.274448 | 0.021 | 0.017 | 1 | 1 | Rpain | 0.004 | Young |
| 0.518672 | 0.040615 | 0.298 | 0.281 | 1 | 1 | Hnrnpab | 0.017 | Young |
| 0.518928 | 0.070582 | 0.022 | 0.018 | 1 | 1 | Cinp | 0.004 | Young |
| 0.51917 | 1.190109 | 0.018 | 0.014 | 1 | 1 | Mgme1 | 0.004 | Young |
| 0.519958 | 0.160072 | 0.038 | 0.032 | 1 | 1 | Ddhd1 | 0.006 | Young |
| 0.520139 | 0.431093 | 0.021 | 0.017 | 1 | 1 | Cisd3 | 0.004 | Young |
| 0.520339 | 0.502018 | 0.021 | 0.017 | 1 | 1 | Cpeb3 | 0.004 | Young |
| 0.520683 | 0.013364 | 0.099 | 0.09 | 1 | 1 | Crlf2 | 0.009 | Young |
| 0.52107 | 0.004034 | 0.057 | 0.05 | 1 | 1 | Mbd6 | 0.007 | Young |
| 0.521809 | 0.000824 | 0.196 | 0.184 | 1 | 1 | Fermt3 | 0.012 | Young |
| 0.522202 | 0.044146 | 0.022 | 0.018 | 1 | 1 | Epor | 0.004 | Young |
| 0.522659 | 0.11905 | 0.088 | 0.08 | 1 | 1 | Sertad2 | 0.008 | Young |
| 0.523886 | 0.203556 | 0.043 | 0.037 | 1 | 1 | Mfsd10 | 0.006 | Young |
| 0.524165 | 0.02307 | 0.047 | 0.041 | 1 | 1 | Scarb1 | 0.006 | Young |
| 0.524565 | 0.703941 | 0.018 | 0.014 | 1 | 1 | Ampd2 | 0.004 | Young |
| 0.5251 | 0.181536 | 0.075 | 0.068 | 1 | 1 | Ints6 | 0.007 | Young |
| 0.525774 | 0.017536 | 0.066 | 0.059 | 1 | 1 | Psmd7 | 0.007 | Young |
| 0.526481 | 0.522294 | 0.032 | 0.027 | 1 | 1 | Nedd1 | 0.005 | Young |
| 0.526514 | 0.3116 | 0.018 | 0.014 | 1 | 1 | Zfp821 | 0.004 | Young |
| 0.527069 | 0.0679 | 0.019 | 0.015 | 1 | 1 | Padi2 | 0.004 | Young |
| 0.527114 | 0.498933 | 0.048 | 0.042 | 1 | 1 | Qars | 0.006 | Young |
| 0.527816 | 0.496871 | 0.018 | 0.014 | 1 | 1 | Hotairm1 | 0.004 | Young |
| 0.528258 | 0.413354 | 0.032 | 0.027 | 1 | 1 | Gadd45gip | 0.005 | Young |
| 0.528484 | 1.044852 | 0.015 | 0.012 | 1 | 1 | Nxpe3 | 0.003 | Young |

| | | | | | | | | |
|---|---|---|---|---|---|---|---|---|
| 0.528541 | 0.297353 | 0.059 | 0.053 | 1 | 1 | Arhgap26 | 0.006 | Young |
| 0.528545 | 0.042645 | 0.021 | 0.017 | 1 | 1 | Tbc1d12 | 0.004 | Young |
| 0.528677 | 0.016883 | 0.013 | 0.017 | 1 | 1 | Brms1l | -0.004 | Young |
| 0.528854 | 0.243316 | 0.041 | 0.036 | 1 | 1 | Isg20l2 | 0.005 | Young |
| 0.529611 | 0.047003 | 0.09 | 0.082 | 1 | 1 | Dhrs7 | 0.008 | Young |
| 0.530079 | 0.443285 | 0.016 | 0.013 | 1 | 1 | Fpgs | 0.003 | Young |
| 0.530616 | 0.085247 | 0.019 | 0.015 | 1 | 1 | Wdcp | 0.004 | Young |
| 0.530859 | 0.030524 | 0.018 | 0.014 | 1 | 1 | Ttc38 | 0.004 | Young |
| 0.530879 | 0.115958 | 0.069 | 0.062 | 1 | 1 | Thoc7 | 0.007 | Young |
| 0.530889 | 0.158389 | 0.062 | 0.055 | 1 | 1 | Cracr2b | 0.007 | Young |
| 0.530898 | 0.25979 | 0.046 | 0.04 | 1 | 1 | Map2k4 | 0.006 | Young |
| 0.531077 | 0.355488 | 0.018 | 0.014 | 1 | 1 | Smg8 | 0.004 | Young |
| 0.531198 | 0.333199 | 0.049 | 0.044 | 1 | 1 | Rbmxl1 | 0.005 | Young |
| 0.531334 | 0.215689 | 0.032 | 0.027 | 1 | 1 | Selenbp1 | 0.005 | Young |
| 0.531444 | 0.654817 | 0.016 | 0.013 | 1 | 1 | Gm26518 | 0.003 | Young |
| 0.531786 | 0.387276 | 0.016 | 0.013 | 1 | 1 | Zc3h3 | 0.003 | Young |
| 0.5325 | 0.035044 | 0.086 | 0.078 | 1 | 1 | Uba6 | 0.008 | Young |
| 0.532664 | 0.116073 | 0.072 | 0.064 | 1 | 1 | Ube2d1 | 0.008 | Young |
| 0.532933 | 0.083108 | 0.044 | 0.039 | 1 | 1 | Ccser2 | 0.005 | Young |
| 0.533531 | 0.070455 | 0.013 | 0.017 | 1 | 1 | Tubb2b | -0.004 | Young |
| 0.533651 | 0.089632 | 0.043 | 0.037 | 1 | 1 | Foxk1 | 0.006 | Young |
| 0.534081 | 0.338508 | 0.064 | 0.058 | 1 | 1 | Tcof1 | 0.006 | Young |
| 0.534179 | 0.368979 | 0.016 | 0.013 | 1 | 1 | Mto1 | 0.003 | Young |
| 0.53436 | 0.198463 | 0.03 | 0.026 | 1 | 1 | Ssna1 | 0.004 | Young |
| 0.534589 | 1.500608 | 0.013 | 0.01 | 1 | 1 | Stradb | 0.003 | Young |
| 0.534726 | 0.100589 | 0.044 | 0.039 | 1 | 1 | Arfip1 | 0.005 | Young |
| 0.534937 | 0.523552 | 0.015 | 0.012 | 1 | 1 | Pdik1l | 0.003 | Young |
| 0.535093 | 0.265181 | 0.016 | 0.013 | 1 | 1 | Gabbr1 | 0.003 | Young |
| 0.535807 | 0.045947 | 0.197 | 0.184 | 1 | 1 | Fam129a | 0.013 | Young |
| 0.535859 | 0.530993 | 0.03 | 0.026 | 1 | 1 | Mob2 | 0.004 | Young |
| 0.536377 | 0.09438 | 0.039 | 0.045 | 1 | 1 | Xpa | -0.006 | Young |
| 0.536464 | 0.199238 | 0.016 | 0.013 | 1 | 1 | Mrpl39 | 0.003 | Young |
| 0.53659 | 0.73987 | 0.029 | 0.024 | 1 | 1 | Gnpat | 0.005 | Young |
| 0.536617 | 0.266132 | 0.015 | 0.012 | 1 | 1 | Usp11 | 0.003 | Young |
| 0.536693 | 0.281677 | 0.016 | 0.013 | 1 | 1 | Rcc1 | 0.003 | Young |
| 0.536693 | 0.238967 | 0.016 | 0.013 | 1 | 1 | Ppil1 | 0.003 | Young |
| 0.537097 | 0.503139 | 0.015 | 0.012 | 1 | 1 | Tatdn3 | 0.003 | Young |
| 0.537165 | 0.067957 | 0.07 | 0.063 | 1 | 1 | Tnpo2 | 0.007 | Young |
| 0.537578 | 0.297951 | 0.015 | 0.012 | 1 | 1 | Zfp777 | 0.003 | Young |
| 0.537723 | 0.024318 | 0.043 | 0.037 | 1 | 1 | Ascc1 | 0.006 | Young |
| 0.537823 | 0.808782 | 0.012 | 0.009 | 1 | 1 | Zfp52 | 0.003 | Young |
| 0.538195 | 0.19941 | 0.03 | 0.026 | 1 | 1 | Zcchc14 | 0.004 | Young |
| 0.538362 | 0.418025 | 0.03 | 0.026 | 1 | 1 | Iffo2 | 0.004 | Young |
| 0.538632 | 0.761074 | 0.012 | 0.009 | 1 | 1 | Ier5l | 0.003 | Young |
| 0.538897 | 0.807459 | 0.013 | 0.01 | 1 | 1 | Iqcc | 0.003 | Young |

| | | | | | | | | |
|---|---|---|---|---|---|---|---|---|
| 0.539171 | 0.756423 | 0.012 | 0.009 | 1 | 1 | Lhpp | 0.003 | Young |
| 0.539365 | 0.331011 | 0.03 | 0.026 | 1 | 1 | Cdkn2aip | 0.004 | Young |
| 0.539474 | 0.043867 | 0.115 | 0.105 | 1 | 1 | Birc2 | 0.01 | Young |
| 0.539533 | 0.121895 | 0.101 | 0.093 | 1 | 1 | Napa | 0.008 | Young |
| 0.540168 | 0.497747 | 0.013 | 0.01 | 1 | 1 | Gpr68 | 0.003 | Young |
| 0.540252 | 0.641443 | 0.012 | 0.009 | 1 | 1 | Mmgt2 | 0.003 | Young |
| 0.541056 | 0.627384 | 0.01 | 0.008 | 1 | 1 | Ecsit | 0.002 | Young |
| 0.541185 | 0.385562 | 0.013 | 0.01 | 1 | 1 | Ube2e2 | 0.003 | Young |
| 0.54143 | 0.064504 | 0.015 | 0.012 | 1 | 1 | Nrg2 | 0.003 | Young |
| 0.54144 | 0.244579 | 0.013 | 0.01 | 1 | 1 | Frat2 | 0.003 | Young |
| 0.541603 | 0.355309 | 0.012 | 0.009 | 1 | 1 | Leng9 | 0.003 | Young |
| 0.541603 | 0.167762 | 0.012 | 0.009 | 1 | 1 | Klrk1 | 0.003 | Young |
| 0.541636 | 0.501058 | 0.01 | 0.008 | 1 | 1 | Cd244 | 0.002 | Young |
| 0.541695 | 0.001597 | 0.013 | 0.01 | 1 | 1 | Abcb10 | 0.003 | Young |
| 0.541874 | 0.58326 | 0.012 | 0.009 | 1 | 1 | 2310040G2 | 0.003 | Young |
| 0.541912 | 0.010075 | 0.015 | 0.012 | 1 | 1 | C87436 | 0.003 | Young |
| 0.542204 | 0.184722 | 0.013 | 0.01 | 1 | 1 | Tecpr2 | 0.003 | Young |
| 0.542636 | 0.39882 | 0.015 | 0.012 | 1 | 1 | Txlng | 0.003 | Young |
| 0.542686 | 0.593586 | 0.012 | 0.009 | 1 | 1 | Rdh13 | 0.003 | Young |
| 0.542713 | 0.467468 | 0.013 | 0.01 | 1 | 1 | Dgka | 0.003 | Young |
| 0.542796 | 0.914549 | 0.01 | 0.008 | 1 | 1 | Polr1b | 0.002 | Young |
| 0.542932 | 0.027438 | 0.139 | 0.129 | 1 | 1 | Ddrgk1 | 0.01 | Young |
| 0.543327 | 0.2626 | 0.092 | 0.085 | 1 | 1 | Gtpbp4 | 0.007 | Young |
| 0.543436 | 0.007626 | 0.043 | 0.037 | 1 | 1 | Cog6 | 0.006 | Young |
| 0.543957 | 0.41367 | 0.01 | 0.008 | 1 | 1 | Clmp | 0.002 | Young |
| 0.544025 | 0.296317 | 0.048 | 0.042 | 1 | 1 | Amn1 | 0.006 | Young |
| 0.545044 | 0.100096 | 0.032 | 0.027 | 1 | 1 | Galnt4 | 0.005 | Young |
| 0.545065 | 0.204727 | 0.009 | 0.012 | 1 | 1 | Hspb1 | -0.003 | Young |
| 0.545668 | 0.190633 | 0.012 | 0.009 | 1 | 1 | Msh3 | 0.003 | Young |
| 0.545668 | 0.085444 | 0.012 | 0.009 | 1 | 1 | Rnaseh1 | 0.003 | Young |
| 0.545766 | 0.121371 | 0.087 | 0.08 | 1 | 1 | BC037034 | 0.007 | Young |
| 0.545907 | 0.232353 | 0.03 | 0.026 | 1 | 1 | Ddx27 | 0.004 | Young |
| 0.54594 | 0.102412 | 0.012 | 0.009 | 1 | 1 | Hemk1 | 0.003 | Young |
| 0.546287 | 0.108289 | 0.013 | 0.01 | 1 | 1 | Surf2 | 0.003 | Young |
| 0.546866 | 0.009949 | 0.01 | 0.008 | 1 | 1 | Kdm4b | 0.002 | Young |
| 0.547054 | 0.172632 | 0.009 | 0.012 | 1 | 1 | Pcca | -0.003 | Young |
| 0.547784 | 0.036836 | 0.081 | 0.073 | 1 | 1 | Zfp217 | 0.008 | Young |
| 0.547894 | 0.188572 | 0.04 | 0.035 | 1 | 1 | Abcf3 | 0.005 | Young |
| 0.548043 | 0.182924 | 0.048 | 0.042 | 1 | 1 | Sap18 | 0.006 | Young |
| 0.548169 | 0.110562 | 0.086 | 0.078 | 1 | 1 | Lin7c | 0.008 | Young |
| 0.548603 | 0.073828 | 0.03 | 0.026 | 1 | 1 | Atr | 0.004 | Young |
| 0.548907 | 0.039254 | 0.01 | 0.008 | 1 | 1 | Ap4b1 | 0.002 | Young |
| 0.549046 | 0.132529 | 0.009 | 0.012 | 1 | 1 | Tti1 | -0.003 | Young |
| 0.549143 | 0.040234 | 0.064 | 0.058 | 1 | 1 | Acsl3 | 0.006 | Young |
| 0.54949 | 0.164148 | 0.01 | 0.008 | 1 | 1 | Rab38 | 0.002 | Young |

| | | | | | | | | |
|---|---|---|---|---|---|---|---|---|
| 0.549902 | 0.078806 | 0.145 | 0.135 | 1 | 1 | Ggnbp2 | 0.01 | Young |
| 0.551017 | 0.19794 | 0.027 | 0.023 | 1 | 1 | Inip | 0.004 | Young |
| 0.551684 | 0.09825 | 0.072 | 0.066 | 1 | 1 | Rftn1 | 0.006 | Young |
| 0.552761 | 0.832729 | 0.024 | 0.021 | 1 | 1 | E130307A1 | 0.003 | Young |
| 0.552931 | 0.037066 | 0.075 | 0.068 | 1 | 1 | Cnot8 | 0.007 | Young |
| 0.553143 | 0.190651 | 0.038 | 0.033 | 1 | 1 | Glt8d1 | 0.005 | Young |
| 0.553781 | 0.213879 | 0.045 | 0.04 | 1 | 1 | Ifih1 | 0.005 | Young |
| 0.55414 | 0.066383 | 0.081 | 0.073 | 1 | 1 | Tmem65 | 0.008 | Young |
| 0.554304 | 0.002071 | 0.059 | 0.053 | 1 | 1 | Rnf139 | 0.006 | Young |
| 0.554458 | 0.008656 | 0.165 | 0.153 | 1 | 1 | Ap1s2 | 0.012 | Young |
| 0.55456 | 0.488721 | 0.035 | 0.031 | 1 | 1 | Mrpl35 | 0.004 | Young |
| 0.555906 | 0.016406 | 0.029 | 0.024 | 1 | 1 | Fam129b | 0.005 | Young |
| 0.556379 | 0.35526 | 0.037 | 0.032 | 1 | 1 | Dcps | 0.005 | Young |
| 0.557365 | 0.091151 | 0.133 | 0.123 | 1 | 1 | Camk2d | 0.01 | Young |
| 0.558448 | 0.345825 | 0.053 | 0.048 | 1 | 1 | Rsrc1 | 0.005 | Young |
| 0.558455 | 0.15427 | 0.037 | 0.032 | 1 | 1 | Lsm14b | 0.005 | Young |
| 0.559308 | 0.010976 | 0.225 | 0.211 | 1 | 1 | Myl12b | 0.014 | Young |
| 0.560657 | 0.301329 | 0.027 | 0.023 | 1 | 1 | Casp2 | 0.004 | Young |
| 0.562451 | 0.032271 | 0.027 | 0.023 | 1 | 1 | Gk | 0.004 | Young |
| 0.564046 | 0.110414 | 0.056 | 0.05 | 1 | 1 | Flot1 | 0.006 | Young |
| 0.56545 | 0.377908 | 0.024 | 0.021 | 1 | 1 | Nectin4 | 0.003 | Young |
| 0.56554 | 0.029616 | 0.038 | 0.033 | 1 | 1 | Ak3 | 0.005 | Young |
| 0.566205 | 0.159551 | 0.026 | 0.022 | 1 | 1 | Ints14 | 0.004 | Young |
| 0.566297 | 0.150783 | 0.023 | 0.027 | 1 | 1 | Hes6 | -0.004 | Young |
| 0.566594 | 0.151783 | 0.024 | 0.021 | 1 | 1 | Dcun1d3 | 0.003 | Young |
| 0.56672 | 0.098843 | 0.064 | 0.058 | 1 | 1 | Ube2w | 0.006 | Young |
| 0.567144 | 0.083446 | 0.047 | 0.041 | 1 | 1 | Trappc1 | 0.006 | Young |
| 0.567506 | 0.167329 | 0.068 | 0.062 | 1 | 1 | Gch1 | 0.006 | Young |
| 0.56779 | 0.31069 | 0.044 | 0.039 | 1 | 1 | Tbcel | 0.005 | Young |
| 0.568076 | 0.268079 | 0.044 | 0.039 | 1 | 1 | Ints1 | 0.005 | Young |
| 0.568841 | 0.240667 | 0.052 | 0.046 | 1 | 1 | Ifi209 | 0.006 | Young |
| 0.569811 | 0.001697 | 0.061 | 0.054 | 1 | 1 | Cd40 | 0.007 | Young |
| 0.570051 | 0.214683 | 0.106 | 0.099 | 1 | 1 | Uso1 | 0.007 | Young |
| 0.570052 | 0.049488 | 0.056 | 0.05 | 1 | 1 | Bccip | 0.006 | Young |
| 0.570599 | 0.014093 | 0.641 | 0.591 | 1 | 1 | Npc2 | 0.05 | Young |
| 0.571013 | 0.38321 | 0.058 | 0.053 | 1 | 1 | Paox | 0.005 | Young |
| 0.571046 | 0.588025 | 0.023 | 0.019 | 1 | 1 | Paqr7 | 0.004 | Young |
| 0.571212 | 0.179531 | 0.052 | 0.046 | 1 | 1 | Tbc1d1 | 0.006 | Young |
| 0.571369 | 0.153799 | 0.024 | 0.021 | 1 | 1 | Coq2 | 0.003 | Young |
| 0.571608 | 0.243075 | 0.05 | 0.045 | 1 | 1 | Rbm15 | 0.005 | Young |
| 0.57213 | 0.351388 | 0.081 | 0.075 | 1 | 1 | Mysm1 | 0.006 | Young |
| 0.572732 | 0.099302 | 0.044 | 0.039 | 1 | 1 | Smurf1 | 0.005 | Young |
| 0.573328 | 0.075866 | 0.069 | 0.063 | 1 | 1 | Zcchc2 | 0.006 | Young |
| 0.573476 | 0.137288 | 0.024 | 0.021 | 1 | 1 | Top3b | 0.003 | Young |
| 0.574562 | 0.174906 | 0.058 | 0.053 | 1 | 1 | Uba5 | 0.005 | Young |

| | | | | | | | | |
|---|---|---|---|---|---|---|---|---|
| 0.576901 | 0.266937 | 0.044 | 0.039 | 1 | 1 | Usp45 | 0.005 | Young |
| 0.57693 | 0.030447 | 0.787 | 0.731 | 1 | 1 | Zeb2 | 0.056 | Young |
| 0.577307 | 0.003947 | 0.131 | 0.122 | 1 | 1 | Mvp | 0.009 | Young |
| 0.579434 | 0.012243 | 0.024 | 0.021 | 1 | 1 | Usp21 | 0.003 | Young |
| 0.579601 | 0.061539 | 0.061 | 0.055 | 1 | 1 | Stim2 | 0.006 | Young |
| 0.581848 | 0.108085 | 0.023 | 0.019 | 1 | 1 | Rbms2 | 0.004 | Young |
| 0.582337 | 0.191579 | 0.027 | 0.031 | 1 | 1 | Pole3 | -0.004 | Young |
| 0.582757 | 0.306814 | 0.021 | 0.018 | 1 | 1 | Gm4951 | 0.003 | Young |
| 0.583409 | 0.000648 | 0.063 | 0.057 | 1 | 1 | Il21r | 0.006 | Young |
| 0.584465 | 0.057628 | 0.075 | 0.068 | 1 | 1 | Clcn3 | 0.007 | Young |
| 0.585427 | 0.065453 | 0.089 | 0.096 | 1 | 1 | Themis2 | -0.007 | Young |
| 0.587153 | 0.155796 | 0.05 | 0.045 | 1 | 1 | Ccdc174 | 0.005 | Young |
| 0.587852 | 0.195293 | 0.058 | 0.053 | 1 | 1 | Manba | 0.005 | Young |
| 0.588451 | 0.222767 | 0.013 | 0.015 | 1 | 1 | St7 | -0.002 | Young |
| 0.589382 | 0.08309 | 0.069 | 0.063 | 1 | 1 | Coro2a | 0.006 | Young |
| 0.589536 | 0.224052 | 0.041 | 0.036 | 1 | 1 | Rnf41 | 0.005 | Young |
| 0.589985 | 0.084946 | 0.021 | 0.018 | 1 | 1 | Zfp68 | 0.003 | Young |
| 0.590405 | 0.290163 | 0.02 | 0.017 | 1 | 1 | Rpap3 | 0.003 | Young |
| 0.590478 | 0.110636 | 0.034 | 0.03 | 1 | 1 | Rwdd4a | 0.004 | Young |
| 0.590835 | 0.194967 | 0.02 | 0.017 | 1 | 1 | Tmem140 | 0.003 | Young |
| 0.590966 | 0.167099 | 0.041 | 0.036 | 1 | 1 | Gpatch2l | 0.005 | Young |
| 0.591136 | 0.048475 | 0.177 | 0.167 | 1 | 1 | Rassf1 | 0.01 | Young |
| 0.591148 | 7.63E-05 | 0.129 | 0.12 | 1 | 1 | mt-Atp8 | 0.009 | Young |
| 0.591294 | 0.231804 | 0.066 | 0.06 | 1 | 1 | Kmt5a | 0.006 | Young |
| 0.592271 | 0.006348 | 0.089 | 0.082 | 1 | 1 | Txnl1 | 0.007 | Young |
| 0.593844 | 0.304155 | 0.02 | 0.017 | 1 | 1 | Ralgapa2 | 0.003 | Young |
| 0.595024 | 0.127697 | 0.024 | 0.028 | 1 | 1 | Hace1 | -0.004 | Young |
| 0.595136 | 0.354667 | 0.02 | 0.017 | 1 | 1 | Hagh | 0.003 | Young |
| 0.595617 | 0.25679 | 0.03 | 0.026 | 1 | 1 | Zbtb2 | 0.004 | Young |
| 0.595993 | 0.164221 | 0.047 | 0.042 | 1 | 1 | Strip1 | 0.005 | Young |
| 0.596274 | 0.235391 | 0.047 | 0.042 | 1 | 1 | Gcn1l1 | 0.005 | Young |
| 0.596777 | 0.10652 | 0.103 | 0.095 | 1 | 1 | Fip1l1 | 0.008 | Young |
| 0.597513 | 0.316341 | 0.064 | 0.059 | 1 | 1 | Rab1b | 0.005 | Young |
| 0.597532 | 0.179994 | 0.018 | 0.022 | 1 | 1 | Eng | -0.004 | Young |
| 0.597608 | 0.095274 | 0.044 | 0.049 | 1 | 1 | Tmem68 | -0.005 | Young |
| 0.597906 | 0.033416 | 0.088 | 0.081 | 1 | 1 | Tbc1d9 | 0.007 | Young |
| 0.598449 | 0.205081 | 0.03 | 0.026 | 1 | 1 | Cyp4v3 | 0.004 | Young |
| 0.599108 | 0.128032 | 0.064 | 0.058 | 1 | 1 | Zdhhc21 | 0.006 | Young |
| 0.599236 | 0.049281 | 0.02 | 0.017 | 1 | 1 | Hectd4 | 0.003 | Young |
| 0.599245 | 0.463291 | 0.018 | 0.015 | 1 | 1 | Phf11d | 0.003 | Young |
| 0.599935 | 0.128887 | 0.01 | 0.013 | 1 | 1 | Smyd2 | -0.003 | Young |
| 0.601568 | 0.448948 | 0.028 | 0.024 | 1 | 1 | Mrpl37 | 0.004 | Young |
| 0.601883 | 0.090807 | 0.01 | 0.013 | 1 | 1 | Egfl7 | -0.003 | Young |
| 0.602167 | 0.256111 | 0.018 | 0.015 | 1 | 1 | Cdk5rap2 | 0.003 | Young |
| 0.60269 | 0.064311 | 0.049 | 0.044 | 1 | 1 | Ppil4 | 0.005 | Young |

| | | | | | | | | |
|---|---|---|---|---|---|---|---|---|
| 0.603063 | 0.160666 | 0.03 | 0.026 | 1 | 1 | Slc27a4 | 0.004 | Young |
| 0.603241 | 0.40286 | 0.03 | 0.026 | 1 | 1 | Mea1 | 0.004 | Young |
| 0.604333 | 0.037976 | 0.078 | 0.071 | 1 | 1 | Vamp4 | 0.007 | Young |
| 0.604486 | 0.123351 | 0.03 | 0.026 | 1 | 1 | Skp2 | 0.004 | Young |
| 0.605069 | 0.026084 | 0.243 | 0.23 | 1 | 1 | Spred1 | 0.013 | Young |
| 0.605547 | 0.342841 | 0.018 | 0.015 | 1 | 1 | Mrps2 | 0.003 | Young |
| 0.605547 | 0.298477 | 0.018 | 0.015 | 1 | 1 | Isca1 | 0.003 | Young |
| 0.605547 | 0.291574 | 0.018 | 0.015 | 1 | 1 | Acvr2b | 0.003 | Young |
| 0.605879 | 0.849744 | 0.017 | 0.014 | 1 | 1 | Pianp | 0.003 | Young |
| 0.605998 | 0.375131 | 0.018 | 0.015 | 1 | 1 | Exosc7 | 0.003 | Young |
| 0.607353 | 0.00529 | 0.016 | 0.019 | 1 | 1 | Cdkal1 | -0.003 | Young |
| 0.607974 | 0.009967 | 0.047 | 0.042 | 1 | 1 | Cdk2ap1 | 0.005 | Young |
| 0.608432 | 0.091338 | 0.028 | 0.032 | 1 | 1 | Exoc7 | -0.004 | Young |
| 0.608765 | 0.18227 | 0.03 | 0.026 | 1 | 1 | Cdyl | 0.004 | Young |
| 0.608918 | 0.257652 | 0.036 | 0.032 | 1 | 1 | Abcg3 | 0.004 | Young |
| 0.609091 | 0.665521 | 0.027 | 0.023 | 1 | 1 | Mpg | 0.004 | Young |
| 0.609604 | 0.270589 | 0.028 | 0.024 | 1 | 1 | Scap | 0.004 | Young |
| 0.609701 | 0.794858 | 0.025 | 0.022 | 1 | 1 | Zfp180 | 0.003 | Young |
| 0.610293 | 0.110196 | 0.018 | 0.015 | 1 | 1 | Snx19 | 0.003 | Young |
| 0.610866 | 0.073972 | 0.098 | 0.091 | 1 | 1 | Eapp | 0.007 | Young |
| 0.611259 | 0.175014 | 0.054 | 0.049 | 1 | 1 | Zmpste24 | 0.005 | Young |
| 0.611648 | 0.669587 | 0.016 | 0.013 | 1 | 1 | Aacs | 0.003 | Young |
| 0.611878 | 0.120888 | 0.018 | 0.015 | 1 | 1 | Mrps10 | 0.003 | Young |
| 0.612637 | 0.800432 | 0.016 | 0.013 | 1 | 1 | Phospho2 | 0.003 | Young |
| 0.613235 | 0.065692 | 0.03 | 0.026 | 1 | 1 | Fkbp1b | 0.004 | Young |
| 0.614061 | 0.268085 | 0.038 | 0.033 | 1 | 1 | Maged1 | 0.005 | Young |
| 0.614742 | 0.714335 | 0.027 | 0.023 | 1 | 1 | Dhx29 | 0.004 | Young |
| 0.615282 | 0.217983 | 0.016 | 0.019 | 1 | 1 | Adprhl2 | -0.003 | Young |
| 0.615399 | 0.030763 | 0.157 | 0.148 | 1 | 1 | Arglu1 | 0.009 | Young |
| 0.615609 | 0.591079 | 0.016 | 0.013 | 1 | 1 | 1500015A( | 0.003 | Young |
| 0.615969 | 0.128728 | 0.038 | 0.033 | 1 | 1 | Idh3a | 0.005 | Young |
| 0.616882 | 0.102662 | 0.046 | 0.041 | 1 | 1 | Lyst | 0.005 | Young |
| 0.617148 | 0.686436 | 0.008 | 0.01 | 1 | 1 | Hps6 | -0.002 | Young |
| 0.617328 | 0.146949 | 0.016 | 0.019 | 1 | 1 | Ppp2r3a | -0.003 | Young |
| 0.617842 | 0.520752 | 0.016 | 0.013 | 1 | 1 | Vps72 | 0.003 | Young |
| 0.618793 | 0.15798 | 0.036 | 0.032 | 1 | 1 | Nagpa | 0.004 | Young |
| 0.620079 | 0.196666 | 0.016 | 0.013 | 1 | 1 | Ift22 | 0.003 | Young |
| 0.620419 | 0.040049 | 0.036 | 0.032 | 1 | 1 | Fam219b | 0.004 | Young |
| 0.620576 | 0.24492 | 0.016 | 0.013 | 1 | 1 | Orc6 | 0.003 | Young |
| 0.621714 | 0.270526 | 0.049 | 0.045 | 1 | 1 | Galnt10 | 0.004 | Young |
| 0.623437 | 0.008448 | 0.039 | 0.035 | 1 | 1 | Tdg | 0.004 | Young |
| 0.623597 | 0.314857 | 0.025 | 0.022 | 1 | 1 | Mid1 | 0.003 | Young |
| 0.623655 | 0.450368 | 0.014 | 0.012 | 1 | 1 | Wdr55 | 0.002 | Young |
| 0.624006 | 0.076474 | 0.058 | 0.053 | 1 | 1 | Mrpl34 | 0.005 | Young |
| 0.624788 | 0.047227 | 0.13 | 0.122 | 1 | 1 | Cnot6 | 0.008 | Young |

| | | | | | | | | |
|---|---|---|---|---|---|---|---|---|
| 0.625942 | 0.481446 | 0.025 | 0.022 | 1 | 1 | Coq5 | 0.003 | Young |
| 0.62596 | 0.180568 | 0.036 | 0.032 | 1 | 1 | Rnaset2a | 0.004 | Young |
| 0.626098 | 0.098595 | 0.02 | 0.023 | 1 | 1 | Alkbh3 | -0.003 | Young |
| 0.62694 | 0.104071 | 0.036 | 0.032 | 1 | 1 | Ocel1 | 0.004 | Young |
| 0.62831 | 0.003638 | 0.016 | 0.013 | 1 | 1 | Trim37 | 0.003 | Young |
| 0.628492 | 0.172206 | 0.02 | 0.023 | 1 | 1 | Pot1b | -0.003 | Young |
| 0.630361 | 0.102663 | 0.043 | 0.039 | 1 | 1 | Slc30a1 | 0.004 | Young |
| 0.630491 | 0.102972 | 0.035 | 0.031 | 1 | 1 | Irgm1 | 0.004 | Young |
| 0.631005 | 0.95495 | 0.013 | 0.01 | 1 | 1 | Nprl2 | 0.003 | Young |
| 0.631014 | 0.215435 | 0.014 | 0.012 | 1 | 1 | Recql5 | 0.002 | Young |
| 0.631057 | 0.065916 | 0.014 | 0.017 | 1 | 1 | Rft1 | -0.003 | Young |
| 0.631951 | 0.113796 | 0.044 | 0.04 | 1 | 1 | Ift52 | 0.004 | Young |
| 0.63212 | 0.649915 | 0.013 | 0.01 | 1 | 1 | Tmem42 | 0.003 | Young |
| 0.632291 | 0.161772 | 0.014 | 0.017 | 1 | 1 | Zfp934 | -0.003 | Young |
| 0.633001 | 0.154196 | 0.025 | 0.022 | 1 | 1 | Ncapg2 | 0.003 | Young |
| 0.633123 | 0.086717 | 0.014 | 0.012 | 1 | 1 | Ercc1 | 0.002 | Young |
| 0.633236 | 0.58503 | 0.013 | 0.01 | 1 | 1 | Pcyt1b | 0.003 | Young |
| 0.633287 | 0.024194 | 0.044 | 0.04 | 1 | 1 | Arl2bp | 0.004 | Young |
| 0.633515 | 0.820522 | 0.013 | 0.01 | 1 | 1 | Hoxb4 | 0.003 | Young |
| 0.633591 | 0.145882 | 0.025 | 0.022 | 1 | 1 | Cxxc1 | 0.003 | Young |
| 0.633649 | 0.029844 | 0.114 | 0.107 | 1 | 1 | Snx30 | 0.007 | Young |
| 0.634423 | 0.052468 | 0.035 | 0.031 | 1 | 1 | Phkb | 0.004 | Young |
| 0.634961 | 0.216918 | 0.085 | 0.08 | 1 | 1 | Psmb5 | 0.005 | Young |
| 0.635165 | 0.151622 | 0.025 | 0.022 | 1 | 1 | Rtl8a | 0.003 | Young |
| 0.635906 | 0.223698 | 0.02 | 0.023 | 1 | 1 | Sik2 | -0.003 | Young |
| 0.63603 | 0.584611 | 0.013 | 0.01 | 1 | 1 | Haus4 | 0.003 | Young |
| 0.636218 | 0.131824 | 0.081 | 0.076 | 1 | 1 | Taf10 | 0.005 | Young |
| 0.637526 | 0.067271 | 0.035 | 0.031 | 1 | 1 | Rce1 | 0.004 | Young |
| 0.637708 | 0.220795 | 0.013 | 0.01 | 1 | 1 | Noa1 | 0.003 | Young |
| 0.638503 | 0.005195 | 0.066 | 0.06 | 1 | 1 | Ppp4r1 | 0.006 | Young |
| 0.639171 | 1.052694 | 0.011 | 0.009 | 1 | 1 | Dbr1 | 0.002 | Young |
| 0.639994 | 0.808581 | 0.022 | 0.019 | 1 | 1 | Agpat2 | 0.003 | Young |
| 0.640229 | 0.137485 | 0.013 | 0.01 | 1 | 1 | Nme4 | 0.003 | Young |
| 0.64051 | 0.256728 | 0.013 | 0.01 | 1 | 1 | 2610021A( | 0.003 | Young |
| 0.64065 | 0.021024 | 0.051 | 0.046 | 1 | 1 | Mrpl11 | 0.005 | Young |
| 0.641247 | 0.026635 | 0.049 | 0.045 | 1 | 1 | Trmt1 | 0.004 | Young |
| 0.641464 | 0.670454 | 0.022 | 0.019 | 1 | 1 | Ap5z1 | 0.003 | Young |
| 0.641672 | 0.05671 | 0.049 | 0.045 | 1 | 1 | Chn2 | 0.004 | Young |
| 0.642456 | 0.367803 | 0.024 | 0.021 | 1 | 1 | Fam110b | 0.003 | Young |
| 0.643355 | 0.593146 | 0.011 | 0.009 | 1 | 1 | Cox18 | 0.002 | Young |
| 0.643578 | 0.440237 | 0.038 | 0.035 | 1 | 1 | Pprc1 | 0.003 | Young |
| 0.643597 | 0.085055 | 0.013 | 0.01 | 1 | 1 | Ccdc191 | 0.003 | Young |
| 0.643654 | 0.623279 | 0.011 | 0.009 | 1 | 1 | Ears2 | 0.002 | Young |
| 0.644494 | 0.356428 | 0.024 | 0.021 | 1 | 1 | 2210016L2 | 0.003 | Young |
| 0.645106 | 0.025796 | 0.024 | 0.021 | 1 | 1 | Dhcr7 | 0.003 | Young |

| | | | | | | | | |
|---|---|---|---|---|---|---|---|---|
| 0.64523 | 0.067603 | 0.041 | 0.037 | 1 | 1 | Zdhhc17 | 0.004 | Young |
| 0.645727 | 0.115684 | 0.032 | 0.028 | 1 | 1 | Icmt | 0.004 | Young |
| 0.645751 | 0.19487 | 0.011 | 0.009 | 1 | 1 | Zfp788 | 0.002 | Young |
| 0.646153 | 0.009545 | 0.018 | 0.021 | 1 | 1 | Arhgef10 | -0.003 | Young |
| 0.64695 | 0.154113 | 0.011 | 0.009 | 1 | 1 | Nlrx1 | 0.002 | Young |
| 0.64727 | 0.083778 | 0.018 | 0.021 | 1 | 1 | Plp2 | -0.003 | Young |
| 0.64815 | 0.197781 | 0.011 | 0.009 | 1 | 1 | Katnb1 | 0.002 | Young |
| 0.64815 | 0.196455 | 0.011 | 0.009 | 1 | 1 | Usp46 | 0.002 | Young |
| 0.64815 | 0.154132 | 0.011 | 0.009 | 1 | 1 | Gstk1 | 0.002 | Young |
| 0.649351 | 0.035398 | 0.011 | 0.009 | 1 | 1 | P2ry10 | 0.002 | Young |
| 0.64943 | 0.964836 | 0.01 | 0.008 | 1 | 1 | Gstz1 | 0.002 | Young |
| 0.649652 | 0.116903 | 0.011 | 0.009 | 1 | 1 | Gm7160 | 0.002 | Young |
| 0.650012 | 0.109634 | 0.024 | 0.021 | 1 | 1 | Pygl | 0.003 | Young |
| 0.651155 | 0.05567 | 0.011 | 0.009 | 1 | 1 | Endog | 0.002 | Young |
| 0.652017 | 0.608836 | 0.01 | 0.008 | 1 | 1 | Nat2 | 0.002 | Young |
| 0.652198 | 0.760975 | 0.029 | 0.026 | 1 | 1 | Xylt1 | 0.003 | Young |
| 0.652358 | 0.129879 | 0.011 | 0.009 | 1 | 1 | Gem | 0.002 | Young |
| 0.652852 | 0.461228 | 0.022 | 0.019 | 1 | 1 | Lancl1 | 0.003 | Young |
| 0.653222 | 0.139697 | 0.033 | 0.03 | 1 | 1 | Mxra7 | 0.003 | Young |
| 0.653635 | 0.35078 | 0.01 | 0.008 | 1 | 1 | Rab3a | 0.002 | Young |
| 0.654171 | 0.52463 | 0.037 | 0.033 | 1 | 1 | Ezh2 | 0.004 | Young |
| 0.654607 | 0.162839 | 0.01 | 0.008 | 1 | 1 | Tfpt | 0.002 | Young |
| 0.65603 | 0.23893 | 0.022 | 0.019 | 1 | 1 | Zbtb43 | 0.003 | Young |
| 0.656553 | 0.179819 | 0.01 | 0.008 | 1 | 1 | Il18bp | 0.002 | Young |
| 0.656686 | 0.031744 | 0.032 | 0.028 | 1 | 1 | Crot | 0.004 | Young |
| 0.65677 | 0.213824 | 0.03 | 0.027 | 1 | 1 | Zfp639 | 0.003 | Young |
| 0.656907 | 0.32488 | 0.018 | 0.021 | 1 | 1 | Ppm1d | -0.003 | Young |
| 0.657686 | 0.075419 | 0.093 | 0.087 | 1 | 1 | Lsm12 | 0.006 | Young |
| 0.657852 | 0.144353 | 0.01 | 0.008 | 1 | 1 | Rtn4ip1 | 0.002 | Young |
| 0.657876 | 0.143079 | 0.045 | 0.041 | 1 | 1 | Rmi1 | 0.004 | Young |
| 0.658042 | 0.338077 | 0.057 | 0.053 | 1 | 1 | Zfp992 | 0.004 | Young |
| 0.658176 | 0.143685 | 0.01 | 0.008 | 1 | 1 | Depdc7 | 0.002 | Young |
| 0.658176 | 0.078002 | 0.01 | 0.008 | 1 | 1 | Tchp | 0.002 | Young |
| 0.659859 | 0.188909 | 0.073 | 0.068 | 1 | 1 | Fundc1 | 0.005 | Young |
| 0.660133 | 0.019113 | 0.053 | 0.049 | 1 | 1 | Aip | 0.004 | Young |
| 0.660497 | 0.091535 | 0.075 | 0.069 | 1 | 1 | Gabpb2 | 0.006 | Young |
| 0.660748 | 0.404919 | 0.029 | 0.026 | 1 | 1 | Phf13 | 0.003 | Young |
| 0.661677 | 0.154814 | 0.03 | 0.027 | 1 | 1 | Prrg2 | 0.003 | Young |
| 0.663634 | 0.52925 | 0.021 | 0.018 | 1 | 1 | Prr5l | 0.003 | Young |
| 0.66389 | 0.067062 | 0.021 | 0.024 | 1 | 1 | Cpsf4 | -0.003 | Young |
| 0.664195 | 0.071454 | 0.073 | 0.068 | 1 | 1 | Cacul1 | 0.005 | Young |
| 0.664516 | 0.208331 | 0.021 | 0.018 | 1 | 1 | Ptrh2 | 0.003 | Young |
| 0.665158 | 0.136216 | 0.038 | 0.042 | 1 | 1 | Tex264 | -0.004 | Young |
| 0.667878 | 0.0906 | 0.03 | 0.027 | 1 | 1 | Prune1 | 0.003 | Young |
| 0.668692 | 0.417148 | 0.01 | 0.012 | 1 | 1 | Dpp4 | -0.002 | Young |

| | | | | | | | | |
|---|---|---|---|---|---|---|---|---|
| 0.668869 | 0.631622 | 0.019 | 0.017 | 1 | 1 | Gm12840 | 0.002 | Young |
| 0.669595 | 0.105534 | 0.021 | 0.018 | 1 | 1 | Alg1 | 0.003 | Young |
| 0.670833 | 0.14577 | 0.057 | 0.053 | 1 | 1 | BC003331 | 0.004 | Young |
| 0.671353 | 0.003472 | 0.03 | 0.027 | 1 | 1 | AU019823 | 0.003 | Young |
| 0.67277 | 0.656575 | 0.019 | 0.017 | 1 | 1 | Tbkbp1 | 0.002 | Young |
| 0.67346 | 0.250101 | 0.019 | 0.017 | 1 | 1 | Lrrc61 | 0.002 | Young |
| 0.674014 | 0.010165 | 0.998 | 0.994 | 1 | 1 | mt-Co1 | 0.004 | Young |
| 0.674784 | 0.132099 | 0.029 | 0.026 | 1 | 1 | Gart | 0.003 | Young |
| 0.675383 | 0.114731 | 0.025 | 0.028 | 1 | 1 | Pxmp4 | -0.003 | Young |
| 0.677416 | 0.167073 | 0.029 | 0.026 | 1 | 1 | B9d2 | 0.003 | Young |
| 0.677833 | 0.022278 | 0.053 | 0.049 | 1 | 1 | Pspc1 | 0.004 | Young |
| 0.678977 | 0.021646 | 0.019 | 0.022 | 1 | 1 | Hddc2 | -0.003 | Young |
| 0.679216 | 0.011607 | 0.019 | 0.017 | 1 | 1 | Wdr70 | 0.002 | Young |
| 0.680693 | 0.243767 | 0.027 | 0.024 | 1 | 1 | Irgm2 | 0.003 | Young |
| 0.681754 | 0.077867 | 0.019 | 0.017 | 1 | 1 | Aifm2 | 0.002 | Young |
| 0.682772 | 0.097093 | 0.102 | 0.096 | 1 | 1 | Mfsd14a | 0.006 | Young |
| 0.683153 | 0.158615 | 0.035 | 0.032 | 1 | 1 | Gars | 0.003 | Young |
| 0.683866 | 0.410186 | 0.018 | 0.015 | 1 | 1 | AI597479 | 0.003 | Young |
| 0.684662 | 0.393949 | 0.025 | 0.028 | 1 | 1 | Poldip2 | -0.003 | Young |
| 0.684748 | 0.046362 | 0.1 | 0.095 | 1 | 1 | Ergic3 | 0.005 | Young |
| 0.685453 | 0.267179 | 0.019 | 0.017 | 1 | 1 | Dlg4 | 0.002 | Young |
| 0.686981 | 0.401767 | 0.032 | 0.03 | 1 | 1 | Prpf4 | 0.002 | Young |
| 0.686995 | 0.418452 | 0.018 | 0.015 | 1 | 1 | Trmt11 | 0.003 | Young |
| 0.687586 | 0.122964 | 0.023 | 0.026 | 1 | 1 | Dhx33 | -0.003 | Young |
| 0.687959 | 0.46081 | 0.018 | 0.015 | 1 | 1 | Zdhhc16 | 0.003 | Young |
| 0.689624 | 0.895558 | 0.016 | 0.014 | 1 | 1 | Hsd17b7 | 0.002 | Young |
| 0.689996 | 0.049527 | 0.013 | 0.015 | 1 | 1 | Slc22a15 | -0.002 | Young |
| 0.690291 | 0.053302 | 0.027 | 0.024 | 1 | 1 | Hsf2 | 0.003 | Young |
| 0.690612 | 0.197883 | 0.018 | 0.015 | 1 | 1 | Alg13 | 0.003 | Young |
| 0.692544 | 0.058281 | 0.035 | 0.032 | 1 | 1 | 1600012H( | 0.003 | Young |
| 0.692637 | 0.177864 | 0.013 | 0.015 | 1 | 1 | Fbxw17 | -0.002 | Young |
| 0.693583 | 0.075617 | 0.064 | 0.059 | 1 | 1 | Ric8a | 0.005 | Young |
| 0.694236 | 0.219413 | 0.018 | 0.015 | 1 | 1 | Mtg1 | 0.003 | Young |
| 0.694328 | 0.219168 | 0.041 | 0.037 | 1 | 1 | Tsfm | 0.004 | Young |
| 0.69447 | 0.649042 | 0.024 | 0.022 | 1 | 1 | Kantr | 0.002 | Young |
| 0.694711 | 0.086756 | 0.023 | 0.026 | 1 | 1 | Isyna1 | -0.003 | Young |
| 0.694895 | 0.02195 | 0.026 | 0.023 | 1 | 1 | Dtx2 | 0.003 | Young |
| 0.69537 | 0.352335 | 0.041 | 0.037 | 1 | 1 | Srebf1 | 0.004 | Young |
| 0.695547 | 0.021965 | 0.013 | 0.015 | 1 | 1 | Tapt1 | -0.002 | Young |
| 0.695727 | 0.139724 | 0.055 | 0.051 | 1 | 1 | Eps15l1 | 0.004 | Young |
| 0.695934 | 0.087077 | 0.049 | 0.045 | 1 | 1 | Tbck | 0.004 | Young |
| 0.69694 | 0.535489 | 0.016 | 0.014 | 1 | 1 | Atm | 0.002 | Young |
| 0.697801 | 0.193411 | 0.059 | 0.055 | 1 | 1 | Slc16a7 | 0.004 | Young |
| 0.700141 | 0.169451 | 0.027 | 0.03 | 1 | 1 | Bop1 | -0.003 | Young |
| 0.700749 | 0.415158 | 0.031 | 0.028 | 1 | 1 | Ikbke | 0.003 | Young |

| | | | | | | | | |
|---|---|---|---|---|---|---|---|---|
| 0.701691 | 0.146263 | 0.024 | 0.022 | 1 | 1 | Prr14l | 0.002 | Young |
| 0.702974 | 1.2351 | 0.015 | 0.013 | 1 | 1 | Pnpla6 | 0.002 | Young |
| 0.703608 | 0.096522 | 0.122 | 0.117 | 1 | 1 | Rmnd5a | 0.005 | Young |
| 0.703904 | 0.117106 | 0.039 | 0.036 | 1 | 1 | Siah1a | 0.003 | Young |
| 0.704031 | 0.148728 | 0.016 | 0.014 | 1 | 1 | 1110012L1 | 0.002 | Young |
| 0.704203 | 0.09283 | 0.041 | 0.037 | 1 | 1 | Nip7 | 0.004 | Young |
| 0.704539 | 0.319862 | 0.016 | 0.014 | 1 | 1 | Vwa8 | 0.002 | Young |
| 0.706135 | 0.088837 | 0.041 | 0.037 | 1 | 1 | Rnf10 | 0.004 | Young |
| 0.706658 | 0.382474 | 0.024 | 0.022 | 1 | 1 | Mcm5 | 0.002 | Young |
| 0.706824 | 0.013367 | 0.016 | 0.014 | 1 | 1 | Tmem241 | 0.002 | Young |
| 0.708449 | 0.038003 | 0.032 | 0.03 | 1 | 1 | Ubl7 | 0.002 | Young |
| 0.708939 | 0.237093 | 0.024 | 0.022 | 1 | 1 | Ptcd2 | 0.002 | Young |
| 0.709169 | 0.04751 | 0.032 | 0.03 | 1 | 1 | Ripor2 | 0.002 | Young |
| 0.70923 | 0.700282 | 0.015 | 0.013 | 1 | 1 | Ddx31 | 0.002 | Young |
| 0.711855 | 0.369094 | 0.023 | 0.021 | 1 | 1 | Gipc1 | 0.002 | Young |
| 0.71199 | 0.044595 | 0.117 | 0.112 | 1 | 1 | Larp4 | 0.005 | Young |
| 0.712591 | 0.111892 | 0.032 | 0.03 | 1 | 1 | Exosc2 | 0.002 | Young |
| 0.712888 | 0.332708 | 0.017 | 0.019 | 1 | 1 | Cep162 | -0.002 | Young |
| 0.713099 | 0.406169 | 0.015 | 0.013 | 1 | 1 | 3110002H1 | 0.002 | Young |
| 0.714473 | 0.014087 | 0.047 | 0.044 | 1 | 1 | Mtf1 | 0.003 | Young |
| 0.714756 | 0.097799 | 0.011 | 0.013 | 1 | 1 | Gm6563 | -0.002 | Young |
| 0.71497 | 0.563017 | 0.015 | 0.013 | 1 | 1 | Eif2b3 | 0.002 | Young |
| 0.715491 | 0.010811 | 0.134 | 0.129 | 1 | 1 | Eif3i | 0.005 | Young |
| 0.715928 | 0.379872 | 0.023 | 0.021 | 1 | 1 | Tmem37 | 0.002 | Young |
| 0.716237 | 0.015261 | 0.089 | 0.084 | 1 | 1 | Dock1 | 0.005 | Young |
| 0.716569 | 0.017611 | 0.031 | 0.028 | 1 | 1 | Mbtps2 | 0.003 | Young |
| 0.716964 | 0.026442 | 0.059 | 0.055 | 1 | 1 | 2310022A1 | 0.004 | Young |
| 0.717645 | 0.012513 | 0.015 | 0.013 | 1 | 1 | Zmat3 | 0.002 | Young |
| 0.717976 | 0.074578 | 0.011 | 0.013 | 1 | 1 | Zfp335 | -0.002 | Young |
| 0.718269 | 0.357931 | 0.011 | 0.013 | 1 | 1 | Apba3 | -0.002 | Young |
| 0.718269 | 0.065356 | 0.011 | 0.013 | 1 | 1 | Bud13 | -0.002 | Young |
| 0.718562 | 0.06133 | 0.011 | 0.013 | 1 | 1 | Slc9a4 | -0.002 | Young |
| 0.719526 | 0.05618 | 0.066 | 0.062 | 1 | 1 | Edc4 | 0.004 | Young |
| 0.720321 | 0.286875 | 0.011 | 0.013 | 1 | 1 | Ptges2 | -0.002 | Young |
| 0.720821 | 0.121458 | 0.031 | 0.028 | 1 | 1 | Agl | 0.003 | Young |
| 0.724106 | 0.767973 | 0.013 | 0.012 | 1 | 1 | Tjap1 | 0.001 | Young |
| 0.724112 | 0.000315 | 0.072 | 0.068 | 1 | 1 | BC004004 | 0.004 | Young |
| 0.725237 | 0.437687 | 0.013 | 0.012 | 1 | 1 | Hdac10 | 0.001 | Young |
| 0.725463 | 0.022457 | 0.2 | 0.193 | 1 | 1 | Ldha | 0.007 | Young |
| 0.725731 | 0.085454 | 0.031 | 0.028 | 1 | 1 | Abhd13 | 0.003 | Young |
| 0.726878 | 0.597685 | 0.021 | 0.019 | 1 | 1 | 4930503L1 | 0.002 | Young |
| 0.727219 | 0.373282 | 0.013 | 0.012 | 1 | 1 | AW554918 | 0.001 | Young |
| 0.727407 | 0.284722 | 0.03 | 0.027 | 1 | 1 | Dhdds | 0.003 | Young |
| 0.728777 | 0.161619 | 0.044 | 0.041 | 1 | 1 | Bin3 | 0.003 | Young |
| 0.728928 | 0.149761 | 0.03 | 0.027 | 1 | 1 | Ifi35 | 0.003 | Young |

| | | | | | | | | |
|---|---|---|---|---|---|---|---|---|
| 0.729386 | 0.177224 | 0.061 | 0.058 | 1 | 1 | Pdha1 | 0.003 | Young |
| 0.729713 | 0.094353 | 0.091 | 0.086 | 1 | 1 | Usp24 | 0.005 | Young |
| 0.730217 | 0.099647 | 0.063 | 0.059 | 1 | 1 | Eif4ebp1 | 0.004 | Young |
| 0.730905 | 0.327758 | 0.013 | 0.012 | 1 | 1 | Phka1 | 0.001 | Young |
| 0.731473 | 0.05544 | 0.013 | 0.012 | 1 | 1 | Ptdss2 | 0.001 | Young |
| 0.731665 | 0.145757 | 0.043 | 0.04 | 1 | 1 | Ate1 | 0.003 | Young |
| 0.731784 | 0.049331 | 0.021 | 0.019 | 1 | 1 | Zc3h12c | 0.002 | Young |
| 0.732041 | 0.138335 | 0.013 | 0.012 | 1 | 1 | Haus7 | 0.001 | Young |
| 0.733046 | 0.089664 | 0.056 | 0.053 | 1 | 1 | Rabggtb | 0.003 | Young |
| 0.733286 | 0.074481 | 0.069 | 0.066 | 1 | 1 | Znfx1 | 0.003 | Young |
| 0.733674 | 0.135983 | 0.056 | 0.053 | 1 | 1 | Zeb2os | 0.003 | Young |
| 0.733813 | 0.118859 | 0.023 | 0.021 | 1 | 1 | Wdyhv1 | 0.002 | Young |
| 0.73457 | 0.206698 | 0.041 | 0.039 | 1 | 1 | Taf3 | 0.002 | Young |
| 0.736771 | 0.818197 | 0.012 | 0.01 | 1 | 1 | Fastkd2 | 0.002 | Young |
| 0.737293 | 0.429732 | 0.028 | 0.026 | 1 | 1 | Brms1 | 0.002 | Young |
| 0.738883 | 0.684618 | 0.012 | 0.01 | 1 | 1 | Qtrt2 | 0.002 | Young |
| 0.739537 | 0.044189 | 0.036 | 0.033 | 1 | 1 | Ints12 | 0.003 | Young |
| 0.74135 | 0.02089 | 0.048 | 0.051 | 1 | 1 | Nedd9 | -0.003 | Young |
| 0.741438 | 0.029503 | 0.049 | 0.046 | 1 | 1 | Tmem245 | 0.003 | Young |
| 0.742737 | 0.087674 | 0.036 | 0.033 | 1 | 1 | Zfp598 | 0.003 | Young |
| 0.74281 | 0.258403 | 0.012 | 0.01 | 1 | 1 | C130050O1 | 0.002 | Young |
| 0.742852 | 0.052928 | 0.03 | 0.027 | 1 | 1 | Zfp991 | 0.003 | Young |
| 0.743113 | 0.606958 | 0.012 | 0.01 | 1 | 1 | Zfp748 | 0.002 | Young |
| 0.743113 | 0.558469 | 0.012 | 0.01 | 1 | 1 | Rhbdd3 | 0.002 | Young |
| 0.743168 | 0.112019 | 0.028 | 0.026 | 1 | 1 | Mtpap | 0.002 | Young |
| 0.743544 | 0.056207 | 0.009 | 0.01 | 1 | 1 | Zfp146 | -0.001 | Young |
| 0.743718 | 0.293204 | 0.012 | 0.01 | 1 | 1 | Gm31718 | 0.002 | Young |
| 0.744928 | 0.068343 | 0.012 | 0.01 | 1 | 1 | Setdb2 | 0.002 | Young |
| 0.746139 | 0.207983 | 0.012 | 0.01 | 1 | 1 | Riox1 | 0.002 | Young |
| 0.746141 | 0.080168 | 0.048 | 0.051 | 1 | 1 | Ebna1bp2 | -0.003 | Young |
| 0.746856 | 0.111644 | 0.009 | 0.01 | 1 | 1 | Tamm41 | -0.001 | Young |
| 0.746857 | 0.199531 | 0.07 | 0.067 | 1 | 1 | Tec | 0.003 | Young |
| 0.747047 | 0.128639 | 0.012 | 0.01 | 1 | 1 | Mettl26 | 0.002 | Young |
| 0.747188 | 0.052847 | 0.009 | 0.01 | 1 | 1 | Lrrc49 | -0.001 | Young |
| 0.747333 | 0.250569 | 0.058 | 0.055 | 1 | 1 | Trim24 | 0.003 | Young |
| 0.74735 | 0.286625 | 0.012 | 0.01 | 1 | 1 | Ptch1 | 0.002 | Young |
| 0.747396 | 0.084009 | 0.026 | 0.028 | 1 | 1 | Atg4d | -0.002 | Young |
| 0.747653 | 0.134986 | 0.012 | 0.01 | 1 | 1 | Rab39 | 0.002 | Young |
| 0.747851 | 0.244761 | 0.009 | 0.01 | 1 | 1 | Purg | -0.001 | Young |
| 0.749169 | 0.031859 | 0.012 | 0.01 | 1 | 1 | Rttn | 0.002 | Young |
| 0.749178 | 0.321517 | 0.009 | 0.01 | 1 | 1 | Creld1 | -0.001 | Young |
| 0.750868 | 0.097445 | 0.02 | 0.018 | 1 | 1 | Habp4 | 0.002 | Young |
| 0.751858 | 0.347912 | 0.045 | 0.042 | 1 | 1 | Matk | 0.003 | Young |
| 0.752175 | 0.001811 | 0.048 | 0.045 | 1 | 1 | Gm28875 | 0.003 | Young |
| 0.753441 | 0.788643 | 0.025 | 0.023 | 1 | 1 | Pip4k2c | 0.002 | Young |

| | | | | | | | | |
|---|---|---|---|---|---|---|---|---|
| 0.754619 | 0.544327 | 0.027 | 0.024 | 1 | 1 | Terf2ip | 0.003 | Young |
| 0.754873 | 0.295528 | 0.052 | 0.049 | 1 | 1 | Sar1b | 0.003 | Young |
| 0.754936 | 0.199753 | 0.026 | 0.028 | 1 | 1 | Scnm1 | -0.002 | Young |
| 0.75648 | 0.019514 | 0.061 | 0.064 | 1 | 1 | Pofut1 | -0.003 | Young |
| 0.756845 | 0.602522 | 0.018 | 0.017 | 1 | 1 | Ccdc112 | 0.001 | Young |
| 0.756994 | 0.208475 | 0.074 | 0.071 | 1 | 1 | Usp47 | 0.003 | Young |
| 0.757245 | 0.581662 | 0.01 | 0.009 | 1 | 1 | Ptchd1 | 0.001 | Young |
| 0.75733 | 0.650063 | 0.018 | 0.017 | 1 | 1 | Abcg2 | 0.001 | Young |
| 0.757543 | 0.498524 | 0.022 | 0.024 | 1 | 1 | Carm1 | -0.002 | Young |
| 0.757648 | 0.151475 | 0.04 | 0.037 | 1 | 1 | Nolc1 | 0.003 | Young |
| 0.75771 | 0.130325 | 0.064 | 0.06 | 1 | 1 | Rnf20 | 0.004 | Young |
| 0.759195 | 0.542859 | 0.01 | 0.009 | 1 | 1 | Cad | 0.001 | Young |
| 0.759515 | 0.223638 | 0.013 | 0.014 | 1 | 1 | Gmeb2 | -0.001 | Young |
| 0.759521 | 0.444306 | 0.01 | 0.009 | 1 | 1 | Pyroxd2 | 0.001 | Young |
| 0.760497 | 0.370319 | 0.01 | 0.009 | 1 | 1 | Fancm | 0.001 | Young |
| 0.761051 | 0.118314 | 0.04 | 0.037 | 1 | 1 | Nr1d2 | 0.003 | Young |
| 0.761148 | 0.196157 | 0.01 | 0.009 | 1 | 1 | F8a | 0.001 | Young |
| 0.762056 | 0.172869 | 0.013 | 0.014 | 1 | 1 | Gpatch1 | -0.001 | Young |
| 0.762613 | 0.25409 | 0.01 | 0.009 | 1 | 1 | Rbm48 | 0.001 | Young |
| 0.763101 | 0.052832 | 0.01 | 0.009 | 1 | 1 | Tle2 | 0.001 | Young |
| 0.763751 | 0.037733 | 0.013 | 0.014 | 1 | 1 | Zfp609 | -0.001 | Young |
| 0.763888 | 0.234951 | 0.018 | 0.017 | 1 | 1 | H2-Ob | 0.001 | Young |
| 0.764079 | 0.117699 | 0.01 | 0.009 | 1 | 1 | Csrp2 | 0.001 | Young |
| 0.76414 | 0.061485 | 0.047 | 0.044 | 1 | 1 | Anapc2 | 0.003 | Young |
| 0.764474 | 0.56251 | 0.025 | 0.023 | 1 | 1 | Tmco3 | 0.002 | Young |
| 0.766191 | 0.015037 | 0.035 | 0.037 | 1 | 1 | Ralgapb | -0.002 | Young |
| 0.767064 | 0.042977 | 0.047 | 0.05 | 1 | 1 | Dapk3 | -0.003 | Young |
| 0.767075 | 0.104286 | 0.047 | 0.044 | 1 | 1 | Cops6 | 0.003 | Young |
| 0.767262 | 0.13601 | 0.05 | 0.048 | 1 | 1 | Mettl23 | 0.002 | Young |
| 0.767539 | 0.051095 | 0.018 | 0.017 | 1 | 1 | Gtpbp1 | 0.001 | Young |
| 0.768269 | 0.198466 | 0.018 | 0.017 | 1 | 1 | Rrp9 | 0.001 | Young |
| 0.769486 | 0.294742 | 0.025 | 0.023 | 1 | 1 | Nbdy | 0.002 | Young |
| 0.771006 | 0.16396 | 0.032 | 0.03 | 1 | 1 | Gatsl2 | 0.002 | Young |
| 0.771047 | 0.081423 | 0.047 | 0.05 | 1 | 1 | Dld | -0.003 | Young |
| 0.771093 | 0.216048 | 0.016 | 0.018 | 1 | 1 | Zfp846 | -0.002 | Young |
| 0.771682 | 0.037357 | 0.018 | 0.017 | 1 | 1 | Gmnn | 0.001 | Young |
| 0.771991 | 0.03516 | 0.125 | 0.121 | 1 | 1 | Pdcd10 | 0.004 | Young |
| 0.774086 | 0.482659 | 0.017 | 0.015 | 1 | 1 | Zw10 | 0.002 | Young |
| 0.774246 | 0.236694 | 0.049 | 0.046 | 1 | 1 | Utp14a | 0.003 | Young |
| 0.774595 | 0.017722 | 0.02 | 0.022 | 1 | 1 | Zfp369 | -0.002 | Young |
| 0.774848 | 0.619349 | 0.017 | 0.015 | 1 | 1 | Tatdn1 | 0.002 | Young |
| 0.77548 | 0.009583 | 0.032 | 0.03 | 1 | 1 | Nup205 | 0.002 | Young |
| 0.776394 | 0.022707 | 0.025 | 0.023 | 1 | 1 | Dis3l2 | 0.002 | Young |
| 0.77722 | 0.003479 | 0.056 | 0.059 | 1 | 1 | Myo18a | -0.003 | Young |
| 0.783141 | 0.028138 | 0.032 | 0.03 | 1 | 1 | Grk5 | 0.002 | Young |

| | | | | | | | | |
|---|---|---|---|---|---|---|---|---|
| 0.783253 | 0.184746 | 0.017 | 0.015 | 1 | 1 | Sh3bp5l | 0.002 | Young |
| 0.783633 | 0.298452 | 0.042 | 0.04 | 1 | 1 | Prpf38a | 0.002 | Young |
| 0.784018 | 0.109061 | 0.061 | 0.058 | 1 | 1 | 1810022K0 | 0.003 | Young |
| 0.784456 | 0.071991 | 0.024 | 0.022 | 1 | 1 | Erh | 0.002 | Young |
| 0.784466 | 0.352038 | 0.083 | 0.081 | 1 | 1 | Flii | 0.002 | Young |
| 0.785298 | 0.078301 | 0.062 | 0.059 | 1 | 1 | Srpk2 | 0.003 | Young |
| 0.78555 | 0.132087 | 0.017 | 0.015 | 1 | 1 | Ppp2r1b | 0.002 | Young |
| 0.786613 | 0.003843 | 0.286 | 0.275 | 1 | 1 | Reep3 | 0.011 | Young |
| 0.787862 | 0.161121 | 0.051 | 0.054 | 1 | 1 | Hlcs | -0.003 | Young |
| 0.789257 | 0.06135 | 0.072 | 0.069 | 1 | 1 | Cenpx | 0.003 | Young |
| 0.792206 | 0.034772 | 0.083 | 0.08 | 1 | 1 | Rcsd1 | 0.003 | Young |
| 0.795497 | 0.166393 | 0.078 | 0.075 | 1 | 1 | Gripap1 | 0.003 | Young |
| 0.795599 | 0.664143 | 0.027 | 0.026 | 1 | 1 | B3galnt2 | 0.001 | Young |
| 0.795746 | 0.387519 | 0.034 | 0.032 | 1 | 1 | Ncoa5 | 0.002 | Young |
| 0.796153 | 0.026413 | 0.01 | 0.012 | 1 | 1 | Wee1 | -0.002 | Young |
| 0.796469 | 0.157874 | 0.01 | 0.012 | 1 | 1 | Cdc42bpg | -0.002 | Young |
| 0.79686 | 0.443069 | 0.022 | 0.021 | 1 | 1 | Ctdp1 | 0.001 | Young |
| 0.797756 | 0.388889 | 0.022 | 0.021 | 1 | 1 | Gpkow | 0.001 | Young |
| 0.798467 | 0.407358 | 0.046 | 0.049 | 1 | 1 | Mkl2 | -0.003 | Young |
| 0.799077 | 0.208215 | 0.035 | 0.033 | 1 | 1 | Kifap3 | 0.002 | Young |
| 0.799326 | 0.337207 | 0.022 | 0.021 | 1 | 1 | Ticam1 | 0.001 | Young |
| 0.800224 | 0.31517 | 0.022 | 0.021 | 1 | 1 | Zfp938 | 0.001 | Young |
| 0.800937 | 0.027005 | 0.021 | 0.023 | 1 | 1 | Nufip1 | -0.002 | Young |
| 0.801246 | 0.206865 | 0.029 | 0.031 | 1 | 1 | Slc9a6 | -0.002 | Young |
| 0.801795 | 0.189718 | 0.022 | 0.021 | 1 | 1 | Xpnpep1 | 0.001 | Young |
| 0.80191 | 0.024565 | 0.016 | 0.014 | 1 | 1 | Pde6d | 0.002 | Young |
| 0.803591 | 0.257971 | 0.022 | 0.021 | 1 | 1 | Zfp329 | 0.001 | Young |
| 0.804028 | 0.056015 | 0.086 | 0.084 | 1 | 1 | Gdap10 | 0.002 | Young |
| 0.804512 | 0.571024 | 0.032 | 0.031 | 1 | 1 | Rab11fip2 | 0.001 | Young |
| 0.806743 | 0.035397 | 0.047 | 0.045 | 1 | 1 | Pex19 | 0.002 | Young |
| 0.80686 | 0.069982 | 0.042 | 0.04 | 1 | 1 | Dhx16 | 0.002 | Young |
| 0.807442 | 0.357016 | 0.014 | 0.015 | 1 | 1 | Ccdc61 | -0.001 | Young |
| 0.807598 | 0.009375 | 0.088 | 0.085 | 1 | 1 | Golga7 | 0.003 | Young |
| 0.810787 | 0.016567 | 0.022 | 0.021 | 1 | 1 | Ptk2 | 0.001 | Young |
| 0.812004 | 0.340757 | 0.021 | 0.019 | 1 | 1 | Ddx19b | 0.002 | Young |
| 0.813453 | 0.744242 | 0.014 | 0.015 | 1 | 1 | Rfc4 | -0.001 | Young |
| 0.81449 | 0.035832 | 0.063 | 0.066 | 1 | 1 | Ncbp2 | -0.003 | Young |
| 0.81604 | 0.13449 | 0.034 | 0.032 | 1 | 1 | Faf1 | 0.002 | Young |
| 0.817866 | 0.147591 | 0.034 | 0.032 | 1 | 1 | Hbegf | 0.002 | Young |
| 0.8184 | 0.209442 | 0.045 | 0.048 | 1 | 1 | Hdac4 | -0.003 | Young |
| 0.819492 | 0.232287 | 0.014 | 0.013 | 1 | 1 | Fbxw9 | 0.001 | Young |
| 0.819694 | 0.034414 | 0.021 | 0.019 | 1 | 1 | Tsta3 | 0.002 | Young |
| 0.820791 | 0.259627 | 0.048 | 0.046 | 1 | 1 | Dtymk | 0.002 | Young |
| 0.820882 | 0.036516 | 0.034 | 0.032 | 1 | 1 | Map3k20 | 0.002 | Young |
| 0.821637 | 0.202353 | 0.027 | 0.026 | 1 | 1 | Sdhaf4 | 0.001 | Young |

| | | | | | | | | |
|---|---|---|---|---|---|---|---|---|
| 0.822609 | 0.237136 | 0.014 | 0.013 | 1 | 1 | Klrb1b | 0.001 | Young |
| 0.822609 | 0.081792 | 0.014 | 0.013 | 1 | 1 | Nr1d1 | 0.001 | Young |
| 0.823742 | 0.085904 | 0.014 | 0.013 | 1 | 1 | Atpaf2 | 0.001 | Young |
| 0.823797 | 0.110403 | 0.032 | 0.031 | 1 | 1 | Btbd10 | 0.001 | Young |
| 0.824593 | 0.040028 | 0.014 | 0.013 | 1 | 1 | Prdm10 | 0.001 | Young |
| 0.826533 | 0.015002 | 0.052 | 0.05 | 1 | 1 | Trip4 | 0.002 | Young |
| 0.826558 | 0.052419 | 0.023 | 0.024 | 1 | 1 | Aen | -0.001 | Young |
| 0.826844 | 0.646528 | 0.019 | 0.018 | 1 | 1 | Stoml1 | 0.001 | Young |
| 0.827438 | 0.12123 | 0.063 | 0.066 | 1 | 1 | Dhdh | -0.003 | Young |
| 0.828059 | 1.154064 | 0.013 | 0.012 | 1 | 1 | Sh3bp4 | 0.001 | Young |
| 0.829291 | 0.185129 | 0.085 | 0.082 | 1 | 1 | Clcn4 | 0.003 | Young |
| 0.830547 | 0.207366 | 0.026 | 0.024 | 1 | 1 | Ppp6r2 | 0.002 | Young |
| 0.830912 | 0.003479 | 0.039 | 0.037 | 1 | 1 | Slc17a5 | 0.002 | Young |
| 0.83095 | 0.014666 | 0.069 | 0.067 | 1 | 1 | Uap1 | 0.002 | Young |
| 0.832432 | 0.274406 | 0.026 | 0.024 | 1 | 1 | Tstd2 | 0.002 | Young |
| 0.833253 | 0.091842 | 0.044 | 0.042 | 1 | 1 | Mier3 | 0.002 | Young |
| 0.833269 | 0.43824 | 0.019 | 0.018 | 1 | 1 | Zbtb24 | 0.001 | Young |
| 0.83327 | 0.235741 | 0.026 | 0.024 | 1 | 1 | Xlr4b | 0.002 | Young |
| 0.833565 | 0.048963 | 0.032 | 0.033 | 1 | 1 | Rcan3 | -0.001 | Young |
| 0.836107 | 0.071046 | 0.068 | 0.066 | 1 | 1 | Plscr3 | 0.002 | Young |
| 0.836906 | 0.064209 | 0.012 | 0.013 | 1 | 1 | Asb1 | -0.001 | Young |
| 0.837507 | 0.086365 | 0.012 | 0.013 | 1 | 1 | Nedd4 | -0.001 | Young |
| 0.837807 | 0.139028 | 0.012 | 0.013 | 1 | 1 | Serpinb9 | -0.001 | Young |
| 0.839948 | 0.071671 | 0.019 | 0.018 | 1 | 1 | Mthfd2l | 0.001 | Young |
| 0.840418 | 0.102126 | 0.108 | 0.105 | 1 | 1 | Plekha2 | 0.003 | Young |
| 0.842008 | 0.125237 | 0.023 | 0.024 | 1 | 1 | Zdhhc23 | -0.001 | Young |
| 0.842015 | 0.320308 | 0.012 | 0.013 | 1 | 1 | Taf8 | -0.001 | Young |
| 0.842037 | 0.166352 | 0.06 | 0.058 | 1 | 1 | Vkorc1 | 0.002 | Young |
| 0.843282 | 0.411666 | 0.016 | 0.017 | 1 | 1 | Pstk | -0.001 | Young |
| 0.843325 | 0.100725 | 0.065 | 0.063 | 1 | 1 | Grk6 | 0.002 | Young |
| 0.843897 | 0.220996 | 0.031 | 0.03 | 1 | 1 | Pitrm1 | 0.001 | Young |
| 0.845201 | 0.053622 | 0.09 | 0.094 | 1 | 1 | Mob1b | -0.004 | Young |
| 0.845854 | 0.030183 | 0.036 | 0.035 | 1 | 1 | Heatr6 | 0.001 | Young |
| 0.847547 | 0.20968 | 0.031 | 0.03 | 1 | 1 | Polr1a | 0.001 | Young |
| 0.850842 | 0.014646 | 0.086 | 0.084 | 1 | 1 | Siva1 | 0.002 | Young |
| 0.85554 | 0.207636 | 0.018 | 0.017 | 1 | 1 | Racgap1 | 0.001 | Young |
| 0.856124 | 0.173354 | 0.028 | 0.03 | 1 | 1 | Gm38399 | -0.002 | Young |
| 0.856383 | 0.186214 | 0.024 | 0.023 | 1 | 1 | Ankrd46 | 0.001 | Young |
| 0.858826 | 0.246046 | 0.011 | 0.01 | 1 | 1 | Cluap1 | 0.001 | Young |
| 0.859594 | 0.465976 | 0.023 | 0.022 | 1 | 1 | Tmem110 | 0.001 | Young |
| 0.859788 | 0.441955 | 0.011 | 0.01 | 1 | 1 | Zc2hc1a | 0.001 | Young |
| 0.861286 | 0.157201 | 0.079 | 0.077 | 1 | 1 | Cacybp | 0.002 | Young |
| 0.861716 | 0.080173 | 0.053 | 0.051 | 1 | 1 | Pafah1b2 | 0.002 | Young |
| 0.862385 | 0.044613 | 0.053 | 0.051 | 1 | 1 | Chrac1 | 0.002 | Young |
| 0.862998 | 0.017662 | 0.011 | 0.01 | 1 | 1 | Jpx | 0.001 | Young |

| | | | | | | | | |
|---|---|---|---|---|---|---|---|---|
| 0.863445 | 0.030987 | 0.057 | 0.055 | 1 | 1 | Wdr41 | 0.002 | Young |
| 0.863752 | 0.110986 | 0.03 | 0.028 | 1 | 1 | Zfp738 | 0.002 | Young |
| 0.864169 | 0.067715 | 0.018 | 0.017 | 1 | 1 | Gm7694 | 0.001 | Young |
| 0.864248 | 0.171431 | 0.021 | 0.022 | 1 | 1 | Zdhhc8 | -0.001 | Young |
| 0.864569 | 0.12266 | 0.041 | 0.04 | 1 | 1 | Atl3 | 0.001 | Young |
| 0.864931 | 0.037276 | 0.017 | 0.018 | 1 | 1 | Cbr3 | -0.001 | Young |
| 0.866077 | 0.351562 | 0.023 | 0.022 | 1 | 1 | Mrpl50 | 0.001 | Young |
| 0.868851 | 0.020069 | 0.058 | 0.057 | 1 | 1 | Yipf1 | 0.001 | Young |
| 0.86891 | 0.066545 | 0.168 | 0.166 | 1 | 1 | Cast | 0.002 | Young |
| 0.869085 | 0.164012 | 0.083 | 0.086 | 1 | 1 | Ipo7 | -0.003 | Young |
| 0.86948 | 0.027803 | 0.03 | 0.028 | 1 | 1 | Fam173b | 0.002 | Young |
| 0.870105 | 0.197539 | 0.023 | 0.022 | 1 | 1 | Btrc | 0.001 | Young |
| 0.870666 | 0.021607 | 0.03 | 0.028 | 1 | 1 | Ddx49 | 0.002 | Young |
| 0.871796 | 0.078473 | 0.017 | 0.018 | 1 | 1 | Tfap4 | -0.001 | Young |
| 0.873799 | 0.052087 | 0.041 | 0.04 | 1 | 1 | Copg2 | 0.001 | Young |
| 0.873828 | 0.58607 | 0.016 | 0.015 | 1 | 1 | Armt1 | 0.001 | Young |
| 0.873962 | 0.032803 | 0.013 | 0.014 | 1 | 1 | Wdr92 | -0.001 | Young |
| 0.874021 | 0.453685 | 0.028 | 0.027 | 1 | 1 | Pacs1 | 0.001 | Young |
| 0.874765 | 0.025811 | 0.072 | 0.071 | 1 | 1 | Ythdf2 | 0.001 | Young |
| 0.876183 | 0.001685 | 0.15 | 0.152 | 1 | 1 | Arcn1 | -0.002 | Young |
| 0.876217 | 0.54858 | 0.016 | 0.015 | 1 | 1 | Asf1a | 0.001 | Young |
| 0.876717 | 0.02678 | 0.03 | 0.031 | 1 | 1 | Ttc32 | -0.001 | Young |
| 0.876833 | 0.140153 | 0.013 | 0.014 | 1 | 1 | Zfp143 | -0.001 | Young |
| 0.8781 | 0.139906 | 0.05 | 0.049 | 1 | 1 | Pgrmc2 | 0.001 | Young |
| 0.878607 | 0.294747 | 0.016 | 0.015 | 1 | 1 | Ndst1 | 0.001 | Young |
| 0.880894 | 0.454928 | 0.021 | 0.021 | 1 | 1 | Clec4d | 0 | Young |
| 0.881295 | 0.178595 | 0.064 | 0.066 | 1 | 1 | Anxa11 | -0.002 | Young |
| 0.881978 | 0.174099 | 0.01 | 0.01 | 1 | 1 | Trim2 | 0 | Young |
| 0.882263 | 0.467297 | 0.032 | 0.031 | 1 | 1 | Lman2l | 0.001 | Young |
| 0.882436 | 0.393513 | 0.013 | 0.014 | 1 | 1 | Cc2d1a | -0.001 | Young |
| 0.882989 | 0.224105 | 0.01 | 0.01 | 1 | 1 | Tmem38a | 0 | Young |
| 0.883085 | 0.925352 | 0.01 | 0.009 | 1 | 1 | Chst15 | 0.001 | Young |
| 0.883085 | 0.413395 | 0.01 | 0.009 | 1 | 1 | Sall2 | 0.001 | Young |
| 0.884523 | 0.075799 | 0.045 | 0.044 | 1 | 1 | Naip2 | 0.001 | Young |
| 0.8848 | 0.187959 | 0.044 | 0.042 | 1 | 1 | Ubap1 | 0.002 | Young |
| 0.885055 | 0.097412 | 0.066 | 0.064 | 1 | 1 | Ppp2r2d | 0.002 | Young |
| 0.885158 | 0.390659 | 0.01 | 0.009 | 1 | 1 | Heg1 | 0.001 | Young |
| 0.887578 | 0.190492 | 0.01 | 0.009 | 1 | 1 | Zfp1 | 0.001 | Young |
| 0.887924 | 0.331703 | 0.01 | 0.009 | 1 | 1 | Mad2l1 | 0.001 | Young |
| 0.887924 | 0.30685 | 0.01 | 0.009 | 1 | 1 | Zfp873 | 0.001 | Young |
| 0.887924 | 0.05401 | 0.01 | 0.009 | 1 | 1 | Rpl36a-ps1 | 0.001 | Young |
| 0.888269 | 0.222115 | 0.01 | 0.009 | 1 | 1 | Tgif2 | 0.001 | Young |
| 0.888388 | 0.076482 | 0.085 | 0.084 | 1 | 1 | Slc25a39 | 0.001 | Young |
| 0.889195 | 0.118968 | 0.112 | 0.116 | 1 | 1 | Ube2s | -0.004 | Young |
| 0.890413 | 1.152959 | 0.01 | 0.01 | 1 | 1 | Cd3eap | 0 | Young |

| | | | | | | | | |
|---|---|---|---|---|---|---|---|---|
| 0.891074 | 0.338367 | 0.037 | 0.036 | 1 | 1 | Slc25a30 | 0.001 | Young |
| 0.893086 | 0.032337 | 0.146 | 0.144 | 1 | 1 | Casp8 | 0.002 | Young |
| 0.893556 | 0.356412 | 0.022 | 0.023 | 1 | 1 | Ebag9 | -0.001 | Young |
| 0.894708 | 0.441746 | 0.015 | 0.014 | 1 | 1 | Exd2 | 0.001 | Young |
| 0.894723 | 0.038209 | 0.144 | 0.141 | 1 | 1 | Sgk3 | 0.003 | Young |
| 0.897779 | 0.103359 | 0.027 | 0.026 | 1 | 1 | Xxylt1 | 0.001 | Young |
| 0.898719 | 0.173362 | 0.027 | 0.026 | 1 | 1 | Maml2 | 0.001 | Young |
| 0.898889 | 0.046898 | 0.032 | 0.031 | 1 | 1 | Naxe | 0.001 | Young |
| 0.900246 | 0.054644 | 0.162 | 0.161 | 1 | 1 | Cd302 | 0.001 | Young |
| 0.900286 | 0.236724 | 0.015 | 0.014 | 1 | 1 | Telo2 | 0.001 | Young |
| 0.902797 | 0.29998 | 0.015 | 0.014 | 1 | 1 | Fam151b | 0.001 | Young |
| 0.90524 | 0.119255 | 0.064 | 0.063 | 1 | 1 | Tk2 | 0.001 | Young |
| 0.906279 | 0.516019 | 0.025 | 0.024 | 1 | 1 | Mrps28 | 0.001 | Young |
| 0.906567 | 0.13359 | 0.015 | 0.014 | 1 | 1 | Zfp866 | 0.001 | Young |
| 0.90668 | 0.392159 | 0.018 | 0.019 | 1 | 1 | Rdh11 | -0.001 | Young |
| 0.907493 | 0.012025 | 0.027 | 0.026 | 1 | 1 | Dhx32 | 0.001 | Young |
| 0.907673 | 0.322546 | 0.02 | 0.019 | 1 | 1 | Cdc16 | 0.001 | Young |
| 0.908423 | 0.403817 | 0.015 | 0.015 | 1 | 1 | Fzd5 | 0 | Young |
| 0.908662 | 0.04577 | 0.015 | 0.014 | 1 | 1 | Pla2g12a | 0.001 | Young |
| 0.910348 | 0.176575 | 0.015 | 0.015 | 1 | 1 | Phlpp2 | 0 | Young |
| 0.91132 | 0.128522 | 0.041 | 0.04 | 1 | 1 | Prr14 | 0.001 | Young |
| 0.911524 | 0.016201 | 0.02 | 0.019 | 1 | 1 | Wdr12 | 0.001 | Young |
| 0.911736 | 0.019275 | 0.015 | 0.014 | 1 | 1 | Hyal2 | 0.001 | Young |
| 0.912323 | 0.014035 | 0.058 | 0.057 | 1 | 1 | Fam53b | 0.001 | Young |
| 0.912408 | 0.096739 | 0.046 | 0.045 | 1 | 1 | Ppp1r7 | 0.001 | Young |
| 0.915577 | 0.530714 | 0.015 | 0.015 | 1 | 1 | Ankrd13d | 0 | Young |
| 0.917037 | 0.167936 | 0.07 | 0.069 | 1 | 1 | Stk24 | 0.001 | Young |
| 0.917405 | 0.053937 | 0.083 | 0.085 | 1 | 1 | Zkscan3 | -0.002 | Young |
| 0.917503 | 0.039501 | 0.011 | 0.012 | 1 | 1 | Kiz | -0.001 | Young |
| 0.917503 | 0.018006 | 0.011 | 0.012 | 1 | 1 | D2hgdh | -0.001 | Young |
| 0.918807 | 0.020085 | 0.038 | 0.039 | 1 | 1 | Iars | -0.001 | Young |
| 0.920706 | 0.158563 | 0.032 | 0.033 | 1 | 1 | Dpm2 | -0.001 | Young |
| 0.921313 | 0.324016 | 0.011 | 0.012 | 1 | 1 | Tmem39b | -0.001 | Young |
| 0.922492 | 0.029985 | 0.032 | 0.033 | 1 | 1 | Pgpep1 | -0.001 | Young |
| 0.923536 | 0.43942 | 0.011 | 0.012 | 1 | 1 | Tgm1 | -0.001 | Young |
| 0.923854 | 0.321946 | 0.011 | 0.012 | 1 | 1 | Ddx20 | -0.001 | Young |
| 0.923973 | 0.599329 | 0.013 | 0.013 | 1 | 1 | Thap1 | 0 | Young |
| 0.924171 | 0.102588 | 0.03 | 0.03 | 1 | 1 | Sart3 | 0 | Young |
| 0.924377 | 0.633676 | 0.018 | 0.018 | 1 | 1 | Apool | 0 | Young |
| 0.925298 | 0.346905 | 0.013 | 0.013 | 1 | 1 | C2cd2 | 0 | Young |
| 0.926917 | 0.316652 | 0.013 | 0.013 | 1 | 1 | Acot7 | 0 | Young |
| 0.927065 | 0.363124 | 0.013 | 0.013 | 1 | 1 | Pef1 | 0 | Young |
| 0.927117 | 0.08042 | 0.03 | 0.03 | 1 | 1 | Lztr1 | 0 | Young |
| 0.927294 | 0.196874 | 0.024 | 0.024 | 1 | 1 | Frs2 | 0 | Young |
| 0.927902 | 0.213347 | 0.02 | 0.021 | 1 | 1 | Tmem242 | -0.001 | Young |

| | | | | | | | | |
|---|---|---|---|---|---|---|---|---|
| 0.928007 | 0.044193 | 0.038 | 0.039 | 1 | 1 | Rnf168 | -0.001 | Young |
| 0.928141 | 0.263202 | 0.024 | 0.023 | 1 | 1 | Prkab1 | 0.001 | Young |
| 0.92839 | 0.203338 | 0.013 | 0.013 | 1 | 1 | Grpel2 | 0 | Young |
| 0.929234 | 0.122784 | 0.038 | 0.039 | 1 | 1 | Gucd1 | -0.001 | Young |
| 0.930048 | 0.19503 | 0.02 | 0.021 | 1 | 1 | Man2b2 | -0.001 | Young |
| 0.930451 | 0.085828 | 0.013 | 0.013 | 1 | 1 | Hmgxb3 | 0 | Young |
| 0.93144 | 0.012505 | 0.03 | 0.03 | 1 | 1 | Tslp | 0 | Young |
| 0.932383 | 0.255088 | 0.018 | 0.018 | 1 | 1 | Elp3 | 0 | Young |
| 0.932702 | 0.047284 | 0.025 | 0.026 | 1 | 1 | Scai | -0.001 | Young |
| 0.934125 | 0.025215 | 0.078 | 0.078 | 1 | 1 | Higd1a | 0 | Young |
| 0.934153 | 0.27788 | 0.029 | 0.028 | 1 | 1 | 9130401M | 0.001 | Young |
| 0.934459 | 0.02746 | 0.034 | 0.035 | 1 | 1 | Rnf34 | -0.001 | Young |
| 0.936769 | 0.154171 | 0.029 | 0.028 | 1 | 1 | Rnf138 | 0.001 | Young |
| 0.93764 | 0.018596 | 0.018 | 0.018 | 1 | 1 | Tuba4a | 0 | Young |
| 0.938987 | 0.069559 | 0.114 | 0.116 | 1 | 1 | Pon2 | -0.002 | Young |
| 0.939002 | 0.115118 | 0.024 | 0.023 | 1 | 1 | Ryk | 0.001 | Young |
| 0.940292 | 0.011305 | 0.029 | 0.028 | 1 | 1 | Trmt2a | 0.001 | Young |
| 0.942109 | 0.131403 | 0.057 | 0.058 | 1 | 1 | Chmp1b | -0.001 | Young |
| 0.94428 | 0.311274 | 0.016 | 0.017 | 1 | 1 | Pde8a | -0.001 | Young |
| 0.944998 | 0.106167 | 0.085 | 0.086 | 1 | 1 | Mknk1 | -0.001 | Young |
| 0.946024 | 0.172785 | 0.032 | 0.032 | 1 | 1 | Vezt | 0 | Young |
| 0.946285 | 0.094107 | 0.034 | 0.033 | 1 | 1 | Pbdc1 | 0.001 | Young |
| 0.94797 | 0.858161 | 0.012 | 0.012 | 1 | 1 | Hist2h2be | 0 | Young |
| 0.948725 | 0.262113 | 0.025 | 0.026 | 1 | 1 | Chst7 | -0.001 | Young |
| 0.948833 | 0.20869 | 0.021 | 0.022 | 1 | 1 | Gm15987 | -0.001 | Young |
| 0.951403 | 0.718351 | 0.012 | 0.012 | 1 | 1 | Spata5 | 0 | Young |
| 0.952335 | 0.244731 | 0.027 | 0.027 | 1 | 1 | Iffo1 | 0 | Young |
| 0.953814 | 0.034199 | 0.013 | 0.013 | 1 | 1 | Rgl3 | 0 | Young |
| 0.955317 | 0.122836 | 0.013 | 0.013 | 1 | 1 | Cactin | 0 | Young |
| 0.955369 | 0.02485 | 0.041 | 0.041 | 1 | 1 | Katnbl1 | 0 | Young |
| 0.956086 | 0.018079 | 0.012 | 0.012 | 1 | 1 | Polr3d | 0 | Young |
| 0.95682 | 0.039517 | 0.013 | 0.013 | 1 | 1 | Atf5 | 0 | Young |
| 0.958022 | 0.319003 | 0.013 | 0.013 | 1 | 1 | Zdhhc12 | 0 | Young |
| 0.958271 | 0.304254 | 0.012 | 0.012 | 1 | 1 | Gm49359 | 0 | Young |
| 0.958271 | 0.072142 | 0.012 | 0.012 | 1 | 1 | 4930522L1 | 0 | Young |
| 0.960086 | 0.154458 | 0.052 | 0.051 | 1 | 1 | Zfp281 | 0.001 | Young |
| 0.961707 | 0.015222 | 0.012 | 0.012 | 1 | 1 | Zfp874a | 0 | Young |
| 0.962032 | 0.050856 | 0.047 | 0.046 | 1 | 1 | Acadvl | 0.001 | Young |
| 0.964396 | 0.059345 | 0.099 | 0.099 | 1 | 1 | Il3ra | 0 | Young |
| 0.966059 | 1.076218 | 0.016 | 0.015 | 1 | 1 | BC002059 | 0.001 | Young |
| 0.96625 | 0.112463 | 0.032 | 0.032 | 1 | 1 | Tdp2 | 0 | Young |
| 0.969519 | 0.00905 | 0.053 | 0.053 | 1 | 1 | Ormdl2 | 0 | Young |
| 0.969858 | 0.427368 | 0.018 | 0.018 | 1 | 1 | Actr5 | 0 | Young |
| 0.971383 | 0.089377 | 0.018 | 0.018 | 1 | 1 | Tdrd3 | 0 | Young |
| 0.972421 | 0.31106 | 0.021 | 0.021 | 1 | 1 | Kif3a | 0 | Young |

| | | | | | | | | |
|---|---|---|---|---|---|---|---|---|
| 0.972715 | 0.140746 | 0.062 | 0.062 | 1 | 1 | Lgals8 | 0 | Young |
| 0.974551 | 0.307617 | 0.021 | 0.021 | 1 | 1 | Nob1 | 0 | Young |
| 0.97497 | 0.038896 | 0.037 | 0.037 | 1 | 1 | Mapk8ip3 | 0 | Young |
| 0.976197 | 0.107684 | 0.031 | 0.031 | 1 | 1 | Apeh | 0 | Young |
| 0.976751 | 0.063426 | 0.037 | 0.037 | 1 | 1 | Nrbf2 | 0 | Young |
| 0.977814 | 0.119796 | 0.026 | 0.026 | 1 | 1 | Gpn1 | 0 | Young |
| 0.977864 | 0.19837 | 0.021 | 0.021 | 1 | 1 | Kctd5 | 0 | Young |
| 0.979048 | 0.183044 | 0.021 | 0.021 | 1 | 1 | Bclaf3 | 0 | Young |
| 0.979758 | 0.206476 | 0.021 | 0.021 | 1 | 1 | Adprm | 0 | Young |
| 0.980346 | 0.211532 | 0.023 | 0.023 | 1 | 1 | Alad | 0 | Young |
| 0.980642 | 0.183791 | 0.044 | 0.044 | 1 | 1 | Dok1 | 0 | Young |
| 0.981191 | 0.001096 | 0.152 | 0.152 | 1 | 1 | Soat1 | 0 | Young |
| 0.982484 | 0.054662 | 0.026 | 0.026 | 1 | 1 | Aatf | 0 | Young |
| 0.982717 | 0.214813 | 0.035 | 0.035 | 1 | 1 | Kifc3 | 0 | Young |
| 0.983164 | 0.040828 | 0.037 | 0.037 | 1 | 1 | Letm1 | 0 | Young |
| 0.983179 | 0.421672 | 0.01 | 0.01 | 1 | 1 | Cnr2 | 0 | Young |
| 0.983308 | 0.021517 | 0.021 | 0.021 | 1 | 1 | Uqcc1 | 0 | Young |
| 0.984321 | 0.039952 | 0.016 | 0.015 | 1 | 1 | Ptcd3 | 0.001 | Young |
| 0.984853 | 0.041886 | 0.014 | 0.014 | 1 | 1 | Fbxl19 | 0 | Young |
| 0.985087 | 0.242613 | 0.033 | 0.033 | 1 | 1 | Usp5 | 0 | Young |
| 0.985202 | 0.050098 | 0.021 | 0.021 | 1 | 1 | Hint3 | 0 | Young |
| 0.98551 | 0.364088 | 0.01 | 0.01 | 1 | 1 | Gpatch3 | 0 | Young |
| 0.986106 | 0.039275 | 0.031 | 0.031 | 1 | 1 | Atad3a | 0 | Young |
| 0.986842 | 0.270061 | 0.01 | 0.01 | 1 | 1 | Glcci1 | 0 | Young |
| 0.988175 | 0.249501 | 0.01 | 0.01 | 1 | 1 | Eif2b1 | 0 | Young |
| 0.988175 | 0.168345 | 0.01 | 0.01 | 1 | 1 | Gin1 | 0 | Young |
| 0.989007 | 0.106536 | 0.01 | 0.01 | 1 | 1 | Amd1 | 0 | Young |
| 0.989227 | 0.074047 | 0.021 | 0.021 | 1 | 1 | Vipas39 | 0 | Young |
| 0.990004 | 0.296117 | 0.023 | 0.023 | 1 | 1 | Rfng | 0 | Young |
| 0.990854 | 0.127883 | 0.014 | 0.014 | 1 | 1 | Zfp579 | 0 | Young |
| 0.990943 | 0.270674 | 0.028 | 0.028 | 1 | 1 | Med11 | 0 | Young |
| 0.991172 | 0.005943 | 0.01 | 0.01 | 1 | 1 | Nmrk1 | 0 | Young |
| 0.991553 | 0.122255 | 0.03 | 0.03 | 1 | 1 | Bag5 | 0 | Young |
| 0.992206 | 0.419369 | 0.046 | 0.046 | 1 | 1 | Mapkap1 | 0 | Young |
| 0.992804 | 0.095652 | 0.078 | 0.078 | 1 | 1 | Usp22 | 0 | Young |
| 0.993865 | 0.053602 | 0.087 | 0.087 | 1 | 1 | Sugt1 | 0 | Young |
| 0.999212 | 0.209048 | 0.038 | 0.039 | 1 | 1 | Ppid | -0.001 | Young |
| 0.999912 | 0.189002 | 0.038 | 0.039 | 1 | 1 | Sord | -0.001 | Young |
| 6.51E-08 | 0.893427 | 0.858 | 0.596 | 0.002021 | 10 | Anxa1 | 0.262 | Old |
| 3.68E-07 | 1.58542 | 0.46 | 0.182 | 0.01143 | 10 | Pros1 | 0.278 | Old |
| 2.20E-06 | 0.929499 | 0.557 | 0.253 | 0.068209 | 10 | Atp6v0d2 | 0.304 | Old |
| 3.09E-06 | 0.637372 | 0.949 | 0.768 | 0.095886 | 10 | Sdc3 | 0.181 | Old |
| 4.14E-06 | 1.205766 | 0.868 | 0.717 | 0.128694 | 10 | Ctsl | 0.151 | Old |
| 6.78E-06 | 0.534408 | 0.949 | 0.869 | 0.210644 | 10 | Anxa2 | 0.08 | Old |
| 9.51E-06 | 0.545052 | 0.985 | 0.939 | 0.295344 | 10 | Lpl | 0.046 | Old |

| | | | | | | | | |
|---|---|---|---|---|---|---|---|---|
| 9.75E-06 | 0.470126 | 0.976 | 0.949 | 0.302704 | 10 | Vim | 0.027 | Old |
| 1.39E-05 | 1.238898 | 0.496 | 0.253 | 0.430336 | 10 | B4galt5 | 0.243 | Old |
| 1.66E-05 | 1.060271 | 0.491 | 0.253 | 0.514963 | 10 | Igf2r | 0.238 | Old |
| 3.02E-05 | 0.647188 | 0.736 | 0.525 | 0.93865 | 10 | Tm6sf1 | 0.211 | Old |
| 4.20E-05 | 5.382208 | 0.186 | 0.02 | 1 | 10 | Cxcl13 | 0.166 | Old |
| 5.00E-05 | 2.267194 | 0.203 | 0.03 | 1 | 10 | Gm28875 | 0.173 | Old |
| 5.55E-05 | 0.493191 | 0.934 | 0.899 | 1 | 10 | Calm2 | 0.035 | Old |
| 5.96E-05 | 1.546898 | 0.254 | 0.061 | 1 | 10 | Abhd17c | 0.193 | Old |
| 6.24E-05 | 0.458173 | 0.848 | 0.717 | 1 | 10 | Lmna | 0.131 | Old |
| 7.48E-05 | 0.643939 | 0.861 | 0.747 | 1 | 10 | Pltp | 0.114 | Old |
| 9.27E-05 | 0.92963 | 0.523 | 0.313 | 1 | 10 | Shtn1 | 0.21 | Old |
| 0.000105 | 1.077526 | 0.633 | 0.455 | 1 | 10 | Clec10a | 0.178 | Old |
| 0.000131 | 0.431826 | 0.941 | 0.808 | 1 | 10 | Capzb | 0.133 | Old |
| 0.000133 | 1.3082 | 0.355 | 0.152 | 1 | 10 | Sik1 | 0.203 | Old |
| 0.000137 | 0.656793 | 0.719 | 0.495 | 1 | 10 | Glg1 | 0.224 | Old |
| 0.000138 | 0.5658 | 0.667 | 0.444 | 1 | 10 | Ifi27l2a | 0.223 | Old |
| 0.000158 | 1.001826 | 0.477 | 0.273 | 1 | 10 | B4galnt1 | 0.204 | Old |
| 0.000165 | 1.102008 | 0.425 | 0.232 | 1 | 10 | Fnbp1l | 0.193 | Old |
| 0.000172 | 0.723123 | 0.452 | 0.232 | 1 | 10 | Tspan31 | 0.22 | Old |
| 0.000191 | 1.772842 | 0.293 | 0.111 | 1 | 10 | Cyp51 | 0.182 | Old |
| 0.000201 | 0.640346 | 0.653 | 0.475 | 1 | 10 | Ppp1r18 | 0.178 | Old |
| 0.000216 | 0.838381 | 0.513 | 0.293 | 1 | 10 | Dusp5 | 0.22 | Old |
| 0.000223 | 0.475965 | 0.888 | 0.778 | 1 | 10 | Dbi | 0.11 | Old |
| 0.000239 | 0.521624 | 0.724 | 0.485 | 1 | 10 | Emp1 | 0.239 | Old |
| 0.000246 | 1.596606 | 0.215 | 0.051 | 1 | 10 | Zc3h4 | 0.164 | Old |
| 0.000266 | 0.933221 | 0.516 | 0.293 | 1 | 10 | Noct | 0.223 | Old |
| 0.000341 | 0.737847 | 0.655 | 0.455 | 1 | 10 | Sppl2a | 0.2 | Old |
| 0.000344 | 0.922601 | 0.477 | 0.263 | 1 | 10 | Dnajc8 | 0.214 | Old |
| 0.000383 | 0.682404 | 0.548 | 0.343 | 1 | 10 | Slc16a10 | 0.205 | Old |
| 0.00039 | 3.134127 | 0.178 | 0.04 | 1 | 10 | Fam198b | 0.138 | Old |
| 0.00039 | 0.978384 | 0.352 | 0.162 | 1 | 10 | Ear2 | 0.19 | Old |
| 0.000413 | 2.52869 | 0.169 | 0.03 | 1 | 10 | Tnip2 | 0.139 | Old |
| 0.000422 | 0.496343 | 0.973 | 0.909 | 1 | 10 | Itm2b | 0.064 | Old |
| 0.000433 | 2.328307 | 0.169 | 0.03 | 1 | 10 | Pccb | 0.139 | Old |
| 0.000434 | 0.994621 | 0.386 | 0.192 | 1 | 10 | Lrp6 | 0.194 | Old |
| 0.000435 | 0.453769 | 0.626 | 0.394 | 1 | 10 | Sh3kbp1 | 0.232 | Old |
| 0.000441 | 0.8884 | 0.565 | 0.384 | 1 | 10 | Dse | 0.181 | Old |
| 0.00046 | 0.354266 | 0.966 | 0.96 | 1 | 10 | Ahnak | 0.006 | Old |
| 0.000489 | 1.138055 | 0.35 | 0.162 | 1 | 10 | Chmp1a | 0.188 | Old |
| 0.000491 | 0.434675 | 0.978 | 0.808 | 1 | 10 | Ccl9 | 0.17 | Old |
| 0.000534 | 1.13159 | 0.281 | 0.111 | 1 | 10 | Chd8 | 0.17 | Old |
| 0.000559 | 0.295599 | 0.985 | 0.939 | 1 | 10 | B2m | 0.046 | Old |
| 0.000681 | 0.381394 | 0.993 | 1 | 1 | 10 | Malat1 | -0.007 | Old |
| 0.000687 | 0.690126 | 0.655 | 0.475 | 1 | 10 | H2-T23 | 0.18 | Old |
| 0.000722 | 1.69946 | 0.164 | 0.03 | 1 | 10 | Gm36161 | 0.134 | Old |

| | | | | | | | | |
|---|---|---|---|---|---|---|---|---|
| 0.000724 | 0.90329 | 0.526 | 0.313 | 1 | 10 | Odc1 | 0.213 | Old |
| 0.000746 | 0.546131 | 0.545 | 0.343 | 1 | 10 | Pycard | 0.202 | Old |
| 0.000769 | 0.372681 | 0.934 | 0.859 | 1 | 10 | Ptp4a2 | 0.075 | Old |
| 0.000783 | 1.009561 | 0.535 | 0.364 | 1 | 10 | Rasa1 | 0.171 | Old |
| 0.000823 | 0.969789 | 0.35 | 0.172 | 1 | 10 | Samm50 | 0.178 | Old |
| 0.000823 | 0.473458 | 0.902 | 0.889 | 1 | 10 | Atp6v1b2 | 0.013 | Old |
| 0.000839 | 1.291606 | 0.237 | 0.081 | 1 | 10 | Mrpl4 | 0.156 | Old |
| 0.000845 | 0.611035 | 0.55 | 0.333 | 1 | 10 | Zdhhc20 | 0.217 | Old |
| 0.000947 | 0.564457 | 0.572 | 0.374 | 1 | 10 | Adcy7 | 0.198 | Old |
| 0.001016 | 1.690643 | 0.225 | 0.081 | 1 | 10 | Cav2 | 0.144 | Old |
| 0.001018 | 0.859443 | 0.372 | 0.182 | 1 | 10 | Dnajb11 | 0.19 | Old |
| 0.001029 | 0.811186 | 0.396 | 0.212 | 1 | 10 | Uhrf1bp1l | 0.184 | Old |
| 0.00103 | 1.109001 | 0.235 | 0.081 | 1 | 10 | Msra | 0.154 | Old |
| 0.001032 | 1.100581 | 0.328 | 0.162 | 1 | 10 | 2210016F1 | 0.166 | Old |
| 0.001054 | 2.517795 | 0.137 | 0.02 | 1 | 10 | Nf2 | 0.117 | Old |
| 0.001058 | 0.733223 | 0.504 | 0.303 | 1 | 10 | Hip1 | 0.201 | Old |
| 0.001087 | 0.680558 | 0.655 | 0.495 | 1 | 10 | Coro1c | 0.16 | Old |
| 0.001092 | 1.134412 | 0.34 | 0.172 | 1 | 10 | Cdk4 | 0.168 | Old |
| 0.001101 | 0.52688 | 0.474 | 0.263 | 1 | 10 | Chmp5 | 0.211 | Old |
| 0.001112 | 2.44931 | 0.137 | 0.02 | 1 | 10 | Vps9d1 | 0.117 | Old |
| 0.001129 | 0.722279 | 0.423 | 0.242 | 1 | 10 | Dennd4c | 0.181 | Old |
| 0.001134 | 0.315092 | 0.77 | 0.596 | 1 | 10 | Atp5k | 0.174 | Old |
| 0.001168 | 0.832392 | 0.633 | 0.485 | 1 | 10 | St8sia4 | 0.148 | Old |
| 0.001181 | 0.329816 | 0.993 | 0.929 | 1 | 10 | Cst3 | 0.064 | Old |
| 0.00119 | 0.786182 | 0.34 | 0.172 | 1 | 10 | C5ar2 | 0.168 | Old |
| 0.001191 | 1.065696 | 0.386 | 0.212 | 1 | 10 | Cep350 | 0.174 | Old |
| 0.001195 | 0.434293 | 0.565 | 0.384 | 1 | 10 | Copa | 0.181 | Old |
| 0.001211 | 0.573429 | 0.636 | 0.444 | 1 | 10 | Plekhb2 | 0.192 | Old |
| 0.001234 | 0.582578 | 0.535 | 0.313 | 1 | 10 | Foxn3 | 0.222 | Old |
| 0.001252 | 1.035543 | 0.386 | 0.212 | 1 | 10 | Bcar3 | 0.174 | Old |
| 0.001266 | 0.698543 | 0.362 | 0.182 | 1 | 10 | Bub3 | 0.18 | Old |
| 0.001287 | 0.726994 | 0.694 | 0.545 | 1 | 10 | Fabp4 | 0.149 | Old |
| 0.001368 | 0.819019 | 0.313 | 0.141 | 1 | 10 | Smpd3 | 0.172 | Old |
| 0.001426 | 0.691422 | 0.631 | 0.455 | 1 | 10 | Foxp1 | 0.176 | Old |
| 0.001495 | 0.360112 | 0.941 | 0.869 | 1 | 10 | Rtn4 | 0.072 | Old |
| 0.001497 | 0.814729 | 0.301 | 0.131 | 1 | 10 | Zdhhc21 | 0.17 | Old |
| 0.001583 | 2.224748 | 0.176 | 0.051 | 1 | 10 | Tusc3 | 0.125 | Old |
| 0.001589 | 0.500984 | 0.802 | 0.697 | 1 | 10 | Adgre1 | 0.105 | Old |
| 0.001648 | 0.460065 | 0.848 | 0.758 | 1 | 10 | 2010107E0 | 0.09 | Old |
| 0.001652 | 0.5707 | 0.741 | 0.586 | 1 | 10 | Evl | 0.155 | Old |
| 0.001708 | 1.270834 | 0.235 | 0.091 | 1 | 10 | Ncbp3 | 0.144 | Old |
| 0.001715 | 1.793835 | 0.191 | 0.061 | 1 | 10 | Eif2ak2 | 0.13 | Old |
| 0.00172 | 3.70661 | 0.11 | 0.01 | 1 | 10 | Net1 | 0.1 | Old |
| 0.001743 | 1.034805 | 0.315 | 0.162 | 1 | 10 | Entpd7 | 0.153 | Old |
| 0.001776 | 0.510392 | 0.692 | 0.545 | 1 | 10 | Sod1 | 0.147 | Old |

| | | | | | | | | |
|---|---|---|---|---|---|---|---|---|
| 0.001821 | 1.144093 | 0.267 | 0.121 | 1 | 10 | Cbfa2t3 | 0.146 | Old |
| 0.001867 | 1.363353 | 0.215 | 0.081 | 1 | 10 | Chm | 0.134 | Old |
| 0.001895 | 0.381919 | 0.836 | 0.717 | 1 | 10 | Lrp1 | 0.119 | Old |
| 0.001912 | 1.338889 | 0.22 | 0.081 | 1 | 10 | Dhrs9 | 0.139 | Old |
| 0.001928 | 4.249969 | 0.09 | 0 | 1 | 10 | Fbxw17 | 0.09 | Old |
| 0.001941 | 1.267713 | 0.235 | 0.091 | 1 | 10 | Vps13b | 0.144 | Old |
| 0.001987 | 0.851779 | 0.433 | 0.273 | 1 | 10 | Osbpl11 | 0.16 | Old |
| 0.002015 | 0.530036 | 0.44 | 0.253 | 1 | 10 | Csk | 0.187 | Old |
| 0.002043 | 0.619751 | 0.328 | 0.162 | 1 | 10 | Gas2l1 | 0.166 | Old |
| 0.002064 | 0.82664 | 0.418 | 0.242 | 1 | 10 | Itsn1 | 0.176 | Old |
| 0.002095 | 1.996903 | 0.159 | 0.04 | 1 | 10 | Exosc9 | 0.119 | Old |
| 0.00213 | 1.185944 | 0.318 | 0.162 | 1 | 10 | Grk3 | 0.156 | Old |
| 0.002149 | 1.835953 | 0.11 | 0.01 | 1 | 10 | Hnmt | 0.1 | Old |
| 0.002165 | 0.232287 | 0.804 | 0.606 | 1 | 10 | Ssr3 | 0.198 | Old |
| 0.00218 | 1.573531 | 0.174 | 0.051 | 1 | 10 | Utp6 | 0.123 | Old |
| 0.00222 | 0.395744 | 0.626 | 0.434 | 1 | 10 | Syngr2 | 0.192 | Old |
| 0.002299 | 0.441656 | 0.687 | 0.485 | 1 | 10 | Rpn2 | 0.202 | Old |
| 0.002351 | 0.323773 | 0.91 | 0.818 | 1 | 10 | Wnk1 | 0.092 | Old |
| 0.002362 | 0.38142 | 0.499 | 0.293 | 1 | 10 | Tcea1 | 0.206 | Old |
| 0.002432 | 0.398312 | 0.462 | 0.273 | 1 | 10 | Lrrc25 | 0.189 | Old |
| 0.002432 | 1.864239 | 0.127 | 0.02 | 1 | 10 | Pip4k2c | 0.107 | Old |
| 0.002447 | 0.64691 | 0.413 | 0.232 | 1 | 10 | Rc3h1 | 0.181 | Old |
| 0.002468 | 1.140085 | 0.293 | 0.141 | 1 | 10 | Arhgap1 | 0.152 | Old |
| 0.002542 | 1.025818 | 0.386 | 0.222 | 1 | 10 | Ptprs | 0.164 | Old |
| 0.002716 | 0.549179 | 0.526 | 0.364 | 1 | 10 | Birc6 | 0.162 | Old |
| 0.002791 | 0.5655 | 0.56 | 0.394 | 1 | 10 | Sec11c | 0.166 | Old |
| 0.002793 | 0.641101 | 0.572 | 0.394 | 1 | 10 | Pepd | 0.178 | Old |
| 0.002824 | 0.562017 | 0.545 | 0.343 | 1 | 10 | Slc27a1 | 0.202 | Old |
| 0.002911 | 0.583943 | 0.333 | 0.162 | 1 | 10 | Yme1l1 | 0.171 | Old |
| 0.002948 | 0.257527 | 0.721 | 0.505 | 1 | 10 | Ap2a2 | 0.216 | Old |
| 0.002986 | 0.422152 | 0.824 | 0.667 | 1 | 10 | Wdr1 | 0.157 | Old |
| 0.002987 | 0.376751 | 0.516 | 0.343 | 1 | 10 | Arhgap9 | 0.173 | Old |
| 0.00299 | 0.854139 | 0.428 | 0.283 | 1 | 10 | Galc | 0.145 | Old |
| 0.003026 | 3.712125 | 0.117 | 0.02 | 1 | 10 | Cd55 | 0.097 | Old |
| 0.003061 | 0.475821 | 0.509 | 0.303 | 1 | 10 | U2af2 | 0.206 | Old |
| 0.003066 | 0.524317 | 0.584 | 0.414 | 1 | 10 | Ier3ip1 | 0.17 | Old |
| 0.003217 | 0.907975 | 0.364 | 0.212 | 1 | 10 | Flii | 0.152 | Old |
| 0.003267 | 0.535493 | 0.567 | 0.404 | 1 | 10 | Mdh1 | 0.163 | Old |
| 0.003305 | 1.755729 | 0.137 | 0.03 | 1 | 10 | C2cd5 | 0.107 | Old |
| 0.003313 | 0.963892 | 0.418 | 0.253 | 1 | 10 | Mmp19 | 0.165 | Old |
| 0.003342 | 0.80581 | 0.413 | 0.253 | 1 | 10 | Nek7 | 0.16 | Old |
| 0.003385 | 0.705883 | 0.345 | 0.182 | 1 | 10 | Ppm1a | 0.163 | Old |
| 0.003406 | 0.546723 | 0.66 | 0.475 | 1 | 10 | Lat2 | 0.185 | Old |
| 0.003411 | 1.510931 | 0.24 | 0.111 | 1 | 10 | Slc38a9 | 0.129 | Old |
| 0.003418 | 0.723122 | 0.663 | 0.495 | 1 | 10 | Klf2 | 0.168 | Old |

| | | | | | | | | |
|---|---|---|---|---|---|---|---|---|
| 0.003552 | 0.77263 | 0.394 | 0.232 | 1 | 10 | Epsti1 | 0.162 | Old |
| 0.003562 | 1.213412 | 0.152 | 0.04 | 1 | 10 | Tmem250- | 0.112 | Old |
| 0.003664 | 0.428853 | 0.677 | 0.485 | 1 | 10 | Abcg1 | 0.192 | Old |
| 0.003713 | 0.746976 | 0.531 | 0.374 | 1 | 10 | Arhgap25 | 0.157 | Old |
| 0.003785 | 0.677909 | 0.572 | 0.394 | 1 | 10 | Etv3 | 0.178 | Old |
| 0.003798 | 0.939014 | 0.264 | 0.121 | 1 | 10 | Gtf2b | 0.143 | Old |
| 0.003878 | 0.350365 | 0.958 | 0.919 | 1 | 10 | Ccl6 | 0.039 | Old |
| 0.003893 | 0.585273 | 0.457 | 0.293 | 1 | 10 | Hadha | 0.164 | Old |
| 0.003899 | 0.238774 | 0.993 | 0.919 | 1 | 10 | Ctss | 0.074 | Old |
| 0.003993 | 0.257549 | 0.621 | 0.434 | 1 | 10 | Mbnl2 | 0.187 | Old |
| 0.004022 | 0.928755 | 0.421 | 0.273 | 1 | 10 | Eif2s3x | 0.148 | Old |
| 0.004089 | 1.177382 | 0.183 | 0.061 | 1 | 10 | Stk25 | 0.122 | Old |
| 0.004098 | 0.67992 | 0.208 | 0.081 | 1 | 10 | Fgl2 | 0.127 | Old |
| 0.004113 | 3.832442 | 0.078 | 0 | 1 | 10 | Atpaf2 | 0.078 | Old |
| 0.004132 | 0.481046 | 0.697 | 0.505 | 1 | 10 | Grk2 | 0.192 | Old |
| 0.004218 | 1.502609 | 0.178 | 0.061 | 1 | 10 | Lipo3 | 0.117 | Old |
| 0.00424 | 0.26769 | 0.963 | 0.899 | 1 | 10 | Plin2 | 0.064 | Old |
| 0.004406 | 0.663782 | 0.274 | 0.131 | 1 | 10 | Mrps25 | 0.143 | Old |
| 0.004419 | 1.803254 | 0.222 | 0.101 | 1 | 10 | Mink1 | 0.121 | Old |
| 0.004447 | 0.81832 | 0.318 | 0.172 | 1 | 10 | Gatad1 | 0.146 | Old |
| 0.004611 | 0.439874 | 0.878 | 0.778 | 1 | 10 | Ptprc | 0.1 | Old |
| 0.004639 | 0.39073 | 0.663 | 0.495 | 1 | 10 | Trip12 | 0.168 | Old |
| 0.00465 | 0.716173 | 0.235 | 0.101 | 1 | 10 | Suclg1 | 0.134 | Old |
| 0.004684 | 1.213437 | 0.276 | 0.141 | 1 | 10 | Ppdpf | 0.135 | Old |
| 0.004776 | 0.463137 | 0.868 | 0.727 | 1 | 10 | Prkcd | 0.141 | Old |
| 0.004818 | 0.288877 | 0.403 | 0.242 | 1 | 10 | Cd200r1 | 0.161 | Old |
| 0.004981 | 1.258935 | 0.188 | 0.071 | 1 | 10 | AI467606 | 0.117 | Old |
| 0.004982 | 0.618799 | 0.496 | 0.333 | 1 | 10 | Rgl1 | 0.163 | Old |
| 0.004993 | 0.42052 | 0.645 | 0.455 | 1 | 10 | 2900097C1 | 0.19 | Old |
| 0.005049 | 0.674 | 0.306 | 0.152 | 1 | 10 | Elp5 | 0.154 | Old |
| 0.00511 | 1.846172 | 0.144 | 0.04 | 1 | 10 | Pitrm1 | 0.104 | Old |
| 0.005146 | 0.776949 | 0.411 | 0.263 | 1 | 10 | Mrpl28 | 0.148 | Old |
| 0.005158 | 0.643984 | 0.364 | 0.202 | 1 | 10 | Sdc1 | 0.162 | Old |
| 0.005173 | 1.52169 | 0.161 | 0.051 | 1 | 10 | Zfp524 | 0.11 | Old |
| 0.005174 | 1.394173 | 0.235 | 0.111 | 1 | 10 | Ranbp10 | 0.124 | Old |
| 0.005272 | 0.458416 | 0.604 | 0.465 | 1 | 10 | Slc37a2 | 0.139 | Old |
| 0.005329 | 0.728315 | 0.372 | 0.212 | 1 | 10 | Agps | 0.16 | Old |
| 0.005375 | 3.746918 | 0.093 | 0.01 | 1 | 10 | Gm26870 | 0.083 | Old |
| 0.005385 | 0.443241 | 0.778 | 0.626 | 1 | 10 | Nceh1 | 0.152 | Old |
| 0.005405 | 0.426773 | 0.592 | 0.404 | 1 | 10 | Cct4 | 0.188 | Old |
| 0.005408 | 0.439287 | 0.775 | 0.626 | 1 | 10 | Prcp | 0.149 | Old |
| 0.005447 | 1.245227 | 0.232 | 0.111 | 1 | 10 | Coro2a | 0.121 | Old |
| 0.005525 | 0.488577 | 0.694 | 0.566 | 1 | 10 | Cndp2 | 0.128 | Old |
| 0.00556 | 3.428689 | 0.073 | 0 | 1 | 10 | Cep70 | 0.073 | Old |
| 0.005599 | 0.768403 | 0.259 | 0.121 | 1 | 10 | Wdr41 | 0.138 | Old |

| | | | | | | | | |
|---|---|---|---|---|---|---|---|---|
| 0.005709 | 0.103118 | 0.479 | 0.273 | 1 | 10 | Gpnmb | 0.206 | Old |
| 0.005722 | 0.842201 | 0.362 | 0.212 | 1 | 10 | Sptbn1 | 0.15 | Old |
| 0.005763 | 0.52796 | 0.677 | 0.545 | 1 | 10 | Tmem256 | 0.132 | Old |
| 0.005776 | 0.618804 | 0.399 | 0.232 | 1 | 10 | Arhgap10 | 0.167 | Old |
| 0.0059 | 0.848095 | 0.355 | 0.202 | 1 | 10 | Ddx54 | 0.153 | Old |
| 0.005934 | 0.726191 | 0.279 | 0.141 | 1 | 10 | Cops5 | 0.138 | Old |
| 0.006051 | 0.529304 | 0.667 | 0.495 | 1 | 10 | Asph | 0.172 | Old |
| 0.006073 | 0.633421 | 0.474 | 0.343 | 1 | 10 | Pla2g15 | 0.131 | Old |
| 0.006176 | 0.372454 | 0.501 | 0.343 | 1 | 10 | 6-Mar | 0.158 | Old |
| 0.006192 | 0.279352 | 0.929 | 0.869 | 1 | 10 | Gm10076 | 0.06 | Old |
| 0.006289 | 0.629299 | 0.443 | 0.293 | 1 | 10 | Rab6a | 0.15 | Old |
| 0.006311 | 1.717729 | 0.181 | 0.071 | 1 | 10 | Akap11 | 0.11 | Old |
| 0.006335 | 0.319799 | 0.848 | 0.727 | 1 | 10 | Dhrs3 | 0.121 | Old |
| 0.006348 | 1.065569 | 0.259 | 0.131 | 1 | 10 | Zmpste24 | 0.128 | Old |
| 0.006378 | 1.685172 | 0.142 | 0.04 | 1 | 10 | Slc10a7 | 0.102 | Old |
| 0.006402 | 1.131394 | 0.159 | 0.051 | 1 | 10 | Asb8 | 0.108 | Old |
| 0.006488 | 1.159281 | 0.249 | 0.121 | 1 | 10 | Bag6 | 0.128 | Old |
| 0.006539 | 0.351134 | 0.841 | 0.717 | 1 | 10 | Aplp2 | 0.124 | Old |
| 0.006539 | 1.217509 | 0.225 | 0.101 | 1 | 10 | Syvn1 | 0.124 | Old |
| 0.006581 | 0.484389 | 0.425 | 0.242 | 1 | 10 | Fam129b | 0.183 | Old |
| 0.006591 | 0.221921 | 0.998 | 0.939 | 1 | 10 | Gpx1 | 0.059 | Old |
| 0.006597 | 0.97721 | 0.337 | 0.202 | 1 | 10 | St6galnac4 | 0.135 | Old |
| 0.006701 | 0.518351 | 0.636 | 0.475 | 1 | 10 | Itpkb | 0.161 | Old |
| 0.006829 | 0.904762 | 0.303 | 0.172 | 1 | 10 | Ntan1 | 0.131 | Old |
| 0.00684 | 0.726927 | 0.406 | 0.263 | 1 | 10 | Asap1 | 0.143 | Old |
| 0.006866 | 1.206551 | 0.198 | 0.081 | 1 | 10 | Ift52 | 0.117 | Old |
| 0.006898 | 0.64359 | 0.389 | 0.232 | 1 | 10 | Tial1 | 0.157 | Old |
| 0.006945 | 0.44019 | 0.709 | 0.525 | 1 | 10 | Gns | 0.184 | Old |
| 0.006973 | 1.487938 | 0.166 | 0.061 | 1 | 10 | Ak3 | 0.105 | Old |
| 0.006993 | 0.514501 | 0.665 | 0.545 | 1 | 10 | Serpinb6a | 0.12 | Old |
| 0.007034 | 1.572191 | 0.181 | 0.071 | 1 | 10 | Gdf3 | 0.11 | Old |
| 0.007106 | 0.484107 | 0.355 | 0.202 | 1 | 10 | Sike1 | 0.153 | Old |
| 0.007135 | 0.45795 | 0.584 | 0.424 | 1 | 10 | Fgd4 | 0.16 | Old |
| 0.007155 | 1.608164 | 0.137 | 0.04 | 1 | 10 | Slc35b3 | 0.097 | Old |
| 0.007357 | 0.668596 | 0.252 | 0.121 | 1 | 10 | Jkamp | 0.131 | Old |
| 0.007423 | 0.608356 | 0.428 | 0.293 | 1 | 10 | Cpt1a | 0.135 | Old |
| 0.007443 | 1.019612 | 0.279 | 0.141 | 1 | 10 | Myo10 | 0.138 | Old |
| 0.007505 | 0.986045 | 0.169 | 0.061 | 1 | 10 | Fubp3 | 0.108 | Old |
| 0.007513 | 3.849878 | 0.068 | 0 | 1 | 10 | Gnb4 | 0.068 | Old |
| 0.007648 | 0.756784 | 0.227 | 0.101 | 1 | 10 | Smad2 | 0.126 | Old |
| 0.007718 | 2.975905 | 0.088 | 0.01 | 1 | 10 | Tmem267 | 0.078 | Old |
| 0.007875 | 2.117449 | 0.12 | 0.03 | 1 | 10 | Ikbkap | 0.09 | Old |
| 0.007907 | 0.69052 | 0.269 | 0.141 | 1 | 10 | Ggta1 | 0.128 | Old |
| 0.00792 | 0.77016 | 0.315 | 0.172 | 1 | 10 | Tcp11l2 | 0.143 | Old |
| 0.007999 | 0.441955 | 0.68 | 0.505 | 1 | 10 | Syk | 0.175 | Old |

| | | | | | | | | |
|---|---|---|---|---|---|---|---|---|
| 0.008069 | 1.286895 | 0.139 | 0.04 | 1 | 10 | Deptor | 0.099 | Old |
| 0.008088 | 0.219488 | 0.587 | 0.394 | 1 | 10 | Ndufa1 | 0.193 | Old |
| 0.008189 | 0.486813 | 0.472 | 0.313 | 1 | 10 | Pitpnc1 | 0.159 | Old |
| 0.008254 | 1.300879 | 0.32 | 0.192 | 1 | 10 | Kitl | 0.128 | Old |
| 0.008315 | 0.699694 | 0.369 | 0.212 | 1 | 10 | Ubr5 | 0.157 | Old |
| 0.008319 | 0.817578 | 0.462 | 0.323 | 1 | 10 | Acbd5 | 0.139 | Old |
| 0.008391 | 0.515761 | 0.482 | 0.313 | 1 | 10 | Pdxk | 0.169 | Old |
| 0.008489 | 0.490366 | 0.567 | 0.394 | 1 | 10 | Fuca1 | 0.173 | Old |
| 0.008497 | 0.87061 | 0.284 | 0.152 | 1 | 10 | Arid1b | 0.132 | Old |
| 0.008561 | 0.66033 | 0.259 | 0.121 | 1 | 10 | Pam | 0.138 | Old |
| 0.008597 | 0.427444 | 0.829 | 0.737 | 1 | 10 | Pld3 | 0.092 | Old |
| 0.008665 | 0.331222 | 0.966 | 0.919 | 1 | 10 | BC005537 | 0.047 | Old |
| 0.008697 | 1.778812 | 0.134 | 0.04 | 1 | 10 | Rtp4 | 0.094 | Old |
| 0.008732 | 3.471453 | 0.066 | 0 | 1 | 10 | Tmed8 | 0.066 | Old |
| 0.008981 | 1.841594 | 0.174 | 0.071 | 1 | 10 | Clasp1 | 0.103 | Old |
| 0.009034 | 0.805769 | 0.323 | 0.182 | 1 | 10 | Eif6 | 0.141 | Old |
| 0.009104 | 0.852771 | 0.311 | 0.182 | 1 | 10 | Mgat2 | 0.129 | Old |
| 0.009174 | 0.836222 | 0.501 | 0.374 | 1 | 10 | Naaa | 0.127 | Old |
| 0.009179 | 0.905459 | 0.279 | 0.141 | 1 | 10 | Fam192a | 0.138 | Old |
| 0.00921 | 0.37908 | 0.521 | 0.343 | 1 | 10 | Ppp1cb | 0.178 | Old |
| 0.009223 | 0.39345 | 0.592 | 0.404 | 1 | 10 | Tgoln1 | 0.188 | Old |
| 0.009235 | 0.65061 | 0.421 | 0.283 | 1 | 10 | Spg21 | 0.138 | Old |
| 0.009273 | 0.409315 | 0.729 | 0.616 | 1 | 10 | Nptn | 0.113 | Old |
| 0.00932 | 0.394024 | 0.377 | 0.212 | 1 | 10 | Tsg101 | 0.165 | Old |
| 0.009453 | 0.349117 | 0.978 | 0.869 | 1 | 10 | mt-Nd3 | 0.109 | Old |
| 0.009579 | 1.332828 | 0.12 | 0.03 | 1 | 10 | Acot2 | 0.09 | Old |
| 0.009638 | 0.988944 | 0.359 | 0.222 | 1 | 10 | Ighm | 0.137 | Old |
| 0.009652 | 0.505197 | 0.599 | 0.444 | 1 | 10 | Xiap | 0.155 | Old |
| 0.009686 | 0.283036 | 0.692 | 0.525 | 1 | 10 | Apbb1ip | 0.167 | Old |
| 0.009702 | 1.076937 | 0.181 | 0.071 | 1 | 10 | Tbc1d17 | 0.11 | Old |
| 0.009707 | 0.630916 | 0.274 | 0.141 | 1 | 10 | Hyou1 | 0.133 | Old |
| 0.009781 | 1.654368 | 0.147 | 0.051 | 1 | 10 | Pigu | 0.096 | Old |
| 0.009786 | 1.121865 | 0.215 | 0.101 | 1 | 10 | Ube3b | 0.114 | Old |
| 0.009847 | 1.56819 | 0.174 | 0.071 | 1 | 10 | Ago4 | 0.103 | Old |
| 0.010065 | 0.738811 | 0.443 | 0.303 | 1 | 10 | Wls | 0.14 | Old |
| 0.010107 | 0.357181 | 0.658 | 0.495 | 1 | 10 | Clec4a3 | 0.163 | Old |
| 0.010186 | 0.560387 | 0.347 | 0.202 | 1 | 10 | Htt | 0.145 | Old |
| 0.010352 | 0.532012 | 0.499 | 0.343 | 1 | 10 | Irf5 | 0.156 | Old |
| 0.01043 | 0.469519 | 0.557 | 0.414 | 1 | 10 | Cdc42se1 | 0.143 | Old |
| 0.010437 | 0.940002 | 0.23 | 0.111 | 1 | 10 | Lpcat1 | 0.119 | Old |
| 0.010529 | 0.357115 | 0.665 | 0.535 | 1 | 10 | Serinc1 | 0.13 | Old |
| 0.010606 | 1.21222 | 0.247 | 0.131 | 1 | 10 | Idi1 | 0.116 | Old |
| 0.01068 | 0.370972 | 0.682 | 0.535 | 1 | 10 | Txndc17 | 0.147 | Old |
| 0.010698 | 0.561358 | 0.474 | 0.323 | 1 | 10 | Lcp2 | 0.151 | Old |
| 0.010731 | 0.481795 | 0.303 | 0.172 | 1 | 10 | Tmem106k | 0.131 | Old |

| | | | | | | | | |
|---|---|---|---|---|---|---|---|---|
| 0.010738 | 1.051524 | 0.252 | 0.131 | 1 | 10 | Steap3 | 0.121 | Old |
| 0.010745 | 0.155196 | 0.998 | 1 | 1 | 10 | mt-Co1 | -0.002 | Old |
| 0.010782 | 0.161146 | 0.445 | 0.293 | 1 | 10 | Qdpr | 0.152 | Old |
| 0.010831 | 1.233972 | 0.188 | 0.081 | 1 | 10 | Ell | 0.107 | Old |
| 0.010896 | 1.938894 | 0.083 | 0.01 | 1 | 10 | Slc13a3 | 0.073 | Old |
| 0.010997 | 0.340064 | 0.509 | 0.333 | 1 | 10 | Tmem165 | 0.176 | Old |
| 0.011031 | 0.592786 | 0.416 | 0.273 | 1 | 10 | Cd37 | 0.143 | Old |
| 0.011032 | 0.640448 | 0.24 | 0.111 | 1 | 10 | Usp6nl | 0.129 | Old |
| 0.011089 | 0.993703 | 0.203 | 0.091 | 1 | 10 | Kdm6a | 0.112 | Old |
| 0.011099 | 0.877467 | 0.244 | 0.121 | 1 | 10 | Trabd | 0.123 | Old |
| 0.011113 | 0.708511 | 0.384 | 0.242 | 1 | 10 | Dynlt3 | 0.142 | Old |
| 0.011199 | 0.800571 | 0.323 | 0.192 | 1 | 10 | Eif2a | 0.131 | Old |
| 0.011303 | 2.585697 | 0.098 | 0.02 | 1 | 10 | Crim1 | 0.078 | Old |
| 0.011319 | 0.495657 | 0.531 | 0.394 | 1 | 10 | Gsk3b | 0.137 | Old |
| 0.011364 | 0.710155 | 0.34 | 0.202 | 1 | 10 | Rbbp4 | 0.138 | Old |
| 0.011573 | 0.731279 | 0.218 | 0.101 | 1 | 10 | Sucla2 | 0.117 | Old |
| 0.011574 | 0.517454 | 0.658 | 0.515 | 1 | 10 | Daglb | 0.143 | Old |
| 0.01168 | 1.202049 | 0.147 | 0.051 | 1 | 10 | Ptgr1 | 0.096 | Old |
| 0.011776 | 2.215069 | 0.098 | 0.02 | 1 | 10 | Hsf2 | 0.078 | Old |
| 0.011794 | 3.521028 | 0.061 | 0 | 1 | 10 | Clybl | 0.061 | Old |
| 0.011794 | 3.313253 | 0.061 | 0 | 1 | 10 | Lyrm9 | 0.061 | Old |
| 0.011794 | 3.120887 | 0.061 | 0 | 1 | 10 | Trim14 | 0.061 | Old |
| 0.011861 | 1.416687 | 0.208 | 0.101 | 1 | 10 | Ctsk | 0.107 | Old |
| 0.011975 | 0.28152 | 0.792 | 0.667 | 1 | 10 | Tmem106a | 0.125 | Old |
| 0.012066 | 1.055399 | 0.289 | 0.172 | 1 | 10 | Stat1 | 0.117 | Old |
| 0.01217 | 2.696185 | 0.095 | 0.02 | 1 | 10 | Reep4 | 0.075 | Old |
| 0.012361 | 0.417693 | 0.68 | 0.515 | 1 | 10 | Xbp1 | 0.165 | Old |
| 0.012427 | 1.870215 | 0.281 | 0.172 | 1 | 10 | Mmp9 | 0.109 | Old |
| 0.012433 | 0.59024 | 0.335 | 0.192 | 1 | 10 | C1d | 0.143 | Old |
| 0.012482 | 0.15586 | 0.995 | 0.96 | 1 | 10 | Laptm5 | 0.035 | Old |
| 0.012512 | 0.477535 | 0.328 | 0.182 | 1 | 10 | Slc25a51 | 0.146 | Old |
| 0.012563 | 0.201131 | 0.998 | 1 | 1 | 10 | Rpl37a | -0.002 | Old |
| 0.012632 | 0.361195 | 0.653 | 0.495 | 1 | 10 | Plekho1 | 0.158 | Old |
| 0.012637 | 0.877161 | 0.315 | 0.192 | 1 | 10 | Rapgef6 | 0.123 | Old |
| 0.012659 | 1.271714 | 0.098 | 0.02 | 1 | 10 | Zfp26 | 0.078 | Old |
| 0.01272 | 0.409805 | 0.501 | 0.354 | 1 | 10 | 2610001J0 | 0.147 | Old |
| 0.012721 | 0.620576 | 0.24 | 0.121 | 1 | 10 | Arhgap39 | 0.119 | Old |
| 0.01278 | 1.040651 | 0.232 | 0.121 | 1 | 10 | Cops6 | 0.111 | Old |
| 0.012806 | 0.366187 | 0.641 | 0.475 | 1 | 10 | Spcs1 | 0.166 | Old |
| 0.0129 | 0.497731 | 0.469 | 0.313 | 1 | 10 | Nono | 0.156 | Old |
| 0.013479 | 0.497472 | 0.301 | 0.162 | 1 | 10 | Gpat4 | 0.139 | Old |
| 0.01366 | 1.476464 | 0.098 | 0.02 | 1 | 10 | Dnajc9 | 0.078 | Old |
| 0.013681 | 1.147078 | 0.142 | 0.051 | 1 | 10 | Ube2l6 | 0.091 | Old |
| 0.013707 | 3.501451 | 0.059 | 0 | 1 | 10 | Tab3 | 0.059 | Old |
| 0.013715 | 0.495073 | 0.386 | 0.242 | 1 | 10 | Bud31 | 0.144 | Old |

| | | | | | | | | |
|---|---|---|---|---|---|---|---|---|
| 0.013732 | 0.555791 | 0.447 | 0.303 | 1 | 10 | Blvra | 0.144 | Old |
| 0.013738 | 0.741646 | 0.306 | 0.172 | 1 | 10 | Atp7a | 0.134 | Old |
| 0.013791 | 1.800493 | 0.139 | 0.051 | 1 | 10 | Prune1 | 0.088 | Old |
| 0.013849 | 0.319083 | 0.753 | 0.586 | 1 | 10 | Cox7a2l | 0.167 | Old |
| 0.013857 | 0.357865 | 0.308 | 0.172 | 1 | 10 | Crebrf | 0.136 | Old |
| 0.013915 | 0.468224 | 0.499 | 0.323 | 1 | 10 | Cd300lb | 0.176 | Old |
| 0.014047 | 0.971766 | 0.377 | 0.253 | 1 | 10 | Galnt7 | 0.124 | Old |
| 0.014174 | 0.237239 | 0.998 | 0.97 | 1 | 10 | mt-Nd2 | 0.028 | Old |
| 0.014348 | 0.573415 | 0.308 | 0.172 | 1 | 10 | Ubqln1 | 0.136 | Old |
| 0.01438 | 1.054135 | 0.183 | 0.081 | 1 | 10 | Riok1 | 0.102 | Old |
| 0.014499 | 1.763929 | 0.125 | 0.04 | 1 | 10 | Wdr7 | 0.085 | Old |
| 0.01453 | 0.288922 | 0.13 | 0.04 | 1 | 10 | Sart3 | 0.09 | Old |
| 0.014613 | 1.943716 | 0.125 | 0.04 | 1 | 10 | Nt5dc2 | 0.085 | Old |
| 0.01471 | 0.244141 | 0.834 | 0.727 | 1 | 10 | Ywhah | 0.107 | Old |
| 0.014738 | 0.538881 | 0.232 | 0.111 | 1 | 10 | Atxn3 | 0.121 | Old |
| 0.014791 | 0.371822 | 0.24 | 0.121 | 1 | 10 | Nudt16l1 | 0.119 | Old |
| 0.014814 | 2.524029 | 0.093 | 0.02 | 1 | 10 | Lclat1 | 0.073 | Old |
| 0.014855 | 1.725481 | 0.11 | 0.03 | 1 | 10 | Aifm1 | 0.08 | Old |
| 0.014969 | 1.741976 | 0.139 | 0.051 | 1 | 10 | Gm5150 | 0.088 | Old |
| 0.014986 | 1.380965 | 0.112 | 0.03 | 1 | 10 | Gucd1 | 0.082 | Old |
| 0.015302 | 0.624472 | 0.443 | 0.303 | 1 | 10 | Pcyt1a | 0.14 | Old |
| 0.015306 | 0.396255 | 0.692 | 0.556 | 1 | 10 | Scp2 | 0.136 | Old |
| 0.015391 | 0.443919 | 0.846 | 0.758 | 1 | 10 | Zfp36l2 | 0.088 | Old |
| 0.015404 | 1.232547 | 0.156 | 0.061 | 1 | 10 | Stat2 | 0.095 | Old |
| 0.015432 | 0.235566 | 0.765 | 0.657 | 1 | 10 | Pdcd6ip | 0.108 | Old |
| 0.015444 | 0.287285 | 0.677 | 0.495 | 1 | 10 | Vwa5a | 0.182 | Old |
| 0.015465 | 0.294303 | 0.614 | 0.444 | 1 | 10 | Rhog | 0.17 | Old |
| 0.015486 | 1.778184 | 0.152 | 0.061 | 1 | 10 | Asxl1 | 0.091 | Old |
| 0.015545 | 0.318256 | 0.841 | 0.717 | 1 | 10 | Cops9 | 0.124 | Old |
| 0.015965 | 0.180617 | 0.993 | 0.98 | 1 | 10 | Rps26 | 0.013 | Old |
| 0.016018 | 0.844056 | 0.183 | 0.081 | 1 | 10 | Casp1 | 0.102 | Old |
| 0.016054 | 0.633392 | 0.259 | 0.141 | 1 | 10 | Csnk1g2 | 0.118 | Old |
| 0.016069 | 0.278479 | 0.912 | 0.818 | 1 | 10 | S100a6 | 0.094 | Old |
| 0.0161 | 1.600009 | 0.152 | 0.061 | 1 | 10 | Nfyc | 0.091 | Old |
| 0.016264 | 0.753563 | 0.112 | 0.03 | 1 | 10 | Rnf214 | 0.082 | Old |
| 0.016295 | 1.064239 | 0.254 | 0.141 | 1 | 10 | Agtrap | 0.113 | Old |
| 0.016319 | 1.123684 | 0.23 | 0.121 | 1 | 10 | Gmps | 0.109 | Old |
| 0.016515 | 0.929483 | 0.267 | 0.152 | 1 | 10 | Lpcat3 | 0.115 | Old |
| 0.016606 | 0.589587 | 0.289 | 0.162 | 1 | 10 | Pip5k1a | 0.127 | Old |
| 0.01671 | 0.363128 | 0.389 | 0.232 | 1 | 10 | Peak1 | 0.157 | Old |
| 0.016831 | 0.540158 | 0.484 | 0.343 | 1 | 10 | Man2a1 | 0.141 | Old |
| 0.016837 | 1.721057 | 0.149 | 0.061 | 1 | 10 | Polr3gl | 0.088 | Old |
| 0.016913 | 0.604877 | 0.391 | 0.263 | 1 | 10 | Baz2b | 0.128 | Old |
| 0.016982 | 0.17428 | 0.526 | 0.323 | 1 | 10 | Adipor1 | 0.203 | Old |
| 0.017058 | 1.182251 | 0.166 | 0.071 | 1 | 10 | Dcun1d1 | 0.095 | Old |

| | | | | | | | | |
|---|---|---|---|---|---|---|---|---|
| 0.017199 | 0.388446 | 0.183 | 0.081 | 1 | 10 | Cnot7 | 0.102 | Old |
| 0.017315 | 0.468246 | 0.445 | 0.293 | 1 | 10 | Cnot1 | 0.152 | Old |
| 0.017316 | 3.421752 | 0.073 | 0.01 | 1 | 10 | Bud13 | 0.063 | Old |
| 0.017347 | 1.073844 | 0.166 | 0.071 | 1 | 10 | Farsa | 0.095 | Old |
| 0.017404 | 1.058238 | 0.281 | 0.172 | 1 | 10 | Fam49a | 0.109 | Old |
| 0.017529 | 0.785922 | 0.355 | 0.222 | 1 | 10 | Ddx50 | 0.133 | Old |
| 0.01754 | 0.440382 | 0.337 | 0.202 | 1 | 10 | Tecr | 0.135 | Old |
| 0.01761 | 0.283646 | 0.401 | 0.263 | 1 | 10 | Cnih1 | 0.138 | Old |
| 0.01766 | 0.601273 | 0.347 | 0.232 | 1 | 10 | Ube2j2 | 0.115 | Old |
| 0.017686 | 0.442346 | 0.413 | 0.263 | 1 | 10 | Mdm2 | 0.15 | Old |
| 0.017739 | 0.446018 | 0.513 | 0.384 | 1 | 10 | Rnh1 | 0.129 | Old |
| 0.017753 | 2.7916 | 0.073 | 0.01 | 1 | 10 | Airn | 0.063 | Old |
| 0.017829 | 1.216855 | 0.237 | 0.131 | 1 | 10 | Bsdc1 | 0.106 | Old |
| 0.017891 | 0.555435 | 0.479 | 0.333 | 1 | 10 | Etfb | 0.146 | Old |
| 0.017982 | 0.385136 | 0.641 | 0.485 | 1 | 10 | Lims1 | 0.156 | Old |
| 0.018065 | 0.612784 | 0.364 | 0.232 | 1 | 10 | Hsd17b4 | 0.132 | Old |
| 0.018169 | 0.612687 | 0.411 | 0.283 | 1 | 10 | Ap1s2 | 0.128 | Old |
| 0.01827 | 0.468876 | 0.675 | 0.566 | 1 | 10 | Jmjd1c | 0.109 | Old |
| 0.018331 | 0.543761 | 0.267 | 0.152 | 1 | 10 | Ptdss1 | 0.115 | Old |
| 0.018332 | 2.192761 | 0.09 | 0.02 | 1 | 10 | Fbxl19 | 0.07 | Old |
| 0.018341 | 0.553075 | 0.421 | 0.303 | 1 | 10 | Rell1 | 0.118 | Old |
| 0.01843 | 0.665541 | 0.254 | 0.141 | 1 | 10 | Plekhf2 | 0.113 | Old |
| 0.018476 | 1.632288 | 0.108 | 0.03 | 1 | 10 | Pcsk7 | 0.078 | Old |
| 0.018517 | 3.495725 | 0.054 | 0 | 1 | 10 | Mill2 | 0.054 | Old |
| 0.018517 | 3.282052 | 0.054 | 0 | 1 | 10 | 1110032A( | 0.054 | Old |
| 0.018517 | 2.972512 | 0.054 | 0 | 1 | 10 | Mgmt | 0.054 | Old |
| 0.018592 | 0.939623 | 0.313 | 0.202 | 1 | 10 | F7 | 0.111 | Old |
| 0.018626 | 1.379016 | 0.2 | 0.101 | 1 | 10 | Ankle2 | 0.099 | Old |
| 0.018791 | 0.220229 | 0.32 | 0.182 | 1 | 10 | Dars | 0.138 | Old |
| 0.019095 | 1.03354 | 0.191 | 0.091 | 1 | 10 | Gnptab | 0.1 | Old |
| 0.019127 | 1.734232 | 0.134 | 0.051 | 1 | 10 | Akap8l | 0.083 | Old |
| 0.019213 | 0.450203 | 0.433 | 0.283 | 1 | 10 | Mcfd2 | 0.15 | Old |
| 0.019217 | 1.996358 | 0.073 | 0.01 | 1 | 10 | Tmem71 | 0.063 | Old |
| 0.019359 | 1.084517 | 0.193 | 0.091 | 1 | 10 | Syne2 | 0.102 | Old |
| 0.019417 | 0.096433 | 0.213 | 0.101 | 1 | 10 | Trnt1 | 0.112 | Old |
| 0.019442 | 0.686804 | 0.225 | 0.111 | 1 | 10 | Trim47 | 0.114 | Old |
| 0.019548 | 0.438754 | 0.142 | 0.051 | 1 | 10 | Cdk20 | 0.091 | Old |
| 0.019622 | 0.421248 | 0.396 | 0.263 | 1 | 10 | Tmem164 | 0.133 | Old |
| 0.01972 | 0.522706 | 0.511 | 0.404 | 1 | 10 | Uhrf2 | 0.107 | Old |
| 0.019844 | 0.415113 | 0.638 | 0.475 | 1 | 10 | Tcirg1 | 0.163 | Old |
| 0.019846 | 0.970798 | 0.205 | 0.101 | 1 | 10 | Ylpm1 | 0.104 | Old |
| 0.019902 | 0.201319 | 0.46 | 0.313 | 1 | 10 | Brd7 | 0.147 | Old |
| 0.019934 | 0.263743 | 0.819 | 0.717 | 1 | 10 | Tkt | 0.102 | Old |
| 0.01995 | 0.177251 | 0.489 | 0.343 | 1 | 10 | Mia2 | 0.146 | Old |
| 0.020071 | 0.157067 | 0.995 | 1 | 1 | 10 | Rpl41 | -0.005 | Old |

| | | | | | | | | |
|---|---|---|---|---|---|---|---|---|
| 0.020077 | 0.435032 | 0.638 | 0.465 | 1 | 10 | 9-Sep | 0.173 | Old |
| 0.020103 | 0.557646 | 0.416 | 0.273 | 1 | 10 | Lamtor3 | 0.143 | Old |
| 0.020205 | 0.954106 | 0.252 | 0.141 | 1 | 10 | Pxk | 0.111 | Old |
| 0.020238 | 0.436603 | 0.677 | 0.556 | 1 | 10 | Myl12a | 0.121 | Old |
| 0.020571 | 0.491015 | 0.484 | 0.333 | 1 | 10 | Snrpb2 | 0.151 | Old |
| 0.020669 | 0.205854 | 0.137 | 0.051 | 1 | 10 | Akr1c13 | 0.086 | Old |
| 0.020676 | 1.157949 | 0.176 | 0.081 | 1 | 10 | Rufy1 | 0.095 | Old |
| 0.020683 | 0.455861 | 0.45 | 0.303 | 1 | 10 | Fli1 | 0.147 | Old |
| 0.020744 | 0.574873 | 0.137 | 0.051 | 1 | 10 | Ctbs | 0.086 | Old |
| 0.020826 | 0.558648 | 0.457 | 0.333 | 1 | 10 | Aco2 | 0.124 | Old |
| 0.020838 | 1.352398 | 0.09 | 0.02 | 1 | 10 | Hsd17b7 | 0.07 | Old |
| 0.020892 | 0.45656 | 0.516 | 0.374 | 1 | 10 | Polr2a | 0.142 | Old |
| 0.020974 | 0.431811 | 0.599 | 0.465 | 1 | 10 | Hpcal1 | 0.134 | Old |
| 0.021121 | 0.350008 | 0.257 | 0.131 | 1 | 10 | Commd1 | 0.126 | Old |
| 0.021208 | 0.739421 | 0.347 | 0.222 | 1 | 10 | Srrt | 0.125 | Old |
| 0.021382 | 0.390645 | 0.386 | 0.253 | 1 | 10 | Retreg2 | 0.133 | Old |
| 0.021513 | 0.37464 | 0.518 | 0.374 | 1 | 10 | Por | 0.144 | Old |
| 0.021521 | 0.285839 | 0.254 | 0.131 | 1 | 10 | Rfc1 | 0.123 | Old |
| 0.021609 | 0.190591 | 0.98 | 0.949 | 1 | 10 | Zeb2 | 0.031 | Old |
| 0.021614 | 1.077924 | 0.122 | 0.04 | 1 | 10 | Map3k5 | 0.082 | Old |
| 0.021951 | 0.93636 | 0.203 | 0.101 | 1 | 10 | Cstf2t | 0.102 | Old |
| 0.022014 | 0.461477 | 0.352 | 0.212 | 1 | 10 | Specc1 | 0.14 | Old |
| 0.022209 | 0.557221 | 0.235 | 0.121 | 1 | 10 | Hfe | 0.114 | Old |
| 0.022486 | 0.537625 | 0.379 | 0.263 | 1 | 10 | Ggh | 0.116 | Old |
| 0.022551 | 0.269983 | 0.352 | 0.212 | 1 | 10 | Agpat5 | 0.14 | Old |
| 0.022554 | 1.108367 | 0.105 | 0.03 | 1 | 10 | P2ry14 | 0.075 | Old |
| 0.022635 | 0.40949 | 0.721 | 0.596 | 1 | 10 | S100a4 | 0.125 | Old |
| 0.022751 | 0.816897 | 0.235 | 0.131 | 1 | 10 | Haus8 | 0.104 | Old |
| 0.022792 | 0.335443 | 0.694 | 0.556 | 1 | 10 | Cd93 | 0.138 | Old |
| 0.022831 | 0.318847 | 0.653 | 0.465 | 1 | 10 | Lamtor1 | 0.188 | Old |
| 0.022833 | 0.522298 | 0.357 | 0.242 | 1 | 10 | Tpcn1 | 0.115 | Old |
| 0.022866 | 0.285362 | 0.318 | 0.192 | 1 | 10 | Stip1 | 0.126 | Old |
| 0.022868 | 0.772035 | 0.225 | 0.121 | 1 | 10 | Chmp2b | 0.104 | Old |
| 0.02287 | 0.721606 | 0.289 | 0.172 | 1 | 10 | Chuk | 0.117 | Old |
| 0.022967 | 1.240225 | 0.132 | 0.051 | 1 | 10 | Usp39 | 0.081 | Old |
| 0.023212 | 0.465986 | 0.579 | 0.444 | 1 | 10 | Uap1l1 | 0.135 | Old |
| 0.023494 | 0.303837 | 0.768 | 0.616 | 1 | 10 | Pdia3 | 0.152 | Old |
| 0.023557 | 1.053555 | 0.171 | 0.081 | 1 | 10 | Phyh | 0.09 | Old |
| 0.023572 | 0.770995 | 0.105 | 0.03 | 1 | 10 | 2300009A( | 0.075 | Old |
| 0.023659 | 0.508835 | 0.46 | 0.323 | 1 | 10 | Emilin2 | 0.137 | Old |
| 0.023739 | 0.207557 | 0.743 | 0.626 | 1 | 10 | Wwp1 | 0.117 | Old |
| 0.023841 | 0.572964 | 0.411 | 0.273 | 1 | 10 | Mitf | 0.138 | Old |
| 0.023933 | 1.312211 | 0.171 | 0.081 | 1 | 10 | Tmem38b | 0.09 | Old |
| 0.024008 | 0.425244 | 0.178 | 0.081 | 1 | 10 | Thra | 0.097 | Old |
| 0.024019 | 1.486636 | 0.156 | 0.071 | 1 | 10 | Ext2 | 0.085 | Old |

| | | | | | | | | |
|---|---|---|---|---|---|---|---|---|
| 0.024042 | 0.502911 | 0.562 | 0.424 | 1 | 10 | Myo1e | 0.138 | Old |
| 0.024085 | 1.562777 | 0.103 | 0.03 | 1 | 10 | Morc2a | 0.073 | Old |
| 0.024403 | 0.492276 | 0.411 | 0.283 | 1 | 10 | Ap3d1 | 0.128 | Old |
| 0.02448 | 0.38183 | 0.191 | 0.091 | 1 | 10 | Foxj3 | 0.1 | Old |
| 0.024525 | 0.960479 | 0.149 | 0.061 | 1 | 10 | Rrp36 | 0.088 | Old |
| 0.024617 | 0.597801 | 0.188 | 0.091 | 1 | 10 | Acot9 | 0.097 | Old |
| 0.024647 | 0.91893 | 0.276 | 0.172 | 1 | 10 | Tec | 0.104 | Old |
| 0.024981 | 0.378047 | 0.357 | 0.242 | 1 | 10 | Ascc3 | 0.115 | Old |
| 0.025002 | 0.602399 | 0.313 | 0.192 | 1 | 10 | Dpep2 | 0.121 | Old |
| 0.02503 | 3.733018 | 0.049 | 0 | 1 | 10 | Serpinb6b | 0.049 | Old |
| 0.02503 | 3.209823 | 0.049 | 0 | 1 | 10 | Dpy19l3 | 0.049 | Old |
| 0.025043 | 1.934092 | 0.086 | 0.02 | 1 | 10 | Mettl21a | 0.066 | Old |
| 0.025095 | 0.187619 | 0.976 | 0.99 | 1 | 10 | Junb | -0.014 | Old |
| 0.025171 | 0.684334 | 0.391 | 0.273 | 1 | 10 | Tmem65 | 0.118 | Old |
| 0.025397 | 1.144857 | 0.156 | 0.071 | 1 | 10 | Nqo2 | 0.085 | Old |
| 0.025505 | 0.177584 | 0.516 | 0.374 | 1 | 10 | Tm9sf2 | 0.142 | Old |
| 0.02562 | 0.205087 | 0.988 | 0.949 | 1 | 10 | H2-K1 | 0.039 | Old |
| 0.025757 | 2.113187 | 0.068 | 0.01 | 1 | 10 | Runx2 | 0.058 | Old |
| 0.025923 | 1.000951 | 0.132 | 0.051 | 1 | 10 | Tmem70 | 0.081 | Old |
| 0.025962 | 1.045442 | 0.183 | 0.091 | 1 | 10 | Arhgap6 | 0.092 | Old |
| 0.026099 | 0.437987 | 0.447 | 0.323 | 1 | 10 | Rap2b | 0.124 | Old |
| 0.026181 | 0.650337 | 0.406 | 0.273 | 1 | 10 | Rasgef1b | 0.133 | Old |
| 0.026276 | 1.662505 | 0.086 | 0.02 | 1 | 10 | Retsat | 0.066 | Old |
| 0.026367 | 0.389386 | 0.562 | 0.444 | 1 | 10 | Sfr1 | 0.118 | Old |
| 0.026394 | 1.437409 | 0.068 | 0.01 | 1 | 10 | Nsun6 | 0.058 | Old |
| 0.026432 | 0.241352 | 0.665 | 0.515 | 1 | 10 | Skap2 | 0.15 | Old |
| 0.026466 | 0.429418 | 0.768 | 0.646 | 1 | 10 | Glul | 0.122 | Old |
| 0.026471 | 0.535369 | 0.281 | 0.172 | 1 | 10 | Ebp | 0.109 | Old |
| 0.02652 | 0.874142 | 0.232 | 0.131 | 1 | 10 | Kif3b | 0.101 | Old |
| 0.026552 | 0.612628 | 0.403 | 0.293 | 1 | 10 | Coro1b | 0.11 | Old |
| 0.026729 | 0.427544 | 0.545 | 0.434 | 1 | 10 | Itpripl2 | 0.111 | Old |
| 0.026861 | 0.774356 | 0.357 | 0.253 | 1 | 10 | Acadl | 0.104 | Old |
| 0.027029 | 0.263114 | 0.352 | 0.222 | 1 | 10 | Stag1 | 0.13 | Old |
| 0.027297 | 0.388954 | 0.577 | 0.434 | 1 | 10 | Selplg | 0.143 | Old |
| 0.027303 | 0.335485 | 0.614 | 0.465 | 1 | 10 | Vps35 | 0.149 | Old |
| 0.027336 | 0.346414 | 0.411 | 0.263 | 1 | 10 | Sbf2 | 0.148 | Old |
| 0.027412 | 0.733541 | 0.279 | 0.162 | 1 | 10 | Smarcad1 | 0.117 | Old |
| 0.027424 | 1.200204 | 0.196 | 0.101 | 1 | 10 | Zdhhc7 | 0.095 | Old |
| 0.027514 | 0.470006 | 0.303 | 0.182 | 1 | 10 | Chp1 | 0.121 | Old |
| 0.027521 | 0.562222 | 0.306 | 0.192 | 1 | 10 | Ube2h | 0.114 | Old |
| 0.027637 | 1.332739 | 0.156 | 0.071 | 1 | 10 | Nt5c2 | 0.085 | Old |
| 0.027656 | 0.437169 | 0.271 | 0.152 | 1 | 10 | Skiv2l | 0.119 | Old |
| 0.027951 | 1.066162 | 0.154 | 0.071 | 1 | 10 | Ephx1 | 0.083 | Old |
| 0.028144 | 0.749315 | 0.161 | 0.071 | 1 | 10 | Trap1 | 0.09 | Old |
| 0.02843 | 1.053592 | 0.171 | 0.081 | 1 | 10 | Zdhhc5 | 0.09 | Old |

| | | | | | | | | |
|---|---|---|---|---|---|---|---|---|
| 0.028462 | 0.21018 | 0.731 | 0.596 | 1 | 10 | Tpr | 0.135 | Old |
| 0.028491 | 1.502178 | 0.127 | 0.051 | 1 | 10 | Traf5 | 0.076 | Old |
| 0.028517 | 0.988435 | 0.249 | 0.152 | 1 | 10 | Pnpla2 | 0.097 | Old |
| 0.028667 | 0.854871 | 0.267 | 0.172 | 1 | 10 | Mogs | 0.095 | Old |
| 0.028674 | 2.421062 | 0.066 | 0.01 | 1 | 10 | Il27ra | 0.056 | Old |
| 0.028897 | 1.587149 | 0.127 | 0.051 | 1 | 10 | Mrs2 | 0.076 | Old |
| 0.028922 | 0.980509 | 0.115 | 0.04 | 1 | 10 | Pdk3 | 0.075 | Old |
| 0.029056 | 0.658115 | 0.271 | 0.162 | 1 | 10 | Lpin1 | 0.109 | Old |
| 0.029061 | 0.764964 | 0.193 | 0.101 | 1 | 10 | Akr1b8 | 0.092 | Old |
| 0.029113 | 3.205194 | 0.046 | 0 | 1 | 10 | Fhdc1 | 0.046 | Old |
| 0.029113 | 3.113556 | 0.046 | 0 | 1 | 10 | Gm42047 | 0.046 | Old |
| 0.029113 | 3.093906 | 0.046 | 0 | 1 | 10 | Rmdn2 | 0.046 | Old |
| 0.029113 | 3.081136 | 0.046 | 0 | 1 | 10 | Zfp930 | 0.046 | Old |
| 0.029113 | 3.001683 | 0.046 | 0 | 1 | 10 | Gm16104 | 0.046 | Old |
| 0.029113 | 2.929919 | 0.046 | 0 | 1 | 10 | Nlrp10 | 0.046 | Old |
| 0.029225 | 0.441851 | 0.557 | 0.444 | 1 | 10 | Gna12 | 0.113 | Old |
| 0.029269 | 1.070072 | 0.144 | 0.061 | 1 | 10 | Traf7 | 0.083 | Old |
| 0.029281 | 1.728704 | 0.098 | 0.03 | 1 | 10 | Terf1 | 0.068 | Old |
| 0.029286 | 0.90657 | 0.156 | 0.071 | 1 | 10 | Gmnn | 0.085 | Old |
| 0.029308 | 1.356952 | 0.127 | 0.051 | 1 | 10 | Hmga1 | 0.076 | Old |
| 0.029331 | 0.42719 | 0.601 | 0.444 | 1 | 10 | Tet3 | 0.157 | Old |
| 0.029345 | 0.168027 | 0.511 | 0.343 | 1 | 10 | Gtf2h5 | 0.168 | Old |
| 0.029437 | 1.682771 | 0.083 | 0.02 | 1 | 10 | Vps72 | 0.063 | Old |
| 0.029532 | 0.208563 | 0.467 | 0.333 | 1 | 10 | Mfhas1 | 0.134 | Old |
| 0.029542 | 0.327828 | 0.565 | 0.434 | 1 | 10 | Idh1 | 0.131 | Old |
| 0.029676 | 0.14217 | 0.971 | 0.909 | 1 | 10 | Npc2 | 0.062 | Old |
| 0.029721 | 0.571331 | 0.381 | 0.253 | 1 | 10 | Cs | 0.128 | Old |
| 0.029774 | 0.387603 | 0.24 | 0.131 | 1 | 10 | Slc39a10 | 0.109 | Old |
| 0.030264 | 0.294186 | 0.535 | 0.404 | 1 | 10 | Tmem59 | 0.131 | Old |
| 0.030442 | 0.528411 | 0.262 | 0.152 | 1 | 10 | Borcs6 | 0.11 | Old |
| 0.030798 | 0.812697 | 0.171 | 0.081 | 1 | 10 | Slfn8 | 0.09 | Old |
| 0.030845 | 0.637106 | 0.23 | 0.131 | 1 | 10 | Slc35c2 | 0.099 | Old |
| 0.030937 | 0.671476 | 0.44 | 0.333 | 1 | 10 | Itch | 0.107 | Old |
| 0.031139 | 0.462595 | 0.418 | 0.293 | 1 | 10 | Huwe1 | 0.125 | Old |
| 0.031214 | 0.115708 | 0.491 | 0.343 | 1 | 10 | Rtf2 | 0.148 | Old |
| 0.031228 | 0.808667 | 0.284 | 0.182 | 1 | 10 | Med1 | 0.102 | Old |
| 0.031253 | 0.751468 | 0.196 | 0.101 | 1 | 10 | AU040320 | 0.095 | Old |
| 0.031376 | 0.288306 | 0.587 | 0.444 | 1 | 10 | Zfp207 | 0.143 | Old |
| 0.031512 | 0.233423 | 0.132 | 0.051 | 1 | 10 | Fam220a | 0.081 | Old |
| 0.031555 | 0.934226 | 0.169 | 0.081 | 1 | 10 | Ulk1 | 0.088 | Old |
| 0.03158 | 0.187987 | 0.677 | 0.545 | 1 | 10 | Dync1i2 | 0.132 | Old |
| 0.031794 | 0.54028 | 0.367 | 0.242 | 1 | 10 | Dip2b | 0.125 | Old |
| 0.031829 | 0.408706 | 0.489 | 0.343 | 1 | 10 | Dock8 | 0.146 | Old |
| 0.031944 | 1.065885 | 0.169 | 0.081 | 1 | 10 | Ap1ar | 0.088 | Old |
| 0.031972 | 0.185686 | 0.227 | 0.121 | 1 | 10 | Rnf128 | 0.106 | Old |

| | | | | | | | | |
|---|---|---|---|---|---|---|---|---|
| 0.03202 | 0.434356 | 0.408 | 0.283 | 1 | 10 | Dennd1a | 0.125 | Old |
| 0.032086 | 0.522157 | 0.496 | 0.364 | 1 | 10 | D17Wsu92 | 0.132 | Old |
| 0.032107 | 0.997536 | 0.178 | 0.091 | 1 | 10 | Pik3r6 | 0.087 | Old |
| 0.032133 | 0.544719 | 0.159 | 0.071 | 1 | 10 | Emc4 | 0.088 | Old |
| 0.032234 | 0.371051 | 0.567 | 0.444 | 1 | 10 | Myo9b | 0.123 | Old |
| 0.032249 | 1.446866 | 0.112 | 0.04 | 1 | 10 | Fam13b | 0.072 | Old |
| 0.032492 | 0.390209 | 0.142 | 0.061 | 1 | 10 | Smarcal1 | 0.081 | Old |
| 0.032655 | 0.616858 | 0.252 | 0.141 | 1 | 10 | Kctd20 | 0.111 | Old |
| 0.032713 | 0.326499 | 0.538 | 0.414 | 1 | 10 | Arhgap18 | 0.124 | Old |
| 0.03288 | 1.0508 | 0.178 | 0.091 | 1 | 10 | Armc1 | 0.087 | Old |
| 0.032906 | 0.08486 | 0.963 | 0.899 | 1 | 10 | Actr3 | 0.064 | Old |
| 0.032961 | 0.425266 | 0.315 | 0.192 | 1 | 10 | Gpr146 | 0.123 | Old |
| 0.032975 | 0.349239 | 0.604 | 0.455 | 1 | 10 | Slc11a1 | 0.149 | Old |
| 0.032992 | 0.596144 | 0.269 | 0.162 | 1 | 10 | Zfp263 | 0.107 | Old |
| 0.033114 | 1.248576 | 0.164 | 0.081 | 1 | 10 | Ms4a8a | 0.083 | Old |
| 0.033187 | 0.598802 | 0.301 | 0.182 | 1 | 10 | Lgals9 | 0.119 | Old |
| 0.033202 | 0.725489 | 0.347 | 0.232 | 1 | 10 | Slc8b1 | 0.115 | Old |
| 0.033264 | 0.466421 | 0.374 | 0.253 | 1 | 10 | Appl1 | 0.121 | Old |
| 0.033297 | 0.478752 | 0.501 | 0.364 | 1 | 10 | Rnf7 | 0.137 | Old |
| 0.033321 | 0.328684 | 0.575 | 0.455 | 1 | 10 | Adam10 | 0.12 | Old |
| 0.033446 | 1.369595 | 0.174 | 0.091 | 1 | 10 | Erlin2 | 0.083 | Old |
| 0.033493 | 0.491089 | 0.083 | 0.02 | 1 | 10 | Haus7 | 0.063 | Old |
| 0.03369 | 0.340778 | 0.631 | 0.505 | 1 | 10 | Dok2 | 0.126 | Old |
| 0.033841 | 1.855336 | 0.081 | 0.02 | 1 | 10 | Pdgfc | 0.061 | Old |
| 0.033873 | 3.37666 | 0.044 | 0 | 1 | 10 | Dmkn | 0.044 | Old |
| 0.033873 | 3.193182 | 0.044 | 0 | 1 | 10 | 2900076A0 | 0.044 | Old |
| 0.033896 | 0.334052 | 0.704 | 0.556 | 1 | 10 | Fyb | 0.148 | Old |
| 0.033945 | 0.188416 | 0.357 | 0.232 | 1 | 10 | Add3 | 0.125 | Old |
| 0.034117 | 0.510878 | 0.421 | 0.303 | 1 | 10 | Adssl1 | 0.118 | Old |
| 0.03435 | 2.310456 | 0.064 | 0.01 | 1 | 10 | Adck5 | 0.054 | Old |
| 0.03461 | 0.451199 | 0.311 | 0.192 | 1 | 10 | Htra2 | 0.119 | Old |
| 0.034692 | 0.534542 | 0.67 | 0.545 | 1 | 10 | Tnip3 | 0.125 | Old |
| 0.034797 | 0.280032 | 0.357 | 0.232 | 1 | 10 | Aurkaip1 | 0.125 | Old |
| 0.034906 | 0.252232 | 0.998 | 1 | 1 | 10 | Tpt1 | -0.002 | Old |
| 0.034935 | 0.289973 | 0.367 | 0.242 | 1 | 10 | Cdk14 | 0.125 | Old |
| 0.034967 | 0.136245 | 0.797 | 0.677 | 1 | 10 | Rgs1 | 0.12 | Old |
| 0.035149 | 0.911861 | 0.193 | 0.101 | 1 | 10 | 9430038I0 | 0.092 | Old |
| 0.03537 | 0.293415 | 0.279 | 0.162 | 1 | 10 | Poldip3 | 0.117 | Old |
| 0.035472 | 0.42597 | 0.112 | 0.04 | 1 | 10 | Camsap1 | 0.072 | Old |
| 0.035938 | 1.213465 | 0.081 | 0.02 | 1 | 10 | Eif2ak4 | 0.061 | Old |
| 0.036068 | 0.222725 | 0.929 | 0.818 | 1 | 10 | Atp6v0b | 0.111 | Old |
| 0.03623 | 0.262746 | 0.484 | 0.343 | 1 | 10 | Ik | 0.141 | Old |
| 0.036264 | 1.297174 | 0.149 | 0.071 | 1 | 10 | Rce1 | 0.078 | Old |
| 0.036476 | 1.113394 | 0.11 | 0.04 | 1 | 10 | Elac1 | 0.07 | Old |
| 0.036678 | 0.613804 | 0.142 | 0.061 | 1 | 10 | D6Wsu163 | 0.081 | Old |

| | | | | | | | | |
|---|---|---|---|---|---|---|---|---|
| 0.036732 | 0.420687 | 0.249 | 0.141 | 1 | 10 | Irf9 | 0.108 | Old |
| 0.036764 | 1.083607 | 0.125 | 0.051 | 1 | 10 | Eya4 | 0.074 | Old |
| 0.036766 | 0.515089 | 0.345 | 0.242 | 1 | 10 | Vps13d | 0.103 | Old |
| 0.036896 | 0.370024 | 0.418 | 0.293 | 1 | 10 | Chic2 | 0.125 | Old |
| 0.037081 | 1.00216 | 0.137 | 0.061 | 1 | 10 | Prep | 0.076 | Old |
| 0.037267 | 0.498611 | 0.377 | 0.263 | 1 | 10 | Lmbrd1 | 0.114 | Old |
| 0.037405 | 0.155796 | 0.166 | 0.081 | 1 | 10 | Nsmce2 | 0.085 | Old |
| 0.037563 | 0.463203 | 0.423 | 0.323 | 1 | 10 | Mfsd12 | 0.1 | Old |
| 0.0376 | 1.003901 | 0.222 | 0.131 | 1 | 10 | Cdk19 | 0.091 | Old |
| 0.037799 | 0.993341 | 0.152 | 0.071 | 1 | 10 | Stard5 | 0.081 | Old |
| 0.037822 | 0.460283 | 0.081 | 0.02 | 1 | 10 | Zdhhc8 | 0.061 | Old |
| 0.037841 | 0.58907 | 0.377 | 0.263 | 1 | 10 | Ndufa8 | 0.114 | Old |
| 0.037936 | 0.192347 | 0.33 | 0.212 | 1 | 10 | Ube2g2 | 0.118 | Old |
| 0.038043 | 0.27107 | 0.513 | 0.374 | 1 | 10 | Ckap4 | 0.139 | Old |
| 0.03812 | 0.778632 | 0.215 | 0.121 | 1 | 10 | Fcf1 | 0.094 | Old |
| 0.038244 | 0.727582 | 0.188 | 0.101 | 1 | 10 | 4930402H2 | 0.087 | Old |
| 0.038325 | 0.6066 | 0.352 | 0.242 | 1 | 10 | Baiap2 | 0.11 | Old |
| 0.038355 | 0.681559 | 0.139 | 0.061 | 1 | 10 | Actr8 | 0.078 | Old |
| 0.038426 | 0.738168 | 0.35 | 0.242 | 1 | 10 | Cyp4f18 | 0.108 | Old |
| 0.038525 | 1.207119 | 0.186 | 0.101 | 1 | 10 | Rnf144b | 0.085 | Old |
| 0.038624 | 0.360963 | 0.242 | 0.141 | 1 | 10 | Ergic2 | 0.101 | Old |
| 0.038841 | 0.527717 | 0.377 | 0.263 | 1 | 10 | Acap2 | 0.114 | Old |
| 0.038931 | 1.144135 | 0.247 | 0.162 | 1 | 10 | Prkch | 0.085 | Old |
| 0.0391 | 0.343928 | 0.328 | 0.212 | 1 | 10 | Akt2 | 0.116 | Old |
| 0.039165 | 0.482071 | 0.555 | 0.444 | 1 | 10 | Rin2 | 0.111 | Old |
| 0.03917 | 0.427476 | 0.421 | 0.293 | 1 | 10 | Psmb10 | 0.128 | Old |
| 0.039229 | 0.549071 | 0.11 | 0.04 | 1 | 10 | Git1 | 0.07 | Old |
| 0.039328 | 0.449212 | 0.244 | 0.141 | 1 | 10 | Grpel1 | 0.103 | Old |
| 0.039382 | 2.034238 | 0.061 | 0.01 | 1 | 10 | Ercc4 | 0.051 | Old |
| 0.039428 | 3.045997 | 0.042 | 0 | 1 | 10 | Ppp2r5b | 0.042 | Old |
| 0.039428 | 2.938516 | 0.042 | 0 | 1 | 10 | Elp4 | 0.042 | Old |
| 0.039428 | 2.833015 | 0.042 | 0 | 1 | 10 | Klf8 | 0.042 | Old |
| 0.039428 | 2.815895 | 0.042 | 0 | 1 | 10 | Anxa9 | 0.042 | Old |
| 0.039428 | 2.583072 | 0.042 | 0 | 1 | 10 | Spdl1 | 0.042 | Old |
| 0.039506 | 0.535457 | 0.293 | 0.192 | 1 | 10 | Pqlc3 | 0.101 | Old |
| 0.039761 | 1.706824 | 0.108 | 0.04 | 1 | 10 | Jpt2 | 0.068 | Old |
| 0.039906 | 1.49265 | 0.108 | 0.04 | 1 | 10 | Dnmt3aos | 0.068 | Old |
| 0.039977 | 0.985495 | 0.149 | 0.071 | 1 | 10 | Pcdh7 | 0.078 | Old |
| 0.040163 | 0.216342 | 0.907 | 0.818 | 1 | 10 | Hexa | 0.089 | Old |
| 0.040307 | 0.318249 | 0.254 | 0.141 | 1 | 10 | Mtmr10 | 0.113 | Old |
| 0.040325 | 1.444623 | 0.134 | 0.061 | 1 | 10 | Adcy3 | 0.073 | Old |
| 0.040392 | 0.44326 | 0.139 | 0.061 | 1 | 10 | Tada1 | 0.078 | Old |
| 0.040542 | 1.75866 | 0.061 | 0.01 | 1 | 10 | Cdc14a | 0.051 | Old |
| 0.040587 | 0.909078 | 0.078 | 0.02 | 1 | 10 | Ttl | 0.058 | Old |
| 0.04063 | 1.005537 | 0.137 | 0.061 | 1 | 10 | Josd2 | 0.076 | Old |

| | | | | | | | | |
|---|---|---|---|---|---|---|---|---|
| 0.040739 | 0.402106 | 0.225 | 0.121 | 1 | 10 | Commd2 | 0.104 | Old |
| 0.04104 | 0.735424 | 0.345 | 0.253 | 1 | 10 | Slc25a39 | 0.092 | Old |
| 0.041067 | 0.438002 | 0.205 | 0.111 | 1 | 10 | Vkorc1 | 0.094 | Old |
| 0.041068 | 1.535538 | 0.132 | 0.061 | 1 | 10 | Bzw2 | 0.071 | Old |
| 0.041098 | 0.5401 | 0.247 | 0.152 | 1 | 10 | Tfeb | 0.095 | Old |
| 0.041293 | 0.890781 | 0.191 | 0.101 | 1 | 10 | Vcl | 0.09 | Old |
| 0.041369 | 2.483924 | 0.076 | 0.02 | 1 | 10 | Xlr4b | 0.056 | Old |
| 0.041688 | 0.182902 | 0.966 | 0.96 | 1 | 10 | Rps15 | 0.006 | Old |
| 0.04171 | 0.431548 | 0.403 | 0.283 | 1 | 10 | mt-Atp8 | 0.12 | Old |
| 0.041833 | 0.471746 | 0.257 | 0.152 | 1 | 10 | Tmem192 | 0.105 | Old |
| 0.041877 | 0.974654 | 0.152 | 0.071 | 1 | 10 | Zfp260 | 0.081 | Old |
| 0.04197 | 0.439033 | 0.169 | 0.081 | 1 | 10 | 2610002M | 0.088 | Old |
| 0.042288 | 0.683348 | 0.186 | 0.101 | 1 | 10 | Sdhaf4 | 0.085 | Old |
| 0.04243 | 0.096421 | 0.462 | 0.333 | 1 | 10 | Dpysl2 | 0.129 | Old |
| 0.042474 | 0.287766 | 0.511 | 0.384 | 1 | 10 | Arcn1 | 0.127 | Old |
| 0.042585 | 0.281088 | 0.704 | 0.586 | 1 | 10 | Gsn | 0.118 | Old |
| 0.042798 | 0.561856 | 0.34 | 0.242 | 1 | 10 | Ptpro | 0.098 | Old |
| 0.042837 | 0.421686 | 0.531 | 0.404 | 1 | 10 | Nadk | 0.127 | Old |
| 0.042846 | 0.176182 | 0.9 | 0.869 | 1 | 10 | Hnrnpk | 0.031 | Old |
| 0.042878 | 0.586228 | 0.369 | 0.263 | 1 | 10 | Txndc9 | 0.106 | Old |
| 0.042953 | 1.20917 | 0.171 | 0.091 | 1 | 10 | Trim12c | 0.08 | Old |
| 0.042972 | 0.647655 | 0.188 | 0.101 | 1 | 10 | Tsc1 | 0.087 | Old |
| 0.043244 | 0.575715 | 0.254 | 0.162 | 1 | 10 | Cetn3 | 0.092 | Old |
| 0.04327 | 1.292696 | 0.12 | 0.051 | 1 | 10 | Trappc12 | 0.069 | Old |
| 0.043303 | 0.13336 | 0.597 | 0.465 | 1 | 10 | Cdk2ap2 | 0.132 | Old |
| 0.043318 | 0.507577 | 0.396 | 0.283 | 1 | 10 | Strn3 | 0.113 | Old |
| 0.043444 | 0.726046 | 0.311 | 0.212 | 1 | 10 | S100a1 | 0.099 | Old |
| 0.043609 | 0.322101 | 0.469 | 0.333 | 1 | 10 | AC149090. | 0.136 | Old |
| 0.043682 | 0.763466 | 0.108 | 0.04 | 1 | 10 | Tbc1d24 | 0.068 | Old |
| 0.044001 | 0.376977 | 0.411 | 0.293 | 1 | 10 | Trim8 | 0.118 | Old |
| 0.044043 | 0.595372 | 0.166 | 0.081 | 1 | 10 | Pfkfb4 | 0.085 | Old |
| 0.044246 | 0.340719 | 0.641 | 0.455 | 1 | 10 | Ccr5 | 0.186 | Old |
| 0.044318 | 0.220145 | 0.663 | 0.505 | 1 | 10 | Atp6v1e1 | 0.158 | Old |
| 0.044393 | 0.431833 | 0.325 | 0.212 | 1 | 10 | Spag7 | 0.113 | Old |
| 0.044541 | 0.688303 | 0.311 | 0.212 | 1 | 10 | Matk | 0.099 | Old |
| 0.044551 | 0.250204 | 0.374 | 0.253 | 1 | 10 | Elmo2 | 0.121 | Old |
| 0.044728 | 0.120914 | 0.998 | 0.99 | 1 | 10 | Rps21 | 0.008 | Old |
| 0.044924 | 0.523365 | 0.269 | 0.162 | 1 | 10 | Cyb5r1 | 0.107 | Old |
| 0.04498 | 1.044601 | 0.144 | 0.071 | 1 | 10 | Cpsf3 | 0.073 | Old |
| 0.045313 | 0.37552 | 0.342 | 0.232 | 1 | 10 | Acly | 0.11 | Old |
| 0.045444 | 1.208469 | 0.076 | 0.02 | 1 | 10 | Unc45a | 0.056 | Old |
| 0.04553 | 0.028511 | 0.203 | 0.111 | 1 | 10 | Tbk1 | 0.092 | Old |
| 0.045538 | 0.468791 | 0.504 | 0.394 | 1 | 10 | Ctdnep1 | 0.11 | Old |
| 0.045562 | 0.270117 | 0.477 | 0.354 | 1 | 10 | Srgap2 | 0.123 | Old |
| 0.045683 | 0.123492 | 0.093 | 0.03 | 1 | 10 | Trim5 | 0.063 | Old |

| | | | | | | | | |
|---|---|---|---|---|---|---|---|---|
| 0.045711 | 0.478462 | 0.281 | 0.172 | 1 | 10 | Slamf8 | 0.109 | Old |
| 0.045796 | 0.335328 | 0.491 | 0.374 | 1 | 10 | Snx9 | 0.117 | Old |
| 0.04592 | 2.953608 | 0.039 | 0 | 1 | 10 | Slc26a6 | 0.039 | Old |
| 0.04592 | 2.66497 | 0.039 | 0 | 1 | 10 | 9030025P2 | 0.039 | Old |
| 0.04592 | 2.60654 | 0.039 | 0 | 1 | 10 | Lmtk2 | 0.039 | Old |
| 0.045969 | 0.383531 | 0.352 | 0.242 | 1 | 10 | Trip11 | 0.11 | Old |
| 0.04599 | 0.342343 | 0.548 | 0.404 | 1 | 10 | Pip4k2a | 0.144 | Old |
| 0.046006 | 0.569396 | 0.306 | 0.202 | 1 | 10 | Szrd1 | 0.104 | Old |
| 0.046194 | 0.154105 | 0.8 | 0.667 | 1 | 10 | Tpd52 | 0.133 | Old |
| 0.04636 | 0.727684 | 0.149 | 0.071 | 1 | 10 | Casp9 | 0.078 | Old |
| 0.046385 | 2.151482 | 0.088 | 0.03 | 1 | 10 | Irf7 | 0.058 | Old |
| 0.046466 | 1.218979 | 0.156 | 0.081 | 1 | 10 | Neurl2 | 0.075 | Old |
| 0.046615 | 0.454424 | 0.367 | 0.263 | 1 | 10 | Mcts1 | 0.104 | Old |
| 0.04666 | 0.518285 | 0.271 | 0.172 | 1 | 10 | Trim35 | 0.099 | Old |
| 0.046789 | 0.20769 | 0.604 | 0.444 | 1 | 10 | Clec2d | 0.16 | Old |
| 0.046936 | 0.996491 | 0.134 | 0.061 | 1 | 10 | Pole3 | 0.073 | Old |
| 0.046955 | 0.144212 | 0.457 | 0.313 | 1 | 10 | Atp6v1h | 0.144 | Old |
| 0.047129 | 1.565887 | 0.09 | 0.03 | 1 | 10 | Ddx28 | 0.06 | Old |
| 0.047159 | 1.560673 | 0.059 | 0.01 | 1 | 10 | Mios | 0.049 | Old |
| 0.047159 | 1.475093 | 0.059 | 0.01 | 1 | 10 | Extl2 | 0.049 | Old |
| 0.04718 | 0.291899 | 0.834 | 0.747 | 1 | 10 | Rbpj | 0.087 | Old |
| 0.047227 | 0.415683 | 0.381 | 0.263 | 1 | 10 | Skp1a | 0.118 | Old |
| 0.047436 | 0.2238 | 0.841 | 0.717 | 1 | 10 | Gnas | 0.124 | Old |
| 0.047558 | 0.226716 | 0.284 | 0.172 | 1 | 10 | Gxylt1 | 0.112 | Old |
| 0.047614 | 1.249654 | 0.059 | 0.01 | 1 | 10 | Nsun3 | 0.049 | Old |
| 0.047646 | 0.692927 | 0.244 | 0.152 | 1 | 10 | Hypk | 0.092 | Old |
| 0.04769 | 1.096714 | 0.134 | 0.061 | 1 | 10 | Rngtt | 0.073 | Old |
| 0.048014 | 0.213528 | 0.912 | 0.798 | 1 | 10 | Uqcrh | 0.114 | Old |
| 0.048058 | 0.13415 | 0.362 | 0.232 | 1 | 10 | Ndufv2 | 0.13 | Old |
| 0.048133 | 0.515639 | 0.32 | 0.222 | 1 | 10 | Mlec | 0.098 | Old |
| 0.048293 | 0.135775 | 0.988 | 0.96 | 1 | 10 | Lgmn | 0.028 | Old |
| 0.048304 | 0.434805 | 0.291 | 0.182 | 1 | 10 | Nagk | 0.109 | Old |
| 0.048328 | 0.367032 | 0.773 | 0.727 | 1 | 10 | Nme1 | 0.046 | Old |
| 0.048478 | 0.64234 | 0.386 | 0.293 | 1 | 10 | Pgap2 | 0.093 | Old |
| 0.048695 | 0.190467 | 0.303 | 0.192 | 1 | 10 | Chd3 | 0.111 | Old |
| 0.048836 | 0.72061 | 0.337 | 0.232 | 1 | 10 | Rsl1d1 | 0.105 | Old |
| 0.048854 | 0.463672 | 0.274 | 0.162 | 1 | 10 | Prpf8 | 0.112 | Old |
| 0.04886 | 0.106792 | 0.988 | 0.97 | 1 | 10 | Anxa5 | 0.018 | Old |
| 0.048912 | 0.657327 | 0.271 | 0.172 | 1 | 10 | Plod1 | 0.099 | Old |
| 0.04894 | 0.486453 | 0.284 | 0.182 | 1 | 10 | Ikbke | 0.102 | Old |
| 0.049037 | 1.243786 | 0.144 | 0.071 | 1 | 10 | Ifi47 | 0.073 | Old |
| 0.049166 | 0.462261 | 0.411 | 0.293 | 1 | 10 | Zcchc11 | 0.118 | Old |
| 0.049188 | 0.336973 | 0.516 | 0.394 | 1 | 10 | Ddi2 | 0.122 | Old |
| 0.049362 | 0.250875 | 0.795 | 0.667 | 1 | 10 | Atp6ap1 | 0.128 | Old |
| 0.049518 | 1.197542 | 0.166 | 0.091 | 1 | 10 | Mob3c | 0.075 | Old |

| | | | | | | | | |
|---|---|---|---|---|---|---|---|---|
| 0.049525 | 0.96184 | 0.12 | 0.051 | 1 | 10 | Bmi1 | 0.069 | Old |
| 0.049541 | 0.443012 | 0.09 | 0.03 | 1 | 10 | Polr3b | 0.06 | Old |
| 0.049547 | 0.773548 | 0.159 | 0.081 | 1 | 10 | Arsa | 0.078 | Old |
| 0.049567 | 0.251554 | 0.672 | 0.556 | 1 | 10 | Sumo2 | 0.116 | Old |
| 0.049854 | 0.295253 | 0.545 | 0.424 | 1 | 10 | Cab39 | 0.121 | Old |
| 0.050209 | 0.463485 | 0.455 | 0.333 | 1 | 10 | Dctn1 | 0.122 | Old |
| 0.050212 | 0.212659 | 0.9 | 0.848 | 1 | 10 | Ywhaz | 0.052 | Old |
| 0.050576 | 0.36837 | 0.665 | 0.576 | 1 | 10 | Lasp1 | 0.089 | Old |
| 0.050867 | 0.689939 | 0.159 | 0.081 | 1 | 10 | Oasl2 | 0.078 | Old |
| 0.050883 | 0.749258 | 0.09 | 0.03 | 1 | 10 | Prr14l | 0.06 | Old |
| 0.050989 | 0.890277 | 0.171 | 0.091 | 1 | 10 | Rexo4 | 0.08 | Old |
| 0.051094 | 0.543056 | 0.161 | 0.081 | 1 | 10 | Dopey2 | 0.08 | Old |
| 0.0511 | 0.323125 | 0.105 | 0.04 | 1 | 10 | Gpkow | 0.065 | Old |
| 0.051223 | 0.389288 | 0.501 | 0.354 | 1 | 10 | Lman2 | 0.147 | Old |
| 0.051312 | 1.829154 | 0.073 | 0.02 | 1 | 10 | Nemp1 | 0.053 | Old |
| 0.051467 | 1.37627 | 0.13 | 0.061 | 1 | 10 | Mbtps2 | 0.069 | Old |
| 0.051503 | 0.198766 | 0.922 | 0.859 | 1 | 10 | Dazap2 | 0.063 | Old |
| 0.051519 | 0.250531 | 0.301 | 0.192 | 1 | 10 | Adprh | 0.109 | Old |
| 0.051542 | 0.570492 | 0.43 | 0.343 | 1 | 10 | Tubb2a | 0.087 | Old |
| 0.051759 | 0.607728 | 0.257 | 0.162 | 1 | 10 | Tspyl1 | 0.095 | Old |
| 0.051786 | 0.537707 | 0.249 | 0.152 | 1 | 10 | Abcd1 | 0.097 | Old |
| 0.051787 | 2.599146 | 0.056 | 0.01 | 1 | 10 | Orc5 | 0.046 | Old |
| 0.052095 | 0.372295 | 0.323 | 0.222 | 1 | 10 | Msmo1 | 0.101 | Old |
| 0.052103 | 0.273165 | 0.34 | 0.222 | 1 | 10 | Ppp2r2a | 0.118 | Old |
| 0.052287 | 0.437494 | 0.306 | 0.202 | 1 | 10 | 4933434E2 | 0.104 | Old |
| 0.052289 | 0.272107 | 0.773 | 0.636 | 1 | 10 | Trim25 | 0.137 | Old |
| 0.052338 | 0.712593 | 0.271 | 0.172 | 1 | 10 | Plp2 | 0.099 | Old |
| 0.052441 | 0.697062 | 0.161 | 0.081 | 1 | 10 | Apeh | 0.08 | Old |
| 0.052563 | 0.88742 | 0.105 | 0.04 | 1 | 10 | Adgrl2 | 0.065 | Old |
| 0.052568 | 0.353899 | 0.291 | 0.192 | 1 | 10 | Actn1 | 0.099 | Old |
| 0.052872 | 0.580143 | 0.227 | 0.131 | 1 | 10 | Irak4 | 0.096 | Old |
| 0.053126 | 0.723238 | 0.257 | 0.162 | 1 | 10 | Araf | 0.095 | Old |
| 0.053129 | 0.242089 | 0.274 | 0.172 | 1 | 10 | Cant1 | 0.102 | Old |
| 0.053169 | 0.156099 | 0.252 | 0.152 | 1 | 10 | Fyn | 0.1 | Old |
| 0.053259 | 0.675293 | 0.134 | 0.061 | 1 | 10 | Decr1 | 0.073 | Old |
| 0.053314 | 1.216815 | 0.088 | 0.03 | 1 | 10 | Fyco1 | 0.058 | Old |
| 0.053515 | 3.035281 | 0.037 | 0 | 1 | 10 | Slfn1 | 0.037 | Old |
| 0.053515 | 2.895117 | 0.037 | 0 | 1 | 10 | Tada2a | 0.037 | Old |
| 0.053515 | 2.894547 | 0.037 | 0 | 1 | 10 | Park2 | 0.037 | Old |
| 0.053515 | 2.641685 | 0.037 | 0 | 1 | 10 | Zrsr1 | 0.037 | Old |
| 0.053515 | 2.625362 | 0.037 | 0 | 1 | 10 | P2ry2 | 0.037 | Old |
| 0.053515 | 2.609495 | 0.037 | 0 | 1 | 10 | Pold2 | 0.037 | Old |
| 0.053751 | 0.574864 | 0.159 | 0.081 | 1 | 10 | Fam53c | 0.078 | Old |
| 0.05379 | 0.259309 | 0.489 | 0.354 | 1 | 10 | Necap2 | 0.135 | Old |
| 0.053852 | 0.498523 | 0.225 | 0.131 | 1 | 10 | Pigx | 0.094 | Old |

| | | | | | | | | |
|---|---|---|---|---|---|---|---|---|
| 0.053985 | 0.911538 | 0.191 | 0.111 | 1 | 10 | Pvr | 0.08 | Old |
| 0.054127 | 0.454825 | 0.269 | 0.172 | 1 | 10 | Ccz1 | 0.097 | Old |
| 0.054153 | 0.729831 | 0.24 | 0.152 | 1 | 10 | Zbtb18 | 0.088 | Old |
| 0.054546 | 0.270748 | 0.513 | 0.384 | 1 | 10 | Mier1 | 0.129 | Old |
| 0.0546 | 0.354451 | 0.289 | 0.182 | 1 | 10 | Rpl7l1 | 0.107 | Old |
| 0.05462 | 1.100667 | 0.181 | 0.101 | 1 | 10 | Akap10 | 0.08 | Old |
| 0.054775 | 0.247914 | 0.306 | 0.202 | 1 | 10 | Cops2 | 0.104 | Old |
| 0.055006 | 0.54209 | 0.462 | 0.364 | 1 | 10 | Hadhb | 0.098 | Old |
| 0.055057 | 0.150435 | 0.325 | 0.212 | 1 | 10 | Nemf | 0.113 | Old |
| 0.055104 | 1.363577 | 0.073 | 0.02 | 1 | 10 | Adora2b | 0.053 | Old |
| 0.055122 | 1.146801 | 0.056 | 0.01 | 1 | 10 | Toe1 | 0.046 | Old |
| 0.055235 | 0.750636 | 0.274 | 0.182 | 1 | 10 | Dpy30 | 0.092 | Old |
| 0.055386 | 1.107993 | 0.056 | 0.01 | 1 | 10 | Uevld | 0.046 | Old |
| 0.0554 | 0.369776 | 0.105 | 0.04 | 1 | 10 | Ngrn | 0.065 | Old |
| 0.055476 | 0.229198 | 0.565 | 0.404 | 1 | 10 | Acer3 | 0.161 | Old |
| 0.055716 | 0.259678 | 0.09 | 0.03 | 1 | 10 | Dgcr8 | 0.06 | Old |
| 0.055898 | 0.122307 | 0.406 | 0.293 | 1 | 10 | Stk38 | 0.113 | Old |
| 0.055992 | 0.451668 | 0.418 | 0.313 | 1 | 10 | Ubald2 | 0.105 | Old |
| 0.056021 | 1.108789 | 0.088 | 0.03 | 1 | 10 | Pstk | 0.058 | Old |
| 0.056041 | 0.848953 | 0.103 | 0.04 | 1 | 10 | Cep290 | 0.063 | Old |
| 0.056109 | 0.70148 | 0.193 | 0.111 | 1 | 10 | mt-Nd6 | 0.082 | Old |
| 0.056171 | 0.820448 | 0.169 | 0.091 | 1 | 10 | Arfip1 | 0.078 | Old |
| 0.056239 | 0.899881 | 0.103 | 0.04 | 1 | 10 | Wrb | 0.063 | Old |
| 0.056246 | 0.781461 | 0.227 | 0.141 | 1 | 10 | C3 | 0.086 | Old |
| 0.056601 | 0.958509 | 0.142 | 0.071 | 1 | 10 | Hspb11 | 0.071 | Old |
| 0.056829 | 0.27836 | 0.198 | 0.111 | 1 | 10 | Tifab | 0.087 | Old |
| 0.05685 | 0.922987 | 0.193 | 0.111 | 1 | 10 | Abca3 | 0.082 | Old |
| 0.056945 | 0.578348 | 0.183 | 0.101 | 1 | 10 | Ccnc | 0.082 | Old |
| 0.056982 | 0.164026 | 0.501 | 0.404 | 1 | 10 | Use1 | 0.097 | Old |
| 0.057075 | 0.350265 | 0.372 | 0.263 | 1 | 10 | Snx4 | 0.109 | Old |
| 0.057143 | 0.317213 | 0.472 | 0.374 | 1 | 10 | Sdf4 | 0.098 | Old |
| 0.05719 | 0.394371 | 0.318 | 0.212 | 1 | 10 | Apc | 0.106 | Old |
| 0.057407 | 0.393336 | 0.208 | 0.121 | 1 | 10 | Pdxdc1 | 0.087 | Old |
| 0.057408 | 0.67617 | 0.267 | 0.172 | 1 | 10 | Tiam1 | 0.095 | Old |
| 0.057447 | 0.456404 | 0.45 | 0.354 | 1 | 10 | Numa1 | 0.096 | Old |
| 0.057547 | 0.221373 | 0.907 | 0.747 | 1 | 10 | mt-Nd4l | 0.16 | Old |
| 0.057904 | 0.31501 | 0.423 | 0.303 | 1 | 10 | Pds5a | 0.12 | Old |
| 0.05824 | 0.444843 | 0.531 | 0.444 | 1 | 10 | Cln8 | 0.087 | Old |
| 0.058295 | 0.385579 | 0.443 | 0.313 | 1 | 10 | Sltm | 0.13 | Old |
| 0.058419 | 0.263293 | 0.582 | 0.434 | 1 | 10 | Ifngr1 | 0.148 | Old |
| 0.058583 | 0.392566 | 0.159 | 0.081 | 1 | 10 | Pign | 0.078 | Old |
| 0.058619 | 1.164788 | 0.088 | 0.03 | 1 | 10 | Tmem88 | 0.058 | Old |
| 0.05866 | 0.620208 | 0.232 | 0.141 | 1 | 10 | Mysm1 | 0.091 | Old |
| 0.058717 | 0.061956 | 0.445 | 0.333 | 1 | 10 | Cpeb4 | 0.112 | Old |
| 0.058719 | 0.261106 | 0.188 | 0.101 | 1 | 10 | Wdr48 | 0.087 | Old |

| | | | | | | | | |
|---|---|---|---|---|---|---|---|---|
| 0.058813 | 0.361228 | 0.394 | 0.283 | 1 | 10 | Dnajc7 | 0.111 | Old |
| 0.058835 | 2.524759 | 0.054 | 0.01 | 1 | 10 | Camk2a | 0.044 | Old |
| 0.058911 | 1.485237 | 0.071 | 0.02 | 1 | 10 | Dexi | 0.051 | Old |
| 0.059029 | 0.818998 | 0.142 | 0.071 | 1 | 10 | Kif1bp | 0.071 | Old |
| 0.059222 | 0.383756 | 0.467 | 0.354 | 1 | 10 | Mxd4 | 0.113 | Old |
| 0.059587 | 0.851366 | 0.254 | 0.172 | 1 | 10 | Rabgef1 | 0.082 | Old |
| 0.059647 | 0.89109 | 0.166 | 0.091 | 1 | 10 | Pold3 | 0.075 | Old |
| 0.059724 | 0.51002 | 0.411 | 0.293 | 1 | 10 | Dmxl1 | 0.118 | Old |
| 0.059746 | 0.67068 | 0.117 | 0.051 | 1 | 10 | Golt1b | 0.066 | Old |
| 0.059746 | 0.60713 | 0.117 | 0.051 | 1 | 10 | Alg10b | 0.066 | Old |
| 0.059865 | 0.229767 | 0.543 | 0.424 | 1 | 10 | Nfam1 | 0.119 | Old |
| 0.059904 | 1.550798 | 0.071 | 0.02 | 1 | 10 | Timmdc1 | 0.051 | Old |
| 0.060276 | 0.551673 | 0.293 | 0.202 | 1 | 10 | Wdfy2 | 0.091 | Old |
| 0.060286 | 0.850737 | 0.086 | 0.03 | 1 | 10 | Nit2 | 0.056 | Old |
| 0.060306 | 0.332453 | 0.369 | 0.273 | 1 | 10 | Ankfy1 | 0.096 | Old |
| 0.060427 | 0.406792 | 0.401 | 0.273 | 1 | 10 | Zranb2 | 0.128 | Old |
| 0.06048 | 1.508014 | 0.112 | 0.051 | 1 | 10 | Mapk8 | 0.061 | Old |
| 0.060537 | 0.465583 | 0.169 | 0.091 | 1 | 10 | Hnrnph3 | 0.078 | Old |
| 0.060714 | 0.653198 | 0.345 | 0.242 | 1 | 10 | Hmgcs1 | 0.103 | Old |
| 0.060788 | 0.298572 | 0.267 | 0.172 | 1 | 10 | Plxna1 | 0.095 | Old |
| 0.060897 | 1.155953 | 0.139 | 0.071 | 1 | 10 | Snx14 | 0.068 | Old |
| 0.061048 | 1.027362 | 0.127 | 0.061 | 1 | 10 | Cox10 | 0.066 | Old |
| 0.061142 | 1.428826 | 0.137 | 0.071 | 1 | 10 | Cyp4f16 | 0.066 | Old |
| 0.061317 | 0.324832 | 0.301 | 0.202 | 1 | 10 | Bmpr2 | 0.099 | Old |
| 0.061379 | 0.272913 | 0.276 | 0.172 | 1 | 10 | Usp22 | 0.104 | Old |
| 0.061476 | 0.262762 | 0.242 | 0.141 | 1 | 10 | Sptlc1 | 0.101 | Old |
| 0.061489 | 1.372608 | 0.112 | 0.051 | 1 | 10 | Cd99l2 | 0.061 | Old |
| 0.061863 | 0.342574 | 0.528 | 0.394 | 1 | 10 | Tmem167 | 0.134 | Old |
| 0.061886 | 0.229289 | 0.421 | 0.313 | 1 | 10 | Mia3 | 0.108 | Old |
| 0.06189 | 0.146279 | 0.934 | 0.929 | 1 | 10 | Ucp2 | 0.005 | Old |
| 0.061919 | 0.600036 | 0.337 | 0.242 | 1 | 10 | Smarcd2 | 0.095 | Old |
| 0.062053 | 0.381178 | 0.169 | 0.091 | 1 | 10 | Stard7 | 0.078 | Old |
| 0.062095 | 1.232988 | 0.1 | 0.04 | 1 | 10 | Fam160b2 | 0.06 | Old |
| 0.062137 | 1.227814 | 0.086 | 0.03 | 1 | 10 | Dolk | 0.056 | Old |
| 0.062415 | 3.136141 | 0.034 | 0 | 1 | 10 | Cacna1a | 0.034 | Old |
| 0.062415 | 2.946735 | 0.034 | 0 | 1 | 10 | Chac2 | 0.034 | Old |
| 0.062415 | 2.894967 | 0.034 | 0 | 1 | 10 | Simc1 | 0.034 | Old |
| 0.062415 | 2.858239 | 0.034 | 0 | 1 | 10 | Ipcef1 | 0.034 | Old |
| 0.062415 | 2.815212 | 0.034 | 0 | 1 | 10 | Myh15 | 0.034 | Old |
| 0.062415 | 2.775124 | 0.034 | 0 | 1 | 10 | Slc25a14 | 0.034 | Old |
| 0.062415 | 2.657014 | 0.034 | 0 | 1 | 10 | Nkrf | 0.034 | Old |
| 0.062415 | 2.593242 | 0.034 | 0 | 1 | 10 | Uros | 0.034 | Old |
| 0.062415 | 2.502474 | 0.034 | 0 | 1 | 10 | Ptcd1 | 0.034 | Old |
| 0.062415 | 2.488004 | 0.034 | 0 | 1 | 10 | Nars2 | 0.034 | Old |
| 0.062415 | 2.476124 | 0.034 | 0 | 1 | 10 | Mx1 | 0.034 | Old |

| | | | | | | | | |
|---|---|---|---|---|---|---|---|---|
| 0.062415 | 2.463769 | 0.034 | 0 | 1 | 10 | Zdhhc1 | 0.034 | Old |
| 0.062415 | 2.145699 | 0.034 | 0 | 1 | 10 | Tmem218 | 0.034 | Old |
| 0.062632 | 0.658662 | 0.504 | 0.424 | 1 | 10 | Ubash3b | 0.08 | Old |
| 0.062706 | 0.194175 | 0.408 | 0.283 | 1 | 10 | Hsd17b11 | 0.125 | Old |
| 0.062783 | 1.253205 | 0.183 | 0.111 | 1 | 10 | Srp68 | 0.072 | Old |
| 0.062864 | 0.654996 | 0.191 | 0.111 | 1 | 10 | Sfxn1 | 0.08 | Old |
| 0.062882 | 0.47332 | 0.271 | 0.182 | 1 | 10 | Setx | 0.089 | Old |
| 0.062915 | 0.407937 | 0.13 | 0.061 | 1 | 10 | Usp40 | 0.069 | Old |
| 0.063018 | 0.511546 | 0.345 | 0.242 | 1 | 10 | Dnajc15 | 0.103 | Old |
| 0.063028 | 0.326534 | 0.506 | 0.394 | 1 | 10 | Mcrip1 | 0.112 | Old |
| 0.063057 | 0.166693 | 0.998 | 1 | 1 | 10 | Psap | -0.002 | Old |
| 0.063167 | 0.649428 | 0.279 | 0.182 | 1 | 10 | Twistnb | 0.097 | Old |
| 0.063228 | 0.640923 | 0.318 | 0.222 | 1 | 10 | Ctdsp1 | 0.096 | Old |
| 0.06329 | 0.196697 | 0.477 | 0.354 | 1 | 10 | Tubb6 | 0.123 | Old |
| 0.06356 | 0.782352 | 0.156 | 0.081 | 1 | 10 | Brap | 0.075 | Old |
| 0.063678 | 0.322763 | 0.45 | 0.333 | 1 | 10 | Mydgf | 0.117 | Old |
| 0.063894 | 0.247355 | 0.682 | 0.515 | 1 | 10 | Vsir | 0.167 | Old |
| 0.064397 | 0.461166 | 0.205 | 0.121 | 1 | 10 | Opa1 | 0.084 | Old |
| 0.064945 | 0.934178 | 0.071 | 0.02 | 1 | 10 | Rem1 | 0.051 | Old |
| 0.065345 | 0.183893 | 0.379 | 0.283 | 1 | 10 | Elmo1 | 0.096 | Old |
| 0.065355 | 0.358446 | 0.267 | 0.172 | 1 | 10 | 1810030O( | 0.095 | Old |
| 0.065558 | 0.899853 | 0.2 | 0.121 | 1 | 10 | Mrpl35 | 0.079 | Old |
| 0.065616 | 0.743137 | 0.071 | 0.02 | 1 | 10 | Dcun1d2 | 0.051 | Old |
| 0.065625 | 0.232854 | 0.878 | 0.798 | 1 | 10 | Lipa | 0.08 | Old |
| 0.065634 | 0.249142 | 0.438 | 0.343 | 1 | 10 | Fem1c | 0.095 | Old |
| 0.065698 | 0.127256 | 0.985 | 0.98 | 1 | 10 | Rps12 | 0.005 | Old |
| 0.065707 | 0.563508 | 0.298 | 0.212 | 1 | 10 | Ccng1 | 0.086 | Old |
| 0.065935 | 0.432795 | 0.482 | 0.384 | 1 | 10 | Pik3r5 | 0.098 | Old |
| 0.065988 | 0.899688 | 0.139 | 0.071 | 1 | 10 | Jagn1 | 0.068 | Old |
| 0.066035 | 0.115249 | 0.993 | 0.99 | 1 | 10 | Rpl38 | 0.003 | Old |
| 0.06605 | 0.243666 | 0.555 | 0.424 | 1 | 10 | Wipf1 | 0.131 | Old |
| 0.066178 | 1.009709 | 0.127 | 0.061 | 1 | 10 | Nup85 | 0.066 | Old |
| 0.066238 | 0.404961 | 0.193 | 0.111 | 1 | 10 | Tubb4b | 0.082 | Old |
| 0.066271 | 0.363077 | 0.196 | 0.111 | 1 | 10 | Slc25a46 | 0.085 | Old |
| 0.06648 | 0.578838 | 0.32 | 0.222 | 1 | 10 | Wbp4 | 0.098 | Old |
| 0.066592 | 0.481112 | 0.345 | 0.253 | 1 | 10 | N4bp2l2 | 0.092 | Old |
| 0.066844 | 0.368941 | 0.301 | 0.202 | 1 | 10 | Mrpl36 | 0.099 | Old |
| 0.066859 | 0.2549 | 0.579 | 0.465 | 1 | 10 | Usp9x | 0.114 | Old |
| 0.066948 | 0.187659 | 0.875 | 0.788 | 1 | 10 | Sh3bgrl | 0.087 | Old |
| 0.066974 | 0.274356 | 0.279 | 0.172 | 1 | 10 | Rab22a | 0.107 | Old |
| 0.067152 | 0.256166 | 0.885 | 0.808 | 1 | 10 | Lgals1 | 0.077 | Old |
| 0.067509 | 2.593049 | 0.051 | 0.01 | 1 | 10 | Rad1 | 0.041 | Old |
| 0.067713 | 0.550175 | 0.1 | 0.04 | 1 | 10 | Aatf | 0.06 | Old |
| 0.067818 | 0.407033 | 0.631 | 0.495 | 1 | 10 | Igf1 | 0.136 | Old |
| 0.067889 | 0.61567 | 0.171 | 0.091 | 1 | 10 | Arfgef2 | 0.08 | Old |

| | | | | | | | | |
|---|---|---|---|---|---|---|---|---|
| 0.067932 | 0.872679 | 0.286 | 0.212 | 1 | 10 | Creld2 | 0.074 | Old |
| 0.068061 | 0.992778 | 0.112 | 0.051 | 1 | 10 | Stx17 | 0.061 | Old |
| 0.068078 | 1.325149 | 0.137 | 0.071 | 1 | 10 | Trim30d | 0.066 | Old |
| 0.068128 | 0.870249 | 0.178 | 0.101 | 1 | 10 | Wfs1 | 0.077 | Old |
| 0.068239 | 0.106543 | 0.225 | 0.131 | 1 | 10 | Gtf3c6 | 0.094 | Old |
| 0.068302 | 1.031326 | 0.098 | 0.04 | 1 | 10 | Atg14 | 0.058 | Old |
| 0.068345 | 0.530207 | 0.318 | 0.222 | 1 | 10 | Lrrc59 | 0.096 | Old |
| 0.068382 | 0.390058 | 0.391 | 0.283 | 1 | 10 | Aup1 | 0.108 | Old |
| 0.068502 | 0.540313 | 0.267 | 0.172 | 1 | 10 | Agap1 | 0.095 | Old |
| 0.068531 | 0.461476 | 0.359 | 0.263 | 1 | 10 | Tmx3 | 0.096 | Old |
| 0.06854 | 1.265586 | 0.098 | 0.04 | 1 | 10 | Dusp10 | 0.058 | Old |
| 0.068712 | 1.112863 | 0.161 | 0.091 | 1 | 10 | Nup205 | 0.07 | Old |
| 0.068763 | 0.234556 | 0.357 | 0.253 | 1 | 10 | Baz1b | 0.104 | Old |
| 0.068918 | 0.256843 | 0.807 | 0.697 | 1 | 10 | Arhgdib | 0.11 | Old |
| 0.068929 | 0.78963 | 0.154 | 0.081 | 1 | 10 | Wdr33 | 0.073 | Old |
| 0.069069 | 0.169083 | 0.995 | 0.98 | 1 | 10 | mt-Nd1 | 0.015 | Old |
| 0.069148 | 2.433074 | 0.051 | 0.01 | 1 | 10 | Msl3l2 | 0.041 | Old |
| 0.069169 | 1.589953 | 0.083 | 0.03 | 1 | 10 | Rdh13 | 0.053 | Old |
| 0.069283 | 0.898013 | 0.152 | 0.081 | 1 | 10 | Fam208b | 0.071 | Old |
| 0.069352 | 0.133852 | 0.756 | 0.586 | 1 | 10 | Trem2 | 0.17 | Old |
| 0.069565 | 0.452798 | 0.235 | 0.141 | 1 | 10 | Plagl2 | 0.094 | Old |
| 0.069685 | 0.286317 | 0.428 | 0.303 | 1 | 10 | Xrn2 | 0.125 | Old |
| 0.069749 | 0.540939 | 0.274 | 0.182 | 1 | 10 | Plekhm1 | 0.092 | Old |
| 0.06981 | 0.985345 | 0.196 | 0.121 | 1 | 10 | Igf2bp2 | 0.075 | Old |
| 0.06995 | 1.36803 | 0.083 | 0.03 | 1 | 10 | Ceacam1 | 0.053 | Old |
| 0.070068 | 0.522425 | 0.196 | 0.111 | 1 | 10 | Trim27 | 0.085 | Old |
| 0.070129 | 0.387582 | 0.347 | 0.242 | 1 | 10 | Was | 0.105 | Old |
| 0.070147 | 2.210266 | 0.051 | 0.01 | 1 | 10 | Prdm15 | 0.041 | Old |
| 0.070198 | 0.623492 | 0.203 | 0.121 | 1 | 10 | Nr2f6 | 0.082 | Old |
| 0.070242 | 0.283304 | 0.533 | 0.434 | 1 | 10 | Lpxn | 0.099 | Old |
| 0.070251 | 1.115779 | 0.125 | 0.061 | 1 | 10 | Hlcs | 0.064 | Old |
| 0.070308 | 0.55104 | 0.323 | 0.232 | 1 | 10 | Plod3 | 0.091 | Old |
| 0.070411 | 0.667568 | 0.24 | 0.152 | 1 | 10 | Atic | 0.088 | Old |
| 0.070421 | 0.252851 | 0.988 | 0.97 | 1 | 10 | Crip1 | 0.018 | Old |
| 0.070576 | 0.628821 | 0.127 | 0.061 | 1 | 10 | Ngly1 | 0.066 | Old |
| 0.070839 | 0.584914 | 0.225 | 0.141 | 1 | 10 | Atp8a1 | 0.084 | Old |
| 0.070945 | 1.055652 | 0.137 | 0.071 | 1 | 10 | Sord | 0.066 | Old |
| 0.071003 | 1.336414 | 0.083 | 0.03 | 1 | 10 | Map3k3 | 0.053 | Old |
| 0.071011 | 0.382212 | 0.259 | 0.172 | 1 | 10 | Dhrs7 | 0.087 | Old |
| 0.07115 | 0.638082 | 0.802 | 0.778 | 1 | 10 | Selenop | 0.024 | Old |
| 0.071228 | 0.290295 | 0.43 | 0.313 | 1 | 10 | Klf10 | 0.117 | Old |
| 0.071301 | 0.34382 | 0.13 | 0.061 | 1 | 10 | Ubtd1 | 0.069 | Old |
| 0.071648 | 0.379239 | 0.511 | 0.434 | 1 | 10 | Sdhc | 0.077 | Old |
| 0.071841 | 0.744354 | 0.139 | 0.071 | 1 | 10 | Diablo | 0.068 | Old |
| 0.071947 | 0.500401 | 0.359 | 0.253 | 1 | 10 | Rasa4 | 0.106 | Old |

| | | | | | | | | |
|---|---|---|---|---|---|---|---|---|
| 0.072119 | 0.919106 | 0.196 | 0.121 | 1 | 10 | BC003965 | 0.075 | Old |
| 0.07218 | 1.873533 | 0.051 | 0.01 | 1 | 10 | Brca1 | 0.041 | Old |
| 0.07218 | 1.663568 | 0.051 | 0.01 | 1 | 10 | Klhl12 | 0.041 | Old |
| 0.07242 | 0.293256 | 0.225 | 0.131 | 1 | 10 | Srf | 0.094 | Old |
| 0.072456 | 0.22862 | 0.271 | 0.172 | 1 | 10 | Rbfa | 0.099 | Old |
| 0.072465 | 0.537491 | 0.303 | 0.212 | 1 | 10 | Vps28 | 0.091 | Old |
| 0.072524 | 1.785546 | 0.051 | 0.01 | 1 | 10 | Zfp709 | 0.041 | Old |
| 0.072534 | 0.955556 | 0.122 | 0.061 | 1 | 10 | Syne3 | 0.061 | Old |
| 0.072554 | 0.364727 | 0.435 | 0.333 | 1 | 10 | Lgals3bp | 0.102 | Old |
| 0.072844 | 0.297048 | 0.645 | 0.525 | 1 | 10 | Dnmt3a | 0.12 | Old |
| 0.072852 | 0.242441 | 0.672 | 0.525 | 1 | 10 | Nsa2 | 0.147 | Old |
| 0.072866 | 2.803994 | 0.032 | 0 | 1 | 10 | Btbd6 | 0.032 | Old |
| 0.072866 | 2.781034 | 0.032 | 0 | 1 | 10 | Colq | 0.032 | Old |
| 0.072866 | 2.697693 | 0.032 | 0 | 1 | 10 | Tmem263 | 0.032 | Old |
| 0.072866 | 2.693412 | 0.032 | 0 | 1 | 10 | Dtwd2 | 0.032 | Old |
| 0.072866 | 2.687676 | 0.032 | 0 | 1 | 10 | Car6 | 0.032 | Old |
| 0.072866 | 2.67933 | 0.032 | 0 | 1 | 10 | Snx33 | 0.032 | Old |
| 0.072866 | 2.651847 | 0.032 | 0 | 1 | 10 | Ifitm5 | 0.032 | Old |
| 0.072866 | 2.615112 | 0.032 | 0 | 1 | 10 | Fkbp7 | 0.032 | Old |
| 0.072866 | 2.586779 | 0.032 | 0 | 1 | 10 | Fndc7 | 0.032 | Old |
| 0.072866 | 2.5788 | 0.032 | 0 | 1 | 10 | Wdr76 | 0.032 | Old |
| 0.072866 | 2.547867 | 0.032 | 0 | 1 | 10 | Acat2 | 0.032 | Old |
| 0.072866 | 2.473127 | 0.032 | 0 | 1 | 10 | Pecam1 | 0.032 | Old |
| 0.072866 | 2.365624 | 0.032 | 0 | 1 | 10 | Lnx2 | 0.032 | Old |
| 0.072877 | 0.943439 | 0.083 | 0.03 | 1 | 10 | Nxt2 | 0.053 | Old |
| 0.072994 | 0.348561 | 0.32 | 0.222 | 1 | 10 | Ppp4r3a | 0.098 | Old |
| 0.073215 | 1.450332 | 0.051 | 0.01 | 1 | 10 | AI597479 | 0.041 | Old |
| 0.073318 | 0.317363 | 0.885 | 0.818 | 1 | 10 | H2afz | 0.067 | Old |
| 0.073392 | 1.155583 | 0.178 | 0.111 | 1 | 10 | Cryl1 | 0.067 | Old |
| 0.073426 | 0.544742 | 0.271 | 0.182 | 1 | 10 | Tomm34 | 0.089 | Old |
| 0.073448 | 0.39025 | 0.487 | 0.384 | 1 | 10 | Map3k1 | 0.103 | Old |
| 0.073458 | 0.452909 | 0.235 | 0.152 | 1 | 10 | Vcpip1 | 0.083 | Old |
| 0.073465 | 0.75298 | 0.296 | 0.212 | 1 | 10 | Ubxn6 | 0.084 | Old |
| 0.073489 | 0.334322 | 0.658 | 0.576 | 1 | 10 | Spag9 | 0.082 | Old |
| 0.073604 | 1.845061 | 0.081 | 0.03 | 1 | 10 | Pi4kb | 0.051 | Old |
| 0.073932 | 0.050079 | 0.614 | 0.444 | 1 | 10 | Acp2 | 0.17 | Old |
| 0.073978 | 0.36794 | 0.447 | 0.323 | 1 | 10 | 1700020I1 | 0.124 | Old |
| 0.073983 | 0.61067 | 0.205 | 0.131 | 1 | 10 | Mroh1 | 0.074 | Old |
| 0.073999 | 0.885843 | 0.125 | 0.061 | 1 | 10 | Tmub2 | 0.064 | Old |
| 0.074022 | 0.652681 | 0.068 | 0.02 | 1 | 10 | Qpctl | 0.048 | Old |
| 0.074244 | 0.139742 | 0.822 | 0.727 | 1 | 10 | Tubb5 | 0.095 | Old |
| 0.074446 | 0.263062 | 0.43 | 0.293 | 1 | 10 | Clint1 | 0.137 | Old |
| 0.074458 | 0.360352 | 0.413 | 0.293 | 1 | 10 | Rex1bd | 0.12 | Old |
| 0.074697 | 1.049822 | 0.149 | 0.081 | 1 | 10 | Arsk | 0.068 | Old |
| 0.075218 | 0.492688 | 0.215 | 0.131 | 1 | 10 | Mapkap1 | 0.084 | Old |

| | | | | | | | | |
|---|---|---|---|---|---|---|---|---|
| 0.075255 | 0.440795 | 0.406 | 0.303 | 1 | 10 | Wac | 0.103 | Old |
| 0.07531 | 0.489097 | 0.645 | 0.576 | 1 | 10 | Atp6v0a1 | 0.069 | Old |
| 0.075347 | 0.494828 | 0.2 | 0.121 | 1 | 10 | Pias4 | 0.079 | Old |
| 0.075408 | 0.137314 | 0.709 | 0.566 | 1 | 10 | Snrpe | 0.143 | Old |
| 0.075487 | 0.312892 | 0.281 | 0.182 | 1 | 10 | Snrpa | 0.099 | Old |
| 0.075535 | 1.059961 | 0.159 | 0.091 | 1 | 10 | Ankrd27 | 0.068 | Old |
| 0.07558 | 0.220867 | 0.565 | 0.434 | 1 | 10 | Mapre2 | 0.131 | Old |
| 0.075581 | 0.29286 | 0.227 | 0.141 | 1 | 10 | Tsnax | 0.086 | Old |
| 0.075784 | 0.061241 | 0.577 | 0.414 | 1 | 10 | Stat3 | 0.163 | Old |
| 0.075796 | 0.096128 | 1 | 1 | 1 | 10 | Actb | 0 | Old |
| 0.075853 | 0.603012 | 0.068 | 0.02 | 1 | 10 | Bcas3 | 0.048 | Old |
| 0.075923 | 0.35821 | 0.753 | 0.687 | 1 | 10 | Rab8b | 0.066 | Old |
| 0.076032 | 0.077026 | 0.293 | 0.192 | 1 | 10 | Psd3 | 0.101 | Old |
| 0.07605 | 0.257625 | 0.804 | 0.677 | 1 | 10 | Tuba1b | 0.127 | Old |
| 0.076197 | 0.429811 | 0.249 | 0.162 | 1 | 10 | Dync1li2 | 0.087 | Old |
| 0.076253 | 0.39556 | 0.345 | 0.242 | 1 | 10 | Fkbp15 | 0.103 | Old |
| 0.076389 | 0.402734 | 0.235 | 0.152 | 1 | 10 | Ckap5 | 0.083 | Old |
| 0.076406 | 0.967874 | 0.112 | 0.051 | 1 | 10 | Ggps1 | 0.061 | Old |
| 0.076415 | 1.376714 | 0.095 | 0.04 | 1 | 10 | Slx4ip | 0.055 | Old |
| 0.076484 | 0.556981 | 0.254 | 0.172 | 1 | 10 | Cpsf7 | 0.082 | Old |
| 0.076628 | 2.085145 | 0.066 | 0.02 | 1 | 10 | Pla2g12a | 0.046 | Old |
| 0.076714 | 0.70004 | 0.21 | 0.131 | 1 | 10 | Acadvl | 0.079 | Old |
| 0.07681 | 0.215526 | 0.308 | 0.202 | 1 | 10 | Smarce1 | 0.106 | Old |
| 0.076861 | 0.013048 | 0.328 | 0.212 | 1 | 10 | Nsun2 | 0.116 | Old |
| 0.076938 | 0.232158 | 0.22 | 0.131 | 1 | 10 | Tmem128 | 0.089 | Old |
| 0.077086 | 0.069041 | 0.687 | 0.576 | 1 | 10 | Akr1b3 | 0.111 | Old |
| 0.077264 | 0.123104 | 0.416 | 0.313 | 1 | 10 | Ly9 | 0.103 | Old |
| 0.077397 | 0.533373 | 0.215 | 0.131 | 1 | 10 | Tpra1 | 0.084 | Old |
| 0.077541 | 0.623192 | 0.098 | 0.04 | 1 | 10 | Hectd3 | 0.058 | Old |
| 0.077616 | 0.253126 | 0.413 | 0.293 | 1 | 10 | 2310022A1 | 0.12 | Old |
| 0.077761 | 0.177639 | 0.631 | 0.515 | 1 | 10 | Arhgap30 | 0.116 | Old |
| 0.077813 | 0.670989 | 0.159 | 0.091 | 1 | 10 | Spg20 | 0.068 | Old |
| 0.078064 | 0.363279 | 0.394 | 0.283 | 1 | 10 | Ids | 0.111 | Old |
| 0.07808 | 0.255263 | 0.227 | 0.141 | 1 | 10 | Aff1 | 0.086 | Old |
| 0.078099 | 0.306087 | 0.452 | 0.354 | 1 | 10 | 7-Mar | 0.098 | Old |
| 0.07826 | 0.136346 | 0.538 | 0.414 | 1 | 10 | 2610507B1 | 0.124 | Old |
| 0.07846 | 1.106888 | 0.11 | 0.051 | 1 | 10 | Pdp1 | 0.059 | Old |
| 0.07846 | 0.846057 | 0.11 | 0.051 | 1 | 10 | Creb3l2 | 0.059 | Old |
| 0.078475 | 0.409241 | 0.425 | 0.323 | 1 | 10 | Pkn1 | 0.102 | Old |
| 0.078568 | 0.407042 | 0.279 | 0.182 | 1 | 10 | Taz | 0.097 | Old |
| 0.078619 | 0.634611 | 0.249 | 0.162 | 1 | 10 | Spopl | 0.087 | Old |
| 0.078681 | 0.791813 | 0.208 | 0.131 | 1 | 10 | Pex2 | 0.077 | Old |
| 0.078749 | 0.236703 | 0.814 | 0.717 | 1 | 10 | S100a10 | 0.097 | Old |
| 0.078749 | 0.269892 | 0.853 | 0.747 | 1 | 10 | Gdi2 | 0.106 | Old |
| 0.078776 | 0.347616 | 0.252 | 0.162 | 1 | 10 | Tcerg1 | 0.09 | Old |

| | | | | | | | | |
|---|---|---|---|---|---|---|---|---|
| 0.078986 | 2.200356 | 0.049 | 0.01 | 1 | 10 | Hspbap1 | 0.039 | Old |
| 0.079197 | 0.097149 | 0.311 | 0.202 | 1 | 10 | Arhgap5 | 0.109 | Old |
| 0.079321 | 0.204451 | 0.359 | 0.263 | 1 | 10 | Chchd1 | 0.096 | Old |
| 0.079795 | 0.523457 | 0.188 | 0.111 | 1 | 10 | Capn7 | 0.077 | Old |
| 0.079832 | 0.293877 | 0.252 | 0.162 | 1 | 10 | Tbc1d5 | 0.09 | Old |
| 0.079901 | 0.226098 | 0.814 | 0.727 | 1 | 10 | Atp6v1a | 0.087 | Old |
| 0.080125 | 2.066579 | 0.049 | 0.01 | 1 | 10 | Hyal1 | 0.039 | Old |
| 0.080166 | 0.560069 | 0.066 | 0.02 | 1 | 10 | Igkc | 0.046 | Old |
| 0.080185 | 1.322327 | 0.081 | 0.03 | 1 | 10 | Ap5z1 | 0.051 | Old |
| 0.080343 | 1.792948 | 0.093 | 0.04 | 1 | 10 | Zbp1 | 0.053 | Old |
| 0.080482 | 1.25978 | 0.081 | 0.03 | 1 | 10 | Ccdc71 | 0.051 | Old |
| 0.080494 | 1.300407 | 0.066 | 0.02 | 1 | 10 | Bphl | 0.046 | Old |
| 0.080613 | 0.215805 | 0.548 | 0.434 | 1 | 10 | Rps6ka3 | 0.114 | Old |
| 0.080702 | 0.641323 | 0.122 | 0.061 | 1 | 10 | Ino80c | 0.061 | Old |
| 0.080748 | 0.111103 | 0.638 | 0.525 | 1 | 10 | Ube2l3 | 0.113 | Old |
| 0.080792 | 0.709123 | 0.159 | 0.091 | 1 | 10 | Zfp646 | 0.068 | Old |
| 0.081051 | 0.411998 | 0.342 | 0.242 | 1 | 10 | Serpinb8 | 0.1 | Old |
| 0.081066 | 0.589078 | 0.174 | 0.101 | 1 | 10 | Dlat | 0.073 | Old |
| 0.081102 | 0.103788 | 0.303 | 0.202 | 1 | 10 | Stom | 0.101 | Old |
| 0.081106 | 0.467818 | 0.232 | 0.152 | 1 | 10 | Vps4a | 0.08 | Old |
| 0.081224 | 0.251593 | 0.394 | 0.273 | 1 | 10 | Acadm | 0.121 | Old |
| 0.081277 | 2.091217 | 0.049 | 0.01 | 1 | 10 | Glmn | 0.039 | Old |
| 0.081296 | 0.613801 | 0.24 | 0.152 | 1 | 10 | Arrdc3 | 0.088 | Old |
| 0.081313 | 0.445102 | 0.22 | 0.141 | 1 | 10 | Insr | 0.079 | Old |
| 0.081437 | 0.422666 | 0.501 | 0.424 | 1 | 10 | Oxct1 | 0.077 | Old |
| 0.081483 | 0.585091 | 0.247 | 0.162 | 1 | 10 | Tgm2 | 0.085 | Old |
| 0.081555 | 0.271142 | 0.425 | 0.313 | 1 | 10 | 2-Mar | 0.112 | Old |
| 0.081679 | 0.6391 | 0.176 | 0.101 | 1 | 10 | Endod1 | 0.075 | Old |
| 0.081809 | 1.309603 | 0.208 | 0.141 | 1 | 10 | Dscr3 | 0.067 | Old |
| 0.081837 | 0.19253 | 0.724 | 0.586 | 1 | 10 | Abi1 | 0.138 | Old |
| 0.081943 | 0.163371 | 0.792 | 0.657 | 1 | 10 | Ifitm2 | 0.135 | Old |
| 0.08204 | 0.380317 | 0.46 | 0.354 | 1 | 10 | Thrap3 | 0.106 | Old |
| 0.082523 | 0.816421 | 0.208 | 0.131 | 1 | 10 | Nae1 | 0.077 | Old |
| 0.082579 | 0.221185 | 0.313 | 0.212 | 1 | 10 | Apaf1 | 0.101 | Old |
| 0.082855 | 0.460995 | 0.401 | 0.303 | 1 | 10 | Olfm1 | 0.098 | Old |
| 0.082859 | 0.381488 | 0.259 | 0.172 | 1 | 10 | Rcor1 | 0.087 | Old |
| 0.082976 | 1.089381 | 0.159 | 0.091 | 1 | 10 | Chid1 | 0.068 | Old |
| 0.083065 | 0.282654 | 0.885 | 0.818 | 1 | 10 | Pim1 | 0.067 | Old |
| 0.083118 | 0.441186 | 0.337 | 0.253 | 1 | 10 | Trim30a | 0.084 | Old |
| 0.083297 | 0.576028 | 0.325 | 0.242 | 1 | 10 | Anxa6 | 0.083 | Old |
| 0.083466 | 0.137697 | 0.125 | 0.061 | 1 | 10 | Smc2 | 0.064 | Old |
| 0.083703 | 0.253805 | 0.452 | 0.333 | 1 | 10 | Herc1 | 0.119 | Old |
| 0.083792 | 0.346472 | 0.267 | 0.172 | 1 | 10 | Copg1 | 0.095 | Old |
| 0.083966 | 0.409623 | 0.269 | 0.172 | 1 | 10 | Idh3b | 0.097 | Old |
| 0.08413 | 0.797047 | 0.134 | 0.071 | 1 | 10 | Rrm1 | 0.063 | Old |

| | | | | | | | | |
|---|---|---|---|---|---|---|---|---|
| 0.084518 | 1.091987 | 0.066 | 0.02 | 1 | 10 | Gtf3c4 | 0.046 | Old |
| 0.08454 | 0.219554 | 0.232 | 0.141 | 1 | 10 | Luc7l | 0.091 | Old |
| 0.08466 | 0.638952 | 0.11 | 0.051 | 1 | 10 | Rgp1 | 0.059 | Old |
| 0.084743 | 0.256386 | 0.242 | 0.152 | 1 | 10 | Crybg1 | 0.09 | Old |
| 0.084802 | 0.172825 | 0.33 | 0.222 | 1 | 10 | Puf60 | 0.108 | Old |
| 0.084814 | 1.384711 | 0.049 | 0.01 | 1 | 10 | Pinx1 | 0.039 | Old |
| 0.084858 | 1.629495 | 0.105 | 0.051 | 1 | 10 | Nelfcd | 0.054 | Old |
| 0.08487 | 0.60152 | 0.267 | 0.182 | 1 | 10 | Dld | 0.085 | Old |
| 0.084929 | 0.237204 | 0.741 | 0.687 | 1 | 10 | Atp5g3 | 0.054 | Old |
| 0.085058 | 0.407403 | 0.232 | 0.152 | 1 | 10 | Xrn1 | 0.08 | Old |
| 0.085163 | 2.661435 | 0.029 | 0 | 1 | 10 | Rbm12b2 | 0.029 | Old |
| 0.085163 | 2.603891 | 0.029 | 0 | 1 | 10 | Zfp438 | 0.029 | Old |
| 0.085163 | 2.567079 | 0.029 | 0 | 1 | 10 | Zfat | 0.029 | Old |
| 0.085163 | 2.564712 | 0.029 | 0 | 1 | 10 | Lyrm1 | 0.029 | Old |
| 0.085163 | 2.476936 | 0.029 | 0 | 1 | 10 | Ceacam19 | 0.029 | Old |
| 0.085163 | 2.391212 | 0.029 | 0 | 1 | 10 | Klra3 | 0.029 | Old |
| 0.085163 | 2.351023 | 0.029 | 0 | 1 | 10 | Slc16a13 | 0.029 | Old |
| 0.085163 | 2.275008 | 0.029 | 0 | 1 | 10 | Tmem150b | 0.029 | Old |
| 0.085163 | 2.255751 | 0.029 | 0 | 1 | 10 | Asb1 | 0.029 | Old |
| 0.085163 | 2.241287 | 0.029 | 0 | 1 | 10 | Acyp2 | 0.029 | Old |
| 0.085163 | 2.144009 | 0.029 | 0 | 1 | 10 | Xlr4a | 0.029 | Old |
| 0.085214 | 1.322044 | 0.049 | 0.01 | 1 | 10 | Ap4e1 | 0.039 | Old |
| 0.085617 | 1.02546 | 0.049 | 0.01 | 1 | 10 | Qtrt2 | 0.039 | Old |
| 0.08572 | 0.347854 | 0.411 | 0.313 | 1 | 10 | Tmem109 | 0.098 | Old |
| 0.085746 | 0.48893 | 0.279 | 0.192 | 1 | 10 | Plaa | 0.087 | Old |
| 0.085754 | 0.842948 | 0.186 | 0.111 | 1 | 10 | Pttg1 | 0.075 | Old |
| 0.085771 | 0.91935 | 0.095 | 0.04 | 1 | 10 | 3110040N1 | 0.055 | Old |
| 0.086157 | 0.705432 | 0.242 | 0.162 | 1 | 10 | Rsl24d1 | 0.08 | Old |
| 0.086338 | 0.505255 | 0.235 | 0.152 | 1 | 10 | Hcfc1 | 0.083 | Old |
| 0.086386 | 0.389082 | 0.35 | 0.263 | 1 | 10 | Cycs | 0.087 | Old |
| 0.086405 | 0.990419 | 0.108 | 0.051 | 1 | 10 | Caap1 | 0.057 | Old |
| 0.086421 | 0.095528 | 0.438 | 0.323 | 1 | 10 | Ago2 | 0.115 | Old |
| 0.086478 | 1.927303 | 0.064 | 0.02 | 1 | 10 | Ip6k2 | 0.044 | Old |
| 0.086509 | 0.359591 | 0.379 | 0.283 | 1 | 10 | Peli1 | 0.096 | Old |
| 0.086536 | 0.560691 | 0.161 | 0.091 | 1 | 10 | Ccdc34 | 0.07 | Old |
| 0.08654 | 0.167334 | 0.467 | 0.374 | 1 | 10 | Hilpda | 0.093 | Old |
| 0.08671 | 0.53969 | 0.164 | 0.091 | 1 | 10 | Rpp25l | 0.073 | Old |
| 0.087015 | 0.315107 | 0.423 | 0.313 | 1 | 10 | Ensa | 0.11 | Old |
| 0.087054 | 1.013409 | 0.132 | 0.071 | 1 | 10 | Msantd4 | 0.061 | Old |
| 0.087141 | 0.32816 | 0.599 | 0.475 | 1 | 10 | Myh9 | 0.124 | Old |
| 0.087161 | 0.245208 | 0.846 | 0.778 | 1 | 10 | Tagln2 | 0.068 | Old |
| 0.08724 | 0.946514 | 0.122 | 0.061 | 1 | 10 | Scnm1 | 0.061 | Old |
| 0.087257 | 0.619128 | 0.183 | 0.111 | 1 | 10 | Mob3a | 0.072 | Old |
| 0.087442 | 0.315208 | 0.55 | 0.465 | 1 | 10 | Papola | 0.085 | Old |
| 0.087904 | 0.118979 | 0.995 | 0.99 | 1 | 10 | mt-Cytb | 0.005 | Old |

| | | | | | | | | |
|---|---|---|---|---|---|---|---|---|
| 0.087906 | 0.684693 | 0.11 | 0.051 | 1 | 10 | Thoc3 | 0.059 | Old |
| 0.088057 | 1.202807 | 0.108 | 0.051 | 1 | 10 | Selenbp1 | 0.057 | Old |
| 0.088098 | 0.273498 | 0.225 | 0.141 | 1 | 10 | Mrpl15 | 0.084 | Old |
| 0.088399 | 0.153159 | 0.355 | 0.263 | 1 | 10 | Anxa11 | 0.092 | Old |
| 0.088578 | 0.270228 | 0.352 | 0.253 | 1 | 10 | Ankrd40 | 0.099 | Old |
| 0.088733 | 1.28262 | 0.105 | 0.051 | 1 | 10 | Poglut1 | 0.054 | Old |
| 0.088987 | 1.659667 | 0.064 | 0.02 | 1 | 10 | Gfod1 | 0.044 | Old |
| 0.089204 | 0.826166 | 0.193 | 0.121 | 1 | 10 | Pmvk | 0.072 | Old |
| 0.089514 | 0.461278 | 0.433 | 0.333 | 1 | 10 | Usp8 | 0.1 | Old |
| 0.08953 | 0.572386 | 0.274 | 0.192 | 1 | 10 | Psme3 | 0.082 | Old |
| 0.089568 | 1.09034 | 0.12 | 0.061 | 1 | 10 | Mbd5 | 0.059 | Old |
| 0.089714 | 1.086773 | 0.064 | 0.02 | 1 | 10 | Zfp426 | 0.044 | Old |
| 0.089853 | 0.995495 | 0.156 | 0.091 | 1 | 10 | Fam43a | 0.065 | Old |
| 0.08987 | 0.086287 | 0.927 | 0.848 | 1 | 10 | Cd63 | 0.079 | Old |
| 0.089914 | 0.55347 | 0.225 | 0.141 | 1 | 10 | Tbc1d22a | 0.084 | Old |
| 0.090194 | 0.826608 | 0.093 | 0.04 | 1 | 10 | Slc46a3 | 0.053 | Old |
| 0.090276 | 0.226639 | 0.377 | 0.273 | 1 | 10 | Gga1 | 0.104 | Old |
| 0.090494 | 0.251576 | 0.257 | 0.162 | 1 | 10 | Frg1 | 0.095 | Old |
| 0.090498 | 1.186218 | 0.093 | 0.04 | 1 | 10 | Leo1 | 0.053 | Old |
| 0.090585 | 0.307748 | 0.134 | 0.071 | 1 | 10 | Ssna1 | 0.063 | Old |
| 0.090894 | 0.154778 | 0.829 | 0.707 | 1 | 10 | Ywhab | 0.122 | Old |
| 0.09093 | 0.034218 | 0.719 | 0.616 | 1 | 10 | Arpc4 | 0.103 | Old |
| 0.090931 | 0.433787 | 0.213 | 0.131 | 1 | 10 | Elovl5 | 0.082 | Old |
| 0.09105 | 0.151513 | 0.895 | 0.848 | 1 | 10 | Cd44 | 0.047 | Old |
| 0.091302 | 0.287212 | 0.455 | 0.323 | 1 | 10 | Paip2 | 0.132 | Old |
| 0.091544 | 0.390984 | 0.455 | 0.354 | 1 | 10 | Hcfc1r1 | 0.101 | Old |
| 0.091613 | 0.661811 | 0.134 | 0.071 | 1 | 10 | Sec24d | 0.063 | Old |
| 0.092297 | 0.37818 | 0.271 | 0.182 | 1 | 10 | Sec13 | 0.089 | Old |
| 0.092537 | 0.141218 | 0.271 | 0.172 | 1 | 10 | Herc2 | 0.099 | Old |
| 0.092556 | 0.882535 | 0.122 | 0.061 | 1 | 10 | Appl2 | 0.061 | Old |
| 0.09265 | 0.35134 | 0.134 | 0.071 | 1 | 10 | Ptger2 | 0.063 | Old |
| 0.092671 | 1.251808 | 0.064 | 0.02 | 1 | 10 | Plpp6 | 0.044 | Old |
| 0.092673 | 0.184946 | 0.809 | 0.697 | 1 | 10 | Celf2 | 0.112 | Old |
| 0.092687 | 0.145934 | 0.978 | 0.97 | 1 | 10 | Rpl39 | 0.008 | Old |
| 0.092755 | 0.107844 | 0.333 | 0.232 | 1 | 10 | Myo9a | 0.101 | Old |
| 0.09282 | 0.983915 | 0.12 | 0.061 | 1 | 10 | Rfwd3 | 0.059 | Old |
| 0.092853 | 1.067769 | 0.078 | 0.03 | 1 | 10 | Kbtbd11 | 0.048 | Old |
| 0.092876 | 2.156181 | 0.046 | 0.01 | 1 | 10 | Fam206a | 0.036 | Old |
| 0.092927 | 0.273662 | 0.597 | 0.505 | 1 | 10 | Gas7 | 0.092 | Old |
| 0.093085 | 0.768562 | 0.169 | 0.101 | 1 | 10 | Smim19 | 0.068 | Old |
| 0.093279 | 0.192701 | 0.139 | 0.071 | 1 | 10 | Pdcd7 | 0.068 | Old |
| 0.093458 | 0.263964 | 0.474 | 0.354 | 1 | 10 | Kdm2a | 0.12 | Old |
| 0.09353 | 0.95944 | 0.078 | 0.03 | 1 | 10 | Cd109 | 0.048 | Old |
| 0.093593 | 0.325432 | 0.457 | 0.364 | 1 | 10 | 1110004F1 | 0.093 | Old |
| 0.094178 | 0.268882 | 0.511 | 0.404 | 1 | 10 | Itgav | 0.107 | Old |

| | | | | | | | | |
|---|---|---|---|---|---|---|---|---|
| 0.094178 | 1.176647 | 0.064 | 0.02 | 1 | 10 | Abcb6 | 0.044 | Old |
| 0.094204 | 0.908575 | 0.176 | 0.111 | 1 | 10 | Mex3c | 0.065 | Old |
| 0.094205 | 2.323333 | 0.046 | 0.01 | 1 | 10 | Fabp3 | 0.036 | Old |
| 0.094205 | 2.169298 | 0.046 | 0.01 | 1 | 10 | Oas3 | 0.036 | Old |
| 0.094219 | 1.24659 | 0.105 | 0.051 | 1 | 10 | Med22 | 0.054 | Old |
| 0.094303 | 1.103897 | 0.166 | 0.101 | 1 | 10 | Fbxw7 | 0.065 | Old |
| 0.094387 | 0.688093 | 0.174 | 0.101 | 1 | 10 | Zmym4 | 0.073 | Old |
| 0.094482 | 0.516191 | 0.225 | 0.141 | 1 | 10 | Socs6 | 0.084 | Old |
| 0.094554 | 1.139702 | 0.078 | 0.03 | 1 | 10 | Uvssa | 0.048 | Old |
| 0.094651 | 2.028373 | 0.046 | 0.01 | 1 | 10 | Mfsd13a | 0.036 | Old |
| 0.094651 | 1.777691 | 0.046 | 0.01 | 1 | 10 | Tldc1 | 0.036 | Old |
| 0.094738 | 0.136001 | 0.758 | 0.667 | 1 | 10 | Ldha | 0.091 | Old |
| 0.094846 | 0.460905 | 0.281 | 0.192 | 1 | 10 | Tsc22d2 | 0.089 | Old |
| 0.094871 | 0.814706 | 0.159 | 0.091 | 1 | 10 | Irgm1 | 0.068 | Old |
| 0.094872 | 0.252433 | 0.787 | 0.727 | 1 | 10 | Pdia6 | 0.06 | Old |
| 0.095049 | 0.228121 | 0.227 | 0.141 | 1 | 10 | Ufl1 | 0.086 | Old |
| 0.095099 | 1.895597 | 0.046 | 0.01 | 1 | 10 | Mfsd4b4 | 0.036 | Old |
| 0.095158 | 0.752168 | 0.237 | 0.162 | 1 | 10 | Man2a2 | 0.075 | Old |
| 0.095317 | 0.675279 | 0.183 | 0.111 | 1 | 10 | Mtf2 | 0.072 | Old |
| 0.095376 | 0.255982 | 0.296 | 0.202 | 1 | 10 | Epb41l1 | 0.094 | Old |
| 0.095482 | 0.342846 | 0.325 | 0.222 | 1 | 10 | Bak1 | 0.103 | Old |
| 0.095714 | 0.659857 | 0.284 | 0.202 | 1 | 10 | Dclre1c | 0.082 | Old |
| 0.096415 | 0.790783 | 0.132 | 0.071 | 1 | 10 | Ryr1 | 0.061 | Old |
| 0.096895 | 0.763294 | 0.325 | 0.242 | 1 | 10 | Dusp2 | 0.083 | Old |
| 0.097158 | 0.173777 | 0.719 | 0.616 | 1 | 10 | Ppp1ca | 0.103 | Old |
| 0.09727 | 0.781185 | 0.13 | 0.071 | 1 | 10 | Pmm2 | 0.059 | Old |
| 0.097503 | 0.271136 | 0.653 | 0.525 | 1 | 10 | Rock2 | 0.128 | Old |
| 0.09768 | 1.034712 | 0.078 | 0.03 | 1 | 10 | Oas2 | 0.048 | Old |
| 0.097822 | 1.244053 | 0.046 | 0.01 | 1 | 10 | 9330160F1 | 0.036 | Old |
| 0.097941 | 0.373512 | 0.496 | 0.404 | 1 | 10 | Ube2r2 | 0.092 | Old |
| 0.09812 | 1.723834 | 0.076 | 0.03 | 1 | 10 | Ptdss2 | 0.046 | Old |
| 0.098286 | 0.387152 | 0.232 | 0.152 | 1 | 10 | Mesd | 0.08 | Old |
| 0.098436 | 0.520253 | 0.311 | 0.232 | 1 | 10 | Tmem219 | 0.079 | Old |
| 0.098496 | 0.545715 | 0.32 | 0.242 | 1 | 10 | Gba | 0.078 | Old |
| 0.098504 | 0.593016 | 0.227 | 0.152 | 1 | 10 | Gskip | 0.075 | Old |
| 0.098589 | 0.719641 | 0.215 | 0.141 | 1 | 10 | Chchd7 | 0.074 | Old |
| 0.098669 | 0.138338 | 0.274 | 0.182 | 1 | 10 | Fam103a1 | 0.092 | Old |
| 0.098746 | 1.178042 | 0.117 | 0.061 | 1 | 10 | Ttc37 | 0.056 | Old |
| 0.09879 | 2.133144 | 0.061 | 0.02 | 1 | 10 | Coq3 | 0.041 | Old |
| 0.098838 | 1.875964 | 0.076 | 0.03 | 1 | 10 | Atm | 0.046 | Old |
| 0.098928 | 1.479155 | 0.103 | 0.051 | 1 | 10 | Aktip | 0.052 | Old |
| 0.098961 | 0.411479 | 0.2 | 0.121 | 1 | 10 | Aaed1 | 0.079 | Old |
| 0.099199 | 1.669784 | 0.076 | 0.03 | 1 | 10 | Scyl3 | 0.046 | Old |
| 0.099282 | 0.777366 | 0.24 | 0.172 | 1 | 10 | Trappc1 | 0.068 | Old |
| 0.099307 | 0.391678 | 0.313 | 0.222 | 1 | 10 | Tia1 | 0.091 | Old |

| | | | | | | | | |
|---|---|---|---|---|---|---|---|---|
| 0.099496 | 0.448774 | 0.242 | 0.162 | 1 | 10 | Tmem184b | 0.08 | Old |
| 0.099587 | 0.197299 | 0.479 | 0.343 | 1 | 10 | Synj1 | 0.136 | Old |
| 0.099672 | 0.922786 | 0.046 | 0.01 | 1 | 10 | Gm26569 | 0.036 | Old |
| 0.099672 | 0.851083 | 0.046 | 0.01 | 1 | 10 | Mvk | 0.036 | Old |
| 0.099674 | 2.824886 | 0.027 | 0 | 1 | 10 | Tgfbr3 | 0.027 | Old |
| 0.099674 | 2.581782 | 0.027 | 0 | 1 | 10 | Strbp | 0.027 | Old |
| 0.099674 | 2.511548 | 0.027 | 0 | 1 | 10 | Gm47507 | 0.027 | Old |
| 0.099674 | 2.509169 | 0.027 | 0 | 1 | 10 | Arv1 | 0.027 | Old |
| 0.099674 | 2.470306 | 0.027 | 0 | 1 | 10 | Ccnb2 | 0.027 | Old |
| 0.099674 | 2.336841 | 0.027 | 0 | 1 | 10 | Pomt1 | 0.027 | Old |
| 0.099674 | 2.251758 | 0.027 | 0 | 1 | 10 | Csprs | 0.027 | Old |
| 0.099674 | 2.250352 | 0.027 | 0 | 1 | 10 | Slc9a7 | 0.027 | Old |
| 0.099674 | 2.24808 | 0.027 | 0 | 1 | 10 | Gm15564 | 0.027 | Old |
| 0.099674 | 2.124513 | 0.027 | 0 | 1 | 10 | Mlkl | 0.027 | Old |
| 0.099674 | 2.043841 | 0.027 | 0 | 1 | 10 | Zfp41 | 0.027 | Old |
| 0.099674 | 2.02531 | 0.027 | 0 | 1 | 10 | Magi2 | 0.027 | Old |
| 0.099677 | 0.397 | 0.347 | 0.263 | 1 | 10 | Il7r | 0.084 | Old |
| 0.099989 | 0.227315 | 0.416 | 0.313 | 1 | 10 | Top2b | 0.103 | Old |
| 0.099991 | 0.410797 | 0.247 | 0.162 | 1 | 10 | Slc25a1 | 0.085 | Old |
| 0.100139 | 0.659752 | 0.046 | 0.01 | 1 | 10 | Cdca8 | 0.036 | Old |
| 0.100139 | 0.335814 | 0.046 | 0.01 | 1 | 10 | Inpp5e | 0.036 | Old |
| 0.100139 | 0.032671 | 0.046 | 0.01 | 1 | 10 | Mtmr4 | 0.036 | Old |
| 0.100423 | 0.626814 | 0.144 | 0.081 | 1 | 10 | Traf3 | 0.063 | Old |
| 0.100437 | 0.319225 | 0.254 | 0.172 | 1 | 10 | Rps27rt | 0.082 | Old |
| 0.100478 | 0.758732 | 0.193 | 0.121 | 1 | 10 | Fcgr4 | 0.072 | Old |
| 0.100498 | 0.587592 | 0.318 | 0.232 | 1 | 10 | Ythdf1 | 0.086 | Old |
| 0.100534 | 0.833457 | 0.22 | 0.152 | 1 | 10 | Mxi1 | 0.068 | Old |
| 0.100658 | 0.681208 | 0.252 | 0.172 | 1 | 10 | Ech1 | 0.08 | Old |
| 0.100811 | 1.855177 | 0.061 | 0.02 | 1 | 10 | Zfp944 | 0.041 | Old |
| 0.100901 | 0.184983 | 0.403 | 0.293 | 1 | 10 | Cd180 | 0.11 | Old |
| 0.100908 | 0.809477 | 0.105 | 0.051 | 1 | 10 | Foxk1 | 0.054 | Old |
| 0.101343 | 1.614237 | 0.112 | 0.061 | 1 | 10 | Wdsub1 | 0.051 | Old |
| 0.101449 | 0.686202 | 0.2 | 0.131 | 1 | 10 | Slc41a2 | 0.069 | Old |
| 0.101534 | 0.673367 | 0.105 | 0.051 | 1 | 10 | Mpg | 0.054 | Old |
| 0.102066 | 0.276993 | 0.406 | 0.323 | 1 | 10 | Mtch1 | 0.083 | Old |
| 0.102302 | 0.55861 | 0.384 | 0.293 | 1 | 10 | Twf1 | 0.091 | Old |
| 0.102329 | 0.357074 | 0.315 | 0.222 | 1 | 10 | Rsf1 | 0.093 | Old |
| 0.102419 | 0.461188 | 0.21 | 0.131 | 1 | 10 | Actl6a | 0.079 | Old |
| 0.102593 | 0.767827 | 0.115 | 0.061 | 1 | 10 | Il16 | 0.054 | Old |
| 0.102654 | 0.257256 | 0.345 | 0.253 | 1 | 10 | Zc3h15 | 0.092 | Old |
| 0.102755 | 0.160739 | 0.355 | 0.253 | 1 | 10 | Tox4 | 0.102 | Old |
| 0.102798 | 0.816905 | 0.181 | 0.111 | 1 | 10 | Klhl28 | 0.07 | Old |
| 0.102986 | 0.563021 | 0.166 | 0.101 | 1 | 10 | Sipa1l1 | 0.065 | Old |
| 0.103105 | 0.828253 | 0.09 | 0.04 | 1 | 10 | Rars2 | 0.05 | Old |
| 0.103525 | 1.168067 | 0.127 | 0.071 | 1 | 10 | Dna2 | 0.056 | Old |

| | | | | | | | | |
|---|---|---|---|---|---|---|---|---|
| 0.103759 | 0.787808 | 0.166 | 0.101 | 1 | 10 | Atg16l1 | 0.065 | Old |
| 0.103797 | 0.907747 | 0.117 | 0.061 | 1 | 10 | Afdn | 0.056 | Old |
| 0.104015 | 0.239255 | 0.609 | 0.505 | 1 | 10 | Ube3a | 0.104 | Old |
| 0.104116 | 0.081209 | 0.134 | 0.071 | 1 | 10 | Ppp1r37 | 0.063 | Old |
| 0.104193 | 0.144933 | 0.844 | 0.737 | 1 | 10 | Rac1 | 0.107 | Old |
| 0.10421 | 0.195676 | 0.183 | 0.111 | 1 | 10 | Cramp1l | 0.072 | Old |
| 0.104258 | 0.124943 | 0.973 | 0.96 | 1 | 10 | H2-D1 | 0.013 | Old |
| 0.104314 | 0.070682 | 0.609 | 0.485 | 1 | 10 | Atrx | 0.124 | Old |
| 0.104456 | 0.570454 | 0.156 | 0.091 | 1 | 10 | Esco1 | 0.065 | Old |
| 0.104524 | 0.30994 | 0.21 | 0.131 | 1 | 10 | Srp54a | 0.079 | Old |
| 0.104532 | 1.485577 | 0.061 | 0.02 | 1 | 10 | Nubpl | 0.041 | Old |
| 0.104532 | 1.279585 | 0.061 | 0.02 | 1 | 10 | Xpnpep3 | 0.041 | Old |
| 0.104532 | 1.207336 | 0.061 | 0.02 | 1 | 10 | Acot1 | 0.041 | Old |
| 0.104667 | 0.424034 | 0.259 | 0.172 | 1 | 10 | Vps11 | 0.087 | Old |
| 0.104713 | 0.638534 | 0.105 | 0.051 | 1 | 10 | Nsmaf | 0.054 | Old |
| 0.104992 | 0.407592 | 0.156 | 0.091 | 1 | 10 | Cnot10 | 0.065 | Old |
| 0.105035 | 0.733663 | 0.105 | 0.051 | 1 | 10 | Fam53a | 0.054 | Old |
| 0.105198 | 0.413889 | 0.213 | 0.131 | 1 | 10 | Gcnt1 | 0.082 | Old |
| 0.1052 | 0.565528 | 0.227 | 0.152 | 1 | 10 | Themis2 | 0.075 | Old |
| 0.105395 | 0.343009 | 0.254 | 0.172 | 1 | 10 | Pcbd2 | 0.082 | Old |
| 0.105749 | 0.260522 | 0.592 | 0.465 | 1 | 10 | Hnrnpc | 0.127 | Old |
| 0.105779 | 0.528043 | 0.33 | 0.242 | 1 | 10 | Msl2 | 0.088 | Old |
| 0.105796 | 1.344733 | 0.061 | 0.02 | 1 | 10 | Wwox | 0.041 | Old |
| 0.105869 | 1.215413 | 0.076 | 0.03 | 1 | 10 | Pcid2 | 0.046 | Old |
| 0.105887 | 0.051198 | 0.594 | 0.495 | 1 | 10 | Alcam | 0.099 | Old |
| 0.105944 | 0.232058 | 0.452 | 0.354 | 1 | 10 | Tmed3 | 0.098 | Old |
| 0.106072 | 0.354618 | 0.655 | 0.596 | 1 | 10 | H2-DMa | 0.059 | Old |
| 0.106105 | 0.548945 | 0.333 | 0.253 | 1 | 10 | Igbp1 | 0.08 | Old |
| 0.10619 | 0.697478 | 0.154 | 0.091 | 1 | 10 | Ccnd3 | 0.063 | Old |
| 0.10622 | 1.353001 | 0.061 | 0.02 | 1 | 10 | Prickle3 | 0.041 | Old |
| 0.106332 | 0.16711 | 0.824 | 0.747 | 1 | 10 | Ndufa4 | 0.077 | Old |
| 0.106527 | 0.590512 | 0.193 | 0.121 | 1 | 10 | Dock1 | 0.072 | Old |
| 0.106657 | 0.568812 | 0.105 | 0.051 | 1 | 10 | Gzf1 | 0.054 | Old |
| 0.107004 | 1.589525 | 0.088 | 0.04 | 1 | 10 | Chkb | 0.048 | Old |
| 0.107019 | 0.082991 | 0.643 | 0.525 | 1 | 10 | Tpm4 | 0.118 | Old |
| 0.107113 | 0.277569 | 0.694 | 0.586 | 1 | 10 | Slfn2 | 0.108 | Old |
| 0.107193 | 0.268555 | 0.205 | 0.131 | 1 | 10 | Fryl | 0.074 | Old |
| 0.107393 | 0.267007 | 0.499 | 0.394 | 1 | 10 | Hnrnpul1 | 0.105 | Old |
| 0.107485 | 0.231775 | 0.117 | 0.061 | 1 | 10 | Rnf170 | 0.056 | Old |
| 0.10751 | 0.674149 | 0.227 | 0.152 | 1 | 10 | Slc36a4 | 0.075 | Old |
| 0.107537 | 0.425026 | 0.222 | 0.152 | 1 | 10 | Zpr1 | 0.07 | Old |
| 0.107558 | 0.80385 | 0.154 | 0.091 | 1 | 10 | Pgap1 | 0.063 | Old |
| 0.107562 | 1.122148 | 0.139 | 0.081 | 1 | 10 | Coq8b | 0.058 | Old |
| 0.107573 | 0.322988 | 0.496 | 0.404 | 1 | 10 | Ppp1r9b | 0.092 | Old |
| 0.10758 | 0.481028 | 0.169 | 0.101 | 1 | 10 | Timm50 | 0.068 | Old |

| | | | | | | | | |
|---|---|---|---|---|---|---|---|---|
| 0.107701 | 1.815897 | 0.044 | 0.01 | 1 | 10 | Snapc4 | 0.034 | Old |
| 0.107824 | 0.514579 | 0.271 | 0.192 | 1 | 10 | Rgs18 | 0.079 | Old |
| 0.107906 | 0.101562 | 0.472 | 0.364 | 1 | 10 | Cpsf6 | 0.108 | Old |
| 0.107923 | 0.324583 | 0.298 | 0.212 | 1 | 10 | Ahnak2 | 0.086 | Old |
| 0.108035 | 0.257699 | 0.399 | 0.303 | 1 | 10 | Bmp2k | 0.096 | Old |
| 0.108109 | 0.641241 | 0.117 | 0.061 | 1 | 10 | Gm38399 | 0.056 | Old |
| 0.108126 | 1.081394 | 0.181 | 0.121 | 1 | 10 | Dgcr2 | 0.06 | Old |
| 0.10824 | 0.173595 | 0.883 | 0.859 | 1 | 10 | Btf3 | 0.024 | Old |
| 0.108242 | 0.521354 | 0.213 | 0.141 | 1 | 10 | Gyg | 0.072 | Old |
| 0.108317 | 0.464582 | 0.264 | 0.182 | 1 | 10 | Drg1 | 0.082 | Old |
| 0.108794 | 1.415819 | 0.088 | 0.04 | 1 | 10 | Tmem206 | 0.048 | Old |
| 0.108803 | 0.283595 | 0.218 | 0.141 | 1 | 10 | Prune2 | 0.077 | Old |
| 0.108821 | 0.030951 | 0.247 | 0.162 | 1 | 10 | Lmbrd2 | 0.085 | Old |
| 0.108913 | 0.051295 | 0.675 | 0.556 | 1 | 10 | Larp4b | 0.119 | Old |
| 0.109022 | 0.276601 | 0.377 | 0.273 | 1 | 10 | Med15 | 0.104 | Old |
| 0.109227 | 1.106647 | 0.061 | 0.02 | 1 | 10 | Ppcs | 0.041 | Old |
| 0.109428 | 0.719767 | 0.115 | 0.061 | 1 | 10 | Urod | 0.054 | Old |
| 0.109496 | 0.38731 | 0.367 | 0.283 | 1 | 10 | Slc4a7 | 0.084 | Old |
| 0.109515 | 0.019476 | 0.374 | 0.283 | 1 | 10 | Ppp1r2 | 0.091 | Old |
| 0.109662 | 1.023566 | 0.061 | 0.02 | 1 | 10 | Zfp65 | 0.041 | Old |
| 0.109662 | 0.362623 | 0.643 | 0.545 | 1 | 10 | Akap13 | 0.098 | Old |
| 0.109714 | 0.26322 | 0.147 | 0.081 | 1 | 10 | Sp3os | 0.066 | Old |
| 0.109747 | 1.842342 | 0.044 | 0.01 | 1 | 10 | Xndc1 | 0.034 | Old |
| 0.109852 | 0.412271 | 0.401 | 0.313 | 1 | 10 | Saraf | 0.088 | Old |
| 0.1099 | 1.054323 | 0.152 | 0.091 | 1 | 10 | Lrrc8a | 0.061 | Old |
| 0.110043 | 0.613614 | 0.142 | 0.081 | 1 | 10 | Daxx | 0.061 | Old |
| 0.110085 | 0.233754 | 0.244 | 0.162 | 1 | 10 | Galns | 0.082 | Old |
| 0.110128 | 0.708019 | 0.23 | 0.162 | 1 | 10 | Plxnc1 | 0.068 | Old |
| 0.110627 | 0.086272 | 0.702 | 0.556 | 1 | 10 | Pfdn5 | 0.146 | Old |
| 0.110647 | 0.172301 | 0.677 | 0.566 | 1 | 10 | Macf1 | 0.111 | Old |
| 0.110764 | 0.840139 | 0.164 | 0.101 | 1 | 10 | Scyl1 | 0.063 | Old |
| 0.111005 | 0.316346 | 0.372 | 0.283 | 1 | 10 | Timm10b | 0.089 | Old |
| 0.111349 | 1.047591 | 0.139 | 0.081 | 1 | 10 | Pik3c3 | 0.058 | Old |
| 0.11136 | 1.076352 | 0.193 | 0.131 | 1 | 10 | Nsmce4a | 0.062 | Old |
| 0.111366 | 0.258664 | 0.582 | 0.444 | 1 | 10 | G3bp2 | 0.138 | Old |
| 0.111392 | 0.459017 | 0.198 | 0.121 | 1 | 10 | Osbp | 0.077 | Old |
| 0.111693 | 0.221284 | 0.599 | 0.485 | 1 | 10 | Ndufb11 | 0.114 | Old |
| 0.11198 | 1.028737 | 0.103 | 0.051 | 1 | 10 | Map4k5 | 0.052 | Old |
| 0.111999 | 0.513428 | 0.401 | 0.313 | 1 | 10 | Cyb5r4 | 0.088 | Old |
| 0.11229 | 0.836908 | 0.09 | 0.04 | 1 | 10 | Nat10 | 0.05 | Old |
| 0.112349 | 1.774916 | 0.044 | 0.01 | 1 | 10 | 2010320M | 0.034 | Old |
| 0.11238 | 0.207967 | 0.567 | 0.475 | 1 | 10 | S100a11 | 0.092 | Old |
| 0.112745 | 0.848506 | 0.061 | 0.02 | 1 | 10 | 10-Sep | 0.041 | Old |
| 0.112875 | 1.597002 | 0.044 | 0.01 | 1 | 10 | Gm29340 | 0.034 | Old |
| 0.11303 | 0.094817 | 0.702 | 0.606 | 1 | 10 | Clec12a | 0.096 | Old |

| | | | | | | | | |
|---|---|---|---|---|---|---|---|---|
| 0.11312 | 0.795647 | 0.166 | 0.101 | 1 | 10 | Slc39a9 | 0.065 | Old |
| 0.113187 | 1.434585 | 0.088 | 0.04 | 1 | 10 | Hmgn5 | 0.048 | Old |
| 0.113189 | 0.808 | 0.2 | 0.131 | 1 | 10 | Hist3h2a | 0.069 | Old |
| 0.11339 | 0.433809 | 0.09 | 0.04 | 1 | 10 | Homer3 | 0.05 | Old |
| 0.113399 | 0.336444 | 0.467 | 0.364 | 1 | 10 | Elavl1 | 0.103 | Old |
| 0.113681 | 0.25417 | 0.677 | 0.576 | 1 | 10 | Rpl27 | 0.101 | Old |
| 0.1137 | 0.300676 | 0.2 | 0.131 | 1 | 10 | Flot2 | 0.069 | Old |
| 0.114025 | 0.38906 | 0.222 | 0.152 | 1 | 10 | Pex19 | 0.07 | Old |
| 0.114107 | 0.79802 | 0.127 | 0.071 | 1 | 10 | Acss1 | 0.056 | Old |
| 0.114499 | 0.190249 | 0.09 | 0.04 | 1 | 10 | Nol9 | 0.05 | Old |
| 0.114514 | 0.189775 | 0.117 | 0.061 | 1 | 10 | Xrcc4 | 0.056 | Old |
| 0.114532 | 0.536312 | 0.254 | 0.182 | 1 | 10 | Fam117b | 0.072 | Old |
| 0.114544 | 0.08012 | 0.587 | 0.444 | 1 | 10 | Clec4a1 | 0.143 | Old |
| 0.114697 | 0.085243 | 0.12 | 0.061 | 1 | 10 | Xpo4 | 0.059 | Old |
| 0.114751 | 1.354887 | 0.073 | 0.03 | 1 | 10 | Slc26a11 | 0.043 | Old |
| 0.115333 | 1.031114 | 0.161 | 0.101 | 1 | 10 | Ttc13 | 0.06 | Old |
| 0.115392 | 0.392834 | 0.242 | 0.162 | 1 | 10 | Glb1 | 0.08 | Old |
| 0.115429 | 1.258638 | 0.1 | 0.051 | 1 | 10 | Cops7b | 0.049 | Old |
| 0.115493 | 0.726438 | 0.164 | 0.101 | 1 | 10 | Eef1akmt1 | 0.063 | Old |
| 0.115646 | 0.369161 | 0.174 | 0.101 | 1 | 10 | Utp18 | 0.073 | Old |
| 0.115665 | 0.33395 | 0.413 | 0.303 | 1 | 10 | Dcun1d5 | 0.11 | Old |
| 0.115939 | 0.080173 | 0.01 | 0.03 | 1 | 10 | Noxo1 | -0.02 | Old |
| 0.116073 | 0.728176 | 0.044 | 0.01 | 1 | 10 | Pdss2 | 0.034 | Old |
| 0.116073 | 0.228867 | 0.044 | 0.01 | 1 | 10 | Zfp429 | 0.034 | Old |
| 0.116437 | 0.261799 | 0.443 | 0.323 | 1 | 10 | Cdk6 | 0.12 | Old |
| 0.116738 | 0.119359 | 0.597 | 0.515 | 1 | 10 | Ppp2r5c | 0.082 | Old |
| 0.116771 | 0.161954 | 0.562 | 0.444 | 1 | 10 | Hectd1 | 0.118 | Old |
| 0.116782 | 0.280468 | 0.418 | 0.323 | 1 | 10 | Zc3hav1 | 0.095 | Old |
| 0.116854 | 3.405302 | 0.024 | 0 | 1 | 10 | Cpeb1 | 0.024 | Old |
| 0.116854 | 2.580814 | 0.024 | 0 | 1 | 10 | Lca5 | 0.024 | Old |
| 0.116854 | 2.504101 | 0.024 | 0 | 1 | 10 | Rab26os | 0.024 | Old |
| 0.116854 | 2.492869 | 0.024 | 0 | 1 | 10 | Kif2c | 0.024 | Old |
| 0.116854 | 2.468711 | 0.024 | 0 | 1 | 10 | Sphk1 | 0.024 | Old |
| 0.116854 | 2.39256 | 0.024 | 0 | 1 | 10 | Mis18bp1 | 0.024 | Old |
| 0.116854 | 2.331502 | 0.024 | 0 | 1 | 10 | Cdk15 | 0.024 | Old |
| 0.116854 | 2.284061 | 0.024 | 0 | 1 | 10 | Enpp4 | 0.024 | Old |
| 0.116854 | 2.235555 | 0.024 | 0 | 1 | 10 | Ttll4 | 0.024 | Old |
| 0.116854 | 2.170614 | 0.024 | 0 | 1 | 10 | Kcnq1 | 0.024 | Old |
| 0.116854 | 2.144507 | 0.024 | 0 | 1 | 10 | Il1rl2 | 0.024 | Old |
| 0.116854 | 2.12011 | 0.024 | 0 | 1 | 10 | C130026I2 | 0.024 | Old |
| 0.116854 | 2.100109 | 0.024 | 0 | 1 | 10 | Hhat | 0.024 | Old |
| 0.116854 | 2.058233 | 0.024 | 0 | 1 | 10 | Kyat1 | 0.024 | Old |
| 0.116854 | 2.053435 | 0.024 | 0 | 1 | 10 | Dennd5b | 0.024 | Old |
| 0.116854 | 1.920495 | 0.024 | 0 | 1 | 10 | Stk39 | 0.024 | Old |
| 0.116854 | 1.870834 | 0.024 | 0 | 1 | 10 | Trmt5 | 0.024 | Old |

| | | | | | | | | |
|---|---|---|---|---|---|---|---|---|
| 0.116854 | 1.844901 | 0.024 | 0 | 1 | 10 | Gm16845 | 0.024 | Old |
| 0.116854 | 1.779317 | 0.024 | 0 | 1 | 10 | 4930555A( | 0.024 | Old |
| 0.116854 | 1.750512 | 0.024 | 0 | 1 | 10 | Klrk1 | 0.024 | Old |
| 0.116899 | 0.446223 | 0.377 | 0.283 | 1 | 10 | Pnpla8 | 0.094 | Old |
| 0.117005 | 0.452637 | 0.178 | 0.111 | 1 | 10 | Lypla1 | 0.067 | Old |
| 0.117103 | 0.231564 | 0.45 | 0.354 | 1 | 10 | Amfr | 0.096 | Old |
| 0.117133 | 0.046385 | 0.587 | 0.444 | 1 | 10 | Vps4b | 0.143 | Old |
| 0.117199 | 0.367705 | 0.164 | 0.101 | 1 | 10 | Nip7 | 0.063 | Old |
| 0.117269 | 0.891335 | 0.149 | 0.091 | 1 | 10 | Mzt1 | 0.058 | Old |
| 0.117269 | 0.868907 | 0.149 | 0.091 | 1 | 10 | Sh3glb2 | 0.058 | Old |
| 0.117291 | 0.197243 | 0.704 | 0.626 | 1 | 10 | Ppp2ca | 0.078 | Old |
| 0.117567 | 0.774341 | 0.103 | 0.051 | 1 | 10 | Cdc42ep2 | 0.052 | Old |
| 0.117567 | 0.74805 | 0.103 | 0.051 | 1 | 10 | Sertad3 | 0.052 | Old |
| 0.1177 | 0.445611 | 0.191 | 0.121 | 1 | 10 | Gm49336 | 0.07 | Old |
| 0.117705 | 0.180233 | 0.905 | 0.838 | 1 | 10 | mt-Nd5 | 0.067 | Old |
| 0.117718 | 0.846352 | 0.088 | 0.04 | 1 | 10 | Dglucy | 0.048 | Old |
| 0.117718 | 0.840015 | 0.088 | 0.04 | 1 | 10 | Ganc | 0.048 | Old |
| 0.117741 | 1.043292 | 0.159 | 0.101 | 1 | 10 | Slc25a17 | 0.058 | Old |
| 0.117785 | 0.697169 | 0.188 | 0.121 | 1 | 10 | Ccdc6 | 0.067 | Old |
| 0.117908 | 1.510679 | 0.059 | 0.02 | 1 | 10 | Rabif | 0.039 | Old |
| 0.117926 | 0.494546 | 0.264 | 0.192 | 1 | 10 | Trappc10 | 0.072 | Old |
| 0.118096 | 0.351502 | 0.293 | 0.202 | 1 | 10 | Herc4 | 0.091 | Old |
| 0.118102 | 0.788456 | 0.088 | 0.04 | 1 | 10 | Uqcc1 | 0.048 | Old |
| 0.118102 | 0.697124 | 0.088 | 0.04 | 1 | 10 | Utp4 | 0.048 | Old |
| 0.118186 | 0.598897 | 0.166 | 0.101 | 1 | 10 | Creb3 | 0.065 | Old |
| 0.11821 | 0.326347 | 0.156 | 0.091 | 1 | 10 | Amn1 | 0.065 | Old |
| 0.118437 | 0.110388 | 0.567 | 0.434 | 1 | 10 | Plxnd1 | 0.133 | Old |
| 0.11926 | 1.120619 | 0.088 | 0.04 | 1 | 10 | Ezh2 | 0.048 | Old |
| 0.119324 | 1.352479 | 0.059 | 0.02 | 1 | 10 | Cd3eap | 0.039 | Old |
| 0.119436 | 0.421821 | 0.306 | 0.222 | 1 | 10 | Hck | 0.084 | Old |
| 0.119486 | 0.056108 | 0.01 | 0.03 | 1 | 10 | Dennd6b | -0.02 | Old |
| 0.119487 | 0.512456 | 0.193 | 0.121 | 1 | 10 | Anapc2 | 0.072 | Old |
| 0.119665 | 0.26941 | 0.721 | 0.636 | 1 | 10 | Apobec1 | 0.085 | Old |
| 0.119684 | 0.229439 | 0.479 | 0.384 | 1 | 10 | Tep1 | 0.095 | Old |
| 0.119909 | 0.227905 | 0.672 | 0.545 | 1 | 10 | Romo1 | 0.127 | Old |
| 0.119928 | 0.375274 | 0.267 | 0.192 | 1 | 10 | Aagab | 0.075 | Old |
| 0.120226 | 1.564294 | 0.086 | 0.04 | 1 | 10 | Plcb1 | 0.046 | Old |
| 0.120367 | 0.296507 | 0.611 | 0.515 | 1 | 10 | 7-Sep | 0.096 | Old |
| 0.120585 | 0.139765 | 0.156 | 0.091 | 1 | 10 | Cabin1 | 0.065 | Old |
| 0.12076 | 0.422799 | 0.391 | 0.303 | 1 | 10 | Rac2 | 0.088 | Old |
| 0.120783 | 0.395692 | 0.267 | 0.182 | 1 | 10 | Crtap | 0.085 | Old |
| 0.120806 | 0.776818 | 0.139 | 0.081 | 1 | 10 | Ssh1 | 0.058 | Old |
| 0.120839 | 0.278935 | 0.396 | 0.303 | 1 | 10 | Acsl4 | 0.093 | Old |
| 0.120886 | 0.152729 | 0.914 | 0.808 | 1 | 10 | Arf5 | 0.106 | Old |
| 0.121222 | 0.685521 | 0.183 | 0.121 | 1 | 10 | Engase | 0.062 | Old |

| | | | | | | | | |
|---|---|---|---|---|---|---|---|---|
| 0.12136 | 0.278237 | 0.658 | 0.545 | 1 | 10 | Slc39a1 | 0.113 | Old |
| 0.121527 | 0.196492 | 0.533 | 0.414 | 1 | 10 | Rab31 | 0.119 | Old |
| 0.12157 | 0.194961 | 0.406 | 0.303 | 1 | 10 | Ist1 | 0.103 | Old |
| 0.12171 | 0.248449 | 0.445 | 0.343 | 1 | 10 | Rufy3 | 0.102 | Old |
| 0.12173 | 0.812707 | 0.115 | 0.061 | 1 | 10 | Numb | 0.054 | Old |
| 0.122133 | 0.586341 | 0.2 | 0.131 | 1 | 10 | Cops8 | 0.069 | Old |
| 0.122171 | 0.621493 | 0.23 | 0.162 | 1 | 10 | Nacc2 | 0.068 | Old |
| 0.122344 | 0.47419 | 0.191 | 0.121 | 1 | 10 | Klhdc4 | 0.07 | Old |
| 0.122391 | 0.454344 | 0.088 | 0.04 | 1 | 10 | Decr2 | 0.048 | Old |
| 0.12275 | 0.1326 | 0.919 | 0.808 | 1 | 10 | Tmbim6 | 0.111 | Old |
| 0.123059 | 1.004374 | 0.1 | 0.051 | 1 | 10 | Etfdh | 0.049 | Old |
| 0.123163 | 1.101687 | 0.059 | 0.02 | 1 | 10 | Tctn3 | 0.039 | Old |
| 0.123775 | 0.234621 | 0.286 | 0.202 | 1 | 10 | Ppp2r2d | 0.084 | Old |
| 0.123803 | 0.21229 | 0.208 | 0.131 | 1 | 10 | Tiprl | 0.077 | Old |
| 0.123805 | 0.044616 | 0.144 | 0.081 | 1 | 10 | Blcap | 0.063 | Old |
| 0.124119 | 0.557453 | 0.269 | 0.192 | 1 | 10 | Micu1 | 0.077 | Old |
| 0.124194 | 0.493561 | 0.254 | 0.182 | 1 | 10 | Manba | 0.072 | Old |
| 0.124369 | 2.374382 | 0.042 | 0.01 | 1 | 10 | Cd2 | 0.032 | Old |
| 0.124451 | 1.380721 | 0.11 | 0.061 | 1 | 10 | Zdhhc17 | 0.049 | Old |
| 0.124512 | 0.580799 | 0.127 | 0.071 | 1 | 10 | Cblb | 0.056 | Old |
| 0.124754 | 0.470042 | 0.406 | 0.313 | 1 | 10 | Itga4 | 0.093 | Old |
| 0.124897 | 0.04244 | 0.281 | 0.192 | 1 | 10 | Slc25a28 | 0.089 | Old |
| 0.124986 | 0.370857 | 0.311 | 0.232 | 1 | 10 | Klc1 | 0.079 | Old |
| 0.125182 | 0.355535 | 0.088 | 0.04 | 1 | 10 | 4921524J1 | 0.048 | Old |
| 0.125356 | 0.016525 | 0.005 | 0.02 | 1 | 10 | Gm48960 | -0.015 | Old |
| 0.125524 | 0.746847 | 0.127 | 0.071 | 1 | 10 | Hsdl2 | 0.056 | Old |
| 0.125541 | 1.911722 | 0.042 | 0.01 | 1 | 10 | Pex12 | 0.032 | Old |
| 0.125834 | 0.569811 | 0.188 | 0.121 | 1 | 10 | Rptor | 0.067 | Old |
| 0.125918 | 0.583404 | 0.161 | 0.101 | 1 | 10 | Ttc5 | 0.06 | Old |
| 0.125955 | 0.164101 | 0.509 | 0.394 | 1 | 10 | Man1c1 | 0.115 | Old |
| 0.12613 | 2.155636 | 0.042 | 0.01 | 1 | 10 | Golm1 | 0.032 | Old |
| 0.12613 | 2.114321 | 0.042 | 0.01 | 1 | 10 | Fdxacb1 | 0.032 | Old |
| 0.126252 | 0.401683 | 0.139 | 0.081 | 1 | 10 | Nit1 | 0.058 | Old |
| 0.126257 | 1.28968 | 0.086 | 0.04 | 1 | 10 | Rhno1 | 0.046 | Old |
| 0.126304 | 0.201041 | 0.557 | 0.444 | 1 | 10 | Tsn | 0.113 | Old |
| 0.126721 | 2.215064 | 0.042 | 0.01 | 1 | 10 | Paip2b | 0.032 | Old |
| 0.126721 | 2.171798 | 0.042 | 0.01 | 1 | 10 | Tdrkh | 0.032 | Old |
| 0.127315 | 2.277232 | 0.042 | 0.01 | 1 | 10 | Ubxn2b | 0.032 | Old |
| 0.127491 | 1.172986 | 0.086 | 0.04 | 1 | 10 | Ccdc122 | 0.046 | Old |
| 0.127561 | 0.75846 | 0.139 | 0.081 | 1 | 10 | Zfp560 | 0.058 | Old |
| 0.127606 | 0.815472 | 0.073 | 0.03 | 1 | 10 | Dnaaf3 | 0.043 | Old |
| 0.127607 | 0.221345 | 0.284 | 0.192 | 1 | 10 | Aida | 0.092 | Old |
| 0.127774 | 0.507374 | 0.213 | 0.141 | 1 | 10 | Polr3c | 0.072 | Old |
| 0.127831 | 0.411053 | 0.396 | 0.313 | 1 | 10 | Pten | 0.083 | Old |
| 0.12791 | 2.149491 | 0.042 | 0.01 | 1 | 10 | Pigg | 0.032 | Old |

| | | | | | | | | |
|---|---|---|---|---|---|---|---|---|
| 0.12791 | 1.903869 | 0.042 | 0.01 | 1 | 10 | Eif2d | 0.032 | Old |
| 0.128211 | 0.414055 | 0.154 | 0.091 | 1 | 10 | Pop7 | 0.063 | Old |
| 0.128331 | 1.901535 | 0.056 | 0.02 | 1 | 10 | Loxl3 | 0.036 | Old |
| 0.128444 | 0.260109 | 0.242 | 0.162 | 1 | 10 | Ilf2 | 0.08 | Old |
| 0.128508 | 1.776042 | 0.042 | 0.01 | 1 | 10 | Tatdn3 | 0.032 | Old |
| 0.128598 | 0.640288 | 0.059 | 0.02 | 1 | 10 | B3gnt5 | 0.039 | Old |
| 0.129024 | 1.119604 | 0.134 | 0.081 | 1 | 10 | Parp9 | 0.053 | Old |
| 0.129102 | 0.502021 | 0.059 | 0.02 | 1 | 10 | Unk | 0.039 | Old |
| 0.129108 | 1.655231 | 0.042 | 0.01 | 1 | 10 | Trdmt1 | 0.032 | Old |
| 0.129297 | 0.508825 | 0.291 | 0.222 | 1 | 10 | Sec22b | 0.069 | Old |
| 0.129346 | 1.575376 | 0.071 | 0.03 | 1 | 10 | Pde4a | 0.041 | Old |
| 0.129417 | 0.134979 | 0.733 | 0.566 | 1 | 10 | Eif5b | 0.167 | Old |
| 0.129432 | 0.134571 | 0.77 | 0.677 | 1 | 10 | Wdr26 | 0.093 | Old |
| 0.129492 | 0.466805 | 0.55 | 0.485 | 1 | 10 | Znrf2 | 0.065 | Old |
| 0.129582 | 0.391134 | 0.237 | 0.162 | 1 | 10 | Adh5 | 0.075 | Old |
| 0.129614 | 0.421442 | 0.227 | 0.152 | 1 | 10 | N4bp2l1 | 0.075 | Old |
| 0.12971 | 1.638923 | 0.042 | 0.01 | 1 | 10 | Slc19a2 | 0.032 | Old |
| 0.129859 | 0.170335 | 0.978 | 0.99 | 1 | 10 | Lcp1 | -0.012 | Old |
| 0.130079 | 0.124823 | 0.743 | 0.646 | 1 | 10 | Itm2c | 0.097 | Old |
| 0.130314 | 1.395618 | 0.042 | 0.01 | 1 | 10 | Cep57l1 | 0.032 | Old |
| 0.130681 | 0.3667 | 0.127 | 0.071 | 1 | 10 | Cdc14b | 0.056 | Old |
| 0.130796 | 0.393707 | 0.337 | 0.253 | 1 | 10 | Cope | 0.084 | Old |
| 0.13083 | 0.125364 | 0.998 | 1 | 1 | 10 | Ctsb | -0.002 | Old |
| 0.130861 | 0.358552 | 0.43 | 0.333 | 1 | 10 | Scarb2 | 0.097 | Old |
| 0.130907 | 0.239391 | 0.271 | 0.192 | 1 | 10 | Mrpl32 | 0.079 | Old |
| 0.131187 | 1.546516 | 0.071 | 0.03 | 1 | 10 | Rnf217 | 0.041 | Old |
| 0.131356 | 0.397611 | 0.269 | 0.192 | 1 | 10 | Rnf213 | 0.077 | Old |
| 0.131508 | 0.392705 | 0.526 | 0.434 | 1 | 10 | Tmem176a | 0.092 | Old |
| 0.131547 | 0.459736 | 0.32 | 0.242 | 1 | 10 | Bmt2 | 0.078 | Old |
| 0.131575 | 0.311927 | 0.487 | 0.384 | 1 | 10 | Psma1 | 0.103 | Old |
| 0.131598 | 0.946709 | 0.112 | 0.061 | 1 | 10 | Dhcr24 | 0.051 | Old |
| 0.13165 | 1.201303 | 0.071 | 0.03 | 1 | 10 | Ccl25 | 0.041 | Old |
| 0.131754 | 0.302386 | 0.384 | 0.303 | 1 | 10 | Med13 | 0.081 | Old |
| 0.131809 | 0.239681 | 0.249 | 0.172 | 1 | 10 | Srpk2 | 0.077 | Old |
| 0.131844 | 0.164821 | 0.33 | 0.232 | 1 | 10 | Ybx3 | 0.098 | Old |
| 0.132521 | 1.198309 | 0.086 | 0.04 | 1 | 10 | Tyw1 | 0.046 | Old |
| 0.132673 | 0.368026 | 0.364 | 0.283 | 1 | 10 | Slc26a2 | 0.081 | Old |
| 0.132898 | 0.138159 | 0.501 | 0.394 | 1 | 10 | Ghitm | 0.107 | Old |
| 0.132994 | 0.179185 | 0.447 | 0.323 | 1 | 10 | Lypla2 | 0.124 | Old |
| 0.133299 | 0.182947 | 0.401 | 0.313 | 1 | 10 | Ube2q1 | 0.088 | Old |
| 0.133368 | 0.96439 | 0.042 | 0.01 | 1 | 10 | Fam102a | 0.032 | Old |
| 0.133368 | 0.92046 | 0.042 | 0.01 | 1 | 10 | A130010J1 | 0.032 | Old |
| 0.133467 | 0.207073 | 0.697 | 0.545 | 1 | 10 | Myof | 0.152 | Old |
| 0.133659 | 0.218271 | 0.308 | 0.222 | 1 | 10 | Kat6a | 0.086 | Old |
| 0.134144 | 0.208885 | 0.215 | 0.141 | 1 | 10 | Sort1 | 0.074 | Old |

| | | | | | | | | |
|---|---|---|---|---|---|---|---|---|
| 0.134367 | 0.265051 | 0.413 | 0.323 | 1 | 10 | Mapk1ip1l | 0.09 | Old |
| 0.134546 | 0.217984 | 0.465 | 0.364 | 1 | 10 | Usp25 | 0.101 | Old |
| 0.134577 | 0.347629 | 0.575 | 0.495 | 1 | 10 | Ap3s1 | 0.08 | Old |
| 0.134661 | 0.951047 | 0.086 | 0.04 | 1 | 10 | Herc3 | 0.046 | Old |
| 0.134855 | 0.167219 | 0.279 | 0.192 | 1 | 10 | Ints6 | 0.087 | Old |
| 0.135205 | 0.488856 | 0.364 | 0.293 | 1 | 10 | Gfpt1 | 0.071 | Old |
| 0.135261 | 0.304549 | 0.284 | 0.202 | 1 | 10 | Tpcn2 | 0.082 | Old |
| 0.135284 | 0.357615 | 0.526 | 0.455 | 1 | 10 | Lbh | 0.071 | Old |
| 0.135403 | 1.346963 | 0.071 | 0.03 | 1 | 10 | Ercc6 | 0.041 | Old |
| 0.135435 | 0.814949 | 0.1 | 0.051 | 1 | 10 | Ints8 | 0.049 | Old |
| 0.135643 | 1.624038 | 0.056 | 0.02 | 1 | 10 | L3mbtl3 | 0.036 | Old |
| 0.135975 | 1.213667 | 0.098 | 0.051 | 1 | 10 | Mrpl22 | 0.047 | Old |
| 0.136004 | 0.139269 | 0.68 | 0.586 | 1 | 10 | Gadd45b | 0.094 | Old |
| 0.136076 | 0.261127 | 0.264 | 0.182 | 1 | 10 | Ireb2 | 0.082 | Old |
| 0.13612 | 0.20717 | 0.408 | 0.313 | 1 | 10 | Pon2 | 0.095 | Old |
| 0.136242 | 0.456533 | 0.149 | 0.091 | 1 | 10 | Gng11 | 0.058 | Old |
| 0.136354 | 1.375513 | 0.071 | 0.03 | 1 | 10 | Glce | 0.041 | Old |
| 0.136382 | 1.014209 | 0.098 | 0.051 | 1 | 10 | Rtl5 | 0.047 | Old |
| 0.13639 | 0.44784 | 0.198 | 0.131 | 1 | 10 | Tmem222 | 0.067 | Old |
| 0.136567 | 0.048252 | 0.513 | 0.384 | 1 | 10 | Rab3il1 | 0.129 | Old |
| 0.136771 | 0.204086 | 0.374 | 0.273 | 1 | 10 | Plekha1 | 0.101 | Old |
| 0.136828 | 0.377102 | 0.086 | 0.04 | 1 | 10 | Pgm3 | 0.046 | Old |
| 0.136844 | 0.435344 | 0.271 | 0.192 | 1 | 10 | Rnf166 | 0.079 | Old |
| 0.136857 | 0.662938 | 0.213 | 0.152 | 1 | 10 | Cdk13 | 0.061 | Old |
| 0.137001 | 0.034542 | 0.215 | 0.141 | 1 | 10 | Apobr | 0.074 | Old |
| 0.137204 | 1.289165 | 0.12 | 0.071 | 1 | 10 | Fbxo4 | 0.049 | Old |
| 0.137278 | 2.919724 | 0.022 | 0 | 1 | 10 | Ifit2 | 0.022 | Old |
| 0.137278 | 2.854316 | 0.022 | 0 | 1 | 10 | Ttn | 0.022 | Old |
| 0.137278 | 2.649554 | 0.022 | 0 | 1 | 10 | E2f2 | 0.022 | Old |
| 0.137278 | 2.54083 | 0.022 | 0 | 1 | 10 | Prkar1b | 0.022 | Old |
| 0.137278 | 2.472765 | 0.022 | 0 | 1 | 10 | Ablim1 | 0.022 | Old |
| 0.137278 | 2.413383 | 0.022 | 0 | 1 | 10 | Tanc1 | 0.022 | Old |
| 0.137278 | 2.288927 | 0.022 | 0 | 1 | 10 | Siah1b | 0.022 | Old |
| 0.137278 | 2.288783 | 0.022 | 0 | 1 | 10 | Gm29650 | 0.022 | Old |
| 0.137278 | 2.272203 | 0.022 | 0 | 1 | 10 | Fitm2 | 0.022 | Old |
| 0.137278 | 2.258583 | 0.022 | 0 | 1 | 10 | Mfsd4a | 0.022 | Old |
| 0.137278 | 2.231127 | 0.022 | 0 | 1 | 10 | Tmem14a | 0.022 | Old |
| 0.137278 | 2.221575 | 0.022 | 0 | 1 | 10 | A630089N( | 0.022 | Old |
| 0.137278 | 2.198016 | 0.022 | 0 | 1 | 10 | Hpn | 0.022 | Old |
| 0.137278 | 2.177144 | 0.022 | 0 | 1 | 10 | Zfp619 | 0.022 | Old |
| 0.137278 | 2.172459 | 0.022 | 0 | 1 | 10 | Ccdc77 | 0.022 | Old |
| 0.137278 | 2.155144 | 0.022 | 0 | 1 | 10 | Tmtc2 | 0.022 | Old |
| 0.137278 | 2.144448 | 0.022 | 0 | 1 | 10 | Armc9 | 0.022 | Old |
| 0.137278 | 2.134346 | 0.022 | 0 | 1 | 10 | Junos | 0.022 | Old |
| 0.137278 | 2.133859 | 0.022 | 0 | 1 | 10 | Slc25a43 | 0.022 | Old |

| | | | | | | | | |
|---|---|---|---|---|---|---|---|---|
| 0.137278 | 2.076287 | 0.022 | 0 | 1 | 10 | Synpo | 0.022 | Old |
| 0.137278 | 2.056503 | 0.022 | 0 | 1 | 10 | Tmem17 | 0.022 | Old |
| 0.137278 | 2.056052 | 0.022 | 0 | 1 | 10 | Satb2 | 0.022 | Old |
| 0.137278 | 2.027936 | 0.022 | 0 | 1 | 10 | Rsph3b | 0.022 | Old |
| 0.137278 | 2.027844 | 0.022 | 0 | 1 | 10 | Als2cl | 0.022 | Old |
| 0.137278 | 2.024184 | 0.022 | 0 | 1 | 10 | Nckap5l | 0.022 | Old |
| 0.137278 | 1.995511 | 0.022 | 0 | 1 | 10 | Magi3 | 0.022 | Old |
| 0.137278 | 1.984316 | 0.022 | 0 | 1 | 10 | Cenpo | 0.022 | Old |
| 0.137278 | 1.957776 | 0.022 | 0 | 1 | 10 | Itpr3 | 0.022 | Old |
| 0.137278 | 1.941914 | 0.022 | 0 | 1 | 10 | Atp5s | 0.022 | Old |
| 0.137278 | 1.89241 | 0.022 | 0 | 1 | 10 | 1700056E2 | 0.022 | Old |
| 0.137278 | 1.889743 | 0.022 | 0 | 1 | 10 | Gpd1 | 0.022 | Old |
| 0.137278 | 1.82317 | 0.022 | 0 | 1 | 10 | A230050P2 | 0.022 | Old |
| 0.137278 | 1.733751 | 0.022 | 0 | 1 | 10 | Dph7 | 0.022 | Old |
| 0.137278 | 1.677173 | 0.022 | 0 | 1 | 10 | Cyp2ab1 | 0.022 | Old |
| 0.137278 | 1.590654 | 0.022 | 0 | 1 | 10 | Rnf24 | 0.022 | Old |
| 0.137536 | 0.083495 | 0.535 | 0.404 | 1 | 10 | Ifnar1 | 0.131 | Old |
| 0.137609 | 0.728297 | 0.098 | 0.051 | 1 | 10 | Ppm1m | 0.047 | Old |
| 0.137692 | 0.766541 | 0.122 | 0.071 | 1 | 10 | Soga1 | 0.051 | Old |
| 0.137969 | 0.362715 | 0.384 | 0.303 | 1 | 10 | Api5 | 0.081 | Old |
| 0.138402 | 0.158491 | 0.844 | 0.737 | 1 | 10 | Tmed10 | 0.107 | Old |
| 0.138559 | 0.6733 | 0.269 | 0.202 | 1 | 10 | Asxl2 | 0.067 | Old |
| 0.13863 | 0.133808 | 0.65 | 0.515 | 1 | 10 | Nckap1l | 0.135 | Old |
| 0.138644 | 0.163435 | 0.758 | 0.677 | 1 | 10 | Nrros | 0.081 | Old |
| 0.138755 | 1.26282 | 0.071 | 0.03 | 1 | 10 | Mad2l2 | 0.041 | Old |
| 0.138844 | 0.655212 | 0.237 | 0.162 | 1 | 10 | Acp5 | 0.075 | Old |
| 0.138952 | 1.343518 | 0.083 | 0.04 | 1 | 10 | Tbp | 0.043 | Old |
| 0.139017 | 0.06823 | 0.623 | 0.505 | 1 | 10 | Plxnb2 | 0.118 | Old |
| 0.139063 | 0.705337 | 0.11 | 0.061 | 1 | 10 | Noc4l | 0.049 | Old |
| 0.139207 | 0.565694 | 0.161 | 0.101 | 1 | 10 | Timm44 | 0.06 | Old |
| 0.139242 | 0.592091 | 0.205 | 0.141 | 1 | 10 | Tceal8 | 0.064 | Old |
| 0.139302 | 0.570724 | 0.152 | 0.091 | 1 | 10 | Acsl3 | 0.061 | Old |
| 0.13937 | 0.109257 | 0.403 | 0.313 | 1 | 10 | Adap2 | 0.09 | Old |
| 0.139417 | 1.355572 | 0.056 | 0.02 | 1 | 10 | Rab13 | 0.036 | Old |
| 0.139461 | 0.199406 | 0.711 | 0.626 | 1 | 10 | Dstn | 0.085 | Old |
| 0.139856 | 0.387333 | 0.2 | 0.131 | 1 | 10 | Gramd1a | 0.069 | Old |
| 0.139931 | 0.442747 | 0.284 | 0.202 | 1 | 10 | Maf1 | 0.082 | Old |
| 0.140046 | 0.285773 | 0.66 | 0.515 | 1 | 10 | Tab2 | 0.145 | Old |
| 0.140331 | 0.646872 | 0.225 | 0.162 | 1 | 10 | Rnf38 | 0.063 | Old |
| 0.14059 | 0.219731 | 0.355 | 0.273 | 1 | 10 | Pnn | 0.082 | Old |
| 0.140698 | 1.01301 | 0.071 | 0.03 | 1 | 10 | Tecpr2 | 0.041 | Old |
| 0.140781 | 0.052757 | 0.548 | 0.455 | 1 | 10 | Malt1 | 0.093 | Old |
| 0.14106 | 0.646208 | 0.056 | 0.02 | 1 | 10 | Tns4 | 0.036 | Old |
| 0.14106 | 0.839486 | 0.056 | 0.02 | 1 | 10 | Lsm11 | 0.036 | Old |
| 0.14161 | 1.161605 | 0.056 | 0.02 | 1 | 10 | Accs | 0.036 | Old |

| | | | | | | | | |
|---|---|---|---|---|---|---|---|---|
| 0.141765 | 0.056095 | 0.379 | 0.293 | 1 | 10 | Ube2s | 0.086 | Old |
| 0.14211 | 0.327278 | 0.315 | 0.232 | 1 | 10 | Ube2f | 0.083 | Old |
| 0.142169 | 0.705522 | 0.071 | 0.03 | 1 | 10 | Mipep | 0.041 | Old |
| 0.142554 | 0.147827 | 0.584 | 0.495 | 1 | 10 | Plec | 0.089 | Old |
| 0.142561 | 1.330377 | 0.083 | 0.04 | 1 | 10 | Tubgcp2 | 0.043 | Old |
| 0.1426 | 0.696072 | 0.098 | 0.051 | 1 | 10 | Ypel2 | 0.047 | Old |
| 0.14264 | 0.922214 | 0.144 | 0.091 | 1 | 10 | Smcr8 | 0.053 | Old |
| 0.142732 | 0.172397 | 0.318 | 0.232 | 1 | 10 | Vps13c | 0.086 | Old |
| 0.142871 | 0.867755 | 0.108 | 0.061 | 1 | 10 | Yod1 | 0.047 | Old |
| 0.143072 | 0.46295 | 0.293 | 0.222 | 1 | 10 | Uba6 | 0.071 | Old |
| 0.143516 | 0.382157 | 0.244 | 0.172 | 1 | 10 | Ktn1 | 0.072 | Old |
| 0.143636 | 0.021837 | 0.995 | 0.98 | 1 | 10 | Iqgap1 | 0.015 | Old |
| 0.143652 | 0.777324 | 0.071 | 0.03 | 1 | 10 | Rbm10 | 0.041 | Old |
| 0.143761 | 0.207341 | 0.303 | 0.222 | 1 | 10 | Tm2d1 | 0.081 | Old |
| 0.14383 | 0.96743 | 0.056 | 0.02 | 1 | 10 | Dusp18 | 0.036 | Old |
| 0.143889 | 0.365018 | 0.176 | 0.111 | 1 | 10 | C1galt1 | 0.065 | Old |
| 0.144295 | 0.574069 | 0.098 | 0.051 | 1 | 10 | Tmem126b | 0.047 | Old |
| 0.144392 | 1.136054 | 0.083 | 0.04 | 1 | 10 | Aptx | 0.043 | Old |
| 0.144564 | 0.53988 | 0.218 | 0.152 | 1 | 10 | Tnfaip1 | 0.066 | Old |
| 0.14457 | 0.460632 | 0.254 | 0.182 | 1 | 10 | Gcc2 | 0.072 | Old |
| 0.144651 | 0.185761 | 0.386 | 0.293 | 1 | 10 | Chmp3 | 0.093 | Old |
| 0.144871 | 0.489333 | 0.125 | 0.071 | 1 | 10 | Aasdhppt | 0.054 | Old |
| 0.144904 | 0.13702 | 0.809 | 0.727 | 1 | 10 | Mrc1 | 0.082 | Old |
| 0.145028 | 0.072193 | 0.518 | 0.414 | 1 | 10 | Copz1 | 0.104 | Old |
| 0.145147 | 0.05005 | 0.071 | 0.03 | 1 | 10 | Zscan21 | 0.041 | Old |
| 0.145266 | 0.386008 | 0.205 | 0.141 | 1 | 10 | Pkib | 0.064 | Old |
| 0.145511 | 0.64444 | 0.056 | 0.02 | 1 | 10 | Rasal2 | 0.036 | Old |
| 0.145572 | 0.677205 | 0.205 | 0.141 | 1 | 10 | Nup50 | 0.064 | Old |
| 0.145928 | 0.12614 | 0.249 | 0.172 | 1 | 10 | D5Ertd579 | 0.077 | Old |
| 0.146075 | 0.246154 | 0.056 | 0.02 | 1 | 10 | Dus2 | 0.036 | Old |
| 0.14615 | 0.805722 | 0.071 | 0.03 | 1 | 10 | Adprm | 0.041 | Old |
| 0.14621 | 0.134423 | 0.579 | 0.455 | 1 | 10 | Il10rb | 0.124 | Old |
| 0.146414 | 1.935159 | 0.039 | 0.01 | 1 | 10 | D3Ertd751 | 0.029 | Old |
| 0.14649 | 0.224587 | 0.487 | 0.404 | 1 | 10 | Stag2 | 0.083 | Old |
| 0.146511 | 0.440938 | 0.247 | 0.172 | 1 | 10 | Dnttip2 | 0.075 | Old |
| 0.146641 | 0.587557 | 0.056 | 0.02 | 1 | 10 | Ccdc91 | 0.036 | Old |
| 0.146643 | 0.983438 | 0.11 | 0.061 | 1 | 10 | Mir17hg | 0.049 | Old |
| 0.146654 | 0.598931 | 0.071 | 0.03 | 1 | 10 | Gstp1 | 0.041 | Old |
| 0.146689 | 0.626324 | 0.134 | 0.081 | 1 | 10 | Btbd10 | 0.053 | Old |
| 0.146779 | 1.10777 | 0.154 | 0.101 | 1 | 10 | Dnpep | 0.053 | Old |
| 0.146799 | 0.246637 | 0.32 | 0.242 | 1 | 10 | Xpr1 | 0.078 | Old |
| 0.147031 | 0.115268 | 0.533 | 0.414 | 1 | 10 | Ralbp1 | 0.119 | Old |
| 0.147082 | 0.338106 | 0.242 | 0.172 | 1 | 10 | Ncoa1 | 0.07 | Old |
| 0.147096 | 2.108085 | 0.039 | 0.01 | 1 | 10 | Acap3 | 0.029 | Old |
| 0.147111 | 0.31642 | 0.205 | 0.141 | 1 | 10 | Zbtb8os | 0.064 | Old |

| | | | | | | | | |
|---|---|---|---|---|---|---|---|---|
| 0.14742 | 0.510093 | 0.205 | 0.141 | 1 | 10 | Srek1ip1 | 0.064 | Old |
| 0.14764 | 1.105921 | 0.083 | 0.04 | 1 | 10 | Trip10 | 0.043 | Old |
| 0.147719 | 0.164085 | 0.484 | 0.384 | 1 | 10 | Bcl10 | 0.1 | Old |
| 0.14778 | 1.842057 | 0.039 | 0.01 | 1 | 10 | Rnf123 | 0.029 | Old |
| 0.147791 | 0.666734 | 0.137 | 0.081 | 1 | 10 | Vezt | 0.056 | Old |
| 0.147805 | 0.619571 | 0.134 | 0.081 | 1 | 10 | Clns1a | 0.053 | Old |
| 0.14787 | 0.565219 | 0.218 | 0.152 | 1 | 10 | Fnta | 0.066 | Old |
| 0.14805 | 0.647962 | 0.249 | 0.182 | 1 | 10 | Ubr2 | 0.067 | Old |
| 0.148217 | 0.197591 | 0.582 | 0.444 | 1 | 10 | Map1lc3b | 0.138 | Old |
| 0.148218 | 0.09277 | 0.271 | 0.192 | 1 | 10 | Rere | 0.079 | Old |
| 0.148764 | 0.181767 | 0.638 | 0.556 | 1 | 10 | H2-DMb1 | 0.082 | Old |
| 0.148804 | 1.359996 | 0.068 | 0.03 | 1 | 10 | Dcun1d3 | 0.038 | Old |
| 0.148804 | 1.276268 | 0.068 | 0.03 | 1 | 10 | Nxpe4 | 0.038 | Old |
| 0.14892 | 0.626256 | 0.056 | 0.02 | 1 | 10 | Zfp846 | 0.036 | Old |
| 0.149146 | 0.306469 | 0.112 | 0.061 | 1 | 10 | Apool | 0.051 | Old |
| 0.149156 | 1.72767 | 0.039 | 0.01 | 1 | 10 | Slc9a4 | 0.029 | Old |
| 0.149196 | 0.19339 | 0.274 | 0.202 | 1 | 10 | Cmc1 | 0.072 | Old |
| 0.14921 | 0.546677 | 0.208 | 0.141 | 1 | 10 | Abcb7 | 0.067 | Old |
| 0.149466 | 0.393563 | 0.438 | 0.343 | 1 | 10 | Ccar1 | 0.095 | Old |
| 0.149494 | 0.517144 | 0.056 | 0.02 | 1 | 10 | Ctps | 0.036 | Old |
| 0.149605 | 0.599629 | 0.347 | 0.293 | 1 | 10 | Mon2 | 0.054 | Old |
| 0.149644 | 0.413343 | 0.347 | 0.263 | 1 | 10 | Secisbp2l | 0.084 | Old |
| 0.149758 | 0.713869 | 0.183 | 0.121 | 1 | 10 | Polr3k | 0.062 | Old |
| 0.149889 | 0.483531 | 0.125 | 0.071 | 1 | 10 | Trim36 | 0.054 | Old |
| 0.149994 | 0.99751 | 0.083 | 0.04 | 1 | 10 | Ndst1 | 0.043 | Old |
| 0.149994 | 0.886103 | 0.083 | 0.04 | 1 | 10 | Tbc1d32 | 0.043 | Old |
| 0.149996 | 0.013354 | 0.222 | 0.141 | 1 | 10 | Dda1 | 0.081 | Old |
| 0.15028 | 0.220988 | 0.125 | 0.071 | 1 | 10 | Cnnm3 | 0.054 | Old |
| 0.150323 | 0.126992 | 0.966 | 0.939 | 1 | 10 | Rpl37 | 0.027 | Old |
| 0.15034 | 0.384942 | 0.335 | 0.253 | 1 | 10 | Lsm12 | 0.082 | Old |
| 0.150363 | 0.421672 | 0.21 | 0.141 | 1 | 10 | Ilkap | 0.069 | Old |
| 0.150388 | 0.068286 | 0.746 | 0.626 | 1 | 10 | Prkar1a | 0.12 | Old |
| 0.15054 | 0.233481 | 0.567 | 0.465 | 1 | 10 | Myadm | 0.102 | Old |
| 0.150885 | 1.439925 | 0.068 | 0.03 | 1 | 10 | Tsr3 | 0.038 | Old |
| 0.150894 | 0.092216 | 0.23 | 0.152 | 1 | 10 | Hibadh | 0.078 | Old |
| 0.150899 | 0.435432 | 0.147 | 0.091 | 1 | 10 | Spryd7 | 0.056 | Old |
| 0.151071 | 1.077162 | 0.12 | 0.071 | 1 | 10 | Dcaf17 | 0.049 | Old |
| 0.151195 | 1.646043 | 0.054 | 0.02 | 1 | 10 | Tor4a | 0.034 | Old |
| 0.151238 | 1.794321 | 0.039 | 0.01 | 1 | 10 | Map2k3os | 0.029 | Old |
| 0.151351 | 0.271386 | 0.785 | 0.697 | 1 | 10 | Rpn1 | 0.088 | Old |
| 0.151559 | 0.10265 | 0.259 | 0.182 | 1 | 10 | Mrpl18 | 0.077 | Old |
| 0.151601 | 0.514022 | 0.149 | 0.091 | 1 | 10 | Ccm2 | 0.058 | Old |
| 0.151674 | 0.074189 | 0.589 | 0.485 | 1 | 10 | Dnajb6 | 0.104 | Old |
| 0.151757 | 0.302489 | 0.484 | 0.394 | 1 | 10 | Klf13 | 0.09 | Old |
| 0.152163 | 0.090246 | 0.301 | 0.212 | 1 | 10 | Syf2 | 0.089 | Old |

| | | | | | | | | |
|---|---|---|---|---|---|---|---|---|
| 0.152176 | 0.301985 | 0.45 | 0.354 | 1 | 10 | Tnks2 | 0.096 | Old |
| 0.152249 | 0.223542 | 0.125 | 0.071 | 1 | 10 | Nostrin | 0.054 | Old |
| 0.152328 | 0.155587 | 0.186 | 0.121 | 1 | 10 | Polr2b | 0.065 | Old |
| 0.152376 | 1.074213 | 0.095 | 0.051 | 1 | 10 | Rtca | 0.044 | Old |
| 0.15251 | 0.173988 | 0.435 | 0.364 | 1 | 10 | Coa5 | 0.071 | Old |
| 0.152621 | 0.297079 | 0.32 | 0.242 | 1 | 10 | R3hdm4 | 0.078 | Old |
| 0.152639 | 1.407024 | 0.039 | 0.01 | 1 | 10 | Trp53i13 | 0.029 | Old |
| 0.152676 | 0.756919 | 0.12 | 0.071 | 1 | 10 | Wrn | 0.049 | Old |
| 0.152754 | 0.008986 | 0.44 | 0.343 | 1 | 10 | Klhl9 | 0.097 | Old |
| 0.152859 | 0.126236 | 0.394 | 0.283 | 1 | 10 | Erbin | 0.111 | Old |
| 0.152867 | 0.379181 | 0.301 | 0.222 | 1 | 10 | Rbm17 | 0.079 | Old |
| 0.152977 | 1.809946 | 0.054 | 0.02 | 1 | 10 | Chst14 | 0.034 | Old |
| 0.15302 | 0.531802 | 0.267 | 0.202 | 1 | 10 | Reps2 | 0.065 | Old |
| 0.153235 | 0.018326 | 0.086 | 0.04 | 1 | 10 | Cdk5 | 0.046 | Old |
| 0.153269 | 0.868214 | 0.11 | 0.061 | 1 | 10 | Fam222b | 0.049 | Old |
| 0.153318 | 0.349097 | 0.396 | 0.323 | 1 | 10 | Fcgr1 | 0.073 | Old |
| 0.153325 | 0.270217 | 0.601 | 0.495 | 1 | 10 | Cct8 | 0.106 | Old |
| 0.153417 | 0.112619 | 0.301 | 0.222 | 1 | 10 | Sdf2l1 | 0.079 | Old |
| 0.153484 | 0.197021 | 0.403 | 0.323 | 1 | 10 | Sdf2 | 0.08 | Old |
| 0.153518 | 1.266444 | 0.068 | 0.03 | 1 | 10 | Slc30a6 | 0.038 | Old |
| 0.153794 | 0.1608 | 0.961 | 0.879 | 1 | 10 | H2-Ab1 | 0.082 | Old |
| 0.153967 | 0.452413 | 0.756 | 0.848 | 1 | 10 | Retnla | -0.092 | Old |
| 0.153969 | 0.435593 | 0.222 | 0.152 | 1 | 10 | Inpp5k | 0.07 | Old |
| 0.154064 | 0.621465 | 0.108 | 0.061 | 1 | 10 | Stk16 | 0.047 | Old |
| 0.154161 | 0.021281 | 0.149 | 0.091 | 1 | 10 | Crnkl1 | 0.058 | Old |
| 0.154165 | 0.353921 | 0.313 | 0.232 | 1 | 10 | Map3k8 | 0.081 | Old |
| 0.154214 | 0.985041 | 0.142 | 0.091 | 1 | 10 | Ltbp3 | 0.051 | Old |
| 0.154427 | 0.553385 | 0.271 | 0.212 | 1 | 10 | Ciita | 0.059 | Old |
| 0.15458 | 1.255631 | 0.068 | 0.03 | 1 | 10 | Pycrl | 0.038 | Old |
| 0.154599 | 0.339978 | 0.281 | 0.202 | 1 | 10 | Rit1 | 0.079 | Old |
| 0.154773 | 0.411727 | 0.323 | 0.253 | 1 | 10 | Cpne3 | 0.07 | Old |
| 0.154961 | 0.312074 | 0.11 | 0.061 | 1 | 10 | Crebzf | 0.049 | Old |
| 0.155198 | 0.28169 | 0.237 | 0.162 | 1 | 10 | Kars | 0.075 | Old |
| 0.155217 | 0.83899 | 0.178 | 0.121 | 1 | 10 | Cep250 | 0.057 | Old |
| 0.155273 | 0.434432 | 0.337 | 0.263 | 1 | 10 | Morc3 | 0.074 | Old |
| 0.155377 | 2.197362 | 0.054 | 0.02 | 1 | 10 | Nudt1 | 0.034 | Old |
| 0.155469 | 0.589701 | 0.039 | 0.01 | 1 | 10 | Mccc2 | 0.029 | Old |
| 0.155469 | 0.363542 | 0.039 | 0.01 | 1 | 10 | BC049352 | 0.029 | Old |
| 0.155649 | 1.223747 | 0.068 | 0.03 | 1 | 10 | Med27 | 0.038 | Old |
| 0.15609 | 0.40631 | 0.235 | 0.162 | 1 | 10 | Tmem127 | 0.073 | Old |
| 0.156183 | 0.641002 | 0.039 | 0.01 | 1 | 10 | Thap1 | 0.029 | Old |
| 0.156183 | 0.554054 | 0.039 | 0.01 | 1 | 10 | Mmab | 0.029 | Old |
| 0.156183 | 0.480333 | 0.039 | 0.01 | 1 | 10 | Il18bp | 0.029 | Old |
| 0.156268 | 0.599632 | 0.252 | 0.192 | 1 | 10 | Fbxl5 | 0.06 | Old |
| 0.156573 | 0.101333 | 0.479 | 0.374 | 1 | 10 | Ostc | 0.105 | Old |

| | | | | | | | | |
|---|---|---|---|---|---|---|---|---|
| 0.156722 | 1.198675 | 0.068 | 0.03 | 1 | 10 | Bub1b | 0.038 | Old |
| 0.156908 | 0.935578 | 0.095 | 0.051 | 1 | 10 | Tbce | 0.044 | Old |
| 0.157196 | 1.33334 | 0.054 | 0.02 | 1 | 10 | Rpap1 | 0.034 | Old |
| 0.157196 | 1.205258 | 0.054 | 0.02 | 1 | 10 | Slc30a4 | 0.034 | Old |
| 0.157227 | 0.594685 | 0.083 | 0.04 | 1 | 10 | Zscan29 | 0.043 | Old |
| 0.157414 | 0.215854 | 0.697 | 0.606 | 1 | 10 | Ap2m1 | 0.091 | Old |
| 0.157812 | 0.206491 | 0.227 | 0.162 | 1 | 10 | Pdcd4 | 0.065 | Old |
| 0.158197 | 0.809277 | 0.132 | 0.081 | 1 | 10 | Zfp277 | 0.051 | Old |
| 0.158309 | 0.324039 | 0.731 | 0.687 | 1 | 10 | Gm26917 | 0.044 | Old |
| 0.158658 | 0.279521 | 0.308 | 0.232 | 1 | 10 | Lars2 | 0.076 | Old |
| 0.158688 | 0.150658 | 0.643 | 0.566 | 1 | 10 | Atp5j2 | 0.077 | Old |
| 0.158795 | 1.278401 | 0.081 | 0.04 | 1 | 10 | Zfp358 | 0.041 | Old |
| 0.159101 | 0.642482 | 0.181 | 0.121 | 1 | 10 | Rbm14 | 0.06 | Old |
| 0.15915 | 0.821405 | 0.142 | 0.091 | 1 | 10 | Pias2 | 0.051 | Old |
| 0.159227 | 0.051424 | 0.597 | 0.525 | 1 | 10 | Atp6v1c1 | 0.072 | Old |
| 0.159268 | 0.132465 | 0.196 | 0.131 | 1 | 10 | Ibtk | 0.065 | Old |
| 0.159432 | 0.818768 | 0.068 | 0.03 | 1 | 10 | D230025D1 | 0.038 | Old |
| 0.159651 | 0.886097 | 0.117 | 0.071 | 1 | 10 | Akna | 0.046 | Old |
| 0.159685 | 0.803421 | 0.108 | 0.061 | 1 | 10 | Vopp1 | 0.047 | Old |
| 0.15996 | 0.170368 | 0.164 | 0.101 | 1 | 10 | Calcrl | 0.063 | Old |
| 0.16008 | 0.566147 | 0.159 | 0.101 | 1 | 10 | Papss1 | 0.058 | Old |
| 0.160274 | 0.181314 | 0.824 | 0.687 | 1 | 10 | Pid1 | 0.137 | Old |
| 0.160374 | 0.372602 | 0.357 | 0.283 | 1 | 10 | Ccnt2 | 0.074 | Old |
| 0.160449 | 0.382472 | 0.269 | 0.202 | 1 | 10 | Magoh | 0.067 | Old |
| 0.160562 | 0.57964 | 0.183 | 0.121 | 1 | 10 | Spr | 0.062 | Old |
| 0.160807 | 1.31753 | 0.081 | 0.04 | 1 | 10 | Ankrd46 | 0.041 | Old |
| 0.160882 | 1.034803 | 0.054 | 0.02 | 1 | 10 | Spata5 | 0.034 | Old |
| 0.160937 | 0.08875 | 0.372 | 0.273 | 1 | 10 | Tnfrsf1a | 0.099 | Old |
| 0.161055 | 0.150661 | 0.526 | 0.414 | 1 | 10 | Nipbl | 0.112 | Old |
| 0.161231 | 0.611043 | 0.169 | 0.111 | 1 | 10 | Slc11a2 | 0.058 | Old |
| 0.161368 | 0.098603 | 0.147 | 0.091 | 1 | 10 | Chchd5 | 0.056 | Old |
| 0.161497 | 0.397693 | 0.193 | 0.131 | 1 | 10 | Golga2 | 0.062 | Old |
| 0.161572 | 0.486336 | 0.215 | 0.141 | 1 | 10 | Stmn1 | 0.074 | Old |
| 0.161682 | 2.607713 | 0.02 | 0 | 1 | 10 | Clec2g | 0.02 | Old |
| 0.161682 | 2.586433 | 0.02 | 0 | 1 | 10 | Rab38 | 0.02 | Old |
| 0.161682 | 2.453829 | 0.02 | 0 | 1 | 10 | Gm43660 | 0.02 | Old |
| 0.161682 | 2.435038 | 0.02 | 0 | 1 | 10 | 2510046G1 | 0.02 | Old |
| 0.161682 | 2.3784 | 0.02 | 0 | 1 | 10 | Tnfaip8l1 | 0.02 | Old |
| 0.161682 | 2.347293 | 0.02 | 0 | 1 | 10 | Gm16638 | 0.02 | Old |
| 0.161682 | 2.290068 | 0.02 | 0 | 1 | 10 | Setmar | 0.02 | Old |
| 0.161682 | 2.248443 | 0.02 | 0 | 1 | 10 | Gm38394 | 0.02 | Old |
| 0.161682 | 2.239801 | 0.02 | 0 | 1 | 10 | Etl4 | 0.02 | Old |
| 0.161682 | 2.221359 | 0.02 | 0 | 1 | 10 | Sirt5 | 0.02 | Old |
| 0.161682 | 2.205894 | 0.02 | 0 | 1 | 10 | Meis1 | 0.02 | Old |
| 0.161682 | 2.203294 | 0.02 | 0 | 1 | 10 | 9930022D1 | 0.02 | Old |

| | | | | | | | | |
|---|---|---|---|---|---|---|---|---|
| 0.161682 | 2.172751 | 0.02 | 0 | 1 | 10 | Ska2 | 0.02 | Old |
| 0.161682 | 2.064646 | 0.02 | 0 | 1 | 10 | Klhl8 | 0.02 | Old |
| 0.161682 | 2.021806 | 0.02 | 0 | 1 | 10 | Gm30025 | 0.02 | Old |
| 0.161682 | 1.988348 | 0.02 | 0 | 1 | 10 | Mdk | 0.02 | Old |
| 0.161682 | 1.977433 | 0.02 | 0 | 1 | 10 | Nuf2 | 0.02 | Old |
| 0.161682 | 1.96668 | 0.02 | 0 | 1 | 10 | Zfp677 | 0.02 | Old |
| 0.161682 | 1.963029 | 0.02 | 0 | 1 | 10 | A830010M | 0.02 | Old |
| 0.161682 | 1.945729 | 0.02 | 0 | 1 | 10 | Tmem129 | 0.02 | Old |
| 0.161682 | 1.936529 | 0.02 | 0 | 1 | 10 | Tmem51os | 0.02 | Old |
| 0.161682 | 1.928573 | 0.02 | 0 | 1 | 10 | Gm45894 | 0.02 | Old |
| 0.161682 | 1.920441 | 0.02 | 0 | 1 | 10 | Elmod3 | 0.02 | Old |
| 0.161682 | 1.900104 | 0.02 | 0 | 1 | 10 | A530032D1 | 0.02 | Old |
| 0.161682 | 1.860558 | 0.02 | 0 | 1 | 10 | Wdpcp | 0.02 | Old |
| 0.161682 | 1.853774 | 0.02 | 0 | 1 | 10 | Cacnb1 | 0.02 | Old |
| 0.161682 | 1.803718 | 0.02 | 0 | 1 | 10 | Top1mt | 0.02 | Old |
| 0.161682 | 1.791762 | 0.02 | 0 | 1 | 10 | Zswim3 | 0.02 | Old |
| 0.161682 | 1.776788 | 0.02 | 0 | 1 | 10 | Zfp51 | 0.02 | Old |
| 0.161682 | 1.776708 | 0.02 | 0 | 1 | 10 | Mief2 | 0.02 | Old |
| 0.161682 | 1.760818 | 0.02 | 0 | 1 | 10 | Aldh1l1 | 0.02 | Old |
| 0.161682 | 1.76057 | 0.02 | 0 | 1 | 10 | Zfp867 | 0.02 | Old |
| 0.161682 | 1.754362 | 0.02 | 0 | 1 | 10 | Gm10269 | 0.02 | Old |
| 0.161682 | 1.753673 | 0.02 | 0 | 1 | 10 | Zfp707 | 0.02 | Old |
| 0.161682 | 1.517404 | 0.02 | 0 | 1 | 10 | Gm14325 | 0.02 | Old |
| 0.161682 | 1.506208 | 0.02 | 0 | 1 | 10 | Gm12089 | 0.02 | Old |
| 0.161682 | 1.501334 | 0.02 | 0 | 1 | 10 | Purg | 0.02 | Old |
| 0.161682 | 1.411156 | 0.02 | 0 | 1 | 10 | Gm15675 | 0.02 | Old |
| 0.161837 | 0.170171 | 0.528 | 0.434 | 1 | 10 | Ppp2r1a | 0.094 | Old |
| 0.16202 | 0.277617 | 0.455 | 0.364 | 1 | 10 | Atp6v1d | 0.091 | Old |
| 0.162202 | 0.407263 | 0.501 | 0.424 | 1 | 10 | Adam8 | 0.077 | Old |
| 0.162323 | 0.089308 | 0.719 | 0.586 | 1 | 10 | Shisa5 | 0.133 | Old |
| 0.16273 | 0.703842 | 0.068 | 0.03 | 1 | 10 | Dcn | 0.038 | Old |
| 0.162959 | 2.064598 | 0.066 | 0.03 | 1 | 10 | Mefv | 0.036 | Old |
| 0.163228 | 0.255286 | 0.482 | 0.384 | 1 | 10 | Wsb1 | 0.098 | Old |
| 0.163286 | 0.087222 | 0.758 | 0.697 | 1 | 10 | Ddx3x | 0.061 | Old |
| 0.163386 | 0.259459 | 0.174 | 0.111 | 1 | 10 | Cox16 | 0.063 | Old |
| 0.16348 | 0.487858 | 0.264 | 0.202 | 1 | 10 | Sec24c | 0.062 | Old |
| 0.163496 | 0.249719 | 0.098 | 0.051 | 1 | 10 | Herc6 | 0.047 | Old |
| 0.163919 | 0.124032 | 0.946 | 0.879 | 1 | 10 | Rps25 | 0.067 | Old |
| 0.164233 | 0.185402 | 0.372 | 0.293 | 1 | 10 | Nudt3 | 0.079 | Old |
| 0.164633 | 0.426141 | 0.054 | 0.02 | 1 | 10 | Unc119b | 0.034 | Old |
| 0.164634 | 0.673052 | 0.178 | 0.121 | 1 | 10 | St3gal4 | 0.057 | Old |
| 0.164956 | 0.375625 | 0.068 | 0.03 | 1 | 10 | Abcb10 | 0.038 | Old |
| 0.164958 | 0.09269 | 0.306 | 0.232 | 1 | 10 | Rbck1 | 0.074 | Old |
| 0.164997 | 0.87069 | 0.154 | 0.101 | 1 | 10 | Smurf1 | 0.053 | Old |
| 0.165101 | 0.025678 | 0.381 | 0.283 | 1 | 10 | Psmd6 | 0.098 | Old |

| | | | | | | | | |
|---|---|---|---|---|---|---|---|---|
| 0.165178 | 0.271577 | 0.237 | 0.162 | 1 | 10 | Ugdh | 0.075 | Old |
| 0.165244 | 0.574177 | 0.274 | 0.212 | 1 | 10 | Mrpl43 | 0.062 | Old |
| 0.165316 | 0.324104 | 0.372 | 0.293 | 1 | 10 | Prpf4b | 0.079 | Old |
| 0.165461 | 0.954262 | 0.105 | 0.061 | 1 | 10 | Setd7 | 0.044 | Old |
| 0.165514 | 0.027379 | 0.188 | 0.121 | 1 | 10 | Oxa1l | 0.067 | Old |
| 0.165735 | 0.076266 | 0.579 | 0.495 | 1 | 10 | Fam46a | 0.084 | Old |
| 0.165898 | 0.786778 | 0.054 | 0.02 | 1 | 10 | Ralgps2 | 0.034 | Old |
| 0.166362 | 0.466283 | 0.213 | 0.152 | 1 | 10 | Kdm4a | 0.061 | Old |
| 0.166458 | 0.13804 | 0.934 | 0.899 | 1 | 10 | Gabarap | 0.035 | Old |
| 0.166555 | 0.480292 | 0.134 | 0.081 | 1 | 10 | Adpgk | 0.053 | Old |
| 0.166756 | 1.053387 | 0.13 | 0.081 | 1 | 10 | Cnot9 | 0.049 | Old |
| 0.16678 | 0.287196 | 0.289 | 0.212 | 1 | 10 | Tbc1d23 | 0.077 | Old |
| 0.166841 | 0.481952 | 0.12 | 0.071 | 1 | 10 | Caml | 0.049 | Old |
| 0.167136 | 1.13326 | 0.093 | 0.051 | 1 | 10 | Gtf2e2 | 0.042 | Old |
| 0.167264 | 0.050259 | 0.191 | 0.121 | 1 | 10 | Aldh16a1 | 0.07 | Old |
| 0.167366 | 0.325831 | 0.213 | 0.152 | 1 | 10 | Cyth3 | 0.061 | Old |
| 0.167482 | 0.206915 | 0.601 | 0.505 | 1 | 10 | Rtn3 | 0.096 | Old |
| 0.167697 | 0.966142 | 0.152 | 0.101 | 1 | 10 | Tmem214 | 0.051 | Old |
| 0.167702 | 0.507072 | 0.213 | 0.152 | 1 | 10 | Ppfibp2 | 0.061 | Old |
| 0.167717 | 0.165103 | 0.369 | 0.283 | 1 | 10 | Vapb | 0.086 | Old |
| 0.167718 | 0.403503 | 0.095 | 0.051 | 1 | 10 | Tirap | 0.044 | Old |
| 0.167808 | 0.615424 | 0.169 | 0.111 | 1 | 10 | Wipf2 | 0.058 | Old |
| 0.167906 | 0.121668 | 0.198 | 0.131 | 1 | 10 | Crybg3 | 0.067 | Old |
| 0.167967 | 0.049993 | 0.276 | 0.202 | 1 | 10 | Dcp2 | 0.074 | Old |
| 0.168095 | 0.618238 | 0.144 | 0.091 | 1 | 10 | Pik3r4 | 0.053 | Old |
| 0.16834 | 0.278351 | 0.068 | 0.03 | 1 | 10 | Jak3 | 0.038 | Old |
| 0.16893 | 0.018248 | 0.33 | 0.242 | 1 | 10 | Paics | 0.088 | Old |
| 0.169002 | 0.124441 | 0.787 | 0.646 | 1 | 10 | Ehd4 | 0.141 | Old |
| 0.169044 | 0.064079 | 0.352 | 0.253 | 1 | 10 | Vti1b | 0.099 | Old |
| 0.16905 | 0.904163 | 0.081 | 0.04 | 1 | 10 | Pym1 | 0.041 | Old |
| 0.169269 | 0.473855 | 0.171 | 0.111 | 1 | 10 | Trappc5 | 0.06 | Old |
| 0.169296 | 1.988862 | 0.037 | 0.01 | 1 | 10 | Eefsec | 0.027 | Old |
| 0.169296 | 1.804319 | 0.037 | 0.01 | 1 | 10 | Gstm4 | 0.027 | Old |
| 0.169479 | 0.463789 | 0.068 | 0.03 | 1 | 10 | Cep68 | 0.038 | Old |
| 0.169671 | 0.035239 | 0.455 | 0.343 | 1 | 10 | Pi4k2a | 0.112 | Old |
| 0.169915 | 0.13123 | 0.11 | 0.061 | 1 | 10 | Ppan | 0.049 | Old |
| 0.170014 | 0.292648 | 0.171 | 0.111 | 1 | 10 | Edc4 | 0.06 | Old |
| 0.170069 | 1.111992 | 0.093 | 0.051 | 1 | 10 | Mrpl44 | 0.042 | Old |
| 0.170111 | 0.134741 | 0.213 | 0.141 | 1 | 10 | Csrp1 | 0.072 | Old |
| 0.170138 | 0.46141 | 0.362 | 0.293 | 1 | 10 | Rab20 | 0.069 | Old |
| 0.170369 | 0.481533 | 0.601 | 0.545 | 1 | 10 | Gda | 0.056 | Old |
| 0.170673 | 0.46987 | 0.235 | 0.172 | 1 | 10 | Pank2 | 0.063 | Old |
| 0.170867 | 2.132642 | 0.037 | 0.01 | 1 | 10 | Map3k10 | 0.027 | Old |
| 0.170867 | 1.823888 | 0.037 | 0.01 | 1 | 10 | Orai2 | 0.027 | Old |
| 0.171044 | 0.146638 | 0.23 | 0.162 | 1 | 10 | Hmgn2 | 0.068 | Old |

| | | | | | | | | |
|---|---|---|---|---|---|---|---|---|
| 0.171191 | 0.529207 | 0.181 | 0.121 | 1 | 10 | G6pdx | 0.06 | Old |
| 0.171221 | 0.170718 | 0.186 | 0.121 | 1 | 10 | Dctn5 | 0.065 | Old |
| 0.171656 | 2.312078 | 0.037 | 0.01 | 1 | 10 | Tmem26 | 0.027 | Old |
| 0.171656 | 1.79493 | 0.037 | 0.01 | 1 | 10 | B930036N1 | 0.027 | Old |
| 0.172396 | 0.100946 | 0.196 | 0.131 | 1 | 10 | Gtf2f1 | 0.065 | Old |
| 0.172448 | 1.486059 | 0.037 | 0.01 | 1 | 10 | Sirpb1a | 0.027 | Old |
| 0.172531 | 0.134996 | 0.247 | 0.182 | 1 | 10 | Anapc1 | 0.065 | Old |
| 0.172543 | 0.472247 | 0.093 | 0.051 | 1 | 10 | Rrn3 | 0.042 | Old |
| 0.172569 | 0.060284 | 0.159 | 0.101 | 1 | 10 | Ppfibp1 | 0.058 | Old |
| 0.172595 | 0.121087 | 0.345 | 0.253 | 1 | 10 | Arpc1a | 0.092 | Old |
| 0.1729 | 0.240644 | 0.663 | 0.606 | 1 | 10 | Uqcrb | 0.057 | Old |
| 0.173041 | 0.065591 | 0.183 | 0.121 | 1 | 10 | Lrch3 | 0.062 | Old |
| 0.173048 | 0.17928 | 0.271 | 0.202 | 1 | 10 | Exoc3 | 0.069 | Old |
| 0.173048 | 0.467628 | 0.271 | 0.202 | 1 | 10 | Baz2a | 0.069 | Old |
| 0.173074 | 0.166146 | 0.758 | 0.646 | 1 | 10 | Csf2ra | 0.112 | Old |
| 0.173307 | 1.741273 | 0.1 | 0.061 | 1 | 10 | Gpatch11 | 0.039 | Old |
| 0.173321 | 1.137581 | 0.105 | 0.061 | 1 | 10 | Hps4 | 0.044 | Old |
| 0.17344 | 0.602407 | 0.2 | 0.141 | 1 | 10 | Stard3 | 0.059 | Old |
| 0.173485 | 0.204942 | 0.374 | 0.293 | 1 | 10 | Cbfb | 0.081 | Old |
| 0.173532 | 0.538219 | 0.154 | 0.101 | 1 | 10 | Nedd9 | 0.053 | Old |
| 0.173869 | 1.658825 | 0.051 | 0.02 | 1 | 10 | Sh3bp5l | 0.031 | Old |
| 0.173882 | 0.352599 | 0.188 | 0.131 | 1 | 10 | Upf3a | 0.057 | Old |
| 0.17404 | 1.911413 | 0.037 | 0.01 | 1 | 10 | Nhej1 | 0.027 | Old |
| 0.17404 | 1.378393 | 0.037 | 0.01 | 1 | 10 | Ggcx | 0.027 | Old |
| 0.174181 | 0.734303 | 0.108 | 0.061 | 1 | 10 | Cep164 | 0.047 | Old |
| 0.174196 | 0.266314 | 0.318 | 0.242 | 1 | 10 | Ndufs4 | 0.076 | Old |
| 0.17454 | 1.76056 | 0.051 | 0.02 | 1 | 10 | Pex14 | 0.031 | Old |
| 0.174815 | 0.201122 | 0.222 | 0.152 | 1 | 10 | Dnajc14 | 0.07 | Old |
| 0.174857 | 0.640368 | 0.318 | 0.263 | 1 | 10 | Adipor2 | 0.055 | Old |
| 0.174959 | 0.204311 | 0.665 | 0.586 | 1 | 10 | Ptpra | 0.079 | Old |
| 0.174966 | 0.118267 | 0.237 | 0.162 | 1 | 10 | Imp4 | 0.075 | Old |
| 0.175037 | 0.205372 | 0.377 | 0.303 | 1 | 10 | 2410015M | 0.074 | Old |
| 0.17511 | 1.28279 | 0.066 | 0.03 | 1 | 10 | Fam26f | 0.036 | Old |
| 0.175136 | 0.187894 | 0.215 | 0.152 | 1 | 10 | Trim33 | 0.063 | Old |
| 0.175222 | 0.367212 | 0.46 | 0.384 | 1 | 10 | Snrpd1 | 0.076 | Old |
| 0.175416 | 1.2082 | 0.115 | 0.071 | 1 | 10 | Mrps2 | 0.044 | Old |
| 0.175562 | 0.534317 | 0.095 | 0.051 | 1 | 10 | Sec22a | 0.044 | Old |
| 0.175562 | 0.008068 | 0.095 | 0.051 | 1 | 10 | Gm45716 | 0.044 | Old |
| 0.175584 | 0.915868 | 0.152 | 0.101 | 1 | 10 | Mapkapk3 | 0.051 | Old |
| 0.175643 | 1.657008 | 0.037 | 0.01 | 1 | 10 | Cnnm2 | 0.027 | Old |
| 0.175643 | 1.485006 | 0.037 | 0.01 | 1 | 10 | Msh6 | 0.027 | Old |
| 0.175643 | 1.461639 | 0.037 | 0.01 | 1 | 10 | Cyp2j6 | 0.027 | Old |
| 0.175714 | 0.143808 | 0.626 | 0.495 | 1 | 10 | Mrpl33 | 0.131 | Old |
| 0.175969 | 0.673981 | 0.178 | 0.121 | 1 | 10 | Akr1b10 | 0.057 | Old |
| 0.175984 | 0.098579 | 0.447 | 0.354 | 1 | 10 | Tmod3 | 0.093 | Old |

| | | | | | | | | |
|---|---|---|---|---|---|---|---|---|
| 0.176079 | 0.145003 | 0.328 | 0.242 | 1 | 10 | Usp14 | 0.086 | Old |
| 0.176277 | 0.214498 | 0.205 | 0.141 | 1 | 10 | Mau2 | 0.064 | Old |
| 0.176449 | 1.361222 | 0.037 | 0.01 | 1 | 10 | Crip2 | 0.027 | Old |
| 0.176566 | 1.646261 | 0.051 | 0.02 | 1 | 10 | Ydjc | 0.031 | Old |
| 0.176566 | 1.541664 | 0.051 | 0.02 | 1 | 10 | Katnal1 | 0.031 | Old |
| 0.176811 | 0.17007 | 0.457 | 0.374 | 1 | 10 | Usp4 | 0.083 | Old |
| 0.177061 | 0.623292 | 0.081 | 0.04 | 1 | 10 | Cds1 | 0.041 | Old |
| 0.177167 | 0.050191 | 0.469 | 0.354 | 1 | 10 | Tmem123 | 0.115 | Old |
| 0.17719 | 0.09679 | 0.98 | 0.98 | 1 | 10 | Rps28 | 0 | Old |
| 0.177257 | 1.426634 | 0.037 | 0.01 | 1 | 10 | Apoo | 0.027 | Old |
| 0.177389 | 0.291979 | 0.335 | 0.253 | 1 | 10 | Trpm7 | 0.082 | Old |
| 0.177435 | 0.582397 | 0.144 | 0.091 | 1 | 10 | Parp8 | 0.053 | Old |
| 0.177709 | 0.436368 | 0.296 | 0.222 | 1 | 10 | Antxr2 | 0.074 | Old |
| 0.177857 | 0.283848 | 0.183 | 0.121 | 1 | 10 | Ddx52 | 0.062 | Old |
| 0.177858 | 0.099566 | 0.604 | 0.475 | 1 | 10 | Rgs10 | 0.129 | Old |
| 0.177915 | 0.025334 | 0.565 | 0.465 | 1 | 10 | Laptm4a | 0.1 | Old |
| 0.177981 | 0.097194 | 0.408 | 0.323 | 1 | 10 | Cbx3 | 0.085 | Old |
| 0.178087 | 0.409872 | 0.147 | 0.091 | 1 | 10 | Nfatc2 | 0.056 | Old |
| 0.178231 | 0.137728 | 0.949 | 0.869 | 1 | 10 | Akr1a1 | 0.08 | Old |
| 0.17824 | 0.422469 | 0.181 | 0.121 | 1 | 10 | Mbd1 | 0.06 | Old |
| 0.178553 | 0.635346 | 0.105 | 0.061 | 1 | 10 | Efl1 | 0.044 | Old |
| 0.178565 | 0.1033 | 0.122 | 0.071 | 1 | 10 | Sharpin | 0.051 | Old |
| 0.178609 | 1.398587 | 0.051 | 0.02 | 1 | 10 | Mfsd8 | 0.031 | Old |
| 0.178698 | 0.227443 | 0.501 | 0.384 | 1 | 10 | Tsc22d3 | 0.117 | Old |
| 0.178703 | 0.562633 | 0.066 | 0.03 | 1 | 10 | Pdlim2 | 0.036 | Old |
| 0.17875 | 0.01727 | 0.274 | 0.192 | 1 | 10 | BC028528 | 0.082 | Old |
| 0.178759 | 0.183178 | 0.249 | 0.172 | 1 | 10 | Cir1 | 0.077 | Old |
| 0.178794 | 0.598888 | 0.21 | 0.152 | 1 | 10 | Exoc2 | 0.058 | Old |
| 0.178962 | 0.18971 | 0.337 | 0.263 | 1 | 10 | Smarcc2 | 0.074 | Old |
| 0.179182 | 0.195314 | 0.286 | 0.212 | 1 | 10 | Tm2d2 | 0.074 | Old |
| 0.179284 | 0.246574 | 0.631 | 0.556 | 1 | 10 | P4hb | 0.075 | Old |
| 0.179307 | 1.0871 | 0.066 | 0.03 | 1 | 10 | Cox15 | 0.036 | Old |
| 0.179331 | 0.437008 | 0.311 | 0.242 | 1 | 10 | Tmem259 | 0.069 | Old |
| 0.179353 | 0.087682 | 0.621 | 0.525 | 1 | 10 | Stxbp3 | 0.096 | Old |
| 0.179485 | 0.0234 | 0.445 | 0.374 | 1 | 10 | Mrps14 | 0.071 | Old |
| 0.179517 | 0.996063 | 0.105 | 0.061 | 1 | 10 | Gphn | 0.044 | Old |
| 0.179522 | 1.351566 | 0.078 | 0.04 | 1 | 10 | Atp8b2 | 0.038 | Old |
| 0.179699 | 0.902264 | 0.037 | 0.01 | 1 | 10 | Cfap45 | 0.027 | Old |
| 0.1803 | 0.164048 | 0.171 | 0.111 | 1 | 10 | Trrap | 0.06 | Old |
| 0.180452 | 0.122521 | 0.186 | 0.121 | 1 | 10 | Plekhg3 | 0.065 | Old |
| 0.180518 | 1.158707 | 0.037 | 0.01 | 1 | 10 | Pms2 | 0.027 | Old |
| 0.180518 | 0.862711 | 0.037 | 0.01 | 1 | 10 | Car9 | 0.027 | Old |
| 0.180518 | 0.528283 | 0.037 | 0.01 | 1 | 10 | Zkscan5 | 0.027 | Old |
| 0.180518 | 0.305975 | 0.037 | 0.01 | 1 | 10 | Nup210 | 0.027 | Old |
| 0.18052 | 0.881336 | 0.066 | 0.03 | 1 | 10 | Wdr55 | 0.036 | Old |

| | | | | | | | | |
|---|---|---|---|---|---|---|---|---|
| 0.18067 | 1.226883 | 0.051 | 0.02 | 1 | 10 | Pold1 | 0.031 | Old |
| 0.180797 | 0.082952 | 0.318 | 0.232 | 1 | 10 | Pcmtd1 | 0.086 | Old |
| 0.181155 | 0.881275 | 0.093 | 0.051 | 1 | 10 | Sh3pxd2a | 0.042 | Old |
| 0.181155 | 0.81825 | 0.093 | 0.051 | 1 | 10 | Cryzl2 | 0.042 | Old |
| 0.181222 | 0.471306 | 0.24 | 0.172 | 1 | 10 | Lsm8 | 0.068 | Old |
| 0.18134 | 0.010332 | 0.037 | 0.01 | 1 | 10 | Pank1 | 0.027 | Old |
| 0.18136 | 1.076047 | 0.051 | 0.02 | 1 | 10 | Supt7l | 0.031 | Old |
| 0.181943 | 0.69089 | 0.105 | 0.061 | 1 | 10 | Nrde2 | 0.044 | Old |
| 0.182047 | 1.136388 | 0.103 | 0.061 | 1 | 10 | Psmg1 | 0.042 | Old |
| 0.182053 | 1.137684 | 0.051 | 0.02 | 1 | 10 | Mcf2l | 0.031 | Old |
| 0.182053 | 1.383845 | 0.051 | 0.02 | 1 | 10 | Hacd1 | 0.031 | Old |
| 0.182173 | 0.638508 | 0.117 | 0.071 | 1 | 10 | Tmem198k | 0.046 | Old |
| 0.182214 | 0.699566 | 0.142 | 0.091 | 1 | 10 | Gm17018 | 0.051 | Old |
| 0.18235 | 0.918147 | 0.066 | 0.03 | 1 | 10 | Eid3 | 0.036 | Old |
| 0.18235 | 0.805866 | 0.066 | 0.03 | 1 | 10 | Alpk1 | 0.036 | Old |
| 0.182748 | 1.122405 | 0.051 | 0.02 | 1 | 10 | Gcdh | 0.031 | Old |
| 0.183293 | 0.424577 | 0.345 | 0.283 | 1 | 10 | Pin4 | 0.062 | Old |
| 0.183448 | 0.43731 | 0.235 | 0.172 | 1 | 10 | BC003331 | 0.063 | Old |
| 0.18354 | 0.100494 | 0.169 | 0.111 | 1 | 10 | Erg28 | 0.058 | Old |
| 0.184143 | 1.164169 | 0.051 | 0.02 | 1 | 10 | Kctd17 | 0.031 | Old |
| 0.184442 | 0.29619 | 0.384 | 0.303 | 1 | 10 | Golph3 | 0.081 | Old |
| 0.184472 | 0.352337 | 0.247 | 0.182 | 1 | 10 | Tars | 0.065 | Old |
| 0.184567 | 0.114112 | 0.205 | 0.141 | 1 | 10 | Sympk | 0.064 | Old |
| 0.184609 | 0.035109 | 0.335 | 0.253 | 1 | 10 | Zfp148 | 0.082 | Old |
| 0.18502 | 0.495448 | 0.291 | 0.222 | 1 | 10 | Dhx38 | 0.069 | Old |
| 0.185115 | 0.443494 | 0.181 | 0.121 | 1 | 10 | Rras | 0.06 | Old |
| 0.185294 | 0.592503 | 0.164 | 0.111 | 1 | 10 | Ubap1 | 0.053 | Old |
| 0.185527 | 0.447074 | 0.178 | 0.121 | 1 | 10 | Pdcl | 0.057 | Old |
| 0.185628 | 0.17157 | 0.32 | 0.242 | 1 | 10 | Ergic3 | 0.078 | Old |
| 0.18564 | 0.096631 | 0.142 | 0.091 | 1 | 10 | Zfp518a | 0.051 | Old |
| 0.185738 | 0.191862 | 0.215 | 0.152 | 1 | 10 | Zufsp | 0.063 | Old |
| 0.185839 | 0.374655 | 0.311 | 0.242 | 1 | 10 | Slc12a7 | 0.069 | Old |
| 0.186033 | 0.105763 | 0.526 | 0.434 | 1 | 10 | Mrps21 | 0.092 | Old |
| 0.186251 | 0.90116 | 0.051 | 0.02 | 1 | 10 | Ice2 | 0.031 | Old |
| 0.186529 | 0.216036 | 0.222 | 0.162 | 1 | 10 | 2810004N: | 0.06 | Old |
| 0.18654 | 0.495865 | 0.325 | 0.263 | 1 | 10 | Parp1 | 0.062 | Old |
| 0.186869 | 0.544705 | 0.2 | 0.141 | 1 | 10 | Utrn | 0.059 | Old |
| 0.187079 | 0.495107 | 0.198 | 0.141 | 1 | 10 | Dapp1 | 0.057 | Old |
| 0.187191 | 1.771452 | 0.064 | 0.03 | 1 | 10 | Ranbp6 | 0.034 | Old |
| 0.187233 | 0.581217 | 0.186 | 0.131 | 1 | 10 | Chrac1 | 0.055 | Old |
| 0.187363 | 0.159904 | 0.244 | 0.182 | 1 | 10 | Lsm7 | 0.062 | Old |
| 0.187372 | 0.041993 | 0.411 | 0.313 | 1 | 10 | Usp34 | 0.098 | Old |
| 0.187814 | 0.453655 | 0.386 | 0.323 | 1 | 10 | C1qbp | 0.063 | Old |
| 0.187941 | 0.387034 | 0.35 | 0.283 | 1 | 10 | Fnip2 | 0.067 | Old |
| 0.188007 | 1.029061 | 0.078 | 0.04 | 1 | 10 | Snapc3 | 0.038 | Old |

| | | | | | | | | |
|---|---|---|---|---|---|---|---|---|
| 0.18802 | 0.077944 | 0.144 | 0.091 | 1 | 10 | Wdr13 | 0.053 | Old |
| 0.188033 | 0.290122 | 0.156 | 0.101 | 1 | 10 | Nelfa | 0.055 | Old |
| 0.188341 | 0.036067 | 0.396 | 0.293 | 1 | 10 | Tprgl | 0.103 | Old |
| 0.188465 | 0.222192 | 0.491 | 0.404 | 1 | 10 | Fgfr1op2 | 0.087 | Old |
| 0.189075 | 0.844925 | 0.09 | 0.051 | 1 | 10 | Mamld1 | 0.039 | Old |
| 0.189166 | 0.183711 | 0.445 | 0.364 | 1 | 10 | Ilk | 0.081 | Old |
| 0.18918 | 0.687258 | 0.066 | 0.03 | 1 | 10 | Rbm45 | 0.036 | Old |
| 0.189225 | 0.66604 | 0.117 | 0.071 | 1 | 10 | Haghl | 0.046 | Old |
| 0.190342 | 0.155287 | 0.298 | 0.222 | 1 | 10 | Usp19 | 0.076 | Old |
| 0.190684 | 0.664649 | 0.174 | 0.121 | 1 | 10 | Mlxip | 0.053 | Old |
| 0.191008 | 0.21563 | 0.565 | 0.455 | 1 | 10 | Psmd13 | 0.11 | Old |
| 0.191027 | 1.570121 | 0.064 | 0.03 | 1 | 10 | Zc3hc1 | 0.034 | Old |
| 0.191035 | 2.707568 | 0.017 | 0 | 1 | 10 | Atp1b1 | 0.017 | Old |
| 0.191035 | 2.443317 | 0.017 | 0 | 1 | 10 | Ccr3 | 0.017 | Old |
| 0.191035 | 2.40575 | 0.017 | 0 | 1 | 10 | Mettl22 | 0.017 | Old |
| 0.191035 | 2.37051 | 0.017 | 0 | 1 | 10 | Zbtb40 | 0.017 | Old |
| 0.191035 | 2.352551 | 0.017 | 0 | 1 | 10 | Il31ra | 0.017 | Old |
| 0.191035 | 2.234658 | 0.017 | 0 | 1 | 10 | Matn2 | 0.017 | Old |
| 0.191035 | 2.204418 | 0.017 | 0 | 1 | 10 | Fam187b | 0.017 | Old |
| 0.191035 | 2.203043 | 0.017 | 0 | 1 | 10 | Gm14403 | 0.017 | Old |
| 0.191035 | 2.150715 | 0.017 | 0 | 1 | 10 | Fam120c | 0.017 | Old |
| 0.191035 | 2.137671 | 0.017 | 0 | 1 | 10 | Gm48066 | 0.017 | Old |
| 0.191035 | 2.127129 | 0.017 | 0 | 1 | 10 | Cep152 | 0.017 | Old |
| 0.191035 | 2.111698 | 0.017 | 0 | 1 | 10 | Htr1b | 0.017 | Old |
| 0.191035 | 2.084408 | 0.017 | 0 | 1 | 10 | Dmwd | 0.017 | Old |
| 0.191035 | 2.07292 | 0.017 | 0 | 1 | 10 | Sgtb | 0.017 | Old |
| 0.191035 | 2.062384 | 0.017 | 0 | 1 | 10 | Setd4 | 0.017 | Old |
| 0.191035 | 2.061094 | 0.017 | 0 | 1 | 10 | Cdyl2 | 0.017 | Old |
| 0.191035 | 2.059428 | 0.017 | 0 | 1 | 10 | Tedc2 | 0.017 | Old |
| 0.191035 | 2.031362 | 0.017 | 0 | 1 | 10 | Map7d3 | 0.017 | Old |
| 0.191035 | 2.031032 | 0.017 | 0 | 1 | 10 | Mettl15 | 0.017 | Old |
| 0.191035 | 2.025486 | 0.017 | 0 | 1 | 10 | Wdr74 | 0.017 | Old |
| 0.191035 | 2.016529 | 0.017 | 0 | 1 | 10 | Serpinb12 | 0.017 | Old |
| 0.191035 | 1.987761 | 0.017 | 0 | 1 | 10 | Zfp532 | 0.017 | Old |
| 0.191035 | 1.955499 | 0.017 | 0 | 1 | 10 | Gm11696 | 0.017 | Old |
| 0.191035 | 1.929158 | 0.017 | 0 | 1 | 10 | Cldn5 | 0.017 | Old |
| 0.191035 | 1.909283 | 0.017 | 0 | 1 | 10 | Gspt2 | 0.017 | Old |
| 0.191035 | 1.894088 | 0.017 | 0 | 1 | 10 | Ppp1r18os | 0.017 | Old |
| 0.191035 | 1.878733 | 0.017 | 0 | 1 | 10 | Hspb7 | 0.017 | Old |
| 0.191035 | 1.87269 | 0.017 | 0 | 1 | 10 | Bhlhb9 | 0.017 | Old |
| 0.191035 | 1.87109 | 0.017 | 0 | 1 | 10 | Tpmt | 0.017 | Old |
| 0.191035 | 1.85357 | 0.017 | 0 | 1 | 10 | Acy1 | 0.017 | Old |
| 0.191035 | 1.847051 | 0.017 | 0 | 1 | 10 | Cldn23 | 0.017 | Old |
| 0.191035 | 1.83239 | 0.017 | 0 | 1 | 10 | Mterf1a | 0.017 | Old |
| 0.191035 | 1.801036 | 0.017 | 0 | 1 | 10 | 6230400D1 | 0.017 | Old |

| | | | | | | | | |
|---|---|---|---|---|---|---|---|---|
| 0.191035 | 1.779501 | 0.017 | 0 | 1 | 10 | Npas4 | 0.017 | Old |
| 0.191035 | 1.770343 | 0.017 | 0 | 1 | 10 | Fam57a | 0.017 | Old |
| 0.191035 | 1.761536 | 0.017 | 0 | 1 | 10 | Rgs12 | 0.017 | Old |
| 0.191035 | 1.753991 | 0.017 | 0 | 1 | 10 | Napepld | 0.017 | Old |
| 0.191035 | 1.689196 | 0.017 | 0 | 1 | 10 | Ssc4d | 0.017 | Old |
| 0.191035 | 1.686832 | 0.017 | 0 | 1 | 10 | 9130230L2 | 0.017 | Old |
| 0.191035 | 1.679663 | 0.017 | 0 | 1 | 10 | Trim6 | 0.017 | Old |
| 0.191035 | 1.663923 | 0.017 | 0 | 1 | 10 | Atp6v0c | 0.017 | Old |
| 0.191035 | 1.653276 | 0.017 | 0 | 1 | 10 | Zfp658 | 0.017 | Old |
| 0.191035 | 1.651558 | 0.017 | 0 | 1 | 10 | Gpx7 | 0.017 | Old |
| 0.191035 | 1.643077 | 0.017 | 0 | 1 | 10 | Gm30329 | 0.017 | Old |
| 0.191035 | 1.625668 | 0.017 | 0 | 1 | 10 | Gstp2 | 0.017 | Old |
| 0.191035 | 1.618254 | 0.017 | 0 | 1 | 10 | Arl6 | 0.017 | Old |
| 0.191035 | 1.616501 | 0.017 | 0 | 1 | 10 | Gm13479 | 0.017 | Old |
| 0.191035 | 1.60857 | 0.017 | 0 | 1 | 10 | Clec1a | 0.017 | Old |
| 0.191035 | 1.559387 | 0.017 | 0 | 1 | 10 | 9130221H1 | 0.017 | Old |
| 0.191035 | 1.55702 | 0.017 | 0 | 1 | 10 | Fam189b | 0.017 | Old |
| 0.191035 | 1.534675 | 0.017 | 0 | 1 | 10 | Taco1 | 0.017 | Old |
| 0.191035 | 1.501882 | 0.017 | 0 | 1 | 10 | Adgrg1 | 0.017 | Old |
| 0.191035 | 1.488273 | 0.017 | 0 | 1 | 10 | A530040E1 | 0.017 | Old |
| 0.191035 | 1.420192 | 0.017 | 0 | 1 | 10 | Mcm10 | 0.017 | Old |
| 0.191035 | 1.402749 | 0.017 | 0 | 1 | 10 | Ss18l1 | 0.017 | Old |
| 0.191035 | 1.400964 | 0.017 | 0 | 1 | 10 | Ccdc62 | 0.017 | Old |
| 0.191035 | 1.397385 | 0.017 | 0 | 1 | 10 | Csrnp2 | 0.017 | Old |
| 0.191035 | 1.396251 | 0.017 | 0 | 1 | 10 | Napb | 0.017 | Old |
| 0.191035 | 1.304993 | 0.017 | 0 | 1 | 10 | Fuk | 0.017 | Old |
| 0.191195 | 0.077908 | 0.533 | 0.455 | 1 | 10 | Smap2 | 0.078 | Old |
| 0.191332 | 0.426408 | 0.279 | 0.212 | 1 | 10 | Bfar | 0.067 | Old |
| 0.191815 | 0.379123 | 0.156 | 0.101 | 1 | 10 | Cd151 | 0.055 | Old |
| 0.192066 | 0.944427 | 0.078 | 0.04 | 1 | 10 | Lrrc61 | 0.038 | Old |
| 0.192068 | 0.016501 | 0.232 | 0.162 | 1 | 10 | Setdb1 | 0.07 | Old |
| 0.192291 | 0.327811 | 0.12 | 0.071 | 1 | 10 | Abhd2 | 0.049 | Old |
| 0.192844 | 0.08932 | 0.032 | 0.061 | 1 | 10 | Fam217b | -0.029 | Old |
| 0.19287 | 0.914513 | 0.09 | 0.051 | 1 | 10 | Dhx58 | 0.039 | Old |
| 0.193584 | 0.227652 | 0.196 | 0.131 | 1 | 10 | Kcnab2 | 0.065 | Old |
| 0.193606 | 0.262135 | 0.367 | 0.283 | 1 | 10 | Pank3 | 0.084 | Old |
| 0.193721 | 0.639367 | 0.154 | 0.101 | 1 | 10 | E2f1 | 0.053 | Old |
| 0.193921 | 0.413224 | 0.188 | 0.131 | 1 | 10 | Hras | 0.057 | Old |
| 0.194267 | 1.19169 | 0.064 | 0.03 | 1 | 10 | Cc2d1a | 0.034 | Old |
| 0.194267 | 0.742404 | 0.064 | 0.03 | 1 | 10 | Zbed3 | 0.034 | Old |
| 0.194336 | 0.609513 | 0.152 | 0.101 | 1 | 10 | Cldnd1 | 0.051 | Old |
| 0.194346 | 0.493521 | 0.213 | 0.152 | 1 | 10 | Naip2 | 0.061 | Old |
| 0.194372 | 0.233644 | 0.557 | 0.465 | 1 | 10 | Atp5o.1 | 0.092 | Old |
| 0.19449 | 0.245955 | 0.484 | 0.414 | 1 | 10 | Tpm1 | 0.07 | Old |
| 0.194656 | 0.189672 | 0.535 | 0.444 | 1 | 10 | Stx7 | 0.091 | Old |

| | | | | | | | | |
|---|---|---|---|---|---|---|---|---|
| 0.195005 | 0.785319 | 0.078 | 0.04 | 1 | 10 | Oas1a | 0.038 | Old |
| 0.19506 | 0.162675 | 0.621 | 0.545 | 1 | 10 | Blnk | 0.076 | Old |
| 0.195482 | 0.216818 | 0.093 | 0.051 | 1 | 10 | Parn | 0.042 | Old |
| 0.195637 | 0.459029 | 0.152 | 0.101 | 1 | 10 | Cfap20 | 0.051 | Old |
| 0.195707 | 0.893596 | 0.103 | 0.061 | 1 | 10 | Pde4dip | 0.042 | Old |
| 0.19583 | 0.33098 | 0.235 | 0.172 | 1 | 10 | Atp9b | 0.063 | Old |
| 0.196006 | 0.145846 | 0.413 | 0.343 | 1 | 10 | Ndufs7 | 0.07 | Old |
| 0.196591 | 0.725441 | 0.127 | 0.081 | 1 | 10 | Elp3 | 0.046 | Old |
| 0.19672 | 0.369965 | 0.09 | 0.051 | 1 | 10 | Rfng | 0.039 | Old |
| 0.1969 | 0.164264 | 0.337 | 0.273 | 1 | 10 | Wtap | 0.064 | Old |
| 0.196907 | 0.648155 | 0.139 | 0.091 | 1 | 10 | Stk19 | 0.048 | Old |
| 0.196981 | 0.060455 | 0.34 | 0.253 | 1 | 10 | Cxcl1 | 0.087 | Old |
| 0.197054 | 0.223307 | 0.303 | 0.222 | 1 | 10 | Snx1 | 0.081 | Old |
| 0.197359 | 0.39997 | 0.284 | 0.212 | 1 | 10 | Tmem120a | 0.072 | Old |
| 0.197385 | 0.084258 | 0.303 | 0.222 | 1 | 10 | Zfr | 0.081 | Old |
| 0.197536 | 0.143955 | 0.127 | 0.081 | 1 | 10 | Tsga10 | 0.046 | Old |
| 0.198382 | 0.278676 | 0.176 | 0.121 | 1 | 10 | Smim20 | 0.055 | Old |
| 0.198643 | 1.845304 | 0.034 | 0.01 | 1 | 10 | Ybey | 0.024 | Old |
| 0.198643 | 1.75052 | 0.034 | 0.01 | 1 | 10 | Fbxo46 | 0.024 | Old |
| 0.198742 | 0.042562 | 0.291 | 0.212 | 1 | 10 | Idh2 | 0.079 | Old |
| 0.198787 | 0.033809 | 0.389 | 0.313 | 1 | 10 | Slc25a4 | 0.076 | Old |
| 0.198861 | 0.235337 | 0.55 | 0.465 | 1 | 10 | Ms4a6c | 0.085 | Old |
| 0.199151 | 0.230355 | 0.237 | 0.172 | 1 | 10 | Phka2 | 0.065 | Old |
| 0.199547 | 0.583322 | 0.156 | 0.101 | 1 | 10 | Spns1 | 0.055 | Old |
| 0.199553 | 1.727061 | 0.034 | 0.01 | 1 | 10 | E2f6 | 0.024 | Old |
| 0.19969 | 0.3235 | 0.318 | 0.253 | 1 | 10 | Inppl1 | 0.065 | Old |
| 0.199751 | 0.824993 | 0.115 | 0.071 | 1 | 10 | Alg1 | 0.044 | Old |
| 0.199859 | 1.966321 | 0.049 | 0.02 | 1 | 10 | Zfp952 | 0.029 | Old |
| 0.200291 | 0.134009 | 0.814 | 0.707 | 1 | 10 | Samhd1 | 0.107 | Old |
| 0.200457 | 0.257049 | 0.445 | 0.374 | 1 | 10 | Ndufb4 | 0.071 | Old |
| 0.200521 | 0.037483 | 0.249 | 0.192 | 1 | 10 | Ccl8 | 0.057 | Old |
| 0.20056 | 0.191559 | 0.22 | 0.152 | 1 | 10 | Kdm2b | 0.068 | Old |
| 0.200617 | 0.105375 | 0.237 | 0.172 | 1 | 10 | Gpr108 | 0.065 | Old |
| 0.200762 | 0.317783 | 0.169 | 0.111 | 1 | 10 | Uimc1 | 0.058 | Old |
| 0.200868 | 1.045541 | 0.064 | 0.03 | 1 | 10 | Dhx35 | 0.034 | Old |
| 0.200935 | 0.097516 | 0.531 | 0.455 | 1 | 10 | Nsf | 0.076 | Old |
| 0.201136 | 0.540192 | 0.208 | 0.152 | 1 | 10 | Arfgap2 | 0.056 | Old |
| 0.201256 | 0.897409 | 0.115 | 0.071 | 1 | 10 | Grk5 | 0.044 | Old |
| 0.201381 | 1.445448 | 0.049 | 0.02 | 1 | 10 | Pter | 0.029 | Old |
| 0.201383 | 1.645322 | 0.034 | 0.01 | 1 | 10 | Ankrd13d | 0.024 | Old |
| 0.20148 | 1.241011 | 0.076 | 0.04 | 1 | 10 | Acad9 | 0.036 | Old |
| 0.201531 | 0.47009 | 0.364 | 0.303 | 1 | 10 | Pgrmc1 | 0.061 | Old |
| 0.201618 | 0.159842 | 0.132 | 0.081 | 1 | 10 | Abcg3 | 0.051 | Old |
| 0.201725 | 0.1286 | 0.535 | 0.424 | 1 | 10 | Pigt | 0.111 | Old |
| 0.201884 | 1.281782 | 0.088 | 0.051 | 1 | 10 | Zfp157 | 0.037 | Old |

| | | | | | | | | |
|---|---|---|---|---|---|---|---|---|
| 0.202108 | 0.599687 | 0.161 | 0.111 | 1 | 10 | Anks1 | 0.05 | Old |
| 0.202146 | 1.691825 | 0.049 | 0.02 | 1 | 10 | Usp30 | 0.029 | Old |
| 0.202226 | 0.334355 | 0.166 | 0.111 | 1 | 10 | Trmt1l | 0.055 | Old |
| 0.202302 | 1.859339 | 0.034 | 0.01 | 1 | 10 | Chst15 | 0.024 | Old |
| 0.202302 | 1.638386 | 0.034 | 0.01 | 1 | 10 | C1ra | 0.024 | Old |
| 0.202302 | 1.63324 | 0.034 | 0.01 | 1 | 10 | Zdhhc23 | 0.024 | Old |
| 0.202302 | 1.461215 | 0.034 | 0.01 | 1 | 10 | Adam22 | 0.024 | Old |
| 0.202768 | 0.736617 | 0.115 | 0.071 | 1 | 10 | Abhd5 | 0.044 | Old |
| 0.202768 | 0.618598 | 0.115 | 0.071 | 1 | 10 | Cyth2 | 0.044 | Old |
| 0.20289 | 0.180767 | 0.78 | 0.636 | 1 | 10 | Gpx4 | 0.144 | Old |
| 0.202925 | 0.420592 | 0.117 | 0.071 | 1 | 10 | Fbxw8 | 0.046 | Old |
| 0.203326 | 1.289762 | 0.076 | 0.04 | 1 | 10 | Phlpp1 | 0.036 | Old |
| 0.20358 | 0.131651 | 0.533 | 0.455 | 1 | 10 | Tbcb | 0.078 | Old |
| 0.20415 | 1.671188 | 0.034 | 0.01 | 1 | 10 | Rbm4 | 0.024 | Old |
| 0.20415 | 1.441238 | 0.034 | 0.01 | 1 | 10 | Ulk3 | 0.024 | Old |
| 0.204257 | 0.358283 | 0.12 | 0.172 | 1 | 10 | Pla2g2d | -0.052 | Old |
| 0.204265 | 0.187473 | 0.335 | 0.253 | 1 | 10 | Slc25a11 | 0.082 | Old |
| 0.20447 | 0.366018 | 0.152 | 0.101 | 1 | 10 | Adar | 0.051 | Old |
| 0.204563 | 1.254473 | 0.076 | 0.04 | 1 | 10 | Ankrd50 | 0.036 | Old |
| 0.204991 | 0.599632 | 0.164 | 0.111 | 1 | 10 | Serpine1 | 0.053 | Old |
| 0.205079 | 1.903821 | 0.034 | 0.01 | 1 | 10 | Mmp13 | 0.024 | Old |
| 0.205079 | 1.455558 | 0.034 | 0.01 | 1 | 10 | Zfp825 | 0.024 | Old |
| 0.205079 | 1.349436 | 0.034 | 0.01 | 1 | 10 | Klhl3 | 0.024 | Old |
| 0.205079 | 1.338669 | 0.034 | 0.01 | 1 | 10 | Zfp408 | 0.024 | Old |
| 0.205079 | 1.304698 | 0.034 | 0.01 | 1 | 10 | Elovl6 | 0.024 | Old |
| 0.205204 | 0.027507 | 0.208 | 0.141 | 1 | 10 | Clasp2 | 0.067 | Old |
| 0.205378 | 0.197012 | 0.702 | 0.606 | 1 | 10 | Vdac2 | 0.096 | Old |
| 0.205395 | 0.21438 | 0.311 | 0.242 | 1 | 10 | Azi2 | 0.069 | Old |
| 0.205586 | 0.821517 | 0.064 | 0.03 | 1 | 10 | Plk4 | 0.034 | Old |
| 0.20576 | 0.612619 | 0.103 | 0.061 | 1 | 10 | Gfm2 | 0.042 | Old |
| 0.205806 | 1.176462 | 0.076 | 0.04 | 1 | 10 | Zfp626 | 0.036 | Old |
| 0.205829 | 0.3614 | 0.262 | 0.192 | 1 | 10 | Rnf11 | 0.07 | Old |
| 0.205989 | 0.579528 | 0.188 | 0.131 | 1 | 10 | Commd10 | 0.057 | Old |
| 0.206011 | 1.278888 | 0.034 | 0.01 | 1 | 10 | Gemin2 | 0.024 | Old |
| 0.206058 | 0.492741 | 0.262 | 0.202 | 1 | 10 | 8-Sep | 0.06 | Old |
| 0.206151 | 1.169255 | 0.112 | 0.071 | 1 | 10 | Siglec1 | 0.041 | Old |
| 0.206294 | 0.388551 | 0.289 | 0.222 | 1 | 10 | Yipf3 | 0.067 | Old |
| 0.206403 | 0.588657 | 0.174 | 0.121 | 1 | 10 | Smim27 | 0.053 | Old |
| 0.206776 | 1.409019 | 0.049 | 0.02 | 1 | 10 | Haus3 | 0.029 | Old |
| 0.206799 | 0.152158 | 0.254 | 0.182 | 1 | 10 | Ten1 | 0.072 | Old |
| 0.206852 | 0.072773 | 0.232 | 0.162 | 1 | 10 | Acvrl1 | 0.07 | Old |
| 0.206945 | 0.972735 | 0.034 | 0.01 | 1 | 10 | Focad | 0.024 | Old |
| 0.207064 | 0.59821 | 0.139 | 0.091 | 1 | 10 | Serhl | 0.048 | Old |
| 0.207676 | 0.340574 | 0.355 | 0.273 | 1 | 10 | Mrpl30 | 0.082 | Old |
| 0.207919 | 0.668138 | 0.264 | 0.212 | 1 | 10 | Pigk | 0.052 | Old |

| | | | | | | | | |
|---|---|---|---|---|---|---|---|---|
| 0.20794 | 0.101281 | 0.342 | 0.253 | 1 | 10 | Tmcc3 | 0.089 | Old |
| 0.208314 | 0.233815 | 0.43 | 0.354 | 1 | 10 | Add1 | 0.076 | Old |
| 0.208336 | 1.10874 | 0.049 | 0.02 | 1 | 10 | Natd1 | 0.029 | Old |
| 0.208336 | 0.916643 | 0.049 | 0.02 | 1 | 10 | Ssh3 | 0.029 | Old |
| 0.208336 | 0.910363 | 0.049 | 0.02 | 1 | 10 | Aspa | 0.029 | Old |
| 0.208443 | 0.180705 | 0.306 | 0.232 | 1 | 10 | Pja2 | 0.074 | Old |
| 0.208824 | 1.050573 | 0.034 | 0.01 | 1 | 10 | Hist2h3c2 | 0.024 | Old |
| 0.208824 | 0.950917 | 0.034 | 0.01 | 1 | 10 | Gm21781 | 0.024 | Old |
| 0.208824 | 0.877063 | 0.034 | 0.01 | 1 | 10 | Tmco4 | 0.024 | Old |
| 0.208824 | 0.744574 | 0.034 | 0.01 | 1 | 10 | Tmem69 | 0.024 | Old |
| 0.208903 | 0.750382 | 0.115 | 0.071 | 1 | 10 | Slc39a3 | 0.044 | Old |
| 0.209004 | 0.771196 | 0.064 | 0.03 | 1 | 10 | Nudt18 | 0.034 | Old |
| 0.209334 | 0.464176 | 0.274 | 0.212 | 1 | 10 | Lhfpl2 | 0.062 | Old |
| 0.20936 | 0.541205 | 0.198 | 0.141 | 1 | 10 | Asl | 0.057 | Old |
| 0.20952 | 0.145524 | 0.751 | 0.636 | 1 | 10 | Clic1 | 0.115 | Old |
| 0.209669 | 0.254226 | 0.477 | 0.384 | 1 | 10 | Folr2 | 0.093 | Old |
| 0.209768 | 0.542029 | 0.034 | 0.01 | 1 | 10 | Pkmyt1 | 0.024 | Old |
| 0.209895 | 0.312693 | 0.345 | 0.273 | 1 | 10 | Srd5a3 | 0.072 | Old |
| 0.210395 | 0.265734 | 0.411 | 0.333 | 1 | 10 | Trappc2l | 0.078 | Old |
| 0.210693 | 0.343469 | 0.049 | 0.02 | 1 | 10 | Spout1 | 0.029 | Old |
| 0.21085 | 0.537041 | 0.139 | 0.091 | 1 | 10 | Dkc1 | 0.048 | Old |
| 0.211075 | 0.755833 | 0.064 | 0.03 | 1 | 10 | Gm46224 | 0.034 | Old |
| 0.211224 | 0.056767 | 0.345 | 0.273 | 1 | 10 | Lamtor5 | 0.072 | Old |
| 0.211548 | 0.931009 | 0.1 | 0.061 | 1 | 10 | Mlycd | 0.039 | Old |
| 0.211653 | 0.52869 | 0.171 | 0.121 | 1 | 10 | Sdhaf2 | 0.05 | Old |
| 0.212089 | 0.462315 | 0.418 | 0.343 | 1 | 10 | Gm26532 | 0.075 | Old |
| 0.212275 | 1.303266 | 0.049 | 0.02 | 1 | 10 | Ndnf | 0.029 | Old |
| 0.212275 | 1.043318 | 0.049 | 0.02 | 1 | 10 | Fabp7 | 0.029 | Old |
| 0.212694 | 0.365799 | 0.13 | 0.081 | 1 | 10 | Trim24 | 0.049 | Old |
| 0.212851 | 0.312444 | 0.103 | 0.061 | 1 | 10 | Pmf1 | 0.042 | Old |
| 0.213053 | 0.231885 | 0.394 | 0.313 | 1 | 10 | Ttc7 | 0.081 | Old |
| 0.21316 | 0.576622 | 0.064 | 0.03 | 1 | 10 | Kbtbd3 | 0.034 | Old |
| 0.213474 | 0.471561 | 0.325 | 0.273 | 1 | 10 | Capn1 | 0.052 | Old |
| 0.213591 | 0.753003 | 0.115 | 0.071 | 1 | 10 | Tmem161k | 0.044 | Old |
| 0.213619 | 0.474289 | 0.127 | 0.081 | 1 | 10 | Stx18 | 0.046 | Old |
| 0.213665 | 0.165143 | 0.807 | 0.737 | 1 | 10 | Cd68 | 0.07 | Old |
| 0.213705 | 0.103341 | 0.406 | 0.323 | 1 | 10 | Ctnnd1 | 0.083 | Old |
| 0.213865 | 1.119934 | 0.049 | 0.02 | 1 | 10 | Armc6 | 0.029 | Old |
| 0.213991 | 0.381612 | 0.303 | 0.232 | 1 | 10 | Zmiz2 | 0.071 | Old |
| 0.214089 | 0.150302 | 0.264 | 0.192 | 1 | 10 | Eif4e2 | 0.072 | Old |
| 0.214116 | 0.305193 | 0.372 | 0.303 | 1 | 10 | Nup98 | 0.069 | Old |
| 0.214134 | 1.022335 | 0.088 | 0.051 | 1 | 10 | Qser1 | 0.037 | Old |
| 0.214586 | 0.258653 | 0.293 | 0.232 | 1 | 10 | Gpalpp1 | 0.061 | Old |
| 0.214606 | 0.240227 | 0.286 | 0.212 | 1 | 10 | Ergic1 | 0.074 | Old |
| 0.215023 | 0.550319 | 0.267 | 0.212 | 1 | 10 | Rnasel | 0.055 | Old |

| | | | | | | | | |
|---|---|---|---|---|---|---|---|---|
| 0.215327 | 0.239473 | 0.23 | 0.172 | 1 | 10 | Scarb1 | 0.058 | Old |
| 0.215915 | 0.453222 | 0.152 | 0.101 | 1 | 10 | Trip4 | 0.051 | Old |
| 0.215947 | 0.961467 | 0.076 | 0.04 | 1 | 10 | Ankmy2 | 0.036 | Old |
| 0.216048 | 0.486915 | 0.154 | 0.101 | 1 | 10 | Atp6v0a2 | 0.053 | Old |
| 0.216127 | 0.918709 | 0.112 | 0.071 | 1 | 10 | Fbxl14 | 0.041 | Old |
| 0.216327 | 0.209926 | 0.222 | 0.162 | 1 | 10 | S1pr2 | 0.06 | Old |
| 0.216513 | 0.228637 | 0.154 | 0.101 | 1 | 10 | Atad3a | 0.053 | Old |
| 0.216586 | 0.986123 | 0.1 | 0.061 | 1 | 10 | Nup155 | 0.039 | Old |
| 0.216586 | 0.968284 | 0.1 | 0.061 | 1 | 10 | Crot | 0.039 | Old |
| 0.21702 | 0.140455 | 0.677 | 0.576 | 1 | 10 | Itgam | 0.101 | Old |
| 0.217071 | 0.675569 | 0.049 | 0.02 | 1 | 10 | Nck2 | 0.029 | Old |
| 0.217229 | 0.295063 | 0.32 | 0.253 | 1 | 10 | 9930111J2 | 0.067 | Old |
| 0.217354 | 0.175103 | 0.633 | 0.556 | 1 | 10 | Ccdc50 | 0.077 | Old |
| 0.217618 | 1.288934 | 0.061 | 0.03 | 1 | 10 | Prkdc | 0.031 | Old |
| 0.217753 | 0.255153 | 0.381 | 0.303 | 1 | 10 | Sgpp1 | 0.078 | Old |
| 0.217878 | 0.113843 | 0.049 | 0.02 | 1 | 10 | Dgke | 0.029 | Old |
| 0.218044 | 0.123528 | 0.284 | 0.222 | 1 | 10 | Jup | 0.062 | Old |
| 0.218083 | 0.266847 | 0.139 | 0.091 | 1 | 10 | Rnaseh2a | 0.048 | Old |
| 0.218258 | 0.45249 | 0.152 | 0.101 | 1 | 10 | Usp45 | 0.051 | Old |
| 0.218458 | 0.333568 | 0.328 | 0.263 | 1 | 10 | Rtf1 | 0.065 | Old |
| 0.218651 | 0.844583 | 0.098 | 0.061 | 1 | 10 | Terf2ip | 0.037 | Old |
| 0.218663 | 0.145668 | 0.35 | 0.273 | 1 | 10 | Usp16 | 0.077 | Old |
| 0.218748 | 0.18658 | 0.577 | 0.515 | 1 | 10 | Evi2a | 0.062 | Old |
| 0.218809 | 0.372871 | 0.112 | 0.071 | 1 | 10 | Myg1 | 0.041 | Old |
| 0.218814 | 0.118272 | 0.667 | 0.576 | 1 | 10 | Cbl | 0.091 | Old |
| 0.218851 | 0.515841 | 0.227 | 0.172 | 1 | 10 | Dhx40 | 0.055 | Old |
| 0.218853 | 0.40688 | 0.22 | 0.162 | 1 | 10 | Naga | 0.058 | Old |
| 0.218881 | 0.102434 | 0.296 | 0.222 | 1 | 10 | Ing4 | 0.074 | Old |
| 0.219043 | 0.227807 | 0.32 | 0.242 | 1 | 10 | Polr2e | 0.078 | Old |
| 0.219061 | 1.476232 | 0.061 | 0.03 | 1 | 10 | Endog | 0.031 | Old |
| 0.219078 | 0.342804 | 0.337 | 0.273 | 1 | 10 | Bag3 | 0.064 | Old |
| 0.219158 | 0.66024 | 0.213 | 0.162 | 1 | 10 | Clcn5 | 0.051 | Old |
| 0.219315 | 0.158827 | 0.264 | 0.192 | 1 | 10 | Mob1b | 0.072 | Old |
| 0.219345 | 0.343463 | 0.399 | 0.343 | 1 | 10 | Slfn5 | 0.056 | Old |
| 0.219421 | 0.707465 | 0.1 | 0.061 | 1 | 10 | Iffo1 | 0.039 | Old |
| 0.219453 | 0.39119 | 0.242 | 0.182 | 1 | 10 | Tor1b | 0.06 | Old |
| 0.220152 | 0.901903 | 0.088 | 0.051 | 1 | 10 | Dock5 | 0.037 | Old |
| 0.220496 | 0.556278 | 0.076 | 0.04 | 1 | 10 | Slc35e2 | 0.036 | Old |
| 0.220596 | 0.6557 | 0.267 | 0.212 | 1 | 10 | Pdlim4 | 0.055 | Old |
| 0.220863 | 0.065102 | 0.707 | 0.636 | 1 | 10 | Ddx6 | 0.071 | Old |
| 0.220894 | 0.186532 | 0.333 | 0.242 | 1 | 10 | Ahcyl2 | 0.091 | Old |
| 0.220934 | 0.157549 | 0.455 | 0.364 | 1 | 10 | Zfand6 | 0.091 | Old |
| 0.221034 | 0.439461 | 0.115 | 0.071 | 1 | 10 | Kif16b | 0.044 | Old |
| 0.221239 | 1.126278 | 0.061 | 0.03 | 1 | 10 | Mul1 | 0.031 | Old |
| 0.221536 | 0.141317 | 0.164 | 0.111 | 1 | 10 | Bcor | 0.053 | Old |

| | | | | | | | | |
|---|---|---|---|---|---|---|---|---|
| 0.221672 | 0.20303 | 0.154 | 0.101 | 1 | 10 | Xpa | 0.053 | Old |
| 0.221777 | 0.0466 | 0.178 | 0.121 | 1 | 10 | Naxd | 0.057 | Old |
| 0.221968 | 1.197771 | 0.061 | 0.03 | 1 | 10 | Gm9493 | 0.031 | Old |
| 0.222013 | 0.13172 | 0.418 | 0.343 | 1 | 10 | Srebf2 | 0.075 | Old |
| 0.222389 | 0.381104 | 0.103 | 0.061 | 1 | 10 | Pef1 | 0.042 | Old |
| 0.222715 | 0.057044 | 0.538 | 0.455 | 1 | 10 | Rtraf | 0.083 | Old |
| 0.222754 | 0.139391 | 0.291 | 0.212 | 1 | 10 | Cdk17 | 0.079 | Old |
| 0.222922 | 0.736791 | 0.203 | 0.152 | 1 | 10 | Slc45a4 | 0.051 | Old |
| 0.222931 | 0.322716 | 0.408 | 0.343 | 1 | 10 | Sbno2 | 0.065 | Old |
| 0.223095 | 0.237578 | 0.724 | 0.677 | 1 | 10 | Abhd12 | 0.047 | Old |
| 0.223191 | 0.159977 | 0.117 | 0.071 | 1 | 10 | Trp53inp1 | 0.046 | Old |
| 0.223254 | 0.541335 | 0.188 | 0.141 | 1 | 10 | Pink1 | 0.047 | Old |
| 0.223281 | 0.057983 | 0.328 | 0.253 | 1 | 10 | Tbc1d10b | 0.075 | Old |
| 0.223451 | 0.272471 | 0.406 | 0.323 | 1 | 10 | Ncoa3 | 0.083 | Old |
| 0.22353 | 0.199379 | 0.103 | 0.061 | 1 | 10 | Pphln1 | 0.042 | Old |
| 0.223851 | 0.009823 | 0.533 | 0.455 | 1 | 10 | Ywhaq | 0.078 | Old |
| 0.223858 | 0.317321 | 0.279 | 0.212 | 1 | 10 | Sft2d1 | 0.067 | Old |
| 0.223912 | 0.359153 | 0.501 | 0.414 | 1 | 10 | Limd1 | 0.087 | Old |
| 0.224436 | 0.403959 | 0.152 | 0.101 | 1 | 10 | Fam126b | 0.051 | Old |
| 0.224903 | 1.179107 | 0.061 | 0.03 | 1 | 10 | Dnal4 | 0.031 | Old |
| 0.225616 | 0.212442 | 0.44 | 0.374 | 1 | 10 | Sdhd | 0.066 | Old |
| 0.225808 | 0.077939 | 0.599 | 0.485 | 1 | 10 | Ndufa3 | 0.114 | Old |
| 0.226077 | 0.498545 | 0.227 | 0.172 | 1 | 10 | Dtx3l | 0.055 | Old |
| 0.226194 | 0.080668 | 0.301 | 0.222 | 1 | 10 | Ypel5 | 0.079 | Old |
| 0.226273 | 0.100451 | 0.584 | 0.495 | 1 | 10 | Cox7b | 0.089 | Old |
| 0.226381 | 1.070215 | 0.061 | 0.03 | 1 | 10 | Tmem185a | 0.031 | Old |
| 0.226381 | 0.825708 | 0.061 | 0.03 | 1 | 10 | Tmem101 | 0.031 | Old |
| 0.226444 | 0.101778 | 0.076 | 0.04 | 1 | 10 | Rundc1 | 0.036 | Old |
| 0.226462 | 0.025012 | 0.269 | 0.202 | 1 | 10 | Sarnp | 0.067 | Old |
| 0.226646 | 2.777134 | 0.015 | 0 | 1 | 10 | Vsig4 | 0.015 | Old |
| 0.226646 | 2.364623 | 0.015 | 0 | 1 | 10 | Zfp455 | 0.015 | Old |
| 0.226646 | 2.090063 | 0.015 | 0 | 1 | 10 | Creb3l4 | 0.015 | Old |
| 0.226646 | 2.02983 | 0.015 | 0 | 1 | 10 | Itgad | 0.015 | Old |
| 0.226646 | 2.027125 | 0.015 | 0 | 1 | 10 | Mettl4 | 0.015 | Old |
| 0.226646 | 2.025446 | 0.015 | 0 | 1 | 10 | Dpep3 | 0.015 | Old |
| 0.226646 | 2.004765 | 0.015 | 0 | 1 | 10 | Zfp458 | 0.015 | Old |
| 0.226646 | 1.962095 | 0.015 | 0 | 1 | 10 | Gm27003 | 0.015 | Old |
| 0.226646 | 1.952822 | 0.015 | 0 | 1 | 10 | Sfn | 0.015 | Old |
| 0.226646 | 1.933365 | 0.015 | 0 | 1 | 10 | 3110009E1 | 0.015 | Old |
| 0.226646 | 1.930822 | 0.015 | 0 | 1 | 10 | Pde7b | 0.015 | Old |
| 0.226646 | 1.906429 | 0.015 | 0 | 1 | 10 | Cyp2s1 | 0.015 | Old |
| 0.226646 | 1.893128 | 0.015 | 0 | 1 | 10 | Col11a2 | 0.015 | Old |
| 0.226646 | 1.887244 | 0.015 | 0 | 1 | 10 | Spata2l | 0.015 | Old |
| 0.226646 | 1.880569 | 0.015 | 0 | 1 | 10 | Swsap1 | 0.015 | Old |
| 0.226646 | 1.875743 | 0.015 | 0 | 1 | 10 | Sgsm3 | 0.015 | Old |

| | | | | | | | | |
|---|---|---|---|---|---|---|---|---|
| 0.226646 | 1.840629 | 0.015 | 0 | 1 | 10 | Bmpr1a | 0.015 | Old |
| 0.226646 | 1.798779 | 0.015 | 0 | 1 | 10 | 9330020H( | 0.015 | Old |
| 0.226646 | 1.793609 | 0.015 | 0 | 1 | 10 | Nme2 | 0.015 | Old |
| 0.226646 | 1.765486 | 0.015 | 0 | 1 | 10 | Gnal | 0.015 | Old |
| 0.226646 | 1.765181 | 0.015 | 0 | 1 | 10 | E2f8 | 0.015 | Old |
| 0.226646 | 1.746339 | 0.015 | 0 | 1 | 10 | Nmb | 0.015 | Old |
| 0.226646 | 1.746052 | 0.015 | 0 | 1 | 10 | Gm10031 | 0.015 | Old |
| 0.226646 | 1.727168 | 0.015 | 0 | 1 | 10 | Rnd1 | 0.015 | Old |
| 0.226646 | 1.719112 | 0.015 | 0 | 1 | 10 | Apoc1 | 0.015 | Old |
| 0.226646 | 1.710974 | 0.015 | 0 | 1 | 10 | Jakmip1 | 0.015 | Old |
| 0.226646 | 1.706681 | 0.015 | 0 | 1 | 10 | Ksr2 | 0.015 | Old |
| 0.226646 | 1.67368 | 0.015 | 0 | 1 | 10 | Ubtd2 | 0.015 | Old |
| 0.226646 | 1.663131 | 0.015 | 0 | 1 | 10 | Ubxn11 | 0.015 | Old |
| 0.226646 | 1.654273 | 0.015 | 0 | 1 | 10 | Ndor1 | 0.015 | Old |
| 0.226646 | 1.635369 | 0.015 | 0 | 1 | 10 | Ttc30a1 | 0.015 | Old |
| 0.226646 | 1.62779 | 0.015 | 0 | 1 | 10 | Zfp523 | 0.015 | Old |
| 0.226646 | 1.622785 | 0.015 | 0 | 1 | 10 | Msrb2 | 0.015 | Old |
| 0.226646 | 1.61904 | 0.015 | 0 | 1 | 10 | Gm7160 | 0.015 | Old |
| 0.226646 | 1.594876 | 0.015 | 0 | 1 | 10 | Gm15726 | 0.015 | Old |
| 0.226646 | 1.59287 | 0.015 | 0 | 1 | 10 | Aoc2 | 0.015 | Old |
| 0.226646 | 1.592685 | 0.015 | 0 | 1 | 10 | Pfkfb2 | 0.015 | Old |
| 0.226646 | 1.590632 | 0.015 | 0 | 1 | 10 | Catip | 0.015 | Old |
| 0.226646 | 1.560614 | 0.015 | 0 | 1 | 10 | Gm12589 | 0.015 | Old |
| 0.226646 | 1.559009 | 0.015 | 0 | 1 | 10 | Pnp2 | 0.015 | Old |
| 0.226646 | 1.556017 | 0.015 | 0 | 1 | 10 | Rsph3a | 0.015 | Old |
| 0.226646 | 1.546569 | 0.015 | 0 | 1 | 10 | Vsig10l | 0.015 | Old |
| 0.226646 | 1.540312 | 0.015 | 0 | 1 | 10 | Fndc10 | 0.015 | Old |
| 0.226646 | 1.53502 | 0.015 | 0 | 1 | 10 | Gm42595 | 0.015 | Old |
| 0.226646 | 1.534643 | 0.015 | 0 | 1 | 10 | Ttll5 | 0.015 | Old |
| 0.226646 | 1.529503 | 0.015 | 0 | 1 | 10 | Zkscan7 | 0.015 | Old |
| 0.226646 | 1.521747 | 0.015 | 0 | 1 | 10 | Mrm3 | 0.015 | Old |
| 0.226646 | 1.514875 | 0.015 | 0 | 1 | 10 | Egfr | 0.015 | Old |
| 0.226646 | 1.504413 | 0.015 | 0 | 1 | 10 | Gm9803 | 0.015 | Old |
| 0.226646 | 1.502583 | 0.015 | 0 | 1 | 10 | Pidd1 | 0.015 | Old |
| 0.226646 | 1.440899 | 0.015 | 0 | 1 | 10 | Cc2d2a | 0.015 | Old |
| 0.226646 | 1.435886 | 0.015 | 0 | 1 | 10 | Spaca9 | 0.015 | Old |
| 0.226646 | 1.422045 | 0.015 | 0 | 1 | 10 | Asf1b | 0.015 | Old |
| 0.226646 | 1.313985 | 0.015 | 0 | 1 | 10 | Mterf1b | 0.015 | Old |
| 0.226646 | 1.231908 | 0.015 | 0 | 1 | 10 | Gvin1 | 0.015 | Old |
| 0.226646 | 1.181633 | 0.015 | 0 | 1 | 10 | L3hypdh | 0.015 | Old |
| 0.226646 | 1.156934 | 0.015 | 0 | 1 | 10 | Spib | 0.015 | Old |
| 0.226646 | 1.057373 | 0.015 | 0 | 1 | 10 | Gm47123 | 0.015 | Old |
| 0.226646 | 1.020194 | 0.015 | 0 | 1 | 10 | Zfp119b | 0.015 | Old |
| 0.226698 | 0.115863 | 0.743 | 0.657 | 1 | 10 | Pak2 | 0.086 | Old |
| 0.226757 | 0.101582 | 0.237 | 0.172 | 1 | 10 | Cc2d1b | 0.065 | Old |

| | | | | | | | | |
|---|---|---|---|---|---|---|---|---|
| 0.227043 | 0.012654 | 0.357 | 0.273 | 1 | 10 | R3hdm2 | 0.084 | Old |
| 0.227193 | 0.055698 | 0.21 | 0.152 | 1 | 10 | Fbxo28 | 0.058 | Old |
| 0.2273 | 0.090008 | 0.457 | 0.384 | 1 | 10 | Rab21 | 0.073 | Old |
| 0.227382 | 1.02838 | 0.098 | 0.061 | 1 | 10 | Nedd1 | 0.037 | Old |
| 0.227464 | 0.114451 | 0.127 | 0.081 | 1 | 10 | Fam160a2 | 0.046 | Old |
| 0.227763 | 0.009206 | 0.526 | 0.434 | 1 | 10 | Atp13a3 | 0.092 | Old |
| 0.227766 | 0.039481 | 0.183 | 0.121 | 1 | 10 | Stoml2 | 0.062 | Old |
| 0.227865 | 1.037108 | 0.061 | 0.03 | 1 | 10 | Ncapd3 | 0.031 | Old |
| 0.227865 | 1.07521 | 0.061 | 0.03 | 1 | 10 | Mettl13 | 0.031 | Old |
| 0.227919 | 1.304303 | 0.073 | 0.04 | 1 | 10 | Dgcr14 | 0.033 | Old |
| 0.227935 | 0.725432 | 0.122 | 0.081 | 1 | 10 | Arhgap21 | 0.041 | Old |
| 0.228039 | 0.16598 | 0.164 | 0.111 | 1 | 10 | Ints12 | 0.053 | Old |
| 0.228062 | 0.003538 | 0.191 | 0.131 | 1 | 10 | Rabggtb | 0.06 | Old |
| 0.228168 | 0.087413 | 0.306 | 0.232 | 1 | 10 | Rab11fip1 | 0.074 | Old |
| 0.228577 | 0.14286 | 0.601 | 0.505 | 1 | 10 | Ndufa7 | 0.096 | Old |
| 0.228662 | 0.470798 | 0.112 | 0.071 | 1 | 10 | Hcfc2 | 0.041 | Old |
| 0.228681 | 0.093193 | 0.599 | 0.525 | 1 | 10 | Trir | 0.074 | Old |
| 0.22878 | 0.573523 | 0.088 | 0.051 | 1 | 10 | Bcl2l13 | 0.037 | Old |
| 0.228932 | 0.49876 | 0.222 | 0.172 | 1 | 10 | H2-T22 | 0.05 | Old |
| 0.22894 | 0.459168 | 0.225 | 0.172 | 1 | 10 | Aggf1 | 0.053 | Old |
| 0.228962 | 0.142689 | 0.178 | 0.121 | 1 | 10 | Nsrp1 | 0.057 | Old |
| 0.229003 | 0.77495 | 0.122 | 0.081 | 1 | 10 | Slc41a3 | 0.041 | Old |
| 0.229419 | 0.352372 | 0.533 | 0.485 | 1 | 10 | Tnfaip2 | 0.048 | Old |
| 0.229635 | 1.864237 | 0.046 | 0.02 | 1 | 10 | Fbf1 | 0.026 | Old |
| 0.22985 | 0.297163 | 0.1 | 0.061 | 1 | 10 | 4930455G( | 0.039 | Old |
| 0.229851 | 0.697683 | 0.1 | 0.061 | 1 | 10 | Pxmp4 | 0.039 | Old |
| 0.229851 | 0.66175 | 0.1 | 0.061 | 1 | 10 | Fkrp | 0.039 | Old |
| 0.230257 | 0.105238 | 0.572 | 0.455 | 1 | 10 | Spcs2 | 0.117 | Old |
| 0.230393 | 0.690507 | 0.166 | 0.121 | 1 | 10 | Tmem106c | 0.045 | Old |
| 0.23044 | 0.606048 | 0.1 | 0.061 | 1 | 10 | Cd79b | 0.039 | Old |
| 0.230448 | 0.256659 | 0.254 | 0.192 | 1 | 10 | Rabl6 | 0.062 | Old |
| 0.230494 | 1.668214 | 0.046 | 0.02 | 1 | 10 | Cacna1f | 0.026 | Old |
| 0.230494 | 1.656251 | 0.046 | 0.02 | 1 | 10 | 1190002N1 | 0.026 | Old |
| 0.230932 | 0.198033 | 0.306 | 0.232 | 1 | 10 | 2-Mar | 0.074 | Old |
| 0.230988 | 0.247465 | 0.227 | 0.172 | 1 | 10 | Inpp5f | 0.055 | Old |
| 0.231055 | 0.211144 | 0.103 | 0.061 | 1 | 10 | Rcan3 | 0.042 | Old |
| 0.23116 | 2.081641 | 0.032 | 0.01 | 1 | 10 | Btbd17 | 0.022 | Old |
| 0.231509 | 0.042489 | 0.02 | 0.04 | 1 | 10 | Mr1 | -0.02 | Old |
| 0.231607 | 0.696289 | 0.061 | 0.03 | 1 | 10 | Fahd1 | 0.031 | Old |
| 0.231622 | 0.431892 | 0.1 | 0.061 | 1 | 10 | Acad8 | 0.039 | Old |
| 0.231667 | 0.403317 | 0.161 | 0.111 | 1 | 10 | Cbx4 | 0.05 | Old |
| 0.231708 | 0.307019 | 0.125 | 0.081 | 1 | 10 | Rpl10-ps3 | 0.044 | Old |
| 0.231919 | 0.537391 | 0.088 | 0.051 | 1 | 10 | Armt1 | 0.037 | Old |
| 0.231969 | 0.442751 | 0.183 | 0.131 | 1 | 10 | Gm6377 | 0.052 | Old |
| 0.23221 | 2.087412 | 0.032 | 0.01 | 1 | 10 | Ffar4 | 0.022 | Old |

| | | | | | | | | |
|---|---|---|---|---|---|---|---|---|
| 0.232693 | 0.234797 | 0.293 | 0.222 | 1 | 10 | Tfrc | 0.071 | Old |
| 0.233057 | 0.299973 | 0.227 | 0.172 | 1 | 10 | Timm10 | 0.055 | Old |
| 0.233273 | 0.183018 | 0.589 | 0.505 | 1 | 10 | Dnajb14 | 0.084 | Old |
| 0.233331 | 0.371844 | 0.205 | 0.152 | 1 | 10 | Retreg3 | 0.053 | Old |
| 0.2334 | 1.13267 | 0.073 | 0.04 | 1 | 10 | Zc3h12d | 0.033 | Old |
| 0.233553 | 0.710358 | 0.134 | 0.091 | 1 | 10 | Dip2c | 0.043 | Old |
| 0.233719 | 0.370648 | 0.284 | 0.222 | 1 | 10 | Rab12 | 0.062 | Old |
| 0.233724 | 0.516767 | 0.196 | 0.141 | 1 | 10 | Cmip | 0.055 | Old |
| 0.233772 | 0.223093 | 0.318 | 0.253 | 1 | 10 | Dcaf8 | 0.065 | Old |
| 0.23427 | 0.23542 | 0.112 | 0.071 | 1 | 10 | Neurl4 | 0.041 | Old |
| 0.234498 | 0.033953 | 0.623 | 0.505 | 1 | 10 | Clec4a2 | 0.118 | Old |
| 0.234587 | 0.210095 | 0.269 | 0.202 | 1 | 10 | Nap1l4 | 0.067 | Old |
| 0.234632 | 0.54369 | 0.061 | 0.03 | 1 | 10 | Mrgbp | 0.031 | Old |
| 0.23464 | 0.039559 | 0.33 | 0.253 | 1 | 10 | Strap | 0.077 | Old |
| 0.234977 | 0.286266 | 0.242 | 0.182 | 1 | 10 | Spata6 | 0.06 | Old |
| 0.235148 | 0.945258 | 0.098 | 0.061 | 1 | 10 | Polr3e | 0.037 | Old |
| 0.23538 | 2.097135 | 0.032 | 0.01 | 1 | 10 | Cdk5r1 | 0.022 | Old |
| 0.23538 | 1.516742 | 0.032 | 0.01 | 1 | 10 | Coq8a | 0.022 | Old |
| 0.23538 | 1.319589 | 0.032 | 0.01 | 1 | 10 | Wdr60 | 0.022 | Old |
| 0.235685 | 0.006595 | 0.2 | 0.141 | 1 | 10 | Fam193a | 0.059 | Old |
| 0.235733 | 0.090033 | 0.408 | 0.313 | 1 | 10 | Etfa | 0.095 | Old |
| 0.235809 | 0.131618 | 0.724 | 0.677 | 1 | 10 | Cmpk1 | 0.047 | Old |
| 0.23589 | 0.051071 | 0.027 | 0.051 | 1 | 10 | Abhd14b | -0.024 | Old |
| 0.235939 | 0.208666 | 0.262 | 0.202 | 1 | 10 | Tnfrsf13b | 0.06 | Old |
| 0.235975 | 0.520257 | 0.227 | 0.172 | 1 | 10 | Pcf11 | 0.055 | Old |
| 0.236155 | 0.752333 | 0.061 | 0.03 | 1 | 10 | Dhx57 | 0.031 | Old |
| 0.236181 | 0.606923 | 0.137 | 0.091 | 1 | 10 | Uckl1 | 0.046 | Old |
| 0.236443 | 1.492663 | 0.032 | 0.01 | 1 | 10 | Nupl2 | 0.022 | Old |
| 0.236593 | 0.17973 | 0.215 | 0.162 | 1 | 10 | Kctd10 | 0.053 | Old |
| 0.23663 | 0.510974 | 0.132 | 0.091 | 1 | 10 | Hdac4 | 0.041 | Old |
| 0.236831 | 0.128822 | 0.802 | 0.788 | 1 | 10 | Atp5f1 | 0.014 | Old |
| 0.237385 | 0.196687 | 0.443 | 0.343 | 1 | 10 | Eif3d | 0.1 | Old |
| 0.237448 | 1.254292 | 0.046 | 0.02 | 1 | 10 | Zcchc14 | 0.026 | Old |
| 0.23751 | 1.518262 | 0.032 | 0.01 | 1 | 10 | Mki67 | 0.022 | Old |
| 0.23751 | 1.455662 | 0.032 | 0.01 | 1 | 10 | Rasal3 | 0.022 | Old |
| 0.237572 | 0.823713 | 0.073 | 0.04 | 1 | 10 | Sprtn | 0.033 | Old |
| 0.237866 | 0.008772 | 0.379 | 0.283 | 1 | 10 | Plaur | 0.096 | Old |
| 0.23814 | 1.262934 | 0.086 | 0.051 | 1 | 10 | Mlh1 | 0.035 | Old |
| 0.238218 | 0.896216 | 0.132 | 0.091 | 1 | 10 | Timm9 | 0.041 | Old |
| 0.238526 | 0.082913 | 0.692 | 0.606 | 1 | 10 | Rnf130 | 0.086 | Old |
| 0.238581 | 1.470527 | 0.032 | 0.01 | 1 | 10 | Ttll1 | 0.022 | Old |
| 0.238581 | 1.415211 | 0.032 | 0.01 | 1 | 10 | 2310040G2 | 0.022 | Old |
| 0.238581 | 1.383796 | 0.032 | 0.01 | 1 | 10 | Ighmbp2 | 0.022 | Old |
| 0.238581 | 1.142285 | 0.032 | 0.01 | 1 | 10 | Chpf | 0.022 | Old |
| 0.238682 | 0.355451 | 0.076 | 0.04 | 1 | 10 | Arel1 | 0.036 | Old |

| | | | | | | | | |
|---|---|---|---|---|---|---|---|---|
| 0.238973 | 0.289343 | 0.171 | 0.121 | 1 | 10 | Abcf2 | 0.05 | Old |
| 0.238981 | 0.191646 | 0.315 | 0.242 | 1 | 10 | Slc25a36 | 0.073 | Old |
| 0.239209 | 1.039884 | 0.046 | 0.02 | 1 | 10 | Eml2 | 0.026 | Old |
| 0.239262 | 0.035465 | 0.33 | 0.263 | 1 | 10 | Trim28 | 0.067 | Old |
| 0.239378 | 0.13332 | 0.325 | 0.263 | 1 | 10 | Csnk2a1 | 0.062 | Old |
| 0.239413 | 0.410201 | 0.1 | 0.061 | 1 | 10 | Slc28a2 | 0.039 | Old |
| 0.239567 | 0.118018 | 0.487 | 0.424 | 1 | 10 | Ctdsp2 | 0.063 | Old |
| 0.239836 | 0.126373 | 0.125 | 0.081 | 1 | 10 | Mob2 | 0.044 | Old |
| 0.239944 | 0.075932 | 0.293 | 0.222 | 1 | 10 | Hmgcr | 0.071 | Old |
| 0.240214 | 0.291806 | 0.641 | 0.576 | 1 | 10 | Ndufb9 | 0.065 | Old |
| 0.240318 | 0.013063 | 0.428 | 0.343 | 1 | 10 | Tfe3 | 0.085 | Old |
| 0.240551 | 0.37781 | 0.112 | 0.071 | 1 | 10 | Elac2 | 0.041 | Old |
| 0.240731 | 1.032294 | 0.032 | 0.01 | 1 | 10 | Mapkapk5 | 0.022 | Old |
| 0.241114 | 0.392733 | 0.191 | 0.141 | 1 | 10 | Secisbp2 | 0.05 | Old |
| 0.241128 | 0.314837 | 0.112 | 0.071 | 1 | 10 | Nhlrc2 | 0.041 | Old |
| 0.241202 | 0.058392 | 0.244 | 0.182 | 1 | 10 | Emc2 | 0.062 | Old |
| 0.241423 | 0.470506 | 0.137 | 0.091 | 1 | 10 | Tmx2 | 0.046 | Old |
| 0.241437 | 0.189896 | 0.347 | 0.273 | 1 | 10 | Coa3 | 0.074 | Old |
| 0.241452 | 0.246741 | 0.455 | 0.374 | 1 | 10 | Bptf | 0.081 | Old |
| 0.241508 | 0.23794 | 0.557 | 0.475 | 1 | 10 | Magt1 | 0.082 | Old |
| 0.24154 | 0.392538 | 0.061 | 0.03 | 1 | 10 | Tbc1d12 | 0.031 | Old |
| 0.241647 | 0.438153 | 0.232 | 0.182 | 1 | 10 | Ube4b | 0.05 | Old |
| 0.241672 | 0.42471 | 0.149 | 0.101 | 1 | 10 | Zfp622 | 0.048 | Old |
| 0.241688 | 0.166513 | 0.391 | 0.333 | 1 | 10 | Cic | 0.058 | Old |
| 0.241797 | 0.935979 | 0.073 | 0.04 | 1 | 10 | Dvl1 | 0.033 | Old |
| 0.241798 | 0.03735 | 0.575 | 0.455 | 1 | 10 | Ptp4a1 | 0.12 | Old |
| 0.241812 | 0.929694 | 0.032 | 0.01 | 1 | 10 | Srr | 0.022 | Old |
| 0.241847 | 0.585894 | 0.1 | 0.061 | 1 | 10 | Siah1a | 0.039 | Old |
| 0.241864 | 0.77144 | 0.098 | 0.061 | 1 | 10 | Erich1 | 0.037 | Old |
| 0.24195 | 0.26476 | 0.677 | 0.636 | 1 | 10 | Mapk3 | 0.041 | Old |
| 0.24206 | 1.928294 | 0.059 | 0.03 | 1 | 10 | Pnpla6 | 0.029 | Old |
| 0.24212 | 0.130298 | 0.939 | 0.828 | 1 | 10 | Arpc5 | 0.111 | Old |
| 0.242285 | 0.217193 | 0.112 | 0.071 | 1 | 10 | Mcu | 0.041 | Old |
| 0.242458 | 0.459507 | 0.1 | 0.061 | 1 | 10 | Prkrip1 | 0.039 | Old |
| 0.24276 | 0.819709 | 0.046 | 0.02 | 1 | 10 | Nop2 | 0.026 | Old |
| 0.243301 | 0.249052 | 0.379 | 0.303 | 1 | 10 | Acbd3 | 0.076 | Old |
| 0.243372 | 0.084957 | 0.782 | 0.677 | 1 | 10 | Tm9sf3 | 0.105 | Old |
| 0.243451 | 0.021116 | 0.645 | 0.576 | 1 | 10 | Nrp1 | 0.069 | Old |
| 0.243654 | 0.995771 | 0.046 | 0.02 | 1 | 10 | Thada | 0.026 | Old |
| 0.243654 | 0.783556 | 0.046 | 0.02 | 1 | 10 | Rsad1 | 0.026 | Old |
| 0.243654 | 0.384828 | 0.046 | 0.02 | 1 | 10 | Ccdc191 | 0.026 | Old |
| 0.243654 | 0.070843 | 0.046 | 0.02 | 1 | 10 | Ccdc152 | 0.026 | Old |
| 0.243659 | 0.364609 | 0.377 | 0.313 | 1 | 10 | Tmem9b | 0.064 | Old |
| 0.243666 | 0.241904 | 0.176 | 0.121 | 1 | 10 | Cdk16 | 0.055 | Old |
| 0.243718 | 0.236303 | 0.152 | 0.101 | 1 | 10 | Chmp6 | 0.051 | Old |

| | | | | | | | | |
|---|---|---|---|---|---|---|---|---|
| 0.243897 | 0.223643 | 0.411 | 0.343 | 1 | 10 | Il10ra | 0.068 | Old |
| 0.243903 | 0.213134 | 0.164 | 0.111 | 1 | 10 | Lcmt1 | 0.053 | Old |
| 0.24396 | 0.048213 | 0.027 | 0.051 | 1 | 10 | Chchd10 | -0.024 | Old |
| 0.243983 | 0.908639 | 0.032 | 0.01 | 1 | 10 | Alg12 | 0.022 | Old |
| 0.243983 | 0.606806 | 0.032 | 0.01 | 1 | 10 | Cbwd1 | 0.022 | Old |
| 0.243983 | 0.267069 | 0.032 | 0.01 | 1 | 10 | Srgap3 | 0.022 | Old |
| 0.243998 | 0.204715 | 0.57 | 0.505 | 1 | 10 | Cct2 | 0.065 | Old |
| 0.244123 | 0.281118 | 0.269 | 0.202 | 1 | 10 | Lrpap1 | 0.067 | Old |
| 0.244194 | 0.339758 | 0.323 | 0.263 | 1 | 10 | Ube2m | 0.06 | Old |
| 0.244443 | 0.373783 | 0.174 | 0.121 | 1 | 10 | Scyl2 | 0.053 | Old |
| 0.244671 | 0.690477 | 0.178 | 0.131 | 1 | 10 | Slc50a1 | 0.047 | Old |
| 0.244717 | 0.131878 | 0.98 | 0.949 | 1 | 10 | Pfn1 | 0.031 | Old |
| 0.245074 | 0.159869 | 0.032 | 0.01 | 1 | 10 | Arap3 | 0.022 | Old |
| 0.245357 | 0.634862 | 0.073 | 0.04 | 1 | 10 | Ptcd2 | 0.033 | Old |
| 0.245447 | 0.856811 | 0.046 | 0.02 | 1 | 10 | Spryd4 | 0.026 | Old |
| 0.245845 | 0.267361 | 0.276 | 0.212 | 1 | 10 | Atg5 | 0.064 | Old |
| 0.24588 | 0.029019 | 0.355 | 0.273 | 1 | 10 | Rcc2 | 0.082 | Old |
| 0.246054 | 0.172655 | 0.557 | 0.475 | 1 | 10 | Psme2 | 0.082 | Old |
| 0.246094 | 0.366109 | 0.088 | 0.051 | 1 | 10 | Stxbp6 | 0.037 | Old |
| 0.246662 | 0.041393 | 0.249 | 0.182 | 1 | 10 | Htatsf1 | 0.067 | Old |
| 0.246833 | 1.339287 | 0.059 | 0.03 | 1 | 10 | Miip | 0.029 | Old |
| 0.24721 | 0.400141 | 0.313 | 0.263 | 1 | 10 | Iah1 | 0.05 | Old |
| 0.247349 | 0.179061 | 0.719 | 0.636 | 1 | 10 | Gusb | 0.083 | Old |
| 0.247386 | 0.383113 | 0.1 | 0.061 | 1 | 10 | Tradd | 0.039 | Old |
| 0.247601 | 0.146195 | 0.125 | 0.081 | 1 | 10 | Rad54l2 | 0.044 | Old |
| 0.247821 | 0.206581 | 0.325 | 0.263 | 1 | 10 | Ocrl | 0.062 | Old |
| 0.248162 | 0.04855 | 0.125 | 0.081 | 1 | 10 | Cdk2 | 0.044 | Old |
| 0.248207 | 0.549805 | 0.159 | 0.111 | 1 | 10 | Phf8 | 0.048 | Old |
| 0.249437 | 0.338973 | 0.137 | 0.091 | 1 | 10 | Prpsap2 | 0.046 | Old |
| 0.249579 | 0.133195 | 0.306 | 0.232 | 1 | 10 | Rabep1 | 0.074 | Old |
| 0.249673 | 0.417002 | 0.21 | 0.162 | 1 | 10 | Vezf1 | 0.048 | Old |
| 0.249752 | 0.001711 | 0.34 | 0.242 | 1 | 10 | Ifi27 | 0.098 | Old |
| 0.249876 | 0.404853 | 0.1 | 0.061 | 1 | 10 | Dnajc12 | 0.039 | Old |
| 0.250971 | 0.080258 | 0.061 | 0.03 | 1 | 10 | Ccar2 | 0.031 | Old |
| 0.251233 | 0.49275 | 0.174 | 0.121 | 1 | 10 | Cog5 | 0.053 | Old |
| 0.251556 | 0.229802 | 0.205 | 0.152 | 1 | 10 | Bcl3 | 0.053 | Old |
| 0.251798 | 0.224481 | 0.046 | 0.02 | 1 | 10 | Pex11g | 0.026 | Old |
| 0.252058 | 0.561513 | 0.22 | 0.172 | 1 | 10 | Impad1 | 0.048 | Old |
| 0.252241 | 0.165583 | 0.408 | 0.313 | 1 | 10 | Snrpd2 | 0.095 | Old |
| 0.252273 | 0.09633 | 0.601 | 0.495 | 1 | 10 | Dnase2a | 0.106 | Old |
| 0.252307 | 0.066619 | 0.433 | 0.343 | 1 | 10 | Ddx3y | 0.09 | Old |
| 0.252392 | 0.470852 | 0.203 | 0.152 | 1 | 10 | Bms1 | 0.051 | Old |
| 0.252715 | 0.428039 | 0.046 | 0.02 | 1 | 10 | Tmem41a | 0.026 | Old |
| 0.252845 | 0.646703 | 0.169 | 0.121 | 1 | 10 | Uap1 | 0.048 | Old |
| 0.253111 | 0.027539 | 0.139 | 0.091 | 1 | 10 | Rbm12 | 0.048 | Old |

| | | | | | | | | |
|---|---|---|---|---|---|---|---|---|
| 0.253311 | 0.201166 | 0.254 | 0.192 | 1 | 10 | St3gal1 | 0.062 | Old |
| 0.253397 | 0.7374 | 0.132 | 0.091 | 1 | 10 | Tceanc2 | 0.041 | Old |
| 0.253691 | 0.008349 | 0.186 | 0.131 | 1 | 10 | Casd1 | 0.055 | Old |
| 0.25378 | 0.442834 | 0.098 | 0.061 | 1 | 10 | Apip | 0.037 | Old |
| 0.253853 | 0.210555 | 0.161 | 0.111 | 1 | 10 | Ints9 | 0.05 | Old |
| 0.254047 | 0.990328 | 0.083 | 0.051 | 1 | 10 | Trmt2b | 0.032 | Old |
| 0.254085 | 0.649409 | 0.12 | 0.081 | 1 | 10 | Plcb3 | 0.039 | Old |
| 0.254085 | 0.169484 | 0.946 | 0.929 | 1 | 10 | Sirpa | 0.017 | Old |
| 0.254178 | 0.125869 | 0.665 | 0.535 | 1 | 10 | Asah1 | 0.13 | Old |
| 0.254231 | 0.379008 | 0.389 | 0.333 | 1 | 10 | Nfic | 0.056 | Old |
| 0.254386 | 0.342892 | 0.183 | 0.131 | 1 | 10 | Pgrmc2 | 0.052 | Old |
| 0.254418 | 0.614471 | 0.098 | 0.061 | 1 | 10 | Ap4m1 | 0.037 | Old |
| 0.254447 | 0.046613 | 0.259 | 0.192 | 1 | 10 | Wdr91 | 0.067 | Old |
| 0.254934 | 1.076721 | 0.059 | 0.03 | 1 | 10 | Paqr4 | 0.029 | Old |
| 0.255112 | 0.50159 | 0.188 | 0.141 | 1 | 10 | Tmem97 | 0.047 | Old |
| 0.255128 | 0.551791 | 0.122 | 0.081 | 1 | 10 | Rwdd4a | 0.041 | Old |
| 0.255205 | 0.148747 | 0.45 | 0.374 | 1 | 10 | Ankrd17 | 0.076 | Old |
| 0.255468 | 0.651996 | 0.086 | 0.051 | 1 | 10 | Ptrh2 | 0.035 | Old |
| 0.255742 | 0.244489 | 0.416 | 0.354 | 1 | 10 | Ppp1cc | 0.062 | Old |
| 0.256512 | 0.17159 | 0.443 | 0.364 | 1 | 10 | Eif3m | 0.079 | Old |
| 0.256631 | 0.119831 | 0.403 | 0.323 | 1 | 10 | Tmem55b | 0.08 | Old |
| 0.257232 | 0.221452 | 0.249 | 0.192 | 1 | 10 | Vps13a | 0.057 | Old |
| 0.257333 | 0.157227 | 0.601 | 0.505 | 1 | 10 | Prpf38b | 0.096 | Old |
| 0.25746 | 0.067114 | 0.308 | 0.242 | 1 | 10 | AB124611 | 0.066 | Old |
| 0.257543 | 0.015225 | 0.364 | 0.283 | 1 | 10 | Snw1 | 0.081 | Old |
| 0.257708 | 0.029785 | 0.435 | 0.343 | 1 | 10 | Psmb7 | 0.092 | Old |
| 0.257858 | 0.241116 | 0.413 | 0.323 | 1 | 10 | Arhgap31 | 0.09 | Old |
| 0.257947 | 0.829412 | 0.164 | 0.121 | 1 | 10 | Thyn1 | 0.043 | Old |
| 0.258091 | 0.318156 | 0.249 | 0.192 | 1 | 10 | Tmem147 | 0.057 | Old |
| 0.258165 | 0.389934 | 0.242 | 0.182 | 1 | 10 | Elk4 | 0.06 | Old |
| 0.258503 | 0.054567 | 0.208 | 0.152 | 1 | 10 | Abcd2 | 0.056 | Old |
| 0.258717 | 0.148999 | 0.215 | 0.162 | 1 | 10 | Cnot2 | 0.053 | Old |
| 0.258753 | 0.62471 | 0.12 | 0.081 | 1 | 10 | Ticam2 | 0.039 | Old |
| 0.258776 | 0.363484 | 0.247 | 0.192 | 1 | 10 | Gpr107 | 0.055 | Old |
| 0.258933 | 0.055491 | 0.499 | 0.414 | 1 | 10 | Scaf11 | 0.085 | Old |
| 0.259053 | 1.308208 | 0.059 | 0.03 | 1 | 10 | Unc13d | 0.029 | Old |
| 0.259077 | 0.100452 | 0.232 | 0.172 | 1 | 10 | Lbr | 0.06 | Old |
| 0.259576 | 0.038451 | 0.125 | 0.081 | 1 | 10 | Psmd5 | 0.044 | Old |
| 0.259594 | 1.129591 | 0.083 | 0.051 | 1 | 10 | Zbed4 | 0.032 | Old |
| 0.259715 | 0.100756 | 0.11 | 0.071 | 1 | 10 | Sdccag8 | 0.039 | Old |
| 0.260135 | 0.100103 | 0.391 | 0.313 | 1 | 10 | Stard3nl | 0.078 | Old |
| 0.260181 | 0.09553 | 0.328 | 0.263 | 1 | 10 | Mrpl34 | 0.065 | Old |
| 0.260503 | 0.059894 | 0.159 | 0.111 | 1 | 10 | Mospd3 | 0.048 | Old |
| 0.260862 | 0.772508 | 0.098 | 0.061 | 1 | 10 | Eif2b2 | 0.037 | Old |
| 0.260967 | 0.546237 | 0.281 | 0.232 | 1 | 10 | Abcc5 | 0.049 | Old |

| | | | | | | | | |
|---|---|---|---|---|---|---|---|---|
| 0.260982 | 0.050208 | 0.198 | 0.141 | 1 | 10 | Paip1 | 0.057 | Old |
| 0.261293 | 0.386247 | 0.191 | 0.141 | 1 | 10 | P2ry10b | 0.05 | Old |
| 0.261546 | 0.957291 | 0.059 | 0.03 | 1 | 10 | Src | 0.029 | Old |
| 0.261555 | 0.323072 | 0.198 | 0.152 | 1 | 10 | Ptgr2 | 0.046 | Old |
| 0.26156 | 0.063248 | 0.164 | 0.111 | 1 | 10 | Vps53 | 0.053 | Old |
| 0.262107 | 0.299701 | 0.303 | 0.242 | 1 | 10 | Tbc1d20 | 0.061 | Old |
| 0.262209 | 0.300589 | 0.073 | 0.04 | 1 | 10 | Hyal2 | 0.033 | Old |
| 0.2623 | 0.077529 | 0.447 | 0.374 | 1 | 10 | Spcs3 | 0.073 | Old |
| 0.262706 | 0.243297 | 0.122 | 0.081 | 1 | 10 | Zcchc8 | 0.041 | Old |
| 0.262776 | 0.187108 | 0.389 | 0.303 | 1 | 10 | Sms | 0.086 | Old |
| 0.263307 | 0.27552 | 0.264 | 0.212 | 1 | 10 | Nans | 0.052 | Old |
| 0.263723 | 0.00903 | 0.249 | 0.192 | 1 | 10 | Nfkbie | 0.057 | Old |
| 0.263792 | 0.506631 | 0.286 | 0.232 | 1 | 10 | Dr1 | 0.054 | Old |
| 0.264056 | 0.539438 | 0.059 | 0.03 | 1 | 10 | Sema4b | 0.029 | Old |
| 0.264073 | 0.548922 | 0.132 | 0.091 | 1 | 10 | Rnf169 | 0.041 | Old |
| 0.264214 | 0.058165 | 0.306 | 0.242 | 1 | 10 | Cyb5b | 0.064 | Old |
| 0.264267 | 0.94642 | 0.071 | 0.04 | 1 | 10 | Tmem94 | 0.031 | Old |
| 0.264281 | 0.066598 | 0.45 | 0.364 | 1 | 10 | Cisd2 | 0.086 | Old |
| 0.264571 | 0.346457 | 0.222 | 0.172 | 1 | 10 | Itfg1 | 0.05 | Old |
| 0.264691 | 1.627815 | 0.044 | 0.02 | 1 | 10 | Trmt10a | 0.024 | Old |
| 0.264785 | 0.548185 | 0.21 | 0.162 | 1 | 10 | Nuak2 | 0.048 | Old |
| 0.265186 | 0.016381 | 0.636 | 0.515 | 1 | 10 | Trf | 0.121 | Old |
| 0.265195 | 0.129374 | 0.156 | 0.111 | 1 | 10 | Ddx58 | 0.045 | Old |
| 0.265237 | 0.073041 | 0.491 | 0.374 | 1 | 10 | 11-Sep | 0.117 | Old |
| 0.265517 | 0.199638 | 0.597 | 0.485 | 1 | 10 | Mpp1 | 0.112 | Old |
| 0.265743 | 0.21922 | 0.205 | 0.152 | 1 | 10 | Hsph1 | 0.053 | Old |
| 0.266126 | 0.908738 | 0.108 | 0.071 | 1 | 10 | Cherp | 0.037 | Old |
| 0.266255 | 0.445446 | 0.122 | 0.081 | 1 | 10 | Ubac1 | 0.041 | Old |
| 0.266255 | 0.026528 | 0.122 | 0.081 | 1 | 10 | Rpusd4 | 0.041 | Old |
| 0.266802 | 0.639 | 0.186 | 0.141 | 1 | 10 | Dap3 | 0.045 | Old |
| 0.267557 | 0.033614 | 0.738 | 0.616 | 1 | 10 | Soat1 | 0.122 | Old |
| 0.267573 | 0.722242 | 0.174 | 0.131 | 1 | 10 | Ddx1 | 0.043 | Old |
| 0.267613 | 1.61651 | 0.044 | 0.02 | 1 | 10 | Cd5l | 0.024 | Old |
| 0.267613 | 1.414349 | 0.044 | 0.02 | 1 | 10 | 5730409E0 | 0.024 | Old |
| 0.267849 | 0.25153 | 0.386 | 0.323 | 1 | 10 | Ndufa10 | 0.063 | Old |
| 0.268575 | 0.323144 | 0.169 | 0.121 | 1 | 10 | Arhgef7 | 0.048 | Old |
| 0.268597 | 0.069388 | 0.401 | 0.313 | 1 | 10 | Mpc2 | 0.088 | Old |
| 0.268714 | 0.148066 | 0.408 | 0.323 | 1 | 10 | Fam168a | 0.085 | Old |
| 0.269008 | 0.002902 | 0.152 | 0.101 | 1 | 10 | Ccnk | 0.051 | Old |
| 0.269083 | 0.159188 | 0.814 | 0.768 | 1 | 10 | Pcbp1 | 0.046 | Old |
| 0.269208 | 0.04622 | 0.511 | 0.424 | 1 | 10 | Pttg1ip | 0.087 | Old |
| 0.269386 | 0.115485 | 0.408 | 0.333 | 1 | 10 | Ube2i | 0.075 | Old |
| 0.269477 | 0.425991 | 0.166 | 0.121 | 1 | 10 | Cd40 | 0.045 | Old |
| 0.26956 | 0.254022 | 0.389 | 0.313 | 1 | 10 | Mib1 | 0.076 | Old |
| 0.269612 | 0.140998 | 0.23 | 0.172 | 1 | 10 | Unc50 | 0.058 | Old |

| | | | | | | | | |
|---|---|---|---|---|---|---|---|---|
| 0.269977 | 0.377125 | 0.059 | 0.03 | 1 | 10 | Fam110a | 0.029 | Old |
| 0.269998 | 0.369804 | 0.159 | 0.111 | 1 | 10 | Lats1 | 0.048 | Old |
| 0.270009 | 0.294628 | 0.379 | 0.313 | 1 | 10 | Fem1b | 0.066 | Old |
| 0.270187 | 1.55805 | 0.068 | 0.04 | 1 | 10 | Pik3r2 | 0.028 | Old |
| 0.27022 | 0.284934 | 0.083 | 0.051 | 1 | 10 | Etfrf1 | 0.032 | Old |
| 0.270358 | 2.651921 | 0.012 | 0 | 1 | 10 | Slc29a2 | 0.012 | Old |
| 0.270358 | 2.649602 | 0.012 | 0 | 1 | 10 | Sox7 | 0.012 | Old |
| 0.270358 | 2.54262 | 0.012 | 0 | 1 | 10 | Mtfp1 | 0.012 | Old |
| 0.270358 | 2.315439 | 0.012 | 0 | 1 | 10 | Tnfrsf26 | 0.012 | Old |
| 0.270358 | 2.308502 | 0.012 | 0 | 1 | 10 | BC147527 | 0.012 | Old |
| 0.270358 | 2.139118 | 0.012 | 0 | 1 | 10 | Tnfaip6 | 0.012 | Old |
| 0.270358 | 2.126239 | 0.012 | 0 | 1 | 10 | Phf19 | 0.012 | Old |
| 0.270358 | 2.079965 | 0.012 | 0 | 1 | 10 | Rftn2 | 0.012 | Old |
| 0.270358 | 2.011593 | 0.012 | 0 | 1 | 10 | Gstp3 | 0.012 | Old |
| 0.270358 | 1.984632 | 0.012 | 0 | 1 | 10 | Cdh6 | 0.012 | Old |
| 0.270358 | 1.952964 | 0.012 | 0 | 1 | 10 | Rad9b | 0.012 | Old |
| 0.270358 | 1.926785 | 0.012 | 0 | 1 | 10 | Agbl3 | 0.012 | Old |
| 0.270358 | 1.924383 | 0.012 | 0 | 1 | 10 | A330069E1 | 0.012 | Old |
| 0.270358 | 1.88438 | 0.012 | 0 | 1 | 10 | Ankrd55 | 0.012 | Old |
| 0.270358 | 1.828586 | 0.012 | 0 | 1 | 10 | Kbtbd8 | 0.012 | Old |
| 0.270358 | 1.81039 | 0.012 | 0 | 1 | 10 | Bend6 | 0.012 | Old |
| 0.270358 | 1.794931 | 0.012 | 0 | 1 | 10 | Gm26699 | 0.012 | Old |
| 0.270358 | 1.794337 | 0.012 | 0 | 1 | 10 | Gm15448 | 0.012 | Old |
| 0.270358 | 1.764166 | 0.012 | 0 | 1 | 10 | Vangl2 | 0.012 | Old |
| 0.270358 | 1.741239 | 0.012 | 0 | 1 | 10 | Cdh1 | 0.012 | Old |
| 0.270358 | 1.715389 | 0.012 | 0 | 1 | 10 | Aqp11 | 0.012 | Old |
| 0.270358 | 1.706375 | 0.012 | 0 | 1 | 10 | Gm12216 | 0.012 | Old |
| 0.270358 | 1.693488 | 0.012 | 0 | 1 | 10 | 4930579G2 | 0.012 | Old |
| 0.270358 | 1.677517 | 0.012 | 0 | 1 | 10 | Slc22a21 | 0.012 | Old |
| 0.270358 | 1.665651 | 0.012 | 0 | 1 | 10 | 1500002C1 | 0.012 | Old |
| 0.270358 | 1.657129 | 0.012 | 0 | 1 | 10 | C1s1 | 0.012 | Old |
| 0.270358 | 1.653308 | 0.012 | 0 | 1 | 10 | Dand5 | 0.012 | Old |
| 0.270358 | 1.651828 | 0.012 | 0 | 1 | 10 | St5 | 0.012 | Old |
| 0.270358 | 1.641777 | 0.012 | 0 | 1 | 10 | E430018J2 | 0.012 | Old |
| 0.270358 | 1.641155 | 0.012 | 0 | 1 | 10 | Zhx2 | 0.012 | Old |
| 0.270358 | 1.640048 | 0.012 | 0 | 1 | 10 | Gabbr2 | 0.012 | Old |
| 0.270358 | 1.627185 | 0.012 | 0 | 1 | 10 | Fads3 | 0.012 | Old |
| 0.270358 | 1.596128 | 0.012 | 0 | 1 | 10 | Mical3 | 0.012 | Old |
| 0.270358 | 1.59577 | 0.012 | 0 | 1 | 10 | Slc22a18 | 0.012 | Old |
| 0.270358 | 1.58605 | 0.012 | 0 | 1 | 10 | Ddias | 0.012 | Old |
| 0.270358 | 1.585157 | 0.012 | 0 | 1 | 10 | Nek8 | 0.012 | Old |
| 0.270358 | 1.578053 | 0.012 | 0 | 1 | 10 | Gimap1 | 0.012 | Old |
| 0.270358 | 1.572399 | 0.012 | 0 | 1 | 10 | G730013B0 | 0.012 | Old |
| 0.270358 | 1.571472 | 0.012 | 0 | 1 | 10 | Slc10a6 | 0.012 | Old |
| 0.270358 | 1.56639 | 0.012 | 0 | 1 | 10 | Frk | 0.012 | Old |

| | | | | | | | | |
|---|---|---|---|---|---|---|---|---|
| 0.270358 | 1.562465 | 0.012 | 0 | 1 | 10 | Klhdc1 | 0.012 | Old |
| 0.270358 | 1.546729 | 0.012 | 0 | 1 | 10 | Zfp764 | 0.012 | Old |
| 0.270358 | 1.537667 | 0.012 | 0 | 1 | 10 | Zbtb7c | 0.012 | Old |
| 0.270358 | 1.52657 | 0.012 | 0 | 1 | 10 | Orc1 | 0.012 | Old |
| 0.270358 | 1.526427 | 0.012 | 0 | 1 | 10 | Tceanc | 0.012 | Old |
| 0.270358 | 1.506327 | 0.012 | 0 | 1 | 10 | Lrrc56 | 0.012 | Old |
| 0.270358 | 1.475993 | 0.012 | 0 | 1 | 10 | Ift81 | 0.012 | Old |
| 0.270358 | 1.473367 | 0.012 | 0 | 1 | 10 | Cacna1b | 0.012 | Old |
| 0.270358 | 1.46488 | 0.012 | 0 | 1 | 10 | 9130019O2 | 0.012 | Old |
| 0.270358 | 1.460205 | 0.012 | 0 | 1 | 10 | 2200002D0 | 0.012 | Old |
| 0.270358 | 1.454966 | 0.012 | 0 | 1 | 10 | Pde2a | 0.012 | Old |
| 0.270358 | 1.44978 | 0.012 | 0 | 1 | 10 | Cd177 | 0.012 | Old |
| 0.270358 | 1.447707 | 0.012 | 0 | 1 | 10 | D830025C0 | 0.012 | Old |
| 0.270358 | 1.432164 | 0.012 | 0 | 1 | 10 | Gm20632 | 0.012 | Old |
| 0.270358 | 1.428999 | 0.012 | 0 | 1 | 10 | Ube2v1 | 0.012 | Old |
| 0.270358 | 1.424178 | 0.012 | 0 | 1 | 10 | Gm45250 | 0.012 | Old |
| 0.270358 | 1.418163 | 0.012 | 0 | 1 | 10 | Olfml2b | 0.012 | Old |
| 0.270358 | 1.418027 | 0.012 | 0 | 1 | 10 | Zfp882 | 0.012 | Old |
| 0.270358 | 1.395025 | 0.012 | 0 | 1 | 10 | Krba1 | 0.012 | Old |
| 0.270358 | 1.393201 | 0.012 | 0 | 1 | 10 | 2210417A0 | 0.012 | Old |
| 0.270358 | 1.38938 | 0.012 | 0 | 1 | 10 | Zfp563 | 0.012 | Old |
| 0.270358 | 1.386812 | 0.012 | 0 | 1 | 10 | Plxna2 | 0.012 | Old |
| 0.270358 | 1.382374 | 0.012 | 0 | 1 | 10 | Arsi | 0.012 | Old |
| 0.270358 | 1.368506 | 0.012 | 0 | 1 | 10 | Col3a1 | 0.012 | Old |
| 0.270358 | 1.363911 | 0.012 | 0 | 1 | 10 | C2 | 0.012 | Old |
| 0.270358 | 1.336445 | 0.012 | 0 | 1 | 10 | Gdpgp1 | 0.012 | Old |
| 0.270358 | 1.33489 | 0.012 | 0 | 1 | 10 | 5430403G1 | 0.012 | Old |
| 0.270358 | 1.320647 | 0.012 | 0 | 1 | 10 | Fam72a | 0.012 | Old |
| 0.270358 | 1.317391 | 0.012 | 0 | 1 | 10 | Usp54 | 0.012 | Old |
| 0.270358 | 1.310003 | 0.012 | 0 | 1 | 10 | Lrrc27 | 0.012 | Old |
| 0.270358 | 1.304826 | 0.012 | 0 | 1 | 10 | Ankrd24 | 0.012 | Old |
| 0.270358 | 1.303422 | 0.012 | 0 | 1 | 10 | 1700030K0 | 0.012 | Old |
| 0.270358 | 1.298137 | 0.012 | 0 | 1 | 10 | Gm16867 | 0.012 | Old |
| 0.270358 | 1.273824 | 0.012 | 0 | 1 | 10 | Angel1 | 0.012 | Old |
| 0.270358 | 1.272986 | 0.012 | 0 | 1 | 10 | A430033K0 | 0.012 | Old |
| 0.270358 | 1.267501 | 0.012 | 0 | 1 | 10 | Myrf | 0.012 | Old |
| 0.270358 | 1.262672 | 0.012 | 0 | 1 | 10 | Serpina3g | 0.012 | Old |
| 0.270358 | 1.262529 | 0.012 | 0 | 1 | 10 | Nphp1 | 0.012 | Old |
| 0.270358 | 1.257382 | 0.012 | 0 | 1 | 10 | Klk8 | 0.012 | Old |
| 0.270358 | 1.254222 | 0.012 | 0 | 1 | 10 | Gm13091 | 0.012 | Old |
| 0.270358 | 1.253832 | 0.012 | 0 | 1 | 10 | Zfp354c | 0.012 | Old |
| 0.270358 | 1.246298 | 0.012 | 0 | 1 | 10 | Cfap126 | 0.012 | Old |
| 0.270358 | 1.232118 | 0.012 | 0 | 1 | 10 | Gm4951 | 0.012 | Old |
| 0.270358 | 1.222321 | 0.012 | 0 | 1 | 10 | Scrn2 | 0.012 | Old |
| 0.270358 | 1.218171 | 0.012 | 0 | 1 | 10 | Melk | 0.012 | Old |

| | | | | | | | | |
|---|---|---|---|---|---|---|---|---|
| 0.270358 | 1.214893 | 0.012 | 0 | 1 | 10 | Spa17 | 0.012 | Old |
| 0.270358 | 1.213952 | 0.012 | 0 | 1 | 10 | Zfp428 | 0.012 | Old |
| 0.270358 | 1.182926 | 0.012 | 0 | 1 | 10 | Lpcat4 | 0.012 | Old |
| 0.270358 | 1.175652 | 0.012 | 0 | 1 | 10 | Gm41307 | 0.012 | Old |
| 0.270358 | 1.172614 | 0.012 | 0 | 1 | 10 | Gm48653 | 0.012 | Old |
| 0.270358 | 1.167259 | 0.012 | 0 | 1 | 10 | Otof | 0.012 | Old |
| 0.270358 | 1.163499 | 0.012 | 0 | 1 | 10 | Gm11772 | 0.012 | Old |
| 0.270358 | 1.14679 | 0.012 | 0 | 1 | 10 | Tiam2 | 0.012 | Old |
| 0.270358 | 1.133016 | 0.012 | 0 | 1 | 10 | Mipol1 | 0.012 | Old |
| 0.270358 | 1.12421 | 0.012 | 0 | 1 | 10 | Gm30246 | 0.012 | Old |
| 0.270358 | 1.121241 | 0.012 | 0 | 1 | 10 | Smim10l2a | 0.012 | Old |
| 0.270358 | 1.103457 | 0.012 | 0 | 1 | 10 | Chtf18 | 0.012 | Old |
| 0.270358 | 1.079447 | 0.012 | 0 | 1 | 10 | Shpk | 0.012 | Old |
| 0.270358 | 1.041112 | 0.012 | 0 | 1 | 10 | Pacrg | 0.012 | Old |
| 0.270358 | 1.034855 | 0.012 | 0 | 1 | 10 | Phf11a | 0.012 | Old |
| 0.270358 | 1.023915 | 0.012 | 0 | 1 | 10 | Raet1d | 0.012 | Old |
| 0.270358 | 0.969 | 0.012 | 0 | 1 | 10 | Cyb5rl | 0.012 | Old |
| 0.270358 | 0.917175 | 0.012 | 0 | 1 | 10 | Slc7a4 | 0.012 | Old |
| 0.270358 | 0.878635 | 0.012 | 0 | 1 | 10 | Gimap6 | 0.012 | Old |
| 0.270591 | 0.080187 | 0.396 | 0.313 | 1 | 10 | Golga7 | 0.083 | Old |
| 0.271272 | 0.146463 | 0.496 | 0.424 | 1 | 10 | Leprot | 0.072 | Old |
| 0.271367 | 0.056255 | 0.359 | 0.303 | 1 | 10 | Trafd1 | 0.056 | Old |
| 0.271474 | 0.052009 | 0.616 | 0.515 | 1 | 10 | S100a13 | 0.101 | Old |
| 0.271548 | 0.431157 | 0.11 | 0.071 | 1 | 10 | Tssc4 | 0.039 | Old |
| 0.271675 | 0.137948 | 0.79 | 0.677 | 1 | 10 | Atp1a1 | 0.113 | Old |
| 0.271734 | 1.855305 | 0.029 | 0.01 | 1 | 10 | Pdk2 | 0.019 | Old |
| 0.271734 | 1.536406 | 0.029 | 0.01 | 1 | 10 | Zfp963 | 0.019 | Old |
| 0.272216 | 0.462381 | 0.289 | 0.242 | 1 | 10 | Elf2 | 0.047 | Old |
| 0.272333 | 0.179908 | 0.721 | 0.586 | 1 | 10 | Tcf4 | 0.135 | Old |
| 0.272404 | 0.10877 | 0.259 | 0.202 | 1 | 10 | Rab3gap2 | 0.057 | Old |
| 0.272414 | 0.280051 | 0.222 | 0.172 | 1 | 10 | Tmem19 | 0.05 | Old |
| 0.272609 | 0.241005 | 0.147 | 0.101 | 1 | 10 | Cipc | 0.046 | Old |
| 0.272686 | 0.071706 | 0.051 | 0.081 | 1 | 10 | Batf | -0.03 | Old |
| 0.272724 | 0.4177 | 0.144 | 0.101 | 1 | 10 | Trim12a | 0.043 | Old |
| 0.272767 | 0.081149 | 0.071 | 0.04 | 1 | 10 | Zfp444 | 0.031 | Old |
| 0.272953 | 1.764747 | 0.029 | 0.01 | 1 | 10 | Bmf | 0.019 | Old |
| 0.272953 | 1.728994 | 0.029 | 0.01 | 1 | 10 | Phka1 | 0.019 | Old |
| 0.272953 | 1.692005 | 0.029 | 0.01 | 1 | 10 | Cep19 | 0.019 | Old |
| 0.272953 | 1.601318 | 0.029 | 0.01 | 1 | 10 | Tmem38a | 0.019 | Old |
| 0.272953 | 1.419818 | 0.029 | 0.01 | 1 | 10 | Gm9844 | 0.019 | Old |
| 0.272953 | 1.4008 | 0.029 | 0.01 | 1 | 10 | Men1 | 0.019 | Old |
| 0.273218 | 1.486462 | 0.093 | 0.061 | 1 | 10 | 0610009B2 | 0.032 | Old |
| 0.273288 | 0.46723 | 0.21 | 0.162 | 1 | 10 | Ppp2r5e | 0.048 | Old |
| 0.273521 | 0.914869 | 0.044 | 0.02 | 1 | 10 | Borcs5 | 0.024 | Old |
| 0.273548 | 0.753879 | 0.071 | 0.04 | 1 | 10 | Fbxo42 | 0.031 | Old |

| | | | | | | | | |
|---|---|---|---|---|---|---|---|---|
| 0.273724 | 0.313571 | 0.274 | 0.212 | 1 | 10 | Slirp | 0.062 | Old |
| 0.273951 | 0.132908 | 0.516 | 0.424 | 1 | 10 | Epn1 | 0.092 | Old |
| 0.274003 | 0.136807 | 0.411 | 0.354 | 1 | 10 | Tex261 | 0.057 | Old |
| 0.274175 | 1.719008 | 0.029 | 0.01 | 1 | 10 | 3300005D( | 0.019 | Old |
| 0.274261 | 0.532248 | 0.059 | 0.03 | 1 | 10 | Entpd5 | 0.029 | Old |
| 0.27447 | 0.582175 | 0.132 | 0.091 | 1 | 10 | Siah2 | 0.041 | Old |
| 0.274723 | 0.114151 | 0.11 | 0.071 | 1 | 10 | Zfp672 | 0.039 | Old |
| 0.27491 | 0.026618 | 0.306 | 0.242 | 1 | 10 | Sec24b | 0.064 | Old |
| 0.275368 | 0.266298 | 0.169 | 0.121 | 1 | 10 | Golga3 | 0.048 | Old |
| 0.275511 | 1.106974 | 0.044 | 0.02 | 1 | 10 | Tacc3 | 0.024 | Old |
| 0.275511 | 0.976948 | 0.044 | 0.02 | 1 | 10 | Sgms2 | 0.024 | Old |
| 0.27552 | 0.265676 | 0.225 | 0.172 | 1 | 10 | Luzp1 | 0.053 | Old |
| 0.275678 | 0.496598 | 0.108 | 0.071 | 1 | 10 | Rrm2b | 0.037 | Old |
| 0.275797 | 0.097854 | 0.149 | 0.101 | 1 | 10 | Prkaa1 | 0.048 | Old |
| 0.275801 | 0.752965 | 0.117 | 0.081 | 1 | 10 | Rnf19a | 0.036 | Old |
| 0.2759 | 0.352408 | 0.335 | 0.283 | 1 | 10 | Papd4 | 0.052 | Old |
| 0.275903 | 0.586409 | 0.071 | 0.04 | 1 | 10 | Rnaseh1 | 0.031 | Old |
| 0.276631 | 1.299472 | 0.029 | 0.01 | 1 | 10 | Brca2 | 0.019 | Old |
| 0.276691 | 0.779743 | 0.071 | 0.04 | 1 | 10 | 2010315B( | 0.031 | Old |
| 0.27672 | 0.022818 | 0.073 | 0.04 | 1 | 10 | Mars | 0.033 | Old |
| 0.276786 | 0.005184 | 0.802 | 0.657 | 1 | 10 | Sub1 | 0.145 | Old |
| 0.276954 | 0.571021 | 0.208 | 0.162 | 1 | 10 | Gtf2i | 0.046 | Old |
| 0.277018 | 0.212706 | 0.235 | 0.182 | 1 | 10 | Tbc1d2b | 0.053 | Old |
| 0.277087 | 0.257861 | 0.147 | 0.101 | 1 | 10 | Ifi35 | 0.046 | Old |
| 0.277148 | 0.219041 | 0.276 | 0.222 | 1 | 10 | Snrnp27 | 0.054 | Old |
| 0.2772 | 0.214147 | 0.274 | 0.212 | 1 | 10 | Rcn2 | 0.062 | Old |
| 0.277311 | 0.369448 | 0.311 | 0.253 | 1 | 10 | Apobec3 | 0.058 | Old |
| 0.277427 | 0.118639 | 0.773 | 0.707 | 1 | 10 | Glud1 | 0.066 | Old |
| 0.27751 | 0.7437 | 0.044 | 0.02 | 1 | 10 | Taf6 | 0.024 | Old |
| 0.27751 | 0.434976 | 0.044 | 0.02 | 1 | 10 | Mettl7a1 | 0.024 | Old |
| 0.27756 | 0.147812 | 0.367 | 0.303 | 1 | 10 | Plin3 | 0.064 | Old |
| 0.277589 | 0.883755 | 0.152 | 0.111 | 1 | 10 | Ptpn23 | 0.041 | Old |
| 0.277605 | 0.53251 | 0.13 | 0.091 | 1 | 10 | Nom1 | 0.039 | Old |
| 0.277864 | 1.293027 | 0.029 | 0.01 | 1 | 10 | Senp8 | 0.019 | Old |
| 0.277864 | 1.071137 | 0.029 | 0.01 | 1 | 10 | Ptch1 | 0.019 | Old |
| 0.277905 | 1.313551 | 0.056 | 0.03 | 1 | 10 | Pigv | 0.026 | Old |
| 0.277997 | 0.403244 | 0.132 | 0.091 | 1 | 10 | Akr1e1 | 0.041 | Old |
| 0.278143 | 0.581287 | 0.152 | 0.111 | 1 | 10 | Zbtb38 | 0.041 | Old |
| 0.2782 | 0.605315 | 0.083 | 0.051 | 1 | 10 | Pkp4 | 0.032 | Old |
| 0.278209 | 0.229861 | 0.418 | 0.354 | 1 | 10 | Tnrc18 | 0.064 | Old |
| 0.278272 | 0.576131 | 0.071 | 0.04 | 1 | 10 | Zbtb6 | 0.031 | Old |
| 0.278327 | 0.504558 | 0.186 | 0.141 | 1 | 10 | Psmd10 | 0.045 | Old |
| 0.27847 | 0.905691 | 0.095 | 0.061 | 1 | 10 | Fem1a | 0.034 | Old |
| 0.27847 | 0.660045 | 0.095 | 0.061 | 1 | 10 | Gt(ROSA)2( | 0.034 | Old |
| 0.278636 | 0.413708 | 0.002 | 0.01 | 1 | 10 | Gm15915 | -0.008 | Old |

| | | | | | | | | |
|---|---|---|---|---|---|---|---|---|
| 0.278636 | 0.263902 | 0.002 | 0.01 | 1 | 10 | Gm43614 | -0.008 | Old |
| 0.278636 | 0.199382 | 0.002 | 0.01 | 1 | 10 | Rpusd2 | -0.008 | Old |
| 0.278636 | 0.085441 | 0.002 | 0.01 | 1 | 10 | Sorbs1 | -0.008 | Old |
| 0.278636 | 0.066886 | 0.002 | 0.01 | 1 | 10 | Samd11 | -0.008 | Old |
| 0.278636 | 0.052187 | 0.002 | 0.01 | 1 | 10 | Gm26764 | -0.008 | Old |
| 0.278636 | 0.047067 | 0.002 | 0.01 | 1 | 10 | Gm9733 | -0.008 | Old |
| 0.278636 | 0.017076 | 0.002 | 0.01 | 1 | 10 | Tbc1d25 | -0.008 | Old |
| 0.278636 | 0.016677 | 0.002 | 0.01 | 1 | 10 | Ccdc157 | -0.008 | Old |
| 0.278697 | 0.035644 | 0.086 | 0.051 | 1 | 10 | Gmppb | 0.035 | Old |
| 0.278795 | 0.677745 | 0.13 | 0.091 | 1 | 10 | Ppid | 0.039 | Old |
| 0.278836 | 0.556861 | 0.186 | 0.141 | 1 | 10 | Ecd | 0.045 | Old |
| 0.279101 | 1.195209 | 0.029 | 0.01 | 1 | 10 | Tmem9 | 0.019 | Old |
| 0.279101 | 1.073738 | 0.029 | 0.01 | 1 | 10 | Plxna3 | 0.019 | Old |
| 0.279101 | 1.03471 | 0.029 | 0.01 | 1 | 10 | Sag | 0.019 | Old |
| 0.279101 | 0.948528 | 0.029 | 0.01 | 1 | 10 | Ncapd2 | 0.019 | Old |
| 0.279519 | 0.809911 | 0.044 | 0.02 | 1 | 10 | 1500015A( | 0.024 | Old |
| 0.279709 | 0.010696 | 0.335 | 0.263 | 1 | 10 | Myo1g | 0.072 | Old |
| 0.280342 | 1.06493 | 0.029 | 0.01 | 1 | 10 | Gm4673 | 0.019 | Old |
| 0.280342 | 0.999995 | 0.029 | 0.01 | 1 | 10 | Hdac8 | 0.019 | Old |
| 0.280342 | 0.971291 | 0.029 | 0.01 | 1 | 10 | Pgap3 | 0.019 | Old |
| 0.280425 | 0.24459 | 0.43 | 0.374 | 1 | 10 | Itgb5 | 0.056 | Old |
| 0.28052 | 0.463714 | 0.2 | 0.152 | 1 | 10 | Atf2 | 0.048 | Old |
| 0.281587 | 0.968231 | 0.029 | 0.01 | 1 | 10 | Map3k12 | 0.019 | Old |
| 0.281587 | 0.886748 | 0.029 | 0.01 | 1 | 10 | Lgi4 | 0.019 | Old |
| 0.281587 | 0.881568 | 0.029 | 0.01 | 1 | 10 | B630019K( | 0.019 | Old |
| 0.281587 | 0.198674 | 0.274 | 0.212 | 1 | 10 | Desi2 | 0.062 | Old |
| 0.281661 | 0.160662 | 0.235 | 0.182 | 1 | 10 | Abraxas2 | 0.053 | Old |
| 0.281861 | 0.117395 | 0.408 | 0.343 | 1 | 10 | Cd2ap | 0.065 | Old |
| 0.282551 | 0.074357 | 0.044 | 0.02 | 1 | 10 | Tspyl4 | 0.024 | Old |
| 0.282835 | 0.721425 | 0.029 | 0.01 | 1 | 10 | Susd1 | 0.019 | Old |
| 0.282835 | 0.668147 | 0.029 | 0.01 | 1 | 10 | Bcorl1 | 0.019 | Old |
| 0.282835 | 0.628432 | 0.029 | 0.01 | 1 | 10 | Ocstamp | 0.019 | Old |
| 0.282984 | 1.213782 | 0.068 | 0.04 | 1 | 10 | Abhd11 | 0.028 | Old |
| 0.283305 | 0.685082 | 0.095 | 0.061 | 1 | 10 | Spsb3 | 0.034 | Old |
| 0.283328 | 0.104502 | 0.44 | 0.364 | 1 | 10 | Mapk6 | 0.076 | Old |
| 0.283574 | 0.284415 | 0.159 | 0.111 | 1 | 10 | Chst11 | 0.048 | Old |
| 0.283608 | 0.159219 | 0.614 | 0.566 | 1 | 10 | Rassf4 | 0.048 | Old |
| 0.283675 | 0.09224 | 0.597 | 0.525 | 1 | 10 | Edem1 | 0.072 | Old |
| 0.283681 | 0.40978 | 0.298 | 0.253 | 1 | 10 | Uvrag | 0.045 | Old |
| 0.283745 | 0.224531 | 0.11 | 0.071 | 1 | 10 | Coq2 | 0.039 | Old |
| 0.283852 | 0.257734 | 0.071 | 0.04 | 1 | 10 | Rpp30 | 0.031 | Old |
| 0.284088 | 0.670725 | 0.029 | 0.01 | 1 | 10 | Tmem132a | 0.019 | Old |
| 0.284088 | 0.560838 | 0.029 | 0.01 | 1 | 10 | Helz2 | 0.019 | Old |
| 0.284088 | 0.485857 | 0.029 | 0.01 | 1 | 10 | Bckdhb | 0.019 | Old |
| 0.284088 | 0.47479 | 0.029 | 0.01 | 1 | 10 | Diaph3 | 0.019 | Old |

| | | | | | | | | |
|---|---|---|---|---|---|---|---|---|
| 0.284088 | 0.430466 | 0.029 | 0.01 | 1 | 10 | Iqcc | 0.019 | Old |
| 0.284088 | 0.410199 | 0.029 | 0.01 | 1 | 10 | Cbx8 | 0.019 | Old |
| 0.284143 | 0.208446 | 0.108 | 0.071 | 1 | 10 | Rnd3 | 0.037 | Old |
| 0.284145 | 0.137315 | 0.751 | 0.687 | 1 | 10 | Jpt1 | 0.064 | Old |
| 0.284801 | 0.474484 | 0.108 | 0.071 | 1 | 10 | Kif13a | 0.037 | Old |
| 0.28507 | 1.081547 | 0.081 | 0.051 | 1 | 10 | Rabgap1l | 0.03 | Old |
| 0.285089 | 0.218898 | 0.257 | 0.202 | 1 | 10 | Rab29 | 0.055 | Old |
| 0.285343 | 0.364507 | 0.029 | 0.01 | 1 | 10 | Pafah2 | 0.019 | Old |
| 0.285343 | 0.207462 | 0.029 | 0.01 | 1 | 10 | A230028O( | 0.019 | Old |
| 0.285605 | 0.499222 | 0.044 | 0.02 | 1 | 10 | Nbeal2 | 0.024 | Old |
| 0.285876 | 0.170609 | 0.181 | 0.131 | 1 | 10 | Kbtbd2 | 0.05 | Old |
| 0.285948 | 1.288233 | 0.056 | 0.03 | 1 | 10 | Pnkp | 0.026 | Old |
| 0.286266 | 0.633509 | 0.071 | 0.04 | 1 | 10 | Mon1b | 0.031 | Old |
| 0.286636 | 0.110808 | 0.213 | 0.152 | 1 | 10 | Hebp1 | 0.061 | Old |
| 0.286783 | 0.367842 | 0.108 | 0.071 | 1 | 10 | Pgm1 | 0.037 | Old |
| 0.286997 | 0.082925 | 0.156 | 0.111 | 1 | 10 | Rnf41 | 0.045 | Old |
| 0.28706 | 0.450948 | 0.186 | 0.141 | 1 | 10 | Stx5a | 0.045 | Old |
| 0.287086 | 0.460885 | 0.083 | 0.051 | 1 | 10 | Usp42 | 0.032 | Old |
| 0.287086 | 0.359909 | 0.083 | 0.051 | 1 | 10 | Gpank1 | 0.032 | Old |
| 0.287337 | 1.132889 | 0.081 | 0.051 | 1 | 10 | Man2c1 | 0.03 | Old |
| 0.287756 | 1.054173 | 0.056 | 0.03 | 1 | 10 | Stn1 | 0.026 | Old |
| 0.287835 | 0.459644 | 0.083 | 0.051 | 1 | 10 | Tpgs2 | 0.032 | Old |
| 0.288131 | 0.077676 | 0.286 | 0.232 | 1 | 10 | Sptlc2 | 0.054 | Old |
| 0.28832 | 0.097833 | 0.262 | 0.192 | 1 | 10 | Plekhm3 | 0.07 | Old |
| 0.288332 | 0.153664 | 0.208 | 0.152 | 1 | 10 | Nf1 | 0.056 | Old |
| 0.288565 | 0.117672 | 0.328 | 0.273 | 1 | 10 | Lrp10 | 0.055 | Old |
| 0.288682 | 0.339546 | 0.044 | 0.02 | 1 | 10 | Syn1 | 0.024 | Old |
| 0.288682 | 0.246006 | 0.044 | 0.02 | 1 | 10 | Slc35d1 | 0.024 | Old |
| 0.288719 | 0.37957 | 0.144 | 0.101 | 1 | 10 | Wdr20 | 0.043 | Old |
| 0.28885 | 0.316064 | 0.472 | 0.404 | 1 | 10 | Cul3 | 0.068 | Old |
| 0.289079 | 0.140849 | 0.147 | 0.101 | 1 | 10 | Pkd2 | 0.046 | Old |
| 0.289572 | 0.907489 | 0.056 | 0.03 | 1 | 10 | Nr2c1 | 0.026 | Old |
| 0.289712 | 0.527443 | 0.044 | 0.02 | 1 | 10 | Mapkbp1 | 0.024 | Old |
| 0.290049 | 0.33249 | 0.2 | 0.152 | 1 | 10 | Stxbp5 | 0.048 | Old |
| 0.290136 | 0.548896 | 0.215 | 0.172 | 1 | 10 | Pip4k2b | 0.043 | Old |
| 0.290379 | 1.080218 | 0.081 | 0.051 | 1 | 10 | Arhgap23 | 0.03 | Old |
| 0.290379 | 1.065416 | 0.081 | 0.051 | 1 | 10 | Tmem175 | 0.03 | Old |
| 0.290483 | 1.164349 | 0.056 | 0.03 | 1 | 10 | Crcp | 0.026 | Old |
| 0.290483 | 1.077811 | 0.056 | 0.03 | 1 | 10 | Chek2 | 0.026 | Old |
| 0.290622 | 0.230961 | 0.645 | 0.586 | 1 | 10 | Psmd2 | 0.059 | Old |
| 0.290745 | 0.288699 | 0.044 | 0.02 | 1 | 10 | Wdr53 | 0.024 | Old |
| 0.290941 | 0.067054 | 1 | 1 | 1 | 10 | mt-Co3 | 0 | Old |
| 0.290985 | 0.122648 | 0.445 | 0.374 | 1 | 10 | Copb1 | 0.071 | Old |
| 0.291029 | 0.121698 | 0.944 | 0.869 | 1 | 10 | Arhgdia | 0.075 | Old |
| 0.291092 | 0.633721 | 0.152 | 0.111 | 1 | 10 | Slc39a11 | 0.041 | Old |

| | | | | | | | | |
|---|---|---|---|---|---|---|---|---|
| 0.291261 | 0.537149 | 0.142 | 0.101 | 1 | 10 | Tspan17 | 0.041 | Old |
| 0.291279 | 0.157272 | 0.609 | 0.556 | 1 | 10 | Actn4 | 0.053 | Old |
| 0.291332 | 0.627522 | 0.161 | 0.121 | 1 | 10 | Tmem181a | 0.04 | Old |
| 0.291396 | 0.239443 | 0.056 | 0.03 | 1 | 10 | Dis3 | 0.026 | Old |
| 0.291814 | 0.11374 | 0.589 | 0.485 | 1 | 10 | Arhgap17 | 0.104 | Old |
| 0.291856 | 0.015241 | 0.337 | 0.283 | 1 | 10 | Ccr1 | 0.054 | Old |
| 0.291948 | 0.10141 | 0.518 | 0.434 | 1 | 10 | Cox14 | 0.084 | Old |
| 0.29201 | 0.626058 | 0.171 | 0.131 | 1 | 10 | Ipo8 | 0.04 | Old |
| 0.292111 | 0.335366 | 0.108 | 0.071 | 1 | 10 | Sik3 | 0.037 | Old |
| 0.292228 | 0.614174 | 0.144 | 0.101 | 1 | 10 | Cavin1 | 0.043 | Old |
| 0.292848 | 0.98136 | 0.068 | 0.04 | 1 | 10 | Lrrc40 | 0.028 | Old |
| 0.294101 | 0.069599 | 0.303 | 0.242 | 1 | 10 | Arrdc1 | 0.061 | Old |
| 0.294223 | 0.145625 | 0.171 | 0.121 | 1 | 10 | Ppa1 | 0.05 | Old |
| 0.294382 | 0.491076 | 0.218 | 0.172 | 1 | 10 | Spg7 | 0.046 | Old |
| 0.294514 | 0.657621 | 0.068 | 0.04 | 1 | 10 | C2cd3 | 0.028 | Old |
| 0.294568 | 0.316028 | 0.095 | 0.061 | 1 | 10 | Zfp414 | 0.034 | Old |
| 0.295022 | 0.228959 | 0.773 | 0.737 | 1 | 10 | Plekho2 | 0.036 | Old |
| 0.295255 | 0.641255 | 0.105 | 0.071 | 1 | 10 | Zfp422 | 0.034 | Old |
| 0.295423 | 0.254152 | 0.235 | 0.182 | 1 | 10 | Mrps12 | 0.053 | Old |
| 0.295519 | 0.234571 | 0.359 | 0.303 | 1 | 10 | Pkig | 0.056 | Old |
| 0.295613 | 0.227332 | 0.462 | 0.404 | 1 | 10 | Bcl2l11 | 0.058 | Old |
| 0.29629 | 0.461209 | 0.188 | 0.141 | 1 | 10 | Prkaca | 0.047 | Old |
| 0.296458 | 0.297809 | 0.67 | 0.606 | 1 | 10 | Dusp3 | 0.064 | Old |
| 0.296559 | 0.392218 | 0.267 | 0.212 | 1 | 10 | Prmt1 | 0.055 | Old |
| 0.29693 | 0.63909 | 0.161 | 0.121 | 1 | 10 | Crtc2 | 0.04 | Old |
| 0.297374 | 0.474901 | 0.117 | 0.081 | 1 | 10 | Rab40c | 0.036 | Old |
| 0.297374 | 0.265339 | 0.21 | 0.162 | 1 | 10 | Triap1 | 0.048 | Old |
| 0.297699 | 0.074509 | 0.337 | 0.253 | 1 | 10 | Lima1 | 0.084 | Old |
| 0.297865 | 0.907196 | 0.068 | 0.04 | 1 | 10 | Fam58b | 0.028 | Old |
| 0.298228 | 0.110248 | 0.428 | 0.374 | 1 | 10 | Ndufa11 | 0.054 | Old |
| 0.298673 | 0.460854 | 0.105 | 0.071 | 1 | 10 | Sfxn2 | 0.034 | Old |
| 0.298768 | 0.65147 | 0.056 | 0.03 | 1 | 10 | Nt5m | 0.026 | Old |
| 0.298794 | 0.265788 | 0.44 | 0.384 | 1 | 10 | Eps8 | 0.056 | Old |
| 0.298896 | 0.214137 | 0.232 | 0.182 | 1 | 10 | Lemd2 | 0.05 | Old |
| 0.299273 | 0.063191 | 0.643 | 0.525 | 1 | 10 | Lnpep | 0.118 | Old |
| 0.299461 | 1.652623 | 0.042 | 0.02 | 1 | 10 | 6330562C2 | 0.022 | Old |
| 0.299748 | 0.200097 | 0.191 | 0.141 | 1 | 10 | Tmem131 | 0.05 | Old |
| 0.29983 | 0.131253 | 0.846 | 0.818 | 1 | 10 | Calr | 0.028 | Old |
| 0.300168 | 0.045962 | 0.178 | 0.131 | 1 | 10 | Auh | 0.047 | Old |
| 0.300631 | 0.52652 | 0.056 | 0.03 | 1 | 10 | Otud1 | 0.026 | Old |
| 0.301008 | 0.443979 | 0.333 | 0.283 | 1 | 10 | Creb1 | 0.05 | Old |
| 0.301197 | 0.786583 | 0.081 | 0.051 | 1 | 10 | Gprc5c | 0.03 | Old |
| 0.301204 | 0.057495 | 0.438 | 0.364 | 1 | 10 | Cdc37 | 0.074 | Old |
| 0.301603 | 0.635144 | 0.149 | 0.111 | 1 | 10 | Taf3 | 0.038 | Old |
| 0.301628 | 1.601416 | 0.042 | 0.02 | 1 | 10 | Ube2c | 0.022 | Old |

| | | | | | | | | |
|---|---|---|---|---|---|---|---|---|
| 0.301669 | 0.128092 | 0.347 | 0.293 | 1 | 10 | Rbm7 | 0.054 | Old |
| 0.30198 | 0.18035 | 0.081 | 0.051 | 1 | 10 | Pop4 | 0.03 | Old |
| 0.302029 | 0.22708 | 0.161 | 0.121 | 1 | 10 | Polr1c | 0.04 | Old |
| 0.30209 | 0.817083 | 0.068 | 0.04 | 1 | 10 | Gm10521 | 0.028 | Old |
| 0.302212 | 0.340515 | 0.284 | 0.242 | 1 | 10 | Gmip | 0.042 | Old |
| 0.302225 | 0.222224 | 0.235 | 0.182 | 1 | 10 | Slc35b1 | 0.053 | Old |
| 0.302255 | 0.228443 | 0.257 | 0.202 | 1 | 10 | Tspan4 | 0.055 | Old |
| 0.302255 | 0.010161 | 0.257 | 0.202 | 1 | 10 | Ogfr | 0.055 | Old |
| 0.302429 | 0.068982 | 0.914 | 0.899 | 1 | 10 | Slc6a6 | 0.015 | Old |
| 0.302501 | 0.494906 | 0.056 | 0.03 | 1 | 10 | Dut | 0.026 | Old |
| 0.302686 | 0.361438 | 0.13 | 0.091 | 1 | 10 | Tsta3 | 0.039 | Old |
| 0.302921 | 0.083333 | 0.132 | 0.091 | 1 | 10 | Fam207a | 0.041 | Old |
| 0.302963 | 0.027029 | 0.411 | 0.354 | 1 | 10 | Klhl24 | 0.057 | Old |
| 0.303016 | 0.740055 | 0.093 | 0.061 | 1 | 10 | Zfp329 | 0.032 | Old |
| 0.303145 | 0.417157 | 0.262 | 0.212 | 1 | 10 | Mrps16 | 0.05 | Old |
| 0.303372 | 0.206276 | 0.323 | 0.263 | 1 | 10 | Arl1 | 0.06 | Old |
| 0.303489 | 0.188052 | 0.645 | 0.586 | 1 | 10 | Rock1 | 0.059 | Old |
| 0.303541 | 0.209208 | 0.156 | 0.111 | 1 | 10 | Osbpl3 | 0.045 | Old |
| 0.303547 | 0.091077 | 0.132 | 0.091 | 1 | 10 | Senp3 | 0.041 | Old |
| 0.30373 | 0.310455 | 0.174 | 0.131 | 1 | 10 | Qpct | 0.043 | Old |
| 0.303791 | 0.853754 | 0.068 | 0.04 | 1 | 10 | Wbp1 | 0.028 | Old |
| 0.30406 | 0.311176 | 0.198 | 0.152 | 1 | 10 | Fbxo3 | 0.046 | Old |
| 0.304064 | 0.080105 | 0.922 | 0.889 | 1 | 10 | Capza2 | 0.033 | Old |
| 0.304185 | 0.353463 | 0.237 | 0.192 | 1 | 10 | Nt5c | 0.045 | Old |
| 0.304378 | 0.266693 | 0.056 | 0.03 | 1 | 10 | Zfand1 | 0.026 | Old |
| 0.30488 | 1.640386 | 0.054 | 0.03 | 1 | 10 | Cep44 | 0.024 | Old |
| 0.304893 | 0.515232 | 0.218 | 0.172 | 1 | 10 | Tmem43 | 0.046 | Old |
| 0.304899 | 1.533257 | 0.042 | 0.02 | 1 | 10 | Tbc1d22b | 0.022 | Old |
| 0.304899 | 1.487286 | 0.042 | 0.02 | 1 | 10 | St7 | 0.022 | Old |
| 0.304899 | 1.189638 | 0.042 | 0.02 | 1 | 10 | Acot11 | 0.022 | Old |
| 0.305164 | 0.067897 | 0.396 | 0.323 | 1 | 10 | Med13l | 0.073 | Old |
| 0.305301 | 0.504177 | 0.139 | 0.101 | 1 | 10 | Pex16 | 0.038 | Old |
| 0.305496 | 0.166383 | 0.411 | 0.333 | 1 | 10 | Bcas2 | 0.078 | Old |
| 0.305764 | 0.063944 | 0.885 | 0.778 | 1 | 10 | Eif4a1 | 0.107 | Old |
| 0.305963 | 0.082566 | 0.093 | 0.061 | 1 | 10 | Fam83f | 0.032 | Old |
| 0.306226 | 0.554293 | 0.127 | 0.091 | 1 | 10 | Pdzd11 | 0.036 | Old |
| 0.306287 | 0.511032 | 0.105 | 0.071 | 1 | 10 | Pja1 | 0.034 | Old |
| 0.306317 | 0.013486 | 0.205 | 0.152 | 1 | 10 | Actr10 | 0.053 | Old |
| 0.30632 | 0.010868 | 0.491 | 0.404 | 1 | 10 | Git2 | 0.087 | Old |
| 0.306354 | 0.375313 | 0.068 | 0.04 | 1 | 10 | Cdkn2aip | 0.028 | Old |
| 0.306467 | 1.051366 | 0.09 | 0.061 | 1 | 10 | Lrrc14 | 0.029 | Old |
| 0.306517 | 0.095506 | 0.411 | 0.333 | 1 | 10 | Prkcsh | 0.078 | Old |
| 0.306858 | 0.507307 | 0.142 | 0.101 | 1 | 10 | Uty | 0.041 | Old |
| 0.306938 | 0.441148 | 0.205 | 0.162 | 1 | 10 | Ubr3 | 0.043 | Old |
| 0.307052 | 0.068313 | 0.665 | 0.566 | 1 | 10 | Ppp3ca | 0.099 | Old |

| | | | | | | | | |
|---|---|---|---|---|---|---|---|---|
| 0.307061 | 0.037049 | 0.357 | 0.283 | 1 | 10 | Ociad1 | 0.074 | Old |
| 0.307086 | 0.209354 | 0.225 | 0.172 | 1 | 10 | Zdhhc3 | 0.053 | Old |
| 0.307092 | 1.425418 | 0.042 | 0.02 | 1 | 10 | Ttbk2 | 0.022 | Old |
| 0.307092 | 1.373952 | 0.042 | 0.02 | 1 | 10 | Sec22c | 0.022 | Old |
| 0.307217 | 0.863679 | 0.125 | 0.091 | 1 | 10 | Ube2w | 0.034 | Old |
| 0.307437 | 0.426751 | 0.259 | 0.212 | 1 | 10 | Hint2 | 0.047 | Old |
| 0.307498 | 0.669001 | 0.081 | 0.051 | 1 | 10 | Med24 | 0.03 | Old |
| 0.307763 | 0.116415 | 0.191 | 0.141 | 1 | 10 | Mrps7 | 0.05 | Old |
| 0.30796 | 0.16024 | 0.394 | 0.323 | 1 | 10 | Arglu1 | 0.071 | Old |
| 0.30799 | 0.238843 | 0.193 | 0.141 | 1 | 10 | Ncln | 0.052 | Old |
| 0.30808 | 0.077962 | 0.35 | 0.283 | 1 | 10 | Cnn2 | 0.067 | Old |
| 0.308081 | 0.210079 | 0.142 | 0.101 | 1 | 10 | Dusp23 | 0.041 | Old |
| 0.308157 | 0.17896 | 0.056 | 0.03 | 1 | 10 | Ap4b1 | 0.026 | Old |
| 0.308193 | 1.543203 | 0.042 | 0.02 | 1 | 10 | Zfp280c | 0.022 | Old |
| 0.308928 | 0.401833 | 0.093 | 0.061 | 1 | 10 | Pou5f2 | 0.032 | Old |
| 0.308974 | 0.356921 | 0.396 | 0.333 | 1 | 10 | Krcc1 | 0.063 | Old |
| 0.309089 | 0.167032 | 0.535 | 0.444 | 1 | 10 | Ddx39b | 0.091 | Old |
| 0.309296 | 1.42856 | 0.042 | 0.02 | 1 | 10 | E130311K1 | 0.022 | Old |
| 0.309672 | 0.679383 | 0.093 | 0.061 | 1 | 10 | Tstd3 | 0.032 | Old |
| 0.309794 | 0.08247 | 0.068 | 0.04 | 1 | 10 | Kctd3 | 0.028 | Old |
| 0.309803 | 0.146571 | 0.244 | 0.192 | 1 | 10 | Sh2b3 | 0.052 | Old |
| 0.310409 | 0.757857 | 0.115 | 0.081 | 1 | 10 | Gpatch2 | 0.034 | Old |
| 0.310681 | 0.72732 | 0.081 | 0.051 | 1 | 10 | 1190007I0 | 0.03 | Old |
| 0.311199 | 0.136186 | 0.105 | 0.071 | 1 | 10 | Drg2 | 0.034 | Old |
| 0.311511 | 1.472117 | 0.042 | 0.02 | 1 | 10 | Phlda3 | 0.022 | Old |
| 0.311669 | 0.144489 | 0.176 | 0.131 | 1 | 10 | 4833420G1 | 0.045 | Old |
| 0.311746 | 0.799062 | 0.137 | 0.101 | 1 | 10 | Coa6 | 0.036 | Old |
| 0.311845 | 0.089019 | 0.252 | 0.192 | 1 | 10 | Mrpl13 | 0.06 | Old |
| 0.311912 | 0.321031 | 0.093 | 0.061 | 1 | 10 | Eif2ak3 | 0.032 | Old |
| 0.311967 | 0.036354 | 0.056 | 0.03 | 1 | 10 | Dclre1b | 0.026 | Old |
| 0.312227 | 0.167742 | 0.13 | 0.091 | 1 | 10 | Ncoa5 | 0.039 | Old |
| 0.312275 | 0.149328 | 0.452 | 0.394 | 1 | 10 | Eif4ebp2 | 0.058 | Old |
| 0.312649 | 0.097983 | 0.873 | 0.758 | 1 | 10 | Tmem50a | 0.115 | Old |
| 0.312677 | 0.140744 | 0.592 | 0.515 | 1 | 10 | Srp14 | 0.077 | Old |
| 0.312708 | 0.056663 | 0.812 | 0.768 | 1 | 10 | Grb2 | 0.044 | Old |
| 0.313055 | 0.173395 | 0.406 | 0.343 | 1 | 10 | Srsf10 | 0.063 | Old |
| 0.313083 | 0.291616 | 0.081 | 0.051 | 1 | 10 | Aamdc | 0.03 | Old |
| 0.31332 | 0.631104 | 0.105 | 0.071 | 1 | 10 | Pus10 | 0.034 | Old |
| 0.313567 | 0.576411 | 0.161 | 0.121 | 1 | 10 | AW112010 | 0.04 | Old |
| 0.313795 | 0.75112 | 0.115 | 0.081 | 1 | 10 | Abcd4 | 0.034 | Old |
| 0.314077 | 0.261033 | 0.846 | 0.758 | 1 | 10 | Serinc3 | 0.088 | Old |
| 0.314262 | 0.370288 | 0.247 | 0.192 | 1 | 10 | Rhoq | 0.055 | Old |
| 0.314737 | 0.424511 | 0.161 | 0.121 | 1 | 10 | Map3k20 | 0.04 | Old |
| 0.314784 | 0.124772 | 0.628 | 0.556 | 1 | 10 | Tbxas1 | 0.072 | Old |
| 0.314844 | 0.183232 | 0.056 | 0.03 | 1 | 10 | E130307A1 | 0.026 | Old |

| | | | | | | | | |
|---|---|---|---|---|---|---|---|---|
| 0.315488 | 0.167837 | 0.533 | 0.455 | 1 | 10 | Ssr1 | 0.078 | Old |
| 0.315755 | 0.127975 | 0.496 | 0.414 | 1 | 10 | Gmfb | 0.082 | Old |
| 0.315971 | 1.427795 | 0.042 | 0.02 | 1 | 10 | Gm7694 | 0.022 | Old |
| 0.316751 | 0.619838 | 0.068 | 0.04 | 1 | 10 | Polr3a | 0.028 | Old |
| 0.316895 | 0.129618 | 0.308 | 0.242 | 1 | 10 | Dennd4b | 0.066 | Old |
| 0.316975 | 2.082513 | 0.027 | 0.01 | 1 | 10 | Nmrk1 | 0.017 | Old |
| 0.316975 | 1.972827 | 0.027 | 0.01 | 1 | 10 | Plekha8 | 0.017 | Old |
| 0.316975 | 1.459739 | 0.027 | 0.01 | 1 | 10 | Znrf3 | 0.017 | Old |
| 0.317016 | 0.061318 | 0.191 | 0.141 | 1 | 10 | Ube2j1 | 0.05 | Old |
| 0.317058 | 1.504904 | 0.066 | 0.04 | 1 | 10 | C1qtnf12 | 0.026 | Old |
| 0.317148 | 0.305901 | 0.357 | 0.313 | 1 | 10 | Ankrd44 | 0.044 | Old |
| 0.317487 | 0.193625 | 0.961 | 0.939 | 1 | 10 | Hsp90b1 | 0.022 | Old |
| 0.317962 | 0.238741 | 0.022 | 0.04 | 1 | 10 | Vsig10 | -0.018 | Old |
| 0.317995 | 0.131797 | 0.619 | 0.545 | 1 | 10 | Ubap2l | 0.074 | Old |
| 0.318216 | 1.134353 | 0.042 | 0.02 | 1 | 10 | Fahd2a | 0.022 | Old |
| 0.318217 | 0.292992 | 0.325 | 0.273 | 1 | 10 | Arl6ip5 | 0.052 | Old |
| 0.318318 | 0.235058 | 0.394 | 0.333 | 1 | 10 | Smg1 | 0.061 | Old |
| 0.31838 | 1.987514 | 0.027 | 0.01 | 1 | 10 | 1700066M | 0.017 | Old |
| 0.31838 | 1.890787 | 0.027 | 0.01 | 1 | 10 | Pxmp2 | 0.017 | Old |
| 0.31838 | 1.74404 | 0.027 | 0.01 | 1 | 10 | Bora | 0.017 | Old |
| 0.31838 | 1.719815 | 0.027 | 0.01 | 1 | 10 | Wdr6 | 0.017 | Old |
| 0.31838 | 1.630543 | 0.027 | 0.01 | 1 | 10 | Vldlr | 0.017 | Old |
| 0.318464 | 0.11945 | 0.511 | 0.465 | 1 | 10 | Frrs1 | 0.046 | Old |
| 0.318491 | 0.084854 | 0.494 | 0.414 | 1 | 10 | Nav1 | 0.08 | Old |
| 0.318673 | 0.01344 | 0.264 | 0.202 | 1 | 10 | Cops4 | 0.062 | Old |
| 0.319343 | 0.964822 | 0.042 | 0.02 | 1 | 10 | Lamc1 | 0.022 | Old |
| 0.319612 | 0.468195 | 0.149 | 0.111 | 1 | 10 | Syne1 | 0.038 | Old |
| 0.319788 | 1.929612 | 0.027 | 0.01 | 1 | 10 | BC035044 | 0.017 | Old |
| 0.319788 | 1.568881 | 0.027 | 0.01 | 1 | 10 | Traf3ip3 | 0.017 | Old |
| 0.319959 | 0.577633 | 0.181 | 0.141 | 1 | 10 | Srebf1 | 0.04 | Old |
| 0.320628 | 0.15182 | 0.161 | 0.121 | 1 | 10 | Pam16 | 0.04 | Old |
| 0.320655 | 0.258076 | 0.13 | 0.091 | 1 | 10 | Klhdc2 | 0.039 | Old |
| 0.32086 | 0.272533 | 0.139 | 0.101 | 1 | 10 | Idh3a | 0.038 | Old |
| 0.321031 | 0.063288 | 0.528 | 0.444 | 1 | 10 | Fam111a | 0.084 | Old |
| 0.321201 | 1.437926 | 0.027 | 0.01 | 1 | 10 | Rhou | 0.017 | Old |
| 0.321201 | 1.349666 | 0.027 | 0.01 | 1 | 10 | Cdc42bpg | 0.017 | Old |
| 0.321201 | 1.345685 | 0.027 | 0.01 | 1 | 10 | Sh3d19 | 0.017 | Old |
| 0.321201 | 1.231251 | 0.027 | 0.01 | 1 | 10 | Bbs9 | 0.017 | Old |
| 0.321201 | 1.12331 | 0.027 | 0.01 | 1 | 10 | Alkbh2 | 0.017 | Old |
| 0.321201 | 1.108643 | 0.027 | 0.01 | 1 | 10 | Angpt2 | 0.017 | Old |
| 0.321201 | 1.033511 | 0.027 | 0.01 | 1 | 10 | Nod2 | 0.017 | Old |
| 0.321309 | 0.134851 | 0.13 | 0.091 | 1 | 10 | Rai1 | 0.039 | Old |
| 0.32137 | 0.647534 | 0.178 | 0.141 | 1 | 10 | Ate1 | 0.037 | Old |
| 0.321588 | 0.035046 | 1 | 1 | 1 | 10 | Tmsb4x | 0 | Old |
| 0.321604 | 0.619256 | 0.042 | 0.02 | 1 | 10 | Panx1 | 0.022 | Old |

| | | | | | | | | |
|---|---|---|---|---|---|---|---|---|
| 0.321661 | 0.946675 | 0.09 | 0.061 | 1 | 10 | Fance | 0.029 | Old |
| 0.321742 | 0.30016 | 0.093 | 0.061 | 1 | 10 | Alkbh8 | 0.032 | Old |
| 0.321798 | 0.251623 | 0.418 | 0.354 | 1 | 10 | Igsf6 | 0.064 | Old |
| 0.321839 | 0.384189 | 0.152 | 0.111 | 1 | 10 | Synj2 | 0.041 | Old |
| 0.322016 | 0.007369 | 0.423 | 0.343 | 1 | 10 | Ppp1r15b | 0.08 | Old |
| 0.322433 | 0.906775 | 0.09 | 0.061 | 1 | 10 | Rnf34 | 0.029 | Old |
| 0.322477 | 1.395368 | 0.054 | 0.03 | 1 | 10 | Ifrd2 | 0.024 | Old |
| 0.322618 | 1.474872 | 0.027 | 0.01 | 1 | 10 | Mdm1 | 0.017 | Old |
| 0.322618 | 1.299326 | 0.027 | 0.01 | 1 | 10 | Zfp472 | 0.017 | Old |
| 0.322618 | 1.03286 | 0.027 | 0.01 | 1 | 10 | 4833418N( | 0.017 | Old |
| 0.322785 | 0.252616 | 0.117 | 0.081 | 1 | 10 | St3gal2 | 0.036 | Old |
| 0.3228 | 0.023855 | 0.944 | 0.859 | 1 | 10 | Canx | 0.085 | Old |
| 0.323073 | 0.099602 | 0.23 | 0.182 | 1 | 10 | Bccip | 0.048 | Old |
| 0.324039 | 1.474633 | 0.027 | 0.01 | 1 | 10 | Car2 | 0.017 | Old |
| 0.324039 | 1.234488 | 0.027 | 0.01 | 1 | 10 | Ugt1a7c | 0.017 | Old |
| 0.324039 | 1.097659 | 0.027 | 0.01 | 1 | 10 | Tfb1m | 0.017 | Old |
| 0.324039 | 0.984708 | 0.027 | 0.01 | 1 | 10 | Cdc6 | 0.017 | Old |
| 0.32445 | 0.253453 | 0.081 | 0.051 | 1 | 10 | Prr3 | 0.03 | Old |
| 0.324767 | 0.661157 | 0.103 | 0.071 | 1 | 10 | Rgs14 | 0.032 | Old |
| 0.324879 | 0.22688 | 0.23 | 0.182 | 1 | 10 | Slmap | 0.048 | Old |
| 0.324918 | 2.150019 | 0.01 | 0 | 1 | 10 | Rnase2a | 0.01 | Old |
| 0.324918 | 2.127926 | 0.01 | 0 | 1 | 10 | Gm8113 | 0.01 | Old |
| 0.324918 | 2.028373 | 0.01 | 0 | 1 | 10 | Runx2os1 | 0.01 | Old |
| 0.324918 | 1.891627 | 0.01 | 0 | 1 | 10 | Slc25a27 | 0.01 | Old |
| 0.324918 | 1.877262 | 0.01 | 0 | 1 | 10 | Adal | 0.01 | Old |
| 0.324918 | 1.867064 | 0.01 | 0 | 1 | 10 | Rab23 | 0.01 | Old |
| 0.324918 | 1.866168 | 0.01 | 0 | 1 | 10 | Kifc5b | 0.01 | Old |
| 0.324918 | 1.856666 | 0.01 | 0 | 1 | 10 | Camk2b | 0.01 | Old |
| 0.324918 | 1.85168 | 0.01 | 0 | 1 | 10 | Il18rap | 0.01 | Old |
| 0.324918 | 1.850959 | 0.01 | 0 | 1 | 10 | Adgra3 | 0.01 | Old |
| 0.324918 | 1.803984 | 0.01 | 0 | 1 | 10 | Zbtb32 | 0.01 | Old |
| 0.324918 | 1.770613 | 0.01 | 0 | 1 | 10 | Gm14548 | 0.01 | Old |
| 0.324918 | 1.763617 | 0.01 | 0 | 1 | 10 | Stpg4 | 0.01 | Old |
| 0.324918 | 1.753061 | 0.01 | 0 | 1 | 10 | Mypop | 0.01 | Old |
| 0.324918 | 1.744784 | 0.01 | 0 | 1 | 10 | Gm16094 | 0.01 | Old |
| 0.324918 | 1.727048 | 0.01 | 0 | 1 | 10 | 9130008F2 | 0.01 | Old |
| 0.324918 | 1.669794 | 0.01 | 0 | 1 | 10 | Dnah12 | 0.01 | Old |
| 0.324918 | 1.651969 | 0.01 | 0 | 1 | 10 | Tal1 | 0.01 | Old |
| 0.324918 | 1.648907 | 0.01 | 0 | 1 | 10 | A530013C2 | 0.01 | Old |
| 0.324918 | 1.618217 | 0.01 | 0 | 1 | 10 | 1700029I1! | 0.01 | Old |
| 0.324918 | 1.612885 | 0.01 | 0 | 1 | 10 | Gm30074 | 0.01 | Old |
| 0.324918 | 1.612676 | 0.01 | 0 | 1 | 10 | Tmem156 | 0.01 | Old |
| 0.324918 | 1.603211 | 0.01 | 0 | 1 | 10 | Tm4sf5 | 0.01 | Old |
| 0.324918 | 1.585525 | 0.01 | 0 | 1 | 10 | Nkapl | 0.01 | Old |
| 0.324918 | 1.579924 | 0.01 | 0 | 1 | 10 | Gm33973 | 0.01 | Old |

| | | | | | | | | |
|---|---|---|---|---|---|---|---|---|
| 0.324918 | 1.572694 | 0.01 | 0 | 1 | 10 | Krtcap3 | 0.01 | Old |
| 0.324918 | 1.56673 | 0.01 | 0 | 1 | 10 | Rab36 | 0.01 | Old |
| 0.324918 | 1.557019 | 0.01 | 0 | 1 | 10 | Tox2 | 0.01 | Old |
| 0.324918 | 1.549605 | 0.01 | 0 | 1 | 10 | Zscan22 | 0.01 | Old |
| 0.324918 | 1.531925 | 0.01 | 0 | 1 | 10 | Gstm2 | 0.01 | Old |
| 0.324918 | 1.531427 | 0.01 | 0 | 1 | 10 | Spred3 | 0.01 | Old |
| 0.324918 | 1.525226 | 0.01 | 0 | 1 | 10 | Rgs5 | 0.01 | Old |
| 0.324918 | 1.512986 | 0.01 | 0 | 1 | 10 | Egln3 | 0.01 | Old |
| 0.324918 | 1.502761 | 0.01 | 0 | 1 | 10 | Atp1a2 | 0.01 | Old |
| 0.324918 | 1.495935 | 0.01 | 0 | 1 | 10 | Zbtb25 | 0.01 | Old |
| 0.324918 | 1.492776 | 0.01 | 0 | 1 | 10 | Dync2li1 | 0.01 | Old |
| 0.324918 | 1.491832 | 0.01 | 0 | 1 | 10 | Rabl2 | 0.01 | Old |
| 0.324918 | 1.49113 | 0.01 | 0 | 1 | 10 | Tesmin | 0.01 | Old |
| 0.324918 | 1.488636 | 0.01 | 0 | 1 | 10 | D630033O | 0.01 | Old |
| 0.324918 | 1.478485 | 0.01 | 0 | 1 | 10 | 1700061G | 0.01 | Old |
| 0.324918 | 1.47177 | 0.01 | 0 | 1 | 10 | Actr3b | 0.01 | Old |
| 0.324918 | 1.468874 | 0.01 | 0 | 1 | 10 | Slc22a5 | 0.01 | Old |
| 0.324918 | 1.462776 | 0.01 | 0 | 1 | 10 | Gm28874 | 0.01 | Old |
| 0.324918 | 1.46079 | 0.01 | 0 | 1 | 10 | Vps25 | 0.01 | Old |
| 0.324918 | 1.458671 | 0.01 | 0 | 1 | 10 | Crisp1 | 0.01 | Old |
| 0.324918 | 1.440591 | 0.01 | 0 | 1 | 10 | Prr15 | 0.01 | Old |
| 0.324918 | 1.43676 | 0.01 | 0 | 1 | 10 | Clic5 | 0.01 | Old |
| 0.324918 | 1.410254 | 0.01 | 0 | 1 | 10 | Cd209e | 0.01 | Old |
| 0.324918 | 1.409276 | 0.01 | 0 | 1 | 10 | 1700019L1 | 0.01 | Old |
| 0.324918 | 1.399599 | 0.01 | 0 | 1 | 10 | Palm3 | 0.01 | Old |
| 0.324918 | 1.399397 | 0.01 | 0 | 1 | 10 | Ankrd42 | 0.01 | Old |
| 0.324918 | 1.399029 | 0.01 | 0 | 1 | 10 | Lgals4 | 0.01 | Old |
| 0.324918 | 1.39593 | 0.01 | 0 | 1 | 10 | Gm14762 | 0.01 | Old |
| 0.324918 | 1.382645 | 0.01 | 0 | 1 | 10 | Cep112 | 0.01 | Old |
| 0.324918 | 1.373007 | 0.01 | 0 | 1 | 10 | Tmem116 | 0.01 | Old |
| 0.324918 | 1.361217 | 0.01 | 0 | 1 | 10 | Camk2n2 | 0.01 | Old |
| 0.324918 | 1.360163 | 0.01 | 0 | 1 | 10 | Kyat3 | 0.01 | Old |
| 0.324918 | 1.348831 | 0.01 | 0 | 1 | 10 | Gm4787 | 0.01 | Old |
| 0.324918 | 1.339847 | 0.01 | 0 | 1 | 10 | AC125149. | 0.01 | Old |
| 0.324918 | 1.332583 | 0.01 | 0 | 1 | 10 | Rad51ap1 | 0.01 | Old |
| 0.324918 | 1.330244 | 0.01 | 0 | 1 | 10 | F830016B0 | 0.01 | Old |
| 0.324918 | 1.313169 | 0.01 | 0 | 1 | 10 | Zfp799 | 0.01 | Old |
| 0.324918 | 1.312881 | 0.01 | 0 | 1 | 10 | Zfp420 | 0.01 | Old |
| 0.324918 | 1.301152 | 0.01 | 0 | 1 | 10 | Dhrs13 | 0.01 | Old |
| 0.324918 | 1.299108 | 0.01 | 0 | 1 | 10 | Gm11496 | 0.01 | Old |
| 0.324918 | 1.294458 | 0.01 | 0 | 1 | 10 | Psrc1 | 0.01 | Old |
| 0.324918 | 1.292392 | 0.01 | 0 | 1 | 10 | Ptgs2os2 | 0.01 | Old |
| 0.324918 | 1.289131 | 0.01 | 0 | 1 | 10 | Rpl7a-ps5 | 0.01 | Old |
| 0.324918 | 1.279222 | 0.01 | 0 | 1 | 10 | Pnma1 | 0.01 | Old |
| 0.324918 | 1.27512 | 0.01 | 0 | 1 | 10 | Htra1 | 0.01 | Old |

| | | | | | | | | |
|---|---|---|---|---|---|---|---|---|
| 0.324918 | 1.253239 | 0.01 | 0 | 1 | 10 | Gm14698 | 0.01 | Old |
| 0.324918 | 1.239095 | 0.01 | 0 | 1 | 10 | Samd10 | 0.01 | Old |
| 0.324918 | 1.231423 | 0.01 | 0 | 1 | 10 | Ly75 | 0.01 | Old |
| 0.324918 | 1.231101 | 0.01 | 0 | 1 | 10 | Rep15 | 0.01 | Old |
| 0.324918 | 1.229825 | 0.01 | 0 | 1 | 10 | Gm29994 | 0.01 | Old |
| 0.324918 | 1.228422 | 0.01 | 0 | 1 | 10 | Angptl6 | 0.01 | Old |
| 0.324918 | 1.225974 | 0.01 | 0 | 1 | 10 | Kifc2 | 0.01 | Old |
| 0.324918 | 1.221137 | 0.01 | 0 | 1 | 10 | Arhgap29 | 0.01 | Old |
| 0.324918 | 1.208606 | 0.01 | 0 | 1 | 10 | Spc24 | 0.01 | Old |
| 0.324918 | 1.208424 | 0.01 | 0 | 1 | 10 | 3110001I2: | 0.01 | Old |
| 0.324918 | 1.187029 | 0.01 | 0 | 1 | 10 | Trim30c | 0.01 | Old |
| 0.324918 | 1.183383 | 0.01 | 0 | 1 | 10 | Zbtb49 | 0.01 | Old |
| 0.324918 | 1.183216 | 0.01 | 0 | 1 | 10 | Prps1l3 | 0.01 | Old |
| 0.324918 | 1.182902 | 0.01 | 0 | 1 | 10 | Srgap1 | 0.01 | Old |
| 0.324918 | 1.182261 | 0.01 | 0 | 1 | 10 | Slfn3 | 0.01 | Old |
| 0.324918 | 1.167497 | 0.01 | 0 | 1 | 10 | Zfp85 | 0.01 | Old |
| 0.324918 | 1.167344 | 0.01 | 0 | 1 | 10 | Wdr54 | 0.01 | Old |
| 0.324918 | 1.166089 | 0.01 | 0 | 1 | 10 | Eaf2 | 0.01 | Old |
| 0.324918 | 1.164623 | 0.01 | 0 | 1 | 10 | Haspin | 0.01 | Old |
| 0.324918 | 1.133025 | 0.01 | 0 | 1 | 10 | Mss51 | 0.01 | Old |
| 0.324918 | 1.128391 | 0.01 | 0 | 1 | 10 | Klhl11 | 0.01 | Old |
| 0.324918 | 1.128194 | 0.01 | 0 | 1 | 10 | Prr7 | 0.01 | Old |
| 0.324918 | 1.125501 | 0.01 | 0 | 1 | 10 | Spns2 | 0.01 | Old |
| 0.324918 | 1.105353 | 0.01 | 0 | 1 | 10 | Gm14461 | 0.01 | Old |
| 0.324918 | 1.092762 | 0.01 | 0 | 1 | 10 | Paqr3 | 0.01 | Old |
| 0.324918 | 1.092695 | 0.01 | 0 | 1 | 10 | Fhit | 0.01 | Old |
| 0.324918 | 1.092351 | 0.01 | 0 | 1 | 10 | Gm20234 | 0.01 | Old |
| 0.324918 | 1.089236 | 0.01 | 0 | 1 | 10 | Tnfsf14 | 0.01 | Old |
| 0.324918 | 1.087041 | 0.01 | 0 | 1 | 10 | Gm42918 | 0.01 | Old |
| 0.324918 | 1.080811 | 0.01 | 0 | 1 | 10 | AC139347. | 0.01 | Old |
| 0.324918 | 1.078179 | 0.01 | 0 | 1 | 10 | Zfp583 | 0.01 | Old |
| 0.324918 | 1.077446 | 0.01 | 0 | 1 | 10 | Gm17354 | 0.01 | Old |
| 0.324918 | 1.053097 | 0.01 | 0 | 1 | 10 | Mdga1 | 0.01 | Old |
| 0.324918 | 1.038557 | 0.01 | 0 | 1 | 10 | Cage1 | 0.01 | Old |
| 0.324918 | 1.031602 | 0.01 | 0 | 1 | 10 | Mmp27 | 0.01 | Old |
| 0.324918 | 1.025023 | 0.01 | 0 | 1 | 10 | B230398E0 | 0.01 | Old |
| 0.324918 | 0.999262 | 0.01 | 0 | 1 | 10 | Fam167b | 0.01 | Old |
| 0.324918 | 0.999026 | 0.01 | 0 | 1 | 10 | D11Wsu47 | 0.01 | Old |
| 0.324918 | 0.972462 | 0.01 | 0 | 1 | 10 | Gm15441 | 0.01 | Old |
| 0.324918 | 0.951752 | 0.01 | 0 | 1 | 10 | Fgf11 | 0.01 | Old |
| 0.324918 | 0.943908 | 0.01 | 0 | 1 | 10 | Gm11131 | 0.01 | Old |
| 0.324918 | 0.937541 | 0.01 | 0 | 1 | 10 | Slc24a5 | 0.01 | Old |
| 0.324918 | 0.929447 | 0.01 | 0 | 1 | 10 | Dusp9 | 0.01 | Old |
| 0.324918 | 0.916465 | 0.01 | 0 | 1 | 10 | Peak1os | 0.01 | Old |
| 0.324918 | 0.905219 | 0.01 | 0 | 1 | 10 | Gm19705 | 0.01 | Old |

| | | | | | | | | |
|---|---|---|---|---|---|---|---|---|
| 0.324918 | 0.902719 | 0.01 | 0 | 1 | 10 | Iqcd | 0.01 | Old |
| 0.324918 | 0.89552 | 0.01 | 0 | 1 | 10 | Gm32856 | 0.01 | Old |
| 0.324918 | 0.886588 | 0.01 | 0 | 1 | 10 | Znrd1as | 0.01 | Old |
| 0.324918 | 0.876716 | 0.01 | 0 | 1 | 10 | Gm47469 | 0.01 | Old |
| 0.324918 | 0.873342 | 0.01 | 0 | 1 | 10 | Chd5 | 0.01 | Old |
| 0.324918 | 0.849715 | 0.01 | 0 | 1 | 10 | Hspa12a | 0.01 | Old |
| 0.324918 | 0.846632 | 0.01 | 0 | 1 | 10 | 1700113A1 | 0.01 | Old |
| 0.324918 | 0.836653 | 0.01 | 0 | 1 | 10 | Zfpm1 | 0.01 | Old |
| 0.324918 | 0.813035 | 0.01 | 0 | 1 | 10 | Ssc5d | 0.01 | Old |
| 0.324918 | 0.811786 | 0.01 | 0 | 1 | 10 | Lag3 | 0.01 | Old |
| 0.324918 | 0.798971 | 0.01 | 0 | 1 | 10 | Ypel1 | 0.01 | Old |
| 0.324918 | 0.79181 | 0.01 | 0 | 1 | 10 | Gm26526 | 0.01 | Old |
| 0.324918 | 0.781549 | 0.01 | 0 | 1 | 10 | Tmsb15b1 | 0.01 | Old |
| 0.324918 | 0.764911 | 0.01 | 0 | 1 | 10 | Gm9774 | 0.01 | Old |
| 0.324918 | 0.744497 | 0.01 | 0 | 1 | 10 | Gm1976 | 0.01 | Old |
| 0.324918 | 0.742908 | 0.01 | 0 | 1 | 10 | Gm14410 | 0.01 | Old |
| 0.324918 | 0.742644 | 0.01 | 0 | 1 | 10 | Gm12355 | 0.01 | Old |
| 0.324918 | 0.699644 | 0.01 | 0 | 1 | 10 | Bcl7a | 0.01 | Old |
| 0.325016 | 0.449728 | 0.042 | 0.02 | 1 | 10 | Zmynd19 | 0.022 | Old |
| 0.325103 | 1.066334 | 0.078 | 0.051 | 1 | 10 | Smim24 | 0.027 | Old |
| 0.325146 | 0.309532 | 0.252 | 0.202 | 1 | 10 | Ndfip1 | 0.05 | Old |
| 0.325464 | 1.349299 | 0.027 | 0.01 | 1 | 10 | Zfp12 | 0.017 | Old |
| 0.325464 | 1.198499 | 0.027 | 0.01 | 1 | 10 | Trim2 | 0.017 | Old |
| 0.325464 | 1.165296 | 0.027 | 0.01 | 1 | 10 | Wdr34 | 0.017 | Old |
| 0.325464 | 0.99212 | 0.027 | 0.01 | 1 | 10 | Gbp9 | 0.017 | Old |
| 0.325774 | 0.102483 | 0.504 | 0.444 | 1 | 10 | Cnot6l | 0.06 | Old |
| 0.325937 | 0.920496 | 0.078 | 0.051 | 1 | 10 | 1110004E0 | 0.027 | Old |
| 0.326883 | 0.552633 | 0.115 | 0.081 | 1 | 10 | Dhx29 | 0.034 | Old |
| 0.326894 | 1.416008 | 0.027 | 0.01 | 1 | 10 | Fastkd3 | 0.017 | Old |
| 0.326894 | 1.210396 | 0.027 | 0.01 | 1 | 10 | Epn2 | 0.017 | Old |
| 0.326894 | 0.940045 | 0.027 | 0.01 | 1 | 10 | Rubcnl | 0.017 | Old |
| 0.326942 | 1.221884 | 0.066 | 0.04 | 1 | 10 | Lilra6 | 0.026 | Old |
| 0.327118 | 0.102097 | 0.093 | 0.061 | 1 | 10 | Magohb | 0.032 | Old |
| 0.327303 | 0.436573 | 0.042 | 0.02 | 1 | 10 | Tprn | 0.022 | Old |
| 0.327403 | 0.064584 | 0.35 | 0.283 | 1 | 10 | Tmx1 | 0.067 | Old |
| 0.32748 | 0.936319 | 0.054 | 0.03 | 1 | 10 | Vps33b | 0.024 | Old |
| 0.327953 | 0.057725 | 0.682 | 0.556 | 1 | 10 | Cd300a | 0.126 | Old |
| 0.328272 | 0.227978 | 0.164 | 0.121 | 1 | 10 | Slc25a30 | 0.043 | Old |
| 0.328327 | 1.064075 | 0.027 | 0.01 | 1 | 10 | Zfp668 | 0.017 | Old |
| 0.328327 | 0.846118 | 0.027 | 0.01 | 1 | 10 | Afap1 | 0.017 | Old |
| 0.328327 | 0.771925 | 0.027 | 0.01 | 1 | 10 | Fn3krp | 0.017 | Old |
| 0.328327 | 0.546906 | 0.027 | 0.01 | 1 | 10 | Cep131 | 0.017 | Old |
| 0.328451 | 0.383726 | 0.042 | 0.02 | 1 | 10 | Gm26853 | 0.022 | Old |
| 0.328754 | 0.188976 | 0.472 | 0.404 | 1 | 10 | Aph1a | 0.068 | Old |
| 0.328995 | 0.493021 | 0.161 | 0.121 | 1 | 10 | Rinl | 0.04 | Old |

| | | | | | | | | |
|---|---|---|---|---|---|---|---|---|
| 0.329017 | 0.191838 | 0.303 | 0.253 | 1 | 10 | Snx12 | 0.05 | Old |
| 0.32944 | 0.367808 | 0.093 | 0.061 | 1 | 10 | Slc25a16 | 0.032 | Old |
| 0.329601 | 0.345911 | 0.042 | 0.02 | 1 | 10 | Ppp1r3e | 0.022 | Old |
| 0.329672 | 0.980452 | 0.066 | 0.04 | 1 | 10 | 1600010M | 0.026 | Old |
| 0.329764 | 0.728417 | 0.027 | 0.01 | 1 | 10 | Gm10053 | 0.017 | Old |
| 0.329764 | 0.678917 | 0.027 | 0.01 | 1 | 10 | Gprc5b | 0.017 | Old |
| 0.33021 | 0.535809 | 0.232 | 0.192 | 1 | 10 | Gemin7 | 0.04 | Old |
| 0.330368 | 0.433159 | 0.21 | 0.162 | 1 | 10 | Tppp3 | 0.048 | Old |
| 0.330618 | 0.278321 | 0.274 | 0.232 | 1 | 10 | Gigyf2 | 0.042 | Old |
| 0.330899 | 0.135444 | 0.269 | 0.222 | 1 | 10 | Anapc5 | 0.047 | Old |
| 0.330993 | 0.057671 | 0.313 | 0.253 | 1 | 10 | Ifi204 | 0.06 | Old |
| 0.331206 | 0.658487 | 0.027 | 0.01 | 1 | 10 | Cptp | 0.017 | Old |
| 0.331206 | 0.472156 | 0.027 | 0.01 | 1 | 10 | Osgepl1 | 0.017 | Old |
| 0.331206 | 0.469935 | 0.027 | 0.01 | 1 | 10 | Zswim1 | 0.017 | Old |
| 0.331206 | 0.286229 | 0.027 | 0.01 | 1 | 10 | Tmlhe | 0.017 | Old |
| 0.331206 | 0.059842 | 0.027 | 0.01 | 1 | 10 | Mylk | 0.017 | Old |
| 0.331319 | 0.000141 | 0.423 | 0.343 | 1 | 10 | Zmynd8 | 0.08 | Old |
| 0.331501 | 0.810942 | 0.066 | 0.04 | 1 | 10 | Rab6b | 0.026 | Old |
| 0.331857 | 0.009092 | 0.638 | 0.576 | 1 | 10 | Atp5b | 0.062 | Old |
| 0.332008 | 0.133217 | 0.452 | 0.364 | 1 | 10 | Egr2 | 0.088 | Old |
| 0.33229 | 0.151916 | 0.232 | 0.192 | 1 | 10 | Bcl6 | 0.04 | Old |
| 0.332498 | 0.391905 | 0.115 | 0.081 | 1 | 10 | Slc35b4 | 0.034 | Old |
| 0.332651 | 0.130612 | 0.027 | 0.01 | 1 | 10 | Rad52 | 0.017 | Old |
| 0.332854 | 0.584326 | 0.171 | 0.131 | 1 | 10 | Acsl1 | 0.04 | Old |
| 0.333372 | 0.271407 | 0.328 | 0.283 | 1 | 10 | Slamf9 | 0.045 | Old |
| 0.333501 | 0.021107 | 0.491 | 0.424 | 1 | 10 | Ndufv3 | 0.067 | Old |
| 0.333733 | 0.232725 | 0.396 | 0.354 | 1 | 10 | Rreb1 | 0.042 | Old |
| 0.334009 | 0.389578 | 0.178 | 0.141 | 1 | 10 | Stx8 | 0.037 | Old |
| 0.334118 | 0.60135 | 0.137 | 0.101 | 1 | 10 | Ctps2 | 0.036 | Old |
| 0.334569 | 0.122289 | 0.054 | 0.03 | 1 | 10 | Rrp1b | 0.024 | Old |
| 0.334797 | 0.189462 | 0.533 | 0.455 | 1 | 10 | Fam120a | 0.078 | Old |
| 0.334867 | 0.140294 | 0.501 | 0.424 | 1 | 10 | Bcap31 | 0.077 | Old |
| 0.335019 | 0.220324 | 0.183 | 0.141 | 1 | 10 | Cln6 | 0.042 | Old |
| 0.335119 | 0.143439 | 0.286 | 0.222 | 1 | 10 | Sf3b4 | 0.064 | Old |
| 0.335196 | 0.896175 | 0.078 | 0.051 | 1 | 10 | Atxn1l | 0.027 | Old |
| 0.335303 | 0.012928 | 0.117 | 0.081 | 1 | 10 | Tsc2 | 0.036 | Old |
| 0.335382 | 0.05201 | 0.156 | 0.111 | 1 | 10 | Stim1 | 0.045 | Old |
| 0.335549 | 0.329703 | 0.345 | 0.293 | 1 | 10 | Pikfyve | 0.052 | Old |
| 0.33559 | 0.882588 | 0.054 | 0.03 | 1 | 10 | Tapbpl | 0.024 | Old |
| 0.33579 | 0.246504 | 0.105 | 0.071 | 1 | 10 | 2900089D1 | 0.034 | Old |
| 0.335852 | 0.293207 | 0.23 | 0.182 | 1 | 10 | Suds3 | 0.048 | Old |
| 0.336203 | 0.055326 | 0.919 | 0.919 | 1 | 10 | Rps17 | 0 | Old |
| 0.336858 | 0.187627 | 0.423 | 0.364 | 1 | 10 | Pdcd10 | 0.059 | Old |
| 0.336999 | 0.359234 | 0.198 | 0.162 | 1 | 10 | Zcchc24 | 0.036 | Old |
| 0.337025 | 0.851508 | 0.066 | 0.04 | 1 | 10 | Ppat | 0.026 | Old |

| | | | | | | | | |
|---|---|---|---|---|---|---|---|---|
| 0.337083 | 0.082169 | 0.142 | 0.101 | 1 | 10 | Zfp397 | 0.041 | Old |
| 0.337404 | 0.222577 | 0.32 | 0.253 | 1 | 10 | Rc3h2 | 0.067 | Old |
| 0.337485 | 0.376185 | 0.318 | 0.273 | 1 | 10 | Wdr89 | 0.045 | Old |
| 0.337637 | 0.608878 | 0.054 | 0.03 | 1 | 10 | 4930430F0 | 0.024 | Old |
| 0.337913 | 0.140536 | 0.127 | 0.091 | 1 | 10 | Rpl27-ps3 | 0.036 | Old |
| 0.338014 | 0.124567 | 0.765 | 0.697 | 1 | 10 | Itgb1 | 0.068 | Old |
| 0.338074 | 0.191097 | 0.237 | 0.182 | 1 | 10 | Clcn3 | 0.055 | Old |
| 0.338554 | 0.064148 | 0.357 | 0.293 | 1 | 10 | Eif4g3 | 0.064 | Old |
| 0.338868 | 0.657641 | 0.103 | 0.071 | 1 | 10 | Ruvbl2 | 0.032 | Old |
| 0.338946 | 0.346125 | 0.149 | 0.111 | 1 | 10 | Insig2 | 0.038 | Old |
| 0.339029 | 0.535377 | 0.191 | 0.152 | 1 | 10 | Epb41l4ao: | 0.039 | Old |
| 0.339366 | 0.082007 | 0.621 | 0.515 | 1 | 10 | Selenof | 0.106 | Old |
| 0.339382 | 0.234823 | 0.176 | 0.131 | 1 | 10 | Cnp | 0.045 | Old |
| 0.339578 | 0.039913 | 0.484 | 0.384 | 1 | 10 | Lmo2 | 0.1 | Old |
| 0.339656 | 0.174107 | 0.577 | 0.505 | 1 | 10 | Atp6v0d1 | 0.072 | Old |
| 0.340436 | 0.140157 | 0.274 | 0.222 | 1 | 10 | Zwint | 0.052 | Old |
| 0.340475 | 0.177801 | 0.215 | 0.172 | 1 | 10 | Grsf1 | 0.043 | Old |
| 0.341756 | 0.417791 | 0.054 | 0.03 | 1 | 10 | Clasrp | 0.024 | Old |
| 0.342414 | 0.285489 | 0.152 | 0.111 | 1 | 10 | Dpp9 | 0.041 | Old |
| 0.342608 | 0.753594 | 0.066 | 0.04 | 1 | 10 | Gorab | 0.026 | Old |
| 0.342608 | 0.600852 | 0.066 | 0.04 | 1 | 10 | Rmnd1 | 0.026 | Old |
| 0.342725 | 0.772954 | 0.1 | 0.071 | 1 | 10 | Bicra | 0.029 | Old |
| 0.342791 | 0.49378 | 0.054 | 0.03 | 1 | 10 | Adamts10 | 0.024 | Old |
| 0.34285 | 0.097919 | 0.081 | 0.051 | 1 | 10 | Zw10 | 0.03 | Old |
| 0.343166 | 0.297803 | 0.183 | 0.141 | 1 | 10 | Ino80e | 0.042 | Old |
| 0.34381 | 0.153888 | 0.616 | 0.545 | 1 | 10 | Lst1 | 0.071 | Old |
| 0.344028 | 0.126059 | 0.958 | 0.919 | 1 | 10 | H2-Aa | 0.039 | Old |
| 0.344253 | 0.109723 | 0.856 | 0.838 | 1 | 10 | Atp5a1 | 0.018 | Old |
| 0.34455 | 0.603342 | 0.09 | 0.061 | 1 | 10 | Rbmx | 0.029 | Old |
| 0.344635 | 0.371126 | 0.308 | 0.263 | 1 | 10 | Xpo1 | 0.045 | Old |
| 0.345488 | 0.328278 | 0.078 | 0.051 | 1 | 10 | Serinc5 | 0.027 | Old |
| 0.345689 | 0.488877 | 0.147 | 0.111 | 1 | 10 | Prpf19 | 0.036 | Old |
| 0.345948 | 0.66759 | 0.174 | 0.141 | 1 | 10 | Pcyox1 | 0.033 | Old |
| 0.346057 | 1.358275 | 0.039 | 0.02 | 1 | 10 | Mapre3 | 0.019 | Old |
| 0.346083 | 0.269073 | 0.115 | 0.081 | 1 | 10 | Fiz1 | 0.034 | Old |
| 0.346791 | 0.011297 | 0.093 | 0.061 | 1 | 10 | Mfn1 | 0.032 | Old |
| 0.347305 | 0.214841 | 0.066 | 0.04 | 1 | 10 | Smad3 | 0.026 | Old |
| 0.347414 | 0.209704 | 0.186 | 0.141 | 1 | 10 | Letm1 | 0.045 | Old |
| 0.347666 | 0.114514 | 0.587 | 0.475 | 1 | 10 | Hnrnpl | 0.112 | Old |
| 0.347994 | 0.202896 | 0.054 | 0.03 | 1 | 10 | Gpatch1 | 0.024 | Old |
| 0.348129 | 0.033316 | 0.117 | 0.081 | 1 | 10 | Nelfb | 0.036 | Old |
| 0.348448 | 0.121547 | 0.213 | 0.162 | 1 | 10 | Dnm1l | 0.051 | Old |
| 0.348904 | 0.064333 | 0.758 | 0.616 | 1 | 10 | 5031439G( | 0.142 | Old |
| 0.348957 | 0.254095 | 0.271 | 0.222 | 1 | 10 | Rela | 0.049 | Old |
| 0.349279 | 0.24324 | 0.433 | 0.364 | 1 | 10 | Nr1h3 | 0.069 | Old |

| | | | | | | | | |
|---|---|---|---|---|---|---|---|---|
| 0.349703 | 0.082276 | 0.623 | 0.525 | 1 | 10 | Ube2d2a | 0.098 | Old |
| 0.35009 | 0.326561 | 0.054 | 0.03 | 1 | 10 | Zfp386 | 0.024 | Old |
| 0.350222 | 0.110011 | 0.291 | 0.242 | 1 | 10 | Ank | 0.049 | Old |
| 0.350307 | 0.173231 | 0.23 | 0.182 | 1 | 10 | Elmsan1 | 0.048 | Old |
| 0.350332 | 0.471567 | 0.169 | 0.131 | 1 | 10 | Zfp593 | 0.038 | Old |
| 0.350445 | 0.430196 | 0.337 | 0.293 | 1 | 10 | Arf2 | 0.044 | Old |
| 0.351257 | 0.733915 | 0.134 | 0.101 | 1 | 10 | Ctso | 0.033 | Old |
| 0.351497 | 0.244801 | 0.311 | 0.263 | 1 | 10 | Stx16 | 0.048 | Old |
| 0.352038 | 0.282604 | 0.152 | 0.111 | 1 | 10 | Gm11808 | 0.041 | Old |
| 0.352043 | 0.633182 | 0.066 | 0.04 | 1 | 10 | Cbr4 | 0.026 | Old |
| 0.352043 | 0.505149 | 0.066 | 0.04 | 1 | 10 | Zfp654 | 0.026 | Old |
| 0.352375 | 0.108477 | 0.65 | 0.556 | 1 | 10 | Tacc1 | 0.094 | Old |
| 0.353117 | 0.097476 | 0.127 | 0.091 | 1 | 10 | Ttc32 | 0.036 | Old |
| 0.353331 | 0.123733 | 0.477 | 0.404 | 1 | 10 | Arid1a | 0.073 | Old |
| 0.353423 | 1.393437 | 0.039 | 0.02 | 1 | 10 | Sirt4 | 0.019 | Old |
| 0.353423 | 1.13476 | 0.039 | 0.02 | 1 | 10 | Atg16l2 | 0.019 | Old |
| 0.353476 | 0.157962 | 0.386 | 0.333 | 1 | 10 | Aldh3b1 | 0.053 | Old |
| 0.353579 | 0.847193 | 0.11 | 0.081 | 1 | 10 | Cgrrf1 | 0.029 | Old |
| 0.353945 | 0.108975 | 0.369 | 0.313 | 1 | 10 | Srp72 | 0.056 | Old |
| 0.354129 | 0.356441 | 0.103 | 0.071 | 1 | 10 | Nup160 | 0.032 | Old |
| 0.354241 | 0.488975 | 0.112 | 0.081 | 1 | 10 | Wdr18 | 0.031 | Old |
| 0.354251 | 0.035455 | 0.308 | 0.253 | 1 | 10 | Plgrkt | 0.055 | Old |
| 0.354305 | 0.022095 | 0.054 | 0.03 | 1 | 10 | Ankrd26 | 0.024 | Old |
| 0.354307 | 0.327713 | 0.161 | 0.121 | 1 | 10 | Gbf1 | 0.04 | Old |
| 0.354577 | 0.288161 | 0.274 | 0.232 | 1 | 10 | Dapk1 | 0.042 | Old |
| 0.354906 | 0.077921 | 0.066 | 0.04 | 1 | 10 | Utp23 | 0.026 | Old |
| 0.355098 | 0.666478 | 0.078 | 0.051 | 1 | 10 | Txlng | 0.027 | Old |
| 0.355188 | 0.34585 | 0.159 | 0.121 | 1 | 10 | Dhdh | 0.038 | Old |
| 0.355212 | 0.043893 | 0.325 | 0.263 | 1 | 10 | Hs6st1 | 0.062 | Old |
| 0.355269 | 0.234145 | 0.125 | 0.091 | 1 | 10 | Nol8 | 0.034 | Old |
| 0.355342 | 0.18164 | 0.249 | 0.202 | 1 | 10 | Flvcr1 | 0.047 | Old |
| 0.355572 | 0.008058 | 0.161 | 0.121 | 1 | 10 | Eif2ak1 | 0.04 | Old |
| 0.355583 | 0.446021 | 0.193 | 0.152 | 1 | 10 | Atp1a3 | 0.041 | Old |
| 0.35602 | 0.18746 | 0.425 | 0.374 | 1 | 10 | Hacd4 | 0.051 | Old |
| 0.356036 | 0.134614 | 0.54 | 0.475 | 1 | 10 | Kdm7a | 0.065 | Old |
| 0.356098 | 0.333937 | 0.411 | 0.364 | 1 | 10 | Herpud1 | 0.047 | Old |
| 0.356621 | 0.162832 | 0.487 | 0.434 | 1 | 10 | Gm16286 | 0.053 | Old |
| 0.356674 | 1.334772 | 0.064 | 0.04 | 1 | 10 | Cdr2 | 0.024 | Old |
| 0.356718 | 0.138681 | 0.134 | 0.101 | 1 | 10 | Exosc8 | 0.033 | Old |
| 0.356788 | 0.03436 | 0.09 | 0.061 | 1 | 10 | Bcl2l12 | 0.029 | Old |
| 0.356822 | 0.377475 | 0.066 | 0.04 | 1 | 10 | Efr3b | 0.026 | Old |
| 0.35691 | 0.146489 | 0.428 | 0.364 | 1 | 10 | Ndufs2 | 0.064 | Old |
| 0.357143 | 1.061311 | 0.039 | 0.02 | 1 | 10 | Hdac6 | 0.019 | Old |
| 0.35737 | 0.294512 | 0.222 | 0.182 | 1 | 10 | Yipf1 | 0.04 | Old |
| 0.357613 | 0.376033 | 0.09 | 0.061 | 1 | 10 | Ncapg2 | 0.029 | Old |

| | | | | | | | | |
|---|---|---|---|---|---|---|---|---|
| 0.357801 | 0.40644 | 0.166 | 0.131 | 1 | 10 | Snx15 | 0.035 | Old |
| 0.357979 | 0.136385 | 0.156 | 0.121 | 1 | 10 | Lztfl1 | 0.035 | Old |
| 0.358705 | 0.005833 | 0.2 | 0.152 | 1 | 10 | Gm26740 | 0.048 | Old |
| 0.35884 | 0.128198 | 0.506 | 0.424 | 1 | 10 | Gm21188 | 0.082 | Old |
| 0.358923 | 0.120983 | 0.071 | 0.101 | 1 | 10 | Snhg20 | -0.03 | Old |
| 0.359128 | 0.124619 | 0.281 | 0.222 | 1 | 10 | Rictor | 0.059 | Old |
| 0.359637 | 0.867698 | 0.039 | 0.02 | 1 | 10 | Fasn | 0.019 | Old |
| 0.359637 | 0.809805 | 0.039 | 0.02 | 1 | 10 | Nmnat1 | 0.019 | Old |
| 0.359965 | 0.122421 | 0.535 | 0.475 | 1 | 10 | Ankrd11 | 0.06 | Old |
| 0.360023 | 0.130264 | 0.161 | 0.121 | 1 | 10 | Lats2 | 0.04 | Old |
| 0.360097 | 0.234928 | 0.09 | 0.061 | 1 | 10 | Lztr1 | 0.029 | Old |
| 0.360302 | 0.436443 | 0.159 | 0.121 | 1 | 10 | S100pbp | 0.038 | Old |
| 0.360541 | 0.546292 | 0.144 | 0.111 | 1 | 10 | Tmem131l | 0.033 | Old |
| 0.360611 | 0.064224 | 0.208 | 0.162 | 1 | 10 | Irs2 | 0.046 | Old |
| 0.360759 | 0.19843 | 0.115 | 0.081 | 1 | 10 | Setd1a | 0.034 | Old |
| 0.360785 | 0.251755 | 0.147 | 0.111 | 1 | 10 | Lnpk | 0.036 | Old |
| 0.360887 | 1.058517 | 0.039 | 0.02 | 1 | 10 | Prepl | 0.019 | Old |
| 0.360887 | 0.991415 | 0.039 | 0.02 | 1 | 10 | Fxn | 0.019 | Old |
| 0.361078 | 0.210306 | 0.247 | 0.202 | 1 | 10 | Madd | 0.045 | Old |
| 0.361195 | 0.342826 | 0.191 | 0.152 | 1 | 10 | Fbrs | 0.039 | Old |
| 0.361506 | 0.407619 | 0.254 | 0.212 | 1 | 10 | Uba5 | 0.042 | Old |
| 0.361522 | 0.457468 | 0.22 | 0.182 | 1 | 10 | Pih1d1 | 0.038 | Old |
| 0.361566 | 0.119033 | 0.523 | 0.465 | 1 | 10 | Zfp706 | 0.058 | Old |
| 0.3618 | 0.30131 | 0.271 | 0.222 | 1 | 10 | Rnf44 | 0.049 | Old |
| 0.362034 | 0.187216 | 0.296 | 0.242 | 1 | 10 | Tbc1d14 | 0.054 | Old |
| 0.362141 | 0.524373 | 0.039 | 0.02 | 1 | 10 | Eva1b | 0.019 | Old |
| 0.362141 | 0.519299 | 0.039 | 0.02 | 1 | 10 | Cep95 | 0.019 | Old |
| 0.362386 | 0.844223 | 0.076 | 0.051 | 1 | 10 | Slc2a8 | 0.025 | Old |
| 0.36241 | 0.118612 | 0.244 | 0.192 | 1 | 10 | Agpat4 | 0.052 | Old |
| 0.362551 | 0.706265 | 0.064 | 0.04 | 1 | 10 | Zbtb34 | 0.024 | Old |
| 0.363522 | 0.542382 | 0.159 | 0.121 | 1 | 10 | Hps5 | 0.038 | Old |
| 0.363581 | 0.050576 | 0.066 | 0.04 | 1 | 10 | Isca1 | 0.026 | Old |
| 0.364168 | 0.402549 | 0.159 | 0.121 | 1 | 10 | Yif1a | 0.038 | Old |
| 0.364656 | 0.595937 | 0.039 | 0.02 | 1 | 10 | Ifitm1 | 0.019 | Old |
| 0.364656 | 0.548484 | 0.039 | 0.02 | 1 | 10 | Il4i1 | 0.019 | Old |
| 0.36501 | 0.060985 | 0.134 | 0.101 | 1 | 10 | Prpsap1 | 0.033 | Old |
| 0.365066 | 0.033687 | 0.103 | 0.071 | 1 | 10 | Dcaf5 | 0.032 | Old |
| 0.365527 | 0.322758 | 0.066 | 0.04 | 1 | 10 | Polr1b | 0.026 | Old |
| 0.365805 | 0.158353 | 0.548 | 0.495 | 1 | 10 | Taok1 | 0.053 | Old |
| 0.365918 | 0.696422 | 0.039 | 0.02 | 1 | 10 | D2hgdh | 0.019 | Old |
| 0.365918 | 0.167632 | 0.039 | 0.02 | 1 | 10 | Nprl3 | 0.019 | Old |
| 0.366102 | 0.327837 | 0.257 | 0.212 | 1 | 10 | Rnf6 | 0.045 | Old |
| 0.366477 | 0.241165 | 0.381 | 0.323 | 1 | 10 | Park7 | 0.058 | Old |
| 0.366833 | 0.10454 | 0.491 | 0.424 | 1 | 10 | Cers6 | 0.067 | Old |
| 0.36696 | 0.088721 | 0.298 | 0.242 | 1 | 10 | Nin | 0.056 | Old |

| | | | | | | | | |
|---|---|---|---|---|---|---|---|---|
| 0.36703 | 0.682648 | 0.142 | 0.111 | 1 | 10 | Mrps34 | 0.031 | Old |
| 0.367443 | 0.281226 | 0.125 | 0.091 | 1 | 10 | Tmem63a | 0.034 | Old |
| 0.367475 | 0.462776 | 0.147 | 0.111 | 1 | 10 | Ythdc2 | 0.036 | Old |
| 0.367674 | 1.11249 | 0.051 | 0.03 | 1 | 10 | Fdxr | 0.021 | Old |
| 0.368315 | 0.417641 | 0.171 | 0.131 | 1 | 10 | Nufip1 | 0.04 | Old |
| 0.36845 | 0.748418 | 0.039 | 0.02 | 1 | 10 | 2700062C0 | 0.019 | Old |
| 0.368489 | 0.120941 | 0.289 | 0.242 | 1 | 10 | Map3k11 | 0.047 | Old |
| 0.368496 | 0.208372 | 0.112 | 0.081 | 1 | 10 | Ankrd10 | 0.031 | Old |
| 0.368559 | 0.474648 | 0.1 | 0.071 | 1 | 10 | 1700025G0 | 0.029 | Old |
| 0.368733 | 0.140951 | 0.391 | 0.333 | 1 | 10 | Dnase1l1 | 0.058 | Old |
| 0.368845 | 0.009315 | 0.548 | 0.465 | 1 | 10 | Npc1 | 0.083 | Old |
| 0.368924 | 0.050791 | 0.998 | 1 | 1 | 10 | Eef1a1 | -0.002 | Old |
| 0.368999 | 0.556752 | 0.122 | 0.091 | 1 | 10 | Tgfbrap1 | 0.031 | Old |
| 0.369484 | 1.032393 | 0.064 | 0.04 | 1 | 10 | Capn15 | 0.024 | Old |
| 0.36972 | 0.523737 | 0.039 | 0.02 | 1 | 10 | Prdm10 | 0.019 | Old |
| 0.369886 | 0.384749 | 0.051 | 0.03 | 1 | 10 | Gadd45a | 0.021 | Old |
| 0.369956 | 0.022802 | 0.526 | 0.455 | 1 | 10 | Azin1 | 0.071 | Old |
| 0.370215 | 0.411387 | 0.264 | 0.222 | 1 | 10 | Arhgap15 | 0.042 | Old |
| 0.37032 | 0.207061 | 0.403 | 0.354 | 1 | 10 | Pold4 | 0.049 | Old |
| 0.370993 | 0.490141 | 0.039 | 0.02 | 1 | 10 | Tigar | 0.019 | Old |
| 0.370993 | 0.411224 | 0.039 | 0.02 | 1 | 10 | Sars2 | 0.019 | Old |
| 0.371496 | 0.775186 | 0.076 | 0.051 | 1 | 10 | Triobp | 0.025 | Old |
| 0.371633 | 0.403241 | 0.127 | 0.091 | 1 | 10 | Ercc3 | 0.036 | Old |
| 0.371949 | 1.801953 | 0.024 | 0.01 | 1 | 10 | Rbm12b1 | 0.014 | Old |
| 0.371949 | 1.633714 | 0.024 | 0.01 | 1 | 10 | Zfp456 | 0.014 | Old |
| 0.372106 | 1.212414 | 0.051 | 0.03 | 1 | 10 | Pomk | 0.021 | Old |
| 0.372137 | 0.172826 | 0.511 | 0.465 | 1 | 10 | Dot1l | 0.046 | Old |
| 0.372268 | 0.551695 | 0.039 | 0.02 | 1 | 10 | BC017158 | 0.019 | Old |
| 0.372305 | 0.496941 | 0.112 | 0.081 | 1 | 10 | Il10 | 0.031 | Old |
| 0.372716 | 0.171576 | 0.134 | 0.101 | 1 | 10 | Snx16 | 0.033 | Old |
| 0.373151 | 0.028411 | 0.839 | 0.768 | 1 | 10 | Cd47 | 0.071 | Old |
| 0.373202 | 0.270881 | 0.347 | 0.293 | 1 | 10 | Znhit1 | 0.054 | Old |
| 0.373219 | 0.946024 | 0.051 | 0.03 | 1 | 10 | Gprasp1 | 0.021 | Old |
| 0.373461 | 0.197299 | 0.284 | 0.232 | 1 | 10 | Cdc26 | 0.052 | Old |
| 0.373572 | 1.651773 | 0.024 | 0.01 | 1 | 10 | Aste1 | 0.014 | Old |
| 0.373572 | 1.551677 | 0.024 | 0.01 | 1 | 10 | Shisa9 | 0.014 | Old |
| 0.373572 | 1.502491 | 0.024 | 0.01 | 1 | 10 | Pkd1l2 | 0.014 | Old |
| 0.373572 | 1.471763 | 0.024 | 0.01 | 1 | 10 | Rarb | 0.014 | Old |
| 0.373572 | 1.209815 | 0.024 | 0.01 | 1 | 10 | Trim68 | 0.014 | Old |
| 0.374334 | 0.857693 | 0.051 | 0.03 | 1 | 10 | Itgb3bp | 0.021 | Old |
| 0.374634 | 0.005776 | 0.966 | 0.96 | 1 | 10 | Tyrobp | 0.006 | Old |
| 0.375018 | 0.472546 | 0.259 | 0.222 | 1 | 10 | Zzef1 | 0.037 | Old |
| 0.3752 | 1.391699 | 0.024 | 0.01 | 1 | 10 | Rgl3 | 0.014 | Old |
| 0.3752 | 1.175427 | 0.024 | 0.01 | 1 | 10 | Zfp566 | 0.014 | Old |
| 0.3752 | 1.058508 | 0.024 | 0.01 | 1 | 10 | Ttc30b | 0.014 | Old |

| | | | | | | | | |
|---|---|---|---|---|---|---|---|---|
| 0.375387 | 0.114194 | 0.281 | 0.232 | 1 | 10 | Ccdc47 | 0.049 | Old |
| 0.375696 | 0.338726 | 0.178 | 0.141 | 1 | 10 | Iars | 0.037 | Old |
| 0.37611 | 0.453683 | 0.039 | 0.02 | 1 | 10 | Zfp810 | 0.019 | Old |
| 0.37611 | 0.180651 | 0.039 | 0.02 | 1 | 10 | Nudt22 | 0.019 | Old |
| 0.376157 | 0.919535 | 0.098 | 0.071 | 1 | 10 | Ccdc97 | 0.027 | Old |
| 0.376354 | 0.542546 | 0.122 | 0.091 | 1 | 10 | Cinp | 0.031 | Old |
| 0.37657 | 0.747935 | 0.051 | 0.03 | 1 | 10 | Gapt | 0.021 | Old |
| 0.376669 | 0.367584 | 0.381 | 0.343 | 1 | 10 | Cfh | 0.038 | Old |
| 0.376832 | 1.205854 | 0.024 | 0.01 | 1 | 10 | 3222401L1 | 0.014 | Old |
| 0.376832 | 1.155476 | 0.024 | 0.01 | 1 | 10 | Cd59a | 0.014 | Old |
| 0.376832 | 1.086731 | 0.024 | 0.01 | 1 | 10 | Tesk2 | 0.014 | Old |
| 0.376836 | 0.204381 | 0.311 | 0.263 | 1 | 10 | Mrps24 | 0.048 | Old |
| 0.377165 | 0.188875 | 0.482 | 0.424 | 1 | 10 | Ranbp2 | 0.058 | Old |
| 0.377397 | 0.10044 | 0.039 | 0.02 | 1 | 10 | Plcxd2 | 0.019 | Old |
| 0.377397 | 0.002997 | 0.039 | 0.02 | 1 | 10 | Hexdc | 0.019 | Old |
| 0.377542 | 0.258243 | 0.753 | 0.697 | 1 | 10 | Aprt | 0.056 | Old |
| 0.377576 | 0.168053 | 0.518 | 0.444 | 1 | 10 | Frmd4b | 0.074 | Old |
| 0.377672 | 0.125478 | 0.134 | 0.101 | 1 | 10 | Tbc1d10a | 0.033 | Old |
| 0.377692 | 0.452999 | 0.051 | 0.03 | 1 | 10 | Cbr3 | 0.021 | Old |
| 0.377956 | 0.665952 | 0.076 | 0.051 | 1 | 10 | Tmem29 | 0.025 | Old |
| 0.377956 | 0.257702 | 0.076 | 0.051 | 1 | 10 | Frs2 | 0.025 | Old |
| 0.378131 | 0.282751 | 0.271 | 0.222 | 1 | 10 | Mbp | 0.049 | Old |
| 0.378468 | 1.080502 | 0.024 | 0.01 | 1 | 10 | Rab3ip | 0.014 | Old |
| 0.378468 | 0.98523 | 0.024 | 0.01 | 1 | 10 | Tmem62 | 0.014 | Old |
| 0.378686 | 0.2322 | 0.039 | 0.02 | 1 | 10 | Zfp1 | 0.019 | Old |
| 0.379428 | 0.138291 | 0.824 | 0.758 | 1 | 10 | Snx5 | 0.066 | Old |
| 0.379733 | 0.226448 | 0.164 | 0.121 | 1 | 10 | Polr1a | 0.043 | Old |
| 0.379782 | 0.162508 | 0.171 | 0.131 | 1 | 10 | Pik3cb | 0.04 | Old |
| 0.37994 | 0.676631 | 0.051 | 0.03 | 1 | 10 | Pcyox1l | 0.021 | Old |
| 0.37994 | 0.624944 | 0.051 | 0.03 | 1 | 10 | Tbc1d16 | 0.021 | Old |
| 0.379976 | 0.022877 | 0.509 | 0.434 | 1 | 10 | Rassf2 | 0.075 | Old |
| 0.379977 | 0.218376 | 0.039 | 0.02 | 1 | 10 | Gna11 | 0.019 | Old |
| 0.380005 | 0.005724 | 0.244 | 0.192 | 1 | 10 | Prdx4 | 0.052 | Old |
| 0.380109 | 1.191298 | 0.024 | 0.01 | 1 | 10 | Chd3os | 0.014 | Old |
| 0.380109 | 1.013025 | 0.024 | 0.01 | 1 | 10 | Xpo5 | 0.014 | Old |
| 0.380417 | 0.692828 | 0.203 | 0.172 | 1 | 10 | Snx24 | 0.031 | Old |
| 0.381068 | 0.314286 | 0.051 | 0.03 | 1 | 10 | Abcg2 | 0.021 | Old |
| 0.381085 | 0.131993 | 0.147 | 0.111 | 1 | 10 | Trappc13 | 0.036 | Old |
| 0.381417 | 0.009735 | 0.35 | 0.293 | 1 | 10 | Cdipt | 0.057 | Old |
| 0.38151 | 0.30096 | 0.1 | 0.071 | 1 | 10 | Zfp362 | 0.029 | Old |
| 0.381679 | 0.494287 | 0.076 | 0.051 | 1 | 10 | Ripk3 | 0.025 | Old |
| 0.381754 | 1.708748 | 0.024 | 0.01 | 1 | 10 | Dmpk | 0.014 | Old |
| 0.381754 | 1.113685 | 0.024 | 0.01 | 1 | 10 | Fgfbp3 | 0.014 | Old |
| 0.381754 | 0.740342 | 0.024 | 0.01 | 1 | 10 | Ankrd13b | 0.014 | Old |
| 0.381754 | 0.682062 | 0.024 | 0.01 | 1 | 10 | Rfxank | 0.014 | Old |

| | | | | | | | | |
|---|---|---|---|---|---|---|---|---|
| 0.38185 | 0.082939 | 0.247 | 0.202 | 1 | 10 | Chka | 0.045 | Old |
| 0.381972 | 0.256984 | 0.225 | 0.182 | 1 | 10 | Ostm1 | 0.043 | Old |
| 0.382654 | 0.612779 | 0.11 | 0.081 | 1 | 10 | Dnttip1 | 0.029 | Old |
| 0.382703 | 0.436081 | 0.132 | 0.101 | 1 | 10 | Pds5b | 0.031 | Old |
| 0.382955 | 0.079081 | 0.23 | 0.182 | 1 | 10 | Brpf1 | 0.048 | Old |
| 0.383036 | 0.123489 | 0.88 | 0.838 | 1 | 10 | Hspa5 | 0.042 | Old |
| 0.383385 | 0.28332 | 0.134 | 0.101 | 1 | 10 | Slc25a20 | 0.033 | Old |
| 0.383404 | 0.868741 | 0.024 | 0.01 | 1 | 10 | Ly6a | 0.014 | Old |
| 0.383404 | 0.86619 | 0.024 | 0.01 | 1 | 10 | Dph2 | 0.014 | Old |
| 0.383404 | 0.762254 | 0.024 | 0.01 | 1 | 10 | Gm31597 | 0.014 | Old |
| 0.383404 | 0.72403 | 0.024 | 0.01 | 1 | 10 | Zfp606 | 0.014 | Old |
| 0.383404 | 0.702612 | 0.024 | 0.01 | 1 | 10 | Tmem143 | 0.014 | Old |
| 0.383404 | 0.609597 | 0.024 | 0.01 | 1 | 10 | Cenpv | 0.014 | Old |
| 0.383549 | 0.554363 | 0.076 | 0.051 | 1 | 10 | Gabpb1 | 0.025 | Old |
| 0.383594 | 0.720946 | 0.064 | 0.04 | 1 | 10 | Lrrc57 | 0.024 | Old |
| 0.383734 | 0.497128 | 0.144 | 0.111 | 1 | 10 | Fdft1 | 0.033 | Old |
| 0.384094 | 0.276859 | 0.333 | 0.293 | 1 | 10 | Mrpl24 | 0.04 | Old |
| 0.384096 | 0.060862 | 0.289 | 0.242 | 1 | 10 | Nfya | 0.047 | Old |
| 0.384462 | 0.467806 | 0.051 | 0.03 | 1 | 10 | Zfp367 | 0.021 | Old |
| 0.384791 | 0.548334 | 0.1 | 0.071 | 1 | 10 | Eipr1 | 0.029 | Old |
| 0.385057 | 0.701358 | 0.024 | 0.01 | 1 | 10 | Tmem221 | 0.014 | Old |
| 0.385057 | 0.67971 | 0.024 | 0.01 | 1 | 10 | Spata13 | 0.014 | Old |
| 0.385057 | 0.652184 | 0.024 | 0.01 | 1 | 10 | Nthl1 | 0.014 | Old |
| 0.385057 | 0.623411 | 0.024 | 0.01 | 1 | 10 | Zc4h2 | 0.014 | Old |
| 0.385524 | 0.040314 | 0.09 | 0.061 | 1 | 10 | Fzd1 | 0.029 | Old |
| 0.385598 | 0.168273 | 0.051 | 0.03 | 1 | 10 | Hirip3 | 0.021 | Old |
| 0.386356 | 0.136799 | 0.279 | 0.232 | 1 | 10 | Chd2 | 0.047 | Old |
| 0.38637 | 0.098854 | 0.264 | 0.222 | 1 | 10 | Lta4h | 0.042 | Old |
| 0.386553 | 0.260417 | 0.215 | 0.172 | 1 | 10 | Tug1 | 0.043 | Old |
| 0.386586 | 0.592553 | 0.11 | 0.081 | 1 | 10 | Sun1 | 0.029 | Old |
| 0.386664 | 0.090092 | 0.152 | 0.192 | 1 | 10 | Tesk1 | -0.04 | Old |
| 0.386715 | 0.50811 | 0.024 | 0.01 | 1 | 10 | 1110002J0 | 0.014 | Old |
| 0.386715 | 0.409582 | 0.024 | 0.01 | 1 | 10 | Mtfmt | 0.014 | Old |
| 0.386715 | 0.277744 | 0.024 | 0.01 | 1 | 10 | Vps51 | 0.014 | Old |
| 0.386715 | 0.01345 | 0.024 | 0.01 | 1 | 10 | Akr1c12 | 0.014 | Old |
| 0.386736 | 0.419826 | 0.051 | 0.03 | 1 | 10 | Crebl2 | 0.021 | Old |
| 0.386812 | 0.194778 | 0.244 | 0.202 | 1 | 10 | Ifi203 | 0.042 | Old |
| 0.387208 | 0.021053 | 0.369 | 0.313 | 1 | 10 | Atg3 | 0.056 | Old |
| 0.387305 | 0.584174 | 0.076 | 0.051 | 1 | 10 | Ppp1r16a | 0.025 | Old |
| 0.387443 | 0.174509 | 0.086 | 0.061 | 1 | 10 | Cog4 | 0.025 | Old |
| 0.387546 | 0.127623 | 0.171 | 0.131 | 1 | 10 | Zbtb1 | 0.04 | Old |
| 0.387686 | 0.974005 | 0.064 | 0.04 | 1 | 10 | Serpinb9 | 0.024 | Old |
| 0.388166 | 0.657836 | 0.11 | 0.081 | 1 | 10 | Dnajc16 | 0.029 | Old |
| 0.388166 | 0.254996 | 0.11 | 0.081 | 1 | 10 | Pot1a | 0.029 | Old |
| 0.388166 | 0.033297 | 0.11 | 0.081 | 1 | 10 | Ifitm6 | 0.029 | Old |

| | | | | | | | | |
|---|---|---|---|---|---|---|---|---|
| 0.388378 | 0.473867 | 0.024 | 0.01 | 1 | 10 | Fgf13 | 0.014 | Old |
| 0.388378 | 0.224551 | 0.024 | 0.01 | 1 | 10 | Bex6 | 0.014 | Old |
| 0.388378 | 0.024336 | 0.024 | 0.01 | 1 | 10 | Slc37a3 | 0.014 | Old |
| 0.388713 | 0.518561 | 0.064 | 0.04 | 1 | 10 | Actr6 | 0.024 | Old |
| 0.389192 | 0.295202 | 0.076 | 0.051 | 1 | 10 | Fam78b | 0.025 | Old |
| 0.38927 | 0.266253 | 0.423 | 0.374 | 1 | 10 | Zbtb20 | 0.049 | Old |
| 0.390276 | 0.074258 | 0.293 | 0.232 | 1 | 10 | Hivep2 | 0.061 | Old |
| 0.390586 | 0.401855 | 0.364 | 0.323 | 1 | 10 | Tle3 | 0.041 | Old |
| 0.390694 | 0.260087 | 0.01 | 0.02 | 1 | 10 | Ch25h | -0.01 | Old |
| 0.390794 | 0.173509 | 0.147 | 0.111 | 1 | 10 | Abcf3 | 0.036 | Old |
| 0.390893 | 0.031813 | 0.364 | 0.293 | 1 | 10 | Mapk14 | 0.071 | Old |
| 0.390962 | 0.047967 | 0.526 | 0.444 | 1 | 10 | Sptssa | 0.082 | Old |
| 0.391002 | 0.805199 | 0.086 | 0.061 | 1 | 10 | Arhgef11 | 0.025 | Old |
| 0.391084 | 0.378258 | 0.076 | 0.051 | 1 | 10 | St8sia6 | 0.025 | Old |
| 0.391307 | 0.22531 | 0.051 | 0.03 | 1 | 10 | Slc37a4 | 0.021 | Old |
| 0.39146 | 0.079933 | 0.235 | 0.192 | 1 | 10 | Pigs | 0.043 | Old |
| 0.391537 | 0.109378 | 0.878 | 0.808 | 1 | 10 | Clta | 0.07 | Old |
| 0.391804 | 0.452786 | 0.064 | 0.04 | 1 | 10 | Mcm2 | 0.024 | Old |
| 0.39192 | 0.099944 | 0.137 | 0.101 | 1 | 10 | Sipa1l3 | 0.036 | Old |
| 0.392263 | 0.08482 | 0.787 | 0.737 | 1 | 10 | Capg | 0.05 | Old |
| 0.392686 | 0.19049 | 0.457 | 0.404 | 1 | 10 | Tnpo3 | 0.053 | Old |
| 0.392699 | 0.03626 | 0.43 | 0.354 | 1 | 10 | Atf6 | 0.076 | Old |
| 0.392789 | 0.775293 | 0.086 | 0.061 | 1 | 10 | Epg5 | 0.025 | Old |
| 0.392837 | 0.403136 | 0.064 | 0.04 | 1 | 10 | Gstcd | 0.024 | Old |
| 0.39285 | 0.525706 | 0.122 | 0.091 | 1 | 10 | Arntl | 0.031 | Old |
| 0.393069 | 0.326306 | 0.1 | 0.071 | 1 | 10 | Ubxn2a | 0.029 | Old |
| 0.39425 | 0.391149 | 0.144 | 0.111 | 1 | 10 | Ptbp2 | 0.033 | Old |
| 0.394495 | 0.354188 | 0.2 | 0.162 | 1 | 10 | Ddx10 | 0.038 | Old |
| 0.394562 | 0.10358 | 0.741 | 0.636 | 1 | 10 | Atp6v0e | 0.105 | Old |
| 0.394665 | 0.143511 | 0.628 | 0.556 | 1 | 10 | Prrc2c | 0.072 | Old |
| 0.39491 | 0.299027 | 0.064 | 0.04 | 1 | 10 | Dynlt1f | 0.024 | Old |
| 0.394915 | 0.787427 | 0.108 | 0.081 | 1 | 10 | Ndufaf3 | 0.027 | Old |
| 0.395133 | 0.319548 | 0.122 | 0.091 | 1 | 10 | Ppfia4 | 0.031 | Old |
| 0.395211 | 0.458742 | 0.01 | 0.02 | 1 | 10 | Arl4d | -0.01 | Old |
| 0.395211 | 0.414899 | 0.01 | 0.02 | 1 | 10 | Jmjd4 | -0.01 | Old |
| 0.395211 | 0.232202 | 0.01 | 0.02 | 1 | 10 | Jchain | -0.01 | Old |
| 0.395948 | 0.177823 | 0.064 | 0.04 | 1 | 10 | Nsun4 | 0.024 | Old |
| 0.396074 | 0.032507 | 0.139 | 0.101 | 1 | 10 | Aga | 0.038 | Old |
| 0.396194 | 0.103042 | 0.281 | 0.232 | 1 | 10 | Dhx36 | 0.049 | Old |
| 0.396467 | 0.210109 | 0.112 | 0.081 | 1 | 10 | Bcl2l2 | 0.031 | Old |
| 0.397114 | 0.312761 | 0.147 | 0.111 | 1 | 10 | Jmjd6 | 0.036 | Old |
| 0.397482 | 0.040926 | 0.01 | 0.02 | 1 | 10 | Fam71f2 | -0.01 | Old |
| 0.397551 | 0.422641 | 0.279 | 0.242 | 1 | 10 | Nsfl1c | 0.037 | Old |
| 0.397752 | 0.314129 | 0.227 | 0.192 | 1 | 10 | Lage3 | 0.035 | Old |
| 0.398145 | 1.418144 | 0.037 | 0.02 | 1 | 10 | Plxdc1 | 0.017 | Old |

| | | | | | | | | |
|---|---|---|---|---|---|---|---|---|
| 0.399063 | 0.033187 | 0.198 | 0.162 | 1 | 10 | Armcx3 | 0.036 | Old |
| 0.399182 | 0.270102 | 0.21 | 0.172 | 1 | 10 | Idnk | 0.038 | Old |
| 0.399184 | 0.013418 | 0.137 | 0.101 | 1 | 10 | Nudt2 | 0.036 | Old |
| 0.399514 | 1.681233 | 0.037 | 0.02 | 1 | 10 | Etfbkmt | 0.017 | Old |
| 0.399514 | 1.368991 | 0.037 | 0.02 | 1 | 10 | Adora2a | 0.017 | Old |
| 0.399591 | 0.379803 | 0.056 | 0.081 | 1 | 10 | Ap3m2 | -0.025 | Old |
| 0.39979 | 0.295553 | 0.21 | 0.172 | 1 | 10 | Tmem205 | 0.038 | Old |
| 0.399895 | 0.265176 | 0.191 | 0.152 | 1 | 10 | Sap30 | 0.039 | Old |
| 0.400119 | 0.060633 | 0.33 | 0.273 | 1 | 10 | Cmklr1 | 0.057 | Old |
| 0.400119 | 0.088306 | 0.064 | 0.04 | 1 | 10 | Fbxw9 | 0.024 | Old |
| 0.400547 | 0.283843 | 0.051 | 0.03 | 1 | 10 | Rmdn1 | 0.021 | Old |
| 0.400826 | 0.211909 | 0.477 | 0.455 | 1 | 10 | Klf3 | 0.022 | Old |
| 0.400886 | 1.568088 | 0.037 | 0.02 | 1 | 10 | Surf6 | 0.017 | Old |
| 0.400968 | 0.035733 | 0.394 | 0.333 | 1 | 10 | Sp100 | 0.061 | Old |
| 0.401298 | 0.04698 | 0.914 | 0.879 | 1 | 10 | Cltc | 0.035 | Old |
| 0.401394 | 0.092134 | 0.936 | 0.889 | 1 | 10 | Rpl36 | 0.047 | Old |
| 0.401986 | 0.114906 | 0.335 | 0.273 | 1 | 10 | Kmt2c | 0.062 | Old |
| 0.402027 | 0.152253 | 0.196 | 0.152 | 1 | 10 | Enox2 | 0.044 | Old |
| 0.40226 | 1.312134 | 0.037 | 0.02 | 1 | 10 | Cd276 | 0.017 | Old |
| 0.402612 | 0.138211 | 0.259 | 0.212 | 1 | 10 | Ide | 0.047 | Old |
| 0.402746 | 0.102358 | 0.269 | 0.222 | 1 | 10 | Eif4ebp1 | 0.047 | Old |
| 0.40277 | 1.476192 | 0.049 | 0.03 | 1 | 10 | Aldh6a1 | 0.019 | Old |
| 0.4028 | 0.135487 | 0.122 | 0.091 | 1 | 10 | Dolpp1 | 0.031 | Old |
| 0.403503 | 0.553306 | 0.144 | 0.111 | 1 | 10 | Tmem37 | 0.033 | Old |
| 0.403774 | 0.079847 | 0.296 | 0.242 | 1 | 10 | Prrc2b | 0.054 | Old |
| 0.403902 | 0.049055 | 0.252 | 0.202 | 1 | 10 | Rmnd5a | 0.05 | Old |
| 0.404176 | 1.039667 | 0.073 | 0.051 | 1 | 10 | Fam136a | 0.022 | Old |
| 0.404279 | 0.098178 | 0.017 | 0.03 | 1 | 10 | Bdh2 | -0.013 | Old |
| 0.404378 | 0.093874 | 0.46 | 0.394 | 1 | 10 | Larp1 | 0.066 | Old |
| 0.404409 | 0.18766 | 0.227 | 0.182 | 1 | 10 | Cdip1 | 0.045 | Old |
| 0.404523 | 0.815937 | 0.086 | 0.061 | 1 | 10 | Kat5 | 0.025 | Old |
| 0.404523 | 0.486805 | 0.086 | 0.061 | 1 | 10 | Pafah1b3 | 0.025 | Old |
| 0.404627 | 0.201599 | 0.098 | 0.071 | 1 | 10 | Ccdc32 | 0.027 | Old |
| 0.404921 | 0.549857 | 0.164 | 0.131 | 1 | 10 | Pdlim1 | 0.033 | Old |
| 0.405018 | 0.991203 | 0.037 | 0.02 | 1 | 10 | Zfp182 | 0.017 | Old |
| 0.405369 | 0.136105 | 0.064 | 0.04 | 1 | 10 | Prmt6 | 0.024 | Old |
| 0.405435 | 0.728348 | 0.086 | 0.061 | 1 | 10 | A530017D2 | 0.025 | Old |
| 0.405452 | 0.195807 | 0.076 | 0.051 | 1 | 10 | Dxo | 0.025 | Old |
| 0.405716 | 0.221077 | 0.386 | 0.343 | 1 | 10 | Naa20 | 0.043 | Old |
| 0.405892 | 0.292229 | 0.122 | 0.091 | 1 | 10 | Ttc4 | 0.031 | Old |
| 0.405921 | 0.149358 | 0.311 | 0.263 | 1 | 10 | Stxbp2 | 0.048 | Old |
| 0.406069 | 0.265456 | 0.017 | 0.03 | 1 | 10 | Kif23 | -0.013 | Old |
| 0.406352 | 1.396671 | 0.049 | 0.03 | 1 | 10 | Gm20604 | 0.019 | Old |
| 0.406525 | 0.424216 | 0.269 | 0.232 | 1 | 10 | Cln3 | 0.037 | Old |
| 0.40755 | 1.133457 | 0.049 | 0.03 | 1 | 10 | Cdpf1 | 0.019 | Old |

| | | | | | | | | |
|---|---|---|---|---|---|---|---|---|
| 0.407683 | 0.112537 | 0.112 | 0.081 | 1 | 10 | Fbxl6 | 0.031 | Old |
| 0.407787 | 1.359672 | 0.037 | 0.02 | 1 | 10 | BC002059 | 0.017 | Old |
| 0.407863 | 0.029009 | 0.017 | 0.03 | 1 | 10 | Gm6133 | -0.013 | Old |
| 0.409007 | 0.351609 | 0.164 | 0.131 | 1 | 10 | Nfx1 | 0.033 | Old |
| 0.409077 | 0.930252 | 0.061 | 0.04 | 1 | 10 | Oasl1 | 0.021 | Old |
| 0.409274 | 0.66594 | 0.142 | 0.111 | 1 | 10 | Rnpc3 | 0.031 | Old |
| 0.409662 | 0.085489 | 0.017 | 0.03 | 1 | 10 | I830077J02 | -0.013 | Old |
| 0.409672 | 0.037228 | 0.523 | 0.475 | 1 | 10 | Ppp2r5a | 0.048 | Old |
| 0.410154 | 0.99123 | 0.061 | 0.04 | 1 | 10 | Zbtb33 | 0.021 | Old |
| 0.410567 | 0.99888 | 0.037 | 0.02 | 1 | 10 | Rnf26 | 0.017 | Old |
| 0.410818 | 0.327883 | 0.303 | 0.263 | 1 | 10 | Col4a3bp | 0.04 | Old |
| 0.410838 | 0.218643 | 0.193 | 0.152 | 1 | 10 | Eif1ad | 0.041 | Old |
| 0.411016 | 0.032948 | 0.418 | 0.333 | 1 | 10 | Pura | 0.085 | Old |
| 0.41127 | 0.098057 | 0.161 | 0.121 | 1 | 10 | Mrto4 | 0.04 | Old |
| 0.411317 | 0.208629 | 0.2 | 0.162 | 1 | 10 | Pisd | 0.038 | Old |
| 0.411711 | 0.076145 | 0.575 | 0.485 | 1 | 10 | H13 | 0.09 | Old |
| 0.411724 | 0.05554 | 0.064 | 0.04 | 1 | 10 | Rhobtb1 | 0.024 | Old |
| 0.411735 | 0.0177 | 0.156 | 0.121 | 1 | 10 | Mta1 | 0.035 | Old |
| 0.411926 | 0.147862 | 0.826 | 0.778 | 1 | 10 | Uqcr11 | 0.048 | Old |
| 0.411961 | 0.931515 | 0.037 | 0.02 | 1 | 10 | Nup37 | 0.017 | Old |
| 0.412169 | 0.965574 | 0.095 | 0.071 | 1 | 10 | 6030458C1 | 0.024 | Old |
| 0.412436 | 0.151177 | 0.156 | 0.121 | 1 | 10 | Senp7 | 0.035 | Old |
| 0.412456 | 0.283468 | 0.301 | 0.263 | 1 | 10 | Samd8 | 0.038 | Old |
| 0.412704 | 0.076441 | 0.971 | 0.97 | 1 | 10 | Rack1 | 0.001 | Old |
| 0.412852 | 0.088944 | 0.237 | 0.192 | 1 | 10 | Tmem141 | 0.045 | Old |
| 0.412886 | 0.58429 | 0.108 | 0.081 | 1 | 10 | Ttc7b | 0.027 | Old |
| 0.412889 | 0.232956 | 0.198 | 0.162 | 1 | 10 | Nifk | 0.036 | Old |
| 0.41292 | 0.387217 | 0.142 | 0.111 | 1 | 10 | Adk | 0.031 | Old |
| 0.413273 | 0.155521 | 0.017 | 0.03 | 1 | 10 | Ccdc106 | -0.013 | Old |
| 0.413343 | 0.260473 | 0.232 | 0.192 | 1 | 10 | Supt6 | 0.04 | Old |
| 0.413652 | 0.625137 | 0.142 | 0.111 | 1 | 10 | Ing5 | 0.031 | Old |
| 0.414039 | 0.709213 | 0.073 | 0.051 | 1 | 10 | Fen1 | 0.022 | Old |
| 0.414351 | 0.083779 | 0.293 | 0.242 | 1 | 10 | Gosr2 | 0.051 | Old |
| 0.414364 | 0.126278 | 0.257 | 0.212 | 1 | 10 | Sesn1 | 0.045 | Old |
| 0.414384 | 0.522039 | 0.142 | 0.111 | 1 | 10 | Dhx8 | 0.031 | Old |
| 0.41471 | 0.105071 | 0.088 | 0.061 | 1 | 10 | Klhl6 | 0.027 | Old |
| 0.414758 | 0.95099 | 0.037 | 0.02 | 1 | 10 | Vcpkmt | 0.017 | Old |
| 0.414758 | 0.410793 | 0.037 | 0.02 | 1 | 10 | 5430416N( | 0.017 | Old |
| 0.414784 | 0.980346 | 0.049 | 0.03 | 1 | 10 | P2ry1 | 0.019 | Old |
| 0.414784 | 0.085266 | 0.049 | 0.03 | 1 | 10 | Fam122b | 0.019 | Old |
| 0.414796 | 0.784745 | 0.095 | 0.071 | 1 | 10 | Nmi | 0.024 | Old |
| 0.414829 | 0.127078 | 0.589 | 0.545 | 1 | 10 | Ccdc88a | 0.044 | Old |
| 0.415175 | 0.331272 | 0.291 | 0.253 | 1 | 10 | Pbx2 | 0.038 | Old |
| 0.415539 | 0.583915 | 0.086 | 0.061 | 1 | 10 | Arfgap3 | 0.025 | Old |
| 0.415569 | 0.782981 | 0.061 | 0.04 | 1 | 10 | Ccdc163 | 0.021 | Old |

| | | | | | | | | |
|---|---|---|---|---|---|---|---|---|
| 0.415993 | 0.093586 | 0.064 | 0.04 | 1 | 10 | Ccdc112 | 0.024 | Old |
| 0.416161 | 0.854173 | 0.037 | 0.02 | 1 | 10 | Nle1 | 0.017 | Old |
| 0.416161 | 0.500703 | 0.037 | 0.02 | 1 | 10 | Zkscan14 | 0.017 | Old |
| 0.416161 | 0.260352 | 0.037 | 0.02 | 1 | 10 | Zfp748 | 0.017 | Old |
| 0.416195 | 0.082418 | 0.137 | 0.101 | 1 | 10 | Nat9 | 0.036 | Old |
| 0.416441 | 0.235845 | 0.147 | 0.111 | 1 | 10 | Rnf121 | 0.036 | Old |
| 0.41653 | 0.071152 | 0.433 | 0.374 | 1 | 10 | Tsc22d1 | 0.059 | Old |
| 0.416801 | 0.069699 | 0.222 | 0.182 | 1 | 10 | Ubqln2 | 0.04 | Old |
| 0.417204 | 0.069275 | 0.323 | 0.273 | 1 | 10 | Mnt | 0.05 | Old |
| 0.417213 | 1.017601 | 0.049 | 0.03 | 1 | 10 | Echdc1 | 0.019 | Old |
| 0.417284 | 0.300379 | 0.22 | 0.182 | 1 | 10 | Lsm3 | 0.038 | Old |
| 0.417566 | 0.738343 | 0.037 | 0.02 | 1 | 10 | Szt2 | 0.017 | Old |
| 0.417566 | 0.441219 | 0.037 | 0.02 | 1 | 10 | Rpl9-ps6 | 0.017 | Old |
| 0.417566 | 0.366261 | 0.037 | 0.02 | 1 | 10 | Cd4 | 0.017 | Old |
| 0.417756 | 0.237967 | 0.218 | 0.182 | 1 | 10 | Nfatc3 | 0.036 | Old |
| 0.417766 | 0.139133 | 0.582 | 0.515 | 1 | 10 | Ptprj | 0.067 | Old |
| 0.418064 | 0.105478 | 0.12 | 0.091 | 1 | 10 | Amz2 | 0.029 | Old |
| 0.418087 | 0.093542 | 0.276 | 0.232 | 1 | 10 | Kansl1 | 0.044 | Old |
| 0.418156 | 0.077775 | 0.076 | 0.051 | 1 | 10 | Zfyve1 | 0.025 | Old |
| 0.418458 | 0.096292 | 0.301 | 0.253 | 1 | 10 | Commd3 | 0.048 | Old |
| 0.418974 | 0.30281 | 0.037 | 0.02 | 1 | 10 | Nif3l1 | 0.017 | Old |
| 0.419023 | 0.592969 | 0.073 | 0.051 | 1 | 10 | Plxna4os1 | 0.022 | Old |
| 0.419649 | 1.013454 | 0.049 | 0.03 | 1 | 10 | Gas8 | 0.019 | Old |
| 0.42043 | 0.100943 | 0.227 | 0.192 | 1 | 10 | Zmym5 | 0.035 | Old |
| 0.420608 | 0.252702 | 0.523 | 0.495 | 1 | 10 | Mdh2 | 0.028 | Old |
| 0.420687 | 0.327666 | 0.296 | 0.253 | 1 | 10 | Nudt21 | 0.043 | Old |
| 0.420902 | 0.160598 | 0.156 | 0.121 | 1 | 10 | Slc25a45 | 0.035 | Old |
| 0.421031 | 0.802349 | 0.147 | 0.121 | 1 | 10 | Hmg20a | 0.026 | Old |
| 0.421083 | 0.009877 | 0.098 | 0.071 | 1 | 10 | Abt1 | 0.027 | Old |
| 0.421805 | 0.067692 | 0.667 | 0.606 | 1 | 10 | Calm3 | 0.061 | Old |
| 0.421863 | 0.078654 | 0.252 | 0.202 | 1 | 10 | Fto | 0.05 | Old |
| 0.42205 | 0.143936 | 0.108 | 0.081 | 1 | 10 | Txnrd2 | 0.027 | Old |
| 0.422094 | 0.681435 | 0.049 | 0.03 | 1 | 10 | Maged2 | 0.019 | Old |
| 0.422094 | 0.330596 | 0.049 | 0.03 | 1 | 10 | Gmpr | 0.019 | Old |
| 0.422118 | 0.00743 | 0.528 | 0.465 | 1 | 10 | Srsf6 | 0.063 | Old |
| 0.422865 | 0.081672 | 0.286 | 0.242 | 1 | 10 | Crat | 0.044 | Old |
| 0.422889 | 0.72457 | 0.108 | 0.081 | 1 | 10 | Prdm4 | 0.027 | Old |
| 0.423216 | 0.687713 | 0.037 | 0.02 | 1 | 10 | Tyw5 | 0.017 | Old |
| 0.424048 | 0.718887 | 0.139 | 0.111 | 1 | 10 | Nucb2 | 0.028 | Old |
| 0.424319 | 0.266404 | 0.061 | 0.04 | 1 | 10 | 2900093K2 | 0.021 | Old |
| 0.424319 | 0.16502 | 0.061 | 0.04 | 1 | 10 | Naa16 | 0.021 | Old |
| 0.424372 | 0.215056 | 0.154 | 0.121 | 1 | 10 | Mcrs1 | 0.033 | Old |
| 0.424547 | 0.609909 | 0.049 | 0.03 | 1 | 10 | Tubgcp6 | 0.019 | Old |
| 0.425291 | 0.168453 | 0.22 | 0.182 | 1 | 10 | Ddhd1 | 0.038 | Old |
| 0.425574 | 0.008315 | 0.11 | 0.081 | 1 | 10 | Brcc3 | 0.029 | Old |

| | | | | | | | | |
|---|---|---|---|---|---|---|---|---|
| 0.425606 | 0.028064 | 0.284 | 0.232 | 1 | 10 | Stam2 | 0.052 | Old |
| 0.425725 | 0.512513 | 0.117 | 0.091 | 1 | 10 | Pskh1 | 0.026 | Old |
| 0.425777 | 0.661437 | 0.049 | 0.03 | 1 | 10 | Tfpt | 0.019 | Old |
| 0.425924 | 0.035472 | 0.328 | 0.263 | 1 | 10 | Ift20 | 0.065 | Old |
| 0.425937 | 0.242958 | 0.274 | 0.232 | 1 | 10 | Stk24 | 0.042 | Old |
| 0.42606 | 0.504107 | 0.073 | 0.051 | 1 | 10 | Trmt13 | 0.022 | Old |
| 0.426153 | 1.027258 | 0.093 | 0.071 | 1 | 10 | Cfap97 | 0.022 | Old |
| 0.426621 | 0.069276 | 0.2 | 0.162 | 1 | 10 | Prpf18 | 0.038 | Old |
| 0.42673 | 0.259339 | 0.086 | 0.061 | 1 | 10 | Prrg2 | 0.025 | Old |
| 0.426855 | 0.162071 | 0.12 | 0.091 | 1 | 10 | Vps45 | 0.029 | Old |
| 0.427008 | 0.130053 | 0.049 | 0.03 | 1 | 10 | Bloc1s3 | 0.019 | Old |
| 0.427497 | 0.0982 | 0.24 | 0.192 | 1 | 10 | Ptpn2 | 0.048 | Old |
| 0.427899 | 0.014087 | 0.134 | 0.101 | 1 | 10 | Foxk2 | 0.033 | Old |
| 0.428083 | 0.55197 | 0.073 | 0.051 | 1 | 10 | Rffl | 0.022 | Old |
| 0.428472 | 0.089377 | 0.306 | 0.263 | 1 | 10 | Lrrfip2 | 0.043 | Old |
| 0.428734 | 0.861879 | 0.061 | 0.04 | 1 | 10 | Iqsec2 | 0.021 | Old |
| 0.429008 | 0.243078 | 0.267 | 0.222 | 1 | 10 | Ttc39b | 0.045 | Old |
| 0.429586 | 0.006545 | 0.455 | 0.404 | 1 | 10 | Phip | 0.051 | Old |
| 0.430499 | 0.165045 | 0.2 | 0.162 | 1 | 10 | Tdrd7 | 0.038 | Old |
| 0.430716 | 0.233532 | 0.049 | 0.03 | 1 | 10 | Lrrc32 | 0.019 | Old |
| 0.431128 | 0.288657 | 0.073 | 0.051 | 1 | 10 | Cpsf4 | 0.022 | Old |
| 0.431313 | 0.24424 | 0.249 | 0.212 | 1 | 10 | Eapp | 0.037 | Old |
| 0.431345 | 0.244413 | 0.132 | 0.101 | 1 | 10 | Xaf1 | 0.031 | Old |
| 0.431455 | 0.030915 | 0.995 | 1 | 1 | 10 | mt-Atp6 | -0.005 | Old |
| 0.432182 | 0.487243 | 0.108 | 0.081 | 1 | 10 | Mtmr14 | 0.027 | Old |
| 0.432617 | 0.381713 | 0.164 | 0.131 | 1 | 10 | Trim26 | 0.033 | Old |
| 0.43321 | 0.025577 | 0.037 | 0.02 | 1 | 10 | Zfp609 | 0.017 | Old |
| 0.433326 | 0.560043 | 0.2 | 0.172 | 1 | 10 | Bin2 | 0.028 | Old |
| 0.433328 | 0.076316 | 0.112 | 0.141 | 1 | 10 | Utp15 | -0.029 | Old |
| 0.433466 | 0.110462 | 0.597 | 0.525 | 1 | 10 | Nipa2 | 0.072 | Old |
| 0.433527 | 0.206885 | 0.144 | 0.111 | 1 | 10 | Crkl | 0.033 | Old |
| 0.433723 | 0.101801 | 0.665 | 0.596 | 1 | 10 | Chmp4b | 0.069 | Old |
| 0.433886 | 0.047144 | 0.237 | 0.192 | 1 | 10 | St6galnac6 | 0.045 | Old |
| 0.433907 | 0.174162 | 0.721 | 0.646 | 1 | 10 | Mfsd1 | 0.075 | Old |
| 0.434252 | 0.109472 | 0.494 | 0.434 | 1 | 10 | Cep170 | 0.06 | Old |
| 0.43433 | 0.771635 | 0.083 | 0.061 | 1 | 10 | Tlr8 | 0.022 | Old |
| 0.435207 | 0.392389 | 0.073 | 0.051 | 1 | 10 | Nbn | 0.022 | Old |
| 0.435207 | 0.229908 | 0.073 | 0.051 | 1 | 10 | Hectd4 | 0.022 | Old |
| 0.435238 | 0.268459 | 0.086 | 0.061 | 1 | 10 | Klf16 | 0.025 | Old |
| 0.435689 | 0.793822 | 0.049 | 0.03 | 1 | 10 | E2f5 | 0.019 | Old |
| 0.436121 | 1.000699 | 0.093 | 0.071 | 1 | 10 | Fra10ac1 | 0.022 | Old |
| 0.436321 | 0.704075 | 0.105 | 0.081 | 1 | 10 | Nsmf | 0.024 | Old |
| 0.436937 | 0.267209 | 0.049 | 0.03 | 1 | 10 | Ripor1 | 0.019 | Old |
| 0.437029 | 0.021452 | 0.098 | 0.071 | 1 | 10 | Ubap2 | 0.027 | Old |
| 0.437057 | 0.016461 | 0.308 | 0.253 | 1 | 10 | Xpo6 | 0.055 | Old |

| | | | | | | | | |
|---|---|---|---|---|---|---|---|---|
| 0.437204 | 1.945921 | 0.022 | 0.01 | 1 | 10 | Timd4 | 0.012 | Old |
| 0.437204 | 1.590201 | 0.022 | 0.01 | 1 | 10 | Arl16 | 0.012 | Old |
| 0.437256 | 0.291356 | 0.073 | 0.051 | 1 | 10 | Kctd11 | 0.022 | Old |
| 0.437745 | 0.317246 | 0.203 | 0.172 | 1 | 10 | Mkl2 | 0.031 | Old |
| 0.43795 | 0.00646 | 0.963 | 0.96 | 1 | 10 | Arpc2 | 0.003 | Old |
| 0.438095 | 0.110516 | 0.086 | 0.061 | 1 | 10 | Zfp84 | 0.025 | Old |
| 0.438188 | 0.318089 | 0.049 | 0.03 | 1 | 10 | Rfc4 | 0.019 | Old |
| 0.438188 | 0.175653 | 0.042 | 0.061 | 1 | 10 | Zbtb24 | -0.019 | Old |
| 0.438282 | 0.199594 | 0.073 | 0.051 | 1 | 10 | Usp21 | 0.022 | Old |
| 0.438297 | 0.057157 | 0.545 | 0.444 | 1 | 10 | Tmco1 | 0.101 | Old |
| 0.438822 | 0.197631 | 0.098 | 0.071 | 1 | 10 | Alg8 | 0.027 | Old |
| 0.438835 | 0.150519 | 0.374 | 0.323 | 1 | 10 | Anapc11 | 0.051 | Old |
| 0.439078 | 1.568443 | 0.022 | 0.01 | 1 | 10 | Gm29291 | 0.012 | Old |
| 0.439078 | 1.36534 | 0.022 | 0.01 | 1 | 10 | Gemin8 | 0.012 | Old |
| 0.439078 | 1.338596 | 0.022 | 0.01 | 1 | 10 | Brat1 | 0.012 | Old |
| 0.439164 | 0.113927 | 0.76 | 0.687 | 1 | 10 | Cyth4 | 0.073 | Old |
| 0.43975 | 0.217394 | 0.425 | 0.394 | 1 | 10 | Crebbp | 0.031 | Old |
| 0.440181 | 0.034517 | 0.408 | 0.343 | 1 | 10 | P2ry6 | 0.065 | Old |
| 0.440191 | 0.188198 | 0.205 | 0.172 | 1 | 10 | 3830406C1 | 0.033 | Old |
| 0.440338 | 0.047449 | 0.073 | 0.051 | 1 | 10 | Mycbp | 0.022 | Old |
| 0.440341 | 0.213086 | 0.186 | 0.152 | 1 | 10 | Ltbr | 0.034 | Old |
| 0.440496 | 0.334515 | 0.142 | 0.111 | 1 | 10 | Prkra | 0.031 | Old |
| 0.440689 | 0.54497 | 0.095 | 0.071 | 1 | 10 | Cyp27a1 | 0.024 | Old |
| 0.440956 | 1.912265 | 0.022 | 0.01 | 1 | 10 | Kcne3 | 0.012 | Old |
| 0.440956 | 1.57275 | 0.022 | 0.01 | 1 | 10 | Cmtr1 | 0.012 | Old |
| 0.440956 | 1.569826 | 0.022 | 0.01 | 1 | 10 | Rfx2 | 0.012 | Old |
| 0.440956 | 1.225495 | 0.022 | 0.01 | 1 | 10 | Otud3 | 0.012 | Old |
| 0.440956 | 1.062361 | 0.022 | 0.01 | 1 | 10 | Rtl6 | 0.012 | Old |
| 0.441333 | 0.412352 | 0.174 | 0.141 | 1 | 10 | Hbs1l | 0.033 | Old |
| 0.4415 | 0.53516 | 0.127 | 0.101 | 1 | 10 | Sorl1 | 0.026 | Old |
| 0.44159 | 0.258955 | 0.32 | 0.283 | 1 | 10 | Ldlrap1 | 0.037 | Old |
| 0.441614 | 1.011207 | 0.093 | 0.071 | 1 | 10 | Ascc1 | 0.022 | Old |
| 0.441656 | 0.216035 | 0.181 | 0.152 | 1 | 10 | Rapgef2 | 0.029 | Old |
| 0.441858 | 0.009265 | 0.372 | 0.303 | 1 | 10 | Mob4 | 0.069 | Old |
| 0.441922 | 0.284178 | 0.086 | 0.061 | 1 | 10 | Tram2 | 0.025 | Old |
| 0.442203 | 0.096844 | 0.22 | 0.182 | 1 | 10 | Slc1a5 | 0.038 | Old |
| 0.44279 | 0.328241 | 0.205 | 0.172 | 1 | 10 | Fam20b | 0.033 | Old |
| 0.442838 | 1.524127 | 0.022 | 0.01 | 1 | 10 | Txnrd3 | 0.012 | Old |
| 0.442838 | 1.468323 | 0.022 | 0.01 | 1 | 10 | B230217C1 | 0.012 | Old |
| 0.442838 | 1.259168 | 0.022 | 0.01 | 1 | 10 | Sirt6 | 0.012 | Old |
| 0.442838 | 1.237645 | 0.022 | 0.01 | 1 | 10 | Gm12689 | 0.012 | Old |
| 0.442838 | 1.015919 | 0.022 | 0.01 | 1 | 10 | Shmt1 | 0.012 | Old |
| 0.44321 | 0.228721 | 0.049 | 0.03 | 1 | 10 | Kantr | 0.019 | Old |
| 0.443266 | 0.233096 | 0.061 | 0.04 | 1 | 10 | Zfp617 | 0.021 | Old |
| 0.443433 | 0.048625 | 0.073 | 0.051 | 1 | 10 | Lmnb1 | 0.022 | Old |

| | | | | | | | | |
|---|---|---|---|---|---|---|---|---|
| 0.443842 | 0.188797 | 0.086 | 0.061 | 1 | 10 | Pnpt1 | 0.025 | Old |
| 0.443995 | 0.719178 | 0.083 | 0.061 | 1 | 10 | Snrnp35 | 0.022 | Old |
| 0.444108 | 0.020963 | 0.45 | 0.374 | 1 | 10 | Klf7 | 0.076 | Old |
| 0.444389 | 0.06131 | 0.188 | 0.152 | 1 | 10 | Slc30a7 | 0.036 | Old |
| 0.444437 | 0.039518 | 0.306 | 0.253 | 1 | 10 | Hcst | 0.053 | Old |
| 0.444438 | 0.16514 | 0.264 | 0.222 | 1 | 10 | Dstyk | 0.042 | Old |
| 0.44447 | 0.066913 | 0.049 | 0.03 | 1 | 10 | Zfp653 | 0.019 | Old |
| 0.444725 | 0.964457 | 0.022 | 0.01 | 1 | 10 | Mfap3l | 0.012 | Old |
| 0.444725 | 0.790692 | 0.022 | 0.01 | 1 | 10 | Ccl27a | 0.012 | Old |
| 0.444888 | 0.207024 | 0.193 | 0.162 | 1 | 10 | Larp7 | 0.031 | Old |
| 0.445 | 0.016751 | 0.609 | 0.525 | 1 | 10 | Tnfaip8 | 0.084 | Old |
| 0.445459 | 0.041486 | 0.406 | 0.343 | 1 | 10 | Psmb8 | 0.063 | Old |
| 0.445866 | 0.733106 | 0.105 | 0.081 | 1 | 10 | Clpp | 0.024 | Old |
| 0.445908 | 0.396836 | 0.108 | 0.081 | 1 | 10 | Ifih1 | 0.027 | Old |
| 0.446044 | 0.244012 | 0.259 | 0.222 | 1 | 10 | Dusp22 | 0.037 | Old |
| 0.44605 | 0.125037 | 0.408 | 0.364 | 1 | 10 | Dram2 | 0.044 | Old |
| 0.446212 | 0.327808 | 0.152 | 0.121 | 1 | 10 | Tmem104 | 0.031 | Old |
| 0.446612 | 0.263685 | 0.291 | 0.253 | 1 | 10 | Slbp | 0.038 | Old |
| 0.446617 | 0.703576 | 0.022 | 0.01 | 1 | 10 | Ccdc51 | 0.012 | Old |
| 0.446617 | 0.664456 | 0.022 | 0.01 | 1 | 10 | Eno3 | 0.012 | Old |
| 0.446676 | 0.208418 | 0.222 | 0.182 | 1 | 10 | Atp11b | 0.04 | Old |
| 0.447592 | 1.366258 | 0.046 | 0.03 | 1 | 10 | Bpgm | 0.016 | Old |
| 0.448513 | 0.870826 | 0.022 | 0.01 | 1 | 10 | Klhl26 | 0.012 | Old |
| 0.448513 | 0.855248 | 0.022 | 0.01 | 1 | 10 | Cdkn1c | 0.012 | Old |
| 0.448513 | 0.814374 | 0.022 | 0.01 | 1 | 10 | Tango6 | 0.012 | Old |
| 0.448513 | 0.724506 | 0.022 | 0.01 | 1 | 10 | Armc2 | 0.012 | Old |
| 0.448513 | 0.706082 | 0.022 | 0.01 | 1 | 10 | Myh10 | 0.012 | Old |
| 0.44902 | 0.275914 | 0.364 | 0.333 | 1 | 10 | Snx8 | 0.031 | Old |
| 0.449165 | 0.1324 | 0.301 | 0.253 | 1 | 10 | Kpna1 | 0.048 | Old |
| 0.449928 | 0.100226 | 0.455 | 0.384 | 1 | 10 | Ggnbp2 | 0.071 | Old |
| 0.450157 | 0.070476 | 0.144 | 0.111 | 1 | 10 | Mcm6 | 0.033 | Old |
| 0.45019 | 0.113094 | 0.2 | 0.162 | 1 | 10 | Angptl4 | 0.038 | Old |
| 0.450414 | 0.89656 | 0.022 | 0.01 | 1 | 10 | Bcs1l | 0.012 | Old |
| 0.450414 | 0.590659 | 0.022 | 0.01 | 1 | 10 | Prdm11 | 0.012 | Old |
| 0.450414 | 0.517474 | 0.022 | 0.01 | 1 | 10 | Rap1gap | 0.012 | Old |
| 0.450414 | 0.481212 | 0.022 | 0.01 | 1 | 10 | Cryba4 | 0.012 | Old |
| 0.450414 | 0.401933 | 0.022 | 0.01 | 1 | 10 | Oas1g | 0.012 | Old |
| 0.450414 | 0.196607 | 0.022 | 0.01 | 1 | 10 | Fam216a | 0.012 | Old |
| 0.451122 | 0.788585 | 0.105 | 0.081 | 1 | 10 | Zdhhc13 | 0.024 | Old |
| 0.451216 | 0.111707 | 0.555 | 0.485 | 1 | 10 | Srsf9 | 0.07 | Old |
| 0.451361 | 0.285108 | 0.315 | 0.283 | 1 | 10 | Ankrd13a | 0.032 | Old |
| 0.452178 | 0.182633 | 0.147 | 0.121 | 1 | 10 | Snrnp25 | 0.026 | Old |
| 0.452319 | 0.437258 | 0.022 | 0.01 | 1 | 10 | Fscn1 | 0.012 | Old |
| 0.452319 | 0.410795 | 0.022 | 0.01 | 1 | 10 | Ddb2 | 0.012 | Old |
| 0.452319 | 0.318696 | 0.022 | 0.01 | 1 | 10 | Hdhd5 | 0.012 | Old |

| | | | | | | | | |
|---|---|---|---|---|---|---|---|---|
| 0.452319 | 0.273892 | 0.022 | 0.01 | 1 | 10 | Gins1 | 0.012 | Old |
| 0.452319 | 0.190381 | 0.022 | 0.01 | 1 | 10 | Prokr1 | 0.012 | Old |
| 0.452319 | 0.164087 | 0.022 | 0.01 | 1 | 10 | Gm26510 | 0.012 | Old |
| 0.452717 | 0.558644 | 0.093 | 0.071 | 1 | 10 | Clip2 | 0.022 | Old |
| 0.45278 | 0.042128 | 0.289 | 0.242 | 1 | 10 | Clcn7 | 0.047 | Old |
| 0.452965 | 0.113237 | 0.176 | 0.141 | 1 | 10 | Inf2 | 0.035 | Old |
| 0.45362 | 0.207315 | 0.154 | 0.121 | 1 | 10 | Mfap3 | 0.033 | Old |
| 0.453652 | 0.287066 | 0.225 | 0.192 | 1 | 10 | Dguok | 0.033 | Old |
| 0.453781 | 0.355594 | 0.083 | 0.061 | 1 | 10 | B4gat1 | 0.022 | Old |
| 0.454094 | 0.046363 | 0.954 | 0.919 | 1 | 10 | Rpl18a | 0.035 | Old |
| 0.455005 | 0.378383 | 0.2 | 0.172 | 1 | 10 | Tctex1d2 | 0.028 | Old |
| 0.455205 | 0.067023 | 0.149 | 0.121 | 1 | 10 | Fut8 | 0.028 | Old |
| 0.455502 | 0.095084 | 0.751 | 0.677 | 1 | 10 | Psmb4 | 0.074 | Old |
| 0.455569 | 0.175642 | 0.516 | 0.465 | 1 | 10 | Tmem176b | 0.051 | Old |
| 0.455611 | 0.191307 | 0.218 | 0.182 | 1 | 10 | Usp24 | 0.036 | Old |
| 0.455614 | 0.200313 | 0.222 | 0.182 | 1 | 10 | Spen | 0.04 | Old |
| 0.455699 | 0.189326 | 0.166 | 0.131 | 1 | 10 | Zbtb21 | 0.035 | Old |
| 0.455753 | 0.549119 | 0.083 | 0.061 | 1 | 10 | Pcnx3 | 0.022 | Old |
| 0.456269 | 0.345559 | 0.117 | 0.091 | 1 | 10 | Slc25a13 | 0.026 | Old |
| 0.456754 | 0.208802 | 0.335 | 0.293 | 1 | 10 | Polr2m | 0.042 | Old |
| 0.45691 | 0.239759 | 0.46 | 0.424 | 1 | 10 | Kdelr1 | 0.036 | Old |
| 0.457241 | 0.302784 | 0.108 | 0.081 | 1 | 10 | Ebag9 | 0.027 | Old |
| 0.457653 | 1.203837 | 0.034 | 0.02 | 1 | 10 | Slx4 | 0.014 | Old |
| 0.457954 | 0.372478 | 0.117 | 0.091 | 1 | 10 | Lsm10 | 0.026 | Old |
| 0.458968 | 0.354203 | 0.081 | 0.061 | 1 | 10 | Mical1 | 0.02 | Old |
| 0.458975 | 0.040755 | 0.381 | 0.323 | 1 | 10 | Smarcc1 | 0.058 | Old |
| 0.459165 | 0.166429 | 0.059 | 0.081 | 1 | 10 | Zfp866 | -0.022 | Old |
| 0.459185 | 0.08939 | 0.164 | 0.131 | 1 | 10 | Mrpl27 | 0.033 | Old |
| 0.459711 | 0.450046 | 0.083 | 0.061 | 1 | 10 | Gm5086 | 0.022 | Old |
| 0.459915 | 0.173745 | 0.164 | 0.131 | 1 | 10 | Gabpa | 0.033 | Old |
| 0.460561 | 0.0168 | 0.939 | 0.98 | 1 | 10 | Fos | -0.041 | Old |
| 0.460969 | 0.307817 | 0.347 | 0.303 | 1 | 10 | Insig1 | 0.044 | Old |
| 0.460976 | 0.807632 | 0.081 | 0.061 | 1 | 10 | Uba7 | 0.02 | Old |
| 0.461269 | 0.180852 | 0.149 | 0.121 | 1 | 10 | Plcl2 | 0.028 | Old |
| 0.461378 | 0.055832 | 0.164 | 0.131 | 1 | 10 | Kmt5c | 0.033 | Old |
| 0.461642 | 0.056866 | 0.108 | 0.081 | 1 | 10 | Idua | 0.027 | Old |
| 0.461834 | 0.172199 | 0.034 | 0.051 | 1 | 10 | Zfp688 | -0.017 | Old |
| 0.462111 | 0.007533 | 0.164 | 0.131 | 1 | 10 | Gatd1 | 0.033 | Old |
| 0.462183 | 0.533472 | 0.117 | 0.091 | 1 | 10 | Mthfsl | 0.026 | Old |
| 0.46224 | 0.047318 | 0.12 | 0.091 | 1 | 10 | Kansl2 | 0.029 | Old |
| 0.463784 | 1.434963 | 0.034 | 0.02 | 1 | 10 | Lrrk2 | 0.014 | Old |
| 0.463837 | 0.12112 | 0.325 | 0.293 | 1 | 10 | Ecm1 | 0.032 | Old |
| 0.46406 | 0.17163 | 0.218 | 0.182 | 1 | 10 | Svil | 0.036 | Old |
| 0.464119 | 0.13772 | 0.154 | 0.121 | 1 | 10 | Crls1 | 0.033 | Old |
| 0.464317 | 0.321371 | 0.149 | 0.121 | 1 | 10 | Nmd3 | 0.028 | Old |

| | | | | | | | | |
|---|---|---|---|---|---|---|---|---|
| 0.464685 | 0.306364 | 0.083 | 0.061 | 1 | 10 | Polr3f | 0.022 | Old |
| 0.464769 | 0.870654 | 0.071 | 0.051 | 1 | 10 | Twsg1 | 0.02 | Old |
| 0.465324 | 1.18693 | 0.034 | 0.02 | 1 | 10 | Cmpk2 | 0.014 | Old |
| 0.465324 | 1.130911 | 0.034 | 0.02 | 1 | 10 | Dhodh | 0.014 | Old |
| 0.465432 | 0.022706 | 0.252 | 0.212 | 1 | 10 | Tgs1 | 0.04 | Old |
| 0.465684 | 0.37974 | 0.083 | 0.061 | 1 | 10 | Hddc2 | 0.022 | Old |
| 0.466016 | 0.920567 | 0.081 | 0.061 | 1 | 10 | Rrnad1 | 0.02 | Old |
| 0.466066 | 0.083706 | 0.235 | 0.192 | 1 | 10 | Pts | 0.043 | Old |
| 0.466245 | 0.026128 | 0.34 | 0.293 | 1 | 10 | Entpd1 | 0.047 | Old |
| 0.466433 | 0.32028 | 0.117 | 0.091 | 1 | 10 | Dcxr | 0.026 | Old |
| 0.466867 | 1.016433 | 0.034 | 0.02 | 1 | 10 | Ccdc66 | 0.014 | Old |
| 0.467262 | 0.225057 | 0.403 | 0.364 | 1 | 10 | Plekhj1 | 0.039 | Old |
| 0.46727 | 0.223867 | 0.474 | 0.424 | 1 | 10 | Far1 | 0.05 | Old |
| 0.467342 | 0.164252 | 0.335 | 0.293 | 1 | 10 | Samd4b | 0.042 | Old |
| 0.467478 | 0.061626 | 0.834 | 0.717 | 1 | 10 | Cox6a1 | 0.117 | Old |
| 0.467661 | 0.136466 | 0.2 | 0.162 | 1 | 10 | Dnmt1 | 0.038 | Old |
| 0.467778 | 0.013678 | 0.474 | 0.424 | 1 | 10 | Capn2 | 0.05 | Old |
| 0.468222 | 0.432818 | 0.127 | 0.101 | 1 | 10 | Plekha3 | 0.026 | Old |
| 0.468478 | 0.425786 | 0.159 | 0.131 | 1 | 10 | Psmg3 | 0.028 | Old |
| 0.469084 | 0.187853 | 0.071 | 0.051 | 1 | 10 | Pde6d | 0.02 | Old |
| 0.469969 | 0.111384 | 0.738 | 0.646 | 1 | 10 | Arf1 | 0.092 | Old |
| 0.471179 | 0.012213 | 0.11 | 0.081 | 1 | 10 | Dppa3 | 0.029 | Old |
| 0.471513 | 0.792865 | 0.034 | 0.02 | 1 | 10 | Phpt1 | 0.014 | Old |
| 0.471591 | 0.417476 | 0.105 | 0.081 | 1 | 10 | Plrg1 | 0.024 | Old |
| 0.47233 | 0.648261 | 0.196 | 0.172 | 1 | 10 | Gid8 | 0.024 | Old |
| 0.472334 | 0.479319 | 0.071 | 0.051 | 1 | 10 | Zfp771 | 0.02 | Old |
| 0.472334 | 0.062878 | 0.071 | 0.051 | 1 | 10 | Ppargc1b | 0.02 | Old |
| 0.472493 | 0.42507 | 0.105 | 0.081 | 1 | 10 | Homer1 | 0.024 | Old |
| 0.472556 | 0.580517 | 0.046 | 0.03 | 1 | 10 | Rac3 | 0.016 | Old |
| 0.473067 | 0.883307 | 0.034 | 0.02 | 1 | 10 | Dnmt3b | 0.014 | Old |
| 0.473067 | 0.753893 | 0.034 | 0.02 | 1 | 10 | Brf2 | 0.014 | Old |
| 0.473124 | 0.922619 | 0.081 | 0.061 | 1 | 10 | Fmo5 | 0.02 | Old |
| 0.473188 | 0.121579 | 0.203 | 0.162 | 1 | 10 | Fh1 | 0.041 | Old |
| 0.473478 | 0.481904 | 0.093 | 0.071 | 1 | 10 | Bcl9 | 0.022 | Old |
| 0.474284 | 0.166476 | 0.196 | 0.162 | 1 | 10 | Cetn2 | 0.034 | Old |
| 0.474624 | 0.800507 | 0.034 | 0.02 | 1 | 10 | Tmem138 | 0.014 | Old |
| 0.474624 | 0.728948 | 0.034 | 0.02 | 1 | 10 | Zhx3 | 0.014 | Old |
| 0.474919 | 0.073434 | 0.301 | 0.253 | 1 | 10 | Zfp395 | 0.048 | Old |
| 0.47514 | 0.257358 | 0.357 | 0.313 | 1 | 10 | Scamp3 | 0.044 | Old |
| 0.475228 | 0.711031 | 0.046 | 0.03 | 1 | 10 | Urb2 | 0.016 | Old |
| 0.475597 | 0.3677 | 0.071 | 0.051 | 1 | 10 | Unkl | 0.02 | Old |
| 0.475873 | 0.075359 | 0.352 | 0.303 | 1 | 10 | Specc1l | 0.049 | Old |
| 0.475885 | 0.221061 | 0.108 | 0.081 | 1 | 10 | Zfp639 | 0.027 | Old |
| 0.475899 | 0.557291 | 0.059 | 0.04 | 1 | 10 | Fastkd2 | 0.019 | Old |
| 0.475899 | 0.318779 | 0.059 | 0.04 | 1 | 10 | Coro2b | 0.019 | Old |

| | | | | | | | | |
|---|---|---|---|---|---|---|---|---|
| 0.476184 | 0.797467 | 0.034 | 0.02 | 1 | 10 | 2610021A( | 0.014 | Old |
| 0.476184 | 0.609648 | 0.034 | 0.02 | 1 | 10 | BC051226 | 0.014 | Old |
| 0.476184 | 0.193371 | 0.034 | 0.02 | 1 | 10 | Lmnb2 | 0.014 | Old |
| 0.476451 | 0.029692 | 0.183 | 0.152 | 1 | 10 | Slc31a2 | 0.031 | Old |
| 0.47651 | 0.151998 | 0.396 | 0.354 | 1 | 10 | Casp8 | 0.042 | Old |
| 0.476567 | 0.621555 | 0.046 | 0.03 | 1 | 10 | 2700097O( | 0.016 | Old |
| 0.476762 | 0.30801 | 0.257 | 0.222 | 1 | 10 | Pik3c2a | 0.035 | Old |
| 0.477014 | 0.459372 | 0.105 | 0.081 | 1 | 10 | Cebpzos | 0.024 | Old |
| 0.477746 | 0.636465 | 0.034 | 0.02 | 1 | 10 | Lig4 | 0.014 | Old |
| 0.477746 | 0.577381 | 0.034 | 0.02 | 1 | 10 | Fastkd5 | 0.014 | Old |
| 0.477746 | 0.031934 | 0.034 | 0.02 | 1 | 10 | Ppp3cc | 0.014 | Old |
| 0.477779 | 0.15373 | 0.071 | 0.051 | 1 | 10 | Mocos | 0.02 | Old |
| 0.477817 | 0.097901 | 0.421 | 0.374 | 1 | 10 | Prelid1 | 0.047 | Old |
| 0.47838 | 0.046146 | 0.663 | 0.586 | 1 | 10 | Spi1 | 0.077 | Old |
| 0.47845 | 0.113319 | 0.43 | 0.374 | 1 | 10 | Uqcrfs1 | 0.056 | Old |
| 0.478772 | 0.065956 | 0.083 | 0.061 | 1 | 10 | B230307C2 | 0.022 | Old |
| 0.478865 | 0.030964 | 0.452 | 0.404 | 1 | 10 | Zmat2 | 0.048 | Old |
| 0.478907 | 0.006716 | 0.465 | 0.404 | 1 | 10 | Snrpg | 0.061 | Old |
| 0.479183 | 0.28047 | 0.169 | 0.141 | 1 | 10 | Ube4a | 0.028 | Old |
| 0.47923 | 0.36998 | 0.093 | 0.071 | 1 | 10 | Man2b2 | 0.022 | Old |
| 0.479384 | 0.233047 | 0.306 | 0.263 | 1 | 10 | Arid5b | 0.043 | Old |
| 0.479488 | 0.524992 | 0.059 | 0.04 | 1 | 10 | Rnase6 | 0.019 | Old |
| 0.480596 | 0.761795 | 0.046 | 0.03 | 1 | 10 | Vill | 0.016 | Old |
| 0.480596 | 0.367249 | 0.046 | 0.03 | 1 | 10 | Myom1 | 0.016 | Old |
| 0.48088 | 0.536203 | 0.034 | 0.02 | 1 | 10 | Plscr4 | 0.014 | Old |
| 0.48088 | 0.275195 | 0.034 | 0.02 | 1 | 10 | Aasdh | 0.014 | Old |
| 0.480995 | 0.218309 | 0.227 | 0.192 | 1 | 10 | Prkd3 | 0.035 | Old |
| 0.481051 | 0.021044 | 0.672 | 0.606 | 1 | 10 | Sec61g | 0.066 | Old |
| 0.481332 | 0.350437 | 0.161 | 0.131 | 1 | 10 | Mrpl45 | 0.03 | Old |
| 0.481658 | 0.247448 | 0.154 | 0.121 | 1 | 10 | Glt8d1 | 0.033 | Old |
| 0.481796 | 0.226868 | 0.572 | 0.515 | 1 | 10 | Selenos | 0.057 | Old |
| 0.481889 | 0.579882 | 0.059 | 0.04 | 1 | 10 | Zfp758 | 0.019 | Old |
| 0.481889 | 0.427563 | 0.059 | 0.04 | 1 | 10 | Chst7 | 0.019 | Old |
| 0.481943 | 0.516443 | 0.046 | 0.03 | 1 | 10 | AU022252 | 0.016 | Old |
| 0.482451 | 0.371211 | 0.034 | 0.02 | 1 | 10 | Spc25 | 0.014 | Old |
| 0.482451 | 0.151391 | 0.034 | 0.02 | 1 | 10 | Dynlt1a | 0.014 | Old |
| 0.482705 | 0.586145 | 0.103 | 0.081 | 1 | 10 | Ttll12 | 0.022 | Old |
| 0.48289 | 0.204483 | 0.325 | 0.283 | 1 | 10 | Fundc2 | 0.042 | Old |
| 0.483258 | 0.285266 | 0.071 | 0.051 | 1 | 10 | Cnnm4 | 0.02 | Old |
| 0.484557 | 0.294839 | 0.222 | 0.192 | 1 | 10 | Rps12-ps3 | 0.03 | Old |
| 0.485019 | 0.369336 | 0.093 | 0.071 | 1 | 10 | Pld2 | 0.022 | Old |
| 0.485144 | 0.058157 | 0.166 | 0.131 | 1 | 10 | Oaz2 | 0.035 | Old |
| 0.485212 | 0.251014 | 0.105 | 0.081 | 1 | 10 | Mrpl16 | 0.024 | Old |
| 0.485519 | 0.073778 | 0.095 | 0.071 | 1 | 10 | Nup133 | 0.024 | Old |
| 0.485792 | 0.178871 | 0.22 | 0.182 | 1 | 10 | Ccdc90b | 0.038 | Old |

| | | | | | | | | |
|---|---|---|---|---|---|---|---|---|
| 0.48582 | 0.242524 | 0.254 | 0.222 | 1 | 10 | Ganab | 0.032 | Old |
| 0.485889 | 0.068033 | 0.164 | 0.131 | 1 | 10 | Faf1 | 0.033 | Old |
| 0.486711 | 0.691608 | 0.059 | 0.04 | 1 | 10 | Inafm2 | 0.019 | Old |
| 0.486925 | 0.127153 | 0.083 | 0.061 | 1 | 10 | Oaf | 0.022 | Old |
| 0.487202 | 0.025415 | 0.968 | 0.98 | 1 | 10 | Cdc42 | -0.012 | Old |
| 0.487352 | 0.392047 | 0.046 | 0.03 | 1 | 10 | Asf1a | 0.016 | Old |
| 0.487712 | 0.105554 | 0.254 | 0.222 | 1 | 10 | Churc1 | 0.032 | Old |
| 0.487798 | 0.149339 | 0.355 | 0.313 | 1 | 10 | Rab18 | 0.042 | Old |
| 0.48792 | 0.076126 | 0.059 | 0.04 | 1 | 10 | Limk1 | 0.019 | Old |
| 0.48804 | 0.148735 | 0.66 | 0.606 | 1 | 10 | Degs1 | 0.054 | Old |
| 0.488372 | 0.097679 | 0.325 | 0.273 | 1 | 10 | Acsl5 | 0.052 | Old |
| 0.488611 | 0.112997 | 0.262 | 0.222 | 1 | 10 | Tlk1 | 0.04 | Old |
| 0.48871 | 0.381213 | 0.046 | 0.03 | 1 | 10 | Zfp874a | 0.016 | Old |
| 0.488763 | 0.123197 | 0.034 | 0.02 | 1 | 10 | Padi2 | 0.014 | Old |
| 0.489321 | 0.378184 | 0.198 | 0.172 | 1 | 10 | Rsrc1 | 0.026 | Old |
| 0.489571 | 0.122523 | 0.653 | 0.596 | 1 | 10 | Psmb1 | 0.057 | Old |
| 0.489835 | 0.020541 | 0.33 | 0.283 | 1 | 10 | Brox | 0.047 | Old |
| 0.490101 | 0.226411 | 0.418 | 0.384 | 1 | 10 | Tnrc6a | 0.034 | Old |
| 0.490166 | 0.097498 | 0.469 | 0.414 | 1 | 10 | Fcho2 | 0.055 | Old |
| 0.490335 | 0.077542 | 0.841 | 0.818 | 1 | 10 | Lamp2 | 0.023 | Old |
| 0.490894 | 0.110712 | 0.631 | 0.545 | 1 | 10 | Scand1 | 0.086 | Old |
| 0.49143 | 0.252664 | 0.046 | 0.03 | 1 | 10 | Sumf2 | 0.016 | Old |
| 0.491532 | 0.132443 | 0.227 | 0.192 | 1 | 10 | Erlec1 | 0.035 | Old |
| 0.491559 | 0.265592 | 0.059 | 0.04 | 1 | 10 | Cactin | 0.019 | Old |
| 0.491559 | 0.199958 | 0.059 | 0.04 | 1 | 10 | Mcm7 | 0.019 | Old |
| 0.49158 | 0.39632 | 0.127 | 0.101 | 1 | 10 | Gpr157 | 0.026 | Old |
| 0.491821 | 0.242676 | 0.093 | 0.071 | 1 | 10 | Slc35e1 | 0.022 | Old |
| 0.491821 | 0.153513 | 0.093 | 0.071 | 1 | 10 | Hjurp | 0.022 | Old |
| 0.491935 | 0.02281 | 0.034 | 0.02 | 1 | 10 | Fam212a | 0.014 | Old |
| 0.491944 | 0.190146 | 0.218 | 0.182 | 1 | 10 | Ppfia1 | 0.036 | Old |
| 0.492096 | 0.233484 | 0.071 | 0.051 | 1 | 10 | Urgcp | 0.02 | Old |
| 0.492231 | 0.051765 | 0.736 | 0.626 | 1 | 10 | Eif3f | 0.11 | Old |
| 0.49293 | 0.191234 | 0.389 | 0.333 | 1 | 10 | Nop53 | 0.056 | Old |
| 0.493279 | 0.681063 | 0.147 | 0.121 | 1 | 10 | Cebpd | 0.026 | Old |
| 0.493324 | 1.463078 | 0.056 | 0.04 | 1 | 10 | Miga2 | 0.016 | Old |
| 0.493346 | 0.187318 | 0.281 | 0.242 | 1 | 10 | Camta2 | 0.039 | Old |
| 0.494159 | 0.243396 | 0.046 | 0.03 | 1 | 10 | Tlcd2 | 0.016 | Old |
| 0.494192 | 0.075735 | 0.095 | 0.071 | 1 | 10 | Sgsh | 0.024 | Old |
| 0.494227 | 0.426618 | 0.159 | 0.131 | 1 | 10 | Cacna1d | 0.028 | Old |
| 0.494318 | 0.028037 | 0.071 | 0.051 | 1 | 10 | Gtf2h4 | 0.02 | Old |
| 0.494564 | 1.300616 | 0.056 | 0.04 | 1 | 10 | Lyrm4 | 0.016 | Old |
| 0.494624 | 0.155412 | 0.149 | 0.121 | 1 | 10 | Naglu | 0.028 | Old |
| 0.494627 | 0.221751 | 0.269 | 0.232 | 1 | 10 | Ncor2 | 0.037 | Old |
| 0.495134 | 0.069016 | 0.995 | 1 | 1 | 10 | mt-Nd4 | -0.005 | Old |
| 0.495527 | 0.330026 | 0.046 | 0.03 | 1 | 10 | Mpst | 0.016 | Old |

| | | | | | | | | |
|---|---|---|---|---|---|---|---|---|
| 0.495639 | 0.085311 | 0.484 | 0.414 | 1 | 10 | Eif3i | 0.07 | Old |
| 0.496279 | 0.249592 | 0.169 | 0.141 | 1 | 10 | Vkorc1l1 | 0.028 | Old |
| 0.497047 | 0.231899 | 0.027 | 0.04 | 1 | 10 | Taf9b | -0.013 | Old |
| 0.49714 | 0.075143 | 0.372 | 0.333 | 1 | 10 | Hipk1 | 0.039 | Old |
| 0.497324 | 0.35507 | 0.193 | 0.162 | 1 | 10 | Sirpb1c | 0.031 | Old |
| 0.497356 | 0.170976 | 0.218 | 0.182 | 1 | 10 | Hmbox1 | 0.036 | Old |
| 0.497941 | 0.795151 | 0.081 | 0.061 | 1 | 10 | Acaca | 0.02 | Old |
| 0.498744 | 0.042638 | 0.897 | 0.879 | 1 | 10 | Mbnl1 | 0.018 | Old |
| 0.498853 | 0.381595 | 0.147 | 0.121 | 1 | 10 | Maml1 | 0.026 | Old |
| 0.498878 | 0.20654 | 0.059 | 0.04 | 1 | 10 | Fam78a | 0.019 | Old |
| 0.498966 | 0.019678 | 0.506 | 0.434 | 1 | 10 | Sypl | 0.072 | Old |
| 0.498997 | 0.300267 | 0.137 | 0.111 | 1 | 10 | Sh3tc1 | 0.026 | Old |
| 0.49955 | 0.106146 | 0.164 | 0.131 | 1 | 10 | Klhl2 | 0.033 | Old |
| 0.499785 | 0.730506 | 0.112 | 0.091 | 1 | 10 | G2e3 | 0.021 | Old |
| 0.50004 | 0.341697 | 0.081 | 0.061 | 1 | 10 | Pnpo | 0.02 | Old |
| 0.500394 | 0.098498 | 0.159 | 0.131 | 1 | 10 | Kdm5c | 0.028 | Old |
| 0.500788 | 0.352329 | 0.156 | 0.131 | 1 | 10 | Usf1 | 0.025 | Old |
| 0.501248 | 0.210272 | 0.174 | 0.141 | 1 | 10 | 1110038F1 | 0.033 | Old |
| 0.501275 | 0.081437 | 0.22 | 0.182 | 1 | 10 | Zbtb4 | 0.038 | Old |
| 0.501627 | 0.278304 | 0.093 | 0.071 | 1 | 10 | Zfp467 | 0.022 | Old |
| 0.501627 | 0.216813 | 0.093 | 0.071 | 1 | 10 | Gm5547 | 0.022 | Old |
| 0.502068 | 0.226569 | 0.225 | 0.192 | 1 | 10 | 4930523C0 | 0.033 | Old |
| 0.502831 | 0.04126 | 0.804 | 0.727 | 1 | 10 | Csf1r | 0.077 | Old |
| 0.502975 | 0.311705 | 0.068 | 0.051 | 1 | 10 | Sema6d | 0.017 | Old |
| 0.503023 | 0.006335 | 0.325 | 0.273 | 1 | 10 | Rb1 | 0.052 | Old |
| 0.503113 | 0.064917 | 0.183 | 0.152 | 1 | 10 | Vac14 | 0.031 | Old |
| 0.50379 | 0.209973 | 0.059 | 0.04 | 1 | 10 | Wdr4 | 0.019 | Old |
| 0.504114 | 0.821929 | 0.068 | 0.051 | 1 | 10 | Nsdhl | 0.017 | Old |
| 0.50429 | 0.48602 | 0.112 | 0.091 | 1 | 10 | Hs1bp3 | 0.021 | Old |
| 0.504326 | 0.3038 | 0.125 | 0.101 | 1 | 10 | Zfp110 | 0.024 | Old |
| 0.504633 | 0.36069 | 0.105 | 0.081 | 1 | 10 | Plekhg2 | 0.024 | Old |
| 0.504791 | 0.077039 | 0.171 | 0.141 | 1 | 10 | Zfp62 | 0.03 | Old |
| 0.504995 | 0.251109 | 0.115 | 0.091 | 1 | 10 | Tnip1 | 0.024 | Old |
| 0.505344 | 0.547389 | 0.134 | 0.111 | 1 | 10 | Fdps | 0.023 | Old |
| 0.505681 | 0.031882 | 0.259 | 0.232 | 1 | 10 | Igsf8 | 0.027 | Old |
| 0.505951 | 0.252363 | 0.347 | 0.323 | 1 | 10 | Cop1 | 0.024 | Old |
| 0.506055 | 0.02332 | 0.35 | 0.303 | 1 | 10 | Impact | 0.047 | Old |
| 0.506397 | 0.605084 | 0.068 | 0.051 | 1 | 10 | Ssbp2 | 0.017 | Old |
| 0.506397 | 0.015004 | 0.068 | 0.051 | 1 | 10 | E4f1 | 0.017 | Old |
| 0.506638 | 0.302086 | 0.247 | 0.222 | 1 | 10 | Dlgap4 | 0.025 | Old |
| 0.50664 | 0.244002 | 0.156 | 0.131 | 1 | 10 | Ppip5k2 | 0.025 | Old |
| 0.506958 | 0.220804 | 0.259 | 0.232 | 1 | 10 | Ptpn22 | 0.027 | Old |
| 0.507049 | 0.126545 | 0.724 | 0.687 | 1 | 10 | Dock2 | 0.037 | Old |
| 0.507212 | 0.08789 | 0.819 | 0.758 | 1 | 10 | Sh3glb1 | 0.061 | Old |
| 0.508876 | 0.039869 | 0.262 | 0.222 | 1 | 10 | Otud5 | 0.04 | Old |

| | | | | | | | | |
|---|---|---|---|---|---|---|---|---|
| 0.510604 | 0.22363 | 0.081 | 0.061 | 1 | 10 | Cks1b | 0.02 | Old |
| 0.511206 | 0.028691 | 0.059 | 0.04 | 1 | 10 | Gtpbp3 | 0.019 | Old |
| 0.511362 | 0.061182 | 0.709 | 0.687 | 1 | 10 | Tpp1 | 0.022 | Old |
| 0.511667 | 0.388491 | 0.081 | 0.061 | 1 | 10 | Bloc1s5 | 0.02 | Old |
| 0.51184 | 0.106683 | 0.567 | 0.525 | 1 | 10 | Rab11b | 0.042 | Old |
| 0.512124 | 0.21004 | 0.125 | 0.101 | 1 | 10 | Irak3 | 0.024 | Old |
| 0.512228 | 0.229688 | 0.181 | 0.152 | 1 | 10 | Thap2 | 0.029 | Old |
| 0.512613 | 0.023497 | 0.477 | 0.414 | 1 | 10 | Irf2 | 0.063 | Old |
| 0.513479 | 0.192751 | 0.191 | 0.162 | 1 | 10 | Rlf | 0.029 | Old |
| 0.513796 | 0.207311 | 0.081 | 0.061 | 1 | 10 | Ggt5 | 0.02 | Old |
| 0.514643 | 0.218266 | 0.318 | 0.293 | 1 | 10 | Cmas | 0.025 | Old |
| 0.514665 | 0.111546 | 0.46 | 0.414 | 1 | 10 | Calu | 0.046 | Old |
| 0.514936 | 1.671738 | 0.02 | 0.01 | 1 | 10 | Zscan20 | 0.01 | Old |
| 0.514936 | 1.561308 | 0.02 | 0.01 | 1 | 10 | Il20rb | 0.01 | Old |
| 0.514936 | 1.553256 | 0.02 | 0.01 | 1 | 10 | 4933433G1 | 0.01 | Old |
| 0.515224 | 0.126249 | 0.352 | 0.303 | 1 | 10 | Sowahc | 0.049 | Old |
| 0.515416 | 1.356968 | 0.044 | 0.03 | 1 | 10 | Rabl3 | 0.014 | Old |
| 0.516092 | 0.088446 | 0.528 | 0.485 | 1 | 10 | Arl8b | 0.043 | Old |
| 0.516344 | 0.082067 | 0.621 | 0.606 | 1 | 10 | Ckb | 0.015 | Old |
| 0.516478 | 0.106339 | 0.303 | 0.263 | 1 | 10 | Btk | 0.04 | Old |
| 0.516941 | 0.42604 | 0.154 | 0.131 | 1 | 10 | Selenoi | 0.023 | Old |
| 0.517134 | 0.031276 | 0.315 | 0.263 | 1 | 10 | Casp4 | 0.052 | Old |
| 0.51763 | 0.123927 | 0.115 | 0.091 | 1 | 10 | Lrrc42 | 0.024 | Old |
| 0.517893 | 0.382875 | 0.068 | 0.051 | 1 | 10 | Rbm15b | 0.017 | Old |
| 0.5179 | 1.218441 | 0.078 | 0.061 | 1 | 10 | Ms4a14 | 0.017 | Old |
| 0.518055 | 0.049955 | 0.78 | 0.737 | 1 | 10 | Eif5a | 0.043 | Old |
| 0.518116 | 0.156839 | 0.472 | 0.444 | 1 | 10 | Elf1 | 0.028 | Old |
| 0.518311 | 0.057849 | 0.958 | 0.939 | 1 | 10 | Rpl5 | 0.019 | Old |
| 0.519256 | 1.421926 | 0.02 | 0.01 | 1 | 10 | Rpusd1 | 0.01 | Old |
| 0.519256 | 1.394673 | 0.02 | 0.01 | 1 | 10 | E130102H2 | 0.01 | Old |
| 0.519256 | 0.883051 | 0.02 | 0.01 | 1 | 10 | Kif22 | 0.01 | Old |
| 0.519698 | 0.971018 | 0.056 | 0.04 | 1 | 10 | Ankrd33b | 0.016 | Old |
| 0.519817 | 0.451781 | 0.137 | 0.111 | 1 | 10 | Zfp322a | 0.026 | Old |
| 0.519963 | 0.07902 | 0.139 | 0.111 | 1 | 10 | Pigm | 0.028 | Old |
| 0.520059 | 0.27706 | 0.267 | 0.232 | 1 | 10 | Herpud2 | 0.035 | Old |
| 0.52066 | 0.09256 | 0.782 | 0.707 | 1 | 10 | Sec61b | 0.075 | Old |
| 0.521116 | 0.129502 | 0.02 | 0.03 | 1 | 10 | C1rl | -0.01 | Old |
| 0.521147 | 1.187592 | 0.044 | 0.03 | 1 | 10 | Tmem202 | 0.014 | Old |
| 0.521147 | 1.136242 | 0.044 | 0.03 | 1 | 10 | Urb1 | 0.014 | Old |
| 0.521242 | 0.293981 | 0.103 | 0.081 | 1 | 10 | Fbxo8 | 0.022 | Old |
| 0.521334 | 0.408319 | 0.21 | 0.182 | 1 | 10 | Oxsr1 | 0.028 | Old |
| 0.521423 | 1.185229 | 0.02 | 0.01 | 1 | 10 | Rad51b | 0.01 | Old |
| 0.521423 | 1.058488 | 0.02 | 0.01 | 1 | 10 | Gm8369 | 0.01 | Old |
| 0.521423 | 1.017619 | 0.02 | 0.01 | 1 | 10 | Limk2 | 0.01 | Old |
| 0.521423 | 0.952344 | 0.02 | 0.01 | 1 | 10 | Tnks1bp1 | 0.01 | Old |

| | | | | | | | | |
|---|---|---|---|---|---|---|---|---|
| 0.521423 | 0.862002 | 0.02 | 0.01 | 1 | 10 | Sdr42e1 | 0.01 | Old |
| 0.521423 | 0.794502 | 0.02 | 0.01 | 1 | 10 | Zfp316 | 0.01 | Old |
| 0.521423 | 0.579352 | 0.02 | 0.01 | 1 | 10 | Tubd1 | 0.01 | Old |
| 0.521502 | 0.09424 | 0.137 | 0.111 | 1 | 10 | Topors | 0.026 | Old |
| 0.521703 | 0.007844 | 0.472 | 0.414 | 1 | 10 | Mindy1 | 0.058 | Old |
| 0.522247 | 0.785558 | 0.056 | 0.04 | 1 | 10 | Rpap2 | 0.016 | Old |
| 0.522643 | 0.371786 | 0.188 | 0.162 | 1 | 10 | Ak6 | 0.026 | Old |
| 0.52277 | 0.114573 | 0.088 | 0.111 | 1 | 10 | Eef2kmt | -0.023 | Old |
| 0.523425 | 0.016057 | 0.286 | 0.242 | 1 | 10 | Man1a2 | 0.044 | Old |
| 0.523595 | 1.164119 | 0.02 | 0.01 | 1 | 10 | Mt3 | 0.01 | Old |
| 0.523595 | 0.765212 | 0.02 | 0.01 | 1 | 10 | Exoc3l4 | 0.01 | Old |
| 0.523595 | 0.723606 | 0.02 | 0.01 | 1 | 10 | Ahrr | 0.01 | Old |
| 0.523595 | 0.673034 | 0.02 | 0.01 | 1 | 10 | Zfp28 | 0.01 | Old |
| 0.523595 | 0.665033 | 0.02 | 0.01 | 1 | 10 | Tha1 | 0.01 | Old |
| 0.523595 | 0.602057 | 0.02 | 0.01 | 1 | 10 | Fam109a | 0.01 | Old |
| 0.523595 | 0.504792 | 0.02 | 0.01 | 1 | 10 | Il15ra | 0.01 | Old |
| 0.523809 | 0.24318 | 0.171 | 0.141 | 1 | 10 | 5-Mar | 0.03 | Old |
| 0.524025 | 0.849992 | 0.044 | 0.03 | 1 | 10 | Pigo | 0.014 | Old |
| 0.524161 | 0.149795 | 0.181 | 0.152 | 1 | 10 | 9930021J0 | 0.029 | Old |
| 0.524231 | 0.007206 | 0.577 | 0.515 | 1 | 10 | Selenot | 0.062 | Old |
| 0.524803 | 0.889407 | 0.056 | 0.04 | 1 | 10 | Smyd3 | 0.016 | Old |
| 0.524809 | 0.176105 | 0.156 | 0.131 | 1 | 10 | D10Wsu10 | 0.025 | Old |
| 0.525085 | 0.306302 | 0.103 | 0.081 | 1 | 10 | Gm31718 | 0.022 | Old |
| 0.525466 | 0.880065 | 0.044 | 0.03 | 1 | 10 | Polr3h | 0.014 | Old |
| 0.525488 | 1.452497 | 0.032 | 0.02 | 1 | 10 | Eps8l1 | 0.012 | Old |
| 0.525532 | 0.146552 | 0.191 | 0.162 | 1 | 10 | Rps6ka4 | 0.029 | Old |
| 0.525771 | 0.515539 | 0.02 | 0.01 | 1 | 10 | Cttn | 0.01 | Old |
| 0.525771 | 0.507087 | 0.02 | 0.01 | 1 | 10 | Prrt1 | 0.01 | Old |
| 0.526404 | 0.042526 | 0.147 | 0.172 | 1 | 10 | Cd33 | -0.025 | Old |
| 0.526815 | 0.721808 | 0.144 | 0.121 | 1 | 10 | St3gal5 | 0.023 | Old |
| 0.52691 | 0.942616 | 0.044 | 0.03 | 1 | 10 | Bvht | 0.014 | Old |
| 0.527364 | 0.60726 | 0.056 | 0.04 | 1 | 10 | Fam84b | 0.016 | Old |
| 0.527632 | 0.014146 | 0.24 | 0.202 | 1 | 10 | C330007P0 | 0.038 | Old |
| 0.5279 | 0.331021 | 0.276 | 0.253 | 1 | 10 | Mbd3 | 0.023 | Old |
| 0.527952 | 1.148591 | 0.02 | 0.01 | 1 | 10 | Erbb3 | 0.01 | Old |
| 0.527952 | 0.739672 | 0.02 | 0.01 | 1 | 10 | Wdr19 | 0.01 | Old |
| 0.527952 | 0.495879 | 0.02 | 0.01 | 1 | 10 | Kbtbd7 | 0.01 | Old |
| 0.527952 | 0.353001 | 0.02 | 0.01 | 1 | 10 | Zfp202 | 0.01 | Old |
| 0.527952 | 0.27261 | 0.02 | 0.01 | 1 | 10 | Kdelc1 | 0.01 | Old |
| 0.527952 | 0.268645 | 0.02 | 0.01 | 1 | 10 | Mrm1 | 0.01 | Old |
| 0.527952 | 0.253396 | 0.02 | 0.01 | 1 | 10 | Hist4h4 | 0.01 | Old |
| 0.527952 | 0.217099 | 0.02 | 0.01 | 1 | 10 | Gm10800 | 0.01 | Old |
| 0.527952 | 0.130047 | 0.02 | 0.01 | 1 | 10 | Lrp4 | 0.01 | Old |
| 0.528319 | 0.075044 | 0.169 | 0.141 | 1 | 10 | Sec16a | 0.028 | Old |
| 0.528354 | 0.065609 | 0.068 | 0.051 | 1 | 10 | Exoc8 | 0.017 | Old |

| | | | | | | | | |
|---|---|---|---|---|---|---|---|---|
| 0.528356 | 0.837504 | 0.044 | 0.03 | 1 | 10 | Tepsin | 0.014 | Old |
| 0.528648 | 0.285416 | 0.056 | 0.04 | 1 | 10 | Fastkd1 | 0.016 | Old |
| 0.52886 | 0.000981 | 0.543 | 0.455 | 1 | 10 | Cast | 0.088 | Old |
| 0.529126 | 0.129534 | 0.198 | 0.172 | 1 | 10 | Med10 | 0.026 | Old |
| 0.529804 | 0.956415 | 0.044 | 0.03 | 1 | 10 | Mtrf1l | 0.014 | Old |
| 0.529804 | 0.314899 | 0.044 | 0.03 | 1 | 10 | Ttc38 | 0.014 | Old |
| 0.529858 | 0.016011 | 0.91 | 0.909 | 1 | 10 | Eif4g2 | 0.001 | Old |
| 0.529908 | 0.238047 | 0.103 | 0.081 | 1 | 10 | Erf | 0.022 | Old |
| 0.53006 | 0.341927 | 0.2 | 0.172 | 1 | 10 | Trmt1 | 0.028 | Old |
| 0.530138 | 0.396193 | 0.02 | 0.01 | 1 | 10 | Ung | 0.01 | Old |
| 0.530138 | 0.356537 | 0.02 | 0.01 | 1 | 10 | Fam20a | 0.01 | Old |
| 0.530138 | 0.162549 | 0.02 | 0.01 | 1 | 10 | Zfp958 | 0.01 | Old |
| 0.530138 | 0.057009 | 0.02 | 0.01 | 1 | 10 | Calml4 | 0.01 | Old |
| 0.530234 | 0.007199 | 0.567 | 0.495 | 1 | 10 | Taok3 | 0.072 | Old |
| 0.531219 | 0.559943 | 0.056 | 0.04 | 1 | 10 | Tuba4a | 0.016 | Old |
| 0.531254 | 0.755429 | 0.044 | 0.03 | 1 | 10 | Zmynd15 | 0.014 | Old |
| 0.532508 | 0.22279 | 0.149 | 0.121 | 1 | 10 | Pygl | 0.028 | Old |
| 0.532813 | 0.152274 | 0.103 | 0.081 | 1 | 10 | Ano10 | 0.022 | Old |
| 0.533115 | 0.55751 | 0.152 | 0.131 | 1 | 10 | Cadm1 | 0.021 | Old |
| 0.533438 | 0.149311 | 0.081 | 0.101 | 1 | 10 | Samd4 | -0.02 | Old |
| 0.533824 | 0.165058 | 0.337 | 0.293 | 1 | 10 | Acaa2 | 0.044 | Old |
| 0.53416 | 0.602853 | 0.044 | 0.03 | 1 | 10 | Cep85l | 0.014 | Old |
| 0.534623 | 0.198562 | 0.09 | 0.071 | 1 | 10 | Lrwd1 | 0.019 | Old |
| 0.534719 | 0.216555 | 0.181 | 0.152 | 1 | 10 | Kcnk13 | 0.029 | Old |
| 0.535472 | 0.291206 | 0.147 | 0.121 | 1 | 10 | Mis12 | 0.026 | Old |
| 0.535722 | 1.161243 | 0.032 | 0.02 | 1 | 10 | Sema6a | 0.012 | Old |
| 0.535722 | 0.709264 | 0.032 | 0.02 | 1 | 10 | 2700038G2 | 0.012 | Old |
| 0.535996 | 0.700026 | 0.139 | 0.121 | 1 | 10 | Hacd3 | 0.018 | Old |
| 0.536564 | 0.149435 | 0.068 | 0.051 | 1 | 10 | Mzt2 | 0.017 | Old |
| 0.536882 | 0.046059 | 0.169 | 0.141 | 1 | 10 | AY036118 | 0.028 | Old |
| 0.537075 | 0.39552 | 0.044 | 0.03 | 1 | 10 | Etaa1 | 0.014 | Old |
| 0.53735 | 0.391834 | 0.112 | 0.091 | 1 | 10 | Rad17 | 0.021 | Old |
| 0.537437 | 0.930866 | 0.032 | 0.02 | 1 | 10 | Gm3448 | 0.012 | Old |
| 0.538608 | 0.03895 | 0.081 | 0.061 | 1 | 10 | Ddx60 | 0.02 | Old |
| 0.538972 | 0.054841 | 0.056 | 0.04 | 1 | 10 | Smg8 | 0.016 | Old |
| 0.539155 | 0.648504 | 0.032 | 0.02 | 1 | 10 | Gm35154 | 0.012 | Old |
| 0.539997 | 0.240232 | 0.044 | 0.03 | 1 | 10 | Batf2 | 0.014 | Old |
| 0.540402 | 0.105685 | 0.384 | 0.333 | 1 | 10 | Elk3 | 0.051 | Old |
| 0.541067 | 0.17891 | 0.408 | 0.374 | 1 | 10 | Sgk3 | 0.034 | Old |
| 0.541245 | 0.21369 | 0.125 | 0.101 | 1 | 10 | Trak2 | 0.024 | Old |
| 0.541461 | 0.378402 | 0.044 | 0.03 | 1 | 10 | Klhl22 | 0.014 | Old |
| 0.542251 | 0.052743 | 0.377 | 0.323 | 1 | 10 | Wsb2 | 0.054 | Old |
| 0.542869 | 0.210271 | 0.056 | 0.04 | 1 | 10 | Irgq | 0.016 | Old |
| 0.543115 | 0.480753 | 0.078 | 0.061 | 1 | 10 | Acvr1 | 0.017 | Old |
| 0.543941 | 0.193798 | 0.672 | 0.626 | 1 | 10 | Fn1 | 0.046 | Old |

| | | | | | | | | |
|---|---|---|---|---|---|---|---|---|
| 0.544081 | 0.110215 | 0.081 | 0.061 | 1 | 10 | Mthfr | 0.02 | Old |
| 0.544326 | 0.747213 | 0.032 | 0.02 | 1 | 10 | Rida | 0.012 | Old |
| 0.544326 | 0.552979 | 0.032 | 0.02 | 1 | 10 | Cox6a2 | 0.012 | Old |
| 0.544326 | 0.483112 | 0.032 | 0.02 | 1 | 10 | Ahdc1 | 0.012 | Old |
| 0.544326 | 0.47906 | 0.032 | 0.02 | 1 | 10 | Ppif | 0.012 | Old |
| 0.544394 | 0.524722 | 0.044 | 0.03 | 1 | 10 | Frat2 | 0.014 | Old |
| 0.544394 | 0.493147 | 0.044 | 0.03 | 1 | 10 | Rtel1 | 0.014 | Old |
| 0.54501 | 0.225456 | 0.557 | 0.525 | 1 | 10 | Jarid2 | 0.032 | Old |
| 0.545136 | 0.119906 | 0.335 | 0.293 | 1 | 10 | Gorasp2 | 0.042 | Old |
| 0.54577 | 0.135336 | 0.372 | 0.343 | 1 | 10 | Supt5 | 0.029 | Old |
| 0.545864 | 0.381019 | 0.044 | 0.03 | 1 | 10 | 2-Sep | 0.014 | Old |
| 0.54603 | 0.505335 | 0.005 | 0.01 | 1 | 10 | Prph | -0.005 | Old |
| 0.54603 | 0.369513 | 0.005 | 0.01 | 1 | 10 | Mycl | -0.005 | Old |
| 0.54603 | 0.190431 | 0.005 | 0.01 | 1 | 10 | Gm12454 | -0.005 | Old |
| 0.54603 | 0.154386 | 0.005 | 0.01 | 1 | 10 | Btnl2 | -0.005 | Old |
| 0.546054 | 0.96735 | 0.032 | 0.02 | 1 | 10 | Jpx | 0.012 | Old |
| 0.546054 | 0.515309 | 0.032 | 0.02 | 1 | 10 | Aaas | 0.012 | Old |
| 0.546054 | 0.42865 | 0.032 | 0.02 | 1 | 10 | Efnb1 | 0.012 | Old |
| 0.546635 | 0.111746 | 0.232 | 0.192 | 1 | 10 | Sertad2 | 0.04 | Old |
| 0.546732 | 0.066064 | 0.45 | 0.404 | 1 | 10 | Cct7 | 0.046 | Old |
| 0.546962 | 0.094302 | 0.335 | 0.293 | 1 | 10 | Mrps15 | 0.042 | Old |
| 0.547044 | 0.433986 | 0.176 | 0.152 | 1 | 10 | Tbpl1 | 0.024 | Old |
| 0.547786 | 0.245773 | 0.032 | 0.02 | 1 | 10 | Pex11a | 0.012 | Old |
| 0.547936 | 0.153484 | 0.012 | 0.02 | 1 | 10 | Psma8 | -0.008 | Old |
| 0.54832 | 0.11695 | 0.159 | 0.131 | 1 | 10 | Ctnnbip1 | 0.028 | Old |
| 0.549521 | 0.476132 | 0.032 | 0.02 | 1 | 10 | Rhobtb2 | 0.012 | Old |
| 0.549521 | 0.248327 | 0.032 | 0.02 | 1 | 10 | Tmc4 | 0.012 | Old |
| 0.549521 | 0.009243 | 0.032 | 0.02 | 1 | 10 | Tbc1d2 | 0.012 | Old |
| 0.549856 | 0.883427 | 0.066 | 0.051 | 1 | 10 | Cntln | 0.015 | Old |
| 0.54986 | 0.610257 | 0.005 | 0.01 | 1 | 10 | Nat8f4 | -0.005 | Old |
| 0.54986 | 0.551928 | 0.005 | 0.01 | 1 | 10 | Serpina3f | -0.005 | Old |
| 0.54986 | 0.550545 | 0.005 | 0.01 | 1 | 10 | Trim46 | -0.005 | Old |
| 0.54986 | 0.511322 | 0.005 | 0.01 | 1 | 10 | Rasl2-9 | -0.005 | Old |
| 0.54986 | 0.494005 | 0.005 | 0.01 | 1 | 10 | Bsn | -0.005 | Old |
| 0.54986 | 0.436003 | 0.005 | 0.01 | 1 | 10 | Fbln2 | -0.005 | Old |
| 0.54986 | 0.37305 | 0.005 | 0.01 | 1 | 10 | Nrg1 | -0.005 | Old |
| 0.54986 | 0.361541 | 0.005 | 0.01 | 1 | 10 | Bloc1s6os | -0.005 | Old |
| 0.54986 | 0.332517 | 0.005 | 0.01 | 1 | 10 | Best1 | -0.005 | Old |
| 0.54986 | 0.32947 | 0.005 | 0.01 | 1 | 10 | Il18r1 | -0.005 | Old |
| 0.54986 | 0.311655 | 0.005 | 0.01 | 1 | 10 | Nxph3 | -0.005 | Old |
| 0.54986 | 0.272084 | 0.005 | 0.01 | 1 | 10 | Fndc4 | -0.005 | Old |
| 0.54986 | 0.26998 | 0.005 | 0.01 | 1 | 10 | Pcyt1b | -0.005 | Old |
| 0.54986 | 0.263552 | 0.005 | 0.01 | 1 | 10 | Nrep | -0.005 | Old |
| 0.54986 | 0.249185 | 0.005 | 0.01 | 1 | 10 | Mttp | -0.005 | Old |
| 0.54986 | 0.1687 | 0.005 | 0.01 | 1 | 10 | Sez6l2 | -0.005 | Old |

| | | | | | | | | |
|---|---|---|---|---|---|---|---|---|
| 0.54986 | 0.161047 | 0.005 | 0.01 | 1 | 10 | Ildr2 | -0.005 | Old |
| 0.54986 | 0.151765 | 0.005 | 0.01 | 1 | 10 | Mfsd9 | -0.005 | Old |
| 0.54986 | 0.118996 | 0.005 | 0.01 | 1 | 10 | Thrsp | -0.005 | Old |
| 0.54986 | 0.112164 | 0.005 | 0.01 | 1 | 10 | Ccna2 | -0.005 | Old |
| 0.54986 | 0.100734 | 0.005 | 0.01 | 1 | 10 | Dsn1 | -0.005 | Old |
| 0.54986 | 0.097574 | 0.005 | 0.01 | 1 | 10 | Tcf15 | -0.005 | Old |
| 0.54986 | 0.066981 | 0.005 | 0.01 | 1 | 10 | Sptbn2 | -0.005 | Old |
| 0.54986 | 0.032461 | 0.005 | 0.01 | 1 | 10 | Wdr35 | -0.005 | Old |
| 0.54986 | 0.005991 | 0.005 | 0.01 | 1 | 10 | Zfp382 | -0.005 | Old |
| 0.54986 | 0.001978 | 0.005 | 0.01 | 1 | 10 | Gm13620 | -0.005 | Old |
| 0.55025 | 0.142005 | 0.169 | 0.141 | 1 | 10 | Sphk2 | 0.028 | Old |
| 0.550456 | 1.07553 | 0.012 | 0.02 | 1 | 10 | Pdk4 | -0.008 | Old |
| 0.550456 | 0.397105 | 0.012 | 0.02 | 1 | 10 | Rita1 | -0.008 | Old |
| 0.550456 | 0.291982 | 0.012 | 0.02 | 1 | 10 | Fut4 | -0.008 | Old |
| 0.550456 | 0.222191 | 0.012 | 0.02 | 1 | 10 | Taco1os | -0.008 | Old |
| 0.550587 | 0.311458 | 0.174 | 0.152 | 1 | 10 | Prorsd1 | 0.022 | Old |
| 0.551245 | 0.644865 | 0.088 | 0.071 | 1 | 10 | Tspyl2 | 0.017 | Old |
| 0.552735 | 0.050317 | 0.875 | 0.808 | 1 | 10 | Irf2bp2 | 0.067 | Old |
| 0.552929 | 0.257747 | 0.174 | 0.152 | 1 | 10 | Kctd12b | 0.022 | Old |
| 0.552944 | 0.212111 | 0.037 | 0.051 | 1 | 10 | Pick1 | -0.014 | Old |
| 0.552982 | 0.598173 | 0.012 | 0.02 | 1 | 10 | Gm13373 | -0.008 | Old |
| 0.552982 | 0.236921 | 0.012 | 0.02 | 1 | 10 | Nfib | -0.008 | Old |
| 0.553205 | 0.280922 | 0.134 | 0.111 | 1 | 10 | Cyp4f13 | 0.023 | Old |
| 0.553214 | 0.366253 | 0.132 | 0.111 | 1 | 10 | Cp | 0.021 | Old |
| 0.554725 | 0.084644 | 0.044 | 0.03 | 1 | 10 | Rfx3 | 0.014 | Old |
| 0.554927 | 0.043926 | 0.12 | 0.141 | 1 | 10 | Eif2b4 | -0.021 | Old |
| 0.555387 | 0.308155 | 0.078 | 0.061 | 1 | 10 | Lix1l | 0.017 | Old |
| 0.555483 | 0.105057 | 0.665 | 0.646 | 1 | 10 | Cfp | 0.019 | Old |
| 0.555831 | 0.189939 | 0.134 | 0.111 | 1 | 10 | Agpat1 | 0.023 | Old |
| 0.556104 | 0.015701 | 0.521 | 0.455 | 1 | 10 | Hnrnph2 | 0.066 | Old |
| 0.556177 | 0.222565 | 0.112 | 0.091 | 1 | 10 | Fkbp1b | 0.021 | Old |
| 0.556294 | 0.077302 | 0.958 | 0.899 | 1 | 10 | Efhd2 | 0.059 | Old |
| 0.556499 | 0.059859 | 0.381 | 0.343 | 1 | 10 | Trpc4ap | 0.038 | Old |
| 0.556707 | 0.341324 | 0.134 | 0.111 | 1 | 10 | Cdk5rap3 | 0.023 | Old |
| 0.556768 | 0.047938 | 0.108 | 0.131 | 1 | 10 | Arl2 | -0.023 | Old |
| 0.556882 | 0.193082 | 0.462 | 0.414 | 1 | 10 | Mrps33 | 0.048 | Old |
| 0.556894 | 0.382867 | 0.122 | 0.101 | 1 | 10 | Gpr137 | 0.021 | Old |
| 0.557673 | 0.479572 | 0.196 | 0.172 | 1 | 10 | Erap1 | 0.024 | Old |
| 0.557694 | 0.140961 | 0.044 | 0.03 | 1 | 10 | Rbm48 | 0.014 | Old |
| 0.557714 | 0.213739 | 0.232 | 0.202 | 1 | 10 | 2310011J0 | 0.03 | Old |
| 0.558758 | 0.214649 | 0.078 | 0.061 | 1 | 10 | St3gal3 | 0.017 | Old |
| 0.558811 | 0.102085 | 0.318 | 0.283 | 1 | 10 | Ndufs3 | 0.035 | Old |
| 0.559539 | 0.013088 | 0.465 | 0.414 | 1 | 10 | Nampt | 0.051 | Old |
| 0.55955 | 0.240769 | 0.144 | 0.121 | 1 | 10 | Casp3 | 0.023 | Old |
| 0.55957 | 0.430417 | 0.066 | 0.051 | 1 | 10 | Ptchd1 | 0.015 | Old |

| | | | | | | | | |
|---|---|---|---|---|---|---|---|---|
| 0.559882 | 0.105066 | 0.191 | 0.162 | 1 | 10 | Tcf20 | 0.029 | Old |
| 0.559962 | 0.286682 | 0.12 | 0.101 | 1 | 10 | Rnf138 | 0.019 | Old |
| 0.55998 | 0.242284 | 0.112 | 0.091 | 1 | 10 | Fut11 | 0.021 | Old |
| 0.560146 | 0.967324 | 0.076 | 0.061 | 1 | 10 | Emilin1 | 0.015 | Old |
| 0.560491 | 0.115811 | 0.535 | 0.485 | 1 | 10 | Gas2l3 | 0.05 | Old |
| 0.560794 | 0.576439 | 0.154 | 0.131 | 1 | 10 | Gle1 | 0.023 | Old |
| 0.560835 | 0.026175 | 0.631 | 0.606 | 1 | 10 | P2rx4 | 0.025 | Old |
| 0.561715 | 0.154356 | 0.09 | 0.071 | 1 | 10 | Trim3 | 0.019 | Old |
| 0.561887 | 0.121364 | 0.112 | 0.091 | 1 | 10 | Sssca1 | 0.021 | Old |
| 0.562138 | 0.398295 | 0.078 | 0.061 | 1 | 10 | Gm15417 | 0.017 | Old |
| 0.562162 | 0.040633 | 0.044 | 0.03 | 1 | 10 | Rap1gap2 | 0.014 | Old |
| 0.562219 | 0.009607 | 0.24 | 0.202 | 1 | 10 | Glod4 | 0.038 | Old |
| 0.56222 | 0.135017 | 0.166 | 0.141 | 1 | 10 | Cwf19l2 | 0.025 | Old |
| 0.562241 | 0.385149 | 0.164 | 0.141 | 1 | 10 | Zmym2 | 0.023 | Old |
| 0.562706 | 0.085553 | 0.384 | 0.333 | 1 | 10 | Ttyh3 | 0.051 | Old |
| 0.563111 | 0.144208 | 0.242 | 0.222 | 1 | 10 | Timm17a | 0.02 | Old |
| 0.563217 | 0.030988 | 0.443 | 0.465 | 1 | 10 | Zfp703 | -0.022 | Old |
| 0.563396 | 0.081997 | 0.425 | 0.364 | 1 | 10 | Txnl1 | 0.061 | Old |
| 0.564286 | 0.085413 | 0.528 | 0.505 | 1 | 10 | Lrrc58 | 0.023 | Old |
| 0.565273 | 0.013518 | 0.699 | 0.616 | 1 | 10 | Ubl5 | 0.083 | Old |
| 0.565506 | 0.017367 | 0.364 | 0.333 | 1 | 10 | Il11ra1 | 0.031 | Old |
| 0.565684 | 0.530463 | 0.066 | 0.051 | 1 | 10 | Gm20696 | 0.015 | Old |
| 0.565696 | 0.023699 | 0.543 | 0.475 | 1 | 10 | Atp5c1 | 0.068 | Old |
| 0.566575 | 0.204066 | 0.21 | 0.182 | 1 | 10 | Tor3a | 0.028 | Old |
| 0.566671 | 0.235505 | 0.222 | 0.192 | 1 | 10 | Fam3c | 0.03 | Old |
| 0.566945 | 0.376503 | 0.152 | 0.131 | 1 | 10 | Gtf3a | 0.021 | Old |
| 0.566988 | 0.786947 | 0.054 | 0.04 | 1 | 10 | Nol10 | 0.014 | Old |
| 0.567036 | 0.006839 | 0.1 | 0.121 | 1 | 10 | Stambp | -0.021 | Old |
| 0.567175 | 0.163219 | 0.482 | 0.424 | 1 | 10 | Pet100 | 0.058 | Old |
| 0.568368 | 0.330825 | 0.267 | 0.242 | 1 | 10 | BC004004 | 0.025 | Old |
| 0.569969 | 0.147404 | 0.611 | 0.545 | 1 | 10 | Mat2a | 0.066 | Old |
| 0.570565 | 0.087159 | 0.2 | 0.172 | 1 | 10 | Atg101 | 0.028 | Old |
| 0.570786 | 0.411671 | 0.171 | 0.152 | 1 | 10 | Dgat1 | 0.019 | Old |
| 0.571055 | 1.105101 | 0.054 | 0.04 | 1 | 10 | Gm20703 | 0.014 | Old |
| 0.571178 | 0.58795 | 0.064 | 0.081 | 1 | 10 | Zranb3 | -0.017 | Old |
| 0.572788 | 0.225762 | 0.164 | 0.141 | 1 | 10 | Itpa | 0.023 | Old |
| 0.572824 | 0.107501 | 0.983 | 0.949 | 1 | 10 | H3f3b | 0.034 | Old |
| 0.572844 | 0.06069 | 0.203 | 0.172 | 1 | 10 | Gsdmd | 0.031 | Old |
| 0.573165 | 0.052771 | 0.11 | 0.091 | 1 | 10 | Vav2 | 0.019 | Old |
| 0.573326 | 0.28104 | 0.218 | 0.192 | 1 | 10 | Tbc1d1 | 0.026 | Old |
| 0.573476 | 0.239039 | 0.078 | 0.061 | 1 | 10 | Tfb2m | 0.017 | Old |
| 0.573487 | 0.017169 | 0.134 | 0.111 | 1 | 10 | Tmem203 | 0.023 | Old |
| 0.573504 | 0.051014 | 0.418 | 0.364 | 1 | 10 | Ppp4r2 | 0.054 | Old |
| 0.574705 | 0.020558 | 0.029 | 0.04 | 1 | 10 | Zfp948 | -0.011 | Old |
| 0.575236 | 0.008726 | 0.249 | 0.222 | 1 | 10 | Elovl1 | 0.027 | Old |

| | | | | | | | | |
|---|---|---|---|---|---|---|---|---|
| 0.575267 | 0.109008 | 0.134 | 0.111 | 1 | 10 | Atg4b | 0.023 | Old |
| 0.575592 | 0.382322 | 0.046 | 0.061 | 1 | 10 | Strada | -0.015 | Old |
| 0.575756 | 0.131576 | 0.078 | 0.061 | 1 | 10 | Gpc1 | 0.017 | Old |
| 0.575969 | 0.094501 | 0.472 | 0.414 | 1 | 10 | Pld4 | 0.058 | Old |
| 0.576125 | 0.207095 | 0.161 | 0.141 | 1 | 10 | Alg5 | 0.02 | Old |
| 0.576284 | 0.14221 | 0.112 | 0.091 | 1 | 10 | Nt5dc3 | 0.021 | Old |
| 0.576377 | 0.08203 | 0.548 | 0.485 | 1 | 10 | Matr3 | 0.063 | Old |
| 0.5765 | 0.584511 | 0.054 | 0.04 | 1 | 10 | Dusp12 | 0.014 | Old |
| 0.576715 | 0.108968 | 0.352 | 0.313 | 1 | 10 | Otulin | 0.039 | Old |
| 0.576798 | 0.115107 | 0.394 | 0.364 | 1 | 10 | Gpr132 | 0.03 | Old |
| 0.576887 | 0.48407 | 0.054 | 0.071 | 1 | 10 | Clpb | -0.017 | Old |
| 0.577219 | 0.436055 | 0.193 | 0.172 | 1 | 10 | Fchsd2 | 0.021 | Old |
| 0.578006 | 0.290282 | 0.066 | 0.051 | 1 | 10 | Gin1 | 0.015 | Old |
| 0.578047 | 0.051168 | 0.355 | 0.323 | 1 | 10 | Vamp3 | 0.032 | Old |
| 0.578147 | 0.286726 | 0.139 | 0.121 | 1 | 10 | Shkbp1 | 0.018 | Old |
| 0.578214 | 0.23891 | 0.408 | 0.364 | 1 | 10 | 4931406C0 | 0.044 | Old |
| 0.578751 | 0.095338 | 0.34 | 0.303 | 1 | 10 | Tmbim4 | 0.037 | Old |
| 0.579243 | 0.217536 | 0.293 | 0.263 | 1 | 10 | Tns1 | 0.03 | Old |
| 0.579406 | 0.028263 | 0.213 | 0.182 | 1 | 10 | Pex6 | 0.031 | Old |
| 0.579467 | 0.18022 | 0.125 | 0.101 | 1 | 10 | Nudt13 | 0.024 | Old |
| 0.57975 | 0.234247 | 0.45 | 0.414 | 1 | 10 | Ddost | 0.036 | Old |
| 0.579981 | 0.449611 | 0.11 | 0.091 | 1 | 10 | Lxn | 0.019 | Old |
| 0.58016 | 0.144833 | 0.55 | 0.525 | 1 | 10 | Cdv3 | 0.025 | Old |
| 0.580475 | 0.031835 | 0.269 | 0.232 | 1 | 10 | Nup88 | 0.037 | Old |
| 0.580599 | 0.704694 | 0.054 | 0.04 | 1 | 10 | Tenm4 | 0.014 | Old |
| 0.580905 | 0.070228 | 0.122 | 0.101 | 1 | 10 | Thoc5 | 0.021 | Old |
| 0.581136 | 0.121119 | 0.186 | 0.162 | 1 | 10 | Zfp800 | 0.024 | Old |
| 0.581177 | 0.04315 | 0.244 | 0.212 | 1 | 10 | Ms4a6b | 0.032 | Old |
| 0.5816 | 0.098569 | 0.029 | 0.04 | 1 | 10 | Pradc1 | -0.011 | Old |
| 0.582472 | 0.024178 | 0.998 | 1 | 1 | 10 | Rps29 | -0.002 | Old |
| 0.582619 | 0.065653 | 0.2 | 0.172 | 1 | 10 | Ssbp1 | 0.028 | Old |
| 0.582668 | 0.369153 | 0.142 | 0.121 | 1 | 10 | Upf2 | 0.021 | Old |
| 0.583541 | 0.137252 | 0.1 | 0.081 | 1 | 10 | Zfp738 | 0.019 | Old |
| 0.584214 | 0.269432 | 0.066 | 0.051 | 1 | 10 | B3galnt1 | 0.015 | Old |
| 0.585063 | 0.223962 | 0.254 | 0.222 | 1 | 10 | Naa38 | 0.032 | Old |
| 0.58509 | 0.09006 | 0.406 | 0.364 | 1 | 10 | Rab11fip5 | 0.042 | Old |
| 0.585504 | 0.489255 | 0.076 | 0.061 | 1 | 10 | Hace1 | 0.015 | Old |
| 0.586338 | 0.011994 | 0.242 | 0.212 | 1 | 10 | Lpgat1 | 0.03 | Old |
| 0.586764 | 1.254494 | 0.042 | 0.03 | 1 | 10 | Abcb4 | 0.012 | Old |
| 0.586901 | 0.39583 | 0.125 | 0.152 | 1 | 10 | Akr7a5 | -0.027 | Old |
| 0.587139 | 0.25438 | 0.227 | 0.202 | 1 | 10 | Atad1 | 0.025 | Old |
| 0.587451 | 0.215738 | 0.122 | 0.101 | 1 | 10 | Proser1 | 0.021 | Old |
| 0.587461 | 0.620063 | 0.054 | 0.04 | 1 | 10 | Pex11b | 0.014 | Old |
| 0.587954 | 0.161984 | 0.066 | 0.051 | 1 | 10 | Rnaset2a | 0.015 | Old |
| 0.588452 | 0.23376 | 0.154 | 0.131 | 1 | 10 | Mrpl19 | 0.023 | Old |

| | | | | | | | | |
|---|---|---|---|---|---|---|---|---|
| 0.588457 | 0.331122 | 0.171 | 0.152 | 1 | 10 | Hk1 | 0.019 | Old |
| 0.588838 | 0.307462 | 0.054 | 0.04 | 1 | 10 | Gtpbp1 | 0.014 | Old |
| 0.589468 | 0.536213 | 0.137 | 0.121 | 1 | 10 | Adgrg6 | 0.016 | Old |
| 0.589656 | 0.041838 | 0.156 | 0.131 | 1 | 10 | Papd5 | 0.025 | Old |
| 0.589677 | 0.313997 | 0.142 | 0.121 | 1 | 10 | Mboat7 | 0.021 | Old |
| 0.589693 | 0.116643 | 0.1 | 0.081 | 1 | 10 | Taf4 | 0.019 | Old |
| 0.589807 | 0.007942 | 0.364 | 0.313 | 1 | 10 | Stx12 | 0.051 | Old |
| 0.590171 | 0.470353 | 0.076 | 0.061 | 1 | 10 | Cars2 | 0.015 | Old |
| 0.590406 | 0.173373 | 0.311 | 0.283 | 1 | 10 | Lactb | 0.028 | Old |
| 0.590495 | 0.128018 | 0.44 | 0.394 | 1 | 10 | Ndufa5 | 0.046 | Old |
| 0.590569 | 0.043159 | 0.252 | 0.222 | 1 | 10 | Slc35a4 | 0.03 | Old |
| 0.591223 | 0.256584 | 0.308 | 0.283 | 1 | 10 | Ccdc71l | 0.025 | Old |
| 0.592402 | 0.131827 | 0.521 | 0.465 | 1 | 10 | Zcchc6 | 0.056 | Old |
| 0.592631 | 0.575776 | 0.108 | 0.091 | 1 | 10 | Prmt7 | 0.017 | Old |
| 0.592978 | 0.382025 | 0.054 | 0.04 | 1 | 10 | Zfp760 | 0.014 | Old |
| 0.593093 | 0.018487 | 0.912 | 0.879 | 1 | 10 | Cotl1 | 0.033 | Old |
| 0.59376 | 0.155024 | 0.029 | 0.04 | 1 | 10 | Piga | -0.011 | Old |
| 0.594006 | 0.560487 | 0.098 | 0.081 | 1 | 10 | Coa4 | 0.017 | Old |
| 0.594968 | 0.117648 | 0.247 | 0.212 | 1 | 10 | Rbm22 | 0.035 | Old |
| 0.595418 | 0.007346 | 0.298 | 0.263 | 1 | 10 | Krit1 | 0.035 | Old |
| 0.595702 | 0.156569 | 0.11 | 0.091 | 1 | 10 | Sgcb | 0.019 | Old |
| 0.596151 | 0.865253 | 0.042 | 0.03 | 1 | 10 | Mfng | 0.012 | Old |
| 0.597131 | 0.379868 | 0.054 | 0.04 | 1 | 10 | Taf8 | 0.014 | Old |
| 0.597131 | 0.3717 | 0.054 | 0.04 | 1 | 10 | Nme7 | 0.014 | Old |
| 0.597549 | 0.11228 | 0.088 | 0.071 | 1 | 10 | 2410089E0 | 0.017 | Old |
| 0.597723 | 0.862347 | 0.042 | 0.03 | 1 | 10 | Flt3l | 0.012 | Old |
| 0.59805 | 0.05264 | 0.518 | 0.495 | 1 | 10 | Runx1 | 0.023 | Old |
| 0.598362 | 0.064593 | 0.443 | 0.404 | 1 | 10 | Fam96a | 0.039 | Old |
| 0.59845 | 0.057147 | 0.176 | 0.152 | 1 | 10 | Nfix | 0.024 | Old |
| 0.598671 | 0.179734 | 0.11 | 0.091 | 1 | 10 | Ppp2r1b | 0.019 | Old |
| 0.599296 | 0.618314 | 0.042 | 0.03 | 1 | 10 | Slc19a1 | 0.012 | Old |
| 0.599743 | 0.022869 | 0.083 | 0.101 | 1 | 10 | Usp36 | -0.018 | Old |
| 0.599907 | 0.209026 | 0.054 | 0.04 | 1 | 10 | Fggy | 0.014 | Old |
| 0.6007 | 0.195419 | 0.408 | 0.374 | 1 | 10 | Cytip | 0.034 | Old |
| 0.600732 | 1.671245 | 0.029 | 0.02 | 1 | 10 | Sh3bgrl2 | 0.009 | Old |
| 0.600761 | 0.410717 | 0.029 | 0.04 | 1 | 10 | Hotairm1 | -0.011 | Old |
| 0.600871 | 0.067189 | 0.042 | 0.03 | 1 | 10 | Btf3l4 | 0.012 | Old |
| 0.601001 | 0.087869 | 0.056 | 0.071 | 1 | 10 | Bnip1 | -0.015 | Old |
| 0.602641 | 0.125119 | 0.11 | 0.091 | 1 | 10 | Rpp14 | 0.019 | Old |
| 0.602936 | 0.444913 | 0.13 | 0.111 | 1 | 10 | Brms1 | 0.019 | Old |
| 0.603235 | 0.085256 | 0.23 | 0.202 | 1 | 10 | Cul5 | 0.028 | Old |
| 0.603625 | 0.104091 | 0.601 | 0.556 | 1 | 10 | Khdrbs1 | 0.045 | Old |
| 0.603995 | 0.013732 | 0.2 | 0.172 | 1 | 10 | Exosc3 | 0.028 | Old |
| 0.604002 | 0.832217 | 0.064 | 0.051 | 1 | 10 | Kctd13 | 0.013 | Old |
| 0.604164 | 0.080278 | 0.435 | 0.384 | 1 | 10 | Ubtf | 0.051 | Old |

| | | | | | | | | |
|---|---|---|---|---|---|---|---|---|
| 0.605023 | 0.066042 | 0.535 | 0.515 | 1 | 10 | Hnrnpa1 | 0.02 | Old |
| 0.605607 | 0.70791 | 0.042 | 0.03 | 1 | 10 | Ap1s3 | 0.012 | Old |
| 0.605607 | 0.589501 | 0.042 | 0.03 | 1 | 10 | Txnl4b | 0.012 | Old |
| 0.605884 | 0.112527 | 0.768 | 0.687 | 1 | 10 | Srsf5 | 0.081 | Old |
| 0.60636 | 1.137606 | 0.029 | 0.02 | 1 | 10 | Fuz | 0.009 | Old |
| 0.606623 | 0.250556 | 0.259 | 0.232 | 1 | 10 | Map3k2 | 0.027 | Old |
| 0.606853 | 0.0452 | 0.907 | 0.859 | 1 | 10 | Eef1b2 | 0.048 | Old |
| 0.607182 | 0.011094 | 0.362 | 0.313 | 1 | 10 | Baz1a | 0.049 | Old |
| 0.607488 | 0.06294 | 0.164 | 0.141 | 1 | 10 | Vps33a | 0.023 | Old |
| 0.607557 | 0.343221 | 0.13 | 0.111 | 1 | 10 | Psd4 | 0.019 | Old |
| 0.607858 | 0.186058 | 0.205 | 0.182 | 1 | 10 | Tmem183a | 0.023 | Old |
| 0.607889 | 0.107377 | 0.186 | 0.162 | 1 | 10 | Tlr4 | 0.024 | Old |
| 0.607904 | 1.618157 | 0.017 | 0.01 | 1 | 10 | Lepr | 0.007 | Old |
| 0.608242 | 1.101284 | 0.029 | 0.02 | 1 | 10 | Gm10642 | 0.009 | Old |
| 0.608242 | 0.952715 | 0.029 | 0.02 | 1 | 10 | Kif3c | 0.009 | Old |
| 0.608269 | 0.0069 | 0.054 | 0.04 | 1 | 10 | Prag1 | 0.014 | Old |
| 0.60828 | 0.415626 | 0.137 | 0.121 | 1 | 10 | Bbx | 0.016 | Old |
| 0.609056 | 0.065789 | 0.188 | 0.162 | 1 | 10 | Get4 | 0.026 | Old |
| 0.609394 | 0.061234 | 0.2 | 0.172 | 1 | 10 | Smim4 | 0.028 | Old |
| 0.609668 | 0.045618 | 0.054 | 0.04 | 1 | 10 | Spryd3 | 0.014 | Old |
| 0.609826 | 0.036231 | 0.658 | 0.606 | 1 | 10 | Bzw1 | 0.052 | Old |
| 0.610125 | 0.940322 | 0.029 | 0.02 | 1 | 10 | Dyrk1b | 0.009 | Old |
| 0.610125 | 0.825768 | 0.029 | 0.02 | 1 | 10 | Knstrn | 0.009 | Old |
| 0.610194 | 0.039653 | 0.076 | 0.061 | 1 | 10 | Eml3 | 0.015 | Old |
| 0.610359 | 0.511855 | 0.042 | 0.03 | 1 | 10 | P3h2 | 0.012 | Old |
| 0.610381 | 1.764295 | 0.017 | 0.01 | 1 | 10 | Enpep | 0.007 | Old |
| 0.610381 | 0.709429 | 0.017 | 0.01 | 1 | 10 | Slc18a1 | 0.007 | Old |
| 0.611681 | 0.11945 | 0.164 | 0.141 | 1 | 10 | Ruvbl1 | 0.023 | Old |
| 0.612012 | 0.846676 | 0.029 | 0.02 | 1 | 10 | Bbs7 | 0.009 | Old |
| 0.612417 | 0.035326 | 0.039 | 0.051 | 1 | 10 | Tslp | -0.012 | Old |
| 0.612791 | 0.007197 | 0.112 | 0.131 | 1 | 10 | Acot13 | -0.019 | Old |
| 0.612863 | 1.295064 | 0.017 | 0.01 | 1 | 10 | Agbl5 | 0.007 | Old |
| 0.612863 | 1.193642 | 0.017 | 0.01 | 1 | 10 | Hgh1 | 0.007 | Old |
| 0.612863 | 1.160725 | 0.017 | 0.01 | 1 | 10 | Heg1 | 0.007 | Old |
| 0.612863 | 0.817412 | 0.017 | 0.01 | 1 | 10 | Zfp951 | 0.007 | Old |
| 0.612863 | 0.801873 | 0.017 | 0.01 | 1 | 10 | Asah2 | 0.007 | Old |
| 0.612863 | 0.719389 | 0.017 | 0.01 | 1 | 10 | Cenpk | 0.007 | Old |
| 0.612875 | 0.252442 | 0.098 | 0.081 | 1 | 10 | Gpn3 | 0.017 | Old |
| 0.613754 | 0.049279 | 0.12 | 0.101 | 1 | 10 | Cox19 | 0.019 | Old |
| 0.613759 | 0.203526 | 0.076 | 0.061 | 1 | 10 | Flnb | 0.015 | Old |
| 0.614722 | 0.079719 | 0.922 | 0.929 | 1 | 10 | Chchd2 | -0.007 | Old |
| 0.614803 | 0.27647 | 0.252 | 0.232 | 1 | 10 | Dapk3 | 0.02 | Old |
| 0.615128 | 0.244812 | 0.042 | 0.03 | 1 | 10 | Mettl17 | 0.012 | Old |
| 0.615349 | 1.505989 | 0.017 | 0.01 | 1 | 10 | Zdhhc24 | 0.007 | Old |
| 0.615349 | 1.068639 | 0.017 | 0.01 | 1 | 10 | Zfp788 | 0.007 | Old |

| | | | | | | | | |
|---|---|---|---|---|---|---|---|---|
| 0.615349 | 0.993556 | 0.017 | 0.01 | 1 | 10 | Fbxo10 | 0.007 | Old |
| 0.615349 | 0.973324 | 0.017 | 0.01 | 1 | 10 | 4931428F0 | 0.007 | Old |
| 0.615349 | 0.62713 | 0.017 | 0.01 | 1 | 10 | Wdr27 | 0.007 | Old |
| 0.615349 | 0.614807 | 0.017 | 0.01 | 1 | 10 | Cmtr2 | 0.007 | Old |
| 0.615349 | 0.592412 | 0.017 | 0.01 | 1 | 10 | Myo1b | 0.007 | Old |
| 0.615349 | 0.58299 | 0.017 | 0.01 | 1 | 10 | Mphosph9 | 0.007 | Old |
| 0.615427 | 0.077218 | 0.171 | 0.152 | 1 | 10 | Wasl | 0.019 | Old |
| 0.615683 | 0.278602 | 0.12 | 0.101 | 1 | 10 | Cep192 | 0.019 | Old |
| 0.615793 | 0.77215 | 0.029 | 0.02 | 1 | 10 | Miga1 | 0.009 | Old |
| 0.616044 | 0.138357 | 0.289 | 0.263 | 1 | 10 | Tsen34 | 0.026 | Old |
| 0.616141 | 0.590987 | 0.076 | 0.061 | 1 | 10 | Supt3 | 0.015 | Old |
| 0.616512 | 0.053979 | 0.433 | 0.384 | 1 | 10 | Fkbp8 | 0.049 | Old |
| 0.617016 | 0.252931 | 0.086 | 0.071 | 1 | 10 | Rnf157 | 0.015 | Old |
| 0.617343 | 0.014036 | 0.523 | 0.465 | 1 | 10 | Psenen | 0.058 | Old |
| 0.617687 | 1.123065 | 0.029 | 0.02 | 1 | 10 | Lrsam1 | 0.009 | Old |
| 0.617687 | 0.805664 | 0.029 | 0.02 | 1 | 10 | Jade3 | 0.009 | Old |
| 0.617687 | 0.593212 | 0.029 | 0.02 | 1 | 10 | Hacl1 | 0.009 | Old |
| 0.617687 | 0.566448 | 0.029 | 0.02 | 1 | 10 | Gm10036 | 0.009 | Old |
| 0.617729 | 0.404746 | 0.108 | 0.091 | 1 | 10 | Zfyve26 | 0.017 | Old |
| 0.61784 | 0.809119 | 0.017 | 0.01 | 1 | 10 | Ptar1 | 0.007 | Old |
| 0.61784 | 0.64073 | 0.017 | 0.01 | 1 | 10 | Gpihbp1 | 0.007 | Old |
| 0.61784 | 0.614856 | 0.017 | 0.01 | 1 | 10 | C920021L1 | 0.007 | Old |
| 0.61784 | 0.601828 | 0.017 | 0.01 | 1 | 10 | Isoc2a | 0.007 | Old |
| 0.61784 | 0.51671 | 0.017 | 0.01 | 1 | 10 | Rnls | 0.007 | Old |
| 0.61784 | 0.421264 | 0.017 | 0.01 | 1 | 10 | AI463229 | 0.007 | Old |
| 0.61784 | 0.417506 | 0.017 | 0.01 | 1 | 10 | Lysmd4 | 0.007 | Old |
| 0.617971 | 0.145544 | 0.171 | 0.192 | 1 | 10 | Etv5 | -0.021 | Old |
| 0.618163 | 0.32622 | 0.098 | 0.081 | 1 | 10 | Paox | 0.017 | Old |
| 0.618178 | 0.366652 | 0.139 | 0.121 | 1 | 10 | Avl9 | 0.018 | Old |
| 0.618698 | 0.143487 | 0.2 | 0.172 | 1 | 10 | Faim | 0.028 | Old |
| 0.619584 | 0.8823 | 0.029 | 0.02 | 1 | 10 | Gm42715 | 0.009 | Old |
| 0.619914 | 0.117168 | 0.042 | 0.03 | 1 | 10 | Mov10 | 0.012 | Old |
| 0.619975 | 0.048799 | 0.315 | 0.283 | 1 | 10 | Heatr5a | 0.032 | Old |
| 0.620056 | 0.319159 | 0.127 | 0.111 | 1 | 10 | Polg | 0.016 | Old |
| 0.620135 | 0.461285 | 0.132 | 0.111 | 1 | 10 | Copg2 | 0.021 | Old |
| 0.620202 | 0.041655 | 0.039 | 0.051 | 1 | 10 | Poc5 | -0.012 | Old |
| 0.620284 | 0.305006 | 0.098 | 0.081 | 1 | 10 | Poc1b | 0.017 | Old |
| 0.620335 | 0.724872 | 0.017 | 0.01 | 1 | 10 | Plpp3 | 0.007 | Old |
| 0.620335 | 0.667097 | 0.017 | 0.01 | 1 | 10 | Dagla | 0.007 | Old |
| 0.620335 | 0.493493 | 0.017 | 0.01 | 1 | 10 | Epor | 0.007 | Old |
| 0.620335 | 0.456635 | 0.017 | 0.01 | 1 | 10 | Zfp652os | 0.007 | Old |
| 0.620335 | 0.377965 | 0.017 | 0.01 | 1 | 10 | Dtnb | 0.007 | Old |
| 0.620335 | 0.259138 | 0.017 | 0.01 | 1 | 10 | Trim16 | 0.007 | Old |
| 0.620335 | 0.190844 | 0.017 | 0.01 | 1 | 10 | Cdiptos | 0.007 | Old |
| 0.621289 | 0.58332 | 0.117 | 0.101 | 1 | 10 | Tomm6 | 0.016 | Old |

| | | | | | | | | |
|---|---|---|---|---|---|---|---|---|
| 0.621483 | 0.324299 | 0.029 | 0.02 | 1 | 10 | Galnt9 | 0.009 | Old |
| 0.621483 | 0.207189 | 0.029 | 0.02 | 1 | 10 | Gm45871 | 0.009 | Old |
| 0.621764 | 0.501352 | 0.039 | 0.051 | 1 | 10 | Sema6b | -0.012 | Old |
| 0.621787 | 0.204457 | 0.108 | 0.091 | 1 | 10 | Rbbp5 | 0.017 | Old |
| 0.621926 | 0.372056 | 0.213 | 0.192 | 1 | 10 | Arhgap12 | 0.021 | Old |
| 0.621938 | 0.476786 | 0.127 | 0.111 | 1 | 10 | Cdk10 | 0.016 | Old |
| 0.622834 | 0.363644 | 0.017 | 0.01 | 1 | 10 | Gm14295 | 0.007 | Old |
| 0.622834 | 0.272925 | 0.017 | 0.01 | 1 | 10 | Vmac | 0.007 | Old |
| 0.622834 | 0.214125 | 0.017 | 0.01 | 1 | 10 | Kctd21 | 0.007 | Old |
| 0.622834 | 0.213577 | 0.017 | 0.01 | 1 | 10 | Gm26624 | 0.007 | Old |
| 0.622834 | 0.19782 | 0.017 | 0.01 | 1 | 10 | Gm11713 | 0.007 | Old |
| 0.622834 | 0.14441 | 0.017 | 0.01 | 1 | 10 | Gm14296 | 0.007 | Old |
| 0.622834 | 0.10317 | 0.017 | 0.01 | 1 | 10 | AC163018. | 0.007 | Old |
| 0.622834 | 0.067611 | 0.017 | 0.01 | 1 | 10 | Zfp324 | 0.007 | Old |
| 0.622834 | 0.005788 | 0.017 | 0.01 | 1 | 10 | Dclk1 | 0.007 | Old |
| 0.622893 | 0.140528 | 0.174 | 0.152 | 1 | 10 | Flrt2 | 0.022 | Old |
| 0.623385 | 0.769389 | 0.029 | 0.02 | 1 | 10 | Spindoc | 0.009 | Old |
| 0.623385 | 0.487504 | 0.029 | 0.02 | 1 | 10 | Gm37168 | 0.009 | Old |
| 0.623385 | 0.371051 | 0.029 | 0.02 | 1 | 10 | Capn5 | 0.009 | Old |
| 0.623385 | 0.309278 | 0.029 | 0.02 | 1 | 10 | Hspa2 | 0.009 | Old |
| 0.623385 | 0.153924 | 0.029 | 0.02 | 1 | 10 | Gm37233 | 0.009 | Old |
| 0.623584 | 0.110046 | 0.296 | 0.263 | 1 | 10 | Golgb1 | 0.033 | Old |
| 0.623706 | 0.103679 | 0.171 | 0.152 | 1 | 10 | Arhgef3 | 0.019 | Old |
| 0.624893 | 0.137324 | 0.039 | 0.051 | 1 | 10 | AI480526 | -0.012 | Old |
| 0.624984 | 0.536629 | 0.095 | 0.081 | 1 | 10 | Bet1l | 0.014 | Old |
| 0.625183 | 0.217426 | 0.164 | 0.141 | 1 | 10 | Xpo7 | 0.023 | Old |
| 0.625289 | 0.318307 | 0.029 | 0.02 | 1 | 10 | Zfhx2 | 0.009 | Old |
| 0.62569 | 0.037098 | 0.11 | 0.091 | 1 | 10 | Mid1 | 0.019 | Old |
| 0.625977 | 0.081896 | 0.183 | 0.162 | 1 | 10 | Ash2l | 0.021 | Old |
| 0.626031 | 0.04485 | 0.164 | 0.141 | 1 | 10 | Cdc42bpb | 0.023 | Old |
| 0.626146 | 0.15538 | 0.355 | 0.323 | 1 | 10 | Slc35f6 | 0.032 | Old |
| 0.626907 | 0.204366 | 0.076 | 0.061 | 1 | 10 | Rbmx2 | 0.015 | Old |
| 0.627731 | 0.090974 | 0.098 | 0.081 | 1 | 10 | Gtpbp10 | 0.017 | Old |
| 0.628108 | 0.304732 | 0.076 | 0.061 | 1 | 10 | Lrch4 | 0.015 | Old |
| 0.62852 | 0.133608 | 0.716 | 0.667 | 1 | 10 | Ogt | 0.049 | Old |
| 0.628653 | 0.123167 | 0.076 | 0.091 | 1 | 10 | D430042O( | -0.015 | Old |
| 0.629054 | 0.262599 | 0.139 | 0.121 | 1 | 10 | Zcchc2 | 0.018 | Old |
| 0.629097 | 0.138308 | 0.421 | 0.384 | 1 | 10 | Rnf4 | 0.037 | Old |
| 0.629105 | 0.356714 | 0.029 | 0.02 | 1 | 10 | Prkd2 | 0.009 | Old |
| 0.629105 | 0.261467 | 0.029 | 0.02 | 1 | 10 | Pou6f1 | 0.009 | Old |
| 0.629233 | 0.24452 | 0.357 | 0.333 | 1 | 10 | Nenf | 0.024 | Old |
| 0.629273 | 0.417176 | 0.159 | 0.141 | 1 | 10 | Agpat3 | 0.018 | Old |
| 0.629899 | 0.306849 | 0.064 | 0.051 | 1 | 10 | Casp7 | 0.013 | Old |
| 0.630392 | 0.015985 | 0.244 | 0.212 | 1 | 10 | Nab1 | 0.032 | Old |
| 0.631143 | 0.063606 | 0.042 | 0.03 | 1 | 10 | Hells | 0.012 | Old |

| | | | | | | | | |
|---|---|---|---|---|---|---|---|---|
| 0.631691 | 0.063152 | 0.66 | 0.626 | 1 | 10 | Naa50 | 0.034 | Old |
| 0.632185 | 0.023934 | 0.198 | 0.172 | 1 | 10 | Emc8 | 0.026 | Old |
| 0.632228 | 0.098824 | 0.022 | 0.03 | 1 | 10 | Zfp763 | -0.008 | Old |
| 0.632746 | 0.069742 | 0.039 | 0.051 | 1 | 10 | Snx25 | -0.012 | Old |
| 0.633026 | 0.051856 | 0.834 | 0.778 | 1 | 10 | Atp5g2 | 0.056 | Old |
| 0.633074 | 0.019017 | 0.098 | 0.081 | 1 | 10 | Brf1 | 0.017 | Old |
| 0.633477 | 0.203472 | 0.271 | 0.253 | 1 | 10 | Atp2c1 | 0.018 | Old |
| 0.633919 | 0.123724 | 0.289 | 0.253 | 1 | 10 | Ugcg | 0.036 | Old |
| 0.634848 | 0.174191 | 0.029 | 0.02 | 1 | 10 | Mcat | 0.009 | Old |
| 0.635066 | 0.052276 | 0.21 | 0.182 | 1 | 10 | Helz | 0.028 | Old |
| 0.635263 | 0.825251 | 0.051 | 0.04 | 1 | 10 | Bid | 0.011 | Old |
| 0.635566 | 0.022116 | 0.949 | 0.899 | 1 | 10 | Rps15a | 0.05 | Old |
| 0.635733 | 0.62322 | 0.112 | 0.101 | 1 | 10 | Mavs | 0.011 | Old |
| 0.636446 | 0.151976 | 0.064 | 0.051 | 1 | 10 | Scai | 0.013 | Old |
| 0.636668 | 0.080777 | 0.724 | 0.657 | 1 | 10 | Hnrnpm | 0.067 | Old |
| 0.636931 | 0.138251 | 0.132 | 0.111 | 1 | 10 | Malsu1 | 0.021 | Old |
| 0.637747 | 0.514303 | 0.073 | 0.061 | 1 | 10 | Creb3l1 | 0.012 | Old |
| 0.638433 | 0.651545 | 0.022 | 0.03 | 1 | 10 | Ccl22 | -0.008 | Old |
| 0.639457 | 0.090042 | 0.748 | 0.747 | 1 | 10 | Vat1 | 0.001 | Old |
| 0.639512 | 0.030704 | 0.264 | 0.232 | 1 | 10 | Ccdc124 | 0.032 | Old |
| 0.640005 | 0.264728 | 0.2 | 0.182 | 1 | 10 | Dicer1 | 0.018 | Old |
| 0.640406 | 0.148636 | 0.171 | 0.152 | 1 | 10 | Dirc2 | 0.019 | Old |
| 0.64108 | 0.649132 | 0.051 | 0.04 | 1 | 10 | Gtf2h3 | 0.011 | Old |
| 0.64108 | 0.627854 | 0.051 | 0.04 | 1 | 10 | Mamdc2 | 0.011 | Old |
| 0.641195 | 0.343874 | 0.095 | 0.081 | 1 | 10 | Trim11 | 0.014 | Old |
| 0.641662 | 0.087674 | 0.098 | 0.081 | 1 | 10 | Inip | 0.017 | Old |
| 0.642148 | 0.032262 | 0.995 | 0.99 | 1 | 10 | Rps24 | 0.005 | Old |
| 0.642583 | 0.580283 | 0.022 | 0.03 | 1 | 10 | Slfn9 | -0.008 | Old |
| 0.642583 | 0.192574 | 0.022 | 0.03 | 1 | 10 | 2610301B2 | -0.008 | Old |
| 0.642739 | 0.154014 | 0.098 | 0.081 | 1 | 10 | R3hcc1l | 0.017 | Old |
| 0.644398 | 0.015138 | 0.24 | 0.212 | 1 | 10 | Mecp2 | 0.028 | Old |
| 0.644458 | 0.009406 | 0.095 | 0.081 | 1 | 10 | Rev1 | 0.014 | Old |
| 0.644584 | 0.18391 | 0.325 | 0.293 | 1 | 10 | Ikzf1 | 0.032 | Old |
| 0.646562 | 0.034561 | 0.086 | 0.071 | 1 | 10 | Mcm3ap | 0.015 | Old |
| 0.646919 | 0.425617 | 0.051 | 0.04 | 1 | 10 | Fam210b | 0.011 | Old |
| 0.646919 | 0.393017 | 0.051 | 0.04 | 1 | 10 | Sugp2 | 0.011 | Old |
| 0.647336 | 0.056047 | 0.264 | 0.232 | 1 | 10 | Cdc73 | 0.032 | Old |
| 0.647636 | 0.196091 | 0.274 | 0.253 | 1 | 10 | Arnt | 0.021 | Old |
| 0.648275 | 0.128852 | 0.399 | 0.374 | 1 | 10 | Pxn | 0.025 | Old |
| 0.648773 | 0.249298 | 0.105 | 0.091 | 1 | 10 | Fzd7 | 0.014 | Old |
| 0.649626 | 0.177679 | 0.064 | 0.051 | 1 | 10 | Ptcd3 | 0.013 | Old |
| 0.649847 | 0.360562 | 0.051 | 0.04 | 1 | 10 | Tmco6 | 0.011 | Old |
| 0.650834 | 0.034649 | 0.144 | 0.162 | 1 | 10 | Med16 | -0.018 | Old |
| 0.651005 | 0.114275 | 0.095 | 0.081 | 1 | 10 | Zdhhc4 | 0.014 | Old |
| 0.651822 | 0.053793 | 0.12 | 0.101 | 1 | 10 | Heatr6 | 0.019 | Old |

| | | | | | | | | |
|---|---|---|---|---|---|---|---|---|
| 0.651822 | 0.033116 | 0.12 | 0.101 | 1 | 10 | Card11 | 0.019 | Old |
| 0.651942 | 0.123254 | 0.176 | 0.152 | 1 | 10 | Smc4 | 0.024 | Old |
| 0.652285 | 0.300884 | 0.379 | 0.354 | 1 | 10 | Spty2d1 | 0.025 | Old |
| 0.653458 | 0.032167 | 0.149 | 0.131 | 1 | 10 | Oard1 | 0.018 | Old |
| 0.653462 | 0.049832 | 0.086 | 0.071 | 1 | 10 | AI987944 | 0.015 | Old |
| 0.653527 | 0.00534 | 0.802 | 0.737 | 1 | 10 | Picalm | 0.065 | Old |
| 0.653602 | 0.288963 | 0.064 | 0.051 | 1 | 10 | AC165953. | 0.013 | Old |
| 0.653916 | 0.210627 | 0.132 | 0.111 | 1 | 10 | Rin3 | 0.021 | Old |
| 0.654155 | 1.071826 | 0.039 | 0.03 | 1 | 10 | Usp46 | 0.009 | Old |
| 0.654249 | 0.523665 | 0.051 | 0.04 | 1 | 10 | Stau2 | 0.011 | Old |
| 0.654598 | 0.000445 | 0.13 | 0.111 | 1 | 10 | Pygo2 | 0.019 | Old |
| 0.654981 | 0.404295 | 0.166 | 0.152 | 1 | 10 | Rnf103 | 0.014 | Old |
| 0.655719 | 0.297992 | 0.051 | 0.04 | 1 | 10 | Setdb2 | 0.011 | Old |
| 0.656176 | 0.24137 | 0.303 | 0.283 | 1 | 10 | Ypel3 | 0.02 | Old |
| 0.656605 | 0.166629 | 0.049 | 0.061 | 1 | 10 | Ccdc28a | -0.012 | Old |
| 0.656668 | 1.054574 | 0.061 | 0.051 | 1 | 10 | Igip | 0.01 | Old |
| 0.657018 | 0.261912 | 0.159 | 0.141 | 1 | 10 | Tmem55a | 0.018 | Old |
| 0.657471 | 0.222935 | 0.073 | 0.061 | 1 | 10 | Lzic | 0.012 | Old |
| 0.657927 | 0.358326 | 0.076 | 0.091 | 1 | 10 | Tpk1 | -0.015 | Old |
| 0.658155 | 0.321504 | 0.127 | 0.111 | 1 | 10 | Gm37494 | 0.016 | Old |
| 0.658663 | 0.121851 | 0.051 | 0.04 | 1 | 10 | Dffa | 0.011 | Old |
| 0.658918 | 0.057126 | 0.064 | 0.051 | 1 | 10 | Slc10a3 | 0.013 | Old |
| 0.659201 | 0.399754 | 0.083 | 0.071 | 1 | 10 | Etv1 | 0.012 | Old |
| 0.660723 | 0.70476 | 0.061 | 0.051 | 1 | 10 | Pan2 | 0.01 | Old |
| 0.660723 | 0.69691 | 0.061 | 0.051 | 1 | 10 | Slc9a3r2 | 0.01 | Old |
| 0.661571 | 0.330459 | 0.137 | 0.121 | 1 | 10 | Dohh | 0.016 | Old |
| 0.661614 | 0.229041 | 0.188 | 0.172 | 1 | 10 | Fam107b | 0.016 | Old |
| 0.661822 | 0.050308 | 0.694 | 0.606 | 1 | 10 | Atp6ap2 | 0.088 | Old |
| 0.661952 | 0.151932 | 0.23 | 0.202 | 1 | 10 | Lrp12 | 0.028 | Old |
| 0.663018 | 0.21409 | 0.218 | 0.202 | 1 | 10 | Commd4 | 0.016 | Old |
| 0.663027 | 0.009049 | 0.711 | 0.667 | 1 | 10 | Ndufa6 | 0.044 | Old |
| 0.663077 | 0.007284 | 0.095 | 0.081 | 1 | 10 | Tacc2 | 0.014 | Old |
| 0.663088 | 0.216726 | 0.051 | 0.04 | 1 | 10 | Pdpr | 0.011 | Old |
| 0.663312 | 0.034099 | 0.318 | 0.293 | 1 | 10 | Hbp1 | 0.025 | Old |
| 0.66423 | 0.341289 | 0.267 | 0.253 | 1 | 10 | Acaa1a | 0.014 | Old |
| 0.664237 | 1.0813 | 0.039 | 0.03 | 1 | 10 | Klhl25 | 0.009 | Old |
| 0.664237 | 0.793688 | 0.039 | 0.03 | 1 | 10 | Haus6 | 0.009 | Old |
| 0.664252 | 0.006223 | 0.059 | 0.071 | 1 | 10 | Atrnl1 | -0.012 | Old |
| 0.66426 | 0.231028 | 0.149 | 0.131 | 1 | 10 | Necap1 | 0.018 | Old |
| 0.664482 | 0.034378 | 0.819 | 0.778 | 1 | 10 | Cox6c | 0.041 | Old |
| 0.664626 | 0.09336 | 0.254 | 0.232 | 1 | 10 | Susd3 | 0.022 | Old |
| 0.665208 | 0.924404 | 0.071 | 0.061 | 1 | 10 | Tada3 | 0.01 | Old |
| 0.666608 | 0.308927 | 0.105 | 0.091 | 1 | 10 | Abi3 | 0.014 | Old |
| 0.666805 | 0.013703 | 0.249 | 0.212 | 1 | 10 | Rnaseh2c | 0.037 | Old |
| 0.667612 | 0.369298 | 0.039 | 0.03 | 1 | 10 | Zbtb16 | 0.009 | Old |

| | | | | | | | | |
|---|---|---|---|---|---|---|---|---|
| 0.667643 | 0.35824 | 0.227 | 0.212 | 1 | 10 | Clock | 0.015 | Old |
| 0.667996 | 0.464488 | 0.115 | 0.101 | 1 | 10 | Tmem184c | 0.014 | Old |
| 0.669302 | 0.902868 | 0.039 | 0.03 | 1 | 10 | Ier5l | 0.009 | Old |
| 0.669582 | 0.011016 | 0.186 | 0.162 | 1 | 10 | Znfx1 | 0.024 | Old |
| 0.669808 | 0.126725 | 0.313 | 0.283 | 1 | 10 | Mrpl51 | 0.03 | Old |
| 0.669891 | 0.032301 | 0.528 | 0.465 | 1 | 10 | Myl12b | 0.063 | Old |
| 0.670491 | 0.341587 | 0.051 | 0.04 | 1 | 10 | Cep128 | 0.011 | Old |
| 0.670491 | 0.06315 | 0.051 | 0.04 | 1 | 10 | Saal1 | 0.011 | Old |
| 0.670546 | 0.051088 | 0.152 | 0.131 | 1 | 10 | Stat5a | 0.021 | Old |
| 0.670888 | 0.212066 | 0.117 | 0.101 | 1 | 10 | Tnfrsf12a | 0.016 | Old |
| 0.670993 | 0.513602 | 0.039 | 0.03 | 1 | 10 | Traf3ip1 | 0.009 | Old |
| 0.671505 | 0.248733 | 0.176 | 0.162 | 1 | 10 | Zfp217 | 0.014 | Old |
| 0.672115 | 0.731887 | 0.083 | 0.071 | 1 | 10 | Gm1673 | 0.012 | Old |
| 0.672686 | 0.591774 | 0.039 | 0.03 | 1 | 10 | Zfp595 | 0.009 | Old |
| 0.672931 | 0.025166 | 0.399 | 0.364 | 1 | 10 | Lsm6 | 0.035 | Old |
| 0.67382 | 0.164236 | 0.13 | 0.111 | 1 | 10 | Ankra2 | 0.019 | Old |
| 0.673975 | 0.10752 | 0.252 | 0.232 | 1 | 10 | Abhd17a | 0.02 | Old |
| 0.674381 | 0.784016 | 0.039 | 0.03 | 1 | 10 | Zfyve28 | 0.009 | Old |
| 0.674381 | 0.259062 | 0.039 | 0.03 | 1 | 10 | Ficd | 0.009 | Old |
| 0.674381 | 0.154124 | 0.039 | 0.03 | 1 | 10 | Clcf1 | 0.009 | Old |
| 0.674474 | 0.351204 | 0.083 | 0.071 | 1 | 10 | Rbm19 | 0.012 | Old |
| 0.675008 | 0.131744 | 0.46 | 0.444 | 1 | 10 | Derl1 | 0.016 | Old |
| 0.675625 | 0.035602 | 0.335 | 0.303 | 1 | 10 | Gngt2 | 0.032 | Old |
| 0.676041 | 0.017458 | 0.149 | 0.131 | 1 | 10 | Bop1 | 0.018 | Old |
| 0.676076 | 0.40963 | 0.156 | 0.141 | 1 | 10 | Ahr | 0.015 | Old |
| 0.677156 | 0.279783 | 0.032 | 0.04 | 1 | 10 | Eif5a2 | -0.008 | Old |
| 0.677775 | 0.423095 | 0.039 | 0.03 | 1 | 10 | Gtf3c5 | 0.009 | Old |
| 0.677775 | 0.338025 | 0.039 | 0.03 | 1 | 10 | Rfx5 | 0.009 | Old |
| 0.677951 | 0.261047 | 0.269 | 0.253 | 1 | 10 | Snrnp200 | 0.016 | Old |
| 0.678675 | 0.092443 | 0.24 | 0.212 | 1 | 10 | Lilrb4a | 0.028 | Old |
| 0.679618 | 0.204571 | 0.364 | 0.343 | 1 | 10 | Zc3h7a | 0.021 | Old |
| 0.68037 | 0.060591 | 0.137 | 0.121 | 1 | 10 | Dnlz | 0.016 | Old |
| 0.680466 | 0.048614 | 0.384 | 0.364 | 1 | 10 | Fam102b | 0.02 | Old |
| 0.680479 | 0.25496 | 0.127 | 0.111 | 1 | 10 | Gpatch2l | 0.016 | Old |
| 0.680626 | 0.382675 | 0.222 | 0.202 | 1 | 10 | Fbxw2 | 0.02 | Old |
| 0.680718 | 0.014594 | 0.381 | 0.394 | 1 | 10 | Rnf19b | -0.013 | Old |
| 0.680744 | 0.091067 | 0.032 | 0.04 | 1 | 10 | Tsen2 | -0.008 | Old |
| 0.681177 | 0.397852 | 0.039 | 0.03 | 1 | 10 | Zfyve9 | 0.009 | Old |
| 0.681177 | 0.162793 | 0.039 | 0.03 | 1 | 10 | Klhl7 | 0.009 | Old |
| 0.682027 | 0.532203 | 0.154 | 0.141 | 1 | 10 | Slc4a1ap | 0.013 | Old |
| 0.682111 | 0.017193 | 0.267 | 0.242 | 1 | 10 | Rnpepl1 | 0.025 | Old |
| 0.68228 | 0.381619 | 0.103 | 0.121 | 1 | 10 | Cwc22 | -0.018 | Old |
| 0.682462 | 0.001244 | 0.357 | 0.313 | 1 | 10 | Nsd3 | 0.044 | Old |
| 0.683664 | 0.0388 | 0.998 | 1 | 1 | 10 | mt-Co2 | -0.002 | Old |
| 0.683888 | 0.330186 | 0.061 | 0.051 | 1 | 10 | Zfyve19 | 0.01 | Old |

| | | | | | | | | |
|---|---|---|---|---|---|---|---|---|
| 0.683888 | 0.269853 | 0.061 | 0.051 | 1 | 10 | Fars2 | 0.01 | Old |
| 0.685343 | 0.007927 | 0.479 | 0.434 | 1 | 10 | Spop | 0.045 | Old |
| 0.685457 | 0.023502 | 0.428 | 0.384 | 1 | 10 | Arhgef2 | 0.044 | Old |
| 0.68629 | 0.094184 | 0.039 | 0.03 | 1 | 10 | Gm16973 | 0.009 | Old |
| 0.686768 | 0.19269 | 0.24 | 0.222 | 1 | 10 | Bcl7b | 0.018 | Old |
| 0.687833 | 0.395766 | 0.105 | 0.091 | 1 | 10 | Lysmd3 | 0.014 | Old |
| 0.68794 | 0.421228 | 0.032 | 0.04 | 1 | 10 | Acvr2b | -0.008 | Old |
| 0.687998 | 0.29049 | 0.039 | 0.03 | 1 | 10 | Stk38l | 0.009 | Old |
| 0.687998 | 0.207004 | 0.039 | 0.03 | 1 | 10 | Pls3 | 0.009 | Old |
| 0.689534 | 0.166874 | 0.271 | 0.253 | 1 | 10 | P2rx7 | 0.018 | Old |
| 0.689664 | 0.390864 | 0.093 | 0.081 | 1 | 10 | Wiz | 0.012 | Old |
| 0.689745 | 0.000233 | 0.521 | 0.455 | 1 | 10 | Spred1 | 0.066 | Old |
| 0.689815 | 0.286178 | 0.059 | 0.071 | 1 | 10 | Serf1 | -0.012 | Old |
| 0.691073 | 0.266082 | 0.083 | 0.071 | 1 | 10 | Tmub1 | 0.012 | Old |
| 0.691377 | 0.081864 | 0.413 | 0.384 | 1 | 10 | Midn | 0.029 | Old |
| 0.692012 | 0.176827 | 0.105 | 0.121 | 1 | 10 | Mtmr2 | -0.016 | Old |
| 0.692484 | 0.219299 | 0.149 | 0.131 | 1 | 10 | Gpr155 | 0.018 | Old |
| 0.693265 | 0.11134 | 0.159 | 0.141 | 1 | 10 | Mcoln1 | 0.018 | Old |
| 0.693559 | 0.111032 | 0.425 | 0.384 | 1 | 10 | Dnajc5 | 0.041 | Old |
| 0.693564 | 0.320604 | 0.147 | 0.131 | 1 | 10 | Cpne2 | 0.016 | Old |
| 0.693743 | 0.084721 | 0.042 | 0.051 | 1 | 10 | 5730480H( | -0.009 | Old |
| 0.69408 | 1.816705 | 0.027 | 0.02 | 1 | 10 | Pclaf | 0.007 | Old |
| 0.69408 | 1.074571 | 0.027 | 0.02 | 1 | 10 | Zfp251 | 0.007 | Old |
| 0.69408 | 0.867295 | 0.027 | 0.02 | 1 | 10 | Alkbh4 | 0.007 | Old |
| 0.69425 | 0.130573 | 0.105 | 0.091 | 1 | 10 | Filip1l | 0.014 | Old |
| 0.695244 | 0.310023 | 0.059 | 0.071 | 1 | 10 | Exd2 | -0.012 | Old |
| 0.695789 | 0.098632 | 0.328 | 0.303 | 1 | 10 | Fam173a | 0.025 | Old |
| 0.696141 | 0.674676 | 0.027 | 0.02 | 1 | 10 | 4632404H1 | 0.007 | Old |
| 0.696275 | 0.022305 | 0.939 | 0.919 | 1 | 10 | Serf2 | 0.02 | Old |
| 0.696452 | 0.23979 | 0.093 | 0.081 | 1 | 10 | Cpox | 0.012 | Old |
| 0.696603 | 0.26882 | 0.059 | 0.071 | 1 | 10 | Atg10 | -0.012 | Old |
| 0.696647 | 0.541977 | 0.142 | 0.131 | 1 | 10 | Mindy3 | 0.011 | Old |
| 0.697557 | 0.335918 | 0.249 | 0.232 | 1 | 10 | Cysltr1 | 0.017 | Old |
| 0.697656 | 0.120071 | 0.061 | 0.051 | 1 | 10 | Ltb4r1 | 0.01 | Old |
| 0.697656 | 0.105586 | 0.061 | 0.051 | 1 | 10 | Rps6ka5 | 0.01 | Old |
| 0.698233 | 0.011256 | 0.078 | 0.091 | 1 | 10 | Maged1 | -0.013 | Old |
| 0.69842 | 0.506619 | 0.015 | 0.02 | 1 | 10 | Zbtb48 | -0.005 | Old |
| 0.69842 | 0.303691 | 0.015 | 0.02 | 1 | 10 | Gm340 | -0.005 | Old |
| 0.69842 | 0.193098 | 0.015 | 0.02 | 1 | 10 | Sfxn5 | -0.005 | Old |
| 0.69842 | 0.064941 | 0.015 | 0.02 | 1 | 10 | Pgm5 | -0.005 | Old |
| 0.698786 | 0.07125 | 0.139 | 0.121 | 1 | 10 | Hic2 | 0.018 | Old |
| 0.699039 | 0.31999 | 0.061 | 0.051 | 1 | 10 | Adgre4 | 0.01 | Old |
| 0.69905 | 0.022621 | 0.311 | 0.283 | 1 | 10 | Wbp1l | 0.028 | Old |
| 0.699141 | 0.133917 | 0.252 | 0.232 | 1 | 10 | Trpv2 | 0.02 | Old |
| 0.700075 | 0.25611 | 0.166 | 0.152 | 1 | 10 | Snhg6 | 0.014 | Old |

| | | | | | | | | |
|---|---|---|---|---|---|---|---|---|
| 0.700114 | 0.330684 | 0.125 | 0.111 | 1 | 10 | Ccpg1 | 0.014 | Old |
| 0.70027 | 1.063005 | 0.027 | 0.02 | 1 | 10 | Zfp931 | 0.007 | Old |
| 0.70027 | 0.984188 | 0.027 | 0.02 | 1 | 10 | Gca | 0.007 | Old |
| 0.70027 | 0.665076 | 0.027 | 0.02 | 1 | 10 | Apex2 | 0.007 | Old |
| 0.70027 | 0.435407 | 0.027 | 0.02 | 1 | 10 | Kiz | 0.007 | Old |
| 0.700688 | 0.162728 | 0.105 | 0.091 | 1 | 10 | Phtf2 | 0.014 | Old |
| 0.701041 | 0.493678 | 0.015 | 0.02 | 1 | 10 | Fat3 | -0.005 | Old |
| 0.701041 | 0.321967 | 0.015 | 0.02 | 1 | 10 | Cd244 | -0.005 | Old |
| 0.701823 | 0.296749 | 0.083 | 0.071 | 1 | 10 | Ipo13 | 0.012 | Old |
| 0.702098 | 0.029553 | 0.127 | 0.111 | 1 | 10 | B9d2 | 0.016 | Old |
| 0.702128 | 0.157743 | 0.093 | 0.081 | 1 | 10 | Mrpl46 | 0.012 | Old |
| 0.702338 | 0.658244 | 0.027 | 0.02 | 1 | 10 | B3gnt3 | 0.007 | Old |
| 0.702338 | 0.55042 | 0.027 | 0.02 | 1 | 10 | Gm43569 | 0.007 | Old |
| 0.702531 | 0.069869 | 0.99 | 1 | 1 | 10 | Rpsa | -0.01 | Old |
| 0.702611 | 0.249399 | 0.22 | 0.202 | 1 | 10 | Kcnk6 | 0.018 | Old |
| 0.703277 | 0.089447 | 0.232 | 0.212 | 1 | 10 | Cdc123 | 0.02 | Old |
| 0.703665 | 0.417837 | 0.015 | 0.02 | 1 | 10 | Amot | -0.005 | Old |
| 0.703665 | 0.321752 | 0.015 | 0.02 | 1 | 10 | Adap2os | -0.005 | Old |
| 0.703665 | 0.307437 | 0.015 | 0.02 | 1 | 10 | Ak8 | -0.005 | Old |
| 0.703665 | 0.269804 | 0.015 | 0.02 | 1 | 10 | Frmd8os | -0.005 | Old |
| 0.703665 | 0.186914 | 0.015 | 0.02 | 1 | 10 | Tspyl3 | -0.005 | Old |
| 0.70411 | 0.149416 | 0.169 | 0.152 | 1 | 10 | Scfd1 | 0.017 | Old |
| 0.704408 | 0.794319 | 0.027 | 0.02 | 1 | 10 | Polrmt | 0.007 | Old |
| 0.704408 | 0.069904 | 0.027 | 0.02 | 1 | 10 | Trim7 | 0.007 | Old |
| 0.704578 | 0.185224 | 0.061 | 0.051 | 1 | 10 | Asb6 | 0.01 | Old |
| 0.705966 | 0.180767 | 0.061 | 0.051 | 1 | 10 | Enc1 | 0.01 | Old |
| 0.705971 | 0.077992 | 0.196 | 0.182 | 1 | 10 | Dram1 | 0.014 | Old |
| 0.706099 | 0.338599 | 0.122 | 0.111 | 1 | 10 | Manbal | 0.011 | Old |
| 0.70648 | 0.34516 | 0.027 | 0.02 | 1 | 10 | Plpp1 | 0.007 | Old |
| 0.708384 | 0.271902 | 0.112 | 0.101 | 1 | 10 | Ubac2 | 0.011 | Old |
| 0.708511 | 0.923222 | 0.059 | 0.051 | 1 | 10 | Mrps31 | 0.008 | Old |
| 0.708555 | 0.5212 | 0.027 | 0.02 | 1 | 10 | Fam185a | 0.007 | Old |
| 0.708555 | 0.098772 | 0.027 | 0.02 | 1 | 10 | Inpp1 | 0.007 | Old |
| 0.708668 | 0.492547 | 0.071 | 0.061 | 1 | 10 | Mettl3 | 0.01 | Old |
| 0.709234 | 0.19795 | 0.242 | 0.222 | 1 | 10 | Dpf2 | 0.02 | Old |
| 0.709537 | 0.15105 | 0.142 | 0.162 | 1 | 10 | Pstpip1 | -0.02 | Old |
| 0.709758 | 0.11639 | 0.166 | 0.152 | 1 | 10 | Ice1 | 0.014 | Old |
| 0.710631 | 0.205894 | 0.027 | 0.02 | 1 | 10 | Upb1 | 0.007 | Old |
| 0.710907 | 0.027896 | 0.369 | 0.333 | 1 | 10 | Piezo1 | 0.036 | Old |
| 0.711005 | 0.500418 | 0.049 | 0.04 | 1 | 10 | Pus3 | 0.009 | Old |
| 0.711037 | 0.032354 | 0.976 | 0.96 | 1 | 10 | Grn | 0.016 | Old |
| 0.711596 | 0.025804 | 0.171 | 0.152 | 1 | 10 | Tdg | 0.019 | Old |
| 0.712386 | 0.029888 | 0.093 | 0.081 | 1 | 10 | Dhrs4 | 0.012 | Old |
| 0.712552 | 0.282646 | 0.049 | 0.04 | 1 | 10 | Nos1ap | 0.009 | Old |
| 0.712647 | 0.065932 | 0.147 | 0.162 | 1 | 10 | Gpr34 | -0.015 | Old |

| | | | | | | | | |
|---|---|---|---|---|---|---|---|---|
| 0.71271 | 0.039941 | 0.027 | 0.02 | 1 | 10 | Ogg1 | 0.007 | Old |
| 0.71271 | 0.039711 | 0.027 | 0.02 | 1 | 10 | Slco3a1 | 0.007 | Old |
| 0.712901 | 0.146688 | 0.213 | 0.192 | 1 | 10 | Smad1 | 0.021 | Old |
| 0.713612 | 0.055513 | 0.108 | 0.121 | 1 | 10 | Surf1 | -0.013 | Old |
| 0.714791 | 0.264689 | 0.027 | 0.02 | 1 | 10 | Zfp873 | 0.007 | Old |
| 0.714791 | 0.041965 | 0.027 | 0.02 | 1 | 10 | C1qtnf6 | 0.007 | Old |
| 0.715395 | 0.109205 | 0.286 | 0.263 | 1 | 10 | Pim3 | 0.023 | Old |
| 0.715951 | 0.384436 | 0.174 | 0.162 | 1 | 10 | Vrk2 | 0.012 | Old |
| 0.717963 | 0.169201 | 0.105 | 0.091 | 1 | 10 | Mb21d1 | 0.014 | Old |
| 0.718037 | 0.308948 | 0.081 | 0.071 | 1 | 10 | Gon7 | 0.01 | Old |
| 0.718291 | 0.0318 | 0.447 | 0.404 | 1 | 10 | Tspan3 | 0.043 | Old |
| 0.718496 | 0.002707 | 0.513 | 0.475 | 1 | 10 | Il6ra | 0.038 | Old |
| 0.71904 | 0.314713 | 0.071 | 0.061 | 1 | 10 | Clmp | 0.01 | Old |
| 0.719254 | 0.134146 | 0.093 | 0.081 | 1 | 10 | Pdp2 | 0.012 | Old |
| 0.719594 | 1.665856 | 0.015 | 0.01 | 1 | 10 | Glis3 | 0.005 | Old |
| 0.719594 | 1.452849 | 0.015 | 0.01 | 1 | 10 | Mtbp | 0.005 | Old |
| 0.719594 | 1.362158 | 0.015 | 0.01 | 1 | 10 | Dock6 | 0.005 | Old |
| 0.719594 | 1.13167 | 0.015 | 0.01 | 1 | 10 | Zfp235 | 0.005 | Old |
| 0.719594 | 1.05321 | 0.015 | 0.01 | 1 | 10 | Eml6 | 0.005 | Old |
| 0.719594 | 1.040232 | 0.015 | 0.01 | 1 | 10 | Gm38251 | 0.005 | Old |
| 0.719594 | 0.99614 | 0.015 | 0.01 | 1 | 10 | Gm36266 | 0.005 | Old |
| 0.719594 | 0.989019 | 0.015 | 0.01 | 1 | 10 | Gm16062 | 0.005 | Old |
| 0.719594 | 0.959621 | 0.015 | 0.01 | 1 | 10 | Rasgrf2 | 0.005 | Old |
| 0.719594 | 0.842785 | 0.015 | 0.01 | 1 | 10 | 9130011E1 | 0.005 | Old |
| 0.719594 | 0.841353 | 0.015 | 0.01 | 1 | 10 | Gpr176 | 0.005 | Old |
| 0.719594 | 0.834339 | 0.015 | 0.01 | 1 | 10 | 1810034E1 | 0.005 | Old |
| 0.719982 | 0.199935 | 0.103 | 0.091 | 1 | 10 | Snx11 | 0.012 | Old |
| 0.721074 | 0.210225 | 0.535 | 0.525 | 1 | 10 | Rnf13 | 0.01 | Old |
| 0.721548 | 0.252396 | 0.088 | 0.101 | 1 | 10 | Tbc1d8 | -0.013 | Old |
| 0.721704 | 0.409546 | 0.081 | 0.071 | 1 | 10 | Dcp1a | 0.01 | Old |
| 0.722423 | 1.268552 | 0.015 | 0.01 | 1 | 10 | 4930430E1 | 0.005 | Old |
| 0.722423 | 0.827869 | 0.015 | 0.01 | 1 | 10 | Rragd | 0.005 | Old |
| 0.722423 | 0.811836 | 0.015 | 0.01 | 1 | 10 | Ttc8 | 0.005 | Old |
| 0.722423 | 0.763408 | 0.015 | 0.01 | 1 | 10 | Xkr8 | 0.005 | Old |
| 0.722423 | 0.726259 | 0.015 | 0.01 | 1 | 10 | Zfp605 | 0.005 | Old |
| 0.722423 | 0.597558 | 0.015 | 0.01 | 1 | 10 | Card6 | 0.005 | Old |
| 0.72296 | 0.057846 | 0.274 | 0.253 | 1 | 10 | Zmynd11 | 0.021 | Old |
| 0.723466 | 0.104673 | 0.169 | 0.152 | 1 | 10 | Tmem173 | 0.017 | Old |
| 0.724366 | 0.104234 | 0.103 | 0.091 | 1 | 10 | Ascc2 | 0.012 | Old |
| 0.724509 | 0.003635 | 0.604 | 0.636 | 1 | 10 | Mapkapk2 | -0.032 | Old |
| 0.725255 | 0.808333 | 0.015 | 0.01 | 1 | 10 | Srd5a1 | 0.005 | Old |
| 0.725255 | 0.773555 | 0.015 | 0.01 | 1 | 10 | Neil2 | 0.005 | Old |
| 0.725255 | 0.740444 | 0.015 | 0.01 | 1 | 10 | Fhad1 | 0.005 | Old |
| 0.725255 | 0.737561 | 0.015 | 0.01 | 1 | 10 | Rab11fip3 | 0.005 | Old |
| 0.725255 | 0.692143 | 0.015 | 0.01 | 1 | 10 | D130020L0 | 0.005 | Old |

| | | | | | | | | |
|---|---|---|---|---|---|---|---|---|
| 0.725255 | 0.661008 | 0.015 | 0.01 | 1 | 10 | Them6 | 0.005 | Old |
| 0.725255 | 0.635422 | 0.015 | 0.01 | 1 | 10 | Htra3 | 0.005 | Old |
| 0.725255 | 0.4075 | 0.015 | 0.01 | 1 | 10 | Acp6 | 0.005 | Old |
| 0.725333 | 0.38103 | 0.191 | 0.182 | 1 | 10 | Trappc3 | 0.009 | Old |
| 0.725704 | 0.102965 | 0.166 | 0.152 | 1 | 10 | Strn | 0.014 | Old |
| 0.726527 | 0.295356 | 0.049 | 0.04 | 1 | 10 | Gramd3 | 0.009 | Old |
| 0.726558 | 0.093578 | 0.156 | 0.141 | 1 | 10 | Sash3 | 0.015 | Old |
| 0.726853 | 0.046321 | 0.071 | 0.061 | 1 | 10 | Megf8 | 0.01 | Old |
| 0.728088 | 0.035459 | 0.274 | 0.253 | 1 | 10 | Paf1 | 0.021 | Old |
| 0.728092 | 0.717081 | 0.015 | 0.01 | 1 | 10 | Zfp623 | 0.005 | Old |
| 0.728092 | 0.677558 | 0.015 | 0.01 | 1 | 10 | Gpr162 | 0.005 | Old |
| 0.728092 | 0.642233 | 0.015 | 0.01 | 1 | 10 | Akap17b | 0.005 | Old |
| 0.728092 | 0.598428 | 0.015 | 0.01 | 1 | 10 | Tspoap1 | 0.005 | Old |
| 0.728092 | 0.552082 | 0.015 | 0.01 | 1 | 10 | Gm16287 | 0.005 | Old |
| 0.728092 | 0.440533 | 0.015 | 0.01 | 1 | 10 | Pnpla1 | 0.005 | Old |
| 0.728092 | 0.401748 | 0.015 | 0.01 | 1 | 10 | Tgif2 | 0.005 | Old |
| 0.728092 | 0.289941 | 0.015 | 0.01 | 1 | 10 | Tmem154 | 0.005 | Old |
| 0.728274 | 0.134088 | 0.051 | 0.061 | 1 | 10 | Cald1 | -0.01 | Old |
| 0.728308 | 0.022385 | 0.213 | 0.232 | 1 | 10 | Tufm | -0.019 | Old |
| 0.728375 | 0.711785 | 0.059 | 0.051 | 1 | 10 | Spice1 | 0.008 | Old |
| 0.728759 | 0.167119 | 0.103 | 0.091 | 1 | 10 | Atr | 0.012 | Old |
| 0.729265 | 0.080244 | 0.154 | 0.141 | 1 | 10 | Napsa | 0.013 | Old |
| 0.730932 | 0.423971 | 0.015 | 0.01 | 1 | 10 | Hal | 0.005 | Old |
| 0.730932 | 0.419691 | 0.015 | 0.01 | 1 | 10 | Ccdc125 | 0.005 | Old |
| 0.730932 | 0.326567 | 0.015 | 0.01 | 1 | 10 | Adipoq | 0.005 | Old |
| 0.730932 | 0.190984 | 0.015 | 0.01 | 1 | 10 | Pex26 | 0.005 | Old |
| 0.730932 | 0.166204 | 0.015 | 0.01 | 1 | 10 | Ccdc166 | 0.005 | Old |
| 0.730958 | 0.120484 | 0.103 | 0.091 | 1 | 10 | Rpa1 | 0.012 | Old |
| 0.731205 | 0.259581 | 0.049 | 0.04 | 1 | 10 | Ksr1 | 0.009 | Old |
| 0.731228 | 0.738224 | 0.059 | 0.051 | 1 | 10 | Fam117a | 0.008 | Old |
| 0.731903 | 0.085595 | 0.088 | 0.101 | 1 | 10 | Ormdl1 | -0.013 | Old |
| 0.73312 | 0.023907 | 0.139 | 0.152 | 1 | 10 | Taf2 | -0.013 | Old |
| 0.733776 | 0.387227 | 0.015 | 0.01 | 1 | 10 | Ispd | 0.005 | Old |
| 0.733776 | 0.08353 | 0.015 | 0.01 | 1 | 10 | Ftx | 0.005 | Old |
| 0.733776 | 0.020425 | 0.015 | 0.01 | 1 | 10 | Sh2d4a | 0.005 | Old |
| 0.734874 | 0.404361 | 0.183 | 0.172 | 1 | 10 | Flot1 | 0.011 | Old |
| 0.73513 | 1.044649 | 0.037 | 0.03 | 1 | 10 | Zfp959 | 0.007 | Old |
| 0.735435 | 0.288464 | 0.122 | 0.111 | 1 | 10 | Cenpc1 | 0.011 | Old |
| 0.736004 | 0.222743 | 0.071 | 0.061 | 1 | 10 | Pyroxd1 | 0.01 | Old |
| 0.736925 | 0.848936 | 0.037 | 0.03 | 1 | 10 | Thumpd2 | 0.007 | Old |
| 0.737542 | 0.382196 | 0.095 | 0.111 | 1 | 10 | Slc33a1 | -0.016 | Old |
| 0.737861 | 0.048281 | 0.139 | 0.152 | 1 | 10 | Ltv1 | -0.013 | Old |
| 0.738293 | 0.062951 | 0.276 | 0.253 | 1 | 10 | Trio | 0.023 | Old |
| 0.738377 | 0.492378 | 0.059 | 0.051 | 1 | 10 | Aldh4a1 | 0.008 | Old |
| 0.73893 | 0.001727 | 0.9 | 0.869 | 1 | 10 | Atp5e | 0.031 | Old |

| | | | | | | | | |
|---|---|---|---|---|---|---|---|---|
| 0.738976 | 0.133958 | 0.23 | 0.212 | 1 | 10 | Ndufa9 | 0.018 | Old |
| 0.739025 | 0.039748 | 0.049 | 0.04 | 1 | 10 | Foxp4 | 0.009 | Old |
| 0.739588 | 0.253053 | 0.132 | 0.121 | 1 | 10 | Zbtb44 | 0.011 | Old |
| 0.739809 | 0.57407 | 0.059 | 0.051 | 1 | 10 | Bloc1s1 | 0.008 | Old |
| 0.739856 | 0.021229 | 0.274 | 0.253 | 1 | 10 | Ube2v2 | 0.021 | Old |
| 0.739937 | 0.014779 | 0.071 | 0.061 | 1 | 10 | Osbpl1a | 0.01 | Old |
| 0.741149 | 0.034286 | 0.093 | 0.081 | 1 | 10 | Mrpl39 | 0.012 | Old |
| 0.741389 | 0.019131 | 0.235 | 0.212 | 1 | 10 | Ss18 | 0.023 | Old |
| 0.74218 | 0.090495 | 0.188 | 0.172 | 1 | 10 | Psmd9 | 0.016 | Old |
| 0.743279 | 0.062589 | 0.897 | 0.909 | 1 | 10 | Rhoa | -0.012 | Old |
| 0.743665 | 0.064181 | 0.134 | 0.121 | 1 | 10 | Polr2h | 0.013 | Old |
| 0.743877 | 0.121037 | 0.496 | 0.485 | 1 | 10 | Tns3 | 0.011 | Old |
| 0.74414 | 0.297862 | 0.024 | 0.03 | 1 | 10 | Arrdc2 | -0.006 | Old |
| 0.744463 | 0.027995 | 0.247 | 0.222 | 1 | 10 | Ddhd2 | 0.025 | Old |
| 0.744572 | 0.136387 | 0.281 | 0.263 | 1 | 10 | Acp1 | 0.018 | Old |
| 0.745076 | 0.167147 | 0.081 | 0.071 | 1 | 10 | Nav2 | 0.01 | Old |
| 0.746055 | 0.023354 | 0.506 | 0.475 | 1 | 10 | Ppig | 0.031 | Old |
| 0.746251 | 0.339115 | 0.024 | 0.03 | 1 | 10 | Palm | -0.006 | Old |
| 0.747731 | 0.756708 | 0.037 | 0.03 | 1 | 10 | B3galt6 | 0.007 | Old |
| 0.747731 | 0.61553 | 0.037 | 0.03 | 1 | 10 | Neil1 | 0.007 | Old |
| 0.747731 | 0.597403 | 0.037 | 0.03 | 1 | 10 | Gm26881 | 0.007 | Old |
| 0.748364 | 0.152431 | 0.024 | 0.03 | 1 | 10 | Gm33103 | -0.006 | Old |
| 0.748364 | 0.092472 | 0.024 | 0.03 | 1 | 10 | Gm5431 | -0.006 | Old |
| 0.748791 | 0.3554 | 0.142 | 0.131 | 1 | 10 | Fbrsl1 | 0.011 | Old |
| 0.749202 | 0.083909 | 0.161 | 0.152 | 1 | 10 | Qrich1 | 0.009 | Old |
| 0.749531 | 0.009201 | 0.164 | 0.152 | 1 | 10 | Atp11c | 0.012 | Old |
| 0.749537 | 0.51015 | 0.037 | 0.03 | 1 | 10 | Tnfsf13os | 0.007 | Old |
| 0.749537 | 0.198703 | 0.037 | 0.03 | 1 | 10 | Macrod1 | 0.007 | Old |
| 0.750435 | 0.001426 | 0.159 | 0.172 | 1 | 10 | Cd2bp2 | -0.013 | Old |
| 0.751443 | 0.060963 | 0.934 | 0.879 | 1 | 10 | Rpl36a | 0.055 | Old |
| 0.752596 | 0.24748 | 0.024 | 0.03 | 1 | 10 | R3hcc1 | -0.006 | Old |
| 0.752744 | 0.061414 | 0.071 | 0.081 | 1 | 10 | Ino80b | -0.01 | Old |
| 0.752817 | 0.202518 | 0.122 | 0.111 | 1 | 10 | Pdrg1 | 0.011 | Old |
| 0.753152 | 0.970319 | 0.037 | 0.03 | 1 | 10 | Csrp2 | 0.007 | Old |
| 0.753198 | 0.500817 | 0.068 | 0.061 | 1 | 10 | Zer1 | 0.007 | Old |
| 0.753748 | 0.061016 | 0.081 | 0.071 | 1 | 10 | Vwa8 | 0.01 | Old |
| 0.75403 | 0.118984 | 0.071 | 0.081 | 1 | 10 | Pag1 | -0.01 | Old |
| 0.754313 | 0.023286 | 0.22 | 0.202 | 1 | 10 | Ormdl2 | 0.018 | Old |
| 0.754344 | 0.065541 | 0.166 | 0.152 | 1 | 10 | Socs4 | 0.014 | Old |
| 0.756303 | 0.23008 | 0.218 | 0.202 | 1 | 10 | Chd9 | 0.016 | Old |
| 0.756774 | 0.453214 | 0.037 | 0.03 | 1 | 10 | Zfp46 | 0.007 | Old |
| 0.756774 | 0.370229 | 0.037 | 0.03 | 1 | 10 | Ankdd1a | 0.007 | Old |
| 0.756774 | 0.219043 | 0.037 | 0.03 | 1 | 10 | 2610044O1 | 0.007 | Old |
| 0.757504 | 0.101825 | 0.584 | 0.545 | 1 | 10 | D8Ertd738 | 0.039 | Old |
| 0.757564 | 0.523451 | 0.176 | 0.172 | 1 | 10 | Nelfe | 0.004 | Old |

| | | | | | | | | |
|---|---|---|---|---|---|---|---|---|
| 0.757956 | 0.428047 | 0.051 | 0.061 | 1 | 10 | Nectin1 | -0.01 | Old |
| 0.758163 | 0.171444 | 0.374 | 0.364 | 1 | 10 | Uso1 | 0.01 | Old |
| 0.758349 | 0.081569 | 0.812 | 0.737 | 1 | 10 | Reep5 | 0.075 | Old |
| 0.758586 | 0.615753 | 0.037 | 0.03 | 1 | 10 | Inpp5b | 0.007 | Old |
| 0.758868 | 0.015372 | 0.154 | 0.141 | 1 | 10 | Il17ra | 0.013 | Old |
| 0.758958 | 0.404471 | 0.024 | 0.03 | 1 | 10 | Pigb | -0.006 | Old |
| 0.760472 | 0.026584 | 0.071 | 0.081 | 1 | 10 | Ofd1 | -0.01 | Old |
| 0.761296 | 0.072858 | 0.601 | 0.535 | 1 | 10 | Eif3e | 0.066 | Old |
| 0.76217 | 0.253934 | 0.1 | 0.091 | 1 | 10 | Phf1 | 0.009 | Old |
| 0.762215 | 0.484829 | 0.037 | 0.03 | 1 | 10 | Cd209b | 0.007 | Old |
| 0.762215 | 0.061654 | 0.037 | 0.03 | 1 | 10 | Zfp275 | 0.007 | Old |
| 0.762234 | 0.042461 | 1 | 0.99 | 1 | 10 | Rplp2 | 0.01 | Old |
| 0.762849 | 0.060169 | 0.059 | 0.051 | 1 | 10 | Pigf | 0.008 | Old |
| 0.76334 | 0.653404 | 0.088 | 0.081 | 1 | 10 | Lmf1 | 0.007 | Old |
| 0.764032 | 0.161881 | 0.037 | 0.03 | 1 | 10 | Ptges3l | 0.007 | Old |
| 0.764825 | 0.037731 | 0.377 | 0.333 | 1 | 10 | Tceal9 | 0.044 | Old |
| 0.765435 | 0.443254 | 0.051 | 0.061 | 1 | 10 | Ints4 | -0.01 | Old |
| 0.765551 | 0.191724 | 0.144 | 0.131 | 1 | 10 | Josd1 | 0.013 | Old |
| 0.765659 | 0.184199 | 0.264 | 0.253 | 1 | 10 | Pdhb | 0.011 | Old |
| 0.765745 | 0.318694 | 0.098 | 0.091 | 1 | 10 | Adcy9 | 0.007 | Old |
| 0.766893 | 0.027422 | 0.306 | 0.293 | 1 | 10 | Usp47 | 0.013 | Old |
| 0.767193 | 0.159849 | 0.059 | 0.051 | 1 | 10 | Tmem110 | 0.008 | Old |
| 0.767669 | 0.062203 | 0.037 | 0.03 | 1 | 10 | Pigz | 0.007 | Old |
| 0.768408 | 0.252766 | 0.1 | 0.111 | 1 | 10 | Kdm5d | -0.011 | Old |
| 0.7693 | 0.083845 | 0.147 | 0.162 | 1 | 10 | Dpp7 | -0.015 | Old |
| 0.769604 | 0.128461 | 0.134 | 0.152 | 1 | 10 | Bad | -0.018 | Old |
| 0.770311 | 0.09639 | 0.134 | 0.121 | 1 | 10 | 2610037D( | 0.013 | Old |
| 0.771188 | 0.140184 | 0.1 | 0.091 | 1 | 10 | Ctu2 | 0.009 | Old |
| 0.771311 | 0.035857 | 0.037 | 0.03 | 1 | 10 | Smug1 | 0.007 | Old |
| 0.772222 | 0.024831 | 0.034 | 0.04 | 1 | 10 | Hdac10 | -0.006 | Old |
| 0.772393 | 0.198731 | 0.122 | 0.111 | 1 | 10 | Cd200r4 | 0.011 | Old |
| 0.772712 | 0.459875 | 0.108 | 0.101 | 1 | 10 | Dcps | 0.007 | Old |
| 0.773134 | 0.137327 | 0.037 | 0.03 | 1 | 10 | Polm | 0.007 | Old |
| 0.773144 | 0.087156 | 0.225 | 0.212 | 1 | 10 | Zrsr2 | 0.013 | Old |
| 0.773428 | 0.032487 | 0.122 | 0.111 | 1 | 10 | Bap1 | 0.011 | Old |
| 0.774296 | 0.208996 | 0.171 | 0.162 | 1 | 10 | Csnk1g1 | 0.009 | Old |
| 0.774742 | 0.071342 | 0.154 | 0.141 | 1 | 10 | Guca1a | 0.013 | Old |
| 0.774912 | 0.756487 | 0.046 | 0.04 | 1 | 10 | Zmym1 | 0.006 | Old |
| 0.774959 | 0.120904 | 0.589 | 0.566 | 1 | 10 | Sgms1 | 0.023 | Old |
| 0.77514 | 0.075872 | 0.276 | 0.263 | 1 | 10 | Zfp445 | 0.013 | Old |
| 0.775515 | 0.033971 | 0.628 | 0.566 | 1 | 10 | Ptpn18 | 0.062 | Old |
| 0.77574 | 0.128023 | 0.09 | 0.081 | 1 | 10 | Pex5 | 0.009 | Old |
| 0.776535 | 0.684855 | 0.046 | 0.04 | 1 | 10 | Nt5dc1 | 0.006 | Old |
| 0.776839 | 0.32089 | 0.1 | 0.091 | 1 | 10 | Sirt1 | 0.009 | Old |
| 0.776927 | 0.001656 | 0.09 | 0.081 | 1 | 10 | Tcp11l1 | 0.009 | Old |

| | | | | | | | | |
|---|---|---|---|---|---|---|---|---|
| 0.777039 | 0.581224 | 0.12 | 0.111 | 1 | 10 | Cd209g | 0.009 | Old |
| 0.777917 | 0.197396 | 0.406 | 0.404 | 1 | 10 | Rer1 | 0.002 | Old |
| 0.778302 | 0.027286 | 0.11 | 0.101 | 1 | 10 | Coprs | 0.009 | Old |
| 0.778412 | 0.011404 | 0.205 | 0.192 | 1 | 10 | Pnpla7 | 0.013 | Old |
| 0.779269 | 0.571048 | 0.108 | 0.101 | 1 | 10 | Prps2 | 0.007 | Old |
| 0.78008 | 0.05902 | 0.196 | 0.182 | 1 | 10 | Sirt7 | 0.014 | Old |
| 0.780152 | 0.243297 | 0.007 | 0.01 | 1 | 10 | Hist1h2ae | -0.003 | Old |
| 0.780152 | 0.184805 | 0.007 | 0.01 | 1 | 10 | Dlgap5 | -0.003 | Old |
| 0.780152 | 0.160984 | 0.007 | 0.01 | 1 | 10 | Sall2 | -0.003 | Old |
| 0.780152 | 0.109952 | 0.007 | 0.01 | 1 | 10 | Grik5 | -0.003 | Old |
| 0.780152 | 0.039346 | 0.007 | 0.01 | 1 | 10 | Eid2 | -0.003 | Old |
| 0.780565 | 0.069191 | 0.335 | 0.313 | 1 | 10 | H2afv | 0.022 | Old |
| 0.781007 | 0.181174 | 0.103 | 0.091 | 1 | 10 | Slc41a1 | 0.012 | Old |
| 0.781183 | 0.138285 | 0.545 | 0.545 | 1 | 10 | Mycbp2 | 0 | Old |
| 0.781409 | 0.235553 | 0.046 | 0.04 | 1 | 10 | Tdrd3 | 0.006 | Old |
| 0.781889 | 1.123476 | 0.066 | 0.061 | 1 | 10 | Lin9 | 0.005 | Old |
| 0.782889 | 0.233048 | 0.23 | 0.222 | 1 | 10 | Btbd7 | 0.008 | Old |
| 0.783036 | 0.682683 | 0.046 | 0.04 | 1 | 10 | Ube2e2 | 0.006 | Old |
| 0.783036 | 0.002372 | 0.044 | 0.051 | 1 | 10 | Spaca6 | -0.007 | Old |
| 0.783709 | 0.012546 | 0.081 | 0.071 | 1 | 10 | Ccdc93 | 0.01 | Old |
| 0.783979 | 0.478302 | 0.007 | 0.01 | 1 | 10 | Zfp105 | -0.003 | Old |
| 0.783979 | 0.327689 | 0.007 | 0.01 | 1 | 10 | Rasip1 | -0.003 | Old |
| 0.783979 | 0.308874 | 0.007 | 0.01 | 1 | 10 | Map3k9 | -0.003 | Old |
| 0.783979 | 0.308412 | 0.007 | 0.01 | 1 | 10 | Bco2 | -0.003 | Old |
| 0.783979 | 0.191425 | 0.007 | 0.01 | 1 | 10 | Gm15050 | -0.003 | Old |
| 0.783979 | 0.185463 | 0.007 | 0.01 | 1 | 10 | Nat8l | -0.003 | Old |
| 0.783979 | 0.18135 | 0.007 | 0.01 | 1 | 10 | Ccl17 | -0.003 | Old |
| 0.783979 | 0.127448 | 0.007 | 0.01 | 1 | 10 | 1700084E1 | -0.003 | Old |
| 0.783979 | 0.125497 | 0.007 | 0.01 | 1 | 10 | Sugct | -0.003 | Old |
| 0.783979 | 0.109804 | 0.007 | 0.01 | 1 | 10 | Garnl3 | -0.003 | Old |
| 0.783979 | 0.094665 | 0.007 | 0.01 | 1 | 10 | Adh1 | -0.003 | Old |
| 0.783979 | 0.094056 | 0.007 | 0.01 | 1 | 10 | Naip1 | -0.003 | Old |
| 0.783979 | 0.039608 | 0.007 | 0.01 | 1 | 10 | AC122413. | -0.003 | Old |
| 0.783979 | 0.038538 | 0.007 | 0.01 | 1 | 10 | Zfp689 | -0.003 | Old |
| 0.783979 | 0.030022 | 0.007 | 0.01 | 1 | 10 | Gen1 | -0.003 | Old |
| 0.783979 | 0.009057 | 0.007 | 0.01 | 1 | 10 | Ccdc15 | -0.003 | Old |
| 0.784588 | 0.003922 | 0.247 | 0.232 | 1 | 10 | Tmem135 | 0.015 | Old |
| 0.784664 | 0.043374 | 0.046 | 0.04 | 1 | 10 | Ano8 | 0.006 | Old |
| 0.785157 | 0.029186 | 0.191 | 0.172 | 1 | 10 | Frmd4a | 0.019 | Old |
| 0.785507 | 0.071431 | 0.068 | 0.061 | 1 | 10 | Mrpl47 | 0.007 | Old |
| 0.786292 | 0.496614 | 0.046 | 0.04 | 1 | 10 | Bclaf3 | 0.006 | Old |
| 0.786471 | 0.125693 | 0.357 | 0.343 | 1 | 10 | Cript | 0.014 | Old |
| 0.786786 | 0.071614 | 0.804 | 0.768 | 1 | 10 | Rab7b | 0.036 | Old |
| 0.786797 | 0.250957 | 0.203 | 0.192 | 1 | 10 | Ssbp3 | 0.011 | Old |
| 0.787386 | 0.083304 | 0.132 | 0.121 | 1 | 10 | Rnf139 | 0.011 | Old |

| | | | | | | | | |
|---|---|---|---|---|---|---|---|---|
| 0.787812 | 0.983496 | 0.007 | 0.01 | 1 | 10 | Tspan2 | -0.003 | Old |
| 0.787812 | 0.792137 | 0.007 | 0.01 | 1 | 10 | Timm23 | -0.003 | Old |
| 0.787812 | 0.733726 | 0.007 | 0.01 | 1 | 10 | Gm15853 | -0.003 | Old |
| 0.787812 | 0.719489 | 0.007 | 0.01 | 1 | 10 | Fam129c | -0.003 | Old |
| 0.787812 | 0.689142 | 0.007 | 0.01 | 1 | 10 | Gm49077 | -0.003 | Old |
| 0.787812 | 0.594938 | 0.007 | 0.01 | 1 | 10 | Rsph9 | -0.003 | Old |
| 0.787812 | 0.534838 | 0.007 | 0.01 | 1 | 10 | Jsrp1 | -0.003 | Old |
| 0.787812 | 0.484318 | 0.007 | 0.01 | 1 | 10 | Sbk2 | -0.003 | Old |
| 0.787812 | 0.477974 | 0.007 | 0.01 | 1 | 10 | Ctla2b | -0.003 | Old |
| 0.787812 | 0.474313 | 0.007 | 0.01 | 1 | 10 | Rrm2 | -0.003 | Old |
| 0.787812 | 0.427883 | 0.007 | 0.01 | 1 | 10 | F5 | -0.003 | Old |
| 0.787812 | 0.387914 | 0.007 | 0.01 | 1 | 10 | Gm4924 | -0.003 | Old |
| 0.787812 | 0.368999 | 0.007 | 0.01 | 1 | 10 | Hmgb3 | -0.003 | Old |
| 0.787812 | 0.359626 | 0.007 | 0.01 | 1 | 10 | Treml1 | -0.003 | Old |
| 0.787812 | 0.314656 | 0.007 | 0.01 | 1 | 10 | Rab19 | -0.003 | Old |
| 0.787812 | 0.289154 | 0.007 | 0.01 | 1 | 10 | Zfand4 | -0.003 | Old |
| 0.787812 | 0.266092 | 0.007 | 0.01 | 1 | 10 | A530076I1 | -0.003 | Old |
| 0.787812 | 0.23774 | 0.007 | 0.01 | 1 | 10 | Acta2 | -0.003 | Old |
| 0.787812 | 0.199359 | 0.007 | 0.01 | 1 | 10 | Tle2 | -0.003 | Old |
| 0.787812 | 0.120104 | 0.007 | 0.01 | 1 | 10 | Nlrp1a | -0.003 | Old |
| 0.787812 | 0.074331 | 0.007 | 0.01 | 1 | 10 | 1700030J2 | -0.003 | Old |
| 0.787922 | 0.340185 | 0.046 | 0.04 | 1 | 10 | Alms1 | 0.006 | Old |
| 0.788467 | 1.26873 | 0.024 | 0.02 | 1 | 10 | Nup35 | 0.004 | Old |
| 0.788555 | 0.57864 | 0.056 | 0.051 | 1 | 10 | Tab1 | 0.005 | Old |
| 0.789556 | 0.212076 | 0.213 | 0.232 | 1 | 10 | Fxyd2 | -0.019 | Old |
| 0.790513 | 0.009498 | 0.034 | 0.04 | 1 | 10 | Zkscan17 | -0.006 | Old |
| 0.7907 | 0.736463 | 0.024 | 0.02 | 1 | 10 | Adat1 | 0.004 | Old |
| 0.7907 | 0.58665 | 0.024 | 0.02 | 1 | 10 | E130308A1 | 0.004 | Old |
| 0.791184 | 0.335425 | 0.046 | 0.04 | 1 | 10 | Ccp110 | 0.006 | Old |
| 0.791779 | 0.08785 | 0.222 | 0.212 | 1 | 10 | Washc3 | 0.01 | Old |
| 0.792348 | 0.266099 | 0.034 | 0.04 | 1 | 10 | Tuft1 | -0.006 | Old |
| 0.792816 | 0.278999 | 0.046 | 0.04 | 1 | 10 | Carf | 0.006 | Old |
| 0.792935 | 0.838707 | 0.024 | 0.02 | 1 | 10 | D930030I0 | 0.004 | Old |
| 0.792935 | 0.613076 | 0.024 | 0.02 | 1 | 10 | Dennd2c | 0.004 | Old |
| 0.793513 | 0.081802 | 0.242 | 0.232 | 1 | 10 | Fam234a | 0.01 | Old |
| 0.793568 | 0.104286 | 0.254 | 0.242 | 1 | 10 | Exoc4 | 0.012 | Old |
| 0.794076 | 0.105444 | 0.161 | 0.152 | 1 | 10 | Smim12 | 0.009 | Old |
| 0.794139 | 0.096205 | 0.264 | 0.283 | 1 | 10 | Sidt2 | -0.019 | Old |
| 0.795002 | 0.07345 | 0.1 | 0.091 | 1 | 10 | Agfg2 | 0.009 | Old |
| 0.795172 | 0.619079 | 0.024 | 0.02 | 1 | 10 | Gm17745 | 0.004 | Old |
| 0.795172 | 0.544799 | 0.024 | 0.02 | 1 | 10 | Ehbp1 | 0.004 | Old |
| 0.796023 | 0.344531 | 0.034 | 0.04 | 1 | 10 | Hemk1 | -0.006 | Old |
| 0.796083 | 0.250386 | 0.046 | 0.04 | 1 | 10 | Actr5 | 0.006 | Old |
| 0.796362 | 0.061973 | 0.068 | 0.061 | 1 | 10 | Foxred1 | 0.007 | Old |
| 0.79724 | 0.045843 | 0.174 | 0.162 | 1 | 10 | Lrif1 | 0.012 | Old |

| | | | | | | | | |
|---|---|---|---|---|---|---|---|---|
| 0.797411 | 0.450795 | 0.024 | 0.02 | 1 | 10 | Ccdc180 | 0.004 | Old |
| 0.798174 | 0.023513 | 0.186 | 0.172 | 1 | 10 | Ccdc186 | 0.014 | Old |
| 0.798341 | 0.022534 | 0.667 | 0.657 | 1 | 10 | Jun | 0.01 | Old |
| 0.798969 | 0.582402 | 0.056 | 0.051 | 1 | 10 | Nlrx1 | 0.005 | Old |
| 0.798991 | 0.015693 | 0.132 | 0.141 | 1 | 10 | Mras | -0.009 | Old |
| 0.799061 | 0.176342 | 0.078 | 0.071 | 1 | 10 | Socs5 | 0.007 | Old |
| 0.799354 | 0.271399 | 0.046 | 0.04 | 1 | 10 | Nmral1 | 0.006 | Old |
| 0.799354 | 0.224946 | 0.046 | 0.04 | 1 | 10 | Rad9a | 0.006 | Old |
| 0.799362 | 0.454692 | 0.088 | 0.081 | 1 | 10 | Ppil3 | 0.007 | Old |
| 0.799526 | 0.009434 | 0.592 | 0.545 | 1 | 10 | Ndufa13 | 0.047 | Old |
| 0.799651 | 0.492566 | 0.024 | 0.02 | 1 | 10 | 5830432E0 | 0.004 | Old |
| 0.80014 | 0.067357 | 0.235 | 0.222 | 1 | 10 | Sav1 | 0.013 | Old |
| 0.800233 | 0.036261 | 0.352 | 0.364 | 1 | 10 | Ddx46 | -0.012 | Old |
| 0.80046 | 0.739558 | 0.056 | 0.051 | 1 | 10 | Snhg9 | 0.005 | Old |
| 0.80046 | 0.557448 | 0.056 | 0.051 | 1 | 10 | Cep85 | 0.005 | Old |
| 0.801224 | 0.427739 | 0.098 | 0.091 | 1 | 10 | Spg11 | 0.007 | Old |
| 0.801843 | 0.0267 | 0.1 | 0.091 | 1 | 10 | Guf1 | 0.009 | Old |
| 0.801892 | 0.692427 | 0.024 | 0.02 | 1 | 10 | Dnah2 | 0.004 | Old |
| 0.801892 | 0.555669 | 0.024 | 0.02 | 1 | 10 | Dcp1b | 0.004 | Old |
| 0.801892 | 0.515952 | 0.024 | 0.02 | 1 | 10 | Ssx2ip | 0.004 | Old |
| 0.801892 | 0.355924 | 0.024 | 0.02 | 1 | 10 | Gm10138 | 0.004 | Old |
| 0.802629 | 0.063713 | 0.046 | 0.04 | 1 | 10 | Trmt61a | 0.006 | Old |
| 0.802989 | 0.276933 | 0.088 | 0.081 | 1 | 10 | Cdk18 | 0.007 | Old |
| 0.803444 | 0.534674 | 0.056 | 0.051 | 1 | 10 | Sp4 | 0.005 | Old |
| 0.803672 | 0.060099 | 0.306 | 0.293 | 1 | 10 | Slc12a6 | 0.013 | Old |
| 0.80468 | 0.30224 | 0.098 | 0.091 | 1 | 10 | Casp6 | 0.007 | Old |
| 0.805907 | 0.241876 | 0.044 | 0.051 | 1 | 10 | Pusl1 | -0.007 | Old |
| 0.806381 | 0.673437 | 0.024 | 0.02 | 1 | 10 | Flt1 | 0.004 | Old |
| 0.806381 | 0.235339 | 0.024 | 0.02 | 1 | 10 | Rnd2 | 0.004 | Old |
| 0.806381 | 0.176065 | 0.024 | 0.02 | 1 | 10 | Gbp2b | 0.004 | Old |
| 0.806381 | 0.016777 | 0.024 | 0.02 | 1 | 10 | Gm14326 | 0.004 | Old |
| 0.80643 | 0.512956 | 0.056 | 0.051 | 1 | 10 | Gm6563 | 0.005 | Old |
| 0.808628 | 0.747287 | 0.024 | 0.02 | 1 | 10 | Thrb | 0.004 | Old |
| 0.808628 | 0.189252 | 0.024 | 0.02 | 1 | 10 | C6 | 0.004 | Old |
| 0.808628 | 0.172176 | 0.024 | 0.02 | 1 | 10 | Tmem220 | 0.004 | Old |
| 0.808628 | 0.107528 | 0.024 | 0.02 | 1 | 10 | Msh3 | 0.004 | Old |
| 0.810249 | 0.014948 | 0.2 | 0.212 | 1 | 10 | Trp53inp2 | -0.012 | Old |
| 0.810256 | 0.084879 | 0.088 | 0.081 | 1 | 10 | Polr3d | 0.007 | Old |
| 0.81083 | 0.013144 | 0.046 | 0.04 | 1 | 10 | Mad1l1 | 0.006 | Old |
| 0.810876 | 0.534878 | 0.024 | 0.02 | 1 | 10 | Gsdme | 0.004 | Old |
| 0.811663 | 0.158963 | 0.054 | 0.061 | 1 | 10 | Nxn | -0.007 | Old |
| 0.811994 | 0.004516 | 0.267 | 0.242 | 1 | 10 | Hipk3 | 0.025 | Old |
| 0.812035 | 0.113462 | 0.152 | 0.162 | 1 | 10 | Jak2 | -0.01 | Old |
| 0.813126 | 0.119804 | 0.024 | 0.02 | 1 | 10 | AC129328. | 0.004 | Old |
| 0.813126 | 0.10536 | 0.024 | 0.02 | 1 | 10 | Zbtb39 | 0.004 | Old |

| | | | | | | | | |
|---|---|---|---|---|---|---|---|---|
| 0.813126 | 0.010283 | 0.024 | 0.02 | 1 | 10 | Tlr5 | 0.004 | Old |
| 0.813753 | 0.218403 | 0.139 | 0.131 | 1 | 10 | Syap1 | 0.008 | Old |
| 0.815073 | 0.297858 | 0.098 | 0.091 | 1 | 10 | Sdhaf1 | 0.007 | Old |
| 0.815281 | 0.05736 | 0.284 | 0.293 | 1 | 10 | Atf7ip | -0.009 | Old |
| 0.816324 | 0.328981 | 0.088 | 0.081 | 1 | 10 | Gtpbp8 | 0.007 | Old |
| 0.816843 | 0.041381 | 0.856 | 0.818 | 1 | 10 | Ptafr | 0.038 | Old |
| 0.816904 | 0.48741 | 0.056 | 0.051 | 1 | 10 | Qtrt1 | 0.005 | Old |
| 0.816904 | 0.456324 | 0.056 | 0.051 | 1 | 10 | Zbtb41 | 0.005 | Old |
| 0.816904 | 0.043042 | 0.056 | 0.051 | 1 | 10 | Mib2 | 0.005 | Old |
| 0.817464 | 0.143507 | 0.093 | 0.101 | 1 | 10 | Helb | -0.008 | Old |
| 0.818272 | 0.124831 | 0.078 | 0.071 | 1 | 10 | Lrig2 | 0.007 | Old |
| 0.818677 | 0.002549 | 0.374 | 0.374 | 1 | 10 | Anpep | 0 | Old |
| 0.818928 | 0.057919 | 0.108 | 0.101 | 1 | 10 | Trp53bp2 | 0.007 | Old |
| 0.819557 | 0.354043 | 0.078 | 0.071 | 1 | 10 | Stx2 | 0.007 | Old |
| 0.819971 | 0.272483 | 0.088 | 0.081 | 1 | 10 | Msto1 | 0.007 | Old |
| 0.820671 | 0.073993 | 0.073 | 0.081 | 1 | 10 | Mpv17 | -0.008 | Old |
| 0.821033 | 0.173684 | 0.171 | 0.182 | 1 | 10 | Fundc1 | -0.011 | Old |
| 0.821179 | 0.134223 | 0.083 | 0.091 | 1 | 10 | Galk1 | -0.008 | Old |
| 0.821404 | 0.364899 | 0.056 | 0.051 | 1 | 10 | Gemin6 | 0.005 | Old |
| 0.821404 | 0.286265 | 0.056 | 0.051 | 1 | 10 | Zbtb5 | 0.005 | Old |
| 0.821517 | 0.240813 | 0.271 | 0.263 | 1 | 10 | Prdm1 | 0.008 | Old |
| 0.82386 | 0.201962 | 0.154 | 0.141 | 1 | 10 | Pknox1 | 0.013 | Old |
| 0.824226 | 0.116072 | 0.11 | 0.101 | 1 | 10 | Pgm2l1 | 0.009 | Old |
| 0.824406 | 0.097772 | 0.056 | 0.051 | 1 | 10 | Mthfsd | 0.005 | Old |
| 0.824475 | 0.168093 | 0.108 | 0.101 | 1 | 10 | Tigd2 | 0.007 | Old |
| 0.824566 | 0.031496 | 0.257 | 0.273 | 1 | 10 | Slc30a5 | -0.016 | Old |
| 0.825136 | 0.031254 | 0.636 | 0.596 | 1 | 10 | Bsg | 0.04 | Old |
| 0.825327 | 0.133162 | 0.11 | 0.101 | 1 | 10 | Arfgap1 | 0.009 | Old |
| 0.825497 | 0.132596 | 0.1 | 0.111 | 1 | 10 | Mrps35 | -0.011 | Old |
| 0.825545 | 0.805598 | 0.034 | 0.03 | 1 | 10 | Gpsm1 | 0.004 | Old |
| 0.825545 | 0.783553 | 0.034 | 0.03 | 1 | 10 | Kdm4b | 0.004 | Old |
| 0.825879 | 0.178654 | 0.073 | 0.081 | 1 | 10 | Rpain | -0.008 | Old |
| 0.826395 | 0.017555 | 0.183 | 0.172 | 1 | 10 | Gm8186 | 0.011 | Old |
| 0.828078 | 0.124239 | 0.193 | 0.192 | 1 | 10 | Usp3 | 0.001 | Old |
| 0.828888 | 0.186512 | 0.066 | 0.061 | 1 | 10 | Ccdc85b | 0.005 | Old |
| 0.829362 | 0.933209 | 0.034 | 0.03 | 1 | 10 | Zfp995 | 0.004 | Old |
| 0.82979 | 0.095472 | 0.073 | 0.081 | 1 | 10 | Prkar2b | -0.008 | Old |
| 0.829962 | 0.0274 | 0.359 | 0.343 | 1 | 10 | Rdx | 0.016 | Old |
| 0.830767 | 0.377764 | 0.017 | 0.02 | 1 | 10 | Rapgef5 | -0.003 | Old |
| 0.831263 | 0.303421 | 0.176 | 0.172 | 1 | 10 | Cdk2ap1 | 0.004 | Old |
| 0.832474 | 0.030397 | 0.298 | 0.283 | 1 | 10 | Mpp6 | 0.015 | Old |
| 0.832663 | 0.15856 | 0.337 | 0.333 | 1 | 10 | Banf1 | 0.004 | Old |
| 0.835094 | 0.286259 | 0.034 | 0.03 | 1 | 10 | Dhrs11 | 0.004 | Old |
| 0.835816 | 0.097778 | 0.088 | 0.081 | 1 | 10 | Irf4 | 0.007 | Old |
| 0.835979 | 0.065874 | 0.017 | 0.02 | 1 | 10 | Mex3b | -0.003 | Old |

| | | | | | | | | |
|---|---|---|---|---|---|---|---|---|
| 0.835979 | 0.023212 | 0.017 | 0.02 | 1 | 10 | Pou5f1 | -0.003 | Old |
| 0.837117 | 0.054484 | 0.098 | 0.091 | 1 | 10 | Cisd3 | 0.007 | Old |
| 0.838588 | 0.08278 | 0.017 | 0.02 | 1 | 10 | 1700003F1 | -0.003 | Old |
| 0.83887 | 0.431423 | 0.147 | 0.141 | 1 | 10 | Rrp7a | 0.006 | Old |
| 0.838939 | 0.26868 | 0.108 | 0.101 | 1 | 10 | Ocel1 | 0.007 | Old |
| 0.839382 | 0.3036 | 0.166 | 0.152 | 1 | 10 | Zfand2a | 0.014 | Old |
| 0.839445 | 0.096024 | 0.098 | 0.091 | 1 | 10 | Hira | 0.007 | Old |
| 0.839458 | 0.020717 | 0.056 | 0.051 | 1 | 10 | Cmss1 | 0.005 | Old |
| 0.839682 | 0.877102 | 0.044 | 0.04 | 1 | 10 | Wdr92 | 0.004 | Old |
| 0.83979 | 0.074719 | 0.298 | 0.323 | 1 | 10 | Pa2g4 | -0.025 | Old |
| 0.840043 | 0.114728 | 0.066 | 0.061 | 1 | 10 | Rfc5 | 0.005 | Old |
| 0.840835 | 0.302099 | 0.034 | 0.03 | 1 | 10 | Zfp628 | 0.004 | Old |
| 0.841199 | 0.41062 | 0.017 | 0.02 | 1 | 10 | Gm37305 | -0.003 | Old |
| 0.841199 | 0.19549 | 0.017 | 0.02 | 1 | 10 | Rab3a | -0.003 | Old |
| 0.841199 | 0.164464 | 0.017 | 0.02 | 1 | 10 | 4930481A1 | -0.003 | Old |
| 0.841199 | 0.022317 | 0.017 | 0.02 | 1 | 10 | Slc18b1 | -0.003 | Old |
| 0.84149 | 0.050939 | 0.142 | 0.131 | 1 | 10 | Socs7 | 0.011 | Old |
| 0.841787 | 0.26843 | 0.147 | 0.141 | 1 | 10 | Ppp1r35 | 0.006 | Old |
| 0.841839 | 0.118094 | 0.23 | 0.242 | 1 | 10 | 1-Mar | -0.012 | Old |
| 0.842103 | 0.027831 | 0.127 | 0.121 | 1 | 10 | Hspa4l | 0.006 | Old |
| 0.843152 | 0.070957 | 0.088 | 0.081 | 1 | 10 | Ring1 | 0.007 | Old |
| 0.843811 | 0.356778 | 0.017 | 0.02 | 1 | 10 | C130050O1 | -0.003 | Old |
| 0.843811 | 0.354281 | 0.017 | 0.02 | 1 | 10 | Cenph | -0.003 | Old |
| 0.843811 | 0.155283 | 0.017 | 0.02 | 1 | 10 | 5830416I19 | -0.003 | Old |
| 0.843811 | 0.086462 | 0.017 | 0.02 | 1 | 10 | A130077B1 | -0.003 | Old |
| 0.844607 | 0.044294 | 0.156 | 0.152 | 1 | 10 | Ccdc25 | 0.004 | Old |
| 0.84509 | 0.114958 | 0.149 | 0.141 | 1 | 10 | Mettl16 | 0.008 | Old |
| 0.845633 | 0.067 | 0.066 | 0.061 | 1 | 10 | Pml | 0.005 | Old |
| 0.8461 | 0.036986 | 0.301 | 0.293 | 1 | 10 | Cdkn1b | 0.008 | Old |
| 0.846425 | 0.327537 | 0.017 | 0.02 | 1 | 10 | Ube2t | -0.003 | Old |
| 0.846425 | 0.176143 | 0.017 | 0.02 | 1 | 10 | 1700010I14 | -0.003 | Old |
| 0.848431 | 0.213201 | 0.066 | 0.061 | 1 | 10 | Ipo4 | 0.005 | Old |
| 0.848502 | 0.064211 | 0.034 | 0.03 | 1 | 10 | Ankrd66 | 0.004 | Old |
| 0.848602 | 0.325527 | 0.147 | 0.141 | 1 | 10 | Cstf3 | 0.006 | Old |
| 0.849564 | 0.299729 | 0.086 | 0.081 | 1 | 10 | Sirpb1b | 0.005 | Old |
| 0.851658 | 0.652053 | 0.017 | 0.02 | 1 | 10 | Zeb1 | -0.003 | Old |
| 0.85195 | 0.013177 | 0.125 | 0.131 | 1 | 10 | Map3k4 | -0.006 | Old |
| 0.852342 | 0.027508 | 0.034 | 0.03 | 1 | 10 | Zfp35 | 0.004 | Old |
| 0.852631 | 0.201311 | 0.066 | 0.061 | 1 | 10 | Pex1 | 0.005 | Old |
| 0.852711 | 0.325491 | 0.027 | 0.03 | 1 | 10 | Aplf | -0.003 | Old |
| 0.852842 | 0.172307 | 0.149 | 0.141 | 1 | 10 | Tspan32 | 0.008 | Old |
| 0.854263 | 0.134314 | 0.034 | 0.03 | 1 | 10 | Trpm2 | 0.004 | Old |
| 0.854408 | 1.089364 | 0.012 | 0.01 | 1 | 10 | 4930524J0 | 0.002 | Old |
| 0.854408 | 1.037005 | 0.012 | 0.01 | 1 | 10 | Trpc4 | 0.002 | Old |
| 0.854408 | 0.949909 | 0.012 | 0.01 | 1 | 10 | Dnm3 | 0.002 | Old |

| | | | | | | | | |
|---|---|---|---|---|---|---|---|---|
| 0.854408 | 0.946481 | 0.012 | 0.01 | 1 | 10 | Tppp | 0.002 | Old |
| 0.854408 | 0.899198 | 0.012 | 0.01 | 1 | 10 | Mast4 | 0.002 | Old |
| 0.854408 | 0.896077 | 0.012 | 0.01 | 1 | 10 | Cysltr2 | 0.002 | Old |
| 0.854408 | 0.786322 | 0.012 | 0.01 | 1 | 10 | Brdt | 0.002 | Old |
| 0.854408 | 0.775521 | 0.012 | 0.01 | 1 | 10 | Rab4a | 0.002 | Old |
| 0.854408 | 0.731268 | 0.012 | 0.01 | 1 | 10 | Rnf122 | 0.002 | Old |
| 0.854408 | 0.710025 | 0.012 | 0.01 | 1 | 10 | Slc9b2 | 0.002 | Old |
| 0.854408 | 0.691947 | 0.012 | 0.01 | 1 | 10 | Gm6710 | 0.002 | Old |
| 0.854408 | 0.619776 | 0.012 | 0.01 | 1 | 10 | Aff3 | 0.002 | Old |
| 0.854646 | 0.052092 | 0.161 | 0.152 | 1 | 10 | Zfyve16 | 0.009 | Old |
| 0.854782 | 0.034784 | 0.149 | 0.141 | 1 | 10 | Myo7a | 0.008 | Old |
| 0.855716 | 0.015182 | 0.078 | 0.071 | 1 | 10 | Mettl14 | 0.007 | Old |
| 0.856179 | 0.234036 | 0.154 | 0.152 | 1 | 10 | Pcnt | 0.002 | Old |
| 0.856229 | 0.326202 | 0.12 | 0.131 | 1 | 10 | Rasgrp4 | -0.011 | Old |
| 0.856632 | 0.580831 | 0.044 | 0.04 | 1 | 10 | Blzf1 | 0.004 | Old |
| 0.857086 | 0.027775 | 0.971 | 0.929 | 1 | 10 | Rpl6 | 0.042 | Old |
| 0.857278 | 0.627316 | 0.073 | 0.081 | 1 | 10 | Lsm14b | -0.008 | Old |
| 0.857608 | 1.23592 | 0.012 | 0.01 | 1 | 10 | Npy | 0.002 | Old |
| 0.857608 | 1.142486 | 0.012 | 0.01 | 1 | 10 | D130017N( | 0.002 | Old |
| 0.857608 | 0.863889 | 0.012 | 0.01 | 1 | 10 | Tpm2 | 0.002 | Old |
| 0.857608 | 0.814875 | 0.012 | 0.01 | 1 | 10 | Arsg | 0.002 | Old |
| 0.857608 | 0.738969 | 0.012 | 0.01 | 1 | 10 | Ccdc17 | 0.002 | Old |
| 0.857608 | 0.732721 | 0.012 | 0.01 | 1 | 10 | Kctd1 | 0.002 | Old |
| 0.857608 | 0.683393 | 0.012 | 0.01 | 1 | 10 | Smkr-ps | 0.002 | Old |
| 0.857608 | 0.672729 | 0.012 | 0.01 | 1 | 10 | Cit | 0.002 | Old |
| 0.857608 | 0.660475 | 0.012 | 0.01 | 1 | 10 | Isg20 | 0.002 | Old |
| 0.857608 | 0.618351 | 0.012 | 0.01 | 1 | 10 | Cd46 | 0.002 | Old |
| 0.857608 | 0.5998 | 0.012 | 0.01 | 1 | 10 | Naprt | 0.002 | Old |
| 0.857608 | 0.338065 | 0.012 | 0.01 | 1 | 10 | Evi5l | 0.002 | Old |
| 0.858311 | 0.059655 | 0.076 | 0.081 | 1 | 10 | Amz1 | -0.005 | Old |
| 0.858419 | 0.025619 | 0.584 | 0.566 | 1 | 10 | Metap2 | 0.018 | Old |
| 0.859046 | 0.118799 | 0.027 | 0.03 | 1 | 10 | Phc1 | -0.003 | Old |
| 0.859762 | 0.434421 | 0.093 | 0.101 | 1 | 10 | Tex2 | -0.008 | Old |
| 0.860031 | 0.080076 | 0.034 | 0.03 | 1 | 10 | Ptrh1 | 0.004 | Old |
| 0.860487 | 0.150894 | 0.269 | 0.263 | 1 | 10 | Sh3bp2 | 0.006 | Old |
| 0.860811 | 0.785145 | 0.012 | 0.01 | 1 | 10 | Tcea3 | 0.002 | Old |
| 0.860811 | 0.719276 | 0.012 | 0.01 | 1 | 10 | Capn10 | 0.002 | Old |
| 0.860811 | 0.620931 | 0.012 | 0.01 | 1 | 10 | Palb2 | 0.002 | Old |
| 0.860811 | 0.545565 | 0.012 | 0.01 | 1 | 10 | Gm35363 | 0.002 | Old |
| 0.860811 | 0.457345 | 0.012 | 0.01 | 1 | 10 | Apold1 | 0.002 | Old |
| 0.860811 | 0.406274 | 0.012 | 0.01 | 1 | 10 | Nynrin | 0.002 | Old |
| 0.860811 | 0.313927 | 0.012 | 0.01 | 1 | 10 | Tmem82 | 0.002 | Old |
| 0.860811 | 0.227305 | 0.012 | 0.01 | 1 | 10 | Dqx1 | 0.002 | Old |
| 0.860811 | 0.080085 | 0.012 | 0.01 | 1 | 10 | Dnajc17 | 0.002 | Old |
| 0.861007 | 0.014773 | 0.352 | 0.343 | 1 | 10 | Cdc5l | 0.009 | Old |

| | | | | | | | | |
|---|---|---|---|---|---|---|---|---|
| 0.861187 | 0.887248 | 0.064 | 0.061 | 1 | 10 | Plpp5 | 0.003 | Old |
| 0.861793 | 0.133942 | 0.037 | 0.04 | 1 | 10 | Stk35 | -0.003 | Old |
| 0.861793 | 0.054416 | 0.037 | 0.04 | 1 | 10 | Gm44751 | -0.003 | Old |
| 0.86186 | 0.111131 | 0.222 | 0.222 | 1 | 10 | Pcgf5 | 0 | Old |
| 0.862168 | 0.039242 | 0.161 | 0.152 | 1 | 10 | Uri1 | 0.009 | Old |
| 0.862946 | 0.103285 | 0.066 | 0.071 | 1 | 10 | Slc25a10 | -0.005 | Old |
| 0.863211 | 0.243924 | 0.086 | 0.081 | 1 | 10 | Prelid2 | 0.005 | Old |
| 0.863877 | 0.244004 | 0.117 | 0.111 | 1 | 10 | Llgl1 | 0.006 | Old |
| 0.864016 | 0.492969 | 0.012 | 0.01 | 1 | 10 | Zfp111 | 0.002 | Old |
| 0.864016 | 0.479239 | 0.012 | 0.01 | 1 | 10 | Lrp8 | 0.002 | Old |
| 0.864016 | 0.290129 | 0.012 | 0.01 | 1 | 10 | Adra1a | 0.002 | Old |
| 0.864016 | 0.277416 | 0.012 | 0.01 | 1 | 10 | Glis2 | 0.002 | Old |
| 0.864016 | 0.211207 | 0.012 | 0.01 | 1 | 10 | Snx32 | 0.002 | Old |
| 0.864016 | 0.148213 | 0.012 | 0.01 | 1 | 10 | 1700099I0 | 0.002 | Old |
| 0.864016 | 0.127533 | 0.012 | 0.01 | 1 | 10 | Angptl2 | 0.002 | Old |
| 0.864016 | 0.052904 | 0.012 | 0.01 | 1 | 10 | Adamtsl4 | 0.002 | Old |
| 0.864016 | 0.025873 | 0.012 | 0.01 | 1 | 10 | Qprt | 0.002 | Old |
| 0.864016 | 0.021555 | 0.012 | 0.01 | 1 | 10 | Cenpm | 0.002 | Old |
| 0.864843 | 0.03042 | 0.227 | 0.232 | 1 | 10 | Sf3a1 | -0.005 | Old |
| 0.865132 | 0.349211 | 0.044 | 0.04 | 1 | 10 | Pcnx4 | 0.004 | Old |
| 0.865391 | 0.103878 | 0.027 | 0.03 | 1 | 10 | Clstn1 | -0.003 | Old |
| 0.865443 | 0.080134 | 0.037 | 0.04 | 1 | 10 | Tubg1 | -0.003 | Old |
| 0.866181 | 0.120073 | 0.289 | 0.273 | 1 | 10 | Tlr13 | 0.016 | Old |
| 0.866529 | 0.280878 | 0.166 | 0.162 | 1 | 10 | Klc4 | 0.004 | Old |
| 0.867223 | 0.052562 | 0.012 | 0.01 | 1 | 10 | Klrb1a | 0.002 | Old |
| 0.867223 | 0.045183 | 0.012 | 0.01 | 1 | 10 | 6430573P0 | 0.002 | Old |
| 0.867223 | 0.035318 | 0.012 | 0.01 | 1 | 10 | Bcl11a | 0.002 | Old |
| 0.867269 | 0.006313 | 0.037 | 0.04 | 1 | 10 | Slc25a33 | -0.003 | Old |
| 0.868115 | 0.287217 | 0.046 | 0.051 | 1 | 10 | Zfp961 | -0.005 | Old |
| 0.868115 | 0.139028 | 0.046 | 0.051 | 1 | 10 | Pilrb1 | -0.005 | Old |
| 0.868193 | 0.052192 | 0.117 | 0.111 | 1 | 10 | Gtf3c1 | 0.006 | Old |
| 0.868294 | 0.302849 | 0.115 | 0.111 | 1 | 10 | Zdhhc16 | 0.004 | Old |
| 0.868426 | 0.200048 | 0.13 | 0.121 | 1 | 10 | Immp1l | 0.009 | Old |
| 0.868862 | 0.158338 | 0.056 | 0.061 | 1 | 10 | Uck1 | -0.005 | Old |
| 0.869221 | 0.223088 | 0.093 | 0.101 | 1 | 10 | Zfp219 | -0.008 | Old |
| 0.869428 | 0.120202 | 0.086 | 0.081 | 1 | 10 | Naf1 | 0.005 | Old |
| 0.869625 | 0.726582 | 0.027 | 0.03 | 1 | 10 | 1110051M | -0.003 | Old |
| 0.86971 | 0.211066 | 0.105 | 0.111 | 1 | 10 | Cdc42ep3 | -0.006 | Old |
| 0.870055 | 0.093616 | 0.22 | 0.212 | 1 | 10 | Sptan1 | 0.008 | Old |
| 0.871943 | 0.108296 | 0.044 | 0.04 | 1 | 10 | Cxcl12 | 0.004 | Old |
| 0.872408 | 0.15402 | 0.076 | 0.071 | 1 | 10 | Mcm9 | 0.005 | Old |
| 0.872683 | 0.011167 | 0.445 | 0.434 | 1 | 10 | Noc2l | 0.011 | Old |
| 0.873018 | 0.225279 | 0.046 | 0.051 | 1 | 10 | Trmt10c | -0.005 | Old |
| 0.873345 | 0.065463 | 0.056 | 0.061 | 1 | 10 | Rabepk | -0.005 | Old |
| 0.873345 | 0.065031 | 0.056 | 0.061 | 1 | 10 | Gm43848 | -0.005 | Old |

| | | | | | | | | |
|---|---|---|---|---|---|---|---|---|
| 0.873863 | 0.51693 | 0.027 | 0.03 | 1 | 10 | H2-T24 | -0.003 | Old |
| 0.874408 | 0.655941 | 0.086 | 0.081 | 1 | 10 | Fdx1 | 0.005 | Old |
| 0.874841 | 0.25852 | 0.056 | 0.061 | 1 | 10 | Hist2h2be | -0.005 | Old |
| 0.875332 | 0.025408 | 0.484 | 0.515 | 1 | 10 | Cd300ld | -0.031 | Old |
| 0.875653 | 0.210523 | 0.086 | 0.081 | 1 | 10 | Rhbdd2 | 0.005 | Old |
| 0.876089 | 0.020706 | 0.594 | 0.556 | 1 | 10 | Elob | 0.038 | Old |
| 0.876409 | 0.348889 | 0.037 | 0.04 | 1 | 10 | Nipal3 | -0.003 | Old |
| 0.877681 | 0.021565 | 0.076 | 0.071 | 1 | 10 | Ccdc167 | 0.005 | Old |
| 0.877704 | 0.006727 | 0.719 | 0.636 | 1 | 10 | Alox5ap | 0.083 | Old |
| 0.877943 | 0.045163 | 0.702 | 0.687 | 1 | 10 | Ier5 | 0.015 | Old |
| 0.878764 | 0.264599 | 0.044 | 0.04 | 1 | 10 | Arid3b | 0.004 | Old |
| 0.879001 | 0.191718 | 0.076 | 0.071 | 1 | 10 | Cass4 | 0.005 | Old |
| 0.879563 | 0.282605 | 0.046 | 0.051 | 1 | 10 | Meis3 | -0.005 | Old |
| 0.880471 | 0.150553 | 0.044 | 0.04 | 1 | 10 | Zfp768 | 0.004 | Old |
| 0.880639 | 0.287456 | 0.086 | 0.081 | 1 | 10 | Atg4c | 0.005 | Old |
| 0.8812 | 0.277925 | 0.046 | 0.051 | 1 | 10 | Cul7 | -0.005 | Old |
| 0.8812 | 0.234249 | 0.046 | 0.051 | 1 | 10 | Fkbp9 | -0.005 | Old |
| 0.881415 | 0.410865 | 0.054 | 0.051 | 1 | 10 | 2610008E1 | 0.003 | Old |
| 0.881989 | 0.082822 | 0.726 | 0.717 | 1 | 10 | Ccnl1 | 0.009 | Old |
| 0.882313 | 0.044833 | 0.159 | 0.152 | 1 | 10 | Mxd1 | 0.007 | Old |
| 0.882793 | 0.204086 | 0.176 | 0.172 | 1 | 10 | Elp2 | 0.004 | Old |
| 0.883734 | 0.465296 | 0.037 | 0.04 | 1 | 10 | Trit1 | -0.003 | Old |
| 0.883734 | 0.273543 | 0.037 | 0.04 | 1 | 10 | Dalrd3 | -0.003 | Old |
| 0.883927 | 0.147923 | 0.095 | 0.101 | 1 | 10 | Carm1 | -0.006 | Old |
| 0.885553 | 0.191885 | 0.125 | 0.121 | 1 | 10 | Rab3d | 0.004 | Old |
| 0.885909 | 0.028471 | 0.932 | 0.899 | 1 | 10 | Rpl3 | 0.033 | Old |
| 0.88611 | 0.002795 | 0.521 | 0.515 | 1 | 10 | Lyn | 0.006 | Old |
| 0.886372 | 0.214222 | 0.259 | 0.253 | 1 | 10 | Slc4a2 | 0.006 | Old |
| 0.886814 | 0.115548 | 0.086 | 0.091 | 1 | 10 | Nuak1 | -0.005 | Old |
| 0.886878 | 0.041556 | 0.086 | 0.081 | 1 | 10 | Cdc42ep4 | 0.005 | Old |
| 0.887397 | 0.067216 | 0.912 | 0.899 | 1 | 10 | Btg1 | 0.013 | Old |
| 0.888575 | 0.116138 | 0.565 | 0.576 | 1 | 10 | Ms4a7 | -0.011 | Old |
| 0.888645 | 0.027757 | 0.995 | 0.99 | 1 | 10 | Rps27a | 0.005 | Old |
| 0.889749 | 0.615746 | 0.064 | 0.061 | 1 | 10 | Leng1 | 0.003 | Old |
| 0.890716 | 0.380177 | 0.054 | 0.051 | 1 | 10 | Spred2 | 0.003 | Old |
| 0.890722 | 0.10764 | 0.044 | 0.04 | 1 | 10 | Ogfod2 | 0.004 | Old |
| 0.892267 | 0.269947 | 0.054 | 0.051 | 1 | 10 | Katnb1 | 0.003 | Old |
| 0.892353 | 0.258798 | 0.095 | 0.091 | 1 | 10 | Tmem161a | 0.004 | Old |
| 0.893 | 0.008645 | 0.311 | 0.293 | 1 | 10 | Cept1 | 0.018 | Old |
| 0.894739 | 0.623471 | 0.037 | 0.04 | 1 | 10 | Srfbp1 | -0.003 | Old |
| 0.894794 | 0.478263 | 0.076 | 0.081 | 1 | 10 | Cdkn2d | -0.005 | Old |
| 0.896455 | 1.059266 | 0.022 | 0.02 | 1 | 10 | Diras2 | 0.002 | Old |
| 0.897566 | 0.065423 | 0.044 | 0.04 | 1 | 10 | Ecsit | 0.004 | Old |
| 0.897831 | 0.039093 | 0.57 | 0.525 | 1 | 10 | Ifi207 | 0.045 | Old |
| 0.89885 | 0.70854 | 0.022 | 0.02 | 1 | 10 | Magee1 | 0.002 | Old |

| | | | | | | | | |
|---|---|---|---|---|---|---|---|---|
| 0.900031 | 0.035955 | 0.054 | 0.051 | 1 | 10 | Tmcc2 | 0.003 | Old |
| 0.900143 | 0.299112 | 0.122 | 0.121 | 1 | 10 | Hgf | 0.001 | Old |
| 0.901087 | 0.276782 | 0.134 | 0.141 | 1 | 10 | Rab9 | -0.007 | Old |
| 0.901246 | 0.695458 | 0.022 | 0.02 | 1 | 10 | Psph | 0.002 | Old |
| 0.901246 | 0.496338 | 0.022 | 0.02 | 1 | 10 | Spata7 | 0.002 | Old |
| 0.901246 | 0.45772 | 0.022 | 0.02 | 1 | 10 | Cercam | 0.002 | Old |
| 0.90126 | 0.276295 | 0.154 | 0.152 | 1 | 10 | Zfp984 | 0.002 | Old |
| 0.901602 | 0.622724 | 0.073 | 0.071 | 1 | 10 | Trp53bp1 | 0.002 | Old |
| 0.901817 | 0.33441 | 0.401 | 0.414 | 1 | 10 | Ly6e | -0.013 | Old |
| 0.902416 | 0.210482 | 0.125 | 0.121 | 1 | 10 | St7l | 0.004 | Old |
| 0.903642 | 0.988054 | 0.022 | 0.02 | 1 | 10 | Krt10 | 0.002 | Old |
| 0.903642 | 0.535594 | 0.022 | 0.02 | 1 | 10 | Itfg2 | 0.002 | Old |
| 0.905262 | 0.369446 | 0.076 | 0.081 | 1 | 10 | Hdac7 | -0.005 | Old |
| 0.905626 | 0.30191 | 0.073 | 0.071 | 1 | 10 | Arl14ep | 0.002 | Old |
| 0.90582 | 0.474293 | 0.046 | 0.051 | 1 | 10 | B230118H( | -0.005 | Old |
| 0.906322 | 0.058361 | 0.364 | 0.343 | 1 | 10 | Hmox2 | 0.021 | Old |
| 0.906418 | 0.043172 | 0.218 | 0.212 | 1 | 10 | Gab2 | 0.006 | Old |
| 0.907025 | 0.113263 | 0.21 | 0.222 | 1 | 10 | 1110008P1 | -0.012 | Old |
| 0.907372 | 0.083216 | 0.222 | 0.222 | 1 | 10 | Lcorl | 0 | Old |
| 0.90831 | 0.366684 | 0.073 | 0.071 | 1 | 10 | Jrkl | 0.002 | Old |
| 0.908913 | 0.027271 | 0.098 | 0.101 | 1 | 10 | Gnpda2 | -0.003 | Old |
| 0.909653 | 0.273153 | 0.073 | 0.071 | 1 | 10 | Borcs8 | 0.002 | Old |
| 0.91011 | 0.011403 | 0.741 | 0.687 | 1 | 10 | Ddx17 | 0.054 | Old |
| 0.910838 | 0.220933 | 0.022 | 0.02 | 1 | 10 | Glb1l | 0.002 | Old |
| 0.91085 | 0.369496 | 0.056 | 0.061 | 1 | 10 | Ints5.1 | -0.005 | Old |
| 0.910917 | 0.039321 | 0.054 | 0.051 | 1 | 10 | Tinf2 | 0.003 | Old |
| 0.912354 | 0.570398 | 0.056 | 0.061 | 1 | 10 | Tfdp2 | -0.005 | Old |
| 0.912644 | 0.00562 | 0.846 | 0.838 | 1 | 10 | Eef2 | 0.008 | Old |
| 0.913004 | 0.076362 | 0.196 | 0.192 | 1 | 10 | Tnfaip8l2 | 0.004 | Old |
| 0.913238 | 0.332238 | 0.022 | 0.02 | 1 | 10 | Usp49 | 0.002 | Old |
| 0.913643 | 0.965662 | 0.032 | 0.03 | 1 | 10 | Clec16a | 0.002 | Old |
| 0.915639 | 0.19204 | 0.022 | 0.02 | 1 | 10 | Ctif | 0.002 | Old |
| 0.915644 | 0.698964 | 0.032 | 0.03 | 1 | 10 | Adck2 | 0.002 | Old |
| 0.916203 | 0.339599 | 0.215 | 0.212 | 1 | 10 | Fam214b | 0.003 | Old |
| 0.916357 | 0.193914 | 0.115 | 0.111 | 1 | 10 | Ginm1 | 0.004 | Old |
| 0.917109 | 0.076112 | 0.098 | 0.101 | 1 | 10 | Grcc10 | -0.003 | Old |
| 0.917476 | 0.022488 | 0.565 | 0.556 | 1 | 10 | Cxcr4 | 0.009 | Old |
| 0.917778 | 0.401686 | 0.086 | 0.091 | 1 | 10 | Ogfod1 | -0.005 | Old |
| 0.918089 | 0.095487 | 0.156 | 0.152 | 1 | 10 | Map1lc3a | 0.004 | Old |
| 0.918352 | 0.154604 | 0.176 | 0.172 | 1 | 10 | Txndc11 | 0.004 | Old |
| 0.91856 | 0.03711 | 0.095 | 0.091 | 1 | 10 | Ammecr1 | 0.004 | Old |
| 0.920443 | 0.54553 | 0.022 | 0.02 | 1 | 10 | Tmem191c | 0.002 | Old |
| 0.920443 | 0.047714 | 0.022 | 0.02 | 1 | 10 | Cpe | 0.002 | Old |
| 0.920745 | 0.099306 | 0.115 | 0.111 | 1 | 10 | Sec23a | 0.004 | Old |
| 0.920905 | 0.001525 | 0.156 | 0.162 | 1 | 10 | Cntrl | -0.006 | Old |

| | | | | | | | | |
|---|---|---|---|---|---|---|---|---|
| 0.921447 | 0.036566 | 0.279 | 0.273 | 1 | 10 | Clip1 | 0.006 | Old |
| 0.921997 | 0.428475 | 0.152 | 0.162 | 1 | 10 | Thbd | -0.01 | Old |
| 0.922608 | 0.03946 | 0.149 | 0.152 | 1 | 10 | BC037034 | -0.003 | Old |
| 0.922679 | 0.006431 | 0.213 | 0.212 | 1 | 10 | Ppil2 | 0.001 | Old |
| 0.922732 | 0.013081 | 0.804 | 0.717 | 1 | 10 | Qk | 0.087 | Old |
| 0.923328 | 0.056515 | 0.166 | 0.172 | 1 | 10 | Fads1 | -0.006 | Old |
| 0.923366 | 0.046332 | 0.271 | 0.263 | 1 | 10 | Rbm6 | 0.008 | Old |
| 0.923652 | 0.984127 | 0.032 | 0.03 | 1 | 10 | Usp11 | 0.002 | Old |
| 0.924264 | 0.071117 | 0.083 | 0.081 | 1 | 10 | Tbcd | 0.002 | Old |
| 0.925619 | 0.838137 | 0.066 | 0.071 | 1 | 10 | Tbc1d4 | -0.005 | Old |
| 0.925655 | 0.533467 | 0.032 | 0.03 | 1 | 10 | Casz1 | 0.002 | Old |
| 0.925724 | 0.139173 | 0.095 | 0.091 | 1 | 10 | Mtpap | 0.004 | Old |
| 0.925748 | 0.109117 | 0.565 | 0.566 | 1 | 10 | Ccr2 | -0.001 | Old |
| 0.92731 | 0.030086 | 0.797 | 0.758 | 1 | 10 | Arpc3 | 0.039 | Old |
| 0.928926 | 0.757495 | 0.071 | 0.071 | 1 | 10 | E130309D( | 0 | Old |
| 0.928979 | 0.197843 | 0.274 | 0.263 | 1 | 10 | Phlda1 | 0.011 | Old |
| 0.929046 | 0.027611 | 0.078 | 0.081 | 1 | 10 | Tef | -0.003 | Old |
| 0.929383 | 0.04407 | 0.33 | 0.323 | 1 | 10 | Bhlhe40 | 0.007 | Old |
| 0.929663 | 0.491981 | 0.032 | 0.03 | 1 | 10 | Zfp729b | 0.002 | Old |
| 0.930089 | 0.082119 | 0.105 | 0.101 | 1 | 10 | Ccdc43 | 0.004 | Old |
| 0.931166 | 0.167185 | 0.073 | 0.071 | 1 | 10 | Ang | 0.002 | Old |
| 0.93122 | 0.047666 | 0.144 | 0.141 | 1 | 10 | Dcaf11 | 0.003 | Old |
| 0.933324 | 0.049611 | 0.858 | 0.879 | 1 | 10 | Emp3 | -0.021 | Old |
| 0.933673 | 0.71623 | 0.032 | 0.03 | 1 | 10 | 4732440D( | 0.002 | Old |
| 0.933931 | 0.418304 | 0.086 | 0.091 | 1 | 10 | Calhm2 | -0.005 | Old |
| 0.934335 | 0.015328 | 0.149 | 0.152 | 1 | 10 | Mcee | -0.003 | Old |
| 0.934381 | 0.633866 | 0.071 | 0.071 | 1 | 10 | Taf1b | 0 | Old |
| 0.934422 | 0.002325 | 0.083 | 0.081 | 1 | 10 | Mkrn2 | 0.002 | Old |
| 0.934603 | 0.000805 | 0.108 | 0.111 | 1 | 10 | B3gnt8 | -0.003 | Old |
| 0.935246 | 0.053664 | 0.059 | 0.061 | 1 | 10 | Rnf113a2 | -0.002 | Old |
| 0.93652 | 0.290722 | 0.042 | 0.04 | 1 | 10 | Zfp212 | 0.002 | Old |
| 0.936553 | 0.095446 | 0.073 | 0.071 | 1 | 10 | Gnl2 | 0.002 | Old |
| 0.936563 | 0.095824 | 0.137 | 0.141 | 1 | 10 | Stim2 | -0.004 | Old |
| 0.939616 | 0.007332 | 0.13 | 0.131 | 1 | 10 | Ctns | -0.001 | Old |
| 0.94004 | 0.427229 | 0.042 | 0.04 | 1 | 10 | Ctdspl | 0.002 | Old |
| 0.940551 | 0.021965 | 0.237 | 0.242 | 1 | 10 | Dpagt1 | -0.005 | Old |
| 0.940572 | 0.187813 | 0.098 | 0.101 | 1 | 10 | Noc3l | -0.003 | Old |
| 0.941203 | 0.480895 | 0.071 | 0.071 | 1 | 10 | Khk | 0 | Old |
| 0.941324 | 0.393652 | 0.142 | 0.141 | 1 | 10 | Pus1 | 0.001 | Old |
| 0.941577 | 0.006493 | 0.049 | 0.051 | 1 | 10 | Igtp | -0.002 | Old |
| 0.941616 | 0.018848 | 0.665 | 0.616 | 1 | 10 | Adam17 | 0.049 | Old |
| 0.942723 | 0.211115 | 0.183 | 0.182 | 1 | 10 | Taf13 | 0.001 | Old |
| 0.942922 | 0.034761 | 0.098 | 0.101 | 1 | 10 | C2cd2l | -0.003 | Old |
| 0.943655 | 0.188516 | 0.137 | 0.141 | 1 | 10 | Cd209f | -0.004 | Old |
| 0.943704 | 0.200133 | 0.032 | 0.03 | 1 | 10 | Gm4070 | 0.002 | Old |

| | | | | | | | | |
|---|---|---|---|---|---|---|---|---|
| 0.944579 | 0.05644 | 0.068 | 0.071 | 1 | 10 | Mfsd7a | -0.003 | Old |
| 0.944971 | 0.115488 | 0.147 | 0.141 | 1 | 10 | Chchd4 | 0.006 | Old |
| 0.945069 | 0.094895 | 0.103 | 0.101 | 1 | 10 | Tpgs1 | 0.002 | Old |
| 0.945712 | 0.073662 | 0.032 | 0.03 | 1 | 10 | Faap24 | 0.002 | Old |
| 0.946224 | 0.17315 | 0.103 | 0.101 | 1 | 10 | Pelp1 | 0.002 | Old |
| 0.946544 | 0.071635 | 0.134 | 0.131 | 1 | 10 | Mapk8ip3 | 0.003 | Old |
| 0.947378 | 0.138153 | 0.103 | 0.101 | 1 | 10 | Kcnj2 | 0.002 | Old |
| 0.947719 | 0.063662 | 0.032 | 0.03 | 1 | 10 | Ankzf1 | 0.002 | Old |
| 0.948154 | 0.044154 | 0.088 | 0.091 | 1 | 10 | Dhcr7 | -0.003 | Old |
| 0.948412 | 0.01789 | 0.43 | 0.424 | 1 | 10 | Srsf11 | 0.006 | Old |
| 0.948598 | 0.267684 | 0.154 | 0.152 | 1 | 10 | Stx3 | 0.002 | Old |
| 0.949083 | 0.405485 | 0.203 | 0.212 | 1 | 10 | Rspry1 | -0.009 | Old |
| 0.949315 | 0.220032 | 0.115 | 0.111 | 1 | 10 | Mitd1 | 0.004 | Old |
| 0.949732 | 0.057611 | 0.631 | 0.626 | 1 | 10 | Tnfrsf1b | 0.005 | Old |
| 0.950606 | 0.518447 | 0.042 | 0.04 | 1 | 10 | Ttc21b | 0.002 | Old |
| 0.950606 | 0.30337 | 0.042 | 0.04 | 1 | 10 | B230354K1 | 0.002 | Old |
| 0.95134 | 0.070829 | 0.726 | 0.646 | 1 | 10 | Rgs2 | 0.08 | Old |
| 0.952224 | 0.144587 | 0.083 | 0.081 | 1 | 10 | Gamt | 0.002 | Old |
| 0.952882 | 0.142353 | 0.068 | 0.071 | 1 | 10 | Gcc1 | -0.003 | Old |
| 0.952882 | 0.094033 | 0.068 | 0.071 | 1 | 10 | Ccdc134 | -0.003 | Old |
| 0.953085 | 0.263416 | 0.059 | 0.061 | 1 | 10 | Gmeb2 | -0.002 | Old |
| 0.953085 | 0.177403 | 0.088 | 0.091 | 1 | 10 | Fbxo9 | -0.003 | Old |
| 0.953786 | 0.072743 | 0.633 | 0.616 | 1 | 10 | Usmg5 | 0.017 | Old |
| 0.954573 | 0.047527 | 0.059 | 0.061 | 1 | 10 | Srcap | -0.002 | Old |
| 0.955122 | 0.078901 | 0.159 | 0.162 | 1 | 10 | Ddx23 | -0.003 | Old |
| 0.956253 | 0.005578 | 0.13 | 0.131 | 1 | 10 | Galnt10 | -0.001 | Old |
| 0.956422 | 0.068383 | 0.029 | 0.03 | 1 | 10 | Rhoj | -0.001 | Old |
| 0.956794 | 0.025607 | 0.983 | 0.97 | 1 | 10 | Rpl23 | 0.013 | Old |
| 0.958019 | 0.103796 | 0.088 | 0.091 | 1 | 10 | Kif3a | -0.003 | Old |
| 0.958204 | 0.015283 | 0.098 | 0.101 | 1 | 10 | Cep63 | -0.003 | Old |
| 0.958586 | 0.344231 | 0.11 | 0.111 | 1 | 10 | Tarbp2 | -0.001 | Old |
| 0.959361 | 0.100476 | 0.039 | 0.04 | 1 | 10 | Phlpp2 | -0.001 | Old |
| 0.959361 | 0.002666 | 0.039 | 0.04 | 1 | 10 | Taf1c | -0.001 | Old |
| 0.959418 | 0.248148 | 0.042 | 0.04 | 1 | 10 | Stk11ip | 0.002 | Old |
| 0.959738 | 0.359494 | 0.186 | 0.192 | 1 | 10 | Evi5 | -0.006 | Old |
| 0.959772 | 0.035897 | 0.032 | 0.03 | 1 | 10 | Cox11 | 0.002 | Old |
| 0.960047 | 0.055612 | 0.093 | 0.091 | 1 | 10 | Atg13 | 0.002 | Old |
| 0.960889 | 0.147 | 0.379 | 0.374 | 1 | 10 | Becn1 | 0.005 | Old |
| 0.961165 | 0.05631 | 0.039 | 0.04 | 1 | 10 | Ick | -0.001 | Old |
| 0.96226 | 0.002307 | 0.934 | 0.909 | 1 | 10 | Rpl28 | 0.025 | Old |
| 0.962643 | 0.065625 | 0.029 | 0.03 | 1 | 10 | Disp1 | -0.001 | Old |
| 0.963326 | 0.219055 | 0.061 | 0.061 | 1 | 10 | Zbtb2 | 0 | Old |
| 0.96385 | 0.150892 | 0.24 | 0.242 | 1 | 10 | Mvb12a | -0.002 | Old |
| 0.964708 | 0.147328 | 0.042 | 0.04 | 1 | 10 | Ogfod3 | 0.002 | Old |
| 0.964775 | 0.121215 | 0.039 | 0.04 | 1 | 10 | 1810024B0 | -0.001 | Old |

| | | | | | | | | |
|---|---|---|---|---|---|---|---|---|
| 0.964935 | 0.503814 | 0.051 | 0.051 | 1 | 10 | Det1 | 0 | Old |
| 0.964935 | 0.197537 | 0.051 | 0.051 | 1 | 10 | Rbm43 | 0 | Old |
| 0.965853 | 0.365643 | 0.02 | 0.02 | 1 | 10 | Iigp1 | 0 | Old |
| 0.965853 | 0.180411 | 0.02 | 0.02 | 1 | 10 | Zfp7 | 0 | Old |
| 0.966258 | 0.325311 | 0.061 | 0.061 | 1 | 10 | Alg11 | 0 | Old |
| 0.966581 | 0.394511 | 0.039 | 0.04 | 1 | 10 | Cd226 | -0.001 | Old |
| 0.966755 | 0.276123 | 0.257 | 0.263 | 1 | 10 | Ptger4 | -0.006 | Old |
| 0.967382 | 0.119668 | 0.142 | 0.141 | 1 | 10 | Man1b1 | 0.001 | Old |
| 0.967523 | 0.086644 | 0.049 | 0.051 | 1 | 10 | Nlrp1b | -0.002 | Old |
| 0.968381 | 0.22133 | 0.02 | 0.02 | 1 | 10 | 1110046J0 | 0 | Old |
| 0.968381 | 0.196134 | 0.02 | 0.02 | 1 | 10 | Slc25a53 | 0 | Old |
| 0.968386 | 0.238723 | 0.039 | 0.04 | 1 | 10 | Ctu1 | -0.001 | Old |
| 0.968951 | 0.015183 | 0.154 | 0.152 | 1 | 10 | Lactb2 | 0.002 | Old |
| 0.969191 | 0.300751 | 0.061 | 0.061 | 1 | 10 | Lsr | 0 | Old |
| 0.969643 | 0.006929 | 0.577 | 0.596 | 1 | 10 | Tra2a | -0.019 | Old |
| 0.969811 | 0.22584 | 0.12 | 0.121 | 1 | 10 | Gab1 | -0.001 | Old |
| 0.97056 | 0.066404 | 0.122 | 0.121 | 1 | 10 | Ap3s2 | 0.001 | Old |
| 0.970769 | 0.11917 | 0.049 | 0.051 | 1 | 10 | D3Ertd254 | -0.002 | Old |
| 0.970909 | 0.278706 | 0.02 | 0.02 | 1 | 10 | P3h1 | 0 | Old |
| 0.970909 | 0.032464 | 0.02 | 0.02 | 1 | 10 | Ercc6l | 0 | Old |
| 0.971308 | 0.117708 | 0.051 | 0.051 | 1 | 10 | Twnk | 0 | Old |
| 0.972683 | 0.193811 | 0.178 | 0.182 | 1 | 10 | Ahctf1 | -0.004 | Old |
| 0.973015 | 0.757213 | 0.029 | 0.03 | 1 | 10 | Gorasp1 | -0.001 | Old |
| 0.973015 | 0.524207 | 0.029 | 0.03 | 1 | 10 | Ikzf2 | -0.001 | Old |
| 0.973015 | 0.418653 | 0.029 | 0.03 | 1 | 10 | AU041133 | -0.001 | Old |
| 0.973015 | 0.201738 | 0.029 | 0.03 | 1 | 10 | Scrn3 | -0.001 | Old |
| 0.973105 | 0.118509 | 0.171 | 0.172 | 1 | 10 | Svbp | -0.001 | Old |
| 0.973438 | 0.684544 | 0.02 | 0.02 | 1 | 10 | Spats2 | 0 | Old |
| 0.973438 | 0.644988 | 0.02 | 0.02 | 1 | 10 | Trip13 | 0 | Old |
| 0.973438 | 0.391315 | 0.02 | 0.02 | 1 | 10 | Ift80 | 0 | Old |
| 0.973438 | 0.265781 | 0.02 | 0.02 | 1 | 10 | Tyms | 0 | Old |
| 0.973438 | 0.099844 | 0.02 | 0.02 | 1 | 10 | Tsacc | 0 | Old |
| 0.973438 | 0.048504 | 0.02 | 0.02 | 1 | 10 | Rad54l | 0 | Old |
| 0.97508 | 0.751717 | 0.01 | 0.01 | 1 | 10 | Prelid3a | 0 | Old |
| 0.97508 | 0.18602 | 0.01 | 0.01 | 1 | 10 | Hist1h1b | 0 | Old |
| 0.97508 | 0.093358 | 0.01 | 0.01 | 1 | 10 | Abcc10 | 0 | Old |
| 0.97508 | 0.078831 | 0.01 | 0.01 | 1 | 10 | Fignl1 | 0 | Old |
| 0.975516 | 0.369586 | 0.156 | 0.162 | 1 | 10 | Errfi1 | -0.006 | Old |
| 0.975543 | 0.066292 | 0.09 | 0.091 | 1 | 10 | Utp14b | -0.001 | Old |
| 0.975707 | 0.206697 | 0.35 | 0.354 | 1 | 10 | Irf1 | -0.004 | Old |
| 0.975967 | 0.190633 | 0.02 | 0.02 | 1 | 10 | Cryz | 0 | Old |
| 0.976524 | 0.15795 | 0.061 | 0.061 | 1 | 10 | Irgm2 | 0 | Old |
| 0.977165 | 0.500814 | 0.029 | 0.03 | 1 | 10 | Pdia5 | -0.001 | Old |
| 0.977821 | 0.080876 | 0.1 | 0.101 | 1 | 10 | Fbxw5 | -0.001 | Old |
| 0.978384 | 0.025203 | 0.924 | 0.899 | 1 | 10 | Mafb | 0.025 | Old |

| | | | | | | | | |
|---|---|---|---|---|---|---|---|---|
| 0.978496 | 0.893978 | 0.02 | 0.02 | 1 | 10 | Cnbd2 | 0 | Old |
| 0.978496 | 0.842964 | 0.02 | 0.02 | 1 | 10 | Mppe1 | 0 | Old |
| 0.978496 | 0.306522 | 0.02 | 0.02 | 1 | 10 | Polg2 | 0 | Old |
| 0.978639 | 0.50392 | 0.01 | 0.01 | 1 | 10 | Pxdc1 | 0 | Old |
| 0.978639 | 0.450723 | 0.01 | 0.01 | 1 | 10 | Rpgr | 0 | Old |
| 0.978639 | 0.445679 | 0.01 | 0.01 | 1 | 10 | Tmem240 | 0 | Old |
| 0.978639 | 0.345625 | 0.01 | 0.01 | 1 | 10 | BC055324 | 0 | Old |
| 0.978639 | 0.319046 | 0.01 | 0.01 | 1 | 10 | E430024I0ɛ | 0 | Old |
| 0.978639 | 0.279053 | 0.01 | 0.01 | 1 | 10 | Pcbp3 | 0 | Old |
| 0.978639 | 0.14698 | 0.01 | 0.01 | 1 | 10 | Sardh | 0 | Old |
| 0.978639 | 0.137961 | 0.01 | 0.01 | 1 | 10 | Enho | 0 | Old |
| 0.978639 | 0.017303 | 0.01 | 0.01 | 1 | 10 | Mrnip | 0 | Old |
| 0.979211 | 0.197274 | 0.09 | 0.091 | 1 | 10 | Strip1 | -0.001 | Old |
| 0.979222 | 0.663815 | 0.039 | 0.04 | 1 | 10 | Phf7 | -0.001 | Old |
| 0.979512 | 0.135041 | 0.12 | 0.121 | 1 | 10 | Ipo11 | -0.001 | Old |
| 0.980189 | 0.294834 | 0.137 | 0.141 | 1 | 10 | Ebi3 | -0.004 | Old |
| 0.980668 | 0.027658 | 0.081 | 0.081 | 1 | 10 | Ddx19b | 0 | Old |
| 0.980812 | 0.527383 | 0.108 | 0.111 | 1 | 10 | Naa40 | -0.003 | Old |
| 0.980925 | 0.140185 | 0.061 | 0.061 | 1 | 10 | Zc3h10 | 0 | Old |
| 0.981025 | 0.727589 | 0.02 | 0.02 | 1 | 10 | Pde8b | 0 | Old |
| 0.981025 | 0.705261 | 0.02 | 0.02 | 1 | 10 | Bach2os | 0 | Old |
| 0.981077 | 0.00826 | 0.232 | 0.232 | 1 | 10 | Zfx | 0 | Old |
| 0.981668 | 0.037282 | 0.12 | 0.121 | 1 | 10 | Nlrc5 | -0.001 | Old |
| 0.981878 | 0.068193 | 0.193 | 0.192 | 1 | 10 | Arrdc4 | 0.001 | Old |
| 0.981956 | 0.270389 | 0.081 | 0.081 | 1 | 10 | Apba1 | 0 | Old |
| 0.982199 | 1.066688 | 0.01 | 0.01 | 1 | 10 | Tceal1 | 0 | Old |
| 0.982199 | 0.81394 | 0.01 | 0.01 | 1 | 10 | N4bp3 | 0 | Old |
| 0.982199 | 0.811898 | 0.01 | 0.01 | 1 | 10 | Hsf2bp | 0 | Old |
| 0.982199 | 0.651884 | 0.01 | 0.01 | 1 | 10 | Mrgpre | 0 | Old |
| 0.982199 | 0.631925 | 0.01 | 0.01 | 1 | 10 | Irak1bp1 | 0 | Old |
| 0.982199 | 0.441923 | 0.01 | 0.01 | 1 | 10 | Clspn | 0 | Old |
| 0.982199 | 0.367769 | 0.01 | 0.01 | 1 | 10 | Il17rd | 0 | Old |
| 0.982199 | 0.358672 | 0.01 | 0.01 | 1 | 10 | Kdr | 0 | Old |
| 0.982199 | 0.324003 | 0.01 | 0.01 | 1 | 10 | Ifi206 | 0 | Old |
| 0.982199 | 0.313894 | 0.01 | 0.01 | 1 | 10 | Pole2 | 0 | Old |
| 0.982199 | 0.285208 | 0.01 | 0.01 | 1 | 10 | Mok | 0 | Old |
| 0.982199 | 0.276489 | 0.01 | 0.01 | 1 | 10 | Zfp59 | 0 | Old |
| 0.982199 | 0.217797 | 0.01 | 0.01 | 1 | 10 | Gm15834 | 0 | Old |
| 0.982199 | 0.183882 | 0.01 | 0.01 | 1 | 10 | Cep72 | 0 | Old |
| 0.982199 | 0.105329 | 0.01 | 0.01 | 1 | 10 | Ccnf | 0 | Old |
| 0.982199 | 0.059266 | 0.01 | 0.01 | 1 | 10 | Ptpn3 | 0 | Old |
| 0.982199 | 0.051965 | 0.01 | 0.01 | 1 | 10 | Gstk1 | 0 | Old |
| 0.982199 | 0.012345 | 0.01 | 0.01 | 1 | 10 | Foxd2 | 0 | Old |
| 0.983186 | 0.081004 | 0.249 | 0.253 | 1 | 10 | Pi4ka | -0.004 | Old |
| 0.983245 | 0.182646 | 0.081 | 0.081 | 1 | 10 | Arhgap22 | 0 | Old |

| | | | | | | | | |
|---|---|---|---|---|---|---|---|---|
| 0.983555 | 0.446631 | 0.02 | 0.02 | 1 | 10 | 1110002L0 | 0 | Old |
| 0.984058 | 0.076126 | 0.051 | 0.051 | 1 | 10 | Apba3 | 0 | Old |
| 0.984369 | 0.395135 | 0.13 | 0.131 | 1 | 10 | Jdp2 | -0.001 | Old |
| 0.985044 | 0.003908 | 0.506 | 0.485 | 1 | 10 | Cct3 | 0.021 | Old |
| 0.985443 | 0.086224 | 0.11 | 0.111 | 1 | 10 | Letmd1 | -0.001 | Old |
| 0.985758 | 0.895675 | 0.01 | 0.01 | 1 | 10 | Pxylp1 | 0 | Old |
| 0.985758 | 0.815956 | 0.01 | 0.01 | 1 | 10 | Ccdc189 | 0 | Old |
| 0.985758 | 0.786542 | 0.01 | 0.01 | 1 | 10 | Amotl2 | 0 | Old |
| 0.985758 | 0.671947 | 0.01 | 0.01 | 1 | 10 | Nemp2 | 0 | Old |
| 0.985758 | 0.665784 | 0.01 | 0.01 | 1 | 10 | Mnd1 | 0 | Old |
| 0.985758 | 0.543174 | 0.01 | 0.01 | 1 | 10 | Cdo1 | 0 | Old |
| 0.985758 | 0.492233 | 0.01 | 0.01 | 1 | 10 | Ajuba | 0 | Old |
| 0.985758 | 0.4394 | 0.01 | 0.01 | 1 | 10 | Gpr19 | 0 | Old |
| 0.985758 | 0.402577 | 0.01 | 0.01 | 1 | 10 | Dnajc28 | 0 | Old |
| 0.985758 | 0.371226 | 0.01 | 0.01 | 1 | 10 | Kmo | 0 | Old |
| 0.985758 | 0.344229 | 0.01 | 0.01 | 1 | 10 | Klc2 | 0 | Old |
| 0.985758 | 0.326958 | 0.01 | 0.01 | 1 | 10 | AL591952.: | 0 | Old |
| 0.986081 | 0.04595 | 0.276 | 0.273 | 1 | 10 | Kdelr2 | 0.003 | Old |
| 0.986677 | 0.275074 | 0.093 | 0.091 | 1 | 10 | Rpa3 | 0.002 | Old |
| 0.987706 | 0.022165 | 0.164 | 0.162 | 1 | 10 | Prpf38a | 0.002 | Old |
| 0.988255 | 0.671749 | 0.039 | 0.04 | 1 | 10 | Mrps27 | -0.001 | Old |
| 0.988405 | 0.003154 | 0.528 | 0.515 | 1 | 10 | Cdc42se2 | 0.013 | Old |
| 0.988615 | 1.188191 | 0.02 | 0.02 | 1 | 10 | Shb | 0 | Old |
| 0.99117 | 0.017683 | 0.149 | 0.152 | 1 | 10 | Cbx5 | -0.003 | Old |
| 0.992266 | 0.465562 | 0.081 | 0.081 | 1 | 10 | Orc4 | 0 | Old |
| 0.993421 | 0.041077 | 0.484 | 0.444 | 1 | 10 | Atxn7l3b | 0.04 | Old |
| 0.993944 | 0.067276 | 0.093 | 0.091 | 1 | 10 | Map3k14 | 0.002 | Old |
| 0.994488 | 0.033052 | 0.262 | 0.263 | 1 | 10 | Dhrs7b | -0.001 | Old |
| 0.997227 | 0.607622 | 0.068 | 0.071 | 1 | 10 | Gbp3 | -0.003 | Old |
| 0.999029 | 0.111604 | 0.154 | 0.152 | 1 | 10 | Sult1a1 | 0.002 | Old |
| 1.99E-12 | 4.302968 | 0.212 | 0.024 | 6.19E-08 | 10 | AA467197 | 0.188 | Young |
| 1.59E-09 | 3.336252 | 0.162 | 0.02 | 4.94E-05 | 10 | Pmepa1 | 0.142 | Young |
| 2.17E-09 | 3.422155 | 0.121 | 0.007 | 6.73E-05 | 10 | Arhgap24 | 0.114 | Young |
| 3.28E-09 | 1.292237 | 0.687 | 0.44 | 0.000102 | 10 | Bcl2a1b | 0.247 | Young |
| 3.26E-08 | 3.995033 | 0.131 | 0.015 | 0.001012 | 10 | Slc15a2 | 0.116 | Young |
| 8.19E-08 | 2.262247 | 0.323 | 0.12 | 0.002543 | 10 | Hspa1b | 0.203 | Young |
| 1.14E-07 | 3.07275 | 0.101 | 0.007 | 0.003539 | 10 | Cd300lf | 0.094 | Young |
| 2.15E-07 | 2.26707 | 0.323 | 0.122 | 0.006667 | 10 | Cx3cr1 | 0.201 | Young |
| 5.69E-07 | 4.333371 | 0.061 | 0 | 0.017681 | 10 | Nt5e | 0.061 | Young |
| 5.79E-07 | 1.289901 | 0.394 | 0.178 | 0.017976 | 10 | Hspa1a | 0.216 | Young |
| 7.14E-07 | 0.824578 | 1 | 1 | 0.02217 | 10 | Fth1 | 0 | Young |
| 8.06E-07 | 3.930708 | 0.091 | 0.007 | 0.025035 | 10 | Ass1 | 0.084 | Young |
| 2.37E-06 | 1.834978 | 0.141 | 0.027 | 0.073635 | 10 | Mtus1 | 0.114 | Young |
| 4.90E-06 | 2.529007 | 0.141 | 0.029 | 0.152215 | 10 | Cxcl14 | 0.112 | Young |
| 8.53E-06 | 3.184419 | 0.071 | 0.005 | 0.264998 | 10 | Acod1 | 0.066 | Young |

| | | | | | | | | |
|---|---|---|---|---|---|---|---|---|
| 8.73E-06 | 2.604981 | 0.192 | 0.056 | 0.271115 | 10 | Slc7a11 | 0.136 | Young |
| 9.48E-06 | 3.566201 | 0.404 | 0.222 | 0.294273 | 10 | Spp1 | 0.182 | Young |
| 9.70E-06 | 2.990794 | 0.172 | 0.046 | 0.301183 | 10 | Clec4n | 0.126 | Young |
| 2.04E-05 | 1.240131 | 0.626 | 0.516 | 0.633283 | 10 | Prdx5 | 0.11 | Young |
| 2.97E-05 | 1.985828 | 0.101 | 0.017 | 0.922454 | 10 | D16Ertd47 | 0.084 | Young |
| 3.22E-05 | 0.956522 | 0.818 | 0.724 | 0.999529 | 10 | Cd14 | 0.094 | Young |
| 3.50E-05 | 1.035782 | 0.616 | 0.462 | 1 | 10 | Id2 | 0.154 | Young |
| 4.54E-05 | 2.78346 | 0.071 | 0.007 | 1 | 10 | Olfml3 | 0.064 | Young |
| 4.93E-05 | 2.757651 | 0.071 | 0.007 | 1 | 10 | Col14a1 | 0.064 | Young |
| 6.05E-05 | 3.54548 | 0.061 | 0.005 | 1 | 10 | Camk2n1 | 0.056 | Young |
| 7.02E-05 | 2.846327 | 0.051 | 0.002 | 1 | 10 | Rapsn | 0.049 | Young |
| 7.73E-05 | 1.70206 | 0.202 | 0.073 | 1 | 10 | Dbf4 | 0.129 | Young |
| 0.000113 | 1.562791 | 0.404 | 0.232 | 1 | 10 | Clec4d | 0.172 | Young |
| 0.000121 | 2.148671 | 0.293 | 0.142 | 1 | 10 | Ccl3 | 0.151 | Young |
| 0.000202 | 1.894349 | 0.071 | 0.01 | 1 | 10 | L3mbtl2 | 0.061 | Young |
| 0.000228 | 1.718281 | 0.212 | 0.088 | 1 | 10 | Cd38 | 0.124 | Young |
| 0.000294 | 1.272015 | 0.343 | 0.186 | 1 | 10 | Ptgs2 | 0.157 | Young |
| 0.000306 | 1.509146 | 0.424 | 0.284 | 1 | 10 | Sgk1 | 0.14 | Young |
| 0.000307 | 2.379394 | 0.081 | 0.015 | 1 | 10 | Slc7a5 | 0.066 | Young |
| 0.000341 | 0.831102 | 0.636 | 0.501 | 1 | 10 | Fcgr2b | 0.135 | Young |
| 0.000371 | 2.343977 | 0.263 | 0.127 | 1 | 10 | Ccl4 | 0.136 | Young |
| 0.000487 | 2.057753 | 0.051 | 0.005 | 1 | 10 | Slc40a1 | 0.046 | Young |
| 0.0006 | 2.591276 | 0.04 | 0.002 | 1 | 10 | Gm26520 | 0.038 | Young |
| 0.000724 | 2.366888 | 0.101 | 0.027 | 1 | 10 | Parvg | 0.074 | Young |
| 0.000816 | 2.222235 | 0.091 | 0.022 | 1 | 10 | Creg2 | 0.069 | Young |
| 0.000851 | 2.199225 | 0.111 | 0.032 | 1 | 10 | Fam46c | 0.079 | Young |
| 0.000895 | 0.11509 | 1 | 1 | 1 | 10 | Gm42418 | 0 | Young |
| 0.001084 | 2.347513 | 0.061 | 0.01 | 1 | 10 | F11r | 0.051 | Young |
| 0.001145 | 1.000514 | 0.465 | 0.325 | 1 | 10 | Tlr2 | 0.14 | Young |
| 0.001191 | 0.295821 | 0.96 | 0.978 | 1 | 10 | Hsp90ab1 | -0.018 | Young |
| 0.00129 | 1.142213 | 0.212 | 0.098 | 1 | 10 | Aoah | 0.114 | Young |
| 0.00138 | 0.511811 | 0.778 | 0.709 | 1 | 10 | Pkm | 0.069 | Young |
| 0.001757 | 1.940609 | 0.081 | 0.02 | 1 | 10 | Als2 | 0.061 | Young |
| 0.00185 | 1.23192 | 0.343 | 0.218 | 1 | 10 | Rap2a | 0.125 | Young |
| 0.001857 | 1.6328 | 0.182 | 0.081 | 1 | 10 | Lpcat2 | 0.101 | Young |
| 0.001869 | 2.932113 | 0.051 | 0.007 | 1 | 10 | 2900026A0 | 0.044 | Young |
| 0.001891 | 1.380083 | 0.182 | 0.078 | 1 | 10 | Rasgrp3 | 0.104 | Young |
| 0.001897 | 0.499997 | 0.768 | 0.716 | 1 | 10 | Srgn | 0.052 | Young |
| 0.002177 | 1.147071 | 0.222 | 0.11 | 1 | 10 | Igf1r | 0.112 | Young |
| 0.002319 | 1.41465 | 0.253 | 0.137 | 1 | 10 | Smg9 | 0.116 | Young |
| 0.00271 | 1.151809 | 0.293 | 0.171 | 1 | 10 | Hps3 | 0.122 | Young |
| 0.003017 | 1.829825 | 0.061 | 0.012 | 1 | 10 | Cfd | 0.049 | Young |
| 0.003063 | 1.644402 | 0.182 | 0.086 | 1 | 10 | Nckipsd | 0.096 | Young |
| 0.003078 | 2.342573 | 0.071 | 0.017 | 1 | 10 | Acss2 | 0.054 | Young |
| 0.003182 | 0.635025 | 0.576 | 0.472 | 1 | 10 | Blvrb | 0.104 | Young |

| | | | | | | | | |
|---|---|---|---|---|---|---|---|---|
| 0.003215 | 2.256393 | 0.071 | 0.017 | 1 | 10 | Hist1h1d | 0.054 | Young |
| 0.003303 | 3.625909 | 0.04 | 0.005 | 1 | 10 | Uchl1 | 0.035 | Young |
| 0.003329 | 2.563696 | 0.071 | 0.017 | 1 | 10 | Gcnt2 | 0.054 | Young |
| 0.003354 | 1.429938 | 0.121 | 0.044 | 1 | 10 | Abcc3 | 0.077 | Young |
| 0.0034 | 1.361078 | 0.636 | 0.545 | 1 | 10 | Hmox1 | 0.091 | Young |
| 0.003436 | 1.397005 | 0.04 | 0.005 | 1 | 10 | Pim2 | 0.035 | Young |
| 0.003446 | 1.550032 | 0.071 | 0.017 | 1 | 10 | Pald1 | 0.054 | Young |
| 0.003483 | 0.619904 | 0.616 | 0.555 | 1 | 10 | Ninj1 | 0.061 | Young |
| 0.003512 | 0.54285 | 0.707 | 0.599 | 1 | 10 | Nfkbiz | 0.108 | Young |
| 0.003744 | 0.933754 | 0.313 | 0.188 | 1 | 10 | Mmp14 | 0.125 | Young |
| 0.003824 | 1.505638 | 0.192 | 0.093 | 1 | 10 | Phactr4 | 0.099 | Young |
| 0.003861 | 0.234722 | 0.98 | 0.983 | 1 | 10 | Rps16 | -0.003 | Young |
| 0.003952 | 1.270927 | 0.273 | 0.166 | 1 | 10 | Ogfrl1 | 0.107 | Young |
| 0.004056 | 3.140434 | 0.02 | 0 | 1 | 10 | Tmeff1 | 0.02 | Young |
| 0.004056 | 2.877276 | 0.02 | 0 | 1 | 10 | Kcnj10 | 0.02 | Young |
| 0.004056 | 2.644062 | 0.02 | 0 | 1 | 10 | Ptges | 0.02 | Young |
| 0.004056 | 2.63774 | 0.02 | 0 | 1 | 10 | Prtn3 | 0.02 | Young |
| 0.004056 | 2.507938 | 0.02 | 0 | 1 | 10 | Gm8251 | 0.02 | Young |
| 0.004056 | 2.471235 | 0.02 | 0 | 1 | 10 | Gm9465 | 0.02 | Young |
| 0.004056 | 2.402892 | 0.02 | 0 | 1 | 10 | Fam89b | 0.02 | Young |
| 0.004056 | 2.356926 | 0.02 | 0 | 1 | 10 | Ica1l | 0.02 | Young |
| 0.004056 | 2.341908 | 0.02 | 0 | 1 | 10 | Grin1 | 0.02 | Young |
| 0.004056 | 2.241944 | 0.02 | 0 | 1 | 10 | Ltb | 0.02 | Young |
| 0.004056 | 2.130564 | 0.02 | 0 | 1 | 10 | Col6a1 | 0.02 | Young |
| 0.004056 | 2.099785 | 0.02 | 0 | 1 | 10 | Kcng2 | 0.02 | Young |
| 0.004056 | 2.088118 | 0.02 | 0 | 1 | 10 | Tmem79 | 0.02 | Young |
| 0.004056 | 2.057724 | 0.02 | 0 | 1 | 10 | Klrd1 | 0.02 | Young |
| 0.004056 | 2.019979 | 0.02 | 0 | 1 | 10 | Gm26912 | 0.02 | Young |
| 0.004056 | 1.888956 | 0.02 | 0 | 1 | 10 | Ccl11 | 0.02 | Young |
| 0.004056 | 1.783056 | 0.02 | 0 | 1 | 10 | Tbx6 | 0.02 | Young |
| 0.004056 | 1.750247 | 0.02 | 0 | 1 | 10 | Tpsb2 | 0.02 | Young |
| 0.004056 | 1.6966 | 0.02 | 0 | 1 | 10 | 3830403N1 | 0.02 | Young |
| 0.004056 | 1.545731 | 0.02 | 0 | 1 | 10 | Hist1h1a | 0.02 | Young |
| 0.004056 | 1.333502 | 0.02 | 0 | 1 | 10 | Bbs1 | 0.02 | Young |
| 0.004116 | 1.672991 | 0.121 | 0.046 | 1 | 10 | Pacs1 | 0.075 | Young |
| 0.004464 | 1.686143 | 0.121 | 0.046 | 1 | 10 | Pdpn | 0.075 | Young |
| 0.004495 | 1.144043 | 0.293 | 0.176 | 1 | 10 | Sntb2 | 0.117 | Young |
| 0.004604 | 1.432198 | 0.162 | 0.073 | 1 | 10 | Rogdi | 0.089 | Young |
| 0.004701 | 1.122618 | 0.192 | 0.093 | 1 | 10 | Creb5 | 0.099 | Young |
| 0.004773 | 1.323766 | 0.182 | 0.088 | 1 | 10 | BC005561 | 0.094 | Young |
| 0.004861 | 4.166698 | 0.03 | 0.002 | 1 | 10 | Tnfsf8 | 0.028 | Young |
| 0.004861 | 3.083944 | 0.03 | 0.002 | 1 | 10 | Zfp341 | 0.028 | Young |
| 0.004861 | 2.970811 | 0.03 | 0.002 | 1 | 10 | H2-Ob | 0.028 | Young |
| 0.004861 | 2.888334 | 0.03 | 0.002 | 1 | 10 | Clec9a | 0.028 | Young |
| 0.004861 | 2.676774 | 0.03 | 0.002 | 1 | 10 | Myl2 | 0.028 | Young |

| | | | | | | | | |
|---|---|---|---|---|---|---|---|---|
| 0.004861 | 2.655196 | 0.03 | 0.002 | 1 | 10 | Fancd2 | 0.028 | Young |
| 0.004936 | 2.394855 | 0.03 | 0.002 | 1 | 10 | Gm26533 | 0.028 | Young |
| 0.004936 | 2.820597 | 0.03 | 0.002 | 1 | 10 | Arg2 | 0.028 | Young |
| 0.004936 | 2.298711 | 0.03 | 0.002 | 1 | 10 | Adarb1 | 0.028 | Young |
| 0.004936 | 2.192271 | 0.03 | 0.002 | 1 | 10 | Trim30b | 0.028 | Young |
| 0.005013 | 1.94214 | 0.03 | 0.002 | 1 | 10 | Gstt1 | 0.028 | Young |
| 0.00518 | 0.512865 | 0.818 | 0.787 | 1 | 10 | Rsrp1 | 0.031 | Young |
| 0.005205 | 0.913946 | 0.424 | 0.293 | 1 | 10 | Gas6 | 0.131 | Young |
| 0.005269 | 1.442068 | 0.172 | 0.081 | 1 | 10 | Rmi1 | 0.091 | Young |
| 0.005317 | 1.290508 | 0.283 | 0.171 | 1 | 10 | Ddx47 | 0.112 | Young |
| 0.005727 | 0.701702 | 0.525 | 0.433 | 1 | 10 | Psmd1 | 0.092 | Young |
| 0.005751 | 1.929952 | 0.131 | 0.056 | 1 | 10 | Dmac2 | 0.075 | Young |
| 0.006114 | 1.07461 | 0.374 | 0.247 | 1 | 10 | Mt2 | 0.127 | Young |
| 0.006177 | 1.661724 | 0.051 | 0.01 | 1 | 10 | Ppbp | 0.041 | Young |
| 0.006221 | 0.792168 | 0.364 | 0.24 | 1 | 10 | Gsto1 | 0.124 | Young |
| 0.006263 | 0.502022 | 0.758 | 0.697 | 1 | 10 | Gapdh | 0.061 | Young |
| 0.00635 | 1.350998 | 0.081 | 0.024 | 1 | 10 | Mettl2 | 0.057 | Young |
| 0.006429 | 1.488383 | 0.212 | 0.112 | 1 | 10 | Ptgs1 | 0.1 | Young |
| 0.006716 | 1.465659 | 0.222 | 0.125 | 1 | 10 | Ltc4s | 0.097 | Young |
| 0.006835 | 1.916224 | 0.061 | 0.015 | 1 | 10 | Zc3h8 | 0.046 | Young |
| 0.007037 | 1.557946 | 0.172 | 0.086 | 1 | 10 | 2510009E0 | 0.086 | Young |
| 0.007073 | 1.581917 | 0.081 | 0.024 | 1 | 10 | Gpr141 | 0.057 | Young |
| 0.007076 | 0.183729 | 0.949 | 0.966 | 1 | 10 | Ppia | -0.017 | Young |
| 0.007078 | 1.738298 | 0.061 | 0.015 | 1 | 10 | Dpysl3 | 0.046 | Young |
| 0.007139 | 1.105106 | 0.061 | 0.015 | 1 | 10 | Pard6a | 0.046 | Young |
| 0.007264 | 1.623729 | 0.061 | 0.015 | 1 | 10 | Gbp5 | 0.046 | Young |
| 0.007302 | 1.508399 | 0.121 | 0.049 | 1 | 10 | Mtfr1 | 0.072 | Young |
| 0.007543 | 1.17632 | 0.303 | 0.193 | 1 | 10 | Thbs1 | 0.11 | Young |
| 0.007587 | 0.7859 | 0.515 | 0.411 | 1 | 10 | Mrpl23 | 0.104 | Young |
| 0.007804 | 1.6624 | 0.131 | 0.056 | 1 | 10 | Rnf31 | 0.075 | Young |
| 0.007965 | 1.500257 | 0.323 | 0.208 | 1 | 10 | Osm | 0.115 | Young |
| 0.008087 | 0.597366 | 0.414 | 0.291 | 1 | 10 | Gatm | 0.123 | Young |
| 0.008294 | 1.205518 | 0.131 | 0.056 | 1 | 10 | Il21r | 0.075 | Young |
| 0.008809 | 0.700171 | 0.394 | 0.276 | 1 | 10 | Coq10b | 0.118 | Young |
| 0.009104 | 1.101537 | 0.364 | 0.262 | 1 | 10 | Eif3g | 0.102 | Young |
| 0.009136 | 0.789624 | 0.374 | 0.254 | 1 | 10 | Cnot4 | 0.12 | Young |
| 0.010326 | 2.068294 | 0.101 | 0.039 | 1 | 10 | Wdyhv1 | 0.062 | Young |
| 0.010326 | 1.931032 | 0.101 | 0.039 | 1 | 10 | Dtx3 | 0.062 | Young |
| 0.010865 | 2.29758 | 0.04 | 0.007 | 1 | 10 | Tulp3 | 0.033 | Young |
| 0.011062 | 1.107045 | 0.313 | 0.205 | 1 | 10 | Il4ra | 0.108 | Young |
| 0.011102 | 2.567171 | 0.04 | 0.007 | 1 | 10 | Mybpc3 | 0.033 | Young |
| 0.011102 | 2.559012 | 0.04 | 0.007 | 1 | 10 | Suv39h2 | 0.033 | Young |
| 0.011102 | 2.503592 | 0.04 | 0.007 | 1 | 10 | Afap1l1 | 0.033 | Young |
| 0.011223 | 2.480926 | 0.04 | 0.007 | 1 | 10 | Fignl2 | 0.033 | Young |
| 0.011223 | 2.216071 | 0.04 | 0.007 | 1 | 10 | Cxxc5 | 0.033 | Young |

| | | | | | | | | |
|---|---|---|---|---|---|---|---|---|
| 0.011422 | 1.934285 | 0.071 | 0.022 | 1 | 10 | Suv39h1 | 0.049 | Young |
| 0.011453 | 0.039905 | 0.141 | 0.271 | 1 | 10 | Tusc1 | -0.13 | Young |
| 0.011467 | 1.754714 | 0.04 | 0.007 | 1 | 10 | Usp2 | 0.033 | Young |
| 0.011467 | 1.509508 | 0.04 | 0.007 | 1 | 10 | Zfp27 | 0.033 | Young |
| 0.011941 | 0.859182 | 0.081 | 0.027 | 1 | 10 | Def6 | 0.054 | Young |
| 0.01197 | 1.373027 | 0.04 | 0.007 | 1 | 10 | Tmem177 | 0.033 | Young |
| 0.012053 | 2.540841 | 0.061 | 0.017 | 1 | 10 | Ap5b1 | 0.044 | Young |
| 0.012272 | 1.14446 | 0.162 | 0.081 | 1 | 10 | Dcaf10 | 0.081 | Young |
| 0.012426 | 0.703415 | 0.535 | 0.44 | 1 | 10 | Ier3 | 0.095 | Young |
| 0.01246 | 1.133942 | 0.152 | 0.073 | 1 | 10 | Akap7 | 0.079 | Young |
| 0.012529 | 0.35848 | 0.899 | 0.861 | 1 | 10 | Cd53 | 0.038 | Young |
| 0.012637 | 2.130904 | 0.061 | 0.017 | 1 | 10 | Hist1h2ap | 0.044 | Young |
| 0.012874 | 0.823117 | 0.465 | 0.377 | 1 | 10 | Kpna4 | 0.088 | Young |
| 0.01304 | 2.020459 | 0.061 | 0.017 | 1 | 10 | Racgap1 | 0.044 | Young |
| 0.013142 | 1.993376 | 0.061 | 0.017 | 1 | 10 | 2700046G( | 0.044 | Young |
| 0.013381 | 3.01987 | 0.051 | 0.012 | 1 | 10 | Ankrd37 | 0.039 | Young |
| 0.01356 | 1.065386 | 0.323 | 0.225 | 1 | 10 | Srek1 | 0.098 | Young |
| 0.013678 | 0.620504 | 0.758 | 0.719 | 1 | 10 | Atf3 | 0.039 | Young |
| 0.013863 | 1.896908 | 0.051 | 0.012 | 1 | 10 | Abtb2 | 0.039 | Young |
| 0.013879 | 2.17552 | 0.061 | 0.017 | 1 | 10 | B3gnt7 | 0.044 | Young |
| 0.013986 | 1.662821 | 0.051 | 0.012 | 1 | 10 | A430046D1 | 0.039 | Young |
| 0.014002 | 1.119161 | 0.162 | 0.083 | 1 | 10 | Tmem243 | 0.079 | Young |
| 0.01421 | 1.231297 | 0.192 | 0.105 | 1 | 10 | Retreg1 | 0.087 | Young |
| 0.015154 | 1.071074 | 0.202 | 0.112 | 1 | 10 | Ntpcr | 0.09 | Young |
| 0.015476 | 0.584436 | 0.697 | 0.619 | 1 | 10 | Usf2 | 0.078 | Young |
| 0.015814 | 0.866291 | 0.212 | 0.12 | 1 | 10 | Hpgd | 0.092 | Young |
| 0.01623 | 1.456778 | 0.192 | 0.11 | 1 | 10 | Rbm15 | 0.082 | Young |
| 0.016307 | 1.168551 | 0.242 | 0.149 | 1 | 10 | Lifr | 0.093 | Young |
| 0.016311 | 0.992637 | 0.111 | 0.046 | 1 | 10 | 1700021F0 | 0.065 | Young |
| 0.01719 | 0.65345 | 0.727 | 0.638 | 1 | 10 | Wfdc17 | 0.089 | Young |
| 0.017536 | 0.831589 | 0.485 | 0.379 | 1 | 10 | Snap23 | 0.106 | Young |
| 0.017735 | 1.308164 | 0.374 | 0.289 | 1 | 10 | Prdx6 | 0.085 | Young |
| 0.017921 | 1.337684 | 0.182 | 0.1 | 1 | 10 | Sephs2 | 0.082 | Young |
| 0.017964 | 0.90721 | 0.303 | 0.21 | 1 | 10 | Xpot | 0.093 | Young |
| 0.018085 | 0.334249 | 0.778 | 0.802 | 1 | 10 | Cox8a | -0.024 | Young |
| 0.018992 | 0.558796 | 0.586 | 0.506 | 1 | 10 | Eef1d | 0.08 | Young |
| 0.019828 | 0.702348 | 0.343 | 0.247 | 1 | 10 | Ube2a | 0.096 | Young |
| 0.020177 | 1.969782 | 0.111 | 0.051 | 1 | 10 | Pgm2 | 0.06 | Young |
| 0.020466 | 1.387135 | 0.162 | 0.086 | 1 | 10 | Tubgcp3 | 0.076 | Young |
| 0.020756 | 1.635824 | 0.101 | 0.044 | 1 | 10 | Pde1b | 0.057 | Young |
| 0.020831 | 0.309012 | 0.859 | 0.853 | 1 | 10 | Cybb | 0.006 | Young |
| 0.021048 | 2.591599 | 0.03 | 0.005 | 1 | 10 | Lgalsl | 0.025 | Young |
| 0.021048 | 1.726391 | 0.03 | 0.005 | 1 | 10 | Fgfrl1 | 0.025 | Young |
| 0.021058 | 0.653247 | 0.384 | 0.274 | 1 | 10 | Sema4d | 0.11 | Young |
| 0.021298 | 2.441654 | 0.03 | 0.005 | 1 | 10 | Htr2b | 0.025 | Young |

| | | | | | | | | |
|---|---|---|---|---|---|---|---|---|
| 0.021298 | 1.873503 | 0.03 | 0.005 | 1 | 10 | Prss46 | 0.025 | Young |
| 0.021806 | 1.382501 | 0.03 | 0.005 | 1 | 10 | Rab27a | 0.025 | Young |
| 0.022058 | 0.368784 | 0.808 | 0.807 | 1 | 10 | App | 0.001 | Young |
| 0.022068 | 0.913081 | 0.273 | 0.183 | 1 | 10 | Srxn1 | 0.09 | Young |
| 0.022176 | 1.206998 | 0.303 | 0.213 | 1 | 10 | Rab32 | 0.09 | Young |
| 0.022267 | 2.355834 | 0.061 | 0.02 | 1 | 10 | Trmo | 0.041 | Young |
| 0.022418 | 0.521672 | 0.424 | 0.328 | 1 | 10 | Cd81 | 0.096 | Young |
| 0.022662 | 0.714982 | 0.091 | 0.037 | 1 | 10 | Ppm1f | 0.054 | Young |
| 0.022723 | 1.368331 | 0.162 | 0.088 | 1 | 10 | Ednrb | 0.074 | Young |
| 0.022793 | 0.0651 | 0.101 | 0.042 | 1 | 10 | Itgal | 0.059 | Young |
| 0.022877 | 0.437615 | 0.747 | 0.699 | 1 | 10 | Rab7 | 0.048 | Young |
| 0.023132 | 0.677055 | 0.202 | 0.117 | 1 | 10 | Fbxo32 | 0.085 | Young |
| 0.023724 | 1.851368 | 0.061 | 0.02 | 1 | 10 | Mxra7 | 0.041 | Young |
| 0.024385 | 0.414227 | 0.778 | 0.753 | 1 | 10 | Txn1 | 0.025 | Young |
| 0.02455 | 0.582704 | 0.596 | 0.518 | 1 | 10 | 9530068E0 | 0.078 | Young |
| 0.024593 | 1.687796 | 0.081 | 0.032 | 1 | 10 | Large1 | 0.049 | Young |
| 0.02579 | 1.21978 | 0.182 | 0.103 | 1 | 10 | Rbm4b | 0.079 | Young |
| 0.025792 | 0.514249 | 0.535 | 0.423 | 1 | 10 | Il1b | 0.112 | Young |
| 0.025845 | 0.235296 | 0.899 | 0.934 | 1 | 10 | Cstb | -0.035 | Young |
| 0.026023 | 0.873069 | 0.192 | 0.112 | 1 | 10 | Pcnx | 0.08 | Young |
| 0.026025 | 1.445213 | 0.162 | 0.088 | 1 | 10 | Smim3 | 0.074 | Young |
| 0.026088 | 0.710489 | 0.515 | 0.44 | 1 | 10 | Txnrd1 | 0.075 | Young |
| 0.026267 | 0.334981 | 0.818 | 0.831 | 1 | 10 | Rpl7a | -0.013 | Young |
| 0.026319 | 1.848956 | 0.051 | 0.015 | 1 | 10 | Inpp4b | 0.036 | Young |
| 0.026573 | 0.825058 | 0.545 | 0.496 | 1 | 10 | Mafg | 0.049 | Young |
| 0.026611 | 1.145556 | 0.141 | 0.073 | 1 | 10 | Rfx7 | 0.068 | Young |
| 0.026831 | 1.845914 | 0.141 | 0.076 | 1 | 10 | Kdsr | 0.065 | Young |
| 0.026931 | 1.061236 | 0.111 | 0.051 | 1 | 10 | Bud23 | 0.06 | Young |
| 0.027124 | 3.232871 | 0.04 | 0.01 | 1 | 10 | Carmil1 | 0.03 | Young |
| 0.027141 | 0.397731 | 0.727 | 0.675 | 1 | 10 | Psma7 | 0.052 | Young |
| 0.027418 | 0.529149 | 0.586 | 0.482 | 1 | 10 | Pdcd5 | 0.104 | Young |
| 0.027435 | 1.308109 | 0.111 | 0.051 | 1 | 10 | Rab2b | 0.06 | Young |
| 0.027619 | 1.881779 | 0.04 | 0.01 | 1 | 10 | Stra6l | 0.03 | Young |
| 0.027786 | 1.267589 | 0.051 | 0.015 | 1 | 10 | Mmaa | 0.036 | Young |
| 0.027993 | 0.274005 | 0.081 | 0.171 | 1 | 10 | Parp2 | -0.09 | Young |
| 0.02818 | 1.134395 | 0.323 | 0.232 | 1 | 10 | Npm3 | 0.091 | Young |
| 0.028877 | 1.377743 | 0.051 | 0.015 | 1 | 10 | Sh2d3c | 0.036 | Young |
| 0.02894 | 2.167535 | 0.101 | 0.046 | 1 | 10 | Iba57 | 0.055 | Young |
| 0.0291 | 0.820507 | 0.051 | 0.015 | 1 | 10 | S100a8 | 0.036 | Young |
| 0.029152 | 1.348325 | 0.04 | 0.01 | 1 | 10 | Per2 | 0.03 | Young |
| 0.029152 | 0.686226 | 0.04 | 0.01 | 1 | 10 | Cxcl9 | 0.03 | Young |
| 0.029228 | 0.54592 | 0.667 | 0.68 | 1 | 10 | Rpl29 | -0.013 | Young |
| 0.029379 | 1.722604 | 0.141 | 0.076 | 1 | 10 | Dzip3 | 0.065 | Young |
| 0.029679 | 0.87657 | 0.04 | 0.01 | 1 | 10 | A630072M | 0.03 | Young |
| 0.029929 | 1.848445 | 0.101 | 0.046 | 1 | 10 | Slc25a15 | 0.055 | Young |

| | | | | | | | | |
|---|---|---|---|---|---|---|---|---|
| 0.030273 | 1.213752 | 0.131 | 0.066 | 1 | 10 | Slc2a1 | 0.065 | Young |
| 0.030298 | 0.217052 | 0.455 | 0.337 | 1 | 10 | Ccl2 | 0.118 | Young |
| 0.030906 | 0.444119 | 0.727 | 0.729 | 1 | 10 | Tgfbi | -0.002 | Young |
| 0.031369 | 0.686292 | 0.535 | 0.482 | 1 | 10 | Tomm5 | 0.053 | Young |
| 0.031539 | 0.804277 | 0.152 | 0.081 | 1 | 10 | Top3b | 0.071 | Young |
| 0.031557 | 0.759009 | 0.394 | 0.303 | 1 | 10 | Rtcb | 0.091 | Young |
| 0.032015 | 1.601425 | 0.071 | 0.027 | 1 | 10 | Acad11 | 0.044 | Young |
| 0.032426 | 0.824048 | 0.202 | 0.125 | 1 | 10 | Zbtb22 | 0.077 | Young |
| 0.032982 | 0.680494 | 0.283 | 0.193 | 1 | 10 | Slc16a6 | 0.09 | Young |
| 0.033319 | 0.561941 | 0.444 | 0.352 | 1 | 10 | Arhgef1 | 0.092 | Young |
| 0.035179 | 0.266135 | 0.838 | 0.792 | 1 | 10 | Hsp90aa1 | 0.046 | Young |
| 0.035955 | 0.275875 | 0.859 | 0.861 | 1 | 10 | Hnrnpf | -0.002 | Young |
| 0.036015 | 0.638254 | 0.384 | 0.298 | 1 | 10 | Uqcc2 | 0.086 | Young |
| 0.036018 | 0.489544 | 0.556 | 0.472 | 1 | 10 | Slc7a8 | 0.084 | Young |
| 0.036294 | 1.49291 | 0.131 | 0.068 | 1 | 10 | Pilra | 0.063 | Young |
| 0.036983 | 0.898765 | 0.202 | 0.125 | 1 | 10 | Csf3r | 0.077 | Young |
| 0.038143 | 0.31296 | 0.949 | 0.954 | 1 | 10 | Cyba | -0.005 | Young |
| 0.038311 | 1.478005 | 0.111 | 0.056 | 1 | 10 | Pwp1 | 0.055 | Young |
| 0.038462 | 2.869487 | 0.02 | 0.002 | 1 | 10 | Rnf144a | 0.018 | Young |
| 0.038462 | 2.610712 | 0.02 | 0.002 | 1 | 10 | Gm10645 | 0.018 | Young |
| 0.038462 | 2.601471 | 0.02 | 0.002 | 1 | 10 | Flt3 | 0.018 | Young |
| 0.038462 | 2.572237 | 0.02 | 0.002 | 1 | 10 | 4930532G1 | 0.018 | Young |
| 0.038462 | 2.365244 | 0.02 | 0.002 | 1 | 10 | Guca1b | 0.018 | Young |
| 0.038462 | 2.049655 | 0.02 | 0.002 | 1 | 10 | Zfp869 | 0.018 | Young |
| 0.038462 | 1.790789 | 0.02 | 0.002 | 1 | 10 | Mageh1 | 0.018 | Young |
| 0.038462 | 1.649995 | 0.02 | 0.002 | 1 | 10 | AC163354. | 0.018 | Young |
| 0.038462 | 1.612015 | 0.02 | 0.002 | 1 | 10 | Gm40117 | 0.018 | Young |
| 0.038462 | 1.157217 | 0.02 | 0.002 | 1 | 10 | Tmem120b | 0.018 | Young |
| 0.038727 | 0.815365 | 0.384 | 0.303 | 1 | 10 | Tgfbr2 | 0.081 | Young |
| 0.038805 | 0.52585 | 0.556 | 0.504 | 1 | 10 | Pgd | 0.052 | Young |
| 0.038836 | 1.079398 | 0.172 | 0.1 | 1 | 10 | Lpar6 | 0.072 | Young |
| 0.039004 | 3.554283 | 0.02 | 0.002 | 1 | 10 | Il1r2 | 0.018 | Young |
| 0.039004 | 2.304866 | 0.02 | 0.002 | 1 | 10 | Triqk | 0.018 | Young |
| 0.039004 | 2.214701 | 0.02 | 0.002 | 1 | 10 | Gm10273 | 0.018 | Young |
| 0.039004 | 1.922394 | 0.02 | 0.002 | 1 | 10 | Txlnb | 0.018 | Young |
| 0.039004 | 1.844255 | 0.02 | 0.002 | 1 | 10 | Mdrl | 0.018 | Young |
| 0.039004 | 1.722897 | 0.02 | 0.002 | 1 | 10 | Mbnl3 | 0.018 | Young |
| 0.039004 | 1.685212 | 0.02 | 0.002 | 1 | 10 | Lzts1 | 0.018 | Young |
| 0.039004 | 1.275727 | 0.02 | 0.002 | 1 | 10 | Nnat | 0.018 | Young |
| 0.039004 | 1.190859 | 0.02 | 0.002 | 1 | 10 | Fmo1 | 0.018 | Young |
| 0.039286 | 1.307401 | 0.232 | 0.156 | 1 | 10 | Ddt | 0.076 | Young |
| 0.03954 | 0.61539 | 0.202 | 0.122 | 1 | 10 | Ethe1 | 0.08 | Young |
| 0.039552 | 1.188467 | 0.02 | 0.002 | 1 | 10 | Nrg2 | 0.018 | Young |
| 0.039552 | 1.050496 | 0.02 | 0.002 | 1 | 10 | Fmo2 | 0.018 | Young |
| 0.039552 | 1.005676 | 0.02 | 0.002 | 1 | 10 | Gm26947 | 0.018 | Young |

| | | | | | | | | |
|---|---|---|---|---|---|---|---|---|
| 0.039552 | 0.933677 | 0.02 | 0.002 | 1 | 10 | Lrfn4 | 0.018 | Young |
| 0.039552 | 0.835804 | 0.02 | 0.002 | 1 | 10 | Syt3 | 0.018 | Young |
| 0.039552 | 0.716484 | 0.02 | 0.002 | 1 | 10 | Plxnb3 | 0.018 | Young |
| 0.039552 | 0.647729 | 0.02 | 0.002 | 1 | 10 | Prr5 | 0.018 | Young |
| 0.041011 | 0.739442 | 0.394 | 0.308 | 1 | 10 | Abcc1 | 0.086 | Young |
| 0.041344 | 0.603816 | 0.525 | 0.46 | 1 | 10 | Skil | 0.065 | Young |
| 0.042062 | 0.483646 | 0.323 | 0.235 | 1 | 10 | Vcam1 | 0.088 | Young |
| 0.042126 | 0.592469 | 0.192 | 0.115 | 1 | 10 | Rsbn1 | 0.077 | Young |
| 0.042519 | 1.052348 | 0.081 | 0.034 | 1 | 10 | Zfp319 | 0.047 | Young |
| 0.042541 | 0.322187 | 1 | 1 | 1 | 10 | Ftl1 | 0 | Young |
| 0.042602 | 3.97445 | 0.01 | 0 | 1 | 10 | Gsta3 | 0.01 | Young |
| 0.042602 | 3.202501 | 0.01 | 0 | 1 | 10 | Slc32a1 | 0.01 | Young |
| 0.042602 | 3.202501 | 0.01 | 0 | 1 | 10 | 1700124L1 | 0.01 | Young |
| 0.042602 | 3.202501 | 0.01 | 0 | 1 | 10 | Gm43964 | 0.01 | Young |
| 0.042602 | 3.202501 | 0.01 | 0 | 1 | 10 | Tmem40 | 0.01 | Young |
| 0.042602 | 3.202501 | 0.01 | 0 | 1 | 10 | Gm26759 | 0.01 | Young |
| 0.042602 | 3.202501 | 0.01 | 0 | 1 | 10 | Rab15 | 0.01 | Young |
| 0.042602 | 3.202501 | 0.01 | 0 | 1 | 10 | Gm4262 | 0.01 | Young |
| 0.042602 | 3.202501 | 0.01 | 0 | 1 | 10 | AC130815. | 0.01 | Young |
| 0.042602 | 3.09391 | 0.01 | 0 | 1 | 10 | D330045A | 0.01 | Young |
| 0.042602 | 3.030146 | 0.01 | 0 | 1 | 10 | Gm44053 | 0.01 | Young |
| 0.042602 | 2.765926 | 0.01 | 0 | 1 | 10 | 4930505N | 0.01 | Young |
| 0.042602 | 2.754676 | 0.01 | 0 | 1 | 10 | Mcoln3 | 0.01 | Young |
| 0.042602 | 2.754676 | 0.01 | 0 | 1 | 10 | Sdr39u1 | 0.01 | Young |
| 0.042602 | 2.71891 | 0.01 | 0 | 1 | 10 | Stx1a | 0.01 | Young |
| 0.042602 | 2.705468 | 0.01 | 0 | 1 | 10 | Gm26523 | 0.01 | Young |
| 0.042602 | 2.58687 | 0.01 | 0 | 1 | 10 | Casp12 | 0.01 | Young |
| 0.042602 | 2.409523 | 0.01 | 0 | 1 | 10 | Fmod | 0.01 | Young |
| 0.042602 | 2.409523 | 0.01 | 0 | 1 | 10 | Notch3 | 0.01 | Young |
| 0.042602 | 2.409523 | 0.01 | 0 | 1 | 10 | Plin4 | 0.01 | Young |
| 0.042602 | 2.36935 | 0.01 | 0 | 1 | 10 | Gm43579 | 0.01 | Young |
| 0.042602 | 2.36935 | 0.01 | 0 | 1 | 10 | Pgbd1 | 0.01 | Young |
| 0.042602 | 2.355529 | 0.01 | 0 | 1 | 10 | Sv2b | 0.01 | Young |
| 0.042602 | 2.355529 | 0.01 | 0 | 1 | 10 | Nkd1 | 0.01 | Young |
| 0.042602 | 2.355529 | 0.01 | 0 | 1 | 10 | Gm16299 | 0.01 | Young |
| 0.042602 | 2.29234 | 0.01 | 0 | 1 | 10 | Mrpl53 | 0.01 | Young |
| 0.042602 | 2.29234 | 0.01 | 0 | 1 | 10 | 4930471E1 | 0.01 | Young |
| 0.042602 | 2.24228 | 0.01 | 0 | 1 | 10 | Atad3aos | 0.01 | Young |
| 0.042602 | 2.24228 | 0.01 | 0 | 1 | 10 | Gm33115 | 0.01 | Young |
| 0.042602 | 2.24228 | 0.01 | 0 | 1 | 10 | 1700020D0 | 0.01 | Young |
| 0.042602 | 2.226442 | 0.01 | 0 | 1 | 10 | Gm13889 | 0.01 | Young |
| 0.042602 | 2.226442 | 0.01 | 0 | 1 | 10 | Slc7a10 | 0.01 | Young |
| 0.042602 | 2.226442 | 0.01 | 0 | 1 | 10 | Tubb4a | 0.01 | Young |
| 0.042602 | 2.203179 | 0.01 | 0 | 1 | 10 | Mettl21c | 0.01 | Young |
| 0.042602 | 2.16876 | 0.01 | 0 | 1 | 10 | Klf15 | 0.01 | Young |

| | | | | | | | | |
|---|---|---|---|---|---|---|---|---|
| 0.042602 | 2.110111 | 0.01 | 0 | 1 | 10 | 9030624G2 | 0.01 | Young |
| 0.042602 | 2.10521 | 0.01 | 0 | 1 | 10 | Lep | 0.01 | Young |
| 0.042602 | 2.099975 | 0.01 | 0 | 1 | 10 | Fgf9 | 0.01 | Young |
| 0.042602 | 2.080393 | 0.01 | 0 | 1 | 10 | Gm42997 | 0.01 | Young |
| 0.042602 | 2.080393 | 0.01 | 0 | 1 | 10 | Gm26786 | 0.01 | Young |
| 0.042602 | 2.058822 | 0.01 | 0 | 1 | 10 | Mettl18 | 0.01 | Young |
| 0.042602 | 2.058822 | 0.01 | 0 | 1 | 10 | 2010013B2 | 0.01 | Young |
| 0.042602 | 2.006452 | 0.01 | 0 | 1 | 10 | Rgs16 | 0.01 | Young |
| 0.042602 | 2.006452 | 0.01 | 0 | 1 | 10 | Gm15228 | 0.01 | Young |
| 0.042602 | 2.006452 | 0.01 | 0 | 1 | 10 | Pde1c | 0.01 | Young |
| 0.042602 | 2.006452 | 0.01 | 0 | 1 | 10 | Osbpl10 | 0.01 | Young |
| 0.042602 | 1.980932 | 0.01 | 0 | 1 | 10 | Tmsb15l | 0.01 | Young |
| 0.042602 | 1.980932 | 0.01 | 0 | 1 | 10 | Lelp1 | 0.01 | Young |
| 0.042602 | 1.980932 | 0.01 | 0 | 1 | 10 | Tppp2 | 0.01 | Young |
| 0.042602 | 1.980932 | 0.01 | 0 | 1 | 10 | St6galnac2 | 0.01 | Young |
| 0.042602 | 1.939217 | 0.01 | 0 | 1 | 10 | Cox4i2 | 0.01 | Young |
| 0.042602 | 1.939217 | 0.01 | 0 | 1 | 10 | Gm16751 | 0.01 | Young |
| 0.042602 | 1.931173 | 0.01 | 0 | 1 | 10 | 4632411P0 | 0.01 | Young |
| 0.042602 | 1.931173 | 0.01 | 0 | 1 | 10 | Prdm5 | 0.01 | Young |
| 0.042602 | 1.931173 | 0.01 | 0 | 1 | 10 | Gm20463 | 0.01 | Young |
| 0.042602 | 1.925853 | 0.01 | 0 | 1 | 10 | Gm45069 | 0.01 | Young |
| 0.042602 | 1.925853 | 0.01 | 0 | 1 | 10 | Gm15879 | 0.01 | Young |
| 0.042602 | 1.910671 | 0.01 | 0 | 1 | 10 | Il1bos | 0.01 | Young |
| 0.042602 | 1.875267 | 0.01 | 0 | 1 | 10 | Fcrlb | 0.01 | Young |
| 0.042602 | 1.875267 | 0.01 | 0 | 1 | 10 | Dok7 | 0.01 | Young |
| 0.042602 | 1.875267 | 0.01 | 0 | 1 | 10 | Tacstd2 | 0.01 | Young |
| 0.042602 | 1.875267 | 0.01 | 0 | 1 | 10 | Slc6a13 | 0.01 | Young |
| 0.042602 | 1.875267 | 0.01 | 0 | 1 | 10 | Edn1 | 0.01 | Young |
| 0.042602 | 1.872775 | 0.01 | 0 | 1 | 10 | Axdnd1 | 0.01 | Young |
| 0.042602 | 1.868914 | 0.01 | 0 | 1 | 10 | Gm32647 | 0.01 | Young |
| 0.042602 | 1.842829 | 0.01 | 0 | 1 | 10 | Lrg1 | 0.01 | Young |
| 0.042602 | 1.830749 | 0.01 | 0 | 1 | 10 | 4921509O0 | 0.01 | Young |
| 0.042602 | 1.828913 | 0.01 | 0 | 1 | 10 | Shcbp1 | 0.01 | Young |
| 0.042602 | 1.828913 | 0.01 | 0 | 1 | 10 | D630036H1 | 0.01 | Young |
| 0.042602 | 1.826559 | 0.01 | 0 | 1 | 10 | 1700067K0 | 0.01 | Young |
| 0.042602 | 1.812576 | 0.01 | 0 | 1 | 10 | A930024E0 | 0.01 | Young |
| 0.042602 | 1.812576 | 0.01 | 0 | 1 | 10 | Rpl3l | 0.01 | Young |
| 0.042602 | 1.794303 | 0.01 | 0 | 1 | 10 | Kcnq5 | 0.01 | Young |
| 0.042602 | 1.794303 | 0.01 | 0 | 1 | 10 | Lalba | 0.01 | Young |
| 0.042602 | 1.691227 | 0.01 | 0 | 1 | 10 | Myl9 | 0.01 | Young |
| 0.042602 | 1.691227 | 0.01 | 0 | 1 | 10 | Ccnj | 0.01 | Young |
| 0.042602 | 1.691227 | 0.01 | 0 | 1 | 10 | Lbx1 | 0.01 | Young |
| 0.042602 | 1.682702 | 0.01 | 0 | 1 | 10 | Kcnd3 | 0.01 | Young |
| 0.042602 | 1.682702 | 0.01 | 0 | 1 | 10 | 5730507C0 | 0.01 | Young |
| 0.042602 | 1.677072 | 0.01 | 0 | 1 | 10 | Crygn | 0.01 | Young |

| | | | | | | | | |
|---|---|---|---|---|---|---|---|---|
| 0.042602 | 1.677072 | 0.01 | 0 | 1 | 10 | Irx3 | 0.01 | Young |
| 0.042602 | 1.677072 | 0.01 | 0 | 1 | 10 | Hoxb5 | 0.01 | Young |
| 0.042602 | 1.672341 | 0.01 | 0 | 1 | 10 | Adra2a | 0.01 | Young |
| 0.042602 | 1.66842 | 0.01 | 0 | 1 | 10 | Hist1h2ac | 0.01 | Young |
| 0.042602 | 1.590795 | 0.01 | 0 | 1 | 10 | Upp1 | 0.01 | Young |
| 0.042602 | 1.580502 | 0.01 | 0 | 1 | 10 | Gm5127 | 0.01 | Young |
| 0.042602 | 1.580502 | 0.01 | 0 | 1 | 10 | Gm49322 | 0.01 | Young |
| 0.042602 | 1.580502 | 0.01 | 0 | 1 | 10 | Apol6 | 0.01 | Young |
| 0.042602 | 1.565889 | 0.01 | 0 | 1 | 10 | AC161165. | 0.01 | Young |
| 0.042602 | 1.565889 | 0.01 | 0 | 1 | 10 | Zfp820 | 0.01 | Young |
| 0.042602 | 1.565889 | 0.01 | 0 | 1 | 10 | Nol4 | 0.01 | Young |
| 0.042602 | 1.526749 | 0.01 | 0 | 1 | 10 | Cecr2 | 0.01 | Young |
| 0.042602 | 1.526749 | 0.01 | 0 | 1 | 10 | Wasf1 | 0.01 | Young |
| 0.042602 | 1.526749 | 0.01 | 0 | 1 | 10 | Neo1 | 0.01 | Young |
| 0.042602 | 1.522028 | 0.01 | 0 | 1 | 10 | St8sia1 | 0.01 | Young |
| 0.042602 | 1.522028 | 0.01 | 0 | 1 | 10 | Sec14l4 | 0.01 | Young |
| 0.042602 | 1.522028 | 0.01 | 0 | 1 | 10 | Aldh3b2 | 0.01 | Young |
| 0.042602 | 1.473577 | 0.01 | 0 | 1 | 10 | Gm26847 | 0.01 | Young |
| 0.042602 | 1.473577 | 0.01 | 0 | 1 | 10 | Pcdhga3 | 0.01 | Young |
| 0.042602 | 1.449643 | 0.01 | 0 | 1 | 10 | Gm13483 | 0.01 | Young |
| 0.042602 | 1.449643 | 0.01 | 0 | 1 | 10 | Capns2 | 0.01 | Young |
| 0.042602 | 1.441696 | 0.01 | 0 | 1 | 10 | 4732496C( | 0.01 | Young |
| 0.042602 | 1.423363 | 0.01 | 0 | 1 | 10 | Gtsf1 | 0.01 | Young |
| 0.042602 | 1.409498 | 0.01 | 0 | 1 | 10 | Rd3 | 0.01 | Young |
| 0.042602 | 1.409498 | 0.01 | 0 | 1 | 10 | Osbp2 | 0.01 | Young |
| 0.042602 | 1.409498 | 0.01 | 0 | 1 | 10 | Mfap4 | 0.01 | Young |
| 0.042602 | 1.409498 | 0.01 | 0 | 1 | 10 | Hspb9 | 0.01 | Young |
| 0.042602 | 1.409498 | 0.01 | 0 | 1 | 10 | Hist1h4m | 0.01 | Young |
| 0.042602 | 1.409498 | 0.01 | 0 | 1 | 10 | Hrasls5 | 0.01 | Young |
| 0.042602 | 1.406701 | 0.01 | 0 | 1 | 10 | Il12rb1 | 0.01 | Young |
| 0.042602 | 1.406701 | 0.01 | 0 | 1 | 10 | Aldh1a7 | 0.01 | Young |
| 0.042602 | 1.376004 | 0.01 | 0 | 1 | 10 | Prss50 | 0.01 | Young |
| 0.042602 | 1.376004 | 0.01 | 0 | 1 | 10 | 9930111J2 | 0.01 | Young |
| 0.042602 | 1.347648 | 0.01 | 0 | 1 | 10 | Ufsp1 | 0.01 | Young |
| 0.042602 | 1.347648 | 0.01 | 0 | 1 | 10 | Slc27a5 | 0.01 | Young |
| 0.042602 | 1.347648 | 0.01 | 0 | 1 | 10 | Myh3 | 0.01 | Young |
| 0.042602 | 1.347648 | 0.01 | 0 | 1 | 10 | Cyp1b1 | 0.01 | Young |
| 0.042602 | 1.335312 | 0.01 | 0 | 1 | 10 | Lpar2 | 0.01 | Young |
| 0.042602 | 1.335312 | 0.01 | 0 | 1 | 10 | Xkr6 | 0.01 | Young |
| 0.042602 | 1.335312 | 0.01 | 0 | 1 | 10 | Gm43951 | 0.01 | Young |
| 0.042602 | 1.335312 | 0.01 | 0 | 1 | 10 | 5-Sep | 0.01 | Young |
| 0.042602 | 1.325603 | 0.01 | 0 | 1 | 10 | Slc36a2 | 0.01 | Young |
| 0.042602 | 1.325409 | 0.01 | 0 | 1 | 10 | Magee2 | 0.01 | Young |
| 0.042602 | 1.325409 | 0.01 | 0 | 1 | 10 | Gm48843 | 0.01 | Young |
| 0.042602 | 1.325409 | 0.01 | 0 | 1 | 10 | 2810457G( | 0.01 | Young |

| | | | | | | | | |
|---|---|---|---|---|---|---|---|---|
| 0.042602 | 1.325409 | 0.01 | 0 | 1 | 10 | Slit3 | 0.01 | Young |
| 0.042602 | 1.322958 | 0.01 | 0 | 1 | 10 | Gm43598 | 0.01 | Young |
| 0.042602 | 1.322958 | 0.01 | 0 | 1 | 10 | Atp6v0e2 | 0.01 | Young |
| 0.042602 | 1.322958 | 0.01 | 0 | 1 | 10 | Cct6b | 0.01 | Young |
| 0.042602 | 1.234128 | 0.01 | 0 | 1 | 10 | Tead2 | 0.01 | Young |
| 0.042602 | 1.234128 | 0.01 | 0 | 1 | 10 | Slc16a9 | 0.01 | Young |
| 0.042602 | 1.234128 | 0.01 | 0 | 1 | 10 | Icam4 | 0.01 | Young |
| 0.042602 | 1.234128 | 0.01 | 0 | 1 | 10 | Gm28727 | 0.01 | Young |
| 0.042602 | 1.227393 | 0.01 | 0 | 1 | 10 | 4930539J0 | 0.01 | Young |
| 0.042602 | 1.227393 | 0.01 | 0 | 1 | 10 | Medag | 0.01 | Young |
| 0.042602 | 1.227393 | 0.01 | 0 | 1 | 10 | Anxa11os | 0.01 | Young |
| 0.042602 | 1.227393 | 0.01 | 0 | 1 | 10 | Zfp651 | 0.01 | Young |
| 0.042602 | 1.227393 | 0.01 | 0 | 1 | 10 | AC102334. | 0.01 | Young |
| 0.042602 | 1.227393 | 0.01 | 0 | 1 | 10 | Pnldc1 | 0.01 | Young |
| 0.042602 | 1.227393 | 0.01 | 0 | 1 | 10 | Emx2 | 0.01 | Young |
| 0.042602 | 1.216843 | 0.01 | 0 | 1 | 10 | Hesx1 | 0.01 | Young |
| 0.042602 | 1.216843 | 0.01 | 0 | 1 | 10 | Gm30918 | 0.01 | Young |
| 0.042602 | 1.216843 | 0.01 | 0 | 1 | 10 | Arhgef28 | 0.01 | Young |
| 0.042602 | 1.216843 | 0.01 | 0 | 1 | 10 | B130046B2 | 0.01 | Young |
| 0.042602 | 1.216089 | 0.01 | 0 | 1 | 10 | Ankrd23 | 0.01 | Young |
| 0.042602 | 1.216089 | 0.01 | 0 | 1 | 10 | Gm10561 | 0.01 | Young |
| 0.042602 | 1.216089 | 0.01 | 0 | 1 | 10 | Cd34 | 0.01 | Young |
| 0.042602 | 1.216089 | 0.01 | 0 | 1 | 10 | Capn3 | 0.01 | Young |
| 0.042602 | 1.216089 | 0.01 | 0 | 1 | 10 | Lmntd2 | 0.01 | Young |
| 0.042602 | 1.216089 | 0.01 | 0 | 1 | 10 | Cmtm8 | 0.01 | Young |
| 0.042602 | 1.216089 | 0.01 | 0 | 1 | 10 | Gm15322 | 0.01 | Young |
| 0.042602 | 1.207754 | 0.01 | 0 | 1 | 10 | Zfp970 | 0.01 | Young |
| 0.042602 | 1.207754 | 0.01 | 0 | 1 | 10 | Kcnj16 | 0.01 | Young |
| 0.042602 | 1.207754 | 0.01 | 0 | 1 | 10 | AC168220. | 0.01 | Young |
| 0.042602 | 1.170204 | 0.01 | 0 | 1 | 10 | Gm43403 | 0.01 | Young |
| 0.042602 | 1.170204 | 0.01 | 0 | 1 | 10 | Tnfsf15 | 0.01 | Young |
| 0.042602 | 1.170204 | 0.01 | 0 | 1 | 10 | Ebf3 | 0.01 | Young |
| 0.042602 | 1.170204 | 0.01 | 0 | 1 | 10 | Grin3b | 0.01 | Young |
| 0.042602 | 1.170204 | 0.01 | 0 | 1 | 10 | Gm19792 | 0.01 | Young |
| 0.042602 | 1.170204 | 0.01 | 0 | 1 | 10 | AL589692.1 | 0.01 | Young |
| 0.042602 | 1.160856 | 0.01 | 0 | 1 | 10 | 4930513N1 | 0.01 | Young |
| 0.042602 | 1.160856 | 0.01 | 0 | 1 | 10 | Gm47237 | 0.01 | Young |
| 0.042602 | 1.155708 | 0.01 | 0 | 1 | 10 | Fam212b | 0.01 | Young |
| 0.042602 | 1.155708 | 0.01 | 0 | 1 | 10 | Ctf1 | 0.01 | Young |
| 0.042602 | 1.155708 | 0.01 | 0 | 1 | 10 | Cfap54 | 0.01 | Young |
| 0.042602 | 1.155708 | 0.01 | 0 | 1 | 10 | Gm17477 | 0.01 | Young |
| 0.042602 | 1.098439 | 0.01 | 0 | 1 | 10 | Tdo2 | 0.01 | Young |
| 0.042602 | 1.098439 | 0.01 | 0 | 1 | 10 | Gm33370 | 0.01 | Young |
| 0.042602 | 1.098439 | 0.01 | 0 | 1 | 10 | Klrb1f | 0.01 | Young |
| 0.042602 | 1.098439 | 0.01 | 0 | 1 | 10 | Gm5464 | 0.01 | Young |

| | | | | | | | | |
|---|---|---|---|---|---|---|---|---|
| 0.042602 | 1.092049 | 0.01 | 0 | 1 | 10 | Mmp23 | 0.01 | Young |
| 0.042602 | 1.092049 | 0.01 | 0 | 1 | 10 | Rin1 | 0.01 | Young |
| 0.042602 | 1.072677 | 0.01 | 0 | 1 | 10 | Gm44204 | 0.01 | Young |
| 0.042602 | 1.072677 | 0.01 | 0 | 1 | 10 | Gm28041 | 0.01 | Young |
| 0.042602 | 1.072677 | 0.01 | 0 | 1 | 10 | Dio3 | 0.01 | Young |
| 0.042602 | 1.072677 | 0.01 | 0 | 1 | 10 | Cst6 | 0.01 | Young |
| 0.042602 | 1.064477 | 0.01 | 0 | 1 | 10 | Gm14137 | 0.01 | Young |
| 0.042602 | 1.064477 | 0.01 | 0 | 1 | 10 | Gm26869 | 0.01 | Young |
| 0.042602 | 1.064477 | 0.01 | 0 | 1 | 10 | Gipc2 | 0.01 | Young |
| 0.042602 | 1.064477 | 0.01 | 0 | 1 | 10 | Vmn1r197 | 0.01 | Young |
| 0.042602 | 1.064477 | 0.01 | 0 | 1 | 10 | Xrcc3 | 0.01 | Young |
| 0.042602 | 1.064477 | 0.01 | 0 | 1 | 10 | 9330136K2 | 0.01 | Young |
| 0.042602 | 1.040643 | 0.01 | 0 | 1 | 10 | Efcab7 | 0.01 | Young |
| 0.042602 | 1.040643 | 0.01 | 0 | 1 | 10 | Rhbdl2 | 0.01 | Young |
| 0.042602 | 1.040643 | 0.01 | 0 | 1 | 10 | Nqo1 | 0.01 | Young |
| 0.042602 | 1.040643 | 0.01 | 0 | 1 | 10 | Mmp8 | 0.01 | Young |
| 0.042602 | 1.040643 | 0.01 | 0 | 1 | 10 | Mboat2 | 0.01 | Young |
| 0.042602 | 1.040643 | 0.01 | 0 | 1 | 10 | A830052D1 | 0.01 | Young |
| 0.042602 | 1.016416 | 0.01 | 0 | 1 | 10 | Slc7a14 | 0.01 | Young |
| 0.042602 | 1.016416 | 0.01 | 0 | 1 | 10 | Fam221a | 0.01 | Young |
| 0.042602 | 1.016416 | 0.01 | 0 | 1 | 10 | Gm47815 | 0.01 | Young |
| 0.042602 | 0.968368 | 0.01 | 0 | 1 | 10 | Ccdc96 | 0.01 | Young |
| 0.042602 | 0.968368 | 0.01 | 0 | 1 | 10 | Clec11a | 0.01 | Young |
| 0.042602 | 0.968368 | 0.01 | 0 | 1 | 10 | Ucp3 | 0.01 | Young |
| 0.042602 | 0.954871 | 0.01 | 0 | 1 | 10 | Slc43a1 | 0.01 | Young |
| 0.042602 | 0.954871 | 0.01 | 0 | 1 | 10 | Gm26899 | 0.01 | Young |
| 0.042602 | 0.954871 | 0.01 | 0 | 1 | 10 | Ripor3 | 0.01 | Young |
| 0.042602 | 0.954871 | 0.01 | 0 | 1 | 10 | Gm32591 | 0.01 | Young |
| 0.042602 | 0.954871 | 0.01 | 0 | 1 | 10 | Olfr472 | 0.01 | Young |
| 0.042602 | 0.954871 | 0.01 | 0 | 1 | 10 | Lrrc14b | 0.01 | Young |
| 0.042602 | 0.954871 | 0.01 | 0 | 1 | 10 | Kdelr3 | 0.01 | Young |
| 0.042602 | 0.954871 | 0.01 | 0 | 1 | 10 | 4632428C0 | 0.01 | Young |
| 0.042602 | 0.954871 | 0.01 | 0 | 1 | 10 | Tigd3 | 0.01 | Young |
| 0.042602 | 0.954871 | 0.01 | 0 | 1 | 10 | Lipm | 0.01 | Young |
| 0.042602 | 0.897144 | 0.01 | 0 | 1 | 10 | Aspm | 0.01 | Young |
| 0.042602 | 0.897144 | 0.01 | 0 | 1 | 10 | Rgsl1 | 0.01 | Young |
| 0.042602 | 0.897144 | 0.01 | 0 | 1 | 10 | Nek2 | 0.01 | Young |
| 0.042602 | 0.897144 | 0.01 | 0 | 1 | 10 | Rarres1 | 0.01 | Young |
| 0.042602 | 0.897144 | 0.01 | 0 | 1 | 10 | Ldlrad1 | 0.01 | Young |
| 0.042602 | 0.897144 | 0.01 | 0 | 1 | 10 | Adamts15 | 0.01 | Young |
| 0.042602 | 0.897144 | 0.01 | 0 | 1 | 10 | Upk1b | 0.01 | Young |
| 0.042602 | 0.897144 | 0.01 | 0 | 1 | 10 | Zfp229 | 0.01 | Young |
| 0.042602 | 0.897144 | 0.01 | 0 | 1 | 10 | Ptcra | 0.01 | Young |
| 0.042602 | 0.897144 | 0.01 | 0 | 1 | 10 | Pcdhga6 | 0.01 | Young |
| 0.042602 | 0.892605 | 0.01 | 0 | 1 | 10 | Rab33a | 0.01 | Young |

| | | | | | | | | |
|---|---|---|---|---|---|---|---|---|
| 0.042602 | 0.892605 | 0.01 | 0 | 1 | 10 | Gm9889 | 0.01 | Young |
| 0.042602 | 0.868838 | 0.01 | 0 | 1 | 10 | Gm26892 | 0.01 | Young |
| 0.042602 | 0.868838 | 0.01 | 0 | 1 | 10 | Grem2 | 0.01 | Young |
| 0.042602 | 0.868838 | 0.01 | 0 | 1 | 10 | Slc52a3 | 0.01 | Young |
| 0.042602 | 0.868838 | 0.01 | 0 | 1 | 10 | Ptger1 | 0.01 | Young |
| 0.042602 | 0.868838 | 0.01 | 0 | 1 | 10 | Ppp1r27 | 0.01 | Young |
| 0.042602 | 0.868838 | 0.01 | 0 | 1 | 10 | Slc12a8 | 0.01 | Young |
| 0.042602 | 0.831195 | 0.01 | 0 | 1 | 10 | Gm13404 | 0.01 | Young |
| 0.042602 | 0.831195 | 0.01 | 0 | 1 | 10 | Gm13814 | 0.01 | Young |
| 0.042602 | 0.831195 | 0.01 | 0 | 1 | 10 | Med12l | 0.01 | Young |
| 0.042602 | 0.831195 | 0.01 | 0 | 1 | 10 | Vps37d | 0.01 | Young |
| 0.042602 | 0.831195 | 0.01 | 0 | 1 | 10 | Ppp1r14a | 0.01 | Young |
| 0.042602 | 0.831195 | 0.01 | 0 | 1 | 10 | Mfsd13b | 0.01 | Young |
| 0.042602 | 0.831195 | 0.01 | 0 | 1 | 10 | Kdm8 | 0.01 | Young |
| 0.042602 | 0.831195 | 0.01 | 0 | 1 | 10 | Gm34934 | 0.01 | Young |
| 0.042602 | 0.831195 | 0.01 | 0 | 1 | 10 | Zkscan4 | 0.01 | Young |
| 0.042602 | 0.831195 | 0.01 | 0 | 1 | 10 | Hist1h3e | 0.01 | Young |
| 0.042602 | 0.831195 | 0.01 | 0 | 1 | 10 | Pcdhgc5 | 0.01 | Young |
| 0.042602 | 0.809981 | 0.01 | 0 | 1 | 10 | Slco5a1 | 0.01 | Young |
| 0.042602 | 0.809981 | 0.01 | 0 | 1 | 10 | Nexmif | 0.01 | Young |
| 0.042602 | 0.809981 | 0.01 | 0 | 1 | 10 | Gm44752 | 0.01 | Young |
| 0.042602 | 0.809981 | 0.01 | 0 | 1 | 10 | Hist1h2bb | 0.01 | Young |
| 0.042602 | 0.809981 | 0.01 | 0 | 1 | 10 | Sh3yl1 | 0.01 | Young |
| 0.042602 | 0.809981 | 0.01 | 0 | 1 | 10 | Syt16 | 0.01 | Young |
| 0.042602 | 0.799898 | 0.01 | 0 | 1 | 10 | Nrarp | 0.01 | Young |
| 0.042602 | 0.799898 | 0.01 | 0 | 1 | 10 | Il12rb2 | 0.01 | Young |
| 0.042602 | 0.799898 | 0.01 | 0 | 1 | 10 | Olfr624 | 0.01 | Young |
| 0.042602 | 0.787617 | 0.01 | 0 | 1 | 10 | Ccdc92 | 0.01 | Young |
| 0.042602 | 0.787617 | 0.01 | 0 | 1 | 10 | Pbp2 | 0.01 | Young |
| 0.042602 | 0.787617 | 0.01 | 0 | 1 | 10 | 4430402I1 | 0.01 | Young |
| 0.042602 | 0.758319 | 0.01 | 0 | 1 | 10 | Cubn | 0.01 | Young |
| 0.042602 | 0.758319 | 0.01 | 0 | 1 | 10 | Gm14252 | 0.01 | Young |
| 0.042602 | 0.758319 | 0.01 | 0 | 1 | 10 | Spin2c | 0.01 | Young |
| 0.042602 | 0.758319 | 0.01 | 0 | 1 | 10 | Gm26546 | 0.01 | Young |
| 0.042602 | 0.758319 | 0.01 | 0 | 1 | 10 | Lrrc63 | 0.01 | Young |
| 0.042602 | 0.758319 | 0.01 | 0 | 1 | 10 | Ephb3 | 0.01 | Young |
| 0.042602 | 0.758319 | 0.01 | 0 | 1 | 10 | Zfp811 | 0.01 | Young |
| 0.042602 | 0.709506 | 0.01 | 0 | 1 | 10 | Mastl | 0.01 | Young |
| 0.042602 | 0.709506 | 0.01 | 0 | 1 | 10 | C8g | 0.01 | Young |
| 0.042602 | 0.709506 | 0.01 | 0 | 1 | 10 | Ttc30a2 | 0.01 | Young |
| 0.042602 | 0.709506 | 0.01 | 0 | 1 | 10 | A730090N | 0.01 | Young |
| 0.042602 | 0.709506 | 0.01 | 0 | 1 | 10 | Cd164l2 | 0.01 | Young |
| 0.042602 | 0.709506 | 0.01 | 0 | 1 | 10 | Msantd1 | 0.01 | Young |
| 0.042602 | 0.709506 | 0.01 | 0 | 1 | 10 | Gprin3 | 0.01 | Young |
| 0.042602 | 0.709506 | 0.01 | 0 | 1 | 10 | Gm44067 | 0.01 | Young |

| | | | | | | | | |
|---|---|---|---|---|---|---|---|---|
| 0.042602 | 0.709506 | 0.01 | 0 | 1 | 10 | D830036C2 | 0.01 | Young |
| 0.042602 | 0.709506 | 0.01 | 0 | 1 | 10 | Klk13 | 0.01 | Young |
| 0.042602 | 0.709506 | 0.01 | 0 | 1 | 10 | Ascl2 | 0.01 | Young |
| 0.042602 | 0.709506 | 0.01 | 0 | 1 | 10 | Pram1 | 0.01 | Young |
| 0.042602 | 0.709506 | 0.01 | 0 | 1 | 10 | Setbp1 | 0.01 | Young |
| 0.042602 | 0.695687 | 0.01 | 0 | 1 | 10 | Sytl5 | 0.01 | Young |
| 0.042602 | 0.695687 | 0.01 | 0 | 1 | 10 | Gm10069 | 0.01 | Young |
| 0.042602 | 0.695687 | 0.01 | 0 | 1 | 10 | Cngb1 | 0.01 | Young |
| 0.042602 | 0.695687 | 0.01 | 0 | 1 | 10 | 4930431P2 | 0.01 | Young |
| 0.042602 | 0.695687 | 0.01 | 0 | 1 | 10 | Mdfi | 0.01 | Young |
| 0.042602 | 0.580592 | 0.01 | 0 | 1 | 10 | Defb25 | 0.01 | Young |
| 0.042602 | 0.580592 | 0.01 | 0 | 1 | 10 | Kcnip4 | 0.01 | Young |
| 0.042602 | 0.580592 | 0.01 | 0 | 1 | 10 | Tjp1 | 0.01 | Young |
| 0.042602 | 0.580592 | 0.01 | 0 | 1 | 10 | Ccdc116 | 0.01 | Young |
| 0.042602 | 0.580592 | 0.01 | 0 | 1 | 10 | Tm4sf19 | 0.01 | Young |
| 0.042602 | 0.580592 | 0.01 | 0 | 1 | 10 | Alpk2 | 0.01 | Young |
| 0.042646 | 0.021793 | 0.101 | 0.193 | 1 | 10 | Pcmtd2 | -0.092 | Young |
| 0.043465 | 0.826726 | 0.242 | 0.164 | 1 | 10 | Dhx30 | 0.078 | Young |
| 0.043714 | 1.758083 | 0.182 | 0.115 | 1 | 10 | Ccnd2 | 0.067 | Young |
| 0.043919 | 0.532538 | 0.525 | 0.472 | 1 | 10 | Ssr4 | 0.053 | Young |
| 0.044055 | 0.509546 | 0.485 | 0.396 | 1 | 10 | Stab1 | 0.089 | Young |
| 0.044779 | 1.161338 | 0.131 | 0.071 | 1 | 10 | Tle1 | 0.06 | Young |
| 0.045322 | 1.054319 | 0.121 | 0.064 | 1 | 10 | Tspan13 | 0.057 | Young |
| 0.045344 | 0.429879 | 0.596 | 0.521 | 1 | 10 | Slc15a3 | 0.075 | Young |
| 0.04581 | 0.931743 | 0.222 | 0.147 | 1 | 10 | Ufd1 | 0.075 | Young |
| 0.046169 | 0.516992 | 0.616 | 0.567 | 1 | 10 | Dnaja1 | 0.049 | Young |
| 0.046262 | 0.71511 | 0.263 | 0.178 | 1 | 10 | Foxj2 | 0.085 | Young |
| 0.046376 | 1.080421 | 0.141 | 0.078 | 1 | 10 | Hsf1 | 0.063 | Young |
| 0.046861 | 0.843688 | 0.131 | 0.071 | 1 | 10 | Nrm | 0.06 | Young |
| 0.046925 | 1.681237 | 0.071 | 0.029 | 1 | 10 | Stoml1 | 0.042 | Young |
| 0.047349 | 0.127921 | 1 | 0.985 | 1 | 10 | Rps3a1 | 0.015 | Young |
| 0.047382 | 0.603483 | 0.354 | 0.259 | 1 | 10 | Cbr2 | 0.095 | Young |
| 0.047442 | 1.458134 | 0.071 | 0.029 | 1 | 10 | Dnajc27 | 0.042 | Young |
| 0.048121 | 0.20944 | 0.98 | 0.99 | 1 | 10 | Eif1 | -0.01 | Young |
| 0.048569 | 0.358389 | 0.616 | 0.577 | 1 | 10 | Nfe2l2 | 0.039 | Young |
| 0.048644 | 0.622176 | 0.485 | 0.418 | 1 | 10 | Phb2 | 0.067 | Young |
| 0.048898 | 1.70564 | 0.051 | 0.017 | 1 | 10 | Lmln | 0.034 | Young |
| 0.048937 | 0.684696 | 0.263 | 0.181 | 1 | 10 | Sae1 | 0.082 | Young |
| 0.049232 | 0.777476 | 0.051 | 0.017 | 1 | 10 | Dedd2 | 0.034 | Young |
| 0.050099 | 1.499865 | 0.071 | 0.029 | 1 | 10 | Ift172 | 0.042 | Young |
| 0.050194 | 0.883459 | 0.101 | 0.049 | 1 | 10 | Gna15 | 0.052 | Young |
| 0.050231 | 0.246519 | 0.99 | 0.978 | 1 | 10 | Hspa8 | 0.012 | Young |
| 0.050937 | 0.957943 | 0.263 | 0.191 | 1 | 10 | Cltb | 0.072 | Young |
| 0.05105 | 1.938191 | 0.091 | 0.044 | 1 | 10 | Kti12 | 0.047 | Young |
| 0.051144 | 0.768687 | 0.091 | 0.042 | 1 | 10 | Zfp568 | 0.049 | Young |

| | | | | | | | | |
|---|---|---|---|---|---|---|---|---|
| 0.051599 | 0.678884 | 0.303 | 0.213 | 1 | 10 | Tfg | 0.09 | Young |
| 0.051752 | 1.503758 | 0.071 | 0.029 | 1 | 10 | Lss | 0.042 | Young |
| 0.052243 | 0.682575 | 0.475 | 0.433 | 1 | 10 | Psmb5 | 0.042 | Young |
| 0.052322 | 1.111335 | 0.051 | 0.017 | 1 | 10 | Zfp949 | 0.034 | Young |
| 0.052322 | 1.040807 | 0.051 | 0.017 | 1 | 10 | Fcrl1 | 0.034 | Young |
| 0.053676 | 1.369009 | 0.202 | 0.137 | 1 | 10 | Nop14 | 0.065 | Young |
| 0.053692 | 1.631692 | 0.101 | 0.051 | 1 | 10 | Tmem18 | 0.05 | Young |
| 0.05422 | 0.66714 | 0.505 | 0.45 | 1 | 10 | Ifngr2 | 0.055 | Young |
| 0.054337 | 0.644557 | 0.525 | 0.469 | 1 | 10 | Eea1 | 0.056 | Young |
| 0.054746 | 0.754184 | 0.394 | 0.328 | 1 | 10 | Camkk2 | 0.066 | Young |
| 0.054872 | 0.936419 | 0.192 | 0.12 | 1 | 10 | Dnajb1 | 0.072 | Young |
| 0.055103 | 0.299806 | 0.646 | 0.535 | 1 | 10 | Cox17 | 0.111 | Young |
| 0.055238 | 0.351441 | 0.778 | 0.765 | 1 | 10 | Atox1 | 0.013 | Young |
| 0.055243 | 1.782863 | 0.081 | 0.037 | 1 | 10 | Zcchc4 | 0.044 | Young |
| 0.055692 | 2.144383 | 0.04 | 0.012 | 1 | 10 | Cxcl10 | 0.028 | Young |
| 0.055692 | 1.707408 | 0.04 | 0.012 | 1 | 10 | Rpp38 | 0.028 | Young |
| 0.055692 | 1.382872 | 0.04 | 0.012 | 1 | 10 | A730063M | 0.028 | Young |
| 0.055703 | 0.892077 | 0.212 | 0.139 | 1 | 10 | Ppil4 | 0.073 | Young |
| 0.055818 | 2.382784 | 0.03 | 0.007 | 1 | 10 | Tmem201 | 0.023 | Young |
| 0.056013 | 0.82127 | 0.152 | 0.088 | 1 | 10 | Qsox1 | 0.064 | Young |
| 0.056055 | 1.385887 | 0.081 | 0.037 | 1 | 10 | P2ry12 | 0.044 | Young |
| 0.05612 | 1.455513 | 0.04 | 0.012 | 1 | 10 | Rcn1 | 0.028 | Young |
| 0.056205 | 1.003249 | 0.364 | 0.286 | 1 | 10 | Tgfbr1 | 0.078 | Young |
| 0.056233 | 0.963481 | 0.121 | 0.066 | 1 | 10 | Kif21b | 0.055 | Young |
| 0.056418 | 1.067423 | 0.283 | 0.208 | 1 | 10 | AU020206 | 0.075 | Young |
| 0.056551 | 1.697386 | 0.04 | 0.012 | 1 | 10 | Atp2b4 | 0.028 | Young |
| 0.056934 | 1.071251 | 0.162 | 0.098 | 1 | 10 | Borcs7 | 0.064 | Young |
| 0.057405 | 1.583863 | 0.03 | 0.007 | 1 | 10 | Cspg4 | 0.023 | Young |
| 0.057405 | 1.461283 | 0.03 | 0.007 | 1 | 10 | Gm15232 | 0.023 | Young |
| 0.057405 | 1.428113 | 0.03 | 0.007 | 1 | 10 | Slc46a1 | 0.023 | Young |
| 0.057908 | 0.797036 | 0.919 | 0.944 | 1 | 10 | Prdx1 | -0.025 | Young |
| 0.057942 | 1.311282 | 0.03 | 0.007 | 1 | 10 | Tkfc | 0.023 | Young |
| 0.057942 | 1.253526 | 0.03 | 0.007 | 1 | 10 | Gm29336 | 0.023 | Young |
| 0.058304 | 0.626688 | 0.04 | 0.012 | 1 | 10 | Numbl | 0.028 | Young |
| 0.058467 | 1.139174 | 0.091 | 0.044 | 1 | 10 | Ldlrad3 | 0.047 | Young |
| 0.058514 | 1.209562 | 0.131 | 0.076 | 1 | 10 | B4galt3 | 0.055 | Young |
| 0.058665 | 1.185246 | 0.101 | 0.051 | 1 | 10 | Pilrb2 | 0.05 | Young |
| 0.058749 | 1.264758 | 0.04 | 0.012 | 1 | 10 | Tmem119 | 0.028 | Young |
| 0.058749 | 0.850888 | 0.04 | 0.012 | 1 | 10 | Sh3bp4 | 0.028 | Young |
| 0.058818 | 0.341614 | 0.737 | 0.741 | 1 | 10 | Hint1 | -0.004 | Young |
| 0.059028 | 1.087192 | 0.03 | 0.007 | 1 | 10 | Spsb1 | 0.023 | Young |
| 0.059578 | 0.902887 | 0.03 | 0.007 | 1 | 10 | A930033H1 | 0.023 | Young |
| 0.059911 | 0.021784 | 0.303 | 0.438 | 1 | 10 | Tnfrsf11a | -0.135 | Young |
| 0.060193 | 0.583238 | 0.535 | 0.484 | 1 | 10 | Neurl3 | 0.051 | Young |
| 0.060267 | 0.953312 | 0.081 | 0.037 | 1 | 10 | Prkce | 0.044 | Young |

| | | | | | | | | |
|---|---|---|---|---|---|---|---|---|
| 0.061364 | 0.250005 | 0.364 | 0.267 | 1 | 10 | Mybbp1a | 0.097 | Young |
| 0.062142 | 0.552066 | 0.323 | 0.24 | 1 | 10 | Bcl7c | 0.083 | Young |
| 0.062168 | 0.61477 | 0.192 | 0.12 | 1 | 10 | Olr1 | 0.072 | Young |
| 0.062776 | 0.77416 | 0.091 | 0.044 | 1 | 10 | Procr | 0.047 | Young |
| 0.062791 | 1.439378 | 0.141 | 0.086 | 1 | 10 | 0610010K1 | 0.055 | Young |
| 0.062973 | 0.794131 | 0.222 | 0.152 | 1 | 10 | Taf5l | 0.07 | Young |
| 0.063474 | 0.013557 | 0.343 | 0.46 | 1 | 10 | Irak1 | -0.117 | Young |
| 0.063503 | 0.733608 | 0.333 | 0.257 | 1 | 10 | St6gal1 | 0.076 | Young |
| 0.063823 | 0.537898 | 0.081 | 0.037 | 1 | 10 | B430306N( | 0.044 | Young |
| 0.064491 | 1.167588 | 0.182 | 0.117 | 1 | 10 | Bcl2a1a | 0.065 | Young |
| 0.064523 | 1.167247 | 0.313 | 0.254 | 1 | 10 | Nek9 | 0.059 | Young |
| 0.064693 | 0.763312 | 0.111 | 0.059 | 1 | 10 | Ghdc | 0.052 | Young |
| 0.064716 | 1.123703 | 0.182 | 0.115 | 1 | 10 | Pycr2 | 0.067 | Young |
| 0.064945 | 1.218489 | 0.111 | 0.059 | 1 | 10 | Havcr2 | 0.052 | Young |
| 0.065509 | 0.294266 | 0.707 | 0.697 | 1 | 10 | Srp9 | 0.01 | Young |
| 0.066015 | 1.213898 | 0.141 | 0.086 | 1 | 10 | Ints11 | 0.055 | Young |
| 0.066181 | 1.413751 | 0.222 | 0.149 | 1 | 10 | Pom121 | 0.073 | Young |
| 0.067267 | 1.523976 | 0.071 | 0.032 | 1 | 10 | Arhgap27 | 0.039 | Young |
| 0.067639 | 0.549013 | 0.091 | 0.044 | 1 | 10 | Rbms2 | 0.047 | Young |
| 0.067699 | 1.051502 | 0.162 | 0.1 | 1 | 10 | Champ1 | 0.062 | Young |
| 0.067969 | 1.384148 | 0.091 | 0.046 | 1 | 10 | Sgf29 | 0.045 | Young |
| 0.068077 | 0.178747 | 0.98 | 0.958 | 1 | 10 | Rps13 | 0.022 | Young |
| 0.06821 | 0.398664 | 0.465 | 0.406 | 1 | 10 | Fam168b | 0.059 | Young |
| 0.069206 | 0.473229 | 0.061 | 0.024 | 1 | 10 | D030056L2 | 0.037 | Young |
| 0.069779 | 0.228613 | 0.909 | 0.907 | 1 | 10 | Rpl14 | 0.002 | Young |
| 0.070411 | 0.062381 | 0.141 | 0.242 | 1 | 10 | Keap1 | -0.101 | Young |
| 0.070569 | 0.281049 | 0.273 | 0.183 | 1 | 10 | Stk40 | 0.09 | Young |
| 0.071054 | 0.573163 | 0.566 | 0.557 | 1 | 10 | Cxcl16 | 0.009 | Young |
| 0.07117 | 0.916715 | 0.313 | 0.235 | 1 | 10 | 1700017B( | 0.078 | Young |
| 0.071397 | 0.725392 | 0.444 | 0.364 | 1 | 10 | Ppp1r15a | 0.08 | Young |
| 0.071544 | 0.828019 | 0.212 | 0.147 | 1 | 10 | Mrpl37 | 0.065 | Young |
| 0.072132 | 0.323423 | 0.545 | 0.489 | 1 | 10 | Abca1 | 0.056 | Young |
| 0.072321 | 0.54955 | 0.374 | 0.301 | 1 | 10 | Psmb2 | 0.073 | Young |
| 0.072596 | 0.895776 | 0.293 | 0.218 | 1 | 10 | Ccl24 | 0.075 | Young |
| 0.072964 | 0.419939 | 0.727 | 0.66 | 1 | 10 | Dab2 | 0.067 | Young |
| 0.073583 | 1.428468 | 0.192 | 0.13 | 1 | 10 | Micall1 | 0.062 | Young |
| 0.073604 | 1.077047 | 0.111 | 0.061 | 1 | 10 | Bhlhe41 | 0.05 | Young |
| 0.074716 | 1.020796 | 0.111 | 0.061 | 1 | 10 | Mir155hg | 0.05 | Young |
| 0.075242 | 0.196923 | 0.949 | 0.924 | 1 | 10 | Rpl26 | 0.025 | Young |
| 0.075967 | 0.195265 | 0.081 | 0.154 | 1 | 10 | Gopc | -0.073 | Young |
| 0.076502 | 1.963646 | 0.051 | 0.02 | 1 | 10 | Cdan1 | 0.031 | Young |
| 0.077163 | 0.647529 | 0.212 | 0.139 | 1 | 10 | Fez2 | 0.073 | Young |
| 0.077609 | 0.912284 | 0.152 | 0.093 | 1 | 10 | Ppp1r8 | 0.059 | Young |
| 0.077727 | 1.46184 | 0.101 | 0.054 | 1 | 10 | Gpr180 | 0.047 | Young |
| 0.077903 | 1.438337 | 0.051 | 0.02 | 1 | 10 | Zbtb10 | 0.031 | Young |

| | | | | | | | | |
|---|---|---|---|---|---|---|---|---|
| 0.077939 | 1.021944 | 0.192 | 0.13 | 1 | 10 | Galnt6 | 0.062 | Young |
| 0.078094 | 1.247742 | 0.152 | 0.093 | 1 | 10 | Lyve1 | 0.059 | Young |
| 0.079126 | 0.333337 | 0.646 | 0.592 | 1 | 10 | Maf | 0.054 | Young |
| 0.07913 | 0.57363 | 0.273 | 0.2 | 1 | 10 | Vps50 | 0.073 | Young |
| 0.079162 | 0.772295 | 0.212 | 0.142 | 1 | 10 | Kdm3a | 0.07 | Young |
| 0.079803 | 1.915644 | 0.051 | 0.02 | 1 | 10 | Ipp | 0.031 | Young |
| 0.0805 | 0.45694 | 0.576 | 0.518 | 1 | 10 | Stx4a | 0.058 | Young |
| 0.080565 | 0.54132 | 0.495 | 0.443 | 1 | 10 | Brd4 | 0.052 | Young |
| 0.081253 | 1.734367 | 0.051 | 0.02 | 1 | 10 | Cep76 | 0.031 | Young |
| 0.081253 | 1.0962 | 0.051 | 0.02 | 1 | 10 | Hist1h1e | 0.031 | Young |
| 0.081495 | 0.412064 | 0.192 | 0.125 | 1 | 10 | Crem | 0.067 | Young |
| 0.08242 | 1.772078 | 0.141 | 0.088 | 1 | 10 | Cd24a | 0.053 | Young |
| 0.082725 | 0.75751 | 0.051 | 0.02 | 1 | 10 | Ddx59 | 0.031 | Young |
| 0.08277 | 0.553259 | 0.152 | 0.09 | 1 | 10 | Tmem8 | 0.062 | Young |
| 0.083471 | 0.496293 | 0.343 | 0.269 | 1 | 10 | Ptk2b | 0.074 | Young |
| 0.083878 | 0.334661 | 0.273 | 0.193 | 1 | 10 | Hivep3 | 0.08 | Young |
| 0.083981 | 0.444986 | 0.394 | 0.315 | 1 | 10 | Tmem208 | 0.079 | Young |
| 0.084588 | 0.167367 | 0.172 | 0.108 | 1 | 10 | Eci2 | 0.064 | Young |
| 0.085741 | 1.814745 | 0.101 | 0.056 | 1 | 10 | Jade2 | 0.045 | Young |
| 0.085863 | 0.16339 | 0.97 | 0.978 | 1 | 10 | Rpl27a | -0.008 | Young |
| 0.085944 | 0.570409 | 0.455 | 0.418 | 1 | 10 | Rsrc2 | 0.037 | Young |
| 0.086233 | 0.29595 | 0.808 | 0.814 | 1 | 10 | Slc25a5 | -0.006 | Young |
| 0.086264 | 1.291389 | 0.131 | 0.078 | 1 | 10 | Apbb2 | 0.053 | Young |
| 0.086425 | 0.329299 | 0.01 | 0.049 | 1 | 10 | Deaf1 | -0.039 | Young |
| 0.086722 | 0.245623 | 0.697 | 0.699 | 1 | 10 | Ubb | -0.002 | Young |
| 0.086828 | 0.894283 | 0.121 | 0.071 | 1 | 10 | Smad7 | 0.05 | Young |
| 0.086928 | 1.286877 | 0.152 | 0.098 | 1 | 10 | Kctd5 | 0.054 | Young |
| 0.088085 | 0.323242 | 0.636 | 0.601 | 1 | 10 | Psmb6 | 0.035 | Young |
| 0.088402 | 0.112884 | 0.98 | 0.993 | 1 | 10 | Rps23 | -0.013 | Young |
| 0.088558 | 1.016246 | 0.131 | 0.078 | 1 | 10 | Ppcdc | 0.053 | Young |
| 0.088576 | 0.560096 | 0.556 | 0.511 | 1 | 10 | Rhoc | 0.045 | Young |
| 0.089001 | 0.954437 | 0.162 | 0.105 | 1 | 10 | Zfp598 | 0.057 | Young |
| 0.08969 | 1.308924 | 0.162 | 0.108 | 1 | 10 | Cdca4 | 0.054 | Young |
| 0.089775 | 1.499211 | 0.091 | 0.049 | 1 | 10 | Mri1 | 0.042 | Young |
| 0.089845 | 1.042824 | 0.121 | 0.071 | 1 | 10 | Alkbh7 | 0.05 | Young |
| 0.090269 | 0.691555 | 0.374 | 0.313 | 1 | 10 | Etnk1 | 0.061 | Young |
| 0.090295 | 0.204154 | 0.859 | 0.902 | 1 | 10 | Rps3 | -0.043 | Young |
| 0.090518 | 0.388861 | 0.616 | 0.601 | 1 | 10 | Lamtor4 | 0.015 | Young |
| 0.090584 | 0.87403 | 0.061 | 0.027 | 1 | 10 | Tmem42 | 0.034 | Young |
| 0.091052 | 0.708265 | 0.061 | 0.027 | 1 | 10 | Areg | 0.034 | Young |
| 0.091081 | 1.234095 | 0.101 | 0.056 | 1 | 10 | Ppip5k1 | 0.045 | Young |
| 0.091094 | 0.192866 | 0.737 | 0.731 | 1 | 10 | Cdkn1a | 0.006 | Young |
| 0.09122 | 1.917043 | 0.091 | 0.049 | 1 | 10 | Runx3 | 0.042 | Young |
| 0.091469 | 0.510235 | 0.657 | 0.65 | 1 | 10 | Plek | 0.007 | Young |
| 0.091556 | 0.274794 | 0.192 | 0.127 | 1 | 10 | Gse1 | 0.065 | Young |

| | | | | | | | | |
|---|---|---|---|---|---|---|---|---|
| 0.091561 | 0.969627 | 0.172 | 0.112 | 1 | 10 | Arsb | 0.06 | Young |
| 0.091585 | 1.403621 | 0.091 | 0.049 | 1 | 10 | Gm20707 | 0.042 | Young |
| 0.092515 | 2.057601 | 0.04 | 0.015 | 1 | 10 | Hist1h4d | 0.025 | Young |
| 0.092698 | 0.876203 | 0.212 | 0.152 | 1 | 10 | Relb | 0.06 | Young |
| 0.092858 | 0.378163 | 0.091 | 0.046 | 1 | 10 | Cdk5rap2 | 0.045 | Young |
| 0.092934 | 0.88433 | 0.162 | 0.105 | 1 | 10 | Akt1s1 | 0.057 | Young |
| 0.092948 | 1.176016 | 0.354 | 0.289 | 1 | 10 | Hist1h2bc | 0.065 | Young |
| 0.092971 | 0.823968 | 0.374 | 0.308 | 1 | 10 | Tcf12 | 0.066 | Young |
| 0.093109 | 0.744638 | 0.172 | 0.112 | 1 | 10 | Pfkl | 0.06 | Young |
| 0.093749 | 0.426314 | 0.141 | 0.086 | 1 | 10 | Ints7 | 0.055 | Young |
| 0.093761 | 0.526631 | 0.485 | 0.44 | 1 | 10 | Polr2k | 0.045 | Young |
| 0.093864 | 0.338146 | 0.727 | 0.694 | 1 | 10 | Aldoa | 0.033 | Young |
| 0.094374 | 1.664362 | 0.04 | 0.015 | 1 | 10 | Uhrf1bp1 | 0.025 | Young |
| 0.095 | 1.976711 | 0.04 | 0.015 | 1 | 10 | Ppp1r13l | 0.025 | Young |
| 0.095 | 1.80828 | 0.04 | 0.015 | 1 | 10 | Izumo4 | 0.025 | Young |
| 0.095667 | 0.073283 | 0.172 | 0.254 | 1 | 10 | Slc30a1 | -0.082 | Young |
| 0.096524 | 0.98004 | 0.232 | 0.166 | 1 | 10 | Prrc1 | 0.066 | Young |
| 0.09662 | 1.524824 | 0.071 | 0.034 | 1 | 10 | Polr1e | 0.037 | Young |
| 0.096899 | 1.515657 | 0.04 | 0.015 | 1 | 10 | Dnaaf5 | 0.025 | Young |
| 0.096967 | 1.003592 | 0.222 | 0.164 | 1 | 10 | Gch1 | 0.058 | Young |
| 0.097391 | 0.634034 | 0.404 | 0.34 | 1 | 10 | Emc6 | 0.064 | Young |
| 0.097667 | 0.175816 | 0.909 | 0.897 | 1 | 10 | Cox7c | 0.012 | Young |
| 0.098136 | 1.110683 | 0.131 | 0.081 | 1 | 10 | Cxxc1 | 0.05 | Young |
| 0.098181 | 1.474164 | 0.04 | 0.015 | 1 | 10 | Wfdc18 | 0.025 | Young |
| 0.098181 | 0.595121 | 0.04 | 0.015 | 1 | 10 | Ift140 | 0.025 | Young |
| 0.098227 | 0.711506 | 0.131 | 0.078 | 1 | 10 | Gtpbp6 | 0.053 | Young |
| 0.098263 | 0.50264 | 0.455 | 0.386 | 1 | 10 | Psmd14 | 0.069 | Young |
| 0.098541 | 0.559362 | 0.424 | 0.367 | 1 | 10 | Psma5 | 0.057 | Young |
| 0.09871 | 0.027864 | 0.091 | 0.156 | 1 | 10 | Fbxo38 | -0.065 | Young |
| 0.098828 | 0.928532 | 0.04 | 0.015 | 1 | 10 | Ccdc114 | 0.025 | Young |
| 0.099175 | 0.961696 | 0.131 | 0.078 | 1 | 10 | Phldb1 | 0.053 | Young |
| 0.099477 | 1.307246 | 0.04 | 0.015 | 1 | 10 | Dennd2d | 0.025 | Young |
| 0.099477 | 0.946125 | 0.04 | 0.015 | 1 | 10 | Zfp442 | 0.025 | Young |
| 0.099696 | 0.733797 | 0.364 | 0.293 | 1 | 10 | Max | 0.071 | Young |
| 0.100411 | 0.885502 | 0.111 | 0.064 | 1 | 10 | Rgmb | 0.047 | Young |
| 0.100412 | 1.112258 | 0.081 | 0.042 | 1 | 10 | Atg2b | 0.039 | Young |
| 0.100787 | 0.868478 | 0.04 | 0.015 | 1 | 10 | Bend3 | 0.025 | Young |
| 0.10113 | 1.32471 | 0.131 | 0.081 | 1 | 10 | Fbxl17 | 0.05 | Young |
| 0.101447 | 0.997957 | 0.04 | 0.015 | 1 | 10 | Tmem186 | 0.025 | Young |
| 0.102777 | 1.844285 | 0.04 | 0.015 | 1 | 10 | Flrt3 | 0.025 | Young |
| 0.103321 | 0.947762 | 0.162 | 0.108 | 1 | 10 | Tmem80 | 0.054 | Young |
| 0.104019 | 1.098769 | 0.121 | 0.073 | 1 | 10 | Hmces | 0.048 | Young |
| 0.10462 | 1.125471 | 0.222 | 0.161 | 1 | 10 | Hvcn1 | 0.061 | Young |
| 0.104681 | 1.001234 | 0.091 | 0.049 | 1 | 10 | Ptgfrn | 0.042 | Young |
| 0.104805 | 0.502023 | 0.232 | 0.166 | 1 | 10 | Zfp992 | 0.066 | Young |

| | | | | | | | | |
|---|---|---|---|---|---|---|---|---|
| 0.105087 | 0.86126 | 0.091 | 0.049 | 1 | 10 | Vdr | 0.042 | Young |
| 0.105509 | 0.565028 | 0.141 | 0.088 | 1 | 10 | Fam219a | 0.053 | Young |
| 0.105599 | 1.414995 | 0.141 | 0.09 | 1 | 10 | Bbc3 | 0.051 | Young |
| 0.105831 | 0.280919 | 0.071 | 0.034 | 1 | 10 | Upp2 | 0.037 | Young |
| 0.105853 | 0.580326 | 0.374 | 0.318 | 1 | 10 | Mrpl14 | 0.056 | Young |
| 0.105961 | 1.013012 | 0.343 | 0.296 | 1 | 10 | Phf14 | 0.047 | Young |
| 0.106146 | 0.805339 | 0.141 | 0.088 | 1 | 10 | Dus1l | 0.053 | Young |
| 0.106515 | 0.264211 | 0.101 | 0.181 | 1 | 10 | Rras2 | -0.08 | Young |
| 0.106774 | 0.782593 | 0.232 | 0.166 | 1 | 10 | Gaa | 0.066 | Young |
| 0.107338 | 0.272483 | 0.818 | 0.792 | 1 | 10 | Arl6ip1 | 0.026 | Young |
| 0.107618 | 0.198043 | 0.111 | 0.064 | 1 | 10 | Clec5a | 0.047 | Young |
| 0.10768 | 1.027548 | 0.273 | 0.222 | 1 | 10 | Lrrc8d | 0.051 | Young |
| 0.107814 | 0.945879 | 0.121 | 0.073 | 1 | 10 | Agpat2 | 0.048 | Young |
| 0.108018 | 0.529551 | 0.374 | 0.306 | 1 | 10 | Zfhx3 | 0.068 | Young |
| 0.109162 | 0.133182 | 0.131 | 0.2 | 1 | 10 | Ric8a | -0.069 | Young |
| 0.109297 | 0.826646 | 0.394 | 0.34 | 1 | 10 | Cited2 | 0.054 | Young |
| 0.109426 | 0.185597 | 0.778 | 0.822 | 1 | 10 | Rpl4 | -0.044 | Young |
| 0.109626 | 0.099889 | 0.091 | 0.164 | 1 | 10 | Wdr3 | -0.073 | Young |
| 0.109898 | 0.190875 | 0.061 | 0.12 | 1 | 10 | Ccser2 | -0.059 | Young |
| 0.10998 | 0.645659 | 0.242 | 0.181 | 1 | 10 | Fmr1 | 0.061 | Young |
| 0.110131 | 0.577183 | 0.394 | 0.325 | 1 | 10 | Krtcap2 | 0.069 | Young |
| 0.110278 | 1.454884 | 0.131 | 0.083 | 1 | 10 | Gbe1 | 0.048 | Young |
| 0.110488 | 0.496586 | 0.091 | 0.049 | 1 | 10 | Snapc2 | 0.042 | Young |
| 0.11085 | 0.757587 | 0.283 | 0.222 | 1 | 10 | Pdgfb | 0.061 | Young |
| 0.110999 | 1.138651 | 0.182 | 0.127 | 1 | 10 | Rassf8 | 0.055 | Young |
| 0.111319 | 0.532322 | 0.818 | 0.831 | 1 | 10 | Mt1 | -0.013 | Young |
| 0.111321 | 0.115446 | 0.929 | 0.941 | 1 | 10 | Rps11 | -0.012 | Young |
| 0.111338 | 0.541067 | 0.091 | 0.049 | 1 | 10 | Cables2 | 0.042 | Young |
| 0.111562 | 0.353174 | 0.535 | 0.489 | 1 | 10 | Stk17b | 0.046 | Young |
| 0.111622 | 2.355437 | 0.03 | 0.01 | 1 | 10 | Hepacam2 | 0.02 | Young |
| 0.111711 | 0.394279 | 0.758 | 0.751 | 1 | 10 | Rnf149 | 0.007 | Young |
| 0.112123 | 0.584304 | 0.434 | 0.386 | 1 | 10 | Zcrb1 | 0.048 | Young |
| 0.112291 | 1.144243 | 0.101 | 0.059 | 1 | 10 | 1110012L1 | 0.042 | Young |
| 0.112475 | 1.984134 | 0.03 | 0.01 | 1 | 10 | Cped1 | 0.02 | Young |
| 0.112475 | 1.838877 | 0.03 | 0.01 | 1 | 10 | Mgp | 0.02 | Young |
| 0.113333 | 2.18224 | 0.03 | 0.01 | 1 | 10 | Fam92a | 0.02 | Young |
| 0.113333 | 2.175503 | 0.03 | 0.01 | 1 | 10 | Cdk1 | 0.02 | Young |
| 0.113333 | 1.766142 | 0.03 | 0.01 | 1 | 10 | Gm10602 | 0.02 | Young |
| 0.113846 | 0.704043 | 0.323 | 0.271 | 1 | 10 | Dcaf12 | 0.052 | Young |
| 0.113937 | 0.197597 | 0.152 | 0.24 | 1 | 10 | Rars | -0.088 | Young |
| 0.114197 | 2.174534 | 0.03 | 0.01 | 1 | 10 | Fut7 | 0.02 | Young |
| 0.114197 | 1.62266 | 0.03 | 0.01 | 1 | 10 | Usp27x | 0.02 | Young |
| 0.114197 | 1.36788 | 0.03 | 0.01 | 1 | 10 | Asrgl1 | 0.02 | Young |
| 0.114848 | 1.174495 | 0.051 | 0.022 | 1 | 10 | Serpinf1 | 0.029 | Young |
| 0.115065 | 2.119926 | 0.03 | 0.01 | 1 | 10 | Colec12 | 0.02 | Young |

| | | | | | | | | |
|---|---|---|---|---|---|---|---|---|
| 0.115457 | 0.804631 | 0.394 | 0.359 | 1 | 10 | Igfbp4 | 0.035 | Young |
| 0.115939 | 1.874181 | 0.03 | 0.01 | 1 | 10 | Ccl5 | 0.02 | Young |
| 0.116818 | 1.401186 | 0.03 | 0.01 | 1 | 10 | Ehd2 | 0.02 | Young |
| 0.116818 | 1.257221 | 0.03 | 0.01 | 1 | 10 | P2ry13 | 0.02 | Young |
| 0.116818 | 1.105476 | 0.03 | 0.01 | 1 | 10 | Hebp2 | 0.02 | Young |
| 0.116955 | 1.180039 | 0.152 | 0.1 | 1 | 10 | Plekhn1 | 0.052 | Young |
| 0.11699 | 1.151157 | 0.111 | 0.066 | 1 | 10 | Ipo9 | 0.045 | Young |
| 0.117438 | 0.939499 | 0.141 | 0.09 | 1 | 10 | Gkap1 | 0.051 | Young |
| 0.118357 | 1.448934 | 0.061 | 0.029 | 1 | 10 | Lppos | 0.032 | Young |
| 0.118373 | 0.533357 | 0.455 | 0.408 | 1 | 10 | 0610012G( | 0.047 | Young |
| 0.118742 | 1.246741 | 0.121 | 0.076 | 1 | 10 | Dhx33 | 0.045 | Young |
| 0.118917 | 0.609995 | 0.101 | 0.056 | 1 | 10 | Cdc23 | 0.045 | Young |
| 0.119004 | 0.856408 | 0.293 | 0.225 | 1 | 10 | Map2k1 | 0.068 | Young |
| 0.119248 | 1.344034 | 0.051 | 0.022 | 1 | 10 | Iqgap3 | 0.029 | Young |
| 0.119353 | 0.728383 | 0.111 | 0.066 | 1 | 10 | Metap1d | 0.045 | Young |
| 0.11974 | 1.005114 | 0.172 | 0.122 | 1 | 10 | Sin3a | 0.05 | Young |
| 0.119749 | 1.123866 | 0.192 | 0.134 | 1 | 10 | Gar1 | 0.058 | Young |
| 0.119837 | 1.192515 | 0.152 | 0.103 | 1 | 10 | Nagpa | 0.049 | Young |
| 0.119872 | 0.63966 | 0.162 | 0.105 | 1 | 10 | Ndst2 | 0.057 | Young |
| 0.120516 | 2.929345 | 0.02 | 0.005 | 1 | 10 | Ctla2a | 0.015 | Young |
| 0.120516 | 2.670992 | 0.02 | 0.005 | 1 | 10 | Gm13391 | 0.015 | Young |
| 0.120516 | 2.381846 | 0.02 | 0.005 | 1 | 10 | Ahcy | 0.015 | Young |
| 0.120516 | 2.363137 | 0.02 | 0.005 | 1 | 10 | Cdc45 | 0.015 | Young |
| 0.120516 | 2.344393 | 0.02 | 0.005 | 1 | 10 | Pcgf2 | 0.015 | Young |
| 0.120516 | 2.326384 | 0.02 | 0.005 | 1 | 10 | Cyp2r1 | 0.015 | Young |
| 0.120516 | 1.984414 | 0.02 | 0.005 | 1 | 10 | Vma21-ps | 0.015 | Young |
| 0.120516 | 1.855798 | 0.02 | 0.005 | 1 | 10 | Cgref1 | 0.015 | Young |
| 0.120516 | 1.847945 | 0.02 | 0.005 | 1 | 10 | 9330104G( | 0.015 | Young |
| 0.12053 | 1.725668 | 0.051 | 0.022 | 1 | 10 | Tut1 | 0.029 | Young |
| 0.120744 | 0.097247 | 0.99 | 1 | 1 | 10 | Rps9 | -0.01 | Young |
| 0.121311 | 0.293816 | 0.879 | 0.844 | 1 | 10 | C1qb | 0.035 | Young |
| 0.121343 | 1.533555 | 0.071 | 0.037 | 1 | 10 | Smpdl3b | 0.034 | Young |
| 0.121462 | 0.562785 | 0.263 | 0.196 | 1 | 10 | Sfswap | 0.067 | Young |
| 0.121712 | 2.424994 | 0.02 | 0.005 | 1 | 10 | Padi3 | 0.015 | Young |
| 0.121712 | 2.347771 | 0.02 | 0.005 | 1 | 10 | Cidec | 0.015 | Young |
| 0.121712 | 1.669159 | 0.02 | 0.005 | 1 | 10 | Aqp1 | 0.015 | Young |
| 0.121712 | 1.653881 | 0.02 | 0.005 | 1 | 10 | Msantd3 | 0.015 | Young |
| 0.121712 | 1.625809 | 0.02 | 0.005 | 1 | 10 | Tmem171 | 0.015 | Young |
| 0.121712 | 1.622716 | 0.02 | 0.005 | 1 | 10 | Ildr1 | 0.015 | Young |
| 0.121712 | 1.523019 | 0.02 | 0.005 | 1 | 10 | Gm6169 | 0.015 | Young |
| 0.121712 | 1.262484 | 0.02 | 0.005 | 1 | 10 | Hist2h2aa1 | 0.015 | Young |
| 0.121712 | 1.183778 | 0.02 | 0.005 | 1 | 10 | Cdkn2a | 0.015 | Young |
| 0.121753 | 0.732229 | 0.111 | 0.066 | 1 | 10 | Rufy2 | 0.045 | Young |
| 0.121822 | 0.82791 | 0.051 | 0.022 | 1 | 10 | Wnt11 | 0.029 | Young |
| 0.121822 | 0.529597 | 0.051 | 0.022 | 1 | 10 | Slc35g1 | 0.029 | Young |

| | | | | | | | | |
|---|---|---|---|---|---|---|---|---|
| 0.122133 | 0.650969 | 0.394 | 0.369 | 1 | 10 | Mpc1 | 0.025 | Young |
| 0.122141 | 0.890766 | 0.121 | 0.076 | 1 | 10 | Sik2 | 0.045 | Young |
| 0.122331 | 0.417652 | 0.212 | 0.149 | 1 | 10 | Thoc6 | 0.063 | Young |
| 0.122917 | 1.403922 | 0.061 | 0.029 | 1 | 10 | Pigh | 0.032 | Young |
| 0.122917 | 2.586718 | 0.02 | 0.005 | 1 | 10 | Med18 | 0.015 | Young |
| 0.122917 | 2.386092 | 0.02 | 0.005 | 1 | 10 | Dlc1 | 0.015 | Young |
| 0.122917 | 1.806578 | 0.02 | 0.005 | 1 | 10 | Nlgn2 | 0.015 | Young |
| 0.122917 | 1.570035 | 0.02 | 0.005 | 1 | 10 | Arhgef40 | 0.015 | Young |
| 0.122917 | 1.309538 | 0.02 | 0.005 | 1 | 10 | Apoc4 | 0.015 | Young |
| 0.122917 | 1.307605 | 0.02 | 0.005 | 1 | 10 | Ly6c2 | 0.015 | Young |
| 0.122917 | 1.228824 | 0.02 | 0.005 | 1 | 10 | Col15a1 | 0.015 | Young |
| 0.122917 | 1.216653 | 0.02 | 0.005 | 1 | 10 | Hist1h4c | 0.015 | Young |
| 0.122917 | 1.204769 | 0.02 | 0.005 | 1 | 10 | Negr1 | 0.015 | Young |
| 0.122917 | 1.099857 | 0.02 | 0.005 | 1 | 10 | Xlr4c | 0.015 | Young |
| 0.122917 | 1.092205 | 0.02 | 0.005 | 1 | 10 | Gm26890 | 0.015 | Young |
| 0.122917 | 1.086371 | 0.02 | 0.005 | 1 | 10 | Tbc1d30 | 0.015 | Young |
| 0.122917 | 0.994041 | 0.02 | 0.005 | 1 | 10 | Zfp647 | 0.015 | Young |
| 0.122917 | 0.939726 | 0.02 | 0.005 | 1 | 10 | Treml2 | 0.015 | Young |
| 0.122917 | 0.645744 | 0.02 | 0.005 | 1 | 10 | Morn2 | 0.015 | Young |
| 0.123496 | 1.408416 | 0.061 | 0.029 | 1 | 10 | Dhfr | 0.032 | Young |
| 0.123679 | 0.167816 | 0.242 | 0.34 | 1 | 10 | Tollip | -0.098 | Young |
| 0.124095 | 0.248916 | 0.232 | 0.159 | 1 | 10 | Ext1 | 0.073 | Young |
| 0.124132 | 1.197399 | 0.02 | 0.005 | 1 | 10 | Gm13710 | 0.015 | Young |
| 0.124132 | 1.146761 | 0.02 | 0.005 | 1 | 10 | Sncg | 0.015 | Young |
| 0.124132 | 0.996884 | 0.02 | 0.005 | 1 | 10 | Npr1 | 0.015 | Young |
| 0.124132 | 0.635074 | 0.02 | 0.005 | 1 | 10 | Abhd6 | 0.015 | Young |
| 0.124221 | 1.393308 | 0.111 | 0.068 | 1 | 10 | Mterf3 | 0.043 | Young |
| 0.124671 | 0.561209 | 0.343 | 0.286 | 1 | 10 | Dtnbp1 | 0.057 | Young |
| 0.125247 | 1.36533 | 0.061 | 0.029 | 1 | 10 | Skint3 | 0.032 | Young |
| 0.125298 | 0.677023 | 0.242 | 0.181 | 1 | 10 | Rfk | 0.061 | Young |
| 0.125356 | 0.759319 | 0.02 | 0.005 | 1 | 10 | Stk26 | 0.015 | Young |
| 0.125837 | 0.64039 | 0.111 | 0.066 | 1 | 10 | Glipr2 | 0.045 | Young |
| 0.125843 | 1.429011 | 0.131 | 0.086 | 1 | 10 | Taf7 | 0.045 | Young |
| 0.126154 | 0.018478 | 0.212 | 0.315 | 1 | 10 | Bbip1 | -0.103 | Young |
| 0.126344 | 0.177384 | 0.071 | 0.132 | 1 | 10 | Maip1 | -0.061 | Young |
| 0.126601 | 1.667675 | 0.081 | 0.044 | 1 | 10 | Kdm1b | 0.037 | Young |
| 0.127513 | 0.215244 | 0.859 | 0.861 | 1 | 10 | Morf4l1 | -0.002 | Young |
| 0.127592 | 0.34465 | 0.131 | 0.081 | 1 | 10 | Ctsf | 0.05 | Young |
| 0.127733 | 0.03468 | 0.061 | 0.115 | 1 | 10 | Brwd1 | -0.054 | Young |
| 0.127748 | 0.182814 | 0.141 | 0.215 | 1 | 10 | Ube3c | -0.074 | Young |
| 0.127815 | 0.721134 | 0.152 | 0.1 | 1 | 10 | Vrk1 | 0.052 | Young |
| 0.128097 | 1.123281 | 0.081 | 0.044 | 1 | 10 | Telo2 | 0.037 | Young |
| 0.1283 | 1.293071 | 0.172 | 0.115 | 1 | 10 | Rgcc | 0.057 | Young |
| 0.128547 | 0.026909 | 0.101 | 0.166 | 1 | 10 | Abhd16a | -0.065 | Young |
| 0.129529 | 0.591525 | 0.162 | 0.108 | 1 | 10 | Nubp2 | 0.054 | Young |

| | | | | | | | | |
|---|---|---|---|---|---|---|---|---|
| 0.129749 | 0.516304 | 0.263 | 0.2 | 1 | 10 | Trappc8 | 0.063 | Young |
| 0.129778 | 0.997691 | 0.586 | 0.606 | 1 | 10 | Sqstm1 | -0.02 | Young |
| 0.130429 | 0.76038 | 0.242 | 0.181 | 1 | 10 | Sacm1l | 0.061 | Young |
| 0.130437 | 0.900366 | 0.071 | 0.037 | 1 | 10 | Cluap1 | 0.034 | Young |
| 0.130911 | 0.589134 | 0.162 | 0.108 | 1 | 10 | Krt80 | 0.054 | Young |
| 0.131224 | 1.005908 | 0.061 | 0.029 | 1 | 10 | Tfcp2 | 0.032 | Young |
| 0.132446 | 1.397852 | 0.061 | 0.029 | 1 | 10 | Tk1 | 0.032 | Young |
| 0.132625 | 0.913365 | 0.283 | 0.232 | 1 | 10 | Ebna1bp2 | 0.051 | Young |
| 0.134455 | 0.193664 | 0.838 | 0.8 | 1 | 10 | Pitpna | 0.038 | Young |
| 0.134563 | 0.285949 | 1 | 0.973 | 1 | 10 | Jund | 0.027 | Young |
| 0.134599 | 0.150661 | 0.869 | 0.836 | 1 | 10 | C1qa | 0.033 | Young |
| 0.135146 | 0.816483 | 0.212 | 0.156 | 1 | 10 | Parp4 | 0.056 | Young |
| 0.135254 | 0.281071 | 0.737 | 0.719 | 1 | 10 | Uqcrq | 0.018 | Young |
| 0.13526 | 0.834489 | 0.081 | 0.044 | 1 | 10 | Rmnd5b | 0.037 | Young |
| 0.135462 | 0.965468 | 0.071 | 0.037 | 1 | 10 | Megf9 | 0.034 | Young |
| 0.135538 | 0.797996 | 0.061 | 0.029 | 1 | 10 | Cenpt | 0.032 | Young |
| 0.135962 | 0.810789 | 0.141 | 0.09 | 1 | 10 | Rcor3 | 0.051 | Young |
| 0.136077 | 0.274211 | 0.263 | 0.2 | 1 | 10 | Arl5c | 0.063 | Young |
| 0.136206 | 0.492096 | 0.515 | 0.518 | 1 | 10 | Psmd8 | -0.003 | Young |
| 0.136613 | 0.965362 | 0.182 | 0.13 | 1 | 10 | Tmem229b | 0.052 | Young |
| 0.136705 | 0.367394 | 0.434 | 0.367 | 1 | 10 | Gpr65 | 0.067 | Young |
| 0.137092 | 1.067807 | 0.091 | 0.051 | 1 | 10 | 1110034G2 | 0.04 | Young |
| 0.137767 | 0.224469 | 0.97 | 0.966 | 1 | 10 | Ctsz | 0.004 | Young |
| 0.13817 | 0.615326 | 0.485 | 0.438 | 1 | 10 | Mindy2 | 0.047 | Young |
| 0.138317 | 0.529018 | 0.071 | 0.037 | 1 | 10 | Tpbgl | 0.034 | Young |
| 0.138578 | 0.472769 | 0.333 | 0.264 | 1 | 10 | Pparg | 0.069 | Young |
| 0.138631 | 0.842517 | 0.333 | 0.279 | 1 | 10 | Mrps17 | 0.054 | Young |
| 0.138729 | 0.375928 | 0.111 | 0.066 | 1 | 10 | Tmem140 | 0.045 | Young |
| 0.139029 | 0.869471 | 0.242 | 0.191 | 1 | 10 | Exoc1 | 0.051 | Young |
| 0.139088 | 0.330246 | 0.485 | 0.418 | 1 | 10 | Rragc | 0.067 | Young |
| 0.139189 | 0.744748 | 0.141 | 0.093 | 1 | 10 | Ikbkg | 0.048 | Young |
| 0.139263 | 0.114754 | 0.051 | 0.103 | 1 | 10 | Sqle | -0.052 | Young |
| 0.139577 | 0.878488 | 0.091 | 0.051 | 1 | 10 | Cnpy4 | 0.04 | Young |
| 0.139654 | 1.01083 | 0.141 | 0.095 | 1 | 10 | Skp2 | 0.046 | Young |
| 0.139908 | 0.984866 | 0.111 | 0.068 | 1 | 10 | Prkab1 | 0.043 | Young |
| 0.141695 | 0.614117 | 0.313 | 0.269 | 1 | 10 | Cd86 | 0.044 | Young |
| 0.14189 | 0.347704 | 0.061 | 0.029 | 1 | 10 | Exo5 | 0.032 | Young |
| 0.142552 | 1.147194 | 0.192 | 0.142 | 1 | 10 | Slc16a3 | 0.05 | Young |
| 0.14311 | 0.115705 | 0.98 | 0.971 | 1 | 10 | Rpl30 | 0.009 | Young |
| 0.14355 | 0.890171 | 0.141 | 0.095 | 1 | 10 | Ptpn9 | 0.046 | Young |
| 0.143572 | 0.65974 | 0.071 | 0.037 | 1 | 10 | Zfp335 | 0.034 | Young |
| 0.143742 | 0.373421 | 0.273 | 0.208 | 1 | 10 | Slc9a9 | 0.065 | Young |
| 0.144338 | 1.05221 | 0.101 | 0.061 | 1 | 10 | Fgd3 | 0.04 | Young |
| 0.144356 | 0.010904 | 0.091 | 0.156 | 1 | 10 | Bckdha | -0.065 | Young |
| 0.144374 | 0.641228 | 0.313 | 0.247 | 1 | 10 | Gsk3a | 0.066 | Young |

| | | | | | | | | |
|---|---|---|---|---|---|---|---|---|
| 0.144708 | 1.663962 | 0.091 | 0.054 | 1 | 10 | Mvb12b | 0.037 | Young |
| 0.144872 | 0.737636 | 0.293 | 0.244 | 1 | 10 | Zhx1 | 0.049 | Young |
| 0.144921 | 0.45346 | 0.081 | 0.044 | 1 | 10 | Sass6 | 0.037 | Young |
| 0.144945 | 0.698667 | 0.434 | 0.401 | 1 | 10 | Vasp | 0.033 | Young |
| 0.144946 | 0.187864 | 0.828 | 0.809 | 1 | 10 | Nfkbia | 0.019 | Young |
| 0.145105 | 0.584306 | 0.162 | 0.11 | 1 | 10 | Mrpl40 | 0.052 | Young |
| 0.145275 | 1.418021 | 0.081 | 0.046 | 1 | 10 | Mpi | 0.035 | Young |
| 0.145654 | 0.402224 | 0.677 | 0.638 | 1 | 10 | Atp1b3 | 0.039 | Young |
| 0.146114 | 0.817626 | 0.232 | 0.178 | 1 | 10 | Scamp5 | 0.054 | Young |
| 0.146341 | 0.44914 | 0.253 | 0.191 | 1 | 10 | Ivns1abp | 0.062 | Young |
| 0.147139 | 2.159758 | 0.04 | 0.017 | 1 | 10 | Crocc | 0.023 | Young |
| 0.147583 | 0.161183 | 0.232 | 0.164 | 1 | 10 | Pex7 | 0.068 | Young |
| 0.147985 | 2.258589 | 0.04 | 0.017 | 1 | 10 | Btbd11 | 0.023 | Young |
| 0.147985 | 2.03337 | 0.04 | 0.017 | 1 | 10 | Gm14005 | 0.023 | Young |
| 0.147985 | 1.541586 | 0.04 | 0.017 | 1 | 10 | Pigw | 0.023 | Young |
| 0.148682 | 0.025074 | 0.03 | 0.071 | 1 | 10 | Prim2 | -0.041 | Young |
| 0.149607 | 0.579775 | 0.313 | 0.264 | 1 | 10 | Pdpk1 | 0.049 | Young |
| 0.149689 | 1.159986 | 0.04 | 0.017 | 1 | 10 | Gm38335 | 0.023 | Young |
| 0.150221 | 0.446473 | 0.384 | 0.328 | 1 | 10 | Nmt1 | 0.056 | Young |
| 0.150546 | 1.48514 | 0.04 | 0.017 | 1 | 10 | Mblac1 | 0.023 | Young |
| 0.150758 | 0.464719 | 0.141 | 0.09 | 1 | 10 | Papd7 | 0.051 | Young |
| 0.151407 | 1.351783 | 0.04 | 0.017 | 1 | 10 | Amacr | 0.023 | Young |
| 0.151596 | 0.434299 | 0.545 | 0.538 | 1 | 10 | Top1 | 0.007 | Young |
| 0.151614 | 0.647095 | 0.141 | 0.093 | 1 | 10 | Pogz | 0.048 | Young |
| 0.152272 | 1.003013 | 0.04 | 0.017 | 1 | 10 | Iqcb1 | 0.023 | Young |
| 0.152324 | 0.558147 | 0.222 | 0.169 | 1 | 10 | Mmp12 | 0.053 | Young |
| 0.152423 | 0.790835 | 0.253 | 0.198 | 1 | 10 | Tmem248 | 0.055 | Young |
| 0.152475 | 1.688149 | 0.141 | 0.103 | 1 | 10 | Rps19bp1 | 0.038 | Young |
| 0.153369 | 0.411249 | 0.283 | 0.22 | 1 | 10 | Rcbtb2 | 0.063 | Young |
| 0.154013 | 1.398271 | 0.04 | 0.017 | 1 | 10 | Pcgf6 | 0.023 | Young |
| 0.154114 | 0.543459 | 0.141 | 0.093 | 1 | 10 | Zbtb17 | 0.048 | Young |
| 0.154126 | 0.321224 | 0.859 | 0.839 | 1 | 10 | Rbm3 | 0.02 | Young |
| 0.154544 | 0.371383 | 0.313 | 0.257 | 1 | 10 | Rsbn1l | 0.056 | Young |
| 0.154765 | 0.064905 | 0.131 | 0.198 | 1 | 10 | Map4k1 | -0.067 | Young |
| 0.154807 | 1.265254 | 0.141 | 0.098 | 1 | 10 | Il1rl1 | 0.043 | Young |
| 0.154889 | 0.819788 | 0.04 | 0.017 | 1 | 10 | Gcat | 0.023 | Young |
| 0.154921 | 0.174409 | 0.111 | 0.174 | 1 | 10 | Tvp23b | -0.063 | Young |
| 0.155765 | 1.110108 | 0.121 | 0.078 | 1 | 10 | Rdh14 | 0.043 | Young |
| 0.155769 | 0.818734 | 0.04 | 0.017 | 1 | 10 | Gm38843 | 0.023 | Young |
| 0.15586 | 1.546908 | 0.081 | 0.046 | 1 | 10 | Anks3 | 0.035 | Young |
| 0.155924 | 0.002778 | 0.283 | 0.369 | 1 | 10 | Ppp6r3 | -0.086 | Young |
| 0.156364 | 0.554426 | 0.313 | 0.252 | 1 | 10 | Mrps36 | 0.061 | Young |
| 0.156489 | 0.272577 | 0.859 | 0.907 | 1 | 10 | Rpl11 | -0.048 | Young |
| 0.15669 | 0.26725 | 0.606 | 0.543 | 1 | 10 | Nfkbid | 0.063 | Young |
| 0.156722 | 0.875845 | 0.091 | 0.054 | 1 | 10 | Rfc3 | 0.037 | Young |

| | | | | | | | | |
|---|---|---|---|---|---|---|---|---|
| 0.158443 | 1.504615 | 0.051 | 0.024 | 1 | 10 | Mettl26 | 0.027 | Young |
| 0.158669 | 0.308821 | 0.354 | 0.286 | 1 | 10 | Cebpz | 0.068 | Young |
| 0.158741 | 1.443478 | 0.081 | 0.046 | 1 | 10 | Egr3 | 0.035 | Young |
| 0.159212 | 1.459154 | 0.051 | 0.024 | 1 | 10 | Gpatch3 | 0.027 | Young |
| 0.159511 | 1.286195 | 0.172 | 0.125 | 1 | 10 | Nol12 | 0.047 | Young |
| 0.159739 | 1.072191 | 0.101 | 0.061 | 1 | 10 | Nes | 0.04 | Young |
| 0.159815 | 1.125089 | 0.101 | 0.064 | 1 | 10 | Cdc25a | 0.037 | Young |
| 0.15992 | 1.011291 | 0.172 | 0.122 | 1 | 10 | Cdkn2aipnl | 0.05 | Young |
| 0.160216 | 1.518181 | 0.061 | 0.032 | 1 | 10 | Zkscan8 | 0.029 | Young |
| 0.160263 | 0.463185 | 0.616 | 0.638 | 1 | 10 | Hdgf | -0.022 | Young |
| 0.160333 | 0.692327 | 0.202 | 0.147 | 1 | 10 | Tmco3 | 0.055 | Young |
| 0.160623 | 0.53948 | 0.162 | 0.11 | 1 | 10 | Dctpp1 | 0.052 | Young |
| 0.161081 | 0.74972 | 0.172 | 0.122 | 1 | 10 | Mrpl58 | 0.05 | Young |
| 0.161145 | 1.083572 | 0.152 | 0.105 | 1 | 10 | Lncpint | 0.047 | Young |
| 0.16143 | 0.380672 | 0.061 | 0.11 | 1 | 10 | Dennd2a | -0.049 | Young |
| 0.161521 | 0.219044 | 0.566 | 0.516 | 1 | 10 | Ndel1 | 0.05 | Young |
| 0.161868 | 0.254827 | 0.101 | 0.161 | 1 | 10 | Vamp7 | -0.06 | Young |
| 0.162318 | 1.50758 | 0.051 | 0.024 | 1 | 10 | Pced1a | 0.027 | Young |
| 0.16232 | 0.582164 | 0.273 | 0.222 | 1 | 10 | Rnf216 | 0.051 | Young |
| 0.162358 | 0.514281 | 0.434 | 0.374 | 1 | 10 | Fxr1 | 0.06 | Young |
| 0.162636 | 0.194808 | 0.081 | 0.139 | 1 | 10 | Trappc11 | -0.058 | Young |
| 0.163102 | 1.323752 | 0.051 | 0.024 | 1 | 10 | Laptm4b | 0.027 | Young |
| 0.16319 | 0.282573 | 0.091 | 0.149 | 1 | 10 | Wdr5 | -0.058 | Young |
| 0.163434 | 1.105868 | 0.081 | 0.046 | 1 | 10 | Stradb | 0.035 | Young |
| 0.164024 | 0.498617 | 0.374 | 0.323 | 1 | 10 | B4galt1 | 0.051 | Young |
| 0.164028 | 1.017104 | 0.071 | 0.039 | 1 | 10 | Mllt1 | 0.032 | Young |
| 0.164623 | 1.249542 | 0.081 | 0.046 | 1 | 10 | Zfp81 | 0.035 | Young |
| 0.164677 | 1.763689 | 0.051 | 0.024 | 1 | 10 | Ccdc84 | 0.027 | Young |
| 0.164677 | 1.535557 | 0.051 | 0.024 | 1 | 10 | Ceacam16 | 0.027 | Young |
| 0.164677 | 1.062428 | 0.051 | 0.024 | 1 | 10 | 2700049A( | 0.027 | Young |
| 0.164937 | 0.405351 | 0.455 | 0.416 | 1 | 10 | Srrm1 | 0.039 | Young |
| 0.165469 | 1.344991 | 0.051 | 0.024 | 1 | 10 | H2-DMb2 | 0.027 | Young |
| 0.165469 | 1.174335 | 0.051 | 0.024 | 1 | 10 | Zfp141 | 0.027 | Young |
| 0.165517 | 0.341329 | 0.091 | 0.054 | 1 | 10 | Ticam1 | 0.037 | Young |
| 0.165685 | 0.470298 | 0.525 | 0.484 | 1 | 10 | Eloc | 0.041 | Young |
| 0.165815 | 1.231287 | 0.061 | 0.032 | 1 | 10 | Zxdb | 0.029 | Young |
| 0.165815 | 0.963783 | 0.061 | 0.032 | 1 | 10 | Cby1 | 0.029 | Young |
| 0.166339 | 0.201894 | 0.646 | 0.577 | 1 | 10 | Egr1 | 0.069 | Young |
| 0.166344 | 0.60915 | 0.384 | 0.345 | 1 | 10 | Glrx3 | 0.039 | Young |
| 0.16661 | 0.829523 | 0.071 | 0.039 | 1 | 10 | Dusp8 | 0.032 | Young |
| 0.167062 | 1.210142 | 0.051 | 0.024 | 1 | 10 | Dbp | 0.027 | Young |
| 0.167233 | 0.688404 | 0.222 | 0.169 | 1 | 10 | Mrps9 | 0.053 | Young |
| 0.167238 | 1.628255 | 0.061 | 0.032 | 1 | 10 | Cnrip1 | 0.029 | Young |
| 0.167261 | 0.748612 | 0.071 | 0.039 | 1 | 10 | Cd320 | 0.032 | Young |
| 0.167633 | 0.811446 | 0.222 | 0.178 | 1 | 10 | Trappc6a | 0.044 | Young |

| | | | | | | | | |
|---|---|---|---|---|---|---|---|---|
| 0.167737 | 0.83955 | 0.242 | 0.188 | 1 | 10 | Nr2c2 | 0.054 | Young |
| 0.167913 | 1.069738 | 0.071 | 0.039 | 1 | 10 | 0610009O2 | 0.032 | Young |
| 0.168382 | 0.317465 | 0.182 | 0.125 | 1 | 10 | AI504432 | 0.057 | Young |
| 0.168567 | 0.880616 | 0.071 | 0.039 | 1 | 10 | Smpd2 | 0.032 | Young |
| 0.169015 | 0.337931 | 0.313 | 0.244 | 1 | 10 | H2-M3 | 0.069 | Young |
| 0.169389 | 1.589637 | 0.061 | 0.032 | 1 | 10 | Gab3 | 0.029 | Young |
| 0.169473 | 1.086193 | 0.051 | 0.024 | 1 | 10 | Cebpe | 0.027 | Young |
| 0.169644 | 0.820747 | 0.222 | 0.171 | 1 | 10 | Dmxl2 | 0.051 | Young |
| 0.170282 | 0.551876 | 0.051 | 0.024 | 1 | 10 | Haus1 | 0.027 | Young |
| 0.170346 | 0.560572 | 0.273 | 0.21 | 1 | 10 | Polr2c | 0.063 | Young |
| 0.170382 | 0.458971 | 0.02 | 0.054 | 1 | 10 | Fbxo31 | -0.034 | Young |
| 0.170809 | 0.990521 | 0.192 | 0.147 | 1 | 10 | Nupr1 | 0.045 | Young |
| 0.171095 | 0.788087 | 0.051 | 0.024 | 1 | 10 | Arhgap35 | 0.027 | Young |
| 0.171095 | 0.539314 | 0.051 | 0.024 | 1 | 10 | 1810055G0 | 0.027 | Young |
| 0.171374 | 0.532546 | 0.212 | 0.159 | 1 | 10 | Bahd1 | 0.053 | Young |
| 0.171743 | 0.131552 | 0.061 | 0.11 | 1 | 10 | 4930453N2 | -0.049 | Young |
| 0.171866 | 1.407646 | 0.071 | 0.039 | 1 | 10 | Eno2 | 0.032 | Young |
| 0.171918 | 0.627135 | 0.414 | 0.386 | 1 | 10 | Arfgef1 | 0.028 | Young |
| 0.171924 | 0.584236 | 0.475 | 0.443 | 1 | 10 | Txn2 | 0.032 | Young |
| 0.172099 | 0.546585 | 0.253 | 0.198 | 1 | 10 | Maff | 0.055 | Young |
| 0.172158 | 0.301285 | 0.131 | 0.086 | 1 | 10 | Dbt | 0.045 | Young |
| 0.172251 | 0.343884 | 0.212 | 0.156 | 1 | 10 | Xpnpep1 | 0.056 | Young |
| 0.172402 | 1.232219 | 0.101 | 0.064 | 1 | 10 | Flywch1 | 0.037 | Young |
| 0.172542 | 0.003697 | 0.374 | 0.457 | 1 | 10 | Cebpa | -0.083 | Young |
| 0.17315 | 0.063856 | 0.475 | 0.56 | 1 | 10 | 1810058I24 | -0.085 | Young |
| 0.174029 | 0.954634 | 0.162 | 0.115 | 1 | 10 | Urm1 | 0.047 | Young |
| 0.174489 | 0.691265 | 0.061 | 0.032 | 1 | 10 | Gatb | 0.029 | Young |
| 0.174895 | 0.332488 | 0.586 | 0.553 | 1 | 10 | Tax1bp1 | 0.033 | Young |
| 0.175661 | 0.961055 | 0.101 | 0.064 | 1 | 10 | Cbx7 | 0.037 | Young |
| 0.176029 | 0.178295 | 0.051 | 0.024 | 1 | 10 | Cish | 0.027 | Young |
| 0.176487 | 0.800169 | 0.495 | 0.462 | 1 | 10 | Esd | 0.033 | Young |
| 0.176819 | 0.340597 | 0.697 | 0.711 | 1 | 10 | Pafah1b1 | -0.014 | Young |
| 0.176969 | 0.071292 | 0.071 | 0.12 | 1 | 10 | Rfx1 | -0.049 | Young |
| 0.177688 | 0.515168 | 0.202 | 0.152 | 1 | 10 | Ebpl | 0.05 | Young |
| 0.177728 | 0.259478 | 0.535 | 0.496 | 1 | 10 | Trps1 | 0.039 | Young |
| 0.17789 | 1.097548 | 0.111 | 0.073 | 1 | 10 | Incenp | 0.038 | Young |
| 0.178587 | 0.214793 | 0.687 | 0.68 | 1 | 10 | Mef2a | 0.007 | Young |
| 0.178697 | 0.378107 | 0.545 | 0.531 | 1 | 10 | Ndufb8 | 0.014 | Young |
| 0.17895 | 1.133107 | 0.061 | 0.032 | 1 | 10 | 1810032O0 | 0.029 | Young |
| 0.178984 | 0.309427 | 0.424 | 0.352 | 1 | 10 | Rab8a | 0.072 | Young |
| 0.180079 | 1.106543 | 0.101 | 0.064 | 1 | 10 | Bod1 | 0.037 | Young |
| 0.180682 | 0.839117 | 0.293 | 0.249 | 1 | 10 | Traf1 | 0.044 | Young |
| 0.180828 | 0.387414 | 0.515 | 0.467 | 1 | 10 | Gapvd1 | 0.048 | Young |
| 0.180919 | 0.635125 | 0.313 | 0.264 | 1 | 10 | Parl | 0.049 | Young |
| 0.181025 | 0.469651 | 0.505 | 0.504 | 1 | 10 | Psmd11 | 0.001 | Young |

| | | | | | | | | |
|---|---|---|---|---|---|---|---|---|
| 0.181127 | 1.253862 | 0.091 | 0.056 | 1 | 10 | Tfpi | 0.035 | Young |
| 0.181161 | 0.462913 | 0.424 | 0.381 | 1 | 10 | Sf3b3 | 0.043 | Young |
| 0.181323 | 0.465298 | 0.081 | 0.046 | 1 | 10 | Them4 | 0.035 | Young |
| 0.181336 | 0.111812 | 0.232 | 0.328 | 1 | 10 | Ago3 | -0.096 | Young |
| 0.18134 | 0.075221 | 0.01 | 0.037 | 1 | 10 | Gstt3 | -0.027 | Young |
| 0.181635 | 0.41148 | 0.343 | 0.286 | 1 | 10 | Fgr | 0.057 | Young |
| 0.181943 | 0.481307 | 0.131 | 0.088 | 1 | 10 | Kat6b | 0.043 | Young |
| 0.182663 | 0.174718 | 0.859 | 0.839 | 1 | 10 | Rpl15 | 0.02 | Young |
| 0.182733 | 0.675498 | 0.061 | 0.032 | 1 | 10 | Zfp276 | 0.029 | Young |
| 0.183233 | 0.483495 | 0.434 | 0.394 | 1 | 10 | Vdac3 | 0.04 | Young |
| 0.183258 | 0.324252 | 0.505 | 0.452 | 1 | 10 | Tnpo1 | 0.053 | Young |
| 0.183445 | 0.30115 | 0.071 | 0.039 | 1 | 10 | Zkscan6 | 0.032 | Young |
| 0.183446 | 1.047016 | 0.101 | 0.064 | 1 | 10 | Slc25a44 | 0.037 | Young |
| 0.183541 | 0.720853 | 0.121 | 0.081 | 1 | 10 | Ift22 | 0.04 | Young |
| 0.18357 | 0.550798 | 0.424 | 0.389 | 1 | 10 | Per1 | 0.035 | Young |
| 0.183604 | 0.845073 | 0.091 | 0.054 | 1 | 10 | Vps37c | 0.037 | Young |
| 0.184038 | 3.492124 | 0.071 | 0.042 | 1 | 10 | Cxcl3 | 0.029 | Young |
| 0.18404 | 0.052674 | 0.556 | 0.692 | 1 | 10 | Anxa4 | -0.136 | Young |
| 0.184052 | 1.025482 | 0.121 | 0.081 | 1 | 10 | Mfge8 | 0.04 | Young |
| 0.184747 | 1.002286 | 0.111 | 0.073 | 1 | 10 | Rpa2 | 0.038 | Young |
| 0.185439 | 1.078953 | 0.141 | 0.1 | 1 | 10 | Atl2 | 0.041 | Young |
| 0.185439 | 1.02384 | 0.141 | 0.1 | 1 | 10 | Dst | 0.041 | Young |
| 0.185851 | 0.36092 | 0.081 | 0.046 | 1 | 10 | Znhit2 | 0.035 | Young |
| 0.186052 | 0.032157 | 0.283 | 0.372 | 1 | 10 | Mff | -0.089 | Young |
| 0.186295 | 0.82683 | 0.172 | 0.125 | 1 | 10 | Cd274 | 0.047 | Young |
| 0.186894 | 1.72608 | 0.03 | 0.012 | 1 | 10 | Platr17 | 0.018 | Young |
| 0.187264 | 0.182582 | 0.333 | 0.425 | 1 | 10 | Eif4e | -0.092 | Young |
| 0.187538 | 0.036305 | 0.263 | 0.372 | 1 | 10 | Cdk11b | -0.109 | Young |
| 0.187788 | 0.251633 | 0.101 | 0.164 | 1 | 10 | C4b | -0.063 | Young |
| 0.188079 | 1.985982 | 0.03 | 0.012 | 1 | 10 | Nudt6 | 0.018 | Young |
| 0.188079 | 1.958051 | 0.03 | 0.012 | 1 | 10 | Csf1 | 0.018 | Young |
| 0.188179 | 0.061824 | 0.04 | 0.081 | 1 | 10 | Mfsd6 | -0.041 | Young |
| 0.188498 | 0.368428 | 0.556 | 0.521 | 1 | 10 | Sf3b5 | 0.035 | Young |
| 0.18878 | 0.454554 | 0.465 | 0.425 | 1 | 10 | Galnt1 | 0.04 | Young |
| 0.189571 | 0.345152 | 0.162 | 0.112 | 1 | 10 | Iqsec1 | 0.05 | Young |
| 0.190282 | 1.932475 | 0.071 | 0.042 | 1 | 10 | Riox1 | 0.029 | Young |
| 0.190369 | 1.154191 | 0.162 | 0.12 | 1 | 10 | Carnmt1 | 0.042 | Young |
| 0.190442 | 0.146262 | 0.03 | 0.066 | 1 | 10 | Scarf1 | -0.036 | Young |
| 0.190464 | 1.498321 | 0.03 | 0.012 | 1 | 10 | Pfkm | 0.018 | Young |
| 0.190464 | 1.455236 | 0.03 | 0.012 | 1 | 10 | Arhgap27o | 0.018 | Young |
| 0.19143 | 0.115251 | 0.96 | 0.971 | 1 | 10 | Rps19 | -0.011 | Young |
| 0.191782 | 0.023772 | 0.152 | 0.222 | 1 | 10 | Selenon | -0.07 | Young |
| 0.191789 | 0.155743 | 0.859 | 0.866 | 1 | 10 | Rps18 | -0.007 | Young |
| 0.192574 | 0.727475 | 0.232 | 0.183 | 1 | 10 | Mrps18a | 0.049 | Young |
| 0.192735 | 0.141319 | 0.97 | 0.983 | 1 | 10 | Rps14 | -0.013 | Young |

| | | | | | | | | |
|---|---|---|---|---|---|---|---|---|
| 0.192872 | 1.51939 | 0.03 | 0.012 | 1 | 10 | 2610035D1 | 0.018 | Young |
| 0.192872 | 1.015524 | 0.03 | 0.012 | 1 | 10 | Gdpd5 | 0.018 | Young |
| 0.192905 | 0.172038 | 0.919 | 0.927 | 1 | 10 | Ddx5 | -0.008 | Young |
| 0.194084 | 0.74755 | 0.03 | 0.012 | 1 | 10 | Cdca3 | 0.018 | Young |
| 0.194265 | 0.686462 | 0.162 | 0.115 | 1 | 10 | Nop9 | 0.047 | Young |
| 0.194487 | 0.517956 | 0.121 | 0.081 | 1 | 10 | Gnpnat1 | 0.04 | Young |
| 0.194726 | 0.016052 | 0.354 | 0.45 | 1 | 10 | Rbm5 | -0.096 | Young |
| 0.194838 | 0.69903 | 0.374 | 0.342 | 1 | 10 | Tulp4 | 0.032 | Young |
| 0.194944 | 0.762564 | 0.152 | 0.108 | 1 | 10 | Med11 | 0.044 | Young |
| 0.195727 | 0.36094 | 0.091 | 0.149 | 1 | 10 | Snrnp40 | -0.058 | Young |
| 0.196028 | 0.089856 | 0.051 | 0.093 | 1 | 10 | Nat6 | -0.042 | Young |
| 0.196525 | 0.777979 | 0.03 | 0.012 | 1 | 10 | Ninl | 0.018 | Young |
| 0.196671 | 0.579902 | 0.273 | 0.222 | 1 | 10 | Rps6kb1 | 0.051 | Young |
| 0.196968 | 1.040293 | 0.141 | 0.1 | 1 | 10 | 2410002F2 | 0.041 | Young |
| 0.197058 | 0.279781 | 0.02 | 0.051 | 1 | 10 | Papolg | -0.031 | Young |
| 0.197064 | 0.512947 | 0.303 | 0.257 | 1 | 10 | Taf1 | 0.046 | Young |
| 0.197097 | 0.666154 | 0.222 | 0.174 | 1 | 10 | Slc35a3 | 0.048 | Young |
| 0.197118 | 0.956593 | 0.081 | 0.049 | 1 | 10 | Zfyve21 | 0.032 | Young |
| 0.197372 | 1.322511 | 0.131 | 0.093 | 1 | 10 | Gadd45g | 0.038 | Young |
| 0.197505 | 0.770579 | 0.354 | 0.308 | 1 | 10 | Eny2 | 0.046 | Young |
| 0.197754 | 1.003874 | 0.03 | 0.012 | 1 | 10 | Rom1 | 0.018 | Young |
| 0.197754 | 0.978276 | 0.03 | 0.012 | 1 | 10 | Zc3h11a | 0.018 | Young |
| 0.198404 | 0.152199 | 0.212 | 0.291 | 1 | 10 | Chtf8 | -0.079 | Young |
| 0.19843 | 0.008536 | 0.505 | 0.582 | 1 | 10 | Rab11a | -0.077 | Young |
| 0.198849 | 1.415534 | 0.071 | 0.042 | 1 | 10 | Sh2b2 | 0.029 | Young |
| 0.198988 | 0.947137 | 0.03 | 0.012 | 1 | 10 | H1fx | 0.018 | Young |
| 0.198988 | 0.705442 | 0.03 | 0.012 | 1 | 10 | Ggnbp1 | 0.018 | Young |
| 0.198988 | 0.695735 | 0.03 | 0.012 | 1 | 10 | B3gnt2 | 0.018 | Young |
| 0.200148 | 0.389682 | 0.232 | 0.176 | 1 | 10 | Ubp1 | 0.056 | Young |
| 0.200229 | 0.274078 | 0.03 | 0.012 | 1 | 10 | Zfp933 | 0.018 | Young |
| 0.200489 | 1.248635 | 0.081 | 0.049 | 1 | 10 | Orc6 | 0.032 | Young |
| 0.20076 | 0.668184 | 0.03 | 0.066 | 1 | 10 | Usp28 | -0.036 | Young |
| 0.200969 | 0.284323 | 0.717 | 0.682 | 1 | 10 | Eif3k | 0.035 | Young |
| 0.201591 | 0.96593 | 0.101 | 0.066 | 1 | 10 | Dph6 | 0.035 | Young |
| 0.20185 | 0.943529 | 0.081 | 0.049 | 1 | 10 | 1600014C1 | 0.032 | Young |
| 0.202003 | 0.120878 | 0.869 | 0.897 | 1 | 10 | Rpl21 | -0.028 | Young |
| 0.202083 | 0.040291 | 0.444 | 0.545 | 1 | 10 | Cmtm3 | -0.101 | Young |
| 0.202495 | 0.500138 | 0.414 | 0.364 | 1 | 10 | Taf1d | 0.05 | Young |
| 0.202726 | 0.409159 | 0.03 | 0.012 | 1 | 10 | Arvcf | 0.018 | Young |
| 0.20292 | 0.256772 | 0.131 | 0.2 | 1 | 10 | Wdr11 | -0.069 | Young |
| 0.203339 | 0.222391 | 0.828 | 0.814 | 1 | 10 | Ndufa2 | 0.014 | Young |
| 0.204081 | 0.16079 | 0.768 | 0.711 | 1 | 10 | Srsf3 | 0.057 | Young |
| 0.20414 | 0.498895 | 0.273 | 0.218 | 1 | 10 | Rchy1 | 0.055 | Young |
| 0.204323 | 0.534739 | 0.343 | 0.296 | 1 | 10 | Sestd1 | 0.047 | Young |
| 0.20435 | 0.187332 | 0.232 | 0.32 | 1 | 10 | Phf20l1 | -0.088 | Young |

| | | | | | | | | |
|---|---|---|---|---|---|---|---|---|
| 0.204947 | 0.123938 | 0.939 | 0.936 | 1 | 10 | Rps2 | 0.003 | Young |
| 0.205787 | 1.204207 | 0.141 | 0.103 | 1 | 10 | Banp | 0.038 | Young |
| 0.205922 | 0.809908 | 0.242 | 0.198 | 1 | 10 | Ppp1r11 | 0.044 | Young |
| 0.207998 | 0.486423 | 0.424 | 0.369 | 1 | 10 | Ell2 | 0.055 | Young |
| 0.208266 | 1.078288 | 0.131 | 0.093 | 1 | 10 | Ctdp1 | 0.038 | Young |
| 0.208308 | 0.304102 | 0.535 | 0.484 | 1 | 10 | 1110008F1 | 0.051 | Young |
| 0.20844 | 0.122564 | 0.121 | 0.183 | 1 | 10 | Mrpl9 | -0.062 | Young |
| 0.208445 | 0.854257 | 0.071 | 0.042 | 1 | 10 | Cenpl | 0.029 | Young |
| 0.208539 | 0.358067 | 0.283 | 0.227 | 1 | 10 | Pnrc2 | 0.056 | Young |
| 0.208582 | 0.629693 | 0.414 | 0.369 | 1 | 10 | Msr1 | 0.045 | Young |
| 0.209663 | 1.615405 | 0.04 | 0.02 | 1 | 10 | Rsl1 | 0.02 | Young |
| 0.209663 | 1.41071 | 0.04 | 0.02 | 1 | 10 | Tnfrsf17 | 0.02 | Young |
| 0.209965 | 0.429168 | 0.222 | 0.174 | 1 | 10 | Abhd17b | 0.048 | Young |
| 0.21009 | 1.223166 | 0.101 | 0.066 | 1 | 10 | Kif19a | 0.035 | Young |
| 0.210243 | 0.61688 | 0.394 | 0.355 | 1 | 10 | Ccnl2 | 0.039 | Young |
| 0.210555 | 0.844118 | 0.141 | 0.1 | 1 | 10 | Thap12 | 0.041 | Young |
| 0.210875 | 0.538789 | 0.263 | 0.22 | 1 | 10 | Dag1 | 0.043 | Young |
| 0.21088 | 0.036119 | 0.343 | 0.443 | 1 | 10 | Srsf1 | -0.1 | Young |
| 0.211043 | 0.350957 | 0.172 | 0.125 | 1 | 10 | Hps1 | 0.047 | Young |
| 0.211331 | 0.482509 | 0.434 | 0.391 | 1 | 10 | Sri | 0.043 | Young |
| 0.212692 | 0.132035 | 0.818 | 0.814 | 1 | 10 | Fus | 0.004 | Young |
| 0.21282 | 1.416587 | 0.051 | 0.027 | 1 | 10 | Lrrc45 | 0.024 | Young |
| 0.213187 | 0.860359 | 0.101 | 0.066 | 1 | 10 | Fbxo6 | 0.035 | Young |
| 0.213187 | 0.853084 | 0.101 | 0.066 | 1 | 10 | Ubxn8 | 0.035 | Young |
| 0.213272 | 0.118323 | 0.051 | 0.09 | 1 | 10 | Tdp2 | -0.039 | Young |
| 0.213542 | 0.795191 | 0.273 | 0.235 | 1 | 10 | Pes1 | 0.038 | Young |
| 0.21373 | 0.262703 | 0.232 | 0.178 | 1 | 10 | Nolc1 | 0.054 | Young |
| 0.213843 | 0.500742 | 0.091 | 0.056 | 1 | 10 | Iffo2 | 0.035 | Young |
| 0.213908 | 1.361875 | 0.04 | 0.02 | 1 | 10 | Osgin1 | 0.02 | Young |
| 0.213927 | 0.280594 | 0.202 | 0.281 | 1 | 10 | Gtpbp2 | -0.079 | Young |
| 0.214395 | 0.611127 | 0.081 | 0.049 | 1 | 10 | Sgsm2 | 0.032 | Young |
| 0.215585 | 0.452233 | 0.293 | 0.244 | 1 | 10 | Cpeb2 | 0.049 | Young |
| 0.215823 | 0.490112 | 0.081 | 0.049 | 1 | 10 | Acsf3 | 0.032 | Young |
| 0.216085 | 0.091573 | 0.889 | 0.958 | 1 | 10 | Rpl34 | -0.069 | Young |
| 0.216292 | 0.559994 | 0.222 | 0.176 | 1 | 10 | Rbm34 | 0.046 | Young |
| 0.216547 | 1.235775 | 0.051 | 0.027 | 1 | 10 | Mtrr | 0.024 | Young |
| 0.216763 | 0.211166 | 0.444 | 0.381 | 1 | 10 | Ubxn4 | 0.063 | Young |
| 0.216972 | 0.553132 | 0.192 | 0.144 | 1 | 10 | Gart | 0.048 | Young |
| 0.217414 | 0.091999 | 0.051 | 0.09 | 1 | 10 | Tmc6 | -0.039 | Young |
| 0.217487 | 1.289647 | 0.051 | 0.027 | 1 | 10 | Gcfc2 | 0.024 | Young |
| 0.217499 | 0.321489 | 0.313 | 0.262 | 1 | 10 | Bdp1 | 0.051 | Young |
| 0.217525 | 0.949038 | 0.061 | 0.034 | 1 | 10 | Wdr47 | 0.027 | Young |
| 0.217643 | 0.01992 | 0.525 | 0.66 | 1 | 10 | Cct5 | -0.135 | Young |
| 0.217673 | 1.062556 | 0.04 | 0.02 | 1 | 10 | Bbs10 | 0.02 | Young |
| 0.218082 | 0.03606 | 0.141 | 0.2 | 1 | 10 | Otud6b | -0.059 | Young |

| | | | | | | | | |
|---|---|---|---|---|---|---|---|---|
| 0.218429 | 1.009672 | 0.051 | 0.027 | 1 | 10 | Cbx6 | 0.024 | Young |
| 0.218608 | 1.228774 | 0.111 | 0.076 | 1 | 10 | Nol4l | 0.035 | Young |
| 0.218822 | 0.340595 | 0.242 | 0.191 | 1 | 10 | Pqbp1 | 0.051 | Young |
| 0.219233 | 0.282879 | 0.091 | 0.056 | 1 | 10 | Aldh7a1 | 0.035 | Young |
| 0.219301 | 1.14841 | 0.04 | 0.02 | 1 | 10 | Cenpq | 0.02 | Young |
| 0.219377 | 0.094707 | 0.111 | 0.174 | 1 | 10 | Sigmar1 | -0.063 | Young |
| 0.219481 | 1.086823 | 0.323 | 0.284 | 1 | 10 | Fam104a | 0.039 | Young |
| 0.219512 | 0.006057 | 0.263 | 0.355 | 1 | 10 | Cul1 | -0.092 | Young |
| 0.219806 | 0.904076 | 0.111 | 0.076 | 1 | 10 | Lin37 | 0.035 | Young |
| 0.219929 | 0.718956 | 0.071 | 0.042 | 1 | 10 | Zfp87 | 0.029 | Young |
| 0.219951 | 0.75362 | 0.152 | 0.11 | 1 | 10 | Gm48065 | 0.042 | Young |
| 0.219959 | 0.229054 | 0.313 | 0.247 | 1 | 10 | Nrip1 | 0.066 | Young |
| 0.220322 | 1.254512 | 0.051 | 0.027 | 1 | 10 | Abcc4 | 0.024 | Young |
| 0.220392 | 0.666844 | 0.04 | 0.02 | 1 | 10 | Cwf19l1 | 0.02 | Young |
| 0.220392 | 0.604436 | 0.04 | 0.02 | 1 | 10 | Atp6v1g2 | 0.02 | Young |
| 0.220696 | 0.484564 | 0.414 | 0.359 | 1 | 10 | Wbp11 | 0.055 | Young |
| 0.220727 | 0.289937 | 0.182 | 0.134 | 1 | 10 | Mphosph8 | 0.048 | Young |
| 0.220747 | 0.306281 | 0.273 | 0.215 | 1 | 10 | Elf4 | 0.058 | Young |
| 0.220956 | 0.557154 | 0.303 | 0.254 | 1 | 10 | Smad4 | 0.049 | Young |
| 0.221486 | 0.691316 | 0.04 | 0.02 | 1 | 10 | Pecr | 0.02 | Young |
| 0.221666 | 0.507045 | 0.566 | 0.548 | 1 | 10 | Tpi1 | 0.018 | Young |
| 0.222173 | 0.495613 | 0.263 | 0.215 | 1 | 10 | Eftud2 | 0.048 | Young |
| 0.22237 | 0.090408 | 0.03 | 0.064 | 1 | 10 | Nudcd1 | -0.034 | Young |
| 0.222435 | 0.248721 | 0.646 | 0.636 | 1 | 10 | Ifitm3 | 0.01 | Young |
| 0.222584 | 0.747314 | 0.04 | 0.02 | 1 | 10 | Tasp1 | 0.02 | Young |
| 0.222584 | 0.471133 | 0.04 | 0.02 | 1 | 10 | Supv3l1 | 0.02 | Young |
| 0.223206 | 0.390768 | 0.253 | 0.203 | 1 | 10 | Coro7 | 0.05 | Young |
| 0.223686 | 0.923324 | 0.04 | 0.02 | 1 | 10 | Hes1 | 0.02 | Young |
| 0.223686 | 0.06654 | 0.04 | 0.02 | 1 | 10 | St18 | 0.02 | Young |
| 0.224191 | 0.054888 | 0.444 | 0.56 | 1 | 10 | Tmem258 | -0.116 | Young |
| 0.224359 | 1.324881 | 0.061 | 0.034 | 1 | 10 | Zfp868 | 0.027 | Young |
| 0.224792 | 0.744409 | 0.04 | 0.02 | 1 | 10 | Ldhb | 0.02 | Young |
| 0.224792 | 0.718524 | 0.04 | 0.02 | 1 | 10 | Letm2 | 0.02 | Young |
| 0.225178 | 0.122773 | 0.949 | 0.983 | 1 | 10 | Rpl17 | -0.034 | Young |
| 0.22555 | 0.790693 | 0.141 | 0.103 | 1 | 10 | Jag1 | 0.038 | Young |
| 0.225591 | 0.493907 | 0.303 | 0.249 | 1 | 10 | Plbd1 | 0.054 | Young |
| 0.226055 | 0.576061 | 0.232 | 0.186 | 1 | 10 | Cyp4v3 | 0.046 | Young |
| 0.226395 | 0.001149 | 0.081 | 0.13 | 1 | 10 | Anapc7 | -0.049 | Young |
| 0.226961 | 1.141824 | 0.061 | 0.034 | 1 | 10 | Ankrd28 | 0.027 | Young |
| 0.227015 | 1.095656 | 0.04 | 0.02 | 1 | 10 | Rbbp9 | 0.02 | Young |
| 0.227102 | 0.461026 | 0.192 | 0.144 | 1 | 10 | Tes | 0.048 | Young |
| 0.227702 | 0.048662 | 0.434 | 0.509 | 1 | 10 | Etf1 | -0.075 | Young |
| 0.228039 | 0.028226 | 0.111 | 0.164 | 1 | 10 | Htatip2 | -0.053 | Young |
| 0.228089 | 0.271289 | 0.414 | 0.359 | 1 | 10 | Uqcrc2 | 0.055 | Young |
| 0.228424 | 0.244909 | 0.808 | 0.866 | 1 | 10 | Rpl10 | -0.058 | Young |

| | | | | | | | | |
|---|---|---|---|---|---|---|---|---|
| 0.228473 | 0.628724 | 0.303 | 0.254 | 1 | 10 | Stard9 | 0.049 | Young |
| 0.228634 | 0.951308 | 0.091 | 0.059 | 1 | 10 | Tex30 | 0.032 | Young |
| 0.228986 | 0.578091 | 0.051 | 0.027 | 1 | 10 | Gm15445 | 0.024 | Young |
| 0.229534 | 0.337845 | 0.455 | 0.401 | 1 | 10 | Dnajc19 | 0.054 | Young |
| 0.229564 | 0.282435 | 0.141 | 0.2 | 1 | 10 | Slu7 | -0.059 | Young |
| 0.229766 | 0.100866 | 0.071 | 0.115 | 1 | 10 | Dusp7 | -0.044 | Young |
| 0.230263 | 0.339809 | 0.798 | 0.778 | 1 | 10 | Csde1 | 0.02 | Young |
| 0.230405 | 0.615403 | 0.212 | 0.169 | 1 | 10 | Vta1 | 0.043 | Young |
| 0.230463 | 0.643877 | 0.061 | 0.034 | 1 | 10 | Usp20 | 0.027 | Young |
| 0.230517 | 0.443422 | 0.465 | 0.418 | 1 | 10 | Leprotl1 | 0.047 | Young |
| 0.230608 | 0.434444 | 0.485 | 0.44 | 1 | 10 | Ftl1-ps1 | 0.045 | Young |
| 0.23079 | 0.152632 | 0.202 | 0.154 | 1 | 10 | Tax1bp3 | 0.048 | Young |
| 0.230944 | 0.642555 | 0.051 | 0.027 | 1 | 10 | Gemin5 | 0.024 | Young |
| 0.231527 | 0.356791 | 0.081 | 0.132 | 1 | 10 | Mdp1 | -0.051 | Young |
| 0.231927 | 0.655397 | 0.051 | 0.027 | 1 | 10 | 9430015G1 | 0.024 | Young |
| 0.232067 | 0.047782 | 0.404 | 0.491 | 1 | 10 | Furin | -0.087 | Young |
| 0.232073 | 0.492934 | 0.364 | 0.323 | 1 | 10 | Alkbh5 | 0.041 | Young |
| 0.232693 | 0.211758 | 0.202 | 0.267 | 1 | 10 | 1810022K0 | -0.065 | Young |
| 0.233042 | 0.150418 | 0.111 | 0.171 | 1 | 10 | Dph5 | -0.06 | Young |
| 0.234506 | 0.050583 | 0.384 | 0.46 | 1 | 10 | Gnpda1 | -0.076 | Young |
| 0.234558 | 1.027896 | 0.111 | 0.076 | 1 | 10 | Adi1 | 0.035 | Young |
| 0.234558 | 0.839735 | 0.111 | 0.076 | 1 | 10 | Rrp12 | 0.035 | Young |
| 0.234659 | 0.244688 | 0.646 | 0.667 | 1 | 10 | Selenok | -0.021 | Young |
| 0.234909 | 0.618824 | 0.091 | 0.059 | 1 | 10 | Slc2a6 | 0.032 | Young |
| 0.235411 | 1.215283 | 0.131 | 0.095 | 1 | 10 | Plekhg5 | 0.036 | Young |
| 0.235613 | 0.752655 | 0.091 | 0.059 | 1 | 10 | Rcn3 | 0.032 | Young |
| 0.235937 | 0.364597 | 0.071 | 0.042 | 1 | 10 | Mterf4 | 0.029 | Young |
| 0.23733 | 0.12398 | 0.212 | 0.284 | 1 | 10 | 2310035C2 | -0.072 | Young |
| 0.237584 | 0.260816 | 0.061 | 0.034 | 1 | 10 | Impa2 | 0.027 | Young |
| 0.237972 | 0.160401 | 0.949 | 0.976 | 1 | 10 | Rpl19 | -0.027 | Young |
| 0.238213 | 0.683412 | 0.313 | 0.271 | 1 | 10 | Agfg1 | 0.042 | Young |
| 0.238981 | 0.434212 | 0.313 | 0.269 | 1 | 10 | Mgl2 | 0.044 | Young |
| 0.239144 | 0.232519 | 0.101 | 0.156 | 1 | 10 | Impdh1 | -0.055 | Young |
| 0.239191 | 0.262468 | 0.768 | 0.78 | 1 | 10 | Hnrnpa0 | -0.012 | Young |
| 0.239219 | 1.44961 | 0.101 | 0.071 | 1 | 10 | Gpr89 | 0.03 | Young |
| 0.239327 | 1.642333 | 0.071 | 0.044 | 1 | 10 | Zbtb45 | 0.027 | Young |
| 0.239375 | 0.040839 | 0.04 | 0.076 | 1 | 10 | Kank3 | -0.036 | Young |
| 0.239847 | 0.359863 | 0.556 | 0.587 | 1 | 10 | Tmem189 | -0.031 | Young |
| 0.240474 | 0.32121 | 0.172 | 0.232 | 1 | 10 | F10 | -0.06 | Young |
| 0.240539 | 2.420094 | 0.02 | 0.007 | 1 | 10 | Zfp607b | 0.013 | Young |
| 0.240539 | 2.190222 | 0.02 | 0.007 | 1 | 10 | Gm26882 | 0.013 | Young |
| 0.240539 | 2.089551 | 0.02 | 0.007 | 1 | 10 | Atp2a3 | 0.013 | Young |
| 0.240539 | 2.019985 | 0.02 | 0.007 | 1 | 10 | Chek1 | 0.013 | Young |
| 0.240539 | 1.85698 | 0.02 | 0.007 | 1 | 10 | Cryab | 0.013 | Young |
| 0.240539 | 1.841013 | 0.02 | 0.007 | 1 | 10 | Homer2 | 0.013 | Young |

| | | | | | | | | |
|---|---|---|---|---|---|---|---|---|
| 0.240539 | 1.319795 | 0.02 | 0.007 | 1 | 10 | Spsb4 | 0.013 | Young |
| 0.240539 | 1.153492 | 0.02 | 0.007 | 1 | 10 | Plppr3 | 0.013 | Young |
| 0.240651 | 0.477494 | 0.242 | 0.193 | 1 | 10 | Hivep1 | 0.049 | Young |
| 0.240771 | 0.024058 | 0.091 | 0.142 | 1 | 10 | Bola1 | -0.051 | Young |
| 0.240957 | 1.260765 | 0.071 | 0.044 | 1 | 10 | Gss | 0.027 | Young |
| 0.240995 | 0.567386 | 0.273 | 0.222 | 1 | 10 | Chordc1 | 0.051 | Young |
| 0.242332 | 2.09608 | 0.02 | 0.007 | 1 | 10 | Rab39 | 0.013 | Young |
| 0.242332 | 2.060544 | 0.02 | 0.007 | 1 | 10 | Phgdh | 0.013 | Young |
| 0.242332 | 1.805899 | 0.02 | 0.007 | 1 | 10 | Treml4 | 0.013 | Young |
| 0.242332 | 1.507301 | 0.02 | 0.007 | 1 | 10 | Cenpn | 0.013 | Young |
| 0.242332 | 1.453688 | 0.02 | 0.007 | 1 | 10 | Adamts5 | 0.013 | Young |
| 0.242332 | 1.371054 | 0.02 | 0.007 | 1 | 10 | Trim59 | 0.013 | Young |
| 0.242332 | 0.861921 | 0.02 | 0.007 | 1 | 10 | Proser3 | 0.013 | Young |
| 0.24243 | 0.262229 | 0.505 | 0.465 | 1 | 10 | Mcub | 0.04 | Young |
| 0.242632 | 0.441823 | 0.556 | 0.533 | 1 | 10 | Mif | 0.023 | Young |
| 0.243079 | 0.462437 | 0.414 | 0.377 | 1 | 10 | Trappc4 | 0.037 | Young |
| 0.243214 | 0.681937 | 0.141 | 0.103 | 1 | 10 | Rcan1 | 0.038 | Young |
| 0.243411 | 0.450137 | 0.717 | 0.729 | 1 | 10 | Basp1 | -0.012 | Young |
| 0.243475 | 0.530792 | 0.192 | 0.147 | 1 | 10 | Zfp740 | 0.045 | Young |
| 0.243896 | 0.267104 | 0.071 | 0.115 | 1 | 10 | Nedd4l | -0.044 | Young |
| 0.244135 | 1.416857 | 0.02 | 0.007 | 1 | 10 | Fam71e1 | 0.013 | Young |
| 0.244135 | 1.308716 | 0.02 | 0.007 | 1 | 10 | Zfp189 | 0.013 | Young |
| 0.244135 | 1.208141 | 0.02 | 0.007 | 1 | 10 | AC162302. | 0.013 | Young |
| 0.244135 | 1.184058 | 0.02 | 0.007 | 1 | 10 | Rad51c | 0.013 | Young |
| 0.244135 | 1.176911 | 0.02 | 0.007 | 1 | 10 | Exo1 | 0.013 | Young |
| 0.244135 | 1.079728 | 0.02 | 0.007 | 1 | 10 | Creb3l3 | 0.013 | Young |
| 0.244135 | 0.966979 | 0.02 | 0.007 | 1 | 10 | Padi4 | 0.013 | Young |
| 0.244135 | 0.94746 | 0.02 | 0.007 | 1 | 10 | Ccdc138 | 0.013 | Young |
| 0.244423 | 0.335638 | 0.626 | 0.665 | 1 | 10 | Gpi1 | -0.039 | Young |
| 0.244454 | 0.709035 | 0.192 | 0.147 | 1 | 10 | Tlr7 | 0.045 | Young |
| 0.245065 | 1.111854 | 0.071 | 0.044 | 1 | 10 | Vwf | 0.027 | Young |
| 0.245074 | 0.064659 | 0.01 | 0.032 | 1 | 10 | Gpt | -0.022 | Young |
| 0.245382 | 0.55065 | 0.121 | 0.181 | 1 | 10 | Ddit3 | -0.06 | Young |
| 0.245636 | 0.368531 | 0.687 | 0.716 | 1 | 10 | Sf3b1 | -0.029 | Young |
| 0.245813 | 0.026272 | 0.182 | 0.247 | 1 | 10 | Snx29 | -0.065 | Young |
| 0.245946 | 2.382642 | 0.02 | 0.007 | 1 | 10 | Hspb1 | 0.013 | Young |
| 0.245946 | 2.084732 | 0.02 | 0.007 | 1 | 10 | Tmem170 | 0.013 | Young |
| 0.245946 | 1.593298 | 0.02 | 0.007 | 1 | 10 | Lbhd1 | 0.013 | Young |
| 0.245946 | 1.218102 | 0.02 | 0.007 | 1 | 10 | Gpat3 | 0.013 | Young |
| 0.245946 | 0.879291 | 0.02 | 0.007 | 1 | 10 | Gsg1 | 0.013 | Young |
| 0.245946 | 0.777619 | 0.02 | 0.007 | 1 | 10 | Pkp3 | 0.013 | Young |
| 0.245946 | 0.704867 | 0.02 | 0.007 | 1 | 10 | Nnt | 0.013 | Young |
| 0.245946 | 0.671817 | 0.02 | 0.007 | 1 | 10 | Lyplal1 | 0.013 | Young |
| 0.245946 | 0.652368 | 0.02 | 0.007 | 1 | 10 | Afp | 0.013 | Young |
| 0.246195 | 0.091069 | 0.374 | 0.479 | 1 | 10 | Camk2d | -0.105 | Young |

| | | | | | | | | |
|---|---|---|---|---|---|---|---|---|
| 0.246211 | 0.46845 | 0.545 | 0.533 | 1 | 10 | Hif1a | 0.012 | Young |
| 0.246659 | 0.650576 | 0.101 | 0.068 | 1 | 10 | Zc3h7b | 0.033 | Young |
| 0.246727 | 0.687738 | 0.152 | 0.112 | 1 | 10 | H2-Q4 | 0.04 | Young |
| 0.246745 | 0.885728 | 0.162 | 0.12 | 1 | 10 | Dnajc11 | 0.042 | Young |
| 0.246766 | 0.79209 | 0.121 | 0.086 | 1 | 10 | Mdc1 | 0.035 | Young |
| 0.247058 | 0.791717 | 0.131 | 0.095 | 1 | 10 | 2310033P0 | 0.036 | Young |
| 0.247285 | 0.071168 | 0.505 | 0.609 | 1 | 10 | Tmem30a | -0.104 | Young |
| 0.247314 | 0.573954 | 0.434 | 0.399 | 1 | 10 | Dock7 | 0.035 | Young |
| 0.24757 | 0.314028 | 0.768 | 0.751 | 1 | 10 | Srrm2 | 0.017 | Young |
| 0.247767 | 0.77591 | 0.02 | 0.007 | 1 | 10 | BC043934 | 0.013 | Young |
| 0.247767 | 0.772954 | 0.02 | 0.007 | 1 | 10 | 2700081O1 | 0.013 | Young |
| 0.247767 | 0.554373 | 0.02 | 0.007 | 1 | 10 | Suox | 0.013 | Young |
| 0.247767 | 0.55349 | 0.02 | 0.007 | 1 | 10 | Frg2f1 | 0.013 | Young |
| 0.247767 | 0.46893 | 0.02 | 0.007 | 1 | 10 | Wdr5b | 0.013 | Young |
| 0.247767 | 0.455845 | 0.02 | 0.007 | 1 | 10 | Fbxo44 | 0.013 | Young |
| 0.247767 | 0.223054 | 0.02 | 0.007 | 1 | 10 | Txk | 0.013 | Young |
| 0.248559 | 1.039211 | 0.111 | 0.078 | 1 | 10 | Prmt9 | 0.033 | Young |
| 0.24857 | 0.721692 | 0.081 | 0.051 | 1 | 10 | Clec2i | 0.03 | Young |
| 0.249598 | 0.465609 | 0.02 | 0.007 | 1 | 10 | Epm2a | 0.013 | Young |
| 0.249598 | 0.225402 | 0.02 | 0.007 | 1 | 10 | Tcea2 | 0.013 | Young |
| 0.249598 | 0.137533 | 0.02 | 0.007 | 1 | 10 | 4932422M | 0.013 | Young |
| 0.250487 | 0.055864 | 0.384 | 0.487 | 1 | 10 | Ikbkb | -0.103 | Young |
| 0.250548 | 0.043964 | 0.283 | 0.362 | 1 | 10 | Cnot6 | -0.079 | Young |
| 0.250554 | 0.076085 | 0.586 | 0.663 | 1 | 10 | Klf4 | -0.077 | Young |
| 0.250653 | 0.316023 | 0.141 | 0.1 | 1 | 10 | Hdgfl2 | 0.041 | Young |
| 0.250752 | 0.232893 | 0.131 | 0.09 | 1 | 10 | Umps | 0.041 | Young |
| 0.250912 | 0.548481 | 0.081 | 0.051 | 1 | 10 | Trim37 | 0.03 | Young |
| 0.251039 | 0.598089 | 0.253 | 0.205 | 1 | 10 | Bcl2a1d | 0.048 | Young |
| 0.251215 | 0.758918 | 0.232 | 0.191 | 1 | 10 | Apopt1 | 0.041 | Young |
| 0.251303 | 0.072357 | 0.131 | 0.186 | 1 | 10 | Tango2 | -0.055 | Young |
| 0.251828 | 0.977946 | 0.131 | 0.095 | 1 | 10 | Abcb1b | 0.036 | Young |
| 0.25215 | 0.464221 | 0.162 | 0.12 | 1 | 10 | Pkd1 | 0.042 | Young |
| 0.252488 | 0.28278 | 0.525 | 0.535 | 1 | 10 | Eif4g1 | -0.01 | Young |
| 0.252532 | 0.541877 | 0.172 | 0.127 | 1 | 10 | Wdr44 | 0.045 | Young |
| 0.25358 | 0.20267 | 0.657 | 0.641 | 1 | 10 | Clk1 | 0.016 | Young |
| 0.25367 | 0.052154 | 0.333 | 0.418 | 1 | 10 | Ak2 | -0.085 | Young |
| 0.25375 | 0.874613 | 0.283 | 0.244 | 1 | 10 | Rbm42 | 0.039 | Young |
| 0.254176 | 0.746375 | 0.384 | 0.347 | 1 | 10 | Akirin2 | 0.037 | Young |
| 0.254275 | 0.893285 | 0.071 | 0.044 | 1 | 10 | Yipf2 | 0.027 | Young |
| 0.254283 | 0.528415 | 0.364 | 0.337 | 1 | 10 | Clptm1 | 0.027 | Young |
| 0.254523 | 0.433288 | 0.475 | 0.46 | 1 | 10 | Stub1 | 0.015 | Young |
| 0.255459 | 0.611192 | 0.263 | 0.215 | 1 | 10 | Glipr1 | 0.048 | Young |
| 0.255943 | 0.256805 | 0.141 | 0.1 | 1 | 10 | Dhx34 | 0.041 | Young |
| 0.256009 | 0.140994 | 0.949 | 0.949 | 1 | 10 | Rpl9 | 0 | Young |
| 0.256225 | 0.53858 | 0.152 | 0.112 | 1 | 10 | Arl4a | 0.04 | Young |

| | | | | | | | | |
|---|---|---|---|---|---|---|---|---|
| 0.256437 | 0.830452 | 0.081 | 0.051 | 1 | 10 | Peli2 | 0.03 | Young |
| 0.256898 | 0.390866 | 0.192 | 0.149 | 1 | 10 | Mepce | 0.043 | Young |
| 0.257413 | 0.255575 | 0.475 | 0.452 | 1 | 10 | Ndufs5 | 0.023 | Young |
| 0.257675 | 0.369827 | 0.141 | 0.103 | 1 | 10 | Mkl1 | 0.038 | Young |
| 0.257931 | 0.307475 | 0.141 | 0.191 | 1 | 10 | Gm34084 | -0.05 | Young |
| 0.25854 | 0.366916 | 0.071 | 0.044 | 1 | 10 | Donson | 0.027 | Young |
| 0.258913 | 0.252584 | 0.475 | 0.443 | 1 | 10 | Edf1 | 0.032 | Young |
| 0.259399 | 0.8004 | 0.071 | 0.044 | 1 | 10 | Bambi | 0.027 | Young |
| 0.259576 | 0.200932 | 0.081 | 0.125 | 1 | 10 | Rab28 | -0.044 | Young |
| 0.260507 | 0.367551 | 0.162 | 0.122 | 1 | 10 | Smox | 0.04 | Young |
| 0.260762 | 0.582872 | 0.434 | 0.406 | 1 | 10 | Exoc5 | 0.028 | Young |
| 0.261268 | 0.437011 | 0.313 | 0.269 | 1 | 10 | Slc23a2 | 0.044 | Young |
| 0.261301 | 0.635817 | 0.293 | 0.249 | 1 | 10 | Csnk1g3 | 0.044 | Young |
| 0.261982 | 0.40989 | 0.091 | 0.059 | 1 | 10 | Cradd | 0.032 | Young |
| 0.263198 | 0.054809 | 0.505 | 0.582 | 1 | 10 | Manf | -0.077 | Young |
| 0.263471 | 0.097579 | 0.061 | 0.098 | 1 | 10 | Bpnt1 | -0.037 | Young |
| 0.26365 | 0.187375 | 0.111 | 0.161 | 1 | 10 | Slc35a2 | -0.05 | Young |
| 0.263723 | 0.629923 | 0.071 | 0.044 | 1 | 10 | Ppil1 | 0.027 | Young |
| 0.264656 | 0.01849 | 0.323 | 0.384 | 1 | 10 | Csnk2a2 | -0.061 | Young |
| 0.264667 | 0.434709 | 0.263 | 0.21 | 1 | 10 | Ophn1 | 0.053 | Young |
| 0.264751 | 1.191968 | 0.111 | 0.081 | 1 | 10 | Tfdp1 | 0.03 | Young |
| 0.26498 | 0.421218 | 0.434 | 0.381 | 1 | 10 | Cux1 | 0.053 | Young |
| 0.265335 | 0.266722 | 0.717 | 0.738 | 1 | 10 | Tomm20 | -0.021 | Young |
| 0.265459 | 0.129292 | 0.909 | 0.922 | 1 | 10 | Rpl12 | -0.013 | Young |
| 0.265593 | 0.74597 | 0.283 | 0.244 | 1 | 10 | Fbxo11 | 0.039 | Young |
| 0.265821 | 0.162029 | 0.172 | 0.125 | 1 | 10 | Ube2g1 | 0.047 | Young |
| 0.265955 | 0.684308 | 0.232 | 0.183 | 1 | 10 | Rai14 | 0.049 | Young |
| 0.266342 | 0.830207 | 0.071 | 0.044 | 1 | 10 | Pwwp2b | 0.027 | Young |
| 0.26637 | 0.721951 | 0.323 | 0.291 | 1 | 10 | Kmt5a | 0.032 | Young |
| 0.266807 | 0.069446 | 0.111 | 0.159 | 1 | 10 | Cracr2b | -0.048 | Young |
| 0.267052 | 0.674307 | 0.222 | 0.181 | 1 | 10 | Tnfsf9 | 0.041 | Young |
| 0.268365 | 0.041725 | 0.556 | 0.672 | 1 | 10 | Gnaq | -0.116 | Young |
| 0.268455 | 0.543972 | 0.121 | 0.086 | 1 | 10 | Ift46 | 0.035 | Young |
| 0.268527 | 0.085947 | 0.253 | 0.328 | 1 | 10 | Rnf111 | -0.075 | Young |
| 0.269645 | 0.145328 | 0.141 | 0.198 | 1 | 10 | Ero1lb | -0.057 | Young |
| 0.269899 | 0.604809 | 0.101 | 0.068 | 1 | 10 | Trmt2a | 0.033 | Young |
| 0.270162 | 0.693495 | 0.091 | 0.061 | 1 | 10 | Asb3 | 0.03 | Young |
| 0.270892 | 1.01576 | 0.182 | 0.149 | 1 | 10 | Cln5 | 0.033 | Young |
| 0.270941 | 0.470456 | 0.354 | 0.315 | 1 | 10 | Uqcc3 | 0.039 | Young |
| 0.271371 | 0.692948 | 0.121 | 0.088 | 1 | 10 | Zfp513 | 0.033 | Young |
| 0.271713 | 0.652951 | 0.253 | 0.213 | 1 | 10 | Inpp4a | 0.04 | Young |
| 0.271864 | 0.499327 | 0.182 | 0.137 | 1 | 10 | Hsd17b10 | 0.045 | Young |
| 0.271948 | 0.380313 | 0.253 | 0.203 | 1 | 10 | Gdap10 | 0.05 | Young |
| 0.272084 | 0.433474 | 0.101 | 0.068 | 1 | 10 | Phtf1 | 0.033 | Young |
| 0.272224 | 0.195273 | 0.051 | 0.086 | 1 | 10 | Xpc | -0.035 | Young |

| | | | | | | | | |
|---|---|---|---|---|---|---|---|---|
| 0.272521 | 0.561241 | 0.071 | 0.044 | 1 | 10 | Dnajc18 | 0.027 | Young |
| 0.272695 | 0.552888 | 0.384 | 0.33 | 1 | 10 | Cacybp | 0.054 | Young |
| 0.273006 | 0.392979 | 0.364 | 0.318 | 1 | 10 | Phf23 | 0.046 | Young |
| 0.273291 | 0.721001 | 0.242 | 0.213 | 1 | 10 | Tbc1d13 | 0.029 | Young |
| 0.273628 | 2.166232 | 0.051 | 0.029 | 1 | 10 | Cdc7 | 0.022 | Young |
| 0.273742 | 0.326895 | 0.505 | 0.457 | 1 | 10 | Nol7 | 0.048 | Young |
| 0.274001 | 0.931619 | 0.121 | 0.088 | 1 | 10 | Mum1 | 0.033 | Young |
| 0.274059 | 0.172175 | 0.081 | 0.122 | 1 | 10 | Tbrg4 | -0.041 | Young |
| 0.27406 | 0.020648 | 0.182 | 0.247 | 1 | 10 | Mpdu1 | -0.065 | Young |
| 0.274439 | 0.799926 | 0.061 | 0.037 | 1 | 10 | Cgnl1 | 0.024 | Young |
| 0.274575 | 0.719389 | 0.273 | 0.235 | 1 | 10 | Prelid3b | 0.038 | Young |
| 0.274786 | 0.374397 | 0.152 | 0.112 | 1 | 10 | Naa60 | 0.04 | Young |
| 0.275273 | 1.076821 | 0.273 | 0.242 | 1 | 10 | Slain2 | 0.031 | Young |
| 0.27531 | 0.100572 | 0.929 | 0.939 | 1 | 10 | Cebpb | -0.01 | Young |
| 0.275527 | 4.87302 | 0.01 | 0.002 | 1 | 10 | Prok2 | 0.008 | Young |
| 0.275527 | 4.442213 | 0.01 | 0.002 | 1 | 10 | Saa3 | 0.008 | Young |
| 0.275527 | 2.666359 | 0.01 | 0.002 | 1 | 10 | Mucl1 | 0.008 | Young |
| 0.275527 | 2.505754 | 0.01 | 0.002 | 1 | 10 | Ctse | 0.008 | Young |
| 0.275527 | 2.474751 | 0.01 | 0.002 | 1 | 10 | Gm46447 | 0.008 | Young |
| 0.275527 | 2.463008 | 0.01 | 0.002 | 1 | 10 | 2010310C0 | 0.008 | Young |
| 0.275527 | 2.36683 | 0.01 | 0.002 | 1 | 10 | Spink10 | 0.008 | Young |
| 0.275527 | 2.288253 | 0.01 | 0.002 | 1 | 10 | Rims3 | 0.008 | Young |
| 0.275527 | 2.197016 | 0.01 | 0.002 | 1 | 10 | Proscos | 0.008 | Young |
| 0.275527 | 2.195831 | 0.01 | 0.002 | 1 | 10 | Gm26737 | 0.008 | Young |
| 0.275527 | 2.174555 | 0.01 | 0.002 | 1 | 10 | Srp54c | 0.008 | Young |
| 0.275527 | 2.171234 | 0.01 | 0.002 | 1 | 10 | Spin4 | 0.008 | Young |
| 0.275527 | 2.150212 | 0.01 | 0.002 | 1 | 10 | Gm45449 | 0.008 | Young |
| 0.275527 | 2.119957 | 0.01 | 0.002 | 1 | 10 | Rab11fip4 | 0.008 | Young |
| 0.275527 | 2.091532 | 0.01 | 0.002 | 1 | 10 | Pir | 0.008 | Young |
| 0.275527 | 2.055168 | 0.01 | 0.002 | 1 | 10 | Enah | 0.008 | Young |
| 0.275527 | 2.039514 | 0.01 | 0.002 | 1 | 10 | Mirt1 | 0.008 | Young |
| 0.275527 | 2.032528 | 0.01 | 0.002 | 1 | 10 | Ptprf | 0.008 | Young |
| 0.275527 | 2.014558 | 0.01 | 0.002 | 1 | 10 | C1qtnf9 | 0.008 | Young |
| 0.275527 | 2.007649 | 0.01 | 0.002 | 1 | 10 | Satb1 | 0.008 | Young |
| 0.275527 | 1.981955 | 0.01 | 0.002 | 1 | 10 | Elavl4 | 0.008 | Young |
| 0.275527 | 1.964886 | 0.01 | 0.002 | 1 | 10 | Prr22 | 0.008 | Young |
| 0.275527 | 1.936013 | 0.01 | 0.002 | 1 | 10 | Kbtbd12 | 0.008 | Young |
| 0.275527 | 1.851217 | 0.01 | 0.002 | 1 | 10 | Gpat2 | 0.008 | Young |
| 0.275527 | 1.849066 | 0.01 | 0.002 | 1 | 10 | Polh | 0.008 | Young |
| 0.275527 | 1.789175 | 0.01 | 0.002 | 1 | 10 | 1700066B1 | 0.008 | Young |
| 0.275527 | 1.779148 | 0.01 | 0.002 | 1 | 10 | Kcnrg | 0.008 | Young |
| 0.275527 | 1.77432 | 0.01 | 0.002 | 1 | 10 | Plat | 0.008 | Young |
| 0.275527 | 1.755777 | 0.01 | 0.002 | 1 | 10 | Gm49339 | 0.008 | Young |
| 0.275527 | 1.752274 | 0.01 | 0.002 | 1 | 10 | Pcna-ps2 | 0.008 | Young |
| 0.275527 | 1.744472 | 0.01 | 0.002 | 1 | 10 | Fbxo5 | 0.008 | Young |

| | | | | | | | | |
|---|---|---|---|---|---|---|---|---|
| 0.275527 | 1.722193 | 0.01 | 0.002 | 1 | 10 | Gm45844 | 0.008 | Young |
| 0.275527 | 1.708795 | 0.01 | 0.002 | 1 | 10 | Dscc1 | 0.008 | Young |
| 0.275527 | 1.695372 | 0.01 | 0.002 | 1 | 10 | Rad51 | 0.008 | Young |
| 0.275527 | 1.689006 | 0.01 | 0.002 | 1 | 10 | AC123724. | 0.008 | Young |
| 0.275527 | 1.616284 | 0.01 | 0.002 | 1 | 10 | A630001O[illegible] | 0.008 | Young |
| 0.275527 | 1.599244 | 0.01 | 0.002 | 1 | 10 | Col27a1 | 0.008 | Young |
| 0.275527 | 1.586895 | 0.01 | 0.002 | 1 | 10 | Fbxl12os | 0.008 | Young |
| 0.275527 | 1.528774 | 0.01 | 0.002 | 1 | 10 | Vegfc | 0.008 | Young |
| 0.275527 | 1.512057 | 0.01 | 0.002 | 1 | 10 | Itgb8 | 0.008 | Young |
| 0.275527 | 1.473138 | 0.01 | 0.002 | 1 | 10 | Apln | 0.008 | Young |
| 0.275527 | 1.463482 | 0.01 | 0.002 | 1 | 10 | Tcap | 0.008 | Young |
| 0.275527 | 1.454836 | 0.01 | 0.002 | 1 | 10 | Gm6598 | 0.008 | Young |
| 0.275527 | 1.44438 | 0.01 | 0.002 | 1 | 10 | Sult1e1 | 0.008 | Young |
| 0.275527 | 1.444028 | 0.01 | 0.002 | 1 | 10 | Gm10522 | 0.008 | Young |
| 0.275527 | 1.437158 | 0.01 | 0.002 | 1 | 10 | Hykk | 0.008 | Young |
| 0.275527 | 1.426337 | 0.01 | 0.002 | 1 | 10 | Gm46440 | 0.008 | Young |
| 0.275527 | 1.414472 | 0.01 | 0.002 | 1 | 10 | Hid1 | 0.008 | Young |
| 0.275527 | 1.389681 | 0.01 | 0.002 | 1 | 10 | Gm14391 | 0.008 | Young |
| 0.275527 | 1.370487 | 0.01 | 0.002 | 1 | 10 | Gne | 0.008 | Young |
| 0.275527 | 1.362674 | 0.01 | 0.002 | 1 | 10 | Zfp457 | 0.008 | Young |
| 0.275527 | 1.35951 | 0.01 | 0.002 | 1 | 10 | Uckl1os | 0.008 | Young |
| 0.275527 | 1.349506 | 0.01 | 0.002 | 1 | 10 | Pi15 | 0.008 | Young |
| 0.275527 | 1.344664 | 0.01 | 0.002 | 1 | 10 | Rbms3 | 0.008 | Young |
| 0.275527 | 1.296079 | 0.01 | 0.002 | 1 | 10 | Fancb | 0.008 | Young |
| 0.275527 | 1.256691 | 0.01 | 0.002 | 1 | 10 | 1700006J1[illegible] | 0.008 | Young |
| 0.275527 | 1.245039 | 0.01 | 0.002 | 1 | 10 | Pard3b | 0.008 | Young |
| 0.275527 | 1.231621 | 0.01 | 0.002 | 1 | 10 | Amigo3 | 0.008 | Young |
| 0.275527 | 1.220369 | 0.01 | 0.002 | 1 | 10 | 1190005I0[illegible] | 0.008 | Young |
| 0.275527 | 1.212339 | 0.01 | 0.002 | 1 | 10 | Soat2 | 0.008 | Young |
| 0.275527 | 1.20975 | 0.01 | 0.002 | 1 | 10 | Gm15972 | 0.008 | Young |
| 0.275527 | 1.206211 | 0.01 | 0.002 | 1 | 10 | Nusap1 | 0.008 | Young |
| 0.275527 | 1.199746 | 0.01 | 0.002 | 1 | 10 | 4933429H1 | 0.008 | Young |
| 0.275527 | 1.163287 | 0.01 | 0.002 | 1 | 10 | Rbpms2 | 0.008 | Young |
| 0.275527 | 1.146594 | 0.01 | 0.002 | 1 | 10 | Mlxipl | 0.008 | Young |
| 0.275527 | 1.131691 | 0.01 | 0.002 | 1 | 10 | Prrg3 | 0.008 | Young |
| 0.275527 | 1.121677 | 0.01 | 0.002 | 1 | 10 | Gm15326 | 0.008 | Young |
| 0.275527 | 1.118443 | 0.01 | 0.002 | 1 | 10 | Sdsl | 0.008 | Young |
| 0.275527 | 1.100129 | 0.01 | 0.002 | 1 | 10 | Muc1 | 0.008 | Young |
| 0.275527 | 1.099954 | 0.01 | 0.002 | 1 | 10 | Gm29666 | 0.008 | Young |
| 0.275527 | 1.093962 | 0.01 | 0.002 | 1 | 10 | A530010L1 | 0.008 | Young |
| 0.275527 | 1.086271 | 0.01 | 0.002 | 1 | 10 | Sptbn4 | 0.008 | Young |
| 0.275527 | 1.082847 | 0.01 | 0.002 | 1 | 10 | Pdzd2 | 0.008 | Young |
| 0.275527 | 1.071131 | 0.01 | 0.002 | 1 | 10 | Col6a3 | 0.008 | Young |
| 0.275527 | 1.06209 | 0.01 | 0.002 | 1 | 10 | Gm26542 | 0.008 | Young |
| 0.275527 | 1.056261 | 0.01 | 0.002 | 1 | 10 | Ccdc146 | 0.008 | Young |

| | | | | | | | | |
|---|---|---|---|---|---|---|---|---|
| 0.275527 | 1.037378 | 0.01 | 0.002 | 1 | 10 | Gm37768 | 0.008 | Young |
| 0.275527 | 1.009325 | 0.01 | 0.002 | 1 | 10 | Gpr50 | 0.008 | Young |
| 0.275527 | 0.987546 | 0.01 | 0.002 | 1 | 10 | Gm32687 | 0.008 | Young |
| 0.275527 | 0.978664 | 0.01 | 0.002 | 1 | 10 | Iglon5 | 0.008 | Young |
| 0.275527 | 0.97555 | 0.01 | 0.002 | 1 | 10 | B230322F0 | 0.008 | Young |
| 0.275527 | 0.974835 | 0.01 | 0.002 | 1 | 10 | Gucy1a1 | 0.008 | Young |
| 0.275527 | 0.955125 | 0.01 | 0.002 | 1 | 10 | Mpzl3 | 0.008 | Young |
| 0.275527 | 0.951896 | 0.01 | 0.002 | 1 | 10 | Sectm1a | 0.008 | Young |
| 0.275527 | 0.937999 | 0.01 | 0.002 | 1 | 10 | Lamb3 | 0.008 | Young |
| 0.275527 | 0.935433 | 0.01 | 0.002 | 1 | 10 | Aarsd1 | 0.008 | Young |
| 0.275527 | 0.933236 | 0.01 | 0.002 | 1 | 10 | Tmeff2 | 0.008 | Young |
| 0.275527 | 0.905201 | 0.01 | 0.002 | 1 | 10 | Tmem91 | 0.008 | Young |
| 0.275527 | 0.904053 | 0.01 | 0.002 | 1 | 10 | Igsf11 | 0.008 | Young |
| 0.275527 | 0.896398 | 0.01 | 0.002 | 1 | 10 | Ccdc14 | 0.008 | Young |
| 0.275527 | 0.880187 | 0.01 | 0.002 | 1 | 10 | Spata25 | 0.008 | Young |
| 0.275527 | 0.878245 | 0.01 | 0.002 | 1 | 10 | 1700064M | 0.008 | Young |
| 0.275527 | 0.842776 | 0.01 | 0.002 | 1 | 10 | Armcx4 | 0.008 | Young |
| 0.275527 | 0.83599 | 0.01 | 0.002 | 1 | 10 | Gm26609 | 0.008 | Young |
| 0.275527 | 0.83583 | 0.01 | 0.002 | 1 | 10 | Bgn | 0.008 | Young |
| 0.275527 | 0.834224 | 0.01 | 0.002 | 1 | 10 | Gm26637 | 0.008 | Young |
| 0.275527 | 0.833981 | 0.01 | 0.002 | 1 | 10 | Ccdc80 | 0.008 | Young |
| 0.275527 | 0.788607 | 0.01 | 0.002 | 1 | 10 | Mgst2 | 0.008 | Young |
| 0.275527 | 0.773636 | 0.01 | 0.002 | 1 | 10 | Hyls1 | 0.008 | Young |
| 0.275527 | 0.766062 | 0.01 | 0.002 | 1 | 10 | Rapgef4 | 0.008 | Young |
| 0.275527 | 0.752289 | 0.01 | 0.002 | 1 | 10 | Bicdl1 | 0.008 | Young |
| 0.275527 | 0.748174 | 0.01 | 0.002 | 1 | 10 | Trim62 | 0.008 | Young |
| 0.275527 | 0.742051 | 0.01 | 0.002 | 1 | 10 | D230017M | 0.008 | Young |
| 0.275527 | 0.738082 | 0.01 | 0.002 | 1 | 10 | Lrrc75a | 0.008 | Young |
| 0.275527 | 0.729138 | 0.01 | 0.002 | 1 | 10 | Nol3 | 0.008 | Young |
| 0.275527 | 0.707047 | 0.01 | 0.002 | 1 | 10 | Gm11520 | 0.008 | Young |
| 0.275527 | 0.704136 | 0.01 | 0.002 | 1 | 10 | Tchh | 0.008 | Young |
| 0.275527 | 0.675254 | 0.01 | 0.002 | 1 | 10 | Hist1h2ab | 0.008 | Young |
| 0.275527 | 0.672209 | 0.01 | 0.002 | 1 | 10 | Col7a1 | 0.008 | Young |
| 0.275527 | 0.669713 | 0.01 | 0.002 | 1 | 10 | Gm31243 | 0.008 | Young |
| 0.275527 | 0.659972 | 0.01 | 0.002 | 1 | 10 | Cdca2 | 0.008 | Young |
| 0.275527 | 0.618239 | 0.01 | 0.002 | 1 | 10 | Hist1h2bj | 0.008 | Young |
| 0.275527 | 0.608079 | 0.01 | 0.002 | 1 | 10 | Zbed5 | 0.008 | Young |
| 0.275527 | 0.608079 | 0.01 | 0.002 | 1 | 10 | 6720489N1 | 0.008 | Young |
| 0.275527 | 0.583164 | 0.01 | 0.002 | 1 | 10 | Mpzl1 | 0.008 | Young |
| 0.275527 | 0.538125 | 0.01 | 0.002 | 1 | 10 | Cplx2 | 0.008 | Young |
| 0.275527 | 0.508195 | 0.01 | 0.002 | 1 | 10 | Itm2a | 0.008 | Young |
| 0.275863 | 0.076404 | 0.172 | 0.232 | 1 | 10 | Tpd52l2 | -0.06 | Young |
| 0.275895 | 0.44127 | 0.192 | 0.152 | 1 | 10 | Gpr160 | 0.04 | Young |
| 0.276387 | 0.901569 | 0.061 | 0.037 | 1 | 10 | Slc12a2 | 0.024 | Young |
| 0.276524 | 0.240034 | 0.101 | 0.147 | 1 | 10 | Prps1 | -0.046 | Young |

| | | | | | | | | |
|---|---|---|---|---|---|---|---|---|
| 0.276634 | 0.342182 | 0.727 | 0.733 | 1 | 10 | C5ar1 | -0.006 | Young |
| 0.27666 | 0.261637 | 0.293 | 0.24 | 1 | 10 | Batf3 | 0.053 | Young |
| 0.276705 | 0.365199 | 0.293 | 0.242 | 1 | 10 | Pqlc1 | 0.051 | Young |
| 0.27683 | 0.950361 | 0.111 | 0.078 | 1 | 10 | Tstd2 | 0.033 | Young |
| 0.277796 | 2.15465 | 0.03 | 0.015 | 1 | 10 | Grasp | 0.015 | Young |
| 0.278 | 0.740021 | 0.162 | 0.125 | 1 | 10 | Aig1 | 0.037 | Young |
| 0.278636 | 0.980721 | 0.01 | 0.002 | 1 | 10 | Nat14 | 0.008 | Young |
| 0.278636 | 0.978649 | 0.01 | 0.002 | 1 | 10 | Gm32200 | 0.008 | Young |
| 0.278636 | 0.859289 | 0.01 | 0.002 | 1 | 10 | Tssk6 | 0.008 | Young |
| 0.278636 | 0.857226 | 0.01 | 0.002 | 1 | 10 | Lrrc39 | 0.008 | Young |
| 0.278636 | 0.82422 | 0.01 | 0.002 | 1 | 10 | Zfp37 | 0.008 | Young |
| 0.278636 | 0.810248 | 0.01 | 0.002 | 1 | 10 | Thsd4 | 0.008 | Young |
| 0.278636 | 0.788456 | 0.01 | 0.002 | 1 | 10 | 1700018L0 | 0.008 | Young |
| 0.278636 | 0.774759 | 0.01 | 0.002 | 1 | 10 | AK157302 | 0.008 | Young |
| 0.278636 | 0.772211 | 0.01 | 0.002 | 1 | 10 | Rab42 | 0.008 | Young |
| 0.278636 | 0.751917 | 0.01 | 0.002 | 1 | 10 | Gm6793 | 0.008 | Young |
| 0.278636 | 0.732287 | 0.01 | 0.002 | 1 | 10 | Mxra8 | 0.008 | Young |
| 0.278636 | 0.730972 | 0.01 | 0.002 | 1 | 10 | Srcin1 | 0.008 | Young |
| 0.278636 | 0.728485 | 0.01 | 0.002 | 1 | 10 | Vmn2r29 | 0.008 | Young |
| 0.278636 | 0.717887 | 0.01 | 0.002 | 1 | 10 | Hspa1l | 0.008 | Young |
| 0.278636 | 0.707736 | 0.01 | 0.002 | 1 | 10 | Gm13830 | 0.008 | Young |
| 0.278636 | 0.703002 | 0.01 | 0.002 | 1 | 10 | Gm16196 | 0.008 | Young |
| 0.278636 | 0.678287 | 0.01 | 0.002 | 1 | 10 | Il1r1 | 0.008 | Young |
| 0.278636 | 0.670596 | 0.01 | 0.002 | 1 | 10 | Comp | 0.008 | Young |
| 0.278636 | 0.669925 | 0.01 | 0.002 | 1 | 10 | Gm16486 | 0.008 | Young |
| 0.278636 | 0.639069 | 0.01 | 0.002 | 1 | 10 | Gfod2 | 0.008 | Young |
| 0.278636 | 0.612193 | 0.01 | 0.002 | 1 | 10 | Hrh2 | 0.008 | Young |
| 0.278636 | 0.609169 | 0.01 | 0.002 | 1 | 10 | Gm20498 | 0.008 | Young |
| 0.278636 | 0.608639 | 0.01 | 0.002 | 1 | 10 | Cygb | 0.008 | Young |
| 0.278636 | 0.588029 | 0.01 | 0.002 | 1 | 10 | Mansc1 | 0.008 | Young |
| 0.278636 | 0.586988 | 0.01 | 0.002 | 1 | 10 | Slc1a4 | 0.008 | Young |
| 0.278636 | 0.583271 | 0.01 | 0.002 | 1 | 10 | Dact1 | 0.008 | Young |
| 0.278636 | 0.563497 | 0.01 | 0.002 | 1 | 10 | Dbn1 | 0.008 | Young |
| 0.278636 | 0.5565 | 0.01 | 0.002 | 1 | 10 | Zfp11 | 0.008 | Young |
| 0.278636 | 0.555547 | 0.01 | 0.002 | 1 | 10 | Edil3 | 0.008 | Young |
| 0.278636 | 0.535302 | 0.01 | 0.002 | 1 | 10 | Cenps | 0.008 | Young |
| 0.278636 | 0.533976 | 0.01 | 0.002 | 1 | 10 | Nup210l | 0.008 | Young |
| 0.278636 | 0.533072 | 0.01 | 0.002 | 1 | 10 | Gm17509 | 0.008 | Young |
| 0.278636 | 0.525232 | 0.01 | 0.002 | 1 | 10 | A430057M | 0.008 | Young |
| 0.278636 | 0.521627 | 0.01 | 0.002 | 1 | 10 | Ankrd34a | 0.008 | Young |
| 0.278636 | 0.520908 | 0.01 | 0.002 | 1 | 10 | Gm13470 | 0.008 | Young |
| 0.278636 | 0.520753 | 0.01 | 0.002 | 1 | 10 | M1ap | 0.008 | Young |
| 0.278636 | 0.519089 | 0.01 | 0.002 | 1 | 10 | Gm9949 | 0.008 | Young |
| 0.278636 | 0.507786 | 0.01 | 0.002 | 1 | 10 | Gm27168 | 0.008 | Young |
| 0.278636 | 0.507532 | 0.01 | 0.002 | 1 | 10 | Hey1 | 0.008 | Young |

| | | | | | | | | |
|---|---|---|---|---|---|---|---|---|
| 0.278636 | 0.505598 | 0.01 | 0.002 | 1 | 10 | Tspan7 | 0.008 | Young |
| 0.278636 | 0.493618 | 0.01 | 0.002 | 1 | 10 | Xntrpc | 0.008 | Young |
| 0.278636 | 0.483312 | 0.01 | 0.002 | 1 | 10 | Gpr4 | 0.008 | Young |
| 0.278636 | 0.479855 | 0.01 | 0.002 | 1 | 10 | B430305J0 | 0.008 | Young |
| 0.278636 | 0.479014 | 0.01 | 0.002 | 1 | 10 | Serac1 | 0.008 | Young |
| 0.278636 | 0.463742 | 0.01 | 0.002 | 1 | 10 | Gng10 | 0.008 | Young |
| 0.278636 | 0.456458 | 0.01 | 0.002 | 1 | 10 | Stard6 | 0.008 | Young |
| 0.278636 | 0.45099 | 0.01 | 0.002 | 1 | 10 | Zfp784 | 0.008 | Young |
| 0.278636 | 0.439843 | 0.01 | 0.002 | 1 | 10 | 1110020A2 | 0.008 | Young |
| 0.278636 | 0.439714 | 0.01 | 0.002 | 1 | 10 | Cyyr1 | 0.008 | Young |
| 0.278636 | 0.438167 | 0.01 | 0.002 | 1 | 10 | Serpinb1b | 0.008 | Young |
| 0.278636 | 0.434067 | 0.01 | 0.002 | 1 | 10 | Gm15964 | 0.008 | Young |
| 0.278636 | 0.421852 | 0.01 | 0.002 | 1 | 10 | Epop | 0.008 | Young |
| 0.278636 | 0.418856 | 0.01 | 0.002 | 1 | 10 | Phactr1 | 0.008 | Young |
| 0.278636 | 0.406495 | 0.01 | 0.002 | 1 | 10 | Galnt14 | 0.008 | Young |
| 0.278636 | 0.393227 | 0.01 | 0.002 | 1 | 10 | Kazald1 | 0.008 | Young |
| 0.278636 | 0.379282 | 0.01 | 0.002 | 1 | 10 | Fzd8 | 0.008 | Young |
| 0.278636 | 0.355658 | 0.01 | 0.002 | 1 | 10 | Gas2 | 0.008 | Young |
| 0.278636 | 0.35512 | 0.01 | 0.002 | 1 | 10 | 4930537H2 | 0.008 | Young |
| 0.278636 | 0.344188 | 0.01 | 0.002 | 1 | 10 | Chst3 | 0.008 | Young |
| 0.278636 | 0.340773 | 0.01 | 0.002 | 1 | 10 | Tcaf1 | 0.008 | Young |
| 0.278636 | 0.327497 | 0.01 | 0.002 | 1 | 10 | Sox12 | 0.008 | Young |
| 0.278636 | 0.327385 | 0.01 | 0.002 | 1 | 10 | Vpreb1 | 0.008 | Young |
| 0.278636 | 0.321615 | 0.01 | 0.002 | 1 | 10 | Hdc | 0.008 | Young |
| 0.278636 | 0.314 | 0.01 | 0.002 | 1 | 10 | Hist1h3a | 0.008 | Young |
| 0.278636 | 0.30598 | 0.01 | 0.002 | 1 | 10 | Rab44 | 0.008 | Young |
| 0.278636 | 0.297304 | 0.01 | 0.002 | 1 | 10 | Syt7 | 0.008 | Young |
| 0.278636 | 0.284507 | 0.01 | 0.002 | 1 | 10 | Prc1 | 0.008 | Young |
| 0.278636 | 0.274843 | 0.01 | 0.002 | 1 | 10 | Sntb1 | 0.008 | Young |
| 0.278636 | 0.261779 | 0.01 | 0.002 | 1 | 10 | Pbk | 0.008 | Young |
| 0.278636 | 0.2596 | 0.01 | 0.002 | 1 | 10 | Rasgrf1 | 0.008 | Young |
| 0.278636 | 0.245599 | 0.01 | 0.002 | 1 | 10 | Lrrc9 | 0.008 | Young |
| 0.278636 | 0.242272 | 0.01 | 0.002 | 1 | 10 | Gm13166 | 0.008 | Young |
| 0.278636 | 0.238461 | 0.01 | 0.002 | 1 | 10 | Fxyd1 | 0.008 | Young |
| 0.278636 | 0.234197 | 0.01 | 0.002 | 1 | 10 | Caly | 0.008 | Young |
| 0.278636 | 0.228871 | 0.01 | 0.002 | 1 | 10 | K230015D0 | 0.008 | Young |
| 0.278636 | 0.217415 | 0.01 | 0.002 | 1 | 10 | 2410004P0 | 0.008 | Young |
| 0.278636 | 0.20883 | 0.01 | 0.002 | 1 | 10 | AA543186 | 0.008 | Young |
| 0.278636 | 0.199177 | 0.01 | 0.002 | 1 | 10 | Prdm16 | 0.008 | Young |
| 0.278636 | 0.192471 | 0.01 | 0.002 | 1 | 10 | Mpz | 0.008 | Young |
| 0.278636 | 0.181484 | 0.01 | 0.002 | 1 | 10 | Tal2 | 0.008 | Young |
| 0.278636 | 0.176631 | 0.01 | 0.002 | 1 | 10 | Enpp6 | 0.008 | Young |
| 0.278636 | 0.167196 | 0.01 | 0.002 | 1 | 10 | Tlcd1 | 0.008 | Young |
| 0.278636 | 0.155777 | 0.01 | 0.002 | 1 | 10 | Gm10762 | 0.008 | Young |
| 0.278636 | 0.146433 | 0.01 | 0.002 | 1 | 10 | Rtkn2 | 0.008 | Young |

| | | | | | | | | |
|---|---|---|---|---|---|---|---|---|
| 0.278636 | 0.129571 | 0.01 | 0.002 | 1 | 10 | Peli3 | 0.008 | Young |
| 0.278636 | 0.127453 | 0.01 | 0.002 | 1 | 10 | Celsr3 | 0.008 | Young |
| 0.278636 | 0.060345 | 0.01 | 0.002 | 1 | 10 | Klhl13 | 0.008 | Young |
| 0.278636 | 0.05726 | 0.01 | 0.002 | 1 | 10 | Ccdc8 | 0.008 | Young |
| 0.278636 | 0.042769 | 0.01 | 0.002 | 1 | 10 | Zfp30 | 0.008 | Young |
| 0.278655 | 0.680372 | 0.414 | 0.416 | 1 | 10 | Msrb1 | -0.002 | Young |
| 0.279099 | 0.870249 | 0.081 | 0.054 | 1 | 10 | Pola1 | 0.027 | Young |
| 0.27912 | 0.561258 | 0.253 | 0.213 | 1 | 10 | Dok1 | 0.04 | Young |
| 0.279326 | 0.530489 | 0.061 | 0.037 | 1 | 10 | Cd300lg | 0.024 | Young |
| 0.279911 | 0.041704 | 0.101 | 0.147 | 1 | 10 | Kpna6 | -0.046 | Young |
| 0.280343 | 0.272499 | 0.404 | 0.352 | 1 | 10 | Pou2f2 | 0.052 | Young |
| 0.281021 | 0.053147 | 0.212 | 0.281 | 1 | 10 | Bcap29 | -0.069 | Young |
| 0.281647 | 0.150861 | 0.152 | 0.208 | 1 | 10 | Ehmt1 | -0.056 | Young |
| 0.282012 | 0.256663 | 0.192 | 0.264 | 1 | 10 | Pbxip1 | -0.072 | Young |
| 0.282255 | 2.189079 | 0.03 | 0.015 | 1 | 10 | Slc2a3 | 0.015 | Young |
| 0.282255 | 1.276029 | 0.03 | 0.015 | 1 | 10 | Trp53rkb | 0.015 | Young |
| 0.282338 | 0.059621 | 0.051 | 0.086 | 1 | 10 | Edc3 | -0.035 | Young |
| 0.282414 | 0.372797 | 0.566 | 0.597 | 1 | 10 | Anxa3 | -0.031 | Young |
| 0.282431 | 1.176098 | 0.081 | 0.054 | 1 | 10 | Hmgxb4 | 0.027 | Young |
| 0.282668 | 0.448823 | 0.313 | 0.271 | 1 | 10 | Sec23b | 0.042 | Young |
| 0.283058 | 0.420033 | 0.212 | 0.166 | 1 | 10 | Pex13 | 0.046 | Young |
| 0.283206 | 0.622782 | 0.162 | 0.125 | 1 | 10 | Tbc1d9 | 0.037 | Young |
| 0.283245 | 0.051082 | 0.303 | 0.384 | 1 | 10 | Yeats4 | -0.081 | Young |
| 0.283404 | 1.139289 | 0.051 | 0.029 | 1 | 10 | Cpne8 | 0.022 | Young |
| 0.283404 | 0.834063 | 0.051 | 0.029 | 1 | 10 | Ccdc181 | 0.022 | Young |
| 0.283753 | 1.565265 | 0.03 | 0.015 | 1 | 10 | Depdc7 | 0.015 | Young |
| 0.283753 | 1.336352 | 0.03 | 0.015 | 1 | 10 | Hr | 0.015 | Young |
| 0.28385 | 0.566009 | 0.03 | 0.059 | 1 | 10 | Atat1 | -0.029 | Young |
| 0.284006 | 0.566271 | 0.374 | 0.337 | 1 | 10 | Tmf1 | 0.037 | Young |
| 0.284207 | 0.464677 | 0.182 | 0.139 | 1 | 10 | Ttc1 | 0.043 | Young |
| 0.284256 | 0.045696 | 0.091 | 0.134 | 1 | 10 | Ermard | -0.043 | Young |
| 0.285129 | 0.600318 | 0.131 | 0.186 | 1 | 10 | Atg12 | -0.055 | Young |
| 0.285255 | 1.241513 | 0.03 | 0.015 | 1 | 10 | Plekhh3 | 0.015 | Young |
| 0.285255 | 1.130973 | 0.03 | 0.015 | 1 | 10 | D130040H: | 0.015 | Young |
| 0.285255 | 0.922056 | 0.03 | 0.015 | 1 | 10 | Cenpw | 0.015 | Young |
| 0.285343 | 1.549959 | 0.04 | 0.022 | 1 | 10 | Mars2 | 0.018 | Young |
| 0.285389 | 0.536107 | 0.172 | 0.132 | 1 | 10 | Ddx49 | 0.04 | Young |
| 0.285608 | 0.904902 | 0.051 | 0.029 | 1 | 10 | Gypc | 0.022 | Young |
| 0.285764 | 0.798672 | 0.091 | 0.061 | 1 | 10 | Golga1 | 0.03 | Young |
| 0.285943 | 1.120586 | 0.091 | 0.064 | 1 | 10 | Fam110b | 0.027 | Young |
| 0.286484 | 0.319413 | 0.687 | 0.65 | 1 | 10 | Cnbp | 0.037 | Young |
| 0.286763 | 1.644169 | 0.03 | 0.015 | 1 | 10 | L1cam | 0.015 | Young |
| 0.286763 | 1.310513 | 0.03 | 0.015 | 1 | 10 | Pde4d | 0.015 | Young |
| 0.286763 | 1.294194 | 0.03 | 0.015 | 1 | 10 | Cd28 | 0.015 | Young |
| 0.286768 | 0.064576 | 0.101 | 0.147 | 1 | 10 | Zfp280d | -0.046 | Young |

| | | | | | | | | |
|---|---|---|---|---|---|---|---|---|
| 0.286824 | 0.362839 | 0.162 | 0.122 | 1 | 10 | E2f4 | 0.04 | Young |
| 0.28724 | 0.73497 | 0.232 | 0.191 | 1 | 10 | Pofut1 | 0.041 | Young |
| 0.287479 | 0.932309 | 0.081 | 0.054 | 1 | 10 | Siglece | 0.027 | Young |
| 0.287824 | 0.718502 | 0.051 | 0.029 | 1 | 10 | Zfp821 | 0.022 | Young |
| 0.287867 | 1.565042 | 0.04 | 0.022 | 1 | 10 | Taf1a | 0.018 | Young |
| 0.288137 | 1.083203 | 0.162 | 0.13 | 1 | 10 | Map4k3 | 0.032 | Young |
| 0.288173 | 0.64895 | 0.242 | 0.198 | 1 | 10 | Cklf | 0.044 | Young |
| 0.288277 | 1.2518 | 0.03 | 0.015 | 1 | 10 | Dus4l | 0.015 | Young |
| 0.289094 | 1.23638 | 0.091 | 0.064 | 1 | 10 | Paqr7 | 0.027 | Young |
| 0.289134 | 1.257271 | 0.04 | 0.022 | 1 | 10 | Arap2 | 0.018 | Young |
| 0.289134 | 0.337476 | 0.04 | 0.022 | 1 | 10 | Lins1 | 0.018 | Young |
| 0.289796 | 1.345799 | 0.03 | 0.015 | 1 | 10 | Stbd1 | 0.015 | Young |
| 0.289796 | 1.25541 | 0.03 | 0.015 | 1 | 10 | Gpr84 | 0.015 | Young |
| 0.289796 | 1.248607 | 0.03 | 0.015 | 1 | 10 | Col4a1 | 0.015 | Young |
| 0.289796 | 0.8482 | 0.03 | 0.015 | 1 | 10 | Tbc1d19 | 0.015 | Young |
| 0.290204 | 0.450658 | 0.535 | 0.526 | 1 | 10 | Zfp91 | 0.009 | Young |
| 0.290372 | 0.132035 | 0.667 | 0.645 | 1 | 10 | Aif1 | 0.022 | Young |
| 0.291169 | 0.701811 | 0.051 | 0.029 | 1 | 10 | Cyb5d2 | 0.022 | Young |
| 0.29132 | 1.121019 | 0.03 | 0.015 | 1 | 10 | Phldb3 | 0.015 | Young |
| 0.29132 | 0.784394 | 0.03 | 0.015 | 1 | 10 | Zfp975 | 0.015 | Young |
| 0.29132 | 0.781608 | 0.03 | 0.015 | 1 | 10 | 4933406I1 | 0.015 | Young |
| 0.29132 | 0.741287 | 0.03 | 0.015 | 1 | 10 | Aurka | 0.015 | Young |
| 0.291679 | 1.079659 | 0.04 | 0.022 | 1 | 10 | Tpx2 | 0.018 | Young |
| 0.291888 | 0.05406 | 0.303 | 0.381 | 1 | 10 | Jtb | -0.078 | Young |
| 0.292162 | 0.561522 | 0.212 | 0.176 | 1 | 10 | Stx6 | 0.036 | Young |
| 0.292201 | 0.163302 | 0.394 | 0.335 | 1 | 10 | Gm10073 | 0.059 | Young |
| 0.29229 | 0.335882 | 0.051 | 0.029 | 1 | 10 | Zfp777 | 0.022 | Young |
| 0.292785 | 0.502522 | 0.505 | 0.511 | 1 | 10 | Ube2b | -0.006 | Young |
| 0.292849 | 1.00696 | 0.03 | 0.015 | 1 | 10 | Ift88 | 0.015 | Young |
| 0.292958 | 0.88386 | 0.04 | 0.022 | 1 | 10 | Armcx5 | 0.018 | Young |
| 0.29328 | 0.973221 | 0.152 | 0.12 | 1 | 10 | Acot7 | 0.032 | Young |
| 0.293858 | 0.166679 | 0.02 | 0.044 | 1 | 10 | Lzts2 | -0.024 | Young |
| 0.29424 | 1.254337 | 0.04 | 0.022 | 1 | 10 | F3 | 0.018 | Young |
| 0.29434 | 0.355428 | 0.061 | 0.037 | 1 | 10 | Zbtb11os1 | 0.024 | Young |
| 0.294384 | 0.540738 | 0.03 | 0.015 | 1 | 10 | C130013H( | 0.015 | Young |
| 0.29567 | 0.547923 | 0.051 | 0.029 | 1 | 10 | Exoc6 | 0.022 | Young |
| 0.296093 | 0.255009 | 0.091 | 0.132 | 1 | 10 | Mrps11 | -0.041 | Young |
| 0.296146 | 0.556306 | 0.192 | 0.154 | 1 | 10 | Slc39a6 | 0.038 | Young |
| 0.296161 | 0.627124 | 0.101 | 0.071 | 1 | 10 | Scn1b | 0.03 | Young |
| 0.296171 | 0.898367 | 0.172 | 0.139 | 1 | 10 | Ccdc127 | 0.033 | Young |
| 0.296192 | 0.370339 | 0.424 | 0.396 | 1 | 10 | Crlf2 | 0.028 | Young |
| 0.296445 | 0.426603 | 0.192 | 0.152 | 1 | 10 | Cnpy3 | 0.04 | Young |
| 0.296496 | 0.344861 | 0.364 | 0.325 | 1 | 10 | Lman1 | 0.039 | Young |
| 0.296765 | 0.642693 | 0.222 | 0.188 | 1 | 10 | Snrpa1 | 0.034 | Young |
| 0.296802 | 0.675112 | 0.051 | 0.029 | 1 | 10 | Zfp809 | 0.022 | Young |

| | | | | | | | | |
|---|---|---|---|---|---|---|---|---|
| 0.29681 | 0.756703 | 0.162 | 0.125 | 1 | 10 | Zdhhc18 | 0.037 | Young |
| 0.296816 | 0.968501 | 0.04 | 0.022 | 1 | 10 | Nicn1 | 0.018 | Young |
| 0.296816 | 0.749697 | 0.04 | 0.022 | 1 | 10 | Zfp503 | 0.018 | Young |
| 0.29686 | 0.002588 | 0.354 | 0.44 | 1 | 10 | Pdcd6 | -0.086 | Young |
| 0.296909 | 0.66588 | 0.263 | 0.225 | 1 | 10 | Psen1 | 0.038 | Young |
| 0.297121 | 0.101049 | 0.152 | 0.213 | 1 | 10 | Babam1 | -0.061 | Young |
| 0.29747 | 0.569015 | 0.03 | 0.015 | 1 | 10 | Kif20b | 0.015 | Young |
| 0.29747 | 0.331563 | 0.03 | 0.015 | 1 | 10 | Poc1a | 0.015 | Young |
| 0.297878 | 0.868493 | 0.091 | 0.064 | 1 | 10 | Thap7 | 0.027 | Young |
| 0.298109 | 0.690079 | 0.04 | 0.022 | 1 | 10 | Mfsd3 | 0.018 | Young |
| 0.298127 | 0.735451 | 0.242 | 0.205 | 1 | 10 | Dusp6 | 0.037 | Young |
| 0.298212 | 0.614548 | 0.343 | 0.308 | 1 | 10 | Abca9 | 0.035 | Young |
| 0.298419 | 0.061034 | 0.465 | 0.594 | 1 | 10 | Tmed9 | -0.129 | Young |
| 0.298447 | 0.238959 | 0.101 | 0.147 | 1 | 10 | Slc5a3 | -0.046 | Young |
| 0.298781 | 0.192813 | 0.273 | 0.357 | 1 | 10 | Zc3h13 | -0.084 | Young |
| 0.299159 | 0.342535 | 0.606 | 0.56 | 1 | 10 | Tardbp | 0.046 | Young |
| 0.29959 | 0.334729 | 0.202 | 0.156 | 1 | 10 | Shc1 | 0.046 | Young |
| 0.300024 | 0.464899 | 0.354 | 0.318 | 1 | 10 | Cyb5a | 0.036 | Young |
| 0.30049 | 0.699512 | 0.232 | 0.196 | 1 | 10 | Gsr | 0.036 | Young |
| 0.300737 | 0.177742 | 0.717 | 0.719 | 1 | 10 | Rab14 | -0.002 | Young |
| 0.301151 | 0.177163 | 0.232 | 0.303 | 1 | 10 | Akirin1 | -0.071 | Young |
| 0.301279 | 0.363093 | 0.566 | 0.555 | 1 | 10 | Snrpf | 0.011 | Young |
| 0.301753 | 0.483918 | 0.313 | 0.276 | 1 | 10 | Atxn10 | 0.037 | Young |
| 0.302012 | 0.657302 | 0.04 | 0.022 | 1 | 10 | Tti1 | 0.018 | Young |
| 0.302133 | 0.401969 | 0.384 | 0.35 | 1 | 10 | Aldh9a1 | 0.034 | Young |
| 0.302508 | 0.47612 | 0.051 | 0.029 | 1 | 10 | Rtn4rl1 | 0.022 | Young |
| 0.302987 | 0.553935 | 0.081 | 0.054 | 1 | 10 | Ccdc28b | 0.027 | Young |
| 0.303089 | 0.708712 | 0.323 | 0.303 | 1 | 10 | Pacsin2 | 0.02 | Young |
| 0.303138 | 0.230284 | 0.485 | 0.438 | 1 | 10 | Aebp2 | 0.047 | Young |
| 0.303248 | 0.15169 | 0.697 | 0.665 | 1 | 10 | Mrpl52 | 0.032 | Young |
| 0.30356 | 1.547339 | 0.091 | 0.064 | 1 | 10 | Traf2 | 0.027 | Young |
| 0.303586 | 0.629163 | 0.222 | 0.186 | 1 | 10 | Zswim8 | 0.036 | Young |
| 0.303628 | 0.075772 | 0.434 | 0.553 | 1 | 10 | Nus1 | -0.119 | Young |
| 0.303658 | 0.317334 | 0.051 | 0.029 | 1 | 10 | Rdh10 | 0.022 | Young |
| 0.304097 | 0.37447 | 0.313 | 0.269 | 1 | 10 | Arrb1 | 0.044 | Young |
| 0.304626 | 0.182124 | 0.879 | 0.841 | 1 | 10 | Rps27l | 0.038 | Young |
| 0.304744 | 0.757891 | 0.081 | 0.054 | 1 | 10 | Wipi1 | 0.027 | Young |
| 0.304932 | 0.038307 | 0.192 | 0.244 | 1 | 10 | Snapin | -0.052 | Young |
| 0.305031 | 0.660182 | 0.162 | 0.127 | 1 | 10 | AW549877 | 0.035 | Young |
| 0.306402 | 0.607478 | 0.293 | 0.267 | 1 | 10 | BC017643 | 0.026 | Young |
| 0.307486 | 1.033695 | 0.152 | 0.12 | 1 | 10 | Bmyc | 0.032 | Young |
| 0.308189 | 0.025277 | 0.162 | 0.215 | 1 | 10 | Ltn1 | -0.053 | Young |
| 0.308592 | 0.340603 | 0.04 | 0.022 | 1 | 10 | Mgll | 0.018 | Young |
| 0.309146 | 0.446579 | 0.424 | 0.399 | 1 | 10 | Tmem234 | 0.025 | Young |
| 0.309344 | 0.57193 | 0.101 | 0.071 | 1 | 10 | 2310039H( | 0.03 | Young |

| | | | | | | | | |
|---|---|---|---|---|---|---|---|---|
| 0.309487 | 0.334133 | 0.162 | 0.12 | 1 | 10 | Smarca2 | 0.042 | Young |
| 0.309902 | 0.057817 | 0.303 | 0.386 | 1 | 10 | Brd1 | -0.083 | Young |
| 0.310039 | 0.061787 | 0.323 | 0.394 | 1 | 10 | Sec31a | -0.071 | Young |
| 0.310058 | 1.485135 | 0.071 | 0.046 | 1 | 10 | Pias3 | 0.025 | Young |
| 0.310634 | 0.975786 | 0.152 | 0.12 | 1 | 10 | Pctp | 0.032 | Young |
| 0.310996 | 0.774787 | 0.192 | 0.159 | 1 | 10 | Ppme1 | 0.033 | Young |
| 0.31107 | 0.585761 | 0.263 | 0.23 | 1 | 10 | Phf5a | 0.033 | Young |
| 0.31175 | 0.411179 | 0.535 | 0.553 | 1 | 10 | Scamp2 | -0.018 | Young |
| 0.312265 | 1.06291 | 0.162 | 0.132 | 1 | 10 | Polr2j | 0.03 | Young |
| 0.312404 | 0.257843 | 0.374 | 0.328 | 1 | 10 | Fam105a | 0.046 | Young |
| 0.312701 | 0.184763 | 0.131 | 0.095 | 1 | 10 | Zfp729a | 0.036 | Young |
| 0.312814 | 0.513344 | 0.222 | 0.181 | 1 | 10 | Dcaf13 | 0.041 | Young |
| 0.312924 | 1.220064 | 0.071 | 0.046 | 1 | 10 | Patz1 | 0.025 | Young |
| 0.312929 | 0.337847 | 0.172 | 0.134 | 1 | 10 | Pgs1 | 0.038 | Young |
| 0.313787 | 0.661958 | 0.111 | 0.081 | 1 | 10 | Exosc7 | 0.03 | Young |
| 0.313852 | 1.021474 | 0.061 | 0.039 | 1 | 10 | Pdik1l | 0.022 | Young |
| 0.314017 | 0.462982 | 0.232 | 0.188 | 1 | 10 | Cdc37l1 | 0.044 | Young |
| 0.314527 | 0.416608 | 0.081 | 0.054 | 1 | 10 | Parp10 | 0.027 | Young |
| 0.315138 | 0.814225 | 0.091 | 0.064 | 1 | 10 | Trpv4 | 0.027 | Young |
| 0.315201 | 0.560373 | 0.313 | 0.279 | 1 | 10 | Rheb | 0.034 | Young |
| 0.315342 | 0.893223 | 0.121 | 0.09 | 1 | 10 | B3galnt2 | 0.031 | Young |
| 0.315942 | 0.351589 | 0.303 | 0.264 | 1 | 10 | Stk10 | 0.039 | Young |
| 0.315976 | 0.502036 | 0.091 | 0.064 | 1 | 10 | Hacd2 | 0.027 | Young |
| 0.31614 | 0.298439 | 0.162 | 0.122 | 1 | 10 | Wwc2 | 0.04 | Young |
| 0.316446 | 1.004359 | 0.232 | 0.191 | 1 | 10 | Gclc | 0.041 | Young |
| 0.317259 | 0.247382 | 0.788 | 0.782 | 1 | 10 | Ncl | 0.006 | Young |
| 0.317574 | 0.541126 | 0.899 | 0.927 | 1 | 10 | Apoe | -0.028 | Young |
| 0.317863 | 0.427447 | 0.384 | 0.357 | 1 | 10 | Emb | 0.027 | Young |
| 0.317885 | 0.723914 | 0.192 | 0.161 | 1 | 10 | Timm8a1 | 0.031 | Young |
| 0.317983 | 0.042386 | 0.293 | 0.364 | 1 | 10 | AI314180 | -0.071 | Young |
| 0.318191 | 0.141093 | 0.838 | 0.792 | 1 | 10 | Atpif1 | 0.046 | Young |
| 0.318226 | 0.362377 | 0.202 | 0.161 | 1 | 10 | Fbxl20 | 0.041 | Young |
| 0.31856 | 0.060771 | 0.303 | 0.381 | 1 | 10 | Dhx9 | -0.078 | Young |
| 0.318826 | 0.569349 | 0.253 | 0.215 | 1 | 10 | Rybp | 0.038 | Young |
| 0.319466 | 0.82584 | 0.172 | 0.139 | 1 | 10 | Virma | 0.033 | Young |
| 0.320074 | 0.814442 | 0.162 | 0.13 | 1 | 10 | Ccl12 | 0.032 | Young |
| 0.320131 | 1.411806 | 0.061 | 0.039 | 1 | 10 | Trim39 | 0.022 | Young |
| 0.320548 | 0.038041 | 0.222 | 0.279 | 1 | 10 | Sft2d2 | -0.057 | Young |
| 0.320781 | 0.972587 | 0.192 | 0.164 | 1 | 10 | Syngr1 | 0.028 | Young |
| 0.320819 | 0.21196 | 0.616 | 0.599 | 1 | 10 | Uqcr10 | 0.017 | Young |
| 0.321626 | 0.041316 | 0.03 | 0.056 | 1 | 10 | Rfesd | -0.026 | Young |
| 0.321834 | 0.143269 | 0.384 | 0.328 | 1 | 10 | Rnf150 | 0.056 | Young |
| 0.321883 | 0.886316 | 0.091 | 0.064 | 1 | 10 | Itpr1 | 0.027 | Young |
| 0.321992 | 0.034276 | 0.051 | 0.081 | 1 | 10 | Nfrkb | -0.03 | Young |
| 0.323264 | 0.840514 | 0.111 | 0.083 | 1 | 10 | Ppih | 0.028 | Young |

| | | | | | | | | |
|---|---|---|---|---|---|---|---|---|
| 0.3233 | 1.291539 | 0.061 | 0.039 | 1 | 10 | Mtx3 | 0.022 | Young |
| 0.323719 | 0.423918 | 0.101 | 0.071 | 1 | 10 | AI606181 | 0.03 | Young |
| 0.324396 | 0.094195 | 0.182 | 0.24 | 1 | 10 | Atxn7l1 | -0.058 | Young |
| 0.324436 | 0.407381 | 0.566 | 0.555 | 1 | 10 | Sec62 | 0.011 | Young |
| 0.324623 | 0.872726 | 0.141 | 0.112 | 1 | 10 | Nxpe5 | 0.029 | Young |
| 0.324784 | 0.101333 | 0.061 | 0.095 | 1 | 10 | Cmbl | -0.034 | Young |
| 0.325357 | 0.122666 | 0.131 | 0.178 | 1 | 10 | Lrrc41 | -0.047 | Young |
| 0.325418 | 0.939251 | 0.141 | 0.11 | 1 | 10 | Sbf1 | 0.031 | Young |
| 0.325607 | 0.070619 | 0.212 | 0.274 | 1 | 10 | Sertad1 | -0.062 | Young |
| 0.325749 | 0.252814 | 0.596 | 0.577 | 1 | 10 | Ppp1r14b | 0.019 | Young |
| 0.325791 | 0.38803 | 0.121 | 0.088 | 1 | 10 | Akip1 | 0.033 | Young |
| 0.326922 | 0.256139 | 0.586 | 0.579 | 1 | 10 | Tra2b | 0.007 | Young |
| 0.326995 | 0.543351 | 0.394 | 0.364 | 1 | 10 | Dctn3 | 0.03 | Young |
| 0.327076 | 0.694751 | 0.182 | 0.149 | 1 | 10 | Ubl4a | 0.033 | Young |
| 0.327211 | 0.089913 | 0.141 | 0.186 | 1 | 10 | Vps37a | -0.045 | Young |
| 0.327303 | 0.060247 | 0.02 | 0.042 | 1 | 10 | C330018D2 | -0.022 | Young |
| 0.327344 | 0.688579 | 0.152 | 0.12 | 1 | 10 | G6pc3 | 0.032 | Young |
| 0.327402 | 0.407467 | 0.212 | 0.171 | 1 | 10 | Sec23ip | 0.041 | Young |
| 0.327515 | 0.459699 | 0.071 | 0.046 | 1 | 10 | Ttc39c | 0.025 | Young |
| 0.327515 | 0.389123 | 0.071 | 0.046 | 1 | 10 | Zc3h6 | 0.025 | Young |
| 0.328066 | 0.199181 | 0.828 | 0.888 | 1 | 10 | Gnb2 | -0.06 | Young |
| 0.328194 | 0.020746 | 0.263 | 0.337 | 1 | 10 | Uqcrc1 | -0.074 | Young |
| 0.328197 | 0.170634 | 0.081 | 0.054 | 1 | 10 | Kifc3 | 0.027 | Young |
| 0.328358 | 0.394496 | 0.303 | 0.269 | 1 | 10 | Pfdn1 | 0.034 | Young |
| 0.32844 | 0.602637 | 0.121 | 0.09 | 1 | 10 | Manea | 0.031 | Young |
| 0.328451 | 0.130765 | 0.02 | 0.042 | 1 | 10 | Nt5c3b | -0.022 | Young |
| 0.328761 | 0.923487 | 0.081 | 0.056 | 1 | 10 | Med6 | 0.025 | Young |
| 0.329493 | 0.376461 | 0.071 | 0.046 | 1 | 10 | Sclt1 | 0.025 | Young |
| 0.329701 | 1.912516 | 0.061 | 0.039 | 1 | 10 | Zfp597 | 0.022 | Young |
| 0.330635 | 0.034874 | 0.343 | 0.445 | 1 | 10 | Rrp1 | -0.102 | Young |
| 0.330967 | 0.248016 | 0.394 | 0.347 | 1 | 10 | Tbl1xr1 | 0.047 | Young |
| 0.331089 | 0.155015 | 0.081 | 0.115 | 1 | 10 | Dhx16 | -0.034 | Young |
| 0.331148 | 0.01699 | 0.343 | 0.44 | 1 | 10 | Usp7 | -0.097 | Young |
| 0.331349 | 0.488985 | 0.202 | 0.166 | 1 | 10 | Usp12 | 0.036 | Young |
| 0.332651 | 0.472656 | 0.01 | 0.027 | 1 | 10 | Esr1 | -0.017 | Young |
| 0.332651 | 0.329571 | 0.01 | 0.027 | 1 | 10 | Gm26756 | -0.017 | Young |
| 0.332651 | 0.239778 | 0.01 | 0.027 | 1 | 10 | C030006K1 | -0.017 | Young |
| 0.332651 | 0.143333 | 0.01 | 0.027 | 1 | 10 | BC052040 | -0.017 | Young |
| 0.333067 | 0.049251 | 0.02 | 0.042 | 1 | 10 | Pola2 | -0.022 | Young |
| 0.333473 | 0.465092 | 0.071 | 0.046 | 1 | 10 | Snx19 | 0.025 | Young |
| 0.334013 | 1.254281 | 0.061 | 0.039 | 1 | 10 | Mag | 0.022 | Young |
| 0.334624 | 0.467326 | 0.465 | 0.406 | 1 | 10 | Ccrl2 | 0.059 | Young |
| 0.334635 | 0.012964 | 0.131 | 0.178 | 1 | 10 | Eif4enif1 | -0.047 | Young |
| 0.335085 | 0.506536 | 0.303 | 0.274 | 1 | 10 | Mospd2 | 0.029 | Young |
| 0.335256 | 0.19055 | 0.323 | 0.274 | 1 | 10 | Dync1li1 | 0.049 | Young |

| | | | | | | | | |
|---|---|---|---|---|---|---|---|---|
| 0.335419 | 0.889943 | 0.121 | 0.093 | 1 | 10 | Isoc1 | 0.028 | Young |
| 0.3361 | 0.633794 | 0.081 | 0.056 | 1 | 10 | Haus4 | 0.025 | Young |
| 0.336183 | 0.627929 | 0.061 | 0.039 | 1 | 10 | Entpd6 | 0.022 | Young |
| 0.336613 | 0.250934 | 0.03 | 0.054 | 1 | 10 | Lrrc20 | -0.024 | Young |
| 0.337104 | 0.142062 | 0.061 | 0.095 | 1 | 10 | Hes6 | -0.034 | Young |
| 0.337483 | 0.214774 | 0.071 | 0.046 | 1 | 10 | Gm16536 | 0.025 | Young |
| 0.338439 | 0.264845 | 0.111 | 0.081 | 1 | 10 | Ccs | 0.03 | Young |
| 0.338551 | 0.55986 | 0.182 | 0.149 | 1 | 10 | Ap4s1 | 0.033 | Young |
| 0.338608 | 0.739592 | 0.212 | 0.181 | 1 | 10 | Gcsh | 0.031 | Young |
| 0.339033 | 0.395948 | 0.162 | 0.127 | 1 | 10 | Fam120b | 0.035 | Young |
| 0.339033 | 0.23925 | 0.101 | 0.142 | 1 | 10 | Ddx41 | -0.041 | Young |
| 0.339562 | 0.005198 | 0.465 | 0.562 | 1 | 10 | Syncrip | -0.097 | Young |
| 0.340467 | 0.046644 | 0.424 | 0.504 | 1 | 10 | Uba1 | -0.08 | Young |
| 0.340509 | 1.193002 | 0.172 | 0.137 | 1 | 10 | Bnip3 | 0.035 | Young |
| 0.340681 | 0.432339 | 0.657 | 0.67 | 1 | 10 | 1810037I1 | -0.013 | Young |
| 0.340772 | 0.580996 | 0.242 | 0.21 | 1 | 10 | Fam114a2 | 0.032 | Young |
| 0.340913 | 0.940566 | 0.091 | 0.064 | 1 | 10 | Nfkbil1 | 0.027 | Young |
| 0.340917 | 0.921138 | 0.222 | 0.188 | 1 | 10 | Slc25a37 | 0.034 | Young |
| 0.340924 | 0.013055 | 0.687 | 0.797 | 1 | 10 | Metrnl | -0.11 | Young |
| 0.340979 | 0.243034 | 0.04 | 0.068 | 1 | 10 | Ccdc117 | -0.028 | Young |
| 0.34104 | 0.01838 | 0.697 | 0.663 | 1 | 10 | Txnip | 0.034 | Young |
| 0.341065 | 0.053502 | 0.232 | 0.296 | 1 | 10 | Csgalnact2 | -0.064 | Young |
| 0.34107 | 0.2728 | 0.576 | 0.55 | 1 | 10 | Ier2 | 0.026 | Young |
| 0.341806 | 1.222097 | 0.051 | 0.032 | 1 | 10 | Zfp932 | 0.019 | Young |
| 0.342033 | 0.409702 | 0.283 | 0.244 | 1 | 10 | Id3 | 0.039 | Young |
| 0.342578 | 0.131509 | 0.909 | 0.924 | 1 | 10 | Rpl32 | -0.015 | Young |
| 0.342654 | 0.201921 | 0.172 | 0.227 | 1 | 10 | Klhdc3 | -0.055 | Young |
| 0.342709 | 0.332443 | 0.091 | 0.13 | 1 | 10 | Usp33 | -0.039 | Young |
| 0.342954 | 0.33319 | 0.131 | 0.176 | 1 | 10 | Bloc1s2 | -0.045 | Young |
| 0.343614 | 0.339788 | 0.263 | 0.215 | 1 | 10 | Hexim1 | 0.048 | Young |
| 0.343715 | 0.076017 | 0.879 | 0.924 | 1 | 10 | Cox4i1 | -0.045 | Young |
| 0.344083 | 0.338793 | 0.374 | 0.35 | 1 | 10 | Dgkz | 0.024 | Young |
| 0.344907 | 0.39419 | 0.182 | 0.147 | 1 | 10 | Actr1b | 0.035 | Young |
| 0.344928 | 0.037413 | 0.141 | 0.183 | 1 | 10 | Dlg1 | -0.042 | Young |
| 0.345199 | 0.637909 | 0.172 | 0.139 | 1 | 10 | Brix1 | 0.033 | Young |
| 0.345728 | 0.056773 | 0.364 | 0.435 | 1 | 10 | Ythdc1 | -0.071 | Young |
| 0.34617 | 0.411114 | 0.202 | 0.161 | 1 | 10 | Ldah | 0.041 | Young |
| 0.346362 | 0.426063 | 0.081 | 0.056 | 1 | 10 | Rint1 | 0.025 | Young |
| 0.346556 | 0.30311 | 0.586 | 0.592 | 1 | 10 | Vapa | -0.006 | Young |
| 0.346815 | 0.445951 | 0.162 | 0.125 | 1 | 10 | Selenoh | 0.037 | Young |
| 0.346841 | 0.311261 | 0.404 | 0.377 | 1 | 10 | Fblim1 | 0.027 | Young |
| 0.347026 | 0.039669 | 0.071 | 0.105 | 1 | 10 | Stx11 | -0.034 | Young |
| 0.347234 | 0.062584 | 0.394 | 0.462 | 1 | 10 | Clptm1l | -0.068 | Young |
| 0.347303 | 0.266885 | 0.111 | 0.156 | 1 | 10 | Pin1 | -0.045 | Young |
| 0.347594 | 0.116688 | 0.061 | 0.093 | 1 | 10 | Tbcc | -0.032 | Young |

| | | | | | | | | |
|---|---|---|---|---|---|---|---|---|
| 0.347842 | 0.065332 | 0.354 | 0.416 | 1 | 10 | Ppp4c | -0.062 | Young |
| 0.348322 | 0.645842 | 0.212 | 0.181 | 1 | 10 | Tanc2 | 0.031 | Young |
| 0.34855 | 0.730418 | 0.444 | 0.445 | 1 | 10 | Marcksl1 | -0.001 | Young |
| 0.348757 | 0.472506 | 0.364 | 0.35 | 1 | 10 | Msl1 | 0.014 | Young |
| 0.348868 | 0.094085 | 0.131 | 0.178 | 1 | 10 | Gpd1l | -0.047 | Young |
| 0.348986 | 0.925136 | 0.091 | 0.066 | 1 | 10 | Uxt | 0.025 | Young |
| 0.349391 | 0.521429 | 0.061 | 0.039 | 1 | 10 | Rnf180 | 0.022 | Young |
| 0.350143 | 0.954694 | 0.081 | 0.056 | 1 | 10 | Med9 | 0.025 | Young |
| 0.350458 | 0.365517 | 0.141 | 0.188 | 1 | 10 | Phb | -0.047 | Young |
| 0.350693 | 0.180475 | 0.091 | 0.064 | 1 | 10 | Pus7 | 0.027 | Young |
| 0.351205 | 0.02649 | 0.071 | 0.1 | 1 | 10 | Rdh11 | -0.029 | Young |
| 0.351425 | 0.617438 | 0.152 | 0.12 | 1 | 10 | S1pr1 | 0.032 | Young |
| 0.351624 | 0.494193 | 0.061 | 0.039 | 1 | 10 | Il1a | 0.022 | Young |
| 0.351826 | 0.23633 | 0.162 | 0.125 | 1 | 10 | Hhex | 0.037 | Young |
| 0.352137 | 0.028183 | 0.485 | 0.445 | 1 | 10 | Ncf1 | 0.04 | Young |
| 0.352193 | 0.3627 | 0.03 | 0.054 | 1 | 10 | Repin1 | -0.024 | Young |
| 0.352528 | 1.123129 | 0.091 | 0.066 | 1 | 10 | Zfp68 | 0.025 | Young |
| 0.352712 | 0.882122 | 0.242 | 0.21 | 1 | 10 | Mbtps1 | 0.032 | Young |
| 0.353136 | 0.671464 | 0.141 | 0.11 | 1 | 10 | Rps6kc1 | 0.031 | Young |
| 0.353417 | 1.223603 | 0.091 | 0.066 | 1 | 10 | 3110002H1 | 0.025 | Young |
| 0.353569 | 0.513466 | 0.253 | 0.215 | 1 | 10 | Xylt1 | 0.038 | Young |
| 0.354249 | 0.020208 | 0.909 | 0.968 | 1 | 10 | Lgals3 | -0.059 | Young |
| 0.354551 | 0.131797 | 0.323 | 0.401 | 1 | 10 | Dap | -0.078 | Young |
| 0.354769 | 0.010745 | 0.303 | 0.364 | 1 | 10 | Nhp2 | -0.061 | Young |
| 0.354906 | 0.572047 | 0.081 | 0.056 | 1 | 10 | 4930503L1 | 0.025 | Young |
| 0.355262 | 0.092073 | 0.202 | 0.259 | 1 | 10 | Atp8b4 | -0.057 | Young |
| 0.355387 | 0.596658 | 0.081 | 0.117 | 1 | 10 | Cog6 | -0.036 | Young |
| 0.355728 | 0.299016 | 0.172 | 0.134 | 1 | 10 | Gga3 | 0.038 | Young |
| 0.35598 | 0.28764 | 0.051 | 0.078 | 1 | 10 | Hmgxb3 | -0.027 | Young |
| 0.356047 | 0.253644 | 0.374 | 0.323 | 1 | 10 | Pkn2 | 0.051 | Young |
| 0.356424 | 0.055889 | 0.03 | 0.054 | 1 | 10 | Gm27010 | -0.024 | Young |
| 0.356576 | 0.010179 | 0.354 | 0.425 | 1 | 10 | Psme4 | -0.071 | Young |
| 0.356822 | 0.097836 | 0.04 | 0.066 | 1 | 10 | Srrd | -0.026 | Young |
| 0.356867 | 0.894378 | 0.192 | 0.161 | 1 | 10 | Thumpd1 | 0.031 | Young |
| 0.35705 | 0.581516 | 0.384 | 0.355 | 1 | 10 | Pfkp | 0.029 | Young |
| 0.357132 | 0.588781 | 0.283 | 0.254 | 1 | 10 | Ccny | 0.029 | Young |
| 0.357617 | 0.485824 | 0.152 | 0.12 | 1 | 10 | Usp10 | 0.032 | Young |
| 0.3578 | 0.668561 | 0.273 | 0.24 | 1 | 10 | Mocs2 | 0.033 | Young |
| 0.358005 | 0.572632 | 0.485 | 0.465 | 1 | 10 | Hdlbp | 0.02 | Young |
| 0.358157 | 0.482037 | 0.545 | 0.553 | 1 | 10 | Pea15a | -0.008 | Young |
| 0.358331 | 0.516125 | 0.131 | 0.1 | 1 | 10 | Tmem199 | 0.031 | Young |
| 0.358548 | 0.027327 | 0.202 | 0.264 | 1 | 10 | Sun2 | -0.062 | Young |
| 0.358634 | 0.556477 | 0.051 | 0.078 | 1 | 10 | Rsad2 | -0.027 | Young |
| 0.358864 | 0.2104 | 0.747 | 0.795 | 1 | 10 | Erp29 | -0.048 | Young |
| 0.359012 | 0.984295 | 0.051 | 0.032 | 1 | 10 | Mblac2 | 0.019 | Young |

| | | | | | | | | |
|---|---|---|---|---|---|---|---|---|
| 0.359536 | 0.442931 | 0.414 | 0.377 | 1 | 10 | Emg1 | 0.037 | Young |
| 0.359544 | 0.074108 | 0.263 | 0.34 | 1 | 10 | Vps41 | -0.077 | Young |
| 0.360168 | 0.199882 | 0.091 | 0.127 | 1 | 10 | Abi2 | -0.036 | Young |
| 0.360189 | 0.363094 | 0.111 | 0.083 | 1 | 10 | Tor2a | 0.028 | Young |
| 0.360674 | 0.71067 | 0.081 | 0.056 | 1 | 10 | Recql5 | 0.025 | Young |
| 0.36121 | 0.704213 | 0.121 | 0.093 | 1 | 10 | Gk | 0.028 | Young |
| 0.361597 | 0.638562 | 0.293 | 0.276 | 1 | 10 | Nudcd3 | 0.017 | Young |
| 0.361955 | 0.368725 | 0.444 | 0.403 | 1 | 10 | Kras | 0.041 | Young |
| 0.362017 | 0.290361 | 0.162 | 0.213 | 1 | 10 | Mrpl2 | -0.051 | Young |
| 0.362072 | 0.005333 | 0.384 | 0.46 | 1 | 10 | Psmc3 | -0.076 | Young |
| 0.362389 | 1.805731 | 0.061 | 0.042 | 1 | 10 | Akap1 | 0.019 | Young |
| 0.362423 | 1.302198 | 0.071 | 0.049 | 1 | 10 | Nup188 | 0.022 | Young |
| 0.362653 | 0.3263 | 0.323 | 0.291 | 1 | 10 | Acat1 | 0.032 | Young |
| 0.362732 | 0.127513 | 0.747 | 0.775 | 1 | 10 | Snrnp70 | -0.028 | Young |
| 0.363165 | 0.600239 | 0.212 | 0.183 | 1 | 10 | Nfxl1 | 0.029 | Young |
| 0.363813 | 1.337059 | 0.04 | 0.024 | 1 | 10 | Epas1 | 0.016 | Young |
| 0.364027 | 0.771393 | 0.051 | 0.032 | 1 | 10 | 3-Mar | 0.019 | Young |
| 0.364065 | 0.062834 | 0.99 | 0.998 | 1 | 10 | Rpl35a | -0.008 | Young |
| 0.364551 | 0.289488 | 0.051 | 0.081 | 1 | 10 | Smyd2 | -0.03 | Young |
| 0.364875 | 1.209557 | 0.091 | 0.068 | 1 | 10 | 2310061I0 | 0.023 | Young |
| 0.365096 | 0.10868 | 0.061 | 0.09 | 1 | 10 | Rora | -0.029 | Young |
| 0.365104 | 0.977733 | 0.091 | 0.066 | 1 | 10 | Fbxl4 | 0.025 | Young |
| 0.365287 | 0.682947 | 0.051 | 0.032 | 1 | 10 | Slc7a6 | 0.019 | Young |
| 0.365814 | 0.214913 | 0.101 | 0.073 | 1 | 10 | 2610524H( | 0.028 | Young |
| 0.366011 | 0.899553 | 0.172 | 0.142 | 1 | 10 | Abl1 | 0.03 | Young |
| 0.366461 | 0.32089 | 0.566 | 0.592 | 1 | 10 | Scpep1 | -0.026 | Young |
| 0.36655 | 0.36895 | 0.051 | 0.032 | 1 | 10 | Dcaf4 | 0.019 | Young |
| 0.366588 | 0.065065 | 0.141 | 0.188 | 1 | 10 | Rbmxl1 | -0.047 | Young |
| 0.36667 | 0.728938 | 0.121 | 0.093 | 1 | 10 | Chp2 | 0.028 | Young |
| 0.366779 | 0.092702 | 0.535 | 0.619 | 1 | 10 | Ndufc1 | -0.084 | Young |
| 0.366919 | 0.441056 | 0.222 | 0.183 | 1 | 10 | Dnajc10 | 0.039 | Young |
| 0.367 | 0.375251 | 0.263 | 0.222 | 1 | 10 | Psmd3 | 0.041 | Young |
| 0.367131 | 0.060637 | 0.162 | 0.205 | 1 | 10 | Zfp385a | -0.043 | Young |
| 0.367176 | 0.823953 | 0.202 | 0.178 | 1 | 10 | Rad50 | 0.024 | Young |
| 0.36735 | 0.387983 | 0.313 | 0.274 | 1 | 10 | Ahcyl1 | 0.039 | Young |
| 0.367377 | 0.368908 | 0.566 | 0.609 | 1 | 10 | Hexb | -0.043 | Young |
| 0.367461 | 0.6413 | 0.596 | 0.582 | 1 | 10 | Cd9 | 0.014 | Young |
| 0.367503 | 0.056508 | 1 | 0.998 | 1 | 10 | Rplp1 | 0.002 | Young |
| 0.367599 | 0.052794 | 0.313 | 0.381 | 1 | 10 | Fndc3a | -0.068 | Young |
| 0.367648 | 0.454167 | 0.172 | 0.134 | 1 | 10 | Cd163 | 0.038 | Young |
| 0.367655 | 0.275816 | 0.424 | 0.396 | 1 | 10 | Sec11a | 0.028 | Young |
| 0.368225 | 0.95547 | 0.101 | 0.076 | 1 | 10 | Mrpl1 | 0.025 | Young |
| 0.36845 | 0.249384 | 0.02 | 0.039 | 1 | 10 | Selenoo | -0.019 | Young |
| 0.368466 | 0.195156 | 0.333 | 0.291 | 1 | 10 | Rap1gds1 | 0.042 | Young |
| 0.368651 | 0.05336 | 0.182 | 0.242 | 1 | 10 | Cenpa | -0.06 | Young |

| | | | | | | | | |
|---|---|---|---|---|---|---|---|---|
| 0.368971 | 0.644697 | 0.121 | 0.095 | 1 | 10 | Ccdc107 | 0.026 | Young |
| 0.370197 | 0.242505 | 0.273 | 0.235 | 1 | 10 | Fyttd1 | 0.038 | Young |
| 0.370277 | 1.257111 | 0.091 | 0.068 | 1 | 10 | Zfp369 | 0.023 | Young |
| 0.370578 | 0.512144 | 0.091 | 0.066 | 1 | 10 | Hist1h4i | 0.025 | Young |
| 0.37081 | 0.880259 | 0.101 | 0.076 | 1 | 10 | Lig1 | 0.025 | Young |
| 0.371685 | 0.066589 | 0.202 | 0.257 | 1 | 10 | Mtm1 | -0.055 | Young |
| 0.3719 | 0.378012 | 0.152 | 0.117 | 1 | 10 | Sh3bp5 | 0.035 | Young |
| 0.372157 | 0.178483 | 0.121 | 0.166 | 1 | 10 | Rnf220 | -0.045 | Young |
| 0.372181 | 0.517232 | 0.121 | 0.093 | 1 | 10 | Sh2b1 | 0.028 | Young |
| 0.372181 | 0.031009 | 0.576 | 0.641 | 1 | 10 | Snrpb | -0.065 | Young |
| 0.372404 | 1.280701 | 0.04 | 0.024 | 1 | 10 | Lcmt2 | 0.016 | Young |
| 0.37286 | 0.685887 | 0.071 | 0.049 | 1 | 10 | Sepsecs | 0.022 | Young |
| 0.372997 | 0.115112 | 0.051 | 0.078 | 1 | 10 | Mthfd2l | -0.027 | Young |
| 0.373187 | 0.386704 | 0.273 | 0.235 | 1 | 10 | Eml4 | 0.038 | Young |
| 0.373212 | 0.039002 | 0.131 | 0.174 | 1 | 10 | Mast3 | -0.043 | Young |
| 0.373388 | 0.106288 | 0.232 | 0.291 | 1 | 10 | Slc25a24 | -0.059 | Young |
| 0.373405 | 0.287981 | 0.232 | 0.191 | 1 | 10 | Nup62 | 0.041 | Young |
| 0.373447 | 0.420874 | 0.293 | 0.262 | 1 | 10 | Ccl7 | 0.031 | Young |
| 0.373546 | 0.130081 | 0.02 | 0.039 | 1 | 10 | Tmem144 | -0.019 | Young |
| 0.373829 | 0.016216 | 0.222 | 0.274 | 1 | 10 | Ehmt2 | -0.052 | Young |
| 0.37397 | 0.789311 | 0.111 | 0.086 | 1 | 10 | Fbxo30 | 0.025 | Young |
| 0.374186 | 0.352707 | 0.051 | 0.032 | 1 | 10 | C030034I2: | 0.019 | Young |
| 0.374714 | 0.783743 | 0.162 | 0.137 | 1 | 10 | Thoc1 | 0.025 | Young |
| 0.374742 | 0.034713 | 0.162 | 0.213 | 1 | 10 | Map1s | -0.051 | Young |
| 0.375005 | 0.442773 | 0.111 | 0.083 | 1 | 10 | Hbegf | 0.028 | Young |
| 0.375069 | 0.483063 | 0.202 | 0.164 | 1 | 10 | Mpnd | 0.038 | Young |
| 0.375116 | 0.085004 | 0.141 | 0.181 | 1 | 10 | Nup93 | -0.04 | Young |
| 0.375264 | 0.168808 | 0.687 | 0.731 | 1 | 10 | Ncf2 | -0.044 | Young |
| 0.375457 | 0.074284 | 0.091 | 0.125 | 1 | 10 | Tjp2 | -0.034 | Young |
| 0.375467 | 0.479154 | 0.293 | 0.254 | 1 | 10 | Gripap1 | 0.039 | Young |
| 0.375551 | 0.013033 | 0.232 | 0.284 | 1 | 10 | Eloa | -0.052 | Young |
| 0.375604 | 0.574134 | 0.162 | 0.134 | 1 | 10 | Lrrk1 | 0.028 | Young |
| 0.375885 | 0.189349 | 0.576 | 0.56 | 1 | 10 | G3bp1 | 0.016 | Young |
| 0.375975 | 0.019162 | 0.566 | 0.633 | 1 | 10 | Prr13 | -0.067 | Young |
| 0.376447 | 0.857709 | 0.111 | 0.086 | 1 | 10 | Vps37b | 0.025 | Young |
| 0.377029 | 0.264257 | 0.475 | 0.457 | 1 | 10 | Hspa4 | 0.018 | Young |
| 0.377029 | 0.433443 | 0.091 | 0.066 | 1 | 10 | Rcc1l | 0.025 | Young |
| 0.37793 | 0.050279 | 0.919 | 0.98 | 1 | 10 | Rpl8 | -0.061 | Young |
| 0.378147 | 0.649017 | 0.071 | 0.049 | 1 | 10 | Snta1 | 0.022 | Young |
| 0.378201 | 1.011446 | 0.04 | 0.024 | 1 | 10 | Zfp874b | 0.016 | Young |
| 0.378653 | 0.055063 | 0.152 | 0.115 | 1 | 10 | Med12 | 0.037 | Young |
| 0.378686 | 0.10496 | 0.02 | 0.039 | 1 | 10 | Gm20300 | -0.019 | Young |
| 0.378949 | 0.280937 | 0.101 | 0.073 | 1 | 10 | Stxbp1 | 0.028 | Young |
| 0.379231 | 0.644412 | 0.162 | 0.132 | 1 | 10 | AU019823 | 0.03 | Young |
| 0.37931 | 0.376643 | 0.525 | 0.487 | 1 | 10 | Sf3b6 | 0.038 | Young |

| | | | | | | | | |
|---|---|---|---|---|---|---|---|---|
| 0.379342 | 0.567211 | 0.121 | 0.093 | 1 | 10 | Vegfb | 0.028 | Young |
| 0.379522 | 0.459168 | 0.222 | 0.191 | 1 | 10 | Itga6 | 0.031 | Young |
| 0.379645 | 0.672075 | 0.131 | 0.103 | 1 | 10 | Zfp991 | 0.028 | Young |
| 0.379659 | 1.049567 | 0.04 | 0.024 | 1 | 10 | Ust | 0.016 | Young |
| 0.380104 | 0.315497 | 0.333 | 0.311 | 1 | 10 | Arhgef10l | 0.022 | Young |
| 0.380105 | 0.569649 | 0.202 | 0.171 | 1 | 10 | Orai3 | 0.031 | Young |
| 0.380511 | 0.066644 | 0.303 | 0.367 | 1 | 10 | Tpp2 | -0.064 | Young |
| 0.380945 | 0.283772 | 0.121 | 0.093 | 1 | 10 | Sdad1 | 0.028 | Young |
| 0.380983 | 0.551163 | 0.141 | 0.112 | 1 | 10 | Bcl2 | 0.029 | Young |
| 0.38112 | 0.959065 | 0.04 | 0.024 | 1 | 10 | Sccpdh | 0.016 | Young |
| 0.38112 | 0.380108 | 0.04 | 0.024 | 1 | 10 | Slc36a3os | 0.016 | Young |
| 0.381325 | 0.157741 | 0.081 | 0.056 | 1 | 10 | Galt | 0.025 | Young |
| 0.381398 | 0.190907 | 0.798 | 0.819 | 1 | 10 | Atp5h | -0.021 | Young |
| 0.38146 | 0.022525 | 0.202 | 0.254 | 1 | 10 | Zfp652 | -0.052 | Young |
| 0.381679 | 1.277258 | 0.091 | 0.066 | 1 | 10 | Cmtm4 | 0.025 | Young |
| 0.381748 | 0.434549 | 0.121 | 0.093 | 1 | 10 | Tlr6 | 0.028 | Young |
| 0.381753 | 1.822411 | 0.02 | 0.01 | 1 | 10 | Rab26 | 0.01 | Young |
| 0.381753 | 1.451048 | 0.02 | 0.01 | 1 | 10 | Zscan18 | 0.01 | Young |
| 0.381765 | 0.003676 | 0.485 | 0.545 | 1 | 10 | Emc7 | -0.06 | Young |
| 0.381827 | 0.05059 | 0.273 | 0.328 | 1 | 10 | 4932438A1 | -0.055 | Young |
| 0.381972 | 0.564875 | 0.242 | 0.21 | 1 | 10 | Pbdc1 | 0.032 | Young |
| 0.382011 | 0.338233 | 0.202 | 0.254 | 1 | 10 | Cfdp1 | -0.052 | Young |
| 0.382023 | 0.267498 | 0.192 | 0.24 | 1 | 10 | Aip | -0.048 | Young |
| 0.382264 | 0.571494 | 0.111 | 0.086 | 1 | 10 | Dusp16 | 0.025 | Young |
| 0.382585 | 1.46117 | 0.04 | 0.024 | 1 | 10 | Rangrf | 0.016 | Young |
| 0.382585 | 1.339365 | 0.04 | 0.024 | 1 | 10 | Tle6 | 0.016 | Young |
| 0.382585 | 0.576641 | 0.04 | 0.024 | 1 | 10 | Dusp19 | 0.016 | Young |
| 0.382602 | 0.569334 | 0.253 | 0.22 | 1 | 10 | Stk11 | 0.033 | Young |
| 0.382718 | 0.336496 | 0.222 | 0.188 | 1 | 10 | Slc38a6 | 0.034 | Young |
| 0.382774 | 0.313819 | 0.737 | 0.714 | 1 | 10 | Tnfaip3 | 0.023 | Young |
| 0.382855 | 0.421567 | 0.444 | 0.45 | 1 | 10 | Ehbp1l1 | -0.006 | Young |
| 0.383178 | 1.73711 | 0.03 | 0.017 | 1 | 10 | Leng9 | 0.013 | Young |
| 0.383178 | 1.607008 | 0.03 | 0.017 | 1 | 10 | Dusp28 | 0.013 | Young |
| 0.383219 | 0.516546 | 0.182 | 0.152 | 1 | 10 | Itga9 | 0.03 | Young |
| 0.383321 | 0.809627 | 0.141 | 0.115 | 1 | 10 | Usp32 | 0.026 | Young |
| 0.383405 | 0.205689 | 0.848 | 0.873 | 1 | 10 | Pcbp2 | -0.025 | Young |
| 0.383667 | 0.61015 | 0.152 | 0.125 | 1 | 10 | Dpm1 | 0.027 | Young |
| 0.383923 | 0.148491 | 0.111 | 0.149 | 1 | 10 | Axin1 | -0.038 | Young |
| 0.383953 | 0.172752 | 0.273 | 0.32 | 1 | 10 | Cpsf2 | -0.047 | Young |
| 0.383976 | 1.54257 | 0.02 | 0.01 | 1 | 10 | Zfp58 | 0.01 | Young |
| 0.384014 | 0.040081 | 0.273 | 0.34 | 1 | 10 | Nbeal1 | -0.067 | Young |
| 0.384053 | 0.678875 | 0.04 | 0.024 | 1 | 10 | Mpv17l | 0.016 | Young |
| 0.38491 | 1.922971 | 0.03 | 0.017 | 1 | 10 | Atad5 | 0.013 | Young |
| 0.385456 | 0.058103 | 0.434 | 0.496 | 1 | 10 | Nop10 | -0.062 | Young |
| 0.385525 | 0.905646 | 0.04 | 0.024 | 1 | 10 | Ifit1 | 0.016 | Young |

| | | | | | | | | |
|---|---|---|---|---|---|---|---|---|
| 0.385572 | 0.187215 | 0.242 | 0.301 | 1 | 10 | Mfsd14a | -0.059 | Young |
| 0.386005 | 0.141826 | 0.131 | 0.174 | 1 | 10 | Phax | -0.043 | Young |
| 0.386125 | 0.232487 | 0.182 | 0.147 | 1 | 10 | Nudt5 | 0.035 | Young |
| 0.386208 | 1.744467 | 0.02 | 0.01 | 1 | 10 | Dchs1 | 0.01 | Young |
| 0.386208 | 1.524346 | 0.02 | 0.01 | 1 | 10 | Cenpe | 0.01 | Young |
| 0.386208 | 1.042456 | 0.02 | 0.01 | 1 | 10 | Zfp287 | 0.01 | Young |
| 0.386208 | 0.701356 | 0.02 | 0.01 | 1 | 10 | Zfp773 | 0.01 | Young |
| 0.386304 | 1.076835 | 0.061 | 0.042 | 1 | 10 | Nod1 | 0.019 | Young |
| 0.386304 | 1.00672 | 0.061 | 0.042 | 1 | 10 | Whamm | 0.019 | Young |
| 0.386311 | 0.33623 | 0.172 | 0.227 | 1 | 10 | Cul2 | -0.055 | Young |
| 0.386337 | 0.100382 | 0.747 | 0.758 | 1 | 10 | Ptms | -0.011 | Young |
| 0.386453 | 0.332808 | 0.283 | 0.237 | 1 | 10 | Sh3bp1 | 0.046 | Young |
| 0.386715 | 0.162005 | 0.01 | 0.024 | 1 | 10 | Tnfrsf25 | -0.014 | Young |
| 0.387 | 1.755735 | 0.04 | 0.024 | 1 | 10 | Gem | 0.016 | Young |
| 0.387 | 0.344814 | 0.04 | 0.024 | 1 | 10 | Icosl | 0.016 | Young |
| 0.387619 | 0.66165 | 0.202 | 0.171 | 1 | 10 | Crtc3 | 0.031 | Young |
| 0.387797 | 0.318975 | 0.455 | 0.455 | 1 | 10 | Got1 | 0 | Young |
| 0.387865 | 0.147268 | 0.101 | 0.073 | 1 | 10 | Mast2 | 0.028 | Young |
| 0.388001 | 0.229509 | 0.152 | 0.196 | 1 | 10 | Ctr9 | -0.044 | Young |
| 0.388153 | 0.126103 | 0.323 | 0.386 | 1 | 10 | Lfng | -0.063 | Young |
| 0.388177 | 0.074808 | 0.101 | 0.139 | 1 | 10 | Shprh | -0.038 | Young |
| 0.388248 | 0.508106 | 0.091 | 0.066 | 1 | 10 | Rbsn | 0.025 | Young |
| 0.388248 | 0.440847 | 0.091 | 0.066 | 1 | 10 | Zc3h12c | 0.025 | Young |
| 0.388378 | 0.530616 | 0.01 | 0.024 | 1 | 10 | Zfp518b | -0.014 | Young |
| 0.388378 | 0.005951 | 0.01 | 0.024 | 1 | 10 | Zfp790 | -0.014 | Young |
| 0.388389 | 1.235081 | 0.03 | 0.017 | 1 | 10 | Dtd1 | 0.013 | Young |
| 0.388389 | 0.923947 | 0.03 | 0.017 | 1 | 10 | 2210408I2 | 0.013 | Young |
| 0.388389 | 0.504639 | 0.03 | 0.017 | 1 | 10 | Birc5 | 0.013 | Young |
| 0.388447 | 2.282744 | 0.02 | 0.01 | 1 | 10 | Gm13205 | 0.01 | Young |
| 0.388447 | 1.720911 | 0.02 | 0.01 | 1 | 10 | Rasgef1a | 0.01 | Young |
| 0.388447 | 1.713157 | 0.02 | 0.01 | 1 | 10 | Zfp112 | 0.01 | Young |
| 0.388447 | 1.591658 | 0.02 | 0.01 | 1 | 10 | Slc39a8 | 0.01 | Young |
| 0.388447 | 1.465987 | 0.02 | 0.01 | 1 | 10 | Lilra5 | 0.01 | Young |
| 0.388447 | 0.831142 | 0.02 | 0.01 | 1 | 10 | 1700008J0 | 0.01 | Young |
| 0.388447 | 0.775831 | 0.02 | 0.01 | 1 | 10 | Ddit4 | 0.01 | Young |
| 0.388447 | 0.727269 | 0.02 | 0.01 | 1 | 10 | Tnfsfm13 | 0.01 | Young |
| 0.388447 | 0.713897 | 0.02 | 0.01 | 1 | 10 | Asgr2 | 0.01 | Young |
| 0.388447 | 0.704457 | 0.02 | 0.01 | 1 | 10 | Ccdc137 | 0.01 | Young |
| 0.388447 | 0.639757 | 0.02 | 0.01 | 1 | 10 | Rilpl1 | 0.01 | Young |
| 0.388447 | 0.596472 | 0.02 | 0.01 | 1 | 10 | AW011738 | 0.01 | Young |
| 0.388447 | 0.565675 | 0.02 | 0.01 | 1 | 10 | Zfp72 | 0.01 | Young |
| 0.388447 | 0.520921 | 0.02 | 0.01 | 1 | 10 | Efnb2 | 0.01 | Young |
| 0.388528 | 0.421241 | 0.222 | 0.191 | 1 | 10 | Trim44 | 0.031 | Young |
| 0.388761 | 0.031382 | 0.242 | 0.303 | 1 | 10 | Abce1 | -0.061 | Young |
| 0.389192 | 1.019332 | 0.091 | 0.066 | 1 | 10 | Cd209a | 0.025 | Young |

| | | | | | | | | |
|---|---|---|---|---|---|---|---|---|
| 0.389487 | 0.484646 | 0.232 | 0.203 | 1 | 10 | Yrdc | 0.029 | Young |
| 0.389645 | 0.199207 | 0.263 | 0.315 | 1 | 10 | Efcab14 | -0.052 | Young |
| 0.389797 | 0.606308 | 0.061 | 0.042 | 1 | 10 | Coa7 | 0.019 | Young |
| 0.39051 | 0.869612 | 0.091 | 0.068 | 1 | 10 | Spidr | 0.023 | Young |
| 0.390529 | 0.117908 | 0.303 | 0.381 | 1 | 10 | Osbpl9 | -0.078 | Young |
| 0.390575 | 0.279452 | 0.141 | 0.11 | 1 | 10 | Gps2 | 0.031 | Young |
| 0.390694 | 1.037196 | 0.02 | 0.01 | 1 | 10 | Haus5 | 0.01 | Young |
| 0.390694 | 0.974575 | 0.02 | 0.01 | 1 | 10 | Etohd2 | 0.01 | Young |
| 0.390694 | 0.771129 | 0.02 | 0.01 | 1 | 10 | Ramp3 | 0.01 | Young |
| 0.390694 | 0.733565 | 0.02 | 0.01 | 1 | 10 | A530064D( | 0.01 | Young |
| 0.390694 | 0.702415 | 0.02 | 0.01 | 1 | 10 | Ppp1r3d | 0.01 | Young |
| 0.390694 | 0.665324 | 0.02 | 0.01 | 1 | 10 | Aldoc | 0.01 | Young |
| 0.390694 | 0.621452 | 0.02 | 0.01 | 1 | 10 | Lrfn1 | 0.01 | Young |
| 0.390694 | 0.495955 | 0.02 | 0.01 | 1 | 10 | Layn | 0.01 | Young |
| 0.390694 | 0.483879 | 0.02 | 0.01 | 1 | 10 | Kif11 | 0.01 | Young |
| 0.390694 | 0.283584 | 0.02 | 0.01 | 1 | 10 | Gm9856 | 0.01 | Young |
| 0.390694 | 0.156354 | 0.02 | 0.01 | 1 | 10 | 2900060B1 | 0.01 | Young |
| 0.39115 | 0.121862 | 0.232 | 0.289 | 1 | 10 | Hmg20b | -0.057 | Young |
| 0.391283 | 0.114127 | 0.354 | 0.43 | 1 | 10 | Napa | -0.076 | Young |
| 0.39139 | 0.476823 | 0.424 | 0.411 | 1 | 10 | Gatad2b | 0.013 | Young |
| 0.391413 | 0.19953 | 0.202 | 0.249 | 1 | 10 | Ccdc22 | -0.047 | Young |
| 0.391445 | 0.280913 | 0.051 | 0.078 | 1 | 10 | Tbccd1 | -0.027 | Young |
| 0.391445 | 0.374369 | 0.04 | 0.024 | 1 | 10 | Tnfsf13b | 0.016 | Young |
| 0.391445 | 0.366441 | 0.04 | 0.024 | 1 | 10 | Mcemp1 | 0.016 | Young |
| 0.391884 | 0.191941 | 0.949 | 0.946 | 1 | 10 | Lyz2 | 0.003 | Young |
| 0.391887 | 1.393819 | 0.03 | 0.017 | 1 | 10 | Ropn1l | 0.013 | Young |
| 0.391887 | 1.167332 | 0.03 | 0.017 | 1 | 10 | Lekr1 | 0.013 | Young |
| 0.391887 | 0.906486 | 0.03 | 0.017 | 1 | 10 | Plekha5 | 0.013 | Young |
| 0.391895 | 0.3934 | 0.101 | 0.076 | 1 | 10 | Slc25a22 | 0.025 | Young |
| 0.392298 | 0.133981 | 0.747 | 0.736 | 1 | 10 | Ctsc | 0.011 | Young |
| 0.392615 | 0.048416 | 0.192 | 0.242 | 1 | 10 | Clk3 | -0.05 | Young |
| 0.39264 | 0.098141 | 0.889 | 0.927 | 1 | 10 | Neat1 | -0.038 | Young |
| 0.392922 | 0.361401 | 0.444 | 0.418 | 1 | 10 | Arl4c | 0.026 | Young |
| 0.392949 | 0.688564 | 0.02 | 0.01 | 1 | 10 | Zfp971 | 0.01 | Young |
| 0.392949 | 0.63769 | 0.02 | 0.01 | 1 | 10 | Hist1h3d | 0.01 | Young |
| 0.392949 | 0.44622 | 0.02 | 0.01 | 1 | 10 | P3h3 | 0.01 | Young |
| 0.392949 | 0.413318 | 0.02 | 0.01 | 1 | 10 | Pigl | 0.01 | Young |
| 0.392949 | 0.331101 | 0.02 | 0.01 | 1 | 10 | Nupr1l | 0.01 | Young |
| 0.392949 | 0.088601 | 0.02 | 0.01 | 1 | 10 | 4933417E1 | 0.01 | Young |
| 0.393196 | 0.441957 | 0.071 | 0.049 | 1 | 10 | Pcca | 0.022 | Young |
| 0.393307 | 0.023728 | 0.374 | 0.452 | 1 | 10 | U2surp | -0.078 | Young |
| 0.393614 | 0.590297 | 0.455 | 0.445 | 1 | 10 | Sbno1 | 0.01 | Young |
| 0.393643 | 1.044973 | 0.03 | 0.017 | 1 | 10 | Ifit3 | 0.013 | Young |
| 0.393643 | 0.856804 | 0.03 | 0.017 | 1 | 10 | Ccdc102a | 0.013 | Young |
| 0.393643 | 0.856005 | 0.03 | 0.017 | 1 | 10 | Efcab2 | 0.013 | Young |

| | | | | | | | | |
|---|---|---|---|---|---|---|---|---|
| 0.393643 | 0.751651 | 0.03 | 0.017 | 1 | 10 | Tmem81 | 0.013 | Young |
| 0.393643 | 0.433937 | 0.03 | 0.017 | 1 | 10 | Tmem67 | 0.013 | Young |
| 0.393691 | 0.681181 | 0.141 | 0.112 | 1 | 10 | Khnyn | 0.029 | Young |
| 0.394147 | 0.349767 | 0.293 | 0.259 | 1 | 10 | Lrrfip1 | 0.034 | Young |
| 0.394426 | 0.390076 | 0.04 | 0.024 | 1 | 10 | Rilp | 0.016 | Young |
| 0.394482 | 0.190426 | 0.141 | 0.188 | 1 | 10 | Sqor | -0.047 | Young |
| 0.394757 | 0.369748 | 0.03 | 0.051 | 1 | 10 | Scimp | -0.021 | Young |
| 0.394816 | 0.035374 | 0.101 | 0.137 | 1 | 10 | 2310001H1 | -0.036 | Young |
| 0.395116 | 0.534123 | 0.061 | 0.09 | 1 | 10 | Tmem64 | -0.029 | Young |
| 0.395183 | 0.091647 | 0.081 | 0.115 | 1 | 10 | Cdadc1 | -0.034 | Young |
| 0.395211 | 0.490584 | 0.02 | 0.01 | 1 | 10 | Dpys | 0.01 | Young |
| 0.395211 | 0.134609 | 0.02 | 0.01 | 1 | 10 | Zfp712 | 0.01 | Young |
| 0.395328 | 0.06668 | 0.586 | 0.577 | 1 | 10 | Rhob | 0.009 | Young |
| 0.395404 | 0.851188 | 0.03 | 0.017 | 1 | 10 | Setd6 | 0.013 | Young |
| 0.395404 | 0.708902 | 0.03 | 0.017 | 1 | 10 | Cdkl2 | 0.013 | Young |
| 0.395567 | 0.372145 | 0.263 | 0.337 | 1 | 10 | Gdi1 | -0.074 | Young |
| 0.395801 | 0.127848 | 0.071 | 0.103 | 1 | 10 | Tars2 | -0.032 | Young |
| 0.395922 | 0.132893 | 0.04 | 0.024 | 1 | 10 | Atp23 | 0.016 | Young |
| 0.396193 | 0.343548 | 0.152 | 0.12 | 1 | 10 | Nr4a3 | 0.032 | Young |
| 0.39663 | 0.094363 | 0.172 | 0.218 | 1 | 10 | Fam204a | -0.046 | Young |
| 0.396947 | 0.506559 | 0.172 | 0.142 | 1 | 10 | Gan | 0.03 | Young |
| 0.396988 | 0.299316 | 0.04 | 0.064 | 1 | 10 | Gm4610 | -0.024 | Young |
| 0.397046 | 0.370945 | 0.253 | 0.222 | 1 | 10 | Tcn2 | 0.031 | Young |
| 0.397105 | 0.093645 | 0.343 | 0.311 | 1 | 10 | Prkcb | 0.032 | Young |
| 0.397169 | 0.770956 | 0.03 | 0.017 | 1 | 10 | Prkca | 0.013 | Young |
| 0.397169 | 0.762437 | 0.03 | 0.017 | 1 | 10 | Fut10 | 0.013 | Young |
| 0.397169 | 0.627945 | 0.03 | 0.017 | 1 | 10 | Gm15446 | 0.013 | Young |
| 0.397169 | 0.28012 | 0.03 | 0.017 | 1 | 10 | Pstpip2 | 0.013 | Young |
| 0.397301 | 0.04884 | 0.354 | 0.438 | 1 | 10 | Gnai3 | -0.084 | Young |
| 0.397424 | 0.299681 | 0.141 | 0.11 | 1 | 10 | Arid5a | 0.031 | Young |
| 0.397568 | 0.52882 | 0.293 | 0.262 | 1 | 10 | Arih2 | 0.031 | Young |
| 0.397931 | 0.728665 | 0.081 | 0.059 | 1 | 10 | Papss2 | 0.022 | Young |
| 0.398027 | 0.096007 | 0.838 | 0.831 | 1 | 10 | C1qc | 0.007 | Young |
| 0.39817 | 0.806763 | 0.131 | 0.105 | 1 | 10 | Zfp994 | 0.026 | Young |
| 0.398279 | 0.143173 | 0.253 | 0.328 | 1 | 10 | Slco2b1 | -0.075 | Young |
| 0.398729 | 0.08731 | 0.202 | 0.244 | 1 | 10 | Myo1c | -0.042 | Young |
| 0.39894 | 2.20734 | 0.03 | 0.017 | 1 | 10 | Inhba | 0.013 | Young |
| 0.39894 | 0.602022 | 0.03 | 0.017 | 1 | 10 | Map3k15 | 0.013 | Young |
| 0.39894 | 0.239972 | 0.03 | 0.017 | 1 | 10 | Tmem158 | 0.013 | Young |
| 0.400451 | 0.116772 | 0.081 | 0.112 | 1 | 10 | Rab10os | -0.031 | Young |
| 0.400583 | 1.59023 | 0.162 | 0.139 | 1 | 10 | Gclm | 0.023 | Young |
| 0.400715 | 1.606366 | 0.03 | 0.017 | 1 | 10 | Gm17586 | 0.013 | Young |
| 0.400973 | 0.415482 | 0.081 | 0.059 | 1 | 10 | Cnot11 | 0.022 | Young |
| 0.401166 | 0.013352 | 0.04 | 0.064 | 1 | 10 | Rcl1 | -0.024 | Young |
| 0.401242 | 0.177161 | 0.586 | 0.587 | 1 | 10 | Adam15 | -0.001 | Young |

| | | | | | | | | |
|---|---|---|---|---|---|---|---|---|
| 0.401362 | 0.745644 | 0.162 | 0.134 | 1 | 10 | Zbtb7b | 0.028 | Young |
| 0.401763 | 0.621538 | 0.111 | 0.086 | 1 | 10 | Rarg | 0.025 | Young |
| 0.4018 | 0.582612 | 0.121 | 0.095 | 1 | 10 | Wdtc1 | 0.026 | Young |
| 0.402116 | 0.232888 | 0.172 | 0.215 | 1 | 10 | Marveld1 | -0.043 | Young |
| 0.402368 | 0.626799 | 0.192 | 0.159 | 1 | 10 | Cspp1 | 0.033 | Young |
| 0.402553 | 0.01749 | 0.051 | 0.076 | 1 | 10 | 2310022B0 | -0.025 | Young |
| 0.402587 | 0.08021 | 0.162 | 0.205 | 1 | 10 | AI413582 | -0.043 | Young |
| 0.403191 | 0.099638 | 0.636 | 0.658 | 1 | 10 | Timm13 | -0.022 | Young |
| 0.403445 | 0.322979 | 0.374 | 0.337 | 1 | 10 | Lsp1 | 0.037 | Young |
| 0.403541 | 0.27253 | 0.758 | 0.804 | 1 | 10 | Mtdh | -0.046 | Young |
| 0.403962 | 0.814984 | 0.061 | 0.042 | 1 | 10 | Dsel | 0.019 | Young |
| 0.404279 | 0.308166 | 0.03 | 0.017 | 1 | 10 | Gm11423 | 0.013 | Young |
| 0.404279 | 0.158751 | 0.03 | 0.017 | 1 | 10 | Zswim9 | 0.013 | Young |
| 0.404413 | 0.30142 | 0.283 | 0.237 | 1 | 10 | Vamp2 | 0.046 | Young |
| 0.404798 | 0.170439 | 0.101 | 0.139 | 1 | 10 | Lemd3 | -0.038 | Young |
| 0.404938 | 0.060292 | 0.111 | 0.144 | 1 | 10 | Itpripl1 | -0.033 | Young |
| 0.405156 | 0.286594 | 0.061 | 0.042 | 1 | 10 | Prr5l | 0.019 | Young |
| 0.405571 | 0.09471 | 0.222 | 0.286 | 1 | 10 | Kmt5b | -0.064 | Young |
| 0.405747 | 0.038184 | 0.111 | 0.083 | 1 | 10 | Cib2 | 0.028 | Young |
| 0.405787 | 0.016019 | 0.465 | 0.518 | 1 | 10 | Ap1b1 | -0.053 | Young |
| 0.405819 | 0.028762 | 0.303 | 0.377 | 1 | 10 | Stat6 | -0.074 | Young |
| 0.405968 | 0.016597 | 0.222 | 0.271 | 1 | 10 | Eif2s3y | -0.049 | Young |
| 0.406069 | 0.328091 | 0.03 | 0.017 | 1 | 10 | Galnt12 | 0.013 | Young |
| 0.406492 | 0.037137 | 0.313 | 0.352 | 1 | 10 | Aph1c | -0.039 | Young |
| 0.406915 | 0.18391 | 0.222 | 0.186 | 1 | 10 | Fam213b | 0.036 | Young |
| 0.407225 | 1.07214 | 0.081 | 0.061 | 1 | 10 | Wdr83 | 0.02 | Young |
| 0.407706 | 0.100058 | 0.939 | 0.963 | 1 | 10 | Ctsd | -0.024 | Young |
| 0.407995 | 0.311962 | 0.253 | 0.218 | 1 | 10 | Msl3 | 0.035 | Young |
| 0.408237 | 0.099022 | 0.071 | 0.1 | 1 | 10 | Edem3 | -0.029 | Young |
| 0.408399 | 0.058336 | 0.343 | 0.389 | 1 | 10 | Golga4 | -0.046 | Young |
| 0.408404 | 1.405785 | 0.121 | 0.095 | 1 | 10 | Lsm1 | 0.026 | Young |
| 0.40909 | 0.636844 | 0.263 | 0.225 | 1 | 10 | Sars | 0.038 | Young |
| 0.409092 | 0.369337 | 0.101 | 0.076 | 1 | 10 | Cmah | 0.025 | Young |
| 0.410168 | 0.123553 | 0.727 | 0.719 | 1 | 10 | Rpl22l1 | 0.008 | Young |
| 0.410306 | 0.369058 | 0.394 | 0.364 | 1 | 10 | Cmtm6 | 0.03 | Young |
| 0.410333 | 0.315433 | 0.172 | 0.139 | 1 | 10 | Cep83 | 0.033 | Young |
| 0.410337 | 0.272809 | 0.717 | 0.758 | 1 | 10 | Ctsh | -0.041 | Young |
| 0.410661 | 0.04448 | 0.04 | 0.064 | 1 | 10 | D17H6S53E | -0.024 | Young |
| 0.411211 | 0.722555 | 0.081 | 0.059 | 1 | 10 | C2cd2 | 0.022 | Young |
| 0.411238 | 0.029146 | 0.101 | 0.134 | 1 | 10 | Usp50 | -0.033 | Young |
| 0.411263 | 0.366323 | 0.232 | 0.2 | 1 | 10 | Slc36a1 | 0.032 | Young |
| 0.411286 | 0.013089 | 0.051 | 0.076 | 1 | 10 | Alad | -0.025 | Young |
| 0.412083 | 0.204839 | 0.485 | 0.452 | 1 | 10 | Anp32a | 0.033 | Young |
| 0.412171 | 0.325333 | 0.202 | 0.169 | 1 | 10 | D10Jhu81e | 0.033 | Young |
| 0.412218 | 0.947106 | 0.071 | 0.051 | 1 | 10 | Top2a | 0.02 | Young |

| | | | | | | | | |
|---|---|---|---|---|---|---|---|---|
| 0.412271 | 0.01334 | 0.212 | 0.259 | 1 | 10 | Ube2d1 | -0.047 | Young |
| 0.412711 | 0.118531 | 0.131 | 0.103 | 1 | 10 | Gdpd1 | 0.028 | Young |
| 0.412788 | 0.01121 | 0.04 | 0.064 | 1 | 10 | Lancl2 | -0.024 | Young |
| 0.412951 | 0.775142 | 0.232 | 0.293 | 1 | 10 | Fbxl3 | -0.061 | Young |
| 0.412981 | 0.113777 | 0.101 | 0.132 | 1 | 10 | Gpr35 | -0.031 | Young |
| 0.413956 | 0.190086 | 0.121 | 0.159 | 1 | 10 | Ap1g2 | -0.038 | Young |
| 0.414107 | 0.094376 | 0.071 | 0.098 | 1 | 10 | Isg20l2 | -0.027 | Young |
| 0.414395 | 0.750845 | 0.141 | 0.112 | 1 | 10 | 1700123O2 | 0.029 | Young |
| 0.414722 | 0.059335 | 0.535 | 0.65 | 1 | 10 | Ndufc2 | -0.115 | Young |
| 0.414991 | 1.251792 | 0.232 | 0.218 | 1 | 10 | Arg1 | 0.014 | Young |
| 0.415212 | 0.237626 | 0.515 | 0.472 | 1 | 10 | Wapl | 0.043 | Young |
| 0.415214 | 1.131742 | 0.051 | 0.034 | 1 | 10 | Ift27 | 0.017 | Young |
| 0.415409 | 0.165032 | 0.222 | 0.188 | 1 | 10 | Arhgap19 | 0.034 | Young |
| 0.415625 | 0.218341 | 0.061 | 0.088 | 1 | 10 | Mcm4 | -0.027 | Young |
| 0.415997 | 0.456839 | 0.061 | 0.042 | 1 | 10 | Rtl8b | 0.019 | Young |
| 0.416239 | 0.62935 | 0.091 | 0.068 | 1 | 10 | Pla2g16 | 0.023 | Young |
| 0.416544 | 1.253117 | 0.051 | 0.034 | 1 | 10 | Slc22a15 | 0.017 | Young |
| 0.417346 | 0.346853 | 0.374 | 0.342 | 1 | 10 | Siva1 | 0.032 | Young |
| 0.417392 | 0.621244 | 0.101 | 0.076 | 1 | 10 | Klhdc10 | 0.025 | Young |
| 0.417638 | 0.473466 | 0.323 | 0.284 | 1 | 10 | Nlrp3 | 0.039 | Young |
| 0.418156 | 0.094331 | 0.051 | 0.076 | 1 | 10 | Ublcp1 | -0.025 | Young |
| 0.418842 | 0.29794 | 0.162 | 0.205 | 1 | 10 | Pofut2 | -0.043 | Young |
| 0.418845 | 0.794133 | 0.152 | 0.127 | 1 | 10 | Bex3 | 0.025 | Young |
| 0.419029 | 0.064258 | 0.96 | 0.954 | 1 | 10 | Marcks | 0.006 | Young |
| 0.419406 | 0.098767 | 0.071 | 0.1 | 1 | 10 | Nt5c3 | -0.029 | Young |
| 0.419419 | 0.281861 | 0.232 | 0.198 | 1 | 10 | Ints6l | 0.034 | Young |
| 0.419524 | 0.163494 | 0.323 | 0.286 | 1 | 10 | Zfc3h1 | 0.037 | Young |
| 0.419755 | 0.28384 | 0.364 | 0.335 | 1 | 10 | Gabarapl2 | 0.029 | Young |
| 0.419861 | 0.573771 | 0.303 | 0.284 | 1 | 10 | Rgs19 | 0.019 | Young |
| 0.419949 | 0.110552 | 0.162 | 0.21 | 1 | 10 | Mea1 | -0.048 | Young |
| 0.41996 | 0.661445 | 0.071 | 0.051 | 1 | 10 | Dnajb2 | 0.02 | Young |
| 0.420282 | 0.139594 | 0.384 | 0.455 | 1 | 10 | Ankrd12 | -0.071 | Young |
| 0.420375 | 0.008411 | 0.081 | 0.108 | 1 | 10 | Eng | -0.027 | Young |
| 0.420532 | 0.277329 | 0.172 | 0.215 | 1 | 10 | Rab3gap1 | -0.043 | Young |
| 0.42087 | 0.369789 | 0.061 | 0.042 | 1 | 10 | Tmem268 | 0.019 | Young |
| 0.42087 | 0.337557 | 0.061 | 0.042 | 1 | 10 | Aven | 0.019 | Young |
| 0.421006 | 1.277306 | 0.091 | 0.071 | 1 | 10 | Ccdc58 | 0.02 | Young |
| 0.421113 | 0.471779 | 0.101 | 0.076 | 1 | 10 | Slc25a19 | 0.025 | Young |
| 0.421124 | 0.238489 | 0.253 | 0.306 | 1 | 10 | Slc44a1 | -0.053 | Young |
| 0.421138 | 0.499725 | 0.061 | 0.088 | 1 | 10 | Trappc9 | -0.027 | Young |
| 0.421211 | 0.358859 | 0.202 | 0.171 | 1 | 10 | Bola3 | 0.031 | Young |
| 0.421276 | 0.092591 | 0.202 | 0.252 | 1 | 10 | Rbm33 | -0.05 | Young |
| 0.421691 | 0.881069 | 0.131 | 0.108 | 1 | 10 | Timm17b | 0.023 | Young |
| 0.422188 | 1.107128 | 0.071 | 0.051 | 1 | 10 | Fbxl18 | 0.02 | Young |
| 0.422286 | 0.152817 | 0.141 | 0.181 | 1 | 10 | Mprip | -0.04 | Young |

| | | | | | | | | |
|---|---|---|---|---|---|---|---|---|
| 0.422352 | 0.02339 | 0.253 | 0.293 | 1 | 10 | Nabp1 | -0.04 | Young |
| 0.423169 | 0.299387 | 0.232 | 0.198 | 1 | 10 | Fbxw4 | 0.034 | Young |
| 0.423319 | 0.429338 | 0.061 | 0.042 | 1 | 10 | Mon1a | 0.019 | Young |
| 0.423633 | 0.400495 | 0.182 | 0.225 | 1 | 10 | Mtfr1l | -0.043 | Young |
| 0.423658 | 0.125269 | 0.697 | 0.711 | 1 | 10 | Brk1 | -0.014 | Young |
| 0.423749 | 0.370085 | 0.071 | 0.1 | 1 | 10 | Sugp1 | -0.029 | Young |
| 0.423829 | 1.372883 | 0.101 | 0.081 | 1 | 10 | Gnl3l | 0.02 | Young |
| 0.424102 | 0.224291 | 0.131 | 0.103 | 1 | 10 | Bloc1s6 | 0.028 | Young |
| 0.424195 | 0.183966 | 0.091 | 0.125 | 1 | 10 | Ermp1 | -0.034 | Young |
| 0.425106 | 0.044691 | 0.354 | 0.418 | 1 | 10 | Washc2 | -0.064 | Young |
| 0.425228 | 0.862364 | 0.172 | 0.147 | 1 | 10 | Tubgcp4 | 0.025 | Young |
| 0.426623 | 0.674085 | 0.222 | 0.2 | 1 | 10 | Klhl5 | 0.022 | Young |
| 0.426908 | 0.212156 | 0.343 | 0.391 | 1 | 10 | Slc38a1 | -0.048 | Young |
| 0.427482 | 0.42268 | 0.02 | 0.037 | 1 | 10 | Fkbpl | -0.017 | Young |
| 0.428242 | 9.39E-05 | 0.03 | 0.049 | 1 | 10 | Bcdin3d | -0.019 | Young |
| 0.428352 | 0.096644 | 0.343 | 0.421 | 1 | 10 | Wbp2 | -0.078 | Young |
| 0.428863 | 0.221597 | 0.424 | 0.386 | 1 | 10 | Ssu72 | 0.038 | Young |
| 0.429107 | 0.675545 | 0.253 | 0.235 | 1 | 10 | Eri1 | 0.018 | Young |
| 0.429331 | 0.021368 | 0.404 | 0.477 | 1 | 10 | Card19 | -0.073 | Young |
| 0.429763 | 0.263196 | 0.081 | 0.11 | 1 | 10 | Zfp512 | -0.029 | Young |
| 0.430113 | 0.250364 | 0.495 | 0.469 | 1 | 10 | Snu13 | 0.026 | Young |
| 0.430128 | 0.00712 | 0.697 | 0.685 | 1 | 10 | Zfp36l1 | 0.012 | Young |
| 0.43027 | 0.589425 | 0.273 | 0.257 | 1 | 10 | Ykt6 | 0.016 | Young |
| 0.430433 | 0.757511 | 0.141 | 0.117 | 1 | 10 | Med7 | 0.024 | Young |
| 0.430797 | 0.020144 | 0.141 | 0.183 | 1 | 10 | Zfp131 | -0.042 | Young |
| 0.430806 | 0.757707 | 0.152 | 0.13 | 1 | 10 | Atad2 | 0.022 | Young |
| 0.431167 | 0.675531 | 0.071 | 0.051 | 1 | 10 | Ppm1d | 0.02 | Young |
| 0.431445 | 0.16859 | 0.061 | 0.086 | 1 | 10 | Kat2a | -0.025 | Young |
| 0.43156 | 0.245835 | 0.626 | 0.623 | 1 | 10 | Axl | 0.003 | Young |
| 0.431657 | 0.444847 | 0.111 | 0.086 | 1 | 10 | Rnf215 | 0.025 | Young |
| 0.431774 | 0.111703 | 0.02 | 0.037 | 1 | 10 | Wars2 | -0.017 | Young |
| 0.432702 | 1.458882 | 0.051 | 0.034 | 1 | 10 | Slc7a2 | 0.017 | Young |
| 0.432917 | 0.149552 | 0.232 | 0.279 | 1 | 10 | Kdm5a | -0.047 | Young |
| 0.433043 | 1.103272 | 0.131 | 0.108 | 1 | 10 | Adrb2 | 0.023 | Young |
| 0.433456 | 0.213848 | 0.071 | 0.098 | 1 | 10 | Arl2bp | -0.027 | Young |
| 0.433456 | 0.596637 | 0.111 | 0.088 | 1 | 10 | Mark4 | 0.023 | Young |
| 0.433627 | 0.207045 | 0.101 | 0.137 | 1 | 10 | Scaf4 | -0.036 | Young |
| 0.433882 | 0.280784 | 0.263 | 0.315 | 1 | 10 | Tnf | -0.052 | Young |
| 0.433934 | 0.577169 | 0.091 | 0.068 | 1 | 10 | Mier2 | 0.023 | Young |
| 0.434347 | 0.219122 | 0.071 | 0.098 | 1 | 10 | Prkci | -0.027 | Young |
| 0.434528 | 0.48342 | 0.091 | 0.125 | 1 | 10 | Lonp2 | -0.034 | Young |
| 0.434529 | 0.102521 | 0.293 | 0.357 | 1 | 10 | Tomm70a | -0.064 | Young |
| 0.434919 | 0.024246 | 0.313 | 0.377 | 1 | 10 | Stt3b | -0.064 | Young |
| 0.434962 | 0.438336 | 0.162 | 0.137 | 1 | 10 | Nde1 | 0.025 | Young |
| 0.435306 | 0.62999 | 0.202 | 0.176 | 1 | 10 | Cog7 | 0.026 | Young |

| | | | | | | | | |
|---|---|---|---|---|---|---|---|---|
| 0.43546 | 0.319426 | 0.141 | 0.115 | 1 | 10 | Enoph1 | 0.026 | Young |
| 0.435829 | 0.093256 | 0.343 | 0.413 | 1 | 10 | Snx20 | -0.07 | Young |
| 0.436317 | 0.404221 | 0.152 | 0.125 | 1 | 10 | Aldh3a2 | 0.027 | Young |
| 0.436452 | 0.525754 | 0.303 | 0.289 | 1 | 10 | Iqgap2 | 0.014 | Young |
| 0.436772 | 0.310043 | 0.222 | 0.191 | 1 | 10 | Supt20 | 0.031 | Young |
| 0.436949 | 0.387207 | 0.212 | 0.178 | 1 | 10 | Nol11 | 0.034 | Young |
| 0.437029 | 0.13151 | 0.071 | 0.098 | 1 | 10 | Slc39a13 | -0.027 | Young |
| 0.437924 | 0.402843 | 0.091 | 0.068 | 1 | 10 | Car3 | 0.023 | Young |
| 0.438375 | 0.20347 | 0.354 | 0.33 | 1 | 10 | Sbds | 0.024 | Young |
| 0.439207 | 0.117409 | 0.141 | 0.181 | 1 | 10 | Tmem11 | -0.04 | Young |
| 0.439277 | 0.780014 | 0.081 | 0.061 | 1 | 10 | Lmbr1l | 0.02 | Young |
| 0.439361 | 0.219991 | 0.475 | 0.457 | 1 | 10 | Csrnp1 | 0.018 | Young |
| 0.439609 | 0.495789 | 0.232 | 0.205 | 1 | 10 | Cdk12 | 0.027 | Young |
| 0.439758 | 0.584976 | 0.293 | 0.259 | 1 | 10 | Cdc40 | 0.034 | Young |
| 0.440338 | 0.12444 | 0.131 | 0.171 | 1 | 10 | Eya3 | -0.04 | Young |
| 0.44042 | 0.265521 | 0.293 | 0.267 | 1 | 10 | Lin7c | 0.026 | Young |
| 0.440523 | 0.189095 | 0.586 | 0.611 | 1 | 10 | Cox5b | -0.025 | Young |
| 0.440694 | 0.267613 | 0.03 | 0.049 | 1 | 10 | Oxnad1 | -0.019 | Young |
| 0.440718 | 0.640532 | 0.212 | 0.186 | 1 | 10 | Parg | 0.026 | Young |
| 0.440898 | 0.364103 | 0.364 | 0.337 | 1 | 10 | Cttnbp2nl | 0.027 | Young |
| 0.440915 | 0.574569 | 0.051 | 0.034 | 1 | 10 | Tmed1 | 0.017 | Young |
| 0.440915 | 0.285392 | 0.051 | 0.034 | 1 | 10 | Ttc28 | 0.017 | Young |
| 0.441382 | 1.079621 | 0.111 | 0.09 | 1 | 10 | Mthfs | 0.021 | Young |
| 0.441434 | 0.216625 | 0.626 | 0.653 | 1 | 10 | Cox7a2 | -0.027 | Young |
| 0.441446 | 0.417271 | 0.192 | 0.164 | 1 | 10 | Irak2 | 0.028 | Young |
| 0.44182 | 0.069677 | 0.838 | 0.885 | 1 | 10 | Ctsa | -0.047 | Young |
| 0.441936 | 0.392467 | 0.091 | 0.068 | 1 | 10 | Pcyt2 | 0.023 | Young |
| 0.441936 | 0.03778 | 0.091 | 0.068 | 1 | 10 | Alox5 | 0.023 | Young |
| 0.443055 | 0.677821 | 0.162 | 0.134 | 1 | 10 | Brwd3 | 0.028 | Young |
| 0.443148 | 0.903526 | 0.162 | 0.137 | 1 | 10 | Hdhd2 | 0.025 | Young |
| 0.44357 | 0.126848 | 0.162 | 0.2 | 1 | 10 | Zc3h14 | -0.038 | Young |
| 0.444263 | 0.284984 | 0.141 | 0.115 | 1 | 10 | Mrps30 | 0.026 | Young |
| 0.444284 | 0.41941 | 0.273 | 0.257 | 1 | 10 | Appbp2 | 0.016 | Young |
| 0.444391 | 0.2324 | 0.283 | 0.337 | 1 | 10 | Pdap1 | -0.054 | Young |
| 0.444704 | 0.06279 | 0.545 | 0.626 | 1 | 10 | Ndufb7 | -0.081 | Young |
| 0.445169 | 0.207363 | 0.222 | 0.269 | 1 | 10 | Gatad2a | -0.047 | Young |
| 0.445316 | 0.351201 | 0.162 | 0.134 | 1 | 10 | Tm2d3 | 0.028 | Young |
| 0.445407 | 0.216396 | 0.131 | 0.174 | 1 | 10 | Slc25a25 | -0.043 | Young |
| 0.445516 | 0.026368 | 0.222 | 0.271 | 1 | 10 | Kif1c | -0.049 | Young |
| 0.44605 | 0.353742 | 0.404 | 0.399 | 1 | 10 | Sar1a | 0.005 | Young |
| 0.446418 | 0.703164 | 0.253 | 0.232 | 1 | 10 | Fuca2 | 0.021 | Young |
| 0.446452 | 0.161812 | 0.152 | 0.186 | 1 | 10 | Aars | -0.034 | Young |
| 0.446522 | 0.450531 | 0.111 | 0.149 | 1 | 10 | Rexo1 | -0.038 | Young |
| 0.446631 | 0.111094 | 0.192 | 0.242 | 1 | 10 | Rnf10 | -0.05 | Young |
| 0.447548 | 1.288973 | 0.091 | 0.071 | 1 | 10 | Pdcd2 | 0.02 | Young |

| | | | | | | | | |
|---|---|---|---|---|---|---|---|---|
| 0.44755 | 0.111259 | 0.404 | 0.472 | 1 | 10 | Psmb3 | -0.068 | Young |
| 0.447777 | 1.002949 | 0.081 | 0.061 | 1 | 10 | Zfp142 | 0.02 | Young |
| 0.447827 | 1.065114 | 0.051 | 0.034 | 1 | 10 | Asb13 | 0.017 | Young |
| 0.447827 | 0.385468 | 0.051 | 0.034 | 1 | 10 | Zfp637 | 0.017 | Young |
| 0.447865 | 0.407761 | 0.323 | 0.315 | 1 | 10 | Dgkd | 0.008 | Young |
| 0.44849 | 0.192674 | 0.121 | 0.095 | 1 | 10 | Mrps18b | 0.026 | Young |
| 0.448742 | 0.419023 | 0.172 | 0.144 | 1 | 10 | Ints1 | 0.028 | Young |
| 0.448784 | 0.015255 | 0.273 | 0.323 | 1 | 10 | Emd | -0.05 | Young |
| 0.448927 | 1.680983 | 0.04 | 0.027 | 1 | 10 | Chaf1a | 0.013 | Young |
| 0.449056 | 0.145328 | 0.232 | 0.289 | 1 | 10 | Rnps1 | -0.057 | Young |
| 0.449236 | 0.107576 | 0.263 | 0.323 | 1 | 10 | 2510039O1 | -0.06 | Young |
| 0.449533 | 0.650173 | 0.03 | 0.049 | 1 | 10 | Fbxl8 | -0.019 | Young |
| 0.449858 | 0.228905 | 0.111 | 0.147 | 1 | 10 | Ttc9c | -0.036 | Young |
| 0.449859 | 0.051698 | 0.141 | 0.178 | 1 | 10 | Zfp950 | -0.037 | Young |
| 0.450206 | 0.470879 | 0.131 | 0.105 | 1 | 10 | Il13ra1 | 0.026 | Young |
| 0.450309 | 0.246535 | 0.141 | 0.117 | 1 | 10 | Las1l | 0.024 | Young |
| 0.450587 | 0.529454 | 0.182 | 0.159 | 1 | 10 | Zfp330 | 0.023 | Young |
| 0.450611 | 0.363843 | 0.071 | 0.051 | 1 | 10 | Eef1akmt2 | 0.02 | Young |
| 0.451149 | 0.435768 | 0.374 | 0.447 | 1 | 10 | Hist1h1c | -0.073 | Young |
| 0.451408 | 0.314342 | 0.222 | 0.196 | 1 | 10 | Ppm1g | 0.026 | Young |
| 0.451653 | 0.242084 | 0.232 | 0.2 | 1 | 10 | Gpr183 | 0.032 | Young |
| 0.452056 | 0.416519 | 0.091 | 0.068 | 1 | 10 | Mex3d | 0.023 | Young |
| 0.452131 | 0.28211 | 0.172 | 0.142 | 1 | 10 | Lcor | 0.03 | Young |
| 0.452157 | 0.312499 | 0.273 | 0.242 | 1 | 10 | Timm8b | 0.031 | Young |
| 0.452323 | 0.097402 | 0.394 | 0.472 | 1 | 10 | Supt16 | -0.078 | Young |
| 0.452796 | 0.055572 | 0.354 | 0.428 | 1 | 10 | Clec4b1 | -0.074 | Young |
| 0.452922 | 0.158526 | 0.121 | 0.152 | 1 | 10 | Reps1 | -0.031 | Young |
| 0.453244 | 0.219838 | 0.343 | 0.306 | 1 | 10 | Eif2s1 | 0.037 | Young |
| 0.4534 | 0.327224 | 0.051 | 0.034 | 1 | 10 | Gpr85 | 0.017 | Young |
| 0.454093 | 0.580547 | 0.071 | 0.051 | 1 | 10 | Tada2b | 0.02 | Young |
| 0.454169 | 0.3966 | 0.212 | 0.183 | 1 | 10 | Got2 | 0.029 | Young |
| 0.454229 | 1.471049 | 0.01 | 0.022 | 1 | 10 | Spint2 | -0.012 | Young |
| 0.454229 | 0.822981 | 0.01 | 0.022 | 1 | 10 | Gatsl3 | -0.012 | Young |
| 0.454229 | 0.740692 | 0.01 | 0.022 | 1 | 10 | Pla2g6 | -0.012 | Young |
| 0.454229 | 0.494216 | 0.01 | 0.022 | 1 | 10 | 1700056N1 | -0.012 | Young |
| 0.454229 | 0.415172 | 0.01 | 0.022 | 1 | 10 | AC163032. | -0.012 | Young |
| 0.454229 | 0.374247 | 0.01 | 0.022 | 1 | 10 | 9930104L0 | -0.012 | Young |
| 0.454229 | 0.099158 | 0.01 | 0.022 | 1 | 10 | Afg1l | -0.012 | Young |
| 0.454229 | 0.01044 | 0.01 | 0.022 | 1 | 10 | 1810009A1 | -0.012 | Young |
| 0.454232 | 0.373034 | 0.071 | 0.098 | 1 | 10 | D730003I1 | -0.027 | Young |
| 0.454628 | 0.816009 | 0.03 | 0.049 | 1 | 10 | Gmds | -0.019 | Young |
| 0.45474 | 0.074137 | 0.697 | 0.636 | 1 | 10 | Anp32b | 0.061 | Young |
| 0.454796 | 0.358646 | 0.141 | 0.115 | 1 | 10 | Gnpat | 0.026 | Young |
| 0.454834 | 0.104606 | 0.545 | 0.645 | 1 | 10 | Mapre1 | -0.1 | Young |
| 0.455447 | 0.192738 | 0.141 | 0.183 | 1 | 10 | Mettl1 | -0.042 | Young |

| | | | | | | | | |
|---|---|---|---|---|---|---|---|---|
| 0.455548 | 0.184306 | 0.414 | 0.491 | 1 | 10 | Nfe2l1 | -0.077 | Young |
| 0.455612 | 0.334427 | 0.141 | 0.115 | 1 | 10 | Chml | 0.026 | Young |
| 0.45607 | 0.54075 | 0.101 | 0.078 | 1 | 10 | Gbp2 | 0.023 | Young |
| 0.45616 | 0.74745 | 0.192 | 0.169 | 1 | 10 | Eed | 0.023 | Young |
| 0.456202 | 0.183336 | 0.051 | 0.034 | 1 | 10 | Kbtbd4 | 0.017 | Young |
| 0.456289 | 0.420277 | 0.111 | 0.086 | 1 | 10 | Pyurf | 0.025 | Young |
| 0.456572 | 0.210322 | 0.364 | 0.335 | 1 | 10 | Med28 | 0.029 | Young |
| 0.456677 | 0.281584 | 0.162 | 0.132 | 1 | 10 | Upf3b | 0.03 | Young |
| 0.456758 | 0.378445 | 0.434 | 0.443 | 1 | 10 | Eif1b | -0.009 | Young |
| 0.45684 | 0.187274 | 0.737 | 0.822 | 1 | 10 | Cd84 | -0.085 | Young |
| 0.45686 | 0.060945 | 0.141 | 0.112 | 1 | 10 | Hopx | 0.029 | Young |
| 0.45717 | 0.23807 | 0.293 | 0.369 | 1 | 10 | Swap70 | -0.076 | Young |
| 0.457954 | 0.723179 | 0.131 | 0.108 | 1 | 10 | Lrrc8c | 0.023 | Young |
| 0.458374 | 0.252757 | 0.172 | 0.144 | 1 | 10 | Mknk1 | 0.028 | Young |
| 0.458629 | 0.473533 | 0.091 | 0.071 | 1 | 10 | Timp1 | 0.02 | Young |
| 0.458699 | 0.094015 | 0.101 | 0.13 | 1 | 10 | Oma1 | -0.029 | Young |
| 0.458728 | 0.588377 | 0.212 | 0.186 | 1 | 10 | Cluh | 0.026 | Young |
| 0.45884 | 0.574918 | 0.232 | 0.203 | 1 | 10 | Egln1 | 0.029 | Young |
| 0.458955 | 0.362072 | 0.606 | 0.604 | 1 | 10 | Srsf2 | 0.002 | Young |
| 0.459266 | 0.105413 | 0.182 | 0.232 | 1 | 10 | Nipsnap3b | -0.05 | Young |
| 0.459309 | 0.203929 | 0.283 | 0.355 | 1 | 10 | Arf6 | -0.072 | Young |
| 0.459376 | 0.205235 | 0.061 | 0.086 | 1 | 10 | Tmtc3 | -0.025 | Young |
| 0.459376 | 0.045929 | 0.061 | 0.086 | 1 | 10 | Gpr68 | -0.025 | Young |
| 0.459403 | 0.393005 | 0.212 | 0.186 | 1 | 10 | Agtpbp1 | 0.026 | Young |
| 0.459617 | 0.958099 | 0.081 | 0.061 | 1 | 10 | Dcakd | 0.02 | Young |
| 0.459617 | 0.736809 | 0.081 | 0.061 | 1 | 10 | Acyp1 | 0.02 | Young |
| 0.459722 | 0.102533 | 0.788 | 0.826 | 1 | 10 | Flna | -0.038 | Young |
| 0.459912 | 1.00662 | 0.04 | 0.027 | 1 | 10 | Slc14a1 | 0.013 | Young |
| 0.460351 | 0.192128 | 0.505 | 0.513 | 1 | 10 | Anxa7 | -0.008 | Young |
| 0.460379 | 0.992365 | 0.071 | 0.054 | 1 | 10 | Eci1 | 0.017 | Young |
| 0.460524 | 0.704933 | 0.182 | 0.156 | 1 | 10 | Exoc6b | 0.026 | Young |
| 0.460576 | 1.400202 | 0.061 | 0.044 | 1 | 10 | Asap2 | 0.017 | Young |
| 0.461029 | 0.217866 | 0.566 | 0.567 | 1 | 10 | Atp2a2 | -0.001 | Young |
| 0.46116 | 0.296085 | 0.485 | 0.526 | 1 | 10 | Rab10 | -0.041 | Young |
| 0.46136 | 0.012043 | 0.273 | 0.333 | 1 | 10 | Lsm14a | -0.06 | Young |
| 0.461494 | 1.285877 | 0.04 | 0.027 | 1 | 10 | Sirt3 | 0.013 | Young |
| 0.461494 | 0.810978 | 0.04 | 0.027 | 1 | 10 | Ncs1 | 0.013 | Young |
| 0.461788 | 0.455327 | 0.081 | 0.061 | 1 | 10 | Sft2d3 | 0.02 | Young |
| 0.462464 | 0.274956 | 0.283 | 0.254 | 1 | 10 | Pmaip1 | 0.029 | Young |
| 0.462547 | 0.068778 | 0.263 | 0.318 | 1 | 10 | Usp48 | -0.055 | Young |
| 0.462635 | 0.133143 | 0.273 | 0.347 | 1 | 10 | Slc29a1 | -0.074 | Young |
| 0.462795 | 0.37778 | 0.545 | 0.579 | 1 | 10 | Pgam1 | -0.034 | Young |
| 0.462981 | 0.32607 | 0.101 | 0.132 | 1 | 10 | Cog8 | -0.031 | Young |
| 0.463061 | 0.033502 | 0.131 | 0.169 | 1 | 10 | Iars2 | -0.038 | Young |
| 0.463079 | 1.375142 | 0.04 | 0.027 | 1 | 10 | Lrmda | 0.013 | Young |

| | | | | | | | | |
|---|---|---|---|---|---|---|---|---|
| 0.463103 | 0.825253 | 0.061 | 0.044 | 1 | 10 | Lrp5 | 0.017 | Young |
| 0.463521 | 0.140443 | 0.192 | 0.235 | 1 | 10 | Nbr1 | -0.043 | Young |
| 0.464003 | 0.770679 | 0.111 | 0.09 | 1 | 10 | Orc2 | 0.021 | Young |
| 0.46425 | 0.408637 | 0.111 | 0.137 | 1 | 10 | Pygb | -0.026 | Young |
| 0.464273 | 1.011239 | 0.101 | 0.081 | 1 | 10 | Ube2o | 0.02 | Young |
| 0.464369 | 1.250066 | 0.061 | 0.044 | 1 | 10 | Zfp574 | 0.017 | Young |
| 0.464369 | 1.106539 | 0.061 | 0.044 | 1 | 10 | Hdac9 | 0.017 | Young |
| 0.464727 | 0.167349 | 0.586 | 0.677 | 1 | 10 | Slc3a2 | -0.091 | Young |
| 0.465303 | 0.567276 | 0.202 | 0.176 | 1 | 10 | Limd2 | 0.026 | Young |
| 0.465637 | 0.673873 | 0.061 | 0.044 | 1 | 10 | Lrrc8b | 0.017 | Young |
| 0.466055 | 0.070799 | 0.141 | 0.178 | 1 | 10 | Mrps23 | -0.037 | Young |
| 0.466148 | 0.486904 | 0.081 | 0.061 | 1 | 10 | Nme6 | 0.02 | Young |
| 0.466257 | 0.500672 | 0.04 | 0.027 | 1 | 10 | B4galt4 | 0.013 | Young |
| 0.466305 | 0.308968 | 0.273 | 0.247 | 1 | 10 | Dtx4 | 0.026 | Young |
| 0.466907 | 0.733151 | 0.061 | 0.044 | 1 | 10 | Mlh3 | 0.017 | Young |
| 0.46699 | 0.144828 | 0.071 | 0.051 | 1 | 10 | Zswim7 | 0.02 | Young |
| 0.467391 | 0.129048 | 0.646 | 0.636 | 1 | 10 | Hspd1 | 0.01 | Young |
| 0.467808 | 0.658093 | 0.091 | 0.071 | 1 | 10 | Adck1 | 0.02 | Young |
| 0.467851 | 1.244667 | 0.04 | 0.027 | 1 | 10 | Serpinb1a | 0.013 | Young |
| 0.468179 | 0.580999 | 0.061 | 0.044 | 1 | 10 | Chchd6 | 0.017 | Young |
| 0.468327 | 0.175932 | 0.192 | 0.24 | 1 | 10 | Fkbp3 | -0.048 | Young |
| 0.469433 | 0.64963 | 0.081 | 0.061 | 1 | 10 | Thap4 | 0.02 | Young |
| 0.469447 | 0.884898 | 0.04 | 0.027 | 1 | 10 | Pus7l | 0.013 | Young |
| 0.470294 | 0.191272 | 0.081 | 0.11 | 1 | 10 | Rif1 | -0.029 | Young |
| 0.470628 | 0.165587 | 0.263 | 0.32 | 1 | 10 | Sp3 | -0.057 | Young |
| 0.470894 | 0.019327 | 0.202 | 0.169 | 1 | 10 | Cd72 | 0.033 | Young |
| 0.470946 | 0.478506 | 0.283 | 0.254 | 1 | 10 | Ndfip2 | 0.029 | Young |
| 0.47135 | 0.382438 | 0.394 | 0.379 | 1 | 10 | Cnppd1 | 0.015 | Young |
| 0.471468 | 0.601019 | 0.172 | 0.149 | 1 | 10 | Sec24a | 0.023 | Young |
| 0.471842 | 0.45284 | 0.051 | 0.073 | 1 | 10 | Wdr70 | -0.022 | Young |
| 0.471989 | 1.081978 | 0.071 | 0.054 | 1 | 10 | 2810402E2 | 0.017 | Young |
| 0.472065 | 0.820979 | 0.121 | 0.1 | 1 | 10 | Lrch1 | 0.021 | Young |
| 0.472115 | 0.348191 | 0.343 | 0.33 | 1 | 10 | Sec63 | 0.013 | Young |
| 0.472165 | 0.23761 | 0.222 | 0.269 | 1 | 10 | Emc3 | -0.047 | Young |
| 0.473189 | 0.301427 | 0.222 | 0.188 | 1 | 10 | Tifa | 0.034 | Young |
| 0.473253 | 0.169885 | 0.475 | 0.474 | 1 | 10 | Hprt | 0.001 | Young |
| 0.473514 | 1.04062 | 0.04 | 0.059 | 1 | 10 | Cav1 | -0.019 | Young |
| 0.473637 | 0.17675 | 0.101 | 0.078 | 1 | 10 | Abhd13 | 0.023 | Young |
| 0.473657 | 0.390885 | 0.182 | 0.156 | 1 | 10 | Ipmk | 0.026 | Young |
| 0.473668 | 0.486414 | 0.283 | 0.257 | 1 | 10 | Synj2bp | 0.026 | Young |
| 0.473685 | 0.101931 | 0.202 | 0.242 | 1 | 10 | Wwp2 | -0.04 | Young |
| 0.473832 | 0.426305 | 0.081 | 0.061 | 1 | 10 | Zcchc10 | 0.02 | Young |
| 0.474255 | 0.576665 | 0.04 | 0.027 | 1 | 10 | B3galt4 | 0.013 | Young |
| 0.474976 | 0.103819 | 0.303 | 0.379 | 1 | 10 | Pcna | -0.076 | Young |
| 0.475003 | 0.196926 | 0.818 | 0.853 | 1 | 10 | Creg1 | -0.035 | Young |

| | | | | | | | | |
|---|---|---|---|---|---|---|---|---|
| 0.475017 | 0.843753 | 0.091 | 0.071 | 1 | 10 | Dus3l | 0.02 | Young |
| 0.47502 | 0.818199 | 0.141 | 0.12 | 1 | 10 | Ppp2r3d | 0.021 | Young |
| 0.47502 | 0.555071 | 0.141 | 0.12 | 1 | 10 | Sumf1 | 0.021 | Young |
| 0.475084 | 0.109755 | 0.747 | 0.729 | 1 | 10 | Mtpn | 0.018 | Young |
| 0.475869 | 0.13693 | 0.141 | 0.183 | 1 | 10 | Dera | -0.042 | Young |
| 0.47591 | 0.109918 | 0.687 | 0.68 | 1 | 10 | Lilr4b | 0.007 | Young |
| 0.475997 | 0.504872 | 0.323 | 0.315 | 1 | 10 | Fam91a1 | 0.008 | Young |
| 0.476041 | 0.755674 | 0.081 | 0.061 | 1 | 10 | Rdm1 | 0.02 | Young |
| 0.476241 | 0.447402 | 0.202 | 0.183 | 1 | 10 | Ccng2 | 0.019 | Young |
| 0.476668 | 0.324814 | 0.091 | 0.12 | 1 | 10 | Rnf126 | -0.029 | Young |
| 0.477039 | 0.683825 | 0.333 | 0.342 | 1 | 10 | Hnrnpr | -0.009 | Young |
| 0.477137 | 0.647585 | 0.061 | 0.044 | 1 | 10 | Rtn1 | 0.017 | Young |
| 0.477787 | 0.848962 | 0.162 | 0.139 | 1 | 10 | Zeb2os | 0.023 | Young |
| 0.477967 | 0.104299 | 0.162 | 0.203 | 1 | 10 | Ddx56 | -0.041 | Young |
| 0.478748 | 0.402671 | 0.253 | 0.301 | 1 | 10 | Zfp644 | -0.048 | Young |
| 0.479682 | 0.109125 | 0.889 | 0.878 | 1 | 10 | Rrbp1 | 0.011 | Young |
| 0.479879 | 0.455609 | 0.626 | 0.633 | 1 | 10 | Pgk1 | -0.007 | Young |
| 0.480023 | 0.601926 | 0.212 | 0.191 | 1 | 10 | Lap3 | 0.021 | Young |
| 0.480094 | 0.971586 | 0.141 | 0.122 | 1 | 10 | Atp11a | 0.019 | Young |
| 0.480179 | 0.074335 | 0.091 | 0.117 | 1 | 10 | Pigq | -0.026 | Young |
| 0.480204 | 0.548062 | 0.091 | 0.071 | 1 | 10 | Tmem159 | 0.02 | Young |
| 0.480465 | 0.08838 | 0.879 | 0.944 | 1 | 10 | H3f3a | -0.065 | Young |
| 0.480708 | 0.531402 | 0.04 | 0.027 | 1 | 10 | 4933427D1 | 0.013 | Young |
| 0.481004 | 1.023966 | 0.061 | 0.044 | 1 | 10 | Gm20186 | 0.017 | Young |
| 0.48216 | 0.280073 | 0.051 | 0.071 | 1 | 10 | Zfp780b | -0.02 | Young |
| 0.482329 | 0.20537 | 0.04 | 0.027 | 1 | 10 | Slc2a9 | 0.013 | Young |
| 0.482451 | 0.152763 | 0.02 | 0.034 | 1 | 10 | Zfp974 | -0.014 | Young |
| 0.482656 | 0.310324 | 0.364 | 0.335 | 1 | 10 | Aimp1 | 0.029 | Young |
| 0.482699 | 0.308951 | 0.081 | 0.061 | 1 | 10 | Ndufaf7 | 0.02 | Young |
| 0.482736 | 0.254177 | 0.495 | 0.496 | 1 | 10 | Prpf40a | -0.001 | Young |
| 0.483268 | 0.427216 | 0.182 | 0.156 | 1 | 10 | Rhot1 | 0.026 | Young |
| 0.483433 | 0.663234 | 0.232 | 0.215 | 1 | 10 | Stk3 | 0.017 | Young |
| 0.483592 | 0.106338 | 0.061 | 0.044 | 1 | 10 | Plscr1 | 0.017 | Young |
| 0.483813 | 0.113234 | 0.798 | 0.802 | 1 | 10 | Cox5a | -0.004 | Young |
| 0.483843 | 0.754169 | 0.152 | 0.13 | 1 | 10 | Anapc10 | 0.022 | Young |
| 0.483952 | 0.007467 | 0.04 | 0.027 | 1 | 10 | Fancc | 0.013 | Young |
| 0.484053 | 0.302294 | 0.192 | 0.249 | 1 | 10 | Orai1 | -0.057 | Young |
| 0.484054 | 1.2208 | 0.051 | 0.037 | 1 | 10 | Rhbdf1 | 0.014 | Young |
| 0.484164 | 0.242036 | 0.152 | 0.193 | 1 | 10 | Eef1e1 | -0.041 | Young |
| 0.484219 | 0.854362 | 0.141 | 0.122 | 1 | 10 | Sipa1l2 | 0.019 | Young |
| 0.484622 | 0.548701 | 0.343 | 0.35 | 1 | 10 | Nrd1 | -0.007 | Young |
| 0.484889 | 0.252559 | 0.061 | 0.044 | 1 | 10 | Zfp715 | 0.017 | Young |
| 0.484934 | 0.213707 | 0.616 | 0.616 | 1 | 10 | Selenow | 0 | Young |
| 0.484977 | 0.180875 | 0.111 | 0.142 | 1 | 10 | Kansl3 | -0.031 | Young |
| 0.485778 | 0.745231 | 0.152 | 0.132 | 1 | 10 | Camsap2 | 0.02 | Young |

| | | | | | | | | |
|---|---|---|---|---|---|---|---|---|
| 0.486262 | 0.431199 | 0.152 | 0.13 | 1 | 10 | Tmem57 | 0.022 | Young |
| 0.486522 | 0.502416 | 0.141 | 0.117 | 1 | 10 | Vps16 | 0.024 | Young |
| 0.486866 | 0.30904 | 0.111 | 0.144 | 1 | 10 | Gmppa | -0.033 | Young |
| 0.487139 | 0.31711 | 0.515 | 0.545 | 1 | 10 | Cers2 | -0.03 | Young |
| 0.48717 | 0.150676 | 0.778 | 0.768 | 1 | 10 | Colgalt1 | 0.01 | Young |
| 0.487448 | 0.114961 | 0.374 | 0.443 | 1 | 10 | Pik3cd | -0.069 | Young |
| 0.487532 | 0.023501 | 0.172 | 0.21 | 1 | 10 | Ing2 | -0.038 | Young |
| 0.487885 | 0.467206 | 0.192 | 0.174 | 1 | 10 | Pde3b | 0.018 | Young |
| 0.488261 | 1.413656 | 0.03 | 0.02 | 1 | 10 | Socs2 | 0.01 | Young |
| 0.48839 | 0.053168 | 0.152 | 0.122 | 1 | 10 | Arhgap11a | 0.03 | Young |
| 0.488512 | 0.767935 | 0.071 | 0.054 | 1 | 10 | Dclre1a | 0.017 | Young |
| 0.488604 | 0.135858 | 0.152 | 0.183 | 1 | 10 | Ezh1 | -0.031 | Young |
| 0.488764 | 0.413165 | 0.374 | 0.362 | 1 | 10 | Lacc1 | 0.012 | Young |
| 0.48879 | 0.455445 | 0.061 | 0.044 | 1 | 10 | Tatdn1 | 0.017 | Young |
| 0.489276 | 0.163291 | 0.212 | 0.252 | 1 | 10 | Bicd2 | -0.04 | Young |
| 0.489321 | 0.586502 | 0.212 | 0.188 | 1 | 10 | Kif2a | 0.024 | Young |
| 0.489441 | 0.13899 | 0.172 | 0.215 | 1 | 10 | Wdr37 | -0.043 | Young |
| 0.489533 | 1.063587 | 0.121 | 0.103 | 1 | 10 | Coasy | 0.018 | Young |
| 0.489535 | 0.105611 | 0.182 | 0.225 | 1 | 10 | Tbrg1 | -0.043 | Young |
| 0.489827 | 0.198601 | 0.687 | 0.716 | 1 | 10 | Litaf | -0.029 | Young |
| 0.490026 | 0.644809 | 0.101 | 0.132 | 1 | 10 | Fcna | -0.031 | Young |
| 0.490163 | 1.965028 | 0.03 | 0.02 | 1 | 10 | Zfp580 | 0.01 | Young |
| 0.49032 | 0.018507 | 0.152 | 0.188 | 1 | 10 | Med17 | -0.036 | Young |
| 0.490347 | 0.018082 | 0.02 | 0.034 | 1 | 10 | A430106G: | -0.014 | Young |
| 0.490438 | 0.002785 | 0.121 | 0.152 | 1 | 10 | Slc35e4 | -0.031 | Young |
| 0.490494 | 0.148342 | 0.192 | 0.232 | 1 | 10 | Preb | -0.04 | Young |
| 0.490669 | 0.376283 | 0.091 | 0.071 | 1 | 10 | Plcb2 | 0.02 | Young |
| 0.490713 | 0.257803 | 0.111 | 0.088 | 1 | 10 | Vps8 | 0.023 | Young |
| 0.49104 | 0.050212 | 0.99 | 0.973 | 1 | 10 | Lamp1 | 0.017 | Young |
| 0.491127 | 0.365002 | 0.384 | 0.357 | 1 | 10 | Ndufaf8 | 0.027 | Young |
| 0.491218 | 0.260727 | 0.172 | 0.213 | 1 | 10 | Ado | -0.041 | Young |
| 0.491231 | 0.065204 | 0.253 | 0.301 | 1 | 10 | Pan3 | -0.048 | Young |
| 0.491292 | 0.184408 | 0.071 | 0.095 | 1 | 10 | Calcoco1 | -0.024 | Young |
| 0.49143 | 0.112488 | 0.03 | 0.046 | 1 | 10 | Aldh18a1 | -0.016 | Young |
| 0.491673 | 1.19058 | 0.091 | 0.073 | 1 | 10 | Lig3 | 0.018 | Young |
| 0.49169 | 0.483632 | 0.192 | 0.171 | 1 | 10 | Chd7 | 0.021 | Young |
| 0.492068 | 1.947382 | 0.03 | 0.02 | 1 | 10 | Rab34 | 0.01 | Young |
| 0.492068 | 1.058719 | 0.03 | 0.02 | 1 | 10 | Vars2 | 0.01 | Young |
| 0.492121 | 0.097265 | 0.232 | 0.284 | 1 | 10 | Mrpl48 | -0.052 | Young |
| 0.49221 | 0.043153 | 0.091 | 0.12 | 1 | 10 | Rpf2 | -0.029 | Young |
| 0.492221 | 0.213062 | 0.485 | 0.479 | 1 | 10 | Ppib | 0.006 | Young |
| 0.492458 | 0.260975 | 0.404 | 0.381 | 1 | 10 | Psen2 | 0.023 | Young |
| 0.492651 | 0.752831 | 0.081 | 0.064 | 1 | 10 | Tpm3-rs7 | 0.017 | Young |
| 0.492709 | 0.269005 | 0.061 | 0.044 | 1 | 10 | Tapt1 | 0.017 | Young |
| 0.492722 | 0.006117 | 0.838 | 0.853 | 1 | 10 | Ptbp3 | -0.015 | Young |

| | | | | | | | | |
|---|---|---|---|---|---|---|---|---|
| 0.492775 | 0.630279 | 0.04 | 0.059 | 1 | 10 | Slc35d2 | -0.019 | Young |
| 0.493088 | 0.474838 | 0.202 | 0.252 | 1 | 10 | Nfu1 | -0.05 | Young |
| 0.493206 | 0.126371 | 0.051 | 0.071 | 1 | 10 | Grwd1 | -0.02 | Young |
| 0.493264 | 0.130346 | 0.091 | 0.117 | 1 | 10 | Lrrcc1 | -0.026 | Young |
| 0.493272 | 0.180048 | 0.101 | 0.127 | 1 | 10 | Naip5 | -0.026 | Young |
| 0.493289 | 0.848911 | 0.071 | 0.054 | 1 | 10 | Zfp64 | 0.017 | Young |
| 0.493389 | 0.082123 | 0.404 | 0.469 | 1 | 10 | Tgif1 | -0.065 | Young |
| 0.493476 | 0.35929 | 0.293 | 0.267 | 1 | 10 | Icam1 | 0.026 | Young |
| 0.493521 | 0.037151 | 0.556 | 0.633 | 1 | 10 | Nisch | -0.077 | Young |
| 0.493525 | 0.224727 | 0.02 | 0.034 | 1 | 10 | Naif1 | -0.014 | Young |
| 0.493525 | 0.20601 | 0.02 | 0.034 | 1 | 10 | Pvt1 | -0.014 | Young |
| 0.493832 | 0.314433 | 0.091 | 0.071 | 1 | 10 | Bag4 | 0.02 | Young |
| 0.493839 | 0.277021 | 0.566 | 0.567 | 1 | 10 | Sdha | -0.001 | Young |
| 0.493901 | 0.417021 | 0.152 | 0.188 | 1 | 10 | Mrpl11 | -0.036 | Young |
| 0.493977 | 1.426961 | 0.03 | 0.02 | 1 | 10 | Dcun1d4 | 0.01 | Young |
| 0.493977 | 1.381027 | 0.03 | 0.02 | 1 | 10 | Steap4 | 0.01 | Young |
| 0.493977 | 1.309139 | 0.03 | 0.02 | 1 | 10 | Caprin2 | 0.01 | Young |
| 0.493977 | 1.026079 | 0.03 | 0.02 | 1 | 10 | Zfp553 | 0.01 | Young |
| 0.494068 | 0.274715 | 0.121 | 0.154 | 1 | 10 | Zfp24 | -0.033 | Young |
| 0.494389 | 0.398086 | 0.424 | 0.428 | 1 | 10 | Smim14 | -0.004 | Young |
| 0.494655 | 0.335895 | 0.192 | 0.237 | 1 | 10 | Nfkb2 | -0.045 | Young |
| 0.495119 | 0.10372 | 0.02 | 0.034 | 1 | 10 | Zfp719 | -0.014 | Young |
| 0.49513 | 0.238122 | 0.242 | 0.303 | 1 | 10 | Ppt2 | -0.061 | Young |
| 0.495205 | 0.170018 | 0.646 | 0.614 | 1 | 10 | Purb | 0.032 | Young |
| 0.495527 | 0.044038 | 0.03 | 0.046 | 1 | 10 | Ttf2 | -0.016 | Young |
| 0.495687 | 0.168292 | 0.071 | 0.054 | 1 | 10 | Armc7 | 0.017 | Young |
| 0.495868 | 0.095374 | 0.859 | 0.873 | 1 | 10 | Ywhae | -0.014 | Young |
| 0.49645 | 0.319808 | 0.141 | 0.178 | 1 | 10 | Faf2 | -0.037 | Young |
| 0.496519 | 0.272199 | 0.172 | 0.147 | 1 | 10 | Slc35f5 | 0.025 | Young |
| 0.496714 | 0.973014 | 0.02 | 0.034 | 1 | 10 | Mypopos | -0.014 | Young |
| 0.496731 | 0.843094 | 0.121 | 0.103 | 1 | 10 | Tbc1d31 | 0.018 | Young |
| 0.497389 | 0.307008 | 0.374 | 0.367 | 1 | 10 | Ptbp1 | 0.007 | Young |
| 0.49778 | 0.113498 | 0.949 | 0.966 | 1 | 10 | Fcer1g | -0.017 | Young |
| 0.497807 | 1.295393 | 0.03 | 0.02 | 1 | 10 | Gnb5 | 0.01 | Young |
| 0.497807 | 0.943529 | 0.03 | 0.02 | 1 | 10 | Cep41 | 0.01 | Young |
| 0.498304 | 0.34367 | 0.242 | 0.218 | 1 | 10 | Hdac5 | 0.024 | Young |
| 0.499276 | 0.064509 | 0.061 | 0.044 | 1 | 10 | Usp53 | 0.017 | Young |
| 0.499296 | 0.458926 | 0.071 | 0.054 | 1 | 10 | Gpx3 | 0.017 | Young |
| 0.499591 | 0.757311 | 0.111 | 0.09 | 1 | 10 | Rnf40 | 0.021 | Young |
| 0.499642 | 0.124035 | 0.03 | 0.046 | 1 | 10 | Mcph1 | -0.016 | Young |
| 0.499642 | 0.05031 | 0.03 | 0.046 | 1 | 10 | Tfec | -0.016 | Young |
| 0.499728 | 1.115017 | 0.03 | 0.02 | 1 | 10 | Gstm5 | 0.01 | Young |
| 0.499811 | 0.45893 | 0.273 | 0.254 | 1 | 10 | 2010111I0 | 0.019 | Young |
| 0.500104 | 0.445719 | 0.04 | 0.059 | 1 | 10 | Gpsm2 | -0.019 | Young |
| 0.500501 | 0.397736 | 0.141 | 0.174 | 1 | 10 | Prpf6 | -0.033 | Young |

| | | | | | | | | |
|---|---|---|---|---|---|---|---|---|
| 0.500557 | 0.388321 | 0.485 | 0.479 | 1 | 10 | Dek | 0.006 | Young |
| 0.500665 | 0.422447 | 0.364 | 0.362 | 1 | 10 | Abcf1 | 0.002 | Young |
| 0.500774 | 0.513626 | 0.283 | 0.357 | 1 | 10 | Ccnd1 | -0.074 | Young |
| 0.500999 | 0.099151 | 0.253 | 0.306 | 1 | 10 | Dennd6a | -0.053 | Young |
| 0.501189 | 1.12414 | 0.051 | 0.037 | 1 | 10 | Plxna4 | 0.014 | Young |
| 0.501443 | 0.437848 | 0.141 | 0.176 | 1 | 10 | Dpm2 | -0.035 | Young |
| 0.501653 | 1.672535 | 0.03 | 0.02 | 1 | 10 | Gm49249 | 0.01 | Young |
| 0.501653 | 0.97322 | 0.03 | 0.02 | 1 | 10 | Zfp469 | 0.01 | Young |
| 0.501836 | 0.340221 | 0.081 | 0.061 | 1 | 10 | Espn | 0.02 | Young |
| 0.501898 | 0.066395 | 0.202 | 0.235 | 1 | 10 | Nploc4 | -0.033 | Young |
| 0.501916 | 0.16435 | 0.061 | 0.044 | 1 | 10 | Mocs1 | 0.017 | Young |
| 0.501942 | 0.389599 | 0.172 | 0.149 | 1 | 10 | Slc7a1 | 0.023 | Young |
| 0.501969 | 0.171471 | 0.071 | 0.095 | 1 | 10 | Ctc1 | -0.024 | Young |
| 0.502068 | 0.567085 | 0.232 | 0.215 | 1 | 10 | Afg3l1 | 0.017 | Young |
| 0.502552 | 0.36241 | 0.152 | 0.13 | 1 | 10 | Pdk1 | 0.022 | Young |
| 0.502594 | 0.256174 | 0.101 | 0.132 | 1 | 10 | Heatr3 | -0.031 | Young |
| 0.502718 | 1.166978 | 0.111 | 0.095 | 1 | 10 | Nr1d2 | 0.016 | Young |
| 0.502734 | 0.460044 | 0.172 | 0.147 | 1 | 10 | Vps54 | 0.025 | Young |
| 0.502805 | 0.523238 | 0.182 | 0.156 | 1 | 10 | Pop5 | 0.026 | Young |
| 0.503093 | 0.736909 | 0.222 | 0.203 | 1 | 10 | Dck | 0.019 | Young |
| 0.503349 | 0.287745 | 0.212 | 0.188 | 1 | 10 | 0610030E2 | 0.024 | Young |
| 0.503582 | 1.780382 | 0.03 | 0.02 | 1 | 10 | Arhgef10 | 0.01 | Young |
| 0.503582 | 1.076202 | 0.03 | 0.02 | 1 | 10 | Slpi | 0.01 | Young |
| 0.503582 | 0.935539 | 0.03 | 0.02 | 1 | 10 | Xlr3b | 0.01 | Young |
| 0.503582 | 0.907444 | 0.03 | 0.02 | 1 | 10 | Abca6 | 0.01 | Young |
| 0.503582 | 0.895485 | 0.03 | 0.02 | 1 | 10 | Tnfrsf23 | 0.01 | Young |
| 0.503657 | 0.195929 | 0.162 | 0.137 | 1 | 10 | Cmc2 | 0.025 | Young |
| 0.503774 | 0.270554 | 0.03 | 0.046 | 1 | 10 | Kptn | -0.016 | Young |
| 0.503825 | 1.176723 | 0.081 | 0.064 | 1 | 10 | Zfp180 | 0.017 | Young |
| 0.503825 | 0.662125 | 0.081 | 0.064 | 1 | 10 | Gdap2 | 0.017 | Young |
| 0.504379 | 0.428383 | 0.111 | 0.09 | 1 | 10 | Myef2 | 0.021 | Young |
| 0.504588 | 0.072866 | 0.071 | 0.093 | 1 | 10 | Rbpms | -0.022 | Young |
| 0.504897 | 0.795169 | 0.121 | 0.103 | 1 | 10 | Naa30 | 0.018 | Young |
| 0.505079 | 0.297972 | 0.081 | 0.108 | 1 | 10 | Taf12 | -0.027 | Young |
| 0.505211 | 0.259366 | 0.242 | 0.215 | 1 | 10 | Ahsa1 | 0.027 | Young |
| 0.505523 | 0.410251 | 0.091 | 0.071 | 1 | 10 | Fas | 0.02 | Young |
| 0.505845 | 0.066003 | 0.111 | 0.139 | 1 | 10 | Mmgt1 | -0.028 | Young |
| 0.506075 | 0.890332 | 0.081 | 0.064 | 1 | 10 | Ptk2 | 0.017 | Young |
| 0.506228 | 0.115415 | 0.818 | 0.851 | 1 | 10 | Unc93b1 | -0.033 | Young |
| 0.506456 | 0.125592 | 0.111 | 0.142 | 1 | 10 | Dnajb4 | -0.031 | Young |
| 0.506585 | 0.202441 | 0.061 | 0.083 | 1 | 10 | Mvd | -0.022 | Young |
| 0.50667 | 0.539442 | 0.152 | 0.13 | 1 | 10 | Chsy1 | 0.022 | Young |
| 0.506919 | 0.251909 | 0.374 | 0.355 | 1 | 10 | Tiparp | 0.019 | Young |
| 0.50743 | 0.131469 | 0.091 | 0.117 | 1 | 10 | Trim34a | -0.026 | Young |
| 0.507451 | 0.693459 | 0.03 | 0.02 | 1 | 10 | Piwil2 | 0.01 | Young |

| | | | | | | | | |
|---|---|---|---|---|---|---|---|---|
| 0.507466 | 0.233227 | 0.434 | 0.408 | 1 | 10 | Atp13a2 | 0.026 | Young |
| 0.507517 | 0.384582 | 0.465 | 0.482 | 1 | 10 | Ep300 | -0.017 | Young |
| 0.507662 | 0.230082 | 0.657 | 0.645 | 1 | 10 | Epb41l2 | 0.012 | Young |
| 0.50777 | 0.772563 | 0.071 | 0.054 | 1 | 10 | Sec61a2 | 0.017 | Young |
| 0.507779 | 0.466336 | 0.141 | 0.117 | 1 | 10 | Tex10 | 0.024 | Young |
| 0.508181 | 0.145297 | 0.384 | 0.46 | 1 | 10 | Senp2 | -0.076 | Young |
| 0.508518 | 0.902471 | 0.141 | 0.122 | 1 | 10 | Thap11 | 0.019 | Young |
| 0.508518 | 0.644877 | 0.141 | 0.122 | 1 | 10 | Nfkbib | 0.019 | Young |
| 0.508714 | 0.017561 | 0.162 | 0.198 | 1 | 10 | Rnf5 | -0.036 | Young |
| 0.508728 | 0.327097 | 0.04 | 0.059 | 1 | 10 | Fzd5 | -0.019 | Young |
| 0.508893 | 0.038787 | 0.626 | 0.689 | 1 | 10 | Capns1 | -0.063 | Young |
| 0.509392 | 0.746282 | 0.03 | 0.02 | 1 | 10 | Aqp9 | 0.01 | Young |
| 0.509894 | 0.612716 | 0.313 | 0.303 | 1 | 10 | Cnpy2 | 0.01 | Young |
| 0.510197 | 0.023233 | 0.929 | 0.968 | 1 | 10 | Rpl23a | -0.039 | Young |
| 0.510604 | 0.714872 | 0.091 | 0.073 | 1 | 10 | Chn2 | 0.018 | Young |
| 0.511097 | 0.164835 | 0.263 | 0.318 | 1 | 10 | Suz12 | -0.055 | Young |
| 0.511331 | 0.736901 | 0.051 | 0.037 | 1 | 10 | Zfp579 | 0.014 | Young |
| 0.511336 | 0.423038 | 0.03 | 0.02 | 1 | 10 | Sesn3 | 0.01 | Young |
| 0.511336 | 0.14685 | 0.03 | 0.02 | 1 | 10 | Cd101 | 0.01 | Young |
| 0.511425 | 0.749554 | 0.071 | 0.054 | 1 | 10 | Pop1 | 0.017 | Young |
| 0.511591 | 0.17636 | 0.424 | 0.401 | 1 | 10 | Emc10 | 0.023 | Young |
| 0.511769 | 0.723868 | 0.192 | 0.169 | 1 | 10 | Slc35a5 | 0.023 | Young |
| 0.511802 | 0.161569 | 0.121 | 0.152 | 1 | 10 | Phf2 | -0.031 | Young |
| 0.512138 | 0.115768 | 0.081 | 0.105 | 1 | 10 | Epm2aip1 | -0.024 | Young |
| 0.512391 | 0.471046 | 0.202 | 0.181 | 1 | 10 | Mak16 | 0.021 | Young |
| 0.512463 | 0.027908 | 0.111 | 0.139 | 1 | 10 | Arl13b | -0.028 | Young |
| 0.512728 | 0.376333 | 0.182 | 0.156 | 1 | 10 | Gm47283 | 0.026 | Young |
| 0.512878 | 0.47836 | 0.141 | 0.12 | 1 | 10 | Mpp5 | 0.021 | Young |
| 0.512905 | 0.066731 | 0.394 | 0.452 | 1 | 10 | Ash1l | -0.058 | Young |
| 0.513285 | 0.492268 | 0.03 | 0.02 | 1 | 10 | Hcar2 | 0.01 | Young |
| 0.513285 | 0.4368 | 0.03 | 0.02 | 1 | 10 | Pex10 | 0.01 | Young |
| 0.513285 | 0.158143 | 0.03 | 0.02 | 1 | 10 | Ms4a4a | 0.01 | Young |
| 0.513294 | 0.328823 | 0.152 | 0.13 | 1 | 10 | Mphosph1 | 0.022 | Young |
| 0.513294 | 0.221061 | 0.242 | 0.284 | 1 | 10 | Tle4 | -0.042 | Young |
| 0.513365 | 1.012827 | 0.121 | 0.105 | 1 | 10 | Ing3 | 0.016 | Young |
| 0.513487 | 0.715685 | 0.03 | 0.046 | 1 | 10 | Tdp1 | -0.016 | Young |
| 0.513494 | 0.272094 | 0.606 | 0.623 | 1 | 10 | Rbx1 | -0.017 | Young |
| 0.514125 | 0.332943 | 0.152 | 0.13 | 1 | 10 | Coq7 | 0.022 | Young |
| 0.514863 | 0.151136 | 0.061 | 0.081 | 1 | 10 | Cdc27 | -0.02 | Young |
| 0.515128 | 0.131844 | 0.343 | 0.391 | 1 | 10 | Phf3 | -0.048 | Young |
| 0.515131 | 1.174933 | 0.081 | 0.064 | 1 | 10 | Wrap73 | 0.017 | Young |
| 0.515131 | 0.61513 | 0.081 | 0.064 | 1 | 10 | Agk | 0.017 | Young |
| 0.515164 | 0.136642 | 0.333 | 0.313 | 1 | 10 | Polr2f | 0.02 | Young |
| 0.515202 | 0.408136 | 0.172 | 0.149 | 1 | 10 | Atrn | 0.023 | Young |
| 0.51522 | 0.190476 | 0.253 | 0.303 | 1 | 10 | Gnl1 | -0.05 | Young |

| | | | | | | | | |
|---|---|---|---|---|---|---|---|---|
| 0.515237 | 0.508223 | 0.03 | 0.02 | 1 | 10 | Plac8 | 0.01 | Young |
| 0.515237 | 0.226182 | 0.03 | 0.02 | 1 | 10 | Mturn | 0.01 | Young |
| 0.515237 | 0.182072 | 0.03 | 0.02 | 1 | 10 | Wdr90 | 0.01 | Young |
| 0.515447 | 0.157396 | 0.434 | 0.494 | 1 | 10 | Grina | -0.06 | Young |
| 0.51571 | 0.511469 | 0.051 | 0.037 | 1 | 10 | Gstt2 | 0.014 | Young |
| 0.51571 | 0.456853 | 0.051 | 0.037 | 1 | 10 | Wdr24 | 0.014 | Young |
| 0.515713 | 1.055308 | 0.071 | 0.056 | 1 | 10 | Ppox | 0.015 | Young |
| 0.516588 | 0.590996 | 0.263 | 0.252 | 1 | 10 | Sema4a | 0.011 | Young |
| 0.51671 | 0.273782 | 0.222 | 0.271 | 1 | 10 | Phc3 | -0.049 | Young |
| 0.516777 | 0.067169 | 0.758 | 0.702 | 1 | 10 | Ran | 0.056 | Young |
| 0.51683 | 0.242837 | 0.919 | 0.885 | 1 | 10 | Eif3a | 0.034 | Young |
| 0.516994 | 0.322204 | 0.192 | 0.169 | 1 | 10 | Mctp1 | 0.023 | Young |
| 0.517019 | 0.472285 | 0.162 | 0.139 | 1 | 10 | Hax1 | 0.023 | Young |
| 0.517105 | 0.143338 | 0.303 | 0.369 | 1 | 10 | Ski | -0.066 | Young |
| 0.517118 | 0.054105 | 0.434 | 0.501 | 1 | 10 | Cdk9 | -0.067 | Young |
| 0.517193 | 1.380849 | 0.03 | 0.02 | 1 | 10 | H2-Q6 | 0.01 | Young |
| 0.517193 | 0.264358 | 0.03 | 0.02 | 1 | 10 | Itga1 | 0.01 | Young |
| 0.517805 | 0.128608 | 0.081 | 0.105 | 1 | 10 | Pogk | -0.024 | Young |
| 0.517828 | 0.127886 | 0.121 | 0.149 | 1 | 10 | Zdhhc6 | -0.028 | Young |
| 0.519358 | 0.510532 | 0.121 | 0.1 | 1 | 10 | Tfam | 0.021 | Young |
| 0.519534 | 0.210806 | 0.071 | 0.093 | 1 | 10 | Eef2k | -0.022 | Young |
| 0.519586 | 0.869897 | 0.121 | 0.103 | 1 | 10 | Riok2 | 0.018 | Young |
| 0.519769 | 0.305464 | 0.141 | 0.12 | 1 | 10 | Zfp608 | 0.021 | Young |
| 0.519806 | 0.950698 | 0.101 | 0.083 | 1 | 10 | Lonrf3 | 0.018 | Young |
| 0.519827 | 0.015364 | 0.293 | 0.328 | 1 | 10 | Wdr45b | -0.035 | Young |
| 0.51986 | 0.801545 | 0.111 | 0.093 | 1 | 10 | Tm7sf3 | 0.018 | Young |
| 0.520039 | 0.333294 | 0.485 | 0.489 | 1 | 10 | Eif3h | -0.004 | Young |
| 0.520112 | 0.331408 | 0.434 | 0.438 | 1 | 10 | Cenpb | -0.004 | Young |
| 0.520395 | 0.157848 | 0.323 | 0.367 | 1 | 10 | Lmo4 | -0.044 | Young |
| 0.520538 | 0.187456 | 0.071 | 0.093 | 1 | 10 | Cks2 | -0.022 | Young |
| 0.520752 | 0.110238 | 0.909 | 0.944 | 1 | 10 | Rpl24 | -0.035 | Young |
| 0.520795 | 0.314397 | 0.212 | 0.257 | 1 | 10 | Cnot3 | -0.045 | Young |
| 0.520819 | 0.739164 | 0.202 | 0.191 | 1 | 10 | Trim56 | 0.011 | Young |
| 0.520944 | 0.504633 | 0.293 | 0.286 | 1 | 10 | Mrpl17 | 0.007 | Young |
| 0.521235 | 0.046572 | 0.222 | 0.259 | 1 | 10 | Aco1 | -0.037 | Young |
| 0.521539 | 0.230473 | 0.162 | 0.2 | 1 | 10 | Tmem245 | -0.038 | Young |
| 0.521637 | 0.112417 | 0.111 | 0.139 | 1 | 10 | Smim26 | -0.028 | Young |
| 0.522049 | 0.023106 | 0.172 | 0.208 | 1 | 10 | Samd9l | -0.036 | Young |
| 0.523068 | 0.635269 | 0.162 | 0.142 | 1 | 10 | Afg3l2 | 0.02 | Young |
| 0.523084 | 0.149936 | 0.03 | 0.02 | 1 | 10 | Tmem150a | 0.01 | Young |
| 0.523084 | 0.091526 | 0.03 | 0.02 | 1 | 10 | E330009J0 | 0.01 | Young |
| 0.523117 | 0.133352 | 0.162 | 0.137 | 1 | 10 | Rprd1b | 0.025 | Young |
| 0.523512 | 0.167576 | 0.121 | 0.149 | 1 | 10 | Nudt9 | -0.028 | Young |
| 0.523551 | 0.007217 | 0.313 | 0.357 | 1 | 10 | Erp44 | -0.044 | Young |
| 0.524293 | 0.309109 | 0.212 | 0.191 | 1 | 10 | Zcchc17 | 0.021 | Young |

| | | | | | | | | |
|---|---|---|---|---|---|---|---|---|
| 0.524389 | 0.066769 | 0.141 | 0.176 | 1 | 10 | Mthfd2 | -0.035 | Young |
| 0.524511 | 0.832511 | 0.091 | 0.073 | 1 | 10 | Pomgnt1 | 0.018 | Young |
| 0.524526 | 0.288054 | 0.051 | 0.037 | 1 | 10 | Pdlim7 | 0.014 | Young |
| 0.524526 | 0.269322 | 0.051 | 0.037 | 1 | 10 | Cnr2 | 0.014 | Young |
| 0.525224 | 0.551309 | 0.212 | 0.193 | 1 | 10 | Rab24 | 0.019 | Young |
| 0.525511 | 0.061045 | 0.364 | 0.406 | 1 | 10 | Rb1cc1 | -0.042 | Young |
| 0.525945 | 0.101268 | 0.242 | 0.286 | 1 | 10 | Ubr4 | -0.044 | Young |
| 0.526055 | 0.128478 | 0.697 | 0.699 | 1 | 10 | St13 | -0.002 | Young |
| 0.52618 | 0.224957 | 0.071 | 0.054 | 1 | 10 | Rpgrip1 | 0.017 | Young |
| 0.526226 | 0.266928 | 0.323 | 0.303 | 1 | 10 | Kxd1 | 0.02 | Young |
| 0.526258 | 0.463849 | 0.313 | 0.303 | 1 | 10 | Gpbp1 | 0.01 | Young |
| 0.526586 | 0.270213 | 0.071 | 0.093 | 1 | 10 | Hscb | -0.022 | Young |
| 0.526646 | 0.788904 | 0.111 | 0.093 | 1 | 10 | Icmt | 0.018 | Young |
| 0.527001 | 0.969259 | 0.081 | 0.066 | 1 | 10 | Tsen15 | 0.015 | Young |
| 0.527188 | 0.168441 | 0.313 | 0.359 | 1 | 10 | R3hdm1 | -0.046 | Young |
| 0.527482 | 0.82758 | 0.051 | 0.037 | 1 | 10 | Nfatc2ip | 0.014 | Young |
| 0.527482 | 0.488682 | 0.051 | 0.037 | 1 | 10 | Abhd18 | 0.014 | Young |
| 0.527597 | 0.071085 | 0.071 | 0.093 | 1 | 10 | Kansl1l | -0.022 | Young |
| 0.527619 | 0.443084 | 0.111 | 0.093 | 1 | 10 | Fam210a | 0.018 | Young |
| 0.528636 | 0.199019 | 0.141 | 0.117 | 1 | 10 | Fam3a | 0.024 | Young |
| 0.528712 | 0.549517 | 0.192 | 0.235 | 1 | 10 | Esyt2 | -0.043 | Young |
| 0.528784 | 0.027161 | 0.808 | 0.822 | 1 | 10 | Serbp1 | -0.014 | Young |
| 0.528963 | 0.261245 | 0.051 | 0.037 | 1 | 10 | Dcaf6 | 0.014 | Young |
| 0.529354 | 0.49727 | 0.242 | 0.227 | 1 | 10 | Acads | 0.015 | Young |
| 0.529508 | 0.039259 | 0.091 | 0.115 | 1 | 10 | Mis18a | -0.024 | Young |
| 0.529868 | 0.091597 | 0.828 | 0.932 | 1 | 10 | Cd36 | -0.104 | Young |
| 0.52991 | 0.901089 | 0.091 | 0.073 | 1 | 10 | Gpatch4 | 0.018 | Young |
| 0.529951 | 0.591095 | 0.202 | 0.181 | 1 | 10 | Ppm1b | 0.021 | Young |
| 0.530171 | 0.338848 | 0.606 | 0.599 | 1 | 10 | Slc43a2 | 0.007 | Young |
| 0.5304 | 0.511496 | 0.242 | 0.215 | 1 | 10 | Pdcd11 | 0.027 | Young |
| 0.530427 | 0.021836 | 0.091 | 0.115 | 1 | 10 | Smagp | -0.024 | Young |
| 0.530457 | 2.02103 | 0.02 | 0.012 | 1 | 10 | Gm48604 | 0.008 | Young |
| 0.530457 | 1.945207 | 0.02 | 0.012 | 1 | 10 | Tm9sf1 | 0.008 | Young |
| 0.530457 | 1.575517 | 0.02 | 0.012 | 1 | 10 | Sigirr | 0.008 | Young |
| 0.530457 | 1.522493 | 0.02 | 0.012 | 1 | 10 | Thnsl1 | 0.008 | Young |
| 0.530457 | 0.985566 | 0.02 | 0.012 | 1 | 10 | Cdk5rap1 | 0.008 | Young |
| 0.53089 | 0.086342 | 0.323 | 0.377 | 1 | 10 | Zyx | -0.054 | Young |
| 0.530914 | 0.058368 | 0.535 | 0.584 | 1 | 10 | Prdx2 | -0.049 | Young |
| 0.530936 | 0.146058 | 0.152 | 0.181 | 1 | 10 | Acox3 | -0.029 | Young |
| 0.530974 | 0.505816 | 0.212 | 0.193 | 1 | 10 | Znrd1 | 0.019 | Young |
| 0.531251 | 0.085439 | 0.253 | 0.298 | 1 | 10 | Ptpn12 | -0.045 | Young |
| 0.531466 | 1.002898 | 0.091 | 0.076 | 1 | 10 | Rnf8 | 0.015 | Young |
| 0.531652 | 0.271318 | 0.071 | 0.093 | 1 | 10 | Pdhx | -0.022 | Young |
| 0.532078 | 0.100608 | 0.061 | 0.081 | 1 | 10 | Rmdn3 | -0.02 | Young |
| 0.532179 | 0.105352 | 0.778 | 0.785 | 1 | 10 | Gltp | -0.007 | Young |

| | | | | | | | | |
|---|---|---|---|---|---|---|---|---|
| 0.532328 | 0.63351 | 0.01 | 0.02 | 1 | 10 | Fsd1l | -0.01 | Young |
| 0.532328 | 0.317824 | 0.01 | 0.02 | 1 | 10 | Jcad | -0.01 | Young |
| 0.532328 | 0.256665 | 0.01 | 0.02 | 1 | 10 | Gm4631 | -0.01 | Young |
| 0.532328 | 0.226906 | 0.01 | 0.02 | 1 | 10 | Amigo1 | -0.01 | Young |
| 0.532328 | 0.210339 | 0.01 | 0.02 | 1 | 10 | Tarbp1 | -0.01 | Young |
| 0.532328 | 0.196773 | 0.01 | 0.02 | 1 | 10 | Tcaim | -0.01 | Young |
| 0.532937 | 2.318861 | 0.02 | 0.012 | 1 | 10 | AC130815. | 0.008 | Young |
| 0.532937 | 1.651323 | 0.02 | 0.012 | 1 | 10 | Egf | 0.008 | Young |
| 0.532937 | 1.404608 | 0.02 | 0.012 | 1 | 10 | Gfi1b | 0.008 | Young |
| 0.532937 | 1.156903 | 0.02 | 0.012 | 1 | 10 | Zfp870 | 0.008 | Young |
| 0.532937 | 0.746909 | 0.02 | 0.012 | 1 | 10 | Ippk | 0.008 | Young |
| 0.533074 | 0.389651 | 0.202 | 0.176 | 1 | 10 | Fam160b1 | 0.026 | Young |
| 0.533638 | 0.659004 | 0.131 | 0.108 | 1 | 10 | AI839979 | 0.023 | Young |
| 0.533708 | 0.192357 | 0.182 | 0.22 | 1 | 10 | Dazap1 | -0.038 | Young |
| 0.533824 | 0.414396 | 0.333 | 0.328 | 1 | 10 | Fam208a | 0.005 | Young |
| 0.53411 | 0.169317 | 0.101 | 0.125 | 1 | 10 | Lyl1 | -0.024 | Young |
| 0.534251 | 0.72533 | 0.061 | 0.081 | 1 | 10 | Zmat3 | -0.02 | Young |
| 0.534251 | 0.147828 | 0.061 | 0.081 | 1 | 10 | Ddx31 | -0.02 | Young |
| 0.534582 | 0.057087 | 0.394 | 0.443 | 1 | 10 | Eprs | -0.049 | Young |
| 0.53466 | 1.178323 | 0.061 | 0.046 | 1 | 10 | Zfp667 | 0.015 | Young |
| 0.53466 | 0.733593 | 0.061 | 0.046 | 1 | 10 | Tmem39b | 0.015 | Young |
| 0.53466 | 0.654935 | 0.061 | 0.046 | 1 | 10 | Trub1 | 0.015 | Young |
| 0.534704 | 0.620956 | 0.071 | 0.093 | 1 | 10 | Exosc1 | -0.022 | Young |
| 0.534784 | 0.103632 | 0.838 | 0.848 | 1 | 10 | Gnb1 | -0.01 | Young |
| 0.535422 | 1.603855 | 0.02 | 0.012 | 1 | 10 | Exog | 0.008 | Young |
| 0.535422 | 1.59345 | 0.02 | 0.012 | 1 | 10 | Pms1 | 0.008 | Young |
| 0.535422 | 0.853565 | 0.02 | 0.012 | 1 | 10 | Ak1 | 0.008 | Young |
| 0.535422 | 0.753637 | 0.02 | 0.012 | 1 | 10 | Cd1d1 | 0.008 | Young |
| 0.535422 | 0.525117 | 0.02 | 0.012 | 1 | 10 | AW146154 | 0.008 | Young |
| 0.53586 | 0.504775 | 0.212 | 0.191 | 1 | 10 | Ccdc88b | 0.021 | Young |
| 0.536122 | 0.809896 | 0.081 | 0.066 | 1 | 10 | 3110082I1 | 0.015 | Young |
| 0.536381 | 0.284922 | 0.04 | 0.056 | 1 | 10 | Npat | -0.016 | Young |
| 0.536427 | 0.688884 | 0.091 | 0.073 | 1 | 10 | Tpst1 | 0.018 | Young |
| 0.536427 | 0.209564 | 0.061 | 0.081 | 1 | 10 | Ears2 | -0.02 | Young |
| 0.536529 | 0.056262 | 0.99 | 0.988 | 1 | 10 | Rps20 | 0.002 | Young |
| 0.536564 | 0.574383 | 0.051 | 0.068 | 1 | 10 | B4galt7 | -0.017 | Young |
| 0.536783 | 0.639239 | 0.152 | 0.137 | 1 | 10 | Extl3 | 0.015 | Young |
| 0.536995 | 0.516565 | 0.182 | 0.164 | 1 | 10 | Scd2 | 0.018 | Young |
| 0.537283 | 0.053485 | 0.222 | 0.259 | 1 | 10 | Ube2e3 | -0.037 | Young |
| 0.537343 | 0.91725 | 0.061 | 0.046 | 1 | 10 | 6-Sep | 0.015 | Young |
| 0.537343 | 0.861302 | 0.061 | 0.046 | 1 | 10 | Invs | 0.015 | Young |
| 0.537343 | 0.483966 | 0.061 | 0.046 | 1 | 10 | Ptges2 | 0.015 | Young |
| 0.537518 | 0.217649 | 0.182 | 0.218 | 1 | 10 | Rabgap1 | -0.036 | Young |
| 0.537818 | 0.359586 | 0.283 | 0.269 | 1 | 10 | Psmb9 | 0.014 | Young |
| 0.537893 | 0.06033 | 0.051 | 0.037 | 1 | 10 | Kat8 | 0.014 | Young |

| | | | | | | | | |
|---|---|---|---|---|---|---|---|---|
| 0.537913 | 0.959192 | 0.02 | 0.012 | 1 | 10 | Zfp775 | 0.008 | Young |
| 0.537913 | 0.687363 | 0.02 | 0.012 | 1 | 10 | Mrvi1 | 0.008 | Young |
| 0.537913 | 0.649821 | 0.02 | 0.012 | 1 | 10 | Fam109b | 0.008 | Young |
| 0.537913 | 0.61057 | 0.02 | 0.012 | 1 | 10 | 5033421B0 | 0.008 | Young |
| 0.537913 | 0.53581 | 0.02 | 0.012 | 1 | 10 | Hist1h2be | 0.008 | Young |
| 0.537917 | 0.830153 | 0.091 | 0.076 | 1 | 10 | Mto1 | 0.015 | Young |
| 0.538057 | 0.022703 | 0.556 | 0.606 | 1 | 10 | Tuba1c | -0.05 | Young |
| 0.538503 | 0.365167 | 0.384 | 0.381 | 1 | 10 | Arpp19 | 0.003 | Young |
| 0.538595 | 0.382891 | 0.091 | 0.12 | 1 | 10 | Ranbp3 | -0.029 | Young |
| 0.538613 | 0.30531 | 0.333 | 0.315 | 1 | 10 | Gabarapl1 | 0.018 | Young |
| 0.538736 | 0.175108 | 0.212 | 0.254 | 1 | 10 | Yipf5 | -0.042 | Young |
| 0.538787 | 0.097955 | 0.071 | 0.093 | 1 | 10 | Fam8a1 | -0.022 | Young |
| 0.539041 | 0.410708 | 0.323 | 0.306 | 1 | 10 | Sp1 | 0.017 | Young |
| 0.539134 | 0.189975 | 0.404 | 0.389 | 1 | 10 | Uhmk1 | 0.015 | Young |
| 0.539389 | 0.342511 | 0.051 | 0.037 | 1 | 10 | Mtg2 | 0.014 | Young |
| 0.539427 | 0.089926 | 0.495 | 0.575 | 1 | 10 | Pum2 | -0.08 | Young |
| 0.539434 | 0.274316 | 0.535 | 0.562 | 1 | 10 | Surf4 | -0.027 | Young |
| 0.539623 | 0.369282 | 0.081 | 0.103 | 1 | 10 | Acvr1b | -0.022 | Young |
| 0.54041 | 0.928473 | 0.02 | 0.012 | 1 | 10 | Cetn4 | 0.008 | Young |
| 0.54041 | 0.897899 | 0.02 | 0.012 | 1 | 10 | Stac3 | 0.008 | Young |
| 0.54041 | 0.639477 | 0.02 | 0.012 | 1 | 10 | Hectd2 | 0.008 | Young |
| 0.54041 | 0.634908 | 0.02 | 0.012 | 1 | 10 | Cd300ld3 | 0.008 | Young |
| 0.540794 | 0.573645 | 0.091 | 0.073 | 1 | 10 | Trim65 | 0.018 | Young |
| 0.541118 | 0.197623 | 0.323 | 0.399 | 1 | 10 | Parvb | -0.076 | Young |
| 0.541128 | 0.414884 | 0.354 | 0.352 | 1 | 10 | Tnfsf12 | 0.002 | Young |
| 0.541641 | 0.230332 | 0.192 | 0.166 | 1 | 10 | Snap29 | 0.026 | Young |
| 0.541869 | 0.121518 | 0.404 | 0.479 | 1 | 10 | Smpdl3a | -0.075 | Young |
| 0.541942 | 0.478146 | 0.152 | 0.13 | 1 | 10 | Csf2rb2 | 0.022 | Young |
| 0.542213 | 1.967201 | 0.01 | 0.005 | 1 | 10 | 2610020C0 | 0.005 | Young |
| 0.542213 | 1.881096 | 0.01 | 0.005 | 1 | 10 | Armcx6 | 0.005 | Young |
| 0.542213 | 1.772689 | 0.01 | 0.005 | 1 | 10 | Jmjd7 | 0.005 | Young |
| 0.542213 | 1.70365 | 0.01 | 0.005 | 1 | 10 | Mindy4 | 0.005 | Young |
| 0.542213 | 1.696421 | 0.01 | 0.005 | 1 | 10 | Gm15952 | 0.005 | Young |
| 0.542213 | 1.639412 | 0.01 | 0.005 | 1 | 10 | Postn | 0.005 | Young |
| 0.542213 | 1.548432 | 0.01 | 0.005 | 1 | 10 | Adgrf5 | 0.005 | Young |
| 0.542213 | 1.523277 | 0.01 | 0.005 | 1 | 10 | Pkn3 | 0.005 | Young |
| 0.542213 | 1.490791 | 0.01 | 0.005 | 1 | 10 | Gm26576 | 0.005 | Young |
| 0.542213 | 1.46923 | 0.01 | 0.005 | 1 | 10 | Gm15283 | 0.005 | Young |
| 0.542213 | 1.439939 | 0.01 | 0.005 | 1 | 10 | Gm26901 | 0.005 | Young |
| 0.542213 | 1.32059 | 0.01 | 0.005 | 1 | 10 | Fam161b | 0.005 | Young |
| 0.542213 | 1.311249 | 0.01 | 0.005 | 1 | 10 | 5930430L0 | 0.005 | Young |
| 0.542213 | 1.266839 | 0.01 | 0.005 | 1 | 10 | Aldh1a3 | 0.005 | Young |
| 0.542213 | 1.244441 | 0.01 | 0.005 | 1 | 10 | Zfp956 | 0.005 | Young |
| 0.542213 | 1.235548 | 0.01 | 0.005 | 1 | 10 | Uggt2 | 0.005 | Young |
| 0.542213 | 1.232671 | 0.01 | 0.005 | 1 | 10 | Dnajb13 | 0.005 | Young |

| | | | | | | | | |
|---|---|---|---|---|---|---|---|---|
| 0.542213 | 1.218654 | 0.01 | 0.005 | 1 | 10 | Scn2b | 0.005 | Young |
| 0.542213 | 1.209614 | 0.01 | 0.005 | 1 | 10 | Atp2b2 | 0.005 | Young |
| 0.542213 | 1.206641 | 0.01 | 0.005 | 1 | 10 | Gm26809 | 0.005 | Young |
| 0.542213 | 1.172767 | 0.01 | 0.005 | 1 | 10 | Bag2 | 0.005 | Young |
| 0.542213 | 1.147824 | 0.01 | 0.005 | 1 | 10 | Il27 | 0.005 | Young |
| 0.542213 | 1.126103 | 0.01 | 0.005 | 1 | 10 | Fam124a | 0.005 | Young |
| 0.542213 | 1.08826 | 0.01 | 0.005 | 1 | 10 | Pkia | 0.005 | Young |
| 0.542213 | 1.05954 | 0.01 | 0.005 | 1 | 10 | Zfp97 | 0.005 | Young |
| 0.542213 | 1.048759 | 0.01 | 0.005 | 1 | 10 | Gm42893 | 0.005 | Young |
| 0.542213 | 1.039603 | 0.01 | 0.005 | 1 | 10 | Carns1 | 0.005 | Young |
| 0.542213 | 0.994813 | 0.01 | 0.005 | 1 | 10 | Ifit3b | 0.005 | Young |
| 0.542213 | 0.922779 | 0.01 | 0.005 | 1 | 10 | Akap2 | 0.005 | Young |
| 0.542213 | 0.887918 | 0.01 | 0.005 | 1 | 10 | Zfp691 | 0.005 | Young |
| 0.542213 | 0.876811 | 0.01 | 0.005 | 1 | 10 | Gm14419 | 0.005 | Young |
| 0.542213 | 0.871444 | 0.01 | 0.005 | 1 | 10 | Colca2 | 0.005 | Young |
| 0.542213 | 0.860976 | 0.01 | 0.005 | 1 | 10 | Atp10a | 0.005 | Young |
| 0.542213 | 0.827084 | 0.01 | 0.005 | 1 | 10 | Efcc1 | 0.005 | Young |
| 0.542213 | 0.823608 | 0.01 | 0.005 | 1 | 10 | Acer2 | 0.005 | Young |
| 0.542213 | 0.776743 | 0.01 | 0.005 | 1 | 10 | 9130604C2 | 0.005 | Young |
| 0.542213 | 0.729468 | 0.01 | 0.005 | 1 | 10 | Zfp449 | 0.005 | Young |
| 0.542213 | 0.722894 | 0.01 | 0.005 | 1 | 10 | Serping1 | 0.005 | Young |
| 0.542213 | 0.714053 | 0.01 | 0.005 | 1 | 10 | Gm32051 | 0.005 | Young |
| 0.542213 | 0.696115 | 0.01 | 0.005 | 1 | 10 | Fam149a | 0.005 | Young |
| 0.542213 | 0.631668 | 0.01 | 0.005 | 1 | 10 | Zwilch | 0.005 | Young |
| 0.542213 | 0.601461 | 0.01 | 0.005 | 1 | 10 | Rnf152 | 0.005 | Young |
| 0.542213 | 0.513613 | 0.01 | 0.005 | 1 | 10 | Ly6g6d | 0.005 | Young |
| 0.542213 | 0.463413 | 0.01 | 0.005 | 1 | 10 | Tedc1 | 0.005 | Young |
| 0.542257 | 0.355354 | 0.172 | 0.149 | 1 | 10 | Zfp592 | 0.023 | Young |
| 0.542359 | 0.621508 | 0.222 | 0.247 | 1 | 10 | Babam2 | -0.025 | Young |
| 0.542728 | 0.729374 | 0.061 | 0.046 | 1 | 10 | Ppp1r13b | 0.015 | Young |
| 0.542913 | 0.755223 | 0.02 | 0.012 | 1 | 10 | Dffb | 0.008 | Young |
| 0.542913 | 0.686004 | 0.02 | 0.012 | 1 | 10 | Ptpdc1 | 0.008 | Young |
| 0.542913 | 0.639213 | 0.02 | 0.012 | 1 | 10 | Rwdd2b | 0.008 | Young |
| 0.542913 | 0.444101 | 0.02 | 0.012 | 1 | 10 | Elp6 | 0.008 | Young |
| 0.542913 | 0.27757 | 0.02 | 0.012 | 1 | 10 | Pcbp4 | 0.008 | Young |
| 0.542913 | 0.137493 | 0.02 | 0.012 | 1 | 10 | Pde5a | 0.008 | Young |
| 0.542984 | 0.478395 | 0.091 | 0.073 | 1 | 10 | Syt11 | 0.018 | Young |
| 0.543401 | 0.711264 | 0.242 | 0.225 | 1 | 10 | Sf3a3 | 0.017 | Young |
| 0.543559 | 0.914633 | 0.202 | 0.188 | 1 | 10 | Clec4e | 0.014 | Young |
| 0.543636 | 0.331398 | 0.354 | 0.33 | 1 | 10 | Ifnar2 | 0.024 | Young |
| 0.544081 | 0.311753 | 0.091 | 0.073 | 1 | 10 | 1500011B0 | 0.018 | Young |
| 0.544559 | 0.42621 | 0.101 | 0.127 | 1 | 10 | Gnptg | -0.026 | Young |
| 0.544625 | 0.63704 | 0.232 | 0.218 | 1 | 10 | Ssr2 | 0.014 | Young |
| 0.545004 | 0.400785 | 0.182 | 0.225 | 1 | 10 | Erlin1 | -0.043 | Young |
| 0.545132 | 0.482835 | 0.081 | 0.064 | 1 | 10 | H2afx | 0.017 | Young |

| | | | | | | | | |
|---|---|---|---|---|---|---|---|---|
| 0.545259 | 0.001082 | 0.485 | 0.555 | 1 | 10 | Arf3 | -0.07 | Young |
| 0.545283 | 0.38151 | 0.131 | 0.112 | 1 | 10 | Uchl3 | 0.019 | Young |
| 0.545422 | 0.517213 | 0.02 | 0.012 | 1 | 10 | Nipsnap1 | 0.008 | Young |
| 0.545422 | 0.438574 | 0.02 | 0.012 | 1 | 10 | Unc5a | 0.008 | Young |
| 0.545422 | 0.426135 | 0.02 | 0.012 | 1 | 10 | Fzd4 | 0.008 | Young |
| 0.545422 | 0.336741 | 0.02 | 0.012 | 1 | 10 | Ccdc171 | 0.008 | Young |
| 0.545422 | 0.318219 | 0.02 | 0.012 | 1 | 10 | Ppm1l | 0.008 | Young |
| 0.545422 | 0.225823 | 0.02 | 0.012 | 1 | 10 | Spata1 | 0.008 | Young |
| 0.545422 | 0.211139 | 0.02 | 0.012 | 1 | 10 | Ccsap | 0.008 | Young |
| 0.545484 | 1.062339 | 0.101 | 0.083 | 1 | 10 | Zfand2b | 0.018 | Young |
| 0.545484 | 0.600919 | 0.101 | 0.083 | 1 | 10 | Vhl | 0.018 | Young |
| 0.54597 | 0.729888 | 0.101 | 0.086 | 1 | 10 | Sp2 | 0.015 | Young |
| 0.54603 | 1.133117 | 0.01 | 0.005 | 1 | 10 | Ighd | 0.005 | Young |
| 0.54603 | 0.91746 | 0.01 | 0.005 | 1 | 10 | Gm16794 | 0.005 | Young |
| 0.54603 | 0.884555 | 0.01 | 0.005 | 1 | 10 | A130051J0 | 0.005 | Young |
| 0.54603 | 0.868398 | 0.01 | 0.005 | 1 | 10 | Kcnc3 | 0.005 | Young |
| 0.54603 | 0.829622 | 0.01 | 0.005 | 1 | 10 | Fap | 0.005 | Young |
| 0.54603 | 0.805676 | 0.01 | 0.005 | 1 | 10 | Ndn | 0.005 | Young |
| 0.54603 | 0.797962 | 0.01 | 0.005 | 1 | 10 | Gm20655 | 0.005 | Young |
| 0.54603 | 0.738783 | 0.01 | 0.005 | 1 | 10 | Ccr7 | 0.005 | Young |
| 0.54603 | 0.711876 | 0.01 | 0.005 | 1 | 10 | Micall2 | 0.005 | Young |
| 0.54603 | 0.709546 | 0.01 | 0.005 | 1 | 10 | Zcwpw1 | 0.005 | Young |
| 0.54603 | 0.691896 | 0.01 | 0.005 | 1 | 10 | Fbxl2 | 0.005 | Young |
| 0.54603 | 0.6557 | 0.01 | 0.005 | 1 | 10 | Gm3336 | 0.005 | Young |
| 0.54603 | 0.654484 | 0.01 | 0.005 | 1 | 10 | Gm26674 | 0.005 | Young |
| 0.54603 | 0.622261 | 0.01 | 0.005 | 1 | 10 | Hist3h2ba | 0.005 | Young |
| 0.54603 | 0.604743 | 0.01 | 0.005 | 1 | 10 | Ctnnd2 | 0.005 | Young |
| 0.54603 | 0.566949 | 0.01 | 0.005 | 1 | 10 | Zfp3 | 0.005 | Young |
| 0.54603 | 0.511086 | 0.01 | 0.005 | 1 | 10 | Ppp1r3f | 0.005 | Young |
| 0.54603 | 0.497012 | 0.01 | 0.005 | 1 | 10 | Lbp | 0.005 | Young |
| 0.54603 | 0.495186 | 0.01 | 0.005 | 1 | 10 | Serpinc1 | 0.005 | Young |
| 0.54603 | 0.463553 | 0.01 | 0.005 | 1 | 10 | Rab4b | 0.005 | Young |
| 0.54603 | 0.440543 | 0.01 | 0.005 | 1 | 10 | Man2c1os | 0.005 | Young |
| 0.54603 | 0.427983 | 0.01 | 0.005 | 1 | 10 | Rpusd3 | 0.005 | Young |
| 0.54603 | 0.426406 | 0.01 | 0.005 | 1 | 10 | Zfp772 | 0.005 | Young |
| 0.54603 | 0.424719 | 0.01 | 0.005 | 1 | 10 | Ndc80 | 0.005 | Young |
| 0.54603 | 0.394733 | 0.01 | 0.005 | 1 | 10 | Omg | 0.005 | Young |
| 0.54603 | 0.393991 | 0.01 | 0.005 | 1 | 10 | Gm38832 | 0.005 | Young |
| 0.54603 | 0.390348 | 0.01 | 0.005 | 1 | 10 | Cadm3 | 0.005 | Young |
| 0.54603 | 0.38766 | 0.01 | 0.005 | 1 | 10 | Trim45 | 0.005 | Young |
| 0.54603 | 0.354514 | 0.01 | 0.005 | 1 | 10 | Slc12a5 | 0.005 | Young |
| 0.54603 | 0.352381 | 0.01 | 0.005 | 1 | 10 | Ttk | 0.005 | Young |
| 0.54603 | 0.338563 | 0.01 | 0.005 | 1 | 10 | Tlr12 | 0.005 | Young |
| 0.54603 | 0.336519 | 0.01 | 0.005 | 1 | 10 | Gng7 | 0.005 | Young |
| 0.54603 | 0.334849 | 0.01 | 0.005 | 1 | 10 | Egfl8 | 0.005 | Young |

| | | | | | | | | |
|---|---|---|---|---|---|---|---|---|
| 0.54603 | 0.319064 | 0.01 | 0.005 | 1 | 10 | Col1a2 | 0.005 | Young |
| 0.54603 | 0.309254 | 0.01 | 0.005 | 1 | 10 | Gm43573 | 0.005 | Young |
| 0.54603 | 0.308564 | 0.01 | 0.005 | 1 | 10 | 2410080I0 | 0.005 | Young |
| 0.54603 | 0.25951 | 0.01 | 0.005 | 1 | 10 | Gm26660 | 0.005 | Young |
| 0.54603 | 0.247992 | 0.01 | 0.005 | 1 | 10 | 5830454E0 | 0.005 | Young |
| 0.54603 | 0.239095 | 0.01 | 0.005 | 1 | 10 | Fermt2 | 0.005 | Young |
| 0.54603 | 0.209136 | 0.01 | 0.005 | 1 | 10 | Gm20633 | 0.005 | Young |
| 0.54603 | 0.183501 | 0.01 | 0.005 | 1 | 10 | Gm15747 | 0.005 | Young |
| 0.54603 | 0.124778 | 0.01 | 0.005 | 1 | 10 | Thbs3 | 0.005 | Young |
| 0.54603 | 0.123344 | 0.01 | 0.005 | 1 | 10 | Anln | 0.005 | Young |
| 0.54603 | 0.10634 | 0.01 | 0.005 | 1 | 10 | Eya1 | 0.005 | Young |
| 0.54603 | 0.09524 | 0.01 | 0.005 | 1 | 10 | Zfp850 | 0.005 | Young |
| 0.54603 | 0.077447 | 0.01 | 0.005 | 1 | 10 | Gm12184 | 0.005 | Young |
| 0.54603 | 0.025684 | 0.01 | 0.005 | 1 | 10 | Gm20522 | 0.005 | Young |
| 0.54603 | 0.021674 | 0.01 | 0.005 | 1 | 10 | Adamts14 | 0.005 | Young |
| 0.546578 | 0.862921 | 0.091 | 0.076 | 1 | 10 | Gm45902 | 0.015 | Young |
| 0.546802 | 0.212081 | 0.838 | 0.861 | 1 | 10 | Cd83 | -0.023 | Young |
| 0.547062 | 0.539614 | 0.212 | 0.196 | 1 | 10 | Ing1 | 0.016 | Young |
| 0.547824 | 0.077823 | 0.182 | 0.22 | 1 | 10 | Snapc5 | -0.038 | Young |
| 0.547915 | 0.231666 | 0.465 | 0.531 | 1 | 10 | Snx2 | -0.066 | Young |
| 0.547936 | 0.008968 | 0.02 | 0.012 | 1 | 10 | Inmt | 0.008 | Young |
| 0.547947 | 0.800845 | 0.141 | 0.125 | 1 | 10 | Usf3 | 0.016 | Young |
| 0.54814 | 0.429618 | 0.061 | 0.046 | 1 | 10 | Bach2 | 0.015 | Young |
| 0.548422 | 0.132522 | 0.273 | 0.244 | 1 | 10 | Idh3g | 0.029 | Young |
| 0.548442 | 0.02257 | 0.081 | 0.103 | 1 | 10 | Nfia | -0.022 | Young |
| 0.548477 | 0.356823 | 0.293 | 0.267 | 1 | 10 | Lyz1 | 0.026 | Young |
| 0.548634 | 0.487983 | 0.343 | 0.33 | 1 | 10 | Smim11 | 0.013 | Young |
| 0.548814 | 0.728748 | 0.071 | 0.056 | 1 | 10 | Zfp146 | 0.015 | Young |
| 0.548814 | 0.679774 | 0.071 | 0.056 | 1 | 10 | Drosha | 0.015 | Young |
| 0.548934 | 0.846545 | 0.111 | 0.095 | 1 | 10 | Tma16 | 0.016 | Young |
| 0.549196 | 1.869943 | 0.04 | 0.029 | 1 | 10 | Mthfd1l | 0.011 | Young |
| 0.549243 | 0.345906 | 0.293 | 0.276 | 1 | 10 | Dnajb9 | 0.017 | Young |
| 0.549497 | 0.53435 | 0.061 | 0.046 | 1 | 10 | Sco1 | 0.015 | Young |
| 0.549497 | 0.076503 | 0.061 | 0.046 | 1 | 10 | Itpkc | 0.015 | Young |
| 0.54986 | 0.404727 | 0.01 | 0.005 | 1 | 10 | Grtp1 | 0.005 | Young |
| 0.54986 | 0.34638 | 0.01 | 0.005 | 1 | 10 | B130006D0 | 0.005 | Young |
| 0.54986 | 0.311332 | 0.01 | 0.005 | 1 | 10 | Dnal1 | 0.005 | Young |
| 0.54986 | 0.283081 | 0.01 | 0.005 | 1 | 10 | Phf21b | 0.005 | Young |
| 0.54986 | 0.252921 | 0.01 | 0.005 | 1 | 10 | Pfkfb1 | 0.005 | Young |
| 0.54986 | 0.231404 | 0.01 | 0.005 | 1 | 10 | Acbd4 | 0.005 | Young |
| 0.54986 | 0.226091 | 0.01 | 0.005 | 1 | 10 | Lyzl4 | 0.005 | Young |
| 0.54986 | 0.221914 | 0.01 | 0.005 | 1 | 10 | Slc8a1 | 0.005 | Young |
| 0.54986 | 0.205486 | 0.01 | 0.005 | 1 | 10 | Gm47428 | 0.005 | Young |
| 0.54986 | 0.205268 | 0.01 | 0.005 | 1 | 10 | Atl1 | 0.005 | Young |
| 0.54986 | 0.2027 | 0.01 | 0.005 | 1 | 10 | AC125351. | 0.005 | Young |

| | | | | | | | | |
|---|---|---|---|---|---|---|---|---|
| 0.54986 | 0.183788 | 0.01 | 0.005 | 1 | 10 | BC030499 | 0.005 | Young |
| 0.54986 | 0.177732 | 0.01 | 0.005 | 1 | 10 | Abcb9 | 0.005 | Young |
| 0.54986 | 0.160205 | 0.01 | 0.005 | 1 | 10 | Prr18 | 0.005 | Young |
| 0.54986 | 0.145333 | 0.01 | 0.005 | 1 | 10 | Jaml | 0.005 | Young |
| 0.54986 | 0.140414 | 0.01 | 0.005 | 1 | 10 | Gm29394 | 0.005 | Young |
| 0.54986 | 0.107557 | 0.01 | 0.005 | 1 | 10 | Rccd1 | 0.005 | Young |
| 0.54986 | 0.103695 | 0.01 | 0.005 | 1 | 10 | Cd69 | 0.005 | Young |
| 0.54986 | 0.096234 | 0.01 | 0.005 | 1 | 10 | Tmem72 | 0.005 | Young |
| 0.54986 | 0.09395 | 0.01 | 0.005 | 1 | 10 | Gm20324 | 0.005 | Young |
| 0.54986 | 0.092071 | 0.01 | 0.005 | 1 | 10 | Gm20939 | 0.005 | Young |
| 0.54986 | 0.073191 | 0.01 | 0.005 | 1 | 10 | Etaa1os | 0.005 | Young |
| 0.54986 | 0.063896 | 0.01 | 0.005 | 1 | 10 | Gm10384 | 0.005 | Young |
| 0.54986 | 0.050597 | 0.01 | 0.005 | 1 | 10 | Lsmem1 | 0.005 | Young |
| 0.54986 | 0.041019 | 0.01 | 0.005 | 1 | 10 | BC022687 | 0.005 | Young |
| 0.54986 | 0.030994 | 0.01 | 0.005 | 1 | 10 | Gm21988 | 0.005 | Young |
| 0.550059 | 0.563422 | 0.071 | 0.056 | 1 | 10 | 2810006K2 | 0.015 | Young |
| 0.550183 | 0.013968 | 0.364 | 0.413 | 1 | 10 | Mrpl54 | -0.049 | Young |
| 0.550243 | 0.265711 | 0.091 | 0.117 | 1 | 10 | Cfap36 | -0.026 | Young |
| 0.550348 | 0.193026 | 0.333 | 0.386 | 1 | 10 | Crk | -0.053 | Young |
| 0.550372 | 0.669914 | 0.343 | 0.347 | 1 | 10 | Dennd4a | -0.004 | Young |
| 0.550384 | 0.293164 | 0.394 | 0.386 | 1 | 10 | Ubn2 | 0.008 | Young |
| 0.550456 | 0.053447 | 0.02 | 0.012 | 1 | 10 | E430024P1 | 0.008 | Young |
| 0.550801 | 0.190185 | 0.768 | 0.817 | 1 | 10 | Cd52 | -0.049 | Young |
| 0.551041 | 0.03875 | 0.141 | 0.169 | 1 | 10 | Chd6 | -0.028 | Young |
| 0.551087 | 0.071349 | 0.323 | 0.364 | 1 | 10 | Neu1 | -0.041 | Young |
| 0.551739 | 0.73457 | 0.101 | 0.083 | 1 | 10 | Fmc1 | 0.018 | Young |
| 0.551914 | 0.043588 | 0.879 | 0.944 | 1 | 10 | Klf6 | -0.065 | Young |
| 0.55217 | 0.050597 | 0.121 | 0.147 | 1 | 10 | Mtx2 | -0.026 | Young |
| 0.552177 | 0.191394 | 0.081 | 0.064 | 1 | 10 | Ric8b | 0.017 | Young |
| 0.55228 | 0.255615 | 0.606 | 0.611 | 1 | 10 | Psma3 | -0.005 | Young |
| 0.552564 | 1.296911 | 0.04 | 0.029 | 1 | 10 | Dnase1l3 | 0.011 | Young |
| 0.552564 | 1.104114 | 0.04 | 0.029 | 1 | 10 | Sbk1 | 0.011 | Young |
| 0.552996 | 0.140248 | 0.616 | 0.67 | 1 | 10 | Diaph1 | -0.054 | Young |
| 0.552997 | 0.573159 | 0.02 | 0.032 | 1 | 10 | Slc5a6 | -0.012 | Young |
| 0.553174 | 0.396981 | 0.051 | 0.068 | 1 | 10 | Syce2 | -0.017 | Young |
| 0.553766 | 0.636276 | 0.354 | 0.347 | 1 | 10 | Hnrnpll | 0.007 | Young |
| 0.554135 | 0.508746 | 0.414 | 0.416 | 1 | 10 | Immt | -0.002 | Young |
| 0.55474 | 0.051436 | 0.02 | 0.032 | 1 | 10 | Mllt3 | -0.012 | Young |
| 0.555287 | 0.48501 | 0.182 | 0.222 | 1 | 10 | Zc3h18 | -0.04 | Young |
| 0.555404 | 0.130593 | 0.071 | 0.09 | 1 | 10 | Tmem87a | -0.019 | Young |
| 0.555552 | 0.214159 | 0.091 | 0.115 | 1 | 10 | Chst12 | -0.024 | Young |
| 0.555943 | 1.597653 | 0.04 | 0.029 | 1 | 10 | Slc25a26 | 0.011 | Young |
| 0.555943 | 1.073353 | 0.04 | 0.029 | 1 | 10 | Gk5 | 0.011 | Young |
| 0.555961 | 0.123739 | 0.04 | 0.056 | 1 | 10 | Elk1 | -0.016 | Young |
| 0.556385 | 0.173539 | 0.121 | 0.147 | 1 | 10 | Rpf1 | -0.026 | Young |

| | | | | | | | | |
|---|---|---|---|---|---|---|---|---|
| 0.556546 | 0.384116 | 0.303 | 0.286 | 1 | 10 | Snd1 | 0.017 | Young |
| 0.557356 | 0.437562 | 0.424 | 0.411 | 1 | 10 | Arrb2 | 0.013 | Young |
| 0.557504 | 0.318975 | 0.071 | 0.09 | 1 | 10 | Bicral | -0.019 | Young |
| 0.557614 | 0.169397 | 0.121 | 0.152 | 1 | 10 | Ccdc174 | -0.031 | Young |
| 0.557964 | 0.487105 | 0.293 | 0.274 | 1 | 10 | Utp11 | 0.019 | Young |
| 0.558053 | 0.318985 | 0.253 | 0.227 | 1 | 10 | Tor1aip2 | 0.026 | Young |
| 0.558064 | 0.274118 | 0.273 | 0.323 | 1 | 10 | Oat | -0.05 | Young |
| 0.558497 | 0.311359 | 0.131 | 0.159 | 1 | 10 | Ddx27 | -0.028 | Young |
| 0.558714 | 0.07555 | 0.212 | 0.257 | 1 | 10 | Rab5b | -0.045 | Young |
| 0.559001 | 0.045846 | 0.768 | 0.831 | 1 | 10 | Vamp8 | -0.063 | Young |
| 0.559036 | 0.223513 | 0.283 | 0.257 | 1 | 10 | Oxr1 | 0.026 | Young |
| 0.559883 | 0.376275 | 0.061 | 0.078 | 1 | 10 | Otud7b | -0.017 | Young |
| 0.559918 | 0.381579 | 0.04 | 0.056 | 1 | 10 | Ddx51 | -0.016 | Young |
| 0.559918 | 0.252021 | 0.04 | 0.056 | 1 | 10 | Trim23 | -0.016 | Young |
| 0.559983 | 0.345373 | 0.02 | 0.032 | 1 | 10 | Ptpn4 | -0.012 | Young |
| 0.559983 | 0.320566 | 0.02 | 0.032 | 1 | 10 | Tcim | -0.012 | Young |
| 0.560353 | 0.339374 | 0.182 | 0.218 | 1 | 10 | Pak1 | -0.036 | Young |
| 0.560382 | 0.349595 | 0.545 | 0.57 | 1 | 10 | Llph | -0.025 | Young |
| 0.560413 | 0.222045 | 0.061 | 0.046 | 1 | 10 | Foxred2 | 0.015 | Young |
| 0.560671 | 1.197729 | 0.051 | 0.039 | 1 | 10 | Chd1l | 0.012 | Young |
| 0.56103 | 0.602885 | 0.04 | 0.029 | 1 | 10 | Prrg1 | 0.011 | Young |
| 0.561196 | 0.121952 | 0.293 | 0.337 | 1 | 10 | Ripk1 | -0.044 | Young |
| 0.561572 | 0.142573 | 0.051 | 0.068 | 1 | 10 | Mbip | -0.017 | Young |
| 0.561715 | 0.237576 | 0.071 | 0.09 | 1 | 10 | Inpp5a | -0.019 | Young |
| 0.561736 | 0.052866 | 0.02 | 0.032 | 1 | 10 | 1110008L1 | -0.012 | Young |
| 0.56181 | 0.06293 | 0.343 | 0.391 | 1 | 10 | Smc6 | -0.048 | Young |
| 0.561878 | 0.74545 | 0.263 | 0.247 | 1 | 10 | Psmg4 | 0.016 | Young |
| 0.562162 | 0.492258 | 0.03 | 0.044 | 1 | 10 | Slc25a40 | -0.014 | Young |
| 0.562303 | 0.465687 | 0.323 | 0.313 | 1 | 10 | Zfp710 | 0.01 | Young |
| 0.562421 | 0.047672 | 0.263 | 0.232 | 1 | 10 | Phf10 | 0.031 | Young |
| 0.562481 | 0.350934 | 0.343 | 0.333 | 1 | 10 | Sys1 | 0.01 | Young |
| 0.562731 | 0.810831 | 0.04 | 0.029 | 1 | 10 | Snap47 | 0.011 | Young |
| 0.563012 | 0.82935 | 0.121 | 0.103 | 1 | 10 | Zfp90 | 0.018 | Young |
| 0.56335 | 0.094611 | 0.273 | 0.311 | 1 | 10 | Ppp1r10 | -0.038 | Young |
| 0.563433 | 0.330727 | 0.101 | 0.127 | 1 | 10 | Arfrp1 | -0.026 | Young |
| 0.563637 | 0.069959 | 0.111 | 0.137 | 1 | 10 | Fam213a | -0.026 | Young |
| 0.563655 | 0.190949 | 0.03 | 0.044 | 1 | 10 | Aacs | -0.014 | Young |
| 0.563655 | 0.13656 | 0.03 | 0.044 | 1 | 10 | Fam35a | -0.014 | Young |
| 0.564354 | 0.195672 | 0.101 | 0.083 | 1 | 10 | Hagh | 0.018 | Young |
| 0.564403 | 0.019106 | 0.566 | 0.663 | 1 | 10 | Capza1 | -0.097 | Young |
| 0.564434 | 0.550989 | 0.04 | 0.029 | 1 | 10 | Poll | 0.011 | Young |
| 0.564534 | 0.45032 | 0.061 | 0.046 | 1 | 10 | Trim32 | 0.015 | Young |
| 0.565001 | 0.339902 | 0.303 | 0.281 | 1 | 10 | Arid4a | 0.022 | Young |
| 0.565057 | 0.134644 | 0.071 | 0.088 | 1 | 10 | Chil3 | -0.017 | Young |
| 0.565158 | 0.437541 | 0.242 | 0.227 | 1 | 10 | Impdh2 | 0.015 | Young |

| | | | | | | | | |
|---|---|---|---|---|---|---|---|---|
| 0.565217 | 0.551747 | 0.182 | 0.164 | 1 | 10 | Itpr2 | 0.018 | Young |
| 0.565225 | 1.262855 | 0.091 | 0.078 | 1 | 10 | Mecr | 0.013 | Young |
| 0.565267 | 0.4539 | 0.081 | 0.105 | 1 | 10 | Prr12 | -0.024 | Young |
| 0.565549 | 0.050536 | 0.394 | 0.443 | 1 | 10 | Eps15 | -0.049 | Young |
| 0.565701 | 0.93794 | 0.101 | 0.086 | 1 | 10 | Msh2 | 0.015 | Young |
| 0.565857 | 0.043886 | 0.576 | 0.587 | 1 | 10 | Rel | -0.011 | Young |
| 0.566087 | 0.116002 | 0.96 | 0.968 | 1 | 10 | Pabpc1 | -0.008 | Young |
| 0.56614 | 0.504025 | 0.04 | 0.029 | 1 | 10 | Dnajc4 | 0.011 | Young |
| 0.566145 | 0.184134 | 0.313 | 0.372 | 1 | 10 | Mknk2 | -0.059 | Young |
| 0.566269 | 0.618547 | 0.111 | 0.137 | 1 | 10 | Med21 | -0.026 | Young |
| 0.566368 | 0.414907 | 0.253 | 0.237 | 1 | 10 | Lyst | 0.016 | Young |
| 0.566414 | 0.172377 | 0.424 | 0.477 | 1 | 10 | Kdm6b | -0.053 | Young |
| 0.566428 | 0.759147 | 0.121 | 0.108 | 1 | 10 | Dock4 | 0.013 | Young |
| 0.567289 | 0.34388 | 0.061 | 0.046 | 1 | 10 | Cnksr3 | 0.015 | Young |
| 0.567529 | 0.037971 | 0.354 | 0.342 | 1 | 10 | Ddrgk1 | 0.012 | Young |
| 0.568049 | 0.425858 | 0.293 | 0.276 | 1 | 10 | Psmd7 | 0.017 | Young |
| 0.568741 | 0.06302 | 0.152 | 0.13 | 1 | 10 | Usp38 | 0.022 | Young |
| 0.568752 | 0.114337 | 0.586 | 0.606 | 1 | 10 | Ewsr1 | -0.02 | Young |
| 0.568891 | 0.319227 | 0.303 | 0.281 | 1 | 10 | Fcrls | 0.022 | Young |
| 0.569222 | 0.425556 | 0.374 | 0.364 | 1 | 10 | Kcmf1 | 0.01 | Young |
| 0.569265 | 0.087189 | 0.384 | 0.445 | 1 | 10 | Rwdd1 | -0.061 | Young |
| 0.569784 | 0.251158 | 0.414 | 0.479 | 1 | 10 | B230219D2 | -0.065 | Young |
| 0.570104 | 0.107063 | 0.232 | 0.276 | 1 | 10 | Trim41 | -0.044 | Young |
| 0.57115 | 0.55428 | 0.03 | 0.044 | 1 | 10 | Zbtb42 | -0.014 | Young |
| 0.571271 | 1.038724 | 0.04 | 0.029 | 1 | 10 | Bcam | 0.011 | Young |
| 0.571271 | 0.763973 | 0.04 | 0.029 | 1 | 10 | Pcolce | 0.011 | Young |
| 0.571271 | 0.586623 | 0.04 | 0.029 | 1 | 10 | Dnajb5 | 0.011 | Young |
| 0.571657 | 0.099308 | 0.131 | 0.159 | 1 | 10 | Fech | -0.028 | Young |
| 0.571665 | 0.258394 | 0.091 | 0.115 | 1 | 10 | Lias | -0.024 | Young |
| 0.571745 | 0.154668 | 0.919 | 0.944 | 1 | 10 | Dusp1 | -0.025 | Young |
| 0.571808 | 0.373939 | 0.121 | 0.149 | 1 | 10 | Mfn2 | -0.028 | Young |
| 0.571859 | 0.889555 | 0.091 | 0.076 | 1 | 10 | Zfp282 | 0.015 | Young |
| 0.571859 | 0.820607 | 0.091 | 0.076 | 1 | 10 | Dedd | 0.015 | Young |
| 0.57205 | 0.197539 | 0.182 | 0.213 | 1 | 10 | Ncoa2 | -0.031 | Young |
| 0.57227 | 0.267762 | 0.515 | 0.489 | 1 | 10 | Ube2n | 0.026 | Young |
| 0.5723 | 0.219038 | 0.081 | 0.103 | 1 | 10 | Rapgef1 | -0.022 | Young |
| 0.572539 | 0.003792 | 0.101 | 0.122 | 1 | 10 | Lonp1 | -0.021 | Young |
| 0.572607 | 0.009076 | 0.636 | 0.699 | 1 | 10 | Fosb | -0.063 | Young |
| 0.572654 | 0.738588 | 0.051 | 0.039 | 1 | 10 | Nr1d1 | 0.012 | Young |
| 0.57268 | 0.4782 | 0.121 | 0.103 | 1 | 10 | Mre11a | 0.018 | Young |
| 0.572783 | 0.110897 | 0.182 | 0.213 | 1 | 10 | Pfdn4 | -0.031 | Young |
| 0.572803 | 0.689403 | 0.111 | 0.095 | 1 | 10 | Ampd3 | 0.016 | Young |
| 0.572841 | 0.29135 | 0.101 | 0.083 | 1 | 10 | Psmf1 | 0.018 | Young |
| 0.572896 | 0.024991 | 0.465 | 0.521 | 1 | 10 | Lamtor2 | -0.056 | Young |
| 0.572987 | 0.650119 | 0.04 | 0.029 | 1 | 10 | Adam33 | 0.011 | Young |

| | | | | | | | | |
|---|---|---|---|---|---|---|---|---|
| 0.573062 | 0.55002 | 0.051 | 0.066 | 1 | 10 | Sufu | -0.015 | Young |
| 0.573065 | 0.139317 | 0.606 | 0.609 | 1 | 10 | Atf4 | -0.003 | Young |
| 0.573078 | 0.052778 | 0.949 | 0.968 | 1 | 10 | Rps4x | -0.019 | Young |
| 0.573283 | 0.298799 | 0.253 | 0.237 | 1 | 10 | Safb | 0.016 | Young |
| 0.573457 | 0.395022 | 0.152 | 0.183 | 1 | 10 | Lrrc47 | -0.031 | Young |
| 0.573636 | 0.003048 | 0.495 | 0.543 | 1 | 10 | Psma2 | -0.048 | Young |
| 0.573664 | 0.543126 | 0.162 | 0.144 | 1 | 10 | Itpk1 | 0.018 | Young |
| 0.57399 | 0.34416 | 0.071 | 0.056 | 1 | 10 | Zfp938 | 0.015 | Young |
| 0.574239 | 0.163164 | 0.333 | 0.306 | 1 | 10 | Dyrk2 | 0.027 | Young |
| 0.574377 | 0.103317 | 0.111 | 0.134 | 1 | 10 | Ralgapb | -0.023 | Young |
| 0.574705 | 0.437539 | 0.04 | 0.029 | 1 | 10 | Btbd3 | 0.011 | Young |
| 0.574705 | 0.260513 | 0.04 | 0.029 | 1 | 10 | Asb4 | 0.011 | Young |
| 0.575177 | 0.127957 | 0.747 | 0.812 | 1 | 10 | Rpl31 | -0.065 | Young |
| 0.575236 | 0.28548 | 0.263 | 0.24 | 1 | 10 | Acbd6 | 0.023 | Young |
| 0.575424 | 0.763558 | 0.141 | 0.125 | 1 | 10 | Ubl7 | 0.016 | Young |
| 0.575754 | 0.340991 | 0.364 | 0.355 | 1 | 10 | Rad23a | 0.009 | Young |
| 0.576425 | 0.089191 | 0.04 | 0.029 | 1 | 10 | Clp1 | 0.011 | Young |
| 0.576479 | 0.21813 | 0.343 | 0.328 | 1 | 10 | Pip5k1c | 0.015 | Young |
| 0.576494 | 0.044321 | 0.707 | 0.748 | 1 | 10 | Nr4a1 | -0.041 | Young |
| 0.576538 | 0.465005 | 0.071 | 0.056 | 1 | 10 | Fam60a | 0.015 | Young |
| 0.57692 | 0.11853 | 0.859 | 0.853 | 1 | 10 | Gng5 | 0.006 | Young |
| 0.577018 | 0.184169 | 0.606 | 0.655 | 1 | 10 | Abracl | -0.049 | Young |
| 0.577061 | 0.682685 | 0.162 | 0.147 | 1 | 10 | Carhsp1 | 0.015 | Young |
| 0.577619 | 0.740824 | 0.131 | 0.117 | 1 | 10 | Klhl21 | 0.014 | Young |
| 0.577678 | 0.019848 | 0.475 | 0.526 | 1 | 10 | B4galt6 | -0.051 | Young |
| 0.57814 | 0.286236 | 0.141 | 0.171 | 1 | 10 | Nabp2 | -0.03 | Young |
| 0.578148 | 0.003673 | 0.04 | 0.029 | 1 | 10 | 1810044D( | 0.011 | Young |
| 0.578358 | 0.047899 | 0.182 | 0.21 | 1 | 10 | Nop58 | -0.028 | Young |
| 0.578412 | 0.006164 | 0.808 | 0.934 | 1 | 10 | H2-Eb1 | -0.126 | Young |
| 0.578438 | 0.101488 | 0.111 | 0.093 | 1 | 10 | Cd82 | 0.018 | Young |
| 0.579195 | 0.003682 | 0.889 | 0.914 | 1 | 10 | Rps6 | -0.025 | Young |
| 0.579307 | 0.236413 | 0.253 | 0.232 | 1 | 10 | Mrps28 | 0.021 | Young |
| 0.579315 | 0.529195 | 0.081 | 0.066 | 1 | 10 | Trpm4 | 0.015 | Young |
| 0.57954 | 0.32753 | 0.253 | 0.237 | 1 | 10 | Tm9sf4 | 0.016 | Young |
| 0.579856 | 0.610023 | 0.111 | 0.095 | 1 | 10 | B3glct | 0.016 | Young |
| 0.580096 | 0.434023 | 0.232 | 0.218 | 1 | 10 | Nasp | 0.014 | Young |
| 0.580504 | 0.561217 | 0.081 | 0.066 | 1 | 10 | Btbd19 | 0.015 | Young |
| 0.580752 | 0.019191 | 0.121 | 0.1 | 1 | 10 | Arl10 | 0.021 | Young |
| 0.580799 | 0.069461 | 0.404 | 0.474 | 1 | 10 | Psmd4 | -0.07 | Young |
| 0.580883 | 0.218199 | 0.182 | 0.213 | 1 | 10 | Ilvbl | -0.031 | Young |
| 0.581082 | 0.27437 | 0.172 | 0.205 | 1 | 10 | Hdac3 | -0.033 | Young |
| 0.581117 | 0.511367 | 0.354 | 0.347 | 1 | 10 | Rad21 | 0.007 | Young |
| 0.581246 | 0.165714 | 0.162 | 0.142 | 1 | 10 | Exosc10 | 0.02 | Young |
| 0.581499 | 0.145632 | 0.343 | 0.384 | 1 | 10 | Csf2rb | -0.041 | Young |
| 0.581721 | 0.912119 | 0.051 | 0.039 | 1 | 10 | Aldh1b1 | 0.012 | Young |

| | | | | | | | | |
|---|---|---|---|---|---|---|---|---|
| 0.581926 | 0.212234 | 0.192 | 0.232 | 1 | 10 | Klf9 | -0.04 | Young |
| 0.582624 | 0.399212 | 0.242 | 0.225 | 1 | 10 | Hars | 0.017 | Young |
| 0.583173 | 0.810969 | 0.131 | 0.117 | 1 | 10 | Dgcr6 | 0.014 | Young |
| 0.583239 | 1.017513 | 0.051 | 0.039 | 1 | 10 | Slamf6 | 0.012 | Young |
| 0.583298 | 0.170331 | 0.162 | 0.193 | 1 | 10 | Micu2 | -0.031 | Young |
| 0.583776 | 0.287054 | 0.455 | 0.44 | 1 | 10 | Cnih4 | 0.015 | Young |
| 0.583823 | 0.316287 | 0.687 | 0.743 | 1 | 10 | Tmsb10 | -0.056 | Young |
| 0.584537 | 0.095255 | 0.232 | 0.269 | 1 | 10 | Tap1 | -0.037 | Young |
| 0.584672 | 0.031788 | 0.889 | 0.91 | 1 | 10 | Ybx1 | -0.021 | Young |
| 0.584758 | 1.100626 | 0.051 | 0.039 | 1 | 10 | Shq1 | 0.012 | Young |
| 0.584927 | 0.222222 | 0.535 | 0.533 | 1 | 10 | Rabac1 | 0.002 | Young |
| 0.585109 | 0.678237 | 0.131 | 0.115 | 1 | 10 | Dcstamp | 0.016 | Young |
| 0.585193 | 0.156414 | 0.263 | 0.306 | 1 | 10 | Ccni | -0.043 | Young |
| 0.585225 | 0.419828 | 0.152 | 0.183 | 1 | 10 | Suco | -0.031 | Young |
| 0.585343 | 0.017847 | 0.939 | 0.966 | 1 | 10 | Rpl7 | -0.027 | Young |
| 0.585514 | 0.194319 | 0.162 | 0.142 | 1 | 10 | Scoc | 0.02 | Young |
| 0.585608 | 0.183465 | 0.697 | 0.731 | 1 | 10 | Fam49b | -0.034 | Young |
| 0.585978 | 0.620154 | 0.121 | 0.105 | 1 | 10 | Sil1 | 0.016 | Young |
| 0.586345 | 0.065434 | 0.172 | 0.205 | 1 | 10 | Tnpo2 | -0.033 | Young |
| 0.586514 | 0.252242 | 0.101 | 0.122 | 1 | 10 | Vps39 | -0.021 | Young |
| 0.586901 | 0.033333 | 0.111 | 0.134 | 1 | 10 | Dctn6 | -0.023 | Young |
| 0.587183 | 0.089687 | 0.222 | 0.257 | 1 | 10 | Ankrd13c | -0.035 | Young |
| 0.587263 | 0.142329 | 0.222 | 0.267 | 1 | 10 | Ppp6r1 | -0.045 | Young |
| 0.588012 | 0.685564 | 0.222 | 0.215 | 1 | 10 | Ssh2 | 0.007 | Young |
| 0.589292 | 0.054895 | 0.121 | 0.103 | 1 | 10 | H6pd | 0.018 | Young |
| 0.589327 | 1.208517 | 0.051 | 0.039 | 1 | 10 | Zmat1 | 0.012 | Young |
| 0.589691 | 0.348027 | 0.303 | 0.355 | 1 | 10 | Plekha2 | -0.052 | Young |
| 0.589967 | 1.144267 | 0.071 | 0.059 | 1 | 10 | Nradd | 0.012 | Young |
| 0.590433 | 0.161954 | 0.374 | 0.355 | 1 | 10 | Ssfa2 | 0.019 | Young |
| 0.590662 | 0.125133 | 0.717 | 0.719 | 1 | 10 | Hspe1 | -0.002 | Young |
| 0.59078 | 0.040878 | 0.485 | 0.535 | 1 | 10 | Rpl13a | -0.05 | Young |
| 0.591093 | 0.028015 | 0.657 | 0.692 | 1 | 10 | Rab5c | -0.035 | Young |
| 0.591207 | 0.331801 | 0.101 | 0.122 | 1 | 10 | Nosip | -0.021 | Young |
| 0.59149 | 0.078762 | 0.475 | 0.56 | 1 | 10 | Pgls | -0.085 | Young |
| 0.591742 | 0.596514 | 0.192 | 0.181 | 1 | 10 | Ldb1 | 0.011 | Young |
| 0.592029 | 0.161766 | 0.212 | 0.259 | 1 | 10 | Eif4a3 | -0.047 | Young |
| 0.592102 | 0.308658 | 0.424 | 0.43 | 1 | 10 | Ralb | -0.006 | Young |
| 0.592244 | 0.795026 | 0.273 | 0.267 | 1 | 10 | Ap3m1 | 0.006 | Young |
| 0.593158 | 0.214719 | 0.263 | 0.303 | 1 | 10 | Tbl1x | -0.04 | Young |
| 0.593199 | 0.472314 | 0.364 | 0.352 | 1 | 10 | Sumo1 | 0.012 | Young |
| 0.593332 | 0.537419 | 0.202 | 0.188 | 1 | 10 | Ss18l2 | 0.014 | Young |
| 0.593419 | 0.009229 | 0.747 | 0.778 | 1 | 10 | Vcp | -0.031 | Young |
| 0.593489 | 0.890334 | 0.141 | 0.125 | 1 | 10 | Phrf1 | 0.016 | Young |
| 0.593497 | 0.298961 | 0.232 | 0.213 | 1 | 10 | Strn4 | 0.019 | Young |
| 0.593788 | 0.403301 | 0.121 | 0.105 | 1 | 10 | Vps26b | 0.016 | Young |

| | | | | | | | | |
|---|---|---|---|---|---|---|---|---|
| 0.593811 | 0.335725 | 0.081 | 0.1 | 1 | 10 | Med20 | -0.019 | Young |
| 0.593913 | 1.052717 | 0.051 | 0.039 | 1 | 10 | Mbd4 | 0.012 | Young |
| 0.593913 | 0.710269 | 0.051 | 0.039 | 1 | 10 | Car5b | 0.012 | Young |
| 0.593913 | 0.709499 | 0.051 | 0.039 | 1 | 10 | Cdkn2c | 0.012 | Young |
| 0.593925 | 0.3428 | 0.152 | 0.183 | 1 | 10 | Usp5 | -0.031 | Young |
| 0.594262 | 0.064299 | 0.071 | 0.088 | 1 | 10 | Cpsf1 | -0.017 | Young |
| 0.594565 | 0.192851 | 0.485 | 0.474 | 1 | 10 | Slc38a2 | 0.011 | Young |
| 0.594595 | 0.483261 | 0.081 | 0.103 | 1 | 10 | Hlx | -0.022 | Young |
| 0.594621 | 0.089955 | 0.101 | 0.12 | 1 | 10 | Parp11 | -0.019 | Young |
| 0.594623 | 0.297554 | 0.394 | 0.406 | 1 | 10 | Copb2 | -0.012 | Young |
| 0.594969 | 0.100334 | 0.737 | 0.79 | 1 | 10 | Sgpl1 | -0.053 | Young |
| 0.595357 | 0.109323 | 0.071 | 0.088 | 1 | 10 | Dhx32 | -0.017 | Young |
| 0.595439 | 0.997553 | 0.152 | 0.139 | 1 | 10 | Utp20 | 0.013 | Young |
| 0.595445 | 0.745269 | 0.051 | 0.039 | 1 | 10 | Trim21 | 0.012 | Young |
| 0.595445 | 0.53433 | 0.051 | 0.039 | 1 | 10 | Sac3d1 | 0.012 | Young |
| 0.595745 | 0.727148 | 0.061 | 0.049 | 1 | 10 | Edrf1 | 0.012 | Young |
| 0.595758 | 0.585556 | 0.182 | 0.164 | 1 | 10 | Flcn | 0.018 | Young |
| 0.595818 | 0.437071 | 0.162 | 0.142 | 1 | 10 | Atmin | 0.02 | Young |
| 0.596277 | 0.244149 | 0.263 | 0.306 | 1 | 10 | Ccnt1 | -0.043 | Young |
| 0.596485 | 0.454734 | 0.101 | 0.083 | 1 | 10 | Tmem39a | 0.018 | Young |
| 0.596503 | 0.029502 | 0.152 | 0.127 | 1 | 10 | 1600012H( | 0.025 | Young |
| 0.596875 | 0.372751 | 0.343 | 0.347 | 1 | 10 | Dpm3 | -0.004 | Young |
| 0.597052 | 0.208643 | 0.717 | 0.76 | 1 | 10 | Luc7l2 | -0.043 | Young |
| 0.597666 | 0.368241 | 0.081 | 0.103 | 1 | 10 | Yars | -0.022 | Young |
| 0.598305 | 0.742936 | 0.141 | 0.13 | 1 | 10 | Rbbp8 | 0.011 | Young |
| 0.598515 | 1.077362 | 0.051 | 0.039 | 1 | 10 | Lysmd1 | 0.012 | Young |
| 0.599177 | 0.360011 | 0.222 | 0.208 | 1 | 10 | Angel2 | 0.014 | Young |
| 0.599278 | 0.187918 | 0.182 | 0.161 | 1 | 10 | Timm22 | 0.021 | Young |
| 0.600332 | 0.130684 | 0.101 | 0.12 | 1 | 10 | Med4 | -0.019 | Young |
| 0.600373 | 0.062875 | 0.131 | 0.154 | 1 | 10 | Cdt1 | -0.023 | Young |
| 0.600762 | 0.287051 | 0.374 | 0.44 | 1 | 10 | Ap1s1 | -0.066 | Young |
| 0.600833 | 0.343066 | 0.354 | 0.423 | 1 | 10 | 2810474O1 | -0.069 | Young |
| 0.601088 | 0.664234 | 0.091 | 0.076 | 1 | 10 | Foxo1 | 0.015 | Young |
| 0.601354 | 0.062941 | 0.333 | 0.311 | 1 | 10 | Chtop | 0.022 | Young |
| 0.601373 | 0.256124 | 0.505 | 0.538 | 1 | 10 | Inpp5d | -0.033 | Young |
| 0.601605 | 0.524833 | 0.091 | 0.115 | 1 | 10 | Slc25a12 | -0.024 | Young |
| 0.602502 | 0.514359 | 0.323 | 0.318 | 1 | 10 | Sppl3 | 0.005 | Young |
| 0.602524 | 0.510445 | 0.222 | 0.213 | 1 | 10 | Slc31a1 | 0.009 | Young |
| 0.602804 | 0.404116 | 0.101 | 0.125 | 1 | 10 | Snhg12 | -0.024 | Young |
| 0.603066 | 0.064006 | 0.212 | 0.244 | 1 | 10 | Ptpn11 | -0.032 | Young |
| 0.603133 | 0.37458 | 0.051 | 0.039 | 1 | 10 | Mcrip2 | 0.012 | Young |
| 0.603402 | 0.068566 | 0.556 | 0.628 | 1 | 10 | Mdfic | -0.072 | Young |
| 0.603523 | 0.162872 | 0.162 | 0.188 | 1 | 10 | Gtf2a2 | -0.026 | Young |
| 0.603545 | 0.609427 | 0.313 | 0.323 | 1 | 10 | Ubn1 | -0.01 | Young |
| 0.603723 | 0.649283 | 0.141 | 0.13 | 1 | 10 | Metap1 | 0.011 | Young |

| | | | | | | | | |
|---|---|---|---|---|---|---|---|---|
| 0.604081 | 0.640095 | 0.061 | 0.049 | 1 | 10 | Gm49359 | 0.012 | Young |
| 0.604081 | 0.074809 | 0.04 | 0.054 | 1 | 10 | Tgds | -0.014 | Young |
| 0.604148 | 0.355844 | 0.071 | 0.088 | 1 | 10 | Ints14 | -0.017 | Young |
| 0.604343 | 0.28098 | 0.111 | 0.095 | 1 | 10 | Kdm4c | 0.016 | Young |
| 0.604535 | 0.25858 | 0.081 | 0.066 | 1 | 10 | A4galt | 0.015 | Young |
| 0.604926 | 0.049796 | 0.152 | 0.176 | 1 | 10 | Cnep1r1 | -0.024 | Young |
| 0.605047 | 0.709378 | 0.182 | 0.174 | 1 | 10 | Tlk2 | 0.008 | Young |
| 0.605047 | 0.212692 | 0.152 | 0.181 | 1 | 10 | Wdr36 | -0.029 | Young |
| 0.605191 | 0.157254 | 0.758 | 0.765 | 1 | 10 | Cox6b1 | -0.007 | Young |
| 0.605317 | 0.207289 | 0.313 | 0.359 | 1 | 10 | Prkacb | -0.046 | Young |
| 0.606202 | 0.288633 | 0.384 | 0.369 | 1 | 10 | Rassf1 | 0.015 | Young |
| 0.606221 | 0.649354 | 0.051 | 0.039 | 1 | 10 | Rnf135 | 0.012 | Young |
| 0.606789 | 0.141414 | 0.414 | 0.487 | 1 | 10 | Mkrn1 | -0.073 | Young |
| 0.606872 | 0.535976 | 0.061 | 0.049 | 1 | 10 | Kremen1 | 0.012 | Young |
| 0.606872 | 0.240738 | 0.061 | 0.049 | 1 | 10 | Phykpl | 0.012 | Young |
| 0.607045 | 0.259703 | 0.525 | 0.496 | 1 | 10 | Tmem14c | 0.029 | Young |
| 0.607222 | 0.017276 | 0.535 | 0.589 | 1 | 10 | Smarca5 | -0.054 | Young |
| 0.607515 | 0.339381 | 0.071 | 0.056 | 1 | 10 | Zfp160 | 0.015 | Young |
| 0.607672 | 1.517635 | 0.03 | 0.022 | 1 | 10 | Ppp1r9a | 0.008 | Young |
| 0.607767 | 0.354784 | 0.051 | 0.039 | 1 | 10 | Alg6 | 0.012 | Young |
| 0.607953 | 0.292959 | 0.131 | 0.161 | 1 | 10 | Memo1 | -0.03 | Young |
| 0.608059 | 0.079915 | 0.556 | 0.55 | 1 | 10 | Ptpre | 0.006 | Young |
| 0.608328 | 0.052532 | 0.313 | 0.286 | 1 | 10 | Nfil3 | 0.027 | Young |
| 0.608456 | 0.063451 | 0.313 | 0.357 | 1 | 10 | Sec14l1 | -0.044 | Young |
| 0.608505 | 0.191431 | 0.212 | 0.193 | 1 | 10 | Hk2 | 0.019 | Young |
| 0.608822 | 0.035887 | 0.071 | 0.056 | 1 | 10 | Mical2 | 0.015 | Young |
| 0.608954 | 0.836871 | 0.121 | 0.108 | 1 | 10 | Nfyb | 0.013 | Young |
| 0.609056 | 0.713327 | 0.192 | 0.181 | 1 | 10 | Fibp | 0.011 | Young |
| 0.609072 | 0.295873 | 0.091 | 0.076 | 1 | 10 | Nadk2 | 0.015 | Young |
| 0.609116 | 0.142349 | 0.121 | 0.103 | 1 | 10 | Gins4 | 0.018 | Young |
| 0.609479 | 0.099712 | 0.192 | 0.225 | 1 | 10 | Casc3 | -0.033 | Young |
| 0.609506 | 0.014395 | 0.091 | 0.073 | 1 | 10 | Zkscan1 | 0.018 | Young |
| 0.609668 | 0.413756 | 0.04 | 0.054 | 1 | 10 | Mettl8 | -0.014 | Young |
| 0.609712 | 0.170249 | 0.313 | 0.357 | 1 | 10 | Srsf7 | -0.044 | Young |
| 0.610072 | 0.646111 | 0.141 | 0.13 | 1 | 10 | Mrps5 | 0.011 | Young |
| 0.61015 | 0.660862 | 0.182 | 0.166 | 1 | 10 | Mtx1 | 0.016 | Young |
| 0.610194 | 0.234221 | 0.061 | 0.076 | 1 | 10 | Haus2 | -0.015 | Young |
| 0.610277 | 0.106301 | 0.172 | 0.198 | 1 | 10 | Anapc15 | -0.026 | Young |
| 0.610411 | 0.232756 | 0.111 | 0.093 | 1 | 10 | Slc7a6os | 0.018 | Young |
| 0.610442 | 0.464805 | 0.192 | 0.23 | 1 | 10 | Prdx3 | -0.038 | Young |
| 0.610622 | 0.26369 | 0.051 | 0.066 | 1 | 10 | Med31 | -0.015 | Young |
| 0.611073 | 0.151237 | 0.242 | 0.215 | 1 | 10 | Atad2b | 0.027 | Young |
| 0.611308 | 0.082247 | 0.101 | 0.125 | 1 | 10 | Cars | -0.024 | Young |
| 0.611638 | 0.109606 | 0.212 | 0.244 | 1 | 10 | Gtf3c2 | -0.032 | Young |
| 0.611735 | 1.072596 | 0.03 | 0.022 | 1 | 10 | Dnm1 | 0.008 | Young |

| | | | | | | | | |
|---|---|---|---|---|---|---|---|---|
| 0.611735 | 0.518183 | 0.03 | 0.022 | 1 | 10 | Klra2 | 0.008 | Young |
| 0.612109 | 0.203125 | 0.545 | 0.56 | 1 | 10 | Ssb | -0.015 | Young |
| 0.612355 | 0.182849 | 0.212 | 0.244 | 1 | 10 | Ddx18 | -0.032 | Young |
| 0.61247 | 0.139459 | 0.04 | 0.054 | 1 | 10 | Trmt11 | -0.014 | Young |
| 0.612993 | 0.692868 | 0.152 | 0.139 | 1 | 10 | Rbm18 | 0.013 | Young |
| 0.613484 | 0.164981 | 0.535 | 0.531 | 1 | 10 | Swi5 | 0.004 | Young |
| 0.613493 | 0.249002 | 0.131 | 0.115 | 1 | 10 | Smarcd1 | 0.016 | Young |
| 0.613547 | 0.053218 | 0.111 | 0.093 | 1 | 10 | 1110059E2 | 0.018 | Young |
| 0.613702 | 0.074185 | 0.354 | 0.406 | 1 | 10 | Rp9 | -0.052 | Young |
| 0.613771 | 1.119566 | 0.03 | 0.022 | 1 | 10 | 8030462N1 | 0.008 | Young |
| 0.613771 | 0.755235 | 0.03 | 0.022 | 1 | 10 | Cables1 | 0.008 | Young |
| 0.613777 | 0.316602 | 0.141 | 0.174 | 1 | 10 | Med30 | -0.033 | Young |
| 0.614156 | 0.07264 | 0.273 | 0.32 | 1 | 10 | Guk1 | -0.047 | Young |
| 0.614269 | 0.242281 | 0.687 | 0.66 | 1 | 10 | Sdc4 | 0.027 | Young |
| 0.614553 | 0.045782 | 0.394 | 0.433 | 1 | 10 | Arhgef6 | -0.039 | Young |
| 0.615515 | 0.170847 | 0.566 | 0.575 | 1 | 10 | Gmfg | -0.009 | Young |
| 0.615526 | 0.254321 | 0.051 | 0.039 | 1 | 10 | Mus81 | 0.012 | Young |
| 0.615526 | 0.181866 | 0.051 | 0.039 | 1 | 10 | Mccc1 | 0.012 | Young |
| 0.615798 | 0.318426 | 0.242 | 0.225 | 1 | 10 | Ankib1 | 0.017 | Young |
| 0.61581 | 1.0179 | 0.03 | 0.022 | 1 | 10 | Primpol | 0.008 | Young |
| 0.61581 | 0.810574 | 0.03 | 0.022 | 1 | 10 | H2-M2 | 0.008 | Young |
| 0.61581 | 0.71993 | 0.03 | 0.022 | 1 | 10 | Nup43 | 0.008 | Young |
| 0.61581 | 0.614886 | 0.03 | 0.022 | 1 | 10 | Mrm2 | 0.008 | Young |
| 0.616282 | 0.445388 | 0.212 | 0.203 | 1 | 10 | Psme2b | 0.009 | Young |
| 0.616301 | 0.182527 | 0.273 | 0.308 | 1 | 10 | Ipo7 | -0.035 | Young |
| 0.616508 | 0.326117 | 0.313 | 0.303 | 1 | 10 | Plk2 | 0.01 | Young |
| 0.616859 | 0.404313 | 0.343 | 0.32 | 1 | 10 | Ube2z | 0.023 | Young |
| 0.617083 | 0.491125 | 0.051 | 0.039 | 1 | 10 | Nudt7 | 0.012 | Young |
| 0.617777 | 0.011212 | 0.444 | 0.496 | 1 | 10 | Psma4 | -0.052 | Young |
| 0.617852 | 1.107685 | 0.03 | 0.022 | 1 | 10 | Wrap53 | 0.008 | Young |
| 0.617852 | 0.749245 | 0.03 | 0.022 | 1 | 10 | Qrsl1 | 0.008 | Young |
| 0.617852 | 0.717184 | 0.03 | 0.022 | 1 | 10 | Chaf1b | 0.008 | Young |
| 0.617852 | 0.515221 | 0.03 | 0.022 | 1 | 10 | Ston2 | 0.008 | Young |
| 0.617852 | 0.419688 | 0.03 | 0.022 | 1 | 10 | Trmu | 0.008 | Young |
| 0.618263 | 0.301872 | 0.051 | 0.066 | 1 | 10 | Mtg1 | -0.015 | Young |
| 0.618783 | 0.514055 | 0.182 | 0.174 | 1 | 10 | Aim2 | 0.008 | Young |
| 0.61891 | 0.157436 | 0.202 | 0.181 | 1 | 10 | Eps15l1 | 0.021 | Young |
| 0.61895 | 0.186475 | 0.576 | 0.638 | 1 | 10 | Cflar | -0.062 | Young |
| 0.619014 | 0.265968 | 0.253 | 0.301 | 1 | 10 | Anapc13 | -0.048 | Young |
| 0.619865 | 0.084564 | 0.192 | 0.225 | 1 | 10 | Brd8 | -0.033 | Young |
| 0.619897 | 0.68941 | 0.03 | 0.022 | 1 | 10 | Eid2b | 0.008 | Young |
| 0.619897 | 0.299013 | 0.03 | 0.022 | 1 | 10 | Eral1 | 0.008 | Young |
| 0.619914 | 0.297348 | 0.03 | 0.042 | 1 | 10 | Acad12 | -0.012 | Young |
| 0.620558 | 0.177997 | 0.091 | 0.076 | 1 | 10 | Dis3l2 | 0.015 | Young |
| 0.620734 | 0.349795 | 0.495 | 0.491 | 1 | 10 | Zbtb7a | 0.004 | Young |

| | | | | | | | | |
|---|---|---|---|---|---|---|---|---|
| 0.620747 | 0.257811 | 0.253 | 0.296 | 1 | 10 | Mark2 | -0.043 | Young |
| 0.620788 | 0.029604 | 0.121 | 0.142 | 1 | 10 | Dubr | -0.021 | Young |
| 0.621191 | 0.172722 | 0.566 | 0.572 | 1 | 10 | Brd2 | -0.006 | Young |
| 0.621345 | 0.564573 | 0.081 | 0.098 | 1 | 10 | Sfi1 | -0.017 | Young |
| 0.621345 | 0.2429 | 0.081 | 0.098 | 1 | 10 | Zadh2 | -0.017 | Young |
| 0.621387 | 0.191766 | 0.101 | 0.086 | 1 | 10 | AU022793 | 0.015 | Young |
| 0.621513 | 0.026785 | 0.03 | 0.042 | 1 | 10 | Rps6kb2 | -0.012 | Young |
| 0.62165 | 0.183636 | 0.505 | 0.509 | 1 | 10 | Dnajc3 | -0.004 | Young |
| 0.621764 | 0.08364 | 0.051 | 0.039 | 1 | 10 | Sdhaf3 | 0.012 | Young |
| 0.621772 | 0.523759 | 0.131 | 0.117 | 1 | 10 | Ino80 | 0.014 | Young |
| 0.621945 | 1.312394 | 0.03 | 0.022 | 1 | 10 | Fpr2 | 0.008 | Young |
| 0.621945 | 1.223161 | 0.03 | 0.022 | 1 | 10 | Tagap1 | 0.008 | Young |
| 0.621972 | 9.76E-06 | 0.172 | 0.198 | 1 | 10 | Atg7 | -0.026 | Young |
| 0.622252 | 0.023019 | 0.222 | 0.254 | 1 | 10 | Vps18 | -0.032 | Young |
| 0.622321 | 0.384651 | 0.061 | 0.049 | 1 | 10 | Arfip2 | 0.012 | Young |
| 0.622321 | 0.165422 | 0.04 | 0.054 | 1 | 10 | Ift43 | -0.014 | Young |
| 0.622344 | 0.802196 | 0.131 | 0.12 | 1 | 10 | Kin | 0.011 | Young |
| 0.622587 | 0.091491 | 0.121 | 0.142 | 1 | 10 | Wdr81 | -0.021 | Young |
| 0.622796 | 0.166654 | 0.131 | 0.154 | 1 | 10 | Ints3 | -0.023 | Young |
| 0.622796 | 0.125378 | 0.131 | 0.154 | 1 | 10 | Arid2 | -0.023 | Young |
| 0.622834 | 0.320071 | 0.01 | 0.017 | 1 | 10 | Cox6b2 | -0.007 | Young |
| 0.622834 | 0.313687 | 0.01 | 0.017 | 1 | 10 | 1700052K1 | -0.007 | Young |
| 0.622834 | 0.30805 | 0.01 | 0.017 | 1 | 10 | C730034F0 | -0.007 | Young |
| 0.622834 | 0.223919 | 0.01 | 0.017 | 1 | 10 | 2510017J1 | -0.007 | Young |
| 0.622834 | 0.105294 | 0.01 | 0.017 | 1 | 10 | 2810454H0 | -0.007 | Young |
| 0.622834 | 0.020082 | 0.01 | 0.017 | 1 | 10 | Wdr78 | -0.007 | Young |
| 0.622836 | 0.404596 | 0.273 | 0.264 | 1 | 10 | Gpbp1l1 | 0.009 | Young |
| 0.622902 | 0.596293 | 0.141 | 0.127 | 1 | 10 | Mphosph6 | 0.014 | Young |
| 0.623336 | 0.253309 | 0.273 | 0.32 | 1 | 10 | Fam172a | -0.047 | Young |
| 0.623385 | 0.739062 | 0.02 | 0.029 | 1 | 10 | Gm47754 | -0.009 | Young |
| 0.623697 | 0.072575 | 0.303 | 0.335 | 1 | 10 | Blmh | -0.032 | Young |
| 0.623779 | 0.098096 | 0.202 | 0.23 | 1 | 10 | Ip6k1 | -0.028 | Young |
| 0.62399 | 0.268564 | 0.141 | 0.166 | 1 | 10 | Eif4e3 | -0.025 | Young |
| 0.623996 | 0.536912 | 0.03 | 0.022 | 1 | 10 | Zfp946 | 0.008 | Young |
| 0.624469 | 0.065374 | 0.232 | 0.269 | 1 | 10 | Zfp292 | -0.037 | Young |
| 0.62457 | 0.654883 | 0.242 | 0.24 | 1 | 10 | Zzz3 | 0.002 | Young |
| 0.624716 | 0.643665 | 0.03 | 0.042 | 1 | 10 | Fadd | -0.012 | Young |
| 0.624738 | 0.062582 | 0.404 | 0.452 | 1 | 10 | Ctnna1 | -0.048 | Young |
| 0.624929 | 0.298915 | 0.182 | 0.161 | 1 | 10 | Mdn1 | 0.021 | Young |
| 0.624952 | 0.302106 | 0.475 | 0.57 | 1 | 10 | Eif5 | -0.095 | Young |
| 0.624991 | 0.256979 | 0.535 | 0.565 | 1 | 10 | Arap1 | -0.03 | Young |
| 0.625338 | 0.857749 | 0.01 | 0.017 | 1 | 10 | Fancm | -0.007 | Young |
| 0.625338 | 0.60481 | 0.01 | 0.017 | 1 | 10 | Uhrf1 | -0.007 | Young |
| 0.625338 | 0.567213 | 0.01 | 0.017 | 1 | 10 | Nptxr | -0.007 | Young |
| 0.625338 | 0.448632 | 0.01 | 0.017 | 1 | 10 | Abhd14a | -0.007 | Young |

| | | | | | | | | |
|---|---|---|---|---|---|---|---|---|
| 0.625338 | 0.430058 | 0.01 | 0.017 | 1 | 10 | Zfp446 | -0.007 | Young |
| 0.625338 | 0.382122 | 0.01 | 0.017 | 1 | 10 | Bbs2 | -0.007 | Young |
| 0.625338 | 0.329155 | 0.01 | 0.017 | 1 | 10 | Zfp60 | -0.007 | Young |
| 0.625338 | 0.190157 | 0.01 | 0.017 | 1 | 10 | Myo19 | -0.007 | Young |
| 0.625338 | 0.139178 | 0.01 | 0.017 | 1 | 10 | Timp3 | -0.007 | Young |
| 0.625338 | 0.127374 | 0.01 | 0.017 | 1 | 10 | Gm26652 | -0.007 | Young |
| 0.625338 | 0.110505 | 0.01 | 0.017 | 1 | 10 | Dctd | -0.007 | Young |
| 0.625338 | 0.088513 | 0.01 | 0.017 | 1 | 10 | Gnat3 | -0.007 | Young |
| 0.625338 | 0.062601 | 0.01 | 0.017 | 1 | 10 | Ccdc94 | -0.007 | Young |
| 0.625338 | 0.048396 | 0.01 | 0.017 | 1 | 10 | Poli | -0.007 | Young |
| 0.62534 | 0.057717 | 0.485 | 0.523 | 1 | 10 | Stap1 | -0.038 | Young |
| 0.625429 | 0.050752 | 0.323 | 0.359 | 1 | 10 | Pde4b | -0.036 | Young |
| 0.625847 | 0.445991 | 0.465 | 0.487 | 1 | 10 | F13a1 | -0.022 | Young |
| 0.625984 | 0.533309 | 0.071 | 0.059 | 1 | 10 | Sergef | 0.012 | Young |
| 0.62605 | 0.972482 | 0.03 | 0.022 | 1 | 10 | Tamm41 | 0.008 | Young |
| 0.62605 | 0.459768 | 0.03 | 0.022 | 1 | 10 | 2810021J2 | 0.008 | Young |
| 0.626069 | 0.089181 | 0.273 | 0.303 | 1 | 10 | Atxn2 | -0.03 | Young |
| 0.626134 | 0.182777 | 0.192 | 0.222 | 1 | 10 | Tpst2 | -0.03 | Young |
| 0.626159 | 0.512639 | 0.081 | 0.1 | 1 | 10 | Spata2 | -0.019 | Young |
| 0.626564 | 0.339008 | 0.061 | 0.049 | 1 | 10 | Bend4 | 0.012 | Young |
| 0.627196 | 0.010739 | 0.02 | 0.029 | 1 | 10 | Ddah2 | -0.009 | Young |
| 0.627281 | 0.03048 | 0.172 | 0.2 | 1 | 10 | Tmem50b | -0.028 | Young |
| 0.627288 | 0.590346 | 0.071 | 0.059 | 1 | 10 | Cyb561d1 | 0.012 | Young |
| 0.627333 | 0.125738 | 0.333 | 0.401 | 1 | 10 | Nucb1 | -0.068 | Young |
| 0.627496 | 0.315552 | 0.121 | 0.105 | 1 | 10 | 0610037L1 | 0.016 | Young |
| 0.627746 | 0.749783 | 0.121 | 0.108 | 1 | 10 | Dnajc25 | 0.013 | Young |
| 0.627849 | 0.595153 | 0.162 | 0.144 | 1 | 10 | Me2 | 0.018 | Young |
| 0.627872 | 0.139239 | 0.354 | 0.403 | 1 | 10 | Gng2 | -0.049 | Young |
| 0.628029 | 0.00687 | 0.051 | 0.039 | 1 | 10 | Atrip | 0.012 | Young |
| 0.628197 | 0.572123 | 0.192 | 0.181 | 1 | 10 | Mcm3 | 0.011 | Young |
| 0.628221 | 0.375828 | 0.212 | 0.198 | 1 | 10 | Rnf20 | 0.014 | Young |
| 0.628373 | 0.811212 | 0.141 | 0.13 | 1 | 10 | Gramd1b | 0.011 | Young |
| 0.628387 | 0.127815 | 0.232 | 0.213 | 1 | 10 | Hgsnat | 0.019 | Young |
| 0.628515 | 0.916887 | 0.051 | 0.066 | 1 | 10 | Shmt2 | -0.015 | Young |
| 0.628579 | 0.604715 | 0.172 | 0.156 | 1 | 10 | Lyrm2 | 0.016 | Young |
| 0.629105 | 0.115049 | 0.02 | 0.029 | 1 | 10 | Rwdd2a | -0.009 | Young |
| 0.6294 | 0.61571 | 0.04 | 0.054 | 1 | 10 | Sh2d1b1 | -0.014 | Young |
| 0.629533 | 0.121079 | 0.03 | 0.042 | 1 | 10 | Pfas | -0.012 | Young |
| 0.629738 | 0.028621 | 0.091 | 0.11 | 1 | 10 | Scly | -0.019 | Young |
| 0.629768 | 0.127435 | 0.131 | 0.154 | 1 | 10 | Stard4 | -0.023 | Young |
| 0.629822 | 0.058851 | 0.172 | 0.198 | 1 | 10 | Cab39l | -0.026 | Young |
| 0.629915 | 0.220431 | 0.586 | 0.638 | 1 | 10 | Eif3c | -0.052 | Young |
| 0.630166 | 0.643779 | 0.03 | 0.022 | 1 | 10 | Cnn3 | 0.008 | Young |
| 0.630326 | 0.43889 | 0.172 | 0.159 | 1 | 10 | Pdgfa | 0.013 | Young |
| 0.630438 | 0.625956 | 0.131 | 0.122 | 1 | 10 | Kifap3 | 0.009 | Young |

| | | | | | | | | |
|---|---|---|---|---|---|---|---|---|
| 0.630819 | 0.129819 | 0.061 | 0.049 | 1 | 10 | Per3 | 0.012 | Young |
| 0.631379 | 0.81853 | 0.091 | 0.078 | 1 | 10 | E2f3 | 0.013 | Young |
| 0.631667 | 0.19508 | 0.525 | 0.513 | 1 | 10 | Sash1 | 0.012 | Young |
| 0.631765 | 0.375136 | 0.091 | 0.11 | 1 | 10 | Prkag2 | -0.019 | Young |
| 0.631913 | 0.21552 | 0.586 | 0.597 | 1 | 10 | Nedd8 | -0.011 | Young |
| 0.631942 | 0.003658 | 0.333 | 0.389 | 1 | 10 | Rbbp6 | -0.056 | Young |
| 0.632189 | 0.262226 | 0.313 | 0.296 | 1 | 10 | Marf1 | 0.017 | Young |
| 0.632197 | 0.285641 | 0.404 | 0.396 | 1 | 10 | Sulf2 | 0.008 | Young |
| 0.632228 | 0.462915 | 0.03 | 0.022 | 1 | 10 | Cnst | 0.008 | Young |
| 0.632228 | 0.094429 | 0.03 | 0.022 | 1 | 10 | Olfr920 | 0.008 | Young |
| 0.632241 | 0.309011 | 0.04 | 0.054 | 1 | 10 | Zfp101 | -0.014 | Young |
| 0.632262 | 0.351233 | 0.101 | 0.122 | 1 | 10 | Vars | -0.021 | Young |
| 0.632432 | 0.250854 | 0.111 | 0.095 | 1 | 10 | Inafm1 | 0.016 | Young |
| 0.632514 | 0.381359 | 0.051 | 0.064 | 1 | 10 | Sppl2b | -0.013 | Young |
| 0.632931 | 0.602232 | 0.02 | 0.029 | 1 | 10 | Bcr | -0.009 | Young |
| 0.633379 | 0.258224 | 0.616 | 0.658 | 1 | 10 | H2afy | -0.042 | Young |
| 0.633663 | 0.258408 | 0.04 | 0.054 | 1 | 10 | Acvr2a | -0.014 | Young |
| 0.634294 | 0.079578 | 0.03 | 0.022 | 1 | 10 | Dnph1 | 0.008 | Young |
| 0.634425 | 0.467711 | 0.212 | 0.198 | 1 | 10 | Zfp384 | 0.014 | Young |
| 0.634497 | 0.361758 | 0.172 | 0.154 | 1 | 10 | Frmd6 | 0.018 | Young |
| 0.634951 | 0.228749 | 0.364 | 0.401 | 1 | 10 | Pacs2 | -0.037 | Young |
| 0.635087 | 0.616082 | 0.04 | 0.054 | 1 | 10 | Dnmbp | -0.014 | Young |
| 0.635087 | 0.463344 | 0.061 | 0.049 | 1 | 10 | Casp2 | 0.012 | Young |
| 0.635134 | 0.247215 | 0.051 | 0.064 | 1 | 10 | Slc20a2 | -0.013 | Young |
| 0.635829 | 0.170799 | 0.091 | 0.11 | 1 | 10 | N4bp2 | -0.019 | Young |
| 0.635901 | 0.215553 | 0.556 | 0.565 | 1 | 10 | Ptpn1 | -0.009 | Young |
| 0.636027 | 0.012291 | 0.152 | 0.178 | 1 | 10 | Txndc15 | -0.026 | Young |
| 0.636512 | 0.366476 | 0.061 | 0.049 | 1 | 10 | Ndufaf6 | 0.012 | Young |
| 0.636685 | 0.380647 | 0.141 | 0.13 | 1 | 10 | Sfmbt1 | 0.011 | Young |
| 0.636767 | 0.734695 | 0.02 | 0.029 | 1 | 10 | AW209491 | -0.009 | Young |
| 0.636767 | 0.537536 | 0.02 | 0.029 | 1 | 10 | Kctd9 | -0.009 | Young |
| 0.636767 | 0.183267 | 0.02 | 0.029 | 1 | 10 | Nudt15 | -0.009 | Young |
| 0.636847 | 0.021 | 0.091 | 0.11 | 1 | 10 | Akt3 | -0.019 | Young |
| 0.637055 | 0.434172 | 0.162 | 0.188 | 1 | 10 | Zfp655 | -0.026 | Young |
| 0.637131 | 0.226171 | 0.273 | 0.315 | 1 | 10 | Commd8 | -0.042 | Young |
| 0.63746 | 0.219486 | 0.424 | 0.487 | 1 | 10 | Nars | -0.063 | Young |
| 0.637611 | 0.903813 | 0.141 | 0.13 | 1 | 10 | Skiv2l2 | 0.011 | Young |
| 0.637781 | 0.019134 | 0.778 | 0.804 | 1 | 10 | Ubl3 | -0.026 | Young |
| 0.637882 | 0.437537 | 0.101 | 0.086 | 1 | 10 | Zfyve27 | 0.015 | Young |
| 0.637953 | 0.309037 | 0.343 | 0.33 | 1 | 10 | Gla | 0.013 | Young |
| 0.638062 | 0.18708 | 0.333 | 0.391 | 1 | 10 | Plbd2 | -0.058 | Young |
| 0.638168 | 0.138915 | 0.04 | 0.051 | 1 | 10 | Ttll3 | -0.011 | Young |
| 0.638331 | 0.185275 | 0.505 | 0.589 | 1 | 10 | Cd300c2 | -0.084 | Young |
| 0.638433 | 0.071859 | 0.03 | 0.022 | 1 | 10 | Gm14286 | 0.008 | Young |
| 0.638688 | 0.581179 | 0.02 | 0.029 | 1 | 10 | Usp18 | -0.009 | Young |

| | | | | | | | | |
|---|---|---|---|---|---|---|---|---|
| 0.638688 | 0.571928 | 0.02 | 0.029 | 1 | 10 | Dync2h1 | -0.009 | Young |
| 0.638703 | 0.516258 | 0.263 | 0.244 | 1 | 10 | Gigyf1 | 0.019 | Young |
| 0.639367 | 0.59679 | 0.061 | 0.049 | 1 | 10 | Diexf | 0.012 | Young |
| 0.639442 | 0.304788 | 0.424 | 0.411 | 1 | 10 | Nxf1 | 0.013 | Young |
| 0.639551 | 0.419939 | 0.152 | 0.137 | 1 | 10 | Traf6 | 0.015 | Young |
| 0.639577 | 0.46824 | 0.101 | 0.088 | 1 | 10 | Gbp7 | 0.013 | Young |
| 0.639645 | 0.017209 | 0.273 | 0.308 | 1 | 10 | Zcchc7 | -0.035 | Young |
| 0.639845 | 0.186993 | 0.343 | 0.335 | 1 | 10 | Ube2e1 | 0.008 | Young |
| 0.640095 | 0.01947 | 0.101 | 0.086 | 1 | 10 | Hook2 | 0.015 | Young |
| 0.640172 | 0.183634 | 0.424 | 0.435 | 1 | 10 | Vps26a | -0.011 | Young |
| 0.640388 | 0.266677 | 0.071 | 0.059 | 1 | 10 | Fam149b | 0.012 | Young |
| 0.640462 | 0.187997 | 0.192 | 0.222 | 1 | 10 | Mrps6 | -0.03 | Young |
| 0.640477 | 0.110188 | 0.182 | 0.21 | 1 | 10 | Fkbp4 | -0.028 | Young |
| 0.640612 | 1.031023 | 0.02 | 0.029 | 1 | 10 | Rbak | -0.009 | Young |
| 0.641046 | 0.094669 | 0.182 | 0.205 | 1 | 10 | Tmem251 | -0.023 | Young |
| 0.641696 | 0.792979 | 0.04 | 0.032 | 1 | 10 | Tbc1d7 | 0.008 | Young |
| 0.64175 | 0.178238 | 0.121 | 0.108 | 1 | 10 | Fam45a | 0.013 | Young |
| 0.641842 | 0.077134 | 0.283 | 0.315 | 1 | 10 | Polr2g | -0.032 | Young |
| 0.642689 | 0.046607 | 0.545 | 0.506 | 1 | 10 | Pla2g7 | 0.039 | Young |
| 0.642893 | 0.079367 | 0.909 | 0.927 | 1 | 10 | Rpl10a | -0.018 | Young |
| 0.643152 | 0.063263 | 0.323 | 0.372 | 1 | 10 | Bin1 | -0.049 | Young |
| 0.643276 | 0.426987 | 0.071 | 0.088 | 1 | 10 | Psip1 | -0.017 | Young |
| 0.64343 | 0.608756 | 0.273 | 0.269 | 1 | 10 | Ddx42 | 0.004 | Young |
| 0.643744 | 0.421721 | 0.162 | 0.191 | 1 | 10 | Ppie | -0.029 | Young |
| 0.643814 | 0.502765 | 0.061 | 0.076 | 1 | 10 | Fancg | -0.015 | Young |
| 0.643877 | 0.612231 | 0.152 | 0.186 | 1 | 10 | Gstm1 | -0.034 | Young |
| 0.644007 | 0.146362 | 0.475 | 0.533 | 1 | 10 | Mlf2 | -0.058 | Young |
| 0.644232 | 0.009046 | 0.556 | 0.623 | 1 | 10 | Ap2s1 | -0.067 | Young |
| 0.644479 | 0.074418 | 0.869 | 0.917 | 1 | 10 | Atp2b1 | -0.048 | Young |
| 0.64469 | 0.184708 | 0.667 | 0.768 | 1 | 10 | Zfand5 | -0.101 | Young |
| 0.644814 | 0.164265 | 0.172 | 0.156 | 1 | 10 | Pfdn6 | 0.016 | Young |
| 0.644903 | 1.182017 | 0.051 | 0.042 | 1 | 10 | Nepro | 0.009 | Young |
| 0.645092 | 0.18365 | 0.061 | 0.049 | 1 | 10 | Klhl15 | 0.012 | Young |
| 0.645207 | 0.570302 | 0.04 | 0.032 | 1 | 10 | Lrrc49 | 0.008 | Young |
| 0.645899 | 0.482469 | 0.152 | 0.137 | 1 | 10 | Scaf1 | 0.015 | Young |
| 0.646078 | 0.130596 | 0.192 | 0.215 | 1 | 10 | Stat5b | -0.023 | Young |
| 0.646208 | 0.162552 | 0.283 | 0.318 | 1 | 10 | Arhgap45 | -0.035 | Young |
| 0.646419 | 0.251424 | 0.172 | 0.205 | 1 | 10 | Katnbl1 | -0.033 | Young |
| 0.646824 | 0.071591 | 0.606 | 0.685 | 1 | 10 | Rbm25 | -0.079 | Young |
| 0.647519 | 0.072833 | 0.323 | 0.357 | 1 | 10 | Smim7 | -0.034 | Young |
| 0.6477 | 0.243277 | 0.121 | 0.105 | 1 | 10 | Dgkh | 0.016 | Young |
| 0.647735 | 0.366682 | 0.293 | 0.342 | 1 | 10 | Dnajc2 | -0.049 | Young |
| 0.648071 | 0.488606 | 0.515 | 0.538 | 1 | 10 | Pmp22 | -0.023 | Young |
| 0.648108 | 0.207342 | 0.333 | 0.379 | 1 | 10 | Sirt2 | -0.046 | Young |
| 0.648382 | 0.161161 | 0.606 | 0.623 | 1 | 10 | Hmgb1 | -0.017 | Young |

| | | | | | | | | |
|---|---|---|---|---|---|---|---|---|
| 0.648637 | 0.162202 | 0.141 | 0.166 | 1 | 10 | Pwwp2a | -0.025 | Young |
| 0.649473 | 0.296706 | 0.101 | 0.088 | 1 | 10 | Pde7a | 0.013 | Young |
| 0.649477 | 0.233526 | 0.232 | 0.264 | 1 | 10 | Wdfy4 | -0.032 | Young |
| 0.649626 | 0.481647 | 0.051 | 0.064 | 1 | 10 | Pwp2 | -0.013 | Young |
| 0.649789 | 0.017734 | 0.222 | 0.249 | 1 | 10 | 8-Mar | -0.027 | Young |
| 0.649872 | 0.600442 | 0.091 | 0.078 | 1 | 10 | Slc35e3 | 0.013 | Young |
| 0.650123 | 0.471633 | 0.242 | 0.237 | 1 | 10 | Nudcd2 | 0.005 | Young |
| 0.650489 | 1.324272 | 0.04 | 0.032 | 1 | 10 | Zfp40 | 0.008 | Young |
| 0.650489 | 0.972523 | 0.04 | 0.032 | 1 | 10 | Ttc12 | 0.008 | Young |
| 0.650584 | 0.529211 | 0.354 | 0.362 | 1 | 10 | Ccdc12 | -0.008 | Young |
| 0.650642 | 0.197963 | 0.263 | 0.303 | 1 | 10 | Cpd | -0.04 | Young |
| 0.650796 | 0.143954 | 0.323 | 0.372 | 1 | 10 | Cuedc2 | -0.049 | Young |
| 0.650821 | 0.312994 | 0.121 | 0.108 | 1 | 10 | Pfkfb3 | 0.013 | Young |
| 0.650861 | 0.908136 | 0.111 | 0.1 | 1 | 10 | Nr2c2ap | 0.011 | Young |
| 0.650907 | 0.591548 | 0.162 | 0.149 | 1 | 10 | Mmd | 0.013 | Young |
| 0.650939 | 0.233737 | 0.354 | 0.391 | 1 | 10 | Pnp | -0.037 | Young |
| 0.651117 | 1.233766 | 0.081 | 0.071 | 1 | 10 | Cul4b | 0.01 | Young |
| 0.651253 | 0.327771 | 0.273 | 0.259 | 1 | 10 | Fmnl1 | 0.014 | Young |
| 0.651361 | 0.225687 | 0.152 | 0.137 | 1 | 10 | Scamp1 | 0.015 | Young |
| 0.651486 | 0.475085 | 0.242 | 0.24 | 1 | 10 | Chmp1b | 0.002 | Young |
| 0.651697 | 0.060542 | 0.505 | 0.484 | 1 | 10 | Mapk1 | 0.021 | Young |
| 0.651856 | 0.575993 | 0.121 | 0.112 | 1 | 10 | Fam118b | 0.009 | Young |
| 0.652276 | 0.285077 | 0.071 | 0.059 | 1 | 10 | Gm45606 | 0.012 | Young |
| 0.652341 | 0.506753 | 0.253 | 0.242 | 1 | 10 | Prpf39 | 0.011 | Young |
| 0.652525 | 0.178025 | 0.111 | 0.095 | 1 | 10 | Orc3 | 0.016 | Young |
| 0.653101 | 0.057518 | 0.505 | 0.567 | 1 | 10 | Prrc2a | -0.062 | Young |
| 0.653482 | 0.168468 | 0.364 | 0.411 | 1 | 10 | Efr3a | -0.047 | Young |
| 0.653551 | 0.762328 | 0.131 | 0.156 | 1 | 10 | Dpy19l1 | -0.025 | Young |
| 0.653602 | 0.239196 | 0.051 | 0.064 | 1 | 10 | Wdcp | -0.013 | Young |
| 0.653767 | 0.272846 | 0.333 | 0.325 | 1 | 10 | Rbm26 | 0.008 | Young |
| 0.653978 | 0.761307 | 0.131 | 0.12 | 1 | 10 | Isca2 | 0.011 | Young |
| 0.653998 | 0.78051 | 0.111 | 0.1 | 1 | 10 | Pgpep1 | 0.011 | Young |
| 0.65402 | 0.974222 | 0.04 | 0.032 | 1 | 10 | Slx1b | 0.008 | Young |
| 0.65402 | 0.786109 | 0.04 | 0.032 | 1 | 10 | 4632415L0 | 0.008 | Young |
| 0.65402 | 0.588947 | 0.04 | 0.032 | 1 | 10 | Sema4c | 0.008 | Young |
| 0.65402 | 0.529664 | 0.04 | 0.032 | 1 | 10 | Ccnyl1 | 0.008 | Young |
| 0.654292 | 0.225694 | 0.172 | 0.156 | 1 | 10 | Usp1 | 0.016 | Young |
| 0.654557 | 0.118206 | 0.192 | 0.22 | 1 | 10 | Ptpn7 | -0.028 | Young |
| 0.654643 | 0.11528 | 0.313 | 0.367 | 1 | 10 | Raly | -0.054 | Young |
| 0.654666 | 0.369226 | 0.263 | 0.254 | 1 | 10 | Tmem86a | 0.009 | Young |
| 0.654929 | 0.173683 | 0.071 | 0.059 | 1 | 10 | Gpn2 | 0.012 | Young |
| 0.655464 | 0.610686 | 0.263 | 0.267 | 1 | 10 | Sde2 | -0.004 | Young |
| 0.655573 | 0.392769 | 0.131 | 0.117 | 1 | 10 | Ap1m1 | 0.014 | Young |
| 0.65569 | 1.212128 | 0.101 | 0.122 | 1 | 10 | Nol6 | -0.021 | Young |
| 0.655947 | 0.065428 | 0.455 | 0.457 | 1 | 10 | Cggbp1 | -0.002 | Young |

| | | | | | | | | |
|---|---|---|---|---|---|---|---|---|
| 0.65652 | 0.679219 | 0.141 | 0.127 | 1 | 10 | Zfp318 | 0.014 | Young |
| 0.656672 | 0.032875 | 0.697 | 0.775 | 1 | 10 | Btg2 | -0.078 | Young |
| 0.657558 | 0.502257 | 0.04 | 0.032 | 1 | 10 | 1700096K1 | 0.008 | Young |
| 0.657588 | 0.518924 | 0.071 | 0.059 | 1 | 10 | Iqce | 0.012 | Young |
| 0.657588 | 0.038902 | 0.051 | 0.064 | 1 | 10 | Recql | -0.013 | Young |
| 0.657649 | 0.053236 | 0.081 | 0.066 | 1 | 10 | Fam122a | 0.015 | Young |
| 0.657764 | 0.240383 | 0.101 | 0.12 | 1 | 10 | Yeats2 | -0.019 | Young |
| 0.657815 | 0.21808 | 0.343 | 0.403 | 1 | 10 | Dlst | -0.06 | Young |
| 0.658677 | 0.034676 | 0.081 | 0.095 | 1 | 10 | Cstf2 | -0.014 | Young |
| 0.658758 | 0.147365 | 0.434 | 0.509 | 1 | 10 | Eif1a | -0.075 | Young |
| 0.659217 | 0.746127 | 0.071 | 0.088 | 1 | 10 | Rcc1 | -0.017 | Young |
| 0.659406 | 0.35164 | 0.091 | 0.11 | 1 | 10 | Osgin2 | -0.019 | Young |
| 0.6596 | 0.120737 | 0.333 | 0.389 | 1 | 10 | Gtpbp4 | -0.056 | Young |
| 0.659776 | 0.162659 | 0.081 | 0.095 | 1 | 10 | Sox4 | -0.014 | Young |
| 0.65993 | 0.12689 | 0.293 | 0.32 | 1 | 10 | Ms4a6d | -0.027 | Young |
| 0.659954 | 0.462295 | 0.081 | 0.068 | 1 | 10 | Cd209d | 0.013 | Young |
| 0.660062 | 0.190598 | 0.697 | 0.778 | 1 | 10 | Chd4 | -0.081 | Young |
| 0.660151 | 0.319229 | 0.152 | 0.178 | 1 | 10 | Pon3 | -0.026 | Young |
| 0.66025 | 0.184872 | 0.071 | 0.059 | 1 | 10 | Snupn | 0.012 | Young |
| 0.660371 | 0.327193 | 0.091 | 0.078 | 1 | 10 | Mtor | 0.013 | Young |
| 0.660542 | 0.379971 | 0.101 | 0.088 | 1 | 10 | Slc9a8 | 0.013 | Young |
| 0.660542 | 0.009899 | 0.101 | 0.088 | 1 | 10 | Naa25 | 0.013 | Young |
| 0.660661 | 0.245028 | 0.162 | 0.144 | 1 | 10 | Cul4a | 0.018 | Young |
| 0.660716 | 0.037545 | 0.182 | 0.208 | 1 | 10 | Itgb1bp1 | -0.026 | Young |
| 0.661369 | 0.054619 | 0.152 | 0.174 | 1 | 10 | Pabpc4 | -0.022 | Young |
| 0.661376 | 0.456905 | 0.202 | 0.196 | 1 | 10 | Cdc34 | 0.006 | Young |
| 0.661583 | 0.409075 | 0.051 | 0.064 | 1 | 10 | Ryk | -0.013 | Young |
| 0.662122 | 0.195038 | 0.293 | 0.274 | 1 | 10 | Glrx2 | 0.019 | Young |
| 0.662187 | 0.051963 | 0.909 | 0.946 | 1 | 10 | Ubc | -0.037 | Young |
| 0.662672 | 0.249982 | 0.394 | 0.457 | 1 | 10 | Plekhm2 | -0.063 | Young |
| 0.662724 | 0.104474 | 0.121 | 0.142 | 1 | 10 | Kri1 | -0.021 | Young |
| 0.662777 | 0.078776 | 0.97 | 0.993 | 1 | 10 | Fau | -0.023 | Young |
| 0.662786 | 0.62176 | 0.162 | 0.152 | 1 | 10 | Yipf6 | 0.01 | Young |
| 0.662881 | 0.253066 | 0.04 | 0.032 | 1 | 10 | Nipa1 | 0.008 | Young |
| 0.662964 | 0.380208 | 0.465 | 0.465 | 1 | 10 | Mkln1 | 0 | Young |
| 0.663088 | 0.600413 | 0.04 | 0.051 | 1 | 10 | Tfip11 | -0.011 | Young |
| 0.663221 | 0.018132 | 0.111 | 0.13 | 1 | 10 | Rrp8 | -0.019 | Young |
| 0.66323 | 0.156909 | 0.525 | 0.66 | 1 | 10 | Eid1 | -0.135 | Young |
| 0.663286 | 0.355596 | 0.131 | 0.156 | 1 | 10 | Mrpl10 | -0.025 | Young |
| 0.663327 | 0.285592 | 0.131 | 0.154 | 1 | 10 | Tmed4 | -0.023 | Young |
| 0.663343 | 0.829538 | 0.081 | 0.098 | 1 | 10 | Narf | -0.017 | Young |
| 0.663343 | 0.399443 | 0.081 | 0.098 | 1 | 10 | Kdelc2 | -0.017 | Young |
| 0.663441 | 0.502614 | 0.141 | 0.132 | 1 | 10 | Zdhhc14 | 0.009 | Young |
| 0.663638 | 0.178613 | 0.273 | 0.264 | 1 | 10 | Mrpl3 | 0.009 | Young |
| 0.663642 | 0.596283 | 0.051 | 0.042 | 1 | 10 | Rrad | 0.009 | Young |

| | | | | | | | | |
|---|---|---|---|---|---|---|---|---|
| 0.663794 | 0.113261 | 0.242 | 0.284 | 1 | 10 | Vamp4 | -0.042 | Young |
| 0.663829 | 0.192476 | 0.384 | 0.389 | 1 | 10 | Tet2 | -0.005 | Young |
| 0.663853 | 0.125385 | 0.293 | 0.328 | 1 | 10 | Adam9 | -0.035 | Young |
| 0.663875 | 0.241197 | 0.111 | 0.098 | 1 | 10 | Fhod1 | 0.013 | Young |
| 0.663886 | 0.135144 | 0.091 | 0.078 | 1 | 10 | Ncdn | 0.013 | Young |
| 0.663973 | 0.49455 | 0.111 | 0.1 | 1 | 10 | Zswim4 | 0.011 | Young |
| 0.664015 | 0.572273 | 0.121 | 0.108 | 1 | 10 | Ciao1 | 0.013 | Young |
| 0.664034 | 0.277052 | 0.091 | 0.108 | 1 | 10 | Slc25a32 | -0.017 | Young |
| 0.664152 | 0.220617 | 0.475 | 0.46 | 1 | 10 | Riok3 | 0.015 | Young |
| 0.66431 | 0.473916 | 0.232 | 0.22 | 1 | 10 | Tatdn2 | 0.012 | Young |
| 0.664592 | 0.485826 | 0.212 | 0.249 | 1 | 10 | Gps1 | -0.037 | Young |
| 0.664792 | 0.190709 | 0.414 | 0.482 | 1 | 10 | Bag1 | -0.068 | Young |
| 0.665029 | 0.168051 | 0.071 | 0.086 | 1 | 10 | Mtap | -0.015 | Young |
| 0.665042 | 0.067263 | 0.253 | 0.232 | 1 | 10 | Ube2q2 | 0.021 | Young |
| 0.665143 | 0.135438 | 0.111 | 0.13 | 1 | 10 | Gpd2 | -0.019 | Young |
| 0.665553 | 0.596178 | 0.111 | 0.1 | 1 | 10 | Mrps10 | 0.011 | Young |
| 0.665768 | 0.590297 | 0.283 | 0.281 | 1 | 10 | Abl2 | 0.002 | Young |
| 0.666178 | 0.307201 | 0.081 | 0.068 | 1 | 10 | Creld1 | 0.013 | Young |
| 0.666244 | 0.093806 | 0.303 | 0.347 | 1 | 10 | Tnrc6c | -0.044 | Young |
| 0.666811 | 0.14 | 0.616 | 0.606 | 1 | 10 | Fkbp1a | 0.01 | Young |
| 0.666821 | 0.085036 | 0.202 | 0.22 | 1 | 10 | Ndufs1 | -0.018 | Young |
| 0.666832 | 0.026265 | 0.919 | 0.912 | 1 | 10 | Rpl35 | 0.007 | Young |
| 0.666902 | 0.408393 | 0.343 | 0.337 | 1 | 10 | Dbnl | 0.006 | Young |
| 0.666925 | 0.199709 | 0.051 | 0.064 | 1 | 10 | Nmnat3 | -0.013 | Young |
| 0.667165 | 0.222977 | 0.475 | 0.487 | 1 | 10 | Bax | -0.012 | Young |
| 0.667352 | 0.318421 | 0.071 | 0.086 | 1 | 10 | Gmcl1 | -0.015 | Young |
| 0.667425 | 0.282132 | 0.081 | 0.068 | 1 | 10 | Cdk7 | 0.013 | Young |
| 0.667791 | 0.347547 | 0.172 | 0.205 | 1 | 10 | Farsb | -0.033 | Young |
| 0.668108 | 0.232847 | 0.273 | 0.315 | 1 | 10 | Arl5a | -0.042 | Young |
| 0.66822 | 0.525192 | 0.04 | 0.032 | 1 | 10 | Sesn2 | 0.008 | Young |
| 0.668553 | 0.351417 | 0.485 | 0.496 | 1 | 10 | Mef2d | -0.011 | Young |
| 0.66856 | 0.221945 | 0.202 | 0.235 | 1 | 10 | Chd1 | -0.033 | Young |
| 0.669414 | 0.147878 | 0.374 | 0.435 | 1 | 10 | Kmt2e | -0.061 | Young |
| 0.669582 | 0.168854 | 0.222 | 0.249 | 1 | 10 | Mta2 | -0.027 | Young |
| 0.669603 | 0.140611 | 0.071 | 0.059 | 1 | 10 | Kctd6 | 0.012 | Young |
| 0.669763 | 0.007522 | 0.212 | 0.193 | 1 | 10 | Hk3 | 0.019 | Young |
| 0.669945 | 0.036511 | 0.354 | 0.381 | 1 | 10 | Crlf3 | -0.027 | Young |
| 0.67041 | 0.168862 | 0.162 | 0.186 | 1 | 10 | Nup153 | -0.024 | Young |
| 0.670414 | 0.417433 | 0.242 | 0.232 | 1 | 10 | Psmc2 | 0.01 | Young |
| 0.670624 | 0.276725 | 0.121 | 0.144 | 1 | 10 | 1110059G1 | -0.023 | Young |
| 0.670978 | 0.214281 | 0.162 | 0.181 | 1 | 10 | Smc5 | -0.019 | Young |
| 0.67126 | 0.117979 | 0.818 | 0.841 | 1 | 10 | Cyfip1 | -0.023 | Young |
| 0.671419 | 0.165223 | 0.384 | 0.389 | 1 | 10 | Fbl | -0.005 | Young |
| 0.672008 | 0.099133 | 0.071 | 0.086 | 1 | 10 | Dnajc24 | -0.015 | Young |
| 0.672141 | 0.083786 | 0.424 | 0.411 | 1 | 10 | Trib1 | 0.013 | Young |

| | | | | | | | | |
|---|---|---|---|---|---|---|---|---|
| 0.672284 | 0.322319 | 0.414 | 0.408 | 1 | 10 | Aftph | 0.006 | Young |
| 0.672423 | 0.142516 | 0.253 | 0.289 | 1 | 10 | Il2rg | -0.036 | Young |
| 0.672781 | 0.372151 | 0.192 | 0.176 | 1 | 10 | Brd3 | 0.016 | Young |
| 0.673092 | 1.240524 | 0.051 | 0.042 | 1 | 10 | Ift57 | 0.009 | Young |
| 0.67336 | 0.462577 | 0.152 | 0.142 | 1 | 10 | Plscr3 | 0.01 | Young |
| 0.673461 | 0.319221 | 0.131 | 0.12 | 1 | 10 | Dok3 | 0.011 | Young |
| 0.673576 | 0.391669 | 0.04 | 0.032 | 1 | 10 | Hibch | 0.008 | Young |
| 0.673576 | 0.129361 | 0.04 | 0.032 | 1 | 10 | Grhpr | 0.008 | Young |
| 0.673676 | 0.459286 | 0.081 | 0.068 | 1 | 10 | Scfd2 | 0.013 | Young |
| 0.673712 | 0.490349 | 0.172 | 0.164 | 1 | 10 | Zfp451 | 0.008 | Young |
| 0.673767 | 0.012346 | 0.202 | 0.22 | 1 | 10 | Slf2 | -0.018 | Young |
| 0.673913 | 0.093045 | 0.101 | 0.117 | 1 | 10 | Kctd2 | -0.016 | Young |
| 0.67397 | 0.231777 | 0.131 | 0.156 | 1 | 10 | Samsn1 | -0.025 | Young |
| 0.674106 | 0.112366 | 0.485 | 0.543 | 1 | 10 | Map4k4 | -0.058 | Young |
| 0.674447 | 0.06324 | 0.202 | 0.222 | 1 | 10 | Tnks | -0.02 | Young |
| 0.674672 | 1.055556 | 0.051 | 0.042 | 1 | 10 | D930016D( | 0.009 | Young |
| 0.674948 | 0.157602 | 0.04 | 0.051 | 1 | 10 | Tmem237 | -0.011 | Young |
| 0.67497 | 0.616152 | 0.051 | 0.064 | 1 | 10 | Ttc27 | -0.013 | Young |
| 0.675523 | 0.011341 | 0.626 | 0.672 | 1 | 10 | Rap1a | -0.046 | Young |
| 0.675757 | 0.170879 | 0.162 | 0.149 | 1 | 10 | Evi2 | 0.013 | Young |
| 0.675885 | 0.753279 | 0.061 | 0.051 | 1 | 10 | Rnaset2b | 0.01 | Young |
| 0.676253 | 0.977005 | 0.051 | 0.042 | 1 | 10 | Ppard | 0.009 | Young |
| 0.676253 | 0.671923 | 0.051 | 0.042 | 1 | 10 | Mettl5 | 0.009 | Young |
| 0.676489 | 0.335087 | 0.081 | 0.098 | 1 | 10 | Txndc12 | -0.017 | Young |
| 0.676747 | 0.009997 | 0.283 | 0.298 | 1 | 10 | Mtch2 | -0.015 | Young |
| 0.676892 | 0.238598 | 0.242 | 0.225 | 1 | 10 | Shoc2 | 0.017 | Young |
| 0.676985 | 0.17427 | 0.172 | 0.203 | 1 | 10 | Spin1 | -0.031 | Young |
| 0.677584 | 2.141953 | 0.02 | 0.015 | 1 | 10 | Trim13 | 0.005 | Young |
| 0.677584 | 1.097219 | 0.02 | 0.015 | 1 | 10 | Tarsl2 | 0.005 | Young |
| 0.677811 | 0.237664 | 0.283 | 0.269 | 1 | 10 | Ccdc115 | 0.014 | Young |
| 0.678745 | 0.03847 | 0.869 | 0.905 | 1 | 10 | Rap1b | -0.036 | Young |
| 0.679013 | 0.281301 | 0.242 | 0.227 | 1 | 10 | Alkbh1 | 0.015 | Young |
| 0.679175 | 0.055651 | 0.919 | 0.966 | 1 | 10 | Sdcbp | -0.047 | Young |
| 0.679629 | 0.184632 | 0.111 | 0.13 | 1 | 10 | Ccdc86 | -0.019 | Young |
| 0.680176 | 1.445265 | 0.02 | 0.015 | 1 | 10 | Lmbr1 | 0.005 | Young |
| 0.680186 | 0.257762 | 0.071 | 0.086 | 1 | 10 | Metrn | -0.015 | Young |
| 0.680744 | 0.336689 | 0.04 | 0.032 | 1 | 10 | Amer1 | 0.008 | Young |
| 0.681211 | 0.01703 | 0.162 | 0.186 | 1 | 10 | Ccnh | -0.024 | Young |
| 0.681823 | 0.395966 | 0.182 | 0.218 | 1 | 10 | Yipf4 | -0.036 | Young |
| 0.681928 | 0.218761 | 0.263 | 0.306 | 1 | 10 | Dcaf7 | -0.043 | Young |
| 0.682144 | 0.308411 | 0.131 | 0.117 | 1 | 10 | Smg6 | 0.014 | Young |
| 0.682156 | 0.488493 | 0.071 | 0.061 | 1 | 10 | Mrrf | 0.01 | Young |
| 0.68226 | 0.06133 | 0.97 | 0.971 | 1 | 10 | Zfp36 | -0.001 | Young |
| 0.682401 | 0.3436 | 0.04 | 0.051 | 1 | 10 | Zmym6 | -0.011 | Young |
| 0.682429 | 0.125598 | 0.354 | 0.408 | 1 | 10 | Adss | -0.054 | Young |

| | | | | | | | | |
|---|---|---|---|---|---|---|---|---|
| 0.682593 | 0.447162 | 0.051 | 0.042 | 1 | 10 | Hoxb4 | 0.009 | Young |
| 0.682766 | 0.216071 | 0.404 | 0.416 | 1 | 10 | Hsd17b12 | -0.012 | Young |
| 0.682771 | 1.779734 | 0.02 | 0.015 | 1 | 10 | Cep89 | 0.005 | Young |
| 0.682771 | 0.909769 | 0.02 | 0.015 | 1 | 10 | Msh5 | 0.005 | Young |
| 0.682771 | 0.856356 | 0.02 | 0.015 | 1 | 10 | Plcl1 | 0.005 | Young |
| 0.682771 | 0.780919 | 0.02 | 0.015 | 1 | 10 | Mettl27 | 0.005 | Young |
| 0.682771 | 0.646123 | 0.02 | 0.015 | 1 | 10 | Gm42031 | 0.005 | Young |
| 0.68288 | 0.22291 | 0.03 | 0.039 | 1 | 10 | Zfp346 | -0.009 | Young |
| 0.683335 | 0.064999 | 0.919 | 0.934 | 1 | 10 | Sh3bgrl3 | -0.015 | Young |
| 0.683527 | 0.038272 | 0.152 | 0.174 | 1 | 10 | Nfatc1 | -0.022 | Young |
| 0.68367 | 0.610599 | 0.343 | 0.357 | 1 | 10 | AC160336. | -0.014 | Young |
| 0.683874 | 0.260592 | 0.212 | 0.249 | 1 | 10 | Psmc4 | -0.037 | Young |
| 0.68418 | 0.745501 | 0.212 | 0.21 | 1 | 10 | Pum3 | 0.002 | Young |
| 0.684352 | 0.597979 | 0.152 | 0.142 | 1 | 10 | 2310009B1 | 0.01 | Young |
| 0.684496 | 0.219033 | 0.394 | 0.44 | 1 | 10 | Gpsm3 | -0.046 | Young |
| 0.684504 | 0.014154 | 0.97 | 0.983 | 1 | 10 | Ptma | -0.013 | Young |
| 0.684985 | 0.119576 | 0.081 | 0.068 | 1 | 10 | Nek4 | 0.013 | Young |
| 0.685247 | 0.456278 | 0.182 | 0.176 | 1 | 10 | Mms19 | 0.006 | Young |
| 0.68537 | 0.890988 | 0.02 | 0.015 | 1 | 10 | Sgce | 0.005 | Young |
| 0.68537 | 0.816952 | 0.02 | 0.015 | 1 | 10 | Ubox5 | 0.005 | Young |
| 0.685542 | 0.23744 | 0.313 | 0.318 | 1 | 10 | Cbx1 | -0.005 | Young |
| 0.685633 | 0.599665 | 0.081 | 0.071 | 1 | 10 | Mrpl49 | 0.01 | Young |
| 0.685633 | 0.553584 | 0.061 | 0.076 | 1 | 10 | F8a | -0.015 | Young |
| 0.685699 | 0.67478 | 0.111 | 0.1 | 1 | 10 | Ctnnbl1 | 0.011 | Young |
| 0.686505 | 0.27858 | 0.303 | 0.352 | 1 | 10 | Bod1l | -0.049 | Young |
| 0.686556 | 0.475571 | 0.212 | 0.244 | 1 | 10 | Med8 | -0.032 | Young |
| 0.686846 | 0.107574 | 0.162 | 0.144 | 1 | 10 | Gon4l | 0.018 | Young |
| 0.687225 | 0.117814 | 0.071 | 0.086 | 1 | 10 | Zfp945 | -0.015 | Young |
| 0.687269 | 0.387145 | 0.434 | 0.516 | 1 | 10 | Hnrnpd | -0.082 | Young |
| 0.687363 | 1.390127 | 0.051 | 0.042 | 1 | 10 | Mpp7 | 0.009 | Young |
| 0.687431 | 0.22702 | 0.343 | 0.337 | 1 | 10 | Mfsd14b | 0.006 | Young |
| 0.687487 | 0.613618 | 0.141 | 0.13 | 1 | 10 | Cfl2 | 0.011 | Young |
| 0.687521 | 0.435257 | 0.232 | 0.222 | 1 | 10 | Chfr | 0.01 | Young |
| 0.687595 | 0.24791 | 0.131 | 0.154 | 1 | 10 | Prkag1 | -0.023 | Young |
| 0.687874 | 0.186078 | 0.283 | 0.315 | 1 | 10 | Otub1 | -0.032 | Young |
| 0.687972 | 1.025841 | 0.02 | 0.015 | 1 | 10 | Hddc3 | 0.005 | Young |
| 0.687972 | 0.857045 | 0.02 | 0.015 | 1 | 10 | Ift122 | 0.005 | Young |
| 0.687972 | 0.81518 | 0.02 | 0.015 | 1 | 10 | Nat8f1 | 0.005 | Young |
| 0.687972 | 0.743878 | 0.02 | 0.015 | 1 | 10 | Klrb1b | 0.005 | Young |
| 0.687972 | 0.536621 | 0.02 | 0.015 | 1 | 10 | Stxbp4 | 0.005 | Young |
| 0.687972 | 0.350773 | 0.02 | 0.015 | 1 | 10 | Taf4b | 0.005 | Young |
| 0.688008 | 0.213253 | 0.051 | 0.061 | 1 | 10 | Mmgt2 | -0.01 | Young |
| 0.688321 | 0.066182 | 0.525 | 0.582 | 1 | 10 | Wasf2 | -0.057 | Young |
| 0.688378 | 0.0355 | 0.111 | 0.13 | 1 | 10 | Mtif2 | -0.019 | Young |
| 0.688381 | 0.158193 | 0.616 | 0.697 | 1 | 10 | Taldo1 | -0.081 | Young |

| | | | | | | | | |
|---|---|---|---|---|---|---|---|---|
| 0.688387 | 0.258346 | 0.04 | 0.051 | 1 | 10 | Def8 | -0.011 | Young |
| 0.688387 | 0.19277 | 0.04 | 0.051 | 1 | 10 | 6720427I0 | -0.011 | Young |
| 0.688522 | 0.434405 | 0.222 | 0.215 | 1 | 10 | 2810403A0 | 0.007 | Young |
| 0.688549 | 0.520624 | 0.182 | 0.174 | 1 | 10 | Dhps | 0.008 | Young |
| 0.688781 | 0.002011 | 0.172 | 0.196 | 1 | 10 | Map2k4 | -0.024 | Young |
| 0.688924 | 0.132931 | 0.141 | 0.127 | 1 | 10 | Cwc25 | 0.014 | Young |
| 0.688998 | 0.133485 | 0.263 | 0.291 | 1 | 10 | Rnf141 | -0.028 | Young |
| 0.689336 | 0.591746 | 0.242 | 0.24 | 1 | 10 | Rbm28 | 0.002 | Young |
| 0.689478 | 0.359408 | 0.273 | 0.308 | 1 | 10 | Rraga | -0.035 | Young |
| 0.689719 | 0.620537 | 0.101 | 0.09 | 1 | 10 | Ptp4a3 | 0.011 | Young |
| 0.69015 | 0.76296 | 0.071 | 0.061 | 1 | 10 | Rasgrp2 | 0.01 | Young |
| 0.690168 | 0.575616 | 0.182 | 0.171 | 1 | 10 | Gcn1l1 | 0.011 | Young |
| 0.690328 | 0.045498 | 0.111 | 0.13 | 1 | 10 | Tmem126a | -0.019 | Young |
| 0.690427 | 0.057234 | 0.232 | 0.257 | 1 | 10 | Gpatch8 | -0.025 | Young |
| 0.69055 | 0.261093 | 0.051 | 0.042 | 1 | 10 | Pank4 | 0.009 | Young |
| 0.690579 | 1.895017 | 0.02 | 0.015 | 1 | 10 | Ank2 | 0.005 | Young |
| 0.690579 | 1.047284 | 0.02 | 0.015 | 1 | 10 | Gm28557 | 0.005 | Young |
| 0.690579 | 0.914423 | 0.02 | 0.015 | 1 | 10 | Kif5c | 0.005 | Young |
| 0.690579 | 0.541278 | 0.02 | 0.015 | 1 | 10 | Helq | 0.005 | Young |
| 0.690579 | 0.536075 | 0.02 | 0.015 | 1 | 10 | Mks1 | 0.005 | Young |
| 0.690579 | 0.24746 | 0.02 | 0.015 | 1 | 10 | Traf3ip2 | 0.005 | Young |
| 0.69062 | 0.62205 | 0.081 | 0.071 | 1 | 10 | 3110062M | 0.01 | Young |
| 0.690665 | 0.171821 | 0.121 | 0.142 | 1 | 10 | Krr1 | -0.021 | Young |
| 0.690896 | 0.202434 | 0.192 | 0.205 | 1 | 10 | Foxo3 | -0.013 | Young |
| 0.691486 | 0.493903 | 0.071 | 0.061 | 1 | 10 | Nup107 | 0.01 | Young |
| 0.691578 | 0.240814 | 0.192 | 0.174 | 1 | 10 | Adap1 | 0.018 | Young |
| 0.692052 | 0.370796 | 0.242 | 0.235 | 1 | 10 | Arid4b | 0.007 | Young |
| 0.692158 | 0.4235 | 0.333 | 0.333 | 1 | 10 | Parp14 | 0 | Young |
| 0.692268 | 0.279402 | 0.141 | 0.13 | 1 | 10 | Tbc1d9b | 0.011 | Young |
| 0.692564 | 0.044599 | 0.061 | 0.073 | 1 | 10 | Cpt2 | -0.012 | Young |
| 0.692595 | 0.063358 | 0.747 | 0.763 | 1 | 10 | Pomp | -0.016 | Young |
| 0.693078 | 0.698158 | 0.101 | 0.09 | 1 | 10 | 1600002K0 | 0.011 | Young |
| 0.693129 | 0.298186 | 0.303 | 0.291 | 1 | 10 | Higd2a | 0.012 | Young |
| 0.693189 | 1.0029 | 0.02 | 0.015 | 1 | 10 | Vamp1 | 0.005 | Young |
| 0.693189 | 0.886224 | 0.02 | 0.015 | 1 | 10 | Gm28151 | 0.005 | Young |
| 0.693189 | 0.521614 | 0.02 | 0.015 | 1 | 10 | Klhl42 | 0.005 | Young |
| 0.693189 | 0.385336 | 0.02 | 0.015 | 1 | 10 | Rpgrip1l | 0.005 | Young |
| 0.693189 | 0.319377 | 0.02 | 0.015 | 1 | 10 | Mtcp1 | 0.005 | Young |
| 0.693189 | 0.286907 | 0.02 | 0.015 | 1 | 10 | Cavin2 | 0.005 | Young |
| 0.693189 | 0.176933 | 0.02 | 0.015 | 1 | 10 | Zfp119a | 0.005 | Young |
| 0.693189 | 0.151629 | 0.02 | 0.015 | 1 | 10 | Uprt | 0.005 | Young |
| 0.69329 | 0.167897 | 0.343 | 0.328 | 1 | 10 | Mgrn1 | 0.015 | Young |
| 0.693522 | 0.120373 | 0.091 | 0.108 | 1 | 10 | Itgb7 | -0.017 | Young |
| 0.693583 | 0.017565 | 0.879 | 0.892 | 1 | 10 | Gnai2 | -0.013 | Young |
| 0.693703 | 0.443658 | 0.202 | 0.188 | 1 | 10 | Edem2 | 0.014 | Young |

| | | | | | | | | |
|---|---|---|---|---|---|---|---|---|
| 0.69373 | 0.162453 | 0.162 | 0.183 | 1 | 10 | Hltf | -0.021 | Young |
| 0.693743 | 0.180423 | 0.051 | 0.042 | 1 | 10 | Eogt | 0.009 | Young |
| 0.694034 | 0.298436 | 0.455 | 0.487 | 1 | 10 | Vdac1 | -0.032 | Young |
| 0.694144 | 0.17329 | 0.192 | 0.22 | 1 | 10 | Hadh | -0.028 | Young |
| 0.694235 | 0.119294 | 0.111 | 0.13 | 1 | 10 | Hs2st1 | -0.019 | Young |
| 0.694248 | 0.079657 | 0.737 | 0.858 | 1 | 10 | C3ar1 | -0.121 | Young |
| 0.694315 | 0.385898 | 0.283 | 0.274 | 1 | 10 | Thoc7 | 0.009 | Young |
| 0.694351 | 0.578126 | 0.354 | 0.347 | 1 | 10 | Mfsd5 | 0.007 | Young |
| 0.694552 | 0.389964 | 0.242 | 0.232 | 1 | 10 | Ythdf2 | 0.01 | Young |
| 0.694736 | 0.042536 | 0.444 | 0.482 | 1 | 10 | Fcgrt | -0.038 | Young |
| 0.695282 | 0.884683 | 0.101 | 0.093 | 1 | 10 | Rubcn | 0.008 | Young |
| 0.695341 | 0.372077 | 0.051 | 0.042 | 1 | 10 | Rbm41 | 0.009 | Young |
| 0.695472 | 0.853488 | 0.091 | 0.081 | 1 | 10 | Gfer | 0.01 | Young |
| 0.695536 | 0.313753 | 0.162 | 0.149 | 1 | 10 | Ppp1r7 | 0.013 | Young |
| 0.695702 | 0.258565 | 0.434 | 0.43 | 1 | 10 | Hpgds | 0.004 | Young |
| 0.695802 | 0.21072 | 0.02 | 0.015 | 1 | 10 | Slc22a4 | 0.005 | Young |
| 0.695802 | 0.084322 | 0.02 | 0.015 | 1 | 10 | Tctn1 | 0.005 | Young |
| 0.695868 | 0.159877 | 0.364 | 0.421 | 1 | 10 | Kif1b | -0.057 | Young |
| 0.696013 | 0.158394 | 0.808 | 0.78 | 1 | 10 | Snx3 | 0.028 | Young |
| 0.696089 | 1.00704 | 0.061 | 0.051 | 1 | 10 | Fancl | 0.01 | Young |
| 0.696365 | 0.444611 | 0.061 | 0.073 | 1 | 10 | Abhd10 | -0.012 | Young |
| 0.696365 | 0.222103 | 0.061 | 0.073 | 1 | 10 | Fam151b | -0.012 | Young |
| 0.696561 | 0.410707 | 0.03 | 0.039 | 1 | 10 | Scd1 | -0.009 | Young |
| 0.696679 | 0.202966 | 0.545 | 0.626 | 1 | 10 | Tmed7 | -0.081 | Young |
| 0.696992 | 0.148407 | 0.202 | 0.188 | 1 | 10 | Fam76b | 0.014 | Young |
| 0.697038 | 0.064047 | 0.091 | 0.078 | 1 | 10 | Rft1 | 0.013 | Young |
| 0.697344 | 0.206736 | 0.162 | 0.149 | 1 | 10 | Senp5 | 0.013 | Young |
| 0.697466 | 0.425311 | 0.111 | 0.1 | 1 | 10 | Ttyh2 | 0.011 | Young |
| 0.697507 | 0.142174 | 0.192 | 0.174 | 1 | 10 | Cyp20a1 | 0.018 | Young |
| 0.697581 | 0.083951 | 0.859 | 0.856 | 1 | 10 | Hnrnpa3 | 0.003 | Young |
| 0.697634 | 0.484995 | 0.121 | 0.11 | 1 | 10 | Csad | 0.011 | Young |
| 0.697651 | 0.499796 | 0.253 | 0.247 | 1 | 10 | Fbxw11 | 0.006 | Young |
| 0.697679 | 0.165891 | 0.515 | 0.501 | 1 | 10 | Fis1 | 0.014 | Young |
| 0.697823 | 0.067851 | 0.626 | 0.665 | 1 | 10 | Hnrnph1 | -0.039 | Young |
| 0.698019 | 0.38402 | 0.566 | 0.641 | 1 | 10 | Zmiz1 | -0.075 | Young |
| 0.698021 | 0.569089 | 0.141 | 0.13 | 1 | 10 | Xylt2 | 0.011 | Young |
| 0.698732 | 0.282602 | 0.626 | 0.653 | 1 | 10 | Hnrnpab | -0.027 | Young |
| 0.699088 | 0.083922 | 0.384 | 0.418 | 1 | 10 | Washc4 | -0.034 | Young |
| 0.699218 | 0.358306 | 0.374 | 0.401 | 1 | 10 | Atxn2l | -0.027 | Young |
| 0.699457 | 0.252376 | 0.152 | 0.137 | 1 | 10 | Tom1 | 0.015 | Young |
| 0.699524 | 0.235298 | 0.152 | 0.176 | 1 | 10 | Wdr82 | -0.024 | Young |
| 0.699856 | 0.157557 | 0.101 | 0.088 | 1 | 10 | Hpse | 0.013 | Young |
| 0.699992 | 0.238227 | 0.313 | 0.357 | 1 | 10 | Stard8 | -0.044 | Young |
| 0.700144 | 0.145206 | 0.051 | 0.042 | 1 | 10 | Tcta | 0.009 | Young |
| 0.700735 | 0.034521 | 0.404 | 0.472 | 1 | 10 | Drap1 | -0.068 | Young |

| | | | | | | | | |
|---|---|---|---|---|---|---|---|---|
| 0.70109 | 0.284998 | 0.111 | 0.13 | 1 | 10 | Med23 | -0.019 | Young |
| 0.701902 | 0.3165 | 0.061 | 0.051 | 1 | 10 | Xrcc6 | 0.01 | Young |
| 0.701956 | 0.092849 | 0.263 | 0.298 | 1 | 10 | Tnrc6b | -0.035 | Young |
| 0.702036 | 0.732217 | 0.111 | 0.103 | 1 | 10 | Pigyl | 0.008 | Young |
| 0.702259 | 0.320907 | 0.364 | 0.355 | 1 | 10 | Snx17 | 0.009 | Young |
| 0.70243 | 0.328404 | 0.273 | 0.315 | 1 | 10 | Nfat5 | -0.042 | Young |
| 0.703024 | 0.411364 | 0.152 | 0.139 | 1 | 10 | Mylip | 0.013 | Young |
| 0.703088 | 0.063564 | 0.131 | 0.117 | 1 | 10 | Heatr1 | 0.014 | Young |
| 0.703169 | 0.118406 | 0.535 | 0.531 | 1 | 10 | Tcp1 | 0.004 | Young |
| 0.703188 | 0.152833 | 0.081 | 0.095 | 1 | 10 | Mtmr9 | -0.014 | Young |
| 0.703192 | 0.111864 | 0.051 | 0.061 | 1 | 10 | Micu3 | -0.01 | Young |
| 0.703257 | 0.137504 | 0.141 | 0.161 | 1 | 10 | Ubqln4 | -0.02 | Young |
| 0.703265 | 0.046906 | 0.101 | 0.088 | 1 | 10 | Klhl20 | 0.013 | Young |
| 0.7034 | 0.009676 | 0.667 | 0.714 | 1 | 10 | Dynll1 | -0.047 | Young |
| 0.703412 | 0.269956 | 0.465 | 0.513 | 1 | 10 | Ogdh | -0.048 | Young |
| 0.703439 | 0.156128 | 0.03 | 0.039 | 1 | 10 | Gm26601 | -0.009 | Young |
| 0.703665 | 0.03048 | 0.02 | 0.015 | 1 | 10 | Gm26669 | 0.005 | Young |
| 0.703865 | 0.240047 | 0.404 | 0.465 | 1 | 10 | Kcnn4 | -0.061 | Young |
| 0.70399 | 0.11028 | 0.061 | 0.073 | 1 | 10 | Fktn | -0.012 | Young |
| 0.704317 | 0.147162 | 0.313 | 0.357 | 1 | 10 | Mgat1 | -0.044 | Young |
| 0.704352 | 0.094992 | 0.323 | 0.379 | 1 | 10 | Os9 | -0.056 | Young |
| 0.704445 | 0.209131 | 0.101 | 0.117 | 1 | 10 | Togaram1 | -0.016 | Young |
| 0.704446 | 0.06898 | 0.384 | 0.44 | 1 | 10 | Susd6 | -0.056 | Young |
| 0.704717 | 0.202067 | 0.172 | 0.156 | 1 | 10 | Adam19 | 0.016 | Young |
| 0.704816 | 0.812845 | 0.061 | 0.051 | 1 | 10 | Erc1 | 0.01 | Young |
| 0.704948 | 0.106417 | 0.222 | 0.24 | 1 | 10 | Raf1 | -0.018 | Young |
| 0.704958 | 0.143435 | 0.051 | 0.042 | 1 | 10 | Wdr73 | 0.009 | Young |
| 0.705044 | 0.353592 | 0.141 | 0.161 | 1 | 10 | Smad5 | -0.02 | Young |
| 0.705107 | 0.000199 | 0.303 | 0.337 | 1 | 10 | Uggt1 | -0.034 | Young |
| 0.705129 | 0.126675 | 0.131 | 0.152 | 1 | 10 | Sh3gl1 | -0.021 | Young |
| 0.705164 | 0.336478 | 0.192 | 0.22 | 1 | 10 | Ftsj3 | -0.028 | Young |
| 0.705661 | 0.65251 | 0.081 | 0.071 | 1 | 10 | Phospho2 | 0.01 | Young |
| 0.706307 | 0.692426 | 0.111 | 0.103 | 1 | 10 | Gosr1 | 0.008 | Young |
| 0.70648 | 0.51401 | 0.02 | 0.027 | 1 | 10 | Rbks | -0.007 | Young |
| 0.706691 | 0.550278 | 0.121 | 0.142 | 1 | 10 | A430005L1 | -0.021 | Young |
| 0.70692 | 0.223499 | 0.081 | 0.071 | 1 | 10 | Bysl | 0.01 | Young |
| 0.707629 | 0.075517 | 0.97 | 0.976 | 1 | 10 | Rps10 | -0.006 | Young |
| 0.707733 | 0.454517 | 0.061 | 0.051 | 1 | 10 | Gm20342 | 0.01 | Young |
| 0.707822 | 0.082021 | 0.071 | 0.083 | 1 | 10 | Pigc | -0.012 | Young |
| 0.707886 | 0.133868 | 0.131 | 0.152 | 1 | 10 | Tcf3 | -0.021 | Young |
| 0.707994 | 0.539737 | 0.172 | 0.161 | 1 | 10 | Atxn7 | 0.011 | Young |
| 0.708175 | 0.618321 | 0.051 | 0.042 | 1 | 10 | Slc25a38 | 0.009 | Young |
| 0.708232 | 0.347783 | 0.162 | 0.149 | 1 | 10 | Hikeshi | 0.013 | Young |
| 0.708398 | 0.470087 | 0.152 | 0.142 | 1 | 10 | Gpaa1 | 0.01 | Young |
| 0.708743 | 0.87706 | 0.051 | 0.061 | 1 | 10 | Acd | -0.01 | Young |

| | | | | | | | | |
|---|---|---|---|---|---|---|---|---|
| 0.708743 | 0.164443 | 0.051 | 0.061 | 1 | 10 | Coq9 | -0.01 | Young |
| 0.709129 | 0.393036 | 0.414 | 0.428 | 1 | 10 | Snrpc | -0.014 | Young |
| 0.70925 | 0.092159 | 0.172 | 0.159 | 1 | 10 | Gm2000 | 0.013 | Young |
| 0.709507 | 0.415264 | 0.424 | 0.44 | 1 | 10 | Ets2 | -0.016 | Young |
| 0.709657 | 0.097835 | 0.293 | 0.279 | 1 | 10 | Fes | 0.014 | Young |
| 0.70981 | 0.303651 | 0.192 | 0.176 | 1 | 10 | Pmpca | 0.016 | Young |
| 0.709836 | 0.380968 | 0.566 | 0.65 | 1 | 10 | Fabp5 | -0.084 | Young |
| 0.709958 | 0.671112 | 0.101 | 0.09 | 1 | 10 | Zbtb37 | 0.011 | Young |
| 0.710134 | 0.254122 | 0.051 | 0.061 | 1 | 10 | Cd22 | -0.01 | Young |
| 0.710287 | 0.775372 | 0.071 | 0.061 | 1 | 10 | Fbxl12 | 0.01 | Young |
| 0.710506 | 0.266074 | 0.152 | 0.139 | 1 | 10 | 9130401M | 0.013 | Young |
| 0.710631 | 1.301991 | 0.02 | 0.027 | 1 | 10 | Gm19951 | -0.007 | Young |
| 0.710631 | 0.599062 | 0.02 | 0.027 | 1 | 10 | Sparc | -0.007 | Young |
| 0.710631 | 0.0505 | 0.02 | 0.027 | 1 | 10 | Rpp40 | -0.007 | Young |
| 0.710655 | 0.671175 | 0.061 | 0.051 | 1 | 10 | Card9 | 0.01 | Young |
| 0.710655 | 0.308996 | 0.061 | 0.051 | 1 | 10 | Trp53rka | 0.01 | Young |
| 0.710701 | 0.574986 | 0.081 | 0.071 | 1 | 10 | Ppm1k | 0.01 | Young |
| 0.711164 | 0.182783 | 0.273 | 0.306 | 1 | 10 | Rlim | -0.033 | Young |
| 0.711211 | 0.420374 | 0.162 | 0.154 | 1 | 10 | Uxs1 | 0.008 | Young |
| 0.711256 | 0.244904 | 0.061 | 0.071 | 1 | 10 | Rab33b | -0.01 | Young |
| 0.711874 | 0.477403 | 0.131 | 0.156 | 1 | 10 | Ago1 | -0.025 | Young |
| 0.712074 | 0.433889 | 0.091 | 0.081 | 1 | 10 | Cela1 | 0.01 | Young |
| 0.712381 | 0.225546 | 0.333 | 0.389 | 1 | 10 | Csnk1d | -0.056 | Young |
| 0.712633 | 0.309757 | 0.071 | 0.083 | 1 | 10 | Fhl3 | -0.012 | Young |
| 0.712694 | 0.139282 | 0.576 | 0.641 | 1 | 10 | Sf3b2 | -0.065 | Young |
| 0.71271 | 1.514043 | 0.03 | 0.024 | 1 | 10 | Arpin | 0.006 | Young |
| 0.71275 | 0.057902 | 0.273 | 0.301 | 1 | 10 | Ino80d | -0.028 | Young |
| 0.712852 | 0.382256 | 0.465 | 0.474 | 1 | 10 | Cebpg | -0.009 | Young |
| 0.712978 | 0.139539 | 0.263 | 0.281 | 1 | 10 | Atf1 | -0.018 | Young |
| 0.713266 | 0.616534 | 0.091 | 0.081 | 1 | 10 | Ndrg1 | 0.01 | Young |
| 0.713332 | 0.214409 | 0.626 | 0.655 | 1 | 10 | Nktr | -0.029 | Young |
| 0.713352 | 0.456467 | 0.101 | 0.09 | 1 | 10 | Thap3 | 0.011 | Young |
| 0.71363 | 0.421355 | 0.242 | 0.232 | 1 | 10 | Arl5b | 0.01 | Young |
| 0.713884 | 0.011638 | 0.99 | 0.998 | 1 | 10 | Actg1 | -0.008 | Young |
| 0.714098 | 0.277537 | 0.141 | 0.127 | 1 | 10 | Ankrd49 | 0.014 | Young |
| 0.714202 | 0.222296 | 0.061 | 0.073 | 1 | 10 | Eif2b3 | -0.012 | Young |
| 0.714623 | 0.102478 | 0.051 | 0.042 | 1 | 10 | AI427809 | 0.009 | Young |
| 0.715044 | 0.305874 | 0.071 | 0.083 | 1 | 10 | Zbtb43 | -0.012 | Young |
| 0.715525 | 0.104877 | 0.152 | 0.169 | 1 | 10 | Zmat5 | -0.017 | Young |
| 0.715534 | 0.889659 | 0.03 | 0.039 | 1 | 10 | Tchp | -0.009 | Young |
| 0.715692 | 0.355341 | 0.071 | 0.061 | 1 | 10 | AL590144.: | 0.01 | Young |
| 0.715723 | 0.00422 | 0.242 | 0.264 | 1 | 10 | Slc9a3r1 | -0.022 | Young |
| 0.715871 | 0.081026 | 0.98 | 0.961 | 1 | 10 | Rpl18 | 0.019 | Young |
| 0.716031 | 0.015463 | 0.374 | 0.406 | 1 | 10 | Npepps | -0.032 | Young |
| 0.716084 | 0.028329 | 0.273 | 0.306 | 1 | 10 | Snx30 | -0.033 | Young |

| | | | | | | | | |
|---|---|---|---|---|---|---|---|---|
| 0.716511 | 0.002591 | 0.737 | 0.736 | 1 | 10 | Sfpq | 0.001 | Young |
| 0.716762 | 0.088789 | 0.384 | 0.408 | 1 | 10 | Psmc1 | -0.024 | Young |
| 0.716874 | 0.678033 | 0.02 | 0.027 | 1 | 10 | Gins2 | -0.007 | Young |
| 0.716879 | 0.895988 | 0.111 | 0.1 | 1 | 10 | Vsig8 | 0.011 | Young |
| 0.717099 | 0.072768 | 0.051 | 0.061 | 1 | 10 | Adsl | -0.01 | Young |
| 0.717099 | 0.062957 | 0.051 | 0.061 | 1 | 10 | Surf2 | -0.01 | Young |
| 0.717271 | 0.273548 | 0.273 | 0.311 | 1 | 10 | Dynll2 | -0.038 | Young |
| 0.71728 | 0.056462 | 0.475 | 0.513 | 1 | 10 | Bclaf1 | -0.038 | Young |
| 0.717457 | 0.126878 | 0.071 | 0.083 | 1 | 10 | Isg15 | -0.012 | Young |
| 0.717633 | 0.681722 | 0.101 | 0.093 | 1 | 10 | Asna1 | 0.008 | Young |
| 0.717886 | 0.381952 | 0.081 | 0.095 | 1 | 10 | Jmy | -0.014 | Young |
| 0.717978 | 0.114029 | 0.061 | 0.051 | 1 | 10 | Zfp942 | 0.01 | Young |
| 0.718046 | 0.363112 | 0.061 | 0.073 | 1 | 10 | Polr2d | -0.012 | Young |
| 0.718403 | 0.19204 | 0.202 | 0.227 | 1 | 10 | Chchd3 | -0.025 | Young |
| 0.718675 | 0.09212 | 0.232 | 0.259 | 1 | 10 | Rest | -0.027 | Young |
| 0.718959 | 0.264642 | 0.02 | 0.027 | 1 | 10 | Sap130 | -0.007 | Young |
| 0.719027 | 0.133409 | 0.172 | 0.159 | 1 | 10 | Zfp120 | 0.013 | Young |
| 0.719199 | 0.093358 | 0.131 | 0.117 | 1 | 10 | Sp140 | 0.014 | Young |
| 0.719341 | 0.60981 | 0.101 | 0.12 | 1 | 10 | Xrcc1 | -0.019 | Young |
| 0.719492 | 0.366531 | 0.303 | 0.301 | 1 | 10 | Socs3 | 0.002 | Young |
| 0.719755 | 0.01084 | 0.071 | 0.061 | 1 | 10 | Timm21 | 0.01 | Young |
| 0.719892 | 0.461428 | 0.051 | 0.061 | 1 | 10 | Dhx37 | -0.01 | Young |
| 0.719903 | 0.099322 | 0.424 | 0.469 | 1 | 10 | Mrpl20 | -0.045 | Young |
| 0.71991 | 0.301601 | 0.323 | 0.308 | 1 | 10 | Foxn2 | 0.015 | Young |
| 0.720367 | 0.230721 | 0.101 | 0.12 | 1 | 10 | Gtf3c3 | -0.019 | Young |
| 0.720816 | 0.071217 | 0.152 | 0.169 | 1 | 10 | Ttpal | -0.017 | Young |
| 0.720818 | 0.337006 | 0.081 | 0.071 | 1 | 10 | Wdr75 | 0.01 | Young |
| 0.720926 | 0.726501 | 0.091 | 0.083 | 1 | 10 | Dtd2 | 0.008 | Young |
| 0.721046 | 0.926962 | 0.03 | 0.024 | 1 | 10 | Tnni2 | 0.006 | Young |
| 0.721046 | 0.221327 | 0.02 | 0.027 | 1 | 10 | Adnp | -0.007 | Young |
| 0.721218 | 0.180546 | 0.091 | 0.105 | 1 | 10 | Kat14 | -0.014 | Young |
| 0.721219 | 0.413555 | 0.283 | 0.279 | 1 | 10 | Rbbp7 | 0.004 | Young |
| 0.721405 | 0.089798 | 0.303 | 0.342 | 1 | 10 | Prkx | -0.039 | Young |
| 0.721428 | 0.04743 | 0.333 | 0.32 | 1 | 10 | Higd1a | 0.013 | Young |
| 0.721561 | 0.172901 | 0.556 | 0.623 | 1 | 10 | Tram1 | -0.067 | Young |
| 0.721743 | 0.459033 | 0.162 | 0.183 | 1 | 10 | Txlna | -0.021 | Young |
| 0.722132 | 0.052877 | 0.919 | 0.951 | 1 | 10 | Arpc1b | -0.032 | Young |
| 0.722185 | 0.308833 | 0.212 | 0.198 | 1 | 10 | Pak1ip1 | 0.014 | Young |
| 0.722582 | 0.532973 | 0.182 | 0.21 | 1 | 10 | Trp53 | -0.028 | Young |
| 0.722679 | 0.18422 | 0.172 | 0.188 | 1 | 10 | Pigp | -0.016 | Young |
| 0.723008 | 0.026434 | 0.263 | 0.303 | 1 | 10 | Nub1 | -0.04 | Young |
| 0.723136 | 0.927962 | 0.02 | 0.027 | 1 | 10 | Siglech | -0.007 | Young |
| 0.723136 | 0.908799 | 0.02 | 0.027 | 1 | 10 | Ace | -0.007 | Young |
| 0.723136 | 0.256576 | 0.02 | 0.027 | 1 | 10 | Syp | -0.007 | Young |
| 0.723136 | 0.115756 | 0.02 | 0.027 | 1 | 10 | Nectin2 | -0.007 | Young |

| | | | | | | | | |
|---|---|---|---|---|---|---|---|---|
| 0.72328 | 0.050729 | 0.96 | 0.946 | 1 | 10 | Rps27 | 0.014 | Young |
| 0.72331 | 0.170976 | 0.384 | 0.455 | 1 | 10 | Taf10 | -0.071 | Young |
| 0.723838 | 0.342428 | 0.323 | 0.323 | 1 | 10 | Btbd1 | 0 | Young |
| 0.723845 | 0.062913 | 0.081 | 0.093 | 1 | 10 | Jade1 | -0.012 | Young |
| 0.724018 | 0.169155 | 0.091 | 0.081 | 1 | 10 | Dtx2 | 0.01 | Young |
| 0.724605 | 0.055604 | 0.121 | 0.11 | 1 | 10 | Coq5 | 0.011 | Young |
| 0.724663 | 0.018602 | 0.364 | 0.403 | 1 | 10 | Ppp4r3b | -0.039 | Young |
| 0.72474 | 0.12936 | 0.737 | 0.79 | 1 | 10 | Hnrnpu | -0.053 | Young |
| 0.724765 | 0.159421 | 0.475 | 0.553 | 1 | 10 | Tor1aip1 | -0.078 | Young |
| 0.725164 | 0.62556 | 0.182 | 0.178 | 1 | 10 | Heca | 0.004 | Young |
| 0.725216 | 0.224895 | 0.091 | 0.081 | 1 | 10 | Tmem2 | 0.01 | Young |
| 0.725227 | 0.998554 | 0.02 | 0.027 | 1 | 10 | Hps6 | -0.007 | Young |
| 0.725227 | 0.951503 | 0.02 | 0.027 | 1 | 10 | Zfp113 | -0.007 | Young |
| 0.725227 | 0.800088 | 0.03 | 0.024 | 1 | 10 | Trmt12 | 0.006 | Young |
| 0.725227 | 0.406838 | 0.02 | 0.027 | 1 | 10 | Acpp | -0.007 | Young |
| 0.725234 | 0.000463 | 0.152 | 0.169 | 1 | 10 | Ctdspl2 | -0.017 | Young |
| 0.725438 | 0.511498 | 0.131 | 0.152 | 1 | 10 | Gtf2f2 | -0.021 | Young |
| 0.725463 | 0.325207 | 0.263 | 0.296 | 1 | 10 | Ppp1r21 | -0.033 | Young |
| 0.725489 | 0.117168 | 0.051 | 0.061 | 1 | 10 | Rtl8a | -0.01 | Young |
| 0.725665 | 0.718302 | 0.091 | 0.083 | 1 | 10 | Wdr45 | 0.008 | Young |
| 0.725682 | 0.921658 | 0.111 | 0.105 | 1 | 10 | Tsr1 | 0.006 | Young |
| 0.725784 | 0.215661 | 0.626 | 0.741 | 1 | 10 | Gm2a | -0.115 | Young |
| 0.72596 | 0.149565 | 0.111 | 0.127 | 1 | 10 | Srprb | -0.016 | Young |
| 0.726119 | 0.06287 | 0.152 | 0.169 | 1 | 10 | Anapc4 | -0.017 | Young |
| 0.726157 | 0.595321 | 0.152 | 0.142 | 1 | 10 | Zbtb11 | 0.01 | Young |
| 0.726314 | 0.111559 | 0.677 | 0.751 | 1 | 10 | Son | -0.074 | Young |
| 0.726397 | 0.01123 | 0.242 | 0.269 | 1 | 10 | Setd1b | -0.027 | Young |
| 0.726853 | 0.478998 | 0.061 | 0.071 | 1 | 10 | Mier3 | -0.01 | Young |
| 0.72691 | 0.170084 | 0.364 | 0.374 | 1 | 10 | Dnajc13 | -0.01 | Young |
| 0.726985 | 0.060112 | 0.586 | 0.643 | 1 | 10 | Cap1 | -0.057 | Young |
| 0.727041 | 0.258074 | 0.141 | 0.13 | 1 | 10 | Arid3a | 0.011 | Young |
| 0.727083 | 0.228414 | 0.364 | 0.369 | 1 | 10 | Ap3b1 | -0.005 | Young |
| 0.727267 | 0.395491 | 0.303 | 0.293 | 1 | 10 | Smu1 | 0.01 | Young |
| 0.72732 | 0.728849 | 0.03 | 0.024 | 1 | 10 | Zscan12 | 0.006 | Young |
| 0.727451 | 0.189033 | 0.323 | 0.325 | 1 | 10 | Ralgds | -0.002 | Young |
| 0.727614 | 0.152209 | 0.303 | 0.333 | 1 | 10 | Mfsd11 | -0.03 | Young |
| 0.727771 | 0.543214 | 0.101 | 0.093 | 1 | 10 | Trappc2 | 0.008 | Young |
| 0.727831 | 0.17196 | 0.071 | 0.081 | 1 | 10 | Rcbtb1 | -0.01 | Young |
| 0.72798 | 0.424157 | 0.242 | 0.24 | 1 | 10 | Dyrk1a | 0.002 | Young |
| 0.728085 | 0.943566 | 0.04 | 0.049 | 1 | 10 | Tbkbp1 | -0.009 | Young |
| 0.728415 | 0.064586 | 0.646 | 0.655 | 1 | 10 | Minos1 | -0.009 | Young |
| 0.728446 | 0.127507 | 0.081 | 0.093 | 1 | 10 | Phf11b | -0.012 | Young |
| 0.728834 | 0.464886 | 0.111 | 0.1 | 1 | 10 | Pprc1 | 0.011 | Young |
| 0.729021 | 0.066664 | 0.192 | 0.176 | 1 | 10 | Mif4gd | 0.016 | Young |
| 0.729066 | 0.210309 | 0.232 | 0.267 | 1 | 10 | Zdhhc9 | -0.035 | Young |

| | | | | | | | | |
|---|---|---|---|---|---|---|---|---|
| 0.729096 | 1.077196 | 0.061 | 0.054 | 1 | 10 | Dgkq | 0.007 | Young |
| 0.729389 | 0.561466 | 0.212 | 0.208 | 1 | 10 | Comtd1 | 0.004 | Young |
| 0.729416 | 0.947429 | 0.03 | 0.024 | 1 | 10 | Mocs3 | 0.006 | Young |
| 0.729416 | 0.920038 | 0.03 | 0.024 | 1 | 10 | Cmc4 | 0.006 | Young |
| 0.729416 | 0.613964 | 0.03 | 0.024 | 1 | 10 | Tsen54 | 0.006 | Young |
| 0.729749 | 1.11042 | 0.081 | 0.073 | 1 | 10 | Rasgrp1 | 0.008 | Young |
| 0.729895 | 0.724615 | 0.051 | 0.044 | 1 | 10 | Parp3 | 0.007 | Young |
| 0.730365 | 0.039997 | 0.303 | 0.333 | 1 | 10 | Il6st | -0.03 | Young |
| 0.730555 | 0.345554 | 0.182 | 0.174 | 1 | 10 | Naa35 | 0.008 | Young |
| 0.730635 | 0.23646 | 0.404 | 0.467 | 1 | 10 | Rassf3 | -0.063 | Young |
| 0.73077 | 0.669425 | 0.061 | 0.071 | 1 | 10 | Fbxo25 | -0.01 | Young |
| 0.73088 | 0.07553 | 0.101 | 0.115 | 1 | 10 | Parp12 | -0.014 | Young |
| 0.730958 | 0.074882 | 0.111 | 0.098 | 1 | 10 | Tubgcp5 | 0.013 | Young |
| 0.73098 | 0.253856 | 0.202 | 0.227 | 1 | 10 | Hdac2 | -0.025 | Young |
| 0.731094 | 0.30212 | 0.222 | 0.215 | 1 | 10 | Zkscan3 | 0.007 | Young |
| 0.731101 | 0.928995 | 0.051 | 0.061 | 1 | 10 | C87436 | -0.01 | Young |
| 0.731218 | 0.23344 | 0.434 | 0.44 | 1 | 10 | Renbp | -0.006 | Young |
| 0.731383 | 0.088474 | 0.222 | 0.213 | 1 | 10 | Rnase4 | 0.009 | Young |
| 0.731488 | 0.785807 | 0.051 | 0.044 | 1 | 10 | Mms22l | 0.007 | Young |
| 0.731513 | 0.610002 | 0.03 | 0.024 | 1 | 10 | Bivm | 0.006 | Young |
| 0.731529 | 0.164278 | 0.253 | 0.281 | 1 | 10 | Ccdc59 | -0.028 | Young |
| 0.731649 | 0.109831 | 0.333 | 0.374 | 1 | 10 | Kpnb1 | -0.041 | Young |
| 0.731765 | 0.049706 | 0.505 | 0.55 | 1 | 10 | Ehd1 | -0.045 | Young |
| 0.731914 | 0.089391 | 0.212 | 0.23 | 1 | 10 | Dock11 | -0.018 | Young |
| 0.732485 | 0.15847 | 0.394 | 0.399 | 1 | 10 | Lpp | -0.005 | Young |
| 0.732688 | 0.123694 | 0.101 | 0.09 | 1 | 10 | Ppwd1 | 0.011 | Young |
| 0.733525 | 0.632493 | 0.081 | 0.073 | 1 | 10 | Ivd | 0.008 | Young |
| 0.733613 | 0.791649 | 0.03 | 0.024 | 1 | 10 | Smyd4 | 0.006 | Young |
| 0.733613 | 0.118808 | 0.03 | 0.024 | 1 | 10 | Spink2 | 0.006 | Young |
| 0.733776 | 0.506777 | 0.01 | 0.015 | 1 | 10 | 9330175E1 | -0.005 | Young |
| 0.733776 | 0.442964 | 0.01 | 0.015 | 1 | 10 | Btd | -0.005 | Young |
| 0.733776 | 0.338629 | 0.01 | 0.015 | 1 | 10 | Rbfox2 | -0.005 | Young |
| 0.733776 | 0.268037 | 0.01 | 0.015 | 1 | 10 | Prodh | -0.005 | Young |
| 0.733776 | 0.060222 | 0.01 | 0.015 | 1 | 10 | Gm16124 | -0.005 | Young |
| 0.733776 | 0.054356 | 0.01 | 0.015 | 1 | 10 | Fam131a | -0.005 | Young |
| 0.733776 | 0.023535 | 0.01 | 0.015 | 1 | 10 | Lmo1 | -0.005 | Young |
| 0.733831 | 0.569265 | 0.101 | 0.09 | 1 | 10 | Zfp511 | 0.011 | Young |
| 0.733981 | 0.353002 | 0.091 | 0.083 | 1 | 10 | Zfp830 | 0.008 | Young |
| 0.734115 | 0.241719 | 0.222 | 0.247 | 1 | 10 | D1Ertd622 | -0.025 | Young |
| 0.734225 | 0.263867 | 0.242 | 0.267 | 1 | 10 | Smndc1 | -0.025 | Young |
| 0.734529 | 0.338311 | 0.192 | 0.183 | 1 | 10 | Ndufv1 | 0.009 | Young |
| 0.73472 | 0.350926 | 0.071 | 0.061 | 1 | 10 | Bank1 | 0.01 | Young |
| 0.735305 | 0.504214 | 0.323 | 0.342 | 1 | 10 | Smc1a | -0.019 | Young |
| 0.735515 | 0.525685 | 0.051 | 0.059 | 1 | 10 | Zc3h3 | -0.008 | Young |
| 0.73557 | 0.325193 | 0.212 | 0.237 | 1 | 10 | P4ha1 | -0.025 | Young |

| | | | | | | | | |
|---|---|---|---|---|---|---|---|---|
| 0.735714 | 1.049413 | 0.03 | 0.024 | 1 | 10 | Hist2h3c1 | 0.006 | Young |
| 0.735714 | 0.146071 | 0.03 | 0.024 | 1 | 10 | Ubiad1 | 0.006 | Young |
| 0.735739 | 0.206071 | 0.525 | 0.55 | 1 | 10 | Hsbp1 | -0.025 | Young |
| 0.735811 | 0.089454 | 0.253 | 0.284 | 1 | 10 | Osgep | -0.031 | Young |
| 0.735894 | 0.046065 | 0.04 | 0.049 | 1 | 10 | Mtrf1 | -0.009 | Young |
| 0.736623 | 1.936577 | 0.01 | 0.015 | 1 | 10 | Gm3435 | -0.005 | Young |
| 0.736623 | 1.760419 | 0.01 | 0.015 | 1 | 10 | Myo1d | -0.005 | Young |
| 0.736623 | 1.684075 | 0.01 | 0.015 | 1 | 10 | Trib3 | -0.005 | Young |
| 0.736623 | 0.882033 | 0.01 | 0.015 | 1 | 10 | Prim1 | -0.005 | Young |
| 0.736623 | 0.821933 | 0.01 | 0.015 | 1 | 10 | Slc25a42 | -0.005 | Young |
| 0.736623 | 0.667226 | 0.01 | 0.015 | 1 | 10 | Pyroxd2 | -0.005 | Young |
| 0.736623 | 0.607567 | 0.01 | 0.015 | 1 | 10 | Gm10605 | -0.005 | Young |
| 0.736623 | 0.598987 | 0.01 | 0.015 | 1 | 10 | Gm11273 | -0.005 | Young |
| 0.736623 | 0.529888 | 0.01 | 0.015 | 1 | 10 | Gm39469 | -0.005 | Young |
| 0.736623 | 0.433024 | 0.01 | 0.015 | 1 | 10 | 1700109H( | -0.005 | Young |
| 0.736623 | 0.432549 | 0.01 | 0.015 | 1 | 10 | Gm26935 | -0.005 | Young |
| 0.736623 | 0.415184 | 0.01 | 0.015 | 1 | 10 | 6430531B1 | -0.005 | Young |
| 0.736623 | 0.382042 | 0.01 | 0.015 | 1 | 10 | Vasn | -0.005 | Young |
| 0.736623 | 0.313093 | 0.01 | 0.015 | 1 | 10 | E030030I0( | -0.005 | Young |
| 0.736623 | 0.071256 | 0.01 | 0.015 | 1 | 10 | Acad10 | -0.005 | Young |
| 0.736677 | 0.446099 | 0.202 | 0.22 | 1 | 10 | Hccs | -0.018 | Young |
| 0.736688 | 0.214405 | 0.364 | 0.403 | 1 | 10 | Nufip2 | -0.039 | Young |
| 0.73692 | 0.424734 | 0.172 | 0.159 | 1 | 10 | Tspan5 | 0.013 | Young |
| 0.737189 | 0.107672 | 0.283 | 0.325 | 1 | 10 | Senp6 | -0.042 | Young |
| 0.7373 | 0.100374 | 0.313 | 0.296 | 1 | 10 | Ap1g1 | 0.017 | Young |
| 0.737567 | 0.240089 | 0.444 | 0.482 | 1 | 10 | Rbm8a | -0.038 | Young |
| 0.737772 | 0.313163 | 0.333 | 0.379 | 1 | 10 | Cat | -0.046 | Young |
| 0.737818 | 0.353385 | 0.03 | 0.024 | 1 | 10 | Ifi213 | 0.006 | Young |
| 0.737818 | 0.281464 | 0.03 | 0.024 | 1 | 10 | Aatk | 0.006 | Young |
| 0.737818 | 0.17841 | 0.03 | 0.024 | 1 | 10 | Gm14305 | 0.006 | Young |
| 0.738096 | 0.288766 | 0.222 | 0.21 | 1 | 10 | Fam53b | 0.012 | Young |
| 0.738201 | 0.20921 | 0.323 | 0.32 | 1 | 10 | Ncoa4 | 0.003 | Young |
| 0.738459 | 0.073134 | 0.162 | 0.152 | 1 | 10 | Nop56 | 0.01 | Young |
| 0.738641 | 0.295547 | 0.081 | 0.071 | 1 | 10 | Faap100 | 0.01 | Young |
| 0.738904 | 0.547768 | 0.141 | 0.132 | 1 | 10 | Ap2a1 | 0.009 | Young |
| 0.738904 | 0.598207 | 0.141 | 0.132 | 1 | 10 | Rhbdf2 | 0.009 | Young |
| 0.739025 | 0.555665 | 0.04 | 0.049 | 1 | 10 | Rtn4ip1 | -0.009 | Young |
| 0.73903 | 0.114178 | 0.596 | 0.597 | 1 | 10 | Ly86 | -0.001 | Young |
| 0.739413 | 0.712464 | 0.111 | 0.132 | 1 | 10 | Abcd3 | -0.021 | Young |
| 0.73947 | 0.954047 | 0.051 | 0.044 | 1 | 10 | Maml2 | 0.007 | Young |
| 0.739588 | 0.132416 | 0.162 | 0.186 | 1 | 10 | Gabpb2 | -0.024 | Young |
| 0.739923 | 0.380684 | 0.03 | 0.024 | 1 | 10 | Med26 | 0.006 | Young |
| 0.739923 | 0.364257 | 0.03 | 0.024 | 1 | 10 | Nsun5 | 0.006 | Young |
| 0.740026 | 0.22027 | 0.354 | 0.425 | 1 | 10 | Mcur1 | -0.071 | Young |
| 0.740142 | 0.42854 | 0.141 | 0.164 | 1 | 10 | Opa3 | -0.023 | Young |

| | | | | | | | | |
|---|---|---|---|---|---|---|---|---|
| 0.740187 | 0.252156 | 0.283 | 0.276 | 1 | 10 | Ankhd1 | 0.007 | Young |
| 0.740593 | 0.144311 | 0.04 | 0.049 | 1 | 10 | Relt | -0.009 | Young |
| 0.740673 | 0.090241 | 0.101 | 0.117 | 1 | 10 | Ankrd16 | -0.016 | Young |
| 0.740701 | 0.654517 | 0.081 | 0.095 | 1 | 10 | Tagap | -0.014 | Young |
| 0.740851 | 0.359818 | 0.141 | 0.132 | 1 | 10 | Samd1 | 0.009 | Young |
| 0.740914 | 0.574039 | 0.212 | 0.213 | 1 | 10 | Tmem41b | -0.001 | Young |
| 0.741065 | 0.964121 | 0.071 | 0.064 | 1 | 10 | Zfp687 | 0.007 | Young |
| 0.741363 | 0.834811 | 0.101 | 0.093 | 1 | 10 | Prpf4 | 0.008 | Young |
| 0.741672 | 0.074077 | 0.162 | 0.181 | 1 | 10 | H2-Q7 | -0.019 | Young |
| 0.741741 | 0.888319 | 0.677 | 0.802 | 1 | 10 | Pf4 | -0.125 | Young |
| 0.741885 | 0.21557 | 0.111 | 0.103 | 1 | 10 | Ndufaf2 | 0.008 | Young |
| 0.741916 | 0.217983 | 0.384 | 0.359 | 1 | 10 | Lair1 | 0.025 | Young |
| 0.742031 | 0.819963 | 0.03 | 0.024 | 1 | 10 | Mapk1ip1 | 0.006 | Young |
| 0.742031 | 0.54437 | 0.03 | 0.024 | 1 | 10 | Mtmr11 | 0.006 | Young |
| 0.742368 | 0.367503 | 0.051 | 0.061 | 1 | 10 | Pcp4l1 | -0.01 | Young |
| 0.743026 | 0.266055 | 0.131 | 0.122 | 1 | 10 | Dhdds | 0.009 | Young |
| 0.743525 | 0.514189 | 0.091 | 0.083 | 1 | 10 | Cog2 | 0.008 | Young |
| 0.743653 | 0.46839 | 0.212 | 0.2 | 1 | 10 | Dhrs1 | 0.012 | Young |
| 0.743877 | 0.729784 | 0.061 | 0.071 | 1 | 10 | Amd1 | -0.01 | Young |
| 0.743877 | 0.120471 | 0.061 | 0.071 | 1 | 10 | Fkbp5 | -0.01 | Young |
| 0.743941 | 0.690948 | 0.121 | 0.142 | 1 | 10 | Brd9 | -0.021 | Young |
| 0.744189 | 0.262314 | 0.455 | 0.477 | 1 | 10 | Ndufb5 | -0.022 | Young |
| 0.744296 | 0.294168 | 0.071 | 0.061 | 1 | 10 | 5430427O1 | 0.01 | Young |
| 0.745006 | 0.322224 | 0.131 | 0.152 | 1 | 10 | Rae1 | -0.021 | Young |
| 0.745107 | 0.024914 | 0.768 | 0.792 | 1 | 10 | Nap1l1 | -0.024 | Young |
| 0.74531 | 0.068332 | 0.091 | 0.103 | 1 | 10 | Fam98c | -0.012 | Young |
| 0.745392 | 0.224818 | 0.152 | 0.174 | 1 | 10 | Nsmce3 | -0.022 | Young |
| 0.745462 | 0.248144 | 0.293 | 0.296 | 1 | 10 | Tmem230 | -0.003 | Young |
| 0.745531 | 0.073149 | 0.838 | 0.897 | 1 | 10 | Mpeg1 | -0.059 | Young |
| 0.745946 | 0.033049 | 0.717 | 0.765 | 1 | 10 | Csnk1a1 | -0.048 | Young |
| 0.745958 | 0.219237 | 0.152 | 0.142 | 1 | 10 | Mtmr12 | 0.01 | Young |
| 0.746378 | 0.476341 | 0.131 | 0.154 | 1 | 10 | Myo18a | -0.023 | Young |
| 0.74641 | 0.267685 | 0.131 | 0.152 | 1 | 10 | Faap20 | -0.021 | Young |
| 0.746732 | 0.087889 | 0.253 | 0.242 | 1 | 10 | Mrpl42 | 0.011 | Young |
| 0.746825 | 0.266266 | 0.293 | 0.32 | 1 | 10 | Ppt1 | -0.027 | Young |
| 0.747485 | 0.006234 | 0.667 | 0.726 | 1 | 10 | Vmp1 | -0.059 | Young |
| 0.747499 | 0.058776 | 0.242 | 0.279 | 1 | 10 | Ubald1 | -0.037 | Young |
| 0.747591 | 0.111319 | 0.101 | 0.09 | 1 | 10 | Rnf168 | 0.011 | Young |
| 0.748097 | 0.130048 | 0.263 | 0.296 | 1 | 10 | Slc35b2 | -0.033 | Young |
| 0.748312 | 0.240153 | 0.131 | 0.149 | 1 | 10 | Minpp1 | -0.018 | Young |
| 0.748451 | 0.022512 | 0.303 | 0.335 | 1 | 10 | Tspan14 | -0.032 | Young |
| 0.748634 | 0.325141 | 0.04 | 0.034 | 1 | 10 | Sorbs3 | 0.006 | Young |
| 0.748656 | 0.087428 | 0.121 | 0.137 | 1 | 10 | Akap8 | -0.016 | Young |
| 0.748678 | 0.135248 | 0.343 | 0.33 | 1 | 10 | Cd302 | 0.013 | Young |
| 0.748983 | 0.019195 | 0.364 | 0.386 | 1 | 10 | Notch2 | -0.022 | Young |

| | | | | | | | | |
|---|---|---|---|---|---|---|---|---|
| 0.749081 | 0.288322 | 0.111 | 0.127 | 1 | 10 | Daam1 | -0.016 | Young |
| 0.74909 | 0.292079 | 0.495 | 0.562 | 1 | 10 | Ubxn1 | -0.067 | Young |
| 0.749732 | 0.35615 | 0.111 | 0.13 | 1 | 10 | Prmt5 | -0.019 | Young |
| 0.749847 | 0.115163 | 0.222 | 0.247 | 1 | 10 | Mmadhc | -0.025 | Young |
| 0.750027 | 0.219188 | 0.606 | 0.645 | 1 | 10 | Tob2 | -0.039 | Young |
| 0.750264 | 0.159831 | 0.313 | 0.342 | 1 | 10 | Glrx5 | -0.029 | Young |
| 0.750357 | 0.320746 | 0.222 | 0.22 | 1 | 10 | Trnau1ap | 0.002 | Young |
| 0.750423 | 0.347187 | 0.162 | 0.152 | 1 | 10 | As3mt | 0.01 | Young |
| 0.75044 | 1.35527 | 0.04 | 0.034 | 1 | 10 | Prpf40b | 0.006 | Young |
| 0.75044 | 1.083324 | 0.04 | 0.034 | 1 | 10 | Tmtc4 | 0.006 | Young |
| 0.75044 | 0.541309 | 0.04 | 0.034 | 1 | 10 | Jmjd8 | 0.006 | Young |
| 0.75046 | 0.441963 | 0.061 | 0.071 | 1 | 10 | 1110065P2 | -0.01 | Young |
| 0.750479 | 0.192214 | 0.03 | 0.024 | 1 | 10 | Usp35 | 0.006 | Young |
| 0.750612 | 0.484657 | 0.121 | 0.137 | 1 | 10 | Terf2 | -0.016 | Young |
| 0.750691 | 0.683755 | 0.051 | 0.044 | 1 | 10 | Mir142hg | 0.007 | Young |
| 0.750691 | 0.304206 | 0.051 | 0.044 | 1 | 10 | Paxx | 0.007 | Young |
| 0.750944 | 0.179829 | 0.091 | 0.1 | 1 | 10 | Hif1an | -0.009 | Young |
| 0.75113 | 0.030559 | 0.848 | 0.929 | 1 | 10 | Rps7 | -0.081 | Young |
| 0.751229 | 0.523261 | 0.081 | 0.073 | 1 | 10 | Adprhl2 | 0.008 | Young |
| 0.751296 | 0.179135 | 0.343 | 0.335 | 1 | 10 | Bcl2l1 | 0.008 | Young |
| 0.751609 | 0.057361 | 0.606 | 0.641 | 1 | 10 | Tmed5 | -0.035 | Young |
| 0.751621 | 0.180535 | 0.384 | 0.425 | 1 | 10 | Snx18 | -0.041 | Young |
| 0.751741 | 0.570883 | 0.131 | 0.125 | 1 | 10 | Cfap43 | 0.006 | Young |
| 0.751778 | 0.128013 | 0.061 | 0.071 | 1 | 10 | Fig4 | -0.01 | Young |
| 0.752166 | 0.168087 | 0.141 | 0.159 | 1 | 10 | Arl11 | -0.018 | Young |
| 0.752248 | 0.87149 | 0.04 | 0.034 | 1 | 10 | 6330403L0 | 0.006 | Young |
| 0.752248 | 0.229019 | 0.04 | 0.034 | 1 | 10 | Plekhg1 | 0.006 | Young |
| 0.752297 | 0.188651 | 0.202 | 0.193 | 1 | 10 | Impa1 | 0.009 | Young |
| 0.752679 | 0.179413 | 0.495 | 0.531 | 1 | 10 | Anp32e | -0.036 | Young |
| 0.752753 | 0.977 | 0.101 | 0.093 | 1 | 10 | Dvl2 | 0.008 | Young |
| 0.752831 | 0.08502 | 0.182 | 0.2 | 1 | 10 | Ubfd1 | -0.018 | Young |
| 0.75298 | 0.479866 | 0.101 | 0.115 | 1 | 10 | Psat1 | -0.014 | Young |
| 0.753075 | 0.836964 | 0.091 | 0.103 | 1 | 10 | Atn1 | -0.012 | Young |
| 0.753103 | 0.272832 | 0.172 | 0.166 | 1 | 10 | Brip1os | 0.006 | Young |
| 0.753108 | 0.393128 | 0.091 | 0.083 | 1 | 10 | Scaper | 0.008 | Young |
| 0.753579 | 0.340529 | 0.121 | 0.139 | 1 | 10 | Gga2 | -0.018 | Young |
| 0.753726 | 0.805146 | 0.172 | 0.164 | 1 | 10 | Gm13212 | 0.008 | Young |
| 0.753751 | 0.322741 | 0.131 | 0.125 | 1 | 10 | 0610010F0 | 0.006 | Young |
| 0.75388 | 0.945428 | 0.061 | 0.054 | 1 | 10 | Adamtsl5 | 0.007 | Young |
| 0.754057 | 0.529905 | 0.04 | 0.034 | 1 | 10 | Nkiras1 | 0.006 | Young |
| 0.754384 | 0.055206 | 0.253 | 0.279 | 1 | 10 | B3gat3 | -0.026 | Young |
| 0.754418 | 0.062895 | 0.061 | 0.071 | 1 | 10 | Xxylt1 | -0.01 | Young |
| 0.754511 | 0.104186 | 0.111 | 0.125 | 1 | 10 | Mrpl38 | -0.014 | Young |
| 0.75472 | 0.885445 | 0.172 | 0.174 | 1 | 10 | Spast | -0.002 | Young |
| 0.754746 | 0.865993 | 0.04 | 0.049 | 1 | 10 | Mtif3 | -0.009 | Young |

| | | | | | | | | |
|---|---|---|---|---|---|---|---|---|
| 0.75513 | 0.163331 | 0.202 | 0.235 | 1 | 10 | Ssbp4 | -0.033 | Young |
| 0.755153 | 0.400354 | 0.202 | 0.23 | 1 | 10 | Nkiras2 | -0.028 | Young |
| 0.755518 | 0.085583 | 0.758 | 0.844 | 1 | 10 | Eef1g | -0.086 | Young |
| 0.755868 | 0.761725 | 0.04 | 0.034 | 1 | 10 | Gm26518 | 0.006 | Young |
| 0.756371 | 0.50641 | 0.263 | 0.274 | 1 | 10 | Utp3 | -0.011 | Young |
| 0.756478 | 0.077425 | 0.131 | 0.142 | 1 | 10 | 1700037H( | -0.011 | Young |
| 0.756667 | 0.112846 | 0.071 | 0.061 | 1 | 10 | Slc35c1 | 0.01 | Young |
| 0.756879 | 0.279639 | 0.333 | 0.364 | 1 | 10 | BC031181 | -0.031 | Young |
| 0.756966 | 0.271016 | 0.101 | 0.112 | 1 | 10 | Slc12a4 | -0.011 | Young |
| 0.757124 | 0.579357 | 0.232 | 0.225 | 1 | 10 | Myd88 | 0.007 | Young |
| 0.757142 | 0.245714 | 0.182 | 0.169 | 1 | 10 | Acadsb | 0.013 | Young |
| 0.757213 | 0.490252 | 0.101 | 0.115 | 1 | 10 | Cryzl1 | -0.014 | Young |
| 0.75768 | 0.449365 | 0.04 | 0.034 | 1 | 10 | Top3a | 0.006 | Young |
| 0.757845 | 0.003608 | 0.818 | 0.87 | 1 | 10 | Naca | -0.052 | Young |
| 0.75789 | 0.206998 | 0.455 | 0.45 | 1 | 10 | Pnisr | 0.005 | Young |
| 0.758301 | 0.071785 | 0.131 | 0.147 | 1 | 10 | Dmtf1 | -0.016 | Young |
| 0.758381 | 0.254885 | 0.455 | 0.474 | 1 | 10 | Ddb1 | -0.019 | Young |
| 0.758585 | 0.16684 | 0.081 | 0.093 | 1 | 10 | Lancl1 | -0.012 | Young |
| 0.759255 | 0.002259 | 0.505 | 0.577 | 1 | 10 | Nras | -0.072 | Young |
| 0.759482 | 0.867428 | 0.04 | 0.049 | 1 | 10 | Btg3 | -0.009 | Young |
| 0.759493 | 0.424303 | 0.04 | 0.034 | 1 | 10 | Zfp629 | 0.006 | Young |
| 0.759525 | 0.416135 | 0.162 | 0.154 | 1 | 10 | Epc2 | 0.008 | Young |
| 0.75972 | 0.17427 | 0.232 | 0.249 | 1 | 10 | Snx13 | -0.017 | Young |
| 0.760123 | 0.841975 | 0.071 | 0.083 | 1 | 10 | Celf4 | -0.012 | Young |
| 0.761029 | 0.409954 | 0.061 | 0.071 | 1 | 10 | Cutc | -0.01 | Young |
| 0.761219 | 0.968642 | 0.061 | 0.054 | 1 | 10 | Il6 | 0.007 | Young |
| 0.761219 | 0.353503 | 0.061 | 0.054 | 1 | 10 | Ercc2 | 0.007 | Young |
| 0.761219 | 0.263119 | 0.061 | 0.054 | 1 | 10 | Rbm38 | 0.007 | Young |
| 0.761524 | 0.365264 | 0.091 | 0.083 | 1 | 10 | Dym | 0.008 | Young |
| 0.76166 | 0.379612 | 0.111 | 0.125 | 1 | 10 | Gmeb1 | -0.014 | Young |
| 0.761965 | 0.375923 | 0.051 | 0.044 | 1 | 10 | Dip2a | 0.007 | Young |
| 0.762148 | 0.223958 | 0.263 | 0.257 | 1 | 10 | Pcm1 | 0.006 | Young |
| 0.762628 | 0.254276 | 0.212 | 0.2 | 1 | 10 | Fgd2 | 0.012 | Young |
| 0.762689 | 0.436991 | 0.061 | 0.054 | 1 | 10 | Ift74 | 0.007 | Young |
| 0.763247 | 0.09654 | 0.626 | 0.709 | 1 | 10 | Tapbp | -0.083 | Young |
| 0.763714 | 0.0605 | 0.313 | 0.308 | 1 | 10 | Derl2 | 0.005 | Young |
| 0.763748 | 0.21513 | 0.222 | 0.247 | 1 | 10 | Hipk2 | -0.025 | Young |
| 0.764032 | 0.331929 | 0.03 | 0.037 | 1 | 10 | Fntb | -0.007 | Young |
| 0.764103 | 0.332773 | 0.283 | 0.303 | 1 | 10 | Gdf15 | -0.02 | Young |
| 0.76416 | 1.003128 | 0.061 | 0.054 | 1 | 10 | Rnf219 | 0.007 | Young |
| 0.764735 | 0.029561 | 0.394 | 0.433 | 1 | 10 | Ncstn | -0.039 | Young |
| 0.764806 | 0.31071 | 0.455 | 0.479 | 1 | 10 | Hook3 | -0.024 | Young |
| 0.764941 | 0.032329 | 0.04 | 0.034 | 1 | 10 | BC024978 | 0.006 | Young |
| 0.765085 | 0.381271 | 0.091 | 0.105 | 1 | 10 | Pde12 | -0.014 | Young |
| 0.765164 | 0.262352 | 0.192 | 0.181 | 1 | 10 | Rxrb | 0.011 | Young |

| | | | | | | | | |
|---|---|---|---|---|---|---|---|---|
| 0.765285 | 0.245289 | 0.141 | 0.134 | 1 | 10 | Topbp1 | 0.007 | Young |
| 0.765343 | 0.679176 | 0.101 | 0.093 | 1 | 10 | Rfxap | 0.008 | Young |
| 0.765462 | 0.200275 | 0.162 | 0.178 | 1 | 10 | Atxn1 | -0.016 | Young |
| 0.765487 | 0.226314 | 0.182 | 0.205 | 1 | 10 | Tob1 | -0.023 | Young |
| 0.765583 | 0.362398 | 0.192 | 0.191 | 1 | 10 | Tor1a | 0.001 | Young |
| 0.765737 | 0.385539 | 0.202 | 0.193 | 1 | 10 | Gramd4 | 0.009 | Young |
| 0.766189 | 0.207134 | 0.071 | 0.081 | 1 | 10 | F630028O1 | -0.01 | Young |
| 0.766247 | 0.144211 | 0.414 | 0.467 | 1 | 10 | Nsd1 | -0.053 | Young |
| 0.766267 | 0.029823 | 0.242 | 0.269 | 1 | 10 | Rasa3 | -0.027 | Young |
| 0.766345 | 0.106659 | 0.141 | 0.13 | 1 | 10 | Hat1 | 0.011 | Young |
| 0.766345 | 0.081729 | 0.121 | 0.134 | 1 | 10 | Cep120 | -0.013 | Young |
| 0.766491 | 0.417645 | 0.101 | 0.093 | 1 | 10 | Rabggta | 0.008 | Young |
| 0.766491 | 0.336285 | 0.101 | 0.093 | 1 | 10 | Trub2 | 0.008 | Young |
| 0.766561 | 0.078952 | 0.414 | 0.457 | 1 | 10 | Cuta | -0.043 | Young |
| 0.766713 | 0.475392 | 0.162 | 0.156 | 1 | 10 | Smim13 | 0.006 | Young |
| 0.766754 | 0.246261 | 0.172 | 0.159 | 1 | 10 | Pspc1 | 0.013 | Young |
| 0.766759 | 0.222887 | 0.04 | 0.034 | 1 | 10 | 1600020E0 | 0.006 | Young |
| 0.766905 | 0.310792 | 0.162 | 0.154 | 1 | 10 | Fam96b | 0.008 | Young |
| 0.767039 | 0.403175 | 0.121 | 0.115 | 1 | 10 | Gipc1 | 0.006 | Young |
| 0.767669 | 0.708374 | 0.03 | 0.037 | 1 | 10 | Cers4 | -0.007 | Young |
| 0.768126 | 0.054742 | 0.505 | 0.562 | 1 | 10 | Wdr83os | -0.057 | Young |
| 0.768171 | 0.093352 | 0.444 | 0.484 | 1 | 10 | Cwc15 | -0.04 | Young |
| 0.768543 | 0.07078 | 0.636 | 0.685 | 1 | 10 | Itgb2 | -0.049 | Young |
| 0.768761 | 0.439235 | 0.091 | 0.083 | 1 | 10 | Golim4 | 0.008 | Young |
| 0.768916 | 0.15435 | 0.364 | 0.389 | 1 | 10 | Gtf2a1 | -0.025 | Young |
| 0.769135 | 0.5986 | 0.222 | 0.222 | 1 | 10 | Birc2 | 0 | Young |
| 0.769473 | 0.223017 | 0.182 | 0.196 | 1 | 10 | Dido1 | -0.014 | Young |
| 0.769561 | 0.295353 | 0.162 | 0.181 | 1 | 10 | Zfp704 | -0.019 | Young |
| 0.769652 | 0.608098 | 0.202 | 0.2 | 1 | 10 | Rpp21 | 0.002 | Young |
| 0.769744 | 0.086651 | 0.111 | 0.12 | 1 | 10 | Mtmr1 | -0.009 | Young |
| 0.769804 | 0.220318 | 0.394 | 0.45 | 1 | 10 | N4bp1 | -0.056 | Young |
| 0.769876 | 0.151212 | 0.141 | 0.156 | 1 | 10 | Fgd6 | -0.015 | Young |
| 0.769995 | 0.025396 | 0.374 | 0.416 | 1 | 10 | Ppp1r12a | -0.042 | Young |
| 0.770054 | 0.353516 | 0.061 | 0.054 | 1 | 10 | Cep135 | 0.007 | Young |
| 0.770093 | 0.236346 | 0.051 | 0.059 | 1 | 10 | 5730455P1 | -0.008 | Young |
| 0.7704 | 0.17188 | 0.04 | 0.034 | 1 | 10 | Icam2 | 0.006 | Young |
| 0.7704 | 0.127915 | 0.04 | 0.034 | 1 | 10 | 1810043G0 | 0.006 | Young |
| 0.770678 | 0.131342 | 0.283 | 0.271 | 1 | 10 | Rgl2 | 0.012 | Young |
| 0.771262 | 0.044457 | 0.98 | 0.971 | 1 | 10 | Cd74 | 0.009 | Young |
| 0.771528 | 0.386507 | 0.141 | 0.159 | 1 | 10 | Clpx | -0.018 | Young |
| 0.772222 | 0.128927 | 0.04 | 0.034 | 1 | 10 | Ms4a4c | 0.006 | Young |
| 0.772222 | 0.083394 | 0.04 | 0.034 | 1 | 10 | Harbi1 | 0.006 | Young |
| 0.772301 | 0.763897 | 0.091 | 0.086 | 1 | 10 | Alkbh6 | 0.005 | Young |
| 0.77242 | 0.30503 | 0.172 | 0.161 | 1 | 10 | Arl6ip4 | 0.011 | Young |
| 0.772429 | 0.347871 | 0.131 | 0.122 | 1 | 10 | Ptrhd1 | 0.009 | Young |

| | | | | | | | | |
|---|---|---|---|---|---|---|---|---|
| 0.77262 | 0.070572 | 0.152 | 0.142 | 1 | 10 | Rhoh | 0.01 | Young |
| 0.772954 | 0.074576 | 0.202 | 0.215 | 1 | 10 | Ramp1 | -0.013 | Young |
| 0.772997 | 0.192567 | 0.051 | 0.059 | 1 | 10 | Eif2b1 | -0.008 | Young |
| 0.772997 | 0.028838 | 0.051 | 0.059 | 1 | 10 | Aph1b | -0.008 | Young |
| 0.773134 | 0.063299 | 0.03 | 0.037 | 1 | 10 | Zbtb26 | -0.007 | Young |
| 0.773271 | 0.137766 | 0.414 | 0.423 | 1 | 10 | Ppp1r12c | -0.009 | Young |
| 0.773672 | 0.288889 | 0.071 | 0.064 | 1 | 10 | Ripk2 | 0.007 | Young |
| 0.774422 | 0.141562 | 0.485 | 0.513 | 1 | 10 | Bst2 | -0.028 | Young |
| 0.774449 | 0.132419 | 0.051 | 0.059 | 1 | 10 | Dcaf1 | -0.008 | Young |
| 0.774912 | 0.813337 | 0.051 | 0.044 | 1 | 10 | 2510002D2 | 0.007 | Young |
| 0.774958 | 0.139994 | 0.03 | 0.037 | 1 | 10 | Slf1 | -0.007 | Young |
| 0.774958 | 0.104213 | 0.03 | 0.037 | 1 | 10 | Tbl2 | -0.007 | Young |
| 0.775277 | 0.012266 | 0.141 | 0.13 | 1 | 10 | Wipi2 | 0.011 | Young |
| 0.77534 | 0.288867 | 0.333 | 0.347 | 1 | 10 | Tank | -0.014 | Young |
| 0.775432 | 0.360916 | 0.121 | 0.115 | 1 | 10 | Mgst3 | 0.006 | Young |
| 0.775792 | 0.001118 | 0.182 | 0.193 | 1 | 10 | Mbd6 | -0.011 | Young |
| 0.775798 | 0.225077 | 0.152 | 0.144 | 1 | 10 | Ndrg3 | 0.008 | Young |
| 0.77606 | 0.40373 | 0.242 | 0.284 | 1 | 10 | Smc3 | -0.042 | Young |
| 0.776133 | 0.263195 | 0.222 | 0.215 | 1 | 10 | Hdac1 | 0.007 | Young |
| 0.776329 | 2.295386 | 0.01 | 0.007 | 1 | 10 | Plcb4 | 0.003 | Young |
| 0.776329 | 1.885428 | 0.01 | 0.007 | 1 | 10 | Ehhadh | 0.003 | Young |
| 0.776329 | 1.7893 | 0.01 | 0.007 | 1 | 10 | 1700001L0 | 0.003 | Young |
| 0.776329 | 1.757664 | 0.01 | 0.007 | 1 | 10 | C920006O1 | 0.003 | Young |
| 0.776329 | 1.751512 | 0.01 | 0.007 | 1 | 10 | Siglecg | 0.003 | Young |
| 0.776329 | 1.488119 | 0.01 | 0.007 | 1 | 10 | Fads6 | 0.003 | Young |
| 0.776329 | 1.479321 | 0.01 | 0.007 | 1 | 10 | Parp16 | 0.003 | Young |
| 0.776329 | 1.411109 | 0.01 | 0.007 | 1 | 10 | C330011M | 0.003 | Young |
| 0.776329 | 1.35939 | 0.01 | 0.007 | 1 | 10 | Map3k6 | 0.003 | Young |
| 0.776329 | 1.324518 | 0.01 | 0.007 | 1 | 10 | Agrn | 0.003 | Young |
| 0.776329 | 1.306163 | 0.01 | 0.007 | 1 | 10 | 9530082P2 | 0.003 | Young |
| 0.776329 | 1.296155 | 0.01 | 0.007 | 1 | 10 | Unc13b | 0.003 | Young |
| 0.776329 | 1.265356 | 0.01 | 0.007 | 1 | 10 | Fanca | 0.003 | Young |
| 0.776329 | 1.248829 | 0.01 | 0.007 | 1 | 10 | Tln2 | 0.003 | Young |
| 0.776329 | 1.180394 | 0.01 | 0.007 | 1 | 10 | Siglecf | 0.003 | Young |
| 0.776329 | 1.153369 | 0.01 | 0.007 | 1 | 10 | Ociad2 | 0.003 | Young |
| 0.776329 | 1.073256 | 0.01 | 0.007 | 1 | 10 | Slc43a3 | 0.003 | Young |
| 0.776329 | 1.010629 | 0.01 | 0.007 | 1 | 10 | Clec3b | 0.003 | Young |
| 0.776329 | 0.986482 | 0.01 | 0.007 | 1 | 10 | Adrb1 | 0.003 | Young |
| 0.776329 | 0.981946 | 0.01 | 0.007 | 1 | 10 | Enkd1 | 0.003 | Young |
| 0.776329 | 0.951906 | 0.01 | 0.007 | 1 | 10 | Ms4a4b | 0.003 | Young |
| 0.776329 | 0.942786 | 0.01 | 0.007 | 1 | 10 | Gja1 | 0.003 | Young |
| 0.776329 | 0.935608 | 0.01 | 0.007 | 1 | 10 | Stox2 | 0.003 | Young |
| 0.776329 | 0.934079 | 0.01 | 0.007 | 1 | 10 | Haao | 0.003 | Young |
| 0.776329 | 0.909008 | 0.01 | 0.007 | 1 | 10 | Zfp273 | 0.003 | Young |
| 0.776329 | 0.870995 | 0.01 | 0.007 | 1 | 10 | Tnfrsf14 | 0.003 | Young |

| | | | | | | | | |
|---|---|---|---|---|---|---|---|---|
| 0.776329 | 0.827751 | 0.01 | 0.007 | 1 | 10 | Gm9796 | 0.003 | Young |
| 0.776329 | 0.786102 | 0.01 | 0.007 | 1 | 10 | Zfp947 | 0.003 | Young |
| 0.776329 | 0.770534 | 0.01 | 0.007 | 1 | 10 | Tjp3 | 0.003 | Young |
| 0.776329 | 0.752266 | 0.01 | 0.007 | 1 | 10 | Fstl1 | 0.003 | Young |
| 0.776329 | 0.724857 | 0.01 | 0.007 | 1 | 10 | Rdh5 | 0.003 | Young |
| 0.776329 | 0.721388 | 0.01 | 0.007 | 1 | 10 | Azin2 | 0.003 | Young |
| 0.776329 | 0.698851 | 0.01 | 0.007 | 1 | 10 | Nsl1 | 0.003 | Young |
| 0.776329 | 0.605806 | 0.01 | 0.007 | 1 | 10 | Neurl1a | 0.003 | Young |
| 0.776329 | 0.592581 | 0.01 | 0.007 | 1 | 10 | Gm48529 | 0.003 | Young |
| 0.776329 | 0.493537 | 0.01 | 0.007 | 1 | 10 | Lix1 | 0.003 | Young |
| 0.776329 | 0.469191 | 0.01 | 0.007 | 1 | 10 | Tonsl | 0.003 | Young |
| 0.776329 | 0.453913 | 0.01 | 0.007 | 1 | 10 | Brip1 | 0.003 | Young |
| 0.776329 | 0.44676 | 0.01 | 0.007 | 1 | 10 | Cxcr3 | 0.003 | Young |
| 0.776329 | 0.427042 | 0.01 | 0.007 | 1 | 10 | Aldh5a1 | 0.003 | Young |
| 0.776329 | 0.330603 | 0.01 | 0.007 | 1 | 10 | B230303O1 | 0.003 | Young |
| 0.776329 | 0.330339 | 0.01 | 0.007 | 1 | 10 | Tst | 0.003 | Young |
| 0.776432 | 0.203597 | 0.505 | 0.538 | 1 | 10 | Mef2c | -0.033 | Young |
| 0.776535 | 0.176936 | 0.051 | 0.044 | 1 | 10 | Pgghg | 0.007 | Young |
| 0.776747 | 0.444564 | 0.081 | 0.073 | 1 | 10 | Usb1 | 0.008 | Young |
| 0.776751 | 0.438358 | 0.131 | 0.149 | 1 | 10 | Rfc2 | -0.018 | Young |
| 0.777082 | 0.03854 | 0.293 | 0.281 | 1 | 10 | Diaph2 | 0.012 | Young |
| 0.777439 | 0.344347 | 0.061 | 0.054 | 1 | 10 | Oraov1 | 0.007 | Young |
| 0.77753 | 0.213311 | 0.303 | 0.342 | 1 | 10 | Usp15 | -0.039 | Young |
| 0.777562 | 0.556459 | 0.172 | 0.171 | 1 | 10 | Fbxo22 | 0.001 | Young |
| 0.777613 | 0.075573 | 0.283 | 0.315 | 1 | 10 | Ufc1 | -0.032 | Young |
| 0.777763 | 0.362549 | 0.162 | 0.156 | 1 | 10 | 2210016L2 | 0.006 | Young |
| 0.778159 | 0.215671 | 0.051 | 0.044 | 1 | 10 | Lipt2 | 0.007 | Young |
| 0.778586 | 0.420628 | 0.121 | 0.115 | 1 | 10 | Abhd4 | 0.006 | Young |
| 0.77861 | 0.362326 | 0.03 | 0.037 | 1 | 10 | Gm15987 | -0.007 | Young |
| 0.77861 | 0.217422 | 0.03 | 0.037 | 1 | 10 | AW554918 | -0.007 | Young |
| 0.778658 | 0.453591 | 0.131 | 0.149 | 1 | 10 | Smn1 | -0.018 | Young |
| 0.778944 | 1.078593 | 0.111 | 0.11 | 1 | 10 | Hint3 | 0.001 | Young |
| 0.779191 | 0.151442 | 0.141 | 0.156 | 1 | 10 | Btaf1 | -0.015 | Young |
| 0.779245 | 0.382094 | 0.222 | 0.22 | 1 | 10 | Npepl1 | 0.002 | Young |
| 0.779301 | 0.016181 | 0.081 | 0.09 | 1 | 10 | Coq10a | -0.009 | Young |
| 0.779555 | 0.596789 | 0.101 | 0.115 | 1 | 10 | Camk2g | -0.014 | Young |
| 0.779613 | 0.052129 | 0.758 | 0.797 | 1 | 10 | Sat1 | -0.039 | Young |
| 0.779643 | 0.119213 | 0.111 | 0.122 | 1 | 10 | Hsdl1 | -0.011 | Young |
| 0.780152 | 1.112092 | 0.01 | 0.007 | 1 | 10 | Acot4 | 0.003 | Young |
| 0.780152 | 0.896372 | 0.01 | 0.007 | 1 | 10 | Tnfrsf9 | 0.003 | Young |
| 0.780152 | 0.619211 | 0.01 | 0.007 | 1 | 10 | 2810414N0 | 0.003 | Young |
| 0.780152 | 0.574187 | 0.01 | 0.007 | 1 | 10 | AC146911. | 0.003 | Young |
| 0.780152 | 0.52459 | 0.01 | 0.007 | 1 | 10 | Wdr62 | 0.003 | Young |
| 0.780152 | 0.503586 | 0.01 | 0.007 | 1 | 10 | Kctd7 | 0.003 | Young |
| 0.780152 | 0.497658 | 0.01 | 0.007 | 1 | 10 | Sparcl1 | 0.003 | Young |

| | | | | | | | | |
|---|---|---|---|---|---|---|---|---|
| 0.780152 | 0.490686 | 0.01 | 0.007 | 1 | 10 | Nos3 | 0.003 | Young |
| 0.780152 | 0.428671 | 0.01 | 0.007 | 1 | 10 | Ifi205 | 0.003 | Young |
| 0.780152 | 0.391055 | 0.01 | 0.007 | 1 | 10 | Hexim2 | 0.003 | Young |
| 0.780152 | 0.380725 | 0.01 | 0.007 | 1 | 10 | 2010007H( | 0.003 | Young |
| 0.780152 | 0.376387 | 0.01 | 0.007 | 1 | 10 | Mrln | 0.003 | Young |
| 0.780152 | 0.329313 | 0.01 | 0.007 | 1 | 10 | Psmc3ip | 0.003 | Young |
| 0.780152 | 0.272362 | 0.01 | 0.007 | 1 | 10 | Gm36839 | 0.003 | Young |
| 0.780152 | 0.261501 | 0.01 | 0.007 | 1 | 10 | Gm19412 | 0.003 | Young |
| 0.780152 | 0.239583 | 0.01 | 0.007 | 1 | 10 | Gm26610 | 0.003 | Young |
| 0.780152 | 0.223568 | 0.01 | 0.007 | 1 | 10 | Igfbp6 | 0.003 | Young |
| 0.780152 | 0.183295 | 0.01 | 0.007 | 1 | 10 | Cerkl | 0.003 | Young |
| 0.780152 | 0.07999 | 0.01 | 0.007 | 1 | 10 | Apol7d | 0.003 | Young |
| 0.780152 | 0.017433 | 0.01 | 0.007 | 1 | 10 | Jag2 | 0.003 | Young |
| 0.78031 | 0.095799 | 0.808 | 0.795 | 1 | 10 | Ifrd1 | 0.013 | Young |
| 0.780624 | 0.023389 | 0.646 | 0.641 | 1 | 10 | Cerk | 0.005 | Young |
| 0.780627 | 0.528085 | 0.091 | 0.105 | 1 | 10 | Ubr1 | -0.014 | Young |
| 0.781088 | 0.013287 | 0.414 | 0.423 | 1 | 10 | Birc3 | -0.009 | Young |
| 0.781889 | 0.24063 | 0.071 | 0.064 | 1 | 10 | Usp31 | 0.007 | Young |
| 0.782476 | 0.272177 | 0.152 | 0.144 | 1 | 10 | Ndufaf4 | 0.008 | Young |
| 0.782633 | 0.463824 | 0.222 | 0.215 | 1 | 10 | Fam162a | 0.007 | Young |
| 0.782814 | 0.00943 | 0.455 | 0.479 | 1 | 10 | Eif3l | -0.024 | Young |
| 0.783112 | 0.371957 | 0.212 | 0.215 | 1 | 10 | Xdh | -0.003 | Young |
| 0.783178 | 0.567692 | 0.212 | 0.213 | 1 | 10 | Sra1 | -0.001 | Young |
| 0.783831 | 0.940889 | 0.121 | 0.117 | 1 | 10 | Mthfd1 | 0.004 | Young |
| 0.783979 | 0.367546 | 0.01 | 0.007 | 1 | 10 | Ptprb | 0.003 | Young |
| 0.783979 | 0.277357 | 0.01 | 0.007 | 1 | 10 | Hk1os | 0.003 | Young |
| 0.783979 | 0.23589 | 0.01 | 0.007 | 1 | 10 | Igf2bp3 | 0.003 | Young |
| 0.783979 | 0.215261 | 0.01 | 0.007 | 1 | 10 | Hspg2 | 0.003 | Young |
| 0.783979 | 0.206578 | 0.01 | 0.007 | 1 | 10 | Ifi214 | 0.003 | Young |
| 0.783979 | 0.114327 | 0.01 | 0.007 | 1 | 10 | 9130023H2 | 0.003 | Young |
| 0.783979 | 0.08469 | 0.01 | 0.007 | 1 | 10 | Pomc | 0.003 | Young |
| 0.783979 | 0.038315 | 0.01 | 0.007 | 1 | 10 | Cd209c | 0.003 | Young |
| 0.783979 | 0.025641 | 0.01 | 0.007 | 1 | 10 | Kntc1 | 0.003 | Young |
| 0.783979 | 0.009109 | 0.01 | 0.007 | 1 | 10 | Hmmr | 0.003 | Young |
| 0.784097 | 0.190279 | 0.03 | 0.037 | 1 | 10 | Tjap1 | -0.007 | Young |
| 0.784242 | 0.081438 | 0.283 | 0.274 | 1 | 10 | Clcn4 | 0.009 | Young |
| 0.784451 | 0.411405 | 0.081 | 0.073 | 1 | 10 | Cog1 | 0.008 | Young |
| 0.784694 | 0.570309 | 0.111 | 0.127 | 1 | 10 | Slc9a6 | -0.016 | Young |
| 0.785395 | 0.56577 | 0.212 | 0.22 | 1 | 10 | Etv6 | -0.008 | Young |
| 0.786007 | 0.325969 | 0.071 | 0.064 | 1 | 10 | Fmnl2 | 0.007 | Young |
| 0.786081 | 0.342284 | 0.162 | 0.156 | 1 | 10 | Tmem60 | 0.006 | Young |
| 0.786329 | 0.951757 | 0.061 | 0.054 | 1 | 10 | Aifm2 | 0.007 | Young |
| 0.786623 | 0.193194 | 0.283 | 0.276 | 1 | 10 | Uba52 | 0.007 | Young |
| 0.787014 | 0.295402 | 0.121 | 0.115 | 1 | 10 | Smurf2 | 0.006 | Young |
| 0.787024 | 0.302622 | 0.081 | 0.073 | 1 | 10 | Ppp1r12b | 0.008 | Young |

| | | | | | | | | |
|---|---|---|---|---|---|---|---|---|
| 0.787082 | 0.206631 | 0.485 | 0.55 | 1 | 10 | Map7d1 | -0.065 | Young |
| 0.78756 | 0.042394 | 0.051 | 0.059 | 1 | 10 | Btbd2 | -0.008 | Young |
| 0.787762 | 0.368596 | 0.03 | 0.037 | 1 | 10 | Map4k2 | -0.007 | Young |
| 0.787762 | 0.322827 | 0.03 | 0.037 | 1 | 10 | C1qtnf1 | -0.007 | Young |
| 0.787812 | 0.190415 | 0.01 | 0.007 | 1 | 10 | Gm44777 | 0.003 | Young |
| 0.787813 | 0.187276 | 0.061 | 0.054 | 1 | 10 | Kank2 | 0.007 | Young |
| 0.788176 | 0.269794 | 0.222 | 0.208 | 1 | 10 | Epb41 | 0.014 | Young |
| 0.788218 | 0.165259 | 0.636 | 0.675 | 1 | 10 | Kif5b | -0.039 | Young |
| 0.788565 | 0.260946 | 0.263 | 0.254 | 1 | 10 | Phc2 | 0.009 | Young |
| 0.78917 | 0.427487 | 0.152 | 0.144 | 1 | 10 | Srsf4 | 0.008 | Young |
| 0.789373 | 0.57785 | 0.091 | 0.083 | 1 | 10 | Rgs3 | 0.008 | Young |
| 0.789552 | 0.02336 | 0.051 | 0.044 | 1 | 10 | Tbc1d8b | 0.007 | Young |
| 0.789646 | 0.310778 | 0.434 | 0.452 | 1 | 10 | Zfp106 | -0.018 | Young |
| 0.790453 | 0.046433 | 0.586 | 0.653 | 1 | 10 | Tspo | -0.067 | Young |
| 0.790636 | 0.343988 | 0.131 | 0.144 | 1 | 10 | Ppp2cb | -0.013 | Young |
| 0.790695 | 0.201557 | 0.303 | 0.335 | 1 | 10 | Gnl3 | -0.032 | Young |
| 0.790784 | 0.211777 | 0.061 | 0.054 | 1 | 10 | Tmem115 | 0.007 | Young |
| 0.79143 | 0.525648 | 0.03 | 0.037 | 1 | 10 | Cry2 | -0.007 | Young |
| 0.792009 | 0.430433 | 0.192 | 0.193 | 1 | 10 | Epc1 | -0.001 | Young |
| 0.792556 | 0.178388 | 0.222 | 0.247 | 1 | 10 | Nck1 | -0.025 | Young |
| 0.792556 | 0.142529 | 0.222 | 0.247 | 1 | 10 | Ranbp9 | -0.025 | Young |
| 0.792575 | 0.10627 | 0.788 | 0.819 | 1 | 10 | Tln1 | -0.031 | Young |
| 0.793495 | 0.378327 | 0.162 | 0.156 | 1 | 10 | Zcchc9 | 0.006 | Young |
| 0.79359 | 0.056133 | 0.081 | 0.09 | 1 | 10 | Ormdl3 | -0.009 | Young |
| 0.793793 | 0.043032 | 0.303 | 0.33 | 1 | 10 | Prkar2a | -0.027 | Young |
| 0.794015 | 0.034412 | 0.141 | 0.132 | 1 | 10 | Lman2l | 0.009 | Young |
| 0.794345 | 0.234733 | 0.232 | 0.252 | 1 | 10 | Phf20 | -0.02 | Young |
| 0.794675 | 0.149675 | 0.192 | 0.183 | 1 | 10 | Sp110 | 0.009 | Young |
| 0.794962 | 0.048107 | 0.212 | 0.205 | 1 | 10 | Eif2b5 | 0.007 | Young |
| 0.795003 | 0.384573 | 0.061 | 0.068 | 1 | 10 | Lipe | -0.007 | Young |
| 0.795104 | 1.245864 | 0.03 | 0.037 | 1 | 10 | Cep78 | -0.007 | Young |
| 0.795265 | 0.064942 | 0.667 | 0.697 | 1 | 10 | Tmed2 | -0.03 | Young |
| 0.79552 | 0.29004 | 0.232 | 0.23 | 1 | 10 | Zfp516 | 0.002 | Young |
| 0.795552 | 0.442654 | 0.162 | 0.183 | 1 | 10 | Pno1 | -0.021 | Young |
| 0.795669 | 0.164265 | 0.374 | 0.403 | 1 | 10 | Esyt1 | -0.029 | Young |
| 0.795855 | 0.285801 | 0.364 | 0.379 | 1 | 10 | Denr | -0.015 | Young |
| 0.796083 | 0.029262 | 0.04 | 0.046 | 1 | 10 | Rhbdd3 | -0.006 | Young |
| 0.796291 | 0.34626 | 0.071 | 0.081 | 1 | 10 | Tbl3 | -0.01 | Young |
| 0.797013 | 0.36799 | 0.111 | 0.127 | 1 | 10 | Fzr1 | -0.016 | Young |
| 0.797174 | 0.782424 | 0.081 | 0.09 | 1 | 10 | Gpt2 | -0.009 | Young |
| 0.797408 | 0.049785 | 0.222 | 0.24 | 1 | 10 | Glrx | -0.018 | Young |
| 0.79755 | 0.671316 | 0.091 | 0.086 | 1 | 10 | Gm26782 | 0.005 | Young |
| 0.797718 | 0.07029 | 0.051 | 0.044 | 1 | 10 | Atg4a | 0.007 | Young |
| 0.797771 | 0.309413 | 0.101 | 0.095 | 1 | 10 | Cep295 | 0.006 | Young |
| 0.797784 | 0.966127 | 0.081 | 0.076 | 1 | 10 | Il15 | 0.005 | Young |

| | | | | | | | | |
|---|---|---|---|---|---|---|---|---|
| 0.797784 | 0.240138 | 0.071 | 0.078 | 1 | 10 | Ccdc88c | -0.007 | Young |
| 0.798224 | 0.021051 | 0.061 | 0.054 | 1 | 10 | Ggct | 0.007 | Young |
| 0.798421 | 0.154656 | 0.495 | 0.562 | 1 | 10 | Psma6 | -0.067 | Young |
| 0.798515 | 0.328601 | 0.404 | 0.423 | 1 | 10 | Glmp | -0.019 | Young |
| 0.799055 | 0.125779 | 0.111 | 0.103 | 1 | 10 | Zdhhc12 | 0.008 | Young |
| 0.799082 | 0.413774 | 0.061 | 0.068 | 1 | 10 | Smpd4 | -0.007 | Young |
| 0.799265 | 0.515378 | 0.051 | 0.059 | 1 | 10 | Cox7a1 | -0.008 | Young |
| 0.799265 | 0.282739 | 0.051 | 0.059 | 1 | 10 | Fbxl15 | -0.008 | Young |
| 0.799415 | 0.08499 | 0.111 | 0.122 | 1 | 10 | Fopnl | -0.011 | Young |
| 0.799589 | 0.107892 | 0.444 | 0.465 | 1 | 10 | Vma21 | -0.021 | Young |
| 0.799769 | 0.327904 | 0.111 | 0.125 | 1 | 10 | Elmod2 | -0.014 | Young |
| 0.799966 | 0.665936 | 0.091 | 0.086 | 1 | 10 | Ece1 | 0.005 | Young |
| 0.800166 | 0.402894 | 0.091 | 0.103 | 1 | 10 | Arhgap26 | -0.012 | Young |
| 0.800184 | 0.019294 | 0.455 | 0.435 | 1 | 10 | Ndufb6 | 0.02 | Young |
| 0.800302 | 0.314482 | 0.222 | 0.22 | 1 | 10 | Pafah1b2 | 0.002 | Young |
| 0.800492 | 0.551079 | 0.192 | 0.196 | 1 | 10 | Dpp3 | -0.004 | Young |
| 0.800595 | 0.073243 | 0.354 | 0.381 | 1 | 10 | Dennd5a | -0.027 | Young |
| 0.800886 | 0.086358 | 0.556 | 0.572 | 1 | 10 | Rbm47 | -0.016 | Young |
| 0.80121 | 0.151624 | 0.444 | 0.494 | 1 | 10 | Snrpd3 | -0.05 | Young |
| 0.801278 | 0.19404 | 0.111 | 0.103 | 1 | 10 | Narfl | 0.008 | Young |
| 0.801822 | 0.750743 | 0.121 | 0.115 | 1 | 10 | Tprkb | 0.006 | Young |
| 0.801822 | 0.225777 | 0.121 | 0.115 | 1 | 10 | Uspl1 | 0.006 | Young |
| 0.801834 | 0.105726 | 0.616 | 0.643 | 1 | 10 | Pebp1 | -0.027 | Young |
| 0.801969 | 0.03072 | 0.667 | 0.677 | 1 | 10 | Pirb | -0.01 | Young |
| 0.802091 | 0.113793 | 0.485 | 0.501 | 1 | 10 | Dynlrb1 | -0.016 | Young |
| 0.802185 | 0.551977 | 0.101 | 0.112 | 1 | 10 | Rab11fip2 | -0.011 | Young |
| 0.802517 | 0.071193 | 0.596 | 0.65 | 1 | 10 | Eif2s2 | -0.054 | Young |
| 0.802526 | 0.2645 | 0.303 | 0.301 | 1 | 10 | Itgax | 0.002 | Young |
| 0.802813 | 0.313319 | 0.283 | 0.271 | 1 | 10 | Ttc3 | 0.012 | Young |
| 0.802892 | 0.715866 | 0.111 | 0.105 | 1 | 10 | Ahsa2 | 0.006 | Young |
| 0.802892 | 0.204129 | 0.111 | 0.105 | 1 | 10 | Sco2 | 0.006 | Young |
| 0.803085 | 0.785636 | 0.152 | 0.154 | 1 | 10 | Pdcd2l | -0.002 | Young |
| 0.803218 | 0.227269 | 0.293 | 0.32 | 1 | 10 | Mrpl12 | -0.027 | Young |
| 0.803525 | 0.503315 | 0.182 | 0.174 | 1 | 10 | Arpc5l | 0.008 | Young |
| 0.804136 | 0.18752 | 0.02 | 0.024 | 1 | 10 | Fam222a | -0.004 | Young |
| 0.804378 | 0.366736 | 0.141 | 0.137 | 1 | 10 | Slc7a7 | 0.004 | Young |
| 0.804415 | 0.029555 | 0.182 | 0.196 | 1 | 10 | Wars | -0.014 | Young |
| 0.804415 | 0.00388 | 0.202 | 0.191 | 1 | 10 | Ngdn | 0.011 | Young |
| 0.80453 | 0.089356 | 0.061 | 0.068 | 1 | 10 | Abcb8 | -0.007 | Young |
| 0.804611 | 0.067135 | 0.212 | 0.225 | 1 | 10 | Polr2l | -0.013 | Young |
| 0.804772 | 0.618294 | 0.152 | 0.149 | 1 | 10 | Nubp1 | 0.003 | Young |
| 0.804775 | 0.452558 | 0.131 | 0.127 | 1 | 10 | N6amt1 | 0.004 | Young |
| 0.8049 | 0.110103 | 0.283 | 0.315 | 1 | 10 | Fip1l1 | -0.032 | Young |
| 0.805135 | 0.533766 | 0.061 | 0.056 | 1 | 10 | Snx7 | 0.005 | Young |
| 0.805522 | 0.487767 | 0.212 | 0.244 | 1 | 10 | Rnf114 | -0.032 | Young |

| | | | | | | | | |
|---|---|---|---|---|---|---|---|---|
| 0.805907 | 1.018329 | 0.04 | 0.046 | 1 | 10 | Gm46430 | -0.006 | Young |
| 0.806227 | 0.370912 | 0.283 | 0.293 | 1 | 10 | Dctn2 | -0.01 | Young |
| 0.806349 | 0.35088 | 0.141 | 0.137 | 1 | 10 | Med25 | 0.004 | Young |
| 0.806381 | 0.164018 | 0.02 | 0.024 | 1 | 10 | Zgpat | -0.004 | Young |
| 0.806381 | 0.128606 | 0.02 | 0.024 | 1 | 10 | Zfp953 | -0.004 | Young |
| 0.806381 | 0.092183 | 0.02 | 0.024 | 1 | 10 | Zfp512b | -0.004 | Young |
| 0.806381 | 0.061897 | 0.02 | 0.024 | 1 | 10 | Praf2 | -0.004 | Young |
| 0.806418 | 0.108895 | 0.283 | 0.301 | 1 | 10 | Cyc1 | -0.018 | Young |
| 0.806557 | 0.087417 | 0.131 | 0.125 | 1 | 10 | Pcgf3 | 0.006 | Young |
| 0.806575 | 0.558149 | 0.071 | 0.066 | 1 | 10 | Tipin | 0.005 | Young |
| 0.806604 | 1.003971 | 0.061 | 0.056 | 1 | 10 | Znhit3 | 0.005 | Young |
| 0.80678 | 0.051304 | 0.586 | 0.594 | 1 | 10 | Atp5d | -0.008 | Young |
| 0.80685 | 0.386776 | 0.081 | 0.093 | 1 | 10 | Gdpd3 | -0.012 | Young |
| 0.806865 | 0.151326 | 0.192 | 0.183 | 1 | 10 | Cib1 | 0.009 | Young |
| 0.806893 | 0.030128 | 0.374 | 0.396 | 1 | 10 | Dusp11 | -0.022 | Young |
| 0.807468 | 0.500826 | 0.101 | 0.115 | 1 | 10 | Pex3 | -0.014 | Young |
| 0.807698 | 0.358355 | 0.323 | 0.372 | 1 | 10 | Wdfy3 | -0.049 | Young |
| 0.808018 | 0.314821 | 0.152 | 0.166 | 1 | 10 | Pnkd | -0.014 | Young |
| 0.808041 | 0.447325 | 0.364 | 0.362 | 1 | 10 | Hp1bp3 | 0.002 | Young |
| 0.808111 | 0.184121 | 0.212 | 0.232 | 1 | 10 | Lgals8 | -0.02 | Young |
| 0.808181 | 0.134562 | 0.313 | 0.311 | 1 | 10 | Kmt2a | 0.002 | Young |
| 0.808597 | 0.340554 | 0.283 | 0.286 | 1 | 10 | Tap2 | -0.003 | Young |
| 0.808628 | 1.051048 | 0.02 | 0.024 | 1 | 10 | Stab2 | -0.004 | Young |
| 0.808628 | 0.128089 | 0.02 | 0.024 | 1 | 10 | Coil | -0.004 | Young |
| 0.808899 | 0.188779 | 0.253 | 0.271 | 1 | 10 | Pik3ca | -0.018 | Young |
| 0.809149 | 0.370931 | 0.081 | 0.09 | 1 | 10 | Ppp2r3c | -0.009 | Young |
| 0.809292 | 0.489697 | 0.071 | 0.078 | 1 | 10 | Ppp6r2 | -0.007 | Young |
| 0.80939 | 0.19621 | 0.343 | 0.335 | 1 | 10 | Nudc | 0.008 | Young |
| 0.809397 | 0.325419 | 0.343 | 0.333 | 1 | 10 | Adgre5 | 0.01 | Young |
| 0.809398 | 0.176964 | 0.303 | 0.333 | 1 | 10 | Gpcpd1 | -0.03 | Young |
| 0.809447 | 0.157019 | 0.121 | 0.112 | 1 | 10 | Exoc7 | 0.009 | Young |
| 0.809482 | 0.257162 | 0.212 | 0.225 | 1 | 10 | Polr2i | -0.013 | Young |
| 0.809486 | 0.304448 | 0.273 | 0.271 | 1 | 10 | Egln2 | 0.002 | Young |
| 0.809732 | 0.022395 | 0.384 | 0.435 | 1 | 10 | Hpf1 | -0.051 | Young |
| 0.809908 | 0.262614 | 0.242 | 0.232 | 1 | 10 | Atf6b | 0.01 | Young |
| 0.810151 | 0.172222 | 0.646 | 0.711 | 1 | 10 | Atp6v1f | -0.065 | Young |
| 0.810427 | 0.108568 | 0.293 | 0.325 | 1 | 10 | Dennd1b | -0.032 | Young |
| 0.810516 | 0.151221 | 0.505 | 0.579 | 1 | 10 | Tcf25 | -0.074 | Young |
| 0.810623 | 0.270168 | 0.111 | 0.105 | 1 | 10 | Phkb | 0.006 | Young |
| 0.81063 | 0.211326 | 0.121 | 0.117 | 1 | 10 | Agl | 0.004 | Young |
| 0.810655 | 0.174523 | 0.424 | 0.467 | 1 | 10 | Arih1 | -0.043 | Young |
| 0.810669 | 0.185042 | 0.525 | 0.523 | 1 | 10 | Sf1 | 0.002 | Young |
| 0.810876 | 0.097873 | 0.02 | 0.024 | 1 | 10 | Marveld2 | -0.004 | Young |
| 0.811451 | 0.286817 | 0.263 | 0.267 | 1 | 10 | Lpin2 | -0.004 | Young |
| 0.811818 | 0.27207 | 0.192 | 0.191 | 1 | 10 | Glo1 | 0.001 | Young |

| | | | | | | | | |
|---|---|---|---|---|---|---|---|---|
| 0.811991 | 0.343784 | 0.303 | 0.301 | 1 | 10 | Mndal | 0.002 | Young |
| 0.812036 | 0.994611 | 0.071 | 0.066 | 1 | 10 | 2410004B1 | 0.005 | Young |
| 0.812132 | 0.196264 | 0.202 | 0.23 | 1 | 10 | Lars | -0.028 | Young |
| 0.812473 | 0.327446 | 0.04 | 0.046 | 1 | 10 | Tmem35b | -0.006 | Young |
| 0.812473 | 0.16386 | 0.04 | 0.046 | 1 | 10 | Dgka | -0.006 | Young |
| 0.812612 | 0.167215 | 0.152 | 0.166 | 1 | 10 | Cops7a | -0.014 | Young |
| 0.812719 | 0.384276 | 0.061 | 0.068 | 1 | 10 | Cep104 | -0.007 | Young |
| 0.812759 | 0.305885 | 0.202 | 0.196 | 1 | 10 | Ly96 | 0.006 | Young |
| 0.812759 | 0.148093 | 0.263 | 0.262 | 1 | 10 | Maz | 0.001 | Young |
| 0.81292 | 0.223272 | 0.303 | 0.32 | 1 | 10 | Rab35 | -0.017 | Young |
| 0.813126 | 0.0238 | 0.02 | 0.024 | 1 | 10 | Plcg1 | -0.004 | Young |
| 0.813616 | 0.300094 | 0.141 | 0.134 | 1 | 10 | Apmap | 0.007 | Young |
| 0.813616 | 0.167825 | 0.141 | 0.134 | 1 | 10 | Cpped1 | 0.007 | Young |
| 0.813628 | 0.260929 | 0.121 | 0.132 | 1 | 10 | Atxn7l3 | -0.011 | Young |
| 0.813643 | 0.050192 | 0.394 | 0.379 | 1 | 10 | Ddx24 | 0.015 | Young |
| 0.813923 | 0.398066 | 0.141 | 0.156 | 1 | 10 | Smg5 | -0.015 | Young |
| 0.813923 | 0.49825 | 0.141 | 0.156 | 1 | 10 | Tcf7l2 | -0.015 | Young |
| 0.814171 | 0.013455 | 0.919 | 0.941 | 1 | 10 | Rps5 | -0.022 | Young |
| 0.814329 | 0.124258 | 0.323 | 0.311 | 1 | 10 | Cr1l | 0.012 | Young |
| 0.814379 | 0.543529 | 0.172 | 0.169 | 1 | 10 | Tecpr1 | 0.003 | Young |
| 0.814503 | 0.603357 | 0.091 | 0.086 | 1 | 10 | Cd80 | 0.005 | Young |
| 0.814558 | 1.111793 | 0.051 | 0.046 | 1 | 10 | Tmod1 | 0.005 | Young |
| 0.814626 | 0.280186 | 0.192 | 0.186 | 1 | 10 | Exosc5 | 0.006 | Young |
| 0.814673 | 0.306623 | 0.313 | 0.311 | 1 | 10 | Zfp638 | 0.002 | Young |
| 0.815092 | 0.126105 | 0.232 | 0.247 | 1 | 10 | Notch1 | -0.015 | Young |
| 0.815153 | 0.923618 | 0.091 | 0.088 | 1 | 10 | Slc16a1 | 0.003 | Young |
| 0.815201 | 0.438873 | 0.152 | 0.147 | 1 | 10 | Cpq | 0.005 | Young |
| 0.815377 | 0.293556 | 0.02 | 0.024 | 1 | 10 | Thop1 | -0.004 | Young |
| 0.815377 | 0.026982 | 0.02 | 0.024 | 1 | 10 | Yars2 | -0.004 | Young |
| 0.815433 | 1.288951 | 0.051 | 0.059 | 1 | 10 | Phf13 | -0.008 | Young |
| 0.815904 | 0.081169 | 0.434 | 0.474 | 1 | 10 | Map2k3 | -0.04 | Young |
| 0.815925 | 0.333887 | 0.242 | 0.269 | 1 | 10 | Mcmbp | -0.027 | Young |
| 0.816021 | 0.009905 | 0.414 | 0.447 | 1 | 10 | Aes | -0.033 | Young |
| 0.81617 | 0.726365 | 0.051 | 0.046 | 1 | 10 | Zfp839 | 0.005 | Young |
| 0.816316 | 0.039536 | 0.152 | 0.161 | 1 | 10 | Naa10 | -0.009 | Young |
| 0.816356 | 0.544558 | 0.081 | 0.09 | 1 | 10 | Zfp865 | -0.009 | Young |
| 0.816525 | 0.905666 | 0.111 | 0.108 | 1 | 10 | Prmt3 | 0.003 | Young |
| 0.816585 | 0.371357 | 0.283 | 0.279 | 1 | 10 | Fam50a | 0.004 | Young |
| 0.81763 | 0.56532 | 0.02 | 0.024 | 1 | 10 | Pdss1 | -0.004 | Young |
| 0.81763 | 0.563503 | 0.02 | 0.024 | 1 | 10 | Trmt44 | -0.004 | Young |
| 0.81763 | 0.121148 | 0.02 | 0.024 | 1 | 10 | L2hgdh | -0.004 | Young |
| 0.81763 | 0.023957 | 0.02 | 0.024 | 1 | 10 | Oit3 | -0.004 | Young |
| 0.817768 | 0.187538 | 0.121 | 0.115 | 1 | 10 | Dnajb12 | 0.006 | Young |
| 0.81777 | 2.191166 | 0.02 | 0.017 | 1 | 10 | Zgrf1 | 0.003 | Young |
| 0.81777 | 1.847533 | 0.02 | 0.017 | 1 | 10 | Dpt | 0.003 | Young |

| | | | | | | | | |
|---|---|---|---|---|---|---|---|---|
| 0.81777 | 1.799734 | 0.02 | 0.017 | 1 | 10 | Zfp955b | 0.003 | Young |
| 0.81777 | 1.432434 | 0.02 | 0.017 | 1 | 10 | Cd200 | 0.003 | Young |
| 0.81777 | 1.363839 | 0.02 | 0.017 | 1 | 10 | BC048403 | 0.003 | Young |
| 0.81777 | 1.094653 | 0.02 | 0.017 | 1 | 10 | Zfp61 | 0.003 | Young |
| 0.817911 | 0.175747 | 0.303 | 0.311 | 1 | 10 | Ncf4 | -0.008 | Young |
| 0.81798 | 0.406079 | 0.253 | 0.244 | 1 | 10 | Atf7 | 0.009 | Young |
| 0.81812 | 0.295221 | 0.212 | 0.237 | 1 | 10 | Mark3 | -0.025 | Young |
| 0.818147 | 0.420104 | 0.091 | 0.086 | 1 | 10 | Qsox2 | 0.005 | Young |
| 0.818215 | 0.197542 | 0.232 | 0.222 | 1 | 10 | Kpna3 | 0.01 | Young |
| 0.818493 | 0.192598 | 0.414 | 0.462 | 1 | 10 | Raph1 | -0.048 | Young |
| 0.818622 | 0.434088 | 0.162 | 0.161 | 1 | 10 | Txnl4a | 0.001 | Young |
| 0.818692 | 0.229225 | 0.131 | 0.122 | 1 | 10 | Mpv17l2 | 0.009 | Young |
| 0.819107 | 0.316991 | 0.152 | 0.161 | 1 | 10 | Casp8ap2 | -0.009 | Young |
| 0.819557 | 0.58175 | 0.081 | 0.076 | 1 | 10 | Larp1b | 0.005 | Young |
| 0.819652 | 0.178786 | 0.323 | 0.359 | 1 | 10 | Ddx21 | -0.036 | Young |
| 0.819856 | 0.588103 | 0.061 | 0.056 | 1 | 10 | Stambpl1 | 0.005 | Young |
| 0.820102 | 0.261465 | 0.545 | 0.57 | 1 | 10 | Tmem160 | -0.025 | Young |
| 0.820222 | 0.65074 | 0.131 | 0.13 | 1 | 10 | Ppp2r5d | 0.001 | Young |
| 0.820365 | 1.025971 | 0.02 | 0.017 | 1 | 10 | Zfp770 | 0.003 | Young |
| 0.820365 | 0.948523 | 0.02 | 0.017 | 1 | 10 | Fsd2 | 0.003 | Young |
| 0.820365 | 0.708126 | 0.02 | 0.017 | 1 | 10 | Copz2 | 0.003 | Young |
| 0.820617 | 0.845406 | 0.202 | 0.205 | 1 | 10 | Rps6ka1 | -0.003 | Young |
| 0.820671 | 0.41069 | 0.081 | 0.073 | 1 | 10 | Cwc27 | 0.008 | Young |
| 0.820673 | 0.668546 | 0.121 | 0.117 | 1 | 10 | Snapc1 | 0.004 | Young |
| 0.820699 | 0.37564 | 0.04 | 0.046 | 1 | 10 | Zfp143 | -0.006 | Young |
| 0.820699 | 0.280959 | 0.04 | 0.046 | 1 | 10 | Efemp2 | -0.006 | Young |
| 0.820854 | 0.301424 | 0.192 | 0.183 | 1 | 10 | Tbck | 0.009 | Young |
| 0.820969 | 0.296885 | 0.162 | 0.159 | 1 | 10 | Nacc1 | 0.003 | Young |
| 0.821095 | 0.177344 | 0.121 | 0.13 | 1 | 10 | Ap5m1 | -0.009 | Young |
| 0.821243 | 0.152012 | 0.394 | 0.452 | 1 | 10 | Snx27 | -0.058 | Young |
| 0.821305 | 0.064639 | 0.091 | 0.1 | 1 | 10 | Rusc2 | -0.009 | Young |
| 0.821366 | 0.324379 | 0.121 | 0.134 | 1 | 10 | Sap30l | -0.013 | Young |
| 0.82158 | 0.480624 | 0.141 | 0.134 | 1 | 10 | Dcaf15 | 0.007 | Young |
| 0.821732 | 0.605757 | 0.121 | 0.117 | 1 | 10 | Bri3bp | 0.004 | Young |
| 0.821746 | 0.149515 | 0.121 | 0.132 | 1 | 10 | Tmem185b | -0.011 | Young |
| 0.821774 | 0.508169 | 0.232 | 0.24 | 1 | 10 | Ppm1h | -0.008 | Young |
| 0.821783 | 0.467407 | 0.364 | 0.396 | 1 | 10 | H1f0 | -0.032 | Young |
| 0.821855 | 0.921672 | 0.04 | 0.037 | 1 | 10 | Spef1 | 0.003 | Young |
| 0.821932 | 0.197817 | 0.576 | 0.645 | 1 | 10 | Atp5j | -0.069 | Young |
| 0.821981 | 0.396722 | 0.121 | 0.137 | 1 | 10 | Med29 | -0.016 | Young |
| 0.822024 | 0.966703 | 0.111 | 0.108 | 1 | 10 | Mcoln2 | 0.003 | Young |
| 0.82214 | 0.970768 | 0.02 | 0.024 | 1 | 10 | Gm12166 | -0.004 | Young |
| 0.822832 | 0.030578 | 0.323 | 0.335 | 1 | 10 | Tmem168 | -0.012 | Young |
| 0.822963 | 1.045592 | 0.02 | 0.017 | 1 | 10 | Tubb2b | 0.003 | Young |
| 0.822963 | 0.803811 | 0.02 | 0.017 | 1 | 10 | Zfp280b | 0.003 | Young |

| | | | | | | | | |
|---|---|---|---|---|---|---|---|---|
| 0.822963 | 0.465774 | 0.02 | 0.017 | 1 | 10 | Uck2 | 0.003 | Young |
| 0.822984 | 1.632964 | 0.071 | 0.066 | 1 | 10 | Espl1 | 0.005 | Young |
| 0.823012 | 0.527909 | 0.091 | 0.086 | 1 | 10 | Tmem63b | 0.005 | Young |
| 0.823041 | 0.23006 | 0.343 | 0.384 | 1 | 10 | Man1a | -0.041 | Young |
| 0.823385 | 0.915273 | 0.131 | 0.132 | 1 | 10 | Mafk | -0.001 | Young |
| 0.823424 | 0.174589 | 0.232 | 0.227 | 1 | 10 | Nsd2 | 0.005 | Young |
| 0.823564 | 0.271265 | 0.364 | 0.384 | 1 | 10 | Pfdn2 | -0.02 | Young |
| 0.824015 | 0.125971 | 0.202 | 0.215 | 1 | 10 | Uchl5 | -0.013 | Young |
| 0.824397 | 1.039887 | 0.02 | 0.024 | 1 | 10 | Tmem86b | -0.004 | Young |
| 0.824397 | 0.733082 | 0.02 | 0.024 | 1 | 10 | Xist | -0.004 | Young |
| 0.824421 | 0.540833 | 0.162 | 0.181 | 1 | 10 | Avpi1 | -0.019 | Young |
| 0.824752 | 0.176898 | 0.111 | 0.125 | 1 | 10 | Mrpl55 | -0.014 | Young |
| 0.82521 | 0.339852 | 0.273 | 0.281 | 1 | 10 | Rnf115 | -0.008 | Young |
| 0.825237 | 0.415564 | 0.172 | 0.188 | 1 | 10 | Smg7 | -0.016 | Young |
| 0.825343 | 0.501146 | 0.313 | 0.335 | 1 | 10 | Alas1 | -0.022 | Young |
| 0.825405 | 0.032145 | 0.121 | 0.134 | 1 | 10 | Slc39a14 | -0.013 | Young |
| 0.825563 | 1.089111 | 0.02 | 0.017 | 1 | 10 | Zfp607a | 0.003 | Young |
| 0.825563 | 0.936644 | 0.02 | 0.017 | 1 | 10 | Fchsd1 | 0.003 | Young |
| 0.825563 | 0.880146 | 0.02 | 0.017 | 1 | 10 | Tnni3 | 0.003 | Young |
| 0.825563 | 0.496297 | 0.02 | 0.017 | 1 | 10 | Gm49368 | 0.003 | Young |
| 0.825563 | 0.181338 | 0.02 | 0.017 | 1 | 10 | Eri2 | 0.003 | Young |
| 0.825574 | 0.252498 | 0.202 | 0.196 | 1 | 10 | Synrg | 0.006 | Young |
| 0.825639 | 0.409756 | 0.232 | 0.264 | 1 | 10 | U2af1 | -0.032 | Young |
| 0.825714 | 0.097297 | 0.162 | 0.154 | 1 | 10 | Mnat1 | 0.008 | Young |
| 0.825776 | 0.565167 | 0.152 | 0.149 | 1 | 10 | Chpf2 | 0.003 | Young |
| 0.825988 | 0.079426 | 0.071 | 0.078 | 1 | 10 | Rusc1 | -0.007 | Young |
| 0.825988 | 0.026814 | 0.071 | 0.078 | 1 | 10 | Pithd1 | -0.007 | Young |
| 0.826656 | 1.259354 | 0.02 | 0.024 | 1 | 10 | Smco4 | -0.004 | Young |
| 0.826656 | 1.209363 | 0.02 | 0.024 | 1 | 10 | Trip6 | -0.004 | Young |
| 0.826829 | 0.313978 | 0.121 | 0.132 | 1 | 10 | Smpd1 | -0.011 | Young |
| 0.827096 | 0.184659 | 0.172 | 0.169 | 1 | 10 | Swt1 | 0.003 | Young |
| 0.82712 | 0.570782 | 0.141 | 0.137 | 1 | 10 | 5031425E2 | 0.004 | Young |
| 0.827277 | 0.84809 | 0.04 | 0.037 | 1 | 10 | Ppp1r3b | 0.003 | Young |
| 0.827294 | 0.589316 | 0.04 | 0.046 | 1 | 10 | Wrnip1 | -0.006 | Young |
| 0.827468 | 0.825218 | 0.03 | 0.027 | 1 | 10 | Svip | 0.003 | Young |
| 0.827531 | 0.030239 | 0.101 | 0.11 | 1 | 10 | Arl3 | -0.009 | Young |
| 0.82757 | 0.613902 | 0.202 | 0.208 | 1 | 10 | Pptc7 | -0.006 | Young |
| 0.827668 | 0.1129 | 0.727 | 0.778 | 1 | 10 | Tma7 | -0.051 | Young |
| 0.827696 | 0.465118 | 0.374 | 0.406 | 1 | 10 | Sod2 | -0.032 | Young |
| 0.827708 | 0.22731 | 0.152 | 0.144 | 1 | 10 | Armc5 | 0.008 | Young |
| 0.827713 | 0.095007 | 0.313 | 0.342 | 1 | 10 | Lrmp | -0.029 | Young |
| 0.827958 | 0.240584 | 0.242 | 0.262 | 1 | 10 | Phf21a | -0.02 | Young |
| 0.828164 | 1.40642 | 0.02 | 0.017 | 1 | 10 | Zfp169 | 0.003 | Young |
| 0.828164 | 0.604731 | 0.02 | 0.017 | 1 | 10 | 6330418K0 | 0.003 | Young |
| 0.828164 | 0.55474 | 0.02 | 0.017 | 1 | 10 | Wdr25 | 0.003 | Young |

| | | | | | | | | |
|---|---|---|---|---|---|---|---|---|
| 0.828164 | 0.303411 | 0.02 | 0.017 | 1 | 10 | 4930522L1 | 0.003 | Young |
| 0.828309 | 0.484291 | 0.182 | 0.186 | 1 | 10 | Nkap | -0.004 | Young |
| 0.828335 | 0.408225 | 0.101 | 0.112 | 1 | 10 | AI837181 | -0.011 | Young |
| 0.828413 | 0.21793 | 0.222 | 0.24 | 1 | 10 | Kmt2d | -0.018 | Young |
| 0.828676 | 0.372689 | 0.131 | 0.147 | 1 | 10 | Tk2 | -0.016 | Young |
| 0.828704 | 0.457963 | 0.394 | 0.401 | 1 | 10 | Srp19 | -0.007 | Young |
| 0.828957 | 0.326726 | 0.162 | 0.156 | 1 | 10 | Pira2 | 0.006 | Young |
| 0.829204 | 0.160393 | 0.152 | 0.166 | 1 | 10 | Btbd9 | -0.014 | Young |
| 0.829262 | 0.238002 | 0.505 | 0.555 | 1 | 10 | Rab2a | -0.05 | Young |
| 0.829466 | 0.244007 | 0.162 | 0.178 | 1 | 10 | Patl1 | -0.016 | Young |
| 0.829566 | 1.290663 | 0.03 | 0.027 | 1 | 10 | Tmem107 | 0.003 | Young |
| 0.830156 | 0.144734 | 0.384 | 0.401 | 1 | 10 | Pdlim5 | -0.017 | Young |
| 0.830592 | 0.684444 | 0.232 | 0.262 | 1 | 10 | Rnmt | -0.03 | Young |
| 0.830767 | 0.564623 | 0.02 | 0.017 | 1 | 10 | Hp | 0.003 | Young |
| 0.830767 | 0.420755 | 0.02 | 0.017 | 1 | 10 | Gm15247 | 0.003 | Young |
| 0.830767 | 0.301303 | 0.02 | 0.017 | 1 | 10 | Cdkl3 | 0.003 | Young |
| 0.830826 | 0.102703 | 0.414 | 0.428 | 1 | 10 | Dnm2 | -0.014 | Young |
| 0.830933 | 0.056095 | 0.081 | 0.088 | 1 | 10 | Prcc | -0.007 | Young |
| 0.831227 | 0.389355 | 0.162 | 0.159 | 1 | 10 | Cops3 | 0.003 | Young |
| 0.831664 | 1.26367 | 0.03 | 0.027 | 1 | 10 | Insl6 | 0.003 | Young |
| 0.83168 | 0.375236 | 0.061 | 0.056 | 1 | 10 | Bcl9l | 0.005 | Young |
| 0.8317 | 0.332652 | 0.111 | 0.105 | 1 | 10 | Gpn1 | 0.006 | Young |
| 0.831878 | 0.102 | 0.848 | 0.892 | 1 | 10 | Npm1 | -0.044 | Young |
| 0.831968 | 0.122313 | 0.354 | 0.347 | 1 | 10 | Kcnq1ot1 | 0.007 | Young |
| 0.83223 | 0.422595 | 0.101 | 0.098 | 1 | 10 | Vgll4 | 0.003 | Young |
| 0.832615 | 0.14758 | 0.192 | 0.21 | 1 | 10 | Galnt11 | -0.018 | Young |
| 0.83316 | 0.749908 | 0.061 | 0.056 | 1 | 10 | Prnp | 0.005 | Young |
| 0.833372 | 1.281467 | 0.02 | 0.017 | 1 | 10 | Nphp3 | 0.003 | Young |
| 0.833372 | 0.349529 | 0.02 | 0.017 | 1 | 10 | Cracr2a | 0.003 | Young |
| 0.833372 | 0.287322 | 0.02 | 0.017 | 1 | 10 | Ugt1a6a | 0.003 | Young |
| 0.833372 | 0.227075 | 0.02 | 0.017 | 1 | 10 | Gm15859 | 0.003 | Young |
| 0.833372 | 0.209689 | 0.02 | 0.017 | 1 | 10 | Nr6a1 | 0.003 | Young |
| 0.833373 | 0.778368 | 0.081 | 0.088 | 1 | 10 | Pak4 | -0.007 | Young |
| 0.833392 | 0.212465 | 0.141 | 0.154 | 1 | 10 | Trak1 | -0.013 | Young |
| 0.833763 | 1.14103 | 0.03 | 0.027 | 1 | 10 | 4931414P1 | 0.003 | Young |
| 0.833763 | 1.02698 | 0.03 | 0.027 | 1 | 10 | Mkks | 0.003 | Young |
| 0.833901 | 0.511701 | 0.04 | 0.046 | 1 | 10 | Mlst8 | -0.006 | Young |
| 0.833932 | 0.01468 | 0.111 | 0.12 | 1 | 10 | Maml3 | -0.009 | Young |
| 0.833936 | 0.232858 | 0.404 | 0.428 | 1 | 10 | Il1rn | -0.024 | Young |
| 0.833958 | 0.219086 | 0.091 | 0.1 | 1 | 10 | Fmnl3 | -0.009 | Young |
| 0.833993 | 0.017749 | 0.586 | 0.638 | 1 | 10 | Bach1 | -0.052 | Young |
| 0.834936 | 0.040766 | 0.051 | 0.056 | 1 | 10 | Dcbld2 | -0.005 | Young |
| 0.835048 | 0.102798 | 0.434 | 0.452 | 1 | 10 | Pdia4 | -0.018 | Young |
| 0.835205 | 0.268872 | 0.091 | 0.086 | 1 | 10 | Nob1 | 0.005 | Young |
| 0.835342 | 0.521218 | 0.071 | 0.066 | 1 | 10 | Dlg4 | 0.005 | Young |

| | | | | | | | | |
|---|---|---|---|---|---|---|---|---|
| 0.835561 | 0.137872 | 0.141 | 0.134 | 1 | 10 | C1galt1c1 | 0.007 | Young |
| 0.835567 | 0.986137 | 0.051 | 0.046 | 1 | 10 | A930007I1 | 0.005 | Young |
| 0.835846 | 0.053083 | 0.141 | 0.152 | 1 | 10 | Klf11 | -0.011 | Young |
| 0.835864 | 1.462427 | 0.03 | 0.027 | 1 | 10 | Ccdc73 | 0.003 | Young |
| 0.835864 | 0.69178 | 0.03 | 0.027 | 1 | 10 | Cenpj | 0.003 | Young |
| 0.8359 | 0.646802 | 0.162 | 0.159 | 1 | 10 | Ncbp1 | 0.003 | Young |
| 0.835979 | 0.905526 | 0.02 | 0.017 | 1 | 10 | Intu | 0.003 | Young |
| 0.835979 | 0.292063 | 0.02 | 0.017 | 1 | 10 | Plag1 | 0.003 | Young |
| 0.835979 | 0.196836 | 0.02 | 0.017 | 1 | 10 | Abhd8 | 0.003 | Young |
| 0.835979 | 0.169313 | 0.02 | 0.017 | 1 | 10 | Fam135a | 0.003 | Young |
| 0.835979 | 0.148683 | 0.02 | 0.017 | 1 | 10 | Prdm9 | 0.003 | Young |
| 0.83602 | 0.203094 | 0.323 | 0.347 | 1 | 10 | Khsrp | -0.024 | Young |
| 0.836152 | 0.137399 | 0.283 | 0.303 | 1 | 10 | Smim10l1 | -0.02 | Young |
| 0.836224 | 0.098165 | 0.455 | 0.474 | 1 | 10 | Sumo3 | -0.019 | Young |
| 0.83636 | 0.402256 | 0.111 | 0.108 | 1 | 10 | Cacfd1 | 0.003 | Young |
| 0.83636 | 0.021816 | 0.101 | 0.11 | 1 | 10 | Lmf2 | -0.009 | Young |
| 0.836492 | 0.878151 | 0.162 | 0.181 | 1 | 10 | Ero1l | -0.019 | Young |
| 0.836669 | 0.006551 | 0.364 | 0.362 | 1 | 10 | Mien1 | 0.002 | Young |
| 0.836912 | 0.259538 | 0.283 | 0.289 | 1 | 10 | Cacul1 | -0.006 | Young |
| 0.837209 | 0.975876 | 0.04 | 0.046 | 1 | 10 | Ercc8 | -0.006 | Young |
| 0.837212 | 0.365542 | 0.152 | 0.152 | 1 | 10 | Pggt1b | 0 | Young |
| 0.837577 | 0.351218 | 0.202 | 0.196 | 1 | 10 | Erh | 0.006 | Young |
| 0.837693 | 0.213633 | 0.515 | 0.545 | 1 | 10 | Eno1 | -0.03 | Young |
| 0.837717 | 0.145402 | 0.444 | 0.518 | 1 | 10 | Cd48 | -0.074 | Young |
| 0.837787 | 0.141741 | 0.232 | 0.254 | 1 | 10 | Fgfr1 | -0.022 | Young |
| 0.837966 | 0.313843 | 0.03 | 0.027 | 1 | 10 | Optn | 0.003 | Young |
| 0.838137 | 0.060803 | 0.374 | 0.399 | 1 | 10 | Srpr | -0.025 | Young |
| 0.838173 | 0.027585 | 0.111 | 0.103 | 1 | 10 | Ttc19 | 0.008 | Young |
| 0.838547 | 0.040425 | 0.273 | 0.262 | 1 | 10 | Ei24 | 0.011 | Young |
| 0.838588 | 0.123451 | 0.02 | 0.017 | 1 | 10 | Lrrc1 | 0.003 | Young |
| 0.838885 | 0.140033 | 0.081 | 0.076 | 1 | 10 | Aen | 0.005 | Young |
| 0.839084 | 0.113795 | 0.798 | 0.841 | 1 | 10 | Rbm39 | -0.043 | Young |
| 0.839473 | 0.413238 | 0.313 | 0.323 | 1 | 10 | Dctn4 | -0.01 | Young |
| 0.839667 | 0.034572 | 0.091 | 0.083 | 1 | 10 | Mad2l1bp | 0.008 | Young |
| 0.839675 | 0.604237 | 0.222 | 0.222 | 1 | 10 | Slc20a1 | 0 | Young |
| 0.839736 | 0.66123 | 0.263 | 0.267 | 1 | 10 | Vbp1 | -0.004 | Young |
| 0.839795 | 0.473799 | 0.263 | 0.271 | 1 | 10 | Dnajc1 | -0.008 | Young |
| 0.839958 | 1.404783 | 0.04 | 0.037 | 1 | 10 | Ripor2 | 0.003 | Young |
| 0.840069 | 0.930222 | 0.03 | 0.027 | 1 | 10 | Rhov | 0.003 | Young |
| 0.840069 | 0.667895 | 0.03 | 0.027 | 1 | 10 | Oxld1 | 0.003 | Young |
| 0.840081 | 0.355682 | 0.121 | 0.132 | 1 | 10 | Psmg2 | -0.011 | Young |
| 0.840245 | 0.21166 | 0.364 | 0.377 | 1 | 10 | Rab5a | -0.013 | Young |
| 0.840393 | 0.167191 | 0.111 | 0.122 | 1 | 10 | Prmt2 | -0.011 | Young |
| 0.840433 | 0.478949 | 0.051 | 0.046 | 1 | 10 | Zmym3 | 0.005 | Young |
| 0.840573 | 0.717837 | 0.061 | 0.056 | 1 | 10 | Commd9 | 0.005 | Young |

| | | | | | | | | |
|---|---|---|---|---|---|---|---|---|
| 0.84059 | 0.037109 | 0.253 | 0.252 | 1 | 10 | Mtmr6 | 0.001 | Young |
| 0.84091 | 0.107585 | 0.545 | 0.57 | 1 | 10 | Rilpl2 | -0.025 | Young |
| 0.841199 | 0.048976 | 0.02 | 0.017 | 1 | 10 | Aurkb | 0.003 | Young |
| 0.841199 | 0.03915 | 0.02 | 0.017 | 1 | 10 | Zbtb46 | 0.003 | Young |
| 0.841484 | 0.381847 | 0.101 | 0.112 | 1 | 10 | Cdc16 | -0.011 | Young |
| 0.841712 | 0.140607 | 0.313 | 0.335 | 1 | 10 | Il3ra | -0.022 | Young |
| 0.841889 | 0.1553 | 0.434 | 0.477 | 1 | 10 | Fnip1 | -0.043 | Young |
| 0.842056 | 1.005147 | 0.051 | 0.046 | 1 | 10 | Ftsj1 | 0.005 | Young |
| 0.842064 | 0.411652 | 0.263 | 0.269 | 1 | 10 | Kat2b | -0.006 | Young |
| 0.842173 | 0.396286 | 0.03 | 0.027 | 1 | 10 | Ccdc69 | 0.003 | Young |
| 0.84248 | 0.059721 | 0.131 | 0.142 | 1 | 10 | Nop16 | -0.011 | Young |
| 0.842768 | 0.162343 | 0.172 | 0.166 | 1 | 10 | Milr1 | 0.006 | Young |
| 0.843589 | 0.452507 | 0.04 | 0.037 | 1 | 10 | Dimt1 | 0.003 | Young |
| 0.84368 | 0.104152 | 0.051 | 0.046 | 1 | 10 | Hus1 | 0.005 | Young |
| 0.843926 | 0.134903 | 0.172 | 0.186 | 1 | 10 | Fam241a | -0.014 | Young |
| 0.844259 | 0.162458 | 0.354 | 0.381 | 1 | 10 | Map4 | -0.027 | Young |
| 0.844293 | 0.700566 | 0.061 | 0.068 | 1 | 10 | St3gal6 | -0.007 | Young |
| 0.84436 | 0.303657 | 0.192 | 0.186 | 1 | 10 | Snrk | 0.006 | Young |
| 0.84437 | 0.198576 | 0.636 | 0.724 | 1 | 10 | Bola2 | -0.088 | Young |
| 0.844627 | 0.202699 | 0.202 | 0.198 | 1 | 10 | Wdr43 | 0.004 | Young |
| 0.845027 | 0.061512 | 0.061 | 0.056 | 1 | 10 | Fbxo21 | 0.005 | Young |
| 0.845106 | 0.230396 | 0.121 | 0.117 | 1 | 10 | Mettl6 | 0.004 | Young |
| 0.845305 | 0.531028 | 0.051 | 0.046 | 1 | 10 | Ddx20 | 0.005 | Young |
| 0.84535 | 0.15208 | 0.081 | 0.076 | 1 | 10 | Scap | 0.005 | Young |
| 0.845406 | 0.457223 | 0.04 | 0.037 | 1 | 10 | Tnnt1 | 0.003 | Young |
| 0.845406 | 0.335991 | 0.04 | 0.037 | 1 | 10 | Agmo | 0.003 | Young |
| 0.845406 | 0.24826 | 0.04 | 0.037 | 1 | 10 | Alyref2 | 0.003 | Young |
| 0.845406 | 0.202143 | 0.04 | 0.037 | 1 | 10 | Zfp53 | 0.003 | Young |
| 0.845495 | 0.60647 | 0.051 | 0.056 | 1 | 10 | Coq6 | -0.005 | Young |
| 0.845498 | 0.410156 | 0.091 | 0.1 | 1 | 10 | Vipas39 | -0.009 | Young |
| 0.84552 | 0.033868 | 0.131 | 0.139 | 1 | 10 | Dopey1 | -0.008 | Young |
| 0.845781 | 0.16939 | 0.545 | 0.601 | 1 | 10 | Nfkb1 | -0.056 | Young |
| 0.845906 | 0.176069 | 0.182 | 0.176 | 1 | 10 | Prpf31 | 0.006 | Young |
| 0.846361 | 0.300297 | 0.071 | 0.066 | 1 | 10 | Yaf2 | 0.005 | Young |
| 0.846361 | 0.287328 | 0.071 | 0.066 | 1 | 10 | Myc | 0.005 | Young |
| 0.846385 | 0.610263 | 0.03 | 0.027 | 1 | 10 | Cntrob | 0.003 | Young |
| 0.846592 | 0.142305 | 0.495 | 0.545 | 1 | 10 | Ost4 | -0.05 | Young |
| 0.846976 | 0.560403 | 0.101 | 0.112 | 1 | 10 | Nudt19 | -0.011 | Young |
| 0.847034 | 0.01827 | 0.172 | 0.181 | 1 | 10 | Ctbp2 | -0.009 | Young |
| 0.847151 | 0.661063 | 0.04 | 0.046 | 1 | 10 | Il18 | -0.006 | Young |
| 0.847434 | 0.139087 | 0.131 | 0.142 | 1 | 10 | Hmbs | -0.011 | Young |
| 0.847438 | 0.810878 | 0.162 | 0.161 | 1 | 10 | Braf | 0.001 | Young |
| 0.847705 | 0.116236 | 0.313 | 0.337 | 1 | 10 | Pdcl3 | -0.024 | Young |
| 0.847748 | 0.056846 | 0.333 | 0.352 | 1 | 10 | Leng8 | -0.019 | Young |
| 0.847767 | 0.036937 | 0.162 | 0.156 | 1 | 10 | Tmx4 | 0.006 | Young |

| | | | | | | | | |
|---|---|---|---|---|---|---|---|---|
| 0.847939 | 0.294423 | 0.081 | 0.076 | 1 | 10 | Med19 | 0.005 | Young |
| 0.847939 | 0.259889 | 0.081 | 0.076 | 1 | 10 | Ints10 | 0.005 | Young |
| 0.848493 | 0.946061 | 0.03 | 0.027 | 1 | 10 | Ehf | 0.003 | Young |
| 0.848493 | 0.286328 | 0.03 | 0.027 | 1 | 10 | Ccdc126 | 0.003 | Young |
| 0.848493 | 0.113066 | 0.03 | 0.027 | 1 | 10 | Arl15 | 0.003 | Young |
| 0.848517 | 0.320365 | 0.051 | 0.056 | 1 | 10 | Brpf3 | -0.005 | Young |
| 0.848736 | 0.189724 | 0.697 | 0.763 | 1 | 10 | Ostf1 | -0.066 | Young |
| 0.849042 | 0.172245 | 0.04 | 0.037 | 1 | 10 | 2410131K1 | 0.003 | Young |
| 0.849042 | 0.023958 | 0.04 | 0.037 | 1 | 10 | Dars2 | 0.003 | Young |
| 0.849727 | 0.33409 | 0.212 | 0.21 | 1 | 10 | Katna1 | 0.002 | Young |
| 0.849793 | 0.084677 | 0.545 | 0.621 | 1 | 10 | Bnip2 | -0.076 | Young |
| 0.849907 | 0.025582 | 0.98 | 0.978 | 1 | 10 | Rpl13 | 0.002 | Young |
| 0.850183 | 0.635787 | 0.051 | 0.046 | 1 | 10 | Gtf2h2 | 0.005 | Young |
| 0.85035 | 0.121664 | 0.303 | 0.323 | 1 | 10 | Taf6l | -0.02 | Young |
| 0.850601 | 0.133221 | 0.03 | 0.027 | 1 | 10 | Uaca | 0.003 | Young |
| 0.850601 | 0.119394 | 0.03 | 0.027 | 1 | 10 | Slc17a9 | 0.003 | Young |
| 0.850836 | 0.114882 | 0.232 | 0.222 | 1 | 10 | Mtmr3 | 0.01 | Young |
| 0.850861 | 0.2931 | 0.04 | 0.037 | 1 | 10 | Fpgt | 0.003 | Young |
| 0.851078 | 0.031736 | 0.404 | 0.396 | 1 | 10 | Sugt1 | 0.008 | Young |
| 0.851107 | 0.464426 | 0.091 | 0.098 | 1 | 10 | Fastk | -0.007 | Young |
| 0.85123 | 0.044632 | 0.061 | 0.066 | 1 | 10 | Ccdc130 | -0.005 | Young |
| 0.851296 | 0.489056 | 0.162 | 0.181 | 1 | 10 | Ccdc82 | -0.019 | Young |
| 0.851528 | 0.086755 | 0.162 | 0.174 | 1 | 10 | Ammecr1l | -0.012 | Young |
| 0.851903 | 0.448015 | 0.182 | 0.183 | 1 | 10 | Bst1 | -0.001 | Young |
| 0.851925 | 0.071325 | 0.515 | 0.555 | 1 | 10 | Ndufs6 | -0.04 | Young |
| 0.852057 | 0.115334 | 0.192 | 0.203 | 1 | 10 | Kif13b | -0.011 | Young |
| 0.852131 | 0.90119 | 0.04 | 0.046 | 1 | 10 | Zfp943 | -0.006 | Young |
| 0.852277 | 0.19003 | 0.283 | 0.308 | 1 | 10 | Mga | -0.025 | Young |
| 0.852397 | 0.039727 | 0.394 | 0.411 | 1 | 10 | Ldlr | -0.017 | Young |
| 0.852406 | 0.065212 | 0.364 | 0.367 | 1 | 10 | Ufm1 | -0.003 | Young |
| 0.852484 | 0.238498 | 0.162 | 0.156 | 1 | 10 | Scrib | 0.006 | Young |
| 0.852681 | 0.607191 | 0.04 | 0.037 | 1 | 10 | Mettl25 | 0.003 | Young |
| 0.852681 | 0.228681 | 0.04 | 0.037 | 1 | 10 | Rabep2 | 0.003 | Young |
| 0.852711 | 0.094505 | 0.03 | 0.027 | 1 | 10 | Gba2 | 0.003 | Young |
| 0.852711 | 0.025776 | 0.03 | 0.027 | 1 | 10 | Tsr2 | 0.003 | Young |
| 0.852842 | 0.0248 | 0.141 | 0.149 | 1 | 10 | Mtr | -0.008 | Young |
| 0.852963 | 0.02512 | 0.212 | 0.218 | 1 | 10 | Nhlrc3 | -0.006 | Young |
| 0.853129 | 0.495136 | 0.131 | 0.147 | 1 | 10 | Golga5 | -0.016 | Young |
| 0.85395 | 0.274482 | 0.061 | 0.056 | 1 | 10 | Fpgs | 0.005 | Young |
| 0.854155 | 0.506187 | 0.121 | 0.12 | 1 | 10 | Slc12a9 | 0.001 | Young |
| 0.854163 | 0.588373 | 0.081 | 0.078 | 1 | 10 | Rev3l | 0.003 | Young |
| 0.854265 | 0.10499 | 0.364 | 0.403 | 1 | 10 | Pik3r1 | -0.039 | Young |
| 0.854381 | 0.172873 | 0.414 | 0.433 | 1 | 10 | Dhx15 | -0.019 | Young |
| 0.854502 | 0.21016 | 0.04 | 0.037 | 1 | 10 | Sbsn | 0.003 | Young |
| 0.854577 | 0.138616 | 0.101 | 0.108 | 1 | 10 | Chmp7 | -0.007 | Young |

| | | | | | | | | |
|---|---|---|---|---|---|---|---|---|
| 0.854822 | 0.154756 | 0.03 | 0.027 | 1 | 10 | Mmachc | 0.003 | Young |
| 0.854992 | 0.077123 | 0.434 | 0.457 | 1 | 10 | Sel1l | -0.023 | Young |
| 0.855066 | 0.352853 | 0.051 | 0.046 | 1 | 10 | Ercc1 | 0.005 | Young |
| 0.855207 | 0.76948 | 0.071 | 0.068 | 1 | 10 | Clcn6 | 0.003 | Young |
| 0.855439 | 0.068396 | 0.061 | 0.056 | 1 | 10 | Dis3l | 0.005 | Young |
| 0.856185 | 0.678178 | 0.03 | 0.034 | 1 | 10 | Tlr3 | -0.004 | Young |
| 0.856229 | 0.71036 | 0.111 | 0.125 | 1 | 10 | Fam173b | -0.014 | Young |
| 0.85624 | 0.133465 | 0.232 | 0.252 | 1 | 10 | Nab2 | -0.02 | Young |
| 0.856324 | 0.412216 | 0.04 | 0.037 | 1 | 10 | Cyfip2 | 0.003 | Young |
| 0.856928 | 0.09851 | 0.061 | 0.056 | 1 | 10 | Alg13 | 0.005 | Young |
| 0.856933 | 0.628638 | 0.03 | 0.027 | 1 | 10 | C77080 | 0.003 | Young |
| 0.857013 | 0.902316 | 0.071 | 0.078 | 1 | 10 | Nxt1 | -0.007 | Young |
| 0.857404 | 0.361082 | 0.172 | 0.176 | 1 | 10 | Tmem68 | -0.004 | Young |
| 0.857424 | 0.006213 | 0.303 | 0.328 | 1 | 10 | Rab1b | -0.025 | Young |
| 0.857534 | 0.342485 | 0.202 | 0.22 | 1 | 10 | Tbc1d15 | -0.018 | Young |
| 0.857694 | 0.094631 | 0.465 | 0.511 | 1 | 10 | Tomm22 | -0.046 | Young |
| 0.858107 | 0.272459 | 0.03 | 0.034 | 1 | 10 | Fam177a | -0.004 | Young |
| 0.85812 | 1.149815 | 0.091 | 0.098 | 1 | 10 | Bmp2 | -0.007 | Young |
| 0.858311 | 0.544967 | 0.071 | 0.078 | 1 | 10 | Ampd2 | -0.007 | Young |
| 0.858311 | 0.135191 | 0.081 | 0.076 | 1 | 10 | Ercc6l2 | 0.005 | Young |
| 0.858364 | 0.121434 | 0.374 | 0.389 | 1 | 10 | Cmtm7 | -0.015 | Young |
| 0.858418 | 1.028787 | 0.061 | 0.056 | 1 | 10 | Styx | 0.005 | Young |
| 0.858562 | 0.18391 | 0.293 | 0.289 | 1 | 10 | Vps36 | 0.004 | Young |
| 0.858811 | 0.221561 | 0.101 | 0.098 | 1 | 10 | Chpt1 | 0.003 | Young |
| 0.858984 | 0.321595 | 0.121 | 0.117 | 1 | 10 | Kdm5b | 0.004 | Young |
| 0.859134 | 0.200547 | 0.212 | 0.232 | 1 | 10 | Washc5 | -0.02 | Young |
| 0.859187 | 0.020999 | 1 | 0.993 | 1 | 10 | Rps8 | 0.007 | Young |
| 0.859584 | 0.239037 | 0.232 | 0.23 | 1 | 10 | Iws1 | 0.002 | Young |
| 0.859614 | 0.374989 | 0.192 | 0.222 | 1 | 10 | Armc8 | -0.03 | Young |
| 0.859829 | 0.353031 | 0.273 | 0.286 | 1 | 10 | Mrps18c | -0.013 | Young |
| 0.859854 | 0.217103 | 0.212 | 0.227 | 1 | 10 | Mgat5 | -0.015 | Young |
| 0.859951 | 0.123548 | 0.354 | 0.35 | 1 | 10 | Sdhb | 0.004 | Young |
| 0.859955 | 0.786945 | 0.051 | 0.046 | 1 | 10 | Cpeb3 | 0.005 | Young |
| 0.859955 | 0.049539 | 0.051 | 0.046 | 1 | 10 | Atf5 | 0.005 | Young |
| 0.85997 | 0.416731 | 0.101 | 0.098 | 1 | 10 | Zfp236 | 0.003 | Young |
| 0.860031 | 0.464004 | 0.03 | 0.034 | 1 | 10 | Spic | -0.004 | Young |
| 0.860031 | 0.053806 | 0.03 | 0.034 | 1 | 10 | Abca2 | -0.004 | Young |
| 0.860095 | 0.129513 | 0.212 | 0.22 | 1 | 10 | Tlr1 | -0.008 | Young |
| 0.860908 | 0.126105 | 0.081 | 0.076 | 1 | 10 | Alg14 | 0.005 | Young |
| 0.860987 | 0.400956 | 0.162 | 0.156 | 1 | 10 | Tmem170b | 0.006 | Young |
| 0.861022 | 0.291255 | 0.182 | 0.174 | 1 | 10 | Prdm2 | 0.008 | Young |
| 0.86103 | 0.15891 | 0.192 | 0.193 | 1 | 10 | Gtf2h1 | -0.001 | Young |
| 0.861044 | 0.198351 | 0.061 | 0.066 | 1 | 10 | Grpel2 | -0.005 | Young |
| 0.86113 | 0.534328 | 0.101 | 0.098 | 1 | 10 | Sos1 | 0.003 | Young |
| 0.86116 | 0.590839 | 0.03 | 0.027 | 1 | 10 | Zbtb12 | 0.003 | Young |

| | | | | | | | | |
|---|---|---|---|---|---|---|---|---|
| 0.86116 | 0.128514 | 0.03 | 0.027 | 1 | 10 | Platr25 | 0.003 | Young |
| 0.861372 | 0.381065 | 0.323 | 0.333 | 1 | 10 | Pcmt1 | -0.01 | Young |
| 0.861442 | 0.092483 | 0.364 | 0.391 | 1 | 10 | Cyb5r3 | -0.027 | Young |
| 0.861648 | 0.398821 | 0.212 | 0.237 | 1 | 10 | Dph3 | -0.025 | Young |
| 0.861658 | 0.242154 | 0.111 | 0.12 | 1 | 10 | BC029722 | -0.009 | Young |
| 0.861831 | 0.285541 | 0.212 | 0.208 | 1 | 10 | Mettl23 | 0.004 | Young |
| 0.861991 | 0.337526 | 0.283 | 0.303 | 1 | 10 | Commd7 | -0.02 | Young |
| 0.862259 | 0.210772 | 0.172 | 0.181 | 1 | 10 | Cnot8 | -0.009 | Young |
| 0.862564 | 0.368259 | 0.232 | 0.252 | 1 | 10 | Map2k7 | -0.02 | Young |
| 0.862968 | 0.842834 | 0.111 | 0.108 | 1 | 10 | Gmpr2 | 0.003 | Young |
| 0.863109 | 0.099098 | 0.152 | 0.161 | 1 | 10 | Elof1 | -0.009 | Young |
| 0.863217 | 0.131451 | 0.051 | 0.046 | 1 | 10 | 5031425F1 | 0.005 | Young |
| 0.863441 | 0.36509 | 0.455 | 0.474 | 1 | 10 | Map2k2 | -0.019 | Young |
| 0.863772 | 0.259259 | 0.343 | 0.367 | 1 | 10 | Rp2 | -0.024 | Young |
| 0.864009 | 0.137954 | 0.081 | 0.088 | 1 | 10 | Zfp954 | -0.007 | Young |
| 0.864016 | 0.065339 | 0.01 | 0.012 | 1 | 10 | Opn3 | -0.002 | Young |
| 0.864016 | 0.006005 | 0.01 | 0.012 | 1 | 10 | Pik3ip1 | -0.002 | Young |
| 0.864131 | 0.324301 | 0.131 | 0.144 | 1 | 10 | Mfap1a | -0.013 | Young |
| 0.864192 | 0.045198 | 0.394 | 0.416 | 1 | 10 | Stt3a | -0.022 | Young |
| 0.864321 | 0.289457 | 0.131 | 0.142 | 1 | 10 | Abtb1 | -0.011 | Young |
| 0.864383 | 0.239217 | 0.061 | 0.056 | 1 | 10 | Cul9 | 0.005 | Young |
| 0.864541 | 0.118352 | 0.222 | 0.244 | 1 | 10 | Ulk2 | -0.022 | Young |
| 0.864849 | 1.015576 | 0.051 | 0.046 | 1 | 10 | Dlg3 | 0.005 | Young |
| 0.864849 | 0.09028 | 0.051 | 0.046 | 1 | 10 | Ppp2r3a | 0.005 | Young |
| 0.865252 | 0.174528 | 0.455 | 0.484 | 1 | 10 | Hmgn1 | -0.029 | Young |
| 0.865317 | 0.284497 | 0.131 | 0.142 | 1 | 10 | Aspscr1 | -0.011 | Young |
| 0.865579 | 0.061918 | 0.838 | 0.858 | 1 | 10 | Mob1a | -0.02 | Young |
| 0.865668 | 0.357005 | 0.253 | 0.257 | 1 | 10 | Tlnrd1 | -0.004 | Young |
| 0.865806 | 1.115246 | 0.03 | 0.034 | 1 | 10 | Polr3g | -0.004 | Young |
| 0.865806 | 0.319028 | 0.03 | 0.034 | 1 | 10 | Nlrc3 | -0.004 | Young |
| 0.865876 | 0.0366 | 0.061 | 0.056 | 1 | 10 | Emc9 | 0.005 | Young |
| 0.866043 | 0.041778 | 0.232 | 0.225 | 1 | 10 | Ndufs8 | 0.007 | Young |
| 0.86619 | 0.220081 | 0.202 | 0.2 | 1 | 10 | Ddx39 | 0.002 | Young |
| 0.866305 | 0.652376 | 0.111 | 0.108 | 1 | 10 | Rnf167 | 0.003 | Young |
| 0.866357 | 0.036501 | 0.131 | 0.127 | 1 | 10 | Phf12 | 0.004 | Young |
| 0.866585 | 0.040966 | 0.677 | 0.68 | 1 | 10 | Smdt1 | -0.003 | Young |
| 0.866807 | 0.280857 | 0.273 | 0.306 | 1 | 10 | Kidins220 | -0.033 | Young |
| 0.86693 | 0.424297 | 0.232 | 0.235 | 1 | 10 | Otud4 | -0.003 | Young |
| 0.867206 | 0.420432 | 0.091 | 0.088 | 1 | 10 | Ubr7 | 0.003 | Young |
| 0.867223 | 0.529188 | 0.01 | 0.012 | 1 | 10 | Cdc42bpa | -0.002 | Young |
| 0.867223 | 0.406773 | 0.01 | 0.012 | 1 | 10 | Gm48420 | -0.002 | Young |
| 0.867223 | 0.380598 | 0.01 | 0.012 | 1 | 10 | Arhgef5 | -0.002 | Young |
| 0.867223 | 0.2636 | 0.01 | 0.012 | 1 | 10 | Gtf2ird1 | -0.002 | Young |
| 0.867223 | 0.254038 | 0.01 | 0.012 | 1 | 10 | Tmem216 | -0.002 | Young |
| 0.867223 | 0.171514 | 0.01 | 0.012 | 1 | 10 | Zcchc3 | -0.002 | Young |

| | | | | | | | | |
|---|---|---|---|---|---|---|---|---|
| 0.867223 | 0.114428 | 0.01 | 0.012 | 1 | 10 | Timeless | -0.002 | Young |
| 0.867223 | 0.091322 | 0.01 | 0.012 | 1 | 10 | Tm7sf2 | -0.002 | Young |
| 0.867223 | 0.059285 | 0.01 | 0.012 | 1 | 10 | Ager | -0.002 | Young |
| 0.867223 | 0.04587 | 0.01 | 0.012 | 1 | 10 | Apbb1 | -0.002 | Young |
| 0.867223 | 0.01706 | 0.01 | 0.012 | 1 | 10 | Mcc | -0.002 | Young |
| 0.867418 | 0.329583 | 0.101 | 0.11 | 1 | 10 | Phkg2 | -0.009 | Young |
| 0.868115 | 0.046382 | 0.051 | 0.046 | 1 | 10 | Nectin4 | 0.005 | Young |
| 0.86814 | 0.135098 | 0.273 | 0.269 | 1 | 10 | Esf1 | 0.004 | Young |
| 0.868223 | 0.199954 | 0.172 | 0.183 | 1 | 10 | Rnf145 | -0.011 | Young |
| 0.868257 | 0.509476 | 0.131 | 0.13 | 1 | 10 | Wdr12 | 0.001 | Young |
| 0.868304 | 0.320276 | 0.131 | 0.142 | 1 | 10 | Nrbf2 | -0.011 | Young |
| 0.868364 | 0.247491 | 0.313 | 0.323 | 1 | 10 | Echs1 | -0.01 | Young |
| 0.868377 | 0.260653 | 0.263 | 0.257 | 1 | 10 | Dpp8 | 0.006 | Young |
| 0.868426 | 0.414779 | 0.131 | 0.127 | 1 | 10 | Mta3 | 0.004 | Young |
| 0.868427 | 0.086211 | 0.424 | 0.443 | 1 | 10 | Eif4a2 | -0.019 | Young |
| 0.868571 | 0.348915 | 0.273 | 0.284 | 1 | 10 | Tnfrsf21 | -0.011 | Young |
| 0.868617 | 0.380334 | 0.121 | 0.117 | 1 | 10 | Ciapin1 | 0.004 | Young |
| 0.868862 | 0.015738 | 0.061 | 0.056 | 1 | 10 | Cad | 0.005 | Young |
| 0.869004 | 0.249538 | 0.444 | 0.447 | 1 | 10 | Slamf7 | -0.003 | Young |
| 0.869487 | 0.40932 | 0.172 | 0.174 | 1 | 10 | Tsfm | -0.002 | Young |
| 0.869571 | 0.525301 | 0.081 | 0.078 | 1 | 10 | Pck2 | 0.003 | Young |
| 0.869643 | 0.316447 | 0.091 | 0.088 | 1 | 10 | Pld1 | 0.003 | Young |
| 0.869749 | 0.023268 | 0.051 | 0.046 | 1 | 10 | Ankrd54 | 0.005 | Young |
| 0.87004 | 0.126446 | 0.444 | 0.443 | 1 | 10 | Ap2b1 | 0.001 | Young |
| 0.870433 | 2.727162 | 0.01 | 0.012 | 1 | 10 | Gm4707 | -0.002 | Young |
| 0.870433 | 1.760151 | 0.01 | 0.012 | 1 | 10 | Gm13431 | -0.002 | Young |
| 0.870433 | 1.670446 | 0.01 | 0.012 | 1 | 10 | 0610040B1 | -0.002 | Young |
| 0.870433 | 1.559282 | 0.01 | 0.012 | 1 | 10 | Gm17655 | -0.002 | Young |
| 0.870433 | 1.086113 | 0.01 | 0.012 | 1 | 10 | Armcx2 | -0.002 | Young |
| 0.870433 | 1.054431 | 0.01 | 0.012 | 1 | 10 | D330050G2 | -0.002 | Young |
| 0.870433 | 0.778039 | 0.01 | 0.012 | 1 | 10 | B3gntl1 | -0.002 | Young |
| 0.870433 | 0.590349 | 0.01 | 0.012 | 1 | 10 | Speg | -0.002 | Young |
| 0.870433 | 0.588447 | 0.01 | 0.012 | 1 | 10 | Sap18b | -0.002 | Young |
| 0.870433 | 0.562433 | 0.01 | 0.012 | 1 | 10 | Macrod2 | -0.002 | Young |
| 0.870433 | 0.513687 | 0.01 | 0.012 | 1 | 10 | Nedd4 | -0.002 | Young |
| 0.870433 | 0.503139 | 0.01 | 0.012 | 1 | 10 | Dgat2 | -0.002 | Young |
| 0.870433 | 0.234231 | 0.01 | 0.012 | 1 | 10 | Fcor | -0.002 | Young |
| 0.870433 | 0.156035 | 0.01 | 0.012 | 1 | 10 | Trnp1 | -0.002 | Young |
| 0.870573 | 0.455008 | 0.182 | 0.183 | 1 | 10 | 6430548M | -0.001 | Young |
| 0.870606 | 0.396031 | 0.343 | 0.369 | 1 | 10 | Aff4 | -0.026 | Young |
| 0.870691 | 0.020053 | 0.848 | 0.856 | 1 | 10 | Bri3 | -0.008 | Young |
| 0.870759 | 0.345852 | 0.111 | 0.108 | 1 | 10 | Prkab2 | 0.003 | Young |
| 0.870976 | 0.15855 | 0.263 | 0.281 | 1 | 10 | Bscl2 | -0.018 | Young |
| 0.871341 | 0.410238 | 0.131 | 0.13 | 1 | 10 | Med14 | 0.001 | Young |
| 0.871349 | 0.178131 | 0.727 | 0.785 | 1 | 10 | Msn | -0.058 | Young |

| | | | | | | | | |
|---|---|---|---|---|---|---|---|---|
| 0.871589 | 0.592162 | 0.03 | 0.034 | 1 | 10 | Amd2 | -0.004 | Young |
| 0.871628 | 0.058089 | 0.303 | 0.298 | 1 | 10 | Irf2bpl | 0.005 | Young |
| 0.871833 | 0.269451 | 0.121 | 0.117 | 1 | 10 | Lin54 | 0.004 | Young |
| 0.871844 | 0.18494 | 0.182 | 0.193 | 1 | 10 | Fbxo18 | -0.011 | Young |
| 0.872595 | 0.391443 | 0.222 | 0.237 | 1 | 10 | Slc6a8 | -0.015 | Young |
| 0.872813 | 0.025304 | 0.838 | 0.863 | 1 | 10 | Ube2d3 | -0.025 | Young |
| 0.873402 | 0.123105 | 0.495 | 0.496 | 1 | 10 | Nr3c1 | -0.001 | Young |
| 0.873442 | 0.342976 | 0.111 | 0.12 | 1 | 10 | Cbll1 | -0.009 | Young |
| 0.873592 | 0.132973 | 0.111 | 0.117 | 1 | 10 | Ttf1 | -0.006 | Young |
| 0.873806 | 0.256223 | 0.364 | 0.411 | 1 | 10 | Mgat4b | -0.047 | Young |
| 0.87453 | 0.091795 | 0.131 | 0.139 | 1 | 10 | Ttc33 | -0.008 | Young |
| 0.875012 | 0.227009 | 0.202 | 0.218 | 1 | 10 | Tmpo | -0.016 | Young |
| 0.875242 | 0.005349 | 0.323 | 0.355 | 1 | 10 | Slc39a7 | -0.032 | Young |
| 0.875523 | 0.249993 | 0.343 | 0.357 | 1 | 10 | 1500011K1 | -0.014 | Young |
| 0.875665 | 0.317071 | 0.404 | 0.435 | 1 | 10 | Rala | -0.031 | Young |
| 0.875953 | 0.244806 | 0.172 | 0.166 | 1 | 10 | Ric1 | 0.006 | Young |
| 0.876531 | 0.270419 | 0.263 | 0.274 | 1 | 10 | Rap2c | -0.011 | Young |
| 0.876835 | 0.065795 | 0.111 | 0.117 | 1 | 10 | Mfap1b | -0.006 | Young |
| 0.877014 | 0.158557 | 0.101 | 0.108 | 1 | 10 | Smim8 | -0.007 | Young |
| 0.877058 | 0.001057 | 0.04 | 0.044 | 1 | 10 | Sacs | -0.004 | Young |
| 0.877425 | 0.22702 | 0.424 | 0.438 | 1 | 10 | Trmt112 | -0.014 | Young |
| 0.877512 | 0.219941 | 0.293 | 0.311 | 1 | 10 | Vav3 | -0.018 | Young |
| 0.877519 | 0.068591 | 0.495 | 0.562 | 1 | 10 | Chmp2a | -0.067 | Young |
| 0.87781 | 0.446809 | 0.343 | 0.377 | 1 | 10 | Eif1ax | -0.034 | Young |
| 0.87782 | 0.239163 | 0.525 | 0.575 | 1 | 10 | Eif4h | -0.05 | Young |
| 0.878129 | 0.157681 | 0.152 | 0.147 | 1 | 10 | Tyk2 | 0.005 | Young |
| 0.878576 | 0.145984 | 0.364 | 0.401 | 1 | 10 | Pias1 | -0.037 | Young |
| 0.879335 | 0.100157 | 0.172 | 0.181 | 1 | 10 | Mllt6 | -0.009 | Young |
| 0.879821 | 0.252776 | 0.121 | 0.13 | 1 | 10 | Dnajc30 | -0.009 | Young |
| 0.879893 | 0.16855 | 0.141 | 0.147 | 1 | 10 | Paxbp1 | -0.006 | Young |
| 0.879942 | 0.128328 | 0.263 | 0.286 | 1 | 10 | Aqr | -0.023 | Young |
| 0.879987 | 0.257121 | 0.222 | 0.227 | 1 | 10 | Slc38a10 | -0.005 | Young |
| 0.880023 | 0.290497 | 0.081 | 0.088 | 1 | 10 | Ppa2 | -0.007 | Young |
| 0.880052 | 0.286674 | 0.141 | 0.137 | 1 | 10 | Selenom | 0.004 | Young |
| 0.880448 | 0.017845 | 0.242 | 0.247 | 1 | 10 | Hspa14 | -0.005 | Young |
| 0.88052 | 0.214368 | 0.424 | 0.465 | 1 | 10 | Fam174a | -0.041 | Young |
| 0.880583 | 0.109717 | 0.121 | 0.127 | 1 | 10 | Mief1 | -0.006 | Young |
| 0.880797 | 0.222689 | 0.202 | 0.2 | 1 | 10 | Map3k7 | 0.002 | Young |
| 0.880891 | 0.108332 | 0.101 | 0.098 | 1 | 10 | Rrp15 | 0.003 | Young |
| 0.880951 | 0.213877 | 0.556 | 0.614 | 1 | 10 | Hcls1 | -0.058 | Young |
| 0.88098 | 0.010527 | 0.071 | 0.076 | 1 | 10 | Fry | -0.005 | Young |
| 0.8812 | 0.550294 | 0.051 | 0.046 | 1 | 10 | Ankrd52 | 0.005 | Young |
| 0.881612 | 0.194962 | 0.394 | 0.403 | 1 | 10 | Smchd1 | -0.009 | Young |
| 0.881657 | 0.194352 | 0.202 | 0.2 | 1 | 10 | Ptov1 | 0.002 | Young |
| 0.881771 | 0.123988 | 0.364 | 0.364 | 1 | 10 | Larp4 | 0 | Young |

| | | | | | | | | |
|---|---|---|---|---|---|---|---|---|
| 0.882143 | 0.549728 | 0.061 | 0.066 | 1 | 10 | Cbfa2t2 | -0.005 | Young |
| 0.882178 | 0.003046 | 0.04 | 0.044 | 1 | 10 | Lrba | -0.004 | Young |
| 0.8825 | 0.252552 | 0.212 | 0.225 | 1 | 10 | Utp14a | -0.013 | Young |
| 0.882545 | 0.055302 | 0.848 | 0.902 | 1 | 10 | Tpm3 | -0.054 | Young |
| 0.882563 | 0.492627 | 0.071 | 0.068 | 1 | 10 | Znhit6 | 0.003 | Young |
| 0.882793 | 0.558856 | 0.152 | 0.169 | 1 | 10 | Zfp871 | -0.017 | Young |
| 0.882916 | 0.079846 | 0.152 | 0.161 | 1 | 10 | Knop1 | -0.009 | Young |
| 0.883039 | 0.124131 | 0.081 | 0.076 | 1 | 10 | Fer | 0.005 | Young |
| 0.883401 | 0.006193 | 0.657 | 0.685 | 1 | 10 | Serp1 | -0.028 | Young |
| 0.883886 | 0.601324 | 0.04 | 0.044 | 1 | 10 | Osbpl7 | -0.004 | Young |
| 0.883934 | 0.335106 | 0.273 | 0.279 | 1 | 10 | Slc15a4 | -0.006 | Young |
| 0.884144 | 0.346287 | 0.101 | 0.11 | 1 | 10 | Tshz1 | -0.009 | Young |
| 0.884158 | 0.622276 | 0.131 | 0.134 | 1 | 10 | Gde1 | -0.003 | Young |
| 0.884281 | 0.638816 | 0.071 | 0.076 | 1 | 10 | Rpl36a-ps1 | -0.005 | Young |
| 0.884289 | 0.561143 | 0.091 | 0.088 | 1 | 10 | Timm29 | 0.003 | Young |
| 0.884338 | 0.196607 | 0.434 | 0.428 | 1 | 10 | Cox20 | 0.006 | Young |
| 0.884601 | 0.068632 | 0.313 | 0.318 | 1 | 10 | Nr4a2 | -0.005 | Young |
| 0.884909 | 0.899648 | 0.061 | 0.059 | 1 | 10 | Mapk7 | 0.002 | Young |
| 0.885004 | 0.608482 | 0.182 | 0.193 | 1 | 10 | Fam133b | -0.011 | Young |
| 0.885108 | 1.102243 | 0.03 | 0.034 | 1 | 10 | Adat2 | -0.004 | Young |
| 0.885108 | 1.066905 | 0.03 | 0.034 | 1 | 10 | Met | -0.004 | Young |
| 0.886277 | 0.217137 | 0.121 | 0.132 | 1 | 10 | Ciz1 | -0.011 | Young |
| 0.887144 | 0.127609 | 0.101 | 0.108 | 1 | 10 | Rasa2 | -0.007 | Young |
| 0.887303 | 0.780888 | 0.04 | 0.044 | 1 | 10 | Ccdc61 | -0.004 | Young |
| 0.887303 | 0.248091 | 0.04 | 0.044 | 1 | 10 | Lrrc28 | -0.004 | Young |
| 0.887303 | 0.010859 | 0.04 | 0.044 | 1 | 10 | Pmm1 | -0.004 | Young |
| 0.888152 | 0.291636 | 0.152 | 0.152 | 1 | 10 | Dpy19l4 | 0 | Young |
| 0.888158 | 0.021125 | 0.626 | 0.663 | 1 | 10 | Celf1 | -0.037 | Young |
| 0.888373 | 0.18787 | 0.485 | 0.511 | 1 | 10 | Tsc22d4 | -0.026 | Young |
| 0.888755 | 0.088586 | 0.545 | 0.567 | 1 | 10 | Arf4 | -0.022 | Young |
| 0.889113 | 0.169627 | 0.343 | 0.374 | 1 | 10 | Gak | -0.031 | Young |
| 0.889196 | 0.627012 | 0.061 | 0.066 | 1 | 10 | Pqlc2 | -0.005 | Young |
| 0.889534 | 0.43838 | 0.333 | 0.345 | 1 | 10 | Mbd2 | -0.012 | Young |
| 0.88997 | 0.29076 | 0.162 | 0.174 | 1 | 10 | Pou2f1 | -0.012 | Young |
| 0.890181 | 0.074844 | 0.606 | 0.626 | 1 | 10 | Myo5a | -0.02 | Young |
| 0.890297 | 0.062158 | 0.303 | 0.323 | 1 | 10 | Setd2 | -0.02 | Young |
| 0.890496 | 0.194935 | 0.444 | 0.462 | 1 | 10 | Fnbp1 | -0.018 | Young |
| 0.890598 | 0.131501 | 0.273 | 0.291 | 1 | 10 | Rbm27 | -0.018 | Young |
| 0.891141 | 0.087786 | 0.263 | 0.274 | 1 | 10 | Tmem5 | -0.011 | Young |
| 0.891158 | 0.342099 | 0.182 | 0.198 | 1 | 10 | Abr | -0.016 | Young |
| 0.891483 | 0.378971 | 0.081 | 0.078 | 1 | 10 | Gtf2e1 | 0.003 | Young |
| 0.89211 | 0.643609 | 0.152 | 0.169 | 1 | 10 | Ccdc9 | -0.017 | Young |
| 0.892376 | 0.355115 | 0.081 | 0.088 | 1 | 10 | Ubxn7 | -0.007 | Young |
| 0.892432 | 0.300269 | 0.04 | 0.044 | 1 | 10 | Nprl2 | -0.004 | Young |
| 0.892432 | 0.134154 | 0.04 | 0.044 | 1 | 10 | Gstz1 | -0.004 | Young |

| | | | | | | | | |
|---|---|---|---|---|---|---|---|---|
| 0.893343 | 0.611364 | 0.091 | 0.098 | 1 | 10 | Nudt14 | -0.007 | Young |
| 0.893534 | 0.107581 | 0.071 | 0.076 | 1 | 10 | Atg4d | -0.005 | Young |
| 0.894076 | 0.69472 | 0.091 | 0.088 | 1 | 10 | Acot8 | 0.003 | Young |
| 0.894076 | 0.503651 | 0.091 | 0.088 | 1 | 10 | Rpia | 0.003 | Young |
| 0.894114 | 0.244681 | 0.414 | 0.469 | 1 | 10 | Tmcc1 | -0.055 | Young |
| 0.894147 | 0.215364 | 0.313 | 0.345 | 1 | 10 | Clk4 | -0.032 | Young |
| 0.894218 | 0.08866 | 0.626 | 0.638 | 1 | 10 | Fosl2 | -0.012 | Young |
| 0.894262 | 0.296404 | 0.242 | 0.24 | 1 | 10 | Akap9 | 0.002 | Young |
| 0.894312 | 0.392169 | 0.212 | 0.215 | 1 | 10 | Ccndbp1 | -0.003 | Young |
| 0.894794 | 0.0534 | 0.081 | 0.076 | 1 | 10 | Pik3ap1 | 0.005 | Young |
| 0.895612 | 0.196877 | 0.141 | 0.147 | 1 | 10 | Palld | -0.006 | Young |
| 0.895844 | 0.3984 | 0.253 | 0.267 | 1 | 10 | Cers5 | -0.014 | Young |
| 0.895921 | 0.311735 | 0.091 | 0.095 | 1 | 10 | Ints13 | -0.004 | Young |
| 0.895996 | 0.320137 | 0.232 | 0.259 | 1 | 10 | Taf15 | -0.027 | Young |
| 0.896652 | 0.408796 | 0.081 | 0.078 | 1 | 10 | Nek1 | 0.003 | Young |
| 0.897088 | 0.264616 | 0.202 | 0.205 | 1 | 10 | Rassf5 | -0.003 | Young |
| 0.897091 | 0.33561 | 0.111 | 0.12 | 1 | 10 | Pgam5 | -0.009 | Young |
| 0.89763 | 0.79991 | 0.121 | 0.117 | 1 | 10 | Zfpl1 | 0.004 | Young |
| 0.897707 | 0.16692 | 0.687 | 0.795 | 1 | 10 | Atp6v1g1 | -0.108 | Young |
| 0.897915 | 0.110908 | 0.455 | 0.472 | 1 | 10 | Ptpa | -0.017 | Young |
| 0.897949 | 1.144621 | 0.051 | 0.049 | 1 | 10 | Parp6 | 0.002 | Young |
| 0.898031 | 0.242792 | 0.212 | 0.218 | 1 | 10 | Tomm40 | -0.006 | Young |
| 0.898196 | 0.219033 | 0.182 | 0.193 | 1 | 10 | 1810013L2 | -0.011 | Young |
| 0.898196 | 0.069055 | 0.182 | 0.193 | 1 | 10 | 4833439L1 | -0.011 | Young |
| 0.898806 | 0.624844 | 0.323 | 0.352 | 1 | 10 | Pcnp | -0.029 | Young |
| 0.898962 | 0.220566 | 0.354 | 0.384 | 1 | 10 | Mgea5 | -0.03 | Young |
| 0.899076 | 0.108091 | 0.182 | 0.193 | 1 | 10 | Ncbp2 | -0.011 | Young |
| 0.899084 | 0.51864 | 0.061 | 0.066 | 1 | 10 | Rnf185 | -0.005 | Young |
| 0.899103 | 0.254868 | 0.182 | 0.191 | 1 | 10 | Zyg11b | -0.009 | Young |
| 0.899195 | 0.069912 | 0.889 | 0.956 | 1 | 10 | Sem1 | -0.067 | Young |
| 0.899236 | 0.175474 | 0.091 | 0.098 | 1 | 10 | Clk2 | -0.007 | Young |
| 0.899334 | 0.369776 | 0.152 | 0.154 | 1 | 10 | Rxra | -0.002 | Young |
| 0.899411 | 0.134841 | 0.121 | 0.127 | 1 | 10 | Rara | -0.006 | Young |
| 0.899558 | 0.512924 | 0.101 | 0.098 | 1 | 10 | Ifi209 | 0.003 | Young |
| 0.899828 | 0.152727 | 0.343 | 0.362 | 1 | 10 | Fam20c | -0.019 | Young |
| 0.900422 | 0.758065 | 0.071 | 0.068 | 1 | 10 | Gys1 | 0.003 | Young |
| 0.900542 | 0.144371 | 0.313 | 0.325 | 1 | 10 | Zc3h12a | -0.012 | Young |
| 0.900602 | 0.343989 | 0.131 | 0.127 | 1 | 10 | Spire1 | 0.004 | Young |
| 0.900727 | 0.303744 | 0.101 | 0.098 | 1 | 10 | Paxip1 | 0.003 | Young |
| 0.900736 | 0.630017 | 0.131 | 0.13 | 1 | 10 | Tmem260 | 0.001 | Young |
| 0.900863 | 0.041533 | 0.121 | 0.117 | 1 | 10 | Kdm3b | 0.004 | Young |
| 0.901154 | 0.669292 | 0.061 | 0.059 | 1 | 10 | Mgme1 | 0.002 | Young |
| 0.901172 | 0.579113 | 0.051 | 0.049 | 1 | 10 | Btrc | 0.002 | Young |
| 0.901213 | 0.378828 | 0.313 | 0.315 | 1 | 10 | Uba2 | -0.002 | Young |
| 0.901602 | 0.198389 | 0.071 | 0.073 | 1 | 10 | Gatsl2 | -0.002 | Young |

| | | | | | | | | |
|---|---|---|---|---|---|---|---|---|
| 0.901692 | 0.027016 | 0.131 | 0.139 | 1 | 10 | Vps52 | -0.008 | Young |
| 0.901769 | 0.971442 | 0.04 | 0.039 | 1 | 10 | Gatc | 0.001 | Young |
| 0.902138 | 0.280248 | 0.263 | 0.284 | 1 | 10 | Actr1a | -0.021 | Young |
| 0.902173 | 0.147983 | 0.677 | 0.711 | 1 | 10 | Tbca | -0.034 | Young |
| 0.902203 | 0.048165 | 0.333 | 0.342 | 1 | 10 | Ythdf3 | -0.009 | Young |
| 0.902257 | 0.51772 | 0.444 | 0.494 | 1 | 10 | Hmgb2 | -0.05 | Young |
| 0.902287 | 0.139891 | 0.131 | 0.13 | 1 | 10 | Wdr77 | 0.001 | Young |
| 0.902705 | 0.455188 | 0.04 | 0.044 | 1 | 10 | Ntmt1 | -0.004 | Young |
| 0.902762 | 0.278882 | 0.283 | 0.293 | 1 | 10 | Atraid | -0.01 | Young |
| 0.902849 | 0.102008 | 0.162 | 0.171 | 1 | 10 | Zfp281 | -0.009 | Young |
| 0.903065 | 0.182732 | 0.414 | 0.433 | 1 | 10 | Ctbp1 | -0.019 | Young |
| 0.903266 | 0.361184 | 0.172 | 0.178 | 1 | 10 | Seh1l | -0.006 | Young |
| 0.903327 | 0.500203 | 0.061 | 0.066 | 1 | 10 | Zfp746 | -0.005 | Young |
| 0.90369 | 0.169027 | 0.535 | 0.575 | 1 | 10 | Psme1 | -0.04 | Young |
| 0.903723 | 0.17347 | 0.212 | 0.21 | 1 | 10 | Lyar | 0.002 | Young |
| 0.903885 | 0.017431 | 0.303 | 0.306 | 1 | 10 | Ndufa12 | -0.003 | Young |
| 0.904056 | 0.138321 | 0.596 | 0.606 | 1 | 10 | Pbrm1 | -0.01 | Young |
| 0.904112 | 0.695282 | 0.061 | 0.059 | 1 | 10 | Vamp5 | 0.002 | Young |
| 0.904509 | 0.356971 | 0.242 | 0.244 | 1 | 10 | Mdm4 | -0.002 | Young |
| 0.904672 | 0.173423 | 0.162 | 0.176 | 1 | 10 | Irf3 | -0.014 | Young |
| 0.904694 | 0.021935 | 0.051 | 0.054 | 1 | 10 | Pramef8 | -0.003 | Young |
| 0.904707 | 0.360136 | 0.162 | 0.169 | 1 | 10 | Pdzd8 | -0.007 | Young |
| 0.904745 | 0.871378 | 0.101 | 0.103 | 1 | 10 | Zfp407 | -0.002 | Young |
| 0.904993 | 0.164271 | 0.212 | 0.232 | 1 | 10 | Napg | -0.02 | Young |
| 0.905404 | 0.22825 | 0.101 | 0.11 | 1 | 10 | Yae1d1 | -0.009 | Young |
| 0.905444 | 0.138756 | 0.091 | 0.095 | 1 | 10 | Asb7 | -0.004 | Young |
| 0.905592 | 0.602365 | 0.061 | 0.059 | 1 | 10 | Wtip | 0.002 | Young |
| 0.90569 | 0.072241 | 0.354 | 0.386 | 1 | 10 | Camk1d | -0.032 | Young |
| 0.905928 | 0.355195 | 0.071 | 0.068 | 1 | 10 | Rprd1a | 0.003 | Young |
| 0.906303 | 0.390671 | 0.212 | 0.24 | 1 | 10 | Cd164 | -0.028 | Young |
| 0.906639 | 0.974266 | 0.121 | 0.125 | 1 | 10 | Nmt2 | -0.004 | Young |
| 0.906687 | 0.331174 | 0.303 | 0.347 | 1 | 10 | Gng12 | -0.044 | Young |
| 0.906929 | 0.040457 | 0.788 | 0.79 | 1 | 10 | Coro1a | -0.002 | Young |
| 0.907328 | 0.319895 | 0.263 | 0.286 | 1 | 10 | Tmem33 | -0.023 | Young |
| 0.907527 | 0.230807 | 0.394 | 0.418 | 1 | 10 | Gspt1 | -0.024 | Young |
| 0.907617 | 0.510977 | 0.131 | 0.134 | 1 | 10 | Thumpd3 | -0.003 | Young |
| 0.907622 | 0.379653 | 0.051 | 0.049 | 1 | 10 | 1300002E1 | 0.002 | Young |
| 0.907729 | 0.085035 | 0.788 | 0.836 | 1 | 10 | Rbms1 | -0.048 | Young |
| 0.907847 | 0.731915 | 0.04 | 0.044 | 1 | 10 | Tmem238 | -0.004 | Young |
| 0.908249 | 0.11092 | 0.141 | 0.137 | 1 | 10 | Rcsd1 | 0.004 | Young |
| 0.908378 | 0.254875 | 0.364 | 0.364 | 1 | 10 | Thoc2 | 0 | Young |
| 0.908589 | 0.19136 | 0.101 | 0.108 | 1 | 10 | Npl | -0.007 | Young |
| 0.908787 | 0.279246 | 0.091 | 0.088 | 1 | 10 | Gid4 | 0.003 | Young |
| 0.909077 | 9.23E-05 | 0.323 | 0.33 | 1 | 10 | 9030624J0 | -0.007 | Young |
| 0.909082 | 0.193963 | 0.444 | 0.484 | 1 | 10 | Dmac1 | -0.04 | Young |

| | | | | | | | | |
|---|---|---|---|---|---|---|---|---|
| 0.909171 | 0.296934 | 0.182 | 0.176 | 1 | 10 | Sla | 0.006 | Young |
| 0.909341 | 0.217588 | 0.141 | 0.142 | 1 | 10 | Taok2 | -0.001 | Young |
| 0.909988 | 0.59217 | 0.222 | 0.252 | 1 | 10 | Pdha1 | -0.03 | Young |
| 0.910127 | 0.020941 | 0.788 | 0.763 | 1 | 10 | Timp2 | 0.025 | Young |
| 0.910339 | 0.219752 | 0.141 | 0.142 | 1 | 10 | Senp1 | -0.001 | Young |
| 0.910646 | 0.196311 | 0.081 | 0.086 | 1 | 10 | Rrp9 | -0.005 | Young |
| 0.91085 | 0.986458 | 0.051 | 0.049 | 1 | 10 | Sephs1 | 0.002 | Young |
| 0.911267 | 0.295397 | 0.061 | 0.064 | 1 | 10 | Slc44a2 | -0.003 | Young |
| 0.911336 | 0.315024 | 0.162 | 0.159 | 1 | 10 | Nsmce1 | 0.003 | Young |
| 0.911415 | 0.066814 | 0.485 | 0.506 | 1 | 10 | Rnpep | -0.021 | Young |
| 0.911827 | 0.228482 | 0.263 | 0.264 | 1 | 10 | Kdm1a | -0.001 | Young |
| 0.911827 | 0.105691 | 0.263 | 0.264 | 1 | 10 | Ctcf | -0.001 | Young |
| 0.912363 | 0.282436 | 0.525 | 0.57 | 1 | 10 | Arl8a | -0.045 | Young |
| 0.912474 | 0.502211 | 0.051 | 0.054 | 1 | 10 | Nme4 | -0.003 | Young |
| 0.912486 | 0.148426 | 0.545 | 0.601 | 1 | 10 | Hspa9 | -0.056 | Young |
| 0.912843 | 0.147368 | 0.343 | 0.35 | 1 | 10 | Ndufb2 | -0.007 | Young |
| 0.913238 | 0.284099 | 0.02 | 0.022 | 1 | 10 | Ldlrad4 | -0.002 | Young |
| 0.913388 | 0.483112 | 0.242 | 0.269 | 1 | 10 | Imp3 | -0.027 | Young |
| 0.913412 | 0.026561 | 0.97 | 0.973 | 1 | 10 | Calm1 | -0.003 | Young |
| 0.91348 | 0.285332 | 0.222 | 0.247 | 1 | 10 | Emsy | -0.025 | Young |
| 0.91354 | 0.095698 | 0.242 | 0.254 | 1 | 10 | Zranb1 | -0.012 | Young |
| 0.913674 | 0.036707 | 0.131 | 0.13 | 1 | 10 | Vrk3 | 0.001 | Young |
| 0.913718 | 0.747735 | 0.111 | 0.117 | 1 | 10 | Ralgapa2 | -0.006 | Young |
| 0.913905 | 0.024194 | 0.212 | 0.21 | 1 | 10 | Ncoa6 | 0.002 | Young |
| 0.914195 | 0.769391 | 0.071 | 0.068 | 1 | 10 | Pde8a | 0.003 | Young |
| 0.914377 | 0.143711 | 0.111 | 0.108 | 1 | 10 | Riox2 | 0.003 | Young |
| 0.914389 | 0.023518 | 0.616 | 0.643 | 1 | 10 | Myo1f | -0.027 | Young |
| 0.914476 | 0.97778 | 0.061 | 0.059 | 1 | 10 | Cstf1 | 0.002 | Young |
| 0.914541 | 0.21332 | 0.232 | 0.247 | 1 | 10 | Dnajc21 | -0.015 | Young |
| 0.914783 | 0.06472 | 0.455 | 0.457 | 1 | 10 | Slc29a3 | -0.002 | Young |
| 0.914925 | 0.811547 | 0.091 | 0.088 | 1 | 10 | Gtdc1 | 0.003 | Young |
| 0.915241 | 0.403234 | 0.101 | 0.105 | 1 | 10 | Eri3 | -0.004 | Young |
| 0.915305 | 0.245869 | 0.141 | 0.149 | 1 | 10 | Nup54 | -0.008 | Young |
| 0.915318 | 0.373389 | 0.141 | 0.147 | 1 | 10 | Zfp664 | -0.006 | Young |
| 0.915574 | 0.313644 | 0.071 | 0.068 | 1 | 10 | Naip6 | 0.003 | Young |
| 0.915694 | 0.418543 | 0.051 | 0.049 | 1 | 10 | Spn | 0.002 | Young |
| 0.915897 | 0.10803 | 0.253 | 0.264 | 1 | 10 | Galk2 | -0.011 | Young |
| 0.915958 | 0.805785 | 0.061 | 0.059 | 1 | 10 | Zfp398 | 0.002 | Young |
| 0.916038 | 0.543702 | 0.111 | 0.122 | 1 | 10 | Fam199x | -0.011 | Young |
| 0.916357 | 0.000338 | 0.111 | 0.115 | 1 | 10 | Exosc4 | -0.004 | Young |
| 0.916377 | 0.26888 | 0.313 | 0.345 | 1 | 10 | Setd3 | -0.032 | Young |
| 0.916502 | 0.409968 | 0.293 | 0.308 | 1 | 10 | Ipo5 | -0.015 | Young |
| 0.916711 | 0.065698 | 0.192 | 0.191 | 1 | 10 | Pianp | 0.001 | Young |
| 0.916792 | 0.140085 | 0.535 | 0.567 | 1 | 10 | Aamp | -0.032 | Young |
| 0.916982 | 0.216573 | 0.111 | 0.117 | 1 | 10 | Nvl | -0.006 | Young |

| | | | | | | | | |
|---|---|---|---|---|---|---|---|---|
| 0.91711 | 0.522446 | 0.121 | 0.12 | 1 | 10 | Plk3 | 0.001 | Young |
| 0.917259 | 0.149785 | 0.101 | 0.1 | 1 | 10 | Umad1 | 0.001 | Young |
| 0.917348 | 0.221887 | 0.212 | 0.225 | 1 | 10 | Tuba1a | -0.013 | Young |
| 0.917368 | 0.194074 | 0.081 | 0.078 | 1 | 10 | Wdfy1 | 0.003 | Young |
| 0.917535 | 0.060621 | 0.303 | 0.32 | 1 | 10 | Ano6 | -0.017 | Young |
| 0.918899 | 0.234429 | 0.192 | 0.2 | 1 | 10 | Rangap1 | -0.008 | Young |
| 0.919133 | 0.391579 | 0.192 | 0.196 | 1 | 10 | Tmem51 | -0.004 | Young |
| 0.91943 | 0.152755 | 0.566 | 0.599 | 1 | 10 | Fcgr3 | -0.033 | Young |
| 0.919685 | 0.508961 | 0.182 | 0.186 | 1 | 10 | Washc1 | -0.004 | Young |
| 0.919685 | 0.343976 | 0.232 | 0.242 | 1 | 10 | 3110043O2 | -0.01 | Young |
| 0.919861 | 0.744775 | 0.04 | 0.044 | 1 | 10 | Car13 | -0.004 | Young |
| 0.920223 | 0.026727 | 0.505 | 0.543 | 1 | 10 | Irf8 | -0.038 | Young |
| 0.920243 | 0.242237 | 0.576 | 0.589 | 1 | 10 | Osbpl8 | -0.013 | Young |
| 0.920349 | 0.418084 | 0.131 | 0.142 | 1 | 10 | Abca7 | -0.011 | Young |
| 0.920443 | 0.435591 | 0.02 | 0.022 | 1 | 10 | Olfr1167 | -0.002 | Young |
| 0.920443 | 0.030056 | 0.02 | 0.022 | 1 | 10 | Gale | -0.002 | Young |
| 0.920458 | 0.60764 | 0.081 | 0.083 | 1 | 10 | Ddx55 | -0.002 | Young |
| 0.920469 | 0.074844 | 0.182 | 0.188 | 1 | 10 | Clcc1 | -0.006 | Young |
| 0.920494 | 0.876216 | 0.091 | 0.098 | 1 | 10 | Zfp410 | -0.007 | Young |
| 0.920624 | 0.009892 | 0.101 | 0.098 | 1 | 10 | Pbx3 | 0.003 | Young |
| 0.921119 | 0.432107 | 0.202 | 0.22 | 1 | 10 | Fbxo33 | -0.018 | Young |
| 0.921174 | 0.006473 | 0.424 | 0.43 | 1 | 10 | Acin1 | -0.006 | Young |
| 0.921373 | 0.130962 | 0.121 | 0.117 | 1 | 10 | Rbl2 | 0.004 | Young |
| 0.921452 | 0.072076 | 0.172 | 0.178 | 1 | 10 | Znrf1 | -0.006 | Young |
| 0.92209 | 0.193958 | 0.101 | 0.105 | 1 | 10 | Pla2g4a | -0.004 | Young |
| 0.922362 | 0.208395 | 0.172 | 0.178 | 1 | 10 | Srpk1 | -0.006 | Young |
| 0.922497 | 0.519597 | 0.121 | 0.125 | 1 | 10 | Suclg2 | -0.004 | Young |
| 0.922634 | 0.025033 | 0.323 | 0.333 | 1 | 10 | Amdhd2 | -0.01 | Young |
| 0.922846 | 0.37279 | 0.02 | 0.022 | 1 | 10 | Tnk2 | -0.002 | Young |
| 0.922846 | 0.3569 | 0.02 | 0.022 | 1 | 10 | 1810011H1 | -0.002 | Young |
| 0.922846 | 0.292033 | 0.02 | 0.022 | 1 | 10 | Endov | -0.002 | Young |
| 0.922846 | 0.169653 | 0.02 | 0.022 | 1 | 10 | Oplah | -0.002 | Young |
| 0.923665 | 0.180066 | 0.566 | 0.609 | 1 | 10 | Ctnnb1 | -0.043 | Young |
| 0.923797 | 0.33297 | 0.202 | 0.218 | 1 | 10 | Smarcb1 | -0.016 | Young |
| 0.924072 | 0.021275 | 0.374 | 0.381 | 1 | 10 | Rexo2 | -0.007 | Young |
| 0.924209 | 0.181149 | 0.061 | 0.064 | 1 | 10 | Adnp2 | -0.003 | Young |
| 0.924405 | 0.402341 | 0.354 | 0.394 | 1 | 10 | Snx6 | -0.04 | Young |
| 0.924686 | 0.329887 | 0.253 | 0.257 | 1 | 10 | Tmem87b | -0.004 | Young |
| 0.924853 | 0.274837 | 0.172 | 0.183 | 1 | 10 | Uba3 | -0.011 | Young |
| 0.924855 | 0.913232 | 0.061 | 0.059 | 1 | 10 | Ercc5 | 0.002 | Young |
| 0.92525 | 0.380803 | 0.02 | 0.022 | 1 | 10 | Frat1 | -0.002 | Young |
| 0.92525 | 0.122398 | 0.02 | 0.022 | 1 | 10 | Zfp692 | -0.002 | Young |
| 0.925369 | 0.156351 | 0.606 | 0.655 | 1 | 10 | Morf4l2 | -0.049 | Young |
| 0.925517 | 0.33414 | 0.101 | 0.105 | 1 | 10 | Ppp5c | -0.004 | Young |
| 0.925531 | 0.471667 | 0.323 | 0.333 | 1 | 10 | Ssrp1 | -0.01 | Young |

| | | | | | | | | |
|---|---|---|---|---|---|---|---|---|
| 0.925694 | 0.035159 | 0.111 | 0.117 | 1 | 10 | Sap30bp | -0.006 | Young |
| 0.926105 | 0.488105 | 0.141 | 0.149 | 1 | 10 | Stau1 | -0.008 | Young |
| 0.926223 | 0.214894 | 0.091 | 0.093 | 1 | 10 | Desi1 | -0.002 | Young |
| 0.926276 | 0.295609 | 0.455 | 0.455 | 1 | 10 | Luc7l3 | 0 | Young |
| 0.926689 | 0.758674 | 0.071 | 0.076 | 1 | 10 | Hars2 | -0.005 | Young |
| 0.926689 | 0.43546 | 0.071 | 0.076 | 1 | 10 | Fbxo34 | -0.005 | Young |
| 0.926858 | 0.19718 | 0.505 | 0.511 | 1 | 10 | Snx10 | -0.006 | Young |
| 0.92692 | 1.126911 | 0.04 | 0.039 | 1 | 10 | Brms1l | 0.001 | Young |
| 0.927008 | 0.229581 | 0.051 | 0.049 | 1 | 10 | Il1rap | 0.002 | Young |
| 0.927026 | 0.37939 | 0.232 | 0.252 | 1 | 10 | Bckdk | -0.02 | Young |
| 0.927119 | 0.147707 | 0.152 | 0.159 | 1 | 10 | Slc16a7 | -0.007 | Young |
| 0.927583 | 0.083628 | 0.374 | 0.399 | 1 | 10 | Akt1 | -0.025 | Young |
| 0.927638 | 0.159952 | 0.434 | 0.44 | 1 | 10 | Hnrnpul2 | -0.006 | Young |
| 0.927655 | 0.374537 | 0.02 | 0.022 | 1 | 10 | Acsf2 | -0.002 | Young |
| 0.927981 | 0.350798 | 0.141 | 0.152 | 1 | 10 | Eepd1 | -0.011 | Young |
| 0.928149 | 0.19203 | 0.172 | 0.181 | 1 | 10 | Mplkip | -0.009 | Young |
| 0.928227 | 0.010186 | 0.859 | 0.919 | 1 | 10 | Atp5l | -0.06 | Young |
| 0.928428 | 0.178248 | 0.111 | 0.115 | 1 | 10 | Ern1 | -0.004 | Young |
| 0.928448 | 0.20568 | 0.091 | 0.088 | 1 | 10 | Xab2 | 0.003 | Young |
| 0.928448 | 0.086515 | 0.091 | 0.088 | 1 | 10 | Snip1 | 0.003 | Young |
| 0.92847 | 0.622229 | 0.152 | 0.149 | 1 | 10 | Rnf181 | 0.003 | Young |
| 0.928489 | 0.344735 | 0.182 | 0.188 | 1 | 10 | Upf1 | -0.006 | Young |
| 0.928807 | 0.09416 | 0.556 | 0.579 | 1 | 10 | Polr1d | -0.023 | Young |
| 0.928922 | 0.121285 | 0.172 | 0.166 | 1 | 10 | Cyhr1 | 0.006 | Young |
| 0.92923 | 0.201584 | 0.131 | 0.13 | 1 | 10 | Nup214 | 0.001 | Young |
| 0.929368 | 0.059152 | 0.172 | 0.183 | 1 | 10 | Plcg2 | -0.011 | Young |
| 0.929374 | 0.145921 | 0.071 | 0.068 | 1 | 10 | Igsf9 | 0.003 | Young |
| 0.929652 | 0.164407 | 0.192 | 0.196 | 1 | 10 | Dtymk | -0.004 | Young |
| 0.930026 | 0.162152 | 0.242 | 0.257 | 1 | 10 | Bcat2 | -0.015 | Young |
| 0.93006 | 0.706138 | 0.02 | 0.022 | 1 | 10 | Aars2 | -0.002 | Young |
| 0.930139 | 0.455118 | 0.192 | 0.208 | 1 | 10 | Cse1l | -0.016 | Young |
| 0.930237 | 0.114582 | 0.374 | 0.396 | 1 | 10 | Slc48a1 | -0.022 | Young |
| 0.930244 | 1.02188 | 0.051 | 0.049 | 1 | 10 | Fam118a | 0.002 | Young |
| 0.930353 | 0.079567 | 0.283 | 0.289 | 1 | 10 | Lsm5 | -0.006 | Young |
| 0.930358 | 0.050431 | 0.869 | 0.929 | 1 | 10 | Slc25a3 | -0.06 | Young |
| 0.930519 | 0.51173 | 0.04 | 0.039 | 1 | 10 | Ulbp1 | 0.001 | Young |
| 0.930519 | 0.430664 | 0.04 | 0.039 | 1 | 10 | Taf5 | 0.001 | Young |
| 0.930755 | 0.474922 | 0.071 | 0.068 | 1 | 10 | Foxo4 | 0.003 | Young |
| 0.93077 | 0.077667 | 0.485 | 0.504 | 1 | 10 | Ptges3 | -0.019 | Young |
| 0.931003 | 0.184637 | 0.333 | 0.345 | 1 | 10 | Ugp2 | -0.012 | Young |
| 0.931223 | 0.552561 | 0.101 | 0.103 | 1 | 10 | Cog3 | -0.002 | Young |
| 0.931317 | 0.724414 | 0.141 | 0.142 | 1 | 10 | Odf2 | -0.001 | Young |
| 0.931862 | 1.07757 | 0.051 | 0.049 | 1 | 10 | Odf2l | 0.002 | Young |
| 0.932344 | 0.360358 | 0.131 | 0.13 | 1 | 10 | Pibf1 | 0.001 | Young |
| 0.932466 | 0.95702 | 0.02 | 0.022 | 1 | 10 | Bbs5 | -0.002 | Young |

| | | | | | | | | |
|---|---|---|---|---|---|---|---|---|
| 0.932942 | 0.169047 | 0.081 | 0.078 | 1 | 10 | Hinfp | 0.003 | Young |
| 0.933017 | 0.572841 | 0.071 | 0.071 | 1 | 10 | Ptgir | 0 | Young |
| 0.933025 | 0.157677 | 0.242 | 0.249 | 1 | 10 | Mllt10 | -0.007 | Young |
| 0.933112 | 0.284748 | 0.152 | 0.154 | 1 | 10 | Lsm2 | -0.002 | Young |
| 0.933237 | 0.508614 | 0.081 | 0.086 | 1 | 10 | Alg2 | -0.005 | Young |
| 0.933237 | 0.392923 | 0.081 | 0.086 | 1 | 10 | Ifi211 | -0.005 | Young |
| 0.933315 | 0.023398 | 0.465 | 0.521 | 1 | 10 | Atp5g1 | -0.056 | Young |
| 0.933481 | 0.955141 | 0.051 | 0.049 | 1 | 10 | Flad1 | 0.002 | Young |
| 0.933481 | 0.135095 | 0.051 | 0.049 | 1 | 10 | Lpar1 | 0.002 | Young |
| 0.933519 | 0.225965 | 0.071 | 0.068 | 1 | 10 | Dvl3 | 0.003 | Young |
| 0.933644 | 0.054522 | 0.202 | 0.21 | 1 | 10 | Bin3 | -0.008 | Young |
| 0.934069 | 0.229417 | 0.121 | 0.127 | 1 | 10 | Poldip2 | -0.006 | Young |
| 0.93412 | 0.240923 | 0.04 | 0.039 | 1 | 10 | 2310057M | 0.001 | Young |
| 0.934251 | 0.053579 | 0.232 | 0.232 | 1 | 10 | Rftn1 | 0 | Young |
| 0.93427 | 0.042065 | 0.869 | 0.888 | 1 | 10 | Actr2 | -0.019 | Young |
| 0.93429 | 0.01217 | 0.061 | 0.064 | 1 | 10 | Camta1 | -0.003 | Young |
| 0.934521 | 0.179898 | 0.162 | 0.169 | 1 | 10 | Fam98b | -0.007 | Young |
| 0.934695 | 0.516252 | 0.141 | 0.154 | 1 | 10 | Bloc1s4 | -0.013 | Young |
| 0.935094 | 0.117669 | 0.374 | 0.379 | 1 | 10 | Iscu | -0.005 | Young |
| 0.935124 | 0.125937 | 0.455 | 0.489 | 1 | 10 | Dock10 | -0.034 | Young |
| 0.935199 | 0.037905 | 0.636 | 0.707 | 1 | 10 | Mtss1 | -0.071 | Young |
| 0.935207 | 0.00232 | 0.121 | 0.125 | 1 | 10 | Golph3l | -0.004 | Young |
| 0.935451 | 0.327037 | 0.212 | 0.222 | 1 | 10 | Sar1b | -0.01 | Young |
| 0.935544 | 0.207865 | 0.343 | 0.372 | 1 | 10 | Stk4 | -0.029 | Young |
| 0.935709 | 0.827674 | 0.03 | 0.029 | 1 | 10 | Tefm | 0.001 | Young |
| 0.935709 | 0.759175 | 0.03 | 0.029 | 1 | 10 | Ovca2 | 0.001 | Young |
| 0.935833 | 0.403688 | 0.343 | 0.362 | 1 | 10 | Alyref | -0.019 | Young |
| 0.935859 | 0.194083 | 0.101 | 0.1 | 1 | 10 | Maoa | 0.001 | Young |
| 0.935885 | 0.138602 | 0.424 | 0.428 | 1 | 10 | Eif3j1 | -0.004 | Young |
| 0.935921 | 0.871474 | 0.04 | 0.039 | 1 | 10 | Zc2hc1a | 0.001 | Young |
| 0.935921 | 0.047151 | 0.04 | 0.039 | 1 | 10 | Cox18 | 0.001 | Young |
| 0.935974 | 0.02502 | 0.293 | 0.311 | 1 | 10 | Cand1 | -0.018 | Young |
| 0.936387 | 0.267365 | 0.162 | 0.169 | 1 | 10 | Cdk8 | -0.007 | Young |
| 0.936786 | 0.260049 | 0.192 | 0.205 | 1 | 10 | Cirbp | -0.013 | Young |
| 0.936867 | 0.161139 | 0.384 | 0.416 | 1 | 10 | Ezr | -0.032 | Young |
| 0.936872 | 0.493101 | 0.232 | 0.244 | 1 | 10 | Rnft1 | -0.012 | Young |
| 0.937188 | 0.494005 | 0.141 | 0.144 | 1 | 10 | Nfs1 | -0.003 | Young |
| 0.937321 | 0.227169 | 0.162 | 0.169 | 1 | 10 | Rprd2 | -0.007 | Young |
| 0.937476 | 0.204962 | 0.364 | 0.408 | 1 | 10 | Psmc5 | -0.044 | Young |
| 0.93748 | 0.339664 | 0.283 | 0.293 | 1 | 10 | Psmd12 | -0.01 | Young |
| 0.937572 | 0.407399 | 0.172 | 0.176 | 1 | 10 | Cds2 | -0.004 | Young |
| 0.93765 | 0.298686 | 0.202 | 0.203 | 1 | 10 | Setd5 | -0.001 | Young |
| 0.937665 | 0.088122 | 0.071 | 0.068 | 1 | 10 | Zbtb14 | 0.003 | Young |
| 0.937723 | 0.31681 | 0.04 | 0.039 | 1 | 10 | Gm12840 | 0.001 | Young |
| 0.937723 | 0.250471 | 0.04 | 0.039 | 1 | 10 | Dgkg | 0.001 | Young |

| | | | | | | | | |
|---|---|---|---|---|---|---|---|---|
| 0.937778 | 0.982268 | 0.03 | 0.029 | 1 | 10 | Disc1 | 0.001 | Young |
| 0.937778 | 0.745135 | 0.03 | 0.029 | 1 | 10 | Dusp4 | 0.001 | Young |
| 0.937778 | 0.689534 | 0.03 | 0.029 | 1 | 10 | Wdhd1 | 0.001 | Young |
| 0.937778 | 0.42864 | 0.03 | 0.029 | 1 | 10 | Zfp52 | 0.001 | Young |
| 0.937992 | 0.366903 | 0.354 | 0.406 | 1 | 10 | Fndc3b | -0.052 | Young |
| 0.938119 | 0.149825 | 0.283 | 0.279 | 1 | 10 | Zfand3 | 0.004 | Young |
| 0.938406 | 0.303891 | 0.242 | 0.252 | 1 | 10 | Itga5 | -0.01 | Young |
| 0.938614 | 0.469383 | 0.061 | 0.064 | 1 | 10 | Pot1b | -0.003 | Young |
| 0.938754 | 0.213752 | 0.242 | 0.257 | 1 | 10 | Ilf3 | -0.015 | Young |
| 0.938907 | 0.291687 | 0.091 | 0.09 | 1 | 10 | Isyna1 | 0.001 | Young |
| 0.939048 | 0.025395 | 0.071 | 0.068 | 1 | 10 | Tom1l2 | 0.003 | Young |
| 0.939192 | 0.027511 | 0.535 | 0.582 | 1 | 10 | Nucks1 | -0.047 | Young |
| 0.939241 | 0.183344 | 0.152 | 0.156 | 1 | 10 | Mfsd10 | -0.004 | Young |
| 0.939248 | 0.23949 | 0.071 | 0.073 | 1 | 10 | Dmap1 | -0.002 | Young |
| 0.939322 | 0.43196 | 0.141 | 0.142 | 1 | 10 | Pi4k2b | -0.001 | Young |
| 0.939958 | 0.371152 | 0.051 | 0.049 | 1 | 10 | Lin52 | 0.002 | Young |
| 0.940591 | 0.083692 | 0.172 | 0.178 | 1 | 10 | Pcif1 | -0.006 | Young |
| 0.940655 | 0.130391 | 0.131 | 0.13 | 1 | 10 | Fam69a | 0.001 | Young |
| 0.941075 | 0.157248 | 0.697 | 0.748 | 1 | 10 | Tgfb1 | -0.051 | Young |
| 0.941237 | 0.127375 | 0.172 | 0.176 | 1 | 10 | Apex1 | -0.004 | Young |
| 0.941326 | 0.23145 | 0.04 | 0.039 | 1 | 10 | Ndufaf5 | 0.001 | Young |
| 0.941764 | 0.347992 | 0.172 | 0.181 | 1 | 10 | Rnf146 | -0.009 | Young |
| 0.941919 | 0.291898 | 0.03 | 0.029 | 1 | 10 | Kpna2 | 0.001 | Young |
| 0.942048 | 0.10832 | 0.081 | 0.083 | 1 | 10 | Oxsm | -0.002 | Young |
| 0.942104 | 0.946581 | 0.051 | 0.054 | 1 | 10 | Heatr5b | -0.003 | Young |
| 0.94224 | 0.262848 | 0.293 | 0.333 | 1 | 10 | Ppp6c | -0.04 | Young |
| 0.942438 | 0.078488 | 0.606 | 0.655 | 1 | 10 | Ywhag | -0.049 | Young |
| 0.94254 | 0.010055 | 0.172 | 0.171 | 1 | 10 | Rnf2 | 0.001 | Young |
| 0.942558 | 0.168691 | 0.182 | 0.191 | 1 | 10 | Acox1 | -0.009 | Young |
| 0.942566 | 0.344717 | 0.091 | 0.09 | 1 | 10 | Zfp266 | 0.001 | Young |
| 0.942732 | 0.06931 | 0.162 | 0.171 | 1 | 10 | Rnf14 | -0.009 | Young |
| 0.942839 | 0.066545 | 0.343 | 0.342 | 1 | 10 | Safb2 | 0.001 | Young |
| 0.942902 | 0.233983 | 0.293 | 0.315 | 1 | 10 | Nrbp1 | -0.022 | Young |
| 0.942939 | 0.382104 | 0.061 | 0.064 | 1 | 10 | Aldh1a2 | -0.003 | Young |
| 0.942939 | 0.332976 | 0.061 | 0.064 | 1 | 10 | Snx21 | -0.003 | Young |
| 0.943128 | 0.087258 | 0.04 | 0.039 | 1 | 10 | Eef1akmt4 | 0.001 | Young |
| 0.943128 | 0.083165 | 0.04 | 0.039 | 1 | 10 | Zfp74 | 0.001 | Young |
| 0.943266 | 0.271805 | 0.192 | 0.2 | 1 | 10 | Slc30a9 | -0.008 | Young |
| 0.943334 | 0.420022 | 0.081 | 0.081 | 1 | 10 | Lonrf1 | 0 | Young |
| 0.943383 | 0.33632 | 0.111 | 0.112 | 1 | 10 | Gfm1 | -0.001 | Young |
| 0.943667 | 0.871081 | 0.051 | 0.054 | 1 | 10 | Tti2 | -0.003 | Young |
| 0.943689 | 0.094692 | 0.121 | 0.125 | 1 | 10 | Map2k5 | -0.004 | Young |
| 0.943704 | 0.514146 | 0.03 | 0.032 | 1 | 10 | Thg1l | -0.002 | Young |
| 0.943704 | 0.02157 | 0.03 | 0.032 | 1 | 10 | Cdc25b | -0.002 | Young |
| 0.943981 | 0.608747 | 0.141 | 0.147 | 1 | 10 | Prr14 | -0.006 | Young |

| | | | | | | | | |
|---|---|---|---|---|---|---|---|---|
| 0.94399 | 1.849358 | 0.03 | 0.029 | 1 | 10 | Asns | 0.001 | Young |
| 0.944162 | 0.086153 | 0.061 | 0.059 | 1 | 10 | Zxdc | 0.002 | Young |
| 0.944337 | 0.036907 | 0.97 | 0.966 | 1 | 10 | Oaz1 | 0.004 | Young |
| 0.945588 | 0.720633 | 0.152 | 0.161 | 1 | 10 | Mut | -0.009 | Young |
| 0.945588 | 0.096916 | 0.162 | 0.159 | 1 | 10 | Ppp4r1 | 0.003 | Young |
| 0.945599 | 0.19144 | 0.111 | 0.112 | 1 | 10 | Tomm40l | -0.001 | Young |
| 0.945712 | 0.425223 | 0.03 | 0.032 | 1 | 10 | Sel1l3 | -0.002 | Young |
| 0.945863 | 0.056811 | 0.081 | 0.083 | 1 | 10 | Crbn | -0.002 | Young |
| 0.946242 | 0.323912 | 0.162 | 0.174 | 1 | 10 | Ralgapa1 | -0.012 | Young |
| 0.946366 | 0.114326 | 0.424 | 0.443 | 1 | 10 | Ndufab1 | -0.019 | Young |
| 0.946438 | 0.116567 | 0.051 | 0.049 | 1 | 10 | Plpp2 | 0.002 | Young |
| 0.947004 | 0.224083 | 0.182 | 0.191 | 1 | 10 | Sf3a2 | -0.009 | Young |
| 0.947135 | 0.404211 | 0.081 | 0.083 | 1 | 10 | Tmem167b | -0.002 | Young |
| 0.947136 | 0.10642 | 0.061 | 0.059 | 1 | 10 | Ankrd39 | 0.002 | Young |
| 0.947205 | 0.079019 | 0.384 | 0.396 | 1 | 10 | Comt | -0.012 | Young |
| 0.947266 | 0.202421 | 0.061 | 0.064 | 1 | 10 | Tspan33 | -0.003 | Young |
| 0.947266 | 0.063054 | 0.061 | 0.064 | 1 | 10 | Mospd1 | -0.003 | Young |
| 0.947864 | 0.299743 | 0.434 | 0.477 | 1 | 10 | Fkbp2 | -0.043 | Young |
| 0.948133 | 0.628535 | 0.03 | 0.029 | 1 | 10 | Tmem241 | 0.001 | Young |
| 0.948133 | 0.470967 | 0.03 | 0.029 | 1 | 10 | Tfap4 | 0.001 | Young |
| 0.948168 | 1.466357 | 0.02 | 0.02 | 1 | 10 | 1700037C1 | 0 | Young |
| 0.948354 | 0.102532 | 0.273 | 0.279 | 1 | 10 | Ola1 | -0.006 | Young |
| 0.94853 | 0.138354 | 0.141 | 0.139 | 1 | 10 | Nudt16 | 0.002 | Young |
| 0.948537 | 0.259193 | 0.04 | 0.039 | 1 | 10 | Cdkal1 | 0.001 | Young |
| 0.948537 | 0.028734 | 0.04 | 0.039 | 1 | 10 | Rnf25 | 0.001 | Young |
| 0.948539 | 0.075365 | 0.515 | 0.531 | 1 | 10 | Ndufb3 | -0.016 | Young |
| 0.948758 | 0.197818 | 0.323 | 0.342 | 1 | 10 | Csnk1e | -0.019 | Young |
| 0.94876 | 0.263313 | 0.192 | 0.193 | 1 | 10 | Meaf6 | -0.001 | Young |
| 0.949284 | 0.126658 | 0.687 | 0.736 | 1 | 10 | Set | -0.049 | Young |
| 0.949336 | 0.150689 | 0.141 | 0.142 | 1 | 10 | Nudt4 | -0.001 | Young |
| 0.949503 | 0.399305 | 0.141 | 0.152 | 1 | 10 | Usp37 | -0.011 | Young |
| 0.949643 | 0.044477 | 0.212 | 0.21 | 1 | 10 | Isy1 | 0.002 | Young |
| 0.94968 | 0.124768 | 0.051 | 0.049 | 1 | 10 | 1110019D1 | 0.002 | Young |
| 0.949727 | 0.389515 | 0.03 | 0.032 | 1 | 10 | Rad18 | -0.002 | Young |
| 0.950096 | 0.118276 | 0.414 | 0.433 | 1 | 10 | Sec61a1 | -0.019 | Young |
| 0.950174 | 0.157028 | 0.313 | 0.325 | 1 | 10 | Fam126a | -0.012 | Young |
| 0.950205 | 0.331445 | 0.03 | 0.029 | 1 | 10 | Crtc1 | 0.001 | Young |
| 0.950205 | 0.14862 | 0.03 | 0.029 | 1 | 10 | Ets1 | 0.001 | Young |
| 0.950304 | 0.341839 | 0.283 | 0.301 | 1 | 10 | Ep400 | -0.018 | Young |
| 0.950304 | 0.332697 | 0.273 | 0.303 | 1 | 10 | Rnf187 | -0.03 | Young |
| 0.950434 | 0.026461 | 0.222 | 0.225 | 1 | 10 | Atg2a | -0.003 | Young |
| 0.950602 | 0.17327 | 0.434 | 0.45 | 1 | 10 | Sin3b | -0.016 | Young |
| 0.950693 | 1.038253 | 0.02 | 0.02 | 1 | 10 | Abraxas1 | 0 | Young |
| 0.951259 | 0.795583 | 0.131 | 0.139 | 1 | 10 | 4931406P1 | -0.008 | Young |
| 0.951736 | 0.020441 | 0.03 | 0.032 | 1 | 10 | Nat2 | -0.002 | Young |

| | | | | | | | | |
|---|---|---|---|---|---|---|---|---|
| 0.952032 | 0.21915 | 0.162 | 0.171 | 1 | 10 | A630001G2 | -0.009 | Young |
| 0.952083 | 0.068488 | 0.232 | 0.242 | 1 | 10 | Adrm1 | -0.01 | Young |
| 0.952123 | 0.176747 | 0.141 | 0.144 | 1 | 10 | Zscan26 | -0.003 | Young |
| 0.952224 | 0.334109 | 0.081 | 0.083 | 1 | 10 | Exosc2 | -0.002 | Young |
| 0.952248 | 0.13836 | 0.253 | 0.269 | 1 | 10 | Arhgef12 | -0.016 | Young |
| 0.952277 | 0.233735 | 0.03 | 0.029 | 1 | 10 | Pkdcc | 0.001 | Young |
| 0.952277 | 0.077452 | 0.03 | 0.029 | 1 | 10 | Dtwd1 | 0.001 | Young |
| 0.95233 | 0.132704 | 0.091 | 0.09 | 1 | 10 | Rnaseh2b | 0.001 | Young |
| 0.952734 | 0.21524 | 0.404 | 0.433 | 1 | 10 | Yy1 | -0.029 | Young |
| 0.952893 | 0.112225 | 0.141 | 0.147 | 1 | 10 | Mosmo | -0.006 | Young |
| 0.953038 | 0.478682 | 0.061 | 0.064 | 1 | 10 | Habp4 | -0.003 | Young |
| 0.953147 | 0.016824 | 0.98 | 0.976 | 1 | 10 | Myl6 | 0.004 | Young |
| 0.953218 | 0.908418 | 0.02 | 0.02 | 1 | 10 | Arhgef18 | 0 | Young |
| 0.953218 | 0.845359 | 0.02 | 0.02 | 1 | 10 | Cdca7l | 0 | Young |
| 0.953218 | 0.674638 | 0.02 | 0.02 | 1 | 10 | Gtf2ird2 | 0 | Young |
| 0.953218 | 0.630724 | 0.02 | 0.02 | 1 | 10 | Zfp934 | 0 | Young |
| 0.953239 | 0.215918 | 0.222 | 0.23 | 1 | 10 | Atp13a1 | -0.008 | Young |
| 0.95357 | 0.637726 | 0.131 | 0.142 | 1 | 10 | Grk6 | -0.011 | Young |
| 0.953613 | 0.307825 | 0.182 | 0.196 | 1 | 10 | Rab43 | -0.014 | Young |
| 0.953884 | 0.062993 | 0.141 | 0.147 | 1 | 10 | Trmt6 | -0.006 | Young |
| 0.953948 | 1.036756 | 0.04 | 0.039 | 1 | 10 | Socs1 | 0.001 | Young |
| 0.953948 | 0.038787 | 0.04 | 0.039 | 1 | 10 | 2810001G2 | 0.001 | Young |
| 0.954378 | 0.557145 | 0.222 | 0.242 | 1 | 10 | Sap18 | -0.02 | Young |
| 0.95447 | 0.426735 | 0.111 | 0.112 | 1 | 10 | Slc27a4 | -0.001 | Young |
| 0.954533 | 0.87198 | 0.061 | 0.061 | 1 | 10 | Phf6 | 0 | Young |
| 0.954637 | 0.075029 | 0.141 | 0.149 | 1 | 10 | Fam214a | -0.008 | Young |
| 0.955502 | 0.230099 | 0.131 | 0.132 | 1 | 10 | Trove2 | -0.001 | Young |
| 0.955622 | 0.128713 | 0.141 | 0.149 | 1 | 10 | Mrps26 | -0.008 | Young |
| 0.955695 | 0.570581 | 0.212 | 0.225 | 1 | 10 | Zfp326 | -0.013 | Young |
| 0.955745 | 0.903366 | 0.02 | 0.02 | 1 | 10 | Slc37a1 | 0 | Young |
| 0.955754 | 0.704916 | 0.03 | 0.032 | 1 | 10 | Mad2l1 | -0.002 | Young |
| 0.955754 | 0.186539 | 0.03 | 0.032 | 1 | 10 | Ada | -0.002 | Young |
| 0.955925 | 0.42134 | 0.061 | 0.064 | 1 | 10 | Tysnd1 | -0.003 | Young |
| 0.956042 | 0.382809 | 0.081 | 0.083 | 1 | 10 | Slc9a1 | -0.002 | Young |
| 0.956263 | 0.288977 | 0.162 | 0.166 | 1 | 10 | Yif1b | -0.004 | Young |
| 0.956422 | 0.149913 | 0.03 | 0.029 | 1 | 10 | Hmgn3 | 0.001 | Young |
| 0.95662 | 0.136347 | 0.212 | 0.213 | 1 | 10 | Pmpcb | -0.001 | Young |
| 0.956762 | 0.298558 | 0.455 | 0.491 | 1 | 10 | Ranbp1 | -0.036 | Young |
| 0.956956 | 0.024615 | 0.556 | 0.582 | 1 | 10 | Nrp2 | -0.026 | Young |
| 0.956987 | 0.050983 | 0.535 | 0.56 | 1 | 10 | Dad1 | -0.025 | Young |
| 0.956996 | 0.42484 | 0.323 | 0.352 | 1 | 10 | Mat2b | -0.029 | Young |
| 0.957035 | 0.007179 | 0.071 | 0.068 | 1 | 10 | Acrbp | 0.003 | Young |
| 0.957279 | 0.072574 | 0.465 | 0.501 | 1 | 10 | Ndufb10 | -0.036 | Young |
| 0.957307 | 0.92845 | 0.071 | 0.076 | 1 | 10 | Mcm5 | -0.005 | Young |
| 0.957521 | 0.218261 | 0.303 | 0.318 | 1 | 10 | Ufsp2 | -0.015 | Young |

| | | | | | | | | |
|---|---|---|---|---|---|---|---|---|
| 0.957756 | 0.33287 | 0.192 | 0.196 | 1 | 10 | CAAA0111ε | -0.004 | Young |
| 0.95784 | 0.148779 | 0.131 | 0.134 | 1 | 10 | Hspa13 | -0.003 | Young |
| 0.957903 | 0.02249 | 0.273 | 0.286 | 1 | 10 | Commd6 | -0.013 | Young |
| 0.958103 | 0.497169 | 0.131 | 0.137 | 1 | 10 | Sc5d | -0.006 | Young |
| 0.958121 | 0.712218 | 0.111 | 0.115 | 1 | 10 | Ambra1 | -0.004 | Young |
| 0.958271 | 1.159436 | 0.02 | 0.02 | 1 | 10 | Homez | 0 | Young |
| 0.958271 | 0.605694 | 0.02 | 0.02 | 1 | 10 | Immp2l | 0 | Young |
| 0.958271 | 0.23921 | 0.02 | 0.02 | 1 | 10 | Gm4285 | 0 | Young |
| 0.958271 | 0.162947 | 0.02 | 0.02 | 1 | 10 | Neurl1b | 0 | Young |
| 0.958271 | 0.13496 | 0.02 | 0.02 | 1 | 10 | Zfp976 | 0 | Young |
| 0.958467 | 0.48632 | 0.202 | 0.218 | 1 | 10 | Mrpl21 | -0.016 | Young |
| 0.958496 | 1.091322 | 0.03 | 0.029 | 1 | 10 | Trmt10b | 0.001 | Young |
| 0.958496 | 0.189256 | 0.03 | 0.029 | 1 | 10 | Spry2 | 0.001 | Young |
| 0.95851 | 0.356661 | 0.455 | 0.499 | 1 | 10 | Bnip3l | -0.044 | Young |
| 0.958565 | 0.363969 | 0.051 | 0.051 | 1 | 10 | Mynn | 0 | Young |
| 0.958604 | 0.032269 | 0.424 | 0.452 | 1 | 10 | Vps29 | -0.028 | Young |
| 0.958824 | 0.061204 | 0.566 | 0.611 | 1 | 10 | Gls | -0.045 | Young |
| 0.958838 | 0.107222 | 0.141 | 0.147 | 1 | 10 | Mlx | -0.006 | Young |
| 0.958908 | 0.442533 | 0.111 | 0.112 | 1 | 10 | Mtf1 | -0.001 | Young |
| 0.959226 | 0.100737 | 0.697 | 0.768 | 1 | 10 | Tomm7 | -0.071 | Young |
| 0.959402 | 0.257345 | 0.424 | 0.479 | 1 | 10 | Mvp | -0.055 | Young |
| 0.959772 | 0.496766 | 0.03 | 0.032 | 1 | 10 | Coq4 | -0.002 | Young |
| 0.959829 | 0.356967 | 0.141 | 0.147 | 1 | 10 | Agap3 | -0.006 | Young |
| 0.95994 | 0.330261 | 0.293 | 0.315 | 1 | 10 | Cenpx | -0.022 | Young |
| 0.960158 | 0.15255 | 0.051 | 0.051 | 1 | 10 | Mrps22 | 0 | Young |
| 0.960257 | 1.020206 | 0.061 | 0.064 | 1 | 10 | Prpf3 | -0.003 | Young |
| 0.960349 | 0.152994 | 0.404 | 0.425 | 1 | 10 | Glyr1 | -0.021 | Young |
| 0.960394 | 0.136935 | 0.192 | 0.196 | 1 | 10 | Wdr61 | -0.004 | Young |
| 0.960666 | 0.185088 | 0.101 | 0.103 | 1 | 10 | Ikzf5 | -0.002 | Young |
| 0.96078 | 0.190047 | 0.242 | 0.252 | 1 | 10 | Mertk | -0.01 | Young |
| 0.960798 | 0.711375 | 0.02 | 0.02 | 1 | 10 | Ralgps1 | 0 | Young |
| 0.960798 | 0.3127 | 0.02 | 0.02 | 1 | 10 | Rttn | 0 | Young |
| 0.960798 | 0.102446 | 0.02 | 0.02 | 1 | 10 | Vcan | 0 | Young |
| 0.960798 | 0.007011 | 0.02 | 0.02 | 1 | 10 | Ptpn21 | 0 | Young |
| 0.960826 | 0.214462 | 0.071 | 0.073 | 1 | 10 | Mllt11 | -0.002 | Young |
| 0.960941 | 0.319617 | 0.303 | 0.333 | 1 | 10 | Naa15 | -0.03 | Young |
| 0.96107 | 0.335866 | 0.323 | 0.357 | 1 | 10 | Vav1 | -0.034 | Young |
| 0.961134 | 0.690107 | 0.081 | 0.083 | 1 | 10 | Siae | -0.002 | Young |
| 0.961259 | 0.283979 | 0.141 | 0.152 | 1 | 10 | Fnbp4 | -0.011 | Young |
| 0.961578 | 0.403902 | 0.192 | 0.2 | 1 | 10 | Asnsd1 | -0.008 | Young |
| 0.961579 | 0.117418 | 0.616 | 0.685 | 1 | 10 | Pnrc1 | -0.069 | Young |
| 0.961782 | 0.316063 | 0.03 | 0.032 | 1 | 10 | Noa1 | -0.002 | Young |
| 0.96195 | 0.091524 | 0.131 | 0.134 | 1 | 10 | Mrpl50 | -0.003 | Young |
| 0.962088 | 0.08021 | 0.414 | 0.425 | 1 | 10 | Twf2 | -0.011 | Young |
| 0.962187 | 0.652179 | 0.131 | 0.137 | 1 | 10 | Tbcel | -0.006 | Young |

| | | | | | | | | |
|---|---|---|---|---|---|---|---|---|
| 0.96242 | 0.017727 | 0.192 | 0.186 | 1 | 10 | Esrra | 0.006 | Young |
| 0.962575 | 0.198584 | 0.071 | 0.068 | 1 | 10 | Dbr1 | 0.003 | Young |
| 0.962643 | 0.253526 | 0.03 | 0.029 | 1 | 10 | C330027CC | 0.001 | Young |
| 0.962822 | 0.406753 | 0.091 | 0.095 | 1 | 10 | Cep57 | -0.004 | Young |
| 0.963094 | 0.452366 | 0.253 | 0.257 | 1 | 10 | Cyth1 | -0.004 | Young |
| 0.963101 | 0.532926 | 0.253 | 0.267 | 1 | 10 | Ncaph2 | -0.014 | Young |
| 0.96313 | 0.25243 | 0.162 | 0.164 | 1 | 10 | Oser1 | -0.002 | Young |
| 0.963145 | 0.518023 | 0.061 | 0.064 | 1 | 10 | Galm | -0.003 | Young |
| 0.963145 | 0.391748 | 0.061 | 0.064 | 1 | 10 | Cbr1 | -0.003 | Young |
| 0.963325 | 0.799647 | 0.02 | 0.02 | 1 | 10 | Sh3rf1 | 0 | Young |
| 0.963325 | 0.123117 | 0.02 | 0.02 | 1 | 10 | Gm29367 | 0 | Young |
| 0.963325 | 0.029822 | 0.02 | 0.02 | 1 | 10 | Fam234b | 0 | Young |
| 0.963325 | 0.0184 | 0.02 | 0.02 | 1 | 10 | Blm | 0 | Young |
| 0.963765 | 0.009696 | 0.293 | 0.296 | 1 | 10 | BC005624 | -0.003 | Young |
| 0.963991 | 0.815565 | 0.051 | 0.054 | 1 | 10 | Nbas | -0.003 | Young |
| 0.964428 | 0.153896 | 0.071 | 0.071 | 1 | 10 | Aimp2 | 0 | Young |
| 0.964599 | 0.230788 | 0.293 | 0.298 | 1 | 10 | Bet1 | -0.005 | Young |
| 0.96461 | 0.191364 | 0.283 | 0.313 | 1 | 10 | Smarca4 | -0.03 | Young |
| 0.964717 | 0.413704 | 0.03 | 0.029 | 1 | 10 | Wdr59 | 0.001 | Young |
| 0.964887 | 0.312439 | 0.091 | 0.093 | 1 | 10 | Pbx1 | -0.002 | Young |
| 0.965103 | 0.011684 | 0.192 | 0.186 | 1 | 10 | Fxr2 | 0.006 | Young |
| 0.965501 | 0.281313 | 0.121 | 0.12 | 1 | 10 | Lsg1 | 0.001 | Young |
| 0.965799 | 0.215697 | 0.303 | 0.311 | 1 | 10 | Ppp3cb | -0.008 | Young |
| 0.965803 | 0.478281 | 0.03 | 0.032 | 1 | 10 | Lhpp | -0.002 | Young |
| 0.965853 | 0.29921 | 0.02 | 0.02 | 1 | 10 | Stac2 | 0 | Young |
| 0.965853 | 0.271556 | 0.02 | 0.02 | 1 | 10 | Pcgf1 | 0 | Young |
| 0.966061 | 0.09815 | 0.131 | 0.134 | 1 | 10 | Nek6 | -0.003 | Young |
| 0.966097 | 0.430884 | 0.091 | 0.093 | 1 | 10 | Ap5s1 | -0.002 | Young |
| 0.966286 | 0.08891 | 0.242 | 0.24 | 1 | 10 | Sdccag3 | 0.002 | Young |
| 0.966399 | 0.211801 | 0.152 | 0.159 | 1 | 10 | Fmn1 | -0.007 | Young |
| 0.966497 | 0.263286 | 0.081 | 0.081 | 1 | 10 | Cyb561d2 | 0 | Young |
| 0.966536 | 0.094526 | 0.586 | 0.633 | 1 | 10 | H2afj | -0.047 | Young |
| 0.966663 | 0.104404 | 0.253 | 0.257 | 1 | 10 | Mgst1 | -0.004 | Young |
| 0.966773 | 0.223942 | 0.172 | 0.174 | 1 | 10 | Rpe | -0.002 | Young |
| 0.966791 | 0.174995 | 0.03 | 0.029 | 1 | 10 | Cbarp | 0.001 | Young |
| 0.966878 | 0.136838 | 0.131 | 0.132 | 1 | 10 | Aak1 | -0.001 | Young |
| 0.966925 | 0.326208 | 0.172 | 0.176 | 1 | 10 | Sart1 | -0.004 | Young |
| 0.96732 | 0.374693 | 0.101 | 0.1 | 1 | 10 | Ttc17 | 0.001 | Young |
| 0.967384 | 0.256071 | 0.414 | 0.462 | 1 | 10 | Lsm4 | -0.048 | Young |
| 0.967431 | 0.011593 | 0.192 | 0.196 | 1 | 10 | Lrpprc | -0.004 | Young |
| 0.967502 | 0.273446 | 0.081 | 0.083 | 1 | 10 | Tmem242 | -0.002 | Young |
| 0.967576 | 1.250484 | 0.071 | 0.073 | 1 | 10 | Nhsl2 | -0.002 | Young |
| 0.967881 | 0.035842 | 0.657 | 0.702 | 1 | 10 | Reep3 | -0.045 | Young |
| 0.967979 | 0.265897 | 0.394 | 0.418 | 1 | 10 | Ppp3r1 | -0.024 | Young |
| 0.968206 | 0.106141 | 0.596 | 0.65 | 1 | 10 | Jak1 | -0.054 | Young |

| | | | | | | | | |
|---|---|---|---|---|---|---|---|---|
| 0.968381 | 0.370159 | 0.02 | 0.02 | 1 | 10 | Smo | 0 | Young |
| 0.968381 | 0.262538 | 0.02 | 0.02 | 1 | 10 | Gm20045 | 0 | Young |
| 0.968381 | 0.174027 | 0.02 | 0.02 | 1 | 10 | Wwc1 | 0 | Young |
| 0.968455 | 0.076483 | 0.707 | 0.782 | 1 | 10 | Clic4 | -0.075 | Young |
| 0.968745 | 0.091263 | 0.131 | 0.13 | 1 | 10 | Nlk | 0.001 | Young |
| 0.968776 | 0.206201 | 0.081 | 0.083 | 1 | 10 | Cdyl | -0.002 | Young |
| 0.969639 | 0.150991 | 0.475 | 0.528 | 1 | 10 | Tmbim1 | -0.053 | Young |
| 0.970049 | 0.11281 | 0.081 | 0.083 | 1 | 10 | Plxdc2 | -0.002 | Young |
| 0.970178 | 0.059108 | 0.636 | 0.663 | 1 | 10 | Dync1h1 | -0.027 | Young |
| 0.970306 | 0.009482 | 0.202 | 0.198 | 1 | 10 | Plpbp | 0.004 | Young |
| 0.970361 | 0.313502 | 0.081 | 0.081 | 1 | 10 | Fbxo45 | 0 | Young |
| 0.970361 | 0.237586 | 0.081 | 0.081 | 1 | 10 | Arhgap4 | 0 | Young |
| 0.970512 | 0.214398 | 0.384 | 0.408 | 1 | 10 | Ttc14 | -0.024 | Young |
| 0.970653 | 0.137587 | 0.091 | 0.09 | 1 | 10 | Irf2bp1 | 0.001 | Young |
| 0.970828 | 0.003665 | 0.131 | 0.13 | 1 | 10 | Dennd1c | 0.001 | Young |
| 0.970909 | 0.17805 | 0.02 | 0.02 | 1 | 10 | Nim1k | 0 | Young |
| 0.970909 | 0.114668 | 0.02 | 0.02 | 1 | 10 | Gm26778 | 0 | Young |
| 0.971308 | 0.572219 | 0.051 | 0.051 | 1 | 10 | Rpap3 | 0 | Young |
| 0.971522 | 2.098641 | 0.01 | 0.01 | 1 | 10 | Gm17259 | 0 | Young |
| 0.971522 | 1.88041 | 0.01 | 0.01 | 1 | 10 | Phf11c | 0 | Young |
| 0.971522 | 1.645759 | 0.01 | 0.01 | 1 | 10 | Rflnb | 0 | Young |
| 0.971522 | 1.504412 | 0.01 | 0.01 | 1 | 10 | Cdh24 | 0 | Young |
| 0.971522 | 1.357869 | 0.01 | 0.01 | 1 | 10 | Bace1 | 0 | Young |
| 0.971522 | 1.258452 | 0.01 | 0.01 | 1 | 10 | Oas1c | 0 | Young |
| 0.971522 | 1.183431 | 0.01 | 0.01 | 1 | 10 | Igf1os | 0 | Young |
| 0.971522 | 1.124271 | 0.01 | 0.01 | 1 | 10 | Dpf3 | 0 | Young |
| 0.971522 | 1.121776 | 0.01 | 0.01 | 1 | 10 | C230037L1 | 0 | Young |
| 0.971522 | 0.932659 | 0.01 | 0.01 | 1 | 10 | Fbln5 | 0 | Young |
| 0.971522 | 0.802416 | 0.01 | 0.01 | 1 | 10 | Pcdhgb4 | 0 | Young |
| 0.971522 | 0.702516 | 0.01 | 0.01 | 1 | 10 | Car7 | 0 | Young |
| 0.971522 | 0.689878 | 0.01 | 0.01 | 1 | 10 | Slc7a13 | 0 | Young |
| 0.971522 | 0.67535 | 0.01 | 0.01 | 1 | 10 | Bahcc1 | 0 | Young |
| 0.971522 | 0.54844 | 0.01 | 0.01 | 1 | 10 | 4930589L2 | 0 | Young |
| 0.971522 | 0.513716 | 0.01 | 0.01 | 1 | 10 | Xlr | 0 | Young |
| 0.971522 | 0.46606 | 0.01 | 0.01 | 1 | 10 | Gm26826 | 0 | Young |
| 0.971522 | 0.464811 | 0.01 | 0.01 | 1 | 10 | Plxnb1 | 0 | Young |
| 0.971522 | 0.462648 | 0.01 | 0.01 | 1 | 10 | Wnt4 | 0 | Young |
| 0.971522 | 0.458372 | 0.01 | 0.01 | 1 | 10 | Kif9 | 0 | Young |
| 0.971522 | 0.428428 | 0.01 | 0.01 | 1 | 10 | Bbs12 | 0 | Young |
| 0.971522 | 0.33388 | 0.01 | 0.01 | 1 | 10 | Gm10603 | 0 | Young |
| 0.971554 | 0.581046 | 0.111 | 0.117 | 1 | 10 | Nbdy | -0.006 | Young |
| 0.971634 | 0.094866 | 0.162 | 0.164 | 1 | 10 | Pitpnm1 | -0.002 | Young |
| 0.971869 | 0.330003 | 0.131 | 0.13 | 1 | 10 | Stam | 0.001 | Young |
| 0.971941 | 0.261835 | 0.222 | 0.232 | 1 | 10 | Sipa1 | -0.01 | Young |
| 0.972033 | 0.14777 | 0.313 | 0.32 | 1 | 10 | Mrpl57 | -0.007 | Young |

| | | | | | | | | |
|---|---|---|---|---|---|---|---|---|
| 0.972124 | 0.707129 | 0.061 | 0.061 | 1 | 10 | Kctd18 | 0 | Young |
| 0.972199 | 0.069785 | 0.606 | 0.636 | 1 | 10 | Gna13 | -0.03 | Young |
| 0.972231 | 0.21677 | 0.333 | 0.359 | 1 | 10 | Sfxn3 | -0.026 | Young |
| 0.972464 | 0.029455 | 0.101 | 0.105 | 1 | 10 | Ndc1 | -0.004 | Young |
| 0.972597 | 1.040694 | 0.081 | 0.083 | 1 | 10 | Mgat4a | -0.002 | Young |
| 0.972807 | 0.329759 | 0.101 | 0.103 | 1 | 10 | Mapk9 | -0.002 | Young |
| 0.972867 | 0.09194 | 0.515 | 0.518 | 1 | 10 | Itsn2 | -0.003 | Young |
| 0.972937 | 0.16081 | 0.081 | 0.081 | 1 | 10 | Ints2 | 0 | Young |
| 0.973269 | 0.324024 | 0.232 | 0.244 | 1 | 10 | Tcof1 | -0.012 | Young |
| 0.973513 | 0.165562 | 0.505 | 0.521 | 1 | 10 | Ncor1 | -0.016 | Young |
| 0.974055 | 0.528897 | 0.141 | 0.144 | 1 | 10 | Tusc2 | -0.003 | Young |
| 0.974367 | 0.017043 | 0.192 | 0.193 | 1 | 10 | Qars | -0.001 | Young |
| 0.974416 | 0.156385 | 0.222 | 0.232 | 1 | 10 | Gars | -0.01 | Young |
| 0.974737 | 0.01091 | 0.919 | 0.961 | 1 | 10 | Rpl22 | -0.042 | Young |
| 0.974963 | 0.219935 | 0.232 | 0.247 | 1 | 10 | Txndc5 | -0.015 | Young |
| 0.97502 | 0.751202 | 0.111 | 0.12 | 1 | 10 | Bag5 | -0.009 | Young |
| 0.97508 | 0.796333 | 0.01 | 0.01 | 1 | 10 | Aldh1a1 | 0 | Young |
| 0.97508 | 0.656409 | 0.01 | 0.01 | 1 | 10 | Gm17146 | 0 | Young |
| 0.97508 | 0.595286 | 0.01 | 0.01 | 1 | 10 | Gabbr1 | 0 | Young |
| 0.97508 | 0.548282 | 0.01 | 0.01 | 1 | 10 | Pcdhga12 | 0 | Young |
| 0.97508 | 0.428339 | 0.01 | 0.01 | 1 | 10 | Ncaph | 0 | Young |
| 0.97508 | 0.427349 | 0.01 | 0.01 | 1 | 10 | Plcd1 | 0 | Young |
| 0.97508 | 0.368233 | 0.01 | 0.01 | 1 | 10 | Fancf | 0 | Young |
| 0.97508 | 0.362186 | 0.01 | 0.01 | 1 | 10 | Zbtb8a | 0 | Young |
| 0.97508 | 0.347558 | 0.01 | 0.01 | 1 | 10 | Rhebl1 | 0 | Young |
| 0.97508 | 0.341818 | 0.01 | 0.01 | 1 | 10 | Tyw3 | 0 | Young |
| 0.97508 | 0.285268 | 0.01 | 0.01 | 1 | 10 | Hspb6 | 0 | Young |
| 0.97508 | 0.278096 | 0.01 | 0.01 | 1 | 10 | Dach1 | 0 | Young |
| 0.97508 | 0.243779 | 0.01 | 0.01 | 1 | 10 | Ciart | 0 | Young |
| 0.97508 | 0.234369 | 0.01 | 0.01 | 1 | 10 | Zscan2 | 0 | Young |
| 0.97508 | 0.23178 | 0.01 | 0.01 | 1 | 10 | Slc25a23 | 0 | Young |
| 0.97508 | 0.224397 | 0.01 | 0.01 | 1 | 10 | Hes7 | 0 | Young |
| 0.97508 | 0.214158 | 0.01 | 0.01 | 1 | 10 | 4930518I1! | 0 | Young |
| 0.97508 | 0.203462 | 0.01 | 0.01 | 1 | 10 | Gm7324 | 0 | Young |
| 0.97508 | 0.200435 | 0.01 | 0.01 | 1 | 10 | Dtl | 0 | Young |
| 0.97508 | 0.052157 | 0.01 | 0.01 | 1 | 10 | Bbs4 | 0 | Young |
| 0.97508 | 0.036438 | 0.01 | 0.01 | 1 | 10 | Eif3j2 | 0 | Young |
| 0.97508 | 0.014482 | 0.01 | 0.01 | 1 | 10 | Ect2 | 0 | Young |
| 0.975219 | 0.364802 | 0.303 | 0.335 | 1 | 10 | Snf8 | -0.032 | Young |
| 0.975235 | 0.217285 | 0.414 | 0.45 | 1 | 10 | Ube2k | -0.036 | Young |
| 0.975516 | 0.483816 | 0.152 | 0.159 | 1 | 10 | Nipsnap2 | -0.007 | Young |
| 0.975629 | 0.138282 | 0.182 | 0.183 | 1 | 10 | Pitpnb | -0.001 | Young |
| 0.975645 | 0.060257 | 0.889 | 0.929 | 1 | 10 | Hnrnpa2b1 | -0.04 | Young |
| 0.97575 | 0.318678 | 0.202 | 0.215 | 1 | 10 | Ncoa7 | -0.013 | Young |
| 0.975905 | 0.259139 | 0.101 | 0.105 | 1 | 10 | Msi2 | -0.004 | Young |

| | | | | | | | | |
|---|---|---|---|---|---|---|---|---|
| 0.976217 | 0.115899 | 0.616 | 0.623 | 1 | 10 | Eif4b | -0.007 | Young |
| 0.976437 | 0.184316 | 0.485 | 0.528 | 1 | 10 | Slk | -0.043 | Young |
| 0.976737 | 0.167635 | 0.071 | 0.071 | 1 | 10 | Nlrc4 | 0 | Young |
| 0.976956 | 0.365653 | 0.152 | 0.159 | 1 | 10 | Fuom | -0.007 | Young |
| 0.97699 | 0.073772 | 0.091 | 0.093 | 1 | 10 | Rbl1 | -0.002 | Young |
| 0.977421 | 0.493728 | 0.212 | 0.225 | 1 | 10 | Anapc16 | -0.013 | Young |
| 0.977577 | 0.108766 | 0.273 | 0.276 | 1 | 10 | Maea | -0.003 | Young |
| 0.977694 | 0.648076 | 0.081 | 0.083 | 1 | 10 | Fam219b | -0.002 | Young |
| 0.977694 | 0.576334 | 0.081 | 0.083 | 1 | 10 | Rhbdd1 | -0.002 | Young |
| 0.978091 | 0.158546 | 0.081 | 0.081 | 1 | 10 | Depdc5 | 0 | Young |
| 0.978105 | 0.252479 | 0.071 | 0.071 | 1 | 10 | Slc52a2 | 0 | Young |
| 0.978563 | 0.880209 | 0.081 | 0.086 | 1 | 10 | Polk | -0.005 | Young |
| 0.978639 | 0.287418 | 0.01 | 0.01 | 1 | 10 | Xrcc2 | 0 | Young |
| 0.978639 | 0.17737 | 0.01 | 0.01 | 1 | 10 | 4930512J1 | 0 | Young |
| 0.978639 | 0.001485 | 0.01 | 0.01 | 1 | 10 | Gm26982 | 0 | Young |
| 0.978698 | 0.275226 | 0.192 | 0.191 | 1 | 10 | Cisd1 | 0.001 | Young |
| 0.97882 | 0.149524 | 0.04 | 0.042 | 1 | 10 | Pomt2 | -0.002 | Young |
| 0.979276 | 0.017895 | 0.051 | 0.051 | 1 | 10 | Alg3 | 0 | Young |
| 0.979504 | 0.079862 | 0.202 | 0.213 | 1 | 10 | Plau | -0.011 | Young |
| 0.979586 | 0.422806 | 0.182 | 0.193 | 1 | 10 | Nln | -0.011 | Young |
| 0.979591 | 0.435336 | 0.313 | 0.333 | 1 | 10 | Pole4 | -0.02 | Young |
| 0.979748 | 0.238333 | 0.101 | 0.103 | 1 | 10 | Mob3b | -0.002 | Young |
| 0.979861 | 0.162263 | 0.212 | 0.222 | 1 | 10 | Hmgcl | -0.01 | Young |
| 0.980313 | 0.096773 | 0.404 | 0.423 | 1 | 10 | Rsu1 | -0.019 | Young |
| 0.980525 | 0.052624 | 0.586 | 0.611 | 1 | 10 | Fubp1 | -0.025 | Young |
| 0.980585 | 0.292059 | 0.04 | 0.042 | 1 | 10 | Isoc2b | -0.002 | Young |
| 0.980607 | 0.46395 | 0.333 | 0.347 | 1 | 10 | Emc1 | -0.014 | Young |
| 0.980668 | 0.445061 | 0.081 | 0.081 | 1 | 10 | Fbxo7 | 0 | Young |
| 0.980674 | 0.192427 | 0.323 | 0.355 | 1 | 10 | Fam32a | -0.032 | Young |
| 0.980676 | 0.238784 | 0.182 | 0.186 | 1 | 10 | Gsap | -0.004 | Young |
| 0.980905 | 0.57173 | 0.101 | 0.103 | 1 | 10 | Rrs1 | -0.002 | Young |
| 0.980925 | 0.480626 | 0.061 | 0.061 | 1 | 10 | Wee1 | 0 | Young |
| 0.981137 | 0.380101 | 0.222 | 0.237 | 1 | 10 | Sgta | -0.015 | Young |
| 0.981137 | 0.186607 | 0.222 | 0.237 | 1 | 10 | Mid1ip1 | -0.015 | Young |
| 0.981585 | 0.152284 | 0.242 | 0.252 | 1 | 10 | Pabpn1 | -0.01 | Young |
| 0.981791 | 0.32593 | 0.242 | 0.262 | 1 | 10 | Cyld | -0.02 | Young |
| 0.982075 | 0.414267 | 0.313 | 0.355 | 1 | 10 | Gpr137b | -0.042 | Young |
| 0.98235 | 0.443503 | 0.04 | 0.042 | 1 | 10 | Ndufaf1 | -0.002 | Young |
| 0.982366 | 0.192255 | 0.384 | 0.394 | 1 | 10 | Camk1 | -0.01 | Young |
| 0.982464 | 0.91111 | 0.051 | 0.051 | 1 | 10 | Srbd1 | 0 | Young |
| 0.982464 | 0.801073 | 0.051 | 0.051 | 1 | 10 | Scmh1 | 0 | Young |
| 0.982556 | 0.05696 | 0.747 | 0.787 | 1 | 10 | M6pr | -0.04 | Young |
| 0.982664 | 0.448015 | 0.192 | 0.205 | 1 | 10 | Ddx19a | -0.013 | Young |
| 0.98276 | 0.388676 | 0.162 | 0.171 | 1 | 10 | Kat7 | -0.009 | Young |
| 0.983204 | 0.026846 | 0.192 | 0.193 | 1 | 10 | Pik3cg | -0.001 | Young |

| | | | | | | | | |
|---|---|---|---|---|---|---|---|---|
| 0.983746 | 0.328065 | 0.222 | 0.225 | 1 | 10 | Tex264 | -0.003 | Young |
| 0.983785 | 0.414269 | 0.071 | 0.073 | 1 | 10 | Ikbip | -0.002 | Young |
| 0.983785 | 0.254264 | 0.071 | 0.073 | 1 | 10 | Grap | -0.002 | Young |
| 0.984058 | 0.419699 | 0.051 | 0.051 | 1 | 10 | Rps6ka2 | 0 | Young |
| 0.98407 | 0.277214 | 0.273 | 0.296 | 1 | 10 | Fam129a | -0.023 | Young |
| 0.984307 | 0.117283 | 0.394 | 0.416 | 1 | 10 | Dnaja2 | -0.022 | Young |
| 0.984565 | 0.429133 | 0.111 | 0.115 | 1 | 10 | Wdr46 | -0.004 | Young |
| 0.98468 | 0.187755 | 0.485 | 0.499 | 1 | 10 | Hnrnpdl | -0.014 | Young |
| 0.985201 | 0.49315 | 0.141 | 0.149 | 1 | 10 | Nrf1 | -0.008 | Young |
| 0.985268 | 0.15545 | 0.162 | 0.161 | 1 | 10 | Nr1h2 | 0.001 | Young |
| 0.985315 | 0.684617 | 0.121 | 0.13 | 1 | 10 | Unc119 | -0.009 | Young |
| 0.985544 | 0.121608 | 0.364 | 0.367 | 1 | 10 | Sh3pxd2b | -0.003 | Young |
| 0.985688 | 0.376665 | 0.131 | 0.137 | 1 | 10 | Vti1a | -0.006 | Young |
| 0.985688 | 0.368089 | 0.131 | 0.137 | 1 | 10 | Srm | -0.006 | Young |
| 0.985693 | 0.039726 | 0.98 | 0.983 | 1 | 10 | Cfl1 | -0.003 | Young |
| 0.985968 | 0.356991 | 0.222 | 0.232 | 1 | 10 | Mettl9 | -0.01 | Young |
| 0.986209 | 0.069966 | 0.727 | 0.77 | 1 | 10 | Fxyd5 | -0.043 | Young |
| 0.98634 | 0.299858 | 0.172 | 0.181 | 1 | 10 | Polb | -0.009 | Young |
| 0.986616 | 0.112136 | 0.121 | 0.122 | 1 | 10 | Gm5617 | -0.001 | Young |
| 0.987189 | 0.302966 | 0.202 | 0.213 | 1 | 10 | Hspbp1 | -0.011 | Young |
| 0.987644 | 0.163742 | 0.04 | 0.042 | 1 | 10 | Zfp317 | -0.002 | Young |
| 0.987658 | 0.092145 | 0.545 | 0.587 | 1 | 10 | Caprin1 | -0.042 | Young |
| 0.987711 | 0.989413 | 0.061 | 0.064 | 1 | 10 | Ahi1 | -0.003 | Young |
| 0.98777 | 0.182464 | 0.091 | 0.09 | 1 | 10 | Osbpl2 | 0.001 | Young |
| 0.987889 | 0.022903 | 0.091 | 0.093 | 1 | 10 | Alkbh3 | -0.002 | Young |
| 0.988053 | 0.017045 | 0.172 | 0.176 | 1 | 10 | Eaf1 | -0.004 | Young |
| 0.988599 | 0.271846 | 0.192 | 0.198 | 1 | 10 | Phactr2 | -0.006 | Young |
| 0.98884 | 0.511295 | 0.051 | 0.051 | 1 | 10 | Cep162 | 0 | Young |
| 0.988935 | 0.036384 | 0.707 | 0.738 | 1 | 10 | Rab1a | -0.031 | Young |
| 0.989051 | 0.550891 | 0.071 | 0.071 | 1 | 10 | Cry1 | 0 | Young |
| 0.9891 | 0.496332 | 0.091 | 0.093 | 1 | 10 | Fgfr1op | -0.002 | Young |
| 0.9891 | 0.397228 | 0.091 | 0.093 | 1 | 10 | AC127341. | -0.002 | Young |
| 0.9891 | 0.171491 | 0.091 | 0.093 | 1 | 10 | Alg9 | -0.002 | Young |
| 0.989121 | 0.50221 | 0.333 | 0.381 | 1 | 10 | Pum1 | -0.048 | Young |
| 0.989147 | 0.148151 | 0.141 | 0.149 | 1 | 10 | Slc38a7 | -0.008 | Young |
| 0.989267 | 0.037574 | 0.152 | 0.152 | 1 | 10 | Gfra2 | 0 | Young |
| 0.989476 | 0.314431 | 0.192 | 0.198 | 1 | 10 | Snrnp48 | -0.006 | Young |
| 0.989833 | 0.268519 | 0.323 | 0.335 | 1 | 10 | Smim15 | -0.012 | Young |
| 0.990001 | 0.381507 | 0.111 | 0.112 | 1 | 10 | Fam114a1 | -0.001 | Young |
| 0.990541 | 0.35659 | 0.071 | 0.073 | 1 | 10 | Atg9a | -0.002 | Young |
| 0.990563 | 0.043768 | 0.929 | 0.954 | 1 | 10 | Mcl1 | -0.025 | Young |
| 0.990594 | 0.091153 | 0.505 | 0.509 | 1 | 10 | Csnk2b | -0.004 | Young |
| 0.990661 | 0.087688 | 0.101 | 0.1 | 1 | 10 | Fam193b | 0.001 | Young |
| 0.990745 | 0.067711 | 0.444 | 0.472 | 1 | 10 | Aldh2 | -0.028 | Young |
| 0.99106 | 0.416897 | 0.162 | 0.166 | 1 | 10 | Naxe | -0.004 | Young |

| | | | | | | | | |
|---|---|---|---|---|---|---|---|---|
| 0.991172 | 0.04849 | 0.97 | 0.968 | 1 | 10 | Rplp0 | 0.002 | Young |
| 0.991174 | 0.490586 | 0.04 | 0.042 | 1 | 10 | Klhl36 | -0.002 | Young |
| 0.991229 | 0.382698 | 0.162 | 0.176 | 1 | 10 | Nomo1 | -0.014 | Young |
| 0.991246 | 0.31969 | 0.111 | 0.117 | 1 | 10 | Klhl18 | -0.006 | Young |
| 0.991665 | 0.174785 | 0.354 | 0.394 | 1 | 10 | Psmc6 | -0.04 | Young |
| 0.992028 | 0.29395 | 0.051 | 0.051 | 1 | 10 | Rad51d | 0 | Young |
| 0.992041 | 0.167803 | 0.747 | 0.785 | 1 | 10 | Cxcl2 | -0.038 | Young |
| 0.992223 | 0.020133 | 0.111 | 0.112 | 1 | 10 | Armc10 | -0.001 | Young |
| 0.992463 | 0.141827 | 0.465 | 0.489 | 1 | 10 | Rpl36al | -0.024 | Young |
| 0.992659 | 0.065299 | 0.545 | 0.553 | 1 | 10 | Fermt3 | -0.008 | Young |
| 0.992799 | 0.181144 | 0.131 | 0.134 | 1 | 10 | Frmd8 | -0.003 | Young |
| 0.993084 | 0.176043 | 0.374 | 0.396 | 1 | 10 | Prex1 | -0.022 | Young |
| 0.993115 | 0.342505 | 0.101 | 0.105 | 1 | 10 | Fam98a | -0.004 | Young |
| 0.9932 | 0.070433 | 0.182 | 0.181 | 1 | 10 | Arl6ip6 | 0.001 | Young |
| 0.993206 | 0.274849 | 0.152 | 0.154 | 1 | 10 | Kmt2b | -0.002 | Young |
| 0.993243 | 0.261325 | 0.152 | 0.156 | 1 | 10 | Atl3 | -0.004 | Young |
| 0.993261 | 0.14961 | 0.505 | 0.543 | 1 | 10 | Ptpn6 | -0.038 | Young |
| 0.993385 | 0.824499 | 0.111 | 0.115 | 1 | 10 | Spsb2 | -0.004 | Young |
| 0.99405 | 0.332564 | 0.303 | 0.311 | 1 | 10 | Fam76a | -0.008 | Young |
| 0.994367 | 0.093211 | 0.354 | 0.372 | 1 | 10 | Mbtd1 | -0.018 | Young |
| 0.994435 | 0.068306 | 0.172 | 0.171 | 1 | 10 | Sos2 | 0.001 | Young |
| 0.994679 | 0.296035 | 0.414 | 0.465 | 1 | 10 | Eif3b | -0.051 | Young |
| 0.994704 | 1.364625 | 0.04 | 0.042 | 1 | 10 | Atpaf1 | -0.002 | Young |
| 0.995556 | 0.126944 | 0.111 | 0.112 | 1 | 10 | Enpp1 | -0.001 | Young |
| 0.995583 | 0.117267 | 0.242 | 0.249 | 1 | 10 | Mrpl41 | -0.007 | Young |
| 0.996143 | 0.285056 | 0.263 | 0.281 | 1 | 10 | Supt4a | -0.018 | Young |
| 0.996366 | 0.239387 | 0.091 | 0.093 | 1 | 10 | Saysd1 | -0.002 | Young |
| 0.996366 | 0.231205 | 0.091 | 0.093 | 1 | 10 | Smyd5 | -0.002 | Young |
| 0.99647 | 0.63077 | 0.04 | 0.042 | 1 | 10 | Msantd2 | -0.002 | Young |
| 0.996527 | 0.156985 | 0.141 | 0.147 | 1 | 10 | Taf11 | -0.006 | Young |
| 0.997111 | 0.199037 | 0.333 | 0.34 | 1 | 10 | Smap1 | -0.007 | Young |
| 0.99712 | 0.500383 | 0.222 | 0.235 | 1 | 10 | Slc17a5 | -0.013 | Young |
| 0.997557 | 0.131773 | 0.768 | 0.829 | 1 | 10 | Kctd12 | -0.061 | Young |
| 0.998095 | 0.2501 | 0.283 | 0.289 | 1 | 10 | Vegfa | -0.006 | Young |
| 0.998528 | 0.432526 | 0.141 | 0.152 | 1 | 10 | Pgp | -0.011 | Young |
| 0.998533 | 0.27863 | 0.061 | 0.061 | 1 | 10 | Galnt4 | 0 | Young |
| 0.998533 | 0.221911 | 0.061 | 0.061 | 1 | 10 | Rhot2 | 0 | Young |
| 0.998533 | 0.100783 | 0.061 | 0.061 | 1 | 10 | Slc35a1 | 0 | Young |
| 0.998533 | 0.078387 | 0.061 | 0.061 | 1 | 10 | Phf11d | 0 | Young |
| 0.998558 | 0.146846 | 0.333 | 0.342 | 1 | 10 | Trappc6b | -0.009 | Young |
| 0.998631 | 0.539589 | 0.071 | 0.071 | 1 | 10 | Tmem209 | 0 | Young |
| 0.99864 | 0.042793 | 0.182 | 0.181 | 1 | 10 | Gadd45gip | 0.001 | Young |
| 0.998649 | 0.530205 | 0.071 | 0.073 | 1 | 10 | Aar2 | -0.002 | Young |
| 0.998897 | 0.709153 | 0.111 | 0.115 | 1 | 10 | Cask | -0.004 | Young |
| 0.999317 | 0.283603 | 0.374 | 0.418 | 1 | 10 | Rad23b | -0.044 | Young |

| | | | | | | | | |
|---|---|---|---|---|---|---|---|---|
| 0.999371 | 0.063701 | 0.566 | 0.614 | 1 | 10 | Man2b1 | -0.048 | Young |
| 0.999382 | 0.164393 | 0.657 | 0.699 | 1 | 10 | Mrfap1 | -0.042 | Young |
| 1.65E-11 | 2.580445 | 0.617 | 0.297 | 5.13E-07 | 11 | Ccl8 | 0.32 | Old |
| 2.70E-10 | 1.555642 | 0.605 | 0.268 | 8.37E-06 | 11 | C3 | 0.337 | Old |
| 1.30E-09 | 2.174883 | 0.395 | 0.109 | 4.03E-05 | 11 | Cxcl13 | 0.286 | Old |
| 3.60E-08 | 1.446693 | 0.508 | 0.21 | 0.001118 | 11 | AC149090. | 0.298 | Old |
| 4.55E-08 | 3.627607 | 0.234 | 0.022 | 0.001411 | 11 | Cd55 | 0.212 | Old |
| 1.70E-07 | 0.892095 | 0.661 | 0.355 | 0.005289 | 11 | Ninj1 | 0.306 | Old |
| 3.64E-07 | 0.700736 | 0.802 | 0.543 | 0.011304 | 11 | Plbd1 | 0.259 | Old |
| 7.87E-07 | 2.157086 | 0.327 | 0.101 | 0.024424 | 11 | Fabp5 | 0.226 | Old |
| 1.07E-06 | 0.583036 | 0.944 | 0.754 | 0.033088 | 11 | Hsp90b1 | 0.19 | Old |
| 1.15E-06 | 0.754047 | 0.794 | 0.594 | 0.035865 | 11 | Cfp | 0.2 | Old |
| 1.19E-06 | 3.599745 | 0.185 | 0.014 | 0.036854 | 11 | Colq | 0.171 | Old |
| 1.75E-06 | 0.811474 | 0.738 | 0.493 | 0.054355 | 11 | Gda | 0.245 | Old |
| 3.49E-06 | 0.867975 | 0.532 | 0.261 | 0.108393 | 11 | Il2rg | 0.271 | Old |
| 3.55E-06 | 0.91815 | 0.516 | 0.239 | 0.110083 | 11 | Tmem14c | 0.277 | Old |
| 4.44E-06 | 0.975068 | 0.786 | 0.58 | 0.137958 | 11 | Clec10a | 0.206 | Old |
| 5.39E-06 | 0.461062 | 0.552 | 0.275 | 0.167309 | 11 | Itgb2 | 0.277 | Old |
| 8.66E-06 | 0.809626 | 0.423 | 0.181 | 0.26884 | 11 | Psmd4 | 0.242 | Old |
| 9.26E-06 | 1.072595 | 0.456 | 0.21 | 0.287595 | 11 | Swi5 | 0.246 | Old |
| 9.48E-06 | 0.755642 | 0.589 | 0.319 | 0.294356 | 11 | Dnajc3 | 0.27 | Old |
| 9.55E-06 | 1.02656 | 0.29 | 0.087 | 0.296421 | 11 | Med1 | 0.203 | Old |
| 1.00E-05 | 0.584535 | 0.879 | 0.688 | 0.311749 | 11 | Tmem176a | 0.191 | Old |
| 1.63E-05 | 0.757107 | 0.544 | 0.268 | 0.507475 | 11 | Mapk3 | 0.276 | Old |
| 1.70E-05 | 1.500903 | 0.278 | 0.087 | 0.527509 | 11 | Hpse | 0.191 | Old |
| 1.87E-05 | 0.81585 | 0.508 | 0.261 | 0.579734 | 11 | Sulf2 | 0.247 | Old |
| 1.90E-05 | 1.090776 | 0.383 | 0.167 | 0.591543 | 11 | Rreb1 | 0.216 | Old |
| 1.91E-05 | 1.777983 | 0.302 | 0.116 | 0.594375 | 11 | Xist | 0.186 | Old |
| 1.94E-05 | 0.484006 | 0.435 | 0.188 | 0.602775 | 11 | Dlst | 0.247 | Old |
| 2.85E-05 | 0.754847 | 0.516 | 0.261 | 0.885298 | 11 | Hpcal1 | 0.255 | Old |
| 2.86E-05 | 0.312696 | 0.323 | 0.109 | 0.887468 | 11 | Slc12a7 | 0.214 | Old |
| 3.09E-05 | 4.666198 | 0.117 | 0 | 0.96098 | 11 | Zfp329 | 0.117 | Old |
| 3.09E-05 | 0.676648 | 0.657 | 0.384 | 0.961012 | 11 | Ctsl | 0.273 | Old |
| 3.63E-05 | 0.999339 | 0.319 | 0.116 | 1 | 11 | Mrpl14 | 0.203 | Old |
| 3.73E-05 | 1.129672 | 0.306 | 0.116 | 1 | 11 | Kidins220 | 0.19 | Old |
| 4.16E-05 | 1.1086 | 0.238 | 0.065 | 1 | 11 | Kif1c | 0.173 | Old |
| 4.23E-05 | 1.483246 | 0.246 | 0.072 | 1 | 11 | Itfg1 | 0.174 | Old |
| 4.62E-05 | 1.157263 | 0.306 | 0.116 | 1 | 11 | Phf20 | 0.19 | Old |
| 4.64E-05 | 1.19644 | 0.355 | 0.152 | 1 | 11 | Mrpl28 | 0.203 | Old |
| 5.04E-05 | 1.592034 | 0.27 | 0.094 | 1 | 11 | Sdf2l1 | 0.176 | Old |
| 5.22E-05 | 2.026445 | 0.181 | 0.036 | 1 | 11 | Stard5 | 0.145 | Old |
| 5.38E-05 | 0.587611 | 0.754 | 0.522 | 1 | 11 | Ptms | 0.232 | Old |
| 6.35E-05 | 0.634785 | 0.552 | 0.304 | 1 | 11 | Psmd2 | 0.248 | Old |
| 6.46E-05 | 1.249267 | 0.31 | 0.123 | 1 | 11 | Marf1 | 0.187 | Old |
| 6.79E-05 | 2.463227 | 0.177 | 0.036 | 1 | 11 | A4galt | 0.141 | Old |

| | | | | | | | | |
|---|---|---|---|---|---|---|---|---|
| 6.84E-05 | 0.681591 | 0.44 | 0.217 | 1 | 11 | Itch | 0.223 | Old |
| 6.93E-05 | 2.419333 | 0.153 | 0.022 | 1 | 11 | Cdca4 | 0.131 | Old |
| 7.05E-05 | 1.088363 | 0.306 | 0.116 | 1 | 11 | Rnf187 | 0.19 | Old |
| 7.12E-05 | 0.870116 | 0.282 | 0.101 | 1 | 11 | Gm36161 | 0.181 | Old |
| 7.28E-05 | 1.612206 | 0.194 | 0.043 | 1 | 11 | Scly | 0.151 | Old |
| 7.30E-05 | 0.752694 | 0.621 | 0.391 | 1 | 11 | AW112010 | 0.23 | Old |
| 7.62E-05 | 0.392877 | 0.452 | 0.21 | 1 | 11 | Uqcrfs1 | 0.242 | Old |
| 7.68E-05 | 1.077219 | 0.302 | 0.116 | 1 | 11 | Dhrs1 | 0.186 | Old |
| 7.69E-05 | 0.440442 | 0.952 | 0.949 | 1 | 11 | H2-Eb1 | 0.003 | Old |
| 7.72E-05 | 1.286911 | 0.319 | 0.13 | 1 | 11 | Cr1l | 0.189 | Old |
| 7.99E-05 | 1.113185 | 0.25 | 0.08 | 1 | 11 | Glb1 | 0.17 | Old |
| 8.11E-05 | 0.762905 | 0.484 | 0.246 | 1 | 11 | Csf2rb | 0.238 | Old |
| 8.17E-05 | 0.281668 | 0.996 | 0.986 | 1 | 11 | B2m | 0.01 | Old |
| 8.20E-05 | 0.796253 | 0.262 | 0.087 | 1 | 11 | Crtc3 | 0.175 | Old |
| 8.33E-05 | 0.720681 | 0.347 | 0.145 | 1 | 11 | Ap3d1 | 0.202 | Old |
| 8.46E-05 | 1.201488 | 0.452 | 0.246 | 1 | 11 | Fxyd2 | 0.206 | Old |
| 8.51E-05 | 0.620814 | 0.806 | 0.558 | 1 | 11 | Cox5a | 0.248 | Old |
| 8.75E-05 | 0.810867 | 0.435 | 0.225 | 1 | 11 | Rab20 | 0.21 | Old |
| 8.75E-05 | 0.596678 | 0.379 | 0.174 | 1 | 11 | Pdxk | 0.205 | Old |
| 9.04E-05 | 1.215284 | 0.278 | 0.101 | 1 | 11 | Tmem208 | 0.177 | Old |
| 9.26E-05 | 0.49623 | 0.544 | 0.29 | 1 | 11 | Nedd8 | 0.254 | Old |
| 9.47E-05 | 0.975946 | 0.282 | 0.101 | 1 | 11 | Suclg1 | 0.181 | Old |
| 0.000101 | 1.507647 | 0.226 | 0.065 | 1 | 11 | Ykt6 | 0.161 | Old |
| 0.000104 | 0.84559 | 0.31 | 0.123 | 1 | 11 | Renbp | 0.187 | Old |
| 0.000109 | 0.946515 | 0.464 | 0.254 | 1 | 11 | Tspan3 | 0.21 | Old |
| 0.000109 | 0.477756 | 0.746 | 0.486 | 1 | 11 | Cstb | 0.26 | Old |
| 0.000113 | 0.564392 | 0.435 | 0.203 | 1 | 11 | Adam10 | 0.232 | Old |
| 0.000118 | 0.909392 | 0.343 | 0.145 | 1 | 11 | Sik1 | 0.198 | Old |
| 0.00012 | 0.470065 | 0.532 | 0.297 | 1 | 11 | Tbxas1 | 0.235 | Old |
| 0.000121 | 0.59316 | 0.681 | 0.428 | 1 | 11 | Cops9 | 0.253 | Old |
| 0.000122 | 1.081497 | 0.185 | 0.043 | 1 | 11 | Ikbke | 0.142 | Old |
| 0.000126 | 1.960697 | 0.194 | 0.051 | 1 | 11 | Nagk | 0.143 | Old |
| 0.000126 | 0.47885 | 0.524 | 0.304 | 1 | 11 | Dusp3 | 0.22 | Old |
| 0.000127 | 0.786926 | 0.423 | 0.217 | 1 | 11 | Cd300c2 | 0.206 | Old |
| 0.000127 | 0.708463 | 0.504 | 0.268 | 1 | 11 | Cox17 | 0.236 | Old |
| 0.000128 | 0.408059 | 0.996 | 0.978 | 1 | 11 | Psap | 0.018 | Old |
| 0.000129 | 0.964359 | 0.335 | 0.145 | 1 | 11 | Fgr | 0.19 | Old |
| 0.00013 | 0.651516 | 0.464 | 0.246 | 1 | 11 | Tmed9 | 0.218 | Old |
| 0.000131 | 0.485322 | 0.407 | 0.188 | 1 | 11 | Tmem51 | 0.219 | Old |
| 0.000134 | 1.704764 | 0.117 | 0.007 | 1 | 11 | Hectd3 | 0.11 | Old |
| 0.000137 | 0.537386 | 0.472 | 0.239 | 1 | 11 | Zfand6 | 0.233 | Old |
| 0.000139 | 0.974834 | 0.306 | 0.123 | 1 | 11 | Psmd6 | 0.183 | Old |
| 0.000142 | 0.83302 | 0.371 | 0.167 | 1 | 11 | Aup1 | 0.204 | Old |
| 0.000145 | 1.296587 | 0.25 | 0.087 | 1 | 11 | Anxa11 | 0.163 | Old |
| 0.000147 | 0.672527 | 0.423 | 0.21 | 1 | 11 | Washc2 | 0.213 | Old |

| | | | | | | | | |
|---|---|---|---|---|---|---|---|---|
| 0.000148 | 0.790001 | 0.419 | 0.217 | 1 | 11 | Atxn2l | 0.202 | Old |
| 0.000149 | 0.569077 | 0.601 | 0.362 | 1 | 11 | Rgl1 | 0.239 | Old |
| 0.000151 | 0.986142 | 0.234 | 0.072 | 1 | 11 | Armcx3 | 0.162 | Old |
| 0.000155 | 2.41229 | 0.129 | 0.014 | 1 | 11 | Nbdy | 0.115 | Old |
| 0.000157 | 0.709697 | 0.685 | 0.442 | 1 | 11 | Snx3 | 0.243 | Old |
| 0.000158 | 0.617614 | 0.46 | 0.239 | 1 | 11 | Vapa | 0.221 | Old |
| 0.000161 | 0.798451 | 0.343 | 0.145 | 1 | 11 | Glrx | 0.198 | Old |
| 0.000164 | 1.118854 | 0.238 | 0.08 | 1 | 11 | Isoc1 | 0.158 | Old |
| 0.000164 | 0.746189 | 0.419 | 0.217 | 1 | 11 | Gfpt1 | 0.202 | Old |
| 0.000165 | 0.674114 | 0.613 | 0.377 | 1 | 11 | Ptpn1 | 0.236 | Old |
| 0.000172 | 0.762473 | 0.544 | 0.326 | 1 | 11 | Vcam1 | 0.218 | Old |
| 0.000176 | 0.627861 | 0.355 | 0.159 | 1 | 11 | Lsm6 | 0.196 | Old |
| 0.000177 | 0.62147 | 0.379 | 0.167 | 1 | 11 | Mospd2 | 0.212 | Old |
| 0.000178 | 1.431467 | 0.246 | 0.087 | 1 | 11 | Galnt7 | 0.159 | Old |
| 0.000179 | 0.76139 | 0.677 | 0.522 | 1 | 11 | Cd63 | 0.155 | Old |
| 0.000184 | 2.191268 | 0.141 | 0.022 | 1 | 11 | Vps52 | 0.119 | Old |
| 0.000186 | 0.983594 | 0.286 | 0.109 | 1 | 11 | Ppp6c | 0.177 | Old |
| 0.000187 | 1.236862 | 0.286 | 0.116 | 1 | 11 | Rheb | 0.17 | Old |
| 0.00019 | 2.703127 | 0.173 | 0.043 | 1 | 11 | Cops3 | 0.13 | Old |
| 0.000192 | 1.100053 | 0.25 | 0.087 | 1 | 11 | Socs4 | 0.163 | Old |
| 0.000195 | 1.913173 | 0.153 | 0.029 | 1 | 11 | Idh3a | 0.124 | Old |
| 0.000199 | 0.538061 | 0.645 | 0.384 | 1 | 11 | Myl12b | 0.261 | Old |
| 0.000212 | 0.783855 | 0.181 | 0.043 | 1 | 11 | Ndufv1 | 0.138 | Old |
| 0.000213 | 0.249395 | 0.286 | 0.109 | 1 | 11 | Serpina3g | 0.177 | Old |
| 0.000214 | 0.769158 | 0.351 | 0.159 | 1 | 11 | Flii | 0.192 | Old |
| 0.00022 | 0.509511 | 0.516 | 0.304 | 1 | 11 | Anxa4 | 0.212 | Old |
| 0.000225 | 1.28994 | 0.427 | 0.254 | 1 | 11 | Mmp9 | 0.173 | Old |
| 0.000225 | 1.872006 | 0.149 | 0.029 | 1 | 11 | Lsm2 | 0.12 | Old |
| 0.000226 | 0.62573 | 0.589 | 0.333 | 1 | 11 | Rp2 | 0.256 | Old |
| 0.00023 | 0.476097 | 0.29 | 0.109 | 1 | 11 | B4galnt1 | 0.181 | Old |
| 0.00023 | 0.953528 | 0.27 | 0.101 | 1 | 11 | Nfu1 | 0.169 | Old |
| 0.000232 | 1.463532 | 0.165 | 0.036 | 1 | 11 | Creb3 | 0.129 | Old |
| 0.000232 | 0.531512 | 0.456 | 0.225 | 1 | 11 | Ppp1r12a | 0.231 | Old |
| 0.000246 | 1.085769 | 0.226 | 0.072 | 1 | 11 | Atf2 | 0.154 | Old |
| 0.000247 | 1.716677 | 0.153 | 0.029 | 1 | 11 | Ero1l | 0.124 | Old |
| 0.000253 | 0.512546 | 0.629 | 0.37 | 1 | 11 | Mfsd1 | 0.259 | Old |
| 0.000255 | 0.678794 | 0.665 | 0.464 | 1 | 11 | Pirb | 0.201 | Old |
| 0.00026 | 0.662259 | 0.512 | 0.283 | 1 | 11 | Sypl | 0.229 | Old |
| 0.00026 | 0.80667 | 0.46 | 0.254 | 1 | 11 | Ppp1cb | 0.206 | Old |
| 0.000262 | 1.484708 | 0.25 | 0.094 | 1 | 11 | Dnajc1 | 0.156 | Old |
| 0.000262 | 0.548655 | 0.395 | 0.181 | 1 | 11 | Ppp4c | 0.214 | Old |
| 0.000267 | 0.618473 | 0.516 | 0.275 | 1 | 11 | Hdlbp | 0.241 | Old |
| 0.000271 | 0.486313 | 0.31 | 0.13 | 1 | 11 | Gnl3 | 0.18 | Old |
| 0.000282 | 0.491223 | 0.524 | 0.275 | 1 | 11 | Map7d1 | 0.249 | Old |
| 0.000284 | 0.607467 | 0.456 | 0.232 | 1 | 11 | Timm13 | 0.224 | Old |

| | | | | | | | | |
|---|---|---|---|---|---|---|---|---|
| 0.000293 | 2.98514 | 0.121 | 0.014 | 1 | 11 | Csprs | 0.107 | Old |
| 0.000302 | 1.81989 | 0.194 | 0.058 | 1 | 11 | Slamf6 | 0.136 | Old |
| 0.000302 | 1.314283 | 0.185 | 0.051 | 1 | 11 | Slc11a2 | 0.134 | Old |
| 0.000308 | 0.696849 | 0.71 | 0.529 | 1 | 11 | Ctsd | 0.181 | Old |
| 0.000309 | 0.560658 | 0.843 | 0.674 | 1 | 11 | Stat1 | 0.169 | Old |
| 0.000321 | 1.643571 | 0.121 | 0.014 | 1 | 11 | Lsg1 | 0.107 | Old |
| 0.000334 | 1.248984 | 0.282 | 0.123 | 1 | 11 | Aimp1 | 0.159 | Old |
| 0.000338 | 2.020652 | 0.169 | 0.043 | 1 | 11 | Atg12 | 0.126 | Old |
| 0.000346 | 0.793966 | 0.246 | 0.087 | 1 | 11 | Btaf1 | 0.159 | Old |
| 0.000352 | 0.463295 | 0.794 | 0.543 | 1 | 11 | Pdia3 | 0.251 | Old |
| 0.000355 | 0.581946 | 0.528 | 0.29 | 1 | 11 | Tmem256 | 0.238 | Old |
| 0.000362 | 0.993394 | 0.25 | 0.094 | 1 | 11 | Actr1a | 0.156 | Old |
| 0.00037 | 1.533825 | 0.286 | 0.13 | 1 | 11 | Nr1h3 | 0.156 | Old |
| 0.00038 | 1.780702 | 0.157 | 0.036 | 1 | 11 | Abhd17c | 0.121 | Old |
| 0.00038 | 0.947921 | 0.185 | 0.051 | 1 | 11 | Atic | 0.134 | Old |
| 0.000384 | 0.439716 | 0.669 | 0.435 | 1 | 11 | Atp6ap2 | 0.234 | Old |
| 0.000386 | 2.208914 | 0.145 | 0.029 | 1 | 11 | Nup214 | 0.116 | Old |
| 0.000397 | 0.404243 | 0.673 | 0.413 | 1 | 11 | Selenok | 0.26 | Old |
| 0.000405 | 0.453716 | 0.855 | 0.703 | 1 | 11 | Pltp | 0.152 | Old |
| 0.000407 | 0.29035 | 0.444 | 0.225 | 1 | 11 | Dennd1a | 0.219 | Old |
| 0.00041 | 0.341151 | 0.968 | 0.964 | 1 | 11 | Itm2b | 0.004 | Old |
| 0.000415 | 0.8246 | 0.278 | 0.116 | 1 | 11 | Pdap1 | 0.162 | Old |
| 0.000418 | 0.947061 | 0.306 | 0.13 | 1 | 11 | Cyb5b | 0.176 | Old |
| 0.000422 | 0.959318 | 0.351 | 0.174 | 1 | 11 | Tmx1 | 0.177 | Old |
| 0.000424 | 0.454265 | 0.5 | 0.275 | 1 | 11 | Lfng | 0.225 | Old |
| 0.000428 | 0.495995 | 0.621 | 0.377 | 1 | 11 | Nipa2 | 0.244 | Old |
| 0.000434 | 0.36231 | 0.286 | 0.123 | 1 | 11 | Ndufv2 | 0.163 | Old |
| 0.000439 | 2.323536 | 0.117 | 0.014 | 1 | 11 | Pitrm1 | 0.103 | Old |
| 0.000444 | 1.582145 | 0.157 | 0.036 | 1 | 11 | Zfp787 | 0.121 | Old |
| 0.000458 | 0.891318 | 0.331 | 0.152 | 1 | 11 | Slc26a2 | 0.179 | Old |
| 0.000475 | 0.980457 | 0.238 | 0.087 | 1 | 11 | Cklf | 0.151 | Old |
| 0.000477 | 2.299511 | 0.141 | 0.029 | 1 | 11 | Thap12 | 0.112 | Old |
| 0.000483 | 0.794309 | 0.327 | 0.152 | 1 | 11 | Uso1 | 0.175 | Old |
| 0.000495 | 2.468882 | 0.101 | 0.007 | 1 | 11 | Arfgap3 | 0.094 | Old |
| 0.000496 | 0.516255 | 0.456 | 0.246 | 1 | 11 | Hnrnpul2 | 0.21 | Old |
| 0.000497 | 3.594058 | 0.113 | 0.014 | 1 | 11 | Art2b | 0.099 | Old |
| 0.0005 | 0.470737 | 0.423 | 0.217 | 1 | 11 | Psmc3 | 0.206 | Old |
| 0.000505 | 0.566939 | 0.343 | 0.159 | 1 | 11 | Anapc11 | 0.184 | Old |
| 0.000511 | 0.852078 | 0.262 | 0.101 | 1 | 11 | Vars | 0.161 | Old |
| 0.000513 | 0.400845 | 0.948 | 0.877 | 1 | 11 | Lamp1 | 0.071 | Old |
| 0.00052 | 0.879667 | 0.266 | 0.109 | 1 | 11 | Anpep | 0.157 | Old |
| 0.000524 | 0.964434 | 0.202 | 0.065 | 1 | 11 | Cabin1 | 0.137 | Old |
| 0.000525 | 0.982198 | 0.218 | 0.072 | 1 | 11 | Ccny | 0.146 | Old |
| 0.000541 | 0.882834 | 0.278 | 0.116 | 1 | 11 | Rabl6 | 0.162 | Old |
| 0.000548 | 0.867129 | 0.238 | 0.087 | 1 | 11 | Ppm1g | 0.151 | Old |

| | | | | | | | | |
|---|---|---|---|---|---|---|---|---|
| 0.000551 | 1.140057 | 0.214 | 0.072 | 1 | 11 | Nupr1 | 0.142 | Old |
| 0.000553 | 2.844112 | 0.113 | 0.014 | 1 | 11 | Lag3 | 0.099 | Old |
| 0.000564 | 1.039974 | 0.294 | 0.13 | 1 | 11 | Ap3b1 | 0.164 | Old |
| 0.000574 | 0.351904 | 0.383 | 0.196 | 1 | 11 | Tmem168 | 0.187 | Old |
| 0.000575 | 1.638354 | 0.165 | 0.043 | 1 | 11 | Arhgap4 | 0.122 | Old |
| 0.000576 | 1.79092 | 0.141 | 0.029 | 1 | 11 | Gdf3 | 0.112 | Old |
| 0.000578 | 0.711775 | 0.246 | 0.094 | 1 | 11 | Lsm3 | 0.152 | Old |
| 0.000585 | 0.782547 | 0.383 | 0.196 | 1 | 11 | Pgam1 | 0.187 | Old |
| 0.000587 | 0.772025 | 0.218 | 0.072 | 1 | 11 | Sptlc1 | 0.146 | Old |
| 0.000595 | 1.444807 | 0.153 | 0.036 | 1 | 11 | Washc5 | 0.117 | Old |
| 0.000606 | 0.978844 | 0.294 | 0.13 | 1 | 11 | Elf2 | 0.164 | Old |
| 0.000616 | 0.485782 | 0.569 | 0.326 | 1 | 11 | Spcs2 | 0.243 | Old |
| 0.000621 | 0.880498 | 0.246 | 0.094 | 1 | 11 | Mrps17 | 0.152 | Old |
| 0.000629 | 3.989871 | 0.081 | 0 | 1 | 11 | Tram2 | 0.081 | Old |
| 0.000634 | 0.321166 | 0.407 | 0.203 | 1 | 11 | Smg1 | 0.204 | Old |
| 0.000641 | 0.295473 | 0.984 | 0.971 | 1 | 11 | H2-Aa | 0.013 | Old |
| 0.000641 | 2.366867 | 0.125 | 0.022 | 1 | 11 | Wdsub1 | 0.103 | Old |
| 0.000646 | 1.076818 | 0.19 | 0.058 | 1 | 11 | Slc25a1 | 0.132 | Old |
| 0.000656 | 0.729773 | 0.214 | 0.072 | 1 | 11 | Ccng1 | 0.142 | Old |
| 0.000662 | 0.739505 | 0.359 | 0.174 | 1 | 11 | Prkcsh | 0.185 | Old |
| 0.000666 | 0.591937 | 0.399 | 0.203 | 1 | 11 | Aco2 | 0.196 | Old |
| 0.000666 | 0.781035 | 0.286 | 0.123 | 1 | 11 | Bcat2 | 0.163 | Old |
| 0.000667 | 0.719873 | 0.226 | 0.08 | 1 | 11 | Pip4k2b | 0.146 | Old |
| 0.000669 | 2.302993 | 0.202 | 0.072 | 1 | 11 | Ly6a | 0.13 | Old |
| 0.000671 | 0.541614 | 0.492 | 0.275 | 1 | 11 | Hspa9 | 0.217 | Old |
| 0.000671 | 0.702211 | 0.411 | 0.225 | 1 | 11 | Sdhd | 0.186 | Old |
| 0.000674 | 0.481701 | 0.355 | 0.181 | 1 | 11 | Cbx3 | 0.174 | Old |
| 0.000682 | 0.497995 | 0.702 | 0.493 | 1 | 11 | Atp5g3 | 0.209 | Old |
| 0.000683 | 1.071636 | 0.113 | 0.014 | 1 | 11 | Tbl3 | 0.099 | Old |
| 0.000684 | 0.482087 | 0.472 | 0.246 | 1 | 11 | Tmem160 | 0.226 | Old |
| 0.000688 | 0.997273 | 0.222 | 0.08 | 1 | 11 | Ahcyl1 | 0.142 | Old |
| 0.000695 | 0.516419 | 0.742 | 0.565 | 1 | 11 | Ctsa | 0.177 | Old |
| 0.000697 | 0.361482 | 0.351 | 0.159 | 1 | 11 | Stt3a | 0.192 | Old |
| 0.000698 | 0.519124 | 0.597 | 0.384 | 1 | 11 | Cd68 | 0.213 | Old |
| 0.000707 | 1.108709 | 0.198 | 0.065 | 1 | 11 | Trp53 | 0.133 | Old |
| 0.000707 | 1.543515 | 0.153 | 0.036 | 1 | 11 | Pmvk | 0.117 | Old |
| 0.00071 | 0.331757 | 0.605 | 0.399 | 1 | 11 | Ccl7 | 0.206 | Old |
| 0.000715 | 1.192089 | 0.23 | 0.087 | 1 | 11 | Hebp1 | 0.143 | Old |
| 0.000728 | 1.380218 | 0.125 | 0.022 | 1 | 11 | Eif2b5 | 0.103 | Old |
| 0.000731 | 0.857061 | 0.391 | 0.217 | 1 | 11 | Sdf2 | 0.174 | Old |
| 0.00075 | 0.223852 | 0.331 | 0.152 | 1 | 11 | Phb2 | 0.179 | Old |
| 0.00075 | 0.943055 | 0.266 | 0.109 | 1 | 11 | Fam213b | 0.157 | Old |
| 0.000766 | 0.697874 | 0.335 | 0.159 | 1 | 11 | Wbp11 | 0.176 | Old |
| 0.000769 | 0.360559 | 0.464 | 0.239 | 1 | 11 | Tank | 0.225 | Old |
| 0.000778 | 0.707885 | 0.315 | 0.145 | 1 | 11 | Kansl1 | 0.17 | Old |

| | | | | | | | | |
|---|---|---|---|---|---|---|---|---|
| 0.000779 | 0.289368 | 0.431 | 0.225 | 1 | 11 | Hsd17b11 | 0.206 | Old |
| 0.000792 | 0.510126 | 0.448 | 0.246 | 1 | 11 | Fis1 | 0.202 | Old |
| 0.000797 | 1.298102 | 0.226 | 0.087 | 1 | 11 | Fam43a | 0.139 | Old |
| 0.000816 | 0.847139 | 0.19 | 0.058 | 1 | 11 | Emc8 | 0.132 | Old |
| 0.000827 | 0.152374 | 0.431 | 0.225 | 1 | 11 | Rragc | 0.206 | Old |
| 0.000833 | 1.520991 | 0.169 | 0.051 | 1 | 11 | BC003965 | 0.118 | Old |
| 0.00084 | 0.614497 | 0.411 | 0.217 | 1 | 11 | 9-Sep | 0.194 | Old |
| 0.000841 | 0.726822 | 0.27 | 0.116 | 1 | 11 | Dusp16 | 0.154 | Old |
| 0.000849 | 0.596443 | 0.569 | 0.37 | 1 | 11 | Folr2 | 0.199 | Old |
| 0.000851 | 1.278056 | 0.137 | 0.029 | 1 | 11 | Apobr | 0.108 | Old |
| 0.000862 | 1.197583 | 0.206 | 0.072 | 1 | 11 | Vps28 | 0.134 | Old |
| 0.000867 | 0.65145 | 0.5 | 0.297 | 1 | 11 | Man2a1 | 0.203 | Old |
| 0.000867 | 0.264172 | 0.427 | 0.217 | 1 | 11 | Psmd11 | 0.21 | Old |
| 0.000868 | 0.556881 | 0.343 | 0.167 | 1 | 11 | Dnajc7 | 0.176 | Old |
| 0.000876 | 4.533627 | 0.077 | 0 | 1 | 11 | Sh2d4a | 0.077 | Old |
| 0.000876 | 4.072613 | 0.077 | 0 | 1 | 11 | Ttc7b | 0.077 | Old |
| 0.000879 | 0.732197 | 0.407 | 0.217 | 1 | 11 | Mif | 0.19 | Old |
| 0.000882 | 0.536528 | 0.77 | 0.543 | 1 | 11 | Son | 0.227 | Old |
| 0.000897 | 0.927171 | 0.298 | 0.145 | 1 | 11 | Apc | 0.153 | Old |
| 0.000903 | 0.499217 | 0.629 | 0.406 | 1 | 11 | 2900097C1 | 0.223 | Old |
| 0.000912 | 1.167054 | 0.161 | 0.043 | 1 | 11 | Fnip2 | 0.118 | Old |
| 0.000929 | 2.732943 | 0.093 | 0.007 | 1 | 11 | Gpr162 | 0.086 | Old |
| 0.000939 | 0.460915 | 0.379 | 0.196 | 1 | 11 | Ralgds | 0.183 | Old |
| 0.000944 | 2.298662 | 0.093 | 0.007 | 1 | 11 | Drg2 | 0.086 | Old |
| 0.000972 | 2.039751 | 0.121 | 0.022 | 1 | 11 | Prcc | 0.099 | Old |
| 0.000976 | 0.42975 | 0.794 | 0.609 | 1 | 11 | Calm2 | 0.185 | Old |
| 0.000979 | 0.626625 | 0.214 | 0.08 | 1 | 11 | Gdf15 | 0.134 | Old |
| 0.000983 | 0.808552 | 0.173 | 0.051 | 1 | 11 | Xpot | 0.122 | Old |
| 0.00099 | 1.168317 | 0.339 | 0.174 | 1 | 11 | Cebpd | 0.165 | Old |
| 0.000991 | 0.493814 | 0.52 | 0.297 | 1 | 11 | Magt1 | 0.223 | Old |
| 0.000998 | 0.732036 | 0.355 | 0.181 | 1 | 11 | Golph3 | 0.174 | Old |
| 0.00101 | 0.621884 | 0.31 | 0.152 | 1 | 11 | Sertad1 | 0.158 | Old |
| 0.001066 | 0.271269 | 0.319 | 0.152 | 1 | 11 | Eloa | 0.167 | Old |
| 0.001079 | 0.78391 | 0.306 | 0.145 | 1 | 11 | Nmt1 | 0.161 | Old |
| 0.001088 | 0.859458 | 0.294 | 0.138 | 1 | 11 | Rhoq | 0.156 | Old |
| 0.001103 | 0.900189 | 0.306 | 0.145 | 1 | 11 | Lrrc8c | 0.161 | Old |
| 0.001114 | 0.563176 | 0.556 | 0.362 | 1 | 11 | Usmg5 | 0.194 | Old |
| 0.001114 | 1.1972 | 0.194 | 0.065 | 1 | 11 | Ints6 | 0.129 | Old |
| 0.001138 | 3.44587 | 0.089 | 0.007 | 1 | 11 | Gpr89 | 0.082 | Old |
| 0.001144 | 2.112644 | 0.117 | 0.022 | 1 | 11 | Naglu | 0.095 | Old |
| 0.001147 | 0.101486 | 0.327 | 0.152 | 1 | 11 | Mydgf | 0.175 | Old |
| 0.00115 | 0.23721 | 0.988 | 0.978 | 1 | 11 | H2-K1 | 0.01 | Old |
| 0.001165 | 0.402941 | 0.488 | 0.283 | 1 | 11 | Eif4ebp2 | 0.205 | Old |
| 0.001179 | 0.587854 | 0.714 | 0.551 | 1 | 11 | Clec4a3 | 0.163 | Old |
| 0.00118 | 1.204079 | 0.133 | 0.029 | 1 | 11 | Arhgef10 | 0.104 | Old |

| | | | | | | | | |
|---|---|---|---|---|---|---|---|---|
| 0.001187 | 0.352111 | 0.629 | 0.391 | 1 | 11 | Asah1 | 0.238 | Old |
| 0.001193 | 1.10811 | 0.226 | 0.087 | 1 | 11 | Srf | 0.139 | Old |
| 0.0012 | 0.442962 | 0.823 | 0.652 | 1 | 11 | Slc25a3 | 0.171 | Old |
| 0.001208 | 0.662244 | 0.423 | 0.239 | 1 | 11 | Coro1c | 0.184 | Old |
| 0.001211 | 0.31488 | 0.903 | 0.761 | 1 | 11 | Anxa5 | 0.142 | Old |
| 0.001216 | 0.245013 | 0.306 | 0.138 | 1 | 11 | Snd1 | 0.168 | Old |
| 0.001227 | 0.259307 | 0.403 | 0.217 | 1 | 11 | Fam26f | 0.186 | Old |
| 0.001239 | 1.029268 | 0.198 | 0.072 | 1 | 11 | Pcbd2 | 0.126 | Old |
| 0.001241 | 1.911724 | 0.129 | 0.029 | 1 | 11 | Gtf3c3 | 0.1 | Old |
| 0.001246 | 1.28956 | 0.165 | 0.051 | 1 | 11 | Dgat1 | 0.114 | Old |
| 0.001256 | 0.808316 | 0.214 | 0.08 | 1 | 11 | Ahctf1 | 0.134 | Old |
| 0.001263 | 0.481722 | 0.46 | 0.254 | 1 | 11 | Polr1d | 0.206 | Old |
| 0.001282 | 1.255834 | 0.226 | 0.094 | 1 | 11 | Fundc2 | 0.132 | Old |
| 0.001285 | 1.944772 | 0.141 | 0.036 | 1 | 11 | Timm50 | 0.105 | Old |
| 0.001287 | 1.239919 | 0.157 | 0.043 | 1 | 11 | Neurl4 | 0.114 | Old |
| 0.001303 | 0.442048 | 0.605 | 0.399 | 1 | 11 | Csf2ra | 0.206 | Old |
| 0.001307 | 2.344133 | 0.089 | 0.007 | 1 | 11 | Endog | 0.082 | Old |
| 0.001307 | 2.069559 | 0.089 | 0.007 | 1 | 11 | Dcaf1 | 0.082 | Old |
| 0.001335 | 1.237534 | 0.129 | 0.029 | 1 | 11 | Tomm40l | 0.1 | Old |
| 0.001338 | 0.453085 | 0.331 | 0.167 | 1 | 11 | Htt | 0.164 | Old |
| 0.001339 | 1.547253 | 0.089 | 0.007 | 1 | 11 | Zfp414 | 0.082 | Old |
| 0.001342 | 0.632208 | 0.621 | 0.435 | 1 | 11 | Atp5k | 0.186 | Old |
| 0.001353 | 0.386686 | 0.399 | 0.203 | 1 | 11 | Ndufs2 | 0.196 | Old |
| 0.00136 | 0.524086 | 0.379 | 0.196 | 1 | 11 | Dnajc19 | 0.183 | Old |
| 0.001365 | 0.375022 | 0.798 | 0.667 | 1 | 11 | Ifitm2 | 0.131 | Old |
| 0.001371 | 0.371831 | 0.839 | 0.616 | 1 | 11 | Arhgdib | 0.223 | Old |
| 0.001385 | 0.539134 | 0.411 | 0.225 | 1 | 11 | Eea1 | 0.186 | Old |
| 0.001386 | 0.40712 | 0.484 | 0.268 | 1 | 11 | Hsbp1 | 0.216 | Old |
| 0.001396 | 0.8465 | 0.335 | 0.174 | 1 | 11 | Hcfc1r1 | 0.161 | Old |
| 0.001402 | 0.549511 | 0.234 | 0.094 | 1 | 11 | Slc7a7 | 0.14 | Old |
| 0.001407 | 0.497662 | 0.742 | 0.543 | 1 | 11 | Clec4a1 | 0.199 | Old |
| 0.001408 | 0.770482 | 0.339 | 0.181 | 1 | 11 | Gpcpd1 | 0.158 | Old |
| 0.00141 | 1.504137 | 0.206 | 0.08 | 1 | 11 | Hk1 | 0.126 | Old |
| 0.001413 | 0.970043 | 0.258 | 0.116 | 1 | 11 | Pigk | 0.142 | Old |
| 0.001415 | 0.422083 | 0.492 | 0.29 | 1 | 11 | Papola | 0.202 | Old |
| 0.001416 | 2.865848 | 0.101 | 0.014 | 1 | 11 | Crebl2 | 0.087 | Old |
| 0.001432 | 0.330066 | 0.81 | 0.58 | 1 | 11 | Rac1 | 0.23 | Old |
| 0.001434 | 0.666922 | 0.419 | 0.239 | 1 | 11 | Map4k4 | 0.18 | Old |
| 0.001443 | 0.483273 | 0.202 | 0.072 | 1 | 11 | Wdr61 | 0.13 | Old |
| 0.001462 | 0.290958 | 0.931 | 0.92 | 1 | 11 | C1qa | 0.011 | Old |
| 0.001463 | 0.45543 | 0.298 | 0.138 | 1 | 11 | Smarca4 | 0.16 | Old |
| 0.001492 | 1.361853 | 0.141 | 0.036 | 1 | 11 | Slc35c1 | 0.105 | Old |
| 0.001524 | 0.533948 | 0.306 | 0.145 | 1 | 11 | Mrpl24 | 0.161 | Old |
| 0.001539 | 0.710522 | 0.141 | 0.036 | 1 | 11 | Sik3 | 0.105 | Old |
| 0.001541 | 0.225146 | 0.395 | 0.203 | 1 | 11 | Setd2 | 0.192 | Old |

| | | | | | | | | |
|---|---|---|---|---|---|---|---|---|
| 0.001548 | 0.414045 | 0.206 | 0.072 | 1 | 11 | Fmr1 | 0.134 | Old |
| 0.00155 | 0.897852 | 0.23 | 0.094 | 1 | 11 | Hvcn1 | 0.136 | Old |
| 0.00156 | 0.682679 | 0.21 | 0.08 | 1 | 11 | Pigs | 0.13 | Old |
| 0.001565 | 0.835716 | 0.282 | 0.138 | 1 | 11 | 2810403A( | 0.144 | Old |
| 0.001572 | 0.538973 | 0.56 | 0.37 | 1 | 11 | Acp2 | 0.19 | Old |
| 0.001573 | 0.412718 | 0.319 | 0.145 | 1 | 11 | Prkar2a | 0.174 | Old |
| 0.00158 | 0.623068 | 0.488 | 0.29 | 1 | 11 | Oxct1 | 0.198 | Old |
| 0.00159 | 0.439748 | 0.375 | 0.203 | 1 | 11 | Arf2 | 0.172 | Old |
| 0.001597 | 0.566041 | 0.161 | 0.051 | 1 | 11 | Arhgap10 | 0.11 | Old |
| 0.001607 | 1.340397 | 0.141 | 0.036 | 1 | 11 | Rcbtb1 | 0.105 | Old |
| 0.001611 | 0.398898 | 0.282 | 0.13 | 1 | 11 | Atg2a | 0.152 | Old |
| 0.001614 | 4.154701 | 0.085 | 0.007 | 1 | 11 | Dmwd | 0.078 | Old |
| 0.001619 | 0.690228 | 0.153 | 0.043 | 1 | 11 | Sec24a | 0.11 | Old |
| 0.001629 | 1.381655 | 0.23 | 0.101 | 1 | 11 | Gngt2 | 0.129 | Old |
| 0.001648 | 1.185446 | 0.226 | 0.094 | 1 | 11 | Cwf19l2 | 0.132 | Old |
| 0.001651 | 0.281081 | 0.339 | 0.159 | 1 | 11 | Slc4a7 | 0.18 | Old |
| 0.001651 | 1.013197 | 0.165 | 0.051 | 1 | 11 | Gaa | 0.114 | Old |
| 0.001654 | 0.766632 | 0.177 | 0.058 | 1 | 11 | Senp5 | 0.119 | Old |
| 0.001678 | 0.204149 | 0.56 | 0.319 | 1 | 11 | G3bp1 | 0.241 | Old |
| 0.001691 | 0.493538 | 0.552 | 0.341 | 1 | 11 | Gm21188 | 0.211 | Old |
| 0.001694 | 0.505324 | 0.379 | 0.196 | 1 | 11 | Atp5g1 | 0.183 | Old |
| 0.001694 | 3.639204 | 0.069 | 0 | 1 | 11 | Klra3 | 0.069 | Old |
| 0.001695 | 2.998282 | 0.085 | 0.007 | 1 | 11 | Irf2bp1 | 0.078 | Old |
| 0.001698 | 0.55397 | 0.355 | 0.188 | 1 | 11 | Larp4 | 0.167 | Old |
| 0.001699 | 0.506642 | 0.282 | 0.13 | 1 | 11 | Anks1 | 0.152 | Old |
| 0.001714 | 3.168985 | 0.133 | 0.036 | 1 | 11 | Mylk | 0.097 | Old |
| 0.001718 | 1.226652 | 0.194 | 0.072 | 1 | 11 | Ndufa9 | 0.122 | Old |
| 0.001735 | 0.716856 | 0.21 | 0.08 | 1 | 11 | Fam98b | 0.13 | Old |
| 0.001737 | 0.798457 | 0.153 | 0.043 | 1 | 11 | Vamp5 | 0.11 | Old |
| 0.001761 | 1.612573 | 0.198 | 0.08 | 1 | 11 | Fam103a1 | 0.118 | Old |
| 0.001763 | 0.418554 | 0.472 | 0.283 | 1 | 11 | Rnasel | 0.189 | Old |
| 0.001795 | 1.610206 | 0.137 | 0.036 | 1 | 11 | Mrpl55 | 0.101 | Old |
| 0.001807 | 0.695735 | 0.286 | 0.138 | 1 | 11 | Inppl1 | 0.148 | Old |
| 0.00181 | 0.415086 | 0.633 | 0.428 | 1 | 11 | Skap2 | 0.205 | Old |
| 0.001817 | 1.338561 | 0.137 | 0.036 | 1 | 11 | H2-Ke6 | 0.101 | Old |
| 0.001817 | 1.213385 | 0.137 | 0.036 | 1 | 11 | Wdr11 | 0.101 | Old |
| 0.001825 | 0.692882 | 0.371 | 0.21 | 1 | 11 | Nfix | 0.161 | Old |
| 0.001837 | 1.776007 | 0.085 | 0.007 | 1 | 11 | Atm | 0.078 | Old |
| 0.001863 | 0.359337 | 0.315 | 0.152 | 1 | 11 | Nacc2 | 0.163 | Old |
| 0.001863 | 1.132769 | 0.161 | 0.051 | 1 | 11 | Snrpa1 | 0.11 | Old |
| 0.001875 | 0.439643 | 0.73 | 0.558 | 1 | 11 | Lst1 | 0.172 | Old |
| 0.001879 | 0.540284 | 0.327 | 0.167 | 1 | 11 | Alyref | 0.16 | Old |
| 0.001916 | 0.306527 | 0.286 | 0.13 | 1 | 11 | Ndufs4 | 0.156 | Old |
| 0.001932 | 0.76145 | 0.302 | 0.152 | 1 | 11 | Amdhd2 | 0.15 | Old |
| 0.001935 | 0.613064 | 0.23 | 0.094 | 1 | 11 | Ndst2 | 0.136 | Old |

| | | | | | | | | |
|---|---|---|---|---|---|---|---|---|
| 0.001946 | 0.861264 | 0.286 | 0.138 | 1 | 11 | Gmip | 0.148 | Old |
| 0.001951 | 1.346784 | 0.149 | 0.043 | 1 | 11 | Atxn7l3 | 0.106 | Old |
| 0.001954 | 0.901181 | 0.23 | 0.094 | 1 | 11 | Cand1 | 0.136 | Old |
| 0.001962 | 0.37899 | 0.544 | 0.333 | 1 | 11 | P4hb | 0.211 | Old |
| 0.001966 | 0.809175 | 0.29 | 0.145 | 1 | 11 | Csnk2a1 | 0.145 | Old |
| 0.001977 | 0.449998 | 0.371 | 0.196 | 1 | 11 | Dclre1c | 0.175 | Old |
| 0.001981 | 0.709282 | 0.48 | 0.319 | 1 | 11 | H2-Q7 | 0.161 | Old |
| 0.001981 | 0.412254 | 0.633 | 0.42 | 1 | 11 | Litaf | 0.213 | Old |
| 0.001982 | 0.333769 | 0.702 | 0.457 | 1 | 11 | G3bp2 | 0.245 | Old |
| 0.001992 | 0.405271 | 0.464 | 0.261 | 1 | 11 | Fam120a | 0.203 | Old |
| 0.002001 | 2.423871 | 0.097 | 0.014 | 1 | 11 | Mrps18b | 0.083 | Old |
| 0.002011 | 0.753875 | 0.173 | 0.058 | 1 | 11 | Ric1 | 0.115 | Old |
| 0.00203 | 2.802224 | 0.097 | 0.014 | 1 | 11 | Mrps11 | 0.083 | Old |
| 0.002045 | 1.399405 | 0.149 | 0.043 | 1 | 11 | Mterf3 | 0.106 | Old |
| 0.002049 | 0.313337 | 0.456 | 0.254 | 1 | 11 | Ms4a6d | 0.202 | Old |
| 0.00205 | 0.437338 | 0.528 | 0.319 | 1 | 11 | D17Wsu92 | 0.209 | Old |
| 0.002054 | 0.596807 | 0.415 | 0.239 | 1 | 11 | Gm26532 | 0.176 | Old |
| 0.002067 | 0.778969 | 0.214 | 0.087 | 1 | 11 | Msmo1 | 0.127 | Old |
| 0.002067 | 1.530528 | 0.137 | 0.036 | 1 | 11 | Ptp4a3 | 0.101 | Old |
| 0.002077 | 1.102684 | 0.214 | 0.087 | 1 | 11 | Rfk | 0.127 | Old |
| 0.002081 | 0.469562 | 0.29 | 0.13 | 1 | 11 | Arrdc4 | 0.16 | Old |
| 0.0021 | 1.186294 | 0.125 | 0.029 | 1 | 11 | Pag1 | 0.096 | Old |
| 0.002105 | 0.492188 | 0.29 | 0.145 | 1 | 11 | Phf23 | 0.145 | Old |
| 0.002106 | 1.793036 | 0.097 | 0.014 | 1 | 11 | Ndufaf2 | 0.083 | Old |
| 0.002106 | 1.713957 | 0.097 | 0.014 | 1 | 11 | Sec24d | 0.083 | Old |
| 0.00211 | 0.476047 | 0.161 | 0.051 | 1 | 11 | Abcg3 | 0.11 | Old |
| 0.002126 | 0.320924 | 0.448 | 0.261 | 1 | 11 | Myof | 0.187 | Old |
| 0.002131 | 0.159497 | 0.528 | 0.319 | 1 | 11 | Fubp1 | 0.209 | Old |
| 0.002144 | 0.415683 | 0.673 | 0.457 | 1 | 11 | Dstn | 0.216 | Old |
| 0.002145 | 0.415983 | 0.673 | 0.464 | 1 | 11 | Sgpl1 | 0.209 | Old |
| 0.002159 | 0.620092 | 0.323 | 0.167 | 1 | 11 | Fam96a | 0.156 | Old |
| 0.002173 | 0.327769 | 0.492 | 0.297 | 1 | 11 | Arhgap15 | 0.195 | Old |
| 0.002185 | 0.379278 | 0.831 | 0.623 | 1 | 11 | Apobec3 | 0.208 | Old |
| 0.002189 | 0.326751 | 0.923 | 0.848 | 1 | 11 | Ly6e | 0.075 | Old |
| 0.002197 | 0.856328 | 0.137 | 0.036 | 1 | 11 | Hmg20a | 0.101 | Old |
| 0.00221 | 0.223898 | 0.532 | 0.333 | 1 | 11 | Gsn | 0.199 | Old |
| 0.002215 | 0.632704 | 0.391 | 0.217 | 1 | 11 | Dip2b | 0.174 | Old |
| 0.002223 | 0.45931 | 0.343 | 0.174 | 1 | 11 | Ndufa5 | 0.169 | Old |
| 0.002224 | 1.090098 | 0.137 | 0.036 | 1 | 11 | Mdn1 | 0.101 | Old |
| 0.00223 | 0.297646 | 0.532 | 0.326 | 1 | 11 | Cct8 | 0.206 | Old |
| 0.002252 | 0.833611 | 0.23 | 0.101 | 1 | 11 | Lrif1 | 0.129 | Old |
| 0.002286 | 1.807194 | 0.165 | 0.058 | 1 | 11 | Dtnbp1 | 0.107 | Old |
| 0.002288 | 0.682967 | 0.492 | 0.319 | 1 | 11 | Eps8 | 0.173 | Old |
| 0.002296 | 0.747945 | 0.125 | 0.029 | 1 | 11 | Jade1 | 0.096 | Old |
| 0.002299 | 0.315932 | 0.911 | 0.804 | 1 | 11 | Pfn1 | 0.107 | Old |

| | | | | | | | | |
|---|---|---|---|---|---|---|---|---|
| 0.002308 | 0.656289 | 0.395 | 0.232 | 1 | 11 | Crk | 0.163 | Old |
| 0.002314 | 0.442739 | 0.157 | 0.051 | 1 | 11 | 1600012H( | 0.106 | Old |
| 0.002319 | 0.327936 | 0.726 | 0.536 | 1 | 11 | Atp5f1 | 0.19 | Old |
| 0.00232 | 0.347083 | 0.407 | 0.217 | 1 | 11 | Tnrc6a | 0.19 | Old |
| 0.002327 | 0.744591 | 0.254 | 0.116 | 1 | 11 | Uqcrc1 | 0.138 | Old |
| 0.002342 | 0.506602 | 0.569 | 0.377 | 1 | 11 | Prex1 | 0.192 | Old |
| 0.002351 | 0.701382 | 0.286 | 0.138 | 1 | 11 | Cript | 0.148 | Old |
| 0.002352 | 0.975813 | 0.198 | 0.08 | 1 | 11 | Slc35a5 | 0.118 | Old |
| 0.002355 | 3.428348 | 0.065 | 0 | 1 | 11 | Mccc1 | 0.065 | Old |
| 0.002356 | 1.625637 | 0.173 | 0.065 | 1 | 11 | Asl | 0.108 | Old |
| 0.002365 | 0.400131 | 0.133 | 0.036 | 1 | 11 | Rpa1 | 0.097 | Old |
| 0.002406 | 1.029014 | 0.343 | 0.188 | 1 | 11 | Eprs | 0.155 | Old |
| 0.00242 | 0.638526 | 0.206 | 0.08 | 1 | 11 | Mrps36 | 0.126 | Old |
| 0.002424 | 1.486918 | 0.133 | 0.036 | 1 | 11 | Pgp | 0.097 | Old |
| 0.002442 | 0.507016 | 0.363 | 0.196 | 1 | 11 | Efcab14 | 0.167 | Old |
| 0.002459 | 2.54994 | 0.081 | 0.007 | 1 | 11 | Spryd3 | 0.074 | Old |
| 0.00247 | 0.676669 | 0.27 | 0.13 | 1 | 11 | 0610012G( | 0.14 | Old |
| 0.002477 | 0.724954 | 0.315 | 0.167 | 1 | 11 | Chd3 | 0.148 | Old |
| 0.002479 | 2.506356 | 0.081 | 0.007 | 1 | 11 | Dmpk | 0.074 | Old |
| 0.002498 | 0.276686 | 0.331 | 0.167 | 1 | 11 | Tpp2 | 0.164 | Old |
| 0.002506 | 0.864152 | 0.145 | 0.043 | 1 | 11 | Gm15987 | 0.102 | Old |
| 0.002507 | 1.050829 | 0.181 | 0.065 | 1 | 11 | Mut | 0.116 | Old |
| 0.002519 | 2.076195 | 0.081 | 0.007 | 1 | 11 | Trim7 | 0.074 | Old |
| 0.002538 | 0.322935 | 0.536 | 0.355 | 1 | 11 | Lipa | 0.181 | Old |
| 0.002539 | 1.713925 | 0.081 | 0.007 | 1 | 11 | Prkar1b | 0.074 | Old |
| 0.002562 | 0.425239 | 0.472 | 0.268 | 1 | 11 | Necap2 | 0.204 | Old |
| 0.002579 | 1.077796 | 0.169 | 0.058 | 1 | 11 | Mfsd10 | 0.111 | Old |
| 0.00258 | 0.949952 | 0.121 | 0.029 | 1 | 11 | Gng11 | 0.092 | Old |
| 0.002596 | 1.815287 | 0.121 | 0.029 | 1 | 11 | Ppp6r2 | 0.092 | Old |
| 0.002616 | 0.719573 | 0.202 | 0.08 | 1 | 11 | Trappc1 | 0.122 | Old |
| 0.002661 | 0.780724 | 0.355 | 0.196 | 1 | 11 | Mib1 | 0.159 | Old |
| 0.00268 | 0.272791 | 0.282 | 0.13 | 1 | 11 | Dhx38 | 0.152 | Old |
| 0.002686 | 0.678569 | 0.48 | 0.312 | 1 | 11 | Spcs1 | 0.168 | Old |
| 0.002687 | 1.044029 | 0.145 | 0.043 | 1 | 11 | Zadh2 | 0.102 | Old |
| 0.002694 | 0.139757 | 0.343 | 0.181 | 1 | 11 | Vps29 | 0.162 | Old |
| 0.002702 | 1.034845 | 0.177 | 0.065 | 1 | 11 | Tmem230 | 0.112 | Old |
| 0.002706 | 1.274714 | 0.165 | 0.058 | 1 | 11 | Cyhr1 | 0.107 | Old |
| 0.002713 | 0.625178 | 0.282 | 0.138 | 1 | 11 | Usp47 | 0.144 | Old |
| 0.002721 | 1.317198 | 0.109 | 0.022 | 1 | 11 | Zfp219 | 0.087 | Old |
| 0.002734 | 0.859326 | 0.145 | 0.043 | 1 | 11 | Nsmce2 | 0.102 | Old |
| 0.002755 | 0.697724 | 0.202 | 0.08 | 1 | 11 | Mocs2 | 0.122 | Old |
| 0.002755 | 1.922045 | 0.129 | 0.036 | 1 | 11 | Eef1e1 | 0.093 | Old |
| 0.002782 | 0.850806 | 0.145 | 0.043 | 1 | 11 | Kri1 | 0.102 | Old |
| 0.002788 | 1.070003 | 0.177 | 0.065 | 1 | 11 | Vta1 | 0.112 | Old |
| 0.00279 | 1.530853 | 0.153 | 0.051 | 1 | 11 | Vrk3 | 0.102 | Old |

| | | | | | | | | |
|---|---|---|---|---|---|---|---|---|
| 0.002798 | 0.323221 | 0.911 | 0.899 | 1 | 11 | C1qb | 0.012 | Old |
| 0.002802 | 0.399695 | 0.863 | 0.732 | 1 | 11 | Lamp2 | 0.131 | Old |
| 0.002808 | 1.215702 | 0.206 | 0.087 | 1 | 11 | Desi2 | 0.119 | Old |
| 0.00282 | 1.408592 | 0.121 | 0.029 | 1 | 11 | Wdr7 | 0.092 | Old |
| 0.002838 | 1.076193 | 0.218 | 0.094 | 1 | 11 | Mrpl12 | 0.124 | Old |
| 0.002841 | 0.409314 | 0.355 | 0.188 | 1 | 11 | Nus1 | 0.167 | Old |
| 0.002859 | 2.33006 | 0.117 | 0.029 | 1 | 11 | Josd2 | 0.088 | Old |
| 0.002864 | 0.413143 | 0.411 | 0.232 | 1 | 11 | Ube2k | 0.179 | Old |
| 0.00287 | 0.476112 | 0.5 | 0.319 | 1 | 11 | Dnmt3a | 0.181 | Old |
| 0.002873 | 0.547073 | 0.351 | 0.181 | 1 | 11 | Smarcd2 | 0.17 | Old |
| 0.002874 | 0.896841 | 0.246 | 0.116 | 1 | 11 | Cdc42ep3 | 0.13 | Old |
| 0.002901 | 0.811533 | 0.169 | 0.058 | 1 | 11 | Gtf2h1 | 0.111 | Old |
| 0.002912 | 1.055105 | 0.206 | 0.087 | 1 | 11 | Dnaaf3 | 0.119 | Old |
| 0.002918 | 0.360608 | 0.504 | 0.297 | 1 | 11 | Uba1 | 0.207 | Old |
| 0.002955 | 0.451224 | 0.246 | 0.109 | 1 | 11 | Hyou1 | 0.137 | Old |
| 0.00297 | 0.52478 | 0.456 | 0.275 | 1 | 11 | Atp2a2 | 0.181 | Old |
| 0.002973 | 0.995231 | 0.157 | 0.051 | 1 | 11 | Tceanc2 | 0.106 | Old |
| 0.002985 | 0.944917 | 0.198 | 0.08 | 1 | 11 | Smim13 | 0.118 | Old |
| 0.003017 | 1.343016 | 0.153 | 0.051 | 1 | 11 | Med11 | 0.102 | Old |
| 0.003018 | 0.375643 | 0.298 | 0.145 | 1 | 11 | Scamp3 | 0.153 | Old |
| 0.003027 | 0.167169 | 1 | 0.993 | 1 | 11 | mt-Atp6 | 0.007 | Old |
| 0.003052 | 0.634132 | 0.294 | 0.145 | 1 | 11 | Tpst2 | 0.149 | Old |
| 0.003056 | 0.434694 | 0.722 | 0.5 | 1 | 11 | Rap1a | 0.222 | Old |
| 0.003076 | 0.005455 | 0.347 | 0.174 | 1 | 11 | Zfp148 | 0.173 | Old |
| 0.003079 | 1.429111 | 0.093 | 0.014 | 1 | 11 | Noc3l | 0.079 | Old |
| 0.003112 | 0.705319 | 0.399 | 0.239 | 1 | 11 | Pdia4 | 0.16 | Old |
| 0.003113 | 1.992925 | 0.105 | 0.022 | 1 | 11 | 1110059E2 | 0.083 | Old |
| 0.003118 | 0.468459 | 0.246 | 0.109 | 1 | 11 | Ipo5 | 0.137 | Old |
| 0.003123 | 0.197299 | 0.157 | 0.051 | 1 | 11 | Hist1h2bc | 0.106 | Old |
| 0.003133 | 0.888769 | 0.266 | 0.13 | 1 | 11 | Aurkaip1 | 0.136 | Old |
| 0.003137 | 3.453149 | 0.077 | 0.007 | 1 | 11 | Manbal | 0.07 | Old |
| 0.003138 | 0.378259 | 0.391 | 0.217 | 1 | 11 | Grk3 | 0.174 | Old |
| 0.003146 | 0.120876 | 0.222 | 0.094 | 1 | 11 | Rab3gap2 | 0.128 | Old |
| 0.003148 | 0.529926 | 0.355 | 0.203 | 1 | 11 | Fam49a | 0.152 | Old |
| 0.003153 | 0.34275 | 0.46 | 0.268 | 1 | 11 | Tmed3 | 0.192 | Old |
| 0.003157 | 0.946353 | 0.262 | 0.13 | 1 | 11 | Med15 | 0.132 | Old |
| 0.003162 | 0.692747 | 0.19 | 0.072 | 1 | 11 | Prkaca | 0.118 | Old |
| 0.003169 | 1.445217 | 0.093 | 0.014 | 1 | 11 | Lancl1 | 0.079 | Old |
| 0.003213 | 0.776766 | 0.165 | 0.058 | 1 | 11 | Tmem120a | 0.107 | Old |
| 0.003232 | 0.283079 | 0.367 | 0.203 | 1 | 11 | Ddb1 | 0.164 | Old |
| 0.003238 | 0.443274 | 0.351 | 0.188 | 1 | 11 | Msrb1 | 0.163 | Old |
| 0.003253 | 0.034963 | 0.323 | 0.167 | 1 | 11 | Smim7 | 0.156 | Old |
| 0.003258 | 0.578347 | 0.19 | 0.072 | 1 | 11 | Dpp3 | 0.118 | Old |
| 0.00326 | 0.623722 | 0.375 | 0.21 | 1 | 11 | 11-Sep | 0.165 | Old |
| 0.003265 | 0.402673 | 0.456 | 0.275 | 1 | 11 | Vps4b | 0.181 | Old |

| | | | | | | | | |
|---|---|---|---|---|---|---|---|---|
| 0.003293 | 1.052399 | 0.25 | 0.123 | 1 | 11 | Relb | 0.127 | Old |
| 0.003344 | 2.804023 | 0.077 | 0.007 | 1 | 11 | Rnf113a2 | 0.07 | Old |
| 0.003347 | 0.977122 | 0.306 | 0.167 | 1 | 11 | Nfkb2 | 0.139 | Old |
| 0.003352 | 1.983401 | 0.105 | 0.022 | 1 | 11 | Mlycd | 0.083 | Old |
| 0.003363 | 0.667554 | 0.294 | 0.152 | 1 | 11 | Dnajc15 | 0.142 | Old |
| 0.003384 | 0.859585 | 0.246 | 0.116 | 1 | 11 | Mrpl13 | 0.13 | Old |
| 0.003393 | 0.255221 | 0.851 | 0.623 | 1 | 11 | App | 0.228 | Old |
| 0.003399 | 0.180664 | 0.359 | 0.188 | 1 | 11 | Osbpl11 | 0.171 | Old |
| 0.003427 | 0.598357 | 0.323 | 0.174 | 1 | 11 | Psmc5 | 0.149 | Old |
| 0.003437 | 1.583566 | 0.149 | 0.051 | 1 | 11 | 1700037H( | 0.098 | Old |
| 0.003446 | 1.140968 | 0.129 | 0.036 | 1 | 11 | Rnf144b | 0.093 | Old |
| 0.003458 | 0.695406 | 0.363 | 0.203 | 1 | 11 | Nsf | 0.16 | Old |
| 0.003464 | 0.128293 | 0.306 | 0.152 | 1 | 11 | Clk4 | 0.154 | Old |
| 0.003491 | 1.259686 | 0.185 | 0.072 | 1 | 11 | 2310022A1 | 0.113 | Old |
| 0.003496 | 0.196982 | 0.133 | 0.036 | 1 | 11 | Trappc11 | 0.097 | Old |
| 0.003497 | 0.850274 | 0.165 | 0.058 | 1 | 11 | Mettl23 | 0.107 | Old |
| 0.003533 | 0.703296 | 0.31 | 0.167 | 1 | 11 | Psmb7 | 0.143 | Old |
| 0.003542 | 0.924858 | 0.141 | 0.043 | 1 | 11 | A430005L1 | 0.098 | Old |
| 0.003551 | 0.412148 | 0.327 | 0.174 | 1 | 11 | Tspan32 | 0.153 | Old |
| 0.003559 | 1.036382 | 0.153 | 0.051 | 1 | 11 | Dnpep | 0.102 | Old |
| 0.003559 | 0.694256 | 0.153 | 0.051 | 1 | 11 | Mphosph8 | 0.102 | Old |
| 0.003589 | 0.578418 | 0.19 | 0.072 | 1 | 11 | Rabgap1 | 0.118 | Old |
| 0.003592 | 0.425936 | 0.677 | 0.464 | 1 | 11 | Ndufa4 | 0.213 | Old |
| 0.003593 | 1.02791 | 0.202 | 0.087 | 1 | 11 | Sar1b | 0.115 | Old |
| 0.003594 | 0.162917 | 0.359 | 0.196 | 1 | 11 | Eif3b | 0.163 | Old |
| 0.003626 | 1.082507 | 0.137 | 0.043 | 1 | 11 | Nhlrc2 | 0.094 | Old |
| 0.003634 | 0.316406 | 0.218 | 0.094 | 1 | 11 | Edem2 | 0.124 | Old |
| 0.003646 | 0.508438 | 0.315 | 0.159 | 1 | 11 | Ndufa12 | 0.156 | Old |
| 0.003662 | 1.495028 | 0.117 | 0.029 | 1 | 11 | Rbm43 | 0.088 | Old |
| 0.003677 | 0.640023 | 0.202 | 0.08 | 1 | 11 | Scarb1 | 0.122 | Old |
| 0.003678 | 0.736663 | 0.153 | 0.051 | 1 | 11 | Pmpca | 0.102 | Old |
| 0.00369 | 0.307964 | 0.605 | 0.42 | 1 | 11 | Rab5c | 0.185 | Old |
| 0.003693 | 0.282967 | 0.423 | 0.225 | 1 | 11 | Pgls | 0.198 | Old |
| 0.003697 | 0.305648 | 0.827 | 0.667 | 1 | 11 | Hexa | 0.16 | Old |
| 0.003702 | 0.807639 | 0.258 | 0.123 | 1 | 11 | Adap1 | 0.135 | Old |
| 0.003718 | 0.262711 | 0.282 | 0.138 | 1 | 11 | Slc25a36 | 0.144 | Old |
| 0.003724 | 1.396901 | 0.129 | 0.036 | 1 | 11 | Gtf3a | 0.093 | Old |
| 0.003726 | 1.08064 | 0.238 | 0.116 | 1 | 11 | Iah1 | 0.122 | Old |
| 0.003731 | 1.097492 | 0.117 | 0.029 | 1 | 11 | Gar1 | 0.088 | Old |
| 0.003732 | 0.743873 | 0.206 | 0.087 | 1 | 11 | 3830406C1 | 0.119 | Old |
| 0.003733 | 0.559918 | 0.339 | 0.181 | 1 | 11 | Arid5b | 0.158 | Old |
| 0.003759 | 0.612771 | 0.29 | 0.152 | 1 | 11 | Usp16 | 0.138 | Old |
| 0.003763 | 0.412679 | 0.262 | 0.123 | 1 | 11 | Gpr108 | 0.139 | Old |
| 0.003817 | 1.289044 | 0.149 | 0.051 | 1 | 11 | Nubp1 | 0.098 | Old |
| 0.003826 | 0.430791 | 0.117 | 0.029 | 1 | 11 | Mief1 | 0.088 | Old |

| | | | | | | | | |
|---|---|---|---|---|---|---|---|---|
| 0.003836 | 1.015627 | 0.141 | 0.043 | 1 | 11 | Pdgfc | 0.098 | Old |
| 0.003843 | 0.684935 | 0.387 | 0.232 | 1 | 11 | Wbp2 | 0.155 | Old |
| 0.003855 | 0.153682 | 1 | 1 | 1 | 11 | Actb | 0 | Old |
| 0.003862 | 0.435989 | 0.234 | 0.101 | 1 | 11 | Erlin1 | 0.133 | Old |
| 0.003876 | 0.084838 | 0.242 | 0.109 | 1 | 11 | Plin3 | 0.133 | Old |
| 0.00388 | 0.322193 | 0.548 | 0.333 | 1 | 11 | Nckap1l | 0.215 | Old |
| 0.003884 | 0.750528 | 0.395 | 0.232 | 1 | 11 | Srpr | 0.163 | Old |
| 0.003887 | 0.283224 | 0.359 | 0.196 | 1 | 11 | Zmynd11 | 0.163 | Old |
| 0.003899 | 0.513465 | 0.238 | 0.109 | 1 | 11 | Irak4 | 0.129 | Old |
| 0.003903 | 0.313426 | 0.948 | 0.862 | 1 | 11 | Grn | 0.086 | Old |
| 0.003919 | 0.28408 | 0.548 | 0.341 | 1 | 11 | Cggbp1 | 0.207 | Old |
| 0.003923 | 1.197428 | 0.117 | 0.029 | 1 | 11 | Rrp7a | 0.088 | Old |
| 0.003959 | 0.603867 | 0.105 | 0.022 | 1 | 11 | Arel1 | 0.083 | Old |
| 0.003963 | 2.101278 | 0.125 | 0.036 | 1 | 11 | Ttc19 | 0.089 | Old |
| 0.003969 | 0.756105 | 0.153 | 0.051 | 1 | 11 | Maml1 | 0.102 | Old |
| 0.003976 | 0.38163 | 0.315 | 0.159 | 1 | 11 | Tmem259 | 0.156 | Old |
| 0.003991 | 0.343544 | 0.375 | 0.203 | 1 | 11 | Tet2 | 0.172 | Old |
| 0.004016 | 0.755226 | 0.254 | 0.123 | 1 | 11 | Commd3 | 0.131 | Old |
| 0.00402 | 1.876566 | 0.157 | 0.058 | 1 | 11 | Tubb4b | 0.099 | Old |
| 0.004036 | 0.935775 | 0.141 | 0.043 | 1 | 11 | Eef1akmt1 | 0.098 | Old |
| 0.004046 | 0.268679 | 0.335 | 0.174 | 1 | 11 | Rab8a | 0.161 | Old |
| 0.004072 | 0.110384 | 0.532 | 0.326 | 1 | 11 | Atp6v1a | 0.206 | Old |
| 0.004083 | 0.421297 | 0.258 | 0.123 | 1 | 11 | Znhit1 | 0.135 | Old |
| 0.004092 | 0.964541 | 0.194 | 0.08 | 1 | 11 | Polr3c | 0.114 | Old |
| 0.004101 | 2.375522 | 0.101 | 0.022 | 1 | 11 | Armc10 | 0.079 | Old |
| 0.004118 | 0.287838 | 0.089 | 0.014 | 1 | 11 | Iqsec1 | 0.075 | Old |
| 0.00414 | 0.531324 | 0.262 | 0.13 | 1 | 11 | Gsap | 0.132 | Old |
| 0.004141 | 0.158375 | 0.347 | 0.196 | 1 | 11 | Pnpla8 | 0.151 | Old |
| 0.004184 | 0.389257 | 0.339 | 0.181 | 1 | 11 | Lrrk1 | 0.158 | Old |
| 0.004192 | 0.091884 | 0.254 | 0.116 | 1 | 11 | Ctdsp1 | 0.138 | Old |
| 0.004195 | 0.072279 | 0.298 | 0.152 | 1 | 11 | Rps27rt | 0.146 | Old |
| 0.004203 | 0.373708 | 0.839 | 0.63 | 1 | 11 | Calr | 0.209 | Old |
| 0.004219 | 0.550016 | 0.351 | 0.188 | 1 | 11 | Twf1 | 0.163 | Old |
| 0.004235 | 1.329652 | 0.149 | 0.051 | 1 | 11 | Abi2 | 0.098 | Old |
| 0.004242 | 0.379671 | 0.359 | 0.196 | 1 | 11 | Edf1 | 0.163 | Old |
| 0.004244 | 1.89967 | 0.113 | 0.029 | 1 | 11 | Zfp362 | 0.084 | Old |
| 0.004269 | 1.513487 | 0.089 | 0.014 | 1 | 11 | Leng1 | 0.075 | Old |
| 0.004273 | 0.489881 | 0.363 | 0.203 | 1 | 11 | Bax | 0.16 | Old |
| 0.004276 | 0.496179 | 0.698 | 0.522 | 1 | 11 | Klf4 | 0.176 | Old |
| 0.004296 | 0.486282 | 0.226 | 0.101 | 1 | 11 | Bcap29 | 0.125 | Old |
| 0.004325 | 0.238144 | 0.56 | 0.341 | 1 | 11 | Selenos | 0.219 | Old |
| 0.00433 | 1.562686 | 0.089 | 0.014 | 1 | 11 | Retsat | 0.075 | Old |
| 0.004332 | 0.329131 | 0.613 | 0.406 | 1 | 11 | Ndufc2 | 0.207 | Old |
| 0.004366 | 0.619165 | 0.141 | 0.043 | 1 | 11 | Jade2 | 0.098 | Old |
| 0.004384 | 2.078706 | 0.101 | 0.022 | 1 | 11 | AC127341. | 0.079 | Old |

| | | | | | | | | |
|---|---|---|---|---|---|---|---|---|
| 0.004391 | 0.643398 | 0.254 | 0.123 | 1 | 11 | Maff | 0.131 | Old |
| 0.004403 | 0.276343 | 0.774 | 0.601 | 1 | 11 | Vamp8 | 0.173 | Old |
| 0.00441 | 0.438077 | 0.496 | 0.304 | 1 | 11 | Sash1 | 0.192 | Old |
| 0.004438 | 3.320561 | 0.073 | 0.007 | 1 | 11 | Tfdp1 | 0.066 | Old |
| 0.004438 | 3.260977 | 0.073 | 0.007 | 1 | 11 | Tfb2m | 0.066 | Old |
| 0.004471 | 0.714233 | 0.149 | 0.051 | 1 | 11 | Pfdn4 | 0.098 | Old |
| 0.004506 | 0.919418 | 0.202 | 0.087 | 1 | 11 | Egln1 | 0.115 | Old |
| 0.00451 | 2.762426 | 0.073 | 0.007 | 1 | 11 | Rbm45 | 0.066 | Old |
| 0.004512 | 0.381361 | 0.746 | 0.536 | 1 | 11 | Pdia6 | 0.21 | Old |
| 0.004531 | 0.477862 | 0.161 | 0.058 | 1 | 11 | 9130401M | 0.103 | Old |
| 0.004568 | 0.77932 | 0.194 | 0.08 | 1 | 11 | Exoc2 | 0.114 | Old |
| 0.004569 | 0.9864 | 0.149 | 0.051 | 1 | 11 | Phf12 | 0.098 | Old |
| 0.004577 | 1.390966 | 0.177 | 0.072 | 1 | 11 | Gfra2 | 0.105 | Old |
| 0.004591 | 0.636465 | 0.282 | 0.145 | 1 | 11 | Ndufs3 | 0.137 | Old |
| 0.004595 | 0.74875 | 0.262 | 0.13 | 1 | 11 | Elmo2 | 0.132 | Old |
| 0.004596 | 0.572697 | 0.214 | 0.094 | 1 | 11 | Stard7 | 0.12 | Old |
| 0.004612 | 0.41554 | 0.415 | 0.239 | 1 | 11 | Camk1 | 0.176 | Old |
| 0.004616 | 0.583901 | 0.403 | 0.232 | 1 | 11 | Ggnbp2 | 0.171 | Old |
| 0.004622 | 0.565287 | 0.234 | 0.109 | 1 | 11 | Rbbp8 | 0.125 | Old |
| 0.004654 | 1.457705 | 0.101 | 0.022 | 1 | 11 | Enoph1 | 0.079 | Old |
| 0.004667 | 0.680163 | 0.254 | 0.123 | 1 | 11 | Blvra | 0.131 | Old |
| 0.00468 | 0.334451 | 0.581 | 0.377 | 1 | 11 | Serinc1 | 0.204 | Old |
| 0.00471 | 0.410366 | 0.565 | 0.399 | 1 | 11 | Syk | 0.166 | Old |
| 0.004727 | 0.60697 | 0.31 | 0.167 | 1 | 11 | Elk4 | 0.143 | Old |
| 0.004741 | 0.857475 | 0.258 | 0.13 | 1 | 11 | Ptk2b | 0.128 | Old |
| 0.004742 | 1.0514 | 0.246 | 0.123 | 1 | 11 | Pdhb | 0.123 | Old |
| 0.004755 | 0.427727 | 0.411 | 0.239 | 1 | 11 | Midn | 0.172 | Old |
| 0.004764 | 1.576501 | 0.125 | 0.036 | 1 | 11 | Scd2 | 0.089 | Old |
| 0.004772 | 0.145362 | 0.516 | 0.312 | 1 | 11 | Wasf2 | 0.204 | Old |
| 0.004779 | 1.529976 | 0.101 | 0.022 | 1 | 11 | Megf8 | 0.079 | Old |
| 0.004788 | 0.304386 | 0.21 | 0.087 | 1 | 11 | Mink1 | 0.123 | Old |
| 0.004805 | 1.956512 | 0.073 | 0.007 | 1 | 11 | Cbfa2t2 | 0.066 | Old |
| 0.004813 | 0.647474 | 0.194 | 0.08 | 1 | 11 | Lrpprc | 0.114 | Old |
| 0.00482 | 1.239054 | 0.125 | 0.036 | 1 | 11 | Pdzd11 | 0.089 | Old |
| 0.004838 | 0.669717 | 0.298 | 0.159 | 1 | 11 | Plbd2 | 0.139 | Old |
| 0.004838 | 1.279684 | 0.113 | 0.029 | 1 | 11 | Actr8 | 0.084 | Old |
| 0.004843 | 1.985958 | 0.073 | 0.007 | 1 | 11 | Abcc4 | 0.066 | Old |
| 0.00486 | 1.007086 | 0.198 | 0.087 | 1 | 11 | Actl6a | 0.111 | Old |
| 0.004868 | 0.720832 | 0.113 | 0.029 | 1 | 11 | Oma1 | 0.084 | Old |
| 0.004875 | 1.108869 | 0.169 | 0.065 | 1 | 11 | Cd4 | 0.104 | Old |
| 0.00488 | 0.181144 | 0.919 | 0.884 | 1 | 11 | Calm1 | 0.035 | Old |
| 0.004899 | 0.457444 | 0.137 | 0.043 | 1 | 11 | Pitpnm1 | 0.094 | Old |
| 0.004902 | 0.307373 | 0.298 | 0.152 | 1 | 11 | Lpcat1 | 0.146 | Old |
| 0.004914 | 0.589759 | 0.347 | 0.203 | 1 | 11 | Plec | 0.144 | Old |
| 0.004915 | 0.841077 | 0.181 | 0.072 | 1 | 11 | Pdcd11 | 0.109 | Old |

| | | | | | | | | |
|---|---|---|---|---|---|---|---|---|
| 0.004932 | 0.673382 | 0.29 | 0.152 | 1 | 11 | Ssfa2 | 0.138 | Old |
| 0.004941 | 0.434882 | 0.48 | 0.304 | 1 | 11 | Surf4 | 0.176 | Old |
| 0.004971 | 1.110075 | 0.101 | 0.022 | 1 | 11 | Prmt7 | 0.079 | Old |
| 0.004986 | 0.369211 | 0.645 | 0.428 | 1 | 11 | Susd6 | 0.217 | Old |
| 0.004989 | 0.469208 | 0.395 | 0.225 | 1 | 11 | B230219D2 | 0.17 | Old |
| 0.00502 | 0.101595 | 0.25 | 0.116 | 1 | 11 | Brox | 0.134 | Old |
| 0.00503 | 0.689848 | 0.379 | 0.225 | 1 | 11 | Huwe1 | 0.154 | Old |
| 0.005076 | 0.558953 | 0.331 | 0.181 | 1 | 11 | Adam9 | 0.15 | Old |
| 0.005083 | 0.822213 | 0.113 | 0.029 | 1 | 11 | Aldh7a1 | 0.084 | Old |
| 0.005084 | 0.816647 | 0.23 | 0.109 | 1 | 11 | Tm2d1 | 0.121 | Old |
| 0.005103 | 0.706792 | 0.101 | 0.022 | 1 | 11 | Anapc7 | 0.079 | Old |
| 0.005107 | 0.52706 | 0.351 | 0.196 | 1 | 11 | Npc1 | 0.155 | Old |
| 0.005123 | 2.619162 | 0.085 | 0.014 | 1 | 11 | Scap | 0.071 | Old |
| 0.005126 | 1.083487 | 0.169 | 0.065 | 1 | 11 | Riok1 | 0.104 | Old |
| 0.005135 | 0.582815 | 0.306 | 0.159 | 1 | 11 | Pan3 | 0.147 | Old |
| 0.005141 | 1.067581 | 0.125 | 0.036 | 1 | 11 | Scarf1 | 0.089 | Old |
| 0.005194 | 0.506779 | 0.391 | 0.225 | 1 | 11 | Was | 0.166 | Old |
| 0.005221 | 0.358877 | 0.399 | 0.232 | 1 | 11 | Pcmtd1 | 0.167 | Old |
| 0.005255 | 0.469614 | 0.319 | 0.167 | 1 | 11 | Vat1 | 0.152 | Old |
| 0.005255 | 1.234736 | 0.169 | 0.065 | 1 | 11 | B4galt3 | 0.104 | Old |
| 0.005308 | 0.596164 | 0.101 | 0.022 | 1 | 11 | C1ra | 0.079 | Old |
| 0.00531 | 2.297824 | 0.085 | 0.014 | 1 | 11 | Kif2c | 0.071 | Old |
| 0.005324 | 0.496628 | 0.125 | 0.036 | 1 | 11 | Uchl3 | 0.089 | Old |
| 0.005344 | 1.207727 | 0.133 | 0.043 | 1 | 11 | Slc25a20 | 0.09 | Old |
| 0.005355 | 0.231523 | 0.379 | 0.217 | 1 | 11 | Tox4 | 0.162 | Old |
| 0.005372 | 0.798744 | 0.113 | 0.029 | 1 | 11 | Stx18 | 0.084 | Old |
| 0.00538 | 0.739995 | 0.177 | 0.072 | 1 | 11 | Pde4dip | 0.105 | Old |
| 0.005387 | 1.148823 | 0.262 | 0.138 | 1 | 11 | Ptpn12 | 0.124 | Old |
| 0.0054 | 1.402367 | 0.109 | 0.029 | 1 | 11 | Cog8 | 0.08 | Old |
| 0.00541 | 0.719725 | 0.202 | 0.087 | 1 | 11 | Atg16l1 | 0.115 | Old |
| 0.005416 | 0.331412 | 0.137 | 0.043 | 1 | 11 | Dcxr | 0.094 | Old |
| 0.00543 | 1.040701 | 0.145 | 0.051 | 1 | 11 | Nat9 | 0.094 | Old |
| 0.005481 | 0.753751 | 0.181 | 0.072 | 1 | 11 | Gmppa | 0.109 | Old |
| 0.005481 | 0.689803 | 0.181 | 0.072 | 1 | 11 | Gemin7 | 0.109 | Old |
| 0.005481 | 0.662646 | 0.319 | 0.181 | 1 | 11 | Ptbp1 | 0.138 | Old |
| 0.005505 | 0.294207 | 0.641 | 0.428 | 1 | 11 | Arf4 | 0.213 | Old |
| 0.005521 | 0.20842 | 0.298 | 0.152 | 1 | 11 | Bscl2 | 0.146 | Old |
| 0.005546 | 0.320409 | 0.427 | 0.246 | 1 | 11 | Mrps21 | 0.181 | Old |
| 0.005551 | 0.151845 | 0.31 | 0.167 | 1 | 11 | Dst | 0.143 | Old |
| 0.005559 | 0.865624 | 0.198 | 0.087 | 1 | 11 | AI413582 | 0.111 | Old |
| 0.005596 | 0.438029 | 0.161 | 0.058 | 1 | 11 | Abr | 0.103 | Old |
| 0.005602 | 0.310568 | 0.383 | 0.217 | 1 | 11 | Ppp3r1 | 0.166 | Old |
| 0.005607 | 1.15748 | 0.145 | 0.051 | 1 | 11 | Atmin | 0.094 | Old |
| 0.00561 | 0.862423 | 0.125 | 0.036 | 1 | 11 | Gm47754 | 0.089 | Old |
| 0.005617 | 0.324063 | 0.298 | 0.152 | 1 | 11 | 4932438A1 | 0.146 | Old |

| | | | | | | | | |
|---|---|---|---|---|---|---|---|---|
| 0.005639 | 2.218935 | 0.109 | 0.029 | 1 | 11 | Sh3glb2 | 0.08 | Old |
| 0.005663 | 1.605431 | 0.085 | 0.014 | 1 | 11 | Cul9 | 0.071 | Old |
| 0.005666 | 0.266465 | 0.31 | 0.159 | 1 | 11 | Pik3r5 | 0.151 | Old |
| 0.005734 | 1.000399 | 0.157 | 0.058 | 1 | 11 | Tmem250- | 0.099 | Old |
| 0.005737 | 0.394281 | 0.266 | 0.138 | 1 | 11 | Capn1 | 0.128 | Old |
| 0.005744 | 0.91275 | 0.254 | 0.13 | 1 | 11 | Rbfa | 0.124 | Old |
| 0.005754 | 0.636378 | 0.48 | 0.319 | 1 | 11 | Lifr | 0.161 | Old |
| 0.005779 | 1.179709 | 0.133 | 0.043 | 1 | 11 | Tlr12 | 0.09 | Old |
| 0.005785 | 1.669083 | 0.085 | 0.014 | 1 | 11 | Camk2a | 0.071 | Old |
| 0.005822 | 0.375827 | 0.274 | 0.138 | 1 | 11 | Plgrkt | 0.136 | Old |
| 0.005824 | 0.525209 | 0.403 | 0.239 | 1 | 11 | Kdelr1 | 0.164 | Old |
| 0.005828 | 0.324982 | 0.282 | 0.145 | 1 | 11 | Rybp | 0.137 | Old |
| 0.005849 | 0.69412 | 0.19 | 0.08 | 1 | 11 | Sart1 | 0.11 | Old |
| 0.005849 | 0.396709 | 0.19 | 0.08 | 1 | 11 | Supt20 | 0.11 | Old |
| 0.005861 | 0.82366 | 0.23 | 0.109 | 1 | 11 | Eif3g | 0.121 | Old |
| 0.005862 | 0.457353 | 0.823 | 0.609 | 1 | 11 | Prkcd | 0.214 | Old |
| 0.005884 | 0.696071 | 0.274 | 0.145 | 1 | 11 | Pcf11 | 0.129 | Old |
| 0.005909 | 0.752857 | 0.085 | 0.014 | 1 | 11 | Ddx55 | 0.071 | Old |
| 0.005919 | 2.035141 | 0.097 | 0.022 | 1 | 11 | Mphosph6 | 0.075 | Old |
| 0.005943 | 0.514821 | 0.125 | 0.036 | 1 | 11 | Mrps26 | 0.089 | Old |
| 0.005946 | 0.970475 | 0.145 | 0.051 | 1 | 11 | Tmem2 | 0.094 | Old |
| 0.00597 | 0.316338 | 0.448 | 0.261 | 1 | 11 | Etf1 | 0.187 | Old |
| 0.00598 | 0.759069 | 0.169 | 0.065 | 1 | 11 | Pop5 | 0.104 | Old |
| 0.005983 | 0.46639 | 0.742 | 0.565 | 1 | 11 | Naaa | 0.177 | Old |
| 0.005987 | 0.812475 | 0.29 | 0.159 | 1 | 11 | Idh3g | 0.131 | Old |
| 0.005992 | 0.435696 | 0.407 | 0.254 | 1 | 11 | Slc11a1 | 0.153 | Old |
| 0.005998 | 0.279176 | 0.383 | 0.217 | 1 | 11 | Ttc7 | 0.166 | Old |
| 0.006029 | 0.414388 | 0.331 | 0.188 | 1 | 11 | Fndc3b | 0.143 | Old |
| 0.006031 | 3.236328 | 0.069 | 0.007 | 1 | 11 | Ndufaf3 | 0.062 | Old |
| 0.006037 | 1.626991 | 0.097 | 0.022 | 1 | 11 | Nol9 | 0.075 | Old |
| 0.006037 | 0.733493 | 0.198 | 0.087 | 1 | 11 | Vps37a | 0.111 | Old |
| 0.006039 | 0.528072 | 0.169 | 0.065 | 1 | 11 | Slc25a10 | 0.104 | Old |
| 0.006042 | 0.58465 | 0.327 | 0.188 | 1 | 11 | Nrbp1 | 0.139 | Old |
| 0.006059 | 0.499388 | 0.21 | 0.094 | 1 | 11 | Cfdp1 | 0.116 | Old |
| 0.006065 | 0.253941 | 0.331 | 0.181 | 1 | 11 | Atp6v1c1 | 0.15 | Old |
| 0.006067 | 0.609202 | 0.157 | 0.058 | 1 | 11 | Rufy1 | 0.099 | Old |
| 0.006115 | 0.498201 | 0.294 | 0.152 | 1 | 11 | Stt3b | 0.142 | Old |
| 0.006128 | 2.082733 | 0.129 | 0.043 | 1 | 11 | Zswim4 | 0.086 | Old |
| 0.006143 | 0.650193 | 0.23 | 0.109 | 1 | 11 | Cers5 | 0.121 | Old |
| 0.00616 | 0.77531 | 0.157 | 0.058 | 1 | 11 | Sphk2 | 0.099 | Old |
| 0.006171 | 1.992893 | 0.185 | 0.087 | 1 | 11 | Tspan13 | 0.098 | Old |
| 0.006178 | 0.482669 | 0.411 | 0.239 | 1 | 11 | Rilpl2 | 0.172 | Old |
| 0.006186 | 1.533057 | 0.109 | 0.029 | 1 | 11 | Gga2 | 0.08 | Old |
| 0.0062 | 0.182866 | 0.319 | 0.167 | 1 | 11 | Ssu72 | 0.152 | Old |
| 0.006204 | 0.221513 | 0.315 | 0.167 | 1 | 11 | Smim10l1 | 0.148 | Old |

| | | | | | | | | |
|---|---|---|---|---|---|---|---|---|
| 0.006234 | 0.26535 | 0.972 | 0.942 | 1 | 11 | Ctsb | 0.03 | Old |
| 0.006282 | 0.311457 | 0.319 | 0.167 | 1 | 11 | Dnajb1 | 0.152 | Old |
| 0.006301 | 1.531013 | 0.109 | 0.029 | 1 | 11 | Tsta3 | 0.08 | Old |
| 0.006303 | 0.257784 | 0.431 | 0.254 | 1 | 11 | Cmklr1 | 0.177 | Old |
| 0.006318 | 0.21778 | 0.302 | 0.159 | 1 | 11 | Cd274 | 0.143 | Old |
| 0.006325 | 4.122719 | 0.052 | 0 | 1 | 11 | Bud13 | 0.052 | Old |
| 0.006325 | 3.375358 | 0.052 | 0 | 1 | 11 | Tsen2 | 0.052 | Old |
| 0.006325 | 3.309047 | 0.052 | 0 | 1 | 11 | Pdcd1lg2 | 0.052 | Old |
| 0.006344 | 0.29582 | 0.722 | 0.522 | 1 | 11 | Pcbp1 | 0.2 | Old |
| 0.006345 | 1.857498 | 0.141 | 0.051 | 1 | 11 | Mrpl19 | 0.09 | Old |
| 0.00635 | 0.994801 | 0.133 | 0.043 | 1 | 11 | Mettl16 | 0.09 | Old |
| 0.006354 | 0.483647 | 0.278 | 0.145 | 1 | 11 | Arhgef12 | 0.133 | Old |
| 0.006375 | 2.383793 | 0.069 | 0.007 | 1 | 11 | Trim37 | 0.062 | Old |
| 0.006378 | 0.508297 | 0.29 | 0.152 | 1 | 11 | Gtpbp4 | 0.138 | Old |
| 0.006409 | 0.240996 | 0.371 | 0.21 | 1 | 11 | Pnn | 0.161 | Old |
| 0.006416 | 0.280955 | 0.315 | 0.167 | 1 | 11 | Brd1 | 0.148 | Old |
| 0.006503 | 0.658273 | 0.202 | 0.087 | 1 | 11 | Smim27 | 0.115 | Old |
| 0.006553 | 0.84214 | 0.198 | 0.087 | 1 | 11 | Tmem68 | 0.111 | Old |
| 0.00658 | 2.134384 | 0.069 | 0.007 | 1 | 11 | Haus4 | 0.062 | Old |
| 0.00658 | 1.912032 | 0.069 | 0.007 | 1 | 11 | Top2a | 0.062 | Old |
| 0.006604 | 2.862223 | 0.081 | 0.014 | 1 | 11 | 0610010K1 | 0.067 | Old |
| 0.00661 | 0.836612 | 0.145 | 0.051 | 1 | 11 | Eipr1 | 0.094 | Old |
| 0.00661 | 0.706808 | 0.145 | 0.051 | 1 | 11 | Akr1b10 | 0.094 | Old |
| 0.006612 | 0.45973 | 0.238 | 0.116 | 1 | 11 | Adrm1 | 0.122 | Old |
| 0.006632 | 2.123501 | 0.069 | 0.007 | 1 | 11 | 3110062M | 0.062 | Old |
| 0.006633 | 0.321746 | 0.81 | 0.587 | 1 | 11 | Prcp | 0.223 | Old |
| 0.006636 | 0.771273 | 0.165 | 0.065 | 1 | 11 | Esrra | 0.1 | Old |
| 0.006681 | 0.332252 | 0.202 | 0.087 | 1 | 11 | Kctd10 | 0.115 | Old |
| 0.006684 | 1.67985 | 0.069 | 0.007 | 1 | 11 | Tor4a | 0.062 | Old |
| 0.006736 | 0.150401 | 0.54 | 0.326 | 1 | 11 | Apbb1ip | 0.214 | Old |
| 0.006737 | 1.261391 | 0.069 | 0.007 | 1 | 11 | Ints14 | 0.062 | Old |
| 0.00675 | 0.667933 | 0.278 | 0.145 | 1 | 11 | Gsdmd | 0.133 | Old |
| 0.006751 | 0.422043 | 0.254 | 0.123 | 1 | 11 | Rab24 | 0.131 | Old |
| 0.006754 | 0.554754 | 0.367 | 0.217 | 1 | 11 | Capn2 | 0.15 | Old |
| 0.006758 | 0.060491 | 0.367 | 0.203 | 1 | 11 | Ptpa | 0.164 | Old |
| 0.006778 | 0.889312 | 0.177 | 0.072 | 1 | 11 | Slc16a7 | 0.105 | Old |
| 0.006798 | 0.336531 | 0.79 | 0.572 | 1 | 11 | Srsf5 | 0.218 | Old |
| 0.006821 | 1.183108 | 0.133 | 0.043 | 1 | 11 | Ldlrap1 | 0.09 | Old |
| 0.006822 | 0.267629 | 0.226 | 0.101 | 1 | 11 | Rspry1 | 0.125 | Old |
| 0.006838 | 0.429491 | 0.282 | 0.145 | 1 | 11 | Vamp4 | 0.137 | Old |
| 0.006849 | 0.736792 | 0.226 | 0.109 | 1 | 11 | Hypk | 0.117 | Old |
| 0.006857 | 0.897793 | 0.145 | 0.051 | 1 | 11 | Xab2 | 0.094 | Old |
| 0.006863 | 0.25962 | 0.109 | 0.029 | 1 | 11 | Nelfcd | 0.08 | Old |
| 0.006864 | 0.454174 | 0.238 | 0.116 | 1 | 11 | Abhd17a | 0.122 | Old |
| 0.00698 | 0.671641 | 0.27 | 0.145 | 1 | 11 | Etv5 | 0.125 | Old |

| | | | | | | | | |
|---|---|---|---|---|---|---|---|---|
| 0.006989 | 0.752595 | 0.153 | 0.058 | 1 | 11 | Cryzl1 | 0.095 | Old |
| 0.006999 | 0.619813 | 0.331 | 0.188 | 1 | 11 | Dram1 | 0.143 | Old |
| 0.007005 | 0.666271 | 0.23 | 0.109 | 1 | 11 | Rit1 | 0.121 | Old |
| 0.00702 | 0.260004 | 0.181 | 0.072 | 1 | 11 | Aldh16a1 | 0.109 | Old |
| 0.007022 | 0.416294 | 0.306 | 0.167 | 1 | 11 | Pik3c2a | 0.139 | Old |
| 0.007028 | 0.509053 | 0.169 | 0.065 | 1 | 11 | Jkamp | 0.104 | Old |
| 0.007079 | 0.999192 | 0.19 | 0.087 | 1 | 11 | Med8 | 0.103 | Old |
| 0.007108 | 0.556217 | 0.165 | 0.065 | 1 | 11 | Ubap1 | 0.1 | Old |
| 0.007151 | 0.517788 | 0.327 | 0.181 | 1 | 11 | Snrpd1 | 0.146 | Old |
| 0.007152 | 0.638688 | 0.121 | 0.036 | 1 | 11 | Parp3 | 0.085 | Old |
| 0.007161 | 0.494577 | 0.194 | 0.087 | 1 | 11 | Slc44a1 | 0.107 | Old |
| 0.007185 | 0.151172 | 0.399 | 0.232 | 1 | 11 | Add1 | 0.167 | Old |
| 0.007192 | 0.649817 | 0.351 | 0.203 | 1 | 11 | R3hdm4 | 0.148 | Old |
| 0.007245 | 2.501069 | 0.081 | 0.014 | 1 | 11 | Jagn1 | 0.067 | Old |
| 0.007248 | 0.770644 | 0.165 | 0.065 | 1 | 11 | E2f4 | 0.1 | Old |
| 0.00729 | 0.995998 | 0.161 | 0.065 | 1 | 11 | Crnkl1 | 0.096 | Old |
| 0.007296 | 2.10668 | 0.081 | 0.014 | 1 | 11 | Pqlc2 | 0.067 | Old |
| 0.007296 | 1.935503 | 0.081 | 0.014 | 1 | 11 | Phlpp1 | 0.067 | Old |
| 0.007331 | 0.201361 | 0.702 | 0.5 | 1 | 11 | Plin2 | 0.202 | Old |
| 0.00736 | 1.356983 | 0.129 | 0.043 | 1 | 11 | Uckl1 | 0.086 | Old |
| 0.007371 | 0.232426 | 0.605 | 0.428 | 1 | 11 | Colgalt1 | 0.177 | Old |
| 0.007397 | 0.285259 | 0.298 | 0.159 | 1 | 11 | Tecr | 0.139 | Old |
| 0.0074 | 2.650558 | 0.081 | 0.014 | 1 | 11 | Kif13a | 0.067 | Old |
| 0.0074 | 2.118768 | 0.081 | 0.014 | 1 | 11 | Riox2 | 0.067 | Old |
| 0.007427 | 0.314072 | 0.427 | 0.261 | 1 | 11 | Mlf2 | 0.166 | Old |
| 0.007429 | 0.309544 | 0.383 | 0.232 | 1 | 11 | Ndufab1 | 0.151 | Old |
| 0.007478 | 0.432021 | 0.246 | 0.123 | 1 | 11 | Sarnp | 0.123 | Old |
| 0.007507 | 0.580586 | 0.238 | 0.123 | 1 | 11 | Fam50a | 0.115 | Old |
| 0.007512 | 0.037453 | 0.343 | 0.181 | 1 | 11 | Pfdn2 | 0.162 | Old |
| 0.007516 | 0.439929 | 0.31 | 0.167 | 1 | 11 | Ctbp1 | 0.143 | Old |
| 0.007521 | 0.237938 | 0.177 | 0.072 | 1 | 11 | Wdr43 | 0.105 | Old |
| 0.007532 | 0.214615 | 0.294 | 0.159 | 1 | 11 | Khsrp | 0.135 | Old |
| 0.007549 | 0.477029 | 0.25 | 0.123 | 1 | 11 | Snrnp200 | 0.127 | Old |
| 0.007617 | 0.511016 | 0.218 | 0.101 | 1 | 11 | Gmps | 0.117 | Old |
| 0.007656 | 0.04194 | 0.31 | 0.167 | 1 | 11 | Ndufb2 | 0.143 | Old |
| 0.007666 | 1.949111 | 0.081 | 0.014 | 1 | 11 | Tbccd1 | 0.067 | Old |
| 0.007668 | 0.425953 | 0.29 | 0.159 | 1 | 11 | Pdcl3 | 0.131 | Old |
| 0.007684 | 0.785026 | 0.173 | 0.072 | 1 | 11 | Avpi1 | 0.101 | Old |
| 0.007721 | 1.895226 | 0.093 | 0.022 | 1 | 11 | Parn | 0.071 | Old |
| 0.007757 | 1.000078 | 0.173 | 0.072 | 1 | 11 | Ddx10 | 0.101 | Old |
| 0.007759 | 1.29698 | 0.117 | 0.036 | 1 | 11 | Rai1 | 0.081 | Old |
| 0.007775 | 1.086844 | 0.129 | 0.043 | 1 | 11 | Tmem268 | 0.086 | Old |
| 0.007813 | 0.207864 | 0.968 | 0.949 | 1 | 11 | H2-Ab1 | 0.019 | Old |
| 0.007818 | 0.297409 | 0.129 | 0.043 | 1 | 11 | Atp11a | 0.086 | Old |
| 0.007822 | 1.672898 | 0.093 | 0.022 | 1 | 11 | Cnnm3 | 0.071 | Old |

| | | | | | | | | |
|---|---|---|---|---|---|---|---|---|
| 0.007838 | 0.487022 | 0.109 | 0.029 | 1 | 11 | Ivd | 0.08 | Old |
| 0.007868 | 0.863346 | 0.173 | 0.072 | 1 | 11 | Erg28 | 0.101 | Old |
| 0.007873 | 1.775187 | 0.093 | 0.022 | 1 | 11 | Slc9a1 | 0.071 | Old |
| 0.007897 | 0.616822 | 0.343 | 0.203 | 1 | 11 | Elk3 | 0.14 | Old |
| 0.007971 | 0.361834 | 0.839 | 0.725 | 1 | 11 | Ctsz | 0.114 | Old |
| 0.007975 | 0.397209 | 0.528 | 0.341 | 1 | 11 | Ldha | 0.187 | Old |
| 0.007976 | 1.696059 | 0.093 | 0.022 | 1 | 11 | Eml3 | 0.071 | Old |
| 0.007993 | 0.064188 | 0.266 | 0.13 | 1 | 11 | Tmem87b | 0.136 | Old |
| 0.007996 | 0.673592 | 0.411 | 0.261 | 1 | 11 | Soat1 | 0.15 | Old |
| 0.008061 | 0.233055 | 0.516 | 0.333 | 1 | 11 | Slamf8 | 0.183 | Old |
| 0.008093 | 0.340459 | 0.347 | 0.196 | 1 | 11 | Acsl5 | 0.151 | Old |
| 0.008095 | 1.473807 | 0.105 | 0.029 | 1 | 11 | Ganc | 0.076 | Old |
| 0.008103 | 0.655665 | 0.355 | 0.21 | 1 | 11 | Wdr45b | 0.145 | Old |
| 0.008188 | 0.331613 | 0.476 | 0.283 | 1 | 11 | Atp6v0d1 | 0.193 | Old |
| 0.008195 | 0.445015 | 0.589 | 0.413 | 1 | 11 | Rpl22l1 | 0.176 | Old |
| 0.00823 | 0.40356 | 0.258 | 0.13 | 1 | 11 | Taz | 0.128 | Old |
| 0.008243 | 0.322585 | 0.157 | 0.058 | 1 | 11 | Tbc1d15 | 0.099 | Old |
| 0.008244 | 1.677965 | 0.105 | 0.029 | 1 | 11 | Rcor3 | 0.076 | Old |
| 0.008265 | 0.183863 | 0.27 | 0.138 | 1 | 11 | Polr2f | 0.132 | Old |
| 0.008288 | 0.243097 | 0.29 | 0.152 | 1 | 11 | St8sia6 | 0.138 | Old |
| 0.008311 | 0.325842 | 0.423 | 0.254 | 1 | 11 | Nfe2l1 | 0.169 | Old |
| 0.008359 | 0.795539 | 0.117 | 0.036 | 1 | 11 | Ocel1 | 0.081 | Old |
| 0.008369 | 0.307546 | 0.339 | 0.188 | 1 | 11 | Ly9 | 0.151 | Old |
| 0.008399 | 0.370336 | 0.419 | 0.246 | 1 | 11 | Casp8 | 0.173 | Old |
| 0.0084 | 1.012303 | 0.093 | 0.022 | 1 | 11 | Alkbh6 | 0.071 | Old |
| 0.008415 | 0.273754 | 0.194 | 0.087 | 1 | 11 | Snap29 | 0.107 | Old |
| 0.00843 | 0.399104 | 0.335 | 0.188 | 1 | 11 | Nceh1 | 0.147 | Old |
| 0.008461 | 1.893152 | 0.113 | 0.036 | 1 | 11 | Cmc2 | 0.077 | Old |
| 0.008481 | 1.921373 | 0.125 | 0.043 | 1 | 11 | Myo7a | 0.082 | Old |
| 0.008488 | 0.304117 | 0.298 | 0.159 | 1 | 11 | Lap3 | 0.139 | Old |
| 0.008497 | 1.322653 | 0.105 | 0.029 | 1 | 11 | E130309D( | 0.076 | Old |
| 0.008501 | 0.789776 | 0.258 | 0.138 | 1 | 11 | Galk2 | 0.12 | Old |
| 0.008512 | 0.174484 | 0.379 | 0.225 | 1 | 11 | Tpi1 | 0.154 | Old |
| 0.008522 | 0.352821 | 0.21 | 0.094 | 1 | 11 | Slc38a6 | 0.116 | Old |
| 0.008525 | 2.525877 | 0.065 | 0.007 | 1 | 11 | Cks1b | 0.058 | Old |
| 0.008543 | 1.344924 | 0.185 | 0.087 | 1 | 11 | Dda1 | 0.098 | Old |
| 0.008545 | 0.346619 | 0.25 | 0.123 | 1 | 11 | Dapk3 | 0.127 | Old |
| 0.008546 | 0.367489 | 0.286 | 0.152 | 1 | 11 | Marveld1 | 0.134 | Old |
| 0.008548 | 1.136594 | 0.105 | 0.029 | 1 | 11 | Nit1 | 0.076 | Old |
| 0.00855 | 0.311338 | 0.298 | 0.159 | 1 | 11 | Atp6v1d | 0.139 | Old |
| 0.008574 | 0.243186 | 0.31 | 0.167 | 1 | 11 | Stx12 | 0.143 | Old |
| 0.008593 | 2.676857 | 0.065 | 0.007 | 1 | 11 | Gm13710 | 0.058 | Old |
| 0.0086 | 0.90409 | 0.117 | 0.036 | 1 | 11 | Polr1c | 0.081 | Old |
| 0.008608 | 0.572959 | 0.141 | 0.051 | 1 | 11 | Ecd | 0.09 | Old |
| 0.008636 | 0.722617 | 0.282 | 0.167 | 1 | 11 | Vps13c | 0.115 | Old |

| | | | | | | | | |
|---|---|---|---|---|---|---|---|---|
| 0.008704 | 1.182725 | 0.105 | 0.029 | 1 | 11 | Fbxl6 | 0.076 | Old |
| 0.008711 | 0.408025 | 0.246 | 0.123 | 1 | 11 | Plcg2 | 0.123 | Old |
| 0.008718 | 0.237716 | 0.153 | 0.058 | 1 | 11 | Anapc15 | 0.095 | Old |
| 0.008741 | 0.426104 | 0.294 | 0.159 | 1 | 11 | Yipf3 | 0.135 | Old |
| 0.008785 | 0.372567 | 0.653 | 0.464 | 1 | 11 | Selenot | 0.189 | Old |
| 0.008799 | 3.340277 | 0.048 | 0 | 1 | 11 | Gm12185 | 0.048 | Old |
| 0.008799 | 3.290567 | 0.048 | 0 | 1 | 11 | Tpk1 | 0.048 | Old |
| 0.008799 | 3.265127 | 0.048 | 0 | 1 | 11 | D3Ertd254 | 0.048 | Old |
| 0.008799 | 3.144587 | 0.048 | 0 | 1 | 11 | Dennd2a | 0.048 | Old |
| 0.008799 | 2.782242 | 0.048 | 0 | 1 | 11 | Twsg1 | 0.048 | Old |
| 0.008799 | 2.746514 | 0.048 | 0 | 1 | 11 | Alg12 | 0.048 | Old |
| 0.008811 | 0.348092 | 0.282 | 0.152 | 1 | 11 | Nudc | 0.13 | Old |
| 0.008829 | 0.199771 | 0.21 | 0.101 | 1 | 11 | Gm8186 | 0.109 | Old |
| 0.008832 | 0.390731 | 0.48 | 0.304 | 1 | 11 | Cd48 | 0.176 | Old |
| 0.008864 | 1.806573 | 0.137 | 0.051 | 1 | 11 | Smg9 | 0.086 | Old |
| 0.008901 | 0.373181 | 0.093 | 0.022 | 1 | 11 | Ints11 | 0.071 | Old |
| 0.008931 | 0.323685 | 0.524 | 0.341 | 1 | 11 | Rbx1 | 0.183 | Old |
| 0.008951 | 0.559678 | 0.302 | 0.174 | 1 | 11 | Rexo2 | 0.128 | Old |
| 0.008959 | 0.272251 | 0.254 | 0.13 | 1 | 11 | Usp48 | 0.124 | Old |
| 0.008977 | 0.361309 | 0.258 | 0.138 | 1 | 11 | Sec23b | 0.12 | Old |
| 0.008987 | 1.016617 | 0.198 | 0.094 | 1 | 11 | Spg7 | 0.104 | Old |
| 0.008997 | 0.062729 | 0.25 | 0.123 | 1 | 11 | Parp8 | 0.127 | Old |
| 0.009012 | 0.364725 | 0.395 | 0.225 | 1 | 11 | Coro1b | 0.17 | Old |
| 0.009023 | 0.236423 | 0.214 | 0.101 | 1 | 11 | Btbd1 | 0.113 | Old |
| 0.009034 | 1.289166 | 0.149 | 0.058 | 1 | 11 | Grpel1 | 0.091 | Old |
| 0.009042 | 0.37269 | 0.984 | 0.964 | 1 | 11 | Ftl1 | 0.02 | Old |
| 0.009078 | 1.00075 | 0.105 | 0.029 | 1 | 11 | Nup85 | 0.076 | Old |
| 0.009078 | 1.917628 | 0.065 | 0.007 | 1 | 11 | Cdkal1 | 0.058 | Old |
| 0.009111 | 0.255223 | 0.141 | 0.051 | 1 | 11 | Gm5547 | 0.09 | Old |
| 0.009125 | 0.299405 | 0.149 | 0.058 | 1 | 11 | Gtf2f1 | 0.091 | Old |
| 0.009149 | 1.796573 | 0.065 | 0.007 | 1 | 11 | Chpf | 0.058 | Old |
| 0.009155 | 1.319442 | 0.125 | 0.043 | 1 | 11 | Prkch | 0.082 | Old |
| 0.009172 | 0.233729 | 0.246 | 0.123 | 1 | 11 | Tmem131l | 0.123 | Old |
| 0.009172 | 0.36187 | 0.488 | 0.326 | 1 | 11 | Rhob | 0.162 | Old |
| 0.009206 | 0.844484 | 0.161 | 0.065 | 1 | 11 | Commd1 | 0.096 | Old |
| 0.009212 | 0.247602 | 0.218 | 0.101 | 1 | 11 | Tnfaip8l2 | 0.117 | Old |
| 0.009258 | 0.658511 | 0.117 | 0.036 | 1 | 11 | Cenpc1 | 0.081 | Old |
| 0.009275 | 0.40057 | 0.359 | 0.21 | 1 | 11 | Map4 | 0.149 | Old |
| 0.009289 | 0.405758 | 0.278 | 0.152 | 1 | 11 | Ppt2 | 0.126 | Old |
| 0.009289 | 0.439964 | 0.258 | 0.13 | 1 | 11 | Cpq | 0.128 | Old |
| 0.009293 | 0.582832 | 0.214 | 0.101 | 1 | 11 | Mrpl34 | 0.113 | Old |
| 0.009293 | 1.429301 | 0.065 | 0.007 | 1 | 11 | Exd2 | 0.058 | Old |
| 0.009293 | 0.689522 | 0.065 | 0.007 | 1 | 11 | Chd1l | 0.058 | Old |
| 0.009295 | 0.438774 | 0.266 | 0.138 | 1 | 11 | Senp2 | 0.128 | Old |
| 0.009306 | 0.191145 | 0.516 | 0.319 | 1 | 11 | Snrpb2 | 0.197 | Old |

| | | | | | | | | |
|---|---|---|---|---|---|---|---|---|
| 0.009311 | 0.968504 | 0.117 | 0.036 | 1 | 11 | Kifap3 | 0.081 | Old |
| 0.009335 | 0.088514 | 0.472 | 0.283 | 1 | 11 | Nme1 | 0.189 | Old |
| 0.009343 | 0.449128 | 0.153 | 0.058 | 1 | 11 | Cpne2 | 0.095 | Old |
| 0.009346 | 0.287584 | 0.819 | 0.71 | 1 | 11 | Aif1 | 0.109 | Old |
| 0.009347 | 0.414213 | 0.52 | 0.348 | 1 | 11 | Ndufa13 | 0.172 | Old |
| 0.009353 | 0.500516 | 0.31 | 0.174 | 1 | 11 | Txnl1 | 0.136 | Old |
| 0.009363 | 0.534465 | 0.117 | 0.036 | 1 | 11 | 2010315BC | 0.081 | Old |
| 0.009377 | 0.336857 | 0.512 | 0.326 | 1 | 11 | Nipbl | 0.186 | Old |
| 0.009384 | 0.099824 | 0.427 | 0.254 | 1 | 11 | Raly | 0.173 | Old |
| 0.009417 | 0.302344 | 0.327 | 0.181 | 1 | 11 | Agps | 0.146 | Old |
| 0.009435 | 0.546914 | 0.298 | 0.167 | 1 | 11 | Idh2 | 0.131 | Old |
| 0.009437 | 0.617045 | 0.19 | 0.087 | 1 | 11 | Dhdh | 0.103 | Old |
| 0.009483 | 1.167491 | 0.137 | 0.051 | 1 | 11 | Thumpd1 | 0.086 | Old |
| 0.009483 | 0.440338 | 0.536 | 0.37 | 1 | 11 | Atp6v1e1 | 0.166 | Old |
| 0.009489 | 0.328462 | 0.093 | 0.022 | 1 | 11 | Mfn1 | 0.071 | Old |
| 0.009504 | 1.260147 | 0.427 | 0.312 | 1 | 11 | Pla2g2d | 0.115 | Old |
| 0.00951 | 1.48662 | 0.125 | 0.043 | 1 | 11 | Csf2rb2 | 0.082 | Old |
| 0.009537 | 0.351815 | 0.508 | 0.341 | 1 | 11 | Sec62 | 0.167 | Old |
| 0.009539 | 0.609397 | 0.298 | 0.167 | 1 | 11 | Cyb5a | 0.131 | Old |
| 0.009545 | 1.015612 | 0.214 | 0.109 | 1 | 11 | Galnt10 | 0.105 | Old |
| 0.009548 | 0.335107 | 0.456 | 0.283 | 1 | 11 | Ghitm | 0.173 | Old |
| 0.009561 | 1.073274 | 0.125 | 0.043 | 1 | 11 | Etfdh | 0.082 | Old |
| 0.009571 | 0.313682 | 0.206 | 0.094 | 1 | 11 | Washc3 | 0.112 | Old |
| 0.00958 | 0.50526 | 0.746 | 0.609 | 1 | 11 | Klf2 | 0.137 | Old |
| 0.00959 | 0.566261 | 0.149 | 0.058 | 1 | 11 | Xpa | 0.091 | Old |
| 0.009601 | 0.741256 | 0.129 | 0.043 | 1 | 11 | Vezt | 0.086 | Old |
| 0.009612 | 0.443002 | 0.387 | 0.239 | 1 | 11 | Sel1l | 0.148 | Old |
| 0.009613 | 0.777317 | 0.266 | 0.145 | 1 | 11 | Wdfy4 | 0.121 | Old |
| 0.009646 | 0.307298 | 0.532 | 0.362 | 1 | 11 | Pgk1 | 0.17 | Old |
| 0.00965 | 0.023231 | 0.298 | 0.159 | 1 | 11 | Slc25a11 | 0.139 | Old |
| 0.009683 | 1.065684 | 0.185 | 0.087 | 1 | 11 | Pdzd8 | 0.098 | Old |
| 0.009705 | 0.267495 | 0.129 | 0.043 | 1 | 11 | BC029722 | 0.086 | Old |
| 0.009725 | 0.565968 | 0.367 | 0.225 | 1 | 11 | Fes | 0.142 | Old |
| 0.00974 | 0.494884 | 0.202 | 0.094 | 1 | 11 | Arhgap1 | 0.108 | Old |
| 0.009804 | 2.219693 | 0.089 | 0.022 | 1 | 11 | Ube2e2 | 0.067 | Old |
| 0.009811 | 0.467835 | 0.383 | 0.232 | 1 | 11 | Cep350 | 0.151 | Old |
| 0.00989 | 0.385637 | 0.161 | 0.065 | 1 | 11 | Stk25 | 0.096 | Old |
| 0.009927 | 0.436147 | 0.629 | 0.435 | 1 | 11 | Psmb9 | 0.194 | Old |
| 0.009942 | 2.078366 | 0.077 | 0.014 | 1 | 11 | Srm | 0.063 | Old |
| 0.009958 | 0.214107 | 0.226 | 0.109 | 1 | 11 | Ttc39b | 0.117 | Old |
| 0.00996 | 0.26788 | 0.407 | 0.246 | 1 | 11 | Rufy3 | 0.161 | Old |
| 0.00996 | 0.060142 | 0.407 | 0.246 | 1 | 11 | Rassf3 | 0.161 | Old |
| 0.009962 | 0.282119 | 0.512 | 0.319 | 1 | 11 | Ssb | 0.193 | Old |
| 0.009963 | 0.77274 | 0.157 | 0.065 | 1 | 11 | Pdxdc1 | 0.092 | Old |
| 0.010011 | 2.672431 | 0.077 | 0.014 | 1 | 11 | Mettl26 | 0.063 | Old |

| | | | | | | | | |
|---|---|---|---|---|---|---|---|---|
| 0.010017 | 0.582455 | 0.169 | 0.072 | 1 | 11 | Nup93 | 0.097 | Old |
| 0.010021 | 0.320604 | 0.181 | 0.08 | 1 | 11 | Stim1 | 0.101 | Old |
| 0.010031 | 0.704751 | 0.274 | 0.152 | 1 | 11 | Psme3 | 0.122 | Old |
| 0.010038 | 0.268348 | 0.254 | 0.123 | 1 | 11 | Ide | 0.131 | Old |
| 0.010078 | 2.273299 | 0.101 | 0.029 | 1 | 11 | 2310022B0 | 0.072 | Old |
| 0.010082 | 2.409411 | 0.077 | 0.014 | 1 | 11 | Rtl8a | 0.063 | Old |
| 0.010082 | 1.705003 | 0.077 | 0.014 | 1 | 11 | Gtf3c4 | 0.063 | Old |
| 0.010095 | 0.270304 | 0.601 | 0.413 | 1 | 11 | Rbm25 | 0.188 | Old |
| 0.010121 | 0.258069 | 0.226 | 0.109 | 1 | 11 | Spopl | 0.117 | Old |
| 0.010124 | 0.373511 | 0.218 | 0.109 | 1 | 11 | Cstf3 | 0.109 | Old |
| 0.010138 | 0.396404 | 0.238 | 0.116 | 1 | 11 | Yipf4 | 0.122 | Old |
| 0.010139 | 1.960145 | 0.101 | 0.029 | 1 | 11 | Rnf19a | 0.072 | Old |
| 0.010149 | 0.601936 | 0.21 | 0.101 | 1 | 11 | Trnt1 | 0.109 | Old |
| 0.01015 | 0.277725 | 0.706 | 0.543 | 1 | 11 | Cd44 | 0.163 | Old |
| 0.010217 | 0.339878 | 0.29 | 0.159 | 1 | 11 | Mknk1 | 0.131 | Old |
| 0.01025 | 0.695378 | 0.254 | 0.138 | 1 | 11 | Mfsd11 | 0.116 | Old |
| 0.010279 | 0.181642 | 0.54 | 0.341 | 1 | 11 | Srgap2 | 0.199 | Old |
| 0.010306 | 0.564135 | 0.19 | 0.087 | 1 | 11 | Pex16 | 0.103 | Old |
| 0.010329 | 0.351685 | 0.173 | 0.072 | 1 | 11 | Cnot7 | 0.101 | Old |
| 0.010353 | 0.308826 | 0.379 | 0.232 | 1 | 11 | Copb1 | 0.147 | Old |
| 0.010378 | 0.23337 | 0.343 | 0.196 | 1 | 11 | Fchsd2 | 0.147 | Old |
| 0.010379 | 0.479312 | 0.266 | 0.138 | 1 | 11 | Hmgcs1 | 0.128 | Old |
| 0.010405 | 0.76369 | 0.137 | 0.051 | 1 | 11 | Tpcn2 | 0.086 | Old |
| 0.010413 | 0.3806 | 0.274 | 0.145 | 1 | 11 | Dennd4b | 0.129 | Old |
| 0.010415 | 0.410411 | 0.456 | 0.29 | 1 | 11 | Dad1 | 0.166 | Old |
| 0.010429 | 0.345517 | 0.165 | 0.072 | 1 | 11 | Mrps16 | 0.093 | Old |
| 0.010458 | 0.864808 | 0.137 | 0.051 | 1 | 11 | Gbp8 | 0.086 | Old |
| 0.010473 | 0.299111 | 0.331 | 0.188 | 1 | 11 | Gpsm3 | 0.143 | Old |
| 0.010478 | 0.830226 | 0.177 | 0.08 | 1 | 11 | Mrpl2 | 0.097 | Old |
| 0.010481 | 0.411828 | 0.181 | 0.08 | 1 | 11 | Vav2 | 0.101 | Old |
| 0.01049 | 0.286135 | 0.883 | 0.652 | 1 | 11 | Cd47 | 0.231 | Old |
| 0.010492 | 0.544946 | 0.282 | 0.152 | 1 | 11 | Rab43 | 0.13 | Old |
| 0.010511 | 1.715092 | 0.101 | 0.029 | 1 | 11 | Adnp2 | 0.072 | Old |
| 0.010524 | 2.063297 | 0.089 | 0.022 | 1 | 11 | Yae1d1 | 0.067 | Old |
| 0.010531 | 0.132175 | 0.274 | 0.145 | 1 | 11 | Ppfia4 | 0.129 | Old |
| 0.010552 | 0.626855 | 0.157 | 0.065 | 1 | 11 | Fam193a | 0.092 | Old |
| 0.01056 | 0.558092 | 0.367 | 0.239 | 1 | 11 | Asap1 | 0.128 | Old |
| 0.010592 | 2.322062 | 0.089 | 0.022 | 1 | 11 | Tgm1 | 0.067 | Old |
| 0.010637 | 1.497373 | 0.101 | 0.029 | 1 | 11 | Map3k20 | 0.072 | Old |
| 0.010701 | 1.992965 | 0.101 | 0.029 | 1 | 11 | Vps8 | 0.072 | Old |
| 0.010721 | 0.309969 | 0.468 | 0.297 | 1 | 11 | Cox14 | 0.171 | Old |
| 0.010741 | 0.241629 | 0.181 | 0.08 | 1 | 11 | Pnpla7 | 0.101 | Old |
| 0.010814 | 0.22798 | 0.359 | 0.21 | 1 | 11 | Cycs | 0.149 | Old |
| 0.010832 | 0.066468 | 0.423 | 0.261 | 1 | 11 | Kif1b | 0.162 | Old |
| 0.010856 | 0.39919 | 0.363 | 0.217 | 1 | 11 | Antxr2 | 0.146 | Old |

| | | | | | | | | |
|---|---|---|---|---|---|---|---|---|
| 0.010878 | 1.059369 | 0.125 | 0.043 | 1 | 11 | Pygb | 0.082 | Old |
| 0.010885 | 0.769117 | 0.077 | 0.014 | 1 | 11 | Fdps | 0.063 | Old |
| 0.010889 | 0.40643 | 0.435 | 0.275 | 1 | 11 | Ankfy1 | 0.16 | Old |
| 0.01092 | 0.272491 | 0.512 | 0.319 | 1 | 11 | Tax1bp1 | 0.193 | Old |
| 0.010937 | 0.52879 | 0.125 | 0.043 | 1 | 11 | Mtmr1 | 0.082 | Old |
| 0.010937 | 0.492546 | 0.125 | 0.043 | 1 | 11 | Zfp991 | 0.082 | Old |
| 0.01094 | 0.535669 | 0.222 | 0.109 | 1 | 11 | Nfya | 0.113 | Old |
| 0.010956 | 0.138229 | 0.19 | 0.087 | 1 | 11 | Snapc1 | 0.103 | Old |
| 0.010983 | 0.516551 | 0.238 | 0.123 | 1 | 11 | Eif2s1 | 0.115 | Old |
| 0.010987 | 0.361513 | 0.242 | 0.123 | 1 | 11 | Fam104a | 0.119 | Old |
| 0.011047 | 0.358607 | 0.29 | 0.159 | 1 | 11 | Trim28 | 0.131 | Old |
| 0.011072 | 0.098336 | 0.319 | 0.174 | 1 | 11 | Gab2 | 0.145 | Old |
| 0.011078 | 0.702895 | 0.21 | 0.101 | 1 | 11 | 1810022K0 | 0.109 | Old |
| 0.011091 | 1.488712 | 0.101 | 0.029 | 1 | 11 | Sh2b1 | 0.072 | Old |
| 0.011097 | 0.542246 | 0.298 | 0.167 | 1 | 11 | Taf6l | 0.131 | Old |
| 0.011103 | 0.526589 | 0.375 | 0.232 | 1 | 11 | Lpp | 0.143 | Old |
| 0.011114 | 1.150519 | 0.077 | 0.014 | 1 | 11 | Recql5 | 0.063 | Old |
| 0.011125 | 0.089335 | 0.335 | 0.181 | 1 | 11 | Mcfd2 | 0.154 | Old |
| 0.011137 | 1.067295 | 0.145 | 0.058 | 1 | 11 | Agpat1 | 0.087 | Old |
| 0.011147 | 1.415232 | 0.089 | 0.022 | 1 | 11 | Usp31 | 0.067 | Old |
| 0.011157 | 1.108964 | 0.101 | 0.029 | 1 | 11 | Tfec | 0.072 | Old |
| 0.011169 | 1.17214 | 0.133 | 0.051 | 1 | 11 | Hps1 | 0.082 | Old |
| 0.011175 | 0.958125 | 0.137 | 0.051 | 1 | 11 | Arl6ip4 | 0.086 | Old |
| 0.011181 | 0.155477 | 0.315 | 0.174 | 1 | 11 | Timm10b | 0.141 | Old |
| 0.011217 | 0.474342 | 0.266 | 0.145 | 1 | 11 | Smg7 | 0.121 | Old |
| 0.011219 | 1.250718 | 0.089 | 0.022 | 1 | 11 | Dph6 | 0.067 | Old |
| 0.01124 | 0.488143 | 0.169 | 0.072 | 1 | 11 | Necap1 | 0.097 | Old |
| 0.011274 | 0.216276 | 0.484 | 0.304 | 1 | 11 | Znrf2 | 0.18 | Old |
| 0.01129 | 1.586648 | 0.089 | 0.022 | 1 | 11 | Cdc23 | 0.067 | Old |
| 0.011296 | 0.510027 | 0.629 | 0.471 | 1 | 11 | Ccr5 | 0.158 | Old |
| 0.01131 | 1.18397 | 0.113 | 0.036 | 1 | 11 | Tirap | 0.077 | Old |
| 0.011335 | 0.47149 | 0.149 | 0.058 | 1 | 11 | Elof1 | 0.091 | Old |
| 0.011358 | 0.757186 | 0.101 | 0.029 | 1 | 11 | Mfsd6 | 0.072 | Old |
| 0.011401 | 3.006651 | 0.06 | 0.007 | 1 | 11 | Il27ra | 0.053 | Old |
| 0.011401 | 2.552629 | 0.06 | 0.007 | 1 | 11 | Usp35 | 0.053 | Old |
| 0.011407 | 0.426532 | 0.488 | 0.304 | 1 | 11 | Tcea1 | 0.184 | Old |
| 0.011418 | 0.422311 | 0.298 | 0.167 | 1 | 11 | Oat | 0.131 | Old |
| 0.011422 | 0.258714 | 0.25 | 0.13 | 1 | 11 | Acaa1a | 0.12 | Old |
| 0.011425 | 0.725285 | 0.077 | 0.014 | 1 | 11 | Dnajc12 | 0.063 | Old |
| 0.011435 | 1.195898 | 0.089 | 0.022 | 1 | 11 | Acod1 | 0.067 | Old |
| 0.011437 | 0.371024 | 0.113 | 0.036 | 1 | 11 | Ccnc | 0.077 | Old |
| 0.011448 | 0.132336 | 0.202 | 0.094 | 1 | 11 | Alas1 | 0.108 | Old |
| 0.011468 | 0.171381 | 0.218 | 0.101 | 1 | 11 | Prpf38a | 0.117 | Old |
| 0.011492 | 0.268341 | 0.976 | 0.891 | 1 | 11 | Ctss | 0.085 | Old |
| 0.011544 | 0.185653 | 0.407 | 0.246 | 1 | 11 | Baz1b | 0.161 | Old |

| | | | | | | | | |
|---|---|---|---|---|---|---|---|---|
| 0.011547 | 0.405492 | 0.548 | 0.355 | 1 | 11 | Ier3ip1 | 0.193 | Old |
| 0.011579 | 0.772966 | 0.137 | 0.051 | 1 | 11 | Lmf2 | 0.086 | Old |
| 0.011581 | 1.136529 | 0.089 | 0.022 | 1 | 11 | Prmt5 | 0.067 | Old |
| 0.011601 | 0.876978 | 0.165 | 0.072 | 1 | 11 | Ssbp1 | 0.093 | Old |
| 0.011631 | 1.288976 | 0.101 | 0.029 | 1 | 11 | AL590144.: | 0.072 | Old |
| 0.011655 | 0.646862 | 0.089 | 0.022 | 1 | 11 | Gnpat | 0.067 | Old |
| 0.011667 | 0.679641 | 0.177 | 0.08 | 1 | 11 | Ilkap | 0.097 | Old |
| 0.011671 | 0.356083 | 0.516 | 0.341 | 1 | 11 | Lmna | 0.175 | Old |
| 0.011719 | 0.926708 | 0.125 | 0.043 | 1 | 11 | Ascc1 | 0.082 | Old |
| 0.011728 | 0.433178 | 0.258 | 0.138 | 1 | 11 | Coa3 | 0.12 | Old |
| 0.011729 | 1.189461 | 0.089 | 0.022 | 1 | 11 | Zfp608 | 0.067 | Old |
| 0.01174 | 0.442279 | 0.262 | 0.138 | 1 | 11 | Hmbox1 | 0.124 | Old |
| 0.011742 | 0.271616 | 0.315 | 0.174 | 1 | 11 | Lman1 | 0.141 | Old |
| 0.01175 | 0.442768 | 0.286 | 0.159 | 1 | 11 | Setd5 | 0.127 | Old |
| 0.011757 | 0.387313 | 0.254 | 0.13 | 1 | 11 | Cyc1 | 0.124 | Old |
| 0.011764 | 2.920109 | 0.06 | 0.007 | 1 | 11 | Lclat1 | 0.053 | Old |
| 0.011771 | 0.435819 | 0.367 | 0.217 | 1 | 11 | Snrpd2 | 0.15 | Old |
| 0.011803 | 0.414155 | 0.407 | 0.246 | 1 | 11 | Ubash3b | 0.161 | Old |
| 0.011839 | 1.419252 | 0.101 | 0.029 | 1 | 11 | Bysl | 0.072 | Old |
| 0.011876 | 0.385986 | 0.137 | 0.051 | 1 | 11 | Tceal8 | 0.086 | Old |
| 0.011879 | 1.398643 | 0.089 | 0.022 | 1 | 11 | Fdx1 | 0.067 | Old |
| 0.011931 | 0.47303 | 0.536 | 0.348 | 1 | 11 | Prrc2a | 0.188 | Old |
| 0.011935 | 0.509003 | 0.254 | 0.138 | 1 | 11 | Mrpl57 | 0.116 | Old |
| 0.01197 | 0.379041 | 0.27 | 0.145 | 1 | 11 | Dhrs7 | 0.125 | Old |
| 0.01199 | 0.499744 | 0.202 | 0.094 | 1 | 11 | Cdc42bpb | 0.108 | Old |
| 0.01201 | 0.144053 | 0.44 | 0.268 | 1 | 11 | Atp5d | 0.172 | Old |
| 0.012023 | 0.440903 | 0.306 | 0.181 | 1 | 11 | Samm50 | 0.125 | Old |
| 0.012111 | 0.480101 | 0.351 | 0.203 | 1 | 11 | Szrd1 | 0.148 | Old |
| 0.012124 | 0.177892 | 0.379 | 0.225 | 1 | 11 | Snu13 | 0.154 | Old |
| 0.012128 | 0.354749 | 0.298 | 0.167 | 1 | 11 | Rnf145 | 0.131 | Old |
| 0.012159 | 0.374456 | 0.351 | 0.21 | 1 | 11 | Mgea5 | 0.141 | Old |
| 0.012164 | 0.520839 | 0.46 | 0.304 | 1 | 11 | Myl12a | 0.156 | Old |
| 0.012191 | 1.603192 | 0.121 | 0.043 | 1 | 11 | Lonp2 | 0.078 | Old |
| 0.012212 | 0.087124 | 0.343 | 0.196 | 1 | 11 | Got1 | 0.147 | Old |
| 0.012232 | 0.221781 | 0.956 | 0.957 | 1 | 11 | H2-D1 | -0.001 | Old |
| 0.012232 | 0.40528 | 0.339 | 0.203 | 1 | 11 | Tbc1d14 | 0.136 | Old |
| 0.012233 | 2.518061 | 0.06 | 0.007 | 1 | 11 | Mgmt | 0.053 | Old |
| 0.012242 | 0.803446 | 0.206 | 0.101 | 1 | 11 | Guca1a | 0.105 | Old |
| 0.012251 | 4.104157 | 0.044 | 0 | 1 | 11 | Gm15056 | 0.044 | Old |
| 0.012284 | 0.078855 | 0.145 | 0.058 | 1 | 11 | Swt1 | 0.087 | Old |
| 0.012304 | 1.430976 | 0.141 | 0.058 | 1 | 11 | Nup205 | 0.083 | Old |
| 0.012309 | 1.060137 | 0.133 | 0.051 | 1 | 11 | Pnpla2 | 0.082 | Old |
| 0.012322 | 0.232637 | 0.383 | 0.217 | 1 | 11 | Arhgap9 | 0.166 | Old |
| 0.012419 | 0.122171 | 0.19 | 0.087 | 1 | 11 | Rabggtb | 0.103 | Old |
| 0.012425 | 2.212807 | 0.06 | 0.007 | 1 | 11 | Osbpl1a | 0.053 | Old |

| | | | | | | | | |
|---|---|---|---|---|---|---|---|---|
| 0.012484 | 1.896758 | 0.109 | 0.036 | 1 | 11 | Dhx29 | 0.073 | Old |
| 0.012493 | 0.148166 | 0.629 | 0.399 | 1 | 11 | Psmb10 | 0.23 | Old |
| 0.01254 | 0.581632 | 0.165 | 0.072 | 1 | 11 | Dctn5 | 0.093 | Old |
| 0.012568 | 0.228928 | 0.319 | 0.181 | 1 | 11 | Psmb2 | 0.138 | Old |
| 0.012574 | 0.195477 | 0.359 | 0.203 | 1 | 11 | Cbfb | 0.156 | Old |
| 0.012626 | 0.462162 | 0.274 | 0.152 | 1 | 11 | Puf60 | 0.122 | Old |
| 0.012633 | 0.257347 | 0.282 | 0.152 | 1 | 11 | Rnf44 | 0.13 | Old |
| 0.012647 | 0.766316 | 0.185 | 0.087 | 1 | 11 | Ctso | 0.098 | Old |
| 0.012655 | 0.892958 | 0.165 | 0.072 | 1 | 11 | Cwc25 | 0.093 | Old |
| 0.012659 | 0.102931 | 0.202 | 0.094 | 1 | 11 | Commd4 | 0.108 | Old |
| 0.012674 | 0.191137 | 0.335 | 0.196 | 1 | 11 | Tomm5 | 0.139 | Old |
| 0.012679 | 0.617474 | 0.137 | 0.051 | 1 | 11 | Exosc9 | 0.086 | Old |
| 0.012726 | 0.397142 | 0.375 | 0.232 | 1 | 11 | Cpt1a | 0.143 | Old |
| 0.012767 | 0.465114 | 0.702 | 0.522 | 1 | 11 | Gnas | 0.18 | Old |
| 0.012772 | 0.237847 | 0.46 | 0.283 | 1 | 11 | Cox7b | 0.177 | Old |
| 0.012782 | 0.033793 | 0.113 | 0.036 | 1 | 11 | Aph1b | 0.077 | Old |
| 0.012786 | 0.424962 | 0.294 | 0.167 | 1 | 11 | Naa38 | 0.127 | Old |
| 0.012788 | 0.345376 | 0.181 | 0.08 | 1 | 11 | Meaf6 | 0.101 | Old |
| 0.012803 | 0.478724 | 0.625 | 0.464 | 1 | 11 | Tkt | 0.161 | Old |
| 0.012817 | 1.61337 | 0.06 | 0.007 | 1 | 11 | Sacs | 0.053 | Old |
| 0.012817 | 1.487271 | 0.06 | 0.007 | 1 | 11 | Cd3eap | 0.053 | Old |
| 0.012817 | 1.368845 | 0.06 | 0.007 | 1 | 11 | Acaca | 0.053 | Old |
| 0.012817 | 1.216845 | 0.06 | 0.007 | 1 | 11 | Zfp68 | 0.053 | Old |
| 0.012817 | 0.172267 | 0.06 | 0.007 | 1 | 11 | Trim2 | 0.053 | Old |
| 0.012833 | 0.622852 | 0.145 | 0.058 | 1 | 11 | Dhx16 | 0.087 | Old |
| 0.01285 | 0.372995 | 0.452 | 0.297 | 1 | 11 | Rac2 | 0.155 | Old |
| 0.012923 | 0.900524 | 0.113 | 0.036 | 1 | 11 | Nectin2 | 0.077 | Old |
| 0.012983 | 1.685626 | 0.109 | 0.036 | 1 | 11 | Urgcp | 0.073 | Old |
| 0.01302 | 0.273898 | 0.464 | 0.297 | 1 | 11 | H2-M3 | 0.167 | Old |
| 0.013028 | 0.486164 | 0.347 | 0.21 | 1 | 11 | Ube2f | 0.137 | Old |
| 0.013063 | 1.341608 | 0.121 | 0.043 | 1 | 11 | Wipi2 | 0.078 | Old |
| 0.013068 | 3.430097 | 0.073 | 0.014 | 1 | 11 | Otud7b | 0.059 | Old |
| 0.013102 | 0.238689 | 0.754 | 0.558 | 1 | 11 | Itm2c | 0.196 | Old |
| 0.013112 | 0.135251 | 0.488 | 0.304 | 1 | 11 | Hspd1 | 0.184 | Old |
| 0.013177 | 0.53923 | 0.218 | 0.109 | 1 | 11 | Acbd6 | 0.109 | Old |
| 0.013215 | 0.511162 | 0.173 | 0.08 | 1 | 11 | Il18 | 0.093 | Old |
| 0.013231 | 0.463512 | 0.157 | 0.065 | 1 | 11 | Polg | 0.092 | Old |
| 0.0133 | 0.22325 | 0.315 | 0.174 | 1 | 11 | Akirin1 | 0.141 | Old |
| 0.013326 | 0.31311 | 0.339 | 0.196 | 1 | 11 | Sec31a | 0.143 | Old |
| 0.013332 | 0.91528 | 0.173 | 0.08 | 1 | 11 | Dpagt1 | 0.093 | Old |
| 0.013423 | 0.361439 | 0.222 | 0.109 | 1 | 11 | Rnaseh2c | 0.113 | Old |
| 0.01348 | 0.388055 | 0.157 | 0.065 | 1 | 11 | Cdk5rap3 | 0.092 | Old |
| 0.013485 | 0.269454 | 0.552 | 0.377 | 1 | 11 | Gusb | 0.175 | Old |
| 0.01352 | 0.268768 | 0.319 | 0.188 | 1 | 11 | Parvb | 0.131 | Old |
| 0.01357 | 0.777016 | 0.173 | 0.08 | 1 | 11 | Trpm2 | 0.093 | Old |

| | | | | | | | | |
|---|---|---|---|---|---|---|---|---|
| 0.013618 | 0.244077 | 0.476 | 0.29 | 1 | 11 | Cct4 | 0.186 | Old |
| 0.013622 | 0.336218 | 0.444 | 0.29 | 1 | 11 | Eif1a | 0.154 | Old |
| 0.013623 | 1.701809 | 0.097 | 0.029 | 1 | 11 | Tbce | 0.068 | Old |
| 0.013635 | 0.299232 | 0.347 | 0.203 | 1 | 11 | Ndufs7 | 0.144 | Old |
| 0.013647 | 0.603745 | 0.31 | 0.181 | 1 | 11 | Hip1 | 0.129 | Old |
| 0.013664 | 0.876297 | 0.153 | 0.065 | 1 | 11 | Casd1 | 0.088 | Old |
| 0.013664 | 1.170269 | 0.153 | 0.065 | 1 | 11 | Ech1 | 0.088 | Old |
| 0.013669 | 0.743688 | 0.165 | 0.072 | 1 | 11 | Nrbf2 | 0.093 | Old |
| 0.013687 | 1.021606 | 0.133 | 0.051 | 1 | 11 | Hdac4 | 0.082 | Old |
| 0.013698 | 0.221131 | 0.306 | 0.167 | 1 | 11 | Ehmt2 | 0.139 | Old |
| 0.013795 | 0.300036 | 0.419 | 0.254 | 1 | 11 | Fam102b | 0.165 | Old |
| 0.013828 | 0.044807 | 0.351 | 0.21 | 1 | 11 | Tns3 | 0.141 | Old |
| 0.013833 | 0.032843 | 0.383 | 0.225 | 1 | 11 | Msl2 | 0.158 | Old |
| 0.01384 | 0.288932 | 0.185 | 0.087 | 1 | 11 | Timm17a | 0.098 | Old |
| 0.013844 | 0.347385 | 0.121 | 0.043 | 1 | 11 | Prpf4 | 0.078 | Old |
| 0.013852 | 0.170598 | 0.343 | 0.203 | 1 | 11 | Fnip1 | 0.14 | Old |
| 0.013872 | 0.245898 | 0.504 | 0.341 | 1 | 11 | Arf3 | 0.163 | Old |
| 0.013899 | 0.640414 | 0.185 | 0.087 | 1 | 11 | Lrrc47 | 0.098 | Old |
| 0.013905 | 0.433836 | 0.27 | 0.145 | 1 | 11 | Pitpnc1 | 0.125 | Old |
| 0.013915 | 0.656786 | 0.234 | 0.123 | 1 | 11 | Raf1 | 0.111 | Old |
| 0.013935 | 0.002271 | 0.298 | 0.159 | 1 | 11 | Trappc2l | 0.139 | Old |
| 0.013993 | 0.527491 | 0.19 | 0.087 | 1 | 11 | Ola1 | 0.103 | Old |
| 0.014016 | 0.191447 | 0.387 | 0.239 | 1 | 11 | Cpne3 | 0.148 | Old |
| 0.014042 | 1.97255 | 0.085 | 0.022 | 1 | 11 | Srprb | 0.063 | Old |
| 0.014046 | 0.182478 | 0.407 | 0.261 | 1 | 11 | Rps6ka3 | 0.146 | Old |
| 0.01405 | 0.31565 | 0.359 | 0.217 | 1 | 11 | Ppp4r2 | 0.142 | Old |
| 0.014068 | 0.231616 | 0.476 | 0.297 | 1 | 11 | Psenen | 0.179 | Old |
| 0.014112 | 0.513563 | 0.19 | 0.087 | 1 | 11 | Lactb | 0.103 | Old |
| 0.014121 | 0.578562 | 0.31 | 0.181 | 1 | 11 | Spag7 | 0.129 | Old |
| 0.014131 | 0.412223 | 0.234 | 0.123 | 1 | 11 | Gpat4 | 0.111 | Old |
| 0.014132 | 0.328921 | 0.367 | 0.225 | 1 | 11 | Tsc22d2 | 0.142 | Old |
| 0.014155 | 0.380038 | 0.294 | 0.167 | 1 | 11 | Acbd5 | 0.127 | Old |
| 0.014177 | 0.31093 | 0.238 | 0.123 | 1 | 11 | Atraid | 0.115 | Old |
| 0.014242 | 0.391868 | 0.173 | 0.08 | 1 | 11 | Virma | 0.093 | Old |
| 0.01427 | 0.6506 | 0.194 | 0.094 | 1 | 11 | Mrpl11 | 0.1 | Old |
| 0.014301 | 0.247549 | 0.266 | 0.145 | 1 | 11 | Gtf3c2 | 0.121 | Old |
| 0.01431 | 0.178976 | 0.706 | 0.58 | 1 | 11 | Ubb | 0.126 | Old |
| 0.014315 | 0.059015 | 0.387 | 0.232 | 1 | 11 | Cenpb | 0.155 | Old |
| 0.014315 | 0.368362 | 0.157 | 0.065 | 1 | 11 | Pom121 | 0.092 | Old |
| 0.014381 | 0.392501 | 0.157 | 0.065 | 1 | 11 | Pias2 | 0.092 | Old |
| 0.014402 | 1.964619 | 0.085 | 0.022 | 1 | 11 | Cep104 | 0.063 | Old |
| 0.014411 | 0.274473 | 0.673 | 0.493 | 1 | 11 | Glul | 0.18 | Old |
| 0.014424 | 0.433058 | 0.355 | 0.217 | 1 | 11 | Myo1g | 0.138 | Old |
| 0.014431 | 0.69784 | 0.173 | 0.08 | 1 | 11 | Gatd1 | 0.093 | Old |
| 0.014459 | 0.60179 | 0.214 | 0.109 | 1 | 11 | Rpe | 0.105 | Old |

| | | | | | | | | |
|---|---|---|---|---|---|---|---|---|
| 0.014464 | 0.453063 | 0.298 | 0.174 | 1 | 11 | Atxn2 | 0.124 | Old |
| 0.014493 | 2.065407 | 0.085 | 0.022 | 1 | 11 | Ly6i | 0.063 | Old |
| 0.014496 | 1.417439 | 0.073 | 0.014 | 1 | 11 | Pam | 0.059 | Old |
| 0.014545 | 0.612838 | 0.145 | 0.058 | 1 | 11 | Apex1 | 0.087 | Old |
| 0.014566 | 0.179609 | 0.21 | 0.101 | 1 | 11 | Cstf2t | 0.109 | Old |
| 0.014584 | 1.842001 | 0.085 | 0.022 | 1 | 11 | Ctnnbl1 | 0.063 | Old |
| 0.014602 | 0.358967 | 0.375 | 0.232 | 1 | 11 | Nucb1 | 0.143 | Old |
| 0.014633 | 1.047586 | 0.161 | 0.072 | 1 | 11 | Akr1c13 | 0.089 | Old |
| 0.01466 | 0.562439 | 0.242 | 0.13 | 1 | 11 | Eapp | 0.112 | Old |
| 0.014665 | 1.090869 | 0.141 | 0.058 | 1 | 11 | Tnfrsf12a | 0.083 | Old |
| 0.014666 | 0.372926 | 0.419 | 0.268 | 1 | 11 | Hectd1 | 0.151 | Old |
| 0.014676 | 1.895212 | 0.085 | 0.022 | 1 | 11 | Ppan | 0.063 | Old |
| 0.014676 | 1.721752 | 0.085 | 0.022 | 1 | 11 | Mrps10 | 0.063 | Old |
| 0.014769 | 1.731415 | 0.085 | 0.022 | 1 | 11 | Caap1 | 0.063 | Old |
| 0.01479 | 0.888188 | 0.153 | 0.065 | 1 | 11 | Dtymk | 0.088 | Old |
| 0.014799 | 0.116316 | 0.472 | 0.297 | 1 | 11 | Myo1f | 0.175 | Old |
| 0.014807 | 0.568831 | 0.214 | 0.109 | 1 | 11 | Upf1 | 0.105 | Old |
| 0.014837 | 0.322526 | 0.306 | 0.181 | 1 | 11 | Ganab | 0.125 | Old |
| 0.014859 | 0.191704 | 0.153 | 0.065 | 1 | 11 | Ppfibp2 | 0.088 | Old |
| 0.014869 | 0.196082 | 0.456 | 0.297 | 1 | 11 | Ubald2 | 0.159 | Old |
| 0.014882 | 0.264394 | 0.21 | 0.109 | 1 | 11 | Ppil4 | 0.101 | Old |
| 0.014897 | 0.703379 | 0.315 | 0.188 | 1 | 11 | Mon2 | 0.127 | Old |
| 0.014899 | 0.743812 | 0.073 | 0.014 | 1 | 11 | Xrcc4 | 0.059 | Old |
| 0.014927 | 0.419289 | 0.375 | 0.239 | 1 | 11 | Pdlim4 | 0.136 | Old |
| 0.014956 | 1.474456 | 0.085 | 0.022 | 1 | 11 | Zfp266 | 0.063 | Old |
| 0.014963 | 0.482432 | 0.262 | 0.152 | 1 | 11 | Stag1 | 0.11 | Old |
| 0.014983 | 0.335258 | 0.177 | 0.08 | 1 | 11 | Fundc1 | 0.097 | Old |
| 0.014984 | 0.38808 | 0.214 | 0.109 | 1 | 11 | Dicer1 | 0.105 | Old |
| 0.015046 | 0.414945 | 0.238 | 0.13 | 1 | 11 | Csgalnact2 | 0.108 | Old |
| 0.015103 | 0.408611 | 0.214 | 0.109 | 1 | 11 | Pik3ca | 0.105 | Old |
| 0.015131 | 0.942715 | 0.185 | 0.087 | 1 | 11 | Rnf146 | 0.098 | Old |
| 0.01515 | 0.376664 | 0.44 | 0.261 | 1 | 11 | Mdh1 | 0.179 | Old |
| 0.015167 | 0.974804 | 0.141 | 0.058 | 1 | 11 | Vopp1 | 0.083 | Old |
| 0.015241 | 0.539268 | 0.448 | 0.297 | 1 | 11 | Mcub | 0.151 | Old |
| 0.015247 | 0.336455 | 0.323 | 0.188 | 1 | 11 | Copb2 | 0.135 | Old |
| 0.015281 | 0.285482 | 0.286 | 0.159 | 1 | 11 | Slc23a2 | 0.127 | Old |
| 0.01529 | 0.340443 | 0.508 | 0.348 | 1 | 11 | Tram1 | 0.16 | Old |
| 0.015321 | 0.514998 | 0.097 | 0.029 | 1 | 11 | Zfp422 | 0.068 | Old |
| 0.015329 | 1.042595 | 0.109 | 0.036 | 1 | 11 | Ticam1 | 0.073 | Old |
| 0.015334 | 0.357082 | 0.242 | 0.13 | 1 | 11 | Fbxo33 | 0.112 | Old |
| 0.015358 | 0.596162 | 0.27 | 0.152 | 1 | 11 | Gch1 | 0.118 | Old |
| 0.015365 | 0.248685 | 0.484 | 0.333 | 1 | 11 | Ralbp1 | 0.151 | Old |
| 0.015375 | 0.144477 | 0.423 | 0.261 | 1 | 11 | Ugcg | 0.162 | Old |
| 0.015387 | 0.799979 | 0.141 | 0.058 | 1 | 11 | Gpr141 | 0.083 | Old |
| 0.015401 | 0.067637 | 0.145 | 0.058 | 1 | 11 | Cd2bp2 | 0.087 | Old |

| | | | | | | | | |
|---|---|---|---|---|---|---|---|---|
| 0.015405 | 0.409272 | 0.198 | 0.094 | 1 | 11 | Sf3a1 | 0.104 | Old |
| 0.015438 | 0.228911 | 0.294 | 0.167 | 1 | 11 | Slc31a1 | 0.127 | Old |
| 0.015582 | 0.408303 | 0.262 | 0.145 | 1 | 11 | Odc1 | 0.117 | Old |
| 0.015591 | 1.270312 | 0.097 | 0.029 | 1 | 11 | Asb3 | 0.068 | Old |
| 0.015591 | 1.027803 | 0.097 | 0.029 | 1 | 11 | Pole3 | 0.068 | Old |
| 0.015613 | 0.116854 | 0.407 | 0.239 | 1 | 11 | Impact | 0.168 | Old |
| 0.015627 | 1.238209 | 0.085 | 0.022 | 1 | 11 | Dvl3 | 0.063 | Old |
| 0.015631 | 0.791947 | 0.194 | 0.094 | 1 | 11 | Flvcr1 | 0.1 | Old |
| 0.015635 | 0.35701 | 0.673 | 0.464 | 1 | 11 | Pak2 | 0.209 | Old |
| 0.015671 | 0.478658 | 0.21 | 0.109 | 1 | 11 | Mindy3 | 0.101 | Old |
| 0.015705 | 0.645919 | 0.153 | 0.065 | 1 | 11 | Dnajc16 | 0.088 | Old |
| 0.015705 | 0.122552 | 0.153 | 0.065 | 1 | 11 | Tmem214 | 0.088 | Old |
| 0.015722 | 0.234867 | 0.379 | 0.232 | 1 | 11 | Ube2s | 0.147 | Old |
| 0.015733 | 1.462071 | 0.137 | 0.058 | 1 | 11 | Ptrhd1 | 0.079 | Old |
| 0.01574 | 0.185445 | 0.31 | 0.181 | 1 | 11 | Lima1 | 0.129 | Old |
| 0.015744 | 0.744371 | 0.173 | 0.08 | 1 | 11 | Fam219a | 0.093 | Old |
| 0.015814 | 0.504009 | 0.133 | 0.051 | 1 | 11 | Ikzf5 | 0.082 | Old |
| 0.015846 | 1.037165 | 0.129 | 0.051 | 1 | 11 | Mrpl10 | 0.078 | Old |
| 0.01589 | 0.660728 | 0.194 | 0.094 | 1 | 11 | Sf3b4 | 0.1 | Old |
| 0.015912 | 0.493566 | 0.141 | 0.058 | 1 | 11 | Pld1 | 0.083 | Old |
| 0.015922 | 0.707388 | 0.085 | 0.022 | 1 | 11 | Pphln1 | 0.063 | Old |
| 0.016007 | 0.579169 | 0.226 | 0.116 | 1 | 11 | Tmem248 | 0.11 | Old |
| 0.01602 | 0.338771 | 0.121 | 0.043 | 1 | 11 | Elmod2 | 0.078 | Old |
| 0.016065 | 0.693919 | 0.141 | 0.058 | 1 | 11 | Crls1 | 0.083 | Old |
| 0.016078 | 0.393607 | 0.351 | 0.21 | 1 | 11 | Mrps14 | 0.141 | Old |
| 0.016143 | 0.957483 | 0.097 | 0.029 | 1 | 11 | Papss2 | 0.068 | Old |
| 0.016174 | 1.722771 | 0.117 | 0.043 | 1 | 11 | Tmx2 | 0.074 | Old |
| 0.016234 | 2.662447 | 0.056 | 0.007 | 1 | 11 | Dgcr14 | 0.049 | Old |
| 0.016234 | 2.505493 | 0.056 | 0.007 | 1 | 11 | Tanc1 | 0.049 | Old |
| 0.016289 | 0.232436 | 0.21 | 0.101 | 1 | 11 | Mbtps1 | 0.109 | Old |
| 0.016306 | 0.617959 | 0.242 | 0.13 | 1 | 11 | Imp3 | 0.112 | Old |
| 0.016328 | 0.641697 | 0.274 | 0.159 | 1 | 11 | Mdm4 | 0.115 | Old |
| 0.016367 | 0.31543 | 0.31 | 0.174 | 1 | 11 | Arl5a | 0.136 | Old |
| 0.0164 | 0.693824 | 0.238 | 0.13 | 1 | 11 | Sun2 | 0.108 | Old |
| 0.016408 | 0.522698 | 0.129 | 0.051 | 1 | 11 | Samd1 | 0.078 | Old |
| 0.016425 | 0.89664 | 0.085 | 0.022 | 1 | 11 | Mag | 0.063 | Old |
| 0.016445 | 0.380141 | 0.198 | 0.094 | 1 | 11 | Pla2g4a | 0.104 | Old |
| 0.016448 | 0.326714 | 0.246 | 0.13 | 1 | 11 | Uba52 | 0.116 | Old |
| 0.016485 | 0.708912 | 0.262 | 0.152 | 1 | 11 | Trim26 | 0.11 | Old |
| 0.016489 | 2.576383 | 0.056 | 0.007 | 1 | 11 | Pinx1 | 0.049 | Old |
| 0.016496 | 0.574876 | 0.286 | 0.167 | 1 | 11 | Kxd1 | 0.119 | Old |
| 0.016521 | 1.059204 | 0.097 | 0.029 | 1 | 11 | Ehf | 0.068 | Old |
| 0.016524 | 0.464829 | 0.121 | 0.043 | 1 | 11 | Thoc3 | 0.078 | Old |
| 0.016527 | 0.762594 | 0.198 | 0.101 | 1 | 11 | Tgfbrap1 | 0.097 | Old |
| 0.0166 | 0.293711 | 0.411 | 0.254 | 1 | 11 | Cmtm3 | 0.157 | Old |

| | | | | | | | | |
|---|---|---|---|---|---|---|---|---|
| 0.01663 | 0.807556 | 0.085 | 0.022 | 1 | 11 | Gzf1 | 0.063 | Old |
| 0.016633 | 0.698292 | 0.266 | 0.145 | 1 | 11 | Map3k11 | 0.121 | Old |
| 0.016633 | 0.360548 | 0.266 | 0.145 | 1 | 11 | Rbbp7 | 0.121 | Old |
| 0.016655 | 0.351887 | 0.391 | 0.246 | 1 | 11 | Sbf2 | 0.145 | Old |
| 0.016669 | 0.318655 | 0.387 | 0.239 | 1 | 11 | Papd4 | 0.148 | Old |
| 0.016723 | 0.171031 | 0.222 | 0.116 | 1 | 11 | Wbp4 | 0.106 | Old |
| 0.016726 | 0.295959 | 0.536 | 0.377 | 1 | 11 | Pepd | 0.159 | Old |
| 0.016734 | 0.66737 | 0.085 | 0.022 | 1 | 11 | Rps6kb2 | 0.063 | Old |
| 0.016739 | 0.76077 | 0.161 | 0.072 | 1 | 11 | Shkbp1 | 0.089 | Old |
| 0.016743 | 0.095801 | 0.444 | 0.29 | 1 | 11 | Sec14l1 | 0.154 | Old |
| 0.016788 | 0.83371 | 0.226 | 0.123 | 1 | 11 | Bst1 | 0.103 | Old |
| 0.016824 | 0.194203 | 0.452 | 0.283 | 1 | 11 | Twf2 | 0.169 | Old |
| 0.016837 | 0.203402 | 0.621 | 0.435 | 1 | 11 | Prpf38b | 0.186 | Old |
| 0.016838 | 0.642514 | 0.085 | 0.022 | 1 | 11 | Cdkn2aip | 0.063 | Old |
| 0.016865 | 0.900126 | 0.19 | 0.094 | 1 | 11 | Sqor | 0.096 | Old |
| 0.016876 | 2.46708 | 0.056 | 0.007 | 1 | 11 | Abcb6 | 0.049 | Old |
| 0.016893 | 0.946154 | 0.185 | 0.094 | 1 | 11 | Ppie | 0.091 | Old |
| 0.0169 | 0.432554 | 0.153 | 0.065 | 1 | 11 | Wdr91 | 0.088 | Old |
| 0.016902 | 0.246884 | 0.585 | 0.406 | 1 | 11 | Gas7 | 0.179 | Old |
| 0.016909 | 0.13652 | 0.109 | 0.036 | 1 | 11 | Eci2 | 0.073 | Old |
| 0.016924 | 0.656208 | 0.262 | 0.145 | 1 | 11 | Fbxo18 | 0.117 | Old |
| 0.016942 | 0.387222 | 0.173 | 0.08 | 1 | 11 | Dscr3 | 0.093 | Old |
| 0.016958 | 1.497448 | 0.234 | 0.138 | 1 | 11 | Cd209g | 0.096 | Old |
| 0.016959 | 0.126472 | 0.399 | 0.254 | 1 | 11 | Tnpo1 | 0.145 | Old |
| 0.016961 | 0.503764 | 0.27 | 0.152 | 1 | 11 | Ankrd13c | 0.118 | Old |
| 0.017007 | 2.07442 | 0.056 | 0.007 | 1 | 11 | Fam234b | 0.049 | Old |
| 0.017018 | 1.878367 | 0.105 | 0.036 | 1 | 11 | Pgm2 | 0.069 | Old |
| 0.017018 | 1.524319 | 0.105 | 0.036 | 1 | 11 | Samd4 | 0.069 | Old |
| 0.017049 | 0.30186 | 0.448 | 0.29 | 1 | 11 | Fgfr1op2 | 0.158 | Old |
| 0.017056 | 0.274906 | 0.315 | 0.188 | 1 | 11 | Qdpr | 0.127 | Old |
| 0.01708 | 3.923292 | 0.04 | 0 | 1 | 11 | Cdh6 | 0.04 | Old |
| 0.01708 | 3.821103 | 0.04 | 0 | 1 | 11 | Eml6 | 0.04 | Old |
| 0.01708 | 3.739873 | 0.04 | 0 | 1 | 11 | Unc5a | 0.04 | Old |
| 0.01708 | 3.729823 | 0.04 | 0 | 1 | 11 | Arhgef5 | 0.04 | Old |
| 0.01708 | 3.462789 | 0.04 | 0 | 1 | 11 | H2-Eb2 | 0.04 | Old |
| 0.01708 | 3.352622 | 0.04 | 0 | 1 | 11 | Dis3 | 0.04 | Old |
| 0.01708 | 3.239207 | 0.04 | 0 | 1 | 11 | Kctd9 | 0.04 | Old |
| 0.01708 | 2.85962 | 0.04 | 0 | 1 | 11 | Echdc1 | 0.04 | Old |
| 0.01708 | 2.715253 | 0.04 | 0 | 1 | 11 | Wrap53 | 0.04 | Old |
| 0.01708 | 2.681827 | 0.04 | 0 | 1 | 11 | Acsf3 | 0.04 | Old |
| 0.017128 | 0.29396 | 0.456 | 0.283 | 1 | 11 | Nup98 | 0.173 | Old |
| 0.017128 | 1.280649 | 0.117 | 0.043 | 1 | 11 | Slc25a19 | 0.074 | Old |
| 0.01714 | 0.565675 | 0.29 | 0.174 | 1 | 11 | Cuedc2 | 0.116 | Old |
| 0.017158 | 0.708447 | 0.21 | 0.109 | 1 | 11 | Fkbp3 | 0.101 | Old |
| 0.017241 | 0.471386 | 0.355 | 0.225 | 1 | 11 | Samd4b | 0.13 | Old |

| | | | | | | | | |
|---|---|---|---|---|---|---|---|---|
| 0.017259 | 0.385608 | 0.085 | 0.022 | 1 | 11 | Parvg | 0.063 | Old |
| 0.017288 | 2.636554 | 0.069 | 0.014 | 1 | 11 | Hook2 | 0.055 | Old |
| 0.0173 | 0.642479 | 0.097 | 0.029 | 1 | 11 | Ppa2 | 0.068 | Old |
| 0.017306 | 0.384303 | 0.121 | 0.043 | 1 | 11 | Hax1 | 0.078 | Old |
| 0.017307 | 0.292836 | 0.165 | 0.072 | 1 | 11 | Erh | 0.093 | Old |
| 0.01731 | 0.528628 | 0.222 | 0.116 | 1 | 11 | Wdr41 | 0.106 | Old |
| 0.017371 | 1.274772 | 0.165 | 0.08 | 1 | 11 | Slc25a24 | 0.085 | Old |
| 0.017394 | 0.178307 | 0.121 | 0.043 | 1 | 11 | B9d2 | 0.078 | Old |
| 0.017396 | 1.48564 | 0.105 | 0.036 | 1 | 11 | Mrps5 | 0.069 | Old |
| 0.017406 | 1.397679 | 0.056 | 0.007 | 1 | 11 | Tmed1 | 0.049 | Old |
| 0.017464 | 0.069356 | 0.411 | 0.246 | 1 | 11 | Kpna4 | 0.165 | Old |
| 0.01749 | 0.174489 | 0.435 | 0.268 | 1 | 11 | Cpsf6 | 0.167 | Old |
| 0.017533 | 0.293696 | 0.173 | 0.08 | 1 | 11 | Agtpbp1 | 0.093 | Old |
| 0.017534 | 0.006085 | 0.444 | 0.275 | 1 | 11 | Syncrip | 0.169 | Old |
| 0.017541 | 1.3291 | 0.056 | 0.007 | 1 | 11 | Arrdc2 | 0.049 | Old |
| 0.017543 | 0.350101 | 0.222 | 0.116 | 1 | 11 | Cln3 | 0.106 | Old |
| 0.017577 | 0.261058 | 0.323 | 0.188 | 1 | 11 | Pank3 | 0.135 | Old |
| 0.01765 | 0.328909 | 0.403 | 0.261 | 1 | 11 | Man1c1 | 0.142 | Old |
| 0.017656 | 0.29172 | 0.302 | 0.181 | 1 | 11 | Wwp2 | 0.121 | Old |
| 0.017676 | 1.183981 | 0.056 | 0.007 | 1 | 11 | Letm2 | 0.049 | Old |
| 0.017738 | 0.231868 | 0.556 | 0.37 | 1 | 11 | Fcho2 | 0.186 | Old |
| 0.017785 | 0.351584 | 0.27 | 0.152 | 1 | 11 | Psmg4 | 0.118 | Old |
| 0.017787 | 0.29452 | 0.512 | 0.348 | 1 | 11 | Ubap2l | 0.164 | Old |
| 0.017806 | 0.756799 | 0.218 | 0.116 | 1 | 11 | Uba7 | 0.102 | Old |
| 0.017814 | 0.270585 | 0.714 | 0.536 | 1 | 11 | Atox1 | 0.178 | Old |
| 0.01784 | 0.299799 | 0.29 | 0.167 | 1 | 11 | Ubr2 | 0.123 | Old |
| 0.017841 | 0.335567 | 0.21 | 0.109 | 1 | 11 | Zyg11b | 0.101 | Old |
| 0.017862 | 0.378375 | 0.234 | 0.123 | 1 | 11 | Plod1 | 0.111 | Old |
| 0.017903 | 0.6859 | 0.177 | 0.087 | 1 | 11 | Sf3a2 | 0.09 | Old |
| 0.017913 | 0.335174 | 0.44 | 0.275 | 1 | 11 | Ifnar1 | 0.165 | Old |
| 0.017914 | 0.33308 | 0.222 | 0.116 | 1 | 11 | Zc3h15 | 0.106 | Old |
| 0.017946 | 0.433408 | 0.117 | 0.043 | 1 | 11 | D10Wsu10 | 0.074 | Old |
| 0.018014 | 0.182434 | 0.323 | 0.181 | 1 | 11 | Smc6 | 0.142 | Old |
| 0.018038 | 1.014407 | 0.117 | 0.043 | 1 | 11 | Mre11a | 0.074 | Old |
| 0.01805 | 0.022788 | 0.427 | 0.275 | 1 | 11 | Trpc4ap | 0.152 | Old |
| 0.018068 | 0.300454 | 0.25 | 0.138 | 1 | 11 | Gpbp1l1 | 0.112 | Old |
| 0.018102 | 0.267155 | 0.52 | 0.348 | 1 | 11 | Bclaf1 | 0.172 | Old |
| 0.018142 | 0.144631 | 0.173 | 0.08 | 1 | 11 | Ctns | 0.093 | Old |
| 0.01819 | 0.146905 | 0.363 | 0.21 | 1 | 11 | Krcc1 | 0.153 | Old |
| 0.018226 | 0.208027 | 0.133 | 0.051 | 1 | 11 | Tmem39a | 0.082 | Old |
| 0.018266 | 0.751082 | 0.169 | 0.08 | 1 | 11 | Cpsf3 | 0.089 | Old |
| 0.018275 | 0.050198 | 0.415 | 0.268 | 1 | 11 | Tmem258 | 0.147 | Old |
| 0.018336 | 0.124424 | 0.395 | 0.246 | 1 | 11 | Lrp6 | 0.149 | Old |
| 0.018355 | 2.612645 | 0.081 | 0.022 | 1 | 11 | Fbxl4 | 0.059 | Old |
| 0.018374 | 0.122721 | 0.468 | 0.297 | 1 | 11 | Sptssa | 0.171 | Old |

| | | | | | | | | |
|---|---|---|---|---|---|---|---|---|
| 0.018381 | 0.479061 | 0.411 | 0.268 | 1 | 11 | Nol7 | 0.143 | Old |
| 0.018381 | 0.691664 | 0.129 | 0.051 | 1 | 11 | Fbxo34 | 0.078 | Old |
| 0.018381 | 0.541232 | 0.129 | 0.051 | 1 | 11 | Trappc13 | 0.078 | Old |
| 0.01839 | 0.53261 | 0.181 | 0.087 | 1 | 11 | Trnau1ap | 0.094 | Old |
| 0.018395 | 0.269055 | 0.391 | 0.246 | 1 | 11 | Zranb2 | 0.145 | Old |
| 0.018398 | 0.122346 | 0.456 | 0.29 | 1 | 11 | Metap2 | 0.166 | Old |
| 0.0184 | 0.935833 | 0.157 | 0.072 | 1 | 11 | Cask | 0.085 | Old |
| 0.018423 | 0.158595 | 0.254 | 0.138 | 1 | 11 | Rictor | 0.116 | Old |
| 0.018472 | 1.811862 | 0.105 | 0.036 | 1 | 11 | Agmo | 0.069 | Old |
| 0.018503 | 0.311484 | 0.238 | 0.123 | 1 | 11 | Nhp2 | 0.115 | Old |
| 0.018509 | 0.98127 | 0.117 | 0.043 | 1 | 11 | Mtf2 | 0.074 | Old |
| 0.018604 | 0.837162 | 0.117 | 0.043 | 1 | 11 | Ormdl1 | 0.074 | Old |
| 0.018615 | 0.181178 | 0.242 | 0.13 | 1 | 11 | Chtf8 | 0.112 | Old |
| 0.018631 | 0.124678 | 0.423 | 0.268 | 1 | 11 | Tmem219 | 0.155 | Old |
| 0.018649 | 0.237357 | 0.77 | 0.551 | 1 | 11 | Arl6ip1 | 0.219 | Old |
| 0.018657 | 0.741951 | 0.206 | 0.109 | 1 | 11 | Smurf1 | 0.097 | Old |
| 0.01867 | 0.287132 | 0.851 | 0.688 | 1 | 11 | Cd53 | 0.163 | Old |
| 0.018676 | 0.211502 | 0.278 | 0.159 | 1 | 11 | Psen1 | 0.119 | Old |
| 0.018683 | 0.266878 | 0.335 | 0.196 | 1 | 11 | Irf9 | 0.139 | Old |
| 0.018702 | 2.30649 | 0.081 | 0.022 | 1 | 11 | Chst2 | 0.059 | Old |
| 0.018726 | 0.237348 | 0.379 | 0.246 | 1 | 11 | Psmb3 | 0.133 | Old |
| 0.018851 | 0.347456 | 0.194 | 0.094 | 1 | 11 | Senp1 | 0.1 | Old |
| 0.018915 | 0.490588 | 0.23 | 0.123 | 1 | 11 | Dctn2 | 0.107 | Old |
| 0.018939 | 0.277271 | 0.673 | 0.5 | 1 | 11 | Zfand5 | 0.173 | Old |
| 0.018967 | 0.080995 | 0.419 | 0.268 | 1 | 11 | Atp13a3 | 0.151 | Old |
| 0.018998 | 0.42497 | 0.302 | 0.181 | 1 | 11 | Pdcd6 | 0.121 | Old |
| 0.019039 | 0.101144 | 0.315 | 0.181 | 1 | 11 | Dcaf8 | 0.134 | Old |
| 0.019083 | 0.119594 | 0.27 | 0.145 | 1 | 11 | Fto | 0.125 | Old |
| 0.019086 | 0.26616 | 0.548 | 0.37 | 1 | 11 | Man1a | 0.178 | Old |
| 0.019095 | 0.473336 | 0.234 | 0.123 | 1 | 11 | Ano6 | 0.111 | Old |
| 0.0191 | 0.078648 | 0.218 | 0.109 | 1 | 11 | Tlnrd1 | 0.109 | Old |
| 0.019101 | 0.348395 | 0.31 | 0.188 | 1 | 11 | Xpr1 | 0.122 | Old |
| 0.019105 | 0.305954 | 0.605 | 0.442 | 1 | 11 | Atp5j2 | 0.163 | Old |
| 0.019158 | 0.836312 | 0.069 | 0.014 | 1 | 11 | Slc22a15 | 0.055 | Old |
| 0.019174 | 0.315207 | 0.48 | 0.304 | 1 | 11 | Smarca5 | 0.176 | Old |
| 0.019309 | 0.518993 | 0.169 | 0.08 | 1 | 11 | Fxr2 | 0.089 | Old |
| 0.01931 | 0.363449 | 0.879 | 0.71 | 1 | 11 | Sat1 | 0.169 | Old |
| 0.019333 | 0.517495 | 0.601 | 0.399 | 1 | 11 | Set | 0.202 | Old |
| 0.01936 | 0.35139 | 0.125 | 0.051 | 1 | 11 | Frs2 | 0.074 | Old |
| 0.019402 | 0.987566 | 0.157 | 0.072 | 1 | 11 | Dnmt1 | 0.085 | Old |
| 0.01941 | 0.318202 | 0.407 | 0.261 | 1 | 11 | Rad23b | 0.146 | Old |
| 0.019451 | 1.219132 | 0.093 | 0.029 | 1 | 11 | Tmem71 | 0.064 | Old |
| 0.019528 | 0.522756 | 0.161 | 0.072 | 1 | 11 | Qars | 0.089 | Old |
| 0.019553 | 1.854111 | 0.069 | 0.014 | 1 | 11 | Gcnt2 | 0.055 | Old |
| 0.019563 | 1.125117 | 0.093 | 0.029 | 1 | 11 | Ppp1r16a | 0.064 | Old |

| | | | | | | | | |
|---|---|---|---|---|---|---|---|---|
| 0.019574 | 0.120016 | 1 | 1 | 1 | 11 | mt-Co1 | 0 | Old |
| 0.019618 | 0.483169 | 0.222 | 0.116 | 1 | 11 | Slc50a1 | 0.106 | Old |
| 0.019701 | 0.11933 | 0.25 | 0.13 | 1 | 11 | Ndufs8 | 0.12 | Old |
| 0.019768 | 0.323268 | 0.565 | 0.391 | 1 | 11 | Eif2s2 | 0.174 | Old |
| 0.019825 | 1.093873 | 0.194 | 0.101 | 1 | 11 | Xpo7 | 0.093 | Old |
| 0.019833 | 1.02936 | 0.157 | 0.072 | 1 | 11 | Irak3 | 0.085 | Old |
| 0.019877 | 0.608757 | 0.234 | 0.13 | 1 | 11 | Hk2 | 0.104 | Old |
| 0.019877 | 0.006192 | 0.669 | 0.478 | 1 | 11 | Arf1 | 0.191 | Old |
| 0.01989 | 0.287544 | 0.452 | 0.297 | 1 | 11 | Cd302 | 0.155 | Old |
| 0.019955 | 0.206747 | 0.565 | 0.384 | 1 | 11 | Rasa4 | 0.181 | Old |
| 0.019982 | 0.252468 | 0.242 | 0.13 | 1 | 11 | Golgb1 | 0.112 | Old |
| 0.019983 | 0.151275 | 0.286 | 0.159 | 1 | 11 | Bdp1 | 0.127 | Old |
| 0.020023 | 0.249943 | 0.351 | 0.217 | 1 | 11 | Arhgef10l | 0.134 | Old |
| 0.020044 | 0.198299 | 0.536 | 0.362 | 1 | 11 | Ndel1 | 0.174 | Old |
| 0.020053 | 0.146707 | 0.262 | 0.145 | 1 | 11 | Ankrd40 | 0.117 | Old |
| 0.02009 | 1.288804 | 0.069 | 0.014 | 1 | 11 | Elk1 | 0.055 | Old |
| 0.020125 | 0.413021 | 0.355 | 0.217 | 1 | 11 | U2surp | 0.138 | Old |
| 0.020143 | 0.747845 | 0.185 | 0.094 | 1 | 11 | Mdp1 | 0.091 | Old |
| 0.020161 | 0.339429 | 0.887 | 0.775 | 1 | 11 | Hspa5 | 0.112 | Old |
| 0.020163 | 0.237132 | 0.343 | 0.21 | 1 | 11 | Ube2e1 | 0.133 | Old |
| 0.020173 | 0.600098 | 0.198 | 0.101 | 1 | 11 | Tmem205 | 0.097 | Old |
| 0.020219 | 0.396652 | 0.23 | 0.123 | 1 | 11 | 2210016F1 | 0.107 | Old |
| 0.02022 | 0.200789 | 0.161 | 0.072 | 1 | 11 | Pccb | 0.089 | Old |
| 0.020225 | 0.15971 | 0.512 | 0.348 | 1 | 11 | Ywhag | 0.164 | Old |
| 0.020273 | 0.165818 | 0.242 | 0.13 | 1 | 11 | Ubqln1 | 0.112 | Old |
| 0.020294 | 0.64327 | 0.117 | 0.043 | 1 | 11 | Mpv17l2 | 0.074 | Old |
| 0.020294 | 0.621892 | 0.117 | 0.043 | 1 | 11 | Dmxl2 | 0.074 | Old |
| 0.020365 | 0.848265 | 0.093 | 0.029 | 1 | 11 | Cib2 | 0.064 | Old |
| 0.020385 | 0.31087 | 0.31 | 0.181 | 1 | 11 | Dnajb11 | 0.129 | Old |
| 0.020397 | 0.588804 | 0.117 | 0.043 | 1 | 11 | Igkc | 0.074 | Old |
| 0.020414 | 0.215464 | 0.911 | 0.833 | 1 | 11 | Ppia | 0.078 | Old |
| 0.020418 | 0.751727 | 0.306 | 0.188 | 1 | 11 | Becn1 | 0.118 | Old |
| 0.02042 | 0.0005 | 0.242 | 0.13 | 1 | 11 | Ssbp4 | 0.112 | Old |
| 0.020491 | 0.588607 | 0.169 | 0.08 | 1 | 11 | Acp5 | 0.089 | Old |
| 0.020529 | 0.184866 | 0.246 | 0.138 | 1 | 11 | Sys1 | 0.108 | Old |
| 0.020668 | 0.306118 | 0.492 | 0.319 | 1 | 11 | Leprot | 0.173 | Old |
| 0.020669 | 0.764439 | 0.278 | 0.174 | 1 | 11 | Tppp3 | 0.104 | Old |
| 0.02069 | 0.706956 | 0.105 | 0.036 | 1 | 11 | Mkrn2 | 0.069 | Old |
| 0.02069 | 0.304055 | 0.105 | 0.036 | 1 | 11 | Lemd3 | 0.069 | Old |
| 0.020718 | 0.775523 | 0.093 | 0.029 | 1 | 11 | Cfb | 0.064 | Old |
| 0.020753 | 0.138906 | 0.161 | 0.072 | 1 | 11 | Dnajc21 | 0.089 | Old |
| 0.020766 | 0.537553 | 0.234 | 0.13 | 1 | 11 | Slc45a4 | 0.104 | Old |
| 0.02078 | 1.086413 | 0.069 | 0.014 | 1 | 11 | Mllt11 | 0.055 | Old |
| 0.020801 | 0.845055 | 0.105 | 0.036 | 1 | 11 | Eif1ad | 0.069 | Old |
| 0.020801 | 0.650616 | 0.105 | 0.036 | 1 | 11 | Txnrd2 | 0.069 | Old |

| | | | | | | | | |
|---|---|---|---|---|---|---|---|---|
| 0.020836 | 1.001322 | 0.093 | 0.029 | 1 | 11 | Slc39a11 | 0.064 | Old |
| 0.020843 | 0.314578 | 0.161 | 0.072 | 1 | 11 | Tex264 | 0.089 | Old |
| 0.020884 | 0.740283 | 0.262 | 0.159 | 1 | 11 | Ddx50 | 0.103 | Old |
| 0.020889 | 0.796113 | 0.125 | 0.051 | 1 | 11 | Cnn3 | 0.074 | Old |
| 0.020907 | 0.083954 | 0.331 | 0.188 | 1 | 11 | Rere | 0.143 | Old |
| 0.020912 | 0.452556 | 0.105 | 0.036 | 1 | 11 | Mast2 | 0.069 | Old |
| 0.020917 | 0.296054 | 0.194 | 0.094 | 1 | 11 | Il11ra1 | 0.1 | Old |
| 0.02096 | 0.535436 | 0.218 | 0.116 | 1 | 11 | Sap18 | 0.102 | Old |
| 0.020972 | 0.427925 | 0.246 | 0.138 | 1 | 11 | Otud4 | 0.108 | Old |
| 0.020982 | 0.203751 | 0.367 | 0.217 | 1 | 11 | Iscu | 0.15 | Old |
| 0.020994 | 0.739869 | 0.157 | 0.072 | 1 | 11 | Bin3 | 0.085 | Old |
| 0.021061 | 0.958623 | 0.069 | 0.014 | 1 | 11 | Atr | 0.055 | Old |
| 0.021074 | 0.354321 | 0.327 | 0.196 | 1 | 11 | Tfrc | 0.131 | Old |
| 0.021086 | 0.917919 | 0.157 | 0.072 | 1 | 11 | Dpy19l4 | 0.085 | Old |
| 0.021115 | 0.229831 | 0.161 | 0.072 | 1 | 11 | Zfp318 | 0.089 | Old |
| 0.021166 | 1.193086 | 0.081 | 0.022 | 1 | 11 | Aar2 | 0.059 | Old |
| 0.021189 | 0.525691 | 0.274 | 0.167 | 1 | 11 | Ms4a8a | 0.107 | Old |
| 0.021241 | 0.249607 | 0.117 | 0.043 | 1 | 11 | Smim26 | 0.074 | Old |
| 0.021248 | 0.389713 | 0.302 | 0.181 | 1 | 11 | Sidt2 | 0.121 | Old |
| 0.021251 | 0.899507 | 0.105 | 0.036 | 1 | 11 | Snx19 | 0.069 | Old |
| 0.021251 | 0.727343 | 0.105 | 0.036 | 1 | 11 | Elp3 | 0.069 | Old |
| 0.021294 | 0.080704 | 0.194 | 0.094 | 1 | 11 | Nsrp1 | 0.1 | Old |
| 0.021296 | 1.279113 | 0.081 | 0.022 | 1 | 11 | Nat6 | 0.059 | Old |
| 0.021356 | 0.177724 | 0.859 | 0.819 | 1 | 11 | Cyba | 0.04 | Old |
| 0.021371 | 0.293126 | 0.302 | 0.174 | 1 | 11 | Rtf1 | 0.128 | Old |
| 0.02141 | 0.197939 | 0.355 | 0.217 | 1 | 11 | mt-Atp8 | 0.138 | Old |
| 0.021414 | 0.775683 | 0.137 | 0.058 | 1 | 11 | Arntl | 0.079 | Old |
| 0.021427 | 1.126268 | 0.081 | 0.022 | 1 | 11 | Smim8 | 0.059 | Old |
| 0.021438 | 0.253318 | 0.093 | 0.029 | 1 | 11 | Ms4a14 | 0.064 | Old |
| 0.021454 | 0.927547 | 0.145 | 0.065 | 1 | 11 | Pex19 | 0.08 | Old |
| 0.021464 | 0.357459 | 0.294 | 0.174 | 1 | 11 | Akirin2 | 0.12 | Old |
| 0.021479 | 0.90539 | 0.105 | 0.036 | 1 | 11 | Golph3l | 0.069 | Old |
| 0.021558 | 0.385282 | 0.27 | 0.159 | 1 | 11 | Irs2 | 0.111 | Old |
| 0.021561 | 1.032466 | 0.093 | 0.029 | 1 | 11 | A530032D[illegible] | 0.064 | Old |
| 0.021561 | 0.989573 | 0.125 | 0.051 | 1 | 11 | Cpped1 | 0.074 | Old |
| 0.021631 | 0.370181 | 0.21 | 0.109 | 1 | 11 | Fkbp4 | 0.101 | Old |
| 0.021691 | 0.975828 | 0.081 | 0.022 | 1 | 11 | Ift22 | 0.059 | Old |
| 0.021742 | 0.83545 | 0.242 | 0.138 | 1 | 11 | Ptpn11 | 0.104 | Old |
| 0.021768 | 1.702473 | 0.101 | 0.036 | 1 | 11 | Rab40c | 0.065 | Old |
| 0.021816 | 0.596123 | 0.137 | 0.058 | 1 | 11 | Scaf1 | 0.079 | Old |
| 0.021816 | 0.582171 | 0.137 | 0.058 | 1 | 11 | Golga5 | 0.079 | Old |
| 0.021819 | 0.150434 | 0.238 | 0.13 | 1 | 11 | Syf2 | 0.108 | Old |
| 0.021828 | 0.533022 | 0.202 | 0.101 | 1 | 11 | Ubac2 | 0.101 | Old |
| 0.021859 | 0.348772 | 0.633 | 0.435 | 1 | 11 | Scp2 | 0.198 | Old |
| 0.021868 | 0.092474 | 0.444 | 0.275 | 1 | 11 | Sirt2 | 0.169 | Old |

| | | | | | | | | |
|---|---|---|---|---|---|---|---|---|
| 0.021878 | 1.046998 | 0.125 | 0.051 | 1 | 11 | Ttc13 | 0.074 | Old |
| 0.02188 | 0.251531 | 0.21 | 0.109 | 1 | 11 | Pafah1b2 | 0.101 | Old |
| 0.021982 | 0.226587 | 0.488 | 0.319 | 1 | 11 | Morf4l2 | 0.169 | Old |
| 0.022037 | 0.226521 | 0.31 | 0.181 | 1 | 11 | Il21r | 0.129 | Old |
| 0.022041 | 1.130647 | 0.113 | 0.043 | 1 | 11 | Hectd4 | 0.07 | Old |
| 0.022056 | 0.949367 | 0.093 | 0.029 | 1 | 11 | Ap5s1 | 0.064 | Old |
| 0.022058 | 0.480368 | 0.105 | 0.036 | 1 | 11 | Rnaseh2b | 0.069 | Old |
| 0.022104 | 0.230425 | 0.694 | 0.486 | 1 | 11 | Pdcd6ip | 0.208 | Old |
| 0.022152 | 0.207174 | 0.274 | 0.152 | 1 | 11 | Cyb5r3 | 0.122 | Old |
| 0.022186 | 0.321971 | 0.274 | 0.159 | 1 | 11 | Setd3 | 0.115 | Old |
| 0.022234 | 3.446885 | 0.052 | 0.007 | 1 | 11 | Sema6a | 0.045 | Old |
| 0.022247 | 0.284387 | 0.435 | 0.29 | 1 | 11 | Ost4 | 0.145 | Old |
| 0.022333 | 0.356474 | 0.339 | 0.21 | 1 | 11 | Uhmk1 | 0.129 | Old |
| 0.02238 | 1.243741 | 0.113 | 0.043 | 1 | 11 | Ptpn23 | 0.07 | Old |
| 0.022414 | 0.93408 | 0.125 | 0.051 | 1 | 11 | Tmem110 | 0.074 | Old |
| 0.02242 | 0.440179 | 0.198 | 0.101 | 1 | 11 | Chd7 | 0.097 | Old |
| 0.022431 | 0.507559 | 0.137 | 0.058 | 1 | 11 | Polr2j | 0.079 | Old |
| 0.022459 | 0.557331 | 0.165 | 0.08 | 1 | 11 | Med30 | 0.085 | Old |
| 0.022466 | 0.201881 | 0.585 | 0.42 | 1 | 11 | Rnf149 | 0.165 | Old |
| 0.022542 | 0.66169 | 0.145 | 0.065 | 1 | 11 | Traf7 | 0.08 | Old |
| 0.022554 | 0.729066 | 0.165 | 0.08 | 1 | 11 | Rsbn1 | 0.085 | Old |
| 0.02258 | 2.45572 | 0.052 | 0.007 | 1 | 11 | 4632415L0 | 0.045 | Old |
| 0.02259 | 0.456151 | 0.335 | 0.21 | 1 | 11 | Cp | 0.125 | Old |
| 0.022623 | 0.62376 | 0.206 | 0.109 | 1 | 11 | Ubqln2 | 0.097 | Old |
| 0.022649 | 0.155324 | 0.25 | 0.138 | 1 | 11 | Tbc1d20 | 0.112 | Old |
| 0.022673 | 0.090151 | 0.335 | 0.196 | 1 | 11 | Adar | 0.139 | Old |
| 0.022695 | 0.229043 | 0.419 | 0.275 | 1 | 11 | Psmd13 | 0.144 | Old |
| 0.022744 | 0.638907 | 0.137 | 0.058 | 1 | 11 | Rgs3 | 0.079 | Old |
| 0.022745 | 0.412437 | 0.165 | 0.08 | 1 | 11 | Cdk16 | 0.085 | Old |
| 0.022755 | 2.984134 | 0.052 | 0.007 | 1 | 11 | Atf5 | 0.045 | Old |
| 0.022811 | 0.374271 | 0.327 | 0.196 | 1 | 11 | Nabp1 | 0.131 | Old |
| 0.022838 | 0.398638 | 0.323 | 0.196 | 1 | 11 | Lamtor3 | 0.127 | Old |
| 0.02284 | 0.328668 | 0.569 | 0.406 | 1 | 11 | Hmgb1 | 0.163 | Old |
| 0.02286 | 0.085273 | 0.29 | 0.167 | 1 | 11 | Gba | 0.123 | Old |
| 0.022902 | 0.215762 | 0.214 | 0.109 | 1 | 11 | Ilf3 | 0.105 | Old |
| 0.022915 | 0.012522 | 0.27 | 0.152 | 1 | 11 | Rsbn1l | 0.118 | Old |
| 0.022918 | 0.27732 | 0.379 | 0.246 | 1 | 11 | Birc2 | 0.133 | Old |
| 0.022931 | 2.862038 | 0.052 | 0.007 | 1 | 11 | Cyp2j6 | 0.045 | Old |
| 0.022955 | 0.690963 | 0.137 | 0.058 | 1 | 11 | Chm | 0.079 | Old |
| 0.022968 | 0.228771 | 0.25 | 0.138 | 1 | 11 | U2af1 | 0.112 | Old |
| 0.023023 | 0.467049 | 0.194 | 0.101 | 1 | 11 | Prpf18 | 0.093 | Old |
| 0.023052 | 0.476556 | 0.145 | 0.065 | 1 | 11 | Pias4 | 0.08 | Old |
| 0.023072 | 1.029336 | 0.113 | 0.043 | 1 | 11 | Dna2 | 0.07 | Old |
| 0.023108 | 2.559763 | 0.052 | 0.007 | 1 | 11 | Adgrg1 | 0.045 | Old |
| 0.023156 | 0.616167 | 0.145 | 0.065 | 1 | 11 | Dopey1 | 0.08 | Old |

| | | | | | | | | |
|---|---|---|---|---|---|---|---|---|
| 0.023161 | 1.733351 | 0.121 | 0.051 | 1 | 11 | C6 | 0.07 | Old |
| 0.023184 | 0.52585 | 0.125 | 0.051 | 1 | 11 | Tusc2 | 0.074 | Old |
| 0.023202 | 0.207958 | 0.323 | 0.196 | 1 | 11 | Uap1l1 | 0.127 | Old |
| 0.023205 | 0.429184 | 0.093 | 0.029 | 1 | 11 | Spon1 | 0.064 | Old |
| 0.023286 | 2.430802 | 0.052 | 0.007 | 1 | 11 | Dbr1 | 0.045 | Old |
| 0.023325 | 0.685076 | 0.165 | 0.08 | 1 | 11 | Rap1gap2 | 0.085 | Old |
| 0.023466 | 2.415634 | 0.052 | 0.007 | 1 | 11 | Treml4 | 0.045 | Old |
| 0.023466 | 2.236336 | 0.052 | 0.007 | 1 | 11 | Utp23 | 0.045 | Old |
| 0.023554 | 0.194623 | 0.363 | 0.225 | 1 | 11 | Eif3l | 0.138 | Old |
| 0.023578 | 0.138267 | 0.427 | 0.268 | 1 | 11 | Srsf9 | 0.159 | Old |
| 0.023597 | 1.53753 | 0.101 | 0.036 | 1 | 11 | Tbcd | 0.065 | Old |
| 0.023609 | 0.089887 | 0.387 | 0.232 | 1 | 11 | Psmb6 | 0.155 | Old |
| 0.023646 | 2.199379 | 0.052 | 0.007 | 1 | 11 | Ric8b | 0.045 | Old |
| 0.023646 | 2.052238 | 0.052 | 0.007 | 1 | 11 | 5730409E0 | 0.045 | Old |
| 0.023663 | 0.069901 | 0.565 | 0.377 | 1 | 11 | Arap1 | 0.188 | Old |
| 0.023663 | 0.93574 | 0.113 | 0.043 | 1 | 11 | Nt5dc3 | 0.07 | Old |
| 0.023688 | 0.202087 | 0.302 | 0.181 | 1 | 11 | Chchd1 | 0.121 | Old |
| 0.023703 | 0.373145 | 0.254 | 0.145 | 1 | 11 | Otulin | 0.109 | Old |
| 0.023706 | 0.470295 | 0.137 | 0.058 | 1 | 11 | C2cd5 | 0.079 | Old |
| 0.023729 | 0.489452 | 0.234 | 0.13 | 1 | 11 | Ing4 | 0.104 | Old |
| 0.023828 | 2.323698 | 0.052 | 0.007 | 1 | 11 | Ttn | 0.045 | Old |
| 0.023828 | 2.17023 | 0.052 | 0.007 | 1 | 11 | Zfp346 | 0.045 | Old |
| 0.023828 | 1.836799 | 0.052 | 0.007 | 1 | 11 | P3h3 | 0.045 | Old |
| 0.023833 | 0.05138 | 0.19 | 0.094 | 1 | 11 | Ranbp9 | 0.096 | Old |
| 0.023834 | 1.17152 | 0.141 | 0.065 | 1 | 11 | Polr2b | 0.076 | Old |
| 0.023855 | 3.652566 | 0.036 | 0 | 1 | 11 | Chchd6 | 0.036 | Old |
| 0.023855 | 3.501448 | 0.036 | 0 | 1 | 11 | Serpinb1a | 0.036 | Old |
| 0.023855 | 3.388974 | 0.036 | 0 | 1 | 11 | Smyd2 | 0.036 | Old |
| 0.023855 | 3.258903 | 0.036 | 0 | 1 | 11 | Armc3 | 0.036 | Old |
| 0.023855 | 2.936444 | 0.036 | 0 | 1 | 11 | Rttn | 0.036 | Old |
| 0.023855 | 2.750082 | 0.036 | 0 | 1 | 11 | Hdc | 0.036 | Old |
| 0.023855 | 2.725874 | 0.036 | 0 | 1 | 11 | 6330403L0 | 0.036 | Old |
| 0.023855 | 2.715964 | 0.036 | 0 | 1 | 11 | Xcr1 | 0.036 | Old |
| 0.023855 | 2.715427 | 0.036 | 0 | 1 | 11 | Cd320 | 0.036 | Old |
| 0.023855 | 2.682658 | 0.036 | 0 | 1 | 11 | Pigh | 0.036 | Old |
| 0.023855 | 2.675223 | 0.036 | 0 | 1 | 11 | 0610040J0 | 0.036 | Old |
| 0.023855 | 2.566961 | 0.036 | 0 | 1 | 11 | 5830432E0 | 0.036 | Old |
| 0.023855 | 2.143437 | 0.036 | 0 | 1 | 11 | Cdk5rap1 | 0.036 | Old |
| 0.023856 | 0.686959 | 0.206 | 0.109 | 1 | 11 | Itga9 | 0.097 | Old |
| 0.023864 | 0.415632 | 0.657 | 0.5 | 1 | 11 | Mapkapk2 | 0.157 | Old |
| 0.023903 | 0.25366 | 0.266 | 0.152 | 1 | 11 | Dock11 | 0.114 | Old |
| 0.023923 | 0.188508 | 0.149 | 0.065 | 1 | 11 | Suclg2 | 0.084 | Old |
| 0.024011 | 1.901166 | 0.052 | 0.007 | 1 | 11 | Tbc1d12 | 0.045 | Old |
| 0.024016 | 0.534039 | 0.286 | 0.174 | 1 | 11 | Emd | 0.112 | Old |
| 0.024047 | 0.255745 | 0.294 | 0.167 | 1 | 11 | Cnih1 | 0.127 | Old |

| | | | | | | | | |
|---|---|---|---|---|---|---|---|---|
| 0.024104 | 0.273251 | 0.258 | 0.145 | 1 | 11 | Dguok | 0.113 | Old |
| 0.024136 | 3.188532 | 0.065 | 0.014 | 1 | 11 | Flrt2 | 0.051 | Old |
| 0.024195 | 1.220866 | 0.052 | 0.007 | 1 | 11 | Lhpp | 0.045 | Old |
| 0.024217 | 0.172956 | 0.286 | 0.167 | 1 | 11 | Mtmr6 | 0.119 | Old |
| 0.024256 | 0.037504 | 0.367 | 0.225 | 1 | 11 | Eif3j1 | 0.142 | Old |
| 0.02432 | 0.1424 | 0.165 | 0.08 | 1 | 11 | Akna | 0.085 | Old |
| 0.024381 | 1.204313 | 0.052 | 0.007 | 1 | 11 | Dnah2 | 0.045 | Old |
| 0.024381 | 0.891539 | 0.052 | 0.007 | 1 | 11 | Rabepk | 0.045 | Old |
| 0.024381 | 0.351494 | 0.052 | 0.007 | 1 | 11 | Nat10 | 0.045 | Old |
| 0.024381 | 0.042661 | 0.052 | 0.007 | 1 | 11 | Mipep | 0.045 | Old |
| 0.024381 | 0.434914 | 0.177 | 0.087 | 1 | 11 | Cog7 | 0.09 | Old |
| 0.024394 | 0.86294 | 0.254 | 0.152 | 1 | 11 | Ss18 | 0.102 | Old |
| 0.024434 | 0.053159 | 0.27 | 0.152 | 1 | 11 | Tcf20 | 0.118 | Old |
| 0.024484 | 1.191689 | 0.141 | 0.065 | 1 | 11 | St3gal5 | 0.076 | Old |
| 0.024521 | 1.595971 | 0.089 | 0.029 | 1 | 11 | Sppl2b | 0.06 | Old |
| 0.024661 | 0.268904 | 0.246 | 0.138 | 1 | 11 | Mpc1 | 0.108 | Old |
| 0.02476 | 1.140565 | 0.113 | 0.043 | 1 | 11 | Rfx5 | 0.07 | Old |
| 0.024784 | 0.115368 | 0.323 | 0.196 | 1 | 11 | Paics | 0.127 | Old |
| 0.024862 | 2.31343 | 0.077 | 0.022 | 1 | 11 | Hspbap1 | 0.055 | Old |
| 0.024912 | 0.506341 | 0.125 | 0.051 | 1 | 11 | Sun1 | 0.074 | Old |
| 0.02492 | 0.317298 | 0.246 | 0.138 | 1 | 11 | Ly96 | 0.108 | Old |
| 0.024937 | 0.488057 | 0.214 | 0.116 | 1 | 11 | Snx1 | 0.098 | Old |
| 0.024983 | 0.266606 | 0.238 | 0.13 | 1 | 11 | Tmem50b | 0.108 | Old |
| 0.025031 | 0.374585 | 0.125 | 0.051 | 1 | 11 | Ccdc97 | 0.074 | Old |
| 0.025087 | 0.380597 | 0.185 | 0.094 | 1 | 11 | Rnf10 | 0.091 | Old |
| 0.02515 | 1.166324 | 0.121 | 0.051 | 1 | 11 | Ppp1r13b | 0.07 | Old |
| 0.025186 | 0.163604 | 0.185 | 0.094 | 1 | 11 | Prelid3b | 0.091 | Old |
| 0.025208 | 0.344615 | 0.29 | 0.174 | 1 | 11 | Rab32 | 0.116 | Old |
| 0.025286 | 0.289421 | 0.319 | 0.196 | 1 | 11 | Vapb | 0.123 | Old |
| 0.025305 | 2.078387 | 0.065 | 0.014 | 1 | 11 | Tada2b | 0.051 | Old |
| 0.025324 | 0.141184 | 0.476 | 0.319 | 1 | 11 | Mkln1 | 0.157 | Old |
| 0.025352 | 0.619479 | 0.165 | 0.08 | 1 | 11 | Gars | 0.085 | Old |
| 0.02537 | 1.101635 | 0.089 | 0.029 | 1 | 11 | Tmem38b | 0.06 | Old |
| 0.025422 | 0.915478 | 0.101 | 0.036 | 1 | 11 | Rps19bp1 | 0.065 | Old |
| 0.025476 | 3.035085 | 0.065 | 0.014 | 1 | 11 | Gm26870 | 0.051 | Old |
| 0.025531 | 0.131109 | 0.508 | 0.333 | 1 | 11 | Plekhb2 | 0.175 | Old |
| 0.025557 | 1.007554 | 0.101 | 0.036 | 1 | 11 | Vegfb | 0.065 | Old |
| 0.025563 | 0.684687 | 0.165 | 0.08 | 1 | 11 | Phf8 | 0.085 | Old |
| 0.025658 | 1.453452 | 0.089 | 0.029 | 1 | 11 | Mtpap | 0.06 | Old |
| 0.025665 | 0.205721 | 0.403 | 0.261 | 1 | 11 | Kras | 0.142 | Old |
| 0.025737 | 0.381008 | 0.27 | 0.152 | 1 | 11 | Stard9 | 0.118 | Old |
| 0.025797 | 1.09598 | 0.133 | 0.058 | 1 | 11 | Stx2 | 0.075 | Old |
| 0.025804 | 1.463201 | 0.089 | 0.029 | 1 | 11 | Tmem185a | 0.06 | Old |
| 0.025821 | 2.057644 | 0.065 | 0.014 | 1 | 11 | Znhit2 | 0.051 | Old |
| 0.025829 | 0.992374 | 0.101 | 0.036 | 1 | 11 | Ap5z1 | 0.065 | Old |

| | | | | | | | | |
|---|---|---|---|---|---|---|---|---|
| 0.025886 | 0.078319 | 0.254 | 0.138 | 1 | 11 | Rara | 0.116 | Old |
| 0.025915 | 0.705032 | 0.133 | 0.058 | 1 | 11 | Myh10 | 0.075 | Old |
| 0.025915 | 0.399773 | 0.133 | 0.058 | 1 | 11 | Decr1 | 0.075 | Old |
| 0.025952 | 1.52765 | 0.129 | 0.058 | 1 | 11 | Rpf1 | 0.071 | Old |
| 0.025962 | 1.571657 | 0.109 | 0.043 | 1 | 11 | Clk2 | 0.066 | Old |
| 0.025967 | 0.359312 | 0.214 | 0.116 | 1 | 11 | Pbx2 | 0.098 | Old |
| 0.026109 | 1.773768 | 0.077 | 0.022 | 1 | 11 | Eogt | 0.055 | Old |
| 0.026114 | 0.405838 | 0.173 | 0.087 | 1 | 11 | Clcc1 | 0.086 | Old |
| 0.026154 | 0.78769 | 0.133 | 0.058 | 1 | 11 | Glt8d1 | 0.075 | Old |
| 0.02616 | 0.458989 | 0.113 | 0.043 | 1 | 11 | Icmt | 0.07 | Old |
| 0.026169 | 2.147513 | 0.065 | 0.014 | 1 | 11 | Hyal2 | 0.051 | Old |
| 0.02618 | 0.46562 | 0.198 | 0.109 | 1 | 11 | Paox | 0.089 | Old |
| 0.026206 | 0.410884 | 0.419 | 0.29 | 1 | 11 | 2-Mar | 0.129 | Old |
| 0.026214 | 0.632804 | 0.145 | 0.065 | 1 | 11 | Smim19 | 0.08 | Old |
| 0.026272 | 0.201955 | 0.254 | 0.145 | 1 | 11 | Acp1 | 0.109 | Old |
| 0.026297 | 0.424004 | 0.226 | 0.13 | 1 | 11 | Zrsr2 | 0.096 | Old |
| 0.026297 | 0.767361 | 0.202 | 0.109 | 1 | 11 | Crkl | 0.093 | Old |
| 0.026313 | 0.453371 | 0.165 | 0.08 | 1 | 11 | Gpr137b | 0.085 | Old |
| 0.026313 | 0.238369 | 0.165 | 0.08 | 1 | 11 | Ints1 | 0.085 | Old |
| 0.02632 | 0.185155 | 0.278 | 0.159 | 1 | 11 | Nudt21 | 0.119 | Old |
| 0.026345 | 1.231812 | 0.065 | 0.014 | 1 | 11 | Atg16l2 | 0.051 | Old |
| 0.026429 | 1.548653 | 0.077 | 0.022 | 1 | 11 | Dhcr7 | 0.055 | Old |
| 0.026482 | 0.364976 | 0.415 | 0.283 | 1 | 11 | Pld3 | 0.132 | Old |
| 0.026502 | 0.372755 | 0.185 | 0.094 | 1 | 11 | Glrx2 | 0.091 | Old |
| 0.026518 | 0.214102 | 0.448 | 0.29 | 1 | 11 | Wapl | 0.158 | Old |
| 0.026522 | 1.555892 | 0.065 | 0.014 | 1 | 11 | 3110002H1 | 0.051 | Old |
| 0.026606 | 0.18338 | 0.185 | 0.094 | 1 | 11 | Fryl | 0.091 | Old |
| 0.026622 | 0.667491 | 0.21 | 0.116 | 1 | 11 | Gbf1 | 0.094 | Old |
| 0.026637 | 0.444721 | 0.133 | 0.058 | 1 | 11 | Pnkd | 0.075 | Old |
| 0.026649 | 0.457797 | 0.218 | 0.116 | 1 | 11 | Rhbdf2 | 0.102 | Old |
| 0.026681 | 0.142141 | 0.403 | 0.254 | 1 | 11 | Cisd2 | 0.149 | Old |
| 0.026697 | 0.193992 | 0.383 | 0.246 | 1 | 11 | Nop53 | 0.137 | Old |
| 0.026725 | 0.668599 | 0.27 | 0.159 | 1 | 11 | Hivep1 | 0.111 | Old |
| 0.026727 | 0.011268 | 0.415 | 0.261 | 1 | 11 | Il4ra | 0.154 | Old |
| 0.026758 | 0.161435 | 0.262 | 0.152 | 1 | 11 | Tlr1 | 0.11 | Old |
| 0.026834 | 0.350118 | 0.306 | 0.196 | 1 | 11 | Eif2s3x | 0.11 | Old |
| 0.026879 | 1.458204 | 0.065 | 0.014 | 1 | 11 | Jaml | 0.051 | Old |
| 0.026879 | 1.071359 | 0.065 | 0.014 | 1 | 11 | Ddx20 | 0.051 | Old |
| 0.026915 | 1.160292 | 0.077 | 0.022 | 1 | 11 | Skp2 | 0.055 | Old |
| 0.026918 | 0.026655 | 0.315 | 0.181 | 1 | 11 | Kcmf1 | 0.134 | Old |
| 0.026944 | 0.226429 | 0.452 | 0.304 | 1 | 11 | Strn3 | 0.148 | Old |
| 0.026968 | 0.806881 | 0.169 | 0.087 | 1 | 11 | Trim44 | 0.082 | Old |
| 0.027059 | 1.619643 | 0.065 | 0.014 | 1 | 11 | Rgp1 | 0.051 | Old |
| 0.027094 | 0.209303 | 0.524 | 0.37 | 1 | 11 | Eif4a2 | 0.154 | Old |
| 0.027198 | 0.619646 | 0.161 | 0.08 | 1 | 11 | Atg4b | 0.081 | Old |

| | | | | | | | | |
|---|---|---|---|---|---|---|---|---|
| 0.027199 | 0.403388 | 0.202 | 0.109 | 1 | 11 | Abhd16a | 0.093 | Old |
| 0.027199 | 0.324258 | 0.206 | 0.109 | 1 | 11 | Ip6k1 | 0.097 | Old |
| 0.027263 | 0.575653 | 0.222 | 0.123 | 1 | 11 | Slc36a1 | 0.099 | Old |
| 0.027296 | 0.186645 | 0.44 | 0.283 | 1 | 11 | Wls | 0.157 | Old |
| 0.027325 | 0.451852 | 0.29 | 0.174 | 1 | 11 | Spty2d1 | 0.116 | Old |
| 0.027378 | 0.183445 | 0.512 | 0.341 | 1 | 11 | Kdm2a | 0.171 | Old |
| 0.027409 | 2.283243 | 0.077 | 0.022 | 1 | 11 | Zbtb17 | 0.055 | Old |
| 0.027409 | 1.377214 | 0.077 | 0.022 | 1 | 11 | Smad7 | 0.055 | Old |
| 0.027416 | 1.245153 | 0.121 | 0.051 | 1 | 11 | Mlx | 0.07 | Old |
| 0.027494 | 0.091532 | 0.327 | 0.196 | 1 | 11 | Rnf7 | 0.131 | Old |
| 0.027512 | 0.867509 | 0.101 | 0.036 | 1 | 11 | Ppard | 0.065 | Old |
| 0.027576 | 1.13394 | 0.077 | 0.022 | 1 | 11 | Hmgn5 | 0.055 | Old |
| 0.027604 | 1.278413 | 0.089 | 0.029 | 1 | 11 | Ncapg2 | 0.06 | Old |
| 0.027606 | 1.024808 | 0.065 | 0.014 | 1 | 11 | Grwd1 | 0.051 | Old |
| 0.027622 | 0.10265 | 0.653 | 0.457 | 1 | 11 | Slc3a2 | 0.196 | Old |
| 0.027626 | 0.606094 | 0.133 | 0.058 | 1 | 11 | Gm17018 | 0.075 | Old |
| 0.027626 | 0.581501 | 0.133 | 0.058 | 1 | 11 | Tusc3 | 0.075 | Old |
| 0.02763 | 0.304614 | 0.597 | 0.428 | 1 | 11 | Tmed2 | 0.169 | Old |
| 0.027657 | 0.29813 | 0.101 | 0.036 | 1 | 11 | Ksr1 | 0.065 | Old |
| 0.027657 | 0.194089 | 0.238 | 0.13 | 1 | 11 | Tbc1d2b | 0.108 | Old |
| 0.027688 | 0.303611 | 0.242 | 0.138 | 1 | 11 | Emc2 | 0.104 | Old |
| 0.027695 | 0.462889 | 0.327 | 0.203 | 1 | 11 | Snx8 | 0.124 | Old |
| 0.027743 | 1.336148 | 0.077 | 0.022 | 1 | 11 | Cog1 | 0.055 | Old |
| 0.027743 | 0.964666 | 0.077 | 0.022 | 1 | 11 | Ccdc167 | 0.055 | Old |
| 0.027786 | 0.117991 | 0.298 | 0.174 | 1 | 11 | Mknk2 | 0.124 | Old |
| 0.02779 | 1.172034 | 0.065 | 0.014 | 1 | 11 | Mad2l2 | 0.051 | Old |
| 0.027797 | 0.263487 | 0.315 | 0.196 | 1 | 11 | Cbfa2t3 | 0.119 | Old |
| 0.027832 | 0.314737 | 0.246 | 0.138 | 1 | 11 | Ppp2r2d | 0.108 | Old |
| 0.027873 | 0.530574 | 0.246 | 0.145 | 1 | 11 | Snx29 | 0.101 | Old |
| 0.027906 | 0.478398 | 0.194 | 0.101 | 1 | 11 | Mmd | 0.093 | Old |
| 0.027912 | 0.084307 | 0.294 | 0.174 | 1 | 11 | Ndufaf8 | 0.12 | Old |
| 0.028036 | 0.031617 | 0.218 | 0.116 | 1 | 11 | Ap3m1 | 0.102 | Old |
| 0.028094 | 0.033421 | 0.101 | 0.036 | 1 | 11 | Rabggta | 0.065 | Old |
| 0.028103 | 0.225406 | 0.367 | 0.232 | 1 | 11 | AC160336. | 0.135 | Old |
| 0.028162 | 1.07167 | 0.065 | 0.014 | 1 | 11 | Aspa | 0.051 | Old |
| 0.028186 | 0.360078 | 0.294 | 0.181 | 1 | 11 | Sft2d2 | 0.113 | Old |
| 0.0282 | 0.208256 | 0.359 | 0.225 | 1 | 11 | Tmem234 | 0.134 | Old |
| 0.028337 | 0.357923 | 0.202 | 0.109 | 1 | 11 | Cnot8 | 0.093 | Old |
| 0.028348 | 0.157112 | 0.46 | 0.29 | 1 | 11 | Pebp1 | 0.17 | Old |
| 0.028386 | 0.492635 | 0.089 | 0.029 | 1 | 11 | Limk1 | 0.06 | Old |
| 0.028422 | 0.257586 | 0.278 | 0.167 | 1 | 11 | Snf8 | 0.111 | Old |
| 0.02844 | 0.080977 | 0.488 | 0.326 | 1 | 11 | Ube2b | 0.162 | Old |
| 0.028467 | 0.684317 | 0.141 | 0.065 | 1 | 11 | Sp2 | 0.076 | Old |
| 0.028537 | 0.563325 | 0.065 | 0.014 | 1 | 11 | Zfp780b | 0.051 | Old |
| 0.028537 | 0.475747 | 0.065 | 0.014 | 1 | 11 | Zbtb34 | 0.051 | Old |

| | | | | | | | | |
|---|---|---|---|---|---|---|---|---|
| 0.028545 | 0.397067 | 0.089 | 0.029 | 1 | 11 | Acvr1b | 0.06 | Old |
| 0.028549 | 0.360339 | 0.214 | 0.116 | 1 | 11 | Clcn3 | 0.098 | Old |
| 0.028575 | 0.257097 | 0.23 | 0.13 | 1 | 11 | Man1a2 | 0.1 | Old |
| 0.028593 | 0.653107 | 0.077 | 0.022 | 1 | 11 | Nudt18 | 0.055 | Old |
| 0.028606 | 0.170751 | 0.145 | 0.065 | 1 | 11 | Foxk1 | 0.08 | Old |
| 0.028647 | 0.697919 | 0.169 | 0.087 | 1 | 11 | Bckdk | 0.082 | Old |
| 0.028668 | 0.230483 | 0.29 | 0.174 | 1 | 11 | Cd37 | 0.116 | Old |
| 0.028677 | 0.261224 | 0.472 | 0.312 | 1 | 11 | Psma4 | 0.16 | Old |
| 0.028727 | 0.302304 | 0.065 | 0.014 | 1 | 11 | Fkbp7 | 0.051 | Old |
| 0.028741 | 0.240502 | 0.198 | 0.101 | 1 | 11 | Arid2 | 0.097 | Old |
| 0.028746 | 0.262808 | 0.431 | 0.283 | 1 | 11 | Igtp | 0.148 | Old |
| 0.028766 | 0.037355 | 0.077 | 0.022 | 1 | 11 | Mllt1 | 0.055 | Old |
| 0.028811 | 0.317785 | 0.153 | 0.072 | 1 | 11 | Gcnt1 | 0.081 | Old |
| 0.028822 | 0.279942 | 0.391 | 0.261 | 1 | 11 | Myo9b | 0.13 | Old |
| 0.028836 | 0.348057 | 0.101 | 0.036 | 1 | 11 | Tstd2 | 0.065 | Old |
| 0.028886 | 0.8249 | 0.242 | 0.152 | 1 | 11 | Reps2 | 0.09 | Old |
| 0.028939 | 0.949301 | 0.077 | 0.022 | 1 | 11 | Cnksr3 | 0.055 | Old |
| 0.028948 | 0.172341 | 0.242 | 0.13 | 1 | 11 | Slc25a45 | 0.112 | Old |
| 0.02897 | 0.532528 | 0.141 | 0.065 | 1 | 11 | Slc25a17 | 0.076 | Old |
| 0.029025 | 0.157141 | 0.089 | 0.029 | 1 | 11 | Lrrcc1 | 0.06 | Old |
| 0.029065 | 0.10359 | 0.714 | 0.514 | 1 | 11 | Wdr1 | 0.2 | Old |
| 0.02907 | 0.482561 | 0.214 | 0.116 | 1 | 11 | Hmgcr | 0.098 | Old |
| 0.029105 | 0.108928 | 0.448 | 0.283 | 1 | 11 | Cab39 | 0.165 | Old |
| 0.02911 | 0.72347 | 0.169 | 0.087 | 1 | 11 | Mrpl4 | 0.082 | Old |
| 0.029125 | 0.382068 | 0.343 | 0.225 | 1 | 11 | Spcs3 | 0.118 | Old |
| 0.029133 | 0.15386 | 0.44 | 0.283 | 1 | 11 | Mkrn1 | 0.157 | Old |
| 0.029173 | 0.451502 | 0.306 | 0.188 | 1 | 11 | BC031181 | 0.118 | Old |
| 0.029208 | 0.586386 | 0.173 | 0.087 | 1 | 11 | A630001G2 | 0.086 | Old |
| 0.02929 | 0.299111 | 0.101 | 0.036 | 1 | 11 | Emilin1 | 0.065 | Old |
| 0.0293 | 0.42451 | 0.133 | 0.058 | 1 | 11 | Gcsh | 0.075 | Old |
| 0.02934 | 0.532571 | 0.181 | 0.094 | 1 | 11 | Bag6 | 0.087 | Old |
| 0.029345 | 0.28882 | 0.399 | 0.268 | 1 | 11 | Slc29a1 | 0.131 | Old |
| 0.029345 | 0.120433 | 0.403 | 0.261 | 1 | 11 | Glmp | 0.142 | Old |
| 0.029394 | 0.782372 | 0.222 | 0.13 | 1 | 11 | Tlr4 | 0.092 | Old |
| 0.029424 | 0.103884 | 0.194 | 0.101 | 1 | 11 | Aig1 | 0.093 | Old |
| 0.029432 | 0.137282 | 0.532 | 0.355 | 1 | 11 | Gpi1 | 0.177 | Old |
| 0.029436 | 0.70306 | 0.121 | 0.051 | 1 | 11 | Brf1 | 0.07 | Old |
| 0.029451 | 0.218311 | 0.278 | 0.167 | 1 | 11 | Kcnk6 | 0.111 | Old |
| 0.029588 | 0.084198 | 0.327 | 0.196 | 1 | 11 | Higd2a | 0.131 | Old |
| 0.029595 | 0.147287 | 0.101 | 0.036 | 1 | 11 | Smim24 | 0.065 | Old |
| 0.029599 | 0.041225 | 0.113 | 0.043 | 1 | 11 | Fem1a | 0.07 | Old |
| 0.029615 | 0.092329 | 0.339 | 0.203 | 1 | 11 | 2410015M | 0.136 | Old |
| 0.029628 | 0.261933 | 0.512 | 0.355 | 1 | 11 | Gna13 | 0.157 | Old |
| 0.029672 | 0.355272 | 0.153 | 0.072 | 1 | 11 | Aph1c | 0.081 | Old |
| 0.029697 | 0.616967 | 0.177 | 0.094 | 1 | 11 | P2ry14 | 0.083 | Old |

| | | | | | | | | |
|---|---|---|---|---|---|---|---|---|
| 0.029749 | 0.220405 | 0.379 | 0.254 | 1 | 11 | Atp6v0a1 | 0.125 | Old |
| 0.029776 | 0.601105 | 0.19 | 0.101 | 1 | 11 | Ing1 | 0.089 | Old |
| 0.029793 | 0.592351 | 0.173 | 0.087 | 1 | 11 | Fam118b | 0.086 | Old |
| 0.029928 | 0.36404 | 0.294 | 0.181 | 1 | 11 | Kdm5a | 0.113 | Old |
| 0.02995 | 0.028332 | 0.198 | 0.101 | 1 | 11 | Gramd1a | 0.097 | Old |
| 0.029978 | 0.177163 | 0.415 | 0.268 | 1 | 11 | Psme2b | 0.147 | Old |
| 0.029999 | 0.335835 | 0.077 | 0.022 | 1 | 11 | Rbm12b2 | 0.055 | Old |
| 0.030007 | 0.379035 | 0.089 | 0.029 | 1 | 11 | Cep295 | 0.06 | Old |
| 0.030048 | 0.570256 | 0.153 | 0.072 | 1 | 11 | Clns1a | 0.081 | Old |
| 0.03012 | 0.134839 | 0.444 | 0.29 | 1 | 11 | Ppt1 | 0.154 | Old |
| 0.030131 | 0.134856 | 0.653 | 0.471 | 1 | 11 | Akap13 | 0.182 | Old |
| 0.030175 | 0.570715 | 0.169 | 0.087 | 1 | 11 | Dock4 | 0.082 | Old |
| 0.030215 | 0.655561 | 0.23 | 0.13 | 1 | 11 | Rbm22 | 0.1 | Old |
| 0.03026 | 0.050566 | 0.145 | 0.065 | 1 | 11 | Cyb5r1 | 0.08 | Old |
| 0.030288 | 0.535463 | 0.19 | 0.101 | 1 | 11 | Rftn1 | 0.089 | Old |
| 0.030289 | 0.17531 | 0.335 | 0.21 | 1 | 11 | Ssrp1 | 0.125 | Old |
| 0.030389 | 0.644424 | 0.161 | 0.08 | 1 | 11 | Slc35f5 | 0.081 | Old |
| 0.030404 | 0.270258 | 0.331 | 0.203 | 1 | 11 | Sav1 | 0.128 | Old |
| 0.030504 | 0.323523 | 0.589 | 0.42 | 1 | 11 | Bsg | 0.169 | Old |
| 0.030566 | 0.70876 | 0.121 | 0.051 | 1 | 11 | Rad17 | 0.07 | Old |
| 0.030596 | 0.183828 | 0.411 | 0.268 | 1 | 11 | Ik | 0.143 | Old |
| 0.03061 | 0.172599 | 0.278 | 0.167 | 1 | 11 | Malt1 | 0.111 | Old |
| 0.030724 | 0.524097 | 0.077 | 0.022 | 1 | 11 | Adprm | 0.055 | Old |
| 0.030746 | 0.913788 | 0.117 | 0.051 | 1 | 11 | Wdr5 | 0.066 | Old |
| 0.03075 | 0.343064 | 0.246 | 0.145 | 1 | 11 | Nsmce4a | 0.101 | Old |
| 0.030774 | 0.254942 | 0.298 | 0.181 | 1 | 11 | Cop1 | 0.117 | Old |
| 0.030797 | 0.1516 | 0.359 | 0.225 | 1 | 11 | Trmt112 | 0.134 | Old |
| 0.030801 | 0.358612 | 0.569 | 0.42 | 1 | 11 | Rpl27 | 0.149 | Old |
| 0.030855 | 0.431741 | 0.315 | 0.203 | 1 | 11 | Abcf1 | 0.112 | Old |
| 0.030874 | 1.677374 | 0.097 | 0.036 | 1 | 11 | Hira | 0.061 | Old |
| 0.030925 | 0.23477 | 0.742 | 0.587 | 1 | 11 | Eif3a | 0.155 | Old |
| 0.031168 | 0.135467 | 0.262 | 0.152 | 1 | 11 | Adam19 | 0.11 | Old |
| 0.031176 | 2.609188 | 0.048 | 0.007 | 1 | 11 | Aacs | 0.041 | Old |
| 0.031226 | 0.230572 | 0.698 | 0.529 | 1 | 11 | Psme1 | 0.169 | Old |
| 0.031228 | 0.416502 | 0.109 | 0.043 | 1 | 11 | Orc3 | 0.066 | Old |
| 0.031285 | 0.186951 | 0.347 | 0.21 | 1 | 11 | B4galt1 | 0.137 | Old |
| 0.031307 | 0.662877 | 0.129 | 0.058 | 1 | 11 | 6430548M | 0.071 | Old |
| 0.031351 | 0.234238 | 0.165 | 0.08 | 1 | 11 | Zfp984 | 0.085 | Old |
| 0.031368 | 0.368531 | 0.294 | 0.181 | 1 | 11 | Chka | 0.113 | Old |
| 0.031412 | 0.098332 | 0.448 | 0.297 | 1 | 11 | Sumo1 | 0.151 | Old |
| 0.031416 | 1.281009 | 0.157 | 0.08 | 1 | 11 | Cisd1 | 0.077 | Old |
| 0.031421 | 0.097903 | 0.202 | 0.109 | 1 | 11 | Hat1 | 0.093 | Old |
| 0.031483 | 0.364161 | 0.31 | 0.196 | 1 | 11 | Poldip3 | 0.114 | Old |
| 0.031571 | 0.355301 | 0.181 | 0.094 | 1 | 11 | Rpl10-ps3 | 0.087 | Old |
| 0.031584 | 0.46748 | 0.121 | 0.051 | 1 | 11 | Usf3 | 0.07 | Old |

| | | | | | | | | |
|---|---|---|---|---|---|---|---|---|
| 0.031642 | 0.356172 | 0.222 | 0.123 | 1 | 11 | Dhx40 | 0.099 | Old |
| 0.031656 | 0.344552 | 0.25 | 0.145 | 1 | 11 | Wars | 0.105 | Old |
| 0.031663 | 0.182559 | 0.496 | 0.333 | 1 | 11 | Ist1 | 0.163 | Old |
| 0.031668 | 0.207815 | 0.657 | 0.478 | 1 | 11 | Atp1a1 | 0.179 | Old |
| 0.031753 | 0.073637 | 0.379 | 0.246 | 1 | 11 | Tmem189 | 0.133 | Old |
| 0.031762 | 0.273227 | 0.738 | 0.529 | 1 | 11 | Rab8b | 0.209 | Old |
| 0.031816 | 0.136603 | 0.556 | 0.377 | 1 | 11 | Ppp1r9b | 0.179 | Old |
| 0.031898 | 2.509504 | 0.048 | 0.007 | 1 | 11 | Tuft1 | 0.041 | Old |
| 0.031941 | 0.088712 | 0.488 | 0.333 | 1 | 11 | Lman2 | 0.155 | Old |
| 0.03202 | 0.954634 | 0.129 | 0.058 | 1 | 11 | Brms1 | 0.071 | Old |
| 0.032036 | 0.441528 | 0.161 | 0.08 | 1 | 11 | Minpp1 | 0.081 | Old |
| 0.032042 | 0.133359 | 0.609 | 0.42 | 1 | 11 | Rpn2 | 0.189 | Old |
| 0.032065 | 0.166292 | 0.31 | 0.196 | 1 | 11 | Slc16a10 | 0.114 | Old |
| 0.032216 | 0.722272 | 0.177 | 0.094 | 1 | 11 | Lta4h | 0.083 | Old |
| 0.032296 | 0.256603 | 0.161 | 0.08 | 1 | 11 | Vps37b | 0.081 | Old |
| 0.03231 | 1.006415 | 0.129 | 0.058 | 1 | 11 | Gm48065 | 0.071 | Old |
| 0.032388 | 1.197284 | 0.117 | 0.051 | 1 | 11 | Krr1 | 0.066 | Old |
| 0.032422 | 0.685518 | 0.218 | 0.123 | 1 | 11 | Chuk | 0.095 | Old |
| 0.032493 | 0.377984 | 0.198 | 0.109 | 1 | 11 | Dcun1d1 | 0.089 | Old |
| 0.032541 | 1.34868 | 0.117 | 0.051 | 1 | 11 | Nup50 | 0.066 | Old |
| 0.032556 | 0.760524 | 0.149 | 0.072 | 1 | 11 | Gatsl2 | 0.077 | Old |
| 0.032558 | 0.537603 | 0.161 | 0.08 | 1 | 11 | Ino80e | 0.081 | Old |
| 0.032607 | 0.323736 | 0.306 | 0.196 | 1 | 11 | Ripk1 | 0.11 | Old |
| 0.032612 | 0.635951 | 0.198 | 0.109 | 1 | 11 | Mindy1 | 0.089 | Old |
| 0.032635 | 2.093069 | 0.048 | 0.007 | 1 | 11 | Sh2b2 | 0.041 | Old |
| 0.032635 | 1.838119 | 0.048 | 0.007 | 1 | 11 | Msh6 | 0.041 | Old |
| 0.032635 | 1.824608 | 0.048 | 0.007 | 1 | 11 | Mars | 0.041 | Old |
| 0.032735 | 0.408656 | 0.262 | 0.152 | 1 | 11 | Rasa3 | 0.11 | Old |
| 0.032776 | 0.757994 | 0.254 | 0.152 | 1 | 11 | Cib1 | 0.102 | Old |
| 0.032795 | 0.415394 | 0.153 | 0.072 | 1 | 11 | Pgap1 | 0.081 | Old |
| 0.032799 | 0.548718 | 0.109 | 0.043 | 1 | 11 | Prps2 | 0.066 | Old |
| 0.032799 | 0.168369 | 0.435 | 0.275 | 1 | 11 | Mef2d | 0.16 | Old |
| 0.032822 | 0.226196 | 0.161 | 0.08 | 1 | 11 | Dhx30 | 0.081 | Old |
| 0.03283 | 0.245399 | 0.149 | 0.072 | 1 | 11 | Dap3 | 0.077 | Old |
| 0.032884 | 2.380083 | 0.048 | 0.007 | 1 | 11 | Usp11 | 0.041 | Old |
| 0.032884 | 2.145537 | 0.048 | 0.007 | 1 | 11 | Prkca | 0.041 | Old |
| 0.032951 | 0.721433 | 0.165 | 0.087 | 1 | 11 | Cdk17 | 0.078 | Old |
| 0.033038 | 1.099908 | 0.097 | 0.036 | 1 | 11 | Rad54l2 | 0.061 | Old |
| 0.033134 | 1.829976 | 0.048 | 0.007 | 1 | 11 | Rad1 | 0.041 | Old |
| 0.033134 | 1.37931 | 0.048 | 0.007 | 1 | 11 | Cpne8 | 0.041 | Old |
| 0.033159 | 1.046942 | 0.117 | 0.051 | 1 | 11 | C130026I2 | 0.066 | Old |
| 0.03321 | 0.943025 | 0.097 | 0.036 | 1 | 11 | Mvd | 0.061 | Old |
| 0.033223 | 0.791468 | 0.177 | 0.094 | 1 | 11 | Rrm1 | 0.083 | Old |
| 0.033309 | 0.08583 | 0.31 | 0.188 | 1 | 11 | Gtpbp2 | 0.122 | Old |
| 0.033349 | 0.978744 | 0.194 | 0.109 | 1 | 11 | Plekhm3 | 0.085 | Old |

| | | | | | | | | |
|---|---|---|---|---|---|---|---|---|
| 0.033354 | 0.367847 | 0.27 | 0.159 | 1 | 11 | Smc3 | 0.111 | Old |
| 0.033372 | 0.208705 | 0.722 | 0.536 | 1 | 11 | Ppp3ca | 0.186 | Old |
| 0.033386 | 1.635496 | 0.048 | 0.007 | 1 | 11 | Osbpl5 | 0.041 | Old |
| 0.033386 | 1.320828 | 0.048 | 0.007 | 1 | 11 | Psph | 0.041 | Old |
| 0.033386 | 1.20397 | 0.048 | 0.007 | 1 | 11 | Ppat | 0.041 | Old |
| 0.033386 | 1.167005 | 0.048 | 0.007 | 1 | 11 | Umps | 0.041 | Old |
| 0.033386 | 1.036582 | 0.048 | 0.007 | 1 | 11 | Ddx28 | 0.041 | Old |
| 0.033386 | 0.942982 | 0.048 | 0.007 | 1 | 11 | Wdr24 | 0.041 | Old |
| 0.033386 | 0.867219 | 0.048 | 0.007 | 1 | 11 | Tmem129 | 0.041 | Old |
| 0.033399 | 3.49041 | 0.032 | 0 | 1 | 11 | Steap4 | 0.032 | Old |
| 0.033399 | 3.228416 | 0.032 | 0 | 1 | 11 | Gm15675 | 0.032 | Old |
| 0.033399 | 3.078246 | 0.032 | 0 | 1 | 11 | Itgb3bp | 0.032 | Old |
| 0.033399 | 3.045176 | 0.032 | 0 | 1 | 11 | Nprl2 | 0.032 | Old |
| 0.033399 | 2.929846 | 0.032 | 0 | 1 | 11 | Zfp456 | 0.032 | Old |
| 0.033399 | 2.879996 | 0.032 | 0 | 1 | 11 | Sectm1a | 0.032 | Old |
| 0.033399 | 2.812256 | 0.032 | 0 | 1 | 11 | Mcrip2 | 0.032 | Old |
| 0.033399 | 2.812067 | 0.032 | 0 | 1 | 11 | Fuz | 0.032 | Old |
| 0.033399 | 2.804179 | 0.032 | 0 | 1 | 11 | Purg | 0.032 | Old |
| 0.033399 | 2.770298 | 0.032 | 0 | 1 | 11 | Nup37 | 0.032 | Old |
| 0.033399 | 2.766431 | 0.032 | 0 | 1 | 11 | Cald1 | 0.032 | Old |
| 0.033399 | 2.708946 | 0.032 | 0 | 1 | 11 | Ndufaf5 | 0.032 | Old |
| 0.033399 | 2.6569 | 0.032 | 0 | 1 | 11 | Neil1 | 0.032 | Old |
| 0.033399 | 2.626323 | 0.032 | 0 | 1 | 11 | Zdhhc24 | 0.032 | Old |
| 0.033399 | 2.60095 | 0.032 | 0 | 1 | 11 | Senp8 | 0.032 | Old |
| 0.033399 | 2.585912 | 0.032 | 0 | 1 | 11 | Mab21l3 | 0.032 | Old |
| 0.033399 | 2.562516 | 0.032 | 0 | 1 | 11 | Jade3 | 0.032 | Old |
| 0.033399 | 2.305977 | 0.032 | 0 | 1 | 11 | Tha1 | 0.032 | Old |
| 0.033399 | 2.271934 | 0.032 | 0 | 1 | 11 | Cbwd1 | 0.032 | Old |
| 0.033399 | 2.250216 | 0.032 | 0 | 1 | 11 | Wdr62 | 0.032 | Old |
| 0.033399 | 2.199036 | 0.032 | 0 | 1 | 11 | Apoo | 0.032 | Old |
| 0.033399 | 1.973581 | 0.032 | 0 | 1 | 11 | Det1 | 0.032 | Old |
| 0.033405 | 0.142558 | 0.641 | 0.464 | 1 | 11 | Vcp | 0.177 | Old |
| 0.033423 | 0.375739 | 0.141 | 0.065 | 1 | 11 | Pggt1b | 0.076 | Old |
| 0.033452 | 0.570774 | 0.222 | 0.13 | 1 | 11 | Vti1b | 0.092 | Old |
| 0.033484 | 0.21094 | 0.593 | 0.42 | 1 | 11 | Stx7 | 0.173 | Old |
| 0.03364 | 1.45007 | 0.048 | 0.007 | 1 | 11 | Cacna1f | 0.041 | Old |
| 0.03364 | 0.933477 | 0.048 | 0.007 | 1 | 11 | Zbtb41 | 0.041 | Old |
| 0.03364 | 0.905413 | 0.048 | 0.007 | 1 | 11 | Ldlrad3 | 0.041 | Old |
| 0.03364 | 0.596934 | 0.048 | 0.007 | 1 | 11 | Nup210 | 0.041 | Old |
| 0.03364 | 0.471552 | 0.048 | 0.007 | 1 | 11 | Carf | 0.041 | Old |
| 0.033723 | 0.179137 | 0.56 | 0.399 | 1 | 11 | Ppp1ca | 0.161 | Old |
| 0.03373 | 0.598731 | 0.097 | 0.036 | 1 | 11 | Fbxl14 | 0.061 | Old |
| 0.033736 | 0.775831 | 0.177 | 0.094 | 1 | 11 | Mtr | 0.083 | Old |
| 0.03376 | 0.046102 | 0.492 | 0.341 | 1 | 11 | 7-Sep | 0.151 | Old |
| 0.033787 | 0.85031 | 0.117 | 0.051 | 1 | 11 | Ebpl | 0.066 | Old |

| | | | | | | | | |
|---|---|---|---|---|---|---|---|---|
| 0.033791 | 1.014607 | 0.129 | 0.058 | 1 | 11 | Rab3d | 0.071 | Old |
| 0.0338 | 0.474831 | 0.238 | 0.138 | 1 | 11 | Psmd3 | 0.1 | Old |
| 0.033886 | 0.182382 | 0.448 | 0.312 | 1 | 11 | Prdx2 | 0.136 | Old |
| 0.033992 | 1.827458 | 0.073 | 0.022 | 1 | 11 | Dus1l | 0.051 | Old |
| 0.034001 | 0.216193 | 0.141 | 0.065 | 1 | 11 | Tnip2 | 0.076 | Old |
| 0.034129 | 0.082804 | 0.294 | 0.174 | 1 | 11 | Erp44 | 0.12 | Old |
| 0.034232 | 0.14518 | 0.726 | 0.565 | 1 | 11 | M6pr | 0.161 | Old |
| 0.034261 | 0.284564 | 0.508 | 0.348 | 1 | 11 | Gapvd1 | 0.16 | Old |
| 0.034264 | 1.043683 | 0.117 | 0.051 | 1 | 11 | Sos1 | 0.066 | Old |
| 0.034264 | 0.819137 | 0.117 | 0.051 | 1 | 11 | Sipa1l3 | 0.066 | Old |
| 0.034271 | 0.124979 | 0.375 | 0.232 | 1 | 11 | Mrpl23 | 0.143 | Old |
| 0.03429 | 0.263828 | 0.258 | 0.152 | 1 | 11 | C5ar2 | 0.106 | Old |
| 0.034332 | 0.27833 | 0.266 | 0.159 | 1 | 11 | Abce1 | 0.107 | Old |
| 0.034506 | 1.668665 | 0.085 | 0.029 | 1 | 11 | Dbt | 0.056 | Old |
| 0.034538 | 0.197804 | 0.194 | 0.101 | 1 | 11 | Cops2 | 0.093 | Old |
| 0.03461 | 1.182611 | 0.073 | 0.022 | 1 | 11 | Tbc1d32 | 0.051 | Old |
| 0.034612 | 0.501621 | 0.097 | 0.036 | 1 | 11 | Akr7a5 | 0.061 | Old |
| 0.034659 | 0.486697 | 0.266 | 0.167 | 1 | 11 | Ddhd1 | 0.099 | Old |
| 0.034666 | 0.199562 | 0.343 | 0.217 | 1 | 11 | Trim41 | 0.126 | Old |
| 0.034667 | 0.884686 | 0.117 | 0.051 | 1 | 11 | Rnf168 | 0.066 | Old |
| 0.034735 | 0.347569 | 0.496 | 0.348 | 1 | 11 | Rnpep | 0.148 | Old |
| 0.034826 | 0.141108 | 0.198 | 0.109 | 1 | 11 | Eif4ebp1 | 0.089 | Old |
| 0.034886 | 0.215991 | 0.153 | 0.072 | 1 | 11 | Zdhhc7 | 0.081 | Old |
| 0.034889 | 1.382204 | 0.085 | 0.029 | 1 | 11 | Pex5 | 0.056 | Old |
| 0.034896 | 0.417478 | 0.468 | 0.355 | 1 | 11 | Epb41l2 | 0.113 | Old |
| 0.03497 | 0.913865 | 0.097 | 0.036 | 1 | 11 | Tmed8 | 0.061 | Old |
| 0.034987 | 0.077462 | 0.226 | 0.123 | 1 | 11 | Gpr107 | 0.103 | Old |
| 0.034999 | 0.415686 | 0.161 | 0.08 | 1 | 11 | Med12 | 0.081 | Old |
| 0.035026 | 1.161833 | 0.073 | 0.022 | 1 | 11 | Zfp821 | 0.051 | Old |
| 0.035026 | 0.971521 | 0.073 | 0.022 | 1 | 11 | Zbed4 | 0.051 | Old |
| 0.035044 | 0.323647 | 0.206 | 0.116 | 1 | 11 | Gigyf2 | 0.09 | Old |
| 0.035082 | 1.105189 | 0.085 | 0.029 | 1 | 11 | Cnot9 | 0.056 | Old |
| 0.035082 | 0.551403 | 0.085 | 0.029 | 1 | 11 | 5730455P1 | 0.056 | Old |
| 0.035135 | 0.247985 | 0.258 | 0.152 | 1 | 11 | Usp22 | 0.106 | Old |
| 0.035197 | 0.6027 | 0.157 | 0.08 | 1 | 11 | Fam20c | 0.077 | Old |
| 0.035233 | 0.309977 | 0.149 | 0.072 | 1 | 11 | Ddx56 | 0.077 | Old |
| 0.035236 | 2.034139 | 0.073 | 0.022 | 1 | 11 | Zcchc10 | 0.051 | Old |
| 0.035335 | 0.345887 | 0.423 | 0.297 | 1 | 11 | Prkx | 0.126 | Old |
| 0.035458 | 1.854843 | 0.06 | 0.014 | 1 | 11 | Btrc | 0.046 | Old |
| 0.035468 | 0.061572 | 0.423 | 0.275 | 1 | 11 | 1110008F1 | 0.148 | Old |
| 0.0355 | 0.233466 | 0.306 | 0.188 | 1 | 11 | Exoc5 | 0.118 | Old |
| 0.035626 | 0.515029 | 0.109 | 0.043 | 1 | 11 | Ikbkg | 0.066 | Old |
| 0.035626 | 0.367147 | 0.109 | 0.043 | 1 | 11 | Tusc1 | 0.066 | Old |
| 0.035668 | 0.403914 | 0.359 | 0.239 | 1 | 11 | Tmem173 | 0.12 | Old |
| 0.035693 | 1.822701 | 0.06 | 0.014 | 1 | 11 | Dusp18 | 0.046 | Old |

| | | | | | | | | |
|---|---|---|---|---|---|---|---|---|
| 0.035707 | 0.215076 | 0.331 | 0.203 | 1 | 11 | Snx17 | 0.128 | Old |
| 0.035851 | 0.589427 | 0.198 | 0.109 | 1 | 11 | Phf5a | 0.089 | Old |
| 0.035873 | 1.190346 | 0.073 | 0.022 | 1 | 11 | Prpf3 | 0.051 | Old |
| 0.035929 | 0.915221 | 0.06 | 0.014 | 1 | 11 | Tmem101 | 0.046 | Old |
| 0.035932 | 0.388278 | 0.27 | 0.167 | 1 | 11 | Stxbp2 | 0.103 | Old |
| 0.036008 | 0.405701 | 0.411 | 0.283 | 1 | 11 | Stard8 | 0.128 | Old |
| 0.036057 | 0.60166 | 0.23 | 0.138 | 1 | 11 | Vps41 | 0.092 | Old |
| 0.036076 | 0.343559 | 0.806 | 0.638 | 1 | 11 | Nrros | 0.168 | Old |
| 0.036126 | 0.438318 | 0.242 | 0.145 | 1 | 11 | Rogdi | 0.097 | Old |
| 0.036144 | 0.42551 | 0.238 | 0.138 | 1 | 11 | Dido1 | 0.1 | Old |
| 0.036166 | 1.058678 | 0.06 | 0.014 | 1 | 11 | Cd22 | 0.046 | Old |
| 0.03632 | 0.730225 | 0.173 | 0.094 | 1 | 11 | Cops8 | 0.079 | Old |
| 0.036333 | 0.069404 | 0.282 | 0.167 | 1 | 11 | Lars2 | 0.115 | Old |
| 0.036405 | 1.676406 | 0.06 | 0.014 | 1 | 11 | Hdhd2 | 0.046 | Old |
| 0.036405 | 1.466158 | 0.06 | 0.014 | 1 | 11 | Dpysl3 | 0.046 | Old |
| 0.036405 | 0.826431 | 0.06 | 0.014 | 1 | 11 | Dexi | 0.046 | Old |
| 0.036449 | 0.000427 | 0.286 | 0.167 | 1 | 11 | Atg5 | 0.119 | Old |
| 0.036532 | 0.225692 | 0.395 | 0.268 | 1 | 11 | Ak2 | 0.127 | Old |
| 0.036536 | 0.042828 | 0.375 | 0.239 | 1 | 11 | Usp8 | 0.136 | Old |
| 0.0366 | 0.18203 | 0.375 | 0.246 | 1 | 11 | Cept1 | 0.129 | Old |
| 0.036621 | 0.289607 | 0.097 | 0.036 | 1 | 11 | Eif2b2 | 0.061 | Old |
| 0.036656 | 0.036071 | 0.5 | 0.348 | 1 | 11 | Acer3 | 0.152 | Old |
| 0.036657 | 1.008882 | 0.085 | 0.029 | 1 | 11 | Arl10 | 0.056 | Old |
| 0.036657 | 0.580887 | 0.085 | 0.029 | 1 | 11 | Camta1 | 0.056 | Old |
| 0.036657 | 0.388969 | 0.085 | 0.029 | 1 | 11 | Usp53 | 0.056 | Old |
| 0.036673 | 0.242723 | 0.371 | 0.239 | 1 | 11 | Rsu1 | 0.132 | Old |
| 0.036736 | 1.18666 | 0.073 | 0.022 | 1 | 11 | Nob1 | 0.051 | Old |
| 0.036736 | 0.934101 | 0.073 | 0.022 | 1 | 11 | Pi4kb | 0.051 | Old |
| 0.036749 | 0.396766 | 0.137 | 0.065 | 1 | 11 | Senp3 | 0.072 | Old |
| 0.036758 | 0.506785 | 0.391 | 0.268 | 1 | 11 | Mapk6 | 0.123 | Old |
| 0.036855 | 0.226866 | 0.234 | 0.13 | 1 | 11 | Coro7 | 0.104 | Old |
| 0.036886 | 1.10875 | 0.06 | 0.014 | 1 | 11 | Nedd4l | 0.046 | Old |
| 0.03691 | 1.119135 | 0.117 | 0.051 | 1 | 11 | Iqsec2 | 0.066 | Old |
| 0.03709 | 0.348698 | 0.129 | 0.058 | 1 | 11 | Chid1 | 0.071 | Old |
| 0.037095 | 0.625445 | 0.254 | 0.152 | 1 | 11 | Ktn1 | 0.102 | Old |
| 0.037346 | 0.367071 | 0.335 | 0.21 | 1 | 11 | Gspt1 | 0.125 | Old |
| 0.037373 | 0.983898 | 0.06 | 0.014 | 1 | 11 | Pdcd2 | 0.046 | Old |
| 0.037418 | 0.073115 | 0.266 | 0.159 | 1 | 11 | Prdx6 | 0.107 | Old |
| 0.037459 | 0.519932 | 0.234 | 0.138 | 1 | 11 | Lmbrd2 | 0.096 | Old |
| 0.037462 | 0.083259 | 0.496 | 0.355 | 1 | 11 | D8Ertd738 | 0.141 | Old |
| 0.037467 | 0.891565 | 0.085 | 0.029 | 1 | 11 | Bet1l | 0.056 | Old |
| 0.037467 | 0.27372 | 0.085 | 0.029 | 1 | 11 | Nudt14 | 0.056 | Old |
| 0.037596 | 0.679513 | 0.262 | 0.167 | 1 | 11 | Fgfr1 | 0.095 | Old |
| 0.037621 | 0.376797 | 0.351 | 0.232 | 1 | 11 | Dnajc8 | 0.119 | Old |
| 0.037699 | 0.11059 | 0.395 | 0.254 | 1 | 11 | Ndufb8 | 0.141 | Old |

| | | | | | | | | |
|---|---|---|---|---|---|---|---|---|
| 0.037713 | 0.290214 | 0.536 | 0.391 | 1 | 11 | Gbp2 | 0.145 | Old |
| 0.03773 | 0.44152 | 0.25 | 0.152 | 1 | 11 | Isy1 | 0.098 | Old |
| 0.037759 | 0.288738 | 0.097 | 0.036 | 1 | 11 | Entpd6 | 0.061 | Old |
| 0.037792 | 0.044271 | 0.19 | 0.101 | 1 | 11 | Fam133b | 0.089 | Old |
| 0.037841 | 1.053894 | 0.073 | 0.022 | 1 | 11 | Slc27a4 | 0.051 | Old |
| 0.037841 | 0.584734 | 0.073 | 0.022 | 1 | 11 | Tfpi | 0.051 | Old |
| 0.037892 | 0.399293 | 0.242 | 0.145 | 1 | 11 | Mrps24 | 0.097 | Old |
| 0.03794 | 0.818408 | 0.149 | 0.072 | 1 | 11 | Phax | 0.077 | Old |
| 0.037971 | 0.550968 | 0.105 | 0.043 | 1 | 11 | Pmf1 | 0.062 | Old |
| 0.037978 | 0.434489 | 0.157 | 0.08 | 1 | 11 | Zmym4 | 0.077 | Old |
| 0.038013 | 0.367514 | 0.246 | 0.145 | 1 | 11 | Kdm1a | 0.101 | Old |
| 0.038062 | 0.436127 | 0.407 | 0.283 | 1 | 11 | Tcp1 | 0.124 | Old |
| 0.038065 | 0.432155 | 0.073 | 0.022 | 1 | 11 | Adam22 | 0.051 | Old |
| 0.038113 | 0.041271 | 0.06 | 0.014 | 1 | 11 | Ip6k3 | 0.046 | Old |
| 0.038144 | 0.242663 | 0.097 | 0.036 | 1 | 11 | Bbc3 | 0.061 | Old |
| 0.038156 | 1.203704 | 0.105 | 0.043 | 1 | 11 | Eef2kmt | 0.062 | Old |
| 0.038214 | 0.574868 | 0.218 | 0.13 | 1 | 11 | Fam53b | 0.088 | Old |
| 0.038286 | 0.197094 | 0.843 | 0.696 | 1 | 11 | Gm10076 | 0.147 | Old |
| 0.038291 | 0.772213 | 0.073 | 0.022 | 1 | 11 | Mmgt1 | 0.051 | Old |
| 0.038311 | 0.10086 | 0.169 | 0.087 | 1 | 11 | Capn7 | 0.082 | Old |
| 0.038343 | 0.196735 | 0.319 | 0.196 | 1 | 11 | Smim11 | 0.123 | Old |
| 0.038379 | 0.208684 | 0.21 | 0.116 | 1 | 11 | Esyt2 | 0.094 | Old |
| 0.038381 | 0.274301 | 0.177 | 0.094 | 1 | 11 | Exoc6b | 0.083 | Old |
| 0.038513 | 0.177696 | 0.21 | 0.116 | 1 | 11 | Fbxw4 | 0.094 | Old |
| 0.038614 | 0.738281 | 0.06 | 0.014 | 1 | 11 | Zmym1 | 0.046 | Old |
| 0.038622 | 0.295068 | 0.298 | 0.181 | 1 | 11 | Retreg2 | 0.117 | Old |
| 0.038636 | 1.663567 | 0.093 | 0.036 | 1 | 11 | Lonp1 | 0.057 | Old |
| 0.038715 | 0.240557 | 0.194 | 0.109 | 1 | 11 | Mrpl17 | 0.085 | Old |
| 0.038715 | 0.908675 | 0.105 | 0.043 | 1 | 11 | Fgd3 | 0.062 | Old |
| 0.038745 | 0.88553 | 0.073 | 0.022 | 1 | 11 | Gfm2 | 0.051 | Old |
| 0.038745 | 0.668242 | 0.073 | 0.022 | 1 | 11 | Gorasp1 | 0.051 | Old |
| 0.038778 | 0.317946 | 0.94 | 0.884 | 1 | 11 | Crip1 | 0.056 | Old |
| 0.038855 | 0.130284 | 0.234 | 0.138 | 1 | 11 | Xpo1 | 0.096 | Old |
| 0.038903 | 0.414092 | 0.383 | 0.254 | 1 | 11 | Cln8 | 0.129 | Old |
| 0.038909 | 0.039085 | 0.343 | 0.217 | 1 | 11 | Mob4 | 0.126 | Old |
| 0.038974 | 1.085677 | 0.073 | 0.022 | 1 | 11 | Vps37c | 0.051 | Old |
| 0.038974 | 0.498613 | 0.073 | 0.022 | 1 | 11 | D230025D1 | 0.051 | Old |
| 0.039123 | 0.349744 | 0.097 | 0.036 | 1 | 11 | Vps72 | 0.061 | Old |
| 0.039198 | 0.142155 | 0.149 | 0.072 | 1 | 11 | Atrnl1 | 0.077 | Old |
| 0.039251 | 0.084602 | 0.274 | 0.167 | 1 | 11 | Polr2g | 0.107 | Old |
| 0.039281 | 1.208503 | 0.105 | 0.043 | 1 | 11 | Nr2c2ap | 0.062 | Old |
| 0.039286 | 0.022928 | 0.125 | 0.058 | 1 | 11 | Tspan17 | 0.067 | Old |
| 0.039469 | 0.515517 | 0.23 | 0.138 | 1 | 11 | D5Ertd579 | 0.092 | Old |
| 0.039577 | 0.053379 | 0.27 | 0.159 | 1 | 11 | Ccdc47 | 0.111 | Old |
| 0.0396 | 0.204278 | 0.923 | 0.826 | 1 | 11 | H3f3a | 0.097 | Old |

| | | | | | | | | |
|---|---|---|---|---|---|---|---|---|
| 0.039616 | 0.224521 | 0.323 | 0.21 | 1 | 11 | Gm6377 | 0.113 | Old |
| 0.039687 | 0.025927 | 0.23 | 0.13 | 1 | 11 | Cdc37l1 | 0.1 | Old |
| 0.039756 | 0.061474 | 0.423 | 0.275 | 1 | 11 | Cast | 0.148 | Old |
| 0.039837 | 0.39884 | 0.137 | 0.065 | 1 | 11 | Trabd | 0.072 | Old |
| 0.039912 | 0.48954 | 0.246 | 0.145 | 1 | 11 | Hibadh | 0.101 | Old |
| 0.039913 | 0.070518 | 0.218 | 0.123 | 1 | 11 | Gabarapl1 | 0.095 | Old |
| 0.040092 | 0.285551 | 0.585 | 0.42 | 1 | 11 | Ptpn6 | 0.165 | Old |
| 0.040092 | 0.128453 | 0.117 | 0.051 | 1 | 11 | Tsga10 | 0.066 | Old |
| 0.040114 | 0.431366 | 0.169 | 0.087 | 1 | 11 | Scn1b | 0.082 | Old |
| 0.040129 | 0.578948 | 0.194 | 0.109 | 1 | 11 | Spin1 | 0.085 | Old |
| 0.040146 | 0.294635 | 0.185 | 0.101 | 1 | 11 | 0610030E2 | 0.084 | Old |
| 0.040224 | 1.02212 | 0.133 | 0.065 | 1 | 11 | Sae1 | 0.068 | Old |
| 0.040256 | 1.142764 | 0.093 | 0.036 | 1 | 11 | Gamt | 0.057 | Old |
| 0.040263 | 0.229163 | 0.569 | 0.413 | 1 | 11 | Atrx | 0.156 | Old |
| 0.040295 | 0.228577 | 0.399 | 0.261 | 1 | 11 | Vcpip1 | 0.138 | Old |
| 0.040328 | 0.129047 | 0.149 | 0.072 | 1 | 11 | Nudt19 | 0.077 | Old |
| 0.040429 | 0.668007 | 0.137 | 0.065 | 1 | 11 | Ago1 | 0.072 | Old |
| 0.040445 | 0.191193 | 0.343 | 0.225 | 1 | 11 | Pdcd5 | 0.118 | Old |
| 0.040463 | 1.209902 | 0.093 | 0.036 | 1 | 11 | Nup155 | 0.057 | Old |
| 0.040483 | 0.103147 | 0.577 | 0.384 | 1 | 11 | Selenof | 0.193 | Old |
| 0.040514 | 0.23225 | 0.242 | 0.145 | 1 | 11 | Myo1e | 0.097 | Old |
| 0.040629 | 1.000724 | 0.105 | 0.043 | 1 | 11 | Rnf167 | 0.062 | Old |
| 0.040646 | 0.110518 | 0.383 | 0.254 | 1 | 11 | Rab6a | 0.129 | Old |
| 0.040665 | 0.162018 | 0.375 | 0.239 | 1 | 11 | Comt | 0.136 | Old |
| 0.040781 | 0.2684 | 0.282 | 0.174 | 1 | 11 | Pabpn1 | 0.108 | Old |
| 0.040834 | 0.749049 | 0.343 | 0.246 | 1 | 11 | Cd209f | 0.097 | Old |
| 0.040846 | 0.20286 | 0.073 | 0.022 | 1 | 11 | Gpatch4 | 0.051 | Old |
| 0.040874 | 1.231935 | 0.125 | 0.058 | 1 | 11 | Cog5 | 0.067 | Old |
| 0.040879 | 1.236535 | 0.093 | 0.036 | 1 | 11 | Acvr2a | 0.057 | Old |
| 0.040959 | 0.804291 | 0.181 | 0.101 | 1 | 11 | Nsd2 | 0.08 | Old |
| 0.041004 | 0.24309 | 0.327 | 0.21 | 1 | 11 | Napa | 0.117 | Old |
| 0.041086 | 0.260604 | 0.266 | 0.167 | 1 | 11 | Fcna | 0.099 | Old |
| 0.041164 | 1.389039 | 0.113 | 0.051 | 1 | 11 | Il4i1 | 0.062 | Old |
| 0.041264 | 1.088991 | 0.133 | 0.065 | 1 | 11 | Dusp23 | 0.068 | Old |
| 0.041372 | 0.376323 | 0.129 | 0.058 | 1 | 11 | Nomo1 | 0.071 | Old |
| 0.041391 | 0.579454 | 0.117 | 0.051 | 1 | 11 | Narf | 0.066 | Old |
| 0.041416 | 0.812877 | 0.125 | 0.058 | 1 | 11 | Mrpl27 | 0.067 | Old |
| 0.041492 | 0.212178 | 0.395 | 0.261 | 1 | 11 | Acap2 | 0.134 | Old |
| 0.041509 | 1.135196 | 0.093 | 0.036 | 1 | 11 | Fam45a | 0.057 | Old |
| 0.041509 | 1.10374 | 0.093 | 0.036 | 1 | 11 | Rbsn | 0.057 | Old |
| 0.041513 | 0.213388 | 0.177 | 0.094 | 1 | 11 | Parp4 | 0.083 | Old |
| 0.041545 | 1.554736 | 0.113 | 0.051 | 1 | 11 | Gucd1 | 0.062 | Old |
| 0.04158 | 0.525662 | 0.117 | 0.051 | 1 | 11 | Fuom | 0.066 | Old |
| 0.041674 | 0.28428 | 0.677 | 0.536 | 1 | 11 | Gns | 0.141 | Old |
| 0.041721 | 1.257789 | 0.093 | 0.036 | 1 | 11 | Ogfod1 | 0.057 | Old |

| | | | | | | | | |
|---|---|---|---|---|---|---|---|---|
| 0.041735 | 0.170908 | 0.323 | 0.21 | 1 | 11 | Ccnl2 | 0.113 | Old |
| 0.041736 | 0.711296 | 0.113 | 0.051 | 1 | 11 | Prmt2 | 0.062 | Old |
| 0.041737 | 0.406705 | 0.218 | 0.13 | 1 | 11 | Usp24 | 0.088 | Old |
| 0.041757 | 0.248071 | 0.157 | 0.08 | 1 | 11 | Micu2 | 0.077 | Old |
| 0.041841 | 0.026573 | 0.371 | 0.232 | 1 | 11 | Leprotl1 | 0.139 | Old |
| 0.041922 | 0.01313 | 0.524 | 0.355 | 1 | 11 | Cd86 | 0.169 | Old |
| 0.041928 | 0.988876 | 0.113 | 0.051 | 1 | 11 | Tfam | 0.062 | Old |
| 0.04195 | 0.16854 | 0.286 | 0.174 | 1 | 11 | Alkbh5 | 0.112 | Old |
| 0.041989 | 0.147809 | 0.27 | 0.159 | 1 | 11 | Gsk3a | 0.111 | Old |
| 0.042035 | 0.305032 | 0.815 | 0.63 | 1 | 11 | Clec2d | 0.185 | Old |
| 0.042048 | 0.084856 | 0.323 | 0.203 | 1 | 11 | Rab18 | 0.12 | Old |
| 0.042075 | 0.109779 | 0.137 | 0.065 | 1 | 11 | Fzr1 | 0.072 | Old |
| 0.042087 | 0.050524 | 0.637 | 0.471 | 1 | 11 | Hnrnpab | 0.166 | Old |
| 0.0421 | 0.869458 | 0.145 | 0.072 | 1 | 11 | Copg1 | 0.073 | Old |
| 0.042165 | 0.27925 | 0.552 | 0.413 | 1 | 11 | Atp5b | 0.139 | Old |
| 0.042167 | 0.14021 | 0.185 | 0.101 | 1 | 11 | Zfp622 | 0.084 | Old |
| 0.042341 | 0.432655 | 0.117 | 0.051 | 1 | 11 | 2210016L2 | 0.066 | Old |
| 0.042354 | 1.73636 | 0.141 | 0.072 | 1 | 11 | Fam220a | 0.069 | Old |
| 0.042364 | 0.192926 | 0.339 | 0.217 | 1 | 11 | Gnpda1 | 0.122 | Old |
| 0.042372 | 0.073632 | 0.56 | 0.399 | 1 | 11 | Ccdc88a | 0.161 | Old |
| 0.04238 | 1.390884 | 0.081 | 0.029 | 1 | 11 | Zfp830 | 0.052 | Old |
| 0.042416 | 0.234057 | 0.722 | 0.543 | 1 | 11 | Sub1 | 0.179 | Old |
| 0.042484 | 0.511372 | 0.194 | 0.109 | 1 | 11 | Tpd52l2 | 0.085 | Old |
| 0.042484 | 0.431979 | 0.181 | 0.101 | 1 | 11 | Kars | 0.08 | Old |
| 0.042528 | 0.058942 | 0.242 | 0.145 | 1 | 11 | Dpp8 | 0.097 | Old |
| 0.042644 | 0.238782 | 0.266 | 0.167 | 1 | 11 | Ncoa1 | 0.099 | Old |
| 0.042749 | 0.175709 | 0.157 | 0.08 | 1 | 11 | Hspbp1 | 0.077 | Old |
| 0.042811 | 0.119415 | 0.992 | 0.978 | 1 | 11 | mt-Cytb | 0.014 | Old |
| 0.04282 | 0.104877 | 0.266 | 0.159 | 1 | 11 | Map2k1 | 0.107 | Old |
| 0.04286 | 0.507924 | 0.185 | 0.101 | 1 | 11 | Zfp326 | 0.084 | Old |
| 0.042889 | 0.371073 | 0.238 | 0.145 | 1 | 11 | Abhd17b | 0.093 | Old |
| 0.04289 | 0.427466 | 0.125 | 0.058 | 1 | 11 | Slc39a13 | 0.067 | Old |
| 0.042891 | 0.119979 | 0.427 | 0.29 | 1 | 11 | Sdhc | 0.137 | Old |
| 0.042938 | 0.118011 | 0.194 | 0.109 | 1 | 11 | Gbp4 | 0.085 | Old |
| 0.043155 | 0.15986 | 0.673 | 0.507 | 1 | 11 | Jak1 | 0.166 | Old |
| 0.04318 | 0.209051 | 0.488 | 0.333 | 1 | 11 | Lrrc58 | 0.155 | Old |
| 0.043224 | 0.408156 | 0.448 | 0.319 | 1 | 11 | Ppp1r15b | 0.129 | Old |
| 0.043377 | 0.277153 | 0.472 | 0.319 | 1 | 11 | Copz1 | 0.153 | Old |
| 0.043425 | 0.182718 | 0.411 | 0.283 | 1 | 11 | Tbcb | 0.128 | Old |
| 0.04343 | 0.009879 | 0.504 | 0.341 | 1 | 11 | Usp9x | 0.163 | Old |
| 0.043527 | 0.212329 | 0.323 | 0.203 | 1 | 11 | Trim56 | 0.12 | Old |
| 0.043645 | 0.461388 | 0.327 | 0.217 | 1 | 11 | Cox20 | 0.11 | Old |
| 0.043707 | 0.0338 | 0.343 | 0.217 | 1 | 11 | Tle4 | 0.126 | Old |
| 0.043729 | 2.286486 | 0.044 | 0.007 | 1 | 11 | Aldh4a1 | 0.037 | Old |
| 0.043729 | 2.089305 | 0.044 | 0.007 | 1 | 11 | Ranbp6 | 0.037 | Old |

| | | | | | | | | |
|---|---|---|---|---|---|---|---|---|
| 0.043739 | 0.247221 | 0.383 | 0.254 | 1 | 11 | Nars | 0.129 | Old |
| 0.043759 | 0.013398 | 0.286 | 0.174 | 1 | 11 | 2510039O1 | 0.112 | Old |
| 0.043762 | 0.229342 | 0.157 | 0.08 | 1 | 11 | Cdk8 | 0.077 | Old |
| 0.043892 | 0.597142 | 0.093 | 0.036 | 1 | 11 | Kdm5b | 0.057 | Old |
| 0.043929 | 0.33497 | 0.125 | 0.058 | 1 | 11 | Pbx1 | 0.067 | Old |
| 0.044061 | 2.444049 | 0.044 | 0.007 | 1 | 11 | Nepro | 0.037 | Old |
| 0.044061 | 2.274932 | 0.044 | 0.007 | 1 | 11 | Ppcs | 0.037 | Old |
| 0.044114 | 1.211974 | 0.093 | 0.036 | 1 | 11 | Eif2b3 | 0.057 | Old |
| 0.044201 | 0.206546 | 0.339 | 0.225 | 1 | 11 | Tubb2a | 0.114 | Old |
| 0.044234 | 0.863828 | 0.153 | 0.08 | 1 | 11 | Cds2 | 0.073 | Old |
| 0.044276 | 0.660391 | 0.105 | 0.043 | 1 | 11 | Yif1a | 0.062 | Old |
| 0.044365 | 0.710953 | 0.161 | 0.087 | 1 | 11 | Ube3b | 0.074 | Old |
| 0.044366 | 2.218166 | 0.056 | 0.014 | 1 | 11 | Pex11b | 0.042 | Old |
| 0.044395 | 2.138781 | 0.044 | 0.007 | 1 | 11 | Gm1720 | 0.037 | Old |
| 0.044395 | 2.134589 | 0.044 | 0.007 | 1 | 11 | Thap4 | 0.037 | Old |
| 0.044402 | 0.257997 | 0.298 | 0.196 | 1 | 11 | Acbd3 | 0.102 | Old |
| 0.044446 | 0.281575 | 0.274 | 0.174 | 1 | 11 | Etfa | 0.1 | Old |
| 0.044486 | 0.665864 | 0.105 | 0.043 | 1 | 11 | Steap3 | 0.062 | Old |
| 0.044621 | 0.180594 | 0.923 | 0.775 | 1 | 11 | Vim | 0.148 | Old |
| 0.044628 | 0.008121 | 0.54 | 0.377 | 1 | 11 | Atp5j | 0.163 | Old |
| 0.044639 | 0.310023 | 0.194 | 0.109 | 1 | 11 | Pisd | 0.085 | Old |
| 0.044731 | 2.156003 | 0.044 | 0.007 | 1 | 11 | Cd1d1 | 0.037 | Old |
| 0.044731 | 2.080743 | 0.044 | 0.007 | 1 | 11 | Oxnad1 | 0.037 | Old |
| 0.044834 | 0.251559 | 0.202 | 0.116 | 1 | 11 | Tmem65 | 0.086 | Old |
| 0.044863 | 0.894869 | 0.181 | 0.101 | 1 | 11 | Znrf1 | 0.08 | Old |
| 0.044865 | 0.057526 | 0.464 | 0.304 | 1 | 11 | Tnrc18 | 0.16 | Old |
| 0.044933 | 0.282483 | 0.153 | 0.08 | 1 | 11 | Naa60 | 0.073 | Old |
| 0.045028 | 0.441898 | 0.21 | 0.123 | 1 | 11 | Pip5k1a | 0.087 | Old |
| 0.045069 | 1.952503 | 0.044 | 0.007 | 1 | 11 | Dppa3 | 0.037 | Old |
| 0.045105 | 0.19814 | 0.351 | 0.225 | 1 | 11 | Pcnp | 0.126 | Old |
| 0.045188 | 0.179019 | 0.181 | 0.101 | 1 | 11 | Ndfip1 | 0.08 | Old |
| 0.045213 | 0.672874 | 0.266 | 0.167 | 1 | 11 | Cnot4 | 0.099 | Old |
| 0.045234 | 1.167401 | 0.081 | 0.029 | 1 | 11 | Arfgap1 | 0.052 | Old |
| 0.045258 | 0.223984 | 0.319 | 0.203 | 1 | 11 | Tor1aip2 | 0.116 | Old |
| 0.045284 | 0.136691 | 0.355 | 0.246 | 1 | 11 | Tnfaip2 | 0.109 | Old |
| 0.045337 | 0.154331 | 0.105 | 0.043 | 1 | 11 | Ss18l2 | 0.062 | Old |
| 0.045409 | 1.775281 | 0.044 | 0.007 | 1 | 11 | Bcorl1 | 0.037 | Old |
| 0.045409 | 1.51228 | 0.044 | 0.007 | 1 | 11 | Zfp934 | 0.037 | Old |
| 0.045409 | 1.455064 | 0.044 | 0.007 | 1 | 11 | Tmtc4 | 0.037 | Old |
| 0.045518 | 0.717793 | 0.113 | 0.051 | 1 | 11 | Slc16a3 | 0.062 | Old |
| 0.045562 | 0.411067 | 0.27 | 0.167 | 1 | 11 | Gpr146 | 0.103 | Old |
| 0.045725 | 0.921144 | 0.081 | 0.029 | 1 | 11 | Ermard | 0.052 | Old |
| 0.045751 | 1.481935 | 0.044 | 0.007 | 1 | 11 | Zfp946 | 0.037 | Old |
| 0.045751 | 1.042891 | 0.044 | 0.007 | 1 | 11 | Hrh1 | 0.037 | Old |
| 0.045767 | 0.118869 | 0.105 | 0.043 | 1 | 11 | Ciapin1 | 0.062 | Old |

| | | | | | | | | |
|---|---|---|---|---|---|---|---|---|
| 0.045779 | 2.655968 | 0.069 | 0.022 | 1 | 11 | Zfp938 | 0.047 | Old |
| 0.045869 | 0.221982 | 0.323 | 0.21 | 1 | 11 | Noc2l | 0.113 | Old |
| 0.045887 | 0.072698 | 0.415 | 0.275 | 1 | 11 | Nav1 | 0.14 | Old |
| 0.04591 | 0.328421 | 0.266 | 0.167 | 1 | 11 | Mark3 | 0.099 | Old |
| 0.045933 | 0.735665 | 0.113 | 0.051 | 1 | 11 | Atg4d | 0.062 | Old |
| 0.045941 | 0.775215 | 0.149 | 0.08 | 1 | 11 | Phactr4 | 0.069 | Old |
| 0.045949 | 0.278283 | 0.165 | 0.087 | 1 | 11 | 4833439L1 | 0.078 | Old |
| 0.045962 | 0.277318 | 0.198 | 0.109 | 1 | 11 | Dnajc14 | 0.089 | Old |
| 0.045996 | 0.036234 | 0.407 | 0.268 | 1 | 11 | Glyr1 | 0.139 | Old |
| 0.046043 | 0.019957 | 0.46 | 0.319 | 1 | 11 | Smap2 | 0.141 | Old |
| 0.046073 | 0.33313 | 0.194 | 0.109 | 1 | 11 | Hars | 0.085 | Old |
| 0.046096 | 1.343424 | 0.044 | 0.007 | 1 | 11 | Kbtbd4 | 0.037 | Old |
| 0.046096 | 1.241098 | 0.044 | 0.007 | 1 | 11 | Tbl2 | 0.037 | Old |
| 0.046157 | 0.482816 | 0.093 | 0.036 | 1 | 11 | Qser1 | 0.057 | Old |
| 0.046182 | 0.247756 | 0.262 | 0.167 | 1 | 11 | Cope | 0.095 | Old |
| 0.046191 | 0.20405 | 0.177 | 0.094 | 1 | 11 | Ten1 | 0.083 | Old |
| 0.04622 | 1.166167 | 0.081 | 0.029 | 1 | 11 | Angptl4 | 0.052 | Old |
| 0.04632 | 1.718251 | 0.069 | 0.022 | 1 | 11 | Jrkl | 0.047 | Old |
| 0.046334 | 0.241983 | 0.448 | 0.312 | 1 | 11 | Atp5o.1 | 0.136 | Old |
| 0.046379 | 0.485467 | 0.145 | 0.072 | 1 | 11 | Zfpl1 | 0.073 | Old |
| 0.046427 | 0.266427 | 0.234 | 0.145 | 1 | 11 | Rab22a | 0.089 | Old |
| 0.046442 | 1.150107 | 0.044 | 0.007 | 1 | 11 | Tnfsf13os | 0.037 | Old |
| 0.046442 | 0.533649 | 0.044 | 0.007 | 1 | 11 | Cdk20 | 0.037 | Old |
| 0.046442 | 0.499717 | 0.044 | 0.007 | 1 | 11 | Ift74 | 0.037 | Old |
| 0.046445 | 2.436553 | 0.056 | 0.014 | 1 | 11 | 5730480H( | 0.042 | Old |
| 0.046627 | 0.212121 | 0.593 | 0.428 | 1 | 11 | Laptm4a | 0.165 | Old |
| 0.046646 | 0.506902 | 0.165 | 0.087 | 1 | 11 | Med17 | 0.078 | Old |
| 0.046658 | 0.203626 | 0.395 | 0.254 | 1 | 11 | Dram2 | 0.141 | Old |
| 0.046732 | 0.239024 | 0.21 | 0.123 | 1 | 11 | Gm49336 | 0.087 | Old |
| 0.046751 | 0.081954 | 0.145 | 0.072 | 1 | 11 | Zdhhc5 | 0.073 | Old |
| 0.046858 | 0.174635 | 0.105 | 0.043 | 1 | 11 | Grcc10 | 0.062 | Old |
| 0.046866 | 1.825444 | 0.069 | 0.022 | 1 | 11 | Gm7694 | 0.047 | Old |
| 0.04687 | 0.093152 | 0.137 | 0.065 | 1 | 11 | Mrps18a | 0.072 | Old |
| 0.04687 | 0.038873 | 0.177 | 0.094 | 1 | 11 | Cntrl | 0.083 | Old |
| 0.046921 | 3.187926 | 0.028 | 0 | 1 | 11 | Pde8b | 0.028 | Old |
| 0.046921 | 2.995433 | 0.028 | 0 | 1 | 11 | Gprc5b | 0.028 | Old |
| 0.046921 | 2.993342 | 0.028 | 0 | 1 | 11 | Zfp810 | 0.028 | Old |
| 0.046921 | 2.954306 | 0.028 | 0 | 1 | 11 | Acad12 | 0.028 | Old |
| 0.046921 | 2.85693 | 0.028 | 0 | 1 | 11 | AC168977. | 0.028 | Old |
| 0.046921 | 2.681538 | 0.028 | 0 | 1 | 11 | Hells | 0.028 | Old |
| 0.046921 | 2.659789 | 0.028 | 0 | 1 | 11 | Zfp953 | 0.028 | Old |
| 0.046921 | 2.640897 | 0.028 | 0 | 1 | 11 | Nat8l | 0.028 | Old |
| 0.046921 | 2.580802 | 0.028 | 0 | 1 | 11 | Pdyn | 0.028 | Old |
| 0.046921 | 2.54309 | 0.028 | 0 | 1 | 11 | Mthfd1l | 0.028 | Old |
| 0.046921 | 2.533747 | 0.028 | 0 | 1 | 11 | B930036N1 | 0.028 | Old |

| | | | | | | | | |
|---|---|---|---|---|---|---|---|---|
| 0.046921 | 2.521136 | 0.028 | 0 | 1 | 11 | Dsel | 0.028 | Old |
| 0.046921 | 2.418606 | 0.028 | 0 | 1 | 11 | Grhpr | 0.028 | Old |
| 0.046921 | 2.307975 | 0.028 | 0 | 1 | 11 | E030030I0 | 0.028 | Old |
| 0.046921 | 2.287388 | 0.028 | 0 | 1 | 11 | Tmem14a | 0.028 | Old |
| 0.046921 | 2.280015 | 0.028 | 0 | 1 | 11 | Plcd1 | 0.028 | Old |
| 0.046921 | 2.254468 | 0.028 | 0 | 1 | 11 | A930033H1 | 0.028 | Old |
| 0.046921 | 2.225687 | 0.028 | 0 | 1 | 11 | Slc7a6 | 0.028 | Old |
| 0.046921 | 2.224493 | 0.028 | 0 | 1 | 11 | E330009J0 | 0.028 | Old |
| 0.046921 | 2.067653 | 0.028 | 0 | 1 | 11 | Gm12089 | 0.028 | Old |
| 0.046921 | 2.063254 | 0.028 | 0 | 1 | 11 | Gale | 0.028 | Old |
| 0.046921 | 1.969402 | 0.028 | 0 | 1 | 11 | Aaas | 0.028 | Old |
| 0.046921 | 1.959699 | 0.028 | 0 | 1 | 11 | Fbxl8 | 0.028 | Old |
| 0.046972 | 1.011473 | 0.081 | 0.029 | 1 | 11 | Gon7 | 0.052 | Old |
| 0.046988 | 0.717918 | 0.101 | 0.043 | 1 | 11 | Prr3 | 0.058 | Old |
| 0.047141 | 2.174005 | 0.069 | 0.022 | 1 | 11 | Cd300lf | 0.047 | Old |
| 0.047173 | 0.336951 | 0.125 | 0.058 | 1 | 11 | Dlat | 0.067 | Old |
| 0.047198 | 0.642613 | 0.133 | 0.065 | 1 | 11 | Iars | 0.068 | Old |
| 0.047213 | 1.176379 | 0.173 | 0.101 | 1 | 11 | Mta1 | 0.072 | Old |
| 0.047251 | 0.52109 | 0.302 | 0.203 | 1 | 11 | Limd2 | 0.099 | Old |
| 0.047285 | 0.21807 | 0.157 | 0.08 | 1 | 11 | St6galnac6 | 0.077 | Old |
| 0.047315 | 0.219667 | 0.145 | 0.072 | 1 | 11 | Faap20 | 0.073 | Old |
| 0.047395 | 0.11144 | 0.661 | 0.449 | 1 | 11 | Sec61b | 0.212 | Old |
| 0.047435 | 0.234461 | 0.25 | 0.152 | 1 | 11 | Dctn3 | 0.098 | Old |
| 0.047628 | 0.362274 | 0.194 | 0.109 | 1 | 11 | Map3k5 | 0.085 | Old |
| 0.047636 | 0.412209 | 0.153 | 0.08 | 1 | 11 | Zmym2 | 0.073 | Old |
| 0.047689 | 0.755833 | 0.254 | 0.159 | 1 | 11 | Smu1 | 0.095 | Old |
| 0.0478 | 0.593452 | 0.093 | 0.036 | 1 | 11 | Tmem199 | 0.057 | Old |
| 0.047837 | 1.119706 | 0.113 | 0.051 | 1 | 11 | Mrpl40 | 0.062 | Old |
| 0.047838 | 0.235165 | 0.56 | 0.413 | 1 | 11 | Atp6v1f | 0.147 | Old |
| 0.047892 | 0.45691 | 0.206 | 0.123 | 1 | 11 | Rnf11 | 0.083 | Old |
| 0.047916 | 0.339627 | 0.198 | 0.109 | 1 | 11 | Cacul1 | 0.089 | Old |
| 0.047937 | 0.270384 | 0.306 | 0.196 | 1 | 11 | Uggt1 | 0.11 | Old |
| 0.04799 | 0.776363 | 0.081 | 0.029 | 1 | 11 | Ankrd49 | 0.052 | Old |
| 0.04799 | 0.514671 | 0.081 | 0.029 | 1 | 11 | Zmym3 | 0.052 | Old |
| 0.048006 | 0.736043 | 0.21 | 0.123 | 1 | 11 | Tmem86a | 0.087 | Old |
| 0.048006 | 0.454647 | 0.21 | 0.123 | 1 | 11 | Psen2 | 0.087 | Old |
| 0.048053 | 0.688627 | 0.113 | 0.051 | 1 | 11 | H2-DMb2 | 0.062 | Old |
| 0.048247 | 0.71428 | 0.081 | 0.029 | 1 | 11 | Dzip3 | 0.052 | Old |
| 0.048255 | 1.32417 | 0.069 | 0.022 | 1 | 11 | Snhg20 | 0.047 | Old |
| 0.048298 | 1.184946 | 0.129 | 0.065 | 1 | 11 | Rexo4 | 0.064 | Old |
| 0.048307 | 0.636786 | 0.008 | 0.036 | 1 | 11 | Spp1 | -0.028 | Old |
| 0.048331 | 0.382122 | 0.339 | 0.217 | 1 | 11 | Agtrap | 0.122 | Old |
| 0.048375 | 0.214507 | 0.488 | 0.333 | 1 | 11 | Rnf13 | 0.155 | Old |
| 0.048459 | 0.370739 | 0.145 | 0.072 | 1 | 11 | Eftud2 | 0.073 | Old |
| 0.048506 | 0.66075 | 0.081 | 0.029 | 1 | 11 | Pop4 | 0.052 | Old |

| | | | | | | | | |
|---|---|---|---|---|---|---|---|---|
| 0.048606 | 1.599345 | 0.056 | 0.014 | 1 | 11 | Gcdh | 0.042 | Old |
| 0.04861 | 0.329013 | 0.125 | 0.058 | 1 | 11 | Utp6 | 0.067 | Old |
| 0.04868 | 0.228233 | 0.202 | 0.116 | 1 | 11 | Ggt5 | 0.086 | Old |
| 0.048705 | 0.500498 | 0.113 | 0.051 | 1 | 11 | Slc39a14 | 0.062 | Old |
| 0.048743 | 0.12467 | 0.206 | 0.116 | 1 | 11 | Tmem170b | 0.09 | Old |
| 0.048766 | 0.576381 | 0.081 | 0.029 | 1 | 11 | Abcd4 | 0.052 | Old |
| 0.04891 | 0.14321 | 0.214 | 0.123 | 1 | 11 | Mbd6 | 0.091 | Old |
| 0.049027 | 0.939498 | 0.081 | 0.029 | 1 | 11 | Mrpl22 | 0.052 | Old |
| 0.04904 | 0.018738 | 0.145 | 0.072 | 1 | 11 | Smagp | 0.073 | Old |
| 0.049063 | 0.408869 | 0.565 | 0.435 | 1 | 11 | Gapdh | 0.13 | Old |
| 0.049104 | 0.677139 | 0.069 | 0.022 | 1 | 11 | Zfp597 | 0.047 | Old |
| 0.049238 | 1.455707 | 0.056 | 0.014 | 1 | 11 | Pstk | 0.042 | Old |
| 0.049301 | 0.104312 | 0.452 | 0.304 | 1 | 11 | Tsc22d4 | 0.148 | Old |
| 0.049447 | 0.312946 | 0.125 | 0.058 | 1 | 11 | Arhgef11 | 0.067 | Old |
| 0.049466 | 0.094032 | 0.302 | 0.196 | 1 | 11 | 1810058I24 | 0.106 | Old |
| 0.049557 | 1.33767 | 0.056 | 0.014 | 1 | 11 | Sccpdh | 0.042 | Old |
| 0.049678 | 0.869906 | 0.069 | 0.022 | 1 | 11 | Igf2r | 0.047 | Old |
| 0.049678 | 0.668507 | 0.069 | 0.022 | 1 | 11 | Aamdc | 0.047 | Old |
| 0.049698 | 0.514107 | 0.206 | 0.123 | 1 | 11 | Ppp2r5e | 0.083 | Old |
| 0.049701 | 0.16801 | 0.29 | 0.188 | 1 | 11 | Rps6ka1 | 0.102 | Old |
| 0.049743 | 0.932344 | 0.101 | 0.043 | 1 | 11 | Ttc17 | 0.058 | Old |
| 0.049794 | 0.495556 | 0.379 | 0.261 | 1 | 11 | Usp15 | 0.118 | Old |
| 0.049878 | 1.535168 | 0.056 | 0.014 | 1 | 11 | Pola2 | 0.042 | Old |
| 0.049878 | 1.421631 | 0.056 | 0.014 | 1 | 11 | Srcap | 0.042 | Old |
| 0.049895 | 0.150249 | 0.165 | 0.087 | 1 | 11 | Chsy1 | 0.078 | Old |
| 0.049895 | 0.042229 | 0.165 | 0.087 | 1 | 11 | Txndc11 | 0.078 | Old |
| 0.049957 | 0.344779 | 0.181 | 0.101 | 1 | 11 | Ilvbl | 0.08 | Old |
| 0.050082 | 0.126708 | 0.081 | 0.029 | 1 | 11 | Pcyt2 | 0.052 | Old |
| 0.05018 | 0.139378 | 0.69 | 0.522 | 1 | 11 | Atpif1 | 0.168 | Old |
| 0.050215 | 1.157873 | 0.101 | 0.043 | 1 | 11 | Src | 0.058 | Old |
| 0.050215 | 0.219489 | 0.173 | 0.094 | 1 | 11 | Lypla1 | 0.079 | Old |
| 0.050249 | 0.159061 | 0.524 | 0.377 | 1 | 11 | Ptpra | 0.147 | Old |
| 0.05038 | 0.317135 | 0.169 | 0.094 | 1 | 11 | Stam2 | 0.075 | Old |
| 0.050461 | 0.646677 | 0.214 | 0.13 | 1 | 11 | Mrpl3 | 0.084 | Old |
| 0.050524 | 1.616829 | 0.056 | 0.014 | 1 | 11 | Ptcd1 | 0.042 | Old |
| 0.050524 | 0.735753 | 0.056 | 0.014 | 1 | 11 | Sorbs3 | 0.042 | Old |
| 0.050573 | 0.626335 | 0.246 | 0.152 | 1 | 11 | Nemf | 0.094 | Old |
| 0.050599 | 0.758582 | 0.141 | 0.072 | 1 | 11 | Zcchc17 | 0.069 | Old |
| 0.050617 | 0.492799 | 0.081 | 0.029 | 1 | 11 | Ap3s2 | 0.052 | Old |
| 0.050622 | 0.131205 | 0.532 | 0.355 | 1 | 11 | Zfp706 | 0.177 | Old |
| 0.050669 | 0.198788 | 0.153 | 0.08 | 1 | 11 | Ube4b | 0.073 | Old |
| 0.050689 | 0.381426 | 0.359 | 0.239 | 1 | 11 | Pgap2 | 0.12 | Old |
| 0.050763 | 0.261498 | 0.218 | 0.13 | 1 | 11 | Maf1 | 0.088 | Old |
| 0.050841 | 0.587497 | 0.069 | 0.022 | 1 | 11 | Mrpl49 | 0.047 | Old |
| 0.05085 | 0.141136 | 0.056 | 0.014 | 1 | 11 | Adck5 | 0.042 | Old |

| | | | | | | | | |
|---|---|---|---|---|---|---|---|---|
| 0.05085 | 0.139813 | 0.056 | 0.014 | 1 | 11 | Tbcc | 0.042 | Old |
| 0.051021 | 0.178791 | 0.754 | 0.601 | 1 | 11 | Bri3 | 0.153 | Old |
| 0.051067 | 0.853654 | 0.121 | 0.058 | 1 | 11 | Hilpda | 0.063 | Old |
| 0.051134 | 0.254439 | 0.173 | 0.094 | 1 | 11 | Nasp | 0.079 | Old |
| 0.051136 | 1.034352 | 0.069 | 0.022 | 1 | 11 | Ttc38 | 0.047 | Old |
| 0.051136 | 0.894611 | 0.069 | 0.022 | 1 | 11 | Rpp30 | 0.047 | Old |
| 0.051136 | 0.641357 | 0.069 | 0.022 | 1 | 11 | St3gal2 | 0.047 | Old |
| 0.051207 | 0.676793 | 0.141 | 0.072 | 1 | 11 | Rapgef1 | 0.069 | Old |
| 0.051228 | 0.313662 | 0.335 | 0.217 | 1 | 11 | Cdipt | 0.118 | Old |
| 0.051411 | 0.868106 | 0.101 | 0.043 | 1 | 11 | Sgsh | 0.058 | Old |
| 0.051561 | 0.34869 | 0.206 | 0.123 | 1 | 11 | Ube2d1 | 0.083 | Old |
| 0.051614 | 0.015349 | 0.246 | 0.145 | 1 | 11 | Ireb2 | 0.101 | Old |
| 0.051653 | 0.836228 | 0.101 | 0.043 | 1 | 11 | Slc39a3 | 0.058 | Old |
| 0.051683 | 0.88334 | 0.169 | 0.094 | 1 | 11 | Wdr44 | 0.075 | Old |
| 0.051699 | 0.299597 | 0.456 | 0.312 | 1 | 11 | Nop10 | 0.144 | Old |
| 0.05171 | 0.77568 | 0.355 | 0.261 | 1 | 11 | Rgs1 | 0.094 | Old |
| 0.051729 | 0.808266 | 0.069 | 0.022 | 1 | 11 | Zfp511 | 0.047 | Old |
| 0.051766 | 0.366684 | 0.218 | 0.13 | 1 | 11 | Slc25a51 | 0.088 | Old |
| 0.051791 | 0.460835 | 0.161 | 0.087 | 1 | 11 | Timm44 | 0.074 | Old |
| 0.051803 | 0.550671 | 0.214 | 0.13 | 1 | 11 | Atp13a1 | 0.084 | Old |
| 0.051838 | 0.671801 | 0.056 | 0.014 | 1 | 11 | Cluap1 | 0.042 | Old |
| 0.051838 | 0.5686 | 0.056 | 0.014 | 1 | 11 | Dtnb | 0.042 | Old |
| 0.051862 | 0.000604 | 0.177 | 0.094 | 1 | 11 | Entpd7 | 0.083 | Old |
| 0.051884 | 0.258791 | 0.23 | 0.138 | 1 | 11 | Hcfc1 | 0.092 | Old |
| 0.051975 | 0.010258 | 0.081 | 0.029 | 1 | 11 | Hbegf | 0.052 | Old |
| 0.052021 | 0.243405 | 0.339 | 0.232 | 1 | 11 | Cdk11b | 0.107 | Old |
| 0.052031 | 0.22843 | 0.847 | 0.812 | 1 | 11 | Tmem176b | 0.035 | Old |
| 0.052076 | 0.189238 | 0.113 | 0.051 | 1 | 11 | Gpkow | 0.062 | Old |
| 0.052126 | 0.408662 | 0.181 | 0.101 | 1 | 11 | Rnf220 | 0.08 | Old |
| 0.052164 | 0.176376 | 0.484 | 0.333 | 1 | 11 | Zfp207 | 0.151 | Old |
| 0.052234 | 0.7621 | 0.141 | 0.072 | 1 | 11 | Pld2 | 0.069 | Old |
| 0.052269 | 0.028651 | 0.56 | 0.399 | 1 | 11 | Tmem59 | 0.161 | Old |
| 0.052348 | 0.439621 | 0.21 | 0.123 | 1 | 11 | Etv1 | 0.087 | Old |
| 0.052495 | 0.24579 | 0.181 | 0.101 | 1 | 11 | Ctbp2 | 0.08 | Old |
| 0.052506 | 0.277901 | 0.056 | 0.014 | 1 | 11 | Prdm15 | 0.042 | Old |
| 0.052506 | 0.164597 | 0.056 | 0.014 | 1 | 11 | Rfc5 | 0.042 | Old |
| 0.052578 | 0.283972 | 0.298 | 0.188 | 1 | 11 | Mff | 0.11 | Old |
| 0.052617 | 0.106775 | 0.399 | 0.275 | 1 | 11 | Ttyh3 | 0.124 | Old |
| 0.052776 | 0.620218 | 0.206 | 0.123 | 1 | 11 | Map3k2 | 0.083 | Old |
| 0.052823 | 0.936876 | 0.089 | 0.036 | 1 | 11 | Dohh | 0.053 | Old |
| 0.052829 | 0.134329 | 0.431 | 0.283 | 1 | 11 | Lamtor2 | 0.148 | Old |
| 0.052913 | 0.144362 | 0.488 | 0.333 | 1 | 11 | Gng2 | 0.155 | Old |
| 0.052933 | 0.2712 | 0.069 | 0.022 | 1 | 11 | Socs5 | 0.047 | Old |
| 0.052955 | 0.412108 | 0.331 | 0.217 | 1 | 11 | 1110004F1 | 0.114 | Old |
| 0.052972 | 0.345923 | 0.278 | 0.181 | 1 | 11 | Klc1 | 0.097 | Old |

| | | | | | | | | |
|---|---|---|---|---|---|---|---|---|
| 0.053016 | 0.148145 | 0.363 | 0.254 | 1 | 11 | Rap2c | 0.109 | Old |
| 0.053041 | 0.164324 | 0.536 | 0.377 | 1 | 11 | Ndufb7 | 0.159 | Old |
| 0.053052 | 0.3363 | 0.149 | 0.08 | 1 | 11 | Fam208b | 0.069 | Old |
| 0.053077 | 0.235068 | 0.19 | 0.109 | 1 | 11 | Ddit3 | 0.081 | Old |
| 0.053098 | 0.575976 | 0.198 | 0.116 | 1 | 11 | Ldah | 0.082 | Old |
| 0.053128 | 0.259538 | 0.347 | 0.225 | 1 | 11 | Cdk6 | 0.122 | Old |
| 0.053129 | 0.224271 | 0.218 | 0.13 | 1 | 11 | Tspan4 | 0.088 | Old |
| 0.053166 | 0.070031 | 0.343 | 0.225 | 1 | 11 | Ndufs5 | 0.118 | Old |
| 0.053252 | 0.241229 | 0.153 | 0.08 | 1 | 11 | Rtl5 | 0.073 | Old |
| 0.053256 | 0.73904 | 0.149 | 0.08 | 1 | 11 | Fyn | 0.069 | Old |
| 0.053256 | 0.650856 | 0.149 | 0.08 | 1 | 11 | Nacc1 | 0.069 | Old |
| 0.053262 | 0.251981 | 0.835 | 0.717 | 1 | 11 | Capzb | 0.118 | Old |
| 0.053282 | 0.220134 | 0.577 | 0.399 | 1 | 11 | Uqcrb | 0.178 | Old |
| 0.053375 | 0.452524 | 0.101 | 0.043 | 1 | 11 | Ccdc107 | 0.058 | Old |
| 0.053375 | 0.088806 | 0.101 | 0.043 | 1 | 11 | Kdsr | 0.058 | Old |
| 0.053381 | 0.159783 | 0.407 | 0.275 | 1 | 11 | Sf3b6 | 0.132 | Old |
| 0.053395 | 0.313721 | 0.21 | 0.123 | 1 | 11 | Smad2 | 0.087 | Old |
| 0.053615 | 0.313251 | 0.133 | 0.065 | 1 | 11 | Rprd1b | 0.068 | Old |
| 0.053616 | 0.434511 | 0.181 | 0.101 | 1 | 11 | Nsmce1 | 0.08 | Old |
| 0.053851 | 0.164714 | 0.069 | 0.022 | 1 | 11 | Uck1 | 0.047 | Old |
| 0.053876 | 0.375931 | 0.101 | 0.043 | 1 | 11 | Phtf2 | 0.058 | Old |
| 0.05391 | 0.190037 | 0.528 | 0.377 | 1 | 11 | Prr13 | 0.151 | Old |
| 0.053948 | 0.045774 | 0.298 | 0.188 | 1 | 11 | Tmf1 | 0.11 | Old |
| 0.054152 | 0.545718 | 0.161 | 0.087 | 1 | 11 | Rcsd1 | 0.074 | Old |
| 0.054374 | 0.327086 | 0.181 | 0.101 | 1 | 11 | Gpatch2 | 0.08 | Old |
| 0.054715 | 0.306466 | 0.133 | 0.065 | 1 | 11 | Vac14 | 0.068 | Old |
| 0.054876 | 0.607077 | 0.218 | 0.13 | 1 | 11 | Cfap43 | 0.088 | Old |
| 0.054913 | 0.125072 | 0.363 | 0.239 | 1 | 11 | Iqgap2 | 0.124 | Old |
| 0.054938 | 0.19088 | 0.133 | 0.065 | 1 | 11 | Oxa1l | 0.068 | Old |
| 0.054959 | 0.409935 | 0.161 | 0.087 | 1 | 11 | Larp7 | 0.074 | Old |
| 0.054966 | 1.116256 | 0.089 | 0.036 | 1 | 11 | Edem3 | 0.053 | Old |
| 0.054966 | 1.316949 | 0.089 | 0.036 | 1 | 11 | Tfip11 | 0.053 | Old |
| 0.055002 | 1.154517 | 0.109 | 0.051 | 1 | 11 | Fra10ac1 | 0.058 | Old |
| 0.055239 | 0.877517 | 0.089 | 0.036 | 1 | 11 | Ikzf2 | 0.053 | Old |
| 0.055354 | 1.58223 | 0.097 | 0.043 | 1 | 11 | Slc7a6os | 0.054 | Old |
| 0.055365 | 0.952699 | 0.129 | 0.065 | 1 | 11 | Zbtb22 | 0.064 | Old |
| 0.055372 | 0.176676 | 0.468 | 0.333 | 1 | 11 | Nras | 0.135 | Old |
| 0.055384 | 0.048604 | 0.133 | 0.065 | 1 | 11 | Mrpl37 | 0.068 | Old |
| 0.055411 | 0.117017 | 0.677 | 0.507 | 1 | 11 | Tma7 | 0.17 | Old |
| 0.055418 | 0.436031 | 0.242 | 0.152 | 1 | 11 | Snx13 | 0.09 | Old |
| 0.055659 | 0.387547 | 0.101 | 0.043 | 1 | 11 | Cdk10 | 0.058 | Old |
| 0.055705 | 0.305601 | 0.605 | 0.457 | 1 | 11 | Pbrm1 | 0.148 | Old |
| 0.055857 | 0.12052 | 0.508 | 0.355 | 1 | 11 | Ndufa3 | 0.153 | Old |
| 0.055913 | 0.266178 | 0.532 | 0.391 | 1 | 11 | Nadk | 0.141 | Old |
| 0.056111 | 0.108758 | 0.319 | 0.196 | 1 | 11 | Uqcc2 | 0.123 | Old |

| | | | | | | | | |
|---|---|---|---|---|---|---|---|---|
| 0.056294 | 0.226462 | 0.141 | 0.072 | 1 | 11 | Taf2 | 0.069 | Old |
| 0.056308 | 0.123645 | 0.181 | 0.101 | 1 | 11 | Stx11 | 0.08 | Old |
| 0.056499 | 0.254771 | 0.246 | 0.152 | 1 | 11 | Sh3bp2 | 0.094 | Old |
| 0.056506 | 1.568478 | 0.077 | 0.029 | 1 | 11 | Rdm1 | 0.048 | Old |
| 0.056621 | 0.702946 | 0.089 | 0.036 | 1 | 11 | Mri1 | 0.053 | Old |
| 0.05702 | 0.05784 | 0.161 | 0.087 | 1 | 11 | Gm13212 | 0.074 | Old |
| 0.057027 | 0.229579 | 0.464 | 0.326 | 1 | 11 | Cdc42se1 | 0.138 | Old |
| 0.05715 | 0.307484 | 0.359 | 0.239 | 1 | 11 | Pros1 | 0.12 | Old |
| 0.057181 | 0.637928 | 0.089 | 0.036 | 1 | 11 | Unc45a | 0.053 | Old |
| 0.057181 | 0.034955 | 0.089 | 0.036 | 1 | 11 | Cars2 | 0.053 | Old |
| 0.057253 | 0.095158 | 0.907 | 0.761 | 1 | 11 | Rap1b | 0.146 | Old |
| 0.057273 | 1.091276 | 0.145 | 0.08 | 1 | 11 | Erlin2 | 0.065 | Old |
| 0.057363 | 0.349185 | 0.198 | 0.116 | 1 | 11 | Trim47 | 0.082 | Old |
| 0.057464 | 1.152909 | 0.089 | 0.036 | 1 | 11 | Zfp954 | 0.053 | Old |
| 0.057496 | 0.215379 | 0.181 | 0.101 | 1 | 11 | Ppfia1 | 0.08 | Old |
| 0.057601 | 0.651484 | 0.202 | 0.123 | 1 | 11 | Sra1 | 0.079 | Old |
| 0.057633 | 0.109627 | 0.984 | 0.964 | 1 | 11 | Ptma | 0.02 | Old |
| 0.057736 | 0.133482 | 0.673 | 0.529 | 1 | 11 | Atp6v1g1 | 0.144 | Old |
| 0.05776 | 0.516551 | 0.121 | 0.058 | 1 | 11 | Mrps34 | 0.063 | Old |
| 0.05776 | 0.316012 | 0.121 | 0.058 | 1 | 11 | Ascc2 | 0.063 | Old |
| 0.057845 | 0.372923 | 0.214 | 0.13 | 1 | 11 | Ckap4 | 0.084 | Old |
| 0.057855 | 0.306143 | 0.141 | 0.072 | 1 | 11 | Mras | 0.069 | Old |
| 0.058077 | 0.029926 | 0.464 | 0.312 | 1 | 11 | Fxr1 | 0.152 | Old |
| 0.058124 | 0.055555 | 0.339 | 0.217 | 1 | 11 | Adcy7 | 0.122 | Old |
| 0.058171 | 0.02817 | 0.173 | 0.094 | 1 | 11 | Sars | 0.079 | Old |
| 0.058262 | 0.269774 | 0.331 | 0.217 | 1 | 11 | Plod3 | 0.114 | Old |
| 0.058272 | 0.206495 | 0.552 | 0.413 | 1 | 11 | Wwp1 | 0.139 | Old |
| 0.058339 | 0.177858 | 0.556 | 0.406 | 1 | 11 | Hif1a | 0.15 | Old |
| 0.058391 | 0.180314 | 0.48 | 0.341 | 1 | 11 | Dtx3l | 0.139 | Old |
| 0.058408 | 0.137939 | 0.315 | 0.203 | 1 | 11 | Gga1 | 0.112 | Old |
| 0.058429 | 1.116016 | 0.117 | 0.058 | 1 | 11 | Nfyb | 0.059 | Old |
| 0.058442 | 0.15764 | 0.528 | 0.37 | 1 | 11 | Nisch | 0.158 | Old |
| 0.058486 | 0.980319 | 0.077 | 0.029 | 1 | 11 | Sertad3 | 0.048 | Old |
| 0.058566 | 0.073185 | 0.504 | 0.348 | 1 | 11 | Ndufb10 | 0.156 | Old |
| 0.058601 | 0.575136 | 0.218 | 0.138 | 1 | 11 | Rcn2 | 0.08 | Old |
| 0.058613 | 0.636782 | 0.129 | 0.065 | 1 | 11 | Prkab2 | 0.064 | Old |
| 0.058678 | 1.145743 | 0.117 | 0.058 | 1 | 11 | Tvp23b | 0.059 | Old |
| 0.058697 | 2.138517 | 0.065 | 0.022 | 1 | 11 | Mtor | 0.043 | Old |
| 0.058746 | 0.433625 | 0.274 | 0.174 | 1 | 11 | Sec24b | 0.1 | Old |
| 0.058833 | 0.228379 | 0.206 | 0.123 | 1 | 11 | Fbxo3 | 0.083 | Old |
| 0.058834 | 1.251552 | 0.097 | 0.043 | 1 | 11 | Nol12 | 0.054 | Old |
| 0.058865 | 0.210587 | 0.194 | 0.116 | 1 | 11 | Kcnq1ot1 | 0.078 | Old |
| 0.058942 | 0.385119 | 0.202 | 0.123 | 1 | 11 | Luc7l | 0.079 | Old |
| 0.05899 | 0.439627 | 0.282 | 0.188 | 1 | 11 | Clock | 0.094 | Old |
| 0.058992 | 0.558491 | 0.141 | 0.072 | 1 | 11 | Scrib | 0.069 | Old |

| | | | | | | | | |
|---|---|---|---|---|---|---|---|---|
| 0.059008 | 0.081477 | 0.327 | 0.21 | 1 | 11 | Rex1bd | 0.117 | Old |
| 0.059038 | 0.098933 | 0.214 | 0.123 | 1 | 11 | Uba6 | 0.091 | Old |
| 0.059107 | 1.201092 | 0.077 | 0.029 | 1 | 11 | Prrg1 | 0.048 | Old |
| 0.059147 | 0.19023 | 0.46 | 0.319 | 1 | 11 | Zfp91 | 0.141 | Old |
| 0.059373 | 0.312697 | 0.101 | 0.043 | 1 | 11 | Fbxo28 | 0.058 | Old |
| 0.059435 | 0.17145 | 0.46 | 0.319 | 1 | 11 | Ppib | 0.141 | Old |
| 0.059548 | 0.643897 | 0.173 | 0.101 | 1 | 11 | Palld | 0.072 | Old |
| 0.05957 | 0.531145 | 0.129 | 0.065 | 1 | 11 | Mrps6 | 0.064 | Old |
| 0.0596 | 0.091734 | 0.387 | 0.261 | 1 | 11 | Blnk | 0.126 | Old |
| 0.059734 | 1.142801 | 0.077 | 0.029 | 1 | 11 | Lmnb1 | 0.048 | Old |
| 0.059959 | 0.288349 | 0.431 | 0.304 | 1 | 11 | Brd4 | 0.127 | Old |
| 0.059973 | 0.429778 | 0.25 | 0.159 | 1 | 11 | Snrnp27 | 0.091 | Old |
| 0.05998 | 0.053886 | 0.121 | 0.058 | 1 | 11 | Gtf2e2 | 0.063 | Old |
| 0.060007 | 0.16804 | 0.286 | 0.181 | 1 | 11 | Srsf10 | 0.105 | Old |
| 0.060014 | 2.487861 | 0.04 | 0.007 | 1 | 11 | Tbc1d16 | 0.033 | Old |
| 0.060014 | 2.200488 | 0.04 | 0.007 | 1 | 11 | Dtd2 | 0.033 | Old |
| 0.060054 | 0.309292 | 0.089 | 0.036 | 1 | 11 | Gpr157 | 0.053 | Old |
| 0.060295 | 0.53686 | 0.129 | 0.065 | 1 | 11 | Abhd4 | 0.064 | Old |
| 0.060465 | 2.666762 | 0.04 | 0.007 | 1 | 11 | Mb21d2 | 0.033 | Old |
| 0.060643 | 0.790931 | 0.089 | 0.036 | 1 | 11 | Adi1 | 0.053 | Old |
| 0.060643 | 0.621488 | 0.089 | 0.036 | 1 | 11 | Pabpc4 | 0.053 | Old |
| 0.060727 | 0.20328 | 0.343 | 0.225 | 1 | 11 | Clptm1 | 0.118 | Old |
| 0.060754 | 0.117793 | 0.339 | 0.217 | 1 | 11 | Igsf6 | 0.122 | Old |
| 0.060808 | 0.203637 | 0.399 | 0.268 | 1 | 11 | Lsm4 | 0.131 | Old |
| 0.060919 | 2.545646 | 0.04 | 0.007 | 1 | 11 | Atp2a3 | 0.033 | Old |
| 0.060919 | 2.026635 | 0.04 | 0.007 | 1 | 11 | Uaca | 0.033 | Old |
| 0.061043 | 0.256759 | 0.504 | 0.377 | 1 | 11 | Mpp1 | 0.127 | Old |
| 0.061088 | 0.47605 | 0.282 | 0.181 | 1 | 11 | Usp34 | 0.101 | Old |
| 0.061153 | 0.60972 | 0.109 | 0.051 | 1 | 11 | Rwdd4a | 0.058 | Old |
| 0.061304 | 0.072967 | 0.319 | 0.21 | 1 | 11 | Bin1 | 0.109 | Old |
| 0.061375 | 1.87761 | 0.04 | 0.007 | 1 | 11 | Coa7 | 0.033 | Old |
| 0.061466 | 0.374352 | 0.371 | 0.268 | 1 | 11 | Egr2 | 0.103 | Old |
| 0.061481 | 0.173777 | 0.431 | 0.297 | 1 | 11 | Eif4g1 | 0.134 | Old |
| 0.061498 | 0.195776 | 0.121 | 0.058 | 1 | 11 | Ddx27 | 0.063 | Old |
| 0.061619 | 0.40443 | 0.169 | 0.094 | 1 | 11 | Adpgk | 0.075 | Old |
| 0.0617 | 0.058531 | 0.415 | 0.275 | 1 | 11 | Ddx58 | 0.14 | Old |
| 0.061835 | 2.371197 | 0.04 | 0.007 | 1 | 11 | Ints5.1 | 0.033 | Old |
| 0.061835 | 2.195126 | 0.04 | 0.007 | 1 | 11 | Slx4 | 0.033 | Old |
| 0.061847 | 0.002888 | 0.286 | 0.188 | 1 | 11 | Tomm70a | 0.098 | Old |
| 0.061861 | 0.250845 | 0.19 | 0.109 | 1 | 11 | Ethe1 | 0.081 | Old |
| 0.061965 | 0.06835 | 0.5 | 0.341 | 1 | 11 | Sdha | 0.159 | Old |
| 0.062057 | 0.234564 | 0.274 | 0.174 | 1 | 11 | Jtb | 0.1 | Old |
| 0.062231 | 0.759029 | 0.109 | 0.051 | 1 | 11 | Rrn3 | 0.058 | Old |
| 0.062243 | 0.178587 | 0.181 | 0.101 | 1 | 11 | Iws1 | 0.08 | Old |
| 0.062247 | 0.108611 | 0.681 | 0.486 | 1 | 11 | Rnf130 | 0.195 | Old |

| | | | | | | | | |
|---|---|---|---|---|---|---|---|---|
| 0.062297 | 2.717039 | 0.04 | 0.007 | 1 | 11 | Slc1a3 | 0.033 | Old |
| 0.062297 | 2.421856 | 0.04 | 0.007 | 1 | 11 | Kyat1 | 0.033 | Old |
| 0.062297 | 1.700001 | 0.04 | 0.007 | 1 | 11 | D130040H2 | 0.033 | Old |
| 0.062382 | 0.10933 | 0.315 | 0.203 | 1 | 11 | Slc15a4 | 0.112 | Old |
| 0.062396 | 2.516041 | 0.052 | 0.014 | 1 | 11 | Lrmda | 0.038 | Old |
| 0.062493 | 0.979336 | 0.097 | 0.043 | 1 | 11 | Pigz | 0.054 | Old |
| 0.062503 | 0.335082 | 0.109 | 0.051 | 1 | 11 | Tshz1 | 0.058 | Old |
| 0.062551 | 0.049268 | 0.238 | 0.152 | 1 | 11 | Pfkfb3 | 0.086 | Old |
| 0.062762 | 1.689309 | 0.04 | 0.007 | 1 | 11 | Aars2 | 0.033 | Old |
| 0.062762 | 1.686489 | 0.04 | 0.007 | 1 | 11 | Usp28 | 0.033 | Old |
| 0.062762 | 1.423454 | 0.04 | 0.007 | 1 | 11 | Ppif | 0.033 | Old |
| 0.062918 | 0.474857 | 0.149 | 0.08 | 1 | 11 | Cep192 | 0.069 | Old |
| 0.062938 | 0.404608 | 0.169 | 0.094 | 1 | 11 | Ppa1 | 0.075 | Old |
| 0.062981 | 1.091118 | 0.145 | 0.08 | 1 | 11 | Rbm4b | 0.065 | Old |
| 0.062983 | 0.388067 | 0.278 | 0.181 | 1 | 11 | Zzef1 | 0.097 | Old |
| 0.063039 | 0.7186 | 0.117 | 0.058 | 1 | 11 | Mcoln2 | 0.059 | Old |
| 0.063046 | 0.445001 | 0.089 | 0.036 | 1 | 11 | Atad3a | 0.053 | Old |
| 0.06305 | 0.07402 | 0.109 | 0.051 | 1 | 11 | Nlk | 0.058 | Old |
| 0.063119 | 0.174157 | 0.19 | 0.109 | 1 | 11 | S100a1 | 0.081 | Old |
| 0.063149 | 0.143924 | 0.27 | 0.174 | 1 | 11 | Sptlc2 | 0.096 | Old |
| 0.063175 | 0.089145 | 0.29 | 0.188 | 1 | 11 | Rbm26 | 0.102 | Old |
| 0.06323 | 1.573842 | 0.04 | 0.007 | 1 | 11 | Gemin5 | 0.033 | Old |
| 0.06323 | 1.39648 | 0.04 | 0.007 | 1 | 11 | Rfc4 | 0.033 | Old |
| 0.06323 | 1.220097 | 0.04 | 0.007 | 1 | 11 | 0610009O2 | 0.033 | Old |
| 0.06323 | 1.145114 | 0.04 | 0.007 | 1 | 11 | Taf5 | 0.033 | Old |
| 0.063248 | 1.766948 | 0.065 | 0.022 | 1 | 11 | Cog2 | 0.043 | Old |
| 0.063298 | 0.395444 | 0.371 | 0.268 | 1 | 11 | Arid4a | 0.103 | Old |
| 0.063312 | 0.529175 | 0.137 | 0.072 | 1 | 11 | Snhg6 | 0.065 | Old |
| 0.063352 | 0.240066 | 0.29 | 0.188 | 1 | 11 | Xdh | 0.102 | Old |
| 0.063387 | 0.27248 | 0.206 | 0.123 | 1 | 11 | Dnase1l1 | 0.083 | Old |
| 0.06344 | 0.160369 | 0.48 | 0.333 | 1 | 11 | Ndufc1 | 0.147 | Old |
| 0.063582 | 0.113298 | 0.315 | 0.203 | 1 | 11 | Nudt3 | 0.112 | Old |
| 0.063592 | 0.014911 | 0.488 | 0.362 | 1 | 11 | Ndufb3 | 0.126 | Old |
| 0.063605 | 2.236601 | 0.052 | 0.014 | 1 | 11 | A530017D2 | 0.038 | Old |
| 0.063605 | 1.869731 | 0.052 | 0.014 | 1 | 11 | Isoc2b | 0.038 | Old |
| 0.063609 | 0.747742 | 0.077 | 0.029 | 1 | 11 | Bzw2 | 0.048 | Old |
| 0.063657 | 0.864314 | 0.097 | 0.043 | 1 | 11 | Tmem87a | 0.054 | Old |
| 0.063701 | 1.387389 | 0.04 | 0.007 | 1 | 11 | Cptp | 0.033 | Old |
| 0.063701 | 1.249802 | 0.04 | 0.007 | 1 | 11 | Abhd14a | 0.033 | Old |
| 0.063701 | 1.216183 | 0.04 | 0.007 | 1 | 11 | Golm1 | 0.033 | Old |
| 0.063751 | 0.024681 | 0.181 | 0.101 | 1 | 11 | Cd300lb | 0.08 | Old |
| 0.063778 | 0.56889 | 0.185 | 0.109 | 1 | 11 | 2310011J0 | 0.076 | Old |
| 0.063879 | 0.244259 | 0.109 | 0.051 | 1 | 11 | Snupn | 0.058 | Old |
| 0.063891 | 0.278951 | 0.23 | 0.145 | 1 | 11 | Gas2l3 | 0.085 | Old |
| 0.06395 | 0.50534 | 0.097 | 0.043 | 1 | 11 | Gstt2 | 0.054 | Old |

| | | | | | | | | |
|---|---|---|---|---|---|---|---|---|
| 0.063958 | 0.094327 | 0.335 | 0.217 | 1 | 11 | H2-Q4 | 0.118 | Old |
| 0.06397 | 0.170356 | 0.266 | 0.167 | 1 | 11 | Aldh3b1 | 0.099 | Old |
| 0.063973 | 0.683304 | 0.065 | 0.022 | 1 | 11 | Wdr70 | 0.043 | Old |
| 0.063993 | 0.329405 | 0.185 | 0.109 | 1 | 11 | Map3k7 | 0.076 | Old |
| 0.064039 | 0.171421 | 0.21 | 0.123 | 1 | 11 | Bet1 | 0.087 | Old |
| 0.064047 | 0.148399 | 0.786 | 0.594 | 1 | 11 | Rbms1 | 0.192 | Old |
| 0.064103 | 0.742971 | 0.117 | 0.058 | 1 | 11 | Gramd1b | 0.059 | Old |
| 0.064104 | 0.325685 | 0.282 | 0.181 | 1 | 11 | 6-Mar | 0.101 | Old |
| 0.064174 | 0.9026 | 0.04 | 0.007 | 1 | 11 | Qrsl1 | 0.033 | Old |
| 0.064174 | 0.852164 | 0.04 | 0.007 | 1 | 11 | Fam78b | 0.033 | Old |
| 0.064174 | 0.721795 | 0.04 | 0.007 | 1 | 11 | Pde4d | 0.033 | Old |
| 0.064174 | 0.334713 | 0.04 | 0.007 | 1 | 11 | Dimt1 | 0.033 | Old |
| 0.064186 | 0.354455 | 0.198 | 0.116 | 1 | 11 | Sgta | 0.082 | Old |
| 0.064272 | 0.340156 | 0.347 | 0.239 | 1 | 11 | Emilin2 | 0.108 | Old |
| 0.064339 | 1.08812 | 0.065 | 0.022 | 1 | 11 | Coprs | 0.043 | Old |
| 0.064506 | 0.36006 | 0.169 | 0.094 | 1 | 11 | Dirc2 | 0.075 | Old |
| 0.064693 | 0.232927 | 0.29 | 0.188 | 1 | 11 | Numa1 | 0.102 | Old |
| 0.064706 | 1.368041 | 0.065 | 0.022 | 1 | 11 | Vldlr | 0.043 | Old |
| 0.06471 | 0.152781 | 0.141 | 0.072 | 1 | 11 | Anapc4 | 0.069 | Old |
| 0.064732 | 0.270038 | 0.448 | 0.312 | 1 | 11 | Morc3 | 0.136 | Old |
| 0.064804 | 0.147011 | 0.165 | 0.094 | 1 | 11 | Faf2 | 0.071 | Old |
| 0.06481 | 0.008006 | 0.266 | 0.174 | 1 | 11 | Fbrsl1 | 0.092 | Old |
| 0.064838 | 0.977541 | 0.097 | 0.043 | 1 | 11 | Gmpr2 | 0.054 | Old |
| 0.064851 | 0.23196 | 0.234 | 0.145 | 1 | 11 | Ncoa4 | 0.089 | Old |
| 0.064862 | 0.167228 | 0.21 | 0.123 | 1 | 11 | Bicd2 | 0.087 | Old |
| 0.06491 | 0.451482 | 0.117 | 0.058 | 1 | 11 | Zfp560 | 0.059 | Old |
| 0.064914 | 0.170968 | 0.177 | 0.101 | 1 | 11 | AU040320 | 0.076 | Old |
| 0.064921 | 0.009972 | 0.476 | 0.333 | 1 | 11 | Vps35 | 0.143 | Old |
| 0.064959 | 0.007534 | 0.169 | 0.094 | 1 | 11 | Gabpa | 0.075 | Old |
| 0.064997 | 0.434179 | 0.109 | 0.051 | 1 | 11 | Ermp1 | 0.058 | Old |
| 0.065065 | 0.119668 | 0.129 | 0.065 | 1 | 11 | Hsd17b10 | 0.064 | Old |
| 0.065074 | 1.165895 | 0.065 | 0.022 | 1 | 11 | Zer1 | 0.043 | Old |
| 0.065405 | 0.463128 | 0.173 | 0.101 | 1 | 11 | Npl | 0.072 | Old |
| 0.065435 | 0.345205 | 0.097 | 0.043 | 1 | 11 | Usp21 | 0.054 | Old |
| 0.065445 | 1.198295 | 0.065 | 0.022 | 1 | 11 | Acap3 | 0.043 | Old |
| 0.065445 | 1.027817 | 0.065 | 0.022 | 1 | 11 | Map4k3 | 0.043 | Old |
| 0.065445 | 0.985301 | 0.065 | 0.022 | 1 | 11 | Fancl | 0.043 | Old |
| 0.065697 | 0.11969 | 0.367 | 0.239 | 1 | 11 | Prelid1 | 0.128 | Old |
| 0.065805 | 0.509401 | 0.137 | 0.072 | 1 | 11 | Abcd3 | 0.065 | Old |
| 0.065935 | 0.176686 | 0.282 | 0.181 | 1 | 11 | Sh2b3 | 0.101 | Old |
| 0.06608 | 0.492846 | 0.052 | 0.014 | 1 | 11 | Bloc1s3 | 0.038 | Old |
| 0.066191 | 1.260443 | 0.065 | 0.022 | 1 | 11 | Dusp12 | 0.043 | Old |
| 0.066191 | 0.526036 | 0.065 | 0.022 | 1 | 11 | Tuba4a | 0.043 | Old |
| 0.066201 | 0.028162 | 0.355 | 0.232 | 1 | 11 | Vma21 | 0.123 | Old |
| 0.066224 | 3.618345 | 0.024 | 0 | 1 | 11 | A630072M | 0.024 | Old |

| | | | | | | | | |
|---|---|---|---|---|---|---|---|---|
| 0.066224 | 3.229049 | 0.024 | 0 | 1 | 11 | Sgsm2 | 0.024 | Old |
| 0.066224 | 3.016499 | 0.024 | 0 | 1 | 11 | Tyw3 | 0.024 | Old |
| 0.066224 | 2.994726 | 0.024 | 0 | 1 | 11 | Stxbp4 | 0.024 | Old |
| 0.066224 | 2.95166 | 0.024 | 0 | 1 | 11 | Hdhd5 | 0.024 | Old |
| 0.066224 | 2.933878 | 0.024 | 0 | 1 | 11 | Nim1k | 0.024 | Old |
| 0.066224 | 2.733617 | 0.024 | 0 | 1 | 11 | Gm48653 | 0.024 | Old |
| 0.066224 | 2.727757 | 0.024 | 0 | 1 | 11 | Tmem38a | 0.024 | Old |
| 0.066224 | 2.668991 | 0.024 | 0 | 1 | 11 | Znrf3 | 0.024 | Old |
| 0.066224 | 2.583119 | 0.024 | 0 | 1 | 11 | Tceanc | 0.024 | Old |
| 0.066224 | 2.5676 | 0.024 | 0 | 1 | 11 | AC139671. | 0.024 | Old |
| 0.066224 | 2.512587 | 0.024 | 0 | 1 | 11 | Ftsj1 | 0.024 | Old |
| 0.066224 | 2.505414 | 0.024 | 0 | 1 | 11 | Ctif | 0.024 | Old |
| 0.066224 | 2.499276 | 0.024 | 0 | 1 | 11 | Rsad1 | 0.024 | Old |
| 0.066224 | 2.496783 | 0.024 | 0 | 1 | 11 | Hsd17b14 | 0.024 | Old |
| 0.066224 | 2.47434 | 0.024 | 0 | 1 | 11 | Exo5 | 0.024 | Old |
| 0.066224 | 2.462926 | 0.024 | 0 | 1 | 11 | Dhodh | 0.024 | Old |
| 0.066224 | 2.36367 | 0.024 | 0 | 1 | 11 | Tedc2 | 0.024 | Old |
| 0.066224 | 2.356898 | 0.024 | 0 | 1 | 11 | Rpp38 | 0.024 | Old |
| 0.066224 | 2.350638 | 0.024 | 0 | 1 | 11 | Samd10 | 0.024 | Old |
| 0.066224 | 2.314945 | 0.024 | 0 | 1 | 11 | 1110019D1 | 0.024 | Old |
| 0.066224 | 2.302568 | 0.024 | 0 | 1 | 11 | Setd4 | 0.024 | Old |
| 0.066224 | 2.256371 | 0.024 | 0 | 1 | 11 | Cfap126 | 0.024 | Old |
| 0.066224 | 2.222149 | 0.024 | 0 | 1 | 11 | Acot11 | 0.024 | Old |
| 0.066224 | 2.220566 | 0.024 | 0 | 1 | 11 | Cdca8 | 0.024 | Old |
| 0.066224 | 2.185263 | 0.024 | 0 | 1 | 11 | Gm12589 | 0.024 | Old |
| 0.066224 | 2.158679 | 0.024 | 0 | 1 | 11 | Gm33973 | 0.024 | Old |
| 0.066224 | 2.139107 | 0.024 | 0 | 1 | 11 | AC125149. | 0.024 | Old |
| 0.066224 | 2.123798 | 0.024 | 0 | 1 | 11 | Ighmbp2 | 0.024 | Old |
| 0.066224 | 2.107912 | 0.024 | 0 | 1 | 11 | Crocc | 0.024 | Old |
| 0.066224 | 2.088978 | 0.024 | 0 | 1 | 11 | Fam135a | 0.024 | Old |
| 0.066224 | 2.081791 | 0.024 | 0 | 1 | 11 | Bmp1 | 0.024 | Old |
| 0.066224 | 2.077039 | 0.024 | 0 | 1 | 11 | Oxld1 | 0.024 | Old |
| 0.066224 | 2.063908 | 0.024 | 0 | 1 | 11 | Frat1 | 0.024 | Old |
| 0.066224 | 2.063765 | 0.024 | 0 | 1 | 11 | Ap4b1 | 0.024 | Old |
| 0.066224 | 2.052793 | 0.024 | 0 | 1 | 11 | 2700046G( | 0.024 | Old |
| 0.066224 | 2.048011 | 0.024 | 0 | 1 | 11 | Cox15 | 0.024 | Old |
| 0.066224 | 2.033533 | 0.024 | 0 | 1 | 11 | Aldh1l1 | 0.024 | Old |
| 0.066224 | 2.00757 | 0.024 | 0 | 1 | 11 | Zswim7 | 0.024 | Old |
| 0.066224 | 1.996719 | 0.024 | 0 | 1 | 11 | Mboat1 | 0.024 | Old |
| 0.066224 | 1.965403 | 0.024 | 0 | 1 | 11 | 1810055G( | 0.024 | Old |
| 0.066224 | 1.936719 | 0.024 | 0 | 1 | 11 | Ift88 | 0.024 | Old |
| 0.066224 | 1.930757 | 0.024 | 0 | 1 | 11 | 1700008J0 | 0.024 | Old |
| 0.066224 | 1.916523 | 0.024 | 0 | 1 | 11 | P3h1 | 0.024 | Old |
| 0.066224 | 1.811179 | 0.024 | 0 | 1 | 11 | Spryd4 | 0.024 | Old |
| 0.066224 | 1.804966 | 0.024 | 0 | 1 | 11 | AI463229 | 0.024 | Old |

| | | | | | | | | |
|---|---|---|---|---|---|---|---|---|
| 0.066224 | 1.79878 | 0.024 | 0 | 1 | 11 | Slc25a35 | 0.024 | Old |
| 0.066224 | 1.780962 | 0.024 | 0 | 1 | 11 | Hist1h2ap | 0.024 | Old |
| 0.066224 | 1.75469 | 0.024 | 0 | 1 | 11 | Pus3 | 0.024 | Old |
| 0.066224 | 1.670642 | 0.024 | 0 | 1 | 11 | Mr1 | 0.024 | Old |
| 0.066224 | 1.627201 | 0.024 | 0 | 1 | 11 | Eid2 | 0.024 | Old |
| 0.066224 | 1.556679 | 0.024 | 0 | 1 | 11 | Dnaaf5 | 0.024 | Old |
| 0.066337 | 0.275513 | 0.169 | 0.094 | 1 | 11 | Trappc6a | 0.075 | Old |
| 0.066372 | 1.846944 | 0.085 | 0.036 | 1 | 11 | Zfp729a | 0.049 | Old |
| 0.06649 | 0.091192 | 0.226 | 0.138 | 1 | 11 | Ccdc71l | 0.088 | Old |
| 0.066618 | 0.275608 | 0.185 | 0.109 | 1 | 11 | Ormdl2 | 0.076 | Old |
| 0.066635 | 0.103971 | 0.129 | 0.065 | 1 | 11 | Cbx7 | 0.064 | Old |
| 0.066783 | 0.102808 | 0.512 | 0.355 | 1 | 11 | Scarb2 | 0.157 | Old |
| 0.066948 | 0.412446 | 0.097 | 0.043 | 1 | 11 | Cdc14a | 0.054 | Old |
| 0.066988 | 0.349991 | 0.222 | 0.138 | 1 | 11 | Plk2 | 0.084 | Old |
| 0.067125 | 0.677777 | 0.145 | 0.08 | 1 | 11 | Sucla2 | 0.065 | Old |
| 0.067165 | 0.194009 | 0.129 | 0.065 | 1 | 11 | Sfi1 | 0.064 | Old |
| 0.067281 | 0.284182 | 0.109 | 0.051 | 1 | 11 | Bad | 0.058 | Old |
| 0.067341 | 0.243435 | 0.077 | 0.029 | 1 | 11 | Rnf126 | 0.048 | Old |
| 0.067342 | 0.208262 | 0.827 | 0.674 | 1 | 11 | Rbm39 | 0.153 | Old |
| 0.067347 | 1.251887 | 0.052 | 0.014 | 1 | 11 | Zfp760 | 0.038 | Old |
| 0.067431 | 0.469909 | 0.129 | 0.065 | 1 | 11 | Usp45 | 0.064 | Old |
| 0.067503 | 0.111195 | 0.169 | 0.094 | 1 | 11 | Crtap | 0.075 | Old |
| 0.067513 | 0.099084 | 0.452 | 0.319 | 1 | 11 | Rab11a | 0.133 | Old |
| 0.067599 | 0.577728 | 0.137 | 0.072 | 1 | 11 | 5031425E2 | 0.065 | Old |
| 0.067622 | 0.163614 | 0.306 | 0.196 | 1 | 11 | Tbl1x | 0.11 | Old |
| 0.067664 | 0.368814 | 0.117 | 0.058 | 1 | 11 | Trmt6 | 0.059 | Old |
| 0.067689 | 0.367145 | 0.077 | 0.029 | 1 | 11 | Usp10 | 0.048 | Old |
| 0.067692 | 0.680855 | 0.173 | 0.101 | 1 | 11 | AW549877 | 0.072 | Old |
| 0.067704 | 1.066211 | 0.065 | 0.022 | 1 | 11 | Nsdhl | 0.043 | Old |
| 0.067723 | 0.266132 | 0.476 | 0.348 | 1 | 11 | Uqcr10 | 0.128 | Old |
| 0.067837 | 0.600577 | 0.165 | 0.094 | 1 | 11 | Tbc1d4 | 0.071 | Old |
| 0.06793 | 0.242349 | 0.476 | 0.341 | 1 | 11 | Degs1 | 0.135 | Old |
| 0.067953 | 0.277212 | 0.157 | 0.087 | 1 | 11 | Spns1 | 0.07 | Old |
| 0.068016 | 0.30442 | 0.387 | 0.283 | 1 | 11 | Pla2g15 | 0.104 | Old |
| 0.068038 | 0.19935 | 0.077 | 0.029 | 1 | 11 | Hes1 | 0.048 | Old |
| 0.068119 | 0.332514 | 0.137 | 0.072 | 1 | 11 | Dok3 | 0.065 | Old |
| 0.068202 | 1.218797 | 0.052 | 0.014 | 1 | 11 | Pwp2 | 0.038 | Old |
| 0.068226 | 0.389125 | 0.117 | 0.058 | 1 | 11 | Kpna6 | 0.059 | Old |
| 0.068235 | 0.171113 | 0.129 | 0.065 | 1 | 11 | Wdr36 | 0.064 | Old |
| 0.068313 | 0.002792 | 0.165 | 0.094 | 1 | 11 | Bms1 | 0.071 | Old |
| 0.068424 | 0.28417 | 0.214 | 0.13 | 1 | 11 | Ubr3 | 0.084 | Old |
| 0.068442 | 0.507962 | 0.157 | 0.087 | 1 | 11 | Ccdc186 | 0.07 | Old |
| 0.06849 | 1.069594 | 0.097 | 0.043 | 1 | 11 | Fas | 0.054 | Old |
| 0.06849 | 0.31997 | 0.097 | 0.043 | 1 | 11 | Tarbp2 | 0.054 | Old |
| 0.068535 | 0.258542 | 0.827 | 0.688 | 1 | 11 | Tagln2 | 0.139 | Old |

| | | | | | | | | |
|---|---|---|---|---|---|---|---|---|
| 0.068598 | 0.060142 | 0.415 | 0.283 | 1 | 11 | Ids | 0.132 | Old |
| 0.068662 | 0.156205 | 0.298 | 0.196 | 1 | 11 | Wdr37 | 0.102 | Old |
| 0.068685 | 0.630051 | 0.105 | 0.051 | 1 | 11 | Mpi | 0.054 | Old |
| 0.068692 | 0.165783 | 0.403 | 0.29 | 1 | 11 | Rpl36al | 0.113 | Old |
| 0.068742 | 0.12085 | 0.109 | 0.051 | 1 | 11 | Ctps2 | 0.058 | Old |
| 0.068791 | 0.217806 | 0.927 | 0.862 | 1 | 11 | Sdc3 | 0.065 | Old |
| 0.068859 | 0.290269 | 0.625 | 0.493 | 1 | 11 | Dock2 | 0.132 | Old |
| 0.068912 | 0.142101 | 0.218 | 0.13 | 1 | 11 | Siah2 | 0.088 | Old |
| 0.068935 | 0.453053 | 0.157 | 0.087 | 1 | 11 | Irf2bpl | 0.07 | Old |
| 0.069048 | 0.147294 | 0.488 | 0.341 | 1 | 11 | Ptges3 | 0.147 | Old |
| 0.069067 | 0.643416 | 0.052 | 0.014 | 1 | 11 | Ap4e1 | 0.038 | Old |
| 0.069099 | 0.120824 | 0.476 | 0.333 | 1 | 11 | Ube3a | 0.143 | Old |
| 0.0691 | 0.005405 | 0.206 | 0.123 | 1 | 11 | Ufl1 | 0.083 | Old |
| 0.069189 | 0.052412 | 0.194 | 0.116 | 1 | 11 | Pigx | 0.078 | Old |
| 0.069343 | 0.339665 | 0.355 | 0.246 | 1 | 11 | Rad21 | 0.109 | Old |
| 0.069423 | 0.19749 | 0.258 | 0.167 | 1 | 11 | Stom | 0.091 | Old |
| 0.069429 | 0.108095 | 0.335 | 0.232 | 1 | 11 | Ddx60 | 0.103 | Old |
| 0.069432 | 0.546105 | 0.137 | 0.072 | 1 | 11 | Kctd2 | 0.065 | Old |
| 0.069432 | 0.472772 | 0.137 | 0.072 | 1 | 11 | 9430038I0: | 0.065 | Old |
| 0.069485 | 0.07533 | 0.694 | 0.522 | 1 | 11 | Rab1a | 0.172 | Old |
| 0.069648 | 0.43592 | 0.117 | 0.058 | 1 | 11 | Rasa2 | 0.059 | Old |
| 0.069693 | 0.27188 | 0.411 | 0.29 | 1 | 11 | Pacs2 | 0.121 | Old |
| 0.069867 | 0.158405 | 0.129 | 0.065 | 1 | 11 | Sntb2 | 0.064 | Old |
| 0.069876 | 0.058693 | 0.923 | 0.862 | 1 | 11 | mt-Nd3 | 0.061 | Old |
| 0.06994 | 0.693455 | 0.052 | 0.014 | 1 | 11 | Ercc2 | 0.038 | Old |
| 0.06994 | 0.504434 | 0.052 | 0.014 | 1 | 11 | Qtrt2 | 0.038 | Old |
| 0.06994 | 0.191675 | 0.052 | 0.014 | 1 | 11 | Plpp5 | 0.038 | Old |
| 0.069965 | 0.584716 | 0.125 | 0.065 | 1 | 11 | Tbck | 0.06 | Old |
| 0.069982 | 0.136391 | 0.206 | 0.123 | 1 | 11 | Dennd4c | 0.083 | Old |
| 0.070128 | 0.321003 | 0.294 | 0.196 | 1 | 11 | Ncf4 | 0.098 | Old |
| 0.070229 | 0.083945 | 0.109 | 0.051 | 1 | 11 | Gnptab | 0.058 | Old |
| 0.070229 | 0.055187 | 0.109 | 0.051 | 1 | 11 | Ccdc85b | 0.058 | Old |
| 0.070356 | 1.151519 | 0.085 | 0.036 | 1 | 11 | Miip | 0.049 | Old |
| 0.07038 | 0.656357 | 0.052 | 0.014 | 1 | 11 | Trim39 | 0.038 | Old |
| 0.070418 | 0.059646 | 0.129 | 0.065 | 1 | 11 | Cdkn2aipnl | 0.064 | Old |
| 0.070522 | 0.967922 | 0.125 | 0.065 | 1 | 11 | Cass4 | 0.06 | Old |
| 0.070554 | 0.102186 | 0.661 | 0.493 | 1 | 11 | Rpn1 | 0.168 | Old |
| 0.070629 | 0.092361 | 0.258 | 0.167 | 1 | 11 | Map2k3 | 0.091 | Old |
| 0.070759 | 0.0498 | 0.331 | 0.217 | 1 | 11 | Rp9 | 0.114 | Old |
| 0.070803 | 0.243577 | 0.117 | 0.058 | 1 | 11 | Rfxap | 0.059 | Old |
| 0.070823 | 0.097779 | 0.052 | 0.014 | 1 | 11 | Plekha8 | 0.038 | Old |
| 0.071017 | 0.039226 | 0.097 | 0.043 | 1 | 11 | Apool | 0.054 | Old |
| 0.071032 | 0.118693 | 0.448 | 0.333 | 1 | 11 | Blvrb | 0.115 | Old |
| 0.071036 | 0.384513 | 0.137 | 0.072 | 1 | 11 | Las1l | 0.065 | Old |
| 0.071038 | 0.516929 | 0.085 | 0.036 | 1 | 11 | Ino80c | 0.049 | Old |

| | | | | | | | | |
|---|---|---|---|---|---|---|---|---|
| 0.071094 | 0.574493 | 0.117 | 0.058 | 1 | 11 | Kdelc2 | 0.059 | Old |
| 0.071109 | 0.927512 | 0.105 | 0.051 | 1 | 11 | Appl2 | 0.054 | Old |
| 0.071314 | 0.703885 | 0.153 | 0.087 | 1 | 11 | Mtf1 | 0.066 | Old |
| 0.071392 | 0.238362 | 0.387 | 0.268 | 1 | 11 | Sbno1 | 0.119 | Old |
| 0.071403 | 0.47751 | 0.21 | 0.13 | 1 | 11 | Tmem141 | 0.08 | Old |
| 0.071491 | 0.52829 | 0.145 | 0.08 | 1 | 11 | Slc35a2 | 0.065 | Old |
| 0.0716 | 0.044227 | 0.149 | 0.08 | 1 | 11 | Mtmr2 | 0.069 | Old |
| 0.071608 | 0.354335 | 0.266 | 0.174 | 1 | 11 | Stim2 | 0.092 | Old |
| 0.07168 | 0.33099 | 0.117 | 0.058 | 1 | 11 | Nubp2 | 0.059 | Old |
| 0.071727 | 0.830829 | 0.085 | 0.036 | 1 | 11 | Galk1 | 0.049 | Old |
| 0.071727 | 0.731987 | 0.085 | 0.036 | 1 | 11 | Chmp6 | 0.049 | Old |
| 0.071767 | 0.292571 | 0.359 | 0.239 | 1 | 11 | Nfil3 | 0.12 | Old |
| 0.071876 | 0.216344 | 0.278 | 0.181 | 1 | 11 | Zfx | 0.097 | Old |
| 0.071916 | 0.090983 | 0.46 | 0.326 | 1 | 11 | Ankrd17 | 0.134 | Old |
| 0.072092 | 0.027041 | 0.129 | 0.065 | 1 | 11 | Tmem60 | 0.064 | Old |
| 0.072194 | 0.313905 | 0.226 | 0.138 | 1 | 11 | B3gat3 | 0.088 | Old |
| 0.072269 | 0.306339 | 0.117 | 0.058 | 1 | 11 | Man2b2 | 0.059 | Old |
| 0.072309 | 0.675617 | 0.097 | 0.043 | 1 | 11 | Dhx34 | 0.054 | Old |
| 0.072396 | 0.56729 | 0.137 | 0.072 | 1 | 11 | Nvl | 0.065 | Old |
| 0.072602 | 0.105205 | 0.177 | 0.101 | 1 | 11 | Cyth3 | 0.076 | Old |
| 0.072635 | 0.444935 | 0.097 | 0.043 | 1 | 11 | Timm8a1 | 0.054 | Old |
| 0.072658 | 0.159962 | 0.129 | 0.065 | 1 | 11 | Zdhhc23 | 0.064 | Old |
| 0.072664 | 0.268104 | 0.109 | 0.051 | 1 | 11 | Rin3 | 0.058 | Old |
| 0.072698 | 0.219823 | 0.315 | 0.21 | 1 | 11 | Dot1l | 0.105 | Old |
| 0.072956 | 0.68624 | 0.177 | 0.109 | 1 | 11 | Nedd9 | 0.068 | Old |
| 0.07299 | 0.353844 | 0.198 | 0.123 | 1 | 11 | Mrpl15 | 0.075 | Old |
| 0.073086 | 0.689989 | 0.145 | 0.08 | 1 | 11 | Ube2g1 | 0.065 | Old |
| 0.073119 | 0.849068 | 0.085 | 0.036 | 1 | 11 | Cd209b | 0.049 | Old |
| 0.073125 | 0.399542 | 0.153 | 0.087 | 1 | 11 | Grk6 | 0.066 | Old |
| 0.073222 | 0.320495 | 0.137 | 0.072 | 1 | 11 | Rngtt | 0.065 | Old |
| 0.073283 | 0.044553 | 0.109 | 0.051 | 1 | 11 | Osgin2 | 0.058 | Old |
| 0.073489 | 0.14567 | 0.52 | 0.384 | 1 | 11 | Cdk2ap2 | 0.136 | Old |
| 0.073499 | 0.30645 | 0.331 | 0.225 | 1 | 11 | Stat6 | 0.106 | Old |
| 0.07376 | 0.175853 | 0.823 | 0.674 | 1 | 11 | Canx | 0.149 | Old |
| 0.073777 | 0.146335 | 0.137 | 0.072 | 1 | 11 | Axin1 | 0.065 | Old |
| 0.074072 | 0.091848 | 0.335 | 0.217 | 1 | 11 | Acly | 0.118 | Old |
| 0.074108 | 0.088466 | 0.359 | 0.254 | 1 | 11 | Eif3d | 0.105 | Old |
| 0.07415 | 0.133301 | 0.633 | 0.478 | 1 | 11 | Pea15a | 0.155 | Old |
| 0.074283 | 0.091312 | 0.097 | 0.043 | 1 | 11 | Abcf2 | 0.054 | Old |
| 0.074327 | 0.083401 | 0.694 | 0.5 | 1 | 11 | Hnrnpa0 | 0.194 | Old |
| 0.074363 | 0.707537 | 0.117 | 0.058 | 1 | 11 | Nostrin | 0.059 | Old |
| 0.074495 | 0.135182 | 0.565 | 0.406 | 1 | 11 | Anxa2 | 0.159 | Old |
| 0.07451 | 0.233215 | 0.278 | 0.181 | 1 | 11 | Rbbp4 | 0.097 | Old |
| 0.074615 | 0.000158 | 0.137 | 0.072 | 1 | 11 | Tinf2 | 0.065 | Old |
| 0.074697 | 0.383116 | 0.173 | 0.101 | 1 | 11 | Mrps12 | 0.072 | Old |

| | | | | | | | | |
|---|---|---|---|---|---|---|---|---|
| 0.074707 | 0.249805 | 0.153 | 0.087 | 1 | 11 | Hmgn2 | 0.066 | Old |
| 0.074736 | 0.17328 | 0.331 | 0.217 | 1 | 11 | Sar1a | 0.114 | Old |
| 0.074777 | 0.1615 | 0.258 | 0.167 | 1 | 11 | Smarcc1 | 0.091 | Old |
| 0.074795 | 0.188429 | 0.601 | 0.449 | 1 | 11 | Syngr2 | 0.152 | Old |
| 0.074891 | 1.041422 | 0.085 | 0.036 | 1 | 11 | Agk | 0.049 | Old |
| 0.07497 | 0.147236 | 0.117 | 0.058 | 1 | 11 | Il15ra | 0.059 | Old |
| 0.07501 | 0.184642 | 0.238 | 0.152 | 1 | 11 | Aldh9a1 | 0.086 | Old |
| 0.075013 | 1.425696 | 0.073 | 0.029 | 1 | 11 | Mcu | 0.044 | Old |
| 0.075118 | 0.305089 | 0.274 | 0.181 | 1 | 11 | Dennd6a | 0.093 | Old |
| 0.075135 | 0.073726 | 0.367 | 0.254 | 1 | 11 | Ppp1r14b | 0.113 | Old |
| 0.075185 | 0.323306 | 0.69 | 0.572 | 1 | 11 | Rpl4 | 0.118 | Old |
| 0.075206 | 0.007735 | 0.29 | 0.196 | 1 | 11 | Plekhj1 | 0.094 | Old |
| 0.075407 | 0.089194 | 0.254 | 0.159 | 1 | 11 | C1qbp | 0.095 | Old |
| 0.075437 | 0.081832 | 0.589 | 0.435 | 1 | 11 | Psme2 | 0.154 | Old |
| 0.07548 | 0.253611 | 0.508 | 0.362 | 1 | 11 | Tmco1 | 0.146 | Old |
| 0.075708 | 0.698144 | 0.125 | 0.065 | 1 | 11 | Sdhaf1 | 0.06 | Old |
| 0.075708 | 0.533728 | 0.125 | 0.065 | 1 | 11 | Hdgfl2 | 0.06 | Old |
| 0.075712 | 0.325669 | 0.202 | 0.123 | 1 | 11 | Htra2 | 0.079 | Old |
| 0.075743 | 0.13075 | 0.379 | 0.268 | 1 | 11 | Rbbp6 | 0.111 | Old |
| 0.075789 | 0.054722 | 0.238 | 0.145 | 1 | 11 | Zdhhc21 | 0.093 | Old |
| 0.075792 | 1.565838 | 0.073 | 0.029 | 1 | 11 | Fam149b | 0.044 | Old |
| 0.075841 | 0.506257 | 0.105 | 0.051 | 1 | 11 | Bnip3 | 0.054 | Old |
| 0.075945 | 0.629493 | 0.161 | 0.094 | 1 | 11 | Agpat3 | 0.067 | Old |
| 0.075971 | 0.534879 | 0.085 | 0.036 | 1 | 11 | Zfyve1 | 0.049 | Old |
| 0.075971 | 0.530781 | 0.085 | 0.036 | 1 | 11 | Gdpd1 | 0.049 | Old |
| 0.075971 | 0.508652 | 0.085 | 0.036 | 1 | 11 | C1rl | 0.049 | Old |
| 0.076051 | 0.058953 | 0.516 | 0.355 | 1 | 11 | Bola2 | 0.161 | Old |
| 0.076166 | 0.588134 | 0.105 | 0.051 | 1 | 11 | Yod1 | 0.054 | Old |
| 0.076266 | 0.450941 | 0.185 | 0.109 | 1 | 11 | Elovl5 | 0.076 | Old |
| 0.076293 | 1.207618 | 0.113 | 0.058 | 1 | 11 | Rinl | 0.055 | Old |
| 0.076303 | 0.706307 | 0.125 | 0.065 | 1 | 11 | Dubr | 0.06 | Old |
| 0.076316 | 0.331204 | 0.137 | 0.072 | 1 | 11 | Nabp2 | 0.065 | Old |
| 0.076368 | 0.140687 | 0.294 | 0.196 | 1 | 11 | Acadm | 0.098 | Old |
| 0.076387 | 0.092981 | 0.548 | 0.406 | 1 | 11 | Cmpk1 | 0.142 | Old |
| 0.076467 | 0.051322 | 0.524 | 0.384 | 1 | 11 | Ddx39b | 0.14 | Old |
| 0.076491 | 0.054181 | 0.27 | 0.167 | 1 | 11 | Rsl1d1 | 0.103 | Old |
| 0.076497 | 0.732777 | 0.133 | 0.072 | 1 | 11 | Cep250 | 0.061 | Old |
| 0.076497 | 0.034287 | 0.415 | 0.297 | 1 | 11 | Fuca1 | 0.118 | Old |
| 0.076607 | 0.203398 | 0.254 | 0.167 | 1 | 11 | Wbp1l | 0.087 | Old |
| 0.076706 | 0.120691 | 0.641 | 0.493 | 1 | 11 | Atp6ap1 | 0.148 | Old |
| 0.076922 | 0.133793 | 0.145 | 0.08 | 1 | 11 | Ino80 | 0.065 | Old |
| 0.077019 | 0.327723 | 0.23 | 0.152 | 1 | 11 | Snx30 | 0.078 | Old |
| 0.077058 | 0.086901 | 0.222 | 0.138 | 1 | 11 | Zdhhc9 | 0.084 | Old |
| 0.077085 | 0.288013 | 0.282 | 0.188 | 1 | 11 | Pcyt1a | 0.094 | Old |
| 0.077273 | 0.545592 | 0.306 | 0.217 | 1 | 11 | D1Ertd622[illegible] | 0.089 | Old |

| | | | | | | | | |
|---|---|---|---|---|---|---|---|---|
| 0.077359 | 0.089381 | 0.258 | 0.167 | 1 | 11 | Hadha | 0.091 | Old |
| 0.077368 | 1.32286 | 0.073 | 0.029 | 1 | 11 | Mgst3 | 0.044 | Old |
| 0.07743 | 0.318367 | 0.085 | 0.036 | 1 | 11 | Creb3l2 | 0.049 | Old |
| 0.07743 | 0.314381 | 0.218 | 0.138 | 1 | 11 | Pgrmc1 | 0.08 | Old |
| 0.077468 | 0.174752 | 0.137 | 0.072 | 1 | 11 | Arfrp1 | 0.065 | Old |
| 0.077504 | 0.244459 | 0.173 | 0.101 | 1 | 11 | Trim33 | 0.072 | Old |
| 0.077553 | 0.939503 | 0.093 | 0.043 | 1 | 11 | Coq7 | 0.05 | Old |
| 0.07778 | 0.103634 | 0.46 | 0.319 | 1 | 11 | Pura | 0.141 | Old |
| 0.077877 | 0.04652 | 0.464 | 0.341 | 1 | 11 | N4bp1 | 0.123 | Old |
| 0.077903 | 0.132936 | 0.5 | 0.377 | 1 | 11 | Vasp | 0.123 | Old |
| 0.078123 | 0.271224 | 0.847 | 0.775 | 1 | 11 | Sirpa | 0.072 | Old |
| 0.078278 | 0.094386 | 0.331 | 0.232 | 1 | 11 | Etfb | 0.099 | Old |
| 0.078341 | 0.053619 | 0.137 | 0.072 | 1 | 11 | Ppid | 0.065 | Old |
| 0.078441 | 0.169813 | 0.165 | 0.094 | 1 | 11 | Tars | 0.071 | Old |
| 0.078522 | 0.837039 | 0.141 | 0.08 | 1 | 11 | Apmap | 0.061 | Old |
| 0.07871 | 0.192114 | 0.165 | 0.094 | 1 | 11 | Stx8 | 0.071 | Old |
| 0.078754 | 0.096219 | 0.573 | 0.42 | 1 | 11 | Adam15 | 0.153 | Old |
| 0.078761 | 0.124122 | 0.226 | 0.138 | 1 | 11 | Rcor1 | 0.088 | Old |
| 0.078783 | 0.30766 | 0.153 | 0.087 | 1 | 11 | Pold3 | 0.066 | Old |
| 0.078864 | 0.065933 | 0.315 | 0.21 | 1 | 11 | Brd7 | 0.105 | Old |
| 0.079062 | 0.161698 | 0.395 | 0.268 | 1 | 11 | Eif3i | 0.127 | Old |
| 0.079068 | 0.086262 | 0.206 | 0.123 | 1 | 11 | Fuca2 | 0.083 | Old |
| 0.07907 | 0.250163 | 0.395 | 0.275 | 1 | 11 | Hspa4 | 0.12 | Old |
| 0.079242 | 0.716448 | 0.177 | 0.109 | 1 | 11 | Pdcd2l | 0.068 | Old |
| 0.079336 | 0.51685 | 0.125 | 0.065 | 1 | 11 | Akap8l | 0.06 | Old |
| 0.07934 | 0.284146 | 0.153 | 0.087 | 1 | 11 | Knop1 | 0.066 | Old |
| 0.079473 | 0.372152 | 0.105 | 0.051 | 1 | 11 | Gm15417 | 0.054 | Old |
| 0.079482 | 0.212035 | 0.315 | 0.21 | 1 | 11 | Mat2b | 0.105 | Old |
| 0.079518 | 0.233669 | 0.165 | 0.094 | 1 | 11 | Nans | 0.071 | Old |
| 0.079591 | 0.215383 | 0.597 | 0.428 | 1 | 11 | Etv3 | 0.169 | Old |
| 0.079605 | 0.567708 | 0.198 | 0.123 | 1 | 11 | Jdp2 | 0.075 | Old |
| 0.080034 | 0.931162 | 0.093 | 0.043 | 1 | 11 | Itpa | 0.05 | Old |
| 0.080101 | 0.415129 | 0.226 | 0.145 | 1 | 11 | 1810030O( | 0.081 | Old |
| 0.080354 | 0.086721 | 0.524 | 0.355 | 1 | 11 | Itsn2 | 0.169 | Old |
| 0.080384 | 0.317698 | 0.351 | 0.232 | 1 | 11 | Frmd4a | 0.119 | Old |
| 0.080417 | 0.335569 | 0.085 | 0.036 | 1 | 11 | Rnf157 | 0.049 | Old |
| 0.080537 | 0.223322 | 0.468 | 0.341 | 1 | 11 | Cd164 | 0.127 | Old |
| 0.0806 | 0.233814 | 0.073 | 0.029 | 1 | 11 | Rtl8b | 0.044 | Old |
| 0.080664 | 0.166338 | 0.173 | 0.101 | 1 | 11 | Ago4 | 0.072 | Old |
| 0.080726 | 0.16877 | 0.677 | 0.522 | 1 | 11 | Emp3 | 0.155 | Old |
| 0.080828 | 0.447625 | 0.105 | 0.051 | 1 | 11 | Ormdl3 | 0.054 | Old |
| 0.080903 | 0.074451 | 0.238 | 0.152 | 1 | 11 | Mmp14 | 0.086 | Old |
| 0.08092 | 0.014618 | 0.145 | 0.08 | 1 | 11 | Akap10 | 0.065 | Old |
| 0.08096 | 0.084878 | 0.379 | 0.261 | 1 | 11 | Idh1 | 0.118 | Old |
| 0.08117 | 0.60397 | 0.105 | 0.051 | 1 | 11 | Asb7 | 0.054 | Old |

| | | | | | | | | |
|---|---|---|---|---|---|---|---|---|
| 0.081399 | 0.28055 | 0.246 | 0.159 | 1 | 11 | Heca | 0.087 | Old |
| 0.081481 | 0.519544 | 0.093 | 0.043 | 1 | 11 | Champ1 | 0.05 | Old |
| 0.081513 | 0.123791 | 0.105 | 0.051 | 1 | 11 | Zfp24 | 0.054 | Old |
| 0.081521 | 0.1292 | 0.528 | 0.399 | 1 | 11 | Hnrnph2 | 0.129 | Old |
| 0.081592 | 0.195396 | 0.726 | 0.543 | 1 | 11 | Fyb | 0.183 | Old |
| 0.08163 | 0.216967 | 0.673 | 0.507 | 1 | 11 | Xbp1 | 0.166 | Old |
| 0.081709 | 0.800809 | 0.121 | 0.065 | 1 | 11 | Ado | 0.056 | Old |
| 0.08176 | 0.14866 | 0.21 | 0.13 | 1 | 11 | Ddx18 | 0.08 | Old |
| 0.082018 | 0.202387 | 0.383 | 0.268 | 1 | 11 | Bptf | 0.115 | Old |
| 0.082223 | 0.188457 | 0.218 | 0.138 | 1 | 11 | Mrpl36 | 0.08 | Old |
| 0.082257 | 0.642185 | 0.073 | 0.029 | 1 | 11 | Ankrd26 | 0.044 | Old |
| 0.082675 | 0.797577 | 0.073 | 0.029 | 1 | 11 | Stk39 | 0.044 | Old |
| 0.082763 | 0.421791 | 0.113 | 0.058 | 1 | 11 | Ctnnbip1 | 0.055 | Old |
| 0.082943 | 0.285388 | 0.125 | 0.065 | 1 | 11 | Urm1 | 0.06 | Old |
| 0.082968 | 0.002188 | 0.242 | 0.152 | 1 | 11 | Cdk19 | 0.09 | Old |
| 0.083033 | 0.115763 | 0.161 | 0.094 | 1 | 11 | Cul4a | 0.067 | Old |
| 0.083039 | 0.034511 | 0.153 | 0.087 | 1 | 11 | Srpk1 | 0.066 | Old |
| 0.083072 | 0.022508 | 0.363 | 0.246 | 1 | 11 | Hp1bp3 | 0.117 | Old |
| 0.083102 | 2.487821 | 0.036 | 0.007 | 1 | 11 | E130311K1 | 0.029 | Old |
| 0.083102 | 2.340979 | 0.036 | 0.007 | 1 | 11 | Ankrd39 | 0.029 | Old |
| 0.083336 | 0.235456 | 0.278 | 0.181 | 1 | 11 | Zc3h12a | 0.097 | Old |
| 0.083426 | 0.457817 | 0.194 | 0.123 | 1 | 11 | Nfatc3 | 0.071 | Old |
| 0.083636 | 1.491961 | 0.06 | 0.022 | 1 | 11 | Pnpo | 0.038 | Old |
| 0.083708 | 0.604566 | 0.133 | 0.072 | 1 | 11 | Hlcs | 0.061 | Old |
| 0.083708 | 0.436508 | 0.133 | 0.072 | 1 | 11 | Ttc1 | 0.061 | Old |
| 0.083719 | 2.333691 | 0.036 | 0.007 | 1 | 11 | Trim59 | 0.029 | Old |
| 0.083719 | 2.172841 | 0.036 | 0.007 | 1 | 11 | Pcyox1l | 0.029 | Old |
| 0.083741 | 0.21424 | 0.367 | 0.254 | 1 | 11 | Bnip3l | 0.113 | Old |
| 0.083768 | 0.106535 | 0.214 | 0.13 | 1 | 11 | Vbp1 | 0.084 | Old |
| 0.083831 | 0.75456 | 0.141 | 0.08 | 1 | 11 | Tdg | 0.061 | Old |
| 0.083863 | 0.129783 | 0.286 | 0.188 | 1 | 11 | Insig1 | 0.098 | Old |
| 0.083885 | 0.332759 | 0.161 | 0.094 | 1 | 11 | AY036118 | 0.067 | Old |
| 0.083928 | 0.088099 | 0.173 | 0.101 | 1 | 11 | Crybg3 | 0.072 | Old |
| 0.083941 | 0.606503 | 0.073 | 0.029 | 1 | 11 | Fars2 | 0.044 | Old |
| 0.084064 | 1.079182 | 0.093 | 0.043 | 1 | 11 | Fmc1 | 0.05 | Old |
| 0.084067 | 0.157438 | 0.125 | 0.065 | 1 | 11 | Snrnp25 | 0.06 | Old |
| 0.084134 | 0.375112 | 0.141 | 0.08 | 1 | 11 | Strn | 0.061 | Old |
| 0.084147 | 0.288759 | 0.29 | 0.196 | 1 | 11 | Eif1ax | 0.094 | Old |
| 0.08417 | 0.485083 | 0.161 | 0.094 | 1 | 11 | Hras | 0.067 | Old |
| 0.084179 | 0.141432 | 0.145 | 0.08 | 1 | 11 | Ptgr2 | 0.065 | Old |
| 0.084249 | 0.28395 | 0.391 | 0.268 | 1 | 11 | Ssr4 | 0.123 | Old |
| 0.084367 | 0.637223 | 0.073 | 0.029 | 1 | 11 | Plekhg1 | 0.044 | Old |
| 0.08438 | 0.10016 | 0.512 | 0.362 | 1 | 11 | Ppp2r5c | 0.15 | Old |
| 0.084451 | 0.135269 | 0.181 | 0.109 | 1 | 11 | Fam120b | 0.072 | Old |
| 0.084467 | 0.280387 | 0.427 | 0.304 | 1 | 11 | Ndufv3 | 0.123 | Old |

| | | | | | | | | |
|---|---|---|---|---|---|---|---|---|
| 0.084571 | 1.710996 | 0.06 | 0.022 | 1 | 11 | Nt5m | 0.038 | Old |
| 0.084571 | 0.933847 | 0.06 | 0.022 | 1 | 11 | Rab13 | 0.038 | Old |
| 0.084652 | 0.251711 | 0.105 | 0.051 | 1 | 11 | Gkap1 | 0.054 | Old |
| 0.084677 | 0.433264 | 0.085 | 0.036 | 1 | 11 | Eci1 | 0.049 | Old |
| 0.084835 | 0.223114 | 0.327 | 0.225 | 1 | 11 | Csnk1e | 0.102 | Old |
| 0.084936 | 0.111934 | 0.758 | 0.587 | 1 | 11 | Sh3bgrl | 0.171 | Old |
| 0.084963 | 2.370053 | 0.036 | 0.007 | 1 | 11 | Rbm12b1 | 0.029 | Old |
| 0.084963 | 1.949951 | 0.036 | 0.007 | 1 | 11 | Tnfsf15 | 0.029 | Old |
| 0.084963 | 1.837871 | 0.036 | 0.007 | 1 | 11 | Rarb | 0.029 | Old |
| 0.084963 | 1.837154 | 0.036 | 0.007 | 1 | 11 | Creb3l1 | 0.029 | Old |
| 0.084977 | 0.192747 | 0.198 | 0.123 | 1 | 11 | Ing2 | 0.075 | Old |
| 0.085042 | 1.256817 | 0.06 | 0.022 | 1 | 11 | Ints10 | 0.038 | Old |
| 0.085042 | 1.128353 | 0.06 | 0.022 | 1 | 11 | Ints8 | 0.038 | Old |
| 0.085042 | 0.872128 | 0.06 | 0.022 | 1 | 11 | Ift140 | 0.038 | Old |
| 0.085087 | 0.062703 | 0.153 | 0.087 | 1 | 11 | Insig2 | 0.066 | Old |
| 0.085105 | 0.06498 | 0.706 | 0.536 | 1 | 11 | Ptprj | 0.17 | Old |
| 0.085141 | 0.288958 | 0.375 | 0.261 | 1 | 11 | Uvrag | 0.114 | Old |
| 0.085191 | 0.470177 | 0.093 | 0.043 | 1 | 11 | Ttll12 | 0.05 | Old |
| 0.085238 | 0.296054 | 0.19 | 0.116 | 1 | 11 | Tpm1 | 0.074 | Old |
| 0.085318 | 2.592216 | 0.048 | 0.014 | 1 | 11 | Xlr4b | 0.034 | Old |
| 0.085319 | 0.066242 | 0.173 | 0.101 | 1 | 11 | Rbmxl1 | 0.072 | Old |
| 0.085591 | 2.144167 | 0.036 | 0.007 | 1 | 11 | Aqp9 | 0.029 | Old |
| 0.085591 | 1.757019 | 0.036 | 0.007 | 1 | 11 | Zfp53 | 0.029 | Old |
| 0.085601 | 0.098014 | 0.508 | 0.37 | 1 | 11 | 2610507B1 | 0.138 | Old |
| 0.085816 | 0.081045 | 0.113 | 0.058 | 1 | 11 | Ints3 | 0.055 | Old |
| 0.085829 | 0.303726 | 0.524 | 0.406 | 1 | 11 | Tbca | 0.118 | Old |
| 0.085913 | 0.266913 | 0.133 | 0.072 | 1 | 11 | Ddx1 | 0.061 | Old |
| 0.085949 | 0.420579 | 0.093 | 0.043 | 1 | 11 | Incenp | 0.05 | Old |
| 0.08599 | 1.046996 | 0.06 | 0.022 | 1 | 11 | Cd200r4 | 0.038 | Old |
| 0.08616 | 0.458642 | 0.113 | 0.058 | 1 | 11 | Gk | 0.055 | Old |
| 0.086223 | 2.248381 | 0.036 | 0.007 | 1 | 11 | Sh3bgrl2 | 0.029 | Old |
| 0.086223 | 2.064826 | 0.036 | 0.007 | 1 | 11 | Ptgfrn | 0.029 | Old |
| 0.086223 | 1.56399 | 0.036 | 0.007 | 1 | 11 | Ffar4 | 0.029 | Old |
| 0.086223 | 1.549974 | 0.036 | 0.007 | 1 | 11 | Akr1c12 | 0.029 | Old |
| 0.086282 | 0.272178 | 0.141 | 0.08 | 1 | 11 | Smim20 | 0.061 | Old |
| 0.086312 | 0.118404 | 0.23 | 0.145 | 1 | 11 | Mid1ip1 | 0.085 | Old |
| 0.086314 | 0.163274 | 0.476 | 0.333 | 1 | 11 | Aamp | 0.143 | Old |
| 0.086373 | 0.06052 | 0.25 | 0.159 | 1 | 11 | Map2k7 | 0.091 | Old |
| 0.086467 | 1.117542 | 0.06 | 0.022 | 1 | 11 | P2ry10b | 0.038 | Old |
| 0.086467 | 0.855906 | 0.06 | 0.022 | 1 | 11 | Ropn1l | 0.038 | Old |
| 0.08652 | 0.353474 | 0.073 | 0.029 | 1 | 11 | Whamm | 0.044 | Old |
| 0.086853 | 0.660478 | 0.113 | 0.058 | 1 | 11 | Eif2b4 | 0.055 | Old |
| 0.086858 | 1.812069 | 0.036 | 0.007 | 1 | 11 | Cmtr2 | 0.029 | Old |
| 0.086858 | 1.58998 | 0.036 | 0.007 | 1 | 11 | Zgpat | 0.029 | Old |
| 0.086858 | 1.545034 | 0.036 | 0.007 | 1 | 11 | Cdc42bpg | 0.029 | Old |

| | | | | | | | | |
|---|---|---|---|---|---|---|---|---|
| 0.086904 | 0.120564 | 0.141 | 0.08 | 1 | 11 | Sgcb | 0.061 | Old |
| 0.086919 | 0.11228 | 0.238 | 0.152 | 1 | 11 | Tmem131 | 0.086 | Old |
| 0.086931 | 1.686366 | 0.048 | 0.014 | 1 | 11 | Ccdc130 | 0.034 | Old |
| 0.086946 | 1.238101 | 0.06 | 0.022 | 1 | 11 | Gpr18 | 0.038 | Old |
| 0.087205 | 0.221578 | 0.476 | 0.348 | 1 | 11 | Tm9sf2 | 0.128 | Old |
| 0.087357 | 0.123705 | 0.214 | 0.13 | 1 | 11 | Zfp217 | 0.084 | Old |
| 0.087378 | 0.15027 | 0.363 | 0.261 | 1 | 11 | Adap2 | 0.102 | Old |
| 0.087394 | 0.181767 | 0.073 | 0.029 | 1 | 11 | Med24 | 0.044 | Old |
| 0.087428 | 0.553133 | 0.06 | 0.022 | 1 | 11 | Pithd1 | 0.038 | Old |
| 0.087474 | 1.680239 | 0.048 | 0.014 | 1 | 11 | Kynu | 0.034 | Old |
| 0.087482 | 1.131984 | 0.093 | 0.043 | 1 | 11 | Ciz1 | 0.05 | Old |
| 0.087482 | 0.436096 | 0.093 | 0.043 | 1 | 11 | Pigc | 0.05 | Old |
| 0.087497 | 1.556351 | 0.036 | 0.007 | 1 | 11 | Lrrc28 | 0.029 | Old |
| 0.087497 | 1.36782 | 0.036 | 0.007 | 1 | 11 | Gm26982 | 0.029 | Old |
| 0.087497 | 1.227068 | 0.036 | 0.007 | 1 | 11 | Cox7a1 | 0.029 | Old |
| 0.087497 | 1.225941 | 0.036 | 0.007 | 1 | 11 | Etaa1 | 0.029 | Old |
| 0.087524 | 0.140795 | 0.105 | 0.051 | 1 | 11 | Agl | 0.054 | Old |
| 0.087806 | 0.31692 | 0.319 | 0.225 | 1 | 11 | Tns1 | 0.094 | Old |
| 0.087911 | 0.938101 | 0.06 | 0.022 | 1 | 11 | Mib2 | 0.038 | Old |
| 0.087915 | 0.233131 | 0.238 | 0.152 | 1 | 11 | 4930523C0 | 0.086 | Old |
| 0.088019 | 1.760061 | 0.048 | 0.014 | 1 | 11 | Ippk | 0.034 | Old |
| 0.088023 | 0.975655 | 0.101 | 0.051 | 1 | 11 | Xxylt1 | 0.05 | Old |
| 0.08814 | 1.23011 | 0.036 | 0.007 | 1 | 11 | Rbm4 | 0.029 | Old |
| 0.08814 | 1.116913 | 0.036 | 0.007 | 1 | 11 | 6330562C2 | 0.029 | Old |
| 0.08814 | 0.644984 | 0.036 | 0.007 | 1 | 11 | Asb6 | 0.029 | Old |
| 0.08814 | 0.641188 | 0.036 | 0.007 | 1 | 11 | Cd209e | 0.029 | Old |
| 0.088163 | 0.095479 | 0.375 | 0.261 | 1 | 11 | Mcur1 | 0.114 | Old |
| 0.088325 | 0.085525 | 0.238 | 0.159 | 1 | 11 | Exoc3 | 0.079 | Old |
| 0.088568 | 1.746116 | 0.048 | 0.014 | 1 | 11 | Frat2 | 0.034 | Old |
| 0.088568 | 1.743428 | 0.048 | 0.014 | 1 | 11 | Acp6 | 0.034 | Old |
| 0.088616 | 0.584533 | 0.185 | 0.116 | 1 | 11 | Dars | 0.069 | Old |
| 0.088734 | 0.32366 | 0.173 | 0.101 | 1 | 11 | Crtc2 | 0.072 | Old |
| 0.088758 | 0.156523 | 0.621 | 0.478 | 1 | 11 | Uqcrq | 0.143 | Old |
| 0.088787 | 1.079152 | 0.036 | 0.007 | 1 | 11 | Zfp930 | 0.029 | Old |
| 0.088787 | 0.881892 | 0.036 | 0.007 | 1 | 11 | Per2 | 0.029 | Old |
| 0.088787 | 0.818451 | 0.036 | 0.007 | 1 | 11 | Ggcx | 0.029 | Old |
| 0.088787 | 0.718398 | 0.036 | 0.007 | 1 | 11 | Exoc3l4 | 0.029 | Old |
| 0.088787 | 0.28144 | 0.036 | 0.007 | 1 | 11 | Kpna2 | 0.029 | Old |
| 0.088795 | 0.166457 | 0.323 | 0.225 | 1 | 11 | Myo9a | 0.098 | Old |
| 0.089089 | 0.06325 | 0.758 | 0.645 | 1 | 11 | Fus | 0.113 | Old |
| 0.089119 | 1.360236 | 0.048 | 0.014 | 1 | 11 | Plk4 | 0.034 | Old |
| 0.089119 | 1.351454 | 0.048 | 0.014 | 1 | 11 | Mical1 | 0.034 | Old |
| 0.089119 | 1.290779 | 0.048 | 0.014 | 1 | 11 | Abhd10 | 0.034 | Old |
| 0.089375 | 0.578473 | 0.06 | 0.022 | 1 | 11 | Fam210a | 0.038 | Old |
| 0.089461 | 0.25666 | 0.367 | 0.261 | 1 | 11 | Abcg1 | 0.106 | Old |

| | | | | | | | | |
|---|---|---|---|---|---|---|---|---|
| 0.089497 | 0.078646 | 0.21 | 0.13 | 1 | 11 | Stk24 | 0.08 | Old |
| 0.089673 | 1.618483 | 0.048 | 0.014 | 1 | 11 | 1810024B0 | 0.034 | Old |
| 0.089673 | 1.343325 | 0.048 | 0.014 | 1 | 11 | Il12rb1 | 0.034 | Old |
| 0.089673 | 1.070233 | 0.048 | 0.014 | 1 | 11 | Clec16a | 0.034 | Old |
| 0.089673 | 1.065655 | 0.048 | 0.014 | 1 | 11 | Tcta | 0.034 | Old |
| 0.089673 | 0.784774 | 0.048 | 0.014 | 1 | 11 | Pnpla1 | 0.034 | Old |
| 0.089781 | 1.752458 | 0.089 | 0.043 | 1 | 11 | Dusp7 | 0.046 | Old |
| 0.089822 | 0.151101 | 0.093 | 0.043 | 1 | 11 | Exoc7 | 0.05 | Old |
| 0.089867 | 0.749834 | 0.06 | 0.022 | 1 | 11 | Mon1a | 0.038 | Old |
| 0.089867 | 0.526473 | 0.06 | 0.022 | 1 | 11 | Taf6 | 0.038 | Old |
| 0.089886 | 0.217339 | 0.698 | 0.551 | 1 | 11 | Reep5 | 0.147 | Old |
| 0.090047 | 0.128613 | 0.206 | 0.13 | 1 | 11 | Tmem5 | 0.076 | Old |
| 0.090189 | 0.006656 | 0.173 | 0.101 | 1 | 11 | Zfp385a | 0.072 | Old |
| 0.09023 | 1.66419 | 0.048 | 0.014 | 1 | 11 | Blzf1 | 0.034 | Old |
| 0.090257 | 0.212118 | 0.387 | 0.268 | 1 | 11 | Tnfrsf1a | 0.119 | Old |
| 0.090313 | 0.042935 | 0.25 | 0.159 | 1 | 11 | Cops4 | 0.091 | Old |
| 0.090314 | 0.168243 | 0.456 | 0.326 | 1 | 11 | Klf13 | 0.13 | Old |
| 0.090353 | 0.534897 | 0.121 | 0.065 | 1 | 11 | Terf2 | 0.056 | Old |
| 0.09057 | 0.97938 | 0.081 | 0.036 | 1 | 11 | Mrpl58 | 0.045 | Old |
| 0.09079 | 1.171254 | 0.048 | 0.014 | 1 | 11 | Trmt12 | 0.034 | Old |
| 0.09079 | 0.804913 | 0.048 | 0.014 | 1 | 11 | Fam58b | 0.034 | Old |
| 0.090925 | 0.122873 | 0.427 | 0.297 | 1 | 11 | Arhgef6 | 0.13 | Old |
| 0.091108 | 0.168344 | 0.673 | 0.486 | 1 | 11 | Hint1 | 0.187 | Old |
| 0.091134 | 0.014865 | 0.331 | 0.217 | 1 | 11 | Mpc2 | 0.114 | Old |
| 0.091262 | 0.076907 | 0.165 | 0.094 | 1 | 11 | Agpat4 | 0.071 | Old |
| 0.091286 | 0.116735 | 0.31 | 0.21 | 1 | 11 | Os9 | 0.1 | Old |
| 0.091352 | 1.171875 | 0.048 | 0.014 | 1 | 11 | Map4k2 | 0.034 | Old |
| 0.091352 | 0.632913 | 0.048 | 0.014 | 1 | 11 | Kptn | 0.034 | Old |
| 0.091357 | 0.214066 | 0.06 | 0.022 | 1 | 11 | Ints7 | 0.038 | Old |
| 0.09141 | 0.398624 | 0.093 | 0.043 | 1 | 11 | Riok2 | 0.05 | Old |
| 0.091453 | 0.158283 | 0.218 | 0.138 | 1 | 11 | Vps13b | 0.08 | Old |
| 0.091562 | 0.464501 | 0.234 | 0.152 | 1 | 11 | Tuba1a | 0.082 | Old |
| 0.091594 | 0.097242 | 0.19 | 0.116 | 1 | 11 | Anapc1 | 0.074 | Old |
| 0.091747 | 0.546409 | 0.121 | 0.065 | 1 | 11 | Spg11 | 0.056 | Old |
| 0.091794 | 0.869531 | 0.101 | 0.051 | 1 | 11 | Cacna1d | 0.05 | Old |
| 0.091852 | 1.035516 | 0.081 | 0.036 | 1 | 11 | Clpp | 0.045 | Old |
| 0.091858 | 0.353034 | 0.06 | 0.022 | 1 | 11 | 2010111I0 | 0.038 | Old |
| 0.091914 | 0.402522 | 0.169 | 0.101 | 1 | 11 | Bin2 | 0.068 | Old |
| 0.091918 | 0.604134 | 0.048 | 0.014 | 1 | 11 | Nck2 | 0.034 | Old |
| 0.091955 | 0.15118 | 0.206 | 0.13 | 1 | 11 | Spen | 0.076 | Old |
| 0.091973 | 0.044884 | 0.427 | 0.304 | 1 | 11 | Map3k1 | 0.123 | Old |
| 0.092178 | 0.503448 | 0.101 | 0.051 | 1 | 11 | Hsdl2 | 0.05 | Old |
| 0.092203 | 0.110982 | 0.21 | 0.13 | 1 | 11 | Suds3 | 0.08 | Old |
| 0.092292 | 0.096262 | 0.512 | 0.377 | 1 | 11 | Lbh | 0.135 | Old |
| 0.092374 | 0.104699 | 0.359 | 0.246 | 1 | 11 | Chtop | 0.113 | Old |

| | | | | | | | | |
|---|---|---|---|---|---|---|---|---|
| 0.092486 | 0.938744 | 0.048 | 0.014 | 1 | 11 | Btbd2 | 0.034 | Old |
| 0.092486 | 0.87535 | 0.048 | 0.014 | 1 | 11 | Ceacam16 | 0.034 | Old |
| 0.092486 | 0.8099 | 0.048 | 0.014 | 1 | 11 | Ift27 | 0.034 | Old |
| 0.092486 | 0.765347 | 0.048 | 0.014 | 1 | 11 | Gstm4 | 0.034 | Old |
| 0.092664 | 0.313096 | 0.141 | 0.08 | 1 | 11 | Tsfm | 0.061 | Old |
| 0.092702 | 0.078838 | 0.282 | 0.188 | 1 | 11 | Maz | 0.094 | Old |
| 0.092762 | 0.326308 | 0.157 | 0.094 | 1 | 11 | Espl1 | 0.063 | Old |
| 0.093074 | 0.618597 | 0.157 | 0.094 | 1 | 11 | Atg101 | 0.063 | Old |
| 0.093322 | 1.206674 | 0.153 | 0.094 | 1 | 11 | Gpr35 | 0.059 | Old |
| 0.093374 | 0.663901 | 0.06 | 0.022 | 1 | 11 | Gys1 | 0.038 | Old |
| 0.093408 | 0.225108 | 0.31 | 0.217 | 1 | 11 | Card19 | 0.093 | Old |
| 0.093411 | 0.059573 | 0.319 | 0.21 | 1 | 11 | Rtcb | 0.109 | Old |
| 0.09363 | 0.774991 | 0.048 | 0.014 | 1 | 11 | Fbxo46 | 0.034 | Old |
| 0.093723 | 0.155833 | 0.161 | 0.094 | 1 | 11 | Snapc5 | 0.067 | Old |
| 0.0938 | 0.027531 | 0.274 | 0.181 | 1 | 11 | Il6st | 0.093 | Old |
| 0.093817 | 0.045126 | 0.238 | 0.152 | 1 | 11 | Adipor2 | 0.086 | Old |
| 0.093874 | 0.215011 | 0.919 | 0.906 | 1 | 11 | Gpx1 | 0.013 | Old |
| 0.093884 | 0.026605 | 0.06 | 0.022 | 1 | 11 | Osbpl7 | 0.038 | Old |
| 0.094023 | 0.782863 | 0.081 | 0.036 | 1 | 11 | Leng9 | 0.045 | Old |
| 0.094078 | 3.790338 | 0.02 | 0 | 1 | 11 | Arg1 | 0.02 | Old |
| 0.094078 | 3.234561 | 0.02 | 0 | 1 | 11 | Plekha4 | 0.02 | Old |
| 0.094078 | 2.797856 | 0.02 | 0 | 1 | 11 | Armcx1 | 0.02 | Old |
| 0.094078 | 2.797189 | 0.02 | 0 | 1 | 11 | Ccdc114 | 0.02 | Old |
| 0.094078 | 2.739477 | 0.02 | 0 | 1 | 11 | Gm16062 | 0.02 | Old |
| 0.094078 | 2.626116 | 0.02 | 0 | 1 | 11 | Nadsyn1 | 0.02 | Old |
| 0.094078 | 2.607202 | 0.02 | 0 | 1 | 11 | Cav1 | 0.02 | Old |
| 0.094078 | 2.592183 | 0.02 | 0 | 1 | 11 | 2210417A( | 0.02 | Old |
| 0.094078 | 2.591828 | 0.02 | 0 | 1 | 11 | Otub2 | 0.02 | Old |
| 0.094078 | 2.569922 | 0.02 | 0 | 1 | 11 | Itfg2 | 0.02 | Old |
| 0.094078 | 2.549605 | 0.02 | 0 | 1 | 11 | Rflnb | 0.02 | Old |
| 0.094078 | 2.538624 | 0.02 | 0 | 1 | 11 | Chic1 | 0.02 | Old |
| 0.094078 | 2.49543 | 0.02 | 0 | 1 | 11 | Map3k15 | 0.02 | Old |
| 0.094078 | 2.494886 | 0.02 | 0 | 1 | 11 | N4bp3 | 0.02 | Old |
| 0.094078 | 2.460286 | 0.02 | 0 | 1 | 11 | Jmjd7 | 0.02 | Old |
| 0.094078 | 2.454524 | 0.02 | 0 | 1 | 11 | Pik3ip1 | 0.02 | Old |
| 0.094078 | 2.445444 | 0.02 | 0 | 1 | 11 | Ovca2 | 0.02 | Old |
| 0.094078 | 2.418963 | 0.02 | 0 | 1 | 11 | Dctd | 0.02 | Old |
| 0.094078 | 2.413921 | 0.02 | 0 | 1 | 11 | Clu | 0.02 | Old |
| 0.094078 | 2.411948 | 0.02 | 0 | 1 | 11 | Rtl6 | 0.02 | Old |
| 0.094078 | 2.39991 | 0.02 | 0 | 1 | 11 | Rnf123 | 0.02 | Old |
| 0.094078 | 2.378268 | 0.02 | 0 | 1 | 11 | Wee1 | 0.02 | Old |
| 0.094078 | 2.375166 | 0.02 | 0 | 1 | 11 | Fxn | 0.02 | Old |
| 0.094078 | 2.362295 | 0.02 | 0 | 1 | 11 | Cdk1 | 0.02 | Old |
| 0.094078 | 2.346352 | 0.02 | 0 | 1 | 11 | 1700037C1 | 0.02 | Old |
| 0.094078 | 2.319724 | 0.02 | 0 | 1 | 11 | Kcnq1 | 0.02 | Old |

| | | | | | | | | |
|---|---|---|---|---|---|---|---|---|
| 0.094078 | 2.317652 | 0.02 | 0 | 1 | 11 | Ddr2 | 0.02 | Old |
| 0.094078 | 2.309903 | 0.02 | 0 | 1 | 11 | Scnn1a | 0.02 | Old |
| 0.094078 | 2.304829 | 0.02 | 0 | 1 | 11 | Gm49085 | 0.02 | Old |
| 0.094078 | 2.2902 | 0.02 | 0 | 1 | 11 | Hist1h2be | 0.02 | Old |
| 0.094078 | 2.242414 | 0.02 | 0 | 1 | 11 | 3222401L1 | 0.02 | Old |
| 0.094078 | 2.21191 | 0.02 | 0 | 1 | 11 | Ptpn13 | 0.02 | Old |
| 0.094078 | 2.180545 | 0.02 | 0 | 1 | 11 | Etl4 | 0.02 | Old |
| 0.094078 | 2.17592 | 0.02 | 0 | 1 | 11 | Zcwpw1 | 0.02 | Old |
| 0.094078 | 2.122783 | 0.02 | 0 | 1 | 11 | Fastkd1 | 0.02 | Old |
| 0.094078 | 2.097111 | 0.02 | 0 | 1 | 11 | Aplf | 0.02 | Old |
| 0.094078 | 2.092991 | 0.02 | 0 | 1 | 11 | Asf1b | 0.02 | Old |
| 0.094078 | 2.075198 | 0.02 | 0 | 1 | 11 | Kctd21 | 0.02 | Old |
| 0.094078 | 2.055496 | 0.02 | 0 | 1 | 11 | Zfp202 | 0.02 | Old |
| 0.094078 | 2.01818 | 0.02 | 0 | 1 | 11 | Otof | 0.02 | Old |
| 0.094078 | 1.999806 | 0.02 | 0 | 1 | 11 | Kif1a | 0.02 | Old |
| 0.094078 | 1.989575 | 0.02 | 0 | 1 | 11 | Zfp366 | 0.02 | Old |
| 0.094078 | 1.977589 | 0.02 | 0 | 1 | 11 | Slc16a13 | 0.02 | Old |
| 0.094078 | 1.976421 | 0.02 | 0 | 1 | 11 | Ulk4 | 0.02 | Old |
| 0.094078 | 1.965189 | 0.02 | 0 | 1 | 11 | Ubox5 | 0.02 | Old |
| 0.094078 | 1.936253 | 0.02 | 0 | 1 | 11 | Tmem143 | 0.02 | Old |
| 0.094078 | 1.932657 | 0.02 | 0 | 1 | 11 | Prdm9 | 0.02 | Old |
| 0.094078 | 1.927994 | 0.02 | 0 | 1 | 11 | Fsd2 | 0.02 | Old |
| 0.094078 | 1.91217 | 0.02 | 0 | 1 | 11 | St14 | 0.02 | Old |
| 0.094078 | 1.889322 | 0.02 | 0 | 1 | 11 | Bvht | 0.02 | Old |
| 0.094078 | 1.862799 | 0.02 | 0 | 1 | 11 | Chchd10 | 0.02 | Old |
| 0.094078 | 1.808927 | 0.02 | 0 | 1 | 11 | Afap1 | 0.02 | Old |
| 0.094078 | 1.807241 | 0.02 | 0 | 1 | 11 | Rbks | 0.02 | Old |
| 0.094078 | 1.799237 | 0.02 | 0 | 1 | 11 | Gprc5a | 0.02 | Old |
| 0.094078 | 1.794484 | 0.02 | 0 | 1 | 11 | Zkscan5 | 0.02 | Old |
| 0.094078 | 1.77784 | 0.02 | 0 | 1 | 11 | Mocos | 0.02 | Old |
| 0.094078 | 1.758493 | 0.02 | 0 | 1 | 11 | Nphp3 | 0.02 | Old |
| 0.094078 | 1.753298 | 0.02 | 0 | 1 | 11 | Fam57a | 0.02 | Old |
| 0.094078 | 1.727041 | 0.02 | 0 | 1 | 11 | Peak1os | 0.02 | Old |
| 0.094078 | 1.66083 | 0.02 | 0 | 1 | 11 | Als2cl | 0.02 | Old |
| 0.094078 | 1.619631 | 0.02 | 0 | 1 | 11 | Lrp4 | 0.02 | Old |
| 0.094078 | 1.612353 | 0.02 | 0 | 1 | 11 | Fignl1 | 0.02 | Old |
| 0.094078 | 1.575489 | 0.02 | 0 | 1 | 11 | Plpp1 | 0.02 | Old |
| 0.094078 | 1.548252 | 0.02 | 0 | 1 | 11 | 3300005D( | 0.02 | Old |
| 0.094226 | 0.431302 | 0.121 | 0.065 | 1 | 11 | Lias | 0.056 | Old |
| 0.094311 | 0.221647 | 0.637 | 0.478 | 1 | 11 | Dync1h1 | 0.159 | Old |
| 0.094335 | 0.048254 | 0.153 | 0.087 | 1 | 11 | Carhsp1 | 0.066 | Old |
| 0.094438 | 0.10581 | 0.262 | 0.174 | 1 | 11 | Dnase1l3 | 0.088 | Old |
| 0.094509 | 1.135838 | 0.101 | 0.051 | 1 | 11 | Actn1 | 0.05 | Old |
| 0.094671 | 0.053208 | 0.56 | 0.406 | 1 | 11 | Rab7 | 0.154 | Old |
| 0.094693 | 0.560495 | 0.109 | 0.058 | 1 | 11 | Ero1lb | 0.051 | Old |

| | | | | | | | | |
|---|---|---|---|---|---|---|---|---|
| 0.094932 | 0.300892 | 0.262 | 0.181 | 1 | 11 | Glrx5 | 0.081 | Old |
| 0.095006 | 0.416478 | 0.149 | 0.087 | 1 | 11 | Manba | 0.062 | Old |
| 0.095115 | 0.240813 | 0.355 | 0.246 | 1 | 11 | Rer1 | 0.109 | Old |
| 0.095203 | 0.057458 | 0.133 | 0.072 | 1 | 11 | Tmub2 | 0.061 | Old |
| 0.095281 | 0.216327 | 0.157 | 0.094 | 1 | 11 | Rasgrp1 | 0.063 | Old |
| 0.095297 | 0.393508 | 0.101 | 0.051 | 1 | 11 | Arl2 | 0.05 | Old |
| 0.095341 | 0.211346 | 0.762 | 0.659 | 1 | 11 | Atp5e | 0.103 | Old |
| 0.095479 | 0.032306 | 0.093 | 0.043 | 1 | 11 | Ubr7 | 0.05 | Old |
| 0.095608 | 0.312729 | 0.161 | 0.094 | 1 | 11 | Brip1os | 0.067 | Old |
| 0.095768 | 0.049063 | 0.319 | 0.217 | 1 | 11 | Galc | 0.102 | Old |
| 0.095788 | 0.29017 | 0.081 | 0.036 | 1 | 11 | Osbpl3 | 0.045 | Old |
| 0.095788 | 0.272618 | 0.081 | 0.036 | 1 | 11 | Metrn | 0.045 | Old |
| 0.095928 | 0.553133 | 0.129 | 0.072 | 1 | 11 | Pik3r4 | 0.057 | Old |
| 0.095966 | 0.067322 | 0.258 | 0.167 | 1 | 11 | Atxn7l1 | 0.091 | Old |
| 0.095988 | 0.317514 | 0.149 | 0.087 | 1 | 11 | Atxn3 | 0.062 | Old |
| 0.096146 | 0.211447 | 0.427 | 0.297 | 1 | 11 | Dmac1 | 0.13 | Old |
| 0.096163 | 0.065156 | 0.141 | 0.08 | 1 | 11 | Zbtb21 | 0.061 | Old |
| 0.096212 | 0.038108 | 0.238 | 0.152 | 1 | 11 | Mgat2 | 0.086 | Old |
| 0.09623 | 0.037435 | 0.399 | 0.283 | 1 | 11 | Rassf1 | 0.116 | Old |
| 0.096234 | 0.635749 | 0.081 | 0.036 | 1 | 11 | Zdhhc8 | 0.045 | Old |
| 0.096611 | 0.212436 | 0.355 | 0.246 | 1 | 11 | Zdhhc20 | 0.109 | Old |
| 0.096633 | 0.495066 | 0.129 | 0.072 | 1 | 11 | Ifi205 | 0.057 | Old |
| 0.096646 | 0.507974 | 0.149 | 0.087 | 1 | 11 | Krt80 | 0.062 | Old |
| 0.096759 | 0.149554 | 0.121 | 0.065 | 1 | 11 | Sin3a | 0.056 | Old |
| 0.097127 | 0.001396 | 0.19 | 0.116 | 1 | 11 | Tm9sf4 | 0.074 | Old |
| 0.097309 | 0.31027 | 0.149 | 0.087 | 1 | 11 | Trak2 | 0.062 | Old |
| 0.097408 | 0.070982 | 0.258 | 0.174 | 1 | 11 | Hipk3 | 0.084 | Old |
| 0.097414 | 0.084391 | 0.113 | 0.058 | 1 | 11 | Pon3 | 0.055 | Old |
| 0.097473 | 0.266466 | 0.367 | 0.261 | 1 | 11 | Ppp6r3 | 0.106 | Old |
| 0.097484 | 0.069302 | 0.31 | 0.217 | 1 | 11 | Slc20a1 | 0.093 | Old |
| 0.097548 | 0.024273 | 0.294 | 0.203 | 1 | 11 | Baz2a | 0.091 | Old |
| 0.097985 | 0.25429 | 0.145 | 0.087 | 1 | 11 | Pi4ka | 0.058 | Old |
| 0.098055 | 0.18297 | 0.129 | 0.072 | 1 | 11 | Pde7a | 0.057 | Old |
| 0.098206 | 0.205966 | 0.383 | 0.275 | 1 | 11 | Sec61a1 | 0.108 | Old |
| 0.098347 | 0.373298 | 0.165 | 0.101 | 1 | 11 | Tnip1 | 0.064 | Old |
| 0.098492 | 0.095513 | 0.274 | 0.181 | 1 | 11 | Kmt5b | 0.093 | Old |
| 0.0985 | 0.333641 | 0.101 | 0.051 | 1 | 11 | Mid1 | 0.05 | Old |
| 0.098726 | 0.235807 | 0.141 | 0.08 | 1 | 11 | Gclc | 0.061 | Old |
| 0.098772 | 0.313717 | 0.129 | 0.072 | 1 | 11 | Exosc3 | 0.057 | Old |
| 0.098814 | 0.416486 | 0.185 | 0.116 | 1 | 11 | Cdk4 | 0.069 | Old |
| 0.098953 | 0.03256 | 0.198 | 0.123 | 1 | 11 | Kdm5c | 0.075 | Old |
| 0.099345 | 0.371991 | 0.121 | 0.065 | 1 | 11 | Kcnk13 | 0.056 | Old |
| 0.099399 | 0.712118 | 0.081 | 0.036 | 1 | 11 | Mrps31 | 0.045 | Old |
| 0.099621 | 0.095054 | 0.206 | 0.13 | 1 | 11 | Rars | 0.076 | Old |
| 0.099794 | 0.144436 | 0.964 | 0.928 | 1 | 11 | Fcer1g | 0.036 | Old |

| | | | | | | | | |
|---|---|---|---|---|---|---|---|---|
| 0.099992 | 0.060321 | 0.73 | 0.58 | 1 | 11 | Gnb1 | 0.15 | Old |
| 0.100115 | 0.14351 | 0.141 | 0.08 | 1 | 11 | Aldh3a2 | 0.061 | Old |
| 0.100465 | 0.218594 | 0.177 | 0.109 | 1 | 11 | Smpd1 | 0.068 | Old |
| 0.100545 | 0.491016 | 0.101 | 0.051 | 1 | 11 | Dcps | 0.05 | Old |
| 0.100584 | 0.7486 | 0.129 | 0.072 | 1 | 11 | Batf | 0.057 | Old |
| 0.100887 | 0.815047 | 0.109 | 0.058 | 1 | 11 | Fbxw5 | 0.051 | Old |
| 0.100939 | 0.011538 | 0.169 | 0.101 | 1 | 11 | Pak1ip1 | 0.068 | Old |
| 0.100958 | 0.3002 | 0.101 | 0.051 | 1 | 11 | Rasgrp2 | 0.05 | Old |
| 0.101098 | 0.249714 | 0.391 | 0.283 | 1 | 11 | Ppp2r1a | 0.108 | Old |
| 0.101227 | 0.29439 | 0.121 | 0.065 | 1 | 11 | Ranbp3 | 0.056 | Old |
| 0.101244 | 0.5087 | 0.081 | 0.036 | 1 | 11 | Lrig2 | 0.045 | Old |
| 0.10125 | 0.195843 | 0.165 | 0.101 | 1 | 11 | Il18bp | 0.064 | Old |
| 0.101293 | 0.096116 | 0.351 | 0.239 | 1 | 11 | Zmat2 | 0.112 | Old |
| 0.101485 | 0.399543 | 0.157 | 0.094 | 1 | 11 | Tufm | 0.063 | Old |
| 0.10153 | 0.153121 | 0.274 | 0.188 | 1 | 11 | Plekhm2 | 0.086 | Old |
| 0.101576 | 0.219607 | 0.165 | 0.101 | 1 | 11 | Zdhhc3 | 0.064 | Old |
| 0.101606 | 0.032083 | 0.121 | 0.065 | 1 | 11 | Uspl1 | 0.056 | Old |
| 0.101875 | 1.104199 | 0.069 | 0.029 | 1 | 11 | Kti12 | 0.04 | Old |
| 0.101875 | 1.103592 | 0.069 | 0.029 | 1 | 11 | Ctdp1 | 0.04 | Old |
| 0.101987 | 0.660932 | 0.202 | 0.138 | 1 | 11 | Cd209d | 0.064 | Old |
| 0.102201 | 0.157238 | 0.766 | 0.63 | 1 | 11 | Akr1a1 | 0.136 | Old |
| 0.102205 | 0.280728 | 0.101 | 0.051 | 1 | 11 | Nt5dc2 | 0.05 | Old |
| 0.102432 | 0.01563 | 0.375 | 0.261 | 1 | 11 | Arl8b | 0.114 | Old |
| 0.102493 | 0.450064 | 0.157 | 0.094 | 1 | 11 | Itpripl1 | 0.063 | Old |
| 0.10259 | 0.23759 | 0.274 | 0.188 | 1 | 11 | Herc4 | 0.086 | Old |
| 0.102621 | 0.162806 | 0.25 | 0.167 | 1 | 11 | Mga | 0.083 | Old |
| 0.102886 | 0.23121 | 0.206 | 0.13 | 1 | 11 | Ubp1 | 0.076 | Old |
| 0.103032 | 0.011896 | 0.44 | 0.312 | 1 | 11 | Tomm22 | 0.128 | Old |
| 0.103043 | 0.234751 | 0.101 | 0.051 | 1 | 11 | Smim12 | 0.05 | Old |
| 0.10315 | 0.017537 | 0.294 | 0.203 | 1 | 11 | Arf6 | 0.091 | Old |
| 0.103155 | 0.169797 | 0.222 | 0.145 | 1 | 11 | Ldb1 | 0.077 | Old |
| 0.103351 | 0.707929 | 0.089 | 0.043 | 1 | 11 | Lsm1 | 0.046 | Old |
| 0.103464 | 0.37288 | 0.101 | 0.051 | 1 | 11 | Cep85l | 0.05 | Old |
| 0.103687 | 0.240295 | 0.262 | 0.188 | 1 | 11 | Tceal9 | 0.074 | Old |
| 0.103696 | 0.586242 | 0.109 | 0.058 | 1 | 11 | Mfsd7a | 0.051 | Old |
| 0.103708 | 0.115172 | 0.492 | 0.362 | 1 | 11 | Rab2a | 0.13 | Old |
| 0.103919 | 0.759297 | 0.069 | 0.029 | 1 | 11 | Kank3 | 0.04 | Old |
| 0.103919 | 0.634651 | 0.069 | 0.029 | 1 | 11 | Atpaf1 | 0.04 | Old |
| 0.103948 | 0.137216 | 0.363 | 0.261 | 1 | 11 | Stk4 | 0.102 | Old |
| 0.104064 | 0.251755 | 0.081 | 0.036 | 1 | 11 | Recql | 0.045 | Old |
| 0.104191 | 0.172847 | 0.157 | 0.094 | 1 | 11 | Psmd10 | 0.063 | Old |
| 0.10421 | 0.223763 | 0.169 | 0.101 | 1 | 11 | Errfi1 | 0.068 | Old |
| 0.104311 | 0.408475 | 0.101 | 0.051 | 1 | 11 | Cfap36 | 0.05 | Old |
| 0.104367 | 0.015404 | 0.298 | 0.203 | 1 | 11 | Ufc1 | 0.095 | Old |
| 0.104435 | 0.786742 | 0.069 | 0.029 | 1 | 11 | Zfyve21 | 0.04 | Old |

| | | | | | | | | |
|---|---|---|---|---|---|---|---|---|
| 0.104533 | 0.103263 | 0.133 | 0.072 | 1 | 11 | Wdr3 | 0.061 | Old |
| 0.10454 | 0.055797 | 0.081 | 0.036 | 1 | 11 | Ndrg1 | 0.045 | Old |
| 0.104633 | 0.341283 | 0.202 | 0.13 | 1 | 11 | Ube2e3 | 0.072 | Old |
| 0.104637 | 0.690472 | 0.177 | 0.116 | 1 | 11 | Plxna1 | 0.061 | Old |
| 0.104662 | 0.177792 | 0.129 | 0.072 | 1 | 11 | Mcrs1 | 0.057 | Old |
| 0.104753 | 0.856829 | 0.117 | 0.065 | 1 | 11 | Apeh | 0.052 | Old |
| 0.104953 | 0.734428 | 0.069 | 0.029 | 1 | 11 | Gtf2h4 | 0.04 | Old |
| 0.104984 | 0.123604 | 0.327 | 0.225 | 1 | 11 | Uqcrc2 | 0.102 | Old |
| 0.105103 | 0.019308 | 0.492 | 0.355 | 1 | 11 | Gdap10 | 0.137 | Old |
| 0.105123 | 0.122335 | 0.431 | 0.304 | 1 | 11 | Ddost | 0.127 | Old |
| 0.105154 | 0.557136 | 0.198 | 0.13 | 1 | 11 | Zfp992 | 0.068 | Old |
| 0.105156 | 0.650274 | 0.089 | 0.043 | 1 | 11 | Dnaja3 | 0.046 | Old |
| 0.105221 | 0.231704 | 0.157 | 0.094 | 1 | 11 | Abl1 | 0.063 | Old |
| 0.105232 | 0.404146 | 0.173 | 0.109 | 1 | 11 | Fbxo4 | 0.064 | Old |
| 0.10525 | 0.057707 | 0.448 | 0.319 | 1 | 11 | Hook3 | 0.129 | Old |
| 0.105404 | 0.062779 | 0.742 | 0.601 | 1 | 11 | Rpl15 | 0.141 | Old |
| 0.105516 | 0.157381 | 0.597 | 0.442 | 1 | 11 | Eif5b | 0.155 | Old |
| 0.105566 | 0.366868 | 0.157 | 0.094 | 1 | 11 | Nom1 | 0.063 | Old |
| 0.105591 | 0.350992 | 0.101 | 0.051 | 1 | 11 | Fbxw8 | 0.05 | Old |
| 0.105591 | 0.075034 | 0.101 | 0.051 | 1 | 11 | Inpp4b | 0.05 | Old |
| 0.105678 | 0.068094 | 0.246 | 0.167 | 1 | 11 | Tnfrsf21 | 0.079 | Old |
| 0.10602 | 0.20755 | 0.101 | 0.051 | 1 | 11 | Cd99l2 | 0.05 | Old |
| 0.106068 | 0.693424 | 0.089 | 0.043 | 1 | 11 | Lrba | 0.046 | Old |
| 0.106296 | 0.07402 | 0.516 | 0.377 | 1 | 11 | Tnfrsf1b | 0.139 | Old |
| 0.106344 | 0.591975 | 0.145 | 0.087 | 1 | 11 | 2310001H1 | 0.058 | Old |
| 0.106443 | 0.048671 | 0.262 | 0.174 | 1 | 11 | Rab1b | 0.088 | Old |
| 0.106935 | 0.04775 | 0.819 | 0.638 | 1 | 11 | Hnrnpf | 0.181 | Old |
| 0.107021 | 0.041518 | 0.444 | 0.312 | 1 | 11 | Mcrip1 | 0.132 | Old |
| 0.107046 | 0.618671 | 0.069 | 0.029 | 1 | 11 | Rnf135 | 0.04 | Old |
| 0.107299 | 0.081252 | 0.222 | 0.145 | 1 | 11 | Cnpy2 | 0.077 | Old |
| 0.107325 | 0.086375 | 0.129 | 0.072 | 1 | 11 | Tbc1d9b | 0.057 | Old |
| 0.107416 | 0.138531 | 0.605 | 0.478 | 1 | 11 | Cndp2 | 0.127 | Old |
| 0.107433 | 0.796331 | 0.137 | 0.08 | 1 | 11 | Bloc1s2 | 0.057 | Old |
| 0.107448 | 0.315298 | 0.089 | 0.043 | 1 | 11 | Vps39 | 0.046 | Old |
| 0.107532 | 0.129759 | 0.609 | 0.442 | 1 | 11 | Prrc2c | 0.167 | Old |
| 0.107582 | 0.202326 | 0.573 | 0.449 | 1 | 11 | Ms4a7 | 0.124 | Old |
| 0.107885 | 2.206836 | 0.056 | 0.022 | 1 | 11 | Fam83f | 0.034 | Old |
| 0.108236 | 0.580649 | 0.109 | 0.058 | 1 | 11 | Ece1 | 0.051 | Old |
| 0.108328 | 0.242343 | 0.629 | 0.464 | 1 | 11 | Rpl29 | 0.165 | Old |
| 0.108711 | 0.172678 | 0.157 | 0.094 | 1 | 11 | Klf11 | 0.063 | Old |
| 0.108732 | 0.036785 | 0.306 | 0.217 | 1 | 11 | Psd3 | 0.089 | Old |
| 0.108826 | 0.104373 | 0.427 | 0.304 | 1 | 11 | Drap1 | 0.123 | Old |
| 0.108853 | 0.239862 | 0.181 | 0.116 | 1 | 11 | Hspa4l | 0.065 | Old |
| 0.108871 | 0.027325 | 0.129 | 0.072 | 1 | 11 | G2e3 | 0.057 | Old |
| 0.108886 | 0.210662 | 0.351 | 0.254 | 1 | 11 | Zfp710 | 0.097 | Old |

| | | | | | | | | |
|---|---|---|---|---|---|---|---|---|
| 0.108933 | 0.499938 | 0.137 | 0.08 | 1 | 11 | Trmt1 | 0.057 | Old |
| 0.108958 | 0.133608 | 0.383 | 0.268 | 1 | 11 | Sec11a | 0.115 | Old |
| 0.109064 | 1.391935 | 0.056 | 0.022 | 1 | 11 | Abhd5 | 0.034 | Old |
| 0.109065 | 0.277476 | 0.157 | 0.094 | 1 | 11 | Ezh1 | 0.063 | Old |
| 0.109172 | 0.807133 | 0.069 | 0.029 | 1 | 11 | Rars2 | 0.04 | Old |
| 0.109309 | 0.062115 | 0.302 | 0.21 | 1 | 11 | Rnf111 | 0.092 | Old |
| 0.109381 | 0.022853 | 0.879 | 0.696 | 1 | 11 | Ms4a6c | 0.183 | Old |
| 0.10953 | 0.647926 | 0.169 | 0.109 | 1 | 11 | Cirbp | 0.06 | Old |
| 0.109558 | 0.019033 | 0.274 | 0.181 | 1 | 11 | Tspyl1 | 0.093 | Old |
| 0.109618 | 0.375122 | 0.169 | 0.101 | 1 | 11 | Map1s | 0.068 | Old |
| 0.109657 | 1.192568 | 0.056 | 0.022 | 1 | 11 | Tmem9 | 0.034 | Old |
| 0.109776 | 0.012003 | 0.157 | 0.094 | 1 | 11 | Tcp11l2 | 0.063 | Old |
| 0.110032 | 0.252181 | 0.222 | 0.145 | 1 | 11 | Aagab | 0.077 | Old |
| 0.11025 | 0.585979 | 0.089 | 0.043 | 1 | 11 | Gps2 | 0.046 | Old |
| 0.110252 | 1.051411 | 0.056 | 0.022 | 1 | 11 | Gm45606 | 0.034 | Old |
| 0.110354 | 0.049489 | 0.585 | 0.457 | 1 | 11 | Gbp7 | 0.128 | Old |
| 0.110592 | 0.035728 | 0.694 | 0.572 | 1 | 11 | Vmp1 | 0.122 | Old |
| 0.110723 | 0.54507 | 0.089 | 0.043 | 1 | 11 | Ralgapa2 | 0.046 | Old |
| 0.110944 | 0.654205 | 0.202 | 0.138 | 1 | 11 | Smad5 | 0.064 | Old |
| 0.110998 | 0.075923 | 0.407 | 0.29 | 1 | 11 | Gatad2b | 0.117 | Old |
| 0.111197 | 0.558459 | 0.089 | 0.043 | 1 | 11 | Klf16 | 0.046 | Old |
| 0.111206 | 0.477771 | 0.109 | 0.058 | 1 | 11 | Fbxo9 | 0.051 | Old |
| 0.111451 | 1.22121 | 0.056 | 0.022 | 1 | 11 | Ifitm5 | 0.034 | Old |
| 0.111451 | 1.207433 | 0.056 | 0.022 | 1 | 11 | Rad9a | 0.034 | Old |
| 0.111607 | 0.105286 | 0.415 | 0.297 | 1 | 11 | Mapre2 | 0.118 | Old |
| 0.111673 | 0.450139 | 0.089 | 0.043 | 1 | 11 | Cdc27 | 0.046 | Old |
| 0.111837 | 0.117049 | 0.774 | 0.652 | 1 | 11 | H2-DMb1 | 0.122 | Old |
| 0.112054 | 0.739698 | 0.056 | 0.022 | 1 | 11 | Hars2 | 0.034 | Old |
| 0.11215 | 0.023612 | 0.089 | 0.043 | 1 | 11 | Ccser2 | 0.046 | Old |
| 0.11216 | 0.044523 | 0.214 | 0.138 | 1 | 11 | Nek9 | 0.076 | Old |
| 0.112214 | 0.026826 | 0.226 | 0.145 | 1 | 11 | Cnot3 | 0.081 | Old |
| 0.112258 | 0.068579 | 0.274 | 0.181 | 1 | 11 | Gm10073 | 0.093 | Old |
| 0.112277 | 0.141313 | 0.194 | 0.123 | 1 | 11 | Casp8ap2 | 0.071 | Old |
| 0.112338 | 0.025662 | 0.254 | 0.167 | 1 | 11 | Atp7a | 0.087 | Old |
| 0.11254 | 0.232221 | 0.181 | 0.116 | 1 | 11 | Crat | 0.065 | Old |
| 0.11266 | 1.029757 | 0.056 | 0.022 | 1 | 11 | Ccdc71 | 0.034 | Old |
| 0.112755 | 0.42776 | 0.153 | 0.094 | 1 | 11 | Ubfd1 | 0.059 | Old |
| 0.112833 | 0.088067 | 0.27 | 0.181 | 1 | 11 | Phf14 | 0.089 | Old |
| 0.112972 | 0.224849 | 0.069 | 0.029 | 1 | 11 | Zfp142 | 0.04 | Old |
| 0.113045 | 0.007998 | 0.569 | 0.435 | 1 | 11 | Sema4a | 0.134 | Old |
| 0.113268 | 1.386662 | 0.056 | 0.022 | 1 | 11 | Eid3 | 0.034 | Old |
| 0.113268 | 1.125373 | 0.056 | 0.022 | 1 | 11 | Ddx51 | 0.034 | Old |
| 0.113319 | 0.181674 | 0.492 | 0.355 | 1 | 11 | Pik3r1 | 0.137 | Old |
| 0.113523 | 0.107234 | 0.069 | 0.029 | 1 | 11 | Ccar2 | 0.04 | Old |
| 0.113751 | 0.281519 | 0.157 | 0.094 | 1 | 11 | Stau1 | 0.063 | Old |

| | | | | | | | | |
|---|---|---|---|---|---|---|---|---|
| 0.113807 | 0.335482 | 0.21 | 0.138 | 1 | 11 | Mtmr12 | 0.072 | Old |
| 0.113969 | 1.860667 | 0.077 | 0.036 | 1 | 11 | Unc13d | 0.041 | Old |
| 0.114134 | 0.670016 | 0.117 | 0.065 | 1 | 11 | Pfdn6 | 0.052 | Old |
| 0.114156 | 0.12354 | 0.492 | 0.355 | 1 | 11 | Eif4b | 0.137 | Old |
| 0.11424 | 0.4807 | 0.109 | 0.058 | 1 | 11 | Hspb11 | 0.051 | Old |
| 0.11429 | 0.172921 | 0.532 | 0.384 | 1 | 11 | Scpep1 | 0.148 | Old |
| 0.114294 | 0.219671 | 0.367 | 0.261 | 1 | 11 | Ankrd12 | 0.106 | Old |
| 0.114312 | 0.017735 | 0.242 | 0.159 | 1 | 11 | Tmem127 | 0.083 | Old |
| 0.114325 | 0.030939 | 0.149 | 0.087 | 1 | 11 | Naxe | 0.062 | Old |
| 0.114492 | 1.176053 | 0.056 | 0.022 | 1 | 11 | Hmga1 | 0.034 | Old |
| 0.114783 | 0.06285 | 0.121 | 0.065 | 1 | 11 | Slc17a5 | 0.056 | Old |
| 0.114848 | 0.122198 | 0.536 | 0.406 | 1 | 11 | Ap2s1 | 0.13 | Old |
| 0.114873 | 0.037004 | 0.496 | 0.37 | 1 | 11 | Cmtm6 | 0.126 | Old |
| 0.115048 | 0.156262 | 0.089 | 0.043 | 1 | 11 | Dvl1 | 0.046 | Old |
| 0.115108 | 0.514526 | 0.056 | 0.022 | 1 | 11 | Epor | 0.034 | Old |
| 0.11519 | 0.139411 | 0.069 | 0.029 | 1 | 11 | Med9 | 0.04 | Old |
| 0.115286 | 0.043167 | 0.23 | 0.152 | 1 | 11 | Noct | 0.078 | Old |
| 0.115494 | 0.270634 | 0.137 | 0.08 | 1 | 11 | Fubp3 | 0.057 | Old |
| 0.115537 | 0.019204 | 0.089 | 0.043 | 1 | 11 | Gtpbp6 | 0.046 | Old |
| 0.115704 | 0.037171 | 0.343 | 0.239 | 1 | 11 | Msr1 | 0.104 | Old |
| 0.11575 | 0.05332 | 0.069 | 0.029 | 1 | 11 | Zfp426 | 0.04 | Old |
| 0.116027 | 0.098655 | 0.089 | 0.043 | 1 | 11 | Eva1b | 0.046 | Old |
| 0.116066 | 0.961696 | 0.077 | 0.036 | 1 | 11 | Pramef8 | 0.041 | Old |
| 0.116126 | 2.056591 | 0.032 | 0.007 | 1 | 11 | Slc35d2 | 0.025 | Old |
| 0.116249 | 0.828981 | 0.125 | 0.072 | 1 | 11 | Poglut1 | 0.053 | Old |
| 0.116348 | 0.830053 | 0.056 | 0.022 | 1 | 11 | D17H6S53I | 0.034 | Old |
| 0.116395 | 0.220235 | 0.472 | 0.355 | 1 | 11 | Rab3il1 | 0.117 | Old |
| 0.116684 | 0.00974 | 0.137 | 0.08 | 1 | 11 | Rnf5 | 0.057 | Old |
| 0.116806 | 0.629106 | 0.097 | 0.051 | 1 | 11 | Rarg | 0.046 | Old |
| 0.116945 | 0.135051 | 0.222 | 0.145 | 1 | 11 | Mif4gd | 0.077 | Old |
| 0.116972 | 0.982945 | 0.056 | 0.022 | 1 | 11 | Dclre1a | 0.034 | Old |
| 0.116972 | 0.627798 | 0.056 | 0.022 | 1 | 11 | Dip2c | 0.034 | Old |
| 0.116972 | 0.267058 | 0.056 | 0.022 | 1 | 11 | 3110040N1 | 0.034 | Old |
| 0.116974 | 1.889268 | 0.032 | 0.007 | 1 | 11 | 2810001G2 | 0.025 | Old |
| 0.116974 | 1.761648 | 0.032 | 0.007 | 1 | 11 | Zfp74 | 0.025 | Old |
| 0.117142 | 0.139598 | 0.774 | 0.616 | 1 | 11 | Ssr3 | 0.158 | Old |
| 0.117272 | 0.632068 | 0.133 | 0.08 | 1 | 11 | Polr3k | 0.053 | Old |
| 0.117343 | 0.071789 | 0.383 | 0.275 | 1 | 11 | Naa50 | 0.108 | Old |
| 0.117599 | 0.773126 | 0.056 | 0.022 | 1 | 11 | Fbxo31 | 0.034 | Old |
| 0.117599 | 0.426142 | 0.056 | 0.022 | 1 | 11 | Cipc | 0.034 | Old |
| 0.117599 | 0.335583 | 0.056 | 0.022 | 1 | 11 | Mgme1 | 0.034 | Old |
| 0.117752 | 0.495757 | 0.097 | 0.051 | 1 | 11 | Fmnl2 | 0.046 | Old |
| 0.117826 | 2.407286 | 0.032 | 0.007 | 1 | 11 | Zbtb10 | 0.025 | Old |
| 0.117826 | 2.294288 | 0.032 | 0.007 | 1 | 11 | Epb41l3 | 0.025 | Old |
| 0.117826 | 2.275179 | 0.032 | 0.007 | 1 | 11 | Nav2 | 0.025 | Old |

| | | | | | | | | |
|---|---|---|---|---|---|---|---|---|
| 0.117826 | 2.062243 | 0.032 | 0.007 | 1 | 11 | Smo | 0.025 | Old |
| 0.117826 | 1.983836 | 0.032 | 0.007 | 1 | 11 | Jchain | 0.025 | Old |
| 0.117826 | 1.935538 | 0.032 | 0.007 | 1 | 11 | Dusp28 | 0.025 | Old |
| 0.117931 | 0.208284 | 0.185 | 0.116 | 1 | 11 | Ergic2 | 0.069 | Old |
| 0.118005 | 0.100355 | 0.089 | 0.043 | 1 | 11 | Smc2 | 0.046 | Old |
| 0.118228 | 0.394528 | 0.056 | 0.022 | 1 | 11 | Amd2 | 0.034 | Old |
| 0.118413 | 0.619453 | 0.117 | 0.065 | 1 | 11 | Zfp518a | 0.052 | Old |
| 0.118504 | 0.419224 | 0.089 | 0.043 | 1 | 11 | Med27 | 0.046 | Old |
| 0.118545 | 1.362072 | 0.044 | 0.014 | 1 | 11 | Znhit3 | 0.03 | Old |
| 0.118684 | 1.971179 | 0.032 | 0.007 | 1 | 11 | Rab38 | 0.025 | Old |
| 0.118684 | 1.779098 | 0.032 | 0.007 | 1 | 11 | Coq6 | 0.025 | Old |
| 0.118785 | 0.259004 | 0.181 | 0.116 | 1 | 11 | Nmi | 0.065 | Old |
| 0.118848 | 0.492752 | 0.117 | 0.065 | 1 | 11 | Psip1 | 0.052 | Old |
| 0.11886 | 0.63398 | 0.056 | 0.022 | 1 | 11 | Btg3 | 0.034 | Old |
| 0.11886 | 0.255723 | 0.056 | 0.022 | 1 | 11 | Tjap1 | 0.034 | Old |
| 0.119091 | 0.526712 | 0.137 | 0.08 | 1 | 11 | Rbm14 | 0.057 | Old |
| 0.119173 | 0.300178 | 0.125 | 0.072 | 1 | 11 | Ak6 | 0.053 | Old |
| 0.119181 | 0.582267 | 0.097 | 0.051 | 1 | 11 | Zfp945 | 0.046 | Old |
| 0.119265 | 1.818613 | 0.044 | 0.014 | 1 | 11 | Tmem238 | 0.03 | Old |
| 0.119265 | 1.227707 | 0.044 | 0.014 | 1 | 11 | Paip2b | 0.03 | Old |
| 0.119267 | 0.933855 | 0.077 | 0.036 | 1 | 11 | Polr3e | 0.041 | Old |
| 0.119281 | 0.194559 | 0.19 | 0.123 | 1 | 11 | Plac8 | 0.067 | Old |
| 0.119493 | 0.121425 | 0.476 | 0.348 | 1 | 11 | Gsk3b | 0.128 | Old |
| 0.119494 | 0.538859 | 0.056 | 0.022 | 1 | 11 | Trmt61a | 0.034 | Old |
| 0.119546 | 2.899125 | 0.032 | 0.007 | 1 | 11 | Prdm10 | 0.025 | Old |
| 0.119546 | 1.997414 | 0.032 | 0.007 | 1 | 11 | Coq4 | 0.025 | Old |
| 0.119546 | 1.770603 | 0.032 | 0.007 | 1 | 11 | Tut1 | 0.025 | Old |
| 0.119546 | 1.613507 | 0.032 | 0.007 | 1 | 11 | Itga2 | 0.025 | Old |
| 0.119546 | 1.455466 | 0.032 | 0.007 | 1 | 11 | Hrh2 | 0.025 | Old |
| 0.119595 | 0.663756 | 0.125 | 0.072 | 1 | 11 | Fbxw7 | 0.053 | Old |
| 0.119807 | 0.734947 | 0.077 | 0.036 | 1 | 11 | Sergef | 0.041 | Old |
| 0.119989 | 1.631411 | 0.044 | 0.014 | 1 | 11 | C130050O1 | 0.03 | Old |
| 0.119989 | 1.539925 | 0.044 | 0.014 | 1 | 11 | Eps8l1 | 0.03 | Old |
| 0.119989 | 1.396249 | 0.044 | 0.014 | 1 | 11 | C330018D2 | 0.03 | Old |
| 0.119989 | 0.7438 | 0.044 | 0.014 | 1 | 11 | Cenpo | 0.03 | Old |
| 0.120017 | 0.302559 | 0.202 | 0.13 | 1 | 11 | Skiv2l | 0.072 | Old |
| 0.120131 | 0.456964 | 0.056 | 0.022 | 1 | 11 | Myc | 0.034 | Old |
| 0.120131 | 0.179909 | 0.056 | 0.022 | 1 | 11 | Gatb | 0.034 | Old |
| 0.120296 | 0.15844 | 0.234 | 0.152 | 1 | 11 | Tlr7 | 0.082 | Old |
| 0.12031 | 0.248611 | 0.137 | 0.08 | 1 | 11 | Coq5 | 0.057 | Old |
| 0.120349 | 0.647405 | 0.077 | 0.036 | 1 | 11 | AC165953. | 0.041 | Old |
| 0.120385 | 0.221494 | 0.714 | 0.551 | 1 | 11 | Ywhab | 0.163 | Old |
| 0.120414 | 1.964261 | 0.032 | 0.007 | 1 | 11 | Zfp568 | 0.025 | Old |
| 0.120414 | 1.675976 | 0.032 | 0.007 | 1 | 11 | Pecam1 | 0.025 | Old |
| 0.120414 | 1.429851 | 0.032 | 0.007 | 1 | 11 | Gcat | 0.025 | Old |

| | | | | | | | | |
|---|---|---|---|---|---|---|---|---|
| 0.120414 | 1.360162 | 0.032 | 0.007 | 1 | 11 | Zfp628 | 0.025 | Old |
| 0.120702 | 0.059869 | 0.202 | 0.13 | 1 | 11 | Cdip1 | 0.072 | Old |
| 0.120716 | 1.761671 | 0.044 | 0.014 | 1 | 11 | Aldh18a1 | 0.03 | Old |
| 0.120716 | 1.76095 | 0.044 | 0.014 | 1 | 11 | Fig4 | 0.03 | Old |
| 0.120716 | 1.154129 | 0.044 | 0.014 | 1 | 11 | AW011738 | 0.03 | Old |
| 0.120716 | 1.014616 | 0.044 | 0.014 | 1 | 11 | Sec61a2 | 0.03 | Old |
| 0.120745 | 0.217515 | 0.496 | 0.37 | 1 | 11 | Rhog | 0.126 | Old |
| 0.120893 | 0.291286 | 0.077 | 0.036 | 1 | 11 | Rcan1 | 0.041 | Old |
| 0.121266 | 0.626876 | 0.185 | 0.123 | 1 | 11 | Gstm1 | 0.062 | Old |
| 0.121286 | 1.626385 | 0.032 | 0.007 | 1 | 11 | Zfp958 | 0.025 | Old |
| 0.121286 | 1.316561 | 0.032 | 0.007 | 1 | 11 | Rida | 0.025 | Old |
| 0.121286 | 1.199009 | 0.032 | 0.007 | 1 | 11 | Plekhh3 | 0.025 | Old |
| 0.121286 | 1.154334 | 0.032 | 0.007 | 1 | 11 | Tmem150a | 0.025 | Old |
| 0.121286 | 0.964264 | 0.032 | 0.007 | 1 | 11 | Iqcc | 0.025 | Old |
| 0.121447 | 1.408132 | 0.044 | 0.014 | 1 | 11 | Pde1b | 0.03 | Old |
| 0.121483 | 0.451995 | 0.117 | 0.065 | 1 | 11 | Kif1bp | 0.052 | Old |
| 0.121614 | 0.248962 | 0.173 | 0.109 | 1 | 11 | Tiprl | 0.064 | Old |
| 0.121699 | 0.086204 | 0.331 | 0.232 | 1 | 11 | Ctnnd1 | 0.099 | Old |
| 0.121826 | 0.254059 | 0.194 | 0.123 | 1 | 11 | Ptprs | 0.071 | Old |
| 0.122009 | 0.267173 | 0.185 | 0.116 | 1 | 11 | Chmp2b | 0.069 | Old |
| 0.122059 | 0.312887 | 0.056 | 0.022 | 1 | 11 | Hcar2 | 0.034 | Old |
| 0.122077 | 0.23301 | 0.435 | 0.319 | 1 | 11 | Dnm2 | 0.116 | Old |
| 0.122163 | 1.076846 | 0.032 | 0.007 | 1 | 11 | Gdf9 | 0.025 | Old |
| 0.122163 | 1.061142 | 0.032 | 0.007 | 1 | 11 | Cabp4 | 0.025 | Old |
| 0.122163 | 1.032082 | 0.032 | 0.007 | 1 | 11 | Hspa2 | 0.025 | Old |
| 0.122163 | 0.794008 | 0.032 | 0.007 | 1 | 11 | Intu | 0.025 | Old |
| 0.122163 | 0.644911 | 0.032 | 0.007 | 1 | 11 | Arfip2 | 0.025 | Old |
| 0.122163 | 0.54778 | 0.032 | 0.007 | 1 | 11 | Trip6 | 0.025 | Old |
| 0.122181 | 1.182713 | 0.044 | 0.014 | 1 | 11 | Ercc4 | 0.03 | Old |
| 0.122181 | 0.981768 | 0.044 | 0.014 | 1 | 11 | Rpgrip1 | 0.03 | Old |
| 0.122181 | 0.915687 | 0.044 | 0.014 | 1 | 11 | Pradc1 | 0.03 | Old |
| 0.122194 | 0.062041 | 0.218 | 0.145 | 1 | 11 | Mtm1 | 0.073 | Old |
| 0.122341 | 0.307647 | 0.109 | 0.058 | 1 | 11 | Mmp19 | 0.051 | Old |
| 0.122371 | 0.656894 | 0.117 | 0.065 | 1 | 11 | Plekha3 | 0.052 | Old |
| 0.122432 | 0.191192 | 0.714 | 0.558 | 1 | 11 | Cox6b1 | 0.156 | Old |
| 0.12245 | 1.407795 | 0.085 | 0.043 | 1 | 11 | Ogfod2 | 0.042 | Old |
| 0.122543 | 0.267285 | 0.105 | 0.058 | 1 | 11 | Alg10b | 0.047 | Old |
| 0.122612 | 0.591975 | 0.133 | 0.08 | 1 | 11 | Prpf31 | 0.053 | Old |
| 0.122706 | 0.190009 | 0.056 | 0.022 | 1 | 11 | Shmt2 | 0.034 | Old |
| 0.122919 | 1.27849 | 0.044 | 0.014 | 1 | 11 | Pex1 | 0.03 | Old |
| 0.122919 | 0.942409 | 0.044 | 0.014 | 1 | 11 | Rnf217 | 0.03 | Old |
| 0.122967 | 0.35524 | 0.379 | 0.283 | 1 | 11 | Thrap3 | 0.096 | Old |
| 0.123046 | 1.015587 | 0.032 | 0.007 | 1 | 11 | Zbtb24 | 0.025 | Old |
| 0.123046 | 1.008536 | 0.032 | 0.007 | 1 | 11 | Klhl15 | 0.025 | Old |
| 0.123046 | 0.717399 | 0.032 | 0.007 | 1 | 11 | Mfsd13a | 0.025 | Old |

| | | | | | | | | |
|---|---|---|---|---|---|---|---|---|
| 0.123046 | 0.672107 | 0.032 | 0.007 | 1 | 11 | Cdyl2 | 0.025 | Old |
| 0.123046 | 0.48557 | 0.032 | 0.007 | 1 | 11 | Timeless | 0.025 | Old |
| 0.123046 | 0.338247 | 0.032 | 0.007 | 1 | 11 | Gstm5 | 0.025 | Old |
| 0.123046 | 0.295915 | 0.032 | 0.007 | 1 | 11 | Fbxo21 | 0.025 | Old |
| 0.123046 | 0.014952 | 0.032 | 0.007 | 1 | 11 | Gna11 | 0.025 | Old |
| 0.123088 | 0.426906 | 0.077 | 0.036 | 1 | 11 | Tmem26 | 0.041 | Old |
| 0.123162 | 0.393268 | 0.149 | 0.094 | 1 | 11 | Cadm1 | 0.055 | Old |
| 0.12345 | 0.485412 | 0.133 | 0.08 | 1 | 11 | Plau | 0.053 | Old |
| 0.123549 | 0.10966 | 0.097 | 0.051 | 1 | 11 | Stxbp1 | 0.046 | Old |
| 0.123597 | 0.063894 | 0.194 | 0.123 | 1 | 11 | Clec4n | 0.071 | Old |
| 0.123642 | 0.235592 | 0.077 | 0.036 | 1 | 11 | Cnot11 | 0.041 | Old |
| 0.12366 | 0.817507 | 0.044 | 0.014 | 1 | 11 | Wrb | 0.03 | Old |
| 0.124183 | 0.161815 | 0.726 | 0.565 | 1 | 11 | Cox6c | 0.161 | Old |
| 0.124404 | 0.915955 | 0.044 | 0.014 | 1 | 11 | Nlrx1 | 0.03 | Old |
| 0.124404 | 0.747186 | 0.044 | 0.014 | 1 | 11 | Nme7 | 0.03 | Old |
| 0.124404 | 0.680779 | 0.044 | 0.014 | 1 | 11 | Ecsit | 0.03 | Old |
| 0.124404 | 0.555046 | 0.044 | 0.014 | 1 | 11 | Tnfsfm13 | 0.03 | Old |
| 0.124615 | 0.772449 | 0.117 | 0.065 | 1 | 11 | Fgfr1op | 0.052 | Old |
| 0.124615 | 0.233219 | 0.117 | 0.065 | 1 | 11 | Mcoln1 | 0.052 | Old |
| 0.124811 | 0.017974 | 0.302 | 0.21 | 1 | 11 | Yy1 | 0.092 | Old |
| 0.125153 | 0.745476 | 0.044 | 0.014 | 1 | 11 | Shb | 0.03 | Old |
| 0.125153 | 0.608326 | 0.044 | 0.014 | 1 | 11 | Ccdc152 | 0.03 | Old |
| 0.125153 | 0.543063 | 0.044 | 0.014 | 1 | 11 | Usp30 | 0.03 | Old |
| 0.125315 | 0.376392 | 0.077 | 0.036 | 1 | 11 | Cep70 | 0.041 | Old |
| 0.125564 | 0.079489 | 0.403 | 0.283 | 1 | 11 | Sri | 0.12 | Old |
| 0.125876 | 0.005523 | 0.601 | 0.478 | 1 | 11 | Herpud1 | 0.123 | Old |
| 0.125904 | 0.487753 | 0.044 | 0.014 | 1 | 11 | Zfp579 | 0.03 | Old |
| 0.126339 | 0.805662 | 0.105 | 0.058 | 1 | 11 | Sirpb1b | 0.047 | Old |
| 0.126347 | 0.108633 | 0.286 | 0.203 | 1 | 11 | Cpeb4 | 0.083 | Old |
| 0.12644 | 0.756953 | 0.077 | 0.036 | 1 | 11 | Slc35a1 | 0.041 | Old |
| 0.12644 | 0.66316 | 0.077 | 0.036 | 1 | 11 | Rora | 0.041 | Old |
| 0.126531 | 0.694508 | 0.097 | 0.051 | 1 | 11 | Dgkh | 0.046 | Old |
| 0.126659 | 0.89681 | 0.044 | 0.014 | 1 | 11 | Trmt11 | 0.03 | Old |
| 0.126659 | 0.356085 | 0.044 | 0.014 | 1 | 11 | Nudt7 | 0.03 | Old |
| 0.126847 | 0.725821 | 0.133 | 0.08 | 1 | 11 | Ak3 | 0.053 | Old |
| 0.126918 | 0.01861 | 0.21 | 0.138 | 1 | 11 | Rras | 0.072 | Old |
| 0.127005 | 0.59179 | 0.077 | 0.036 | 1 | 11 | Oxsm | 0.041 | Old |
| 0.127157 | 0.645555 | 0.177 | 0.116 | 1 | 11 | Phf2 | 0.061 | Old |
| 0.127175 | 0.149542 | 0.23 | 0.152 | 1 | 11 | Emc6 | 0.078 | Old |
| 0.127213 | 0.206187 | 0.395 | 0.297 | 1 | 11 | Anxa3 | 0.098 | Old |
| 0.127349 | 0.201718 | 0.117 | 0.065 | 1 | 11 | Carm1 | 0.052 | Old |
| 0.127572 | 0.55717 | 0.077 | 0.036 | 1 | 11 | Hmgxb3 | 0.041 | Old |
| 0.12764 | 0.970094 | 0.113 | 0.065 | 1 | 11 | Fam46c | 0.048 | Old |
| 0.127708 | 0.538001 | 0.133 | 0.08 | 1 | 11 | Prkra | 0.053 | Old |
| 0.127778 | 0.182591 | 0.476 | 0.355 | 1 | 11 | Mbnl2 | 0.121 | Old |

| | | | | | | | | |
|---|---|---|---|---|---|---|---|---|
| 0.127796 | 0.496906 | 0.065 | 0.029 | 1 | 11 | Rab2b | 0.036 | Old |
| 0.127844 | 0.45609 | 0.125 | 0.072 | 1 | 11 | Dctn6 | 0.053 | Old |
| 0.12829 | 0.222188 | 0.125 | 0.072 | 1 | 11 | Rab28 | 0.053 | Old |
| 0.128356 | 0.172289 | 0.423 | 0.319 | 1 | 11 | Sh3bp1 | 0.104 | Old |
| 0.128375 | 0.089783 | 0.468 | 0.348 | 1 | 11 | Pias1 | 0.12 | Old |
| 0.128412 | 0.074471 | 0.492 | 0.377 | 1 | 11 | Psma3 | 0.115 | Old |
| 0.128738 | 0.40152 | 0.125 | 0.072 | 1 | 11 | Odf2 | 0.053 | Old |
| 0.128758 | 0.179375 | 0.105 | 0.058 | 1 | 11 | Slc46a3 | 0.047 | Old |
| 0.129105 | 0.082972 | 0.258 | 0.174 | 1 | 11 | Appbp2 | 0.084 | Old |
| 0.129162 | 0.433599 | 0.206 | 0.138 | 1 | 11 | Uchl5 | 0.068 | Old |
| 0.129571 | 0.159264 | 0.27 | 0.188 | 1 | 11 | Pts | 0.082 | Old |
| 0.129736 | 0.64629 | 0.105 | 0.058 | 1 | 11 | P2ry13 | 0.047 | Old |
| 0.129752 | 0.32316 | 0.198 | 0.13 | 1 | 11 | Tor1b | 0.068 | Old |
| 0.13008 | 0.078038 | 0.097 | 0.051 | 1 | 11 | Gpatch2l | 0.046 | Old |
| 0.130215 | 0.062964 | 0.871 | 0.761 | 1 | 11 | Gabarap | 0.11 | Old |
| 0.130252 | 0.053469 | 0.258 | 0.174 | 1 | 11 | Polr2m | 0.084 | Old |
| 0.130296 | 0.879582 | 0.065 | 0.029 | 1 | 11 | Prkar2b | 0.036 | Old |
| 0.130317 | 0.523499 | 0.133 | 0.08 | 1 | 11 | Dph5 | 0.053 | Old |
| 0.130331 | 0.301409 | 0.161 | 0.101 | 1 | 11 | Mafk | 0.06 | Old |
| 0.130405 | 0.12998 | 0.286 | 0.196 | 1 | 11 | Mgrn1 | 0.09 | Old |
| 0.130439 | 0.016566 | 0.077 | 0.036 | 1 | 11 | Rcan3 | 0.041 | Old |
| 0.130597 | 0.147093 | 0.117 | 0.065 | 1 | 11 | Rbms2 | 0.052 | Old |
| 0.130767 | 0.085285 | 0.274 | 0.188 | 1 | 11 | Cebpz | 0.086 | Old |
| 0.130831 | 0.051029 | 0.484 | 0.362 | 1 | 11 | Epn1 | 0.122 | Old |
| 0.130926 | 1.21035 | 0.065 | 0.029 | 1 | 11 | Rai14 | 0.036 | Old |
| 0.131068 | 0.171707 | 0.331 | 0.232 | 1 | 11 | Il1rn | 0.099 | Old |
| 0.131627 | 0.000715 | 0.343 | 0.239 | 1 | 11 | Zmynd8 | 0.104 | Old |
| 0.131903 | 0.201363 | 0.125 | 0.072 | 1 | 11 | Slc4a1ap | 0.053 | Old |
| 0.132195 | 1.067112 | 0.065 | 0.029 | 1 | 11 | Rnf25 | 0.036 | Old |
| 0.132348 | 0.488978 | 0.161 | 0.101 | 1 | 11 | Pmpcb | 0.06 | Old |
| 0.132356 | 0.082152 | 0.625 | 0.478 | 1 | 11 | Rock2 | 0.147 | Old |
| 0.132379 | 0.214646 | 0.25 | 0.174 | 1 | 11 | Ugp2 | 0.076 | Old |
| 0.132552 | 0.610865 | 0.149 | 0.094 | 1 | 11 | Apopt1 | 0.055 | Old |
| 0.132678 | 0.686347 | 0.185 | 0.123 | 1 | 11 | Abca7 | 0.062 | Old |
| 0.132706 | 0.868358 | 0.121 | 0.072 | 1 | 11 | Vcl | 0.049 | Old |
| 0.13357 | 0.117551 | 0.161 | 0.101 | 1 | 11 | Mrpl41 | 0.06 | Old |
| 0.133704 | 0.108495 | 0.105 | 0.058 | 1 | 11 | Zfp236 | 0.047 | Old |
| 0.134097 | 0.105214 | 0.556 | 0.413 | 1 | 11 | Capza1 | 0.143 | Old |
| 0.134116 | 1.006364 | 0.065 | 0.029 | 1 | 11 | Cox18 | 0.036 | Old |
| 0.134222 | 0.032214 | 0.278 | 0.188 | 1 | 11 | Ergic3 | 0.09 | Old |
| 0.134229 | 0.111665 | 0.097 | 0.051 | 1 | 11 | Kctd12b | 0.046 | Old |
| 0.134518 | 0.133538 | 0.379 | 0.275 | 1 | 11 | Dcp2 | 0.104 | Old |
| 0.134791 | 0.001462 | 0.31 | 0.225 | 1 | 11 | Ypel3 | 0.085 | Old |
| 0.134909 | 3.518883 | 0.016 | 0 | 1 | 11 | Ablim1 | 0.016 | Old |
| 0.134909 | 2.98299 | 0.016 | 0 | 1 | 11 | Trem3 | 0.016 | Old |

| | | | | | | | | |
|---|---|---|---|---|---|---|---|---|
| 0.134909 | 2.770208 | 0.016 | 0 | 1 | 11 | Zfp61 | 0.016 | Old |
| 0.134909 | 2.717775 | 0.016 | 0 | 1 | 11 | Trim13 | 0.016 | Old |
| 0.134909 | 2.664027 | 0.016 | 0 | 1 | 11 | Homer2 | 0.016 | Old |
| 0.134909 | 2.599143 | 0.016 | 0 | 1 | 11 | Nthl1 | 0.016 | Old |
| 0.134909 | 2.58137 | 0.016 | 0 | 1 | 11 | Matn2 | 0.016 | Old |
| 0.134909 | 2.466792 | 0.016 | 0 | 1 | 11 | Ankrd33b | 0.016 | Old |
| 0.134909 | 2.430661 | 0.016 | 0 | 1 | 11 | Hoxb7 | 0.016 | Old |
| 0.134909 | 2.429706 | 0.016 | 0 | 1 | 11 | Slc16a12 | 0.016 | Old |
| 0.134909 | 2.414707 | 0.016 | 0 | 1 | 11 | Atad3aos | 0.016 | Old |
| 0.134909 | 2.356505 | 0.016 | 0 | 1 | 11 | Tspyl4 | 0.016 | Old |
| 0.134909 | 2.317088 | 0.016 | 0 | 1 | 11 | Bend3 | 0.016 | Old |
| 0.134909 | 2.256774 | 0.016 | 0 | 1 | 11 | Tlr9 | 0.016 | Old |
| 0.134909 | 2.218874 | 0.016 | 0 | 1 | 11 | Klhl8 | 0.016 | Old |
| 0.134909 | 2.199198 | 0.016 | 0 | 1 | 11 | Slc25a14 | 0.016 | Old |
| 0.134909 | 2.194662 | 0.016 | 0 | 1 | 11 | Gm35339 | 0.016 | Old |
| 0.134909 | 2.188299 | 0.016 | 0 | 1 | 11 | Kdr | 0.016 | Old |
| 0.134909 | 2.178907 | 0.016 | 0 | 1 | 11 | L3hypdh | 0.016 | Old |
| 0.134909 | 2.138691 | 0.016 | 0 | 1 | 11 | Akr1c19 | 0.016 | Old |
| 0.134909 | 2.118758 | 0.016 | 0 | 1 | 11 | Lrrc63 | 0.016 | Old |
| 0.134909 | 2.105558 | 0.016 | 0 | 1 | 11 | Magi1 | 0.016 | Old |
| 0.134909 | 2.094809 | 0.016 | 0 | 1 | 11 | Prss23 | 0.016 | Old |
| 0.134909 | 2.080532 | 0.016 | 0 | 1 | 11 | Hyal1 | 0.016 | Old |
| 0.134909 | 2.077465 | 0.016 | 0 | 1 | 11 | Zfp503 | 0.016 | Old |
| 0.134909 | 2.069928 | 0.016 | 0 | 1 | 11 | Klf8 | 0.016 | Old |
| 0.134909 | 2.066799 | 0.016 | 0 | 1 | 11 | Rita1 | 0.016 | Old |
| 0.134909 | 2.063743 | 0.016 | 0 | 1 | 11 | Taco1os | 0.016 | Old |
| 0.134909 | 2.060041 | 0.016 | 0 | 1 | 11 | Cxcr3 | 0.016 | Old |
| 0.134909 | 2.057039 | 0.016 | 0 | 1 | 11 | AI480526 | 0.016 | Old |
| 0.134909 | 2.029734 | 0.016 | 0 | 1 | 11 | Gm10521 | 0.016 | Old |
| 0.134909 | 2.012167 | 0.016 | 0 | 1 | 11 | Srgap3 | 0.016 | Old |
| 0.134909 | 2.009708 | 0.016 | 0 | 1 | 11 | Lrp8 | 0.016 | Old |
| 0.134909 | 2.001491 | 0.016 | 0 | 1 | 11 | Ldhb | 0.016 | Old |
| 0.134909 | 1.994114 | 0.016 | 0 | 1 | 11 | Gm34342 | 0.016 | Old |
| 0.134909 | 1.99308 | 0.016 | 0 | 1 | 11 | 2810405F1 | 0.016 | Old |
| 0.134909 | 1.981255 | 0.016 | 0 | 1 | 11 | Gm19951 | 0.016 | Old |
| 0.134909 | 1.975567 | 0.016 | 0 | 1 | 11 | Cracr2a | 0.016 | Old |
| 0.134909 | 1.945446 | 0.016 | 0 | 1 | 11 | Gemin8 | 0.016 | Old |
| 0.134909 | 1.919007 | 0.016 | 0 | 1 | 11 | Slc22a5 | 0.016 | Old |
| 0.134909 | 1.91593 | 0.016 | 0 | 1 | 11 | Pou5f1 | 0.016 | Old |
| 0.134909 | 1.906519 | 0.016 | 0 | 1 | 11 | Fbxo32 | 0.016 | Old |
| 0.134909 | 1.89345 | 0.016 | 0 | 1 | 11 | Creb3l3 | 0.016 | Old |
| 0.134909 | 1.881195 | 0.016 | 0 | 1 | 11 | AU022793 | 0.016 | Old |
| 0.134909 | 1.868863 | 0.016 | 0 | 1 | 11 | Bbs7 | 0.016 | Old |
| 0.134909 | 1.854339 | 0.016 | 0 | 1 | 11 | Slc7a4 | 0.016 | Old |
| 0.134909 | 1.838695 | 0.016 | 0 | 1 | 11 | Ddb2 | 0.016 | Old |

| | | | | | | | | |
|---|---|---|---|---|---|---|---|---|
| 0.134909 | 1.833455 | 0.016 | 0 | 1 | 11 | Gm4631 | 0.016 | Old |
| 0.134909 | 1.800957 | 0.016 | 0 | 1 | 11 | Zfp882 | 0.016 | Old |
| 0.134909 | 1.786973 | 0.016 | 0 | 1 | 11 | Map3k9 | 0.016 | Old |
| 0.134909 | 1.773954 | 0.016 | 0 | 1 | 11 | Prox2 | 0.016 | Old |
| 0.134909 | 1.751538 | 0.016 | 0 | 1 | 11 | Fbxo44 | 0.016 | Old |
| 0.134909 | 1.750335 | 0.016 | 0 | 1 | 11 | Rad51c | 0.016 | Old |
| 0.134909 | 1.749407 | 0.016 | 0 | 1 | 11 | Fgf11 | 0.016 | Old |
| 0.134909 | 1.744619 | 0.016 | 0 | 1 | 11 | Plxdc1 | 0.016 | Old |
| 0.134909 | 1.732234 | 0.016 | 0 | 1 | 11 | Pex11g | 0.016 | Old |
| 0.134909 | 1.687304 | 0.016 | 0 | 1 | 11 | 4933424M | 0.016 | Old |
| 0.134909 | 1.671959 | 0.016 | 0 | 1 | 11 | Zfp773 | 0.016 | Old |
| 0.134909 | 1.665879 | 0.016 | 0 | 1 | 11 | Tmem95 | 0.016 | Old |
| 0.134909 | 1.660063 | 0.016 | 0 | 1 | 11 | Rmdn2 | 0.016 | Old |
| 0.134909 | 1.653468 | 0.016 | 0 | 1 | 11 | Egf | 0.016 | Old |
| 0.134909 | 1.641469 | 0.016 | 0 | 1 | 11 | Gk5 | 0.016 | Old |
| 0.134909 | 1.639766 | 0.016 | 0 | 1 | 11 | Ssx2ip | 0.016 | Old |
| 0.134909 | 1.639177 | 0.016 | 0 | 1 | 11 | Nme2 | 0.016 | Old |
| 0.134909 | 1.634334 | 0.016 | 0 | 1 | 11 | Soat2 | 0.016 | Old |
| 0.134909 | 1.608789 | 0.016 | 0 | 1 | 11 | Gm15446 | 0.016 | Old |
| 0.134909 | 1.602259 | 0.016 | 0 | 1 | 11 | Nmrk1 | 0.016 | Old |
| 0.134909 | 1.597273 | 0.016 | 0 | 1 | 11 | Fcrl1 | 0.016 | Old |
| 0.134909 | 1.570806 | 0.016 | 0 | 1 | 11 | Lmbr1 | 0.016 | Old |
| 0.134909 | 1.53749 | 0.016 | 0 | 1 | 11 | Car7 | 0.016 | Old |
| 0.134909 | 1.520861 | 0.016 | 0 | 1 | 11 | Vsig10l | 0.016 | Old |
| 0.134909 | 1.519744 | 0.016 | 0 | 1 | 11 | Slfn3 | 0.016 | Old |
| 0.134909 | 1.517752 | 0.016 | 0 | 1 | 11 | Tnni2 | 0.016 | Old |
| 0.134909 | 1.498695 | 0.016 | 0 | 1 | 11 | Ube2v1 | 0.016 | Old |
| 0.134909 | 1.477862 | 0.016 | 0 | 1 | 11 | Mettl22 | 0.016 | Old |
| 0.134909 | 1.475155 | 0.016 | 0 | 1 | 11 | BC052040 | 0.016 | Old |
| 0.134909 | 1.452826 | 0.016 | 0 | 1 | 11 | Gm11110 | 0.016 | Old |
| 0.134909 | 1.446853 | 0.016 | 0 | 1 | 11 | Tyms | 0.016 | Old |
| 0.134909 | 1.433918 | 0.016 | 0 | 1 | 11 | 5830416I1 | 0.016 | Old |
| 0.134909 | 1.428051 | 0.016 | 0 | 1 | 11 | Zfp784 | 0.016 | Old |
| 0.134909 | 1.412419 | 0.016 | 0 | 1 | 11 | Adora3 | 0.016 | Old |
| 0.134909 | 1.403526 | 0.016 | 0 | 1 | 11 | Uprt | 0.016 | Old |
| 0.134909 | 1.39525 | 0.016 | 0 | 1 | 11 | Atxn7l2 | 0.016 | Old |
| 0.134909 | 1.381002 | 0.016 | 0 | 1 | 11 | Zfp763 | 0.016 | Old |
| 0.134909 | 1.365434 | 0.016 | 0 | 1 | 11 | Fut7 | 0.016 | Old |
| 0.134909 | 1.364187 | 0.016 | 0 | 1 | 11 | Disp1 | 0.016 | Old |
| 0.134909 | 1.36279 | 0.016 | 0 | 1 | 11 | Gm45871 | 0.016 | Old |
| 0.134909 | 1.346387 | 0.016 | 0 | 1 | 11 | Gm42722 | 0.016 | Old |
| 0.134909 | 1.336293 | 0.016 | 0 | 1 | 11 | Pigl | 0.016 | Old |
| 0.134909 | 1.313435 | 0.016 | 0 | 1 | 11 | Kcnk5 | 0.016 | Old |
| 0.134909 | 1.303677 | 0.016 | 0 | 1 | 11 | 1700006J1 | 0.016 | Old |
| 0.134909 | 1.298031 | 0.016 | 0 | 1 | 11 | Cdca5 | 0.016 | Old |

| | | | | | | | | |
|---|---|---|---|---|---|---|---|---|
| 0.134909 | 1.29309 | 0.016 | 0 | 1 | 11 | Zfp420 | 0.016 | Old |
| 0.134909 | 1.291242 | 0.016 | 0 | 1 | 11 | Creg2 | 0.016 | Old |
| 0.134909 | 1.282614 | 0.016 | 0 | 1 | 11 | 9130023H2 | 0.016 | Old |
| 0.134909 | 1.271531 | 0.016 | 0 | 1 | 11 | Ispd | 0.016 | Old |
| 0.134909 | 1.258263 | 0.016 | 0 | 1 | 11 | Rdh5 | 0.016 | Old |
| 0.134909 | 1.247927 | 0.016 | 0 | 1 | 11 | Gm10031 | 0.016 | Old |
| 0.134909 | 1.223239 | 0.016 | 0 | 1 | 11 | Spag1 | 0.016 | Old |
| 0.134909 | 1.220659 | 0.016 | 0 | 1 | 11 | Ccpg1os | 0.016 | Old |
| 0.134909 | 1.213183 | 0.016 | 0 | 1 | 11 | Tsen54 | 0.016 | Old |
| 0.134909 | 1.20923 | 0.016 | 0 | 1 | 11 | P2rx5 | 0.016 | Old |
| 0.134909 | 1.207677 | 0.016 | 0 | 1 | 11 | Anxa9 | 0.016 | Old |
| 0.134909 | 1.13959 | 0.016 | 0 | 1 | 11 | Guca1b | 0.016 | Old |
| 0.134909 | 1.122757 | 0.016 | 0 | 1 | 11 | Lrp11 | 0.016 | Old |
| 0.135274 | 0.460414 | 0.173 | 0.116 | 1 | 11 | Zfyve16 | 0.057 | Old |
| 0.13551 | 0.903884 | 0.121 | 0.072 | 1 | 11 | Tmem8 | 0.049 | Old |
| 0.135552 | 0.001209 | 0.21 | 0.138 | 1 | 11 | Alkbh1 | 0.072 | Old |
| 0.135557 | 0.210034 | 0.278 | 0.203 | 1 | 11 | Micu1 | 0.075 | Old |
| 0.135723 | 0.413527 | 0.105 | 0.058 | 1 | 11 | Ddt | 0.047 | Old |
| 0.135757 | 0.075367 | 0.387 | 0.283 | 1 | 11 | Llph | 0.104 | Old |
| 0.13658 | 0.028525 | 0.145 | 0.087 | 1 | 11 | Slc39a9 | 0.058 | Old |
| 0.137142 | 0.253826 | 0.677 | 0.565 | 1 | 11 | Gltp | 0.112 | Old |
| 0.137392 | 2.111524 | 0.052 | 0.022 | 1 | 11 | Piga | 0.03 | Old |
| 0.137523 | 0.208359 | 0.609 | 0.486 | 1 | 11 | 1810037I1 | 0.123 | Old |
| 0.137765 | 0.236348 | 0.105 | 0.058 | 1 | 11 | Ppfibp1 | 0.047 | Old |
| 0.137801 | 0.30769 | 0.153 | 0.094 | 1 | 11 | Plekhm1 | 0.059 | Old |
| 0.137983 | 0.033765 | 0.56 | 0.413 | 1 | 11 | Inpp5d | 0.147 | Old |
| 0.138022 | 0.737735 | 0.065 | 0.029 | 1 | 11 | Spred2 | 0.036 | Old |
| 0.138068 | 0.02495 | 0.202 | 0.13 | 1 | 11 | Ltbr | 0.072 | Old |
| 0.138103 | 0.031 | 0.395 | 0.283 | 1 | 11 | Eif2ak2 | 0.112 | Old |
| 0.13828 | 0.360988 | 0.105 | 0.058 | 1 | 11 | Plcb3 | 0.047 | Old |
| 0.138681 | 0.939019 | 0.065 | 0.029 | 1 | 11 | Armc7 | 0.036 | Old |
| 0.138795 | 0.115195 | 0.105 | 0.058 | 1 | 11 | Mfap1a | 0.047 | Old |
| 0.138876 | 0.54936 | 0.085 | 0.043 | 1 | 11 | Gphn | 0.042 | Old |
| 0.139049 | 0.125116 | 0.649 | 0.522 | 1 | 11 | Psmb4 | 0.127 | Old |
| 0.139451 | 0.683157 | 0.085 | 0.043 | 1 | 11 | Bcl9l | 0.042 | Old |
| 0.139856 | 0.265024 | 0.206 | 0.138 | 1 | 11 | Ubxn6 | 0.068 | Old |
| 0.140007 | 0.756113 | 0.065 | 0.029 | 1 | 11 | C2cd2 | 0.036 | Old |
| 0.140007 | 0.58605 | 0.065 | 0.029 | 1 | 11 | Golga1 | 0.036 | Old |
| 0.140007 | 0.42597 | 0.065 | 0.029 | 1 | 11 | Zfp961 | 0.036 | Old |
| 0.140287 | 0.665426 | 0.121 | 0.072 | 1 | 11 | Ergic1 | 0.049 | Old |
| 0.140334 | 1.667527 | 0.052 | 0.022 | 1 | 11 | Ice2 | 0.03 | Old |
| 0.140387 | 0.009373 | 0.444 | 0.319 | 1 | 11 | Hmox2 | 0.125 | Old |
| 0.140528 | 0.410578 | 0.169 | 0.109 | 1 | 11 | Mrpl43 | 0.06 | Old |
| 0.140599 | 0.107684 | 0.544 | 0.413 | 1 | 11 | Dnajb6 | 0.131 | Old |
| 0.140885 | 0.005144 | 0.359 | 0.254 | 1 | 11 | Rlim | 0.105 | Old |

| | | | | | | | | |
|---|---|---|---|---|---|---|---|---|
| 0.140887 | 0.148539 | 0.294 | 0.21 | 1 | 11 | Tulp4 | 0.084 | Old |
| 0.140944 | 0.193227 | 0.169 | 0.109 | 1 | 11 | Lage3 | 0.06 | Old |
| 0.140964 | 0.309359 | 0.141 | 0.087 | 1 | 11 | Pbdc1 | 0.054 | Old |
| 0.14107 | 0.161869 | 0.532 | 0.42 | 1 | 11 | Ap2m1 | 0.112 | Old |
| 0.141189 | 0.398277 | 0.085 | 0.043 | 1 | 11 | Sfxn1 | 0.042 | Old |
| 0.141531 | 0.244655 | 0.27 | 0.188 | 1 | 11 | Mdm2 | 0.082 | Old |
| 0.142245 | 0.206257 | 0.125 | 0.072 | 1 | 11 | Vps16 | 0.053 | Old |
| 0.142356 | 0.289961 | 0.085 | 0.043 | 1 | 11 | Cebpzos | 0.042 | Old |
| 0.142391 | 0.008119 | 0.181 | 0.116 | 1 | 11 | Hdac2 | 0.065 | Old |
| 0.142573 | 0.102504 | 0.133 | 0.08 | 1 | 11 | Dpp9 | 0.053 | Old |
| 0.142636 | 0.006414 | 0.198 | 0.13 | 1 | 11 | Kdm6a | 0.068 | Old |
| 0.142934 | 0.214 | 0.375 | 0.29 | 1 | 11 | Anxa1 | 0.085 | Old |
| 0.143004 | 0.142379 | 0.181 | 0.116 | 1 | 11 | Spr | 0.065 | Old |
| 0.143024 | 0.149571 | 0.153 | 0.094 | 1 | 11 | Fgd6 | 0.059 | Old |
| 0.143324 | 0.702324 | 0.052 | 0.022 | 1 | 11 | Vill | 0.03 | Old |
| 0.143365 | 0.303961 | 0.065 | 0.029 | 1 | 11 | Bag4 | 0.036 | Old |
| 0.143365 | 0.264067 | 0.065 | 0.029 | 1 | 11 | Mynn | 0.036 | Old |
| 0.143531 | 0.206416 | 0.085 | 0.043 | 1 | 11 | Pam16 | 0.042 | Old |
| 0.143747 | 0.151741 | 0.113 | 0.065 | 1 | 11 | Pign | 0.048 | Old |
| 0.143751 | 0.172429 | 0.762 | 0.645 | 1 | 11 | Fosb | 0.117 | Old |
| 0.143813 | 0.080998 | 0.319 | 0.232 | 1 | 11 | Calu | 0.087 | Old |
| 0.143842 | 0.105651 | 0.448 | 0.326 | 1 | 11 | Cox5b | 0.122 | Old |
| 0.143934 | 0.413626 | 0.157 | 0.101 | 1 | 11 | Chp2 | 0.056 | Old |
| 0.144214 | 0.149133 | 0.198 | 0.13 | 1 | 11 | Sirpb1c | 0.068 | Old |
| 0.144504 | 0.012222 | 0.504 | 0.384 | 1 | 11 | Lgals9 | 0.12 | Old |
| 0.144713 | 0.426962 | 0.085 | 0.043 | 1 | 11 | Nup133 | 0.042 | Old |
| 0.144827 | 0.070371 | 0.258 | 0.174 | 1 | 11 | Zfp644 | 0.084 | Old |
| 0.144838 | 1.413716 | 0.052 | 0.022 | 1 | 11 | Phka1 | 0.03 | Old |
| 0.1451 | 0.026216 | 0.246 | 0.167 | 1 | 11 | Tbc1d23 | 0.079 | Old |
| 0.145286 | 0.084999 | 0.113 | 0.065 | 1 | 11 | Rpp25l | 0.048 | Old |
| 0.14541 | 0.281395 | 0.065 | 0.029 | 1 | 11 | Mapkbp1 | 0.036 | Old |
| 0.145903 | 0.132046 | 0.085 | 0.043 | 1 | 11 | Slc10a7 | 0.042 | Old |
| 0.146134 | 0.298166 | 0.194 | 0.13 | 1 | 11 | Fbxo11 | 0.064 | Old |
| 0.146318 | 0.093232 | 0.113 | 0.065 | 1 | 11 | D6Wsu163 | 0.048 | Old |
| 0.146363 | 0.882026 | 0.052 | 0.022 | 1 | 11 | Gemin6 | 0.03 | Old |
| 0.146719 | 1.040699 | 0.073 | 0.036 | 1 | 11 | Ino80b | 0.037 | Old |
| 0.146722 | 0.087931 | 0.371 | 0.268 | 1 | 11 | Irak1 | 0.103 | Old |
| 0.146749 | 0.17691 | 0.185 | 0.123 | 1 | 11 | Naga | 0.062 | Old |
| 0.146931 | 0.059936 | 0.145 | 0.087 | 1 | 11 | Nmt2 | 0.058 | Old |
| 0.14698 | 0.233026 | 0.149 | 0.094 | 1 | 11 | Mrpl21 | 0.055 | Old |
| 0.147131 | 1.356183 | 0.052 | 0.022 | 1 | 11 | Dis3l | 0.03 | Old |
| 0.147131 | 1.318391 | 0.052 | 0.022 | 1 | 11 | Suv39h1 | 0.03 | Old |
| 0.147152 | 0.145142 | 0.161 | 0.101 | 1 | 11 | Utrn | 0.06 | Old |
| 0.14733 | 0.054145 | 0.133 | 0.08 | 1 | 11 | Ampd3 | 0.053 | Old |
| 0.147335 | 0.314581 | 0.569 | 0.442 | 1 | 11 | Gpx4 | 0.127 | Old |

| | | | | | | | | |
|---|---|---|---|---|---|---|---|---|
| 0.147476 | 0.150823 | 0.065 | 0.029 | 1 | 11 | L3mbtl3 | 0.036 | Old |
| 0.14749 | 0.186441 | 0.431 | 0.333 | 1 | 11 | Raph1 | 0.098 | Old |
| 0.147508 | 0.271002 | 0.165 | 0.109 | 1 | 11 | Trpv2 | 0.056 | Old |
| 0.14768 | 0.269935 | 0.21 | 0.145 | 1 | 11 | Stk3 | 0.065 | Old |
| 0.148017 | 0.069873 | 0.206 | 0.138 | 1 | 11 | Zfp950 | 0.068 | Old |
| 0.148296 | 0.151203 | 0.133 | 0.08 | 1 | 11 | Selenon | 0.053 | Old |
| 0.148414 | 0.084932 | 0.206 | 0.138 | 1 | 11 | Wdr82 | 0.068 | Old |
| 0.148445 | 0.013422 | 0.657 | 0.522 | 1 | 11 | Birc3 | 0.135 | Old |
| 0.148611 | 0.236986 | 0.185 | 0.123 | 1 | 11 | Plekhg3 | 0.062 | Old |
| 0.148626 | 0.161108 | 0.899 | 0.797 | 1 | 11 | H2-T23 | 0.102 | Old |
| 0.148674 | 0.984156 | 0.052 | 0.022 | 1 | 11 | Lrrc14 | 0.03 | Old |
| 0.148674 | 0.707942 | 0.052 | 0.022 | 1 | 11 | Atat1 | 0.03 | Old |
| 0.148876 | 0.014795 | 0.661 | 0.507 | 1 | 11 | Atp6v1b2 | 0.154 | Old |
| 0.148887 | 0.515053 | 0.093 | 0.051 | 1 | 11 | Urod | 0.042 | Old |
| 0.149152 | 0.094965 | 0.992 | 0.978 | 1 | 11 | mt-Nd4 | 0.014 | Old |
| 0.149289 | 0.312469 | 0.141 | 0.087 | 1 | 11 | Acox1 | 0.054 | Old |
| 0.14932 | 0.922939 | 0.073 | 0.036 | 1 | 11 | Ndst1 | 0.037 | Old |
| 0.149325 | 0.035508 | 0.242 | 0.167 | 1 | 11 | H2-Q6 | 0.075 | Old |
| 0.149444 | 0.089068 | 0.665 | 0.493 | 1 | 11 | Selenow | 0.172 | Old |
| 0.149451 | 0.834888 | 0.052 | 0.022 | 1 | 11 | Atp1a3 | 0.03 | Old |
| 0.149763 | 0.081614 | 0.141 | 0.087 | 1 | 11 | Tmem183a | 0.054 | Old |
| 0.149972 | 0.136605 | 0.105 | 0.058 | 1 | 11 | Serpinb9 | 0.047 | Old |
| 0.149976 | 0.29856 | 0.073 | 0.036 | 1 | 11 | Donson | 0.037 | Old |
| 0.149977 | 0.219209 | 0.113 | 0.065 | 1 | 11 | Pnpt1 | 0.048 | Old |
| 0.150106 | 0.217202 | 0.653 | 0.529 | 1 | 11 | Gm2a | 0.124 | Old |
| 0.150126 | 0.205109 | 0.085 | 0.043 | 1 | 11 | B230118H( | 0.042 | Old |
| 0.150231 | 1.178005 | 0.052 | 0.022 | 1 | 11 | Adora2a | 0.03 | Old |
| 0.150231 | 0.30656 | 0.052 | 0.022 | 1 | 11 | H2afx | 0.03 | Old |
| 0.150334 | 0.006531 | 0.169 | 0.109 | 1 | 11 | Srpk2 | 0.06 | Old |
| 0.150418 | 0.161837 | 0.746 | 0.594 | 1 | 11 | Grb2 | 0.152 | Old |
| 0.150626 | 0.459779 | 0.093 | 0.051 | 1 | 11 | Ankrd10 | 0.042 | Old |
| 0.150634 | 0.394732 | 0.073 | 0.036 | 1 | 11 | Maml3 | 0.037 | Old |
| 0.150917 | 0.03607 | 0.294 | 0.21 | 1 | 11 | Mfsd14a | 0.084 | Old |
| 0.151013 | 1.532224 | 0.052 | 0.022 | 1 | 11 | Ghdc | 0.03 | Old |
| 0.151013 | 0.785716 | 0.052 | 0.022 | 1 | 11 | Ccdc181 | 0.03 | Old |
| 0.151043 | 0.180678 | 0.407 | 0.312 | 1 | 11 | Herc6 | 0.095 | Old |
| 0.151172 | 0.060647 | 0.984 | 0.964 | 1 | 11 | Rpl41 | 0.02 | Old |
| 0.151294 | 1.158524 | 0.073 | 0.036 | 1 | 11 | Mrs2 | 0.037 | Old |
| 0.151541 | 0.106703 | 0.399 | 0.283 | 1 | 11 | Gnai3 | 0.116 | Old |
| 0.151799 | 2.302688 | 0.052 | 0.022 | 1 | 11 | Dpt | 0.03 | Old |
| 0.151956 | 1.201364 | 0.073 | 0.036 | 1 | 11 | Tsr1 | 0.037 | Old |
| 0.152189 | 0.236828 | 0.25 | 0.181 | 1 | 11 | Nr4a3 | 0.069 | Old |
| 0.152588 | 0.955162 | 0.052 | 0.022 | 1 | 11 | Tlr8 | 0.03 | Old |
| 0.152588 | 0.752023 | 0.052 | 0.022 | 1 | 11 | Eef1akmt4 | 0.03 | Old |
| 0.15262 | 0.992908 | 0.073 | 0.036 | 1 | 11 | Dock5 | 0.037 | Old |

| | | | | | | | | |
|---|---|---|---|---|---|---|---|---|
| 0.152631 | 0.159773 | 0.714 | 0.587 | 1 | 11 | Eif4a1 | 0.127 | Old |
| 0.152631 | 0.301964 | 0.137 | 0.087 | 1 | 11 | Nolc1 | 0.05 | Old |
| 0.152898 | 0.646008 | 0.169 | 0.116 | 1 | 11 | Plekhg5 | 0.053 | Old |
| 0.153015 | 0.018243 | 0.609 | 0.464 | 1 | 11 | Pip4k2a | 0.145 | Old |
| 0.153167 | 0.144949 | 0.113 | 0.065 | 1 | 11 | Mzt1 | 0.048 | Old |
| 0.153238 | 0.290423 | 0.165 | 0.109 | 1 | 11 | Hjurp | 0.056 | Old |
| 0.153267 | 0.392576 | 0.129 | 0.08 | 1 | 11 | Eif2ak1 | 0.049 | Old |
| 0.153287 | 0.819586 | 0.073 | 0.036 | 1 | 11 | 4930503L1 | 0.037 | Old |
| 0.153287 | 0.405476 | 0.073 | 0.036 | 1 | 11 | Tmem80 | 0.037 | Old |
| 0.153559 | 0.08802 | 0.093 | 0.051 | 1 | 11 | Med23 | 0.042 | Old |
| 0.153905 | 0.214956 | 0.214 | 0.145 | 1 | 11 | Zzz3 | 0.069 | Old |
| 0.153956 | 0.214801 | 0.073 | 0.036 | 1 | 11 | Mycbp | 0.037 | Old |
| 0.154134 | 0.459033 | 0.165 | 0.109 | 1 | 11 | Ube3c | 0.056 | Old |
| 0.154151 | 0.633193 | 0.093 | 0.051 | 1 | 11 | Zfp771 | 0.042 | Old |
| 0.154176 | 0.573359 | 0.052 | 0.022 | 1 | 11 | Mlh3 | 0.03 | Old |
| 0.154572 | 0.15476 | 0.734 | 0.616 | 1 | 11 | Cd81 | 0.118 | Old |
| 0.154627 | 0.607825 | 0.073 | 0.036 | 1 | 11 | Spidr | 0.037 | Old |
| 0.154681 | 0.073216 | 0.952 | 0.913 | 1 | 11 | Cfl1 | 0.039 | Old |
| 0.154974 | 0.78896 | 0.052 | 0.022 | 1 | 11 | Slc2a8 | 0.03 | Old |
| 0.154974 | 0.216522 | 0.052 | 0.022 | 1 | 11 | Csrp2 | 0.03 | Old |
| 0.15513 | 0.061097 | 0.46 | 0.341 | 1 | 11 | Washc4 | 0.119 | Old |
| 0.15534 | 0.060101 | 0.093 | 0.051 | 1 | 11 | Plcb2 | 0.042 | Old |
| 0.155585 | 0.662386 | 0.101 | 0.058 | 1 | 11 | Zdhhc6 | 0.043 | Old |
| 0.155748 | 0.049208 | 0.714 | 0.551 | 1 | 11 | Eef1g | 0.163 | Old |
| 0.155775 | 0.771977 | 0.052 | 0.022 | 1 | 11 | Ttc37 | 0.03 | Old |
| 0.15579 | 0.434599 | 0.129 | 0.08 | 1 | 11 | Tmem63b | 0.049 | Old |
| 0.155911 | 0.130786 | 0.335 | 0.246 | 1 | 11 | Hexim1 | 0.089 | Old |
| 0.155944 | 0.633142 | 0.109 | 0.065 | 1 | 11 | Fam219b | 0.044 | Old |
| 0.155976 | 0.644256 | 0.073 | 0.036 | 1 | 11 | Lin37 | 0.037 | Old |
| 0.156144 | 0.007785 | 0.262 | 0.181 | 1 | 11 | Thoc7 | 0.081 | Old |
| 0.15616 | 0.074639 | 0.306 | 0.217 | 1 | 11 | Zc3h13 | 0.089 | Old |
| 0.15625 | 0.183596 | 0.198 | 0.138 | 1 | 11 | Cenpx | 0.06 | Old |
| 0.156415 | 0.129244 | 0.25 | 0.174 | 1 | 11 | Itgb5 | 0.076 | Old |
| 0.15658 | 0.257047 | 0.052 | 0.022 | 1 | 11 | Timm21 | 0.03 | Old |
| 0.156731 | 0.638354 | 0.101 | 0.058 | 1 | 11 | Acsl3 | 0.043 | Old |
| 0.156797 | 0.226413 | 0.387 | 0.29 | 1 | 11 | Fem1c | 0.097 | Old |
| 0.156808 | 0.472052 | 0.129 | 0.08 | 1 | 11 | Sart3 | 0.049 | Old |
| 0.157306 | 0.944693 | 0.101 | 0.058 | 1 | 11 | Lsm10 | 0.043 | Old |
| 0.157334 | 0.580628 | 0.073 | 0.036 | 1 | 11 | Abtb1 | 0.037 | Old |
| 0.15738 | 0.645348 | 0.153 | 0.101 | 1 | 11 | Shprh | 0.052 | Old |
| 0.157388 | 0.311655 | 0.052 | 0.022 | 1 | 11 | Mbd4 | 0.03 | Old |
| 0.157534 | 0.989261 | 0.133 | 0.087 | 1 | 11 | Ncbp2 | 0.046 | Old |
| 0.157979 | 0.062607 | 0.399 | 0.283 | 1 | 11 | Srsf11 | 0.116 | Old |
| 0.158199 | 0.60646 | 0.052 | 0.022 | 1 | 11 | Gm20604 | 0.03 | Old |
| 0.158199 | 0.241477 | 0.052 | 0.022 | 1 | 11 | Def8 | 0.03 | Old |

| | | | | | | | | |
|---|---|---|---|---|---|---|---|---|
| 0.158207 | 0.113127 | 0.149 | 0.094 | 1 | 11 | Ttc9c | 0.055 | Old |
| 0.158689 | 0.227145 | 0.149 | 0.094 | 1 | 11 | Bcor | 0.055 | Old |
| 0.158948 | 0.16541 | 0.093 | 0.051 | 1 | 11 | Git1 | 0.042 | Old |
| 0.15899 | 0.708262 | 0.177 | 0.123 | 1 | 11 | Adgre4 | 0.054 | Old |
| 0.159005 | 0.249583 | 0.19 | 0.13 | 1 | 11 | Bhlhe40 | 0.06 | Old |
| 0.159013 | 0.583587 | 0.052 | 0.022 | 1 | 11 | Cry2 | 0.03 | Old |
| 0.159042 | 0.364337 | 0.101 | 0.058 | 1 | 11 | Pdlim2 | 0.043 | Old |
| 0.159089 | 0.280614 | 0.121 | 0.072 | 1 | 11 | Gpr137 | 0.049 | Old |
| 0.159285 | 0.02466 | 0.242 | 0.167 | 1 | 11 | Ntan1 | 0.075 | Old |
| 0.159541 | 0.456987 | 0.137 | 0.087 | 1 | 11 | Skiv2l2 | 0.05 | Old |
| 0.159623 | 0.496585 | 0.121 | 0.072 | 1 | 11 | Tigd2 | 0.049 | Old |
| 0.159732 | 0.213141 | 0.222 | 0.152 | 1 | 11 | Gas2l1 | 0.07 | Old |
| 0.160165 | 0.2047 | 0.185 | 0.123 | 1 | 11 | Ppil2 | 0.062 | Old |
| 0.16019 | 0.170786 | 0.685 | 0.507 | 1 | 11 | Uqcr11 | 0.178 | Old |
| 0.160296 | 1.480887 | 0.04 | 0.014 | 1 | 11 | Ppp1r12b | 0.026 | Old |
| 0.160296 | 1.0668 | 0.04 | 0.014 | 1 | 11 | Zfp667 | 0.026 | Old |
| 0.160469 | 0.027246 | 0.234 | 0.159 | 1 | 11 | Nap1l4 | 0.075 | Old |
| 0.160767 | 0.414179 | 0.073 | 0.036 | 1 | 11 | Papolg | 0.037 | Old |
| 0.160896 | 0.121028 | 0.379 | 0.283 | 1 | 11 | Rnf114 | 0.096 | Old |
| 0.161231 | 0.4352 | 0.121 | 0.072 | 1 | 11 | Zfp407 | 0.049 | Old |
| 0.161241 | 1.611006 | 0.04 | 0.014 | 1 | 11 | Gm12216 | 0.026 | Old |
| 0.161241 | 1.584414 | 0.04 | 0.014 | 1 | 11 | Fam117a | 0.026 | Old |
| 0.161275 | 0.038617 | 0.48 | 0.348 | 1 | 11 | Dync1i2 | 0.132 | Old |
| 0.161389 | 0.307128 | 0.093 | 0.051 | 1 | 11 | Gdap2 | 0.042 | Old |
| 0.161474 | 0.112681 | 0.052 | 0.022 | 1 | 11 | Hp | 0.03 | Old |
| 0.161474 | 0.030613 | 0.052 | 0.022 | 1 | 11 | Gtpbp8 | 0.03 | Old |
| 0.161906 | 0.674845 | 0.117 | 0.072 | 1 | 11 | Banp | 0.045 | Old |
| 0.16197 | 0.07077 | 0.335 | 0.246 | 1 | 11 | Map3k8 | 0.089 | Old |
| 0.16219 | 1.478249 | 0.04 | 0.014 | 1 | 11 | Ndc1 | 0.026 | Old |
| 0.16219 | 1.198448 | 0.04 | 0.014 | 1 | 11 | Adamtsl5 | 0.026 | Old |
| 0.16219 | 0.906946 | 0.04 | 0.014 | 1 | 11 | Ndufaf7 | 0.026 | Old |
| 0.162298 | 0.029703 | 0.423 | 0.319 | 1 | 11 | Specc1l | 0.104 | Old |
| 0.162491 | 0.212216 | 0.181 | 0.123 | 1 | 11 | Pold4 | 0.058 | Old |
| 0.162622 | 1.924248 | 0.028 | 0.007 | 1 | 11 | Zfp873 | 0.021 | Old |
| 0.162622 | 1.719978 | 0.028 | 0.007 | 1 | 11 | Dhrs11 | 0.021 | Old |
| 0.162924 | 1.440974 | 0.06 | 0.029 | 1 | 11 | Scd1 | 0.031 | Old |
| 0.163143 | 1.312925 | 0.04 | 0.014 | 1 | 11 | Abhd11 | 0.026 | Old |
| 0.163148 | 0.488419 | 0.101 | 0.058 | 1 | 11 | Hinfp | 0.043 | Old |
| 0.163181 | 0.958889 | 0.081 | 0.043 | 1 | 11 | Trim30b | 0.038 | Old |
| 0.16334 | 0.024887 | 0.254 | 0.181 | 1 | 11 | Safb | 0.073 | Old |
| 0.163384 | 0.176078 | 0.492 | 0.377 | 1 | 11 | Zmiz1 | 0.115 | Old |
| 0.163394 | 0.433091 | 0.121 | 0.072 | 1 | 11 | Acot9 | 0.049 | Old |
| 0.163631 | 0.603875 | 0.161 | 0.109 | 1 | 11 | Fam214b | 0.052 | Old |
| 0.163691 | 1.684861 | 0.06 | 0.029 | 1 | 11 | Myo10 | 0.031 | Old |
| 0.163741 | 0.496698 | 0.101 | 0.058 | 1 | 11 | Metap1 | 0.043 | Old |

| | | | | | | | | |
|---|---|---|---|---|---|---|---|---|
| 0.163762 | 0.20311 | 0.762 | 0.609 | 1 | 11 | Fam49b | 0.153 | Old |
| 0.163786 | 2.26079 | 0.028 | 0.007 | 1 | 11 | Ptpn4 | 0.021 | Old |
| 0.163786 | 2.11461 | 0.028 | 0.007 | 1 | 11 | Ugt1a7c | 0.021 | Old |
| 0.163786 | 2.08041 | 0.028 | 0.007 | 1 | 11 | Cd244 | 0.021 | Old |
| 0.163786 | 1.97296 | 0.028 | 0.007 | 1 | 11 | Dpy19l3 | 0.021 | Old |
| 0.163802 | 0.00342 | 0.484 | 0.362 | 1 | 11 | Txndc17 | 0.122 | Old |
| 0.163909 | 0.131278 | 0.319 | 0.232 | 1 | 11 | Golga4 | 0.087 | Old |
| 0.164094 | 0.052477 | 0.343 | 0.254 | 1 | 11 | Rbm47 | 0.089 | Old |
| 0.164101 | 0.996105 | 0.04 | 0.014 | 1 | 11 | Smpd4 | 0.026 | Old |
| 0.16446 | 1.099171 | 0.06 | 0.029 | 1 | 11 | Glce | 0.031 | Old |
| 0.164507 | 0.801696 | 0.081 | 0.043 | 1 | 11 | Klc4 | 0.038 | Old |
| 0.164619 | 0.130695 | 0.157 | 0.101 | 1 | 11 | Itgb1bp1 | 0.056 | Old |
| 0.164742 | 0.048183 | 0.113 | 0.065 | 1 | 11 | Sdccag8 | 0.048 | Old |
| 0.164956 | 2.207123 | 0.028 | 0.007 | 1 | 11 | Ndufaf6 | 0.021 | Old |
| 0.164956 | 2.18324 | 0.028 | 0.007 | 1 | 11 | Bbs2 | 0.021 | Old |
| 0.164956 | 1.983417 | 0.028 | 0.007 | 1 | 11 | Zfp111 | 0.021 | Old |
| 0.164956 | 1.889446 | 0.028 | 0.007 | 1 | 11 | Klhl12 | 0.021 | Old |
| 0.164956 | 1.711116 | 0.028 | 0.007 | 1 | 11 | Platr25 | 0.021 | Old |
| 0.164956 | 1.579806 | 0.028 | 0.007 | 1 | 11 | H2-Q10 | 0.021 | Old |
| 0.165062 | 1.065811 | 0.04 | 0.014 | 1 | 11 | Tacc3 | 0.026 | Old |
| 0.165062 | 0.717176 | 0.04 | 0.014 | 1 | 11 | Cnnm2 | 0.026 | Old |
| 0.165062 | 0.649732 | 0.04 | 0.014 | 1 | 11 | Cactin | 0.026 | Old |
| 0.165062 | 0.545115 | 0.04 | 0.014 | 1 | 11 | Ada | 0.026 | Old |
| 0.165142 | 0.195358 | 0.19 | 0.13 | 1 | 11 | Pfdn1 | 0.06 | Old |
| 0.165205 | 0.206897 | 0.726 | 0.638 | 1 | 11 | Atp5a1 | 0.088 | Old |
| 0.165669 | 0.152632 | 0.492 | 0.362 | 1 | 11 | Aes | 0.13 | Old |
| 0.165761 | 0.034307 | 0.54 | 0.42 | 1 | 11 | Stxbp3 | 0.12 | Old |
| 0.165841 | 0.928094 | 0.081 | 0.043 | 1 | 11 | Cdk5rap2 | 0.038 | Old |
| 0.166016 | 0.044668 | 0.504 | 0.384 | 1 | 11 | Cd300ld | 0.12 | Old |
| 0.166028 | 0.822396 | 0.04 | 0.014 | 1 | 11 | Chst15 | 0.026 | Old |
| 0.166028 | 0.800238 | 0.04 | 0.014 | 1 | 11 | Haus2 | 0.026 | Old |
| 0.166028 | 0.74474 | 0.04 | 0.014 | 1 | 11 | Spout1 | 0.026 | Old |
| 0.166028 | 0.719363 | 0.04 | 0.014 | 1 | 11 | Pgap3 | 0.026 | Old |
| 0.166028 | 0.035972 | 0.04 | 0.014 | 1 | 11 | Bmpr1a | 0.026 | Old |
| 0.166032 | 0.079696 | 0.141 | 0.087 | 1 | 11 | Flot1 | 0.054 | Old |
| 0.166046 | 0.305162 | 0.177 | 0.116 | 1 | 11 | Armc1 | 0.061 | Old |
| 0.166085 | 0.05665 | 0.597 | 0.457 | 1 | 11 | Plxnb2 | 0.14 | Old |
| 0.16613 | 0.026202 | 0.121 | 0.072 | 1 | 11 | Heatr6 | 0.049 | Old |
| 0.166133 | 2.149025 | 0.028 | 0.007 | 1 | 11 | Tcaim | 0.021 | Old |
| 0.166133 | 1.753633 | 0.028 | 0.007 | 1 | 11 | Amigo3 | 0.021 | Old |
| 0.166133 | 1.688516 | 0.028 | 0.007 | 1 | 11 | Rilp | 0.021 | Old |
| 0.166133 | 1.675894 | 0.028 | 0.007 | 1 | 11 | Mpp7 | 0.021 | Old |
| 0.166133 | 1.576581 | 0.028 | 0.007 | 1 | 11 | Enpep | 0.021 | Old |
| 0.166133 | 1.473428 | 0.028 | 0.007 | 1 | 11 | Slc25a26 | 0.021 | Old |
| 0.166133 | 1.460009 | 0.028 | 0.007 | 1 | 11 | 2900093K2 | 0.021 | Old |

| | | | | | | | | |
|---|---|---|---|---|---|---|---|---|
| 0.166133 | 1.438014 | 0.028 | 0.007 | 1 | 11 | Gm17586 | 0.021 | Old |
| 0.166133 | 1.378172 | 0.028 | 0.007 | 1 | 11 | Slc43a3 | 0.021 | Old |
| 0.166294 | 0.145901 | 0.149 | 0.094 | 1 | 11 | Lrch1 | 0.055 | Old |
| 0.166378 | 0.411058 | 0.073 | 0.036 | 1 | 11 | Commd9 | 0.037 | Old |
| 0.166378 | 0.156248 | 0.073 | 0.036 | 1 | 11 | Dnttip1 | 0.037 | Old |
| 0.166512 | 0.796268 | 0.081 | 0.043 | 1 | 11 | 1600002K0 | 0.038 | Old |
| 0.166544 | 0.12816 | 0.23 | 0.159 | 1 | 11 | Rnf139 | 0.071 | Old |
| 0.166656 | 0.040451 | 0.323 | 0.232 | 1 | 11 | Ythdf3 | 0.091 | Old |
| 0.166731 | 0.138497 | 0.101 | 0.058 | 1 | 11 | Malsu1 | 0.043 | Old |
| 0.166998 | 0.474838 | 0.04 | 0.014 | 1 | 11 | Trmt10b | 0.026 | Old |
| 0.167014 | 0.009209 | 0.548 | 0.413 | 1 | 11 | Nrp2 | 0.135 | Old |
| 0.167139 | 0.179921 | 0.839 | 0.746 | 1 | 11 | Uqcrh | 0.093 | Old |
| 0.167316 | 3.550737 | 0.028 | 0.007 | 1 | 11 | Gpnmb | 0.021 | Old |
| 0.167316 | 1.636588 | 0.028 | 0.007 | 1 | 11 | Prepl | 0.021 | Old |
| 0.167316 | 1.608587 | 0.028 | 0.007 | 1 | 11 | Gorab | 0.021 | Old |
| 0.167316 | 1.555876 | 0.028 | 0.007 | 1 | 11 | Srr | 0.021 | Old |
| 0.167316 | 1.450552 | 0.028 | 0.007 | 1 | 11 | Cox11 | 0.021 | Old |
| 0.167316 | 1.307435 | 0.028 | 0.007 | 1 | 11 | D2hgdh | 0.021 | Old |
| 0.167316 | 1.18912 | 0.028 | 0.007 | 1 | 11 | Zfp606 | 0.021 | Old |
| 0.167334 | 0.257696 | 0.101 | 0.058 | 1 | 11 | Cpox | 0.043 | Old |
| 0.167589 | 0.479546 | 0.177 | 0.123 | 1 | 11 | Kif3b | 0.054 | Old |
| 0.167787 | 0.254892 | 0.121 | 0.072 | 1 | 11 | Gpr155 | 0.049 | Old |
| 0.167858 | 0.308854 | 0.081 | 0.043 | 1 | 11 | Mpg | 0.038 | Old |
| 0.167938 | 0.314612 | 0.101 | 0.058 | 1 | 11 | Togaram1 | 0.043 | Old |
| 0.167944 | 0.484933 | 0.169 | 0.116 | 1 | 11 | Shoc2 | 0.053 | Old |
| 0.167972 | 0.667854 | 0.04 | 0.014 | 1 | 11 | Snn | 0.026 | Old |
| 0.167972 | 0.66626 | 0.04 | 0.014 | 1 | 11 | Zfand1 | 0.026 | Old |
| 0.167972 | 0.543339 | 0.04 | 0.014 | 1 | 11 | Cnr2 | 0.026 | Old |
| 0.167972 | 0.413022 | 0.04 | 0.014 | 1 | 11 | Glb1l | 0.026 | Old |
| 0.167972 | 0.400916 | 0.04 | 0.014 | 1 | 11 | Kbtbd3 | 0.026 | Old |
| 0.167972 | 0.224046 | 0.04 | 0.014 | 1 | 11 | Gmds | 0.026 | Old |
| 0.168408 | 0.483472 | 0.109 | 0.065 | 1 | 11 | Bola3 | 0.044 | Old |
| 0.168426 | 0.178987 | 0.214 | 0.152 | 1 | 11 | Uap1 | 0.062 | Old |
| 0.168505 | 1.639245 | 0.028 | 0.007 | 1 | 11 | Lrrk2 | 0.021 | Old |
| 0.168505 | 1.434511 | 0.028 | 0.007 | 1 | 11 | Tctn1 | 0.021 | Old |
| 0.168505 | 1.391809 | 0.028 | 0.007 | 1 | 11 | Fancm | 0.021 | Old |
| 0.168505 | 1.267598 | 0.028 | 0.007 | 1 | 11 | Yars2 | 0.021 | Old |
| 0.168505 | 1.239618 | 0.028 | 0.007 | 1 | 11 | Armc6 | 0.021 | Old |
| 0.168505 | 1.231006 | 0.028 | 0.007 | 1 | 11 | Mtcp1 | 0.021 | Old |
| 0.168505 | 1.229781 | 0.028 | 0.007 | 1 | 11 | Tprn | 0.021 | Old |
| 0.168505 | 1.222174 | 0.028 | 0.007 | 1 | 11 | Cacna1a | 0.021 | Old |
| 0.168505 | 1.184148 | 0.028 | 0.007 | 1 | 11 | Zfp995 | 0.021 | Old |
| 0.168505 | 1.155252 | 0.028 | 0.007 | 1 | 11 | Casp12 | 0.021 | Old |
| 0.168505 | 1.048395 | 0.028 | 0.007 | 1 | 11 | Sema7a | 0.021 | Old |
| 0.168505 | 0.962739 | 0.028 | 0.007 | 1 | 11 | Tbc1d7 | 0.021 | Old |

| | | | | | | | | |
|---|---|---|---|---|---|---|---|---|
| 0.168951 | 0.69088 | 0.04 | 0.014 | 1 | 11 | Zfp319 | 0.026 | Old |
| 0.168951 | 0.610581 | 0.04 | 0.014 | 1 | 11 | Mxra7 | 0.026 | Old |
| 0.168951 | 0.570532 | 0.04 | 0.014 | 1 | 11 | Zfp668 | 0.026 | Old |
| 0.168951 | 0.362806 | 0.04 | 0.014 | 1 | 11 | Gm37233 | 0.026 | Old |
| 0.169012 | 0.16431 | 0.157 | 0.101 | 1 | 11 | Gps1 | 0.056 | Old |
| 0.169152 | 0.292906 | 0.101 | 0.058 | 1 | 11 | Cryzl2 | 0.043 | Old |
| 0.169421 | 0.30752 | 0.129 | 0.08 | 1 | 11 | Zbtb1 | 0.049 | Old |
| 0.169492 | 0.297512 | 0.153 | 0.101 | 1 | 11 | Smc5 | 0.052 | Old |
| 0.169701 | 1.244644 | 0.028 | 0.007 | 1 | 11 | Zfhx2 | 0.021 | Old |
| 0.169701 | 1.157409 | 0.028 | 0.007 | 1 | 11 | Tnfsf14 | 0.021 | Old |
| 0.169701 | 0.995885 | 0.028 | 0.007 | 1 | 11 | Rmdn1 | 0.021 | Old |
| 0.169701 | 0.946296 | 0.028 | 0.007 | 1 | 11 | 4632404H1 | 0.021 | Old |
| 0.169701 | 0.847637 | 0.028 | 0.007 | 1 | 11 | Aatk | 0.021 | Old |
| 0.169701 | 0.726403 | 0.028 | 0.007 | 1 | 11 | Snap47 | 0.021 | Old |
| 0.169701 | 0.7169 | 0.028 | 0.007 | 1 | 11 | Gm48604 | 0.021 | Old |
| 0.169701 | 0.523532 | 0.028 | 0.007 | 1 | 11 | Tmc4 | 0.021 | Old |
| 0.169701 | 0.504097 | 0.028 | 0.007 | 1 | 11 | 2-Sep | 0.021 | Old |
| 0.169701 | 0.503482 | 0.028 | 0.007 | 1 | 11 | Mkks | 0.021 | Old |
| 0.169701 | 0.374293 | 0.028 | 0.007 | 1 | 11 | Ss18l1 | 0.021 | Old |
| 0.169894 | 0.838794 | 0.081 | 0.043 | 1 | 11 | Gpr68 | 0.038 | Old |
| 0.169934 | 0.44835 | 0.04 | 0.014 | 1 | 11 | Cnbd2 | 0.026 | Old |
| 0.169934 | 0.274369 | 0.04 | 0.014 | 1 | 11 | Synpo | 0.026 | Old |
| 0.169959 | 0.127289 | 0.073 | 0.036 | 1 | 11 | Alad | 0.037 | Old |
| 0.169959 | 0.042517 | 0.073 | 0.036 | 1 | 11 | Slc16a1 | 0.037 | Old |
| 0.170016 | 0.027638 | 0.121 | 0.072 | 1 | 11 | Noc4l | 0.049 | Old |
| 0.170373 | 0.186229 | 0.101 | 0.058 | 1 | 11 | Pparg | 0.043 | Old |
| 0.170468 | 0.004321 | 0.605 | 0.449 | 1 | 11 | Ube2d2a | 0.156 | Old |
| 0.170472 | 0.267622 | 0.165 | 0.109 | 1 | 11 | Itpr2 | 0.056 | Old |
| 0.170576 | 0.359857 | 0.081 | 0.043 | 1 | 11 | Inafm1 | 0.038 | Old |
| 0.17061 | 0.45719 | 0.137 | 0.087 | 1 | 11 | Srebf1 | 0.05 | Old |
| 0.170682 | 0.108887 | 0.073 | 0.036 | 1 | 11 | Rpp14 | 0.037 | Old |
| 0.170903 | 0.966751 | 0.028 | 0.007 | 1 | 11 | Cttn | 0.021 | Old |
| 0.170903 | 0.876271 | 0.028 | 0.007 | 1 | 11 | Spib | 0.021 | Old |
| 0.170903 | 0.750505 | 0.028 | 0.007 | 1 | 11 | E2f2 | 0.021 | Old |
| 0.170903 | 0.662438 | 0.028 | 0.007 | 1 | 11 | St5 | 0.021 | Old |
| 0.170903 | 0.654732 | 0.028 | 0.007 | 1 | 11 | Pnp2 | 0.021 | Old |
| 0.170903 | 0.431711 | 0.028 | 0.007 | 1 | 11 | Setd6 | 0.021 | Old |
| 0.170903 | 0.394175 | 0.028 | 0.007 | 1 | 11 | Cep128 | 0.021 | Old |
| 0.170903 | 0.345391 | 0.028 | 0.007 | 1 | 11 | Ankrd9 | 0.021 | Old |
| 0.170903 | 0.32755 | 0.028 | 0.007 | 1 | 11 | Slc9a4 | 0.021 | Old |
| 0.170903 | 0.313217 | 0.028 | 0.007 | 1 | 11 | Snapc4 | 0.021 | Old |
| 0.170903 | 0.289909 | 0.028 | 0.007 | 1 | 11 | Gm28557 | 0.021 | Old |
| 0.170903 | 0.208841 | 0.028 | 0.007 | 1 | 11 | Nynrin | 0.021 | Old |
| 0.170903 | 0.176152 | 0.028 | 0.007 | 1 | 11 | Lzts2 | 0.021 | Old |
| 0.170903 | 0.175517 | 0.028 | 0.007 | 1 | 11 | Syn1 | 0.021 | Old |

| | | | | | | | | |
|---|---|---|---|---|---|---|---|---|
| 0.170903 | 0.092408 | 0.028 | 0.007 | 1 | 11 | Spata1 | 0.021 | Old |
| 0.170903 | 0.037214 | 0.028 | 0.007 | 1 | 11 | Ubiad1 | 0.021 | Old |
| 0.170921 | 0.104103 | 0.04 | 0.014 | 1 | 11 | Nudt22 | 0.026 | Old |
| 0.170969 | 0.131574 | 0.403 | 0.304 | 1 | 11 | Ppp1r12c | 0.099 | Old |
| 0.171071 | 0.0366 | 0.988 | 0.964 | 1 | 11 | Actg1 | 0.024 | Old |
| 0.171375 | 0.069048 | 0.532 | 0.406 | 1 | 11 | Peli1 | 0.126 | Old |
| 0.171778 | 0.133623 | 0.375 | 0.283 | 1 | 11 | Vdac3 | 0.092 | Old |
| 0.171947 | 0.155233 | 0.456 | 0.341 | 1 | 11 | Eloc | 0.115 | Old |
| 0.172199 | 0.060946 | 0.266 | 0.188 | 1 | 11 | Gosr2 | 0.078 | Old |
| 0.172268 | 0.137227 | 0.121 | 0.072 | 1 | 11 | Foxk2 | 0.049 | Old |
| 0.172305 | 0.687533 | 0.06 | 0.029 | 1 | 11 | Gnb4 | 0.031 | Old |
| 0.172527 | 0.430369 | 0.109 | 0.065 | 1 | 11 | Fbxo7 | 0.044 | Old |
| 0.172527 | 0.177149 | 0.109 | 0.065 | 1 | 11 | Lztfl1 | 0.044 | Old |
| 0.173015 | 0.048912 | 0.448 | 0.341 | 1 | 11 | Olfm1 | 0.107 | Old |
| 0.173105 | 1.306378 | 0.06 | 0.029 | 1 | 11 | Ercc5 | 0.031 | Old |
| 0.173907 | 1.005185 | 0.06 | 0.029 | 1 | 11 | Cables2 | 0.031 | Old |
| 0.174302 | 0.404218 | 0.161 | 0.109 | 1 | 11 | Commd2 | 0.052 | Old |
| 0.174315 | 0.335748 | 0.109 | 0.065 | 1 | 11 | Lin54 | 0.044 | Old |
| 0.174343 | 0.064803 | 0.129 | 0.08 | 1 | 11 | Sorl1 | 0.049 | Old |
| 0.174414 | 0.060645 | 0.621 | 0.486 | 1 | 11 | Nap1l1 | 0.135 | Old |
| 0.174528 | 0.500384 | 0.153 | 0.101 | 1 | 11 | Mrpl16 | 0.052 | Old |
| 0.174713 | 0.534134 | 0.06 | 0.029 | 1 | 11 | Gnpnat1 | 0.031 | Old |
| 0.174835 | 0.575306 | 0.117 | 0.072 | 1 | 11 | Kcnab2 | 0.045 | Old |
| 0.175063 | 0.020038 | 0.613 | 0.464 | 1 | 11 | Nrp1 | 0.149 | Old |
| 0.175126 | 0.069247 | 0.996 | 1 | 1 | 11 | mt-Co2 | -0.004 | Old |
| 0.175297 | 0.283942 | 0.161 | 0.109 | 1 | 11 | Mrpl35 | 0.052 | Old |
| 0.175521 | 0.544182 | 0.06 | 0.029 | 1 | 11 | Pank1 | 0.031 | Old |
| 0.175618 | 0.006379 | 0.5 | 0.37 | 1 | 11 | Anp32a | 0.13 | Old |
| 0.175679 | 0.247159 | 0.907 | 0.891 | 1 | 11 | Selenop | 0.016 | Old |
| 0.175868 | 0.611714 | 0.145 | 0.094 | 1 | 11 | Ate1 | 0.051 | Old |
| 0.175946 | 0.086439 | 0.101 | 0.058 | 1 | 11 | Klhl2 | 0.043 | Old |
| 0.17602 | 0.21563 | 0.157 | 0.101 | 1 | 11 | Ncoa5 | 0.056 | Old |
| 0.176153 | 0.871213 | 0.089 | 0.051 | 1 | 11 | Borcs7 | 0.038 | Old |
| 0.176331 | 0.828994 | 0.06 | 0.029 | 1 | 11 | Zfp160 | 0.031 | Old |
| 0.176331 | 0.452933 | 0.06 | 0.029 | 1 | 11 | Rrp9 | 0.031 | Old |
| 0.176574 | 0.167801 | 0.153 | 0.101 | 1 | 11 | Ngly1 | 0.052 | Old |
| 0.176812 | 1.261539 | 0.081 | 0.043 | 1 | 11 | Dgcr8 | 0.038 | Old |
| 0.177145 | 0.167827 | 0.06 | 0.029 | 1 | 11 | Sesn3 | 0.031 | Old |
| 0.177145 | 0.167548 | 0.06 | 0.029 | 1 | 11 | Gm26782 | 0.031 | Old |
| 0.17716 | 0.312991 | 0.117 | 0.072 | 1 | 11 | Tmem181a | 0.045 | Old |
| 0.177682 | 0.297029 | 0.129 | 0.08 | 1 | 11 | Abhd2 | 0.049 | Old |
| 0.177849 | 0.124136 | 0.137 | 0.087 | 1 | 11 | Anapc2 | 0.05 | Old |
| 0.177961 | 0.388782 | 0.06 | 0.029 | 1 | 11 | Cbr3 | 0.031 | Old |
| 0.178415 | 1.555043 | 0.069 | 0.036 | 1 | 11 | Tpgs2 | 0.033 | Old |
| 0.178415 | 1.498048 | 0.069 | 0.036 | 1 | 11 | Crbn | 0.033 | Old |

| | | | | | | | | |
|---|---|---|---|---|---|---|---|---|
| 0.17878 | 0.69996 | 0.06 | 0.029 | 1 | 11 | Zscan21 | 0.031 | Old |
| 0.178792 | 0.053625 | 0.27 | 0.188 | 1 | 11 | Fyttd1 | 0.082 | Old |
| 0.179151 | 0.247503 | 0.109 | 0.065 | 1 | 11 | Cdc42ep2 | 0.044 | Old |
| 0.179151 | 0.102102 | 0.109 | 0.065 | 1 | 11 | Exoc6 | 0.044 | Old |
| 0.179228 | 0.257495 | 0.181 | 0.123 | 1 | 11 | Snrpa | 0.058 | Old |
| 0.179504 | 0.350432 | 0.089 | 0.051 | 1 | 11 | Ctu2 | 0.038 | Old |
| 0.179602 | 0.109545 | 0.06 | 0.029 | 1 | 11 | St7 | 0.031 | Old |
| 0.179662 | 0.143935 | 0.673 | 0.529 | 1 | 11 | Ncf2 | 0.144 | Old |
| 0.17982 | 0.103617 | 0.21 | 0.145 | 1 | 11 | Rxra | 0.065 | Old |
| 0.179843 | 0.276745 | 0.298 | 0.232 | 1 | 11 | Kitl | 0.066 | Old |
| 0.180275 | 0.061416 | 0.21 | 0.145 | 1 | 11 | Usp14 | 0.065 | Old |
| 0.180502 | 0.060493 | 0.343 | 0.254 | 1 | 11 | Nfat5 | 0.089 | Old |
| 0.180858 | 0.68001 | 0.089 | 0.051 | 1 | 11 | Mthfsl | 0.038 | Old |
| 0.180973 | 0.07276 | 0.419 | 0.297 | 1 | 11 | Mdh2 | 0.122 | Old |
| 0.181255 | 0.472675 | 0.06 | 0.029 | 1 | 11 | Tlcd2 | 0.031 | Old |
| 0.181566 | 0.007109 | 0.262 | 0.181 | 1 | 11 | Gng12 | 0.081 | Old |
| 0.182085 | 0.46064 | 0.06 | 0.029 | 1 | 11 | Slc44a2 | 0.031 | Old |
| 0.182476 | 0.499062 | 0.117 | 0.072 | 1 | 11 | Farsb | 0.045 | Old |
| 0.182495 | 0.315139 | 0.081 | 0.043 | 1 | 11 | Ano10 | 0.038 | Old |
| 0.182577 | 0.066215 | 0.645 | 0.478 | 1 | 11 | Brk1 | 0.167 | Old |
| 0.182665 | 0.045548 | 0.298 | 0.217 | 1 | 11 | Pfkp | 0.081 | Old |
| 0.182843 | 0.333146 | 0.109 | 0.065 | 1 | 11 | Dhdds | 0.044 | Old |
| 0.182918 | 0.086861 | 0.06 | 0.029 | 1 | 11 | Cep290 | 0.031 | Old |
| 0.182937 | 0.215691 | 0.875 | 0.79 | 1 | 11 | Neat1 | 0.085 | Old |
| 0.183215 | 0.645352 | 0.081 | 0.043 | 1 | 11 | BC147527 | 0.038 | Old |
| 0.183242 | 0.135753 | 0.157 | 0.101 | 1 | 11 | Sil1 | 0.056 | Old |
| 0.183754 | 0.261108 | 0.06 | 0.029 | 1 | 11 | Dhx37 | 0.031 | Old |
| 0.183929 | 0.301865 | 0.161 | 0.109 | 1 | 11 | Ilf2 | 0.052 | Old |
| 0.183937 | 0.085824 | 0.081 | 0.043 | 1 | 11 | Abi3 | 0.038 | Old |
| 0.184485 | 0.007822 | 0.194 | 0.13 | 1 | 11 | Ehmt1 | 0.064 | Old |
| 0.184593 | 0.285852 | 0.06 | 0.029 | 1 | 11 | Tbrg4 | 0.031 | Old |
| 0.185368 | 0.143471 | 0.782 | 0.616 | 1 | 11 | Erp29 | 0.166 | Old |
| 0.185597 | 0.630814 | 0.169 | 0.116 | 1 | 11 | Igsf8 | 0.053 | Old |
| 0.185728 | 0.07312 | 0.831 | 0.667 | 1 | 11 | Ctsh | 0.164 | Old |
| 0.186279 | 0.522374 | 0.06 | 0.029 | 1 | 11 | Flt1 | 0.031 | Old |
| 0.186279 | 0.264291 | 0.06 | 0.029 | 1 | 11 | Cep135 | 0.031 | Old |
| 0.186404 | 0.128128 | 0.157 | 0.101 | 1 | 11 | Slc4a2 | 0.056 | Old |
| 0.186408 | 0.038085 | 0.427 | 0.326 | 1 | 11 | Chmp2a | 0.101 | Old |
| 0.187044 | 0.388549 | 0.089 | 0.051 | 1 | 11 | Zcchc8 | 0.038 | Old |
| 0.187204 | 0.003247 | 0.323 | 0.232 | 1 | 11 | Srp72 | 0.091 | Old |
| 0.187252 | 1.084863 | 0.097 | 0.058 | 1 | 11 | Pja1 | 0.039 | Old |
| 0.187292 | 0.104905 | 0.77 | 0.638 | 1 | 11 | Srrm2 | 0.132 | Old |
| 0.187605 | 0.12449 | 0.153 | 0.101 | 1 | 11 | Cspp1 | 0.052 | Old |
| 0.187764 | 0.918343 | 0.069 | 0.036 | 1 | 11 | St3gal3 | 0.033 | Old |
| 0.187764 | 0.857877 | 0.069 | 0.036 | 1 | 11 | Prkdc | 0.033 | Old |

| | | | | | | | | |
|---|---|---|---|---|---|---|---|---|
| 0.187852 | 0.167647 | 0.109 | 0.065 | 1 | 11 | Utp15 | 0.044 | Old |
| 0.187853 | 0.038689 | 0.137 | 0.087 | 1 | 11 | Glipr2 | 0.05 | Old |
| 0.187862 | 0.01341 | 0.359 | 0.254 | 1 | 11 | Nlrc5 | 0.105 | Old |
| 0.187896 | 0.163059 | 0.23 | 0.167 | 1 | 11 | Mb21d1 | 0.063 | Old |
| 0.187977 | 0.313816 | 0.06 | 0.029 | 1 | 11 | Isyna1 | 0.031 | Old |
| 0.187977 | 0.22679 | 0.06 | 0.029 | 1 | 11 | Dffa | 0.031 | Old |
| 0.18844 | 0.086986 | 0.089 | 0.051 | 1 | 11 | Taf12 | 0.038 | Old |
| 0.188687 | 0.270531 | 0.254 | 0.188 | 1 | 11 | R3hdm2 | 0.066 | Old |
| 0.18873 | 0.397668 | 0.133 | 0.087 | 1 | 11 | Rbm15 | 0.046 | Old |
| 0.189881 | 0.374367 | 0.133 | 0.087 | 1 | 11 | Trim65 | 0.046 | Old |
| 0.19032 | 0.182694 | 0.472 | 0.355 | 1 | 11 | Ube2l3 | 0.117 | Old |
| 0.190362 | 0.396846 | 0.117 | 0.072 | 1 | 11 | Ttc32 | 0.045 | Old |
| 0.190395 | 0.022474 | 0.109 | 0.065 | 1 | 11 | Cd226 | 0.044 | Old |
| 0.190735 | 0.827589 | 0.157 | 0.109 | 1 | 11 | Fut11 | 0.048 | Old |
| 0.190805 | 0.136007 | 0.484 | 0.37 | 1 | 11 | Anp32b | 0.114 | Old |
| 0.191026 | 0.06617 | 0.274 | 0.203 | 1 | 11 | Pim3 | 0.071 | Old |
| 0.191765 | 0.769306 | 0.069 | 0.036 | 1 | 11 | Gm28875 | 0.033 | Old |
| 0.191786 | 0.085599 | 0.843 | 0.746 | 1 | 11 | Rpl36 | 0.097 | Old |
| 0.191939 | 0.044838 | 0.153 | 0.101 | 1 | 11 | Tcf3 | 0.052 | Old |
| 0.191963 | 0.388031 | 0.089 | 0.051 | 1 | 11 | Oard1 | 0.038 | Old |
| 0.191963 | 0.350515 | 0.089 | 0.051 | 1 | 11 | Inpp5a | 0.038 | Old |
| 0.192126 | 0.086434 | 0.508 | 0.399 | 1 | 11 | Slc39a1 | 0.109 | Old |
| 0.19227 | 0.142319 | 0.06 | 0.029 | 1 | 11 | Ccdc58 | 0.031 | Old |
| 0.192459 | 0.071211 | 0.528 | 0.406 | 1 | 11 | Spop | 0.122 | Old |
| 0.192598 | 0.132088 | 0.282 | 0.21 | 1 | 11 | Cnppd1 | 0.072 | Old |
| 0.192673 | 0.55827 | 0.089 | 0.051 | 1 | 11 | Mrpl39 | 0.038 | Old |
| 0.192673 | 0.211159 | 0.089 | 0.051 | 1 | 11 | Comtd1 | 0.038 | Old |
| 0.192866 | 0.32343 | 0.625 | 0.493 | 1 | 11 | Foxp1 | 0.132 | Old |
| 0.193227 | 0.037703 | 0.222 | 0.152 | 1 | 11 | Tmem9b | 0.07 | Old |
| 0.193516 | 0.023358 | 0.081 | 0.043 | 1 | 11 | Maml2 | 0.038 | Old |
| 0.19354 | 0.69575 | 0.121 | 0.08 | 1 | 11 | Kmo | 0.041 | Old |
| 0.193583 | 0.467727 | 0.153 | 0.101 | 1 | 11 | Ear2 | 0.052 | Old |
| 0.194374 | 0.919976 | 0.048 | 0.022 | 1 | 11 | Rpl36a-ps1 | 0.026 | Old |
| 0.19501 | 1.019714 | 0.069 | 0.036 | 1 | 11 | Spint2 | 0.033 | Old |
| 0.195353 | 0.988371 | 0.048 | 0.022 | 1 | 11 | Rabif | 0.026 | Old |
| 0.195947 | 0.091194 | 0.685 | 0.558 | 1 | 11 | Plekho2 | 0.127 | Old |
| 0.196181 | 0.104135 | 0.181 | 0.123 | 1 | 11 | Eed | 0.058 | Old |
| 0.196266 | 2.786116 | 0.012 | 0 | 1 | 11 | Dpyd | 0.012 | Old |
| 0.196266 | 2.770389 | 0.012 | 0 | 1 | 11 | Gm11464 | 0.012 | Old |
| 0.196266 | 2.762086 | 0.012 | 0 | 1 | 11 | Adamdec1 | 0.012 | Old |
| 0.196266 | 2.73129 | 0.012 | 0 | 1 | 11 | Cndp1 | 0.012 | Old |
| 0.196266 | 2.653812 | 0.012 | 0 | 1 | 11 | Dmtn | 0.012 | Old |
| 0.196266 | 2.636649 | 0.012 | 0 | 1 | 11 | Nanos1 | 0.012 | Old |
| 0.196266 | 2.619893 | 0.012 | 0 | 1 | 11 | 4931428F0 | 0.012 | Old |
| 0.196266 | 2.574903 | 0.012 | 0 | 1 | 11 | Dand5 | 0.012 | Old |

| | | | | | | | | |
|---|---|---|---|---|---|---|---|---|
| 0.196266 | 2.555203 | 0.012 | 0 | 1 | 11 | Arntl2 | 0.012 | Old |
| 0.196266 | 2.5548 | 0.012 | 0 | 1 | 11 | Scarf2 | 0.012 | Old |
| 0.196266 | 2.485801 | 0.012 | 0 | 1 | 11 | Col16a1 | 0.012 | Old |
| 0.196266 | 2.444141 | 0.012 | 0 | 1 | 11 | Rab34 | 0.012 | Old |
| 0.196266 | 2.331866 | 0.012 | 0 | 1 | 11 | Apol9b | 0.012 | Old |
| 0.196266 | 2.249481 | 0.012 | 0 | 1 | 11 | Tns2 | 0.012 | Old |
| 0.196266 | 2.219943 | 0.012 | 0 | 1 | 11 | Gm47438 | 0.012 | Old |
| 0.196266 | 2.206004 | 0.012 | 0 | 1 | 11 | Eno2 | 0.012 | Old |
| 0.196266 | 2.181358 | 0.012 | 0 | 1 | 11 | Btc | 0.012 | Old |
| 0.196266 | 2.169433 | 0.012 | 0 | 1 | 11 | Dab2ip | 0.012 | Old |
| 0.196266 | 2.166062 | 0.012 | 0 | 1 | 11 | Cd209c | 0.012 | Old |
| 0.196266 | 2.131299 | 0.012 | 0 | 1 | 11 | Gfi1b | 0.012 | Old |
| 0.196266 | 2.127945 | 0.012 | 0 | 1 | 11 | Mmp23 | 0.012 | Old |
| 0.196266 | 2.11503 | 0.012 | 0 | 1 | 11 | Nos3 | 0.012 | Old |
| 0.196266 | 2.087966 | 0.012 | 0 | 1 | 11 | 1110046J0 | 0.012 | Old |
| 0.196266 | 2.086406 | 0.012 | 0 | 1 | 11 | Tsku | 0.012 | Old |
| 0.196266 | 2.073791 | 0.012 | 0 | 1 | 11 | Bicdl1 | 0.012 | Old |
| 0.196266 | 2.046261 | 0.012 | 0 | 1 | 11 | Nos2 | 0.012 | Old |
| 0.196266 | 2.03365 | 0.012 | 0 | 1 | 11 | Hspb6 | 0.012 | Old |
| 0.196266 | 2.018454 | 0.012 | 0 | 1 | 11 | Crisp1 | 0.012 | Old |
| 0.196266 | 2.016891 | 0.012 | 0 | 1 | 11 | Adat1 | 0.012 | Old |
| 0.196266 | 2.006499 | 0.012 | 0 | 1 | 11 | Gm7967 | 0.012 | Old |
| 0.196266 | 2.006004 | 0.012 | 0 | 1 | 11 | Phf24 | 0.012 | Old |
| 0.196266 | 1.983114 | 0.012 | 0 | 1 | 11 | Gm28874 | 0.012 | Old |
| 0.196266 | 1.979662 | 0.012 | 0 | 1 | 11 | Ppm1l | 0.012 | Old |
| 0.196266 | 1.964634 | 0.012 | 0 | 1 | 11 | Xndc1 | 0.012 | Old |
| 0.196266 | 1.961458 | 0.012 | 0 | 1 | 11 | 1810034E1 | 0.012 | Old |
| 0.196266 | 1.959372 | 0.012 | 0 | 1 | 11 | Nup210l | 0.012 | Old |
| 0.196266 | 1.947771 | 0.012 | 0 | 1 | 11 | Dnph1 | 0.012 | Old |
| 0.196266 | 1.947024 | 0.012 | 0 | 1 | 11 | Cygb | 0.012 | Old |
| 0.196266 | 1.923887 | 0.012 | 0 | 1 | 11 | Gm17092 | 0.012 | Old |
| 0.196266 | 1.912297 | 0.012 | 0 | 1 | 11 | Slc6a9 | 0.012 | Old |
| 0.196266 | 1.889055 | 0.012 | 0 | 1 | 11 | Rad54l | 0.012 | Old |
| 0.196266 | 1.8885 | 0.012 | 0 | 1 | 11 | Trmu | 0.012 | Old |
| 0.196266 | 1.861902 | 0.012 | 0 | 1 | 11 | Slc16a9 | 0.012 | Old |
| 0.196266 | 1.853789 | 0.012 | 0 | 1 | 11 | Zfp3 | 0.012 | Old |
| 0.196266 | 1.847913 | 0.012 | 0 | 1 | 11 | Chaf1b | 0.012 | Old |
| 0.196266 | 1.847099 | 0.012 | 0 | 1 | 11 | 5430403G1 | 0.012 | Old |
| 0.196266 | 1.841566 | 0.012 | 0 | 1 | 11 | 1200007C1 | 0.012 | Old |
| 0.196266 | 1.814347 | 0.012 | 0 | 1 | 11 | Dbndd2 | 0.012 | Old |
| 0.196266 | 1.773647 | 0.012 | 0 | 1 | 11 | Wdr74 | 0.012 | Old |
| 0.196266 | 1.769622 | 0.012 | 0 | 1 | 11 | Hsf4 | 0.012 | Old |
| 0.196266 | 1.760393 | 0.012 | 0 | 1 | 11 | Gm16124 | 0.012 | Old |
| 0.196266 | 1.760072 | 0.012 | 0 | 1 | 11 | 9930111J2 | 0.012 | Old |
| 0.196266 | 1.759069 | 0.012 | 0 | 1 | 11 | Tubg1 | 0.012 | Old |

| | | | | | | | | |
|---|---|---|---|---|---|---|---|---|
| 0.196266 | 1.756304 | 0.012 | 0 | 1 | 11 | Dus4l | 0.012 | Old |
| 0.196266 | 1.753811 | 0.012 | 0 | 1 | 11 | Diaph3 | 0.012 | Old |
| 0.196266 | 1.748128 | 0.012 | 0 | 1 | 11 | Mnd1 | 0.012 | Old |
| 0.196266 | 1.737968 | 0.012 | 0 | 1 | 11 | Clic5 | 0.012 | Old |
| 0.196266 | 1.732507 | 0.012 | 0 | 1 | 11 | Hoxb3 | 0.012 | Old |
| 0.196266 | 1.711007 | 0.012 | 0 | 1 | 11 | Cenpn | 0.012 | Old |
| 0.196266 | 1.700932 | 0.012 | 0 | 1 | 11 | Ak8 | 0.012 | Old |
| 0.196266 | 1.697803 | 0.012 | 0 | 1 | 11 | Cdk5r1 | 0.012 | Old |
| 0.196266 | 1.689373 | 0.012 | 0 | 1 | 11 | Recql4 | 0.012 | Old |
| 0.196266 | 1.682954 | 0.012 | 0 | 1 | 11 | Piwil2 | 0.012 | Old |
| 0.196266 | 1.668816 | 0.012 | 0 | 1 | 11 | Tcaf1 | 0.012 | Old |
| 0.196266 | 1.654139 | 0.012 | 0 | 1 | 11 | Zbtb7c | 0.012 | Old |
| 0.196266 | 1.649898 | 0.012 | 0 | 1 | 11 | Gm20513 | 0.012 | Old |
| 0.196266 | 1.629898 | 0.012 | 0 | 1 | 11 | Iglc2 | 0.012 | Old |
| 0.196266 | 1.625009 | 0.012 | 0 | 1 | 11 | Adgrl3 | 0.012 | Old |
| 0.196266 | 1.616138 | 0.012 | 0 | 1 | 11 | Pard6a | 0.012 | Old |
| 0.196266 | 1.609659 | 0.012 | 0 | 1 | 11 | Eri2 | 0.012 | Old |
| 0.196266 | 1.607744 | 0.012 | 0 | 1 | 11 | Tmem204 | 0.012 | Old |
| 0.196266 | 1.603119 | 0.012 | 0 | 1 | 11 | Efemp1 | 0.012 | Old |
| 0.196266 | 1.599131 | 0.012 | 0 | 1 | 11 | Rtn2 | 0.012 | Old |
| 0.196266 | 1.595979 | 0.012 | 0 | 1 | 11 | Zbtb8a | 0.012 | Old |
| 0.196266 | 1.594877 | 0.012 | 0 | 1 | 11 | Hykk | 0.012 | Old |
| 0.196266 | 1.591671 | 0.012 | 0 | 1 | 11 | Prim1 | 0.012 | Old |
| 0.196266 | 1.560198 | 0.012 | 0 | 1 | 11 | Gm20632 | 0.012 | Old |
| 0.196266 | 1.553813 | 0.012 | 0 | 1 | 11 | Zfp383 | 0.012 | Old |
| 0.196266 | 1.549966 | 0.012 | 0 | 1 | 11 | Txlnb | 0.012 | Old |
| 0.196266 | 1.54079 | 0.012 | 0 | 1 | 11 | Srd5a1 | 0.012 | Old |
| 0.196266 | 1.538505 | 0.012 | 0 | 1 | 11 | Magi2 | 0.012 | Old |
| 0.196266 | 1.538489 | 0.012 | 0 | 1 | 11 | Fgf2 | 0.012 | Old |
| 0.196266 | 1.516764 | 0.012 | 0 | 1 | 11 | Ndst4 | 0.012 | Old |
| 0.196266 | 1.514675 | 0.012 | 0 | 1 | 11 | Ppp1r3f | 0.012 | Old |
| 0.196266 | 1.511658 | 0.012 | 0 | 1 | 11 | Rad51 | 0.012 | Old |
| 0.196266 | 1.488857 | 0.012 | 0 | 1 | 11 | Rnf128 | 0.012 | Old |
| 0.196266 | 1.484016 | 0.012 | 0 | 1 | 11 | Dync2li1 | 0.012 | Old |
| 0.196266 | 1.480859 | 0.012 | 0 | 1 | 11 | Suv39h2 | 0.012 | Old |
| 0.196266 | 1.468181 | 0.012 | 0 | 1 | 11 | Pcdhgc4 | 0.012 | Old |
| 0.196266 | 1.461303 | 0.012 | 0 | 1 | 11 | Gm47815 | 0.012 | Old |
| 0.196266 | 1.456171 | 0.012 | 0 | 1 | 11 | Gm37305 | 0.012 | Old |
| 0.196266 | 1.455833 | 0.012 | 0 | 1 | 11 | Mbnl3 | 0.012 | Old |
| 0.196266 | 1.449946 | 0.012 | 0 | 1 | 11 | Ablim3 | 0.012 | Old |
| 0.196266 | 1.431772 | 0.012 | 0 | 1 | 11 | Colca2 | 0.012 | Old |
| 0.196266 | 1.398897 | 0.012 | 0 | 1 | 11 | Dclk2 | 0.012 | Old |
| 0.196266 | 1.395222 | 0.012 | 0 | 1 | 11 | Dennd6b | 0.012 | Old |
| 0.196266 | 1.393468 | 0.012 | 0 | 1 | 11 | Icos | 0.012 | Old |
| 0.196266 | 1.393309 | 0.012 | 0 | 1 | 11 | Mipol1 | 0.012 | Old |

| | | | | | | | | |
|---|---|---|---|---|---|---|---|---|
| 0.196266 | 1.388705 | 0.012 | 0 | 1 | 11 | Swsap1 | 0.012 | Old |
| 0.196266 | 1.385372 | 0.012 | 0 | 1 | 11 | Sparcl1 | 0.012 | Old |
| 0.196266 | 1.377701 | 0.012 | 0 | 1 | 11 | Fabp3 | 0.012 | Old |
| 0.196266 | 1.374243 | 0.012 | 0 | 1 | 11 | Vgll3 | 0.012 | Old |
| 0.196266 | 1.366735 | 0.012 | 0 | 1 | 11 | Ckap2l | 0.012 | Old |
| 0.196266 | 1.36218 | 0.012 | 0 | 1 | 11 | Gm42595 | 0.012 | Old |
| 0.196266 | 1.318653 | 0.012 | 0 | 1 | 11 | Gdpd5 | 0.012 | Old |
| 0.196266 | 1.306135 | 0.012 | 0 | 1 | 11 | 2210408I2 | 0.012 | Old |
| 0.196266 | 1.297551 | 0.012 | 0 | 1 | 11 | Ccdc157 | 0.012 | Old |
| 0.196266 | 1.289573 | 0.012 | 0 | 1 | 11 | Bex6 | 0.012 | Old |
| 0.196266 | 1.287281 | 0.012 | 0 | 1 | 11 | Cdc42ep5 | 0.012 | Old |
| 0.196266 | 1.275872 | 0.012 | 0 | 1 | 11 | Dclk1 | 0.012 | Old |
| 0.196266 | 1.272606 | 0.012 | 0 | 1 | 11 | 1500002C1 | 0.012 | Old |
| 0.196266 | 1.270721 | 0.012 | 0 | 1 | 11 | Ltbp2 | 0.012 | Old |
| 0.196266 | 1.26813 | 0.012 | 0 | 1 | 11 | 1700003F1 | 0.012 | Old |
| 0.196266 | 1.245039 | 0.012 | 0 | 1 | 11 | Rab19 | 0.012 | Old |
| 0.196266 | 1.226964 | 0.012 | 0 | 1 | 11 | Galnt9 | 0.012 | Old |
| 0.196266 | 1.226797 | 0.012 | 0 | 1 | 11 | Cit | 0.012 | Old |
| 0.196266 | 1.220778 | 0.012 | 0 | 1 | 11 | Serpinb6b | 0.012 | Old |
| 0.196266 | 1.21471 | 0.012 | 0 | 1 | 11 | Pcdhb14 | 0.012 | Old |
| 0.196266 | 1.20794 | 0.012 | 0 | 1 | 11 | Cyp2r1 | 0.012 | Old |
| 0.196266 | 1.193207 | 0.012 | 0 | 1 | 11 | Rgs12 | 0.012 | Old |
| 0.196266 | 1.192859 | 0.012 | 0 | 1 | 11 | Gm16287 | 0.012 | Old |
| 0.196266 | 1.188319 | 0.012 | 0 | 1 | 11 | Txnrd3 | 0.012 | Old |
| 0.196266 | 1.183228 | 0.012 | 0 | 1 | 11 | Slc22a4 | 0.012 | Old |
| 0.196266 | 1.17819 | 0.012 | 0 | 1 | 11 | Zfp607b | 0.012 | Old |
| 0.196266 | 1.174263 | 0.012 | 0 | 1 | 11 | Gm45351 | 0.012 | Old |
| 0.196266 | 1.169352 | 0.012 | 0 | 1 | 11 | Kiss1r | 0.012 | Old |
| 0.196266 | 1.163568 | 0.012 | 0 | 1 | 11 | 4930430E1 | 0.012 | Old |
| 0.196266 | 1.147788 | 0.012 | 0 | 1 | 11 | Cenpk | 0.012 | Old |
| 0.196266 | 1.146762 | 0.012 | 0 | 1 | 11 | Pcdhb22 | 0.012 | Old |
| 0.196266 | 1.139908 | 0.012 | 0 | 1 | 11 | Tox2 | 0.012 | Old |
| 0.196266 | 1.134136 | 0.012 | 0 | 1 | 11 | CAAA0114 | 0.012 | Old |
| 0.196266 | 1.132389 | 0.012 | 0 | 1 | 11 | Prodh | 0.012 | Old |
| 0.196266 | 1.119973 | 0.012 | 0 | 1 | 11 | Gm17745 | 0.012 | Old |
| 0.196266 | 1.110437 | 0.012 | 0 | 1 | 11 | Hdhd3 | 0.012 | Old |
| 0.196266 | 1.091452 | 0.012 | 0 | 1 | 11 | Bbs9 | 0.012 | Old |
| 0.196266 | 1.077717 | 0.012 | 0 | 1 | 11 | A930024E0 | 0.012 | Old |
| 0.196266 | 1.063855 | 0.012 | 0 | 1 | 11 | Galnt15 | 0.012 | Old |
| 0.196266 | 1.062913 | 0.012 | 0 | 1 | 11 | Gm20655 | 0.012 | Old |
| 0.196266 | 1.057834 | 0.012 | 0 | 1 | 11 | 1110035H1 | 0.012 | Old |
| 0.196266 | 1.041577 | 0.012 | 0 | 1 | 11 | Gm10642 | 0.012 | Old |
| 0.196266 | 1.040728 | 0.012 | 0 | 1 | 11 | Rwdd2a | 0.012 | Old |
| 0.196266 | 1.034378 | 0.012 | 0 | 1 | 11 | Capns2 | 0.012 | Old |
| 0.196266 | 1.023849 | 0.012 | 0 | 1 | 11 | Gng7 | 0.012 | Old |

| | | | | | | | | |
|---|---|---|---|---|---|---|---|---|
| 0.196266 | 1.013675 | 0.012 | 0 | 1 | 11 | 3110009E1 | 0.012 | Old |
| 0.196266 | 1.009674 | 0.012 | 0 | 1 | 11 | M1ap | 0.012 | Old |
| 0.196266 | 1.004944 | 0.012 | 0 | 1 | 11 | Phldb3 | 0.012 | Old |
| 0.196266 | 1.004706 | 0.012 | 0 | 1 | 11 | Mpzl3 | 0.012 | Old |
| 0.196266 | 0.996304 | 0.012 | 0 | 1 | 11 | Slc39a8 | 0.012 | Old |
| 0.196266 | 0.992425 | 0.012 | 0 | 1 | 11 | Wdr19 | 0.012 | Old |
| 0.196266 | 0.978642 | 0.012 | 0 | 1 | 11 | Rab11fip3 | 0.012 | Old |
| 0.196266 | 0.954452 | 0.012 | 0 | 1 | 11 | 1700010I1 | 0.012 | Old |
| 0.196266 | 0.951627 | 0.012 | 0 | 1 | 11 | Rwdd2b | 0.012 | Old |
| 0.196266 | 0.949551 | 0.012 | 0 | 1 | 11 | Pde2a | 0.012 | Old |
| 0.196266 | 0.942583 | 0.012 | 0 | 1 | 11 | Thbs3 | 0.012 | Old |
| 0.196266 | 0.942073 | 0.012 | 0 | 1 | 11 | Marveld2 | 0.012 | Old |
| 0.196266 | 0.93729 | 0.012 | 0 | 1 | 11 | D11Wsu47 | 0.012 | Old |
| 0.196266 | 0.901853 | 0.012 | 0 | 1 | 11 | Dqx1 | 0.012 | Old |
| 0.196266 | 0.89861 | 0.012 | 0 | 1 | 11 | Tfb1m | 0.012 | Old |
| 0.196266 | 0.884822 | 0.012 | 0 | 1 | 11 | Myo15 | 0.012 | Old |
| 0.196266 | 0.87722 | 0.012 | 0 | 1 | 11 | Zfp382 | 0.012 | Old |
| 0.196266 | 0.854757 | 0.012 | 0 | 1 | 11 | Hmga1b | 0.012 | Old |
| 0.196266 | 0.834125 | 0.012 | 0 | 1 | 11 | Haus5 | 0.012 | Old |
| 0.196266 | 0.810362 | 0.012 | 0 | 1 | 11 | Ankrd37 | 0.012 | Old |
| 0.196266 | 0.808651 | 0.012 | 0 | 1 | 11 | Gm29666 | 0.012 | Old |
| 0.196266 | 0.735807 | 0.012 | 0 | 1 | 11 | Gm34961 | 0.012 | Old |
| 0.196336 | 0.494843 | 0.048 | 0.022 | 1 | 11 | Crtc1 | 0.026 | Old |
| 0.196782 | 0.012458 | 0.617 | 0.478 | 1 | 11 | Man2b1 | 0.139 | Old |
| 0.197291 | 0.118902 | 0.742 | 0.587 | 1 | 11 | Msn | 0.155 | Old |
| 0.197322 | 0.508661 | 0.048 | 0.022 | 1 | 11 | Fam206a | 0.026 | Old |
| 0.197322 | 0.477804 | 0.048 | 0.022 | 1 | 11 | Chst14 | 0.026 | Old |
| 0.197435 | 0.753337 | 0.097 | 0.058 | 1 | 11 | Brd9 | 0.039 | Old |
| 0.197632 | 0.071016 | 0.335 | 0.254 | 1 | 11 | Rab5a | 0.081 | Old |
| 0.197694 | 0.065545 | 0.657 | 0.529 | 1 | 11 | Grk2 | 0.128 | Old |
| 0.197703 | 0.451199 | 0.185 | 0.13 | 1 | 11 | Vav3 | 0.055 | Old |
| 0.19808 | 0.451736 | 0.177 | 0.123 | 1 | 11 | Slmap | 0.054 | Old |
| 0.198312 | 1.217179 | 0.048 | 0.022 | 1 | 11 | Nxpe4 | 0.026 | Old |
| 0.198312 | 0.85764 | 0.048 | 0.022 | 1 | 11 | Card9 | 0.026 | Old |
| 0.198312 | 0.519009 | 0.048 | 0.022 | 1 | 11 | Dcun1d4 | 0.026 | Old |
| 0.198823 | 0.184268 | 0.097 | 0.058 | 1 | 11 | Mapk8ip3 | 0.039 | Old |
| 0.199123 | 0.855639 | 0.069 | 0.036 | 1 | 11 | Gbe1 | 0.033 | Old |
| 0.199305 | 0.656404 | 0.048 | 0.022 | 1 | 11 | Tysnd1 | 0.026 | Old |
| 0.199585 | 1.361761 | 0.085 | 0.051 | 1 | 11 | Mettl14 | 0.034 | Old |
| 0.199634 | 0.298225 | 0.363 | 0.275 | 1 | 11 | Aebp2 | 0.088 | Old |
| 0.199922 | 0.134189 | 0.161 | 0.109 | 1 | 11 | Pcyox1 | 0.052 | Old |
| 0.200031 | 0.111709 | 0.387 | 0.297 | 1 | 11 | Arglu1 | 0.09 | Old |
| 0.200303 | 0.68699 | 0.048 | 0.022 | 1 | 11 | Akap1 | 0.026 | Old |
| 0.200303 | 0.238063 | 0.048 | 0.022 | 1 | 11 | Lrrc61 | 0.026 | Old |
| 0.200303 | 0.069543 | 0.048 | 0.022 | 1 | 11 | Wdr55 | 0.026 | Old |

| | | | | | | | | |
|---|---|---|---|---|---|---|---|---|
| 0.200511 | 0.22629 | 0.202 | 0.145 | 1 | 11 | C1d | 0.057 | Old |
| 0.200898 | 0.001056 | 0.165 | 0.109 | 1 | 11 | Tpcn1 | 0.056 | Old |
| 0.201256 | 0.008945 | 0.254 | 0.181 | 1 | 11 | Srebf2 | 0.073 | Old |
| 0.201359 | 0.116636 | 0.56 | 0.442 | 1 | 11 | St13 | 0.118 | Old |
| 0.201621 | 0.449083 | 0.069 | 0.036 | 1 | 11 | Fam53a | 0.033 | Old |
| 0.202254 | 0.896142 | 0.077 | 0.043 | 1 | 11 | Slc12a2 | 0.034 | Old |
| 0.20226 | 0.876752 | 0.113 | 0.072 | 1 | 11 | Lyar | 0.041 | Old |
| 0.202308 | 0.540563 | 0.048 | 0.022 | 1 | 11 | Rtn4ip1 | 0.026 | Old |
| 0.202324 | 0.109731 | 0.097 | 0.058 | 1 | 11 | Zfp513 | 0.039 | Old |
| 0.202459 | 0.671511 | 0.069 | 0.036 | 1 | 11 | Polr3f | 0.033 | Old |
| 0.202526 | 0.196134 | 0.153 | 0.101 | 1 | 11 | S100pbp | 0.052 | Old |
| 0.202893 | 0.490666 | 0.133 | 0.087 | 1 | 11 | Slc7a1 | 0.046 | Old |
| 0.203299 | 0.037553 | 0.069 | 0.036 | 1 | 11 | Wdtc1 | 0.033 | Old |
| 0.203316 | 0.921144 | 0.048 | 0.022 | 1 | 11 | Sh3d19 | 0.026 | Old |
| 0.203316 | 0.870954 | 0.048 | 0.022 | 1 | 11 | Ccdc28b | 0.026 | Old |
| 0.203316 | 0.542962 | 0.048 | 0.022 | 1 | 11 | Arap3 | 0.026 | Old |
| 0.203621 | 0.254474 | 0.145 | 0.094 | 1 | 11 | Camk2g | 0.051 | Old |
| 0.203736 | 0.391632 | 0.097 | 0.058 | 1 | 11 | Ttc5 | 0.039 | Old |
| 0.204136 | 0.016142 | 0.25 | 0.181 | 1 | 11 | Zcrb1 | 0.069 | Old |
| 0.204141 | 0.562159 | 0.105 | 0.065 | 1 | 11 | Txndc12 | 0.04 | Old |
| 0.204196 | 0.017687 | 0.548 | 0.428 | 1 | 11 | Dock10 | 0.12 | Old |
| 0.204204 | 0.065635 | 0.323 | 0.246 | 1 | 11 | Prpf4b | 0.077 | Old |
| 0.20426 | 0.082386 | 0.56 | 0.42 | 1 | 11 | Lamtor4 | 0.14 | Old |
| 0.204327 | 0.242277 | 0.048 | 0.022 | 1 | 11 | Casp2 | 0.026 | Old |
| 0.204327 | 0.089475 | 0.048 | 0.022 | 1 | 11 | Aptx | 0.026 | Old |
| 0.204446 | 0.289711 | 0.097 | 0.058 | 1 | 11 | Selenbp1 | 0.039 | Old |
| 0.204923 | 0.363723 | 0.117 | 0.072 | 1 | 11 | Tgm2 | 0.045 | Old |
| 0.204987 | 0.22545 | 0.069 | 0.036 | 1 | 11 | R3hcc1l | 0.033 | Old |
| 0.205343 | 0.084697 | 0.048 | 0.022 | 1 | 11 | Rrs1 | 0.026 | Old |
| 0.205378 | 0.027648 | 0.653 | 0.507 | 1 | 11 | Tpr | 0.146 | Old |
| 0.205382 | 0.081218 | 0.145 | 0.094 | 1 | 11 | Rexo1 | 0.051 | Old |
| 0.205869 | 0.359403 | 0.097 | 0.058 | 1 | 11 | Upf3b | 0.039 | Old |
| 0.205938 | 0.394332 | 0.133 | 0.087 | 1 | 11 | Ppp2r3d | 0.046 | Old |
| 0.206518 | 0.031488 | 0.718 | 0.565 | 1 | 11 | Dbi | 0.153 | Old |
| 0.206685 | 0.642207 | 0.069 | 0.036 | 1 | 11 | Gipc1 | 0.033 | Old |
| 0.206748 | 0.044751 | 0.125 | 0.08 | 1 | 11 | Prrc1 | 0.045 | Old |
| 0.207384 | 0.144242 | 0.048 | 0.022 | 1 | 11 | Lbx2 | 0.026 | Old |
| 0.207538 | 0.118012 | 0.069 | 0.036 | 1 | 11 | Chkb | 0.033 | Old |
| 0.207566 | 0.280913 | 0.105 | 0.065 | 1 | 11 | Ap4m1 | 0.04 | Old |
| 0.207725 | 2.564191 | 0.036 | 0.014 | 1 | 11 | Macrod1 | 0.022 | Old |
| 0.208394 | 0.93182 | 0.069 | 0.036 | 1 | 11 | Tbp | 0.033 | Old |
| 0.208399 | 0.253413 | 0.133 | 0.087 | 1 | 11 | Ctr9 | 0.046 | Old |
| 0.208779 | 0.126404 | 0.98 | 0.971 | 1 | 11 | Cd74 | 0.009 | Old |
| 0.208997 | 0.073255 | 0.915 | 0.92 | 1 | 11 | C1qc | -0.005 | Old |
| 0.20903 | 0.010605 | 0.315 | 0.239 | 1 | 11 | Cmpk2 | 0.076 | Old |

| | | | | | | | | |
|---|---|---|---|---|---|---|---|---|
| 0.209403 | 0.119052 | 0.294 | 0.225 | 1 | 11 | Irgm2 | 0.069 | Old |
| 0.209441 | 0.092741 | 0.048 | 0.022 | 1 | 11 | Numb | 0.026 | Old |
| 0.209459 | 0.370771 | 0.097 | 0.058 | 1 | 11 | Timm29 | 0.039 | Old |
| 0.209591 | 0.024919 | 0.29 | 0.217 | 1 | 11 | Slc38a1 | 0.073 | Old |
| 0.210118 | 2.299627 | 0.036 | 0.014 | 1 | 11 | 2700049A( | 0.022 | Old |
| 0.210118 | 1.267163 | 0.036 | 0.014 | 1 | 11 | Nudt1 | 0.022 | Old |
| 0.210183 | 0.084904 | 0.097 | 0.058 | 1 | 11 | Traf6 | 0.039 | Old |
| 0.21056 | 0.193356 | 0.141 | 0.094 | 1 | 11 | Trip4 | 0.047 | Old |
| 0.210562 | 0.004537 | 0.161 | 0.109 | 1 | 11 | Scfd1 | 0.052 | Old |
| 0.210865 | 0.013505 | 0.214 | 0.152 | 1 | 11 | Tmem128 | 0.062 | Old |
| 0.211076 | 0.641966 | 0.077 | 0.043 | 1 | 11 | Dnajb12 | 0.034 | Old |
| 0.211322 | 1.959371 | 0.036 | 0.014 | 1 | 11 | Gm26881 | 0.022 | Old |
| 0.211341 | 0.087179 | 0.444 | 0.341 | 1 | 11 | Snx20 | 0.103 | Old |
| 0.211372 | 0.165195 | 0.859 | 0.703 | 1 | 11 | Zfp36l2 | 0.156 | Old |
| 0.211731 | 0.213444 | 0.105 | 0.065 | 1 | 11 | AU019823 | 0.04 | Old |
| 0.212268 | 0.127885 | 0.423 | 0.326 | 1 | 11 | Ssr1 | 0.097 | Old |
| 0.21234 | 0.128869 | 0.169 | 0.116 | 1 | 11 | Bank1 | 0.053 | Old |
| 0.212346 | 0.375381 | 0.222 | 0.167 | 1 | 11 | Rsf1 | 0.055 | Old |
| 0.212364 | 0.329365 | 0.097 | 0.058 | 1 | 11 | Ipo13 | 0.039 | Old |
| 0.212709 | 0.571181 | 0.077 | 0.043 | 1 | 11 | Mrto4 | 0.034 | Old |
| 0.21271 | 0.059051 | 0.488 | 0.391 | 1 | 11 | Tcirg1 | 0.097 | Old |
| 0.212753 | 0.105227 | 0.206 | 0.145 | 1 | 11 | Trappc10 | 0.061 | Old |
| 0.213475 | 0.874032 | 0.056 | 0.029 | 1 | 11 | Plpp2 | 0.027 | Old |
| 0.213529 | 0.713794 | 0.077 | 0.043 | 1 | 11 | Ppm1d | 0.034 | Old |
| 0.21356 | 0.379525 | 0.113 | 0.072 | 1 | 11 | Usp5 | 0.041 | Old |
| 0.213581 | 0.012097 | 0.069 | 0.036 | 1 | 11 | Nsun3 | 0.033 | Old |
| 0.213836 | 0.198842 | 0.105 | 0.065 | 1 | 11 | Ltv1 | 0.04 | Old |
| 0.214213 | 0.160595 | 0.141 | 0.094 | 1 | 11 | Pstpip1 | 0.047 | Old |
| 0.214964 | 1.4502 | 0.036 | 0.014 | 1 | 11 | Ankzf1 | 0.022 | Old |
| 0.215176 | 0.721831 | 0.077 | 0.043 | 1 | 11 | Epg5 | 0.034 | Old |
| 0.215298 | 0.1323 | 0.097 | 0.058 | 1 | 11 | Fech | 0.039 | Old |
| 0.215458 | 0.077275 | 0.625 | 0.478 | 1 | 11 | Ndufa6 | 0.147 | Old |
| 0.216187 | 1.697397 | 0.036 | 0.014 | 1 | 11 | Nudcd1 | 0.022 | Old |
| 0.216187 | 1.465032 | 0.036 | 0.014 | 1 | 11 | Unc119b | 0.022 | Old |
| 0.21686 | 0.112615 | 0.371 | 0.283 | 1 | 11 | Lrrc25 | 0.088 | Old |
| 0.216907 | 0.395886 | 0.161 | 0.109 | 1 | 11 | Cops5 | 0.052 | Old |
| 0.21706 | 0.002171 | 0.125 | 0.08 | 1 | 11 | Umad1 | 0.045 | Old |
| 0.217335 | 0.619206 | 0.056 | 0.029 | 1 | 11 | 1300002E1 | 0.027 | Old |
| 0.218003 | 0.058722 | 0.218 | 0.159 | 1 | 11 | Tomm34 | 0.059 | Old |
| 0.218308 | 1.056153 | 0.056 | 0.029 | 1 | 11 | Il1rap | 0.027 | Old |
| 0.218646 | 0.276641 | 0.21 | 0.152 | 1 | 11 | Slirp | 0.058 | Old |
| 0.21865 | 1.404714 | 0.036 | 0.014 | 1 | 11 | Krt10 | 0.022 | Old |
| 0.21865 | 1.184448 | 0.036 | 0.014 | 1 | 11 | Rabl3 | 0.022 | Old |
| 0.21865 | 1.175963 | 0.036 | 0.014 | 1 | 11 | Nup35 | 0.022 | Old |
| 0.21865 | 0.661732 | 0.036 | 0.014 | 1 | 11 | Vars2 | 0.022 | Old |

| | | | | | | | | |
|---|---|---|---|---|---|---|---|---|
| 0.218664 | 0.368199 | 0.149 | 0.101 | 1 | 11 | Mrps25 | 0.048 | Old |
| 0.218702 | 0.102697 | 0.681 | 0.558 | 1 | 11 | Gmfg | 0.123 | Old |
| 0.218765 | 0.778398 | 0.085 | 0.051 | 1 | 11 | Rab11fip2 | 0.034 | Old |
| 0.21907 | 1.141702 | 0.093 | 0.058 | 1 | 11 | Triobp | 0.035 | Old |
| 0.219101 | 0.102915 | 0.29 | 0.217 | 1 | 11 | Rdx | 0.073 | Old |
| 0.219116 | 0.120165 | 0.403 | 0.326 | 1 | 11 | Serpinb6a | 0.077 | Old |
| 0.219284 | 1.004571 | 0.056 | 0.029 | 1 | 11 | Dhcr24 | 0.027 | Old |
| 0.219284 | 0.890473 | 0.056 | 0.029 | 1 | 11 | Dars2 | 0.027 | Old |
| 0.219522 | 0.127022 | 0.105 | 0.065 | 1 | 11 | Fbxo6 | 0.04 | Old |
| 0.219734 | 0.347185 | 0.133 | 0.087 | 1 | 11 | Ubr1 | 0.046 | Old |
| 0.219844 | 0.09001 | 0.992 | 0.964 | 1 | 11 | mt-Nd1 | 0.028 | Old |
| 0.219866 | 0.025936 | 0.222 | 0.159 | 1 | 11 | Bccip | 0.063 | Old |
| 0.219889 | 0.931634 | 0.036 | 0.014 | 1 | 11 | Pomt2 | 0.022 | Old |
| 0.219889 | 0.846518 | 0.036 | 0.014 | 1 | 11 | Thg1l | 0.022 | Old |
| 0.219889 | 0.64353 | 0.036 | 0.014 | 1 | 11 | C030034I2 | 0.022 | Old |
| 0.219889 | 0.036262 | 0.036 | 0.014 | 1 | 11 | Cenpv | 0.022 | Old |
| 0.220172 | 0.307036 | 0.077 | 0.043 | 1 | 11 | Cdk7 | 0.034 | Old |
| 0.221013 | 0.134301 | 0.077 | 0.043 | 1 | 11 | Ubac1 | 0.034 | Old |
| 0.221133 | 1.796991 | 0.036 | 0.014 | 1 | 11 | Smyd4 | 0.022 | Old |
| 0.221133 | 0.728999 | 0.036 | 0.014 | 1 | 11 | Tmem150b | 0.022 | Old |
| 0.221133 | 0.51842 | 0.036 | 0.014 | 1 | 11 | Wbp1 | 0.022 | Old |
| 0.221245 | 1.383407 | 0.056 | 0.029 | 1 | 11 | Ccdc191 | 0.027 | Old |
| 0.221245 | 0.689018 | 0.056 | 0.029 | 1 | 11 | Tmem39b | 0.027 | Old |
| 0.221682 | 0.192092 | 0.105 | 0.065 | 1 | 11 | Yeats2 | 0.04 | Old |
| 0.221856 | 0.942853 | 0.077 | 0.043 | 1 | 11 | Rabgap1l | 0.034 | Old |
| 0.221856 | 0.442273 | 0.077 | 0.043 | 1 | 11 | 2310039H0 | 0.034 | Old |
| 0.221953 | 0.314125 | 0.085 | 0.051 | 1 | 11 | Alg2 | 0.034 | Old |
| 0.222382 | 0.858632 | 0.036 | 0.014 | 1 | 11 | Sgce | 0.022 | Old |
| 0.222382 | 0.814811 | 0.036 | 0.014 | 1 | 11 | Dcaf6 | 0.022 | Old |
| 0.222415 | 0.026294 | 0.492 | 0.384 | 1 | 11 | Ctdsp2 | 0.108 | Old |
| 0.222702 | 0.63179 | 0.077 | 0.043 | 1 | 11 | Rtca | 0.034 | Old |
| 0.222797 | 0.1266 | 0.371 | 0.283 | 1 | 11 | Clec4b1 | 0.088 | Old |
| 0.222945 | 0.473711 | 0.129 | 0.087 | 1 | 11 | Pogz | 0.042 | Old |
| 0.223237 | 0.162555 | 0.794 | 0.681 | 1 | 11 | mt-Nd4l | 0.113 | Old |
| 0.223636 | 0.561373 | 0.036 | 0.014 | 1 | 11 | Fam210b | 0.022 | Old |
| 0.223636 | 0.452243 | 0.036 | 0.014 | 1 | 11 | Zfp874b | 0.022 | Old |
| 0.223636 | 0.443879 | 0.036 | 0.014 | 1 | 11 | Cacnb1 | 0.022 | Old |
| 0.223636 | 0.332585 | 0.036 | 0.014 | 1 | 11 | Zfp87 | 0.022 | Old |
| 0.223858 | 0.450003 | 0.105 | 0.065 | 1 | 11 | Ipo11 | 0.04 | Old |
| 0.224211 | 0.913707 | 0.056 | 0.029 | 1 | 11 | Sco1 | 0.027 | Old |
| 0.224775 | 0.379129 | 0.226 | 0.167 | 1 | 11 | Cnpy3 | 0.059 | Old |
| 0.224895 | 0.884617 | 0.036 | 0.014 | 1 | 11 | 2310040G2 | 0.022 | Old |
| 0.224895 | 0.5106 | 0.036 | 0.014 | 1 | 11 | Dtd1 | 0.022 | Old |
| 0.224895 | 0.477854 | 0.036 | 0.014 | 1 | 11 | Gpt2 | 0.022 | Old |
| 0.224895 | 0.349609 | 0.036 | 0.014 | 1 | 11 | 2900076A0 | 0.022 | Old |

| | | | | | | | | |
|---|---|---|---|---|---|---|---|---|
| 0.224895 | 0.344386 | 0.036 | 0.014 | 1 | 11 | Hexdc | 0.022 | Old |
| 0.224895 | 0.305705 | 0.036 | 0.014 | 1 | 11 | Rnf122 | 0.022 | Old |
| 0.225309 | 0.539214 | 0.113 | 0.072 | 1 | 11 | Fyco1 | 0.041 | Old |
| 0.225971 | 0.075839 | 0.875 | 0.812 | 1 | 11 | Rplp0 | 0.063 | Old |
| 0.22616 | 0.654997 | 0.036 | 0.014 | 1 | 11 | Stk11ip | 0.022 | Old |
| 0.226787 | 0.091596 | 0.48 | 0.362 | 1 | 11 | Tcf25 | 0.118 | Old |
| 0.226797 | 0.483317 | 0.085 | 0.051 | 1 | 11 | Gpn2 | 0.034 | Old |
| 0.226963 | 0.261373 | 0.077 | 0.043 | 1 | 11 | Med6 | 0.034 | Old |
| 0.227126 | 0.157006 | 0.746 | 0.623 | 1 | 11 | Eef2 | 0.123 | Old |
| 0.227429 | 0.312439 | 0.036 | 0.014 | 1 | 11 | Zfp944 | 0.022 | Old |
| 0.227429 | 0.244141 | 0.036 | 0.014 | 1 | 11 | Traf3ip1 | 0.022 | Old |
| 0.228077 | 0.072867 | 0.468 | 0.348 | 1 | 11 | Mvp | 0.12 | Old |
| 0.228209 | 0.389412 | 0.056 | 0.029 | 1 | 11 | Fbf1 | 0.027 | Old |
| 0.228373 | 2.80173 | 0.024 | 0.007 | 1 | 11 | Cd300e | 0.017 | Old |
| 0.228373 | 2.062894 | 0.024 | 0.007 | 1 | 11 | Gm42715 | 0.017 | Old |
| 0.228373 | 1.872901 | 0.024 | 0.007 | 1 | 11 | Sncg | 0.017 | Old |
| 0.228373 | 1.674581 | 0.024 | 0.007 | 1 | 11 | Slc2a9 | 0.017 | Old |
| 0.228667 | 0.017316 | 0.681 | 0.514 | 1 | 11 | Ywhah | 0.167 | Old |
| 0.228703 | 0.378821 | 0.036 | 0.014 | 1 | 11 | Tsix | 0.022 | Old |
| 0.229216 | 0.218841 | 0.056 | 0.029 | 1 | 11 | Hap1 | 0.027 | Old |
| 0.229451 | 0.050791 | 0.629 | 0.522 | 1 | 11 | Tmsb10 | 0.107 | Old |
| 0.229875 | 0.053002 | 0.185 | 0.13 | 1 | 11 | Spata6 | 0.055 | Old |
| 0.229968 | 1.915099 | 0.024 | 0.007 | 1 | 11 | BC017158 | 0.017 | Old |
| 0.229968 | 1.606112 | 0.024 | 0.007 | 1 | 11 | Gm9844 | 0.017 | Old |
| 0.229968 | 1.454496 | 0.024 | 0.007 | 1 | 11 | Prim2 | 0.017 | Old |
| 0.229968 | 1.330323 | 0.024 | 0.007 | 1 | 11 | Zfp442 | 0.017 | Old |
| 0.229968 | 1.285735 | 0.024 | 0.007 | 1 | 11 | Tspyl3 | 0.017 | Old |
| 0.229968 | 1.256224 | 0.024 | 0.007 | 1 | 11 | Pusl1 | 0.017 | Old |
| 0.229968 | 1.238609 | 0.024 | 0.007 | 1 | 11 | Zfp747 | 0.017 | Old |
| 0.229999 | 0.135714 | 0.718 | 0.609 | 1 | 11 | Ehd4 | 0.109 | Old |
| 0.230227 | 0.261343 | 0.056 | 0.029 | 1 | 11 | Arhgap27 | 0.027 | Old |
| 0.230508 | 0.222401 | 0.137 | 0.094 | 1 | 11 | Arl6ip6 | 0.043 | Old |
| 0.230716 | 0.067371 | 0.508 | 0.406 | 1 | 11 | Capns1 | 0.102 | Old |
| 0.231282 | 0.233336 | 0.077 | 0.043 | 1 | 11 | Mrps35 | 0.034 | Old |
| 0.231328 | 0.259576 | 0.121 | 0.08 | 1 | 11 | Alg5 | 0.041 | Old |
| 0.23157 | 2.186947 | 0.024 | 0.007 | 1 | 11 | Ube2t | 0.017 | Old |
| 0.23157 | 2.085335 | 0.024 | 0.007 | 1 | 11 | Hhat | 0.017 | Old |
| 0.23157 | 1.878729 | 0.024 | 0.007 | 1 | 11 | Tstd1 | 0.017 | Old |
| 0.23157 | 1.582734 | 0.024 | 0.007 | 1 | 11 | Mtrr | 0.017 | Old |
| 0.23157 | 1.182577 | 0.024 | 0.007 | 1 | 11 | Runx2 | 0.017 | Old |
| 0.23157 | 1.144039 | 0.024 | 0.007 | 1 | 11 | Fndc4 | 0.017 | Old |
| 0.231965 | 0.05896 | 0.105 | 0.065 | 1 | 11 | 2310033P( | 0.04 | Old |
| 0.232022 | 0.376154 | 0.121 | 0.08 | 1 | 11 | Zfp512 | 0.041 | Old |
| 0.232052 | 0.879337 | 0.065 | 0.036 | 1 | 11 | Helb | 0.029 | Old |
| 0.232176 | 0.250908 | 0.157 | 0.109 | 1 | 11 | Fcf1 | 0.048 | Old |

| | | | | | | | | |
|---|---|---|---|---|---|---|---|---|
| 0.232291 | 0.120956 | 0.516 | 0.406 | 1 | 11 | Myadm | 0.11 | Old |
| 0.232428 | 0.125991 | 0.98 | 0.935 | 1 | 11 | Dusp1 | 0.045 | Old |
| 0.233153 | 0.178254 | 0.113 | 0.072 | 1 | 11 | Rps6kc1 | 0.041 | Old |
| 0.233181 | 1.77504 | 0.024 | 0.007 | 1 | 11 | Poc1a | 0.017 | Old |
| 0.233181 | 1.656275 | 0.024 | 0.007 | 1 | 11 | Cep78 | 0.017 | Old |
| 0.233181 | 1.461788 | 0.024 | 0.007 | 1 | 11 | Dyrk1b | 0.017 | Old |
| 0.233181 | 1.431261 | 0.024 | 0.007 | 1 | 11 | Snx32 | 0.017 | Old |
| 0.233181 | 1.391814 | 0.024 | 0.007 | 1 | 11 | AW209491 | 0.017 | Old |
| 0.233181 | 1.122007 | 0.024 | 0.007 | 1 | 11 | Gm26778 | 0.017 | Old |
| 0.233181 | 0.98864 | 0.024 | 0.007 | 1 | 11 | Hectd2 | 0.017 | Old |
| 0.233524 | 0.031928 | 0.399 | 0.297 | 1 | 11 | Zc3h7a | 0.102 | Old |
| 0.234432 | 0.281346 | 0.177 | 0.13 | 1 | 11 | Phf10 | 0.047 | Old |
| 0.234752 | 0.003989 | 0.516 | 0.391 | 1 | 11 | Calm3 | 0.125 | Old |
| 0.234799 | 1.749996 | 0.024 | 0.007 | 1 | 11 | Pcnx4 | 0.017 | Old |
| 0.234799 | 1.512812 | 0.024 | 0.007 | 1 | 11 | 9030025P2 | 0.017 | Old |
| 0.234799 | 1.351377 | 0.024 | 0.007 | 1 | 11 | A430033K( | 0.017 | Old |
| 0.234799 | 1.225462 | 0.024 | 0.007 | 1 | 11 | Poll | 0.017 | Old |
| 0.234799 | 1.207613 | 0.024 | 0.007 | 1 | 11 | Gm31597 | 0.017 | Old |
| 0.234799 | 1.189266 | 0.024 | 0.007 | 1 | 11 | Prpf40b | 0.017 | Old |
| 0.234799 | 1.182056 | 0.024 | 0.007 | 1 | 11 | 2510046G1 | 0.017 | Old |
| 0.234799 | 1.074431 | 0.024 | 0.007 | 1 | 11 | Syp | 0.017 | Old |
| 0.234799 | 1.066748 | 0.024 | 0.007 | 1 | 11 | Tarbp1 | 0.017 | Old |
| 0.234799 | 0.789086 | 0.024 | 0.007 | 1 | 11 | Fam109b | 0.017 | Old |
| 0.235327 | 0.278878 | 0.056 | 0.029 | 1 | 11 | Palm | 0.027 | Old |
| 0.235327 | 0.034429 | 0.056 | 0.029 | 1 | 11 | Adprhl2 | 0.027 | Old |
| 0.235327 | 0.00878 | 0.056 | 0.029 | 1 | 11 | Mon1b | 0.027 | Old |
| 0.235389 | 0.684933 | 0.093 | 0.058 | 1 | 11 | Ints12 | 0.035 | Old |
| 0.235846 | 0.14276 | 0.681 | 0.543 | 1 | 11 | Ddx3x | 0.138 | Old |
| 0.235859 | 0.745717 | 0.065 | 0.036 | 1 | 11 | Gna15 | 0.029 | Old |
| 0.236356 | 0.536229 | 0.056 | 0.029 | 1 | 11 | Prune1 | 0.027 | Old |
| 0.236425 | 0.869523 | 0.024 | 0.007 | 1 | 11 | Paqr9 | 0.017 | Old |
| 0.236425 | 0.753644 | 0.024 | 0.007 | 1 | 11 | Rbp1 | 0.017 | Old |
| 0.236425 | 0.569276 | 0.024 | 0.007 | 1 | 11 | Gm43796 | 0.017 | Old |
| 0.236425 | 0.079072 | 0.024 | 0.007 | 1 | 11 | Cby1 | 0.017 | Old |
| 0.23677 | 0.183557 | 0.847 | 0.717 | 1 | 11 | Tapbp | 0.13 | Old |
| 0.236925 | 0.066752 | 0.121 | 0.08 | 1 | 11 | Pin1 | 0.041 | Old |
| 0.237389 | 0.295301 | 0.056 | 0.029 | 1 | 11 | Zfp358 | 0.027 | Old |
| 0.237389 | 0.257666 | 0.056 | 0.029 | 1 | 11 | Slc25a16 | 0.027 | Old |
| 0.237541 | 0.076634 | 0.181 | 0.13 | 1 | 11 | Bcl7c | 0.051 | Old |
| 0.237778 | 0.597702 | 0.065 | 0.036 | 1 | 11 | 6030458C1 | 0.029 | Old |
| 0.238059 | 0.670389 | 0.024 | 0.007 | 1 | 11 | Dlg3 | 0.017 | Old |
| 0.238059 | 0.569477 | 0.024 | 0.007 | 1 | 11 | Trp53rkb | 0.017 | Old |
| 0.238059 | 0.522469 | 0.024 | 0.007 | 1 | 11 | Rfx2 | 0.017 | Old |
| 0.238059 | 0.492499 | 0.024 | 0.007 | 1 | 11 | Xrcc6 | 0.017 | Old |
| 0.238059 | 0.438515 | 0.024 | 0.007 | 1 | 11 | Zfp446 | 0.017 | Old |

| | | | | | | | | |
|---|---|---|---|---|---|---|---|---|
| 0.238059 | 0.433176 | 0.024 | 0.007 | 1 | 11 | Ighd | 0.017 | Old |
| 0.238059 | 0.360428 | 0.024 | 0.007 | 1 | 11 | Ccdc17 | 0.017 | Old |
| 0.238059 | 0.343975 | 0.024 | 0.007 | 1 | 11 | Pms2 | 0.017 | Old |
| 0.238059 | 0.302246 | 0.024 | 0.007 | 1 | 11 | Slc19a1 | 0.017 | Old |
| 0.238059 | 0.17182 | 0.024 | 0.007 | 1 | 11 | Cpeb1 | 0.017 | Old |
| 0.238059 | 0.107177 | 0.024 | 0.007 | 1 | 11 | Zfp408 | 0.017 | Old |
| 0.238059 | 0.05848 | 0.024 | 0.007 | 1 | 11 | Tm9sf1 | 0.017 | Old |
| 0.238059 | 0.018736 | 0.024 | 0.007 | 1 | 11 | Hdac6 | 0.017 | Old |
| 0.239709 | 1.181537 | 0.065 | 0.036 | 1 | 11 | Dolk | 0.029 | Old |
| 0.239738 | 0.01623 | 0.141 | 0.094 | 1 | 11 | Lpcat3 | 0.047 | Old |
| 0.239805 | 0.115848 | 0.242 | 0.181 | 1 | 11 | Tbc1d5 | 0.061 | Old |
| 0.240082 | 0.731483 | 0.085 | 0.051 | 1 | 11 | Plscr1 | 0.034 | Old |
| 0.240158 | 0.047326 | 0.976 | 0.986 | 1 | 11 | mt-Nd2 | -0.01 | Old |
| 0.240356 | 0.468143 | 0.161 | 0.116 | 1 | 11 | Ssbp3 | 0.045 | Old |
| 0.240678 | 1.209549 | 0.065 | 0.036 | 1 | 11 | Ust | 0.029 | Old |
| 0.241015 | 0.412728 | 0.093 | 0.058 | 1 | 11 | Bmi1 | 0.035 | Old |
| 0.241396 | 0.058021 | 0.165 | 0.116 | 1 | 11 | Zbtb44 | 0.049 | Old |
| 0.24165 | 1.196458 | 0.065 | 0.036 | 1 | 11 | Nfatc2ip | 0.029 | Old |
| 0.24165 | 0.683971 | 0.065 | 0.036 | 1 | 11 | Asgr2 | 0.029 | Old |
| 0.242625 | 1.006086 | 0.065 | 0.036 | 1 | 11 | Gab3 | 0.029 | Old |
| 0.243015 | 0.056326 | 0.742 | 0.587 | 1 | 11 | Slc25a5 | 0.155 | Old |
| 0.243217 | 0.080567 | 0.129 | 0.087 | 1 | 11 | Gab1 | 0.042 | Old |
| 0.243403 | 0.225231 | 0.27 | 0.203 | 1 | 11 | Eif2a | 0.067 | Old |
| 0.243482 | 0.21571 | 0.161 | 0.116 | 1 | 11 | Flot2 | 0.045 | Old |
| 0.243773 | 0.530248 | 0.145 | 0.101 | 1 | 11 | Ddx39 | 0.044 | Old |
| 0.243941 | 0.310597 | 0.343 | 0.268 | 1 | 11 | Phf3 | 0.075 | Old |
| 0.244517 | 0.429909 | 0.149 | 0.101 | 1 | 11 | Mea1 | 0.048 | Old |
| 0.244542 | 0.380083 | 0.101 | 0.065 | 1 | 11 | Rnaseh2a | 0.036 | Old |
| 0.244583 | 0.010227 | 0.722 | 0.58 | 1 | 11 | Pld4 | 0.142 | Old |
| 0.244583 | 0.178713 | 0.065 | 0.036 | 1 | 11 | Tmem175 | 0.029 | Old |
| 0.245013 | 0.048296 | 0.798 | 0.601 | 1 | 11 | Atp6v0b | 0.197 | Old |
| 0.245567 | 0.350012 | 0.065 | 0.036 | 1 | 11 | Ubap2 | 0.029 | Old |
| 0.245912 | 0.236775 | 0.093 | 0.058 | 1 | 11 | Pdp2 | 0.035 | Old |
| 0.245975 | 0.027674 | 0.19 | 0.13 | 1 | 11 | Gtf2a2 | 0.06 | Old |
| 0.246063 | 0.142454 | 0.085 | 0.051 | 1 | 11 | Epm2aip1 | 0.034 | Old |
| 0.246441 | 0.011075 | 0.141 | 0.094 | 1 | 11 | Pgrmc2 | 0.047 | Old |
| 0.246503 | 0.067929 | 0.629 | 0.5 | 1 | 11 | Scamp2 | 0.129 | Old |
| 0.246505 | 0.025777 | 0.996 | 1 | 1 | 11 | mt-Co3 | -0.004 | Old |
| 0.246552 | 0.996545 | 0.065 | 0.036 | 1 | 11 | Borcs5 | 0.029 | Old |
| 0.246689 | 0.024203 | 0.395 | 0.297 | 1 | 11 | Vamp3 | 0.098 | Old |
| 0.247648 | 0.086174 | 0.165 | 0.116 | 1 | 11 | 1810013L2 | 0.049 | Old |
| 0.247668 | 0.01506 | 0.976 | 0.978 | 1 | 11 | Rps26 | -0.002 | Old |
| 0.248489 | 0.51623 | 0.101 | 0.065 | 1 | 11 | Cpsf4 | 0.036 | Old |
| 0.248985 | 0.134197 | 0.21 | 0.152 | 1 | 11 | Akap9 | 0.058 | Old |
| 0.249065 | 0.637069 | 0.109 | 0.072 | 1 | 11 | Hsdl1 | 0.037 | Old |

| | | | | | | | | |
|---|---|---|---|---|---|---|---|---|
| 0.249327 | 1.363121 | 0.044 | 0.022 | 1 | 11 | Pcgf1 | 0.022 | Old |
| 0.249708 | 0.320272 | 0.153 | 0.109 | 1 | 11 | Klhl6 | 0.044 | Old |
| 0.250536 | 1.291175 | 0.044 | 0.022 | 1 | 11 | 3110001I2: | 0.022 | Old |
| 0.250536 | 0.898918 | 0.044 | 0.022 | 1 | 11 | 6-Sep | 0.022 | Old |
| 0.25186 | 0.058048 | 0.157 | 0.109 | 1 | 11 | Tmem140 | 0.048 | Old |
| 0.252046 | 0.136706 | 0.262 | 0.196 | 1 | 11 | Socs6 | 0.066 | Old |
| 0.252527 | 0.351299 | 0.065 | 0.036 | 1 | 11 | Heg1 | 0.029 | Old |
| 0.253366 | 0.301781 | 0.198 | 0.145 | 1 | 11 | Nhlrc3 | 0.053 | Old |
| 0.253388 | 0.123246 | 0.093 | 0.058 | 1 | 11 | Dhps | 0.035 | Old |
| 0.253757 | 0.163369 | 0.871 | 0.754 | 1 | 11 | Mob1a | 0.117 | Old |
| 0.25378 | 0.112762 | 0.181 | 0.13 | 1 | 11 | Glipr1 | 0.051 | Old |
| 0.253813 | 0.046954 | 0.145 | 0.101 | 1 | 11 | Nudcd2 | 0.044 | Old |
| 0.253996 | 0.142322 | 0.306 | 0.232 | 1 | 11 | Phc3 | 0.074 | Old |
| 0.254064 | 0.858882 | 0.073 | 0.043 | 1 | 11 | Ripor2 | 0.03 | Old |
| 0.25419 | 1.411905 | 0.044 | 0.022 | 1 | 11 | Ip6k2 | 0.022 | Old |
| 0.25419 | 0.70088 | 0.044 | 0.022 | 1 | 11 | Wdcp | 0.022 | Old |
| 0.254207 | 0.123383 | 0.327 | 0.254 | 1 | 11 | Nfic | 0.073 | Old |
| 0.254229 | 0.058316 | 0.093 | 0.058 | 1 | 11 | Uvssa | 0.035 | Old |
| 0.25454 | 0.313076 | 0.065 | 0.036 | 1 | 11 | Tmem35b | 0.029 | Old |
| 0.255298 | 0.141099 | 0.641 | 0.514 | 1 | 11 | Trim25 | 0.127 | Old |
| 0.255416 | 0.718155 | 0.044 | 0.022 | 1 | 11 | 2900052N( | 0.022 | Old |
| 0.255495 | 0.054298 | 0.254 | 0.188 | 1 | 11 | Ino80d | 0.066 | Old |
| 0.255551 | 0.491646 | 0.065 | 0.036 | 1 | 11 | Hes6 | 0.029 | Old |
| 0.256484 | 0.001261 | 0.206 | 0.145 | 1 | 11 | Oxr1 | 0.061 | Old |
| 0.256565 | 0.228452 | 0.065 | 0.036 | 1 | 11 | Ift172 | 0.029 | Old |
| 0.256646 | 1.18129 | 0.044 | 0.022 | 1 | 11 | Rrnad1 | 0.022 | Old |
| 0.256646 | 1.109391 | 0.044 | 0.022 | 1 | 11 | Ciart | 0.022 | Old |
| 0.2568 | 0.627375 | 0.109 | 0.072 | 1 | 11 | Iffo2 | 0.037 | Old |
| 0.256916 | 0.722557 | 0.073 | 0.043 | 1 | 11 | Mettl21a | 0.03 | Old |
| 0.256979 | 0.279732 | 0.153 | 0.109 | 1 | 11 | Kbtbd11 | 0.044 | Old |
| 0.257582 | 0.359859 | 0.065 | 0.036 | 1 | 11 | A530040E1 | 0.029 | Old |
| 0.25788 | 0.664047 | 0.044 | 0.022 | 1 | 11 | Zcchc4 | 0.022 | Old |
| 0.258821 | 0.075837 | 0.54 | 0.457 | 1 | 11 | Gas6 | 0.083 | Old |
| 0.259119 | 1.082633 | 0.044 | 0.022 | 1 | 11 | 2300009A( | 0.022 | Old |
| 0.259119 | 0.669128 | 0.044 | 0.022 | 1 | 11 | Plcl1 | 0.022 | Old |
| 0.259612 | 0.051002 | 0.129 | 0.087 | 1 | 11 | Psmd9 | 0.042 | Old |
| 0.259628 | 0.347065 | 0.145 | 0.101 | 1 | 11 | Lpar1 | 0.044 | Old |
| 0.260158 | 0.479587 | 0.149 | 0.109 | 1 | 11 | Stard4 | 0.04 | Old |
| 0.260361 | 0.570059 | 0.044 | 0.022 | 1 | 11 | Creld1 | 0.022 | Old |
| 0.260361 | 0.126806 | 0.044 | 0.022 | 1 | 11 | Endov | 0.022 | Old |
| 0.260648 | 0.293027 | 0.065 | 0.036 | 1 | 11 | Ifitm1 | 0.029 | Old |
| 0.260752 | 0.83207 | 0.073 | 0.043 | 1 | 11 | Mefv | 0.03 | Old |
| 0.261021 | 0.567754 | 0.093 | 0.058 | 1 | 11 | Ankrd16 | 0.035 | Old |
| 0.261166 | 0.045442 | 0.137 | 0.094 | 1 | 11 | Tada1 | 0.043 | Old |
| 0.261418 | 0.346289 | 0.101 | 0.065 | 1 | 11 | Tmem126a | 0.036 | Old |

| | | | | | | | | |
|---|---|---|---|---|---|---|---|---|
| 0.261608 | 0.379117 | 0.044 | 0.022 | 1 | 11 | Zfp709 | 0.022 | Old |
| 0.261608 | 0.249969 | 0.044 | 0.022 | 1 | 11 | Acyp1 | 0.022 | Old |
| 0.261676 | 0.309461 | 0.065 | 0.036 | 1 | 11 | Enpp1 | 0.029 | Old |
| 0.262678 | 0.163944 | 0.347 | 0.268 | 1 | 11 | Aprt | 0.079 | Old |
| 0.262706 | 0.056651 | 0.065 | 0.036 | 1 | 11 | Pdhx | 0.029 | Old |
| 0.262859 | 0.52508 | 0.044 | 0.022 | 1 | 11 | Cry1 | 0.022 | Old |
| 0.262859 | 0.390657 | 0.044 | 0.022 | 1 | 11 | Mrrf | 0.022 | Old |
| 0.262859 | 0.345614 | 0.044 | 0.022 | 1 | 11 | Zfp182 | 0.022 | Old |
| 0.263656 | 0.444141 | 0.073 | 0.043 | 1 | 11 | Sap30bp | 0.03 | Old |
| 0.263835 | 0.345365 | 0.234 | 0.181 | 1 | 11 | Atp2c1 | 0.053 | Old |
| 0.264672 | 0.234795 | 0.29 | 0.217 | 1 | 11 | Slc39a7 | 0.073 | Old |
| 0.264776 | 0.106974 | 0.065 | 0.036 | 1 | 11 | Tnni3 | 0.029 | Old |
| 0.264776 | 0.07445 | 0.065 | 0.036 | 1 | 11 | Zw10 | 0.029 | Old |
| 0.26533 | 0.188718 | 0.093 | 0.058 | 1 | 11 | Acd | 0.035 | Old |
| 0.265372 | 0.427792 | 0.044 | 0.022 | 1 | 11 | Tnfrsf25 | 0.022 | Old |
| 0.265673 | 0.034738 | 0.512 | 0.406 | 1 | 11 | Dynll1 | 0.106 | Old |
| 0.266636 | 1.108011 | 0.044 | 0.022 | 1 | 11 | Zbtb45 | 0.022 | Old |
| 0.266636 | 0.326868 | 0.044 | 0.022 | 1 | 11 | Bco2 | 0.022 | Old |
| 0.266636 | 0.272454 | 0.044 | 0.022 | 1 | 11 | Tmem67 | 0.022 | Old |
| 0.266662 | 0.014649 | 0.004 | 0.014 | 1 | 11 | Fam221b | -0.01 | Old |
| 0.268251 | 1.557887 | 0.052 | 0.029 | 1 | 11 | Wwox | 0.023 | Old |
| 0.268825 | 0.46264 | 0.19 | 0.138 | 1 | 11 | Ythdf2 | 0.052 | Old |
| 0.268856 | 0.199617 | 0.125 | 0.087 | 1 | 11 | C1qtnf1 | 0.038 | Old |
| 0.268979 | 0.838834 | 0.081 | 0.051 | 1 | 11 | Reep4 | 0.03 | Old |
| 0.269174 | 0.294311 | 0.044 | 0.022 | 1 | 11 | 1700021F0 | 0.022 | Old |
| 0.269174 | 0.012186 | 0.044 | 0.022 | 1 | 11 | Wdr75 | 0.022 | Old |
| 0.269336 | 0.137992 | 0.242 | 0.181 | 1 | 11 | Esyt1 | 0.061 | Old |
| 0.269788 | 0.001273 | 0.597 | 0.464 | 1 | 11 | Creg1 | 0.133 | Old |
| 0.27045 | 0.019674 | 0.044 | 0.022 | 1 | 11 | Mamdc2 | 0.022 | Old |
| 0.270572 | 0.407472 | 0.101 | 0.065 | 1 | 11 | Zbtb37 | 0.036 | Old |
| 0.270573 | 1.154716 | 0.052 | 0.029 | 1 | 11 | Mrpl44 | 0.023 | Old |
| 0.271729 | 0.127893 | 0.044 | 0.022 | 1 | 11 | Havcr2 | 0.022 | Old |
| 0.271952 | 0.204145 | 0.109 | 0.072 | 1 | 11 | Nmd3 | 0.037 | Old |
| 0.272118 | 0.002305 | 0.323 | 0.254 | 1 | 11 | Mfsd14b | 0.069 | Old |
| 0.2725 | 0.343809 | 0.073 | 0.043 | 1 | 11 | Fzd5 | 0.03 | Old |
| 0.2725 | 0.255551 | 0.073 | 0.043 | 1 | 11 | Cep57 | 0.03 | Old |
| 0.272637 | 0.099952 | 0.766 | 0.674 | 1 | 11 | Naca | 0.092 | Old |
| 0.273108 | 0.96354 | 0.089 | 0.058 | 1 | 11 | Pgm1 | 0.031 | Old |
| 0.273309 | 0.011778 | 0.79 | 0.681 | 1 | 11 | Clta | 0.109 | Old |
| 0.273383 | 0.056151 | 0.536 | 0.406 | 1 | 11 | Hnrnpc | 0.13 | Old |
| 0.274174 | 0.096899 | 0.125 | 0.087 | 1 | 11 | Arhgap11a | 0.038 | Old |
| 0.274493 | 0.283866 | 0.073 | 0.043 | 1 | 11 | Eepd1 | 0.03 | Old |
| 0.274493 | 0.052479 | 0.524 | 0.413 | 1 | 11 | Smchd1 | 0.111 | Old |
| 0.274804 | 0.323807 | 0.101 | 0.065 | 1 | 11 | Smn1 | 0.036 | Old |
| 0.27556 | 0.699158 | 0.081 | 0.051 | 1 | 11 | Rev1 | 0.03 | Old |

| | | | | | | | | |
|---|---|---|---|---|---|---|---|---|
| 0.27556 | 0.688648 | 0.081 | 0.051 | 1 | 11 | Tubgcp2 | 0.03 | Old |
| 0.275809 | 0.993176 | 0.089 | 0.058 | 1 | 11 | Exosc8 | 0.031 | Old |
| 0.275863 | 0.325449 | 0.202 | 0.152 | 1 | 11 | Lpgat1 | 0.05 | Old |
| 0.276221 | 0.043047 | 0.298 | 0.232 | 1 | 11 | Trim12a | 0.066 | Old |
| 0.276509 | 0.647991 | 0.081 | 0.051 | 1 | 11 | Pde12 | 0.03 | Old |
| 0.277365 | 0.096449 | 0.101 | 0.065 | 1 | 11 | Mrpl1 | 0.036 | Old |
| 0.2775 | 0.064143 | 0.073 | 0.043 | 1 | 11 | Polr3b | 0.03 | Old |
| 0.278413 | 0.181548 | 0.081 | 0.051 | 1 | 11 | Trim3 | 0.03 | Old |
| 0.278497 | 0.075495 | 0.863 | 0.71 | 1 | 11 | Tpm3 | 0.153 | Old |
| 0.278807 | 0.922634 | 0.052 | 0.029 | 1 | 11 | Tdrd3 | 0.023 | Old |
| 0.278807 | 0.362132 | 0.052 | 0.029 | 1 | 11 | Rrp12 | 0.023 | Old |
| 0.279146 | 0.082578 | 0.452 | 0.348 | 1 | 11 | Ikzf1 | 0.104 | Old |
| 0.280327 | 0.492484 | 0.081 | 0.051 | 1 | 11 | Nsmf | 0.03 | Old |
| 0.281468 | 0.020489 | 0.242 | 0.181 | 1 | 11 | Usp12 | 0.061 | Old |
| 0.281774 | 0.098649 | 0.411 | 0.326 | 1 | 11 | Mgat1 | 0.085 | Old |
| 0.282157 | 0.268737 | 0.185 | 0.138 | 1 | 11 | Anapc16 | 0.047 | Old |
| 0.282388 | 0.947666 | 0.052 | 0.029 | 1 | 11 | Gpm6b | 0.023 | Old |
| 0.282674 | 0.185639 | 0.125 | 0.087 | 1 | 11 | Gpd1l | 0.038 | Old |
| 0.282676 | 0.050585 | 0.117 | 0.08 | 1 | 11 | Ulk1 | 0.037 | Old |
| 0.283576 | 0.021565 | 0.649 | 0.493 | 1 | 11 | Synj1 | 0.156 | Old |
| 0.283582 | 0.008946 | 0.194 | 0.145 | 1 | 11 | Pqlc1 | 0.049 | Old |
| 0.283588 | 0.69194 | 0.052 | 0.029 | 1 | 11 | Ttbk2 | 0.023 | Old |
| 0.283588 | 0.344447 | 0.052 | 0.029 | 1 | 11 | Alkbh7 | 0.023 | Old |
| 0.283953 | 0.535466 | 0.149 | 0.109 | 1 | 11 | Ccdc25 | 0.04 | Old |
| 0.284091 | 0.102293 | 0.435 | 0.341 | 1 | 11 | Pycard | 0.094 | Old |
| 0.284792 | 0.518342 | 0.052 | 0.029 | 1 | 11 | Ppp2r3a | 0.023 | Old |
| 0.285331 | 0.182002 | 0.161 | 0.116 | 1 | 11 | Dlg1 | 0.045 | Old |
| 0.285528 | 0.080752 | 0.222 | 0.159 | 1 | 11 | Itgav | 0.063 | Old |
| 0.285869 | 0.527064 | 0.089 | 0.058 | 1 | 11 | Prr14l | 0.031 | Old |
| 0.285925 | 1.366328 | 0.032 | 0.014 | 1 | 11 | Pex12 | 0.018 | Old |
| 0.286121 | 0.081773 | 0.081 | 0.051 | 1 | 11 | Hint3 | 0.03 | Old |
| 0.28721 | 0.943493 | 0.052 | 0.029 | 1 | 11 | E130102H2 | 0.023 | Old |
| 0.28721 | 0.711455 | 0.052 | 0.029 | 1 | 11 | Sephs2 | 0.023 | Old |
| 0.287705 | 0.031112 | 0.649 | 0.536 | 1 | 11 | Pomp | 0.113 | Old |
| 0.28785 | 0.216568 | 0.097 | 0.065 | 1 | 11 | Zfp282 | 0.032 | Old |
| 0.288637 | 0.26658 | 0.165 | 0.123 | 1 | 11 | Eps15l1 | 0.042 | Old |
| 0.288781 | 0.01214 | 0.347 | 0.268 | 1 | 11 | Polr2k | 0.079 | Old |
| 0.28881 | 0.041085 | 0.629 | 0.522 | 1 | 11 | Rnf213 | 0.107 | Old |
| 0.289588 | 0.75299 | 0.089 | 0.058 | 1 | 11 | Pik3cb | 0.031 | Old |
| 0.28976 | 0.335921 | 0.125 | 0.087 | 1 | 11 | Golga2 | 0.038 | Old |
| 0.290141 | 0.086665 | 0.411 | 0.326 | 1 | 11 | Cd300a | 0.085 | Old |
| 0.290523 | 0.354567 | 0.089 | 0.058 | 1 | 11 | Sfmbt1 | 0.031 | Old |
| 0.290864 | 0.540779 | 0.052 | 0.029 | 1 | 11 | Sema6b | 0.023 | Old |
| 0.291295 | 0.083714 | 0.927 | 0.833 | 1 | 11 | Gnai2 | 0.094 | Old |
| 0.29135 | 0.356774 | 0.125 | 0.087 | 1 | 11 | Ubxn2a | 0.038 | Old |

| | | | | | | | | |
|---|---|---|---|---|---|---|---|---|
| 0.291459 | 0.681912 | 0.089 | 0.058 | 1 | 11 | Scnm1 | 0.031 | Old |
| 0.291461 | 0.015805 | 0.569 | 0.435 | 1 | 11 | Mapre1 | 0.134 | Old |
| 0.292218 | 1.405413 | 0.032 | 0.014 | 1 | 11 | Sap18b | 0.018 | Old |
| 0.292218 | 1.012788 | 0.032 | 0.014 | 1 | 11 | Gm15564 | 0.018 | Old |
| 0.292218 | 0.933898 | 0.032 | 0.014 | 1 | 11 | Coil | 0.018 | Old |
| 0.293316 | 0.522486 | 0.052 | 0.029 | 1 | 11 | F10 | 0.023 | Old |
| 0.293316 | 0.184592 | 0.052 | 0.029 | 1 | 11 | Pym1 | 0.023 | Old |
| 0.293806 | 1.218524 | 0.032 | 0.014 | 1 | 11 | Prickle3 | 0.018 | Old |
| 0.293806 | 1.140934 | 0.032 | 0.014 | 1 | 11 | Tfap4 | 0.018 | Old |
| 0.293806 | 0.913271 | 0.032 | 0.014 | 1 | 11 | Nek1 | 0.018 | Old |
| 0.293806 | 0.659372 | 0.032 | 0.014 | 1 | 11 | Zbtb39 | 0.018 | Old |
| 0.294117 | 0.12268 | 0.117 | 0.08 | 1 | 11 | Dpp7 | 0.037 | Old |
| 0.294964 | 0.151628 | 0.081 | 0.051 | 1 | 11 | Tgtp2 | 0.03 | Old |
| 0.295399 | 0.973493 | 0.032 | 0.014 | 1 | 11 | 4930518I1! | 0.018 | Old |
| 0.295399 | 0.951583 | 0.032 | 0.014 | 1 | 11 | Rcn1 | 0.018 | Old |
| 0.295399 | 0.519044 | 0.032 | 0.014 | 1 | 11 | Urb1 | 0.018 | Old |
| 0.295399 | 0.185227 | 0.032 | 0.014 | 1 | 11 | Drosha | 0.018 | Old |
| 0.295655 | 0.887482 | 0.06 | 0.036 | 1 | 11 | Tmem184c | 0.024 | Old |
| 0.295655 | 0.814568 | 0.06 | 0.036 | 1 | 11 | Snx25 | 0.024 | Old |
| 0.295655 | 0.380711 | 0.06 | 0.036 | 1 | 11 | Sirpb1a | 0.024 | Old |
| 0.295783 | 0.012795 | 0.052 | 0.029 | 1 | 11 | Gm13822 | 0.023 | Old |
| 0.295938 | 0.472149 | 0.097 | 0.065 | 1 | 11 | Gprc5c | 0.032 | Old |
| 0.295951 | 0.153682 | 0.169 | 0.123 | 1 | 11 | Tbc1d1 | 0.046 | Old |
| 0.296599 | 0.154589 | 0.133 | 0.094 | 1 | 11 | Nckipsd | 0.039 | Old |
| 0.296999 | 0.762023 | 0.032 | 0.014 | 1 | 11 | Fam177a | 0.018 | Old |
| 0.296999 | 0.502336 | 0.032 | 0.014 | 1 | 11 | Ccdc91 | 0.018 | Old |
| 0.297022 | 0.297546 | 0.052 | 0.029 | 1 | 11 | Nrg2 | 0.023 | Old |
| 0.297022 | 0.276969 | 0.052 | 0.029 | 1 | 11 | Dynlt1f | 0.023 | Old |
| 0.297485 | 0.146851 | 0.302 | 0.232 | 1 | 11 | Arid5a | 0.07 | Old |
| 0.297949 | 1.127297 | 0.06 | 0.036 | 1 | 11 | Ptrh2 | 0.024 | Old |
| 0.298604 | 0.888178 | 0.032 | 0.014 | 1 | 11 | Wdhd1 | 0.018 | Old |
| 0.298604 | 0.825954 | 0.032 | 0.014 | 1 | 11 | Tarm1 | 0.018 | Old |
| 0.298604 | 0.586213 | 0.032 | 0.014 | 1 | 11 | Zfp868 | 0.018 | Old |
| 0.298604 | 0.534261 | 0.032 | 0.014 | 1 | 11 | Rnf24 | 0.018 | Old |
| 0.298604 | 0.506142 | 0.032 | 0.014 | 1 | 11 | Nme6 | 0.018 | Old |
| 0.298604 | 0.251939 | 0.032 | 0.014 | 1 | 11 | Arhgap35 | 0.018 | Old |
| 0.298604 | 0.012352 | 0.032 | 0.014 | 1 | 11 | Spaca9 | 0.018 | Old |
| 0.29945 | 0.32937 | 0.141 | 0.101 | 1 | 11 | Rad50 | 0.04 | Old |
| 0.299509 | 0.465769 | 0.052 | 0.029 | 1 | 11 | Nek6 | 0.023 | Old |
| 0.299509 | 0.150384 | 0.052 | 0.029 | 1 | 11 | Fasn | 0.023 | Old |
| 0.300215 | 0.539879 | 0.032 | 0.014 | 1 | 11 | C920006O1 | 0.018 | Old |
| 0.300215 | 0.277187 | 0.032 | 0.014 | 1 | 11 | Fads6 | 0.018 | Old |
| 0.300215 | 0.268936 | 0.032 | 0.014 | 1 | 11 | Gm17435 | 0.018 | Old |
| 0.300215 | 0.168314 | 0.032 | 0.014 | 1 | 11 | Zmym6 | 0.018 | Old |
| 0.300215 | 0.128414 | 0.032 | 0.014 | 1 | 11 | Apex2 | 0.018 | Old |

| | | | | | | | | |
|---|---|---|---|---|---|---|---|---|
| 0.300215 | 0.12044 | 0.032 | 0.014 | 1 | 11 | Evi5l | 0.018 | Old |
| 0.301414 | 0.612024 | 0.06 | 0.036 | 1 | 11 | Bud23 | 0.024 | Old |
| 0.301414 | 0.471018 | 0.06 | 0.036 | 1 | 11 | Myg1 | 0.024 | Old |
| 0.301831 | 0.390371 | 0.032 | 0.014 | 1 | 11 | Gm26890 | 0.018 | Old |
| 0.301831 | 0.304837 | 0.032 | 0.014 | 1 | 11 | Tmod1 | 0.018 | Old |
| 0.30201 | 0.224094 | 0.052 | 0.029 | 1 | 11 | Cmss1 | 0.023 | Old |
| 0.302976 | 0.382213 | 0.081 | 0.051 | 1 | 11 | Sssca1 | 0.03 | Old |
| 0.303454 | 0.824861 | 0.032 | 0.014 | 1 | 11 | Nxpe3 | 0.018 | Old |
| 0.303454 | 0.143812 | 0.032 | 0.014 | 1 | 11 | Gtf2h2 | 0.018 | Old |
| 0.303454 | 0.107494 | 0.032 | 0.014 | 1 | 11 | Tmem171 | 0.018 | Old |
| 0.303454 | 0.065203 | 0.032 | 0.014 | 1 | 11 | Tubd1 | 0.018 | Old |
| 0.303701 | 0.079066 | 0.133 | 0.094 | 1 | 11 | Vti1a | 0.039 | Old |
| 0.303738 | 0.673681 | 0.06 | 0.036 | 1 | 11 | Ccdc88c | 0.024 | Old |
| 0.303823 | 0.138162 | 0.089 | 0.058 | 1 | 11 | Fbxo8 | 0.031 | Old |
| 0.303834 | 0.126847 | 0.319 | 0.254 | 1 | 11 | Cerk | 0.065 | Old |
| 0.304713 | 0.020284 | 0.694 | 0.609 | 1 | 11 | Tpd52 | 0.085 | Old |
| 0.304788 | 0.129123 | 0.089 | 0.058 | 1 | 11 | Nr1d2 | 0.031 | Old |
| 0.304791 | 0.036096 | 0.831 | 0.754 | 1 | 11 | Rrbp1 | 0.077 | Old |
| 0.304804 | 0.371737 | 0.173 | 0.13 | 1 | 11 | Pptc7 | 0.043 | Old |
| 0.304905 | 0.739819 | 0.06 | 0.036 | 1 | 11 | Ltbp3 | 0.024 | Old |
| 0.305082 | 0.206883 | 0.032 | 0.014 | 1 | 11 | Gm20045 | 0.018 | Old |
| 0.305082 | 0.039858 | 0.032 | 0.014 | 1 | 11 | Adcy3 | 0.018 | Old |
| 0.305755 | 0.253007 | 0.089 | 0.058 | 1 | 11 | Galns | 0.031 | Old |
| 0.305755 | 0.133984 | 0.089 | 0.058 | 1 | 11 | Sco2 | 0.031 | Old |
| 0.305788 | 0.075474 | 0.052 | 0.029 | 1 | 11 | Smyd5 | 0.023 | Old |
| 0.305855 | 0.731653 | 0.121 | 0.087 | 1 | 11 | Ccl5 | 0.034 | Old |
| 0.306017 | 0.20842 | 0.081 | 0.051 | 1 | 11 | Chst12 | 0.03 | Old |
| 0.306039 | 0.388427 | 0.105 | 0.072 | 1 | 11 | Fam3a | 0.033 | Old |
| 0.306126 | 0.12293 | 0.286 | 0.225 | 1 | 11 | Sltm | 0.061 | Old |
| 0.306716 | 0.114577 | 0.032 | 0.014 | 1 | 11 | Sag | 0.018 | Old |
| 0.306755 | 0.055749 | 0.125 | 0.087 | 1 | 11 | Tmem19 | 0.038 | Old |
| 0.306807 | 0.594355 | 0.113 | 0.08 | 1 | 11 | Brix1 | 0.033 | Old |
| 0.307297 | 0.240564 | 0.423 | 0.355 | 1 | 11 | C4b | 0.068 | Old |
| 0.307669 | 0.263208 | 0.113 | 0.08 | 1 | 11 | Camsap2 | 0.033 | Old |
| 0.308423 | 0.253854 | 0.06 | 0.036 | 1 | 11 | Zfp994 | 0.024 | Old |
| 0.309302 | 0.088024 | 0.133 | 0.094 | 1 | 11 | Got2 | 0.039 | Old |
| 0.309781 | 0.018596 | 0.194 | 0.145 | 1 | 11 | Klhl5 | 0.049 | Old |
| 0.309801 | 0.110142 | 0.21 | 0.159 | 1 | 11 | Oasl1 | 0.051 | Old |
| 0.309944 | 0.020421 | 0.149 | 0.109 | 1 | 11 | Sdccag3 | 0.04 | Old |
| 0.310124 | 0.005485 | 0.464 | 0.348 | 1 | 11 | Fermt3 | 0.116 | Old |
| 0.310783 | 0.350292 | 0.06 | 0.036 | 1 | 11 | Rhbdf1 | 0.024 | Old |
| 0.311481 | 0.094231 | 0.165 | 0.123 | 1 | 11 | Ptpn7 | 0.042 | Old |
| 0.311835 | 0.096104 | 0.218 | 0.167 | 1 | 11 | Mgat4a | 0.051 | Old |
| 0.311967 | 0.814473 | 0.06 | 0.036 | 1 | 11 | Tmem29 | 0.024 | Old |
| 0.312429 | 0.072987 | 0.77 | 0.652 | 1 | 11 | Timp2 | 0.118 | Old |

| | | | | | | | | |
|---|---|---|---|---|---|---|---|---|
| 0.313155 | 0.065943 | 0.06 | 0.036 | 1 | 11 | Mbip | 0.024 | Old |
| 0.313227 | 0.154193 | 0.105 | 0.072 | 1 | 11 | Hspa13 | 0.033 | Old |
| 0.314772 | 0.337502 | 0.145 | 0.109 | 1 | 11 | Tbpl1 | 0.036 | Old |
| 0.315539 | 0.203515 | 0.06 | 0.036 | 1 | 11 | Miga1 | 0.024 | Old |
| 0.315867 | 0.096528 | 0.565 | 0.457 | 1 | 11 | Hspe1 | 0.108 | Old |
| 0.316098 | 0.004236 | 0.149 | 0.109 | 1 | 11 | Pigp | 0.04 | Old |
| 0.316735 | 0.40574 | 0.06 | 0.036 | 1 | 11 | Ttc39c | 0.024 | Old |
| 0.316932 | 0.0521 | 0.774 | 0.616 | 1 | 11 | Tuba1b | 0.158 | Old |
| 0.317892 | 0.655311 | 0.069 | 0.043 | 1 | 11 | Stk35 | 0.026 | Old |
| 0.317892 | 0.648599 | 0.069 | 0.043 | 1 | 11 | Cish | 0.026 | Old |
| 0.317934 | 0.119005 | 0.06 | 0.036 | 1 | 11 | Rbmx2 | 0.024 | Old |
| 0.318141 | 0.391506 | 0.113 | 0.08 | 1 | 11 | Gpx3 | 0.033 | Old |
| 0.3185 | 0.436959 | 0.121 | 0.087 | 1 | 11 | Tmem161b | 0.034 | Old |
| 0.319194 | 0.033997 | 0.097 | 0.065 | 1 | 11 | Lrrc42 | 0.032 | Old |
| 0.320143 | 0.231416 | 0.21 | 0.159 | 1 | 11 | Smc4 | 0.051 | Old |
| 0.320342 | 0.417822 | 0.06 | 0.036 | 1 | 11 | Yaf2 | 0.024 | Old |
| 0.320553 | 0.109265 | 0.698 | 0.58 | 1 | 11 | Cox8a | 0.118 | Old |
| 0.320879 | 0.093009 | 0.895 | 0.761 | 1 | 11 | Serf2 | 0.134 | Old |
| 0.32155 | 0.395139 | 0.06 | 0.036 | 1 | 11 | Polr3d | 0.024 | Old |
| 0.32155 | 0.141345 | 0.06 | 0.036 | 1 | 11 | Fzd7 | 0.024 | Old |
| 0.321857 | 0.162505 | 0.722 | 0.587 | 1 | 11 | Sfpq | 0.135 | Old |
| 0.321862 | 2.393006 | 0.02 | 0.007 | 1 | 11 | Hes7 | 0.013 | Old |
| 0.321862 | 2.257746 | 0.02 | 0.007 | 1 | 11 | Gm28707 | 0.013 | Old |
| 0.321862 | 2.019881 | 0.02 | 0.007 | 1 | 11 | Zfp974 | 0.013 | Old |
| 0.321862 | 1.550977 | 0.02 | 0.007 | 1 | 11 | Ccdc137 | 0.013 | Old |
| 0.322762 | 0.222598 | 0.06 | 0.036 | 1 | 11 | Zc3hc1 | 0.024 | Old |
| 0.322994 | 1.023755 | 0.04 | 0.022 | 1 | 11 | Mpst | 0.018 | Old |
| 0.324037 | 2.195508 | 0.02 | 0.007 | 1 | 11 | Rxrg | 0.013 | Old |
| 0.324037 | 1.970887 | 0.02 | 0.007 | 1 | 11 | Hepacam2 | 0.013 | Old |
| 0.324037 | 1.6618 | 0.02 | 0.007 | 1 | 11 | Gm13431 | 0.013 | Old |
| 0.324037 | 1.622846 | 0.02 | 0.007 | 1 | 11 | Slc7a13 | 0.013 | Old |
| 0.324037 | 1.477925 | 0.02 | 0.007 | 1 | 11 | Slc7a5 | 0.013 | Old |
| 0.324037 | 1.448901 | 0.02 | 0.007 | 1 | 11 | Gm26853 | 0.013 | Old |
| 0.324037 | 1.325816 | 0.02 | 0.007 | 1 | 11 | Pmm1 | 0.013 | Old |
| 0.324037 | 1.202438 | 0.02 | 0.007 | 1 | 11 | Bahcc1 | 0.013 | Old |
| 0.324037 | 1.12159 | 0.02 | 0.007 | 1 | 11 | Katnal1 | 0.013 | Old |
| 0.324356 | 0.378034 | 0.113 | 0.08 | 1 | 11 | Klrk1 | 0.033 | Old |
| 0.324488 | 1.500458 | 0.04 | 0.022 | 1 | 11 | Ier5l | 0.018 | Old |
| 0.324949 | 0.315731 | 0.097 | 0.065 | 1 | 11 | Ranbp10 | 0.032 | Old |
| 0.325142 | 0.165702 | 0.105 | 0.072 | 1 | 11 | Hacd3 | 0.033 | Old |
| 0.325193 | 0.192702 | 0.06 | 0.036 | 1 | 11 | Rfc3 | 0.024 | Old |
| 0.325193 | 0.156499 | 0.06 | 0.036 | 1 | 11 | Zfp942 | 0.024 | Old |
| 0.325473 | 0.099266 | 0.198 | 0.152 | 1 | 11 | Acsl1 | 0.046 | Old |
| 0.325825 | 0.660568 | 0.069 | 0.043 | 1 | 11 | Gmeb2 | 0.026 | Old |
| 0.326221 | 1.715588 | 0.02 | 0.007 | 1 | 11 | Afap1l1 | 0.013 | Old |

| | | | | | | | | |
|---|---|---|---|---|---|---|---|---|
| 0.326221 | 1.488155 | 0.02 | 0.007 | 1 | 11 | Prps1l3 | 0.013 | Old |
| 0.326221 | 1.430202 | 0.02 | 0.007 | 1 | 11 | Rnf144a | 0.013 | Old |
| 0.326221 | 1.398425 | 0.02 | 0.007 | 1 | 11 | Rhou | 0.013 | Old |
| 0.326221 | 1.262755 | 0.02 | 0.007 | 1 | 11 | Zfp688 | 0.013 | Old |
| 0.326221 | 0.7643 | 0.02 | 0.007 | 1 | 11 | E2f8 | 0.013 | Old |
| 0.326413 | 0.129153 | 0.06 | 0.036 | 1 | 11 | Kctd18 | 0.024 | Old |
| 0.326413 | 0.081119 | 0.06 | 0.036 | 1 | 11 | Fam122a | 0.024 | Old |
| 0.326413 | 0.07828 | 0.06 | 0.036 | 1 | 11 | Men1 | 0.024 | Old |
| 0.327435 | 0.530511 | 0.077 | 0.051 | 1 | 11 | Rrp36 | 0.026 | Old |
| 0.327472 | 0.056823 | 0.548 | 0.449 | 1 | 11 | Ctnnb1 | 0.099 | Old |
| 0.328415 | 1.642138 | 0.02 | 0.007 | 1 | 11 | Fabp1 | 0.013 | Old |
| 0.328415 | 1.516757 | 0.02 | 0.007 | 1 | 11 | Prnp | 0.013 | Old |
| 0.328415 | 1.276543 | 0.02 | 0.007 | 1 | 11 | Ftx | 0.013 | Old |
| 0.328415 | 1.1255 | 0.02 | 0.007 | 1 | 11 | Megf9 | 0.013 | Old |
| 0.328415 | 0.951515 | 0.02 | 0.007 | 1 | 11 | 1110002L0 | 0.013 | Old |
| 0.328415 | 0.843562 | 0.02 | 0.007 | 1 | 11 | 4930522L1 | 0.013 | Old |
| 0.328415 | 0.711901 | 0.02 | 0.007 | 1 | 11 | Pomt1 | 0.013 | Old |
| 0.328415 | 0.690398 | 0.02 | 0.007 | 1 | 11 | Adam23 | 0.013 | Old |
| 0.328415 | 0.644436 | 0.02 | 0.007 | 1 | 11 | Armc9 | 0.013 | Old |
| 0.328415 | 0.56377 | 0.02 | 0.007 | 1 | 11 | Hist1h4d | 0.013 | Old |
| 0.328415 | 0.554388 | 0.02 | 0.007 | 1 | 11 | Immp2l | 0.013 | Old |
| 0.32886 | 0.541693 | 0.121 | 0.087 | 1 | 11 | Mta3 | 0.034 | Old |
| 0.328995 | 1.091676 | 0.04 | 0.022 | 1 | 11 | Mapkapk5 | 0.018 | Old |
| 0.329604 | 0.433001 | 0.077 | 0.051 | 1 | 11 | Ap3m2 | 0.026 | Old |
| 0.329679 | 0.061989 | 0.496 | 0.384 | 1 | 11 | Neurl3 | 0.112 | Old |
| 0.329879 | 0.213277 | 0.181 | 0.138 | 1 | 11 | Sos2 | 0.043 | Old |
| 0.330415 | 0.687916 | 0.069 | 0.043 | 1 | 11 | Dnajc9 | 0.026 | Old |
| 0.330506 | 0.56197 | 0.04 | 0.022 | 1 | 11 | Cenpl | 0.018 | Old |
| 0.330506 | 0.41663 | 0.04 | 0.022 | 1 | 11 | Polg2 | 0.018 | Old |
| 0.330618 | 1.436424 | 0.02 | 0.007 | 1 | 11 | Tgfbr3 | 0.013 | Old |
| 0.330618 | 0.943775 | 0.02 | 0.007 | 1 | 11 | Tmem69 | 0.013 | Old |
| 0.330618 | 0.888057 | 0.02 | 0.007 | 1 | 11 | Odf2l | 0.013 | Old |
| 0.330618 | 0.857212 | 0.02 | 0.007 | 1 | 11 | Retn | 0.013 | Old |
| 0.330618 | 0.798771 | 0.02 | 0.007 | 1 | 11 | Ubtd2 | 0.013 | Old |
| 0.330618 | 0.758545 | 0.02 | 0.007 | 1 | 11 | Bora | 0.013 | Old |
| 0.330618 | 0.680174 | 0.02 | 0.007 | 1 | 11 | Thnsl1 | 0.013 | Old |
| 0.330618 | 0.646084 | 0.02 | 0.007 | 1 | 11 | Pyroxd2 | 0.013 | Old |
| 0.330618 | 0.645544 | 0.02 | 0.007 | 1 | 11 | Trim62 | 0.013 | Old |
| 0.330618 | 0.638708 | 0.02 | 0.007 | 1 | 11 | Zfp607a | 0.013 | Old |
| 0.330618 | 0.53659 | 0.02 | 0.007 | 1 | 11 | Gstk1 | 0.013 | Old |
| 0.330618 | 0.518307 | 0.02 | 0.007 | 1 | 11 | Maged2 | 0.013 | Old |
| 0.330618 | 0.467605 | 0.02 | 0.007 | 1 | 11 | Tubb2b | 0.013 | Old |
| 0.330618 | 0.270259 | 0.02 | 0.007 | 1 | 11 | AW146154 | 0.013 | Old |
| 0.330618 | 0.219138 | 0.02 | 0.007 | 1 | 11 | Gm38394 | 0.013 | Old |
| 0.330771 | 0.010989 | 0.097 | 0.065 | 1 | 11 | AI606181 | 0.032 | Old |

| | | | | | | | | |
|---|---|---|---|---|---|---|---|---|
| 0.331081 | 0.298023 | 0.198 | 0.152 | 1 | 11 | Rprd2 | 0.046 | Old |
| 0.331569 | 0.242879 | 0.069 | 0.043 | 1 | 11 | Chchd4 | 0.026 | Old |
| 0.331742 | 0.086157 | 0.351 | 0.297 | 1 | 11 | Zfp703 | 0.054 | Old |
| 0.332022 | 0.726942 | 0.04 | 0.022 | 1 | 11 | Cdc25a | 0.018 | Old |
| 0.332463 | 0.282416 | 0.113 | 0.08 | 1 | 11 | Zdhhc14 | 0.033 | Old |
| 0.332479 | 0.102314 | 0.464 | 0.377 | 1 | 11 | Rab31 | 0.087 | Old |
| 0.332538 | 0.051359 | 0.5 | 0.391 | 1 | 11 | Lamtor1 | 0.109 | Old |
| 0.332831 | 0.519549 | 0.02 | 0.007 | 1 | 11 | D030056L2 | 0.013 | Old |
| 0.332831 | 0.457238 | 0.02 | 0.007 | 1 | 11 | Polr3g | 0.013 | Old |
| 0.332831 | 0.343213 | 0.02 | 0.007 | 1 | 11 | Hdac8 | 0.013 | Old |
| 0.332831 | 0.227895 | 0.02 | 0.007 | 1 | 11 | A330069E1 | 0.013 | Old |
| 0.332831 | 0.134741 | 0.02 | 0.007 | 1 | 11 | Fastkd3 | 0.013 | Old |
| 0.332831 | 0.119311 | 0.02 | 0.007 | 1 | 11 | Cadm3 | 0.013 | Old |
| 0.332831 | 0.075321 | 0.02 | 0.007 | 1 | 11 | Dennd2d | 0.013 | Old |
| 0.333542 | 0.541338 | 0.04 | 0.022 | 1 | 11 | Gm26518 | 0.018 | Old |
| 0.333542 | 0.494516 | 0.04 | 0.022 | 1 | 11 | Gm20696 | 0.018 | Old |
| 0.333953 | 0.166573 | 0.165 | 0.123 | 1 | 11 | Slc27a1 | 0.042 | Old |
| 0.334715 | 0.10656 | 0.157 | 0.116 | 1 | 11 | Clasp1 | 0.041 | Old |
| 0.335067 | 0.74892 | 0.04 | 0.022 | 1 | 11 | Tab1 | 0.018 | Old |
| 0.335067 | 0.684183 | 0.04 | 0.022 | 1 | 11 | Tapt1 | 0.018 | Old |
| 0.335067 | 0.639899 | 0.04 | 0.022 | 1 | 11 | Racgap1 | 0.018 | Old |
| 0.336211 | 0.143932 | 0.069 | 0.043 | 1 | 11 | Rhbdd1 | 0.026 | Old |
| 0.336596 | 0.807832 | 0.04 | 0.022 | 1 | 11 | Btf3l4 | 0.018 | Old |
| 0.336596 | 0.754285 | 0.04 | 0.022 | 1 | 11 | Lmtk2 | 0.018 | Old |
| 0.336917 | 0.160956 | 0.702 | 0.572 | 1 | 11 | 2010107E0 | 0.13 | Old |
| 0.337378 | 0.056978 | 0.069 | 0.043 | 1 | 11 | Zfp746 | 0.026 | Old |
| 0.337993 | 0.041031 | 0.21 | 0.159 | 1 | 11 | Rpl7l1 | 0.051 | Old |
| 0.33813 | 0.72038 | 0.04 | 0.022 | 1 | 11 | Mansc1 | 0.018 | Old |
| 0.33813 | 0.547206 | 0.04 | 0.022 | 1 | 11 | Aste1 | 0.018 | Old |
| 0.33813 | 0.35861 | 0.04 | 0.022 | 1 | 11 | Ccdc77 | 0.018 | Old |
| 0.33813 | 0.116866 | 0.04 | 0.022 | 1 | 11 | Lmln | 0.018 | Old |
| 0.338707 | 0.040549 | 0.181 | 0.138 | 1 | 11 | Rbm33 | 0.043 | Old |
| 0.339668 | 0.553163 | 0.04 | 0.022 | 1 | 11 | Trit1 | 0.018 | Old |
| 0.339668 | 0.458251 | 0.04 | 0.022 | 1 | 11 | Usp2 | 0.018 | Old |
| 0.339668 | 0.312789 | 0.04 | 0.022 | 1 | 11 | Bach2 | 0.018 | Old |
| 0.339668 | 0.228066 | 0.04 | 0.022 | 1 | 11 | Ahrr | 0.018 | Old |
| 0.340381 | 0.071158 | 0.649 | 0.558 | 1 | 11 | Tomm7 | 0.091 | Old |
| 0.341211 | 0.651487 | 0.04 | 0.022 | 1 | 11 | Wdr76 | 0.018 | Old |
| 0.341736 | 0.481465 | 0.085 | 0.058 | 1 | 11 | Slc41a2 | 0.027 | Old |
| 0.341978 | 0.317925 | 0.145 | 0.109 | 1 | 11 | Rnd3 | 0.036 | Old |
| 0.342112 | 0.267514 | 0.121 | 0.087 | 1 | 11 | Wwc2 | 0.034 | Old |
| 0.342758 | 0.395913 | 0.04 | 0.022 | 1 | 11 | Eral1 | 0.018 | Old |
| 0.343473 | 0.046294 | 0.113 | 0.08 | 1 | 11 | Abcf3 | 0.033 | Old |
| 0.343592 | 0.097967 | 0.306 | 0.254 | 1 | 11 | Smim15 | 0.052 | Old |
| 0.34431 | 0.127428 | 0.04 | 0.022 | 1 | 11 | Synj2 | 0.018 | Old |

| | | | | | | | | |
|---|---|---|---|---|---|---|---|---|
| 0.345621 | 0.504087 | 0.069 | 0.043 | 1 | 11 | Fiz1 | 0.026 | Old |
| 0.345866 | 0.042987 | 0.04 | 0.022 | 1 | 11 | Slc12a4 | 0.018 | Old |
| 0.345947 | 0.414716 | 0.218 | 0.174 | 1 | 11 | Eif4a3 | 0.044 | Old |
| 0.345983 | 0.540224 | 0.085 | 0.058 | 1 | 11 | Nsun6 | 0.027 | Old |
| 0.347091 | 0.133791 | 0.242 | 0.188 | 1 | 11 | Ube2h | 0.054 | Old |
| 0.347427 | 0.325395 | 0.04 | 0.022 | 1 | 11 | Mtfr1 | 0.018 | Old |
| 0.347427 | 0.256765 | 0.04 | 0.022 | 1 | 11 | Zfp472 | 0.018 | Old |
| 0.347427 | 0.094268 | 0.04 | 0.022 | 1 | 11 | Cyfip2 | 0.018 | Old |
| 0.348119 | 0.456765 | 0.085 | 0.058 | 1 | 11 | Bcl2l12 | 0.027 | Old |
| 0.348389 | 0.193489 | 0.282 | 0.225 | 1 | 11 | Rnf115 | 0.057 | Old |
| 0.348711 | 0.15928 | 0.25 | 0.203 | 1 | 11 | Yipf1 | 0.047 | Old |
| 0.348992 | 0.176244 | 0.04 | 0.022 | 1 | 11 | Katnb1 | 0.018 | Old |
| 0.348992 | 0.116537 | 0.04 | 0.022 | 1 | 11 | Pald1 | 0.018 | Old |
| 0.349419 | 1.067833 | 0.048 | 0.029 | 1 | 11 | Atg4c | 0.019 | Old |
| 0.349542 | 0.258632 | 0.077 | 0.051 | 1 | 11 | Pxmp4 | 0.026 | Old |
| 0.349542 | 0.163805 | 0.077 | 0.051 | 1 | 11 | Mpv17 | 0.026 | Old |
| 0.35025 | 0.056055 | 0.73 | 0.616 | 1 | 11 | Aplp2 | 0.114 | Old |
| 0.350284 | 0.06057 | 0.137 | 0.101 | 1 | 11 | Zc3h14 | 0.036 | Old |
| 0.351118 | 0.227278 | 0.161 | 0.123 | 1 | 11 | Mcemp1 | 0.038 | Old |
| 0.351804 | 0.281884 | 0.077 | 0.051 | 1 | 11 | Cdr2 | 0.026 | Old |
| 0.351928 | 0.014296 | 0.145 | 0.109 | 1 | 11 | Braf | 0.036 | Old |
| 0.353694 | 0.913417 | 0.048 | 0.029 | 1 | 11 | Pdik1l | 0.019 | Old |
| 0.354646 | 0.348519 | 0.093 | 0.065 | 1 | 11 | Dpep2 | 0.028 | Old |
| 0.355286 | 0.135319 | 0.145 | 0.109 | 1 | 11 | Cir1 | 0.036 | Old |
| 0.35567 | 0.019324 | 0.133 | 0.094 | 1 | 11 | Topbp1 | 0.039 | Old |
| 0.356562 | 0.998772 | 0.048 | 0.029 | 1 | 11 | Ptges2 | 0.019 | Old |
| 0.35672 | 0.429711 | 0.093 | 0.065 | 1 | 11 | Pkp4 | 0.028 | Old |
| 0.357498 | 0.219133 | 0.077 | 0.051 | 1 | 11 | Atg9a | 0.026 | Old |
| 0.358001 | 0.745921 | 0.048 | 0.029 | 1 | 11 | Pdpr | 0.019 | Old |
| 0.358381 | 0.550374 | 0.157 | 0.123 | 1 | 11 | Egr3 | 0.034 | Old |
| 0.358577 | 0.031294 | 0.327 | 0.261 | 1 | 11 | Uhrf2 | 0.066 | Old |
| 0.358644 | 0.324021 | 0.077 | 0.051 | 1 | 11 | Panx1 | 0.026 | Old |
| 0.358687 | 0.051483 | 0.367 | 0.29 | 1 | 11 | Pikfyve | 0.077 | Old |
| 0.358718 | 0.477805 | 0.246 | 0.203 | 1 | 11 | Ell2 | 0.043 | Old |
| 0.359561 | 0.070635 | 0.153 | 0.116 | 1 | 11 | Alox5 | 0.037 | Old |
| 0.360891 | 0.007129 | 0.048 | 0.029 | 1 | 11 | Hpn | 0.019 | Old |
| 0.362342 | 0.356553 | 0.048 | 0.029 | 1 | 11 | Trmt10a | 0.019 | Old |
| 0.362406 | 0.113736 | 0.137 | 0.101 | 1 | 11 | Aspscr1 | 0.036 | Old |
| 0.363251 | 0.405501 | 0.077 | 0.051 | 1 | 11 | Mettl1 | 0.026 | Old |
| 0.363346 | 0.076037 | 0.238 | 0.188 | 1 | 11 | Rbm27 | 0.05 | Old |
| 0.364871 | 0.086068 | 0.129 | 0.094 | 1 | 11 | Itpr1 | 0.035 | Old |
| 0.365254 | 0.80152 | 0.048 | 0.029 | 1 | 11 | Mettl8 | 0.019 | Old |
| 0.365616 | 0.038681 | 0.569 | 0.471 | 1 | 11 | Itgam | 0.098 | Old |
| 0.366145 | 0.286642 | 0.093 | 0.065 | 1 | 11 | Isca1 | 0.028 | Old |
| 0.366221 | 0.132508 | 0.153 | 0.116 | 1 | 11 | Napsa | 0.037 | Old |

| | | | | | | | | |
|---|---|---|---|---|---|---|---|---|
| 0.366715 | 1.029527 | 0.048 | 0.029 | 1 | 11 | 1110032A( | 0.019 | Old |
| 0.367718 | 0.476435 | 0.085 | 0.058 | 1 | 11 | Cbx6 | 0.027 | Old |
| 0.367718 | 0.470796 | 0.085 | 0.058 | 1 | 11 | Igip | 0.027 | Old |
| 0.36818 | 0.476438 | 0.048 | 0.029 | 1 | 11 | Hotairm1 | 0.019 | Old |
| 0.36818 | 0.396064 | 0.048 | 0.029 | 1 | 11 | Ptger2 | 0.019 | Old |
| 0.36826 | 0.445962 | 0.093 | 0.065 | 1 | 11 | Prkag2 | 0.028 | Old |
| 0.36882 | 0.085887 | 0.544 | 0.449 | 1 | 11 | Ranbp2 | 0.095 | Old |
| 0.369337 | 0.168723 | 0.161 | 0.123 | 1 | 11 | Mthfr | 0.038 | Old |
| 0.369649 | 0.155288 | 0.048 | 0.029 | 1 | 11 | Mospd1 | 0.019 | Old |
| 0.371016 | 0.087181 | 0.665 | 0.536 | 1 | 11 | Rps27l | 0.129 | Old |
| 0.371121 | 0.469081 | 0.048 | 0.029 | 1 | 11 | Zfp595 | 0.019 | Old |
| 0.371222 | 0.094121 | 0.137 | 0.101 | 1 | 11 | Cyp20a1 | 0.036 | Old |
| 0.372597 | 0.3865 | 0.048 | 0.029 | 1 | 11 | Slx4ip | 0.019 | Old |
| 0.372597 | 0.242368 | 0.048 | 0.029 | 1 | 11 | Gm4707 | 0.019 | Old |
| 0.372597 | 0.11647 | 0.048 | 0.029 | 1 | 11 | Rnf219 | 0.019 | Old |
| 0.372807 | 0.130464 | 0.177 | 0.138 | 1 | 11 | Rsl24d1 | 0.039 | Old |
| 0.373582 | 0.205852 | 0.093 | 0.065 | 1 | 11 | Pdk3 | 0.028 | Old |
| 0.374077 | 0.011745 | 0.048 | 0.029 | 1 | 11 | Ttll3 | 0.019 | Old |
| 0.374318 | 0.021502 | 0.702 | 0.58 | 1 | 11 | Serbp1 | 0.122 | Old |
| 0.374832 | 0.827927 | 0.056 | 0.036 | 1 | 11 | Kank2 | 0.02 | Old |
| 0.374832 | 0.660359 | 0.056 | 0.036 | 1 | 11 | Ankrd13d | 0.02 | Old |
| 0.375503 | 0.212637 | 0.71 | 0.587 | 1 | 11 | Hnrnpu | 0.123 | Old |
| 0.37556 | 0.02072 | 0.048 | 0.029 | 1 | 11 | Dbf4 | 0.019 | Old |
| 0.375784 | 0.293503 | 0.109 | 0.08 | 1 | 11 | Kmt5c | 0.029 | Old |
| 0.377116 | 0.012239 | 0.815 | 0.681 | 1 | 11 | Picalm | 0.134 | Old |
| 0.377531 | 0.141832 | 0.774 | 0.667 | 1 | 11 | Ifi207 | 0.107 | Old |
| 0.377571 | 0.842706 | 0.056 | 0.036 | 1 | 11 | Stambp | 0.02 | Old |
| 0.377829 | 0.031696 | 0.399 | 0.312 | 1 | 11 | Arhgap18 | 0.087 | Old |
| 0.37893 | 0.036659 | 0.371 | 0.297 | 1 | 11 | Tsc22d1 | 0.074 | Old |
| 0.379187 | 0.00831 | 0.177 | 0.138 | 1 | 11 | AI467606 | 0.039 | Old |
| 0.379229 | 0.178523 | 0.169 | 0.13 | 1 | 11 | Ctsf | 0.039 | Old |
| 0.38003 | 0.173583 | 0.093 | 0.065 | 1 | 11 | Psmg3 | 0.028 | Old |
| 0.380322 | 0.592235 | 0.056 | 0.036 | 1 | 11 | Fam213a | 0.02 | Old |
| 0.380545 | 0.00862 | 0.161 | 0.123 | 1 | 11 | Pvr | 0.038 | Old |
| 0.381529 | 0.016257 | 0.048 | 0.029 | 1 | 11 | Mthfd1 | 0.019 | Old |
| 0.381703 | 0.252405 | 0.056 | 0.036 | 1 | 11 | Atg4a | 0.02 | Old |
| 0.381834 | 0.062889 | 0.891 | 0.804 | 1 | 11 | Hnrnpa2b1 | 0.087 | Old |
| 0.383151 | 1.615061 | 0.028 | 0.014 | 1 | 11 | Pigf | 0.014 | Old |
| 0.383151 | 1.600664 | 0.028 | 0.014 | 1 | 11 | Gatsl3 | 0.014 | Old |
| 0.383151 | 1.477123 | 0.028 | 0.014 | 1 | 11 | Nle1 | 0.014 | Old |
| 0.384473 | 0.377549 | 0.056 | 0.036 | 1 | 11 | Spsb2 | 0.02 | Old |
| 0.385572 | 0.316995 | 0.117 | 0.087 | 1 | 11 | Amn1 | 0.03 | Old |
| 0.386546 | 0.08122 | 0.093 | 0.065 | 1 | 11 | Ssna1 | 0.028 | Old |
| 0.387136 | 1.767093 | 0.028 | 0.014 | 1 | 11 | Zfp971 | 0.014 | Old |
| 0.387136 | 0.943901 | 0.028 | 0.014 | 1 | 11 | Mterf4 | 0.014 | Old |

| | | | | | | | | |
|---|---|---|---|---|---|---|---|---|
| 0.387211 | 0.059313 | 0.149 | 0.116 | 1 | 11 | Nop56 | 0.033 | Old |
| 0.387255 | 0.249557 | 0.056 | 0.036 | 1 | 11 | Zmat3 | 0.02 | Old |
| 0.387841 | 0.853714 | 0.065 | 0.043 | 1 | 11 | Gm20707 | 0.022 | Old |
| 0.387841 | 0.746273 | 0.065 | 0.043 | 1 | 11 | Txnl4a | 0.022 | Old |
| 0.38878 | 0.043716 | 0.895 | 0.833 | 1 | 11 | Mrc1 | 0.062 | Old |
| 0.389138 | 1.203299 | 0.028 | 0.014 | 1 | 11 | Meis3 | 0.014 | Old |
| 0.389138 | 1.128166 | 0.028 | 0.014 | 1 | 11 | Wdr6 | 0.014 | Old |
| 0.389138 | 1.11029 | 0.028 | 0.014 | 1 | 11 | Snapc2 | 0.014 | Old |
| 0.389138 | 1.076337 | 0.028 | 0.014 | 1 | 11 | Zmat1 | 0.014 | Old |
| 0.389145 | 1.210417 | 0.065 | 0.043 | 1 | 11 | Nit2 | 0.022 | Old |
| 0.38924 | 0.252828 | 0.214 | 0.167 | 1 | 11 | Mbd3 | 0.047 | Old |
| 0.389297 | 0.069726 | 0.101 | 0.072 | 1 | 11 | Arfgef2 | 0.029 | Old |
| 0.390049 | 0.387349 | 0.056 | 0.036 | 1 | 11 | Akip1 | 0.02 | Old |
| 0.390049 | 0.065443 | 0.056 | 0.036 | 1 | 11 | Mlh1 | 0.02 | Old |
| 0.390451 | 0.527405 | 0.065 | 0.043 | 1 | 11 | Pot1b | 0.022 | Old |
| 0.390464 | 0.158451 | 0.117 | 0.087 | 1 | 11 | Dpm1 | 0.03 | Old |
| 0.390853 | 0.077137 | 0.109 | 0.08 | 1 | 11 | Pot1a | 0.029 | Old |
| 0.391147 | 1.140575 | 0.028 | 0.014 | 1 | 11 | Ccdc84 | 0.014 | Old |
| 0.391147 | 1.067112 | 0.028 | 0.014 | 1 | 11 | Zfp335 | 0.014 | Old |
| 0.391147 | 0.912799 | 0.028 | 0.014 | 1 | 11 | Fahd2a | 0.014 | Old |
| 0.391451 | 0.344083 | 0.056 | 0.036 | 1 | 11 | Wrnip1 | 0.02 | Old |
| 0.391451 | 0.061246 | 0.056 | 0.036 | 1 | 11 | Zfand2a | 0.02 | Old |
| 0.391751 | 0.164527 | 0.181 | 0.145 | 1 | 11 | Sde2 | 0.036 | Old |
| 0.392444 | 0.110265 | 0.254 | 0.203 | 1 | 11 | Mrps18c | 0.051 | Old |
| 0.393161 | 1.195011 | 0.028 | 0.014 | 1 | 11 | Ankrd66 | 0.014 | Old |
| 0.393161 | 0.666837 | 0.028 | 0.014 | 1 | 11 | Zfp948 | 0.014 | Old |
| 0.393161 | 0.159181 | 0.028 | 0.014 | 1 | 11 | Spaca6 | 0.014 | Old |
| 0.393198 | 0.378782 | 0.133 | 0.101 | 1 | 11 | Trim27 | 0.032 | Old |
| 0.394234 | 0.056258 | 0.093 | 0.065 | 1 | 11 | B3galnt1 | 0.028 | Old |
| 0.394264 | 0.542852 | 0.056 | 0.036 | 1 | 11 | Gtpbp10 | 0.02 | Old |
| 0.394264 | 0.445436 | 0.056 | 0.036 | 1 | 11 | Efl1 | 0.02 | Old |
| 0.394264 | 0.379915 | 0.056 | 0.036 | 1 | 11 | Vwa8 | 0.02 | Old |
| 0.394384 | 0.341282 | 0.065 | 0.043 | 1 | 11 | 1110012L1 | 0.022 | Old |
| 0.394647 | 1.069444 | 0.073 | 0.051 | 1 | 11 | Rdh14 | 0.022 | Old |
| 0.395182 | 0.645319 | 0.028 | 0.014 | 1 | 11 | Mageh1 | 0.014 | Old |
| 0.395182 | 0.489086 | 0.028 | 0.014 | 1 | 11 | Adgrl2 | 0.014 | Old |
| 0.396013 | 0.089681 | 0.681 | 0.63 | 1 | 11 | Trf | 0.051 | Old |
| 0.396851 | 0.035572 | 0.198 | 0.152 | 1 | 11 | Foxj2 | 0.046 | Old |
| 0.39702 | 0.785345 | 0.065 | 0.043 | 1 | 11 | Oraov1 | 0.022 | Old |
| 0.397209 | 1.258209 | 0.028 | 0.014 | 1 | 11 | Ahdc1 | 0.014 | Old |
| 0.397209 | 0.751018 | 0.028 | 0.014 | 1 | 11 | Lcmt2 | 0.014 | Old |
| 0.397209 | 0.589614 | 0.028 | 0.014 | 1 | 11 | Ick | 0.014 | Old |
| 0.398342 | 0.415633 | 0.065 | 0.043 | 1 | 11 | Rufy2 | 0.022 | Old |
| 0.398506 | 0.113296 | 0.056 | 0.036 | 1 | 11 | Mblac2 | 0.02 | Old |
| 0.398758 | 0.041281 | 0.185 | 0.145 | 1 | 11 | Lin7c | 0.04 | Old |

| | | | | | | | | |
|---|---|---|---|---|---|---|---|---|
| 0.398926 | 0.01614 | 0.125 | 0.094 | 1 | 11 | Zbtb8os | 0.031 | Old |
| 0.399243 | 0.86003 | 0.028 | 0.014 | 1 | 11 | Zfp287 | 0.014 | Old |
| 0.399243 | 0.694771 | 0.028 | 0.014 | 1 | 11 | Eml2 | 0.014 | Old |
| 0.399243 | 0.495229 | 0.028 | 0.014 | 1 | 11 | Mex3b | 0.014 | Old |
| 0.399243 | 0.46506 | 0.028 | 0.014 | 1 | 11 | Mtbp | 0.014 | Old |
| 0.399243 | 0.437418 | 0.028 | 0.014 | 1 | 11 | Ttc21b | 0.014 | Old |
| 0.399243 | 0.394171 | 0.028 | 0.014 | 1 | 11 | Car9 | 0.014 | Old |
| 0.399243 | 0.388507 | 0.028 | 0.014 | 1 | 11 | A130010J1 | 0.014 | Old |
| 0.399243 | 0.297566 | 0.028 | 0.014 | 1 | 11 | Zfp251 | 0.014 | Old |
| 0.399243 | 0.164213 | 0.028 | 0.014 | 1 | 11 | Ralgps1 | 0.014 | Old |
| 0.40098 | 0.0962 | 0.347 | 0.275 | 1 | 11 | Rap2b | 0.072 | Old |
| 0.401283 | 0.173105 | 0.028 | 0.014 | 1 | 11 | Selenoo | 0.014 | Old |
| 0.401283 | 0.039205 | 0.028 | 0.014 | 1 | 11 | Mgp | 0.014 | Old |
| 0.403329 | 0.148427 | 0.028 | 0.014 | 1 | 11 | Nat2 | 0.014 | Old |
| 0.403329 | 0.087539 | 0.028 | 0.014 | 1 | 11 | Armcx5 | 0.014 | Old |
| 0.403329 | 0.083058 | 0.028 | 0.014 | 1 | 11 | Gpr85 | 0.014 | Old |
| 0.403329 | 0.023471 | 0.028 | 0.014 | 1 | 11 | Cox6a2 | 0.014 | Old |
| 0.403329 | 0.013462 | 0.028 | 0.014 | 1 | 11 | Slc9a3r2 | 0.014 | Old |
| 0.403379 | 0.465801 | 0.073 | 0.051 | 1 | 11 | Tpgs1 | 0.022 | Old |
| 0.404208 | 0.093896 | 0.153 | 0.116 | 1 | 11 | Ube4a | 0.037 | Old |
| 0.405091 | 0.220943 | 0.298 | 0.254 | 1 | 11 | Dr1 | 0.044 | Old |
| 0.405381 | 0.150362 | 0.028 | 0.014 | 1 | 11 | Mcat | 0.014 | Old |
| 0.405895 | 0.285792 | 0.073 | 0.051 | 1 | 11 | Kifc3 | 0.022 | Old |
| 0.407377 | 0.117302 | 0.117 | 0.087 | 1 | 11 | Tagap | 0.03 | Old |
| 0.408027 | 0.562936 | 0.016 | 0.029 | 1 | 11 | C2 | -0.013 | Old |
| 0.40842 | 0.401099 | 0.073 | 0.051 | 1 | 11 | 4930453N2 | 0.022 | Old |
| 0.410358 | 0.224148 | 0.065 | 0.043 | 1 | 11 | Pwp1 | 0.022 | Old |
| 0.412268 | 0.529605 | 0.121 | 0.094 | 1 | 11 | Pspc1 | 0.027 | Old |
| 0.413057 | 0.188318 | 0.065 | 0.043 | 1 | 11 | Gid4 | 0.022 | Old |
| 0.413869 | 0.018509 | 0.827 | 0.703 | 1 | 11 | Sh3bgrl3 | 0.124 | Old |
| 0.413906 | 0.076752 | 0.278 | 0.232 | 1 | 11 | Slfn1 | 0.046 | Old |
| 0.414774 | 0.182697 | 0.073 | 0.051 | 1 | 11 | Ptgr1 | 0.022 | Old |
| 0.415184 | 0.054615 | 0.835 | 0.761 | 1 | 11 | Ucp2 | 0.074 | Old |
| 0.415767 | 0.102757 | 0.065 | 0.043 | 1 | 11 | Usp42 | 0.022 | Old |
| 0.415767 | 0.097334 | 0.065 | 0.043 | 1 | 11 | Pyroxd1 | 0.022 | Old |
| 0.415767 | 0.005994 | 0.065 | 0.043 | 1 | 11 | Sema4b | 0.022 | Old |
| 0.415854 | 1.503964 | 0.036 | 0.022 | 1 | 11 | Bpgm | 0.014 | Old |
| 0.416052 | 0.60286 | 0.073 | 0.051 | 1 | 11 | Prkce | 0.022 | Old |
| 0.416828 | 0.225733 | 0.133 | 0.101 | 1 | 11 | Uxs1 | 0.032 | Old |
| 0.41753 | 0.093363 | 0.331 | 0.283 | 1 | 11 | Eno1 | 0.048 | Old |
| 0.417611 | 0.185842 | 0.206 | 0.167 | 1 | 11 | Egln2 | 0.039 | Old |
| 0.419485 | 0.07414 | 0.125 | 0.094 | 1 | 11 | Tmem88 | 0.031 | Old |
| 0.419491 | 0.943131 | 0.036 | 0.022 | 1 | 11 | Uhrf1bp1 | 0.014 | Old |
| 0.419966 | 0.054516 | 0.165 | 0.13 | 1 | 11 | Mysm1 | 0.035 | Old |
| 0.420069 | 0.004714 | 0.718 | 0.58 | 1 | 11 | Tspo | 0.138 | Old |

| | | | | | | | | |
|---|---|---|---|---|---|---|---|---|
| 0.420649 | 0.353363 | 0.113 | 0.087 | 1 | 11 | Sc5d | 0.026 | Old |
| 0.420841 | 0.107759 | 0.948 | 0.884 | 1 | 11 | Myl6 | 0.064 | Old |
| 0.422472 | 0.266945 | 0.125 | 0.094 | 1 | 11 | Chmp7 | 0.031 | Old |
| 0.423148 | 1.095088 | 0.036 | 0.022 | 1 | 11 | Gadd45a | 0.014 | Old |
| 0.423332 | 0.794252 | 0.097 | 0.072 | 1 | 11 | Phkg2 | 0.025 | Old |
| 0.423771 | 0.584203 | 0.113 | 0.087 | 1 | 11 | Akap7 | 0.026 | Old |
| 0.424292 | 0.444403 | 0.121 | 0.094 | 1 | 11 | Kdm4c | 0.027 | Old |
| 0.425577 | 0.113128 | 0.089 | 0.065 | 1 | 11 | Slc9a6 | 0.024 | Old |
| 0.426422 | 0.025527 | 0.359 | 0.29 | 1 | 11 | Gtf2h5 | 0.069 | Old |
| 0.42672 | 0.449996 | 0.105 | 0.08 | 1 | 11 | Tom1 | 0.025 | Old |
| 0.427926 | 0.010674 | 0.806 | 0.674 | 1 | 11 | Morf4l1 | 0.132 | Old |
| 0.427944 | 0.225401 | 0.081 | 0.058 | 1 | 11 | Tcp11l1 | 0.023 | Old |
| 0.427944 | 0.18762 | 0.081 | 0.058 | 1 | 11 | Sigmar1 | 0.023 | Old |
| 0.427944 | 0.051938 | 0.081 | 0.058 | 1 | 11 | Prdm4 | 0.023 | Old |
| 0.428517 | 0.161963 | 0.27 | 0.225 | 1 | 11 | Prrc2b | 0.045 | Old |
| 0.428667 | 0.871522 | 0.036 | 0.022 | 1 | 11 | 1110065P2 | 0.014 | Old |
| 0.430416 | 0.181498 | 0.081 | 0.058 | 1 | 11 | Wdr77 | 0.023 | Old |
| 0.430461 | 0.06103 | 0.206 | 0.167 | 1 | 11 | Smad4 | 0.039 | Old |
| 0.430516 | 0.943085 | 0.036 | 0.022 | 1 | 11 | Ydjc | 0.014 | Old |
| 0.430748 | 0.008443 | 0.157 | 0.123 | 1 | 11 | Pogk | 0.034 | Old |
| 0.432162 | 0.262471 | 0.105 | 0.08 | 1 | 11 | Phf11a | 0.025 | Old |
| 0.43237 | 0.647985 | 0.036 | 0.022 | 1 | 11 | Vamp1 | 0.014 | Old |
| 0.43237 | 0.51797 | 0.036 | 0.022 | 1 | 11 | Ctdspl | 0.014 | Old |
| 0.43237 | 0.000686 | 0.036 | 0.022 | 1 | 11 | Gemin2 | 0.014 | Old |
| 0.433153 | 0.004065 | 0.681 | 0.565 | 1 | 11 | Lyn | 0.116 | Old |
| 0.433154 | 0.012372 | 0.452 | 0.377 | 1 | 11 | Itpkb | 0.075 | Old |
| 0.434228 | 0.709065 | 0.036 | 0.022 | 1 | 11 | Tnfrsf17 | 0.014 | Old |
| 0.434228 | 0.493713 | 0.036 | 0.022 | 1 | 11 | Tfpt | 0.014 | Old |
| 0.434228 | 0.389297 | 0.036 | 0.022 | 1 | 11 | Gtpbp3 | 0.014 | Old |
| 0.436091 | 0.435529 | 0.036 | 0.022 | 1 | 11 | Rhbdd3 | 0.014 | Old |
| 0.437199 | 0.009946 | 0.226 | 0.181 | 1 | 11 | Pcmt1 | 0.045 | Old |
| 0.437959 | 0.420919 | 0.036 | 0.022 | 1 | 11 | B3galt6 | 0.014 | Old |
| 0.439297 | 0.034685 | 0.996 | 0.986 | 1 | 11 | Fth1 | 0.01 | Old |
| 0.440383 | 0.106991 | 0.081 | 0.058 | 1 | 11 | Casp6 | 0.023 | Old |
| 0.441707 | 0.49778 | 0.036 | 0.022 | 1 | 11 | Adsl | 0.014 | Old |
| 0.441707 | 0.345822 | 0.036 | 0.022 | 1 | 11 | Iqcb1 | 0.014 | Old |
| 0.441707 | 0.170311 | 0.036 | 0.022 | 1 | 11 | Apba3 | 0.014 | Old |
| 0.441707 | 0.035187 | 0.036 | 0.022 | 1 | 11 | 4930555A0 | 0.014 | Old |
| 0.441713 | 0.435872 | 0.113 | 0.087 | 1 | 11 | Phb | 0.026 | Old |
| 0.441873 | 0.103094 | 0.028 | 0.043 | 1 | 11 | Enc1 | -0.015 | Old |
| 0.443163 | 0.153109 | 0.105 | 0.08 | 1 | 11 | Rbmx | 0.025 | Old |
| 0.443236 | 0.003439 | 0.141 | 0.109 | 1 | 11 | Mtmr10 | 0.032 | Old |
| 0.443852 | 0.070894 | 0.113 | 0.087 | 1 | 11 | Upf2 | 0.026 | Old |
| 0.444287 | 0.007148 | 0.472 | 0.399 | 1 | 11 | B4galt6 | 0.073 | Old |
| 0.444525 | 0.015071 | 0.444 | 0.377 | 1 | 11 | Cbr2 | 0.067 | Old |

| | | | | | | | | |
|---|---|---|---|---|---|---|---|---|
| 0.444588 | 0.03232 | 0.089 | 0.065 | 1 | 11 | Sik2 | 0.024 | Old |
| 0.445286 | 0.67071 | 0.044 | 0.029 | 1 | 11 | Coro2b | 0.015 | Old |
| 0.446679 | 0.16781 | 0.081 | 0.058 | 1 | 11 | Llgl1 | 0.023 | Old |
| 0.446998 | 0.037297 | 0.089 | 0.065 | 1 | 11 | Syap1 | 0.024 | Old |
| 0.450388 | 1.056106 | 0.044 | 0.029 | 1 | 11 | Zfp609 | 0.015 | Old |
| 0.450388 | 0.762559 | 0.044 | 0.029 | 1 | 11 | 9930104L0 | 0.015 | Old |
| 0.451837 | 0.099504 | 0.867 | 0.812 | 1 | 11 | Ifitm3 | 0.055 | Old |
| 0.452108 | 0.043476 | 0.19 | 0.152 | 1 | 11 | Utp3 | 0.038 | Old |
| 0.45221 | 0.230982 | 0.19 | 0.159 | 1 | 11 | Cxcl12 | 0.031 | Old |
| 0.454109 | 0.025351 | 0.073 | 0.051 | 1 | 11 | Jmy | 0.022 | Old |
| 0.455662 | 2.148185 | 0.016 | 0.007 | 1 | 11 | Kyat3 | 0.009 | Old |
| 0.455662 | 2.002687 | 0.016 | 0.007 | 1 | 11 | Trp53i11 | 0.009 | Old |
| 0.455662 | 1.631126 | 0.016 | 0.007 | 1 | 11 | Nmb | 0.009 | Old |
| 0.455662 | 1.46177 | 0.016 | 0.007 | 1 | 11 | Asrgl1 | 0.009 | Old |
| 0.455662 | 1.388032 | 0.016 | 0.007 | 1 | 11 | Fam216a | 0.009 | Old |
| 0.455662 | 0.967695 | 0.016 | 0.007 | 1 | 11 | Lmnb2 | 0.009 | Old |
| 0.455662 | 0.897931 | 0.016 | 0.007 | 1 | 11 | Zfp949 | 0.009 | Old |
| 0.455662 | 0.788725 | 0.016 | 0.007 | 1 | 11 | 2700081O1 | 0.009 | Old |
| 0.455662 | 0.723071 | 0.016 | 0.007 | 1 | 11 | C1qtnf12 | 0.009 | Old |
| 0.456044 | 0.074417 | 0.903 | 0.855 | 1 | 11 | Mafb | 0.048 | Old |
| 0.456937 | 0.00794 | 0.141 | 0.109 | 1 | 11 | H2-T24 | 0.032 | Old |
| 0.457241 | 0.563576 | 0.044 | 0.029 | 1 | 11 | Stn1 | 0.015 | Old |
| 0.457765 | 0.83077 | 0.052 | 0.036 | 1 | 11 | Adamts10 | 0.016 | Old |
| 0.458029 | 0.088139 | 0.504 | 0.435 | 1 | 11 | Socs3 | 0.069 | Old |
| 0.458603 | 2.915738 | 0.016 | 0.007 | 1 | 11 | Dock9 | 0.009 | Old |
| 0.458603 | 2.253503 | 0.016 | 0.007 | 1 | 11 | Srpx | 0.009 | Old |
| 0.458603 | 1.966673 | 0.016 | 0.007 | 1 | 11 | Gm16867 | 0.009 | Old |
| 0.458603 | 1.715703 | 0.016 | 0.007 | 1 | 11 | Cep72 | 0.009 | Old |
| 0.458603 | 1.567506 | 0.016 | 0.007 | 1 | 11 | Mks1 | 0.009 | Old |
| 0.458603 | 1.355752 | 0.016 | 0.007 | 1 | 11 | Rad52 | 0.009 | Old |
| 0.458603 | 1.269655 | 0.016 | 0.007 | 1 | 11 | Siah1b | 0.009 | Old |
| 0.458603 | 1.216403 | 0.016 | 0.007 | 1 | 11 | Pced1b | 0.009 | Old |
| 0.458603 | 1.199362 | 0.016 | 0.007 | 1 | 11 | Pms1 | 0.009 | Old |
| 0.458603 | 1.157541 | 0.016 | 0.007 | 1 | 11 | Ttc12 | 0.009 | Old |
| 0.458603 | 1.148688 | 0.016 | 0.007 | 1 | 11 | Gm15706 | 0.009 | Old |
| 0.458603 | 1.14866 | 0.016 | 0.007 | 1 | 11 | Nptxr | 0.009 | Old |
| 0.458603 | 0.806201 | 0.016 | 0.007 | 1 | 11 | Fam92a | 0.009 | Old |
| 0.458603 | 0.805432 | 0.016 | 0.007 | 1 | 11 | Car13 | 0.009 | Old |
| 0.458819 | 0.050951 | 0.536 | 0.435 | 1 | 11 | Arhgap17 | 0.101 | Old |
| 0.458832 | 0.087196 | 0.105 | 0.08 | 1 | 11 | Gpn1 | 0.025 | Old |
| 0.458964 | 0.87188 | 0.044 | 0.029 | 1 | 11 | Rcl1 | 0.015 | Old |
| 0.458964 | 0.553386 | 0.044 | 0.029 | 1 | 11 | Rasal2 | 0.015 | Old |
| 0.459345 | 1.23661 | 0.052 | 0.036 | 1 | 11 | Ruvbl2 | 0.016 | Old |
| 0.459555 | 0.042556 | 0.782 | 0.659 | 1 | 11 | Ywhae | 0.123 | Old |
| 0.459832 | 0.051098 | 0.427 | 0.362 | 1 | 11 | Far1 | 0.065 | Old |

| | | | | | | | | |
|---|---|---|---|---|---|---|---|---|
| 0.460393 | 0.252868 | 0.133 | 0.101 | 1 | 11 | 1700123O2 | 0.032 | Old |
| 0.46069 | 0.710717 | 0.044 | 0.029 | 1 | 11 | Dpp4 | 0.015 | Old |
| 0.46069 | 0.431902 | 0.044 | 0.029 | 1 | 11 | Nr1d1 | 0.015 | Old |
| 0.460927 | 0.975886 | 0.052 | 0.036 | 1 | 11 | Trmt10c | 0.016 | Old |
| 0.461294 | 0.055279 | 0.129 | 0.101 | 1 | 11 | Lyst | 0.028 | Old |
| 0.461555 | 1.344416 | 0.016 | 0.007 | 1 | 11 | Tmem81 | 0.009 | Old |
| 0.461555 | 1.284227 | 0.016 | 0.007 | 1 | 11 | Tex14 | 0.009 | Old |
| 0.461555 | 1.125216 | 0.016 | 0.007 | 1 | 11 | Gm17025 | 0.009 | Old |
| 0.461555 | 1.010164 | 0.016 | 0.007 | 1 | 11 | Stx1b | 0.009 | Old |
| 0.461555 | 0.991549 | 0.016 | 0.007 | 1 | 11 | Stab2 | 0.009 | Old |
| 0.461555 | 0.93754 | 0.016 | 0.007 | 1 | 11 | Magee1 | 0.009 | Old |
| 0.461555 | 0.878618 | 0.016 | 0.007 | 1 | 11 | Crybb1 | 0.009 | Old |
| 0.461555 | 0.805117 | 0.016 | 0.007 | 1 | 11 | Pla2g6 | 0.009 | Old |
| 0.461555 | 0.779189 | 0.016 | 0.007 | 1 | 11 | Hyls1 | 0.009 | Old |
| 0.461555 | 0.770167 | 0.016 | 0.007 | 1 | 11 | Gas8 | 0.009 | Old |
| 0.461555 | 0.741157 | 0.016 | 0.007 | 1 | 11 | Cox6b2 | 0.009 | Old |
| 0.461555 | 0.737439 | 0.016 | 0.007 | 1 | 11 | Galnt6 | 0.009 | Old |
| 0.461555 | 0.705874 | 0.016 | 0.007 | 1 | 11 | Gm13479 | 0.009 | Old |
| 0.461555 | 0.702267 | 0.016 | 0.007 | 1 | 11 | Cgnl1 | 0.009 | Old |
| 0.461555 | 0.558133 | 0.016 | 0.007 | 1 | 11 | B3gnt2 | 0.009 | Old |
| 0.462194 | 0.15233 | 0.153 | 0.123 | 1 | 11 | Zfp131 | 0.03 | Old |
| 0.462419 | 0.692701 | 0.044 | 0.029 | 1 | 11 | Zfp865 | 0.015 | Old |
| 0.462419 | 0.433926 | 0.044 | 0.029 | 1 | 11 | Hdac7 | 0.015 | Old |
| 0.462419 | 0.389678 | 0.044 | 0.029 | 1 | 11 | Zfp212 | 0.015 | Old |
| 0.462419 | 0.311721 | 0.044 | 0.029 | 1 | 11 | Pef1 | 0.015 | Old |
| 0.462419 | 0.275666 | 0.044 | 0.029 | 1 | 11 | Gm13373 | 0.015 | Old |
| 0.462449 | 0.054248 | 0.097 | 0.072 | 1 | 11 | Sp3os | 0.025 | Old |
| 0.463095 | 0.138646 | 0.641 | 0.514 | 1 | 11 | Atf4 | 0.127 | Old |
| 0.463694 | 0.033851 | 0.585 | 0.5 | 1 | 11 | Pnp | 0.085 | Old |
| 0.464153 | 0.738267 | 0.044 | 0.029 | 1 | 11 | Tsen15 | 0.015 | Old |
| 0.464517 | 1.369347 | 0.016 | 0.007 | 1 | 11 | Palb2 | 0.009 | Old |
| 0.464517 | 1.238109 | 0.016 | 0.007 | 1 | 11 | Ddr1 | 0.009 | Old |
| 0.464517 | 1.054724 | 0.016 | 0.007 | 1 | 11 | Pkmyt1 | 0.009 | Old |
| 0.464517 | 0.987724 | 0.016 | 0.007 | 1 | 11 | Frs3 | 0.009 | Old |
| 0.464517 | 0.845925 | 0.016 | 0.007 | 1 | 11 | Fancf | 0.009 | Old |
| 0.464517 | 0.76108 | 0.016 | 0.007 | 1 | 11 | Prelid3a | 0.009 | Old |
| 0.464517 | 0.667137 | 0.016 | 0.007 | 1 | 11 | Gm36839 | 0.009 | Old |
| 0.464517 | 0.596258 | 0.016 | 0.007 | 1 | 11 | Rgcc | 0.009 | Old |
| 0.464517 | 0.591696 | 0.016 | 0.007 | 1 | 11 | Nhej1 | 0.009 | Old |
| 0.464517 | 0.519028 | 0.016 | 0.007 | 1 | 11 | Acss2 | 0.009 | Old |
| 0.464517 | 0.498071 | 0.016 | 0.007 | 1 | 11 | 6330418K0 | 0.009 | Old |
| 0.464517 | 0.433926 | 0.016 | 0.007 | 1 | 11 | Kcna2 | 0.009 | Old |
| 0.464517 | 0.430663 | 0.016 | 0.007 | 1 | 11 | Focad | 0.009 | Old |
| 0.464517 | 0.427084 | 0.016 | 0.007 | 1 | 11 | Ccdc180 | 0.009 | Old |
| 0.464517 | 0.414518 | 0.016 | 0.007 | 1 | 11 | Gm26522 | 0.009 | Old |

| | | | | | | | | |
|---|---|---|---|---|---|---|---|---|
| 0.464517 | 0.401159 | 0.016 | 0.007 | 1 | 11 | Arsg | 0.009 | Old |
| 0.464517 | 0.328722 | 0.016 | 0.007 | 1 | 11 | Gm6133 | 0.009 | Old |
| 0.464517 | 0.311991 | 0.016 | 0.007 | 1 | 11 | Poli | 0.009 | Old |
| 0.464517 | 0.311509 | 0.016 | 0.007 | 1 | 11 | Ube2c | 0.009 | Old |
| 0.464517 | 0.072816 | 0.016 | 0.007 | 1 | 11 | Sac3d1 | 0.009 | Old |
| 0.46589 | 0.622943 | 0.044 | 0.029 | 1 | 11 | F11r | 0.015 | Old |
| 0.466282 | 0.347365 | 0.012 | 0.022 | 1 | 11 | AC163032. | -0.01 | Old |
| 0.466569 | 0.045199 | 0.532 | 0.442 | 1 | 11 | Mtss1 | 0.09 | Old |
| 0.46749 | 0.335989 | 0.016 | 0.007 | 1 | 11 | Fancc | 0.009 | Old |
| 0.46749 | 0.287622 | 0.016 | 0.007 | 1 | 11 | Tkfc | 0.009 | Old |
| 0.46749 | 0.285298 | 0.016 | 0.007 | 1 | 11 | Fam13a | 0.009 | Old |
| 0.46749 | 0.270944 | 0.016 | 0.007 | 1 | 11 | Lins1 | 0.009 | Old |
| 0.46749 | 0.256424 | 0.016 | 0.007 | 1 | 11 | Pafah2 | 0.009 | Old |
| 0.46749 | 0.251462 | 0.016 | 0.007 | 1 | 11 | Gm21887 | 0.009 | Old |
| 0.46749 | 0.241407 | 0.016 | 0.007 | 1 | 11 | Zfp316 | 0.009 | Old |
| 0.46749 | 0.206647 | 0.016 | 0.007 | 1 | 11 | F5 | 0.009 | Old |
| 0.46749 | 0.177897 | 0.016 | 0.007 | 1 | 11 | Enpp2 | 0.009 | Old |
| 0.46749 | 0.131299 | 0.016 | 0.007 | 1 | 11 | Akap17b | 0.009 | Old |
| 0.46749 | 0.124868 | 0.016 | 0.007 | 1 | 11 | Pclaf | 0.009 | Old |
| 0.46749 | 0.12424 | 0.016 | 0.007 | 1 | 11 | Gne | 0.009 | Old |
| 0.46749 | 0.088198 | 0.016 | 0.007 | 1 | 11 | Armc2 | 0.009 | Old |
| 0.46749 | 0.075226 | 0.016 | 0.007 | 1 | 11 | Bbs5 | 0.009 | Old |
| 0.46749 | 0.060342 | 0.016 | 0.007 | 1 | 11 | Slc4a8 | 0.009 | Old |
| 0.46763 | 0.128182 | 0.044 | 0.029 | 1 | 11 | Zfp143 | 0.015 | Old |
| 0.46763 | 0.049276 | 0.044 | 0.029 | 1 | 11 | Zfp369 | 0.015 | Old |
| 0.469374 | 0.049476 | 0.044 | 0.029 | 1 | 11 | Mettl17 | 0.015 | Old |
| 0.469508 | 0.036689 | 0.206 | 0.167 | 1 | 11 | Gnl1 | 0.039 | Old |
| 0.470949 | 0.55855 | 0.069 | 0.051 | 1 | 11 | Ndufaf4 | 0.018 | Old |
| 0.473157 | 0.051427 | 0.915 | 0.819 | 1 | 11 | Pabpc1 | 0.096 | Old |
| 0.473654 | 0.417724 | 0.105 | 0.08 | 1 | 11 | Atxn1l | 0.025 | Old |
| 0.475523 | 0.007699 | 0.097 | 0.072 | 1 | 11 | Fastk | 0.025 | Old |
| 0.480153 | 0.0521 | 0.052 | 0.036 | 1 | 11 | Inpp5b | 0.016 | Old |
| 0.480337 | 0.007851 | 0.883 | 0.826 | 1 | 11 | Unc93b1 | 0.057 | Old |
| 0.482217 | 0.567844 | 0.069 | 0.051 | 1 | 11 | B3glct | 0.018 | Old |
| 0.484239 | 0.112267 | 0.129 | 0.101 | 1 | 11 | Arl4a | 0.028 | Old |
| 0.485027 | 0.256485 | 0.052 | 0.036 | 1 | 11 | Zfp574 | 0.016 | Old |
| 0.486657 | 0.077382 | 0.052 | 0.036 | 1 | 11 | Erich1 | 0.016 | Old |
| 0.486816 | 0.061093 | 0.315 | 0.275 | 1 | 11 | Ddx3y | 0.04 | Old |
| 0.487383 | 0.149118 | 0.202 | 0.167 | 1 | 11 | Sec13 | 0.035 | Old |
| 0.487804 | 0.132782 | 0.06 | 0.043 | 1 | 11 | Guf1 | 0.017 | Old |
| 0.489333 | 0.537091 | 0.069 | 0.051 | 1 | 11 | Mpp5 | 0.018 | Old |
| 0.491567 | 0.011828 | 0.052 | 0.036 | 1 | 11 | Ppih | 0.016 | Old |
| 0.492372 | 0.012209 | 0.06 | 0.043 | 1 | 11 | Gm16536 | 0.017 | Old |
| 0.493542 | 0.204226 | 0.306 | 0.261 | 1 | 11 | Saraf | 0.045 | Old |
| 0.4939 | 0.459727 | 0.06 | 0.043 | 1 | 11 | Cul4b | 0.017 | Old |

| | | | | | | | | |
|---|---|---|---|---|---|---|---|---|
| 0.4939 | 0.133255 | 0.06 | 0.043 | 1 | 11 | Flt3l | 0.017 | Old |
| 0.495066 | 0.424981 | 0.069 | 0.051 | 1 | 11 | Pdp1 | 0.018 | Old |
| 0.495195 | 0.321065 | 0.093 | 0.072 | 1 | 11 | Tspan5 | 0.021 | Old |
| 0.495431 | 0.210332 | 0.06 | 0.043 | 1 | 11 | Trim14 | 0.017 | Old |
| 0.495661 | 0.245679 | 0.077 | 0.058 | 1 | 11 | Bri3bp | 0.019 | Old |
| 0.496505 | 0.044191 | 0.069 | 0.051 | 1 | 11 | Siah1a | 0.018 | Old |
| 0.497023 | 0.299314 | 0.077 | 0.058 | 1 | 11 | Cdkn2d | 0.019 | Old |
| 0.497425 | 0.013441 | 0.266 | 0.217 | 1 | 11 | Pml | 0.049 | Old |
| 0.497904 | 0.018643 | 0.165 | 0.138 | 1 | 11 | Deptor | 0.027 | Old |
| 0.497946 | 0.27287 | 0.069 | 0.051 | 1 | 11 | Cep63 | 0.018 | Old |
| 0.499755 | 0.123915 | 0.077 | 0.058 | 1 | 11 | Dnajc24 | 0.019 | Old |
| 0.499944 | 0.424008 | 0.085 | 0.065 | 1 | 11 | Mill2 | 0.02 | Old |
| 0.501155 | 0.011096 | 0.512 | 0.435 | 1 | 11 | Pmaip1 | 0.077 | Old |
| 0.502043 | 0.090434 | 0.145 | 0.116 | 1 | 11 | Rras2 | 0.029 | Old |
| 0.504747 | 0.138057 | 0.185 | 0.152 | 1 | 11 | Homer1 | 0.033 | Old |
| 0.504776 | 0.206341 | 0.125 | 0.101 | 1 | 11 | Pilrb2 | 0.024 | Old |
| 0.506127 | 0.310133 | 0.343 | 0.312 | 1 | 11 | Cd38 | 0.031 | Old |
| 0.506619 | 0.008622 | 0.077 | 0.058 | 1 | 11 | Map4k1 | 0.019 | Old |
| 0.506783 | 0.240595 | 0.117 | 0.094 | 1 | 11 | 5430427O1 | 0.023 | Old |
| 0.506883 | 0.005434 | 0.206 | 0.167 | 1 | 11 | Pbx3 | 0.039 | Old |
| 0.50824 | 0.121301 | 0.109 | 0.087 | 1 | 11 | Stx17 | 0.022 | Old |
| 0.509556 | 0.019117 | 0.194 | 0.159 | 1 | 11 | Ccdc115 | 0.035 | Old |
| 0.510012 | 0.071749 | 0.024 | 0.036 | 1 | 11 | Gimap6 | -0.012 | Old |
| 0.510744 | 0.290793 | 0.19 | 0.159 | 1 | 11 | Kpna1 | 0.031 | Old |
| 0.510761 | 2.183229 | 0.024 | 0.014 | 1 | 11 | Nfib | 0.01 | Old |
| 0.510761 | 0.062978 | 0.077 | 0.058 | 1 | 11 | Lrp12 | 0.019 | Old |
| 0.510902 | 0.309787 | 0.734 | 0.71 | 1 | 11 | Egr1 | 0.024 | Old |
| 0.513243 | 1.239321 | 0.024 | 0.014 | 1 | 11 | Mrvi1 | 0.01 | Old |
| 0.513243 | 0.995839 | 0.024 | 0.014 | 1 | 11 | Capn10 | 0.01 | Old |
| 0.518226 | 1.481295 | 0.024 | 0.014 | 1 | 11 | Zfp677 | 0.01 | Old |
| 0.520289 | 0.120507 | 0.331 | 0.283 | 1 | 11 | S100a11 | 0.048 | Old |
| 0.520727 | 0.535868 | 0.024 | 0.014 | 1 | 11 | Pold2 | 0.01 | Old |
| 0.520727 | 0.418741 | 0.024 | 0.014 | 1 | 11 | Fut10 | 0.01 | Old |
| 0.523234 | 0.950198 | 0.024 | 0.014 | 1 | 11 | Slc27a3 | 0.01 | Old |
| 0.523234 | 0.873722 | 0.024 | 0.014 | 1 | 11 | Sh3rf1 | 0.01 | Old |
| 0.523234 | 0.750983 | 0.024 | 0.014 | 1 | 11 | Tada2a | 0.01 | Old |
| 0.523234 | 0.666083 | 0.024 | 0.014 | 1 | 11 | Bub1b | 0.01 | Old |
| 0.523234 | 0.631915 | 0.024 | 0.014 | 1 | 11 | Atp23 | 0.01 | Old |
| 0.523234 | 0.614271 | 0.024 | 0.014 | 1 | 11 | Zfp40 | 0.01 | Old |
| 0.523234 | 0.592526 | 0.024 | 0.014 | 1 | 11 | Tcim | 0.01 | Old |
| 0.523234 | 0.515279 | 0.024 | 0.014 | 1 | 11 | 4933427D1 | 0.01 | Old |
| 0.523837 | 0.026445 | 0.117 | 0.094 | 1 | 11 | Usp6nl | 0.023 | Old |
| 0.525024 | 0.309086 | 0.085 | 0.065 | 1 | 11 | Selenoi | 0.02 | Old |
| 0.525748 | 1.507728 | 0.024 | 0.014 | 1 | 11 | Ccdc138 | 0.01 | Old |
| 0.525748 | 0.67908 | 0.024 | 0.014 | 1 | 11 | Rasgrf2 | 0.01 | Old |

| | | | | | | | | |
|---|---|---|---|---|---|---|---|---|
| 0.525748 | 0.59806 | 0.024 | 0.014 | 1 | 11 | Rhobtb2 | 0.01 | Old |
| 0.525748 | 0.305564 | 0.024 | 0.014 | 1 | 11 | Exog | 0.01 | Old |
| 0.525748 | 0.267581 | 0.024 | 0.014 | 1 | 11 | Mtmr11 | 0.01 | Old |
| 0.526451 | 0.035378 | 0.544 | 0.464 | 1 | 11 | Trafd1 | 0.08 | Old |
| 0.527504 | 0.004214 | 0.077 | 0.058 | 1 | 11 | Espn | 0.019 | Old |
| 0.528267 | 1.002788 | 0.024 | 0.014 | 1 | 11 | Gdpgp1 | 0.01 | Old |
| 0.528267 | 0.245153 | 0.024 | 0.014 | 1 | 11 | Tmem201 | 0.01 | Old |
| 0.528267 | 0.243826 | 0.024 | 0.014 | 1 | 11 | Gm7160 | 0.01 | Old |
| 0.528267 | 0.235219 | 0.024 | 0.014 | 1 | 11 | Ogfod3 | 0.01 | Old |
| 0.528267 | 0.219854 | 0.024 | 0.014 | 1 | 11 | Clec1a | 0.01 | Old |
| 0.528267 | 0.096119 | 0.024 | 0.014 | 1 | 11 | Nsun5 | 0.01 | Old |
| 0.528912 | 0.00699 | 0.077 | 0.058 | 1 | 11 | Camsap1 | 0.019 | Old |
| 0.529042 | 0.282668 | 0.085 | 0.065 | 1 | 11 | Polk | 0.02 | Old |
| 0.529042 | 0.11066 | 0.085 | 0.065 | 1 | 11 | Fkbp1b | 0.02 | Old |
| 0.529301 | 0.033875 | 0.093 | 0.072 | 1 | 11 | Engase | 0.021 | Old |
| 0.530321 | 0.065726 | 0.077 | 0.058 | 1 | 11 | Scyl3 | 0.019 | Old |
| 0.530793 | 0.645549 | 0.024 | 0.014 | 1 | 11 | Harbi1 | 0.01 | Old |
| 0.530793 | 0.463805 | 0.024 | 0.014 | 1 | 11 | Ifrd2 | 0.01 | Old |
| 0.530793 | 0.365001 | 0.024 | 0.014 | 1 | 11 | Trub1 | 0.01 | Old |
| 0.531697 | 0.011335 | 0.069 | 0.051 | 1 | 11 | Tmem18 | 0.018 | Old |
| 0.532006 | 0.018311 | 0.891 | 0.783 | 1 | 11 | Cox4i1 | 0.108 | Old |
| 0.533048 | 0.117998 | 0.109 | 0.087 | 1 | 11 | Pira2 | 0.022 | Old |
| 0.533325 | 0.307736 | 0.024 | 0.014 | 1 | 11 | Pcolce | 0.01 | Old |
| 0.533325 | 0.171783 | 0.024 | 0.014 | 1 | 11 | Gm11696 | 0.01 | Old |
| 0.533325 | 0.074744 | 0.024 | 0.014 | 1 | 11 | Gstt1 | 0.01 | Old |
| 0.533994 | 0.035323 | 0.766 | 0.688 | 1 | 11 | Hnrnpa3 | 0.078 | Old |
| 0.534507 | 0.005415 | 0.637 | 0.536 | 1 | 11 | Ighm | 0.101 | Old |
| 0.534562 | 0.019939 | 0.077 | 0.058 | 1 | 11 | Kctd6 | 0.019 | Old |
| 0.534768 | 0.045579 | 0.101 | 0.08 | 1 | 11 | Mapkapk3 | 0.021 | Old |
| 0.543942 | 0.405776 | 0.153 | 0.13 | 1 | 11 | Mt2 | 0.023 | Old |
| 0.546722 | 0.923407 | 0.032 | 0.022 | 1 | 11 | 1110002J0 | 0.01 | Old |
| 0.546722 | 0.737211 | 0.032 | 0.022 | 1 | 11 | Riox1 | 0.01 | Old |
| 0.547547 | 0.070727 | 0.194 | 0.159 | 1 | 11 | Eif4e2 | 0.035 | Old |
| 0.548228 | 0.022012 | 0.694 | 0.659 | 1 | 11 | Atf3 | 0.035 | Old |
| 0.548922 | 0.505904 | 0.032 | 0.022 | 1 | 11 | Pomk | 0.01 | Old |
| 0.551126 | 1.022456 | 0.032 | 0.022 | 1 | 11 | Rint1 | 0.01 | Old |
| 0.551126 | 0.93192 | 0.032 | 0.022 | 1 | 11 | Ankrd54 | 0.01 | Old |
| 0.551395 | 0.062064 | 0.758 | 0.717 | 1 | 11 | Cox7c | 0.041 | Old |
| 0.552372 | 0.066171 | 0.008 | 0.014 | 1 | 11 | Gm47232 | -0.006 | Old |
| 0.553334 | 0.552702 | 0.032 | 0.022 | 1 | 11 | Snx33 | 0.01 | Old |
| 0.553334 | 0.530707 | 0.032 | 0.022 | 1 | 11 | Fance | 0.01 | Old |
| 0.553334 | 0.450771 | 0.032 | 0.022 | 1 | 11 | Susd1 | 0.01 | Old |
| 0.553334 | 0.223235 | 0.032 | 0.022 | 1 | 11 | Nkiras1 | 0.01 | Old |
| 0.555546 | 0.539402 | 0.032 | 0.022 | 1 | 11 | Nubpl | 0.01 | Old |
| 0.556006 | 0.411364 | 0.008 | 0.014 | 1 | 11 | Gm17089 | -0.006 | Old |

| | | | | | | | | |
|---|---|---|---|---|---|---|---|---|
| 0.556909 | 0.081943 | 0.238 | 0.203 | 1 | 11 | Phf11d | 0.035 | Old |
| 0.557763 | 0.571561 | 0.032 | 0.022 | 1 | 11 | Ssh3 | 0.01 | Old |
| 0.557763 | 0.25107 | 0.032 | 0.022 | 1 | 11 | Rab3ip | 0.01 | Old |
| 0.557763 | 0.081357 | 0.032 | 0.022 | 1 | 11 | Mus81 | 0.01 | Old |
| 0.559653 | 1.211061 | 0.008 | 0.014 | 1 | 11 | Dennd3 | -0.006 | Old |
| 0.559653 | 0.688702 | 0.008 | 0.014 | 1 | 11 | Nipsnap1 | -0.006 | Old |
| 0.559653 | 0.432581 | 0.008 | 0.014 | 1 | 11 | Edn1 | -0.006 | Old |
| 0.559653 | 0.13553 | 0.008 | 0.014 | 1 | 11 | Srfbp1 | -0.006 | Old |
| 0.559984 | 0.377739 | 0.032 | 0.022 | 1 | 11 | Nmnat1 | 0.01 | Old |
| 0.559984 | 0.075258 | 0.032 | 0.022 | 1 | 11 | Rsl1 | 0.01 | Old |
| 0.561771 | 0.173353 | 0.044 | 0.058 | 1 | 11 | Prkd2 | -0.014 | Old |
| 0.56221 | 0.337758 | 0.032 | 0.022 | 1 | 11 | Ndnf | 0.01 | Old |
| 0.56221 | 0.278409 | 0.032 | 0.022 | 1 | 11 | Inpp5e | 0.01 | Old |
| 0.56221 | 0.176478 | 0.032 | 0.022 | 1 | 11 | Dedd2 | 0.01 | Old |
| 0.56296 | 0.045977 | 0.835 | 0.717 | 1 | 11 | Rpl36a | 0.118 | Old |
| 0.563627 | 0.56248 | 0.04 | 0.029 | 1 | 11 | AC163018. | 0.011 | Old |
| 0.564439 | 0.305292 | 0.032 | 0.022 | 1 | 11 | Ptgs2os2 | 0.01 | Old |
| 0.564439 | 0.211026 | 0.032 | 0.022 | 1 | 11 | Gm49359 | 0.01 | Old |
| 0.564439 | 0.014733 | 0.032 | 0.022 | 1 | 11 | Gvin1 | 0.01 | Old |
| 0.564874 | 0.043977 | 0.754 | 0.638 | 1 | 11 | Txnip | 0.116 | Old |
| 0.565454 | 0.185938 | 0.125 | 0.101 | 1 | 11 | Rnpc3 | 0.024 | Old |
| 0.565613 | 0.583598 | 0.04 | 0.029 | 1 | 11 | Gin1 | 0.011 | Old |
| 0.566673 | 0.157465 | 0.032 | 0.022 | 1 | 11 | Cep152 | 0.01 | Old |
| 0.567603 | 1.095766 | 0.04 | 0.029 | 1 | 11 | Kif19a | 0.011 | Old |
| 0.569019 | 0.083359 | 0.786 | 0.732 | 1 | 11 | Cd52 | 0.054 | Old |
| 0.570795 | 0.314321 | 0.113 | 0.094 | 1 | 11 | Nde1 | 0.019 | Old |
| 0.571593 | 0.299865 | 0.04 | 0.029 | 1 | 11 | Coq3 | 0.011 | Old |
| 0.571593 | 0.033755 | 0.04 | 0.029 | 1 | 11 | A930007I1 | 0.011 | Old |
| 0.573033 | 0.495162 | 0.048 | 0.036 | 1 | 11 | Rfxank | 0.012 | Old |
| 0.573592 | 0.081076 | 0.032 | 0.043 | 1 | 11 | Dclre1b | -0.011 | Old |
| 0.573887 | 0.064063 | 0.089 | 0.072 | 1 | 11 | Plekhg2 | 0.017 | Old |
| 0.574857 | 0.810269 | 0.048 | 0.036 | 1 | 11 | Alg13 | 0.012 | Old |
| 0.576276 | 0.403966 | 0.056 | 0.043 | 1 | 11 | Polr1a | 0.013 | Old |
| 0.576684 | 0.579489 | 0.048 | 0.036 | 1 | 11 | Rnaseh1 | 0.012 | Old |
| 0.579611 | 0.060032 | 0.04 | 0.029 | 1 | 11 | Zfp788 | 0.011 | Old |
| 0.58004 | 0.365712 | 0.105 | 0.087 | 1 | 11 | Ankrd27 | 0.018 | Old |
| 0.58205 | 0.356199 | 0.089 | 0.072 | 1 | 11 | Psmd5 | 0.017 | Old |
| 0.583143 | 0.245429 | 0.073 | 0.058 | 1 | 11 | Fam160b2 | 0.015 | Old |
| 0.58364 | 0.204181 | 0.04 | 0.029 | 1 | 11 | Erc1 | 0.011 | Old |
| 0.584018 | 0.331484 | 0.048 | 0.036 | 1 | 11 | Parp6 | 0.012 | Old |
| 0.584171 | 0.187038 | 0.113 | 0.094 | 1 | 11 | Naip5 | 0.019 | Old |
| 0.584762 | 0.437808 | 0.056 | 0.043 | 1 | 11 | Actr5 | 0.013 | Old |
| 0.585414 | 0.052952 | 0.903 | 0.819 | 1 | 11 | Actr3 | 0.084 | Old |
| 0.587701 | 0.989403 | 0.048 | 0.036 | 1 | 11 | 0610009B2 | 0.012 | Old |
| 0.588112 | 0.154993 | 0.754 | 0.717 | 1 | 11 | Retnla | 0.037 | Old |

| | | | | | | | | |
|---|---|---|---|---|---|---|---|---|
| 0.589071 | 0.312675 | 0.113 | 0.094 | 1 | 11 | Chml | 0.019 | Old |
| 0.591737 | 0.01009 | 0.04 | 0.029 | 1 | 11 | Zfp101 | 0.011 | Old |
| 0.592026 | 0.414595 | 0.137 | 0.116 | 1 | 11 | Sass6 | 0.021 | Old |
| 0.592801 | 0.165405 | 0.065 | 0.051 | 1 | 11 | Chil3 | 0.014 | Old |
| 0.593307 | 0.281188 | 0.056 | 0.043 | 1 | 11 | Smyd3 | 0.013 | Old |
| 0.593713 | 0.335963 | 0.073 | 0.058 | 1 | 11 | Fam222b | 0.015 | Old |
| 0.593716 | 0.335327 | 0.02 | 0.029 | 1 | 11 | Anln | -0.009 | Old |
| 0.594396 | 0.250581 | 0.089 | 0.072 | 1 | 11 | Mlkl | 0.017 | Old |
| 0.595099 | 0.165287 | 0.048 | 0.036 | 1 | 11 | Tstd3 | 0.012 | Old |
| 0.59523 | 0.242462 | 0.073 | 0.058 | 1 | 11 | Cdk5 | 0.015 | Old |
| 0.59523 | 0.17621 | 0.073 | 0.058 | 1 | 11 | Spryd7 | 0.015 | Old |
| 0.596084 | 0.090536 | 0.081 | 0.065 | 1 | 11 | Ccdc112 | 0.016 | Old |
| 0.598766 | 0.004385 | 0.02 | 0.029 | 1 | 11 | Tcea3 | -0.009 | Old |
| 0.600849 | 0.170366 | 0.065 | 0.051 | 1 | 11 | Carnmt1 | 0.014 | Old |
| 0.601103 | 0.520566 | 0.181 | 0.159 | 1 | 11 | Clec4e | 0.022 | Old |
| 0.601907 | 0.033813 | 0.056 | 0.043 | 1 | 11 | Nadk2 | 0.013 | Old |
| 0.60254 | 0.2503 | 0.048 | 0.036 | 1 | 11 | Fahd1 | 0.012 | Old |
| 0.60358 | 0.028382 | 0.907 | 0.819 | 1 | 11 | Rpl5 | 0.088 | Old |
| 0.604119 | 0.043541 | 0.327 | 0.29 | 1 | 11 | Hivep2 | 0.037 | Old |
| 0.606377 | 1.032319 | 0.02 | 0.029 | 1 | 11 | Paqr4 | -0.009 | Old |
| 0.606607 | 0.118394 | 0.121 | 0.101 | 1 | 11 | Atp11c | 0.02 | Old |
| 0.607095 | 0.102989 | 0.056 | 0.043 | 1 | 11 | Supt7l | 0.013 | Old |
| 0.607095 | 0.082129 | 0.056 | 0.043 | 1 | 11 | Tma16 | 0.013 | Old |
| 0.608828 | 0.120077 | 0.056 | 0.043 | 1 | 11 | Atg2b | 0.013 | Old |
| 0.608945 | 0.068281 | 0.065 | 0.051 | 1 | 11 | Zc3h12c | 0.014 | Old |
| 0.60926 | 0.078694 | 0.899 | 0.841 | 1 | 11 | Ddx5 | 0.058 | Old |
| 0.611039 | 0.03777 | 0.089 | 0.072 | 1 | 11 | Gnpda2 | 0.017 | Old |
| 0.61306 | 0.296702 | 0.298 | 0.268 | 1 | 11 | Id3 | 0.03 | Old |
| 0.613525 | 0.073313 | 0.081 | 0.065 | 1 | 11 | Cgrrf1 | 0.016 | Old |
| 0.615492 | 0.04262 | 0.052 | 0.065 | 1 | 11 | Naip6 | -0.013 | Old |
| 0.620858 | 0.159487 | 0.081 | 0.065 | 1 | 11 | Vegfa | 0.016 | Old |
| 0.62356 | 0.081299 | 0.145 | 0.123 | 1 | 11 | Setdb1 | 0.022 | Old |
| 0.629482 | 0.025795 | 0.887 | 0.783 | 1 | 11 | Sem1 | 0.104 | Old |
| 0.634857 | 0.042182 | 0.145 | 0.123 | 1 | 11 | Spg20 | 0.022 | Old |
| 0.642076 | 0.025148 | 0.089 | 0.072 | 1 | 11 | Ipo9 | 0.017 | Old |
| 0.644165 | 0.0066 | 0.19 | 0.167 | 1 | 11 | Csf3r | 0.023 | Old |
| 0.645702 | 0.108233 | 0.794 | 0.804 | 1 | 11 | Ccl6 | -0.01 | Old |
| 0.64851 | 1.844674 | 0.012 | 0.007 | 1 | 11 | Fn3krp | 0.005 | Old |
| 0.64851 | 1.612294 | 0.012 | 0.007 | 1 | 11 | Isoc2a | 0.005 | Old |
| 0.64851 | 1.521591 | 0.012 | 0.007 | 1 | 11 | Aurkb | 0.005 | Old |
| 0.64851 | 1.401512 | 0.012 | 0.007 | 1 | 11 | Osgepl1 | 0.005 | Old |
| 0.64851 | 1.400276 | 0.012 | 0.007 | 1 | 11 | Ccno | 0.005 | Old |
| 0.64851 | 1.397966 | 0.012 | 0.007 | 1 | 11 | Frk | 0.005 | Old |
| 0.64851 | 1.222288 | 0.012 | 0.007 | 1 | 11 | Xlr3b | 0.005 | Old |
| 0.64851 | 1.187297 | 0.012 | 0.007 | 1 | 11 | Tmem202 | 0.005 | Old |

| | | | | | | | | |
|---|---|---|---|---|---|---|---|---|
| 0.64851 | 1.132398 | 0.012 | 0.007 | 1 | 11 | Plppr3 | 0.005 | Old |
| 0.64851 | 1.106751 | 0.012 | 0.007 | 1 | 11 | Ppef2 | 0.005 | Old |
| 0.64851 | 0.967852 | 0.012 | 0.007 | 1 | 11 | Zfp119b | 0.005 | Old |
| 0.64851 | 0.955276 | 0.012 | 0.007 | 1 | 11 | Ctla2a | 0.005 | Old |
| 0.64851 | 0.938837 | 0.012 | 0.007 | 1 | 11 | Mad2l1 | 0.005 | Old |
| 0.64851 | 0.920034 | 0.012 | 0.007 | 1 | 11 | Fcho1 | 0.005 | Old |
| 0.64851 | 0.919578 | 0.012 | 0.007 | 1 | 11 | Tmem82 | 0.005 | Old |
| 0.64851 | 0.789485 | 0.012 | 0.007 | 1 | 11 | Ccdc51 | 0.005 | Old |
| 0.64851 | 0.758247 | 0.012 | 0.007 | 1 | 11 | Mrm3 | 0.005 | Old |
| 0.64851 | 0.729306 | 0.012 | 0.007 | 1 | 11 | Htr1b | 0.005 | Old |
| 0.64851 | 0.651038 | 0.012 | 0.007 | 1 | 11 | 1700056N1 | 0.005 | Old |
| 0.64851 | 0.449474 | 0.012 | 0.007 | 1 | 11 | Plxnb3 | 0.005 | Old |
| 0.648857 | 0.030014 | 0.819 | 0.725 | 1 | 11 | Rhoa | 0.094 | Old |
| 0.652418 | 1.839757 | 0.012 | 0.007 | 1 | 11 | Mmp2 | 0.005 | Old |
| 0.652418 | 1.543402 | 0.012 | 0.007 | 1 | 11 | C030034L1 | 0.005 | Old |
| 0.652418 | 1.432518 | 0.012 | 0.007 | 1 | 11 | Gm13391 | 0.005 | Old |
| 0.652418 | 1.247332 | 0.012 | 0.007 | 1 | 11 | Gm20517 | 0.005 | Old |
| 0.652418 | 1.18873 | 0.012 | 0.007 | 1 | 11 | Acot1 | 0.005 | Old |
| 0.652418 | 1.158611 | 0.012 | 0.007 | 1 | 11 | E230032D2 | 0.005 | Old |
| 0.652418 | 1.06969 | 0.012 | 0.007 | 1 | 11 | 1700030K0 | 0.005 | Old |
| 0.652418 | 1.009917 | 0.012 | 0.007 | 1 | 11 | Mcm8 | 0.005 | Old |
| 0.652418 | 0.974907 | 0.012 | 0.007 | 1 | 11 | Kif22 | 0.005 | Old |
| 0.652418 | 0.967568 | 0.012 | 0.007 | 1 | 11 | Ccr9 | 0.005 | Old |
| 0.652418 | 0.871195 | 0.012 | 0.007 | 1 | 11 | Zfp870 | 0.005 | Old |
| 0.652418 | 0.824688 | 0.012 | 0.007 | 1 | 11 | Fignl2 | 0.005 | Old |
| 0.652418 | 0.798305 | 0.012 | 0.007 | 1 | 11 | Spc25 | 0.005 | Old |
| 0.652418 | 0.778746 | 0.012 | 0.007 | 1 | 11 | Rpl39l | 0.005 | Old |
| 0.652418 | 0.765497 | 0.012 | 0.007 | 1 | 11 | Ccdc125 | 0.005 | Old |
| 0.652418 | 0.750568 | 0.012 | 0.007 | 1 | 11 | 9130011E1 | 0.005 | Old |
| 0.652418 | 0.708604 | 0.012 | 0.007 | 1 | 11 | Il2 | 0.005 | Old |
| 0.652418 | 0.665293 | 0.012 | 0.007 | 1 | 11 | Ubxn11 | 0.005 | Old |
| 0.652418 | 0.657956 | 0.012 | 0.007 | 1 | 11 | Zfp748 | 0.005 | Old |
| 0.652418 | 0.616111 | 0.012 | 0.007 | 1 | 11 | Siglecg | 0.005 | Old |
| 0.652418 | 0.59679 | 0.012 | 0.007 | 1 | 11 | Rftn2 | 0.005 | Old |
| 0.652418 | 0.51098 | 0.012 | 0.007 | 1 | 11 | Zfp933 | 0.005 | Old |
| 0.652418 | 0.462954 | 0.012 | 0.007 | 1 | 11 | Dnajc17 | 0.005 | Old |
| 0.652418 | 0.388629 | 0.012 | 0.007 | 1 | 11 | Cage1 | 0.005 | Old |
| 0.652418 | 0.222585 | 0.012 | 0.007 | 1 | 11 | Igfbp7 | 0.005 | Old |
| 0.652616 | 0.043744 | 0.891 | 0.862 | 1 | 11 | Ahnak | 0.029 | Old |
| 0.653886 | 0.021172 | 0.105 | 0.087 | 1 | 11 | Smg6 | 0.018 | Old |
| 0.655361 | 0.197712 | 0.101 | 0.087 | 1 | 11 | Gypc | 0.014 | Old |
| 0.65539 | 0.080911 | 0.109 | 0.094 | 1 | 11 | Anapc10 | 0.015 | Old |
| 0.656335 | 2.075247 | 0.012 | 0.007 | 1 | 11 | Lif | 0.005 | Old |
| 0.656335 | 1.17961 | 0.012 | 0.007 | 1 | 11 | Eaf2 | 0.005 | Old |
| 0.656335 | 0.679989 | 0.012 | 0.007 | 1 | 11 | Fam185a | 0.005 | Old |

| | | | | | | | | |
|---|---|---|---|---|---|---|---|---|
| 0.656335 | 0.657151 | 0.012 | 0.007 | 1 | 11 | Plcg1 | 0.005 | Old |
| 0.656335 | 0.471294 | 0.012 | 0.007 | 1 | 11 | Cavin2 | 0.005 | Old |
| 0.656335 | 0.422717 | 0.012 | 0.007 | 1 | 11 | Lysmd2 | 0.005 | Old |
| 0.656335 | 0.397576 | 0.012 | 0.007 | 1 | 11 | Gm16638 | 0.005 | Old |
| 0.656335 | 0.363643 | 0.012 | 0.007 | 1 | 11 | P2ry1 | 0.005 | Old |
| 0.656335 | 0.332583 | 0.012 | 0.007 | 1 | 11 | Grik5 | 0.005 | Old |
| 0.656335 | 0.3281 | 0.012 | 0.007 | 1 | 11 | 9130221H1 | 0.005 | Old |
| 0.656335 | 0.304341 | 0.012 | 0.007 | 1 | 11 | 3-Sep | 0.005 | Old |
| 0.656335 | 0.288166 | 0.012 | 0.007 | 1 | 11 | Mrm2 | 0.005 | Old |
| 0.656335 | 0.27718 | 0.012 | 0.007 | 1 | 11 | Col4a1 | 0.005 | Old |
| 0.656335 | 0.236606 | 0.012 | 0.007 | 1 | 11 | Gm12166 | 0.005 | Old |
| 0.656335 | 0.230686 | 0.012 | 0.007 | 1 | 11 | Nmnat3 | 0.005 | Old |
| 0.656335 | 0.229828 | 0.012 | 0.007 | 1 | 11 | Rsph3b | 0.005 | Old |
| 0.656335 | 0.220629 | 0.012 | 0.007 | 1 | 11 | Phf21b | 0.005 | Old |
| 0.656335 | 0.220597 | 0.012 | 0.007 | 1 | 11 | A530013C2 | 0.005 | Old |
| 0.656335 | 0.173823 | 0.012 | 0.007 | 1 | 11 | Ccdc94 | 0.005 | Old |
| 0.656335 | 0.164542 | 0.012 | 0.007 | 1 | 11 | Pcbp4 | 0.005 | Old |
| 0.656335 | 0.152544 | 0.012 | 0.007 | 1 | 11 | Stbd1 | 0.005 | Old |
| 0.656335 | 0.143439 | 0.012 | 0.007 | 1 | 11 | B230217C1 | 0.005 | Old |
| 0.656335 | 0.11905 | 0.012 | 0.007 | 1 | 11 | Bhlhe41 | 0.005 | Old |
| 0.656335 | 0.091789 | 0.012 | 0.007 | 1 | 11 | Zfp692 | 0.005 | Old |
| 0.656335 | 0.081502 | 0.012 | 0.007 | 1 | 11 | Gm29291 | 0.005 | Old |
| 0.656335 | 0.077654 | 0.012 | 0.007 | 1 | 11 | Nphp1 | 0.005 | Old |
| 0.656335 | 0.06806 | 0.012 | 0.007 | 1 | 11 | Jag1 | 0.005 | Old |
| 0.656335 | 0.049712 | 0.012 | 0.007 | 1 | 11 | Tchp | 0.005 | Old |
| 0.657286 | 0.012898 | 0.556 | 0.478 | 1 | 11 | Eif3e | 0.078 | Old |
| 0.660261 | 0.139849 | 0.012 | 0.007 | 1 | 11 | Gm3435 | 0.005 | Old |
| 0.660261 | 0.089784 | 0.012 | 0.007 | 1 | 11 | Shpk | 0.005 | Old |
| 0.660261 | 0.089601 | 0.012 | 0.007 | 1 | 11 | Kbtbd8 | 0.005 | Old |
| 0.660261 | 0.036331 | 0.012 | 0.007 | 1 | 11 | Gm4117 | 0.005 | Old |
| 0.660261 | 0.021403 | 0.012 | 0.007 | 1 | 11 | Cenps | 0.005 | Old |
| 0.661269 | 0.998078 | 0.077 | 0.065 | 1 | 11 | Slc35e4 | 0.012 | Old |
| 0.667019 | 0.051976 | 0.815 | 0.71 | 1 | 11 | Ywhaz | 0.105 | Old |
| 0.677278 | 1.004716 | 0.044 | 0.036 | 1 | 11 | Ccdc69 | 0.008 | Old |
| 0.678102 | 0.131241 | 0.085 | 0.072 | 1 | 11 | Rrp15 | 0.013 | Old |
| 0.680657 | 0.765492 | 0.028 | 0.036 | 1 | 11 | Qsox2 | -0.008 | Old |
| 0.681444 | 0.01899 | 0.093 | 0.08 | 1 | 11 | Rpia | 0.013 | Old |
| 0.681744 | 0.240103 | 0.052 | 0.043 | 1 | 11 | Fam98a | 0.009 | Old |
| 0.681985 | 1.329951 | 0.02 | 0.014 | 1 | 11 | Cd177 | 0.006 | Old |
| 0.681985 | 1.238168 | 0.02 | 0.014 | 1 | 11 | Zfp931 | 0.006 | Old |
| 0.681985 | 1.062646 | 0.02 | 0.014 | 1 | 11 | Zfp580 | 0.006 | Old |
| 0.685008 | 1.576559 | 0.02 | 0.014 | 1 | 11 | Nemp2 | 0.006 | Old |
| 0.685008 | 0.924883 | 0.02 | 0.014 | 1 | 11 | F7 | 0.006 | Old |
| 0.685008 | 0.761291 | 0.02 | 0.014 | 1 | 11 | Wdr34 | 0.006 | Old |
| 0.685996 | 0.099298 | 0.077 | 0.065 | 1 | 11 | Ttyh2 | 0.012 | Old |

| | | | | | | | | |
|---|---|---|---|---|---|---|---|---|
| 0.688037 | 0.906799 | 0.02 | 0.014 | 1 | 11 | Asap2 | 0.006 | Old |
| 0.688037 | 0.728526 | 0.02 | 0.014 | 1 | 11 | Zfp7 | 0.006 | Old |
| 0.688037 | 0.379591 | 0.02 | 0.014 | 1 | 11 | Gm15247 | 0.006 | Old |
| 0.688037 | 0.154017 | 0.02 | 0.014 | 1 | 11 | Aldoc | 0.006 | Old |
| 0.688037 | 0.007899 | 0.02 | 0.014 | 1 | 11 | Itpr3 | 0.006 | Old |
| 0.68824 | 0.939745 | 0.028 | 0.022 | 1 | 11 | Nars2 | 0.006 | Old |
| 0.689419 | 0.511419 | 0.044 | 0.036 | 1 | 11 | Pde7b | 0.008 | Old |
| 0.690788 | 0.729812 | 0.028 | 0.022 | 1 | 11 | Fam167b | 0.006 | Old |
| 0.690852 | 0.023234 | 0.109 | 0.094 | 1 | 11 | Larp1b | 0.015 | Old |
| 0.691071 | 0.925941 | 0.02 | 0.014 | 1 | 11 | 1810011H1 | 0.006 | Old |
| 0.691071 | 0.613966 | 0.02 | 0.014 | 1 | 11 | Shmt1 | 0.006 | Old |
| 0.691071 | 0.442387 | 0.02 | 0.014 | 1 | 11 | Glmn | 0.006 | Old |
| 0.691071 | 0.358327 | 0.02 | 0.014 | 1 | 11 | Suox | 0.006 | Old |
| 0.691071 | 0.216447 | 0.02 | 0.014 | 1 | 11 | Vangl2 | 0.006 | Old |
| 0.691071 | 0.208562 | 0.02 | 0.014 | 1 | 11 | Prr5 | 0.006 | Old |
| 0.691071 | 0.184633 | 0.02 | 0.014 | 1 | 11 | Efcab2 | 0.006 | Old |
| 0.691081 | 0.350679 | 0.052 | 0.043 | 1 | 11 | Mcm7 | 0.009 | Old |
| 0.692071 | 0.056651 | 0.069 | 0.058 | 1 | 11 | Gpatch1 | 0.011 | Old |
| 0.693484 | 0.667198 | 0.044 | 0.036 | 1 | 11 | Tsr2 | 0.008 | Old |
| 0.69411 | 1.529911 | 0.02 | 0.014 | 1 | 11 | Zfp707 | 0.006 | Old |
| 0.69411 | 0.578714 | 0.02 | 0.014 | 1 | 11 | Ptar1 | 0.006 | Old |
| 0.69411 | 0.469282 | 0.02 | 0.014 | 1 | 11 | Dtwd1 | 0.006 | Old |
| 0.69411 | 0.36949 | 0.02 | 0.014 | 1 | 11 | Chd3os | 0.006 | Old |
| 0.69411 | 0.277454 | 0.02 | 0.014 | 1 | 11 | Ntmt1 | 0.006 | Old |
| 0.69411 | 0.262422 | 0.02 | 0.014 | 1 | 11 | Lyrm1 | 0.006 | Old |
| 0.69411 | 0.26105 | 0.02 | 0.014 | 1 | 11 | Praf2 | 0.006 | Old |
| 0.69411 | 0.071178 | 0.02 | 0.014 | 1 | 11 | Zfp119a | 0.006 | Old |
| 0.697154 | 0.79083 | 0.02 | 0.014 | 1 | 11 | Marco | 0.006 | Old |
| 0.697154 | 0.30373 | 0.02 | 0.014 | 1 | 11 | Zfp658 | 0.006 | Old |
| 0.697154 | 0.203491 | 0.02 | 0.014 | 1 | 11 | Cfap45 | 0.006 | Old |
| 0.697154 | 0.163363 | 0.02 | 0.014 | 1 | 11 | Gm30025 | 0.006 | Old |
| 0.697154 | 0.142259 | 0.02 | 0.014 | 1 | 11 | Elovl6 | 0.006 | Old |
| 0.697154 | 0.129993 | 0.02 | 0.014 | 1 | 11 | Mtrf1 | 0.006 | Old |
| 0.697154 | 0.117039 | 0.02 | 0.014 | 1 | 11 | Zkscan7 | 0.006 | Old |
| 0.697154 | 0.116435 | 0.02 | 0.014 | 1 | 11 | 4930532G1 | 0.006 | Old |
| 0.697154 | 0.038555 | 0.02 | 0.014 | 1 | 11 | Lca5 | 0.006 | Old |
| 0.697558 | 0.641868 | 0.044 | 0.036 | 1 | 11 | Ube2o | 0.008 | Old |
| 0.697558 | 0.30327 | 0.044 | 0.036 | 1 | 11 | Taf8 | 0.008 | Old |
| 0.697558 | 0.252422 | 0.044 | 0.036 | 1 | 11 | Tyw5 | 0.008 | Old |
| 0.697776 | 0.996954 | 0.036 | 0.029 | 1 | 11 | Eefsec | 0.007 | Old |
| 0.697776 | 0.491059 | 0.036 | 0.029 | 1 | 11 | Rbbp9 | 0.007 | Old |
| 0.697776 | 0.159266 | 0.036 | 0.029 | 1 | 11 | BC002059 | 0.007 | Old |
| 0.697812 | 0.553284 | 0.113 | 0.101 | 1 | 11 | Atl3 | 0.012 | Old |
| 0.698453 | 0.206469 | 0.028 | 0.022 | 1 | 11 | Kiz | 0.006 | Old |
| 0.698489 | 0.052031 | 0.077 | 0.065 | 1 | 11 | Exosc7 | 0.012 | Old |

| | | | | | | | | |
|---|---|---|---|---|---|---|---|---|
| 0.699375 | 0.225174 | 0.06 | 0.051 | 1 | 11 | Ripor1 | 0.009 | Old |
| 0.699867 | 0.043651 | 0.988 | 0.993 | 1 | 11 | Lyz2 | -0.005 | Old |
| 0.700202 | 0.183747 | 0.016 | 0.022 | 1 | 11 | Tesk2 | -0.006 | Old |
| 0.700202 | 0.036744 | 0.016 | 0.022 | 1 | 11 | Slc46a1 | -0.006 | Old |
| 0.700358 | 0.024295 | 0.815 | 0.754 | 1 | 11 | Rtn4 | 0.061 | Old |
| 0.701016 | 1.222727 | 0.028 | 0.022 | 1 | 11 | Ccr7 | 0.006 | Old |
| 0.701016 | 0.488154 | 0.028 | 0.022 | 1 | 11 | Tnk2 | 0.006 | Old |
| 0.701016 | 0.216155 | 0.028 | 0.022 | 1 | 11 | 4930430F0 | 0.006 | Old |
| 0.701016 | 0.157708 | 0.028 | 0.022 | 1 | 11 | Gm41307 | 0.006 | Old |
| 0.701016 | 0.039303 | 0.028 | 0.022 | 1 | 11 | Cnst | 0.006 | Old |
| 0.701728 | 0.437802 | 0.016 | 0.022 | 1 | 11 | Tmlhe | -0.006 | Old |
| 0.702289 | 0.309868 | 0.036 | 0.029 | 1 | 11 | Efemp2 | 0.007 | Old |
| 0.703256 | 0.148622 | 0.016 | 0.022 | 1 | 11 | Fam189b | -0.006 | Old |
| 0.703581 | 0.901395 | 0.028 | 0.022 | 1 | 11 | Zfp768 | 0.006 | Old |
| 0.703581 | 0.50164 | 0.028 | 0.022 | 1 | 11 | 1700096K1 | 0.006 | Old |
| 0.703581 | 0.448444 | 0.028 | 0.022 | 1 | 11 | Peli2 | 0.006 | Old |
| 0.703581 | 0.31988 | 0.028 | 0.022 | 1 | 11 | Zkscan17 | 0.006 | Old |
| 0.703581 | 0.245384 | 0.028 | 0.022 | 1 | 11 | Gm47675 | 0.006 | Old |
| 0.704232 | 0.041597 | 0.052 | 0.043 | 1 | 11 | Zbtb43 | 0.009 | Old |
| 0.704767 | 0.008858 | 0.077 | 0.065 | 1 | 11 | Dnajc25 | 0.012 | Old |
| 0.705082 | 0.101739 | 0.802 | 0.696 | 1 | 11 | Arpc5 | 0.106 | Old |
| 0.705732 | 0.23682 | 0.044 | 0.036 | 1 | 11 | Paqr7 | 0.008 | Old |
| 0.70615 | 0.377364 | 0.028 | 0.022 | 1 | 11 | Fam136a | 0.006 | Old |
| 0.70615 | 0.038308 | 0.028 | 0.022 | 1 | 11 | Large1 | 0.006 | Old |
| 0.706412 | 0.122118 | 0.085 | 0.072 | 1 | 11 | Emc4 | 0.013 | Old |
| 0.706812 | 0.236198 | 0.036 | 0.029 | 1 | 11 | Slc37a1 | 0.007 | Old |
| 0.707781 | 0.135592 | 0.044 | 0.036 | 1 | 11 | Bnip1 | 0.008 | Old |
| 0.708006 | 0.032772 | 0.052 | 0.043 | 1 | 11 | Itpkc | 0.009 | Old |
| 0.708167 | 0.160703 | 0.06 | 0.051 | 1 | 11 | Tmem94 | 0.009 | Old |
| 0.708722 | 1.605738 | 0.028 | 0.022 | 1 | 11 | Grap2 | 0.006 | Old |
| 0.708722 | 0.246328 | 0.028 | 0.022 | 1 | 11 | Rdh13 | 0.006 | Old |
| 0.708722 | 0.030919 | 0.028 | 0.022 | 1 | 11 | Card11 | 0.006 | Old |
| 0.711298 | 0.060207 | 0.028 | 0.022 | 1 | 11 | Mmp12 | 0.006 | Old |
| 0.711345 | 0.419058 | 0.036 | 0.029 | 1 | 11 | 2310057M | 0.007 | Old |
| 0.711926 | 0.22173 | 0.069 | 0.058 | 1 | 11 | Supt3 | 0.011 | Old |
| 0.71376 | 0.043843 | 0.77 | 0.688 | 1 | 11 | Ifi27l2a | 0.082 | Old |
| 0.713876 | 0.071221 | 0.028 | 0.022 | 1 | 11 | Ccp110 | 0.006 | Old |
| 0.71394 | 0.700745 | 0.044 | 0.036 | 1 | 11 | Fancg | 0.008 | Old |
| 0.714713 | 0.186853 | 0.048 | 0.058 | 1 | 11 | Zxdc | -0.01 | Old |
| 0.715575 | 0.005034 | 0.052 | 0.043 | 1 | 11 | Mtx3 | 0.009 | Old |
| 0.715889 | 0.064956 | 0.036 | 0.029 | 1 | 11 | Bivm | 0.007 | Old |
| 0.715889 | 0.024683 | 0.036 | 0.029 | 1 | 11 | Bcl11a | 0.007 | Old |
| 0.718164 | 0.078491 | 0.036 | 0.029 | 1 | 11 | Spats2 | 0.007 | Old |
| 0.719479 | 0.068211 | 0.54 | 0.507 | 1 | 11 | Cfh | 0.033 | Old |
| 0.720542 | 0.052587 | 0.06 | 0.051 | 1 | 11 | Cd24a | 0.009 | Old |

| | | | | | | | | |
|---|---|---|---|---|---|---|---|---|
| 0.722723 | 0.348631 | 0.036 | 0.029 | 1 | 11 | Neurl1b | 0.007 | Old |
| 0.722723 | 0.100394 | 0.036 | 0.029 | 1 | 11 | Polr3a | 0.007 | Old |
| 0.724247 | 0.04202 | 0.044 | 0.036 | 1 | 11 | Gm12840 | 0.008 | Old |
| 0.730445 | 0.068724 | 0.056 | 0.065 | 1 | 11 | Scaper | -0.009 | Old |
| 0.735674 | 0.019731 | 0.597 | 0.514 | 1 | 11 | Hnrnpm | 0.083 | Old |
| 0.738179 | 0.099033 | 0.077 | 0.087 | 1 | 11 | Fkrp | -0.01 | Old |
| 0.743396 | 0.01543 | 0.399 | 0.362 | 1 | 11 | Tiparp | 0.037 | Old |
| 0.743899 | 0.090742 | 0.105 | 0.094 | 1 | 11 | Tlr6 | 0.011 | Old |
| 0.747495 | 0.033835 | 0.754 | 0.696 | 1 | 11 | Cotl1 | 0.058 | Old |
| 0.766775 | 0.002634 | 0.27 | 0.239 | 1 | 11 | Higd1a | 0.031 | Old |
| 0.778816 | 0.31154 | 0.113 | 0.101 | 1 | 11 | Haus8 | 0.012 | Old |
| 0.784618 | 0.039241 | 0.181 | 0.159 | 1 | 11 | Taok2 | 0.022 | Old |
| 0.794611 | 0.860825 | 0.056 | 0.051 | 1 | 11 | Crcp | 0.005 | Old |
| 0.798003 | 0.123324 | 0.758 | 0.659 | 1 | 11 | Tcf4 | 0.099 | Old |
| 0.798611 | 0.008252 | 0.952 | 0.884 | 1 | 11 | Rps19 | 0.068 | Old |
| 0.805521 | 0.628918 | 0.048 | 0.043 | 1 | 11 | Zfp141 | 0.005 | Old |
| 0.807714 | 0.249474 | 0.056 | 0.051 | 1 | 11 | Klrb1b | 0.005 | Old |
| 0.810382 | 0.124865 | 0.065 | 0.058 | 1 | 11 | Gm4070 | 0.007 | Old |
| 0.810382 | 0.033893 | 0.065 | 0.058 | 1 | 11 | Zbtb46 | 0.007 | Old |
| 0.81604 | 0.715823 | 0.109 | 0.116 | 1 | 11 | Gt(ROSA)2( | -0.007 | Old |
| 0.819669 | 0.307976 | 0.048 | 0.043 | 1 | 11 | Gm5150 | 0.005 | Old |
| 0.820978 | 0.147284 | 0.065 | 0.058 | 1 | 11 | Fam13b | 0.007 | Old |
| 0.821695 | 0.087358 | 0.048 | 0.043 | 1 | 11 | Copz2 | 0.005 | Old |
| 0.821695 | 0.08287 | 0.048 | 0.043 | 1 | 11 | Iqce | 0.005 | Old |
| 0.827144 | 0.044851 | 0.081 | 0.072 | 1 | 11 | 1600010M | 0.009 | Old |
| 0.828304 | 0.010787 | 0.911 | 0.754 | 1 | 11 | Sdcbp | 0.157 | Old |
| 0.829811 | 0.299684 | 0.048 | 0.043 | 1 | 11 | Eif2ak4 | 0.005 | Old |
| 0.829985 | 0.143321 | 0.032 | 0.036 | 1 | 11 | Slc30a6 | -0.004 | Old |
| 0.831201 | 0.114881 | 0.415 | 0.377 | 1 | 11 | Fcgr4 | 0.038 | Old |
| 0.833876 | 0.002242 | 0.048 | 0.043 | 1 | 11 | Prmt9 | 0.005 | Old |
| 0.836007 | 0.046195 | 0.117 | 0.109 | 1 | 11 | Unc119 | 0.008 | Old |
| 0.843961 | 0.074106 | 0.012 | 0.014 | 1 | 11 | Pstpip2 | -0.002 | Old |
| 0.843961 | 0.054406 | 0.012 | 0.014 | 1 | 11 | Epas1 | -0.002 | Old |
| 0.843961 | 0.00295 | 0.012 | 0.014 | 1 | 11 | Taf4b | -0.002 | Old |
| 0.844261 | 0.090722 | 0.032 | 0.036 | 1 | 11 | Rcc1 | -0.004 | Old |
| 0.845626 | 0.999289 | 0.032 | 0.029 | 1 | 11 | Hdac10 | 0.003 | Old |
| 0.847766 | 0.659816 | 0.012 | 0.014 | 1 | 11 | Zfp341 | -0.002 | Old |
| 0.847766 | 0.44097 | 0.012 | 0.014 | 1 | 11 | Unc13a | -0.002 | Old |
| 0.847766 | 0.377705 | 0.012 | 0.014 | 1 | 11 | Kif20b | -0.002 | Old |
| 0.847766 | 0.109695 | 0.012 | 0.014 | 1 | 11 | Adamtsl4 | -0.002 | Old |
| 0.847766 | 0.061188 | 0.012 | 0.014 | 1 | 11 | Ift122 | -0.002 | Old |
| 0.847766 | 0.007535 | 0.012 | 0.014 | 1 | 11 | D830025C( | -0.002 | Old |
| 0.847828 | 0.246977 | 0.04 | 0.036 | 1 | 11 | Prmt6 | 0.004 | Old |
| 0.848105 | 0.831091 | 0.032 | 0.029 | 1 | 11 | Negr1 | 0.003 | Old |
| 0.850056 | 0.014414 | 0.04 | 0.036 | 1 | 11 | Herc3 | 0.004 | Old |

| | | | | | | | | |
|---|---|---|---|---|---|---|---|---|
| 0.850586 | 0.651339 | 0.032 | 0.029 | 1 | 11 | Fastkd5 | 0.003 | Old |
| 0.851575 | 0.110837 | 0.012 | 0.014 | 1 | 11 | Nsl1 | -0.002 | Old |
| 0.853068 | 0.333564 | 0.032 | 0.029 | 1 | 11 | Gm42047 | 0.003 | Old |
| 0.85356 | 0.21593 | 0.117 | 0.109 | 1 | 11 | Gm2000 | 0.008 | Old |
| 0.855387 | 1.299031 | 0.012 | 0.014 | 1 | 11 | Cryz | -0.002 | Old |
| 0.855387 | 0.531202 | 0.012 | 0.014 | 1 | 11 | Barx2 | -0.002 | Old |
| 0.85687 | 0.24576 | 0.278 | 0.261 | 1 | 11 | Fabp4 | 0.017 | Old |
| 0.858037 | 0.488625 | 0.032 | 0.029 | 1 | 11 | Tmem154 | 0.003 | Old |
| 0.858588 | 0.589053 | 0.032 | 0.036 | 1 | 11 | Slc7a11 | -0.004 | Old |
| 0.860524 | 0.460962 | 0.032 | 0.029 | 1 | 11 | Uck2 | 0.003 | Old |
| 0.860524 | 0.13007 | 0.032 | 0.029 | 1 | 11 | Tmem126b | 0.003 | Old |
| 0.860524 | 0.088747 | 0.032 | 0.029 | 1 | 11 | Snx21 | 0.003 | Old |
| 0.863012 | 0.253578 | 0.032 | 0.029 | 1 | 11 | Zfp637 | 0.003 | Old |
| 0.86324 | 0.002392 | 0.657 | 0.558 | 1 | 11 | Clk1 | 0.099 | Old |
| 0.865501 | 0.126231 | 0.032 | 0.029 | 1 | 11 | Hemk1 | 0.003 | Old |
| 0.870484 | 0.135568 | 0.032 | 0.029 | 1 | 11 | Gm20703 | 0.003 | Old |
| 0.871211 | 0.493005 | 0.069 | 0.065 | 1 | 11 | Ppp1r9a | 0.004 | Old |
| 0.871274 | 0.625331 | 0.024 | 0.022 | 1 | 11 | Rab23 | 0.002 | Old |
| 0.871274 | 0.355919 | 0.024 | 0.022 | 1 | 11 | Mamld1 | 0.002 | Old |
| 0.872977 | 0.38189 | 0.032 | 0.029 | 1 | 11 | Sdhaf3 | 0.003 | Old |
| 0.872977 | 0.067462 | 0.032 | 0.029 | 1 | 11 | Rpap3 | 0.003 | Old |
| 0.874142 | 0.423486 | 0.024 | 0.022 | 1 | 11 | Zfp874a | 0.002 | Old |
| 0.874142 | 0.224745 | 0.024 | 0.022 | 1 | 11 | Bbs12 | 0.002 | Old |
| 0.874475 | 0.105911 | 0.31 | 0.319 | 1 | 11 | Oas3 | -0.009 | Old |
| 0.879884 | 0.652526 | 0.024 | 0.022 | 1 | 11 | Impa2 | 0.002 | Old |
| 0.882757 | 0.057312 | 0.024 | 0.022 | 1 | 11 | Cad | 0.002 | Old |
| 0.885632 | 0.609108 | 0.024 | 0.022 | 1 | 11 | Tspan7 | 0.002 | Old |
| 0.888713 | 0.877709 | 0.016 | 0.014 | 1 | 11 | Map7d3 | 0.002 | Old |
| 0.888713 | 0.481013 | 0.016 | 0.014 | 1 | 11 | Bcl7a | 0.002 | Old |
| 0.891696 | 0.277777 | 0.109 | 0.101 | 1 | 11 | Adam8 | 0.008 | Old |
| 0.892224 | 1.007383 | 0.016 | 0.014 | 1 | 11 | Ttc30a1 | 0.002 | Old |
| 0.892328 | 0.088639 | 0.085 | 0.08 | 1 | 11 | Slc2a6 | 0.005 | Old |
| 0.895737 | 0.898661 | 0.016 | 0.014 | 1 | 11 | Acpp | 0.002 | Old |
| 0.895737 | 0.59069 | 0.016 | 0.014 | 1 | 11 | Ogg1 | 0.002 | Old |
| 0.895737 | 0.51905 | 0.016 | 0.014 | 1 | 11 | Slc25a43 | 0.002 | Old |
| 0.895737 | 0.377388 | 0.016 | 0.014 | 1 | 11 | Polr1b | 0.002 | Old |
| 0.895737 | 0.358981 | 0.016 | 0.014 | 1 | 11 | 8030462N1 | 0.002 | Old |
| 0.898898 | 0.030888 | 0.101 | 0.094 | 1 | 11 | Cfap20 | 0.007 | Old |
| 0.899252 | 2.029883 | 0.016 | 0.014 | 1 | 11 | Cysltr2 | 0.002 | Old |
| 0.899252 | 0.521306 | 0.016 | 0.014 | 1 | 11 | Gpc1 | 0.002 | Old |
| 0.899252 | 0.447026 | 0.016 | 0.014 | 1 | 11 | Chaf1a | 0.002 | Old |
| 0.899252 | 0.437734 | 0.016 | 0.014 | 1 | 11 | Zbtb3 | 0.002 | Old |
| 0.899252 | 0.396796 | 0.016 | 0.014 | 1 | 11 | Armcx6 | 0.002 | Old |
| 0.899252 | 0.299096 | 0.016 | 0.014 | 1 | 11 | D930030I0 | 0.002 | Old |
| 0.899252 | 0.284307 | 0.016 | 0.014 | 1 | 11 | Card6 | 0.002 | Old |

| | | | | | | | | |
|---|---|---|---|---|---|---|---|---|
| 0.899252 | 0.219796 | 0.016 | 0.014 | 1 | 11 | Glcci1 | 0.002 | Old |
| 0.899252 | 0.076717 | 0.016 | 0.014 | 1 | 11 | Fam198b | 0.002 | Old |
| 0.902769 | 0.899153 | 0.016 | 0.014 | 1 | 11 | Lum | 0.002 | Old |
| 0.902769 | 0.309853 | 0.016 | 0.014 | 1 | 11 | Kcne3 | 0.002 | Old |
| 0.902769 | 0.209137 | 0.016 | 0.014 | 1 | 11 | 1810009A1 | 0.002 | Old |
| 0.902769 | 0.198036 | 0.016 | 0.014 | 1 | 11 | Homez | 0.002 | Old |
| 0.902769 | 0.074787 | 0.016 | 0.014 | 1 | 11 | Opn3 | 0.002 | Old |
| 0.902769 | 0.032892 | 0.016 | 0.014 | 1 | 11 | Pdia5 | 0.002 | Old |
| 0.904444 | 0.00033 | 0.895 | 0.841 | 1 | 11 | Tmbim6 | 0.054 | Old |
| 0.906288 | 0.097243 | 0.016 | 0.014 | 1 | 11 | 1700029I1 | 0.002 | Old |
| 0.906288 | 0.056326 | 0.016 | 0.014 | 1 | 11 | C1s1 | 0.002 | Old |
| 0.910935 | 0.01673 | 0.02 | 0.022 | 1 | 11 | Crip2 | -0.002 | Old |
| 0.913994 | 0.636925 | 0.02 | 0.022 | 1 | 11 | Elavl4 | -0.002 | Old |
| 0.917055 | 0.090319 | 0.02 | 0.022 | 1 | 11 | Amotl2 | -0.002 | Old |
| 0.917587 | 0.043191 | 0.105 | 0.101 | 1 | 11 | C1galt1c1 | 0.004 | Old |
| 0.918091 | 0.018551 | 0.048 | 0.051 | 1 | 11 | Bend4 | -0.003 | Old |
| 0.920116 | 0.200281 | 0.02 | 0.022 | 1 | 11 | Aff3 | -0.002 | Old |
| 0.920529 | 0.066412 | 0.161 | 0.159 | 1 | 11 | Klhdc3 | 0.002 | Old |
| 0.923179 | 1.656155 | 0.02 | 0.022 | 1 | 11 | Platr17 | -0.002 | Old |
| 0.926243 | 0.216923 | 0.02 | 0.022 | 1 | 11 | Cetn4 | -0.002 | Old |
| 0.929308 | 0.628532 | 0.02 | 0.022 | 1 | 11 | Trmo | -0.002 | Old |
| 0.930669 | 0.164402 | 0.06 | 0.058 | 1 | 11 | Snhg9 | 0.002 | Old |
| 0.941306 | 0.202145 | 0.044 | 0.043 | 1 | 11 | Mettl7a1 | 0.001 | Old |
| 0.944982 | 0.015132 | 0.984 | 0.957 | 1 | 11 | Jund | 0.027 | Old |
| 0.948373 | 0.485131 | 0.048 | 0.051 | 1 | 11 | Myom1 | -0.003 | Old |
| 0.953915 | 0.511258 | 0.028 | 0.029 | 1 | 11 | Chek2 | -0.001 | Old |
| 0.954096 | 0.594306 | 0.044 | 0.043 | 1 | 11 | Ank2 | 0.001 | Old |
| 0.956546 | 0.010044 | 0.028 | 0.029 | 1 | 11 | Atpaf2 | -0.001 | Old |
| 0.961809 | 0.200271 | 0.028 | 0.029 | 1 | 11 | Gstp3 | -0.001 | Old |
| 0.962629 | 0.489381 | 0.044 | 0.043 | 1 | 11 | Ddah2 | 0.001 | Old |
| 0.96442 | 0.132633 | 0.052 | 0.051 | 1 | 11 | Psmf1 | 0.001 | Old |
| 0.965041 | 0.298821 | 0.056 | 0.058 | 1 | 11 | Tab3 | -0.002 | Old |
| 0.974212 | 0.363641 | 0.036 | 0.036 | 1 | 11 | Ms4a4b | 0 | Old |
| 0.980398 | 0.335277 | 0.101 | 0.101 | 1 | 11 | Cxcl1 | 0 | Old |
| 0.9924 | 0.159682 | 0.073 | 0.072 | 1 | 11 | 1500011B0 | 0.001 | Old |
| 0.99531 | 0.598064 | 0.036 | 0.036 | 1 | 11 | Slc25a13 | 0 | Old |
| 4.37E-07 | 0.527491 | 0.971 | 0.972 | 0.013575 | 11 | Marcks | -0.001 | Young |
| 4.55E-07 | 5.405029 | 0.123 | 0.008 | 0.014144 | 11 | Pmepa1 | 0.115 | Young |
| 1.73E-06 | 0.864389 | 0.891 | 0.851 | 0.053575 | 11 | Cd83 | 0.04 | Young |
| 2.56E-06 | 1.569188 | 0.493 | 0.323 | 0.079482 | 11 | Gatm | 0.17 | Young |
| 0.000285 | 4.63397 | 0.065 | 0.004 | 1 | 11 | Col14a1 | 0.061 | Young |
| 0.000286 | 0.962374 | 0.616 | 0.532 | 1 | 11 | Axl | 0.084 | Young |
| 0.000317 | 1.296795 | 0.493 | 0.355 | 1 | 11 | Hspa1b | 0.138 | Young |
| 0.000398 | 1.468367 | 0.261 | 0.129 | 1 | 11 | Ccnd2 | 0.132 | Young |
| 0.000429 | 0.743032 | 0.659 | 0.528 | 1 | 11 | Il1b | 0.131 | Young |

| | | | | | | | | |
|---|---|---|---|---|---|---|---|---|
| 0.000503 | 0.940862 | 0.587 | 0.48 | 1 | 11 | Skil | 0.107 | Young |
| 0.000567 | 1.89866 | 0.21 | 0.093 | 1 | 11 | Arsb | 0.117 | Young |
| 0.000642 | 0.31308 | 0.986 | 0.984 | 1 | 11 | Rps9 | 0.002 | Young |
| 0.000803 | 2.584568 | 0.058 | 0.004 | 1 | 11 | Tln2 | 0.054 | Young |
| 0.000819 | 1.640069 | 0.312 | 0.19 | 1 | 11 | Hpgd | 0.122 | Young |
| 0.000957 | 4.084735 | 0.043 | 0 | 1 | 11 | Tnfsf8 | 0.043 | Young |
| 0.00126 | 0.374345 | 0.942 | 0.972 | 1 | 11 | Rps14 | -0.03 | Young |
| 0.001271 | 0.26877 | 0.964 | 0.976 | 1 | 11 | Rps23 | -0.012 | Young |
| 0.001316 | 0.253498 | 0.971 | 0.984 | 1 | 11 | Rps24 | -0.013 | Young |
| 0.002503 | 0.340539 | 0.935 | 0.935 | 1 | 11 | Rps16 | 0 | Young |
| 0.00302 | 0.033308 | 0.167 | 0.335 | 1 | 11 | Klhl24 | -0.168 | Young |
| 0.003262 | 2.223393 | 0.058 | 0.008 | 1 | 11 | Gm26601 | 0.05 | Young |
| 0.003309 | 0.375853 | 0.986 | 0.972 | 1 | 11 | Cst3 | 0.014 | Young |
| 0.003449 | 1.579701 | 0.058 | 0.008 | 1 | 11 | Cfd | 0.05 | Young |
| 0.003461 | 0.499237 | 0.942 | 0.903 | 1 | 11 | Apoe | 0.039 | Young |
| 0.003538 | 0.535672 | 0.065 | 0.181 | 1 | 11 | Zswim8 | -0.116 | Young |
| 0.003545 | 0.176519 | 1 | 1 | 1 | 11 | Gm42418 | 0 | Young |
| 0.003733 | 0.22283 | 0.08 | 0.198 | 1 | 11 | Mrpl42 | -0.118 | Young |
| 0.003767 | 1.548172 | 0.254 | 0.153 | 1 | 11 | 1700017B0 | 0.101 | Young |
| 0.003863 | 0.805281 | 0.594 | 0.548 | 1 | 11 | Mef2c | 0.046 | Young |
| 0.00436 | 0.706901 | 0.58 | 0.504 | 1 | 11 | Rassf4 | 0.076 | Young |
| 0.004371 | 0.028679 | 0.08 | 0.198 | 1 | 11 | Mgst1 | -0.118 | Young |
| 0.004513 | 0.160952 | 0.964 | 0.976 | 1 | 11 | Rps12 | -0.012 | Young |
| 0.004564 | 0.509313 | 0.833 | 0.863 | 1 | 11 | Nfkbiz | -0.03 | Young |
| 0.004607 | 0.9031 | 0.609 | 0.536 | 1 | 11 | Cd36 | 0.073 | Young |
| 0.005042 | 1.000724 | 0.42 | 0.29 | 1 | 11 | Ifit2 | 0.13 | Young |
| 0.005129 | 0.709782 | 0.638 | 0.544 | 1 | 11 | Cxcl2 | 0.094 | Young |
| 0.006169 | 0.461865 | 0.109 | 0.23 | 1 | 11 | Mogs | -0.121 | Young |
| 0.007045 | 0.199597 | 0.986 | 0.988 | 1 | 11 | Rpl35a | -0.002 | Young |
| 0.007169 | 3.470716 | 0.029 | 0 | 1 | 11 | Il2ra | 0.029 | Young |
| 0.007169 | 3.352491 | 0.029 | 0 | 1 | 11 | Mttp | 0.029 | Young |
| 0.007688 | 1.529276 | 0.297 | 0.198 | 1 | 11 | Tnfsf9 | 0.099 | Young |
| 0.007817 | 1.101102 | 0.391 | 0.294 | 1 | 11 | Cx3cr1 | 0.097 | Young |
| 0.007967 | 0.188868 | 0.065 | 0.169 | 1 | 11 | Mapk8 | -0.104 | Young |
| 0.008147 | 0.382522 | 0.514 | 0.411 | 1 | 11 | Hspa1a | 0.103 | Young |
| 0.008679 | 0.255916 | 0.558 | 0.782 | 1 | 11 | Prdx1 | -0.224 | Young |
| 0.009149 | 2.868766 | 0.08 | 0.024 | 1 | 11 | Sparc | 0.056 | Young |
| 0.009287 | 0.344876 | 0.891 | 0.887 | 1 | 11 | Rpl26 | 0.004 | Young |
| 0.009507 | 0.282612 | 0.116 | 0.242 | 1 | 11 | Batf2 | -0.126 | Young |
| 0.00966 | 0.056076 | 0.13 | 0.266 | 1 | 11 | Ap1g1 | -0.136 | Young |
| 0.00969 | 0.274338 | 0.957 | 0.976 | 1 | 11 | Fau | -0.019 | Young |
| 0.009745 | 0.228319 | 0.993 | 0.988 | 1 | 11 | Rplp2 | 0.005 | Young |
| 0.009878 | 1.079542 | 0.522 | 0.472 | 1 | 11 | Hexb | 0.05 | Young |
| 0.010282 | 0.607033 | 0.645 | 0.605 | 1 | 11 | Cd14 | 0.04 | Young |
| 0.011337 | 0.656111 | 0.558 | 0.508 | 1 | 11 | Mycbp2 | 0.05 | Young |

| | | | | | | | | |
|---|---|---|---|---|---|---|---|---|
| 0.012185 | 0.08533 | 0.254 | 0.407 | 1 | 11 | Smpdl3a | -0.153 | Young |
| 0.012274 | 0.165254 | 0.181 | 0.331 | 1 | 11 | Tspan14 | -0.15 | Young |
| 0.013118 | 0.065457 | 0.239 | 0.395 | 1 | 11 | Rbm5 | -0.156 | Young |
| 0.013365 | 0.896958 | 0.145 | 0.069 | 1 | 11 | Chn2 | 0.076 | Young |
| 0.013866 | 0.590641 | 0.732 | 0.718 | 1 | 11 | Nr4a1 | 0.014 | Young |
| 0.014546 | 0.235172 | 0.949 | 0.968 | 1 | 11 | Rpsa | -0.019 | Young |
| 0.014917 | 0.887281 | 0.036 | 0.004 | 1 | 11 | Socs2 | 0.032 | Young |
| 0.015021 | 0.150875 | 0.072 | 0.165 | 1 | 11 | Zscan26 | -0.093 | Young |
| 0.015397 | 0.013951 | 0.188 | 0.339 | 1 | 11 | Ggh | -0.151 | Young |
| 0.015983 | 0.469869 | 0.08 | 0.177 | 1 | 11 | Ndufs1 | -0.097 | Young |
| 0.016323 | 0.155389 | 0.022 | 0.085 | 1 | 11 | Tjp2 | -0.063 | Young |
| 0.016333 | 0.056101 | 0.181 | 0.31 | 1 | 11 | Hsd17b4 | -0.129 | Young |
| 0.016711 | 0.321369 | 0.109 | 0.222 | 1 | 11 | Traf3 | -0.113 | Young |
| 0.017306 | 0.287708 | 0.043 | 0.121 | 1 | 11 | Lrch4 | -0.078 | Young |
| 0.017699 | 0.060119 | 0.123 | 0.238 | 1 | 11 | Slf2 | -0.115 | Young |
| 0.018006 | 0.729241 | 0.572 | 0.56 | 1 | 11 | Runx1 | 0.012 | Young |
| 0.01839 | 0.012078 | 0.087 | 0.181 | 1 | 11 | Washc1 | -0.094 | Young |
| 0.018478 | 0.246292 | 0.152 | 0.278 | 1 | 11 | Erap1 | -0.126 | Young |
| 0.018601 | 0.153197 | 0.058 | 0.141 | 1 | 11 | Ssh1 | -0.083 | Young |
| 0.018835 | 1.380868 | 0.297 | 0.218 | 1 | 11 | Dhx58 | 0.079 | Young |
| 0.01899 | 0.241841 | 0.196 | 0.351 | 1 | 11 | Rad23a | -0.155 | Young |
| 0.019276 | 0.099328 | 0.13 | 0.254 | 1 | 11 | Lrrc59 | -0.124 | Young |
| 0.019351 | 0.056735 | 0.145 | 0.27 | 1 | 11 | Ddx54 | -0.125 | Young |
| 0.019766 | 0.115794 | 0.13 | 0.242 | 1 | 11 | Aco1 | -0.112 | Young |
| 0.019789 | 0.416298 | 0.804 | 0.835 | 1 | 11 | Nfkbia | -0.031 | Young |
| 0.020081 | 6.306705 | 0.022 | 0 | 1 | 11 | Ccl17 | 0.022 | Young |
| 0.020081 | 4.579369 | 0.022 | 0 | 1 | 11 | Gm30122 | 0.022 | Young |
| 0.020081 | 4.218125 | 0.022 | 0 | 1 | 11 | Il12b | 0.022 | Young |
| 0.020081 | 3.380182 | 0.022 | 0 | 1 | 11 | Apoc1 | 0.022 | Young |
| 0.020081 | 3.217108 | 0.022 | 0 | 1 | 11 | Xlr4a | 0.022 | Young |
| 0.020081 | 3.102617 | 0.022 | 0 | 1 | 11 | Cdk18 | 0.022 | Young |
| 0.020081 | 2.987648 | 0.022 | 0 | 1 | 11 | Rapgef5 | 0.022 | Young |
| 0.020081 | 2.868782 | 0.022 | 0 | 1 | 11 | Gm5113 | 0.022 | Young |
| 0.020081 | 2.502501 | 0.022 | 0 | 1 | 11 | Tagln | 0.022 | Young |
| 0.020081 | 2.417539 | 0.022 | 0 | 1 | 11 | Lgr4 | 0.022 | Young |
| 0.020081 | 2.416832 | 0.022 | 0 | 1 | 11 | Ppfia3 | 0.022 | Young |
| 0.020081 | 1.988285 | 0.022 | 0 | 1 | 11 | Bbs1 | 0.022 | Young |
| 0.020081 | 1.916569 | 0.022 | 0 | 1 | 11 | L1cam | 0.022 | Young |
| 0.020081 | 1.855399 | 0.022 | 0 | 1 | 11 | Proser3 | 0.022 | Young |
| 0.020081 | 1.648089 | 0.022 | 0 | 1 | 11 | Il1rl2 | 0.022 | Young |
| 0.020081 | 1.588186 | 0.022 | 0 | 1 | 11 | Klrb1a | 0.022 | Young |
| 0.020127 | 1.512043 | 0.043 | 0.008 | 1 | 11 | Cxxc5 | 0.035 | Young |
| 0.020872 | 0.306366 | 0.899 | 0.903 | 1 | 11 | Rpl17 | -0.004 | Young |
| 0.021243 | 0.260006 | 0.188 | 0.331 | 1 | 11 | Pja2 | -0.143 | Young |
| 0.021593 | 0.159557 | 0.036 | 0.105 | 1 | 11 | Ppp1r8 | -0.069 | Young |

| | | | | | | | | |
|---|---|---|---|---|---|---|---|---|
| 0.021674 | 0.215309 | 0.043 | 0.117 | 1 | 11 | 0610010F0 | -0.074 | Young |
| 0.022029 | 1.896781 | 0.051 | 0.012 | 1 | 11 | Xlr | 0.039 | Young |
| 0.022081 | 0.056093 | 0.239 | 0.399 | 1 | 11 | Birc6 | -0.16 | Young |
| 0.022093 | 0.085104 | 0.022 | 0.081 | 1 | 11 | Rmnd1 | -0.059 | Young |
| 0.022224 | 0.291614 | 0.913 | 0.919 | 1 | 11 | Lgmn | -0.006 | Young |
| 0.022519 | 0.296165 | 0.804 | 0.794 | 1 | 11 | Rpl10a | 0.01 | Young |
| 0.022542 | 0.14263 | 1 | 1 | 1 | 11 | Tmsb4x | 0 | Young |
| 0.022974 | 0.023294 | 0.254 | 0.399 | 1 | 11 | Sf3b5 | -0.145 | Young |
| 0.023472 | 0.028105 | 0.384 | 0.569 | 1 | 11 | Tpp1 | -0.185 | Young |
| 0.024544 | 0.523028 | 0.029 | 0.093 | 1 | 11 | Lrp5 | -0.064 | Young |
| 0.024863 | 0.174116 | 0.109 | 0.214 | 1 | 11 | Magoh | -0.105 | Young |
| 0.025153 | 0.367987 | 0.072 | 0.161 | 1 | 11 | Acads | -0.089 | Young |
| 0.025165 | 0.196424 | 0.123 | 0.238 | 1 | 11 | Ccz1 | -0.115 | Young |
| 0.025438 | 0.18294 | 0.065 | 0.149 | 1 | 11 | Cnot10 | -0.084 | Young |
| 0.025589 | 0.082224 | 0.072 | 0.157 | 1 | 11 | Auh | -0.085 | Young |
| 0.025692 | 0.151683 | 0.145 | 0.262 | 1 | 11 | Pole4 | -0.117 | Young |
| 0.025757 | 0.129278 | 0.065 | 0.145 | 1 | 11 | Tctex1d2 | -0.08 | Young |
| 0.026477 | 0.118398 | 0.203 | 0.339 | 1 | 11 | Cct3 | -0.136 | Young |
| 0.027405 | 1.073276 | 0.536 | 0.512 | 1 | 11 | Abca1 | 0.024 | Young |
| 0.027611 | 0.112387 | 0.072 | 0.157 | 1 | 11 | Zfp330 | -0.085 | Young |
| 0.027961 | 0.07928 | 0.072 | 0.157 | 1 | 11 | Pdcl | -0.085 | Young |
| 0.02861 | 0.094482 | 0.13 | 0.234 | 1 | 11 | Dgkz | -0.104 | Young |
| 0.02881 | 0.096844 | 0.225 | 0.375 | 1 | 11 | Ube2q1 | -0.15 | Young |
| 0.028906 | 0.099937 | 0.058 | 0.133 | 1 | 11 | Hs1bp3 | -0.075 | Young |
| 0.028944 | 0.222665 | 0.087 | 0.177 | 1 | 11 | Cd82 | -0.09 | Young |
| 0.02897 | 0.037937 | 0.065 | 0.141 | 1 | 11 | Zfp260 | -0.076 | Young |
| 0.0291 | 0.185396 | 0.362 | 0.544 | 1 | 11 | Slamf7 | -0.182 | Young |
| 0.029841 | 0.434651 | 0.739 | 0.762 | 1 | 11 | Pcbp2 | -0.023 | Young |
| 0.029999 | 0.361397 | 0.768 | 0.827 | 1 | 11 | Eif4g2 | -0.059 | Young |
| 0.030571 | 0.160946 | 0.159 | 0.278 | 1 | 11 | Slc35f6 | -0.119 | Young |
| 0.030725 | 1.110306 | 0.152 | 0.085 | 1 | 11 | Adam33 | 0.067 | Young |
| 0.030777 | 0.053943 | 0.116 | 0.218 | 1 | 11 | Rnps1 | -0.102 | Young |
| 0.03085 | 0.059645 | 0.08 | 0.165 | 1 | 11 | Nploc4 | -0.085 | Young |
| 0.031369 | 0.556671 | 0.551 | 0.532 | 1 | 11 | Rel | 0.019 | Young |
| 0.03156 | 0.562324 | 0.37 | 0.274 | 1 | 11 | Ifit3 | 0.096 | Young |
| 0.031581 | 0.009172 | 0.21 | 0.339 | 1 | 11 | Rwdd1 | -0.129 | Young |
| 0.031626 | 0.125589 | 0.196 | 0.327 | 1 | 11 | Atp8a1 | -0.131 | Young |
| 0.032029 | 0.094366 | 0.116 | 0.214 | 1 | 11 | Ccdc82 | -0.098 | Young |
| 0.032293 | 0.235441 | 0.065 | 0.141 | 1 | 11 | Ift52 | -0.076 | Young |
| 0.032427 | 0.069326 | 0.42 | 0.577 | 1 | 11 | Srsf2 | -0.157 | Young |
| 0.032618 | 2.501242 | 0.08 | 0.032 | 1 | 11 | Trp53rka | 0.048 | Young |
| 0.032898 | 1.09871 | 0.094 | 0.04 | 1 | 11 | Timd4 | 0.054 | Young |
| 0.033187 | 0.106964 | 0.159 | 0.274 | 1 | 11 | 8-Sep | -0.115 | Young |
| 0.034749 | 0.016394 | 0.116 | 0.214 | 1 | 11 | Mrps9 | -0.098 | Young |
| 0.035846 | 1.643552 | 0.08 | 0.032 | 1 | 11 | Accs | 0.048 | Young |

| | | | | | | | | |
|---|---|---|---|---|---|---|---|---|
| 0.035934 | 1.569301 | 0.109 | 0.052 | 1 | 11 | Mir155hg | 0.057 | Young |
| 0.036131 | 0.00145 | 0.109 | 0.206 | 1 | 11 | Lsm7 | -0.097 | Young |
| 0.036649 | 0.368886 | 0.109 | 0.206 | 1 | 11 | Chfr | -0.097 | Young |
| 0.036771 | 0.100088 | 0.203 | 0.331 | 1 | 11 | Pip5k1c | -0.128 | Young |
| 0.036906 | 0.004091 | 0.065 | 0.137 | 1 | 11 | Pigq | -0.072 | Young |
| 0.036935 | 0.137035 | 0.275 | 0.435 | 1 | 11 | Copa | -0.16 | Young |
| 0.037296 | 3.062024 | 0.029 | 0.004 | 1 | 11 | Gm43660 | 0.025 | Young |
| 0.037444 | 0.310109 | 0.225 | 0.363 | 1 | 11 | Dnajc5 | -0.138 | Young |
| 0.037517 | 1.208562 | 0.13 | 0.069 | 1 | 11 | Qpct | 0.061 | Young |
| 0.037644 | 0.550643 | 0.138 | 0.246 | 1 | 11 | Frg1 | -0.108 | Young |
| 0.037742 | 3.202701 | 0.029 | 0.004 | 1 | 11 | E430024P1 | 0.025 | Young |
| 0.037742 | 1.950956 | 0.029 | 0.004 | 1 | 11 | Sigirr | 0.025 | Young |
| 0.037792 | 0.146776 | 0.101 | 0.19 | 1 | 11 | Ammecr1l | -0.089 | Young |
| 0.038192 | 0.330558 | 0.225 | 0.363 | 1 | 11 | Ilk | -0.138 | Young |
| 0.038192 | 2.232117 | 0.029 | 0.004 | 1 | 11 | Frg2f1 | 0.025 | Young |
| 0.038379 | 0.039236 | 0.13 | 0.23 | 1 | 11 | Mlxip | -0.1 | Young |
| 0.038612 | 0.141185 | 0.21 | 0.343 | 1 | 11 | Atg3 | -0.133 | Young |
| 0.038647 | 1.425224 | 0.029 | 0.004 | 1 | 11 | Abhd6 | 0.025 | Young |
| 0.038723 | 0.014434 | 0.29 | 0.431 | 1 | 11 | Ddx24 | -0.141 | Young |
| 0.038768 | 0.001434 | 0.13 | 0.23 | 1 | 11 | Nin | -0.1 | Young |
| 0.039198 | 0.172787 | 0.072 | 0.149 | 1 | 11 | Atg7 | -0.077 | Young |
| 0.039941 | 0.049649 | 0.21 | 0.339 | 1 | 11 | Trip11 | -0.129 | Young |
| 0.040815 | 2.279417 | 0.065 | 0.024 | 1 | 11 | Plxna4 | 0.041 | Young |
| 0.041085 | 0.021875 | 0.203 | 0.323 | 1 | 11 | Senp6 | -0.12 | Young |
| 0.04203 | 0.086538 | 0.239 | 0.375 | 1 | 11 | Eif1b | -0.136 | Young |
| 0.042437 | 0.125378 | 0.116 | 0.21 | 1 | 11 | Mospd3 | -0.094 | Young |
| 0.042478 | 0.054688 | 0.333 | 0.488 | 1 | 11 | Tap2 | -0.155 | Young |
| 0.042563 | 0.145485 | 0.181 | 0.302 | 1 | 11 | Srd5a3 | -0.121 | Young |
| 0.042756 | 0.061136 | 0.087 | 0.165 | 1 | 11 | Rrad | -0.078 | Young |
| 0.042762 | 0.17731 | 0.978 | 0.988 | 1 | 11 | Rpl37a | -0.01 | Young |
| 0.042814 | 0.571973 | 0.514 | 0.476 | 1 | 11 | Lacc1 | 0.038 | Young |
| 0.042853 | 0.007237 | 0.109 | 0.198 | 1 | 11 | Tnfaip1 | -0.089 | Young |
| 0.043198 | 0.373842 | 0.058 | 0.129 | 1 | 11 | Cln5 | -0.071 | Young |
| 0.043586 | 0.04729 | 0.167 | 0.274 | 1 | 11 | Mxd1 | -0.107 | Young |
| 0.043902 | 0.007794 | 0.558 | 0.73 | 1 | 11 | Ifrd1 | -0.172 | Young |
| 0.044254 | 2.520836 | 0.058 | 0.02 | 1 | 11 | A530064D( | 0.038 | Young |
| 0.044544 | 0.076846 | 0.246 | 0.391 | 1 | 11 | Supt16 | -0.145 | Young |
| 0.044776 | 0.416927 | 0.123 | 0.218 | 1 | 11 | Guk1 | -0.095 | Young |
| 0.044785 | 0.043011 | 0.167 | 0.278 | 1 | 11 | C1galt1 | -0.111 | Young |
| 0.045076 | 0.021549 | 0.21 | 0.347 | 1 | 11 | Tmem164 | -0.137 | Young |
| 0.045228 | 0.018832 | 0.123 | 0.218 | 1 | 11 | Rbm28 | -0.095 | Young |
| 0.045283 | 0.156308 | 0.413 | 0.593 | 1 | 11 | Atp6v0e | -0.18 | Young |
| 0.045337 | 0.087613 | 0.043 | 0.105 | 1 | 11 | Akr1e1 | -0.062 | Young |
| 0.04543 | 0.08711 | 0.232 | 0.371 | 1 | 11 | Anxa6 | -0.139 | Young |
| 0.045447 | 0.557204 | 0.087 | 0.169 | 1 | 11 | Zfp384 | -0.082 | Young |

| | | | | | | | | |
|---|---|---|---|---|---|---|---|---|
| 0.045524 | 0.002719 | 0.333 | 0.468 | 1 | 11 | Tep1 | -0.135 | Young |
| 0.046016 | 3.607916 | 0.036 | 0.008 | 1 | 11 | Rrm2 | 0.028 | Young |
| 0.046042 | 0.028282 | 0.254 | 0.395 | 1 | 11 | Pkn1 | -0.141 | Young |
| 0.046299 | 0.117106 | 0.167 | 0.278 | 1 | 11 | Rnf6 | -0.111 | Young |
| 0.046467 | 1.941765 | 0.036 | 0.008 | 1 | 11 | Mycl | 0.028 | Young |
| 0.046921 | 2.439324 | 0.036 | 0.008 | 1 | 11 | Grasp | 0.028 | Young |
| 0.047117 | 0.263308 | 0.138 | 0.234 | 1 | 11 | Nt5c | -0.096 | Young |
| 0.047353 | 0.282738 | 0.087 | 0.169 | 1 | 11 | Mpdu1 | -0.082 | Young |
| 0.047496 | 0.300072 | 0.906 | 0.923 | 1 | 11 | Rps27 | -0.017 | Young |
| 0.047842 | 1.621233 | 0.036 | 0.008 | 1 | 11 | Apold1 | 0.028 | Young |
| 0.048218 | 0.068434 | 0.232 | 0.375 | 1 | 11 | Fkbp2 | -0.143 | Young |
| 0.048238 | 0.710616 | 0.529 | 0.524 | 1 | 11 | Tmcc1 | 0.005 | Young |
| 0.04852 | 0.015111 | 0.036 | 0.093 | 1 | 11 | Mecr | -0.057 | Young |
| 0.048572 | 0.338147 | 0.768 | 0.786 | 1 | 11 | Slfn5 | -0.018 | Young |
| 0.04869 | 0.180908 | 0.971 | 0.98 | 1 | 11 | Rps20 | -0.009 | Young |
| 0.048745 | 0.092929 | 0.312 | 0.456 | 1 | 11 | Ndufa1 | -0.144 | Young |
| 0.048995 | 0.316319 | 0.217 | 0.343 | 1 | 11 | Phc2 | -0.126 | Young |
| 0.049004 | 0.327302 | 0.036 | 0.093 | 1 | 11 | Ap2a1 | -0.057 | Young |
| 0.049197 | 0.015724 | 0.181 | 0.294 | 1 | 11 | Sla | -0.113 | Young |
| 0.049427 | 0.510684 | 0.406 | 0.331 | 1 | 11 | Lsp1 | 0.075 | Young |
| 0.049734 | 0.706159 | 0.435 | 0.387 | 1 | 11 | St8sia4 | 0.048 | Young |
| 0.050065 | 0.298151 | 0.239 | 0.383 | 1 | 11 | Dhx15 | -0.144 | Young |
| 0.050089 | 0.116627 | 0.08 | 0.153 | 1 | 11 | Klhl28 | -0.073 | Young |
| 0.050321 | 1.875933 | 0.043 | 0.012 | 1 | 11 | H2-Ob | 0.031 | Young |
| 0.050487 | 0.008131 | 0.319 | 0.452 | 1 | 11 | Cxcr4 | -0.133 | Young |
| 0.050725 | 0.945287 | 0.196 | 0.125 | 1 | 11 | Crim1 | 0.071 | Young |
| 0.050953 | 0.521641 | 0.138 | 0.242 | 1 | 11 | Plekha1 | -0.104 | Young |
| 0.051122 | 0.054953 | 0.094 | 0.177 | 1 | 11 | Gcc2 | -0.083 | Young |
| 0.051219 | 0.020056 | 0.072 | 0.145 | 1 | 11 | Parg | -0.073 | Young |
| 0.051297 | 0.101012 | 0.268 | 0.395 | 1 | 11 | BC017643 | -0.127 | Young |
| 0.051507 | 0.118714 | 0.014 | 0.056 | 1 | 11 | 2810006K2 | -0.042 | Young |
| 0.051641 | 0.017799 | 0.217 | 0.339 | 1 | 11 | Csnk1d | -0.122 | Young |
| 0.051838 | 0.218779 | 0.014 | 0.056 | 1 | 11 | Fbxo42 | -0.042 | Young |
| 0.051914 | 0.47261 | 0.203 | 0.327 | 1 | 11 | Cnot1 | -0.124 | Young |
| 0.052143 | 0.113334 | 0.123 | 0.218 | 1 | 11 | Ppm1b | -0.095 | Young |
| 0.052183 | 0.737045 | 0.058 | 0.02 | 1 | 11 | Bcdin3d | 0.038 | Young |
| 0.052378 | 0.006952 | 0.167 | 0.274 | 1 | 11 | Ncaph2 | -0.107 | Young |
| 0.052717 | 0.1534 | 0.377 | 0.56 | 1 | 11 | Bnip2 | -0.183 | Young |
| 0.052773 | 0.185863 | 0.051 | 0.113 | 1 | 11 | Heatr1 | -0.062 | Young |
| 0.052893 | 0.239188 | 0.913 | 0.956 | 1 | 11 | Rps4x | -0.043 | Young |
| 0.052894 | 0.047651 | 0.239 | 0.367 | 1 | 11 | Ifnar2 | -0.128 | Young |
| 0.053132 | 0.037018 | 0.239 | 0.375 | 1 | 11 | Srsf7 | -0.136 | Young |
| 0.05359 | 0.237995 | 0.058 | 0.125 | 1 | 11 | Pdgfa | -0.067 | Young |
| 0.053775 | 0.0486 | 0.203 | 0.319 | 1 | 11 | Hs6st1 | -0.116 | Young |
| 0.054075 | 1.044432 | 0.312 | 0.258 | 1 | 11 | Dyrk2 | 0.054 | Young |

| | | | | | | | | |
|---|---|---|---|---|---|---|---|---|
| 0.054375 | 0.104972 | 0.094 | 0.169 | 1 | 11 | Phyh | -0.075 | Young |
| 0.054747 | 0.106939 | 0.072 | 0.145 | 1 | 11 | Ggta1 | -0.073 | Young |
| 0.055103 | 0.108265 | 0.08 | 0.153 | 1 | 11 | Spata2 | -0.073 | Young |
| 0.05522 | 0.144138 | 0.159 | 0.262 | 1 | 11 | Atf7 | -0.103 | Young |
| 0.055397 | 0.176775 | 0.051 | 0.113 | 1 | 11 | Sbf1 | -0.062 | Young |
| 0.055397 | 0.03634 | 0.051 | 0.113 | 1 | 11 | Ncbp1 | -0.062 | Young |
| 0.055732 | 0.043679 | 0.08 | 0.153 | 1 | 11 | Zfp280d | -0.073 | Young |
| 0.056395 | 0.270063 | 0.087 | 0.161 | 1 | 11 | Dpy30 | -0.074 | Young |
| 0.056395 | 0.127301 | 0.174 | 0.282 | 1 | 11 | Hfe | -0.108 | Young |
| 0.057 | 0.16788 | 0.203 | 0.319 | 1 | 11 | Supt5 | -0.116 | Young |
| 0.057301 | 0.033305 | 0.239 | 0.359 | 1 | 11 | Snrpc | -0.12 | Young |
| 0.057727 | 0.045115 | 0.312 | 0.468 | 1 | 11 | Psma6 | -0.156 | Young |
| 0.057757 | 0.004411 | 0.043 | 0.101 | 1 | 11 | Rfx1 | -0.058 | Young |
| 0.05784 | 0.650689 | 0.507 | 0.476 | 1 | 11 | Tlr2 | 0.031 | Young |
| 0.058009 | 0.210251 | 0.087 | 0.165 | 1 | 11 | Qrich1 | -0.078 | Young |
| 0.058162 | 4.1284 | 0.014 | 0 | 1 | 11 | Gm6297 | 0.014 | Young |
| 0.058162 | 4.062335 | 0.014 | 0 | 1 | 11 | Slc2a3 | 0.014 | Young |
| 0.058162 | 3.688479 | 0.014 | 0 | 1 | 11 | Prag1 | 0.014 | Young |
| 0.058162 | 2.999699 | 0.014 | 0 | 1 | 11 | Lbx1 | 0.014 | Young |
| 0.058162 | 2.997861 | 0.014 | 0 | 1 | 11 | Ildr2 | 0.014 | Young |
| 0.058162 | 2.956289 | 0.014 | 0 | 1 | 11 | Gm16894 | 0.014 | Young |
| 0.058162 | 2.943479 | 0.014 | 0 | 1 | 11 | Mettl15 | 0.014 | Young |
| 0.058162 | 2.845852 | 0.014 | 0 | 1 | 11 | 4933406I1 | 0.014 | Young |
| 0.058162 | 2.829735 | 0.014 | 0 | 1 | 11 | Ephb3 | 0.014 | Young |
| 0.058162 | 2.779524 | 0.014 | 0 | 1 | 11 | Fgf13 | 0.014 | Young |
| 0.058162 | 2.771927 | 0.014 | 0 | 1 | 11 | Itm2a | 0.014 | Young |
| 0.058162 | 2.742961 | 0.014 | 0 | 1 | 11 | Fxyd6 | 0.014 | Young |
| 0.058162 | 2.735482 | 0.014 | 0 | 1 | 11 | Gfod2 | 0.014 | Young |
| 0.058162 | 2.666268 | 0.014 | 0 | 1 | 11 | Hmgcs2 | 0.014 | Young |
| 0.058162 | 2.659161 | 0.014 | 0 | 1 | 11 | Tbx21 | 0.014 | Young |
| 0.058162 | 2.591281 | 0.014 | 0 | 1 | 11 | Gm15728 | 0.014 | Young |
| 0.058162 | 2.572414 | 0.014 | 0 | 1 | 11 | Mucl1 | 0.014 | Young |
| 0.058162 | 2.568245 | 0.014 | 0 | 1 | 11 | Gm48545 | 0.014 | Young |
| 0.058162 | 2.501443 | 0.014 | 0 | 1 | 11 | Ly6g6d | 0.014 | Young |
| 0.058162 | 2.500885 | 0.014 | 0 | 1 | 11 | Rhof | 0.014 | Young |
| 0.058162 | 2.500885 | 0.014 | 0 | 1 | 11 | Agbl3 | 0.014 | Young |
| 0.058162 | 2.483909 | 0.014 | 0 | 1 | 11 | H1fx | 0.014 | Young |
| 0.058162 | 2.399899 | 0.014 | 0 | 1 | 11 | Gm31763 | 0.014 | Young |
| 0.058162 | 2.396552 | 0.014 | 0 | 1 | 11 | Alb | 0.014 | Young |
| 0.058162 | 2.376003 | 0.014 | 0 | 1 | 11 | Hist1h4c | 0.014 | Young |
| 0.058162 | 2.358785 | 0.014 | 0 | 1 | 11 | Ebf1 | 0.014 | Young |
| 0.058162 | 2.309093 | 0.014 | 0 | 1 | 11 | Pcdhga1 | 0.014 | Young |
| 0.058162 | 2.303486 | 0.014 | 0 | 1 | 11 | Hs3st3a1 | 0.014 | Young |
| 0.058162 | 2.301371 | 0.014 | 0 | 1 | 11 | Smpd5 | 0.014 | Young |
| 0.058162 | 2.283897 | 0.014 | 0 | 1 | 11 | Zfyve28 | 0.014 | Young |

| | | | | | | | | |
|---|---|---|---|---|---|---|---|---|
| 0.058162 | 2.266517 | 0.014 | 0 | 1 | 11 | Cyp2s1 | 0.014 | Young |
| 0.058162 | 2.219323 | 0.014 | 0 | 1 | 11 | Tgtp1 | 0.014 | Young |
| 0.058162 | 2.120913 | 0.014 | 0 | 1 | 11 | Sgo2a | 0.014 | Young |
| 0.058162 | 2.117547 | 0.014 | 0 | 1 | 11 | Clec1b | 0.014 | Young |
| 0.058162 | 2.094682 | 0.014 | 0 | 1 | 11 | Ets1 | 0.014 | Young |
| 0.058162 | 2.051482 | 0.014 | 0 | 1 | 11 | Fbn1 | 0.014 | Young |
| 0.058162 | 2.038334 | 0.014 | 0 | 1 | 11 | Gm5914 | 0.014 | Young |
| 0.058162 | 1.844873 | 0.014 | 0 | 1 | 11 | Akr1c18 | 0.014 | Young |
| 0.058162 | 1.834317 | 0.014 | 0 | 1 | 11 | Bfsp1 | 0.014 | Young |
| 0.058162 | 1.819213 | 0.014 | 0 | 1 | 11 | Whrn | 0.014 | Young |
| 0.058162 | 1.758013 | 0.014 | 0 | 1 | 11 | Apbb2 | 0.014 | Young |
| 0.058162 | 1.747719 | 0.014 | 0 | 1 | 11 | Abca14 | 0.014 | Young |
| 0.058162 | 1.713236 | 0.014 | 0 | 1 | 11 | Spin4 | 0.014 | Young |
| 0.058162 | 1.694962 | 0.014 | 0 | 1 | 11 | Gm42726 | 0.014 | Young |
| 0.058162 | 1.692396 | 0.014 | 0 | 1 | 11 | Gm5141 | 0.014 | Young |
| 0.058162 | 1.535555 | 0.014 | 0 | 1 | 11 | Slc17a8 | 0.014 | Young |
| 0.058162 | 1.53293 | 0.014 | 0 | 1 | 11 | Gm17227 | 0.014 | Young |
| 0.058162 | 1.509165 | 0.014 | 0 | 1 | 11 | Zfp799 | 0.014 | Young |
| 0.058162 | 1.462689 | 0.014 | 0 | 1 | 11 | Kazald1 | 0.014 | Young |
| 0.058162 | 1.459373 | 0.014 | 0 | 1 | 11 | Aspn | 0.014 | Young |
| 0.058162 | 1.395947 | 0.014 | 0 | 1 | 11 | Slc4a11 | 0.014 | Young |
| 0.058162 | 1.357069 | 0.014 | 0 | 1 | 11 | Spry1 | 0.014 | Young |
| 0.058162 | 1.351545 | 0.014 | 0 | 1 | 11 | Cdca3 | 0.014 | Young |
| 0.058162 | 1.279861 | 0.014 | 0 | 1 | 11 | Mst1r | 0.014 | Young |
| 0.058162 | 1.206612 | 0.014 | 0 | 1 | 11 | Dlc1 | 0.014 | Young |
| 0.058289 | 0.085052 | 0.304 | 0.476 | 1 | 11 | Ndufb5 | -0.172 | Young |
| 0.058783 | 1.311096 | 0.123 | 0.069 | 1 | 11 | Frmd6 | 0.054 | Young |
| 0.058975 | 1.509398 | 0.08 | 0.036 | 1 | 11 | 2510009E0 | 0.044 | Young |
| 0.059133 | 0.057637 | 0.275 | 0.415 | 1 | 11 | Sec11c | -0.14 | Young |
| 0.059161 | 0.048256 | 0.109 | 0.198 | 1 | 11 | Rab3gap1 | -0.089 | Young |
| 0.059598 | 0.056246 | 0.297 | 0.444 | 1 | 11 | Limd1 | -0.147 | Young |
| 0.060022 | 0.094307 | 0.065 | 0.133 | 1 | 11 | Med21 | -0.068 | Young |
| 0.060743 | 0.06746 | 0.232 | 0.355 | 1 | 11 | Bcl3 | -0.123 | Young |
| 0.060891 | 0.013731 | 0.087 | 0.161 | 1 | 11 | Fads1 | -0.074 | Young |
| 0.061226 | 0.126736 | 0.123 | 0.21 | 1 | 11 | Nod1 | -0.087 | Young |
| 0.061382 | 0.03302 | 0.203 | 0.319 | 1 | 11 | Smarcc2 | -0.116 | Young |
| 0.061545 | 0.206912 | 0.13 | 0.23 | 1 | 11 | Pin4 | -0.1 | Young |
| 0.061657 | 0.058785 | 0.261 | 0.403 | 1 | 11 | Psmd8 | -0.142 | Young |
| 0.06179 | 0.353794 | 0.754 | 0.802 | 1 | 11 | Mpeg1 | -0.048 | Young |
| 0.061997 | 0.006585 | 0.181 | 0.286 | 1 | 11 | Rbm17 | -0.105 | Young |
| 0.062454 | 0.013296 | 0.087 | 0.161 | 1 | 11 | Serhl | -0.074 | Young |
| 0.062667 | 0.281685 | 0.13 | 0.226 | 1 | 11 | Pank2 | -0.096 | Young |
| 0.062764 | 0.055809 | 0.428 | 0.625 | 1 | 11 | Ppp2ca | -0.197 | Young |
| 0.063079 | 0.259595 | 0.109 | 0.194 | 1 | 11 | Vps36 | -0.085 | Young |
| 0.063284 | 0.173561 | 0.442 | 0.593 | 1 | 11 | Snrpb | -0.151 | Young |

| | | | | | | | | |
|---|---|---|---|---|---|---|---|---|
| 0.063462 | 0.291751 | 0.855 | 0.859 | 1 | 11 | Rpl34 | -0.004 | Young |
| 0.063536 | 0.18772 | 0.13 | 0.218 | 1 | 11 | Ccnd3 | -0.088 | Young |
| 0.063795 | 0.82666 | 0.239 | 0.173 | 1 | 11 | Zfp395 | 0.066 | Young |
| 0.063941 | 0.075822 | 0.029 | 0.077 | 1 | 11 | Loxl3 | -0.048 | Young |
| 0.064148 | 0.25194 | 0.181 | 0.29 | 1 | 11 | Spg21 | -0.109 | Young |
| 0.064174 | 0.007218 | 0.007 | 0.04 | 1 | 11 | Ccdc61 | -0.033 | Young |
| 0.064533 | 0.039846 | 0.225 | 0.343 | 1 | 11 | Fam126a | -0.118 | Young |
| 0.064982 | 0.227532 | 0.138 | 0.23 | 1 | 11 | Enox2 | -0.092 | Young |
| 0.065289 | 0.198441 | 0.101 | 0.181 | 1 | 11 | Lysmd3 | -0.08 | Young |
| 0.065479 | 0.243384 | 0.152 | 0.25 | 1 | 11 | Foxn2 | -0.098 | Young |
| 0.066035 | 0.588721 | 0.08 | 0.149 | 1 | 11 | Ddx23 | -0.069 | Young |
| 0.066045 | 0.171896 | 0.174 | 0.278 | 1 | 11 | Trio | -0.104 | Young |
| 0.066302 | 0.104962 | 0.101 | 0.185 | 1 | 11 | Txndc15 | -0.084 | Young |
| 0.066519 | 0.100724 | 0.058 | 0.121 | 1 | 11 | Wdr48 | -0.063 | Young |
| 0.066797 | 0.648675 | 0.036 | 0.089 | 1 | 11 | Fbxl18 | -0.053 | Young |
| 0.067118 | 0.512588 | 0.036 | 0.089 | 1 | 11 | Pdk1 | -0.053 | Young |
| 0.067207 | 0.16266 | 0.913 | 0.935 | 1 | 11 | Rpl39 | -0.022 | Young |
| 0.067658 | 0.048305 | 0.254 | 0.387 | 1 | 11 | Pds5a | -0.133 | Young |
| 0.067711 | 0.024616 | 0.159 | 0.25 | 1 | 11 | Plagl2 | -0.091 | Young |
| 0.06785 | 0.060301 | 0.116 | 0.194 | 1 | 11 | Lats2 | -0.078 | Young |
| 0.068768 | 0.070801 | 0.138 | 0.226 | 1 | 11 | Csnk1g2 | -0.088 | Young |
| 0.068787 | 0.245347 | 0.239 | 0.367 | 1 | 11 | Tex261 | -0.128 | Young |
| 0.069117 | 0.152008 | 0.188 | 0.294 | 1 | 11 | Emc3 | -0.106 | Young |
| 0.069312 | 0.119214 | 0.167 | 0.27 | 1 | 11 | Sec22b | -0.103 | Young |
| 0.06978 | 0.226331 | 0.087 | 0.161 | 1 | 11 | Exosc5 | -0.074 | Young |
| 0.070297 | 0.039595 | 0.312 | 0.452 | 1 | 11 | Pigt | -0.14 | Young |
| 0.07053 | 0.173235 | 0.051 | 0.109 | 1 | 11 | Smad3 | -0.058 | Young |
| 0.070648 | 0.180444 | 0.123 | 0.21 | 1 | 11 | Prdm2 | -0.087 | Young |
| 0.070679 | 0.002345 | 0.159 | 0.246 | 1 | 11 | Tifab | -0.087 | Young |
| 0.070823 | 0.421337 | 0.014 | 0.052 | 1 | 11 | Cela1 | -0.038 | Young |
| 0.071271 | 0.269914 | 0.101 | 0.181 | 1 | 11 | Usp32 | -0.08 | Young |
| 0.071367 | 0.296256 | 0.246 | 0.379 | 1 | 11 | Rcc2 | -0.133 | Young |
| 0.071473 | 0.857336 | 0.116 | 0.065 | 1 | 11 | Unkl | 0.051 | Young |
| 0.07148 | 0.041281 | 0.094 | 0.165 | 1 | 11 | Acadvl | -0.071 | Young |
| 0.071809 | 0.03859 | 0.297 | 0.452 | 1 | 11 | Ensa | -0.155 | Young |
| 0.072145 | 0.061821 | 0.203 | 0.315 | 1 | 11 | Ddrgk1 | -0.112 | Young |
| 0.072215 | 0.634725 | 0.217 | 0.149 | 1 | 11 | Eif2s3y | 0.068 | Young |
| 0.072269 | 0.12249 | 0.058 | 0.117 | 1 | 11 | Nfx1 | -0.059 | Young |
| 0.072286 | 0.322396 | 0.652 | 0.609 | 1 | 11 | Cxcl16 | 0.043 | Young |
| 0.072337 | 0.112499 | 0.094 | 0.169 | 1 | 11 | Suco | -0.075 | Young |
| 0.073645 | 1.714534 | 0.145 | 0.093 | 1 | 11 | Dopey2 | 0.052 | Young |
| 0.073655 | 2.922343 | 0.065 | 0.028 | 1 | 11 | Mmp13 | 0.037 | Young |
| 0.073759 | 0.22913 | 0.058 | 0.117 | 1 | 11 | Cherp | -0.059 | Young |
| 0.073951 | 0.041997 | 0.043 | 0.097 | 1 | 11 | Dhx33 | -0.054 | Young |
| 0.07422 | 0.603026 | 0.123 | 0.069 | 1 | 11 | Cd9 | 0.054 | Young |

| | | | | | | | | |
|---|---|---|---|---|---|---|---|---|
| 0.074253 | 0.029788 | 0.094 | 0.165 | 1 | 11 | Ylpm1 | -0.071 | Young |
| 0.074395 | 0.057232 | 0.196 | 0.302 | 1 | 11 | Vamp2 | -0.106 | Young |
| 0.074561 | 0.015875 | 0.239 | 0.363 | 1 | 11 | Dock7 | -0.124 | Young |
| 0.074716 | 0.203932 | 0.181 | 0.282 | 1 | 11 | Herc2 | -0.101 | Young |
| 0.074763 | 1.010568 | 0.261 | 0.198 | 1 | 11 | Cd72 | 0.063 | Young |
| 0.074778 | 0.308232 | 0.138 | 0.238 | 1 | 11 | Eif6 | -0.1 | Young |
| 0.074807 | 0.053128 | 0.087 | 0.157 | 1 | 11 | Sema6d | -0.07 | Young |
| 0.074874 | 0.205298 | 0.196 | 0.298 | 1 | 11 | Anapc13 | -0.102 | Young |
| 0.075306 | 0.218946 | 0.138 | 0.226 | 1 | 11 | Atf6b | -0.088 | Young |
| 0.075432 | 0.242669 | 0.058 | 0.121 | 1 | 11 | Mroh1 | -0.063 | Young |
| 0.075888 | 0.008307 | 0.058 | 0.117 | 1 | 11 | Wdr20 | -0.059 | Young |
| 0.07662 | 2.7036 | 0.101 | 0.056 | 1 | 11 | Itga6 | 0.045 | Young |
| 0.076666 | 0.053894 | 0.094 | 0.169 | 1 | 11 | Ddx47 | -0.075 | Young |
| 0.076774 | 0.043173 | 0.341 | 0.492 | 1 | 11 | Zbtb7a | -0.151 | Young |
| 0.077101 | 0.035033 | 0.297 | 0.431 | 1 | 11 | Stag2 | -0.134 | Young |
| 0.077293 | 0.426378 | 0.739 | 0.827 | 1 | 11 | Marcksl1 | -0.088 | Young |
| 0.07733 | 0.038892 | 0.283 | 0.411 | 1 | 11 | Eif4g3 | -0.128 | Young |
| 0.077642 | 0.00967 | 0.094 | 0.165 | 1 | 11 | Dld | -0.071 | Young |
| 0.07778 | 0.02305 | 0.355 | 0.504 | 1 | 11 | Caprin1 | -0.149 | Young |
| 0.077808 | 0.288828 | 0.065 | 0.125 | 1 | 11 | Prpsap1 | -0.06 | Young |
| 0.078214 | 0.046415 | 0.167 | 0.27 | 1 | 11 | Glrx3 | -0.103 | Young |
| 0.078539 | 0.120846 | 0.036 | 0.085 | 1 | 11 | Clp1 | -0.049 | Young |
| 0.078589 | 0.047324 | 0.225 | 0.331 | 1 | 11 | Zfp638 | -0.106 | Young |
| 0.078669 | 0.15683 | 0.232 | 0.363 | 1 | 11 | Rbm8a | -0.131 | Young |
| 0.078695 | 0.054181 | 0.159 | 0.258 | 1 | 11 | Adprh | -0.099 | Young |
| 0.078729 | 0.241098 | 0.754 | 0.798 | 1 | 11 | Rps3 | -0.044 | Young |
| 0.078901 | 0.535174 | 0.08 | 0.145 | 1 | 11 | Selenoh | -0.065 | Young |
| 0.079312 | 0.199433 | 0.891 | 0.903 | 1 | 11 | Rpl22 | -0.012 | Young |
| 0.079337 | 0.905763 | 0.254 | 0.194 | 1 | 11 | Mx1 | 0.06 | Young |
| 0.079349 | 0.084357 | 0.275 | 0.391 | 1 | 11 | Arih1 | -0.116 | Young |
| 0.079738 | 0.377197 | 0.71 | 0.782 | 1 | 11 | Sh3glb1 | -0.072 | Young |
| 0.080115 | 2.276813 | 0.058 | 0.024 | 1 | 11 | Ppcdc | 0.034 | Young |
| 0.080482 | 0.2318 | 0.891 | 0.911 | 1 | 11 | Rpl30 | -0.02 | Young |
| 0.080703 | 0.072745 | 0.384 | 0.532 | 1 | 11 | Notch2 | -0.148 | Young |
| 0.080985 | 0.233361 | 0.167 | 0.262 | 1 | 11 | Casp1 | -0.095 | Young |
| 0.081444 | 0.574892 | 0.239 | 0.355 | 1 | 11 | Cxcl9 | -0.116 | Young |
| 0.081877 | 0.163041 | 0.159 | 0.25 | 1 | 11 | Cdk14 | -0.091 | Young |
| 0.081965 | 0.085692 | 0.181 | 0.282 | 1 | 11 | Csnk1g3 | -0.101 | Young |
| 0.082132 | 0.142677 | 0.116 | 0.198 | 1 | 11 | Aqr | -0.082 | Young |
| 0.082305 | 0.227334 | 0.884 | 0.887 | 1 | 11 | Capza2 | -0.003 | Young |
| 0.082549 | 0.08352 | 0.051 | 0.105 | 1 | 11 | Mgat5 | -0.054 | Young |
| 0.08258 | 0.400722 | 0.043 | 0.093 | 1 | 11 | Alpk1 | -0.05 | Young |
| 0.082719 | 0.753631 | 0.036 | 0.085 | 1 | 11 | Gde1 | -0.049 | Young |
| 0.082911 | 0.27409 | 0.754 | 0.794 | 1 | 11 | Rpl11 | -0.04 | Young |
| 0.083132 | 0.41318 | 0.072 | 0.137 | 1 | 11 | Psd4 | -0.065 | Young |

| | | | | | | | | |
|---|---|---|---|---|---|---|---|---|
| 0.083548 | 0.254581 | 0.21 | 0.323 | 1 | 11 | Mrps33 | -0.113 | Young |
| 0.083634 | 0.0663 | 0.123 | 0.202 | 1 | 11 | Commd6 | -0.079 | Young |
| 0.084071 | 0.044902 | 0.116 | 0.194 | 1 | 11 | Ccdc6 | -0.078 | Young |
| 0.08428 | 0.185934 | 0.174 | 0.27 | 1 | 11 | Synj2bp | -0.096 | Young |
| 0.084298 | 0.081697 | 0.051 | 0.105 | 1 | 11 | Med7 | -0.054 | Young |
| 0.084463 | 0.173485 | 0.29 | 0.403 | 1 | 11 | Coa5 | -0.113 | Young |
| 0.084726 | 0.054373 | 0.261 | 0.387 | 1 | 11 | Itsn1 | -0.126 | Young |
| 0.085086 | 0.2396 | 0.08 | 0.145 | 1 | 11 | Ipo8 | -0.065 | Young |
| 0.085191 | 0.290783 | 0.819 | 0.859 | 1 | 11 | Btg2 | -0.04 | Young |
| 0.085438 | 0.252705 | 0.913 | 0.923 | 1 | 11 | Rpl23a | -0.01 | Young |
| 0.085595 | 0.113959 | 0.072 | 0.133 | 1 | 11 | Sdhaf2 | -0.061 | Young |
| 0.085806 | 0.318377 | 0.217 | 0.331 | 1 | 11 | Kmt2c | -0.114 | Young |
| 0.085947 | 0.036679 | 0.312 | 0.435 | 1 | 11 | Cers6 | -0.123 | Young |
| 0.086682 | 0.420055 | 0.065 | 0.125 | 1 | 11 | Sdhaf4 | -0.06 | Young |
| 0.087012 | 1.606689 | 0.101 | 0.056 | 1 | 11 | Tyw1 | 0.045 | Young |
| 0.087497 | 0.040183 | 0.391 | 0.52 | 1 | 11 | Hdgf | -0.129 | Young |
| 0.087888 | 0.407546 | 0.051 | 0.105 | 1 | 11 | Tomm6 | -0.054 | Young |
| 0.087914 | 0.078779 | 0.145 | 0.234 | 1 | 11 | 2-Mar | -0.089 | Young |
| 0.088005 | 1.335918 | 0.13 | 0.081 | 1 | 11 | Smpdl3b | 0.049 | Young |
| 0.088235 | 0.98939 | 0.384 | 0.375 | 1 | 11 | Cnot6l | 0.009 | Young |
| 0.088607 | 0.146896 | 0.138 | 0.226 | 1 | 11 | Mtmr3 | -0.088 | Young |
| 0.088687 | 0.207501 | 0.232 | 0.339 | 1 | 11 | Taf10 | -0.107 | Young |
| 0.088787 | 0.355822 | 0.007 | 0.036 | 1 | 11 | Il1a | -0.029 | Young |
| 0.088787 | 0.102909 | 0.007 | 0.036 | 1 | 11 | Lnx2 | -0.029 | Young |
| 0.088957 | 0.095469 | 0.101 | 0.169 | 1 | 11 | Oas1a | -0.068 | Young |
| 0.088978 | 0.503586 | 0.065 | 0.121 | 1 | 11 | Selenom | -0.056 | Young |
| 0.089087 | 0.047622 | 0.159 | 0.246 | 1 | 11 | Zcchc7 | -0.087 | Young |
| 0.089151 | 0.032592 | 0.217 | 0.319 | 1 | 11 | Pi4k2a | -0.102 | Young |
| 0.089423 | 0.506865 | 0.08 | 0.145 | 1 | 11 | Tlr13 | -0.065 | Young |
| 0.089423 | 0.086632 | 0.08 | 0.145 | 1 | 11 | Nudt5 | -0.065 | Young |
| 0.089595 | 0.417409 | 0.087 | 0.157 | 1 | 11 | Aak1 | -0.07 | Young |
| 0.089653 | 0.184025 | 0.254 | 0.379 | 1 | 11 | Arpp19 | -0.125 | Young |
| 0.089703 | 0.052108 | 0.065 | 0.125 | 1 | 11 | Arsa | -0.06 | Young |
| 0.089706 | 0.342207 | 0.167 | 0.266 | 1 | 11 | Yme1l1 | -0.099 | Young |
| 0.090043 | 0.353136 | 0.065 | 0.125 | 1 | 11 | Rnf181 | -0.06 | Young |
| 0.090433 | 0.17283 | 0.261 | 0.379 | 1 | 11 | Tubb6 | -0.118 | Young |
| 0.090508 | 0.068915 | 0.029 | 0.073 | 1 | 11 | Gmppb | -0.044 | Young |
| 0.090595 | 0.249039 | 0.152 | 0.242 | 1 | 11 | Acvrl1 | -0.09 | Young |
| 0.090834 | 0.00384 | 0.355 | 0.508 | 1 | 11 | Rab11b | -0.153 | Young |
| 0.090915 | 2.25575 | 0.051 | 0.02 | 1 | 11 | Deaf1 | 0.031 | Young |
| 0.091072 | 0.97871 | 0.138 | 0.085 | 1 | 11 | Lman2l | 0.053 | Young |
| 0.091167 | 0.121935 | 0.087 | 0.153 | 1 | 11 | Il17ra | -0.066 | Young |
| 0.091357 | 1.03179 | 0.022 | 0.06 | 1 | 11 | Aimp2 | -0.038 | Young |
| 0.091521 | 0.25081 | 0.884 | 0.915 | 1 | 11 | Rpl8 | -0.031 | Young |
| 0.091522 | 2.19123 | 0.051 | 0.02 | 1 | 11 | Fam84b | 0.031 | Young |

| | | | | | | | | |
|---|---|---|---|---|---|---|---|---|
| 0.091522 | 2.090989 | 0.051 | 0.02 | 1 | 11 | Micu3 | 0.031 | Young |
| 0.091615 | 0.55338 | 0.101 | 0.169 | 1 | 11 | Arfip1 | -0.068 | Young |
| 0.091784 | 0.241112 | 0.246 | 0.363 | 1 | 11 | Top2b | -0.117 | Young |
| 0.091796 | 0.940817 | 0.145 | 0.093 | 1 | 11 | Sh3bp5 | 0.052 | Young |
| 0.091858 | 0.29791 | 0.022 | 0.06 | 1 | 11 | Ttf2 | -0.038 | Young |
| 0.092256 | 0.179788 | 0.181 | 0.278 | 1 | 11 | Rasgrp4 | -0.097 | Young |
| 0.093175 | 1.05801 | 0.37 | 0.331 | 1 | 11 | Stab1 | 0.039 | Young |
| 0.093594 | 0.104861 | 0.268 | 0.407 | 1 | 11 | Rab11fip5 | -0.139 | Young |
| 0.093765 | 0.284603 | 0.855 | 0.911 | 1 | 11 | Cebpb | -0.056 | Young |
| 0.094925 | 0.057389 | 0.261 | 0.395 | 1 | 11 | Baz2b | -0.134 | Young |
| 0.095004 | 0.04843 | 0.109 | 0.181 | 1 | 11 | Rnf2 | -0.072 | Young |
| 0.095038 | 0.249607 | 0.181 | 0.286 | 1 | 11 | Nudcd3 | -0.105 | Young |
| 0.095166 | 0.35581 | 0.116 | 0.194 | 1 | 11 | Babam1 | -0.078 | Young |
| 0.095215 | 0.29579 | 0.145 | 0.234 | 1 | 11 | Commd7 | -0.089 | Young |
| 0.095368 | 0.00118 | 0.014 | 0.048 | 1 | 11 | Wfdc18 | -0.034 | Young |
| 0.095409 | 0.125536 | 0.174 | 0.27 | 1 | 11 | Lsm12 | -0.096 | Young |
| 0.095871 | 0.241314 | 0.261 | 0.367 | 1 | 11 | Med13 | -0.106 | Young |
| 0.096686 | 0.005722 | 0.275 | 0.391 | 1 | 11 | Msl1 | -0.116 | Young |
| 0.096727 | 0.160764 | 0.043 | 0.093 | 1 | 11 | Ercc6l2 | -0.05 | Young |
| 0.097349 | 0.058798 | 0.159 | 0.246 | 1 | 11 | Prpf8 | -0.087 | Young |
| 0.097493 | 0.072381 | 0.138 | 0.218 | 1 | 11 | BC028528 | -0.08 | Young |
| 0.09752 | 0.131572 | 0.116 | 0.194 | 1 | 11 | Abl2 | -0.078 | Young |
| 0.097541 | 0.383643 | 0.072 | 0.137 | 1 | 11 | Gtf3c1 | -0.065 | Young |
| 0.097551 | 2.345256 | 0.072 | 0.036 | 1 | 11 | Mthfs | 0.036 | Young |
| 0.098043 | 0.853367 | 0.167 | 0.254 | 1 | 11 | P4ha1 | -0.087 | Young |
| 0.098309 | 0.20343 | 0.254 | 0.375 | 1 | 11 | Vav1 | -0.121 | Young |
| 0.098445 | 0.042749 | 0.145 | 0.23 | 1 | 11 | Zc3h18 | -0.085 | Young |
| 0.098466 | 0.260327 | 0.21 | 0.31 | 1 | 11 | Atf7ip | -0.1 | Young |
| 0.098591 | 0.994083 | 0.268 | 0.206 | 1 | 11 | Ccl3 | 0.062 | Young |
| 0.098758 | 0.024475 | 0.572 | 0.722 | 1 | 11 | Arpc3 | -0.15 | Young |
| 0.099002 | 0.273444 | 0.145 | 0.23 | 1 | 11 | Unc50 | -0.085 | Young |
| 0.099009 | 3.137012 | 0.022 | 0.004 | 1 | 11 | Acbd4 | 0.018 | Young |
| 0.099009 | 2.777894 | 0.022 | 0.004 | 1 | 11 | Vnn3 | 0.018 | Young |
| 0.099009 | 2.545962 | 0.022 | 0.004 | 1 | 11 | Nipal3 | 0.018 | Young |
| 0.099009 | 2.352422 | 0.022 | 0.004 | 1 | 11 | C030006K1 | 0.018 | Young |
| 0.099009 | 2.285714 | 0.022 | 0.004 | 1 | 11 | Xkr8 | 0.018 | Young |
| 0.099009 | 2.100785 | 0.022 | 0.004 | 1 | 11 | Krtcap3 | 0.018 | Young |
| 0.099009 | 1.812224 | 0.022 | 0.004 | 1 | 11 | Zc4h2 | 0.018 | Young |
| 0.099016 | 0.056843 | 0.101 | 0.169 | 1 | 11 | Elp5 | -0.068 | Young |
| 0.099232 | 0.046202 | 0.116 | 0.19 | 1 | 11 | CAAA0111 | -0.074 | Young |
| 0.099723 | 0.193235 | 0.058 | 0.113 | 1 | 11 | Pop7 | -0.055 | Young |
| 0.099858 | 0.176973 | 0.036 | 0.081 | 1 | 11 | Zc3h3 | -0.045 | Young |
| 0.100124 | 2.138069 | 0.022 | 0.004 | 1 | 11 | Prdm16 | 0.018 | Young |
| 0.100124 | 2.027316 | 0.022 | 0.004 | 1 | 11 | Timp3 | 0.018 | Young |
| 0.100703 | 0.001985 | 0.21 | 0.306 | 1 | 11 | Srp19 | -0.096 | Young |

| | | | | | | | | |
|---|---|---|---|---|---|---|---|---|
| 0.100883 | 0.24432 | 0.486 | 0.448 | 1 | 11 | Cited2 | 0.038 | Young |
| 0.101249 | 2.172781 | 0.022 | 0.004 | 1 | 11 | F3 | 0.018 | Young |
| 0.101249 | 2.132182 | 0.022 | 0.004 | 1 | 11 | 9130019O2 | 0.018 | Young |
| 0.101249 | 1.94687 | 0.022 | 0.004 | 1 | 11 | Mmp8 | 0.018 | Young |
| 0.101249 | 1.695062 | 0.022 | 0.004 | 1 | 11 | Mical2 | 0.018 | Young |
| 0.101249 | 1.414705 | 0.022 | 0.004 | 1 | 11 | Npr1 | 0.018 | Young |
| 0.101249 | 1.193925 | 0.022 | 0.004 | 1 | 11 | Fam120aos | 0.018 | Young |
| 0.101286 | 0.029231 | 0.058 | 0.113 | 1 | 11 | Btbd9 | -0.055 | Young |
| 0.101417 | 0.424736 | 0.043 | 0.093 | 1 | 11 | Pkdcc | -0.05 | Young |
| 0.101927 | 0.244028 | 0.181 | 0.278 | 1 | 11 | Myo1c | -0.097 | Young |
| 0.101987 | 0.120095 | 0.065 | 0.121 | 1 | 11 | Sipa1l1 | -0.056 | Young |
| 0.102012 | 0.347742 | 0.609 | 0.649 | 1 | 11 | Ogt | -0.04 | Young |
| 0.102179 | 2.440896 | 0.043 | 0.016 | 1 | 11 | Nup107 | 0.027 | Young |
| 0.102179 | 2.187079 | 0.043 | 0.016 | 1 | 11 | Zfp770 | 0.027 | Young |
| 0.102385 | 1.091934 | 0.022 | 0.004 | 1 | 11 | Smim1 | 0.018 | Young |
| 0.102385 | 0.943757 | 0.022 | 0.004 | 1 | 11 | Gm9938 | 0.018 | Young |
| 0.102385 | 0.913746 | 0.022 | 0.004 | 1 | 11 | Rapsn | 0.018 | Young |
| 0.102385 | 0.86587 | 0.022 | 0.004 | 1 | 11 | Tshz3 | 0.018 | Young |
| 0.102646 | 1.48835 | 0.094 | 0.052 | 1 | 11 | Plxdc2 | 0.042 | Young |
| 0.10337 | 0.332476 | 0.203 | 0.315 | 1 | 11 | Psmb5 | -0.112 | Young |
| 0.103563 | 0.072151 | 0.159 | 0.25 | 1 | 11 | Siva1 | -0.091 | Young |
| 0.104031 | 0.077004 | 0.174 | 0.266 | 1 | 11 | Ddx46 | -0.092 | Young |
| 0.10419 | 1.496404 | 0.109 | 0.065 | 1 | 11 | Lilra5 | 0.044 | Young |
| 0.104197 | 0.261304 | 0.08 | 0.141 | 1 | 11 | Rnaset2a | -0.061 | Young |
| 0.1043 | 0.003189 | 0.159 | 0.242 | 1 | 11 | Inpp5k | -0.083 | Young |
| 0.104736 | 0.400422 | 0.051 | 0.101 | 1 | 11 | Zc3h7b | -0.05 | Young |
| 0.105074 | 0.37142 | 0.065 | 0.121 | 1 | 11 | Serpina3f | -0.056 | Young |
| 0.105115 | 1.090838 | 0.043 | 0.016 | 1 | 11 | Knstrn | 0.027 | Young |
| 0.105124 | 0.033856 | 0.152 | 0.238 | 1 | 11 | Chchd7 | -0.086 | Young |
| 0.105373 | 0.272768 | 0.13 | 0.214 | 1 | 11 | Fam192a | -0.084 | Young |
| 0.106057 | 2.420537 | 0.036 | 0.012 | 1 | 11 | E2f6 | 0.024 | Young |
| 0.106146 | 0.861078 | 0.42 | 0.403 | 1 | 11 | Bcl2a1b | 0.017 | Young |
| 0.10675 | 0.02054 | 0.428 | 0.556 | 1 | 11 | Sec61g | -0.128 | Young |
| 0.106867 | 0.089694 | 0.094 | 0.161 | 1 | 11 | S1pr1 | -0.067 | Young |
| 0.106894 | 2.800846 | 0.036 | 0.012 | 1 | 11 | Atp6v0c | 0.024 | Young |
| 0.106894 | 2.132543 | 0.036 | 0.012 | 1 | 11 | Dus2 | 0.024 | Young |
| 0.107091 | 0.210406 | 0.29 | 0.427 | 1 | 11 | Snrpd3 | -0.137 | Young |
| 0.107225 | 0.066068 | 0.101 | 0.169 | 1 | 11 | Cpeb2 | -0.068 | Young |
| 0.107367 | 0.179915 | 0.957 | 0.952 | 1 | 11 | Serinc3 | 0.005 | Young |
| 0.107547 | 0.03407 | 0.203 | 0.298 | 1 | 11 | Appl1 | -0.095 | Young |
| 0.107736 | 2.012699 | 0.036 | 0.012 | 1 | 11 | Ttc8 | 0.024 | Young |
| 0.107736 | 1.661459 | 0.036 | 0.012 | 1 | 11 | Gm15964 | 0.024 | Young |
| 0.107783 | 0.074853 | 0.268 | 0.363 | 1 | 11 | Hps3 | -0.095 | Young |
| 0.10792 | 0.477955 | 0.036 | 0.081 | 1 | 11 | Lxn | -0.045 | Young |
| 0.107985 | 0.199915 | 0.377 | 0.516 | 1 | 11 | Myo5a | -0.139 | Young |

| | | | | | | | | |
|---|---|---|---|---|---|---|---|---|
| 0.107986 | 0.168461 | 0.188 | 0.282 | 1 | 11 | 2310035C2 | -0.094 | Young |
| 0.108296 | 2.96483 | 0.029 | 0.008 | 1 | 11 | Bcl2l14 | 0.021 | Young |
| 0.108296 | 2.940033 | 0.029 | 0.008 | 1 | 11 | 2610035D1 | 0.021 | Young |
| 0.108296 | 2.660836 | 0.029 | 0.008 | 1 | 11 | Cdkn1c | 0.021 | Young |
| 0.108296 | 2.225893 | 0.029 | 0.008 | 1 | 11 | Zfp825 | 0.021 | Young |
| 0.108582 | 3.607207 | 0.036 | 0.012 | 1 | 11 | Hspb1 | 0.024 | Young |
| 0.108637 | 0.541259 | 0.029 | 0.069 | 1 | 11 | Tmub1 | -0.04 | Young |
| 0.108848 | 0.165205 | 0.246 | 0.363 | 1 | 11 | Stub1 | -0.117 | Young |
| 0.109058 | 0.501096 | 0.13 | 0.214 | 1 | 11 | Elovl1 | -0.084 | Young |
| 0.109145 | 0.041197 | 0.312 | 0.427 | 1 | 11 | Sgpp1 | -0.115 | Young |
| 0.109172 | 0.279629 | 0.217 | 0.323 | 1 | 11 | 2610001J0 | -0.106 | Young |
| 0.109276 | 2.863475 | 0.029 | 0.008 | 1 | 11 | Atp2b4 | 0.021 | Young |
| 0.109276 | 2.155049 | 0.029 | 0.008 | 1 | 11 | Map10 | 0.021 | Young |
| 0.109805 | 0.105937 | 0.159 | 0.242 | 1 | 11 | Slc25a28 | -0.083 | Young |
| 0.109844 | 0.137176 | 0.065 | 0.121 | 1 | 11 | Rnf40 | -0.056 | Young |
| 0.110062 | 0.643673 | 0.145 | 0.226 | 1 | 11 | Derl2 | -0.081 | Young |
| 0.110262 | 2.822998 | 0.029 | 0.008 | 1 | 11 | Egfl7 | 0.021 | Young |
| 0.11043 | 1.243414 | 0.217 | 0.169 | 1 | 11 | Hdac1 | 0.048 | Young |
| 0.110431 | 0.086528 | 0.123 | 0.198 | 1 | 11 | Mpnd | -0.075 | Young |
| 0.110836 | 0.021589 | 0.051 | 0.101 | 1 | 11 | Fopnl | -0.05 | Young |
| 0.111079 | 1.118905 | 0.072 | 0.036 | 1 | 11 | Acad8 | 0.036 | Young |
| 0.111079 | 0.986873 | 0.072 | 0.036 | 1 | 11 | Fhl3 | 0.036 | Young |
| 0.111154 | 1.396375 | 0.036 | 0.012 | 1 | 11 | Fdxacb1 | 0.024 | Young |
| 0.111199 | 0.044323 | 0.203 | 0.294 | 1 | 11 | Rab35 | -0.091 | Young |
| 0.111279 | 0.095287 | 0.348 | 0.468 | 1 | 11 | Dok2 | -0.12 | Young |
| 0.111512 | 0.354118 | 0.304 | 0.435 | 1 | 11 | Cd40 | -0.131 | Young |
| 0.111557 | 0.285095 | 0.254 | 0.363 | 1 | 11 | Ncoa3 | -0.109 | Young |
| 0.111889 | 0.423626 | 0.087 | 0.153 | 1 | 11 | Ints9 | -0.066 | Young |
| 0.112022 | 1.150372 | 0.036 | 0.012 | 1 | 11 | Fads3 | 0.024 | Young |
| 0.112054 | 2.468912 | 0.065 | 0.032 | 1 | 11 | Mcph1 | 0.033 | Young |
| 0.112292 | 0.289337 | 0.065 | 0.121 | 1 | 11 | Zfp639 | -0.056 | Young |
| 0.112369 | 0.097301 | 0.159 | 0.246 | 1 | 11 | Snx4 | -0.087 | Young |
| 0.112405 | 0.0174 | 0.138 | 0.218 | 1 | 11 | Por | -0.08 | Young |
| 0.113118 | 0.306818 | 0.065 | 0.121 | 1 | 11 | Ccdc134 | -0.056 | Young |
| 0.113872 | 0.040648 | 0.333 | 0.456 | 1 | 11 | Sumo3 | -0.123 | Young |
| 0.114044 | 0.16842 | 0.225 | 0.323 | 1 | 11 | Dhx9 | -0.098 | Young |
| 0.114085 | 0.082561 | 0.145 | 0.226 | 1 | 11 | Emsy | -0.081 | Young |
| 0.114111 | 0.126204 | 0.123 | 0.202 | 1 | 11 | Pih1d1 | -0.079 | Young |
| 0.114604 | 0.028843 | 0.072 | 0.125 | 1 | 11 | Bloc1s4 | -0.053 | Young |
| 0.114632 | 0.248278 | 0.029 | 0.069 | 1 | 11 | Acad9 | -0.04 | Young |
| 0.114773 | 0.344356 | 0.123 | 0.202 | 1 | 11 | Smg5 | -0.079 | Young |
| 0.114776 | 0.30091 | 0.138 | 0.21 | 1 | 11 | Ulk2 | -0.072 | Young |
| 0.115018 | 0.368439 | 0.667 | 0.673 | 1 | 11 | Ifi203 | -0.006 | Young |
| 0.115202 | 0.30769 | 0.065 | 0.121 | 1 | 11 | Ppp2cb | -0.056 | Young |
| 0.11556 | 0.131142 | 0.058 | 0.109 | 1 | 11 | Rmdn3 | -0.051 | Young |

| | | | | | | | | |
|---|---|---|---|---|---|---|---|---|
| 0.11557 | 0.256286 | 0.21 | 0.319 | 1 | 11 | Usp19 | -0.109 | Young |
| 0.115675 | 0.276294 | 0.196 | 0.294 | 1 | 11 | Suz12 | -0.098 | Young |
| 0.115966 | 0.087505 | 0.094 | 0.157 | 1 | 11 | Vamp7 | -0.063 | Young |
| 0.116003 | 0.302285 | 0.058 | 0.109 | 1 | 11 | Zfyve26 | -0.051 | Young |
| 0.116028 | 0.0746 | 0.116 | 0.19 | 1 | 11 | Ticam2 | -0.074 | Young |
| 0.116045 | 0.028655 | 0.072 | 0.129 | 1 | 11 | Gm5617 | -0.057 | Young |
| 0.116339 | 0.254671 | 0.094 | 0.157 | 1 | 11 | Sdad1 | -0.063 | Young |
| 0.116447 | 0.221626 | 0.058 | 0.109 | 1 | 11 | Ttc33 | -0.051 | Young |
| 0.1165 | 0.236092 | 0.109 | 0.177 | 1 | 11 | Fbrs | -0.068 | Young |
| 0.116854 | 0.133394 | 0.109 | 0.177 | 1 | 11 | Vps33a | -0.068 | Young |
| 0.11686 | 0.869599 | 0.087 | 0.048 | 1 | 11 | Hacd2 | 0.039 | Young |
| 0.116892 | 0.324194 | 0.058 | 0.109 | 1 | 11 | Med25 | -0.051 | Young |
| 0.117024 | 0.035615 | 0.116 | 0.181 | 1 | 11 | Yrdc | -0.065 | Young |
| 0.117073 | 0.004911 | 0.188 | 0.282 | 1 | 11 | Fam32a | -0.094 | Young |
| 0.117245 | 0.217995 | 0.159 | 0.246 | 1 | 11 | 4933434E2 | -0.087 | Young |
| 0.117317 | 0.035561 | 0.065 | 0.121 | 1 | 11 | Diablo | -0.056 | Young |
| 0.117404 | 0.135164 | 0.239 | 0.347 | 1 | 11 | Apaf1 | -0.108 | Young |
| 0.117414 | 0.313194 | 0.138 | 0.222 | 1 | 11 | Psmd7 | -0.084 | Young |
| 0.117824 | 0.021024 | 0.21 | 0.31 | 1 | 11 | Sppl3 | -0.1 | Young |
| 0.117838 | 0.108188 | 0.094 | 0.157 | 1 | 11 | Acadsb | -0.063 | Young |
| 0.118025 | 0.244957 | 0.094 | 0.161 | 1 | 11 | Hdac5 | -0.067 | Young |
| 0.118126 | 0.189809 | 0.203 | 0.298 | 1 | 11 | Tmbim4 | -0.095 | Young |
| 0.118171 | 0.069636 | 0.297 | 0.444 | 1 | 11 | Rpl13a | -0.147 | Young |
| 0.118319 | 0.10029 | 0.188 | 0.278 | 1 | 11 | Wtap | -0.09 | Young |
| 0.118639 | 0.18981 | 0.109 | 0.177 | 1 | 11 | Tbc1d22a | -0.068 | Young |
| 0.11886 | 0.126796 | 0.022 | 0.056 | 1 | 11 | Klhl22 | -0.034 | Young |
| 0.119004 | 0.264722 | 0.043 | 0.089 | 1 | 11 | Ap1g2 | -0.046 | Young |
| 0.119004 | 0.153821 | 0.043 | 0.089 | 1 | 11 | Rhot2 | -0.046 | Young |
| 0.119029 | 0.156864 | 0.21 | 0.306 | 1 | 11 | Rc3h2 | -0.096 | Young |
| 0.11913 | 0.313109 | 0.196 | 0.29 | 1 | 11 | Mcmbp | -0.094 | Young |
| 0.119706 | 0.67056 | 0.087 | 0.149 | 1 | 11 | Tfeb | -0.062 | Young |
| 0.119944 | 0.011312 | 0.181 | 0.27 | 1 | 11 | Gorasp2 | -0.089 | Young |
| 0.120098 | 0.946851 | 0.159 | 0.109 | 1 | 11 | Pygo2 | 0.05 | Young |
| 0.120131 | 0.06001 | 0.022 | 0.056 | 1 | 11 | Mad2l1bp | -0.034 | Young |
| 0.120131 | 0.014437 | 0.022 | 0.056 | 1 | 11 | Kbtbd7 | -0.034 | Young |
| 0.120591 | 0.184988 | 0.254 | 0.363 | 1 | 11 | Dpy19l1 | -0.109 | Young |
| 0.120629 | 0.000717 | 0.138 | 0.214 | 1 | 11 | Polr2i | -0.076 | Young |
| 0.121041 | 0.166987 | 0.065 | 0.117 | 1 | 11 | Snx15 | -0.052 | Young |
| 0.121647 | 0.110762 | 0.159 | 0.246 | 1 | 11 | Dock1 | -0.087 | Young |
| 0.121826 | 0.442192 | 0.123 | 0.194 | 1 | 11 | Sfswap | -0.071 | Young |
| 0.122362 | 0.250827 | 0.08 | 0.137 | 1 | 11 | Calhm2 | -0.057 | Young |
| 0.122706 | 0.996668 | 0.065 | 0.032 | 1 | 11 | Rusc2 | 0.033 | Young |
| 0.122706 | 0.089855 | 0.022 | 0.056 | 1 | 11 | Spice1 | -0.034 | Young |
| 0.122706 | 0.001733 | 0.022 | 0.056 | 1 | 11 | Decr2 | -0.034 | Young |
| 0.122804 | 0.193655 | 0.058 | 0.109 | 1 | 11 | Ddx41 | -0.051 | Young |

| | | | | | | | | |
|---|---|---|---|---|---|---|---|---|
| 0.122993 | 0.0835 | 0.428 | 0.593 | 1 | 11 | Cd84 | -0.165 | Young |
| 0.123025 | 0.295006 | 0.203 | 0.306 | 1 | 11 | Mfsd5 | -0.103 | Young |
| 0.123046 | 0.15554 | 0.007 | 0.032 | 1 | 11 | Zfp41 | -0.025 | Young |
| 0.123046 | 0.127386 | 0.007 | 0.032 | 1 | 11 | Amacr | -0.025 | Young |
| 0.123065 | 0.190803 | 0.043 | 0.089 | 1 | 11 | Rcc1l | -0.046 | Young |
| 0.123107 | 0.296564 | 0.203 | 0.31 | 1 | 11 | Tsen34 | -0.107 | Young |
| 0.123206 | 0.224705 | 0.094 | 0.157 | 1 | 11 | Asnsd1 | -0.063 | Young |
| 0.1233 | 0.01888 | 0.196 | 0.29 | 1 | 11 | Park7 | -0.094 | Young |
| 0.12366 | 0.01494 | 0.014 | 0.044 | 1 | 11 | Snx7 | -0.03 | Young |
| 0.123697 | 0.219912 | 0.899 | 0.944 | 1 | 11 | Rps10 | -0.045 | Young |
| 0.123841 | 0.211759 | 0.109 | 0.173 | 1 | 11 | Hadh | -0.064 | Young |
| 0.123991 | 0.001365 | 0.246 | 0.347 | 1 | 11 | Emc10 | -0.101 | Young |
| 0.124097 | 0.183768 | 0.043 | 0.089 | 1 | 11 | Esr1 | -0.046 | Young |
| 0.124423 | 0.037349 | 0.101 | 0.161 | 1 | 11 | Mrps7 | -0.06 | Young |
| 0.12487 | 0.046921 | 0.145 | 0.222 | 1 | 11 | Abcc1 | -0.077 | Young |
| 0.124878 | 0.454105 | 0.072 | 0.129 | 1 | 11 | Nufip1 | -0.057 | Young |
| 0.125245 | 0.172791 | 0.174 | 0.262 | 1 | 11 | Pon2 | -0.088 | Young |
| 0.125395 | 0.01676 | 0.167 | 0.25 | 1 | 11 | Tfg | -0.083 | Young |
| 0.125722 | 0.227003 | 0.109 | 0.173 | 1 | 11 | Trpv4 | -0.064 | Young |
| 0.125876 | 0.045784 | 0.464 | 0.609 | 1 | 11 | Arpc4 | -0.145 | Young |
| 0.12596 | 0.144292 | 0.094 | 0.157 | 1 | 11 | Aggf1 | -0.063 | Young |
| 0.126147 | 0.245064 | 0.21 | 0.31 | 1 | 11 | Ttc3 | -0.1 | Young |
| 0.126315 | 0.827745 | 0.304 | 0.25 | 1 | 11 | Dusp6 | 0.054 | Young |
| 0.126358 | 0.073132 | 0.094 | 0.157 | 1 | 11 | Ccdc22 | -0.063 | Young |
| 0.126659 | 0.031736 | 0.014 | 0.044 | 1 | 11 | Depdc7 | -0.03 | Young |
| 0.126744 | 0.28181 | 0.109 | 0.177 | 1 | 11 | Tmem147 | -0.068 | Young |
| 0.127316 | 1.309195 | 0.065 | 0.032 | 1 | 11 | Nup188 | 0.033 | Young |
| 0.127459 | 0.286528 | 0.275 | 0.391 | 1 | 11 | Nsd1 | -0.116 | Young |
| 0.127496 | 0.071069 | 0.072 | 0.129 | 1 | 11 | Sec23a | -0.057 | Young |
| 0.127502 | 0.125123 | 0.109 | 0.177 | 1 | 11 | Mybbp1a | -0.068 | Young |
| 0.127702 | 0.056274 | 0.304 | 0.423 | 1 | 11 | Rnf141 | -0.119 | Young |
| 0.128042 | 0.508072 | 0.051 | 0.097 | 1 | 11 | Paxip1 | -0.046 | Young |
| 0.12818 | 0.781458 | 0.014 | 0.044 | 1 | 11 | Bphl | -0.03 | Young |
| 0.12818 | 0.590418 | 0.014 | 0.044 | 1 | 11 | Sirt3 | -0.03 | Young |
| 0.12818 | 0.117298 | 0.014 | 0.044 | 1 | 11 | Scn2b | -0.03 | Young |
| 0.128229 | 0.182592 | 0.167 | 0.254 | 1 | 11 | St3gal1 | -0.087 | Young |
| 0.128466 | 0.304115 | 0.058 | 0.109 | 1 | 11 | Ptk2 | -0.051 | Young |
| 0.128953 | 0.72549 | 0.5 | 0.516 | 1 | 11 | Ppp1r18 | -0.016 | Young |
| 0.129086 | 0.048745 | 0.167 | 0.246 | 1 | 11 | Mien1 | -0.079 | Young |
| 0.129283 | 0.411986 | 0.116 | 0.19 | 1 | 11 | Pex6 | -0.074 | Young |
| 0.129333 | 1.71268 | 0.065 | 0.032 | 1 | 11 | Zfp719 | 0.033 | Young |
| 0.129388 | 0.29974 | 0.21 | 0.319 | 1 | 11 | Gm4951 | -0.109 | Young |
| 0.129396 | 0.191744 | 0.123 | 0.194 | 1 | 11 | Zmym5 | -0.071 | Young |
| 0.129668 | 0.24047 | 0.942 | 0.964 | 1 | 11 | Klf6 | -0.022 | Young |
| 0.13008 | 0.124662 | 0.051 | 0.097 | 1 | 11 | Brpf3 | -0.046 | Young |

| | | | | | | | | |
|---|---|---|---|---|---|---|---|---|
| 0.13026 | 0.157733 | 0.159 | 0.238 | 1 | 11 | Gatad1 | -0.079 | Young |
| 0.130439 | 0.568321 | 0.036 | 0.077 | 1 | 11 | Cwc22 | -0.041 | Young |
| 0.130505 | 0.296511 | 0.761 | 0.77 | 1 | 11 | Rpl35 | -0.009 | Young |
| 0.131176 | 0.206887 | 0.159 | 0.246 | 1 | 11 | Arid1b | -0.087 | Young |
| 0.131436 | 0.064585 | 0.145 | 0.222 | 1 | 11 | Tnfsf12 | -0.077 | Young |
| 0.131512 | 0.460194 | 0.043 | 0.089 | 1 | 11 | Med4 | -0.046 | Young |
| 0.131613 | 0.134861 | 0.094 | 0.157 | 1 | 11 | Stx6 | -0.063 | Young |
| 0.131954 | 0.190828 | 0.072 | 0.129 | 1 | 11 | Rbm38 | -0.057 | Young |
| 0.132113 | 0.481476 | 0.043 | 0.085 | 1 | 11 | Phf13 | -0.042 | Young |
| 0.132205 | 0.689104 | 0.058 | 0.105 | 1 | 11 | Cks2 | -0.047 | Young |
| 0.132405 | 0.166477 | 0.283 | 0.403 | 1 | 11 | Tmem109 | -0.12 | Young |
| 0.132482 | 0.021571 | 0.239 | 0.339 | 1 | 11 | Hsd17b12 | -0.1 | Young |
| 0.132482 | 0.184606 | 0.065 | 0.117 | 1 | 11 | Tbc1d31 | -0.052 | Young |
| 0.132686 | 0.250607 | 0.087 | 0.145 | 1 | 11 | Pex2 | -0.058 | Young |
| 0.13286 | 0.420333 | 0.072 | 0.129 | 1 | 11 | Ambra1 | -0.057 | Young |
| 0.133833 | 0.577295 | 0.058 | 0.109 | 1 | 11 | Cdyl | -0.051 | Young |
| 0.133922 | 0.30061 | 0.196 | 0.286 | 1 | 11 | Dynlt3 | -0.09 | Young |
| 0.134102 | 0.050131 | 0.507 | 0.694 | 1 | 11 | Serp1 | -0.187 | Young |
| 0.134262 | 0.200425 | 0.109 | 0.173 | 1 | 11 | Snx12 | -0.064 | Young |
| 0.134512 | 0.465334 | 0.094 | 0.157 | 1 | 11 | Rbm18 | -0.063 | Young |
| 0.134781 | 0.041406 | 0.464 | 0.621 | 1 | 11 | H2afy | -0.157 | Young |
| 0.134783 | 0.282857 | 0.826 | 0.879 | 1 | 11 | Zfp36l1 | -0.053 | Young |
| 0.134868 | 0.399016 | 0.065 | 0.117 | 1 | 11 | Slc25a30 | -0.052 | Young |
| 0.135097 | 0.07552 | 0.261 | 0.367 | 1 | 11 | Klhl9 | -0.106 | Young |
| 0.135993 | 0.309363 | 0.254 | 0.371 | 1 | 11 | Ppp1r21 | -0.117 | Young |
| 0.136327 | 0.062461 | 0.159 | 0.246 | 1 | 11 | Cdc40 | -0.087 | Young |
| 0.13688 | 0.10012 | 0.29 | 0.399 | 1 | 11 | Ago2 | -0.109 | Young |
| 0.136913 | 0.037878 | 0.167 | 0.246 | 1 | 11 | Cpsf7 | -0.079 | Young |
| 0.137053 | 0.090849 | 0.312 | 0.431 | 1 | 11 | Hnrnpul1 | -0.119 | Young |
| 0.137522 | 0.152599 | 0.036 | 0.077 | 1 | 11 | Letmd1 | -0.041 | Young |
| 0.137729 | 0.357097 | 0.239 | 0.347 | 1 | 11 | Pum1 | -0.108 | Young |
| 0.137797 | 0.682315 | 0.37 | 0.331 | 1 | 11 | Camkk2 | 0.039 | Young |
| 0.138068 | 0.192278 | 0.181 | 0.27 | 1 | 11 | Denr | -0.089 | Young |
| 0.138125 | 0.080899 | 0.13 | 0.206 | 1 | 11 | Pdha1 | -0.076 | Young |
| 0.138125 | 0.283143 | 0.036 | 0.077 | 1 | 11 | Wdyhv1 | -0.041 | Young |
| 0.138199 | 1.491017 | 0.058 | 0.028 | 1 | 11 | Serinc5 | 0.03 | Young |
| 0.138201 | 0.672326 | 0.391 | 0.343 | 1 | 11 | Fcrls | 0.048 | Young |
| 0.139264 | 0.004745 | 0.181 | 0.262 | 1 | 11 | Tcof1 | -0.081 | Young |
| 0.139559 | 0.04969 | 0.217 | 0.315 | 1 | 11 | Pkn2 | -0.098 | Young |
| 0.140518 | 1.076339 | 0.138 | 0.093 | 1 | 11 | Tes | 0.045 | Young |
| 0.141009 | 0.060285 | 0.087 | 0.145 | 1 | 11 | Gyg | -0.058 | Young |
| 0.141664 | 0.054562 | 0.123 | 0.19 | 1 | 11 | Plpbp | -0.067 | Young |
| 0.141836 | 0.699943 | 0.362 | 0.315 | 1 | 11 | Cysltr1 | 0.047 | Young |
| 0.14207 | 0.048116 | 0.254 | 0.363 | 1 | 11 | Zmiz2 | -0.109 | Young |
| 0.142114 | 0.302118 | 0.181 | 0.266 | 1 | 11 | Nsfl1c | -0.085 | Young |

| | | | | | | | | |
|---|---|---|---|---|---|---|---|---|
| 0.142143 | 0.307885 | 0.094 | 0.153 | 1 | 11 | Fnta | -0.059 | Young |
| 0.142285 | 0.485212 | 0.051 | 0.097 | 1 | 11 | Timm9 | -0.046 | Young |
| 0.142288 | 0.069159 | 0.225 | 0.315 | 1 | 11 | Plekha2 | -0.09 | Young |
| 0.142735 | 0.113108 | 0.246 | 0.343 | 1 | 11 | Rb1cc1 | -0.097 | Young |
| 0.142763 | 0.117133 | 0.261 | 0.371 | 1 | 11 | Hadhb | -0.11 | Young |
| 0.142923 | 0.00894 | 0.138 | 0.206 | 1 | 11 | Creld2 | -0.068 | Young |
| 0.143292 | 0.0191 | 0.377 | 0.516 | 1 | 11 | Matr3 | -0.139 | Young |
| 0.143502 | 0.146698 | 0.246 | 0.351 | 1 | 11 | Ncstn | -0.105 | Young |
| 0.144058 | 0.233252 | 0.188 | 0.29 | 1 | 11 | Zfp800 | -0.102 | Young |
| 0.14416 | 0.168659 | 0.217 | 0.306 | 1 | 11 | Fam91a1 | -0.089 | Young |
| 0.144282 | 0.304943 | 0.116 | 0.185 | 1 | 11 | Fem1b | -0.069 | Young |
| 0.144808 | 2.325093 | 0.087 | 0.052 | 1 | 11 | Mtmr9 | 0.035 | Young |
| 0.145514 | 0.043054 | 0.428 | 0.577 | 1 | 11 | Mrpl52 | -0.149 | Young |
| 0.145609 | 0.325481 | 0.051 | 0.097 | 1 | 11 | Tmem70 | -0.046 | Young |
| 0.145704 | 0.146311 | 0.826 | 0.806 | 1 | 11 | Rps2 | 0.02 | Young |
| 0.14589 | 0.068988 | 0.188 | 0.274 | 1 | 11 | Vrk2 | -0.086 | Young |
| 0.146173 | 0.080707 | 0.058 | 0.105 | 1 | 11 | Fam173b | -0.047 | Young |
| 0.146454 | 0.100324 | 0.594 | 0.722 | 1 | 11 | Wfdc17 | -0.128 | Young |
| 0.146731 | 0.368713 | 0.051 | 0.097 | 1 | 11 | Mtmr4 | -0.046 | Young |
| 0.146738 | 0.246651 | 0.862 | 0.903 | 1 | 11 | Pim1 | -0.041 | Young |
| 0.146897 | 0.08523 | 0.145 | 0.222 | 1 | 11 | Plaa | -0.077 | Young |
| 0.146915 | 0.157226 | 0.123 | 0.19 | 1 | 11 | Uimc1 | -0.067 | Young |
| 0.147017 | 0.080216 | 1 | 1 | 1 | 11 | Rps29 | 0 | Young |
| 0.147394 | 0.161587 | 0.087 | 0.145 | 1 | 11 | Lipe | -0.058 | Young |
| 0.147406 | 0.0456 | 0.174 | 0.254 | 1 | 11 | Atad2b | -0.08 | Young |
| 0.147644 | 0.559787 | 0.072 | 0.125 | 1 | 11 | Usb1 | -0.053 | Young |
| 0.147665 | 0.486499 | 0.167 | 0.246 | 1 | 11 | Gripap1 | -0.079 | Young |
| 0.148304 | 0.137997 | 0.043 | 0.085 | 1 | 11 | Hic2 | -0.042 | Young |
| 0.148324 | 0.40136 | 0.087 | 0.145 | 1 | 11 | Ccpg1 | -0.058 | Young |
| 0.148346 | 0.002819 | 0.087 | 0.141 | 1 | 11 | Tatdn2 | -0.054 | Young |
| 0.148349 | 0.995101 | 0.167 | 0.121 | 1 | 11 | Ctdspl2 | 0.046 | Young |
| 0.149029 | 0.236164 | 0.13 | 0.198 | 1 | 11 | Idi1 | -0.068 | Young |
| 0.149487 | 0.003282 | 0.319 | 0.431 | 1 | 11 | Dnase2a | -0.112 | Young |
| 0.149565 | 0.11405 | 0.029 | 0.065 | 1 | 11 | 4921524J1 | -0.036 | Young |
| 0.149565 | 0.099881 | 0.029 | 0.065 | 1 | 11 | Arl14ep | -0.036 | Young |
| 0.149626 | 0.09231 | 0.109 | 0.173 | 1 | 11 | Erlec1 | -0.064 | Young |
| 0.149787 | 0.193681 | 0.268 | 0.391 | 1 | 11 | 7-Mar | -0.123 | Young |
| 0.149963 | 0.073281 | 0.152 | 0.226 | 1 | 11 | Clcn7 | -0.074 | Young |
| 0.150346 | 0.236506 | 0.819 | 0.867 | 1 | 11 | Rps6 | -0.048 | Young |
| 0.150636 | 0.296572 | 0.819 | 0.774 | 1 | 11 | Ccl9 | 0.045 | Young |
| 0.150736 | 0.285377 | 0.043 | 0.085 | 1 | 11 | Acot13 | -0.042 | Young |
| 0.15097 | 0.364525 | 0.029 | 0.065 | 1 | 11 | Scai | -0.036 | Young |
| 0.15097 | 0.05631 | 0.029 | 0.065 | 1 | 11 | Wdr53 | -0.036 | Young |
| 0.150971 | 0.009369 | 0.196 | 0.278 | 1 | 11 | Cyth1 | -0.082 | Young |
| 0.151013 | 0.199336 | 0.196 | 0.274 | 1 | 11 | Mrpl30 | -0.078 | Young |

| | | | | | | | | |
|---|---|---|---|---|---|---|---|---|
| 0.151072 | 0.465892 | 0.058 | 0.105 | 1 | 11 | Pycr2 | -0.047 | Young |
| 0.151507 | 0.107829 | 0.145 | 0.214 | 1 | 11 | Rabgef1 | -0.069 | Young |
| 0.151566 | 0.306521 | 0.065 | 0.113 | 1 | 11 | Pigyl | -0.048 | Young |
| 0.151901 | 0.012545 | 0.13 | 0.202 | 1 | 11 | Nub1 | -0.072 | Young |
| 0.152177 | 0.583358 | 0.058 | 0.105 | 1 | 11 | Akt1s1 | -0.047 | Young |
| 0.152304 | 0.112497 | 0.167 | 0.25 | 1 | 11 | Abcd1 | -0.083 | Young |
| 0.152588 | 0.448406 | 0.022 | 0.052 | 1 | 11 | Sesn2 | -0.03 | Young |
| 0.152684 | 0.020837 | 0.152 | 0.226 | 1 | 11 | Ext1 | -0.074 | Young |
| 0.152983 | 0.106657 | 0.094 | 0.149 | 1 | 11 | Ebna1bp2 | -0.055 | Young |
| 0.153076 | 0.447859 | 0.152 | 0.226 | 1 | 11 | Ndufa10 | -0.074 | Young |
| 0.153097 | 1.638104 | 0.072 | 0.04 | 1 | 11 | Ppp2r5d | 0.032 | Young |
| 0.153167 | 0.16083 | 0.065 | 0.113 | 1 | 11 | Tspan33 | -0.048 | Young |
| 0.153188 | 0.339471 | 0.101 | 0.161 | 1 | 11 | Vps4a | -0.06 | Young |
| 0.153198 | 0.116111 | 0.043 | 0.085 | 1 | 11 | Narfl | -0.042 | Young |
| 0.1533 | 0.152829 | 0.928 | 0.94 | 1 | 11 | Rpl13 | -0.012 | Young |
| 0.153334 | 0.147533 | 0.145 | 0.218 | 1 | 11 | Nuak1 | -0.073 | Young |
| 0.153608 | 2.244375 | 0.051 | 0.024 | 1 | 11 | Plcb1 | 0.027 | Young |
| 0.154016 | 0.22832 | 0.159 | 0.238 | 1 | 11 | Angel2 | -0.079 | Young |
| 0.154244 | 0.022369 | 0.167 | 0.242 | 1 | 11 | Ebp | -0.075 | Young |
| 0.154488 | 2.144597 | 0.051 | 0.024 | 1 | 11 | Zc3h8 | 0.027 | Young |
| 0.154677 | 0.172912 | 0.717 | 0.738 | 1 | 11 | Rbpj | -0.021 | Young |
| 0.154782 | 0.921931 | 0.123 | 0.081 | 1 | 11 | Pi4k2b | 0.042 | Young |
| 0.154824 | 0.096498 | 0.464 | 0.661 | 1 | 11 | 5031439G( | -0.197 | Young |
| 0.154854 | 0.083285 | 0.123 | 0.19 | 1 | 11 | Ibtk | -0.067 | Young |
| 0.155372 | 1.820762 | 0.051 | 0.024 | 1 | 11 | Ttc28 | 0.027 | Young |
| 0.155652 | 0.041918 | 0.232 | 0.323 | 1 | 11 | Esd | -0.091 | Young |
| 0.155691 | 0.107541 | 0.043 | 0.085 | 1 | 11 | Trmt2a | -0.042 | Young |
| 0.155769 | 0.050778 | 0.196 | 0.27 | 1 | 11 | Ccr1 | -0.074 | Young |
| 0.156094 | 0.30069 | 0.058 | 0.105 | 1 | 11 | Isg20l2 | -0.047 | Young |
| 0.156094 | 0.111757 | 0.058 | 0.105 | 1 | 11 | Patz1 | -0.047 | Young |
| 0.156153 | 0.206631 | 0.152 | 0.23 | 1 | 11 | Dpf2 | -0.078 | Young |
| 0.156155 | 0.102128 | 0.239 | 0.351 | 1 | 11 | Phf21a | -0.112 | Young |
| 0.15629 | 0.113065 | 0.246 | 0.359 | 1 | 11 | Stx16 | -0.113 | Young |
| 0.156319 | 0.58101 | 0.043 | 0.085 | 1 | 11 | Slc25a38 | -0.042 | Young |
| 0.156336 | 0.018885 | 0.152 | 0.218 | 1 | 11 | Cnp | -0.066 | Young |
| 0.156439 | 0.010639 | 0.159 | 0.23 | 1 | 11 | Ddx42 | -0.071 | Young |
| 0.157247 | 0.151477 | 0.094 | 0.149 | 1 | 11 | Ext2 | -0.055 | Young |
| 0.157782 | 0.019455 | 0.138 | 0.206 | 1 | 11 | Ccm2 | -0.068 | Young |
| 0.158046 | 1.751306 | 0.051 | 0.024 | 1 | 11 | Zc3h6 | 0.027 | Young |
| 0.15814 | 0.090272 | 0.297 | 0.399 | 1 | 11 | Chmp5 | -0.102 | Young |
| 0.158199 | 1.082089 | 0.022 | 0.052 | 1 | 11 | Extl2 | -0.03 | Young |
| 0.158727 | 0.253119 | 0.123 | 0.19 | 1 | 11 | Slain2 | -0.067 | Young |
| 0.158913 | 0.235435 | 0.167 | 0.254 | 1 | 11 | Cant1 | -0.087 | Young |
| 0.158974 | 0.02105 | 0.087 | 0.141 | 1 | 11 | Gan | -0.054 | Young |
| 0.159556 | 0.155943 | 0.051 | 0.093 | 1 | 11 | Gm43848 | -0.042 | Young |

| | | | | | | | | |
|---|---|---|---|---|---|---|---|---|
| 0.159598 | 0.062747 | 0.246 | 0.355 | 1 | 11 | Azi2 | -0.109 | Young |
| 0.15983 | 0.304574 | 0.022 | 0.052 | 1 | 11 | Otud1 | -0.03 | Young |
| 0.160651 | 0.014603 | 0.022 | 0.052 | 1 | 11 | Tmem161a | -0.03 | Young |
| 0.160686 | 0.213572 | 0.232 | 0.319 | 1 | 11 | Ndufb6 | -0.087 | Young |
| 0.160772 | 0.081978 | 0.116 | 0.181 | 1 | 11 | Mrpl18 | -0.065 | Young |
| 0.160785 | 0.255023 | 0.732 | 0.762 | 1 | 11 | Ier5 | -0.03 | Young |
| 0.160813 | 0.192784 | 0.065 | 0.113 | 1 | 11 | Cenpa | -0.048 | Young |
| 0.16094 | 0.304008 | 0.174 | 0.254 | 1 | 11 | Ufsp2 | -0.08 | Young |
| 0.161023 | 0.18439 | 0.283 | 0.403 | 1 | 11 | Snx6 | -0.12 | Young |
| 0.161217 | 0.43996 | 0.138 | 0.21 | 1 | 11 | Acaa2 | -0.072 | Young |
| 0.161414 | 0.239992 | 0.297 | 0.411 | 1 | 11 | Nufip2 | -0.114 | Young |
| 0.161666 | 1.740396 | 0.051 | 0.024 | 1 | 11 | Trmt44 | 0.027 | Young |
| 0.161789 | 0.375196 | 0.123 | 0.19 | 1 | 11 | Epc1 | -0.067 | Young |
| 0.162003 | 0.166631 | 0.051 | 0.093 | 1 | 11 | Trappc2 | -0.042 | Young |
| 0.162157 | 0.614573 | 0.036 | 0.073 | 1 | 11 | Fggy | -0.037 | Young |
| 0.162231 | 0.01446 | 0.181 | 0.262 | 1 | 11 | Luzp1 | -0.081 | Young |
| 0.162255 | 0.228031 | 0.217 | 0.306 | 1 | 11 | Psmd1 | -0.089 | Young |
| 0.162345 | 0.040033 | 0.196 | 0.278 | 1 | 11 | Xrn1 | -0.082 | Young |
| 0.162488 | 0.151156 | 0.065 | 0.113 | 1 | 11 | Bid | -0.048 | Young |
| 0.163154 | 0.04017 | 0.145 | 0.21 | 1 | 11 | Slc35a4 | -0.065 | Young |
| 0.163318 | 0.052767 | 0.203 | 0.29 | 1 | 11 | Srrt | -0.087 | Young |
| 0.163716 | 0.222658 | 0.123 | 0.185 | 1 | 11 | Insr | -0.062 | Young |
| 0.16379 | 0.163696 | 0.145 | 0.214 | 1 | 11 | Cdc123 | -0.069 | Young |
| 0.163841 | 0.305396 | 0.188 | 0.274 | 1 | 11 | Mta2 | -0.086 | Young |
| 0.163858 | 0.459347 | 0.051 | 0.093 | 1 | 11 | Gpank1 | -0.042 | Young |
| 0.163892 | 0.459328 | 0.08 | 0.133 | 1 | 11 | Fam193b | -0.053 | Young |
| 0.163927 | 0.217409 | 0.116 | 0.181 | 1 | 11 | Uhrf1bp1l | -0.065 | Young |
| 0.163939 | 0.056809 | 0.072 | 0.121 | 1 | 11 | Gtdc1 | -0.049 | Young |
| 0.164075 | 0.061672 | 0.623 | 0.81 | 1 | 11 | Plek | -0.187 | Young |
| 0.164092 | 0.035026 | 0.094 | 0.145 | 1 | 11 | Hs2st1 | -0.051 | Young |
| 0.164164 | 0.484544 | 0.217 | 0.319 | 1 | 11 | Thoc2 | -0.102 | Young |
| 0.164454 | 0.026013 | 0.116 | 0.173 | 1 | 11 | Adssl1 | -0.057 | Young |
| 0.164702 | 0.190422 | 0.261 | 0.371 | 1 | 11 | Cul3 | -0.11 | Young |
| 0.16499 | 0.108963 | 0.174 | 0.25 | 1 | 11 | Rnft1 | -0.076 | Young |
| 0.165573 | 0.08754 | 0.13 | 0.194 | 1 | 11 | Dyrk1a | -0.064 | Young |
| 0.165792 | 0.185215 | 0.123 | 0.19 | 1 | 11 | Utp14a | -0.067 | Young |
| 0.165898 | 0.148086 | 0.174 | 0.258 | 1 | 11 | Dnajc2 | -0.084 | Young |
| 0.166523 | 0.516324 | 0.167 | 0.238 | 1 | 11 | Tmem229b | -0.071 | Young |
| 0.166545 | 0.234249 | 0.087 | 0.141 | 1 | 11 | Gsto1 | -0.054 | Young |
| 0.166592 | 0.238757 | 0.217 | 0.323 | 1 | 11 | Atp13a2 | -0.106 | Young |
| 0.166709 | 1.282482 | 0.123 | 0.085 | 1 | 11 | Pus1 | 0.038 | Young |
| 0.166724 | 0.005077 | 0.471 | 0.601 | 1 | 11 | Ehd1 | -0.13 | Young |
| 0.166856 | 0.213618 | 0.152 | 0.226 | 1 | 11 | Idh3b | -0.074 | Young |
| 0.166922 | 0.127436 | 0.884 | 0.887 | 1 | 11 | Rps11 | -0.003 | Young |
| 0.166962 | 0.081641 | 0.391 | 0.516 | 1 | 11 | Cd2ap | -0.125 | Young |

| | | | | | | | | |
|---|---|---|---|---|---|---|---|---|
| 0.166998 | 1.044757 | 0.014 | 0.04 | 1 | 11 | Msh3 | -0.026 | Young |
| 0.167028 | 0.092566 | 0.232 | 0.331 | 1 | 11 | Txnrd1 | -0.099 | Young |
| 0.16709 | 0.769725 | 0.036 | 0.073 | 1 | 11 | Hist1h4i | -0.037 | Young |
| 0.16709 | 0.059845 | 0.036 | 0.073 | 1 | 11 | Zdhhc16 | -0.037 | Young |
| 0.167164 | 0.655315 | 0.5 | 0.508 | 1 | 11 | Rgs2 | -0.008 | Young |
| 0.167689 | 0.066234 | 0.196 | 0.278 | 1 | 11 | Smndc1 | -0.082 | Young |
| 0.167792 | 0.640413 | 0.174 | 0.25 | 1 | 11 | Setx | -0.076 | Young |
| 0.167819 | 0.125555 | 0.978 | 0.972 | 1 | 11 | Rps3a1 | 0.006 | Young |
| 0.168047 | 0.001282 | 0.529 | 0.694 | 1 | 11 | Cyth4 | -0.165 | Young |
| 0.168673 | 0.127457 | 0.232 | 0.327 | 1 | 11 | Sdhb | -0.095 | Young |
| 0.16888 | 0.351666 | 0.051 | 0.093 | 1 | 11 | Dhx8 | -0.042 | Young |
| 0.168951 | 0.561731 | 0.014 | 0.04 | 1 | 11 | Aifm2 | -0.026 | Young |
| 0.169009 | 0.054766 | 0.239 | 0.331 | 1 | 11 | Commd8 | -0.092 | Young |
| 0.169091 | 0.128509 | 0.181 | 0.262 | 1 | 11 | Hcst | -0.081 | Young |
| 0.169189 | 0.080788 | 0.167 | 0.238 | 1 | 11 | Filip1l | -0.071 | Young |
| 0.169235 | 0.912744 | 0.341 | 0.323 | 1 | 11 | Arhgap31 | 0.018 | Young |
| 0.16983 | 0.045511 | 0.341 | 0.46 | 1 | 11 | Tet3 | -0.119 | Young |
| 0.169934 | 0.415714 | 0.014 | 0.04 | 1 | 11 | Acsf2 | -0.026 | Young |
| 0.169948 | 0.178076 | 0.174 | 0.246 | 1 | 11 | Hnrnpll | -0.072 | Young |
| 0.170016 | 0.226543 | 0.072 | 0.121 | 1 | 11 | Golim4 | -0.049 | Young |
| 0.170156 | 0.100315 | 0.949 | 0.952 | 1 | 11 | Mcl1 | -0.003 | Young |
| 0.170207 | 0.204039 | 0.145 | 0.214 | 1 | 11 | Col4a3bp | -0.069 | Young |
| 0.170268 | 0.215163 | 0.319 | 0.456 | 1 | 11 | Ube2r2 | -0.137 | Young |
| 0.170691 | 0.281985 | 0.087 | 0.141 | 1 | 11 | Nkiras2 | -0.054 | Young |
| 0.170857 | 0.049383 | 0.5 | 0.621 | 1 | 11 | Fosl2 | -0.121 | Young |
| 0.170903 | 0.647052 | 0.007 | 0.028 | 1 | 11 | Klhl26 | -0.021 | Young |
| 0.170903 | 0.251614 | 0.007 | 0.028 | 1 | 11 | Lipt2 | -0.021 | Young |
| 0.170903 | 0.231733 | 0.007 | 0.028 | 1 | 11 | Zfp146 | -0.021 | Young |
| 0.170903 | 0.013843 | 0.007 | 0.028 | 1 | 11 | Tmem119 | -0.021 | Young |
| 0.170921 | 0.038402 | 0.014 | 0.04 | 1 | 11 | Lrrc20 | -0.026 | Young |
| 0.171018 | 0.543081 | 0.109 | 0.169 | 1 | 11 | Hsph1 | -0.06 | Young |
| 0.171089 | 0.021797 | 0.094 | 0.149 | 1 | 11 | Snrnp40 | -0.055 | Young |
| 0.171408 | 0.257575 | 0.036 | 0.073 | 1 | 11 | Fkbp9 | -0.037 | Young |
| 0.171703 | 0.301 | 0.072 | 0.121 | 1 | 11 | Clec2i | -0.049 | Young |
| 0.171819 | 0.037524 | 0.384 | 0.524 | 1 | 11 | Larp1 | -0.14 | Young |
| 0.171943 | 0.34457 | 0.051 | 0.024 | 1 | 11 | Rab39 | 0.027 | Young |
| 0.172361 | 0.087311 | 0.275 | 0.387 | 1 | 11 | 1700020I1 | -0.112 | Young |
| 0.172414 | 0.172974 | 0.159 | 0.23 | 1 | 11 | Synrg | -0.071 | Young |
| 0.17244 | 1.311304 | 0.087 | 0.052 | 1 | 11 | Cnrip1 | 0.035 | Young |
| 0.172723 | 0.184179 | 0.283 | 0.391 | 1 | 11 | Stard3nl | -0.108 | Young |
| 0.172919 | 0.199882 | 0.116 | 0.173 | 1 | 11 | Ccdc59 | -0.057 | Young |
| 0.172919 | 0.428245 | 0.174 | 0.254 | 1 | 11 | Rgs19 | -0.08 | Young |
| 0.173079 | 0.122845 | 0.116 | 0.177 | 1 | 11 | Brpf1 | -0.061 | Young |
| 0.173123 | 0.124282 | 0.312 | 0.444 | 1 | 11 | Per1 | -0.132 | Young |
| 0.173355 | 0.217434 | 0.899 | 0.931 | 1 | 11 | Cybb | -0.032 | Young |

| | | | | | | | | |
|---|---|---|---|---|---|---|---|---|
| 0.173405 | 0.390947 | 0.065 | 0.113 | 1 | 11 | Nsmce3 | -0.048 | Young |
| 0.173442 | 0.25209 | 0.109 | 0.169 | 1 | 11 | Dok1 | -0.06 | Young |
| 0.173515 | 0.150898 | 0.152 | 0.218 | 1 | 11 | Scamp1 | -0.066 | Young |
| 0.173629 | 0.076514 | 0.406 | 0.532 | 1 | 11 | Abracl | -0.126 | Young |
| 0.173654 | 0.19831 | 0.094 | 0.149 | 1 | 11 | Tbrg1 | -0.055 | Young |
| 0.173896 | 0.551835 | 0.167 | 0.242 | 1 | 11 | Ints6l | -0.075 | Young |
| 0.173974 | 0.117282 | 0.413 | 0.556 | 1 | 11 | Ran | -0.143 | Young |
| 0.174136 | 0.037697 | 0.522 | 0.653 | 1 | 11 | Fgl2 | -0.131 | Young |
| 0.174736 | 0.590663 | 0.167 | 0.242 | 1 | 11 | Zfc3h1 | -0.075 | Young |
| 0.174744 | 0.148889 | 0.174 | 0.25 | 1 | 11 | Ubr5 | -0.076 | Young |
| 0.174822 | 0.189359 | 0.964 | 0.988 | 1 | 11 | Rps27a | -0.024 | Young |
| 0.174915 | 0.529033 | 0.087 | 0.141 | 1 | 11 | Ctbs | -0.054 | Young |
| 0.175222 | 2.235306 | 0.043 | 0.02 | 1 | 11 | Szt2 | 0.023 | Young |
| 0.175665 | 0.170944 | 0.123 | 0.185 | 1 | 11 | Pitpnb | -0.062 | Young |
| 0.17571 | 0.340646 | 0.254 | 0.347 | 1 | 11 | Lmo4 | -0.093 | Young |
| 0.175808 | 0.002266 | 0.283 | 0.379 | 1 | 11 | Nfkbie | -0.096 | Young |
| 0.17589 | 0.012005 | 0.167 | 0.238 | 1 | 11 | Phactr2 | -0.071 | Young |
| 0.175946 | 0.018897 | 0.058 | 0.101 | 1 | 11 | Dnlz | -0.043 | Young |
| 0.176039 | 0.003709 | 0.457 | 0.585 | 1 | 11 | Ubl5 | -0.128 | Young |
| 0.17605 | 0.173147 | 0.261 | 0.367 | 1 | 11 | Ccar1 | -0.106 | Young |
| 0.176315 | 0.127304 | 0.167 | 0.238 | 1 | 11 | R3hdm1 | -0.071 | Young |
| 0.176579 | 0.426291 | 0.181 | 0.262 | 1 | 11 | Exoc4 | -0.081 | Young |
| 0.176872 | 0.0843 | 0.152 | 0.222 | 1 | 11 | Slc28a2 | -0.07 | Young |
| 0.176878 | 0.015692 | 0.109 | 0.169 | 1 | 11 | Ptdss1 | -0.06 | Young |
| 0.177038 | 0.516332 | 0.101 | 0.157 | 1 | 11 | Psmc2 | -0.056 | Young |
| 0.177818 | 0.015683 | 0.094 | 0.149 | 1 | 11 | Plxnc1 | -0.055 | Young |
| 0.177834 | 0.041308 | 0.058 | 0.101 | 1 | 11 | Slc33a1 | -0.043 | Young |
| 0.178061 | 0.28448 | 0.101 | 0.157 | 1 | 11 | Wdr81 | -0.056 | Young |
| 0.178178 | 0.184842 | 0.283 | 0.399 | 1 | 11 | Pet100 | -0.116 | Young |
| 0.178991 | 0.319676 | 0.123 | 0.185 | 1 | 11 | Fam96b | -0.062 | Young |
| 0.179968 | 0.095191 | 0.225 | 0.315 | 1 | 11 | Sptbn1 | -0.09 | Young |
| 0.18012 | 0.195963 | 0.101 | 0.157 | 1 | 11 | Gtf3c6 | -0.056 | Young |
| 0.180587 | 0.149753 | 0.087 | 0.137 | 1 | 11 | Naa35 | -0.05 | Young |
| 0.181201 | 0.097021 | 0.261 | 0.367 | 1 | 11 | Ubtf | -0.106 | Young |
| 0.181308 | 0.208195 | 0.152 | 0.222 | 1 | 11 | Osbp | -0.07 | Young |
| 0.181543 | 0.048457 | 0.406 | 0.54 | 1 | 11 | Macf1 | -0.134 | Young |
| 0.181763 | 0.190558 | 0.333 | 0.44 | 1 | 11 | Pnisr | -0.107 | Young |
| 0.182086 | 0.755924 | 0.072 | 0.121 | 1 | 11 | Fbxl17 | -0.049 | Young |
| 0.182156 | 0.205634 | 0.384 | 0.536 | 1 | 11 | Galnt1 | -0.152 | Young |
| 0.182211 | 0.112408 | 0.196 | 0.274 | 1 | 11 | Asxl2 | -0.078 | Young |
| 0.182297 | 0.116443 | 0.058 | 0.101 | 1 | 11 | Cep83 | -0.043 | Young |
| 0.182315 | 0.490955 | 0.457 | 0.444 | 1 | 11 | Slamf9 | 0.013 | Young |
| 0.182495 | 0.175861 | 0.043 | 0.081 | 1 | 11 | Kat14 | -0.038 | Young |
| 0.182781 | 0.10327 | 0.08 | 0.129 | 1 | 11 | Klhdc4 | -0.049 | Young |
| 0.182781 | 0.045118 | 0.08 | 0.129 | 1 | 11 | Pofut1 | -0.049 | Young |

| | | | | | | | | |
|---|---|---|---|---|---|---|---|---|
| 0.182845 | 0.023363 | 0.188 | 0.266 | 1 | 11 | Tbc1d13 | -0.078 | Young |
| 0.182918 | 0.345535 | 0.029 | 0.06 | 1 | 11 | Wfs1 | -0.031 | Young |
| 0.182918 | 0.074608 | 0.029 | 0.06 | 1 | 11 | Rbm19 | -0.031 | Young |
| 0.182961 | 0.229952 | 0.312 | 0.427 | 1 | 11 | Derl1 | -0.115 | Young |
| 0.183075 | 1.412542 | 0.065 | 0.036 | 1 | 11 | Zfp629 | 0.029 | Young |
| 0.183928 | 0.145252 | 0.08 | 0.129 | 1 | 11 | Fblim1 | -0.049 | Young |
| 0.18395 | 0.293367 | 0.21 | 0.29 | 1 | 11 | Bbip1 | -0.08 | Young |
| 0.18427 | 0.009365 | 0.507 | 0.677 | 1 | 11 | Nktr | -0.17 | Young |
| 0.184282 | 0.030162 | 0.196 | 0.274 | 1 | 11 | Btbd7 | -0.078 | Young |
| 0.184424 | 0.054574 | 0.094 | 0.145 | 1 | 11 | Tmem55a | -0.051 | Young |
| 0.184593 | 0.052678 | 0.029 | 0.06 | 1 | 11 | Flad1 | -0.031 | Young |
| 0.184661 | 0.510408 | 0.043 | 0.081 | 1 | 11 | Rprd1a | -0.038 | Young |
| 0.184875 | 1.278271 | 0.043 | 0.02 | 1 | 11 | Gm14548 | 0.023 | Young |
| 0.184875 | 0.931594 | 0.043 | 0.02 | 1 | 11 | Repin1 | 0.023 | Young |
| 0.184916 | 0.133216 | 0.152 | 0.222 | 1 | 11 | Wsb2 | -0.07 | Young |
| 0.185329 | 0.81197 | 0.399 | 0.387 | 1 | 11 | Tmcc3 | 0.012 | Young |
| 0.185388 | 0.26027 | 0.043 | 0.081 | 1 | 11 | Cdc16 | -0.038 | Young |
| 0.185388 | 0.08586 | 0.043 | 0.081 | 1 | 11 | Atg13 | -0.038 | Young |
| 0.185732 | 0.22475 | 0.138 | 0.206 | 1 | 11 | Slc38a10 | -0.068 | Young |
| 0.185962 | 0.395425 | 0.065 | 0.109 | 1 | 11 | Rnf14 | -0.044 | Young |
| 0.185998 | 0.476397 | 0.109 | 0.165 | 1 | 11 | Sptan1 | -0.056 | Young |
| 0.186279 | 0.008776 | 0.029 | 0.06 | 1 | 11 | Rundc1 | -0.031 | Young |
| 0.186719 | 1.292305 | 0.203 | 0.165 | 1 | 11 | Bcl2l1 | 0.038 | Young |
| 0.187073 | 1.42967 | 0.043 | 0.02 | 1 | 11 | Gabbr1 | 0.023 | Young |
| 0.187221 | 0.205438 | 0.065 | 0.109 | 1 | 11 | Klhl20 | -0.044 | Young |
| 0.187397 | 1.195277 | 0.065 | 0.036 | 1 | 11 | Gpatch11 | 0.029 | Young |
| 0.188143 | 1.158214 | 0.159 | 0.121 | 1 | 11 | Rnf38 | 0.038 | Young |
| 0.188157 | 0.151176 | 0.058 | 0.101 | 1 | 11 | Pprc1 | -0.043 | Young |
| 0.188215 | 0.047064 | 0.159 | 0.23 | 1 | 11 | Fam234a | -0.071 | Young |
| 0.188954 | 0.098953 | 0.275 | 0.387 | 1 | 11 | Ppp1cc | -0.112 | Young |
| 0.188996 | 0.324103 | 0.652 | 0.673 | 1 | 11 | Tnfaip3 | -0.021 | Young |
| 0.189686 | 0.203402 | 0.029 | 0.06 | 1 | 11 | Tlr5 | -0.031 | Young |
| 0.189686 | 0.098551 | 0.029 | 0.06 | 1 | 11 | Gm5431 | -0.031 | Young |
| 0.189692 | 0.152721 | 0.188 | 0.27 | 1 | 11 | St6galnac4 | -0.082 | Young |
| 0.189821 | 0.083904 | 0.13 | 0.194 | 1 | 11 | Fam114a2 | -0.064 | Young |
| 0.190141 | 0.423943 | 0.058 | 0.101 | 1 | 11 | Dis3l2 | -0.043 | Young |
| 0.190143 | 0.074898 | 0.101 | 0.157 | 1 | 11 | Nipsnap3b | -0.056 | Young |
| 0.19053 | 0.247258 | 0.043 | 0.081 | 1 | 11 | Pde6d | -0.038 | Young |
| 0.19053 | 0.158629 | 0.043 | 0.081 | 1 | 11 | Gm45902 | -0.038 | Young |
| 0.190548 | 0.226188 | 0.051 | 0.089 | 1 | 11 | Ublcp1 | -0.038 | Young |
| 0.190699 | 0.217548 | 0.42 | 0.56 | 1 | 11 | Dek | -0.14 | Young |
| 0.190764 | 0.095062 | 0.986 | 0.992 | 1 | 11 | Eef1a1 | -0.006 | Young |
| 0.190901 | 0.127618 | 0.043 | 0.081 | 1 | 11 | Dusp10 | -0.038 | Young |
| 0.191177 | 0.096635 | 0.203 | 0.282 | 1 | 11 | Stat5b | -0.079 | Young |
| 0.191278 | 0.227893 | 0.087 | 0.137 | 1 | 11 | Nf1 | -0.05 | Young |

| | | | | | | | | |
|---|---|---|---|---|---|---|---|---|
| 0.191389 | 0.477339 | 0.174 | 0.258 | 1 | 11 | Chmp1a | -0.084 | Young |
| 0.192096 | 0.062377 | 0.406 | 0.544 | 1 | 11 | Smdt1 | -0.138 | Young |
| 0.19232 | 0.119538 | 0.333 | 0.456 | 1 | 11 | Cwc15 | -0.123 | Young |
| 0.192343 | 0.958863 | 0.123 | 0.085 | 1 | 11 | Plxna4os1 | 0.038 | Young |
| 0.192556 | 0.196243 | 0.304 | 0.423 | 1 | 11 | Usp7 | -0.119 | Young |
| 0.192787 | 0.324887 | 0.297 | 0.435 | 1 | 11 | Pten | -0.138 | Young |
| 0.192838 | 0.205718 | 0.072 | 0.117 | 1 | 11 | Mum1 | -0.045 | Young |
| 0.192876 | 0.10906 | 0.283 | 0.379 | 1 | 11 | Ccnt2 | -0.096 | Young |
| 0.193003 | 0.448269 | 0.138 | 0.097 | 1 | 11 | Arhgap26 | 0.041 | Young |
| 0.193022 | 0.215146 | 0.203 | 0.286 | 1 | 11 | Rc3h1 | -0.083 | Young |
| 0.193138 | 1.065132 | 0.029 | 0.06 | 1 | 11 | Cars | -0.031 | Young |
| 0.193385 | 0.015473 | 0.051 | 0.089 | 1 | 11 | Bola1 | -0.038 | Young |
| 0.193411 | 0.012035 | 0.116 | 0.173 | 1 | 11 | Faim | -0.057 | Young |
| 0.193913 | 0.185637 | 0.341 | 0.48 | 1 | 11 | Tmed7 | -0.139 | Young |
| 0.194008 | 0.578533 | 0.029 | 0.06 | 1 | 11 | Galt | -0.031 | Young |
| 0.194235 | 0.088031 | 0.145 | 0.206 | 1 | 11 | Mrps15 | -0.061 | Young |
| 0.194392 | 0.486754 | 0.217 | 0.319 | 1 | 11 | Ccni | -0.102 | Young |
| 0.194484 | 0.461476 | 0.101 | 0.157 | 1 | 11 | Abraxas2 | -0.056 | Young |
| 0.194822 | 0.609786 | 0.109 | 0.169 | 1 | 11 | Slc25a25 | -0.06 | Young |
| 0.194995 | 0.067922 | 0.138 | 0.198 | 1 | 11 | Chmp1b | -0.06 | Young |
| 0.195484 | 0.541742 | 0.457 | 0.444 | 1 | 11 | Arl4c | 0.013 | Young |
| 0.195489 | 0.139931 | 0.138 | 0.198 | 1 | 11 | Mtfr1l | -0.06 | Young |
| 0.195502 | 0.212925 | 0.116 | 0.173 | 1 | 11 | Gsr | -0.057 | Young |
| 0.195602 | 0.074212 | 0.275 | 0.367 | 1 | 11 | Kat2b | -0.092 | Young |
| 0.195959 | 0.260531 | 0.312 | 0.427 | 1 | 11 | Ldlr | -0.115 | Young |
| 0.196277 | 0.305498 | 0.167 | 0.242 | 1 | 11 | Hist3h2a | -0.075 | Young |
| 0.19649 | 1.782439 | 0.101 | 0.069 | 1 | 11 | Mir17hg | 0.032 | Young |
| 0.196647 | 0.862074 | 0.036 | 0.069 | 1 | 11 | Edc3 | -0.033 | Young |
| 0.19667 | 1.171293 | 0.232 | 0.198 | 1 | 11 | Chd8 | 0.034 | Young |
| 0.196797 | 0.119495 | 0.08 | 0.125 | 1 | 11 | Pdrg1 | -0.045 | Young |
| 0.1977 | 0.021976 | 0.051 | 0.089 | 1 | 11 | Prkaa1 | -0.038 | Young |
| 0.198165 | 0.689625 | 0.399 | 0.395 | 1 | 11 | Pde4b | 0.004 | Young |
| 0.198289 | 0.511182 | 0.13 | 0.194 | 1 | 11 | Msl3 | -0.064 | Young |
| 0.198525 | 0.301672 | 0.13 | 0.198 | 1 | 11 | Ythdc2 | -0.068 | Young |
| 0.198725 | 0.396467 | 0.08 | 0.129 | 1 | 11 | Zbtb11 | -0.049 | Young |
| 0.199388 | 0.210505 | 0.138 | 0.202 | 1 | 11 | Dstyk | -0.064 | Young |
| 0.199534 | 0.73261 | 0.13 | 0.198 | 1 | 11 | Snx24 | -0.068 | Young |
| 0.200155 | 0.377951 | 0.065 | 0.109 | 1 | 11 | Gmcl1 | -0.044 | Young |
| 0.20023 | 0.00514 | 0.529 | 0.669 | 1 | 11 | Frmd4b | -0.14 | Young |
| 0.200336 | 0.337933 | 0.123 | 0.181 | 1 | 11 | Epb41l1 | -0.058 | Young |
| 0.200567 | 2.319539 | 0.036 | 0.016 | 1 | 11 | Surf6 | 0.02 | Young |
| 0.200567 | 1.652223 | 0.036 | 0.016 | 1 | 11 | Cdc6 | 0.02 | Young |
| 0.200965 | 0.238924 | 0.754 | 0.815 | 1 | 11 | Rpl12 | -0.061 | Young |
| 0.200973 | 0.198662 | 0.123 | 0.185 | 1 | 11 | Cmip | -0.062 | Young |
| 0.201175 | 0.155763 | 0.877 | 0.899 | 1 | 11 | Rps25 | -0.022 | Young |

| | | | | | | | | |
|---|---|---|---|---|---|---|---|---|
| 0.201303 | 0.034188 | 0.022 | 0.048 | 1 | 11 | Zfyve9 | -0.026 | Young |
| 0.201384 | 0.813865 | 0.123 | 0.181 | 1 | 11 | Mcm3 | -0.058 | Young |
| 0.201755 | 0.085874 | 0.486 | 0.641 | 1 | 11 | Gls | -0.155 | Young |
| 0.201851 | 1.857216 | 0.036 | 0.016 | 1 | 11 | Lrrc49 | 0.02 | Young |
| 0.201851 | 1.784938 | 0.036 | 0.016 | 1 | 11 | Tfdp2 | 0.02 | Young |
| 0.201958 | 0.270339 | 0.101 | 0.153 | 1 | 11 | Irf3 | -0.052 | Young |
| 0.202334 | 0.316496 | 0.072 | 0.117 | 1 | 11 | Cetn2 | -0.045 | Young |
| 0.202847 | 0.225222 | 0.174 | 0.25 | 1 | 11 | Rps6kb1 | -0.076 | Young |
| 0.203141 | 2.316754 | 0.036 | 0.016 | 1 | 11 | Hist1h1e | 0.02 | Young |
| 0.203141 | 1.881004 | 0.036 | 0.016 | 1 | 11 | Oaf | 0.02 | Young |
| 0.203141 | 1.861635 | 0.036 | 0.016 | 1 | 11 | Ybey | 0.02 | Young |
| 0.203141 | 1.399171 | 0.036 | 0.016 | 1 | 11 | Cdkn2c | 0.02 | Young |
| 0.203316 | 0.017584 | 0.022 | 0.048 | 1 | 11 | Ccdc163 | -0.026 | Young |
| 0.203782 | 0.578259 | 0.5 | 0.532 | 1 | 11 | Nfam1 | -0.032 | Young |
| 0.204168 | 0.073335 | 0.203 | 0.27 | 1 | 11 | Ivns1abp | -0.067 | Young |
| 0.204244 | 0.09753 | 0.326 | 0.435 | 1 | 11 | Pttg1ip | -0.109 | Young |
| 0.204327 | 0.042351 | 0.022 | 0.048 | 1 | 11 | Clpb | -0.026 | Young |
| 0.204379 | 0.718477 | 0.406 | 0.387 | 1 | 11 | Foxn3 | 0.019 | Young |
| 0.204436 | 1.850528 | 0.036 | 0.016 | 1 | 11 | Elp4 | 0.02 | Young |
| 0.205232 | 0.114725 | 0.217 | 0.298 | 1 | 11 | Mrpl20 | -0.081 | Young |
| 0.205385 | 0.186612 | 0.239 | 0.335 | 1 | 11 | Ccnt1 | -0.096 | Young |
| 0.205618 | 0.215668 | 0.254 | 0.355 | 1 | 11 | Pdlim5 | -0.101 | Young |
| 0.205738 | 2.065873 | 0.036 | 0.016 | 1 | 11 | Naprt | 0.02 | Young |
| 0.205738 | 1.7314 | 0.036 | 0.016 | 1 | 11 | Zfp959 | 0.02 | Young |
| 0.205738 | 1.62865 | 0.036 | 0.016 | 1 | 11 | Mmaa | 0.02 | Young |
| 0.205738 | 1.520509 | 0.036 | 0.016 | 1 | 11 | Lekr1 | 0.02 | Young |
| 0.206362 | 0.744629 | 0.022 | 0.048 | 1 | 11 | Clip2 | -0.026 | Young |
| 0.206522 | 1.095131 | 0.116 | 0.081 | 1 | 11 | Surf1 | 0.035 | Young |
| 0.206528 | 0.794348 | 0.123 | 0.085 | 1 | 11 | Stxbp6 | 0.038 | Young |
| 0.206751 | 0.199886 | 0.116 | 0.173 | 1 | 11 | Adh5 | -0.057 | Young |
| 0.20689 | 0.011676 | 0.145 | 0.21 | 1 | 11 | Ube2j2 | -0.065 | Young |
| 0.207321 | 0.317132 | 0.297 | 0.415 | 1 | 11 | Cdc42se2 | -0.118 | Young |
| 0.207936 | 0.013453 | 0.196 | 0.278 | 1 | 11 | Csnk2a2 | -0.082 | Young |
| 0.208186 | 0.215242 | 0.246 | 0.339 | 1 | 11 | Csnk2b | -0.093 | Young |
| 0.20836 | 0.964178 | 0.036 | 0.016 | 1 | 11 | Iba57 | 0.02 | Young |
| 0.208411 | 0.388849 | 0.022 | 0.048 | 1 | 11 | Zxdb | -0.026 | Young |
| 0.20872 | 0.236875 | 0.717 | 0.79 | 1 | 11 | Ube2d3 | -0.073 | Young |
| 0.208852 | 0.039293 | 0.072 | 0.117 | 1 | 11 | Pdcd7 | -0.045 | Young |
| 0.208938 | 0.175188 | 0.094 | 0.145 | 1 | 11 | Cmas | -0.051 | Young |
| 0.209423 | 0.218454 | 0.109 | 0.161 | 1 | 11 | Mrpl38 | -0.052 | Young |
| 0.209441 | 0.100493 | 0.022 | 0.048 | 1 | 11 | Ccdc28a | -0.026 | Young |
| 0.209458 | 0.031008 | 0.457 | 0.621 | 1 | 11 | Ifngr1 | -0.164 | Young |
| 0.209468 | 0.248631 | 0.261 | 0.367 | 1 | 11 | Fam208a | -0.106 | Young |
| 0.209548 | 0.30185 | 0.051 | 0.089 | 1 | 11 | Alg14 | -0.038 | Young |
| 0.209679 | 1.47716 | 0.036 | 0.016 | 1 | 11 | Tpbgl | 0.02 | Young |

| | | | | | | | | |
|---|---|---|---|---|---|---|---|---|
| 0.209837 | 0.02268 | 0.181 | 0.254 | 1 | 11 | Slc9a3r1 | -0.073 | Young |
| 0.209856 | 0.732018 | 0.123 | 0.085 | 1 | 11 | 1600014C1 | 0.038 | Young |
| 0.209932 | 0.351061 | 0.08 | 0.125 | 1 | 11 | Gga3 | -0.045 | Young |
| 0.21006 | 0.479563 | 0.196 | 0.274 | 1 | 11 | Cacybp | -0.078 | Young |
| 0.210112 | 0.087032 | 0.036 | 0.069 | 1 | 11 | Nop16 | -0.033 | Young |
| 0.210253 | 0.130579 | 0.377 | 0.512 | 1 | 11 | Cct5 | -0.135 | Young |
| 0.210612 | 0.747685 | 0.217 | 0.177 | 1 | 11 | Emp1 | 0.04 | Young |
| 0.210976 | 0.320069 | 0.036 | 0.069 | 1 | 11 | Wdr46 | -0.033 | Young |
| 0.211005 | 1.296493 | 0.036 | 0.016 | 1 | 11 | Ptch1 | 0.02 | Young |
| 0.211076 | 0.413474 | 0.043 | 0.077 | 1 | 11 | Vipas39 | -0.034 | Young |
| 0.211133 | 0.155601 | 0.109 | 0.161 | 1 | 11 | Sacm1l | -0.052 | Young |
| 0.211512 | 0.679792 | 0.022 | 0.048 | 1 | 11 | Klhl7 | -0.026 | Young |
| 0.211559 | 0.610565 | 0.087 | 0.137 | 1 | 11 | Polr3gl | -0.05 | Young |
| 0.211592 | 0.757824 | 0.384 | 0.399 | 1 | 11 | Eif3h | -0.015 | Young |
| 0.211806 | 0.137343 | 0.261 | 0.355 | 1 | 11 | Zcchc24 | -0.094 | Young |
| 0.212038 | 0.281732 | 0.152 | 0.218 | 1 | 11 | Pak1 | -0.066 | Young |
| 0.21206 | 0.081328 | 0.268 | 0.359 | 1 | 11 | Slc7a8 | -0.091 | Young |
| 0.212337 | 0.779459 | 0.036 | 0.016 | 1 | 11 | Zfp235 | 0.02 | Young |
| 0.212343 | 0.887269 | 0.065 | 0.109 | 1 | 11 | Snx11 | -0.044 | Young |
| 0.212364 | 0.651437 | 0.058 | 0.097 | 1 | 11 | St7l | -0.039 | Young |
| 0.212595 | 0.110682 | 0.522 | 0.694 | 1 | 11 | Il6ra | -0.172 | Young |
| 0.212676 | 0.384802 | 0.181 | 0.254 | 1 | 11 | Snx27 | -0.073 | Young |
| 0.21271 | 0.21437 | 0.036 | 0.069 | 1 | 11 | Srrd | -0.033 | Young |
| 0.213106 | 0.172023 | 0.319 | 0.46 | 1 | 11 | Fli1 | -0.141 | Young |
| 0.213134 | 0.010954 | 0.42 | 0.556 | 1 | 11 | Gnaq | -0.136 | Young |
| 0.213581 | 0.205743 | 0.036 | 0.069 | 1 | 11 | Mcm5 | -0.033 | Young |
| 0.213581 | 0.105807 | 0.036 | 0.069 | 1 | 11 | Arhgap6 | -0.033 | Young |
| 0.214352 | 0.12106 | 0.043 | 0.077 | 1 | 11 | Fkbp5 | -0.034 | Young |
| 0.215293 | 0.85959 | 0.145 | 0.105 | 1 | 11 | Ramp1 | 0.04 | Young |
| 0.215394 | 0.206708 | 0.275 | 0.375 | 1 | 11 | Traf1 | -0.1 | Young |
| 0.215661 | 0.343608 | 0.051 | 0.089 | 1 | 11 | Mettl6 | -0.038 | Young |
| 0.216005 | 0.428961 | 0.348 | 0.472 | 1 | 11 | Mdfic | -0.124 | Young |
| 0.216407 | 1.263133 | 0.058 | 0.032 | 1 | 11 | Gbp6 | 0.026 | Young |
| 0.216434 | 0.521603 | 0.051 | 0.089 | 1 | 11 | Bambi | -0.038 | Young |
| 0.216502 | 0.072598 | 0.377 | 0.508 | 1 | 11 | Rabac1 | -0.131 | Young |
| 0.216511 | 0.079741 | 0.355 | 0.46 | 1 | 11 | Elob | -0.105 | Young |
| 0.216663 | 0.171457 | 0.08 | 0.048 | 1 | 11 | Fry | 0.032 | Young |
| 0.217377 | 0.054364 | 0.065 | 0.105 | 1 | 11 | Slc13a3 | -0.04 | Young |
| 0.217436 | 0.091376 | 0.116 | 0.169 | 1 | 11 | 1110059G1 | -0.053 | Young |
| 0.217543 | 0.017262 | 0.362 | 0.484 | 1 | 11 | Tpm4 | -0.122 | Young |
| 0.217616 | 0.010402 | 0.348 | 0.452 | 1 | 11 | Gna12 | -0.104 | Young |
| 0.218805 | 0.093031 | 0.065 | 0.105 | 1 | 11 | Matk | -0.04 | Young |
| 0.21974 | 0.412714 | 0.536 | 0.573 | 1 | 11 | Khdrbs1 | -0.037 | Young |
| 0.21981 | 0.985764 | 0.174 | 0.133 | 1 | 11 | Bmp2 | 0.041 | Young |
| 0.219939 | 0.147753 | 0.232 | 0.315 | 1 | 11 | Tcn2 | -0.083 | Young |

| | | | | | | | | |
|---|---|---|---|---|---|---|---|---|
| 0.220329 | 0.346302 | 0.051 | 0.089 | 1 | 11 | 5031425F1 | -0.038 | Young |
| 0.220425 | 0.069702 | 0.109 | 0.161 | 1 | 11 | Sec16a | -0.052 | Young |
| 0.220486 | 0.207979 | 0.101 | 0.149 | 1 | 11 | Sumf1 | -0.048 | Young |
| 0.220626 | 0.089208 | 0.217 | 0.302 | 1 | 11 | Nbeal1 | -0.085 | Young |
| 0.220691 | 0.151031 | 0.246 | 0.331 | 1 | 11 | Vdac1 | -0.085 | Young |
| 0.221252 | 0.151173 | 0.058 | 0.097 | 1 | 11 | Traf2 | -0.039 | Young |
| 0.221607 | 0.112835 | 0.109 | 0.161 | 1 | 11 | Csnk1g1 | -0.052 | Young |
| 0.221651 | 0.587652 | 0.072 | 0.117 | 1 | 11 | Bicra | -0.045 | Young |
| 0.221856 | 0.5244 | 0.043 | 0.077 | 1 | 11 | Dym | -0.034 | Young |
| 0.222199 | 0.472012 | 0.109 | 0.161 | 1 | 11 | Fam162a | -0.052 | Young |
| 0.222284 | 0.296001 | 0.094 | 0.141 | 1 | 11 | Taf11 | -0.047 | Young |
| 0.222616 | 0.084032 | 0.319 | 0.431 | 1 | 11 | Lmo2 | -0.112 | Young |
| 0.222728 | 1.078795 | 0.072 | 0.044 | 1 | 11 | Exoc8 | 0.028 | Young |
| 0.222796 | 0.034953 | 0.319 | 0.415 | 1 | 11 | Cnn2 | -0.096 | Young |
| 0.22302 | 0.661927 | 0.08 | 0.125 | 1 | 11 | Ing5 | -0.045 | Young |
| 0.223351 | 0.068351 | 0.225 | 0.31 | 1 | 11 | Nck1 | -0.085 | Young |
| 0.223636 | 2.21615 | 0.014 | 0.036 | 1 | 11 | Syne3 | -0.022 | Young |
| 0.223663 | 0.99216 | 0.058 | 0.032 | 1 | 11 | Nol4l | 0.026 | Young |
| 0.223864 | 0.038288 | 0.58 | 0.75 | 1 | 11 | Ap2a2 | -0.17 | Young |
| 0.224005 | 0.01511 | 0.196 | 0.258 | 1 | 11 | Ankhd1 | -0.062 | Young |
| 0.224232 | 0.831972 | 0.036 | 0.069 | 1 | 11 | Cpsf1 | -0.033 | Young |
| 0.224421 | 0.129127 | 0.891 | 0.863 | 1 | 11 | Iqgap1 | 0.028 | Young |
| 0.224463 | 0.089428 | 0.08 | 0.121 | 1 | 11 | Mcm6 | -0.041 | Young |
| 0.224535 | 0.00122 | 0.188 | 0.258 | 1 | 11 | Hpf1 | -0.07 | Young |
| 0.224586 | 0.158182 | 0.065 | 0.105 | 1 | 11 | Nol8 | -0.04 | Young |
| 0.224895 | 0.156114 | 0.014 | 0.036 | 1 | 11 | Tmem115 | -0.022 | Young |
| 0.22515 | 0.036534 | 0.232 | 0.315 | 1 | 11 | Inpp4a | -0.083 | Young |
| 0.225208 | 2.456718 | 0.029 | 0.012 | 1 | 11 | Nicn1 | 0.017 | Young |
| 0.225478 | 0.24435 | 0.145 | 0.202 | 1 | 11 | Rap2a | -0.057 | Young |
| 0.22616 | 0.709295 | 0.014 | 0.036 | 1 | 11 | Abhd18 | -0.022 | Young |
| 0.22616 | 0.085846 | 0.014 | 0.036 | 1 | 11 | Adck2 | -0.022 | Young |
| 0.22619 | 0.059551 | 0.13 | 0.185 | 1 | 11 | Casp3 | -0.055 | Young |
| 0.226225 | 0.293822 | 0.246 | 0.339 | 1 | 11 | Wdfy3 | -0.093 | Young |
| 0.226782 | 0.464041 | 0.065 | 0.105 | 1 | 11 | Commd10 | -0.04 | Young |
| 0.226786 | 3.07583 | 0.029 | 0.012 | 1 | 11 | Sfxn5 | 0.017 | Young |
| 0.226786 | 2.192095 | 0.029 | 0.012 | 1 | 11 | Tmtc2 | 0.017 | Young |
| 0.226804 | 0.108246 | 0.841 | 0.851 | 1 | 11 | Rpl3 | -0.01 | Young |
| 0.227083 | 0.233439 | 0.804 | 0.863 | 1 | 11 | BC005537 | -0.059 | Young |
| 0.227207 | 0.043247 | 0.123 | 0.177 | 1 | 11 | Rbpms | -0.054 | Young |
| 0.227379 | 0.338986 | 0.681 | 0.75 | 1 | 11 | Sppl2a | -0.069 | Young |
| 0.227517 | 0.225325 | 0.065 | 0.105 | 1 | 11 | Isca2 | -0.04 | Young |
| 0.227592 | 0.004791 | 0.145 | 0.206 | 1 | 11 | Smarca2 | -0.061 | Young |
| 0.228187 | 0.231822 | 0.109 | 0.161 | 1 | 11 | Klhl18 | -0.052 | Young |
| 0.228373 | 1.981257 | 0.029 | 0.012 | 1 | 11 | Ptrh1 | 0.017 | Young |
| 0.228373 | 1.958715 | 0.029 | 0.012 | 1 | 11 | Wnt11 | 0.017 | Young |

| | | | | | | | | |
|---|---|---|---|---|---|---|---|---|
| 0.228703 | 0.391836 | 0.014 | 0.036 | 1 | 11 | Gm16104 | -0.022 | Young |
| 0.228703 | 0.093045 | 0.014 | 0.036 | 1 | 11 | Ppp3cc | -0.022 | Young |
| 0.228913 | 0.62687 | 0.123 | 0.181 | 1 | 11 | Dcaf13 | -0.058 | Young |
| 0.229207 | 1.050973 | 0.145 | 0.109 | 1 | 11 | Nup54 | 0.036 | Young |
| 0.229314 | 1.187108 | 0.123 | 0.089 | 1 | 11 | Pgm2l1 | 0.034 | Young |
| 0.229405 | 0.287274 | 0.246 | 0.351 | 1 | 11 | Cuta | -0.105 | Young |
| 0.229629 | 0.100055 | 0.442 | 0.589 | 1 | 11 | Tgoln1 | -0.147 | Young |
| 0.229761 | 0.041674 | 0.101 | 0.149 | 1 | 11 | Brwd3 | -0.048 | Young |
| 0.22978 | 0.289016 | 0.08 | 0.125 | 1 | 11 | Mavs | -0.045 | Young |
| 0.229968 | 2.145411 | 0.029 | 0.012 | 1 | 11 | Arhgef40 | 0.017 | Young |
| 0.229968 | 1.761348 | 0.029 | 0.012 | 1 | 11 | Zfp764 | 0.017 | Young |
| 0.229968 | 1.386321 | 0.029 | 0.012 | 1 | 11 | Rarres2 | 0.017 | Young |
| 0.229982 | 0.725963 | 0.014 | 0.036 | 1 | 11 | Mapre3 | -0.022 | Young |
| 0.229982 | 0.552899 | 0.014 | 0.036 | 1 | 11 | Tsr3 | -0.022 | Young |
| 0.229982 | 0.459686 | 0.014 | 0.036 | 1 | 11 | Rbak | -0.022 | Young |
| 0.230018 | 0.259283 | 0.196 | 0.278 | 1 | 11 | Lrp10 | -0.082 | Young |
| 0.230044 | 0.029451 | 0.232 | 0.319 | 1 | 11 | Dctn4 | -0.087 | Young |
| 0.230404 | 0.423727 | 0.058 | 0.097 | 1 | 11 | Ttf1 | -0.039 | Young |
| 0.230891 | 0.208822 | 0.051 | 0.085 | 1 | 11 | Aifm1 | -0.034 | Young |
| 0.230917 | 0.073238 | 0.203 | 0.27 | 1 | 11 | Sec63 | -0.067 | Young |
| 0.230973 | 0.040836 | 0.457 | 0.613 | 1 | 11 | Shisa5 | -0.156 | Young |
| 0.231505 | 1.283698 | 0.138 | 0.105 | 1 | 11 | Tmem63a | 0.033 | Young |
| 0.23157 | 0.832035 | 0.029 | 0.012 | 1 | 11 | Capn5 | 0.017 | Young |
| 0.232123 | 0.259045 | 0.275 | 0.371 | 1 | 11 | Neu1 | -0.096 | Young |
| 0.232282 | 0.187589 | 0.217 | 0.302 | 1 | 11 | Cat | -0.085 | Young |
| 0.232452 | 0.124924 | 0.13 | 0.185 | 1 | 11 | Slc30a7 | -0.055 | Young |
| 0.232867 | 0.030715 | 0.246 | 0.331 | 1 | 11 | Ankrd13a | -0.085 | Young |
| 0.232954 | 0.507222 | 0.152 | 0.226 | 1 | 11 | Kmt5a | -0.074 | Young |
| 0.233181 | 1.883528 | 0.029 | 0.012 | 1 | 11 | Ramp3 | 0.017 | Young |
| 0.233181 | 0.814139 | 0.029 | 0.012 | 1 | 11 | Chrna5 | 0.017 | Young |
| 0.233461 | 0.149536 | 0.065 | 0.105 | 1 | 11 | Abcd2 | -0.04 | Young |
| 0.233493 | 0.816712 | 0.268 | 0.242 | 1 | 11 | Rnf150 | 0.026 | Young |
| 0.233559 | 1.273151 | 0.13 | 0.097 | 1 | 11 | AU020206 | 0.033 | Young |
| 0.233614 | 0.330418 | 0.268 | 0.367 | 1 | 11 | Tmbim1 | -0.099 | Young |
| 0.233812 | 0.753457 | 0.312 | 0.29 | 1 | 11 | Ypel5 | 0.022 | Young |
| 0.234799 | 1.128426 | 0.029 | 0.012 | 1 | 11 | Zswim3 | 0.017 | Young |
| 0.234799 | 1.115022 | 0.029 | 0.012 | 1 | 11 | Acat2 | 0.017 | Young |
| 0.234847 | 0.119205 | 0.877 | 0.883 | 1 | 11 | Rps5 | -0.006 | Young |
| 0.235037 | 0.221248 | 0.051 | 0.085 | 1 | 11 | Orai2 | -0.034 | Young |
| 0.23508 | 0.362172 | 0.058 | 0.097 | 1 | 11 | Immp1l | -0.039 | Young |
| 0.235129 | 1.860423 | 0.051 | 0.028 | 1 | 11 | Cox19 | 0.023 | Young |
| 0.235146 | 0.36223 | 0.312 | 0.431 | 1 | 11 | Anxa7 | -0.119 | Young |
| 0.235327 | 0.243039 | 0.029 | 0.056 | 1 | 11 | Fam217b | -0.027 | Young |
| 0.235327 | 0.110675 | 0.029 | 0.056 | 1 | 11 | Yars | -0.027 | Young |
| 0.235389 | 0.641747 | 0.058 | 0.093 | 1 | 11 | Arhgap21 | -0.035 | Young |

| | | | | | | | | |
|---|---|---|---|---|---|---|---|---|
| 0.235463 | 0.190877 | 0.275 | 0.375 | 1 | 11 | Fgd4 | -0.1 | Young |
| 0.235498 | 0.143836 | 0.884 | 0.94 | 1 | 11 | Rpl18a | -0.056 | Young |
| 0.235571 | 0.14643 | 0.297 | 0.395 | 1 | 11 | Dynlrb1 | -0.098 | Young |
| 0.235659 | 0.094528 | 0.043 | 0.077 | 1 | 11 | Cacfd1 | -0.034 | Young |
| 0.235677 | 0.055287 | 0.196 | 0.266 | 1 | 11 | Piezo1 | -0.07 | Young |
| 0.236041 | 0.156516 | 0.312 | 0.415 | 1 | 11 | Evl | -0.103 | Young |
| 0.236541 | 0.909666 | 0.043 | 0.077 | 1 | 11 | Stk16 | -0.034 | Young |
| 0.236541 | 0.599835 | 0.043 | 0.077 | 1 | 11 | Cep164 | -0.034 | Young |
| 0.236541 | 0.547599 | 0.043 | 0.077 | 1 | 11 | Pgam5 | -0.034 | Young |
| 0.236541 | 0.330409 | 0.043 | 0.077 | 1 | 11 | Gnl2 | -0.034 | Young |
| 0.236855 | 0.163875 | 0.355 | 0.46 | 1 | 11 | Mia2 | -0.105 | Young |
| 0.236913 | 1.216879 | 0.152 | 0.117 | 1 | 11 | Ice1 | 0.035 | Young |
| 0.237169 | 0.047981 | 0.471 | 0.581 | 1 | 11 | Fnbp1 | -0.11 | Young |
| 0.23729 | 0.000396 | 0.341 | 0.456 | 1 | 11 | Wdr83os | -0.115 | Young |
| 0.237443 | 0.192779 | 0.138 | 0.202 | 1 | 11 | Nsmaf | -0.064 | Young |
| 0.237474 | 0.441298 | 0.159 | 0.226 | 1 | 11 | Gpalpp1 | -0.067 | Young |
| 0.237567 | 0.664477 | 0.109 | 0.165 | 1 | 11 | Zfp62 | -0.056 | Young |
| 0.237729 | 0.816602 | 0.304 | 0.274 | 1 | 11 | Mxd4 | 0.03 | Young |
| 0.238059 | 1.01213 | 0.007 | 0.024 | 1 | 11 | Abraxas1 | -0.017 | Young |
| 0.238059 | 0.559153 | 0.007 | 0.024 | 1 | 11 | Tmem177 | -0.017 | Young |
| 0.238059 | 0.437875 | 0.007 | 0.024 | 1 | 11 | Mgll | -0.017 | Young |
| 0.238059 | 0.307232 | 0.007 | 0.024 | 1 | 11 | Mvk | -0.017 | Young |
| 0.238059 | 0.30346 | 0.007 | 0.024 | 1 | 11 | Gstt3 | -0.017 | Young |
| 0.238059 | 0.243323 | 0.007 | 0.024 | 1 | 11 | Zbtb25 | -0.017 | Young |
| 0.238059 | 0.109879 | 0.007 | 0.024 | 1 | 11 | Nup43 | -0.017 | Young |
| 0.238059 | 0.07666 | 0.007 | 0.024 | 1 | 11 | Gtf3c5 | -0.017 | Young |
| 0.238059 | 0.009893 | 0.007 | 0.024 | 1 | 11 | Rangrf | -0.017 | Young |
| 0.238059 | 0.005378 | 0.007 | 0.024 | 1 | 11 | Zdhhc1 | -0.017 | Young |
| 0.23807 | 0.727755 | 0.138 | 0.198 | 1 | 11 | Srp54a | -0.06 | Young |
| 0.238086 | 0.456881 | 0.116 | 0.169 | 1 | 11 | Cln6 | -0.053 | Young |
| 0.238313 | 0.184617 | 0.043 | 0.077 | 1 | 11 | Zbtb14 | -0.034 | Young |
| 0.238559 | 0.04147 | 0.493 | 0.645 | 1 | 11 | Ostf1 | -0.152 | Young |
| 0.238751 | 0.2607 | 0.065 | 0.105 | 1 | 11 | Slc25a33 | -0.04 | Young |
| 0.238783 | 0.497597 | 0.08 | 0.125 | 1 | 11 | Zbtb2 | -0.045 | Young |
| 0.238809 | 0.039763 | 0.536 | 0.677 | 1 | 11 | Bach1 | -0.141 | Young |
| 0.238815 | 0.189096 | 0.493 | 0.597 | 1 | 11 | Lgals3 | -0.104 | Young |
| 0.238989 | 0.17284 | 0.428 | 0.573 | 1 | 11 | Klf3 | -0.145 | Young |
| 0.239544 | 0.53467 | 0.362 | 0.347 | 1 | 11 | Cebpg | 0.015 | Young |
| 0.239611 | 0.164133 | 0.145 | 0.202 | 1 | 11 | Taf1 | -0.057 | Young |
| 0.239966 | 0.138109 | 0.957 | 0.984 | 1 | 11 | H3f3b | -0.027 | Young |
| 0.24001 | 0.306678 | 0.13 | 0.185 | 1 | 11 | Vps13d | -0.055 | Young |
| 0.240081 | 0.466895 | 0.13 | 0.19 | 1 | 11 | Nr1h2 | -0.06 | Young |
| 0.240082 | 0.075617 | 0.051 | 0.085 | 1 | 11 | H2-M2 | -0.034 | Young |
| 0.24028 | 0.22057 | 0.087 | 0.133 | 1 | 11 | Ppp1r7 | -0.046 | Young |
| 0.240348 | 0.026343 | 0.239 | 0.331 | 1 | 11 | Fbxl5 | -0.092 | Young |

| | | | | | | | | |
|---|---|---|---|---|---|---|---|---|
| 0.240394 | 0.208726 | 0.232 | 0.323 | 1 | 11 | Ubn1 | -0.091 | Young |
| 0.240441 | 0.118318 | 0.087 | 0.129 | 1 | 11 | Mrpl45 | -0.042 | Young |
| 0.240507 | 0.183276 | 0.029 | 0.056 | 1 | 11 | Trp53bp1 | -0.027 | Young |
| 0.240507 | 0.128051 | 0.029 | 0.056 | 1 | 11 | Ddx31 | -0.027 | Young |
| 0.240514 | 0.242873 | 0.217 | 0.29 | 1 | 11 | Abcc5 | -0.073 | Young |
| 0.241044 | 0.480279 | 0.065 | 0.105 | 1 | 11 | Vps45 | -0.04 | Young |
| 0.241065 | 0.05542 | 0.362 | 0.472 | 1 | 11 | Tmod3 | -0.11 | Young |
| 0.241066 | 0.266784 | 0.406 | 0.532 | 1 | 11 | Tardbp | -0.126 | Young |
| 0.241194 | 0.080522 | 0.355 | 0.464 | 1 | 11 | Slc43a2 | -0.109 | Young |
| 0.241352 | 0.519841 | 0.029 | 0.012 | 1 | 11 | Bckdhb | 0.017 | Young |
| 0.241904 | 0.088307 | 0.101 | 0.149 | 1 | 11 | Vps50 | -0.048 | Young |
| 0.241934 | 0.712595 | 0.268 | 0.242 | 1 | 11 | Ifi206 | 0.026 | Young |
| 0.242119 | 0.043192 | 0.283 | 0.383 | 1 | 11 | Arhgef1 | -0.1 | Young |
| 0.242394 | 0.608088 | 0.109 | 0.161 | 1 | 11 | Usp38 | -0.052 | Young |
| 0.242395 | 0.092136 | 0.167 | 0.23 | 1 | 11 | Vps11 | -0.063 | Young |
| 0.242403 | 0.157157 | 0.094 | 0.141 | 1 | 11 | Drg1 | -0.047 | Young |
| 0.242601 | 0.076594 | 0.029 | 0.056 | 1 | 11 | Scfd2 | -0.027 | Young |
| 0.242657 | 0.111776 | 0.181 | 0.25 | 1 | 11 | Casc3 | -0.069 | Young |
| 0.242674 | 0.184977 | 0.196 | 0.262 | 1 | 11 | Ociad1 | -0.066 | Young |
| 0.24268 | 0.248418 | 0.188 | 0.258 | 1 | 11 | Jup | -0.07 | Young |
| 0.243644 | 0.026745 | 0.406 | 0.492 | 1 | 11 | Pla2g7 | -0.086 | Young |
| 0.243861 | 0.31896 | 0.101 | 0.149 | 1 | 11 | Nelfe | -0.048 | Young |
| 0.243969 | 0.125109 | 0.239 | 0.323 | 1 | 11 | Sipa1 | -0.084 | Young |
| 0.244498 | 0.212809 | 0.181 | 0.25 | 1 | 11 | Ipo7 | -0.069 | Young |
| 0.245173 | 0.080245 | 0.101 | 0.149 | 1 | 11 | Zbtb38 | -0.048 | Young |
| 0.24558 | 0.204506 | 0.152 | 0.218 | 1 | 11 | Rfc1 | -0.066 | Young |
| 0.245722 | 1.472434 | 0.051 | 0.028 | 1 | 11 | Usp20 | 0.023 | Young |
| 0.245766 | 1.27676 | 0.029 | 0.056 | 1 | 11 | Med19 | -0.027 | Young |
| 0.245766 | 0.362261 | 0.029 | 0.056 | 1 | 11 | Dmap1 | -0.027 | Young |
| 0.245912 | 0.200914 | 0.058 | 0.093 | 1 | 11 | Desi1 | -0.035 | Young |
| 0.24637 | 0.483129 | 0.188 | 0.27 | 1 | 11 | Xpo6 | -0.082 | Young |
| 0.246584 | 0.192743 | 0.159 | 0.222 | 1 | 11 | Atf1 | -0.063 | Young |
| 0.246602 | 0.25157 | 0.717 | 0.81 | 1 | 11 | Arf5 | -0.093 | Young |
| 0.246719 | 0.321275 | 0.152 | 0.214 | 1 | 11 | Kdm2b | -0.062 | Young |
| 0.246839 | 0.025104 | 0.109 | 0.161 | 1 | 11 | Ppdpf | -0.052 | Young |
| 0.24692 | 1.859504 | 0.051 | 0.028 | 1 | 11 | Taf1c | 0.023 | Young |
| 0.246957 | 0.204693 | 0.775 | 0.815 | 1 | 11 | Rps17 | -0.04 | Young |
| 0.246988 | 0.560981 | 0.478 | 0.484 | 1 | 11 | Akr1b3 | -0.006 | Young |
| 0.246998 | 0.147571 | 0.21 | 0.274 | 1 | 11 | AI314180 | -0.064 | Young |
| 0.247232 | 0.419563 | 0.203 | 0.282 | 1 | 11 | Blmh | -0.079 | Young |
| 0.247421 | 0.955883 | 0.152 | 0.117 | 1 | 11 | Proser1 | 0.035 | Young |
| 0.247892 | 0.981708 | 0.029 | 0.056 | 1 | 11 | Psat1 | -0.027 | Young |
| 0.248214 | 0.381298 | 0.167 | 0.234 | 1 | 11 | Heatr5a | -0.067 | Young |
| 0.248224 | 0.0092 | 0.051 | 0.085 | 1 | 11 | Ggct | -0.034 | Young |
| 0.248406 | 0.127959 | 0.08 | 0.121 | 1 | 11 | Eef2k | -0.041 | Young |

| | | | | | | | | |
|---|---|---|---|---|---|---|---|---|
| 0.248447 | 0.026437 | 0.435 | 0.56 | 1 | 11 | Cox7a2 | -0.125 | Young |
| 0.248723 | 0.499321 | 0.21 | 0.294 | 1 | 11 | Golga7 | -0.084 | Young |
| 0.248958 | 0.791628 | 0.217 | 0.181 | 1 | 11 | Ltc4s | 0.036 | Young |
| 0.249032 | 0.122194 | 0.957 | 0.984 | 1 | 11 | Junb | -0.027 | Young |
| 0.249327 | 2.125521 | 0.051 | 0.028 | 1 | 11 | Ccl22 | 0.023 | Young |
| 0.24972 | 0.068819 | 0.623 | 0.694 | 1 | 11 | Mt1 | -0.071 | Young |
| 0.249841 | 0.276073 | 0.094 | 0.141 | 1 | 11 | Ltn1 | -0.047 | Young |
| 0.250058 | 0.580552 | 0.043 | 0.077 | 1 | 11 | Capn15 | -0.034 | Young |
| 0.250103 | 0.256244 | 0.29 | 0.379 | 1 | 11 | Pkig | -0.089 | Young |
| 0.250396 | 2.875253 | 0.022 | 0.008 | 1 | 11 | Brf2 | 0.014 | Young |
| 0.250396 | 2.242779 | 0.022 | 0.008 | 1 | 11 | Abhd15 | 0.014 | Young |
| 0.250396 | 2.236873 | 0.022 | 0.008 | 1 | 11 | Armcx2 | 0.014 | Young |
| 0.250396 | 2.173989 | 0.022 | 0.008 | 1 | 11 | Sirt5 | 0.014 | Young |
| 0.250396 | 1.90094 | 0.022 | 0.008 | 1 | 11 | Gm9725 | 0.014 | Young |
| 0.250396 | 1.892581 | 0.022 | 0.008 | 1 | 11 | Morn2 | 0.014 | Young |
| 0.250486 | 0.192854 | 0.283 | 0.371 | 1 | 11 | Mob3c | -0.088 | Young |
| 0.250644 | 0.110455 | 0.406 | 0.532 | 1 | 11 | Psmb1 | -0.126 | Young |
| 0.250652 | 0.226143 | 0.225 | 0.302 | 1 | 11 | Ezr | -0.077 | Young |
| 0.250787 | 0.00539 | 0.167 | 0.226 | 1 | 11 | Oaz2 | -0.059 | Young |
| 0.250832 | 0.570396 | 0.486 | 0.508 | 1 | 11 | Ncf1 | -0.022 | Young |
| 0.251192 | 0.103595 | 0.993 | 0.984 | 1 | 11 | Rps8 | 0.009 | Young |
| 0.251365 | 0.206739 | 0.13 | 0.185 | 1 | 11 | Snapin | -0.055 | Young |
| 0.251605 | 1.573694 | 0.065 | 0.04 | 1 | 11 | Mki67 | 0.025 | Young |
| 0.251695 | 0.535754 | 0.072 | 0.113 | 1 | 11 | Tbc1d10a | -0.041 | Young |
| 0.251695 | 0.057191 | 0.072 | 0.113 | 1 | 11 | Mdc1 | -0.041 | Young |
| 0.25175 | 1.379315 | 0.051 | 0.028 | 1 | 11 | Tti2 | 0.023 | Young |
| 0.251949 | 0.464053 | 0.065 | 0.105 | 1 | 11 | Bcl2l13 | -0.04 | Young |
| 0.252147 | 0.711499 | 0.051 | 0.085 | 1 | 11 | Dglucy | -0.034 | Young |
| 0.252342 | 0.43117 | 0.297 | 0.403 | 1 | 11 | Trim8 | -0.106 | Young |
| 0.252382 | 0.42772 | 0.493 | 0.516 | 1 | 11 | Nr3c1 | -0.023 | Young |
| 0.252405 | 1.664096 | 0.022 | 0.008 | 1 | 11 | Agrn | 0.014 | Young |
| 0.252405 | 1.525973 | 0.022 | 0.008 | 1 | 11 | Atad5 | 0.014 | Young |
| 0.252405 | 1.247331 | 0.022 | 0.008 | 1 | 11 | Pdlim7 | 0.014 | Young |
| 0.252474 | 0.193549 | 0.101 | 0.149 | 1 | 11 | Usf1 | -0.048 | Young |
| 0.252527 | 0.652071 | 0.036 | 0.065 | 1 | 11 | Bloc1s1 | -0.029 | Young |
| 0.252987 | 0.022643 | 0.341 | 0.435 | 1 | 11 | Stx4a | -0.094 | Young |
| 0.253817 | 0.844059 | 0.101 | 0.149 | 1 | 11 | Dcaf15 | -0.048 | Young |
| 0.254229 | 0.095361 | 0.058 | 0.093 | 1 | 11 | Rpa3 | -0.035 | Young |
| 0.254412 | 0.237038 | 0.304 | 0.415 | 1 | 11 | Bcl10 | -0.111 | Young |
| 0.254424 | 3.124704 | 0.022 | 0.008 | 1 | 11 | Pole2 | 0.014 | Young |
| 0.254424 | 2.215913 | 0.022 | 0.008 | 1 | 11 | 1700056E2 | 0.014 | Young |
| 0.254424 | 1.708056 | 0.022 | 0.008 | 1 | 11 | Slc6a8 | 0.014 | Young |
| 0.254424 | 1.64481 | 0.022 | 0.008 | 1 | 11 | Ccdc171 | 0.014 | Young |
| 0.254424 | 1.46033 | 0.022 | 0.008 | 1 | 11 | Tmem42 | 0.014 | Young |
| 0.254424 | 1.371622 | 0.022 | 0.008 | 1 | 11 | Dynlt1a | 0.014 | Young |

| | | | | | | | | |
|---|---|---|---|---|---|---|---|---|
| 0.254424 | 1.251187 | 0.022 | 0.008 | 1 | 11 | C330027C( | 0.014 | Young |
| 0.254424 | 1.193476 | 0.022 | 0.008 | 1 | 11 | Rnf152 | 0.014 | Young |
| 0.254424 | 0.817915 | 0.022 | 0.008 | 1 | 11 | Zhx3 | 0.014 | Young |
| 0.255396 | 0.355409 | 0.587 | 0.629 | 1 | 11 | Eif3f | -0.042 | Young |
| 0.255551 | 0.316683 | 0.036 | 0.065 | 1 | 11 | Sqle | -0.029 | Young |
| 0.255671 | 0.577398 | 0.051 | 0.085 | 1 | 11 | Haus3 | -0.034 | Young |
| 0.255698 | 0.769805 | 0.21 | 0.298 | 1 | 11 | Lrmp | -0.088 | Young |
| 0.25571 | 0.635574 | 0.312 | 0.278 | 1 | 11 | Dusp2 | 0.034 | Young |
| 0.2558 | 0.034151 | 0.109 | 0.157 | 1 | 11 | Parp2 | -0.048 | Young |
| 0.256142 | 0.358893 | 0.188 | 0.258 | 1 | 11 | C330007P( | -0.07 | Young |
| 0.256445 | 0.052139 | 0.384 | 0.504 | 1 | 11 | Eps15 | -0.12 | Young |
| 0.256456 | 1.301033 | 0.022 | 0.008 | 1 | 11 | H2-Oa | 0.014 | Young |
| 0.256456 | 1.001567 | 0.022 | 0.008 | 1 | 11 | Klhl13 | 0.014 | Young |
| 0.256456 | 0.975339 | 0.022 | 0.008 | 1 | 11 | Bace1 | 0.014 | Young |
| 0.256456 | 0.887697 | 0.022 | 0.008 | 1 | 11 | Rccd1 | 0.014 | Young |
| 0.256591 | 0.027066 | 0.116 | 0.165 | 1 | 11 | Ddx52 | -0.049 | Young |
| 0.256726 | 0.716442 | 0.065 | 0.105 | 1 | 11 | Foxred1 | -0.04 | Young |
| 0.257658 | 0.363531 | 0.174 | 0.238 | 1 | 11 | Banf1 | -0.064 | Young |
| 0.257995 | 0.238419 | 0.261 | 0.347 | 1 | 11 | Tmpo | -0.086 | Young |
| 0.258336 | 0.381073 | 0.051 | 0.085 | 1 | 11 | Mcm3ap | -0.034 | Young |
| 0.258336 | 0.081662 | 0.051 | 0.085 | 1 | 11 | Tmc6 | -0.034 | Young |
| 0.258377 | 0.263493 | 0.188 | 0.262 | 1 | 11 | Arnt | -0.074 | Young |
| 0.258498 | 1.008124 | 0.022 | 0.008 | 1 | 11 | Cd28 | 0.014 | Young |
| 0.258498 | 0.454279 | 0.022 | 0.008 | 1 | 11 | Qprt | 0.014 | Young |
| 0.258498 | 0.390572 | 0.022 | 0.008 | 1 | 11 | Cep112 | 0.014 | Young |
| 0.258566 | 0.040442 | 0.283 | 0.363 | 1 | 11 | Cdkn1b | -0.08 | Young |
| 0.25866 | 0.331282 | 0.319 | 0.444 | 1 | 11 | Dennd1b | -0.125 | Young |
| 0.25903 | 1.470688 | 0.065 | 0.04 | 1 | 11 | Stmn1 | 0.025 | Young |
| 0.25903 | 1.055299 | 0.065 | 0.04 | 1 | 11 | Mthfd2l | 0.025 | Young |
| 0.259092 | 0.005557 | 0.797 | 0.774 | 1 | 11 | Jun | 0.023 | Young |
| 0.259116 | 0.095506 | 0.109 | 0.157 | 1 | 11 | Lats1 | -0.048 | Young |
| 0.259687 | 0.061594 | 0.486 | 0.597 | 1 | 11 | Stat3 | -0.111 | Young |
| 0.259746 | 0.206384 | 0.094 | 0.137 | 1 | 11 | Vps53 | -0.043 | Young |
| 0.259772 | 0.07864 | 0.29 | 0.375 | 1 | 11 | Ap1s2 | -0.085 | Young |
| 0.260052 | 0.085048 | 0.471 | 0.613 | 1 | 11 | Cnbp | -0.142 | Young |
| 0.260067 | 0.233217 | 0.196 | 0.274 | 1 | 11 | Peak1 | -0.078 | Young |
| 0.260104 | 1.415056 | 0.065 | 0.04 | 1 | 11 | Pcid2 | 0.025 | Young |
| 0.260201 | 0.679307 | 0.348 | 0.331 | 1 | 11 | Elmo1 | 0.017 | Young |
| 0.260307 | 1.477726 | 0.101 | 0.073 | 1 | 11 | Pno1 | 0.028 | Young |
| 0.260345 | 0.10371 | 0.37 | 0.484 | 1 | 11 | Ankrd11 | -0.114 | Young |
| 0.26038 | 0.169809 | 0.312 | 0.435 | 1 | 11 | Pxn | -0.123 | Young |
| 0.260552 | 0.445848 | 0.022 | 0.008 | 1 | 11 | Ly6c1 | 0.014 | Young |
| 0.260552 | 0.427685 | 0.022 | 0.008 | 1 | 11 | Cenpe | 0.014 | Young |
| 0.260648 | 0.115825 | 0.036 | 0.065 | 1 | 11 | Zcchc14 | -0.029 | Young |
| 0.260755 | 0.415043 | 0.065 | 0.105 | 1 | 11 | Glo1 | -0.04 | Young |

| | | | | | | | | |
|---|---|---|---|---|---|---|---|---|
| 0.261235 | 1.126827 | 0.08 | 0.052 | 1 | 11 | Pcnx3 | 0.028 | Young |
| 0.261271 | 0.130163 | 0.217 | 0.29 | 1 | 11 | Trappc4 | -0.073 | Young |
| 0.26131 | 3.510711 | 0.014 | 0.004 | 1 | 11 | Dennd5b | 0.01 | Young |
| 0.26131 | 3.137262 | 0.014 | 0.004 | 1 | 11 | AC113595. | 0.01 | Young |
| 0.26131 | 3.055717 | 0.014 | 0.004 | 1 | 11 | Slc25a27 | 0.01 | Young |
| 0.26131 | 2.901612 | 0.014 | 0.004 | 1 | 11 | Tmem8b | 0.01 | Young |
| 0.26131 | 2.774232 | 0.014 | 0.004 | 1 | 11 | Tcf15 | 0.01 | Young |
| 0.26131 | 2.711836 | 0.014 | 0.004 | 1 | 11 | Fgf9 | 0.01 | Young |
| 0.26131 | 2.691346 | 0.014 | 0.004 | 1 | 11 | Slc7a2 | 0.01 | Young |
| 0.26131 | 2.527051 | 0.014 | 0.004 | 1 | 11 | Zfp512b | 0.01 | Young |
| 0.26131 | 2.481173 | 0.014 | 0.004 | 1 | 11 | Zfp324 | 0.01 | Young |
| 0.26131 | 2.323421 | 0.014 | 0.004 | 1 | 11 | Mrnip | 0.01 | Young |
| 0.26131 | 2.316236 | 0.014 | 0.004 | 1 | 11 | Prkcg | 0.01 | Young |
| 0.26131 | 2.304383 | 0.014 | 0.004 | 1 | 11 | Cxcl14 | 0.01 | Young |
| 0.26131 | 2.273357 | 0.014 | 0.004 | 1 | 11 | Bcl6b | 0.01 | Young |
| 0.26131 | 2.250764 | 0.014 | 0.004 | 1 | 11 | Camk2n1 | 0.01 | Young |
| 0.26131 | 2.155448 | 0.014 | 0.004 | 1 | 11 | Mss51 | 0.01 | Young |
| 0.26131 | 2.140472 | 0.014 | 0.004 | 1 | 11 | 2810457G( | 0.01 | Young |
| 0.26131 | 2.139823 | 0.014 | 0.004 | 1 | 11 | Ankrd24 | 0.01 | Young |
| 0.26131 | 2.038378 | 0.014 | 0.004 | 1 | 11 | Klrg2 | 0.01 | Young |
| 0.26131 | 1.988781 | 0.014 | 0.004 | 1 | 11 | Jcad | 0.01 | Young |
| 0.26131 | 1.959241 | 0.014 | 0.004 | 1 | 11 | Samd14 | 0.01 | Young |
| 0.26131 | 1.840769 | 0.014 | 0.004 | 1 | 11 | Stk26 | 0.01 | Young |
| 0.26131 | 1.837563 | 0.014 | 0.004 | 1 | 11 | Gnai1 | 0.01 | Young |
| 0.26131 | 1.803032 | 0.014 | 0.004 | 1 | 11 | Zfp30 | 0.01 | Young |
| 0.26131 | 1.799848 | 0.014 | 0.004 | 1 | 11 | Mars2 | 0.01 | Young |
| 0.26131 | 1.749933 | 0.014 | 0.004 | 1 | 11 | Tgfb3 | 0.01 | Young |
| 0.26131 | 1.706982 | 0.014 | 0.004 | 1 | 11 | Aldh1a1 | 0.01 | Young |
| 0.26131 | 1.70506 | 0.014 | 0.004 | 1 | 11 | Adamts2 | 0.01 | Young |
| 0.26131 | 1.699886 | 0.014 | 0.004 | 1 | 11 | Gm42670 | 0.01 | Young |
| 0.26131 | 1.655015 | 0.014 | 0.004 | 1 | 11 | Nbl1 | 0.01 | Young |
| 0.26131 | 1.556961 | 0.014 | 0.004 | 1 | 11 | Hist1h4h | 0.01 | Young |
| 0.26131 | 1.544795 | 0.014 | 0.004 | 1 | 11 | Olfml3 | 0.01 | Young |
| 0.26131 | 1.53485 | 0.014 | 0.004 | 1 | 11 | Fgfbp3 | 0.01 | Young |
| 0.26131 | 1.4961 | 0.014 | 0.004 | 1 | 11 | Irak1bp1 | 0.01 | Young |
| 0.26131 | 1.390852 | 0.014 | 0.004 | 1 | 11 | Wdpcp | 0.01 | Young |
| 0.26131 | 1.291204 | 0.014 | 0.004 | 1 | 11 | Efna2 | 0.01 | Young |
| 0.26131 | 1.154728 | 0.014 | 0.004 | 1 | 11 | Ltb | 0.01 | Young |
| 0.26131 | 1.078752 | 0.014 | 0.004 | 1 | 11 | Efnb2 | 0.01 | Young |
| 0.26131 | 0.860421 | 0.014 | 0.004 | 1 | 11 | Eif3j2 | 0.01 | Young |
| 0.26131 | 0.836578 | 0.014 | 0.004 | 1 | 11 | Spats2l | 0.01 | Young |
| 0.262013 | 0.290252 | 0.428 | 0.387 | 1 | 11 | S100a4 | 0.041 | Young |
| 0.262526 | 0.204235 | 0.391 | 0.536 | 1 | 11 | Glg1 | -0.145 | Young |
| 0.262538 | 0.301738 | 0.08 | 0.121 | 1 | 11 | Spic | -0.041 | Young |
| 0.263136 | 0.008837 | 0.507 | 0.677 | 1 | 11 | Pafah1b1 | -0.17 | Young |

| | | | | | | | | |
|---|---|---|---|---|---|---|---|---|
| 0.263376 | 0.300993 | 0.225 | 0.31 | 1 | 11 | Bmt2 | -0.085 | Young |
| 0.263977 | 2.099192 | 0.014 | 0.004 | 1 | 11 | Tnfrsf10b | 0.01 | Young |
| 0.263977 | 1.986424 | 0.014 | 0.004 | 1 | 11 | Arvcf | 0.01 | Young |
| 0.263977 | 1.985274 | 0.014 | 0.004 | 1 | 11 | Zfp937 | 0.01 | Young |
| 0.263977 | 1.729833 | 0.014 | 0.004 | 1 | 11 | Tbc1d19 | 0.01 | Young |
| 0.263977 | 1.61081 | 0.014 | 0.004 | 1 | 11 | B3galt2 | 0.01 | Young |
| 0.263977 | 1.538201 | 0.014 | 0.004 | 1 | 11 | Sv2b | 0.01 | Young |
| 0.263977 | 1.516449 | 0.014 | 0.004 | 1 | 11 | Mdga1 | 0.01 | Young |
| 0.263977 | 1.235135 | 0.014 | 0.004 | 1 | 11 | Prc1 | 0.01 | Young |
| 0.263977 | 1.192085 | 0.014 | 0.004 | 1 | 11 | Satb1 | 0.01 | Young |
| 0.263977 | 1.186385 | 0.014 | 0.004 | 1 | 11 | Kcnd1 | 0.01 | Young |
| 0.263977 | 1.149932 | 0.014 | 0.004 | 1 | 11 | Thsd1 | 0.01 | Young |
| 0.263977 | 1.146347 | 0.014 | 0.004 | 1 | 11 | Synpo2 | 0.01 | Young |
| 0.263977 | 1.10313 | 0.014 | 0.004 | 1 | 11 | Nlrc3 | 0.01 | Young |
| 0.263977 | 1.09493 | 0.014 | 0.004 | 1 | 11 | Tigd5 | 0.01 | Young |
| 0.263977 | 1.068975 | 0.014 | 0.004 | 1 | 11 | Gm29562 | 0.01 | Young |
| 0.263993 | 0.160169 | 0.37 | 0.456 | 1 | 11 | Id2 | -0.086 | Young |
| 0.264132 | 0.261756 | 0.232 | 0.31 | 1 | 11 | Krtcap2 | -0.078 | Young |
| 0.264359 | 0.127869 | 0.848 | 0.867 | 1 | 11 | Ptprc | -0.019 | Young |
| 0.264587 | 0.045417 | 0.362 | 0.468 | 1 | 11 | Arl8a | -0.106 | Young |
| 0.264671 | 0.402683 | 0.565 | 0.629 | 1 | 11 | Rbm3 | -0.064 | Young |
| 0.265035 | 0.063022 | 0.391 | 0.496 | 1 | 11 | Manf | -0.105 | Young |
| 0.26533 | 0.248144 | 0.058 | 0.093 | 1 | 11 | Car3 | -0.035 | Young |
| 0.265815 | 0.55038 | 0.036 | 0.065 | 1 | 11 | Pias3 | -0.029 | Young |
| 0.265815 | 0.071241 | 0.036 | 0.065 | 1 | 11 | Fmo5 | -0.029 | Young |
| 0.265895 | 0.079971 | 0.239 | 0.31 | 1 | 11 | Creb5 | -0.071 | Young |
| 0.266662 | 1.269171 | 0.014 | 0.004 | 1 | 11 | Ccdc173 | 0.01 | Young |
| 0.266662 | 0.996053 | 0.014 | 0.004 | 1 | 11 | E130208F1 | 0.01 | Young |
| 0.266662 | 0.818799 | 0.014 | 0.004 | 1 | 11 | Lrrc32 | 0.01 | Young |
| 0.266662 | 0.550984 | 0.014 | 0.004 | 1 | 11 | Clec4g | 0.01 | Young |
| 0.266662 | 0.516863 | 0.014 | 0.004 | 1 | 11 | Cd200 | 0.01 | Young |
| 0.266662 | 0.498052 | 0.014 | 0.004 | 1 | 11 | Klrd1 | 0.01 | Young |
| 0.266662 | 0.456807 | 0.014 | 0.004 | 1 | 11 | Hacd1 | 0.01 | Young |
| 0.266662 | 0.192751 | 0.014 | 0.004 | 1 | 11 | Cd46 | 0.01 | Young |
| 0.26678 | 0.703002 | 0.283 | 0.254 | 1 | 11 | Ssh2 | 0.029 | Young |
| 0.266857 | 0.487475 | 0.036 | 0.065 | 1 | 11 | Surf2 | -0.029 | Young |
| 0.266988 | 1.598141 | 0.145 | 0.117 | 1 | 11 | Trappc5 | 0.028 | Young |
| 0.267114 | 0.072029 | 0.08 | 0.121 | 1 | 11 | Slc41a3 | -0.041 | Young |
| 0.26721 | 0.2228 | 0.333 | 0.431 | 1 | 11 | Igf1 | -0.098 | Young |
| 0.267233 | 0.043629 | 0.348 | 0.46 | 1 | 11 | Hmgb2 | -0.112 | Young |
| 0.267633 | 0.294913 | 0.181 | 0.242 | 1 | 11 | Strap | -0.061 | Young |
| 0.267701 | 0.845964 | 0.065 | 0.04 | 1 | 11 | 3110082I1 | 0.025 | Young |
| 0.267882 | 0.407258 | 0.08 | 0.121 | 1 | 11 | B3galnt2 | -0.041 | Young |
| 0.267884 | 0.272632 | 0.109 | 0.161 | 1 | 11 | Tk2 | -0.052 | Young |
| 0.268139 | 0.160013 | 0.13 | 0.185 | 1 | 11 | Mfsd12 | -0.055 | Young |

| | | | | | | | | |
|---|---|---|---|---|---|---|---|---|
| 0.268176 | 0.092952 | 0.116 | 0.161 | 1 | 11 | Fh1 | -0.045 | Young |
| 0.26818 | 0.080112 | 0.362 | 0.435 | 1 | 11 | Zyx | -0.073 | Young |
| 0.268313 | 0.013187 | 0.536 | 0.681 | 1 | 11 | Trps1 | -0.145 | Young |
| 0.268346 | 0.138554 | 0.101 | 0.145 | 1 | 11 | Rxrb | -0.044 | Young |
| 0.268716 | 0.38185 | 0.152 | 0.218 | 1 | 11 | Rnpepl1 | -0.066 | Young |
| 0.268949 | 0.501538 | 0.036 | 0.065 | 1 | 11 | Amd1 | -0.029 | Young |
| 0.268949 | 0.288041 | 0.036 | 0.065 | 1 | 11 | Amz1 | -0.029 | Young |
| 0.269144 | 0.175974 | 0.181 | 0.254 | 1 | 11 | Stk40 | -0.073 | Young |
| 0.26953 | 0.108808 | 0.043 | 0.073 | 1 | 11 | B4gat1 | -0.03 | Young |
| 0.269687 | 0.287599 | 0.058 | 0.093 | 1 | 11 | Fam207a | -0.035 | Young |
| 0.269714 | 0.133826 | 0.587 | 0.54 | 1 | 11 | Ccr2 | 0.047 | Young |
| 0.26973 | 0.75638 | 0.065 | 0.101 | 1 | 11 | Pcgf3 | -0.036 | Young |
| 0.269918 | 0.570368 | 0.109 | 0.161 | 1 | 11 | Msi2 | -0.052 | Young |
| 0.269999 | 0.234789 | 0.036 | 0.065 | 1 | 11 | Ripk3 | -0.029 | Young |
| 0.270328 | 0.43747 | 0.101 | 0.149 | 1 | 11 | Wdr13 | -0.048 | Young |
| 0.270703 | 0.051872 | 0.246 | 0.343 | 1 | 11 | Safb2 | -0.097 | Young |
| 0.270858 | 0.202124 | 0.261 | 0.351 | 1 | 11 | Med13l | -0.09 | Young |
| 0.271262 | 0.123114 | 0.145 | 0.202 | 1 | 11 | Tollip | -0.057 | Young |
| 0.271388 | 0.311198 | 0.167 | 0.234 | 1 | 11 | Yipf5 | -0.067 | Young |
| 0.271407 | 0.132968 | 0.167 | 0.226 | 1 | 11 | Nfia | -0.059 | Young |
| 0.271414 | 0.096235 | 0.065 | 0.101 | 1 | 11 | Nudt13 | -0.036 | Young |
| 0.271444 | 0.580872 | 0.058 | 0.093 | 1 | 11 | Hmbs | -0.035 | Young |
| 0.271507 | 0.069995 | 0.043 | 0.073 | 1 | 11 | Zfyve19 | -0.03 | Young |
| 0.271727 | 0.264491 | 0.681 | 0.726 | 1 | 11 | H2afz | -0.045 | Young |
| 0.271729 | 0.436098 | 0.022 | 0.044 | 1 | 11 | 2310061I0 | -0.022 | Young |
| 0.271729 | 0.403258 | 0.022 | 0.044 | 1 | 11 | Arhgap22 | -0.022 | Young |
| 0.272123 | 0.340865 | 0.319 | 0.419 | 1 | 11 | Fam129a | -0.1 | Young |
| 0.272325 | 0.338426 | 0.058 | 0.093 | 1 | 11 | Gfm1 | -0.035 | Young |
| 0.273113 | 1.038076 | 0.123 | 0.093 | 1 | 11 | Hikeshi | 0.03 | Young |
| 0.273451 | 0.418763 | 0.094 | 0.137 | 1 | 11 | Ccdc86 | -0.043 | Young |
| 0.273691 | 0.234896 | 0.319 | 0.444 | 1 | 11 | Phf11b | -0.125 | Young |
| 0.273721 | 0.096757 | 0.087 | 0.129 | 1 | 11 | Naa25 | -0.042 | Young |
| 0.273864 | 0.112589 | 0.42 | 0.548 | 1 | 11 | Eid1 | -0.128 | Young |
| 0.273954 | 0.10122 | 0.065 | 0.101 | 1 | 11 | Tex10 | -0.036 | Young |
| 0.274147 | 0.117669 | 0.493 | 0.605 | 1 | 11 | Lgals3bp | -0.112 | Young |
| 0.27491 | 0.147107 | 0.377 | 0.508 | 1 | 11 | H2afj | -0.131 | Young |
| 0.274951 | 0.09523 | 0.116 | 0.161 | 1 | 11 | Pcif1 | -0.045 | Young |
| 0.275218 | 0.186282 | 0.399 | 0.52 | 1 | 11 | Vwa5a | -0.121 | Young |
| 0.275243 | 0.213975 | 0.145 | 0.206 | 1 | 11 | Ptpn22 | -0.061 | Young |
| 0.275313 | 0.441944 | 0.145 | 0.198 | 1 | 11 | Hspa14 | -0.053 | Young |
| 0.27545 | 0.825353 | 0.065 | 0.04 | 1 | 11 | Zeb1 | 0.025 | Young |
| 0.275593 | 0.633052 | 0.022 | 0.044 | 1 | 11 | Ulk3 | -0.022 | Young |
| 0.275909 | 0.405356 | 0.174 | 0.238 | 1 | 11 | Mrpl51 | -0.064 | Young |
| 0.275958 | 0.275392 | 0.13 | 0.181 | 1 | 11 | Rab29 | -0.051 | Young |
| 0.276017 | 0.042546 | 0.638 | 0.774 | 1 | 11 | Srgn | -0.136 | Young |

| | | | | | | | | |
|---|---|---|---|---|---|---|---|---|
| 0.276087 | 1.429117 | 0.174 | 0.149 | 1 | 11 | Mprip | 0.025 | Young |
| 0.27651 | 0.056445 | 0.065 | 0.101 | 1 | 11 | Nrde2 | -0.036 | Young |
| 0.276809 | 0.291505 | 0.333 | 0.444 | 1 | 11 | Nsd3 | -0.111 | Young |
| 0.276856 | 0.218244 | 0.145 | 0.202 | 1 | 11 | Timm8b | -0.057 | Young |
| 0.276889 | 1.726526 | 0.022 | 0.044 | 1 | 11 | Kdm1b | -0.022 | Young |
| 0.277012 | 0.013705 | 0.152 | 0.206 | 1 | 11 | Chrac1 | -0.054 | Young |
| 0.277138 | 0.056114 | 0.094 | 0.137 | 1 | 11 | Slc25a46 | -0.043 | Young |
| 0.277308 | 0.364707 | 0.203 | 0.266 | 1 | 11 | Trpm7 | -0.063 | Young |
| 0.277375 | 1.778645 | 0.072 | 0.048 | 1 | 11 | Rft1 | 0.024 | Young |
| 0.2775 | 0.069336 | 0.043 | 0.073 | 1 | 11 | Gabpb1 | -0.03 | Young |
| 0.277673 | 0.462884 | 0.362 | 0.468 | 1 | 11 | Gpr65 | -0.106 | Young |
| 0.277866 | 0.342929 | 0.116 | 0.165 | 1 | 11 | Slc37a2 | -0.049 | Young |
| 0.277867 | 0.08091 | 0.08 | 0.117 | 1 | 11 | Slc38a7 | -0.037 | Young |
| 0.278174 | 0.334885 | 0.304 | 0.411 | 1 | 11 | Ifih1 | -0.107 | Young |
| 0.278223 | 2.330109 | 0.043 | 0.024 | 1 | 11 | Ms4a4a | 0.019 | Young |
| 0.27846 | 0.180672 | 0.159 | 0.218 | 1 | 11 | Supt4a | -0.059 | Young |
| 0.278501 | 0.581978 | 0.036 | 0.065 | 1 | 11 | Cisd3 | -0.029 | Young |
| 0.278516 | 0.043774 | 0.188 | 0.25 | 1 | 11 | Sesn1 | -0.062 | Young |
| 0.278549 | 0.265034 | 0.116 | 0.165 | 1 | 11 | Hnrnph3 | -0.049 | Young |
| 0.27925 | 0.025958 | 0.688 | 0.794 | 1 | 11 | Arhgdia | -0.106 | Young |
| 0.279517 | 0.00183 | 0.043 | 0.073 | 1 | 11 | Dnajb5 | -0.03 | Young |
| 0.279577 | 0.79806 | 0.036 | 0.065 | 1 | 11 | Dip2a | -0.029 | Young |
| 0.279713 | 0.697106 | 0.239 | 0.21 | 1 | 11 | Fam3c | 0.029 | Young |
| 0.280167 | 0.059555 | 0.072 | 0.109 | 1 | 11 | Tmco3 | -0.037 | Young |
| 0.280344 | 0.356268 | 0.058 | 0.093 | 1 | 11 | Pmm2 | -0.035 | Young |
| 0.280655 | 0.733253 | 0.036 | 0.065 | 1 | 11 | Actr6 | -0.029 | Young |
| 0.281039 | 1.596821 | 0.043 | 0.024 | 1 | 11 | Pls3 | 0.019 | Young |
| 0.281169 | 0.334313 | 0.08 | 0.121 | 1 | 11 | Wrn | -0.041 | Young |
| 0.28118 | 0.232105 | 0.094 | 0.141 | 1 | 11 | Avl9 | -0.047 | Young |
| 0.281295 | 0.394336 | 0.116 | 0.165 | 1 | 11 | Dag1 | -0.049 | Young |
| 0.281545 | 0.1391 | 0.043 | 0.073 | 1 | 11 | E2f5 | -0.03 | Young |
| 0.281567 | 0.096384 | 0.21 | 0.278 | 1 | 11 | Mfhas1 | -0.068 | Young |
| 0.28179 | 0.091224 | 0.174 | 0.238 | 1 | 11 | Ostm1 | -0.064 | Young |
| 0.281791 | 0.346345 | 0.167 | 0.23 | 1 | 11 | Atp6v0a2 | -0.063 | Young |
| 0.281871 | 0.222015 | 0.326 | 0.423 | 1 | 11 | Ash1l | -0.097 | Young |
| 0.281978 | 0.547235 | 0.181 | 0.254 | 1 | 11 | Ufm1 | -0.073 | Young |
| 0.281983 | 0.206952 | 0.384 | 0.48 | 1 | 11 | Phip | -0.096 | Young |
| 0.282454 | 1.355858 | 0.043 | 0.024 | 1 | 11 | Eif2b1 | 0.019 | Young |
| 0.282536 | 0.086715 | 0.065 | 0.101 | 1 | 11 | Ikbip | -0.036 | Young |
| 0.284518 | 0.333877 | 0.196 | 0.262 | 1 | 11 | Smarce1 | -0.066 | Young |
| 0.284605 | 0.073205 | 0.043 | 0.073 | 1 | 11 | Gstp1 | -0.03 | Young |
| 0.28501 | 0.193587 | 0.348 | 0.46 | 1 | 11 | Prkcb | -0.112 | Young |
| 0.285033 | 0.157333 | 0.319 | 0.427 | 1 | 11 | Efr3a | -0.108 | Young |
| 0.285224 | 0.407174 | 0.087 | 0.129 | 1 | 11 | Usp33 | -0.042 | Young |
| 0.285299 | 1.246584 | 0.043 | 0.024 | 1 | 11 | Kif5c | 0.019 | Young |

| | | | | | | | | |
|---|---|---|---|---|---|---|---|---|
| 0.285368 | 0.147222 | 0.094 | 0.137 | 1 | 11 | Cyth2 | -0.043 | Young |
| 0.286051 | 0.433425 | 0.283 | 0.391 | 1 | 11 | Cdk9 | -0.108 | Young |
| 0.286102 | 0.305533 | 0.174 | 0.234 | 1 | 11 | Churc1 | -0.06 | Young |
| 0.286145 | 0.624059 | 0.116 | 0.165 | 1 | 11 | Magohb | -0.049 | Young |
| 0.286392 | 0.258083 | 0.167 | 0.226 | 1 | 11 | Pdpk1 | -0.059 | Young |
| 0.286704 | 0.024387 | 0.174 | 0.234 | 1 | 11 | Araf | -0.06 | Young |
| 0.286729 | 1.28868 | 0.043 | 0.024 | 1 | 11 | Egfr | 0.019 | Young |
| 0.286782 | 0.329869 | 0.087 | 0.129 | 1 | 11 | Ash2l | -0.042 | Young |
| 0.286903 | 0.072388 | 0.362 | 0.472 | 1 | 11 | Snrpf | -0.11 | Young |
| 0.287123 | 0.149152 | 0.297 | 0.387 | 1 | 11 | Aff4 | -0.09 | Young |
| 0.287132 | 0.226216 | 0.232 | 0.319 | 1 | 11 | Rtf2 | -0.087 | Young |
| 0.287355 | 1.181159 | 0.203 | 0.177 | 1 | 11 | Ebi3 | 0.026 | Young |
| 0.287406 | 0.027834 | 0.377 | 0.48 | 1 | 11 | Dck | -0.103 | Young |
| 0.287865 | 0.321835 | 0.13 | 0.181 | 1 | 11 | Ccndbp1 | -0.051 | Young |
| 0.288164 | 1.416417 | 0.043 | 0.024 | 1 | 11 | Gm8369 | 0.019 | Young |
| 0.288175 | 0.979057 | 0.13 | 0.101 | 1 | 11 | Tango2 | 0.029 | Young |
| 0.288651 | 0.194929 | 0.065 | 0.101 | 1 | 11 | Sash3 | -0.036 | Young |
| 0.288692 | 0.209037 | 0.188 | 0.254 | 1 | 11 | Pcna | -0.066 | Young |
| 0.288719 | 0.224592 | 0.043 | 0.073 | 1 | 11 | Prep | -0.03 | Young |
| 0.288739 | 0.048236 | 0.268 | 0.351 | 1 | 11 | Fkbp8 | -0.083 | Young |
| 0.288753 | 0.036454 | 0.239 | 0.315 | 1 | 11 | Klf10 | -0.076 | Young |
| 0.288793 | 0.17529 | 0.406 | 0.532 | 1 | 11 | Trip12 | -0.126 | Young |
| 0.288926 | 1.05456 | 0.217 | 0.19 | 1 | 11 | Samsn1 | 0.027 | Young |
| 0.288941 | 0.446692 | 0.116 | 0.165 | 1 | 11 | Irak2 | -0.049 | Young |
| 0.289213 | 0.52933 | 0.522 | 0.536 | 1 | 11 | Ier3 | -0.014 | Young |
| 0.289261 | 0.200105 | 0.268 | 0.351 | 1 | 11 | Tnpo3 | -0.083 | Young |
| 0.289341 | 0.161817 | 0.101 | 0.145 | 1 | 11 | Lrrc41 | -0.044 | Young |
| 0.289363 | 1.400235 | 0.13 | 0.101 | 1 | 11 | mt-Nd6 | 0.029 | Young |
| 0.289712 | 0.248074 | 0.101 | 0.145 | 1 | 11 | Atxn10 | -0.044 | Young |
| 0.289723 | 0.409801 | 0.384 | 0.528 | 1 | 11 | Ccnd1 | -0.144 | Young |
| 0.289754 | 0.244797 | 0.043 | 0.073 | 1 | 11 | Sh3bp4 | -0.03 | Young |
| 0.290814 | 0.931959 | 0.08 | 0.121 | 1 | 11 | Stoml2 | -0.041 | Young |
| 0.2909 | 0.574083 | 0.167 | 0.23 | 1 | 11 | BC004004 | -0.063 | Young |
| 0.291047 | 0.85172 | 0.043 | 0.024 | 1 | 11 | Tmem186 | 0.019 | Young |
| 0.29105 | 0.036919 | 0.145 | 0.194 | 1 | 11 | Dusp5 | -0.049 | Young |
| 0.291098 | 0.241104 | 0.072 | 0.109 | 1 | 11 | Tecpr1 | -0.037 | Young |
| 0.291233 | 0.351869 | 0.13 | 0.181 | 1 | 11 | Lars | -0.051 | Young |
| 0.291473 | 0.153577 | 0.261 | 0.335 | 1 | 11 | B4galt5 | -0.074 | Young |
| 0.291831 | 0.006368 | 0.043 | 0.073 | 1 | 11 | Zc3h10 | -0.03 | Young |
| 0.29195 | 0.086624 | 0.072 | 0.109 | 1 | 11 | Cluh | -0.037 | Young |
| 0.29228 | 0.523666 | 0.087 | 0.129 | 1 | 11 | Rnf31 | -0.042 | Young |
| 0.292321 | 0.064837 | 0.101 | 0.145 | 1 | 11 | Gid8 | -0.044 | Young |
| 0.292396 | 0.310198 | 0.116 | 0.161 | 1 | 11 | Cox16 | -0.045 | Young |
| 0.292496 | 1.287953 | 0.043 | 0.024 | 1 | 11 | 2700062C0 | 0.019 | Young |
| 0.293053 | 0.056235 | 0.928 | 0.944 | 1 | 11 | Zeb2 | -0.016 | Young |

| | | | | | | | | |
|---|---|---|---|---|---|---|---|---|
| 0.293073 | 0.419124 | 0.065 | 0.101 | 1 | 11 | Bcl2l2 | -0.036 | Young |
| 0.29329 | 0.344959 | 0.08 | 0.117 | 1 | 11 | Cops7a | -0.037 | Young |
| 0.29366 | 0.216921 | 0.072 | 0.109 | 1 | 11 | Zfp704 | -0.037 | Young |
| 0.293946 | 0.303887 | 0.13 | 0.181 | 1 | 11 | Tomm40 | -0.051 | Young |
| 0.29395 | 0.937936 | 0.043 | 0.024 | 1 | 11 | Pdk2 | 0.019 | Young |
| 0.294046 | 0.113279 | 0.268 | 0.347 | 1 | 11 | Ftl1-ps1 | -0.079 | Young |
| 0.294575 | 0.013219 | 0.159 | 0.214 | 1 | 11 | Echs1 | -0.055 | Young |
| 0.294647 | 0.23313 | 0.254 | 0.331 | 1 | 11 | Immt | -0.077 | Young |
| 0.294696 | 0.195269 | 0.232 | 0.302 | 1 | 11 | Lamtor5 | -0.07 | Young |
| 0.295306 | 0.206044 | 0.094 | 0.137 | 1 | 11 | Ftsj3 | -0.043 | Young |
| 0.295575 | 0.08253 | 0.319 | 0.419 | 1 | 11 | Hnrnpl | -0.1 | Young |
| 0.295748 | 0.244218 | 0.065 | 0.101 | 1 | 11 | Usp36 | -0.036 | Young |
| 0.295783 | 0.039877 | 0.029 | 0.052 | 1 | 11 | Elac2 | -0.023 | Young |
| 0.29608 | 0.58769 | 0.094 | 0.137 | 1 | 11 | Rubcn | -0.043 | Young |
| 0.296648 | 1.345751 | 0.094 | 0.069 | 1 | 11 | Arl2bp | 0.025 | Young |
| 0.296817 | 0.16011 | 0.261 | 0.339 | 1 | 11 | Ube2i | -0.078 | Young |
| 0.296873 | 0.894912 | 0.043 | 0.024 | 1 | 11 | Atp6v0d2 | 0.019 | Young |
| 0.296953 | 1.063876 | 0.087 | 0.06 | 1 | 11 | Cracr2b | 0.027 | Young |
| 0.297022 | 0.609782 | 0.029 | 0.052 | 1 | 11 | Fastkd2 | -0.023 | Young |
| 0.297057 | 0.013296 | 0.406 | 0.496 | 1 | 11 | Tmem123 | -0.09 | Young |
| 0.297068 | 0.411955 | 0.043 | 0.073 | 1 | 11 | Fam160a2 | -0.03 | Young |
| 0.297249 | 0.085226 | 0.406 | 0.532 | 1 | 11 | Aftph | -0.126 | Young |
| 0.297345 | 0.053395 | 0.587 | 0.734 | 1 | 11 | Tor1aip1 | -0.147 | Young |
| 0.297411 | 0.219563 | 0.116 | 0.161 | 1 | 11 | Armc5 | -0.045 | Young |
| 0.297964 | 0.397401 | 0.072 | 0.109 | 1 | 11 | Xpo4 | -0.037 | Young |
| 0.298074 | 0.304683 | 0.058 | 0.089 | 1 | 11 | Siae | -0.031 | Young |
| 0.298277 | 0.107825 | 0.08 | 0.117 | 1 | 11 | Brwd1 | -0.037 | Young |
| 0.298343 | 0.621776 | 0.101 | 0.145 | 1 | 11 | Bahd1 | -0.044 | Young |
| 0.29847 | 0.171409 | 0.906 | 0.923 | 1 | 11 | Rps15 | -0.017 | Young |
| 0.298494 | 0.624522 | 0.428 | 0.44 | 1 | 11 | Wipf1 | -0.012 | Young |
| 0.298604 | 0.272433 | 0.014 | 0.032 | 1 | 11 | Cul7 | -0.018 | Young |
| 0.298604 | 0.17808 | 0.014 | 0.032 | 1 | 11 | Chac2 | -0.018 | Young |
| 0.298605 | 0.001559 | 0.232 | 0.302 | 1 | 11 | Il3ra | -0.07 | Young |
| 0.29877 | 0.00136 | 0.123 | 0.169 | 1 | 11 | Sec23ip | -0.046 | Young |
| 0.299175 | 0.343204 | 0.333 | 0.444 | 1 | 11 | Paip2 | -0.111 | Young |
| 0.299851 | 0.420759 | 0.754 | 0.778 | 1 | 11 | Btg1 | -0.024 | Young |
| 0.299975 | 0.304082 | 0.188 | 0.254 | 1 | 11 | Sugt1 | -0.066 | Young |
| 0.300051 | 0.481125 | 0.239 | 0.335 | 1 | 11 | Tprgl | -0.096 | Young |
| 0.300215 | 0.117192 | 0.014 | 0.032 | 1 | 11 | Phykpl | -0.018 | Young |
| 0.300215 | 0.059401 | 0.014 | 0.032 | 1 | 11 | Trim68 | -0.018 | Young |
| 0.300939 | 0.268944 | 0.058 | 0.089 | 1 | 11 | Irf4 | -0.031 | Young |
| 0.301096 | 0.707342 | 0.413 | 0.407 | 1 | 11 | Asph | 0.006 | Young |
| 0.301298 | 0.100745 | 0.152 | 0.214 | 1 | 11 | Cdk13 | -0.062 | Young |
| 0.301437 | 0.039243 | 0.37 | 0.468 | 1 | 11 | Ogdh | -0.098 | Young |
| 0.301606 | 0.833777 | 0.159 | 0.129 | 1 | 11 | Slfn9 | 0.03 | Young |

| | | | | | | | | |
|---|---|---|---|---|---|---|---|---|
| 0.301755 | 0.971356 | 0.145 | 0.117 | 1 | 11 | Fam107b | 0.028 | Young |
| 0.301777 | 0.346455 | 0.493 | 0.484 | 1 | 11 | Tgfbr2 | 0.009 | Young |
| 0.301831 | 0.224042 | 0.014 | 0.032 | 1 | 11 | Abca2 | -0.018 | Young |
| 0.30201 | 1.207409 | 0.058 | 0.036 | 1 | 11 | Wdr12 | 0.022 | Young |
| 0.302061 | 1.141137 | 0.167 | 0.141 | 1 | 11 | Stk19 | 0.026 | Young |
| 0.302183 | 0.48052 | 0.138 | 0.19 | 1 | 11 | Jmjd6 | -0.052 | Young |
| 0.302673 | 0.076486 | 0.406 | 0.52 | 1 | 11 | 9530068E0 | -0.114 | Young |
| 0.302719 | 0.906592 | 0.225 | 0.198 | 1 | 11 | Mbp | 0.027 | Young |
| 0.302976 | 0.487115 | 0.051 | 0.081 | 1 | 11 | 2410089E0 | -0.03 | Young |
| 0.303266 | 1.503363 | 0.058 | 0.036 | 1 | 11 | Trim23 | 0.022 | Young |
| 0.303303 | 1.301138 | 0.072 | 0.048 | 1 | 11 | Tprkb | 0.024 | Young |
| 0.303454 | 0.342999 | 0.014 | 0.032 | 1 | 11 | Xpo5 | -0.018 | Young |
| 0.303454 | 0.170012 | 0.014 | 0.032 | 1 | 11 | Pola1 | -0.018 | Young |
| 0.30348 | 0.019152 | 0.174 | 0.23 | 1 | 11 | Polr2c | -0.056 | Young |
| 0.303571 | 0.219788 | 0.304 | 0.387 | 1 | 11 | Zfp106 | -0.083 | Young |
| 0.304461 | 0.036908 | 0.123 | 0.169 | 1 | 11 | Trim5 | -0.046 | Young |
| 0.304656 | 0.156848 | 0.123 | 0.173 | 1 | 11 | Cnot2 | -0.05 | Young |
| 0.304734 | 0.219554 | 0.471 | 0.435 | 1 | 11 | Iigp1 | 0.036 | Young |
| 0.304827 | 0.749391 | 0.101 | 0.073 | 1 | 11 | Tmem260 | 0.028 | Young |
| 0.304853 | 0.759465 | 0.08 | 0.121 | 1 | 11 | Zmynd15 | -0.041 | Young |
| 0.305001 | 0.314563 | 0.051 | 0.081 | 1 | 11 | Tecpr2 | -0.03 | Young |
| 0.305015 | 0.154112 | 0.08 | 0.117 | 1 | 11 | Rnf103 | -0.037 | Young |
| 0.305082 | 0.334074 | 0.014 | 0.032 | 1 | 11 | Polrmt | -0.018 | Young |
| 0.305082 | 0.158094 | 0.014 | 0.032 | 1 | 11 | Casz1 | -0.018 | Young |
| 0.305678 | 0.02655 | 0.42 | 0.516 | 1 | 11 | Xiap | -0.096 | Young |
| 0.305755 | 0.126022 | 0.442 | 0.581 | 1 | 11 | Scand1 | -0.139 | Young |
| 0.305779 | 0.26331 | 0.217 | 0.294 | 1 | 11 | Usp4 | -0.077 | Young |
| 0.305788 | 0.220387 | 0.029 | 0.052 | 1 | 11 | Aven | -0.023 | Young |
| 0.305818 | 0.473309 | 0.072 | 0.109 | 1 | 11 | Dolpp1 | -0.037 | Young |
| 0.30606 | 0.113381 | 0.848 | 0.907 | 1 | 11 | Arpc2 | -0.059 | Young |
| 0.306092 | 0.227201 | 0.174 | 0.234 | 1 | 11 | Cetn3 | -0.06 | Young |
| 0.306141 | 0.165106 | 0.116 | 0.161 | 1 | 11 | Letm1 | -0.045 | Young |
| 0.30639 | 0.24298 | 0.246 | 0.331 | 1 | 11 | Akt2 | -0.085 | Young |
| 0.306614 | 0.372741 | 0.123 | 0.169 | 1 | 11 | Ube2w | -0.046 | Young |
| 0.306651 | 1.211218 | 0.043 | 0.073 | 1 | 11 | Nbn | -0.03 | Young |
| 0.306716 | 0.957624 | 0.014 | 0.032 | 1 | 11 | Gcfc2 | -0.018 | Young |
| 0.306716 | 0.929599 | 0.014 | 0.032 | 1 | 11 | BC049352 | -0.018 | Young |
| 0.306716 | 0.913911 | 0.014 | 0.032 | 1 | 11 | Gatc | -0.018 | Young |
| 0.306716 | 0.529477 | 0.014 | 0.032 | 1 | 11 | Unk | -0.018 | Young |
| 0.306716 | 0.380406 | 0.014 | 0.032 | 1 | 11 | Hmgxb4 | -0.018 | Young |
| 0.306716 | 0.217843 | 0.014 | 0.032 | 1 | 11 | Nkrf | -0.018 | Young |
| 0.307018 | 1.007958 | 0.181 | 0.149 | 1 | 11 | Arhgap5 | 0.032 | Young |
| 0.307036 | 0.249767 | 0.051 | 0.081 | 1 | 11 | Ebag9 | -0.03 | Young |
| 0.307038 | 0.768423 | 0.188 | 0.157 | 1 | 11 | Kctd20 | 0.031 | Young |
| 0.307305 | 0.210211 | 0.203 | 0.262 | 1 | 11 | Cyb5r4 | -0.059 | Young |

| | | | | | | | | |
|---|---|---|---|---|---|---|---|---|
| 0.307531 | 0.269011 | 0.101 | 0.145 | 1 | 11 | Procr | -0.044 | Young |
| 0.307581 | 0.855883 | 0.087 | 0.125 | 1 | 11 | Dera | -0.038 | Young |
| 0.308217 | 0.220865 | 0.174 | 0.23 | 1 | 11 | Lsm5 | -0.056 | Young |
| 0.308742 | 0.311539 | 0.101 | 0.141 | 1 | 11 | Nucb2 | -0.04 | Young |
| 0.309079 | 0.078673 | 0.051 | 0.081 | 1 | 11 | Med22 | -0.03 | Young |
| 0.30942 | 0.133623 | 0.159 | 0.214 | 1 | 11 | Rnf216 | -0.055 | Young |
| 0.309577 | 0.353278 | 0.167 | 0.226 | 1 | 11 | Arih2 | -0.059 | Young |
| 0.309724 | 0.462106 | 0.138 | 0.19 | 1 | 11 | Slc35b1 | -0.052 | Young |
| 0.309765 | 0.92835 | 0.167 | 0.137 | 1 | 11 | Ank | 0.03 | Young |
| 0.309828 | 0.131891 | 0.116 | 0.161 | 1 | 11 | Klf9 | -0.045 | Young |
| 0.310104 | 0.098302 | 0.051 | 0.081 | 1 | 11 | Sell | -0.03 | Young |
| 0.310259 | 0.143497 | 0.978 | 0.98 | 1 | 11 | Rps21 | -0.002 | Young |
| 0.310498 | 0.062907 | 0.203 | 0.27 | 1 | 11 | Ahsa1 | -0.067 | Young |
| 0.310874 | 0.850943 | 0.058 | 0.036 | 1 | 11 | Wtip | 0.022 | Young |
| 0.310932 | 0.018909 | 0.196 | 0.25 | 1 | 11 | Zfp516 | -0.054 | Young |
| 0.310992 | 0.476922 | 0.08 | 0.117 | 1 | 11 | Yif1b | -0.037 | Young |
| 0.310992 | 0.307819 | 0.08 | 0.117 | 1 | 11 | 4930402H2 | -0.037 | Young |
| 0.311414 | 0.593606 | 0.101 | 0.145 | 1 | 11 | Tpra1 | -0.044 | Young |
| 0.311771 | 0.198304 | 0.217 | 0.282 | 1 | 11 | Skp1a | -0.065 | Young |
| 0.312085 | 0.018835 | 0.116 | 0.157 | 1 | 11 | Nifk | -0.041 | Young |
| 0.312295 | 0.051662 | 0.312 | 0.407 | 1 | 11 | Arhgef2 | -0.095 | Young |
| 0.312416 | 0.220686 | 0.232 | 0.298 | 1 | 11 | Ndufa11 | -0.066 | Young |
| 0.312583 | 0.016249 | 0.058 | 0.089 | 1 | 11 | Ddx49 | -0.031 | Young |
| 0.312907 | 0.388428 | 0.203 | 0.27 | 1 | 11 | 1500011K1 | -0.067 | Young |
| 0.313193 | 0.085526 | 0.051 | 0.081 | 1 | 11 | Lztr1 | -0.03 | Young |
| 0.313292 | 0.158699 | 0.21 | 0.278 | 1 | 11 | Agfg1 | -0.068 | Young |
| 0.313429 | 1.565511 | 0.072 | 0.048 | 1 | 11 | 2900089D1 | 0.024 | Young |
| 0.313429 | 0.737744 | 0.072 | 0.048 | 1 | 11 | Prmt3 | 0.024 | Young |
| 0.313826 | 0.403236 | 0.188 | 0.25 | 1 | 11 | Parp1 | -0.062 | Young |
| 0.314073 | 0.030335 | 0.442 | 0.54 | 1 | 11 | Lgals1 | -0.098 | Young |
| 0.314438 | 0.626674 | 0.21 | 0.286 | 1 | 11 | Rmnd5a | -0.076 | Young |
| 0.314531 | 0.869381 | 0.043 | 0.069 | 1 | 11 | Kat6b | -0.026 | Young |
| 0.314531 | 0.308853 | 0.043 | 0.069 | 1 | 11 | Akt3 | -0.026 | Young |
| 0.314552 | 0.055363 | 0.058 | 0.089 | 1 | 11 | Tle1 | -0.031 | Young |
| 0.314567 | 0.24467 | 0.326 | 0.423 | 1 | 11 | Herc1 | -0.097 | Young |
| 0.314724 | 0.774864 | 0.029 | 0.052 | 1 | 11 | Sort1 | -0.023 | Young |
| 0.314978 | 1.371243 | 0.13 | 0.105 | 1 | 11 | Ripk2 | 0.025 | Young |
| 0.315309 | 0.149129 | 0.08 | 0.117 | 1 | 11 | Rpl27-ps3 | -0.037 | Young |
| 0.315916 | 0.198612 | 0.13 | 0.177 | 1 | 11 | Lcor | -0.047 | Young |
| 0.316059 | 0.228625 | 0.123 | 0.169 | 1 | 11 | Evi2 | -0.046 | Young |
| 0.316128 | 0.187565 | 0.261 | 0.347 | 1 | 11 | Cdv3 | -0.086 | Young |
| 0.316301 | 1.150795 | 0.087 | 0.06 | 1 | 11 | Tmem37 | 0.027 | Young |
| 0.316604 | 1.189229 | 0.159 | 0.133 | 1 | 11 | Brd3 | 0.026 | Young |
| 0.317047 | 0.142627 | 0.08 | 0.117 | 1 | 11 | Esco1 | -0.037 | Young |
| 0.317236 | 0.307735 | 0.159 | 0.218 | 1 | 11 | Ecm1 | -0.059 | Young |

| | | | | | | | | |
|---|---|---|---|---|---|---|---|---|
| 0.317514 | 0.945721 | 0.232 | 0.198 | 1 | 11 | Pdgfb | 0.034 | Young |
| 0.318045 | 1.483581 | 0.08 | 0.056 | 1 | 11 | Btbd10 | 0.024 | Young |
| 0.318223 | 0.96345 | 0.145 | 0.117 | 1 | 11 | Ccng2 | 0.028 | Young |
| 0.318241 | 0.186299 | 0.094 | 0.133 | 1 | 11 | Sfxn2 | -0.039 | Young |
| 0.318242 | 0.070636 | 0.065 | 0.097 | 1 | 11 | Ppargc1b | -0.032 | Young |
| 0.318385 | 1.395246 | 0.051 | 0.081 | 1 | 11 | Rnaset2b | -0.03 | Young |
| 0.318515 | 0.100381 | 0.058 | 0.089 | 1 | 11 | Sharpin | -0.031 | Young |
| 0.318552 | 0.127358 | 0.37 | 0.5 | 1 | 11 | Arid1a | -0.13 | Young |
| 0.318564 | 0.176501 | 0.159 | 0.218 | 1 | 11 | Tbk1 | -0.059 | Young |
| 0.318694 | 0.031954 | 0.855 | 0.919 | 1 | 11 | Hsp90ab1 | -0.064 | Young |
| 0.31923 | 0.343736 | 0.159 | 0.218 | 1 | 11 | Naa20 | -0.059 | Young |
| 0.319292 | 0.169062 | 0.196 | 0.258 | 1 | 11 | Gm34084 | -0.062 | Young |
| 0.319584 | 0.391307 | 0.543 | 0.569 | 1 | 11 | Gadd45b | -0.026 | Young |
| 0.319977 | 0.426712 | 0.167 | 0.226 | 1 | 11 | Gpatch8 | -0.059 | Young |
| 0.320272 | 0.295079 | 0.145 | 0.202 | 1 | 11 | Abca3 | -0.057 | Young |
| 0.32044 | 0.416589 | 0.109 | 0.153 | 1 | 11 | Zfp664 | -0.044 | Young |
| 0.320541 | 0.163832 | 0.08 | 0.117 | 1 | 11 | Zfp90 | -0.037 | Young |
| 0.320567 | 0.232903 | 0.13 | 0.181 | 1 | 11 | Ahr | -0.051 | Young |
| 0.320916 | 0.530275 | 0.145 | 0.198 | 1 | 11 | Ifi27 | -0.053 | Young |
| 0.320972 | 0.366178 | 0.13 | 0.177 | 1 | 11 | Nudt4 | -0.047 | Young |
| 0.321069 | 1.278069 | 0.123 | 0.097 | 1 | 11 | Pde3b | 0.026 | Young |
| 0.321211 | 0.786496 | 0.058 | 0.036 | 1 | 11 | Ypel2 | 0.022 | Young |
| 0.32155 | 0.366981 | 0.036 | 0.06 | 1 | 11 | Tom1l2 | -0.024 | Young |
| 0.321636 | 0.270256 | 0.217 | 0.294 | 1 | 11 | Slc25a37 | -0.077 | Young |
| 0.321979 | 0.148832 | 0.152 | 0.206 | 1 | 11 | Zbtb18 | -0.054 | Young |
| 0.322023 | 0.40205 | 0.167 | 0.23 | 1 | 11 | Ccdc88b | -0.063 | Young |
| 0.322528 | 2.000461 | 0.036 | 0.02 | 1 | 11 | Slc25a40 | 0.016 | Young |
| 0.322579 | 0.877592 | 0.051 | 0.081 | 1 | 11 | Wdr18 | -0.03 | Young |
| 0.322729 | 0.144053 | 0.275 | 0.351 | 1 | 11 | Kmt2a | -0.076 | Young |
| 0.32277 | 0.306402 | 0.109 | 0.153 | 1 | 11 | Fam160b1 | -0.044 | Young |
| 0.32366 | 0.31665 | 0.246 | 0.327 | 1 | 11 | Nab1 | -0.081 | Young |
| 0.323993 | 0.176922 | 0.268 | 0.343 | 1 | 11 | Zufsp | -0.075 | Young |
| 0.324329 | 0.476938 | 0.109 | 0.153 | 1 | 11 | Ugdh | -0.044 | Young |
| 0.324341 | 0.061334 | 0.145 | 0.194 | 1 | 11 | Esf1 | -0.049 | Young |
| 0.32452 | 0.298114 | 0.058 | 0.089 | 1 | 11 | Usp39 | -0.031 | Young |
| 0.32452 | 0.227932 | 0.058 | 0.089 | 1 | 11 | Mrpl46 | -0.031 | Young |
| 0.32452 | 0.095133 | 0.058 | 0.089 | 1 | 11 | Bpnt1 | -0.031 | Young |
| 0.324949 | 0.19036 | 0.065 | 0.097 | 1 | 11 | Zdhhc18 | -0.032 | Young |
| 0.325313 | 0.250926 | 0.188 | 0.254 | 1 | 11 | Txndc9 | -0.066 | Young |
| 0.32548 | 1.133861 | 0.152 | 0.125 | 1 | 11 | Khnyn | 0.027 | Young |
| 0.325867 | 0.134204 | 0.217 | 0.282 | 1 | 11 | Cul1 | -0.065 | Young |
| 0.325923 | 2.137285 | 0.036 | 0.02 | 1 | 11 | Kcnj2 | 0.016 | Young |
| 0.325923 | 1.627193 | 0.036 | 0.02 | 1 | 11 | Mcc | 0.016 | Young |
| 0.326103 | 0.567296 | 0.072 | 0.048 | 1 | 11 | Yipf2 | 0.024 | Young |
| 0.326506 | 0.3679 | 0.145 | 0.198 | 1 | 11 | Smarcb1 | -0.053 | Young |

| | | | | | | | | |
|---|---|---|---|---|---|---|---|---|
| 0.326507 | 0.586275 | 0.072 | 0.109 | 1 | 11 | Med20 | -0.037 | Young |
| 0.326972 | 0.400229 | 0.087 | 0.125 | 1 | 11 | Trp53inp2 | -0.038 | Young |
| 0.327104 | 0.348275 | 0.123 | 0.165 | 1 | 11 | Cul2 | -0.042 | Young |
| 0.32763 | 1.739962 | 0.036 | 0.02 | 1 | 11 | Cbr4 | 0.016 | Young |
| 0.327636 | 0.552014 | 0.036 | 0.06 | 1 | 11 | Ppm1f | -0.024 | Young |
| 0.328073 | 0.127518 | 0.196 | 0.258 | 1 | 11 | Fbxl3 | -0.062 | Young |
| 0.3283 | 0.116509 | 0.174 | 0.226 | 1 | 11 | Arl1 | -0.052 | Young |
| 0.328862 | 0.446987 | 0.036 | 0.06 | 1 | 11 | Cep85 | -0.024 | Young |
| 0.328881 | 0.415866 | 0.101 | 0.145 | 1 | 11 | Bicral | -0.044 | Young |
| 0.328938 | 0.586809 | 0.051 | 0.081 | 1 | 11 | Slc41a1 | -0.03 | Young |
| 0.328955 | 0.286354 | 0.326 | 0.427 | 1 | 11 | Mapk1 | -0.101 | Young |
| 0.329138 | 0.200676 | 0.514 | 0.677 | 1 | 11 | Nfkb1 | -0.163 | Young |
| 0.329263 | 0.193977 | 0.167 | 0.226 | 1 | 11 | Dap | -0.059 | Young |
| 0.329286 | 0.131574 | 0.152 | 0.202 | 1 | 11 | Twistnb | -0.05 | Young |
| 0.329445 | 0.396881 | 0.464 | 0.46 | 1 | 11 | Isg15 | 0.004 | Young |
| 0.32958 | 0.322052 | 0.058 | 0.089 | 1 | 11 | Zfp467 | -0.031 | Young |
| 0.329785 | 0.13451 | 0.21 | 0.27 | 1 | 11 | Ppp4r3a | -0.06 | Young |
| 0.330271 | 1.347062 | 0.109 | 0.085 | 1 | 11 | Gtf2f2 | 0.024 | Young |
| 0.330502 | 0.418855 | 0.13 | 0.177 | 1 | 11 | Usp3 | -0.047 | Young |
| 0.330682 | 0.510852 | 0.254 | 0.347 | 1 | 11 | Tuba1c | -0.093 | Young |
| 0.330771 | 0.312614 | 0.065 | 0.097 | 1 | 11 | Nfyc | -0.032 | Young |
| 0.33106 | 2.414447 | 0.036 | 0.02 | 1 | 11 | Rab26os | 0.016 | Young |
| 0.331157 | 0.783797 | 0.145 | 0.117 | 1 | 11 | Atl2 | 0.028 | Young |
| 0.331279 | 0.315602 | 0.138 | 0.19 | 1 | 11 | Opa1 | -0.052 | Young |
| 0.33179 | 0.025057 | 0.116 | 0.161 | 1 | 11 | Fpr2 | -0.045 | Young |
| 0.332462 | 0.318512 | 0.283 | 0.375 | 1 | 11 | Eif3m | -0.092 | Young |
| 0.332641 | 0.438042 | 0.058 | 0.089 | 1 | 11 | Tmem159 | -0.031 | Young |
| 0.332783 | 1.497555 | 0.036 | 0.02 | 1 | 11 | Micall1 | 0.016 | Young |
| 0.332783 | 1.290844 | 0.036 | 0.02 | 1 | 11 | Zfp46 | 0.016 | Young |
| 0.332831 | 1.237508 | 0.007 | 0.02 | 1 | 11 | Mocs3 | -0.013 | Young |
| 0.332831 | 1.00621 | 0.007 | 0.02 | 1 | 11 | Mppe1 | -0.013 | Young |
| 0.332831 | 0.872656 | 0.007 | 0.02 | 1 | 11 | Slc6a1 | -0.013 | Young |
| 0.332831 | 0.765172 | 0.007 | 0.02 | 1 | 11 | Tk1 | -0.013 | Young |
| 0.332831 | 0.660718 | 0.007 | 0.02 | 1 | 11 | 1110051M | -0.013 | Young |
| 0.332831 | 0.639017 | 0.007 | 0.02 | 1 | 11 | Abcc10 | -0.013 | Young |
| 0.332831 | 0.598413 | 0.007 | 0.02 | 1 | 11 | Phf19 | -0.013 | Young |
| 0.332831 | 0.471222 | 0.007 | 0.02 | 1 | 11 | Cebpe | -0.013 | Young |
| 0.332831 | 0.430926 | 0.007 | 0.02 | 1 | 11 | Adgrg5 | -0.013 | Young |
| 0.332831 | 0.421818 | 0.007 | 0.02 | 1 | 11 | Pxmp2 | -0.013 | Young |
| 0.332831 | 0.316291 | 0.007 | 0.02 | 1 | 11 | Fpgs | -0.013 | Young |
| 0.332831 | 0.314384 | 0.007 | 0.02 | 1 | 11 | Ttll5 | -0.013 | Young |
| 0.332831 | 0.235926 | 0.007 | 0.02 | 1 | 11 | BC030499 | -0.013 | Young |
| 0.332831 | 0.221369 | 0.007 | 0.02 | 1 | 11 | Mdm1 | -0.013 | Young |
| 0.332831 | 0.215331 | 0.007 | 0.02 | 1 | 11 | Napb | -0.013 | Young |
| 0.332831 | 0.199982 | 0.007 | 0.02 | 1 | 11 | 1110004E0 | -0.013 | Young |

| | | | | | | | | |
|---|---|---|---|---|---|---|---|---|
| 0.332831 | 0.137314 | 0.007 | 0.02 | 1 | 11 | Upp1 | -0.013 | Young |
| 0.332831 | 0.084483 | 0.007 | 0.02 | 1 | 11 | Rab3a | -0.013 | Young |
| 0.332831 | 0.077184 | 0.007 | 0.02 | 1 | 11 | Rpp40 | -0.013 | Young |
| 0.332831 | 0.07028 | 0.007 | 0.02 | 1 | 11 | Azin2 | -0.013 | Young |
| 0.332831 | 0.067655 | 0.007 | 0.02 | 1 | 11 | 4931414P1 | -0.013 | Young |
| 0.332831 | 0.021942 | 0.007 | 0.02 | 1 | 11 | Ahnak2 | -0.013 | Young |
| 0.332858 | 0.383844 | 0.145 | 0.194 | 1 | 11 | Lpin1 | -0.049 | Young |
| 0.333853 | 0.241099 | 0.333 | 0.444 | 1 | 11 | Psma2 | -0.111 | Young |
| 0.334348 | 0.091824 | 0.283 | 0.363 | 1 | 11 | Gm16286 | -0.08 | Young |
| 0.334513 | 1.79163 | 0.036 | 0.02 | 1 | 11 | Zbtb42 | 0.016 | Young |
| 0.334791 | 0.529842 | 0.159 | 0.218 | 1 | 11 | Mllt6 | -0.059 | Young |
| 0.335037 | 0.404141 | 0.036 | 0.06 | 1 | 11 | Slc37a4 | -0.024 | Young |
| 0.335082 | 0.215219 | 0.101 | 0.137 | 1 | 11 | Ppp1r35 | -0.036 | Young |
| 0.335597 | 0.667149 | 0.174 | 0.234 | 1 | 11 | Nfatc1 | -0.06 | Young |
| 0.335795 | 0.172291 | 0.899 | 0.944 | 1 | 11 | Npc2 | -0.045 | Young |
| 0.335823 | 0.023344 | 0.167 | 0.214 | 1 | 11 | Mlec | -0.047 | Young |
| 0.335845 | 0.090236 | 0.109 | 0.149 | 1 | 11 | Fam204a | -0.04 | Young |
| 0.336054 | 0.005229 | 0.29 | 0.359 | 1 | 11 | Camk2d | -0.069 | Young |
| 0.336248 | 1.185951 | 0.036 | 0.02 | 1 | 11 | Mtfmt | 0.016 | Young |
| 0.336365 | 0.586641 | 0.188 | 0.262 | 1 | 11 | Mtch2 | -0.074 | Young |
| 0.336743 | 0.547974 | 0.217 | 0.181 | 1 | 11 | Arrdc3 | 0.036 | Young |
| 0.336943 | 0.327136 | 0.283 | 0.367 | 1 | 11 | Mbtd1 | -0.084 | Young |
| 0.337179 | 0.068913 | 0.384 | 0.488 | 1 | 11 | Nucks1 | -0.104 | Young |
| 0.337528 | 0.773919 | 0.036 | 0.06 | 1 | 11 | Ipo4 | -0.024 | Young |
| 0.337648 | 0.045588 | 0.065 | 0.097 | 1 | 11 | Gosr1 | -0.032 | Young |
| 0.338816 | 0.275791 | 0.058 | 0.089 | 1 | 11 | Msantd4 | -0.031 | Young |
| 0.339252 | 0.431739 | 0.072 | 0.105 | 1 | 11 | Chchd5 | -0.033 | Young |
| 0.339347 | 0.187117 | 0.884 | 0.935 | 1 | 11 | Rps13 | -0.051 | Young |
| 0.339392 | 0.042937 | 0.159 | 0.21 | 1 | 11 | Tapbpl | -0.051 | Young |
| 0.339736 | 0.918126 | 0.036 | 0.02 | 1 | 11 | Fktn | 0.016 | Young |
| 0.339853 | 0.974624 | 0.058 | 0.089 | 1 | 11 | Vkorc1l1 | -0.031 | Young |
| 0.340026 | 0.251686 | 0.174 | 0.23 | 1 | 11 | Eny2 | -0.056 | Young |
| 0.340031 | 0.570891 | 0.036 | 0.06 | 1 | 11 | Uxt | -0.024 | Young |
| 0.340202 | 1.394478 | 0.051 | 0.032 | 1 | 11 | Kif23 | 0.019 | Young |
| 0.340336 | 0.083612 | 0.377 | 0.492 | 1 | 11 | Mrpl33 | -0.115 | Young |
| 0.340353 | 0.067829 | 0.297 | 0.375 | 1 | 11 | Rell1 | -0.078 | Young |
| 0.340644 | 0.014727 | 0.37 | 0.484 | 1 | 11 | S100a13 | -0.114 | Young |
| 0.3407 | 1.050633 | 0.152 | 0.125 | 1 | 11 | AI504432 | 0.027 | Young |
| 0.341111 | 0.079328 | 0.225 | 0.29 | 1 | 11 | Madd | -0.065 | Young |
| 0.341118 | 0.545312 | 0.08 | 0.117 | 1 | 11 | Idua | -0.037 | Young |
| 0.341347 | 0.063361 | 0.536 | 0.694 | 1 | 11 | Nampt | -0.158 | Young |
| 0.341431 | 0.735573 | 0.167 | 0.23 | 1 | 11 | Slc25a39 | -0.063 | Young |
| 0.341625 | 0.191082 | 0.217 | 0.282 | 1 | 11 | Clasp2 | -0.065 | Young |
| 0.341881 | 0.060244 | 0.493 | 0.613 | 1 | 11 | Lcp2 | -0.12 | Young |
| 0.341932 | 0.683715 | 0.058 | 0.089 | 1 | 11 | Nol6 | -0.031 | Young |

| | | | | | | | | |
|---|---|---|---|---|---|---|---|---|
| 0.342223 | 0.164915 | 0.116 | 0.157 | 1 | 11 | Klra2 | -0.041 | Young |
| 0.342545 | 1.419927 | 0.065 | 0.044 | 1 | 11 | Fmnl3 | 0.021 | Young |
| 0.34261 | 0.447364 | 0.094 | 0.133 | 1 | 11 | Tmem104 | -0.039 | Young |
| 0.34261 | 0.270332 | 0.094 | 0.133 | 1 | 11 | Sh3pxd2a | -0.039 | Young |
| 0.34269 | 0.593335 | 0.087 | 0.125 | 1 | 11 | Me2 | -0.038 | Young |
| 0.342758 | 0.192774 | 0.022 | 0.04 | 1 | 11 | Zfp715 | -0.018 | Young |
| 0.343089 | 1.531739 | 0.051 | 0.032 | 1 | 11 | Tmem242 | 0.019 | Young |
| 0.343226 | 0.144419 | 0.297 | 0.371 | 1 | 11 | Lypla2 | -0.074 | Young |
| 0.343247 | 0.666621 | 0.036 | 0.02 | 1 | 11 | Sumf2 | 0.016 | Young |
| 0.34349 | 0.419178 | 0.196 | 0.266 | 1 | 11 | Clcn4 | -0.07 | Young |
| 0.343807 | 1.183743 | 0.036 | 0.06 | 1 | 11 | Cdc42ep4 | -0.024 | Young |
| 0.344009 | 0.010483 | 0.312 | 0.407 | 1 | 11 | Batf3 | -0.095 | Young |
| 0.344017 | 0.231817 | 0.268 | 0.347 | 1 | 11 | Mbd2 | -0.079 | Young |
| 0.344019 | 0.78206 | 0.058 | 0.089 | 1 | 11 | Endod1 | -0.031 | Young |
| 0.34431 | 0.411398 | 0.022 | 0.04 | 1 | 11 | Vps51 | -0.018 | Young |
| 0.345062 | 0.055979 | 0.558 | 0.706 | 1 | 11 | Fcgr3 | -0.148 | Young |
| 0.345356 | 0.268124 | 0.087 | 0.125 | 1 | 11 | Slc30a9 | -0.038 | Young |
| 0.345416 | 0.440238 | 0.116 | 0.157 | 1 | 11 | Yipf6 | -0.041 | Young |
| 0.345866 | 1.337264 | 0.022 | 0.04 | 1 | 11 | Trdmt1 | -0.018 | Young |
| 0.345866 | 0.116052 | 0.022 | 0.04 | 1 | 11 | Dgkg | -0.018 | Young |
| 0.34596 | 0.181953 | 0.159 | 0.206 | 1 | 11 | Epc2 | -0.047 | Young |
| 0.346035 | 0.203568 | 0.217 | 0.282 | 1 | 11 | Cs | -0.065 | Young |
| 0.346605 | 0.062665 | 0.087 | 0.121 | 1 | 11 | Eaf1 | -0.034 | Young |
| 0.346624 | 0.434095 | 0.065 | 0.097 | 1 | 11 | Ubtd1 | -0.032 | Young |
| 0.346632 | 0.506423 | 0.138 | 0.185 | 1 | 11 | Fbxw2 | -0.047 | Young |
| 0.346632 | 0.330144 | 0.138 | 0.185 | 1 | 11 | Smarcad1 | -0.047 | Young |
| 0.346732 | 0.425967 | 0.196 | 0.266 | 1 | 11 | Otub1 | -0.07 | Young |
| 0.346935 | 0.640579 | 0.072 | 0.105 | 1 | 11 | Caml | -0.033 | Young |
| 0.346954 | 0.330214 | 0.312 | 0.403 | 1 | 11 | Adipor1 | -0.091 | Young |
| 0.347019 | 0.544299 | 0.116 | 0.157 | 1 | 11 | Vrk1 | -0.041 | Young |
| 0.347448 | 1.60965 | 0.051 | 0.032 | 1 | 11 | Zfp157 | 0.019 | Young |
| 0.34761 | 1.841957 | 0.065 | 0.044 | 1 | 11 | Def6 | 0.021 | Young |
| 0.347825 | 0.244972 | 0.326 | 0.415 | 1 | 11 | Dusp11 | -0.089 | Young |
| 0.347999 | 0.164332 | 0.268 | 0.355 | 1 | 11 | Map2k2 | -0.087 | Young |
| 0.347999 | 0.034271 | 0.043 | 0.069 | 1 | 11 | B430306N( | -0.026 | Young |
| 0.348102 | 0.137776 | 0.406 | 0.52 | 1 | 11 | Romo1 | -0.114 | Young |
| 0.348441 | 0.398901 | 0.138 | 0.181 | 1 | 11 | Ssr2 | -0.043 | Young |
| 0.348628 | 0.861414 | 0.159 | 0.133 | 1 | 11 | Trem2 | 0.026 | Young |
| 0.348759 | 0.464393 | 0.145 | 0.198 | 1 | 11 | Maea | -0.053 | Young |
| 0.348908 | 1.474538 | 0.051 | 0.032 | 1 | 11 | Thap7 | 0.019 | Young |
| 0.348992 | 0.391934 | 0.022 | 0.04 | 1 | 11 | Pgm3 | -0.018 | Young |
| 0.349046 | 0.529545 | 0.167 | 0.226 | 1 | 11 | Zfr | -0.059 | Young |
| 0.349119 | 0.064781 | 0.239 | 0.31 | 1 | 11 | Tgs1 | -0.071 | Young |
| 0.349197 | 0.889726 | 0.181 | 0.157 | 1 | 11 | Ralgps2 | 0.024 | Young |
| 0.34934 | 0.359019 | 0.188 | 0.254 | 1 | 11 | Fam117b | -0.066 | Young |

| | | | | | | | | |
|---|---|---|---|---|---|---|---|---|
| 0.349629 | 0.810913 | 0.123 | 0.097 | 1 | 11 | Orc4 | 0.026 | Young |
| 0.349633 | 0.182699 | 0.138 | 0.185 | 1 | 11 | Mesd | -0.047 | Young |
| 0.350333 | 0.2547 | 0.775 | 0.851 | 1 | 11 | Csf1r | -0.076 | Young |
| 0.350373 | 1.566147 | 0.051 | 0.032 | 1 | 11 | Ofd1 | 0.019 | Young |
| 0.350449 | 0.143555 | 0.355 | 0.448 | 1 | 11 | Ogfr | -0.093 | Young |
| 0.350561 | 0.113569 | 0.022 | 0.04 | 1 | 11 | Cep57l1 | -0.018 | Young |
| 0.350561 | 0.07916 | 0.022 | 0.04 | 1 | 11 | Bclaf3 | -0.018 | Young |
| 0.350561 | 0.028383 | 0.022 | 0.04 | 1 | 11 | Gpatch3 | -0.018 | Young |
| 0.351187 | 0.423964 | 0.101 | 0.141 | 1 | 11 | Pum3 | -0.04 | Young |
| 0.3519 | 0.425383 | 0.145 | 0.185 | 1 | 11 | Gla | -0.04 | Young |
| 0.352135 | 0.644593 | 0.022 | 0.04 | 1 | 11 | Rhox8 | -0.018 | Young |
| 0.352135 | 0.510161 | 0.022 | 0.04 | 1 | 11 | Heatr5b | -0.018 | Young |
| 0.352135 | 0.214857 | 0.022 | 0.04 | 1 | 11 | Gm6563 | -0.018 | Young |
| 0.352135 | 0.16149 | 0.022 | 0.04 | 1 | 11 | Dbp | -0.018 | Young |
| 0.352135 | 0.141056 | 0.022 | 0.04 | 1 | 11 | Tmem237 | -0.018 | Young |
| 0.352135 | 0.106315 | 0.022 | 0.04 | 1 | 11 | Tmco4 | -0.018 | Young |
| 0.35215 | 0.230672 | 0.159 | 0.21 | 1 | 11 | Gabarapl2 | -0.051 | Young |
| 0.35223 | 0.079348 | 0.232 | 0.298 | 1 | 11 | Polb | -0.066 | Young |
| 0.352412 | 0.491393 | 0.13 | 0.177 | 1 | 11 | Slc38a9 | -0.047 | Young |
| 0.352609 | 0.184222 | 0.167 | 0.218 | 1 | 11 | Prpf39 | -0.051 | Young |
| 0.353314 | 1.259246 | 0.051 | 0.032 | 1 | 11 | 1810032O( | 0.019 | Young |
| 0.353356 | 0.501304 | 0.123 | 0.169 | 1 | 11 | Arhgap19 | -0.046 | Young |
| 0.353477 | 0.4472 | 0.217 | 0.286 | 1 | 11 | Igbp1 | -0.069 | Young |
| 0.35374 | 0.373601 | 0.152 | 0.206 | 1 | 11 | Ndfip2 | -0.054 | Young |
| 0.353991 | 0.13318 | 0.043 | 0.069 | 1 | 11 | Srxn1 | -0.026 | Young |
| 0.354085 | 0.165193 | 0.232 | 0.294 | 1 | 11 | Gtf2a1 | -0.062 | Young |
| 0.354125 | 0.005062 | 0.094 | 0.129 | 1 | 11 | Ppm1m | -0.035 | Young |
| 0.354907 | 0.512514 | 0.261 | 0.347 | 1 | 11 | Cyld | -0.086 | Young |
| 0.355197 | 0.338508 | 0.043 | 0.069 | 1 | 11 | Kremen1 | -0.026 | Young |
| 0.355197 | 0.249575 | 0.043 | 0.069 | 1 | 11 | Mob2 | -0.026 | Young |
| 0.355243 | 1.642657 | 0.123 | 0.101 | 1 | 11 | Kin | 0.022 | Young |
| 0.355283 | 0.101773 | 0.138 | 0.181 | 1 | 11 | Mvb12a | -0.043 | Young |
| 0.355296 | 1.145901 | 0.065 | 0.044 | 1 | 11 | B3gnt8 | 0.021 | Young |
| 0.35571 | 0.565164 | 0.072 | 0.105 | 1 | 11 | Extl3 | -0.033 | Young |
| 0.356141 | 0.764667 | 0.13 | 0.173 | 1 | 11 | Cd209a | -0.043 | Young |
| 0.356396 | 0.201737 | 0.377 | 0.484 | 1 | 11 | Srsf6 | -0.107 | Young |
| 0.356619 | 0.312567 | 0.087 | 0.121 | 1 | 11 | Mrpl9 | -0.034 | Young |
| 0.356884 | 0.713671 | 0.022 | 0.04 | 1 | 11 | Rpap2 | -0.018 | Young |
| 0.356931 | 0.595575 | 0.5 | 0.532 | 1 | 11 | Ms4a6b | -0.032 | Young |
| 0.357356 | 0.090142 | 0.29 | 0.379 | 1 | 11 | Pdcd10 | -0.089 | Young |
| 0.357619 | 0.116161 | 0.681 | 0.738 | 1 | 11 | Tln1 | -0.057 | Young |
| 0.357829 | 0.45959 | 0.181 | 0.242 | 1 | 11 | Hbp1 | -0.061 | Young |
| 0.358047 | 0.182166 | 0.101 | 0.137 | 1 | 11 | Edc4 | -0.036 | Young |
| 0.358361 | 0.391793 | 0.493 | 0.524 | 1 | 11 | Spi1 | -0.031 | Young |
| 0.35846 | 0.426723 | 0.087 | 0.121 | 1 | 11 | Atxn1 | -0.034 | Young |

| | | | | | | | | |
|---|---|---|---|---|---|---|---|---|
| 0.358512 | 0.420973 | 0.08 | 0.113 | 1 | 11 | Rpf2 | -0.033 | Young |
| 0.358644 | 0.047451 | 0.051 | 0.077 | 1 | 11 | Ifi214 | -0.026 | Young |
| 0.358788 | 0.620911 | 0.116 | 0.161 | 1 | 11 | Copg2 | -0.045 | Young |
| 0.358916 | 0.602003 | 0.101 | 0.137 | 1 | 11 | Rhot1 | -0.036 | Young |
| 0.359165 | 0.048129 | 0.232 | 0.298 | 1 | 11 | Tdrd7 | -0.066 | Young |
| 0.359465 | 0.931367 | 0.08 | 0.113 | 1 | 11 | E2f1 | -0.033 | Young |
| 0.359757 | 0.243198 | 0.159 | 0.214 | 1 | 11 | Stx5a | -0.055 | Young |
| 0.360267 | 0.166682 | 0.138 | 0.185 | 1 | 11 | Zfp281 | -0.047 | Young |
| 0.360418 | 0.575198 | 0.319 | 0.294 | 1 | 11 | Rnase4 | 0.025 | Young |
| 0.360877 | 0.752442 | 0.08 | 0.056 | 1 | 11 | Tbkbp1 | 0.024 | Young |
| 0.361049 | 1.313196 | 0.087 | 0.065 | 1 | 11 | Skint3 | 0.022 | Young |
| 0.361138 | 0.756864 | 0.152 | 0.21 | 1 | 11 | BC037034 | -0.058 | Young |
| 0.361221 | 0.019643 | 0.899 | 0.935 | 1 | 11 | Rack1 | -0.036 | Young |
| 0.361267 | 0.104546 | 0.094 | 0.129 | 1 | 11 | F630028O1 | -0.035 | Young |
| 0.361664 | 0.267516 | 0.116 | 0.157 | 1 | 11 | Syne2 | -0.041 | Young |
| 0.361685 | 0.150929 | 0.152 | 0.202 | 1 | 11 | Tlk1 | -0.05 | Young |
| 0.362097 | 0.567461 | 0.123 | 0.169 | 1 | 11 | Ptbp2 | -0.046 | Young |
| 0.362351 | 0.291633 | 0.101 | 0.141 | 1 | 11 | Pqbp1 | -0.04 | Young |
| 0.363295 | 0.402645 | 0.08 | 0.113 | 1 | 11 | Zfp110 | -0.033 | Young |
| 0.363295 | 0.138794 | 0.08 | 0.113 | 1 | 11 | Ccdc127 | -0.033 | Young |
| 0.363304 | 0.136144 | 0.058 | 0.085 | 1 | 11 | Coq8b | -0.027 | Young |
| 0.363644 | 0.26776 | 0.312 | 0.407 | 1 | 11 | Ndufs6 | -0.095 | Young |
| 0.363655 | 0.356677 | 0.094 | 0.133 | 1 | 11 | Crebzf | -0.039 | Young |
| 0.364531 | 0.301033 | 0.152 | 0.198 | 1 | 11 | Ncoa2 | -0.046 | Young |
| 0.364549 | 0.443993 | 0.145 | 0.194 | 1 | 11 | Fam76b | -0.049 | Young |
| 0.364625 | 0.277016 | 0.283 | 0.371 | 1 | 11 | Xaf1 | -0.088 | Young |
| 0.364671 | 0.512976 | 0.275 | 0.371 | 1 | 11 | Ccdc12 | -0.096 | Young |
| 0.364812 | 0.978343 | 0.203 | 0.181 | 1 | 11 | Hltf | 0.022 | Young |
| 0.364874 | 0.307356 | 0.471 | 0.476 | 1 | 11 | Lrp1 | -0.005 | Young |
| 0.364938 | 0.496396 | 0.043 | 0.069 | 1 | 11 | Psmg2 | -0.026 | Young |
| 0.365233 | 0.57069 | 0.051 | 0.032 | 1 | 11 | Xpc | 0.019 | Young |
| 0.36538 | 0.221784 | 0.167 | 0.222 | 1 | 11 | Aim2 | -0.055 | Young |
| 0.3655 | 0.86133 | 0.123 | 0.165 | 1 | 11 | Hist1h1c | -0.042 | Young |
| 0.365568 | 0.070177 | 0.051 | 0.077 | 1 | 11 | Rce1 | -0.026 | Young |
| 0.365578 | 0.381081 | 0.623 | 0.665 | 1 | 11 | Slfn2 | -0.042 | Young |
| 0.36574 | 0.131219 | 0.167 | 0.222 | 1 | 11 | Tmem106b | -0.055 | Young |
| 0.366169 | 0.311527 | 0.283 | 0.367 | 1 | 11 | H2-T22 | -0.084 | Young |
| 0.366611 | 0.02033 | 0.058 | 0.085 | 1 | 11 | Cdadc1 | -0.027 | Young |
| 0.36673 | 0.374833 | 0.051 | 0.077 | 1 | 11 | Ankrd46 | -0.026 | Young |
| 0.36673 | 0.182911 | 0.051 | 0.077 | 1 | 11 | Maged1 | -0.026 | Young |
| 0.366761 | 0.071963 | 0.457 | 0.573 | 1 | 11 | Kif5b | -0.116 | Young |
| 0.367109 | 1.147005 | 0.065 | 0.097 | 1 | 11 | Cdc14b | -0.032 | Young |
| 0.367346 | 0.64755 | 0.109 | 0.149 | 1 | 11 | Fam69a | -0.04 | Young |
| 0.367894 | 0.375896 | 0.051 | 0.077 | 1 | 11 | 1810043G0 | -0.026 | Young |
| 0.368564 | 0.103402 | 0.101 | 0.137 | 1 | 11 | Pigm | -0.036 | Young |

| | | | | | | | | |
|---|---|---|---|---|---|---|---|---|
| 0.368827 | 0.144413 | 0.138 | 0.181 | 1 | 11 | 8-Mar | -0.043 | Young |
| 0.369448 | 0.157069 | 0.254 | 0.323 | 1 | 11 | Slc35b2 | -0.069 | Young |
| 0.369541 | 0.132443 | 0.138 | 0.185 | 1 | 11 | Atp9b | -0.047 | Young |
| 0.369607 | 0.159849 | 0.348 | 0.431 | 1 | 11 | Fam168b | -0.083 | Young |
| 0.369649 | 0.023138 | 0.029 | 0.048 | 1 | 11 | Pick1 | -0.019 | Young |
| 0.369652 | 0.771847 | 0.341 | 0.343 | 1 | 11 | Arl5b | -0.002 | Young |
| 0.369773 | 0.38133 | 0.601 | 0.645 | 1 | 11 | Ncl | -0.044 | Young |
| 0.370191 | 0.06147 | 0.217 | 0.278 | 1 | 11 | Sp1 | -0.061 | Young |
| 0.37023 | 0.330159 | 0.051 | 0.077 | 1 | 11 | Abt1 | -0.026 | Young |
| 0.370771 | 0.191592 | 0.268 | 0.347 | 1 | 11 | Psma1 | -0.079 | Young |
| 0.370822 | 0.28588 | 0.42 | 0.532 | 1 | 11 | Apobec1 | -0.112 | Young |
| 0.371265 | 0.136321 | 0.116 | 0.153 | 1 | 11 | Gon4l | -0.037 | Young |
| 0.371286 | 0.720931 | 0.051 | 0.032 | 1 | 11 | Ccl25 | 0.019 | Young |
| 0.371494 | 0.096172 | 0.312 | 0.395 | 1 | 11 | Dnajc13 | -0.083 | Young |
| 0.371604 | 0.060215 | 0.145 | 0.19 | 1 | 11 | Paxbp1 | -0.045 | Young |
| 0.371967 | 0.27092 | 0.261 | 0.347 | 1 | 11 | P2ry6 | -0.086 | Young |
| 0.372111 | 0.289155 | 0.101 | 0.137 | 1 | 11 | Get4 | -0.036 | Young |
| 0.372716 | 0.204229 | 0.304 | 0.391 | 1 | 11 | Prkacb | -0.087 | Young |
| 0.372851 | 0.2704 | 0.101 | 0.141 | 1 | 11 | Nae1 | -0.04 | Young |
| 0.372874 | 0.311156 | 0.674 | 0.77 | 1 | 11 | Tmem50a | -0.096 | Young |
| 0.372981 | 1.110863 | 0.08 | 0.113 | 1 | 11 | Gadd45gip | -0.033 | Young |
| 0.373063 | 0.038281 | 0.094 | 0.129 | 1 | 11 | Snhg12 | -0.035 | Young |
| 0.373149 | 0.930619 | 0.181 | 0.157 | 1 | 11 | Htatsf1 | 0.024 | Young |
| 0.37367 | 0.074067 | 0.152 | 0.202 | 1 | 11 | Ube2j1 | -0.05 | Young |
| 0.374077 | 0.409837 | 0.029 | 0.048 | 1 | 11 | 1190007I0 | -0.019 | Young |
| 0.374077 | 0.158199 | 0.029 | 0.048 | 1 | 11 | Pold1 | -0.019 | Young |
| 0.374173 | 0.093833 | 0.29 | 0.371 | 1 | 11 | Daglb | -0.081 | Young |
| 0.374211 | 0.375763 | 0.232 | 0.302 | 1 | 11 | N4bp2l2 | -0.07 | Young |
| 0.374224 | 0.437958 | 0.174 | 0.23 | 1 | 11 | Cdk12 | -0.056 | Young |
| 0.374245 | 0.06385 | 0.225 | 0.278 | 1 | 11 | Cebpa | -0.053 | Young |
| 0.374653 | 0.171971 | 0.116 | 0.153 | 1 | 11 | Mkl1 | -0.037 | Young |
| 0.374729 | 0.622454 | 0.138 | 0.181 | 1 | 11 | Nup88 | -0.043 | Young |
| 0.374876 | 0.348629 | 0.145 | 0.194 | 1 | 11 | Parp10 | -0.049 | Young |
| 0.374899 | 0.055715 | 0.094 | 0.129 | 1 | 11 | Mrps28 | -0.035 | Young |
| 0.375522 | 0.112333 | 0.058 | 0.085 | 1 | 11 | Tpm3-rs7 | -0.027 | Young |
| 0.37556 | 1.524086 | 0.029 | 0.048 | 1 | 11 | Syce2 | -0.019 | Young |
| 0.37641 | 0.182241 | 0.167 | 0.218 | 1 | 11 | Bcl7b | -0.051 | Young |
| 0.376441 | 0.102321 | 0.145 | 0.185 | 1 | 11 | Celf4 | -0.04 | Young |
| 0.376645 | 0.513185 | 0.058 | 0.085 | 1 | 11 | Memo1 | -0.027 | Young |
| 0.376645 | 0.004496 | 0.058 | 0.085 | 1 | 11 | Zfp410 | -0.027 | Young |
| 0.376731 | 0.086531 | 0.239 | 0.302 | 1 | 11 | Fam173a | -0.063 | Young |
| 0.376752 | 0.307858 | 0.304 | 0.387 | 1 | 11 | Arfgef1 | -0.083 | Young |
| 0.376852 | 0.018768 | 0.428 | 0.532 | 1 | 11 | Nono | -0.104 | Young |
| 0.377047 | 0.142646 | 0.029 | 0.048 | 1 | 11 | Wdr83 | -0.019 | Young |
| 0.377464 | 0.364828 | 0.529 | 0.565 | 1 | 11 | Tmem30a | -0.036 | Young |

| | | | | | | | | |
|---|---|---|---|---|---|---|---|---|
| 0.378278 | 0.436589 | 0.283 | 0.25 | 1 | 11 | Ms4a4c | 0.033 | Young |
| 0.378588 | 0.553816 | 0.094 | 0.129 | 1 | 11 | Kdm3b | -0.035 | Young |
| 0.378871 | 0.073677 | 0.123 | 0.161 | 1 | 11 | Lrrc8d | -0.038 | Young |
| 0.378898 | 0.248086 | 0.058 | 0.085 | 1 | 11 | Gfer | -0.027 | Young |
| 0.379139 | 0.445246 | 0.268 | 0.351 | 1 | 11 | Taf15 | -0.083 | Young |
| 0.379257 | 0.53336 | 0.13 | 0.173 | 1 | 11 | Agpat5 | -0.043 | Young |
| 0.379271 | 0.186064 | 0.101 | 0.137 | 1 | 11 | Hopx | -0.036 | Young |
| 0.380032 | 0.51935 | 0.029 | 0.048 | 1 | 11 | Slc35d1 | -0.019 | Young |
| 0.38005 | 0.378766 | 0.087 | 0.121 | 1 | 11 | 1110038F1 | -0.034 | Young |
| 0.380073 | 0.192668 | 0.275 | 0.347 | 1 | 11 | Swap70 | -0.072 | Young |
| 0.380207 | 2.325721 | 0.029 | 0.016 | 1 | 11 | Serpine1 | 0.013 | Young |
| 0.380207 | 1.526939 | 0.029 | 0.016 | 1 | 11 | E130308A1 | 0.013 | Young |
| 0.38094 | 0.240522 | 0.145 | 0.19 | 1 | 11 | Txlna | -0.045 | Young |
| 0.381178 | 0.822897 | 0.225 | 0.206 | 1 | 11 | Tanc2 | 0.019 | Young |
| 0.381638 | 0.041041 | 0.333 | 0.399 | 1 | 11 | Tspan31 | -0.066 | Young |
| 0.381897 | 0.051175 | 0.355 | 0.456 | 1 | 11 | Edem1 | -0.101 | Young |
| 0.381965 | 0.532812 | 0.087 | 0.121 | 1 | 11 | Tesk1 | -0.034 | Young |
| 0.381977 | 0.207491 | 0.101 | 0.137 | 1 | 11 | Arpc5l | -0.036 | Young |
| 0.382162 | 0.335467 | 0.297 | 0.375 | 1 | 11 | AB124611 | -0.078 | Young |
| 0.382292 | 0.526064 | 0.058 | 0.085 | 1 | 11 | Zfp654 | -0.027 | Young |
| 0.382298 | 0.632773 | 0.094 | 0.129 | 1 | 11 | Cbx5 | -0.035 | Young |
| 0.382305 | 2.216404 | 0.029 | 0.016 | 1 | 11 | Tmem144 | 0.013 | Young |
| 0.382884 | 0.468635 | 0.326 | 0.31 | 1 | 11 | Cd93 | 0.016 | Young |
| 0.382884 | 0.330677 | 0.268 | 0.343 | 1 | 11 | Eif4e | -0.075 | Young |
| 0.383016 | 0.1389 | 0.783 | 0.831 | 1 | 11 | Hnrnpk | -0.048 | Young |
| 0.383031 | 0.419172 | 0.029 | 0.048 | 1 | 11 | Als2 | -0.019 | Young |
| 0.383724 | 1.052748 | 0.159 | 0.137 | 1 | 11 | BC035044 | 0.022 | Young |
| 0.384056 | 0.199143 | 0.072 | 0.101 | 1 | 11 | Zfp398 | -0.029 | Young |
| 0.384057 | 0.244992 | 0.21 | 0.27 | 1 | 11 | Ubr4 | -0.06 | Young |
| 0.38441 | 1.717777 | 0.029 | 0.016 | 1 | 11 | Ctu1 | 0.013 | Young |
| 0.384629 | 0.427706 | 0.42 | 0.435 | 1 | 11 | Stat2 | -0.015 | Young |
| 0.384873 | 0.902072 | 0.167 | 0.141 | 1 | 11 | Lrrfip2 | 0.026 | Young |
| 0.385252 | 0.166515 | 0.384 | 0.512 | 1 | 11 | Cep170 | -0.128 | Young |
| 0.38552 | 0.428679 | 0.109 | 0.149 | 1 | 11 | 4931406P1 | -0.04 | Young |
| 0.38598 | 0.286738 | 0.391 | 0.5 | 1 | 11 | Ap1b1 | -0.109 | Young |
| 0.386044 | 0.686538 | 0.029 | 0.048 | 1 | 11 | C87436 | -0.019 | Young |
| 0.386522 | 1.895264 | 0.029 | 0.016 | 1 | 11 | Rilpl1 | 0.013 | Young |
| 0.386522 | 1.734125 | 0.029 | 0.016 | 1 | 11 | Nudt6 | 0.013 | Young |
| 0.387195 | 0.265893 | 0.072 | 0.101 | 1 | 11 | Apba1 | -0.029 | Young |
| 0.387284 | 0.228348 | 0.123 | 0.161 | 1 | 11 | Sestd1 | -0.038 | Young |
| 0.387355 | 0.894759 | 0.159 | 0.218 | 1 | 11 | Utp11 | -0.059 | Young |
| 0.387444 | 0.347008 | 0.152 | 0.202 | 1 | 11 | Sirt7 | -0.05 | Young |
| 0.387556 | 0.365709 | 0.029 | 0.048 | 1 | 11 | Trub2 | -0.019 | Young |
| 0.38765 | 0.940398 | 0.138 | 0.113 | 1 | 11 | Mboat7 | 0.025 | Young |
| 0.38799 | 0.198032 | 0.188 | 0.246 | 1 | 11 | Rps12-ps3 | -0.058 | Young |

| | | | | | | | | |
|---|---|---|---|---|---|---|---|---|
| 0.388038 | 0.558842 | 0.051 | 0.077 | 1 | 11 | Fer | -0.026 | Young |
| 0.388195 | 0.167895 | 0.862 | 0.911 | 1 | 11 | Mbnl1 | -0.049 | Young |
| 0.38832 | 1.087056 | 0.181 | 0.157 | 1 | 11 | Parl | 0.024 | Young |
| 0.38864 | 1.564035 | 0.029 | 0.016 | 1 | 11 | Pex26 | 0.013 | Young |
| 0.38864 | 1.431879 | 0.029 | 0.016 | 1 | 11 | Colec12 | 0.013 | Young |
| 0.38864 | 1.318561 | 0.029 | 0.016 | 1 | 11 | 1700066M | 0.013 | Young |
| 0.38864 | 1.243523 | 0.029 | 0.016 | 1 | 11 | Gpr84 | 0.013 | Young |
| 0.38898 | 0.297656 | 0.123 | 0.161 | 1 | 11 | Tmem243 | -0.038 | Young |
| 0.389136 | 0.432558 | 0.058 | 0.085 | 1 | 11 | Thyn1 | -0.027 | Young |
| 0.389208 | 0.434721 | 0.138 | 0.109 | 1 | 11 | Tm2d3 | 0.029 | Young |
| 0.38947 | 0.102556 | 0.942 | 0.935 | 1 | 11 | Rpl27a | 0.007 | Young |
| 0.389497 | 0.207279 | 0.123 | 0.165 | 1 | 11 | Thoc6 | -0.042 | Young |
| 0.38995 | 0.17465 | 0.732 | 0.758 | 1 | 11 | Btf3 | -0.026 | Young |
| 0.390048 | 1.178598 | 0.297 | 0.282 | 1 | 11 | Ccl24 | 0.015 | Young |
| 0.39075 | 0.695456 | 0.181 | 0.157 | 1 | 11 | Zmpste24 | 0.024 | Young |
| 0.390766 | 1.451928 | 0.029 | 0.016 | 1 | 11 | Ctnnd2 | 0.013 | Young |
| 0.390766 | 1.312936 | 0.029 | 0.016 | 1 | 11 | Mrps22 | 0.013 | Young |
| 0.390766 | 1.064239 | 0.029 | 0.016 | 1 | 11 | Rnf180 | 0.013 | Young |
| 0.390766 | 0.943471 | 0.029 | 0.016 | 1 | 11 | Wdr4 | 0.013 | Young |
| 0.391075 | 0.078717 | 0.283 | 0.343 | 1 | 11 | Cd180 | -0.06 | Young |
| 0.391155 | 0.44997 | 0.275 | 0.363 | 1 | 11 | Cdc37 | -0.088 | Young |
| 0.391294 | 1.075778 | 0.072 | 0.052 | 1 | 11 | Tbc1d24 | 0.02 | Young |
| 0.391447 | 0.055025 | 0.087 | 0.117 | 1 | 11 | Cyp4v3 | -0.03 | Young |
| 0.392033 | 0.276419 | 0.123 | 0.165 | 1 | 11 | Dgcr2 | -0.042 | Young |
| 0.392115 | 0.864801 | 0.029 | 0.048 | 1 | 11 | Spata5 | -0.019 | Young |
| 0.392374 | 0.428552 | 0.138 | 0.181 | 1 | 11 | Pink1 | -0.043 | Young |
| 0.392462 | 0.587314 | 0.072 | 0.101 | 1 | 11 | Prkci | -0.029 | Young |
| 0.392586 | 0.215731 | 0.058 | 0.085 | 1 | 11 | Trim36 | -0.027 | Young |
| 0.392899 | 1.237848 | 0.029 | 0.016 | 1 | 11 | Slc1a2 | 0.013 | Young |
| 0.393027 | 0.233686 | 0.239 | 0.306 | 1 | 11 | Zfp445 | -0.067 | Young |
| 0.393198 | 0.802871 | 0.138 | 0.113 | 1 | 11 | AI837181 | 0.025 | Young |
| 0.39391 | 0.025159 | 0.145 | 0.185 | 1 | 11 | Actr10 | -0.04 | Young |
| 0.394178 | 0.362079 | 0.232 | 0.306 | 1 | 11 | Yeats4 | -0.074 | Young |
| 0.394264 | 0.037524 | 0.036 | 0.056 | 1 | 11 | Mllt3 | -0.02 | Young |
| 0.394496 | 0.328197 | 0.116 | 0.153 | 1 | 11 | Pds5b | -0.037 | Young |
| 0.394819 | 0.283741 | 0.138 | 0.181 | 1 | 11 | Uba5 | -0.043 | Young |
| 0.394896 | 0.486897 | 0.058 | 0.085 | 1 | 11 | Cxxc1 | -0.027 | Young |
| 0.395039 | 1.202 | 0.029 | 0.016 | 1 | 11 | Mfsd3 | 0.013 | Young |
| 0.395039 | 0.400359 | 0.029 | 0.016 | 1 | 11 | Hibch | 0.013 | Young |
| 0.395171 | 1.163782 | 0.029 | 0.048 | 1 | 11 | Tti1 | -0.019 | Young |
| 0.395335 | 0.07472 | 0.109 | 0.145 | 1 | 11 | Tec | -0.036 | Young |
| 0.396054 | 0.238462 | 0.058 | 0.085 | 1 | 11 | Rffl | -0.027 | Young |
| 0.396181 | 0.119721 | 0.181 | 0.234 | 1 | 11 | Ptov1 | -0.053 | Young |
| 0.396384 | 0.050541 | 0.087 | 0.117 | 1 | 11 | Papd5 | -0.03 | Young |
| 0.396705 | 1.084395 | 0.029 | 0.048 | 1 | 11 | Dalrd3 | -0.019 | Young |

| | | | | | | | | |
|---|---|---|---|---|---|---|---|---|
| 0.397098 | 0.091006 | 0.507 | 0.617 | 1 | 11 | Fcgrt | -0.11 | Young |
| 0.397242 | 0.334424 | 0.283 | 0.367 | 1 | 11 | Psme4 | -0.084 | Young |
| 0.397294 | 0.031719 | 0.58 | 0.573 | 1 | 11 | S100a6 | 0.007 | Young |
| 0.397621 | 0.236469 | 0.145 | 0.19 | 1 | 11 | Zc3h4 | -0.045 | Young |
| 0.398037 | 1.492794 | 0.109 | 0.089 | 1 | 11 | Serpinf1 | 0.02 | Young |
| 0.398462 | 0.356396 | 0.101 | 0.137 | 1 | 11 | Smurf2 | -0.036 | Young |
| 0.398506 | 0.297433 | 0.036 | 0.056 | 1 | 11 | Zfp64 | -0.02 | Young |
| 0.398657 | 0.122214 | 0.217 | 0.278 | 1 | 11 | Wdfy2 | -0.061 | Young |
| 0.398669 | 0.284547 | 0.065 | 0.093 | 1 | 11 | Tars2 | -0.028 | Young |
| 0.399341 | 0.427846 | 0.029 | 0.016 | 1 | 11 | Enho | 0.013 | Young |
| 0.399341 | 0.356827 | 0.029 | 0.016 | 1 | 11 | Zfp438 | 0.013 | Young |
| 0.399341 | 0.342255 | 0.029 | 0.016 | 1 | 11 | Pxdc1 | 0.013 | Young |
| 0.399343 | 0.209058 | 0.21 | 0.274 | 1 | 11 | Lsm14a | -0.064 | Young |
| 0.399532 | 0.211187 | 0.659 | 0.641 | 1 | 11 | Dab2 | 0.018 | Young |
| 0.399705 | 0.908294 | 0.203 | 0.181 | 1 | 11 | Fam20b | 0.022 | Young |
| 0.399926 | 0.110522 | 0.036 | 0.056 | 1 | 11 | Haus7 | -0.02 | Young |
| 0.400068 | 0.197916 | 0.08 | 0.109 | 1 | 11 | 4930455G( | -0.029 | Young |
| 0.400211 | 0.0523 | 0.174 | 0.222 | 1 | 11 | Elf4 | -0.048 | Young |
| 0.400304 | 0.664476 | 0.362 | 0.375 | 1 | 11 | Snap23 | -0.013 | Young |
| 0.400686 | 0.364823 | 0.5 | 0.577 | 1 | 11 | Srp9 | -0.077 | Young |
| 0.401056 | 0.204611 | 0.304 | 0.395 | 1 | 11 | Ube2n | -0.091 | Young |
| 0.401283 | 0.165146 | 0.014 | 0.028 | 1 | 11 | Lmo1 | -0.014 | Young |
| 0.401422 | 1.375093 | 0.043 | 0.028 | 1 | 11 | Tubgcp6 | 0.015 | Young |
| 0.401502 | 0.586308 | 0.029 | 0.016 | 1 | 11 | Olfr920 | 0.013 | Young |
| 0.401502 | 0.333666 | 0.029 | 0.016 | 1 | 11 | Spink2 | 0.013 | Young |
| 0.401502 | 0.313373 | 0.029 | 0.016 | 1 | 11 | Klhl42 | 0.013 | Young |
| 0.402076 | 0.348477 | 0.145 | 0.194 | 1 | 11 | Tnks | -0.049 | Young |
| 0.402144 | 0.239492 | 0.181 | 0.23 | 1 | 11 | Tmem33 | -0.049 | Young |
| 0.40224 | 1.120159 | 0.058 | 0.04 | 1 | 11 | Osbpl2 | 0.018 | Young |
| 0.402426 | 0.280379 | 0.261 | 0.339 | 1 | 11 | Nenf | -0.078 | Young |
| 0.403036 | 0.303002 | 0.232 | 0.294 | 1 | 11 | Ppp6r1 | -0.062 | Young |
| 0.403126 | 0.15158 | 0.072 | 0.101 | 1 | 11 | BC005561 | -0.029 | Young |
| 0.403311 | 0.033864 | 0.616 | 0.746 | 1 | 11 | Ccl2 | -0.13 | Young |
| 0.403329 | 0.257313 | 0.014 | 0.028 | 1 | 11 | Abcb8 | -0.014 | Young |
| 0.403329 | 0.188955 | 0.014 | 0.028 | 1 | 11 | Cdc25b | -0.014 | Young |
| 0.403329 | 0.165536 | 0.014 | 0.028 | 1 | 11 | Btbd19 | -0.014 | Young |
| 0.403329 | 0.073838 | 0.014 | 0.028 | 1 | 11 | Tmem218 | -0.014 | Young |
| 0.403603 | 0.199252 | 0.37 | 0.472 | 1 | 11 | Map1lc3b | -0.102 | Young |
| 0.404065 | 0.201633 | 0.094 | 0.129 | 1 | 11 | Isg20 | -0.035 | Young |
| 0.404202 | 0.252085 | 0.072 | 0.101 | 1 | 11 | Ubl4a | -0.029 | Young |
| 0.404262 | 0.078868 | 0.522 | 0.649 | 1 | 11 | Clic1 | -0.127 | Young |
| 0.40439 | 0.385332 | 0.109 | 0.145 | 1 | 11 | Kmt2b | -0.036 | Young |
| 0.404522 | 1.221552 | 0.101 | 0.081 | 1 | 11 | Utp20 | 0.02 | Young |
| 0.404531 | 0.323239 | 0.225 | 0.294 | 1 | 11 | Sp140 | -0.069 | Young |
| 0.404651 | 0.482043 | 0.188 | 0.246 | 1 | 11 | Nuak2 | -0.058 | Young |

| | | | | | | | | |
|---|---|---|---|---|---|---|---|---|
| 0.404798 | 0.017453 | 0.145 | 0.181 | 1 | 11 | Itgax | -0.036 | Young |
| 0.405173 | 0.464077 | 0.058 | 0.04 | 1 | 11 | Sdc1 | 0.018 | Young |
| 0.405179 | 0.091457 | 0.957 | 0.976 | 1 | 11 | Zfp36 | -0.019 | Young |
| 0.405221 | 0.137072 | 0.246 | 0.306 | 1 | 11 | Armc8 | -0.06 | Young |
| 0.405381 | 1.042973 | 0.014 | 0.028 | 1 | 11 | Zfp518b | -0.014 | Young |
| 0.405381 | 0.99821 | 0.014 | 0.028 | 1 | 11 | Vdr | -0.014 | Young |
| 0.405381 | 0.760465 | 0.014 | 0.028 | 1 | 11 | 9330175E1 | -0.014 | Young |
| 0.405381 | 0.505701 | 0.014 | 0.028 | 1 | 11 | Serf1 | -0.014 | Young |
| 0.405381 | 0.339506 | 0.014 | 0.028 | 1 | 11 | Elmod3 | -0.014 | Young |
| 0.405381 | 0.298035 | 0.014 | 0.028 | 1 | 11 | Zmynd19 | -0.014 | Young |
| 0.405519 | 0.287441 | 0.138 | 0.181 | 1 | 11 | Slc35c2 | -0.043 | Young |
| 0.405637 | 0.996605 | 0.036 | 0.056 | 1 | 11 | Ano8 | -0.02 | Young |
| 0.405911 | 0.140645 | 0.319 | 0.407 | 1 | 11 | Icam1 | -0.088 | Young |
| 0.406055 | 0.203144 | 0.667 | 0.746 | 1 | 11 | Rpl7a | -0.079 | Young |
| 0.406211 | 0.223908 | 0.138 | 0.177 | 1 | 11 | Tnpo2 | -0.039 | Young |
| 0.406949 | 0.954413 | 0.109 | 0.149 | 1 | 11 | Rbm12 | -0.04 | Young |
| 0.407439 | 1.921099 | 0.014 | 0.028 | 1 | 11 | Tgds | -0.014 | Young |
| 0.407439 | 0.662807 | 0.014 | 0.028 | 1 | 11 | Tepsin | -0.014 | Young |
| 0.407439 | 0.612017 | 0.014 | 0.028 | 1 | 11 | Rabep2 | -0.014 | Young |
| 0.407439 | 0.569226 | 0.014 | 0.028 | 1 | 11 | Nudt15 | -0.014 | Young |
| 0.407439 | 0.433893 | 0.014 | 0.028 | 1 | 11 | Cmtr1 | -0.014 | Young |
| 0.407439 | 0.373615 | 0.014 | 0.028 | 1 | 11 | Cd300ld3 | -0.014 | Young |
| 0.407439 | 0.16595 | 0.014 | 0.028 | 1 | 11 | C1qtnf6 | -0.014 | Young |
| 0.407439 | 0.069708 | 0.014 | 0.028 | 1 | 11 | Pfas | -0.014 | Young |
| 0.407736 | 0.506099 | 0.145 | 0.194 | 1 | 11 | Dennd1c | -0.049 | Young |
| 0.407947 | 0.108153 | 0.196 | 0.25 | 1 | 11 | Eif4e3 | -0.054 | Young |
| 0.408226 | 0.035383 | 0.101 | 0.133 | 1 | 11 | Cnep1r1 | -0.032 | Young |
| 0.408292 | 1.594774 | 0.043 | 0.028 | 1 | 11 | Sh3bp5l | 0.015 | Young |
| 0.408292 | 1.249377 | 0.043 | 0.028 | 1 | 11 | Tctn3 | 0.015 | Young |
| 0.408523 | 0.422873 | 0.072 | 0.101 | 1 | 11 | Rfx7 | -0.029 | Young |
| 0.408639 | 0.149282 | 0.094 | 0.125 | 1 | 11 | Tbc1d17 | -0.031 | Young |
| 0.408929 | 0.748375 | 0.058 | 0.085 | 1 | 11 | Rdh11 | -0.027 | Young |
| 0.408955 | 0.504591 | 0.196 | 0.165 | 1 | 11 | Lyve1 | 0.031 | Young |
| 0.409596 | 0.288949 | 0.254 | 0.319 | 1 | 11 | Dnajb9 | -0.065 | Young |
| 0.409952 | 0.805966 | 0.036 | 0.056 | 1 | 11 | 2410004B1 | -0.02 | Young |
| 0.410244 | 0.081889 | 0.13 | 0.169 | 1 | 11 | Dnajc10 | -0.039 | Young |
| 0.410406 | 0.315051 | 0.087 | 0.117 | 1 | 11 | Fam126b | -0.03 | Young |
| 0.410406 | 0.244522 | 0.087 | 0.117 | 1 | 11 | Erf | -0.03 | Young |
| 0.410588 | 0.951721 | 0.152 | 0.125 | 1 | 11 | Gbp9 | 0.027 | Young |
| 0.410834 | 0.660451 | 0.094 | 0.129 | 1 | 11 | Plk3 | -0.035 | Young |
| 0.41146 | 0.34452 | 0.188 | 0.242 | 1 | 11 | Pnrc2 | -0.054 | Young |
| 0.411666 | 0.190439 | 0.341 | 0.435 | 1 | 11 | Wac | -0.094 | Young |
| 0.412116 | 0.080108 | 0.326 | 0.407 | 1 | 11 | Zcchc11 | -0.081 | Young |
| 0.412557 | 0.510827 | 0.08 | 0.109 | 1 | 11 | Eng | -0.029 | Young |
| 0.412843 | 0.680994 | 0.036 | 0.056 | 1 | 11 | Cdt1 | -0.02 | Young |

| | | | | | | | | |
|---|---|---|---|---|---|---|---|---|
| 0.413221 | 0.234725 | 0.116 | 0.157 | 1 | 11 | Itpk1 | -0.041 | Young |
| 0.413499 | 0.004051 | 0.051 | 0.073 | 1 | 11 | Tada3 | -0.022 | Young |
| 0.413887 | 0.386808 | 0.159 | 0.206 | 1 | 11 | Rab5b | -0.047 | Young |
| 0.413962 | 0.406611 | 0.072 | 0.101 | 1 | 11 | N4bp2 | -0.029 | Young |
| 0.414411 | 0.035545 | 0.043 | 0.065 | 1 | 11 | Casp9 | -0.022 | Young |
| 0.414548 | 0.228698 | 0.13 | 0.169 | 1 | 11 | Cab39l | -0.039 | Young |
| 0.414918 | 0.106979 | 0.957 | 0.984 | 1 | 11 | Laptm5 | -0.027 | Young |
| 0.414975 | 0.133823 | 0.616 | 0.685 | 1 | 11 | Ndufa2 | -0.069 | Young |
| 0.41512 | 0.125086 | 0.174 | 0.218 | 1 | 11 | Cdc73 | -0.044 | Young |
| 0.415176 | 0.082455 | 0.145 | 0.185 | 1 | 11 | Siglec1 | -0.04 | Young |
| 0.415767 | 0.259065 | 0.043 | 0.065 | 1 | 11 | Calcoco1 | -0.022 | Young |
| 0.416002 | 0.202309 | 0.21 | 0.27 | 1 | 11 | Atp11b | -0.06 | Young |
| 0.416209 | 0.194901 | 0.152 | 0.194 | 1 | 11 | Oser1 | -0.042 | Young |
| 0.416329 | 0.09387 | 0.217 | 0.27 | 1 | 11 | Chp1 | -0.053 | Young |
| 0.416979 | 1.647836 | 0.043 | 0.028 | 1 | 11 | Aldh6a1 | 0.015 | Young |
| 0.418663 | 0.165368 | 0.036 | 0.056 | 1 | 11 | Slc2a1 | -0.02 | Young |
| 0.418706 | 0.240763 | 0.283 | 0.367 | 1 | 11 | Mtch1 | -0.084 | Young |
| 0.418865 | 0.35676 | 0.283 | 0.355 | 1 | 11 | Stk38 | -0.072 | Young |
| 0.418957 | 0.089501 | 0.391 | 0.48 | 1 | 11 | Zbtb20 | -0.089 | Young |
| 0.418992 | 0.543628 | 0.065 | 0.093 | 1 | 11 | Exosc2 | -0.028 | Young |
| 0.419359 | 0.598978 | 0.109 | 0.085 | 1 | 11 | Dusp8 | 0.024 | Young |
| 0.419512 | 0.087982 | 0.341 | 0.411 | 1 | 11 | Hprt | -0.07 | Young |
| 0.419947 | 0.102711 | 0.08 | 0.109 | 1 | 11 | Cblb | -0.029 | Young |
| 0.4206 | 0.313719 | 0.087 | 0.117 | 1 | 11 | Vhl | -0.03 | Young |
| 0.420723 | 0.374128 | 0.196 | 0.266 | 1 | 11 | Rbm42 | -0.07 | Young |
| 0.420847 | 0.802859 | 0.058 | 0.085 | 1 | 11 | Pctp | -0.027 | Young |
| 0.420962 | 0.510252 | 0.152 | 0.198 | 1 | 11 | Epb41 | -0.046 | Young |
| 0.420997 | 0.210548 | 0.109 | 0.145 | 1 | 11 | Napg | -0.036 | Young |
| 0.421001 | 0.386485 | 0.428 | 0.456 | 1 | 11 | Slc15a3 | -0.028 | Young |
| 0.42138 | 1.130013 | 0.08 | 0.06 | 1 | 11 | Coa6 | 0.02 | Young |
| 0.421591 | 0.454793 | 0.036 | 0.056 | 1 | 11 | Rpap1 | -0.02 | Young |
| 0.421678 | 0.694431 | 0.145 | 0.194 | 1 | 11 | Nsun2 | -0.049 | Young |
| 0.422183 | 0.401057 | 0.116 | 0.149 | 1 | 11 | Tmem57 | -0.033 | Young |
| 0.422242 | 1.089118 | 0.043 | 0.028 | 1 | 11 | Znhit6 | 0.015 | Young |
| 0.422435 | 0.32165 | 0.232 | 0.294 | 1 | 11 | Mertk | -0.062 | Young |
| 0.422435 | 0.17659 | 0.304 | 0.391 | 1 | 11 | Max | -0.087 | Young |
| 0.422537 | 1.463315 | 0.138 | 0.117 | 1 | 11 | Arid3a | 0.021 | Young |
| 0.422821 | 0.756902 | 0.355 | 0.383 | 1 | 11 | Mia3 | -0.028 | Young |
| 0.422924 | 0.176128 | 0.457 | 0.448 | 1 | 11 | Pou2f2 | 0.009 | Young |
| 0.423138 | 0.384726 | 0.08 | 0.109 | 1 | 11 | Prkab1 | -0.029 | Young |
| 0.42314 | 0.700476 | 0.116 | 0.153 | 1 | 11 | Nxpe5 | -0.037 | Young |
| 0.42347 | 0.414616 | 0.094 | 0.125 | 1 | 11 | Prpf19 | -0.031 | Young |
| 0.423475 | 1.266774 | 0.109 | 0.089 | 1 | 11 | Cntln | 0.02 | Young |
| 0.423958 | 0.323527 | 0.043 | 0.065 | 1 | 11 | Ttc4 | -0.022 | Young |
| 0.423979 | 0.470093 | 0.101 | 0.137 | 1 | 11 | Mcee | -0.036 | Young |

| | | | | | | | | |
|---|---|---|---|---|---|---|---|---|
| 0.424006 | 0.657054 | 0.043 | 0.028 | 1 | 11 | Cdc42bpa | 0.015 | Young |
| 0.424054 | 0.171904 | 0.116 | 0.153 | 1 | 11 | Arhgap39 | -0.037 | Young |
| 0.42453 | 0.397248 | 0.036 | 0.056 | 1 | 11 | Zfp943 | -0.02 | Young |
| 0.424553 | 0.826674 | 0.058 | 0.04 | 1 | 11 | Arl15 | 0.018 | Young |
| 0.425213 | 0.380048 | 0.159 | 0.21 | 1 | 11 | Clk3 | -0.051 | Young |
| 0.425273 | 0.548303 | 0.08 | 0.109 | 1 | 11 | Dcaf5 | -0.029 | Young |
| 0.425904 | 0.231891 | 0.319 | 0.415 | 1 | 11 | Ttc14 | -0.096 | Young |
| 0.426349 | 0.183131 | 0.565 | 0.54 | 1 | 11 | Mgl2 | 0.025 | Young |
| 0.426556 | 1.112748 | 0.254 | 0.242 | 1 | 11 | Bub3 | 0.012 | Young |
| 0.42671 | 0.220425 | 0.043 | 0.065 | 1 | 11 | Rpa2 | -0.022 | Young |
| 0.42674 | 0.364079 | 0.159 | 0.206 | 1 | 11 | Lemd2 | -0.047 | Young |
| 0.427189 | 0.388516 | 0.217 | 0.282 | 1 | 11 | Slc8b1 | -0.065 | Young |
| 0.427582 | 0.916986 | 0.058 | 0.04 | 1 | 11 | Faap100 | 0.018 | Young |
| 0.427582 | 0.709012 | 0.058 | 0.04 | 1 | 11 | Trpm4 | 0.018 | Young |
| 0.427693 | 0.328941 | 0.203 | 0.266 | 1 | 11 | Arhgef3 | -0.063 | Young |
| 0.427748 | 0.129023 | 0.145 | 0.185 | 1 | 11 | Hmg20b | -0.04 | Young |
| 0.427839 | 0.540394 | 0.101 | 0.137 | 1 | 11 | Srp68 | -0.036 | Young |
| 0.427998 | 0.078202 | 0.746 | 0.819 | 1 | 11 | Atp5l | -0.073 | Young |
| 0.428643 | 0.401671 | 0.116 | 0.153 | 1 | 11 | Tob1 | -0.037 | Young |
| 0.428667 | 0.126341 | 0.022 | 0.036 | 1 | 11 | Ass1 | -0.014 | Young |
| 0.429165 | 0.186979 | 0.319 | 0.407 | 1 | 11 | Hnrnpdl | -0.088 | Young |
| 0.429285 | 0.102997 | 0.377 | 0.492 | 1 | 11 | Larp4b | -0.115 | Young |
| 0.429321 | 0.622995 | 0.043 | 0.028 | 1 | 11 | Zfp932 | 0.015 | Young |
| 0.429471 | 0.236951 | 0.043 | 0.065 | 1 | 11 | Mfsd4a | -0.022 | Young |
| 0.42949 | 0.199925 | 0.094 | 0.125 | 1 | 11 | Gart | -0.031 | Young |
| 0.429849 | 0.114925 | 0.217 | 0.278 | 1 | 11 | Zfp263 | -0.061 | Young |
| 0.42996 | 0.453715 | 0.065 | 0.093 | 1 | 11 | Thap11 | -0.028 | Young |
| 0.429977 | 0.640855 | 0.072 | 0.101 | 1 | 11 | Mak16 | -0.029 | Young |
| 0.430014 | 0.642221 | 0.196 | 0.173 | 1 | 11 | Rnase6 | 0.023 | Young |
| 0.430285 | 0.165382 | 0.065 | 0.089 | 1 | 11 | Itgb7 | -0.024 | Young |
| 0.430362 | 0.26718 | 0.37 | 0.468 | 1 | 11 | Dbnl | -0.098 | Young |
| 0.430414 | 0.820896 | 0.116 | 0.157 | 1 | 11 | Mrps23 | -0.041 | Young |
| 0.430434 | 0.31915 | 0.225 | 0.278 | 1 | 11 | Osgep | -0.053 | Young |
| 0.430535 | 1.079441 | 0.072 | 0.101 | 1 | 11 | Dpm2 | -0.029 | Young |
| 0.430558 | 0.198371 | 0.13 | 0.165 | 1 | 11 | Tlk2 | -0.035 | Young |
| 0.430713 | 0.278196 | 0.42 | 0.54 | 1 | 11 | Usf2 | -0.12 | Young |
| 0.430748 | 0.610542 | 0.101 | 0.137 | 1 | 11 | Trappc12 | -0.036 | Young |
| 0.430842 | 0.258062 | 0.152 | 0.198 | 1 | 11 | Chordc1 | -0.046 | Young |
| 0.430856 | 0.697688 | 0.043 | 0.065 | 1 | 11 | Dgcr6 | -0.022 | Young |
| 0.430856 | 0.259995 | 0.043 | 0.065 | 1 | 11 | Clcn6 | -0.022 | Young |
| 0.431045 | 0.518937 | 0.246 | 0.319 | 1 | 11 | Vps26a | -0.073 | Young |
| 0.431223 | 0.001042 | 0.072 | 0.097 | 1 | 11 | Mfap1b | -0.025 | Young |
| 0.431531 | 0.164266 | 0.159 | 0.202 | 1 | 11 | Zbtb4 | -0.043 | Young |
| 0.431716 | 0.244693 | 0.08 | 0.109 | 1 | 11 | Sap30l | -0.029 | Young |
| 0.432244 | 0.595004 | 0.043 | 0.065 | 1 | 11 | Smg8 | -0.022 | Young |

| | | | | | | | | |
|---|---|---|---|---|---|---|---|---|
| 0.433267 | 0.719763 | 0.333 | 0.315 | 1 | 11 | Kat6a | 0.018 | Young |
| 0.43353 | 0.211769 | 0.094 | 0.125 | 1 | 11 | Soga1 | -0.031 | Young |
| 0.433634 | 0.446977 | 0.043 | 0.065 | 1 | 11 | Clmp | -0.022 | Young |
| 0.433657 | 0.587183 | 0.174 | 0.23 | 1 | 11 | Csrp1 | -0.056 | Young |
| 0.433867 | 0.369938 | 0.232 | 0.298 | 1 | 11 | Bmpr2 | -0.066 | Young |
| 0.434362 | 0.185371 | 0.304 | 0.363 | 1 | 11 | Rab21 | -0.059 | Young |
| 0.434393 | 0.150987 | 0.123 | 0.157 | 1 | 11 | Foxo3 | -0.034 | Young |
| 0.434554 | 0.067315 | 0.109 | 0.141 | 1 | 11 | D10Jhu81e | -0.032 | Young |
| 0.434643 | 0.189794 | 0.101 | 0.137 | 1 | 11 | Pqlc3 | -0.036 | Young |
| 0.43465 | 0.12812 | 0.442 | 0.56 | 1 | 11 | Sbno2 | -0.118 | Young |
| 0.435026 | 0.242844 | 0.043 | 0.065 | 1 | 11 | Cyb561d2 | -0.022 | Young |
| 0.435383 | 0.619646 | 0.058 | 0.081 | 1 | 11 | Poldip2 | -0.023 | Young |
| 0.435514 | 0.595868 | 0.109 | 0.141 | 1 | 11 | Hhex | -0.032 | Young |
| 0.435683 | 0.975478 | 0.21 | 0.194 | 1 | 11 | St3gal6 | 0.016 | Young |
| 0.436011 | 0.341676 | 0.123 | 0.161 | 1 | 11 | Rlf | -0.038 | Young |
| 0.436421 | 0.922783 | 0.043 | 0.065 | 1 | 11 | Rhno1 | -0.022 | Young |
| 0.436602 | 0.501064 | 0.217 | 0.282 | 1 | 11 | Slc31a2 | -0.065 | Young |
| 0.43705 | 0.532543 | 0.152 | 0.129 | 1 | 11 | As3mt | 0.023 | Young |
| 0.437503 | 0.431443 | 0.152 | 0.198 | 1 | 11 | Tcerg1 | -0.046 | Young |
| 0.437605 | 0.501 | 0.196 | 0.238 | 1 | 11 | Cytip | -0.042 | Young |
| 0.437665 | 1.275473 | 0.065 | 0.048 | 1 | 11 | Cpeb3 | 0.017 | Young |
| 0.437819 | 0.504242 | 0.043 | 0.065 | 1 | 11 | AI839979 | -0.022 | Young |
| 0.437959 | 0.793565 | 0.022 | 0.036 | 1 | 11 | Gm2814 | -0.014 | Young |
| 0.437959 | 0.105972 | 0.022 | 0.036 | 1 | 11 | Dhrs4 | -0.014 | Young |
| 0.437959 | 0.003628 | 0.022 | 0.036 | 1 | 11 | Cbarp | -0.014 | Young |
| 0.43815 | 0.583619 | 0.217 | 0.27 | 1 | 11 | Stip1 | -0.053 | Young |
| 0.438559 | 0.922243 | 0.101 | 0.137 | 1 | 11 | Myo18a | -0.036 | Young |
| 0.438612 | 0.489534 | 0.094 | 0.125 | 1 | 11 | Rfwd3 | -0.031 | Young |
| 0.438935 | 0.354942 | 0.232 | 0.302 | 1 | 11 | Sbds | -0.07 | Young |
| 0.439293 | 0.105432 | 0.457 | 0.565 | 1 | 11 | Sod1 | -0.108 | Young |
| 0.43932 | 0.301858 | 0.087 | 0.117 | 1 | 11 | Pgpep1 | -0.03 | Young |
| 0.439447 | 0.27622 | 0.051 | 0.073 | 1 | 11 | Pou5f2 | -0.022 | Young |
| 0.439626 | 0.583273 | 0.13 | 0.173 | 1 | 11 | Slc39a10 | -0.043 | Young |
| 0.440622 | 0.461573 | 0.043 | 0.065 | 1 | 11 | G6pc3 | -0.022 | Young |
| 0.440622 | 0.418918 | 0.043 | 0.065 | 1 | 11 | Tmem198b | -0.022 | Young |
| 0.440661 | 0.551085 | 0.072 | 0.101 | 1 | 11 | Trim21 | -0.029 | Young |
| 0.440752 | 0.479206 | 0.312 | 0.411 | 1 | 11 | Lasp1 | -0.099 | Young |
| 0.441313 | 0.318383 | 0.319 | 0.427 | 1 | 11 | Ctdnep1 | -0.108 | Young |
| 0.44145 | 0.030008 | 0.384 | 0.476 | 1 | 11 | Sgk3 | -0.092 | Young |
| 0.441707 | 0.025305 | 0.022 | 0.036 | 1 | 11 | Zfp429 | -0.014 | Young |
| 0.441707 | 0.01826 | 0.022 | 0.036 | 1 | 11 | Ttc27 | -0.014 | Young |
| 0.441769 | 0.249605 | 0.254 | 0.323 | 1 | 11 | Myd88 | -0.069 | Young |
| 0.441885 | 0.572331 | 0.275 | 0.254 | 1 | 11 | Ifi213 | 0.021 | Young |
| 0.442135 | 0.87125 | 0.21 | 0.194 | 1 | 11 | Uqcc3 | 0.016 | Young |
| 0.442808 | 0.450411 | 0.283 | 0.27 | 1 | 11 | Osbpl8 | 0.013 | Young |

| | | | | | | | | |
|---|---|---|---|---|---|---|---|---|
| 0.443589 | 0.577877 | 0.022 | 0.036 | 1 | 11 | Efnb1 | -0.014 | Young |
| 0.443589 | 0.447702 | 0.022 | 0.036 | 1 | 11 | Rrp1b | -0.014 | Young |
| 0.443648 | 0.602413 | 0.116 | 0.149 | 1 | 11 | Tmem106c | -0.033 | Young |
| 0.443683 | 0.019604 | 0.043 | 0.028 | 1 | 11 | Ptdss2 | 0.015 | Young |
| 0.444025 | 0.196398 | 0.348 | 0.427 | 1 | 11 | Tfe3 | -0.079 | Young |
| 0.444155 | 0.582461 | 0.058 | 0.081 | 1 | 11 | Fscn1 | -0.023 | Young |
| 0.444155 | 0.26488 | 0.058 | 0.081 | 1 | 11 | Smarcd1 | -0.023 | Young |
| 0.444256 | 0.420854 | 0.167 | 0.214 | 1 | 11 | Aff1 | -0.047 | Young |
| 0.444515 | 0.394378 | 0.13 | 0.169 | 1 | 11 | Mettl9 | -0.039 | Young |
| 0.444893 | 0.446514 | 0.333 | 0.327 | 1 | 11 | Stap1 | 0.006 | Young |
| 0.4454 | 0.296578 | 0.116 | 0.153 | 1 | 11 | Trappc8 | -0.037 | Young |
| 0.44574 | 0.557916 | 0.138 | 0.177 | 1 | 11 | Zfp740 | -0.039 | Young |
| 0.445963 | 0.27036 | 0.217 | 0.278 | 1 | 11 | Rala | -0.061 | Young |
| 0.446078 | 0.139502 | 0.051 | 0.073 | 1 | 11 | Plekhn1 | -0.022 | Young |
| 0.446292 | 1.238148 | 0.065 | 0.048 | 1 | 11 | Trmt13 | 0.017 | Young |
| 0.446418 | 0.319128 | 0.08 | 0.109 | 1 | 11 | Stk38l | -0.029 | Young |
| 0.446461 | 0.072915 | 0.42 | 0.536 | 1 | 11 | Sf3b2 | -0.116 | Young |
| 0.447213 | 1.011265 | 0.181 | 0.161 | 1 | 11 | Vezf1 | 0.02 | Young |
| 0.447365 | 0.923239 | 0.022 | 0.036 | 1 | 11 | Zfp444 | -0.014 | Young |
| 0.448206 | 0.014552 | 0.065 | 0.089 | 1 | 11 | Msra | -0.024 | Young |
| 0.448295 | 0.135679 | 0.551 | 0.702 | 1 | 11 | Csnk1a1 | -0.151 | Young |
| 0.448385 | 0.188311 | 0.297 | 0.367 | 1 | 11 | Dgkd | -0.07 | Young |
| 0.448746 | 0.553593 | 0.051 | 0.073 | 1 | 11 | Cwc27 | -0.022 | Young |
| 0.448929 | 0.390507 | 0.217 | 0.278 | 1 | 11 | Arpc1a | -0.061 | Young |
| 0.448936 | 0.825568 | 0.174 | 0.153 | 1 | 11 | Smox | 0.021 | Young |
| 0.449163 | 0.305488 | 0.341 | 0.44 | 1 | 11 | Gbp3 | -0.099 | Young |
| 0.44926 | 0.600749 | 0.022 | 0.036 | 1 | 11 | Dcun1d2 | -0.014 | Young |
| 0.44926 | 0.027914 | 0.022 | 0.036 | 1 | 11 | Csf1 | -0.014 | Young |
| 0.449382 | 0.199543 | 0.138 | 0.173 | 1 | 11 | Kdm4a | -0.035 | Young |
| 0.449744 | 0.118654 | 0.167 | 0.206 | 1 | 11 | Pttg1 | -0.039 | Young |
| 0.450459 | 0.092326 | 0.884 | 0.919 | 1 | 11 | Oaz1 | -0.035 | Young |
| 0.450478 | 0.039292 | 0.587 | 0.706 | 1 | 11 | Cox6a1 | -0.119 | Young |
| 0.450481 | 0.182031 | 0.058 | 0.081 | 1 | 11 | Kat2a | -0.023 | Young |
| 0.450643 | 0.361522 | 0.232 | 0.29 | 1 | 11 | Kpnb1 | -0.058 | Young |
| 0.450786 | 0.723645 | 0.094 | 0.129 | 1 | 11 | Ckap5 | -0.035 | Young |
| 0.451159 | 0.491341 | 0.022 | 0.036 | 1 | 11 | Zfp84 | -0.014 | Young |
| 0.451159 | 0.13857 | 0.022 | 0.036 | 1 | 11 | Ptgir | -0.014 | Young |
| 0.451258 | 0.001465 | 0.116 | 0.149 | 1 | 11 | Cds1 | -0.033 | Young |
| 0.451286 | 0.210214 | 0.13 | 0.165 | 1 | 11 | D16Ertd47 | -0.035 | Young |
| 0.45158 | 0.024398 | 0.42 | 0.508 | 1 | 11 | Aldoa | -0.088 | Young |
| 0.452467 | 0.089926 | 0.087 | 0.113 | 1 | 11 | Mov10 | -0.026 | Young |
| 0.452801 | 2.529664 | 0.022 | 0.012 | 1 | 11 | Zfp951 | 0.01 | Young |
| 0.452801 | 2.155241 | 0.022 | 0.012 | 1 | 11 | Ipp | 0.01 | Young |
| 0.452801 | 2.151358 | 0.022 | 0.012 | 1 | 11 | Gm26759 | 0.01 | Young |
| 0.452801 | 2.099248 | 0.022 | 0.012 | 1 | 11 | Plcxd2 | 0.01 | Young |

| | | | | | | | | |
|---|---|---|---|---|---|---|---|---|
| 0.452801 | 1.695683 | 0.022 | 0.012 | 1 | 11 | Fpr1 | 0.01 | Young |
| 0.452801 | 1.58467 | 0.022 | 0.012 | 1 | 11 | Neil2 | 0.01 | Young |
| 0.452801 | 1.470977 | 0.022 | 0.012 | 1 | 11 | Meis1 | 0.01 | Young |
| 0.45302 | 0.315471 | 0.094 | 0.125 | 1 | 11 | 9930021J0 | -0.031 | Young |
| 0.453063 | 0.722393 | 0.022 | 0.036 | 1 | 11 | Lrrc8b | -0.014 | Young |
| 0.453063 | 0.595271 | 0.022 | 0.036 | 1 | 11 | Pde8a | -0.014 | Young |
| 0.453063 | 0.541302 | 0.022 | 0.036 | 1 | 11 | 4933433G1 | -0.014 | Young |
| 0.453063 | 0.519781 | 0.022 | 0.036 | 1 | 11 | Amer1 | -0.014 | Young |
| 0.453063 | 0.500289 | 0.022 | 0.036 | 1 | 11 | Tmem107 | -0.014 | Young |
| 0.453063 | 0.495005 | 0.022 | 0.036 | 1 | 11 | Supv3l1 | -0.014 | Young |
| 0.45406 | 0.719272 | 0.094 | 0.125 | 1 | 11 | Coasy | -0.031 | Young |
| 0.454075 | 0.265164 | 0.616 | 0.669 | 1 | 11 | Mndal | -0.053 | Young |
| 0.454302 | 0.468625 | 0.058 | 0.081 | 1 | 11 | Tspyl2 | -0.023 | Young |
| 0.454851 | 0.421157 | 0.159 | 0.202 | 1 | 11 | Dazap1 | -0.043 | Young |
| 0.455112 | 1.666215 | 0.051 | 0.036 | 1 | 11 | Clybl | 0.015 | Young |
| 0.455264 | 1.169202 | 0.145 | 0.129 | 1 | 11 | Gtf2b | 0.016 | Young |
| 0.45548 | 1.918166 | 0.022 | 0.012 | 1 | 11 | Rpgr | 0.01 | Young |
| 0.45548 | 1.367831 | 0.022 | 0.012 | 1 | 11 | Svip | 0.01 | Young |
| 0.45548 | 1.291503 | 0.022 | 0.012 | 1 | 11 | Gm14286 | 0.01 | Young |
| 0.455492 | 0.460643 | 0.094 | 0.073 | 1 | 11 | Tnfsf10 | 0.021 | Young |
| 0.456393 | 0.511038 | 0.217 | 0.278 | 1 | 11 | Phlda1 | -0.061 | Young |
| 0.456454 | 0.317333 | 0.203 | 0.258 | 1 | 11 | Tnfrsf13b | -0.055 | Young |
| 0.457744 | 1.361388 | 0.101 | 0.081 | 1 | 11 | Ginm1 | 0.02 | Young |
| 0.457759 | 1.485837 | 0.072 | 0.056 | 1 | 11 | Cd300lg | 0.016 | Young |
| 0.457986 | 0.640082 | 0.312 | 0.319 | 1 | 11 | Ubxn4 | -0.007 | Young |
| 0.458047 | 0.809455 | 0.08 | 0.109 | 1 | 11 | Naxd | -0.029 | Young |
| 0.458167 | 2.11007 | 0.022 | 0.012 | 1 | 11 | 9430015G1 | 0.01 | Young |
| 0.458167 | 1.641091 | 0.022 | 0.012 | 1 | 11 | Ptpdc1 | 0.01 | Young |
| 0.458167 | 1.38305 | 0.022 | 0.012 | 1 | 11 | Zgrf1 | 0.01 | Young |
| 0.458258 | 0.862819 | 0.167 | 0.214 | 1 | 11 | 3110043O2 | -0.047 | Young |
| 0.458441 | 0.347887 | 0.239 | 0.298 | 1 | 11 | Oas2 | -0.059 | Young |
| 0.458687 | 0.375001 | 0.384 | 0.504 | 1 | 11 | Luc7l3 | -0.12 | Young |
| 0.459463 | 0.364726 | 0.558 | 0.605 | 1 | 11 | Wdr26 | -0.047 | Young |
| 0.459774 | 0.569218 | 0.181 | 0.157 | 1 | 11 | Helz | 0.024 | Young |
| 0.459903 | 0.362381 | 0.109 | 0.141 | 1 | 11 | Ralgapb | -0.032 | Young |
| 0.460393 | 0.602506 | 0.101 | 0.133 | 1 | 11 | Mosmo | -0.032 | Young |
| 0.460863 | 2.001211 | 0.022 | 0.012 | 1 | 11 | Gpt | 0.01 | Young |
| 0.460863 | 1.792941 | 0.022 | 0.012 | 1 | 11 | Slc22a17 | 0.01 | Young |
| 0.460863 | 1.441301 | 0.022 | 0.012 | 1 | 11 | Zrsr1 | 0.01 | Young |
| 0.460863 | 1.140431 | 0.022 | 0.012 | 1 | 11 | Dync2h1 | 0.01 | Young |
| 0.460863 | 0.892963 | 0.022 | 0.012 | 1 | 11 | Gm13713 | 0.01 | Young |
| 0.460863 | 0.715068 | 0.022 | 0.012 | 1 | 11 | Insl6 | 0.01 | Young |
| 0.46109 | 0.515419 | 0.203 | 0.258 | 1 | 11 | Tmem165 | -0.055 | Young |
| 0.461192 | 0.946782 | 0.21 | 0.198 | 1 | 11 | Katna1 | 0.012 | Young |
| 0.461412 | 0.494551 | 0.101 | 0.133 | 1 | 11 | Cdk2ap1 | -0.032 | Young |

| | | | | | | | | |
|---|---|---|---|---|---|---|---|---|
| 0.461633 | 0.2767 | 0.507 | 0.556 | 1 | 11 | Snrpe | -0.049 | Young |
| 0.461902 | 1.363511 | 0.072 | 0.056 | 1 | 11 | Kantr | 0.016 | Young |
| 0.462054 | 0.260019 | 0.312 | 0.399 | 1 | 11 | Nek7 | -0.087 | Young |
| 0.462093 | 0.748636 | 0.232 | 0.294 | 1 | 11 | Tsg101 | -0.062 | Young |
| 0.462157 | 0.178681 | 0.449 | 0.548 | 1 | 11 | Fcgr1 | -0.099 | Young |
| 0.462431 | 0.026844 | 0.174 | 0.218 | 1 | 11 | Impad1 | -0.044 | Young |
| 0.462562 | 0.218339 | 0.152 | 0.194 | 1 | 11 | Srek1 | -0.042 | Young |
| 0.462904 | 0.283498 | 0.225 | 0.278 | 1 | 11 | Krit1 | -0.053 | Young |
| 0.463024 | 0.287505 | 0.493 | 0.508 | 1 | 11 | Eif3k | -0.015 | Young |
| 0.463367 | 0.140873 | 0.087 | 0.113 | 1 | 11 | Impdh1 | -0.026 | Young |
| 0.463473 | 1.255235 | 0.051 | 0.036 | 1 | 11 | Vcan | 0.015 | Young |
| 0.463481 | 0.036111 | 0.435 | 0.512 | 1 | 11 | Ptp4a1 | -0.077 | Young |
| 0.463568 | 1.06327 | 0.022 | 0.012 | 1 | 11 | Rtn4rl1 | 0.01 | Young |
| 0.463568 | 0.942385 | 0.022 | 0.012 | 1 | 11 | Zscan20 | 0.01 | Young |
| 0.463568 | 0.591061 | 0.022 | 0.012 | 1 | 11 | Tasp1 | 0.01 | Young |
| 0.463568 | 0.322815 | 0.022 | 0.012 | 1 | 11 | Ncapd2 | 0.01 | Young |
| 0.463568 | 0.315073 | 0.022 | 0.012 | 1 | 11 | A730063M | 0.01 | Young |
| 0.463568 | 0.102959 | 0.022 | 0.012 | 1 | 11 | Flt3 | 0.01 | Young |
| 0.46358 | 0.83352 | 0.051 | 0.073 | 1 | 11 | Man2c1 | -0.022 | Young |
| 0.463832 | 1.01098 | 0.065 | 0.048 | 1 | 11 | Metap1d | 0.017 | Young |
| 0.463832 | 0.654355 | 0.065 | 0.048 | 1 | 11 | Bod1 | 0.017 | Young |
| 0.463935 | 0.644125 | 0.246 | 0.23 | 1 | 11 | Trim34a | 0.016 | Young |
| 0.464076 | 0.141613 | 0.065 | 0.089 | 1 | 11 | Fam129b | -0.024 | Young |
| 0.464205 | 0.373133 | 0.145 | 0.19 | 1 | 11 | Pfkfb4 | -0.045 | Young |
| 0.464517 | 0.07052 | 0.007 | 0.016 | 1 | 11 | Arhgef18 | -0.009 | Young |
| 0.464517 | 0.00238 | 0.007 | 0.016 | 1 | 11 | Saa3 | -0.009 | Young |
| 0.464564 | 0.011822 | 0.254 | 0.302 | 1 | 11 | Notch1 | -0.048 | Young |
| 0.464578 | 0.400796 | 0.058 | 0.081 | 1 | 11 | Golt1b | -0.023 | Young |
| 0.464816 | 0.650472 | 0.355 | 0.371 | 1 | 11 | Dnaja2 | -0.016 | Young |
| 0.465126 | 0.764797 | 0.196 | 0.177 | 1 | 11 | Ddhd2 | 0.019 | Young |
| 0.465214 | 0.964482 | 0.109 | 0.145 | 1 | 11 | Zbtb7b | -0.036 | Young |
| 0.465565 | 0.841727 | 0.116 | 0.097 | 1 | 11 | Zmat5 | 0.019 | Young |
| 0.465724 | 0.29484 | 0.377 | 0.48 | 1 | 11 | H13 | -0.103 | Young |
| 0.465745 | 0.047633 | 0.406 | 0.508 | 1 | 11 | Atxn7l3b | -0.102 | Young |
| 0.46589 | 0.151079 | 0.029 | 0.044 | 1 | 11 | Wipi1 | -0.015 | Young |
| 0.46589 | 0.120869 | 0.029 | 0.044 | 1 | 11 | Lig3 | -0.015 | Young |
| 0.466282 | 0.14596 | 0.022 | 0.012 | 1 | 11 | Galnt12 | 0.01 | Young |
| 0.466866 | 0.08378 | 0.109 | 0.141 | 1 | 11 | Runx3 | -0.032 | Young |
| 0.46749 | 1.546006 | 0.007 | 0.016 | 1 | 11 | Nnt | -0.009 | Young |
| 0.46749 | 0.945438 | 0.007 | 0.016 | 1 | 11 | Gm14005 | -0.009 | Young |
| 0.46749 | 0.842003 | 0.007 | 0.016 | 1 | 11 | 4833445I0 | -0.009 | Young |
| 0.46749 | 0.838208 | 0.007 | 0.016 | 1 | 11 | Lysmd1 | -0.009 | Young |
| 0.46749 | 0.791525 | 0.007 | 0.016 | 1 | 11 | Chst1 | -0.009 | Young |
| 0.46749 | 0.791372 | 0.007 | 0.016 | 1 | 11 | Fam222a | -0.009 | Young |
| 0.46749 | 0.771621 | 0.007 | 0.016 | 1 | 11 | Gm10791 | -0.009 | Young |

| | | | | | | | | |
|---|---|---|---|---|---|---|---|---|
| 0.46749 | 0.620173 | 0.007 | 0.016 | 1 | 11 | Zscan22 | -0.009 | Young |
| 0.46749 | 0.594466 | 0.007 | 0.016 | 1 | 11 | Nid2 | -0.009 | Young |
| 0.46749 | 0.589308 | 0.007 | 0.016 | 1 | 11 | Phlda3 | -0.009 | Young |
| 0.46749 | 0.553232 | 0.007 | 0.016 | 1 | 11 | Junos | -0.009 | Young |
| 0.46749 | 0.541625 | 0.007 | 0.016 | 1 | 11 | Zfp975 | -0.009 | Young |
| 0.46749 | 0.517165 | 0.007 | 0.016 | 1 | 11 | Camkmt | -0.009 | Young |
| 0.46749 | 0.48857 | 0.007 | 0.016 | 1 | 11 | Clec2g | -0.009 | Young |
| 0.46749 | 0.418549 | 0.007 | 0.016 | 1 | 11 | Gm9856 | -0.009 | Young |
| 0.46749 | 0.407231 | 0.007 | 0.016 | 1 | 11 | Gm13091 | -0.009 | Young |
| 0.46749 | 0.394079 | 0.007 | 0.016 | 1 | 11 | Gm11131 | -0.009 | Young |
| 0.46749 | 0.346367 | 0.007 | 0.016 | 1 | 11 | Id1 | -0.009 | Young |
| 0.46749 | 0.344957 | 0.007 | 0.016 | 1 | 11 | Gm26802 | -0.009 | Young |
| 0.46749 | 0.336728 | 0.007 | 0.016 | 1 | 11 | Fitm2 | -0.009 | Young |
| 0.46749 | 0.327613 | 0.007 | 0.016 | 1 | 11 | Zfp867 | -0.009 | Young |
| 0.46749 | 0.286685 | 0.007 | 0.016 | 1 | 11 | Tigar | -0.009 | Young |
| 0.46749 | 0.276682 | 0.007 | 0.016 | 1 | 11 | 2410131K1 | -0.009 | Young |
| 0.46749 | 0.260115 | 0.007 | 0.016 | 1 | 11 | Cntrob | -0.009 | Young |
| 0.46749 | 0.219201 | 0.007 | 0.016 | 1 | 11 | Ajuba | -0.009 | Young |
| 0.46749 | 0.217076 | 0.007 | 0.016 | 1 | 11 | Cep89 | -0.009 | Young |
| 0.46749 | 0.204314 | 0.007 | 0.016 | 1 | 11 | Pcyt1b | -0.009 | Young |
| 0.46749 | 0.103997 | 0.007 | 0.016 | 1 | 11 | Cyb5rl | -0.009 | Young |
| 0.46749 | 0.102927 | 0.007 | 0.016 | 1 | 11 | Gm14326 | -0.009 | Young |
| 0.46749 | 0.098246 | 0.007 | 0.016 | 1 | 11 | Gm15050 | -0.009 | Young |
| 0.46749 | 0.079314 | 0.007 | 0.016 | 1 | 11 | Ddit4 | -0.009 | Young |
| 0.46749 | 0.077743 | 0.007 | 0.016 | 1 | 11 | Lrsam1 | -0.009 | Young |
| 0.46749 | 0.048578 | 0.007 | 0.016 | 1 | 11 | Tmem120b | -0.009 | Young |
| 0.46749 | 0.034226 | 0.007 | 0.016 | 1 | 11 | Upb1 | -0.009 | Young |
| 0.46763 | 0.433856 | 0.029 | 0.044 | 1 | 11 | Med31 | -0.015 | Young |
| 0.467782 | 0.331218 | 0.065 | 0.089 | 1 | 11 | Poc1b | -0.024 | Young |
| 0.46781 | 0.035379 | 0.478 | 0.589 | 1 | 11 | Tmed5 | -0.111 | Young |
| 0.467821 | 0.424361 | 0.254 | 0.319 | 1 | 11 | Setd1b | -0.065 | Young |
| 0.468211 | 0.217648 | 0.558 | 0.613 | 1 | 11 | Itgb1 | -0.055 | Young |
| 0.468237 | 0.413366 | 0.174 | 0.218 | 1 | 11 | Polr2l | -0.044 | Young |
| 0.468775 | 0.105321 | 0.232 | 0.294 | 1 | 11 | Diaph2 | -0.062 | Young |
| 0.469005 | 1.694926 | 0.022 | 0.012 | 1 | 11 | Tldc1 | 0.01 | Young |
| 0.469005 | 0.177534 | 0.022 | 0.012 | 1 | 11 | Zfp605 | 0.01 | Young |
| 0.469021 | 0.422542 | 0.065 | 0.089 | 1 | 11 | Ptcd2 | -0.024 | Young |
| 0.469066 | 0.467042 | 0.109 | 0.089 | 1 | 11 | Brap | 0.02 | Young |
| 0.469177 | 0.721783 | 0.109 | 0.145 | 1 | 11 | Tubgcp5 | -0.036 | Young |
| 0.469281 | 0.613879 | 0.116 | 0.153 | 1 | 11 | Il16 | -0.037 | Young |
| 0.469292 | 0.013777 | 0.768 | 0.871 | 1 | 11 | Arpc1b | -0.103 | Young |
| 0.469368 | 0.534521 | 0.087 | 0.117 | 1 | 11 | Nol11 | -0.03 | Young |
| 0.469374 | 0.096678 | 0.029 | 0.044 | 1 | 11 | Dnajc27 | -0.015 | Young |
| 0.470103 | 0.222509 | 0.341 | 0.423 | 1 | 11 | Ikbkb | -0.082 | Young |
| 0.470157 | 0.600673 | 0.196 | 0.246 | 1 | 11 | Dusp22 | -0.05 | Young |

| | | | | | | | | |
|---|---|---|---|---|---|---|---|---|
| 0.470261 | 0.338585 | 0.065 | 0.089 | 1 | 11 | Strip1 | -0.024 | Young |
| 0.470334 | 0.568127 | 0.188 | 0.234 | 1 | 11 | Camta2 | -0.046 | Young |
| 0.470512 | 0.618908 | 0.087 | 0.069 | 1 | 11 | Smim3 | 0.018 | Young |
| 0.470545 | 0.435365 | 0.217 | 0.27 | 1 | 11 | Lilrb4a | -0.053 | Young |
| 0.470638 | 0.545372 | 0.101 | 0.133 | 1 | 11 | Elp2 | -0.032 | Young |
| 0.471248 | 0.084881 | 0.413 | 0.496 | 1 | 11 | Jarid2 | -0.083 | Young |
| 0.471736 | 0.210028 | 0.022 | 0.012 | 1 | 11 | Fam20a | 0.01 | Young |
| 0.47174 | 0.239725 | 0.152 | 0.19 | 1 | 11 | Herpud2 | -0.038 | Young |
| 0.471932 | 0.430031 | 0.094 | 0.125 | 1 | 11 | Frmd8 | -0.031 | Young |
| 0.471937 | 0.404217 | 0.072 | 0.097 | 1 | 11 | Ubxn7 | -0.025 | Young |
| 0.47242 | 0.700361 | 0.362 | 0.359 | 1 | 11 | Ccl4 | 0.003 | Young |
| 0.472673 | 0.300003 | 0.254 | 0.315 | 1 | 11 | Tnrc6b | -0.061 | Young |
| 0.472749 | 0.404787 | 0.065 | 0.089 | 1 | 11 | Aen | -0.024 | Young |
| 0.472873 | 0.026002 | 0.029 | 0.044 | 1 | 11 | Ttll4 | -0.015 | Young |
| 0.473131 | 0.43092 | 0.072 | 0.097 | 1 | 11 | Cd151 | -0.025 | Young |
| 0.473505 | 0.325462 | 0.13 | 0.165 | 1 | 11 | Tax1bp3 | -0.035 | Young |
| 0.474 | 0.415864 | 0.167 | 0.21 | 1 | 11 | St6gal1 | -0.043 | Young |
| 0.474238 | 1.108738 | 0.065 | 0.048 | 1 | 11 | Cavin1 | 0.017 | Young |
| 0.474272 | 0.26796 | 0.464 | 0.492 | 1 | 11 | Kdm7a | -0.028 | Young |
| 0.474334 | 0.395214 | 0.261 | 0.339 | 1 | 11 | Klf7 | -0.078 | Young |
| 0.474413 | 0.7147 | 0.116 | 0.097 | 1 | 11 | Lnpk | 0.019 | Young |
| 0.474628 | 0.34791 | 0.029 | 0.044 | 1 | 11 | Cc2d1a | -0.015 | Young |
| 0.474983 | 0.610376 | 0.058 | 0.081 | 1 | 11 | Mettl5 | -0.023 | Young |
| 0.47545 | 0.20405 | 0.181 | 0.218 | 1 | 11 | Paip1 | -0.037 | Young |
| 0.475651 | 1.166853 | 0.087 | 0.069 | 1 | 11 | Tns4 | 0.018 | Young |
| 0.475671 | 0.383723 | 0.239 | 0.306 | 1 | 11 | Uba2 | -0.067 | Young |
| 0.475699 | 0.014772 | 0.145 | 0.185 | 1 | 11 | Tcf7l2 | -0.04 | Young |
| 0.476148 | 0.949354 | 0.167 | 0.149 | 1 | 11 | Ppip5k2 | 0.018 | Young |
| 0.476387 | 0.512745 | 0.029 | 0.044 | 1 | 11 | Top3a | -0.015 | Young |
| 0.476387 | 0.44472 | 0.029 | 0.044 | 1 | 11 | R3hcc1 | -0.015 | Young |
| 0.476418 | 0.537073 | 0.181 | 0.226 | 1 | 11 | Arhgap25 | -0.045 | Young |
| 0.476493 | 1.087618 | 0.065 | 0.089 | 1 | 11 | Pak4 | -0.024 | Young |
| 0.476639 | 0.007487 | 0.087 | 0.113 | 1 | 11 | Aars | -0.026 | Young |
| 0.477028 | 0.755656 | 0.051 | 0.036 | 1 | 11 | Ahi1 | 0.015 | Young |
| 0.477262 | 0.629873 | 0.094 | 0.125 | 1 | 11 | Atn1 | -0.031 | Young |
| 0.477926 | 0.671304 | 0.109 | 0.141 | 1 | 11 | Tbc1d8 | -0.032 | Young |
| 0.478149 | 0.190538 | 0.029 | 0.044 | 1 | 11 | Slc25a15 | -0.015 | Young |
| 0.47868 | 1.168693 | 0.072 | 0.056 | 1 | 11 | Tex2 | 0.016 | Young |
| 0.478726 | 1.077417 | 0.036 | 0.024 | 1 | 11 | Ankrd52 | 0.012 | Young |
| 0.478874 | 0.377697 | 0.362 | 0.464 | 1 | 11 | Ifngr2 | -0.102 | Young |
| 0.478998 | 0.037314 | 0.065 | 0.089 | 1 | 11 | Ercc3 | -0.024 | Young |
| 0.479236 | 0.043469 | 0.268 | 0.331 | 1 | 11 | Sod2 | -0.063 | Young |
| 0.479242 | 0.414054 | 0.341 | 0.427 | 1 | 11 | Leng8 | -0.086 | Young |
| 0.47953 | 0.265978 | 0.304 | 0.395 | 1 | 11 | Mapk14 | -0.091 | Young |
| 0.479621 | 0.172513 | 0.862 | 0.887 | 1 | 11 | Rpl18 | -0.025 | Young |

| | | | | | | | | |
|---|---|---|---|---|---|---|---|---|
| 0.480778 | 1.409995 | 0.036 | 0.024 | 1 | 11 | 2610008E1 | 0.012 | Young |
| 0.480778 | 1.359707 | 0.036 | 0.024 | 1 | 11 | Ercc6 | 0.012 | Young |
| 0.480778 | 1.359108 | 0.036 | 0.024 | 1 | 11 | Vmac | 0.012 | Young |
| 0.480778 | 1.214545 | 0.036 | 0.024 | 1 | 11 | Mms22l | 0.012 | Young |
| 0.48113 | 0.070466 | 0.174 | 0.214 | 1 | 11 | Rtn1 | -0.04 | Young |
| 0.481683 | 0.995107 | 0.029 | 0.044 | 1 | 11 | Rmnd5b | -0.015 | Young |
| 0.481744 | 0.331716 | 0.08 | 0.105 | 1 | 11 | Naa30 | -0.025 | Young |
| 0.482309 | 0.739357 | 0.159 | 0.141 | 1 | 11 | Mepce | 0.018 | Young |
| 0.482769 | 0.564892 | 0.065 | 0.089 | 1 | 11 | Nop14 | -0.024 | Young |
| 0.482836 | 1.316836 | 0.036 | 0.024 | 1 | 11 | Nes | 0.012 | Young |
| 0.482926 | 1.453788 | 0.072 | 0.056 | 1 | 11 | Zfand2b | 0.016 | Young |
| 0.482926 | 1.066259 | 0.072 | 0.056 | 1 | 11 | Sgf29 | 0.016 | Young |
| 0.483048 | 0.195882 | 0.094 | 0.121 | 1 | 11 | Josd1 | -0.027 | Young |
| 0.483116 | 0.206667 | 0.37 | 0.456 | 1 | 11 | Abhd12 | -0.086 | Young |
| 0.483332 | 0.136435 | 0.717 | 0.831 | 1 | 11 | Maf | -0.114 | Young |
| 0.483404 | 0.396764 | 0.188 | 0.234 | 1 | 11 | Polr2e | -0.046 | Young |
| 0.483456 | 0.5009 | 0.029 | 0.044 | 1 | 11 | Fbxl15 | -0.015 | Young |
| 0.483581 | 0.327206 | 0.348 | 0.452 | 1 | 11 | Slc29a3 | -0.104 | Young |
| 0.483886 | 0.70639 | 0.051 | 0.036 | 1 | 11 | Gm1673 | 0.015 | Young |
| 0.484029 | 0.589132 | 0.065 | 0.089 | 1 | 11 | Inpp5f | -0.024 | Young |
| 0.484407 | 0.116038 | 0.754 | 0.859 | 1 | 11 | Actr2 | -0.105 | Young |
| 0.484512 | 0.491078 | 0.522 | 0.589 | 1 | 11 | Vsir | -0.067 | Young |
| 0.484898 | 1.467761 | 0.036 | 0.024 | 1 | 11 | Tgif2 | 0.012 | Young |
| 0.484898 | 1.392164 | 0.036 | 0.024 | 1 | 11 | Nedd4 | 0.012 | Young |
| 0.484898 | 1.110474 | 0.036 | 0.024 | 1 | 11 | Phldb1 | 0.012 | Young |
| 0.484898 | 0.75321 | 0.036 | 0.024 | 1 | 11 | Zbtb26 | 0.012 | Young |
| 0.485405 | 0.248324 | 0.116 | 0.149 | 1 | 11 | Tmem251 | -0.033 | Young |
| 0.486657 | 0.036137 | 0.036 | 0.052 | 1 | 11 | Ryr1 | -0.016 | Young |
| 0.486965 | 1.022209 | 0.036 | 0.024 | 1 | 11 | B4galt4 | 0.012 | Young |
| 0.487012 | 0.611742 | 0.029 | 0.044 | 1 | 11 | Gm9733 | -0.015 | Young |
| 0.487282 | 1.249291 | 0.101 | 0.085 | 1 | 11 | Uty | 0.016 | Young |
| 0.487335 | 1.007769 | 0.051 | 0.036 | 1 | 11 | Dcstamp | 0.015 | Young |
| 0.487407 | 0.54018 | 0.094 | 0.121 | 1 | 11 | Phf1 | -0.027 | Young |
| 0.487568 | 0.644298 | 0.08 | 0.105 | 1 | 11 | Arl3 | -0.025 | Young |
| 0.487568 | 0.545156 | 0.08 | 0.105 | 1 | 11 | E4f1 | -0.025 | Young |
| 0.487583 | 0.314357 | 0.072 | 0.097 | 1 | 11 | Pcsk7 | -0.025 | Young |
| 0.487947 | 0.098673 | 0.275 | 0.343 | 1 | 11 | Tbl1xr1 | -0.068 | Young |
| 0.488098 | 0.433488 | 0.391 | 0.403 | 1 | 11 | Ddi2 | -0.012 | Young |
| 0.488737 | 0.642048 | 0.08 | 0.105 | 1 | 11 | Rnf121 | -0.025 | Young |
| 0.489036 | 0.859506 | 0.036 | 0.024 | 1 | 11 | Diras2 | 0.012 | Young |
| 0.489402 | 0.065745 | 0.891 | 0.927 | 1 | 11 | Samhd1 | -0.036 | Young |
| 0.489601 | 0.47626 | 0.13 | 0.165 | 1 | 11 | Hgsnat | -0.035 | Young |
| 0.489907 | 0.532957 | 0.152 | 0.194 | 1 | 11 | Cbx4 | -0.042 | Young |
| 0.489968 | 0.355611 | 0.362 | 0.448 | 1 | 11 | Ywhaq | -0.086 | Young |
| 0.490445 | 0.601724 | 0.101 | 0.133 | 1 | 11 | Patl1 | -0.032 | Young |

| | | | | | | | | |
|---|---|---|---|---|---|---|---|---|
| 0.490461 | 0.240878 | 0.341 | 0.419 | 1 | 11 | Parp9 | -0.078 | Young |
| 0.490664 | 0.20251 | 0.239 | 0.29 | 1 | 11 | Chd1 | -0.051 | Young |
| 0.490867 | 1.129312 | 0.283 | 0.286 | 1 | 11 | Tnf | -0.003 | Young |
| 0.490903 | 0.738393 | 0.152 | 0.202 | 1 | 11 | Ncln | -0.05 | Young |
| 0.491113 | 0.596925 | 0.036 | 0.024 | 1 | 11 | Kcnj10 | 0.012 | Young |
| 0.491187 | 0.020025 | 0.174 | 0.206 | 1 | 11 | Ube2g2 | -0.032 | Young |
| 0.491223 | 0.706281 | 0.261 | 0.254 | 1 | 11 | Arrdc1 | 0.007 | Young |
| 0.491976 | 0.272496 | 0.123 | 0.161 | 1 | 11 | Naip2 | -0.038 | Young |
| 0.492247 | 0.214116 | 0.13 | 0.161 | 1 | 11 | Zfp871 | -0.031 | Young |
| 0.492534 | 0.534812 | 0.051 | 0.036 | 1 | 11 | Ccnyl1 | 0.015 | Young |
| 0.492534 | 0.503341 | 0.051 | 0.036 | 1 | 11 | Mypopos | 0.015 | Young |
| 0.492912 | 0.965985 | 0.072 | 0.056 | 1 | 11 | Aktip | 0.016 | Young |
| 0.493013 | 0.678908 | 0.181 | 0.234 | 1 | 11 | Babam2 | -0.053 | Young |
| 0.493194 | 0.648034 | 0.036 | 0.024 | 1 | 11 | Dtx3 | 0.012 | Young |
| 0.493364 | 0.000176 | 0.312 | 0.379 | 1 | 11 | Abca9 | -0.067 | Young |
| 0.493439 | 0.08257 | 0.435 | 0.544 | 1 | 11 | Dpysl2 | -0.109 | Young |
| 0.493904 | 0.675603 | 0.188 | 0.242 | 1 | 11 | Prune2 | -0.054 | Young |
| 0.493986 | 0.176369 | 0.094 | 0.121 | 1 | 11 | Gm11808 | -0.027 | Young |
| 0.494166 | 1.477072 | 0.029 | 0.044 | 1 | 11 | Acot2 | -0.015 | Young |
| 0.494406 | 0.08122 | 0.384 | 0.448 | 1 | 11 | Nrip1 | -0.064 | Young |
| 0.494854 | 0.394458 | 0.036 | 0.052 | 1 | 11 | Hsd17b7 | -0.016 | Young |
| 0.494897 | 0.49067 | 0.072 | 0.097 | 1 | 11 | Mphosph1 | -0.025 | Young |
| 0.495182 | 0.393812 | 0.261 | 0.331 | 1 | 11 | Akt1 | -0.07 | Young |
| 0.49528 | 0.629548 | 0.036 | 0.024 | 1 | 11 | Pcgf6 | 0.012 | Young |
| 0.495528 | 0.160312 | 0.225 | 0.274 | 1 | 11 | Psmc1 | -0.049 | Young |
| 0.497371 | 0.707061 | 0.036 | 0.024 | 1 | 11 | Gem | 0.012 | Young |
| 0.498082 | 0.141388 | 0.101 | 0.129 | 1 | 11 | Phka2 | -0.028 | Young |
| 0.498154 | 0.855 | 0.036 | 0.052 | 1 | 11 | Cbr1 | -0.016 | Young |
| 0.498523 | 0.547098 | 0.377 | 0.375 | 1 | 11 | Ifit1 | 0.002 | Young |
| 0.498576 | 0.814198 | 0.072 | 0.097 | 1 | 11 | Nosip | -0.025 | Young |
| 0.49932 | 0.393309 | 0.109 | 0.137 | 1 | 11 | Setd7 | -0.028 | Young |
| 0.499328 | 0.478989 | 0.08 | 0.105 | 1 | 11 | Slc1a5 | -0.025 | Young |
| 0.499432 | 1.165067 | 0.138 | 0.125 | 1 | 11 | Reps1 | 0.013 | Young |
| 0.499466 | 0.218801 | 0.036 | 0.024 | 1 | 11 | Cenpq | 0.012 | Young |
| 0.499805 | 0.371665 | 0.072 | 0.097 | 1 | 11 | Nectin4 | -0.025 | Young |
| 0.499808 | 1.010146 | 0.036 | 0.052 | 1 | 11 | Agpat2 | -0.016 | Young |
| 0.499808 | 0.232607 | 0.036 | 0.052 | 1 | 11 | Aldh1b1 | -0.016 | Young |
| 0.499905 | 0.324862 | 0.167 | 0.21 | 1 | 11 | Il15 | -0.043 | Young |
| 0.500112 | 0.832322 | 0.072 | 0.056 | 1 | 11 | BC051226 | 0.016 | Young |
| 0.500366 | 1.067016 | 0.138 | 0.121 | 1 | 11 | Tmem245 | 0.017 | Young |
| 0.501263 | 0.428276 | 0.051 | 0.036 | 1 | 11 | Zc2hc1a | 0.015 | Young |
| 0.501465 | 0.469108 | 0.036 | 0.052 | 1 | 11 | Inafm2 | -0.016 | Young |
| 0.50153 | 0.623715 | 0.275 | 0.266 | 1 | 11 | Rela | 0.009 | Young |
| 0.501698 | 0.833041 | 0.08 | 0.105 | 1 | 11 | Lsr | -0.025 | Young |
| 0.501776 | 0.368043 | 0.196 | 0.238 | 1 | 11 | Ybx3 | -0.042 | Young |

| | | | | | | | | |
|---|---|---|---|---|---|---|---|---|
| 0.502015 | 0.607365 | 0.297 | 0.302 | 1 | 11 | Atp6v1h | -0.005 | Young |
| 0.502269 | 0.503402 | 0.072 | 0.097 | 1 | 11 | Ell | -0.025 | Young |
| 0.502334 | 0.954477 | 0.094 | 0.077 | 1 | 11 | Bap1 | 0.017 | Young |
| 0.502917 | 0.288598 | 0.21 | 0.262 | 1 | 11 | Bcas2 | -0.052 | Young |
| 0.503074 | 0.514031 | 0.13 | 0.165 | 1 | 11 | Nop58 | -0.035 | Young |
| 0.50312 | 0.497771 | 0.043 | 0.06 | 1 | 11 | Zbtb5 | -0.017 | Young |
| 0.50341 | 0.018727 | 0.761 | 0.835 | 1 | 11 | Rps7 | -0.074 | Young |
| 0.503462 | 0.192539 | 0.101 | 0.129 | 1 | 11 | Pofut2 | -0.028 | Young |
| 0.50351 | 0.222263 | 0.109 | 0.137 | 1 | 11 | Glod4 | -0.028 | Young |
| 0.503586 | 0.582818 | 0.109 | 0.141 | 1 | 11 | Chpf2 | -0.032 | Young |
| 0.503701 | 0.341431 | 0.565 | 0.641 | 1 | 11 | Lnpep | -0.076 | Young |
| 0.503794 | 0.125591 | 0.623 | 0.746 | 1 | 11 | Ubl3 | -0.123 | Young |
| 0.504208 | 1.061168 | 0.167 | 0.153 | 1 | 11 | Mctp1 | 0.014 | Young |
| 0.504613 | 0.061428 | 0.993 | 1 | 1 | 11 | Tpt1 | -0.007 | Young |
| 0.504665 | 0.209623 | 0.043 | 0.06 | 1 | 11 | Dcakd | -0.017 | Young |
| 0.504787 | 0.145366 | 0.036 | 0.052 | 1 | 11 | Dgkq | -0.016 | Young |
| 0.505 | 0.478568 | 0.297 | 0.375 | 1 | 11 | Erbin | -0.078 | Young |
| 0.505265 | 0.330191 | 0.08 | 0.105 | 1 | 11 | Nf2 | -0.025 | Young |
| 0.506196 | 1.171617 | 0.116 | 0.101 | 1 | 11 | Clec5a | 0.015 | Young |
| 0.506481 | 1.592875 | 0.087 | 0.073 | 1 | 11 | Kat5 | 0.014 | Young |
| 0.506549 | 0.14949 | 0.159 | 0.198 | 1 | 11 | Txndc5 | -0.039 | Young |
| 0.506944 | 0.26304 | 0.174 | 0.218 | 1 | 11 | Slc48a1 | -0.044 | Young |
| 0.507045 | 0.103351 | 0.384 | 0.472 | 1 | 11 | Ythdc1 | -0.088 | Young |
| 0.507956 | 0.157649 | 0.21 | 0.25 | 1 | 11 | Ube2l6 | -0.04 | Young |
| 0.508172 | 0.179071 | 0.413 | 0.52 | 1 | 11 | Itpripl2 | -0.107 | Young |
| 0.508204 | 0.297628 | 0.123 | 0.153 | 1 | 11 | Nrf1 | -0.03 | Young |
| 0.50882 | 0.074444 | 0.906 | 0.952 | 1 | 11 | Rpl7 | -0.046 | Young |
| 0.509162 | 0.685882 | 0.203 | 0.194 | 1 | 11 | Hk3 | 0.009 | Young |
| 0.509257 | 0.028202 | 0.159 | 0.198 | 1 | 11 | Sertad2 | -0.039 | Young |
| 0.509317 | 0.001845 | 0.043 | 0.06 | 1 | 11 | Stam | -0.017 | Young |
| 0.509378 | 0.265738 | 0.058 | 0.077 | 1 | 11 | Zfp738 | -0.019 | Young |
| 0.509508 | 0.711059 | 0.377 | 0.391 | 1 | 11 | Tle3 | -0.014 | Young |
| 0.50959 | 0.274253 | 0.174 | 0.214 | 1 | 11 | Ppm1h | -0.04 | Young |
| 0.509752 | 0.872398 | 0.138 | 0.117 | 1 | 11 | Thbs1 | 0.021 | Young |
| 0.509793 | 0.77653 | 0.036 | 0.052 | 1 | 11 | Saal1 | -0.016 | Young |
| 0.509793 | 0.59673 | 0.036 | 0.052 | 1 | 11 | Per3 | -0.016 | Young |
| 0.510765 | 0.188444 | 0.13 | 0.161 | 1 | 11 | Gtf2i | -0.031 | Young |
| 0.510884 | 0.479515 | 0.181 | 0.23 | 1 | 11 | Tor1a | -0.049 | Young |
| 0.511282 | 0.570287 | 0.116 | 0.145 | 1 | 11 | Prkag1 | -0.029 | Young |
| 0.511376 | 0.332443 | 0.21 | 0.258 | 1 | 11 | Ube2m | -0.048 | Young |
| 0.511467 | 0.732726 | 0.036 | 0.052 | 1 | 11 | Btbd3 | -0.016 | Young |
| 0.511572 | 0.322713 | 0.435 | 0.472 | 1 | 11 | Eif5 | -0.037 | Young |
| 0.511958 | 0.394057 | 0.333 | 0.431 | 1 | 11 | Fndc3a | -0.098 | Young |
| 0.511984 | 0.461989 | 0.283 | 0.274 | 1 | 11 | Usp18 | 0.009 | Young |
| 0.512655 | 0.400823 | 0.348 | 0.343 | 1 | 11 | Fn1 | 0.005 | Young |

| | | | | | | | | |
|---|---|---|---|---|---|---|---|---|
| 0.51286 | 0.264768 | 0.138 | 0.169 | 1 | 11 | Kcnn4 | -0.031 | Young |
| 0.513305 | 0.216816 | 0.138 | 0.173 | 1 | 11 | BC003331 | -0.035 | Young |
| 0.513699 | 0.45859 | 0.116 | 0.149 | 1 | 11 | Prdx3 | -0.033 | Young |
| 0.513796 | 0.285872 | 0.13 | 0.165 | 1 | 11 | Dmtf1 | -0.035 | Young |
| 0.51399 | 0.405266 | 0.043 | 0.06 | 1 | 11 | Cstf1 | -0.017 | Young |
| 0.515252 | 0.272363 | 0.399 | 0.524 | 1 | 11 | Wsb1 | -0.125 | Young |
| 0.515334 | 0.714739 | 0.188 | 0.173 | 1 | 11 | Gpr34 | 0.015 | Young |
| 0.515681 | 0.237711 | 0.283 | 0.355 | 1 | 11 | Snrpg | -0.072 | Young |
| 0.515709 | 0.125282 | 0.065 | 0.085 | 1 | 11 | Rnf170 | -0.02 | Young |
| 0.515867 | 0.084962 | 0.572 | 0.69 | 1 | 11 | Tmem106a | -0.118 | Young |
| 0.516146 | 0.635341 | 0.072 | 0.056 | 1 | 11 | Setd1a | 0.016 | Young |
| 0.517035 | 0.110951 | 0.065 | 0.085 | 1 | 11 | Rassf8 | -0.02 | Young |
| 0.517118 | 0.671402 | 0.043 | 0.06 | 1 | 11 | Fbxo45 | -0.017 | Young |
| 0.517187 | 0.90974 | 0.072 | 0.097 | 1 | 11 | Scyl1 | -0.025 | Young |
| 0.517248 | 0.528445 | 0.08 | 0.105 | 1 | 11 | Rrm2b | -0.025 | Young |
| 0.517957 | 0.069904 | 0.101 | 0.125 | 1 | 11 | Stat5a | -0.024 | Young |
| 0.518171 | 0.851094 | 0.181 | 0.161 | 1 | 11 | Dapk1 | 0.02 | Young |
| 0.518282 | 0.232986 | 0.471 | 0.605 | 1 | 11 | Mtpn | -0.134 | Young |
| 0.518454 | 0.685512 | 0.08 | 0.105 | 1 | 11 | Mbtps2 | -0.025 | Young |
| 0.518712 | 0.396715 | 0.63 | 0.69 | 1 | 11 | Gm26917 | -0.06 | Young |
| 0.519246 | 0.360703 | 0.464 | 0.504 | 1 | 11 | Tnfaip8 | -0.04 | Young |
| 0.519691 | 0.186003 | 0.065 | 0.085 | 1 | 11 | Ciao1 | -0.02 | Young |
| 0.519827 | 0.231902 | 0.051 | 0.069 | 1 | 11 | Sugp1 | -0.018 | Young |
| 0.520256 | 1.474021 | 0.043 | 0.06 | 1 | 11 | Nt5dc1 | -0.017 | Young |
| 0.520256 | 0.304257 | 0.043 | 0.06 | 1 | 11 | Lin9 | -0.017 | Young |
| 0.520883 | 0.318009 | 0.101 | 0.129 | 1 | 11 | Phkb | -0.028 | Young |
| 0.521422 | 1.149912 | 0.058 | 0.044 | 1 | 11 | Ncdn | 0.014 | Young |
| 0.521432 | 0.084063 | 0.833 | 0.907 | 1 | 11 | Rpl6 | -0.074 | Young |
| 0.521892 | 0.996448 | 0.08 | 0.065 | 1 | 11 | Slc14a1 | 0.015 | Young |
| 0.522129 | 0.451575 | 0.283 | 0.274 | 1 | 11 | Parp12 | 0.009 | Young |
| 0.522466 | 0.172909 | 0.348 | 0.427 | 1 | 11 | Arcn1 | -0.079 | Young |
| 0.522516 | 0.17542 | 0.275 | 0.343 | 1 | 11 | Acsl4 | -0.068 | Young |
| 0.522593 | 0.567192 | 0.159 | 0.141 | 1 | 11 | Hccs | 0.018 | Young |
| 0.522657 | 0.416296 | 0.13 | 0.165 | 1 | 11 | Mmadhc | -0.035 | Young |
| 0.522726 | 0.251136 | 0.37 | 0.472 | 1 | 11 | Ndufb11 | -0.102 | Young |
| 0.523404 | 1.022765 | 0.043 | 0.06 | 1 | 11 | Lrrc40 | -0.017 | Young |
| 0.523469 | 0.593597 | 0.072 | 0.097 | 1 | 11 | Flnb | -0.025 | Young |
| 0.523602 | 0.348743 | 0.094 | 0.121 | 1 | 11 | Exoc1 | -0.027 | Young |
| 0.523933 | 0.279504 | 0.283 | 0.347 | 1 | 11 | Osbpl9 | -0.064 | Young |
| 0.524033 | 0.366017 | 0.174 | 0.218 | 1 | 11 | Prmt1 | -0.044 | Young |
| 0.524508 | 0.640284 | 0.08 | 0.105 | 1 | 11 | Rbl1 | -0.025 | Young |
| 0.524694 | 0.15274 | 0.058 | 0.077 | 1 | 11 | Aasdhppt | -0.019 | Young |
| 0.524727 | 1.447453 | 0.058 | 0.044 | 1 | 11 | Disc1 | 0.014 | Young |
| 0.525748 | 0.278763 | 0.014 | 0.024 | 1 | 11 | Ncaph | -0.01 | Young |
| 0.52639 | 0.41493 | 0.101 | 0.129 | 1 | 11 | Fbxo22 | -0.028 | Young |

| | | | | | | | | |
|---|---|---|---|---|---|---|---|---|
| 0.526411 | 0.017089 | 0.377 | 0.44 | 1 | 11 | Atp5c1 | -0.063 | Young |
| 0.526561 | 0.989796 | 0.043 | 0.06 | 1 | 11 | Mcm2 | -0.017 | Young |
| 0.526561 | 0.603516 | 0.043 | 0.06 | 1 | 11 | Slc37a3 | -0.017 | Young |
| 0.52723 | 0.825816 | 0.051 | 0.069 | 1 | 11 | Nedd1 | -0.018 | Young |
| 0.52723 | 0.060047 | 0.051 | 0.069 | 1 | 11 | Rusc1 | -0.018 | Young |
| 0.52749 | 0.293588 | 0.181 | 0.226 | 1 | 11 | Mpp6 | -0.045 | Young |
| 0.527504 | 0.432529 | 0.058 | 0.077 | 1 | 11 | Prrg2 | -0.019 | Young |
| 0.527504 | 0.319085 | 0.058 | 0.077 | 1 | 11 | Neurl2 | -0.019 | Young |
| 0.527504 | 0.032948 | 0.058 | 0.077 | 1 | 11 | Rps6ka5 | -0.019 | Young |
| 0.528017 | 0.085282 | 0.072 | 0.093 | 1 | 11 | Gclm | -0.021 | Young |
| 0.528043 | 0.425958 | 0.167 | 0.149 | 1 | 11 | Cmah | 0.018 | Young |
| 0.52811 | 0.606749 | 0.13 | 0.165 | 1 | 11 | Cops6 | -0.035 | Young |
| 0.528267 | 0.422462 | 0.014 | 0.024 | 1 | 11 | Rbm48 | -0.01 | Young |
| 0.528267 | 0.069428 | 0.014 | 0.024 | 1 | 11 | Mrps27 | -0.01 | Young |
| 0.528638 | 0.433005 | 0.13 | 0.161 | 1 | 11 | Rfc2 | -0.031 | Young |
| 0.528717 | 0.337615 | 0.051 | 0.069 | 1 | 11 | Cox10 | -0.018 | Young |
| 0.528818 | 0.160268 | 0.725 | 0.806 | 1 | 11 | Ybx1 | -0.081 | Young |
| 0.528995 | 0.543445 | 0.638 | 0.69 | 1 | 11 | Sdc4 | -0.052 | Young |
| 0.52906 | 0.231369 | 0.297 | 0.371 | 1 | 11 | Bcap31 | -0.074 | Young |
| 0.529436 | 0.216041 | 0.536 | 0.629 | 1 | 11 | Tab2 | -0.093 | Young |
| 0.529628 | 0.54791 | 0.196 | 0.238 | 1 | 11 | Lrrfip1 | -0.042 | Young |
| 0.529704 | 0.966056 | 0.058 | 0.044 | 1 | 11 | Snip1 | 0.014 | Young |
| 0.529704 | 0.738803 | 0.058 | 0.044 | 1 | 11 | Ryk | 0.014 | Young |
| 0.529857 | 0.016757 | 0.971 | 0.996 | 1 | 11 | Rplp1 | -0.025 | Young |
| 0.530168 | 0.089915 | 0.217 | 0.266 | 1 | 11 | Bak1 | -0.049 | Young |
| 0.530206 | 0.327862 | 0.051 | 0.069 | 1 | 11 | Mrpl47 | -0.018 | Young |
| 0.53028 | 0.838277 | 0.196 | 0.181 | 1 | 11 | Ift20 | 0.015 | Young |
| 0.530697 | 0.957424 | 0.188 | 0.177 | 1 | 11 | Ccdc124 | 0.011 | Young |
| 0.530793 | 0.157545 | 0.014 | 0.024 | 1 | 11 | Nradd | -0.01 | Young |
| 0.531465 | 0.276917 | 0.246 | 0.302 | 1 | 11 | Rrp1 | -0.056 | Young |
| 0.532311 | 1.898002 | 0.043 | 0.032 | 1 | 11 | Optn | 0.011 | Young |
| 0.532359 | 0.266332 | 0.471 | 0.577 | 1 | 11 | Cbl | -0.106 | Young |
| 0.533325 | 0.808756 | 0.014 | 0.024 | 1 | 11 | Dcp1b | -0.01 | Young |
| 0.533325 | 0.477583 | 0.014 | 0.024 | 1 | 11 | Gm26510 | -0.01 | Young |
| 0.533325 | 0.217081 | 0.014 | 0.024 | 1 | 11 | Ppip5k1 | -0.01 | Young |
| 0.533325 | 0.201726 | 0.014 | 0.024 | 1 | 11 | 1110008L1 | -0.01 | Young |
| 0.533476 | 0.13322 | 0.638 | 0.738 | 1 | 11 | Alox5ap | -0.1 | Young |
| 0.533656 | 0.393132 | 0.21 | 0.258 | 1 | 11 | Slc25a4 | -0.048 | Young |
| 0.534423 | 0.399834 | 0.065 | 0.085 | 1 | 11 | Zfp322a | -0.02 | Young |
| 0.534497 | 1.002528 | 0.043 | 0.06 | 1 | 11 | Atg10 | -0.017 | Young |
| 0.534562 | 0.709594 | 0.08 | 0.065 | 1 | 11 | Bcl9 | 0.015 | Young |
| 0.535863 | 0.749199 | 0.014 | 0.024 | 1 | 11 | Mpv17l | -0.01 | Young |
| 0.535863 | 0.627563 | 0.014 | 0.024 | 1 | 11 | Ift43 | -0.01 | Young |
| 0.535863 | 0.563704 | 0.014 | 0.024 | 1 | 11 | Rnd1 | -0.01 | Young |
| 0.535863 | 0.4105 | 0.014 | 0.024 | 1 | 11 | Telo2 | -0.01 | Young |

| | | | | | | | | |
|---|---|---|---|---|---|---|---|---|
| 0.535979 | 0.671797 | 0.08 | 0.065 | 1 | 11 | Zfp277 | 0.015 | Young |
| 0.536177 | 2.401547 | 0.043 | 0.032 | 1 | 11 | Txlng | 0.011 | Young |
| 0.536177 | 1.704821 | 0.043 | 0.032 | 1 | 11 | Edrf1 | 0.011 | Young |
| 0.536177 | 1.702412 | 0.043 | 0.032 | 1 | 11 | Gm4841 | 0.011 | Young |
| 0.536347 | 0.385253 | 0.116 | 0.145 | 1 | 11 | Mob3a | -0.029 | Young |
| 0.536347 | 0.329022 | 0.116 | 0.145 | 1 | 11 | Kif13b | -0.029 | Young |
| 0.536377 | 0.722496 | 0.058 | 0.044 | 1 | 11 | Msantd2 | 0.014 | Young |
| 0.536705 | 0.094213 | 0.623 | 0.766 | 1 | 11 | Ddx17 | -0.143 | Young |
| 0.536729 | 0.882257 | 0.174 | 0.222 | 1 | 11 | Pa2g4 | -0.048 | Young |
| 0.537124 | 0.429122 | 0.065 | 0.085 | 1 | 11 | Tipin | -0.02 | Young |
| 0.537218 | 0.405903 | 0.159 | 0.194 | 1 | 11 | Rchy1 | -0.035 | Young |
| 0.538407 | 2.184248 | 0.014 | 0.024 | 1 | 11 | 2610021A( | -0.01 | Young |
| 0.538407 | 1.400401 | 0.014 | 0.024 | 1 | 11 | Zfp553 | -0.01 | Young |
| 0.538407 | 0.523241 | 0.014 | 0.024 | 1 | 11 | Carmil1 | -0.01 | Young |
| 0.538407 | 0.036646 | 0.014 | 0.024 | 1 | 11 | Arpin | -0.01 | Young |
| 0.538477 | 0.286672 | 0.065 | 0.085 | 1 | 11 | Fdft1 | -0.02 | Young |
| 0.538781 | 0.491286 | 0.123 | 0.153 | 1 | 11 | Srsf4 | -0.03 | Young |
| 0.538909 | 0.462834 | 0.449 | 0.492 | 1 | 11 | Rassf2 | -0.043 | Young |
| 0.539288 | 0.909932 | 0.043 | 0.06 | 1 | 11 | Sec22a | -0.017 | Young |
| 0.539327 | 0.342861 | 0.116 | 0.141 | 1 | 11 | Ipmk | -0.025 | Young |
| 0.539327 | 0.137323 | 0.116 | 0.141 | 1 | 11 | Ccnh | -0.025 | Young |
| 0.539434 | 0.199511 | 0.659 | 0.762 | 1 | 11 | Npm1 | -0.103 | Young |
| 0.539631 | 0.13518 | 0.072 | 0.093 | 1 | 11 | Dtx2 | -0.021 | Young |
| 0.539832 | 0.180931 | 0.065 | 0.085 | 1 | 11 | Mbd5 | -0.02 | Young |
| 0.539976 | 0.443421 | 0.188 | 0.169 | 1 | 11 | St3gal4 | 0.019 | Young |
| 0.540057 | 1.815669 | 0.043 | 0.032 | 1 | 11 | Zdhhc12 | 0.011 | Young |
| 0.540247 | 0.234631 | 0.087 | 0.109 | 1 | 11 | Zfp598 | -0.022 | Young |
| 0.540275 | 0.371558 | 0.428 | 0.456 | 1 | 11 | Ckb | -0.028 | Young |
| 0.540888 | 0.185492 | 0.333 | 0.407 | 1 | 11 | Mapk1ip1l | -0.074 | Young |
| 0.541175 | 0.113697 | 0.196 | 0.234 | 1 | 11 | Rbm7 | -0.038 | Young |
| 0.542194 | 0.41081 | 0.051 | 0.069 | 1 | 11 | Kctd3 | -0.018 | Young |
| 0.542347 | 0.144391 | 0.094 | 0.117 | 1 | 11 | Agap3 | -0.023 | Young |
| 0.54254 | 0.201053 | 0.312 | 0.375 | 1 | 11 | Snx9 | -0.063 | Young |
| 0.543005 | 0.150774 | 0.384 | 0.468 | 1 | 11 | Fkbp1a | -0.084 | Young |
| 0.543119 | 0.063738 | 0.326 | 0.383 | 1 | 11 | Creb1 | -0.057 | Young |
| 0.543175 | 0.6482 | 0.29 | 0.294 | 1 | 11 | Fmnl1 | -0.004 | Young |
| 0.543617 | 0.496603 | 0.181 | 0.222 | 1 | 11 | Lrpap1 | -0.041 | Young |
| 0.544231 | 0.383064 | 0.217 | 0.266 | 1 | 11 | Ptpn2 | -0.049 | Young |
| 0.544522 | 0.435789 | 0.058 | 0.077 | 1 | 11 | Zfp280c | -0.019 | Young |
| 0.544835 | 0.258415 | 0.58 | 0.698 | 1 | 11 | Cdkn1a | -0.118 | Young |
| 0.545137 | 3.330175 | 0.014 | 0.008 | 1 | 11 | Pfkfb1 | 0.006 | Young |
| 0.545137 | 2.718688 | 0.014 | 0.008 | 1 | 11 | S100a16 | 0.006 | Young |
| 0.545137 | 2.713979 | 0.014 | 0.008 | 1 | 11 | Fam212b | 0.006 | Young |
| 0.545137 | 2.585936 | 0.014 | 0.008 | 1 | 11 | Rgs5 | 0.006 | Young |
| 0.545137 | 2.350491 | 0.014 | 0.008 | 1 | 11 | Rhebl1 | 0.006 | Young |

| | | | | | | | | |
|---|---|---|---|---|---|---|---|---|
| 0.545137 | 2.339756 | 0.014 | 0.008 | 1 | 11 | Klhl11 | 0.006 | Young |
| 0.545137 | 2.231625 | 0.014 | 0.008 | 1 | 11 | Sardh | 0.006 | Young |
| 0.545137 | 1.971855 | 0.014 | 0.008 | 1 | 11 | Mmab | 0.006 | Young |
| 0.545137 | 1.918801 | 0.014 | 0.008 | 1 | 11 | Tulp3 | 0.006 | Young |
| 0.545137 | 1.895813 | 0.014 | 0.008 | 1 | 11 | Arap2 | 0.006 | Young |
| 0.545137 | 1.836332 | 0.014 | 0.008 | 1 | 11 | Farp2 | 0.006 | Young |
| 0.545137 | 1.73792 | 0.014 | 0.008 | 1 | 11 | Gm9484 | 0.006 | Young |
| 0.545137 | 1.631638 | 0.014 | 0.008 | 1 | 11 | Lhfpl2 | 0.006 | Young |
| 0.545137 | 1.607991 | 0.014 | 0.008 | 1 | 11 | Them4 | 0.006 | Young |
| 0.545137 | 1.573136 | 0.014 | 0.008 | 1 | 11 | Aurka | 0.006 | Young |
| 0.545137 | 1.445384 | 0.014 | 0.008 | 1 | 11 | Kif21a | 0.006 | Young |
| 0.545137 | 1.391804 | 0.014 | 0.008 | 1 | 11 | Ldlrad4 | 0.006 | Young |
| 0.545137 | 1.32231 | 0.014 | 0.008 | 1 | 11 | Gm12184 | 0.006 | Young |
| 0.545137 | 1.25318 | 0.014 | 0.008 | 1 | 11 | Bag2 | 0.006 | Young |
| 0.545137 | 1.014968 | 0.014 | 0.008 | 1 | 11 | Slc36a3os | 0.006 | Young |
| 0.545137 | 0.809175 | 0.014 | 0.008 | 1 | 11 | Slc17a9 | 0.006 | Young |
| 0.545137 | 0.584641 | 0.014 | 0.008 | 1 | 11 | A830010M | 0.006 | Young |
| 0.545304 | 0.064227 | 0.217 | 0.258 | 1 | 11 | Trim35 | -0.041 | Young |
| 0.545668 | 0.854236 | 0.174 | 0.222 | 1 | 11 | Ncoa6 | -0.048 | Young |
| 0.545903 | 0.407823 | 0.116 | 0.145 | 1 | 11 | Fam199x | -0.029 | Young |
| 0.546004 | 0.027178 | 0.449 | 0.52 | 1 | 11 | Lilr4b | -0.071 | Young |
| 0.547017 | 0.530726 | 0.159 | 0.198 | 1 | 11 | Znrd1 | -0.039 | Young |
| 0.547385 | 0.627378 | 0.058 | 0.077 | 1 | 11 | Nxt1 | -0.019 | Young |
| 0.547608 | 0.491735 | 0.167 | 0.206 | 1 | 11 | Mnt | -0.039 | Young |
| 0.54799 | 0.282512 | 0.319 | 0.315 | 1 | 11 | Sgms1 | 0.004 | Young |
| 0.547997 | 0.851059 | 0.087 | 0.073 | 1 | 11 | Ch25h | 0.014 | Young |
| 0.54801 | 0.264004 | 0.188 | 0.23 | 1 | 11 | Aip | -0.042 | Young |
| 0.548197 | 0.673094 | 0.094 | 0.117 | 1 | 11 | Arsk | -0.023 | Young |
| 0.548748 | 2.574603 | 0.014 | 0.008 | 1 | 11 | Mtus2 | 0.006 | Young |
| 0.548748 | 2.500576 | 0.014 | 0.008 | 1 | 11 | Tpx2 | 0.006 | Young |
| 0.548748 | 2.298575 | 0.014 | 0.008 | 1 | 11 | Gm45894 | 0.006 | Young |
| 0.548748 | 2.171415 | 0.014 | 0.008 | 1 | 11 | Slpi | 0.006 | Young |
| 0.548748 | 2.026338 | 0.014 | 0.008 | 1 | 11 | Slc39a12 | 0.006 | Young |
| 0.548748 | 1.862365 | 0.014 | 0.008 | 1 | 11 | Zfp14 | 0.006 | Young |
| 0.548748 | 1.603305 | 0.014 | 0.008 | 1 | 11 | 2200002D( | 0.006 | Young |
| 0.548748 | 1.587483 | 0.014 | 0.008 | 1 | 11 | Fermt2 | 0.006 | Young |
| 0.548748 | 1.579634 | 0.014 | 0.008 | 1 | 11 | Pkp3 | 0.006 | Young |
| 0.548748 | 1.32384 | 0.014 | 0.008 | 1 | 11 | 3010003L2 | 0.006 | Young |
| 0.548748 | 1.294994 | 0.014 | 0.008 | 1 | 11 | Rnls | 0.006 | Young |
| 0.548748 | 1.269002 | 0.014 | 0.008 | 1 | 11 | Gm40773 | 0.006 | Young |
| 0.548748 | 1.037539 | 0.014 | 0.008 | 1 | 11 | Spsb1 | 0.006 | Young |
| 0.548748 | 1.028408 | 0.014 | 0.008 | 1 | 11 | Gpr21 | 0.006 | Young |
| 0.548748 | 0.926663 | 0.014 | 0.008 | 1 | 11 | Spred3 | 0.006 | Young |
| 0.548748 | 0.919928 | 0.014 | 0.008 | 1 | 11 | Zfp85 | 0.006 | Young |
| 0.548748 | 0.879273 | 0.014 | 0.008 | 1 | 11 | Ceacam19 | 0.006 | Young |

| | | | | | | | | |
|---|---|---|---|---|---|---|---|---|
| 0.548748 | 0.73126 | 0.014 | 0.008 | 1 | 11 | Cdh5 | 0.006 | Young |
| 0.548752 | 0.354223 | 0.072 | 0.093 | 1 | 11 | Sord | -0.021 | Young |
| 0.548842 | 0.203578 | 0.319 | 0.395 | 1 | 11 | Nxf1 | -0.076 | Young |
| 0.549364 | 0.593322 | 0.065 | 0.085 | 1 | 11 | Ppil3 | -0.02 | Young |
| 0.549848 | 0.462107 | 0.058 | 0.044 | 1 | 11 | Gm20342 | 0.014 | Young |
| 0.550543 | 0.664551 | 0.268 | 0.274 | 1 | 11 | Cttnbp2nl | -0.006 | Young |
| 0.551126 | 0.408304 | 0.022 | 0.032 | 1 | 11 | Akr1b8 | -0.01 | Young |
| 0.55178 | 0.243059 | 0.043 | 0.032 | 1 | 11 | Polr2d | 0.011 | Young |
| 0.552081 | 1.498941 | 0.065 | 0.052 | 1 | 11 | D930016D( | 0.013 | Young |
| 0.552097 | 0.417712 | 0.159 | 0.141 | 1 | 11 | Tmem222 | 0.018 | Young |
| 0.552102 | 0.243514 | 0.065 | 0.085 | 1 | 11 | Ring1 | -0.02 | Young |
| 0.552229 | 0.620931 | 0.138 | 0.169 | 1 | 11 | Rhobtb1 | -0.031 | Young |
| 0.552273 | 0.185564 | 0.239 | 0.294 | 1 | 11 | Nfxl1 | -0.055 | Young |
| 0.552292 | 0.522361 | 0.08 | 0.101 | 1 | 11 | Elac1 | -0.021 | Young |
| 0.552292 | 0.090796 | 0.08 | 0.101 | 1 | 11 | Gpaa1 | -0.021 | Young |
| 0.552372 | 2.302596 | 0.014 | 0.008 | 1 | 11 | Vsig4 | 0.006 | Young |
| 0.552372 | 1.622694 | 0.014 | 0.008 | 1 | 11 | Med12l | 0.006 | Young |
| 0.552372 | 0.945007 | 0.014 | 0.008 | 1 | 11 | Taf1a | 0.006 | Young |
| 0.552372 | 0.785685 | 0.014 | 0.008 | 1 | 11 | Thsd4 | 0.006 | Young |
| 0.552372 | 0.759627 | 0.014 | 0.008 | 1 | 11 | Rasip1 | 0.006 | Young |
| 0.552372 | 0.675323 | 0.014 | 0.008 | 1 | 11 | Kif18a | 0.006 | Young |
| 0.552372 | 0.659776 | 0.014 | 0.008 | 1 | 11 | Cep170b | 0.006 | Young |
| 0.552372 | 0.572223 | 0.014 | 0.008 | 1 | 11 | Gm28151 | 0.006 | Young |
| 0.552372 | 0.55375 | 0.014 | 0.008 | 1 | 11 | Pde5a | 0.006 | Young |
| 0.552372 | 0.52594 | 0.014 | 0.008 | 1 | 11 | Kif9 | 0.006 | Young |
| 0.552372 | 0.519844 | 0.014 | 0.008 | 1 | 11 | Slc25a42 | 0.006 | Young |
| 0.552372 | 0.448738 | 0.014 | 0.008 | 1 | 11 | Zbtb48 | 0.006 | Young |
| 0.552372 | 0.434078 | 0.014 | 0.008 | 1 | 11 | Gm12689 | 0.006 | Young |
| 0.552372 | 0.432118 | 0.014 | 0.008 | 1 | 11 | Oplah | 0.006 | Young |
| 0.552372 | 0.409659 | 0.014 | 0.008 | 1 | 11 | Cped1 | 0.006 | Young |
| 0.552372 | 0.369752 | 0.014 | 0.008 | 1 | 11 | Enkd1 | 0.006 | Young |
| 0.552372 | 0.362617 | 0.014 | 0.008 | 1 | 11 | Prtn3 | 0.006 | Young |
| 0.552372 | 0.350871 | 0.014 | 0.008 | 1 | 11 | Mgst2 | 0.006 | Young |
| 0.552372 | 0.300313 | 0.014 | 0.008 | 1 | 11 | Cdkl3 | 0.006 | Young |
| 0.552372 | 0.221837 | 0.014 | 0.008 | 1 | 11 | Ttll1 | 0.006 | Young |
| 0.552372 | 0.220633 | 0.014 | 0.008 | 1 | 11 | Ctsk | 0.006 | Young |
| 0.552372 | 0.190555 | 0.014 | 0.008 | 1 | 11 | Sall2 | 0.006 | Young |
| 0.552372 | 0.074207 | 0.014 | 0.008 | 1 | 11 | Ttc25 | 0.006 | Young |
| 0.552372 | 0.041248 | 0.014 | 0.008 | 1 | 11 | Fbxo10 | 0.006 | Young |
| 0.552392 | 0.187984 | 0.058 | 0.044 | 1 | 11 | Depdc5 | 0.014 | Young |
| 0.552936 | 0.831542 | 0.094 | 0.121 | 1 | 11 | Aga | -0.027 | Young |
| 0.55358 | 0.438941 | 0.13 | 0.117 | 1 | 11 | Retreg1 | 0.013 | Young |
| 0.553746 | 1.55885 | 0.043 | 0.032 | 1 | 11 | Thap1 | 0.011 | Young |
| 0.553784 | 0.78583 | 0.152 | 0.141 | 1 | 11 | Upf3a | 0.011 | Young |
| 0.553833 | 0.063442 | 0.746 | 0.831 | 1 | 11 | H2-DMa | -0.085 | Young |

| | | | | | | | | |
|---|---|---|---|---|---|---|---|---|
| 0.553902 | 0.353343 | 0.312 | 0.395 | 1 | 11 | Ap2b1 | -0.083 | Young |
| 0.553957 | 0.059917 | 0.101 | 0.125 | 1 | 11 | Taf3 | -0.024 | Young |
| 0.554396 | 0.23304 | 0.116 | 0.141 | 1 | 11 | Nup62 | -0.025 | Young |
| 0.554802 | 0.265814 | 0.319 | 0.302 | 1 | 11 | Sema4d | 0.017 | Young |
| 0.555538 | 0.040475 | 0.109 | 0.133 | 1 | 11 | Hdac9 | -0.024 | Young |
| 0.555538 | 0.002742 | 0.109 | 0.133 | 1 | 11 | Pknox1 | -0.024 | Young |
| 0.555543 | 0.300554 | 0.116 | 0.145 | 1 | 11 | Impa1 | -0.029 | Young |
| 0.555644 | 0.401278 | 0.486 | 0.504 | 1 | 11 | Oasl2 | -0.018 | Young |
| 0.555715 | 0.811631 | 0.043 | 0.032 | 1 | 11 | Mccc2 | 0.011 | Young |
| 0.556006 | 0.574736 | 0.014 | 0.008 | 1 | 11 | Gm16740 | 0.006 | Young |
| 0.556006 | 0.420255 | 0.014 | 0.008 | 1 | 11 | C920021L1 | 0.006 | Young |
| 0.556006 | 0.323509 | 0.014 | 0.008 | 1 | 11 | Igfbp6 | 0.006 | Young |
| 0.556006 | 0.181157 | 0.014 | 0.008 | 1 | 11 | Spata2l | 0.006 | Young |
| 0.556006 | 0.176448 | 0.014 | 0.008 | 1 | 11 | Zfp661 | 0.006 | Young |
| 0.556009 | 0.377057 | 0.087 | 0.109 | 1 | 11 | Hif1an | -0.022 | Young |
| 0.556222 | 0.391338 | 0.065 | 0.085 | 1 | 11 | Nrm | -0.02 | Young |
| 0.556222 | 0.241376 | 0.065 | 0.085 | 1 | 11 | Relt | -0.02 | Young |
| 0.55644 | 0.243386 | 0.094 | 0.117 | 1 | 11 | Ern1 | -0.023 | Young |
| 0.55644 | 0.121962 | 0.094 | 0.117 | 1 | 11 | Usp50 | -0.023 | Young |
| 0.557015 | 0.333575 | 0.37 | 0.476 | 1 | 11 | Bmp2k | -0.106 | Young |
| 0.557035 | 0.090954 | 0.457 | 0.536 | 1 | 11 | Irgm1 | -0.079 | Young |
| 0.557306 | 0.046868 | 0.942 | 0.96 | 1 | 11 | Ctsc | -0.018 | Young |
| 0.55761 | 0.005164 | 0.826 | 0.907 | 1 | 11 | Lcp1 | -0.081 | Young |
| 0.557767 | 0.181233 | 0.109 | 0.133 | 1 | 11 | Ifi44 | -0.024 | Young |
| 0.558026 | 0.524868 | 0.225 | 0.278 | 1 | 11 | Rbm6 | -0.053 | Young |
| 0.558331 | 0.086394 | 0.297 | 0.351 | 1 | 11 | Furin | -0.054 | Young |
| 0.558418 | 0.161958 | 0.312 | 0.383 | 1 | 11 | Sin3b | -0.071 | Young |
| 0.558454 | 0.544567 | 0.087 | 0.109 | 1 | 11 | Thra | -0.022 | Young |
| 0.558505 | 0.190682 | 0.399 | 0.488 | 1 | 11 | Tor3a | -0.089 | Young |
| 0.558517 | 0.70668 | 0.109 | 0.137 | 1 | 11 | Cbll1 | -0.028 | Young |
| 0.558867 | 0.731351 | 0.362 | 0.375 | 1 | 11 | Cmtm7 | -0.013 | Young |
| 0.558926 | 1.823001 | 0.029 | 0.02 | 1 | 11 | B3gnt5 | 0.009 | Young |
| 0.559287 | 0.22952 | 0.232 | 0.286 | 1 | 11 | Btk | -0.054 | Young |
| 0.559653 | 0.121767 | 0.014 | 0.008 | 1 | 11 | 4930570G1 | 0.006 | Young |
| 0.559653 | 0.005943 | 0.014 | 0.008 | 1 | 11 | Ctla2b | 0.006 | Young |
| 0.559696 | 0.289897 | 0.486 | 0.536 | 1 | 11 | Fam174a | -0.05 | Young |
| 0.559863 | 0.580362 | 0.377 | 0.407 | 1 | 11 | Camk1d | -0.03 | Young |
| 0.559954 | 0.732386 | 0.094 | 0.121 | 1 | 11 | Snx14 | -0.027 | Young |
| 0.559984 | 0.23835 | 0.022 | 0.032 | 1 | 11 | Naif1 | -0.01 | Young |
| 0.560415 | 0.664514 | 0.138 | 0.173 | 1 | 11 | Gse1 | -0.035 | Young |
| 0.560628 | 0.111521 | 0.152 | 0.181 | 1 | 11 | Man2a2 | -0.029 | Young |
| 0.560732 | 0.918892 | 0.109 | 0.137 | 1 | 11 | Asb8 | -0.028 | Young |
| 0.561378 | 1.746975 | 0.029 | 0.02 | 1 | 11 | Tdp1 | 0.009 | Young |
| 0.561912 | 0.393944 | 0.072 | 0.093 | 1 | 11 | Anks3 | -0.021 | Young |
| 0.56221 | 0.973872 | 0.022 | 0.032 | 1 | 11 | Zbed3 | -0.01 | Young |

| | | | | | | | | |
|---|---|---|---|---|---|---|---|---|
| 0.56221 | 0.693517 | 0.022 | 0.032 | 1 | 11 | Gm38843 | -0.01 | Young |
| 0.56221 | 0.65031 | 0.022 | 0.032 | 1 | 11 | Eif5a2 | -0.01 | Young |
| 0.56221 | 0.216479 | 0.022 | 0.032 | 1 | 11 | Zfp276 | -0.01 | Young |
| 0.562672 | 0.790401 | 0.225 | 0.218 | 1 | 11 | Mecp2 | 0.007 | Young |
| 0.563223 | 1.314759 | 0.065 | 0.052 | 1 | 11 | Sema4c | 0.013 | Young |
| 0.563704 | 0.232404 | 0.08 | 0.101 | 1 | 11 | Ing3 | -0.021 | Young |
| 0.563723 | 0.971992 | 0.13 | 0.117 | 1 | 11 | Prpf6 | 0.013 | Young |
| 0.564439 | 0.342222 | 0.022 | 0.032 | 1 | 11 | Olfr56 | -0.01 | Young |
| 0.564439 | 0.294367 | 0.022 | 0.032 | 1 | 11 | Gm43936 | -0.01 | Young |
| 0.564439 | 0.268906 | 0.022 | 0.032 | 1 | 11 | Map3k10 | -0.01 | Young |
| 0.564439 | 0.131474 | 0.022 | 0.032 | 1 | 11 | Gm26669 | -0.01 | Young |
| 0.564465 | 0.156526 | 0.449 | 0.556 | 1 | 11 | Fam111a | -0.107 | Young |
| 0.564589 | 0.317317 | 0.087 | 0.109 | 1 | 11 | Stx3 | -0.022 | Young |
| 0.564616 | 0.263435 | 0.261 | 0.302 | 1 | 11 | Emb | -0.041 | Young |
| 0.564949 | 0.419983 | 0.123 | 0.153 | 1 | 11 | Mms19 | -0.03 | Young |
| 0.565602 | 0.233389 | 0.203 | 0.242 | 1 | 11 | Sult1a1 | -0.039 | Young |
| 0.565934 | 0.147739 | 0.094 | 0.117 | 1 | 11 | Eya3 | -0.023 | Young |
| 0.566288 | 1.004016 | 0.109 | 0.137 | 1 | 11 | Mplkip | -0.028 | Young |
| 0.566297 | 2.158179 | 0.029 | 0.02 | 1 | 11 | Zfp523 | 0.009 | Young |
| 0.566297 | 1.43358 | 0.029 | 0.02 | 1 | 11 | Urb2 | 0.009 | Young |
| 0.566297 | 1.428216 | 0.029 | 0.02 | 1 | 11 | Fam72a | 0.009 | Young |
| 0.566347 | 0.253432 | 0.355 | 0.452 | 1 | 11 | Sdf4 | -0.097 | Young |
| 0.566381 | 0.394598 | 0.203 | 0.246 | 1 | 11 | Rab12 | -0.043 | Young |
| 0.566673 | 0.085771 | 0.022 | 0.032 | 1 | 11 | Pcca | -0.01 | Young |
| 0.567554 | 0.585084 | 0.196 | 0.234 | 1 | 11 | Tm2d2 | -0.038 | Young |
| 0.567603 | 0.033802 | 0.029 | 0.04 | 1 | 11 | Cep44 | -0.011 | Young |
| 0.568031 | 0.550407 | 0.065 | 0.052 | 1 | 11 | Ikbkap | 0.013 | Young |
| 0.568043 | 1.06845 | 0.101 | 0.129 | 1 | 11 | Ap1ar | -0.028 | Young |
| 0.568285 | 0.534397 | 0.087 | 0.109 | 1 | 11 | 2810004N2 | -0.022 | Young |
| 0.568764 | 1.747571 | 0.029 | 0.02 | 1 | 11 | Gm16973 | 0.009 | Young |
| 0.568764 | 1.518218 | 0.029 | 0.02 | 1 | 11 | 9330160F1 | 0.009 | Young |
| 0.568764 | 1.364361 | 0.029 | 0.02 | 1 | 11 | Glis2 | 0.009 | Young |
| 0.568764 | 1.230654 | 0.029 | 0.02 | 1 | 11 | Lyrm4 | 0.009 | Young |
| 0.568764 | 1.125613 | 0.029 | 0.02 | 1 | 11 | Pop1 | 0.009 | Young |
| 0.568764 | 1.108996 | 0.029 | 0.02 | 1 | 11 | 2810454H0 | 0.009 | Young |
| 0.568764 | 0.93121 | 0.029 | 0.02 | 1 | 11 | Itga1 | 0.009 | Young |
| 0.568912 | 0.342046 | 0.022 | 0.032 | 1 | 11 | Gstz1 | -0.01 | Young |
| 0.568912 | 0.281969 | 0.022 | 0.032 | 1 | 11 | Timmdc1 | -0.01 | Young |
| 0.568912 | 0.275532 | 0.022 | 0.032 | 1 | 11 | Aplnr | -0.01 | Young |
| 0.568912 | 0.172465 | 0.022 | 0.032 | 1 | 11 | Ncapd3 | -0.01 | Young |
| 0.568912 | 0.065418 | 0.022 | 0.032 | 1 | 11 | Nop2 | -0.01 | Young |
| 0.568979 | 0.166131 | 0.109 | 0.133 | 1 | 11 | Bex3 | -0.024 | Young |
| 0.569334 | 0.183605 | 0.196 | 0.234 | 1 | 11 | Arl6ip5 | -0.038 | Young |
| 0.569476 | 0.711191 | 0.138 | 0.173 | 1 | 11 | Scamp5 | -0.035 | Young |
| 0.56952 | 0.166789 | 0.145 | 0.173 | 1 | 11 | Zfp592 | -0.028 | Young |

| | | | | | | | | |
|---|---|---|---|---|---|---|---|---|
| 0.569686 | 0.289174 | 0.101 | 0.085 | 1 | 11 | Zfyve27 | 0.016 | Young |
| 0.569864 | 0.139888 | 0.275 | 0.335 | 1 | 11 | Med28 | -0.06 | Young |
| 0.570029 | 0.988307 | 0.174 | 0.161 | 1 | 11 | Igf1r | 0.013 | Young |
| 0.570084 | 0.436781 | 0.101 | 0.125 | 1 | 11 | 2610002M | -0.024 | Young |
| 0.570247 | 0.182191 | 0.442 | 0.536 | 1 | 11 | Rtn3 | -0.094 | Young |
| 0.570344 | 0.674827 | 0.413 | 0.456 | 1 | 11 | Ptger4 | -0.043 | Young |
| 0.570444 | 1.268447 | 0.145 | 0.137 | 1 | 11 | Acox3 | 0.008 | Young |
| 0.571154 | 0.310686 | 0.022 | 0.032 | 1 | 11 | Tmem86b | -0.01 | Young |
| 0.571237 | 1.625031 | 0.029 | 0.02 | 1 | 11 | Fam102a | 0.009 | Young |
| 0.571237 | 1.158112 | 0.029 | 0.02 | 1 | 11 | Slc5a6 | 0.009 | Young |
| 0.571237 | 1.066721 | 0.029 | 0.02 | 1 | 11 | Lppos | 0.009 | Young |
| 0.573098 | 0.224129 | 0.283 | 0.339 | 1 | 11 | Smc1a | -0.056 | Young |
| 0.573169 | 0.263573 | 0.638 | 0.758 | 1 | 11 | Fam46a | -0.12 | Young |
| 0.5733 | 0.874167 | 0.101 | 0.089 | 1 | 11 | Mfn2 | 0.012 | Young |
| 0.573308 | 0.419214 | 0.159 | 0.194 | 1 | 11 | Arhgap12 | -0.035 | Young |
| 0.5734 | 0.721453 | 0.022 | 0.032 | 1 | 11 | BC024978 | -0.01 | Young |
| 0.5734 | 0.549297 | 0.022 | 0.032 | 1 | 11 | Gba2 | -0.01 | Young |
| 0.57348 | 0.674465 | 0.058 | 0.077 | 1 | 11 | Ccs | -0.019 | Young |
| 0.573592 | 0.323711 | 0.043 | 0.032 | 1 | 11 | Rbm41 | 0.011 | Young |
| 0.573714 | 0.976055 | 0.029 | 0.02 | 1 | 11 | Btbd6 | 0.009 | Young |
| 0.573714 | 0.973801 | 0.029 | 0.02 | 1 | 11 | Ptchd1 | 0.009 | Young |
| 0.573714 | 0.658567 | 0.029 | 0.02 | 1 | 11 | Ift57 | 0.009 | Young |
| 0.573942 | 0.787982 | 0.08 | 0.101 | 1 | 11 | Dcaf11 | -0.021 | Young |
| 0.573942 | 0.265676 | 0.08 | 0.101 | 1 | 11 | Manea | -0.021 | Young |
| 0.573942 | 0.255332 | 0.08 | 0.101 | 1 | 11 | Prr5l | -0.021 | Young |
| 0.574475 | 0.866394 | 0.145 | 0.181 | 1 | 11 | Hdac3 | -0.036 | Young |
| 0.574516 | 0.946132 | 0.152 | 0.137 | 1 | 11 | Pilra | 0.015 | Young |
| 0.574585 | 0.917391 | 0.101 | 0.089 | 1 | 11 | Cdk2 | 0.012 | Young |
| 0.574625 | 0.491566 | 0.109 | 0.133 | 1 | 11 | Clpx | -0.024 | Young |
| 0.574952 | 0.384234 | 0.188 | 0.173 | 1 | 11 | Osm | 0.015 | Young |
| 0.575595 | 0.20743 | 0.029 | 0.04 | 1 | 11 | Gm20300 | -0.011 | Young |
| 0.575651 | 1.399618 | 0.022 | 0.032 | 1 | 11 | Hist2h2be | -0.01 | Young |
| 0.575651 | 0.69891 | 0.022 | 0.032 | 1 | 11 | Haus1 | -0.01 | Young |
| 0.575651 | 0.623455 | 0.022 | 0.032 | 1 | 11 | Zkscan8 | -0.01 | Young |
| 0.575757 | 0.384543 | 0.109 | 0.133 | 1 | 11 | Trove2 | -0.024 | Young |
| 0.576055 | 0.527386 | 0.37 | 0.391 | 1 | 11 | Tgif1 | -0.021 | Young |
| 0.576197 | 1.203772 | 0.029 | 0.02 | 1 | 11 | Zfp52 | 0.009 | Young |
| 0.576197 | 1.064273 | 0.029 | 0.02 | 1 | 11 | Acad11 | 0.009 | Young |
| 0.576197 | 0.546024 | 0.029 | 0.02 | 1 | 11 | D3Ertd751 | 0.009 | Young |
| 0.576562 | 0.423293 | 0.072 | 0.093 | 1 | 11 | Hbs1l | -0.021 | Young |
| 0.577344 | 0.016615 | 0.478 | 0.589 | 1 | 11 | Dse | -0.111 | Young |
| 0.577437 | 0.642585 | 0.275 | 0.351 | 1 | 11 | Aoah | -0.076 | Young |
| 0.577452 | 0.283245 | 0.188 | 0.234 | 1 | 11 | Tug1 | -0.046 | Young |
| 0.577602 | 0.128197 | 0.029 | 0.04 | 1 | 11 | Lamc1 | -0.011 | Young |
| 0.577906 | 0.456916 | 0.022 | 0.032 | 1 | 11 | Abtb2 | -0.01 | Young |

| | | | | | | | | |
|---|---|---|---|---|---|---|---|---|
| 0.578039 | 0.491136 | 0.145 | 0.177 | 1 | 11 | Eif4enif1 | -0.032 | Young |
| 0.578591 | 0.4542 | 0.138 | 0.169 | 1 | 11 | Ncor2 | -0.031 | Young |
| 0.578685 | 0.734023 | 0.029 | 0.02 | 1 | 11 | Tnks1bp1 | 0.009 | Young |
| 0.578685 | 0.598291 | 0.029 | 0.02 | 1 | 11 | Rpl9-ps6 | 0.009 | Young |
| 0.578685 | 0.432764 | 0.029 | 0.02 | 1 | 11 | Smpd3 | 0.009 | Young |
| 0.579103 | 0.499687 | 0.138 | 0.169 | 1 | 11 | Fnbp4 | -0.031 | Young |
| 0.579358 | 0.382932 | 0.08 | 0.065 | 1 | 11 | Slc30a4 | 0.015 | Young |
| 0.579566 | 0.588961 | 0.101 | 0.129 | 1 | 11 | Fbxo38 | -0.028 | Young |
| 0.579611 | 0.693318 | 0.029 | 0.04 | 1 | 11 | Pigg | -0.011 | Young |
| 0.580029 | 0.459797 | 0.051 | 0.065 | 1 | 11 | 2410002F2 | -0.014 | Young |
| 0.58008 | 0.439417 | 0.159 | 0.194 | 1 | 11 | Kbtbd2 | -0.035 | Young |
| 0.580129 | 0.270152 | 0.246 | 0.294 | 1 | 11 | Kmt2d | -0.048 | Young |
| 0.580165 | 1.689035 | 0.022 | 0.032 | 1 | 11 | Lss | -0.01 | Young |
| 0.580165 | 0.953158 | 0.022 | 0.032 | 1 | 11 | Invs | -0.01 | Young |
| 0.581177 | 0.811981 | 0.029 | 0.02 | 1 | 11 | Nckap5l | 0.009 | Young |
| 0.581624 | 1.116008 | 0.029 | 0.04 | 1 | 11 | Rfx3 | -0.011 | Young |
| 0.581749 | 0.173886 | 0.283 | 0.339 | 1 | 11 | 5-Mar | -0.056 | Young |
| 0.581805 | 0.474652 | 0.116 | 0.141 | 1 | 11 | Ankra2 | -0.025 | Young |
| 0.581934 | 1.045865 | 0.094 | 0.081 | 1 | 11 | Calcrl | 0.013 | Young |
| 0.58218 | 1.946827 | 0.051 | 0.04 | 1 | 11 | Ppox | 0.011 | Young |
| 0.582676 | 0.501648 | 0.138 | 0.165 | 1 | 11 | Susd3 | -0.027 | Young |
| 0.583675 | 0.670951 | 0.029 | 0.02 | 1 | 11 | Sars2 | 0.009 | Young |
| 0.583712 | 0.168655 | 0.138 | 0.165 | 1 | 11 | Pou2f1 | -0.027 | Young |
| 0.584198 | 0.837752 | 0.065 | 0.052 | 1 | 11 | Etfrf1 | 0.013 | Young |
| 0.584245 | 0.428049 | 0.138 | 0.121 | 1 | 11 | Igsf9 | 0.017 | Young |
| 0.584628 | 1.260968 | 0.072 | 0.093 | 1 | 11 | F830016B0 | -0.021 | Young |
| 0.584628 | 0.718265 | 0.094 | 0.081 | 1 | 11 | Afdn | 0.013 | Young |
| 0.585511 | 0.090468 | 0.362 | 0.435 | 1 | 11 | Themis2 | -0.073 | Young |
| 0.58566 | 0.196588 | 0.029 | 0.04 | 1 | 11 | Entpd5 | -0.011 | Young |
| 0.585686 | 0.721729 | 0.188 | 0.173 | 1 | 11 | Adrb2 | 0.015 | Young |
| 0.585858 | 1.996735 | 0.051 | 0.04 | 1 | 11 | Nrg1 | 0.011 | Young |
| 0.585858 | 0.550117 | 0.036 | 0.048 | 1 | 11 | Brat1 | -0.012 | Young |
| 0.585912 | 0.128345 | 0.457 | 0.569 | 1 | 11 | Hcls1 | -0.112 | Young |
| 0.585978 | 0.882525 | 0.094 | 0.081 | 1 | 11 | Abhd13 | 0.013 | Young |
| 0.586178 | 0.539411 | 0.029 | 0.02 | 1 | 11 | Gins2 | 0.009 | Young |
| 0.586178 | 0.301109 | 0.029 | 0.02 | 1 | 11 | Kit | 0.009 | Young |
| 0.586214 | 1.070464 | 0.101 | 0.089 | 1 | 11 | Icosl | 0.012 | Young |
| 0.587502 | 0.51041 | 0.167 | 0.206 | 1 | 11 | Impdh2 | -0.039 | Young |
| 0.58787 | 0.254298 | 0.746 | 0.806 | 1 | 11 | Ifi204 | -0.06 | Young |
| 0.587997 | 0.450178 | 0.051 | 0.065 | 1 | 11 | Gtf2e1 | -0.014 | Young |
| 0.588282 | 0.537049 | 0.109 | 0.133 | 1 | 11 | Zwint | -0.024 | Young |
| 0.588686 | 0.196361 | 0.029 | 0.02 | 1 | 11 | Cwf19l1 | 0.009 | Young |
| 0.588686 | 0.138229 | 0.029 | 0.02 | 1 | 11 | Olr1 | 0.009 | Young |
| 0.589547 | 0.022764 | 0.036 | 0.048 | 1 | 11 | Pnkp | -0.012 | Young |
| 0.589708 | 1.140595 | 0.029 | 0.04 | 1 | 11 | Snrnp35 | -0.011 | Young |

| | | | | | | | | |
|---|---|---|---|---|---|---|---|---|
| 0.589708 | 0.700434 | 0.029 | 0.04 | 1 | 11 | Tbc1d30 | -0.011 | Young |
| 0.58984 | 0.565037 | 0.254 | 0.306 | 1 | 11 | Acat1 | -0.052 | Young |
| 0.589996 | 0.264283 | 0.196 | 0.234 | 1 | 11 | Otud5 | -0.038 | Young |
| 0.590429 | 0.598365 | 0.13 | 0.161 | 1 | 11 | Chchd3 | -0.031 | Young |
| 0.590708 | 0.093127 | 0.949 | 0.972 | 1 | 11 | Rpl38 | -0.023 | Young |
| 0.591395 | 1.51592 | 0.051 | 0.04 | 1 | 11 | Zc3h12d | 0.011 | Young |
| 0.591584 | 0.718097 | 0.116 | 0.145 | 1 | 11 | Scyl2 | -0.029 | Young |
| 0.591737 | 0.380484 | 0.029 | 0.04 | 1 | 11 | Ercc8 | -0.011 | Young |
| 0.591737 | 0.330623 | 0.029 | 0.04 | 1 | 11 | Zfp653 | -0.011 | Young |
| 0.59262 | 0.460579 | 0.239 | 0.294 | 1 | 11 | Chic2 | -0.055 | Young |
| 0.592795 | 0.853401 | 0.152 | 0.137 | 1 | 11 | Nudt9 | 0.015 | Young |
| 0.592795 | 0.46934 | 0.152 | 0.137 | 1 | 11 | Mier3 | 0.015 | Young |
| 0.592868 | 0.573933 | 0.109 | 0.133 | 1 | 11 | Cpd | -0.024 | Young |
| 0.593076 | 0.22542 | 0.138 | 0.165 | 1 | 11 | Rnmt | -0.027 | Young |
| 0.593246 | 0.454122 | 0.036 | 0.048 | 1 | 11 | Irgq | -0.012 | Young |
| 0.593307 | 0.177765 | 0.043 | 0.056 | 1 | 11 | Atg14 | -0.013 | Young |
| 0.59377 | 0.776474 | 0.029 | 0.04 | 1 | 11 | Nif3l1 | -0.011 | Young |
| 0.59377 | 0.604972 | 0.029 | 0.04 | 1 | 11 | 1600020E0 | -0.011 | Young |
| 0.59377 | 0.377727 | 0.029 | 0.04 | 1 | 11 | Atrip | -0.011 | Young |
| 0.59377 | 0.06241 | 0.029 | 0.04 | 1 | 11 | Trim32 | -0.011 | Young |
| 0.594284 | 0.270222 | 0.174 | 0.21 | 1 | 11 | Mex3c | -0.036 | Young |
| 0.59468 | 0.92803 | 0.08 | 0.101 | 1 | 11 | Tef | -0.021 | Young |
| 0.595099 | 0.713965 | 0.036 | 0.048 | 1 | 11 | Pomgnt1 | -0.012 | Young |
| 0.595099 | 0.701807 | 0.036 | 0.048 | 1 | 11 | Alyref2 | -0.012 | Young |
| 0.595785 | 0.036692 | 0.768 | 0.815 | 1 | 11 | mt-Nd5 | -0.047 | Young |
| 0.595805 | 0.843918 | 0.029 | 0.04 | 1 | 11 | Cdpf1 | -0.011 | Young |
| 0.595805 | 0.202542 | 0.029 | 0.04 | 1 | 11 | Kif3a | -0.011 | Young |
| 0.59598 | 0.162138 | 0.203 | 0.238 | 1 | 11 | Fam76a | -0.035 | Young |
| 0.596037 | 0.138753 | 0.391 | 0.5 | 1 | 11 | Srp14 | -0.109 | Young |
| 0.596093 | 0.519219 | 0.094 | 0.117 | 1 | 11 | Farsa | -0.023 | Young |
| 0.596192 | 0.861468 | 0.232 | 0.234 | 1 | 11 | Tnrc6c | -0.002 | Young |
| 0.596224 | 0.385296 | 0.13 | 0.157 | 1 | 11 | Preb | -0.027 | Young |
| 0.596304 | 0.196189 | 0.116 | 0.141 | 1 | 11 | Smcr8 | -0.025 | Young |
| 0.596353 | 0.237189 | 0.196 | 0.234 | 1 | 11 | Kdelr2 | -0.038 | Young |
| 0.596357 | 0.370596 | 0.283 | 0.347 | 1 | 11 | Emc7 | -0.064 | Young |
| 0.596619 | 0.028261 | 0.486 | 0.581 | 1 | 11 | Slc38a2 | -0.095 | Young |
| 0.596702 | 0.024869 | 0.609 | 0.734 | 1 | 11 | Cyfip1 | -0.125 | Young |
| 0.596749 | 0.280486 | 0.058 | 0.073 | 1 | 11 | Bloc1s5 | -0.015 | Young |
| 0.596955 | 0.390926 | 0.036 | 0.048 | 1 | 11 | Cpt2 | -0.012 | Young |
| 0.596985 | 0.342045 | 0.21 | 0.254 | 1 | 11 | Dcaf12 | -0.044 | Young |
| 0.597018 | 0.48816 | 0.167 | 0.202 | 1 | 11 | Plekhf2 | -0.035 | Young |
| 0.597117 | 0.074045 | 0.609 | 0.746 | 1 | 11 | Pitpna | -0.137 | Young |
| 0.597132 | 0.051603 | 0.217 | 0.254 | 1 | 11 | Taf1d | -0.037 | Young |
| 0.597429 | 0.507458 | 0.188 | 0.234 | 1 | 11 | Rnh1 | -0.046 | Young |
| 0.597844 | 0.490637 | 0.029 | 0.04 | 1 | 11 | Brms1l | -0.011 | Young |

| | | | | | | | | |
|---|---|---|---|---|---|---|---|---|
| 0.597951 | 1.301816 | 0.101 | 0.089 | 1 | 11 | Polr2h | 0.012 | Young |
| 0.598042 | 0.92794 | 0.174 | 0.218 | 1 | 11 | Acadl | -0.044 | Young |
| 0.598148 | 0.147646 | 0.543 | 0.629 | 1 | 11 | Psma7 | -0.086 | Young |
| 0.598778 | 0.422602 | 0.493 | 0.548 | 1 | 11 | Ptpre | -0.055 | Young |
| 0.598814 | 0.079614 | 0.036 | 0.048 | 1 | 11 | Gprasp1 | -0.012 | Young |
| 0.599119 | 0.252346 | 0.297 | 0.367 | 1 | 11 | Ski | -0.07 | Young |
| 0.599235 | 0.026339 | 0.051 | 0.065 | 1 | 11 | D430042O( | -0.014 | Young |
| 0.599265 | 1.072911 | 0.072 | 0.06 | 1 | 11 | Lmbr1l | 0.012 | Young |
| 0.599681 | 0.097353 | 0.833 | 0.915 | 1 | 11 | Rpl9 | -0.082 | Young |
| 0.599917 | 1.315936 | 0.08 | 0.101 | 1 | 11 | Ggps1 | -0.021 | Young |
| 0.600183 | 0.041599 | 0.043 | 0.056 | 1 | 11 | Hagh | -0.013 | Young |
| 0.600483 | 0.504785 | 0.507 | 0.56 | 1 | 11 | Usp25 | -0.053 | Young |
| 0.600676 | 0.508558 | 0.036 | 0.048 | 1 | 11 | Lilra6 | -0.012 | Young |
| 0.600676 | 0.466192 | 0.036 | 0.048 | 1 | 11 | Xpnpep3 | -0.012 | Young |
| 0.600676 | 0.2083 | 0.036 | 0.048 | 1 | 11 | Plpp6 | -0.012 | Young |
| 0.600733 | 0.510109 | 0.159 | 0.198 | 1 | 11 | Vps54 | -0.039 | Young |
| 0.600996 | 0.198917 | 0.326 | 0.383 | 1 | 11 | Irf5 | -0.057 | Young |
| 0.60103 | 0.234279 | 0.225 | 0.266 | 1 | 11 | Chmp3 | -0.041 | Young |
| 0.601802 | 1.048777 | 0.123 | 0.113 | 1 | 11 | Rbbp5 | 0.01 | Young |
| 0.603167 | 0.105105 | 0.42 | 0.508 | 1 | 11 | Eif4h | -0.088 | Young |
| 0.603347 | 0.490713 | 0.261 | 0.335 | 1 | 11 | Orai1 | -0.074 | Young |
| 0.603634 | 0.643251 | 0.043 | 0.056 | 1 | 11 | Rfng | -0.013 | Young |
| 0.603651 | 0.845123 | 0.072 | 0.093 | 1 | 11 | Lyrm2 | -0.021 | Young |
| 0.603716 | 0.442706 | 0.225 | 0.274 | 1 | 11 | Bcl2a1d | -0.049 | Young |
| 0.60388 | 0.382214 | 0.08 | 0.097 | 1 | 11 | Rnf138 | -0.017 | Young |
| 0.603936 | 0.09306 | 0.268 | 0.319 | 1 | 11 | Riok3 | -0.051 | Young |
| 0.603979 | 0.86491 | 0.029 | 0.04 | 1 | 11 | Tamm41 | -0.011 | Young |
| 0.604081 | 0.007255 | 0.051 | 0.065 | 1 | 11 | Gss | -0.014 | Young |
| 0.604406 | 1.256562 | 0.051 | 0.04 | 1 | 11 | A230050P2 | 0.011 | Young |
| 0.604406 | 0.400008 | 0.036 | 0.048 | 1 | 11 | Dnajc4 | -0.012 | Young |
| 0.604767 | 0.364372 | 0.174 | 0.21 | 1 | 11 | Rnf166 | -0.036 | Young |
| 0.604773 | 0.223538 | 0.167 | 0.202 | 1 | 11 | Frrs1 | -0.035 | Young |
| 0.605263 | 0.042587 | 0.261 | 0.298 | 1 | 11 | Fbl | -0.037 | Young |
| 0.605452 | 0.643735 | 0.225 | 0.282 | 1 | 11 | Gxylt1 | -0.057 | Young |
| 0.605773 | 1.289759 | 0.159 | 0.149 | 1 | 11 | Psmc4 | 0.01 | Young |
| 0.606113 | 0.263838 | 0.399 | 0.484 | 1 | 11 | Ndufa7 | -0.085 | Young |
| 0.606255 | 0.229479 | 0.377 | 0.448 | 1 | 11 | Clint1 | -0.071 | Young |
| 0.606276 | 1.109076 | 0.051 | 0.04 | 1 | 11 | 2700097O( | 0.011 | Young |
| 0.606744 | 0.154673 | 0.21 | 0.254 | 1 | 11 | Gak | -0.044 | Young |
| 0.607041 | 0.825947 | 0.072 | 0.06 | 1 | 11 | Dkc1 | 0.012 | Young |
| 0.607458 | 0.358825 | 0.152 | 0.181 | 1 | 11 | Epb41l4ao: | -0.029 | Young |
| 0.608084 | 1.206476 | 0.029 | 0.04 | 1 | 11 | Zfp35 | -0.011 | Young |
| 0.608113 | 0.684156 | 0.377 | 0.415 | 1 | 11 | Dock8 | -0.038 | Young |
| 0.608148 | 0.491885 | 0.036 | 0.048 | 1 | 11 | E2f3 | -0.012 | Young |
| 0.608768 | 0.328615 | 0.239 | 0.286 | 1 | 11 | H1f0 | -0.047 | Young |

| | | | | | | | | |
|---|---|---|---|---|---|---|---|---|
| 0.608828 | 0.161233 | 0.043 | 0.056 | 1 | 11 | Asah2 | -0.013 | Young |
| 0.609021 | 0.223068 | 0.384 | 0.472 | 1 | 11 | Ndufb9 | -0.088 | Young |
| 0.61052 | 0.500687 | 0.174 | 0.21 | 1 | 11 | Ube2a | -0.036 | Young |
| 0.610603 | 0.179373 | 0.065 | 0.081 | 1 | 11 | Aldh1a2 | -0.016 | Young |
| 0.611024 | 0.628225 | 0.13 | 0.117 | 1 | 11 | Gm37494 | 0.013 | Young |
| 0.61127 | 0.473923 | 0.341 | 0.415 | 1 | 11 | Acin1 | -0.074 | Young |
| 0.611899 | 1.400326 | 0.036 | 0.048 | 1 | 11 | Slc35e2 | -0.012 | Young |
| 0.611899 | 0.458674 | 0.036 | 0.048 | 1 | 11 | Il27 | -0.012 | Young |
| 0.612063 | 0.167177 | 0.261 | 0.306 | 1 | 11 | Gpbp1 | -0.045 | Young |
| 0.612199 | 0.321456 | 0.319 | 0.387 | 1 | 11 | Plxnd1 | -0.068 | Young |
| 0.612677 | 1.444746 | 0.116 | 0.105 | 1 | 11 | Timm10 | 0.011 | Young |
| 0.61282 | 0.898839 | 0.109 | 0.137 | 1 | 11 | Rbm34 | -0.028 | Young |
| 0.612897 | 1.342453 | 0.087 | 0.077 | 1 | 11 | Gnptg | 0.01 | Young |
| 0.613097 | 0.126019 | 0.435 | 0.512 | 1 | 11 | S100a10 | -0.077 | Young |
| 0.613518 | 0.67273 | 0.087 | 0.109 | 1 | 11 | Pde4a | -0.022 | Young |
| 0.613762 | 0.077197 | 0.268 | 0.327 | 1 | 11 | Etv6 | -0.059 | Young |
| 0.613778 | 1.002954 | 0.051 | 0.04 | 1 | 11 | Slc26a11 | 0.011 | Young |
| 0.614042 | 0.165094 | 0.043 | 0.056 | 1 | 11 | Asna1 | -0.013 | Young |
| 0.614215 | 0.778496 | 0.152 | 0.185 | 1 | 11 | Sfxn3 | -0.033 | Young |
| 0.614869 | 0.43021 | 0.109 | 0.133 | 1 | 11 | Ccdc34 | -0.024 | Young |
| 0.615376 | 0.605646 | 0.268 | 0.258 | 1 | 11 | Tmx3 | 0.01 | Young |
| 0.615661 | 1.185745 | 0.051 | 0.04 | 1 | 11 | Adck1 | 0.011 | Young |
| 0.61577 | 0.494203 | 0.094 | 0.117 | 1 | 11 | Fhod1 | -0.023 | Young |
| 0.615784 | 0.26336 | 0.043 | 0.056 | 1 | 11 | Rnf8 | -0.013 | Young |
| 0.616431 | 0.565255 | 0.072 | 0.06 | 1 | 11 | Mto1 | 0.012 | Young |
| 0.616647 | 0.298622 | 0.116 | 0.141 | 1 | 11 | Slc5a3 | -0.025 | Young |
| 0.616815 | 0.39752 | 0.529 | 0.605 | 1 | 11 | Tacc1 | -0.076 | Young |
| 0.616911 | 0.038774 | 0.109 | 0.129 | 1 | 11 | Rangap1 | -0.02 | Young |
| 0.61726 | 0.083893 | 0.928 | 0.98 | 1 | 11 | Rpl23 | -0.052 | Young |
| 0.617394 | 1.665163 | 0.08 | 0.069 | 1 | 11 | Cd69 | 0.011 | Young |
| 0.617529 | 0.389902 | 0.043 | 0.056 | 1 | 11 | Pacs1 | -0.013 | Young |
| 0.617545 | 0.813014 | 0.036 | 0.048 | 1 | 11 | Qtrt1 | -0.012 | Young |
| 0.618202 | 0.000593 | 0.891 | 0.944 | 1 | 11 | Rpl37 | -0.053 | Young |
| 0.618641 | 0.722885 | 0.087 | 0.109 | 1 | 11 | Uba3 | -0.022 | Young |
| 0.618818 | 0.373482 | 0.217 | 0.25 | 1 | 11 | Slc30a1 | -0.033 | Young |
| 0.618865 | 0.848027 | 0.196 | 0.246 | 1 | 11 | Lrch3 | -0.05 | Young |
| 0.618896 | 0.391573 | 0.232 | 0.274 | 1 | 11 | Slc30a5 | -0.042 | Young |
| 0.61918 | 0.11046 | 0.203 | 0.238 | 1 | 11 | Tmem184b | -0.035 | Young |
| 0.619225 | 0.07983 | 0.435 | 0.512 | 1 | 11 | Pkm | -0.077 | Young |
| 0.619275 | 0.956985 | 0.043 | 0.056 | 1 | 11 | Poc5 | -0.013 | Young |
| 0.619432 | 0.718379 | 0.036 | 0.048 | 1 | 11 | Wrap73 | -0.012 | Young |
| 0.619432 | 0.658358 | 0.036 | 0.048 | 1 | 11 | Slc25a32 | -0.012 | Young |
| 0.619575 | 0.607215 | 0.072 | 0.06 | 1 | 11 | Ammecr1 | 0.012 | Young |
| 0.619825 | 0.190152 | 0.609 | 0.738 | 1 | 11 | Qk | -0.129 | Young |
| 0.620445 | 0.422787 | 0.159 | 0.19 | 1 | 11 | Tsnax | -0.031 | Young |

| | | | | | | | | |
|---|---|---|---|---|---|---|---|---|
| 0.620459 | 0.932184 | 0.232 | 0.234 | 1 | 11 | Rasa1 | -0.002 | Young |
| 0.621024 | 0.32145 | 0.043 | 0.056 | 1 | 11 | Dnajc30 | -0.013 | Young |
| 0.621192 | 0.103244 | 0.246 | 0.282 | 1 | 11 | Hpgds | -0.036 | Young |
| 0.621209 | 1.495672 | 0.138 | 0.129 | 1 | 11 | Mapk9 | 0.009 | Young |
| 0.621253 | 1.042256 | 0.123 | 0.153 | 1 | 11 | Akap8 | -0.03 | Young |
| 0.621322 | 0.762897 | 0.036 | 0.048 | 1 | 11 | Cd276 | -0.012 | Young |
| 0.621601 | 0.41768 | 0.217 | 0.258 | 1 | 11 | BC005624 | -0.041 | Young |
| 0.621632 | 1.109544 | 0.203 | 0.206 | 1 | 11 | Lpar6 | -0.003 | Young |
| 0.621978 | 0.731055 | 0.094 | 0.117 | 1 | 11 | Nop9 | -0.023 | Young |
| 0.622666 | 0.373187 | 0.08 | 0.097 | 1 | 11 | Slc36a4 | -0.017 | Young |
| 0.622925 | 0.825133 | 0.174 | 0.214 | 1 | 11 | Pbxip1 | -0.04 | Young |
| 0.622956 | 0.302332 | 0.217 | 0.262 | 1 | 11 | Ago3 | -0.045 | Young |
| 0.623149 | 0.851141 | 0.138 | 0.173 | 1 | 11 | Sf3a3 | -0.035 | Young |
| 0.623624 | 0.115322 | 0.087 | 0.105 | 1 | 11 | Vps26b | -0.018 | Young |
| 0.623639 | 1.90267 | 0.065 | 0.056 | 1 | 11 | Thoc5 | 0.009 | Young |
| 0.623639 | 0.69043 | 0.051 | 0.065 | 1 | 11 | Cnpy4 | -0.014 | Young |
| 0.624104 | 0.040133 | 0.558 | 0.661 | 1 | 11 | Snx5 | -0.103 | Young |
| 0.624383 | 0.061353 | 0.058 | 0.073 | 1 | 11 | Inip | -0.015 | Young |
| 0.624386 | 0.462908 | 0.152 | 0.185 | 1 | 11 | Npepl1 | -0.033 | Young |
| 0.62464 | 0.818901 | 0.116 | 0.141 | 1 | 11 | Chpt1 | -0.025 | Young |
| 0.624927 | 0.101421 | 0.087 | 0.105 | 1 | 11 | Lsm14b | -0.018 | Young |
| 0.625065 | 0.813271 | 0.072 | 0.089 | 1 | 11 | Trim11 | -0.017 | Young |
| 0.625065 | 0.318324 | 0.072 | 0.089 | 1 | 11 | Xylt2 | -0.017 | Young |
| 0.625109 | 0.898396 | 0.036 | 0.048 | 1 | 11 | Habp4 | -0.012 | Young |
| 0.625181 | 0.111863 | 0.413 | 0.492 | 1 | 11 | Ep300 | -0.079 | Young |
| 0.625293 | 0.137105 | 0.449 | 0.54 | 1 | 11 | Spred1 | -0.091 | Young |
| 0.625645 | 0.353365 | 0.558 | 0.613 | 1 | 11 | Tra2a | -0.055 | Young |
| 0.626259 | 0.515872 | 0.486 | 0.56 | 1 | 11 | Lpl | -0.074 | Young |
| 0.626493 | 0.453228 | 0.159 | 0.19 | 1 | 11 | Supt6 | -0.031 | Young |
| 0.626925 | 0.394402 | 0.051 | 0.065 | 1 | 11 | Net1 | -0.014 | Young |
| 0.627259 | 0.057962 | 0.514 | 0.629 | 1 | 11 | Nfe2l2 | -0.115 | Young |
| 0.62822 | 1.007575 | 0.123 | 0.113 | 1 | 11 | Nr2c2 | 0.01 | Young |
| 0.62839 | 0.445751 | 0.239 | 0.29 | 1 | 11 | Cnot6 | -0.051 | Young |
| 0.628477 | 0.082267 | 0.101 | 0.121 | 1 | 11 | Foxj3 | -0.02 | Young |
| 0.628934 | 0.23252 | 0.094 | 0.113 | 1 | 11 | Dnajc11 | -0.019 | Young |
| 0.629043 | 0.46768 | 0.058 | 0.073 | 1 | 11 | Acot8 | -0.015 | Young |
| 0.6293 | 0.216105 | 0.072 | 0.089 | 1 | 11 | Borcs8 | -0.017 | Young |
| 0.629554 | 0.501669 | 0.268 | 0.331 | 1 | 11 | Fkbp15 | -0.063 | Young |
| 0.6298 | 0.433452 | 0.043 | 0.056 | 1 | 11 | Fam110a | -0.013 | Young |
| 0.630106 | 0.355267 | 0.217 | 0.254 | 1 | 11 | Dmxl1 | -0.037 | Young |
| 0.630151 | 1.146405 | 0.087 | 0.105 | 1 | 11 | Alg1 | -0.018 | Young |
| 0.630333 | 0.148593 | 0.449 | 0.544 | 1 | 11 | Rab10 | -0.095 | Young |
| 0.630363 | 1.55125 | 0.036 | 0.028 | 1 | 11 | Lin52 | 0.008 | Young |
| 0.630714 | 0.195693 | 0.072 | 0.089 | 1 | 11 | Ly6c2 | -0.017 | Young |
| 0.631073 | 0.179472 | 0.384 | 0.468 | 1 | 11 | Scaf11 | -0.084 | Young |

| | | | | | | | | |
|---|---|---|---|---|---|---|---|---|
| 0.631315 | 0.189869 | 0.819 | 0.843 | 1 | 11 | Rpl32 | -0.024 | Young |
| 0.631525 | 0.136225 | 0.536 | 0.601 | 1 | 11 | Clec12a | -0.065 | Young |
| 0.631814 | 0.049009 | 0.239 | 0.278 | 1 | 11 | Crybg1 | -0.039 | Young |
| 0.632293 | 0.120929 | 0.217 | 0.242 | 1 | 11 | Ptgs1 | -0.025 | Young |
| 0.632571 | 0.156416 | 0.138 | 0.161 | 1 | 11 | Trmt1l | -0.023 | Young |
| 0.632614 | 1.536937 | 0.036 | 0.028 | 1 | 11 | Rfesd | 0.008 | Young |
| 0.633079 | 1.095181 | 0.13 | 0.121 | 1 | 11 | Trrap | 0.009 | Young |
| 0.633212 | 0.219104 | 0.63 | 0.738 | 1 | 11 | Clic4 | -0.108 | Young |
| 0.633546 | 0.588879 | 0.072 | 0.089 | 1 | 11 | Hmces | -0.017 | Young |
| 0.634611 | 0.21904 | 0.101 | 0.121 | 1 | 11 | Zcchc9 | -0.02 | Young |
| 0.634619 | 1.470301 | 0.036 | 0.048 | 1 | 11 | Lzic | -0.012 | Young |
| 0.634869 | 1.53441 | 0.036 | 0.028 | 1 | 11 | Rasal3 | 0.008 | Young |
| 0.634965 | 0.867748 | 0.072 | 0.089 | 1 | 11 | Acvr1 | -0.017 | Young |
| 0.635091 | 0.508998 | 0.043 | 0.056 | 1 | 11 | Thrb | -0.013 | Young |
| 0.635278 | 0.673493 | 0.058 | 0.073 | 1 | 11 | Zbtb6 | -0.015 | Young |
| 0.6354 | 0.528904 | 0.072 | 0.06 | 1 | 11 | Slc25a44 | 0.012 | Young |
| 0.636384 | 0.416654 | 0.072 | 0.089 | 1 | 11 | Tmtc3 | -0.017 | Young |
| 0.636384 | 0.181397 | 0.072 | 0.089 | 1 | 11 | Rab10os | -0.017 | Young |
| 0.636671 | 0.673586 | 0.275 | 0.278 | 1 | 11 | Bcl6 | -0.003 | Young |
| 0.638485 | 0.493856 | 0.051 | 0.065 | 1 | 11 | Sclt1 | -0.014 | Young |
| 0.63861 | 0.375541 | 0.065 | 0.081 | 1 | 11 | Gcc1 | -0.016 | Young |
| 0.638967 | 0.287091 | 0.196 | 0.238 | 1 | 11 | Tbc1d10b | -0.042 | Young |
| 0.639338 | 0.114981 | 0.087 | 0.105 | 1 | 11 | 0610037L1 | -0.018 | Young |
| 0.639598 | 0.529638 | 0.333 | 0.347 | 1 | 11 | Serpinb8 | -0.014 | Young |
| 0.639864 | 0.14314 | 0.717 | 0.746 | 1 | 11 | Adgre1 | -0.029 | Young |
| 0.639971 | 0.183047 | 0.058 | 0.073 | 1 | 11 | Tubgcp3 | -0.015 | Young |
| 0.640135 | 0.089186 | 0.333 | 0.391 | 1 | 11 | Tnks2 | -0.058 | Young |
| 0.640257 | 0.602545 | 0.145 | 0.177 | 1 | 11 | Pla2g16 | -0.032 | Young |
| 0.640651 | 0.733136 | 0.072 | 0.089 | 1 | 11 | Agfg2 | -0.017 | Young |
| 0.640702 | 0.236426 | 0.109 | 0.129 | 1 | 11 | Csad | -0.02 | Young |
| 0.640763 | 0.27416 | 0.63 | 0.726 | 1 | 11 | Pnrc1 | -0.096 | Young |
| 0.641392 | 0.27858 | 0.116 | 0.137 | 1 | 11 | Naa10 | -0.021 | Young |
| 0.641445 | 0.086321 | 0.08 | 0.069 | 1 | 11 | Rpusd4 | 0.011 | Young |
| 0.641531 | 0.105943 | 0.594 | 0.69 | 1 | 11 | Rab14 | -0.096 | Young |
| 0.641614 | 0.144393 | 0.13 | 0.153 | 1 | 11 | Eri1 | -0.023 | Young |
| 0.641654 | 1.520636 | 0.036 | 0.028 | 1 | 11 | Cmc4 | 0.008 | Young |
| 0.641654 | 1.200515 | 0.036 | 0.028 | 1 | 11 | Wars2 | 0.008 | Young |
| 0.641778 | 0.077745 | 0.072 | 0.06 | 1 | 11 | Ints13 | 0.012 | Young |
| 0.641899 | 0.397858 | 0.384 | 0.48 | 1 | 11 | Sqstm1 | -0.096 | Young |
| 0.642529 | 0.235952 | 0.348 | 0.407 | 1 | 11 | Kmt2e | -0.059 | Young |
| 0.642821 | 0.114624 | 0.159 | 0.185 | 1 | 11 | Pdlim1 | -0.026 | Young |
| 0.642943 | 0.676616 | 0.297 | 0.302 | 1 | 11 | Nlrp3 | -0.005 | Young |
| 0.642998 | 0.775953 | 0.13 | 0.157 | 1 | 11 | Slc35a3 | -0.027 | Young |
| 0.64307 | 0.824301 | 0.08 | 0.097 | 1 | 11 | Bckdha | -0.017 | Young |
| 0.643094 | 0.212943 | 0.362 | 0.44 | 1 | 11 | Elavl1 | -0.078 | Young |

| | | | | | | | | |
|---|---|---|---|---|---|---|---|---|
| 0.643108 | 0.837884 | 0.058 | 0.073 | 1 | 11 | Rad51d | -0.015 | Young |
| 0.643108 | 0.029908 | 0.058 | 0.073 | 1 | 11 | Eri3 | -0.015 | Young |
| 0.643923 | 1.257831 | 0.036 | 0.028 | 1 | 11 | Cenpt | 0.008 | Young |
| 0.644233 | 0.438387 | 0.116 | 0.141 | 1 | 11 | Lipo3 | -0.025 | Young |
| 0.64521 | 0.651415 | 0.152 | 0.145 | 1 | 11 | Mrpl32 | 0.007 | Young |
| 0.645728 | 0.545006 | 0.058 | 0.048 | 1 | 11 | Trip10 | 0.01 | Young |
| 0.645834 | 0.774264 | 0.087 | 0.109 | 1 | 11 | Pibf1 | -0.022 | Young |
| 0.646028 | 0.831541 | 0.174 | 0.169 | 1 | 11 | Psmd12 | 0.005 | Young |
| 0.646195 | 1.562113 | 0.036 | 0.028 | 1 | 11 | Styx | 0.008 | Young |
| 0.646572 | 0.482949 | 0.37 | 0.391 | 1 | 11 | Trim30d | -0.021 | Young |
| 0.646656 | 0.446963 | 0.138 | 0.125 | 1 | 11 | Ifit3b | 0.013 | Young |
| 0.64671 | 0.276266 | 0.109 | 0.129 | 1 | 11 | Uri1 | -0.02 | Young |
| 0.647033 | 0.183653 | 0.145 | 0.169 | 1 | 11 | Cc2d1b | -0.024 | Young |
| 0.647184 | 0.376323 | 0.08 | 0.097 | 1 | 11 | H6pd | -0.017 | Young |
| 0.647334 | 0.864579 | 0.087 | 0.077 | 1 | 11 | Slc15a2 | 0.01 | Young |
| 0.647508 | 1.199327 | 0.043 | 0.056 | 1 | 11 | Qpctl | -0.013 | Young |
| 0.648404 | 0.363793 | 0.5 | 0.625 | 1 | 11 | Il10ra | -0.125 | Young |
| 0.648526 | 0.42667 | 0.123 | 0.145 | 1 | 11 | Shc1 | -0.022 | Young |
| 0.648668 | 0.354207 | 0.507 | 0.56 | 1 | 11 | Jmjd1c | -0.053 | Young |
| 0.64904 | 0.536589 | 0.08 | 0.069 | 1 | 11 | Clec4d | 0.011 | Young |
| 0.650562 | 0.226419 | 0.065 | 0.081 | 1 | 11 | Maip1 | -0.016 | Young |
| 0.650748 | 0.616809 | 0.036 | 0.028 | 1 | 11 | Zbtb40 | 0.008 | Young |
| 0.651173 | 0.297637 | 0.333 | 0.403 | 1 | 11 | Use1 | -0.07 | Young |
| 0.651231 | 0.258695 | 0.087 | 0.105 | 1 | 11 | Htatip2 | -0.018 | Young |
| 0.652557 | 0.594783 | 0.058 | 0.073 | 1 | 11 | Cinp | -0.015 | Young |
| 0.652726 | 0.803024 | 0.181 | 0.226 | 1 | 11 | Prdm1 | -0.045 | Young |
| 0.65303 | 1.070746 | 0.036 | 0.028 | 1 | 11 | Tppp | 0.008 | Young |
| 0.653631 | 0.070996 | 0.319 | 0.363 | 1 | 11 | Ccrl2 | -0.044 | Young |
| 0.653679 | 0.481369 | 0.362 | 0.383 | 1 | 11 | Polr2a | -0.021 | Young |
| 0.65409 | 0.740107 | 0.13 | 0.157 | 1 | 11 | Ubl7 | -0.027 | Young |
| 0.654291 | 0.878525 | 0.116 | 0.137 | 1 | 11 | Iars2 | -0.021 | Young |
| 0.654648 | 0.710132 | 0.058 | 0.048 | 1 | 11 | Fmn1 | 0.01 | Young |
| 0.654725 | 0.228848 | 0.297 | 0.347 | 1 | 11 | Cyp4f18 | -0.05 | Young |
| 0.65496 | 0.559114 | 0.072 | 0.089 | 1 | 11 | Apip | -0.017 | Young |
| 0.655124 | 0.732321 | 0.123 | 0.113 | 1 | 11 | Slc25a12 | 0.01 | Young |
| 0.655314 | 0.988257 | 0.036 | 0.028 | 1 | 11 | Plscr4 | 0.008 | Young |
| 0.655361 | 0.989301 | 0.101 | 0.093 | 1 | 11 | Cog4 | 0.008 | Young |
| 0.656335 | 0.285955 | 0.007 | 0.012 | 1 | 11 | Nipa1 | -0.005 | Young |
| 0.656335 | 0.05549 | 0.007 | 0.012 | 1 | 11 | Kdm8 | -0.005 | Young |
| 0.656459 | 0.38661 | 0.239 | 0.286 | 1 | 11 | Ndufa8 | -0.047 | Young |
| 0.656629 | 0.394778 | 0.094 | 0.081 | 1 | 11 | Socs1 | 0.013 | Young |
| 0.657603 | 0.347798 | 0.036 | 0.028 | 1 | 11 | Car5b | 0.008 | Young |
| 0.657603 | 0.234824 | 0.036 | 0.028 | 1 | 11 | Pycrl | 0.008 | Young |
| 0.657707 | 0.536497 | 0.181 | 0.214 | 1 | 11 | Fam172a | -0.033 | Young |
| 0.658637 | 0.052377 | 0.601 | 0.714 | 1 | 11 | Atp5h | -0.113 | Young |

| | | | | | | | | |
|---|---|---|---|---|---|---|---|---|
| 0.658988 | 0.142057 | 0.087 | 0.077 | 1 | 11 | Myef2 | 0.01 | Young |
| 0.659803 | 0.221773 | 0.428 | 0.552 | 1 | 11 | Prdx5 | -0.124 | Young |
| 0.659969 | 0.167658 | 0.797 | 0.843 | 1 | 11 | Wnk1 | -0.046 | Young |
| 0.660261 | 1.825121 | 0.007 | 0.012 | 1 | 11 | Plpp3 | -0.005 | Young |
| 0.660261 | 1.758789 | 0.007 | 0.012 | 1 | 11 | Nr6a1 | -0.005 | Young |
| 0.660261 | 1.457886 | 0.007 | 0.012 | 1 | 11 | Ramp2 | -0.005 | Young |
| 0.660261 | 1.443425 | 0.007 | 0.012 | 1 | 11 | Gnb1l | -0.005 | Young |
| 0.660261 | 1.326079 | 0.007 | 0.012 | 1 | 11 | Snhg18 | -0.005 | Young |
| 0.660261 | 1.290155 | 0.007 | 0.012 | 1 | 11 | Msh5 | -0.005 | Young |
| 0.660261 | 1.283015 | 0.007 | 0.012 | 1 | 11 | Fanca | -0.005 | Young |
| 0.660261 | 1.277939 | 0.007 | 0.012 | 1 | 11 | Cdca7l | -0.005 | Young |
| 0.660261 | 1.181912 | 0.007 | 0.012 | 1 | 11 | Zfp963 | -0.005 | Young |
| 0.660261 | 1.099555 | 0.007 | 0.012 | 1 | 11 | Nlgn2 | -0.005 | Young |
| 0.660261 | 1.094226 | 0.007 | 0.012 | 1 | 11 | Garnl3 | -0.005 | Young |
| 0.660261 | 1.060756 | 0.007 | 0.012 | 1 | 11 | Vsig8 | -0.005 | Young |
| 0.660261 | 1.053927 | 0.007 | 0.012 | 1 | 11 | Brdt | -0.005 | Young |
| 0.660261 | 1.051252 | 0.007 | 0.012 | 1 | 11 | Adat2 | -0.005 | Young |
| 0.660261 | 0.99386 | 0.007 | 0.012 | 1 | 11 | B3gnt7 | -0.005 | Young |
| 0.660261 | 0.97253 | 0.007 | 0.012 | 1 | 11 | Tmem216 | -0.005 | Young |
| 0.660261 | 0.958818 | 0.007 | 0.012 | 1 | 11 | Atp10a | -0.005 | Young |
| 0.660261 | 0.948199 | 0.007 | 0.012 | 1 | 11 | Cdca7 | -0.005 | Young |
| 0.660261 | 0.91325 | 0.007 | 0.012 | 1 | 11 | Hist1h1d | -0.005 | Young |
| 0.660261 | 0.868956 | 0.007 | 0.012 | 1 | 11 | Gm9796 | -0.005 | Young |
| 0.660261 | 0.860573 | 0.007 | 0.012 | 1 | 11 | 9930022D1 | -0.005 | Young |
| 0.660261 | 0.849255 | 0.007 | 0.012 | 1 | 11 | Mphosph9 | -0.005 | Young |
| 0.660261 | 0.849097 | 0.007 | 0.012 | 1 | 11 | BC048403 | -0.005 | Young |
| 0.660261 | 0.839979 | 0.007 | 0.012 | 1 | 11 | St18 | -0.005 | Young |
| 0.660261 | 0.802711 | 0.007 | 0.012 | 1 | 11 | Mterf1a | -0.005 | Young |
| 0.660261 | 0.801938 | 0.007 | 0.012 | 1 | 11 | Ggact | -0.005 | Young |
| 0.660261 | 0.749009 | 0.007 | 0.012 | 1 | 11 | Tjp3 | -0.005 | Young |
| 0.660261 | 0.70608 | 0.007 | 0.012 | 1 | 11 | Fam131a | -0.005 | Young |
| 0.660261 | 0.660154 | 0.007 | 0.012 | 1 | 11 | Trmt5 | -0.005 | Young |
| 0.660261 | 0.649857 | 0.007 | 0.012 | 1 | 11 | Cd200r3 | -0.005 | Young |
| 0.660261 | 0.611435 | 0.007 | 0.012 | 1 | 11 | Trip13 | -0.005 | Young |
| 0.660261 | 0.607938 | 0.007 | 0.012 | 1 | 11 | Gm17354 | -0.005 | Young |
| 0.660261 | 0.547001 | 0.007 | 0.012 | 1 | 11 | Brca2 | -0.005 | Young |
| 0.660261 | 0.53955 | 0.007 | 0.012 | 1 | 11 | Gm38079 | -0.005 | Young |
| 0.660261 | 0.468373 | 0.007 | 0.012 | 1 | 11 | Catip | -0.005 | Young |
| 0.660261 | 0.461522 | 0.007 | 0.012 | 1 | 11 | Gm14296 | -0.005 | Young |
| 0.660261 | 0.435689 | 0.007 | 0.012 | 1 | 11 | Sox4 | -0.005 | Young |
| 0.660261 | 0.427954 | 0.007 | 0.012 | 1 | 11 | Fam161a | -0.005 | Young |
| 0.660261 | 0.423982 | 0.007 | 0.012 | 1 | 11 | Ptges3l | -0.005 | Young |
| 0.660261 | 0.394594 | 0.007 | 0.012 | 1 | 11 | Rsph3a | -0.005 | Young |
| 0.660261 | 0.369105 | 0.007 | 0.012 | 1 | 11 | Dph7 | -0.005 | Young |
| 0.660261 | 0.35581 | 0.007 | 0.012 | 1 | 11 | Ugt1a6a | -0.005 | Young |

| | | | | | | | | |
|---|---|---|---|---|---|---|---|---|
| 0.660261 | 0.352572 | 0.007 | 0.012 | 1 | 11 | 6230400D1 | -0.005 | Young |
| 0.660261 | 0.344312 | 0.007 | 0.012 | 1 | 11 | Elp6 | -0.005 | Young |
| 0.660261 | 0.333762 | 0.007 | 0.012 | 1 | 11 | Zfp647 | -0.005 | Young |
| 0.660261 | 0.322505 | 0.007 | 0.012 | 1 | 11 | AI427809 | -0.005 | Young |
| 0.660261 | 0.315497 | 0.007 | 0.012 | 1 | 11 | Hist1h2ae | -0.005 | Young |
| 0.660261 | 0.236838 | 0.007 | 0.012 | 1 | 11 | Usp27x | -0.005 | Young |
| 0.660261 | 0.209407 | 0.007 | 0.012 | 1 | 11 | Gm31410 | -0.005 | Young |
| 0.660261 | 0.173725 | 0.007 | 0.012 | 1 | 11 | Alkbh4 | -0.005 | Young |
| 0.660261 | 0.169201 | 0.007 | 0.012 | 1 | 11 | Stox2 | -0.005 | Young |
| 0.660261 | 0.062164 | 0.007 | 0.012 | 1 | 11 | Mis18bp1 | -0.005 | Young |
| 0.660261 | 0.059883 | 0.007 | 0.012 | 1 | 11 | 6530409C1 | -0.005 | Young |
| 0.660261 | 0.045416 | 0.007 | 0.012 | 1 | 11 | Gm13205 | -0.005 | Young |
| 0.660261 | 0.027423 | 0.007 | 0.012 | 1 | 11 | Zfp976 | -0.005 | Young |
| 0.660557 | 0.34491 | 0.138 | 0.129 | 1 | 11 | Pex7 | 0.009 | Young |
| 0.660783 | 0.79825 | 0.152 | 0.145 | 1 | 11 | Syvn1 | 0.007 | Young |
| 0.660831 | 0.296903 | 0.123 | 0.149 | 1 | 11 | Itga5 | -0.026 | Young |
| 0.660995 | 0.110843 | 0.174 | 0.202 | 1 | 11 | Ubald1 | -0.028 | Young |
| 0.661221 | 0.589946 | 0.109 | 0.129 | 1 | 11 | Tssc4 | -0.02 | Young |
| 0.661271 | 0.373507 | 0.246 | 0.302 | 1 | 11 | Arl5c | -0.056 | Young |
| 0.662436 | 0.457611 | 0.109 | 0.129 | 1 | 11 | Tmem41b | -0.02 | Young |
| 0.662485 | 0.644164 | 0.196 | 0.226 | 1 | 11 | Dhx36 | -0.03 | Young |
| 0.663342 | 0.323081 | 0.232 | 0.278 | 1 | 11 | Ocrl | -0.046 | Young |
| 0.663618 | 1.161811 | 0.058 | 0.048 | 1 | 11 | Fut4 | 0.01 | Young |
| 0.663652 | 0.34508 | 0.109 | 0.129 | 1 | 11 | Nudt16 | -0.02 | Young |
| 0.664405 | 0.385154 | 0.239 | 0.286 | 1 | 11 | Atxn7 | -0.047 | Young |
| 0.664486 | 0.944469 | 0.094 | 0.085 | 1 | 11 | Hcfc2 | 0.009 | Young |
| 0.664486 | 0.42766 | 0.036 | 0.028 | 1 | 11 | Trp53i13 | 0.008 | Young |
| 0.66489 | 0.136411 | 0.435 | 0.452 | 1 | 11 | Crebbp | -0.017 | Young |
| 0.665254 | 0.751559 | 0.145 | 0.173 | 1 | 11 | Zeb2os | -0.028 | Young |
| 0.665418 | 0.540247 | 0.058 | 0.048 | 1 | 11 | Mtus1 | 0.01 | Young |
| 0.665583 | 0.056944 | 0.551 | 0.637 | 1 | 11 | Psmb8 | -0.086 | Young |
| 0.665629 | 1.096893 | 0.065 | 0.081 | 1 | 11 | Ufd1 | -0.016 | Young |
| 0.666087 | 0.26253 | 0.109 | 0.129 | 1 | 11 | Rab9 | -0.02 | Young |
| 0.666108 | 0.301775 | 0.116 | 0.137 | 1 | 11 | Prpsap2 | -0.021 | Young |
| 0.666523 | 0.548945 | 0.08 | 0.097 | 1 | 11 | Pip4k2c | -0.017 | Young |
| 0.666787 | 0.6388 | 0.036 | 0.028 | 1 | 11 | Ctps | 0.008 | Young |
| 0.666787 | 0.482779 | 0.036 | 0.028 | 1 | 11 | Ccdc66 | 0.008 | Young |
| 0.666787 | 0.338757 | 0.036 | 0.028 | 1 | 11 | Acvr2b | 0.008 | Young |
| 0.667227 | 0.713701 | 0.087 | 0.105 | 1 | 11 | Faf1 | -0.018 | Young |
| 0.668376 | 0.407913 | 0.42 | 0.464 | 1 | 11 | Zc3hav1 | -0.044 | Young |
| 0.668574 | 0.404426 | 0.145 | 0.169 | 1 | 11 | Cramp1l | -0.024 | Young |
| 0.669371 | 0.513839 | 0.239 | 0.242 | 1 | 11 | Bod1l | -0.003 | Young |
| 0.669396 | 0.662301 | 0.072 | 0.089 | 1 | 11 | Atrn | -0.017 | Young |
| 0.669646 | 0.462832 | 0.29 | 0.347 | 1 | 11 | Phf20l1 | -0.057 | Young |
| 0.670335 | 0.592282 | 0.203 | 0.246 | 1 | 11 | Idnk | -0.043 | Young |

| | | | | | | | | |
|---|---|---|---|---|---|---|---|---|
| 0.670346 | 0.159654 | 0.268 | 0.306 | 1 | 11 | Ppp3cb | -0.038 | Young |
| 0.670648 | 0.506709 | 0.297 | 0.359 | 1 | 11 | Sh3kbp1 | -0.062 | Young |
| 0.670859 | 0.236521 | 0.116 | 0.137 | 1 | 11 | Rsrc1 | -0.021 | Young |
| 0.670967 | 0.400251 | 0.109 | 0.129 | 1 | 11 | Taf5l | -0.02 | Young |
| 0.671233 | 0.469925 | 0.167 | 0.198 | 1 | 11 | 4931406CC | -0.031 | Young |
| 0.671923 | 0.268013 | 0.181 | 0.21 | 1 | 11 | Brd8 | -0.029 | Young |
| 0.673687 | 1.072886 | 0.109 | 0.101 | 1 | 11 | Taf7 | 0.008 | Young |
| 0.673938 | 1.05707 | 0.087 | 0.105 | 1 | 11 | Stoml1 | -0.018 | Young |
| 0.6741 | 0.437308 | 0.174 | 0.202 | 1 | 11 | Adss | -0.028 | Young |
| 0.675713 | 0.863523 | 0.101 | 0.121 | 1 | 11 | Nip7 | -0.02 | Young |
| 0.675831 | 0.215582 | 0.181 | 0.206 | 1 | 11 | Lpcat2 | -0.025 | Young |
| 0.675952 | 1.897317 | 0.022 | 0.016 | 1 | 11 | Dcaf4 | 0.006 | Young |
| 0.675952 | 1.704313 | 0.022 | 0.016 | 1 | 11 | Thrsp | 0.006 | Young |
| 0.675952 | 1.679282 | 0.022 | 0.016 | 1 | 11 | Lancl2 | 0.006 | Young |
| 0.675952 | 1.665544 | 0.022 | 0.016 | 1 | 11 | Ears2 | 0.006 | Young |
| 0.675952 | 1.644265 | 0.022 | 0.016 | 1 | 11 | B3galt4 | 0.006 | Young |
| 0.675952 | 1.584022 | 0.022 | 0.016 | 1 | 11 | Zscan12 | 0.006 | Young |
| 0.675952 | 1.396222 | 0.022 | 0.016 | 1 | 11 | Cenpw | 0.006 | Young |
| 0.675952 | 1.303993 | 0.022 | 0.016 | 1 | 11 | Mrm1 | 0.006 | Young |
| 0.675952 | 0.871661 | 0.022 | 0.016 | 1 | 11 | Patj | 0.006 | Young |
| 0.676661 | 1.18416 | 0.072 | 0.089 | 1 | 11 | Tmem97 | -0.017 | Young |
| 0.677139 | 0.033311 | 0.174 | 0.202 | 1 | 11 | Sft2d1 | -0.028 | Young |
| 0.677268 | 0.311701 | 0.145 | 0.169 | 1 | 11 | Oxsr1 | -0.024 | Young |
| 0.677821 | 0.56255 | 0.297 | 0.31 | 1 | 11 | Daxx | -0.013 | Young |
| 0.67818 | 0.282134 | 0.312 | 0.363 | 1 | 11 | Hck | -0.051 | Young |
| 0.678243 | 0.541552 | 0.246 | 0.246 | 1 | 11 | 9030624J0 | 0 | Young |
| 0.678338 | 0.641377 | 0.036 | 0.028 | 1 | 11 | Usp46 | 0.008 | Young |
| 0.67866 | 0.199438 | 0.174 | 0.202 | 1 | 11 | Rhoc | -0.028 | Young |
| 0.678966 | 1.268307 | 0.022 | 0.016 | 1 | 11 | Phf11c | 0.006 | Young |
| 0.679265 | 0.39961 | 0.174 | 0.206 | 1 | 11 | Bsdc1 | -0.032 | Young |
| 0.679958 | 0.524589 | 0.181 | 0.218 | 1 | 11 | Arid4b | -0.037 | Young |
| 0.681985 | 1.272552 | 0.022 | 0.016 | 1 | 11 | Igfbp5 | 0.006 | Young |
| 0.681985 | 1.195587 | 0.022 | 0.016 | 1 | 11 | Wdr90 | 0.006 | Young |
| 0.681985 | 1.08335 | 0.022 | 0.016 | 1 | 11 | Il7r | 0.006 | Young |
| 0.681985 | 1.007189 | 0.022 | 0.016 | 1 | 11 | Tmem138 | 0.006 | Young |
| 0.682123 | 0.609965 | 0.145 | 0.137 | 1 | 11 | Ntpcr | 0.008 | Young |
| 0.682494 | 1.184255 | 0.072 | 0.089 | 1 | 11 | Hsf2 | -0.017 | Young |
| 0.683061 | 0.402485 | 0.159 | 0.19 | 1 | 11 | Cmc1 | -0.031 | Young |
| 0.683879 | 0.734878 | 0.065 | 0.056 | 1 | 11 | Pik3ap1 | 0.009 | Young |
| 0.684192 | 0.263432 | 0.543 | 0.573 | 1 | 11 | Flna | -0.03 | Young |
| 0.684303 | 0.021893 | 0.029 | 0.036 | 1 | 11 | Gm20186 | -0.007 | Young |
| 0.685008 | 1.848801 | 0.022 | 0.016 | 1 | 11 | Zfp213 | 0.006 | Young |
| 0.685008 | 1.223021 | 0.022 | 0.016 | 1 | 11 | Dffb | 0.006 | Young |
| 0.685008 | 1.023328 | 0.022 | 0.016 | 1 | 11 | Cers4 | 0.006 | Young |
| 0.685008 | 0.985963 | 0.022 | 0.016 | 1 | 11 | Hps6 | 0.006 | Young |

| | | | | | | | | |
|---|---|---|---|---|---|---|---|---|
| 0.685008 | 0.928369 | 0.022 | 0.016 | 1 | 11 | Sec22c | 0.006 | Young |
| 0.685008 | 0.862056 | 0.022 | 0.016 | 1 | 11 | Oas1c | 0.006 | Young |
| 0.685008 | 0.748691 | 0.022 | 0.016 | 1 | 11 | Acy1 | 0.006 | Young |
| 0.685348 | 0.138746 | 0.413 | 0.504 | 1 | 11 | Adam17 | -0.091 | Young |
| 0.686077 | 0.284029 | 0.304 | 0.351 | 1 | 11 | Anp32e | -0.047 | Young |
| 0.686606 | 0.883632 | 0.101 | 0.093 | 1 | 11 | Traf5 | 0.008 | Young |
| 0.686884 | 0.595569 | 0.145 | 0.173 | 1 | 11 | S1pr2 | -0.028 | Young |
| 0.687134 | 0.921514 | 0.094 | 0.085 | 1 | 11 | Fbxw17 | 0.009 | Young |
| 0.687553 | 0.028828 | 0.065 | 0.077 | 1 | 11 | AI987944 | -0.012 | Young |
| 0.68794 | 0.590621 | 0.362 | 0.391 | 1 | 11 | Amfr | -0.029 | Young |
| 0.687978 | 0.871926 | 0.101 | 0.093 | 1 | 11 | Blcap | 0.008 | Young |
| 0.687978 | 0.073398 | 0.087 | 0.101 | 1 | 11 | Gm45716 | -0.014 | Young |
| 0.688037 | 1.43369 | 0.022 | 0.016 | 1 | 11 | Znrd1as | 0.006 | Young |
| 0.688037 | 0.780678 | 0.022 | 0.016 | 1 | 11 | Gm10053 | 0.006 | Young |
| 0.688037 | 0.696534 | 0.022 | 0.016 | 1 | 11 | B3gntl1 | 0.006 | Young |
| 0.688037 | 0.512104 | 0.022 | 0.016 | 1 | 11 | Col3a1 | 0.006 | Young |
| 0.688037 | 0.274111 | 0.022 | 0.016 | 1 | 11 | 2610301B2 | 0.006 | Young |
| 0.688075 | 0.168442 | 0.232 | 0.266 | 1 | 11 | Crebrf | -0.034 | Young |
| 0.688174 | 0.626507 | 0.239 | 0.238 | 1 | 11 | Tpst1 | 0.001 | Young |
| 0.688684 | 0.711183 | 0.232 | 0.282 | 1 | 11 | Rb1 | -0.05 | Young |
| 0.689706 | 0.5007 | 0.261 | 0.27 | 1 | 11 | Slc9a9 | -0.009 | Young |
| 0.689985 | 0.65871 | 0.094 | 0.085 | 1 | 11 | Nqo2 | 0.009 | Young |
| 0.690174 | 0.158132 | 0.13 | 0.153 | 1 | 11 | Arfgap2 | -0.023 | Young |
| 0.690829 | 0.833413 | 0.123 | 0.149 | 1 | 11 | Hint2 | -0.026 | Young |
| 0.691071 | 0.820748 | 0.022 | 0.016 | 1 | 11 | Ppp1r3d | 0.006 | Young |
| 0.691071 | 0.789708 | 0.022 | 0.016 | 1 | 11 | Gm26756 | 0.006 | Young |
| 0.691071 | 0.688514 | 0.022 | 0.016 | 1 | 11 | Lrfn1 | 0.006 | Young |
| 0.691071 | 0.582106 | 0.022 | 0.016 | 1 | 11 | Galm | 0.006 | Young |
| 0.691071 | 0.369012 | 0.022 | 0.016 | 1 | 11 | Rad18 | 0.006 | Young |
| 0.691071 | 0.046595 | 0.014 | 0.02 | 1 | 11 | Hmgn3 | -0.006 | Young |
| 0.691458 | 0.528478 | 0.072 | 0.085 | 1 | 11 | Thap2 | -0.013 | Young |
| 0.691506 | 0.744108 | 0.087 | 0.105 | 1 | 11 | Mtx1 | -0.018 | Young |
| 0.691774 | 0.388207 | 0.101 | 0.117 | 1 | 11 | Tmem209 | -0.016 | Young |
| 0.692071 | 0.063894 | 0.058 | 0.069 | 1 | 11 | Mlst8 | -0.011 | Young |
| 0.692466 | 0.042493 | 0.13 | 0.153 | 1 | 11 | Spast | -0.023 | Young |
| 0.693122 | 0.578548 | 0.225 | 0.27 | 1 | 11 | Bbx | -0.045 | Young |
| 0.693378 | 0.212855 | 0.355 | 0.411 | 1 | 11 | Coq10b | -0.056 | Young |
| 0.69365 | 0.255831 | 0.355 | 0.419 | 1 | 11 | Cct2 | -0.064 | Young |
| 0.694004 | 0.088843 | 0.891 | 0.879 | 1 | 11 | Tyrobp | 0.012 | Young |
| 0.69411 | 0.911339 | 0.022 | 0.016 | 1 | 11 | Sbk1 | 0.006 | Young |
| 0.69411 | 0.79965 | 0.022 | 0.016 | 1 | 11 | Sirt4 | 0.006 | Young |
| 0.69411 | 0.566986 | 0.022 | 0.016 | 1 | 11 | Mios | 0.006 | Young |
| 0.69411 | 0.319324 | 0.014 | 0.02 | 1 | 11 | Dnm1 | -0.006 | Young |
| 0.69411 | 0.269277 | 0.022 | 0.016 | 1 | 11 | Gm49339 | 0.006 | Young |
| 0.69411 | 0.184682 | 0.014 | 0.02 | 1 | 11 | Slc22a18 | -0.006 | Young |

| | | | | | | | | |
|---|---|---|---|---|---|---|---|---|
| 0.69411 | 0.142693 | 0.022 | 0.016 | 1 | 11 | Zfp1 | 0.006 | Young |
| 0.694537 | 1.234363 | 0.08 | 0.097 | 1 | 11 | Afg3l2 | -0.017 | Young |
| 0.695027 | 0.754817 | 0.217 | 0.226 | 1 | 11 | Mgat4b | -0.009 | Young |
| 0.6951 | 0.299758 | 0.543 | 0.641 | 1 | 11 | Kdm6b | -0.098 | Young |
| 0.69561 | 0.820777 | 0.145 | 0.173 | 1 | 11 | Atad1 | -0.028 | Young |
| 0.695753 | 1.274088 | 0.072 | 0.065 | 1 | 11 | Dedd | 0.007 | Young |
| 0.695753 | 0.801666 | 0.072 | 0.065 | 1 | 11 | Pelp1 | 0.007 | Young |
| 0.695845 | 0.74064 | 0.065 | 0.056 | 1 | 11 | Prelid2 | 0.009 | Young |
| 0.69588 | 1.827406 | 0.043 | 0.036 | 1 | 11 | Rdh10 | 0.007 | Young |
| 0.695885 | 0.389972 | 0.486 | 0.569 | 1 | 11 | Abi1 | -0.083 | Young |
| 0.695886 | 0.201257 | 0.145 | 0.169 | 1 | 11 | Ngdn | -0.024 | Young |
| 0.696166 | 0.743656 | 0.109 | 0.101 | 1 | 11 | Tmx4 | 0.008 | Young |
| 0.696972 | 0.168144 | 0.181 | 0.21 | 1 | 11 | N4bp2l1 | -0.029 | Young |
| 0.697154 | 0.572922 | 0.014 | 0.02 | 1 | 11 | Gm340 | -0.006 | Young |
| 0.697154 | 0.417962 | 0.014 | 0.02 | 1 | 11 | Gm16094 | -0.006 | Young |
| 0.697154 | 0.388387 | 0.022 | 0.016 | 1 | 11 | Icam2 | 0.006 | Young |
| 0.697154 | 0.257818 | 0.014 | 0.02 | 1 | 11 | Igf2bp2 | -0.006 | Young |
| 0.697154 | 0.235298 | 0.014 | 0.02 | 1 | 11 | Pecr | -0.006 | Young |
| 0.697154 | 0.063944 | 0.022 | 0.016 | 1 | 11 | Dnal4 | 0.006 | Young |
| 0.697332 | 0.645787 | 0.232 | 0.23 | 1 | 11 | Zcchc2 | 0.002 | Young |
| 0.698363 | 0.539458 | 0.239 | 0.286 | 1 | 11 | Sh3pxd2b | -0.047 | Young |
| 0.698407 | 0.398727 | 0.145 | 0.137 | 1 | 11 | Slc25a22 | 0.008 | Young |
| 0.698607 | 0.721087 | 0.159 | 0.153 | 1 | 11 | Wdr33 | 0.006 | Young |
| 0.698677 | 0.70783 | 0.014 | 0.02 | 1 | 11 | Cenpj | -0.006 | Young |
| 0.698711 | 0.02875 | 0.667 | 0.798 | 1 | 11 | Ptbp3 | -0.131 | Young |
| 0.698803 | 0.411979 | 0.471 | 0.528 | 1 | 11 | Pum2 | -0.057 | Young |
| 0.699027 | 0.705968 | 0.167 | 0.198 | 1 | 11 | Vps18 | -0.031 | Young |
| 0.700202 | 0.743396 | 0.014 | 0.02 | 1 | 11 | 0610040B1 | -0.006 | Young |
| 0.700202 | 0.524665 | 0.014 | 0.02 | 1 | 11 | Ptpn21 | -0.006 | Young |
| 0.700202 | 0.438623 | 0.014 | 0.02 | 1 | 11 | Tnfrsf14 | -0.006 | Young |
| 0.700202 | 0.434912 | 0.014 | 0.02 | 1 | 11 | Alkbh2 | -0.006 | Young |
| 0.700202 | 0.381732 | 0.014 | 0.02 | 1 | 11 | Ubd | -0.006 | Young |
| 0.700202 | 0.311251 | 0.014 | 0.02 | 1 | 11 | Fuk | -0.006 | Young |
| 0.700202 | 0.216152 | 0.022 | 0.016 | 1 | 11 | Mettl2 | 0.006 | Young |
| 0.700202 | 0.196022 | 0.014 | 0.02 | 1 | 11 | Polr3h | -0.006 | Young |
| 0.700202 | 0.109401 | 0.014 | 0.02 | 1 | 11 | Fbxw9 | -0.006 | Young |
| 0.700284 | 0.798959 | 0.138 | 0.165 | 1 | 11 | Plscr3 | -0.027 | Young |
| 0.701495 | 0.516695 | 0.152 | 0.145 | 1 | 11 | Fbxl20 | 0.007 | Young |
| 0.70186 | 0.425111 | 0.203 | 0.242 | 1 | 11 | Dcaf7 | -0.039 | Young |
| 0.701947 | 0.326319 | 0.442 | 0.536 | 1 | 11 | Arrb2 | -0.094 | Young |
| 0.70196 | 0.569347 | 0.355 | 0.383 | 1 | 11 | Git2 | -0.028 | Young |
| 0.701998 | 0.364537 | 0.116 | 0.133 | 1 | 11 | Ccdc9 | -0.017 | Young |
| 0.702179 | 0.619281 | 0.312 | 0.363 | 1 | 11 | Ifi211 | -0.051 | Young |
| 0.702289 | 0.36532 | 0.029 | 0.036 | 1 | 11 | Ptprm | -0.007 | Young |
| 0.702627 | 0.68082 | 0.094 | 0.113 | 1 | 11 | Fibp | -0.019 | Young |

| | | | | | | | | |
|---|---|---|---|---|---|---|---|---|
| 0.703046 | 0.57324 | 0.159 | 0.19 | 1 | 11 | Lsm8 | -0.031 | Young |
| 0.703256 | 0.985535 | 0.014 | 0.02 | 1 | 11 | Dhfr | -0.006 | Young |
| 0.703256 | 0.959276 | 0.014 | 0.02 | 1 | 11 | 2610524H( | -0.006 | Young |
| 0.703256 | 0.736847 | 0.014 | 0.02 | 1 | 11 | 2700038G2 | -0.006 | Young |
| 0.703256 | 0.385222 | 0.014 | 0.02 | 1 | 11 | Strbp | -0.006 | Young |
| 0.703378 | 0.085122 | 0.101 | 0.117 | 1 | 11 | Helz2 | -0.016 | Young |
| 0.703933 | 0.386573 | 0.275 | 0.29 | 1 | 11 | Il13ra1 | -0.015 | Young |
| 0.704263 | 0.400968 | 0.21 | 0.238 | 1 | 11 | Fbxw11 | -0.028 | Young |
| 0.704304 | 1.157977 | 0.043 | 0.036 | 1 | 11 | Mmgt2 | 0.007 | Young |
| 0.704304 | 0.6822 | 0.043 | 0.036 | 1 | 11 | Fam212a | 0.007 | Young |
| 0.704307 | 0.56026 | 0.304 | 0.375 | 1 | 11 | Pgd | -0.071 | Young |
| 0.704439 | 0.387765 | 0.065 | 0.056 | 1 | 11 | Orc5 | 0.009 | Young |
| 0.70444 | 0.40592 | 0.116 | 0.133 | 1 | 11 | Mis12 | -0.017 | Young |
| 0.704668 | 0.656918 | 0.275 | 0.335 | 1 | 11 | Bcl2l11 | -0.06 | Young |
| 0.704738 | 0.845684 | 0.123 | 0.117 | 1 | 11 | Cd80 | 0.006 | Young |
| 0.705063 | 0.224232 | 0.529 | 0.593 | 1 | 11 | Tm9sf3 | -0.064 | Young |
| 0.70594 | 0.605918 | 0.203 | 0.246 | 1 | 11 | Vps13a | -0.043 | Young |
| 0.705945 | 0.084285 | 0.428 | 0.492 | 1 | 11 | Casp4 | -0.064 | Young |
| 0.706233 | 0.957221 | 0.101 | 0.121 | 1 | 11 | Gcn1l1 | -0.02 | Young |
| 0.706314 | 1.995967 | 0.014 | 0.02 | 1 | 11 | Usp49 | -0.006 | Young |
| 0.706314 | 1.718335 | 0.014 | 0.02 | 1 | 11 | Map2k3os | -0.006 | Young |
| 0.706314 | 1.534257 | 0.014 | 0.02 | 1 | 11 | Alg3 | -0.006 | Young |
| 0.706314 | 1.512729 | 0.014 | 0.02 | 1 | 11 | Smco4 | -0.006 | Young |
| 0.706314 | 1.393578 | 0.014 | 0.02 | 1 | 11 | Gm15232 | -0.006 | Young |
| 0.706314 | 1.2287 | 0.014 | 0.02 | 1 | 11 | Cd5l | -0.006 | Young |
| 0.706314 | 1.020891 | 0.014 | 0.02 | 1 | 11 | Pex10 | -0.006 | Young |
| 0.706314 | 0.9753 | 0.014 | 0.02 | 1 | 11 | Them6 | -0.006 | Young |
| 0.706314 | 0.781997 | 0.014 | 0.02 | 1 | 11 | Zfp777 | -0.006 | Young |
| 0.706314 | 0.707748 | 0.014 | 0.02 | 1 | 11 | Rem1 | -0.006 | Young |
| 0.706314 | 0.655963 | 0.014 | 0.02 | 1 | 11 | Blm | -0.006 | Young |
| 0.706314 | 0.604868 | 0.014 | 0.02 | 1 | 11 | Msl3l2 | -0.006 | Young |
| 0.706314 | 0.532714 | 0.014 | 0.02 | 1 | 11 | Pigb | -0.006 | Young |
| 0.706314 | 0.36341 | 0.014 | 0.02 | 1 | 11 | Sap130 | -0.006 | Young |
| 0.706339 | 0.146321 | 0.065 | 0.077 | 1 | 11 | Pgghg | -0.012 | Young |
| 0.706351 | 0.297783 | 0.5 | 0.544 | 1 | 11 | Rab7b | -0.044 | Young |
| 0.706406 | 0.159453 | 0.051 | 0.06 | 1 | 11 | Mex3d | -0.009 | Young |
| 0.706714 | 0.046191 | 0.225 | 0.262 | 1 | 11 | Keap1 | -0.037 | Young |
| 0.706726 | 0.70076 | 0.138 | 0.161 | 1 | 11 | Enpp4 | -0.023 | Young |
| 0.707121 | 1.227579 | 0.072 | 0.065 | 1 | 11 | Nxt2 | 0.007 | Young |
| 0.707218 | 0.314546 | 0.464 | 0.5 | 1 | 11 | Zfhx3 | -0.036 | Young |
| 0.70728 | 0.119323 | 0.087 | 0.101 | 1 | 11 | Trap1 | -0.014 | Young |
| 0.707331 | 0.277751 | 0.391 | 0.448 | 1 | 11 | Rin2 | -0.057 | Young |
| 0.707524 | 0.18511 | 0.493 | 0.585 | 1 | 11 | Dhrs3 | -0.092 | Young |
| 0.708006 | 0.109288 | 0.043 | 0.052 | 1 | 11 | Rev3l | -0.009 | Young |
| 0.70853 | 0.649552 | 0.043 | 0.036 | 1 | 11 | Lsm11 | 0.007 | Young |

| | | | | | | | | |
|---|---|---|---|---|---|---|---|---|
| 0.708557 | 0.4161 | 0.101 | 0.117 | 1 | 11 | Dph3 | -0.016 | Young |
| 0.708625 | 0.210515 | 0.08 | 0.093 | 1 | 11 | Tmem64 | -0.013 | Young |
| 0.708722 | 1.390157 | 0.022 | 0.028 | 1 | 11 | Dgke | -0.006 | Young |
| 0.708722 | 0.326806 | 0.022 | 0.028 | 1 | 11 | Plppr4 | -0.006 | Young |
| 0.708722 | 0.065746 | 0.022 | 0.028 | 1 | 11 | Paxx | -0.006 | Young |
| 0.70929 | 0.395819 | 0.196 | 0.194 | 1 | 11 | Alcam | 0.002 | Young |
| 0.709516 | 0.651653 | 0.152 | 0.145 | 1 | 11 | Med10 | 0.007 | Young |
| 0.709795 | 0.750696 | 0.138 | 0.133 | 1 | 11 | Rcbtb2 | 0.005 | Young |
| 0.70993 | 0.730466 | 0.051 | 0.06 | 1 | 11 | Alkbh8 | -0.009 | Young |
| 0.710054 | 0.475835 | 0.087 | 0.101 | 1 | 11 | Bop1 | -0.014 | Young |
| 0.710436 | 0.017848 | 0.174 | 0.198 | 1 | 11 | Rnf20 | -0.024 | Young |
| 0.710647 | 0.822438 | 0.043 | 0.036 | 1 | 11 | Kctd11 | 0.007 | Young |
| 0.710857 | 0.213653 | 0.094 | 0.109 | 1 | 11 | Pcnt | -0.015 | Young |
| 0.711252 | 0.498363 | 0.138 | 0.161 | 1 | 11 | Txndc16 | -0.023 | Young |
| 0.711298 | 0.872153 | 0.022 | 0.028 | 1 | 11 | Spef1 | -0.006 | Young |
| 0.711298 | 0.392688 | 0.022 | 0.028 | 1 | 11 | Wdr60 | -0.006 | Young |
| 0.711298 | 0.124792 | 0.022 | 0.028 | 1 | 11 | Neurl1a | -0.006 | Young |
| 0.711298 | 0.108997 | 0.022 | 0.028 | 1 | 11 | Bcr | -0.006 | Young |
| 0.711417 | 0.136846 | 0.493 | 0.601 | 1 | 11 | Top1 | -0.108 | Young |
| 0.711442 | 0.819312 | 0.087 | 0.101 | 1 | 11 | Vgll4 | -0.014 | Young |
| 0.711787 | 0.501816 | 0.043 | 0.052 | 1 | 11 | Dcun1d3 | -0.009 | Young |
| 0.711885 | 0.131531 | 0.036 | 0.044 | 1 | 11 | Mzt2 | -0.008 | Young |
| 0.712183 | 0.084944 | 0.196 | 0.226 | 1 | 11 | Eml4 | -0.03 | Young |
| 0.712765 | 1.018762 | 0.043 | 0.036 | 1 | 11 | Nbeal2 | 0.007 | Young |
| 0.713425 | 1.44949 | 0.087 | 0.081 | 1 | 11 | Cmtm4 | 0.006 | Young |
| 0.713461 | 0.221235 | 0.051 | 0.06 | 1 | 11 | Gm38399 | -0.009 | Young |
| 0.713564 | 0.170036 | 0.326 | 0.383 | 1 | 11 | Snx18 | -0.057 | Young |
| 0.713616 | 0.08534 | 0.029 | 0.036 | 1 | 11 | Asf1a | -0.007 | Young |
| 0.713616 | 0.004956 | 0.029 | 0.036 | 1 | 11 | Gm31718 | -0.007 | Young |
| 0.713876 | 0.617829 | 0.022 | 0.028 | 1 | 11 | Mmp25 | -0.006 | Young |
| 0.713876 | 0.369386 | 0.022 | 0.028 | 1 | 11 | Kcng2 | -0.006 | Young |
| 0.713876 | 0.044748 | 0.022 | 0.028 | 1 | 11 | Cyb5d2 | -0.006 | Young |
| 0.714221 | 0.196201 | 0.065 | 0.077 | 1 | 11 | Ceacam1 | -0.012 | Young |
| 0.71424 | 0.416274 | 0.116 | 0.133 | 1 | 11 | Coq2 | -0.017 | Young |
| 0.714253 | 0.720197 | 0.348 | 0.415 | 1 | 11 | Cxcl10 | -0.067 | Young |
| 0.714396 | 0.131302 | 0.08 | 0.093 | 1 | 11 | 2310009B1 | -0.013 | Young |
| 0.714607 | 0.279333 | 0.254 | 0.302 | 1 | 11 | Ppp1r10 | -0.048 | Young |
| 0.714814 | 0.301022 | 0.109 | 0.125 | 1 | 11 | Cep120 | -0.016 | Young |
| 0.715229 | 0.147677 | 0.051 | 0.06 | 1 | 11 | Zfp180 | -0.009 | Young |
| 0.715237 | 0.502685 | 0.312 | 0.343 | 1 | 11 | Xrn2 | -0.031 | Young |
| 0.715254 | 0.382947 | 0.058 | 0.069 | 1 | 11 | Dlg4 | -0.011 | Young |
| 0.715575 | 0.27709 | 0.043 | 0.052 | 1 | 11 | Stambpl1 | -0.009 | Young |
| 0.715575 | 0.125469 | 0.043 | 0.052 | 1 | 11 | Calml4 | -0.009 | Young |
| 0.715607 | 0.479956 | 0.152 | 0.177 | 1 | 11 | Ncbp3 | -0.025 | Young |
| 0.715739 | 0.125256 | 0.188 | 0.181 | 1 | 11 | Itgal | 0.007 | Young |

| | | | | | | | | |
|---|---|---|---|---|---|---|---|---|
| 0.715801 | 0.198127 | 0.065 | 0.077 | 1 | 11 | Dcaf17 | -0.012 | Young |
| 0.715889 | 0.033797 | 0.029 | 0.036 | 1 | 11 | Zfp386 | -0.007 | Young |
| 0.715947 | 0.28588 | 0.239 | 0.274 | 1 | 11 | Mob1b | -0.035 | Young |
| 0.715997 | 0.870181 | 0.036 | 0.044 | 1 | 11 | Gm10138 | -0.008 | Young |
| 0.716074 | 0.080878 | 0.884 | 0.935 | 1 | 11 | Cdc42 | -0.051 | Young |
| 0.716459 | 1.17678 | 0.022 | 0.028 | 1 | 11 | Zfp619 | -0.006 | Young |
| 0.716459 | 0.768465 | 0.022 | 0.028 | 1 | 11 | Gm9493 | -0.006 | Young |
| 0.716459 | 0.550678 | 0.022 | 0.028 | 1 | 11 | Sirt6 | -0.006 | Young |
| 0.716459 | 0.358155 | 0.022 | 0.028 | 1 | 11 | Eid2b | -0.006 | Young |
| 0.716459 | 0.307821 | 0.022 | 0.028 | 1 | 11 | Pnpla6 | -0.006 | Young |
| 0.716459 | 0.225493 | 0.022 | 0.028 | 1 | 11 | Vcpkmt | -0.006 | Young |
| 0.716919 | 0.569646 | 0.072 | 0.065 | 1 | 11 | Fam8a1 | 0.007 | Young |
| 0.717472 | 0.113444 | 0.043 | 0.052 | 1 | 11 | Uqcc1 | -0.009 | Young |
| 0.717621 | 0.148108 | 0.674 | 0.794 | 1 | 11 | Slc6a6 | -0.12 | Young |
| 0.717789 | 0.533219 | 0.152 | 0.177 | 1 | 11 | Cul5 | -0.025 | Young |
| 0.71799 | 0.261889 | 0.471 | 0.573 | 1 | 11 | Ets2 | -0.102 | Young |
| 0.718057 | 0.368455 | 0.036 | 0.044 | 1 | 11 | Tradd | -0.008 | Young |
| 0.718164 | 0.671153 | 0.029 | 0.036 | 1 | 11 | Tmem51os | -0.007 | Young |
| 0.718588 | 0.089008 | 0.058 | 0.069 | 1 | 11 | Gm27010 | -0.011 | Young |
| 0.718769 | 0.72214 | 0.051 | 0.06 | 1 | 11 | Cfap97 | -0.009 | Young |
| 0.719044 | 0.92599 | 0.022 | 0.028 | 1 | 11 | Ift80 | -0.006 | Young |
| 0.719044 | 0.854957 | 0.022 | 0.028 | 1 | 11 | 2010320M | -0.006 | Young |
| 0.719044 | 0.41478 | 0.022 | 0.028 | 1 | 11 | Slc8a1 | -0.006 | Young |
| 0.719044 | 0.109726 | 0.022 | 0.028 | 1 | 11 | L3mbtl2 | -0.006 | Young |
| 0.719404 | 0.476751 | 0.232 | 0.23 | 1 | 11 | Sp3 | 0.002 | Young |
| 0.719477 | 0.560913 | 0.152 | 0.181 | 1 | 11 | Srek1ip1 | -0.029 | Young |
| 0.719477 | 0.867396 | 0.174 | 0.169 | 1 | 11 | Pgs1 | 0.005 | Young |
| 0.719598 | 0.484902 | 0.406 | 0.452 | 1 | 11 | Sfr1 | -0.046 | Young |
| 0.71987 | 0.955816 | 0.123 | 0.117 | 1 | 11 | Vps9d1 | 0.006 | Young |
| 0.71987 | 0.650812 | 0.123 | 0.117 | 1 | 11 | Wasl | 0.006 | Young |
| 0.720195 | 0.704332 | 0.072 | 0.065 | 1 | 11 | Mettl13 | 0.007 | Young |
| 0.720442 | 1.083929 | 0.029 | 0.036 | 1 | 11 | Ccdc126 | -0.007 | Young |
| 0.720442 | 1.083734 | 0.029 | 0.036 | 1 | 11 | Mrps2 | -0.007 | Young |
| 0.720442 | 0.600592 | 0.029 | 0.036 | 1 | 11 | Zkscan6 | -0.007 | Young |
| 0.720442 | 0.442215 | 0.029 | 0.036 | 1 | 11 | AU022252 | -0.007 | Young |
| 0.720542 | 0.403115 | 0.051 | 0.06 | 1 | 11 | Ipcef1 | -0.009 | Young |
| 0.720548 | 0.225992 | 0.065 | 0.077 | 1 | 11 | Gtpbp1 | -0.012 | Young |
| 0.720805 | 0.039699 | 0.529 | 0.597 | 1 | 11 | Fxyd5 | -0.068 | Young |
| 0.720958 | 0.626699 | 0.08 | 0.073 | 1 | 11 | Snta1 | 0.007 | Young |
| 0.721185 | 0.439904 | 0.101 | 0.093 | 1 | 11 | Bloc1s6 | 0.008 | Young |
| 0.721479 | 0.352826 | 0.072 | 0.085 | 1 | 11 | Klhl21 | -0.013 | Young |
| 0.721632 | 0.945753 | 0.022 | 0.028 | 1 | 11 | Miga2 | -0.006 | Young |
| 0.721632 | 0.307887 | 0.022 | 0.028 | 1 | 11 | Hr | -0.006 | Young |
| 0.721633 | 0.427296 | 0.08 | 0.093 | 1 | 11 | Brcc3 | -0.013 | Young |
| 0.721706 | 0.679741 | 0.13 | 0.121 | 1 | 11 | Abcb1b | 0.009 | Young |

| | | | | | | | | |
|---|---|---|---|---|---|---|---|---|
| 0.72174 | 0.355253 | 0.065 | 0.056 | 1 | 11 | Cnnm4 | 0.009 | Young |
| 0.721762 | 0.758851 | 0.232 | 0.242 | 1 | 11 | Ncoa7 | -0.01 | Young |
| 0.722181 | 0.505587 | 0.036 | 0.044 | 1 | 11 | Aasdh | -0.008 | Young |
| 0.722247 | 0.242815 | 0.341 | 0.415 | 1 | 11 | Rtraf | -0.074 | Young |
| 0.722316 | 0.404128 | 0.051 | 0.06 | 1 | 11 | Gm35154 | -0.009 | Young |
| 0.722316 | 0.110708 | 0.051 | 0.06 | 1 | 11 | Fen1 | -0.009 | Young |
| 0.722394 | 0.068444 | 0.239 | 0.274 | 1 | 11 | Crem | -0.035 | Young |
| 0.722559 | 0.082765 | 0.993 | 0.984 | 1 | 11 | Malat1 | 0.009 | Young |
| 0.722723 | 1.371944 | 0.029 | 0.036 | 1 | 11 | Fam122b | -0.007 | Young |
| 0.722723 | 1.12546 | 0.029 | 0.036 | 1 | 11 | Phc1 | -0.007 | Young |
| 0.722723 | 0.690708 | 0.029 | 0.036 | 1 | 11 | 1500015A( | -0.007 | Young |
| 0.722723 | 0.190949 | 0.029 | 0.036 | 1 | 11 | Ppp2r1b | -0.007 | Young |
| 0.722814 | 0.338016 | 0.13 | 0.149 | 1 | 11 | Imp4 | -0.019 | Young |
| 0.723173 | 0.179584 | 0.043 | 0.052 | 1 | 11 | Alg6 | -0.009 | Young |
| 0.723285 | 0.578904 | 0.159 | 0.181 | 1 | 11 | Pwwp2a | -0.022 | Young |
| 0.723345 | 0.830811 | 0.159 | 0.153 | 1 | 11 | Chd6 | 0.006 | Young |
| 0.723391 | 0.466311 | 0.043 | 0.036 | 1 | 11 | Primpol | 0.007 | Young |
| 0.723391 | 0.310965 | 0.043 | 0.036 | 1 | 11 | B3gnt3 | 0.007 | Young |
| 0.723391 | 0.108443 | 0.043 | 0.036 | 1 | 11 | Tdrkh | 0.007 | Young |
| 0.723391 | 0.091244 | 0.043 | 0.036 | 1 | 11 | Dennd2c | 0.007 | Young |
| 0.72349 | 0.21244 | 0.471 | 0.577 | 1 | 11 | 9930111J2 | -0.106 | Young |
| 0.723559 | 0.3282 | 0.297 | 0.298 | 1 | 11 | Ascc3 | -0.001 | Young |
| 0.723666 | 0.199242 | 0.42 | 0.496 | 1 | 11 | Ncor1 | -0.076 | Young |
| 0.723808 | 0.895793 | 0.159 | 0.185 | 1 | 11 | Ankib1 | -0.026 | Young |
| 0.724079 | 1.199382 | 0.08 | 0.073 | 1 | 11 | Thoc1 | 0.007 | Young |
| 0.724224 | 0.572501 | 0.022 | 0.028 | 1 | 11 | Met | -0.006 | Young |
| 0.724224 | 0.538172 | 0.022 | 0.028 | 1 | 11 | Nod2 | -0.006 | Young |
| 0.724247 | 0.956642 | 0.036 | 0.044 | 1 | 11 | Pan2 | -0.008 | Young |
| 0.724391 | 0.289616 | 0.406 | 0.508 | 1 | 11 | Gpr132 | -0.102 | Young |
| 0.724523 | 0.181429 | 0.63 | 0.718 | 1 | 11 | Pid1 | -0.088 | Young |
| 0.724889 | 0.693897 | 0.138 | 0.161 | 1 | 11 | Zfp451 | -0.023 | Young |
| 0.725112 | 0.00496 | 0.435 | 0.496 | 1 | 11 | Ciita | -0.061 | Young |
| 0.725306 | 0.523016 | 0.065 | 0.077 | 1 | 11 | Heatr3 | -0.012 | Young |
| 0.725402 | 0.389472 | 0.37 | 0.448 | 1 | 11 | Dennd5a | -0.078 | Young |
| 0.725655 | 0.39729 | 0.094 | 0.109 | 1 | 11 | Ralgapa1 | -0.015 | Young |
| 0.725703 | 0.26878 | 0.493 | 0.597 | 1 | 11 | Csrnp1 | -0.104 | Young |
| 0.726144 | 0.307175 | 0.572 | 0.661 | 1 | 11 | Irf8 | -0.089 | Young |
| 0.726314 | 0.549586 | 0.036 | 0.044 | 1 | 11 | Fndc7 | -0.008 | Young |
| 0.726314 | 0.071425 | 0.036 | 0.044 | 1 | 11 | Fbxl19 | -0.008 | Young |
| 0.726555 | 0.138934 | 0.116 | 0.133 | 1 | 11 | Secisbp2 | -0.017 | Young |
| 0.726773 | 0.525782 | 0.087 | 0.101 | 1 | 11 | Phf6 | -0.014 | Young |
| 0.726773 | 0.188707 | 0.087 | 0.101 | 1 | 11 | Ahsa2 | -0.014 | Young |
| 0.726812 | 0.511538 | 0.152 | 0.145 | 1 | 11 | Baiap2 | 0.007 | Young |
| 0.726819 | 0.920494 | 0.022 | 0.028 | 1 | 11 | Thumpd2 | -0.006 | Young |
| 0.726894 | 0.80886 | 0.065 | 0.077 | 1 | 11 | Nelfb | -0.012 | Young |

| | | | | | | | | |
|---|---|---|---|---|---|---|---|---|
| 0.726947 | 0.212943 | 0.058 | 0.069 | 1 | 11 | Cradd | -0.011 | Young |
| 0.726982 | 0.148996 | 0.043 | 0.052 | 1 | 11 | Fam35a | -0.009 | Young |
| 0.726999 | 1.225688 | 0.051 | 0.044 | 1 | 11 | Hnmt | 0.007 | Young |
| 0.727057 | 0.123892 | 0.732 | 0.823 | 1 | 11 | Tgfbi | -0.091 | Young |
| 0.72717 | 0.812974 | 0.138 | 0.161 | 1 | 11 | Wipf2 | -0.023 | Young |
| 0.727292 | 0.530116 | 0.029 | 0.036 | 1 | 11 | Utp4 | -0.007 | Young |
| 0.727292 | 0.458959 | 0.029 | 0.036 | 1 | 11 | Pdss2 | -0.007 | Young |
| 0.727292 | 0.395504 | 0.029 | 0.036 | 1 | 11 | Rpain | -0.007 | Young |
| 0.727292 | 0.218468 | 0.029 | 0.036 | 1 | 11 | Pigw | -0.007 | Young |
| 0.727343 | 0.151725 | 0.514 | 0.617 | 1 | 11 | Celf1 | -0.103 | Young |
| 0.727537 | 0.30454 | 0.072 | 0.085 | 1 | 11 | Gpn3 | -0.013 | Young |
| 0.727646 | 0.146521 | 0.051 | 0.06 | 1 | 11 | Nlrc4 | -0.009 | Young |
| 0.728357 | 0.455362 | 0.094 | 0.109 | 1 | 11 | Prr12 | -0.015 | Young |
| 0.728622 | 0.437205 | 0.058 | 0.069 | 1 | 11 | Tslp | -0.011 | Young |
| 0.729054 | 0.067374 | 0.072 | 0.085 | 1 | 11 | Nfs1 | -0.013 | Young |
| 0.729197 | 0.071134 | 0.174 | 0.198 | 1 | 11 | Fam241a | -0.024 | Young |
| 0.729417 | 1.494613 | 0.022 | 0.028 | 1 | 11 | Phpt1 | -0.006 | Young |
| 0.729426 | 0.408511 | 0.051 | 0.06 | 1 | 11 | Ubqln4 | -0.009 | Young |
| 0.72958 | 0.98914 | 0.029 | 0.036 | 1 | 11 | Kdm4b | -0.007 | Young |
| 0.72958 | 0.172588 | 0.029 | 0.036 | 1 | 11 | Kctd17 | -0.007 | Young |
| 0.729723 | 0.227171 | 0.797 | 0.899 | 1 | 11 | Ly86 | -0.102 | Young |
| 0.72994 | 0.013962 | 0.572 | 0.698 | 1 | 11 | Sp100 | -0.126 | Young |
| 0.729977 | 0.37216 | 0.261 | 0.31 | 1 | 11 | Ndufb4 | -0.049 | Young |
| 0.730075 | 0.378044 | 0.065 | 0.077 | 1 | 11 | Khk | -0.012 | Young |
| 0.730198 | 0.371874 | 0.123 | 0.141 | 1 | 11 | Mapkap1 | -0.018 | Young |
| 0.730455 | 0.284388 | 0.036 | 0.044 | 1 | 11 | Diexf | -0.008 | Young |
| 0.730797 | 0.504242 | 0.043 | 0.052 | 1 | 11 | Bcar3 | -0.009 | Young |
| 0.730797 | 0.238801 | 0.043 | 0.052 | 1 | 11 | Dhx35 | -0.009 | Young |
| 0.730974 | 0.41753 | 0.087 | 0.101 | 1 | 11 | Actr1b | -0.014 | Young |
| 0.730974 | 0.371698 | 0.087 | 0.101 | 1 | 11 | Pfkl | -0.014 | Young |
| 0.731015 | 0.786469 | 0.225 | 0.234 | 1 | 11 | Ap1s1 | -0.009 | Young |
| 0.731062 | 0.492117 | 0.094 | 0.109 | 1 | 11 | Smarcal1 | -0.015 | Young |
| 0.731721 | 0.448885 | 0.196 | 0.23 | 1 | 11 | Psmd14 | -0.034 | Young |
| 0.73193 | 0.123781 | 0.043 | 0.036 | 1 | 11 | Ccdc122 | 0.007 | Young |
| 0.731977 | 0.147865 | 0.058 | 0.069 | 1 | 11 | Wdfy1 | -0.011 | Young |
| 0.732376 | 0.492126 | 0.087 | 0.101 | 1 | 11 | Prorsd1 | -0.014 | Young |
| 0.732431 | 0.092291 | 0.254 | 0.282 | 1 | 11 | Anapc5 | -0.028 | Young |
| 0.732528 | 0.414969 | 0.036 | 0.044 | 1 | 11 | Txnl4b | -0.008 | Young |
| 0.733323 | 0.733271 | 0.225 | 0.23 | 1 | 11 | Mllt10 | -0.005 | Young |
| 0.733818 | 0.498743 | 0.123 | 0.141 | 1 | 11 | Papss1 | -0.018 | Young |
| 0.734079 | 0.583612 | 0.188 | 0.185 | 1 | 11 | Borcs6 | 0.003 | Young |
| 0.734604 | 0.739136 | 0.036 | 0.044 | 1 | 11 | Pced1a | -0.008 | Young |
| 0.734604 | 0.669832 | 0.036 | 0.044 | 1 | 11 | AI597479 | -0.008 | Young |
| 0.734604 | 0.428151 | 0.036 | 0.044 | 1 | 11 | Slc10a3 | -0.008 | Young |
| 0.734604 | 0.362681 | 0.036 | 0.044 | 1 | 11 | Cutc | -0.008 | Young |

| | | | | | | | | |
|---|---|---|---|---|---|---|---|---|
| 0.73462 | 1.432138 | 0.043 | 0.052 | 1 | 11 | Clasrp | -0.009 | Young |
| 0.73462 | 0.251829 | 0.043 | 0.052 | 1 | 11 | Ccdc32 | -0.009 | Young |
| 0.734774 | 0.536259 | 0.051 | 0.06 | 1 | 11 | Ankmy2 | -0.009 | Young |
| 0.734837 | 0.956358 | 0.051 | 0.044 | 1 | 11 | Fam214a | 0.007 | Young |
| 0.734854 | 0.454052 | 0.065 | 0.077 | 1 | 11 | Pskh1 | -0.012 | Young |
| 0.734854 | 0.285522 | 0.065 | 0.077 | 1 | 11 | Nos1ap | -0.012 | Young |
| 0.734899 | 0.504329 | 0.232 | 0.23 | 1 | 11 | Paf1 | 0.002 | Young |
| 0.735113 | 0.830822 | 0.123 | 0.117 | 1 | 11 | Zpr1 | 0.006 | Young |
| 0.735219 | 1.162474 | 0.13 | 0.125 | 1 | 11 | Tmem43 | 0.005 | Young |
| 0.73564 | 0.246149 | 0.268 | 0.31 | 1 | 11 | Api5 | -0.042 | Young |
| 0.736459 | 0.505223 | 0.116 | 0.133 | 1 | 11 | Ccdc90b | -0.017 | Young |
| 0.736483 | 0.731076 | 0.094 | 0.109 | 1 | 11 | Ppp5c | -0.015 | Young |
| 0.736681 | 1.403886 | 0.036 | 0.044 | 1 | 11 | Thop1 | -0.008 | Young |
| 0.737165 | 0.85534 | 0.181 | 0.177 | 1 | 11 | Cyp51 | 0.004 | Young |
| 0.737701 | 0.558388 | 0.116 | 0.133 | 1 | 11 | Gskip | -0.017 | Young |
| 0.738045 | 0.248472 | 0.065 | 0.077 | 1 | 11 | Dcp1a | -0.012 | Young |
| 0.738356 | 0.630527 | 0.043 | 0.036 | 1 | 11 | Vps33b | 0.007 | Young |
| 0.738755 | 1.014465 | 0.029 | 0.036 | 1 | 11 | Mfsd4b4 | -0.007 | Young |
| 0.739198 | 0.386801 | 0.094 | 0.109 | 1 | 11 | Eif2ak3 | -0.015 | Young |
| 0.739387 | 0.51449 | 0.254 | 0.298 | 1 | 11 | Rassf5 | -0.044 | Young |
| 0.739399 | 0.06998 | 0.087 | 0.101 | 1 | 11 | Trim30c | -0.014 | Young |
| 0.739958 | 0.751458 | 0.072 | 0.065 | 1 | 11 | Nectin1 | 0.007 | Young |
| 0.740135 | 1.085741 | 0.051 | 0.06 | 1 | 11 | Casp7 | -0.009 | Young |
| 0.740135 | 0.299632 | 0.051 | 0.06 | 1 | 11 | Rrp8 | -0.009 | Young |
| 0.740197 | 0.100881 | 0.616 | 0.673 | 1 | 11 | Gng5 | -0.057 | Young |
| 0.740366 | 0.510363 | 0.043 | 0.052 | 1 | 11 | Zfp729b | -0.009 | Young |
| 0.740519 | 0.153381 | 0.399 | 0.46 | 1 | 11 | Spag9 | -0.061 | Young |
| 0.740743 | 0.269023 | 0.174 | 0.198 | 1 | 11 | Lyl1 | -0.024 | Young |
| 0.741055 | 1.093195 | 0.029 | 0.036 | 1 | 11 | Tmem206 | -0.007 | Young |
| 0.741242 | 0.539037 | 0.065 | 0.077 | 1 | 11 | Gnl3l | -0.012 | Young |
| 0.741242 | 0.383401 | 0.065 | 0.077 | 1 | 11 | Rnf185 | -0.012 | Young |
| 0.741242 | 0.280083 | 0.065 | 0.077 | 1 | 11 | Mtap | -0.012 | Young |
| 0.741495 | 0.09162 | 0.109 | 0.125 | 1 | 11 | Spn | -0.016 | Young |
| 0.742284 | 1.412216 | 0.043 | 0.052 | 1 | 11 | Dut | -0.009 | Young |
| 0.742284 | 0.826032 | 0.043 | 0.052 | 1 | 11 | Psmg1 | -0.009 | Young |
| 0.742284 | 0.782644 | 0.043 | 0.052 | 1 | 11 | Gmnn | -0.009 | Young |
| 0.742544 | 0.029688 | 0.101 | 0.117 | 1 | 11 | Mob3b | -0.016 | Young |
| 0.742611 | 0.058663 | 0.949 | 0.972 | 1 | 11 | Hspa8 | -0.023 | Young |
| 0.742663 | 0.172657 | 0.145 | 0.161 | 1 | 11 | Seh1l | -0.016 | Young |
| 0.742672 | 0.530464 | 0.116 | 0.133 | 1 | 11 | Slu7 | -0.017 | Young |
| 0.74281 | 1.271752 | 0.058 | 0.052 | 1 | 11 | Zdhhc13 | 0.006 | Young |
| 0.742992 | 0.277701 | 0.304 | 0.347 | 1 | 11 | Psma5 | -0.043 | Young |
| 0.743098 | 0.493655 | 0.225 | 0.23 | 1 | 11 | Trappc6b | -0.005 | Young |
| 0.743196 | 1.230071 | 0.109 | 0.129 | 1 | 11 | Tbcel | -0.02 | Young |
| 0.743277 | 0.30928 | 0.094 | 0.109 | 1 | 11 | Spire1 | -0.015 | Young |

| | | | | | | | | |
|---|---|---|---|---|---|---|---|---|
| 0.743327 | 0.368844 | 0.188 | 0.214 | 1 | 11 | Evi5 | -0.026 | Young |
| 0.743716 | 0.694144 | 0.051 | 0.06 | 1 | 11 | 2510002D2 | -0.009 | Young |
| 0.74376 | 1.319503 | 0.058 | 0.069 | 1 | 11 | Thbd | -0.011 | Young |
| 0.744443 | 0.669101 | 0.065 | 0.077 | 1 | 11 | Mrps30 | -0.012 | Young |
| 0.744702 | 0.26157 | 0.181 | 0.177 | 1 | 11 | Plaur | 0.004 | Young |
| 0.744957 | 0.35338 | 0.08 | 0.093 | 1 | 11 | Xrcc1 | -0.013 | Young |
| 0.745461 | 0.072216 | 0.725 | 0.77 | 1 | 11 | Gnb2 | -0.045 | Young |
| 0.74562 | 0.41225 | 0.493 | 0.621 | 1 | 11 | Chmp4b | -0.128 | Young |
| 0.746047 | 0.774785 | 0.08 | 0.073 | 1 | 11 | Pigu | 0.007 | Young |
| 0.746047 | 0.64146 | 0.08 | 0.073 | 1 | 11 | Phlpp2 | 0.007 | Young |
| 0.746126 | 0.708048 | 0.043 | 0.052 | 1 | 11 | Sephs1 | -0.009 | Young |
| 0.746326 | 0.336511 | 0.348 | 0.411 | 1 | 11 | Srrm1 | -0.063 | Young |
| 0.746379 | 1.862264 | 0.029 | 0.024 | 1 | 11 | Nt5c3b | 0.005 | Young |
| 0.746379 | 1.397181 | 0.029 | 0.024 | 1 | 11 | Gtf2h3 | 0.005 | Young |
| 0.747096 | 0.833785 | 0.036 | 0.044 | 1 | 11 | Npat | -0.008 | Young |
| 0.747138 | 0.748659 | 0.058 | 0.069 | 1 | 11 | Armt1 | -0.011 | Young |
| 0.747303 | 0.595967 | 0.051 | 0.06 | 1 | 11 | Dnajb2 | -0.009 | Young |
| 0.747648 | 0.827108 | 0.065 | 0.077 | 1 | 11 | Naa40 | -0.012 | Young |
| 0.747687 | 0.382186 | 0.225 | 0.258 | 1 | 11 | Zfp652 | -0.033 | Young |
| 0.747856 | 0.507556 | 0.087 | 0.101 | 1 | 11 | Prps1 | -0.014 | Young |
| 0.74805 | 0.846838 | 0.043 | 0.052 | 1 | 11 | Bag5 | -0.009 | Young |
| 0.750502 | 0.097105 | 0.58 | 0.694 | 1 | 11 | Cox7a2l | -0.114 | Young |
| 0.750925 | 0.304085 | 0.522 | 0.641 | 1 | 11 | Dnaja1 | -0.119 | Young |
| 0.750947 | 0.354648 | 0.384 | 0.456 | 1 | 11 | Bag1 | -0.072 | Young |
| 0.750955 | 0.2417 | 0.435 | 0.504 | 1 | 11 | Vdac2 | -0.069 | Young |
| 0.751339 | 1.194399 | 0.203 | 0.202 | 1 | 11 | Gadd45g | 0.001 | Young |
| 0.751456 | 0.959939 | 0.109 | 0.101 | 1 | 11 | Rae1 | 0.008 | Young |
| 0.751624 | 2.333882 | 0.029 | 0.024 | 1 | 11 | Gstcd | 0.005 | Young |
| 0.751624 | 1.870357 | 0.029 | 0.024 | 1 | 11 | Bcs1l | 0.005 | Young |
| 0.751624 | 1.503479 | 0.029 | 0.024 | 1 | 11 | Arv1 | 0.005 | Young |
| 0.751624 | 1.476003 | 0.029 | 0.024 | 1 | 11 | Klhl25 | 0.005 | Young |
| 0.752254 | 0.972992 | 0.13 | 0.129 | 1 | 11 | Ppp4r1 | 0.001 | Young |
| 0.752461 | 0.065825 | 0.217 | 0.238 | 1 | 11 | Parp11 | -0.021 | Young |
| 0.752488 | 0.92444 | 0.254 | 0.27 | 1 | 11 | Ahcyl2 | -0.016 | Young |
| 0.752561 | 0.279531 | 0.297 | 0.343 | 1 | 11 | Dctn1 | -0.046 | Young |
| 0.753052 | 0.43187 | 0.319 | 0.343 | 1 | 11 | Ppp4r3b | -0.024 | Young |
| 0.753604 | 0.361647 | 0.326 | 0.387 | 1 | 11 | Hnrnpa1 | -0.061 | Young |
| 0.753765 | 0.51688 | 0.08 | 0.093 | 1 | 11 | Snapc3 | -0.013 | Young |
| 0.75425 | 1.328572 | 0.029 | 0.024 | 1 | 11 | Phactr1 | 0.005 | Young |
| 0.75425 | 0.71761 | 0.029 | 0.024 | 1 | 11 | Atp5s | 0.005 | Young |
| 0.754269 | 0.235525 | 0.522 | 0.597 | 1 | 11 | C5ar1 | -0.075 | Young |
| 0.754313 | 0.690494 | 0.123 | 0.117 | 1 | 11 | Thumpd3 | 0.006 | Young |
| 0.755398 | 0.290384 | 0.268 | 0.274 | 1 | 11 | Sf3b3 | -0.006 | Young |
| 0.755461 | 1.302594 | 0.036 | 0.044 | 1 | 11 | N6amt1 | -0.008 | Young |
| 0.755557 | 0.378324 | 0.109 | 0.101 | 1 | 11 | Zfp397 | 0.008 | Young |

| | | | | | | | | |
|---|---|---|---|---|---|---|---|---|
| 0.755599 | 1.03646 | 0.123 | 0.117 | 1 | 11 | Nipsnap2 | 0.006 | Young |
| 0.756293 | 0.95067 | 0.051 | 0.06 | 1 | 11 | Zfp626 | -0.009 | Young |
| 0.756293 | 0.725475 | 0.051 | 0.06 | 1 | 11 | Ifit1bl1 | -0.009 | Young |
| 0.756398 | 0.832417 | 0.116 | 0.133 | 1 | 11 | Mtx2 | -0.017 | Young |
| 0.756531 | 0.485709 | 0.051 | 0.044 | 1 | 11 | Fam110b | 0.007 | Young |
| 0.756561 | 0.984497 | 0.072 | 0.065 | 1 | 11 | Bmf | 0.007 | Young |
| 0.756561 | 0.449574 | 0.072 | 0.065 | 1 | 11 | Dctpp1 | 0.007 | Young |
| 0.756709 | 0.717453 | 0.08 | 0.093 | 1 | 11 | Rbm15b | -0.013 | Young |
| 0.756754 | 0.015101 | 0.123 | 0.137 | 1 | 11 | Retreg3 | -0.014 | Young |
| 0.756879 | 2.271023 | 0.029 | 0.024 | 1 | 11 | AA467197 | 0.005 | Young |
| 0.756879 | 1.502884 | 0.029 | 0.024 | 1 | 11 | Lrrc45 | 0.005 | Young |
| 0.756879 | 1.269171 | 0.029 | 0.024 | 1 | 11 | Cd2 | 0.005 | Young |
| 0.756879 | 1.146654 | 0.029 | 0.024 | 1 | 11 | Slc40a1 | 0.005 | Young |
| 0.756879 | 1.132491 | 0.029 | 0.024 | 1 | 11 | Tango6 | 0.005 | Young |
| 0.756879 | 0.899657 | 0.029 | 0.024 | 1 | 11 | Gm47242 | 0.005 | Young |
| 0.757122 | 0.153667 | 0.514 | 0.524 | 1 | 11 | Igfbp4 | -0.01 | Young |
| 0.757557 | 0.963771 | 0.036 | 0.044 | 1 | 11 | Bcas3 | -0.008 | Young |
| 0.75761 | 0.9364 | 0.167 | 0.165 | 1 | 11 | Hmgcl | 0.002 | Young |
| 0.75769 | 1.62448 | 0.043 | 0.052 | 1 | 11 | Pyurf | -0.009 | Young |
| 0.758095 | 0.770721 | 0.051 | 0.06 | 1 | 11 | Slc35b4 | -0.009 | Young |
| 0.758513 | 0.778384 | 0.051 | 0.044 | 1 | 11 | Nme4 | 0.007 | Young |
| 0.758514 | 0.168147 | 0.587 | 0.685 | 1 | 11 | Srsf3 | -0.098 | Young |
| 0.758913 | 0.384635 | 0.174 | 0.202 | 1 | 11 | Ctcf | -0.028 | Young |
| 0.759326 | 0.567122 | 0.123 | 0.141 | 1 | 11 | Rnf169 | -0.018 | Young |
| 0.759487 | 0.609226 | 0.13 | 0.149 | 1 | 11 | Siglece | -0.019 | Young |
| 0.759512 | 1.438212 | 0.029 | 0.024 | 1 | 11 | Galnt3 | 0.005 | Young |
| 0.759622 | 1.305849 | 0.043 | 0.052 | 1 | 11 | C2cd2l | -0.009 | Young |
| 0.759664 | 0.694567 | 0.094 | 0.109 | 1 | 11 | Rmi1 | -0.015 | Young |
| 0.760598 | 0.435363 | 0.087 | 0.101 | 1 | 11 | Syne1 | -0.014 | Young |
| 0.761012 | 0.1561 | 0.558 | 0.593 | 1 | 11 | Atp1b3 | -0.035 | Young |
| 0.761117 | 0.333433 | 0.159 | 0.177 | 1 | 11 | Kif2a | -0.018 | Young |
| 0.761446 | 0.708126 | 0.145 | 0.165 | 1 | 11 | Rbl2 | -0.02 | Young |
| 0.761703 | 0.984859 | 0.051 | 0.06 | 1 | 11 | Mul1 | -0.009 | Young |
| 0.762146 | 0.890663 | 0.029 | 0.024 | 1 | 11 | Rgl3 | 0.005 | Young |
| 0.762146 | 0.855101 | 0.029 | 0.024 | 1 | 11 | Simc1 | 0.005 | Young |
| 0.762588 | 0.356247 | 0.145 | 0.165 | 1 | 11 | Mast3 | -0.02 | Young |
| 0.763135 | 0.718142 | 0.21 | 0.218 | 1 | 11 | Pik3r6 | -0.008 | Young |
| 0.763354 | 0.67926 | 0.254 | 0.266 | 1 | 11 | Arhgap45 | -0.012 | Young |
| 0.763399 | 0.145241 | 0.486 | 0.556 | 1 | 11 | Rnf19b | -0.07 | Young |
| 0.763918 | 0.919438 | 0.13 | 0.125 | 1 | 11 | Il6 | 0.005 | Young |
| 0.764293 | 0.525024 | 0.254 | 0.266 | 1 | 11 | Cpsf2 | -0.012 | Young |
| 0.764468 | 0.772393 | 0.051 | 0.044 | 1 | 11 | Dxo | 0.007 | Young |
| 0.764468 | 0.563985 | 0.051 | 0.044 | 1 | 11 | Zfp839 | 0.007 | Young |
| 0.764809 | 0.236547 | 0.283 | 0.331 | 1 | 11 | Bud31 | -0.048 | Young |
| 0.76486 | 0.056643 | 0.101 | 0.093 | 1 | 11 | Kansl2 | 0.008 | Young |

| | | | | | | | | |
|---|---|---|---|---|---|---|---|---|
| 0.765072 | 0.911283 | 0.188 | 0.19 | 1 | 11 | Zranb1 | -0.002 | Young |
| 0.765173 | 0.976639 | 0.116 | 0.133 | 1 | 11 | Hps5 | -0.017 | Young |
| 0.765316 | 0.915527 | 0.051 | 0.06 | 1 | 11 | Cops7b | -0.009 | Young |
| 0.765439 | 0.911269 | 0.138 | 0.133 | 1 | 11 | Arl13b | 0.005 | Young |
| 0.766078 | 0.328398 | 0.13 | 0.145 | 1 | 11 | Cdc34 | -0.015 | Young |
| 0.766427 | 0.510478 | 0.152 | 0.149 | 1 | 11 | Mthfd2 | 0.003 | Young |
| 0.766457 | 0.34913 | 0.051 | 0.044 | 1 | 11 | Haus6 | 0.007 | Young |
| 0.766462 | 0.174762 | 0.79 | 0.883 | 1 | 11 | Dazap2 | -0.093 | Young |
| 0.76665 | 0.635681 | 0.174 | 0.169 | 1 | 11 | Dlgap4 | 0.005 | Young |
| 0.766731 | 1.182391 | 0.058 | 0.052 | 1 | 11 | Rhbdd2 | 0.006 | Young |
| 0.76708 | 0.848451 | 0.203 | 0.214 | 1 | 11 | Clip1 | -0.011 | Young |
| 0.767425 | 1.32976 | 0.029 | 0.024 | 1 | 11 | Mtif3 | 0.005 | Young |
| 0.767425 | 1.087568 | 0.029 | 0.024 | 1 | 11 | Lyrm9 | 0.005 | Young |
| 0.76758 | 0.098971 | 0.725 | 0.823 | 1 | 11 | Rpl14 | -0.098 | Young |
| 0.767583 | 1.005824 | 0.101 | 0.097 | 1 | 11 | Lncpint | 0.004 | Young |
| 0.767996 | 0.58367 | 0.268 | 0.282 | 1 | 11 | Elmsan1 | -0.014 | Young |
| 0.768816 | 0.875257 | 0.167 | 0.165 | 1 | 11 | Svbp | 0.002 | Young |
| 0.769114 | 0.513029 | 0.138 | 0.133 | 1 | 11 | Snrnp48 | 0.005 | Young |
| 0.770068 | 1.051657 | 0.029 | 0.024 | 1 | 11 | Slc52a2 | 0.005 | Young |
| 0.770068 | 0.660876 | 0.029 | 0.024 | 1 | 11 | Ubxn2b | 0.005 | Young |
| 0.770068 | 0.517428 | 0.029 | 0.024 | 1 | 11 | Flrt3 | 0.005 | Young |
| 0.770423 | 0.008869 | 0.732 | 0.778 | 1 | 11 | Rpl10 | -0.046 | Young |
| 0.770432 | 1.154333 | 0.058 | 0.052 | 1 | 11 | Jpx | 0.006 | Young |
| 0.770432 | 1.124133 | 0.058 | 0.052 | 1 | 11 | Pafah1b3 | 0.006 | Young |
| 0.770438 | 1.245767 | 0.051 | 0.044 | 1 | 11 | Pank4 | 0.007 | Young |
| 0.770998 | 0.111622 | 0.377 | 0.423 | 1 | 11 | Csk | -0.046 | Young |
| 0.771409 | 0.380707 | 0.08 | 0.073 | 1 | 11 | Gpr183 | 0.007 | Young |
| 0.771503 | 0.01366 | 0.616 | 0.714 | 1 | 11 | Epsti1 | -0.098 | Young |
| 0.771518 | 0.522635 | 0.254 | 0.302 | 1 | 11 | Tial1 | -0.048 | Young |
| 0.772037 | 0.866279 | 0.13 | 0.149 | 1 | 11 | Man1b1 | -0.019 | Young |
| 0.772712 | 1.476895 | 0.065 | 0.06 | 1 | 11 | Msh2 | 0.005 | Young |
| 0.772714 | 0.700868 | 0.029 | 0.024 | 1 | 11 | Helq | 0.005 | Young |
| 0.772714 | 0.282347 | 0.029 | 0.024 | 1 | 11 | Mier2 | 0.005 | Young |
| 0.772976 | 0.126543 | 0.551 | 0.613 | 1 | 11 | Mtdh | -0.062 | Young |
| 0.773036 | 0.67033 | 0.101 | 0.117 | 1 | 11 | Atp8b4 | -0.016 | Young |
| 0.773666 | 0.604095 | 0.109 | 0.125 | 1 | 11 | 1700025G( | -0.016 | Young |
| 0.775 | 0.087007 | 0.174 | 0.161 | 1 | 11 | Nhsl2 | 0.013 | Young |
| 0.775324 | 0.539011 | 0.246 | 0.286 | 1 | 11 | Cic | -0.04 | Young |
| 0.775771 | 0.449982 | 0.094 | 0.105 | 1 | 11 | Adcy9 | -0.011 | Young |
| 0.775993 | 1.154156 | 0.058 | 0.052 | 1 | 11 | Tex30 | 0.006 | Young |
| 0.776116 | 0.278764 | 0.101 | 0.113 | 1 | 11 | Agap1 | -0.012 | Young |
| 0.776194 | 0.901758 | 0.152 | 0.153 | 1 | 11 | Sms | -0.001 | Young |
| 0.776674 | 0.785354 | 0.087 | 0.081 | 1 | 11 | Rnf215 | 0.006 | Young |
| 0.77674 | 0.322382 | 0.399 | 0.484 | 1 | 11 | Diaph1 | -0.085 | Young |
| 0.777041 | 0.633107 | 0.101 | 0.117 | 1 | 11 | Nt5c2 | -0.016 | Young |

| | | | | | | | | |
|---|---|---|---|---|---|---|---|---|
| 0.777272 | 0.401056 | 0.203 | 0.23 | 1 | 11 | Gm26740 | -0.027 | Young |
| 0.777555 | 0.898555 | 0.109 | 0.125 | 1 | 11 | Med16 | -0.016 | Young |
| 0.777863 | 0.809638 | 0.109 | 0.105 | 1 | 11 | Med29 | 0.004 | Young |
| 0.778013 | 0.553707 | 0.029 | 0.024 | 1 | 11 | Adnp | 0.005 | Young |
| 0.778013 | 0.269812 | 0.029 | 0.024 | 1 | 11 | Gm11772 | 0.005 | Young |
| 0.778013 | 0.081844 | 0.029 | 0.024 | 1 | 11 | Coa4 | 0.005 | Young |
| 0.77836 | 0.288192 | 0.116 | 0.129 | 1 | 11 | Gle1 | -0.013 | Young |
| 0.778987 | 0.912505 | 0.254 | 0.274 | 1 | 11 | Hnrnpr | -0.02 | Young |
| 0.779216 | 0.208851 | 0.442 | 0.532 | 1 | 11 | Irf2 | -0.09 | Young |
| 0.779623 | 0.245997 | 0.225 | 0.25 | 1 | 11 | Smap1 | -0.025 | Young |
| 0.779871 | 0.65525 | 0.275 | 0.323 | 1 | 11 | Smim14 | -0.048 | Young |
| 0.782736 | 0.611342 | 0.167 | 0.185 | 1 | 11 | Ppp1r11 | -0.018 | Young |
| 0.782784 | 0.849416 | 0.072 | 0.085 | 1 | 11 | Ctc1 | -0.013 | Young |
| 0.783323 | 0.426749 | 0.029 | 0.024 | 1 | 11 | Cdan1 | 0.005 | Young |
| 0.783459 | 0.006072 | 0.116 | 0.129 | 1 | 11 | Ccnk | -0.013 | Young |
| 0.783516 | 1.350409 | 0.152 | 0.157 | 1 | 11 | Gabpb2 | -0.005 | Young |
| 0.783728 | 0.188985 | 0.116 | 0.109 | 1 | 11 | Scoc | 0.007 | Young |
| 0.78448 | 0.669047 | 0.246 | 0.262 | 1 | 11 | Nrd1 | -0.016 | Young |
| 0.784561 | 0.128476 | 0.5 | 0.573 | 1 | 11 | Etnk1 | -0.073 | Young |
| 0.784882 | 0.449724 | 0.196 | 0.19 | 1 | 11 | Fut8 | 0.006 | Young |
| 0.784918 | 0.353757 | 0.167 | 0.161 | 1 | 11 | Nfkbib | 0.006 | Young |
| 0.785067 | 0.557304 | 0.116 | 0.109 | 1 | 11 | Rgs14 | 0.007 | Young |
| 0.785982 | 0.046162 | 0.029 | 0.024 | 1 | 11 | Ttl | 0.005 | Young |
| 0.786529 | 0.978386 | 0.188 | 0.198 | 1 | 11 | Stard3 | -0.01 | Young |
| 0.78661 | 0.039939 | 0.667 | 0.786 | 1 | 11 | Gdi2 | -0.119 | Young |
| 0.786648 | 0.668664 | 0.109 | 0.125 | 1 | 11 | Nagpa | -0.016 | Young |
| 0.786845 | 0.462867 | 0.152 | 0.173 | 1 | 11 | Prkd3 | -0.021 | Young |
| 0.787056 | 0.375655 | 0.275 | 0.31 | 1 | 11 | Cnih4 | -0.035 | Young |
| 0.788346 | 0.242339 | 0.522 | 0.577 | 1 | 11 | Mrfap1 | -0.055 | Young |
| 0.78876 | 0.034163 | 0.08 | 0.089 | 1 | 11 | Exosc4 | -0.009 | Young |
| 0.789554 | 0.293837 | 0.261 | 0.266 | 1 | 11 | Add3 | -0.005 | Young |
| 0.790404 | 0.509107 | 0.341 | 0.335 | 1 | 11 | Ptgs2 | 0.006 | Young |
| 0.790779 | 0.340965 | 0.297 | 0.355 | 1 | 11 | Rnf4 | -0.058 | Young |
| 0.79128 | 0.131175 | 0.616 | 0.718 | 1 | 11 | Ptpn18 | -0.102 | Young |
| 0.792293 | 0.713835 | 0.486 | 0.548 | 1 | 11 | Ccl12 | -0.062 | Young |
| 0.792848 | 0.399002 | 0.449 | 0.52 | 1 | 11 | Cap1 | -0.071 | Young |
| 0.792997 | 0.155064 | 0.261 | 0.286 | 1 | 11 | Lmbrd1 | -0.025 | Young |
| 0.793189 | 1.665224 | 0.058 | 0.069 | 1 | 11 | Orc2 | -0.011 | Young |
| 0.793286 | 0.282178 | 0.348 | 0.367 | 1 | 11 | Nr4a2 | -0.019 | Young |
| 0.793804 | 0.41085 | 0.232 | 0.274 | 1 | 11 | Rraga | -0.042 | Young |
| 0.795042 | 0.483784 | 0.087 | 0.081 | 1 | 11 | Kctd5 | 0.006 | Young |
| 0.79517 | 0.224508 | 0.152 | 0.169 | 1 | 11 | Gpr160 | -0.017 | Young |
| 0.795325 | 1.060817 | 0.072 | 0.069 | 1 | 11 | Syt11 | 0.003 | Young |
| 0.795759 | 0.284259 | 0.551 | 0.637 | 1 | 11 | Nfkbid | -0.086 | Young |
| 0.79607 | 0.496103 | 0.304 | 0.327 | 1 | 11 | Snw1 | -0.023 | Young |

| | | | | | | | | |
|---|---|---|---|---|---|---|---|---|
| 0.796303 | 1.35993 | 0.152 | 0.157 | 1 | 11 | Ube2v2 | -0.005 | Young |
| 0.796838 | 0.494711 | 0.174 | 0.173 | 1 | 11 | Papd7 | 0.001 | Young |
| 0.797984 | 0.379728 | 0.333 | 0.379 | 1 | 11 | Azin1 | -0.046 | Young |
| 0.79834 | 0.818531 | 0.072 | 0.085 | 1 | 11 | Cog6 | -0.013 | Young |
| 0.798806 | 0.0586 | 0.594 | 0.698 | 1 | 11 | Jpt1 | -0.104 | Young |
| 0.798815 | 0.428084 | 0.348 | 0.423 | 1 | 11 | Prpf40a | -0.075 | Young |
| 0.798865 | 0.805085 | 0.152 | 0.177 | 1 | 11 | Kpna3 | -0.025 | Young |
| 0.799116 | 0.653756 | 0.196 | 0.23 | 1 | 11 | Chd2 | -0.034 | Young |
| 0.799238 | 0.443318 | 0.304 | 0.371 | 1 | 11 | Atf6 | -0.067 | Young |
| 0.799253 | 1.430275 | 0.036 | 0.032 | 1 | 11 | Smpd2 | 0.004 | Young |
| 0.799468 | 0.367437 | 0.275 | 0.335 | 1 | 11 | Tmem167 | -0.06 | Young |
| 0.79959 | 1.245782 | 0.152 | 0.153 | 1 | 11 | Svil | -0.001 | Young |
| 0.800219 | 0.447866 | 0.058 | 0.052 | 1 | 11 | Siglech | 0.006 | Young |
| 0.800232 | 0.0743 | 0.572 | 0.661 | 1 | 11 | Ddx6 | -0.089 | Young |
| 0.800699 | 0.541658 | 0.225 | 0.258 | 1 | 11 | Bfar | -0.033 | Young |
| 0.801086 | 0.901128 | 0.29 | 0.319 | 1 | 11 | Gbp5 | -0.029 | Young |
| 0.801579 | 0.032792 | 0.058 | 0.065 | 1 | 11 | 1110008P1 | -0.007 | Young |
| 0.80239 | 0.409089 | 0.08 | 0.089 | 1 | 11 | Tsc2 | -0.009 | Young |
| 0.802649 | 0.171789 | 0.08 | 0.073 | 1 | 11 | Otud6b | 0.007 | Young |
| 0.803867 | 0.321138 | 0.297 | 0.319 | 1 | 11 | Ostc | -0.022 | Young |
| 0.803944 | 0.269049 | 0.116 | 0.129 | 1 | 11 | Lcmt1 | -0.013 | Young |
| 0.803962 | 1.462758 | 0.036 | 0.032 | 1 | 11 | Pwwp2b | 0.004 | Young |
| 0.803964 | 0.248304 | 0.058 | 0.052 | 1 | 11 | Alg11 | 0.006 | Young |
| 0.803971 | 0.50671 | 0.145 | 0.141 | 1 | 11 | Grsf1 | 0.004 | Young |
| 0.804303 | 0.419462 | 0.188 | 0.214 | 1 | 11 | Zkscan3 | -0.026 | Young |
| 0.804479 | 0.272812 | 0.522 | 0.629 | 1 | 11 | Stk17b | -0.107 | Young |
| 0.805063 | 0.538124 | 0.384 | 0.484 | 1 | 11 | Crlf3 | -0.1 | Young |
| 0.805306 | 0.904716 | 0.072 | 0.069 | 1 | 11 | Ston2 | 0.003 | Young |
| 0.805627 | 0.16814 | 0.391 | 0.448 | 1 | 11 | Sowahc | -0.057 | Young |
| 0.806077 | 0.598838 | 0.391 | 0.508 | 1 | 11 | Znfx1 | -0.117 | Young |
| 0.806231 | 1.057261 | 0.109 | 0.125 | 1 | 11 | Daam1 | -0.016 | Young |
| 0.806292 | 0.030679 | 0.79 | 0.855 | 1 | 11 | Rpl24 | -0.065 | Young |
| 0.806319 | 1.046675 | 0.036 | 0.032 | 1 | 11 | Dnmbp | 0.004 | Young |
| 0.806858 | 0.677793 | 0.138 | 0.153 | 1 | 11 | Med14 | -0.015 | Young |
| 0.806917 | 0.770883 | 0.196 | 0.226 | 1 | 11 | Clcn5 | -0.03 | Young |
| 0.807697 | 0.374522 | 0.109 | 0.121 | 1 | 11 | Trappc9 | -0.012 | Young |
| 0.807865 | 0.133147 | 0.493 | 0.581 | 1 | 11 | Minos1 | -0.088 | Young |
| 0.807905 | 0.214762 | 0.312 | 0.363 | 1 | 11 | Ap3s1 | -0.051 | Young |
| 0.807945 | 0.186406 | 0.645 | 0.694 | 1 | 11 | Cflar | -0.049 | Young |
| 0.808432 | 0.557322 | 0.087 | 0.097 | 1 | 11 | Sirt1 | -0.01 | Young |
| 0.808467 | 0.198455 | 0.08 | 0.089 | 1 | 11 | Map3k3 | -0.009 | Young |
| 0.808678 | 1.41306 | 0.036 | 0.032 | 1 | 11 | Vwf | 0.004 | Young |
| 0.809108 | 0.557339 | 0.159 | 0.181 | 1 | 11 | Hlx | -0.022 | Young |
| 0.809324 | 0.50383 | 0.181 | 0.206 | 1 | 11 | Orai3 | -0.025 | Young |
| 0.809501 | 0.882229 | 0.065 | 0.06 | 1 | 11 | Zkscan1 | 0.005 | Young |

| | | | | | | | | |
|---|---|---|---|---|---|---|---|---|
| 0.809958 | 0.623167 | 0.123 | 0.121 | 1 | 11 | P3h2 | 0.002 | Young |
| 0.810197 | 0.400076 | 0.507 | 0.593 | 1 | 11 | Evi2a | -0.086 | Young |
| 0.811039 | 1.14471 | 0.036 | 0.032 | 1 | 11 | Tlr11 | 0.004 | Young |
| 0.811347 | 0.420654 | 0.174 | 0.177 | 1 | 11 | 2610037D0 | -0.003 | Young |
| 0.811424 | 0.222521 | 0.268 | 0.306 | 1 | 11 | P2rx7 | -0.038 | Young |
| 0.812398 | 0.433594 | 0.225 | 0.258 | 1 | 11 | Naa15 | -0.033 | Young |
| 0.813028 | 0.069854 | 0.065 | 0.06 | 1 | 11 | Mcm4 | 0.005 | Young |
| 0.813334 | 0.911584 | 0.101 | 0.117 | 1 | 11 | Pex13 | -0.016 | Young |
| 0.813401 | 0.894258 | 0.036 | 0.032 | 1 | 11 | Mtrf1l | 0.004 | Young |
| 0.813524 | 0.196774 | 0.087 | 0.081 | 1 | 11 | Prkrip1 | 0.006 | Young |
| 0.814388 | 0.340362 | 0.072 | 0.081 | 1 | 11 | Inf2 | -0.009 | Young |
| 0.814388 | 0.147617 | 0.072 | 0.081 | 1 | 11 | Mrpl50 | -0.009 | Young |
| 0.815766 | 0.828349 | 0.036 | 0.032 | 1 | 11 | Flywch1 | 0.004 | Young |
| 0.816099 | 0.522959 | 0.268 | 0.31 | 1 | 11 | Fnbp1l | -0.042 | Young |
| 0.816559 | 0.58966 | 0.065 | 0.06 | 1 | 11 | Sepsecs | 0.005 | Young |
| 0.816559 | 0.304213 | 0.065 | 0.06 | 1 | 11 | Pkib | 0.005 | Young |
| 0.816991 | 0.361894 | 0.065 | 0.073 | 1 | 11 | Coq10a | -0.008 | Young |
| 0.816991 | 0.272472 | 0.065 | 0.073 | 1 | 11 | Gramd4 | -0.008 | Young |
| 0.817106 | 0.189658 | 0.058 | 0.052 | 1 | 11 | Naf1 | 0.006 | Young |
| 0.818116 | 0.203863 | 0.507 | 0.621 | 1 | 11 | Pfdn5 | -0.114 | Young |
| 0.81825 | 0.689195 | 0.101 | 0.113 | 1 | 11 | Tmem185b | -0.012 | Young |
| 0.818335 | 0.258648 | 0.428 | 0.492 | 1 | 11 | Tomm20 | -0.064 | Young |
| 0.818461 | 0.810102 | 0.167 | 0.19 | 1 | 11 | Mau2 | -0.023 | Young |
| 0.818988 | 0.765142 | 0.051 | 0.056 | 1 | 11 | Nfkbil1 | -0.005 | Young |
| 0.819308 | 0.228135 | 0.42 | 0.492 | 1 | 11 | Snx10 | -0.072 | Young |
| 0.820337 | 0.692411 | 0.072 | 0.069 | 1 | 11 | Terf1 | 0.003 | Young |
| 0.820499 | 0.828246 | 0.036 | 0.032 | 1 | 11 | Zfp113 | 0.004 | Young |
| 0.82074 | 0.6374 | 0.094 | 0.105 | 1 | 11 | Tm7sf3 | -0.011 | Young |
| 0.820771 | 0.103821 | 0.645 | 0.718 | 1 | 11 | Parp14 | -0.073 | Young |
| 0.821174 | 0.069272 | 0.638 | 0.702 | 1 | 11 | Atp5g2 | -0.064 | Young |
| 0.821218 | 0.39697 | 0.109 | 0.117 | 1 | 11 | Ephx1 | -0.008 | Young |
| 0.821297 | 0.980778 | 0.174 | 0.181 | 1 | 11 | P2ry12 | -0.007 | Young |
| 0.821471 | 0.439176 | 0.159 | 0.177 | 1 | 11 | Ei24 | -0.018 | Young |
| 0.821766 | 0.617894 | 0.181 | 0.181 | 1 | 11 | Gigyf1 | 0 | Young |
| 0.822173 | 0.0852 | 0.536 | 0.637 | 1 | 11 | Tm6sf1 | -0.101 | Young |
| 0.822355 | 0.194495 | 0.072 | 0.081 | 1 | 11 | Mtif2 | -0.009 | Young |
| 0.822358 | 0.37372 | 0.101 | 0.113 | 1 | 11 | Rpp21 | -0.012 | Young |
| 0.822826 | 0.484165 | 0.5 | 0.516 | 1 | 11 | Rsad2 | -0.016 | Young |
| 0.822868 | 0.476653 | 0.036 | 0.032 | 1 | 11 | Zfp758 | 0.004 | Young |
| 0.823572 | 0.327356 | 0.094 | 0.105 | 1 | 11 | Rptor | -0.011 | Young |
| 0.823687 | 0.121221 | 0.065 | 0.073 | 1 | 11 | 1190002N1 | -0.008 | Young |
| 0.824039 | 0.46602 | 0.355 | 0.395 | 1 | 11 | Ppig | -0.04 | Young |
| 0.824522 | 0.125398 | 0.507 | 0.629 | 1 | 11 | Reep3 | -0.122 | Young |
| 0.824552 | 0.502766 | 0.087 | 0.097 | 1 | 11 | Map2k5 | -0.01 | Young |
| 0.824989 | 0.005531 | 0.094 | 0.105 | 1 | 11 | Cyp4f13 | -0.011 | Young |

| | | | | | | | | |
|---|---|---|---|---|---|---|---|---|
| 0.825239 | 0.678728 | 0.036 | 0.032 | 1 | 11 | Toe1 | 0.004 | Young |
| 0.825239 | 0.464975 | 0.036 | 0.032 | 1 | 11 | Spata13 | 0.004 | Young |
| 0.825313 | 0.416392 | 0.094 | 0.089 | 1 | 11 | Fam60a | 0.005 | Young |
| 0.825317 | 0.090136 | 0.406 | 0.472 | 1 | 11 | Sp110 | -0.066 | Young |
| 0.825721 | 0.611536 | 0.254 | 0.29 | 1 | 11 | Dynll2 | -0.036 | Young |
| 0.825751 | 0.016202 | 0.043 | 0.048 | 1 | 11 | AW554918 | -0.005 | Young |
| 0.825965 | 0.679462 | 0.275 | 0.339 | 1 | 11 | Ogfrl1 | -0.064 | Young |
| 0.826323 | 0.308114 | 0.428 | 0.484 | 1 | 11 | Lims1 | -0.056 | Young |
| 0.826404 | 0.574866 | 0.362 | 0.415 | 1 | 11 | Ubn2 | -0.053 | Young |
| 0.82656 | 1.964674 | 0.043 | 0.04 | 1 | 11 | Mvb12b | 0.003 | Young |
| 0.827174 | 0.356664 | 0.065 | 0.06 | 1 | 11 | Morc2a | 0.005 | Young |
| 0.827505 | 0.158023 | 0.167 | 0.181 | 1 | 11 | Cldnd1 | -0.014 | Young |
| 0.827954 | 0.01424 | 0.681 | 0.786 | 1 | 11 | Celf2 | -0.105 | Young |
| 0.82806 | 0.33199 | 0.058 | 0.065 | 1 | 11 | Zfp524 | -0.007 | Young |
| 0.82806 | 0.286901 | 0.058 | 0.065 | 1 | 11 | Trmt2b | -0.007 | Young |
| 0.828495 | 0.466509 | 0.174 | 0.198 | 1 | 11 | Fam168a | -0.024 | Young |
| 0.828934 | 0.286228 | 0.109 | 0.121 | 1 | 11 | Zfp655 | -0.012 | Young |
| 0.829319 | 0.702279 | 0.138 | 0.137 | 1 | 11 | Galnt11 | 0.001 | Young |
| 0.82955 | 0.569801 | 0.304 | 0.331 | 1 | 11 | Lbr | -0.027 | Young |
| 0.82968 | 0.385874 | 0.145 | 0.161 | 1 | 11 | Nbr1 | -0.016 | Young |
| 0.829917 | 0.334316 | 0.304 | 0.347 | 1 | 11 | Secisbp2l | -0.043 | Young |
| 0.829985 | 0.333552 | 0.036 | 0.032 | 1 | 11 | Cbx8 | 0.004 | Young |
| 0.83034 | 0.342321 | 0.072 | 0.081 | 1 | 11 | Tmem167b | -0.009 | Young |
| 0.830793 | 0.911195 | 0.145 | 0.145 | 1 | 11 | Tgfbr1 | 0 | Young |
| 0.830857 | 1.37171 | 0.043 | 0.04 | 1 | 11 | Kat8 | 0.003 | Young |
| 0.830996 | 0.771005 | 0.181 | 0.185 | 1 | 11 | Mitd1 | -0.004 | Young |
| 0.83132 | 0.046505 | 0.225 | 0.222 | 1 | 11 | Lyz1 | 0.003 | Young |
| 0.831568 | 0.300664 | 0.109 | 0.105 | 1 | 11 | Ddx19a | 0.004 | Young |
| 0.831606 | 0.305996 | 0.058 | 0.065 | 1 | 11 | Qsox1 | -0.007 | Young |
| 0.832361 | 0.785647 | 0.036 | 0.032 | 1 | 11 | Tmem241 | 0.004 | Young |
| 0.832568 | 2.80577 | 0.014 | 0.012 | 1 | 11 | Spdl1 | 0.002 | Young |
| 0.832568 | 2.085374 | 0.014 | 0.012 | 1 | 11 | Cep41 | 0.002 | Young |
| 0.832568 | 1.969151 | 0.014 | 0.012 | 1 | 11 | Pou6f1 | 0.002 | Young |
| 0.832568 | 1.703039 | 0.014 | 0.012 | 1 | 11 | Cidec | 0.002 | Young |
| 0.832568 | 1.689363 | 0.014 | 0.012 | 1 | 11 | Gm9530 | 0.002 | Young |
| 0.832568 | 1.591329 | 0.014 | 0.012 | 1 | 11 | Gm30978 | 0.002 | Young |
| 0.832568 | 1.54405 | 0.014 | 0.012 | 1 | 11 | Chst11 | 0.002 | Young |
| 0.832568 | 1.441129 | 0.014 | 0.012 | 1 | 11 | Satb2 | 0.002 | Young |
| 0.832568 | 1.30788 | 0.014 | 0.012 | 1 | 11 | Nat8f1 | 0.002 | Young |
| 0.832568 | 1.158122 | 0.014 | 0.012 | 1 | 11 | Tiam2 | 0.002 | Young |
| 0.832568 | 0.802291 | 0.014 | 0.012 | 1 | 11 | Timp1 | 0.002 | Young |
| 0.832887 | 0.782614 | 0.08 | 0.089 | 1 | 11 | C2cd3 | -0.009 | Young |
| 0.833007 | 1.214252 | 0.043 | 0.04 | 1 | 11 | Zfp617 | 0.003 | Young |
| 0.833178 | 0.726487 | 0.145 | 0.141 | 1 | 11 | Il10 | 0.004 | Young |
| 0.833237 | 0.835282 | 0.152 | 0.153 | 1 | 11 | Senp7 | -0.001 | Young |

| | | | | | | | | |
|---|---|---|---|---|---|---|---|---|
| 0.833376 | 0.462754 | 0.087 | 0.097 | 1 | 11 | Slc9a8 | -0.01 | Young |
| 0.833876 | 0.158149 | 0.043 | 0.048 | 1 | 11 | Nek4 | -0.005 | Young |
| 0.834079 | 0.215519 | 0.051 | 0.056 | 1 | 11 | Cep162 | -0.005 | Young |
| 0.834087 | 0.514516 | 0.123 | 0.137 | 1 | 11 | Tmem135 | -0.014 | Young |
| 0.834087 | 0.385639 | 0.123 | 0.137 | 1 | 11 | Fam53c | -0.014 | Young |
| 0.835435 | 0.760731 | 0.065 | 0.073 | 1 | 11 | Scmh1 | -0.008 | Young |
| 0.835435 | 0.662578 | 0.065 | 0.073 | 1 | 11 | Fam98c | -0.008 | Young |
| 0.835435 | 0.394934 | 0.065 | 0.073 | 1 | 11 | Atp8b2 | -0.008 | Young |
| 0.83551 | 0.089549 | 0.725 | 0.734 | 1 | 11 | Pf4 | -0.009 | Young |
| 0.835735 | 0.145899 | 0.101 | 0.109 | 1 | 11 | Kat7 | -0.008 | Young |
| 0.836167 | 0.421351 | 0.399 | 0.488 | 1 | 11 | Grina | -0.089 | Young |
| 0.836251 | 0.399189 | 0.138 | 0.153 | 1 | 11 | Usp37 | -0.015 | Young |
| 0.83634 | 0.050026 | 0.92 | 0.948 | 1 | 11 | Eif1 | -0.028 | Young |
| 0.836343 | 1.01101 | 0.101 | 0.101 | 1 | 11 | Tmed4 | 0 | Young |
| 0.836362 | 1.884276 | 0.014 | 0.012 | 1 | 11 | Asb4 | 0.002 | Young |
| 0.836362 | 1.146926 | 0.014 | 0.012 | 1 | 11 | Sgsm3 | 0.002 | Young |
| 0.836362 | 1.118315 | 0.014 | 0.012 | 1 | 11 | C130013H( | 0.002 | Young |
| 0.836362 | 1.094964 | 0.014 | 0.012 | 1 | 11 | Gm38251 | 0.002 | Young |
| 0.836362 | 1.056334 | 0.014 | 0.012 | 1 | 11 | Wdr25 | 0.002 | Young |
| 0.836362 | 1.052519 | 0.014 | 0.012 | 1 | 11 | Gm14305 | 0.002 | Young |
| 0.836362 | 1.038759 | 0.014 | 0.012 | 1 | 11 | Taf9b | 0.002 | Young |
| 0.836362 | 0.987649 | 0.014 | 0.012 | 1 | 11 | Numbl | 0.002 | Young |
| 0.836362 | 0.978531 | 0.014 | 0.012 | 1 | 11 | Dlgap5 | 0.002 | Young |
| 0.836362 | 0.940629 | 0.014 | 0.012 | 1 | 11 | 1700030J2 | 0.002 | Young |
| 0.836362 | 0.864286 | 0.014 | 0.012 | 1 | 11 | Tceal1 | 0.002 | Young |
| 0.836362 | 0.855026 | 0.014 | 0.012 | 1 | 11 | Smtn | 0.002 | Young |
| 0.836362 | 0.752014 | 0.014 | 0.012 | 1 | 11 | Tm4sf5 | 0.002 | Young |
| 0.836362 | 0.719864 | 0.014 | 0.012 | 1 | 11 | Spata24 | 0.002 | Young |
| 0.836362 | 0.672648 | 0.014 | 0.012 | 1 | 11 | Gm49368 | 0.002 | Young |
| 0.836362 | 0.646572 | 0.014 | 0.012 | 1 | 11 | Lockd | 0.002 | Young |
| 0.836362 | 0.556811 | 0.014 | 0.012 | 1 | 11 | Cep19 | 0.002 | Young |
| 0.836362 | 0.469391 | 0.014 | 0.012 | 1 | 11 | 5430416N( | 0.002 | Young |
| 0.836932 | 0.844976 | 0.058 | 0.065 | 1 | 11 | Tor2a | -0.007 | Young |
| 0.836932 | 0.74012 | 0.058 | 0.065 | 1 | 11 | Ints2 | -0.007 | Young |
| 0.837116 | 0.745876 | 0.036 | 0.032 | 1 | 11 | Tenm4 | 0.004 | Young |
| 0.837311 | 1.18187 | 0.043 | 0.04 | 1 | 11 | Tatdn3 | 0.003 | Young |
| 0.83746 | 0.621993 | 0.101 | 0.113 | 1 | 11 | Map3k4 | -0.012 | Young |
| 0.837485 | 0.590345 | 0.08 | 0.089 | 1 | 11 | Sipa1l2 | -0.009 | Young |
| 0.837946 | 0.392605 | 0.043 | 0.048 | 1 | 11 | Sp4 | -0.005 | Young |
| 0.838343 | 1.101511 | 0.072 | 0.081 | 1 | 11 | Cep68 | -0.009 | Young |
| 0.838451 | 0.090073 | 0.333 | 0.375 | 1 | 11 | Samd9l | -0.042 | Young |
| 0.838798 | 0.521325 | 0.065 | 0.073 | 1 | 11 | Nemp1 | -0.008 | Young |
| 0.838988 | 0.517909 | 0.225 | 0.262 | 1 | 11 | Hipk2 | -0.037 | Young |
| 0.839019 | 0.398881 | 0.08 | 0.089 | 1 | 11 | Mis18a | -0.009 | Young |
| 0.839136 | 0.137835 | 0.746 | 0.786 | 1 | 11 | Rsrp1 | -0.04 | Young |

| | | | | | | | | |
|---|---|---|---|---|---|---|---|---|
| 0.839724 | 0.534146 | 0.159 | 0.173 | 1 | 11 | Lcorl | -0.014 | Young |
| 0.839754 | 0.078953 | 0.051 | 0.056 | 1 | 11 | B230307C2 | -0.005 | Young |
| 0.840098 | 0.577123 | 0.159 | 0.161 | 1 | 11 | Ric8a | -0.002 | Young |
| 0.840159 | 1.545183 | 0.014 | 0.012 | 1 | 11 | Mettl25 | 0.002 | Young |
| 0.840159 | 0.781009 | 0.014 | 0.012 | 1 | 11 | Zswim9 | 0.002 | Young |
| 0.840159 | 0.687264 | 0.014 | 0.012 | 1 | 11 | Zswim1 | 0.002 | Young |
| 0.840159 | 0.672802 | 0.014 | 0.012 | 1 | 11 | 1-Sep | 0.002 | Young |
| 0.840159 | 0.633213 | 0.014 | 0.012 | 1 | 11 | Cldn10 | 0.002 | Young |
| 0.840159 | 0.449709 | 0.014 | 0.012 | 1 | 11 | Sphk1 | 0.002 | Young |
| 0.840159 | 0.436065 | 0.014 | 0.012 | 1 | 11 | Zkscan4 | 0.002 | Young |
| 0.840159 | 0.399872 | 0.014 | 0.012 | 1 | 11 | Art3 | 0.002 | Young |
| 0.840159 | 0.229148 | 0.014 | 0.012 | 1 | 11 | Gsdme | 0.002 | Young |
| 0.840159 | 0.163392 | 0.014 | 0.012 | 1 | 11 | Map3k12 | 0.002 | Young |
| 0.840159 | 0.09201 | 0.014 | 0.012 | 1 | 11 | Gm29336 | 0.002 | Young |
| 0.840481 | 0.237494 | 0.065 | 0.073 | 1 | 11 | Hsf1 | -0.008 | Young |
| 0.841047 | 0.304121 | 0.355 | 0.395 | 1 | 11 | Capg | -0.04 | Young |
| 0.841109 | 0.268426 | 0.138 | 0.149 | 1 | 11 | Cltb | -0.011 | Young |
| 0.841549 | 0.066266 | 0.551 | 0.581 | 1 | 11 | Tob2 | -0.03 | Young |
| 0.84162 | 0.762367 | 0.043 | 0.04 | 1 | 11 | Kctd13 | 0.003 | Young |
| 0.84162 | 0.360927 | 0.043 | 0.04 | 1 | 11 | Mad1l1 | 0.003 | Young |
| 0.842019 | 0.140413 | 0.043 | 0.048 | 1 | 11 | Alkbh3 | -0.005 | Young |
| 0.84202 | 0.178512 | 0.565 | 0.617 | 1 | 11 | Snx2 | -0.052 | Young |
| 0.842088 | 0.322236 | 0.08 | 0.089 | 1 | 11 | Rnf214 | -0.009 | Young |
| 0.842164 | 0.559005 | 0.065 | 0.073 | 1 | 11 | Hscb | -0.008 | Young |
| 0.842265 | 0.328288 | 0.058 | 0.065 | 1 | 11 | Leo1 | -0.007 | Young |
| 0.842419 | 0.808203 | 0.174 | 0.181 | 1 | 11 | Strn4 | -0.007 | Young |
| 0.842697 | 0.571025 | 0.297 | 0.319 | 1 | 11 | Entpd1 | -0.022 | Young |
| 0.843153 | 0.586308 | 0.072 | 0.081 | 1 | 11 | Saysd1 | -0.009 | Young |
| 0.843377 | 0.035948 | 0.036 | 0.04 | 1 | 11 | Zkscan14 | -0.004 | Young |
| 0.843961 | 1.225361 | 0.014 | 0.012 | 1 | 11 | Nxn | 0.002 | Young |
| 0.843961 | 1.166031 | 0.014 | 0.012 | 1 | 11 | 1700052K1 | 0.002 | Young |
| 0.843961 | 0.635949 | 0.014 | 0.012 | 1 | 11 | Arhgap27o | 0.002 | Young |
| 0.843961 | 0.504423 | 0.014 | 0.012 | 1 | 11 | Lig4 | 0.002 | Young |
| 0.843961 | 0.487614 | 0.014 | 0.012 | 1 | 11 | Gm5086 | 0.002 | Young |
| 0.843961 | 0.384296 | 0.014 | 0.012 | 1 | 11 | Slc26a6 | 0.002 | Young |
| 0.843961 | 0.378155 | 0.014 | 0.012 | 1 | 11 | Slc10a6 | 0.002 | Young |
| 0.843961 | 0.276782 | 0.014 | 0.012 | 1 | 11 | Arsi | 0.002 | Young |
| 0.843961 | 0.197239 | 0.014 | 0.012 | 1 | 11 | Cacna1b | 0.002 | Young |
| 0.843961 | 0.067805 | 0.014 | 0.012 | 1 | 11 | Stat4 | 0.002 | Young |
| 0.844149 | 0.879845 | 0.145 | 0.145 | 1 | 11 | Prdx4 | 0 | Young |
| 0.844435 | 1.347227 | 0.094 | 0.093 | 1 | 11 | Hps4 | 0.001 | Young |
| 0.844657 | 0.994688 | 0.123 | 0.125 | 1 | 11 | Nln | -0.002 | Young |
| 0.845934 | 0.84984 | 0.043 | 0.04 | 1 | 11 | Lrwd1 | 0.003 | Young |
| 0.846307 | 1.281934 | 0.094 | 0.105 | 1 | 11 | Ppp1r37 | -0.011 | Young |
| 0.846362 | 1.156307 | 0.072 | 0.081 | 1 | 11 | Klhdc10 | -0.009 | Young |

| | | | | | | | | |
|---|---|---|---|---|---|---|---|---|
| 0.846362 | 0.56065 | 0.072 | 0.081 | 1 | 11 | Slc35e3 | -0.009 | Young |
| 0.846543 | 0.439424 | 0.167 | 0.181 | 1 | 11 | Pcmtd2 | -0.014 | Young |
| 0.847219 | 0.998719 | 0.072 | 0.069 | 1 | 11 | 6720427I0 | 0.003 | Young |
| 0.847277 | 0.368628 | 0.333 | 0.399 | 1 | 11 | Aldh2 | -0.066 | Young |
| 0.847283 | 0.073782 | 0.507 | 0.565 | 1 | 11 | Metrnl | -0.058 | Young |
| 0.847345 | 0.374635 | 0.413 | 0.484 | 1 | 11 | Ppp1r15a | -0.071 | Young |
| 0.847424 | 0.599984 | 0.268 | 0.29 | 1 | 11 | Lpxn | -0.022 | Young |
| 0.847766 | 0.156097 | 0.014 | 0.012 | 1 | 11 | L2hgdh | 0.002 | Young |
| 0.847766 | 0.011591 | 0.014 | 0.012 | 1 | 11 | Mturn | 0.002 | Young |
| 0.847766 | 0.004297 | 0.014 | 0.012 | 1 | 11 | Jmjd8 | 0.002 | Young |
| 0.848094 | 0.527077 | 0.254 | 0.262 | 1 | 11 | Cd33 | -0.008 | Young |
| 0.848365 | 0.435459 | 0.268 | 0.298 | 1 | 11 | Pacsin2 | -0.03 | Young |
| 0.848646 | 0.390978 | 0.312 | 0.339 | 1 | 11 | Hipk1 | -0.027 | Young |
| 0.848789 | 0.253464 | 0.167 | 0.181 | 1 | 11 | Dync1li2 | -0.014 | Young |
| 0.849113 | 0.739588 | 0.167 | 0.185 | 1 | 11 | Bcl2a1a | -0.018 | Young |
| 0.849215 | 0.028837 | 0.13 | 0.141 | 1 | 11 | Asxl1 | -0.011 | Young |
| 0.849231 | 1.135691 | 0.051 | 0.056 | 1 | 11 | Zfp687 | -0.005 | Young |
| 0.849388 | 0.677749 | 0.058 | 0.065 | 1 | 11 | Ssbp2 | -0.007 | Young |
| 0.849454 | 0.751389 | 0.181 | 0.19 | 1 | 11 | Sap30 | -0.009 | Young |
| 0.849679 | 0.568908 | 0.304 | 0.335 | 1 | 11 | Lat2 | -0.031 | Young |
| 0.850056 | 0.208418 | 0.036 | 0.04 | 1 | 11 | Ppil1 | -0.004 | Young |
| 0.850584 | 0.664556 | 0.094 | 0.105 | 1 | 11 | Nr2f6 | -0.011 | Young |
| 0.850858 | 0.266877 | 0.362 | 0.403 | 1 | 11 | Ankrd44 | -0.041 | Young |
| 0.850931 | 0.272768 | 0.413 | 0.444 | 1 | 11 | Ehbp1l1 | -0.031 | Young |
| 0.851058 | 0.093219 | 0.616 | 0.742 | 1 | 11 | Ptafr | -0.126 | Young |
| 0.851129 | 0.701907 | 0.051 | 0.056 | 1 | 11 | Hddc2 | -0.005 | Young |
| 0.851129 | 0.241304 | 0.051 | 0.056 | 1 | 11 | Tdp2 | -0.005 | Young |
| 0.851672 | 0.077585 | 0.471 | 0.552 | 1 | 11 | Nptn | -0.081 | Young |
| 0.851857 | 0.685956 | 0.188 | 0.198 | 1 | 11 | Ankle2 | -0.01 | Young |
| 0.85217 | 0.28046 | 0.457 | 0.552 | 1 | 11 | Prkar1a | -0.095 | Young |
| 0.852222 | 0.703007 | 0.043 | 0.048 | 1 | 11 | Arid3b | -0.005 | Young |
| 0.852463 | 0.922516 | 0.123 | 0.125 | 1 | 11 | Mtmr14 | -0.002 | Young |
| 0.852622 | 0.66295 | 0.101 | 0.113 | 1 | 11 | Crot | -0.012 | Young |
| 0.852729 | 0.185477 | 0.616 | 0.726 | 1 | 11 | Ier2 | -0.11 | Young |
| 0.852954 | 0.744785 | 0.058 | 0.065 | 1 | 11 | Cyb561d1 | -0.007 | Young |
| 0.852954 | 0.223224 | 0.058 | 0.065 | 1 | 11 | Mcm9 | -0.007 | Young |
| 0.853663 | 0.312096 | 0.326 | 0.363 | 1 | 11 | Ubxn1 | -0.037 | Young |
| 0.85394 | 1.213217 | 0.051 | 0.048 | 1 | 11 | Pik3r2 | 0.003 | Young |
| 0.854003 | 0.721706 | 0.101 | 0.113 | 1 | 11 | Fbxo30 | -0.012 | Young |
| 0.854265 | 0.40844 | 0.043 | 0.048 | 1 | 11 | Apol7c | -0.005 | Young |
| 0.854389 | 1.375997 | 0.08 | 0.089 | 1 | 11 | Ttpal | -0.009 | Young |
| 0.854514 | 0.122315 | 0.036 | 0.04 | 1 | 11 | Smug1 | -0.004 | Young |
| 0.854575 | 0.865073 | 0.043 | 0.04 | 1 | 11 | Ppp1r3b | 0.003 | Young |
| 0.85549 | 0.719068 | 0.159 | 0.177 | 1 | 11 | Acss1 | -0.018 | Young |
| 0.855526 | 1.255549 | 0.087 | 0.097 | 1 | 11 | Utp14b | -0.01 | Young |

| | | | | | | | | |
|---|---|---|---|---|---|---|---|---|
| 0.855526 | 0.179004 | 0.087 | 0.097 | 1 | 11 | Taf4 | -0.01 | Young |
| 0.855658 | 0.763803 | 0.065 | 0.073 | 1 | 11 | Nbas | -0.008 | Young |
| 0.855871 | 0.241646 | 0.246 | 0.27 | 1 | 11 | Rapgef6 | -0.024 | Young |
| 0.85598 | 0.449959 | 0.348 | 0.375 | 1 | 11 | Actn4 | -0.027 | Young |
| 0.856738 | 0.655627 | 0.043 | 0.04 | 1 | 11 | Zfp81 | 0.003 | Young |
| 0.856745 | 0.051363 | 0.036 | 0.04 | 1 | 11 | Pex11a | -0.004 | Young |
| 0.856828 | 0.47716 | 0.051 | 0.056 | 1 | 11 | Pigv | -0.005 | Young |
| 0.856879 | 0.456925 | 0.123 | 0.137 | 1 | 11 | Gm46224 | -0.014 | Young |
| 0.857007 | 0.678009 | 0.094 | 0.093 | 1 | 11 | Iffo1 | 0.001 | Young |
| 0.857436 | 0.166508 | 0.616 | 0.685 | 1 | 11 | Mat2a | -0.069 | Young |
| 0.85747 | 0.860564 | 0.08 | 0.089 | 1 | 11 | Slc35b3 | -0.009 | Young |
| 0.858422 | 0.976235 | 0.08 | 0.077 | 1 | 11 | Tmem203 | 0.003 | Young |
| 0.858422 | 0.320461 | 0.08 | 0.077 | 1 | 11 | Fam114a1 | 0.003 | Young |
| 0.859153 | 0.760149 | 0.094 | 0.105 | 1 | 11 | Cd79b | -0.011 | Young |
| 0.859229 | 0.731793 | 0.094 | 0.093 | 1 | 11 | Mir142hg | 0.001 | Young |
| 0.859339 | 0.399306 | 0.37 | 0.448 | 1 | 11 | Hacd4 | -0.078 | Young |
| 0.859813 | 0.626462 | 0.138 | 0.153 | 1 | 11 | Dnajb4 | -0.015 | Young |
| 0.85992 | 1.275169 | 0.051 | 0.048 | 1 | 11 | Naa16 | 0.003 | Young |
| 0.860147 | 0.001734 | 0.123 | 0.133 | 1 | 11 | Ap4s1 | -0.01 | Young |
| 0.860524 | 0.492817 | 0.029 | 0.032 | 1 | 11 | Gpsm2 | -0.003 | Young |
| 0.860729 | 0.743719 | 0.072 | 0.069 | 1 | 11 | Fbxo25 | 0.003 | Young |
| 0.860729 | 0.357688 | 0.072 | 0.069 | 1 | 11 | Ngrn | 0.003 | Young |
| 0.861211 | 0.775235 | 0.036 | 0.04 | 1 | 11 | Zbtb12 | -0.004 | Young |
| 0.861211 | 0.568319 | 0.036 | 0.04 | 1 | 11 | Dhx32 | -0.004 | Young |
| 0.861211 | 0.394717 | 0.036 | 0.04 | 1 | 11 | Thada | -0.004 | Young |
| 0.861211 | 0.213815 | 0.036 | 0.04 | 1 | 11 | Zfp275 | -0.004 | Young |
| 0.861587 | 0.661984 | 0.145 | 0.161 | 1 | 11 | Map1lc3a | -0.016 | Young |
| 0.861881 | 0.263069 | 0.058 | 0.065 | 1 | 11 | 1110034G2 | -0.007 | Young |
| 0.86215 | 1.421023 | 0.043 | 0.04 | 1 | 11 | Abhd14b | 0.003 | Young |
| 0.862535 | 0.783569 | 0.051 | 0.056 | 1 | 11 | Dcaf10 | -0.005 | Young |
| 0.862535 | 0.623572 | 0.051 | 0.056 | 1 | 11 | Sft2d3 | -0.005 | Young |
| 0.862949 | 0.250639 | 0.536 | 0.637 | 1 | 11 | Tgfb1 | -0.101 | Young |
| 0.863232 | 0.726167 | 0.13 | 0.133 | 1 | 11 | Akap11 | -0.003 | Young |
| 0.863449 | 0.581316 | 0.138 | 0.153 | 1 | 11 | Afg3l1 | -0.015 | Young |
| 0.86434 | 0.509093 | 0.333 | 0.391 | 1 | 11 | Mindy2 | -0.058 | Young |
| 0.864344 | 1.003519 | 0.21 | 0.246 | 1 | 11 | Gdi1 | -0.036 | Young |
| 0.864344 | 0.487317 | 0.225 | 0.238 | 1 | 11 | Ddx21 | -0.013 | Young |
| 0.864497 | 0.817109 | 0.043 | 0.048 | 1 | 11 | Dnajc18 | -0.005 | Young |
| 0.864497 | 0.804291 | 0.043 | 0.048 | 1 | 11 | Abcg2 | -0.005 | Young |
| 0.864708 | 0.135906 | 0.471 | 0.577 | 1 | 11 | Eif5a | -0.106 | Young |
| 0.86494 | 0.624862 | 0.217 | 0.25 | 1 | 11 | Sike1 | -0.033 | Young |
| 0.865501 | 0.090548 | 0.029 | 0.032 | 1 | 11 | Treml2 | -0.003 | Young |
| 0.865641 | 0.069227 | 0.08 | 0.085 | 1 | 11 | Lmf1 | -0.005 | Young |
| 0.86568 | 0.552128 | 0.036 | 0.04 | 1 | 11 | Nprl3 | -0.004 | Young |
| 0.866547 | 1.025724 | 0.043 | 0.048 | 1 | 11 | Trp53inp1 | -0.005 | Young |

| | | | | | | | | |
|---|---|---|---|---|---|---|---|---|
| 0.866547 | 1.000244 | 0.043 | 0.048 | 1 | 11 | Xylt1 | -0.005 | Young |
| 0.866726 | 0.762787 | 0.087 | 0.085 | 1 | 11 | Rab6b | 0.002 | Young |
| 0.867023 | 0.300012 | 0.232 | 0.258 | 1 | 11 | Fip1l1 | -0.026 | Young |
| 0.867286 | 0.987298 | 0.072 | 0.081 | 1 | 11 | Thap3 | -0.009 | Young |
| 0.8677 | 0.047391 | 0.623 | 0.633 | 1 | 11 | F13a1 | -0.01 | Young |
| 0.867738 | 0.671318 | 0.094 | 0.105 | 1 | 11 | Ankrd50 | -0.011 | Young |
| 0.867755 | 0.411958 | 0.13 | 0.145 | 1 | 11 | Mfap3 | -0.015 | Young |
| 0.868249 | 0.984847 | 0.051 | 0.056 | 1 | 11 | Zbtb16 | -0.005 | Young |
| 0.868376 | 1.210295 | 0.138 | 0.141 | 1 | 11 | Bmyc | -0.003 | Young |
| 0.868469 | 0.349424 | 0.109 | 0.117 | 1 | 11 | Triap1 | -0.008 | Young |
| 0.8689 | 0.827246 | 0.072 | 0.081 | 1 | 11 | Jak3 | -0.009 | Young |
| 0.869133 | 0.576405 | 0.29 | 0.339 | 1 | 11 | Crlf2 | -0.049 | Young |
| 0.869194 | 0.782488 | 0.065 | 0.073 | 1 | 11 | Grap | -0.008 | Young |
| 0.869605 | 0.252586 | 0.159 | 0.157 | 1 | 11 | Smim4 | 0.002 | Young |
| 0.869738 | 0.732985 | 0.043 | 0.04 | 1 | 11 | Tmcc2 | 0.003 | Young |
| 0.869821 | 0.512719 | 0.174 | 0.181 | 1 | 11 | Mylip | -0.007 | Young |
| 0.870155 | 1.007767 | 0.036 | 0.04 | 1 | 11 | Slfn4 | -0.004 | Young |
| 0.870267 | 0.486566 | 0.181 | 0.202 | 1 | 11 | Rgl2 | -0.021 | Young |
| 0.870746 | 0.308892 | 0.428 | 0.48 | 1 | 11 | Sf1 | -0.052 | Young |
| 0.871161 | 0.079035 | 0.558 | 0.669 | 1 | 11 | Zcchc6 | -0.111 | Young |
| 0.871171 | 0.600969 | 0.384 | 0.468 | 1 | 11 | Tsc22d3 | -0.084 | Young |
| 0.871361 | 1.287371 | 0.08 | 0.089 | 1 | 11 | Kansl1l | -0.009 | Young |
| 0.871361 | 0.691956 | 0.08 | 0.089 | 1 | 11 | Pex3 | -0.009 | Young |
| 0.871667 | 1.162392 | 0.145 | 0.153 | 1 | 11 | Gm47283 | -0.008 | Young |
| 0.872036 | 0.600979 | 0.094 | 0.105 | 1 | 11 | Tubgcp4 | -0.011 | Young |
| 0.872393 | 0.650271 | 0.036 | 0.04 | 1 | 11 | Mettl3 | -0.004 | Young |
| 0.872533 | 0.452322 | 0.268 | 0.306 | 1 | 11 | Cdc5l | -0.038 | Young |
| 0.872584 | 0.198365 | 0.072 | 0.069 | 1 | 11 | Kif16b | 0.003 | Young |
| 0.872967 | 0.385692 | 0.13 | 0.141 | 1 | 11 | Rif1 | -0.011 | Young |
| 0.872977 | 0.811922 | 0.029 | 0.032 | 1 | 11 | Gm29340 | -0.003 | Young |
| 0.872977 | 0.115259 | 0.029 | 0.032 | 1 | 11 | Twnk | -0.003 | Young |
| 0.873436 | 0.232361 | 0.101 | 0.109 | 1 | 11 | Scaf4 | -0.008 | Young |
| 0.873623 | 0.66597 | 0.13 | 0.129 | 1 | 11 | Cyp4f16 | 0.001 | Young |
| 0.873904 | 1.435373 | 0.051 | 0.048 | 1 | 11 | Cables1 | 0.003 | Young |
| 0.874079 | 1.016847 | 0.109 | 0.109 | 1 | 11 | Arhgef7 | 0 | Young |
| 0.874079 | 0.815833 | 0.109 | 0.109 | 1 | 11 | Tmem11 | 0 | Young |
| 0.874241 | 0.147308 | 0.63 | 0.706 | 1 | 11 | Fcgr2b | -0.076 | Young |
| 0.874633 | 0.936365 | 0.036 | 0.04 | 1 | 11 | Ulbp1 | -0.004 | Young |
| 0.874752 | 1.483574 | 0.043 | 0.048 | 1 | 11 | Arhgap23 | -0.005 | Young |
| 0.874752 | 0.949891 | 0.043 | 0.048 | 1 | 11 | B230354K1 | -0.005 | Young |
| 0.875343 | 0.403944 | 0.63 | 0.734 | 1 | 11 | Hsp90aa1 | -0.104 | Young |
| 0.875905 | 0.743624 | 0.051 | 0.048 | 1 | 11 | Pus10 | 0.003 | Young |
| 0.876806 | 1.657503 | 0.043 | 0.048 | 1 | 11 | Mark4 | -0.005 | Young |
| 0.876806 | 1.324807 | 0.043 | 0.048 | 1 | 11 | Sufu | -0.005 | Young |
| 0.876844 | 0.449438 | 0.355 | 0.407 | 1 | 11 | Txn2 | -0.052 | Young |

| | | | | | | | | |
|---|---|---|---|---|---|---|---|---|
| 0.877827 | 0.184457 | 0.174 | 0.185 | 1 | 11 | Snrk | -0.011 | Young |
| 0.877907 | 1.098738 | 0.051 | 0.048 | 1 | 11 | Traf3ip3 | 0.003 | Young |
| 0.877926 | 0.213272 | 0.174 | 0.173 | 1 | 11 | Coro2a | 0.001 | Young |
| 0.877968 | 0.432725 | 0.029 | 0.032 | 1 | 11 | Dusp19 | -0.003 | Young |
| 0.878079 | 0.345874 | 0.507 | 0.585 | 1 | 11 | P2rx4 | -0.078 | Young |
| 0.87832 | 0.547066 | 0.152 | 0.169 | 1 | 11 | Dhrs7b | -0.017 | Young |
| 0.878592 | 0.658724 | 0.072 | 0.081 | 1 | 11 | Wiz | -0.009 | Young |
| 0.878691 | 0.560527 | 0.159 | 0.161 | 1 | 11 | Gatad2a | -0.002 | Young |
| 0.879008 | 1.248474 | 0.116 | 0.133 | 1 | 11 | Map2k4 | -0.017 | Young |
| 0.879013 | 0.224535 | 0.384 | 0.415 | 1 | 11 | Rsrc2 | -0.031 | Young |
| 0.879095 | 1.08269 | 0.08 | 0.089 | 1 | 11 | Phrf1 | -0.009 | Young |
| 0.879115 | 0.740092 | 0.036 | 0.04 | 1 | 11 | Orc6 | -0.004 | Young |
| 0.879115 | 0.726177 | 0.036 | 0.04 | 1 | 11 | Gpr180 | -0.004 | Young |
| 0.879749 | 1.049912 | 0.101 | 0.109 | 1 | 11 | Il1rl1 | -0.008 | Young |
| 0.879846 | 0.177305 | 0.312 | 0.331 | 1 | 11 | Zfand3 | -0.019 | Young |
| 0.880022 | 0.604353 | 0.094 | 0.093 | 1 | 11 | Zfp646 | 0.001 | Young |
| 0.880465 | 1.485641 | 0.029 | 0.032 | 1 | 11 | Ddx19b | -0.003 | Young |
| 0.880465 | 0.999452 | 0.029 | 0.032 | 1 | 11 | Phf7 | -0.003 | Young |
| 0.880465 | 0.855591 | 0.029 | 0.032 | 1 | 11 | Map4k5 | -0.003 | Young |
| 0.880465 | 0.5258 | 0.029 | 0.032 | 1 | 11 | Ptcd3 | -0.003 | Young |
| 0.880465 | 0.503514 | 0.029 | 0.032 | 1 | 11 | Hgf | -0.003 | Young |
| 0.880465 | 0.478872 | 0.029 | 0.032 | 1 | 11 | Clstn1 | -0.003 | Young |
| 0.880598 | 0.952254 | 0.043 | 0.04 | 1 | 11 | Adora2b | 0.003 | Young |
| 0.881341 | 0.351606 | 0.13 | 0.129 | 1 | 11 | Ednrb | 0.001 | Young |
| 0.881358 | 0.977773 | 0.036 | 0.04 | 1 | 11 | Gm44751 | -0.004 | Young |
| 0.881511 | 1.003852 | 0.094 | 0.093 | 1 | 11 | Pygl | 0.001 | Young |
| 0.882217 | 0.782974 | 0.188 | 0.198 | 1 | 11 | Mitf | -0.01 | Young |
| 0.882529 | 0.495397 | 0.21 | 0.21 | 1 | 11 | Gbp2b | 0 | Young |
| 0.882757 | 0.488019 | 0.022 | 0.024 | 1 | 11 | Clcf1 | -0.002 | Young |
| 0.882757 | 0.27446 | 0.022 | 0.024 | 1 | 11 | Rnf26 | -0.002 | Young |
| 0.882757 | 0.102148 | 0.022 | 0.024 | 1 | 11 | Mapk1ip1 | -0.002 | Young |
| 0.882963 | 0.638765 | 0.029 | 0.032 | 1 | 11 | Coq9 | -0.003 | Young |
| 0.883009 | 0.135651 | 0.326 | 0.323 | 1 | 11 | Tnip3 | 0.003 | Young |
| 0.883341 | 0.421204 | 0.384 | 0.444 | 1 | 11 | Eef1d | -0.06 | Young |
| 0.883602 | 0.868921 | 0.036 | 0.04 | 1 | 11 | Slf1 | -0.004 | Young |
| 0.883652 | 0.590387 | 0.326 | 0.371 | 1 | 11 | Srsf1 | -0.045 | Young |
| 0.883703 | 0.133231 | 0.601 | 0.685 | 1 | 11 | Snrnp70 | -0.084 | Young |
| 0.883806 | 1.375029 | 0.065 | 0.065 | 1 | 11 | Ift46 | 0 | Young |
| 0.883978 | 0.353677 | 0.152 | 0.165 | 1 | 11 | Map3k14 | -0.013 | Young |
| 0.884094 | 0.234079 | 0.565 | 0.633 | 1 | 11 | Tap1 | -0.068 | Young |
| 0.884156 | 0.700977 | 0.152 | 0.169 | 1 | 11 | Cyp27a1 | -0.017 | Young |
| 0.884322 | 0.464282 | 0.37 | 0.411 | 1 | 11 | U2af2 | -0.041 | Young |
| 0.884782 | 0.52119 | 0.333 | 0.371 | 1 | 11 | Rap1gds1 | -0.038 | Young |
| 0.885028 | 0.785334 | 0.043 | 0.048 | 1 | 11 | Mfsd8 | -0.005 | Young |
| 0.885462 | 1.412716 | 0.029 | 0.032 | 1 | 11 | Dgka | -0.003 | Young |

| | | | | | | | | |
|---|---|---|---|---|---|---|---|---|
| 0.885462 | 1.060603 | 0.029 | 0.032 | 1 | 11 | Alms1 | -0.003 | Young |
| 0.885632 | 1.117889 | 0.022 | 0.024 | 1 | 11 | Slx1b | -0.002 | Young |
| 0.885632 | 0.418719 | 0.022 | 0.024 | 1 | 11 | Gm46430 | -0.002 | Young |
| 0.885632 | 0.083331 | 0.022 | 0.024 | 1 | 11 | Atp6v1g2 | -0.002 | Young |
| 0.885632 | 0.012821 | 0.022 | 0.024 | 1 | 11 | Cst7 | -0.002 | Young |
| 0.88674 | 0.410498 | 0.377 | 0.44 | 1 | 11 | Fgd2 | -0.063 | Young |
| 0.886882 | 0.058478 | 0.783 | 0.839 | 1 | 11 | Rpl28 | -0.056 | Young |
| 0.887452 | 0.924006 | 0.123 | 0.137 | 1 | 11 | Trappc3 | -0.014 | Young |
| 0.887452 | 0.861238 | 0.123 | 0.137 | 1 | 11 | Abcb7 | -0.014 | Young |
| 0.887469 | 0.697195 | 0.094 | 0.093 | 1 | 11 | Zfp120 | 0.001 | Young |
| 0.887606 | 0.618699 | 0.326 | 0.371 | 1 | 11 | Aph1a | -0.045 | Young |
| 0.887928 | 0.786371 | 0.051 | 0.048 | 1 | 11 | Ubxn8 | 0.003 | Young |
| 0.887963 | 0.837473 | 0.029 | 0.032 | 1 | 11 | Ppp1r3e | -0.003 | Young |
| 0.887963 | 0.712659 | 0.029 | 0.032 | 1 | 11 | Cep95 | -0.003 | Young |
| 0.888508 | 0.508999 | 0.022 | 0.024 | 1 | 11 | Rps6ka2 | -0.002 | Young |
| 0.888508 | 0.503074 | 0.022 | 0.024 | 1 | 11 | Slc18b1 | -0.002 | Young |
| 0.888749 | 0.538229 | 0.254 | 0.27 | 1 | 11 | Tnfrsf11a | -0.016 | Young |
| 0.889068 | 0.788303 | 0.123 | 0.125 | 1 | 11 | Sh3tc1 | -0.002 | Young |
| 0.889287 | 0.961924 | 0.109 | 0.121 | 1 | 11 | Gpd2 | -0.012 | Young |
| 0.889359 | 0.089879 | 0.739 | 0.823 | 1 | 11 | Rpl21 | -0.084 | Young |
| 0.890464 | 0.985823 | 0.029 | 0.032 | 1 | 11 | Kdelc1 | -0.003 | Young |
| 0.890464 | 0.844994 | 0.029 | 0.032 | 1 | 11 | Stradb | -0.003 | Young |
| 0.890464 | 0.617613 | 0.029 | 0.032 | 1 | 11 | Abcc3 | -0.003 | Young |
| 0.890669 | 0.170026 | 0.493 | 0.544 | 1 | 11 | Pik3cd | -0.051 | Young |
| 0.890745 | 0.604321 | 0.333 | 0.383 | 1 | 11 | Rasgef1b | -0.05 | Young |
| 0.890843 | 0.874827 | 0.232 | 0.254 | 1 | 11 | Slbp | -0.022 | Young |
| 0.890944 | 0.117004 | 0.572 | 0.698 | 1 | 11 | Glud1 | -0.126 | Young |
| 0.890953 | 0.400979 | 0.261 | 0.274 | 1 | 11 | Ranbp1 | -0.013 | Young |
| 0.891173 | 0.390915 | 0.478 | 0.589 | 1 | 11 | Trib1 | -0.111 | Young |
| 0.891386 | 0.969728 | 0.022 | 0.024 | 1 | 11 | Pvt1 | -0.002 | Young |
| 0.891386 | 0.918274 | 0.022 | 0.024 | 1 | 11 | Uevld | -0.002 | Young |
| 0.891386 | 0.814614 | 0.022 | 0.024 | 1 | 11 | Rhoj | -0.002 | Young |
| 0.891386 | 0.432565 | 0.022 | 0.024 | 1 | 11 | Rab27a | -0.002 | Young |
| 0.891386 | 0.365774 | 0.022 | 0.024 | 1 | 11 | Afg1l | -0.002 | Young |
| 0.891645 | 1.449587 | 0.058 | 0.056 | 1 | 11 | Galnt4 | 0.002 | Young |
| 0.891645 | 0.871493 | 0.058 | 0.056 | 1 | 11 | Terf2ip | 0.002 | Young |
| 0.891731 | 0.215081 | 0.428 | 0.52 | 1 | 11 | Selplg | -0.092 | Young |
| 0.89216 | 0.553822 | 0.283 | 0.323 | 1 | 11 | Pcgf5 | -0.04 | Young |
| 0.892565 | 0.95979 | 0.13 | 0.133 | 1 | 11 | Ptpn9 | -0.003 | Young |
| 0.892585 | 1.491161 | 0.036 | 0.04 | 1 | 11 | Tatdn1 | -0.004 | Young |
| 0.893085 | 1.739382 | 0.051 | 0.056 | 1 | 11 | Eef1akmt2 | -0.005 | Young |
| 0.893434 | 0.894995 | 0.094 | 0.093 | 1 | 11 | Stxbp5 | 0.001 | Young |
| 0.893527 | 0.429732 | 0.268 | 0.294 | 1 | 11 | Emc1 | -0.026 | Young |
| 0.893529 | 0.275985 | 0.377 | 0.431 | 1 | 11 | Ralb | -0.054 | Young |
| 0.893748 | 0.157187 | 0.645 | 0.738 | 1 | 11 | Ccnl1 | -0.093 | Young |

| | | | | | | | | |
|---|---|---|---|---|---|---|---|---|
| 0.89415 | 0.460587 | 0.203 | 0.214 | 1 | 11 | Cdc26 | -0.011 | Young |
| 0.894265 | 0.677822 | 0.022 | 0.024 | 1 | 11 | Adrb1 | -0.002 | Young |
| 0.894265 | 0.556931 | 0.022 | 0.024 | 1 | 11 | Tfcp2 | -0.002 | Young |
| 0.894265 | 0.551386 | 0.022 | 0.024 | 1 | 11 | Kif3c | -0.002 | Young |
| 0.894265 | 0.497499 | 0.022 | 0.024 | 1 | 11 | B4galt7 | -0.002 | Young |
| 0.894265 | 0.247902 | 0.022 | 0.024 | 1 | 11 | Tbc1d22b | -0.002 | Young |
| 0.894265 | 0.229552 | 0.022 | 0.024 | 1 | 11 | Abhd8 | -0.002 | Young |
| 0.89429 | 0.005298 | 0.746 | 0.823 | 1 | 11 | Eef1b2 | -0.077 | Young |
| 0.894384 | 1.102462 | 0.065 | 0.065 | 1 | 11 | Ccdc43 | 0 | Young |
| 0.894688 | 0.728561 | 0.116 | 0.117 | 1 | 11 | Pkd1 | -0.001 | Young |
| 0.894833 | 1.001151 | 0.036 | 0.04 | 1 | 11 | Hoxb4 | -0.004 | Young |
| 0.89516 | 0.536183 | 0.246 | 0.286 | 1 | 11 | Arrb1 | -0.04 | Young |
| 0.895211 | 0.219487 | 0.493 | 0.585 | 1 | 11 | Chd4 | -0.092 | Young |
| 0.895323 | 1.242589 | 0.043 | 0.048 | 1 | 11 | Ercc1 | -0.005 | Young |
| 0.895394 | 0.246134 | 0.362 | 0.415 | 1 | 11 | Tsn | -0.053 | Young |
| 0.895633 | 0.648391 | 0.123 | 0.125 | 1 | 11 | Cd200r1 | -0.002 | Young |
| 0.895661 | 1.644119 | 0.022 | 0.02 | 1 | 11 | A430046D1 | 0.002 | Young |
| 0.895661 | 1.594868 | 0.022 | 0.02 | 1 | 11 | E130307A1 | 0.002 | Young |
| 0.895661 | 1.554378 | 0.022 | 0.02 | 1 | 11 | A130077B1 | 0.002 | Young |
| 0.895661 | 1.428279 | 0.022 | 0.02 | 1 | 11 | Tnfaip8l1 | 0.002 | Young |
| 0.895661 | 1.386441 | 0.022 | 0.02 | 1 | 11 | Scrn3 | 0.002 | Young |
| 0.895661 | 1.317767 | 0.022 | 0.02 | 1 | 11 | Hist2h2aa1 | 0.002 | Young |
| 0.895661 | 1.087842 | 0.022 | 0.02 | 1 | 11 | Med26 | 0.002 | Young |
| 0.895661 | 1.050725 | 0.022 | 0.02 | 1 | 11 | Zfp809 | 0.002 | Young |
| 0.895958 | 1.134795 | 0.058 | 0.065 | 1 | 11 | Dtx4 | -0.007 | Young |
| 0.895984 | 1.133943 | 0.094 | 0.101 | 1 | 11 | Zfp593 | -0.007 | Young |
| 0.896149 | 1.043142 | 0.065 | 0.065 | 1 | 11 | Fpgt | 0 | Young |
| 0.896151 | 1.153201 | 0.08 | 0.089 | 1 | 11 | Exosc10 | -0.009 | Young |
| 0.896165 | 0.074131 | 0.362 | 0.387 | 1 | 11 | Pmp22 | -0.025 | Young |
| 0.896946 | 0.57581 | 0.123 | 0.125 | 1 | 11 | Ube2q2 | -0.002 | Young |
| 0.897051 | 0.247402 | 0.08 | 0.085 | 1 | 11 | Dcn | -0.005 | Young |
| 0.897145 | 0.945046 | 0.022 | 0.024 | 1 | 11 | Wdr59 | -0.002 | Young |
| 0.897145 | 0.937366 | 0.022 | 0.024 | 1 | 11 | Zfp790 | -0.002 | Young |
| 0.897145 | 0.814224 | 0.022 | 0.024 | 1 | 11 | Fzd1 | -0.002 | Young |
| 0.897145 | 0.706556 | 0.022 | 0.024 | 1 | 11 | Tmem220 | -0.002 | Young |
| 0.897145 | 0.622082 | 0.022 | 0.024 | 1 | 11 | Btd | -0.002 | Young |
| 0.897145 | 0.514291 | 0.022 | 0.024 | 1 | 11 | Asb2 | -0.002 | Young |
| 0.897381 | 0.645055 | 0.174 | 0.181 | 1 | 11 | Lgals8 | -0.007 | Young |
| 0.897391 | 1.29635 | 0.116 | 0.117 | 1 | 11 | Utp18 | -0.001 | Young |
| 0.897503 | 0.029131 | 0.71 | 0.802 | 1 | 11 | Ptp4a2 | -0.092 | Young |
| 0.898713 | 1.097589 | 0.022 | 0.02 | 1 | 11 | Fdxr | 0.002 | Young |
| 0.898713 | 1.02571 | 0.022 | 0.02 | 1 | 11 | Wdr92 | 0.002 | Young |
| 0.89954 | 0.243103 | 0.507 | 0.637 | 1 | 11 | Txn1 | -0.13 | Young |
| 0.899557 | 0.971498 | 0.058 | 0.065 | 1 | 11 | Phospho2 | -0.007 | Young |
| 0.899632 | 0.351677 | 0.464 | 0.552 | 1 | 11 | Plekho1 | -0.088 | Young |

| | | | | | | | | |
|---|---|---|---|---|---|---|---|---|
| 0.8997 | 0.154044 | 0.471 | 0.52 | 1 | 11 | Eif3c | -0.049 | Young |
| 0.899827 | 0.633867 | 0.138 | 0.149 | 1 | 11 | Trak1 | -0.011 | Young |
| 0.899976 | 0.562234 | 0.051 | 0.048 | 1 | 11 | Spsb3 | 0.003 | Young |
| 0.900027 | 1.334688 | 0.022 | 0.024 | 1 | 11 | Arl6 | -0.002 | Young |
| 0.900027 | 1.250334 | 0.022 | 0.024 | 1 | 11 | Tmem132a | -0.002 | Young |
| 0.900027 | 0.790648 | 0.022 | 0.024 | 1 | 11 | Tbc1d8b | -0.002 | Young |
| 0.900291 | 0.851459 | 0.152 | 0.157 | 1 | 11 | Ifi208 | -0.005 | Young |
| 0.900482 | 1.809945 | 0.029 | 0.032 | 1 | 11 | Areg | -0.003 | Young |
| 0.900623 | 0.232698 | 0.413 | 0.476 | 1 | 11 | Mier1 | -0.063 | Young |
| 0.901016 | 0.691572 | 0.058 | 0.056 | 1 | 11 | Rubcnl | 0.002 | Young |
| 0.901016 | 0.409686 | 0.058 | 0.056 | 1 | 11 | Dus3l | 0.002 | Young |
| 0.901202 | 0.965552 | 0.101 | 0.113 | 1 | 11 | Gmeb1 | -0.012 | Young |
| 0.901451 | 1.206441 | 0.072 | 0.073 | 1 | 11 | Nmral1 | -0.001 | Young |
| 0.901766 | 1.404076 | 0.022 | 0.02 | 1 | 11 | Pus7 | 0.002 | Young |
| 0.901766 | 1.365194 | 0.022 | 0.02 | 1 | 11 | Gm4673 | 0.002 | Young |
| 0.901766 | 0.94749 | 0.022 | 0.02 | 1 | 11 | Hirip3 | 0.002 | Young |
| 0.901766 | 0.857554 | 0.022 | 0.02 | 1 | 11 | Prr7 | 0.002 | Young |
| 0.901766 | 0.845809 | 0.022 | 0.02 | 1 | 11 | Pigo | 0.002 | Young |
| 0.901766 | 0.76432 | 0.022 | 0.02 | 1 | 11 | Pi16 | 0.002 | Young |
| 0.901766 | 0.762016 | 0.022 | 0.02 | 1 | 11 | Cep131 | 0.002 | Young |
| 0.901766 | 0.658862 | 0.022 | 0.02 | 1 | 11 | Ap5b1 | 0.002 | Young |
| 0.901871 | 0.215947 | 0.377 | 0.44 | 1 | 11 | Elf1 | -0.063 | Young |
| 0.902224 | 0.650328 | 0.101 | 0.101 | 1 | 11 | Chst7 | 0 | Young |
| 0.902566 | 0.071898 | 0.326 | 0.347 | 1 | 11 | Adgre5 | -0.021 | Young |
| 0.902769 | 0.95021 | 0.014 | 0.016 | 1 | 11 | Zfp12 | -0.002 | Young |
| 0.902769 | 0.420577 | 0.014 | 0.016 | 1 | 11 | Gm28187 | -0.002 | Young |
| 0.902769 | 0.042408 | 0.014 | 0.016 | 1 | 11 | Tshz2 | -0.002 | Young |
| 0.902911 | 1.916134 | 0.022 | 0.024 | 1 | 11 | Gm48066 | -0.002 | Young |
| 0.902911 | 1.559251 | 0.022 | 0.024 | 1 | 11 | Klhl36 | -0.002 | Young |
| 0.902911 | 1.296895 | 0.022 | 0.024 | 1 | 11 | Stau2 | -0.002 | Young |
| 0.903401 | 0.064121 | 0.935 | 0.972 | 1 | 11 | Rps28 | -0.037 | Young |
| 0.903572 | 0.132798 | 0.572 | 0.661 | 1 | 11 | Mef2a | -0.089 | Young |
| 0.903997 | 0.430132 | 0.051 | 0.048 | 1 | 11 | Sh2d1b1 | 0.003 | Young |
| 0.904821 | 1.870838 | 0.022 | 0.02 | 1 | 11 | 4930599N2 | 0.002 | Young |
| 0.904821 | 1.651107 | 0.022 | 0.02 | 1 | 11 | Zfp691 | 0.002 | Young |
| 0.904821 | 1.608141 | 0.022 | 0.02 | 1 | 11 | Tal1 | 0.002 | Young |
| 0.904821 | 1.596042 | 0.022 | 0.02 | 1 | 11 | Gm3448 | 0.002 | Young |
| 0.904821 | 1.564685 | 0.022 | 0.02 | 1 | 11 | Trim6 | 0.002 | Young |
| 0.904821 | 1.38543 | 0.022 | 0.02 | 1 | 11 | Uros | 0.002 | Young |
| 0.904821 | 1.214286 | 0.022 | 0.02 | 1 | 11 | Zhx2 | 0.002 | Young |
| 0.904821 | 0.96218 | 0.022 | 0.02 | 1 | 11 | Arl4d | 0.002 | Young |
| 0.90493 | 0.497684 | 0.087 | 0.093 | 1 | 11 | Timm22 | -0.006 | Young |
| 0.905783 | 0.435846 | 0.101 | 0.109 | 1 | 11 | Cog3 | -0.008 | Young |
| 0.905795 | 1.117406 | 0.022 | 0.024 | 1 | 11 | Bend6 | -0.002 | Young |
| 0.906053 | 0.10066 | 0.688 | 0.782 | 1 | 11 | Chchd2 | -0.094 | Young |

| | | | | | | | | |
|---|---|---|---|---|---|---|---|---|
| 0.906288 | 0.623749 | 0.014 | 0.016 | 1 | 11 | Ska2 | -0.002 | Young |
| 0.906288 | 0.612595 | 0.014 | 0.016 | 1 | 11 | Eya4 | -0.002 | Young |
| 0.906288 | 0.523154 | 0.014 | 0.016 | 1 | 11 | Tm7sf2 | -0.002 | Young |
| 0.906288 | 0.478059 | 0.014 | 0.016 | 1 | 11 | Zbtb11os1 | -0.002 | Young |
| 0.906288 | 0.441368 | 0.014 | 0.016 | 1 | 11 | Brip1 | -0.002 | Young |
| 0.906288 | 0.250748 | 0.014 | 0.016 | 1 | 11 | 1700019L1 | -0.002 | Young |
| 0.906288 | 0.188989 | 0.014 | 0.016 | 1 | 11 | Mblac1 | -0.002 | Young |
| 0.906288 | 0.175459 | 0.014 | 0.016 | 1 | 11 | Gm26825 | -0.002 | Young |
| 0.906288 | 0.145515 | 0.014 | 0.016 | 1 | 11 | 4732440D( | -0.002 | Young |
| 0.906288 | 0.05886 | 0.014 | 0.016 | 1 | 11 | Zfp60 | -0.002 | Young |
| 0.906288 | 0.051696 | 0.014 | 0.016 | 1 | 11 | Btbd17 | -0.002 | Young |
| 0.906771 | 0.767617 | 0.254 | 0.302 | 1 | 11 | Ythdf1 | -0.048 | Young |
| 0.906898 | 0.679129 | 0.283 | 0.323 | 1 | 11 | Dpm3 | -0.04 | Young |
| 0.907466 | 0.566119 | 0.123 | 0.125 | 1 | 11 | Topors | -0.002 | Young |
| 0.90758 | 0.526235 | 0.167 | 0.173 | 1 | 11 | Tiam1 | -0.006 | Young |
| 0.907877 | 1.87474 | 0.022 | 0.02 | 1 | 11 | Zfp280b | 0.002 | Young |
| 0.907877 | 1.21854 | 0.022 | 0.02 | 1 | 11 | Spindoc | 0.002 | Young |
| 0.907877 | 0.766059 | 0.022 | 0.02 | 1 | 11 | Zfat | 0.002 | Young |
| 0.907877 | 0.649367 | 0.022 | 0.02 | 1 | 11 | Htra3 | 0.002 | Young |
| 0.907877 | 0.627685 | 0.022 | 0.02 | 1 | 11 | Pdss1 | 0.002 | Young |
| 0.907877 | 0.406669 | 0.022 | 0.02 | 1 | 11 | 1810044D( | 0.002 | Young |
| 0.908239 | 0.189371 | 0.5 | 0.56 | 1 | 11 | Rock1 | -0.06 | Young |
| 0.908523 | 1.416408 | 0.058 | 0.056 | 1 | 11 | Tnfsf13b | 0.002 | Young |
| 0.90894 | 0.043532 | 0.123 | 0.129 | 1 | 11 | Sh3gl1 | -0.006 | Young |
| 0.909723 | 0.420206 | 0.413 | 0.464 | 1 | 11 | Ewsr1 | -0.051 | Young |
| 0.909809 | 1.246817 | 0.014 | 0.016 | 1 | 11 | Col1a2 | -0.002 | Young |
| 0.909809 | 1.13288 | 0.014 | 0.016 | 1 | 11 | Ccdc73 | -0.002 | Young |
| 0.909809 | 1.007342 | 0.014 | 0.016 | 1 | 11 | Gm10036 | -0.002 | Young |
| 0.909809 | 0.963806 | 0.014 | 0.016 | 1 | 11 | Slc25a53 | -0.002 | Young |
| 0.909809 | 0.748903 | 0.014 | 0.016 | 1 | 11 | Gm15859 | -0.002 | Young |
| 0.909809 | 0.723735 | 0.014 | 0.016 | 1 | 11 | Gm43672 | -0.002 | Young |
| 0.909809 | 0.691138 | 0.014 | 0.016 | 1 | 11 | Otud3 | -0.002 | Young |
| 0.909809 | 0.671543 | 0.014 | 0.016 | 1 | 11 | Hus1 | -0.002 | Young |
| 0.909809 | 0.650807 | 0.014 | 0.016 | 1 | 11 | Magi3 | -0.002 | Young |
| 0.909809 | 0.619918 | 0.014 | 0.016 | 1 | 11 | Ehbp1 | -0.002 | Young |
| 0.909809 | 0.562581 | 0.014 | 0.016 | 1 | 11 | Gpr150 | -0.002 | Young |
| 0.909809 | 0.545069 | 0.014 | 0.016 | 1 | 11 | Adap2os | -0.002 | Young |
| 0.909809 | 0.527371 | 0.014 | 0.016 | 1 | 11 | Amigo1 | -0.002 | Young |
| 0.909809 | 0.450124 | 0.014 | 0.016 | 1 | 11 | Faap24 | -0.002 | Young |
| 0.909809 | 0.353684 | 0.014 | 0.016 | 1 | 11 | 4833418N( | -0.002 | Young |
| 0.909809 | 0.333552 | 0.014 | 0.016 | 1 | 11 | B630019K( | -0.002 | Young |
| 0.909809 | 0.256423 | 0.014 | 0.016 | 1 | 11 | Slc35g1 | -0.002 | Young |
| 0.909809 | 0.208723 | 0.014 | 0.016 | 1 | 11 | Gpsm1 | -0.002 | Young |
| 0.909809 | 0.149434 | 0.014 | 0.016 | 1 | 11 | Bgn | -0.002 | Young |
| 0.909809 | 0.133578 | 0.014 | 0.016 | 1 | 11 | Gm15445 | -0.002 | Young |

| | | | | | | | | |
|---|---|---|---|---|---|---|---|---|
| 0.909809 | 0.100175 | 0.014 | 0.016 | 1 | 11 | Fadd | -0.002 | Young |
| 0.909809 | 0.090029 | 0.014 | 0.016 | 1 | 11 | 9530082P2 | -0.002 | Young |
| 0.909809 | 0.089271 | 0.014 | 0.016 | 1 | 11 | Gm826 | -0.002 | Young |
| 0.910034 | 0.565666 | 0.051 | 0.048 | 1 | 11 | Alg9 | 0.003 | Young |
| 0.910034 | 0.543798 | 0.051 | 0.048 | 1 | 11 | Wdr45 | 0.003 | Young |
| 0.910224 | 0.689752 | 0.217 | 0.25 | 1 | 11 | Zfp292 | -0.033 | Young |
| 0.910401 | 0.552668 | 0.058 | 0.056 | 1 | 11 | Fam78a | 0.002 | Young |
| 0.910704 | 0.476608 | 0.268 | 0.31 | 1 | 11 | Tifa | -0.042 | Young |
| 0.910935 | 2.292504 | 0.022 | 0.02 | 1 | 11 | Inpp1 | 0.002 | Young |
| 0.910935 | 0.235611 | 0.022 | 0.02 | 1 | 11 | Upp2 | 0.002 | Young |
| 0.910988 | 0.370721 | 0.087 | 0.093 | 1 | 11 | Pilrb1 | -0.006 | Young |
| 0.911036 | 0.134722 | 0.065 | 0.069 | 1 | 11 | Haghl | -0.004 | Young |
| 0.911135 | 0.400618 | 0.514 | 0.625 | 1 | 11 | C3ar1 | -0.111 | Young |
| 0.911237 | 0.158681 | 0.13 | 0.137 | 1 | 11 | Usp1 | -0.007 | Young |
| 0.912688 | 0.515791 | 0.29 | 0.339 | 1 | 11 | Clptm1l | -0.049 | Young |
| 0.912774 | 0.075058 | 0.065 | 0.069 | 1 | 11 | Kdm5d | -0.004 | Young |
| 0.912821 | 0.251807 | 0.08 | 0.085 | 1 | 11 | Adk | -0.005 | Young |
| 0.913332 | 1.335634 | 0.014 | 0.016 | 1 | 11 | Slc25a23 | -0.002 | Young |
| 0.913332 | 1.099955 | 0.014 | 0.016 | 1 | 11 | Wnt4 | -0.002 | Young |
| 0.913332 | 1.037399 | 0.014 | 0.016 | 1 | 11 | Top1mt | -0.002 | Young |
| 0.913332 | 0.946678 | 0.014 | 0.016 | 1 | 11 | Plekha5 | -0.002 | Young |
| 0.913332 | 0.905475 | 0.014 | 0.016 | 1 | 11 | Cdkn2a | -0.002 | Young |
| 0.913332 | 0.874511 | 0.014 | 0.016 | 1 | 11 | Prdm11 | -0.002 | Young |
| 0.913332 | 0.820696 | 0.014 | 0.016 | 1 | 11 | Polr1e | -0.002 | Young |
| 0.913332 | 0.726058 | 0.014 | 0.016 | 1 | 11 | Polm | -0.002 | Young |
| 0.913332 | 0.649622 | 0.014 | 0.016 | 1 | 11 | Caprin2 | -0.002 | Young |
| 0.913332 | 0.593366 | 0.014 | 0.016 | 1 | 11 | Trim16 | -0.002 | Young |
| 0.913332 | 0.542981 | 0.014 | 0.016 | 1 | 11 | Rpgrip1l | -0.002 | Young |
| 0.913332 | 0.475058 | 0.014 | 0.016 | 1 | 11 | Arhgap29 | -0.002 | Young |
| 0.913332 | 0.451694 | 0.014 | 0.016 | 1 | 11 | Pter | -0.002 | Young |
| 0.913625 | 0.387254 | 0.203 | 0.218 | 1 | 11 | Smad1 | -0.015 | Young |
| 0.913825 | 0.863805 | 0.065 | 0.065 | 1 | 11 | Taf1b | 0 | Young |
| 0.913994 | 1.427614 | 0.022 | 0.02 | 1 | 11 | Tefm | 0.002 | Young |
| 0.913994 | 1.008123 | 0.022 | 0.02 | 1 | 11 | Olfr1167 | 0.002 | Young |
| 0.913994 | 0.775332 | 0.022 | 0.02 | 1 | 11 | Ficd | 0.002 | Young |
| 0.913994 | 0.730496 | 0.022 | 0.02 | 1 | 11 | Natd1 | 0.002 | Young |
| 0.913994 | 0.403558 | 0.022 | 0.02 | 1 | 11 | Mcm10 | 0.002 | Young |
| 0.913994 | 0.362196 | 0.022 | 0.02 | 1 | 11 | Mast4 | 0.002 | Young |
| 0.913994 | 0.280985 | 0.022 | 0.02 | 1 | 11 | Fkbpl | 0.002 | Young |
| 0.913994 | 0.170746 | 0.022 | 0.02 | 1 | 11 | Park2 | 0.002 | Young |
| 0.9144 | 0.357038 | 0.08 | 0.085 | 1 | 11 | Ccdc117 | -0.005 | Young |
| 0.914421 | 1.322169 | 0.087 | 0.089 | 1 | 11 | Ap5m1 | -0.002 | Young |
| 0.915008 | 0.00735 | 0.826 | 0.895 | 1 | 11 | Cltc | -0.069 | Young |
| 0.915167 | 0.481005 | 0.283 | 0.306 | 1 | 11 | Psmc6 | -0.023 | Young |
| 0.916654 | 0.61026 | 0.196 | 0.21 | 1 | 11 | Slc12a9 | -0.014 | Young |

| | | | | | | | | |
|---|---|---|---|---|---|---|---|---|
| 0.916856 | 1.748118 | 0.014 | 0.016 | 1 | 11 | Jmjd4 | -0.002 | Young |
| 0.916856 | 1.245076 | 0.014 | 0.016 | 1 | 11 | Zfp623 | -0.002 | Young |
| 0.916856 | 1.182194 | 0.014 | 0.016 | 1 | 11 | Klhdc8b | -0.002 | Young |
| 0.916856 | 1.168326 | 0.014 | 0.016 | 1 | 11 | Plxna3 | -0.002 | Young |
| 0.916856 | 0.728746 | 0.014 | 0.016 | 1 | 11 | Paqr3 | -0.002 | Young |
| 0.916856 | 0.690367 | 0.014 | 0.016 | 1 | 11 | Nupl2 | -0.002 | Young |
| 0.916856 | 0.586805 | 0.014 | 0.016 | 1 | 11 | Cdc7 | -0.002 | Young |
| 0.917926 | 0.135851 | 0.087 | 0.085 | 1 | 11 | Gopc | 0.002 | Young |
| 0.918077 | 0.917408 | 0.123 | 0.129 | 1 | 11 | Kansl3 | -0.006 | Young |
| 0.918219 | 0.46747 | 0.188 | 0.198 | 1 | 11 | Stk10 | -0.01 | Young |
| 0.918268 | 0.467124 | 0.145 | 0.157 | 1 | 11 | Taf13 | -0.012 | Young |
| 0.918569 | 0.010714 | 0.087 | 0.093 | 1 | 11 | Grk5 | -0.006 | Young |
| 0.919202 | 0.245645 | 0.435 | 0.512 | 1 | 11 | Taok1 | -0.077 | Young |
| 0.919333 | 0.578331 | 0.094 | 0.101 | 1 | 11 | Zdhhc17 | -0.007 | Young |
| 0.919912 | 0.885147 | 0.072 | 0.073 | 1 | 11 | Lig1 | -0.001 | Young |
| 0.920116 | 0.621124 | 0.022 | 0.02 | 1 | 11 | Map2k6 | 0.002 | Young |
| 0.920116 | 0.403945 | 0.022 | 0.02 | 1 | 11 | Msrb2 | 0.002 | Young |
| 0.920116 | 0.322201 | 0.022 | 0.02 | 1 | 11 | B130006D( | 0.002 | Young |
| 0.920908 | 0.932285 | 0.065 | 0.065 | 1 | 11 | Plrg1 | 0 | Young |
| 0.920908 | 0.908321 | 0.065 | 0.065 | 1 | 11 | Nr2c1 | 0 | Young |
| 0.920908 | 0.63783 | 0.065 | 0.065 | 1 | 11 | Cav2 | 0 | Young |
| 0.921103 | 0.733357 | 0.167 | 0.185 | 1 | 11 | Nudt16l1 | -0.018 | Young |
| 0.921415 | 0.040901 | 0.841 | 0.871 | 1 | 11 | Ubc | -0.03 | Young |
| 0.922256 | 0.89442 | 0.094 | 0.101 | 1 | 11 | Rcn3 | -0.007 | Young |
| 0.922679 | 1.206716 | 0.065 | 0.065 | 1 | 11 | Clec9a | 0 | Young |
| 0.923114 | 0.314438 | 0.217 | 0.234 | 1 | 11 | Nt5c3 | -0.017 | Young |
| 0.923179 | 0.659318 | 0.022 | 0.02 | 1 | 11 | Osgin1 | 0.002 | Young |
| 0.923179 | 0.313494 | 0.022 | 0.02 | 1 | 11 | Gm44067 | 0.002 | Young |
| 0.923303 | 0.842967 | 0.123 | 0.129 | 1 | 11 | Lactb2 | -0.006 | Young |
| 0.924557 | 0.534722 | 0.232 | 0.258 | 1 | 11 | Rabep1 | -0.026 | Young |
| 0.924946 | 0.047047 | 0.065 | 0.069 | 1 | 11 | Gins4 | -0.004 | Young |
| 0.925196 | 0.396391 | 0.275 | 0.315 | 1 | 11 | Npepps | -0.04 | Young |
| 0.92572 | 1.27437 | 0.109 | 0.113 | 1 | 11 | Bcl2 | -0.004 | Young |
| 0.925777 | 0.215445 | 0.377 | 0.435 | 1 | 11 | Cers2 | -0.058 | Young |
| 0.92588 | 0.575996 | 0.188 | 0.198 | 1 | 11 | Sec24c | -0.01 | Young |
| 0.925952 | 0.421775 | 0.377 | 0.427 | 1 | 11 | Zbp1 | -0.05 | Young |
| 0.92625 | 0.465084 | 0.514 | 0.585 | 1 | 11 | Irf1 | -0.071 | Young |
| 0.926294 | 0.494063 | 0.399 | 0.456 | 1 | 11 | Trir | -0.057 | Young |
| 0.926719 | 0.324114 | 0.152 | 0.153 | 1 | 11 | Ppm1k | -0.001 | Young |
| 0.926719 | 0.239901 | 0.152 | 0.153 | 1 | 11 | Samd8 | -0.001 | Young |
| 0.92693 | 0.129806 | 0.268 | 0.29 | 1 | 11 | Ppp1r2 | -0.022 | Young |
| 0.926957 | 0.049755 | 0.551 | 0.649 | 1 | 11 | Ifi47 | -0.098 | Young |
| 0.927129 | 0.403269 | 0.333 | 0.363 | 1 | 11 | Rtp4 | -0.03 | Young |
| 0.927813 | 1.183058 | 0.072 | 0.077 | 1 | 11 | Zfp672 | -0.005 | Young |
| 0.927819 | 0.175704 | 0.246 | 0.254 | 1 | 11 | Slco2b1 | -0.008 | Young |

| | | | | | | | | |
|---|---|---|---|---|---|---|---|---|
| 0.927997 | 0.554649 | 0.065 | 0.065 | 1 | 11 | Gca | 0 | Young |
| 0.928663 | 0.414401 | 0.413 | 0.492 | 1 | 11 | Shtn1 | -0.079 | Young |
| 0.928876 | 1.28441 | 0.167 | 0.19 | 1 | 11 | Rest | -0.023 | Young |
| 0.929208 | 0.248677 | 0.058 | 0.056 | 1 | 11 | Ltb4r1 | 0.002 | Young |
| 0.929405 | 0.360654 | 0.37 | 0.419 | 1 | 11 | Ppp2r5a | -0.049 | Young |
| 0.92971 | 0.307686 | 0.217 | 0.226 | 1 | 11 | Ifitm6 | -0.009 | Young |
| 0.930065 | 0.439773 | 0.239 | 0.262 | 1 | 11 | Pxk | -0.023 | Young |
| 0.930218 | 0.078044 | 0.601 | 0.726 | 1 | 11 | Trim30a | -0.125 | Young |
| 0.930649 | 0.42374 | 0.138 | 0.149 | 1 | 11 | Plcl2 | -0.011 | Young |
| 0.930789 | 0.683108 | 0.196 | 0.206 | 1 | 11 | Rgs18 | -0.01 | Young |
| 0.931091 | 0.355058 | 0.058 | 0.056 | 1 | 11 | Rasgrp3 | 0.002 | Young |
| 0.931262 | 0.636792 | 0.116 | 0.117 | 1 | 11 | Specc1 | -0.001 | Young |
| 0.931747 | 0.437027 | 0.13 | 0.129 | 1 | 11 | Adgrg6 | 0.001 | Young |
| 0.932513 | 0.253972 | 0.058 | 0.06 | 1 | 11 | Mocs1 | -0.002 | Young |
| 0.932637 | 0.323129 | 0.406 | 0.472 | 1 | 11 | Taldo1 | -0.066 | Young |
| 0.93289 | 1.731724 | 0.029 | 0.028 | 1 | 11 | Wdr73 | 0.001 | Young |
| 0.93289 | 1.426812 | 0.029 | 0.028 | 1 | 11 | Pus7l | 0.001 | Young |
| 0.933622 | 0.293583 | 0.536 | 0.661 | 1 | 11 | Ccdc50 | -0.125 | Young |
| 0.933652 | 1.015527 | 0.065 | 0.069 | 1 | 11 | Acot7 | -0.004 | Young |
| 0.933831 | 0.55423 | 0.123 | 0.125 | 1 | 11 | Tsc1 | -0.002 | Young |
| 0.934016 | 0.311325 | 0.203 | 0.222 | 1 | 11 | Dapp1 | -0.019 | Young |
| 0.93459 | 0.526168 | 0.138 | 0.141 | 1 | 11 | Cryl1 | -0.003 | Young |
| 0.934899 | 0.396045 | 0.13 | 0.133 | 1 | 11 | Aaed1 | -0.003 | Young |
| 0.935218 | 0.150718 | 0.413 | 0.472 | 1 | 11 | Slk | -0.059 | Young |
| 0.935232 | 0.51548 | 0.167 | 0.173 | 1 | 11 | Nup153 | -0.006 | Young |
| 0.935912 | 0.681558 | 0.152 | 0.165 | 1 | 11 | Ophn1 | -0.013 | Young |
| 0.936092 | 0.68829 | 0.072 | 0.077 | 1 | 11 | Jpt2 | -0.005 | Young |
| 0.936224 | 0.389144 | 0.217 | 0.23 | 1 | 11 | Jak2 | -0.013 | Young |
| 0.936537 | 0.495312 | 0.08 | 0.085 | 1 | 11 | Rab33b | -0.005 | Young |
| 0.936541 | 0.048048 | 0.703 | 0.778 | 1 | 11 | Rps18 | -0.075 | Young |
| 0.936568 | 0.701 | 0.159 | 0.173 | 1 | 11 | Golga3 | -0.014 | Young |
| 0.936733 | 1.022683 | 0.072 | 0.073 | 1 | 11 | Klhdc2 | -0.001 | Young |
| 0.937011 | 0.034314 | 0.464 | 0.536 | 1 | 11 | Taok3 | -0.072 | Young |
| 0.937123 | 0.505053 | 0.333 | 0.383 | 1 | 11 | 1-Mar | -0.05 | Young |
| 0.937203 | 0.341379 | 0.377 | 0.435 | 1 | 11 | Ifi209 | -0.058 | Young |
| 0.937404 | 0.072022 | 0.928 | 0.96 | 1 | 11 | Fos | -0.032 | Young |
| 0.938054 | 0.717924 | 0.116 | 0.117 | 1 | 11 | Rgmb | -0.001 | Young |
| 0.938054 | 0.537423 | 0.116 | 0.117 | 1 | 11 | Usp40 | -0.001 | Young |
| 0.938125 | 0.541029 | 0.109 | 0.117 | 1 | 11 | Kdm3a | -0.008 | Young |
| 0.938125 | 0.454187 | 0.109 | 0.117 | 1 | 11 | Mxi1 | -0.008 | Young |
| 0.938416 | 0.702431 | 0.072 | 0.073 | 1 | 11 | Amz2 | -0.001 | Young |
| 0.938486 | 0.741896 | 0.29 | 0.323 | 1 | 11 | Aida | -0.033 | Young |
| 0.938732 | 0.167881 | 0.783 | 0.835 | 1 | 11 | Efhd2 | -0.052 | Young |
| 0.939441 | 0.426771 | 0.435 | 0.484 | 1 | 11 | Itga4 | -0.049 | Young |
| 0.93953 | 0.89809 | 0.203 | 0.218 | 1 | 11 | Zhx1 | -0.015 | Young |

| | | | | | | | | |
|---|---|---|---|---|---|---|---|---|
| 0.939737 | 0.130596 | 0.688 | 0.73 | 1 | 11 | Bst2 | -0.042 | Young |
| 0.939894 | 0.242669 | 0.058 | 0.06 | 1 | 11 | I830077J02 | -0.002 | Young |
| 0.94034 | 0.194498 | 0.428 | 0.492 | 1 | 11 | Brd2 | -0.064 | Young |
| 0.940769 | 1.111497 | 0.029 | 0.028 | 1 | 11 | Cdkl2 | 0.001 | Young |
| 0.940803 | 0.46288 | 0.304 | 0.351 | 1 | 11 | Hnrnpd | -0.047 | Young |
| 0.941619 | 0.593885 | 0.261 | 0.282 | 1 | 11 | Emg1 | -0.021 | Young |
| 0.94174 | 0.237229 | 0.058 | 0.06 | 1 | 11 | Pck2 | -0.002 | Young |
| 0.941756 | 0.492884 | 0.123 | 0.125 | 1 | 11 | 4833420G1 | -0.002 | Young |
| 0.942177 | 0.312978 | 0.203 | 0.218 | 1 | 11 | Dnttip2 | -0.015 | Young |
| 0.942192 | 0.747945 | 0.065 | 0.065 | 1 | 11 | Kif21b | 0 | Young |
| 0.942257 | 0.72548 | 0.319 | 0.359 | 1 | 11 | Ctnna1 | -0.04 | Young |
| 0.942366 | 0.38613 | 0.065 | 0.069 | 1 | 11 | Foxp4 | -0.004 | Young |
| 0.94272 | 0.583522 | 0.072 | 0.077 | 1 | 11 | Ppwd1 | -0.005 | Young |
| 0.942804 | 0.43908 | 0.181 | 0.194 | 1 | 11 | Fez2 | -0.013 | Young |
| 0.943238 | 0.317569 | 0.145 | 0.153 | 1 | 11 | Ccdc174 | -0.008 | Young |
| 0.943333 | 0.887748 | 0.101 | 0.105 | 1 | 11 | Mnat1 | -0.004 | Young |
| 0.943397 | 1.777173 | 0.029 | 0.028 | 1 | 11 | Dcbld2 | 0.001 | Young |
| 0.943397 | 0.908849 | 0.029 | 0.028 | 1 | 11 | Epop | 0.001 | Young |
| 0.943397 | 0.786711 | 0.029 | 0.028 | 1 | 11 | Cdiptos | 0.001 | Young |
| 0.943571 | 0.771474 | 0.225 | 0.254 | 1 | 11 | Rapgef2 | -0.029 | Young |
| 0.943967 | 0.587665 | 0.065 | 0.065 | 1 | 11 | 2810402E2 | 0 | Young |
| 0.944448 | 1.000009 | 0.08 | 0.081 | 1 | 11 | Alg8 | -0.001 | Young |
| 0.945021 | 0.581505 | 0.181 | 0.194 | 1 | 11 | Milr1 | -0.013 | Young |
| 0.945154 | 0.569198 | 0.072 | 0.073 | 1 | 11 | Top3b | -0.001 | Young |
| 0.946026 | 0.929146 | 0.029 | 0.028 | 1 | 11 | Izumo4 | 0.001 | Young |
| 0.946026 | 0.723005 | 0.029 | 0.028 | 1 | 11 | Emc9 | 0.001 | Young |
| 0.946026 | 0.400844 | 0.029 | 0.028 | 1 | 11 | Tacc2 | 0.001 | Young |
| 0.946362 | 0.770837 | 0.109 | 0.117 | 1 | 11 | Pcnx | -0.008 | Young |
| 0.946652 | 0.250572 | 0.051 | 0.052 | 1 | 11 | Foxo4 | -0.001 | Young |
| 0.947173 | 0.169915 | 0.565 | 0.685 | 1 | 11 | Hnrnph1 | -0.12 | Young |
| 0.947257 | 0.272738 | 0.188 | 0.206 | 1 | 11 | Tbc1d9 | -0.018 | Young |
| 0.94728 | 0.501303 | 0.058 | 0.06 | 1 | 11 | Rbm10 | -0.002 | Young |
| 0.947356 | 0.971834 | 0.159 | 0.177 | 1 | 11 | Tyk2 | -0.018 | Young |
| 0.947547 | 0.392871 | 0.442 | 0.552 | 1 | 11 | Bzw1 | -0.11 | Young |
| 0.947598 | 0.12152 | 0.841 | 0.915 | 1 | 11 | Kctd12 | -0.074 | Young |
| 0.94838 | 0.444416 | 0.362 | 0.431 | 1 | 11 | Dnajb14 | -0.069 | Young |
| 0.948387 | 0.799544 | 0.239 | 0.262 | 1 | 11 | Dcun1d5 | -0.023 | Young |
| 0.948405 | 0.351477 | 0.138 | 0.141 | 1 | 11 | Mbd1 | -0.003 | Young |
| 0.948453 | 0.488731 | 0.246 | 0.27 | 1 | 11 | Ep400 | -0.024 | Young |
| 0.948648 | 0.428834 | 0.326 | 0.375 | 1 | 11 | 2810474O1 | -0.049 | Young |
| 0.948655 | 0.916523 | 0.029 | 0.028 | 1 | 11 | Tmco6 | 0.001 | Young |
| 0.948655 | 0.894596 | 0.029 | 0.028 | 1 | 11 | Mfng | 0.001 | Young |
| 0.948655 | 0.460593 | 0.029 | 0.028 | 1 | 11 | Ddx59 | 0.001 | Young |
| 0.948695 | 0.776913 | 0.152 | 0.169 | 1 | 11 | Tmem192 | -0.017 | Young |
| 0.949296 | 0.636207 | 0.065 | 0.065 | 1 | 11 | Ang | 0 | Young |

| | | | | | | | | |
|---|---|---|---|---|---|---|---|---|
| 0.95021 | 0.711476 | 0.072 | 0.073 | 1 | 11 | Fam118a | -0.001 | Young |
| 0.950442 | 0.2058 | 0.254 | 0.278 | 1 | 11 | Tcf12 | -0.024 | Young |
| 0.950791 | 1.726087 | 0.036 | 0.036 | 1 | 11 | F8a | 0 | Young |
| 0.951073 | 0.619517 | 0.065 | 0.065 | 1 | 11 | Zscan29 | 0 | Young |
| 0.951285 | 0.881335 | 0.029 | 0.028 | 1 | 11 | Etfbkmt | 0.001 | Young |
| 0.951285 | 0.820102 | 0.029 | 0.028 | 1 | 11 | Phtf1 | 0.001 | Young |
| 0.951285 | 0.252971 | 0.029 | 0.028 | 1 | 11 | Laptm4b | 0.001 | Young |
| 0.951358 | 0.05841 | 0.819 | 0.883 | 1 | 11 | Rps15a | -0.064 | Young |
| 0.951415 | 0.788578 | 0.261 | 0.286 | 1 | 11 | Lpin2 | -0.025 | Young |
| 0.951649 | 1.628156 | 0.109 | 0.121 | 1 | 11 | Tlr3 | -0.012 | Young |
| 0.951784 | 1.01338 | 0.196 | 0.214 | 1 | 11 | Pik3cg | -0.018 | Young |
| 0.951856 | 1.146308 | 0.109 | 0.117 | 1 | 11 | Ndrg3 | -0.008 | Young |
| 0.95225 | 0.427711 | 0.225 | 0.246 | 1 | 11 | Ifi35 | -0.021 | Young |
| 0.952268 | 0.231464 | 0.478 | 0.556 | 1 | 11 | Rgs10 | -0.078 | Young |
| 0.952831 | 0.580519 | 0.065 | 0.069 | 1 | 11 | Snx16 | -0.004 | Young |
| 0.952983 | 0.400102 | 0.413 | 0.468 | 1 | 11 | Il10rb | -0.055 | Young |
| 0.953059 | 0.244499 | 0.297 | 0.327 | 1 | 11 | Mafg | -0.03 | Young |
| 0.953313 | 0.906386 | 0.13 | 0.137 | 1 | 11 | G6pdx | -0.007 | Young |
| 0.953327 | 0.686471 | 0.145 | 0.157 | 1 | 11 | Opa3 | -0.012 | Young |
| 0.953362 | 0.840516 | 0.101 | 0.105 | 1 | 11 | Atad2 | -0.004 | Young |
| 0.953581 | 0.392797 | 0.072 | 0.073 | 1 | 11 | Trp53bp2 | -0.001 | Young |
| 0.953915 | 0.441752 | 0.029 | 0.028 | 1 | 11 | Tmem229a | 0.001 | Young |
| 0.954065 | 0.640463 | 0.275 | 0.327 | 1 | 11 | Fam105a | -0.052 | Young |
| 0.954627 | 0.388355 | 0.065 | 0.065 | 1 | 11 | Zbtb33 | 0 | Young |
| 0.954671 | 0.55037 | 0.058 | 0.06 | 1 | 11 | Nfrkb | -0.002 | Young |
| 0.955103 | 0.392543 | 0.203 | 0.222 | 1 | 11 | Rab11fip1 | -0.019 | Young |
| 0.95523 | 0.546618 | 0.116 | 0.125 | 1 | 11 | Sympk | -0.009 | Young |
| 0.955319 | 0.759607 | 0.225 | 0.258 | 1 | 11 | Cux1 | -0.033 | Young |
| 0.955706 | 0.635689 | 0.08 | 0.081 | 1 | 11 | Foxred2 | -0.001 | Young |
| 0.955979 | 0.421093 | 0.232 | 0.246 | 1 | 11 | Mcts1 | -0.014 | Young |
| 0.955992 | 0.59036 | 0.087 | 0.089 | 1 | 11 | Aatf | -0.002 | Young |
| 0.956321 | 0.655517 | 0.065 | 0.069 | 1 | 11 | Zfp26 | -0.004 | Young |
| 0.956332 | 0.488909 | 0.145 | 0.149 | 1 | 11 | Nab2 | -0.004 | Young |
| 0.956521 | 0.196092 | 0.051 | 0.052 | 1 | 11 | Wdr47 | -0.001 | Young |
| 0.956546 | 0.529462 | 0.029 | 0.028 | 1 | 11 | Ndufaf1 | 0.001 | Young |
| 0.956546 | 0.506301 | 0.029 | 0.028 | 1 | 11 | Pla2g12a | 0.001 | Young |
| 0.956546 | 0.033753 | 0.029 | 0.028 | 1 | 11 | Mtg2 | 0.001 | Young |
| 0.956862 | 0.13838 | 0.478 | 0.536 | 1 | 11 | Clec4a2 | -0.058 | Young |
| 0.957723 | 0.72641 | 0.246 | 0.282 | 1 | 11 | Chd9 | -0.036 | Young |
| 0.957827 | 0.508449 | 0.239 | 0.25 | 1 | 11 | Ube2z | -0.011 | Young |
| 0.958495 | 0.250161 | 0.051 | 0.052 | 1 | 11 | Sprtn | -0.001 | Young |
| 0.958742 | 0.751985 | 0.196 | 0.214 | 1 | 11 | Scimp | -0.018 | Young |
| 0.95902 | 0.152107 | 0.406 | 0.431 | 1 | 11 | Basp1 | -0.025 | Young |
| 0.959077 | 0.73138 | 0.087 | 0.089 | 1 | 11 | Xpnpep1 | -0.002 | Young |
| 0.959177 | 0.467301 | 0.029 | 0.028 | 1 | 11 | 2610044O1 | 0.001 | Young |

| | | | | | | | | |
|---|---|---|---|---|---|---|---|---|
| 0.959177 | 0.211433 | 0.029 | 0.028 | 1 | 11 | Shq1 | 0.001 | Young |
| 0.959371 | 0.100598 | 0.094 | 0.093 | 1 | 11 | Pkd2 | 0.001 | Young |
| 0.959383 | 0.756774 | 0.145 | 0.157 | 1 | 11 | Flcn | -0.012 | Young |
| 0.959968 | 0.308369 | 0.123 | 0.129 | 1 | 11 | Katnbl1 | -0.006 | Young |
| 0.960657 | 0.371072 | 0.138 | 0.145 | 1 | 11 | Rnf34 | -0.007 | Young |
| 0.961379 | 0.429791 | 0.312 | 0.339 | 1 | 11 | Cd163 | -0.027 | Young |
| 0.961606 | 0.923378 | 0.13 | 0.145 | 1 | 11 | Prr14 | -0.015 | Young |
| 0.961809 | 0.649106 | 0.029 | 0.028 | 1 | 11 | Pfkm | 0.001 | Young |
| 0.961809 | 0.476661 | 0.029 | 0.028 | 1 | 11 | Limk2 | 0.001 | Young |
| 0.962065 | 0.756484 | 0.058 | 0.06 | 1 | 11 | 10-Sep | -0.002 | Young |
| 0.962445 | 0.35464 | 0.051 | 0.052 | 1 | 11 | Epn2 | -0.001 | Young |
| 0.962497 | 1.164857 | 0.036 | 0.036 | 1 | 11 | Cstf2 | 0 | Young |
| 0.962658 | 0.654359 | 0.087 | 0.093 | 1 | 11 | Pik3c3 | -0.006 | Young |
| 0.963167 | 0.840083 | 0.109 | 0.113 | 1 | 11 | Nkap | -0.004 | Young |
| 0.963529 | 0.987664 | 0.239 | 0.27 | 1 | 11 | Slc12a6 | -0.031 | Young |
| 0.963915 | 0.794577 | 0.058 | 0.06 | 1 | 11 | Zfp367 | -0.002 | Young |
| 0.963915 | 0.667188 | 0.058 | 0.06 | 1 | 11 | Zfp866 | -0.002 | Young |
| 0.964181 | 0.643292 | 0.087 | 0.093 | 1 | 11 | Ccdc93 | -0.006 | Young |
| 0.9644 | 0.374156 | 0.254 | 0.274 | 1 | 11 | Ppm1a | -0.02 | Young |
| 0.964441 | 0.318537 | 0.029 | 0.028 | 1 | 11 | Rac3 | 0.001 | Young |
| 0.964441 | 0.091055 | 0.029 | 0.028 | 1 | 11 | Noa1 | 0.001 | Young |
| 0.964476 | 0.304044 | 0.275 | 0.298 | 1 | 11 | Ppp2r2a | -0.023 | Young |
| 0.964887 | 0.203044 | 0.109 | 0.109 | 1 | 11 | Slc35e1 | 0 | Young |
| 0.965317 | 0.611542 | 0.261 | 0.302 | 1 | 11 | Cbx1 | -0.041 | Young |
| 0.965764 | 0.595814 | 0.058 | 0.06 | 1 | 11 | Ezh2 | -0.002 | Young |
| 0.966396 | 0.471402 | 0.051 | 0.052 | 1 | 11 | Dmac2 | -0.001 | Young |
| 0.967374 | 0.439682 | 0.13 | 0.137 | 1 | 11 | Vkorc1 | -0.007 | Young |
| 0.967614 | 0.876697 | 0.058 | 0.06 | 1 | 11 | Asb13 | -0.002 | Young |
| 0.967846 | 0.135735 | 0.514 | 0.585 | 1 | 11 | Arhgap30 | -0.071 | Young |
| 0.967865 | 0.724861 | 0.145 | 0.157 | 1 | 11 | Dnm1l | -0.012 | Young |
| 0.968544 | 0.754001 | 0.065 | 0.069 | 1 | 11 | Lonrf1 | -0.004 | Young |
| 0.968668 | 0.809024 | 0.181 | 0.202 | 1 | 11 | Cse1l | -0.021 | Young |
| 0.968751 | 0.93684 | 0.087 | 0.093 | 1 | 11 | Arl11 | -0.006 | Young |
| 0.968763 | 0.220529 | 0.072 | 0.073 | 1 | 11 | Sugp2 | -0.001 | Young |
| 0.969425 | 0.337893 | 0.362 | 0.411 | 1 | 11 | Dennd4a | -0.049 | Young |
| 0.969463 | 1.256945 | 0.058 | 0.06 | 1 | 11 | Gapt | -0.002 | Young |
| 0.969525 | 1.795998 | 0.036 | 0.036 | 1 | 11 | Acrbp | 0 | Young |
| 0.969525 | 1.099975 | 0.036 | 0.036 | 1 | 11 | Fbxl12 | 0 | Young |
| 0.969525 | 1.055741 | 0.036 | 0.036 | 1 | 11 | Zfp846 | 0 | Young |
| 0.969525 | 0.913312 | 0.036 | 0.036 | 1 | 11 | Mapk7 | 0 | Young |
| 0.969706 | 0.223861 | 0.029 | 0.028 | 1 | 11 | Tmem221 | 0.001 | Young |
| 0.969777 | 0.055678 | 0.58 | 0.702 | 1 | 11 | Csde1 | -0.122 | Young |
| 0.969831 | 0.61908 | 0.08 | 0.085 | 1 | 11 | Cfl2 | -0.005 | Young |
| 0.969881 | 0.458169 | 0.087 | 0.089 | 1 | 11 | Strada | -0.002 | Young |
| 0.970291 | 0.641512 | 0.065 | 0.069 | 1 | 11 | Lix1l | -0.004 | Young |

| | | | | | | | | |
|---|---|---|---|---|---|---|---|---|
| 0.971869 | 0.365408 | 0.036 | 0.036 | 1 | 11 | Lrrc57 | 0 | Young |
| 0.971892 | 0.548982 | 0.138 | 0.145 | 1 | 11 | Scaf8 | -0.007 | Young |
| 0.972323 | 0.311569 | 0.051 | 0.052 | 1 | 11 | Nol10 | -0.001 | Young |
| 0.972969 | 0.502711 | 0.087 | 0.089 | 1 | 11 | Slc39a6 | -0.002 | Young |
| 0.973246 | 0.149941 | 0.384 | 0.431 | 1 | 11 | Ptpro | -0.047 | Young |
| 0.973302 | 0.461585 | 0.043 | 0.044 | 1 | 11 | Padi2 | -0.001 | Young |
| 0.973623 | 0.129424 | 0.413 | 0.452 | 1 | 11 | Slfn8 | -0.039 | Young |
| 0.973988 | 0.447091 | 0.203 | 0.218 | 1 | 11 | Stk11 | -0.015 | Young |
| 0.974253 | 0.576784 | 0.275 | 0.315 | 1 | 11 | H2afv | -0.04 | Young |
| 0.974931 | 0.276794 | 0.203 | 0.214 | 1 | 11 | Tia1 | -0.011 | Young |
| 0.975437 | 0.136412 | 0.043 | 0.044 | 1 | 11 | Ints4 | -0.001 | Young |
| 0.97597 | 0.509976 | 0.065 | 0.065 | 1 | 11 | Zdhhc4 | 0 | Young |
| 0.975971 | 0.220564 | 0.457 | 0.512 | 1 | 11 | Hmox1 | -0.055 | Young |
| 0.976105 | 0.810562 | 0.181 | 0.194 | 1 | 11 | Mrpl54 | -0.013 | Young |
| 0.97618 | 0.576448 | 0.145 | 0.153 | 1 | 11 | Slc16a6 | -0.008 | Young |
| 0.976241 | 0.224101 | 0.572 | 0.665 | 1 | 11 | Tubb5 | -0.093 | Young |
| 0.976276 | 0.428665 | 0.051 | 0.052 | 1 | 11 | Nlrp1b | -0.001 | Young |
| 0.976496 | 0.638658 | 0.094 | 0.101 | 1 | 11 | Ruvbl1 | -0.007 | Young |
| 0.976556 | 1.053702 | 0.036 | 0.036 | 1 | 11 | Grpel2 | 0 | Young |
| 0.976556 | 0.386437 | 0.036 | 0.036 | 1 | 11 | Homer3 | 0 | Young |
| 0.976636 | 0.405853 | 0.08 | 0.081 | 1 | 11 | Setdb2 | -0.001 | Young |
| 0.976719 | 0.415503 | 0.377 | 0.444 | 1 | 11 | Tmem55b | -0.067 | Young |
| 0.977138 | 0.431803 | 0.159 | 0.169 | 1 | 11 | Pes1 | -0.01 | Young |
| 0.977203 | 0.517602 | 0.072 | 0.073 | 1 | 11 | Rad51b | -0.001 | Young |
| 0.978487 | 0.338371 | 0.457 | 0.524 | 1 | 11 | Myh9 | -0.067 | Young |
| 0.978899 | 1.095194 | 0.036 | 0.036 | 1 | 11 | Msto1 | 0 | Young |
| 0.978899 | 0.630548 | 0.036 | 0.036 | 1 | 11 | Slc20a2 | 0 | Young |
| 0.978997 | 0.85796 | 0.181 | 0.198 | 1 | 11 | Npm3 | -0.017 | Young |
| 0.979146 | 0.412882 | 0.087 | 0.089 | 1 | 11 | Trim24 | -0.002 | Young |
| 0.979357 | 0.907839 | 0.109 | 0.117 | 1 | 11 | Ppp2r3c | -0.008 | Young |
| 0.979708 | 1.461466 | 0.043 | 0.044 | 1 | 11 | Pex14 | -0.001 | Young |
| 0.979708 | 0.379344 | 0.043 | 0.044 | 1 | 11 | Mfge8 | -0.001 | Young |
| 0.980229 | 1.087849 | 0.051 | 0.052 | 1 | 11 | Oas1g | -0.001 | Young |
| 0.980412 | 0.965384 | 0.225 | 0.258 | 1 | 11 | Nfatc2 | -0.033 | Young |
| 0.981244 | 0.179684 | 0.036 | 0.036 | 1 | 11 | Eif2d | 0 | Young |
| 0.981386 | 0.063155 | 0.906 | 0.899 | 1 | 11 | Rpl19 | 0.007 | Young |
| 0.981444 | 0.541421 | 0.13 | 0.137 | 1 | 11 | Hivep3 | -0.007 | Young |
| 0.981839 | 0.332901 | 0.333 | 0.371 | 1 | 11 | Gmfb | -0.038 | Young |
| 0.981843 | 1.070562 | 0.043 | 0.044 | 1 | 11 | Nsun4 | -0.001 | Young |
| 0.981843 | 0.308142 | 0.043 | 0.044 | 1 | 11 | Exosc1 | -0.001 | Young |
| 0.982521 | 0.577224 | 0.065 | 0.069 | 1 | 11 | Gramd3 | -0.004 | Young |
| 0.982706 | 0.148346 | 0.652 | 0.746 | 1 | 11 | Rpl31 | -0.094 | Young |
| 0.982872 | 0.235043 | 0.217 | 0.23 | 1 | 11 | Dync1li1 | -0.013 | Young |
| 0.982957 | 0.321399 | 0.188 | 0.198 | 1 | 11 | Mrpl48 | -0.01 | Young |
| 0.983514 | 0.235802 | 0.174 | 0.181 | 1 | 11 | Rhoh | -0.007 | Young |

| | | | | | | | | |
|---|---|---|---|---|---|---|---|---|
| 0.983588 | 1.164568 | 0.036 | 0.036 | 1 | 11 | Gm21781 | 0 | Young |
| 0.983588 | 0.572292 | 0.036 | 0.036 | 1 | 11 | Fam151b | 0 | Young |
| 0.983588 | 0.32335 | 0.036 | 0.036 | 1 | 11 | Asb1 | 0 | Young |
| 0.983979 | 0.569068 | 0.043 | 0.044 | 1 | 11 | Bcam | -0.001 | Young |
| 0.983979 | 0.053925 | 0.043 | 0.044 | 1 | 11 | Zfp952 | -0.001 | Young |
| 0.984183 | 1.220923 | 0.051 | 0.052 | 1 | 11 | Ampd2 | -0.001 | Young |
| 0.984183 | 0.695458 | 0.051 | 0.052 | 1 | 11 | Gdpd3 | -0.001 | Young |
| 0.984269 | 1.552049 | 0.065 | 0.069 | 1 | 11 | Dvl2 | -0.004 | Young |
| 0.98476 | 0.349164 | 0.384 | 0.44 | 1 | 11 | Irf7 | -0.056 | Young |
| 0.984953 | 0.437908 | 0.362 | 0.415 | 1 | 11 | Trim12c | -0.053 | Young |
| 0.985403 | 0.679384 | 0.21 | 0.23 | 1 | 11 | Mark2 | -0.02 | Young |
| 0.985707 | 1.262205 | 0.08 | 0.085 | 1 | 11 | Rnf41 | -0.005 | Young |
| 0.98616 | 0.849012 | 0.051 | 0.052 | 1 | 11 | Nup160 | -0.001 | Young |
| 0.987334 | 0.106045 | 0.072 | 0.073 | 1 | 11 | Foxo1 | -0.001 | Young |
| 0.987681 | 0.337559 | 0.5 | 0.589 | 1 | 11 | Nsa2 | -0.089 | Young |
| 0.988137 | 1.023255 | 0.051 | 0.052 | 1 | 11 | Nudt2 | -0.001 | Young |
| 0.988251 | 0.764423 | 0.043 | 0.044 | 1 | 11 | Mthfsd | -0.001 | Young |
| 0.988277 | 0.510795 | 0.036 | 0.036 | 1 | 11 | Zfp317 | 0 | Young |
| 0.988784 | 0.160596 | 0.493 | 0.556 | 1 | 11 | Lair1 | -0.063 | Young |
| 0.989283 | 0.858557 | 0.217 | 0.246 | 1 | 11 | Rbck1 | -0.029 | Young |
| 0.989525 | 0.099017 | 0.08 | 0.081 | 1 | 11 | Hace1 | -0.001 | Young |
| 0.990387 | 0.754422 | 0.043 | 0.044 | 1 | 11 | Mrgbp | -0.001 | Young |
| 0.990387 | 0.261118 | 0.043 | 0.044 | 1 | 11 | Srbd1 | -0.001 | Young |
| 0.9905 | 0.033244 | 0.855 | 0.907 | 1 | 11 | Atp2b1 | -0.052 | Young |
| 0.990621 | 1.510051 | 0.036 | 0.036 | 1 | 11 | Ccdc102a | 0 | Young |
| 0.990621 | 0.017914 | 0.036 | 0.036 | 1 | 11 | Gm42031 | 0 | Young |
| 0.99075 | 0.665019 | 0.188 | 0.206 | 1 | 11 | Sgk1 | -0.018 | Young |
| 0.991058 | 0.492276 | 0.181 | 0.19 | 1 | 11 | Bag3 | -0.009 | Young |
| 0.991148 | 0.409673 | 0.159 | 0.165 | 1 | 11 | Slco3a1 | -0.006 | Young |
| 0.991546 | 0.545537 | 0.319 | 0.371 | 1 | 11 | Baz1a | -0.052 | Young |
| 0.992059 | 1.192826 | 0.08 | 0.085 | 1 | 11 | Ap1m1 | -0.005 | Young |
| 0.992091 | 0.601711 | 0.051 | 0.052 | 1 | 11 | Zfp65 | -0.001 | Young |
| 0.992222 | 0.214817 | 0.587 | 0.669 | 1 | 11 | Luc7l2 | -0.082 | Young |
| 0.992276 | 0.069749 | 0.609 | 0.718 | 1 | 11 | Tmed10 | -0.109 | Young |
| 0.992311 | 0.58123 | 0.304 | 0.351 | 1 | 11 | Cct7 | -0.047 | Young |
| 0.992926 | 0.532079 | 0.188 | 0.206 | 1 | 11 | Pcm1 | -0.018 | Young |
| 0.993048 | 0.502057 | 0.087 | 0.089 | 1 | 11 | Ppme1 | -0.002 | Young |
| 0.993539 | 0.845166 | 0.101 | 0.105 | 1 | 11 | Maoa | -0.004 | Young |
| 0.994091 | 0.527014 | 0.123 | 0.129 | 1 | 11 | Pdcd4 | -0.006 | Young |
| 0.994476 | 0.209625 | 0.514 | 0.617 | 1 | 11 | Sumo2 | -0.103 | Young |
| 0.994949 | 0.059278 | 0.587 | 0.677 | 1 | 11 | Sf3b1 | -0.09 | Young |
| 0.99531 | 1.029251 | 0.036 | 0.036 | 1 | 11 | Mtg1 | 0 | Young |
| 0.99542 | 0.531804 | 0.297 | 0.335 | 1 | 11 | Hmgn1 | -0.038 | Young |
| 0.995971 | 0.548949 | 0.08 | 0.081 | 1 | 11 | Mkl2 | -0.001 | Young |
| 0.996191 | 0.366777 | 0.196 | 0.202 | 1 | 11 | Wdr89 | -0.006 | Young |

| | | | | | | | | |
|---|---|---|---|---|---|---|---|---|
| 0.996504 | 1.428265 | 0.065 | 0.069 | 1 | 11 | Timm17b | -0.004 | Young |
| 0.996796 | 0.945951 | 0.043 | 0.044 | 1 | 11 | Plp2 | -0.001 | Young |
| 0.997614 | 0.270173 | 0.514 | 0.577 | 1 | 11 | Tra2b | -0.063 | Young |
| 0.997717 | 0.076021 | 0.804 | 0.859 | 1 | 11 | Irf2bp2 | -0.055 | Young |
| 0.998023 | 0.746675 | 0.051 | 0.052 | 1 | 11 | Dhx57 | -0.001 | Young |
| 0.99808 | 0.414357 | 0.13 | 0.137 | 1 | 11 | Socs7 | -0.007 | Young |
| 0.998932 | 0.061345 | 0.043 | 0.044 | 1 | 11 | Pianp | -0.001 | Young |
| 0.999214 | 0.156723 | 0.551 | 0.617 | 1 | 11 | Purb | -0.066 | Young |
| 0.999325 | 1.024436 | 0.116 | 0.121 | 1 | 11 | Lrrc8a | -0.005 | Young |
| 0.999343 | 0.617096 | 0.123 | 0.129 | 1 | 11 | Rps6ka4 | -0.006 | Young |
| 0.000596 | 3.120419 | 0.119 | 0.015 | 1 | 12 | Zfp637 | 0.104 | Old |
| 0.00064 | 1.123835 | 0.356 | 0.182 | 1 | 12 | Ndufb2 | 0.174 | Old |
| 0.001413 | 1.640158 | 0.229 | 0.088 | 1 | 12 | Akr1b3 | 0.141 | Old |
| 0.002251 | 0.779764 | 0.441 | 0.248 | 1 | 12 | Nap1l5 | 0.193 | Old |
| 0.002261 | 1.2566 | 0.254 | 0.109 | 1 | 12 | Tmem147 | 0.145 | Old |
| 0.002562 | 0.860157 | 0.534 | 0.365 | 1 | 12 | Atp5b | 0.169 | Old |
| 0.003685 | 2.123652 | 0.136 | 0.036 | 1 | 12 | Appl1 | 0.1 | Old |
| 0.004685 | 3.497418 | 0.076 | 0.007 | 1 | 12 | Cpsf2 | 0.069 | Old |
| 0.005039 | 1.361486 | 0.178 | 0.066 | 1 | 12 | Mrpl55 | 0.112 | Old |
| 0.006623 | 1.970638 | 0.136 | 0.044 | 1 | 12 | Eml2 | 0.092 | Old |
| 0.007763 | 6.411522 | 0.051 | 0 | 1 | 12 | Kif26b | 0.051 | Old |
| 0.007763 | 6.31591 | 0.051 | 0 | 1 | 12 | Sympk | 0.051 | Old |
| 0.007763 | 6.304463 | 0.051 | 0 | 1 | 12 | Cspp1 | 0.051 | Old |
| 0.007763 | 6.207585 | 0.051 | 0 | 1 | 12 | Arl8b | 0.051 | Old |
| 0.007763 | 6.167105 | 0.051 | 0 | 1 | 12 | Naa15 | 0.051 | Old |
| 0.008438 | 1.152906 | 0.246 | 0.124 | 1 | 12 | Pfdn1 | 0.122 | Old |
| 0.009557 | 2.848285 | 0.068 | 0.007 | 1 | 12 | Cndp2 | 0.061 | Old |
| 0.009712 | 0.718775 | 0.364 | 0.219 | 1 | 12 | Gsk3b | 0.145 | Old |
| 0.009857 | 2.87418 | 0.068 | 0.007 | 1 | 12 | Usp5 | 0.061 | Old |
| 0.009857 | 2.856028 | 0.068 | 0.007 | 1 | 12 | Mark2 | 0.061 | Old |
| 0.009857 | 2.61669 | 0.068 | 0.007 | 1 | 12 | Exoc5 | 0.061 | Old |
| 0.010994 | 2.539977 | 0.093 | 0.022 | 1 | 12 | Tmem60 | 0.071 | Old |
| 0.01258 | 2.170734 | 0.093 | 0.022 | 1 | 12 | Apip | 0.071 | Old |
| 0.013876 | 0.456642 | 0.644 | 0.555 | 1 | 12 | Atp5h | 0.089 | Old |
| 0.014489 | 2.390905 | 0.102 | 0.029 | 1 | 12 | Ntpcr | 0.073 | Old |
| 0.014965 | 2.539946 | 0.076 | 0.015 | 1 | 12 | Lnpep | 0.061 | Old |
| 0.015267 | 2.611185 | 0.076 | 0.015 | 1 | 12 | Plk2 | 0.061 | Old |
| 0.015314 | 6.235678 | 0.042 | 0 | 1 | 12 | Gm17586 | 0.042 | Old |
| 0.015314 | 6.136796 | 0.042 | 0 | 1 | 12 | Thop1 | 0.042 | Old |
| 0.015314 | 6.116626 | 0.042 | 0 | 1 | 12 | Ptpn3 | 0.042 | Old |
| 0.015314 | 6.074822 | 0.042 | 0 | 1 | 12 | Smyd3 | 0.042 | Old |
| 0.015314 | 6.064247 | 0.042 | 0 | 1 | 12 | Wdr45 | 0.042 | Old |
| 0.015314 | 6.046112 | 0.042 | 0 | 1 | 12 | Rrm2b | 0.042 | Old |
| 0.015314 | 6.029354 | 0.042 | 0 | 1 | 12 | Zpr1 | 0.042 | Old |
| 0.015314 | 5.938327 | 0.042 | 0 | 1 | 12 | 1700096K1 | 0.042 | Old |

| | | | | | | | | |
|---|---|---|---|---|---|---|---|---|
| 0.015314 | 5.929119 | 0.042 | 0 | 1 | 12 | Klhl13 | 0.042 | Old |
| 0.015314 | 5.897547 | 0.042 | 0 | 1 | 12 | Car11 | 0.042 | Old |
| 0.015314 | 5.895732 | 0.042 | 0 | 1 | 12 | Nbas | 0.042 | Old |
| 0.015314 | 5.872065 | 0.042 | 0 | 1 | 12 | Asxl1 | 0.042 | Old |
| 0.015314 | 5.845031 | 0.042 | 0 | 1 | 12 | Chst7 | 0.042 | Old |
| 0.015314 | 5.760691 | 0.042 | 0 | 1 | 12 | Myh9 | 0.042 | Old |
| 0.015314 | 5.533321 | 0.042 | 0 | 1 | 12 | Cnpy3 | 0.042 | Old |
| 0.015314 | 5.517911 | 0.042 | 0 | 1 | 12 | Usf2 | 0.042 | Old |
| 0.015476 | 1.592531 | 0.136 | 0.051 | 1 | 12 | Ech1 | 0.085 | Old |
| 0.0156 | 1.481467 | 0.127 | 0.044 | 1 | 12 | Dnajc7 | 0.083 | Old |
| 0.015669 | 1.941027 | 0.102 | 0.029 | 1 | 12 | Strap | 0.073 | Old |
| 0.015995 | 2.295694 | 0.076 | 0.015 | 1 | 12 | Tnfaip1 | 0.061 | Old |
| 0.016941 | 3.455673 | 0.059 | 0.007 | 1 | 12 | Rras2 | 0.052 | Old |
| 0.017086 | 2.126866 | 0.076 | 0.015 | 1 | 12 | H13 | 0.061 | Old |
| 0.017468 | 3.120052 | 0.059 | 0.007 | 1 | 12 | Erich1 | 0.052 | Old |
| 0.017468 | 3.031459 | 0.059 | 0.007 | 1 | 12 | Bcat2 | 0.052 | Old |
| 0.017507 | 0.875312 | 0.229 | 0.117 | 1 | 12 | Eef1e1 | 0.112 | Old |
| 0.017621 | 1.117123 | 0.161 | 0.066 | 1 | 12 | Celf6 | 0.095 | Old |
| 0.017736 | 2.956095 | 0.059 | 0.007 | 1 | 12 | Rnf217 | 0.052 | Old |
| 0.018009 | 2.892244 | 0.059 | 0.007 | 1 | 12 | Camk2b | 0.052 | Old |
| 0.018285 | 2.791958 | 0.059 | 0.007 | 1 | 12 | Ecsit | 0.052 | Old |
| 0.018384 | 1.064394 | 0.161 | 0.066 | 1 | 12 | Usp34 | 0.095 | Old |
| 0.018906 | 1.410022 | 0.136 | 0.051 | 1 | 12 | Mrpl32 | 0.085 | Old |
| 0.019306 | 1.645778 | 0.102 | 0.029 | 1 | 12 | Cmip | 0.073 | Old |
| 0.020053 | 2.493683 | 0.085 | 0.022 | 1 | 12 | BC004004 | 0.063 | Old |
| 0.020053 | 2.373612 | 0.085 | 0.022 | 1 | 12 | Sar1a | 0.063 | Old |
| 0.02016 | 1.39991 | 0.102 | 0.029 | 1 | 12 | Eif5b | 0.073 | Old |
| 0.021505 | 1.611492 | 0.11 | 0.036 | 1 | 12 | Csnk1g3 | 0.074 | Old |
| 0.021797 | 1.10391 | 0.203 | 0.102 | 1 | 12 | Marcksl1 | 0.101 | Old |
| 0.022337 | 1.386477 | 0.153 | 0.066 | 1 | 12 | Gm10076 | 0.087 | Old |
| 0.022833 | 1.930025 | 0.085 | 0.022 | 1 | 12 | Copz1 | 0.063 | Old |
| 0.022833 | 2.024877 | 0.085 | 0.022 | 1 | 12 | Tmem126a | 0.063 | Old |
| 0.023014 | 0.312098 | 0.856 | 0.774 | 1 | 12 | Hint1 | 0.082 | Old |
| 0.023198 | 1.616467 | 0.119 | 0.044 | 1 | 12 | Mgat4b | 0.075 | Old |
| 0.023647 | 1.886354 | 0.085 | 0.022 | 1 | 12 | Tug1 | 0.063 | Old |
| 0.024063 | 1.829498 | 0.085 | 0.022 | 1 | 12 | Tmem11 | 0.063 | Old |
| 0.024486 | 1.94454 | 0.085 | 0.022 | 1 | 12 | Rfc2 | 0.063 | Old |
| 0.024772 | 1.811879 | 0.085 | 0.022 | 1 | 12 | Lsm3 | 0.063 | Old |
| 0.024772 | 1.744762 | 0.085 | 0.022 | 1 | 12 | Ccdc106 | 0.063 | Old |
| 0.026364 | 0.693576 | 0.356 | 0.219 | 1 | 12 | Coa3 | 0.137 | Old |
| 0.027192 | 1.157916 | 0.161 | 0.073 | 1 | 12 | Cox14 | 0.088 | Old |
| 0.027769 | 0.727564 | 0.356 | 0.241 | 1 | 12 | Pomp | 0.115 | Old |
| 0.027995 | 0.949316 | 0.22 | 0.117 | 1 | 12 | Mtx2 | 0.103 | Old |
| 0.028362 | 1.778816 | 0.093 | 0.029 | 1 | 12 | Degs1 | 0.064 | Old |
| 0.028897 | 2.27537 | 0.068 | 0.015 | 1 | 12 | Pdcd2 | 0.053 | Old |

| | | | | | | | | |
|---|---|---|---|---|---|---|---|---|
| 0.029412 | 0.43409 | 0.686 | 0.577 | 1 | 12 | Cfl1 | 0.109 | Old |
| 0.02959 | 1.610534 | 0.093 | 0.029 | 1 | 12 | Sarnp | 0.064 | Old |
| 0.03003 | 2.081383 | 0.068 | 0.015 | 1 | 12 | Serpinf1 | 0.053 | Old |
| 0.030146 | 0.418944 | 0.534 | 0.431 | 1 | 12 | Aes | 0.103 | Old |
| 0.030479 | 5.881427 | 0.034 | 0 | 1 | 12 | Slc38a9 | 0.034 | Old |
| 0.030479 | 5.876875 | 0.034 | 0 | 1 | 12 | Efr3a | 0.034 | Old |
| 0.030479 | 5.865028 | 0.034 | 0 | 1 | 12 | Cpox | 0.034 | Old |
| 0.030479 | 5.848524 | 0.034 | 0 | 1 | 12 | Tmem115 | 0.034 | Old |
| 0.030479 | 5.81494 | 0.034 | 0 | 1 | 12 | Smim15 | 0.034 | Old |
| 0.030479 | 5.795366 | 0.034 | 0 | 1 | 12 | Tmem41b | 0.034 | Old |
| 0.030479 | 5.767788 | 0.034 | 0 | 1 | 12 | Myef2 | 0.034 | Old |
| 0.030479 | 5.74899 | 0.034 | 0 | 1 | 12 | Il6st | 0.034 | Old |
| 0.030479 | 5.74408 | 0.034 | 0 | 1 | 12 | Agps | 0.034 | Old |
| 0.030479 | 5.736298 | 0.034 | 0 | 1 | 12 | Mkl1 | 0.034 | Old |
| 0.030479 | 5.728508 | 0.034 | 0 | 1 | 12 | Asb1 | 0.034 | Old |
| 0.030479 | 5.701011 | 0.034 | 0 | 1 | 12 | E2f4 | 0.034 | Old |
| 0.030479 | 5.697284 | 0.034 | 0 | 1 | 12 | Rfc4 | 0.034 | Old |
| 0.030479 | 5.678717 | 0.034 | 0 | 1 | 12 | Odf2 | 0.034 | Old |
| 0.030479 | 5.670938 | 0.034 | 0 | 1 | 12 | Syt16 | 0.034 | Old |
| 0.030479 | 5.667466 | 0.034 | 0 | 1 | 12 | Xrcc1 | 0.034 | Old |
| 0.030479 | 5.657678 | 0.034 | 0 | 1 | 12 | Atp7a | 0.034 | Old |
| 0.030479 | 5.654212 | 0.034 | 0 | 1 | 12 | Secisbp2 | 0.034 | Old |
| 0.030479 | 5.64147 | 0.034 | 0 | 1 | 12 | Chd9 | 0.034 | Old |
| 0.030479 | 5.59835 | 0.034 | 0 | 1 | 12 | Fbxo11 | 0.034 | Old |
| 0.030479 | 5.5919 | 0.034 | 0 | 1 | 12 | Adam22 | 0.034 | Old |
| 0.030479 | 5.586431 | 0.034 | 0 | 1 | 12 | Zfyve9 | 0.034 | Old |
| 0.030479 | 5.58352 | 0.034 | 0 | 1 | 12 | Tssc4 | 0.034 | Old |
| 0.030479 | 5.567996 | 0.034 | 0 | 1 | 12 | Poldip2 | 0.034 | Old |
| 0.030479 | 5.56773 | 0.034 | 0 | 1 | 12 | Gskip | 0.034 | Old |
| 0.030479 | 5.557252 | 0.034 | 0 | 1 | 12 | Btg1 | 0.034 | Old |
| 0.030479 | 5.550314 | 0.034 | 0 | 1 | 12 | Kpna1 | 0.034 | Old |
| 0.030479 | 5.550088 | 0.034 | 0 | 1 | 12 | Prkcb | 0.034 | Old |
| 0.030479 | 5.549182 | 0.034 | 0 | 1 | 12 | Cdc42bpb | 0.034 | Old |
| 0.030479 | 5.538104 | 0.034 | 0 | 1 | 12 | Zfp672 | 0.034 | Old |
| 0.030479 | 5.517725 | 0.034 | 0 | 1 | 12 | Iars | 0.034 | Old |
| 0.030479 | 5.5153 | 0.034 | 0 | 1 | 12 | Lypla1 | 0.034 | Old |
| 0.030479 | 5.495113 | 0.034 | 0 | 1 | 12 | Zfp950 | 0.034 | Old |
| 0.030479 | 5.482075 | 0.034 | 0 | 1 | 12 | Rab3gap2 | 0.034 | Old |
| 0.030479 | 5.376768 | 0.034 | 0 | 1 | 12 | Cd36 | 0.034 | Old |
| 0.030479 | 5.269731 | 0.034 | 0 | 1 | 12 | Ube2l6 | 0.034 | Old |
| 0.030479 | 5.239329 | 0.034 | 0 | 1 | 12 | Cbx4 | 0.034 | Old |
| 0.030479 | 5.20923 | 0.034 | 0 | 1 | 12 | Tmem203 | 0.034 | Old |
| 0.030479 | 5.182894 | 0.034 | 0 | 1 | 12 | Cxcl2 | 0.034 | Old |
| 0.030479 | 5.00923 | 0.034 | 0 | 1 | 12 | Psme3 | 0.034 | Old |
| 0.031599 | 1.960179 | 0.068 | 0.015 | 1 | 12 | Calb1 | 0.053 | Old |

| | | | | | | | | |
|---|---|---|---|---|---|---|---|---|
| 0.032045 | 0.567484 | 0.288 | 0.161 | 1 | 12 | Ndufv2 | 0.127 | Old |
| 0.032948 | 3.07081 | 0.051 | 0.007 | 1 | 12 | Gid8 | 0.044 | Old |
| 0.032948 | 2.987685 | 0.051 | 0.007 | 1 | 12 | Crem | 0.044 | Old |
| 0.033291 | 1.766341 | 0.102 | 0.036 | 1 | 12 | Cmbl | 0.066 | Old |
| 0.033445 | 3.141007 | 0.051 | 0.007 | 1 | 12 | Smu1 | 0.044 | Old |
| 0.033445 | 2.941632 | 0.051 | 0.007 | 1 | 12 | Neat1 | 0.044 | Old |
| 0.033551 | 1.426997 | 0.093 | 0.029 | 1 | 12 | Srsf7 | 0.064 | Old |
| 0.033949 | 2.771744 | 0.051 | 0.007 | 1 | 12 | Coq5 | 0.044 | Old |
| 0.034975 | 2.455402 | 0.051 | 0.007 | 1 | 12 | Sgpl1 | 0.044 | Old |
| 0.034975 | 2.350784 | 0.051 | 0.007 | 1 | 12 | Tmed5 | 0.044 | Old |
| 0.035498 | 2.587459 | 0.051 | 0.007 | 1 | 12 | Ddr1 | 0.044 | Old |
| 0.035498 | 2.394032 | 0.051 | 0.007 | 1 | 12 | Gga2 | 0.044 | Old |
| 0.036028 | 2.27931 | 0.051 | 0.007 | 1 | 12 | Ilvbl | 0.044 | Old |
| 0.036028 | 1.786934 | 0.051 | 0.007 | 1 | 12 | Nifk | 0.044 | Old |
| 0.036028 | 1.687179 | 0.051 | 0.007 | 1 | 12 | Ttc19 | 0.044 | Old |
| 0.036543 | 0.993985 | 0.186 | 0.095 | 1 | 12 | Acat1 | 0.091 | Old |
| 0.03656 | 2.177295 | 0.076 | 0.022 | 1 | 12 | Atad1 | 0.054 | Old |
| 0.036693 | 1.064776 | 0.195 | 0.102 | 1 | 12 | Cmpk1 | 0.093 | Old |
| 0.036979 | 2.037355 | 0.076 | 0.022 | 1 | 12 | Cntn1 | 0.054 | Old |
| 0.039132 | 1.894135 | 0.076 | 0.022 | 1 | 12 | Bud31 | 0.054 | Old |
| 0.040269 | 0.867795 | 0.186 | 0.095 | 1 | 12 | Nisch | 0.091 | Old |
| 0.040467 | 0.552095 | 0.263 | 0.153 | 1 | 12 | Dynlt3 | 0.11 | Old |
| 0.041573 | 0.913127 | 0.22 | 0.124 | 1 | 12 | Hmgcs1 | 0.096 | Old |
| 0.042089 | 1.715182 | 0.076 | 0.022 | 1 | 12 | Fam69b | 0.054 | Old |
| 0.042325 | 1.710467 | 0.076 | 0.022 | 1 | 12 | Ap1ar | 0.054 | Old |
| 0.042798 | 1.648006 | 0.076 | 0.022 | 1 | 12 | Apopt1 | 0.054 | Old |
| 0.043759 | 1.671179 | 0.076 | 0.022 | 1 | 12 | Lysmd2 | 0.054 | Old |
| 0.043759 | 1.640412 | 0.076 | 0.022 | 1 | 12 | Ptpn1 | 0.054 | Old |
| 0.04684 | 1.271597 | 0.119 | 0.051 | 1 | 12 | Cnih1 | 0.068 | Old |
| 0.046858 | 0.861326 | 0.229 | 0.139 | 1 | 12 | Emc10 | 0.09 | Old |
| 0.048195 | 1.759734 | 0.085 | 0.029 | 1 | 12 | Necap1 | 0.056 | Old |
| 0.048507 | 0.987838 | 0.178 | 0.095 | 1 | 12 | Rhoa | 0.083 | Old |
| 0.049966 | 2.659297 | 0.059 | 0.015 | 1 | 12 | Znhit6 | 0.044 | Old |
| 0.049966 | 2.483935 | 0.059 | 0.015 | 1 | 12 | Otub1 | 0.044 | Old |
| 0.051452 | 0.877713 | 0.186 | 0.102 | 1 | 12 | Cidea | 0.084 | Old |
| 0.051636 | 0.859348 | 0.195 | 0.109 | 1 | 12 | Dgcr6 | 0.086 | Old |
| 0.05182 | 1.141536 | 0.153 | 0.08 | 1 | 12 | Soga3 | 0.073 | Old |
| 0.05184 | 1.092997 | 0.169 | 0.095 | 1 | 12 | Etfb | 0.074 | Old |
| 0.051986 | 0.550811 | 0.288 | 0.19 | 1 | 12 | Tmem50a | 0.098 | Old |
| 0.052508 | 2.116417 | 0.059 | 0.015 | 1 | 12 | Grpel2 | 0.044 | Old |
| 0.052508 | 2.073942 | 0.059 | 0.015 | 1 | 12 | Gpatch4 | 0.044 | Old |
| 0.052846 | 0.907817 | 0.22 | 0.139 | 1 | 12 | Pmvk | 0.081 | Old |
| 0.05316 | 2.270113 | 0.059 | 0.015 | 1 | 12 | Zfp36 | 0.044 | Old |
| 0.053644 | 0.172383 | 0.915 | 0.934 | 1 | 12 | Slc25a4 | -0.019 | Old |
| 0.053819 | 2.065116 | 0.059 | 0.015 | 1 | 12 | Csnk1g2 | 0.044 | Old |

| | | | | | | | | |
|---|---|---|---|---|---|---|---|---|
| 0.053819 | 1.972859 | 0.059 | 0.015 | 1 | 12 | Irak1bp1 | 0.044 | Old |
| 0.05412 | 1.213431 | 0.127 | 0.058 | 1 | 12 | Rerg | 0.069 | Old |
| 0.054128 | 0.838088 | 0.212 | 0.124 | 1 | 12 | Alcam | 0.088 | Old |
| 0.054484 | 1.918923 | 0.059 | 0.015 | 1 | 12 | Appbp2 | 0.044 | Old |
| 0.055325 | 1.351624 | 0.144 | 0.073 | 1 | 12 | Mrpl43 | 0.071 | Old |
| 0.055794 | 0.722859 | 0.237 | 0.146 | 1 | 12 | Cetn3 | 0.091 | Old |
| 0.055836 | 1.770017 | 0.059 | 0.015 | 1 | 12 | Lmbrd2 | 0.044 | Old |
| 0.056295 | 1.18702 | 0.127 | 0.058 | 1 | 12 | Aip | 0.069 | Old |
| 0.057096 | 1.034551 | 0.136 | 0.066 | 1 | 12 | Retreg1 | 0.07 | Old |
| 0.057194 | 0.313506 | 0.619 | 0.482 | 1 | 12 | Rpl35a | 0.137 | Old |
| 0.057215 | 1.840328 | 0.059 | 0.015 | 1 | 12 | Desi2 | 0.044 | Old |
| 0.057547 | 0.489829 | 0.424 | 0.299 | 1 | 12 | Dctn2 | 0.125 | Old |
| 0.059317 | 1.439606 | 0.093 | 0.036 | 1 | 12 | Sdf4 | 0.057 | Old |
| 0.059338 | 1.598599 | 0.059 | 0.015 | 1 | 12 | Herc1 | 0.044 | Old |
| 0.059338 | 1.331856 | 0.059 | 0.015 | 1 | 12 | Ehd4 | 0.044 | Old |
| 0.059714 | 1.182775 | 0.085 | 0.029 | 1 | 12 | Ppp2r2a | 0.056 | Old |
| 0.06004 | 0.273666 | 0.89 | 0.774 | 1 | 12 | Rps24 | 0.116 | Old |
| 0.061535 | 1.365932 | 0.093 | 0.036 | 1 | 12 | Hdac2 | 0.057 | Old |
| 0.061611 | 5.619823 | 0.025 | 0 | 1 | 12 | Xpo6 | 0.025 | Old |
| 0.061611 | 5.619563 | 0.025 | 0 | 1 | 12 | Cyld | 0.025 | Old |
| 0.061611 | 5.60939 | 0.025 | 0 | 1 | 12 | Slc2a1 | 0.025 | Old |
| 0.061611 | 5.573104 | 0.025 | 0 | 1 | 12 | Stk32a | 0.025 | Old |
| 0.061611 | 5.544068 | 0.025 | 0 | 1 | 12 | Rgs3 | 0.025 | Old |
| 0.061611 | 5.464645 | 0.025 | 0 | 1 | 12 | Uxt | 0.025 | Old |
| 0.061611 | 5.455883 | 0.025 | 0 | 1 | 12 | Zc3h13 | 0.025 | Old |
| 0.061611 | 5.450158 | 0.025 | 0 | 1 | 12 | Isyna1 | 0.025 | Old |
| 0.061611 | 5.449344 | 0.025 | 0 | 1 | 12 | Ncoa4 | 0.025 | Old |
| 0.061611 | 5.426046 | 0.025 | 0 | 1 | 12 | Adgra1 | 0.025 | Old |
| 0.061611 | 5.404356 | 0.025 | 0 | 1 | 12 | Rdh11 | 0.025 | Old |
| 0.061611 | 5.402538 | 0.025 | 0 | 1 | 12 | Dok1 | 0.025 | Old |
| 0.061611 | 5.392691 | 0.025 | 0 | 1 | 12 | Fzd5 | 0.025 | Old |
| 0.061611 | 5.391127 | 0.025 | 0 | 1 | 12 | Uts2b | 0.025 | Old |
| 0.061611 | 5.39043 | 0.025 | 0 | 1 | 12 | Lypla2 | 0.025 | Old |
| 0.061611 | 5.386041 | 0.025 | 0 | 1 | 12 | Slc35c2 | 0.025 | Old |
| 0.061611 | 5.383997 | 0.025 | 0 | 1 | 12 | Dpm1 | 0.025 | Old |
| 0.061611 | 5.382867 | 0.025 | 0 | 1 | 12 | Vps45 | 0.025 | Old |
| 0.061611 | 5.376496 | 0.025 | 0 | 1 | 12 | Pex5l | 0.025 | Old |
| 0.061611 | 5.374521 | 0.025 | 0 | 1 | 12 | Nrf1 | 0.025 | Old |
| 0.061611 | 5.364349 | 0.025 | 0 | 1 | 12 | Nfkbie | 0.025 | Old |
| 0.061611 | 5.363431 | 0.025 | 0 | 1 | 12 | 8-Sep | 0.025 | Old |
| 0.061611 | 5.350556 | 0.025 | 0 | 1 | 12 | Jade1 | 0.025 | Old |
| 0.061611 | 5.346772 | 0.025 | 0 | 1 | 12 | Ankrd13a | 0.025 | Old |
| 0.061611 | 5.328777 | 0.025 | 0 | 1 | 12 | Pik3cb | 0.025 | Old |
| 0.061611 | 5.321089 | 0.025 | 0 | 1 | 12 | Golim4 | 0.025 | Old |
| 0.061611 | 5.320897 | 0.025 | 0 | 1 | 12 | Abhd6 | 0.025 | Old |

| | | | | | | | | |
|---|---|---|---|---|---|---|---|---|
| 0.061611 | 5.320624 | 0.025 | 0 | 1 | 12 | Cd2bp2 | 0.025 | Old |
| 0.061611 | 5.316526 | 0.025 | 0 | 1 | 12 | Tmem145 | 0.025 | Old |
| 0.061611 | 5.314567 | 0.025 | 0 | 1 | 12 | Prmt8 | 0.025 | Old |
| 0.061611 | 5.310074 | 0.025 | 0 | 1 | 12 | L3mbtl2 | 0.025 | Old |
| 0.061611 | 5.309167 | 0.025 | 0 | 1 | 12 | Pip4k2c | 0.025 | Old |
| 0.061611 | 5.303793 | 0.025 | 0 | 1 | 12 | Dffa | 0.025 | Old |
| 0.061611 | 5.295622 | 0.025 | 0 | 1 | 12 | Pde4b | 0.025 | Old |
| 0.061611 | 5.29457 | 0.025 | 0 | 1 | 12 | Grhpr | 0.025 | Old |
| 0.061611 | 5.292422 | 0.025 | 0 | 1 | 12 | Smox | 0.025 | Old |
| 0.061611 | 5.290537 | 0.025 | 0 | 1 | 12 | Fgfbp3 | 0.025 | Old |
| 0.061611 | 5.287426 | 0.025 | 0 | 1 | 12 | Zfp532 | 0.025 | Old |
| 0.061611 | 5.282759 | 0.025 | 0 | 1 | 12 | Adgrb1 | 0.025 | Old |
| 0.061611 | 5.275714 | 0.025 | 0 | 1 | 12 | Gm2464 | 0.025 | Old |
| 0.061611 | 5.270903 | 0.025 | 0 | 1 | 12 | Esyt1 | 0.025 | Old |
| 0.061611 | 5.253767 | 0.025 | 0 | 1 | 12 | Slc20a1 | 0.025 | Old |
| 0.061611 | 5.227636 | 0.025 | 0 | 1 | 12 | Ogfod1 | 0.025 | Old |
| 0.061611 | 5.226879 | 0.025 | 0 | 1 | 12 | Pgap1 | 0.025 | Old |
| 0.061611 | 5.223266 | 0.025 | 0 | 1 | 12 | Mfsd14b | 0.025 | Old |
| 0.061611 | 5.221172 | 0.025 | 0 | 1 | 12 | Ddx49 | 0.025 | Old |
| 0.061611 | 5.219993 | 0.025 | 0 | 1 | 12 | Pcna | 0.025 | Old |
| 0.061611 | 5.20678 | 0.025 | 0 | 1 | 12 | Serping1 | 0.025 | Old |
| 0.061611 | 5.158788 | 0.025 | 0 | 1 | 12 | Rwdd4a | 0.025 | Old |
| 0.061611 | 5.156174 | 0.025 | 0 | 1 | 12 | Tmem97 | 0.025 | Old |
| 0.061611 | 5.152375 | 0.025 | 0 | 1 | 12 | Syne2 | 0.025 | Old |
| 0.061611 | 5.141377 | 0.025 | 0 | 1 | 12 | Snapc1 | 0.025 | Old |
| 0.061611 | 5.140164 | 0.025 | 0 | 1 | 12 | Trim26 | 0.025 | Old |
| 0.061611 | 5.123834 | 0.025 | 0 | 1 | 12 | Cables2 | 0.025 | Old |
| 0.061611 | 5.114618 | 0.025 | 0 | 1 | 12 | Ube2j1 | 0.025 | Old |
| 0.061611 | 5.114113 | 0.025 | 0 | 1 | 12 | Cln5 | 0.025 | Old |
| 0.061611 | 5.107465 | 0.025 | 0 | 1 | 12 | Slf2 | 0.025 | Old |
| 0.061611 | 5.086277 | 0.025 | 0 | 1 | 12 | Sspn | 0.025 | Old |
| 0.061611 | 5.073151 | 0.025 | 0 | 1 | 12 | Tcte2 | 0.025 | Old |
| 0.061611 | 5.070009 | 0.025 | 0 | 1 | 12 | Msto1 | 0.025 | Old |
| 0.061611 | 5.062361 | 0.025 | 0 | 1 | 12 | Prkx | 0.025 | Old |
| 0.061611 | 5.05661 | 0.025 | 0 | 1 | 12 | C330007P( | 0.025 | Old |
| 0.061611 | 5.051716 | 0.025 | 0 | 1 | 12 | Crebbp | 0.025 | Old |
| 0.061611 | 5.049394 | 0.025 | 0 | 1 | 12 | Sos1 | 0.025 | Old |
| 0.061611 | 5.041258 | 0.025 | 0 | 1 | 12 | Kctd15 | 0.025 | Old |
| 0.061611 | 5.026054 | 0.025 | 0 | 1 | 12 | Fance | 0.025 | Old |
| 0.061611 | 5.024801 | 0.025 | 0 | 1 | 12 | Rbm28 | 0.025 | Old |
| 0.061611 | 5.019125 | 0.025 | 0 | 1 | 12 | Tmbim1 | 0.025 | Old |
| 0.061611 | 5.016988 | 0.025 | 0 | 1 | 12 | Eno3 | 0.025 | Old |
| 0.061611 | 5.007845 | 0.025 | 0 | 1 | 12 | Cul4a | 0.025 | Old |
| 0.061611 | 4.99305 | 0.025 | 0 | 1 | 12 | Fam149a | 0.025 | Old |
| 0.061611 | 4.991687 | 0.025 | 0 | 1 | 12 | Metap1 | 0.025 | Old |

| | | | | | | | | |
|---|---|---|---|---|---|---|---|---|
| 0.061611 | 4.975093 | 0.025 | 0 | 1 | 12 | Ctcf | 0.025 | Old |
| 0.061611 | 4.927682 | 0.025 | 0 | 1 | 12 | Gm15417 | 0.025 | Old |
| 0.061611 | 4.892696 | 0.025 | 0 | 1 | 12 | Arhgap17 | 0.025 | Old |
| 0.061611 | 4.869733 | 0.025 | 0 | 1 | 12 | Cenpq | 0.025 | Old |
| 0.061611 | 4.847929 | 0.025 | 0 | 1 | 12 | Med14 | 0.025 | Old |
| 0.061611 | 4.840385 | 0.025 | 0 | 1 | 12 | Vamp8 | 0.025 | Old |
| 0.061611 | 4.828633 | 0.025 | 0 | 1 | 12 | Pias1 | 0.025 | Old |
| 0.061611 | 4.828598 | 0.025 | 0 | 1 | 12 | Rbm15b | 0.025 | Old |
| 0.061611 | 4.824888 | 0.025 | 0 | 1 | 12 | Lgals3 | 0.025 | Old |
| 0.061611 | 4.814076 | 0.025 | 0 | 1 | 12 | Homer1 | 0.025 | Old |
| 0.061611 | 4.767438 | 0.025 | 0 | 1 | 12 | Ctdsp2 | 0.025 | Old |
| 0.061611 | 4.745678 | 0.025 | 0 | 1 | 12 | Slc9a9 | 0.025 | Old |
| 0.061611 | 4.724865 | 0.025 | 0 | 1 | 12 | Slc39a13 | 0.025 | Old |
| 0.061611 | 4.716414 | 0.025 | 0 | 1 | 12 | Nrp2 | 0.025 | Old |
| 0.061611 | 4.693134 | 0.025 | 0 | 1 | 12 | Erp44 | 0.025 | Old |
| 0.061611 | 4.651059 | 0.025 | 0 | 1 | 12 | Itsn2 | 0.025 | Old |
| 0.061611 | 4.591495 | 0.025 | 0 | 1 | 12 | Twnk | 0.025 | Old |
| 0.061611 | 4.569075 | 0.025 | 0 | 1 | 12 | Bst2 | 0.025 | Old |
| 0.061611 | 4.506601 | 0.025 | 0 | 1 | 12 | Kmt2c | 0.025 | Old |
| 0.061611 | 4.478347 | 0.025 | 0 | 1 | 12 | Ciz1 | 0.025 | Old |
| 0.061611 | 4.383871 | 0.025 | 0 | 1 | 12 | Cenpw | 0.025 | Old |
| 0.063151 | 1.041616 | 0.153 | 0.08 | 1 | 12 | Bub3 | 0.073 | Old |
| 0.063745 | 0.873121 | 0.169 | 0.095 | 1 | 12 | Canx | 0.074 | Old |
| 0.06409 | 3.219496 | 0.042 | 0.007 | 1 | 12 | Lxn | 0.035 | Old |
| 0.06409 | 2.985115 | 0.042 | 0.007 | 1 | 12 | Acsl6 | 0.035 | Old |
| 0.06409 | 2.892965 | 0.042 | 0.007 | 1 | 12 | D10Jhu81e | 0.035 | Old |
| 0.06409 | 2.865805 | 0.042 | 0.007 | 1 | 12 | Nmnat2 | 0.035 | Old |
| 0.06409 | 2.799005 | 0.042 | 0.007 | 1 | 12 | Mecp2 | 0.035 | Old |
| 0.06409 | 2.725477 | 0.042 | 0.007 | 1 | 12 | Zbtb25 | 0.035 | Old |
| 0.06409 | 2.710174 | 0.042 | 0.007 | 1 | 12 | Picalm | 0.035 | Old |
| 0.065027 | 2.84701 | 0.042 | 0.007 | 1 | 12 | Rab3gap1 | 0.035 | Old |
| 0.065027 | 2.795625 | 0.042 | 0.007 | 1 | 12 | Sec22b | 0.035 | Old |
| 0.065027 | 2.771385 | 0.042 | 0.007 | 1 | 12 | Pum2 | 0.035 | Old |
| 0.065027 | 2.75336 | 0.042 | 0.007 | 1 | 12 | Pbdc1 | 0.035 | Old |
| 0.065027 | 2.709678 | 0.042 | 0.007 | 1 | 12 | Oard1 | 0.035 | Old |
| 0.065027 | 2.665649 | 0.042 | 0.007 | 1 | 12 | Mettl1 | 0.035 | Old |
| 0.065027 | 2.535161 | 0.042 | 0.007 | 1 | 12 | 2510039O1 | 0.035 | Old |
| 0.065027 | 2.389131 | 0.042 | 0.007 | 1 | 12 | Gtf2f1 | 0.035 | Old |
| 0.065976 | 2.59298 | 0.042 | 0.007 | 1 | 12 | Ganab | 0.035 | Old |
| 0.065976 | 2.546574 | 0.042 | 0.007 | 1 | 12 | Ap3d1 | 0.035 | Old |
| 0.065976 | 2.523587 | 0.042 | 0.007 | 1 | 12 | Fzd3 | 0.035 | Old |
| 0.065976 | 2.485661 | 0.042 | 0.007 | 1 | 12 | Aptx | 0.035 | Old |
| 0.065976 | 2.45088 | 0.042 | 0.007 | 1 | 12 | 4933434E2 | 0.035 | Old |
| 0.066077 | 0.944234 | 0.169 | 0.095 | 1 | 12 | Stxbp1 | 0.074 | Old |
| 0.066225 | 0.47143 | 0.314 | 0.226 | 1 | 12 | App | 0.088 | Old |

| | | | | | | | | |
|---|---|---|---|---|---|---|---|---|
| 0.066741 | 0.879816 | 0.144 | 0.073 | 1 | 12 | Fam104a | 0.071 | Old |
| 0.066937 | 2.454332 | 0.042 | 0.007 | 1 | 12 | Hltf | 0.035 | Old |
| 0.066937 | 2.442782 | 0.042 | 0.007 | 1 | 12 | Cox10 | 0.035 | Old |
| 0.066937 | 2.422604 | 0.042 | 0.007 | 1 | 12 | Stx4a | 0.035 | Old |
| 0.066937 | 2.355402 | 0.042 | 0.007 | 1 | 12 | Rc3h1 | 0.035 | Old |
| 0.066937 | 2.345178 | 0.042 | 0.007 | 1 | 12 | Msi2 | 0.035 | Old |
| 0.066937 | 2.322552 | 0.042 | 0.007 | 1 | 12 | Trim67 | 0.035 | Old |
| 0.066937 | 2.298821 | 0.042 | 0.007 | 1 | 12 | Phc2 | 0.035 | Old |
| 0.066937 | 2.251851 | 0.042 | 0.007 | 1 | 12 | Unc80 | 0.035 | Old |
| 0.067022 | 0.856705 | 0.161 | 0.088 | 1 | 12 | Ap1s1 | 0.073 | Old |
| 0.067105 | 1.9702 | 0.068 | 0.022 | 1 | 12 | Fbl | 0.046 | Old |
| 0.067382 | 1.178695 | 0.093 | 0.036 | 1 | 12 | Tbrg1 | 0.057 | Old |
| 0.067459 | 0.404788 | 0.517 | 0.431 | 1 | 12 | Rpl39 | 0.086 | Old |
| 0.067909 | 2.121289 | 0.042 | 0.007 | 1 | 12 | Zyx | 0.035 | Old |
| 0.067909 | 2.118836 | 0.042 | 0.007 | 1 | 12 | Ip6k1 | 0.035 | Old |
| 0.067909 | 2.014684 | 0.042 | 0.007 | 1 | 12 | Hdac11 | 0.035 | Old |
| 0.067909 | 1.984158 | 0.042 | 0.007 | 1 | 12 | Synj1 | 0.035 | Old |
| 0.068814 | 1.275967 | 0.102 | 0.044 | 1 | 12 | Plekhb2 | 0.058 | Old |
| 0.068892 | 2.234462 | 0.042 | 0.007 | 1 | 12 | Hsd17b11 | 0.035 | Old |
| 0.068892 | 1.999624 | 0.042 | 0.007 | 1 | 12 | Usp47 | 0.035 | Old |
| 0.068892 | 1.886741 | 0.042 | 0.007 | 1 | 12 | Trip11 | 0.035 | Old |
| 0.069301 | 1.812761 | 0.068 | 0.022 | 1 | 12 | Cs | 0.046 | Old |
| 0.070045 | 1.75726 | 0.068 | 0.022 | 1 | 12 | Arl4c | 0.046 | Old |
| 0.070797 | 1.656754 | 0.068 | 0.022 | 1 | 12 | Elovl6 | 0.046 | Old |
| 0.070904 | 1.200047 | 0.119 | 0.058 | 1 | 12 | Hars | 0.061 | Old |
| 0.071151 | 0.423755 | 0.475 | 0.372 | 1 | 12 | Btf3 | 0.103 | Old |
| 0.071554 | 1.706236 | 0.068 | 0.022 | 1 | 12 | Nktr | 0.046 | Old |
| 0.072319 | 1.610395 | 0.068 | 0.022 | 1 | 12 | Hnrnph2 | 0.046 | Old |
| 0.072783 | 0.807219 | 0.263 | 0.175 | 1 | 12 | Vapa | 0.088 | Old |
| 0.07309 | 1.60564 | 0.068 | 0.022 | 1 | 12 | Srpr | 0.046 | Old |
| 0.073188 | 1.315111 | 0.11 | 0.051 | 1 | 12 | Eno2 | 0.059 | Old |
| 0.073597 | 0.351819 | 0.644 | 0.577 | 1 | 12 | Rps10 | 0.067 | Old |
| 0.073658 | 0.591191 | 0.212 | 0.124 | 1 | 12 | Pja2 | 0.088 | Old |
| 0.073868 | 1.595728 | 0.068 | 0.022 | 1 | 12 | 4930402H2 | 0.046 | Old |
| 0.073868 | 1.535246 | 0.068 | 0.022 | 1 | 12 | Rdh14 | 0.046 | Old |
| 0.074664 | 1.133899 | 0.11 | 0.051 | 1 | 12 | Ubl3 | 0.059 | Old |
| 0.075447 | 0.283587 | 0.559 | 0.46 | 1 | 12 | Rpl10a | 0.099 | Old |
| 0.076164 | 1.153585 | 0.11 | 0.051 | 1 | 12 | Vta1 | 0.059 | Old |
| 0.076242 | 1.435908 | 0.068 | 0.022 | 1 | 12 | Sike1 | 0.046 | Old |
| 0.077002 | 0.898621 | 0.093 | 0.036 | 1 | 12 | Larp4 | 0.057 | Old |
| 0.077047 | 1.422513 | 0.068 | 0.022 | 1 | 12 | Islr2 | 0.046 | Old |
| 0.078615 | 1.279497 | 0.11 | 0.051 | 1 | 12 | Gripap1 | 0.059 | Old |
| 0.078678 | 1.523192 | 0.068 | 0.022 | 1 | 12 | Trim28 | 0.046 | Old |
| 0.079898 | 1.682141 | 0.076 | 0.029 | 1 | 12 | Sdc3 | 0.047 | Old |
| 0.081937 | 1.177511 | 0.127 | 0.066 | 1 | 12 | Phf5a | 0.061 | Old |

| | | | | | | | | |
|---|---|---|---|---|---|---|---|---|
| 0.082678 | 0.994473 | 0.161 | 0.095 | 1 | 12 | Ube3a | 0.066 | Old |
| 0.083023 | 1.617524 | 0.076 | 0.029 | 1 | 12 | Cisd2 | 0.047 | Old |
| 0.083056 | 1.158394 | 0.136 | 0.073 | 1 | 12 | Ndfip1 | 0.063 | Old |
| 0.083534 | 0.814006 | 0.186 | 0.109 | 1 | 12 | Scand1 | 0.077 | Old |
| 0.086351 | 0.84993 | 0.153 | 0.088 | 1 | 12 | Tspan31 | 0.065 | Old |
| 0.087068 | 1.414942 | 0.076 | 0.029 | 1 | 12 | Capza1 | 0.047 | Old |
| 0.089237 | 0.990279 | 0.119 | 0.058 | 1 | 12 | Mtch2 | 0.061 | Old |
| 0.08957 | 1.297578 | 0.076 | 0.029 | 1 | 12 | Smpd1 | 0.047 | Old |
| 0.090417 | 1.33989 | 0.076 | 0.029 | 1 | 12 | R3hdm2 | 0.047 | Old |
| 0.090623 | 0.607131 | 0.271 | 0.19 | 1 | 12 | Zfand5 | 0.081 | Old |
| 0.090843 | 1.11803 | 0.076 | 0.029 | 1 | 12 | Upf3b | 0.047 | Old |
| 0.091467 | 0.500915 | 0.237 | 0.153 | 1 | 12 | 2410015M | 0.084 | Old |
| 0.091839 | 1.008323 | 0.127 | 0.066 | 1 | 12 | Maged1 | 0.061 | Old |
| 0.09213 | 1.293669 | 0.076 | 0.029 | 1 | 12 | Srrm1 | 0.047 | Old |
| 0.092183 | 0.421041 | 0.576 | 0.526 | 1 | 12 | Cd9 | 0.05 | Old |
| 0.092239 | 0.986286 | 0.119 | 0.058 | 1 | 12 | Ktn1 | 0.061 | Old |
| 0.092345 | 2.488058 | 0.051 | 0.015 | 1 | 12 | Pole4 | 0.036 | Old |
| 0.092345 | 2.243711 | 0.051 | 0.015 | 1 | 12 | Capn10 | 0.036 | Old |
| 0.092834 | 0.466633 | 0.381 | 0.307 | 1 | 12 | Rps17 | 0.074 | Old |
| 0.093442 | 2.167013 | 0.051 | 0.015 | 1 | 12 | Lnpk | 0.036 | Old |
| 0.093442 | 2.154309 | 0.051 | 0.015 | 1 | 12 | Tmem184c | 0.036 | Old |
| 0.093442 | 2.13425 | 0.051 | 0.015 | 1 | 12 | Ctu2 | 0.036 | Old |
| 0.093988 | 0.137002 | 1 | 0.985 | 1 | 12 | Stmn2 | 0.015 | Old |
| 0.093993 | 1.969929 | 0.051 | 0.015 | 1 | 12 | Dhx15 | 0.036 | Old |
| 0.094143 | 0.480915 | 0.373 | 0.27 | 1 | 12 | Selenof | 0.103 | Old |
| 0.094549 | 2.29183 | 0.051 | 0.015 | 1 | 12 | Mosmo | 0.036 | Old |
| 0.094549 | 2.230174 | 0.051 | 0.015 | 1 | 12 | Mfsd11 | 0.036 | Old |
| 0.094549 | 1.830286 | 0.051 | 0.015 | 1 | 12 | Ift27 | 0.036 | Old |
| 0.094574 | 0.483567 | 0.314 | 0.219 | 1 | 12 | mt-Nd4l | 0.095 | Old |
| 0.095666 | 2.047711 | 0.051 | 0.015 | 1 | 12 | Ckap5 | 0.036 | Old |
| 0.095704 | 0.211323 | 0.831 | 0.759 | 1 | 12 | S100a10 | 0.072 | Old |
| 0.096793 | 1.791259 | 0.051 | 0.015 | 1 | 12 | Suz12 | 0.036 | Old |
| 0.096794 | 1.935689 | 0.051 | 0.015 | 1 | 12 | Trappc2 | 0.036 | Old |
| 0.096794 | 1.743672 | 0.051 | 0.015 | 1 | 12 | Knop1 | 0.036 | Old |
| 0.097426 | 1.195926 | 0.076 | 0.029 | 1 | 12 | Spns1 | 0.047 | Old |
| 0.097426 | 0.865403 | 0.076 | 0.029 | 1 | 12 | Hmbox1 | 0.047 | Old |
| 0.097932 | 1.799894 | 0.051 | 0.015 | 1 | 12 | Macrod2 | 0.036 | Old |
| 0.097932 | 1.653864 | 0.051 | 0.015 | 1 | 12 | Rad21 | 0.036 | Old |
| 0.099082 | 1.935718 | 0.051 | 0.015 | 1 | 12 | Tbc1d7 | 0.036 | Old |
| 0.099082 | 1.775603 | 0.051 | 0.015 | 1 | 12 | Gm10419 | 0.036 | Old |
| 0.100242 | 1.862884 | 0.051 | 0.015 | 1 | 12 | Smim4 | 0.036 | Old |
| 0.100242 | 1.778966 | 0.051 | 0.015 | 1 | 12 | Tmem150c | 0.036 | Old |
| 0.100644 | 0.85589 | 0.119 | 0.058 | 1 | 12 | Mcts1 | 0.061 | Old |
| 0.10109 | 1.044597 | 0.076 | 0.029 | 1 | 12 | Bcap29 | 0.047 | Old |
| 0.101368 | 1.344326 | 0.093 | 0.044 | 1 | 12 | Zcchc11 | 0.049 | Old |

| | | | | | | | | |
|---|---|---|---|---|---|---|---|---|
| 0.102972 | 1.272236 | 0.085 | 0.036 | 1 | 12 | Fbll1 | 0.049 | Old |
| 0.10302 | 1.379678 | 0.093 | 0.044 | 1 | 12 | Nipsnap3b | 0.049 | Old |
| 0.103207 | 0.813102 | 0.119 | 0.058 | 1 | 12 | Atp6v1a | 0.061 | Old |
| 0.103564 | 0.331809 | 0.432 | 0.343 | 1 | 12 | Edf1 | 0.089 | Old |
| 0.103786 | 1.586149 | 0.051 | 0.015 | 1 | 12 | Nudt4 | 0.036 | Old |
| 0.103786 | 1.366259 | 0.051 | 0.015 | 1 | 12 | Cmss1 | 0.036 | Old |
| 0.10453 | 0.445407 | 0.322 | 0.234 | 1 | 12 | Ube2s | 0.088 | Old |
| 0.105644 | 0.885108 | 0.085 | 0.036 | 1 | 12 | Zmat2 | 0.049 | Old |
| 0.106205 | 1.401073 | 0.051 | 0.015 | 1 | 12 | Flrt1 | 0.036 | Old |
| 0.106205 | 1.138264 | 0.051 | 0.015 | 1 | 12 | Dmwd | 0.036 | Old |
| 0.106391 | 1.356115 | 0.093 | 0.044 | 1 | 12 | Fh1 | 0.049 | Old |
| 0.107456 | 1.147011 | 0.085 | 0.036 | 1 | 12 | Hnrnpul2 | 0.049 | Old |
| 0.108568 | 0.452281 | 0.407 | 0.321 | 1 | 12 | Ndufc2 | 0.086 | Old |
| 0.109292 | 1.023447 | 0.085 | 0.036 | 1 | 12 | Gyg | 0.049 | Old |
| 0.109978 | 0.727093 | 0.271 | 0.197 | 1 | 12 | Eif3h | 0.074 | Old |
| 0.112367 | 0.783753 | 0.212 | 0.146 | 1 | 12 | Fkbp4 | 0.066 | Old |
| 0.11248 | 0.752598 | 0.153 | 0.088 | 1 | 12 | Tro | 0.065 | Old |
| 0.112498 | 0.924828 | 0.093 | 0.044 | 1 | 12 | Auh | 0.049 | Old |
| 0.113574 | 2.043399 | 0.059 | 0.022 | 1 | 12 | Cdc42bpa | 0.037 | Old |
| 0.114293 | 1.273416 | 0.093 | 0.044 | 1 | 12 | Smyd2 | 0.049 | Old |
| 0.114293 | 1.107034 | 0.093 | 0.044 | 1 | 12 | Znrf1 | 0.049 | Old |
| 0.115903 | 2.119047 | 0.059 | 0.022 | 1 | 12 | 2310039H( | 0.037 | Old |
| 0.115915 | 0.817628 | 0.085 | 0.036 | 1 | 12 | Usp15 | 0.049 | Old |
| 0.117081 | 1.868025 | 0.059 | 0.022 | 1 | 12 | Usp14 | 0.037 | Old |
| 0.117287 | 1.083124 | 0.11 | 0.058 | 1 | 12 | Ssna1 | 0.052 | Old |
| 0.119467 | 1.754858 | 0.059 | 0.022 | 1 | 12 | Prpf19 | 0.037 | Old |
| 0.119467 | 1.584556 | 0.059 | 0.022 | 1 | 12 | Creg2 | 0.037 | Old |
| 0.120213 | 0.602249 | 0.263 | 0.19 | 1 | 12 | Tcf25 | 0.073 | Old |
| 0.120526 | 0.217178 | 0.873 | 0.818 | 1 | 12 | Ptma | 0.055 | Old |
| 0.120658 | 0.536957 | 0.288 | 0.219 | 1 | 12 | Ndufv3 | 0.069 | Old |
| 0.120668 | 1.094675 | 0.11 | 0.058 | 1 | 12 | Anapc11 | 0.052 | Old |
| 0.120674 | 1.467843 | 0.059 | 0.022 | 1 | 12 | Pak2 | 0.037 | Old |
| 0.12189 | 1.574253 | 0.059 | 0.022 | 1 | 12 | Timm44 | 0.037 | Old |
| 0.122173 | 0.903997 | 0.093 | 0.044 | 1 | 12 | Arglu1 | 0.049 | Old |
| 0.12265 | 0.965602 | 0.093 | 0.044 | 1 | 12 | Dnajb14 | 0.049 | Old |
| 0.123388 | 0.455525 | 0.186 | 0.117 | 1 | 12 | Dst | 0.069 | Old |
| 0.123733 | 0.339605 | 0.415 | 0.314 | 1 | 12 | Arl8a | 0.101 | Old |
| 0.124352 | 1.527442 | 0.059 | 0.022 | 1 | 12 | Stx1b | 0.037 | Old |
| 0.124352 | 1.466195 | 0.059 | 0.022 | 1 | 12 | Ngdn | 0.037 | Old |
| 0.124352 | 1.419037 | 0.059 | 0.022 | 1 | 12 | Cdk9 | 0.037 | Old |
| 0.124376 | 0.683266 | 0.237 | 0.161 | 1 | 12 | Cd24a | 0.076 | Old |
| 0.124571 | 0.959973 | 0.093 | 0.044 | 1 | 12 | Bnip3 | 0.049 | Old |
| 0.124773 | 3.068807 | 0.034 | 0.007 | 1 | 12 | Ntrk1 | 0.027 | Old |
| 0.124773 | 3.004022 | 0.034 | 0.007 | 1 | 12 | Wdr77 | 0.027 | Old |
| 0.124773 | 2.734789 | 0.034 | 0.007 | 1 | 12 | Wdr11 | 0.027 | Old |

| | | | | | | | | |
|---|---|---|---|---|---|---|---|---|
| 0.124773 | 2.699641 | 0.034 | 0.007 | 1 | 12 | Tsen54 | 0.027 | Old |
| 0.124773 | 2.674791 | 0.034 | 0.007 | 1 | 12 | Arvcf | 0.027 | Old |
| 0.124773 | 2.665487 | 0.034 | 0.007 | 1 | 12 | Rmdn1 | 0.027 | Old |
| 0.124773 | 2.659352 | 0.034 | 0.007 | 1 | 12 | Minpp1 | 0.027 | Old |
| 0.124773 | 2.61006 | 0.034 | 0.007 | 1 | 12 | Rilpl1 | 0.027 | Old |
| 0.124773 | 2.379081 | 0.034 | 0.007 | 1 | 12 | Itga3 | 0.027 | Old |
| 0.124773 | 2.278485 | 0.034 | 0.007 | 1 | 12 | Usp48 | 0.027 | Old |
| 0.124949 | 0.57763 | 0.203 | 0.131 | 1 | 12 | Tkt | 0.072 | Old |
| 0.125421 | 0.537269 | 0.203 | 0.124 | 1 | 12 | Rrp1 | 0.079 | Old |
| 0.125536 | 0.611399 | 0.246 | 0.175 | 1 | 12 | Timm8b | 0.071 | Old |
| 0.125597 | 1.498992 | 0.059 | 0.022 | 1 | 12 | Bbs4 | 0.037 | Old |
| 0.125597 | 1.483755 | 0.059 | 0.022 | 1 | 12 | Usp8 | 0.037 | Old |
| 0.125597 | 1.438042 | 0.059 | 0.022 | 1 | 12 | Ginm1 | 0.037 | Old |
| 0.125597 | 1.430091 | 0.059 | 0.022 | 1 | 12 | Kctd6 | 0.037 | Old |
| 0.125597 | 1.399976 | 0.059 | 0.022 | 1 | 12 | Dnajc30 | 0.037 | Old |
| 0.125631 | 0.999458 | 0.102 | 0.051 | 1 | 12 | Cib2 | 0.051 | Old |
| 0.125631 | 0.977866 | 0.102 | 0.051 | 1 | 12 | Pyurf | 0.051 | Old |
| 0.12598 | 0.310172 | 0.322 | 0.234 | 1 | 12 | Atp5k | 0.088 | Old |
| 0.126377 | 0.648986 | 0.212 | 0.139 | 1 | 12 | Psmd12 | 0.073 | Old |
| 0.126524 | 2.535877 | 0.034 | 0.007 | 1 | 12 | Sidt2 | 0.027 | Old |
| 0.126524 | 2.510038 | 0.034 | 0.007 | 1 | 12 | Nfkbil1 | 0.027 | Old |
| 0.126524 | 2.464228 | 0.034 | 0.007 | 1 | 12 | Zcchc7 | 0.027 | Old |
| 0.126524 | 2.357504 | 0.034 | 0.007 | 1 | 12 | Tor1aip1 | 0.027 | Old |
| 0.126524 | 2.355676 | 0.034 | 0.007 | 1 | 12 | Lrrn2 | 0.027 | Old |
| 0.126524 | 2.266776 | 0.034 | 0.007 | 1 | 12 | Zdhhc2 | 0.027 | Old |
| 0.126524 | 2.238496 | 0.034 | 0.007 | 1 | 12 | Htatip2 | 0.027 | Old |
| 0.126524 | 2.231025 | 0.034 | 0.007 | 1 | 12 | Osgepl1 | 0.027 | Old |
| 0.126524 | 2.214806 | 0.034 | 0.007 | 1 | 12 | Rab22a | 0.027 | Old |
| 0.126524 | 2.205158 | 0.034 | 0.007 | 1 | 12 | Clgn | 0.027 | Old |
| 0.126524 | 2.203278 | 0.034 | 0.007 | 1 | 12 | Sat2 | 0.027 | Old |
| 0.126524 | 2.202149 | 0.034 | 0.007 | 1 | 12 | Slc35f5 | 0.027 | Old |
| 0.126524 | 2.193967 | 0.034 | 0.007 | 1 | 12 | Med11 | 0.027 | Old |
| 0.126524 | 2.169328 | 0.034 | 0.007 | 1 | 12 | Sf3b4 | 0.027 | Old |
| 0.126524 | 2.145674 | 0.034 | 0.007 | 1 | 12 | Med15 | 0.027 | Old |
| 0.126524 | 2.075195 | 0.034 | 0.007 | 1 | 12 | Nfs1 | 0.027 | Old |
| 0.126524 | 2.021552 | 0.034 | 0.007 | 1 | 12 | Snx5 | 0.027 | Old |
| 0.126557 | 0.957724 | 0.102 | 0.051 | 1 | 12 | Eif1a | 0.051 | Old |
| 0.126557 | 1.03009 | 0.102 | 0.051 | 1 | 12 | Hp1bp3 | 0.051 | Old |
| 0.126852 | 1.457181 | 0.059 | 0.022 | 1 | 12 | Kctd17 | 0.037 | Old |
| 0.127213 | 0.954497 | 0.11 | 0.058 | 1 | 12 | Nudt19 | 0.052 | Old |
| 0.127405 | 2.027209 | 0.034 | 0.007 | 1 | 12 | Atrnl1 | 0.027 | Old |
| 0.128198 | 5.382123 | 0.017 | 0 | 1 | 12 | Fam19a2 | 0.017 | Old |
| 0.128198 | 5.196085 | 0.017 | 0 | 1 | 12 | Sertad1 | 0.017 | Old |
| 0.128198 | 5.075804 | 0.017 | 0 | 1 | 12 | Ebf4 | 0.017 | Old |
| 0.128198 | 5.068956 | 0.017 | 0 | 1 | 12 | Zfp263 | 0.017 | Old |

| | | | | | | | | |
|---|---|---|---|---|---|---|---|---|
| 0.128198 | 5.027938 | 0.017 | 0 | 1 | 12 | Gm26890 | 0.017 | Old |
| 0.128198 | 5.02256 | 0.017 | 0 | 1 | 12 | Apc2 | 0.017 | Old |
| 0.128198 | 5.018257 | 0.017 | 0 | 1 | 12 | Cr1l | 0.017 | Old |
| 0.128198 | 5.015432 | 0.017 | 0 | 1 | 12 | Adm | 0.017 | Old |
| 0.128198 | 5.009995 | 0.017 | 0 | 1 | 12 | 5033430I1! | 0.017 | Old |
| 0.128198 | 5.008426 | 0.017 | 0 | 1 | 12 | Arhgef18 | 0.017 | Old |
| 0.128198 | 5.008426 | 0.017 | 0 | 1 | 12 | Zfp948 | 0.017 | Old |
| 0.128198 | 5.002065 | 0.017 | 0 | 1 | 12 | Pcdha11.1 | 0.017 | Old |
| 0.128198 | 4.991853 | 0.017 | 0 | 1 | 12 | Map2k5 | 0.017 | Old |
| 0.128198 | 4.988285 | 0.017 | 0 | 1 | 12 | Tmem150a | 0.017 | Old |
| 0.128198 | 4.984285 | 0.017 | 0 | 1 | 12 | Gabpb1 | 0.017 | Old |
| 0.128198 | 4.980528 | 0.017 | 0 | 1 | 12 | Hhip | 0.017 | Old |
| 0.128198 | 4.977038 | 0.017 | 0 | 1 | 12 | Gm11808 | 0.017 | Old |
| 0.128198 | 4.975186 | 0.017 | 0 | 1 | 12 | Polrmt | 0.017 | Old |
| 0.128198 | 4.969147 | 0.017 | 0 | 1 | 12 | Mtrf1 | 0.017 | Old |
| 0.128198 | 4.967921 | 0.017 | 0 | 1 | 12 | Gm10767 | 0.017 | Old |
| 0.128198 | 4.967422 | 0.017 | 0 | 1 | 12 | Hecw1 | 0.017 | Old |
| 0.128198 | 4.964905 | 0.017 | 0 | 1 | 12 | Gclc | 0.017 | Old |
| 0.128198 | 4.946802 | 0.017 | 0 | 1 | 12 | Asnsd1 | 0.017 | Old |
| 0.128198 | 4.937212 | 0.017 | 0 | 1 | 12 | Gbp7 | 0.017 | Old |
| 0.128198 | 4.934054 | 0.017 | 0 | 1 | 12 | Zfp513 | 0.017 | Old |
| 0.128198 | 4.932661 | 0.017 | 0 | 1 | 12 | Rec114 | 0.017 | Old |
| 0.128198 | 4.927282 | 0.017 | 0 | 1 | 12 | Bend7 | 0.017 | Old |
| 0.128198 | 4.91929 | 0.017 | 0 | 1 | 12 | Krt1 | 0.017 | Old |
| 0.128198 | 4.915228 | 0.017 | 0 | 1 | 12 | Cenpl | 0.017 | Old |
| 0.128198 | 4.912599 | 0.017 | 0 | 1 | 12 | Tepsin | 0.017 | Old |
| 0.128198 | 4.910909 | 0.017 | 0 | 1 | 12 | Pogz | 0.017 | Old |
| 0.128198 | 4.9104 | 0.017 | 0 | 1 | 12 | Ddx21 | 0.017 | Old |
| 0.128198 | 4.908039 | 0.017 | 0 | 1 | 12 | Pogk | 0.017 | Old |
| 0.128198 | 4.901874 | 0.017 | 0 | 1 | 12 | Kat8 | 0.017 | Old |
| 0.128198 | 4.900155 | 0.017 | 0 | 1 | 12 | Sdc1 | 0.017 | Old |
| 0.128198 | 4.900038 | 0.017 | 0 | 1 | 12 | Pla2g15 | 0.017 | Old |
| 0.128198 | 4.894911 | 0.017 | 0 | 1 | 12 | C130073E2 | 0.017 | Old |
| 0.128198 | 4.891004 | 0.017 | 0 | 1 | 12 | Syt14 | 0.017 | Old |
| 0.128198 | 4.889235 | 0.017 | 0 | 1 | 12 | Dbr1 | 0.017 | Old |
| 0.128198 | 4.88588 | 0.017 | 0 | 1 | 12 | Mpv17 | 0.017 | Old |
| 0.128198 | 4.88234 | 0.017 | 0 | 1 | 12 | Dcaf6 | 0.017 | Old |
| 0.128198 | 4.881043 | 0.017 | 0 | 1 | 12 | Blm | 0.017 | Old |
| 0.128198 | 4.877696 | 0.017 | 0 | 1 | 12 | Ednrb | 0.017 | Old |
| 0.128198 | 4.876936 | 0.017 | 0 | 1 | 12 | Tmem248 | 0.017 | Old |
| 0.128198 | 4.872974 | 0.017 | 0 | 1 | 12 | Ptprg | 0.017 | Old |
| 0.128198 | 4.87272 | 0.017 | 0 | 1 | 12 | Gm17092 | 0.017 | Old |
| 0.128198 | 4.866644 | 0.017 | 0 | 1 | 12 | Papd7 | 0.017 | Old |
| 0.128198 | 4.8653 | 0.017 | 0 | 1 | 12 | Diexf | 0.017 | Old |
| 0.128198 | 4.864454 | 0.017 | 0 | 1 | 12 | Begain | 0.017 | Old |

| | | | | | | | | |
|---|---|---|---|---|---|---|---|---|
| 0.128198 | 4.863829 | 0.017 | 0 | 1 | 12 | Akip1 | 0.017 | Old |
| 0.128198 | 4.862175 | 0.017 | 0 | 1 | 12 | Pus10 | 0.017 | Old |
| 0.128198 | 4.857662 | 0.017 | 0 | 1 | 12 | Umps | 0.017 | Old |
| 0.128198 | 4.856278 | 0.017 | 0 | 1 | 12 | Ooep | 0.017 | Old |
| 0.128198 | 4.852991 | 0.017 | 0 | 1 | 12 | Sptlc1 | 0.017 | Old |
| 0.128198 | 4.849204 | 0.017 | 0 | 1 | 12 | Rgs12 | 0.017 | Old |
| 0.128198 | 4.848915 | 0.017 | 0 | 1 | 12 | Gpr158 | 0.017 | Old |
| 0.128198 | 4.848622 | 0.017 | 0 | 1 | 12 | Samd4b | 0.017 | Old |
| 0.128198 | 4.847723 | 0.017 | 0 | 1 | 12 | Hsbp1l1 | 0.017 | Old |
| 0.128198 | 4.83698 | 0.017 | 0 | 1 | 12 | Ppt1 | 0.017 | Old |
| 0.128198 | 4.835094 | 0.017 | 0 | 1 | 12 | D130017N( | 0.017 | Old |
| 0.128198 | 4.829104 | 0.017 | 0 | 1 | 12 | Syce2 | 0.017 | Old |
| 0.128198 | 4.829087 | 0.017 | 0 | 1 | 12 | Dchs1 | 0.017 | Old |
| 0.128198 | 4.826783 | 0.017 | 0 | 1 | 12 | Eif4ebp2 | 0.017 | Old |
| 0.128198 | 4.82634 | 0.017 | 0 | 1 | 12 | Nmral1 | 0.017 | Old |
| 0.128198 | 4.818158 | 0.017 | 0 | 1 | 12 | Sowahc | 0.017 | Old |
| 0.128198 | 4.812795 | 0.017 | 0 | 1 | 12 | Gle1 | 0.017 | Old |
| 0.128198 | 4.81188 | 0.017 | 0 | 1 | 12 | Ccdc166 | 0.017 | Old |
| 0.128198 | 4.811434 | 0.017 | 0 | 1 | 12 | Exoc1 | 0.017 | Old |
| 0.128198 | 4.807236 | 0.017 | 0 | 1 | 12 | Spred2 | 0.017 | Old |
| 0.128198 | 4.807214 | 0.017 | 0 | 1 | 12 | Mef2c | 0.017 | Old |
| 0.128198 | 4.801508 | 0.017 | 0 | 1 | 12 | Phtf2 | 0.017 | Old |
| 0.128198 | 4.800011 | 0.017 | 0 | 1 | 12 | Zfp846 | 0.017 | Old |
| 0.128198 | 4.797747 | 0.017 | 0 | 1 | 12 | Rarb | 0.017 | Old |
| 0.128198 | 4.797148 | 0.017 | 0 | 1 | 12 | Cox16 | 0.017 | Old |
| 0.128198 | 4.795219 | 0.017 | 0 | 1 | 12 | Pcdh10 | 0.017 | Old |
| 0.128198 | 4.795213 | 0.017 | 0 | 1 | 12 | Rad9b | 0.017 | Old |
| 0.128198 | 4.787993 | 0.017 | 0 | 1 | 12 | Ikbip | 0.017 | Old |
| 0.128198 | 4.786188 | 0.017 | 0 | 1 | 12 | Smg8 | 0.017 | Old |
| 0.128198 | 4.78604 | 0.017 | 0 | 1 | 12 | Gm48551 | 0.017 | Old |
| 0.128198 | 4.785861 | 0.017 | 0 | 1 | 12 | Scn11a | 0.017 | Old |
| 0.128198 | 4.781948 | 0.017 | 0 | 1 | 12 | Ogfod2 | 0.017 | Old |
| 0.128198 | 4.781948 | 0.017 | 0 | 1 | 12 | Zfp426 | 0.017 | Old |
| 0.128198 | 4.778636 | 0.017 | 0 | 1 | 12 | Nsmaf | 0.017 | Old |
| 0.128198 | 4.776272 | 0.017 | 0 | 1 | 12 | 2310068J1 | 0.017 | Old |
| 0.128198 | 4.772704 | 0.017 | 0 | 1 | 12 | Dhx34 | 0.017 | Old |
| 0.128198 | 4.771586 | 0.017 | 0 | 1 | 12 | Vps52 | 0.017 | Old |
| 0.128198 | 4.771586 | 0.017 | 0 | 1 | 12 | Ttc39c | 0.017 | Old |
| 0.128198 | 4.77 | 0.017 | 0 | 1 | 12 | Zfp692 | 0.017 | Old |
| 0.128198 | 4.769448 | 0.017 | 0 | 1 | 12 | Dis3 | 0.017 | Old |
| 0.128198 | 4.762862 | 0.017 | 0 | 1 | 12 | Rab30 | 0.017 | Old |
| 0.128198 | 4.762618 | 0.017 | 0 | 1 | 12 | Mtag2 | 0.017 | Old |
| 0.128198 | 4.761145 | 0.017 | 0 | 1 | 12 | Kdm2b | 0.017 | Old |
| 0.128198 | 4.760748 | 0.017 | 0 | 1 | 12 | Mut | 0.017 | Old |
| 0.128198 | 4.759521 | 0.017 | 0 | 1 | 12 | Cgrrf1 | 0.017 | Old |

| | | | | | | | | |
|---|---|---|---|---|---|---|---|---|
| 0.128198 | 4.759179 | 0.017 | 0 | 1 | 12 | Rhof | 0.017 | Old |
| 0.128198 | 4.757405 | 0.017 | 0 | 1 | 12 | Zfp362 | 0.017 | Old |
| 0.128198 | 4.756942 | 0.017 | 0 | 1 | 12 | Fam102a | 0.017 | Old |
| 0.128198 | 4.756432 | 0.017 | 0 | 1 | 12 | Dtx3l | 0.017 | Old |
| 0.128198 | 4.755547 | 0.017 | 0 | 1 | 12 | Lrrc75a | 0.017 | Old |
| 0.128198 | 4.751277 | 0.017 | 0 | 1 | 12 | Gm27030 | 0.017 | Old |
| 0.128198 | 4.74981 | 0.017 | 0 | 1 | 12 | Dmxl2 | 0.017 | Old |
| 0.128198 | 4.744557 | 0.017 | 0 | 1 | 12 | Mtmr2 | 0.017 | Old |
| 0.128198 | 4.742082 | 0.017 | 0 | 1 | 12 | Ftsj3 | 0.017 | Old |
| 0.128198 | 4.742041 | 0.017 | 0 | 1 | 12 | Prkar2b | 0.017 | Old |
| 0.128198 | 4.741939 | 0.017 | 0 | 1 | 12 | Spata2l | 0.017 | Old |
| 0.128198 | 4.74094 | 0.017 | 0 | 1 | 12 | Golm1 | 0.017 | Old |
| 0.128198 | 4.738851 | 0.017 | 0 | 1 | 12 | Fitm2 | 0.017 | Old |
| 0.128198 | 4.738029 | 0.017 | 0 | 1 | 12 | Haus1 | 0.017 | Old |
| 0.128198 | 4.737856 | 0.017 | 0 | 1 | 12 | Mbtps2 | 0.017 | Old |
| 0.128198 | 4.735458 | 0.017 | 0 | 1 | 12 | Coro1b | 0.017 | Old |
| 0.128198 | 4.73459 | 0.017 | 0 | 1 | 12 | Nek4 | 0.017 | Old |
| 0.128198 | 4.734571 | 0.017 | 0 | 1 | 12 | 2810001G2 | 0.017 | Old |
| 0.128198 | 4.734068 | 0.017 | 0 | 1 | 12 | Tob2 | 0.017 | Old |
| 0.128198 | 4.731338 | 0.017 | 0 | 1 | 12 | Hdac7 | 0.017 | Old |
| 0.128198 | 4.730567 | 0.017 | 0 | 1 | 12 | Uvrag | 0.017 | Old |
| 0.128198 | 4.729914 | 0.017 | 0 | 1 | 12 | Kcnc3 | 0.017 | Old |
| 0.128198 | 4.723647 | 0.017 | 0 | 1 | 12 | Pecr | 0.017 | Old |
| 0.128198 | 4.721901 | 0.017 | 0 | 1 | 12 | Gins2 | 0.017 | Old |
| 0.128198 | 4.721895 | 0.017 | 0 | 1 | 12 | Fam20c | 0.017 | Old |
| 0.128198 | 4.718348 | 0.017 | 0 | 1 | 12 | Rasl10b | 0.017 | Old |
| 0.128198 | 4.707626 | 0.017 | 0 | 1 | 12 | 5-Sep | 0.017 | Old |
| 0.128198 | 4.704955 | 0.017 | 0 | 1 | 12 | Tkfc | 0.017 | Old |
| 0.128198 | 4.704547 | 0.017 | 0 | 1 | 12 | Xpo7 | 0.017 | Old |
| 0.128198 | 4.702523 | 0.017 | 0 | 1 | 12 | Fuz | 0.017 | Old |
| 0.128198 | 4.700346 | 0.017 | 0 | 1 | 12 | Perp | 0.017 | Old |
| 0.128198 | 4.698353 | 0.017 | 0 | 1 | 12 | Igf2r | 0.017 | Old |
| 0.128198 | 4.696689 | 0.017 | 0 | 1 | 12 | Alyref2 | 0.017 | Old |
| 0.128198 | 4.696689 | 0.017 | 0 | 1 | 12 | Rnmt | 0.017 | Old |
| 0.128198 | 4.696524 | 0.017 | 0 | 1 | 12 | Cox7a1 | 0.017 | Old |
| 0.128198 | 4.692744 | 0.017 | 0 | 1 | 12 | Ccz1 | 0.017 | Old |
| 0.128198 | 4.691875 | 0.017 | 0 | 1 | 12 | Mboat1 | 0.017 | Old |
| 0.128198 | 4.688915 | 0.017 | 0 | 1 | 12 | Arrdc1 | 0.017 | Old |
| 0.128198 | 4.68796 | 0.017 | 0 | 1 | 12 | Paqr4 | 0.017 | Old |
| 0.128198 | 4.686355 | 0.017 | 0 | 1 | 12 | Dmd | 0.017 | Old |
| 0.128198 | 4.683531 | 0.017 | 0 | 1 | 12 | Aebp2 | 0.017 | Old |
| 0.128198 | 4.682249 | 0.017 | 0 | 1 | 12 | N4bp2 | 0.017 | Old |
| 0.128198 | 4.679721 | 0.017 | 0 | 1 | 12 | Ltbr | 0.017 | Old |
| 0.128198 | 4.677156 | 0.017 | 0 | 1 | 12 | Lpl | 0.017 | Old |
| 0.128198 | 4.675644 | 0.017 | 0 | 1 | 12 | Tmpo | 0.017 | Old |

| | | | | | | | | |
|---|---|---|---|---|---|---|---|---|
| 0.128198 | 4.664221 | 0.017 | 0 | 1 | 12 | Tcea2 | 0.017 | Old |
| 0.128198 | 4.662442 | 0.017 | 0 | 1 | 12 | Fgf14 | 0.017 | Old |
| 0.128198 | 4.658308 | 0.017 | 0 | 1 | 12 | Asb13 | 0.017 | Old |
| 0.128198 | 4.653628 | 0.017 | 0 | 1 | 12 | Eral1 | 0.017 | Old |
| 0.128198 | 4.652021 | 0.017 | 0 | 1 | 12 | Gm26881 | 0.017 | Old |
| 0.128198 | 4.645915 | 0.017 | 0 | 1 | 12 | Kctd18 | 0.017 | Old |
| 0.128198 | 4.64588 | 0.017 | 0 | 1 | 12 | Spg11 | 0.017 | Old |
| 0.128198 | 4.644093 | 0.017 | 0 | 1 | 12 | Psmb9 | 0.017 | Old |
| 0.128198 | 4.644002 | 0.017 | 0 | 1 | 12 | D6Wsu163 | 0.017 | Old |
| 0.128198 | 4.643414 | 0.017 | 0 | 1 | 12 | Ampd2 | 0.017 | Old |
| 0.128198 | 4.640966 | 0.017 | 0 | 1 | 12 | Slc1a4 | 0.017 | Old |
| 0.128198 | 4.640953 | 0.017 | 0 | 1 | 12 | Ints9 | 0.017 | Old |
| 0.128198 | 4.637728 | 0.017 | 0 | 1 | 12 | 5730409E0 | 0.017 | Old |
| 0.128198 | 4.636456 | 0.017 | 0 | 1 | 12 | Zfp512b | 0.017 | Old |
| 0.128198 | 4.636078 | 0.017 | 0 | 1 | 12 | 2810403A0 | 0.017 | Old |
| 0.128198 | 4.63583 | 0.017 | 0 | 1 | 12 | Fbxo8 | 0.017 | Old |
| 0.128198 | 4.626274 | 0.017 | 0 | 1 | 12 | Tnpo3 | 0.017 | Old |
| 0.128198 | 4.62529 | 0.017 | 0 | 1 | 12 | Tnip3 | 0.017 | Old |
| 0.128198 | 4.623149 | 0.017 | 0 | 1 | 12 | Champ1 | 0.017 | Old |
| 0.128198 | 4.61804 | 0.017 | 0 | 1 | 12 | Gm26888 | 0.017 | Old |
| 0.128198 | 4.617887 | 0.017 | 0 | 1 | 12 | Mccc1 | 0.017 | Old |
| 0.128198 | 4.615606 | 0.017 | 0 | 1 | 12 | Omg | 0.017 | Old |
| 0.128198 | 4.614182 | 0.017 | 0 | 1 | 12 | Cnnm3 | 0.017 | Old |
| 0.128198 | 4.610947 | 0.017 | 0 | 1 | 12 | Sntb2 | 0.017 | Old |
| 0.128198 | 4.609383 | 0.017 | 0 | 1 | 12 | Hdac1 | 0.017 | Old |
| 0.128198 | 4.606973 | 0.017 | 0 | 1 | 12 | Slitrk5 | 0.017 | Old |
| 0.128198 | 4.604732 | 0.017 | 0 | 1 | 12 | Zfp113 | 0.017 | Old |
| 0.128198 | 4.599919 | 0.017 | 0 | 1 | 12 | Atp11a | 0.017 | Old |
| 0.128198 | 4.593625 | 0.017 | 0 | 1 | 12 | Camk4 | 0.017 | Old |
| 0.128198 | 4.593484 | 0.017 | 0 | 1 | 12 | Gtf2h2 | 0.017 | Old |
| 0.128198 | 4.592424 | 0.017 | 0 | 1 | 12 | Zkscan4 | 0.017 | Old |
| 0.128198 | 4.590955 | 0.017 | 0 | 1 | 12 | Nsun5 | 0.017 | Old |
| 0.128198 | 4.586311 | 0.017 | 0 | 1 | 12 | Prrg2 | 0.017 | Old |
| 0.128198 | 4.583875 | 0.017 | 0 | 1 | 12 | Lca5 | 0.017 | Old |
| 0.128198 | 4.583875 | 0.017 | 0 | 1 | 12 | Adcy5 | 0.017 | Old |
| 0.128198 | 4.583035 | 0.017 | 0 | 1 | 12 | Ddhd1 | 0.017 | Old |
| 0.128198 | 4.582979 | 0.017 | 0 | 1 | 12 | Peak1 | 0.017 | Old |
| 0.128198 | 4.581113 | 0.017 | 0 | 1 | 12 | Gtf3c3 | 0.017 | Old |
| 0.128198 | 4.578792 | 0.017 | 0 | 1 | 12 | Tpmt | 0.017 | Old |
| 0.128198 | 4.573636 | 0.017 | 0 | 1 | 12 | Mtmr9 | 0.017 | Old |
| 0.128198 | 4.568163 | 0.017 | 0 | 1 | 12 | Mier2 | 0.017 | Old |
| 0.128198 | 4.566186 | 0.017 | 0 | 1 | 12 | Unc45a | 0.017 | Old |
| 0.128198 | 4.560688 | 0.017 | 0 | 1 | 12 | 3110009E1 | 0.017 | Old |
| 0.128198 | 4.554029 | 0.017 | 0 | 1 | 12 | Ptpn23 | 0.017 | Old |
| 0.128198 | 4.553845 | 0.017 | 0 | 1 | 12 | Agrn | 0.017 | Old |

| | | | | | | | | |
|---|---|---|---|---|---|---|---|---|
| 0.128198 | 4.550426 | 0.017 | 0 | 1 | 12 | Mto1 | 0.017 | Old |
| 0.128198 | 4.547618 | 0.017 | 0 | 1 | 12 | Pdzd4 | 0.017 | Old |
| 0.128198 | 4.54595 | 0.017 | 0 | 1 | 12 | Jrkl | 0.017 | Old |
| 0.128198 | 4.535989 | 0.017 | 0 | 1 | 12 | Rnf103 | 0.017 | Old |
| 0.128198 | 4.526575 | 0.017 | 0 | 1 | 12 | Tmem130 | 0.017 | Old |
| 0.128198 | 4.524052 | 0.017 | 0 | 1 | 12 | Mtmr11 | 0.017 | Old |
| 0.128198 | 4.513086 | 0.017 | 0 | 1 | 12 | Smg5 | 0.017 | Old |
| 0.128198 | 4.512973 | 0.017 | 0 | 1 | 12 | Pabpc4 | 0.017 | Old |
| 0.128198 | 4.508182 | 0.017 | 0 | 1 | 12 | Nova2 | 0.017 | Old |
| 0.128198 | 4.50656 | 0.017 | 0 | 1 | 12 | Alg3 | 0.017 | Old |
| 0.128198 | 4.504965 | 0.017 | 0 | 1 | 12 | St8sia4 | 0.017 | Old |
| 0.128198 | 4.502646 | 0.017 | 0 | 1 | 12 | Rfx1 | 0.017 | Old |
| 0.128198 | 4.502362 | 0.017 | 0 | 1 | 12 | Wdr3 | 0.017 | Old |
| 0.128198 | 4.495923 | 0.017 | 0 | 1 | 12 | Tbx2 | 0.017 | Old |
| 0.128198 | 4.489956 | 0.017 | 0 | 1 | 12 | Syngr2 | 0.017 | Old |
| 0.128198 | 4.486348 | 0.017 | 0 | 1 | 12 | Tfcp2 | 0.017 | Old |
| 0.128198 | 4.483282 | 0.017 | 0 | 1 | 12 | Ccdc74a | 0.017 | Old |
| 0.128198 | 4.482736 | 0.017 | 0 | 1 | 12 | Fjx1 | 0.017 | Old |
| 0.128198 | 4.482736 | 0.017 | 0 | 1 | 12 | Abhd2 | 0.017 | Old |
| 0.128198 | 4.476095 | 0.017 | 0 | 1 | 12 | Gm5113 | 0.017 | Old |
| 0.128198 | 4.473603 | 0.017 | 0 | 1 | 12 | Gpr162 | 0.017 | Old |
| 0.128198 | 4.47288 | 0.017 | 0 | 1 | 12 | Wdr89 | 0.017 | Old |
| 0.128198 | 4.469262 | 0.017 | 0 | 1 | 12 | Orai2 | 0.017 | Old |
| 0.128198 | 4.468994 | 0.017 | 0 | 1 | 12 | Mfsd3 | 0.017 | Old |
| 0.128198 | 4.468036 | 0.017 | 0 | 1 | 12 | Tsr1 | 0.017 | Old |
| 0.128198 | 4.463131 | 0.017 | 0 | 1 | 12 | Trip10 | 0.017 | Old |
| 0.128198 | 4.46007 | 0.017 | 0 | 1 | 12 | Mmp9 | 0.017 | Old |
| 0.128198 | 4.450261 | 0.017 | 0 | 1 | 12 | Plbd1 | 0.017 | Old |
| 0.128198 | 4.438499 | 0.017 | 0 | 1 | 12 | Ptprr | 0.017 | Old |
| 0.128198 | 4.43699 | 0.017 | 0 | 1 | 12 | Gm17197 | 0.017 | Old |
| 0.128198 | 4.415375 | 0.017 | 0 | 1 | 12 | Haus4 | 0.017 | Old |
| 0.128198 | 4.407272 | 0.017 | 0 | 1 | 12 | Lcor | 0.017 | Old |
| 0.128198 | 4.397725 | 0.017 | 0 | 1 | 12 | Tti2 | 0.017 | Old |
| 0.128198 | 4.376049 | 0.017 | 0 | 1 | 12 | Snupn | 0.017 | Old |
| 0.128198 | 4.369058 | 0.017 | 0 | 1 | 12 | Jag2 | 0.017 | Old |
| 0.128198 | 4.360818 | 0.017 | 0 | 1 | 12 | Rgs7 | 0.017 | Old |
| 0.128198 | 4.345377 | 0.017 | 0 | 1 | 12 | Osbpl5 | 0.017 | Old |
| 0.128198 | 4.340655 | 0.017 | 0 | 1 | 12 | 2610044O1 | 0.017 | Old |
| 0.128198 | 4.336254 | 0.017 | 0 | 1 | 12 | Srek1ip1 | 0.017 | Old |
| 0.128198 | 4.329271 | 0.017 | 0 | 1 | 12 | Rpusd3 | 0.017 | Old |
| 0.128198 | 4.320172 | 0.017 | 0 | 1 | 12 | Prrxl1 | 0.017 | Old |
| 0.128198 | 4.319782 | 0.017 | 0 | 1 | 12 | Uhrf1bp1l | 0.017 | Old |
| 0.128198 | 4.310723 | 0.017 | 0 | 1 | 12 | Sp1 | 0.017 | Old |
| 0.128198 | 4.309703 | 0.017 | 0 | 1 | 12 | Sall2 | 0.017 | Old |
| 0.128198 | 4.308207 | 0.017 | 0 | 1 | 12 | Osbpl8 | 0.017 | Old |

| | | | | | | | | |
|---|---|---|---|---|---|---|---|---|
| 0.128198 | 4.308207 | 0.017 | 0 | 1 | 12 | Rnf130 | 0.017 | Old |
| 0.128198 | 4.277015 | 0.017 | 0 | 1 | 12 | Fxyd5 | 0.017 | Old |
| 0.128198 | 4.271065 | 0.017 | 0 | 1 | 12 | Gss | 0.017 | Old |
| 0.128198 | 4.255853 | 0.017 | 0 | 1 | 12 | Palld | 0.017 | Old |
| 0.128198 | 4.252198 | 0.017 | 0 | 1 | 12 | Susd6 | 0.017 | Old |
| 0.128198 | 4.242013 | 0.017 | 0 | 1 | 12 | Phc1 | 0.017 | Old |
| 0.128198 | 4.23744 | 0.017 | 0 | 1 | 12 | Tank | 0.017 | Old |
| 0.128198 | 4.226094 | 0.017 | 0 | 1 | 12 | Lrrc25 | 0.017 | Old |
| 0.128198 | 4.199913 | 0.017 | 0 | 1 | 12 | Shank1 | 0.017 | Old |
| 0.128198 | 4.192774 | 0.017 | 0 | 1 | 12 | Arrdc3 | 0.017 | Old |
| 0.128198 | 4.191178 | 0.017 | 0 | 1 | 12 | Usp32 | 0.017 | Old |
| 0.128198 | 4.179054 | 0.017 | 0 | 1 | 12 | Safb2 | 0.017 | Old |
| 0.128198 | 4.176683 | 0.017 | 0 | 1 | 12 | Ncoa3 | 0.017 | Old |
| 0.128198 | 4.16173 | 0.017 | 0 | 1 | 12 | Ogg1 | 0.017 | Old |
| 0.128198 | 4.160541 | 0.017 | 0 | 1 | 12 | Ep300 | 0.017 | Old |
| 0.128198 | 4.158371 | 0.017 | 0 | 1 | 12 | Axin1 | 0.017 | Old |
| 0.128198 | 4.139511 | 0.017 | 0 | 1 | 12 | Sypl | 0.017 | Old |
| 0.128198 | 4.127128 | 0.017 | 0 | 1 | 12 | Sgk3 | 0.017 | Old |
| 0.128198 | 4.126113 | 0.017 | 0 | 1 | 12 | Mybbp1a | 0.017 | Old |
| 0.128198 | 4.118954 | 0.017 | 0 | 1 | 12 | Rnf213 | 0.017 | Old |
| 0.128198 | 4.118488 | 0.017 | 0 | 1 | 12 | C5ar1 | 0.017 | Old |
| 0.128198 | 4.112867 | 0.017 | 0 | 1 | 12 | Gatb | 0.017 | Old |
| 0.128198 | 4.094402 | 0.017 | 0 | 1 | 12 | Mdga2 | 0.017 | Old |
| 0.128198 | 4.093699 | 0.017 | 0 | 1 | 12 | Cybb | 0.017 | Old |
| 0.128198 | 4.085774 | 0.017 | 0 | 1 | 12 | Selenbp1 | 0.017 | Old |
| 0.128198 | 4.081001 | 0.017 | 0 | 1 | 12 | Cd38 | 0.017 | Old |
| 0.128198 | 4.07708 | 0.017 | 0 | 1 | 12 | Ms4a6c | 0.017 | Old |
| 0.128198 | 4.055828 | 0.017 | 0 | 1 | 12 | Tnfsfm13 | 0.017 | Old |
| 0.128198 | 4.053917 | 0.017 | 0 | 1 | 12 | Usp19 | 0.017 | Old |
| 0.128198 | 4.033137 | 0.017 | 0 | 1 | 12 | Zfp952 | 0.017 | Old |
| 0.128198 | 4.023832 | 0.017 | 0 | 1 | 12 | Furin | 0.017 | Old |
| 0.128198 | 4.020117 | 0.017 | 0 | 1 | 12 | Rhobtb1 | 0.017 | Old |
| 0.128198 | 4.01095 | 0.017 | 0 | 1 | 12 | Zfp740 | 0.017 | Old |
| 0.128198 | 3.992488 | 0.017 | 0 | 1 | 12 | P4ha1 | 0.017 | Old |
| 0.128198 | 3.967075 | 0.017 | 0 | 1 | 12 | Alox5ap | 0.017 | Old |
| 0.128198 | 3.954101 | 0.017 | 0 | 1 | 12 | Smarcal1 | 0.017 | Old |
| 0.128198 | 3.954101 | 0.017 | 0 | 1 | 12 | Ifi27l2a | 0.017 | Old |
| 0.128198 | 3.945507 | 0.017 | 0 | 1 | 12 | Ncf1 | 0.017 | Old |
| 0.128198 | 3.943798 | 0.017 | 0 | 1 | 12 | Phlda1 | 0.017 | Old |
| 0.128198 | 3.926864 | 0.017 | 0 | 1 | 12 | Ss18 | 0.017 | Old |
| 0.128198 | 3.905933 | 0.017 | 0 | 1 | 12 | Ranbp2 | 0.017 | Old |
| 0.128198 | 3.898738 | 0.017 | 0 | 1 | 12 | Pdzd8 | 0.017 | Old |
| 0.128198 | 3.892237 | 0.017 | 0 | 1 | 12 | Gpr146 | 0.017 | Old |
| 0.128198 | 3.880966 | 0.017 | 0 | 1 | 12 | 6030458C1 | 0.017 | Old |
| 0.128198 | 3.880966 | 0.017 | 0 | 1 | 12 | Phf1 | 0.017 | Old |

| | | | | | | | | |
|---|---|---|---|---|---|---|---|---|
| 0.128198 | 3.852576 | 0.017 | 0 | 1 | 12 | Foxred1 | 0.017 | Old |
| 0.128198 | 3.851024 | 0.017 | 0 | 1 | 12 | Spata2 | 0.017 | Old |
| 0.128198 | 3.841065 | 0.017 | 0 | 1 | 12 | Sirpa | 0.017 | Old |
| 0.128198 | 3.840225 | 0.017 | 0 | 1 | 12 | Npepl1 | 0.017 | Old |
| 0.128198 | 3.840225 | 0.017 | 0 | 1 | 12 | Gk5 | 0.017 | Old |
| 0.128198 | 3.840225 | 0.017 | 0 | 1 | 12 | Retreg3 | 0.017 | Old |
| 0.128198 | 3.81426 | 0.017 | 0 | 1 | 12 | Sash1 | 0.017 | Old |
| 0.128198 | 3.802636 | 0.017 | 0 | 1 | 12 | Lyn | 0.017 | Old |
| 0.128198 | 3.802636 | 0.017 | 0 | 1 | 12 | Guf1 | 0.017 | Old |
| 0.128198 | 3.733079 | 0.017 | 0 | 1 | 12 | Lrrk1 | 0.017 | Old |
| 0.128198 | 3.733079 | 0.017 | 0 | 1 | 12 | Mfhas1 | 0.017 | Old |
| 0.128198 | 3.733079 | 0.017 | 0 | 1 | 12 | Mcm5 | 0.017 | Old |
| 0.128198 | 3.716816 | 0.017 | 0 | 1 | 12 | Prkcd | 0.017 | Old |
| 0.128198 | 3.716816 | 0.017 | 0 | 1 | 12 | F13a1 | 0.017 | Old |
| 0.128198 | 3.712795 | 0.017 | 0 | 1 | 12 | Baz2a | 0.017 | Old |
| 0.128198 | 3.712795 | 0.017 | 0 | 1 | 12 | Pdcd11 | 0.017 | Old |
| 0.128198 | 3.703481 | 0.017 | 0 | 1 | 12 | Osbpl2 | 0.017 | Old |
| 0.128198 | 3.703481 | 0.017 | 0 | 1 | 12 | Bank1 | 0.017 | Old |
| 0.128198 | 3.690339 | 0.017 | 0 | 1 | 12 | Daglb | 0.017 | Old |
| 0.128198 | 3.690339 | 0.017 | 0 | 1 | 12 | Ccdc97 | 0.017 | Old |
| 0.128198 | 3.664616 | 0.017 | 0 | 1 | 12 | Nup214 | 0.017 | Old |
| 0.128198 | 3.611697 | 0.017 | 0 | 1 | 12 | Zfp703 | 0.017 | Old |
| 0.128198 | 3.610504 | 0.017 | 0 | 1 | 12 | Itsn1 | 0.017 | Old |
| 0.128198 | 3.468878 | 0.017 | 0 | 1 | 12 | Tnfaip3 | 0.017 | Old |
| 0.128198 | 3.450873 | 0.017 | 0 | 1 | 12 | Dennd4c | 0.017 | Old |
| 0.128198 | 3.433252 | 0.017 | 0 | 1 | 12 | Sox10 | 0.017 | Old |
| 0.128294 | 2.304033 | 0.034 | 0.007 | 1 | 12 | S100pbp | 0.027 | Old |
| 0.128294 | 2.206027 | 0.034 | 0.007 | 1 | 12 | Mrps28 | 0.027 | Old |
| 0.128294 | 2.183823 | 0.034 | 0.007 | 1 | 12 | Krt222 | 0.027 | Old |
| 0.128294 | 2.174529 | 0.034 | 0.007 | 1 | 12 | 0610009O2 | 0.027 | Old |
| 0.128294 | 2.160347 | 0.034 | 0.007 | 1 | 12 | Bin3 | 0.027 | Old |
| 0.128294 | 2.134564 | 0.034 | 0.007 | 1 | 12 | Prkca | 0.027 | Old |
| 0.128294 | 2.119927 | 0.034 | 0.007 | 1 | 12 | 4930452B0 | 0.027 | Old |
| 0.128294 | 2.119879 | 0.034 | 0.007 | 1 | 12 | Rims2 | 0.027 | Old |
| 0.128294 | 2.103366 | 0.034 | 0.007 | 1 | 12 | Lclat1 | 0.027 | Old |
| 0.128294 | 2.051006 | 0.034 | 0.007 | 1 | 12 | Crcp | 0.027 | Old |
| 0.128294 | 1.978486 | 0.034 | 0.007 | 1 | 12 | Carnmt1 | 0.027 | Old |
| 0.128294 | 1.941218 | 0.034 | 0.007 | 1 | 12 | Sec22a | 0.027 | Old |
| 0.128294 | 1.803963 | 0.034 | 0.007 | 1 | 12 | Dnmt1 | 0.027 | Old |
| 0.129481 | 1.010716 | 0.136 | 0.08 | 1 | 12 | Mrps26 | 0.056 | Old |
| 0.130083 | 1.981354 | 0.034 | 0.007 | 1 | 12 | Xpo1 | 0.027 | Old |
| 0.130083 | 1.954995 | 0.034 | 0.007 | 1 | 12 | Rnf152 | 0.027 | Old |
| 0.130083 | 1.954711 | 0.034 | 0.007 | 1 | 12 | A330102I1 | 0.027 | Old |
| 0.130083 | 1.946038 | 0.034 | 0.007 | 1 | 12 | Rrp36 | 0.027 | Old |
| 0.130083 | 1.943944 | 0.034 | 0.007 | 1 | 12 | Thoc5 | 0.027 | Old |

| | | | | | | | | |
|---|---|---|---|---|---|---|---|---|
| 0.130083 | 1.934307 | 0.034 | 0.007 | 1 | 12 | Rabl2 | 0.027 | Old |
| 0.130083 | 1.890557 | 0.034 | 0.007 | 1 | 12 | Sec13 | 0.027 | Old |
| 0.130083 | 1.865648 | 0.034 | 0.007 | 1 | 12 | Mesd | 0.027 | Old |
| 0.130083 | 1.831513 | 0.034 | 0.007 | 1 | 12 | Cacna1b | 0.027 | Old |
| 0.130083 | 1.723543 | 0.034 | 0.007 | 1 | 12 | Atcay | 0.027 | Old |
| 0.130083 | 1.714129 | 0.034 | 0.007 | 1 | 12 | As3mt | 0.027 | Old |
| 0.130083 | 1.706638 | 0.034 | 0.007 | 1 | 12 | Nab1 | 0.027 | Old |
| 0.130083 | 1.677477 | 0.034 | 0.007 | 1 | 12 | Agpat1 | 0.027 | Old |
| 0.130083 | 1.595299 | 0.034 | 0.007 | 1 | 12 | Mgat2 | 0.027 | Old |
| 0.130083 | 1.553217 | 0.034 | 0.007 | 1 | 12 | Rnpep | 0.027 | Old |
| 0.13036 | 1.033008 | 0.11 | 0.058 | 1 | 12 | Commd7 | 0.052 | Old |
| 0.130675 | 1.30568 | 0.059 | 0.022 | 1 | 12 | Eif2b2 | 0.037 | Old |
| 0.131677 | 0.860693 | 0.153 | 0.095 | 1 | 12 | Nsfl1c | 0.058 | Old |
| 0.131892 | 2.081806 | 0.034 | 0.007 | 1 | 12 | Sec24c | 0.027 | Old |
| 0.131892 | 1.987247 | 0.034 | 0.007 | 1 | 12 | Slc25a36 | 0.027 | Old |
| 0.131892 | 1.951634 | 0.034 | 0.007 | 1 | 12 | Rnaseh2a | 0.027 | Old |
| 0.131892 | 1.925999 | 0.034 | 0.007 | 1 | 12 | Rbm5 | 0.027 | Old |
| 0.131892 | 1.836425 | 0.034 | 0.007 | 1 | 12 | Slc6a17 | 0.027 | Old |
| 0.131892 | 1.783036 | 0.034 | 0.007 | 1 | 12 | Slc44a2 | 0.027 | Old |
| 0.131892 | 1.772138 | 0.034 | 0.007 | 1 | 12 | Tbl1x | 0.027 | Old |
| 0.131892 | 1.688401 | 0.034 | 0.007 | 1 | 12 | Vps37a | 0.027 | Old |
| 0.131892 | 1.669634 | 0.034 | 0.007 | 1 | 12 | Rtkn | 0.027 | Old |
| 0.131892 | 1.655821 | 0.034 | 0.007 | 1 | 12 | Ccl24 | 0.027 | Old |
| 0.131892 | 1.65259 | 0.034 | 0.007 | 1 | 12 | Cnot11 | 0.027 | Old |
| 0.131892 | 1.595872 | 0.034 | 0.007 | 1 | 12 | Nampt | 0.027 | Old |
| 0.131892 | 1.558483 | 0.034 | 0.007 | 1 | 12 | Psd3 | 0.027 | Old |
| 0.131892 | 1.527883 | 0.034 | 0.007 | 1 | 12 | Vgll4 | 0.027 | Old |
| 0.131892 | 1.447473 | 0.034 | 0.007 | 1 | 12 | Ptpro | 0.027 | Old |
| 0.132181 | 0.595208 | 0.322 | 0.248 | 1 | 12 | Kif5c | 0.074 | Old |
| 0.132645 | 0.930604 | 0.11 | 0.058 | 1 | 12 | Tcaf1 | 0.052 | Old |
| 0.133273 | 0.777903 | 0.059 | 0.022 | 1 | 12 | Bin1 | 0.037 | Old |
| 0.133418 | 0.633592 | 0.263 | 0.197 | 1 | 12 | Eif3c | 0.066 | Old |
| 0.133425 | 0.511719 | 0.246 | 0.175 | 1 | 12 | Vps29 | 0.071 | Old |
| 0.134495 | 0.957856 | 0.11 | 0.058 | 1 | 12 | Slc48a1 | 0.052 | Old |
| 0.134587 | 1.001358 | 0.059 | 0.022 | 1 | 12 | 2610507B1 | 0.037 | Old |
| 0.135701 | 1.442372 | 0.068 | 0.029 | 1 | 12 | Wasl | 0.039 | Old |
| 0.136839 | 0.678055 | 0.254 | 0.182 | 1 | 12 | Gnb1 | 0.072 | Old |
| 0.137403 | 0.989162 | 0.119 | 0.066 | 1 | 12 | H2-Ab1 | 0.053 | Old |
| 0.138152 | 1.52201 | 0.068 | 0.029 | 1 | 12 | Mark3 | 0.039 | Old |
| 0.140164 | 0.928925 | 0.11 | 0.058 | 1 | 12 | Ube2r2 | 0.052 | Old |
| 0.14049 | 0.395863 | 0.364 | 0.277 | 1 | 12 | Rabac1 | 0.087 | Old |
| 0.140636 | 1.376114 | 0.068 | 0.029 | 1 | 12 | Pde10a | 0.039 | Old |
| 0.14115 | 0.833069 | 0.153 | 0.095 | 1 | 12 | Arpc1b | 0.058 | Old |
| 0.141532 | 0.953237 | 0.119 | 0.066 | 1 | 12 | Yaf2 | 0.053 | Old |
| 0.141601 | 0.613565 | 0.237 | 0.168 | 1 | 12 | Aprt | 0.069 | Old |

| | | | | | | | | |
|---|---|---|---|---|---|---|---|---|
| 0.143154 | 1.33843 | 0.068 | 0.029 | 1 | 12 | Zeb2 | 0.039 | Old |
| 0.143154 | 1.309072 | 0.068 | 0.029 | 1 | 12 | Nop58 | 0.039 | Old |
| 0.14339 | 0.292501 | 0.475 | 0.387 | 1 | 12 | Rpl17 | 0.088 | Old |
| 0.145357 | 0.407867 | 0.483 | 0.416 | 1 | 12 | Rps6 | 0.067 | Old |
| 0.145512 | 0.77059 | 0.136 | 0.08 | 1 | 12 | Cuta | 0.056 | Old |
| 0.146254 | 0.370689 | 0.475 | 0.387 | 1 | 12 | Vdac3 | 0.088 | Old |
| 0.146335 | 0.416601 | 0.203 | 0.131 | 1 | 12 | Psmc2 | 0.072 | Old |
| 0.146913 | 0.63096 | 0.153 | 0.095 | 1 | 12 | Metap2 | 0.058 | Old |
| 0.148294 | 1.20609 | 0.068 | 0.029 | 1 | 12 | Lancl1 | 0.039 | Old |
| 0.149045 | 1.334045 | 0.076 | 0.036 | 1 | 12 | Gadd45gip | 0.04 | Old |
| 0.150043 | 0.223565 | 0.746 | 0.708 | 1 | 12 | Hspa8 | 0.038 | Old |
| 0.15018 | 0.667811 | 0.127 | 0.073 | 1 | 12 | Grpel1 | 0.054 | Old |
| 0.150857 | 0.415041 | 0.331 | 0.248 | 1 | 12 | Cox7a2l | 0.083 | Old |
| 0.150917 | 1.148636 | 0.068 | 0.029 | 1 | 12 | Pop7 | 0.039 | Old |
| 0.150917 | 0.817852 | 0.068 | 0.029 | 1 | 12 | Acsl5 | 0.039 | Old |
| 0.151491 | 1.24889 | 0.076 | 0.036 | 1 | 12 | Tmem176a | 0.04 | Old |
| 0.151884 | 0.261231 | 0.644 | 0.584 | 1 | 12 | Rpl26 | 0.06 | Old |
| 0.152241 | 1.084074 | 0.068 | 0.029 | 1 | 12 | Afdn | 0.039 | Old |
| 0.153278 | 1.319799 | 0.085 | 0.044 | 1 | 12 | Dut | 0.041 | Old |
| 0.153574 | 1.093347 | 0.068 | 0.029 | 1 | 12 | Gtf3a | 0.039 | Old |
| 0.154191 | 0.668951 | 0.229 | 0.168 | 1 | 12 | Snx10 | 0.061 | Old |
| 0.155642 | 0.545497 | 0.288 | 0.226 | 1 | 12 | Pcmt1 | 0.062 | Old |
| 0.156267 | 1.043928 | 0.068 | 0.029 | 1 | 12 | Pcdh9 | 0.039 | Old |
| 0.157045 | 0.790629 | 0.186 | 0.131 | 1 | 12 | Tmed10 | 0.055 | Old |
| 0.157214 | 0.510462 | 0.229 | 0.153 | 1 | 12 | Spcs2 | 0.076 | Old |
| 0.157327 | 0.612313 | 0.136 | 0.08 | 1 | 12 | Rabep1 | 0.056 | Old |
| 0.158996 | 0.968329 | 0.068 | 0.029 | 1 | 12 | Smim26 | 0.039 | Old |
| 0.162061 | 0.537308 | 0.136 | 0.08 | 1 | 12 | Pin1 | 0.056 | Old |
| 0.162823 | 1.102659 | 0.085 | 0.044 | 1 | 12 | Rsrc1 | 0.041 | Old |
| 0.163008 | 0.559672 | 0.22 | 0.153 | 1 | 12 | Psmb7 | 0.067 | Old |
| 0.163008 | 1.018634 | 0.169 | 0.117 | 1 | 12 | Cd74 | 0.052 | Old |
| 0.163156 | 0.44003 | 0.068 | 0.029 | 1 | 12 | Dbn1 | 0.039 | Old |
| 0.164181 | 1.074099 | 0.076 | 0.036 | 1 | 12 | Epn1 | 0.04 | Old |
| 0.165542 | 0.324564 | 0.246 | 0.175 | 1 | 12 | Gm1673 | 0.071 | Old |
| 0.166516 | 1.080873 | 0.085 | 0.044 | 1 | 12 | Smc6 | 0.041 | Old |
| 0.166812 | 1.02559 | 0.076 | 0.036 | 1 | 12 | Chmp1a | 0.04 | Old |
| 0.167391 | 0.515856 | 0.195 | 0.131 | 1 | 12 | Mrpl12 | 0.064 | Old |
| 0.167474 | 0.943889 | 0.076 | 0.036 | 1 | 12 | 2610001J0 | 0.04 | Old |
| 0.168139 | 1.04084 | 0.076 | 0.036 | 1 | 12 | Ldah | 0.04 | Old |
| 0.168139 | 0.963134 | 0.076 | 0.036 | 1 | 12 | Raly | 0.04 | Old |
| 0.168299 | 2.266185 | 0.042 | 0.015 | 1 | 12 | Egr1 | 0.027 | Old |
| 0.168299 | 1.931254 | 0.042 | 0.015 | 1 | 12 | Stk19 | 0.027 | Old |
| 0.168762 | 0.625419 | 0.127 | 0.073 | 1 | 12 | Cbr1 | 0.054 | Old |
| 0.170165 | 1.84242 | 0.042 | 0.015 | 1 | 12 | Rnf6 | 0.027 | Old |
| 0.170272 | 0.745775 | 0.068 | 0.029 | 1 | 12 | Id3 | 0.039 | Old |

| | | | | | | | | |
|---|---|---|---|---|---|---|---|---|
| 0.171586 | 1.049964 | 0.093 | 0.051 | 1 | 12 | Cbarp | 0.042 | Old |
| 0.172047 | 1.790326 | 0.042 | 0.015 | 1 | 12 | Dlat | 0.027 | Old |
| 0.172047 | 1.619831 | 0.042 | 0.015 | 1 | 12 | Akap7 | 0.027 | Old |
| 0.172513 | 0.625017 | 0.203 | 0.139 | 1 | 12 | Pmm1 | 0.064 | Old |
| 0.17281 | 1.095281 | 0.085 | 0.044 | 1 | 12 | Idnk | 0.041 | Old |
| 0.17281 | 0.99724 | 0.085 | 0.044 | 1 | 12 | Srsf10 | 0.041 | Old |
| 0.173629 | 1.190225 | 0.102 | 0.058 | 1 | 12 | Fuom | 0.044 | Old |
| 0.173866 | 0.401639 | 0.254 | 0.19 | 1 | 12 | Tlx2 | 0.064 | Old |
| 0.173944 | 1.866552 | 0.042 | 0.015 | 1 | 12 | Scamp3 | 0.027 | Old |
| 0.173944 | 1.556002 | 0.042 | 0.015 | 1 | 12 | Rftn1 | 0.027 | Old |
| 0.175856 | 1.826435 | 0.042 | 0.015 | 1 | 12 | BC005561 | 0.027 | Old |
| 0.175856 | 1.808443 | 0.042 | 0.015 | 1 | 12 | Prune1 | 0.027 | Old |
| 0.175856 | 1.659902 | 0.042 | 0.015 | 1 | 12 | Faf1 | 0.027 | Old |
| 0.175856 | 1.642117 | 0.042 | 0.015 | 1 | 12 | Pid1 | 0.027 | Old |
| 0.175856 | 1.419208 | 0.042 | 0.015 | 1 | 12 | Sf3a3 | 0.027 | Old |
| 0.176159 | 0.639003 | 0.195 | 0.131 | 1 | 12 | Zcchc17 | 0.064 | Old |
| 0.176271 | 0.795404 | 0.076 | 0.036 | 1 | 12 | Unc119 | 0.04 | Old |
| 0.176271 | 0.728937 | 0.076 | 0.036 | 1 | 12 | Mt1 | 0.04 | Old |
| 0.177175 | 0.297299 | 0.508 | 0.416 | 1 | 12 | Fkbp1a | 0.092 | Old |
| 0.177784 | 1.643587 | 0.042 | 0.015 | 1 | 12 | Endog | 0.027 | Old |
| 0.177784 | 1.544794 | 0.042 | 0.015 | 1 | 12 | Ggt7 | 0.027 | Old |
| 0.177784 | 1.521294 | 0.042 | 0.015 | 1 | 12 | Tnrc6b | 0.027 | Old |
| 0.177784 | 1.493949 | 0.042 | 0.015 | 1 | 12 | Mcts2 | 0.027 | Old |
| 0.177784 | 1.427908 | 0.042 | 0.015 | 1 | 12 | Cks1b | 0.027 | Old |
| 0.177784 | 1.337477 | 0.042 | 0.015 | 1 | 12 | Usp10 | 0.027 | Old |
| 0.177784 | 1.319837 | 0.042 | 0.015 | 1 | 12 | Bbx | 0.027 | Old |
| 0.178753 | 1.491375 | 0.042 | 0.015 | 1 | 12 | Pex19 | 0.027 | Old |
| 0.17928 | 0.963292 | 0.085 | 0.044 | 1 | 12 | Mfsd6 | 0.041 | Old |
| 0.179651 | 0.685538 | 0.153 | 0.102 | 1 | 12 | Tomm7 | 0.051 | Old |
| 0.179728 | 1.560062 | 0.042 | 0.015 | 1 | 12 | Ly6e | 0.027 | Old |
| 0.179728 | 1.558084 | 0.042 | 0.015 | 1 | 12 | Snhg12 | 0.027 | Old |
| 0.179728 | 1.520142 | 0.042 | 0.015 | 1 | 12 | Dync2li1 | 0.027 | Old |
| 0.179728 | 1.470927 | 0.042 | 0.015 | 1 | 12 | Myh10 | 0.027 | Old |
| 0.179728 | 1.43303 | 0.042 | 0.015 | 1 | 12 | 1700123O2 | 0.027 | Old |
| 0.179728 | 1.408138 | 0.042 | 0.015 | 1 | 12 | Nubp2 | 0.027 | Old |
| 0.179728 | 1.407177 | 0.042 | 0.015 | 1 | 12 | Slc38a10 | 0.027 | Old |
| 0.179728 | 1.355607 | 0.042 | 0.015 | 1 | 12 | St6galnac5 | 0.027 | Old |
| 0.179728 | 1.325796 | 0.042 | 0.015 | 1 | 12 | Fbxw9 | 0.027 | Old |
| 0.179728 | 1.325492 | 0.042 | 0.015 | 1 | 12 | Ctsa | 0.027 | Old |
| 0.179728 | 1.299179 | 0.042 | 0.015 | 1 | 12 | Aldh9a1 | 0.027 | Old |
| 0.179728 | 1.177356 | 0.042 | 0.015 | 1 | 12 | Vezf1 | 0.027 | Old |
| 0.179728 | 0.734567 | 0.042 | 0.015 | 1 | 12 | Tango2 | 0.027 | Old |
| 0.181104 | 0.607519 | 0.161 | 0.109 | 1 | 12 | Cops9 | 0.052 | Old |
| 0.18122 | 0.456484 | 0.237 | 0.168 | 1 | 12 | Tmem59 | 0.069 | Old |
| 0.18149 | 1.093814 | 0.119 | 0.073 | 1 | 12 | Rgs4 | 0.046 | Old |

| | | | | | | | | |
|---|---|---|---|---|---|---|---|---|
| 0.181688 | 1.398774 | 0.042 | 0.015 | 1 | 12 | Ppat | 0.027 | Old |
| 0.181688 | 1.215448 | 0.042 | 0.015 | 1 | 12 | Tom1 | 0.027 | Old |
| 0.181688 | 1.203686 | 0.042 | 0.015 | 1 | 12 | Tm9sf3 | 0.027 | Old |
| 0.181688 | 1.078834 | 0.042 | 0.015 | 1 | 12 | Smn1 | 0.027 | Old |
| 0.181752 | 0.310203 | 0.297 | 0.219 | 1 | 12 | Hsp90b1 | 0.078 | Old |
| 0.183466 | 0.591441 | 0.161 | 0.109 | 1 | 12 | Dusp26 | 0.052 | Old |
| 0.183663 | 1.2339 | 0.042 | 0.015 | 1 | 12 | Scn2a | 0.027 | Old |
| 0.183663 | 1.229828 | 0.042 | 0.015 | 1 | 12 | Smad1 | 0.027 | Old |
| 0.183663 | 1.082898 | 0.042 | 0.015 | 1 | 12 | Rnf5 | 0.027 | Old |
| 0.183663 | 0.963405 | 0.042 | 0.015 | 1 | 12 | Ethe1 | 0.027 | Old |
| 0.184828 | 0.457347 | 0.271 | 0.204 | 1 | 12 | Arhgdia | 0.067 | Old |
| 0.185364 | 0.812278 | 0.119 | 0.073 | 1 | 12 | Cyhr1 | 0.046 | Old |
| 0.185654 | 1.186385 | 0.042 | 0.015 | 1 | 12 | Cog4 | 0.027 | Old |
| 0.185654 | 1.045464 | 0.042 | 0.015 | 1 | 12 | Trio | 0.027 | Old |
| 0.185744 | 0.607619 | 0.178 | 0.117 | 1 | 12 | Cct3 | 0.061 | Old |
| 0.185923 | 1.033702 | 0.119 | 0.073 | 1 | 12 | H2-Aa | 0.046 | Old |
| 0.185929 | 0.860145 | 0.085 | 0.044 | 1 | 12 | Cops8 | 0.041 | Old |
| 0.186623 | 0.248454 | 0.254 | 0.175 | 1 | 12 | Cacybp | 0.079 | Old |
| 0.1869 | 0.410968 | 0.483 | 0.438 | 1 | 12 | Rpl28 | 0.045 | Old |
| 0.186909 | 0.897074 | 0.127 | 0.08 | 1 | 12 | Idh3a | 0.047 | Old |
| 0.187662 | 1.222667 | 0.042 | 0.015 | 1 | 12 | Hmgn2 | 0.027 | Old |
| 0.187662 | 1.14307 | 0.042 | 0.015 | 1 | 12 | Kcnb1 | 0.027 | Old |
| 0.187662 | 1.10541 | 0.042 | 0.015 | 1 | 12 | Tor1a | 0.027 | Old |
| 0.187662 | 0.740066 | 0.042 | 0.015 | 1 | 12 | H1f0 | 0.027 | Old |
| 0.187662 | 0.725779 | 0.042 | 0.015 | 1 | 12 | Bdh2 | 0.027 | Old |
| 0.187798 | 0.521812 | 0.263 | 0.204 | 1 | 12 | Psma5 | 0.059 | Old |
| 0.188509 | 0.382022 | 0.229 | 0.161 | 1 | 12 | Tsn | 0.068 | Old |
| 0.190005 | 0.811063 | 0.085 | 0.044 | 1 | 12 | Emb | 0.041 | Old |
| 0.191215 | 0.563945 | 0.254 | 0.19 | 1 | 12 | Stub1 | 0.064 | Old |
| 0.191378 | 0.788045 | 0.085 | 0.044 | 1 | 12 | Morn2 | 0.041 | Old |
| 0.193757 | 0.803335 | 0.144 | 0.095 | 1 | 12 | Ccnd1 | 0.049 | Old |
| 0.194146 | 0.722157 | 0.085 | 0.044 | 1 | 12 | Zmynd11 | 0.041 | Old |
| 0.195541 | 0.615165 | 0.085 | 0.044 | 1 | 12 | Egln1 | 0.041 | Old |
| 0.197773 | 0.737633 | 0.136 | 0.088 | 1 | 12 | Cdv3 | 0.048 | Old |
| 0.198236 | 0.809885 | 0.093 | 0.051 | 1 | 12 | Cyb561d2 | 0.042 | Old |
| 0.199577 | 0.819206 | 0.093 | 0.051 | 1 | 12 | Magoh | 0.042 | Old |
| 0.199577 | 0.743266 | 0.093 | 0.051 | 1 | 12 | Snrnp25 | 0.042 | Old |
| 0.199855 | 0.259447 | 0.627 | 0.518 | 1 | 12 | Rpl24 | 0.109 | Old |
| 0.201383 | 0.689622 | 0.127 | 0.08 | 1 | 12 | Napb | 0.047 | Old |
| 0.202039 | 1.40417 | 0.051 | 0.022 | 1 | 12 | Gtpbp4 | 0.029 | Old |
| 0.203929 | 1.347494 | 0.051 | 0.022 | 1 | 12 | Ppp2r3c | 0.029 | Old |
| 0.205832 | 1.867441 | 0.051 | 0.022 | 1 | 12 | Selenop | 0.029 | Old |
| 0.205832 | 1.46944 | 0.051 | 0.022 | 1 | 12 | Cux1 | 0.029 | Old |
| 0.205832 | 1.407788 | 0.051 | 0.022 | 1 | 12 | Rad23b | 0.029 | Old |
| 0.205832 | 1.363982 | 0.051 | 0.022 | 1 | 12 | Cop1 | 0.029 | Old |

| | | | | | | | | |
|---|---|---|---|---|---|---|---|---|
| 0.207748 | 1.423588 | 0.051 | 0.022 | 1 | 12 | H2-Eb1 | 0.029 | Old |
| 0.207748 | 1.356175 | 0.051 | 0.022 | 1 | 12 | Ublcp1 | 0.029 | Old |
| 0.207748 | 1.344861 | 0.051 | 0.022 | 1 | 12 | 7-Mar | 0.029 | Old |
| 0.207748 | 1.246927 | 0.051 | 0.022 | 1 | 12 | Cdc42se2 | 0.029 | Old |
| 0.208923 | 0.626857 | 0.102 | 0.058 | 1 | 12 | Garnl3 | 0.044 | Old |
| 0.209676 | 1.271898 | 0.051 | 0.022 | 1 | 12 | Arhgap35 | 0.029 | Old |
| 0.209676 | 1.195354 | 0.051 | 0.022 | 1 | 12 | Emc3 | 0.029 | Old |
| 0.210653 | 0.330257 | 0.432 | 0.35 | 1 | 12 | Ndufa13 | 0.082 | Old |
| 0.210915 | 0.715597 | 0.102 | 0.058 | 1 | 12 | Gpx1 | 0.044 | Old |
| 0.211618 | 1.451818 | 0.051 | 0.022 | 1 | 12 | Rabl6 | 0.029 | Old |
| 0.211618 | 1.256314 | 0.051 | 0.022 | 1 | 12 | Abcd3 | 0.029 | Old |
| 0.212703 | 0.742599 | 0.186 | 0.139 | 1 | 12 | Cd63 | 0.047 | Old |
| 0.213572 | 1.304663 | 0.051 | 0.022 | 1 | 12 | Agpat4 | 0.029 | Old |
| 0.213572 | 1.265361 | 0.051 | 0.022 | 1 | 12 | Ypel5 | 0.029 | Old |
| 0.213572 | 1.210758 | 0.051 | 0.022 | 1 | 12 | Upp2 | 0.029 | Old |
| 0.213572 | 1.183458 | 0.051 | 0.022 | 1 | 12 | Babam2 | 0.029 | Old |
| 0.213572 | 1.108702 | 0.051 | 0.022 | 1 | 12 | Lmbrd1 | 0.029 | Old |
| 0.213572 | 1.019482 | 0.051 | 0.022 | 1 | 12 | Nufip2 | 0.029 | Old |
| 0.213747 | 0.180413 | 0.864 | 0.891 | 1 | 12 | Stmn3 | -0.027 | Old |
| 0.214214 | 0.68502 | 0.119 | 0.073 | 1 | 12 | Cfap36 | 0.046 | Old |
| 0.214496 | 0.475191 | 0.271 | 0.204 | 1 | 12 | Sub1 | 0.067 | Old |
| 0.214554 | 1.251408 | 0.051 | 0.022 | 1 | 12 | Dcaf11 | 0.029 | Old |
| 0.21494 | 0.627672 | 0.102 | 0.058 | 1 | 12 | Fam213b | 0.044 | Old |
| 0.215157 | 0.748958 | 0.186 | 0.139 | 1 | 12 | 7-Sep | 0.047 | Old |
| 0.21554 | 1.129525 | 0.051 | 0.022 | 1 | 12 | Rnf8 | 0.029 | Old |
| 0.215812 | 0.577769 | 0.195 | 0.139 | 1 | 12 | Lmo1 | 0.056 | Old |
| 0.21752 | 1.33782 | 0.051 | 0.022 | 1 | 12 | Rcn2 | 0.029 | Old |
| 0.21752 | 1.082967 | 0.051 | 0.022 | 1 | 12 | Pja1 | 0.029 | Old |
| 0.21752 | 1.063431 | 0.051 | 0.022 | 1 | 12 | Rnf149 | 0.029 | Old |
| 0.21752 | 1.013565 | 0.051 | 0.022 | 1 | 12 | Ebag9 | 0.029 | Old |
| 0.21752 | 0.854025 | 0.051 | 0.022 | 1 | 12 | Msrb1 | 0.029 | Old |
| 0.219407 | 0.633556 | 0.186 | 0.131 | 1 | 12 | Mrpl57 | 0.055 | Old |
| 0.219514 | 0.938437 | 0.051 | 0.022 | 1 | 12 | Nrcam | 0.029 | Old |
| 0.219514 | 0.914279 | 0.051 | 0.022 | 1 | 12 | Frg1 | 0.029 | Old |
| 0.219683 | 0.671303 | 0.11 | 0.066 | 1 | 12 | Ss18l2 | 0.044 | Old |
| 0.220025 | 0.225677 | 0.398 | 0.314 | 1 | 12 | Vdac2 | 0.084 | Old |
| 0.224633 | 0.413742 | 0.203 | 0.146 | 1 | 12 | Ube2v2 | 0.057 | Old |
| 0.225442 | 0.270167 | 0.449 | 0.394 | 1 | 12 | Pfn2 | 0.055 | Old |
| 0.225573 | 0.453959 | 0.051 | 0.022 | 1 | 12 | Ahsa2 | 0.029 | Old |
| 0.225846 | 0.29256 | 0.356 | 0.285 | 1 | 12 | Tagln2 | 0.071 | Old |
| 0.226525 | 1.504446 | 0.059 | 0.029 | 1 | 12 | Lias | 0.03 | Old |
| 0.226525 | 1.303633 | 0.059 | 0.029 | 1 | 12 | Ccser2 | 0.03 | Old |
| 0.226525 | 1.254288 | 0.059 | 0.029 | 1 | 12 | Dctn4 | 0.03 | Old |
| 0.227619 | 0.995156 | 0.051 | 0.022 | 1 | 12 | Kpnb1 | 0.029 | Old |
| 0.227619 | 0.870567 | 0.051 | 0.022 | 1 | 12 | Ccdc115 | 0.029 | Old |

| | | | | | | | | |
|---|---|---|---|---|---|---|---|---|
| 0.228389 | 1.489344 | 0.059 | 0.029 | 1 | 12 | 9130401M | 0.03 | Old |
| 0.228389 | 1.460115 | 0.059 | 0.029 | 1 | 12 | Cnep1r1 | 0.03 | Old |
| 0.229724 | 0.656667 | 0.229 | 0.182 | 1 | 12 | Psap | 0.047 | Old |
| 0.231751 | 0.689982 | 0.051 | 0.022 | 1 | 12 | Nipa2 | 0.029 | Old |
| 0.232125 | 0.492655 | 0.203 | 0.146 | 1 | 12 | Cct5 | 0.057 | Old |
| 0.232147 | 1.454233 | 0.059 | 0.029 | 1 | 12 | Cnih4 | 0.03 | Old |
| 0.232147 | 1.156929 | 0.059 | 0.029 | 1 | 12 | Rap2a | 0.03 | Old |
| 0.232147 | 1.14386 | 0.059 | 0.029 | 1 | 12 | Zfand3 | 0.03 | Old |
| 0.232147 | 1.099608 | 0.059 | 0.029 | 1 | 12 | Trabd | 0.03 | Old |
| 0.232383 | 0.363303 | 0.195 | 0.139 | 1 | 12 | Zcrb1 | 0.056 | Old |
| 0.23381 | 1.398332 | 0.068 | 0.036 | 1 | 12 | Nop56 | 0.032 | Old |
| 0.23381 | 1.260045 | 0.068 | 0.036 | 1 | 12 | Hnrnpl | 0.032 | Old |
| 0.234043 | 1.12542 | 0.059 | 0.029 | 1 | 12 | Tex264 | 0.03 | Old |
| 0.23595 | 1.115491 | 0.059 | 0.029 | 1 | 12 | Cyp20a1 | 0.03 | Old |
| 0.23626 | 0.603191 | 0.136 | 0.088 | 1 | 12 | Mrpl28 | 0.048 | Old |
| 0.236305 | 0.826534 | 0.169 | 0.124 | 1 | 12 | Hspb1 | 0.045 | Old |
| 0.237731 | 0.364637 | 0.203 | 0.146 | 1 | 12 | Mrps24 | 0.057 | Old |
| 0.237868 | 1.432874 | 0.059 | 0.029 | 1 | 12 | Idh3g | 0.03 | Old |
| 0.237868 | 1.095805 | 0.059 | 0.029 | 1 | 12 | Akt1s1 | 0.03 | Old |
| 0.237868 | 1.091467 | 0.059 | 0.029 | 1 | 12 | Pgam5 | 0.03 | Old |
| 0.237868 | 1.001524 | 0.059 | 0.029 | 1 | 12 | Pgd | 0.03 | Old |
| 0.238082 | 0.409149 | 0.22 | 0.161 | 1 | 12 | Serinc1 | 0.059 | Old |
| 0.239285 | 0.362806 | 0.483 | 0.453 | 1 | 12 | Bsg | 0.03 | Old |
| 0.239399 | 0.486987 | 0.178 | 0.124 | 1 | 12 | Brk1 | 0.054 | Old |
| 0.239796 | 0.751889 | 0.059 | 0.029 | 1 | 12 | Rpn2 | 0.03 | Old |
| 0.240086 | 0.12926 | 0.458 | 0.358 | 1 | 12 | Rpl38 | 0.1 | Old |
| 0.240904 | 1.186171 | 0.068 | 0.036 | 1 | 12 | Acads | 0.032 | Old |
| 0.24159 | 0.367499 | 0.297 | 0.226 | 1 | 12 | Prdx5 | 0.071 | Old |
| 0.241736 | 0.928878 | 0.059 | 0.029 | 1 | 12 | Msl1 | 0.03 | Old |
| 0.241736 | 0.861083 | 0.059 | 0.029 | 1 | 12 | Snrpb2 | 0.03 | Old |
| 0.241736 | 0.848485 | 0.059 | 0.029 | 1 | 12 | Amfr | 0.03 | Old |
| 0.241736 | 0.801335 | 0.059 | 0.029 | 1 | 12 | Fam136a | 0.03 | Old |
| 0.242087 | 1.094354 | 0.102 | 0.066 | 1 | 12 | Pdlim2 | 0.036 | Old |
| 0.242153 | 0.36408 | 0.237 | 0.175 | 1 | 12 | Thra | 0.062 | Old |
| 0.243227 | 2.337495 | 0.025 | 0.007 | 1 | 12 | Snph | 0.018 | Old |
| 0.243229 | 2.574578 | 0.025 | 0.007 | 1 | 12 | Tnfrsf12a | 0.018 | Old |
| 0.243229 | 2.537831 | 0.025 | 0.007 | 1 | 12 | Grik5 | 0.018 | Old |
| 0.243229 | 2.483584 | 0.025 | 0.007 | 1 | 12 | Caap1 | 0.018 | Old |
| 0.243229 | 2.342945 | 0.025 | 0.007 | 1 | 12 | Mettl16 | 0.018 | Old |
| 0.243229 | 2.333603 | 0.025 | 0.007 | 1 | 12 | Gtf2f2 | 0.018 | Old |
| 0.243229 | 2.329385 | 0.025 | 0.007 | 1 | 12 | H2afx | 0.018 | Old |
| 0.243229 | 2.289517 | 0.025 | 0.007 | 1 | 12 | Kcna6 | 0.018 | Old |
| 0.243229 | 2.27952 | 0.025 | 0.007 | 1 | 12 | Hid1 | 0.018 | Old |
| 0.243229 | 2.259069 | 0.025 | 0.007 | 1 | 12 | Zfp60 | 0.018 | Old |
| 0.243229 | 2.234087 | 0.025 | 0.007 | 1 | 12 | Dnajc27 | 0.018 | Old |

| | | | | | | | | |
|---|---|---|---|---|---|---|---|---|
| 0.243229 | 2.232585 | 0.025 | 0.007 | 1 | 12 | 1810030O( | 0.018 | Old |
| 0.243229 | 2.18778 | 0.025 | 0.007 | 1 | 12 | Gpr107 | 0.018 | Old |
| 0.243229 | 2.178214 | 0.025 | 0.007 | 1 | 12 | Lrwd1 | 0.018 | Old |
| 0.243229 | 2.157552 | 0.025 | 0.007 | 1 | 12 | Lrrc57 | 0.018 | Old |
| 0.243229 | 2.137458 | 0.025 | 0.007 | 1 | 12 | Ttyh3 | 0.018 | Old |
| 0.243229 | 2.067091 | 0.025 | 0.007 | 1 | 12 | Dtx2 | 0.018 | Old |
| 0.243229 | 2.048154 | 0.025 | 0.007 | 1 | 12 | Asf1a | 0.018 | Old |
| 0.243229 | 2.031926 | 0.025 | 0.007 | 1 | 12 | Dxo | 0.018 | Old |
| 0.243229 | 2.019794 | 0.025 | 0.007 | 1 | 12 | Ttc8 | 0.018 | Old |
| 0.243229 | 1.994817 | 0.025 | 0.007 | 1 | 12 | Pir | 0.018 | Old |
| 0.243229 | 1.991282 | 0.025 | 0.007 | 1 | 12 | Agfg2 | 0.018 | Old |
| 0.243229 | 1.990809 | 0.025 | 0.007 | 1 | 12 | Mysm1 | 0.018 | Old |
| 0.243229 | 1.982717 | 0.025 | 0.007 | 1 | 12 | Zmym2 | 0.018 | Old |
| 0.243229 | 1.959296 | 0.025 | 0.007 | 1 | 12 | Gm20604 | 0.018 | Old |
| 0.243229 | 1.955355 | 0.025 | 0.007 | 1 | 12 | Id2 | 0.018 | Old |
| 0.243229 | 1.910977 | 0.025 | 0.007 | 1 | 12 | Cbx1 | 0.018 | Old |
| 0.243229 | 1.905117 | 0.025 | 0.007 | 1 | 12 | Mnt | 0.018 | Old |
| 0.243687 | 0.863183 | 0.059 | 0.029 | 1 | 12 | Taok1 | 0.03 | Old |
| 0.243687 | 0.739405 | 0.059 | 0.029 | 1 | 12 | Azin1 | 0.03 | Old |
| 0.244431 | 0.493001 | 0.28 | 0.219 | 1 | 12 | Cct7 | 0.061 | Old |
| 0.244828 | 1.979997 | 0.025 | 0.007 | 1 | 12 | Ttc7b | 0.018 | Old |
| 0.245649 | 0.880739 | 0.059 | 0.029 | 1 | 12 | Nacad | 0.03 | Old |
| 0.246322 | 1.044893 | 0.068 | 0.036 | 1 | 12 | Cntfr | 0.032 | Old |
| 0.246439 | 2.186686 | 0.025 | 0.007 | 1 | 12 | Lsg1 | 0.018 | Old |
| 0.246439 | 2.154535 | 0.025 | 0.007 | 1 | 12 | Nr3c1 | 0.018 | Old |
| 0.246439 | 2.076775 | 0.025 | 0.007 | 1 | 12 | St6galnac6 | 0.018 | Old |
| 0.246439 | 2.049222 | 0.025 | 0.007 | 1 | 12 | Hectd4 | 0.018 | Old |
| 0.246439 | 2.017946 | 0.025 | 0.007 | 1 | 12 | Gfod1 | 0.018 | Old |
| 0.246439 | 1.974418 | 0.025 | 0.007 | 1 | 12 | Fam173b | 0.018 | Old |
| 0.246439 | 1.969397 | 0.025 | 0.007 | 1 | 12 | Zfp445 | 0.018 | Old |
| 0.246439 | 1.960143 | 0.025 | 0.007 | 1 | 12 | Ppp2cb | 0.018 | Old |
| 0.246439 | 1.940194 | 0.025 | 0.007 | 1 | 12 | Reep3 | 0.018 | Old |
| 0.246439 | 1.929457 | 0.025 | 0.007 | 1 | 12 | Unc93b1 | 0.018 | Old |
| 0.246439 | 1.902549 | 0.025 | 0.007 | 1 | 12 | Dpagt1 | 0.018 | Old |
| 0.246439 | 1.891809 | 0.025 | 0.007 | 1 | 12 | Cept1 | 0.018 | Old |
| 0.246439 | 1.875672 | 0.025 | 0.007 | 1 | 12 | Cd3eap | 0.018 | Old |
| 0.246439 | 1.862733 | 0.025 | 0.007 | 1 | 12 | Dpf2 | 0.018 | Old |
| 0.246439 | 1.852685 | 0.025 | 0.007 | 1 | 12 | Gm10762 | 0.018 | Old |
| 0.246439 | 1.839272 | 0.025 | 0.007 | 1 | 12 | Pgpep1 | 0.018 | Old |
| 0.246439 | 1.82114 | 0.025 | 0.007 | 1 | 12 | Smarcd2 | 0.018 | Old |
| 0.246439 | 1.817364 | 0.025 | 0.007 | 1 | 12 | Ccng1 | 0.018 | Old |
| 0.246439 | 1.817235 | 0.025 | 0.007 | 1 | 12 | Armcx4 | 0.018 | Old |
| 0.246439 | 1.809966 | 0.025 | 0.007 | 1 | 12 | Whamm | 0.018 | Old |
| 0.246439 | 1.778915 | 0.025 | 0.007 | 1 | 12 | Kcnh1 | 0.018 | Old |
| 0.246439 | 1.774193 | 0.025 | 0.007 | 1 | 12 | Dtx1 | 0.018 | Old |

| | | | | | | | | |
|---|---|---|---|---|---|---|---|---|
| 0.246439 | 1.761831 | 0.025 | 0.007 | 1 | 12 | Sugp1 | 0.018 | Old |
| 0.246439 | 1.755099 | 0.025 | 0.007 | 1 | 12 | Glyr1 | 0.018 | Old |
| 0.246439 | 1.74887 | 0.025 | 0.007 | 1 | 12 | Gm20300 | 0.018 | Old |
| 0.246439 | 1.737446 | 0.025 | 0.007 | 1 | 12 | Slc38a7 | 0.018 | Old |
| 0.246439 | 1.733859 | 0.025 | 0.007 | 1 | 12 | Ptprt | 0.018 | Old |
| 0.246439 | 1.726243 | 0.025 | 0.007 | 1 | 12 | Ggct | 0.018 | Old |
| 0.246439 | 1.718761 | 0.025 | 0.007 | 1 | 12 | Dbndd2 | 0.018 | Old |
| 0.246439 | 1.710229 | 0.025 | 0.007 | 1 | 12 | Zfp523 | 0.018 | Old |
| 0.246439 | 1.701521 | 0.025 | 0.007 | 1 | 12 | Stk11 | 0.018 | Old |
| 0.246439 | 1.700016 | 0.025 | 0.007 | 1 | 12 | Mettl2 | 0.018 | Old |
| 0.246439 | 1.686727 | 0.025 | 0.007 | 1 | 12 | Fam171a1 | 0.018 | Old |
| 0.246439 | 1.685546 | 0.025 | 0.007 | 1 | 12 | Rnf123 | 0.018 | Old |
| 0.246439 | 1.685411 | 0.025 | 0.007 | 1 | 12 | Rnf19b | 0.018 | Old |
| 0.246439 | 1.682475 | 0.025 | 0.007 | 1 | 12 | Uimc1 | 0.018 | Old |
| 0.246439 | 1.661075 | 0.025 | 0.007 | 1 | 12 | Fam199x | 0.018 | Old |
| 0.246439 | 1.634006 | 0.025 | 0.007 | 1 | 12 | Lancl2 | 0.018 | Old |
| 0.246439 | 1.60958 | 0.025 | 0.007 | 1 | 12 | Rbm4b | 0.018 | Old |
| 0.246439 | 1.603063 | 0.025 | 0.007 | 1 | 12 | Dyrk2 | 0.018 | Old |
| 0.246439 | 1.601758 | 0.025 | 0.007 | 1 | 12 | Rab11fip5 | 0.018 | Old |
| 0.246439 | 1.57249 | 0.025 | 0.007 | 1 | 12 | Rcan1 | 0.018 | Old |
| 0.246439 | 1.561189 | 0.025 | 0.007 | 1 | 12 | Celf1 | 0.018 | Old |
| 0.246439 | 1.556483 | 0.025 | 0.007 | 1 | 12 | 2310057M | 0.018 | Old |
| 0.247296 | 0.340727 | 0.254 | 0.197 | 1 | 12 | Cfl2 | 0.057 | Old |
| 0.247621 | 0.950566 | 0.059 | 0.029 | 1 | 12 | Tbx20 | 0.03 | Old |
| 0.247621 | 0.831597 | 0.059 | 0.029 | 1 | 12 | Pcmtd1 | 0.03 | Old |
| 0.247621 | 0.747528 | 0.059 | 0.029 | 1 | 12 | Fam103a1 | 0.03 | Old |
| 0.249678 | 1.981471 | 0.025 | 0.007 | 1 | 12 | Pitrm1 | 0.018 | Old |
| 0.249678 | 1.84765 | 0.025 | 0.007 | 1 | 12 | Zc3hc1 | 0.018 | Old |
| 0.249678 | 1.794369 | 0.025 | 0.007 | 1 | 12 | Mrps9 | 0.018 | Old |
| 0.249678 | 1.784528 | 0.025 | 0.007 | 1 | 12 | Slc25a37 | 0.018 | Old |
| 0.249678 | 1.779874 | 0.025 | 0.007 | 1 | 12 | Gmcl1 | 0.018 | Old |
| 0.249678 | 1.752304 | 0.025 | 0.007 | 1 | 12 | Gins4 | 0.018 | Old |
| 0.249678 | 1.712715 | 0.025 | 0.007 | 1 | 12 | Osbp | 0.018 | Old |
| 0.249678 | 1.709881 | 0.025 | 0.007 | 1 | 12 | Pwp1 | 0.018 | Old |
| 0.249678 | 1.688364 | 0.025 | 0.007 | 1 | 12 | Rpp25l | 0.018 | Old |
| 0.249678 | 1.683016 | 0.025 | 0.007 | 1 | 12 | Slc25a19 | 0.018 | Old |
| 0.249678 | 1.67642 | 0.025 | 0.007 | 1 | 12 | Zfp414 | 0.018 | Old |
| 0.249678 | 1.666656 | 0.025 | 0.007 | 1 | 12 | Wdr5 | 0.018 | Old |
| 0.249678 | 1.665287 | 0.025 | 0.007 | 1 | 12 | Exosc1 | 0.018 | Old |
| 0.249678 | 1.652016 | 0.025 | 0.007 | 1 | 12 | Rpp30 | 0.018 | Old |
| 0.249678 | 1.650387 | 0.025 | 0.007 | 1 | 12 | Cd209g | 0.018 | Old |
| 0.249678 | 1.64751 | 0.025 | 0.007 | 1 | 12 | Fgf1 | 0.018 | Old |
| 0.249678 | 1.621803 | 0.025 | 0.007 | 1 | 12 | 3110040N1 | 0.018 | Old |
| 0.249678 | 1.606627 | 0.025 | 0.007 | 1 | 12 | Ankrd28 | 0.018 | Old |
| 0.249678 | 1.586682 | 0.025 | 0.007 | 1 | 12 | Rab35 | 0.018 | Old |

| | | | | | | | | |
|---|---|---|---|---|---|---|---|---|
| 0.249678 | 1.582442 | 0.025 | 0.007 | 1 | 12 | 5730455P1 | 0.018 | Old |
| 0.249678 | 1.580612 | 0.025 | 0.007 | 1 | 12 | Arrb1 | 0.018 | Old |
| 0.249678 | 1.57992 | 0.025 | 0.007 | 1 | 12 | Cfh | 0.018 | Old |
| 0.249678 | 1.5768 | 0.025 | 0.007 | 1 | 12 | S100a16 | 0.018 | Old |
| 0.249678 | 1.567125 | 0.025 | 0.007 | 1 | 12 | Cyp4f13 | 0.018 | Old |
| 0.249678 | 1.564136 | 0.025 | 0.007 | 1 | 12 | Zfp704 | 0.018 | Old |
| 0.249678 | 1.552062 | 0.025 | 0.007 | 1 | 12 | Rps6ka3 | 0.018 | Old |
| 0.249678 | 1.528686 | 0.025 | 0.007 | 1 | 12 | Cnnm4 | 0.018 | Old |
| 0.249678 | 1.503453 | 0.025 | 0.007 | 1 | 12 | Lmf1 | 0.018 | Old |
| 0.249678 | 1.463514 | 0.025 | 0.007 | 1 | 12 | Nyap1 | 0.018 | Old |
| 0.249678 | 1.44433 | 0.025 | 0.007 | 1 | 12 | Qk | 0.018 | Old |
| 0.249678 | 1.419982 | 0.025 | 0.007 | 1 | 12 | Ireb2 | 0.018 | Old |
| 0.249678 | 1.365653 | 0.025 | 0.007 | 1 | 12 | Zfp292 | 0.018 | Old |
| 0.249678 | 1.329355 | 0.025 | 0.007 | 1 | 12 | Litaf | 0.018 | Old |
| 0.249678 | 1.323686 | 0.025 | 0.007 | 1 | 12 | Ipo7 | 0.018 | Old |
| 0.249678 | 1.321395 | 0.025 | 0.007 | 1 | 12 | Pno1 | 0.018 | Old |
| 0.249678 | 1.304831 | 0.025 | 0.007 | 1 | 12 | Man2b1 | 0.018 | Old |
| 0.249678 | 1.236616 | 0.025 | 0.007 | 1 | 12 | Abl2 | 0.018 | Old |
| 0.249678 | 1.182641 | 0.025 | 0.007 | 1 | 12 | Trp53bp1 | 0.018 | Old |
| 0.249678 | 1.100374 | 0.025 | 0.007 | 1 | 12 | Ier5 | 0.018 | Old |
| 0.249678 | 1.08909 | 0.025 | 0.007 | 1 | 12 | Letm2 | 0.018 | Old |
| 0.249678 | 0.9527 | 0.025 | 0.007 | 1 | 12 | Arhgap1 | 0.018 | Old |
| 0.249777 | 0.227101 | 0.466 | 0.409 | 1 | 12 | Rps14 | 0.057 | Old |
| 0.249982 | 0.981927 | 0.068 | 0.036 | 1 | 12 | Ash2l | 0.032 | Old |
| 0.250928 | 0.500041 | 0.229 | 0.175 | 1 | 12 | Cfdp1 | 0.054 | Old |
| 0.251826 | 0.980098 | 0.068 | 0.036 | 1 | 12 | Uhmk1 | 0.032 | Old |
| 0.251826 | 0.876003 | 0.068 | 0.036 | 1 | 12 | Rnf10 | 0.032 | Old |
| 0.252403 | 0.750104 | 0.136 | 0.095 | 1 | 12 | Fam171b | 0.041 | Old |
| 0.252613 | 1.083917 | 0.085 | 0.051 | 1 | 12 | Tusc2 | 0.034 | Old |
| 0.252948 | 1.628901 | 0.025 | 0.007 | 1 | 12 | Pex14 | 0.018 | Old |
| 0.252948 | 1.560076 | 0.025 | 0.007 | 1 | 12 | Gm5617 | 0.018 | Old |
| 0.252948 | 1.48229 | 0.025 | 0.007 | 1 | 12 | Kcnc1 | 0.018 | Old |
| 0.252948 | 1.480224 | 0.025 | 0.007 | 1 | 12 | AW549877 | 0.018 | Old |
| 0.252948 | 1.469047 | 0.025 | 0.007 | 1 | 12 | B4galnt4 | 0.018 | Old |
| 0.252948 | 1.461417 | 0.025 | 0.007 | 1 | 12 | Tmem184b | 0.018 | Old |
| 0.252948 | 1.456622 | 0.025 | 0.007 | 1 | 12 | Map3k7 | 0.018 | Old |
| 0.252948 | 1.441261 | 0.025 | 0.007 | 1 | 12 | Pnpo | 0.018 | Old |
| 0.252948 | 1.44092 | 0.025 | 0.007 | 1 | 12 | Ccdc58 | 0.018 | Old |
| 0.252948 | 1.433629 | 0.025 | 0.007 | 1 | 12 | Rims4 | 0.018 | Old |
| 0.252948 | 1.419204 | 0.025 | 0.007 | 1 | 12 | Ccnl1 | 0.018 | Old |
| 0.252948 | 1.416857 | 0.025 | 0.007 | 1 | 12 | Zdhhc16 | 0.018 | Old |
| 0.252948 | 1.408224 | 0.025 | 0.007 | 1 | 12 | Lcp1 | 0.018 | Old |
| 0.252948 | 1.393908 | 0.025 | 0.007 | 1 | 12 | Wdtc1 | 0.018 | Old |
| 0.252948 | 1.358887 | 0.025 | 0.007 | 1 | 12 | Fcer1g | 0.018 | Old |
| 0.252948 | 1.333363 | 0.025 | 0.007 | 1 | 12 | Svip | 0.018 | Old |

| | | | | | | | | |
|---|---|---|---|---|---|---|---|---|
| 0.252948 | 1.310857 | 0.025 | 0.007 | 1 | 12 | Clec10a | 0.018 | Old |
| 0.252948 | 1.238057 | 0.025 | 0.007 | 1 | 12 | Usp16 | 0.018 | Old |
| 0.252948 | 1.19781 | 0.025 | 0.007 | 1 | 12 | Casp9 | 0.018 | Old |
| 0.252948 | 1.195727 | 0.025 | 0.007 | 1 | 12 | 1110046J0 | 0.018 | Old |
| 0.252948 | 1.182642 | 0.025 | 0.007 | 1 | 12 | Tmem38b | 0.018 | Old |
| 0.252948 | 1.154299 | 0.025 | 0.007 | 1 | 12 | Nip7 | 0.018 | Old |
| 0.252948 | 1.144826 | 0.025 | 0.007 | 1 | 12 | Ngly1 | 0.018 | Old |
| 0.252948 | 1.140267 | 0.025 | 0.007 | 1 | 12 | Dlx1 | 0.018 | Old |
| 0.252948 | 1.113331 | 0.025 | 0.007 | 1 | 12 | Amacr | 0.018 | Old |
| 0.252948 | 1.10114 | 0.025 | 0.007 | 1 | 12 | Lox | 0.018 | Old |
| 0.252948 | 1.070923 | 0.025 | 0.007 | 1 | 12 | Taok3 | 0.018 | Old |
| 0.252948 | 1.056353 | 0.025 | 0.007 | 1 | 12 | Osbpl1a | 0.018 | Old |
| 0.252948 | 1.042047 | 0.025 | 0.007 | 1 | 12 | Vps51 | 0.018 | Old |
| 0.252948 | 0.948453 | 0.025 | 0.007 | 1 | 12 | Skap2 | 0.018 | Old |
| 0.252948 | 0.945724 | 0.025 | 0.007 | 1 | 12 | Samhd1 | 0.018 | Old |
| 0.252948 | 0.941899 | 0.025 | 0.007 | 1 | 12 | Rrp9 | 0.018 | Old |
| 0.252948 | 0.938483 | 0.025 | 0.007 | 1 | 12 | Ccnt2 | 0.018 | Old |
| 0.252948 | 0.936548 | 0.025 | 0.007 | 1 | 12 | Lrrc41 | 0.018 | Old |
| 0.252948 | 0.870249 | 0.025 | 0.007 | 1 | 12 | Wsb1 | 0.018 | Old |
| 0.252948 | 0.768015 | 0.025 | 0.007 | 1 | 12 | Paqr7 | 0.018 | Old |
| 0.252948 | 0.724278 | 0.025 | 0.007 | 1 | 12 | Trove2 | 0.018 | Old |
| 0.252948 | 0.713277 | 0.025 | 0.007 | 1 | 12 | Ptprc | 0.018 | Old |
| 0.253679 | 0.877309 | 0.068 | 0.036 | 1 | 12 | Fubp1 | 0.032 | Old |
| 0.254241 | 0.638579 | 0.153 | 0.109 | 1 | 12 | Dync1i2 | 0.044 | Old |
| 0.2543 | 0.92088 | 0.076 | 0.044 | 1 | 12 | Gnb5 | 0.032 | Old |
| 0.255542 | 0.975762 | 0.068 | 0.036 | 1 | 12 | Atp6v0a1 | 0.032 | Old |
| 0.25717 | 0.278921 | 0.263 | 0.197 | 1 | 12 | Dnajb6 | 0.066 | Old |
| 0.257414 | 0.909143 | 0.068 | 0.036 | 1 | 12 | Akt1 | 0.032 | Old |
| 0.257414 | 0.810513 | 0.068 | 0.036 | 1 | 12 | Eef1d | 0.032 | Old |
| 0.257517 | 0.350446 | 0.398 | 0.35 | 1 | 12 | Ran | 0.048 | Old |
| 0.258354 | 0.874881 | 0.068 | 0.036 | 1 | 12 | Ascc1 | 0.032 | Old |
| 0.259296 | 0.921513 | 0.068 | 0.036 | 1 | 12 | Pvt1 | 0.032 | Old |
| 0.259296 | 0.646661 | 0.068 | 0.036 | 1 | 12 | Rrbp1 | 0.032 | Old |
| 0.260774 | 0.308788 | 0.466 | 0.409 | 1 | 12 | Cox7a2 | 0.057 | Old |
| 0.260852 | 0.498432 | 0.195 | 0.139 | 1 | 12 | Aig1 | 0.056 | Old |
| 0.260871 | 0.974667 | 0.085 | 0.051 | 1 | 12 | Sin3b | 0.034 | Old |
| 0.261188 | 0.917718 | 0.068 | 0.036 | 1 | 12 | Cstb | 0.032 | Old |
| 0.261742 | 0.573962 | 0.059 | 0.029 | 1 | 12 | Ctps2 | 0.03 | Old |
| 0.262545 | 0.89154 | 0.085 | 0.051 | 1 | 12 | Naa38 | 0.034 | Old |
| 0.262808 | 0.789935 | 0.102 | 0.066 | 1 | 12 | Naxe | 0.036 | Old |
| 0.26288 | 0.108098 | 0.992 | 0.985 | 1 | 12 | Tuba1a | 0.007 | Old |
| 0.263089 | 0.748092 | 0.068 | 0.036 | 1 | 12 | Bpnt1 | 0.032 | Old |
| 0.263089 | 0.670144 | 0.068 | 0.036 | 1 | 12 | Arf3 | 0.032 | Old |
| 0.263102 | 0.905036 | 0.076 | 0.044 | 1 | 12 | Rab5c | 0.032 | Old |
| 0.266682 | 0.901517 | 0.076 | 0.044 | 1 | 12 | Me1 | 0.032 | Old |

| | | | | | | | | |
|---|---|---|---|---|---|---|---|---|
| 0.266919 | 0.859569 | 0.068 | 0.036 | 1 | 12 | Ttl | 0.032 | Old |
| 0.267882 | 0.705178 | 0.068 | 0.036 | 1 | 12 | Ppp2r5c | 0.032 | Old |
| 0.268484 | 0.838464 | 0.076 | 0.044 | 1 | 12 | Hnrnpdl | 0.032 | Old |
| 0.268849 | 0.767233 | 0.068 | 0.036 | 1 | 12 | Vma21 | 0.032 | Old |
| 0.272037 | 0.4967 | 0.186 | 0.139 | 1 | 12 | Mphosph8 | 0.047 | Old |
| 0.272736 | 0.487255 | 0.068 | 0.036 | 1 | 12 | Rab39b | 0.032 | Old |
| 0.273151 | 0.334262 | 0.22 | 0.168 | 1 | 12 | Snx3 | 0.052 | Old |
| 0.27394 | 0.797214 | 0.076 | 0.044 | 1 | 12 | Dgkb | 0.032 | Old |
| 0.273967 | 0.791724 | 0.093 | 0.058 | 1 | 12 | Arl2bp | 0.035 | Old |
| 0.274469 | 0.976656 | 0.085 | 0.051 | 1 | 12 | Sap18 | 0.034 | Old |
| 0.274694 | 0.704524 | 0.068 | 0.036 | 1 | 12 | Farsb | 0.032 | Old |
| 0.275505 | 0.666797 | 0.136 | 0.095 | 1 | 12 | Ntan1 | 0.041 | Old |
| 0.276202 | 0.973277 | 0.085 | 0.051 | 1 | 12 | Xbp1 | 0.034 | Old |
| 0.276435 | 0.719876 | 0.093 | 0.058 | 1 | 12 | Psmc1 | 0.035 | Old |
| 0.277071 | 0.88966 | 0.085 | 0.051 | 1 | 12 | Dap3 | 0.034 | Old |
| 0.277357 | 0.205979 | 0.737 | 0.642 | 1 | 12 | Atp1b1 | 0.095 | Old |
| 0.277619 | 0.686369 | 0.076 | 0.044 | 1 | 12 | Ninj1 | 0.032 | Old |
| 0.278338 | 0.756415 | 0.102 | 0.066 | 1 | 12 | Sh3glb2 | 0.036 | Old |
| 0.279776 | 0.739172 | 0.119 | 0.08 | 1 | 12 | Top1 | 0.039 | Old |
| 0.280386 | 0.598932 | 0.11 | 0.073 | 1 | 12 | Hdhd2 | 0.037 | Old |
| 0.282623 | 0.673092 | 0.068 | 0.036 | 1 | 12 | Ilf2 | 0.032 | Old |
| 0.283091 | 0.734719 | 0.093 | 0.058 | 1 | 12 | Polb | 0.035 | Old |
| 0.283278 | 0.14767 | 0.847 | 0.81 | 1 | 12 | Calm2 | 0.037 | Old |
| 0.284629 | 0.509471 | 0.068 | 0.036 | 1 | 12 | Cdk5rap3 | 0.032 | Old |
| 0.286333 | 0.576653 | 0.102 | 0.066 | 1 | 12 | Prr13 | 0.036 | Old |
| 0.286539 | 0.682727 | 0.11 | 0.073 | 1 | 12 | Ndufa6 | 0.037 | Old |
| 0.287363 | 0.408204 | 0.195 | 0.146 | 1 | 12 | Sfr1 | 0.049 | Old |
| 0.287614 | 0.305031 | 0.305 | 0.241 | 1 | 12 | Snap25 | 0.064 | Old |
| 0.288385 | 0.361697 | 0.322 | 0.27 | 1 | 12 | Tomm20 | 0.052 | Old |
| 0.289649 | 0.444829 | 0.11 | 0.073 | 1 | 12 | Paics | 0.037 | Old |
| 0.289809 | 0.53914 | 0.076 | 0.044 | 1 | 12 | Dnm3 | 0.032 | Old |
| 0.289933 | 0.371739 | 0.28 | 0.219 | 1 | 12 | D8Ertd738[illegible] | 0.061 | Old |
| 0.291239 | 0.776376 | 0.127 | 0.088 | 1 | 12 | Chgb | 0.039 | Old |
| 0.292672 | 0.626165 | 0.076 | 0.044 | 1 | 12 | Sars | 0.032 | Old |
| 0.294283 | 0.549663 | 0.195 | 0.153 | 1 | 12 | Psmd4 | 0.042 | Old |
| 0.294358 | 0.74873 | 0.11 | 0.073 | 1 | 12 | Atl1 | 0.037 | Old |
| 0.294591 | 0.642152 | 0.076 | 0.044 | 1 | 12 | Vamp7 | 0.032 | Old |
| 0.294998 | 0.607504 | 0.093 | 0.058 | 1 | 12 | Cgref1 | 0.035 | Old |
| 0.295324 | 0.443403 | 0.178 | 0.139 | 1 | 12 | Snf8 | 0.039 | Old |
| 0.295759 | 0.579261 | 0.085 | 0.051 | 1 | 12 | Tmed4 | 0.034 | Old |
| 0.296835 | 0.38837 | 0.322 | 0.277 | 1 | 12 | Oxct1 | 0.045 | Old |
| 0.296925 | 0.378046 | 0.203 | 0.153 | 1 | 12 | Rpl22 | 0.05 | Old |
| 0.297631 | 0.300781 | 0.339 | 0.277 | 1 | 12 | Mgst3 | 0.062 | Old |
| 0.298565 | 0.372921 | 0.144 | 0.102 | 1 | 12 | Sec62 | 0.042 | Old |
| 0.29961 | 0.116456 | 0.915 | 0.927 | 1 | 12 | Tubb2b | -0.012 | Old |

| | | | | | | | | |
|---|---|---|---|---|---|---|---|---|
| 0.300946 | 1.826026 | 0.034 | 0.015 | 1 | 12 | Rmnd5b | 0.019 | Old |
| 0.300946 | 1.715195 | 0.034 | 0.015 | 1 | 12 | Irs2 | 0.019 | Old |
| 0.300946 | 1.622407 | 0.034 | 0.015 | 1 | 12 | Ppcs | 0.019 | Old |
| 0.300946 | 1.548843 | 0.034 | 0.015 | 1 | 12 | Ankrd24 | 0.019 | Old |
| 0.300946 | 1.541879 | 0.034 | 0.015 | 1 | 12 | Tra2b | 0.019 | Old |
| 0.300946 | 1.518731 | 0.034 | 0.015 | 1 | 12 | Tmem141 | 0.019 | Old |
| 0.300946 | 1.472341 | 0.034 | 0.015 | 1 | 12 | Nln | 0.019 | Old |
| 0.300946 | 1.434736 | 0.034 | 0.015 | 1 | 12 | Ddx3x | 0.019 | Old |
| 0.300946 | 1.430851 | 0.034 | 0.015 | 1 | 12 | 9330158H( | 0.019 | Old |
| 0.300993 | 0.343682 | 0.28 | 0.226 | 1 | 12 | Pgk1 | 0.054 | Old |
| 0.301563 | 0.412006 | 0.127 | 0.088 | 1 | 12 | Mxra7 | 0.039 | Old |
| 0.301849 | 0.215406 | 0.22 | 0.161 | 1 | 12 | Cyb5a | 0.059 | Old |
| 0.30195 | 0.612325 | 0.093 | 0.058 | 1 | 12 | Acyp2 | 0.035 | Old |
| 0.3029 | 0.50863 | 0.136 | 0.095 | 1 | 12 | Tcf4 | 0.041 | Old |
| 0.303094 | 0.674095 | 0.085 | 0.051 | 1 | 12 | Ralgps2 | 0.034 | Old |
| 0.303987 | 1.840471 | 0.034 | 0.015 | 1 | 12 | 1110059E2 | 0.019 | Old |
| 0.303987 | 1.719483 | 0.034 | 0.015 | 1 | 12 | Trmt6 | 0.019 | Old |
| 0.303987 | 1.640589 | 0.034 | 0.015 | 1 | 12 | Gm26722 | 0.019 | Old |
| 0.303987 | 1.589397 | 0.034 | 0.015 | 1 | 12 | Kin | 0.019 | Old |
| 0.303987 | 1.567281 | 0.034 | 0.015 | 1 | 12 | Gnl2 | 0.019 | Old |
| 0.303987 | 1.553271 | 0.034 | 0.015 | 1 | 12 | Dus3l | 0.019 | Old |
| 0.303987 | 1.451343 | 0.034 | 0.015 | 1 | 12 | Dnm2 | 0.019 | Old |
| 0.303987 | 1.409591 | 0.034 | 0.015 | 1 | 12 | Phf10 | 0.019 | Old |
| 0.303987 | 1.374248 | 0.034 | 0.015 | 1 | 12 | Rfc1 | 0.019 | Old |
| 0.304078 | 0.139391 | 0.907 | 0.854 | 1 | 12 | Sncg | 0.053 | Old |
| 0.304618 | 0.161294 | 0.169 | 0.117 | 1 | 12 | Vapb | 0.052 | Old |
| 0.304789 | 0.203199 | 0.229 | 0.175 | 1 | 12 | Sec61b | 0.054 | Old |
| 0.304948 | 0.522582 | 0.085 | 0.051 | 1 | 12 | Ccni | 0.034 | Old |
| 0.305918 | 0.370881 | 0.186 | 0.139 | 1 | 12 | Pdap1 | 0.047 | Old |
| 0.307048 | 1.617049 | 0.034 | 0.015 | 1 | 12 | Zranb1 | 0.019 | Old |
| 0.307048 | 1.564877 | 0.034 | 0.015 | 1 | 12 | Ahcyl1 | 0.019 | Old |
| 0.307048 | 1.526157 | 0.034 | 0.015 | 1 | 12 | Slit3 | 0.019 | Old |
| 0.307048 | 1.47213 | 0.034 | 0.015 | 1 | 12 | Memo1 | 0.019 | Old |
| 0.307048 | 1.45119 | 0.034 | 0.015 | 1 | 12 | Arl5a | 0.019 | Old |
| 0.307048 | 1.449436 | 0.034 | 0.015 | 1 | 12 | Chka | 0.019 | Old |
| 0.307048 | 1.443836 | 0.034 | 0.015 | 1 | 12 | Vcl | 0.019 | Old |
| 0.307048 | 1.393527 | 0.034 | 0.015 | 1 | 12 | Camk2n1 | 0.019 | Old |
| 0.307048 | 1.282217 | 0.034 | 0.015 | 1 | 12 | Gpx3 | 0.019 | Old |
| 0.307048 | 1.266606 | 0.034 | 0.015 | 1 | 12 | Fnbp1l | 0.019 | Old |
| 0.307048 | 1.212919 | 0.034 | 0.015 | 1 | 12 | Lgi2 | 0.019 | Old |
| 0.307235 | 0.510037 | 0.093 | 0.058 | 1 | 12 | Larp1 | 0.035 | Old |
| 0.307839 | 0.550181 | 0.102 | 0.066 | 1 | 12 | Cnpy2 | 0.036 | Old |
| 0.31013 | 1.43775 | 0.034 | 0.015 | 1 | 12 | Kcna4 | 0.019 | Old |
| 0.31013 | 1.351932 | 0.034 | 0.015 | 1 | 12 | Tysnd1 | 0.019 | Old |
| 0.31013 | 1.339512 | 0.034 | 0.015 | 1 | 12 | Abat | 0.019 | Old |

| | | | | | | | | |
|---|---|---|---|---|---|---|---|---|
| 0.31013 | 1.322748 | 0.034 | 0.015 | 1 | 12 | Cux2 | 0.019 | Old |
| 0.31013 | 1.297735 | 0.034 | 0.015 | 1 | 12 | Nsd1 | 0.019 | Old |
| 0.31013 | 1.268187 | 0.034 | 0.015 | 1 | 12 | Dlgap3 | 0.019 | Old |
| 0.31013 | 1.241774 | 0.034 | 0.015 | 1 | 12 | Nipal3 | 0.019 | Old |
| 0.31013 | 1.241649 | 0.034 | 0.015 | 1 | 12 | Utp18 | 0.019 | Old |
| 0.31013 | 1.214958 | 0.034 | 0.015 | 1 | 12 | Akap17b | 0.019 | Old |
| 0.31013 | 1.178234 | 0.034 | 0.015 | 1 | 12 | Mapk1ip1l | 0.019 | Old |
| 0.31013 | 1.045673 | 0.034 | 0.015 | 1 | 12 | Dusp11 | 0.019 | Old |
| 0.312057 | 0.577069 | 0.11 | 0.073 | 1 | 12 | Akain1 | 0.037 | Old |
| 0.312094 | 0.62029 | 0.102 | 0.066 | 1 | 12 | Kcmf1 | 0.036 | Old |
| 0.313231 | 1.423576 | 0.034 | 0.015 | 1 | 12 | Coro2a | 0.019 | Old |
| 0.313231 | 1.289612 | 0.034 | 0.015 | 1 | 12 | Fads1 | 0.019 | Old |
| 0.313231 | 1.249365 | 0.034 | 0.015 | 1 | 12 | Opa1 | 0.019 | Old |
| 0.313231 | 1.241674 | 0.034 | 0.015 | 1 | 12 | 1700037H( | 0.019 | Old |
| 0.313231 | 1.219609 | 0.034 | 0.015 | 1 | 12 | Mphosph6 | 0.019 | Old |
| 0.313231 | 1.199792 | 0.034 | 0.015 | 1 | 12 | Cisd3 | 0.019 | Old |
| 0.313231 | 1.194033 | 0.034 | 0.015 | 1 | 12 | Dcaf12l1 | 0.019 | Old |
| 0.313231 | 1.18638 | 0.034 | 0.015 | 1 | 12 | Ice1 | 0.019 | Old |
| 0.313231 | 1.176154 | 0.034 | 0.015 | 1 | 12 | Mrpl45 | 0.019 | Old |
| 0.313231 | 1.169905 | 0.034 | 0.015 | 1 | 12 | Pum3 | 0.019 | Old |
| 0.313231 | 1.169449 | 0.034 | 0.015 | 1 | 12 | Ptpn12 | 0.019 | Old |
| 0.313231 | 1.154709 | 0.034 | 0.015 | 1 | 12 | Fbxo44 | 0.019 | Old |
| 0.313231 | 1.136855 | 0.034 | 0.015 | 1 | 12 | Clec2d | 0.019 | Old |
| 0.313231 | 1.130579 | 0.034 | 0.015 | 1 | 12 | Bptf | 0.019 | Old |
| 0.313231 | 1.112645 | 0.034 | 0.015 | 1 | 12 | Pkia | 0.019 | Old |
| 0.313231 | 1.101471 | 0.034 | 0.015 | 1 | 12 | Ssrp1 | 0.019 | Old |
| 0.313231 | 1.084203 | 0.034 | 0.015 | 1 | 12 | Wwp1 | 0.019 | Old |
| 0.313231 | 1.082469 | 0.034 | 0.015 | 1 | 12 | Rnf41 | 0.019 | Old |
| 0.313231 | 1.082378 | 0.034 | 0.015 | 1 | 12 | Usp3 | 0.019 | Old |
| 0.313231 | 1.052331 | 0.034 | 0.015 | 1 | 12 | Ppp2r5d | 0.019 | Old |
| 0.313231 | 1.037194 | 0.034 | 0.015 | 1 | 12 | Ppp1r12c | 0.019 | Old |
| 0.313231 | 1.031734 | 0.034 | 0.015 | 1 | 12 | Nasp | 0.019 | Old |
| 0.313231 | 1.003142 | 0.034 | 0.015 | 1 | 12 | Tmem245 | 0.019 | Old |
| 0.313231 | 0.98734 | 0.034 | 0.015 | 1 | 12 | Qrich1 | 0.019 | Old |
| 0.313231 | 0.907347 | 0.034 | 0.015 | 1 | 12 | Washc2 | 0.019 | Old |
| 0.313231 | 0.810001 | 0.034 | 0.015 | 1 | 12 | Commd8 | 0.019 | Old |
| 0.31327 | 0.477308 | 0.186 | 0.146 | 1 | 12 | Polr2j | 0.04 | Old |
| 0.314789 | 1.112111 | 0.034 | 0.015 | 1 | 12 | Jazf1 | 0.019 | Old |
| 0.314937 | 0.47907 | 0.119 | 0.08 | 1 | 12 | Ghitm | 0.039 | Old |
| 0.316179 | 0.38344 | 0.093 | 0.058 | 1 | 12 | Sltm | 0.035 | Old |
| 0.316352 | 0.955953 | 0.034 | 0.015 | 1 | 12 | Spata24 | 0.019 | Old |
| 0.316353 | 1.429275 | 0.034 | 0.015 | 1 | 12 | Tubgcp5 | 0.019 | Old |
| 0.316353 | 1.251752 | 0.034 | 0.015 | 1 | 12 | Gmds | 0.019 | Old |
| 0.316353 | 1.133211 | 0.034 | 0.015 | 1 | 12 | Wdr53 | 0.019 | Old |
| 0.316353 | 1.105201 | 0.034 | 0.015 | 1 | 12 | Ankrd13d | 0.019 | Old |

| | | | | | | | | |
|---|---|---|---|---|---|---|---|---|
| 0.316353 | 1.077853 | 0.034 | 0.015 | 1 | 12 | Mtus2 | 0.019 | Old |
| 0.316353 | 1.065351 | 0.034 | 0.015 | 1 | 12 | Dkk3 | 0.019 | Old |
| 0.316353 | 1.056522 | 0.034 | 0.015 | 1 | 12 | Irak1 | 0.019 | Old |
| 0.316353 | 1.04198 | 0.034 | 0.015 | 1 | 12 | Defb42 | 0.019 | Old |
| 0.316353 | 1.031286 | 0.034 | 0.015 | 1 | 12 | Dnajc6 | 0.019 | Old |
| 0.316353 | 1.018927 | 0.034 | 0.015 | 1 | 12 | Fam189b | 0.019 | Old |
| 0.316353 | 1.015 | 0.034 | 0.015 | 1 | 12 | Erg28 | 0.019 | Old |
| 0.316353 | 0.999385 | 0.034 | 0.015 | 1 | 12 | Pibf1 | 0.019 | Old |
| 0.316353 | 0.974505 | 0.034 | 0.015 | 1 | 12 | Wac | 0.019 | Old |
| 0.316353 | 0.897796 | 0.034 | 0.015 | 1 | 12 | Xkr6 | 0.019 | Old |
| 0.316353 | 0.879558 | 0.034 | 0.015 | 1 | 12 | Alg5 | 0.019 | Old |
| 0.316353 | 0.803608 | 0.034 | 0.015 | 1 | 12 | Lman2 | 0.019 | Old |
| 0.316353 | 0.702785 | 0.034 | 0.015 | 1 | 12 | Mib1 | 0.019 | Old |
| 0.316353 | 0.555233 | 0.034 | 0.015 | 1 | 12 | Wnk1 | 0.019 | Old |
| 0.317061 | 0.414715 | 0.144 | 0.102 | 1 | 12 | 1110008F1 | 0.042 | Old |
| 0.317921 | 1.158786 | 0.034 | 0.015 | 1 | 12 | Rabgap1l | 0.019 | Old |
| 0.318116 | 0.514971 | 0.102 | 0.066 | 1 | 12 | Gemin7 | 0.036 | Old |
| 0.318126 | 0.451654 | 0.119 | 0.08 | 1 | 12 | Map2 | 0.039 | Old |
| 0.319496 | 1.677548 | 0.034 | 0.015 | 1 | 12 | C1qa | 0.019 | Old |
| 0.319496 | 1.049816 | 0.034 | 0.015 | 1 | 12 | Zfp9 | 0.019 | Old |
| 0.319496 | 0.988938 | 0.034 | 0.015 | 1 | 12 | B3galt6 | 0.019 | Old |
| 0.319496 | 0.955315 | 0.034 | 0.015 | 1 | 12 | Arl6 | 0.019 | Old |
| 0.319496 | 0.94557 | 0.034 | 0.015 | 1 | 12 | Lrrc75b | 0.019 | Old |
| 0.319496 | 0.931794 | 0.034 | 0.015 | 1 | 12 | Zmym3 | 0.019 | Old |
| 0.319496 | 0.920568 | 0.034 | 0.015 | 1 | 12 | Pigs | 0.019 | Old |
| 0.319496 | 0.800683 | 0.034 | 0.015 | 1 | 12 | Gab2 | 0.019 | Old |
| 0.319496 | 0.76785 | 0.034 | 0.015 | 1 | 12 | Impad1 | 0.019 | Old |
| 0.319496 | 0.582946 | 0.034 | 0.015 | 1 | 12 | Sostdc1 | 0.019 | Old |
| 0.319978 | 0.162891 | 0.525 | 0.453 | 1 | 12 | 6330403K0 | 0.072 | Old |
| 0.320922 | 0.515176 | 0.186 | 0.146 | 1 | 12 | Dynll2 | 0.04 | Old |
| 0.322659 | 1.011214 | 0.034 | 0.015 | 1 | 12 | Katnal1 | 0.019 | Old |
| 0.322659 | 0.928762 | 0.034 | 0.015 | 1 | 12 | Cog8 | 0.019 | Old |
| 0.322659 | 0.921577 | 0.034 | 0.015 | 1 | 12 | Snx18 | 0.019 | Old |
| 0.322659 | 0.834558 | 0.034 | 0.015 | 1 | 12 | Elp6 | 0.019 | Old |
| 0.322659 | 0.801013 | 0.034 | 0.015 | 1 | 12 | Sharpin | 0.019 | Old |
| 0.322659 | 0.548362 | 0.034 | 0.015 | 1 | 12 | Slc6a6 | 0.019 | Old |
| 0.325842 | 0.809218 | 0.034 | 0.015 | 1 | 12 | Zfp639 | 0.019 | Old |
| 0.325842 | 0.683397 | 0.034 | 0.015 | 1 | 12 | Tmem189 | 0.019 | Old |
| 0.325842 | 0.557172 | 0.034 | 0.015 | 1 | 12 | P2rx4 | 0.019 | Old |
| 0.325842 | 0.434666 | 0.034 | 0.015 | 1 | 12 | Ube2o | 0.019 | Old |
| 0.327087 | 0.132434 | 0.754 | 0.686 | 1 | 12 | Oaz1 | 0.068 | Old |
| 0.32784 | 0.34017 | 0.297 | 0.248 | 1 | 12 | Ube2b | 0.049 | Old |
| 0.328464 | 0.163516 | 0.805 | 0.854 | 1 | 12 | Nefl | -0.049 | Old |
| 0.330714 | 1.399837 | 0.042 | 0.022 | 1 | 12 | Srrm3 | 0.02 | Old |
| 0.330935 | 0.28706 | 0.153 | 0.109 | 1 | 12 | 1110065P2 | 0.044 | Old |

| | | | | | | | | |
|---|---|---|---|---|---|---|---|---|
| 0.33176 | 0.168589 | 0.508 | 0.453 | 1 | 12 | Cox6c | 0.055 | Old |
| 0.332736 | 0.294347 | 0.322 | 0.27 | 1 | 12 | Minos1 | 0.052 | Old |
| 0.333525 | 1.400178 | 0.042 | 0.022 | 1 | 12 | Lage3 | 0.02 | Old |
| 0.334602 | 0.547064 | 0.186 | 0.146 | 1 | 12 | Ufc1 | 0.04 | Old |
| 0.334662 | 0.61463 | 0.136 | 0.102 | 1 | 12 | Mrpl34 | 0.034 | Old |
| 0.334935 | 1.386074 | 0.042 | 0.022 | 1 | 12 | Pdcd7 | 0.02 | Old |
| 0.336269 | 0.146606 | 0.831 | 0.752 | 1 | 12 | Tubb2a | 0.079 | Old |
| 0.336351 | 1.368129 | 0.042 | 0.022 | 1 | 12 | Vps53 | 0.02 | Old |
| 0.336351 | 1.319346 | 0.042 | 0.022 | 1 | 12 | Alkbh7 | 0.02 | Old |
| 0.336351 | 1.297571 | 0.042 | 0.022 | 1 | 12 | Tril | 0.02 | Old |
| 0.339193 | 1.285288 | 0.042 | 0.022 | 1 | 12 | Ehd1 | 0.02 | Old |
| 0.339193 | 1.246541 | 0.042 | 0.022 | 1 | 12 | Mlycd | 0.02 | Old |
| 0.339193 | 1.191018 | 0.042 | 0.022 | 1 | 12 | Efna5 | 0.02 | Old |
| 0.339193 | 1.074047 | 0.042 | 0.022 | 1 | 12 | Disp2 | 0.02 | Old |
| 0.339512 | 0.217462 | 0.542 | 0.496 | 1 | 12 | Rps7 | 0.046 | Old |
| 0.341143 | 0.174431 | 0.983 | 0.942 | 1 | 12 | Actg1 | 0.041 | Old |
| 0.34205 | 1.205299 | 0.042 | 0.022 | 1 | 12 | Scx | 0.02 | Old |
| 0.34205 | 1.17481 | 0.042 | 0.022 | 1 | 12 | Zfp617 | 0.02 | Old |
| 0.34205 | 1.16951 | 0.042 | 0.022 | 1 | 12 | Ptrh2 | 0.02 | Old |
| 0.34205 | 1.158819 | 0.042 | 0.022 | 1 | 12 | Nsun2 | 0.02 | Old |
| 0.34205 | 1.127739 | 0.042 | 0.022 | 1 | 12 | Scnm1 | 0.02 | Old |
| 0.34205 | 1.079155 | 0.042 | 0.022 | 1 | 12 | U2af2 | 0.02 | Old |
| 0.34205 | 1.072025 | 0.042 | 0.022 | 1 | 12 | Zfp318 | 0.02 | Old |
| 0.34205 | 1.020427 | 0.042 | 0.022 | 1 | 12 | Utp15 | 0.02 | Old |
| 0.344193 | 0.264286 | 0.466 | 0.409 | 1 | 12 | Rpl22l1 | 0.057 | Old |
| 0.344922 | 1.425233 | 0.042 | 0.022 | 1 | 12 | Pak1 | 0.02 | Old |
| 0.344922 | 1.193709 | 0.042 | 0.022 | 1 | 12 | Acta1 | 0.02 | Old |
| 0.344922 | 1.077934 | 0.042 | 0.022 | 1 | 12 | Slc30a9 | 0.02 | Old |
| 0.344922 | 1.046925 | 0.042 | 0.022 | 1 | 12 | Pef1 | 0.02 | Old |
| 0.344922 | 1.039955 | 0.042 | 0.022 | 1 | 12 | Cd2ap | 0.02 | Old |
| 0.344922 | 0.917986 | 0.042 | 0.022 | 1 | 12 | Gstz1 | 0.02 | Old |
| 0.344922 | 0.864045 | 0.042 | 0.022 | 1 | 12 | Decr1 | 0.02 | Old |
| 0.344998 | 1.749255 | 0.051 | 0.029 | 1 | 12 | Nmt1 | 0.022 | Old |
| 0.346363 | 0.77256 | 0.042 | 0.022 | 1 | 12 | Ehmt2 | 0.02 | Old |
| 0.347809 | 1.01187 | 0.042 | 0.022 | 1 | 12 | Ddx50 | 0.02 | Old |
| 0.34781 | 1.197451 | 0.042 | 0.022 | 1 | 12 | Mmgt2 | 0.02 | Old |
| 0.34781 | 1.115771 | 0.042 | 0.022 | 1 | 12 | Sdhaf2 | 0.02 | Old |
| 0.34781 | 1.028068 | 0.042 | 0.022 | 1 | 12 | Rfesd | 0.02 | Old |
| 0.34781 | 0.831795 | 0.042 | 0.022 | 1 | 12 | Nt5dc3 | 0.02 | Old |
| 0.34781 | 0.817606 | 0.042 | 0.022 | 1 | 12 | Cryzl1 | 0.02 | Old |
| 0.348623 | 0.340919 | 0.119 | 0.08 | 1 | 12 | Chic2 | 0.039 | Old |
| 0.349023 | 0.114121 | 0.492 | 0.431 | 1 | 12 | Rpl36 | 0.061 | Old |
| 0.349273 | 0.361079 | 0.22 | 0.175 | 1 | 12 | Guk1 | 0.045 | Old |
| 0.350414 | 0.896202 | 0.085 | 0.058 | 1 | 12 | Eri3 | 0.027 | Old |
| 0.350713 | 0.976015 | 0.042 | 0.022 | 1 | 12 | Rusc1 | 0.02 | Old |

| | | | | | | | | |
|---|---|---|---|---|---|---|---|---|
| 0.350713 | 0.962504 | 0.042 | 0.022 | 1 | 12 | Zfp207 | 0.02 | Old |
| 0.350713 | 0.921418 | 0.042 | 0.022 | 1 | 12 | Map3k10 | 0.02 | Old |
| 0.350713 | 0.888143 | 0.042 | 0.022 | 1 | 12 | Apc | 0.02 | Old |
| 0.350713 | 0.781618 | 0.042 | 0.022 | 1 | 12 | Dhcr7 | 0.02 | Old |
| 0.35224 | 0.783412 | 0.127 | 0.095 | 1 | 12 | Znhit1 | 0.032 | Old |
| 0.353233 | 0.295381 | 0.331 | 0.299 | 1 | 12 | Rnf187 | 0.032 | Old |
| 0.353632 | 1.061367 | 0.042 | 0.022 | 1 | 12 | Tmem109 | 0.02 | Old |
| 0.353632 | 0.918725 | 0.042 | 0.022 | 1 | 12 | Timp2 | 0.02 | Old |
| 0.353632 | 0.915293 | 0.042 | 0.022 | 1 | 12 | Mrpl46 | 0.02 | Old |
| 0.353632 | 0.885549 | 0.042 | 0.022 | 1 | 12 | Rmnd1 | 0.02 | Old |
| 0.353632 | 0.830588 | 0.042 | 0.022 | 1 | 12 | Arfgef1 | 0.02 | Old |
| 0.353632 | 0.804396 | 0.042 | 0.022 | 1 | 12 | Dido1 | 0.02 | Old |
| 0.353632 | 0.638707 | 0.042 | 0.022 | 1 | 12 | Blvrb | 0.02 | Old |
| 0.354629 | 0.45385 | 0.229 | 0.19 | 1 | 12 | Atp6v0d1 | 0.039 | Old |
| 0.355443 | 1.097387 | 0.051 | 0.029 | 1 | 12 | Dlg4 | 0.022 | Old |
| 0.356566 | 0.937074 | 0.042 | 0.022 | 1 | 12 | Ddx10 | 0.02 | Old |
| 0.356566 | 0.858114 | 0.042 | 0.022 | 1 | 12 | Itgb1bp1 | 0.02 | Old |
| 0.356566 | 0.850737 | 0.042 | 0.022 | 1 | 12 | Pnma2 | 0.02 | Old |
| 0.356566 | 0.838389 | 0.042 | 0.022 | 1 | 12 | Copb1 | 0.02 | Old |
| 0.356566 | 0.825317 | 0.042 | 0.022 | 1 | 12 | Nrg1 | 0.02 | Old |
| 0.356566 | 0.809955 | 0.042 | 0.022 | 1 | 12 | Arhgap21 | 0.02 | Old |
| 0.356566 | 0.65397 | 0.042 | 0.022 | 1 | 12 | Stk39 | 0.02 | Old |
| 0.358086 | 1.165533 | 0.051 | 0.029 | 1 | 12 | Lonp1 | 0.022 | Old |
| 0.358086 | 1.077943 | 0.051 | 0.029 | 1 | 12 | 5031439G( | 0.022 | Old |
| 0.359383 | 1.029231 | 0.059 | 0.036 | 1 | 12 | Fstl5 | 0.023 | Old |
| 0.359515 | 0.834954 | 0.042 | 0.022 | 1 | 12 | Ccdc34 | 0.02 | Old |
| 0.359515 | 0.787944 | 0.042 | 0.022 | 1 | 12 | Tomm34 | 0.02 | Old |
| 0.359515 | 0.454693 | 0.042 | 0.022 | 1 | 12 | Sbno1 | 0.02 | Old |
| 0.359515 | 0.407517 | 0.042 | 0.022 | 1 | 12 | Add3 | 0.02 | Old |
| 0.36019 | 1.080973 | 0.093 | 0.066 | 1 | 12 | Eif4h | 0.027 | Old |
| 0.360428 | 0.67053 | 0.136 | 0.102 | 1 | 12 | Psmd13 | 0.034 | Old |
| 0.36074 | 0.897016 | 0.051 | 0.029 | 1 | 12 | Rnf4 | 0.022 | Old |
| 0.361342 | 0.429928 | 0.203 | 0.168 | 1 | 12 | Pam16 | 0.035 | Old |
| 0.361821 | 0.957855 | 0.059 | 0.036 | 1 | 12 | Ddx56 | 0.023 | Old |
| 0.362479 | 0.713723 | 0.042 | 0.022 | 1 | 12 | Ubfd1 | 0.02 | Old |
| 0.362479 | 0.676553 | 0.042 | 0.022 | 1 | 12 | Dhrs7b | 0.02 | Old |
| 0.362479 | 0.6754 | 0.042 | 0.022 | 1 | 12 | Rnf157 | 0.02 | Old |
| 0.362479 | 0.564681 | 0.042 | 0.022 | 1 | 12 | Nipbl | 0.02 | Old |
| 0.363407 | 1.071532 | 0.051 | 0.029 | 1 | 12 | Epdr1 | 0.022 | Old |
| 0.363407 | 1.022563 | 0.051 | 0.029 | 1 | 12 | Shf | 0.022 | Old |
| 0.363407 | 0.989231 | 0.051 | 0.029 | 1 | 12 | Gfra4 | 0.022 | Old |
| 0.363407 | 0.877229 | 0.051 | 0.029 | 1 | 12 | Nup88 | 0.022 | Old |
| 0.363687 | 0.488499 | 0.144 | 0.109 | 1 | 12 | Phyh | 0.035 | Old |
| 0.364204 | 0.238164 | 0.381 | 0.328 | 1 | 12 | Fdps | 0.053 | Old |
| 0.364247 | 0.267776 | 0.28 | 0.234 | 1 | 12 | Timm13 | 0.046 | Old |

| | | | | | | | | |
|---|---|---|---|---|---|---|---|---|
| 0.365459 | 0.73884 | 0.042 | 0.022 | 1 | 12 | Dixdc1 | 0.02 | Old |
| 0.365459 | 0.668459 | 0.042 | 0.022 | 1 | 12 | Hddc2 | 0.02 | Old |
| 0.365459 | 0.393766 | 0.042 | 0.022 | 1 | 12 | Mrpl3 | 0.02 | Old |
| 0.366086 | 0.831226 | 0.051 | 0.029 | 1 | 12 | Serp1 | 0.022 | Old |
| 0.366086 | 0.812476 | 0.051 | 0.029 | 1 | 12 | Lyrm2 | 0.022 | Old |
| 0.366276 | 0.230306 | 0.254 | 0.204 | 1 | 12 | 1110008P1 | 0.05 | Old |
| 0.366728 | 1.080239 | 0.059 | 0.036 | 1 | 12 | Kdm1a | 0.023 | Old |
| 0.366904 | 0.158843 | 0.364 | 0.307 | 1 | 12 | Pfn1 | 0.057 | Old |
| 0.368778 | 0.792529 | 0.051 | 0.029 | 1 | 12 | Map7 | 0.022 | Old |
| 0.371482 | 0.992616 | 0.051 | 0.029 | 1 | 12 | Gpatch11 | 0.022 | Old |
| 0.371482 | 0.836642 | 0.051 | 0.029 | 1 | 12 | Bdh1 | 0.022 | Old |
| 0.371482 | 0.825713 | 0.051 | 0.029 | 1 | 12 | Wdr70 | 0.022 | Old |
| 0.371482 | 0.804876 | 0.051 | 0.029 | 1 | 12 | Irf2bpl | 0.022 | Old |
| 0.371702 | 0.937907 | 0.076 | 0.051 | 1 | 12 | Jakmip1 | 0.025 | Old |
| 0.372191 | 0.196943 | 0.542 | 0.518 | 1 | 12 | Nsg1 | 0.024 | Old |
| 0.372564 | 0.734041 | 0.102 | 0.073 | 1 | 12 | Mpnd | 0.029 | Old |
| 0.372837 | 0.852495 | 0.051 | 0.029 | 1 | 12 | Mrto4 | 0.022 | Old |
| 0.373055 | 0.854768 | 0.068 | 0.044 | 1 | 12 | Ptrhd1 | 0.024 | Old |
| 0.373735 | 0.196298 | 0.576 | 0.511 | 1 | 12 | Pebp1 | 0.065 | Old |
| 0.374198 | 0.843029 | 0.051 | 0.029 | 1 | 12 | Nelfe | 0.022 | Old |
| 0.374198 | 0.776612 | 0.051 | 0.029 | 1 | 12 | R3hcc1 | 0.022 | Old |
| 0.374198 | 0.636632 | 0.051 | 0.029 | 1 | 12 | Gsto1 | 0.022 | Old |
| 0.374198 | 0.408965 | 0.051 | 0.029 | 1 | 12 | 1810043G( | 0.022 | Old |
| 0.374489 | 0.244037 | 0.042 | 0.022 | 1 | 12 | Sfxn3 | 0.02 | Old |
| 0.374785 | 0.114762 | 0.492 | 0.453 | 1 | 12 | Rtn4 | 0.039 | Old |
| 0.375374 | 0.873039 | 0.068 | 0.044 | 1 | 12 | Mapk8ip2 | 0.024 | Old |
| 0.375999 | 0.227114 | 0.161 | 0.117 | 1 | 12 | Set | 0.044 | Old |
| 0.376666 | 1.100475 | 0.059 | 0.036 | 1 | 12 | Gipc1 | 0.023 | Old |
| 0.376666 | 1.036022 | 0.059 | 0.036 | 1 | 12 | Rab11fip2 | 0.023 | Old |
| 0.376926 | 0.759711 | 0.051 | 0.029 | 1 | 12 | Cdk2ap2 | 0.022 | Old |
| 0.377015 | 0.229413 | 0.153 | 0.117 | 1 | 12 | Ddt | 0.036 | Old |
| 0.377702 | 0.788652 | 0.068 | 0.044 | 1 | 12 | Ubn1 | 0.024 | Old |
| 0.378244 | 0.925782 | 0.076 | 0.051 | 1 | 12 | Zfp771 | 0.025 | Old |
| 0.379151 | 0.275719 | 0.381 | 0.328 | 1 | 12 | Mt3 | 0.053 | Old |
| 0.379176 | 0.72618 | 0.059 | 0.036 | 1 | 12 | Nolc1 | 0.023 | Old |
| 0.379666 | 0.818921 | 0.051 | 0.029 | 1 | 12 | Tigd2 | 0.022 | Old |
| 0.379666 | 0.768785 | 0.051 | 0.029 | 1 | 12 | Smim19 | 0.022 | Old |
| 0.380039 | 0.892354 | 0.068 | 0.044 | 1 | 12 | Nradd | 0.024 | Old |
| 0.382419 | 0.650022 | 0.051 | 0.029 | 1 | 12 | Angptl6 | 0.022 | Old |
| 0.382964 | 0.725754 | 0.102 | 0.073 | 1 | 12 | Selenoh | 0.029 | Old |
| 0.383139 | 0.217985 | 0.22 | 0.175 | 1 | 12 | Arl6ip1 | 0.045 | Old |
| 0.384136 | 0.205069 | 0.178 | 0.139 | 1 | 12 | Serf1 | 0.039 | Old |
| 0.38439 | 0.232594 | 0.254 | 0.204 | 1 | 12 | Vcp | 0.05 | Old |
| 0.384739 | 0.481942 | 0.068 | 0.044 | 1 | 12 | Alg2 | 0.024 | Old |
| 0.38657 | 0.339516 | 0.136 | 0.102 | 1 | 12 | Rpl27 | 0.034 | Old |

| | | | | | | | | |
|---|---|---|---|---|---|---|---|---|
| 0.386768 | 0.738183 | 0.059 | 0.036 | 1 | 12 | Nans | 0.023 | Old |
| 0.387386 | 0.464829 | 0.127 | 0.095 | 1 | 12 | Dnm1l | 0.032 | Old |
| 0.387754 | 0.074422 | 0.898 | 0.825 | 1 | 12 | Chchd2 | 0.073 | Old |
| 0.387961 | 0.646363 | 0.051 | 0.029 | 1 | 12 | Mettl22 | 0.022 | Old |
| 0.388042 | 0.697868 | 0.059 | 0.036 | 1 | 12 | Snap91 | 0.023 | Old |
| 0.389304 | 0.669625 | 0.076 | 0.051 | 1 | 12 | Nxn | 0.025 | Old |
| 0.389319 | 0.93628 | 0.059 | 0.036 | 1 | 12 | Ngrn | 0.023 | Old |
| 0.389319 | 0.74227 | 0.059 | 0.036 | 1 | 12 | Sh3gl3 | 0.023 | Old |
| 0.389319 | 0.678945 | 0.059 | 0.036 | 1 | 12 | Adam19 | 0.023 | Old |
| 0.389319 | 0.659121 | 0.059 | 0.036 | 1 | 12 | Smim11 | 0.023 | Old |
| 0.389474 | 0.77236 | 0.068 | 0.044 | 1 | 12 | Nol12 | 0.024 | Old |
| 0.389475 | 0.820447 | 0.068 | 0.044 | 1 | 12 | AC160336. | 0.024 | Old |
| 0.390859 | 0.371108 | 0.144 | 0.109 | 1 | 12 | Arl2 | 0.035 | Old |
| 0.391539 | 0.617225 | 0.076 | 0.051 | 1 | 12 | Mtdh | 0.025 | Old |
| 0.391765 | 0.384709 | 0.161 | 0.124 | 1 | 12 | Fabp3 | 0.037 | Old |
| 0.391958 | 0.587731 | 0.119 | 0.088 | 1 | 12 | Acot13 | 0.031 | Old |
| 0.392271 | 0.370325 | 0.178 | 0.139 | 1 | 12 | Cystm1 | 0.039 | Old |
| 0.392527 | 0.318489 | 0.381 | 0.35 | 1 | 12 | Gabarapl2 | 0.031 | Old |
| 0.392659 | 0.530565 | 0.076 | 0.051 | 1 | 12 | Sncb | 0.025 | Old |
| 0.393532 | 0.103363 | 0.5 | 0.438 | 1 | 12 | Pkm | 0.062 | Old |
| 0.394246 | 0.84231 | 0.068 | 0.044 | 1 | 12 | Chchd4 | 0.024 | Old |
| 0.394246 | 0.645133 | 0.068 | 0.044 | 1 | 12 | Nbl1 | 0.024 | Old |
| 0.396365 | 0.542643 | 0.051 | 0.029 | 1 | 12 | Fam8a1 | 0.022 | Old |
| 0.396365 | 0.417519 | 0.051 | 0.029 | 1 | 12 | Dnajc21 | 0.022 | Old |
| 0.396458 | 0.481635 | 0.119 | 0.088 | 1 | 12 | Npc2 | 0.031 | Old |
| 0.398292 | 0.668593 | 0.076 | 0.051 | 1 | 12 | Unc50 | 0.025 | Old |
| 0.398328 | 0.527874 | 0.059 | 0.036 | 1 | 12 | Slc35g2 | 0.023 | Old |
| 0.398794 | 0.545246 | 0.093 | 0.066 | 1 | 12 | Pls3 | 0.027 | Old |
| 0.398957 | 0.163248 | 0.398 | 0.343 | 1 | 12 | Fkbp3 | 0.055 | Old |
| 0.399052 | 0.545236 | 0.068 | 0.044 | 1 | 12 | Slc39a6 | 0.024 | Old |
| 0.399424 | 0.673005 | 0.076 | 0.051 | 1 | 12 | Sdha | 0.025 | Old |
| 0.399561 | 0.451447 | 0.127 | 0.095 | 1 | 12 | Bex4 | 0.032 | Old |
| 0.399626 | 0.632578 | 0.059 | 0.036 | 1 | 12 | Mboat7 | 0.023 | Old |
| 0.399626 | 0.613831 | 0.059 | 0.036 | 1 | 12 | Mcam | 0.023 | Old |
| 0.399711 | 0.25634 | 0.508 | 0.496 | 1 | 12 | Ywhae | 0.012 | Old |
| 0.400361 | 0.468455 | 0.102 | 0.073 | 1 | 12 | Ptp4a1 | 0.029 | Old |
| 0.400558 | 0.67558 | 0.076 | 0.051 | 1 | 12 | Amn1 | 0.025 | Old |
| 0.400809 | 0.390152 | 0.153 | 0.117 | 1 | 12 | Smim14 | 0.036 | Old |
| 0.401468 | 0.726668 | 0.068 | 0.044 | 1 | 12 | Pnisr | 0.024 | Old |
| 0.401468 | 0.624749 | 0.068 | 0.044 | 1 | 12 | Gfra2 | 0.024 | Old |
| 0.401468 | 0.557629 | 0.068 | 0.044 | 1 | 12 | Sema6d | 0.024 | Old |
| 0.402029 | 0.520551 | 0.051 | 0.029 | 1 | 12 | Polr3k | 0.022 | Old |
| 0.402181 | 0.415914 | 0.212 | 0.182 | 1 | 12 | Acot7 | 0.03 | Old |
| 0.402228 | 0.576303 | 0.059 | 0.036 | 1 | 12 | Mrpl2 | 0.023 | Old |
| 0.403893 | 0.628741 | 0.068 | 0.044 | 1 | 12 | Trappc10 | 0.024 | Old |

| | | | | | | | | |
|---|---|---|---|---|---|---|---|---|
| 0.403998 | 0.40178 | 0.237 | 0.204 | 1 | 12 | Cct2 | 0.033 | Old |
| 0.404236 | 0.709385 | 0.085 | 0.058 | 1 | 12 | Aco2 | 0.027 | Old |
| 0.404236 | 0.528532 | 0.085 | 0.058 | 1 | 12 | Necab1 | 0.027 | Old |
| 0.404895 | 0.40508 | 0.144 | 0.109 | 1 | 12 | Rab10 | 0.035 | Old |
| 0.405114 | 0.279865 | 0.076 | 0.051 | 1 | 12 | BC005624 | 0.025 | Old |
| 0.405856 | 0.177469 | 0.161 | 0.117 | 1 | 12 | Gdi2 | 0.044 | Old |
| 0.406258 | 0.564889 | 0.076 | 0.051 | 1 | 12 | Ptpa | 0.025 | Old |
| 0.406327 | 0.624398 | 0.068 | 0.044 | 1 | 12 | Spsb3 | 0.024 | Old |
| 0.407011 | 0.48744 | 0.093 | 0.066 | 1 | 12 | Timm22 | 0.027 | Old |
| 0.407424 | 0.649225 | 0.161 | 0.131 | 1 | 12 | Supt4a | 0.03 | Old |
| 0.407462 | 0.583751 | 0.059 | 0.036 | 1 | 12 | Aspscr1 | 0.023 | Old |
| 0.40774 | 0.211959 | 0.051 | 0.029 | 1 | 12 | Nudt2 | 0.022 | Old |
| 0.407921 | 0.340628 | 0.22 | 0.182 | 1 | 12 | Ap2m1 | 0.038 | Old |
| 0.408571 | 0.559105 | 0.085 | 0.058 | 1 | 12 | Nsmce4a | 0.027 | Old |
| 0.40864 | 0.475176 | 0.136 | 0.102 | 1 | 12 | Hsd17b10 | 0.034 | Old |
| 0.408769 | 0.625875 | 0.068 | 0.044 | 1 | 12 | Prpf38a | 0.024 | Old |
| 0.408769 | 0.513346 | 0.068 | 0.044 | 1 | 12 | Mcee | 0.024 | Old |
| 0.408777 | 0.473096 | 0.059 | 0.036 | 1 | 12 | Cxxc5 | 0.023 | Old |
| 0.409701 | 0.605063 | 0.076 | 0.051 | 1 | 12 | Parl | 0.025 | Old |
| 0.410094 | 0.523626 | 0.059 | 0.036 | 1 | 12 | Aida | 0.023 | Old |
| 0.410749 | 0.587426 | 0.085 | 0.058 | 1 | 12 | Bola2 | 0.027 | Old |
| 0.410761 | 0.14291 | 0.144 | 0.109 | 1 | 12 | Sqstm1 | 0.035 | Old |
| 0.411021 | 0.42705 | 0.271 | 0.226 | 1 | 12 | Dstn | 0.045 | Old |
| 0.411063 | 0.438124 | 0.119 | 0.088 | 1 | 12 | Txndc9 | 0.031 | Old |
| 0.411395 | 0.287571 | 0.153 | 0.117 | 1 | 12 | Rtl8b | 0.036 | Old |
| 0.412006 | 0.581141 | 0.076 | 0.051 | 1 | 12 | Ykt6 | 0.025 | Old |
| 0.412057 | 0.278269 | 0.331 | 0.285 | 1 | 12 | Tma7 | 0.046 | Old |
| 0.412123 | 0.252983 | 0.381 | 0.343 | 1 | 12 | Akr1a1 | 0.038 | Old |
| 0.41228 | 0.179164 | 0.186 | 0.146 | 1 | 12 | Fermt2 | 0.04 | Old |
| 0.412538 | 0.17854 | 0.737 | 0.708 | 1 | 12 | Selenow | 0.029 | Old |
| 0.412934 | 0.45993 | 0.085 | 0.058 | 1 | 12 | Kctd12 | 0.027 | Old |
| 0.413087 | 0.35571 | 0.212 | 0.175 | 1 | 12 | Esd | 0.037 | Old |
| 0.41368 | 0.55562 | 0.068 | 0.044 | 1 | 12 | Rars | 0.024 | Old |
| 0.41368 | 0.54143 | 0.068 | 0.044 | 1 | 12 | Prnp | 0.024 | Old |
| 0.414319 | 0.43721 | 0.076 | 0.051 | 1 | 12 | Pink1 | 0.025 | Old |
| 0.416217 | 0.473687 | 0.102 | 0.073 | 1 | 12 | Phb2 | 0.029 | Old |
| 0.416639 | 0.519166 | 0.076 | 0.051 | 1 | 12 | Chtop | 0.025 | Old |
| 0.417163 | 0.381183 | 0.153 | 0.117 | 1 | 12 | Elavl4 | 0.036 | Old |
| 0.417297 | 0.389154 | 0.195 | 0.161 | 1 | 12 | Uqcrc1 | 0.034 | Old |
| 0.417488 | 0.155656 | 0.525 | 0.489 | 1 | 12 | Rpl37a | 0.036 | Old |
| 0.418214 | 0.274955 | 0.169 | 0.131 | 1 | 12 | Higd1a | 0.038 | Old |
| 0.418846 | 0.299412 | 0.178 | 0.139 | 1 | 12 | Ndufs2 | 0.039 | Old |
| 0.418967 | 0.654731 | 0.076 | 0.051 | 1 | 12 | Ufm1 | 0.025 | Old |
| 0.419162 | 0.155324 | 0.746 | 0.708 | 1 | 12 | Dynll1 | 0.038 | Old |
| 0.41953 | 0.550378 | 0.085 | 0.058 | 1 | 12 | Tmem258 | 0.027 | Old |

| | | | | | | | | |
|---|---|---|---|---|---|---|---|---|
| 0.41953 | 0.448638 | 0.085 | 0.058 | 1 | 12 | Hypk | 0.027 | Old |
| 0.41955 | 0.397178 | 0.161 | 0.124 | 1 | 12 | Mrps21 | 0.037 | Old |
| 0.421112 | 0.491519 | 0.068 | 0.044 | 1 | 12 | Uri1 | 0.024 | Old |
| 0.422686 | 0.470904 | 0.093 | 0.066 | 1 | 12 | Chmp2a | 0.027 | Old |
| 0.423268 | 0.385542 | 0.102 | 0.073 | 1 | 12 | Fgf13 | 0.029 | Old |
| 0.423646 | 0.40544 | 0.076 | 0.051 | 1 | 12 | Srsf6 | 0.025 | Old |
| 0.425996 | 0.503035 | 0.076 | 0.051 | 1 | 12 | Cacna2d1 | 0.025 | Old |
| 0.4261 | 0.076724 | 0.059 | 0.036 | 1 | 12 | Mrps25 | 0.023 | Old |
| 0.42611 | 0.328379 | 0.068 | 0.044 | 1 | 12 | Fam168b | 0.024 | Old |
| 0.428355 | 0.53401 | 0.076 | 0.051 | 1 | 12 | Mrpl15 | 0.025 | Old |
| 0.428355 | 0.334574 | 0.076 | 0.051 | 1 | 12 | Dner | 0.025 | Old |
| 0.428421 | 0.579621 | 0.085 | 0.058 | 1 | 12 | Dusp1 | 0.027 | Old |
| 0.428579 | 0.433584 | 0.11 | 0.08 | 1 | 12 | Pfdn6 | 0.03 | Old |
| 0.429616 | 0.464884 | 0.169 | 0.139 | 1 | 12 | Ndufb3 | 0.03 | Old |
| 0.431526 | 0.370313 | 0.11 | 0.08 | 1 | 12 | Jkamp | 0.03 | Old |
| 0.433094 | 0.429873 | 0.076 | 0.051 | 1 | 12 | Paxx | 0.025 | Old |
| 0.43333 | 0.591275 | 0.093 | 0.066 | 1 | 12 | Anapc16 | 0.027 | Old |
| 0.433562 | 0.383088 | 0.119 | 0.088 | 1 | 12 | Fos | 0.031 | Old |
| 0.43449 | 0.44351 | 0.102 | 0.073 | 1 | 12 | Dnmt3a | 0.029 | Old |
| 0.43516 | 0.357472 | 0.085 | 0.058 | 1 | 12 | Dhx36 | 0.027 | Old |
| 0.435469 | 0.207001 | 0.119 | 0.088 | 1 | 12 | Aimp1 | 0.031 | Old |
| 0.438877 | 0.232232 | 0.347 | 0.292 | 1 | 12 | Nrsn1 | 0.055 | Old |
| 0.438955 | 0.016505 | 0.195 | 0.161 | 1 | 12 | Bex1 | 0.034 | Old |
| 0.441155 | 0.406102 | 0.127 | 0.095 | 1 | 12 | Chchd1 | 0.032 | Old |
| 0.441173 | 0.22102 | 0.458 | 0.423 | 1 | 12 | Mdh1 | 0.035 | Old |
| 0.441309 | 0.329611 | 0.068 | 0.044 | 1 | 12 | Atic | 0.024 | Old |
| 0.441309 | 0.285359 | 0.068 | 0.044 | 1 | 12 | Ago2 | 0.024 | Old |
| 0.442664 | 0.467859 | 0.076 | 0.051 | 1 | 12 | Vps35 | 0.025 | Old |
| 0.44609 | 0.184123 | 0.458 | 0.431 | 1 | 12 | Tecr | 0.027 | Old |
| 0.446281 | 0.068406 | 0.246 | 0.197 | 1 | 12 | Psmd7 | 0.049 | Old |
| 0.446913 | 0.265278 | 0.186 | 0.146 | 1 | 12 | Pdia3 | 0.04 | Old |
| 0.450058 | 0.310425 | 0.203 | 0.175 | 1 | 12 | Gna11 | 0.028 | Old |
| 0.451062 | 0.227879 | 0.271 | 0.234 | 1 | 12 | Psma2 | 0.037 | Old |
| 0.452874 | 0.513119 | 0.093 | 0.066 | 1 | 12 | Ntrk3 | 0.027 | Old |
| 0.453132 | 0.21007 | 0.136 | 0.102 | 1 | 12 | Rab7 | 0.034 | Old |
| 0.453427 | 0.036354 | 0.085 | 0.058 | 1 | 12 | Upf3a | 0.027 | Old |
| 0.454012 | 0.142503 | 0.534 | 0.489 | 1 | 12 | Phox2a | 0.045 | Old |
| 0.45625 | 0.385489 | 0.161 | 0.131 | 1 | 12 | 2210016L2 | 0.03 | Old |
| 0.459726 | 0.233138 | 0.237 | 0.204 | 1 | 12 | Calm3 | 0.033 | Old |
| 0.461823 | 0.536505 | 0.153 | 0.124 | 1 | 12 | Srsf2 | 0.029 | Old |
| 0.462669 | 0.182914 | 0.763 | 0.766 | 1 | 12 | Rps27a | -0.003 | Old |
| 0.462721 | 0.142196 | 0.085 | 0.058 | 1 | 12 | Gtf2b | 0.027 | Old |
| 0.463541 | 0.213962 | 0.407 | 0.372 | 1 | 12 | Gpx4 | 0.035 | Old |
| 0.464298 | 0.202257 | 0.263 | 0.219 | 1 | 12 | Saraf | 0.044 | Old |
| 0.464427 | 0.258573 | 0.153 | 0.124 | 1 | 12 | Hmgn3 | 0.029 | Old |

| | | | | | | | | |
|---|---|---|---|---|---|---|---|---|
| 0.469422 | 0.077319 | 0.568 | 0.489 | 1 | 12 | Mtch1 | 0.079 | Old |
| 0.47416 | 2.094138 | 0.017 | 0.007 | 1 | 12 | Gm8797 | 0.01 | Old |
| 0.474162 | 2.452794 | 0.017 | 0.007 | 1 | 12 | Stxbp5l | 0.01 | Old |
| 0.474162 | 2.17497 | 0.017 | 0.007 | 1 | 12 | Cab39l | 0.01 | Old |
| 0.474162 | 2.06035 | 0.017 | 0.007 | 1 | 12 | Smarcc2 | 0.01 | Old |
| 0.474162 | 2.04892 | 0.017 | 0.007 | 1 | 12 | Cdc42ep3 | 0.01 | Old |
| 0.474162 | 2.015305 | 0.017 | 0.007 | 1 | 12 | Tlk2 | 0.01 | Old |
| 0.474162 | 1.925627 | 0.017 | 0.007 | 1 | 12 | Rhobtb3 | 0.01 | Old |
| 0.474162 | 1.873164 | 0.017 | 0.007 | 1 | 12 | Wdr54 | 0.01 | Old |
| 0.474162 | 1.8528 | 0.017 | 0.007 | 1 | 12 | Fbxo34 | 0.01 | Old |
| 0.474162 | 1.802564 | 0.017 | 0.007 | 1 | 12 | Gns | 0.01 | Old |
| 0.474162 | 1.781412 | 0.017 | 0.007 | 1 | 12 | Uros | 0.01 | Old |
| 0.474162 | 1.770013 | 0.017 | 0.007 | 1 | 12 | Ccdc43 | 0.01 | Old |
| 0.474162 | 1.748333 | 0.017 | 0.007 | 1 | 12 | Npat | 0.01 | Old |
| 0.474162 | 1.74727 | 0.017 | 0.007 | 1 | 12 | Gem | 0.01 | Old |
| 0.474162 | 1.746785 | 0.017 | 0.007 | 1 | 12 | Hmx1 | 0.01 | Old |
| 0.474162 | 1.745703 | 0.017 | 0.007 | 1 | 12 | Saxo2 | 0.01 | Old |
| 0.474162 | 1.736583 | 0.017 | 0.007 | 1 | 12 | Atp8b3 | 0.01 | Old |
| 0.474162 | 1.73164 | 0.017 | 0.007 | 1 | 12 | Sgtb | 0.01 | Old |
| 0.474162 | 1.720333 | 0.017 | 0.007 | 1 | 12 | Ncapd3 | 0.01 | Old |
| 0.474162 | 1.703706 | 0.017 | 0.007 | 1 | 12 | Usf3 | 0.01 | Old |
| 0.474162 | 1.690591 | 0.017 | 0.007 | 1 | 12 | Gm16099 | 0.01 | Old |
| 0.474162 | 1.67946 | 0.017 | 0.007 | 1 | 12 | Rybp | 0.01 | Old |
| 0.474162 | 1.667888 | 0.017 | 0.007 | 1 | 12 | Samd12 | 0.01 | Old |
| 0.474162 | 1.664538 | 0.017 | 0.007 | 1 | 12 | Ankrd54 | 0.01 | Old |
| 0.474162 | 1.657979 | 0.017 | 0.007 | 1 | 12 | Stap2 | 0.01 | Old |
| 0.474162 | 1.648418 | 0.017 | 0.007 | 1 | 12 | Zfp81 | 0.01 | Old |
| 0.474162 | 1.638207 | 0.017 | 0.007 | 1 | 12 | Hivep3 | 0.01 | Old |
| 0.474162 | 1.624028 | 0.017 | 0.007 | 1 | 12 | Mbtd1 | 0.01 | Old |
| 0.474162 | 1.622225 | 0.017 | 0.007 | 1 | 12 | Vps11 | 0.01 | Old |
| 0.474162 | 1.616335 | 0.017 | 0.007 | 1 | 12 | Dcaf13 | 0.01 | Old |
| 0.474162 | 1.610021 | 0.017 | 0.007 | 1 | 12 | Cmc4 | 0.01 | Old |
| 0.474162 | 1.60951 | 0.017 | 0.007 | 1 | 12 | Itgb3bp | 0.01 | Old |
| 0.474162 | 1.590637 | 0.017 | 0.007 | 1 | 12 | Cfp | 0.01 | Old |
| 0.474162 | 1.580332 | 0.017 | 0.007 | 1 | 12 | Lrp3 | 0.01 | Old |
| 0.474162 | 1.577499 | 0.017 | 0.007 | 1 | 12 | Txlna | 0.01 | Old |
| 0.474162 | 1.574859 | 0.017 | 0.007 | 1 | 12 | Surf6 | 0.01 | Old |
| 0.474162 | 1.572487 | 0.017 | 0.007 | 1 | 12 | Wdr41 | 0.01 | Old |
| 0.474162 | 1.569582 | 0.017 | 0.007 | 1 | 12 | Vac14 | 0.01 | Old |
| 0.474162 | 1.562568 | 0.017 | 0.007 | 1 | 12 | Otulin | 0.01 | Old |
| 0.474162 | 1.559208 | 0.017 | 0.007 | 1 | 12 | Golga3 | 0.01 | Old |
| 0.474162 | 1.558399 | 0.017 | 0.007 | 1 | 12 | Snip1 | 0.01 | Old |
| 0.474162 | 1.553734 | 0.017 | 0.007 | 1 | 12 | Pcmtd2 | 0.01 | Old |
| 0.474162 | 1.548843 | 0.017 | 0.007 | 1 | 12 | Zfyve27 | 0.01 | Old |
| 0.474162 | 1.547539 | 0.017 | 0.007 | 1 | 12 | Hat1 | 0.01 | Old |

| | | | | | | | | |
|---|---|---|---|---|---|---|---|---|
| 0.474162 | 1.546638 | 0.017 | 0.007 | 1 | 12 | C1qc | 0.01 | Old |
| 0.474162 | 1.545702 | 0.017 | 0.007 | 1 | 12 | Sdsl | 0.01 | Old |
| 0.474162 | 1.536931 | 0.017 | 0.007 | 1 | 12 | Pdxdc1 | 0.01 | Old |
| 0.474162 | 1.535162 | 0.017 | 0.007 | 1 | 12 | Fam135a | 0.01 | Old |
| 0.474162 | 1.535116 | 0.017 | 0.007 | 1 | 12 | Lilr4b | 0.01 | Old |
| 0.474162 | 1.525607 | 0.017 | 0.007 | 1 | 12 | Acox1 | 0.01 | Old |
| 0.474162 | 1.521063 | 0.017 | 0.007 | 1 | 12 | Tnpo1 | 0.01 | Old |
| 0.474162 | 1.511179 | 0.017 | 0.007 | 1 | 12 | Rabl3 | 0.01 | Old |
| 0.474162 | 1.509657 | 0.017 | 0.007 | 1 | 12 | Galnt13 | 0.01 | Old |
| 0.474162 | 1.504176 | 0.017 | 0.007 | 1 | 12 | Zscan26 | 0.01 | Old |
| 0.474162 | 1.503774 | 0.017 | 0.007 | 1 | 12 | Stard4 | 0.01 | Old |
| 0.474162 | 1.5016 | 0.017 | 0.007 | 1 | 12 | Cap2 | 0.01 | Old |
| 0.474162 | 1.487664 | 0.017 | 0.007 | 1 | 12 | 4931428F0 | 0.01 | Old |
| 0.474162 | 1.481942 | 0.017 | 0.007 | 1 | 12 | Smc5 | 0.01 | Old |
| 0.474162 | 1.481031 | 0.017 | 0.007 | 1 | 12 | Mau2 | 0.01 | Old |
| 0.474162 | 1.47123 | 0.017 | 0.007 | 1 | 12 | Gpat4 | 0.01 | Old |
| 0.474162 | 1.468174 | 0.017 | 0.007 | 1 | 12 | Ncoa2 | 0.01 | Old |
| 0.474162 | 1.467393 | 0.017 | 0.007 | 1 | 12 | Gm6563 | 0.01 | Old |
| 0.474162 | 1.465436 | 0.017 | 0.007 | 1 | 12 | Ciao1 | 0.01 | Old |
| 0.474162 | 1.460446 | 0.017 | 0.007 | 1 | 12 | Tob1 | 0.01 | Old |
| 0.474162 | 1.457279 | 0.017 | 0.007 | 1 | 12 | Cela1 | 0.01 | Old |
| 0.474162 | 1.45127 | 0.017 | 0.007 | 1 | 12 | Tceal5 | 0.01 | Old |
| 0.474162 | 1.445467 | 0.017 | 0.007 | 1 | 12 | Wdr60 | 0.01 | Old |
| 0.474162 | 1.445436 | 0.017 | 0.007 | 1 | 12 | Mettl6 | 0.01 | Old |
| 0.474162 | 1.443917 | 0.017 | 0.007 | 1 | 12 | Snx31 | 0.01 | Old |
| 0.474162 | 1.432946 | 0.017 | 0.007 | 1 | 12 | Metrn | 0.01 | Old |
| 0.474162 | 1.428434 | 0.017 | 0.007 | 1 | 12 | Fmnl1 | 0.01 | Old |
| 0.474162 | 1.427454 | 0.017 | 0.007 | 1 | 12 | Dapk1 | 0.01 | Old |
| 0.474162 | 1.421263 | 0.017 | 0.007 | 1 | 12 | Adat2 | 0.01 | Old |
| 0.474162 | 1.41953 | 0.017 | 0.007 | 1 | 12 | Mirg | 0.01 | Old |
| 0.474162 | 1.405649 | 0.017 | 0.007 | 1 | 12 | Rassf7 | 0.01 | Old |
| 0.474162 | 1.400522 | 0.017 | 0.007 | 1 | 12 | Rec8 | 0.01 | Old |
| 0.474162 | 1.400387 | 0.017 | 0.007 | 1 | 12 | Cwc22 | 0.01 | Old |
| 0.474162 | 1.399762 | 0.017 | 0.007 | 1 | 12 | Grin1 | 0.01 | Old |
| 0.474162 | 1.39822 | 0.017 | 0.007 | 1 | 12 | Tm6sf1 | 0.01 | Old |
| 0.474162 | 1.389544 | 0.017 | 0.007 | 1 | 12 | Rbbp8 | 0.01 | Old |
| 0.474162 | 1.388684 | 0.017 | 0.007 | 1 | 12 | Stam | 0.01 | Old |
| 0.474162 | 1.387971 | 0.017 | 0.007 | 1 | 12 | Aar2 | 0.01 | Old |
| 0.474162 | 1.387431 | 0.017 | 0.007 | 1 | 12 | Znrd1as | 0.01 | Old |
| 0.474162 | 1.385866 | 0.017 | 0.007 | 1 | 12 | Zfp707 | 0.01 | Old |
| 0.474162 | 1.383575 | 0.017 | 0.007 | 1 | 12 | Bag4 | 0.01 | Old |
| 0.474162 | 1.382783 | 0.017 | 0.007 | 1 | 12 | Maneal | 0.01 | Old |
| 0.474162 | 1.380414 | 0.017 | 0.007 | 1 | 12 | Oraov1 | 0.01 | Old |
| 0.474162 | 1.371181 | 0.017 | 0.007 | 1 | 12 | Bgn | 0.01 | Old |
| 0.474162 | 1.369309 | 0.017 | 0.007 | 1 | 12 | Zcchc3 | 0.01 | Old |

| | | | | | | | | |
|---|---|---|---|---|---|---|---|---|
| 0.474162 | 1.364068 | 0.017 | 0.007 | 1 | 12 | Kndc1 | 0.01 | Old |
| 0.474162 | 1.363705 | 0.017 | 0.007 | 1 | 12 | Zc4h2 | 0.01 | Old |
| 0.474162 | 1.362177 | 0.017 | 0.007 | 1 | 12 | Polk | 0.01 | Old |
| 0.474162 | 1.359807 | 0.017 | 0.007 | 1 | 12 | Fry | 0.01 | Old |
| 0.474162 | 1.340748 | 0.017 | 0.007 | 1 | 12 | Sobp | 0.01 | Old |
| 0.474162 | 1.336955 | 0.017 | 0.007 | 1 | 12 | Rab28 | 0.01 | Old |
| 0.474162 | 1.330313 | 0.017 | 0.007 | 1 | 12 | Hook2 | 0.01 | Old |
| 0.474162 | 1.325464 | 0.017 | 0.007 | 1 | 12 | Josd1 | 0.01 | Old |
| 0.474162 | 1.324782 | 0.017 | 0.007 | 1 | 12 | Tfe3 | 0.01 | Old |
| 0.474162 | 1.31817 | 0.017 | 0.007 | 1 | 12 | Pgbd5 | 0.01 | Old |
| 0.474162 | 1.313962 | 0.017 | 0.007 | 1 | 12 | Kiz | 0.01 | Old |
| 0.474162 | 1.307759 | 0.017 | 0.007 | 1 | 12 | Faim2 | 0.01 | Old |
| 0.474162 | 1.304416 | 0.017 | 0.007 | 1 | 12 | Zmiz2 | 0.01 | Old |
| 0.474162 | 1.302035 | 0.017 | 0.007 | 1 | 12 | Kdsr | 0.01 | Old |
| 0.474162 | 1.297831 | 0.017 | 0.007 | 1 | 12 | 4931406C0 | 0.01 | Old |
| 0.474162 | 1.297167 | 0.017 | 0.007 | 1 | 12 | Vamp1 | 0.01 | Old |
| 0.474162 | 1.287976 | 0.017 | 0.007 | 1 | 12 | Dync2h1 | 0.01 | Old |
| 0.474162 | 1.287598 | 0.017 | 0.007 | 1 | 12 | Fmn1 | 0.01 | Old |
| 0.474162 | 1.287598 | 0.017 | 0.007 | 1 | 12 | Uckl1 | 0.01 | Old |
| 0.474162 | 1.282397 | 0.017 | 0.007 | 1 | 12 | Pptc7 | 0.01 | Old |
| 0.474162 | 1.272006 | 0.017 | 0.007 | 1 | 12 | Clstn3 | 0.01 | Old |
| 0.474162 | 1.267218 | 0.017 | 0.007 | 1 | 12 | Clk3 | 0.01 | Old |
| 0.474162 | 1.260859 | 0.017 | 0.007 | 1 | 12 | Nup43 | 0.01 | Old |
| 0.474162 | 1.26 | 0.017 | 0.007 | 1 | 12 | Arnt | 0.01 | Old |
| 0.474162 | 1.252939 | 0.017 | 0.007 | 1 | 12 | Vps37d | 0.01 | Old |
| 0.474162 | 1.241382 | 0.017 | 0.007 | 1 | 12 | Cul7 | 0.01 | Old |
| 0.474162 | 1.233832 | 0.017 | 0.007 | 1 | 12 | Zfp37 | 0.01 | Old |
| 0.474162 | 1.231376 | 0.017 | 0.007 | 1 | 12 | Klhl28 | 0.01 | Old |
| 0.474162 | 1.2212 | 0.017 | 0.007 | 1 | 12 | Rsph9 | 0.01 | Old |
| 0.474162 | 1.188859 | 0.017 | 0.007 | 1 | 12 | Cacna2d3 | 0.01 | Old |
| 0.474162 | 1.168092 | 0.017 | 0.007 | 1 | 12 | Lyrm7 | 0.01 | Old |
| 0.476149 | 0.400988 | 0.136 | 0.109 | 1 | 12 | Psme2 | 0.027 | Old |
| 0.476388 | 0.097913 | 0.28 | 0.241 | 1 | 12 | 2010111I0 | 0.039 | Old |
| 0.47698 | 1.476449 | 0.017 | 0.007 | 1 | 12 | Ddx47 | 0.01 | Old |
| 0.477741 | 0.345665 | 0.195 | 0.168 | 1 | 12 | Ndufs8 | 0.027 | Old |
| 0.479811 | 1.43732 | 0.017 | 0.007 | 1 | 12 | Rbm15 | 0.01 | Old |
| 0.479811 | 1.433196 | 0.017 | 0.007 | 1 | 12 | Crnkl1 | 0.01 | Old |
| 0.479811 | 1.416051 | 0.017 | 0.007 | 1 | 12 | Rap1gap2 | 0.01 | Old |
| 0.479811 | 1.399414 | 0.017 | 0.007 | 1 | 12 | Rhou | 0.01 | Old |
| 0.479811 | 1.386035 | 0.017 | 0.007 | 1 | 12 | Traf4 | 0.01 | Old |
| 0.479811 | 1.382569 | 0.017 | 0.007 | 1 | 12 | Rest | 0.01 | Old |
| 0.479811 | 1.367946 | 0.017 | 0.007 | 1 | 12 | Ino80e | 0.01 | Old |
| 0.479811 | 1.365092 | 0.017 | 0.007 | 1 | 12 | Npr2 | 0.01 | Old |
| 0.479811 | 1.36232 | 0.017 | 0.007 | 1 | 12 | E2f5 | 0.01 | Old |
| 0.479811 | 1.347773 | 0.017 | 0.007 | 1 | 12 | Crtc3 | 0.01 | Old |

| | | | | | | | | |
|---|---|---|---|---|---|---|---|---|
| 0.479811 | 1.334871 | 0.017 | 0.007 | 1 | 12 | Elp4 | 0.01 | Old |
| 0.479811 | 1.326149 | 0.017 | 0.007 | 1 | 12 | Nop9 | 0.01 | Old |
| 0.479811 | 1.320936 | 0.017 | 0.007 | 1 | 12 | Abcf3 | 0.01 | Old |
| 0.479811 | 1.319422 | 0.017 | 0.007 | 1 | 12 | Lgalsl | 0.01 | Old |
| 0.479811 | 1.304894 | 0.017 | 0.007 | 1 | 12 | Zfp512 | 0.01 | Old |
| 0.479811 | 1.304026 | 0.017 | 0.007 | 1 | 12 | Mbd1 | 0.01 | Old |
| 0.479811 | 1.293577 | 0.017 | 0.007 | 1 | 12 | Ppip5k2 | 0.01 | Old |
| 0.479811 | 1.282165 | 0.017 | 0.007 | 1 | 12 | Pik3cd | 0.01 | Old |
| 0.479811 | 1.26824 | 0.017 | 0.007 | 1 | 12 | 2700062C0 | 0.01 | Old |
| 0.479811 | 1.266411 | 0.017 | 0.007 | 1 | 12 | Wnk3 | 0.01 | Old |
| 0.479811 | 1.257859 | 0.017 | 0.007 | 1 | 12 | Rnd3 | 0.01 | Old |
| 0.479811 | 1.257207 | 0.017 | 0.007 | 1 | 12 | Kcnb2 | 0.01 | Old |
| 0.479811 | 1.255175 | 0.017 | 0.007 | 1 | 12 | Pkn2 | 0.01 | Old |
| 0.479811 | 1.254469 | 0.017 | 0.007 | 1 | 12 | Wdr7 | 0.01 | Old |
| 0.479811 | 1.248931 | 0.017 | 0.007 | 1 | 12 | Sdc4 | 0.01 | Old |
| 0.479811 | 1.242561 | 0.017 | 0.007 | 1 | 12 | Gps2 | 0.01 | Old |
| 0.479811 | 1.235233 | 0.017 | 0.007 | 1 | 12 | Tmem8b | 0.01 | Old |
| 0.479811 | 1.233444 | 0.017 | 0.007 | 1 | 12 | Fkbp9 | 0.01 | Old |
| 0.479811 | 1.231133 | 0.017 | 0.007 | 1 | 12 | Cnot3 | 0.01 | Old |
| 0.479811 | 1.228549 | 0.017 | 0.007 | 1 | 12 | Diaph1 | 0.01 | Old |
| 0.479811 | 1.227706 | 0.017 | 0.007 | 1 | 12 | Gpr135 | 0.01 | Old |
| 0.479811 | 1.22568 | 0.017 | 0.007 | 1 | 12 | Myo1b | 0.01 | Old |
| 0.479811 | 1.220606 | 0.017 | 0.007 | 1 | 12 | Dennd6b | 0.01 | Old |
| 0.479811 | 1.2193 | 0.017 | 0.007 | 1 | 12 | Nbeal1 | 0.01 | Old |
| 0.479811 | 1.218307 | 0.017 | 0.007 | 1 | 12 | Ccl2 | 0.01 | Old |
| 0.479811 | 1.214106 | 0.017 | 0.007 | 1 | 12 | Atg4a | 0.01 | Old |
| 0.479811 | 1.21338 | 0.017 | 0.007 | 1 | 12 | Psen1 | 0.01 | Old |
| 0.479811 | 1.211816 | 0.017 | 0.007 | 1 | 12 | Zfp871 | 0.01 | Old |
| 0.479811 | 1.198384 | 0.017 | 0.007 | 1 | 12 | Stox2 | 0.01 | Old |
| 0.479811 | 1.198201 | 0.017 | 0.007 | 1 | 12 | Exoc2 | 0.01 | Old |
| 0.479811 | 1.196219 | 0.017 | 0.007 | 1 | 12 | Lrrc51 | 0.01 | Old |
| 0.479811 | 1.185659 | 0.017 | 0.007 | 1 | 12 | Usb1 | 0.01 | Old |
| 0.479811 | 1.181143 | 0.017 | 0.007 | 1 | 12 | Trub1 | 0.01 | Old |
| 0.479811 | 1.171095 | 0.017 | 0.007 | 1 | 12 | Ormdl3 | 0.01 | Old |
| 0.479811 | 1.170193 | 0.017 | 0.007 | 1 | 12 | Rexo1 | 0.01 | Old |
| 0.479811 | 1.157844 | 0.017 | 0.007 | 1 | 12 | Nfe2l2 | 0.01 | Old |
| 0.479811 | 1.15211 | 0.017 | 0.007 | 1 | 12 | Iglon5 | 0.01 | Old |
| 0.479811 | 1.150483 | 0.017 | 0.007 | 1 | 12 | Dusp6 | 0.01 | Old |
| 0.479811 | 1.146523 | 0.017 | 0.007 | 1 | 12 | Mctp1 | 0.01 | Old |
| 0.479811 | 1.145544 | 0.017 | 0.007 | 1 | 12 | Ints11 | 0.01 | Old |
| 0.479811 | 1.144863 | 0.017 | 0.007 | 1 | 12 | Wdr59 | 0.01 | Old |
| 0.479811 | 1.144096 | 0.017 | 0.007 | 1 | 12 | Tatdn3 | 0.01 | Old |
| 0.479811 | 1.138296 | 0.017 | 0.007 | 1 | 12 | Tfb2m | 0.01 | Old |
| 0.479811 | 1.130882 | 0.017 | 0.007 | 1 | 12 | 2810429I0 | 0.01 | Old |
| 0.479811 | 1.129336 | 0.017 | 0.007 | 1 | 12 | Abr | 0.01 | Old |

| | | | | | | | | |
|---|---|---|---|---|---|---|---|---|
| 0.479811 | 1.126769 | 0.017 | 0.007 | 1 | 12 | Hmgxb3 | 0.01 | Old |
| 0.479811 | 1.124155 | 0.017 | 0.007 | 1 | 12 | Gm14419 | 0.01 | Old |
| 0.479811 | 1.122719 | 0.017 | 0.007 | 1 | 12 | 4933431E2 | 0.01 | Old |
| 0.479811 | 1.121468 | 0.017 | 0.007 | 1 | 12 | Chd8 | 0.01 | Old |
| 0.479811 | 1.11563 | 0.017 | 0.007 | 1 | 12 | Senp8 | 0.01 | Old |
| 0.479811 | 1.115407 | 0.017 | 0.007 | 1 | 12 | Hey1 | 0.01 | Old |
| 0.479811 | 1.114729 | 0.017 | 0.007 | 1 | 12 | Amer1 | 0.01 | Old |
| 0.479811 | 1.109084 | 0.017 | 0.007 | 1 | 12 | Shkbp1 | 0.01 | Old |
| 0.479811 | 1.105148 | 0.017 | 0.007 | 1 | 12 | Phf8 | 0.01 | Old |
| 0.479811 | 1.083305 | 0.017 | 0.007 | 1 | 12 | Pithd1 | 0.01 | Old |
| 0.479811 | 1.082963 | 0.017 | 0.007 | 1 | 12 | Nrde2 | 0.01 | Old |
| 0.479811 | 1.082012 | 0.017 | 0.007 | 1 | 12 | Stat3 | 0.01 | Old |
| 0.479811 | 1.081737 | 0.017 | 0.007 | 1 | 12 | Smarcad1 | 0.01 | Old |
| 0.479811 | 1.077125 | 0.017 | 0.007 | 1 | 12 | Zfx | 0.01 | Old |
| 0.479811 | 1.07664 | 0.017 | 0.007 | 1 | 12 | Gigyf2 | 0.01 | Old |
| 0.479811 | 1.067866 | 0.017 | 0.007 | 1 | 12 | D230017M | 0.01 | Old |
| 0.479811 | 1.064872 | 0.017 | 0.007 | 1 | 12 | Tmem251 | 0.01 | Old |
| 0.479811 | 1.059703 | 0.017 | 0.007 | 1 | 12 | D5Ertd579 | 0.01 | Old |
| 0.479811 | 1.052772 | 0.017 | 0.007 | 1 | 12 | Tbc1d12 | 0.01 | Old |
| 0.479811 | 1.052077 | 0.017 | 0.007 | 1 | 12 | Psmd5 | 0.01 | Old |
| 0.479811 | 1.049038 | 0.017 | 0.007 | 1 | 12 | Slc35f3 | 0.01 | Old |
| 0.479811 | 1.047993 | 0.017 | 0.007 | 1 | 12 | Maf | 0.01 | Old |
| 0.479811 | 1.047235 | 0.017 | 0.007 | 1 | 12 | Dis3l2 | 0.01 | Old |
| 0.479811 | 1.044191 | 0.017 | 0.007 | 1 | 12 | Platr25 | 0.01 | Old |
| 0.479811 | 1.043977 | 0.017 | 0.007 | 1 | 12 | AU022252 | 0.01 | Old |
| 0.479811 | 1.041725 | 0.017 | 0.007 | 1 | 12 | Crebzf | 0.01 | Old |
| 0.479811 | 1.039959 | 0.017 | 0.007 | 1 | 12 | Pou2f2 | 0.01 | Old |
| 0.479811 | 1.037216 | 0.017 | 0.007 | 1 | 12 | Ivns1abp | 0.01 | Old |
| 0.479811 | 1.034709 | 0.017 | 0.007 | 1 | 12 | Ascl1 | 0.01 | Old |
| 0.479811 | 1.034235 | 0.017 | 0.007 | 1 | 12 | Tubg2 | 0.01 | Old |
| 0.479811 | 1.017213 | 0.017 | 0.007 | 1 | 12 | Tgfbi | 0.01 | Old |
| 0.479811 | 1.011483 | 0.017 | 0.007 | 1 | 12 | 2210008F0 | 0.01 | Old |
| 0.479811 | 1.008918 | 0.017 | 0.007 | 1 | 12 | Zdhhc6 | 0.01 | Old |
| 0.479811 | 1.006036 | 0.017 | 0.007 | 1 | 12 | Ctsc | 0.01 | Old |
| 0.479811 | 1.005101 | 0.017 | 0.007 | 1 | 12 | Edem2 | 0.01 | Old |
| 0.479811 | 1.004316 | 0.017 | 0.007 | 1 | 12 | Zfp608 | 0.01 | Old |
| 0.479811 | 1.001801 | 0.017 | 0.007 | 1 | 12 | Sirt1 | 0.01 | Old |
| 0.479811 | 0.9963 | 0.017 | 0.007 | 1 | 12 | Ccdc22 | 0.01 | Old |
| 0.479811 | 0.994707 | 0.017 | 0.007 | 1 | 12 | Itch | 0.01 | Old |
| 0.479811 | 0.989029 | 0.017 | 0.007 | 1 | 12 | Tmem91 | 0.01 | Old |
| 0.479811 | 0.975295 | 0.017 | 0.007 | 1 | 12 | Ano10 | 0.01 | Old |
| 0.479811 | 0.963815 | 0.017 | 0.007 | 1 | 12 | Fam219b | 0.01 | Old |
| 0.479811 | 0.940944 | 0.017 | 0.007 | 1 | 12 | A4galt | 0.01 | Old |
| 0.479811 | 0.919776 | 0.017 | 0.007 | 1 | 12 | Col4a3bp | 0.01 | Old |
| 0.479811 | 0.918105 | 0.017 | 0.007 | 1 | 12 | Lsm2 | 0.01 | Old |

| | | | | | | | | |
|---|---|---|---|---|---|---|---|---|
| 0.479811 | 0.910301 | 0.017 | 0.007 | 1 | 12 | Cstf1 | 0.01 | Old |
| 0.479811 | 0.908657 | 0.017 | 0.007 | 1 | 12 | Dtd2 | 0.01 | Old |
| 0.479811 | 0.900052 | 0.017 | 0.007 | 1 | 12 | Plcg1 | 0.01 | Old |
| 0.479811 | 0.886879 | 0.017 | 0.007 | 1 | 12 | Clock | 0.01 | Old |
| 0.479811 | 0.857161 | 0.017 | 0.007 | 1 | 12 | Itpa | 0.01 | Old |
| 0.479811 | 0.834042 | 0.017 | 0.007 | 1 | 12 | Lrp10 | 0.01 | Old |
| 0.479811 | 0.799252 | 0.017 | 0.007 | 1 | 12 | Map2k7 | 0.01 | Old |
| 0.479811 | 0.762392 | 0.017 | 0.007 | 1 | 12 | Gfm2 | 0.01 | Old |
| 0.479811 | 0.694145 | 0.017 | 0.007 | 1 | 12 | Zdhhc20 | 0.01 | Old |
| 0.479811 | 0.599582 | 0.017 | 0.007 | 1 | 12 | Ppwd1 | 0.01 | Old |
| 0.479811 | 0.598687 | 0.017 | 0.007 | 1 | 12 | Smpd4 | 0.01 | Old |
| 0.479811 | 0.58732 | 0.017 | 0.007 | 1 | 12 | Rab8b | 0.01 | Old |
| 0.479811 | 0.581374 | 0.017 | 0.007 | 1 | 12 | Ipo13 | 0.01 | Old |
| 0.479811 | 0.580402 | 0.017 | 0.007 | 1 | 12 | Epc2 | 0.01 | Old |
| 0.479811 | 0.552642 | 0.017 | 0.007 | 1 | 12 | Ptdss1 | 0.01 | Old |
| 0.479811 | 0.531929 | 0.017 | 0.007 | 1 | 12 | Sf3b3 | 0.01 | Old |
| 0.479811 | 0.531722 | 0.017 | 0.007 | 1 | 12 | Fnbp1 | 0.01 | Old |
| 0.479811 | 0.48921 | 0.017 | 0.007 | 1 | 12 | Cpeb3 | 0.01 | Old |
| 0.479811 | 0.45627 | 0.017 | 0.007 | 1 | 12 | Nceh1 | 0.01 | Old |
| 0.479811 | 0.410402 | 0.017 | 0.007 | 1 | 12 | Mtm1 | 0.01 | Old |
| 0.479811 | 0.386522 | 0.017 | 0.007 | 1 | 12 | Wwp2 | 0.01 | Old |
| 0.480317 | 0.083997 | 0.212 | 0.175 | 1 | 12 | Sh3glb1 | 0.037 | Old |
| 0.48098 | 0.304395 | 0.161 | 0.131 | 1 | 12 | Tmem179 | 0.03 | Old |
| 0.482647 | 0.985345 | 0.017 | 0.007 | 1 | 12 | Kmt2b | 0.01 | Old |
| 0.482672 | 0.313473 | 0.136 | 0.109 | 1 | 12 | Ttll7 | 0.027 | Old |
| 0.485084 | 0.312155 | 0.127 | 0.102 | 1 | 12 | Pcbp1 | 0.025 | Old |
| 0.485402 | 0.303678 | 0.178 | 0.146 | 1 | 12 | Zfp706 | 0.032 | Old |
| 0.485497 | 1.129202 | 0.017 | 0.007 | 1 | 12 | Eml3 | 0.01 | Old |
| 0.485497 | 1.078076 | 0.017 | 0.007 | 1 | 12 | Csnk1g1 | 0.01 | Old |
| 0.485497 | 1.048473 | 0.017 | 0.007 | 1 | 12 | Sec24b | 0.01 | Old |
| 0.485497 | 1.030003 | 0.017 | 0.007 | 1 | 12 | Cers5 | 0.01 | Old |
| 0.485497 | 1.022745 | 0.017 | 0.007 | 1 | 12 | Pank1 | 0.01 | Old |
| 0.485497 | 1.020433 | 0.017 | 0.007 | 1 | 12 | Stab1 | 0.01 | Old |
| 0.485497 | 1.018853 | 0.017 | 0.007 | 1 | 12 | Taf3 | 0.01 | Old |
| 0.485497 | 1.006587 | 0.017 | 0.007 | 1 | 12 | Med4 | 0.01 | Old |
| 0.485497 | 1.006058 | 0.017 | 0.007 | 1 | 12 | Tpp1 | 0.01 | Old |
| 0.485497 | 1.003938 | 0.017 | 0.007 | 1 | 12 | Ramp1 | 0.01 | Old |
| 0.485497 | 0.994536 | 0.017 | 0.007 | 1 | 12 | Neurl1b | 0.01 | Old |
| 0.485497 | 0.984531 | 0.017 | 0.007 | 1 | 12 | Yme1l1 | 0.01 | Old |
| 0.485497 | 0.965401 | 0.017 | 0.007 | 1 | 12 | Qsox1 | 0.01 | Old |
| 0.485497 | 0.961375 | 0.017 | 0.007 | 1 | 12 | Dynlt1f | 0.01 | Old |
| 0.485497 | 0.960013 | 0.017 | 0.007 | 1 | 12 | Avl9 | 0.01 | Old |
| 0.485497 | 0.949117 | 0.017 | 0.007 | 1 | 12 | Ttc39b | 0.01 | Old |
| 0.485497 | 0.948459 | 0.017 | 0.007 | 1 | 12 | Tmcc2 | 0.01 | Old |
| 0.485497 | 0.946959 | 0.017 | 0.007 | 1 | 12 | Kdm6a | 0.01 | Old |

| | | | | | | | | |
|---|---|---|---|---|---|---|---|---|
| 0.485497 | 0.944637 | 0.017 | 0.007 | 1 | 12 | Kank3 | 0.01 | Old |
| 0.485497 | 0.944385 | 0.017 | 0.007 | 1 | 12 | Capn5 | 0.01 | Old |
| 0.485497 | 0.932474 | 0.017 | 0.007 | 1 | 12 | Xrcc4 | 0.01 | Old |
| 0.485497 | 0.931424 | 0.017 | 0.007 | 1 | 12 | Zbtb18 | 0.01 | Old |
| 0.485497 | 0.929503 | 0.017 | 0.007 | 1 | 12 | Suco | 0.01 | Old |
| 0.485497 | 0.928862 | 0.017 | 0.007 | 1 | 12 | Cited1 | 0.01 | Old |
| 0.485497 | 0.927638 | 0.017 | 0.007 | 1 | 12 | Setdb1 | 0.01 | Old |
| 0.485497 | 0.927616 | 0.017 | 0.007 | 1 | 12 | Ldlrad3 | 0.01 | Old |
| 0.485497 | 0.905933 | 0.017 | 0.007 | 1 | 12 | Igsf9b | 0.01 | Old |
| 0.485497 | 0.894646 | 0.017 | 0.007 | 1 | 12 | Zfp40 | 0.01 | Old |
| 0.485497 | 0.884723 | 0.017 | 0.007 | 1 | 12 | Gdpd5 | 0.01 | Old |
| 0.485497 | 0.88178 | 0.017 | 0.007 | 1 | 12 | Haus2 | 0.01 | Old |
| 0.485497 | 0.881751 | 0.017 | 0.007 | 1 | 12 | Ccne2 | 0.01 | Old |
| 0.485497 | 0.88112 | 0.017 | 0.007 | 1 | 12 | Siah1b | 0.01 | Old |
| 0.485497 | 0.880665 | 0.017 | 0.007 | 1 | 12 | Dsel | 0.01 | Old |
| 0.485497 | 0.876295 | 0.017 | 0.007 | 1 | 12 | Mapk11 | 0.01 | Old |
| 0.485497 | 0.867625 | 0.017 | 0.007 | 1 | 12 | Zfp868 | 0.01 | Old |
| 0.485497 | 0.86311 | 0.017 | 0.007 | 1 | 12 | Mrm3 | 0.01 | Old |
| 0.485497 | 0.850851 | 0.017 | 0.007 | 1 | 12 | P3h4 | 0.01 | Old |
| 0.485497 | 0.848251 | 0.017 | 0.007 | 1 | 12 | Ap5z1 | 0.01 | Old |
| 0.485497 | 0.847713 | 0.017 | 0.007 | 1 | 12 | Mrpl37 | 0.01 | Old |
| 0.485497 | 0.846344 | 0.017 | 0.007 | 1 | 12 | Arhgap23 | 0.01 | Old |
| 0.485497 | 0.843405 | 0.017 | 0.007 | 1 | 12 | Strn | 0.01 | Old |
| 0.485497 | 0.839041 | 0.017 | 0.007 | 1 | 12 | Slc35a2 | 0.01 | Old |
| 0.485497 | 0.838564 | 0.017 | 0.007 | 1 | 12 | Rnf32 | 0.01 | Old |
| 0.485497 | 0.828213 | 0.017 | 0.007 | 1 | 12 | Sp3os | 0.01 | Old |
| 0.485497 | 0.823941 | 0.017 | 0.007 | 1 | 12 | Plxdc2 | 0.01 | Old |
| 0.485497 | 0.81501 | 0.017 | 0.007 | 1 | 12 | Il13ra1 | 0.01 | Old |
| 0.485497 | 0.813467 | 0.017 | 0.007 | 1 | 12 | Gtf3c2 | 0.01 | Old |
| 0.485497 | 0.807406 | 0.017 | 0.007 | 1 | 12 | Agl | 0.01 | Old |
| 0.485497 | 0.801105 | 0.017 | 0.007 | 1 | 12 | Naa60 | 0.01 | Old |
| 0.485497 | 0.800886 | 0.017 | 0.007 | 1 | 12 | Kbtbd7 | 0.01 | Old |
| 0.485497 | 0.798841 | 0.017 | 0.007 | 1 | 12 | Rasl11b | 0.01 | Old |
| 0.485497 | 0.791355 | 0.017 | 0.007 | 1 | 12 | Heatr6 | 0.01 | Old |
| 0.485497 | 0.791265 | 0.017 | 0.007 | 1 | 12 | A830010M | 0.01 | Old |
| 0.485497 | 0.788219 | 0.017 | 0.007 | 1 | 12 | Relb | 0.01 | Old |
| 0.485497 | 0.779546 | 0.017 | 0.007 | 1 | 12 | Fam169a | 0.01 | Old |
| 0.485497 | 0.753531 | 0.017 | 0.007 | 1 | 12 | Fkbpl | 0.01 | Old |
| 0.485497 | 0.749686 | 0.017 | 0.007 | 1 | 12 | Stk38l | 0.01 | Old |
| 0.485497 | 0.740234 | 0.017 | 0.007 | 1 | 12 | Gm15446 | 0.01 | Old |
| 0.485497 | 0.737861 | 0.017 | 0.007 | 1 | 12 | Tctn1 | 0.01 | Old |
| 0.485497 | 0.733509 | 0.017 | 0.007 | 1 | 12 | Heca | 0.01 | Old |
| 0.485497 | 0.727727 | 0.017 | 0.007 | 1 | 12 | Fcho2 | 0.01 | Old |
| 0.485497 | 0.726199 | 0.017 | 0.007 | 1 | 12 | Uck1 | 0.01 | Old |
| 0.485497 | 0.719955 | 0.017 | 0.007 | 1 | 12 | Sppl2b | 0.01 | Old |

| | | | | | | | | |
|---|---|---|---|---|---|---|---|---|
| 0.485497 | 0.718571 | 0.017 | 0.007 | 1 | 12 | Man1b1 | 0.01 | Old |
| 0.485497 | 0.716614 | 0.017 | 0.007 | 1 | 12 | 2310001H1 | 0.01 | Old |
| 0.485497 | 0.716336 | 0.017 | 0.007 | 1 | 12 | Rbm7 | 0.01 | Old |
| 0.485497 | 0.713764 | 0.017 | 0.007 | 1 | 12 | Pianp | 0.01 | Old |
| 0.485497 | 0.713547 | 0.017 | 0.007 | 1 | 12 | Hmces | 0.01 | Old |
| 0.485497 | 0.701507 | 0.017 | 0.007 | 1 | 12 | Fabp4 | 0.01 | Old |
| 0.485497 | 0.678606 | 0.017 | 0.007 | 1 | 12 | Ano8 | 0.01 | Old |
| 0.485497 | 0.673632 | 0.017 | 0.007 | 1 | 12 | Spire2 | 0.01 | Old |
| 0.485497 | 0.66653 | 0.017 | 0.007 | 1 | 12 | Dcaf15 | 0.01 | Old |
| 0.485497 | 0.630968 | 0.017 | 0.007 | 1 | 12 | Bud23 | 0.01 | Old |
| 0.485497 | 0.624211 | 0.017 | 0.007 | 1 | 12 | Anapc1 | 0.01 | Old |
| 0.485497 | 0.622948 | 0.017 | 0.007 | 1 | 12 | Apeh | 0.01 | Old |
| 0.485497 | 0.620217 | 0.017 | 0.007 | 1 | 12 | Washc4 | 0.01 | Old |
| 0.485497 | 0.60166 | 0.017 | 0.007 | 1 | 12 | Trmt12 | 0.01 | Old |
| 0.485497 | 0.588305 | 0.017 | 0.007 | 1 | 12 | Gm14325 | 0.01 | Old |
| 0.485497 | 0.581421 | 0.017 | 0.007 | 1 | 12 | Ppp2r3d | 0.01 | Old |
| 0.485497 | 0.574378 | 0.017 | 0.007 | 1 | 12 | Mex3c | 0.01 | Old |
| 0.485497 | 0.568543 | 0.017 | 0.007 | 1 | 12 | Parn | 0.01 | Old |
| 0.485497 | 0.560239 | 0.017 | 0.007 | 1 | 12 | Arfgap3 | 0.01 | Old |
| 0.485497 | 0.540711 | 0.017 | 0.007 | 1 | 12 | Baz1b | 0.01 | Old |
| 0.485497 | 0.539979 | 0.017 | 0.007 | 1 | 12 | Edem1 | 0.01 | Old |
| 0.485497 | 0.530144 | 0.017 | 0.007 | 1 | 12 | 1700113A1 | 0.01 | Old |
| 0.485497 | 0.520361 | 0.017 | 0.007 | 1 | 12 | Dhrs4 | 0.01 | Old |
| 0.485497 | 0.372485 | 0.017 | 0.007 | 1 | 12 | Cog1 | 0.01 | Old |
| 0.485497 | 0.353826 | 0.017 | 0.007 | 1 | 12 | Phf23 | 0.01 | Old |
| 0.485497 | 0.279094 | 0.017 | 0.007 | 1 | 12 | Igfbp4 | 0.01 | Old |
| 0.485497 | 0.273892 | 0.017 | 0.007 | 1 | 12 | Vps13a | 0.01 | Old |
| 0.485497 | 0.261992 | 0.017 | 0.007 | 1 | 12 | Anks1 | 0.01 | Old |
| 0.485497 | 0.255795 | 0.017 | 0.007 | 1 | 12 | Pigf | 0.01 | Old |
| 0.485497 | 0.240617 | 0.017 | 0.007 | 1 | 12 | Tcf20 | 0.01 | Old |
| 0.485497 | 0.221785 | 0.017 | 0.007 | 1 | 12 | Med1 | 0.01 | Old |
| 0.485497 | 0.194598 | 0.017 | 0.007 | 1 | 12 | Ppp1r9b | 0.01 | Old |
| 0.485497 | 0.188956 | 0.017 | 0.007 | 1 | 12 | Aif1 | 0.01 | Old |
| 0.485497 | 0.167699 | 0.017 | 0.007 | 1 | 12 | Ccl8 | 0.01 | Old |
| 0.485497 | 0.166486 | 0.017 | 0.007 | 1 | 12 | Sumf1 | 0.01 | Old |
| 0.485497 | 0.161302 | 0.017 | 0.007 | 1 | 12 | Cwf19l2 | 0.01 | Old |
| 0.485497 | 0.086446 | 0.017 | 0.007 | 1 | 12 | Fam91a1 | 0.01 | Old |
| 0.485636 | 0.474551 | 0.229 | 0.197 | 1 | 12 | 0610012G0 | 0.032 | Old |
| 0.485993 | 0.605354 | 0.119 | 0.095 | 1 | 12 | Pcbp2 | 0.024 | Old |
| 0.487058 | 0.175406 | 0.161 | 0.131 | 1 | 12 | Wbp4 | 0.03 | Old |
| 0.487376 | 0.189183 | 0.186 | 0.153 | 1 | 12 | Tpgs2 | 0.033 | Old |
| 0.489751 | 0.250559 | 0.322 | 0.285 | 1 | 12 | Ass1 | 0.037 | Old |
| 0.49022 | 0.245043 | 0.203 | 0.168 | 1 | 12 | Arpc5 | 0.035 | Old |
| 0.490655 | 0.219046 | 0.186 | 0.161 | 1 | 12 | Pitpna | 0.025 | Old |
| 0.491232 | 0.184728 | 0.195 | 0.168 | 1 | 12 | Igfbp7 | 0.027 | Old |

| | | | | | | | | |
|---|---|---|---|---|---|---|---|---|
| 0.491469 | 0.057482 | 0.195 | 0.153 | 1 | 12 | Rbm39 | 0.042 | Old |
| 0.492931 | 0.272614 | 0.322 | 0.292 | 1 | 12 | Ptov1 | 0.03 | Old |
| 0.493022 | 0.385451 | 0.136 | 0.109 | 1 | 12 | Nckap1 | 0.027 | Old |
| 0.493089 | 0.033691 | 0.212 | 0.168 | 1 | 12 | Snrpe | 0.044 | Old |
| 0.493329 | 0.220758 | 0.373 | 0.336 | 1 | 12 | Naca | 0.037 | Old |
| 0.493769 | 0.317751 | 0.178 | 0.146 | 1 | 12 | 1810058I24 | 0.032 | Old |
| 0.495092 | 0.133781 | 0.297 | 0.255 | 1 | 12 | Eef1g | 0.042 | Old |
| 0.497916 | 0.103434 | 0.407 | 0.365 | 1 | 12 | Olfm1 | 0.042 | Old |
| 0.502892 | 0.532863 | 0.102 | 0.08 | 1 | 12 | Dnajc8 | 0.022 | Old |
| 0.502893 | 0.550099 | 0.102 | 0.08 | 1 | 12 | Ccdc88a | 0.022 | Old |
| 0.504978 | 0.357616 | 0.203 | 0.175 | 1 | 12 | Ndufa12 | 0.028 | Old |
| 0.50711 | 0.463944 | 0.11 | 0.088 | 1 | 12 | Gnl3l | 0.022 | Old |
| 0.50873 | 0.350649 | 0.186 | 0.161 | 1 | 12 | C1qbp | 0.025 | Old |
| 0.510544 | 0.38214 | 0.127 | 0.102 | 1 | 12 | Mrpl17 | 0.025 | Old |
| 0.51065 | 0.122613 | 0.449 | 0.394 | 1 | 12 | Sh3bgrl3 | 0.055 | Old |
| 0.51347 | 0.227863 | 0.11 | 0.088 | 1 | 12 | Tmem208 | 0.022 | Old |
| 0.51347 | 0.412284 | 0.11 | 0.088 | 1 | 12 | Nap1l1 | 0.022 | Old |
| 0.513878 | 0.42048 | 0.102 | 0.08 | 1 | 12 | Fmc1 | 0.022 | Old |
| 0.514522 | 0.374834 | 0.127 | 0.102 | 1 | 12 | Zfp91 | 0.025 | Old |
| 0.515941 | 0.365187 | 0.144 | 0.117 | 1 | 12 | Cltc | 0.027 | Old |
| 0.516401 | 0.609788 | 0.085 | 0.066 | 1 | 12 | Cnrip1 | 0.019 | Old |
| 0.518487 | 0.391607 | 0.119 | 0.095 | 1 | 12 | Id1 | 0.024 | Old |
| 0.520266 | 0.37195 | 0.093 | 0.073 | 1 | 12 | Rtf2 | 0.02 | Old |
| 0.521542 | 0.834344 | 0.068 | 0.051 | 1 | 12 | Utp3 | 0.017 | Old |
| 0.521568 | 0.235706 | 0.246 | 0.204 | 1 | 12 | Mien1 | 0.042 | Old |
| 0.523392 | 1.563311 | 0.025 | 0.015 | 1 | 12 | Aaed1 | 0.01 | Old |
| 0.523392 | 1.416596 | 0.025 | 0.015 | 1 | 12 | Slc9b2 | 0.01 | Old |
| 0.523392 | 1.403045 | 0.025 | 0.015 | 1 | 12 | Nrip3 | 0.01 | Old |
| 0.523392 | 1.3717 | 0.025 | 0.015 | 1 | 12 | Mpv17l | 0.01 | Old |
| 0.523392 | 1.333403 | 0.025 | 0.015 | 1 | 12 | Gm17484 | 0.01 | Old |
| 0.523392 | 1.327169 | 0.025 | 0.015 | 1 | 12 | Gtf3c1 | 0.01 | Old |
| 0.523392 | 1.316167 | 0.025 | 0.015 | 1 | 12 | Hps4 | 0.01 | Old |
| 0.523392 | 1.250621 | 0.025 | 0.015 | 1 | 12 | Naa10 | 0.01 | Old |
| 0.523392 | 1.234409 | 0.025 | 0.015 | 1 | 12 | Upb1 | 0.01 | Old |
| 0.523392 | 1.209044 | 0.025 | 0.015 | 1 | 12 | Surf2 | 0.01 | Old |
| 0.523392 | 1.184344 | 0.025 | 0.015 | 1 | 12 | Laptm5 | 0.01 | Old |
| 0.523392 | 1.167193 | 0.025 | 0.015 | 1 | 12 | Cnot6l | 0.01 | Old |
| 0.523392 | 1.139671 | 0.025 | 0.015 | 1 | 12 | Zfp622 | 0.01 | Old |
| 0.523392 | 1.124368 | 0.025 | 0.015 | 1 | 12 | Tead1 | 0.01 | Old |
| 0.523392 | 1.113617 | 0.025 | 0.015 | 1 | 12 | Klhl9 | 0.01 | Old |
| 0.523392 | 1.113157 | 0.025 | 0.015 | 1 | 12 | Med27 | 0.01 | Old |
| 0.523392 | 1.082451 | 0.025 | 0.015 | 1 | 12 | Dhx29 | 0.01 | Old |
| 0.523392 | 1.057191 | 0.025 | 0.015 | 1 | 12 | Ube2t | 0.01 | Old |
| 0.523392 | 1.051944 | 0.025 | 0.015 | 1 | 12 | Gale | 0.01 | Old |
| 0.523392 | 1.050217 | 0.025 | 0.015 | 1 | 12 | Nosip | 0.01 | Old |

| | | | | | | | | |
|---|---|---|---|---|---|---|---|---|
| 0.523392 | 1.009594 | 0.025 | 0.015 | 1 | 12 | Tvp23b | 0.01 | Old |
| 0.523392 | 0.981757 | 0.025 | 0.015 | 1 | 12 | Mafb | 0.01 | Old |
| 0.523735 | 0.676664 | 0.127 | 0.109 | 1 | 12 | Ppp1r2 | 0.018 | Old |
| 0.523793 | 0.251383 | 0.136 | 0.109 | 1 | 12 | Mcrip1 | 0.027 | Old |
| 0.524702 | 0.368394 | 0.119 | 0.095 | 1 | 12 | Tpd52 | 0.024 | Old |
| 0.526105 | 0.014832 | 0.076 | 0.102 | 1 | 12 | Ptprn | -0.026 | Old |
| 0.526139 | 0.207626 | 0.339 | 0.299 | 1 | 12 | Ndufb4 | 0.04 | Old |
| 0.527328 | 0.186693 | 0.729 | 0.708 | 1 | 12 | Cox5a | 0.021 | Old |
| 0.528016 | 1.082324 | 0.025 | 0.015 | 1 | 12 | Glipr2 | 0.01 | Old |
| 0.528018 | 1.642359 | 0.025 | 0.015 | 1 | 12 | Syt2 | 0.01 | Old |
| 0.528018 | 1.275616 | 0.025 | 0.015 | 1 | 12 | Ankrd10 | 0.01 | Old |
| 0.528018 | 1.212371 | 0.025 | 0.015 | 1 | 12 | Sos2 | 0.01 | Old |
| 0.528018 | 1.208816 | 0.025 | 0.015 | 1 | 12 | Rpap3 | 0.01 | Old |
| 0.528018 | 1.194061 | 0.025 | 0.015 | 1 | 12 | Trak2 | 0.01 | Old |
| 0.528018 | 1.141082 | 0.025 | 0.015 | 1 | 12 | Ankrd39 | 0.01 | Old |
| 0.528018 | 1.12478 | 0.025 | 0.015 | 1 | 12 | Zcchc10 | 0.01 | Old |
| 0.528018 | 1.025544 | 0.025 | 0.015 | 1 | 12 | Pdss2 | 0.01 | Old |
| 0.528018 | 1.013967 | 0.025 | 0.015 | 1 | 12 | Acbd4 | 0.01 | Old |
| 0.528018 | 1.007588 | 0.025 | 0.015 | 1 | 12 | Mtap | 0.01 | Old |
| 0.528018 | 0.976095 | 0.025 | 0.015 | 1 | 12 | Dnajc14 | 0.01 | Old |
| 0.528018 | 0.970464 | 0.025 | 0.015 | 1 | 12 | Coq6 | 0.01 | Old |
| 0.528018 | 0.948299 | 0.025 | 0.015 | 1 | 12 | Eapp | 0.01 | Old |
| 0.528018 | 0.946678 | 0.025 | 0.015 | 1 | 12 | Rprd1b | 0.01 | Old |
| 0.528018 | 0.890439 | 0.025 | 0.015 | 1 | 12 | Irf2 | 0.01 | Old |
| 0.528018 | 0.824889 | 0.025 | 0.015 | 1 | 12 | Zfp358 | 0.01 | Old |
| 0.528353 | 0.119003 | 0.153 | 0.124 | 1 | 12 | Mlf2 | 0.029 | Old |
| 0.52841 | 0.495094 | 0.093 | 0.073 | 1 | 12 | Wdr6 | 0.02 | Old |
| 0.529181 | 0.108169 | 0.237 | 0.212 | 1 | 12 | Dbi | 0.025 | Old |
| 0.529714 | 0.653891 | 0.068 | 0.051 | 1 | 12 | Nt5m | 0.017 | Old |
| 0.529785 | 0.438193 | 0.085 | 0.066 | 1 | 12 | Vegfb | 0.019 | Old |
| 0.530337 | 0.877483 | 0.025 | 0.015 | 1 | 12 | A430005L1 | 0.01 | Old |
| 0.53062 | 0.105745 | 0.559 | 0.489 | 1 | 12 | Rps18 | 0.07 | Old |
| 0.531671 | 0.958548 | 0.059 | 0.044 | 1 | 12 | Exoc7 | 0.015 | Old |
| 0.532452 | 0.607574 | 0.068 | 0.051 | 1 | 12 | Tra2a | 0.017 | Old |
| 0.532664 | 1.63044 | 0.025 | 0.015 | 1 | 12 | Limd2 | 0.01 | Old |
| 0.532664 | 1.22159 | 0.025 | 0.015 | 1 | 12 | Lrrc49 | 0.01 | Old |
| 0.532664 | 1.186878 | 0.025 | 0.015 | 1 | 12 | Scpep1 | 0.01 | Old |
| 0.532664 | 1.105078 | 0.025 | 0.015 | 1 | 12 | Phpt1 | 0.01 | Old |
| 0.532664 | 1.0755 | 0.025 | 0.015 | 1 | 12 | Gstm7 | 0.01 | Old |
| 0.532664 | 1.039659 | 0.025 | 0.015 | 1 | 12 | Ank | 0.01 | Old |
| 0.532664 | 1.039608 | 0.025 | 0.015 | 1 | 12 | Kpna4 | 0.01 | Old |
| 0.532664 | 1.031065 | 0.025 | 0.015 | 1 | 12 | Yipf3 | 0.01 | Old |
| 0.532664 | 0.943567 | 0.025 | 0.015 | 1 | 12 | Camk2g | 0.01 | Old |
| 0.532664 | 0.93777 | 0.025 | 0.015 | 1 | 12 | Trim2 | 0.01 | Old |
| 0.532664 | 0.935622 | 0.025 | 0.015 | 1 | 12 | Pdcd2l | 0.01 | Old |

| | | | | | | | | |
|---|---|---|---|---|---|---|---|---|
| 0.532664 | 0.931071 | 0.025 | 0.015 | 1 | 12 | Grb14 | 0.01 | Old |
| 0.532664 | 0.928011 | 0.025 | 0.015 | 1 | 12 | Lemd2 | 0.01 | Old |
| 0.532664 | 0.926481 | 0.025 | 0.015 | 1 | 12 | Arhgap10 | 0.01 | Old |
| 0.532664 | 0.924837 | 0.025 | 0.015 | 1 | 12 | Hacd2 | 0.01 | Old |
| 0.532664 | 0.922993 | 0.025 | 0.015 | 1 | 12 | Ptgr2 | 0.01 | Old |
| 0.532664 | 0.92048 | 0.025 | 0.015 | 1 | 12 | Apoa2 | 0.01 | Old |
| 0.532664 | 0.911423 | 0.025 | 0.015 | 1 | 12 | Rps6kb2 | 0.01 | Old |
| 0.532664 | 0.910171 | 0.025 | 0.015 | 1 | 12 | Sdhaf3 | 0.01 | Old |
| 0.532664 | 0.907643 | 0.025 | 0.015 | 1 | 12 | Atg9a | 0.01 | Old |
| 0.532664 | 0.907002 | 0.025 | 0.015 | 1 | 12 | Adgrl3 | 0.01 | Old |
| 0.532664 | 0.902852 | 0.025 | 0.015 | 1 | 12 | Gm11837 | 0.01 | Old |
| 0.532664 | 0.900361 | 0.025 | 0.015 | 1 | 12 | 2700081O1 | 0.01 | Old |
| 0.532664 | 0.896065 | 0.025 | 0.015 | 1 | 12 | Tab2 | 0.01 | Old |
| 0.532664 | 0.88607 | 0.025 | 0.015 | 1 | 12 | Ski | 0.01 | Old |
| 0.532664 | 0.883217 | 0.025 | 0.015 | 1 | 12 | Pex13 | 0.01 | Old |
| 0.532664 | 0.882355 | 0.025 | 0.015 | 1 | 12 | Cuedc1 | 0.01 | Old |
| 0.532664 | 0.872852 | 0.025 | 0.015 | 1 | 12 | Ift80 | 0.01 | Old |
| 0.532664 | 0.870537 | 0.025 | 0.015 | 1 | 12 | Mtg1 | 0.01 | Old |
| 0.532664 | 0.867278 | 0.025 | 0.015 | 1 | 12 | Ammecr1l | 0.01 | Old |
| 0.532664 | 0.831806 | 0.025 | 0.015 | 1 | 12 | Hspa1b | 0.01 | Old |
| 0.532664 | 0.813429 | 0.025 | 0.015 | 1 | 12 | Kcnk10 | 0.01 | Old |
| 0.532664 | 0.798935 | 0.025 | 0.015 | 1 | 12 | Flvcr1 | 0.01 | Old |
| 0.532664 | 0.757363 | 0.025 | 0.015 | 1 | 12 | Cyba | 0.01 | Old |
| 0.532664 | 0.750268 | 0.025 | 0.015 | 1 | 12 | Txnip | 0.01 | Old |
| 0.532664 | 0.719671 | 0.025 | 0.015 | 1 | 12 | C1galt1c1 | 0.01 | Old |
| 0.532664 | 0.578376 | 0.025 | 0.015 | 1 | 12 | Pcgf3 | 0.01 | Old |
| 0.533795 | 0.73226 | 0.076 | 0.058 | 1 | 12 | Inafm1 | 0.018 | Old |
| 0.534198 | 1.118943 | 0.051 | 0.036 | 1 | 12 | Hpf1 | 0.015 | Old |
| 0.534607 | 0.899104 | 0.059 | 0.044 | 1 | 12 | Timm8a1 | 0.015 | Old |
| 0.534994 | 0.789781 | 0.025 | 0.015 | 1 | 12 | Grn | 0.01 | Old |
| 0.535895 | 0.101401 | 0.551 | 0.54 | 1 | 12 | Nme1 | 0.011 | Old |
| 0.53662 | 0.412066 | 0.093 | 0.073 | 1 | 12 | Caprin1 | 0.02 | Old |
| 0.537331 | 1.246637 | 0.025 | 0.015 | 1 | 12 | St8sia2 | 0.01 | Old |
| 0.537331 | 1.230419 | 0.025 | 0.015 | 1 | 12 | Kif26a | 0.01 | Old |
| 0.537331 | 0.969683 | 0.025 | 0.015 | 1 | 12 | Fam13b | 0.01 | Old |
| 0.537331 | 0.895715 | 0.025 | 0.015 | 1 | 12 | Dpysl4 | 0.01 | Old |
| 0.537331 | 0.890822 | 0.025 | 0.015 | 1 | 12 | Fam149b | 0.01 | Old |
| 0.537331 | 0.89082 | 0.025 | 0.015 | 1 | 12 | Nup133 | 0.01 | Old |
| 0.537331 | 0.884238 | 0.025 | 0.015 | 1 | 12 | Tbc1d17 | 0.01 | Old |
| 0.537331 | 0.874838 | 0.025 | 0.015 | 1 | 12 | Pkn1 | 0.01 | Old |
| 0.537331 | 0.859335 | 0.025 | 0.015 | 1 | 12 | Usp20 | 0.01 | Old |
| 0.537331 | 0.854319 | 0.025 | 0.015 | 1 | 12 | Slc7a14 | 0.01 | Old |
| 0.537331 | 0.840658 | 0.025 | 0.015 | 1 | 12 | Wrb | 0.01 | Old |
| 0.537331 | 0.832808 | 0.025 | 0.015 | 1 | 12 | Jakmip2 | 0.01 | Old |
| 0.537331 | 0.830069 | 0.025 | 0.015 | 1 | 12 | Ogfod3 | 0.01 | Old |

| | | | | | | | | |
|---|---|---|---|---|---|---|---|---|
| 0.537331 | 0.823059 | 0.025 | 0.015 | 1 | 12 | Gtf3c6 | 0.01 | Old |
| 0.537331 | 0.820386 | 0.025 | 0.015 | 1 | 12 | Gnal | 0.01 | Old |
| 0.537331 | 0.800027 | 0.025 | 0.015 | 1 | 12 | Rragb | 0.01 | Old |
| 0.537331 | 0.799577 | 0.025 | 0.015 | 1 | 12 | Atg16l1 | 0.01 | Old |
| 0.537331 | 0.788984 | 0.025 | 0.015 | 1 | 12 | Tyw1 | 0.01 | Old |
| 0.537331 | 0.768045 | 0.025 | 0.015 | 1 | 12 | Nr2f6 | 0.01 | Old |
| 0.537331 | 0.758907 | 0.025 | 0.015 | 1 | 12 | Aggf1 | 0.01 | Old |
| 0.537331 | 0.749158 | 0.025 | 0.015 | 1 | 12 | Taf9b | 0.01 | Old |
| 0.537331 | 0.748241 | 0.025 | 0.015 | 1 | 12 | Lonp2 | 0.01 | Old |
| 0.537331 | 0.736154 | 0.025 | 0.015 | 1 | 12 | Mbd5 | 0.01 | Old |
| 0.537331 | 0.729289 | 0.025 | 0.015 | 1 | 12 | Znhit2 | 0.01 | Old |
| 0.537331 | 0.725957 | 0.025 | 0.015 | 1 | 12 | Rhot1 | 0.01 | Old |
| 0.537331 | 0.725761 | 0.025 | 0.015 | 1 | 12 | C530008M | 0.01 | Old |
| 0.537331 | 0.718194 | 0.025 | 0.015 | 1 | 12 | Chd2 | 0.01 | Old |
| 0.537331 | 0.713599 | 0.025 | 0.015 | 1 | 12 | 9-Mar | 0.01 | Old |
| 0.537331 | 0.705629 | 0.025 | 0.015 | 1 | 12 | Slc25a16 | 0.01 | Old |
| 0.537331 | 0.69243 | 0.025 | 0.015 | 1 | 12 | Trp53rka | 0.01 | Old |
| 0.537331 | 0.690882 | 0.025 | 0.015 | 1 | 12 | Trip12 | 0.01 | Old |
| 0.537331 | 0.689624 | 0.025 | 0.015 | 1 | 12 | Xiap | 0.01 | Old |
| 0.537331 | 0.689533 | 0.025 | 0.015 | 1 | 12 | Preb | 0.01 | Old |
| 0.537331 | 0.689243 | 0.025 | 0.015 | 1 | 12 | Mast2 | 0.01 | Old |
| 0.537331 | 0.681279 | 0.025 | 0.015 | 1 | 12 | Cdk2ap1 | 0.01 | Old |
| 0.537331 | 0.677257 | 0.025 | 0.015 | 1 | 12 | Sh3kbp1 | 0.01 | Old |
| 0.537331 | 0.670237 | 0.025 | 0.015 | 1 | 12 | Slc25a51 | 0.01 | Old |
| 0.537331 | 0.621514 | 0.025 | 0.015 | 1 | 12 | Golga4 | 0.01 | Old |
| 0.537331 | 0.568307 | 0.025 | 0.015 | 1 | 12 | Stx16 | 0.01 | Old |
| 0.537331 | 0.490921 | 0.025 | 0.015 | 1 | 12 | Glrb | 0.01 | Old |
| 0.537331 | 0.43453 | 0.025 | 0.015 | 1 | 12 | Ap1b1 | 0.01 | Old |
| 0.537331 | 0.025368 | 0.025 | 0.015 | 1 | 12 | Pkib | 0.01 | Old |
| 0.537385 | 1.168447 | 0.051 | 0.036 | 1 | 12 | Polr2c | 0.015 | Old |
| 0.537385 | 1.136418 | 0.051 | 0.036 | 1 | 12 | Tmem108 | 0.015 | Old |
| 0.537385 | 0.856273 | 0.051 | 0.036 | 1 | 12 | Slc29a4 | 0.015 | Old |
| 0.537465 | 1.236998 | 0.034 | 0.022 | 1 | 12 | Nptxr | 0.012 | Old |
| 0.537465 | 1.092846 | 0.034 | 0.022 | 1 | 12 | Rgs10 | 0.012 | Old |
| 0.537685 | 0.446091 | 0.076 | 0.058 | 1 | 12 | Egln2 | 0.018 | Old |
| 0.53795 | 0.521597 | 0.068 | 0.051 | 1 | 12 | Anapc10 | 0.017 | Old |
| 0.539523 | 0.292624 | 0.237 | 0.204 | 1 | 12 | Clic1 | 0.033 | Old |
| 0.53954 | 0.310742 | 0.136 | 0.109 | 1 | 12 | Caly | 0.027 | Old |
| 0.539624 | 0.22621 | 0.169 | 0.146 | 1 | 12 | Klf7 | 0.023 | Old |
| 0.539968 | 0.086304 | 0.017 | 0.029 | 1 | 12 | Mul1 | -0.012 | Old |
| 0.540581 | 0.148918 | 0.034 | 0.051 | 1 | 12 | Hspa4l | -0.017 | Old |
| 0.540767 | 0.409437 | 0.127 | 0.102 | 1 | 12 | Dync1h1 | 0.025 | Old |
| 0.540867 | 0.465623 | 0.085 | 0.066 | 1 | 12 | Vps36 | 0.019 | Old |
| 0.541441 | 1.098376 | 0.034 | 0.022 | 1 | 12 | Kidins220 | 0.012 | Old |
| 0.541441 | 0.94628 | 0.034 | 0.022 | 1 | 12 | Adss | 0.012 | Old |

| | | | | | | | | |
|---|---|---|---|---|---|---|---|---|
| 0.542015 | 0.335713 | 0.178 | 0.146 | 1 | 12 | Eif4a1 | 0.032 | Old |
| 0.542018 | 0.87916 | 0.025 | 0.015 | 1 | 12 | Zfp444 | 0.01 | Old |
| 0.542018 | 0.870266 | 0.025 | 0.015 | 1 | 12 | Dhrs7 | 0.01 | Old |
| 0.542018 | 0.765503 | 0.025 | 0.015 | 1 | 12 | Tnfsf12 | 0.01 | Old |
| 0.542018 | 0.748609 | 0.025 | 0.015 | 1 | 12 | Sord | 0.01 | Old |
| 0.542018 | 0.724136 | 0.025 | 0.015 | 1 | 12 | 9330159F1 | 0.01 | Old |
| 0.542018 | 0.714413 | 0.025 | 0.015 | 1 | 12 | 4833420G1 | 0.01 | Old |
| 0.542018 | 0.70929 | 0.025 | 0.015 | 1 | 12 | G6pc3 | 0.01 | Old |
| 0.542018 | 0.701159 | 0.025 | 0.015 | 1 | 12 | Dnajc17 | 0.01 | Old |
| 0.542018 | 0.688706 | 0.025 | 0.015 | 1 | 12 | Exosc7 | 0.01 | Old |
| 0.542018 | 0.674594 | 0.025 | 0.015 | 1 | 12 | Armc8 | 0.01 | Old |
| 0.542018 | 0.664947 | 0.025 | 0.015 | 1 | 12 | Brd7 | 0.01 | Old |
| 0.542018 | 0.642807 | 0.025 | 0.015 | 1 | 12 | Ntng1 | 0.01 | Old |
| 0.542018 | 0.639571 | 0.025 | 0.015 | 1 | 12 | Med17 | 0.01 | Old |
| 0.542018 | 0.625263 | 0.025 | 0.015 | 1 | 12 | Uqcc1 | 0.01 | Old |
| 0.542018 | 0.61982 | 0.025 | 0.015 | 1 | 12 | Prepl | 0.01 | Old |
| 0.542018 | 0.609745 | 0.025 | 0.015 | 1 | 12 | Mbtps1 | 0.01 | Old |
| 0.542018 | 0.606592 | 0.025 | 0.015 | 1 | 12 | Srsf4 | 0.01 | Old |
| 0.542018 | 0.599849 | 0.025 | 0.015 | 1 | 12 | Gpn1 | 0.01 | Old |
| 0.542018 | 0.590143 | 0.025 | 0.015 | 1 | 12 | Rnf167 | 0.01 | Old |
| 0.542018 | 0.582486 | 0.025 | 0.015 | 1 | 12 | Ccdc112 | 0.01 | Old |
| 0.542018 | 0.559503 | 0.025 | 0.015 | 1 | 12 | Rnf19a | 0.01 | Old |
| 0.542018 | 0.552955 | 0.025 | 0.015 | 1 | 12 | Txlng | 0.01 | Old |
| 0.542018 | 0.548863 | 0.025 | 0.015 | 1 | 12 | Gigyf1 | 0.01 | Old |
| 0.542018 | 0.510195 | 0.025 | 0.015 | 1 | 12 | Sacs | 0.01 | Old |
| 0.542018 | 0.506776 | 0.025 | 0.015 | 1 | 12 | Nomo1 | 0.01 | Old |
| 0.542018 | 0.454044 | 0.025 | 0.015 | 1 | 12 | Irf2bp2 | 0.01 | Old |
| 0.542018 | 0.409311 | 0.025 | 0.015 | 1 | 12 | Map1s | 0.01 | Old |
| 0.542018 | 0.404133 | 0.025 | 0.015 | 1 | 12 | Tubb6 | 0.01 | Old |
| 0.542018 | 0.395198 | 0.025 | 0.015 | 1 | 12 | Tuba4a | 0.01 | Old |
| 0.542018 | 0.302971 | 0.025 | 0.015 | 1 | 12 | Ythdf1 | 0.01 | Old |
| 0.542018 | 0.289579 | 0.025 | 0.015 | 1 | 12 | Ctbp2 | 0.01 | Old |
| 0.542018 | 0.289316 | 0.025 | 0.015 | 1 | 12 | Nhsl2 | 0.01 | Old |
| 0.542018 | 0.284486 | 0.025 | 0.015 | 1 | 12 | Bdp1 | 0.01 | Old |
| 0.542018 | 0.27729 | 0.025 | 0.015 | 1 | 12 | Htt | 0.01 | Old |
| 0.542018 | 0.271511 | 0.025 | 0.015 | 1 | 12 | Sppl2a | 0.01 | Old |
| 0.542018 | 0.261152 | 0.025 | 0.015 | 1 | 12 | Ube4a | 0.01 | Old |
| 0.542018 | 0.210778 | 0.025 | 0.015 | 1 | 12 | Kpna6 | 0.01 | Old |
| 0.542018 | 0.205867 | 0.025 | 0.015 | 1 | 12 | Scyl2 | 0.01 | Old |
| 0.542018 | 0.141151 | 0.025 | 0.015 | 1 | 12 | Dars | 0.01 | Old |
| 0.542018 | 0.105738 | 0.025 | 0.015 | 1 | 12 | Tmub2 | 0.01 | Old |
| 0.542518 | 0.296699 | 0.136 | 0.109 | 1 | 12 | Zfp580 | 0.027 | Old |
| 0.543683 | 0.824993 | 0.042 | 0.029 | 1 | 12 | Mprip | 0.013 | Old |
| 0.543786 | 0.870682 | 0.051 | 0.036 | 1 | 12 | Comtd1 | 0.015 | Old |
| 0.543786 | 0.825565 | 0.051 | 0.036 | 1 | 12 | Nudcd3 | 0.015 | Old |

| | | | | | | | | |
|---|---|---|---|---|---|---|---|---|
| 0.545432 | 1.101635 | 0.034 | 0.022 | 1 | 12 | Plppr3 | 0.012 | Old |
| 0.545432 | 1.060343 | 0.034 | 0.022 | 1 | 12 | Washc3 | 0.012 | Old |
| 0.545432 | 1.045869 | 0.034 | 0.022 | 1 | 12 | Adck1 | 0.012 | Old |
| 0.545432 | 0.987764 | 0.034 | 0.022 | 1 | 12 | Cavin3 | 0.012 | Old |
| 0.545432 | 0.905863 | 0.034 | 0.022 | 1 | 12 | Fam122a | 0.012 | Old |
| 0.545432 | 0.845567 | 0.034 | 0.022 | 1 | 12 | Tpst2 | 0.012 | Old |
| 0.54608 | 0.366898 | 0.093 | 0.073 | 1 | 12 | Pigp | 0.02 | Old |
| 0.546108 | 0.462711 | 0.161 | 0.146 | 1 | 12 | 1110004F1 | 0.015 | Old |
| 0.546249 | 0.573663 | 0.068 | 0.051 | 1 | 12 | Fam98c | 0.017 | Old |
| 0.546336 | 0.317612 | 0.195 | 0.168 | 1 | 12 | Fam174a | 0.027 | Old |
| 0.54643 | 0.645404 | 0.059 | 0.044 | 1 | 12 | Asphd2 | 0.015 | Old |
| 0.546726 | 0.623115 | 0.025 | 0.015 | 1 | 12 | Ubr7 | 0.01 | Old |
| 0.546726 | 0.602965 | 0.025 | 0.015 | 1 | 12 | Slc4a3 | 0.01 | Old |
| 0.546726 | 0.569784 | 0.025 | 0.015 | 1 | 12 | Fam76a | 0.01 | Old |
| 0.546726 | 0.556722 | 0.025 | 0.015 | 1 | 12 | Slc7a3 | 0.01 | Old |
| 0.546726 | 0.528814 | 0.025 | 0.015 | 1 | 12 | Dck | 0.01 | Old |
| 0.546726 | 0.470748 | 0.025 | 0.015 | 1 | 12 | Cdc42ep2 | 0.01 | Old |
| 0.546726 | 0.430289 | 0.025 | 0.015 | 1 | 12 | Tcerg1 | 0.01 | Old |
| 0.546726 | 0.419323 | 0.025 | 0.015 | 1 | 12 | Hist3h2a | 0.01 | Old |
| 0.546726 | 0.312279 | 0.025 | 0.015 | 1 | 12 | Adam15 | 0.01 | Old |
| 0.546726 | 0.299599 | 0.025 | 0.015 | 1 | 12 | Fbxo18 | 0.01 | Old |
| 0.546726 | 0.227072 | 0.025 | 0.015 | 1 | 12 | Gas2l1 | 0.01 | Old |
| 0.546726 | 0.218808 | 0.025 | 0.015 | 1 | 12 | Ppp1r37 | 0.01 | Old |
| 0.546726 | 0.099095 | 0.025 | 0.015 | 1 | 12 | Asah1 | 0.01 | Old |
| 0.546726 | 0.004628 | 0.025 | 0.015 | 1 | 12 | Trim25 | 0.01 | Old |
| 0.547224 | 0.890683 | 0.042 | 0.029 | 1 | 12 | Hmg20b | 0.013 | Old |
| 0.547224 | 0.747634 | 0.042 | 0.029 | 1 | 12 | Uchl5 | 0.013 | Old |
| 0.547501 | 0.323253 | 0.136 | 0.109 | 1 | 12 | Fam122b | 0.027 | Old |
| 0.547568 | 0.331031 | 0.102 | 0.08 | 1 | 12 | Yif1b | 0.022 | Old |
| 0.547814 | 0.177612 | 0.119 | 0.095 | 1 | 12 | Mrps17 | 0.024 | Old |
| 0.549405 | 0.667718 | 0.059 | 0.044 | 1 | 12 | Impact | 0.015 | Old |
| 0.549405 | 0.603677 | 0.059 | 0.044 | 1 | 12 | Mat2a | 0.015 | Old |
| 0.549438 | 1.006536 | 0.034 | 0.022 | 1 | 12 | Kxd1 | 0.012 | Old |
| 0.549438 | 0.75916 | 0.034 | 0.022 | 1 | 12 | Csnk2a1 | 0.012 | Old |
| 0.549438 | 0.74615 | 0.034 | 0.022 | 1 | 12 | Morc3 | 0.012 | Old |
| 0.549438 | 0.636244 | 0.034 | 0.022 | 1 | 12 | Ddx39 | 0.012 | Old |
| 0.549565 | 0.370122 | 0.085 | 0.066 | 1 | 12 | Mrpl48 | 0.019 | Old |
| 0.550225 | 0.779956 | 0.051 | 0.036 | 1 | 12 | Sgpp2 | 0.015 | Old |
| 0.550775 | 0.874743 | 0.042 | 0.029 | 1 | 12 | Trim37 | 0.013 | Old |
| 0.550775 | 0.853845 | 0.042 | 0.029 | 1 | 12 | Rnaseh2c | 0.013 | Old |
| 0.550775 | 0.761172 | 0.042 | 0.029 | 1 | 12 | Max | 0.013 | Old |
| 0.550775 | 0.753407 | 0.042 | 0.029 | 1 | 12 | Gdap1l1 | 0.013 | Old |
| 0.550896 | 0.59557 | 0.059 | 0.044 | 1 | 12 | Med19 | 0.015 | Old |
| 0.551454 | 0.526521 | 0.025 | 0.015 | 1 | 12 | Slc25a46 | 0.01 | Old |
| 0.551454 | 0.475891 | 0.025 | 0.015 | 1 | 12 | Mogs | 0.01 | Old |

| | | | | | | | | |
|---|---|---|---|---|---|---|---|---|
| 0.551454 | 0.45987 | 0.025 | 0.015 | 1 | 12 | Ccm2 | 0.01 | Old |
| 0.551454 | 0.395579 | 0.025 | 0.015 | 1 | 12 | Brd3 | 0.01 | Old |
| 0.551454 | 0.381432 | 0.025 | 0.015 | 1 | 12 | Wdr44 | 0.01 | Old |
| 0.551454 | 0.350005 | 0.025 | 0.015 | 1 | 12 | Ino80d | 0.01 | Old |
| 0.551454 | 0.31695 | 0.025 | 0.015 | 1 | 12 | Smim10l2a | 0.01 | Old |
| 0.551454 | 0.315286 | 0.025 | 0.015 | 1 | 12 | Dnajc4 | 0.01 | Old |
| 0.551454 | 0.211795 | 0.025 | 0.015 | 1 | 12 | Noc4l | 0.01 | Old |
| 0.551454 | 0.048085 | 0.025 | 0.015 | 1 | 12 | Tex30 | 0.01 | Old |
| 0.552389 | 0.601793 | 0.059 | 0.044 | 1 | 12 | Cebpz | 0.015 | Old |
| 0.553457 | 0.618569 | 0.034 | 0.022 | 1 | 12 | Ptpn11 | 0.012 | Old |
| 0.553458 | 0.876405 | 0.034 | 0.022 | 1 | 12 | Dmtn | 0.012 | Old |
| 0.553458 | 0.82317 | 0.034 | 0.022 | 1 | 12 | Pkp3 | 0.012 | Old |
| 0.553458 | 0.799535 | 0.034 | 0.022 | 1 | 12 | 9530068E0 | 0.012 | Old |
| 0.553458 | 0.798691 | 0.034 | 0.022 | 1 | 12 | Tmem35a | 0.012 | Old |
| 0.553458 | 0.78647 | 0.034 | 0.022 | 1 | 12 | Clec2l | 0.012 | Old |
| 0.553458 | 0.731984 | 0.034 | 0.022 | 1 | 12 | Nek7 | 0.012 | Old |
| 0.553458 | 0.713315 | 0.034 | 0.022 | 1 | 12 | Mplkip | 0.012 | Old |
| 0.553458 | 0.648828 | 0.034 | 0.022 | 1 | 12 | Atf2 | 0.012 | Old |
| 0.553458 | 0.612626 | 0.034 | 0.022 | 1 | 12 | Fuca1 | 0.012 | Old |
| 0.553458 | 0.601016 | 0.034 | 0.022 | 1 | 12 | Iqsec3 | 0.012 | Old |
| 0.553459 | 0.701843 | 0.051 | 0.036 | 1 | 12 | Clcn4 | 0.015 | Old |
| 0.553459 | 0.533129 | 0.051 | 0.036 | 1 | 12 | Shc3 | 0.015 | Old |
| 0.554338 | 0.924574 | 0.042 | 0.029 | 1 | 12 | Wdr47 | 0.013 | Old |
| 0.554338 | 0.900421 | 0.042 | 0.029 | 1 | 12 | Acaa2 | 0.013 | Old |
| 0.554338 | 0.678018 | 0.042 | 0.029 | 1 | 12 | Ppp1r9a | 0.013 | Old |
| 0.554709 | 0.493647 | 0.076 | 0.058 | 1 | 12 | Creld1 | 0.018 | Old |
| 0.555329 | 0.161268 | 0.407 | 0.38 | 1 | 12 | Rplp1 | 0.027 | Old |
| 0.555603 | 0.406137 | 0.254 | 0.234 | 1 | 12 | Pcp4 | 0.02 | Old |
| 0.555621 | 0.434309 | 0.093 | 0.073 | 1 | 12 | Tomm70a | 0.02 | Old |
| 0.556702 | 0.695863 | 0.051 | 0.036 | 1 | 12 | Haghl | 0.015 | Old |
| 0.556702 | 0.669381 | 0.051 | 0.036 | 1 | 12 | Mrps5 | 0.015 | Old |
| 0.557351 | 0.380139 | 0.076 | 0.058 | 1 | 12 | Exoc3 | 0.018 | Old |
| 0.557492 | 0.845932 | 0.034 | 0.022 | 1 | 12 | Haus8 | 0.012 | Old |
| 0.557492 | 0.824773 | 0.034 | 0.022 | 1 | 12 | Fars2 | 0.012 | Old |
| 0.557492 | 0.765536 | 0.034 | 0.022 | 1 | 12 | Ufsp1 | 0.012 | Old |
| 0.557492 | 0.70154 | 0.034 | 0.022 | 1 | 12 | Sdf2l1 | 0.012 | Old |
| 0.557492 | 0.699037 | 0.034 | 0.022 | 1 | 12 | Zfp397 | 0.012 | Old |
| 0.557492 | 0.691184 | 0.034 | 0.022 | 1 | 12 | Chadl | 0.012 | Old |
| 0.557492 | 0.679563 | 0.034 | 0.022 | 1 | 12 | Fam3a | 0.012 | Old |
| 0.557492 | 0.631193 | 0.034 | 0.022 | 1 | 12 | Atg10 | 0.012 | Old |
| 0.557492 | 0.613151 | 0.034 | 0.022 | 1 | 12 | Smndc1 | 0.012 | Old |
| 0.557492 | 0.593671 | 0.034 | 0.022 | 1 | 12 | Ube2g1 | 0.012 | Old |
| 0.557492 | 0.491183 | 0.034 | 0.022 | 1 | 12 | Rchy1 | 0.012 | Old |
| 0.557492 | 0.460092 | 0.034 | 0.022 | 1 | 12 | Rfk | 0.012 | Old |
| 0.557492 | 0.401293 | 0.034 | 0.022 | 1 | 12 | Msra | 0.012 | Old |

| | | | | | | | | |
|---|---|---|---|---|---|---|---|---|
| 0.557492 | 0.395447 | 0.034 | 0.022 | 1 | 12 | Usp7 | 0.012 | Old |
| 0.557912 | 0.680134 | 0.042 | 0.029 | 1 | 12 | Tnks2 | 0.013 | Old |
| 0.557912 | 0.776884 | 0.042 | 0.029 | 1 | 12 | Sv2a | 0.013 | Old |
| 0.557912 | 0.609062 | 0.042 | 0.029 | 1 | 12 | Nucb1 | 0.013 | Old |
| 0.557912 | 0.601985 | 0.042 | 0.029 | 1 | 12 | Cdc5l | 0.013 | Old |
| 0.55838 | 0.472729 | 0.059 | 0.044 | 1 | 12 | Gstm1 | 0.015 | Old |
| 0.559082 | 0.177205 | 0.22 | 0.19 | 1 | 12 | Srrm2 | 0.03 | Old |
| 0.559514 | 0.58683 | 0.034 | 0.022 | 1 | 12 | Rexo4 | 0.012 | Old |
| 0.559747 | 0.045579 | 1 | 1 | 1 | 12 | Tmsb4x | 0 | Old |
| 0.559954 | 0.623577 | 0.051 | 0.036 | 1 | 12 | Mrps11 | 0.015 | Old |
| 0.559954 | 0.593957 | 0.051 | 0.036 | 1 | 12 | Il11ra1 | 0.015 | Old |
| 0.560223 | 0.515215 | 0.068 | 0.051 | 1 | 12 | Cebpb | 0.017 | Old |
| 0.561498 | 0.713029 | 0.042 | 0.029 | 1 | 12 | Mkln1 | 0.013 | Old |
| 0.561498 | 0.669609 | 0.042 | 0.029 | 1 | 12 | Rab4a | 0.013 | Old |
| 0.561498 | 0.637117 | 0.042 | 0.029 | 1 | 12 | Cul2 | 0.013 | Old |
| 0.561498 | 0.620354 | 0.042 | 0.029 | 1 | 12 | Fam192a | 0.013 | Old |
| 0.561498 | 0.60665 | 0.042 | 0.029 | 1 | 12 | Rnf168 | 0.013 | Old |
| 0.561498 | 0.591786 | 0.042 | 0.029 | 1 | 12 | BC003965 | 0.013 | Old |
| 0.561498 | 0.57393 | 0.042 | 0.029 | 1 | 12 | Exosc8 | 0.013 | Old |
| 0.561498 | 0.473222 | 0.042 | 0.029 | 1 | 12 | Bcl7b | 0.013 | Old |
| 0.561531 | 0.297327 | 0.246 | 0.226 | 1 | 12 | Rheb | 0.02 | Old |
| 0.561541 | 0.631443 | 0.034 | 0.022 | 1 | 12 | Trim27 | 0.012 | Old |
| 0.561541 | 0.602724 | 0.034 | 0.022 | 1 | 12 | Nprl2 | 0.012 | Old |
| 0.561541 | 0.586045 | 0.034 | 0.022 | 1 | 12 | Ezr | 0.012 | Old |
| 0.561541 | 0.571486 | 0.034 | 0.022 | 1 | 12 | Steap3 | 0.012 | Old |
| 0.561541 | 0.571227 | 0.034 | 0.022 | 1 | 12 | Cast | 0.012 | Old |
| 0.561541 | 0.374793 | 0.034 | 0.022 | 1 | 12 | Csnk1d | 0.012 | Old |
| 0.561541 | 0.03727 | 0.034 | 0.022 | 1 | 12 | Supt16 | 0.012 | Old |
| 0.561626 | 0.385124 | 0.093 | 0.073 | 1 | 12 | Bzw2 | 0.02 | Old |
| 0.561626 | 0.280232 | 0.093 | 0.073 | 1 | 12 | Gdap1 | 0.02 | Old |
| 0.563038 | 0.465467 | 0.068 | 0.051 | 1 | 12 | Armc1 | 0.017 | Old |
| 0.563215 | 0.542294 | 0.051 | 0.036 | 1 | 12 | Ift57 | 0.015 | Old |
| 0.563294 | 0.638311 | 0.042 | 0.029 | 1 | 12 | Ppp3cb | 0.013 | Old |
| 0.563371 | 0.379911 | 0.085 | 0.066 | 1 | 12 | Kdelr1 | 0.019 | Old |
| 0.563464 | 0.359937 | 0.212 | 0.19 | 1 | 12 | Drap1 | 0.022 | Old |
| 0.564152 | 0.151089 | 0.737 | 0.686 | 1 | 12 | Syn2 | 0.051 | Old |
| 0.56432 | 0.215554 | 0.153 | 0.124 | 1 | 12 | Sod1 | 0.029 | Old |
| 0.564622 | 0.325991 | 0.246 | 0.226 | 1 | 12 | Srsf3 | 0.02 | Old |
| 0.565094 | 0.731317 | 0.042 | 0.029 | 1 | 12 | Syt9 | 0.013 | Old |
| 0.565094 | 0.65644 | 0.042 | 0.029 | 1 | 12 | Eif2b5 | 0.013 | Old |
| 0.565094 | 0.641872 | 0.042 | 0.029 | 1 | 12 | Bcl2 | 0.013 | Old |
| 0.565094 | 0.615361 | 0.042 | 0.029 | 1 | 12 | Fam57b | 0.013 | Old |
| 0.565094 | 0.608019 | 0.042 | 0.029 | 1 | 12 | Ptn | 0.013 | Old |
| 0.565094 | 0.524175 | 0.042 | 0.029 | 1 | 12 | Ddx25 | 0.013 | Old |
| 0.565094 | 0.520589 | 0.042 | 0.029 | 1 | 12 | Fndc4 | 0.013 | Old |

| | | | | | | | | |
|---|---|---|---|---|---|---|---|---|
| 0.565094 | 0.487412 | 0.042 | 0.029 | 1 | 12 | Baz2b | 0.013 | Old |
| 0.565285 | 0.348556 | 0.144 | 0.124 | 1 | 12 | Emc2 | 0.02 | Old |
| 0.565603 | 0.645554 | 0.034 | 0.022 | 1 | 12 | Pex11b | 0.012 | Old |
| 0.565603 | 0.609494 | 0.034 | 0.022 | 1 | 12 | Mpp1 | 0.012 | Old |
| 0.565603 | 0.602266 | 0.034 | 0.022 | 1 | 12 | Fbxl5 | 0.012 | Old |
| 0.565603 | 0.500369 | 0.034 | 0.022 | 1 | 12 | AI314180 | 0.012 | Old |
| 0.565603 | 0.470708 | 0.034 | 0.022 | 1 | 12 | Fdx1 | 0.012 | Old |
| 0.565603 | 0.313037 | 0.034 | 0.022 | 1 | 12 | Tpm4 | 0.012 | Old |
| 0.56586 | 0.265076 | 0.068 | 0.051 | 1 | 12 | Ssbp1 | 0.017 | Old |
| 0.566183 | 0.149211 | 0.508 | 0.445 | 1 | 12 | Arhgdig | 0.063 | Old |
| 0.566485 | 0.533193 | 0.051 | 0.036 | 1 | 12 | Dnpep | 0.015 | Old |
| 0.566485 | 0.51747 | 0.051 | 0.036 | 1 | 12 | Gm45716 | 0.015 | Old |
| 0.566485 | 0.413746 | 0.051 | 0.036 | 1 | 12 | Pnp | 0.015 | Old |
| 0.567426 | 0.361833 | 0.059 | 0.044 | 1 | 12 | Pigt | 0.015 | Old |
| 0.567426 | 0.503143 | 0.059 | 0.044 | 1 | 12 | Acbd5 | 0.015 | Old |
| 0.567426 | 0.451233 | 0.059 | 0.044 | 1 | 12 | Atg5 | 0.015 | Old |
| 0.567981 | 0.295618 | 0.076 | 0.058 | 1 | 12 | Retreg2 | 0.018 | Old |
| 0.567981 | 0.189045 | 0.076 | 0.058 | 1 | 12 | Nfib | 0.018 | Old |
| 0.568421 | 0.457971 | 0.161 | 0.139 | 1 | 12 | Vat1l | 0.022 | Old |
| 0.568433 | 0.30525 | 0.085 | 0.066 | 1 | 12 | Snapin | 0.019 | Old |
| 0.568701 | 0.611258 | 0.042 | 0.029 | 1 | 12 | Phf3 | 0.013 | Old |
| 0.568701 | 0.623065 | 0.042 | 0.029 | 1 | 12 | Dnttip1 | 0.013 | Old |
| 0.568701 | 0.606296 | 0.042 | 0.029 | 1 | 12 | Eef1akmt2 | 0.013 | Old |
| 0.568701 | 0.554222 | 0.042 | 0.029 | 1 | 12 | Dpp3 | 0.013 | Old |
| 0.568701 | 0.486018 | 0.042 | 0.029 | 1 | 12 | Ipo5 | 0.013 | Old |
| 0.569142 | 0.077359 | 0.356 | 0.321 | 1 | 12 | Rbx1 | 0.035 | Old |
| 0.569679 | 0.559723 | 0.034 | 0.022 | 1 | 12 | Metap1d | 0.012 | Old |
| 0.569679 | 0.482794 | 0.034 | 0.022 | 1 | 12 | Ccdc25 | 0.012 | Old |
| 0.56968 | 0.595387 | 0.034 | 0.022 | 1 | 12 | Tmem41a | 0.012 | Old |
| 0.56968 | 0.497686 | 0.034 | 0.022 | 1 | 12 | Rsf1 | 0.012 | Old |
| 0.56968 | 0.436558 | 0.034 | 0.022 | 1 | 12 | Rasa1 | 0.012 | Old |
| 0.56968 | 0.425282 | 0.034 | 0.022 | 1 | 12 | Aasdhppt | 0.012 | Old |
| 0.56968 | 0.419572 | 0.034 | 0.022 | 1 | 12 | Ogdh | 0.012 | Old |
| 0.56968 | 0.402192 | 0.034 | 0.022 | 1 | 12 | Epn2 | 0.012 | Old |
| 0.56968 | 0.185125 | 0.034 | 0.022 | 1 | 12 | Ids | 0.012 | Old |
| 0.569765 | 0.512347 | 0.051 | 0.036 | 1 | 12 | Coq10a | 0.015 | Old |
| 0.569765 | 0.505588 | 0.051 | 0.036 | 1 | 12 | Ddhd2 | 0.015 | Old |
| 0.570457 | 0.492938 | 0.059 | 0.044 | 1 | 12 | Orc6 | 0.015 | Old |
| 0.570619 | 0.206188 | 0.102 | 0.08 | 1 | 12 | Mrpl52 | 0.022 | Old |
| 0.571407 | 0.448125 | 0.051 | 0.036 | 1 | 12 | Ndufs1 | 0.015 | Old |
| 0.571525 | 0.474586 | 0.068 | 0.051 | 1 | 12 | Bloc1s2 | 0.017 | Old |
| 0.571525 | 0.341936 | 0.068 | 0.051 | 1 | 12 | Acadm | 0.017 | Old |
| 0.571525 | 0.303544 | 0.068 | 0.051 | 1 | 12 | Ppp2r2d | 0.017 | Old |
| 0.571525 | 0.29376 | 0.068 | 0.051 | 1 | 12 | Oat | 0.017 | Old |
| 0.572182 | 0.349644 | 0.263 | 0.248 | 1 | 12 | Sult4a1 | 0.015 | Old |

| | | | | | | | | |
|---|---|---|---|---|---|---|---|---|
| 0.57232 | 0.581953 | 0.042 | 0.029 | 1 | 12 | Tgs1 | 0.013 | Old |
| 0.57232 | 0.522555 | 0.042 | 0.029 | 1 | 12 | Crbn | 0.013 | Old |
| 0.57232 | 0.415009 | 0.042 | 0.029 | 1 | 12 | Ucp2 | 0.013 | Old |
| 0.573053 | 0.504616 | 0.051 | 0.036 | 1 | 12 | Herc3 | 0.015 | Old |
| 0.573053 | 0.226447 | 0.051 | 0.036 | 1 | 12 | Ccdc174 | 0.015 | Old |
| 0.573495 | 0.568392 | 0.059 | 0.044 | 1 | 12 | Khdrbs3 | 0.015 | Old |
| 0.573516 | 0.200235 | 0.085 | 0.066 | 1 | 12 | Dync1li1 | 0.019 | Old |
| 0.573771 | 0.506887 | 0.034 | 0.022 | 1 | 12 | Mthfd2l | 0.012 | Old |
| 0.573771 | 0.493899 | 0.034 | 0.022 | 1 | 12 | Gnl1 | 0.012 | Old |
| 0.573771 | 0.468106 | 0.034 | 0.022 | 1 | 12 | Smarcd1 | 0.012 | Old |
| 0.573771 | 0.421724 | 0.034 | 0.022 | 1 | 12 | Rgs7bp | 0.012 | Old |
| 0.573771 | 0.406827 | 0.034 | 0.022 | 1 | 12 | Lrrc4b | 0.012 | Old |
| 0.573771 | 0.406196 | 0.034 | 0.022 | 1 | 12 | Wipi2 | 0.012 | Old |
| 0.573771 | 0.2218 | 0.034 | 0.022 | 1 | 12 | Otud5 | 0.012 | Old |
| 0.573771 | 0.216941 | 0.034 | 0.022 | 1 | 12 | Exosc5 | 0.012 | Old |
| 0.573771 | 0.155605 | 0.034 | 0.022 | 1 | 12 | Snx32 | 0.012 | Old |
| 0.573771 | 0.147606 | 0.034 | 0.022 | 1 | 12 | Brd9 | 0.012 | Old |
| 0.574367 | 0.24994 | 0.068 | 0.051 | 1 | 12 | Esf1 | 0.017 | Old |
| 0.574946 | 0.266663 | 0.093 | 0.073 | 1 | 12 | Jph4 | 0.02 | Old |
| 0.575897 | 0.152728 | 0.11 | 0.088 | 1 | 12 | Chrna3 | 0.022 | Old |
| 0.575949 | 0.473894 | 0.042 | 0.029 | 1 | 12 | Epb41l1 | 0.013 | Old |
| 0.575949 | 0.324055 | 0.042 | 0.029 | 1 | 12 | Wdr26 | 0.013 | Old |
| 0.576065 | 0.167905 | 0.085 | 0.066 | 1 | 12 | Tpm3 | 0.019 | Old |
| 0.576233 | 0.32017 | 0.178 | 0.153 | 1 | 12 | Hand2 | 0.025 | Old |
| 0.576541 | 0.425732 | 0.059 | 0.044 | 1 | 12 | Amdhd2 | 0.015 | Old |
| 0.576542 | 0.45217 | 0.059 | 0.044 | 1 | 12 | Tial1 | 0.015 | Old |
| 0.576542 | 0.399374 | 0.059 | 0.044 | 1 | 12 | 2900097C1 | 0.015 | Old |
| 0.57736 | 0.509112 | 0.076 | 0.058 | 1 | 12 | Luc7l | 0.018 | Old |
| 0.577372 | 0.160988 | 0.347 | 0.343 | 1 | 12 | Dctn3 | 0.004 | Old |
| 0.577383 | 0.200635 | 0.093 | 0.073 | 1 | 12 | Uba2 | 0.02 | Old |
| 0.577876 | 0.372434 | 0.034 | 0.022 | 1 | 12 | C1qtnf12 | 0.012 | Old |
| 0.577876 | 0.34501 | 0.034 | 0.022 | 1 | 12 | Erlec1 | 0.012 | Old |
| 0.577876 | 0.235001 | 0.034 | 0.022 | 1 | 12 | Qpct | 0.012 | Old |
| 0.577876 | 0.142892 | 0.034 | 0.022 | 1 | 12 | Mvp | 0.012 | Old |
| 0.577876 | 0.026462 | 0.034 | 0.022 | 1 | 12 | Txndc12 | 0.012 | Old |
| 0.578067 | 0.404722 | 0.059 | 0.044 | 1 | 12 | Ap1s2 | 0.015 | Old |
| 0.578178 | 0.026953 | 0.127 | 0.102 | 1 | 12 | Slc3a2 | 0.025 | Old |
| 0.578396 | 0.133003 | 0.508 | 0.482 | 1 | 12 | Atp5e | 0.026 | Old |
| 0.578508 | 0.175419 | 0.203 | 0.175 | 1 | 12 | Ppp2ca | 0.028 | Old |
| 0.579589 | 0.529016 | 0.042 | 0.029 | 1 | 12 | Agfg1 | 0.013 | Old |
| 0.579589 | 0.478821 | 0.042 | 0.029 | 1 | 12 | Tmem159 | 0.013 | Old |
| 0.579589 | 0.43154 | 0.042 | 0.029 | 1 | 12 | Plscr3 | 0.013 | Old |
| 0.579589 | 0.411797 | 0.042 | 0.029 | 1 | 12 | Ube2f | 0.013 | Old |
| 0.579589 | 0.388957 | 0.042 | 0.029 | 1 | 12 | Tmem128 | 0.013 | Old |
| 0.579589 | 0.332788 | 0.042 | 0.029 | 1 | 12 | Etfa | 0.013 | Old |

| | | | | | | | | |
|---|---|---|---|---|---|---|---|---|
| 0.579595 | 0.435609 | 0.059 | 0.044 | 1 | 12 | Nipsnap1 | 0.015 | Old |
| 0.579595 | 0.445865 | 0.059 | 0.044 | 1 | 12 | Ube2e2 | 0.015 | Old |
| 0.579595 | 0.439077 | 0.059 | 0.044 | 1 | 12 | Wwc2 | 0.015 | Old |
| 0.579657 | 0.28622 | 0.051 | 0.036 | 1 | 12 | Slc25a17 | 0.015 | Old |
| 0.580053 | 0.224743 | 0.076 | 0.058 | 1 | 12 | Dnajc2 | 0.018 | Old |
| 0.580071 | 0.403947 | 0.068 | 0.051 | 1 | 12 | Trappc5 | 0.017 | Old |
| 0.58154 | 0.019442 | 0.11 | 0.088 | 1 | 12 | Podxl2 | 0.022 | Old |
| 0.581994 | 0.010114 | 0.034 | 0.022 | 1 | 12 | Aatk | 0.012 | Old |
| 0.582657 | 0.286252 | 0.059 | 0.044 | 1 | 12 | Dph6 | 0.015 | Old |
| 0.582972 | 0.273744 | 0.051 | 0.036 | 1 | 12 | Ttc14 | 0.015 | Old |
| 0.58324 | 0.31862 | 0.042 | 0.029 | 1 | 12 | Psen2 | 0.013 | Old |
| 0.583468 | 0.086042 | 0.127 | 0.102 | 1 | 12 | Slc25a11 | 0.025 | Old |
| 0.584667 | 0.19911 | 0.102 | 0.08 | 1 | 12 | Abhd17a | 0.022 | Old |
| 0.585273 | 0.161829 | 0.407 | 0.416 | 1 | 12 | Arf5 | -0.009 | Old |
| 0.585394 | 0.091806 | 0.398 | 0.372 | 1 | 12 | Gpi1 | 0.026 | Old |
| 0.586284 | 0.406654 | 0.153 | 0.131 | 1 | 12 | Arl3 | 0.022 | Old |
| 0.586296 | 0.369991 | 0.051 | 0.036 | 1 | 12 | Clasp2 | 0.015 | Old |
| 0.586296 | 0.219905 | 0.051 | 0.036 | 1 | 12 | Prkcsh | 0.015 | Old |
| 0.586902 | 0.424429 | 0.042 | 0.029 | 1 | 12 | Spast | 0.013 | Old |
| 0.586902 | 0.287852 | 0.042 | 0.029 | 1 | 12 | Mrpl39 | 0.013 | Old |
| 0.586902 | 0.220154 | 0.042 | 0.029 | 1 | 12 | Clstn1 | 0.013 | Old |
| 0.587168 | 0.337548 | 0.136 | 0.117 | 1 | 12 | Ydjc | 0.019 | Old |
| 0.588205 | 0.01173 | 0.102 | 0.08 | 1 | 12 | Pts | 0.022 | Old |
| 0.588684 | 0.305998 | 0.169 | 0.146 | 1 | 12 | Srp19 | 0.023 | Old |
| 0.588736 | 0.32941 | 0.042 | 0.029 | 1 | 12 | Iah1 | 0.013 | Old |
| 0.589629 | 0.360144 | 0.051 | 0.036 | 1 | 12 | Mrpl49 | 0.015 | Old |
| 0.589629 | 0.355658 | 0.051 | 0.036 | 1 | 12 | Nsd2 | 0.015 | Old |
| 0.589629 | 0.169837 | 0.051 | 0.036 | 1 | 12 | Ogt | 0.015 | Old |
| 0.58964 | 0.199814 | 0.373 | 0.365 | 1 | 12 | Rpl29 | 0.008 | Old |
| 0.589644 | 0.255662 | 0.093 | 0.073 | 1 | 12 | Acbd6 | 0.02 | Old |
| 0.590574 | 0.465179 | 0.042 | 0.029 | 1 | 12 | Ndufaf1 | 0.013 | Old |
| 0.590574 | 0.216588 | 0.042 | 0.029 | 1 | 12 | Selenot | 0.013 | Old |
| 0.590574 | 0.052767 | 0.042 | 0.029 | 1 | 12 | Tpp2 | 0.013 | Old |
| 0.591275 | 0.087228 | 0.246 | 0.219 | 1 | 12 | Nop10 | 0.027 | Old |
| 0.591885 | 0.374658 | 0.059 | 0.044 | 1 | 12 | Shisa5 | 0.015 | Old |
| 0.591885 | 0.337426 | 0.059 | 0.044 | 1 | 12 | Tex261 | 0.015 | Old |
| 0.592971 | 0.488851 | 0.051 | 0.036 | 1 | 12 | Rit2 | 0.015 | Old |
| 0.592971 | 0.17086 | 0.051 | 0.036 | 1 | 12 | Ccnc | 0.015 | Old |
| 0.594043 | 0.031327 | 0.983 | 0.978 | 1 | 12 | Ppia | 0.005 | Old |
| 0.594257 | 0.343275 | 0.042 | 0.029 | 1 | 12 | Pycrl | 0.013 | Old |
| 0.594257 | 0.045932 | 0.042 | 0.029 | 1 | 12 | Yipf5 | 0.013 | Old |
| 0.594311 | 0.337909 | 0.127 | 0.109 | 1 | 12 | Snrpb | 0.018 | Old |
| 0.594448 | 0.210549 | 0.068 | 0.051 | 1 | 12 | Sqle | 0.017 | Old |
| 0.595186 | 0.044076 | 0.11 | 0.088 | 1 | 12 | Gmfb | 0.022 | Old |
| 0.596321 | 0.295114 | 0.051 | 0.036 | 1 | 12 | Arid5b | 0.015 | Old |

| | | | | | | | | |
|---|---|---|---|---|---|---|---|---|
| 0.596321 | 0.14632 | 0.051 | 0.036 | 1 | 12 | Tardbp | 0.015 | Old |
| 0.597342 | 0.22506 | 0.068 | 0.051 | 1 | 12 | Cotl1 | 0.017 | Old |
| 0.597609 | 0.31185 | 0.229 | 0.204 | 1 | 12 | Vamp2 | 0.025 | Old |
| 0.598075 | 0.146511 | 0.059 | 0.044 | 1 | 12 | Nipa1 | 0.015 | Old |
| 0.598963 | 0.495044 | 0.169 | 0.153 | 1 | 12 | Ndrg4 | 0.016 | Old |
| 0.59968 | 0.255082 | 0.051 | 0.036 | 1 | 12 | Htatsf1 | 0.015 | Old |
| 0.600543 | 0.023957 | 0.992 | 0.978 | 1 | 12 | Hsp90ab1 | 0.014 | Old |
| 0.600762 | 0.173049 | 0.178 | 0.153 | 1 | 12 | Ube2l3 | 0.025 | Old |
| 0.602064 | 0.06379 | 0.085 | 0.109 | 1 | 12 | Gnai1 | -0.024 | Old |
| 0.603588 | 0.107001 | 0.263 | 0.226 | 1 | 12 | Cntn5 | 0.037 | Old |
| 0.604156 | 0.062367 | 0.822 | 0.81 | 1 | 12 | Atpif1 | 0.012 | Old |
| 0.604294 | 0.205196 | 0.059 | 0.044 | 1 | 12 | Sys1 | 0.015 | Old |
| 0.604294 | 0.037166 | 0.059 | 0.044 | 1 | 12 | Clns1a | 0.015 | Old |
| 0.604525 | 0.259779 | 0.169 | 0.153 | 1 | 12 | Rufy3 | 0.016 | Old |
| 0.606084 | 0.18784 | 0.322 | 0.285 | 1 | 12 | Atxn7l3b | 0.037 | Old |
| 0.606423 | 0.258198 | 0.051 | 0.036 | 1 | 12 | Chmp6 | 0.015 | Old |
| 0.60797 | 0.400671 | 0.119 | 0.102 | 1 | 12 | Use1 | 0.017 | Old |
| 0.609307 | 0.307326 | 0.424 | 0.431 | 1 | 12 | Ctsb | -0.007 | Old |
| 0.609965 | 0.056353 | 0.178 | 0.153 | 1 | 12 | Ppa1 | 0.025 | Old |
| 0.610446 | 0.089014 | 0.068 | 0.051 | 1 | 12 | Arpc4 | 0.017 | Old |
| 0.610553 | 0.30676 | 0.314 | 0.299 | 1 | 12 | Kif21a | 0.015 | Old |
| 0.610631 | 0.051107 | 0.407 | 0.445 | 1 | 12 | H3f3b | -0.038 | Old |
| 0.61067 | 0.204868 | 0.178 | 0.161 | 1 | 12 | Krtcap2 | 0.017 | Old |
| 0.613985 | 0.297351 | 0.144 | 0.124 | 1 | 12 | Kif5b | 0.02 | Old |
| 0.616818 | 0.043588 | 0.042 | 0.058 | 1 | 12 | Cul5 | -0.016 | Old |
| 0.618987 | 0.603355 | 0.119 | 0.102 | 1 | 12 | Mrpl27 | 0.017 | Old |
| 0.623187 | 0.379725 | 0.203 | 0.19 | 1 | 12 | Psma4 | 0.013 | Old |
| 0.6235 | 0.11951 | 0.246 | 0.219 | 1 | 12 | Resp18 | 0.027 | Old |
| 0.625728 | 0.177449 | 0.025 | 0.036 | 1 | 12 | Myo9a | -0.011 | Old |
| 0.628783 | 0.084509 | 0.432 | 0.511 | 1 | 12 | Eef1b2 | -0.079 | Old |
| 0.629534 | 0.285022 | 0.127 | 0.109 | 1 | 12 | Denr | 0.018 | Old |
| 0.631386 | 0.068571 | 0.695 | 0.708 | 1 | 12 | Fau | -0.013 | Old |
| 0.632322 | 0.35578 | 0.119 | 0.102 | 1 | 12 | Sap30l | 0.017 | Old |
| 0.635632 | 0.551851 | 0.093 | 0.08 | 1 | 12 | Cops4 | 0.013 | Old |
| 0.637486 | 0.014075 | 0.229 | 0.263 | 1 | 12 | Cisd1 | -0.034 | Old |
| 0.642619 | 0.171133 | 0.263 | 0.241 | 1 | 12 | Eloc | 0.022 | Old |
| 0.643324 | 0.300878 | 0.136 | 0.117 | 1 | 12 | Hist3h2ba | 0.019 | Old |
| 0.6449 | 0.374368 | 0.161 | 0.146 | 1 | 12 | Gng5 | 0.015 | Old |
| 0.647125 | 0.116557 | 0.576 | 0.555 | 1 | 12 | Psma7 | 0.021 | Old |
| 0.648887 | 0.07566 | 0.195 | 0.226 | 1 | 12 | Txnl1 | -0.031 | Old |
| 0.653233 | 0.071026 | 0.169 | 0.146 | 1 | 12 | Qdpr | 0.023 | Old |
| 0.653903 | 0.337479 | 0.203 | 0.19 | 1 | 12 | Rpl23a | 0.013 | Old |
| 0.654545 | 0.009767 | 0.068 | 0.088 | 1 | 12 | Cacnb3 | -0.02 | Old |
| 0.660268 | 0.258427 | 0.144 | 0.124 | 1 | 12 | Syt4 | 0.02 | Old |
| 0.661647 | 0.930585 | 0.008 | 0.015 | 1 | 12 | Ppp6r2 | -0.007 | Old |

| | | | | | | | | |
|---|---|---|---|---|---|---|---|---|
| 0.661647 | 0.82201 | 0.008 | 0.015 | 1 | 12 | Cradd | -0.007 | Old |
| 0.661647 | 0.35465 | 0.008 | 0.015 | 1 | 12 | Cpq | -0.007 | Old |
| 0.661647 | 0.30634 | 0.008 | 0.015 | 1 | 12 | Prpf8 | -0.007 | Old |
| 0.661647 | 0.264763 | 0.008 | 0.015 | 1 | 12 | Mettl27 | -0.007 | Old |
| 0.661647 | 0.171827 | 0.008 | 0.015 | 1 | 12 | Nell2 | -0.007 | Old |
| 0.661647 | 0.130593 | 0.008 | 0.015 | 1 | 12 | Pih1d1 | -0.007 | Old |
| 0.661647 | 0.128003 | 0.008 | 0.015 | 1 | 12 | Nrsn2 | -0.007 | Old |
| 0.661647 | 0.100578 | 0.008 | 0.015 | 1 | 12 | Supt5 | -0.007 | Old |
| 0.661647 | 0.065505 | 0.008 | 0.015 | 1 | 12 | Crkl | -0.007 | Old |
| 0.662575 | 0.181158 | 0.051 | 0.066 | 1 | 12 | 1700025G( | -0.015 | Old |
| 0.66276 | 0.387522 | 0.119 | 0.102 | 1 | 12 | Hacd1 | 0.017 | Old |
| 0.663774 | 0.000925 | 0.153 | 0.182 | 1 | 12 | Fam162a | -0.029 | Old |
| 0.665868 | 0.128524 | 0.551 | 0.555 | 1 | 12 | Gng3 | -0.004 | Old |
| 0.667782 | 0.429781 | 0.085 | 0.073 | 1 | 12 | Sccpdh | 0.012 | Old |
| 0.668161 | 0.000325 | 0.373 | 0.431 | 1 | 12 | Ppp2r2b | -0.058 | Old |
| 0.668482 | 0.505814 | 0.161 | 0.146 | 1 | 12 | Grina | 0.015 | Old |
| 0.671774 | 0.374961 | 0.085 | 0.073 | 1 | 12 | Cyfip2 | 0.012 | Old |
| 0.67217 | 0.221976 | 0.11 | 0.095 | 1 | 12 | Eif3m | 0.015 | Old |
| 0.674231 | 0.134861 | 0.178 | 0.161 | 1 | 12 | Ubxn6 | 0.017 | Old |
| 0.675138 | 0.149364 | 0.186 | 0.175 | 1 | 12 | Kif1a | 0.011 | Old |
| 0.67657 | 0.047889 | 0.314 | 0.292 | 1 | 12 | Uqcrfs1 | 0.022 | Old |
| 0.676809 | 0.260524 | 0.102 | 0.088 | 1 | 12 | Tiprl | 0.014 | Old |
| 0.678604 | 0.475216 | 0.093 | 0.08 | 1 | 12 | Nars | 0.013 | Old |
| 0.680161 | 0.272134 | 0.407 | 0.409 | 1 | 12 | Tmem176k | -0.002 | Old |
| 0.68092 | 0.314044 | 0.22 | 0.204 | 1 | 12 | Ypel3 | 0.016 | Old |
| 0.681124 | 0.304387 | 0.085 | 0.073 | 1 | 12 | Dnajc12 | 0.012 | Old |
| 0.686672 | 0.247937 | 0.102 | 0.088 | 1 | 12 | Scn9a | 0.014 | Old |
| 0.686762 | 0.255695 | 0.195 | 0.182 | 1 | 12 | Serp2 | 0.013 | Old |
| 0.687245 | 0.461491 | 0.186 | 0.175 | 1 | 12 | Gnao1 | 0.011 | Old |
| 0.688875 | 0.240225 | 0.093 | 0.08 | 1 | 12 | Snrpf | 0.013 | Old |
| 0.689177 | 0.424648 | 0.085 | 0.073 | 1 | 12 | Tm2d2 | 0.012 | Old |
| 0.689177 | 0.363127 | 0.085 | 0.073 | 1 | 12 | Nap1l4 | 0.012 | Old |
| 0.690002 | 0.244898 | 0.11 | 0.095 | 1 | 12 | Acyp1 | 0.015 | Old |
| 0.69145 | 0.218064 | 0.119 | 0.102 | 1 | 12 | Lgmn | 0.017 | Old |
| 0.69187 | 0.172413 | 0.085 | 0.073 | 1 | 12 | Them4 | 0.012 | Old |
| 0.697266 | 0.361357 | 0.085 | 0.073 | 1 | 12 | Ebpl | 0.012 | Old |
| 0.697266 | 0.275407 | 0.085 | 0.073 | 1 | 12 | Taf13 | 0.012 | Old |
| 0.698959 | 0.098026 | 0.153 | 0.131 | 1 | 12 | Purb | 0.022 | Old |
| 0.69997 | 0.337273 | 0.085 | 0.073 | 1 | 12 | Fbxw7 | 0.012 | Old |
| 0.700377 | 0.093249 | 0.246 | 0.27 | 1 | 12 | Ndufb5 | -0.024 | Old |
| 0.701566 | 0.19862 | 0.102 | 0.088 | 1 | 12 | Gdpd1 | 0.014 | Old |
| 0.701796 | 0.242226 | 0.093 | 0.08 | 1 | 12 | B2m | 0.013 | Old |
| 0.704038 | 0.326971 | 0.305 | 0.307 | 1 | 12 | Isl1 | -0.002 | Old |
| 0.70439 | 0.334606 | 0.093 | 0.08 | 1 | 12 | Matr3 | 0.013 | Old |
| 0.704763 | 0.023592 | 0.203 | 0.182 | 1 | 12 | Mrpl54 | 0.021 | Old |

| | | | | | | | | |
|---|---|---|---|---|---|---|---|---|
| 0.706044 | 0.249672 | 0.144 | 0.131 | 1 | 12 | Prkacb | 0.013 | Old |
| 0.706121 | 0.148391 | 0.263 | 0.255 | 1 | 12 | Paip2 | 0.008 | Old |
| 0.706761 | 0.263486 | 0.381 | 0.394 | 1 | 12 | Ndufab1 | -0.013 | Old |
| 0.706988 | 0.388742 | 0.093 | 0.08 | 1 | 12 | Naa20 | 0.013 | Old |
| 0.707217 | 0.147628 | 0.246 | 0.226 | 1 | 12 | Dpysl2 | 0.02 | Old |
| 0.709057 | 0.217225 | 0.102 | 0.088 | 1 | 12 | Ppfibp2 | 0.014 | Old |
| 0.709219 | 0.302703 | 0.144 | 0.131 | 1 | 12 | Snca | 0.013 | Old |
| 0.70977 | 0.068333 | 0.034 | 0.044 | 1 | 12 | Cdc37l1 | -0.01 | Old |
| 0.710462 | 0.124077 | 0.644 | 0.642 | 1 | 12 | Rps11 | 0.002 | Old |
| 0.710872 | 0.418785 | 0.068 | 0.058 | 1 | 12 | Drg1 | 0.01 | Old |
| 0.711219 | 0.049086 | 0.119 | 0.102 | 1 | 12 | Pdcl3 | 0.017 | Old |
| 0.712194 | 0.288674 | 0.093 | 0.08 | 1 | 12 | Ctsz | 0.013 | Old |
| 0.713892 | 0.603647 | 0.068 | 0.058 | 1 | 12 | Cdipt | 0.01 | Old |
| 0.714048 | 0.138014 | 0.11 | 0.095 | 1 | 12 | Cmas | 0.015 | Old |
| 0.714803 | 0.113081 | 0.093 | 0.08 | 1 | 12 | Mrps14 | 0.013 | Old |
| 0.715483 | 0.606156 | 0.059 | 0.051 | 1 | 12 | Eif1ad | 0.008 | Old |
| 0.716273 | 0.334312 | 0.085 | 0.073 | 1 | 12 | Rsrc2 | 0.012 | Old |
| 0.716824 | 0.151558 | 0.542 | 0.555 | 1 | 12 | Fxyd7 | -0.013 | Old |
| 0.716917 | 0.442887 | 0.068 | 0.058 | 1 | 12 | Hsd17b12 | 0.01 | Old |
| 0.717058 | 0.322676 | 0.076 | 0.066 | 1 | 12 | Csde1 | 0.01 | Old |
| 0.717956 | 0.136437 | 0.322 | 0.307 | 1 | 12 | Map1lc3b | 0.015 | Old |
| 0.718913 | 0.001526 | 0.602 | 0.657 | 1 | 12 | Rps9 | -0.055 | Old |
| 0.719003 | 0.197954 | 0.085 | 0.073 | 1 | 12 | Anp32a | 0.012 | Old |
| 0.719179 | 0.097317 | 0.195 | 0.182 | 1 | 12 | Cct8 | 0.013 | Old |
| 0.719925 | 0.342896 | 0.076 | 0.066 | 1 | 12 | Vbp1 | 0.01 | Old |
| 0.719925 | 0.269274 | 0.076 | 0.066 | 1 | 12 | Syt5 | 0.01 | Old |
| 0.720029 | 0.211996 | 0.093 | 0.08 | 1 | 12 | Phax | 0.013 | Old |
| 0.720825 | 0.003893 | 0.585 | 0.562 | 1 | 12 | Cox5b | 0.023 | Old |
| 0.721383 | 0.124585 | 0.203 | 0.182 | 1 | 12 | Phactr1 | 0.021 | Old |
| 0.722085 | 0.253276 | 0.339 | 0.343 | 1 | 12 | Uqcrb | -0.004 | Old |
| 0.722533 | 0.081948 | 0.093 | 0.109 | 1 | 12 | Rraga | -0.016 | Old |
| 0.722789 | 0.257865 | 0.136 | 0.124 | 1 | 12 | Adrm1 | 0.012 | Old |
| 0.725167 | 0.535824 | 0.059 | 0.051 | 1 | 12 | Hdgfl2 | 0.008 | Old |
| 0.725672 | 0.094445 | 0.076 | 0.066 | 1 | 12 | Tgfb1 | 0.01 | Old |
| 0.725843 | 0.092756 | 0.085 | 0.073 | 1 | 12 | Atg12 | 0.012 | Old |
| 0.726017 | 0.511244 | 0.068 | 0.058 | 1 | 12 | Dusp3 | 0.01 | Old |
| 0.727537 | 0.451992 | 0.068 | 0.058 | 1 | 12 | Lyz2 | 0.01 | Old |
| 0.728706 | 0.057885 | 0.034 | 0.044 | 1 | 12 | Rtf1 | -0.01 | Old |
| 0.728837 | 0.250974 | 0.169 | 0.161 | 1 | 12 | Atp6v1d | 0.008 | Old |
| 0.728891 | 0.065416 | 0.627 | 0.599 | 1 | 12 | Basp1 | 0.028 | Old |
| 0.729059 | 0.375998 | 0.068 | 0.058 | 1 | 12 | Chordc1 | 0.01 | Old |
| 0.730608 | 0.377623 | 0.034 | 0.044 | 1 | 12 | Bbc3 | -0.01 | Old |
| 0.732105 | 0.135881 | 0.068 | 0.058 | 1 | 12 | Scp2 | 0.01 | Old |
| 0.732656 | 0.100187 | 0.136 | 0.124 | 1 | 12 | Lsm6 | 0.012 | Old |
| 0.734034 | 0.031369 | 0.195 | 0.219 | 1 | 12 | C1d | -0.024 | Old |

| | | | | | | | | |
|---|---|---|---|---|---|---|---|---|
| 0.734065 | 0.014515 | 0.966 | 0.956 | 1 | 12 | Calm1 | 0.01 | Old |
| 0.73432 | 0.288911 | 0.076 | 0.066 | 1 | 12 | Mrps18b | 0.01 | Old |
| 0.735004 | 0.037948 | 0.068 | 0.08 | 1 | 12 | Arpp19 | -0.012 | Old |
| 0.735155 | 0.275729 | 0.068 | 0.058 | 1 | 12 | Prelid3b | 0.01 | Old |
| 0.735156 | 0.458049 | 0.068 | 0.058 | 1 | 12 | Commd4 | 0.01 | Old |
| 0.735781 | 0.012538 | 0.153 | 0.175 | 1 | 12 | Cdkn1a | -0.022 | Old |
| 0.737211 | 0.241979 | 0.076 | 0.066 | 1 | 12 | Rnf208 | 0.01 | Old |
| 0.738126 | 0.156833 | 0.347 | 0.343 | 1 | 12 | Cox7b | 0.004 | Old |
| 0.738148 | 0.447956 | 0.059 | 0.051 | 1 | 12 | Rnf126 | 0.008 | Old |
| 0.73821 | 0.348121 | 0.068 | 0.058 | 1 | 12 | Slirp | 0.01 | Old |
| 0.738211 | 0.013757 | 0.085 | 0.073 | 1 | 12 | Tmed9 | 0.012 | Old |
| 0.738633 | 0.115764 | 0.331 | 0.314 | 1 | 12 | Cox6a1 | 0.017 | Old |
| 0.740385 | 0.06316 | 0.466 | 0.453 | 1 | 12 | mt-Nd1 | 0.013 | Old |
| 0.740693 | 0.054527 | 0.246 | 0.226 | 1 | 12 | Kif1b | 0.02 | Old |
| 0.74087 | 0.103827 | 0.364 | 0.372 | 1 | 12 | Psmb5 | -0.008 | Old |
| 0.741405 | 0.74462 | 0.059 | 0.051 | 1 | 12 | Banf1 | 0.008 | Old |
| 0.741405 | 0.442169 | 0.059 | 0.051 | 1 | 12 | Map2k1 | 0.008 | Old |
| 0.742969 | 0.128733 | 0.229 | 0.212 | 1 | 12 | Arpc5l | 0.017 | Old |
| 0.743002 | 0.14715 | 0.076 | 0.066 | 1 | 12 | Eif4e | 0.01 | Old |
| 0.743002 | 0.143449 | 0.076 | 0.066 | 1 | 12 | Pbrm1 | 0.01 | Old |
| 0.743767 | 0.079502 | 0.703 | 0.708 | 1 | 12 | Rpl15 | -0.005 | Old |
| 0.744154 | 0.039722 | 0.178 | 0.161 | 1 | 12 | Cltb | 0.017 | Old |
| 0.744277 | 0.066308 | 0.915 | 0.912 | 1 | 12 | Cartpt | 0.003 | Old |
| 0.744775 | 0.30838 | 0.136 | 0.124 | 1 | 12 | Mrpl33 | 0.012 | Old |
| 0.74483 | 0.191212 | 0.22 | 0.212 | 1 | 12 | Psmb4 | 0.008 | Old |
| 0.745295 | 0.326817 | 0.144 | 0.139 | 1 | 12 | Psmc4 | 0.005 | Old |
| 0.745904 | 0.153698 | 0.076 | 0.066 | 1 | 12 | Senp6 | 0.01 | Old |
| 0.747421 | 0.423981 | 0.11 | 0.102 | 1 | 12 | Rab3b | 0.008 | Old |
| 0.747933 | 0.547961 | 0.059 | 0.051 | 1 | 12 | Junb | 0.008 | Old |
| 0.747933 | 0.413608 | 0.059 | 0.051 | 1 | 12 | Rangap1 | 0.008 | Old |
| 0.748977 | 0.113928 | 0.585 | 0.547 | 1 | 12 | Rpl21 | 0.038 | Old |
| 0.75047 | 0.221278 | 0.068 | 0.058 | 1 | 12 | Hikeshi | 0.01 | Old |
| 0.750569 | 0.453523 | 0.051 | 0.044 | 1 | 12 | Rbfox1 | 0.007 | Old |
| 0.751204 | 0.279918 | 0.059 | 0.051 | 1 | 12 | Chd3os | 0.008 | Old |
| 0.753173 | 0.073425 | 0.076 | 0.066 | 1 | 12 | Osgep | 0.01 | Old |
| 0.753545 | 0.453973 | 0.068 | 0.058 | 1 | 12 | Ncoa7 | 0.01 | Old |
| 0.754217 | 0.101138 | 0.314 | 0.299 | 1 | 12 | Vdac1 | 0.015 | Old |
| 0.754479 | 0.262685 | 0.059 | 0.051 | 1 | 12 | Nsd3 | 0.008 | Old |
| 0.75463 | 0.125001 | 0.076 | 0.066 | 1 | 12 | Snrnp40 | 0.01 | Old |
| 0.754965 | 0.164448 | 0.364 | 0.343 | 1 | 12 | Reep5 | 0.021 | Old |
| 0.756177 | 0.138782 | 0.127 | 0.117 | 1 | 12 | Sod2 | 0.01 | Old |
| 0.756624 | 0.241325 | 0.068 | 0.058 | 1 | 12 | Tsnax | 0.01 | Old |
| 0.75771 | 0.073713 | 0.881 | 0.876 | 1 | 12 | Rpl13 | 0.005 | Old |
| 0.757759 | 0.355457 | 0.059 | 0.051 | 1 | 12 | Bola3 | 0.008 | Old |
| 0.757759 | 0.321563 | 0.059 | 0.051 | 1 | 12 | Lamp2 | 0.008 | Old |

| | | | | | | | | |
|---|---|---|---|---|---|---|---|---|
| 0.758777 | 0.075615 | 0.203 | 0.19 | 1 | 12 | Rab3c | 0.013 | Old |
| 0.759529 | 0.337475 | 0.11 | 0.102 | 1 | 12 | Suclg1 | 0.008 | Old |
| 0.75968 | 0.348479 | 0.102 | 0.117 | 1 | 12 | Psmc3 | -0.015 | Old |
| 0.759696 | 0.134472 | 0.161 | 0.146 | 1 | 12 | Fam89a | 0.015 | Old |
| 0.759707 | 0.258941 | 0.068 | 0.058 | 1 | 12 | Snx15 | 0.01 | Old |
| 0.761153 | 0.399398 | 0.051 | 0.044 | 1 | 12 | Rap1gds1 | 0.007 | Old |
| 0.762793 | 0.266174 | 0.068 | 0.058 | 1 | 12 | Bbip1 | 0.01 | Old |
| 0.762793 | 0.219612 | 0.068 | 0.058 | 1 | 12 | Meis1 | 0.01 | Old |
| 0.762793 | 0.160228 | 0.068 | 0.058 | 1 | 12 | Map6 | 0.01 | Old |
| 0.762954 | 0.17045 | 0.305 | 0.307 | 1 | 12 | Zwint | -0.002 | Old |
| 0.764331 | 0.311934 | 0.059 | 0.051 | 1 | 12 | Smim20 | 0.008 | Old |
| 0.764331 | 0.150495 | 0.059 | 0.051 | 1 | 12 | Rbbp4 | 0.008 | Old |
| 0.764691 | 0.322732 | 0.051 | 0.044 | 1 | 12 | AI413582 | 0.007 | Old |
| 0.764691 | 0.648353 | 0.051 | 0.044 | 1 | 12 | Gadd45g | 0.007 | Old |
| 0.764691 | 0.497283 | 0.051 | 0.044 | 1 | 12 | Nfkbib | 0.007 | Old |
| 0.765884 | 0.238101 | 0.068 | 0.058 | 1 | 12 | Nudt17 | 0.01 | Old |
| 0.766313 | 0.145921 | 0.076 | 0.066 | 1 | 12 | Cep164 | 0.01 | Old |
| 0.766313 | 0.131497 | 0.076 | 0.066 | 1 | 12 | Hmx3 | 0.01 | Old |
| 0.766803 | 0.283793 | 0.119 | 0.109 | 1 | 12 | Bend5 | 0.01 | Old |
| 0.767624 | 0.257724 | 0.059 | 0.051 | 1 | 12 | Yipf1 | 0.008 | Old |
| 0.768234 | 0.374252 | 0.051 | 0.044 | 1 | 12 | Atp2c1 | 0.007 | Old |
| 0.768234 | 0.242159 | 0.051 | 0.044 | 1 | 12 | Mvb12a | 0.007 | Old |
| 0.768318 | 0.002005 | 0.415 | 0.46 | 1 | 12 | Rab6b | -0.045 | Old |
| 0.768978 | 0.151351 | 0.068 | 0.058 | 1 | 12 | Leprot | 0.01 | Old |
| 0.769067 | 0.009981 | 0.068 | 0.08 | 1 | 12 | Celf3 | -0.012 | Old |
| 0.769163 | 0.180302 | 0.119 | 0.109 | 1 | 12 | Pafah1b2 | 0.01 | Old |
| 0.770921 | 0.258898 | 0.059 | 0.051 | 1 | 12 | Ppp2r2c | 0.008 | Old |
| 0.770921 | 0.199375 | 0.059 | 0.051 | 1 | 12 | Coasy | 0.008 | Old |
| 0.771782 | 0.436764 | 0.051 | 0.044 | 1 | 12 | Lrrc58 | 0.007 | Old |
| 0.772076 | 0.11975 | 0.068 | 0.058 | 1 | 12 | Glud1 | 0.01 | Old |
| 0.772076 | 0.099496 | 0.068 | 0.058 | 1 | 12 | Etnk1 | 0.01 | Old |
| 0.773558 | 0.063926 | 0.042 | 0.051 | 1 | 12 | Igsf8 | -0.009 | Old |
| 0.774222 | 0.127268 | 0.059 | 0.051 | 1 | 12 | Eif4g3 | 0.008 | Old |
| 0.774222 | 0.126103 | 0.059 | 0.051 | 1 | 12 | Btbd1 | 0.008 | Old |
| 0.774313 | 0.094126 | 0.085 | 0.073 | 1 | 12 | Timm10b | 0.012 | Old |
| 0.77477 | 0.34759 | 0.093 | 0.088 | 1 | 12 | Lsm1 | 0.005 | Old |
| 0.774812 | 0.647927 | 0.042 | 0.036 | 1 | 12 | Usp12 | 0.006 | Old |
| 0.774812 | 0.627927 | 0.042 | 0.036 | 1 | 12 | Micu3 | 0.006 | Old |
| 0.775177 | 0.074914 | 0.068 | 0.058 | 1 | 12 | H2afv | 0.01 | Old |
| 0.775335 | 0.371217 | 0.051 | 0.044 | 1 | 12 | Impa1 | 0.007 | Old |
| 0.775335 | 0.346698 | 0.051 | 0.044 | 1 | 12 | Kmt2e | 0.007 | Old |
| 0.775335 | 0.315974 | 0.051 | 0.044 | 1 | 12 | Plin2 | 0.007 | Old |
| 0.775335 | 0.232013 | 0.051 | 0.044 | 1 | 12 | Mettl9 | 0.007 | Old |
| 0.777112 | 0.296253 | 0.051 | 0.044 | 1 | 12 | Ptger3 | 0.007 | Old |
| 0.777808 | 0.044048 | 0.076 | 0.088 | 1 | 12 | Mrpl18 | -0.012 | Old |

| | | | | | | | | |
|---|---|---|---|---|---|---|---|---|
| 0.778693 | 0.481371 | 0.042 | 0.036 | 1 | 12 | Ccdc136 | 0.006 | Old |
| 0.778892 | 0.555588 | 0.051 | 0.044 | 1 | 12 | Acbd3 | 0.007 | Old |
| 0.778892 | 0.275298 | 0.051 | 0.044 | 1 | 12 | Ttll1 | 0.007 | Old |
| 0.778892 | 0.039623 | 0.042 | 0.051 | 1 | 12 | Rgs19 | -0.009 | Old |
| 0.779427 | 0.092084 | 0.017 | 0.022 | 1 | 12 | Irgq | -0.005 | Old |
| 0.780273 | 0.011288 | 0.441 | 0.423 | 1 | 12 | Psmb6 | 0.018 | Old |
| 0.780836 | 0.180034 | 0.059 | 0.051 | 1 | 12 | Eml5 | 0.008 | Old |
| 0.780836 | 0.080641 | 0.059 | 0.051 | 1 | 12 | Rtcb | 0.008 | Old |
| 0.78139 | 0.025008 | 0.068 | 0.058 | 1 | 12 | Cox19 | 0.01 | Old |
| 0.781434 | 0.192848 | 0.415 | 0.431 | 1 | 12 | H3f3a | -0.016 | Old |
| 0.782021 | 0.067572 | 0.212 | 0.234 | 1 | 12 | Fkbp2 | -0.022 | Old |
| 0.782454 | 0.243526 | 0.051 | 0.044 | 1 | 12 | Nsf | 0.007 | Old |
| 0.782492 | 0.181386 | 0.059 | 0.051 | 1 | 12 | Ndrg3 | 0.008 | Old |
| 0.78258 | 0.531505 | 0.042 | 0.036 | 1 | 12 | Mpped2 | 0.006 | Old |
| 0.78258 | 0.50301 | 0.042 | 0.036 | 1 | 12 | 1110032A( | 0.006 | Old |
| 0.78258 | 0.424032 | 0.042 | 0.036 | 1 | 12 | Myt1l | 0.006 | Old |
| 0.78258 | 0.414255 | 0.042 | 0.036 | 1 | 12 | Nrbf2 | 0.006 | Old |
| 0.782937 | 0.109373 | 0.169 | 0.161 | 1 | 12 | Kifap3 | 0.008 | Old |
| 0.783614 | 0.132251 | 0.161 | 0.146 | 1 | 12 | Mrpl42 | 0.015 | Old |
| 0.784149 | 0.414035 | 0.059 | 0.051 | 1 | 12 | Ankrd46 | 0.008 | Old |
| 0.784149 | 0.182583 | 0.059 | 0.051 | 1 | 12 | Prkra | 0.008 | Old |
| 0.784643 | 0.004742 | 0.144 | 0.131 | 1 | 12 | Tmem256 | 0.013 | Old |
| 0.784872 | 0.424134 | 0.017 | 0.022 | 1 | 12 | Areg | -0.005 | Old |
| 0.786471 | 0.49222 | 0.042 | 0.036 | 1 | 12 | Mdp1 | 0.006 | Old |
| 0.786471 | 0.466815 | 0.042 | 0.036 | 1 | 12 | Supt3 | 0.006 | Old |
| 0.786528 | 0.116565 | 0.432 | 0.416 | 1 | 12 | Ndufb8 | 0.016 | Old |
| 0.787466 | 0.201049 | 0.059 | 0.051 | 1 | 12 | Mrpl24 | 0.008 | Old |
| 0.788784 | 0.009343 | 0.076 | 0.088 | 1 | 12 | Txnrd1 | -0.012 | Old |
| 0.789592 | 0.236993 | 0.051 | 0.044 | 1 | 12 | Glg1 | 0.007 | Old |
| 0.790101 | 0.135729 | 0.229 | 0.219 | 1 | 12 | Nedd4 | 0.01 | Old |
| 0.790327 | 0.399317 | 0.017 | 0.022 | 1 | 12 | Borcs8 | -0.005 | Old |
| 0.790327 | 0.279086 | 0.017 | 0.022 | 1 | 12 | Ccl7 | -0.005 | Old |
| 0.790327 | 0.107661 | 0.017 | 0.022 | 1 | 12 | Zfp180 | -0.005 | Old |
| 0.790327 | 0.106656 | 0.017 | 0.022 | 1 | 12 | Tmem219 | -0.005 | Old |
| 0.790327 | 0.05299 | 0.017 | 0.022 | 1 | 12 | Atp5s | -0.005 | Old |
| 0.790327 | 0.04069 | 0.017 | 0.022 | 1 | 12 | Dalrd3 | -0.005 | Old |
| 0.790327 | 0.02555 | 0.017 | 0.022 | 1 | 12 | Rspry1 | -0.005 | Old |
| 0.790369 | 0.481651 | 0.042 | 0.036 | 1 | 12 | Zfp644 | 0.006 | Old |
| 0.790369 | 0.476499 | 0.042 | 0.036 | 1 | 12 | Syt17 | 0.006 | Old |
| 0.790369 | 0.406922 | 0.042 | 0.036 | 1 | 12 | Mcl1 | 0.006 | Old |
| 0.790369 | 0.262396 | 0.042 | 0.036 | 1 | 12 | Lin37 | 0.006 | Old |
| 0.791378 | 0.072887 | 0.051 | 0.044 | 1 | 12 | Hdac3 | 0.007 | Old |
| 0.791991 | 0.36573 | 0.136 | 0.131 | 1 | 12 | Tmod2 | 0.005 | Old |
| 0.792319 | 0.522698 | 0.042 | 0.036 | 1 | 12 | Mtg2 | 0.006 | Old |
| 0.793167 | 0.294311 | 0.051 | 0.044 | 1 | 12 | Mbd3 | 0.007 | Old |

| | | | | | | | | |
|---|---|---|---|---|---|---|---|---|
| 0.794271 | 0.580392 | 0.042 | 0.036 | 1 | 12 | Mrpl44 | 0.006 | Old |
| 0.794271 | 0.439856 | 0.042 | 0.036 | 1 | 12 | Hint3 | 0.006 | Old |
| 0.794271 | 0.406321 | 0.042 | 0.036 | 1 | 12 | Thy1 | 0.006 | Old |
| 0.794271 | 0.400539 | 0.042 | 0.036 | 1 | 12 | Galt | 0.006 | Old |
| 0.794271 | 0.202799 | 0.042 | 0.036 | 1 | 12 | Tle4 | 0.006 | Old |
| 0.794525 | 0.312013 | 0.093 | 0.088 | 1 | 12 | Sptan1 | 0.005 | Old |
| 0.795472 | 0.095117 | 0.805 | 0.774 | 1 | 12 | Eef1a1 | 0.031 | Old |
| 0.795793 | 0.216151 | 0.017 | 0.022 | 1 | 12 | Bysl | -0.005 | Old |
| 0.795793 | 0.213482 | 0.017 | 0.022 | 1 | 12 | Msantd3 | -0.005 | Old |
| 0.795793 | 0.183382 | 0.017 | 0.022 | 1 | 12 | Hjurp | -0.005 | Old |
| 0.795793 | 0.182463 | 0.017 | 0.022 | 1 | 12 | Dtx3 | -0.005 | Old |
| 0.795793 | 0.158627 | 0.017 | 0.022 | 1 | 12 | Fam168a | -0.005 | Old |
| 0.795793 | 0.132848 | 0.017 | 0.022 | 1 | 12 | Efl1 | -0.005 | Old |
| 0.795793 | 0.091186 | 0.017 | 0.022 | 1 | 12 | Rnf146 | -0.005 | Old |
| 0.795793 | 0.05621 | 0.017 | 0.022 | 1 | 12 | Arhgap36 | -0.005 | Old |
| 0.795793 | 0.050274 | 0.017 | 0.022 | 1 | 12 | Atxn3 | -0.005 | Old |
| 0.796747 | 0.17867 | 0.051 | 0.044 | 1 | 12 | Carhsp1 | 0.007 | Old |
| 0.79744 | 0.143112 | 0.059 | 0.051 | 1 | 12 | Slc2a3 | 0.008 | Old |
| 0.797517 | 0.719817 | 0.034 | 0.029 | 1 | 12 | Spred1 | 0.005 | Old |
| 0.797517 | 0.690032 | 0.034 | 0.029 | 1 | 12 | Taf12 | 0.005 | Old |
| 0.797904 | 0.153787 | 0.254 | 0.248 | 1 | 12 | Arpc1a | 0.006 | Old |
| 0.798179 | 0.387385 | 0.042 | 0.036 | 1 | 12 | Timm17b | 0.006 | Old |
| 0.798179 | 0.296052 | 0.042 | 0.036 | 1 | 12 | Ap3s2 | 0.006 | Old |
| 0.798179 | 0.287457 | 0.042 | 0.036 | 1 | 12 | Tmem14a | 0.006 | Old |
| 0.799699 | 0.040669 | 0.347 | 0.321 | 1 | 12 | Mpc2 | 0.026 | Old |
| 0.802052 | 0.072211 | 0.331 | 0.336 | 1 | 12 | Mdh2 | -0.005 | Old |
| 0.802091 | 0.621975 | 0.042 | 0.036 | 1 | 12 | Ddb1 | 0.006 | Old |
| 0.802091 | 0.409379 | 0.042 | 0.036 | 1 | 12 | Gm2990 | 0.006 | Old |
| 0.802091 | 0.35254 | 0.042 | 0.036 | 1 | 12 | Gpr108 | 0.006 | Old |
| 0.802091 | 0.346581 | 0.042 | 0.036 | 1 | 12 | Ppm1g | 0.006 | Old |
| 0.802091 | 0.327577 | 0.042 | 0.036 | 1 | 12 | Tmem33 | 0.006 | Old |
| 0.802091 | 0.324394 | 0.042 | 0.036 | 1 | 12 | Fhl1 | 0.006 | Old |
| 0.802091 | 0.308943 | 0.042 | 0.036 | 1 | 12 | Spint2 | 0.006 | Old |
| 0.802091 | 0.307452 | 0.042 | 0.036 | 1 | 12 | AI837181 | 0.006 | Old |
| 0.802091 | 0.265731 | 0.042 | 0.036 | 1 | 12 | Hmbs | 0.006 | Old |
| 0.802091 | 0.16608 | 0.042 | 0.036 | 1 | 12 | Atp11b | 0.006 | Old |
| 0.80361 | 0.083635 | 0.119 | 0.109 | 1 | 12 | Dclk1 | 0.01 | Old |
| 0.803919 | 0.294572 | 0.051 | 0.044 | 1 | 12 | Glul | 0.007 | Old |
| 0.803919 | 0.118501 | 0.051 | 0.044 | 1 | 12 | Ythdc1 | 0.007 | Old |
| 0.803919 | 0.108746 | 0.051 | 0.044 | 1 | 12 | Mat2b | 0.007 | Old |
| 0.804104 | 0.116717 | 0.11 | 0.124 | 1 | 12 | Cript | -0.014 | Old |
| 0.805385 | 0.20246 | 0.297 | 0.292 | 1 | 12 | Morf4l1 | 0.005 | Old |
| 0.806009 | 0.261765 | 0.042 | 0.036 | 1 | 12 | Rcan3 | 0.006 | Old |
| 0.806009 | 0.223521 | 0.042 | 0.036 | 1 | 12 | Chrac1 | 0.006 | Old |
| 0.806009 | 0.182458 | 0.042 | 0.036 | 1 | 12 | Golga7b | 0.006 | Old |

| | | | | | | | | |
|---|---|---|---|---|---|---|---|---|
| 0.806009 | 0.082202 | 0.042 | 0.036 | 1 | 12 | Cap1 | 0.006 | Old |
| 0.806044 | 0.298068 | 0.212 | 0.212 | 1 | 12 | Eef2 | 0 | Old |
| 0.806237 | 0.825322 | 0.034 | 0.029 | 1 | 12 | Ifngr2 | 0.005 | Old |
| 0.806237 | 0.527201 | 0.034 | 0.029 | 1 | 12 | Srek1 | 0.005 | Old |
| 0.807419 | 0.173237 | 0.203 | 0.197 | 1 | 12 | Ssu72 | 0.006 | Old |
| 0.807512 | 0.18594 | 0.051 | 0.044 | 1 | 12 | Ufd1 | 0.007 | Old |
| 0.807512 | 0.129369 | 0.051 | 0.044 | 1 | 12 | Kif1bp | 0.007 | Old |
| 0.807512 | 0.083996 | 0.051 | 0.044 | 1 | 12 | Mex3d | 0.007 | Old |
| 0.809931 | 0.278068 | 0.042 | 0.036 | 1 | 12 | Xrn2 | 0.006 | Old |
| 0.809931 | 0.230687 | 0.042 | 0.036 | 1 | 12 | Ccs | 0.006 | Old |
| 0.809931 | 0.229759 | 0.042 | 0.036 | 1 | 12 | Aimp2 | 0.006 | Old |
| 0.809931 | 0.154535 | 0.042 | 0.036 | 1 | 12 | 2310009B1 | 0.006 | Old |
| 0.809931 | 0.077663 | 0.042 | 0.036 | 1 | 12 | Auts2 | 0.006 | Old |
| 0.810605 | 0.641692 | 0.034 | 0.029 | 1 | 12 | Fbxo25 | 0.005 | Old |
| 0.810605 | 0.543954 | 0.034 | 0.029 | 1 | 12 | Srxn1 | 0.005 | Old |
| 0.810605 | 0.51031 | 0.034 | 0.029 | 1 | 12 | Scamp1 | 0.005 | Old |
| 0.810605 | 0.45512 | 0.034 | 0.029 | 1 | 12 | Kpna3 | 0.005 | Old |
| 0.813859 | 0.281397 | 0.042 | 0.036 | 1 | 12 | Drg2 | 0.006 | Old |
| 0.813859 | 0.227857 | 0.042 | 0.036 | 1 | 12 | Trib2 | 0.006 | Old |
| 0.814718 | 0.141916 | 0.11 | 0.124 | 1 | 12 | Camta1 | -0.014 | Old |
| 0.81498 | 0.73632 | 0.034 | 0.029 | 1 | 12 | Bag6 | 0.005 | Old |
| 0.81498 | 0.406584 | 0.034 | 0.029 | 1 | 12 | 1500011K1 | 0.005 | Old |
| 0.817183 | 0.275447 | 0.11 | 0.102 | 1 | 12 | Ppdpf | 0.008 | Old |
| 0.817791 | 0.130972 | 0.042 | 0.036 | 1 | 12 | Syngr1 | 0.006 | Old |
| 0.81936 | 0.43344 | 0.034 | 0.029 | 1 | 12 | Fam96a | 0.005 | Old |
| 0.81936 | 0.387568 | 0.034 | 0.029 | 1 | 12 | Smad7 | 0.005 | Old |
| 0.81936 | 0.380277 | 0.034 | 0.029 | 1 | 12 | Fbxw5 | 0.005 | Old |
| 0.81936 | 0.322079 | 0.034 | 0.029 | 1 | 12 | Ifrd1 | 0.005 | Old |
| 0.81936 | 0.312632 | 0.034 | 0.029 | 1 | 12 | R3hdm4 | 0.005 | Old |
| 0.81936 | 0.292319 | 0.034 | 0.029 | 1 | 12 | 2310011J0 | 0.005 | Old |
| 0.81936 | 0.245523 | 0.034 | 0.029 | 1 | 12 | Zfp131 | 0.005 | Old |
| 0.819758 | 0.129485 | 0.042 | 0.036 | 1 | 12 | Plp2 | 0.006 | Old |
| 0.820839 | 0.142805 | 0.051 | 0.058 | 1 | 12 | Stx1a | -0.007 | Old |
| 0.821151 | 0.304618 | 0.102 | 0.095 | 1 | 12 | Pten | 0.007 | Old |
| 0.821666 | 0.208757 | 0.153 | 0.146 | 1 | 12 | Morf4l2 | 0.007 | Old |
| 0.821727 | 0.23926 | 0.042 | 0.036 | 1 | 12 | Ubap2l | 0.006 | Old |
| 0.821727 | 0.145852 | 0.042 | 0.036 | 1 | 12 | Arl16 | 0.006 | Old |
| 0.821727 | 0.12648 | 0.042 | 0.036 | 1 | 12 | Gpr27 | 0.006 | Old |
| 0.822194 | 0.170621 | 0.424 | 0.445 | 1 | 12 | Atp5j | -0.021 | Old |
| 0.822593 | 0.062069 | 0.28 | 0.263 | 1 | 12 | Tspan3 | 0.017 | Old |
| 0.823565 | 0.193553 | 0.178 | 0.175 | 1 | 12 | Bmyc | 0.003 | Old |
| 0.823746 | 0.479927 | 0.034 | 0.029 | 1 | 12 | Btbd2 | 0.005 | Old |
| 0.823746 | 0.394977 | 0.034 | 0.029 | 1 | 12 | Lyar | 0.005 | Old |
| 0.823746 | 0.392238 | 0.034 | 0.029 | 1 | 12 | Hdac5 | 0.005 | Old |
| 0.823746 | 0.347071 | 0.034 | 0.029 | 1 | 12 | Brd1 | 0.005 | Old |

| | | | | | | | | |
|---|---|---|---|---|---|---|---|---|
| 0.823746 | 0.34086 | 0.034 | 0.029 | 1 | 12 | Dicer1 | 0.005 | Old |
| 0.823746 | 0.335024 | 0.034 | 0.029 | 1 | 12 | Pik3ca | 0.005 | Old |
| 0.823746 | 0.305674 | 0.034 | 0.029 | 1 | 12 | Rabif | 0.005 | Old |
| 0.823746 | 0.271248 | 0.034 | 0.029 | 1 | 12 | Clptm1 | 0.005 | Old |
| 0.823746 | 0.191245 | 0.034 | 0.029 | 1 | 12 | Ahsa1 | 0.005 | Old |
| 0.823746 | 0.184419 | 0.034 | 0.029 | 1 | 12 | Daam1 | 0.005 | Old |
| 0.825069 | 0.226664 | 0.093 | 0.088 | 1 | 12 | Pdrg1 | 0.005 | Old |
| 0.826403 | 0.2243 | 0.093 | 0.088 | 1 | 12 | Hspb8 | 0.005 | Old |
| 0.826648 | 0.079601 | 0.127 | 0.117 | 1 | 12 | S100a13 | 0.01 | Old |
| 0.828138 | 0.57063 | 0.034 | 0.029 | 1 | 12 | Tfrc | 0.005 | Old |
| 0.828138 | 0.367733 | 0.034 | 0.029 | 1 | 12 | Mgea5 | 0.005 | Old |
| 0.828138 | 0.317187 | 0.034 | 0.029 | 1 | 12 | 11-Sep | 0.005 | Old |
| 0.828138 | 0.309728 | 0.034 | 0.029 | 1 | 12 | Ssbp4 | 0.005 | Old |
| 0.828138 | 0.296816 | 0.034 | 0.029 | 1 | 12 | Gfod2 | 0.005 | Old |
| 0.828138 | 0.278069 | 0.034 | 0.029 | 1 | 12 | Zdhhc18 | 0.005 | Old |
| 0.828138 | 0.262953 | 0.034 | 0.029 | 1 | 12 | N4bp1 | 0.005 | Old |
| 0.828138 | 0.254709 | 0.034 | 0.029 | 1 | 12 | Trappc3 | 0.005 | Old |
| 0.828138 | 0.241175 | 0.034 | 0.029 | 1 | 12 | Ndufaf8 | 0.005 | Old |
| 0.828138 | 0.239265 | 0.034 | 0.029 | 1 | 12 | Hmx2 | 0.005 | Old |
| 0.828138 | 0.235662 | 0.034 | 0.029 | 1 | 12 | Tub | 0.005 | Old |
| 0.828138 | 0.185099 | 0.034 | 0.029 | 1 | 12 | Gon4l | 0.005 | Old |
| 0.828138 | 0.180016 | 0.034 | 0.029 | 1 | 12 | Cetn2 | 0.005 | Old |
| 0.829614 | 0.169252 | 0.042 | 0.036 | 1 | 12 | Parp1 | 0.006 | Old |
| 0.830918 | 0.330645 | 0.051 | 0.058 | 1 | 12 | Atp9a | -0.007 | Old |
| 0.831034 | 0.14207 | 0.178 | 0.168 | 1 | 12 | Capza2 | 0.01 | Old |
| 0.831693 | 0.13484 | 0.161 | 0.153 | 1 | 12 | Skp1a | 0.008 | Old |
| 0.831836 | 0.051404 | 0.136 | 0.146 | 1 | 12 | Puf60 | -0.01 | Old |
| 0.832228 | 0.466751 | 0.136 | 0.131 | 1 | 12 | Retnla | 0.005 | Old |
| 0.832535 | 0.273239 | 0.034 | 0.029 | 1 | 12 | Dlg2 | 0.005 | Old |
| 0.832535 | 0.272238 | 0.034 | 0.029 | 1 | 12 | Pgs1 | 0.005 | Old |
| 0.832535 | 0.264605 | 0.034 | 0.029 | 1 | 12 | Copa | 0.005 | Old |
| 0.832535 | 0.252816 | 0.034 | 0.029 | 1 | 12 | Ccnl2 | 0.005 | Old |
| 0.832535 | 0.250155 | 0.034 | 0.029 | 1 | 12 | Anapc4 | 0.005 | Old |
| 0.832535 | 0.24715 | 0.034 | 0.029 | 1 | 12 | Gas8 | 0.005 | Old |
| 0.832535 | 0.231723 | 0.034 | 0.029 | 1 | 12 | Mapk9 | 0.005 | Old |
| 0.832535 | 0.219938 | 0.034 | 0.029 | 1 | 12 | Bcl7c | 0.005 | Old |
| 0.832535 | 0.186002 | 0.034 | 0.029 | 1 | 12 | Zbtb38 | 0.005 | Old |
| 0.832535 | 0.142824 | 0.034 | 0.029 | 1 | 12 | Tmod3 | 0.005 | Old |
| 0.832535 | 0.13365 | 0.034 | 0.029 | 1 | 12 | Alkbh6 | 0.005 | Old |
| 0.832535 | 0.1029 | 0.034 | 0.029 | 1 | 12 | Ngb | 0.005 | Old |
| 0.832535 | 0.067832 | 0.034 | 0.029 | 1 | 12 | Ube2w | 0.005 | Old |
| 0.83277 | 0.168356 | 0.28 | 0.263 | 1 | 12 | Ube2m | 0.017 | Old |
| 0.833065 | 0.797083 | 0.025 | 0.022 | 1 | 12 | Hspa13 | 0.003 | Old |
| 0.833065 | 0.685107 | 0.025 | 0.022 | 1 | 12 | Mfsd10 | 0.003 | Old |
| 0.833065 | 0.673611 | 0.025 | 0.022 | 1 | 12 | Pdpk1 | 0.003 | Old |

| | | | | | | | | |
|---|---|---|---|---|---|---|---|---|
| 0.833065 | 0.616618 | 0.025 | 0.022 | 1 | 12 | Scn1a | 0.003 | Old |
| 0.833065 | 0.512502 | 0.025 | 0.022 | 1 | 12 | Naa30 | 0.003 | Old |
| 0.833065 | 0.499918 | 0.025 | 0.022 | 1 | 12 | Adck5 | 0.003 | Old |
| 0.833065 | 0.488716 | 0.025 | 0.022 | 1 | 12 | Gpr89 | 0.003 | Old |
| 0.833065 | 0.472408 | 0.025 | 0.022 | 1 | 12 | Golph3 | 0.003 | Old |
| 0.833065 | 0.448648 | 0.025 | 0.022 | 1 | 12 | Bloc1s5 | 0.003 | Old |
| 0.833065 | 0.364557 | 0.025 | 0.022 | 1 | 12 | Arpp21 | 0.003 | Old |
| 0.833365 | 0.10127 | 0.22 | 0.219 | 1 | 12 | Trir | 0.001 | Old |
| 0.834735 | 0.2077 | 0.034 | 0.029 | 1 | 12 | Far1 | 0.005 | Old |
| 0.835517 | 0.082402 | 0.788 | 0.825 | 1 | 12 | Fth1 | -0.037 | Old |
| 0.836937 | 0.119811 | 0.034 | 0.029 | 1 | 12 | Taf7 | 0.005 | Old |
| 0.836937 | 0.229648 | 0.034 | 0.029 | 1 | 12 | Tmcc3 | 0.005 | Old |
| 0.837274 | 0.404169 | 0.085 | 0.08 | 1 | 12 | Nefh | 0.005 | Old |
| 0.837479 | 0.074069 | 0.314 | 0.321 | 1 | 12 | Erp29 | -0.007 | Old |
| 0.837652 | 0.085058 | 0.051 | 0.058 | 1 | 12 | BC005537 | -0.007 | Old |
| 0.838129 | 0.909218 | 0.025 | 0.022 | 1 | 12 | 4-Sep | 0.003 | Old |
| 0.838129 | 0.740684 | 0.025 | 0.022 | 1 | 12 | Tmem74 | 0.003 | Old |
| 0.838129 | 0.69869 | 0.025 | 0.022 | 1 | 12 | Rabgap1 | 0.003 | Old |
| 0.838129 | 0.652294 | 0.025 | 0.022 | 1 | 12 | Tmx3 | 0.003 | Old |
| 0.838129 | 0.650954 | 0.025 | 0.022 | 1 | 12 | Mapk8ip3 | 0.003 | Old |
| 0.838129 | 0.575476 | 0.025 | 0.022 | 1 | 12 | Asb3 | 0.003 | Old |
| 0.838129 | 0.55319 | 0.025 | 0.022 | 1 | 12 | Abca7 | 0.003 | Old |
| 0.838129 | 0.479133 | 0.025 | 0.022 | 1 | 12 | Med7 | 0.003 | Old |
| 0.838129 | 0.475585 | 0.025 | 0.022 | 1 | 12 | Mypop | 0.003 | Old |
| 0.838129 | 0.45791 | 0.025 | 0.022 | 1 | 12 | Pds5b | 0.003 | Old |
| 0.838129 | 0.390237 | 0.025 | 0.022 | 1 | 12 | Ogfr | 0.003 | Old |
| 0.841165 | 0.056383 | 0.568 | 0.577 | 1 | 12 | Rps19 | -0.009 | Old |
| 0.841344 | 0.145372 | 0.034 | 0.029 | 1 | 12 | Slc39a10 | 0.005 | Old |
| 0.841344 | 0.076458 | 0.034 | 0.029 | 1 | 12 | Mpdu1 | 0.005 | Old |
| 0.841344 | 0.071858 | 0.034 | 0.029 | 1 | 12 | Spg21 | 0.005 | Old |
| 0.842711 | 0.037996 | 0.051 | 0.058 | 1 | 12 | Gm10605 | -0.007 | Old |
| 0.8432 | 0.668766 | 0.025 | 0.022 | 1 | 12 | Narf | 0.003 | Old |
| 0.8432 | 0.614315 | 0.025 | 0.022 | 1 | 12 | Rc3h2 | 0.003 | Old |
| 0.8432 | 0.60573 | 0.025 | 0.022 | 1 | 12 | Dnajb4 | 0.003 | Old |
| 0.8432 | 0.597359 | 0.025 | 0.022 | 1 | 12 | Ptp4a3 | 0.003 | Old |
| 0.8432 | 0.494886 | 0.025 | 0.022 | 1 | 12 | Bcl9 | 0.003 | Old |
| 0.8432 | 0.487084 | 0.025 | 0.022 | 1 | 12 | Phf24 | 0.003 | Old |
| 0.8432 | 0.476732 | 0.025 | 0.022 | 1 | 12 | Senp5 | 0.003 | Old |
| 0.8432 | 0.454662 | 0.025 | 0.022 | 1 | 12 | Grk3 | 0.003 | Old |
| 0.8432 | 0.427198 | 0.025 | 0.022 | 1 | 12 | Cpne3 | 0.003 | Old |
| 0.8432 | 0.426263 | 0.025 | 0.022 | 1 | 12 | Ebna1bp2 | 0.003 | Old |
| 0.8432 | 0.409143 | 0.025 | 0.022 | 1 | 12 | Sec14l1 | 0.003 | Old |
| 0.8432 | 0.390865 | 0.025 | 0.022 | 1 | 12 | Samd10 | 0.003 | Old |
| 0.8432 | 0.370616 | 0.025 | 0.022 | 1 | 12 | Stk32c | 0.003 | Old |
| 0.8432 | 0.369142 | 0.025 | 0.022 | 1 | 12 | Nudt22 | 0.003 | Old |

| | | | | | | | | |
|---|---|---|---|---|---|---|---|---|
| 0.8432 | 0.357922 | 0.025 | 0.022 | 1 | 12 | H2afy2 | 0.003 | Old |
| 0.8432 | 0.268671 | 0.025 | 0.022 | 1 | 12 | Dnajb1 | 0.003 | Old |
| 0.844472 | 0.044132 | 0.347 | 0.328 | 1 | 12 | Uqcr11 | 0.019 | Old |
| 0.844536 | 0.092908 | 0.11 | 0.102 | 1 | 12 | Prdx6 | 0.008 | Old |
| 0.845738 | 0.332659 | 0.025 | 0.022 | 1 | 12 | Dpy19l4 | 0.003 | Old |
| 0.845757 | 0.098377 | 0.034 | 0.029 | 1 | 12 | Ubr3 | 0.005 | Old |
| 0.845757 | 0.075752 | 0.034 | 0.029 | 1 | 12 | Krcc1 | 0.005 | Old |
| 0.845757 | 0.06536 | 0.034 | 0.029 | 1 | 12 | Izumo4 | 0.005 | Old |
| 0.845757 | 0.017002 | 0.034 | 0.029 | 1 | 12 | Snrpg | 0.005 | Old |
| 0.845757 | 0.002041 | 0.034 | 0.029 | 1 | 12 | 8-Mar | 0.005 | Old |
| 0.848278 | 0.79032 | 0.025 | 0.022 | 1 | 12 | Plekha5 | 0.003 | Old |
| 0.848278 | 0.652607 | 0.025 | 0.022 | 1 | 12 | Rsl24d1 | 0.003 | Old |
| 0.848278 | 0.52934 | 0.025 | 0.022 | 1 | 12 | Ccdc12 | 0.003 | Old |
| 0.848278 | 0.523037 | 0.025 | 0.022 | 1 | 12 | Tmed8 | 0.003 | Old |
| 0.848278 | 0.466429 | 0.025 | 0.022 | 1 | 12 | Hps3 | 0.003 | Old |
| 0.848278 | 0.461859 | 0.025 | 0.022 | 1 | 12 | Zfp612 | 0.003 | Old |
| 0.848278 | 0.444869 | 0.025 | 0.022 | 1 | 12 | Cep120 | 0.003 | Old |
| 0.848278 | 0.441095 | 0.025 | 0.022 | 1 | 12 | Maml3 | 0.003 | Old |
| 0.848278 | 0.438009 | 0.025 | 0.022 | 1 | 12 | Rbm11 | 0.003 | Old |
| 0.848278 | 0.435191 | 0.025 | 0.022 | 1 | 12 | Uba6 | 0.003 | Old |
| 0.848278 | 0.433507 | 0.025 | 0.022 | 1 | 12 | Odc1 | 0.003 | Old |
| 0.848278 | 0.410201 | 0.025 | 0.022 | 1 | 12 | Actl6a | 0.003 | Old |
| 0.848278 | 0.356747 | 0.025 | 0.022 | 1 | 12 | Lztr1 | 0.003 | Old |
| 0.848278 | 0.356617 | 0.025 | 0.022 | 1 | 12 | Rapgef6 | 0.003 | Old |
| 0.848278 | 0.356531 | 0.025 | 0.022 | 1 | 12 | Prr3 | 0.003 | Old |
| 0.848278 | 0.346428 | 0.025 | 0.022 | 1 | 12 | Ctdnep1 | 0.003 | Old |
| 0.848278 | 0.311874 | 0.025 | 0.022 | 1 | 12 | Zfp688 | 0.003 | Old |
| 0.848278 | 0.295653 | 0.025 | 0.022 | 1 | 12 | Fbxw8 | 0.003 | Old |
| 0.848278 | 0.276183 | 0.025 | 0.022 | 1 | 12 | Nup54 | 0.003 | Old |
| 0.848278 | 0.268547 | 0.025 | 0.022 | 1 | 12 | 1110002L0 | 0.003 | Old |
| 0.848278 | 0.218094 | 0.025 | 0.022 | 1 | 12 | Il1b | 0.003 | Old |
| 0.848278 | 0.209776 | 0.025 | 0.022 | 1 | 12 | Slmap | 0.003 | Old |
| 0.848278 | 0.171045 | 0.025 | 0.022 | 1 | 12 | Ikbkb | 0.003 | Old |
| 0.848278 | 0.127268 | 0.025 | 0.022 | 1 | 12 | Nfkbiz | 0.003 | Old |
| 0.850174 | 0.080499 | 0.034 | 0.029 | 1 | 12 | Ppil3 | 0.005 | Old |
| 0.850174 | 0.027621 | 0.034 | 0.029 | 1 | 12 | Rnf215 | 0.005 | Old |
| 0.850174 | 0.001692 | 0.034 | 0.029 | 1 | 12 | Glrx | 0.005 | Old |
| 0.850818 | 0.243485 | 0.025 | 0.022 | 1 | 12 | Ablim2 | 0.003 | Old |
| 0.853361 | 0.664995 | 0.025 | 0.022 | 1 | 12 | Camsap2 | 0.003 | Old |
| 0.853361 | 0.360538 | 0.025 | 0.022 | 1 | 12 | 9930021J0 | 0.003 | Old |
| 0.853361 | 0.3219 | 0.025 | 0.022 | 1 | 12 | Pccb | 0.003 | Old |
| 0.853361 | 0.3134 | 0.025 | 0.022 | 1 | 12 | Dnaja3 | 0.003 | Old |
| 0.853361 | 0.300411 | 0.025 | 0.022 | 1 | 12 | Ndufaf6 | 0.003 | Old |
| 0.853361 | 0.294783 | 0.025 | 0.022 | 1 | 12 | Ppm1b | 0.003 | Old |
| 0.853361 | 0.293367 | 0.025 | 0.022 | 1 | 12 | Colq | 0.003 | Old |

| | | | | | | | | |
|---|---|---|---|---|---|---|---|---|
| 0.853361 | 0.282787 | 0.025 | 0.022 | 1 | 12 | Phospho2 | 0.003 | Old |
| 0.853361 | 0.277156 | 0.025 | 0.022 | 1 | 12 | Ubqln1 | 0.003 | Old |
| 0.853361 | 0.258942 | 0.025 | 0.022 | 1 | 12 | Pqlc1 | 0.003 | Old |
| 0.853361 | 0.254633 | 0.025 | 0.022 | 1 | 12 | Necab2 | 0.003 | Old |
| 0.853361 | 0.253108 | 0.025 | 0.022 | 1 | 12 | Nvl | 0.003 | Old |
| 0.853361 | 0.245721 | 0.025 | 0.022 | 1 | 12 | Hsdl1 | 0.003 | Old |
| 0.853361 | 0.234889 | 0.025 | 0.022 | 1 | 12 | Trpv2 | 0.003 | Old |
| 0.853361 | 0.22878 | 0.025 | 0.022 | 1 | 12 | Cdc40 | 0.003 | Old |
| 0.853361 | 0.211464 | 0.025 | 0.022 | 1 | 12 | Sde2 | 0.003 | Old |
| 0.853361 | 0.17118 | 0.025 | 0.022 | 1 | 12 | Cluh | 0.003 | Old |
| 0.853361 | 0.168975 | 0.025 | 0.022 | 1 | 12 | Med22 | 0.003 | Old |
| 0.853361 | 0.166703 | 0.025 | 0.022 | 1 | 12 | Chid1 | 0.003 | Old |
| 0.853361 | 0.155872 | 0.025 | 0.022 | 1 | 12 | Flii | 0.003 | Old |
| 0.853361 | 0.136639 | 0.025 | 0.022 | 1 | 12 | Cpeb1 | 0.003 | Old |
| 0.853361 | 0.02276 | 0.025 | 0.022 | 1 | 12 | Elovl5 | 0.003 | Old |
| 0.853449 | 0.14672 | 0.432 | 0.431 | 1 | 12 | Vim | 0.001 | Old |
| 0.854535 | 0.169282 | 0.093 | 0.088 | 1 | 12 | Polr2m | 0.005 | Old |
| 0.855905 | 0.261411 | 0.025 | 0.022 | 1 | 12 | Magt1 | 0.003 | Old |
| 0.856773 | 0.093553 | 0.025 | 0.029 | 1 | 12 | Larp4b | -0.004 | Old |
| 0.857319 | 0.130681 | 0.229 | 0.219 | 1 | 12 | Trappc4 | 0.01 | Old |
| 0.857727 | 0.243366 | 0.169 | 0.161 | 1 | 12 | Mycbp2 | 0.008 | Old |
| 0.858451 | 0.318316 | 0.025 | 0.022 | 1 | 12 | Arih1 | 0.003 | Old |
| 0.858451 | 0.293946 | 0.025 | 0.022 | 1 | 12 | Poglut1 | 0.003 | Old |
| 0.858451 | 0.247922 | 0.025 | 0.022 | 1 | 12 | Fam193b | 0.003 | Old |
| 0.858451 | 0.241793 | 0.025 | 0.022 | 1 | 12 | Fam219a | 0.003 | Old |
| 0.858451 | 0.199108 | 0.025 | 0.022 | 1 | 12 | Prcp | 0.003 | Old |
| 0.858451 | 0.1813 | 0.025 | 0.022 | 1 | 12 | Naa25 | 0.003 | Old |
| 0.858451 | 0.173829 | 0.025 | 0.022 | 1 | 12 | Ntrk2 | 0.003 | Old |
| 0.858451 | 0.163722 | 0.025 | 0.022 | 1 | 12 | Zmat3 | 0.003 | Old |
| 0.858451 | 0.139926 | 0.025 | 0.022 | 1 | 12 | Abhd14a | 0.003 | Old |
| 0.858451 | 0.139053 | 0.025 | 0.022 | 1 | 12 | Nat8l | 0.003 | Old |
| 0.858451 | 0.138434 | 0.025 | 0.022 | 1 | 12 | Aldh7a1 | 0.003 | Old |
| 0.858451 | 0.132761 | 0.025 | 0.022 | 1 | 12 | Mab21l1 | 0.003 | Old |
| 0.858451 | 0.098048 | 0.025 | 0.022 | 1 | 12 | Lrrfip2 | 0.003 | Old |
| 0.858451 | 0.077742 | 0.025 | 0.022 | 1 | 12 | Fam213a | 0.003 | Old |
| 0.858451 | 0.064843 | 0.025 | 0.022 | 1 | 12 | Cwc27 | 0.003 | Old |
| 0.859853 | 0.072328 | 0.102 | 0.095 | 1 | 12 | Rbbp7 | 0.007 | Old |
| 0.860955 | 0.022583 | 0.22 | 0.212 | 1 | 12 | Mapk3 | 0.008 | Old |
| 0.861063 | 0.126322 | 0.085 | 0.08 | 1 | 12 | Sf3b1 | 0.005 | Old |
| 0.863403 | 0.254091 | 0.076 | 0.073 | 1 | 12 | Tbx3 | 0.003 | Old |
| 0.863547 | 0.500668 | 0.025 | 0.022 | 1 | 12 | Saysd1 | 0.003 | Old |
| 0.863547 | 0.205813 | 0.025 | 0.022 | 1 | 12 | Pusl1 | 0.003 | Old |
| 0.863547 | 0.160375 | 0.025 | 0.022 | 1 | 12 | Ica1 | 0.003 | Old |
| 0.863547 | 0.15523 | 0.025 | 0.022 | 1 | 12 | Lin7b | 0.003 | Old |
| 0.863547 | 0.153482 | 0.025 | 0.022 | 1 | 12 | Herpud1 | 0.003 | Old |

| | | | | | | | | |
|---|---|---|---|---|---|---|---|---|
| 0.863547 | 0.139306 | 0.025 | 0.022 | 1 | 12 | Mrs2 | 0.003 | Old |
| 0.863547 | 0.110447 | 0.025 | 0.022 | 1 | 12 | Otud6b | 0.003 | Old |
| 0.863547 | 0.101852 | 0.025 | 0.022 | 1 | 12 | Hes1 | 0.003 | Old |
| 0.863547 | 0.079706 | 0.025 | 0.022 | 1 | 12 | Ggact | 0.003 | Old |
| 0.863547 | 0.044613 | 0.025 | 0.022 | 1 | 12 | Eif4b | 0.003 | Old |
| 0.863547 | 0.035575 | 0.025 | 0.022 | 1 | 12 | Bvht | 0.003 | Old |
| 0.863547 | 0.009086 | 0.025 | 0.022 | 1 | 12 | Tmem106c | 0.003 | Old |
| 0.86504 | 0.026894 | 0.102 | 0.095 | 1 | 12 | Tgfb1i1 | 0.007 | Old |
| 0.866228 | 0.0653 | 0.025 | 0.029 | 1 | 12 | Snx16 | -0.004 | Old |
| 0.866228 | 0.03744 | 0.025 | 0.029 | 1 | 12 | Trpc4ap | -0.004 | Old |
| 0.866228 | 0.029685 | 0.025 | 0.029 | 1 | 12 | Surf4 | -0.004 | Old |
| 0.866228 | 0.013952 | 0.025 | 0.029 | 1 | 12 | Frmd4b | -0.004 | Old |
| 0.866485 | 0.111835 | 0.059 | 0.066 | 1 | 12 | Zfr | -0.007 | Old |
| 0.867999 | 0.117352 | 0.085 | 0.095 | 1 | 12 | Wbp2 | -0.01 | Old |
| 0.868085 | 0.081436 | 0.085 | 0.08 | 1 | 12 | Bnip3l | 0.005 | Old |
| 0.868649 | 0.136353 | 0.025 | 0.022 | 1 | 12 | Ubl4a | 0.003 | Old |
| 0.868649 | 0.003065 | 0.025 | 0.022 | 1 | 12 | Rras | 0.003 | Old |
| 0.868935 | 0.092222 | 0.093 | 0.102 | 1 | 12 | Praf2 | -0.009 | Old |
| 0.869288 | 0.335241 | 0.076 | 0.073 | 1 | 12 | Cbx6 | 0.003 | Old |
| 0.870234 | 0.00719 | 0.102 | 0.095 | 1 | 12 | Cadm1 | 0.007 | Old |
| 0.870963 | 0.02449 | 0.025 | 0.029 | 1 | 12 | Abce1 | -0.004 | Old |
| 0.870963 | 0.12831 | 0.025 | 0.029 | 1 | 12 | Nelfb | -0.004 | Old |
| 0.870963 | 0.053199 | 0.025 | 0.029 | 1 | 12 | Phb | -0.004 | Old |
| 0.871159 | 1.18105 | 0.017 | 0.015 | 1 | 12 | Rpp14 | 0.002 | Old |
| 0.871159 | 1.064887 | 0.017 | 0.015 | 1 | 12 | Sumf2 | 0.002 | Old |
| 0.871159 | 1.017751 | 0.017 | 0.015 | 1 | 12 | Nyap2 | 0.002 | Old |
| 0.871159 | 0.948983 | 0.017 | 0.015 | 1 | 12 | Tbc1d19 | 0.002 | Old |
| 0.871159 | 0.885191 | 0.017 | 0.015 | 1 | 12 | Mast1 | 0.002 | Old |
| 0.871159 | 0.876719 | 0.017 | 0.015 | 1 | 12 | Gramd1b | 0.002 | Old |
| 0.871159 | 0.846323 | 0.017 | 0.015 | 1 | 12 | Fam210a | 0.002 | Old |
| 0.871159 | 0.843551 | 0.017 | 0.015 | 1 | 12 | Mfsd1 | 0.002 | Old |
| 0.871159 | 0.834101 | 0.017 | 0.015 | 1 | 12 | Braf | 0.002 | Old |
| 0.871159 | 0.832539 | 0.017 | 0.015 | 1 | 12 | Rela | 0.002 | Old |
| 0.871159 | 0.820601 | 0.017 | 0.015 | 1 | 12 | Elf2 | 0.002 | Old |
| 0.871159 | 0.791239 | 0.017 | 0.015 | 1 | 12 | Gmpr2 | 0.002 | Old |
| 0.871159 | 0.790112 | 0.017 | 0.015 | 1 | 12 | Kat2a | 0.002 | Old |
| 0.871159 | 0.78196 | 0.017 | 0.015 | 1 | 12 | Tada2b | 0.002 | Old |
| 0.871159 | 0.746599 | 0.017 | 0.015 | 1 | 12 | Elk3 | 0.002 | Old |
| 0.871159 | 0.738856 | 0.017 | 0.015 | 1 | 12 | Cacng4 | 0.002 | Old |
| 0.871159 | 0.732995 | 0.017 | 0.015 | 1 | 12 | Ralgapb | 0.002 | Old |
| 0.871159 | 0.725633 | 0.017 | 0.015 | 1 | 12 | Taf1 | 0.002 | Old |
| 0.871159 | 0.664215 | 0.017 | 0.015 | 1 | 12 | Bhlhb9 | 0.002 | Old |
| 0.871159 | 0.652901 | 0.017 | 0.015 | 1 | 12 | Dgkd | 0.002 | Old |
| 0.871159 | 0.617282 | 0.017 | 0.015 | 1 | 12 | Enpp4 | 0.002 | Old |
| 0.871159 | 0.616795 | 0.017 | 0.015 | 1 | 12 | Rnf34 | 0.002 | Old |

| | | | | | | | | |
|---|---|---|---|---|---|---|---|---|
| 0.871159 | 0.606731 | 0.017 | 0.015 | 1 | 12 | Bmpr1b | 0.002 | Old |
| 0.871159 | 0.606054 | 0.017 | 0.015 | 1 | 12 | Tbcd | 0.002 | Old |
| 0.871159 | 0.597723 | 0.017 | 0.015 | 1 | 12 | Eci1 | 0.002 | Old |
| 0.871159 | 0.595298 | 0.017 | 0.015 | 1 | 12 | Tsta3 | 0.002 | Old |
| 0.871159 | 0.591876 | 0.017 | 0.015 | 1 | 12 | Taz | 0.002 | Old |
| 0.871159 | 0.573894 | 0.017 | 0.015 | 1 | 12 | Nrxn3 | 0.002 | Old |
| 0.871159 | 0.559887 | 0.017 | 0.015 | 1 | 12 | Smg6 | 0.002 | Old |
| 0.871159 | 0.556317 | 0.017 | 0.015 | 1 | 12 | Ppp1r10 | 0.002 | Old |
| 0.871159 | 0.544632 | 0.017 | 0.015 | 1 | 12 | Eipr1 | 0.002 | Old |
| 0.871159 | 0.537971 | 0.017 | 0.015 | 1 | 12 | Tmem63b | 0.002 | Old |
| 0.871159 | 0.53336 | 0.017 | 0.015 | 1 | 12 | Rlim | 0.002 | Old |
| 0.871159 | 0.523259 | 0.017 | 0.015 | 1 | 12 | Zfp945 | 0.002 | Old |
| 0.871159 | 0.52171 | 0.017 | 0.015 | 1 | 12 | Tecpr1 | 0.002 | Old |
| 0.871159 | 0.512919 | 0.017 | 0.015 | 1 | 12 | Pik3ip1 | 0.002 | Old |
| 0.871159 | 0.50404 | 0.017 | 0.015 | 1 | 12 | Tmem214 | 0.002 | Old |
| 0.871159 | 0.502351 | 0.017 | 0.015 | 1 | 12 | Alg14 | 0.002 | Old |
| 0.871159 | 0.496932 | 0.017 | 0.015 | 1 | 12 | Adcyap1r1 | 0.002 | Old |
| 0.871159 | 0.495284 | 0.017 | 0.015 | 1 | 12 | Asap2 | 0.002 | Old |
| 0.871159 | 0.49266 | 0.017 | 0.015 | 1 | 12 | Srrd | 0.002 | Old |
| 0.871159 | 0.489047 | 0.017 | 0.015 | 1 | 12 | Zkscan1 | 0.002 | Old |
| 0.871159 | 0.487297 | 0.017 | 0.015 | 1 | 12 | Nsmf | 0.002 | Old |
| 0.871159 | 0.486618 | 0.017 | 0.015 | 1 | 12 | Ide | 0.002 | Old |
| 0.871159 | 0.468274 | 0.017 | 0.015 | 1 | 12 | Ppp2r5e | 0.002 | Old |
| 0.871159 | 0.466295 | 0.017 | 0.015 | 1 | 12 | Ormdl1 | 0.002 | Old |
| 0.871159 | 0.440496 | 0.017 | 0.015 | 1 | 12 | Susd2 | 0.002 | Old |
| 0.871159 | 0.438666 | 0.017 | 0.015 | 1 | 12 | Trmt10a | 0.002 | Old |
| 0.871159 | 0.437078 | 0.017 | 0.015 | 1 | 12 | Emc1 | 0.002 | Old |
| 0.871159 | 0.424457 | 0.017 | 0.015 | 1 | 12 | Rcor3 | 0.002 | Old |
| 0.871159 | 0.421324 | 0.017 | 0.015 | 1 | 12 | Orc5 | 0.002 | Old |
| 0.871159 | 0.417835 | 0.017 | 0.015 | 1 | 12 | Atad3a | 0.002 | Old |
| 0.871159 | 0.376553 | 0.017 | 0.015 | 1 | 12 | Telo2 | 0.002 | Old |
| 0.871159 | 0.353144 | 0.017 | 0.015 | 1 | 12 | Cbr4 | 0.002 | Old |
| 0.871159 | 0.32976 | 0.017 | 0.015 | 1 | 12 | Snhg6 | 0.002 | Old |
| 0.871159 | 0.317078 | 0.017 | 0.015 | 1 | 12 | Msh2 | 0.002 | Old |
| 0.871159 | 0.296864 | 0.017 | 0.015 | 1 | 12 | Zfp61 | 0.002 | Old |
| 0.872045 | 0.119246 | 0.093 | 0.088 | 1 | 12 | Ppp2r1a | 0.005 | Old |
| 0.872633 | 0.111992 | 0.144 | 0.139 | 1 | 12 | H2afj | 0.005 | Old |
| 0.872863 | 0.011029 | 0.059 | 0.066 | 1 | 12 | Enah | -0.007 | Old |
| 0.873332 | 0.040513 | 0.025 | 0.029 | 1 | 12 | Slc17a5 | -0.004 | Old |
| 0.873706 | 0.232145 | 0.076 | 0.073 | 1 | 12 | Cdk16 | 0.003 | Old |
| 0.874274 | 0.402484 | 0.017 | 0.015 | 1 | 12 | 2310035C2 | 0.002 | Old |
| 0.874745 | 0.032687 | 0.093 | 0.088 | 1 | 12 | Eif6 | 0.005 | Old |
| 0.874745 | 0.154104 | 0.093 | 0.088 | 1 | 12 | Cops3 | 0.005 | Old |
| 0.875179 | 0.354433 | 0.076 | 0.073 | 1 | 12 | Tmem205 | 0.003 | Old |
| 0.875703 | 0.259289 | 0.025 | 0.029 | 1 | 12 | Urod | -0.004 | Old |

| | | | | | | | | |
|---|---|---|---|---|---|---|---|---|
| 0.875703 | 0.11433 | 0.025 | 0.029 | 1 | 12 | Yars2 | -0.004 | Old |
| 0.875703 | 0.100055 | 0.025 | 0.029 | 1 | 12 | Sdr39u1 | -0.004 | Old |
| 0.875703 | 0.068996 | 0.025 | 0.029 | 1 | 12 | Tmem192 | -0.004 | Old |
| 0.875703 | 0.012752 | 0.025 | 0.029 | 1 | 12 | Sprn | -0.004 | Old |
| 0.877102 | 0.033007 | 0.102 | 0.109 | 1 | 12 | U2af1 | -0.007 | Old |
| 0.877393 | 1.070272 | 0.017 | 0.015 | 1 | 12 | Mpzl1 | 0.002 | Old |
| 0.877393 | 0.815826 | 0.017 | 0.015 | 1 | 12 | Cald1 | 0.002 | Old |
| 0.877393 | 0.635029 | 0.017 | 0.015 | 1 | 12 | Mospd1 | 0.002 | Old |
| 0.877393 | 0.611441 | 0.017 | 0.015 | 1 | 12 | Ythdf3 | 0.002 | Old |
| 0.877393 | 0.568206 | 0.017 | 0.015 | 1 | 12 | Proser2 | 0.002 | Old |
| 0.877393 | 0.55392 | 0.017 | 0.015 | 1 | 12 | Ctr9 | 0.002 | Old |
| 0.877393 | 0.5487 | 0.017 | 0.015 | 1 | 12 | Slc25a14 | 0.002 | Old |
| 0.877393 | 0.538506 | 0.017 | 0.015 | 1 | 12 | Coq4 | 0.002 | Old |
| 0.877393 | 0.496943 | 0.017 | 0.015 | 1 | 12 | Mri1 | 0.002 | Old |
| 0.877393 | 0.496617 | 0.017 | 0.015 | 1 | 12 | Xaf1 | 0.002 | Old |
| 0.877393 | 0.474829 | 0.017 | 0.015 | 1 | 12 | Cxcl14 | 0.002 | Old |
| 0.877393 | 0.469142 | 0.017 | 0.015 | 1 | 12 | Gpx7 | 0.002 | Old |
| 0.877393 | 0.465371 | 0.017 | 0.015 | 1 | 12 | Aacs | 0.002 | Old |
| 0.877393 | 0.43913 | 0.017 | 0.015 | 1 | 12 | Cep350 | 0.002 | Old |
| 0.877393 | 0.419554 | 0.017 | 0.015 | 1 | 12 | Plgrkt | 0.002 | Old |
| 0.877393 | 0.416829 | 0.017 | 0.015 | 1 | 12 | Cpt1c | 0.002 | Old |
| 0.877393 | 0.415121 | 0.017 | 0.015 | 1 | 12 | Amz2 | 0.002 | Old |
| 0.877393 | 0.413118 | 0.017 | 0.015 | 1 | 12 | Socs5 | 0.002 | Old |
| 0.877393 | 0.410586 | 0.017 | 0.015 | 1 | 12 | Rpusd4 | 0.002 | Old |
| 0.877393 | 0.388059 | 0.017 | 0.015 | 1 | 12 | Ankrd13b | 0.002 | Old |
| 0.877393 | 0.372112 | 0.017 | 0.015 | 1 | 12 | Orai3 | 0.002 | Old |
| 0.877393 | 0.370714 | 0.017 | 0.015 | 1 | 12 | Rpn1 | 0.002 | Old |
| 0.877393 | 0.341698 | 0.017 | 0.015 | 1 | 12 | Csrnp3 | 0.002 | Old |
| 0.877393 | 0.338294 | 0.017 | 0.015 | 1 | 12 | Hadh | 0.002 | Old |
| 0.877393 | 0.337954 | 0.017 | 0.015 | 1 | 12 | Gm14418 | 0.002 | Old |
| 0.877393 | 0.335326 | 0.017 | 0.015 | 1 | 12 | Sema3c | 0.002 | Old |
| 0.877393 | 0.329629 | 0.017 | 0.015 | 1 | 12 | Tmem110 | 0.002 | Old |
| 0.877393 | 0.316263 | 0.017 | 0.015 | 1 | 12 | Eif5a2 | 0.002 | Old |
| 0.877393 | 0.31607 | 0.017 | 0.015 | 1 | 12 | Thap11 | 0.002 | Old |
| 0.877393 | 0.314311 | 0.017 | 0.015 | 1 | 12 | Hexb | 0.002 | Old |
| 0.877393 | 0.294259 | 0.017 | 0.015 | 1 | 12 | Tmem191c | 0.002 | Old |
| 0.877393 | 0.291177 | 0.017 | 0.015 | 1 | 12 | Chpf | 0.002 | Old |
| 0.877393 | 0.287626 | 0.017 | 0.015 | 1 | 12 | Dcps | 0.002 | Old |
| 0.877393 | 0.269133 | 0.017 | 0.015 | 1 | 12 | Stambpl1 | 0.002 | Old |
| 0.877393 | 0.261948 | 0.017 | 0.015 | 1 | 12 | Acsl4 | 0.002 | Old |
| 0.877393 | 0.260213 | 0.017 | 0.015 | 1 | 12 | Fam155a | 0.002 | Old |
| 0.877393 | 0.247224 | 0.017 | 0.015 | 1 | 12 | Gm9493 | 0.002 | Old |
| 0.878128 | 0.245183 | 0.076 | 0.073 | 1 | 12 | Palm | 0.003 | Old |
| 0.880447 | 0.348512 | 0.025 | 0.029 | 1 | 12 | Tex2 | -0.004 | Old |
| 0.880447 | 0.29198 | 0.025 | 0.029 | 1 | 12 | Pim1 | -0.004 | Old |

| | | | | | | | | |
|---|---|---|---|---|---|---|---|---|
| 0.880447 | 0.289847 | 0.025 | 0.029 | 1 | 12 | Rangrf | -0.004 | Old |
| 0.880447 | 0.25364 | 0.025 | 0.029 | 1 | 12 | Zbtb7a | -0.004 | Old |
| 0.880447 | 0.2265 | 0.025 | 0.029 | 1 | 12 | Mtor | -0.004 | Old |
| 0.880447 | 0.190259 | 0.025 | 0.029 | 1 | 12 | Stx2 | -0.004 | Old |
| 0.880447 | 0.089154 | 0.025 | 0.029 | 1 | 12 | Sgf29 | -0.004 | Old |
| 0.880447 | 0.068592 | 0.025 | 0.029 | 1 | 12 | Eif4ebp1 | -0.004 | Old |
| 0.880512 | 0.39854 | 0.017 | 0.015 | 1 | 12 | Alg1 | 0.002 | Old |
| 0.880757 | 0.079313 | 0.237 | 0.234 | 1 | 12 | Eid1 | 0.003 | Old |
| 0.881078 | 0.22301 | 0.076 | 0.073 | 1 | 12 | Ak2 | 0.003 | Old |
| 0.881078 | 0.287904 | 0.076 | 0.073 | 1 | 12 | Mrpl50 | 0.003 | Old |
| 0.882446 | 0.158107 | 0.059 | 0.066 | 1 | 12 | Scg3 | -0.007 | Old |
| 0.882565 | 0.00689 | 0.237 | 0.234 | 1 | 12 | Ttc9b | 0.003 | Old |
| 0.88328 | 0.026369 | 0.331 | 0.314 | 1 | 12 | Capns1 | 0.017 | Old |
| 0.883634 | 0.47089 | 0.017 | 0.015 | 1 | 12 | Ormdl2 | 0.002 | Old |
| 0.883634 | 0.470313 | 0.017 | 0.015 | 1 | 12 | Emd | 0.002 | Old |
| 0.883634 | 0.461327 | 0.017 | 0.015 | 1 | 12 | Nf2 | 0.002 | Old |
| 0.883634 | 0.444844 | 0.017 | 0.015 | 1 | 12 | Hivep1 | 0.002 | Old |
| 0.883634 | 0.402825 | 0.017 | 0.015 | 1 | 12 | Atf3 | 0.002 | Old |
| 0.883634 | 0.397277 | 0.017 | 0.015 | 1 | 12 | Wdr48 | 0.002 | Old |
| 0.883634 | 0.385982 | 0.017 | 0.015 | 1 | 12 | Cluap1 | 0.002 | Old |
| 0.883634 | 0.365304 | 0.017 | 0.015 | 1 | 12 | Snx25 | 0.002 | Old |
| 0.883634 | 0.363262 | 0.017 | 0.015 | 1 | 12 | Tuba8 | 0.002 | Old |
| 0.883634 | 0.351176 | 0.017 | 0.015 | 1 | 12 | Swt1 | 0.002 | Old |
| 0.883634 | 0.347293 | 0.017 | 0.015 | 1 | 12 | Ehmt1 | 0.002 | Old |
| 0.883634 | 0.343167 | 0.017 | 0.015 | 1 | 12 | Ghdc | 0.002 | Old |
| 0.883634 | 0.330719 | 0.017 | 0.015 | 1 | 12 | Coil | 0.002 | Old |
| 0.883634 | 0.31492 | 0.017 | 0.015 | 1 | 12 | Kars | 0.002 | Old |
| 0.883634 | 0.310933 | 0.017 | 0.015 | 1 | 12 | Zfp407 | 0.002 | Old |
| 0.883634 | 0.306497 | 0.017 | 0.015 | 1 | 12 | Sema4f | 0.002 | Old |
| 0.883634 | 0.303572 | 0.017 | 0.015 | 1 | 12 | Tmem161a | 0.002 | Old |
| 0.883634 | 0.2993 | 0.017 | 0.015 | 1 | 12 | Endod1 | 0.002 | Old |
| 0.883634 | 0.290337 | 0.017 | 0.015 | 1 | 12 | Ero1l | 0.002 | Old |
| 0.883634 | 0.287712 | 0.017 | 0.015 | 1 | 12 | Ppm1h | 0.002 | Old |
| 0.883634 | 0.285692 | 0.017 | 0.015 | 1 | 12 | Ist1 | 0.002 | Old |
| 0.883634 | 0.28492 | 0.017 | 0.015 | 1 | 12 | Selenon | 0.002 | Old |
| 0.883634 | 0.28355 | 0.017 | 0.015 | 1 | 12 | Trrap | 0.002 | Old |
| 0.883634 | 0.283417 | 0.017 | 0.015 | 1 | 12 | Tmem165 | 0.002 | Old |
| 0.883634 | 0.280694 | 0.017 | 0.015 | 1 | 12 | Hist1h2bc | 0.002 | Old |
| 0.883634 | 0.27732 | 0.017 | 0.015 | 1 | 12 | Ptpn4 | 0.002 | Old |
| 0.883634 | 0.264557 | 0.017 | 0.015 | 1 | 12 | Akt2 | 0.002 | Old |
| 0.883634 | 0.260239 | 0.017 | 0.015 | 1 | 12 | Riox1 | 0.002 | Old |
| 0.883634 | 0.255684 | 0.017 | 0.015 | 1 | 12 | Fxn | 0.002 | Old |
| 0.883634 | 0.251737 | 0.017 | 0.015 | 1 | 12 | Gm2000 | 0.002 | Old |
| 0.883634 | 0.250908 | 0.017 | 0.015 | 1 | 12 | Wdr43 | 0.002 | Old |
| 0.883634 | 0.249073 | 0.017 | 0.015 | 1 | 12 | Nup93 | 0.002 | Old |

| | | | | | | | | |
|---|---|---|---|---|---|---|---|---|
| 0.883634 | 0.245281 | 0.017 | 0.015 | 1 | 12 | Tbc1d16 | 0.002 | Old |
| 0.883634 | 0.245031 | 0.017 | 0.015 | 1 | 12 | Smim1 | 0.002 | Old |
| 0.883634 | 0.244359 | 0.017 | 0.015 | 1 | 12 | Vps9d1 | 0.002 | Old |
| 0.883634 | 0.243125 | 0.017 | 0.015 | 1 | 12 | F2rl2 | 0.002 | Old |
| 0.883634 | 0.238209 | 0.017 | 0.015 | 1 | 12 | Akr1e1 | 0.002 | Old |
| 0.883634 | 0.23348 | 0.017 | 0.015 | 1 | 12 | Wdr61 | 0.002 | Old |
| 0.883634 | 0.233034 | 0.017 | 0.015 | 1 | 12 | Ndufaf5 | 0.002 | Old |
| 0.883634 | 0.231133 | 0.017 | 0.015 | 1 | 12 | Atpaf2 | 0.002 | Old |
| 0.883634 | 0.226156 | 0.017 | 0.015 | 1 | 12 | Mtmr14 | 0.002 | Old |
| 0.883634 | 0.224611 | 0.017 | 0.015 | 1 | 12 | Pinx1 | 0.002 | Old |
| 0.883634 | 0.222776 | 0.017 | 0.015 | 1 | 12 | Pcgf6 | 0.002 | Old |
| 0.883634 | 0.22164 | 0.017 | 0.015 | 1 | 12 | Nudt11 | 0.002 | Old |
| 0.883634 | 0.219758 | 0.017 | 0.015 | 1 | 12 | Aga | 0.002 | Old |
| 0.883634 | 0.218732 | 0.017 | 0.015 | 1 | 12 | Prtn3 | 0.002 | Old |
| 0.883634 | 0.218606 | 0.017 | 0.015 | 1 | 12 | Tradd | 0.002 | Old |
| 0.883634 | 0.218317 | 0.017 | 0.015 | 1 | 12 | Armcx5 | 0.002 | Old |
| 0.883634 | 0.208442 | 0.017 | 0.015 | 1 | 12 | Ccdc167 | 0.002 | Old |
| 0.883634 | 0.206166 | 0.017 | 0.015 | 1 | 12 | Mboat2 | 0.002 | Old |
| 0.883634 | 0.197553 | 0.017 | 0.015 | 1 | 12 | Atf6b | 0.002 | Old |
| 0.883634 | 0.196288 | 0.017 | 0.015 | 1 | 12 | Rgmb | 0.002 | Old |
| 0.883634 | 0.196186 | 0.017 | 0.015 | 1 | 12 | Gzf1 | 0.002 | Old |
| 0.883634 | 0.192201 | 0.017 | 0.015 | 1 | 12 | Tbc1d15 | 0.002 | Old |
| 0.883634 | 0.189678 | 0.017 | 0.015 | 1 | 12 | Hbs1l | 0.002 | Old |
| 0.883634 | 0.18788 | 0.017 | 0.015 | 1 | 12 | Ralyl | 0.002 | Old |
| 0.883634 | 0.186258 | 0.017 | 0.015 | 1 | 12 | C230004F1 | 0.002 | Old |
| 0.883634 | 0.185431 | 0.017 | 0.015 | 1 | 12 | Ppie | 0.002 | Old |
| 0.883634 | 0.184378 | 0.017 | 0.015 | 1 | 12 | Adam23 | 0.002 | Old |
| 0.883634 | 0.172516 | 0.017 | 0.015 | 1 | 12 | Dok5 | 0.002 | Old |
| 0.883634 | 0.17161 | 0.017 | 0.015 | 1 | 12 | Arhgap33 | 0.002 | Old |
| 0.883634 | 0.166771 | 0.017 | 0.015 | 1 | 12 | St8sia3 | 0.002 | Old |
| 0.883634 | 0.165047 | 0.017 | 0.015 | 1 | 12 | Tatdn1 | 0.002 | Old |
| 0.883634 | 0.164747 | 0.017 | 0.015 | 1 | 12 | Tmem175 | 0.002 | Old |
| 0.883634 | 0.16269 | 0.017 | 0.015 | 1 | 12 | Pnkp | 0.002 | Old |
| 0.883634 | 0.161514 | 0.017 | 0.015 | 1 | 12 | Tshz2 | 0.002 | Old |
| 0.883634 | 0.160752 | 0.017 | 0.015 | 1 | 12 | Asl | 0.002 | Old |
| 0.883634 | 0.145715 | 0.017 | 0.015 | 1 | 12 | Ylpm1 | 0.002 | Old |
| 0.883634 | 0.134852 | 0.017 | 0.015 | 1 | 12 | Sap30 | 0.002 | Old |
| 0.883634 | 0.124534 | 0.017 | 0.015 | 1 | 12 | Eif4enif1 | 0.002 | Old |
| 0.883634 | 0.116343 | 0.017 | 0.015 | 1 | 12 | Rif1 | 0.002 | Old |
| 0.883634 | 0.110159 | 0.017 | 0.015 | 1 | 12 | Bcas3 | 0.002 | Old |
| 0.883634 | 0.085611 | 0.017 | 0.015 | 1 | 12 | Arrb2 | 0.002 | Old |
| 0.883634 | 0.081326 | 0.017 | 0.015 | 1 | 12 | Elovl7 | 0.002 | Old |
| 0.883634 | 0.069224 | 0.017 | 0.015 | 1 | 12 | Mllt3 | 0.002 | Old |
| 0.883634 | 0.062212 | 0.017 | 0.015 | 1 | 12 | Lss | 0.002 | Old |
| 0.883634 | 0.059279 | 0.017 | 0.015 | 1 | 12 | Gsk3a | 0.002 | Old |

| | | | | | | | | |
|---|---|---|---|---|---|---|---|---|
| 0.883634 | 0.055626 | 0.017 | 0.015 | 1 | 12 | Bcl2l1 | 0.002 | Old |
| 0.883634 | 0.039955 | 0.017 | 0.015 | 1 | 12 | Pemt | 0.002 | Old |
| 0.883634 | 0.024402 | 0.017 | 0.015 | 1 | 12 | Nudt1 | 0.002 | Old |
| 0.883634 | 0.002937 | 0.017 | 0.015 | 1 | 12 | Iws1 | 0.002 | Old |
| 0.885195 | 0.318912 | 0.025 | 0.029 | 1 | 12 | Mettl23 | -0.004 | Old |
| 0.885195 | 0.152699 | 0.025 | 0.029 | 1 | 12 | Abcc5 | -0.004 | Old |
| 0.885644 | 0.179147 | 0.059 | 0.066 | 1 | 12 | Stoml2 | -0.007 | Old |
| 0.888265 | 0.072411 | 0.093 | 0.088 | 1 | 12 | Rab11a | 0.005 | Old |
| 0.888963 | 0.31466 | 0.11 | 0.109 | 1 | 12 | Slc10a4 | 0.001 | Old |
| 0.889883 | 0.183693 | 0.017 | 0.015 | 1 | 12 | Armc2 | 0.002 | Old |
| 0.889883 | 0.138964 | 0.017 | 0.015 | 1 | 12 | Mob1b | 0.002 | Old |
| 0.889883 | 0.13431 | 0.017 | 0.015 | 1 | 12 | Pwwp2a | 0.002 | Old |
| 0.889883 | 0.120927 | 0.017 | 0.015 | 1 | 12 | Timm29 | 0.002 | Old |
| 0.889883 | 0.113306 | 0.017 | 0.015 | 1 | 12 | Tmed1 | 0.002 | Old |
| 0.889883 | 0.10978 | 0.017 | 0.015 | 1 | 12 | Sh2b1 | 0.002 | Old |
| 0.889883 | 0.104519 | 0.017 | 0.015 | 1 | 12 | Pcnx3 | 0.002 | Old |
| 0.889883 | 0.092673 | 0.017 | 0.015 | 1 | 12 | Dkk1 | 0.002 | Old |
| 0.889883 | 0.087032 | 0.017 | 0.015 | 1 | 12 | Rida | 0.002 | Old |
| 0.889883 | 0.086629 | 0.017 | 0.015 | 1 | 12 | Vps50 | 0.002 | Old |
| 0.889883 | 0.07843 | 0.017 | 0.015 | 1 | 12 | Dcbld2 | 0.002 | Old |
| 0.889883 | 0.076352 | 0.017 | 0.015 | 1 | 12 | Nr2c2 | 0.002 | Old |
| 0.889883 | 0.075697 | 0.017 | 0.015 | 1 | 12 | Lyst | 0.002 | Old |
| 0.889883 | 0.072756 | 0.017 | 0.015 | 1 | 12 | Coq10b | 0.002 | Old |
| 0.889883 | 0.065667 | 0.017 | 0.015 | 1 | 12 | Prdm4 | 0.002 | Old |
| 0.889883 | 0.064278 | 0.017 | 0.015 | 1 | 12 | Fam58b | 0.002 | Old |
| 0.889883 | 0.062908 | 0.017 | 0.015 | 1 | 12 | Sirt7 | 0.002 | Old |
| 0.889883 | 0.054694 | 0.017 | 0.015 | 1 | 12 | Ldlr | 0.002 | Old |
| 0.889883 | 0.037119 | 0.017 | 0.015 | 1 | 12 | Cks2 | 0.002 | Old |
| 0.889883 | 0.035111 | 0.017 | 0.015 | 1 | 12 | Brd8 | 0.002 | Old |
| 0.889883 | 0.031637 | 0.017 | 0.015 | 1 | 12 | Pdp1 | 0.002 | Old |
| 0.889883 | 0.017108 | 0.017 | 0.015 | 1 | 12 | Gpbp1l1 | 0.002 | Old |
| 0.891063 | 0.145549 | 0.102 | 0.095 | 1 | 12 | Fam92a | 0.007 | Old |
| 0.892034 | 0.138661 | 0.085 | 0.08 | 1 | 12 | Slc25a1 | 0.005 | Old |
| 0.892206 | 0.168445 | 0.102 | 0.109 | 1 | 12 | Mpp6 | -0.007 | Old |
| 0.892894 | 0.247012 | 0.076 | 0.073 | 1 | 12 | Atp6ap1 | 0.003 | Old |
| 0.892894 | 0.169448 | 0.076 | 0.073 | 1 | 12 | Isca2 | 0.003 | Old |
| 0.89301 | 0.078706 | 0.017 | 0.015 | 1 | 12 | Srgap2 | 0.002 | Old |
| 0.894677 | 0.012698 | 0.119 | 0.117 | 1 | 12 | Mrpl11 | 0.002 | Old |
| 0.894728 | 0.163077 | 0.102 | 0.109 | 1 | 12 | Tmem30a | -0.007 | Old |
| 0.895042 | 0.079814 | 0.068 | 0.066 | 1 | 12 | Ggnbp2 | 0.002 | Old |
| 0.896139 | 0.079257 | 0.017 | 0.015 | 1 | 12 | Lgals8 | 0.002 | Old |
| 0.896139 | 0.06917 | 0.017 | 0.015 | 1 | 12 | Arhgef12 | 0.002 | Old |
| 0.896139 | 0.032597 | 0.017 | 0.015 | 1 | 12 | Get4 | 0.002 | Old |
| 0.896139 | 0.025572 | 0.017 | 0.015 | 1 | 12 | Spred3 | 0.002 | Old |
| 0.896139 | 0.012599 | 0.017 | 0.015 | 1 | 12 | Pold4 | 0.002 | Old |

| | | | | | | | | |
|---|---|---|---|---|---|---|---|---|
| 0.897035 | 0.034135 | 0.305 | 0.307 | 1 | 12 | Rps21 | -0.002 | Old |
| 0.898812 | 0.204091 | 0.076 | 0.073 | 1 | 12 | Mrps10 | 0.003 | Old |
| 0.899104 | 0.062408 | 0.093 | 0.088 | 1 | 12 | Fyttd1 | 0.005 | Old |
| 0.900814 | 0.009248 | 0.068 | 0.073 | 1 | 12 | Dph3 | -0.005 | Old |
| 0.900814 | 0.001206 | 0.068 | 0.073 | 1 | 12 | Ahnak2 | -0.005 | Old |
| 0.901275 | 0.307234 | 0.068 | 0.066 | 1 | 12 | Nt5c3b | 0.002 | Old |
| 0.901365 | 0.037476 | 0.136 | 0.131 | 1 | 12 | H1fx | 0.005 | Old |
| 0.901511 | 0.046638 | 0.161 | 0.153 | 1 | 12 | Cope | 0.008 | Old |
| 0.901662 | 0.228725 | 0.059 | 0.066 | 1 | 12 | Acadvl | -0.007 | Old |
| 0.90204 | 0.024628 | 0.212 | 0.219 | 1 | 12 | Spock3 | -0.007 | Old |
| 0.902613 | 0.139973 | 0.11 | 0.109 | 1 | 12 | Gprasp1 | 0.001 | Old |
| 0.907051 | 0.127827 | 0.144 | 0.146 | 1 | 12 | Sf3b6 | -0.002 | Old |
| 0.907089 | 0.284738 | 0.136 | 0.139 | 1 | 12 | Uqcc2 | -0.003 | Old |
| 0.908821 | 0.010768 | 0.034 | 0.036 | 1 | 12 | Pdlim5 | -0.002 | Old |
| 0.909178 | 0.12547 | 0.144 | 0.153 | 1 | 12 | Rab11b | -0.009 | Old |
| 0.910636 | 0.211906 | 0.068 | 0.066 | 1 | 12 | Strbp | 0.002 | Old |
| 0.910663 | 0.097502 | 0.398 | 0.409 | 1 | 12 | Ptms | -0.011 | Old |
| 0.913237 | 0.062209 | 0.161 | 0.153 | 1 | 12 | Vat1 | 0.008 | Old |
| 0.91367 | 0.210999 | 0.102 | 0.109 | 1 | 12 | Ddx39b | -0.007 | Old |
| 0.913701 | 0.073045 | 0.449 | 0.474 | 1 | 12 | Uqcrh | -0.025 | Old |
| 0.913759 | 0.193235 | 0.068 | 0.066 | 1 | 12 | Mir670hg | 0.002 | Old |
| 0.914516 | 0.196116 | 0.246 | 0.255 | 1 | 12 | Nbdy | -0.009 | Old |
| 0.915485 | 0.13516 | 0.271 | 0.263 | 1 | 12 | Eif4g2 | 0.008 | Old |
| 0.916883 | 0.145386 | 0.068 | 0.066 | 1 | 12 | Atg3 | 0.002 | Old |
| 0.917271 | 0.143204 | 0.034 | 0.036 | 1 | 12 | Cpne7 | -0.002 | Old |
| 0.918783 | 0.000229 | 0.11 | 0.109 | 1 | 12 | Mrps34 | 0.001 | Old |
| 0.919987 | 0.122935 | 0.076 | 0.08 | 1 | 12 | Srsf5 | -0.004 | Old |
| 0.920172 | 0.576061 | 0.059 | 0.058 | 1 | 12 | Kdm5b | 0.001 | Old |
| 0.920172 | 0.402992 | 0.059 | 0.058 | 1 | 12 | Nt5dc2 | 0.001 | Old |
| 0.920744 | 0.168685 | 0.195 | 0.197 | 1 | 12 | Lamtor4 | -0.002 | Old |
| 0.921499 | 0.037823 | 0.034 | 0.036 | 1 | 12 | Atp2b4 | -0.002 | Old |
| 0.9221 | 0.032659 | 0.669 | 0.693 | 1 | 12 | Mllt11 | -0.024 | Old |
| 0.922534 | 0.037801 | 0.076 | 0.073 | 1 | 12 | Araf | 0.003 | Old |
| 0.922534 | 0.00224 | 0.076 | 0.073 | 1 | 12 | Ctnnb1 | 0.003 | Old |
| 0.92573 | 0.121249 | 0.034 | 0.036 | 1 | 12 | Apex1 | -0.002 | Old |
| 0.92573 | 0.103265 | 0.034 | 0.036 | 1 | 12 | Vkorc1 | -0.002 | Old |
| 0.92573 | 0.029964 | 0.034 | 0.036 | 1 | 12 | Fndc3a | -0.002 | Old |
| 0.92573 | 0.010446 | 0.034 | 0.036 | 1 | 12 | Iqcb1 | -0.002 | Old |
| 0.928193 | 0.233119 | 0.068 | 0.073 | 1 | 12 | Ccdc50 | -0.005 | Old |
| 0.929963 | 0.130867 | 0.034 | 0.036 | 1 | 12 | Acaca | -0.002 | Old |
| 0.929963 | 0.099578 | 0.034 | 0.036 | 1 | 12 | Itfg1 | -0.002 | Old |
| 0.9323 | 0.14757 | 0.11 | 0.117 | 1 | 12 | Gsn | -0.007 | Old |
| 0.932524 | 0.079699 | 0.068 | 0.066 | 1 | 12 | Serinc3 | 0.002 | Old |
| 0.932524 | 0.002335 | 0.068 | 0.066 | 1 | 12 | Coq2 | 0.002 | Old |
| 0.935655 | 0.049119 | 0.068 | 0.066 | 1 | 12 | Cbx5 | 0.002 | Old |

| | | | | | | | | |
|---|---|---|---|---|---|---|---|---|
| 0.935984 | 0.189411 | 0.11 | 0.117 | 1 | 12 | Chga | -0.007 | Old |
| 0.937557 | 0.003278 | 0.153 | 0.146 | 1 | 12 | G3bp2 | 0.007 | Old |
| 0.93774 | 0.003347 | 0.212 | 0.219 | 1 | 12 | Pafah1b3 | -0.007 | Old |
| 0.938434 | 0.238011 | 0.034 | 0.036 | 1 | 12 | Ric3 | -0.002 | Old |
| 0.938434 | 0.192654 | 0.034 | 0.036 | 1 | 12 | Gtf2i | -0.002 | Old |
| 0.938434 | 0.187607 | 0.034 | 0.036 | 1 | 12 | Idh2 | -0.002 | Old |
| 0.938434 | 0.169337 | 0.034 | 0.036 | 1 | 12 | Psmg3 | -0.002 | Old |
| 0.938434 | 0.167666 | 0.034 | 0.036 | 1 | 12 | Rbck1 | -0.002 | Old |
| 0.938434 | 0.162849 | 0.034 | 0.036 | 1 | 12 | Limk2 | -0.002 | Old |
| 0.938434 | 0.1176 | 0.034 | 0.036 | 1 | 12 | Tm2d3 | -0.002 | Old |
| 0.939257 | 0.143353 | 0.102 | 0.102 | 1 | 12 | Llph | 0 | Old |
| 0.940085 | 0.205115 | 0.059 | 0.058 | 1 | 12 | Tmem59l | 0.001 | Old |
| 0.94031 | 0.082246 | 0.076 | 0.08 | 1 | 12 | Actl6b | -0.004 | Old |
| 0.940654 | 0.091875 | 0.636 | 0.606 | 1 | 12 | Slc25a3 | 0.03 | Old |
| 0.942672 | 0.181139 | 0.034 | 0.036 | 1 | 12 | Syncrip | -0.002 | Old |
| 0.943216 | 0.32808 | 0.076 | 0.08 | 1 | 12 | Fhod3 | -0.004 | Old |
| 0.945056 | 0.178729 | 0.068 | 0.066 | 1 | 12 | Dmac1 | 0.002 | Old |
| 0.946732 | 0.283344 | 0.059 | 0.058 | 1 | 12 | Tmbim6 | 0.001 | Old |
| 0.948191 | 0.020868 | 0.068 | 0.066 | 1 | 12 | Cyth2 | 0.002 | Old |
| 0.948717 | 0.079498 | 0.11 | 0.109 | 1 | 12 | Tmem246 | 0.001 | Old |
| 0.948932 | 0.093972 | 0.136 | 0.139 | 1 | 12 | Cd200 | -0.003 | Old |
| 0.949935 | 0.103777 | 0.178 | 0.175 | 1 | 12 | Ppp1r11 | 0.003 | Old |
| 0.951374 | 0.005889 | 0.737 | 0.745 | 1 | 12 | Pcsk1n | -0.008 | Old |
| 0.951941 | 0.089928 | 0.076 | 0.08 | 1 | 12 | Zc2hc1a | -0.004 | Old |
| 0.952164 | 0.172157 | 0.102 | 0.102 | 1 | 12 | Rit1 | 0 | Old |
| 0.953366 | 0.131211 | 0.246 | 0.248 | 1 | 12 | Psmb2 | -0.002 | Old |
| 0.953382 | 0.323704 | 0.059 | 0.058 | 1 | 12 | Dpm2 | 0.001 | Old |
| 0.955565 | 0.015157 | 0.186 | 0.197 | 1 | 12 | Ranbp1 | -0.011 | Old |
| 0.956032 | 0.015563 | 0.068 | 0.066 | 1 | 12 | Apbb1 | 0.002 | Old |
| 0.956551 | 0.018691 | 0.195 | 0.19 | 1 | 12 | Dnajc15 | 0.005 | Old |
| 0.956709 | 0.177657 | 0.059 | 0.058 | 1 | 12 | Slc36a1 | 0.001 | Old |
| 0.957716 | 0.13716 | 0.169 | 0.175 | 1 | 12 | Sec61g | -0.006 | Old |
| 0.959562 | 0.049724 | 0.085 | 0.088 | 1 | 12 | Fam49b | -0.003 | Old |
| 0.959936 | 0.025932 | 1 | 1 | 1 | 12 | Gnas | 0 | Old |
| 0.960036 | 0.400388 | 0.059 | 0.058 | 1 | 12 | Tbcc | 0.001 | Old |
| 0.960036 | 0.117715 | 0.059 | 0.058 | 1 | 12 | Pdxp | 0.001 | Old |
| 0.960091 | 0.097439 | 0.144 | 0.146 | 1 | 12 | Itgb1 | -0.002 | Old |
| 0.961214 | 0.030666 | 0.11 | 0.109 | 1 | 12 | Comt | 0.001 | Old |
| 0.962333 | 0.017074 | 0.314 | 0.328 | 1 | 12 | Chchd10 | -0.014 | Old |
| 0.962586 | 0.284854 | 0.051 | 0.051 | 1 | 12 | Plec | 0 | Old |
| 0.965274 | 0.305165 | 0.042 | 0.044 | 1 | 12 | Anxa7 | -0.002 | Old |
| 0.965274 | 0.083586 | 0.042 | 0.044 | 1 | 12 | Tia1 | -0.002 | Old |
| 0.965274 | 0.079681 | 0.042 | 0.044 | 1 | 12 | Dlc1 | -0.002 | Old |
| 0.965274 | 0.075939 | 0.042 | 0.044 | 1 | 12 | Sac3d1 | -0.002 | Old |
| 0.965274 | 0.004988 | 0.042 | 0.044 | 1 | 12 | Pnrc1 | -0.002 | Old |

| | | | | | | | | |
|---|---|---|---|---|---|---|---|---|
| 0.965274 | 0.003409 | 0.042 | 0.044 | 1 | 12 | Cdpf1 | -0.002 | Old |
| 0.966147 | 0.595398 | 0.051 | 0.051 | 1 | 12 | Akap11 | 0 | Old |
| 0.966215 | 0.126339 | 0.11 | 0.109 | 1 | 12 | Tsg101 | 0.001 | Old |
| 0.966215 | 0.115295 | 0.11 | 0.109 | 1 | 12 | Flot2 | 0.001 | Old |
| 0.96913 | 0.073174 | 0.042 | 0.044 | 1 | 12 | Rbm18 | -0.002 | Old |
| 0.969708 | 0.301348 | 0.051 | 0.051 | 1 | 12 | Habp4 | 0 | Old |
| 0.971282 | 0.036201 | 0.119 | 0.124 | 1 | 12 | Napa | -0.005 | Old |
| 0.972347 | 0.06445 | 0.542 | 0.584 | 1 | 12 | Jund | -0.042 | Old |
| 0.972987 | 0.150827 | 0.042 | 0.044 | 1 | 12 | Arid4b | -0.002 | Old |
| 0.972987 | 0.046356 | 0.042 | 0.044 | 1 | 12 | Dnm1 | -0.002 | Old |
| 0.973948 | 0.201373 | 0.195 | 0.212 | 1 | 12 | Atox1 | -0.017 | Old |
| 0.975231 | 0.123241 | 0.076 | 0.08 | 1 | 12 | Fibp | -0.004 | Old |
| 0.976845 | 0.039102 | 0.042 | 0.044 | 1 | 12 | Cyb5r4 | -0.002 | Old |
| 0.976845 | 0.164368 | 0.042 | 0.044 | 1 | 12 | Ncdn | -0.002 | Old |
| 0.976845 | 0.075996 | 0.042 | 0.044 | 1 | 12 | Dll3 | -0.002 | Old |
| 0.976845 | 0.030683 | 0.042 | 0.044 | 1 | 12 | Wtap | -0.002 | Old |
| 0.978011 | 0.041493 | 0.102 | 0.102 | 1 | 12 | Hnrnpd | 0 | Old |
| 0.978426 | 0.060572 | 0.136 | 0.139 | 1 | 12 | Kif3a | -0.003 | Old |
| 0.978509 | 0.069466 | 0.28 | 0.277 | 1 | 12 | Usmg5 | 0.003 | Old |
| 0.980243 | 0.136731 | 0.186 | 0.197 | 1 | 12 | Atp1a1 | -0.011 | Old |
| 0.980703 | 0.151033 | 0.042 | 0.044 | 1 | 12 | Efhd2 | -0.002 | Old |
| 0.980703 | 0.137288 | 0.042 | 0.044 | 1 | 12 | Chn2 | -0.002 | Old |
| 0.980979 | 0.038521 | 0.958 | 0.927 | 1 | 12 | mt-Cytb | 0.031 | Old |
| 0.981104 | 0.02715 | 0.5 | 0.496 | 1 | 12 | Txn1 | 0.004 | Old |
| 0.981526 | 0.059887 | 0.449 | 0.474 | 1 | 12 | Tubb4b | -0.025 | Old |
| 0.98167 | 0.023332 | 0.271 | 0.263 | 1 | 12 | Ncald | 0.008 | Old |
| 0.984562 | 0.12355 | 0.042 | 0.044 | 1 | 12 | Rhbdd2 | -0.002 | Old |
| 0.984562 | 0.113762 | 0.042 | 0.044 | 1 | 12 | Psmd10 | -0.002 | Old |
| 0.984761 | 0.043689 | 0.627 | 0.693 | 1 | 12 | Rpl41 | -0.066 | Old |
| 0.986835 | 0.146487 | 0.119 | 0.124 | 1 | 12 | Mff | -0.005 | Old |
| 0.987859 | 0.02881 | 0.119 | 0.117 | 1 | 12 | Hagh | 0.002 | Old |
| 0.988421 | 0.231442 | 0.042 | 0.044 | 1 | 12 | Atg101 | -0.002 | Old |
| 0.988421 | 0.227536 | 0.042 | 0.044 | 1 | 12 | Mtx1 | -0.002 | Old |
| 0.988421 | 0.200382 | 0.042 | 0.044 | 1 | 12 | Insig1 | -0.002 | Old |
| 0.989876 | 0.081177 | 0.288 | 0.299 | 1 | 12 | Ahi1 | -0.011 | Old |
| 0.991089 | 0.243095 | 0.051 | 0.051 | 1 | 12 | Med31 | 0 | Old |
| 0.992281 | 0.273925 | 0.042 | 0.044 | 1 | 12 | Snrpc | -0.002 | Old |
| 0.992498 | 0.287409 | 0.212 | 0.226 | 1 | 12 | Pdcd5 | -0.014 | Old |
| 0.992564 | 0.145821 | 0.153 | 0.168 | 1 | 12 | Parva | -0.015 | Old |
| 0.993295 | 0.012344 | 0.093 | 0.095 | 1 | 12 | Hnrnpa2b1 | -0.002 | Old |
| 0.994176 | 0.10011 | 0.127 | 0.131 | 1 | 12 | Anapc5 | -0.004 | Old |
| 0.994653 | 0.07993 | 0.051 | 0.051 | 1 | 12 | Tmem55a | 0 | Old |
| 0.994653 | 0.050788 | 0.051 | 0.051 | 1 | 12 | Sergef | 0 | Old |
| 0.99614 | 0.255801 | 0.042 | 0.044 | 1 | 12 | Hebp2 | -0.002 | Old |
| 0.99614 | 0.344907 | 0.042 | 0.044 | 1 | 12 | Ate1 | -0.002 | Old |

| | | | | | | | | |
|---|---|---|---|---|---|---|---|---|
| 0.996668 | 0.066374 | 0.059 | 0.058 | 1 | 12 | Flywch2 | 0.001 | Old |
| 0.998218 | 0.085807 | 0.051 | 0.051 | 1 | 12 | Cnot4 | 0 | Old |
| 0.998218 | 0.066449 | 0.051 | 0.051 | 1 | 12 | Tram1l1 | 0 | Old |
| 0.998218 | 0.04671 | 0.051 | 0.051 | 1 | 12 | 1110051M | 0 | Old |
| 0.998605 | 0.194316 | 0.085 | 0.088 | 1 | 12 | Pnkd | -0.003 | Old |
| 0.001297 | 1.522956 | 0.197 | 0.059 | 1 | 12 | Anp32b | 0.138 | Young |
| 0.002252 | 0.890995 | 0.496 | 0.314 | 1 | 12 | Uqcrq | 0.182 | Young |
| 0.002837 | 6.715484 | 0.073 | 0 | 1 | 12 | Rpain | 0.073 | Young |
| 0.002962 | 1.391615 | 0.226 | 0.085 | 1 | 12 | H2afy | 0.141 | Young |
| 0.003792 | 1.465189 | 0.197 | 0.068 | 1 | 12 | Ache | 0.129 | Young |
| 0.004685 | 2.901681 | 0.088 | 0.008 | 1 | 12 | Add2 | 0.08 | Young |
| 0.005088 | 3.054176 | 0.102 | 0.017 | 1 | 12 | Mrpl36 | 0.085 | Young |
| 0.005518 | 2.241966 | 0.117 | 0.025 | 1 | 12 | Pla2g12a | 0.092 | Young |
| 0.005553 | 2.352863 | 0.102 | 0.017 | 1 | 12 | Herc2 | 0.085 | Young |
| 0.006019 | 1.849946 | 0.131 | 0.034 | 1 | 12 | Agtpbp1 | 0.097 | Young |
| 0.006585 | 4.602091 | 0.08 | 0.008 | 1 | 12 | Cers2 | 0.072 | Young |
| 0.006624 | 0.831134 | 0.365 | 0.203 | 1 | 12 | Rala | 0.162 | Young |
| 0.006772 | 3.333365 | 0.08 | 0.008 | 1 | 12 | Lsm5 | 0.072 | Young |
| 0.007653 | 1.423982 | 0.226 | 0.102 | 1 | 12 | Acp1 | 0.124 | Young |
| 0.007723 | 1.785837 | 0.139 | 0.042 | 1 | 12 | Evl | 0.097 | Young |
| 0.008172 | 1.593288 | 0.153 | 0.051 | 1 | 12 | Slc22a17 | 0.102 | Young |
| 0.009659 | 1.280309 | 0.124 | 0.034 | 1 | 12 | Trim44 | 0.09 | Young |
| 0.011096 | 3.238471 | 0.073 | 0.008 | 1 | 12 | Dcaf7 | 0.065 | Young |
| 0.011482 | 1.583514 | 0.146 | 0.051 | 1 | 12 | Dtd1 | 0.095 | Young |
| 0.011565 | 3.073314 | 0.073 | 0.008 | 1 | 12 | Ppp1cc | 0.065 | Young |
| 0.011696 | 1.563228 | 0.146 | 0.051 | 1 | 12 | Mrpl13 | 0.095 | Young |
| 0.011725 | 3.030891 | 0.073 | 0.008 | 1 | 12 | Map4k4 | 0.065 | Young |
| 0.012705 | 1.481716 | 0.19 | 0.085 | 1 | 12 | Emc7 | 0.105 | Young |
| 0.012794 | 0.972983 | 0.292 | 0.169 | 1 | 12 | 2010107E0 | 0.123 | Young |
| 0.013078 | 6.411063 | 0.051 | 0 | 1 | 12 | Gphn | 0.051 | Young |
| 0.013078 | 6.381994 | 0.051 | 0 | 1 | 12 | Dirc2 | 0.051 | Young |
| 0.013078 | 6.214907 | 0.051 | 0 | 1 | 12 | Zmpste24 | 0.051 | Young |
| 0.013078 | 6.196915 | 0.051 | 0 | 1 | 12 | Arhgap26 | 0.051 | Young |
| 0.013078 | 6.183275 | 0.051 | 0 | 1 | 12 | Bend6 | 0.051 | Young |
| 0.013078 | 6.152598 | 0.051 | 0 | 1 | 12 | Polr3g | 0.051 | Young |
| 0.013078 | 6.115805 | 0.051 | 0 | 1 | 12 | Arhgef4 | 0.051 | Young |
| 0.013078 | 6.007203 | 0.051 | 0 | 1 | 12 | Tmem42 | 0.051 | Young |
| 0.013106 | 0.357298 | 0.759 | 0.636 | 1 | 12 | mt-Nd4 | 0.123 | Young |
| 0.014359 | 0.854539 | 0.285 | 0.153 | 1 | 12 | Ctsl | 0.132 | Young |
| 0.014401 | 1.811727 | 0.139 | 0.051 | 1 | 12 | Ybx3 | 0.088 | Young |
| 0.01517 | 1.194374 | 0.182 | 0.076 | 1 | 12 | Cpe | 0.106 | Young |
| 0.016203 | 1.500789 | 0.161 | 0.068 | 1 | 12 | Ube2d3 | 0.093 | Young |
| 0.016584 | 1.129008 | 0.241 | 0.127 | 1 | 12 | Mrpl51 | 0.114 | Young |
| 0.017107 | 1.798475 | 0.139 | 0.051 | 1 | 12 | Vamp3 | 0.088 | Young |
| 0.018171 | 3.489863 | 0.066 | 0.008 | 1 | 12 | Cdk5 | 0.058 | Young |

| | | | | | | | | |
|---|---|---|---|---|---|---|---|---|
| 0.018421 | 3.316244 | 0.066 | 0.008 | 1 | 12 | 2810004N2 | 0.058 | Young |
| 0.018421 | 3.290141 | 0.066 | 0.008 | 1 | 12 | Pttg1ip | 0.058 | Young |
| 0.01945 | 2.872718 | 0.066 | 0.008 | 1 | 12 | Ogfrl1 | 0.058 | Young |
| 0.019715 | 2.859771 | 0.066 | 0.008 | 1 | 12 | Acaa1a | 0.058 | Young |
| 0.019902 | 2.136927 | 0.109 | 0.034 | 1 | 12 | Etf1 | 0.075 | Young |
| 0.02071 | 1.752722 | 0.109 | 0.034 | 1 | 12 | Acat2 | 0.075 | Young |
| 0.020916 | 2.098471 | 0.109 | 0.034 | 1 | 12 | St13 | 0.075 | Young |
| 0.021124 | 1.741483 | 0.109 | 0.034 | 1 | 12 | Huwe1 | 0.075 | Young |
| 0.021502 | 2.079737 | 0.095 | 0.025 | 1 | 12 | Nr2c2ap | 0.07 | Young |
| 0.021758 | 1.389644 | 0.168 | 0.076 | 1 | 12 | Bcap31 | 0.092 | Young |
| 0.021874 | 6.250058 | 0.044 | 0 | 1 | 12 | Gm14296 | 0.044 | Young |
| 0.021874 | 6.189635 | 0.044 | 0 | 1 | 12 | Unc5c | 0.044 | Young |
| 0.021874 | 6.181779 | 0.044 | 0 | 1 | 12 | Adipor2 | 0.044 | Young |
| 0.021874 | 6.118121 | 0.044 | 0 | 1 | 12 | Galk1 | 0.044 | Young |
| 0.021874 | 6.097655 | 0.044 | 0 | 1 | 12 | B630019K0 | 0.044 | Young |
| 0.021874 | 6.09586 | 0.044 | 0 | 1 | 12 | Nufip1 | 0.044 | Young |
| 0.021874 | 6.031031 | 0.044 | 0 | 1 | 12 | Fbxl15 | 0.044 | Young |
| 0.021874 | 5.984665 | 0.044 | 0 | 1 | 12 | Arcn1 | 0.044 | Young |
| 0.021874 | 5.972335 | 0.044 | 0 | 1 | 12 | Fam50a | 0.044 | Young |
| 0.021874 | 5.916586 | 0.044 | 0 | 1 | 12 | Ipo8 | 0.044 | Young |
| 0.021874 | 5.884168 | 0.044 | 0 | 1 | 12 | Tdrd3 | 0.044 | Young |
| 0.021874 | 5.86061 | 0.044 | 0 | 1 | 12 | Pde4dip | 0.044 | Young |
| 0.021874 | 5.855835 | 0.044 | 0 | 1 | 12 | Sdad1 | 0.044 | Young |
| 0.021874 | 5.825985 | 0.044 | 0 | 1 | 12 | Kif3c | 0.044 | Young |
| 0.021874 | 5.719799 | 0.044 | 0 | 1 | 12 | Pigx | 0.044 | Young |
| 0.022304 | 2.20284 | 0.08 | 0.017 | 1 | 12 | Pdcd6 | 0.063 | Young |
| 0.022366 | 1.276159 | 0.161 | 0.068 | 1 | 12 | Rtl8a | 0.093 | Young |
| 0.026531 | 0.536405 | 0.555 | 0.424 | 1 | 12 | Rpl37 | 0.131 | Young |
| 0.028451 | 1.018089 | 0.168 | 0.076 | 1 | 12 | Sri | 0.092 | Young |
| 0.029204 | 0.66639 | 0.401 | 0.28 | 1 | 12 | Ubc | 0.121 | Young |
| 0.029721 | 1.192412 | 0.197 | 0.102 | 1 | 12 | Snu13 | 0.095 | Young |
| 0.029814 | 0.376909 | 0.715 | 0.653 | 1 | 12 | Snhg11 | 0.062 | Young |
| 0.030159 | 3.37625 | 0.058 | 0.008 | 1 | 12 | Prpf4b | 0.05 | Young |
| 0.031396 | 2.792578 | 0.058 | 0.008 | 1 | 12 | Cox18 | 0.05 | Young |
| 0.031396 | 2.639854 | 0.058 | 0.008 | 1 | 12 | Insig2 | 0.05 | Young |
| 0.031427 | 0.482848 | 0.372 | 0.246 | 1 | 12 | Lix1 | 0.126 | Young |
| 0.031818 | 2.906239 | 0.058 | 0.008 | 1 | 12 | Pcnp | 0.05 | Young |
| 0.032245 | 2.726529 | 0.058 | 0.008 | 1 | 12 | Gopc | 0.05 | Young |
| 0.032245 | 2.638455 | 0.058 | 0.008 | 1 | 12 | Wasf1 | 0.05 | Young |
| 0.032763 | 0.704891 | 0.35 | 0.229 | 1 | 12 | Tpi1 | 0.121 | Young |
| 0.033113 | 2.315343 | 0.058 | 0.008 | 1 | 12 | Nudt16l1 | 0.05 | Young |
| 0.033214 | 1.194146 | 0.131 | 0.051 | 1 | 12 | Mydgf | 0.08 | Young |
| 0.033356 | 2.265789 | 0.073 | 0.017 | 1 | 12 | Kansl2 | 0.056 | Young |
| 0.033555 | 2.332814 | 0.058 | 0.008 | 1 | 12 | Azin2 | 0.05 | Young |
| 0.035199 | 0.85893 | 0.248 | 0.144 | 1 | 12 | Ndufs4 | 0.104 | Young |

| | | | | | | | | |
|---|---|---|---|---|---|---|---|---|
| 0.036146 | 2.130714 | 0.073 | 0.017 | 1 | 12 | Yif1a | 0.056 | Young |
| 0.03679 | 6.083477 | 0.036 | 0 | 1 | 12 | Safb | 0.036 | Young |
| 0.03679 | 6.051216 | 0.036 | 0 | 1 | 12 | Man1a2 | 0.036 | Young |
| 0.03679 | 6.012421 | 0.036 | 0 | 1 | 12 | Cnot2 | 0.036 | Young |
| 0.03679 | 6.005648 | 0.036 | 0 | 1 | 12 | Rnf219 | 0.036 | Young |
| 0.03679 | 5.961171 | 0.036 | 0 | 1 | 12 | Tmem120a | 0.036 | Young |
| 0.03679 | 5.93766 | 0.036 | 0 | 1 | 12 | Plpp5 | 0.036 | Young |
| 0.03679 | 5.920819 | 0.036 | 0 | 1 | 12 | Celf5 | 0.036 | Young |
| 0.03679 | 5.91395 | 0.036 | 0 | 1 | 12 | Kcnc4 | 0.036 | Young |
| 0.03679 | 5.902062 | 0.036 | 0 | 1 | 12 | Fam222b | 0.036 | Young |
| 0.03679 | 5.831598 | 0.036 | 0 | 1 | 12 | Pdss1 | 0.036 | Young |
| 0.03679 | 5.822332 | 0.036 | 0 | 1 | 12 | 2010320M | 0.036 | Young |
| 0.03679 | 5.798357 | 0.036 | 0 | 1 | 12 | Fsd1l | 0.036 | Young |
| 0.03679 | 5.798081 | 0.036 | 0 | 1 | 12 | Nfkbia | 0.036 | Young |
| 0.03679 | 5.782682 | 0.036 | 0 | 1 | 12 | Stambp | 0.036 | Young |
| 0.03679 | 5.750332 | 0.036 | 0 | 1 | 12 | Cipc | 0.036 | Young |
| 0.03679 | 5.740183 | 0.036 | 0 | 1 | 12 | Lipa | 0.036 | Young |
| 0.03679 | 5.723573 | 0.036 | 0 | 1 | 12 | Mrps31 | 0.036 | Young |
| 0.03679 | 5.697972 | 0.036 | 0 | 1 | 12 | Dstyk | 0.036 | Young |
| 0.03679 | 5.66276 | 0.036 | 0 | 1 | 12 | Rnd2 | 0.036 | Young |
| 0.03679 | 5.659087 | 0.036 | 0 | 1 | 12 | Cd59a | 0.036 | Young |
| 0.03679 | 5.657258 | 0.036 | 0 | 1 | 12 | 5730480H( | 0.036 | Young |
| 0.03679 | 5.656265 | 0.036 | 0 | 1 | 12 | Rbp1 | 0.036 | Young |
| 0.03679 | 5.638357 | 0.036 | 0 | 1 | 12 | Lrrc3b | 0.036 | Young |
| 0.03679 | 5.627608 | 0.036 | 0 | 1 | 12 | Scn2b | 0.036 | Young |
| 0.03679 | 5.614161 | 0.036 | 0 | 1 | 12 | Creb5 | 0.036 | Young |
| 0.03679 | 5.612575 | 0.036 | 0 | 1 | 12 | Zfhx2 | 0.036 | Young |
| 0.03679 | 5.608941 | 0.036 | 0 | 1 | 12 | Lrrtm1 | 0.036 | Young |
| 0.03679 | 5.560005 | 0.036 | 0 | 1 | 12 | Plekhm3 | 0.036 | Young |
| 0.036811 | 1.713378 | 0.088 | 0.025 | 1 | 12 | Xist | 0.063 | Young |
| 0.036827 | 0.161919 | 0.978 | 0.975 | 1 | 12 | Ubb | 0.003 | Young |
| 0.037514 | 1.322401 | 0.146 | 0.068 | 1 | 12 | Gm16286 | 0.078 | Young |
| 0.037567 | 0.939567 | 0.175 | 0.085 | 1 | 12 | Psenen | 0.09 | Young |
| 0.038639 | 0.902674 | 0.175 | 0.085 | 1 | 12 | Rex1bd | 0.09 | Young |
| 0.038866 | 1.212941 | 0.139 | 0.059 | 1 | 12 | Prrc2b | 0.08 | Young |
| 0.039481 | 0.552524 | 0.423 | 0.28 | 1 | 12 | Mif | 0.143 | Young |
| 0.039904 | 0.484225 | 0.599 | 0.492 | 1 | 12 | Rps16 | 0.107 | Young |
| 0.041113 | 1.448861 | 0.124 | 0.051 | 1 | 12 | Tln1 | 0.073 | Young |
| 0.041146 | 1.215774 | 0.102 | 0.034 | 1 | 12 | Ahnak | 0.068 | Young |
| 0.04246 | 0.878505 | 0.255 | 0.153 | 1 | 12 | Scg5 | 0.102 | Young |
| 0.042823 | 1.599991 | 0.109 | 0.042 | 1 | 12 | Maoa | 0.067 | Young |
| 0.045947 | 0.985892 | 0.139 | 0.059 | 1 | 12 | Psmd2 | 0.08 | Young |
| 0.046479 | 1.045797 | 0.139 | 0.059 | 1 | 12 | Nt5c | 0.08 | Young |
| 0.047353 | 0.353946 | 0.788 | 0.678 | 1 | 12 | Rpl11 | 0.11 | Young |
| 0.047364 | 0.993838 | 0.161 | 0.076 | 1 | 12 | Chchd7 | 0.085 | Young |

| | | | | | | | | |
|---|---|---|---|---|---|---|---|---|
| 0.047385 | 2.064489 | 0.095 | 0.034 | 1 | 12 | Srsf11 | 0.061 | Young |
| 0.047558 | 0.949435 | 0.146 | 0.068 | 1 | 12 | Pcbp4 | 0.078 | Young |
| 0.047826 | 1.754731 | 0.095 | 0.034 | 1 | 12 | Rab1a | 0.061 | Young |
| 0.048496 | 0.692572 | 0.27 | 0.169 | 1 | 12 | Ubl5 | 0.101 | Young |
| 0.04938 | 0.310219 | 0.511 | 0.415 | 1 | 12 | Atp5d | 0.096 | Young |
| 0.04943 | 2.097336 | 0.08 | 0.025 | 1 | 12 | Ttc1 | 0.055 | Young |
| 0.049628 | 1.576008 | 0.095 | 0.034 | 1 | 12 | Cdc37 | 0.061 | Young |
| 0.05006 | 3.16287 | 0.051 | 0.008 | 1 | 12 | Ddx54 | 0.043 | Young |
| 0.050723 | 2.65854 | 0.051 | 0.008 | 1 | 12 | Pex7 | 0.043 | Young |
| 0.050947 | 1.879771 | 0.08 | 0.025 | 1 | 12 | Setd3 | 0.055 | Young |
| 0.051017 | 1.498935 | 0.095 | 0.034 | 1 | 12 | Prdx3 | 0.061 | Young |
| 0.051396 | 0.909114 | 0.124 | 0.051 | 1 | 12 | Gnpda2 | 0.073 | Young |
| 0.051592 | 0.246249 | 0.73 | 0.602 | 1 | 12 | Map1lc3a | 0.128 | Young |
| 0.05196 | 1.449629 | 0.095 | 0.034 | 1 | 12 | Zrsr2 | 0.061 | Young |
| 0.052072 | 2.771112 | 0.051 | 0.008 | 1 | 12 | Lonrf2 | 0.043 | Young |
| 0.052208 | 0.990544 | 0.109 | 0.042 | 1 | 12 | Aldoc | 0.067 | Young |
| 0.052624 | 0.659222 | 0.277 | 0.169 | 1 | 12 | Ndufs6 | 0.108 | Young |
| 0.053127 | 0.60265 | 0.423 | 0.322 | 1 | 12 | Rplp2 | 0.101 | Young |
| 0.053291 | 2.166319 | 0.066 | 0.017 | 1 | 12 | Synpr | 0.049 | Young |
| 0.053451 | 2.363738 | 0.051 | 0.008 | 1 | 12 | Meis3 | 0.043 | Young |
| 0.053451 | 2.380421 | 0.051 | 0.008 | 1 | 12 | Yars | 0.043 | Young |
| 0.053451 | 2.335562 | 0.051 | 0.008 | 1 | 12 | Papola | 0.043 | Young |
| 0.053563 | 1.935346 | 0.08 | 0.025 | 1 | 12 | Tmem9 | 0.055 | Young |
| 0.054152 | 2.211049 | 0.051 | 0.008 | 1 | 12 | Sbf1 | 0.043 | Young |
| 0.054152 | 2.164334 | 0.051 | 0.008 | 1 | 12 | Rnf13 | 0.043 | Young |
| 0.05474 | 0.437977 | 0.701 | 0.627 | 1 | 12 | Rpl9 | 0.074 | Young |
| 0.05486 | 2.137874 | 0.051 | 0.008 | 1 | 12 | Dnal4 | 0.043 | Young |
| 0.055087 | 0.947022 | 0.168 | 0.085 | 1 | 12 | Hmgb3 | 0.083 | Young |
| 0.055106 | 2.218352 | 0.066 | 0.017 | 1 | 12 | Blmh | 0.049 | Young |
| 0.05629 | 1.788783 | 0.08 | 0.025 | 1 | 12 | Nkiras1 | 0.055 | Young |
| 0.056344 | 2.013165 | 0.066 | 0.017 | 1 | 12 | Taf11 | 0.049 | Young |
| 0.05636 | 0.472327 | 0.263 | 0.153 | 1 | 12 | Rab6a | 0.11 | Young |
| 0.05639 | 0.468184 | 0.321 | 0.203 | 1 | 12 | Rab2a | 0.118 | Young |
| 0.056849 | 1.714626 | 0.08 | 0.025 | 1 | 12 | Gtf2e2 | 0.055 | Young |
| 0.056972 | 2.155217 | 0.066 | 0.017 | 1 | 12 | Ppme1 | 0.049 | Young |
| 0.057312 | 1.083428 | 0.131 | 0.059 | 1 | 12 | Ccl27a | 0.072 | Young |
| 0.057605 | 1.687177 | 0.066 | 0.017 | 1 | 12 | Keap1 | 0.049 | Young |
| 0.05889 | 1.80821 | 0.066 | 0.017 | 1 | 12 | Gmps | 0.049 | Young |
| 0.05889 | 1.781006 | 0.066 | 0.017 | 1 | 12 | Ppp3r1 | 0.049 | Young |
| 0.05889 | 1.764461 | 0.066 | 0.017 | 1 | 12 | BC051226 | 0.049 | Young |
| 0.059131 | 1.498031 | 0.08 | 0.025 | 1 | 12 | Ank3 | 0.055 | Young |
| 0.059541 | 1.697297 | 0.066 | 0.017 | 1 | 12 | Rae1 | 0.049 | Young |
| 0.061281 | 1.006249 | 0.161 | 0.085 | 1 | 12 | Vgf | 0.076 | Young |
| 0.061774 | 1.150196 | 0.197 | 0.119 | 1 | 12 | Tspo | 0.078 | Young |
| 0.062106 | 1.045047 | 0.161 | 0.085 | 1 | 12 | Dnajc19 | 0.076 | Young |

| | | | | | | | | |
|---|---|---|---|---|---|---|---|---|
| 0.062107 | 1.203305 | 0.117 | 0.051 | 1 | 12 | Sgta | 0.066 | Young |
| 0.062438 | 5.838339 | 0.029 | 0 | 1 | 12 | Dbh | 0.029 | Young |
| 0.062438 | 5.667731 | 0.029 | 0 | 1 | 12 | Brsk1 | 0.029 | Young |
| 0.062438 | 5.660478 | 0.029 | 0 | 1 | 12 | Zmynd19 | 0.029 | Young |
| 0.062438 | 5.645156 | 0.029 | 0 | 1 | 12 | Ccl25 | 0.029 | Young |
| 0.062438 | 5.629525 | 0.029 | 0 | 1 | 12 | Fam208a | 0.029 | Young |
| 0.062438 | 5.600147 | 0.029 | 0 | 1 | 12 | Unc13a | 0.029 | Young |
| 0.062438 | 5.592655 | 0.029 | 0 | 1 | 12 | Slc25a10 | 0.029 | Young |
| 0.062438 | 5.587642 | 0.029 | 0 | 1 | 12 | Ppp1r12b | 0.029 | Young |
| 0.062438 | 5.565015 | 0.029 | 0 | 1 | 12 | Ptbp2 | 0.029 | Young |
| 0.062438 | 5.554269 | 0.029 | 0 | 1 | 12 | Crebl2 | 0.029 | Young |
| 0.062438 | 5.543098 | 0.029 | 0 | 1 | 12 | Mrpl47 | 0.029 | Young |
| 0.062438 | 5.533821 | 0.029 | 0 | 1 | 12 | Ptges2 | 0.029 | Young |
| 0.062438 | 5.527625 | 0.029 | 0 | 1 | 12 | Slc4a8 | 0.029 | Young |
| 0.062438 | 5.521121 | 0.029 | 0 | 1 | 12 | Mthfsl | 0.029 | Young |
| 0.062438 | 5.517367 | 0.029 | 0 | 1 | 12 | Srsf9 | 0.029 | Young |
| 0.062438 | 5.517106 | 0.029 | 0 | 1 | 12 | Cep68 | 0.029 | Young |
| 0.062438 | 5.508134 | 0.029 | 0 | 1 | 12 | Zzef1 | 0.029 | Young |
| 0.062438 | 5.504891 | 0.029 | 0 | 1 | 12 | Flt1 | 0.029 | Young |
| 0.062438 | 5.501766 | 0.029 | 0 | 1 | 12 | Kbtbd4 | 0.029 | Young |
| 0.062438 | 5.490915 | 0.029 | 0 | 1 | 12 | Dnajc18 | 0.029 | Young |
| 0.062438 | 5.490776 | 0.029 | 0 | 1 | 12 | Exosc3 | 0.029 | Young |
| 0.062438 | 5.471175 | 0.029 | 0 | 1 | 12 | Gpr19 | 0.029 | Young |
| 0.062438 | 5.469288 | 0.029 | 0 | 1 | 12 | Trafd1 | 0.029 | Young |
| 0.062438 | 5.456483 | 0.029 | 0 | 1 | 12 | Phactr2 | 0.029 | Young |
| 0.062438 | 5.454016 | 0.029 | 0 | 1 | 12 | Mynn | 0.029 | Young |
| 0.062438 | 5.44529 | 0.029 | 0 | 1 | 12 | Ints12 | 0.029 | Young |
| 0.062438 | 5.441265 | 0.029 | 0 | 1 | 12 | Nudt16 | 0.029 | Young |
| 0.062438 | 5.422591 | 0.029 | 0 | 1 | 12 | Lnx1 | 0.029 | Young |
| 0.062438 | 5.41566 | 0.029 | 0 | 1 | 12 | Chchd5 | 0.029 | Young |
| 0.062438 | 5.402588 | 0.029 | 0 | 1 | 12 | Ppfia3 | 0.029 | Young |
| 0.062438 | 5.400253 | 0.029 | 0 | 1 | 12 | 1810041L1 | 0.029 | Young |
| 0.062438 | 5.390072 | 0.029 | 0 | 1 | 12 | Sptbn2 | 0.029 | Young |
| 0.062438 | 5.379832 | 0.029 | 0 | 1 | 12 | Pknox1 | 0.029 | Young |
| 0.062438 | 5.376649 | 0.029 | 0 | 1 | 12 | Mpst | 0.029 | Young |
| 0.062438 | 5.376502 | 0.029 | 0 | 1 | 12 | Pisd | 0.029 | Young |
| 0.062438 | 5.375731 | 0.029 | 0 | 1 | 12 | Tmem50b | 0.029 | Young |
| 0.062438 | 5.37462 | 0.029 | 0 | 1 | 12 | Ncaph2 | 0.029 | Young |
| 0.062438 | 5.351594 | 0.029 | 0 | 1 | 12 | Cep290 | 0.029 | Young |
| 0.062438 | 5.346468 | 0.029 | 0 | 1 | 12 | Ccdc86 | 0.029 | Young |
| 0.062438 | 5.336923 | 0.029 | 0 | 1 | 12 | Tmem135 | 0.029 | Young |
| 0.062438 | 5.334601 | 0.029 | 0 | 1 | 12 | Tcp11l1 | 0.029 | Young |
| 0.062438 | 5.330281 | 0.029 | 0 | 1 | 12 | Trnt1 | 0.029 | Young |
| 0.062438 | 5.316947 | 0.029 | 0 | 1 | 12 | Btbd6 | 0.029 | Young |
| 0.062438 | 5.310628 | 0.029 | 0 | 1 | 12 | Stk38 | 0.029 | Young |

| | | | | | | | | |
|---|---|---|---|---|---|---|---|---|
| 0.062438 | 5.306669 | 0.029 | 0 | 1 | 12 | Hnrnpul1 | 0.029 | Young |
| 0.062438 | 5.306311 | 0.029 | 0 | 1 | 12 | Setd2 | 0.029 | Young |
| 0.062438 | 5.295447 | 0.029 | 0 | 1 | 12 | Rpp38 | 0.029 | Young |
| 0.062438 | 5.290574 | 0.029 | 0 | 1 | 12 | Akap6 | 0.029 | Young |
| 0.062438 | 5.269519 | 0.029 | 0 | 1 | 12 | Rasa2 | 0.029 | Young |
| 0.062438 | 5.227273 | 0.029 | 0 | 1 | 12 | Mark4 | 0.029 | Young |
| 0.062438 | 5.223621 | 0.029 | 0 | 1 | 12 | Ackr1 | 0.029 | Young |
| 0.062438 | 5.206161 | 0.029 | 0 | 1 | 12 | Tnfrsf21 | 0.029 | Young |
| 0.062438 | 5.203679 | 0.029 | 0 | 1 | 12 | Chrnb2 | 0.029 | Young |
| 0.062438 | 5.148698 | 0.029 | 0 | 1 | 12 | 9330182L0 | 0.029 | Young |
| 0.062438 | 5.126422 | 0.029 | 0 | 1 | 12 | Fam193a | 0.029 | Young |
| 0.062438 | 5.123304 | 0.029 | 0 | 1 | 12 | Zfp655 | 0.029 | Young |
| 0.062438 | 5.114739 | 0.029 | 0 | 1 | 12 | Zfp652os | 0.029 | Young |
| 0.062438 | 5.04159 | 0.029 | 0 | 1 | 12 | Zfp148 | 0.029 | Young |
| 0.062697 | 1.05691 | 0.08 | 0.025 | 1 | 12 | Yeats4 | 0.055 | Young |
| 0.062698 | 0.984255 | 0.131 | 0.059 | 1 | 12 | Got2 | 0.072 | Young |
| 0.063924 | 1.147138 | 0.08 | 0.025 | 1 | 12 | Rgs9 | 0.055 | Young |
| 0.063926 | 0.445439 | 0.423 | 0.322 | 1 | 12 | Rbm3 | 0.101 | Young |
| 0.065096 | 0.957933 | 0.175 | 0.093 | 1 | 12 | Sms | 0.082 | Young |
| 0.067085 | 0.742432 | 0.219 | 0.127 | 1 | 12 | Ube2e1 | 0.092 | Young |
| 0.06826 | 1.313412 | 0.102 | 0.042 | 1 | 12 | Trappc1 | 0.06 | Young |
| 0.069942 | 0.997496 | 0.161 | 0.085 | 1 | 12 | Dpy30 | 0.076 | Young |
| 0.071459 | 0.363024 | 0.401 | 0.271 | 1 | 12 | Ndufa5 | 0.13 | Young |
| 0.072003 | 0.972217 | 0.117 | 0.051 | 1 | 12 | Spr | 0.066 | Young |
| 0.072236 | 0.469915 | 0.365 | 0.263 | 1 | 12 | Ndufb7 | 0.102 | Young |
| 0.073679 | 0.961378 | 0.117 | 0.051 | 1 | 12 | 2900011O( | 0.066 | Young |
| 0.07404 | 0.639293 | 0.277 | 0.186 | 1 | 12 | Npdc1 | 0.091 | Young |
| 0.074244 | 0.907332 | 0.117 | 0.051 | 1 | 12 | Actr10 | 0.066 | Young |
| 0.07813 | 0.65361 | 0.255 | 0.161 | 1 | 12 | Cops6 | 0.094 | Young |
| 0.079132 | 1.884086 | 0.073 | 0.025 | 1 | 12 | Rab1b | 0.048 | Young |
| 0.08039 | 1.28416 | 0.109 | 0.051 | 1 | 12 | Borcs7 | 0.058 | Young |
| 0.082233 | 1.676139 | 0.073 | 0.025 | 1 | 12 | Eif2b3 | 0.048 | Young |
| 0.08316 | 4.600799 | 0.044 | 0.008 | 1 | 12 | Zfyve19 | 0.036 | Young |
| 0.08316 | 4.423676 | 0.044 | 0.008 | 1 | 12 | Larp1b | 0.036 | Young |
| 0.08316 | 3.057706 | 0.044 | 0.008 | 1 | 12 | B4galt6 | 0.036 | Young |
| 0.08316 | 2.763195 | 0.044 | 0.008 | 1 | 12 | Rsbn1 | 0.036 | Young |
| 0.083891 | 0.642397 | 0.292 | 0.195 | 1 | 12 | Rtn3 | 0.097 | Young |
| 0.084235 | 2.766824 | 0.044 | 0.008 | 1 | 12 | Sec31a | 0.036 | Young |
| 0.084235 | 2.602488 | 0.044 | 0.008 | 1 | 12 | Siah1a | 0.036 | Young |
| 0.084235 | 2.588517 | 0.044 | 0.008 | 1 | 12 | Rbbp6 | 0.036 | Young |
| 0.084235 | 2.549732 | 0.044 | 0.008 | 1 | 12 | Tmf1 | 0.036 | Young |
| 0.085322 | 2.631858 | 0.044 | 0.008 | 1 | 12 | Arhgap28 | 0.036 | Young |
| 0.085322 | 2.542432 | 0.044 | 0.008 | 1 | 12 | Dnajb9 | 0.036 | Young |
| 0.085322 | 2.458988 | 0.044 | 0.008 | 1 | 12 | Thrap3 | 0.036 | Young |
| 0.085322 | 2.435311 | 0.044 | 0.008 | 1 | 12 | Xpnpep1 | 0.036 | Young |

| | | | | | | | | |
|---|---|---|---|---|---|---|---|---|
| 0.085322 | 2.310268 | 0.044 | 0.008 | 1 | 12 | Pgap2 | 0.036 | Young |
| 0.085459 | 0.991209 | 0.109 | 0.051 | 1 | 12 | Cers4 | 0.058 | Young |
| 0.08568 | 0.810941 | 0.212 | 0.127 | 1 | 12 | Coq7 | 0.085 | Young |
| 0.086247 | 1.586086 | 0.073 | 0.025 | 1 | 12 | BC003331 | 0.048 | Young |
| 0.086247 | 1.460023 | 0.073 | 0.025 | 1 | 12 | Fam204a | 0.048 | Young |
| 0.086532 | 1.874682 | 0.058 | 0.017 | 1 | 12 | Clvs1 | 0.041 | Young |
| 0.087068 | 1.504556 | 0.073 | 0.025 | 1 | 12 | Peg3 | 0.048 | Young |
| 0.087068 | 1.446003 | 0.073 | 0.025 | 1 | 12 | Naa50 | 0.048 | Young |
| 0.08715 | 0.4556 | 0.431 | 0.322 | 1 | 12 | Gnb2 | 0.109 | Young |
| 0.087169 | 1.12069 | 0.131 | 0.068 | 1 | 12 | Pdia6 | 0.063 | Young |
| 0.087466 | 1.890699 | 0.058 | 0.017 | 1 | 12 | Mob2 | 0.041 | Young |
| 0.08753 | 2.360289 | 0.044 | 0.008 | 1 | 12 | Cryab | 0.036 | Young |
| 0.08753 | 2.285559 | 0.044 | 0.008 | 1 | 12 | Shoc2 | 0.036 | Young |
| 0.08753 | 2.131869 | 0.044 | 0.008 | 1 | 12 | 3632451O( | 0.036 | Young |
| 0.08753 | 2.027885 | 0.044 | 0.008 | 1 | 12 | Mccc2 | 0.036 | Young |
| 0.08753 | 2.024631 | 0.044 | 0.008 | 1 | 12 | Tbc1d9b | 0.036 | Young |
| 0.087896 | 1.465038 | 0.073 | 0.025 | 1 | 12 | Uba5 | 0.048 | Young |
| 0.088408 | 1.940733 | 0.058 | 0.017 | 1 | 12 | Utp11 | 0.041 | Young |
| 0.088651 | 2.231696 | 0.044 | 0.008 | 1 | 12 | Gmpr | 0.036 | Young |
| 0.088651 | 1.997023 | 0.044 | 0.008 | 1 | 12 | Pdgfa | 0.036 | Young |
| 0.088651 | 1.977884 | 0.044 | 0.008 | 1 | 12 | Zfp664 | 0.036 | Young |
| 0.088651 | 1.90273 | 0.044 | 0.008 | 1 | 12 | Cd83 | 0.036 | Young |
| 0.088651 | 1.883162 | 0.044 | 0.008 | 1 | 12 | Wdr37 | 0.036 | Young |
| 0.08873 | 1.554576 | 0.073 | 0.025 | 1 | 12 | Adgre1 | 0.048 | Young |
| 0.08873 | 1.390553 | 0.073 | 0.025 | 1 | 12 | Gng12 | 0.048 | Young |
| 0.08957 | 1.304942 | 0.073 | 0.025 | 1 | 12 | Nudcd2 | 0.048 | Young |
| 0.089782 | 1.992938 | 0.044 | 0.008 | 1 | 12 | Snrnp48 | 0.036 | Young |
| 0.089783 | 2.030896 | 0.044 | 0.008 | 1 | 12 | Farsa | 0.036 | Young |
| 0.089783 | 1.782616 | 0.044 | 0.008 | 1 | 12 | Emp3 | 0.036 | Young |
| 0.090317 | 1.684143 | 0.058 | 0.017 | 1 | 12 | Riok3 | 0.041 | Young |
| 0.091271 | 1.068769 | 0.073 | 0.025 | 1 | 12 | Mturn | 0.048 | Young |
| 0.091284 | 1.638593 | 0.058 | 0.017 | 1 | 12 | Bfar | 0.041 | Young |
| 0.091772 | 0.796867 | 0.175 | 0.102 | 1 | 12 | Sugt1 | 0.073 | Young |
| 0.092534 | 0.68461 | 0.285 | 0.195 | 1 | 12 | Arpc3 | 0.09 | Young |
| 0.092997 | 1.26266 | 0.073 | 0.025 | 1 | 12 | Actr1b | 0.048 | Young |
| 0.094164 | 1.032127 | 0.139 | 0.076 | 1 | 12 | Pfkp | 0.063 | Young |
| 0.094234 | 1.404766 | 0.058 | 0.017 | 1 | 12 | Triap1 | 0.041 | Young |
| 0.095234 | 0.855029 | 0.058 | 0.017 | 1 | 12 | Add1 | 0.041 | Young |
| 0.095378 | 0.293456 | 0.445 | 0.314 | 1 | 12 | Syt1 | 0.131 | Young |
| 0.096242 | 1.354777 | 0.058 | 0.017 | 1 | 12 | Nrn1 | 0.041 | Young |
| 0.096242 | 1.096124 | 0.058 | 0.017 | 1 | 12 | Ebp | 0.041 | Young |
| 0.097853 | 0.471116 | 0.307 | 0.203 | 1 | 12 | Dad1 | 0.104 | Young |
| 0.098286 | 1.369115 | 0.058 | 0.017 | 1 | 12 | Fbxo28 | 0.041 | Young |
| 0.098758 | 1.237264 | 0.095 | 0.042 | 1 | 12 | Gucy1b1 | 0.053 | Young |
| 0.100037 | 0.699835 | 0.219 | 0.136 | 1 | 12 | Polr1d | 0.083 | Young |

| | | | | | | | | |
|---|---|---|---|---|---|---|---|---|
| 0.10113 | 1.233779 | 0.095 | 0.042 | 1 | 12 | Pnck | 0.053 | Young |
| 0.102416 | 0.415594 | 0.307 | 0.212 | 1 | 12 | Mrps33 | 0.095 | Young |
| 0.102854 | 1.024409 | 0.117 | 0.059 | 1 | 12 | Yipf4 | 0.058 | Young |
| 0.103282 | 0.823849 | 0.161 | 0.093 | 1 | 12 | Ssr2 | 0.068 | Young |
| 0.105041 | 1.219892 | 0.117 | 0.059 | 1 | 12 | Iscu | 0.058 | Young |
| 0.105057 | 0.216642 | 0.869 | 0.847 | 1 | 12 | Map1b | 0.022 | Young |
| 0.105184 | 1.080993 | 0.095 | 0.042 | 1 | 12 | Hdgfl3 | 0.053 | Young |
| 0.107564 | 5.364604 | 0.022 | 0 | 1 | 12 | Pigw | 0.022 | Young |
| 0.107564 | 5.341447 | 0.022 | 0 | 1 | 12 | Ppcdc | 0.022 | Young |
| 0.107564 | 5.336288 | 0.022 | 0 | 1 | 12 | Kmt5b | 0.022 | Young |
| 0.107564 | 5.314074 | 0.022 | 0 | 1 | 12 | Srsf1 | 0.022 | Young |
| 0.107564 | 5.313845 | 0.022 | 0 | 1 | 12 | Rell2 | 0.022 | Young |
| 0.107564 | 5.307731 | 0.022 | 0 | 1 | 12 | Atp6v0a2 | 0.022 | Young |
| 0.107564 | 5.297156 | 0.022 | 0 | 1 | 12 | Fam126b | 0.022 | Young |
| 0.107564 | 5.291937 | 0.022 | 0 | 1 | 12 | Trmt10c | 0.022 | Young |
| 0.107564 | 5.290256 | 0.022 | 0 | 1 | 12 | Fastkd2 | 0.022 | Young |
| 0.107564 | 5.283015 | 0.022 | 0 | 1 | 12 | Ralgapa2 | 0.022 | Young |
| 0.107564 | 5.270994 | 0.022 | 0 | 1 | 12 | Lipt2 | 0.022 | Young |
| 0.107564 | 5.264123 | 0.022 | 0 | 1 | 12 | Adgrb2 | 0.022 | Young |
| 0.107564 | 5.255959 | 0.022 | 0 | 1 | 12 | Cst12 | 0.022 | Young |
| 0.107564 | 5.243882 | 0.022 | 0 | 1 | 12 | Scamp5 | 0.022 | Young |
| 0.107564 | 5.238906 | 0.022 | 0 | 1 | 12 | Lrch3 | 0.022 | Young |
| 0.107564 | 5.231732 | 0.022 | 0 | 1 | 12 | Mipol1 | 0.022 | Young |
| 0.107564 | 5.222952 | 0.022 | 0 | 1 | 12 | Tle1 | 0.022 | Young |
| 0.107564 | 5.211684 | 0.022 | 0 | 1 | 12 | Chm | 0.022 | Young |
| 0.107564 | 5.211066 | 0.022 | 0 | 1 | 12 | Rad54l2 | 0.022 | Young |
| 0.107564 | 5.206114 | 0.022 | 0 | 1 | 12 | Iqsec1 | 0.022 | Young |
| 0.107564 | 5.204332 | 0.022 | 0 | 1 | 12 | Neu1 | 0.022 | Young |
| 0.107564 | 5.201306 | 0.022 | 0 | 1 | 12 | Strip1 | 0.022 | Young |
| 0.107564 | 5.196968 | 0.022 | 0 | 1 | 12 | Cdk17 | 0.022 | Young |
| 0.107564 | 5.195935 | 0.022 | 0 | 1 | 12 | Xkr4 | 0.022 | Young |
| 0.107564 | 5.193026 | 0.022 | 0 | 1 | 12 | Ift46 | 0.022 | Young |
| 0.107564 | 5.189223 | 0.022 | 0 | 1 | 12 | Fam129b | 0.022 | Young |
| 0.107564 | 5.17685 | 0.022 | 0 | 1 | 12 | Rnf216 | 0.022 | Young |
| 0.107564 | 5.176654 | 0.022 | 0 | 1 | 12 | Slc25a26 | 0.022 | Young |
| 0.107564 | 5.171392 | 0.022 | 0 | 1 | 12 | Hccs | 0.022 | Young |
| 0.107564 | 5.168129 | 0.022 | 0 | 1 | 12 | Nkapl | 0.022 | Young |
| 0.107564 | 5.167678 | 0.022 | 0 | 1 | 12 | Ubxn7 | 0.022 | Young |
| 0.107564 | 5.164465 | 0.022 | 0 | 1 | 12 | Mmgt1 | 0.022 | Young |
| 0.107564 | 5.163941 | 0.022 | 0 | 1 | 12 | Coq9 | 0.022 | Young |
| 0.107564 | 5.160088 | 0.022 | 0 | 1 | 12 | Lysmd4 | 0.022 | Young |
| 0.107564 | 5.159049 | 0.022 | 0 | 1 | 12 | Otud4 | 0.022 | Young |
| 0.107564 | 5.149419 | 0.022 | 0 | 1 | 12 | Phgdh | 0.022 | Young |
| 0.107564 | 5.142658 | 0.022 | 0 | 1 | 12 | Pex3 | 0.022 | Young |
| 0.107564 | 5.139978 | 0.022 | 0 | 1 | 12 | 5033428I2 | 0.022 | Young |

| | | | | | | | | |
|---|---|---|---|---|---|---|---|---|
| 0.107564 | 5.135841 | 0.022 | 0 | 1 | 12 | Dhx57 | 0.022 | Young |
| 0.107564 | 5.133794 | 0.022 | 0 | 1 | 12 | Rbm43 | 0.022 | Young |
| 0.107564 | 5.132813 | 0.022 | 0 | 1 | 12 | Gm11549 | 0.022 | Young |
| 0.107564 | 5.129244 | 0.022 | 0 | 1 | 12 | Atg2b | 0.022 | Young |
| 0.107564 | 5.127885 | 0.022 | 0 | 1 | 12 | Sparcl1 | 0.022 | Young |
| 0.107564 | 5.123708 | 0.022 | 0 | 1 | 12 | Pex6 | 0.022 | Young |
| 0.107564 | 5.11792 | 0.022 | 0 | 1 | 12 | Pold3 | 0.022 | Young |
| 0.107564 | 5.113279 | 0.022 | 0 | 1 | 12 | Rcbtb2 | 0.022 | Young |
| 0.107564 | 5.112869 | 0.022 | 0 | 1 | 12 | Ssh2 | 0.022 | Young |
| 0.107564 | 5.112847 | 0.022 | 0 | 1 | 12 | 1600002K0 | 0.022 | Young |
| 0.107564 | 5.112305 | 0.022 | 0 | 1 | 12 | Sf3a1 | 0.022 | Young |
| 0.107564 | 5.102564 | 0.022 | 0 | 1 | 12 | C1qtnf7 | 0.022 | Young |
| 0.107564 | 5.100328 | 0.022 | 0 | 1 | 12 | Tvp23a | 0.022 | Young |
| 0.107564 | 5.098849 | 0.022 | 0 | 1 | 12 | Mtif3 | 0.022 | Young |
| 0.107564 | 5.092513 | 0.022 | 0 | 1 | 12 | Ctdspl2 | 0.022 | Young |
| 0.107564 | 5.087578 | 0.022 | 0 | 1 | 12 | Zfp760 | 0.022 | Young |
| 0.107564 | 5.086293 | 0.022 | 0 | 1 | 12 | Qars | 0.022 | Young |
| 0.107564 | 5.083466 | 0.022 | 0 | 1 | 12 | Brms1l | 0.022 | Young |
| 0.107564 | 5.075372 | 0.022 | 0 | 1 | 12 | Rcc1l | 0.022 | Young |
| 0.107564 | 5.0749 | 0.022 | 0 | 1 | 12 | B3glct | 0.022 | Young |
| 0.107564 | 5.074175 | 0.022 | 0 | 1 | 12 | Btbd3 | 0.022 | Young |
| 0.107564 | 5.071286 | 0.022 | 0 | 1 | 12 | Taf1b | 0.022 | Young |
| 0.107564 | 5.071034 | 0.022 | 0 | 1 | 12 | Rce1 | 0.022 | Young |
| 0.107564 | 5.070884 | 0.022 | 0 | 1 | 12 | Cenpt | 0.022 | Young |
| 0.107564 | 5.068218 | 0.022 | 0 | 1 | 12 | Mcm6 | 0.022 | Young |
| 0.107564 | 5.064764 | 0.022 | 0 | 1 | 12 | Rbfox3 | 0.022 | Young |
| 0.107564 | 5.064534 | 0.022 | 0 | 1 | 12 | Zbtb37 | 0.022 | Young |
| 0.107564 | 5.062936 | 0.022 | 0 | 1 | 12 | Nphp1 | 0.022 | Young |
| 0.107564 | 5.055508 | 0.022 | 0 | 1 | 12 | 2900055J2 | 0.022 | Young |
| 0.107564 | 5.051063 | 0.022 | 0 | 1 | 12 | Ptpn2 | 0.022 | Young |
| 0.107564 | 5.044634 | 0.022 | 0 | 1 | 12 | Oxsr1 | 0.022 | Young |
| 0.107564 | 5.042545 | 0.022 | 0 | 1 | 12 | Plscr4 | 0.022 | Young |
| 0.107564 | 5.041126 | 0.022 | 0 | 1 | 12 | Fam234a | 0.022 | Young |
| 0.107564 | 5.038369 | 0.022 | 0 | 1 | 12 | Rmdn3 | 0.022 | Young |
| 0.107564 | 5.037097 | 0.022 | 0 | 1 | 12 | Rhno1 | 0.022 | Young |
| 0.107564 | 5.036581 | 0.022 | 0 | 1 | 12 | Cstf2 | 0.022 | Young |
| 0.107564 | 5.034867 | 0.022 | 0 | 1 | 12 | Ift122 | 0.022 | Young |
| 0.107564 | 5.03285 | 0.022 | 0 | 1 | 12 | Cntnap1 | 0.022 | Young |
| 0.107564 | 5.032677 | 0.022 | 0 | 1 | 12 | Hip1 | 0.022 | Young |
| 0.107564 | 5.03071 | 0.022 | 0 | 1 | 12 | Kcnmb4 | 0.022 | Young |
| 0.107564 | 5.030526 | 0.022 | 0 | 1 | 12 | Serpinh1 | 0.022 | Young |
| 0.107564 | 5.028521 | 0.022 | 0 | 1 | 12 | Mapk12 | 0.022 | Young |
| 0.107564 | 5.021786 | 0.022 | 0 | 1 | 12 | Gabbr2 | 0.022 | Young |
| 0.107564 | 5.020397 | 0.022 | 0 | 1 | 12 | Ercc4 | 0.022 | Young |
| 0.107564 | 5.018255 | 0.022 | 0 | 1 | 12 | Rcor1 | 0.022 | Young |

| | | | | | | | | |
|---|---|---|---|---|---|---|---|---|
| 0.107564 | 5.016322 | 0.022 | 0 | 1 | 12 | Smug1 | 0.022 | Young |
| 0.107564 | 5.012442 | 0.022 | 0 | 1 | 12 | Lyz1 | 0.022 | Young |
| 0.107564 | 5.009192 | 0.022 | 0 | 1 | 12 | Gm49336 | 0.022 | Young |
| 0.107564 | 5.003368 | 0.022 | 0 | 1 | 12 | Syap1 | 0.022 | Young |
| 0.107564 | 4.999408 | 0.022 | 0 | 1 | 12 | Camta2 | 0.022 | Young |
| 0.107564 | 4.997291 | 0.022 | 0 | 1 | 12 | Ctnnd1 | 0.022 | Young |
| 0.107564 | 4.995221 | 0.022 | 0 | 1 | 12 | Tacc1 | 0.022 | Young |
| 0.107564 | 4.983881 | 0.022 | 0 | 1 | 12 | Zc3h7b | 0.022 | Young |
| 0.107564 | 4.982495 | 0.022 | 0 | 1 | 12 | Ulk2 | 0.022 | Young |
| 0.107564 | 4.979823 | 0.022 | 0 | 1 | 12 | Zdhhc22 | 0.022 | Young |
| 0.107564 | 4.976925 | 0.022 | 0 | 1 | 12 | Zfp597 | 0.022 | Young |
| 0.107564 | 4.971193 | 0.022 | 0 | 1 | 12 | Afg3l1 | 0.022 | Young |
| 0.107564 | 4.968436 | 0.022 | 0 | 1 | 12 | Map2k4 | 0.022 | Young |
| 0.107564 | 4.966022 | 0.022 | 0 | 1 | 12 | Gm26614 | 0.022 | Young |
| 0.107564 | 4.96566 | 0.022 | 0 | 1 | 12 | Sync | 0.022 | Young |
| 0.107564 | 4.9648 | 0.022 | 0 | 1 | 12 | Erlin2 | 0.022 | Young |
| 0.107564 | 4.955525 | 0.022 | 0 | 1 | 12 | Kbtbd3 | 0.022 | Young |
| 0.107564 | 4.942594 | 0.022 | 0 | 1 | 12 | Cpeb2 | 0.022 | Young |
| 0.107564 | 4.941166 | 0.022 | 0 | 1 | 12 | Borcs5 | 0.022 | Young |
| 0.107564 | 4.941124 | 0.022 | 0 | 1 | 12 | Nxt1 | 0.022 | Young |
| 0.107564 | 4.93767 | 0.022 | 0 | 1 | 12 | Mtcl1 | 0.022 | Young |
| 0.107564 | 4.935113 | 0.022 | 0 | 1 | 12 | Dgka | 0.022 | Young |
| 0.107564 | 4.933703 | 0.022 | 0 | 1 | 12 | Isy1 | 0.022 | Young |
| 0.107564 | 4.932712 | 0.022 | 0 | 1 | 12 | Magohb | 0.022 | Young |
| 0.107564 | 4.926596 | 0.022 | 0 | 1 | 12 | Bet1 | 0.022 | Young |
| 0.107564 | 4.924127 | 0.022 | 0 | 1 | 12 | Ryk | 0.022 | Young |
| 0.107564 | 4.917802 | 0.022 | 0 | 1 | 12 | Gtf2a1 | 0.022 | Young |
| 0.107564 | 4.916024 | 0.022 | 0 | 1 | 12 | Igfbp5 | 0.022 | Young |
| 0.107564 | 4.90968 | 0.022 | 0 | 1 | 12 | Cbwd1 | 0.022 | Young |
| 0.107564 | 4.903854 | 0.022 | 0 | 1 | 12 | Cdc23 | 0.022 | Young |
| 0.107564 | 4.898397 | 0.022 | 0 | 1 | 12 | Scube1 | 0.022 | Young |
| 0.107564 | 4.893438 | 0.022 | 0 | 1 | 12 | Thsd7a | 0.022 | Young |
| 0.107564 | 4.891967 | 0.022 | 0 | 1 | 12 | Sulf2 | 0.022 | Young |
| 0.107564 | 4.888022 | 0.022 | 0 | 1 | 12 | Fam49a | 0.022 | Young |
| 0.107564 | 4.885505 | 0.022 | 0 | 1 | 12 | Atrip | 0.022 | Young |
| 0.107564 | 4.866899 | 0.022 | 0 | 1 | 12 | Heatr3 | 0.022 | Young |
| 0.107564 | 4.866564 | 0.022 | 0 | 1 | 12 | Nol11 | 0.022 | Young |
| 0.107564 | 4.85787 | 0.022 | 0 | 1 | 12 | Pcdha5 | 0.022 | Young |
| 0.107564 | 4.855508 | 0.022 | 0 | 1 | 12 | Dyrk1a | 0.022 | Young |
| 0.107564 | 4.844011 | 0.022 | 0 | 1 | 12 | Tyw5 | 0.022 | Young |
| 0.107564 | 4.838222 | 0.022 | 0 | 1 | 12 | Hipk2 | 0.022 | Young |
| 0.107564 | 4.834842 | 0.022 | 0 | 1 | 12 | Parp2 | 0.022 | Young |
| 0.107564 | 4.826836 | 0.022 | 0 | 1 | 12 | Rnf150 | 0.022 | Young |
| 0.107564 | 4.822274 | 0.022 | 0 | 1 | 12 | Chst12 | 0.022 | Young |
| 0.107564 | 4.807041 | 0.022 | 0 | 1 | 12 | Rab33b | 0.022 | Young |

| | | | | | | | | |
|---|---|---|---|---|---|---|---|---|
| 0.107564 | 4.800321 | 0.022 | 0 | 1 | 12 | Igf1r | 0.022 | Young |
| 0.107564 | 4.799825 | 0.022 | 0 | 1 | 12 | Creb3l1 | 0.022 | Young |
| 0.107564 | 4.789363 | 0.022 | 0 | 1 | 12 | Urgcp | 0.022 | Young |
| 0.107564 | 4.772457 | 0.022 | 0 | 1 | 12 | Mthfsd | 0.022 | Young |
| 0.107564 | 4.771389 | 0.022 | 0 | 1 | 12 | Camk1d | 0.022 | Young |
| 0.107564 | 4.710319 | 0.022 | 0 | 1 | 12 | Neo1 | 0.022 | Young |
| 0.107564 | 4.67942 | 0.022 | 0 | 1 | 12 | Sort1 | 0.022 | Young |
| 0.107564 | 4.671813 | 0.022 | 0 | 1 | 12 | Mpg | 0.022 | Young |
| 0.107564 | 4.670579 | 0.022 | 0 | 1 | 12 | E130309D( | 0.022 | Young |
| 0.107564 | 4.660383 | 0.022 | 0 | 1 | 12 | Dusp23 | 0.022 | Young |
| 0.107564 | 4.657346 | 0.022 | 0 | 1 | 12 | Apool | 0.022 | Young |
| 0.107752 | 0.657097 | 0.131 | 0.068 | 1 | 12 | Nxf1 | 0.063 | Young |
| 0.108145 | 0.616715 | 0.27 | 0.195 | 1 | 12 | Smap1 | 0.075 | Young |
| 0.108413 | 1.242675 | 0.102 | 0.051 | 1 | 12 | Prrc2c | 0.051 | Young |
| 0.108519 | 1.079652 | 0.095 | 0.042 | 1 | 12 | Rbms1 | 0.053 | Young |
| 0.109905 | 1.075407 | 0.117 | 0.059 | 1 | 12 | Mrpl40 | 0.058 | Young |
| 0.110443 | 0.608213 | 0.285 | 0.195 | 1 | 12 | Rab14 | 0.09 | Young |
| 0.111793 | 0.791593 | 0.182 | 0.11 | 1 | 12 | Bzw1 | 0.072 | Young |
| 0.112987 | 0.98809 | 0.117 | 0.059 | 1 | 12 | Maged2 | 0.058 | Young |
| 0.113645 | 0.397043 | 0.496 | 0.407 | 1 | 12 | Tppp3 | 0.089 | Young |
| 0.113992 | 1.275943 | 0.08 | 0.034 | 1 | 12 | Gnaq | 0.046 | Young |
| 0.115059 | 1.149358 | 0.124 | 0.068 | 1 | 12 | Rhob | 0.056 | Young |
| 0.116414 | 0.136957 | 0.993 | 0.983 | 1 | 12 | mt-Atp6 | 0.01 | Young |
| 0.117111 | 1.119399 | 0.102 | 0.051 | 1 | 12 | Sumo3 | 0.051 | Young |
| 0.117672 | 0.991956 | 0.095 | 0.042 | 1 | 12 | Yae1d1 | 0.053 | Young |
| 0.118523 | 1.032653 | 0.124 | 0.068 | 1 | 12 | Commd3 | 0.056 | Young |
| 0.119261 | 1.229004 | 0.102 | 0.051 | 1 | 12 | Ociad1 | 0.051 | Young |
| 0.120987 | 0.897058 | 0.117 | 0.059 | 1 | 12 | Evi5 | 0.058 | Young |
| 0.12184 | 1.314146 | 0.08 | 0.034 | 1 | 12 | Gsr | 0.046 | Young |
| 0.122821 | 1.577866 | 0.066 | 0.025 | 1 | 12 | Rnf115 | 0.041 | Young |
| 0.123183 | 0.420183 | 0.394 | 0.305 | 1 | 12 | Pfdn5 | 0.089 | Young |
| 0.123427 | 0.612773 | 0.234 | 0.161 | 1 | 12 | Aplp1 | 0.073 | Young |
| 0.123891 | 0.74712 | 0.117 | 0.059 | 1 | 12 | Sra1 | 0.058 | Young |
| 0.123952 | 1.564924 | 0.066 | 0.025 | 1 | 12 | Ablim1 | 0.041 | Young |
| 0.123952 | 1.544065 | 0.066 | 0.025 | 1 | 12 | Mtfr1l | 0.041 | Young |
| 0.12444 | 0.375379 | 0.474 | 0.39 | 1 | 12 | Eif5a | 0.084 | Young |
| 0.125073 | 0.564743 | 0.212 | 0.136 | 1 | 12 | Zc3h15 | 0.076 | Young |
| 0.12509 | 1.684159 | 0.066 | 0.025 | 1 | 12 | Paip2b | 0.041 | Young |
| 0.12509 | 1.594631 | 0.066 | 0.025 | 1 | 12 | Nudt10 | 0.041 | Young |
| 0.125107 | 1.072937 | 0.131 | 0.076 | 1 | 12 | Mrps7 | 0.055 | Young |
| 0.12598 | 0.493805 | 0.314 | 0.229 | 1 | 12 | Ndufb11 | 0.085 | Young |
| 0.12739 | 1.420736 | 0.066 | 0.025 | 1 | 12 | Lars | 0.041 | Young |
| 0.12739 | 1.410018 | 0.066 | 0.025 | 1 | 12 | Cnp | 0.041 | Young |
| 0.127701 | 0.838444 | 0.139 | 0.076 | 1 | 12 | Dhps | 0.063 | Young |
| 0.129595 | 0.322268 | 0.511 | 0.432 | 1 | 12 | Rps15a | 0.079 | Young |

| | | | | | | | | |
|---|---|---|---|---|---|---|---|---|
| 0.129723 | 1.402286 | 0.066 | 0.025 | 1 | 12 | Gm47283 | 0.041 | Young |
| 0.12987 | 0.210477 | 0.81 | 0.754 | 1 | 12 | Uchl1 | 0.056 | Young |
| 0.130014 | 0.968815 | 0.102 | 0.051 | 1 | 12 | mt-Atp8 | 0.051 | Young |
| 0.130772 | 0.719271 | 0.146 | 0.085 | 1 | 12 | Diablo | 0.061 | Young |
| 0.130891 | 0.511136 | 0.197 | 0.127 | 1 | 12 | Ndufa1 | 0.07 | Young |
| 0.130903 | 1.475988 | 0.066 | 0.025 | 1 | 12 | Cul3 | 0.041 | Young |
| 0.130903 | 1.312343 | 0.066 | 0.025 | 1 | 12 | Vps41 | 0.041 | Young |
| 0.131166 | 0.68741 | 0.08 | 0.034 | 1 | 12 | Apoo | 0.046 | Young |
| 0.13155 | 0.46699 | 0.321 | 0.246 | 1 | 12 | Higd2a | 0.075 | Young |
| 0.13209 | 1.32367 | 0.066 | 0.025 | 1 | 12 | Lyrm9 | 0.041 | Young |
| 0.132372 | 2.743186 | 0.051 | 0.017 | 1 | 12 | Zbtb4 | 0.034 | Young |
| 0.133115 | 0.781884 | 0.146 | 0.085 | 1 | 12 | Emc4 | 0.061 | Young |
| 0.133151 | 0.322048 | 0.46 | 0.339 | 1 | 12 | Eif1b | 0.121 | Young |
| 0.134484 | 0.683065 | 0.124 | 0.068 | 1 | 12 | Nfe2l1 | 0.056 | Young |
| 0.134489 | 1.506977 | 0.066 | 0.025 | 1 | 12 | Rbm42 | 0.041 | Young |
| 0.134489 | 1.034353 | 0.066 | 0.025 | 1 | 12 | Ncam2 | 0.041 | Young |
| 0.134694 | 0.838965 | 0.146 | 0.085 | 1 | 12 | Ubxn4 | 0.061 | Young |
| 0.13471 | 0.728281 | 0.102 | 0.051 | 1 | 12 | Idh3b | 0.051 | Young |
| 0.135134 | 1.891975 | 0.051 | 0.017 | 1 | 12 | Rufy2 | 0.034 | Young |
| 0.135417 | 0.603605 | 0.321 | 0.246 | 1 | 12 | Ank2 | 0.075 | Young |
| 0.136531 | 2.411145 | 0.051 | 0.017 | 1 | 12 | 2410002F2 | 0.034 | Young |
| 0.136531 | 1.877764 | 0.051 | 0.017 | 1 | 12 | Nbea | 0.034 | Young |
| 0.136531 | 1.837508 | 0.051 | 0.017 | 1 | 12 | Fez2 | 0.034 | Young |
| 0.13794 | 2.200905 | 0.051 | 0.017 | 1 | 12 | Sptlc2 | 0.034 | Young |
| 0.138369 | 4.29867 | 0.036 | 0.008 | 1 | 12 | Sqor | 0.028 | Young |
| 0.138369 | 4.099825 | 0.036 | 0.008 | 1 | 12 | Ccnd3 | 0.028 | Young |
| 0.138369 | 4.074813 | 0.036 | 0.008 | 1 | 12 | Casd1 | 0.028 | Young |
| 0.138369 | 3.988545 | 0.036 | 0.008 | 1 | 12 | Psmg2 | 0.028 | Young |
| 0.138369 | 2.516931 | 0.036 | 0.008 | 1 | 12 | Tfpt | 0.028 | Young |
| 0.138369 | 2.486294 | 0.036 | 0.008 | 1 | 12 | Mgat4a | 0.028 | Young |
| 0.138369 | 2.473228 | 0.036 | 0.008 | 1 | 12 | Slc7a1 | 0.028 | Young |
| 0.138369 | 2.462746 | 0.036 | 0.008 | 1 | 12 | Chuk | 0.028 | Young |
| 0.138369 | 2.333699 | 0.036 | 0.008 | 1 | 12 | Stag2 | 0.028 | Young |
| 0.138369 | 2.309473 | 0.036 | 0.008 | 1 | 12 | Fto | 0.028 | Young |
| 0.138369 | 2.154074 | 0.036 | 0.008 | 1 | 12 | Erbin | 0.028 | Young |
| 0.140103 | 2.591906 | 0.036 | 0.008 | 1 | 12 | Bcam | 0.028 | Young |
| 0.140103 | 2.352003 | 0.036 | 0.008 | 1 | 12 | Dlst | 0.028 | Young |
| 0.140103 | 2.326306 | 0.036 | 0.008 | 1 | 12 | Nudt18 | 0.028 | Young |
| 0.140103 | 2.290863 | 0.036 | 0.008 | 1 | 12 | Cep83 | 0.028 | Young |
| 0.140103 | 2.265493 | 0.036 | 0.008 | 1 | 12 | Clip3 | 0.028 | Young |
| 0.140103 | 2.256489 | 0.036 | 0.008 | 1 | 12 | Blzf1 | 0.028 | Young |
| 0.140103 | 2.211922 | 0.036 | 0.008 | 1 | 12 | Rhot2 | 0.028 | Young |
| 0.140103 | 2.205296 | 0.036 | 0.008 | 1 | 12 | Scyl1 | 0.028 | Young |
| 0.140103 | 2.092896 | 0.036 | 0.008 | 1 | 12 | Rbsn | 0.028 | Young |
| 0.140103 | 2.028237 | 0.036 | 0.008 | 1 | 12 | Dcaf12 | 0.028 | Young |

| | | | | | | | | |
|---|---|---|---|---|---|---|---|---|
| 0.14051 | 1.11491 | 0.109 | 0.059 | 1 | 12 | Nptn | 0.05 | Young |
| 0.14079 | 1.577237 | 0.051 | 0.017 | 1 | 12 | Ctdspl | 0.034 | Young |
| 0.141455 | 0.390692 | 0.307 | 0.22 | 1 | 12 | Tmem234 | 0.087 | Young |
| 0.141854 | 2.208872 | 0.036 | 0.008 | 1 | 12 | Ttc4 | 0.028 | Young |
| 0.141854 | 2.160568 | 0.036 | 0.008 | 1 | 12 | Bclaf1 | 0.028 | Young |
| 0.141854 | 2.088848 | 0.036 | 0.008 | 1 | 12 | Gusb | 0.028 | Young |
| 0.141854 | 1.927084 | 0.036 | 0.008 | 1 | 12 | Naa35 | 0.028 | Young |
| 0.142232 | 1.520181 | 0.051 | 0.017 | 1 | 12 | Plcl1 | 0.034 | Young |
| 0.142233 | 1.692984 | 0.051 | 0.017 | 1 | 12 | Pak3 | 0.034 | Young |
| 0.142233 | 1.616745 | 0.051 | 0.017 | 1 | 12 | Hnrnph1 | 0.034 | Young |
| 0.142233 | 1.545196 | 0.051 | 0.017 | 1 | 12 | Capn7 | 0.034 | Young |
| 0.142233 | 1.499272 | 0.051 | 0.017 | 1 | 12 | Pnpla8 | 0.034 | Young |
| 0.14329 | 0.768667 | 0.124 | 0.068 | 1 | 12 | Txn2 | 0.056 | Young |
| 0.143622 | 2.035096 | 0.036 | 0.008 | 1 | 12 | Sft2d3 | 0.028 | Young |
| 0.143622 | 1.984773 | 0.036 | 0.008 | 1 | 12 | Klc2 | 0.028 | Young |
| 0.143622 | 1.97652 | 0.036 | 0.008 | 1 | 12 | Adgrb3 | 0.028 | Young |
| 0.143622 | 1.93743 | 0.036 | 0.008 | 1 | 12 | Haus7 | 0.028 | Young |
| 0.143622 | 1.922892 | 0.036 | 0.008 | 1 | 12 | Slk | 0.028 | Young |
| 0.143622 | 1.914401 | 0.036 | 0.008 | 1 | 12 | Ppm1l | 0.028 | Young |
| 0.143622 | 1.91165 | 0.036 | 0.008 | 1 | 12 | Rad50 | 0.028 | Young |
| 0.143622 | 1.857568 | 0.036 | 0.008 | 1 | 12 | Csk | 0.028 | Young |
| 0.143622 | 1.842424 | 0.036 | 0.008 | 1 | 12 | Tmem242 | 0.028 | Young |
| 0.143622 | 1.834949 | 0.036 | 0.008 | 1 | 12 | Tmem263 | 0.028 | Young |
| 0.143622 | 1.816053 | 0.036 | 0.008 | 1 | 12 | Gtf2ird1 | 0.028 | Young |
| 0.143686 | 1.534607 | 0.051 | 0.017 | 1 | 12 | Arid4a | 0.034 | Young |
| 0.143686 | 1.470251 | 0.051 | 0.017 | 1 | 12 | Spats2 | 0.034 | Young |
| 0.143686 | 1.44677 | 0.051 | 0.017 | 1 | 12 | Map7d1 | 0.034 | Young |
| 0.143686 | 0.846479 | 0.051 | 0.017 | 1 | 12 | Sdhaf1 | 0.034 | Young |
| 0.144597 | 0.637073 | 0.168 | 0.102 | 1 | 12 | Chmp5 | 0.066 | Young |
| 0.145151 | 1.568869 | 0.051 | 0.017 | 1 | 12 | B9d2 | 0.034 | Young |
| 0.145151 | 1.548851 | 0.051 | 0.017 | 1 | 12 | Phactr3 | 0.034 | Young |
| 0.145151 | 1.455566 | 0.051 | 0.017 | 1 | 12 | Wdr13 | 0.034 | Young |
| 0.145151 | 1.396801 | 0.051 | 0.017 | 1 | 12 | Sirt2 | 0.034 | Young |
| 0.145151 | 1.346402 | 0.051 | 0.017 | 1 | 12 | Tor1aip2 | 0.034 | Young |
| 0.145406 | 2.084502 | 0.036 | 0.008 | 1 | 12 | Fahd1 | 0.028 | Young |
| 0.145406 | 1.948812 | 0.036 | 0.008 | 1 | 12 | Tmem70 | 0.028 | Young |
| 0.145406 | 1.936356 | 0.036 | 0.008 | 1 | 12 | Dtnbp1 | 0.028 | Young |
| 0.145406 | 1.857701 | 0.036 | 0.008 | 1 | 12 | 4931406P1 | 0.028 | Young |
| 0.145406 | 1.760705 | 0.036 | 0.008 | 1 | 12 | Cby1 | 0.028 | Young |
| 0.145406 | 1.667134 | 0.036 | 0.008 | 1 | 12 | Tnni1 | 0.028 | Young |
| 0.145406 | 1.614986 | 0.036 | 0.008 | 1 | 12 | Rxfp1 | 0.028 | Young |
| 0.145406 | 1.613795 | 0.036 | 0.008 | 1 | 12 | Ifi35 | 0.028 | Young |
| 0.145406 | 1.585767 | 0.036 | 0.008 | 1 | 12 | Lactb | 0.028 | Young |
| 0.146176 | 1.100758 | 0.088 | 0.042 | 1 | 12 | Magee1 | 0.046 | Young |
| 0.146177 | 0.772193 | 0.088 | 0.042 | 1 | 12 | Tpd52l2 | 0.046 | Young |

| | | | | | | | | |
|---|---|---|---|---|---|---|---|---|
| 0.146628 | 1.390885 | 0.051 | 0.017 | 1 | 12 | Smarca4 | 0.034 | Young |
| 0.146628 | 1.34735 | 0.051 | 0.017 | 1 | 12 | Pphln1 | 0.034 | Young |
| 0.146695 | 0.861671 | 0.153 | 0.093 | 1 | 12 | Polr2k | 0.06 | Young |
| 0.147208 | 1.819568 | 0.036 | 0.008 | 1 | 12 | Rock1 | 0.028 | Young |
| 0.147208 | 1.757793 | 0.036 | 0.008 | 1 | 12 | Faf2 | 0.028 | Young |
| 0.147208 | 1.691043 | 0.036 | 0.008 | 1 | 12 | Htra2 | 0.028 | Young |
| 0.147208 | 1.638752 | 0.036 | 0.008 | 1 | 12 | Eif2ak1 | 0.028 | Young |
| 0.147208 | 1.638105 | 0.036 | 0.008 | 1 | 12 | Mapk8ip1 | 0.028 | Young |
| 0.147208 | 1.633883 | 0.036 | 0.008 | 1 | 12 | Eif2a | 0.028 | Young |
| 0.147208 | 1.493897 | 0.036 | 0.008 | 1 | 12 | Fzr1 | 0.028 | Young |
| 0.147208 | 1.436549 | 0.036 | 0.008 | 1 | 12 | Ablim3 | 0.028 | Young |
| 0.147208 | 1.397838 | 0.036 | 0.008 | 1 | 12 | Desi1 | 0.028 | Young |
| 0.147208 | 1.384439 | 0.036 | 0.008 | 1 | 12 | Ccnh | 0.028 | Young |
| 0.14748 | 0.358366 | 0.343 | 0.254 | 1 | 12 | Tbca | 0.089 | Young |
| 0.147782 | 1.263294 | 0.095 | 0.051 | 1 | 12 | Cggbp1 | 0.044 | Young |
| 0.147996 | 0.585024 | 0.219 | 0.144 | 1 | 12 | Rps28 | 0.075 | Young |
| 0.148116 | 1.360853 | 0.051 | 0.017 | 1 | 12 | Papss1 | 0.034 | Young |
| 0.148116 | 1.356738 | 0.051 | 0.017 | 1 | 12 | Madd | 0.034 | Young |
| 0.148116 | 1.217121 | 0.051 | 0.017 | 1 | 12 | Ndufaf2 | 0.034 | Young |
| 0.149183 | 1.260911 | 0.109 | 0.059 | 1 | 12 | Rbm25 | 0.05 | Young |
| 0.149615 | 1.361876 | 0.051 | 0.017 | 1 | 12 | Btg2 | 0.034 | Young |
| 0.149615 | 1.031486 | 0.051 | 0.017 | 1 | 12 | Stoml1 | 0.034 | Young |
| 0.150326 | 0.326085 | 0.489 | 0.398 | 1 | 12 | Rpl34 | 0.091 | Young |
| 0.151126 | 1.165992 | 0.051 | 0.017 | 1 | 12 | Polr2d | 0.034 | Young |
| 0.15116 | 0.962109 | 0.088 | 0.042 | 1 | 12 | Atp1a3 | 0.046 | Young |
| 0.151782 | 0.548128 | 0.168 | 0.102 | 1 | 12 | Pfdn4 | 0.066 | Young |
| 0.1526 | 0.254891 | 0.555 | 0.458 | 1 | 12 | Rpl18a | 0.097 | Young |
| 0.152782 | 0.443602 | 0.299 | 0.212 | 1 | 12 | Scoc | 0.087 | Young |
| 0.153166 | 0.928436 | 0.109 | 0.059 | 1 | 12 | Srrm4 | 0.05 | Young |
| 0.153166 | 0.877939 | 0.109 | 0.059 | 1 | 12 | Fundc2 | 0.05 | Young |
| 0.154175 | 0.724256 | 0.109 | 0.059 | 1 | 12 | Mafg | 0.05 | Young |
| 0.154518 | 0.388884 | 0.38 | 0.314 | 1 | 12 | Csrp1 | 0.066 | Young |
| 0.155697 | 0.895822 | 0.109 | 0.059 | 1 | 12 | Stk16 | 0.05 | Young |
| 0.156251 | 0.611387 | 0.175 | 0.11 | 1 | 12 | Ube2d2a | 0.065 | Young |
| 0.156847 | 0.90424 | 0.088 | 0.042 | 1 | 12 | 11-Mar | 0.046 | Young |
| 0.157028 | 0.46324 | 0.35 | 0.254 | 1 | 12 | Ndufs7 | 0.096 | Young |
| 0.157327 | 0.830818 | 0.131 | 0.076 | 1 | 12 | Mrpl21 | 0.055 | Young |
| 0.158259 | 0.952369 | 0.109 | 0.059 | 1 | 12 | Leprotl1 | 0.05 | Young |
| 0.159641 | 0.772274 | 0.168 | 0.11 | 1 | 12 | Tspyl4 | 0.058 | Young |
| 0.159751 | 0.88561 | 0.088 | 0.042 | 1 | 12 | Atrx | 0.046 | Young |
| 0.160293 | 1.640738 | 0.073 | 0.034 | 1 | 12 | Tpgs1 | 0.039 | Young |
| 0.160535 | 0.270548 | 0.672 | 0.576 | 1 | 12 | Rpl32 | 0.096 | Young |
| 0.163582 | 0.344463 | 0.504 | 0.407 | 1 | 12 | Cdc42 | 0.097 | Young |
| 0.16568 | 0.740604 | 0.088 | 0.042 | 1 | 12 | Pnn | 0.046 | Young |
| 0.166151 | 1.260972 | 0.073 | 0.034 | 1 | 12 | Cplx2 | 0.039 | Young |

| | | | | | | | | |
|---|---|---|---|---|---|---|---|---|
| 0.166812 | 1.17445 | 0.073 | 0.034 | 1 | 12 | Cbx3 | 0.039 | Young |
| 0.166985 | 1.073386 | 0.095 | 0.051 | 1 | 12 | Ndufaf3 | 0.044 | Young |
| 0.16739 | 0.826517 | 0.131 | 0.076 | 1 | 12 | Akirin2 | 0.055 | Young |
| 0.168661 | 0.927594 | 0.117 | 0.068 | 1 | 12 | Mapre3 | 0.049 | Young |
| 0.17075 | 0.884233 | 0.117 | 0.068 | 1 | 12 | Prdx4 | 0.049 | Young |
| 0.172859 | 0.882791 | 0.117 | 0.068 | 1 | 12 | Arl1 | 0.049 | Young |
| 0.173137 | 0.461346 | 0.175 | 0.11 | 1 | 12 | Gabarapl1 | 0.065 | Young |
| 0.173529 | 1.234839 | 0.073 | 0.034 | 1 | 12 | Fbxw2 | 0.039 | Young |
| 0.173529 | 1.137804 | 0.073 | 0.034 | 1 | 12 | Med30 | 0.039 | Young |
| 0.173696 | 0.47253 | 0.263 | 0.186 | 1 | 12 | Ndufc1 | 0.077 | Young |
| 0.17517 | 1.017528 | 0.095 | 0.051 | 1 | 12 | Caml | 0.044 | Young |
| 0.175363 | 0.391053 | 0.547 | 0.466 | 1 | 12 | Atp5c1 | 0.081 | Young |
| 0.176724 | 0.591355 | 0.219 | 0.161 | 1 | 12 | Prelid1 | 0.058 | Young |
| 0.177135 | 0.862216 | 0.117 | 0.068 | 1 | 12 | Sparc | 0.049 | Young |
| 0.177532 | 0.421278 | 0.401 | 0.331 | 1 | 12 | Mrfap1 | 0.07 | Young |
| 0.178768 | 0.941989 | 0.095 | 0.051 | 1 | 12 | Pcbd2 | 0.044 | Young |
| 0.178783 | 0.714601 | 0.109 | 0.059 | 1 | 12 | Ppfia2 | 0.05 | Young |
| 0.179197 | 0.23422 | 0.409 | 0.331 | 1 | 12 | Txndc17 | 0.078 | Young |
| 0.179902 | 0.478798 | 0.431 | 0.347 | 1 | 12 | Slc18a3 | 0.084 | Young |
| 0.180346 | 0.985635 | 0.131 | 0.085 | 1 | 12 | Rnh1 | 0.046 | Young |
| 0.181652 | 0.362102 | 0.358 | 0.28 | 1 | 12 | Ywhab | 0.078 | Young |
| 0.181853 | 0.86179 | 0.073 | 0.034 | 1 | 12 | Reep2 | 0.039 | Young |
| 0.182959 | 0.55725 | 0.234 | 0.169 | 1 | 12 | Taldo1 | 0.065 | Young |
| 0.183269 | 0.997148 | 0.073 | 0.034 | 1 | 12 | Fbxo22 | 0.039 | Young |
| 0.183422 | 1.489907 | 0.058 | 0.025 | 1 | 12 | Cmc1 | 0.033 | Young |
| 0.183576 | 0.231486 | 0.613 | 0.568 | 1 | 12 | Myl6 | 0.045 | Young |
| 0.18452 | 0.844188 | 0.153 | 0.102 | 1 | 12 | Ctbp1 | 0.051 | Young |
| 0.184693 | 0.945038 | 0.073 | 0.034 | 1 | 12 | Mindy3 | 0.039 | Young |
| 0.184693 | 0.827869 | 0.073 | 0.034 | 1 | 12 | Ssbp2 | 0.039 | Young |
| 0.18523 | 0.422759 | 0.234 | 0.161 | 1 | 12 | Mrpl41 | 0.073 | Young |
| 0.186125 | 0.931605 | 0.073 | 0.034 | 1 | 12 | Pttg1 | 0.039 | Young |
| 0.187565 | 0.838908 | 0.073 | 0.034 | 1 | 12 | Rtn2 | 0.039 | Young |
| 0.188155 | 0.44423 | 0.248 | 0.178 | 1 | 12 | Cnbp | 0.07 | Young |
| 0.188432 | 0.204592 | 0.62 | 0.568 | 1 | 12 | Cox6b1 | 0.052 | Young |
| 0.189877 | 1.557142 | 0.058 | 0.025 | 1 | 12 | Sat1 | 0.033 | Young |
| 0.190362 | 4.571351 | 0.015 | 0 | 1 | 12 | Rnf113a1 | 0.015 | Young |
| 0.190366 | 4.85416 | 0.015 | 0 | 1 | 12 | Ubxn2b | 0.015 | Young |
| 0.190366 | 4.836551 | 0.015 | 0 | 1 | 12 | Tfr2 | 0.015 | Young |
| 0.190366 | 4.811735 | 0.015 | 0 | 1 | 12 | B230217C1 | 0.015 | Young |
| 0.190366 | 4.807766 | 0.015 | 0 | 1 | 12 | Rpf1 | 0.015 | Young |
| 0.190366 | 4.801411 | 0.015 | 0 | 1 | 12 | Hcn2 | 0.015 | Young |
| 0.190366 | 4.799432 | 0.015 | 0 | 1 | 12 | Zcchc9 | 0.015 | Young |
| 0.190366 | 4.799156 | 0.015 | 0 | 1 | 12 | Ispd | 0.015 | Young |
| 0.190366 | 4.793772 | 0.015 | 0 | 1 | 12 | Spink10 | 0.015 | Young |
| 0.190366 | 4.790914 | 0.015 | 0 | 1 | 12 | Foxp1 | 0.015 | Young |

| | | | | | | | | |
|---|---|---|---|---|---|---|---|---|
| 0.190366 | 4.784934 | 0.015 | 0 | 1 | 12 | Scai | 0.015 | Young |
| 0.190366 | 4.781217 | 0.015 | 0 | 1 | 12 | Plxna1 | 0.015 | Young |
| 0.190366 | 4.77892 | 0.015 | 0 | 1 | 12 | Nkiras2 | 0.015 | Young |
| 0.190366 | 4.776086 | 0.015 | 0 | 1 | 12 | Dleu7 | 0.015 | Young |
| 0.190366 | 4.774836 | 0.015 | 0 | 1 | 12 | Ccpg1 | 0.015 | Young |
| 0.190366 | 4.770902 | 0.015 | 0 | 1 | 12 | Aoah | 0.015 | Young |
| 0.190366 | 4.769546 | 0.015 | 0 | 1 | 12 | Snap23 | 0.015 | Young |
| 0.190366 | 4.768683 | 0.015 | 0 | 1 | 12 | Mcmbp | 0.015 | Young |
| 0.190366 | 4.766185 | 0.015 | 0 | 1 | 12 | Fktn | 0.015 | Young |
| 0.190366 | 4.765187 | 0.015 | 0 | 1 | 12 | Mthfd1l | 0.015 | Young |
| 0.190366 | 4.762437 | 0.015 | 0 | 1 | 12 | Taok2 | 0.015 | Young |
| 0.190366 | 4.759829 | 0.015 | 0 | 1 | 12 | Scfd2 | 0.015 | Young |
| 0.190366 | 4.756717 | 0.015 | 0 | 1 | 12 | Incenp | 0.015 | Young |
| 0.190366 | 4.75426 | 0.015 | 0 | 1 | 12 | Arl6ip6 | 0.015 | Young |
| 0.190366 | 4.753227 | 0.015 | 0 | 1 | 12 | Zfp866 | 0.015 | Young |
| 0.190366 | 4.751774 | 0.015 | 0 | 1 | 12 | Mfap2 | 0.015 | Young |
| 0.190366 | 4.750675 | 0.015 | 0 | 1 | 12 | A830005F2 | 0.015 | Young |
| 0.190366 | 4.748489 | 0.015 | 0 | 1 | 12 | Ermard | 0.015 | Young |
| 0.190366 | 4.747848 | 0.015 | 0 | 1 | 12 | Il17rd | 0.015 | Young |
| 0.190366 | 4.742712 | 0.015 | 0 | 1 | 12 | Cilp2 | 0.015 | Young |
| 0.190366 | 4.741806 | 0.015 | 0 | 1 | 12 | Rad17 | 0.015 | Young |
| 0.190366 | 4.741788 | 0.015 | 0 | 1 | 12 | Ptk2 | 0.015 | Young |
| 0.190366 | 4.741557 | 0.015 | 0 | 1 | 12 | Myo1d | 0.015 | Young |
| 0.190366 | 4.737028 | 0.015 | 0 | 1 | 12 | Slc8a3 | 0.015 | Young |
| 0.190366 | 4.734699 | 0.015 | 0 | 1 | 12 | C1ql1 | 0.015 | Young |
| 0.190366 | 4.734217 | 0.015 | 0 | 1 | 12 | Cdk13 | 0.015 | Young |
| 0.190366 | 4.733784 | 0.015 | 0 | 1 | 12 | Ccnt1 | 0.015 | Young |
| 0.190366 | 4.733067 | 0.015 | 0 | 1 | 12 | C2cd5 | 0.015 | Young |
| 0.190366 | 4.724891 | 0.015 | 0 | 1 | 12 | Timp1 | 0.015 | Young |
| 0.190366 | 4.724704 | 0.015 | 0 | 1 | 12 | Zfp7 | 0.015 | Young |
| 0.190366 | 4.719645 | 0.015 | 0 | 1 | 12 | Ubxn8 | 0.015 | Young |
| 0.190366 | 4.71648 | 0.015 | 0 | 1 | 12 | Tmem181a | 0.015 | Young |
| 0.190366 | 4.713428 | 0.015 | 0 | 1 | 12 | Agk | 0.015 | Young |
| 0.190366 | 4.711029 | 0.015 | 0 | 1 | 12 | Vasn | 0.015 | Young |
| 0.190366 | 4.708763 | 0.015 | 0 | 1 | 12 | A230006K0 | 0.015 | Young |
| 0.190366 | 4.703564 | 0.015 | 0 | 1 | 12 | Parp8 | 0.015 | Young |
| 0.190366 | 4.698378 | 0.015 | 0 | 1 | 12 | Tmem54 | 0.015 | Young |
| 0.190366 | 4.692421 | 0.015 | 0 | 1 | 12 | St3gal5 | 0.015 | Young |
| 0.190366 | 4.691289 | 0.015 | 0 | 1 | 12 | Zc3h18 | 0.015 | Young |
| 0.190366 | 4.690934 | 0.015 | 0 | 1 | 12 | Gm15418 | 0.015 | Young |
| 0.190366 | 4.690849 | 0.015 | 0 | 1 | 12 | Stard3 | 0.015 | Young |
| 0.190366 | 4.6871 | 0.015 | 0 | 1 | 12 | Gm10053 | 0.015 | Young |
| 0.190366 | 4.682863 | 0.015 | 0 | 1 | 12 | Samd1 | 0.015 | Young |
| 0.190366 | 4.680742 | 0.015 | 0 | 1 | 12 | Ppil4 | 0.015 | Young |
| 0.190366 | 4.678143 | 0.015 | 0 | 1 | 12 | Metrnl | 0.015 | Young |

| | | | | | | | | |
|---|---|---|---|---|---|---|---|---|
| 0.190366 | 4.675053 | 0.015 | 0 | 1 | 12 | Rusc2 | 0.015 | Young |
| 0.190366 | 4.673362 | 0.015 | 0 | 1 | 12 | Ptcd3 | 0.015 | Young |
| 0.190366 | 4.672277 | 0.015 | 0 | 1 | 12 | Fam126a | 0.015 | Young |
| 0.190366 | 4.671408 | 0.015 | 0 | 1 | 12 | Dtnb | 0.015 | Young |
| 0.190366 | 4.668427 | 0.015 | 0 | 1 | 12 | Lhpp | 0.015 | Young |
| 0.190366 | 4.667967 | 0.015 | 0 | 1 | 12 | Fam131b | 0.015 | Young |
| 0.190366 | 4.660077 | 0.015 | 0 | 1 | 12 | Noa1 | 0.015 | Young |
| 0.190366 | 4.659455 | 0.015 | 0 | 1 | 12 | Osbpl3 | 0.015 | Young |
| 0.190366 | 4.659019 | 0.015 | 0 | 1 | 12 | Tsga10 | 0.015 | Young |
| 0.190366 | 4.657687 | 0.015 | 0 | 1 | 12 | Porcn | 0.015 | Young |
| 0.190366 | 4.655403 | 0.015 | 0 | 1 | 12 | Bcl10 | 0.015 | Young |
| 0.190366 | 4.655187 | 0.015 | 0 | 1 | 12 | Nlgn2 | 0.015 | Young |
| 0.190366 | 4.654735 | 0.015 | 0 | 1 | 12 | Pcyox1l | 0.015 | Young |
| 0.190366 | 4.654312 | 0.015 | 0 | 1 | 12 | Atg13 | 0.015 | Young |
| 0.190366 | 4.651134 | 0.015 | 0 | 1 | 12 | Rasal2 | 0.015 | Young |
| 0.190366 | 4.648759 | 0.015 | 0 | 1 | 12 | C330011M | 0.015 | Young |
| 0.190366 | 4.645016 | 0.015 | 0 | 1 | 12 | Rft1 | 0.015 | Young |
| 0.190366 | 4.641929 | 0.015 | 0 | 1 | 12 | Fermt3 | 0.015 | Young |
| 0.190366 | 4.640996 | 0.015 | 0 | 1 | 12 | Nlrx1 | 0.015 | Young |
| 0.190366 | 4.639981 | 0.015 | 0 | 1 | 12 | Dera | 0.015 | Young |
| 0.190366 | 4.639981 | 0.015 | 0 | 1 | 12 | Sphk2 | 0.015 | Young |
| 0.190366 | 4.638636 | 0.015 | 0 | 1 | 12 | Cdh18 | 0.015 | Young |
| 0.190366 | 4.637286 | 0.015 | 0 | 1 | 12 | Mxd1 | 0.015 | Young |
| 0.190366 | 4.636209 | 0.015 | 0 | 1 | 12 | Rarres1 | 0.015 | Young |
| 0.190366 | 4.635647 | 0.015 | 0 | 1 | 12 | Dgkh | 0.015 | Young |
| 0.190366 | 4.631515 | 0.015 | 0 | 1 | 12 | Arhgef15 | 0.015 | Young |
| 0.190366 | 4.63138 | 0.015 | 0 | 1 | 12 | Mta3 | 0.015 | Young |
| 0.190366 | 4.628469 | 0.015 | 0 | 1 | 12 | Zgpat | 0.015 | Young |
| 0.190366 | 4.627675 | 0.015 | 0 | 1 | 12 | Zfp943 | 0.015 | Young |
| 0.190366 | 4.623634 | 0.015 | 0 | 1 | 12 | Bbs2 | 0.015 | Young |
| 0.190366 | 4.62272 | 0.015 | 0 | 1 | 12 | Il1rl1 | 0.015 | Young |
| 0.190366 | 4.621847 | 0.015 | 0 | 1 | 12 | Arfgap2 | 0.015 | Young |
| 0.190366 | 4.621401 | 0.015 | 0 | 1 | 12 | Sec24a | 0.015 | Young |
| 0.190366 | 4.61987 | 0.015 | 0 | 1 | 12 | 9030624J0 | 0.015 | Young |
| 0.190366 | 4.617234 | 0.015 | 0 | 1 | 12 | Psmf1 | 0.015 | Young |
| 0.190366 | 4.615506 | 0.015 | 0 | 1 | 12 | Zfp275 | 0.015 | Young |
| 0.190366 | 4.612121 | 0.015 | 0 | 1 | 12 | Hdac9 | 0.015 | Young |
| 0.190366 | 4.611586 | 0.015 | 0 | 1 | 12 | Gfer | 0.015 | Young |
| 0.190366 | 4.611458 | 0.015 | 0 | 1 | 12 | Junos | 0.015 | Young |
| 0.190366 | 4.608526 | 0.015 | 0 | 1 | 12 | Inppl1 | 0.015 | Young |
| 0.190366 | 4.607467 | 0.015 | 0 | 1 | 12 | Tti1 | 0.015 | Young |
| 0.190366 | 4.606179 | 0.015 | 0 | 1 | 12 | Fli1 | 0.015 | Young |
| 0.190366 | 4.605305 | 0.015 | 0 | 1 | 12 | Rasl11a | 0.015 | Young |
| 0.190366 | 4.604527 | 0.015 | 0 | 1 | 12 | Kat14 | 0.015 | Young |
| 0.190366 | 4.603679 | 0.015 | 0 | 1 | 12 | Rcc2 | 0.015 | Young |

| | | | | | | | | |
|---|---|---|---|---|---|---|---|---|
| 0.190366 | 4.600164 | 0.015 | 0 | 1 | 12 | Neurl2 | 0.015 | Young |
| 0.190366 | 4.599192 | 0.015 | 0 | 1 | 12 | Foxj3 | 0.015 | Young |
| 0.190366 | 4.599192 | 0.015 | 0 | 1 | 12 | Abca5 | 0.015 | Young |
| 0.190366 | 4.596135 | 0.015 | 0 | 1 | 12 | Mdn1 | 0.015 | Young |
| 0.190366 | 4.596095 | 0.015 | 0 | 1 | 12 | Akr1b8 | 0.015 | Young |
| 0.190366 | 4.59609 | 0.015 | 0 | 1 | 12 | Rad51d | 0.015 | Young |
| 0.190366 | 4.595299 | 0.015 | 0 | 1 | 12 | Trappc8 | 0.015 | Young |
| 0.190366 | 4.595157 | 0.015 | 0 | 1 | 12 | Noct | 0.015 | Young |
| 0.190366 | 4.593543 | 0.015 | 0 | 1 | 12 | Atxn7 | 0.015 | Young |
| 0.190366 | 4.591631 | 0.015 | 0 | 1 | 12 | Chrna7 | 0.015 | Young |
| 0.190366 | 4.591506 | 0.015 | 0 | 1 | 12 | Vegfa | 0.015 | Young |
| 0.190366 | 4.58781 | 0.015 | 0 | 1 | 12 | Mfap3l | 0.015 | Young |
| 0.190366 | 4.586975 | 0.015 | 0 | 1 | 12 | Mmp15 | 0.015 | Young |
| 0.190366 | 4.586975 | 0.015 | 0 | 1 | 12 | Trim11 | 0.015 | Young |
| 0.190366 | 4.585834 | 0.015 | 0 | 1 | 12 | Ncln | 0.015 | Young |
| 0.190366 | 4.585834 | 0.015 | 0 | 1 | 12 | Hmgxb4 | 0.015 | Young |
| 0.190366 | 4.582261 | 0.015 | 0 | 1 | 12 | 1190005I0 | 0.015 | Young |
| 0.190366 | 4.579664 | 0.015 | 0 | 1 | 12 | Opa3 | 0.015 | Young |
| 0.190366 | 4.578493 | 0.015 | 0 | 1 | 12 | Serinc5 | 0.015 | Young |
| 0.190366 | 4.578002 | 0.015 | 0 | 1 | 12 | Eef2k | 0.015 | Young |
| 0.190366 | 4.577737 | 0.015 | 0 | 1 | 12 | E2f6 | 0.015 | Young |
| 0.190366 | 4.576605 | 0.015 | 0 | 1 | 12 | Ttll12 | 0.015 | Young |
| 0.190366 | 4.576033 | 0.015 | 0 | 1 | 12 | Chml | 0.015 | Young |
| 0.190366 | 4.575921 | 0.015 | 0 | 1 | 12 | Pcdhb7 | 0.015 | Young |
| 0.190366 | 4.568917 | 0.015 | 0 | 1 | 12 | Tbc1d30 | 0.015 | Young |
| 0.190366 | 4.567319 | 0.015 | 0 | 1 | 12 | Dnaja4 | 0.015 | Young |
| 0.190366 | 4.566305 | 0.015 | 0 | 1 | 12 | Rpa2 | 0.015 | Young |
| 0.190366 | 4.564594 | 0.015 | 0 | 1 | 12 | Ppp1r3c | 0.015 | Young |
| 0.190366 | 4.559202 | 0.015 | 0 | 1 | 12 | Kcns3 | 0.015 | Young |
| 0.190366 | 4.559194 | 0.015 | 0 | 1 | 12 | Gm34006 | 0.015 | Young |
| 0.190366 | 4.555785 | 0.015 | 0 | 1 | 12 | Prmt3 | 0.015 | Young |
| 0.190366 | 4.555315 | 0.015 | 0 | 1 | 12 | Tox4 | 0.015 | Young |
| 0.190366 | 4.554694 | 0.015 | 0 | 1 | 12 | Etnk2 | 0.015 | Young |
| 0.190366 | 4.55452 | 0.015 | 0 | 1 | 12 | Plxnb2 | 0.015 | Young |
| 0.190366 | 4.55219 | 0.015 | 0 | 1 | 12 | Nol9 | 0.015 | Young |
| 0.190366 | 4.548624 | 0.015 | 0 | 1 | 12 | Lrfn3 | 0.015 | Young |
| 0.190366 | 4.547848 | 0.015 | 0 | 1 | 12 | Ttc21b | 0.015 | Young |
| 0.190366 | 4.547411 | 0.015 | 0 | 1 | 12 | Dnajc1 | 0.015 | Young |
| 0.190366 | 4.545059 | 0.015 | 0 | 1 | 12 | Cdc25a | 0.015 | Young |
| 0.190366 | 4.544782 | 0.015 | 0 | 1 | 12 | Rab9b | 0.015 | Young |
| 0.190366 | 4.544527 | 0.015 | 0 | 1 | 12 | Polr1a | 0.015 | Young |
| 0.190366 | 4.543382 | 0.015 | 0 | 1 | 12 | Ypel4 | 0.015 | Young |
| 0.190366 | 4.543355 | 0.015 | 0 | 1 | 12 | Brwd1 | 0.015 | Young |
| 0.190366 | 4.542649 | 0.015 | 0 | 1 | 12 | Ipmk | 0.015 | Young |
| 0.190366 | 4.5422 | 0.015 | 0 | 1 | 12 | Smad5 | 0.015 | Young |

| | | | | | | | | |
|---|---|---|---|---|---|---|---|---|
| 0.190366 | 4.541892 | 0.015 | 0 | 1 | 12 | Cacna1c | 0.015 | Young |
| 0.190366 | 4.541494 | 0.015 | 0 | 1 | 12 | Kcnn2 | 0.015 | Young |
| 0.190366 | 4.540305 | 0.015 | 0 | 1 | 12 | Zfp711 | 0.015 | Young |
| 0.190366 | 4.538479 | 0.015 | 0 | 1 | 12 | C030014I2: | 0.015 | Young |
| 0.190366 | 4.537461 | 0.015 | 0 | 1 | 12 | Map2k6 | 0.015 | Young |
| 0.190366 | 4.535088 | 0.015 | 0 | 1 | 12 | Mpp2 | 0.015 | Young |
| 0.190366 | 4.530506 | 0.015 | 0 | 1 | 12 | Tcaim | 0.015 | Young |
| 0.190366 | 4.5293 | 0.015 | 0 | 1 | 12 | Spin2c | 0.015 | Young |
| 0.190366 | 4.526008 | 0.015 | 0 | 1 | 12 | Zbtb46 | 0.015 | Young |
| 0.190366 | 4.523948 | 0.015 | 0 | 1 | 12 | Zfp940 | 0.015 | Young |
| 0.190366 | 4.522699 | 0.015 | 0 | 1 | 12 | Ifit1bl2 | 0.015 | Young |
| 0.190366 | 4.521336 | 0.015 | 0 | 1 | 12 | Plpp3 | 0.015 | Young |
| 0.190366 | 4.520908 | 0.015 | 0 | 1 | 12 | Alg10b | 0.015 | Young |
| 0.190366 | 4.519447 | 0.015 | 0 | 1 | 12 | Mtss1l | 0.015 | Young |
| 0.190366 | 4.518476 | 0.015 | 0 | 1 | 12 | Sipa1 | 0.015 | Young |
| 0.190366 | 4.515686 | 0.015 | 0 | 1 | 12 | Gid4 | 0.015 | Young |
| 0.190366 | 4.514547 | 0.015 | 0 | 1 | 12 | Chodl | 0.015 | Young |
| 0.190366 | 4.513572 | 0.015 | 0 | 1 | 12 | Hcn4 | 0.015 | Young |
| 0.190366 | 4.50844 | 0.015 | 0 | 1 | 12 | Chd1l | 0.015 | Young |
| 0.190366 | 4.504337 | 0.015 | 0 | 1 | 12 | Zfp248 | 0.015 | Young |
| 0.190366 | 4.5017 | 0.015 | 0 | 1 | 12 | Acsl1 | 0.015 | Young |
| 0.190366 | 4.501416 | 0.015 | 0 | 1 | 12 | Zfp784 | 0.015 | Young |
| 0.190366 | 4.498001 | 0.015 | 0 | 1 | 12 | Pcnx | 0.015 | Young |
| 0.190366 | 4.497478 | 0.015 | 0 | 1 | 12 | Gadd45b | 0.015 | Young |
| 0.190366 | 4.496963 | 0.015 | 0 | 1 | 12 | Rpap2 | 0.015 | Young |
| 0.190366 | 4.494399 | 0.015 | 0 | 1 | 12 | Cc2d1a | 0.015 | Young |
| 0.190366 | 4.493778 | 0.015 | 0 | 1 | 12 | Cenpf | 0.015 | Young |
| 0.190366 | 4.488226 | 0.015 | 0 | 1 | 12 | Mgl2 | 0.015 | Young |
| 0.190366 | 4.487464 | 0.015 | 0 | 1 | 12 | Cep44 | 0.015 | Young |
| 0.190366 | 4.484771 | 0.015 | 0 | 1 | 12 | Rab36 | 0.015 | Young |
| 0.190366 | 4.48333 | 0.015 | 0 | 1 | 12 | Lmo2 | 0.015 | Young |
| 0.190366 | 4.48333 | 0.015 | 0 | 1 | 12 | Zc3h8 | 0.015 | Young |
| 0.190366 | 4.48025 | 0.015 | 0 | 1 | 12 | Myo16 | 0.015 | Young |
| 0.190366 | 4.480227 | 0.015 | 0 | 1 | 12 | Wipf2 | 0.015 | Young |
| 0.190366 | 4.479209 | 0.015 | 0 | 1 | 12 | 4930539J0 | 0.015 | Young |
| 0.190366 | 4.479093 | 0.015 | 0 | 1 | 12 | Prickle2 | 0.015 | Young |
| 0.190366 | 4.478423 | 0.015 | 0 | 1 | 12 | Slc9a6 | 0.015 | Young |
| 0.190366 | 4.477449 | 0.015 | 0 | 1 | 12 | Mfsd13a | 0.015 | Young |
| 0.190366 | 4.473022 | 0.015 | 0 | 1 | 12 | Daxx | 0.015 | Young |
| 0.190366 | 4.470085 | 0.015 | 0 | 1 | 12 | Rassf3 | 0.015 | Young |
| 0.190366 | 4.468875 | 0.015 | 0 | 1 | 12 | Topbp1 | 0.015 | Young |
| 0.190366 | 4.467846 | 0.015 | 0 | 1 | 12 | Dot1l | 0.015 | Young |
| 0.190366 | 4.467642 | 0.015 | 0 | 1 | 12 | Fbxo16 | 0.015 | Young |
| 0.190366 | 4.466429 | 0.015 | 0 | 1 | 12 | Usp1 | 0.015 | Young |
| 0.190366 | 4.464469 | 0.015 | 0 | 1 | 12 | Tmem100 | 0.015 | Young |

| | | | | | | | | |
|---|---|---|---|---|---|---|---|---|
| 0.190366 | 4.46231 | 0.015 | 0 | 1 | 12 | Rapgef2 | 0.015 | Young |
| 0.190366 | 4.459819 | 0.015 | 0 | 1 | 12 | Etfbkmt | 0.015 | Young |
| 0.190366 | 4.459695 | 0.015 | 0 | 1 | 12 | Stx3 | 0.015 | Young |
| 0.190366 | 4.458812 | 0.015 | 0 | 1 | 12 | Ehbp1 | 0.015 | Young |
| 0.190366 | 4.458728 | 0.015 | 0 | 1 | 12 | Traf7 | 0.015 | Young |
| 0.190366 | 4.458372 | 0.015 | 0 | 1 | 12 | Med20 | 0.015 | Young |
| 0.190366 | 4.458276 | 0.015 | 0 | 1 | 12 | Igdcc4 | 0.015 | Young |
| 0.190366 | 4.457314 | 0.015 | 0 | 1 | 12 | Enoph1 | 0.015 | Young |
| 0.190366 | 4.456742 | 0.015 | 0 | 1 | 12 | Rgp1 | 0.015 | Young |
| 0.190366 | 4.456742 | 0.015 | 0 | 1 | 12 | Pcdh1 | 0.015 | Young |
| 0.190366 | 4.453938 | 0.015 | 0 | 1 | 12 | Spire1 | 0.015 | Young |
| 0.190366 | 4.4521 | 0.015 | 0 | 1 | 12 | Banp | 0.015 | Young |
| 0.190366 | 4.451489 | 0.015 | 0 | 1 | 12 | Zfp2 | 0.015 | Young |
| 0.190366 | 4.450104 | 0.015 | 0 | 1 | 12 | Dcun1d2 | 0.015 | Young |
| 0.190366 | 4.449424 | 0.015 | 0 | 1 | 12 | Pex11a | 0.015 | Young |
| 0.190366 | 4.448629 | 0.015 | 0 | 1 | 12 | Zfp768 | 0.015 | Young |
| 0.190366 | 4.445619 | 0.015 | 0 | 1 | 12 | Lins1 | 0.015 | Young |
| 0.190366 | 4.445569 | 0.015 | 0 | 1 | 12 | Gm10138 | 0.015 | Young |
| 0.190366 | 4.445194 | 0.015 | 0 | 1 | 12 | Ncstn | 0.015 | Young |
| 0.190366 | 4.445106 | 0.015 | 0 | 1 | 12 | Fam98a | 0.015 | Young |
| 0.190366 | 4.440234 | 0.015 | 0 | 1 | 12 | Tmtc3 | 0.015 | Young |
| 0.190366 | 4.439557 | 0.015 | 0 | 1 | 12 | Brf1 | 0.015 | Young |
| 0.190366 | 4.436586 | 0.015 | 0 | 1 | 12 | Slc25a23 | 0.015 | Young |
| 0.190366 | 4.432421 | 0.015 | 0 | 1 | 12 | Slc29a3 | 0.015 | Young |
| 0.190366 | 4.432316 | 0.015 | 0 | 1 | 12 | Papolg | 0.015 | Young |
| 0.190366 | 4.430569 | 0.015 | 0 | 1 | 12 | Rhog | 0.015 | Young |
| 0.190366 | 4.429509 | 0.015 | 0 | 1 | 12 | Pick1 | 0.015 | Young |
| 0.190366 | 4.414987 | 0.015 | 0 | 1 | 12 | Usp36 | 0.015 | Young |
| 0.190366 | 4.41436 | 0.015 | 0 | 1 | 12 | Stt3b | 0.015 | Young |
| 0.190366 | 4.414255 | 0.015 | 0 | 1 | 12 | Prps1l3 | 0.015 | Young |
| 0.190366 | 4.413501 | 0.015 | 0 | 1 | 12 | Ankrd16 | 0.015 | Young |
| 0.190366 | 4.413265 | 0.015 | 0 | 1 | 12 | Trim13 | 0.015 | Young |
| 0.190366 | 4.411998 | 0.015 | 0 | 1 | 12 | Pcsk2 | 0.015 | Young |
| 0.190366 | 4.41123 | 0.015 | 0 | 1 | 12 | Edn3 | 0.015 | Young |
| 0.190366 | 4.410998 | 0.015 | 0 | 1 | 12 | Scd1 | 0.015 | Young |
| 0.190366 | 4.409003 | 0.015 | 0 | 1 | 12 | Ttf2 | 0.015 | Young |
| 0.190366 | 4.40861 | 0.015 | 0 | 1 | 12 | Hrasls | 0.015 | Young |
| 0.190366 | 4.408371 | 0.015 | 0 | 1 | 12 | Uhrf2 | 0.015 | Young |
| 0.190366 | 4.406042 | 0.015 | 0 | 1 | 12 | Ptbp1 | 0.015 | Young |
| 0.190366 | 4.405571 | 0.015 | 0 | 1 | 12 | Slc35e4 | 0.015 | Young |
| 0.190366 | 4.403289 | 0.015 | 0 | 1 | 12 | S100a4 | 0.015 | Young |
| 0.190366 | 4.402611 | 0.015 | 0 | 1 | 12 | Zfp536 | 0.015 | Young |
| 0.190366 | 4.401898 | 0.015 | 0 | 1 | 12 | Gprin1 | 0.015 | Young |
| 0.190366 | 4.399823 | 0.015 | 0 | 1 | 12 | Pigh | 0.015 | Young |
| 0.190366 | 4.399715 | 0.015 | 0 | 1 | 12 | Gm12258 | 0.015 | Young |

| | | | | | | | | |
|---|---|---|---|---|---|---|---|---|
| 0.190366 | 4.395838 | 0.015 | 0 | 1 | 12 | Ddx55 | 0.015 | Young |
| 0.190366 | 4.395368 | 0.015 | 0 | 1 | 12 | Prkaa1 | 0.015 | Young |
| 0.190366 | 4.394089 | 0.015 | 0 | 1 | 12 | Gm16701 | 0.015 | Young |
| 0.190366 | 4.394089 | 0.015 | 0 | 1 | 12 | 2010015M | 0.015 | Young |
| 0.190366 | 4.393861 | 0.015 | 0 | 1 | 12 | Cep72 | 0.015 | Young |
| 0.190366 | 4.392777 | 0.015 | 0 | 1 | 12 | Rbm14 | 0.015 | Young |
| 0.190366 | 4.390969 | 0.015 | 0 | 1 | 12 | Ncan | 0.015 | Young |
| 0.190366 | 4.390322 | 0.015 | 0 | 1 | 12 | Ube3b | 0.015 | Young |
| 0.190366 | 4.388238 | 0.015 | 0 | 1 | 12 | Myo7a | 0.015 | Young |
| 0.190366 | 4.385354 | 0.015 | 0 | 1 | 12 | Ostm1 | 0.015 | Young |
| 0.190366 | 4.385335 | 0.015 | 0 | 1 | 12 | Gal | 0.015 | Young |
| 0.190366 | 4.384939 | 0.015 | 0 | 1 | 12 | Nkrf | 0.015 | Young |
| 0.190366 | 4.380013 | 0.015 | 0 | 1 | 12 | Fkbp5 | 0.015 | Young |
| 0.190366 | 4.379318 | 0.015 | 0 | 1 | 12 | Tsr2 | 0.015 | Young |
| 0.190366 | 4.378471 | 0.015 | 0 | 1 | 12 | Bloc1s1 | 0.015 | Young |
| 0.190366 | 4.377249 | 0.015 | 0 | 1 | 12 | Gm30648 | 0.015 | Young |
| 0.190366 | 4.376054 | 0.015 | 0 | 1 | 12 | Agbl3 | 0.015 | Young |
| 0.190366 | 4.37561 | 0.015 | 0 | 1 | 12 | Zfp28 | 0.015 | Young |
| 0.190366 | 4.375445 | 0.015 | 0 | 1 | 12 | Polr3d | 0.015 | Young |
| 0.190366 | 4.372128 | 0.015 | 0 | 1 | 12 | 6030419C1 | 0.015 | Young |
| 0.190366 | 4.370399 | 0.015 | 0 | 1 | 12 | Lonrf1 | 0.015 | Young |
| 0.190366 | 4.368775 | 0.015 | 0 | 1 | 12 | Slc39a3 | 0.015 | Young |
| 0.190366 | 4.368216 | 0.015 | 0 | 1 | 12 | Gm42517 | 0.015 | Young |
| 0.190366 | 4.367185 | 0.015 | 0 | 1 | 12 | Thrsp | 0.015 | Young |
| 0.190366 | 4.365078 | 0.015 | 0 | 1 | 12 | Cenpk | 0.015 | Young |
| 0.190366 | 4.364659 | 0.015 | 0 | 1 | 12 | Gm10602 | 0.015 | Young |
| 0.190366 | 4.364659 | 0.015 | 0 | 1 | 12 | Nrep | 0.015 | Young |
| 0.190366 | 4.362411 | 0.015 | 0 | 1 | 12 | Slc16a7 | 0.015 | Young |
| 0.190366 | 4.361429 | 0.015 | 0 | 1 | 12 | Tbck | 0.015 | Young |
| 0.190366 | 4.361082 | 0.015 | 0 | 1 | 12 | Fer | 0.015 | Young |
| 0.190366 | 4.360383 | 0.015 | 0 | 1 | 12 | Wfs1 | 0.015 | Young |
| 0.190366 | 4.359209 | 0.015 | 0 | 1 | 12 | Zfp367 | 0.015 | Young |
| 0.190366 | 4.358597 | 0.015 | 0 | 1 | 12 | Mlh1 | 0.015 | Young |
| 0.190366 | 4.355602 | 0.015 | 0 | 1 | 12 | Rilpl2 | 0.015 | Young |
| 0.190366 | 4.355596 | 0.015 | 0 | 1 | 12 | Cnksr2 | 0.015 | Young |
| 0.190366 | 4.351259 | 0.015 | 0 | 1 | 12 | Zfp282 | 0.015 | Young |
| 0.190366 | 4.346582 | 0.015 | 0 | 1 | 12 | Entpd7 | 0.015 | Young |
| 0.190366 | 4.343533 | 0.015 | 0 | 1 | 12 | 4930481A1 | 0.015 | Young |
| 0.190366 | 4.342513 | 0.015 | 0 | 1 | 12 | Btrc | 0.015 | Young |
| 0.190366 | 4.3423 | 0.015 | 0 | 1 | 12 | Epb41l4aos | 0.015 | Young |
| 0.190366 | 4.33954 | 0.015 | 0 | 1 | 12 | Anapc15 | 0.015 | Young |
| 0.190366 | 4.33954 | 0.015 | 0 | 1 | 12 | Creld2 | 0.015 | Young |
| 0.190366 | 4.33917 | 0.015 | 0 | 1 | 12 | BC022687 | 0.015 | Young |
| 0.190366 | 4.337625 | 0.015 | 0 | 1 | 12 | Mdm2 | 0.015 | Young |
| 0.190366 | 4.331655 | 0.015 | 0 | 1 | 12 | Polr3a | 0.015 | Young |

| | | | | | | | | |
|---|---|---|---|---|---|---|---|---|
| 0.190366 | 4.331505 | 0.015 | 0 | 1 | 12 | Ccne1 | 0.015 | Young |
| 0.190366 | 4.327697 | 0.015 | 0 | 1 | 12 | Bcor | 0.015 | Young |
| 0.190366 | 4.323601 | 0.015 | 0 | 1 | 12 | Ociad2 | 0.015 | Young |
| 0.190366 | 4.32344 | 0.015 | 0 | 1 | 12 | Clic4 | 0.015 | Young |
| 0.190366 | 4.320464 | 0.015 | 0 | 1 | 12 | Ino80 | 0.015 | Young |
| 0.190366 | 4.320174 | 0.015 | 0 | 1 | 12 | Tmem168 | 0.015 | Young |
| 0.190366 | 4.320052 | 0.015 | 0 | 1 | 12 | Pcnx2 | 0.015 | Young |
| 0.190366 | 4.319338 | 0.015 | 0 | 1 | 12 | Oxnad1 | 0.015 | Young |
| 0.190366 | 4.318301 | 0.015 | 0 | 1 | 12 | Gm10069 | 0.015 | Young |
| 0.190366 | 4.315829 | 0.015 | 0 | 1 | 12 | Casp6 | 0.015 | Young |
| 0.190366 | 4.313869 | 0.015 | 0 | 1 | 12 | Prpf6 | 0.015 | Young |
| 0.190366 | 4.313195 | 0.015 | 0 | 1 | 12 | 1700086O( | 0.015 | Young |
| 0.190366 | 4.313091 | 0.015 | 0 | 1 | 12 | Atad5 | 0.015 | Young |
| 0.190366 | 4.304907 | 0.015 | 0 | 1 | 12 | Nexmif | 0.015 | Young |
| 0.190366 | 4.304515 | 0.015 | 0 | 1 | 12 | Afg1l | 0.015 | Young |
| 0.190366 | 4.303738 | 0.015 | 0 | 1 | 12 | Slc25a28 | 0.015 | Young |
| 0.190366 | 4.300125 | 0.015 | 0 | 1 | 12 | Gm6169 | 0.015 | Young |
| 0.190366 | 4.299106 | 0.015 | 0 | 1 | 12 | Lyrm4 | 0.015 | Young |
| 0.190366 | 4.295175 | 0.015 | 0 | 1 | 12 | Adcyap1 | 0.015 | Young |
| 0.190366 | 4.287678 | 0.015 | 0 | 1 | 12 | Lats1 | 0.015 | Young |
| 0.190366 | 4.285629 | 0.015 | 0 | 1 | 12 | Nr4a1 | 0.015 | Young |
| 0.190366 | 4.285574 | 0.015 | 0 | 1 | 12 | Slc10a7 | 0.015 | Young |
| 0.190366 | 4.28463 | 0.015 | 0 | 1 | 12 | Ctnnbl1 | 0.015 | Young |
| 0.190366 | 4.281573 | 0.015 | 0 | 1 | 12 | Hotairm1 | 0.015 | Young |
| 0.190366 | 4.28068 | 0.015 | 0 | 1 | 12 | Fkbp15 | 0.015 | Young |
| 0.190366 | 4.278651 | 0.015 | 0 | 1 | 12 | Gorasp1 | 0.015 | Young |
| 0.190366 | 4.273645 | 0.015 | 0 | 1 | 12 | Rabggta | 0.015 | Young |
| 0.190366 | 4.269673 | 0.015 | 0 | 1 | 12 | Cytip | 0.015 | Young |
| 0.190366 | 4.269446 | 0.015 | 0 | 1 | 12 | Hoxb2 | 0.015 | Young |
| 0.190366 | 4.266431 | 0.015 | 0 | 1 | 12 | Ints10 | 0.015 | Young |
| 0.190366 | 4.26305 | 0.015 | 0 | 1 | 12 | Mapk1ip1 | 0.015 | Young |
| 0.190366 | 4.262417 | 0.015 | 0 | 1 | 12 | Nek9 | 0.015 | Young |
| 0.190366 | 4.261687 | 0.015 | 0 | 1 | 12 | A830092H1 | 0.015 | Young |
| 0.190366 | 4.257199 | 0.015 | 0 | 1 | 12 | Cep126 | 0.015 | Young |
| 0.190366 | 4.254772 | 0.015 | 0 | 1 | 12 | Abcb9 | 0.015 | Young |
| 0.190366 | 4.250309 | 0.015 | 0 | 1 | 12 | Myc | 0.015 | Young |
| 0.190366 | 4.25 | 0.015 | 0 | 1 | 12 | Mkln1os | 0.015 | Young |
| 0.190366 | 4.248826 | 0.015 | 0 | 1 | 12 | Ascc3 | 0.015 | Young |
| 0.190366 | 4.248151 | 0.015 | 0 | 1 | 12 | Myl4 | 0.015 | Young |
| 0.190366 | 4.247893 | 0.015 | 0 | 1 | 12 | Rpl9-ps6 | 0.015 | Young |
| 0.190366 | 4.246203 | 0.015 | 0 | 1 | 12 | Negr1 | 0.015 | Young |
| 0.190366 | 4.245951 | 0.015 | 0 | 1 | 12 | Sel1l3 | 0.015 | Young |
| 0.190366 | 4.243742 | 0.015 | 0 | 1 | 12 | Phka2 | 0.015 | Young |
| 0.190366 | 4.240527 | 0.015 | 0 | 1 | 12 | Zrsr1 | 0.015 | Young |
| 0.190366 | 4.236251 | 0.015 | 0 | 1 | 12 | Spc24 | 0.015 | Young |

| | | | | | | | | |
|---|---|---|---|---|---|---|---|---|
| 0.190366 | 4.234798 | 0.015 | 0 | 1 | 12 | Zfc3h1 | 0.015 | Young |
| 0.190366 | 4.233977 | 0.015 | 0 | 1 | 12 | Trpm2 | 0.015 | Young |
| 0.190366 | 4.232056 | 0.015 | 0 | 1 | 12 | Hif1an | 0.015 | Young |
| 0.190366 | 4.229489 | 0.015 | 0 | 1 | 12 | Kptn | 0.015 | Young |
| 0.190366 | 4.227253 | 0.015 | 0 | 1 | 12 | Filip1l | 0.015 | Young |
| 0.190366 | 4.220903 | 0.015 | 0 | 1 | 12 | Ptpn6 | 0.015 | Young |
| 0.190366 | 4.218826 | 0.015 | 0 | 1 | 12 | Abt1 | 0.015 | Young |
| 0.190366 | 4.217994 | 0.015 | 0 | 1 | 12 | Zfp277 | 0.015 | Young |
| 0.190366 | 4.216298 | 0.015 | 0 | 1 | 12 | Mier3 | 0.015 | Young |
| 0.190366 | 4.213659 | 0.015 | 0 | 1 | 12 | Qpctl | 0.015 | Young |
| 0.190366 | 4.207719 | 0.015 | 0 | 1 | 12 | Ap5m1 | 0.015 | Young |
| 0.190366 | 4.203704 | 0.015 | 0 | 1 | 12 | Camsap3 | 0.015 | Young |
| 0.190366 | 4.202951 | 0.015 | 0 | 1 | 12 | Utp14b | 0.015 | Young |
| 0.190366 | 4.190836 | 0.015 | 0 | 1 | 12 | Sarm1 | 0.015 | Young |
| 0.190366 | 4.186122 | 0.015 | 0 | 1 | 12 | Cnppd1 | 0.015 | Young |
| 0.190366 | 4.174569 | 0.015 | 0 | 1 | 12 | Oser1 | 0.015 | Young |
| 0.190366 | 4.164428 | 0.015 | 0 | 1 | 12 | Ppm1d | 0.015 | Young |
| 0.190366 | 4.163565 | 0.015 | 0 | 1 | 12 | Rhov | 0.015 | Young |
| 0.190366 | 4.163424 | 0.015 | 0 | 1 | 12 | Tm7sf2 | 0.015 | Young |
| 0.190366 | 4.157337 | 0.015 | 0 | 1 | 12 | Mns1 | 0.015 | Young |
| 0.190366 | 4.152813 | 0.015 | 0 | 1 | 12 | Thap7 | 0.015 | Young |
| 0.190366 | 4.150714 | 0.015 | 0 | 1 | 12 | Samd5 | 0.015 | Young |
| 0.190366 | 4.150107 | 0.015 | 0 | 1 | 12 | Pcyox1 | 0.015 | Young |
| 0.190366 | 4.147773 | 0.015 | 0 | 1 | 12 | Spata5 | 0.015 | Young |
| 0.190366 | 4.147773 | 0.015 | 0 | 1 | 12 | C2cd2l | 0.015 | Young |
| 0.190366 | 4.14565 | 0.015 | 0 | 1 | 12 | Atf1 | 0.015 | Young |
| 0.190366 | 4.144739 | 0.015 | 0 | 1 | 12 | Primpol | 0.015 | Young |
| 0.190366 | 4.13295 | 0.015 | 0 | 1 | 12 | Paqr8 | 0.015 | Young |
| 0.190366 | 4.132622 | 0.015 | 0 | 1 | 12 | Pcdh20 | 0.015 | Young |
| 0.190366 | 4.132547 | 0.015 | 0 | 1 | 12 | Oxld1 | 0.015 | Young |
| 0.190366 | 4.128133 | 0.015 | 0 | 1 | 12 | Mfap1a | 0.015 | Young |
| 0.190366 | 4.128047 | 0.015 | 0 | 1 | 12 | Gm14322 | 0.015 | Young |
| 0.190366 | 4.122238 | 0.015 | 0 | 1 | 12 | Mkrn2 | 0.015 | Young |
| 0.190366 | 4.120032 | 0.015 | 0 | 1 | 12 | Kdm5c | 0.015 | Young |
| 0.190366 | 4.120032 | 0.015 | 0 | 1 | 12 | Taf2 | 0.015 | Young |
| 0.190366 | 4.119919 | 0.015 | 0 | 1 | 12 | Mnat1 | 0.015 | Young |
| 0.190366 | 4.116683 | 0.015 | 0 | 1 | 12 | Tufm | 0.015 | Young |
| 0.190366 | 4.114954 | 0.015 | 0 | 1 | 12 | Uggt2 | 0.015 | Young |
| 0.190366 | 4.113619 | 0.015 | 0 | 1 | 12 | Zcchc2 | 0.015 | Young |
| 0.190366 | 4.110949 | 0.015 | 0 | 1 | 12 | Banf2os | 0.015 | Young |
| 0.190366 | 4.101465 | 0.015 | 0 | 1 | 12 | Zbtb7c | 0.015 | Young |
| 0.190366 | 4.098658 | 0.015 | 0 | 1 | 12 | Supt20 | 0.015 | Young |
| 0.190366 | 4.095691 | 0.015 | 0 | 1 | 12 | Slc35b2 | 0.015 | Young |
| 0.190366 | 4.093598 | 0.015 | 0 | 1 | 12 | 1110034G2 | 0.015 | Young |
| 0.190366 | 4.068092 | 0.015 | 0 | 1 | 12 | Tspan5 | 0.015 | Young |

| | | | | | | | | |
|---|---|---|---|---|---|---|---|---|
| 0.190366 | 4.067651 | 0.015 | 0 | 1 | 12 | Faah | 0.015 | Young |
| 0.190366 | 4.066136 | 0.015 | 0 | 1 | 12 | Bclaf3 | 0.015 | Young |
| 0.190366 | 4.065975 | 0.015 | 0 | 1 | 12 | Khsrp | 0.015 | Young |
| 0.190366 | 4.052607 | 0.015 | 0 | 1 | 12 | Parvb | 0.015 | Young |
| 0.190366 | 4.051675 | 0.015 | 0 | 1 | 12 | Pde11a | 0.015 | Young |
| 0.190366 | 4.043738 | 0.015 | 0 | 1 | 12 | Icam5 | 0.015 | Young |
| 0.190366 | 4.042721 | 0.015 | 0 | 1 | 12 | 1700088E0 | 0.015 | Young |
| 0.190366 | 4.042324 | 0.015 | 0 | 1 | 12 | Prr14l | 0.015 | Young |
| 0.190366 | 4.039909 | 0.015 | 0 | 1 | 12 | Aak1 | 0.015 | Young |
| 0.190366 | 4.026696 | 0.015 | 0 | 1 | 12 | Rlf | 0.015 | Young |
| 0.190366 | 4.017092 | 0.015 | 0 | 1 | 12 | Hsd17b4 | 0.015 | Young |
| 0.190366 | 4.015488 | 0.015 | 0 | 1 | 12 | Abca8a | 0.015 | Young |
| 0.190366 | 4.004927 | 0.015 | 0 | 1 | 12 | Pdhx | 0.015 | Young |
| 0.190366 | 3.99864 | 0.015 | 0 | 1 | 12 | Cd1d1 | 0.015 | Young |
| 0.190366 | 3.986181 | 0.015 | 0 | 1 | 12 | Eif2s3y | 0.015 | Young |
| 0.190366 | 3.981036 | 0.015 | 0 | 1 | 12 | Lyrm1 | 0.015 | Young |
| 0.190366 | 3.970801 | 0.015 | 0 | 1 | 12 | Mterf3 | 0.015 | Young |
| 0.190366 | 3.965018 | 0.015 | 0 | 1 | 12 | Mdrl | 0.015 | Young |
| 0.190366 | 3.964874 | 0.015 | 0 | 1 | 12 | Gpm6b | 0.015 | Young |
| 0.190366 | 3.937166 | 0.015 | 0 | 1 | 12 | Rflnb | 0.015 | Young |
| 0.190366 | 3.872021 | 0.015 | 0 | 1 | 12 | Immp2l | 0.015 | Young |
| 0.190366 | 3.757623 | 0.015 | 0 | 1 | 12 | Men1 | 0.015 | Young |
| 0.190471 | 0.879332 | 0.073 | 0.034 | 1 | 12 | Slc35b1 | 0.039 | Young |
| 0.191577 | 0.771801 | 0.117 | 0.068 | 1 | 12 | Tmeff2 | 0.049 | Young |
| 0.191862 | 0.37688 | 0.496 | 0.424 | 1 | 12 | Ndufs5 | 0.072 | Young |
| 0.192433 | 0.86812 | 0.095 | 0.051 | 1 | 12 | Jagn1 | 0.044 | Young |
| 0.192433 | 0.779794 | 0.095 | 0.051 | 1 | 12 | Rabggtb | 0.044 | Young |
| 0.193166 | 1.381801 | 0.058 | 0.025 | 1 | 12 | Cttn | 0.033 | Young |
| 0.193409 | 0.705522 | 0.073 | 0.034 | 1 | 12 | Smap2 | 0.039 | Young |
| 0.19429 | 0.587885 | 0.168 | 0.11 | 1 | 12 | Tpm1 | 0.058 | Young |
| 0.194827 | 1.490497 | 0.058 | 0.025 | 1 | 12 | Synj2bp | 0.033 | Young |
| 0.194827 | 1.306183 | 0.058 | 0.025 | 1 | 12 | Grsf1 | 0.033 | Young |
| 0.194827 | 1.287748 | 0.058 | 0.025 | 1 | 12 | Nap1l3 | 0.033 | Young |
| 0.194827 | 1.265067 | 0.058 | 0.025 | 1 | 12 | Cmc2 | 0.033 | Young |
| 0.194827 | 1.054359 | 0.058 | 0.025 | 1 | 12 | Rab27b | 0.033 | Young |
| 0.194842 | 0.909407 | 0.08 | 0.042 | 1 | 12 | Fxr1 | 0.038 | Young |
| 0.195163 | 0.710546 | 0.124 | 0.076 | 1 | 12 | Pde6d | 0.048 | Young |
| 0.195541 | 1.311839 | 0.08 | 0.042 | 1 | 12 | Eif3j1 | 0.038 | Young |
| 0.195722 | 0.781508 | 0.124 | 0.076 | 1 | 12 | Tsc22d1 | 0.048 | Young |
| 0.195814 | 0.482935 | 0.248 | 0.178 | 1 | 12 | Snrpd2 | 0.07 | Young |
| 0.196393 | 0.543683 | 0.27 | 0.212 | 1 | 12 | Atp6v1b2 | 0.058 | Young |
| 0.198179 | 1.164054 | 0.058 | 0.025 | 1 | 12 | Thumpd1 | 0.033 | Young |
| 0.199061 | 1.201984 | 0.08 | 0.042 | 1 | 12 | Palmd | 0.038 | Young |
| 0.199871 | 1.126944 | 0.058 | 0.025 | 1 | 12 | Bpgm | 0.033 | Young |
| 0.201573 | 0.986967 | 0.058 | 0.025 | 1 | 12 | Gaa | 0.033 | Young |

| | | | | | | | | |
|---|---|---|---|---|---|---|---|---|
| 0.201573 | 1.14113 | 0.058 | 0.025 | 1 | 12 | Cyb561 | 0.033 | Young |
| 0.201573 | 0.979074 | 0.058 | 0.025 | 1 | 12 | Vmp1 | 0.033 | Young |
| 0.201717 | 0.704943 | 0.212 | 0.153 | 1 | 12 | Eif5 | 0.059 | Young |
| 0.202428 | 1.094292 | 0.058 | 0.025 | 1 | 12 | Hacd3 | 0.033 | Young |
| 0.202842 | 0.730486 | 0.095 | 0.051 | 1 | 12 | Svbp | 0.044 | Young |
| 0.203286 | 1.064277 | 0.058 | 0.025 | 1 | 12 | Ddx46 | 0.033 | Young |
| 0.203286 | 0.895125 | 0.058 | 0.025 | 1 | 12 | Polr2h | 0.033 | Young |
| 0.204836 | 0.867339 | 0.124 | 0.076 | 1 | 12 | Hadhb | 0.048 | Young |
| 0.204914 | 0.745702 | 0.168 | 0.119 | 1 | 12 | Hprt | 0.049 | Young |
| 0.206666 | 0.419663 | 0.197 | 0.127 | 1 | 12 | Stx7 | 0.07 | Young |
| 0.206744 | 0.962889 | 0.058 | 0.025 | 1 | 12 | Tctex1d2 | 0.033 | Young |
| 0.206744 | 0.875241 | 0.058 | 0.025 | 1 | 12 | Asna1 | 0.033 | Young |
| 0.206967 | 1.112293 | 0.08 | 0.042 | 1 | 12 | Rbm17 | 0.038 | Young |
| 0.207006 | 0.267528 | 0.701 | 0.669 | 1 | 12 | Hsp90aa1 | 0.032 | Young |
| 0.207135 | 0.278745 | 0.321 | 0.246 | 1 | 12 | Rack1 | 0.075 | Young |
| 0.207696 | 1.007029 | 0.08 | 0.042 | 1 | 12 | B3gat3 | 0.038 | Young |
| 0.208 | 1.025679 | 0.102 | 0.059 | 1 | 12 | 5-Mar | 0.043 | Young |
| 0.208079 | 0.404294 | 0.182 | 0.119 | 1 | 12 | Zranb2 | 0.063 | Young |
| 0.208091 | 0.636687 | 0.19 | 0.136 | 1 | 12 | Cct4 | 0.054 | Young |
| 0.208489 | 0.952018 | 0.058 | 0.025 | 1 | 12 | Ccdc28b | 0.033 | Young |
| 0.210746 | 0.294277 | 0.686 | 0.585 | 1 | 12 | Ywhah | 0.101 | Young |
| 0.211675 | 1.761132 | 0.044 | 0.017 | 1 | 12 | Ik | 0.027 | Young |
| 0.212115 | 0.989333 | 0.08 | 0.042 | 1 | 12 | Adipor1 | 0.038 | Young |
| 0.212858 | 0.963044 | 0.08 | 0.042 | 1 | 12 | Fsd1 | 0.038 | Young |
| 0.213384 | 0.419178 | 0.226 | 0.161 | 1 | 12 | Rsrp1 | 0.065 | Young |
| 0.213744 | 1.56261 | 0.044 | 0.017 | 1 | 12 | Wdr75 | 0.027 | Young |
| 0.213787 | 0.739103 | 0.058 | 0.025 | 1 | 12 | Ppil1 | 0.033 | Young |
| 0.21435 | 0.872808 | 0.08 | 0.042 | 1 | 12 | Cntnap5a | 0.038 | Young |
| 0.21486 | 0.131997 | 0.664 | 0.602 | 1 | 12 | Rps3a1 | 0.062 | Young |
| 0.215574 | 0.782129 | 0.058 | 0.025 | 1 | 12 | Dusp8 | 0.033 | Young |
| 0.215829 | 2.003964 | 0.044 | 0.017 | 1 | 12 | Pmpca | 0.027 | Young |
| 0.215829 | 1.545357 | 0.044 | 0.017 | 1 | 12 | Epb41l3 | 0.027 | Young |
| 0.216289 | 0.669527 | 0.153 | 0.102 | 1 | 12 | Rap1b | 0.051 | Young |
| 0.217927 | 1.434594 | 0.044 | 0.017 | 1 | 12 | Dync1i1 | 0.027 | Young |
| 0.217927 | 1.448971 | 0.044 | 0.017 | 1 | 12 | Cdc34 | 0.027 | Young |
| 0.217927 | 1.433621 | 0.044 | 0.017 | 1 | 12 | Gm9885 | 0.027 | Young |
| 0.219567 | 0.358187 | 0.234 | 0.169 | 1 | 12 | Srp9 | 0.065 | Young |
| 0.220041 | 1.726791 | 0.044 | 0.017 | 1 | 12 | Ostf1 | 0.027 | Young |
| 0.220041 | 1.568352 | 0.044 | 0.017 | 1 | 12 | Polr2l | 0.027 | Young |
| 0.220041 | 1.387769 | 0.044 | 0.017 | 1 | 12 | Gpr22 | 0.027 | Young |
| 0.220041 | 1.330412 | 0.044 | 0.017 | 1 | 12 | Wtip | 0.027 | Young |
| 0.220244 | 0.620081 | 0.19 | 0.136 | 1 | 12 | Ddx5 | 0.054 | Young |
| 0.220391 | 0.862212 | 0.08 | 0.042 | 1 | 12 | Chn1 | 0.038 | Young |
| 0.220391 | 0.819343 | 0.08 | 0.042 | 1 | 12 | Dnajc5 | 0.038 | Young |
| 0.220554 | 0.54432 | 0.212 | 0.153 | 1 | 12 | Tesc | 0.059 | Young |

| | | | | | | | | |
|---|---|---|---|---|---|---|---|---|
| 0.221941 | 0.609737 | 0.182 | 0.127 | 1 | 12 | Celf4 | 0.055 | Young |
| 0.222168 | 1.261796 | 0.044 | 0.017 | 1 | 12 | Rtca | 0.027 | Young |
| 0.222169 | 1.429378 | 0.044 | 0.017 | 1 | 12 | Zfp830 | 0.027 | Young |
| 0.222169 | 1.329675 | 0.044 | 0.017 | 1 | 12 | Dazap2 | 0.027 | Young |
| 0.222169 | 1.32449 | 0.044 | 0.017 | 1 | 12 | Zfp451 | 0.027 | Young |
| 0.222169 | 1.313657 | 0.044 | 0.017 | 1 | 12 | Zdhhc3 | 0.027 | Young |
| 0.222169 | 1.313465 | 0.044 | 0.017 | 1 | 12 | Mfn2 | 0.027 | Young |
| 0.222169 | 1.30438 | 0.044 | 0.017 | 1 | 12 | Bop1 | 0.027 | Young |
| 0.222169 | 1.289542 | 0.044 | 0.017 | 1 | 12 | Cebpg | 0.027 | Young |
| 0.222169 | 1.249776 | 0.044 | 0.017 | 1 | 12 | Tmco3 | 0.027 | Young |
| 0.223457 | 0.799404 | 0.08 | 0.042 | 1 | 12 | Rbfa | 0.038 | Young |
| 0.224312 | 1.346546 | 0.044 | 0.017 | 1 | 12 | Ssx2ip | 0.027 | Young |
| 0.224312 | 1.286407 | 0.044 | 0.017 | 1 | 12 | Tmem5 | 0.027 | Young |
| 0.224312 | 1.224626 | 0.044 | 0.017 | 1 | 12 | Mef2a | 0.027 | Young |
| 0.22647 | 1.407654 | 0.044 | 0.017 | 1 | 12 | Fip1l1 | 0.027 | Young |
| 0.22647 | 1.201817 | 0.044 | 0.017 | 1 | 12 | Vps26b | 0.027 | Young |
| 0.22647 | 1.138739 | 0.044 | 0.017 | 1 | 12 | Zfp386 | 0.027 | Young |
| 0.228112 | 0.753958 | 0.08 | 0.042 | 1 | 12 | Akap12 | 0.038 | Young |
| 0.228588 | 1.205876 | 0.066 | 0.034 | 1 | 12 | Rps19bp1 | 0.032 | Young |
| 0.228642 | 1.2281 | 0.044 | 0.017 | 1 | 12 | Mospd3 | 0.027 | Young |
| 0.22942 | 0.434648 | 0.35 | 0.28 | 1 | 12 | Rpl31 | 0.07 | Young |
| 0.230319 | 1.307058 | 0.066 | 0.034 | 1 | 12 | Ncs1 | 0.032 | Young |
| 0.230817 | 4.256386 | 0.029 | 0.008 | 1 | 12 | Ube2z | 0.021 | Young |
| 0.230817 | 4.047607 | 0.029 | 0.008 | 1 | 12 | Gm40841 | 0.021 | Young |
| 0.230817 | 3.986394 | 0.029 | 0.008 | 1 | 12 | Cers6 | 0.021 | Young |
| 0.230817 | 3.891048 | 0.029 | 0.008 | 1 | 12 | Slc7a8 | 0.021 | Young |
| 0.230817 | 3.787064 | 0.029 | 0.008 | 1 | 12 | Zfp36l1 | 0.021 | Young |
| 0.230817 | 3.296562 | 0.029 | 0.008 | 1 | 12 | Cln8 | 0.021 | Young |
| 0.230817 | 2.440803 | 0.029 | 0.008 | 1 | 12 | Zeb1 | 0.021 | Young |
| 0.230817 | 2.404146 | 0.029 | 0.008 | 1 | 12 | Nfyb | 0.021 | Young |
| 0.230817 | 2.327985 | 0.029 | 0.008 | 1 | 12 | Polr3gl | 0.021 | Young |
| 0.230817 | 2.283948 | 0.029 | 0.008 | 1 | 12 | Fgfr1 | 0.021 | Young |
| 0.230817 | 2.256156 | 0.029 | 0.008 | 1 | 12 | Atg7 | 0.021 | Young |
| 0.230817 | 2.211846 | 0.029 | 0.008 | 1 | 12 | Myo5a | 0.021 | Young |
| 0.230817 | 2.14057 | 0.029 | 0.008 | 1 | 12 | Zfp266 | 0.021 | Young |
| 0.230817 | 2.128564 | 0.029 | 0.008 | 1 | 12 | Ypel1 | 0.021 | Young |
| 0.230817 | 2.124781 | 0.029 | 0.008 | 1 | 12 | Ankrd6 | 0.021 | Young |
| 0.230817 | 2.122369 | 0.029 | 0.008 | 1 | 12 | Rom1 | 0.021 | Young |
| 0.230817 | 2.118669 | 0.029 | 0.008 | 1 | 12 | Pip5k1c | 0.021 | Young |
| 0.230817 | 2.11145 | 0.029 | 0.008 | 1 | 12 | Otud7b | 0.021 | Young |
| 0.230817 | 1.863442 | 0.029 | 0.008 | 1 | 12 | Scg2 | 0.021 | Young |
| 0.230817 | 1.861834 | 0.029 | 0.008 | 1 | 12 | Trappc9 | 0.021 | Young |
| 0.230817 | 1.823321 | 0.029 | 0.008 | 1 | 12 | Impdh2 | 0.021 | Young |
| 0.230829 | 1.041291 | 0.044 | 0.017 | 1 | 12 | Rpf2 | 0.027 | Young |
| 0.231927 | 0.971974 | 0.044 | 0.017 | 1 | 12 | Ap2b1 | 0.027 | Young |

| | | | | | | | | |
|---|---|---|---|---|---|---|---|---|
| 0.23206 | 1.534475 | 0.066 | 0.034 | 1 | 12 | Ubald2 | 0.032 | Young |
| 0.232522 | 0.237469 | 0.153 | 0.093 | 1 | 12 | Hmgcr | 0.06 | Young |
| 0.233001 | 0.47273 | 0.153 | 0.102 | 1 | 12 | Mrpl14 | 0.051 | Young |
| 0.23303 | 1.022341 | 0.044 | 0.017 | 1 | 12 | 2510002D2 | 0.027 | Young |
| 0.233588 | 2.113304 | 0.029 | 0.008 | 1 | 12 | Adora2a | 0.021 | Young |
| 0.233588 | 2.071446 | 0.029 | 0.008 | 1 | 12 | Skiv2l2 | 0.021 | Young |
| 0.233588 | 2.062832 | 0.029 | 0.008 | 1 | 12 | Wbp1 | 0.021 | Young |
| 0.233588 | 2.059171 | 0.029 | 0.008 | 1 | 12 | Ubr4 | 0.021 | Young |
| 0.233588 | 2.027671 | 0.029 | 0.008 | 1 | 12 | Lrrc3 | 0.021 | Young |
| 0.233588 | 2.000812 | 0.029 | 0.008 | 1 | 12 | Srd5a1 | 0.021 | Young |
| 0.233588 | 1.960219 | 0.029 | 0.008 | 1 | 12 | Prps2 | 0.021 | Young |
| 0.233588 | 1.941941 | 0.029 | 0.008 | 1 | 12 | Thap12 | 0.021 | Young |
| 0.233588 | 1.928022 | 0.029 | 0.008 | 1 | 12 | Setd5 | 0.021 | Young |
| 0.233588 | 1.926032 | 0.029 | 0.008 | 1 | 12 | Aifm1 | 0.021 | Young |
| 0.233588 | 1.839315 | 0.029 | 0.008 | 1 | 12 | Gpsm3 | 0.021 | Young |
| 0.233588 | 1.810148 | 0.029 | 0.008 | 1 | 12 | Rcan2 | 0.021 | Young |
| 0.233588 | 1.757387 | 0.029 | 0.008 | 1 | 12 | Coq3 | 0.021 | Young |
| 0.234403 | 0.65529 | 0.131 | 0.085 | 1 | 12 | Nek1 | 0.046 | Young |
| 0.235248 | 0.926717 | 0.044 | 0.017 | 1 | 12 | Dnajc3 | 0.027 | Young |
| 0.235391 | 0.3508 | 0.387 | 0.322 | 1 | 12 | Hsbp1 | 0.065 | Young |
| 0.236383 | 1.945672 | 0.029 | 0.008 | 1 | 12 | Cdkn2aipnl | 0.021 | Young |
| 0.236383 | 1.940482 | 0.029 | 0.008 | 1 | 12 | Dhrs13 | 0.021 | Young |
| 0.236383 | 1.92415 | 0.029 | 0.008 | 1 | 12 | Ralgapa1 | 0.021 | Young |
| 0.236383 | 1.863625 | 0.029 | 0.008 | 1 | 12 | Leo1 | 0.021 | Young |
| 0.236383 | 1.850573 | 0.029 | 0.008 | 1 | 12 | Zfand2a | 0.021 | Young |
| 0.236383 | 1.849403 | 0.029 | 0.008 | 1 | 12 | Phf6 | 0.021 | Young |
| 0.236383 | 1.826332 | 0.029 | 0.008 | 1 | 12 | Acot8 | 0.021 | Young |
| 0.236383 | 1.816991 | 0.029 | 0.008 | 1 | 12 | Tet3 | 0.021 | Young |
| 0.236383 | 1.798501 | 0.029 | 0.008 | 1 | 12 | Ptdss2 | 0.021 | Young |
| 0.236383 | 1.759081 | 0.029 | 0.008 | 1 | 12 | Slc31a2 | 0.021 | Young |
| 0.236383 | 1.745695 | 0.029 | 0.008 | 1 | 12 | Tcp11l2 | 0.021 | Young |
| 0.236383 | 1.709838 | 0.029 | 0.008 | 1 | 12 | Bmp3 | 0.021 | Young |
| 0.236383 | 1.703744 | 0.029 | 0.008 | 1 | 12 | Lrrc24 | 0.021 | Young |
| 0.236383 | 1.699558 | 0.029 | 0.008 | 1 | 12 | Phkg2 | 0.021 | Young |
| 0.236383 | 1.69107 | 0.029 | 0.008 | 1 | 12 | Paox | 0.021 | Young |
| 0.236383 | 1.685285 | 0.029 | 0.008 | 1 | 12 | Nbr1 | 0.021 | Young |
| 0.236383 | 1.679555 | 0.029 | 0.008 | 1 | 12 | Pcyt1a | 0.021 | Young |
| 0.236383 | 1.672473 | 0.029 | 0.008 | 1 | 12 | Rmnd5a | 0.021 | Young |
| 0.236383 | 1.653726 | 0.029 | 0.008 | 1 | 12 | Tram1 | 0.021 | Young |
| 0.236383 | 1.629915 | 0.029 | 0.008 | 1 | 12 | Spsb2 | 0.021 | Young |
| 0.236383 | 1.627568 | 0.029 | 0.008 | 1 | 12 | Orc2 | 0.021 | Young |
| 0.236383 | 1.596049 | 0.029 | 0.008 | 1 | 12 | Gpatch2l | 0.021 | Young |
| 0.236383 | 1.575831 | 0.029 | 0.008 | 1 | 12 | Igsf3 | 0.021 | Young |
| 0.236383 | 1.53426 | 0.029 | 0.008 | 1 | 12 | Tbce | 0.021 | Young |
| 0.236383 | 1.477366 | 0.029 | 0.008 | 1 | 12 | St6galnac4 | 0.021 | Young |

| | | | | | | | | |
|---|---|---|---|---|---|---|---|---|
| 0.237784 | 0.817641 | 0.109 | 0.068 | 1 | 12 | Lhfpl5 | 0.041 | Young |
| 0.239201 | 1.690274 | 0.029 | 0.008 | 1 | 12 | Map3k2 | 0.021 | Young |
| 0.239201 | 1.641918 | 0.029 | 0.008 | 1 | 12 | Mum1 | 0.021 | Young |
| 0.239201 | 1.603155 | 0.029 | 0.008 | 1 | 12 | Polr3h | 0.021 | Young |
| 0.239201 | 1.601888 | 0.029 | 0.008 | 1 | 12 | Hook3 | 0.021 | Young |
| 0.239201 | 1.592197 | 0.029 | 0.008 | 1 | 12 | Evi5l | 0.021 | Young |
| 0.239201 | 1.579441 | 0.029 | 0.008 | 1 | 12 | Tcf3 | 0.021 | Young |
| 0.239201 | 1.565206 | 0.029 | 0.008 | 1 | 12 | AC154509. | 0.021 | Young |
| 0.239201 | 1.561422 | 0.029 | 0.008 | 1 | 12 | Adcy1 | 0.021 | Young |
| 0.239201 | 1.545688 | 0.029 | 0.008 | 1 | 12 | Lrrc73 | 0.021 | Young |
| 0.239201 | 1.541451 | 0.029 | 0.008 | 1 | 12 | D030056L2 | 0.021 | Young |
| 0.239201 | 1.510126 | 0.029 | 0.008 | 1 | 12 | Clip1 | 0.021 | Young |
| 0.239201 | 1.490759 | 0.029 | 0.008 | 1 | 12 | Rnf220 | 0.021 | Young |
| 0.239201 | 1.485179 | 0.029 | 0.008 | 1 | 12 | Os9 | 0.021 | Young |
| 0.239201 | 1.484606 | 0.029 | 0.008 | 1 | 12 | Tmem121 | 0.021 | Young |
| 0.239201 | 1.481517 | 0.029 | 0.008 | 1 | 12 | Zfp593 | 0.021 | Young |
| 0.239201 | 1.479851 | 0.029 | 0.008 | 1 | 12 | Creb3 | 0.021 | Young |
| 0.239201 | 1.4452 | 0.029 | 0.008 | 1 | 12 | Trim16 | 0.021 | Young |
| 0.239201 | 1.443264 | 0.029 | 0.008 | 1 | 12 | Atpaf1 | 0.021 | Young |
| 0.239201 | 1.399386 | 0.029 | 0.008 | 1 | 12 | Cabp1 | 0.021 | Young |
| 0.239201 | 1.39897 | 0.029 | 0.008 | 1 | 12 | Ppp4c | 0.021 | Young |
| 0.239201 | 1.3814 | 0.029 | 0.008 | 1 | 12 | Sh3gl1 | 0.021 | Young |
| 0.239201 | 1.324061 | 0.029 | 0.008 | 1 | 12 | Ift52 | 0.021 | Young |
| 0.239201 | 1.17991 | 0.029 | 0.008 | 1 | 12 | Uap1l1 | 0.021 | Young |
| 0.240521 | 0.45952 | 0.153 | 0.102 | 1 | 12 | Mrps16 | 0.051 | Young |
| 0.240958 | 0.166137 | 0.533 | 0.449 | 1 | 12 | Pgam1 | 0.084 | Young |
| 0.240994 | 0.376345 | 0.394 | 0.347 | 1 | 12 | Psma3 | 0.047 | Young |
| 0.241253 | 0.581528 | 0.175 | 0.127 | 1 | 12 | Lamtor5 | 0.048 | Young |
| 0.242043 | 1.554274 | 0.029 | 0.008 | 1 | 12 | Pard3 | 0.021 | Young |
| 0.242043 | 1.529277 | 0.029 | 0.008 | 1 | 12 | Bloc1s6 | 0.021 | Young |
| 0.242043 | 1.481231 | 0.029 | 0.008 | 1 | 12 | Gpc1 | 0.021 | Young |
| 0.242043 | 1.476798 | 0.029 | 0.008 | 1 | 12 | Dcun1d4 | 0.021 | Young |
| 0.242043 | 1.450495 | 0.029 | 0.008 | 1 | 12 | Smc2 | 0.021 | Young |
| 0.242043 | 1.438754 | 0.029 | 0.008 | 1 | 12 | Cpne5 | 0.021 | Young |
| 0.242043 | 1.389401 | 0.029 | 0.008 | 1 | 12 | Trpc6 | 0.021 | Young |
| 0.242043 | 1.383006 | 0.029 | 0.008 | 1 | 12 | Fmr1 | 0.021 | Young |
| 0.242043 | 1.382334 | 0.029 | 0.008 | 1 | 12 | Wdr36 | 0.021 | Young |
| 0.242043 | 1.368302 | 0.029 | 0.008 | 1 | 12 | Ppp6r3 | 0.021 | Young |
| 0.242043 | 1.358395 | 0.029 | 0.008 | 1 | 12 | Ncbp1 | 0.021 | Young |
| 0.242043 | 1.352961 | 0.029 | 0.008 | 1 | 12 | Clybl | 0.021 | Young |
| 0.242043 | 1.351608 | 0.029 | 0.008 | 1 | 12 | 1700020I1 | 0.021 | Young |
| 0.242043 | 1.349395 | 0.029 | 0.008 | 1 | 12 | Lrrc45 | 0.021 | Young |
| 0.242043 | 1.303549 | 0.029 | 0.008 | 1 | 12 | Abhd10 | 0.021 | Young |
| 0.242043 | 1.291882 | 0.029 | 0.008 | 1 | 12 | Prps1 | 0.021 | Young |
| 0.242043 | 1.247568 | 0.029 | 0.008 | 1 | 12 | Atp23 | 0.021 | Young |

| | | | | | | | | |
|---|---|---|---|---|---|---|---|---|
| 0.242043 | 1.237063 | 0.029 | 0.008 | 1 | 12 | Rprd2 | 0.021 | Young |
| 0.242043 | 1.231641 | 0.029 | 0.008 | 1 | 12 | Ssr3 | 0.021 | Young |
| 0.242043 | 1.220217 | 0.029 | 0.008 | 1 | 12 | Rnf138 | 0.021 | Young |
| 0.242043 | 1.218643 | 0.029 | 0.008 | 1 | 12 | Nit1 | 0.021 | Young |
| 0.242043 | 1.218501 | 0.029 | 0.008 | 1 | 12 | Svop | 0.021 | Young |
| 0.242043 | 1.125077 | 0.029 | 0.008 | 1 | 12 | Elavl1 | 0.021 | Young |
| 0.242043 | 1.122993 | 0.029 | 0.008 | 1 | 12 | Zhx1 | 0.021 | Young |
| 0.242043 | 1.082494 | 0.029 | 0.008 | 1 | 12 | Pbx2 | 0.021 | Young |
| 0.242043 | 1.078237 | 0.029 | 0.008 | 1 | 12 | Mcoln1 | 0.021 | Young |
| 0.242043 | 1.050225 | 0.029 | 0.008 | 1 | 12 | Pstk | 0.021 | Young |
| 0.242043 | 0.818553 | 0.029 | 0.008 | 1 | 12 | Gxylt2 | 0.021 | Young |
| 0.242598 | 0.3597 | 0.146 | 0.093 | 1 | 12 | Lamtor1 | 0.053 | Young |
| 0.242641 | 0.459024 | 0.234 | 0.169 | 1 | 12 | Cend1 | 0.065 | Young |
| 0.242869 | 0.606824 | 0.153 | 0.102 | 1 | 12 | Idi1 | 0.051 | Young |
| 0.243313 | 0.295547 | 0.46 | 0.398 | 1 | 12 | Srp14 | 0.062 | Young |
| 0.243791 | 0.428454 | 0.131 | 0.085 | 1 | 12 | Tcp1 | 0.046 | Young |
| 0.24381 | 0.517285 | 0.146 | 0.093 | 1 | 12 | Shisa4 | 0.053 | Young |
| 0.244426 | 0.7316 | 0.131 | 0.085 | 1 | 12 | Chmp2b | 0.046 | Young |
| 0.244507 | 1.03789 | 0.066 | 0.034 | 1 | 12 | Ankra2 | 0.032 | Young |
| 0.244657 | 0.737397 | 0.168 | 0.119 | 1 | 12 | Ncl | 0.049 | Young |
| 0.246171 | 0.278913 | 0.285 | 0.22 | 1 | 12 | Ap2s1 | 0.065 | Young |
| 0.246322 | 0.98326 | 0.066 | 0.034 | 1 | 12 | Nrd1 | 0.032 | Young |
| 0.246322 | 0.979125 | 0.066 | 0.034 | 1 | 12 | Wdr1 | 0.032 | Young |
| 0.247156 | 0.49981 | 0.102 | 0.059 | 1 | 12 | Casc4 | 0.043 | Young |
| 0.247596 | 0.816067 | 0.088 | 0.051 | 1 | 12 | Tulp4 | 0.037 | Young |
| 0.248147 | 1.100453 | 0.066 | 0.034 | 1 | 12 | Copb2 | 0.032 | Young |
| 0.248147 | 0.948864 | 0.066 | 0.034 | 1 | 12 | Atp6v1h | 0.032 | Young |
| 0.248905 | 0.607963 | 0.131 | 0.085 | 1 | 12 | Slc25a39 | 0.046 | Young |
| 0.249561 | 0.768087 | 0.109 | 0.068 | 1 | 12 | Sssca1 | 0.041 | Young |
| 0.249981 | 0.974446 | 0.066 | 0.034 | 1 | 12 | Gng4 | 0.032 | Young |
| 0.249982 | 0.671231 | 0.066 | 0.034 | 1 | 12 | Serpine2 | 0.032 | Young |
| 0.250267 | 0.582268 | 0.109 | 0.068 | 1 | 12 | Spag7 | 0.041 | Young |
| 0.251826 | 1.015984 | 0.066 | 0.034 | 1 | 12 | Dnajb11 | 0.032 | Young |
| 0.251826 | 0.88206 | 0.066 | 0.034 | 1 | 12 | Rab15 | 0.032 | Young |
| 0.251895 | 0.512702 | 0.161 | 0.11 | 1 | 12 | Fkbp8 | 0.051 | Young |
| 0.252332 | 0.96329 | 0.088 | 0.051 | 1 | 12 | Hdlbp | 0.037 | Young |
| 0.252661 | 0.268575 | 0.299 | 0.237 | 1 | 12 | Sptbn1 | 0.062 | Young |
| 0.252894 | 0.313587 | 0.387 | 0.339 | 1 | 12 | Anxa5 | 0.048 | Young |
| 0.254204 | 0.349525 | 0.19 | 0.136 | 1 | 12 | Tusc3 | 0.054 | Young |
| 0.254527 | 0.682507 | 0.109 | 0.068 | 1 | 12 | Mrpl23 | 0.041 | Young |
| 0.255119 | 0.493678 | 0.197 | 0.144 | 1 | 12 | Psmc6 | 0.053 | Young |
| 0.255542 | 0.888634 | 0.066 | 0.034 | 1 | 12 | Eif2s1 | 0.032 | Young |
| 0.25626 | 0.824483 | 0.117 | 0.076 | 1 | 12 | Immt | 0.041 | Young |
| 0.256496 | 0.694475 | 0.146 | 0.102 | 1 | 12 | Actr3 | 0.044 | Young |
| 0.257079 | 0.426147 | 0.124 | 0.076 | 1 | 12 | Rp9 | 0.048 | Young |

| | | | | | | | | |
|---|---|---|---|---|---|---|---|---|
| 0.257414 | 1.043833 | 0.066 | 0.034 | 1 | 12 | Zfand6 | 0.032 | Young |
| 0.257739 | 0.716081 | 0.146 | 0.102 | 1 | 12 | Gars | 0.044 | Young |
| 0.258827 | 0.369986 | 0.307 | 0.254 | 1 | 12 | Rps15 | 0.053 | Young |
| 0.259035 | 0.681241 | 0.117 | 0.076 | 1 | 12 | Lims1 | 0.041 | Young |
| 0.259089 | 0.596042 | 0.234 | 0.178 | 1 | 12 | Ssb | 0.056 | Young |
| 0.259296 | 0.959514 | 0.066 | 0.034 | 1 | 12 | Aamdc | 0.032 | Young |
| 0.26021 | 0.613843 | 0.168 | 0.119 | 1 | 12 | Hnrnpa0 | 0.049 | Young |
| 0.260218 | 0.459742 | 0.182 | 0.127 | 1 | 12 | Syp | 0.055 | Young |
| 0.260365 | 0.741117 | 0.088 | 0.051 | 1 | 12 | Hes6 | 0.037 | Young |
| 0.262393 | 0.422435 | 0.248 | 0.195 | 1 | 12 | Uqcrc2 | 0.053 | Young |
| 0.263089 | 0.799421 | 0.066 | 0.034 | 1 | 12 | 6-Sep | 0.032 | Young |
| 0.263089 | 0.755816 | 0.066 | 0.034 | 1 | 12 | Srebf2 | 0.032 | Young |
| 0.263208 | 0.511549 | 0.175 | 0.127 | 1 | 12 | Ncam1 | 0.048 | Young |
| 0.266063 | 0.775227 | 0.117 | 0.076 | 1 | 12 | Ndufa9 | 0.041 | Young |
| 0.267625 | 0.163428 | 0.263 | 0.195 | 1 | 12 | Cyc1 | 0.068 | Young |
| 0.268495 | 0.599927 | 0.153 | 0.102 | 1 | 12 | Mrpl30 | 0.051 | Young |
| 0.268911 | 0.69892 | 0.117 | 0.076 | 1 | 12 | Bax | 0.041 | Young |
| 0.27246 | 0.329104 | 0.394 | 0.339 | 1 | 12 | Ywhaq | 0.055 | Young |
| 0.272736 | 0.580716 | 0.066 | 0.034 | 1 | 12 | Elovl4 | 0.032 | Young |
| 0.272736 | 0.53272 | 0.066 | 0.034 | 1 | 12 | Laptm4b | 0.032 | Young |
| 0.273435 | 0.165917 | 0.482 | 0.398 | 1 | 12 | Gabarap | 0.084 | Young |
| 0.273624 | 0.45735 | 0.204 | 0.153 | 1 | 12 | Klhdc2 | 0.051 | Young |
| 0.274694 | 0.630686 | 0.066 | 0.034 | 1 | 12 | Commd9 | 0.032 | Young |
| 0.276791 | 0.40384 | 0.153 | 0.102 | 1 | 12 | Oaz2 | 0.051 | Young |
| 0.277178 | 0.748024 | 0.095 | 0.059 | 1 | 12 | Nsmce3 | 0.036 | Young |
| 0.278607 | 0.337116 | 0.219 | 0.161 | 1 | 12 | Timm17a | 0.058 | Young |
| 0.278655 | 0.716362 | 0.088 | 0.051 | 1 | 12 | Ric8a | 0.037 | Young |
| 0.279192 | 0.451299 | 0.175 | 0.127 | 1 | 12 | Eif4a2 | 0.048 | Young |
| 0.279675 | 1.568344 | 0.051 | 0.025 | 1 | 12 | Plekhj1 | 0.026 | Young |
| 0.28036 | 0.710101 | 0.088 | 0.051 | 1 | 12 | Ubac1 | 0.037 | Young |
| 0.28042 | 0.529514 | 0.095 | 0.059 | 1 | 12 | Actr1a | 0.036 | Young |
| 0.283296 | 0.708026 | 0.124 | 0.085 | 1 | 12 | Ppig | 0.039 | Young |
| 0.283791 | 0.540582 | 0.088 | 0.051 | 1 | 12 | L1cam | 0.037 | Young |
| 0.284204 | 1.432316 | 0.051 | 0.025 | 1 | 12 | Sdhc | 0.026 | Young |
| 0.284204 | 1.2995 | 0.051 | 0.025 | 1 | 12 | Sbds | 0.026 | Young |
| 0.284782 | 0.282436 | 0.285 | 0.22 | 1 | 12 | Ndufa2 | 0.065 | Young |
| 0.284947 | 0.71251 | 0.117 | 0.076 | 1 | 12 | Ndfip2 | 0.041 | Young |
| 0.285776 | 0.49794 | 0.212 | 0.161 | 1 | 12 | Cirbp | 0.051 | Young |
| 0.28646 | 0.523385 | 0.394 | 0.339 | 1 | 12 | Pafah1b1 | 0.055 | Young |
| 0.286476 | 0.336744 | 0.255 | 0.186 | 1 | 12 | Eif3f | 0.069 | Young |
| 0.286487 | 1.094635 | 0.051 | 0.025 | 1 | 12 | Erh | 0.026 | Young |
| 0.28728 | 0.576783 | 0.263 | 0.212 | 1 | 12 | Hmgb1 | 0.051 | Young |
| 0.288859 | 0.855375 | 0.073 | 0.042 | 1 | 12 | Fbxl2 | 0.031 | Young |
| 0.290788 | 0.448684 | 0.168 | 0.119 | 1 | 12 | Csnk1e | 0.049 | Young |
| 0.29109 | 1.088375 | 0.051 | 0.025 | 1 | 12 | Slc50a1 | 0.026 | Young |

| | | | | | | | | |
|---|---|---|---|---|---|---|---|---|
| 0.29109 | 1.080715 | 0.051 | 0.025 | 1 | 12 | Slc38a2 | 0.026 | Young |
| 0.29109 | 1.039292 | 0.051 | 0.025 | 1 | 12 | Nop16 | 0.026 | Young |
| 0.29109 | 0.938671 | 0.051 | 0.025 | 1 | 12 | Csrp2 | 0.026 | Young |
| 0.29109 | 0.834025 | 0.051 | 0.025 | 1 | 12 | Ugcg | 0.026 | Young |
| 0.292672 | 1.040305 | 0.073 | 0.042 | 1 | 12 | Cebpzos | 0.031 | Young |
| 0.293569 | 0.426269 | 0.109 | 0.068 | 1 | 12 | Tmed3 | 0.041 | Young |
| 0.295279 | 0.18312 | 0.19 | 0.136 | 1 | 12 | Pam | 0.054 | Young |
| 0.295554 | 0.931692 | 0.073 | 0.042 | 1 | 12 | Spock2 | 0.031 | Young |
| 0.295743 | 0.985467 | 0.051 | 0.025 | 1 | 12 | Selenos | 0.026 | Young |
| 0.295938 | 0.376682 | 0.109 | 0.068 | 1 | 12 | Txnl4a | 0.041 | Young |
| 0.296018 | 0.499992 | 0.204 | 0.161 | 1 | 12 | Glrx3 | 0.043 | Young |
| 0.29652 | 0.867528 | 0.073 | 0.042 | 1 | 12 | Cab39 | 0.031 | Young |
| 0.296913 | 0.931223 | 0.051 | 0.025 | 1 | 12 | Rab9 | 0.026 | Young |
| 0.298088 | 1.001793 | 0.051 | 0.025 | 1 | 12 | Vps4a | 0.026 | Young |
| 0.298088 | 0.989109 | 0.051 | 0.025 | 1 | 12 | Ak6 | 0.026 | Young |
| 0.298456 | 0.692011 | 0.073 | 0.042 | 1 | 12 | Tenm2 | 0.031 | Young |
| 0.300445 | 1.023652 | 0.051 | 0.025 | 1 | 12 | Exosc9 | 0.026 | Young |
| 0.300445 | 0.995976 | 0.051 | 0.025 | 1 | 12 | Cacfd1 | 0.026 | Young |
| 0.300445 | 0.929595 | 0.051 | 0.025 | 1 | 12 | Armcx3 | 0.026 | Young |
| 0.300445 | 0.90093 | 0.051 | 0.025 | 1 | 12 | Plxna4 | 0.026 | Young |
| 0.300445 | 0.819065 | 0.051 | 0.025 | 1 | 12 | Gorasp2 | 0.026 | Young |
| 0.301362 | 0.747966 | 0.139 | 0.102 | 1 | 12 | Crmp1 | 0.037 | Young |
| 0.302815 | 0.96601 | 0.051 | 0.025 | 1 | 12 | Pet100 | 0.026 | Young |
| 0.305197 | 0.854343 | 0.051 | 0.025 | 1 | 12 | Lpgat1 | 0.026 | Young |
| 0.305197 | 0.847201 | 0.051 | 0.025 | 1 | 12 | Ppp6c | 0.026 | Young |
| 0.305197 | 0.831969 | 0.051 | 0.025 | 1 | 12 | Fam102b | 0.026 | Young |
| 0.305197 | 0.718025 | 0.051 | 0.025 | 1 | 12 | Ak1 | 0.026 | Young |
| 0.305197 | 0.352014 | 0.051 | 0.025 | 1 | 12 | Gria2 | 0.026 | Young |
| 0.305559 | 0.784741 | 0.095 | 0.059 | 1 | 12 | Srpk2 | 0.036 | Young |
| 0.307592 | 0.735308 | 0.051 | 0.025 | 1 | 12 | Pdzd11 | 0.026 | Young |
| 0.307838 | 0.497001 | 0.117 | 0.076 | 1 | 12 | Sar1b | 0.041 | Young |
| 0.309999 | 0.682757 | 0.051 | 0.025 | 1 | 12 | Zfp326 | 0.026 | Young |
| 0.310251 | 0.692646 | 0.073 | 0.042 | 1 | 12 | Tceal3 | 0.031 | Young |
| 0.31211 | 0.590944 | 0.241 | 0.195 | 1 | 12 | Rpl5 | 0.046 | Young |
| 0.312246 | 0.718601 | 0.073 | 0.042 | 1 | 12 | Rgs17 | 0.031 | Young |
| 0.312246 | 0.560001 | 0.073 | 0.042 | 1 | 12 | Ccdc32 | 0.031 | Young |
| 0.312508 | 0.662248 | 0.095 | 0.059 | 1 | 12 | Ddrgk1 | 0.036 | Young |
| 0.313724 | 0.816127 | 0.168 | 0.127 | 1 | 12 | Chl1 | 0.041 | Young |
| 0.314324 | 0.867724 | 0.08 | 0.051 | 1 | 12 | Syf2 | 0.029 | Young |
| 0.314423 | 0.803764 | 0.124 | 0.085 | 1 | 12 | Spop | 0.039 | Young |
| 0.317787 | 0.812343 | 0.139 | 0.102 | 1 | 12 | Kif5a | 0.037 | Young |
| 0.318127 | 0.917711 | 0.08 | 0.051 | 1 | 12 | Mrpl38 | 0.029 | Young |
| 0.319296 | 0.655557 | 0.073 | 0.042 | 1 | 12 | Smc1a | 0.031 | Young |
| 0.319335 | 0.408335 | 0.263 | 0.212 | 1 | 12 | Mocs2 | 0.051 | Young |
| 0.31975 | 0.16925 | 0.051 | 0.025 | 1 | 12 | Erc1 | 0.026 | Young |

| | | | | | | | | |
|---|---|---|---|---|---|---|---|---|
| 0.320089 | 0.2713 | 0.445 | 0.398 | 1 | 12 | S100a6 | 0.047 | Young |
| 0.321787 | 1.709829 | 0.058 | 0.034 | 1 | 12 | Hspb11 | 0.024 | Young |
| 0.323219 | 0.164012 | 0.715 | 0.703 | 1 | 12 | Rpsa | 0.012 | Young |
| 0.325495 | 0.728263 | 0.109 | 0.076 | 1 | 12 | Fscn1 | 0.033 | Young |
| 0.326668 | 0.533174 | 0.102 | 0.068 | 1 | 12 | Churc1 | 0.034 | Young |
| 0.326715 | 0.476852 | 0.095 | 0.059 | 1 | 12 | Maea | 0.036 | Young |
| 0.328263 | 2.470182 | 0.036 | 0.017 | 1 | 12 | Gatm | 0.019 | Young |
| 0.328263 | 2.301415 | 0.036 | 0.017 | 1 | 12 | Pabpn1 | 0.019 | Young |
| 0.328263 | 1.993556 | 0.036 | 0.017 | 1 | 12 | Cpeb4 | 0.019 | Young |
| 0.328263 | 1.62064 | 0.036 | 0.017 | 1 | 12 | Sh3gl2 | 0.019 | Young |
| 0.328263 | 1.479423 | 0.036 | 0.017 | 1 | 12 | Rab33a | 0.019 | Young |
| 0.328263 | 1.372151 | 0.036 | 0.017 | 1 | 12 | Cep19 | 0.019 | Young |
| 0.328396 | 0.61262 | 0.102 | 0.068 | 1 | 12 | Mapk1 | 0.034 | Young |
| 0.329593 | 0.476167 | 0.146 | 0.102 | 1 | 12 | Ostc | 0.044 | Young |
| 0.330936 | 1.379032 | 0.058 | 0.034 | 1 | 12 | Slc39a7 | 0.024 | Young |
| 0.331249 | 1.91417 | 0.036 | 0.017 | 1 | 12 | Trnau1ap | 0.019 | Young |
| 0.331249 | 1.680339 | 0.036 | 0.017 | 1 | 12 | Sh3bgrl | 0.019 | Young |
| 0.331249 | 1.385006 | 0.036 | 0.017 | 1 | 12 | Sorl1 | 0.019 | Young |
| 0.331674 | 0.894636 | 0.08 | 0.051 | 1 | 12 | Ncor1 | 0.029 | Young |
| 0.331817 | 0.107814 | 0.708 | 0.636 | 1 | 12 | Rpl8 | 0.072 | Young |
| 0.332108 | 0.416516 | 0.372 | 0.314 | 1 | 12 | Ttc3 | 0.058 | Young |
| 0.332993 | 0.652463 | 0.109 | 0.076 | 1 | 12 | Elavl3 | 0.033 | Young |
| 0.334252 | 1.858589 | 0.036 | 0.017 | 1 | 12 | Plekho1 | 0.019 | Young |
| 0.334252 | 1.429157 | 0.036 | 0.017 | 1 | 12 | Fstl1 | 0.019 | Young |
| 0.334252 | 1.396116 | 0.036 | 0.017 | 1 | 12 | Dnajc9 | 0.019 | Young |
| 0.334252 | 1.387994 | 0.036 | 0.017 | 1 | 12 | Fra10ac1 | 0.019 | Young |
| 0.334252 | 1.38614 | 0.036 | 0.017 | 1 | 12 | Srsf12 | 0.019 | Young |
| 0.334252 | 1.226062 | 0.036 | 0.017 | 1 | 12 | Mtmr6 | 0.019 | Young |
| 0.334252 | 1.137579 | 0.036 | 0.017 | 1 | 12 | Cfap97 | 0.019 | Young |
| 0.335804 | 0.380957 | 0.095 | 0.059 | 1 | 12 | Prpsap1 | 0.036 | Young |
| 0.337272 | 1.911844 | 0.036 | 0.017 | 1 | 12 | Siva1 | 0.019 | Young |
| 0.337272 | 1.677715 | 0.036 | 0.017 | 1 | 12 | Smpdl3a | 0.019 | Young |
| 0.337272 | 1.522211 | 0.036 | 0.017 | 1 | 12 | Hax1 | 0.019 | Young |
| 0.337272 | 1.413042 | 0.036 | 0.017 | 1 | 12 | Twistnb | 0.019 | Young |
| 0.337272 | 1.286657 | 0.036 | 0.017 | 1 | 12 | Sc5d | 0.019 | Young |
| 0.337272 | 1.279522 | 0.036 | 0.017 | 1 | 12 | Ica1l | 0.019 | Young |
| 0.337272 | 1.279387 | 0.036 | 0.017 | 1 | 12 | Erc2 | 0.019 | Young |
| 0.337272 | 1.223166 | 0.036 | 0.017 | 1 | 12 | Zfpm1 | 0.019 | Young |
| 0.337272 | 1.205416 | 0.036 | 0.017 | 1 | 12 | Mrps22 | 0.019 | Young |
| 0.337272 | 1.204181 | 0.036 | 0.017 | 1 | 12 | Zmat5 | 0.019 | Young |
| 0.337592 | 0.831193 | 0.08 | 0.051 | 1 | 12 | Akirin1 | 0.029 | Young |
| 0.337651 | 0.435883 | 0.212 | 0.161 | 1 | 12 | Camk2n2 | 0.051 | Young |
| 0.338245 | 0.435647 | 0.131 | 0.093 | 1 | 12 | Tspan13 | 0.038 | Young |
| 0.339594 | 0.503449 | 0.124 | 0.085 | 1 | 12 | Polr2f | 0.039 | Young |
| 0.340173 | 0.518074 | 0.212 | 0.161 | 1 | 12 | Cdkn2d | 0.051 | Young |

| | | | | | | | | |
|---|---|---|---|---|---|---|---|---|
| 0.340229 | 0.10983 | 0.745 | 0.695 | 1 | 12 | Gapdh | 0.05 | Young |
| 0.34031 | 1.507653 | 0.036 | 0.017 | 1 | 12 | Shd | 0.019 | Young |
| 0.34031 | 1.221115 | 0.036 | 0.017 | 1 | 12 | Psd | 0.019 | Young |
| 0.34031 | 1.1749 | 0.036 | 0.017 | 1 | 12 | Pxk | 0.019 | Young |
| 0.34031 | 1.156138 | 0.036 | 0.017 | 1 | 12 | Zfp938 | 0.019 | Young |
| 0.34031 | 1.085257 | 0.036 | 0.017 | 1 | 12 | Asb8 | 0.019 | Young |
| 0.34031 | 1.059961 | 0.036 | 0.017 | 1 | 12 | Prcc | 0.019 | Young |
| 0.34031 | 1.054172 | 0.036 | 0.017 | 1 | 12 | Mcf2l | 0.019 | Young |
| 0.340654 | 0.568098 | 0.102 | 0.068 | 1 | 12 | Ddc | 0.034 | Young |
| 0.340781 | 0.268914 | 0.431 | 0.356 | 1 | 12 | Jpt1 | 0.075 | Young |
| 0.341834 | 1.522579 | 0.036 | 0.017 | 1 | 12 | Lrpprc | 0.019 | Young |
| 0.342101 | 0.176678 | 0.62 | 0.542 | 1 | 12 | Rpl14 | 0.078 | Young |
| 0.342309 | 0.82004 | 0.109 | 0.076 | 1 | 12 | Tpr | 0.033 | Young |
| 0.342607 | 0.995241 | 0.058 | 0.034 | 1 | 12 | Snrk | 0.024 | Young |
| 0.343365 | 1.494932 | 0.036 | 0.017 | 1 | 12 | Gart | 0.019 | Young |
| 0.343365 | 1.185858 | 0.036 | 0.017 | 1 | 12 | Pole3 | 0.019 | Young |
| 0.343365 | 1.168268 | 0.036 | 0.017 | 1 | 12 | Stx18 | 0.019 | Young |
| 0.343365 | 1.088327 | 0.036 | 0.017 | 1 | 12 | Dubr | 0.019 | Young |
| 0.343365 | 1.084837 | 0.036 | 0.017 | 1 | 12 | Pcyt1b | 0.019 | Young |
| 0.343365 | 1.033239 | 0.036 | 0.017 | 1 | 12 | Ubap2 | 0.019 | Young |
| 0.343365 | 1.026348 | 0.036 | 0.017 | 1 | 12 | Krt10 | 0.019 | Young |
| 0.343365 | 1.017484 | 0.036 | 0.017 | 1 | 12 | Tprkb | 0.019 | Young |
| 0.343365 | 1.013942 | 0.036 | 0.017 | 1 | 12 | Gpr137 | 0.019 | Young |
| 0.343365 | 1.003361 | 0.036 | 0.017 | 1 | 12 | Serpini1 | 0.019 | Young |
| 0.343365 | 0.987984 | 0.036 | 0.017 | 1 | 12 | Gpr85 | 0.019 | Young |
| 0.343365 | 0.975058 | 0.036 | 0.017 | 1 | 12 | Arxes2 | 0.019 | Young |
| 0.343365 | 0.959034 | 0.036 | 0.017 | 1 | 12 | Phf20 | 0.019 | Young |
| 0.343365 | 0.958895 | 0.036 | 0.017 | 1 | 12 | Hscb | 0.019 | Young |
| 0.343578 | 0.668741 | 0.08 | 0.051 | 1 | 12 | Arfgef3 | 0.029 | Young |
| 0.34414 | 0.067994 | 0.971 | 0.907 | 1 | 12 | Rtn1 | 0.064 | Young |
| 0.344973 | 1.119949 | 0.058 | 0.034 | 1 | 12 | Prpf40a | 0.024 | Young |
| 0.344973 | 0.990242 | 0.058 | 0.034 | 1 | 12 | Sec23a | 0.024 | Young |
| 0.345997 | 0.574963 | 0.102 | 0.068 | 1 | 12 | Rnf11 | 0.034 | Young |
| 0.346438 | 1.25602 | 0.036 | 0.017 | 1 | 12 | Dgkg | 0.019 | Young |
| 0.346438 | 1.233642 | 0.036 | 0.017 | 1 | 12 | Zfp579 | 0.019 | Young |
| 0.346438 | 1.227803 | 0.036 | 0.017 | 1 | 12 | Ier3 | 0.019 | Young |
| 0.346438 | 1.184813 | 0.036 | 0.017 | 1 | 12 | Usp31 | 0.019 | Young |
| 0.346438 | 1.061052 | 0.036 | 0.017 | 1 | 12 | AC149090. | 0.019 | Young |
| 0.346438 | 1.03797 | 0.036 | 0.017 | 1 | 12 | Ilk | 0.019 | Young |
| 0.346438 | 0.997584 | 0.036 | 0.017 | 1 | 12 | Aars | 0.019 | Young |
| 0.346438 | 0.982504 | 0.036 | 0.017 | 1 | 12 | Mars | 0.019 | Young |
| 0.347349 | 0.889507 | 0.058 | 0.034 | 1 | 12 | Spryd7 | 0.024 | Young |
| 0.347606 | 0.896168 | 0.08 | 0.051 | 1 | 12 | Rbfox2 | 0.029 | Young |
| 0.349358 | 0.235613 | 0.321 | 0.263 | 1 | 12 | Ywhaz | 0.058 | Young |
| 0.349528 | 0.99115 | 0.036 | 0.017 | 1 | 12 | Kcnip4 | 0.019 | Young |

| | | | | | | | | |
|---|---|---|---|---|---|---|---|---|
| 0.349528 | 0.982155 | 0.036 | 0.017 | 1 | 12 | Derl1 | 0.019 | Young |
| 0.349528 | 0.912862 | 0.036 | 0.017 | 1 | 12 | Gstp1 | 0.019 | Young |
| 0.349528 | 0.834293 | 0.036 | 0.017 | 1 | 12 | Mettl5 | 0.019 | Young |
| 0.349528 | 0.70535 | 0.036 | 0.017 | 1 | 12 | Pigyl | 0.019 | Young |
| 0.349528 | 0.699238 | 0.036 | 0.017 | 1 | 12 | Slc38a1 | 0.019 | Young |
| 0.349735 | 1.067321 | 0.058 | 0.034 | 1 | 12 | F2r | 0.024 | Young |
| 0.349735 | 0.900676 | 0.058 | 0.034 | 1 | 12 | 3-Sep | 0.024 | Young |
| 0.350458 | 0.180278 | 0.482 | 0.432 | 1 | 12 | Rps29 | 0.05 | Young |
| 0.351079 | 0.909423 | 0.036 | 0.017 | 1 | 12 | Ino80b | 0.019 | Young |
| 0.351392 | 0.321513 | 0.102 | 0.068 | 1 | 12 | Prr15 | 0.034 | Young |
| 0.352131 | 1.047881 | 0.058 | 0.034 | 1 | 12 | Rbmxl1 | 0.024 | Young |
| 0.352131 | 0.806914 | 0.058 | 0.034 | 1 | 12 | Azi2 | 0.024 | Young |
| 0.352636 | 0.992924 | 0.036 | 0.017 | 1 | 12 | Mark1 | 0.019 | Young |
| 0.352636 | 0.944105 | 0.036 | 0.017 | 1 | 12 | Lrrn3 | 0.019 | Young |
| 0.352636 | 0.930193 | 0.036 | 0.017 | 1 | 12 | Snx27 | 0.019 | Young |
| 0.352636 | 0.89001 | 0.036 | 0.017 | 1 | 12 | Susd4 | 0.019 | Young |
| 0.352636 | 0.864503 | 0.036 | 0.017 | 1 | 12 | Mmd | 0.019 | Young |
| 0.352636 | 0.860688 | 0.036 | 0.017 | 1 | 12 | Zfp24 | 0.019 | Young |
| 0.352636 | 0.843906 | 0.036 | 0.017 | 1 | 12 | Aff2 | 0.019 | Young |
| 0.352636 | 0.835846 | 0.036 | 0.017 | 1 | 12 | Lman1 | 0.019 | Young |
| 0.352636 | 0.818845 | 0.036 | 0.017 | 1 | 12 | Ubr5 | 0.019 | Young |
| 0.352636 | 0.800318 | 0.036 | 0.017 | 1 | 12 | Arid1a | 0.019 | Young |
| 0.352636 | 0.761605 | 0.036 | 0.017 | 1 | 12 | Kdelr2 | 0.019 | Young |
| 0.352636 | 0.758629 | 0.036 | 0.017 | 1 | 12 | Thap3 | 0.019 | Young |
| 0.353449 | 0.135558 | 0.409 | 0.364 | 1 | 12 | Htr3a | 0.045 | Young |
| 0.353704 | 0.592789 | 0.08 | 0.051 | 1 | 12 | Tmem38a | 0.029 | Young |
| 0.354058 | 0.499724 | 0.131 | 0.093 | 1 | 12 | Prmt2 | 0.038 | Young |
| 0.354403 | 0.766465 | 0.109 | 0.076 | 1 | 12 | Tmbim4 | 0.033 | Young |
| 0.354538 | 0.802746 | 0.058 | 0.034 | 1 | 12 | Poldip3 | 0.024 | Young |
| 0.354538 | 0.7614 | 0.058 | 0.034 | 1 | 12 | Tmem237 | 0.024 | Young |
| 0.355761 | 0.785218 | 0.036 | 0.017 | 1 | 12 | Rpl7l1 | 0.019 | Young |
| 0.355761 | 0.74225 | 0.036 | 0.017 | 1 | 12 | Cdk12 | 0.019 | Young |
| 0.355761 | 0.620263 | 0.036 | 0.017 | 1 | 12 | Dap | 0.019 | Young |
| 0.355761 | 0.598192 | 0.036 | 0.017 | 1 | 12 | Lrrc8b | 0.019 | Young |
| 0.356956 | 1.09381 | 0.058 | 0.034 | 1 | 12 | Rap1a | 0.024 | Young |
| 0.357846 | 0.22425 | 0.168 | 0.127 | 1 | 12 | Fam173a | 0.041 | Young |
| 0.358084 | 0.397379 | 0.19 | 0.144 | 1 | 12 | Pgp | 0.046 | Young |
| 0.358903 | 0.719017 | 0.036 | 0.017 | 1 | 12 | Smarcc1 | 0.019 | Young |
| 0.358903 | 0.685688 | 0.036 | 0.017 | 1 | 12 | Ranbp6 | 0.019 | Young |
| 0.358903 | 0.676855 | 0.036 | 0.017 | 1 | 12 | Arfrp1 | 0.019 | Young |
| 0.359383 | 0.80592 | 0.058 | 0.034 | 1 | 12 | Ccdc59 | 0.024 | Young |
| 0.359869 | 0.583862 | 0.08 | 0.051 | 1 | 12 | Mrpl22 | 0.029 | Young |
| 0.360502 | 0.465213 | 0.102 | 0.068 | 1 | 12 | Txndc15 | 0.034 | Young |
| 0.361421 | 0.454355 | 0.102 | 0.068 | 1 | 12 | Malsu1 | 0.034 | Young |
| 0.361421 | 0.331078 | 0.102 | 0.068 | 1 | 12 | Smim7 | 0.034 | Young |

| | | | | | | | | |
|---|---|---|---|---|---|---|---|---|
| 0.361821 | 0.862741 | 0.058 | 0.034 | 1 | 12 | Fam117b | 0.024 | Young |
| 0.361821 | 0.744823 | 0.058 | 0.034 | 1 | 12 | Chd4 | 0.024 | Young |
| 0.361821 | 0.711456 | 0.058 | 0.034 | 1 | 12 | Ube2i | 0.024 | Young |
| 0.362029 | 0.340512 | 0.255 | 0.212 | 1 | 12 | Ywhag | 0.043 | Young |
| 0.362449 | 0.831943 | 0.088 | 0.059 | 1 | 12 | Msmo1 | 0.029 | Young |
| 0.363436 | 0.864357 | 0.088 | 0.059 | 1 | 12 | Sdf2 | 0.029 | Young |
| 0.363926 | 0.359765 | 0.153 | 0.11 | 1 | 12 | Rac3 | 0.043 | Young |
| 0.36427 | 0.737561 | 0.058 | 0.034 | 1 | 12 | Lrrc8c | 0.024 | Young |
| 0.36427 | 0.699745 | 0.058 | 0.034 | 1 | 12 | Arhgap22 | 0.024 | Young |
| 0.36427 | 0.650985 | 0.058 | 0.034 | 1 | 12 | Tubg1 | 0.024 | Young |
| 0.364582 | 0.470315 | 0.131 | 0.093 | 1 | 12 | Kif2a | 0.038 | Young |
| 0.366519 | 0.352658 | 0.343 | 0.305 | 1 | 12 | Sdhb | 0.038 | Young |
| 0.367398 | 0.751008 | 0.088 | 0.059 | 1 | 12 | Cox20 | 0.029 | Young |
| 0.367715 | 0.306346 | 0.124 | 0.085 | 1 | 12 | Arhgef28 | 0.039 | Young |
| 0.367961 | 0.696767 | 0.058 | 0.034 | 1 | 12 | Zdhhc12 | 0.024 | Young |
| 0.368017 | 0.407411 | 0.212 | 0.169 | 1 | 12 | Mrps12 | 0.043 | Young |
| 0.368194 | 0.215621 | 0.08 | 0.051 | 1 | 12 | Tspan6 | 0.029 | Young |
| 0.369013 | 1.144946 | 0.095 | 0.068 | 1 | 12 | Atf4 | 0.027 | Young |
| 0.369197 | 0.664837 | 0.058 | 0.034 | 1 | 12 | Rfxap | 0.024 | Young |
| 0.369197 | 0.649889 | 0.058 | 0.034 | 1 | 12 | Zfp560 | 0.024 | Young |
| 0.370294 | 0.243541 | 0.08 | 0.051 | 1 | 12 | Nsa2 | 0.029 | Young |
| 0.370343 | 0.557753 | 0.109 | 0.076 | 1 | 12 | Dcun1d5 | 0.033 | Young |
| 0.371853 | 0.414324 | 0.219 | 0.169 | 1 | 12 | Ssr4 | 0.05 | Young |
| 0.372118 | 0.53791 | 0.139 | 0.102 | 1 | 12 | Thoc7 | 0.037 | Young |
| 0.374515 | 0.570982 | 0.08 | 0.051 | 1 | 12 | Nicn1 | 0.029 | Young |
| 0.375341 | 0.429069 | 0.139 | 0.102 | 1 | 12 | Elof1 | 0.037 | Young |
| 0.375562 | 0.531198 | 0.146 | 0.11 | 1 | 12 | Ier3ip1 | 0.036 | Young |
| 0.375754 | 0.379878 | 0.109 | 0.076 | 1 | 12 | Dlgap1 | 0.033 | Young |
| 0.375754 | 0.581427 | 0.109 | 0.076 | 1 | 12 | Sec11a | 0.033 | Young |
| 0.376637 | 0.442403 | 0.08 | 0.051 | 1 | 12 | Rhoc | 0.029 | Young |
| 0.376665 | 0.553209 | 0.058 | 0.034 | 1 | 12 | Map9 | 0.024 | Young |
| 0.376935 | 0.275957 | 0.321 | 0.28 | 1 | 12 | Nenf | 0.041 | Young |
| 0.377421 | 0.650394 | 0.088 | 0.059 | 1 | 12 | Atp8a1 | 0.029 | Young |
| 0.377677 | 0.12851 | 0.65 | 0.602 | 1 | 12 | Rps3 | 0.048 | Young |
| 0.378089 | 0.256107 | 0.496 | 0.458 | 1 | 12 | Uqcr10 | 0.038 | Young |
| 0.379176 | 0.588165 | 0.058 | 0.034 | 1 | 12 | Slbp | 0.024 | Young |
| 0.379389 | 0.583501 | 0.109 | 0.076 | 1 | 12 | Dnaja2 | 0.033 | Young |
| 0.38046 | 0.700521 | 0.088 | 0.059 | 1 | 12 | Snapc5 | 0.029 | Young |
| 0.380741 | 0.13602 | 0.416 | 0.356 | 1 | 12 | Ndufb9 | 0.06 | Young |
| 0.381697 | 0.356533 | 0.058 | 0.034 | 1 | 12 | Emc8 | 0.024 | Young |
| 0.383048 | 0.270843 | 0.08 | 0.051 | 1 | 12 | Sae1 | 0.029 | Young |
| 0.383234 | 0.238214 | 0.292 | 0.237 | 1 | 12 | Rtraf | 0.055 | Young |
| 0.383455 | 0.081034 | 0.613 | 0.534 | 1 | 12 | Rps23 | 0.079 | Young |
| 0.384227 | 0.472788 | 0.058 | 0.034 | 1 | 12 | Snx2 | 0.024 | Young |
| 0.384739 | 1.042692 | 0.066 | 0.042 | 1 | 12 | Commd6 | 0.024 | Young |

| | | | | | | | | |
|---|---|---|---|---|---|---|---|---|
| 0.38556 | 0.471569 | 0.088 | 0.059 | 1 | 12 | Dhrs1 | 0.029 | Young |
| 0.386317 | 3.845531 | 0.022 | 0.008 | 1 | 12 | Tipin | 0.014 | Young |
| 0.386317 | 3.632148 | 0.022 | 0.008 | 1 | 12 | Noc2l | 0.014 | Young |
| 0.386317 | 3.542181 | 0.022 | 0.008 | 1 | 12 | Cpt1a | 0.014 | Young |
| 0.386317 | 3.478723 | 0.022 | 0.008 | 1 | 12 | Hyou1 | 0.014 | Young |
| 0.386317 | 3.434851 | 0.022 | 0.008 | 1 | 12 | Galnt7 | 0.014 | Young |
| 0.386317 | 3.354919 | 0.022 | 0.008 | 1 | 12 | Sh3bgrl2 | 0.014 | Young |
| 0.386317 | 3.312834 | 0.022 | 0.008 | 1 | 12 | Mthfd1 | 0.014 | Young |
| 0.386317 | 3.302867 | 0.022 | 0.008 | 1 | 12 | Sec61a1 | 0.014 | Young |
| 0.386317 | 3.292975 | 0.022 | 0.008 | 1 | 12 | Gm10073 | 0.014 | Young |
| 0.386317 | 3.118006 | 0.022 | 0.008 | 1 | 12 | Scamp2 | 0.014 | Young |
| 0.386317 | 2.71007 | 0.022 | 0.008 | 1 | 12 | Numa1 | 0.014 | Young |
| 0.386317 | 2.679738 | 0.022 | 0.008 | 1 | 12 | Galnt1 | 0.014 | Young |
| 0.386317 | 2.625129 | 0.022 | 0.008 | 1 | 12 | Mapk14 | 0.014 | Young |
| 0.386317 | 2.616803 | 0.022 | 0.008 | 1 | 12 | Klhdc10 | 0.014 | Young |
| 0.386317 | 2.256403 | 0.022 | 0.008 | 1 | 12 | Papd4 | 0.014 | Young |
| 0.386317 | 1.949749 | 0.022 | 0.008 | 1 | 12 | Prpf31 | 0.014 | Young |
| 0.386317 | 1.927969 | 0.022 | 0.008 | 1 | 12 | Stx5a | 0.014 | Young |
| 0.386317 | 1.918914 | 0.022 | 0.008 | 1 | 12 | Fam13c | 0.014 | Young |
| 0.386317 | 1.866732 | 0.022 | 0.008 | 1 | 12 | Pspc1 | 0.014 | Young |
| 0.386317 | 1.854199 | 0.022 | 0.008 | 1 | 12 | Mbd2 | 0.014 | Young |
| 0.386317 | 1.824946 | 0.022 | 0.008 | 1 | 12 | Ppih | 0.014 | Young |
| 0.386317 | 1.801917 | 0.022 | 0.008 | 1 | 12 | Cdr2 | 0.014 | Young |
| 0.386317 | 1.773674 | 0.022 | 0.008 | 1 | 12 | Gt(ROSA)2( | 0.014 | Young |
| 0.386317 | 1.768842 | 0.022 | 0.008 | 1 | 12 | Cox17 | 0.014 | Young |
| 0.386317 | 1.746585 | 0.022 | 0.008 | 1 | 12 | Ccdc71 | 0.014 | Young |
| 0.386317 | 1.724019 | 0.022 | 0.008 | 1 | 12 | Pias2 | 0.014 | Young |
| 0.386317 | 1.720414 | 0.022 | 0.008 | 1 | 12 | Fign | 0.014 | Young |
| 0.386317 | 1.709694 | 0.022 | 0.008 | 1 | 12 | Snrnp200 | 0.014 | Young |
| 0.386317 | 1.701558 | 0.022 | 0.008 | 1 | 12 | Nalcn | 0.014 | Young |
| 0.386317 | 1.695253 | 0.022 | 0.008 | 1 | 12 | Pnpla2 | 0.014 | Young |
| 0.386317 | 1.671786 | 0.022 | 0.008 | 1 | 12 | Gmppb | 0.014 | Young |
| 0.386317 | 1.65846 | 0.022 | 0.008 | 1 | 12 | Cdk5r1 | 0.014 | Young |
| 0.386317 | 1.647186 | 0.022 | 0.008 | 1 | 12 | Slf1 | 0.014 | Young |
| 0.386317 | 1.634137 | 0.022 | 0.008 | 1 | 12 | Gan | 0.014 | Young |
| 0.386317 | 1.629272 | 0.022 | 0.008 | 1 | 12 | Ptpdc1 | 0.014 | Young |
| 0.386317 | 1.621389 | 0.022 | 0.008 | 1 | 12 | Pomgnt1 | 0.014 | Young |
| 0.386317 | 1.603526 | 0.022 | 0.008 | 1 | 12 | Mfsd14a | 0.014 | Young |
| 0.386317 | 1.579286 | 0.022 | 0.008 | 1 | 12 | Usp46 | 0.014 | Young |
| 0.386317 | 1.565005 | 0.022 | 0.008 | 1 | 12 | Vezt | 0.014 | Young |
| 0.386317 | 1.542523 | 0.022 | 0.008 | 1 | 12 | Trak1 | 0.014 | Young |
| 0.386317 | 1.533734 | 0.022 | 0.008 | 1 | 12 | Dr1 | 0.014 | Young |
| 0.386317 | 1.522004 | 0.022 | 0.008 | 1 | 12 | Sh3pxd2a | 0.014 | Young |
| 0.386317 | 1.509829 | 0.022 | 0.008 | 1 | 12 | Arih2 | 0.014 | Young |
| 0.386317 | 1.475795 | 0.022 | 0.008 | 1 | 12 | Alad | 0.014 | Young |

| | | | | | | | | |
|---|---|---|---|---|---|---|---|---|
| 0.386317 | 1.471832 | 0.022 | 0.008 | 1 | 12 | Sugp2 | 0.014 | Young |
| 0.386317 | 1.461257 | 0.022 | 0.008 | 1 | 12 | Zfp30 | 0.014 | Young |
| 0.386317 | 1.436621 | 0.022 | 0.008 | 1 | 12 | Mrps27 | 0.014 | Young |
| 0.386317 | 1.433497 | 0.022 | 0.008 | 1 | 12 | Itga6 | 0.014 | Young |
| 0.386317 | 1.429157 | 0.022 | 0.008 | 1 | 12 | Tstd3 | 0.014 | Young |
| 0.386317 | 1.42193 | 0.022 | 0.008 | 1 | 12 | Pex2 | 0.014 | Young |
| 0.386722 | 0.421808 | 0.109 | 0.076 | 1 | 12 | 9530059O1 | 0.033 | Young |
| 0.386961 | 0.485414 | 0.234 | 0.195 | 1 | 12 | Gng2 | 0.039 | Young |
| 0.388223 | 0.185402 | 0.745 | 0.746 | 1 | 12 | Lgals1 | -0.001 | Young |
| 0.388405 | 0.491193 | 0.139 | 0.102 | 1 | 12 | Nagk | 0.037 | Young |
| 0.388569 | 0.502178 | 0.109 | 0.076 | 1 | 12 | Rsu1 | 0.033 | Young |
| 0.389475 | 0.862 | 0.066 | 0.042 | 1 | 12 | Rad23a | 0.024 | Young |
| 0.389669 | 0.496613 | 0.088 | 0.059 | 1 | 12 | Coprs | 0.029 | Young |
| 0.389669 | 0.37409 | 0.088 | 0.059 | 1 | 12 | Atp6ap2 | 0.029 | Young |
| 0.390599 | 0.308829 | 0.197 | 0.153 | 1 | 12 | Rps27 | 0.044 | Young |
| 0.39067 | 2.179864 | 0.022 | 0.008 | 1 | 12 | Slc24a5 | 0.014 | Young |
| 0.39067 | 1.98857 | 0.022 | 0.008 | 1 | 12 | Spout1 | 0.014 | Young |
| 0.39067 | 1.85829 | 0.022 | 0.008 | 1 | 12 | Iffo1 | 0.014 | Young |
| 0.39067 | 1.712732 | 0.022 | 0.008 | 1 | 12 | Mib2 | 0.014 | Young |
| 0.39067 | 1.650489 | 0.022 | 0.008 | 1 | 12 | Nuak1 | 0.014 | Young |
| 0.39067 | 1.592103 | 0.022 | 0.008 | 1 | 12 | Tfam | 0.014 | Young |
| 0.39067 | 1.574718 | 0.022 | 0.008 | 1 | 12 | Ranbp9 | 0.014 | Young |
| 0.39067 | 1.557618 | 0.022 | 0.008 | 1 | 12 | Rngtt | 0.014 | Young |
| 0.39067 | 1.553122 | 0.022 | 0.008 | 1 | 12 | Bnip1 | 0.014 | Young |
| 0.39067 | 1.534542 | 0.022 | 0.008 | 1 | 12 | 4932438A1 | 0.014 | Young |
| 0.39067 | 1.507984 | 0.022 | 0.008 | 1 | 12 | Esco1 | 0.014 | Young |
| 0.39067 | 1.482769 | 0.022 | 0.008 | 1 | 12 | Tfip11 | 0.014 | Young |
| 0.39067 | 1.459539 | 0.022 | 0.008 | 1 | 12 | Tspyl1 | 0.014 | Young |
| 0.39067 | 1.453865 | 0.022 | 0.008 | 1 | 12 | Pnrc2 | 0.014 | Young |
| 0.39067 | 1.450136 | 0.022 | 0.008 | 1 | 12 | Msh3 | 0.014 | Young |
| 0.39067 | 1.429643 | 0.022 | 0.008 | 1 | 12 | Adprm | 0.014 | Young |
| 0.39067 | 1.429636 | 0.022 | 0.008 | 1 | 12 | Zfp618 | 0.014 | Young |
| 0.39067 | 1.424657 | 0.022 | 0.008 | 1 | 12 | 2810006K2 | 0.014 | Young |
| 0.39067 | 1.417786 | 0.022 | 0.008 | 1 | 12 | AU019823 | 0.014 | Young |
| 0.39067 | 1.415198 | 0.022 | 0.008 | 1 | 12 | Tbc1d10b | 0.014 | Young |
| 0.39067 | 1.405507 | 0.022 | 0.008 | 1 | 12 | Bckdhb | 0.014 | Young |
| 0.39067 | 1.396679 | 0.022 | 0.008 | 1 | 12 | Dnajc25 | 0.014 | Young |
| 0.39067 | 1.394867 | 0.022 | 0.008 | 1 | 12 | Pik3r3 | 0.014 | Young |
| 0.39067 | 1.392248 | 0.022 | 0.008 | 1 | 12 | Chfr | 0.014 | Young |
| 0.39067 | 1.386723 | 0.022 | 0.008 | 1 | 12 | Ube4b | 0.014 | Young |
| 0.39067 | 1.384115 | 0.022 | 0.008 | 1 | 12 | Selenoi | 0.014 | Young |
| 0.39067 | 1.376975 | 0.022 | 0.008 | 1 | 12 | Olfr78 | 0.014 | Young |
| 0.39067 | 1.372184 | 0.022 | 0.008 | 1 | 12 | Mta1 | 0.014 | Young |
| 0.39067 | 1.356732 | 0.022 | 0.008 | 1 | 12 | Tceal1 | 0.014 | Young |
| 0.39067 | 1.349081 | 0.022 | 0.008 | 1 | 12 | Dock3 | 0.014 | Young |

| | | | | | | | | |
|---|---|---|---|---|---|---|---|---|
| 0.39067 | 1.347103 | 0.022 | 0.008 | 1 | 12 | Tgfbrap1 | 0.014 | Young |
| 0.39067 | 1.34488 | 0.022 | 0.008 | 1 | 12 | Orai1 | 0.014 | Young |
| 0.39067 | 1.335893 | 0.022 | 0.008 | 1 | 12 | 6-Mar | 0.014 | Young |
| 0.39067 | 1.335264 | 0.022 | 0.008 | 1 | 12 | Btbd9 | 0.014 | Young |
| 0.39067 | 1.325012 | 0.022 | 0.008 | 1 | 12 | Tceal6 | 0.014 | Young |
| 0.39067 | 1.317808 | 0.022 | 0.008 | 1 | 12 | Abhd17b | 0.014 | Young |
| 0.39067 | 1.309955 | 0.022 | 0.008 | 1 | 12 | Leng8 | 0.014 | Young |
| 0.39067 | 1.274056 | 0.022 | 0.008 | 1 | 12 | Ddx18 | 0.014 | Young |
| 0.39067 | 1.262333 | 0.022 | 0.008 | 1 | 12 | Hibch | 0.014 | Young |
| 0.39067 | 1.251127 | 0.022 | 0.008 | 1 | 12 | Tmsb15b1 | 0.014 | Young |
| 0.39067 | 1.222809 | 0.022 | 0.008 | 1 | 12 | Zfp365 | 0.014 | Young |
| 0.39067 | 1.15495 | 0.022 | 0.008 | 1 | 12 | Nectin2 | 0.014 | Young |
| 0.39067 | 1.109688 | 0.022 | 0.008 | 1 | 12 | Tgfbr1 | 0.014 | Young |
| 0.391714 | 0.22947 | 0.139 | 0.102 | 1 | 12 | Eif1ax | 0.037 | Young |
| 0.392855 | 1.397372 | 0.022 | 0.008 | 1 | 12 | Mrpl19 | 0.014 | Young |
| 0.393282 | 0.655498 | 0.131 | 0.102 | 1 | 12 | Tceal9 | 0.029 | Young |
| 0.393737 | 0.140451 | 0.482 | 0.424 | 1 | 12 | Atp6v0b | 0.058 | Young |
| 0.394151 | 0.340954 | 0.321 | 0.28 | 1 | 12 | Crip2 | 0.041 | Young |
| 0.394289 | 0.074296 | 0.781 | 0.678 | 1 | 12 | Rps4x | 0.103 | Young |
| 0.395052 | 1.443761 | 0.022 | 0.008 | 1 | 12 | 5033406O( | 0.014 | Young |
| 0.395052 | 1.432673 | 0.022 | 0.008 | 1 | 12 | Lrrc42 | 0.014 | Young |
| 0.395052 | 1.361687 | 0.022 | 0.008 | 1 | 12 | B4galt2 | 0.014 | Young |
| 0.395052 | 1.354804 | 0.022 | 0.008 | 1 | 12 | Smarca1 | 0.014 | Young |
| 0.395052 | 1.326831 | 0.022 | 0.008 | 1 | 12 | Usp50 | 0.014 | Young |
| 0.395052 | 1.317293 | 0.022 | 0.008 | 1 | 12 | Lsm11 | 0.014 | Young |
| 0.395052 | 1.308168 | 0.022 | 0.008 | 1 | 12 | Rab11fip4 | 0.014 | Young |
| 0.395052 | 1.301498 | 0.022 | 0.008 | 1 | 12 | Actr6 | 0.014 | Young |
| 0.395052 | 1.288791 | 0.022 | 0.008 | 1 | 12 | Spryd4 | 0.014 | Young |
| 0.395052 | 1.283356 | 0.022 | 0.008 | 1 | 12 | Tada3 | 0.014 | Young |
| 0.395052 | 1.272557 | 0.022 | 0.008 | 1 | 12 | Magi2 | 0.014 | Young |
| 0.395052 | 1.268297 | 0.022 | 0.008 | 1 | 12 | Kcnk2 | 0.014 | Young |
| 0.395052 | 1.256862 | 0.022 | 0.008 | 1 | 12 | Bmi1 | 0.014 | Young |
| 0.395052 | 1.237143 | 0.022 | 0.008 | 1 | 12 | Gnptab | 0.014 | Young |
| 0.395052 | 1.237129 | 0.022 | 0.008 | 1 | 12 | Med25 | 0.014 | Young |
| 0.395052 | 1.216049 | 0.022 | 0.008 | 1 | 12 | Tspyl2 | 0.014 | Young |
| 0.395052 | 1.215097 | 0.022 | 0.008 | 1 | 12 | Tbcel | 0.014 | Young |
| 0.395052 | 1.208951 | 0.022 | 0.008 | 1 | 12 | Arhgef1 | 0.014 | Young |
| 0.395052 | 1.205621 | 0.022 | 0.008 | 1 | 12 | Dscr3 | 0.014 | Young |
| 0.395052 | 1.190938 | 0.022 | 0.008 | 1 | 12 | Numbl | 0.014 | Young |
| 0.395052 | 1.181948 | 0.022 | 0.008 | 1 | 12 | Trim3 | 0.014 | Young |
| 0.395052 | 1.178152 | 0.022 | 0.008 | 1 | 12 | Rbpj | 0.014 | Young |
| 0.395052 | 1.174 | 0.022 | 0.008 | 1 | 12 | F8a | 0.014 | Young |
| 0.395052 | 1.164659 | 0.022 | 0.008 | 1 | 12 | Snx8 | 0.014 | Young |
| 0.395052 | 1.157454 | 0.022 | 0.008 | 1 | 12 | Efnb2 | 0.014 | Young |
| 0.395052 | 1.151801 | 0.022 | 0.008 | 1 | 12 | Pank2 | 0.014 | Young |

| | | | | | | | | |
|---|---|---|---|---|---|---|---|---|
| 0.395052 | 1.138575 | 0.022 | 0.008 | 1 | 12 | Jmjd6 | 0.014 | Young |
| 0.395052 | 1.132774 | 0.022 | 0.008 | 1 | 12 | Dusp19 | 0.014 | Young |
| 0.395052 | 1.126288 | 0.022 | 0.008 | 1 | 12 | Slc2a8 | 0.014 | Young |
| 0.395052 | 1.116855 | 0.022 | 0.008 | 1 | 12 | Stx17 | 0.014 | Young |
| 0.395052 | 1.113815 | 0.022 | 0.008 | 1 | 12 | Coa4 | 0.014 | Young |
| 0.395052 | 1.1113 | 0.022 | 0.008 | 1 | 12 | Tmem255b | 0.014 | Young |
| 0.395052 | 1.108692 | 0.022 | 0.008 | 1 | 12 | Adamts5 | 0.014 | Young |
| 0.395052 | 1.084306 | 0.022 | 0.008 | 1 | 12 | Arhgap12 | 0.014 | Young |
| 0.395052 | 1.068502 | 0.022 | 0.008 | 1 | 12 | Cdk5r2 | 0.014 | Young |
| 0.395052 | 1.067277 | 0.022 | 0.008 | 1 | 12 | Sart1 | 0.014 | Young |
| 0.395052 | 1.046491 | 0.022 | 0.008 | 1 | 12 | Clcn6 | 0.014 | Young |
| 0.395052 | 1.044011 | 0.022 | 0.008 | 1 | 12 | Ankmy2 | 0.014 | Young |
| 0.395052 | 1.039099 | 0.022 | 0.008 | 1 | 12 | Bet1l | 0.014 | Young |
| 0.395052 | 1.029106 | 0.022 | 0.008 | 1 | 12 | Snx12 | 0.014 | Young |
| 0.395052 | 1.028509 | 0.022 | 0.008 | 1 | 12 | Siah2 | 0.014 | Young |
| 0.395052 | 1.020801 | 0.022 | 0.008 | 1 | 12 | Mum1l1 | 0.014 | Young |
| 0.395052 | 1.020227 | 0.022 | 0.008 | 1 | 12 | Rita1 | 0.014 | Young |
| 0.395052 | 1.016769 | 0.022 | 0.008 | 1 | 12 | Smim18 | 0.014 | Young |
| 0.395052 | 1.007028 | 0.022 | 0.008 | 1 | 12 | Vps26a | 0.014 | Young |
| 0.395052 | 0.999023 | 0.022 | 0.008 | 1 | 12 | Sco2 | 0.014 | Young |
| 0.395052 | 0.988909 | 0.022 | 0.008 | 1 | 12 | Zkscan3 | 0.014 | Young |
| 0.395052 | 0.968062 | 0.022 | 0.008 | 1 | 12 | Polr1c | 0.014 | Young |
| 0.395052 | 0.956529 | 0.022 | 0.008 | 1 | 12 | Nol8 | 0.014 | Young |
| 0.395052 | 0.939441 | 0.022 | 0.008 | 1 | 12 | Optn | 0.014 | Young |
| 0.395052 | 0.929836 | 0.022 | 0.008 | 1 | 12 | Arrdc4 | 0.014 | Young |
| 0.395052 | 0.927713 | 0.022 | 0.008 | 1 | 12 | Slc44a1 | 0.014 | Young |
| 0.395052 | 0.866454 | 0.022 | 0.008 | 1 | 12 | Ubl7 | 0.014 | Young |
| 0.396067 | 0.196207 | 0.08 | 0.051 | 1 | 12 | Fgfr1op2 | 0.029 | Young |
| 0.396261 | 0.890912 | 0.095 | 0.068 | 1 | 12 | Lrpap1 | 0.027 | Young |
| 0.396452 | 0.285456 | 0.555 | 0.508 | 1 | 12 | Smdt1 | 0.047 | Young |
| 0.398342 | 0.175039 | 0.401 | 0.356 | 1 | 12 | Rpl4 | 0.045 | Young |
| 0.399455 | 0.45148 | 0.153 | 0.119 | 1 | 12 | Psma1 | 0.034 | Young |
| 0.399461 | 1.178453 | 0.022 | 0.008 | 1 | 12 | Sp3 | 0.014 | Young |
| 0.399463 | 1.211314 | 0.022 | 0.008 | 1 | 12 | Creg1 | 0.014 | Young |
| 0.399463 | 1.198189 | 0.022 | 0.008 | 1 | 12 | Sf1 | 0.014 | Young |
| 0.399463 | 1.170953 | 0.022 | 0.008 | 1 | 12 | Zmynd8 | 0.014 | Young |
| 0.399463 | 1.15133 | 0.022 | 0.008 | 1 | 12 | Fasn | 0.014 | Young |
| 0.399463 | 1.151203 | 0.022 | 0.008 | 1 | 12 | Mtif2 | 0.014 | Young |
| 0.399463 | 1.137756 | 0.022 | 0.008 | 1 | 12 | Wdr34 | 0.014 | Young |
| 0.399463 | 1.131664 | 0.022 | 0.008 | 1 | 12 | Uso1 | 0.014 | Young |
| 0.399463 | 1.127146 | 0.022 | 0.008 | 1 | 12 | Suclg2 | 0.014 | Young |
| 0.399463 | 1.118928 | 0.022 | 0.008 | 1 | 12 | Polr2b | 0.014 | Young |
| 0.399463 | 1.106356 | 0.022 | 0.008 | 1 | 12 | Chtf8 | 0.014 | Young |
| 0.399463 | 1.095138 | 0.022 | 0.008 | 1 | 12 | Bcl7a | 0.014 | Young |
| 0.399463 | 1.08599 | 0.022 | 0.008 | 1 | 12 | Pes1 | 0.014 | Young |

| | | | | | | | | |
|---|---|---|---|---|---|---|---|---|
| 0.399463 | 1.079586 | 0.022 | 0.008 | 1 | 12 | Mrnip | 0.014 | Young |
| 0.399463 | 1.078205 | 0.022 | 0.008 | 1 | 12 | Pcdh17 | 0.014 | Young |
| 0.399463 | 1.074609 | 0.022 | 0.008 | 1 | 12 | Fdxr | 0.014 | Young |
| 0.399463 | 1.059438 | 0.022 | 0.008 | 1 | 12 | Fbxw11 | 0.014 | Young |
| 0.399463 | 1.014578 | 0.022 | 0.008 | 1 | 12 | Tmem216 | 0.014 | Young |
| 0.399463 | 1.005414 | 0.022 | 0.008 | 1 | 12 | Diras1 | 0.014 | Young |
| 0.399463 | 0.976234 | 0.022 | 0.008 | 1 | 12 | Nfx1 | 0.014 | Young |
| 0.399463 | 0.956773 | 0.022 | 0.008 | 1 | 12 | Iars2 | 0.014 | Young |
| 0.399463 | 0.951036 | 0.022 | 0.008 | 1 | 12 | Arhgef40 | 0.014 | Young |
| 0.399463 | 0.949233 | 0.022 | 0.008 | 1 | 12 | Chpt1 | 0.014 | Young |
| 0.399463 | 0.944164 | 0.022 | 0.008 | 1 | 12 | Mpdz | 0.014 | Young |
| 0.399463 | 0.938174 | 0.022 | 0.008 | 1 | 12 | Hist1h1e | 0.014 | Young |
| 0.399463 | 0.931775 | 0.022 | 0.008 | 1 | 12 | Rps6kb1 | 0.014 | Young |
| 0.399463 | 0.923475 | 0.022 | 0.008 | 1 | 12 | Pim3 | 0.014 | Young |
| 0.399463 | 0.918924 | 0.022 | 0.008 | 1 | 12 | Ftsj1 | 0.014 | Young |
| 0.399463 | 0.90493 | 0.022 | 0.008 | 1 | 12 | Nsrp1 | 0.014 | Young |
| 0.399463 | 0.900032 | 0.022 | 0.008 | 1 | 12 | Jph3 | 0.014 | Young |
| 0.399463 | 0.895836 | 0.022 | 0.008 | 1 | 12 | Fpgs | 0.014 | Young |
| 0.399463 | 0.893531 | 0.022 | 0.008 | 1 | 12 | Cdkl5 | 0.014 | Young |
| 0.399463 | 0.892241 | 0.022 | 0.008 | 1 | 12 | Adar | 0.014 | Young |
| 0.399463 | 0.885336 | 0.022 | 0.008 | 1 | 12 | Cenpv | 0.014 | Young |
| 0.399463 | 0.875672 | 0.022 | 0.008 | 1 | 12 | Spg20 | 0.014 | Young |
| 0.399463 | 0.87438 | 0.022 | 0.008 | 1 | 12 | Golph3l | 0.014 | Young |
| 0.399463 | 0.844237 | 0.022 | 0.008 | 1 | 12 | Casc3 | 0.014 | Young |
| 0.399463 | 0.829666 | 0.022 | 0.008 | 1 | 12 | Cog7 | 0.014 | Young |
| 0.399463 | 0.82274 | 0.022 | 0.008 | 1 | 12 | Bri3bp | 0.014 | Young |
| 0.399463 | 0.793669 | 0.022 | 0.008 | 1 | 12 | H2-T22 | 0.014 | Young |
| 0.399463 | 0.786967 | 0.022 | 0.008 | 1 | 12 | 5031425E2 | 0.014 | Young |
| 0.399463 | 0.780307 | 0.022 | 0.008 | 1 | 12 | Zdhhc13 | 0.014 | Young |
| 0.399463 | 0.777467 | 0.022 | 0.008 | 1 | 12 | Ctnnd2 | 0.014 | Young |
| 0.399463 | 0.713169 | 0.022 | 0.008 | 1 | 12 | Snx13 | 0.014 | Young |
| 0.399463 | 0.709839 | 0.022 | 0.008 | 1 | 12 | Prima1 | 0.014 | Young |
| 0.399463 | 0.675326 | 0.022 | 0.008 | 1 | 12 | Homer2 | 0.014 | Young |
| 0.399463 | 0.637815 | 0.022 | 0.008 | 1 | 12 | Ctnna2 | 0.014 | Young |
| 0.399463 | 0.620264 | 0.022 | 0.008 | 1 | 12 | Gatad2a | 0.014 | Young |
| 0.399463 | 0.584645 | 0.022 | 0.008 | 1 | 12 | Commd10 | 0.014 | Young |
| 0.399463 | 0.561219 | 0.022 | 0.008 | 1 | 12 | Dnajb2 | 0.014 | Young |
| 0.399463 | 0.559348 | 0.022 | 0.008 | 1 | 12 | Cntn2 | 0.014 | Young |
| 0.399463 | 0.343569 | 0.022 | 0.008 | 1 | 12 | Zswim7 | 0.014 | Young |
| 0.399626 | 0.296057 | 0.058 | 0.034 | 1 | 12 | Abhd12 | 0.024 | Young |
| 0.399762 | 0.404703 | 0.109 | 0.076 | 1 | 12 | Tmem55b | 0.033 | Young |
| 0.401199 | 0.454485 | 0.197 | 0.161 | 1 | 12 | Ptges3 | 0.036 | Young |
| 0.401332 | 0.309627 | 0.372 | 0.356 | 1 | 12 | Atp6v1f | 0.016 | Young |
| 0.401898 | 0.494956 | 0.117 | 0.085 | 1 | 12 | Adprh | 0.032 | Young |
| 0.402984 | 1.639371 | 0.044 | 0.025 | 1 | 12 | Dgcr2 | 0.019 | Young |

| | | | | | | | | |
|---|---|---|---|---|---|---|---|---|
| 0.40426 | 0.532592 | 0.088 | 0.059 | 1 | 12 | Nrxn1 | 0.029 | Young |
| 0.405549 | 0.469912 | 0.117 | 0.085 | 1 | 12 | Hint2 | 0.032 | Young |
| 0.406327 | 0.791717 | 0.066 | 0.042 | 1 | 12 | Gls | 0.024 | Young |
| 0.406327 | 0.468046 | 0.066 | 0.042 | 1 | 12 | Sfpq | 0.024 | Young |
| 0.411221 | 0.702578 | 0.066 | 0.042 | 1 | 12 | Vamp4 | 0.024 | Young |
| 0.411221 | 0.653111 | 0.066 | 0.042 | 1 | 12 | Ddx6 | 0.024 | Young |
| 0.412363 | 0.532737 | 0.095 | 0.068 | 1 | 12 | Atp6v1g2 | 0.027 | Young |
| 0.412912 | 0.521599 | 0.117 | 0.085 | 1 | 12 | Cwc15 | 0.032 | Young |
| 0.41368 | 0.940636 | 0.066 | 0.042 | 1 | 12 | Ppp1r7 | 0.024 | Young |
| 0.41368 | 0.651451 | 0.066 | 0.042 | 1 | 12 | Faim | 0.024 | Young |
| 0.41368 | 0.650518 | 0.066 | 0.042 | 1 | 12 | Kcnq3 | 0.024 | Young |
| 0.41488 | 0.470795 | 0.088 | 0.059 | 1 | 12 | Ina | 0.029 | Young |
| 0.41488 | 0.538395 | 0.088 | 0.059 | 1 | 12 | Mtpn | 0.029 | Young |
| 0.415074 | 1.055908 | 0.044 | 0.025 | 1 | 12 | Abi2 | 0.019 | Young |
| 0.416149 | 0.949878 | 0.066 | 0.042 | 1 | 12 | Tnrc6c | 0.024 | Young |
| 0.416149 | 0.63238 | 0.066 | 0.042 | 1 | 12 | Cdkl2 | 0.024 | Young |
| 0.416448 | 0.534025 | 0.095 | 0.068 | 1 | 12 | Pak1ip1 | 0.027 | Young |
| 0.41691 | 0.344097 | 0.109 | 0.076 | 1 | 12 | Marcks | 0.033 | Young |
| 0.41813 | 1.349246 | 0.044 | 0.025 | 1 | 12 | Dapk3 | 0.019 | Young |
| 0.41813 | 1.284003 | 0.044 | 0.025 | 1 | 12 | Mlx | 0.019 | Young |
| 0.41813 | 1.173752 | 0.044 | 0.025 | 1 | 12 | Fam133b | 0.019 | Young |
| 0.41813 | 1.00349 | 0.044 | 0.025 | 1 | 12 | Tnik | 0.019 | Young |
| 0.418161 | 0.260117 | 0.54 | 0.517 | 1 | 12 | Prdx2 | 0.023 | Young |
| 0.418514 | 0.351662 | 0.204 | 0.169 | 1 | 12 | mt-Nd5 | 0.035 | Young |
| 0.418842 | 0.356322 | 0.109 | 0.076 | 1 | 12 | Arl4a | 0.033 | Young |
| 0.419004 | 0.234339 | 0.168 | 0.127 | 1 | 12 | 1810022K0 | 0.041 | Young |
| 0.419173 | 0.427374 | 0.088 | 0.059 | 1 | 12 | Ddx42 | 0.029 | Young |
| 0.419173 | 0.222286 | 0.088 | 0.059 | 1 | 12 | U2surp | 0.029 | Young |
| 0.421112 | 0.363991 | 0.066 | 0.042 | 1 | 12 | Hnrnpc | 0.024 | Young |
| 0.421199 | 1.374915 | 0.044 | 0.025 | 1 | 12 | Glod4 | 0.019 | Young |
| 0.421199 | 0.974995 | 0.044 | 0.025 | 1 | 12 | Ttbk2 | 0.019 | Young |
| 0.421199 | 0.9481 | 0.044 | 0.025 | 1 | 12 | Ewsr1 | 0.019 | Young |
| 0.421199 | 0.929495 | 0.044 | 0.025 | 1 | 12 | Tesk1 | 0.019 | Young |
| 0.421199 | 0.873001 | 0.044 | 0.025 | 1 | 12 | Slc31a1 | 0.019 | Young |
| 0.424281 | 1.210458 | 0.044 | 0.025 | 1 | 12 | Zfp931 | 0.019 | Young |
| 0.424759 | 0.14849 | 0.591 | 0.534 | 1 | 12 | Myl1 | 0.057 | Young |
| 0.424857 | 0.354188 | 0.066 | 0.042 | 1 | 12 | Aamp | 0.024 | Young |
| 0.426765 | 0.548613 | 0.095 | 0.068 | 1 | 12 | Ift20 | 0.027 | Young |
| 0.427376 | 1.05074 | 0.044 | 0.025 | 1 | 12 | Mgll | 0.019 | Young |
| 0.427376 | 0.928415 | 0.044 | 0.025 | 1 | 12 | Gnai3 | 0.019 | Young |
| 0.427376 | 0.906475 | 0.044 | 0.025 | 1 | 12 | Fam114a2 | 0.019 | Young |
| 0.427376 | 0.753426 | 0.044 | 0.025 | 1 | 12 | 2610524H0 | 0.019 | Young |
| 0.428591 | 0.307032 | 0.299 | 0.246 | 1 | 12 | Dynlrb1 | 0.053 | Young |
| 0.428622 | 0.554553 | 0.066 | 0.042 | 1 | 12 | Mrps35 | 0.024 | Young |
| 0.429654 | 0.490537 | 0.182 | 0.153 | 1 | 12 | Gnai2 | 0.029 | Young |

| | | | | | | | | |
|---|---|---|---|---|---|---|---|---|
| 0.429764 | 0.319448 | 0.117 | 0.085 | 1 | 12 | Laptm4a | 0.032 | Young |
| 0.430021 | 0.507197 | 0.088 | 0.059 | 1 | 12 | Mrps18c | 0.029 | Young |
| 0.430484 | 0.638732 | 0.044 | 0.025 | 1 | 12 | Fam69a | 0.019 | Young |
| 0.430484 | 0.868365 | 0.044 | 0.025 | 1 | 12 | Prkcz | 0.019 | Young |
| 0.430484 | 0.763054 | 0.044 | 0.025 | 1 | 12 | Zdhhc17 | 0.019 | Young |
| 0.430484 | 0.761811 | 0.044 | 0.025 | 1 | 12 | Ufsp2 | 0.019 | Young |
| 0.430484 | 0.754968 | 0.044 | 0.025 | 1 | 12 | Pcsk1 | 0.019 | Young |
| 0.430484 | 0.748587 | 0.044 | 0.025 | 1 | 12 | CAAA0111ɛ | 0.019 | Young |
| 0.431142 | 0.545119 | 0.066 | 0.042 | 1 | 12 | Rnps1 | 0.024 | Young |
| 0.431142 | 0.483766 | 0.066 | 0.042 | 1 | 12 | Sirt3 | 0.024 | Young |
| 0.432065 | 0.413114 | 0.153 | 0.119 | 1 | 12 | Ap3s1 | 0.034 | Young |
| 0.43271 | 0.428273 | 0.182 | 0.153 | 1 | 12 | Gstm5 | 0.029 | Young |
| 0.432797 | 0.284878 | 0.365 | 0.331 | 1 | 12 | Atp5a1 | 0.034 | Young |
| 0.433606 | 0.946467 | 0.044 | 0.025 | 1 | 12 | Uap1 | 0.019 | Young |
| 0.433606 | 0.752612 | 0.044 | 0.025 | 1 | 12 | Igbp1 | 0.019 | Young |
| 0.433606 | 0.74006 | 0.044 | 0.025 | 1 | 12 | Cst6 | 0.019 | Young |
| 0.433606 | 0.718502 | 0.044 | 0.025 | 1 | 12 | Strn3 | 0.019 | Young |
| 0.433606 | 0.684463 | 0.044 | 0.025 | 1 | 12 | Spata7 | 0.019 | Young |
| 0.433606 | 0.664163 | 0.044 | 0.025 | 1 | 12 | Sgcb | 0.019 | Young |
| 0.433606 | 0.175363 | 0.044 | 0.025 | 1 | 12 | Mical3 | 0.019 | Young |
| 0.434074 | 0.635449 | 0.095 | 0.068 | 1 | 12 | 1500011B( | 0.027 | Young |
| 0.434588 | 0.170718 | 0.474 | 0.39 | 1 | 12 | Rpl7 | 0.084 | Young |
| 0.435125 | 0.58953 | 0.095 | 0.068 | 1 | 12 | Ppp3ca | 0.027 | Young |
| 0.435475 | 0.689445 | 0.073 | 0.051 | 1 | 12 | Nsmce1 | 0.022 | Young |
| 0.436209 | 0.639733 | 0.066 | 0.042 | 1 | 12 | Fam216a | 0.024 | Young |
| 0.436209 | 0.093934 | 0.066 | 0.042 | 1 | 12 | Prkaca | 0.024 | Young |
| 0.43734 | 0.400786 | 0.139 | 0.11 | 1 | 12 | Sf3b2 | 0.029 | Young |
| 0.43748 | 0.413092 | 0.066 | 0.042 | 1 | 12 | Ppp2r5b | 0.024 | Young |
| 0.439297 | 0.240508 | 0.124 | 0.093 | 1 | 12 | Slc25a33 | 0.031 | Young |
| 0.439888 | 0.688783 | 0.044 | 0.025 | 1 | 12 | Mvd | 0.019 | Young |
| 0.439888 | 0.597271 | 0.044 | 0.025 | 1 | 12 | Ubxn2a | 0.019 | Young |
| 0.439888 | 0.547444 | 0.044 | 0.025 | 1 | 12 | Agpat5 | 0.019 | Young |
| 0.440688 | 0.296234 | 0.234 | 0.203 | 1 | 12 | Serf2 | 0.031 | Young |
| 0.441308 | 0.442391 | 0.131 | 0.102 | 1 | 12 | Dnlz | 0.029 | Young |
| 0.441309 | 0.440329 | 0.066 | 0.042 | 1 | 12 | Stau2 | 0.024 | Young |
| 0.442413 | 0.094551 | 0.27 | 0.212 | 1 | 12 | Flot1 | 0.058 | Young |
| 0.442664 | 0.865585 | 0.073 | 0.051 | 1 | 12 | Srm | 0.022 | Young |
| 0.443048 | 0.615097 | 0.044 | 0.025 | 1 | 12 | Ube2q1 | 0.019 | Young |
| 0.443578 | 0.439864 | 0.095 | 0.068 | 1 | 12 | Epha5 | 0.027 | Young |
| 0.444572 | 0.34606 | 0.161 | 0.127 | 1 | 12 | Ngfr | 0.034 | Young |
| 0.44604 | 0.427273 | 0.117 | 0.085 | 1 | 12 | Ten1 | 0.032 | Young |
| 0.446661 | 0.378079 | 0.234 | 0.195 | 1 | 12 | Atxn10 | 0.039 | Young |
| 0.447888 | 0.353203 | 0.139 | 0.11 | 1 | 12 | Hdgf | 0.029 | Young |
| 0.448948 | 0.40814 | 0.117 | 0.085 | 1 | 12 | Pdcd10 | 0.032 | Young |
| 0.449408 | 0.543061 | 0.044 | 0.025 | 1 | 12 | Suds3 | 0.019 | Young |

| | | | | | | | | |
|---|---|---|---|---|---|---|---|---|
| 0.449408 | 0.459102 | 0.044 | 0.025 | 1 | 12 | Xpa | 0.019 | Young |
| 0.449475 | 0.489739 | 0.131 | 0.102 | 1 | 12 | Jak1 | 0.029 | Young |
| 0.450547 | 0.506094 | 0.168 | 0.136 | 1 | 12 | Phlda3 | 0.032 | Young |
| 0.450893 | 0.397896 | 0.117 | 0.085 | 1 | 12 | Reep1 | 0.032 | Young |
| 0.450893 | 0.357862 | 0.117 | 0.085 | 1 | 12 | Tmem167 | 0.032 | Young |
| 0.451135 | 0.75868 | 0.073 | 0.051 | 1 | 12 | Hspbp1 | 0.022 | Young |
| 0.452125 | 0.457382 | 0.095 | 0.068 | 1 | 12 | Mrpl35 | 0.027 | Young |
| 0.452607 | 0.507535 | 0.044 | 0.025 | 1 | 12 | Orc4 | 0.019 | Young |
| 0.452607 | 0.426693 | 0.044 | 0.025 | 1 | 12 | Slc9a3r2 | 0.019 | Young |
| 0.452607 | 0.365363 | 0.044 | 0.025 | 1 | 12 | Nop53 | 0.019 | Young |
| 0.453018 | 0.49333 | 0.197 | 0.169 | 1 | 12 | Tln2 | 0.028 | Young |
| 0.453371 | 0.2777 | 0.35 | 0.305 | 1 | 12 | Rpl7a | 0.045 | Young |
| 0.454793 | 0.745094 | 0.073 | 0.051 | 1 | 12 | Ap4s1 | 0.022 | Young |
| 0.455776 | 0.0648 | 0.117 | 0.085 | 1 | 12 | Tubb4a | 0.032 | Young |
| 0.455819 | 0.342517 | 0.044 | 0.025 | 1 | 12 | Acap3 | 0.019 | Young |
| 0.456433 | 0.434415 | 0.095 | 0.068 | 1 | 12 | Asph | 0.027 | Young |
| 0.457242 | 0.606616 | 0.073 | 0.051 | 1 | 12 | Samd14 | 0.022 | Young |
| 0.459698 | 0.733769 | 0.073 | 0.051 | 1 | 12 | Sigmar1 | 0.022 | Young |
| 0.460033 | 0.263929 | 0.474 | 0.441 | 1 | 12 | Prdx1 | 0.033 | Young |
| 0.460764 | 0.450777 | 0.095 | 0.068 | 1 | 12 | Commd1 | 0.027 | Young |
| 0.460764 | 0.398799 | 0.095 | 0.068 | 1 | 12 | Cadps | 0.027 | Young |
| 0.462161 | 0.496033 | 0.073 | 0.051 | 1 | 12 | Synrg | 0.022 | Young |
| 0.463077 | 0.535192 | 0.139 | 0.11 | 1 | 12 | Gclm | 0.029 | Young |
| 0.463303 | 0.265237 | 0.372 | 0.339 | 1 | 12 | Selenom | 0.033 | Young |
| 0.463395 | 0.503907 | 0.073 | 0.051 | 1 | 12 | Atraid | 0.022 | Young |
| 0.463806 | 0.347856 | 0.124 | 0.093 | 1 | 12 | Mrps36 | 0.031 | Young |
| 0.464631 | 0.240665 | 0.073 | 0.051 | 1 | 12 | Nt5c3 | 0.022 | Young |
| 0.464667 | 0.320561 | 0.168 | 0.127 | 1 | 12 | Ppp1r1a | 0.041 | Young |
| 0.467109 | 0.681898 | 0.073 | 0.051 | 1 | 12 | Prkar1b | 0.022 | Young |
| 0.467109 | 0.605267 | 0.073 | 0.051 | 1 | 12 | Jtb | 0.022 | Young |
| 0.467109 | 0.509884 | 0.073 | 0.051 | 1 | 12 | Mthfd2 | 0.022 | Young |
| 0.468074 | 0.904763 | 0.051 | 0.034 | 1 | 12 | Thoc3 | 0.017 | Young |
| 0.468409 | 0.067597 | 0.358 | 0.288 | 1 | 12 | Rpl12 | 0.07 | Young |
| 0.468821 | 0.590218 | 0.109 | 0.085 | 1 | 12 | Zcchc12 | 0.024 | Young |
| 0.46956 | 1.021041 | 0.051 | 0.034 | 1 | 12 | Cdk11b | 0.017 | Young |
| 0.469594 | 0.644828 | 0.073 | 0.051 | 1 | 12 | Smarca5 | 0.022 | Young |
| 0.469594 | 0.529311 | 0.073 | 0.051 | 1 | 12 | Atp6v1c1 | 0.022 | Young |
| 0.469642 | 0.447109 | 0.102 | 0.076 | 1 | 12 | Sfxn1 | 0.026 | Young |
| 0.470034 | 0.157278 | 0.401 | 0.331 | 1 | 12 | Ndufa11 | 0.07 | Young |
| 0.47169 | 0.285825 | 0.095 | 0.068 | 1 | 12 | Golga7 | 0.027 | Young |
| 0.472511 | 0.236763 | 0.358 | 0.322 | 1 | 12 | Swi5 | 0.036 | Young |
| 0.472604 | 0.234345 | 0.131 | 0.102 | 1 | 12 | Sptssa | 0.029 | Young |
| 0.475523 | 0.907597 | 0.051 | 0.034 | 1 | 12 | N6amt1 | 0.017 | Young |
| 0.476962 | 0.544622 | 0.109 | 0.085 | 1 | 12 | Nav1 | 0.024 | Young |
| 0.477042 | 0.441356 | 0.102 | 0.076 | 1 | 12 | Cplx1 | 0.026 | Young |

| | | | | | | | | |
|---|---|---|---|---|---|---|---|---|
| 0.477093 | 0.520908 | 0.073 | 0.051 | 1 | 12 | Tmem29 | 0.022 | Young |
| 0.478521 | 0.834105 | 0.051 | 0.034 | 1 | 12 | Pcdh7 | 0.017 | Young |
| 0.478864 | 0.095431 | 0.766 | 0.754 | 1 | 12 | Stmn1 | 0.012 | Young |
| 0.479607 | 0.489234 | 0.073 | 0.051 | 1 | 12 | Tmem199 | 0.022 | Young |
| 0.47967 | 0.264273 | 0.445 | 0.415 | 1 | 12 | Park7 | 0.03 | Young |
| 0.480023 | 1.075858 | 0.051 | 0.034 | 1 | 12 | Washc1 | 0.017 | Young |
| 0.480387 | 0.364692 | 0.139 | 0.11 | 1 | 12 | Stk25 | 0.029 | Young |
| 0.480535 | 0.294258 | 0.095 | 0.068 | 1 | 12 | Samm50 | 0.027 | Young |
| 0.481529 | 0.750641 | 0.051 | 0.034 | 1 | 12 | Ccdc127 | 0.017 | Young |
| 0.482847 | 0.499461 | 0.153 | 0.127 | 1 | 12 | Ubxn1 | 0.026 | Young |
| 0.484657 | 0.488735 | 0.073 | 0.051 | 1 | 12 | Atp2a2 | 0.022 | Young |
| 0.485497 | 0.103943 | 0.007 | 0.017 | 1 | 12 | Dennd5a | -0.01 | Young |
| 0.485497 | 0.06333 | 0.007 | 0.017 | 1 | 12 | Sec63 | -0.01 | Young |
| 0.485497 | 0.062493 | 0.007 | 0.017 | 1 | 12 | Prkab2 | -0.01 | Young |
| 0.485497 | 0.05443 | 0.007 | 0.017 | 1 | 12 | Col4a2 | -0.01 | Young |
| 0.485497 | 0.036765 | 0.007 | 0.017 | 1 | 12 | Smurf2 | -0.01 | Young |
| 0.487192 | 0.42323 | 0.073 | 0.051 | 1 | 12 | Thyn1 | 0.022 | Young |
| 0.487575 | 0.942618 | 0.051 | 0.034 | 1 | 12 | Spryd3 | 0.017 | Young |
| 0.487575 | 0.888049 | 0.051 | 0.034 | 1 | 12 | Snrpa | 0.017 | Young |
| 0.487575 | 0.841122 | 0.051 | 0.034 | 1 | 12 | Fxyd2 | 0.017 | Young |
| 0.487614 | 0.610172 | 0.08 | 0.059 | 1 | 12 | Pirt | 0.021 | Young |
| 0.490613 | 0.778482 | 0.051 | 0.034 | 1 | 12 | D430019H: | 0.017 | Young |
| 0.490613 | 0.622853 | 0.051 | 0.034 | 1 | 12 | Nudt3 | 0.017 | Young |
| 0.490954 | 0.316787 | 0.102 | 0.076 | 1 | 12 | Smim12 | 0.026 | Young |
| 0.491229 | 0.790338 | 0.08 | 0.059 | 1 | 12 | Eif4a3 | 0.021 | Young |
| 0.492136 | 0.695825 | 0.051 | 0.034 | 1 | 12 | Zfp330 | 0.017 | Young |
| 0.492285 | 0.328054 | 0.073 | 0.051 | 1 | 12 | Mvk | 0.022 | Young |
| 0.492546 | 0.281327 | 0.255 | 0.22 | 1 | 12 | Pcbd1 | 0.035 | Young |
| 0.493364 | 0.417568 | 0.139 | 0.11 | 1 | 12 | Arl6ip5 | 0.029 | Young |
| 0.493646 | 0.697182 | 0.08 | 0.059 | 1 | 12 | Hspa4 | 0.021 | Young |
| 0.493662 | 0.816471 | 0.051 | 0.034 | 1 | 12 | Aen | 0.017 | Young |
| 0.493662 | 0.786909 | 0.051 | 0.034 | 1 | 12 | Dpm3 | 0.017 | Young |
| 0.493662 | 0.612945 | 0.051 | 0.034 | 1 | 12 | Lamtor3 | 0.017 | Young |
| 0.494449 | 0.268882 | 0.131 | 0.102 | 1 | 12 | Glo1 | 0.029 | Young |
| 0.49607 | 0.618622 | 0.08 | 0.059 | 1 | 12 | Ankrd11 | 0.021 | Young |
| 0.49672 | 0.809905 | 0.051 | 0.034 | 1 | 12 | Ap2a1 | 0.017 | Young |
| 0.49672 | 0.634358 | 0.051 | 0.034 | 1 | 12 | Ttc33 | 0.017 | Young |
| 0.497406 | 0.27551 | 0.073 | 0.051 | 1 | 12 | Gkap1 | 0.022 | Young |
| 0.499621 | 0.323293 | 0.102 | 0.076 | 1 | 12 | Elp5 | 0.026 | Young |
| 0.499789 | 0.652464 | 0.051 | 0.034 | 1 | 12 | Pycr2 | 0.017 | Young |
| 0.499789 | 0.57675 | 0.051 | 0.034 | 1 | 12 | Tm4sf1 | 0.017 | Young |
| 0.499977 | 0.343011 | 0.073 | 0.051 | 1 | 12 | Rab24 | 0.022 | Young |
| 0.501643 | 0.289034 | 0.212 | 0.178 | 1 | 12 | Csnk1a1 | 0.034 | Young |
| 0.50247 | 0.091897 | 0.898 | 0.864 | 1 | 12 | mt-Co3 | 0.034 | Young |
| 0.502684 | 0.735915 | 0.088 | 0.068 | 1 | 12 | Arl6ip4 | 0.02 | Young |

| | | | | | | | | |
|---|---|---|---|---|---|---|---|---|
| 0.502868 | 0.629771 | 0.051 | 0.034 | 1 | 12 | Gm26782 | 0.017 | Young |
| 0.502868 | 0.613371 | 0.051 | 0.034 | 1 | 12 | Rpl36al | 0.017 | Young |
| 0.502868 | 0.566229 | 0.051 | 0.034 | 1 | 12 | Lrrc59 | 0.017 | Young |
| 0.50338 | 0.67643 | 0.08 | 0.059 | 1 | 12 | Ppp1cb | 0.021 | Young |
| 0.503847 | 0.261052 | 0.073 | 0.051 | 1 | 12 | Hspa2 | 0.022 | Young |
| 0.504134 | 0.296696 | 0.117 | 0.093 | 1 | 12 | Actr2 | 0.024 | Young |
| 0.504603 | 0.599192 | 0.08 | 0.059 | 1 | 12 | Man2a1 | 0.021 | Young |
| 0.504958 | 0.258664 | 0.124 | 0.093 | 1 | 12 | Nol7 | 0.031 | Young |
| 0.504999 | 0.384456 | 0.109 | 0.085 | 1 | 12 | Mob4 | 0.024 | Young |
| 0.505085 | 0.403656 | 0.131 | 0.102 | 1 | 12 | Psmd14 | 0.029 | Young |
| 0.505115 | 0.07385 | 0.869 | 0.881 | 1 | 12 | Prph | -0.012 | Young |
| 0.505323 | 0.324298 | 0.168 | 0.136 | 1 | 12 | Lamp1 | 0.032 | Young |
| 0.505957 | 0.709152 | 0.051 | 0.034 | 1 | 12 | Dcaf8 | 0.017 | Young |
| 0.505957 | 0.536926 | 0.051 | 0.034 | 1 | 12 | Phf14 | 0.017 | Young |
| 0.506214 | 1.921483 | 0.029 | 0.017 | 1 | 12 | Cyfip1 | 0.012 | Young |
| 0.506214 | 1.756571 | 0.029 | 0.017 | 1 | 12 | Pdia4 | 0.012 | Young |
| 0.506214 | 1.626804 | 0.029 | 0.017 | 1 | 12 | Smchd1 | 0.012 | Young |
| 0.506214 | 1.53675 | 0.029 | 0.017 | 1 | 12 | Gtf2h1 | 0.012 | Young |
| 0.506214 | 1.335666 | 0.029 | 0.017 | 1 | 12 | Rbmx2 | 0.012 | Young |
| 0.506214 | 1.295818 | 0.029 | 0.017 | 1 | 12 | Gnpnat1 | 0.012 | Young |
| 0.506214 | 1.261633 | 0.029 | 0.017 | 1 | 12 | Atxn7l3 | 0.012 | Young |
| 0.506214 | 1.08033 | 0.029 | 0.017 | 1 | 12 | Pygb | 0.012 | Young |
| 0.508284 | 0.492985 | 0.08 | 0.059 | 1 | 12 | Hnrnpf | 0.021 | Young |
| 0.509055 | 0.50129 | 0.051 | 0.034 | 1 | 12 | Msrb2 | 0.017 | Young |
| 0.509137 | 0.093399 | 0.577 | 0.5 | 1 | 12 | Elob | 0.077 | Young |
| 0.509741 | 0.303677 | 0.182 | 0.153 | 1 | 12 | Map2k2 | 0.029 | Young |
| 0.510332 | 0.214246 | 0.073 | 0.051 | 1 | 12 | Mkrn1 | 0.022 | Young |
| 0.510372 | 1.739239 | 0.029 | 0.017 | 1 | 12 | Ddx17 | 0.012 | Young |
| 0.510372 | 1.539753 | 0.029 | 0.017 | 1 | 12 | Socs4 | 0.012 | Young |
| 0.510372 | 1.508841 | 0.029 | 0.017 | 1 | 12 | 2210016F1 | 0.012 | Young |
| 0.510372 | 1.252395 | 0.029 | 0.017 | 1 | 12 | Ppp1r35 | 0.012 | Young |
| 0.510372 | 1.135987 | 0.029 | 0.017 | 1 | 12 | Gna12 | 0.012 | Young |
| 0.510372 | 1.112561 | 0.029 | 0.017 | 1 | 12 | Prkrip1 | 0.012 | Young |
| 0.510372 | 1.07611 | 0.029 | 0.017 | 1 | 12 | Camsap1 | 0.012 | Young |
| 0.510372 | 1.068881 | 0.029 | 0.017 | 1 | 12 | Taf1d | 0.012 | Young |
| 0.510372 | 1.059304 | 0.029 | 0.017 | 1 | 12 | Ugp2 | 0.012 | Young |
| 0.510372 | 0.94289 | 0.029 | 0.017 | 1 | 12 | Srgap3 | 0.012 | Young |
| 0.512164 | 0.076669 | 0.051 | 0.034 | 1 | 12 | Med9 | 0.017 | Young |
| 0.512407 | 0.433046 | 0.109 | 0.085 | 1 | 12 | Lsm7 | 0.024 | Young |
| 0.513683 | 0.004535 | 0.65 | 0.602 | 1 | 12 | Ybx1 | 0.048 | Young |
| 0.514079 | 0.150113 | 0.774 | 0.712 | 1 | 12 | Rpl19 | 0.062 | Young |
| 0.514548 | 1.497294 | 0.029 | 0.017 | 1 | 12 | Abracl | 0.012 | Young |
| 0.514548 | 1.455763 | 0.029 | 0.017 | 1 | 12 | P4hb | 0.012 | Young |
| 0.514548 | 1.38566 | 0.029 | 0.017 | 1 | 12 | Slu7 | 0.012 | Young |
| 0.514548 | 1.094391 | 0.029 | 0.017 | 1 | 12 | Micu1 | 0.012 | Young |

| | | | | | | | | |
|---|---|---|---|---|---|---|---|---|
| 0.514548 | 1.047867 | 0.029 | 0.017 | 1 | 12 | Ctps | 0.012 | Young |
| 0.514548 | 1.042267 | 0.029 | 0.017 | 1 | 12 | Immp1l | 0.012 | Young |
| 0.514548 | 1.027596 | 0.029 | 0.017 | 1 | 12 | Topors | 0.012 | Young |
| 0.514548 | 0.992923 | 0.029 | 0.017 | 1 | 12 | Gpr150 | 0.012 | Young |
| 0.514548 | 0.98437 | 0.029 | 0.017 | 1 | 12 | Slc25a22 | 0.012 | Young |
| 0.514548 | 0.940943 | 0.029 | 0.017 | 1 | 12 | Arpin | 0.012 | Young |
| 0.514548 | 0.882461 | 0.029 | 0.017 | 1 | 12 | Aup1 | 0.012 | Young |
| 0.514548 | 0.863193 | 0.029 | 0.017 | 1 | 12 | Gcc2 | 0.012 | Young |
| 0.514548 | 0.821642 | 0.029 | 0.017 | 1 | 12 | Kat7 | 0.012 | Young |
| 0.514548 | 0.748975 | 0.029 | 0.017 | 1 | 12 | Cdkn1b | 0.012 | Young |
| 0.514796 | 0.352191 | 0.146 | 0.119 | 1 | 12 | Capzb | 0.027 | Young |
| 0.514983 | 0.315453 | 0.102 | 0.076 | 1 | 12 | Ptpra | 0.026 | Young |
| 0.515283 | 0.584031 | 0.051 | 0.034 | 1 | 12 | Capn2 | 0.017 | Young |
| 0.515283 | 0.369244 | 0.051 | 0.034 | 1 | 12 | Smim8 | 0.017 | Young |
| 0.516666 | 0.344778 | 0.109 | 0.085 | 1 | 12 | Polr2g | 0.024 | Young |
| 0.516845 | 0.472965 | 0.051 | 0.034 | 1 | 12 | 2300009A( | 0.017 | Young |
| 0.516845 | 0.443769 | 0.051 | 0.034 | 1 | 12 | Tusc1 | 0.017 | Young |
| 0.517514 | 0.165843 | 0.708 | 0.695 | 1 | 12 | Meg3 | 0.013 | Young |
| 0.517991 | 0.408381 | 0.234 | 0.203 | 1 | 12 | Plcb4 | 0.031 | Young |
| 0.518411 | 0.444279 | 0.051 | 0.034 | 1 | 12 | Chd6 | 0.017 | Young |
| 0.518741 | 1.456681 | 0.029 | 0.017 | 1 | 12 | Mterf4 | 0.012 | Young |
| 0.518741 | 1.412869 | 0.029 | 0.017 | 1 | 12 | Tmem80 | 0.012 | Young |
| 0.518741 | 1.289644 | 0.029 | 0.017 | 1 | 12 | Prkag1 | 0.012 | Young |
| 0.518741 | 1.230666 | 0.029 | 0.017 | 1 | 12 | Elmo2 | 0.012 | Young |
| 0.518741 | 1.105504 | 0.029 | 0.017 | 1 | 12 | Adra2c | 0.012 | Young |
| 0.518741 | 0.973637 | 0.029 | 0.017 | 1 | 12 | M6pr | 0.012 | Young |
| 0.518741 | 0.944216 | 0.029 | 0.017 | 1 | 12 | Kif3b | 0.012 | Young |
| 0.518741 | 0.938545 | 0.029 | 0.017 | 1 | 12 | 3830406C1 | 0.012 | Young |
| 0.518741 | 0.934777 | 0.029 | 0.017 | 1 | 12 | Plbd2 | 0.012 | Young |
| 0.518741 | 0.894502 | 0.029 | 0.017 | 1 | 12 | Zfp322a | 0.012 | Young |
| 0.518741 | 0.872844 | 0.029 | 0.017 | 1 | 12 | Hist1h1c | 0.012 | Young |
| 0.518741 | 0.822362 | 0.029 | 0.017 | 1 | 12 | Pbx1 | 0.012 | Young |
| 0.518741 | 0.820733 | 0.029 | 0.017 | 1 | 12 | Trim24 | 0.012 | Young |
| 0.518741 | 0.796544 | 0.029 | 0.017 | 1 | 12 | Shtn1 | 0.012 | Young |
| 0.518741 | 0.790185 | 0.029 | 0.017 | 1 | 12 | Gm13889 | 0.012 | Young |
| 0.519119 | 0.553361 | 0.088 | 0.068 | 1 | 12 | Snrnp27 | 0.02 | Young |
| 0.519978 | 0.319072 | 0.051 | 0.034 | 1 | 12 | Deaf1 | 0.017 | Young |
| 0.520456 | 0.343662 | 0.204 | 0.178 | 1 | 12 | Aurkaip1 | 0.026 | Young |
| 0.520554 | 0.550232 | 0.117 | 0.093 | 1 | 12 | Stard3nl | 0.024 | Young |
| 0.521549 | 0.337392 | 0.051 | 0.034 | 1 | 12 | Tmem230 | 0.017 | Young |
| 0.522951 | 1.224197 | 0.029 | 0.017 | 1 | 12 | Tubgcp2 | 0.012 | Young |
| 0.522951 | 1.196411 | 0.029 | 0.017 | 1 | 12 | Tnks | 0.012 | Young |
| 0.522951 | 1.066487 | 0.029 | 0.017 | 1 | 12 | Mtss1 | 0.012 | Young |
| 0.522951 | 1.055701 | 0.029 | 0.017 | 1 | 12 | Tfdp1 | 0.012 | Young |
| 0.522951 | 0.966502 | 0.029 | 0.017 | 1 | 12 | H2-Ke6 | 0.012 | Young |

| | | | | | | | | |
|---|---|---|---|---|---|---|---|---|
| 0.522951 | 0.963082 | 0.029 | 0.017 | 1 | 12 | Uspl1 | 0.012 | Young |
| 0.522951 | 0.929792 | 0.029 | 0.017 | 1 | 12 | Sppl3 | 0.012 | Young |
| 0.522951 | 0.917853 | 0.029 | 0.017 | 1 | 12 | Stau1 | 0.012 | Young |
| 0.522951 | 0.905448 | 0.029 | 0.017 | 1 | 12 | Ehd3 | 0.012 | Young |
| 0.522951 | 0.870217 | 0.029 | 0.017 | 1 | 12 | Pmpcb | 0.012 | Young |
| 0.522951 | 0.858842 | 0.029 | 0.017 | 1 | 12 | 2410131K1 | 0.012 | Young |
| 0.522951 | 0.819518 | 0.029 | 0.017 | 1 | 12 | Eif2b4 | 0.012 | Young |
| 0.522951 | 0.817623 | 0.029 | 0.017 | 1 | 12 | Gamt | 0.012 | Young |
| 0.522951 | 0.805934 | 0.029 | 0.017 | 1 | 12 | Dnajc11 | 0.012 | Young |
| 0.522951 | 0.802092 | 0.029 | 0.017 | 1 | 12 | Bckdk | 0.012 | Young |
| 0.522951 | 0.796173 | 0.029 | 0.017 | 1 | 12 | Rcl1 | 0.012 | Young |
| 0.522951 | 0.786983 | 0.029 | 0.017 | 1 | 12 | Cdh2 | 0.012 | Young |
| 0.522951 | 0.748459 | 0.029 | 0.017 | 1 | 12 | Cyb5r1 | 0.012 | Young |
| 0.522951 | 0.74168 | 0.029 | 0.017 | 1 | 12 | Snx4 | 0.012 | Young |
| 0.522951 | 0.731563 | 0.029 | 0.017 | 1 | 12 | Upf1 | 0.012 | Young |
| 0.522951 | 0.712647 | 0.029 | 0.017 | 1 | 12 | Nap1l2 | 0.012 | Young |
| 0.522951 | 0.708067 | 0.029 | 0.017 | 1 | 12 | Kat5 | 0.012 | Young |
| 0.522951 | 0.689192 | 0.029 | 0.017 | 1 | 12 | Lysmd1 | 0.012 | Young |
| 0.522951 | 0.683506 | 0.029 | 0.017 | 1 | 12 | Cbfb | 0.012 | Young |
| 0.522951 | 0.533378 | 0.029 | 0.017 | 1 | 12 | Timm50 | 0.012 | Young |
| 0.522951 | 0.317921 | 0.029 | 0.017 | 1 | 12 | Spsb4 | 0.012 | Young |
| 0.524162 | 0.40454 | 0.109 | 0.085 | 1 | 12 | Rbm8a | 0.024 | Young |
| 0.524697 | 0.210594 | 0.051 | 0.034 | 1 | 12 | Borcs6 | 0.017 | Young |
| 0.52718 | 0.773476 | 0.029 | 0.017 | 1 | 12 | Dctn5 | 0.012 | Young |
| 0.52718 | 0.761405 | 0.029 | 0.017 | 1 | 12 | Zfp821 | 0.012 | Young |
| 0.52718 | 0.739959 | 0.029 | 0.017 | 1 | 12 | Wdfy3 | 0.012 | Young |
| 0.52718 | 0.702553 | 0.029 | 0.017 | 1 | 12 | Ppp1r12a | 0.012 | Young |
| 0.52718 | 0.685899 | 0.029 | 0.017 | 1 | 12 | Cinp | 0.012 | Young |
| 0.52718 | 0.65455 | 0.029 | 0.017 | 1 | 12 | Nub1 | 0.012 | Young |
| 0.52718 | 0.621322 | 0.029 | 0.017 | 1 | 12 | Txnrd2 | 0.012 | Young |
| 0.52718 | 0.61858 | 0.029 | 0.017 | 1 | 12 | Plpp7 | 0.012 | Young |
| 0.52718 | 0.603211 | 0.029 | 0.017 | 1 | 12 | Nom1 | 0.012 | Young |
| 0.52718 | 0.590965 | 0.029 | 0.017 | 1 | 12 | Nat14 | 0.012 | Young |
| 0.52718 | 0.581296 | 0.029 | 0.017 | 1 | 12 | Zfp804a | 0.012 | Young |
| 0.52718 | 0.570289 | 0.029 | 0.017 | 1 | 12 | Tmem151a | 0.012 | Young |
| 0.52718 | 0.565826 | 0.029 | 0.017 | 1 | 12 | Pard6a | 0.012 | Young |
| 0.52718 | 0.515482 | 0.029 | 0.017 | 1 | 12 | Gm2a | 0.012 | Young |
| 0.52718 | 0.464219 | 0.029 | 0.017 | 1 | 12 | Kctd20 | 0.012 | Young |
| 0.52718 | 0.27272 | 0.029 | 0.017 | 1 | 12 | Ccl9 | 0.012 | Young |
| 0.527854 | 0.379466 | 0.051 | 0.034 | 1 | 12 | Med8 | 0.017 | Young |
| 0.52815 | 0.581955 | 0.08 | 0.059 | 1 | 12 | Med28 | 0.021 | Young |
| 0.52815 | 0.426784 | 0.08 | 0.059 | 1 | 12 | Sv2c | 0.021 | Young |
| 0.52815 | 0.380665 | 0.08 | 0.059 | 1 | 12 | Chp1 | 0.021 | Young |
| 0.529859 | 0.352933 | 0.248 | 0.229 | 1 | 12 | Snap47 | 0.019 | Young |
| 0.531022 | 0.292546 | 0.051 | 0.034 | 1 | 12 | Ppp4r2 | 0.017 | Young |

| | | | | | | | | |
|---|---|---|---|---|---|---|---|---|
| 0.531022 | 0.273675 | 0.051 | 0.034 | 1 | 12 | Dab2ip | 0.017 | Young |
| 0.531425 | 0.700318 | 0.029 | 0.017 | 1 | 12 | Acsl3 | 0.012 | Young |
| 0.531425 | 0.623111 | 0.029 | 0.017 | 1 | 12 | Hspa12a | 0.012 | Young |
| 0.531425 | 0.56584 | 0.029 | 0.017 | 1 | 12 | Dact3 | 0.012 | Young |
| 0.531425 | 0.521448 | 0.029 | 0.017 | 1 | 12 | Extl2 | 0.012 | Young |
| 0.531425 | 0.47131 | 0.029 | 0.017 | 1 | 12 | Snw1 | 0.012 | Young |
| 0.531425 | 0.468451 | 0.029 | 0.017 | 1 | 12 | Bag2 | 0.012 | Young |
| 0.531425 | 0.454186 | 0.029 | 0.017 | 1 | 12 | Prrg3 | 0.012 | Young |
| 0.531425 | 0.439602 | 0.029 | 0.017 | 1 | 12 | Msantd4 | 0.012 | Young |
| 0.531425 | 0.431681 | 0.029 | 0.017 | 1 | 12 | Thoc6 | 0.012 | Young |
| 0.532171 | 0.057643 | 0.701 | 0.61 | 1 | 12 | Dpysl3 | 0.091 | Young |
| 0.532836 | 0.237676 | 0.102 | 0.076 | 1 | 12 | Ldhb | 0.026 | Young |
| 0.534692 | 0.26556 | 0.131 | 0.102 | 1 | 12 | Ret | 0.029 | Young |
| 0.535148 | 0.292357 | 0.153 | 0.127 | 1 | 12 | Map4 | 0.026 | Young |
| 0.535688 | 0.466712 | 0.029 | 0.017 | 1 | 12 | Vps4b | 0.012 | Young |
| 0.535688 | 0.392909 | 0.029 | 0.017 | 1 | 12 | Ppt2 | 0.012 | Young |
| 0.535688 | 0.386835 | 0.029 | 0.017 | 1 | 12 | Cdk10 | 0.012 | Young |
| 0.535688 | 0.343686 | 0.029 | 0.017 | 1 | 12 | Wbp11 | 0.012 | Young |
| 0.535688 | 0.341612 | 0.029 | 0.017 | 1 | 12 | Tnr | 0.012 | Young |
| 0.535688 | 0.337133 | 0.029 | 0.017 | 1 | 12 | Nrbp1 | 0.012 | Young |
| 0.536054 | 0.503005 | 0.109 | 0.085 | 1 | 12 | Vstm2l | 0.024 | Young |
| 0.536258 | 0.329734 | 0.161 | 0.136 | 1 | 12 | Mapre1 | 0.025 | Young |
| 0.537347 | 0.027721 | 0.102 | 0.076 | 1 | 12 | Ddost | 0.026 | Young |
| 0.53823 | 0.29509 | 0.109 | 0.085 | 1 | 12 | Slco3a1 | 0.024 | Young |
| 0.53894 | 0.332707 | 0.197 | 0.169 | 1 | 12 | Ddx24 | 0.028 | Young |
| 0.539746 | 0.493837 | 0.124 | 0.102 | 1 | 12 | Snx6 | 0.022 | Young |
| 0.539967 | 0.404275 | 0.029 | 0.017 | 1 | 12 | Stac | 0.012 | Young |
| 0.539967 | 0.274321 | 0.029 | 0.017 | 1 | 12 | Them6 | 0.012 | Young |
| 0.539968 | 0.554746 | 0.029 | 0.017 | 1 | 12 | AW209491 | 0.012 | Young |
| 0.539968 | 0.506861 | 0.029 | 0.017 | 1 | 12 | Gon7 | 0.012 | Young |
| 0.539968 | 0.410612 | 0.029 | 0.017 | 1 | 12 | Frmd3 | 0.012 | Young |
| 0.539968 | 0.402777 | 0.029 | 0.017 | 1 | 12 | Smarcd3 | 0.012 | Young |
| 0.539968 | 0.306229 | 0.029 | 0.017 | 1 | 12 | Armc6 | 0.012 | Young |
| 0.539968 | 0.247294 | 0.029 | 0.017 | 1 | 12 | Chmp7 | 0.012 | Young |
| 0.539968 | 0.246668 | 0.029 | 0.017 | 1 | 12 | Lars2 | 0.012 | Young |
| 0.539968 | 0.219406 | 0.029 | 0.017 | 1 | 12 | Senp2 | 0.012 | Young |
| 0.539968 | 0.145756 | 0.029 | 0.017 | 1 | 12 | Mrc1 | 0.012 | Young |
| 0.540502 | 0.668833 | 0.058 | 0.042 | 1 | 12 | Kctd13 | 0.016 | Young |
| 0.540581 | 0.090876 | 0.051 | 0.034 | 1 | 12 | Surf1 | 0.017 | Young |
| 0.541175 | 0.2889 | 0.146 | 0.119 | 1 | 12 | Tmem14c | 0.027 | Young |
| 0.541732 | 0.070275 | 0.752 | 0.746 | 1 | 12 | Cox8a | 0.006 | Young |
| 0.542018 | 0.132922 | 0.015 | 0.025 | 1 | 12 | Nf1 | -0.01 | Young |
| 0.542517 | 0.179816 | 0.117 | 0.093 | 1 | 12 | Eny2 | 0.024 | Young |
| 0.543462 | 0.59082 | 0.058 | 0.042 | 1 | 12 | Nae1 | 0.016 | Young |
| 0.54589 | 0.428391 | 0.124 | 0.102 | 1 | 12 | Mcub | 0.022 | Young |

| | | | | | | | | |
|---|---|---|---|---|---|---|---|---|
| 0.546726 | 0.007204 | 0.015 | 0.025 | 1 | 12 | Zfhx3 | -0.01 | Young |
| 0.546914 | 0.053812 | 0.124 | 0.093 | 1 | 12 | Th | 0.031 | Young |
| 0.547126 | 0.281154 | 0.08 | 0.059 | 1 | 12 | Klf13 | 0.021 | Young |
| 0.549405 | 0.713681 | 0.058 | 0.042 | 1 | 12 | Asrgl1 | 0.016 | Young |
| 0.549405 | 0.553975 | 0.058 | 0.042 | 1 | 12 | Atp13a2 | 0.016 | Young |
| 0.550637 | 0.246002 | 0.168 | 0.136 | 1 | 12 | Taf10 | 0.032 | Young |
| 0.550852 | 0.154034 | 0.35 | 0.331 | 1 | 12 | Ckb | 0.019 | Young |
| 0.551454 | 0.134276 | 0.015 | 0.025 | 1 | 12 | Actn4 | -0.01 | Young |
| 0.551454 | 0.114725 | 0.015 | 0.025 | 1 | 12 | Fam3c | -0.01 | Young |
| 0.552389 | 0.524514 | 0.058 | 0.042 | 1 | 12 | Chd5 | 0.016 | Young |
| 0.552803 | 0.544943 | 0.088 | 0.068 | 1 | 12 | Tspan7 | 0.02 | Young |
| 0.55351 | 0.438006 | 0.131 | 0.11 | 1 | 12 | Napg | 0.021 | Young |
| 0.55359 | 0.173392 | 0.109 | 0.085 | 1 | 12 | Gps1 | 0.024 | Young |
| 0.555249 | 0.461874 | 0.088 | 0.068 | 1 | 12 | Pkig | 0.02 | Young |
| 0.555271 | 0.578112 | 0.117 | 0.093 | 1 | 12 | Apoe | 0.024 | Young |
| 0.555381 | 0.693731 | 0.058 | 0.042 | 1 | 12 | Scaf11 | 0.016 | Young |
| 0.555381 | 0.539692 | 0.058 | 0.042 | 1 | 12 | Faap20 | 0.016 | Young |
| 0.55558 | 0.193652 | 0.146 | 0.119 | 1 | 12 | Rac1 | 0.027 | Young |
| 0.556219 | 0.291361 | 0.175 | 0.144 | 1 | 12 | Trmt112 | 0.031 | Young |
| 0.556302 | 0.145813 | 0.518 | 0.492 | 1 | 12 | Rab3a | 0.026 | Young |
| 0.556362 | 0.27977 | 0.248 | 0.237 | 1 | 12 | Psmc5 | 0.011 | Young |
| 0.557701 | 0.368066 | 0.088 | 0.068 | 1 | 12 | Nrxn2 | 0.02 | Young |
| 0.558214 | 0.266802 | 0.161 | 0.136 | 1 | 12 | Cuedc2 | 0.025 | Young |
| 0.55838 | 0.520527 | 0.058 | 0.042 | 1 | 12 | Rsl1d1 | 0.016 | Young |
| 0.55838 | 0.487412 | 0.058 | 0.042 | 1 | 12 | 1500009L1 | 0.016 | Young |
| 0.561388 | 0.509218 | 0.058 | 0.042 | 1 | 12 | Myl12a | 0.016 | Young |
| 0.561882 | 0.113177 | 0.321 | 0.271 | 1 | 12 | Ndufa7 | 0.05 | Young |
| 0.56218 | 0.225342 | 0.314 | 0.288 | 1 | 12 | Rnf7 | 0.026 | Young |
| 0.564746 | 0.180902 | 0.27 | 0.22 | 1 | 12 | Eif3k | 0.05 | Young |
| 0.566168 | 0.476389 | 0.139 | 0.119 | 1 | 12 | Pdlim7 | 0.02 | Young |
| 0.566726 | 0.030561 | 0.19 | 0.153 | 1 | 12 | Nucks1 | 0.037 | Young |
| 0.567426 | 0.448775 | 0.058 | 0.042 | 1 | 12 | Mapre2 | 0.016 | Young |
| 0.567426 | 0.385424 | 0.058 | 0.042 | 1 | 12 | Fem1b | 0.016 | Young |
| 0.569033 | 0.171596 | 0.08 | 0.059 | 1 | 12 | Uchl3 | 0.021 | Young |
| 0.570037 | 0.342486 | 0.088 | 0.068 | 1 | 12 | Echs1 | 0.02 | Young |
| 0.570105 | 0.711097 | 0.066 | 0.051 | 1 | 12 | Ap2a2 | 0.015 | Young |
| 0.570457 | 0.493688 | 0.058 | 0.042 | 1 | 12 | 2410004B1 | 0.016 | Young |
| 0.570457 | 0.441003 | 0.058 | 0.042 | 1 | 12 | Rab5a | 0.016 | Young |
| 0.570457 | 0.384902 | 0.058 | 0.042 | 1 | 12 | Crlf2 | 0.016 | Young |
| 0.570709 | 0.558335 | 0.131 | 0.11 | 1 | 12 | Ddah1 | 0.021 | Young |
| 0.570783 | 0.198778 | 0.234 | 0.195 | 1 | 12 | Rexo2 | 0.039 | Young |
| 0.57338 | 0.379387 | 0.212 | 0.186 | 1 | 12 | Zcchc18 | 0.026 | Young |
| 0.573495 | 0.378346 | 0.058 | 0.042 | 1 | 12 | Mapk10 | 0.016 | Young |
| 0.573495 | 0.318711 | 0.058 | 0.042 | 1 | 12 | Aktip | 0.016 | Young |
| 0.573586 | 0.248861 | 0.117 | 0.093 | 1 | 12 | Elmo1 | 0.024 | Young |

| | | | | | | | | |
|---|---|---|---|---|---|---|---|---|
| 0.575017 | 0.376677 | 0.058 | 0.042 | 1 | 12 | Rrp7a | 0.016 | Young |
| 0.57546 | 0.237177 | 0.182 | 0.153 | 1 | 12 | Ctnnbip1 | 0.029 | Young |
| 0.576254 | 0.279979 | 0.088 | 0.068 | 1 | 12 | Fam171a2 | 0.02 | Young |
| 0.576842 | 0.609558 | 0.102 | 0.085 | 1 | 12 | Lcmt1 | 0.017 | Young |
| 0.579595 | 0.461999 | 0.058 | 0.042 | 1 | 12 | Mrps23 | 0.016 | Young |
| 0.579595 | 0.348722 | 0.058 | 0.042 | 1 | 12 | Birc6 | 0.016 | Young |
| 0.579595 | 0.331045 | 0.058 | 0.042 | 1 | 12 | Riiad1 | 0.016 | Young |
| 0.579595 | 0.313183 | 0.058 | 0.042 | 1 | 12 | Fnta | 0.016 | Young |
| 0.579933 | 0.210169 | 0.182 | 0.161 | 1 | 12 | Emc6 | 0.021 | Young |
| 0.579999 | 0.422101 | 0.088 | 0.068 | 1 | 12 | Map1a | 0.02 | Young |
| 0.580056 | 0.484454 | 0.161 | 0.144 | 1 | 12 | Lsm4 | 0.017 | Young |
| 0.581994 | 0.164556 | 0.022 | 0.034 | 1 | 12 | Zcchc6 | -0.012 | Young |
| 0.582547 | 0.009246 | 0.182 | 0.144 | 1 | 12 | Fam96b | 0.038 | Young |
| 0.582657 | 0.314684 | 0.058 | 0.042 | 1 | 12 | Ap3b2 | 0.016 | Young |
| 0.582657 | 0.282754 | 0.058 | 0.042 | 1 | 12 | Hnrnpr | 0.016 | Young |
| 0.585411 | 0.349614 | 0.095 | 0.076 | 1 | 12 | Nhp2 | 0.019 | Young |
| 0.585725 | 0.401091 | 0.058 | 0.042 | 1 | 12 | Rnf181 | 0.016 | Young |
| 0.586126 | 0.316012 | 0.022 | 0.034 | 1 | 12 | Rab8a | -0.012 | Young |
| 0.586126 | 0.108263 | 0.022 | 0.034 | 1 | 12 | Cenpb | -0.012 | Young |
| 0.586126 | 0.071793 | 0.022 | 0.034 | 1 | 12 | Ifitm2 | -0.012 | Young |
| 0.586348 | 1.528635 | 0.036 | 0.025 | 1 | 12 | Brix1 | 0.011 | Young |
| 0.586348 | 1.06454 | 0.036 | 0.025 | 1 | 12 | Pf4 | 0.011 | Young |
| 0.586484 | 0.292619 | 0.241 | 0.212 | 1 | 12 | Bri3 | 0.029 | Young |
| 0.586812 | 1.075707 | 0.073 | 0.059 | 1 | 12 | Pik3r1 | 0.014 | Young |
| 0.587717 | 0.262393 | 0.124 | 0.102 | 1 | 12 | Ube2a | 0.022 | Young |
| 0.588461 | 0.070785 | 0.139 | 0.169 | 1 | 12 | Ly6h | -0.03 | Young |
| 0.588461 | 0.04609 | 0.168 | 0.136 | 1 | 12 | Cyb5r3 | 0.032 | Young |
| 0.589192 | 0.245181 | 0.131 | 0.11 | 1 | 12 | Tmed2 | 0.021 | Young |
| 0.591885 | 0.292278 | 0.058 | 0.042 | 1 | 12 | Tsfm | 0.016 | Young |
| 0.591885 | 0.271846 | 0.058 | 0.042 | 1 | 12 | Tmem47 | 0.016 | Young |
| 0.591885 | 0.214058 | 0.058 | 0.042 | 1 | 12 | Naxd | 0.016 | Young |
| 0.591885 | 0.121432 | 0.058 | 0.042 | 1 | 12 | Clu | 0.016 | Young |
| 0.593334 | 0.097442 | 0.628 | 0.585 | 1 | 12 | Cox7c | 0.043 | Young |
| 0.594049 | 0.072134 | 0.153 | 0.127 | 1 | 12 | Nudc | 0.026 | Young |
| 0.594132 | 1.230604 | 0.036 | 0.025 | 1 | 12 | Tm9sf2 | 0.011 | Young |
| 0.594132 | 1.147024 | 0.036 | 0.025 | 1 | 12 | Hexa | 0.011 | Young |
| 0.594132 | 0.829705 | 0.036 | 0.025 | 1 | 12 | Eif4e3 | 0.011 | Young |
| 0.594132 | 0.708067 | 0.036 | 0.025 | 1 | 12 | Nr1h2 | 0.011 | Young |
| 0.594132 | 0.680011 | 0.036 | 0.025 | 1 | 12 | Crtac1 | 0.011 | Young |
| 0.594327 | 0.061775 | 0.234 | 0.195 | 1 | 12 | Arpc2 | 0.039 | Young |
| 0.594448 | 0.649704 | 0.066 | 0.051 | 1 | 12 | Mbp | 0.015 | Young |
| 0.594448 | 0.533124 | 0.066 | 0.051 | 1 | 12 | Cops2 | 0.015 | Young |
| 0.594508 | 0.461581 | 0.146 | 0.127 | 1 | 12 | Rer1 | 0.019 | Young |
| 0.594977 | 0.328279 | 0.058 | 0.042 | 1 | 12 | Mras | 0.016 | Young |
| 0.594977 | 0.205587 | 0.058 | 0.042 | 1 | 12 | Mblac2 | 0.016 | Young |

| | | | | | | | | |
|---|---|---|---|---|---|---|---|---|
| 0.595494 | 0.029158 | 0.095 | 0.119 | 1 | 12 | Scrn1 | -0.024 | Young |
| 0.597943 | 0.13996 | 0.635 | 0.585 | 1 | 12 | Rpl18 | 0.05 | Young |
| 0.598041 | 0.860037 | 0.036 | 0.025 | 1 | 12 | Abraxas2 | 0.011 | Young |
| 0.598041 | 0.669119 | 0.036 | 0.025 | 1 | 12 | Ttc5 | 0.011 | Young |
| 0.598431 | 0.287474 | 0.226 | 0.203 | 1 | 12 | Syngr3 | 0.023 | Young |
| 0.598704 | 0.388341 | 0.095 | 0.076 | 1 | 12 | Bean1 | 0.019 | Young |
| 0.598812 | 0.193894 | 0.117 | 0.093 | 1 | 12 | Hmox2 | 0.024 | Young |
| 0.598882 | 0.12803 | 0.547 | 0.492 | 1 | 12 | Rpl30 | 0.055 | Young |
| 0.599474 | 0.274095 | 0.124 | 0.102 | 1 | 12 | Hnrnpa1 | 0.022 | Young |
| 0.599928 | 0.273134 | 0.204 | 0.186 | 1 | 12 | Nsg2 | 0.018 | Young |
| 0.600075 | 0.246537 | 0.226 | 0.203 | 1 | 12 | Slc25a5 | 0.023 | Young |
| 0.600438 | 0.06282 | 0.058 | 0.076 | 1 | 12 | Hspa9 | -0.018 | Young |
| 0.600461 | 0.107454 | 0.642 | 0.644 | 1 | 12 | Atp5g1 | -0.002 | Young |
| 0.601181 | 0.057121 | 0.058 | 0.042 | 1 | 12 | Chchd3 | 0.016 | Young |
| 0.601425 | 0.37486 | 0.146 | 0.127 | 1 | 12 | Aplp2 | 0.019 | Young |
| 0.601962 | 1.118518 | 0.036 | 0.025 | 1 | 12 | Hadha | 0.011 | Young |
| 0.602198 | 0.237176 | 0.255 | 0.22 | 1 | 12 | BC031181 | 0.035 | Young |
| 0.602437 | 0.128478 | 0.161 | 0.136 | 1 | 12 | Csnk2b | 0.025 | Young |
| 0.603927 | 0.723206 | 0.036 | 0.025 | 1 | 12 | Sema4d | 0.011 | Young |
| 0.604294 | 0.174606 | 0.058 | 0.042 | 1 | 12 | Dbnl | 0.016 | Young |
| 0.604294 | 0.080295 | 0.058 | 0.042 | 1 | 12 | Mtmr7 | 0.016 | Young |
| 0.604514 | 0.010692 | 0.073 | 0.093 | 1 | 12 | Maz | -0.02 | Young |
| 0.60479 | 0.116787 | 0.095 | 0.076 | 1 | 12 | Eif3g | 0.019 | Young |
| 0.605367 | 0.212453 | 0.029 | 0.042 | 1 | 12 | Pigk | -0.013 | Young |
| 0.605865 | 0.52493 | 0.102 | 0.085 | 1 | 12 | Stx12 | 0.017 | Young |
| 0.605895 | 0.973866 | 0.036 | 0.025 | 1 | 12 | Zfp638 | 0.011 | Young |
| 0.605895 | 0.882036 | 0.036 | 0.025 | 1 | 12 | Rab5b | 0.011 | Young |
| 0.605895 | 0.856151 | 0.036 | 0.025 | 1 | 12 | Brox | 0.011 | Young |
| 0.605895 | 0.774284 | 0.036 | 0.025 | 1 | 12 | Ier2 | 0.011 | Young |
| 0.605895 | 0.679664 | 0.036 | 0.025 | 1 | 12 | Itfg2 | 0.011 | Young |
| 0.605895 | 0.656201 | 0.036 | 0.025 | 1 | 12 | Zdhhc21 | 0.011 | Young |
| 0.605895 | 0.535009 | 0.036 | 0.025 | 1 | 12 | Pitpnb | 0.011 | Young |
| 0.605932 | 0.204243 | 0.124 | 0.102 | 1 | 12 | Tomm22 | 0.022 | Young |
| 0.607197 | 0.241282 | 0.175 | 0.153 | 1 | 12 | Ccdc124 | 0.022 | Young |
| 0.607523 | 0.677246 | 0.066 | 0.051 | 1 | 12 | B4gat1 | 0.015 | Young |
| 0.607807 | 0.149141 | 0.088 | 0.068 | 1 | 12 | Ube2j2 | 0.02 | Young |
| 0.608361 | 0.086762 | 0.321 | 0.288 | 1 | 12 | Rpl35 | 0.033 | Young |
| 0.608377 | 0.271171 | 0.139 | 0.119 | 1 | 12 | Pop5 | 0.02 | Young |
| 0.609839 | 0.904675 | 0.036 | 0.025 | 1 | 12 | Cul1 | 0.011 | Young |
| 0.609839 | 0.855674 | 0.036 | 0.025 | 1 | 12 | Ubr2 | 0.011 | Young |
| 0.609839 | 0.695438 | 0.036 | 0.025 | 1 | 12 | Nemf | 0.011 | Young |
| 0.609839 | 0.627385 | 0.036 | 0.025 | 1 | 12 | Ube2q2 | 0.011 | Young |
| 0.609839 | 0.612684 | 0.036 | 0.025 | 1 | 12 | Dek | 0.011 | Young |
| 0.609839 | 0.541265 | 0.036 | 0.025 | 1 | 12 | Phyhipl | 0.011 | Young |
| 0.609839 | 0.529426 | 0.036 | 0.025 | 1 | 12 | Bicd1 | 0.011 | Young |

| | | | | | | | | |
|---|---|---|---|---|---|---|---|---|
| 0.609839 | 0.523751 | 0.036 | 0.025 | 1 | 12 | Filip1 | 0.011 | Young |
| 0.609913 | 0.205224 | 0.255 | 0.229 | 1 | 12 | Ube2n | 0.026 | Young |
| 0.609935 | 0.165718 | 0.117 | 0.093 | 1 | 12 | Cd151 | 0.024 | Young |
| 0.610446 | 0.39031 | 0.066 | 0.051 | 1 | 12 | Gabbr1 | 0.015 | Young |
| 0.611266 | 0.65461 | 0.117 | 0.102 | 1 | 12 | Calr | 0.015 | Young |
| 0.611911 | 0.427566 | 0.066 | 0.051 | 1 | 12 | Dtymk | 0.015 | Young |
| 0.612131 | 0.171737 | 0.095 | 0.076 | 1 | 12 | Chmp4b | 0.019 | Young |
| 0.6132 | 0.016454 | 0.036 | 0.051 | 1 | 12 | Gna13 | -0.015 | Young |
| 0.613795 | 0.894037 | 0.036 | 0.025 | 1 | 12 | 2310033P0 | 0.011 | Young |
| 0.613795 | 0.719096 | 0.036 | 0.025 | 1 | 12 | Blvra | 0.011 | Young |
| 0.613795 | 0.619931 | 0.036 | 0.025 | 1 | 12 | Sbsn | 0.011 | Young |
| 0.613795 | 0.547321 | 0.036 | 0.025 | 1 | 12 | Dazap1 | 0.011 | Young |
| 0.613795 | 0.544185 | 0.036 | 0.025 | 1 | 12 | Armcx2 | 0.011 | Young |
| 0.613795 | 0.539101 | 0.036 | 0.025 | 1 | 12 | Cir1 | 0.011 | Young |
| 0.613795 | 0.52566 | 0.036 | 0.025 | 1 | 12 | Ankrd13c | 0.011 | Young |
| 0.613795 | 0.487925 | 0.036 | 0.025 | 1 | 12 | Nsmce2 | 0.011 | Young |
| 0.614209 | 0.183517 | 0.088 | 0.068 | 1 | 12 | Atp6v0e | 0.02 | Young |
| 0.614843 | 0.412916 | 0.066 | 0.051 | 1 | 12 | Fam220a | 0.015 | Young |
| 0.614843 | 0.472 | 0.066 | 0.051 | 1 | 12 | Mcrs1 | 0.015 | Young |
| 0.615776 | 0.441286 | 0.036 | 0.025 | 1 | 12 | 1110059G1 | 0.011 | Young |
| 0.615816 | 0.036912 | 0.095 | 0.076 | 1 | 12 | Polr2i | 0.019 | Young |
| 0.616366 | 0.285026 | 0.131 | 0.11 | 1 | 12 | Tsen34 | 0.021 | Young |
| 0.617758 | 0.174233 | 0.715 | 0.653 | 1 | 12 | Rps8 | 0.062 | Young |
| 0.617761 | 0.741286 | 0.036 | 0.025 | 1 | 12 | Lsm8 | 0.011 | Young |
| 0.617761 | 0.717661 | 0.036 | 0.025 | 1 | 12 | G3bp1 | 0.011 | Young |
| 0.617761 | 0.694712 | 0.036 | 0.025 | 1 | 12 | D10Wsu10 | 0.011 | Young |
| 0.617761 | 0.413993 | 0.036 | 0.025 | 1 | 12 | Dpp6 | 0.011 | Young |
| 0.617782 | 0.300264 | 0.066 | 0.051 | 1 | 12 | Dhx9 | 0.015 | Young |
| 0.617782 | 0.407967 | 0.066 | 0.051 | 1 | 12 | Eif3b | 0.015 | Young |
| 0.617782 | 0.347944 | 0.066 | 0.051 | 1 | 12 | Frmd4a | 0.015 | Young |
| 0.618073 | 0.211565 | 0.197 | 0.169 | 1 | 12 | Trnp1 | 0.028 | Young |
| 0.618279 | 0.478115 | 0.095 | 0.076 | 1 | 12 | Camk2d | 0.019 | Young |
| 0.61829 | 0.068359 | 0.073 | 0.093 | 1 | 12 | Sucla2 | -0.02 | Young |
| 0.618792 | 0.10099 | 0.292 | 0.271 | 1 | 12 | Rps26 | 0.021 | Young |
| 0.620727 | 0.500747 | 0.066 | 0.051 | 1 | 12 | Arhgap5 | 0.015 | Young |
| 0.621739 | 0.66313 | 0.036 | 0.025 | 1 | 12 | Stag1 | 0.011 | Young |
| 0.621739 | 0.481419 | 0.036 | 0.025 | 1 | 12 | Ccdc47 | 0.011 | Young |
| 0.621739 | 0.450738 | 0.036 | 0.025 | 1 | 12 | Csdc2 | 0.011 | Young |
| 0.621739 | 0.450551 | 0.036 | 0.025 | 1 | 12 | Kcnq1ot1 | 0.011 | Young |
| 0.621739 | 0.442116 | 0.036 | 0.025 | 1 | 12 | Ruvbl1 | 0.011 | Young |
| 0.621739 | 0.429474 | 0.036 | 0.025 | 1 | 12 | Parp6 | 0.011 | Young |
| 0.621739 | 0.414497 | 0.036 | 0.025 | 1 | 12 | Dhx32 | 0.011 | Young |
| 0.621739 | 0.389782 | 0.036 | 0.025 | 1 | 12 | Scn1b | 0.011 | Young |
| 0.621739 | 0.388104 | 0.036 | 0.025 | 1 | 12 | Etaa1 | 0.011 | Young |
| 0.621739 | 0.368521 | 0.036 | 0.025 | 1 | 12 | Coa7 | 0.011 | Young |

| | | | | | | | | |
|---|---|---|---|---|---|---|---|---|
| 0.621739 | 0.365685 | 0.036 | 0.025 | 1 | 12 | Bccip | 0.011 | Young |
| 0.621739 | 0.319645 | 0.036 | 0.025 | 1 | 12 | Zfp329 | 0.011 | Young |
| 0.622686 | 0.330265 | 0.241 | 0.22 | 1 | 12 | Atp2b1 | 0.021 | Young |
| 0.623122 | 0.116865 | 0.044 | 0.059 | 1 | 12 | Dhcr24 | -0.015 | Young |
| 0.623678 | 0.423906 | 0.066 | 0.051 | 1 | 12 | Spag9 | 0.015 | Young |
| 0.623678 | 0.344804 | 0.066 | 0.051 | 1 | 12 | 0610010F0 | 0.015 | Young |
| 0.623678 | 0.260913 | 0.066 | 0.051 | 1 | 12 | Fus | 0.015 | Young |
| 0.625728 | 0.681746 | 0.036 | 0.025 | 1 | 12 | Psmg4 | 0.011 | Young |
| 0.625728 | 0.40472 | 0.036 | 0.025 | 1 | 12 | Anp32e | 0.011 | Young |
| 0.625728 | 0.384397 | 0.036 | 0.025 | 1 | 12 | Josd2 | 0.011 | Young |
| 0.625728 | 0.240462 | 0.036 | 0.025 | 1 | 12 | Lpin2 | 0.011 | Young |
| 0.625728 | 0.236083 | 0.036 | 0.025 | 1 | 12 | 4833439L1 | 0.011 | Young |
| 0.625728 | 0.232727 | 0.036 | 0.025 | 1 | 12 | Dlgap4 | 0.011 | Young |
| 0.625728 | 0.128515 | 0.036 | 0.025 | 1 | 12 | Tbpl1 | 0.011 | Young |
| 0.625795 | 0.445708 | 0.204 | 0.186 | 1 | 12 | Ctxn1 | 0.018 | Young |
| 0.629271 | 0.064946 | 0.985 | 0.975 | 1 | 12 | mt-Co2 | 0.01 | Young |
| 0.629573 | 0.328318 | 0.153 | 0.136 | 1 | 12 | Dda1 | 0.017 | Young |
| 0.629727 | 0.325719 | 0.036 | 0.025 | 1 | 12 | Kmt2a | 0.011 | Young |
| 0.629728 | 0.347081 | 0.036 | 0.025 | 1 | 12 | Tmem259 | 0.011 | Young |
| 0.629728 | 0.289125 | 0.036 | 0.025 | 1 | 12 | Gm42372 | 0.011 | Young |
| 0.630133 | 0.468436 | 0.109 | 0.093 | 1 | 12 | Ptp4a2 | 0.016 | Young |
| 0.630907 | 0.403431 | 0.073 | 0.059 | 1 | 12 | Fabp7 | 0.014 | Young |
| 0.631285 | 0.322226 | 0.109 | 0.093 | 1 | 12 | Asap1 | 0.016 | Young |
| 0.631337 | 0.184621 | 0.467 | 0.449 | 1 | 12 | Atp5f1 | 0.018 | Young |
| 0.631677 | 0.094246 | 0.453 | 0.381 | 1 | 12 | Psmb1 | 0.072 | Young |
| 0.631932 | 0.518719 | 0.102 | 0.085 | 1 | 12 | Cfap20 | 0.017 | Young |
| 0.633739 | 0.102833 | 0.036 | 0.025 | 1 | 12 | Adgrl1 | 0.011 | Young |
| 0.635541 | 0.293457 | 0.066 | 0.051 | 1 | 12 | Top2b | 0.015 | Young |
| 0.635633 | 0.103038 | 0.095 | 0.076 | 1 | 12 | Cdc123 | 0.019 | Young |
| 0.637761 | 0.311854 | 0.036 | 0.025 | 1 | 12 | Dexi | 0.011 | Young |
| 0.637761 | 0.277986 | 0.036 | 0.025 | 1 | 12 | Aph1a | 0.011 | Young |
| 0.637761 | 0.214461 | 0.036 | 0.025 | 1 | 12 | Wasf2 | 0.011 | Young |
| 0.637761 | 0.020693 | 0.036 | 0.025 | 1 | 12 | Clint1 | 0.011 | Young |
| 0.638747 | 0.09063 | 0.146 | 0.169 | 1 | 12 | Hnrnpk | -0.023 | Young |
| 0.639329 | 0.436879 | 0.073 | 0.059 | 1 | 12 | Pdha1 | 0.014 | Young |
| 0.639689 | 0.093168 | 0.182 | 0.161 | 1 | 12 | Crip1 | 0.021 | Young |
| 0.639826 | 0.196068 | 0.175 | 0.153 | 1 | 12 | Ola1 | 0.022 | Young |
| 0.641087 | 0.125306 | 0.409 | 0.347 | 1 | 12 | Clta | 0.062 | Young |
| 0.641454 | 0.087605 | 0.745 | 0.695 | 1 | 12 | Ndufa4 | 0.05 | Young |
| 0.641508 | 0.242717 | 0.066 | 0.051 | 1 | 12 | 4921524J1 | 0.015 | Young |
| 0.641698 | 0.205919 | 0.109 | 0.093 | 1 | 12 | Tceal8 | 0.016 | Young |
| 0.641793 | 0.158499 | 0.036 | 0.025 | 1 | 12 | Dgkz | 0.011 | Young |
| 0.643144 | 1.190647 | 0.044 | 0.034 | 1 | 12 | Reps1 | 0.01 | Young |
| 0.644501 | 0.181729 | 0.066 | 0.051 | 1 | 12 | Adap1 | 0.015 | Young |
| 0.644501 | 0.128104 | 0.066 | 0.051 | 1 | 12 | Itm2c | 0.015 | Young |

| | | | | | | | | |
|---|---|---|---|---|---|---|---|---|
| 0.644624 | 0.100438 | 0.365 | 0.331 | 1 | 12 | Elavl2 | 0.034 | Young |
| 0.645446 | 0.019762 | 0.117 | 0.136 | 1 | 12 | Lsm12 | -0.019 | Young |
| 0.64564 | 0.081768 | 0.095 | 0.076 | 1 | 12 | Vti1b | 0.019 | Young |
| 0.645836 | 0.058357 | 0.036 | 0.025 | 1 | 12 | Impdh1 | 0.011 | Young |
| 0.646306 | 0.028194 | 0.27 | 0.246 | 1 | 12 | Spcs1 | 0.024 | Young |
| 0.647499 | 0.143464 | 0.066 | 0.051 | 1 | 12 | Rab3d | 0.015 | Young |
| 0.64848 | 3.24023 | 0.015 | 0.008 | 1 | 12 | Trpc1 | 0.007 | Young |
| 0.64848 | 3.236167 | 0.015 | 0.008 | 1 | 12 | Atxn2l | 0.007 | Young |
| 0.64848 | 3.213005 | 0.015 | 0.008 | 1 | 12 | Rnasel | 0.007 | Young |
| 0.64848 | 3.146559 | 0.015 | 0.008 | 1 | 12 | Sephs2 | 0.007 | Young |
| 0.64848 | 3.120247 | 0.015 | 0.008 | 1 | 12 | Dip2b | 0.007 | Young |
| 0.64848 | 3.095847 | 0.015 | 0.008 | 1 | 12 | Ring1 | 0.007 | Young |
| 0.64848 | 3.071403 | 0.015 | 0.008 | 1 | 12 | Eftud2 | 0.007 | Young |
| 0.64848 | 3.059767 | 0.015 | 0.008 | 1 | 12 | Slc30a1 | 0.007 | Young |
| 0.64848 | 3.040453 | 0.015 | 0.008 | 1 | 12 | Stt3a | 0.007 | Young |
| 0.64848 | 3.03613 | 0.015 | 0.008 | 1 | 12 | Glb1 | 0.007 | Young |
| 0.64848 | 2.984448 | 0.015 | 0.008 | 1 | 12 | Epb41l2 | 0.007 | Young |
| 0.64848 | 2.952694 | 0.015 | 0.008 | 1 | 12 | Fut8 | 0.007 | Young |
| 0.64848 | 2.926069 | 0.015 | 0.008 | 1 | 12 | Arhgap31 | 0.007 | Young |
| 0.64848 | 2.917218 | 0.015 | 0.008 | 1 | 12 | Rrp12 | 0.007 | Young |
| 0.64848 | 2.82944 | 0.015 | 0.008 | 1 | 12 | Rb1 | 0.007 | Young |
| 0.64848 | 2.785394 | 0.015 | 0.008 | 1 | 12 | Usp21 | 0.007 | Young |
| 0.64848 | 2.742841 | 0.015 | 0.008 | 1 | 12 | Anks3 | 0.007 | Young |
| 0.64848 | 2.726902 | 0.015 | 0.008 | 1 | 12 | Bak1 | 0.007 | Young |
| 0.64848 | 2.71225 | 0.015 | 0.008 | 1 | 12 | Hmgcl | 0.007 | Young |
| 0.64848 | 2.711301 | 0.015 | 0.008 | 1 | 12 | 1190007I0 | 0.007 | Young |
| 0.64848 | 2.648962 | 0.015 | 0.008 | 1 | 12 | Arhgef2 | 0.007 | Young |
| 0.64848 | 2.638464 | 0.015 | 0.008 | 1 | 12 | Adgrl2 | 0.007 | Young |
| 0.64848 | 2.606687 | 0.015 | 0.008 | 1 | 12 | Rbm33 | 0.007 | Young |
| 0.64848 | 2.592616 | 0.015 | 0.008 | 1 | 12 | Klf11 | 0.007 | Young |
| 0.64848 | 2.583988 | 0.015 | 0.008 | 1 | 12 | Tdrd7 | 0.007 | Young |
| 0.64848 | 2.489385 | 0.015 | 0.008 | 1 | 12 | Nfkb1 | 0.007 | Young |
| 0.64848 | 2.406682 | 0.015 | 0.008 | 1 | 12 | Gpn2 | 0.007 | Young |
| 0.64848 | 2.108521 | 0.015 | 0.008 | 1 | 12 | Dnajc13 | 0.007 | Young |
| 0.64848 | 1.876234 | 0.015 | 0.008 | 1 | 12 | Ankfy1 | 0.007 | Young |
| 0.64848 | 1.868651 | 0.015 | 0.008 | 1 | 12 | Ap3m2 | 0.007 | Young |
| 0.64848 | 1.85461 | 0.015 | 0.008 | 1 | 12 | Trp53 | 0.007 | Young |
| 0.64848 | 1.74868 | 0.015 | 0.008 | 1 | 12 | Med16 | 0.007 | Young |
| 0.64848 | 1.727474 | 0.015 | 0.008 | 1 | 12 | Fam241a | 0.007 | Young |
| 0.64848 | 1.600885 | 0.015 | 0.008 | 1 | 12 | Gda | 0.007 | Young |
| 0.64848 | 1.559564 | 0.015 | 0.008 | 1 | 12 | Oasl2 | 0.007 | Young |
| 0.64848 | 1.528066 | 0.015 | 0.008 | 1 | 12 | Ppp3cc | 0.007 | Young |
| 0.64848 | 1.492528 | 0.015 | 0.008 | 1 | 12 | Lrrc8a | 0.007 | Young |
| 0.64848 | 1.476784 | 0.015 | 0.008 | 1 | 12 | Ddx27 | 0.007 | Young |
| 0.64848 | 1.471785 | 0.015 | 0.008 | 1 | 12 | Armc5 | 0.007 | Young |

| | | | | | | | | |
|---|---|---|---|---|---|---|---|---|
| 0.64848 | 1.455234 | 0.015 | 0.008 | 1 | 12 | Mcm4 | 0.007 | Young |
| 0.64848 | 1.450717 | 0.015 | 0.008 | 1 | 12 | Carmil3 | 0.007 | Young |
| 0.64848 | 1.447212 | 0.015 | 0.008 | 1 | 12 | Tox2 | 0.007 | Young |
| 0.64848 | 1.432452 | 0.015 | 0.008 | 1 | 12 | Plp1 | 0.007 | Young |
| 0.64848 | 1.417785 | 0.015 | 0.008 | 1 | 12 | Itgam | 0.007 | Young |
| 0.64848 | 1.392192 | 0.015 | 0.008 | 1 | 12 | 4732440D( | 0.007 | Young |
| 0.64848 | 1.375589 | 0.015 | 0.008 | 1 | 12 | Mrps30 | 0.007 | Young |
| 0.64848 | 1.355675 | 0.015 | 0.008 | 1 | 12 | Acadsb | 0.007 | Young |
| 0.64848 | 1.325901 | 0.015 | 0.008 | 1 | 12 | Hsdl2 | 0.007 | Young |
| 0.64848 | 1.267459 | 0.015 | 0.008 | 1 | 12 | Dpp8 | 0.007 | Young |
| 0.64848 | 1.265805 | 0.015 | 0.008 | 1 | 12 | Crtap | 0.007 | Young |
| 0.64848 | 1.25 | 0.015 | 0.008 | 1 | 12 | Zfp511 | 0.007 | Young |
| 0.64848 | 1.249562 | 0.015 | 0.008 | 1 | 12 | Fam234b | 0.007 | Young |
| 0.64848 | 1.240299 | 0.015 | 0.008 | 1 | 12 | Ino80c | 0.007 | Young |
| 0.64848 | 1.226562 | 0.015 | 0.008 | 1 | 12 | Ap5s1 | 0.007 | Young |
| 0.64848 | 1.22271 | 0.015 | 0.008 | 1 | 12 | Leng1 | 0.007 | Young |
| 0.64848 | 1.216541 | 0.015 | 0.008 | 1 | 12 | Slc22a15 | 0.007 | Young |
| 0.64848 | 1.190507 | 0.015 | 0.008 | 1 | 12 | Fam98b | 0.007 | Young |
| 0.64848 | 1.164149 | 0.015 | 0.008 | 1 | 12 | Anapc7 | 0.007 | Young |
| 0.64848 | 1.162599 | 0.015 | 0.008 | 1 | 12 | Limk1 | 0.007 | Young |
| 0.64848 | 1.148083 | 0.015 | 0.008 | 1 | 12 | Kansl3 | 0.007 | Young |
| 0.64848 | 1.147192 | 0.015 | 0.008 | 1 | 12 | Gosr2 | 0.007 | Young |
| 0.64848 | 1.129697 | 0.015 | 0.008 | 1 | 12 | Pcnx4 | 0.007 | Young |
| 0.64848 | 1.129332 | 0.015 | 0.008 | 1 | 12 | Apbb2 | 0.007 | Young |
| 0.64848 | 1.126376 | 0.015 | 0.008 | 1 | 12 | Tmcc1 | 0.007 | Young |
| 0.64848 | 1.122063 | 0.015 | 0.008 | 1 | 12 | Tmem62 | 0.007 | Young |
| 0.64848 | 1.119697 | 0.015 | 0.008 | 1 | 12 | Cnpy4 | 0.007 | Young |
| 0.64848 | 1.116272 | 0.015 | 0.008 | 1 | 12 | Cblb | 0.007 | Young |
| 0.64848 | 1.105182 | 0.015 | 0.008 | 1 | 12 | Nlk | 0.007 | Young |
| 0.64848 | 1.098334 | 0.015 | 0.008 | 1 | 12 | Lgi1 | 0.007 | Young |
| 0.64848 | 1.086785 | 0.015 | 0.008 | 1 | 12 | Hmg20a | 0.007 | Young |
| 0.64848 | 1.080637 | 0.015 | 0.008 | 1 | 12 | Hdac6 | 0.007 | Young |
| 0.64848 | 1.079951 | 0.015 | 0.008 | 1 | 12 | Ecd | 0.007 | Young |
| 0.64848 | 1.062971 | 0.015 | 0.008 | 1 | 12 | Adcy3 | 0.007 | Young |
| 0.64848 | 1.052263 | 0.015 | 0.008 | 1 | 12 | Foxj2 | 0.007 | Young |
| 0.64848 | 1.043291 | 0.015 | 0.008 | 1 | 12 | Klhl8 | 0.007 | Young |
| 0.64848 | 1.042107 | 0.015 | 0.008 | 1 | 12 | Atxn7l1 | 0.007 | Young |
| 0.64848 | 1.036295 | 0.015 | 0.008 | 1 | 12 | Rims3 | 0.007 | Young |
| 0.64848 | 1.02688 | 0.015 | 0.008 | 1 | 12 | Fnbp4 | 0.007 | Young |
| 0.64848 | 1.023651 | 0.015 | 0.008 | 1 | 12 | Mxd4 | 0.007 | Young |
| 0.64848 | 1.023303 | 0.015 | 0.008 | 1 | 12 | Pus7 | 0.007 | Young |
| 0.64848 | 1.017948 | 0.015 | 0.008 | 1 | 12 | Brms1 | 0.007 | Young |
| 0.64848 | 1.005478 | 0.015 | 0.008 | 1 | 12 | Gm16499.: | 0.007 | Young |
| 0.64848 | 0.991747 | 0.015 | 0.008 | 1 | 12 | Gak | 0.007 | Young |
| 0.64848 | 0.986828 | 0.015 | 0.008 | 1 | 12 | Ccng2 | 0.007 | Young |

| | | | | | | | | |
|---|---|---|---|---|---|---|---|---|
| 0.64848 | 0.986382 | 0.015 | 0.008 | 1 | 12 | Abcb8 | 0.007 | Young |
| 0.64848 | 0.979644 | 0.015 | 0.008 | 1 | 12 | Npy2r | 0.007 | Young |
| 0.64848 | 0.975196 | 0.015 | 0.008 | 1 | 12 | Enpp5 | 0.007 | Young |
| 0.64848 | 0.969883 | 0.015 | 0.008 | 1 | 12 | Arid1b | 0.007 | Young |
| 0.64848 | 0.96313 | 0.015 | 0.008 | 1 | 12 | Eif2d | 0.007 | Young |
| 0.64848 | 0.96263 | 0.015 | 0.008 | 1 | 12 | Elac2 | 0.007 | Young |
| 0.64848 | 0.955371 | 0.015 | 0.008 | 1 | 12 | Zbtb10 | 0.007 | Young |
| 0.64848 | 0.952624 | 0.015 | 0.008 | 1 | 12 | Taf6 | 0.007 | Young |
| 0.64848 | 0.940566 | 0.015 | 0.008 | 1 | 12 | Tmem86a | 0.007 | Young |
| 0.64848 | 0.936653 | 0.015 | 0.008 | 1 | 12 | Tmem238 | 0.007 | Young |
| 0.64848 | 0.936119 | 0.015 | 0.008 | 1 | 12 | Mtus1 | 0.007 | Young |
| 0.64848 | 0.934626 | 0.015 | 0.008 | 1 | 12 | Casz1 | 0.007 | Young |
| 0.64848 | 0.931138 | 0.015 | 0.008 | 1 | 12 | Letmd1 | 0.007 | Young |
| 0.64848 | 0.927059 | 0.015 | 0.008 | 1 | 12 | Pdk2 | 0.007 | Young |
| 0.64848 | 0.924867 | 0.015 | 0.008 | 1 | 12 | Xrcc6 | 0.007 | Young |
| 0.64848 | 0.922267 | 0.015 | 0.008 | 1 | 12 | Nfat5 | 0.007 | Young |
| 0.64848 | 0.920664 | 0.015 | 0.008 | 1 | 12 | Tstd2 | 0.007 | Young |
| 0.64848 | 0.916143 | 0.015 | 0.008 | 1 | 12 | Cobl | 0.007 | Young |
| 0.64848 | 0.914631 | 0.015 | 0.008 | 1 | 12 | Cog5 | 0.007 | Young |
| 0.64848 | 0.900625 | 0.015 | 0.008 | 1 | 12 | Zfp518a | 0.007 | Young |
| 0.64848 | 0.89614 | 0.015 | 0.008 | 1 | 12 | Mt2 | 0.007 | Young |
| 0.64848 | 0.892386 | 0.015 | 0.008 | 1 | 12 | Gpn3 | 0.007 | Young |
| 0.64848 | 0.888469 | 0.015 | 0.008 | 1 | 12 | Zzz3 | 0.007 | Young |
| 0.64848 | 0.883166 | 0.015 | 0.008 | 1 | 12 | Seh1l | 0.007 | Young |
| 0.64848 | 0.869373 | 0.015 | 0.008 | 1 | 12 | Purg | 0.007 | Young |
| 0.64848 | 0.856973 | 0.015 | 0.008 | 1 | 12 | Pla2g6 | 0.007 | Young |
| 0.64848 | 0.856769 | 0.015 | 0.008 | 1 | 12 | Zfp64 | 0.007 | Young |
| 0.64848 | 0.851088 | 0.015 | 0.008 | 1 | 12 | Scaper | 0.007 | Young |
| 0.64848 | 0.850234 | 0.015 | 0.008 | 1 | 12 | Dgki | 0.007 | Young |
| 0.64848 | 0.846363 | 0.015 | 0.008 | 1 | 12 | Fbxl7 | 0.007 | Young |
| 0.64848 | 0.845184 | 0.015 | 0.008 | 1 | 12 | Cav3 | 0.007 | Young |
| 0.64848 | 0.844539 | 0.015 | 0.008 | 1 | 12 | Ezh1 | 0.007 | Young |
| 0.64848 | 0.843114 | 0.015 | 0.008 | 1 | 12 | Stradb | 0.007 | Young |
| 0.64848 | 0.840595 | 0.015 | 0.008 | 1 | 12 | Rrp1b | 0.007 | Young |
| 0.64848 | 0.837498 | 0.015 | 0.008 | 1 | 12 | Tollip | 0.007 | Young |
| 0.64848 | 0.833115 | 0.015 | 0.008 | 1 | 12 | Qser1 | 0.007 | Young |
| 0.64848 | 0.831877 | 0.015 | 0.008 | 1 | 12 | Dag1 | 0.007 | Young |
| 0.64848 | 0.791499 | 0.015 | 0.008 | 1 | 12 | Rnaseh2b | 0.007 | Young |
| 0.648551 | 0.09716 | 0.197 | 0.169 | 1 | 12 | Ndufv1 | 0.028 | Young |
| 0.650411 | 0.0479 | 0.175 | 0.203 | 1 | 12 | Bcas2 | -0.028 | Young |
| 0.650449 | 0.203955 | 0.153 | 0.136 | 1 | 12 | Inpp5f | 0.017 | Young |
| 0.650503 | 0.206886 | 0.066 | 0.051 | 1 | 12 | Anapc13 | 0.015 | Young |
| 0.65176 | 0.923426 | 0.015 | 0.008 | 1 | 12 | Wapl | 0.007 | Young |
| 0.652007 | 0.205803 | 0.066 | 0.051 | 1 | 12 | Gtpbp6 | 0.015 | Young |
| 0.653353 | 0.215054 | 0.255 | 0.229 | 1 | 12 | Rpl36a | 0.026 | Young |

| | | | | | | | | |
|---|---|---|---|---|---|---|---|---|
| 0.653462 | 0.103878 | 0.73 | 0.661 | 1 | 12 | Rps20 | 0.069 | Young |
| 0.654662 | 0.135947 | 0.212 | 0.186 | 1 | 12 | Pfdn2 | 0.026 | Young |
| 0.65505 | 1.131498 | 0.015 | 0.008 | 1 | 12 | Grcc10 | 0.007 | Young |
| 0.65505 | 1.119725 | 0.015 | 0.008 | 1 | 12 | Man1a | 0.007 | Young |
| 0.65505 | 1.112635 | 0.015 | 0.008 | 1 | 12 | Slc52a2 | 0.007 | Young |
| 0.65505 | 1.078895 | 0.015 | 0.008 | 1 | 12 | Aldh3b1 | 0.007 | Young |
| 0.65505 | 1.070495 | 0.015 | 0.008 | 1 | 12 | Fbxl3 | 0.007 | Young |
| 0.65505 | 1.021969 | 0.015 | 0.008 | 1 | 12 | Cpne4 | 0.007 | Young |
| 0.65505 | 1.010109 | 0.015 | 0.008 | 1 | 12 | Nmt2 | 0.007 | Young |
| 0.65505 | 0.977134 | 0.015 | 0.008 | 1 | 12 | Atl3 | 0.007 | Young |
| 0.65505 | 0.974817 | 0.015 | 0.008 | 1 | 12 | Dynlt1a | 0.007 | Young |
| 0.65505 | 0.9472 | 0.015 | 0.008 | 1 | 12 | Exoc6b | 0.007 | Young |
| 0.65505 | 0.946976 | 0.015 | 0.008 | 1 | 12 | Socs7 | 0.007 | Young |
| 0.65505 | 0.944828 | 0.015 | 0.008 | 1 | 12 | Dcaf4 | 0.007 | Young |
| 0.65505 | 0.90829 | 0.015 | 0.008 | 1 | 12 | Klhl24 | 0.007 | Young |
| 0.65505 | 0.908014 | 0.015 | 0.008 | 1 | 12 | Adprhl2 | 0.007 | Young |
| 0.65505 | 0.904634 | 0.015 | 0.008 | 1 | 12 | Cit | 0.007 | Young |
| 0.65505 | 0.902685 | 0.015 | 0.008 | 1 | 12 | Brf2 | 0.007 | Young |
| 0.65505 | 0.89669 | 0.015 | 0.008 | 1 | 12 | Cherp | 0.007 | Young |
| 0.65505 | 0.883981 | 0.015 | 0.008 | 1 | 12 | Scaf8 | 0.007 | Young |
| 0.65505 | 0.877601 | 0.015 | 0.008 | 1 | 12 | Ccdc82 | 0.007 | Young |
| 0.65505 | 0.868155 | 0.015 | 0.008 | 1 | 12 | Cd320 | 0.007 | Young |
| 0.65505 | 0.860822 | 0.015 | 0.008 | 1 | 12 | Ado | 0.007 | Young |
| 0.65505 | 0.854821 | 0.015 | 0.008 | 1 | 12 | Tmem218 | 0.007 | Young |
| 0.65505 | 0.849818 | 0.015 | 0.008 | 1 | 12 | Sgms1 | 0.007 | Young |
| 0.65505 | 0.847202 | 0.015 | 0.008 | 1 | 12 | Uba52 | 0.007 | Young |
| 0.65505 | 0.843181 | 0.015 | 0.008 | 1 | 12 | Ergic1 | 0.007 | Young |
| 0.65505 | 0.833901 | 0.015 | 0.008 | 1 | 12 | Gemin2 | 0.007 | Young |
| 0.65505 | 0.820332 | 0.015 | 0.008 | 1 | 12 | Chd1 | 0.007 | Young |
| 0.65505 | 0.819512 | 0.015 | 0.008 | 1 | 12 | BC017643 | 0.007 | Young |
| 0.65505 | 0.818709 | 0.015 | 0.008 | 1 | 12 | Vipas39 | 0.007 | Young |
| 0.65505 | 0.817662 | 0.015 | 0.008 | 1 | 12 | Cd164 | 0.007 | Young |
| 0.65505 | 0.817223 | 0.015 | 0.008 | 1 | 12 | Nol4l | 0.007 | Young |
| 0.65505 | 0.817132 | 0.015 | 0.008 | 1 | 12 | Wars | 0.007 | Young |
| 0.65505 | 0.81066 | 0.015 | 0.008 | 1 | 12 | Chat | 0.007 | Young |
| 0.65505 | 0.802277 | 0.015 | 0.008 | 1 | 12 | Slc27a4 | 0.007 | Young |
| 0.65505 | 0.79727 | 0.015 | 0.008 | 1 | 12 | Ptprd | 0.007 | Young |
| 0.65505 | 0.796612 | 0.015 | 0.008 | 1 | 12 | Inpp5k | 0.007 | Young |
| 0.65505 | 0.789486 | 0.015 | 0.008 | 1 | 12 | Syt3 | 0.007 | Young |
| 0.65505 | 0.789354 | 0.015 | 0.008 | 1 | 12 | Lman2l | 0.007 | Young |
| 0.65505 | 0.784208 | 0.015 | 0.008 | 1 | 12 | Hunk | 0.007 | Young |
| 0.65505 | 0.779754 | 0.015 | 0.008 | 1 | 12 | Fundc1 | 0.007 | Young |
| 0.65505 | 0.778217 | 0.015 | 0.008 | 1 | 12 | Nlgn1 | 0.007 | Young |
| 0.65505 | 0.777804 | 0.015 | 0.008 | 1 | 12 | Dusp28 | 0.007 | Young |
| 0.65505 | 0.769637 | 0.015 | 0.008 | 1 | 12 | Ppef1 | 0.007 | Young |

| | | | | | | | | |
|---|---|---|---|---|---|---|---|---|
| 0.65505 | 0.76778 | 0.015 | 0.008 | 1 | 12 | Pck2 | 0.007 | Young |
| 0.65505 | 0.766269 | 0.015 | 0.008 | 1 | 12 | Zbtb26 | 0.007 | Young |
| 0.65505 | 0.765443 | 0.015 | 0.008 | 1 | 12 | Rab10os | 0.007 | Young |
| 0.65505 | 0.765346 | 0.015 | 0.008 | 1 | 12 | Ubald1 | 0.007 | Young |
| 0.65505 | 0.765148 | 0.015 | 0.008 | 1 | 12 | Rtl6 | 0.007 | Young |
| 0.65505 | 0.759439 | 0.015 | 0.008 | 1 | 12 | 0610010K1 | 0.007 | Young |
| 0.65505 | 0.757999 | 0.015 | 0.008 | 1 | 12 | Ocel1 | 0.007 | Young |
| 0.65505 | 0.756149 | 0.015 | 0.008 | 1 | 12 | Adsl | 0.007 | Young |
| 0.65505 | 0.755945 | 0.015 | 0.008 | 1 | 12 | Rad52 | 0.007 | Young |
| 0.65505 | 0.754626 | 0.015 | 0.008 | 1 | 12 | Cbl | 0.007 | Young |
| 0.65505 | 0.747756 | 0.015 | 0.008 | 1 | 12 | Tmem136 | 0.007 | Young |
| 0.65505 | 0.747295 | 0.015 | 0.008 | 1 | 12 | Rgs2 | 0.007 | Young |
| 0.65505 | 0.736301 | 0.015 | 0.008 | 1 | 12 | Tceanc2 | 0.007 | Young |
| 0.65505 | 0.734346 | 0.015 | 0.008 | 1 | 12 | Ipo4 | 0.007 | Young |
| 0.65505 | 0.734255 | 0.015 | 0.008 | 1 | 12 | Rbm27 | 0.007 | Young |
| 0.65505 | 0.733775 | 0.015 | 0.008 | 1 | 12 | Sec61a2 | 0.007 | Young |
| 0.65505 | 0.732331 | 0.015 | 0.008 | 1 | 12 | Pgr | 0.007 | Young |
| 0.65505 | 0.730582 | 0.015 | 0.008 | 1 | 12 | Maip1 | 0.007 | Young |
| 0.65505 | 0.728284 | 0.015 | 0.008 | 1 | 12 | Lpcat3 | 0.007 | Young |
| 0.65505 | 0.728261 | 0.015 | 0.008 | 1 | 12 | Capn15 | 0.007 | Young |
| 0.65505 | 0.725799 | 0.015 | 0.008 | 1 | 12 | Pmm2 | 0.007 | Young |
| 0.65505 | 0.722787 | 0.015 | 0.008 | 1 | 12 | Dennd4a | 0.007 | Young |
| 0.65505 | 0.720947 | 0.015 | 0.008 | 1 | 12 | Clcn5 | 0.007 | Young |
| 0.65505 | 0.712648 | 0.015 | 0.008 | 1 | 12 | Fndc5 | 0.007 | Young |
| 0.65505 | 0.710909 | 0.015 | 0.008 | 1 | 12 | Ryr2 | 0.007 | Young |
| 0.65505 | 0.710441 | 0.015 | 0.008 | 1 | 12 | Dusp18 | 0.007 | Young |
| 0.65505 | 0.705306 | 0.015 | 0.008 | 1 | 12 | Qrsl1 | 0.007 | Young |
| 0.65505 | 0.705293 | 0.015 | 0.008 | 1 | 12 | mt-Nd6 | 0.007 | Young |
| 0.65505 | 0.701854 | 0.015 | 0.008 | 1 | 12 | Hsf1 | 0.007 | Young |
| 0.65505 | 0.701034 | 0.015 | 0.008 | 1 | 12 | Brinp1 | 0.007 | Young |
| 0.65505 | 0.697633 | 0.015 | 0.008 | 1 | 12 | Ksr2 | 0.007 | Young |
| 0.65505 | 0.695308 | 0.015 | 0.008 | 1 | 12 | Cthrc1 | 0.007 | Young |
| 0.65505 | 0.695028 | 0.015 | 0.008 | 1 | 12 | Shc2 | 0.007 | Young |
| 0.65505 | 0.68096 | 0.015 | 0.008 | 1 | 12 | Chac2 | 0.007 | Young |
| 0.65505 | 0.672071 | 0.015 | 0.008 | 1 | 12 | Mgat3 | 0.007 | Young |
| 0.65505 | 0.671021 | 0.015 | 0.008 | 1 | 12 | Afg3l2 | 0.007 | Young |
| 0.65505 | 0.669392 | 0.015 | 0.008 | 1 | 12 | Skiv2l | 0.007 | Young |
| 0.65505 | 0.669228 | 0.015 | 0.008 | 1 | 12 | Thap2 | 0.007 | Young |
| 0.65505 | 0.669067 | 0.015 | 0.008 | 1 | 12 | Cadm2 | 0.007 | Young |
| 0.65505 | 0.667529 | 0.015 | 0.008 | 1 | 12 | Ptrh1 | 0.007 | Young |
| 0.65505 | 0.663381 | 0.015 | 0.008 | 1 | 12 | Atrn | 0.007 | Young |
| 0.65505 | 0.65945 | 0.015 | 0.008 | 1 | 12 | Tmem198 | 0.007 | Young |
| 0.65505 | 0.657679 | 0.015 | 0.008 | 1 | 12 | Gm14305 | 0.007 | Young |
| 0.65505 | 0.655947 | 0.015 | 0.008 | 1 | 12 | Rbks | 0.007 | Young |
| 0.65505 | 0.652703 | 0.015 | 0.008 | 1 | 12 | Hexim1 | 0.007 | Young |

| | | | | | | | | |
|---|---|---|---|---|---|---|---|---|
| 0.65505 | 0.635906 | 0.015 | 0.008 | 1 | 12 | Tm7sf3 | 0.007 | Young |
| 0.65505 | 0.621326 | 0.015 | 0.008 | 1 | 12 | Zfp968 | 0.007 | Young |
| 0.65505 | 0.6209 | 0.015 | 0.008 | 1 | 12 | Asb6 | 0.007 | Young |
| 0.65505 | 0.617122 | 0.015 | 0.008 | 1 | 12 | Rptor | 0.007 | Young |
| 0.65505 | 0.615022 | 0.015 | 0.008 | 1 | 12 | Wdr20 | 0.007 | Young |
| 0.65505 | 0.612507 | 0.015 | 0.008 | 1 | 12 | Pkp4 | 0.007 | Young |
| 0.65505 | 0.610721 | 0.015 | 0.008 | 1 | 12 | Fam229b | 0.007 | Young |
| 0.65505 | 0.610051 | 0.015 | 0.008 | 1 | 12 | Rapgef1 | 0.007 | Young |
| 0.65505 | 0.608803 | 0.015 | 0.008 | 1 | 12 | Txndc5 | 0.007 | Young |
| 0.65505 | 0.597713 | 0.015 | 0.008 | 1 | 12 | Coq8a | 0.007 | Young |
| 0.65505 | 0.594405 | 0.015 | 0.008 | 1 | 12 | Thnsl1 | 0.007 | Young |
| 0.65505 | 0.593235 | 0.015 | 0.008 | 1 | 12 | Timmdc1 | 0.007 | Young |
| 0.65505 | 0.589286 | 0.015 | 0.008 | 1 | 12 | Sirt5 | 0.007 | Young |
| 0.65505 | 0.559513 | 0.015 | 0.008 | 1 | 12 | Hspbap1 | 0.007 | Young |
| 0.65505 | 0.537271 | 0.015 | 0.008 | 1 | 12 | Tspan18 | 0.007 | Young |
| 0.65505 | 0.535228 | 0.015 | 0.008 | 1 | 12 | Grwd1 | 0.007 | Young |
| 0.65505 | 0.530923 | 0.015 | 0.008 | 1 | 12 | Armcx1 | 0.007 | Young |
| 0.65505 | 0.51984 | 0.015 | 0.008 | 1 | 12 | Ddx23 | 0.007 | Young |
| 0.65505 | 0.519385 | 0.015 | 0.008 | 1 | 12 | Wdr74 | 0.007 | Young |
| 0.65505 | 0.516769 | 0.015 | 0.008 | 1 | 12 | Slc25a27 | 0.007 | Young |
| 0.65505 | 0.491771 | 0.015 | 0.008 | 1 | 12 | Hes5 | 0.007 | Young |
| 0.65505 | 0.487959 | 0.015 | 0.008 | 1 | 12 | B230307C2 | 0.007 | Young |
| 0.65505 | 0.484804 | 0.015 | 0.008 | 1 | 12 | Prrc2a | 0.007 | Young |
| 0.65505 | 0.480823 | 0.015 | 0.008 | 1 | 12 | Nfxl1 | 0.007 | Young |
| 0.65505 | 0.47851 | 0.015 | 0.008 | 1 | 12 | Eif2s3x | 0.007 | Young |
| 0.65505 | 0.478349 | 0.015 | 0.008 | 1 | 12 | Stxbp5 | 0.007 | Young |
| 0.65505 | 0.461381 | 0.015 | 0.008 | 1 | 12 | Myg1 | 0.007 | Young |
| 0.65505 | 0.44134 | 0.015 | 0.008 | 1 | 12 | Spats2l | 0.007 | Young |
| 0.65505 | 0.437994 | 0.015 | 0.008 | 1 | 12 | Acvr2b | 0.007 | Young |
| 0.65505 | 0.322038 | 0.015 | 0.008 | 1 | 12 | Chst15 | 0.007 | Young |
| 0.655997 | 0.205782 | 0.321 | 0.28 | 1 | 12 | Rps25 | 0.041 | Young |
| 0.656314 | 0.316499 | 0.073 | 0.059 | 1 | 12 | Ccar1 | 0.014 | Young |
| 0.657716 | 0.180864 | 0.307 | 0.28 | 1 | 12 | Gtf2h5 | 0.027 | Young |
| 0.657727 | 0.983877 | 0.044 | 0.034 | 1 | 12 | Mest | 0.01 | Young |
| 0.657727 | 0.666027 | 0.044 | 0.034 | 1 | 12 | Gpatch8 | 0.01 | Young |
| 0.657727 | 0.509575 | 0.044 | 0.034 | 1 | 12 | Clptm1l | 0.01 | Young |
| 0.657737 | 0.300123 | 0.073 | 0.059 | 1 | 12 | Babam1 | 0.014 | Young |
| 0.658344 | 0.458926 | 0.015 | 0.008 | 1 | 12 | H2-DMa | 0.007 | Young |
| 0.658819 | 0.164176 | 0.423 | 0.407 | 1 | 12 | Snrpn | 0.016 | Young |
| 0.66037 | 0.277129 | 0.139 | 0.119 | 1 | 12 | Snrpd1 | 0.02 | Young |
| 0.661394 | 0.817113 | 0.044 | 0.034 | 1 | 12 | H2-K1 | 0.01 | Young |
| 0.661646 | 0.707229 | 0.015 | 0.008 | 1 | 12 | Gstk1 | 0.007 | Young |
| 0.661646 | 0.68777 | 0.015 | 0.008 | 1 | 12 | A430046D1 | 0.007 | Young |
| 0.661647 | 0.682985 | 0.015 | 0.008 | 1 | 12 | Fam217b | 0.007 | Young |
| 0.661647 | 0.679335 | 0.015 | 0.008 | 1 | 12 | Lrrc47 | 0.007 | Young |

| | | | | | | | | |
|---|---|---|---|---|---|---|---|---|
| 0.661647 | 0.671271 | 0.015 | 0.008 | 1 | 12 | Sbk1 | 0.007 | Young |
| 0.661647 | 0.661797 | 0.015 | 0.008 | 1 | 12 | Mga | 0.007 | Young |
| 0.661647 | 0.653912 | 0.015 | 0.008 | 1 | 12 | Vrk3 | 0.007 | Young |
| 0.661647 | 0.6449 | 0.015 | 0.008 | 1 | 12 | Upf2 | 0.007 | Young |
| 0.661647 | 0.640579 | 0.015 | 0.008 | 1 | 12 | Cadm4 | 0.007 | Young |
| 0.661647 | 0.639942 | 0.015 | 0.008 | 1 | 12 | Fut11 | 0.007 | Young |
| 0.661647 | 0.631647 | 0.015 | 0.008 | 1 | 12 | Vps37b | 0.007 | Young |
| 0.661647 | 0.629504 | 0.015 | 0.008 | 1 | 12 | Tmem74b | 0.007 | Young |
| 0.661647 | 0.617653 | 0.015 | 0.008 | 1 | 12 | Cib1 | 0.007 | Young |
| 0.661647 | 0.613692 | 0.015 | 0.008 | 1 | 12 | Rnf138rt1 | 0.007 | Young |
| 0.661647 | 0.609179 | 0.015 | 0.008 | 1 | 12 | Wdr92 | 0.007 | Young |
| 0.661647 | 0.595839 | 0.015 | 0.008 | 1 | 12 | Usp39 | 0.007 | Young |
| 0.661647 | 0.594828 | 0.015 | 0.008 | 1 | 12 | Gys1 | 0.007 | Young |
| 0.661647 | 0.592621 | 0.015 | 0.008 | 1 | 12 | Nup85 | 0.007 | Young |
| 0.661647 | 0.59141 | 0.015 | 0.008 | 1 | 12 | Dhfr | 0.007 | Young |
| 0.661647 | 0.589473 | 0.015 | 0.008 | 1 | 12 | Actr3b | 0.007 | Young |
| 0.661647 | 0.572502 | 0.015 | 0.008 | 1 | 12 | Prkdc | 0.007 | Young |
| 0.661647 | 0.571893 | 0.015 | 0.008 | 1 | 12 | Hpcal4 | 0.007 | Young |
| 0.661647 | 0.563299 | 0.015 | 0.008 | 1 | 12 | Tars2 | 0.007 | Young |
| 0.661647 | 0.560078 | 0.015 | 0.008 | 1 | 12 | BC037034 | 0.007 | Young |
| 0.661647 | 0.558675 | 0.015 | 0.008 | 1 | 12 | Atp8b2 | 0.007 | Young |
| 0.661647 | 0.552271 | 0.015 | 0.008 | 1 | 12 | Pcgf1 | 0.007 | Young |
| 0.661647 | 0.551539 | 0.015 | 0.008 | 1 | 12 | Sel1l | 0.007 | Young |
| 0.661647 | 0.54876 | 0.015 | 0.008 | 1 | 12 | Brd4 | 0.007 | Young |
| 0.661647 | 0.54633 | 0.015 | 0.008 | 1 | 12 | Tmem56 | 0.007 | Young |
| 0.661647 | 0.543572 | 0.015 | 0.008 | 1 | 12 | Abca3 | 0.007 | Young |
| 0.661647 | 0.538951 | 0.015 | 0.008 | 1 | 12 | Crtc1 | 0.007 | Young |
| 0.661647 | 0.538573 | 0.015 | 0.008 | 1 | 12 | Phf20l1 | 0.007 | Young |
| 0.661647 | 0.537013 | 0.015 | 0.008 | 1 | 12 | 3110043O2 | 0.007 | Young |
| 0.661647 | 0.535789 | 0.015 | 0.008 | 1 | 12 | Fbxw4 | 0.007 | Young |
| 0.661647 | 0.533396 | 0.015 | 0.008 | 1 | 12 | Faap100 | 0.007 | Young |
| 0.661647 | 0.530515 | 0.015 | 0.008 | 1 | 12 | Mcph1 | 0.007 | Young |
| 0.661647 | 0.525659 | 0.015 | 0.008 | 1 | 12 | Ythdf2 | 0.007 | Young |
| 0.661647 | 0.521051 | 0.015 | 0.008 | 1 | 12 | Agap1 | 0.007 | Young |
| 0.661647 | 0.520298 | 0.015 | 0.008 | 1 | 12 | Fbf1 | 0.007 | Young |
| 0.661647 | 0.519007 | 0.015 | 0.008 | 1 | 12 | Jmy | 0.007 | Young |
| 0.661647 | 0.516252 | 0.015 | 0.008 | 1 | 12 | Hs3st5 | 0.007 | Young |
| 0.661647 | 0.510015 | 0.015 | 0.008 | 1 | 12 | Ccdc40 | 0.007 | Young |
| 0.661647 | 0.507698 | 0.015 | 0.008 | 1 | 12 | Sox11 | 0.007 | Young |
| 0.661647 | 0.507588 | 0.015 | 0.008 | 1 | 12 | Acvr1 | 0.007 | Young |
| 0.661647 | 0.505798 | 0.015 | 0.008 | 1 | 12 | Zfp932 | 0.007 | Young |
| 0.661647 | 0.504402 | 0.015 | 0.008 | 1 | 12 | Pitpnm1 | 0.007 | Young |
| 0.661647 | 0.504017 | 0.015 | 0.008 | 1 | 12 | Gmeb1 | 0.007 | Young |
| 0.661647 | 0.50382 | 0.015 | 0.008 | 1 | 12 | Atf5 | 0.007 | Young |
| 0.661647 | 0.497023 | 0.015 | 0.008 | 1 | 12 | Rims1 | 0.007 | Young |

| | | | | | | | | |
|---|---|---|---|---|---|---|---|---|
| 0.661647 | 0.483258 | 0.015 | 0.008 | 1 | 12 | Sestd1 | 0.007 | Young |
| 0.661647 | 0.474105 | 0.015 | 0.008 | 1 | 12 | Eva1c | 0.007 | Young |
| 0.661647 | 0.473546 | 0.015 | 0.008 | 1 | 12 | Ercc3 | 0.007 | Young |
| 0.661647 | 0.472409 | 0.015 | 0.008 | 1 | 12 | Snn | 0.007 | Young |
| 0.661647 | 0.468631 | 0.015 | 0.008 | 1 | 12 | Pi4ka | 0.007 | Young |
| 0.661647 | 0.46771 | 0.015 | 0.008 | 1 | 12 | Ankrd26 | 0.007 | Young |
| 0.661647 | 0.461868 | 0.015 | 0.008 | 1 | 12 | Ahcyl2 | 0.007 | Young |
| 0.661647 | 0.459344 | 0.015 | 0.008 | 1 | 12 | Tdrp | 0.007 | Young |
| 0.661647 | 0.456259 | 0.015 | 0.008 | 1 | 12 | Hip1r | 0.007 | Young |
| 0.661647 | 0.453971 | 0.015 | 0.008 | 1 | 12 | Ammecr1 | 0.007 | Young |
| 0.661647 | 0.449879 | 0.015 | 0.008 | 1 | 12 | Actr8 | 0.007 | Young |
| 0.661647 | 0.449459 | 0.015 | 0.008 | 1 | 12 | Etl4 | 0.007 | Young |
| 0.661647 | 0.447647 | 0.015 | 0.008 | 1 | 12 | Gm27217 | 0.007 | Young |
| 0.661647 | 0.447437 | 0.015 | 0.008 | 1 | 12 | Entpd3 | 0.007 | Young |
| 0.661647 | 0.443667 | 0.015 | 0.008 | 1 | 12 | Bcan | 0.007 | Young |
| 0.661647 | 0.440817 | 0.015 | 0.008 | 1 | 12 | Zfp758 | 0.007 | Young |
| 0.661647 | 0.435782 | 0.015 | 0.008 | 1 | 12 | Snd1 | 0.007 | Young |
| 0.661647 | 0.421571 | 0.015 | 0.008 | 1 | 12 | Prep | 0.007 | Young |
| 0.661647 | 0.420641 | 0.015 | 0.008 | 1 | 12 | Mak16 | 0.007 | Young |
| 0.661647 | 0.417265 | 0.015 | 0.008 | 1 | 12 | Hs2st1 | 0.007 | Young |
| 0.661647 | 0.400533 | 0.015 | 0.008 | 1 | 12 | Taco1os | 0.007 | Young |
| 0.661647 | 0.396995 | 0.015 | 0.008 | 1 | 12 | Tspan2 | 0.007 | Young |
| 0.661647 | 0.396 | 0.015 | 0.008 | 1 | 12 | Prkci | 0.007 | Young |
| 0.661647 | 0.39558 | 0.015 | 0.008 | 1 | 12 | Wnk2 | 0.007 | Young |
| 0.661647 | 0.388251 | 0.015 | 0.008 | 1 | 12 | Fbxo45 | 0.007 | Young |
| 0.661647 | 0.384656 | 0.015 | 0.008 | 1 | 12 | Hexdc | 0.007 | Young |
| 0.661647 | 0.378623 | 0.015 | 0.008 | 1 | 12 | Mpeg1 | 0.007 | Young |
| 0.661647 | 0.377826 | 0.015 | 0.008 | 1 | 12 | Traf3 | 0.007 | Young |
| 0.661647 | 0.376881 | 0.015 | 0.008 | 1 | 12 | Ube2g2 | 0.007 | Young |
| 0.661647 | 0.376613 | 0.015 | 0.008 | 1 | 12 | Ankrd44 | 0.007 | Young |
| 0.661647 | 0.364011 | 0.015 | 0.008 | 1 | 12 | Nfe2l3 | 0.007 | Young |
| 0.661647 | 0.363097 | 0.015 | 0.008 | 1 | 12 | Pnpt1 | 0.007 | Young |
| 0.661647 | 0.360652 | 0.015 | 0.008 | 1 | 12 | Abcb10 | 0.007 | Young |
| 0.661647 | 0.348066 | 0.015 | 0.008 | 1 | 12 | Nectin3 | 0.007 | Young |
| 0.661647 | 0.340633 | 0.015 | 0.008 | 1 | 12 | Crls1 | 0.007 | Young |
| 0.661647 | 0.337083 | 0.015 | 0.008 | 1 | 12 | Gria3 | 0.007 | Young |
| 0.661647 | 0.332905 | 0.015 | 0.008 | 1 | 12 | Qtrt1 | 0.007 | Young |
| 0.661647 | 0.318866 | 0.015 | 0.008 | 1 | 12 | Bcl2l13 | 0.007 | Young |
| 0.661647 | 0.318369 | 0.015 | 0.008 | 1 | 12 | Zdbf2 | 0.007 | Young |
| 0.661647 | 0.305273 | 0.015 | 0.008 | 1 | 12 | Wdpcp | 0.007 | Young |
| 0.661647 | 0.302518 | 0.015 | 0.008 | 1 | 12 | 4930430F0 | 0.007 | Young |
| 0.661647 | 0.302323 | 0.015 | 0.008 | 1 | 12 | Sec23ip | 0.007 | Young |
| 0.661647 | 0.300982 | 0.015 | 0.008 | 1 | 12 | Ddx41 | 0.007 | Young |
| 0.661647 | 0.29921 | 0.015 | 0.008 | 1 | 12 | Fam105a | 0.007 | Young |
| 0.661647 | 0.293568 | 0.015 | 0.008 | 1 | 12 | Trpm7 | 0.007 | Young |

| | | | | | | | | |
|---|---|---|---|---|---|---|---|---|
| 0.661647 | 0.288036 | 0.015 | 0.008 | 1 | 12 | Pdcl | 0.007 | Young |
| 0.661647 | 0.275559 | 0.015 | 0.008 | 1 | 12 | Usp25 | 0.007 | Young |
| 0.661647 | 0.275129 | 0.015 | 0.008 | 1 | 12 | Mitd1 | 0.007 | Young |
| 0.661647 | 0.267196 | 0.015 | 0.008 | 1 | 12 | Dpp10 | 0.007 | Young |
| 0.661647 | 0.263023 | 0.015 | 0.008 | 1 | 12 | Rbm34 | 0.007 | Young |
| 0.661647 | 0.256469 | 0.015 | 0.008 | 1 | 12 | Rnf112 | 0.007 | Young |
| 0.661647 | 0.254997 | 0.015 | 0.008 | 1 | 12 | Vwa5a | 0.007 | Young |
| 0.661647 | 0.247626 | 0.015 | 0.008 | 1 | 12 | Khk | 0.007 | Young |
| 0.661647 | 0.245733 | 0.015 | 0.008 | 1 | 12 | Foxo3 | 0.007 | Young |
| 0.661647 | 0.242189 | 0.015 | 0.008 | 1 | 12 | Atg4b | 0.007 | Young |
| 0.661647 | 0.220319 | 0.015 | 0.008 | 1 | 12 | Sox12 | 0.007 | Young |
| 0.661647 | 0.203022 | 0.015 | 0.008 | 1 | 12 | Ddit3 | 0.007 | Young |
| 0.661647 | 0.191689 | 0.015 | 0.008 | 1 | 12 | Ankrd37 | 0.007 | Young |
| 0.661647 | 0.190654 | 0.015 | 0.008 | 1 | 12 | Ccl6 | 0.007 | Young |
| 0.661647 | 0.1873 | 0.015 | 0.008 | 1 | 12 | Dnajb5 | 0.007 | Young |
| 0.661647 | 0.177356 | 0.015 | 0.008 | 1 | 12 | B4galnt1 | 0.007 | Young |
| 0.661647 | 0.17567 | 0.015 | 0.008 | 1 | 12 | Kmt5a | 0.007 | Young |
| 0.661647 | 0.172773 | 0.015 | 0.008 | 1 | 12 | Clasrp | 0.007 | Young |
| 0.661647 | 0.161868 | 0.015 | 0.008 | 1 | 12 | Dnph1 | 0.007 | Young |
| 0.661647 | 0.141135 | 0.015 | 0.008 | 1 | 12 | Klf10 | 0.007 | Young |
| 0.661647 | 0.138178 | 0.015 | 0.008 | 1 | 12 | Hnrnph3 | 0.007 | Young |
| 0.661647 | 0.136634 | 0.015 | 0.008 | 1 | 12 | Tmem126b | 0.007 | Young |
| 0.661647 | 0.115944 | 0.015 | 0.008 | 1 | 12 | Gabrg2 | 0.007 | Young |
| 0.661647 | 0.089245 | 0.015 | 0.008 | 1 | 12 | Tab1 | 0.007 | Young |
| 0.661647 | 0.078802 | 0.015 | 0.008 | 1 | 12 | Vps16 | 0.007 | Young |
| 0.661647 | 0.072302 | 0.015 | 0.008 | 1 | 12 | Tcf7l2 | 0.007 | Young |
| 0.661647 | 0.052288 | 0.015 | 0.008 | 1 | 12 | B9d1 | 0.007 | Young |
| 0.661647 | 0.041882 | 0.015 | 0.008 | 1 | 12 | Elovl1 | 0.007 | Young |
| 0.661647 | 0.026041 | 0.015 | 0.008 | 1 | 12 | Gabrg3 | 0.007 | Young |
| 0.662016 | 0.388408 | 0.073 | 0.059 | 1 | 12 | Coro1a | 0.014 | Young |
| 0.662575 | 0.114174 | 0.066 | 0.051 | 1 | 12 | Polr2e | 0.015 | Young |
| 0.664853 | 0.327976 | 0.285 | 0.28 | 1 | 12 | Rps12 | 0.005 | Young |
| 0.666443 | 0.221662 | 0.175 | 0.153 | 1 | 12 | Stmn4 | 0.022 | Young |
| 0.667481 | 0.13463 | 0.803 | 0.831 | 1 | 12 | Eif1 | -0.028 | Young |
| 0.668751 | 0.798658 | 0.044 | 0.034 | 1 | 12 | Twf1 | 0.01 | Young |
| 0.668751 | 0.73743 | 0.044 | 0.034 | 1 | 12 | Lztfl1 | 0.01 | Young |
| 0.668751 | 0.710909 | 0.044 | 0.034 | 1 | 12 | Spin1 | 0.01 | Young |
| 0.668751 | 0.560645 | 0.044 | 0.034 | 1 | 12 | Tacc2 | 0.01 | Young |
| 0.668751 | 0.484164 | 0.044 | 0.034 | 1 | 12 | Pgrmc2 | 0.01 | Young |
| 0.669944 | 0.237029 | 0.372 | 0.322 | 1 | 12 | Tspan17 | 0.05 | Young |
| 0.671872 | 0.037734 | 0.81 | 0.754 | 1 | 12 | Fabp5 | 0.056 | Young |
| 0.672044 | 0.323871 | 0.139 | 0.119 | 1 | 12 | Mab21l2 | 0.02 | Young |
| 0.672442 | 0.861253 | 0.044 | 0.034 | 1 | 12 | Dld | 0.01 | Young |
| 0.672442 | 0.796155 | 0.044 | 0.034 | 1 | 12 | Slc35a1 | 0.01 | Young |
| 0.672442 | 0.55352 | 0.044 | 0.034 | 1 | 12 | Urm1 | 0.01 | Young |

| | | | | | | | | |
|---|---|---|---|---|---|---|---|---|
| 0.672442 | 0.540784 | 0.044 | 0.034 | 1 | 12 | Nit2 | 0.01 | Young |
| 0.672574 | 0.243864 | 0.234 | 0.212 | 1 | 12 | Atp6v1e1 | 0.022 | Young |
| 0.673124 | 0.157492 | 0.102 | 0.085 | 1 | 12 | Dctn6 | 0.017 | Young |
| 0.674491 | 0.293795 | 0.124 | 0.11 | 1 | 12 | Ralbp1 | 0.014 | Young |
| 0.674537 | 0.192867 | 0.109 | 0.093 | 1 | 12 | Sdhd | 0.016 | Young |
| 0.675075 | 0.052749 | 0.139 | 0.161 | 1 | 12 | Chd3 | -0.022 | Young |
| 0.676121 | 0.064424 | 0.584 | 0.542 | 1 | 12 | Fxyd6 | 0.042 | Young |
| 0.67614 | 0.745799 | 0.044 | 0.034 | 1 | 12 | Ndel1 | 0.01 | Young |
| 0.67614 | 0.443417 | 0.044 | 0.034 | 1 | 12 | Pcyt2 | 0.01 | Young |
| 0.67614 | 0.434755 | 0.044 | 0.034 | 1 | 12 | Ankrd40 | 0.01 | Young |
| 0.676665 | 0.501345 | 0.08 | 0.068 | 1 | 12 | Trim35 | 0.012 | Young |
| 0.677621 | 0.290486 | 0.117 | 0.102 | 1 | 12 | Chchd6 | 0.015 | Young |
| 0.679403 | 0.338452 | 0.08 | 0.068 | 1 | 12 | Ddx1 | 0.012 | Young |
| 0.679847 | 0.954907 | 0.044 | 0.034 | 1 | 12 | Tfg | 0.01 | Young |
| 0.682145 | 0.420076 | 0.08 | 0.068 | 1 | 12 | Tax1bp1 | 0.012 | Young |
| 0.68356 | 0.366633 | 0.044 | 0.034 | 1 | 12 | Brd2 | 0.01 | Young |
| 0.683561 | 0.549885 | 0.044 | 0.034 | 1 | 12 | Supt6 | 0.01 | Young |
| 0.683561 | 0.434256 | 0.044 | 0.034 | 1 | 12 | Tmem65 | 0.01 | Young |
| 0.683561 | 0.234799 | 0.044 | 0.034 | 1 | 12 | Tm9sf4 | 0.01 | Young |
| 0.68357 | 0.361183 | 0.073 | 0.059 | 1 | 12 | Ubtf | 0.014 | Young |
| 0.683645 | 0.017971 | 0.555 | 0.585 | 1 | 12 | mt-Nd2 | -0.03 | Young |
| 0.683961 | 0.17963 | 0.146 | 0.127 | 1 | 12 | Eif3e | 0.019 | Young |
| 0.684201 | 0.04597 | 0.102 | 0.085 | 1 | 12 | Camk1 | 0.017 | Young |
| 0.68445 | 0.102999 | 0.635 | 0.602 | 1 | 12 | Rplp0 | 0.033 | Young |
| 0.685484 | 0.020238 | 0.234 | 0.254 | 1 | 12 | Pgls | -0.02 | Young |
| 0.687283 | 0.568508 | 0.044 | 0.034 | 1 | 12 | Ufl1 | 0.01 | Young |
| 0.687283 | 0.399476 | 0.044 | 0.034 | 1 | 12 | Prkar2a | 0.01 | Young |
| 0.687283 | 0.380001 | 0.044 | 0.034 | 1 | 12 | Npepps | 0.01 | Young |
| 0.687283 | 0.365863 | 0.044 | 0.034 | 1 | 12 | Zfand2b | 0.01 | Young |
| 0.687283 | 0.345036 | 0.044 | 0.034 | 1 | 12 | Msl3 | 0.01 | Young |
| 0.687283 | 0.339193 | 0.044 | 0.034 | 1 | 12 | Rab2b | 0.01 | Young |
| 0.687283 | 0.306031 | 0.044 | 0.034 | 1 | 12 | Alyref | 0.01 | Young |
| 0.687641 | 0.332103 | 0.08 | 0.068 | 1 | 12 | Ccdc92 | 0.012 | Young |
| 0.689453 | 0.085425 | 0.387 | 0.347 | 1 | 12 | Nefm | 0.04 | Young |
| 0.689536 | 0.108636 | 0.328 | 0.297 | 1 | 12 | Hspe1 | 0.031 | Young |
| 0.690396 | 0.317841 | 0.08 | 0.068 | 1 | 12 | Snx17 | 0.012 | Young |
| 0.690396 | 0.303602 | 0.08 | 0.068 | 1 | 12 | Lrp11 | 0.012 | Young |
| 0.691012 | 0.631967 | 0.044 | 0.034 | 1 | 12 | Tcta | 0.01 | Young |
| 0.691012 | 0.44201 | 0.044 | 0.034 | 1 | 12 | 1110038F1 | 0.01 | Young |
| 0.691012 | 0.32574 | 0.044 | 0.034 | 1 | 12 | Sh3bp5 | 0.01 | Young |
| 0.691012 | 0.323514 | 0.044 | 0.034 | 1 | 12 | Cenpx | 0.01 | Young |
| 0.691012 | 0.310918 | 0.044 | 0.034 | 1 | 12 | Gnptg | 0.01 | Young |
| 0.691012 | 0.296578 | 0.044 | 0.034 | 1 | 12 | Rb1cc1 | 0.01 | Young |
| 0.691012 | 0.284015 | 0.044 | 0.034 | 1 | 12 | Psmd9 | 0.01 | Young |
| 0.691012 | 0.160465 | 0.044 | 0.034 | 1 | 12 | Cdc26 | 0.01 | Young |

| | | | | | | | | |
|---|---|---|---|---|---|---|---|---|
| 0.691394 | 0.234022 | 0.161 | 0.144 | 1 | 12 | Ppib | 0.017 | Young |
| 0.692231 | 0.18371 | 0.161 | 0.136 | 1 | 12 | Psmd6 | 0.025 | Young |
| 0.694749 | 0.365095 | 0.044 | 0.034 | 1 | 12 | Uba1 | 0.01 | Young |
| 0.694955 | 0.691375 | 0.051 | 0.042 | 1 | 12 | Epm2aip1 | 0.009 | Young |
| 0.694983 | 0.123031 | 0.416 | 0.407 | 1 | 12 | Sumo2 | 0.009 | Young |
| 0.696629 | 0.170133 | 0.073 | 0.059 | 1 | 12 | Pop4 | 0.014 | Young |
| 0.698493 | 0.289567 | 0.044 | 0.034 | 1 | 12 | S100a1 | 0.01 | Young |
| 0.698493 | 0.278385 | 0.044 | 0.034 | 1 | 12 | Gcsh | 0.01 | Young |
| 0.698684 | 0.289948 | 0.08 | 0.068 | 1 | 12 | Ube2k | 0.012 | Young |
| 0.698684 | 0.400502 | 0.08 | 0.068 | 1 | 12 | Nkap | 0.012 | Young |
| 0.698824 | 0.041819 | 0.358 | 0.322 | 1 | 12 | S100a11 | 0.036 | Young |
| 0.698892 | 0.015115 | 0.54 | 0.517 | 1 | 12 | Atp5l | 0.023 | Young |
| 0.700361 | 0.158912 | 0.161 | 0.136 | 1 | 12 | Pcbp3 | 0.025 | Young |
| 0.70113 | 0.033821 | 0.226 | 0.195 | 1 | 12 | Fbxo2 | 0.031 | Young |
| 0.701385 | 0.086612 | 0.394 | 0.356 | 1 | 12 | Ldha | 0.038 | Young |
| 0.701455 | 0.281728 | 0.08 | 0.068 | 1 | 12 | Tsc22d3 | 0.012 | Young |
| 0.701891 | 0.129455 | 0.117 | 0.102 | 1 | 12 | Rnf14 | 0.015 | Young |
| 0.702245 | 0.196609 | 0.044 | 0.034 | 1 | 12 | C1qtnf4 | 0.01 | Young |
| 0.702245 | 0.169004 | 0.044 | 0.034 | 1 | 12 | Plpp2 | 0.01 | Young |
| 0.702245 | 0.162922 | 0.044 | 0.034 | 1 | 12 | Ndufaf7 | 0.01 | Young |
| 0.702245 | 0.115357 | 0.044 | 0.034 | 1 | 12 | Klf9 | 0.01 | Young |
| 0.702245 | 0.098674 | 0.044 | 0.034 | 1 | 12 | Gnaz | 0.01 | Young |
| 0.702925 | 0.209548 | 0.197 | 0.186 | 1 | 12 | Eno1 | 0.011 | Young |
| 0.703191 | 0.034797 | 0.109 | 0.093 | 1 | 12 | Glrx2 | 0.016 | Young |
| 0.704154 | 0.054553 | 0.445 | 0.398 | 1 | 12 | Klc1 | 0.047 | Young |
| 0.704466 | 0.205363 | 0.285 | 0.271 | 1 | 12 | Bag1 | 0.014 | Young |
| 0.705022 | 0.022954 | 0.62 | 0.593 | 1 | 12 | Rpl27a | 0.027 | Young |
| 0.705386 | 0.109698 | 0.073 | 0.059 | 1 | 12 | Ensa | 0.014 | Young |
| 0.706004 | 0.193505 | 0.044 | 0.034 | 1 | 12 | Ccny | 0.01 | Young |
| 0.707626 | 0.351903 | 0.088 | 0.076 | 1 | 12 | Scd2 | 0.012 | Young |
| 0.70977 | 0.198216 | 0.044 | 0.034 | 1 | 12 | Mzt2 | 0.01 | Young |
| 0.70977 | 0.136622 | 0.044 | 0.034 | 1 | 12 | Ift22 | 0.01 | Young |
| 0.70977 | 0.135614 | 0.044 | 0.034 | 1 | 12 | Scarb2 | 0.01 | Young |
| 0.709779 | 0.113247 | 0.073 | 0.059 | 1 | 12 | Fastk | 0.014 | Young |
| 0.70979 | 0.372574 | 0.08 | 0.068 | 1 | 12 | Srpk1 | 0.012 | Young |
| 0.709791 | 0.19797 | 0.08 | 0.068 | 1 | 12 | Mrpl16 | 0.012 | Young |
| 0.710285 | 0.412666 | 0.088 | 0.076 | 1 | 12 | Rpp21 | 0.012 | Young |
| 0.711246 | 0.118418 | 0.073 | 0.059 | 1 | 12 | Ppm1a | 0.014 | Young |
| 0.711321 | 0.147267 | 0.234 | 0.212 | 1 | 12 | Serpinb6a | 0.022 | Young |
| 0.712577 | 0.33838 | 0.08 | 0.068 | 1 | 12 | Mrps6 | 0.012 | Young |
| 0.712587 | 0.338535 | 0.095 | 0.085 | 1 | 12 | Mrpl58 | 0.01 | Young |
| 0.712646 | 0.164395 | 0.19 | 0.169 | 1 | 12 | Nfic | 0.021 | Young |
| 0.713543 | 0.106478 | 0.044 | 0.034 | 1 | 12 | Zyg11b | 0.01 | Young |
| 0.714048 | 0.152985 | 0.109 | 0.093 | 1 | 12 | Manbal | 0.016 | Young |
| 0.715367 | 0.276085 | 0.08 | 0.068 | 1 | 12 | Pacsin1 | 0.012 | Young |

| | | | | | | | | |
|---|---|---|---|---|---|---|---|---|
| 0.715367 | 0.239525 | 0.08 | 0.068 | 1 | 12 | Eif3d | 0.012 | Young |
| 0.715367 | 0.181022 | 0.08 | 0.068 | 1 | 12 | Mlec | 0.012 | Young |
| 0.717324 | 0.067603 | 0.044 | 0.034 | 1 | 12 | Dusp22 | 0.01 | Young |
| 0.71737 | 0.533222 | 0.051 | 0.042 | 1 | 12 | Lmo4 | 0.009 | Young |
| 0.718652 | 0.003811 | 0.255 | 0.288 | 1 | 12 | Tuba1b | -0.033 | Young |
| 0.719105 | 0.515368 | 0.051 | 0.042 | 1 | 12 | Cnot7 | 0.009 | Young |
| 0.71956 | 0.306638 | 0.08 | 0.068 | 1 | 12 | Abcf1 | 0.012 | Young |
| 0.720952 | 0.384465 | 0.088 | 0.076 | 1 | 12 | Eif4e2 | 0.012 | Young |
| 0.720959 | 0.294666 | 0.08 | 0.068 | 1 | 12 | Smc3 | 0.012 | Young |
| 0.722579 | 0.292806 | 0.051 | 0.042 | 1 | 12 | Commd2 | 0.009 | Young |
| 0.725744 | 0.051818 | 0.971 | 0.949 | 1 | 12 | mt-Co1 | 0.022 | Young |
| 0.726058 | 0.537368 | 0.051 | 0.042 | 1 | 12 | Atp1b3 | 0.009 | Young |
| 0.728405 | 0.585573 | 0.058 | 0.051 | 1 | 12 | Tmem183a | 0.007 | Young |
| 0.729252 | 0.065984 | 0.788 | 0.746 | 1 | 12 | Actb | 0.042 | Young |
| 0.72976 | 0.082644 | 0.19 | 0.169 | 1 | 12 | Psma6 | 0.021 | Young |
| 0.731648 | 0.750747 | 0.058 | 0.051 | 1 | 12 | Eif4g1 | 0.007 | Young |
| 0.731674 | 0.388383 | 0.088 | 0.076 | 1 | 12 | Srp72 | 0.012 | Young |
| 0.732514 | 0.011856 | 0.044 | 0.034 | 1 | 12 | Tsen15 | 0.01 | Young |
| 0.733034 | 0.358587 | 0.051 | 0.042 | 1 | 12 | Pcm1 | 0.009 | Young |
| 0.738148 | 0.501865 | 0.058 | 0.051 | 1 | 12 | 5330434G( | 0.007 | Young |
| 0.739589 | 0.065138 | 0.073 | 0.085 | 1 | 12 | Asns | -0.012 | Young |
| 0.740258 | 0.111694 | 0.168 | 0.153 | 1 | 12 | Ppp1r14b | 0.015 | Young |
| 0.740647 | 0.137417 | 0.08 | 0.068 | 1 | 12 | Ndufa3 | 0.012 | Young |
| 0.741586 | 0.181493 | 0.401 | 0.39 | 1 | 12 | Nnat | 0.011 | Young |
| 0.741944 | 0.127992 | 0.555 | 0.568 | 1 | 12 | Bex3 | -0.013 | Young |
| 0.742002 | 0.12774 | 0.766 | 0.754 | 1 | 12 | Npy | 0.012 | Young |
| 0.743474 | 0.108306 | 0.08 | 0.068 | 1 | 12 | Exosc4 | 0.012 | Young |
| 0.743539 | 0.28826 | 0.051 | 0.042 | 1 | 12 | Mrpl9 | 0.009 | Young |
| 0.74354 | 0.198146 | 0.124 | 0.11 | 1 | 12 | Mrps18a | 0.014 | Young |
| 0.743541 | 0.21804 | 0.109 | 0.127 | 1 | 12 | Psme1 | -0.018 | Young |
| 0.744201 | 0.105958 | 0.117 | 0.102 | 1 | 12 | Sf3b5 | 0.015 | Young |
| 0.745151 | 0.219042 | 0.088 | 0.076 | 1 | 12 | Ost4 | 0.012 | Young |
| 0.74588 | 0.040008 | 0.139 | 0.119 | 1 | 12 | Nabp2 | 0.02 | Young |
| 0.745944 | 0.290655 | 0.131 | 0.119 | 1 | 12 | Gde1 | 0.012 | Young |
| 0.746037 | 0.001593 | 0.526 | 0.585 | 1 | 12 | Anxa2 | -0.059 | Young |
| 0.747051 | 0.23663 | 0.051 | 0.042 | 1 | 12 | Ei24 | 0.009 | Young |
| 0.747929 | 0.324704 | 0.139 | 0.127 | 1 | 12 | Eid2 | 0.012 | Young |
| 0.748658 | 0.300668 | 0.095 | 0.085 | 1 | 12 | Hk1 | 0.01 | Young |
| 0.749957 | 0.267611 | 0.095 | 0.085 | 1 | 12 | Wdr83os | 0.01 | Young |
| 0.750569 | 0.246169 | 0.051 | 0.042 | 1 | 12 | Apba1 | 0.009 | Young |
| 0.750569 | 0.208561 | 0.051 | 0.042 | 1 | 12 | Tdrkh | 0.009 | Young |
| 0.751047 | 0.403391 | 0.109 | 0.102 | 1 | 12 | Gm17018 | 0.007 | Young |
| 0.754092 | 0.279255 | 0.051 | 0.042 | 1 | 12 | Pepd | 0.009 | Young |
| 0.754092 | 0.221692 | 0.051 | 0.042 | 1 | 12 | Sdccag3 | 0.009 | Young |
| 0.757414 | 0.005901 | 0.168 | 0.153 | 1 | 12 | Tomm5 | 0.015 | Young |

| | | | | | | | | |
|---|---|---|---|---|---|---|---|---|
| 0.75762 | 0.338531 | 0.051 | 0.042 | 1 | 12 | Pi4k2a | 0.009 | Young |
| 0.758021 | 0.196748 | 0.175 | 0.161 | 1 | 12 | Rwdd1 | 0.014 | Young |
| 0.7587 | 0.04667 | 0.153 | 0.136 | 1 | 12 | Zfp428 | 0.017 | Young |
| 0.760506 | 0.063439 | 0.08 | 0.068 | 1 | 12 | Ckmt1 | 0.012 | Young |
| 0.761043 | 0.592764 | 0.058 | 0.051 | 1 | 12 | Abhd8 | 0.007 | Young |
| 0.761153 | 0.354253 | 0.051 | 0.042 | 1 | 12 | Gabrb3 | 0.009 | Young |
| 0.761153 | 0.189188 | 0.051 | 0.042 | 1 | 12 | Nop14 | 0.009 | Young |
| 0.761681 | 0.331692 | 0.095 | 0.085 | 1 | 12 | Ergic3 | 0.01 | Young |
| 0.761681 | 0.242569 | 0.095 | 0.085 | 1 | 12 | Rab18 | 0.01 | Young |
| 0.761681 | 0.092067 | 0.095 | 0.085 | 1 | 12 | Msn | 0.01 | Young |
| 0.763158 | 1.793608 | 0.022 | 0.017 | 1 | 12 | Lifr | 0.005 | Young |
| 0.763158 | 1.6871 | 0.022 | 0.017 | 1 | 12 | Pus1 | 0.005 | Young |
| 0.763158 | 1.329047 | 0.022 | 0.017 | 1 | 12 | Xpr1 | 0.005 | Young |
| 0.763158 | 1.322958 | 0.022 | 0.017 | 1 | 12 | Kdm5a | 0.005 | Young |
| 0.763158 | 1.271489 | 0.022 | 0.017 | 1 | 12 | Oxr1 | 0.005 | Young |
| 0.763158 | 0.806865 | 0.022 | 0.017 | 1 | 12 | Zc3h14 | 0.005 | Young |
| 0.763158 | 0.766905 | 0.022 | 0.017 | 1 | 12 | Fbxl17 | 0.005 | Young |
| 0.763158 | 0.762241 | 0.022 | 0.017 | 1 | 12 | Epb41l4a | 0.005 | Young |
| 0.763158 | 0.757828 | 0.022 | 0.017 | 1 | 12 | Plpbp | 0.005 | Young |
| 0.763158 | 0.755897 | 0.022 | 0.017 | 1 | 12 | Rprd1a | 0.005 | Young |
| 0.763158 | 0.68191 | 0.022 | 0.017 | 1 | 12 | Ptcd2 | 0.005 | Young |
| 0.763158 | 0.660575 | 0.022 | 0.017 | 1 | 12 | Colgalt1 | 0.005 | Young |
| 0.763158 | 0.627266 | 0.022 | 0.017 | 1 | 12 | Sh3bp5l | 0.005 | Young |
| 0.763158 | 0.491381 | 0.022 | 0.017 | 1 | 12 | Senp7 | 0.005 | Young |
| 0.764691 | 0.122193 | 0.051 | 0.042 | 1 | 12 | Bscl2 | 0.009 | Young |
| 0.76621 | 0.208519 | 0.08 | 0.068 | 1 | 12 | Uqcc3 | 0.012 | Young |
| 0.766451 | 0.325235 | 0.102 | 0.093 | 1 | 12 | B230219D2 | 0.009 | Young |
| 0.76804 | 0.267944 | 0.109 | 0.102 | 1 | 12 | Smim13 | 0.007 | Young |
| 0.768234 | 0.163585 | 0.051 | 0.042 | 1 | 12 | Syn1 | 0.009 | Young |
| 0.768234 | 0.103953 | 0.051 | 0.042 | 1 | 12 | Ilkap | 0.009 | Young |
| 0.768234 | 0.098048 | 0.051 | 0.042 | 1 | 12 | Abhd11 | 0.009 | Young |
| 0.768435 | 0.102848 | 0.336 | 0.322 | 1 | 12 | Psmb3 | 0.014 | Young |
| 0.768569 | 1.335253 | 0.022 | 0.017 | 1 | 12 | Acap2 | 0.005 | Young |
| 0.768569 | 1.231848 | 0.022 | 0.017 | 1 | 12 | Flrt2 | 0.005 | Young |
| 0.768569 | 1.198973 | 0.022 | 0.017 | 1 | 12 | Rnf2 | 0.005 | Young |
| 0.768569 | 0.889583 | 0.022 | 0.017 | 1 | 12 | Khdrbs2 | 0.005 | Young |
| 0.768569 | 0.880049 | 0.022 | 0.017 | 1 | 12 | Cyp2u1 | 0.005 | Young |
| 0.768569 | 0.813174 | 0.022 | 0.017 | 1 | 12 | Tbc1d10a | 0.005 | Young |
| 0.768569 | 0.790288 | 0.022 | 0.017 | 1 | 12 | Mindy1 | 0.005 | Young |
| 0.768569 | 0.746262 | 0.022 | 0.017 | 1 | 12 | Golga2 | 0.005 | Young |
| 0.768569 | 0.731059 | 0.022 | 0.017 | 1 | 12 | Cep170b | 0.005 | Young |
| 0.768569 | 0.721647 | 0.022 | 0.017 | 1 | 12 | Ppp1r15a | 0.005 | Young |
| 0.768569 | 0.631293 | 0.022 | 0.017 | 1 | 12 | Lrfn1 | 0.005 | Young |
| 0.768569 | 0.62244 | 0.022 | 0.017 | 1 | 12 | Arhgef25 | 0.005 | Young |
| 0.768569 | 0.611458 | 0.022 | 0.017 | 1 | 12 | Trip4 | 0.005 | Young |

| | | | | | | | | |
|---|---|---|---|---|---|---|---|---|
| 0.768569 | 0.608172 | 0.022 | 0.017 | 1 | 12 | Pias4 | 0.005 | Young |
| 0.768569 | 0.582914 | 0.022 | 0.017 | 1 | 12 | Clasp1 | 0.005 | Young |
| 0.768569 | 0.56516 | 0.022 | 0.017 | 1 | 12 | Mapkap1 | 0.005 | Young |
| 0.768569 | 0.561761 | 0.022 | 0.017 | 1 | 12 | Trappc13 | 0.005 | Young |
| 0.768569 | 0.535989 | 0.022 | 0.017 | 1 | 12 | Cdip1 | 0.005 | Young |
| 0.768569 | 0.513788 | 0.022 | 0.017 | 1 | 12 | AW551984 | 0.005 | Young |
| 0.768569 | 0.510462 | 0.022 | 0.017 | 1 | 12 | Wipi1 | 0.005 | Young |
| 0.768569 | 0.483018 | 0.022 | 0.017 | 1 | 12 | Fahd2a | 0.005 | Young |
| 0.768569 | 0.476564 | 0.022 | 0.017 | 1 | 12 | Uxs1 | 0.005 | Young |
| 0.768569 | 0.470658 | 0.022 | 0.017 | 1 | 12 | Coro2b | 0.005 | Young |
| 0.768569 | 0.465851 | 0.022 | 0.017 | 1 | 12 | Lin7c | 0.005 | Young |
| 0.768978 | 0.00306 | 0.058 | 0.068 | 1 | 12 | Tmco1 | -0.01 | Young |
| 0.769604 | 0.180756 | 0.088 | 0.076 | 1 | 12 | Hnrnpa3 | 0.012 | Young |
| 0.772175 | 0.061332 | 0.066 | 0.076 | 1 | 12 | Sdcbp | -0.01 | Young |
| 0.773802 | 0.141035 | 0.088 | 0.102 | 1 | 12 | Imp3 | -0.014 | Young |
| 0.773992 | 0.547361 | 0.022 | 0.017 | 1 | 12 | Syne1 | 0.005 | Young |
| 0.773993 | 1.29792 | 0.022 | 0.017 | 1 | 12 | Api5 | 0.005 | Young |
| 0.773993 | 1.070795 | 0.022 | 0.017 | 1 | 12 | Iqgap1 | 0.005 | Young |
| 0.773993 | 1.040003 | 0.022 | 0.017 | 1 | 12 | Smad2 | 0.005 | Young |
| 0.773993 | 1.005359 | 0.022 | 0.017 | 1 | 12 | Csf2ra | 0.005 | Young |
| 0.773993 | 0.969522 | 0.022 | 0.017 | 1 | 12 | Hs6st1 | 0.005 | Young |
| 0.773993 | 0.9409 | 0.022 | 0.017 | 1 | 12 | Acp2 | 0.005 | Young |
| 0.773993 | 0.925275 | 0.022 | 0.017 | 1 | 12 | Luzp1 | 0.005 | Young |
| 0.773993 | 0.736723 | 0.022 | 0.017 | 1 | 12 | Anapc2 | 0.005 | Young |
| 0.773993 | 0.71759 | 0.022 | 0.017 | 1 | 12 | Pqbp1 | 0.005 | Young |
| 0.773993 | 0.696303 | 0.022 | 0.017 | 1 | 12 | Hsd17b7 | 0.005 | Young |
| 0.773993 | 0.68241 | 0.022 | 0.017 | 1 | 12 | Rragd | 0.005 | Young |
| 0.773993 | 0.648108 | 0.022 | 0.017 | 1 | 12 | 0610037L1 | 0.005 | Young |
| 0.773993 | 0.628701 | 0.022 | 0.017 | 1 | 12 | Srcin1 | 0.005 | Young |
| 0.773993 | 0.627651 | 0.022 | 0.017 | 1 | 12 | Mrrf | 0.005 | Young |
| 0.773993 | 0.6156 | 0.022 | 0.017 | 1 | 12 | Ttc9 | 0.005 | Young |
| 0.773993 | 0.607032 | 0.022 | 0.017 | 1 | 12 | Peli1 | 0.005 | Young |
| 0.773993 | 0.602607 | 0.022 | 0.017 | 1 | 12 | Cds2 | 0.005 | Young |
| 0.773993 | 0.582183 | 0.022 | 0.017 | 1 | 12 | Ift43 | 0.005 | Young |
| 0.773993 | 0.582043 | 0.022 | 0.017 | 1 | 12 | Rpl13a | 0.005 | Young |
| 0.773993 | 0.574917 | 0.022 | 0.017 | 1 | 12 | Gas6 | 0.005 | Young |
| 0.773993 | 0.571754 | 0.022 | 0.017 | 1 | 12 | Clpb | 0.005 | Young |
| 0.773993 | 0.559641 | 0.022 | 0.017 | 1 | 12 | Mvb12b | 0.005 | Young |
| 0.773993 | 0.540169 | 0.022 | 0.017 | 1 | 12 | Scn3a | 0.005 | Young |
| 0.773993 | 0.490052 | 0.022 | 0.017 | 1 | 12 | Brap | 0.005 | Young |
| 0.773993 | 0.489986 | 0.022 | 0.017 | 1 | 12 | Gm6710 | 0.005 | Young |
| 0.773993 | 0.479888 | 0.022 | 0.017 | 1 | 12 | Spg7 | 0.005 | Young |
| 0.773993 | 0.468979 | 0.022 | 0.017 | 1 | 12 | Pltp | 0.005 | Young |
| 0.773993 | 0.462558 | 0.022 | 0.017 | 1 | 12 | Coa6 | 0.005 | Young |
| 0.773993 | 0.459227 | 0.022 | 0.017 | 1 | 12 | Calu | 0.005 | Young |

| | | | | | | | | |
|---|---|---|---|---|---|---|---|---|
| 0.773993 | 0.452128 | 0.022 | 0.017 | 1 | 12 | Wdr55 | 0.005 | Young |
| 0.773993 | 0.447699 | 0.022 | 0.017 | 1 | 12 | Psmg1 | 0.005 | Young |
| 0.773993 | 0.446103 | 0.022 | 0.017 | 1 | 12 | Eml4 | 0.005 | Young |
| 0.773993 | 0.434034 | 0.022 | 0.017 | 1 | 12 | Pacs1 | 0.005 | Young |
| 0.773993 | 0.430888 | 0.022 | 0.017 | 1 | 12 | Krit1 | 0.005 | Young |
| 0.773993 | 0.412412 | 0.022 | 0.017 | 1 | 12 | Las1l | 0.005 | Young |
| 0.773993 | 0.410583 | 0.022 | 0.017 | 1 | 12 | Cabin1 | 0.005 | Young |
| 0.773993 | 0.406806 | 0.022 | 0.017 | 1 | 12 | Thoc1 | 0.005 | Young |
| 0.773993 | 0.403626 | 0.022 | 0.017 | 1 | 12 | Arid2 | 0.005 | Young |
| 0.773993 | 0.403173 | 0.022 | 0.017 | 1 | 12 | Osbpl9 | 0.005 | Young |
| 0.773993 | 0.393788 | 0.022 | 0.017 | 1 | 12 | Snx11 | 0.005 | Young |
| 0.773993 | 0.389239 | 0.022 | 0.017 | 1 | 12 | Gatad2b | 0.005 | Young |
| 0.773993 | 0.386017 | 0.022 | 0.017 | 1 | 12 | Bace2 | 0.005 | Young |
| 0.773993 | 0.376325 | 0.022 | 0.017 | 1 | 12 | Camkk2 | 0.005 | Young |
| 0.773993 | 0.364079 | 0.022 | 0.017 | 1 | 12 | Fem1a | 0.005 | Young |
| 0.773993 | 0.363639 | 0.022 | 0.017 | 1 | 12 | Cpsf6 | 0.005 | Young |
| 0.773993 | 0.36123 | 0.022 | 0.017 | 1 | 12 | N4bp2l1 | 0.005 | Young |
| 0.773993 | 0.351629 | 0.022 | 0.017 | 1 | 12 | Pom121 | 0.005 | Young |
| 0.773993 | 0.300327 | 0.022 | 0.017 | 1 | 12 | Tbc1d9 | 0.005 | Young |
| 0.774222 | 0.460218 | 0.058 | 0.051 | 1 | 12 | Cnn3 | 0.007 | Young |
| 0.774771 | 0.054935 | 0.095 | 0.085 | 1 | 12 | Nol3 | 0.01 | Young |
| 0.775177 | 0.017757 | 0.058 | 0.068 | 1 | 12 | Mpv17l2 | -0.01 | Young |
| 0.777396 | 0.152057 | 0.095 | 0.085 | 1 | 12 | Mea1 | 0.01 | Young |
| 0.777515 | 0.076332 | 0.533 | 0.542 | 1 | 12 | Cox4i1 | -0.009 | Young |
| 0.777527 | 0.410743 | 0.058 | 0.051 | 1 | 12 | Egfl7 | 0.007 | Young |
| 0.777545 | 0.29662 | 0.197 | 0.195 | 1 | 12 | Ciapin1 | 0.002 | Young |
| 0.779091 | 0.365232 | 0.124 | 0.119 | 1 | 12 | Hpcal1 | 0.005 | Young |
| 0.779427 | 0.852001 | 0.022 | 0.017 | 1 | 12 | Mecr | 0.005 | Young |
| 0.779427 | 0.813914 | 0.022 | 0.017 | 1 | 12 | Mtmr4 | 0.005 | Young |
| 0.779427 | 0.758646 | 0.022 | 0.017 | 1 | 12 | Rufy1 | 0.005 | Young |
| 0.779427 | 0.756934 | 0.022 | 0.017 | 1 | 12 | Akap8l | 0.005 | Young |
| 0.779427 | 0.706278 | 0.022 | 0.017 | 1 | 12 | Scaf1 | 0.005 | Young |
| 0.779427 | 0.573985 | 0.022 | 0.017 | 1 | 12 | Galk2 | 0.005 | Young |
| 0.779427 | 0.538518 | 0.022 | 0.017 | 1 | 12 | Plppr4 | 0.005 | Young |
| 0.779427 | 0.501546 | 0.022 | 0.017 | 1 | 12 | Fgf12 | 0.005 | Young |
| 0.779427 | 0.498269 | 0.022 | 0.017 | 1 | 12 | Synm | 0.005 | Young |
| 0.779427 | 0.482765 | 0.022 | 0.017 | 1 | 12 | Prkag2 | 0.005 | Young |
| 0.779427 | 0.479943 | 0.022 | 0.017 | 1 | 12 | Abi1 | 0.005 | Young |
| 0.779427 | 0.465555 | 0.022 | 0.017 | 1 | 12 | Pold2 | 0.005 | Young |
| 0.779427 | 0.462719 | 0.022 | 0.017 | 1 | 12 | Rnf170 | 0.005 | Young |
| 0.779427 | 0.462049 | 0.022 | 0.017 | 1 | 12 | Grk6 | 0.005 | Young |
| 0.779427 | 0.44891 | 0.022 | 0.017 | 1 | 12 | Gatd1 | 0.005 | Young |
| 0.779427 | 0.435043 | 0.022 | 0.017 | 1 | 12 | Fam241b | 0.005 | Young |
| 0.779427 | 0.431005 | 0.022 | 0.017 | 1 | 12 | Kcnma1 | 0.005 | Young |
| 0.779427 | 0.410538 | 0.022 | 0.017 | 1 | 12 | D1Ertd622 | 0.005 | Young |

| | | | | | | | | |
|---|---|---|---|---|---|---|---|---|
| 0.779427 | 0.403253 | 0.022 | 0.017 | 1 | 12 | Kctd5 | 0.005 | Young |
| 0.779427 | 0.397053 | 0.022 | 0.017 | 1 | 12 | Cdk8 | 0.005 | Young |
| 0.779427 | 0.395761 | 0.022 | 0.017 | 1 | 12 | Slc8a1 | 0.005 | Young |
| 0.779427 | 0.38913 | 0.022 | 0.017 | 1 | 12 | Rgs11 | 0.005 | Young |
| 0.779427 | 0.384929 | 0.022 | 0.017 | 1 | 12 | Wdr46 | 0.005 | Young |
| 0.779427 | 0.380786 | 0.022 | 0.017 | 1 | 12 | Fbxl6 | 0.005 | Young |
| 0.779427 | 0.367439 | 0.022 | 0.017 | 1 | 12 | Slc29a1 | 0.005 | Young |
| 0.779427 | 0.359377 | 0.022 | 0.017 | 1 | 12 | Sdc2 | 0.005 | Young |
| 0.779427 | 0.346471 | 0.022 | 0.017 | 1 | 12 | Nelfcd | 0.005 | Young |
| 0.779427 | 0.345035 | 0.022 | 0.017 | 1 | 12 | Gtpbp8 | 0.005 | Young |
| 0.779427 | 0.341551 | 0.022 | 0.017 | 1 | 12 | Nck1 | 0.005 | Young |
| 0.779427 | 0.334426 | 0.022 | 0.017 | 1 | 12 | Zfp422 | 0.005 | Young |
| 0.779427 | 0.315478 | 0.022 | 0.017 | 1 | 12 | Dusp12 | 0.005 | Young |
| 0.779427 | 0.292853 | 0.022 | 0.017 | 1 | 12 | Lrrc4c | 0.005 | Young |
| 0.779427 | 0.288647 | 0.022 | 0.017 | 1 | 12 | Sh3yl1 | 0.005 | Young |
| 0.779427 | 0.284011 | 0.022 | 0.017 | 1 | 12 | Sesn3 | 0.005 | Young |
| 0.779427 | 0.282418 | 0.022 | 0.017 | 1 | 12 | Ccdc85b | 0.005 | Young |
| 0.779427 | 0.280637 | 0.022 | 0.017 | 1 | 12 | Abcb6 | 0.005 | Young |
| 0.779427 | 0.276018 | 0.022 | 0.017 | 1 | 12 | Ephx1 | 0.005 | Young |
| 0.779427 | 0.262013 | 0.022 | 0.017 | 1 | 12 | Mtf2 | 0.005 | Young |
| 0.779427 | 0.218916 | 0.022 | 0.017 | 1 | 12 | Layn | 0.005 | Young |
| 0.779427 | 0.188369 | 0.022 | 0.017 | 1 | 12 | B3galt1 | 0.005 | Young |
| 0.780618 | 0.031133 | 0.905 | 0.915 | 1 | 12 | Tubb3 | -0.01 | Young |
| 0.780836 | 0.317885 | 0.058 | 0.051 | 1 | 12 | Rprm | 0.007 | Young |
| 0.782147 | 0.25807 | 0.022 | 0.017 | 1 | 12 | Stk24 | 0.005 | Young |
| 0.78392 | 0.371448 | 0.109 | 0.102 | 1 | 12 | Mbnl2 | 0.007 | Young |
| 0.784147 | 0.281442 | 0.102 | 0.093 | 1 | 12 | Snrnp70 | 0.009 | Young |
| 0.784149 | 0.228448 | 0.058 | 0.051 | 1 | 12 | Arxes1 | 0.007 | Young |
| 0.784503 | 0.03513 | 0.058 | 0.068 | 1 | 12 | Macf1 | -0.01 | Young |
| 0.784872 | 0.380584 | 0.022 | 0.017 | 1 | 12 | Cep295 | 0.005 | Young |
| 0.784872 | 0.351516 | 0.022 | 0.017 | 1 | 12 | Capn1 | 0.005 | Young |
| 0.784872 | 0.34662 | 0.022 | 0.017 | 1 | 12 | Nfrkb | 0.005 | Young |
| 0.784872 | 0.33989 | 0.022 | 0.017 | 1 | 12 | Trmt10b | 0.005 | Young |
| 0.784872 | 0.323049 | 0.022 | 0.017 | 1 | 12 | Dimt1 | 0.005 | Young |
| 0.784872 | 0.300362 | 0.022 | 0.017 | 1 | 12 | Eid2b | 0.005 | Young |
| 0.784872 | 0.294215 | 0.022 | 0.017 | 1 | 12 | Nup35 | 0.005 | Young |
| 0.784872 | 0.287 | 0.022 | 0.017 | 1 | 12 | Nubp1 | 0.005 | Young |
| 0.784872 | 0.285096 | 0.022 | 0.017 | 1 | 12 | Ythdc2 | 0.005 | Young |
| 0.784872 | 0.267672 | 0.022 | 0.017 | 1 | 12 | 4930453N2 | 0.005 | Young |
| 0.784872 | 0.266325 | 0.022 | 0.017 | 1 | 12 | Mtf1 | 0.005 | Young |
| 0.784872 | 0.263977 | 0.022 | 0.017 | 1 | 12 | Rhbdd3 | 0.005 | Young |
| 0.784872 | 0.261598 | 0.022 | 0.017 | 1 | 12 | Epc1 | 0.005 | Young |
| 0.784872 | 0.254342 | 0.022 | 0.017 | 1 | 12 | Thoc2 | 0.005 | Young |
| 0.784872 | 0.249782 | 0.022 | 0.017 | 1 | 12 | Scfd1 | 0.005 | Young |
| 0.784872 | 0.243881 | 0.022 | 0.017 | 1 | 12 | Bace1 | 0.005 | Young |

| | | | | | | | | |
|---|---|---|---|---|---|---|---|---|
| 0.784872 | 0.229068 | 0.022 | 0.017 | 1 | 12 | Hspa14 | 0.005 | Young |
| 0.784872 | 0.227985 | 0.022 | 0.017 | 1 | 12 | Frat2 | 0.005 | Young |
| 0.784872 | 0.221627 | 0.022 | 0.017 | 1 | 12 | Hoxc4 | 0.005 | Young |
| 0.784872 | 0.214385 | 0.022 | 0.017 | 1 | 12 | Xpc | 0.005 | Young |
| 0.784872 | 0.208798 | 0.022 | 0.017 | 1 | 12 | Ecel1 | 0.005 | Young |
| 0.784872 | 0.208109 | 0.022 | 0.017 | 1 | 12 | Atxn2 | 0.005 | Young |
| 0.784872 | 0.203797 | 0.022 | 0.017 | 1 | 12 | Lpcat1 | 0.005 | Young |
| 0.784872 | 0.199749 | 0.022 | 0.017 | 1 | 12 | Rab3ip | 0.005 | Young |
| 0.784872 | 0.184625 | 0.022 | 0.017 | 1 | 12 | Ccdc184 | 0.005 | Young |
| 0.784872 | 0.184137 | 0.022 | 0.017 | 1 | 12 | Gpaa1 | 0.005 | Young |
| 0.784872 | 0.171127 | 0.022 | 0.017 | 1 | 12 | Cic | 0.005 | Young |
| 0.784872 | 0.155088 | 0.022 | 0.017 | 1 | 12 | Abhd5 | 0.005 | Young |
| 0.784872 | 0.154289 | 0.022 | 0.017 | 1 | 12 | Hibadh | 0.005 | Young |
| 0.784872 | 0.148872 | 0.022 | 0.017 | 1 | 12 | Dusp2 | 0.005 | Young |
| 0.784872 | 0.144931 | 0.022 | 0.017 | 1 | 12 | Cdr2l | 0.005 | Young |
| 0.784872 | 0.131639 | 0.022 | 0.017 | 1 | 12 | Arf6 | 0.005 | Young |
| 0.784872 | 0.130792 | 0.022 | 0.017 | 1 | 12 | Jarid2 | 0.005 | Young |
| 0.784872 | 0.130017 | 0.022 | 0.017 | 1 | 12 | Fndc10 | 0.005 | Young |
| 0.784872 | 0.119818 | 0.022 | 0.017 | 1 | 12 | Casp8ap2 | 0.005 | Young |
| 0.784872 | 0.116217 | 0.022 | 0.017 | 1 | 12 | Lrrtm3 | 0.005 | Young |
| 0.784872 | 0.103635 | 0.022 | 0.017 | 1 | 12 | Rnf165 | 0.005 | Young |
| 0.784872 | 0.071147 | 0.022 | 0.017 | 1 | 12 | Tax1bp3 | 0.005 | Young |
| 0.784872 | 0.043615 | 0.022 | 0.017 | 1 | 12 | Slc16a3 | 0.005 | Young |
| 0.784872 | 0.025325 | 0.022 | 0.017 | 1 | 12 | Arhgef7 | 0.005 | Young |
| 0.785423 | 0.065598 | 0.701 | 0.729 | 1 | 12 | Rps5 | -0.028 | Young |
| 0.785807 | 0.413041 | 0.058 | 0.051 | 1 | 12 | Agap3 | 0.007 | Young |
| 0.785807 | 0.297648 | 0.058 | 0.051 | 1 | 12 | Tanc2 | 0.007 | Young |
| 0.785807 | 0.139147 | 0.051 | 0.059 | 1 | 12 | Usp9x | -0.008 | Young |
| 0.786075 | 0.192712 | 0.146 | 0.136 | 1 | 12 | Trappc2l | 0.01 | Young |
| 0.787941 | 0.098342 | 0.328 | 0.314 | 1 | 12 | Atp6v1g1 | 0.014 | Young |
| 0.788117 | 0.265196 | 0.117 | 0.11 | 1 | 12 | Bad | 0.007 | Young |
| 0.790327 | 0.231341 | 0.022 | 0.017 | 1 | 12 | Cerk | 0.005 | Young |
| 0.790327 | 0.222299 | 0.022 | 0.017 | 1 | 12 | Gbf1 | 0.005 | Young |
| 0.790327 | 0.220877 | 0.022 | 0.017 | 1 | 12 | Plppr2 | 0.005 | Young |
| 0.790327 | 0.179448 | 0.022 | 0.017 | 1 | 12 | Spry1 | 0.005 | Young |
| 0.790327 | 0.172272 | 0.022 | 0.017 | 1 | 12 | Brsk2 | 0.005 | Young |
| 0.790327 | 0.154632 | 0.022 | 0.017 | 1 | 12 | Lsm10 | 0.005 | Young |
| 0.790327 | 0.153863 | 0.022 | 0.017 | 1 | 12 | Oxa1l | 0.005 | Young |
| 0.790327 | 0.138271 | 0.022 | 0.017 | 1 | 12 | Tbc1d8 | 0.005 | Young |
| 0.790327 | 0.135741 | 0.022 | 0.017 | 1 | 12 | Sgsm3 | 0.005 | Young |
| 0.790327 | 0.12384 | 0.022 | 0.017 | 1 | 12 | Slc41a2 | 0.005 | Young |
| 0.790327 | 0.075138 | 0.022 | 0.017 | 1 | 12 | Fam53a | 0.005 | Young |
| 0.790327 | 0.068056 | 0.022 | 0.017 | 1 | 12 | Cd99l2 | 0.005 | Young |
| 0.790327 | 0.040488 | 0.022 | 0.017 | 1 | 12 | Tspyl3 | 0.005 | Young |
| 0.790327 | 0.030255 | 0.022 | 0.017 | 1 | 12 | Lingo1 | 0.005 | Young |

| | | | | | | | | |
|---|---|---|---|---|---|---|---|---|
| 0.790404 | 0.285269 | 0.146 | 0.136 | 1 | 12 | Gtf2a2 | 0.01 | Young |
| 0.790787 | 0.387124 | 0.058 | 0.051 | 1 | 12 | Mid1ip1 | 0.007 | Young |
| 0.790787 | 0.191109 | 0.058 | 0.051 | 1 | 12 | Ptprs | 0.007 | Young |
| 0.790964 | 0.105558 | 0.35 | 0.339 | 1 | 12 | Hras | 0.011 | Young |
| 0.791247 | 0.055725 | 0.635 | 0.627 | 1 | 12 | Cst3 | 0.008 | Young |
| 0.792233 | 0.097865 | 0.219 | 0.203 | 1 | 12 | Tprgl | 0.016 | Young |
| 0.793736 | 0.033904 | 0.73 | 0.669 | 1 | 12 | Aldoa | 0.061 | Young |
| 0.79431 | 0.225029 | 0.102 | 0.093 | 1 | 12 | Hmgn1 | 0.009 | Young |
| 0.795793 | 0.101253 | 0.022 | 0.017 | 1 | 12 | Wdr33 | 0.005 | Young |
| 0.795793 | 0.059934 | 0.022 | 0.017 | 1 | 12 | Rtl5 | 0.005 | Young |
| 0.795793 | 0.050707 | 0.022 | 0.017 | 1 | 12 | Dhdds | 0.005 | Young |
| 0.795793 | 0.046549 | 0.022 | 0.017 | 1 | 12 | 1600020E0 | 0.005 | Young |
| 0.795793 | 0.00391 | 0.022 | 0.017 | 1 | 12 | Fosb | 0.005 | Young |
| 0.795847 | 0.302351 | 0.095 | 0.085 | 1 | 12 | Psmd1 | 0.01 | Young |
| 0.796747 | 0.065421 | 0.044 | 0.051 | 1 | 12 | Mindy2 | -0.007 | Young |
| 0.79744 | 0.312287 | 0.058 | 0.051 | 1 | 12 | Ing4 | 0.007 | Young |
| 0.797562 | 0.23278 | 0.234 | 0.237 | 1 | 12 | Hcfc1r1 | -0.003 | Young |
| 0.797568 | 0.021618 | 0.365 | 0.398 | 1 | 12 | Socs2 | -0.033 | Young |
| 0.800114 | 0.22068 | 0.066 | 0.059 | 1 | 12 | Copg1 | 0.007 | Young |
| 0.800619 | 0.048846 | 0.124 | 0.136 | 1 | 12 | Olfm2 | -0.012 | Young |
| 0.800772 | 0.40497 | 0.058 | 0.051 | 1 | 12 | Flywch1 | 0.007 | Young |
| 0.801075 | 0.056802 | 0.825 | 0.805 | 1 | 12 | Tmsb10 | 0.02 | Young |
| 0.8022 | 0.006451 | 0.131 | 0.144 | 1 | 12 | Cldnd1 | -0.013 | Young |
| 0.80244 | 0.1507 | 0.058 | 0.051 | 1 | 12 | Bag5 | 0.007 | Young |
| 0.802899 | 0.012575 | 0.343 | 0.314 | 1 | 12 | H2afz | 0.029 | Young |
| 0.803245 | 0.306454 | 0.066 | 0.059 | 1 | 12 | Med29 | 0.007 | Young |
| 0.803246 | 0.380298 | 0.066 | 0.059 | 1 | 12 | Clpp | 0.007 | Young |
| 0.804108 | 0.049262 | 0.058 | 0.051 | 1 | 12 | Parm1 | 0.007 | Young |
| 0.807151 | 0.09753 | 0.131 | 0.119 | 1 | 12 | Grb2 | 0.012 | Young |
| 0.807447 | 0.317389 | 0.058 | 0.051 | 1 | 12 | Clcn3 | 0.007 | Young |
| 0.807447 | 0.268749 | 0.058 | 0.051 | 1 | 12 | Nsdhl | 0.007 | Young |
| 0.807447 | 0.258828 | 0.058 | 0.051 | 1 | 12 | Atp2b2 | 0.007 | Young |
| 0.807447 | 0.080115 | 0.058 | 0.051 | 1 | 12 | Tm2d1 | 0.007 | Young |
| 0.80962 | 0.135627 | 0.102 | 0.093 | 1 | 12 | Snrpd3 | 0.009 | Young |
| 0.809944 | 0.012478 | 0.365 | 0.347 | 1 | 12 | Dnaja1 | 0.018 | Young |
| 0.810513 | 0.109536 | 0.263 | 0.263 | 1 | 12 | Sem1 | 0 | Young |
| 0.811847 | 0.014836 | 0.533 | 0.534 | 1 | 12 | Isoc1 | -0.001 | Young |
| 0.812179 | 0.011772 | 0.102 | 0.093 | 1 | 12 | Ube2e3 | 0.009 | Young |
| 0.81266 | 0.352218 | 0.058 | 0.068 | 1 | 12 | Mettl26 | -0.01 | Young |
| 0.813235 | 0.169273 | 0.146 | 0.136 | 1 | 12 | Arf4 | 0.01 | Young |
| 0.813554 | 0.299022 | 0.073 | 0.068 | 1 | 12 | Bod1 | 0.005 | Young |
| 0.813555 | 0.021576 | 0.204 | 0.186 | 1 | 12 | Phox2b | 0.018 | Young |
| 0.813859 | 0.020334 | 0.036 | 0.042 | 1 | 12 | Prmt7 | -0.006 | Young |
| 0.814191 | 0.052204 | 0.153 | 0.144 | 1 | 12 | Klf6 | 0.009 | Young |
| 0.815405 | 0.071701 | 0.569 | 0.542 | 1 | 12 | Mpc1 | 0.027 | Young |

| | | | | | | | | |
|---|---|---|---|---|---|---|---|---|
| 0.815805 | 0.257757 | 0.066 | 0.059 | 1 | 12 | Pabpc1 | 0.007 | Young |
| 0.816977 | 0.135731 | 0.394 | 0.373 | 1 | 12 | Npm1 | 0.021 | Young |
| 0.817486 | 0.185087 | 0.058 | 0.051 | 1 | 12 | Golgb1 | 0.007 | Young |
| 0.817486 | 0.046121 | 0.051 | 0.059 | 1 | 12 | Ckap4 | -0.008 | Young |
| 0.817791 | 0.016078 | 0.036 | 0.042 | 1 | 12 | Ggh | -0.006 | Young |
| 0.818313 | 0.05094 | 0.044 | 0.051 | 1 | 12 | Coro1c | -0.007 | Young |
| 0.818598 | 0.014963 | 0.139 | 0.153 | 1 | 12 | A730017C2 | -0.014 | Young |
| 0.81956 | 0.050328 | 0.204 | 0.212 | 1 | 12 | Eif2s2 | -0.008 | Young |
| 0.820759 | 0.006178 | 0.139 | 0.153 | 1 | 12 | Son | -0.014 | Young |
| 0.820839 | 0.019892 | 0.058 | 0.051 | 1 | 12 | Dctn1 | 0.007 | Young |
| 0.821571 | 0.078269 | 0.182 | 0.169 | 1 | 12 | Sumo1 | 0.013 | Young |
| 0.821921 | 0.07906 | 0.044 | 0.051 | 1 | 12 | Zfhx4 | -0.007 | Young |
| 0.822102 | 0.359159 | 0.066 | 0.059 | 1 | 12 | Rdx | 0.007 | Young |
| 0.824988 | 0.184607 | 0.175 | 0.169 | 1 | 12 | A830018L1 | 0.006 | Young |
| 0.825256 | 0.318019 | 0.066 | 0.059 | 1 | 12 | Usp22 | 0.007 | Young |
| 0.825479 | 0.599988 | 0.073 | 0.068 | 1 | 12 | Sgip1 | 0.005 | Young |
| 0.825668 | 0.095126 | 0.036 | 0.042 | 1 | 12 | Ilf3 | -0.006 | Young |
| 0.825668 | 0.078018 | 0.036 | 0.042 | 1 | 12 | Gpbp1 | -0.006 | Young |
| 0.827392 | 0.076093 | 0.613 | 0.559 | 1 | 12 | Rps2 | 0.054 | Young |
| 0.827555 | 0.087015 | 0.051 | 0.059 | 1 | 12 | Cadm3 | -0.008 | Young |
| 0.827574 | 0.017472 | 0.102 | 0.093 | 1 | 12 | Gata3 | 0.009 | Young |
| 0.828028 | 0.041264 | 0.182 | 0.195 | 1 | 12 | Eif3i | -0.013 | Young |
| 0.828138 | 0.083254 | 0.029 | 0.034 | 1 | 12 | Ube2d1 | -0.005 | Young |
| 0.828138 | 0.051913 | 0.029 | 0.034 | 1 | 12 | Adk | -0.005 | Young |
| 0.828138 | 0.044405 | 0.029 | 0.034 | 1 | 12 | Ubn2 | -0.005 | Young |
| 0.828536 | 0.732222 | 0.029 | 0.025 | 1 | 12 | Osbpl6 | 0.004 | Young |
| 0.829149 | 0.168845 | 0.044 | 0.051 | 1 | 12 | Cxxc4 | -0.007 | Young |
| 0.829149 | 0.028068 | 0.044 | 0.051 | 1 | 12 | Tsr3 | -0.007 | Young |
| 0.829158 | 0.010477 | 0.307 | 0.347 | 1 | 12 | Ndufb10 | -0.04 | Young |
| 0.82949 | 0.023781 | 0.474 | 0.517 | 1 | 12 | Rps13 | -0.043 | Young |
| 0.83066 | 0.067311 | 0.35 | 0.339 | 1 | 12 | Fkbp1b | 0.011 | Young |
| 0.831443 | 0.128858 | 0.35 | 0.339 | 1 | 12 | Fis1 | 0.011 | Young |
| 0.83157 | 0.261723 | 0.066 | 0.059 | 1 | 12 | Smarce1 | 0.007 | Young |
| 0.832535 | 0.081976 | 0.029 | 0.034 | 1 | 12 | Zmym4 | -0.005 | Young |
| 0.833228 | 0.838946 | 0.029 | 0.025 | 1 | 12 | Zfp68 | 0.004 | Young |
| 0.833563 | 0.047829 | 0.036 | 0.042 | 1 | 12 | Smpd3 | -0.006 | Young |
| 0.834141 | 0.038618 | 0.518 | 0.5 | 1 | 12 | Atp5j2 | 0.018 | Young |
| 0.834308 | 0.176524 | 0.292 | 0.28 | 1 | 12 | Pmp22 | 0.012 | Young |
| 0.834449 | 0.130725 | 0.073 | 0.068 | 1 | 12 | Stard10 | 0.005 | Young |
| 0.835211 | 0.108972 | 0.197 | 0.186 | 1 | 12 | Trappc6b | 0.011 | Young |
| 0.835296 | 0.102703 | 0.102 | 0.093 | 1 | 12 | Rundc3a | 0.009 | Young |
| 0.835479 | 0.018623 | 0.299 | 0.314 | 1 | 12 | Cd81 | -0.015 | Young |
| 0.836391 | 0.604711 | 0.044 | 0.051 | 1 | 12 | Chmp3 | -0.007 | Young |
| 0.836391 | 0.162031 | 0.044 | 0.051 | 1 | 12 | Ssbp3 | -0.007 | Young |
| 0.837056 | 0.119395 | 0.109 | 0.102 | 1 | 12 | Sec11c | 0.007 | Young |

| | | | | | | | | |
|---|---|---|---|---|---|---|---|---|
| 0.837517 | 0.154687 | 0.036 | 0.042 | 1 | 12 | Fcf1 | -0.006 | Young |
| 0.837896 | 0.104386 | 0.066 | 0.059 | 1 | 12 | Akap9 | 0.007 | Young |
| 0.837925 | 0.879945 | 0.029 | 0.025 | 1 | 12 | Dock7 | 0.004 | Young |
| 0.837925 | 0.863345 | 0.029 | 0.025 | 1 | 12 | Trmt1 | 0.004 | Young |
| 0.837925 | 0.709068 | 0.029 | 0.025 | 1 | 12 | Tor1b | 0.004 | Young |
| 0.837925 | 0.515182 | 0.029 | 0.025 | 1 | 12 | Nras | 0.004 | Young |
| 0.837925 | 0.507114 | 0.029 | 0.025 | 1 | 12 | Trim32 | 0.004 | Young |
| 0.837925 | 0.492044 | 0.029 | 0.025 | 1 | 12 | Galnt11 | 0.004 | Young |
| 0.837925 | 0.46836 | 0.029 | 0.025 | 1 | 12 | Ift74 | 0.004 | Young |
| 0.83838 | 0.102445 | 0.19 | 0.178 | 1 | 12 | Lamtor2 | 0.012 | Young |
| 0.840696 | 0.08091 | 0.117 | 0.127 | 1 | 12 | Psmd11 | -0.01 | Young |
| 0.841344 | 0.258317 | 0.029 | 0.034 | 1 | 12 | Cyth3 | -0.005 | Young |
| 0.841344 | 0.116882 | 0.029 | 0.034 | 1 | 12 | Ankrd17 | -0.005 | Young |
| 0.841344 | 0.037359 | 0.029 | 0.034 | 1 | 12 | Pgm2 | -0.005 | Young |
| 0.842206 | 0.275931 | 0.08 | 0.076 | 1 | 12 | Lsamp | 0.004 | Young |
| 0.842629 | 0.850265 | 0.029 | 0.025 | 1 | 12 | Slc25a12 | 0.004 | Young |
| 0.842629 | 0.756329 | 0.029 | 0.025 | 1 | 12 | Stx6 | 0.004 | Young |
| 0.842629 | 0.730407 | 0.029 | 0.025 | 1 | 12 | Cyb5b | 0.004 | Young |
| 0.842629 | 0.684129 | 0.029 | 0.025 | 1 | 12 | Coa5 | 0.004 | Young |
| 0.842629 | 0.525626 | 0.029 | 0.025 | 1 | 12 | Ubash3b | 0.004 | Young |
| 0.842629 | 0.510369 | 0.029 | 0.025 | 1 | 12 | Mrpl10 | 0.004 | Young |
| 0.842629 | 0.424453 | 0.029 | 0.025 | 1 | 12 | Fads3 | 0.004 | Young |
| 0.842629 | 0.423612 | 0.029 | 0.025 | 1 | 12 | Sez6l2 | 0.004 | Young |
| 0.842629 | 0.395761 | 0.029 | 0.025 | 1 | 12 | Pcif1 | 0.004 | Young |
| 0.842629 | 0.385127 | 0.029 | 0.025 | 1 | 12 | Mzt1 | 0.004 | Young |
| 0.842629 | 0.353365 | 0.029 | 0.025 | 1 | 12 | Wdr18 | 0.004 | Young |
| 0.842629 | 0.340626 | 0.029 | 0.025 | 1 | 12 | Nme6 | 0.004 | Young |
| 0.842629 | 0.281964 | 0.029 | 0.025 | 1 | 12 | Satb1 | 0.004 | Young |
| 0.842629 | 0.252457 | 0.029 | 0.025 | 1 | 12 | Stard7 | 0.004 | Young |
| 0.842629 | 0.231067 | 0.029 | 0.025 | 1 | 12 | Lhfpl4 | 0.004 | Young |
| 0.84344 | 0.36434 | 0.073 | 0.068 | 1 | 12 | Stip1 | 0.005 | Young |
| 0.8451 | 0.051744 | 0.307 | 0.314 | 1 | 12 | Ndufa8 | -0.007 | Young |
| 0.845437 | 0.236308 | 0.036 | 0.042 | 1 | 12 | Pea15a | -0.006 | Young |
| 0.845581 | 0.20549 | 0.19 | 0.186 | 1 | 12 | Hspd1 | 0.004 | Young |
| 0.845757 | 0.215627 | 0.029 | 0.034 | 1 | 12 | Tlk1 | -0.005 | Young |
| 0.845757 | 0.176645 | 0.029 | 0.034 | 1 | 12 | Rere | -0.005 | Young |
| 0.845757 | 0.066365 | 0.029 | 0.034 | 1 | 12 | C1qb | -0.005 | Young |
| 0.846795 | 0.079203 | 0.139 | 0.153 | 1 | 12 | Psip1 | -0.014 | Young |
| 0.847338 | 0.721629 | 0.029 | 0.025 | 1 | 12 | Znrf2 | 0.004 | Young |
| 0.847338 | 0.680596 | 0.029 | 0.025 | 1 | 12 | Fbxo3 | 0.004 | Young |
| 0.847338 | 0.612455 | 0.029 | 0.025 | 1 | 12 | Plekha1 | 0.004 | Young |
| 0.847338 | 0.599974 | 0.029 | 0.025 | 1 | 12 | Ehbp1l1 | 0.004 | Young |
| 0.847338 | 0.460086 | 0.029 | 0.025 | 1 | 12 | Panx1 | 0.004 | Young |
| 0.847338 | 0.4535 | 0.029 | 0.025 | 1 | 12 | Ikbkap | 0.004 | Young |
| 0.847338 | 0.434827 | 0.029 | 0.025 | 1 | 12 | Hnrnpm | 0.004 | Young |

| | | | | | | | | |
|---|---|---|---|---|---|---|---|---|
| 0.847338 | 0.328025 | 0.029 | 0.025 | 1 | 12 | B3galnt1 | 0.004 | Young |
| 0.847338 | 0.308933 | 0.029 | 0.025 | 1 | 12 | Hsph1 | 0.004 | Young |
| 0.847944 | 0.18774 | 0.073 | 0.068 | 1 | 12 | Tmx4 | 0.005 | Young |
| 0.849322 | 0.103917 | 0.117 | 0.11 | 1 | 12 | Tmeff1 | 0.007 | Young |
| 0.849446 | 0.239791 | 0.073 | 0.068 | 1 | 12 | Chrm2 | 0.005 | Young |
| 0.850174 | 0.261603 | 0.029 | 0.034 | 1 | 12 | Cstf3 | -0.005 | Young |
| 0.850174 | 0.243288 | 0.029 | 0.034 | 1 | 12 | Gramd1a | -0.005 | Young |
| 0.850246 | 0.057123 | 0.168 | 0.178 | 1 | 12 | Cd47 | -0.01 | Young |
| 0.850578 | 0.096526 | 0.066 | 0.059 | 1 | 12 | Smarca2 | 0.007 | Young |
| 0.850908 | 0.0796 | 0.226 | 0.22 | 1 | 12 | Arf1 | 0.006 | Young |
| 0.851093 | 0.135728 | 0.153 | 0.169 | 1 | 12 | mt-Nd3 | -0.016 | Young |
| 0.851185 | 0.01022 | 0.161 | 0.144 | 1 | 12 | Tomm40 | 0.017 | Young |
| 0.852053 | 0.919963 | 0.029 | 0.025 | 1 | 12 | Rrs1 | 0.004 | Young |
| 0.852053 | 0.63234 | 0.029 | 0.025 | 1 | 12 | Tmem9b | 0.004 | Young |
| 0.852053 | 0.535654 | 0.029 | 0.025 | 1 | 12 | Zc3h7a | 0.004 | Young |
| 0.852053 | 0.511152 | 0.029 | 0.025 | 1 | 12 | Usf1 | 0.004 | Young |
| 0.852053 | 0.504711 | 0.029 | 0.025 | 1 | 12 | Yy1 | 0.004 | Young |
| 0.852053 | 0.43231 | 0.029 | 0.025 | 1 | 12 | Nmd3 | 0.004 | Young |
| 0.852053 | 0.313168 | 0.029 | 0.025 | 1 | 12 | Dcakd | 0.004 | Young |
| 0.852053 | 0.309742 | 0.029 | 0.025 | 1 | 12 | Fam110b | 0.004 | Young |
| 0.852053 | 0.279945 | 0.029 | 0.025 | 1 | 12 | Acer3 | 0.004 | Young |
| 0.852053 | 0.243636 | 0.029 | 0.025 | 1 | 12 | Myo18a | 0.004 | Young |
| 0.852053 | 0.203029 | 0.029 | 0.025 | 1 | 12 | Dcaf5 | 0.004 | Young |
| 0.852053 | 0.182232 | 0.029 | 0.025 | 1 | 12 | Tom1l2 | 0.004 | Young |
| 0.852053 | 0.099864 | 0.029 | 0.025 | 1 | 12 | Clk1 | 0.004 | Young |
| 0.852404 | 0.008707 | 0.409 | 0.432 | 1 | 12 | Nedd8 | -0.023 | Young |
| 0.852555 | 0.088955 | 0.168 | 0.161 | 1 | 12 | Ndufb6 | 0.007 | Young |
| 0.853361 | 0.102007 | 0.022 | 0.025 | 1 | 12 | Adrb2 | -0.003 | Young |
| 0.853361 | 0.010784 | 0.022 | 0.025 | 1 | 12 | Zfp260 | -0.003 | Young |
| 0.853755 | 0.006337 | 0.066 | 0.059 | 1 | 12 | Abhd16a | 0.007 | Young |
| 0.853984 | 0.104038 | 0.197 | 0.203 | 1 | 12 | Ppp1ca | -0.006 | Young |
| 0.85453 | 0.224254 | 0.109 | 0.102 | 1 | 12 | Rbms3 | 0.007 | Young |
| 0.854595 | 0.032943 | 0.029 | 0.034 | 1 | 12 | Gspt1 | -0.005 | Young |
| 0.854596 | 0.333301 | 0.029 | 0.034 | 1 | 12 | Aldh2 | -0.005 | Young |
| 0.854596 | 0.134074 | 0.029 | 0.034 | 1 | 12 | Pmepa1 | -0.005 | Young |
| 0.854596 | 0.010921 | 0.029 | 0.034 | 1 | 12 | Polr2a | -0.005 | Young |
| 0.855461 | 0.178875 | 0.073 | 0.068 | 1 | 12 | Smim10l1 | 0.005 | Young |
| 0.855515 | 0.155089 | 0.234 | 0.22 | 1 | 12 | Ndufa10 | 0.014 | Young |
| 0.856773 | 0.473097 | 0.029 | 0.025 | 1 | 12 | Kcnk3 | 0.004 | Young |
| 0.856773 | 0.455968 | 0.029 | 0.025 | 1 | 12 | Med10 | 0.004 | Young |
| 0.856773 | 0.424287 | 0.029 | 0.025 | 1 | 12 | Snhg20 | 0.004 | Young |
| 0.856773 | 0.351339 | 0.029 | 0.025 | 1 | 12 | Nt5c2 | 0.004 | Young |
| 0.856773 | 0.336034 | 0.029 | 0.025 | 1 | 12 | Eea1 | 0.004 | Young |
| 0.856773 | 0.291129 | 0.029 | 0.025 | 1 | 12 | Pex16 | 0.004 | Young |
| 0.856773 | 0.279395 | 0.029 | 0.025 | 1 | 12 | Tbc1d14 | 0.004 | Young |

| | | | | | | | | |
|---|---|---|---|---|---|---|---|---|
| 0.856773 | 0.273934 | 0.029 | 0.025 | 1 | 12 | Cacna2d2 | 0.004 | Young |
| 0.856773 | 0.247339 | 0.029 | 0.025 | 1 | 12 | Wdr83 | 0.004 | Young |
| 0.856773 | 0.233116 | 0.029 | 0.025 | 1 | 12 | Fbxo21 | 0.004 | Young |
| 0.856773 | 0.22768 | 0.029 | 0.025 | 1 | 12 | Nus1 | 0.004 | Young |
| 0.856773 | 0.199981 | 0.029 | 0.025 | 1 | 12 | Nob1 | 0.004 | Young |
| 0.856773 | 0.188109 | 0.029 | 0.025 | 1 | 12 | Mapk8 | 0.004 | Young |
| 0.856773 | 0.169117 | 0.029 | 0.025 | 1 | 12 | Ankib1 | 0.004 | Young |
| 0.856773 | 0.14851 | 0.029 | 0.025 | 1 | 12 | Cdc27 | 0.004 | Young |
| 0.856773 | 0.131563 | 0.029 | 0.025 | 1 | 12 | Brd3os | 0.004 | Young |
| 0.856773 | 0.111046 | 0.029 | 0.025 | 1 | 12 | Asic2 | 0.004 | Young |
| 0.856773 | 0.097513 | 0.029 | 0.025 | 1 | 12 | Acot9 | 0.004 | Young |
| 0.856808 | 0.321521 | 0.029 | 0.034 | 1 | 12 | Ntmt1 | -0.005 | Young |
| 0.858451 | 0.234801 | 0.022 | 0.025 | 1 | 12 | Cand1 | -0.003 | Young |
| 0.858451 | 0.22692 | 0.022 | 0.025 | 1 | 12 | Pld3 | -0.003 | Young |
| 0.858451 | 0.186363 | 0.022 | 0.025 | 1 | 12 | Smc4 | -0.003 | Young |
| 0.858451 | 0.142302 | 0.022 | 0.025 | 1 | 12 | Lap3 | -0.003 | Young |
| 0.858451 | 0.085995 | 0.022 | 0.025 | 1 | 12 | Slc35a4 | -0.003 | Young |
| 0.858451 | 0.070836 | 0.022 | 0.025 | 1 | 12 | Tapbp | -0.003 | Young |
| 0.858451 | 0.062781 | 0.022 | 0.025 | 1 | 12 | Arfip2 | -0.003 | Young |
| 0.859022 | 0.4927 | 0.029 | 0.034 | 1 | 12 | Cat | -0.005 | Young |
| 0.859022 | 0.359316 | 0.029 | 0.034 | 1 | 12 | Atp13a3 | -0.005 | Young |
| 0.859022 | 0.332292 | 0.029 | 0.034 | 1 | 12 | Rogdi | -0.005 | Young |
| 0.859022 | 0.297064 | 0.029 | 0.034 | 1 | 12 | N4bp2l2 | -0.005 | Young |
| 0.859022 | 0.134116 | 0.029 | 0.034 | 1 | 12 | Ppil2 | -0.005 | Young |
| 0.859022 | 0.034256 | 0.029 | 0.034 | 1 | 12 | Mphosph1 | -0.005 | Young |
| 0.859537 | 0.039161 | 0.109 | 0.102 | 1 | 12 | Idh1 | 0.007 | Young |
| 0.860023 | 0.033728 | 0.46 | 0.458 | 1 | 12 | Selenok | 0.002 | Young |
| 0.860115 | 0.004094 | 0.066 | 0.059 | 1 | 12 | Ubqln2 | 0.007 | Young |
| 0.861324 | 0.249795 | 0.036 | 0.042 | 1 | 12 | D17Wsu92 | -0.006 | Young |
| 0.861434 | 0.1418 | 0.153 | 0.144 | 1 | 12 | Sox4 | 0.009 | Young |
| 0.861497 | 0.047412 | 0.029 | 0.025 | 1 | 12 | Ncor2 | 0.004 | Young |
| 0.861498 | 0.163087 | 0.029 | 0.025 | 1 | 12 | Lbh | 0.004 | Young |
| 0.861498 | 0.541182 | 0.029 | 0.025 | 1 | 12 | Ap1g1 | 0.004 | Young |
| 0.861498 | 0.247728 | 0.029 | 0.025 | 1 | 12 | Glmn | 0.004 | Young |
| 0.861498 | 0.201808 | 0.029 | 0.025 | 1 | 12 | Acd | 0.004 | Young |
| 0.861498 | 0.179921 | 0.029 | 0.025 | 1 | 12 | Mbnl1 | 0.004 | Young |
| 0.861498 | 0.136565 | 0.029 | 0.025 | 1 | 12 | Fam174b | 0.004 | Young |
| 0.861498 | 0.122643 | 0.029 | 0.025 | 1 | 12 | Gprasp2 | 0.004 | Young |
| 0.861498 | 0.106891 | 0.029 | 0.025 | 1 | 12 | Trappc6a | 0.004 | Young |
| 0.861498 | 0.082597 | 0.029 | 0.025 | 1 | 12 | Fam32a | 0.004 | Young |
| 0.861498 | 0.077605 | 0.029 | 0.025 | 1 | 12 | 1110012L1 | 0.004 | Young |
| 0.861498 | 0.073168 | 0.029 | 0.025 | 1 | 12 | Larp7 | 0.004 | Young |
| 0.861498 | 0.068941 | 0.029 | 0.025 | 1 | 12 | Ocrl | 0.004 | Young |
| 0.861498 | 0.039092 | 0.029 | 0.025 | 1 | 12 | Zfp467 | 0.004 | Young |
| 0.861498 | 0.029351 | 0.029 | 0.025 | 1 | 12 | Apmap | 0.004 | Young |

| | | | | | | | | |
|---|---|---|---|---|---|---|---|---|
| 0.861498 | 0.024948 | 0.029 | 0.025 | 1 | 12 | Htr3b | 0.004 | Young |
| 0.861498 | 0.006722 | 0.029 | 0.025 | 1 | 12 | Alas1 | 0.004 | Young |
| 0.863042 | 0.124135 | 0.124 | 0.119 | 1 | 12 | Lmna | 0.005 | Young |
| 0.863547 | 0.303993 | 0.022 | 0.025 | 1 | 12 | Rars2 | -0.003 | Young |
| 0.863547 | 0.25957 | 0.022 | 0.025 | 1 | 12 | Zfand1 | -0.003 | Young |
| 0.863547 | 0.221156 | 0.022 | 0.025 | 1 | 12 | Gnl3 | -0.003 | Young |
| 0.863547 | 0.191278 | 0.022 | 0.025 | 1 | 12 | Arf2 | -0.003 | Young |
| 0.863547 | 0.138282 | 0.022 | 0.025 | 1 | 12 | Vps72 | -0.003 | Young |
| 0.863547 | 0.10076 | 0.022 | 0.025 | 1 | 12 | Ralb | -0.003 | Young |
| 0.863547 | 0.018154 | 0.022 | 0.025 | 1 | 12 | Zfpl1 | -0.003 | Young |
| 0.863547 | 0.011722 | 0.022 | 0.025 | 1 | 12 | Gse1 | -0.003 | Young |
| 0.863547 | 0.011158 | 0.022 | 0.025 | 1 | 12 | Zfp106 | -0.003 | Young |
| 0.863547 | 0.004437 | 0.022 | 0.025 | 1 | 12 | Dnajc10 | -0.003 | Young |
| 0.863547 | 0.000943 | 0.022 | 0.025 | 1 | 12 | Trim68 | -0.003 | Young |
| 0.863862 | 0.292763 | 0.029 | 0.025 | 1 | 12 | Sacm1l | 0.004 | Young |
| 0.866228 | 0.269902 | 0.029 | 0.025 | 1 | 12 | Yrdc | 0.004 | Young |
| 0.866228 | 0.214761 | 0.029 | 0.025 | 1 | 12 | Myadm | 0.004 | Young |
| 0.866228 | 0.135993 | 0.029 | 0.025 | 1 | 12 | Alkbh5 | 0.004 | Young |
| 0.866228 | 0.114844 | 0.029 | 0.025 | 1 | 12 | Klhdc8b | 0.004 | Young |
| 0.866228 | 0.09376 | 0.029 | 0.025 | 1 | 12 | Foxn3 | 0.004 | Young |
| 0.866228 | 0.083926 | 0.029 | 0.025 | 1 | 12 | Ltv1 | 0.004 | Young |
| 0.866228 | 0.081513 | 0.029 | 0.025 | 1 | 12 | 6430548M | 0.004 | Young |
| 0.866228 | 0.023611 | 0.029 | 0.025 | 1 | 12 | Dkc1 | 0.004 | Young |
| 0.866228 | 0.021499 | 0.029 | 0.025 | 1 | 12 | Mospd2 | 0.004 | Young |
| 0.867511 | 0.091153 | 0.175 | 0.169 | 1 | 12 | Fez1 | 0.006 | Young |
| 0.867888 | 0.358114 | 0.029 | 0.034 | 1 | 12 | Prpf38b | -0.005 | Young |
| 0.867888 | 0.094132 | 0.029 | 0.034 | 1 | 12 | Hebp1 | -0.005 | Young |
| 0.868121 | 0.139198 | 0.19 | 0.186 | 1 | 12 | Kras | 0.004 | Young |
| 0.868595 | 0.072112 | 0.029 | 0.025 | 1 | 12 | Eya1 | 0.004 | Young |
| 0.868648 | 0.540633 | 0.022 | 0.025 | 1 | 12 | Tnfrsf1a | -0.003 | Young |
| 0.868649 | 0.423167 | 0.022 | 0.025 | 1 | 12 | Aftph | -0.003 | Young |
| 0.868649 | 0.419442 | 0.022 | 0.025 | 1 | 12 | Ecm1 | -0.003 | Young |
| 0.868649 | 0.380249 | 0.022 | 0.025 | 1 | 12 | Pitpnc1 | -0.003 | Young |
| 0.868649 | 0.326695 | 0.022 | 0.025 | 1 | 12 | Zmiz1 | -0.003 | Young |
| 0.868649 | 0.031365 | 0.022 | 0.025 | 1 | 12 | Gpr45 | -0.003 | Young |
| 0.869566 | 0.040937 | 0.109 | 0.102 | 1 | 12 | Timm9 | 0.007 | Young |
| 0.869566 | 0.05507 | 0.109 | 0.102 | 1 | 12 | Gnb4 | 0.007 | Young |
| 0.870533 | 0.021512 | 0.073 | 0.068 | 1 | 12 | Ergic2 | 0.005 | Young |
| 0.870963 | 0.115155 | 0.029 | 0.025 | 1 | 12 | Eif2b1 | 0.004 | Young |
| 0.870963 | 0.000784 | 0.029 | 0.025 | 1 | 12 | Mdm4 | 0.004 | Young |
| 0.873554 | 0.133374 | 0.073 | 0.068 | 1 | 12 | Pdhb | 0.005 | Young |
| 0.873756 | 0.454424 | 0.022 | 0.025 | 1 | 12 | Xab2 | -0.003 | Young |
| 0.873756 | 0.431747 | 0.022 | 0.025 | 1 | 12 | Thumpd3 | -0.003 | Young |
| 0.873756 | 0.418263 | 0.022 | 0.025 | 1 | 12 | Dnttip2 | -0.003 | Young |
| 0.873756 | 0.020447 | 0.022 | 0.025 | 1 | 12 | Slc6a15 | -0.003 | Young |

| | | | | | | | | |
|---|---|---|---|---|---|---|---|---|
| 0.873756 | 0.018757 | 0.022 | 0.025 | 1 | 12 | Pnmal2 | -0.003 | Young |
| 0.874412 | 0.212887 | 0.153 | 0.144 | 1 | 12 | Ndufs3 | 0.009 | Young |
| 0.875703 | 0.052677 | 0.029 | 0.025 | 1 | 12 | Cacna1a | 0.004 | Young |
| 0.875703 | 0.012475 | 0.029 | 0.025 | 1 | 12 | Terf2ip | 0.004 | Young |
| 0.878197 | 0.052109 | 0.971 | 0.992 | 1 | 12 | Malat1 | -0.021 | Young |
| 0.878804 | 0.005803 | 0.182 | 0.169 | 1 | 12 | Serbp1 | 0.013 | Young |
| 0.878867 | 0.086846 | 0.022 | 0.025 | 1 | 12 | 2610002M | -0.003 | Young |
| 0.878868 | 0.646519 | 0.022 | 0.025 | 1 | 12 | Itpripl2 | -0.003 | Young |
| 0.878868 | 0.540262 | 0.022 | 0.025 | 1 | 12 | Ccdc6 | -0.003 | Young |
| 0.878868 | 0.206671 | 0.022 | 0.025 | 1 | 12 | Ube3c | -0.003 | Young |
| 0.878868 | 0.19355 | 0.022 | 0.025 | 1 | 12 | Ptprf | -0.003 | Young |
| 0.878868 | 0.159383 | 0.022 | 0.025 | 1 | 12 | Slc45a1 | -0.003 | Young |
| 0.878868 | 0.027123 | 0.022 | 0.025 | 1 | 12 | Zfp651 | -0.003 | Young |
| 0.879512 | 0.254363 | 0.08 | 0.076 | 1 | 12 | Pank3 | 0.004 | Young |
| 0.87983 | 0.041941 | 0.146 | 0.153 | 1 | 12 | Card19 | -0.007 | Young |
| 0.881112 | 0.095927 | 0.073 | 0.068 | 1 | 12 | Cygb | 0.005 | Young |
| 0.882394 | 0.312754 | 0.08 | 0.076 | 1 | 12 | Plpp1 | 0.004 | Young |
| 0.883537 | 0.698863 | 0.036 | 0.034 | 1 | 12 | Cflar | 0.002 | Young |
| 0.883537 | 0.391264 | 0.036 | 0.034 | 1 | 12 | Dpysl5 | 0.002 | Young |
| 0.883634 | 0.440281 | 0.015 | 0.017 | 1 | 12 | Necap2 | -0.002 | Young |
| 0.883634 | 0.411191 | 0.015 | 0.017 | 1 | 12 | Dennd1a | -0.002 | Young |
| 0.883634 | 0.287957 | 0.015 | 0.017 | 1 | 12 | Ubap1 | -0.002 | Young |
| 0.883634 | 0.225794 | 0.015 | 0.017 | 1 | 12 | Flna | -0.002 | Young |
| 0.883634 | 0.198115 | 0.015 | 0.017 | 1 | 12 | Taf6l | -0.002 | Young |
| 0.883634 | 0.100668 | 0.015 | 0.017 | 1 | 12 | Sec23b | -0.002 | Young |
| 0.883634 | 0.082688 | 0.015 | 0.017 | 1 | 12 | Ppp1r14c | -0.002 | Young |
| 0.883634 | 0.00651 | 0.015 | 0.017 | 1 | 12 | Tdg | -0.002 | Young |
| 0.883634 | 0.001541 | 0.015 | 0.017 | 1 | 12 | Tmem87b | -0.002 | Young |
| 0.884545 | 0.054924 | 0.095 | 0.102 | 1 | 12 | Gatad1 | -0.007 | Young |
| 0.884561 | 0.057915 | 0.117 | 0.11 | 1 | 12 | Ctsd | 0.007 | Young |
| 0.884619 | 0.173458 | 0.088 | 0.085 | 1 | 12 | Dohh | 0.003 | Young |
| 0.887 | 0.037904 | 0.117 | 0.11 | 1 | 12 | Fst | 0.007 | Young |
| 0.887744 | 0.730871 | 0.036 | 0.034 | 1 | 12 | R3hdm1 | 0.002 | Young |
| 0.887744 | 0.35428 | 0.036 | 0.034 | 1 | 12 | Dhx30 | 0.002 | Young |
| 0.888265 | 0.008832 | 0.088 | 0.093 | 1 | 12 | Hnrnpu | -0.005 | Young |
| 0.888426 | 0.075002 | 0.102 | 0.11 | 1 | 12 | Fbxo9 | -0.008 | Young |
| 0.888499 | 0.031519 | 0.38 | 0.347 | 1 | 12 | Ndn | 0.033 | Young |
| 0.888682 | 0.109523 | 0.073 | 0.068 | 1 | 12 | Adh5 | 0.005 | Young |
| 0.889883 | 1.127778 | 0.015 | 0.017 | 1 | 12 | Gm26917 | -0.002 | Young |
| 0.889883 | 0.796951 | 0.015 | 0.017 | 1 | 12 | Mef2d | -0.002 | Young |
| 0.889883 | 0.648013 | 0.015 | 0.017 | 1 | 12 | Cd14 | -0.002 | Young |
| 0.889883 | 0.583138 | 0.015 | 0.017 | 1 | 12 | Ddx52 | -0.002 | Young |
| 0.889883 | 0.557586 | 0.015 | 0.017 | 1 | 12 | Cpsf7 | -0.002 | Young |
| 0.889883 | 0.533915 | 0.015 | 0.017 | 1 | 12 | Gltp | -0.002 | Young |
| 0.889883 | 0.491942 | 0.015 | 0.017 | 1 | 12 | Fam120a | -0.002 | Young |

| | | | | | | | | |
|---|---|---|---|---|---|---|---|---|
| 0.889883 | 0.453153 | 0.015 | 0.017 | 1 | 12 | Map2k3 | -0.002 | Young |
| 0.889883 | 0.283462 | 0.015 | 0.017 | 1 | 12 | Lasp1 | -0.002 | Young |
| 0.889883 | 0.244965 | 0.015 | 0.017 | 1 | 12 | Mob1a | -0.002 | Young |
| 0.889883 | 0.154075 | 0.015 | 0.017 | 1 | 12 | Utp14a | -0.002 | Young |
| 0.889883 | 0.152199 | 0.015 | 0.017 | 1 | 12 | Nadk | -0.002 | Young |
| 0.889883 | 0.132564 | 0.015 | 0.017 | 1 | 12 | Arfgef2 | -0.002 | Young |
| 0.889883 | 0.131308 | 0.015 | 0.017 | 1 | 12 | Rnf44 | -0.002 | Young |
| 0.889883 | 0.130316 | 0.015 | 0.017 | 1 | 12 | Mucl1 | -0.002 | Young |
| 0.889883 | 0.121493 | 0.015 | 0.017 | 1 | 12 | Srrt | -0.002 | Young |
| 0.889883 | 0.103099 | 0.015 | 0.017 | 1 | 12 | Cdadc1 | -0.002 | Young |
| 0.889883 | 0.096273 | 0.015 | 0.017 | 1 | 12 | Zfp937 | -0.002 | Young |
| 0.889883 | 0.089598 | 0.015 | 0.017 | 1 | 12 | Lrrc20 | -0.002 | Young |
| 0.889883 | 0.071703 | 0.015 | 0.017 | 1 | 12 | Prkaa2 | -0.002 | Young |
| 0.889883 | 0.062416 | 0.015 | 0.017 | 1 | 12 | Mettl3 | -0.002 | Young |
| 0.889883 | 0.038952 | 0.015 | 0.017 | 1 | 12 | Def8 | -0.002 | Young |
| 0.889883 | 0.038781 | 0.015 | 0.017 | 1 | 12 | Clmp | -0.002 | Young |
| 0.889883 | 0.034789 | 0.015 | 0.017 | 1 | 12 | Kcnk1 | -0.002 | Young |
| 0.889883 | 0.020304 | 0.015 | 0.017 | 1 | 12 | Abcg2 | -0.002 | Young |
| 0.889883 | 0.001018 | 0.015 | 0.017 | 1 | 12 | Fam172a | -0.002 | Young |
| 0.891953 | 0.450359 | 0.036 | 0.034 | 1 | 12 | Ptpn9 | 0.002 | Young |
| 0.894177 | 0.134037 | 0.314 | 0.305 | 1 | 12 | Tagln3 | 0.009 | Young |
| 0.896138 | 0.344251 | 0.015 | 0.017 | 1 | 12 | 3110002H1 | -0.002 | Young |
| 0.896138 | 0.268153 | 0.015 | 0.017 | 1 | 12 | Mtrf1l | -0.002 | Young |
| 0.896139 | 1.252466 | 0.015 | 0.017 | 1 | 12 | Rabepk | -0.002 | Young |
| 0.896139 | 1.061421 | 0.015 | 0.017 | 1 | 12 | Mia3 | -0.002 | Young |
| 0.896139 | 1.046578 | 0.015 | 0.017 | 1 | 12 | 0610030E2 | -0.002 | Young |
| 0.896139 | 0.960038 | 0.015 | 0.017 | 1 | 12 | Gpr180 | -0.002 | Young |
| 0.896139 | 0.95644 | 0.015 | 0.017 | 1 | 12 | Exoc4 | -0.002 | Young |
| 0.896139 | 0.94453 | 0.015 | 0.017 | 1 | 12 | Tyrobp | -0.002 | Young |
| 0.896139 | 0.874039 | 0.015 | 0.017 | 1 | 12 | Bok | -0.002 | Young |
| 0.896139 | 0.868519 | 0.015 | 0.017 | 1 | 12 | Spata6 | -0.002 | Young |
| 0.896139 | 0.802762 | 0.015 | 0.017 | 1 | 12 | Rrp8 | -0.002 | Young |
| 0.896139 | 0.781595 | 0.015 | 0.017 | 1 | 12 | Eps8 | -0.002 | Young |
| 0.896139 | 0.761677 | 0.015 | 0.017 | 1 | 12 | Rbm22 | -0.002 | Young |
| 0.896139 | 0.750765 | 0.015 | 0.017 | 1 | 12 | Elp2 | -0.002 | Young |
| 0.896139 | 0.732472 | 0.015 | 0.017 | 1 | 12 | Brcc3 | -0.002 | Young |
| 0.896139 | 0.723055 | 0.015 | 0.017 | 1 | 12 | Tbl3 | -0.002 | Young |
| 0.896139 | 0.717005 | 0.015 | 0.017 | 1 | 12 | Dhx40 | -0.002 | Young |
| 0.896139 | 0.716715 | 0.015 | 0.017 | 1 | 12 | Jmjd1c | -0.002 | Young |
| 0.896139 | 0.685132 | 0.015 | 0.017 | 1 | 12 | Pnoc | -0.002 | Young |
| 0.896139 | 0.605546 | 0.015 | 0.017 | 1 | 12 | Zmym5 | -0.002 | Young |
| 0.896139 | 0.578923 | 0.015 | 0.017 | 1 | 12 | Lama4 | -0.002 | Young |
| 0.896139 | 0.468042 | 0.015 | 0.017 | 1 | 12 | Nab2 | -0.002 | Young |
| 0.896139 | 0.43358 | 0.015 | 0.017 | 1 | 12 | Arhgdib | -0.002 | Young |
| 0.896139 | 0.431264 | 0.015 | 0.017 | 1 | 12 | Smg7 | -0.002 | Young |

| | | | | | | | | |
|---|---|---|---|---|---|---|---|---|
| 0.896139 | 0.387641 | 0.015 | 0.017 | 1 | 12 | Cdk7 | -0.002 | Young |
| 0.896139 | 0.329737 | 0.015 | 0.017 | 1 | 12 | Ak3 | -0.002 | Young |
| 0.896139 | 0.301044 | 0.015 | 0.017 | 1 | 12 | Ccdc91 | -0.002 | Young |
| 0.896139 | 0.284872 | 0.015 | 0.017 | 1 | 12 | Abtb2 | -0.002 | Young |
| 0.896139 | 0.27644 | 0.015 | 0.017 | 1 | 12 | Ctss | -0.002 | Young |
| 0.896139 | 0.26967 | 0.015 | 0.017 | 1 | 12 | Dbndd1 | -0.002 | Young |
| 0.896139 | 0.25775 | 0.015 | 0.017 | 1 | 12 | Chpf2 | -0.002 | Young |
| 0.896139 | 0.249537 | 0.015 | 0.017 | 1 | 12 | Umad1 | -0.002 | Young |
| 0.896139 | 0.241527 | 0.015 | 0.017 | 1 | 12 | Rab40c | -0.002 | Young |
| 0.896139 | 0.24065 | 0.015 | 0.017 | 1 | 12 | Riok2 | -0.002 | Young |
| 0.896139 | 0.233777 | 0.015 | 0.017 | 1 | 12 | Tmtc1 | -0.002 | Young |
| 0.896139 | 0.202608 | 0.015 | 0.017 | 1 | 12 | Kti12 | -0.002 | Young |
| 0.896139 | 0.201862 | 0.015 | 0.017 | 1 | 12 | Nudt14 | -0.002 | Young |
| 0.896139 | 0.173912 | 0.015 | 0.017 | 1 | 12 | Pnma3 | -0.002 | Young |
| 0.896139 | 0.158283 | 0.015 | 0.017 | 1 | 12 | Dus1l | -0.002 | Young |
| 0.896139 | 0.156515 | 0.015 | 0.017 | 1 | 12 | Micu2 | -0.002 | Young |
| 0.896139 | 0.146598 | 0.015 | 0.017 | 1 | 12 | Rrn3 | -0.002 | Young |
| 0.896139 | 0.142285 | 0.015 | 0.017 | 1 | 12 | Ctxn2 | -0.002 | Young |
| 0.896139 | 0.130392 | 0.015 | 0.017 | 1 | 12 | Gpatch2 | -0.002 | Young |
| 0.896139 | 0.124753 | 0.015 | 0.017 | 1 | 12 | Pdcd4 | -0.002 | Young |
| 0.896139 | 0.124039 | 0.015 | 0.017 | 1 | 12 | Gcnt2 | -0.002 | Young |
| 0.896139 | 0.117524 | 0.015 | 0.017 | 1 | 12 | Stim1 | -0.002 | Young |
| 0.896139 | 0.093722 | 0.015 | 0.017 | 1 | 12 | Akt3 | -0.002 | Young |
| 0.896139 | 0.091622 | 0.015 | 0.017 | 1 | 12 | Phip | -0.002 | Young |
| 0.896139 | 0.086949 | 0.015 | 0.017 | 1 | 12 | Aagab | -0.002 | Young |
| 0.896139 | 0.070901 | 0.015 | 0.017 | 1 | 12 | Hap1 | -0.002 | Young |
| 0.896139 | 0.066702 | 0.015 | 0.017 | 1 | 12 | Fyn | -0.002 | Young |
| 0.896139 | 0.059078 | 0.015 | 0.017 | 1 | 12 | Fam207a | -0.002 | Young |
| 0.896139 | 0.051366 | 0.015 | 0.017 | 1 | 12 | D730003I1 | -0.002 | Young |
| 0.896139 | 0.041012 | 0.015 | 0.017 | 1 | 12 | Lrp8 | -0.002 | Young |
| 0.896139 | 0.039863 | 0.015 | 0.017 | 1 | 12 | Frs2 | -0.002 | Young |
| 0.896139 | 0.010226 | 0.015 | 0.017 | 1 | 12 | Cmpk2 | -0.002 | Young |
| 0.896166 | 0.487039 | 0.036 | 0.034 | 1 | 12 | Ctsf | 0.002 | Young |
| 0.896166 | 0.260854 | 0.036 | 0.034 | 1 | 12 | Ankrd29 | 0.002 | Young |
| 0.896393 | 0.020068 | 0.088 | 0.093 | 1 | 12 | Eprs | -0.005 | Young |
| 0.897991 | 0.075239 | 0.117 | 0.11 | 1 | 12 | Pin4 | 0.007 | Young |
| 0.898512 | 0.007603 | 0.109 | 0.102 | 1 | 12 | Map7d2 | 0.007 | Young |
| 0.899717 | 0.186427 | 0.08 | 0.076 | 1 | 12 | Anxa6 | 0.004 | Young |
| 0.899717 | 0.005418 | 0.08 | 0.076 | 1 | 12 | Ncbp2 | 0.004 | Young |
| 0.900382 | 0.405958 | 0.036 | 0.034 | 1 | 12 | Abcf2 | 0.002 | Young |
| 0.900382 | 0.350731 | 0.036 | 0.034 | 1 | 12 | Efcab14 | 0.002 | Young |
| 0.900502 | 0.032369 | 0.62 | 0.636 | 1 | 12 | Ftl1 | -0.016 | Young |
| 0.900854 | 0.150485 | 0.197 | 0.212 | 1 | 12 | Rps27l | -0.015 | Young |
| 0.902609 | 0.001459 | 0.08 | 0.076 | 1 | 12 | Gm34030 | 0.004 | Young |
| 0.903742 | 0.694295 | 0.044 | 0.042 | 1 | 12 | Pdcd6ip | 0.002 | Young |

| | | | | | | | | |
|---|---|---|---|---|---|---|---|---|
| 0.9046 | 0.260682 | 0.036 | 0.034 | 1 | 12 | Ccndbp1 | 0.002 | Young |
| 0.9046 | 0.229768 | 0.036 | 0.034 | 1 | 12 | 1110004E0 | 0.002 | Young |
| 0.9046 | 0.2262 | 0.036 | 0.034 | 1 | 12 | Cptp | 0.002 | Young |
| 0.9046 | 0.213173 | 0.036 | 0.034 | 1 | 12 | Zbtb8os | 0.002 | Young |
| 0.9046 | 0.165513 | 0.036 | 0.034 | 1 | 12 | Mageh1 | 0.002 | Young |
| 0.9046 | 0.084486 | 0.036 | 0.034 | 1 | 12 | Kctd2 | 0.002 | Young |
| 0.90591 | 0.049828 | 0.153 | 0.144 | 1 | 12 | Cops5 | 0.009 | Young |
| 0.906344 | 0.305252 | 0.095 | 0.093 | 1 | 12 | Eef1akmt1 | 0.002 | Young |
| 0.90671 | 0.227565 | 0.036 | 0.034 | 1 | 12 | Tmem64 | 0.002 | Young |
| 0.908821 | 0.285279 | 0.036 | 0.034 | 1 | 12 | Ptbp3 | 0.002 | Young |
| 0.908821 | 0.202863 | 0.036 | 0.034 | 1 | 12 | Scn3b | 0.002 | Young |
| 0.908821 | 0.196966 | 0.036 | 0.034 | 1 | 12 | Letm1 | 0.002 | Young |
| 0.909519 | 0.287586 | 0.088 | 0.085 | 1 | 12 | Manf | 0.003 | Young |
| 0.91145 | 0.135494 | 0.117 | 0.11 | 1 | 12 | Got1 | 0.007 | Young |
| 0.912906 | 0.063443 | 0.277 | 0.271 | 1 | 12 | Rpl10 | 0.006 | Young |
| 0.913045 | 0.18391 | 0.036 | 0.034 | 1 | 12 | Snx7 | 0.002 | Young |
| 0.913045 | 0.505166 | 0.036 | 0.034 | 1 | 12 | Tgoln1 | 0.002 | Young |
| 0.913045 | 0.144366 | 0.036 | 0.034 | 1 | 12 | Apba2 | 0.002 | Young |
| 0.913045 | 0.034339 | 0.036 | 0.034 | 1 | 12 | Bloc1s4 | 0.002 | Young |
| 0.913454 | 0.049419 | 0.65 | 0.661 | 1 | 12 | Rpl3 | -0.011 | Young |
| 0.915157 | 0.055153 | 0.036 | 0.034 | 1 | 12 | Dzip3 | 0.002 | Young |
| 0.915355 | 0.080673 | 0.555 | 0.602 | 1 | 12 | Bex2 | -0.047 | Young |
| 0.9162 | 0.086104 | 0.109 | 0.102 | 1 | 12 | Tmx2 | 0.007 | Young |
| 0.916596 | 0.017554 | 0.073 | 0.076 | 1 | 12 | Emg1 | -0.003 | Young |
| 0.917203 | 0.090462 | 0.095 | 0.102 | 1 | 12 | Ifi27 | -0.007 | Young |
| 0.917271 | 0.210024 | 0.036 | 0.034 | 1 | 12 | Srgap1 | 0.002 | Young |
| 0.917271 | 0.153075 | 0.036 | 0.034 | 1 | 12 | Snrpa1 | 0.002 | Young |
| 0.917271 | 0.137339 | 0.036 | 0.034 | 1 | 12 | Paip1 | 0.002 | Young |
| 0.917271 | 0.115382 | 0.036 | 0.034 | 1 | 12 | Mrpl4 | 0.002 | Young |
| 0.917271 | 0.113991 | 0.036 | 0.034 | 1 | 12 | Tsc22d4 | 0.002 | Young |
| 0.917271 | 0.068591 | 0.036 | 0.034 | 1 | 12 | Ap1m1 | 0.002 | Young |
| 0.917271 | 0.035631 | 0.036 | 0.034 | 1 | 12 | Id4 | 0.002 | Young |
| 0.917271 | 0.00554 | 0.036 | 0.034 | 1 | 12 | Ano6 | 0.002 | Young |
| 0.918118 | 0.044467 | 0.088 | 0.093 | 1 | 12 | Eif3l | -0.005 | Young |
| 0.91834 | 0.191742 | 0.095 | 0.093 | 1 | 12 | Bod1l | 0.002 | Young |
| 0.919565 | 0.016957 | 0.073 | 0.076 | 1 | 12 | Ppid | -0.003 | Young |
| 0.920289 | 0.023087 | 0.263 | 0.246 | 1 | 12 | Slc5a7 | 0.017 | Young |
| 0.920616 | 0.151833 | 0.088 | 0.085 | 1 | 12 | Med21 | 0.003 | Young |
| 0.921499 | 0.207723 | 0.036 | 0.034 | 1 | 12 | Cep170 | 0.002 | Young |
| 0.921499 | 0.093212 | 0.036 | 0.034 | 1 | 12 | Cnst | 0.002 | Young |
| 0.921499 | 0.088638 | 0.036 | 0.034 | 1 | 12 | Parg | 0.002 | Young |
| 0.921499 | 0.01697 | 0.036 | 0.034 | 1 | 12 | Zbtb20 | 0.002 | Young |
| 0.921499 | 0.00322 | 0.036 | 0.034 | 1 | 12 | Zfp787 | 0.002 | Young |
| 0.921562 | 0.0402 | 0.591 | 0.619 | 1 | 12 | Gm42418 | -0.028 | Young |
| 0.922327 | 0.012032 | 0.204 | 0.195 | 1 | 12 | Vps28 | 0.009 | Young |

| | | | | | | | | |
|---|---|---|---|---|---|---|---|---|
| 0.922926 | 0.373021 | 0.044 | 0.042 | 1 | 12 | Cdk4 | 0.002 | Young |
| 0.923106 | 0.149505 | 0.175 | 0.186 | 1 | 12 | Atp6v0e2 | -0.011 | Young |
| 0.923171 | 0.082436 | 0.08 | 0.085 | 1 | 12 | Luc7l3 | -0.005 | Young |
| 0.92573 | 0.018558 | 0.036 | 0.034 | 1 | 12 | Stx8 | 0.002 | Young |
| 0.92573 | 0.034582 | 0.036 | 0.034 | 1 | 12 | Plaa | 0.002 | Young |
| 0.925789 | 0.023801 | 0.08 | 0.076 | 1 | 12 | B230118H( | 0.004 | Young |
| 0.926281 | 0.03507 | 0.088 | 0.093 | 1 | 12 | 0610009B2 | -0.005 | Young |
| 0.926491 | 0.216754 | 0.175 | 0.178 | 1 | 12 | 1810037I1 | -0.003 | Young |
| 0.926769 | 0.446244 | 0.044 | 0.042 | 1 | 12 | Hpca | 0.002 | Young |
| 0.926769 | 0.286834 | 0.044 | 0.042 | 1 | 12 | Ercc1 | 0.002 | Young |
| 0.927684 | 0.282129 | 0.095 | 0.093 | 1 | 12 | Plcxd3 | 0.002 | Young |
| 0.928461 | 0.260267 | 0.117 | 0.119 | 1 | 12 | Nfu1 | -0.002 | Young |
| 0.928477 | 0.196811 | 0.073 | 0.076 | 1 | 12 | Khdrbs1 | -0.003 | Young |
| 0.929963 | 0.094588 | 0.036 | 0.034 | 1 | 12 | Ssr1 | 0.002 | Young |
| 0.929963 | 0.162169 | 0.036 | 0.034 | 1 | 12 | Tmx1 | 0.002 | Young |
| 0.930613 | 0.296815 | 0.044 | 0.042 | 1 | 12 | Syndig1 | 0.002 | Young |
| 0.930613 | 0.180216 | 0.044 | 0.042 | 1 | 12 | Tmod1 | 0.002 | Young |
| 0.930999 | 0.026391 | 0.153 | 0.144 | 1 | 12 | Eef1a2 | 0.009 | Young |
| 0.93208 | 0.060814 | 0.036 | 0.034 | 1 | 12 | Psme4 | 0.002 | Young |
| 0.93413 | 0.570655 | 0.051 | 0.051 | 1 | 12 | Ube2h | 0 | Young |
| 0.935106 | 0.193941 | 0.292 | 0.297 | 1 | 12 | Mapt | -0.005 | Young |
| 0.936315 | 0.009032 | 0.036 | 0.034 | 1 | 12 | Rragc | 0.002 | Young |
| 0.938228 | 0.014761 | 0.92 | 0.924 | 1 | 12 | Tubb5 | -0.004 | Young |
| 0.938306 | 0.430795 | 0.044 | 0.042 | 1 | 12 | Btbd10 | 0.002 | Young |
| 0.938306 | 0.358045 | 0.044 | 0.042 | 1 | 12 | Raph1 | 0.002 | Young |
| 0.938306 | 0.285303 | 0.044 | 0.042 | 1 | 12 | Prr7 | 0.002 | Young |
| 0.938306 | 0.219931 | 0.044 | 0.042 | 1 | 12 | Cnr1 | 0.002 | Young |
| 0.938787 | 0.135067 | 0.08 | 0.085 | 1 | 12 | Cyp51 | -0.005 | Young |
| 0.940373 | 0.124376 | 0.073 | 0.076 | 1 | 12 | Hectd1 | -0.003 | Young |
| 0.940774 | 0.003931 | 0.343 | 0.347 | 1 | 12 | Mrpl20 | -0.004 | Young |
| 0.941861 | 0.090014 | 0.073 | 0.076 | 1 | 12 | Meaf6 | -0.003 | Young |
| 0.942155 | 0.144714 | 0.044 | 0.042 | 1 | 12 | Hnrnpab | 0.002 | Young |
| 0.942155 | 0.172336 | 0.044 | 0.042 | 1 | 12 | Akr7a5 | 0.002 | Young |
| 0.942155 | 0.064058 | 0.044 | 0.042 | 1 | 12 | Tmem57 | 0.002 | Young |
| 0.94408 | 0.146598 | 0.044 | 0.042 | 1 | 12 | Dguok | 0.002 | Young |
| 0.946005 | 0.373035 | 0.044 | 0.042 | 1 | 12 | Wdr12 | 0.002 | Young |
| 0.946005 | 0.312709 | 0.044 | 0.042 | 1 | 12 | Mia2 | 0.002 | Young |
| 0.946005 | 0.103775 | 0.044 | 0.042 | 1 | 12 | Nipsnap2 | 0.002 | Young |
| 0.946722 | 0.069132 | 0.088 | 0.093 | 1 | 12 | Eif3a | -0.005 | Young |
| 0.947028 | 0.034156 | 0.088 | 0.085 | 1 | 12 | Ppa2 | 0.003 | Young |
| 0.947276 | 0.039023 | 0.642 | 0.661 | 1 | 12 | Itm2b | -0.019 | Young |
| 0.949857 | 0.195085 | 0.044 | 0.042 | 1 | 12 | Rnf214 | 0.002 | Young |
| 0.949857 | 0.095178 | 0.044 | 0.042 | 1 | 12 | Rsbn1l | 0.002 | Young |
| 0.949857 | 0.085929 | 0.044 | 0.042 | 1 | 12 | Rab34 | 0.002 | Young |
| 0.951316 | 0.224234 | 0.095 | 0.102 | 1 | 12 | Wsb2 | -0.007 | Young |

| | | | | | | | | |
|---|---|---|---|---|---|---|---|---|
| 0.953709 | 0.233061 | 0.044 | 0.042 | 1 | 12 | Akap13 | 0.002 | Young |
| 0.953709 | 0.152823 | 0.044 | 0.042 | 1 | 12 | Dctpp1 | 0.002 | Young |
| 0.955733 | 0.005999 | 0.27 | 0.271 | 1 | 12 | Psmd8 | -0.001 | Young |
| 0.957476 | 0.186996 | 0.131 | 0.136 | 1 | 12 | Pgrmc1 | -0.005 | Young |
| 0.957563 | 0.116666 | 0.044 | 0.042 | 1 | 12 | Csnk2a2 | 0.002 | Young |
| 0.957563 | 0.022881 | 0.044 | 0.042 | 1 | 12 | Ggps1 | 0.002 | Young |
| 0.957563 | 0.016672 | 0.044 | 0.042 | 1 | 12 | Znrd1 | 0.002 | Young |
| 0.958447 | 0.088467 | 0.095 | 0.093 | 1 | 12 | Hspa5 | 0.002 | Young |
| 0.959025 | 0.287557 | 0.051 | 0.051 | 1 | 12 | Snx21 | 0 | Young |
| 0.960143 | 0.15249 | 0.175 | 0.178 | 1 | 12 | Pura | -0.003 | Young |
| 0.961418 | 0.119907 | 0.044 | 0.042 | 1 | 12 | Eps15 | 0.002 | Young |
| 0.961853 | 0.476502 | 0.102 | 0.11 | 1 | 12 | Gdi1 | -0.008 | Young |
| 0.962308 | 0.021819 | 0.066 | 0.068 | 1 | 12 | Gch1 | -0.002 | Young |
| 0.962308 | 0.06284 | 0.066 | 0.068 | 1 | 12 | Cnih2 | -0.002 | Young |
| 0.963346 | 0.020229 | 0.044 | 0.042 | 1 | 12 | Rhoq | 0.002 | Young |
| 0.963746 | 0.013803 | 0.752 | 0.763 | 1 | 12 | Gap43 | -0.011 | Young |
| 0.963805 | 0.087918 | 0.095 | 0.093 | 1 | 12 | Ndrg2 | 0.002 | Young |
| 0.965274 | 0.580556 | 0.044 | 0.042 | 1 | 12 | Bmpr2 | 0.002 | Young |
| 0.965274 | 0.011268 | 0.044 | 0.042 | 1 | 12 | Pum1 | 0.002 | Young |
| 0.966679 | 0.155053 | 0.168 | 0.169 | 1 | 12 | Cited2 | -0.001 | Young |
| 0.967227 | 0.082777 | 0.117 | 0.119 | 1 | 12 | Mrps15 | -0.002 | Young |
| 0.969237 | 0.066566 | 0.745 | 0.788 | 1 | 12 | Tpt1 | -0.043 | Young |
| 0.969708 | 0.410514 | 0.051 | 0.051 | 1 | 12 | Bag3 | 0 | Young |
| 0.970156 | 0.11222 | 0.066 | 0.068 | 1 | 12 | Becn1 | -0.002 | Young |
| 0.972372 | 0.107112 | 0.438 | 0.449 | 1 | 12 | Myl12b | -0.011 | Young |
| 0.972987 | 0.117915 | 0.044 | 0.042 | 1 | 12 | Znhit3 | 0.002 | Young |
| 0.973271 | 0.045381 | 0.051 | 0.051 | 1 | 12 | Sdhaf4 | 0 | Young |
| 0.973351 | 0.155735 | 0.058 | 0.059 | 1 | 12 | Pa2g4 | -0.001 | Young |
| 0.974867 | 0.060173 | 0.066 | 0.068 | 1 | 12 | Ralgds | -0.002 | Young |
| 0.975016 | 0.043075 | 0.058 | 0.059 | 1 | 12 | Ndufaf4 | -0.001 | Young |
| 0.976222 | 0.225137 | 0.109 | 0.11 | 1 | 12 | Pgm2l1 | -0.001 | Young |
| 0.980012 | 0.227465 | 0.058 | 0.059 | 1 | 12 | Ccdc107 | -0.001 | Young |
| 0.980012 | 0.177743 | 0.058 | 0.059 | 1 | 12 | Eci2 | -0.001 | Young |
| 0.980012 | 0.126806 | 0.058 | 0.059 | 1 | 12 | Cops7a | -0.001 | Young |
| 0.980397 | 0.192288 | 0.051 | 0.051 | 1 | 12 | Hspa1a | 0 | Young |
| 0.980397 | 0.166856 | 0.051 | 0.051 | 1 | 12 | Agpat3 | 0 | Young |
| 0.980397 | 0.079404 | 0.051 | 0.051 | 1 | 12 | Psmb10 | 0 | Young |
| 0.980626 | 0.012262 | 0.453 | 0.441 | 1 | 12 | Atp5o.1 | 0.012 | Young |
| 0.981203 | 0.141928 | 0.423 | 0.441 | 1 | 12 | Atp5g3 | -0.018 | Young |
| 0.982186 | 0.077681 | 0.336 | 0.331 | 1 | 12 | Tbcb | 0.005 | Young |
| 0.982847 | 0.087662 | 0.737 | 0.72 | 1 | 12 | Rpl23 | 0.017 | Young |
| 0.983005 | 0.206248 | 0.161 | 0.161 | 1 | 12 | Tmem160 | 0 | Young |
| 0.983365 | 0.065087 | 0.146 | 0.144 | 1 | 12 | Jun | 0.002 | Young |
| 0.98396 | 0.268759 | 0.051 | 0.051 | 1 | 12 | Rock2 | 0 | Young |
| 0.98396 | 0.026682 | 0.051 | 0.051 | 1 | 12 | Tmem158 | 0 | Young |

| | | | | | | | | |
|---|---|---|---|---|---|---|---|---|
| 0.98396 | 0.023893 | 0.051 | 0.051 | 1 | 12 | Dync1li2 | 0 | Young |
| 0.984474 | 0.059392 | 0.146 | 0.144 | 1 | 12 | Fdft1 | 0.002 | Young |
| 0.984477 | 0.031801 | 0.102 | 0.102 | 1 | 12 | Timm10 | 0 | Young |
| 0.985525 | 0.100377 | 0.204 | 0.203 | 1 | 12 | Acadl | 0.001 | Young |
| 0.986674 | 0.162324 | 0.058 | 0.059 | 1 | 12 | Ankrd12 | -0.001 | Young |
| 0.986674 | 0.064338 | 0.058 | 0.059 | 1 | 12 | Psmd3 | -0.001 | Young |
| 0.986674 | 0.038649 | 0.058 | 0.059 | 1 | 12 | Smarcb1 | -0.001 | Young |
| 0.987524 | 0.153681 | 0.051 | 0.051 | 1 | 12 | Bola1 | 0 | Young |
| 0.987715 | 0.026705 | 0.263 | 0.263 | 1 | 12 | Prkar1a | 0 | Young |
| 0.987821 | 0.08106 | 0.409 | 0.415 | 1 | 12 | Atp5g2 | -0.006 | Young |
| 0.987931 | 0.014437 | 0.095 | 0.093 | 1 | 12 | Luc7l2 | 0.002 | Young |
| 0.989579 | 0.057096 | 0.27 | 0.271 | 1 | 12 | Cycs | -0.001 | Young |
| 0.990005 | 0.038521 | 0.058 | 0.059 | 1 | 12 | Tmem222 | -0.001 | Young |
| 0.990613 | 0.061605 | 0.095 | 0.093 | 1 | 12 | Enho | 0.002 | Young |
| 0.991089 | 0.007866 | 0.051 | 0.051 | 1 | 12 | Neurl1a | 0 | Young |
| 0.991128 | 0.016204 | 0.146 | 0.144 | 1 | 12 | Ddah2 | 0.002 | Young |
| 0.992102 | 0.111458 | 0.263 | 0.263 | 1 | 12 | Romo1 | 0 | Young |
| 0.992145 | 0.17111 | 0.066 | 0.068 | 1 | 12 | H2-D1 | -0.002 | Young |
| 0.992145 | 0.120474 | 0.066 | 0.068 | 1 | 12 | Prune2 | -0.002 | Young |
| 0.99249 | 0.061921 | 0.109 | 0.11 | 1 | 12 | 1700021F0 | -0.001 | Young |
| 0.993337 | 0.34546 | 0.058 | 0.059 | 1 | 12 | Acin1 | -0.001 | Young |
| 0.993337 | 0.234103 | 0.058 | 0.059 | 1 | 12 | Nudt21 | -0.001 | Young |
| 0.993337 | 0.070046 | 0.058 | 0.059 | 1 | 12 | Rab21 | -0.001 | Young |
| 0.994653 | 0.030551 | 0.051 | 0.051 | 1 | 12 | Avpi1 | 0 | Young |
| 0.994653 | 0.013598 | 0.051 | 0.051 | 1 | 12 | Pclo | 0 | Young |
| 0.994689 | 0.066406 | 0.161 | 0.161 | 1 | 12 | Gucy1a1 | 0 | Young |
| 0.994933 | 0.037829 | 0.182 | 0.178 | 1 | 12 | Prmt1 | 0.004 | Young |
| 0.995287 | 0.275518 | 0.066 | 0.068 | 1 | 12 | Fopnl | -0.002 | Young |
| 0.995287 | 0.192507 | 0.066 | 0.068 | 1 | 12 | Nudt9 | -0.002 | Young |
| 0.996249 | 0.080005 | 0.219 | 0.22 | 1 | 12 | Syt11 | -0.001 | Young |
| 0.996539 | 0.022782 | 0.657 | 0.678 | 1 | 12 | Rpl6 | -0.021 | Young |
| 0.996668 | 0.115349 | 0.058 | 0.059 | 1 | 12 | Tcea1 | -0.001 | Young |
| 0.999069 | 0.046205 | 0.226 | 0.22 | 1 | 12 | Glrx5 | 0.006 | Young |
| 7.08E-17 | 3.360285 | 0.598 | 0.079 | 2.20E-12 | 13 | Cd55 | 0.519 | Old |
| 4.41E-16 | 3.146026 | 0.73 | 0.289 | 1.37E-11 | 13 | C3 | 0.441 | Old |
| 6.11E-16 | 0.612817 | 1 | 1 | 1.90E-11 | 13 | B2m | 0 | Old |
| 6.29E-16 | 3.300269 | 0.697 | 0.202 | 1.95E-11 | 13 | Cxcl13 | 0.495 | Old |
| 8.31E-11 | 1.547529 | 0.852 | 0.746 | 2.58E-06 | 13 | Mmp9 | 0.106 | Old |
| 2.05E-10 | 4.511625 | 0.303 | 0 | 6.37E-06 | 13 | Colq | 0.303 | Old |
| 1.70E-09 | 1.001185 | 0.959 | 0.956 | 5.29E-05 | 13 | Ly6e | 0.003 | Old |
| 1.24E-08 | 0.390568 | 1 | 1 | 0.000384 | 13 | H2-K1 | 0 | Old |
| 2.64E-08 | 0.737853 | 0.992 | 0.974 | 0.00082 | 13 | Cstb | 0.018 | Old |
| 9.03E-08 | 0.402045 | 1 | 1 | 0.002803 | 13 | Ctsz | 0 | Old |
| 1.51E-07 | 0.249663 | 1 | 1 | 0.004676 | 13 | mt-Co1 | 0 | Old |
| 7.78E-07 | 0.404024 | 1 | 1 | 0.024156 | 13 | mt-Cytb | 0 | Old |

| | | | | | | | | |
|---|---|---|---|---|---|---|---|---|
| 1.20E-06 | 0.30856 | 1 | 1 | 0.037389 | 13 | mt-Co3 | 0 | Old |
| 1.38E-06 | 0.48597 | 1 | 1 | 0.04277 | 13 | Gda | 0 | Old |
| 1.41E-06 | 0.604163 | 0.893 | 0.746 | 0.043751 | 13 | Ppp4c | 0.147 | Old |
| 1.94E-06 | 0.409773 | 1 | 1 | 0.060245 | 13 | Ctss | 0 | Old |
| 2.05E-06 | 1.748243 | 0.336 | 0.079 | 0.063695 | 13 | C6 | 0.257 | Old |
| 2.23E-06 | 0.815355 | 0.893 | 0.64 | 0.069138 | 13 | Pdxk | 0.253 | Old |
| 2.39E-06 | 0.259853 | 1 | 1 | 0.074108 | 13 | Psap | 0 | Old |
| 2.57E-06 | 3.806365 | 0.197 | 0.009 | 0.079774 | 13 | Ttn | 0.188 | Old |
| 2.57E-06 | 0.360436 | 1 | 1 | 0.079872 | 13 | Sirpa | 0 | Old |
| 2.73E-06 | 0.724211 | 0.91 | 0.754 | 0.08483 | 13 | Cd300a | 0.156 | Old |
| 3.38E-06 | 0.674217 | 0.975 | 0.939 | 0.104983 | 13 | Lmna | 0.036 | Old |
| 4.67E-06 | 0.904722 | 0.951 | 0.921 | 0.145156 | 13 | Anxa1 | 0.03 | Old |
| 4.69E-06 | 0.333933 | 1 | 1 | 0.145563 | 13 | Sdc3 | 0 | Old |
| 4.82E-06 | 0.443655 | 1 | 1 | 0.149597 | 13 | Cd44 | 0 | Old |
| 5.04E-06 | 2.695117 | 0.246 | 0.035 | 0.15643 | 13 | Xist | 0.211 | Old |
| 6.56E-06 | 0.993639 | 0.836 | 0.728 | 0.203592 | 13 | Ccl8 | 0.108 | Old |
| 7.74E-06 | 0.632938 | 0.918 | 0.868 | 0.240405 | 13 | Rgl1 | 0.05 | Old |
| 7.98E-06 | 0.395611 | 1 | 1 | 0.247788 | 13 | mt-Nd3 | 0 | Old |
| 8.34E-06 | 0.463745 | 1 | 1 | 0.259069 | 13 | Ifi27l2a | 0 | Old |
| 8.78E-06 | 0.801146 | 0.82 | 0.64 | 0.272623 | 13 | Il2rg | 0.18 | Old |
| 1.35E-05 | 1.677532 | 0.377 | 0.132 | 0.418222 | 13 | Slc41a2 | 0.245 | Old |
| 1.38E-05 | 1.618713 | 0.361 | 0.114 | 0.428453 | 13 | Gng11 | 0.247 | Old |
| 1.40E-05 | 2.516416 | 0.197 | 0.018 | 0.434328 | 13 | Kif2c | 0.179 | Old |
| 1.66E-05 | 1.5944 | 0.738 | 0.5 | 0.515916 | 13 | Fabp5 | 0.238 | Old |
| 1.80E-05 | 2.791812 | 0.189 | 0.018 | 0.55929 | 13 | Dmpk | 0.171 | Old |
| 1.86E-05 | 1.323786 | 0.459 | 0.202 | 0.57881 | 13 | Entpd7 | 0.257 | Old |
| 1.97E-05 | 0.708088 | 0.869 | 0.789 | 0.612042 | 13 | Tcirg1 | 0.08 | Old |
| 2.44E-05 | 2.701481 | 0.262 | 0.061 | 0.759027 | 13 | Gdf3 | 0.201 | Old |
| 2.49E-05 | 0.454338 | 0.984 | 0.974 | 0.772885 | 13 | Clec4a3 | 0.01 | Old |
| 2.77E-05 | 1.442872 | 0.402 | 0.158 | 0.860432 | 13 | Oas1a | 0.244 | Old |
| 2.99E-05 | 2.842299 | 0.164 | 0.009 | 0.927518 | 13 | Vldlr | 0.155 | Old |
| 3.44E-05 | 0.664934 | 0.885 | 0.632 | 1 | 13 | Pi4k2a | 0.253 | Old |
| 3.50E-05 | 0.78509 | 0.779 | 0.596 | 1 | 13 | Plod3 | 0.183 | Old |
| 3.57E-05 | 0.578341 | 0.984 | 1 | 1 | 13 | Plin2 | -0.016 | Old |
| 3.71E-05 | 0.932373 | 0.689 | 0.447 | 1 | 13 | Etv5 | 0.242 | Old |
| 4.11E-05 | 0.284426 | 1 | 0.991 | 1 | 13 | Sdcbp | 0.009 | Old |
| 4.45E-05 | 0.54532 | 0.91 | 0.781 | 1 | 13 | Ap2b1 | 0.129 | Old |
| 4.50E-05 | 1.403722 | 0.475 | 0.246 | 1 | 13 | Hpse | 0.229 | Old |
| 5.07E-05 | 2.411898 | 0.279 | 0.079 | 1 | 13 | Atp6v0d2 | 0.2 | Old |
| 5.13E-05 | 0.357448 | 1 | 0.991 | 1 | 13 | Cyfip1 | 0.009 | Old |
| 5.21E-05 | 0.465281 | 0.992 | 0.982 | 1 | 13 | Ldha | 0.01 | Old |
| 5.54E-05 | 1.938489 | 0.254 | 0.061 | 1 | 13 | Mcoln2 | 0.193 | Old |
| 6.01E-05 | 2.724768 | 0.189 | 0.026 | 1 | 13 | Camk2a | 0.163 | Old |
| 6.53E-05 | 0.673632 | 0.836 | 0.64 | 1 | 13 | Ap1s1 | 0.196 | Old |
| 6.82E-05 | 0.834392 | 0.705 | 0.465 | 1 | 13 | Snx1 | 0.24 | Old |

| | | | | | | | | |
|---|---|---|---|---|---|---|---|---|
| 7.17E-05 | 0.450416 | 0.934 | 0.904 | 1 | 13 | Larp1 | 0.03 | Old |
| 7.21E-05 | 2.16777 | 0.172 | 0.018 | 1 | 13 | Spint2 | 0.154 | Old |
| 7.35E-05 | 0.62909 | 0.779 | 0.535 | 1 | 13 | Vcam1 | 0.244 | Old |
| 7.50E-05 | 2.201751 | 0.189 | 0.026 | 1 | 13 | Sema6a | 0.163 | Old |
| 7.67E-05 | 0.333286 | 0.992 | 1 | 1 | 13 | Tuba1b | -0.008 | Old |
| 8.51E-05 | 0.406046 | 1 | 1 | 1 | 13 | Ctsb | 0 | Old |
| 8.73E-05 | 0.508653 | 0.967 | 0.851 | 1 | 13 | Slc39a1 | 0.116 | Old |
| 9.06E-05 | 1.10351 | 0.598 | 0.342 | 1 | 13 | Gm36161 | 0.256 | Old |
| 9.66E-05 | 0.295277 | 1 | 0.991 | 1 | 13 | Prkcd | 0.009 | Old |
| 9.66E-05 | 0.634413 | 0.893 | 0.912 | 1 | 13 | Apobec3 | -0.019 | Old |
| 0.000103 | 1.026956 | 0.582 | 0.342 | 1 | 13 | Ly9 | 0.24 | Old |
| 0.000115 | 3.085496 | 0.123 | 0 | 1 | 13 | Sh2d4a | 0.123 | Old |
| 0.000116 | 1.136868 | 0.713 | 0.526 | 1 | 13 | Odc1 | 0.187 | Old |
| 0.00012 | 0.342981 | 0.992 | 0.982 | 1 | 13 | Rpn1 | 0.01 | Old |
| 0.000124 | 0.830751 | 0.672 | 0.439 | 1 | 13 | Slc16a10 | 0.233 | Old |
| 0.000149 | 1.154164 | 0.492 | 0.263 | 1 | 13 | Ifi47 | 0.229 | Old |
| 0.000161 | 0.561695 | 1 | 1 | 1 | 13 | Ctsd | 0 | Old |
| 0.000161 | 2.288021 | 0.238 | 0.061 | 1 | 13 | F7 | 0.177 | Old |
| 0.000161 | 2.334193 | 0.139 | 0.009 | 1 | 13 | Purg | 0.13 | Old |
| 0.000215 | 0.275819 | 1 | 1 | 1 | 13 | mt-Nd4 | 0 | Old |
| 0.000225 | 0.3251 | 1 | 1 | 1 | 13 | mt-Atp6 | 0 | Old |
| 0.000235 | 1.844292 | 0.189 | 0.035 | 1 | 13 | Dmwd | 0.154 | Old |
| 0.000245 | 1.053134 | 0.451 | 0.219 | 1 | 13 | Stard5 | 0.232 | Old |
| 0.000248 | 0.281321 | 1 | 1 | 1 | 13 | 2010107E0 | 0 | Old |
| 0.000251 | 1.150661 | 0.475 | 0.237 | 1 | 13 | Ikbke | 0.238 | Old |
| 0.000275 | 0.280021 | 1 | 1 | 1 | 13 | Akr1a1 | 0 | Old |
| 0.000281 | 0.253475 | 1 | 1 | 1 | 13 | H2-D1 | 0 | Old |
| 0.000307 | 0.326563 | 0.992 | 1 | 1 | 13 | Plekho2 | -0.008 | Old |
| 0.000314 | 0.350396 | 1 | 1 | 1 | 13 | Vim | 0 | Old |
| 0.000319 | 2.117898 | 0.131 | 0.009 | 1 | 13 | Csprs | 0.122 | Old |
| 0.000328 | 0.509052 | 0.795 | 0.64 | 1 | 13 | Adam9 | 0.155 | Old |
| 0.000329 | 0.555872 | 0.91 | 0.816 | 1 | 13 | Anxa4 | 0.094 | Old |
| 0.000342 | 0.248262 | 1 | 1 | 1 | 13 | Lamp1 | 0 | Old |
| 0.000362 | 0.490756 | 0.951 | 0.86 | 1 | 13 | Hspa9 | 0.091 | Old |
| 0.000369 | 1.874618 | 0.27 | 0.096 | 1 | 13 | Actn1 | 0.174 | Old |
| 0.000375 | 0.981792 | 0.5 | 0.272 | 1 | 13 | St8sia6 | 0.228 | Old |
| 0.000385 | 0.393193 | 0.992 | 0.965 | 1 | 13 | Pdia6 | 0.027 | Old |
| 0.000395 | 0.671357 | 0.803 | 0.57 | 1 | 13 | Otulin | 0.233 | Old |
| 0.000397 | 0.368028 | 0.984 | 0.965 | 1 | 13 | Atp5g3 | 0.019 | Old |
| 0.000398 | 0.702978 | 0.705 | 0.482 | 1 | 13 | Mindy1 | 0.223 | Old |
| 0.000405 | 0.338005 | 1 | 1 | 1 | 13 | Cox6a1 | 0 | Old |
| 0.00041 | 1.087634 | 0.484 | 0.289 | 1 | 13 | Ddx58 | 0.195 | Old |
| 0.000413 | 1.886426 | 0.164 | 0.026 | 1 | 13 | Sh3bgrl2 | 0.138 | Old |
| 0.000413 | 0.660626 | 0.623 | 0.377 | 1 | 13 | Msmo1 | 0.246 | Old |
| 0.000417 | 0.222101 | 1 | 1 | 1 | 13 | Tmbim6 | 0 | Old |

| | | | | | | | | |
|---|---|---|---|---|---|---|---|---|
| 0.000422 | 0.478737 | 0.926 | 0.789 | 1 | 13 | Actn4 | 0.137 | Old |
| 0.000425 | 1.136316 | 0.41 | 0.193 | 1 | 13 | Pex16 | 0.217 | Old |
| 0.000426 | 0.449813 | 0.967 | 0.956 | 1 | 13 | Clec2d | 0.011 | Old |
| 0.000436 | 1.121712 | 0.246 | 0.07 | 1 | 13 | Dhcr7 | 0.176 | Old |
| 0.000468 | 0.242353 | 0.992 | 1 | 1 | 13 | Lamp2 | -0.008 | Old |
| 0.000471 | 0.322505 | 0.975 | 0.965 | 1 | 13 | Dync1h1 | 0.01 | Old |
| 0.000485 | 0.279942 | 1 | 1 | 1 | 13 | Cd53 | 0 | Old |
| 0.000517 | 0.640244 | 0.844 | 0.728 | 1 | 13 | Tspan3 | 0.116 | Old |
| 0.000543 | 0.855561 | 0.451 | 0.228 | 1 | 13 | Il21r | 0.223 | Old |
| 0.000547 | 0.430558 | 0.975 | 0.895 | 1 | 13 | Dnmt3a | 0.08 | Old |
| 0.000551 | 0.598331 | 0.984 | 0.991 | 1 | 13 | Lpl | -0.007 | Old |
| 0.000555 | 0.268487 | 0.984 | 0.982 | 1 | 13 | Capns1 | 0.002 | Old |
| 0.00056 | 2.02021 | 0.123 | 0.009 | 1 | 13 | Gm15675 | 0.114 | Old |
| 0.000563 | 0.245367 | 1 | 1 | 1 | 13 | Cfp | 0 | Old |
| 0.000575 | 1.536394 | 0.221 | 0.061 | 1 | 13 | Dock5 | 0.16 | Old |
| 0.000605 | 1.31682 | 0.287 | 0.105 | 1 | 13 | Recql | 0.182 | Old |
| 0.000617 | 0.53909 | 0.861 | 0.746 | 1 | 13 | Slfn5 | 0.115 | Old |
| 0.000637 | 1.538948 | 0.139 | 0.018 | 1 | 13 | Rbm12b2 | 0.121 | Old |
| 0.000647 | 1.721653 | 0.189 | 0.044 | 1 | 13 | Erc1 | 0.145 | Old |
| 0.000654 | 1.982302 | 0.156 | 0.026 | 1 | 13 | Adgrg1 | 0.13 | Old |
| 0.000681 | 1.244815 | 0.393 | 0.202 | 1 | 13 | Igf2r | 0.191 | Old |
| 0.000683 | 0.457969 | 0.992 | 0.956 | 1 | 13 | Trim25 | 0.036 | Old |
| 0.000693 | 0.481509 | 0.836 | 0.737 | 1 | 13 | Psmc1 | 0.099 | Old |
| 0.000693 | 1.036251 | 0.377 | 0.167 | 1 | 13 | Abcd3 | 0.21 | Old |
| 0.000697 | 0.358595 | 0.975 | 0.974 | 1 | 13 | Atp5k | 0.001 | Old |
| 0.000715 | 0.519589 | 0.836 | 0.623 | 1 | 13 | Yipf3 | 0.213 | Old |
| 0.000733 | 0.687473 | 0.721 | 0.544 | 1 | 13 | Gas2l1 | 0.177 | Old |
| 0.000774 | 0.290192 | 0.992 | 0.974 | 1 | 13 | Rab7 | 0.018 | Old |
| 0.000781 | 1.415037 | 0.279 | 0.105 | 1 | 13 | Cd4 | 0.174 | Old |
| 0.000787 | 0.237583 | 1 | 1 | 1 | 13 | Ccl9 | 0 | Old |
| 0.000809 | 0.262093 | 1 | 1 | 1 | 13 | Anxa2 | 0 | Old |
| 0.000839 | 0.528664 | 0.811 | 0.684 | 1 | 13 | Alyref | 0.127 | Old |
| 0.000855 | 0.284873 | 0.975 | 0.965 | 1 | 13 | Cops9 | 0.01 | Old |
| 0.000897 | 1.50525 | 0.213 | 0.061 | 1 | 13 | Mlh1 | 0.152 | Old |
| 0.00091 | 0.468716 | 0.975 | 0.982 | 1 | 13 | Clec10a | -0.007 | Old |
| 0.000913 | 1.157964 | 0.377 | 0.184 | 1 | 13 | Oas2 | 0.193 | Old |
| 0.000938 | 1.241425 | 0.27 | 0.096 | 1 | 13 | Sgcb | 0.174 | Old |
| 0.001004 | 0.459017 | 0.943 | 0.991 | 1 | 13 | Cd63 | -0.048 | Old |
| 0.001018 | 1.588896 | 0.221 | 0.07 | 1 | 13 | Pi16 | 0.151 | Old |
| 0.001057 | 0.963532 | 0.492 | 0.289 | 1 | 13 | Fam20c | 0.203 | Old |
| 0.001065 | 2.351368 | 0.09 | 0 | 1 | 13 | Cdh6 | 0.09 | Old |
| 0.00114 | 0.411539 | 0.959 | 0.939 | 1 | 13 | Clec4a1 | 0.02 | Old |
| 0.001168 | 0.196395 | 1 | 1 | 1 | 13 | mt-Co2 | 0 | Old |
| 0.001181 | 1.105306 | 0.344 | 0.149 | 1 | 13 | Nxt1 | 0.195 | Old |
| 0.001184 | 0.635722 | 0.672 | 0.456 | 1 | 13 | 2310022A1 | 0.216 | Old |

| | | | | | | | | |
|---|---|---|---|---|---|---|---|---|
| 0.001185 | 1.361588 | 0.287 | 0.114 | 1 | 13 | Mylk | 0.173 | Old |
| 0.001213 | 0.732516 | 0.533 | 0.307 | 1 | 13 | Slc39a9 | 0.226 | Old |
| 0.001221 | 0.673985 | 0.557 | 0.316 | 1 | 13 | Ldlrap1 | 0.241 | Old |
| 0.001224 | 0.331725 | 0.992 | 0.982 | 1 | 13 | Ssr3 | 0.01 | Old |
| 0.001272 | 0.634463 | 0.77 | 0.588 | 1 | 13 | Acaa1a | 0.182 | Old |
| 0.001316 | 1.445171 | 0.18 | 0.044 | 1 | 13 | Kyat1 | 0.136 | Old |
| 0.001389 | 2.124575 | 0.107 | 0.009 | 1 | 13 | Amacr | 0.098 | Old |
| 0.001417 | 0.236961 | 1 | 0.974 | 1 | 13 | Pcbp1 | 0.026 | Old |
| 0.001418 | 0.686444 | 0.656 | 0.491 | 1 | 13 | Pdlim4 | 0.165 | Old |
| 0.001426 | 0.163574 | 1 | 1 | 1 | 13 | Rps26 | 0 | Old |
| 0.001464 | 0.688736 | 0.672 | 0.509 | 1 | 13 | Renbp | 0.163 | Old |
| 0.001476 | 0.937309 | 0.393 | 0.193 | 1 | 13 | Dnaaf3 | 0.2 | Old |
| 0.001513 | 0.344564 | 1 | 0.991 | 1 | 13 | Pkm | 0.009 | Old |
| 0.001541 | 1.264523 | 0.467 | 0.289 | 1 | 13 | Fcgr4 | 0.178 | Old |
| 0.001574 | 0.291459 | 0.984 | 1 | 1 | 13 | Tapbp | -0.016 | Old |
| 0.001615 | 0.694513 | 0.73 | 0.544 | 1 | 13 | Nfkb2 | 0.186 | Old |
| 0.001642 | 0.618836 | 0.787 | 0.588 | 1 | 13 | Lgals3bp | 0.199 | Old |
| 0.001657 | 1.831916 | 0.107 | 0.009 | 1 | 13 | Gm15446 | 0.098 | Old |
| 0.001724 | 0.322393 | 0.984 | 0.921 | 1 | 13 | Gns | 0.063 | Old |
| 0.001758 | 0.702153 | 0.615 | 0.465 | 1 | 13 | Elovl1 | 0.15 | Old |
| 0.001765 | 0.341694 | 0.934 | 0.895 | 1 | 13 | Sdha | 0.039 | Old |
| 0.00179 | 1.200316 | 0.287 | 0.114 | 1 | 13 | 2010111I0 | 0.173 | Old |
| 0.001849 | 2.128575 | 0.082 | 0 | 1 | 13 | Osbpl5 | 0.082 | Old |
| 0.001849 | 1.867195 | 0.082 | 0 | 1 | 13 | Spsb4 | 0.082 | Old |
| 0.001946 | 0.44529 | 0.713 | 0.5 | 1 | 13 | Pank3 | 0.213 | Old |
| 0.001949 | 1.371283 | 0.213 | 0.07 | 1 | 13 | Inpp5e | 0.143 | Old |
| 0.00195 | 0.200145 | 1 | 1 | 1 | 13 | mt-Nd1 | 0 | Old |
| 0.001978 | 1.137022 | 0.254 | 0.096 | 1 | 13 | Oxsm | 0.158 | Old |
| 0.002005 | 0.625736 | 0.689 | 0.526 | 1 | 13 | Cdk4 | 0.163 | Old |
| 0.002034 | 1.128183 | 0.336 | 0.167 | 1 | 13 | Riok2 | 0.169 | Old |
| 0.00204 | 0.218179 | 1 | 1 | 1 | 13 | Ccl6 | 0 | Old |
| 0.002062 | 1.615953 | 0.139 | 0.026 | 1 | 13 | Cacna1a | 0.113 | Old |
| 0.002095 | 1.188294 | 0.254 | 0.096 | 1 | 13 | Hmgxb4 | 0.158 | Old |
| 0.00212 | 0.310108 | 1 | 1 | 1 | 13 | Calr | 0 | Old |
| 0.002122 | 0.607818 | 0.689 | 0.535 | 1 | 13 | Uap1l1 | 0.154 | Old |
| 0.002296 | 0.711973 | 0.574 | 0.36 | 1 | 13 | Tomm40 | 0.214 | Old |
| 0.002314 | 0.533575 | 0.787 | 0.675 | 1 | 13 | Gorasp2 | 0.112 | Old |
| 0.002317 | 0.386017 | 0.975 | 0.93 | 1 | 13 | Sbno2 | 0.045 | Old |
| 0.002331 | 0.855833 | 0.467 | 0.289 | 1 | 13 | Rpf2 | 0.178 | Old |
| 0.002438 | 0.376981 | 0.992 | 0.93 | 1 | 13 | Tmem106a | 0.062 | Old |
| 0.002478 | 0.641067 | 0.656 | 0.447 | 1 | 13 | Wsb2 | 0.209 | Old |
| 0.002494 | 0.514566 | 0.918 | 0.921 | 1 | 13 | Pgam1 | -0.003 | Old |
| 0.002516 | 0.262761 | 1 | 1 | 1 | 13 | Cotl1 | 0 | Old |
| 0.002666 | 0.540074 | 0.705 | 0.518 | 1 | 13 | Ndufa9 | 0.187 | Old |
| 0.002684 | 1.378375 | 0.18 | 0.053 | 1 | 13 | Jpx | 0.127 | Old |

| | | | | | | | | |
|---|---|---|---|---|---|---|---|---|
| 0.002731 | 0.245026 | 1 | 1 | 1 | 13 | Neat1 | 0 | Old |
| 0.002765 | 0.71078 | 0.598 | 0.386 | 1 | 13 | Csrp1 | 0.212 | Old |
| 0.002793 | 0.45361 | 0.754 | 0.623 | 1 | 13 | Gak | 0.131 | Old |
| 0.0028 | 0.307516 | 0.992 | 0.965 | 1 | 13 | Rpl36al | 0.027 | Old |
| 0.002843 | 0.925396 | 0.377 | 0.193 | 1 | 13 | B3glct | 0.184 | Old |
| 0.002849 | 1.389106 | 0.311 | 0.149 | 1 | 13 | Nr1h3 | 0.162 | Old |
| 0.002856 | 1.810338 | 0.148 | 0.035 | 1 | 13 | Trim2 | 0.113 | Old |
| 0.002867 | 0.581723 | 0.967 | 0.904 | 1 | 13 | Gm26917 | 0.063 | Old |
| 0.002888 | 0.332399 | 0.984 | 0.982 | 1 | 13 | Plbd1 | 0.002 | Old |
| 0.002888 | 0.296839 | 0.984 | 0.991 | 1 | 13 | Litaf | -0.007 | Old |
| 0.002945 | 0.894059 | 0.369 | 0.202 | 1 | 13 | Rnaset2b | 0.167 | Old |
| 0.00298 | 1.150023 | 0.279 | 0.123 | 1 | 13 | Cstf1 | 0.156 | Old |
| 0.003028 | 0.956664 | 0.443 | 0.246 | 1 | 13 | Slc35f5 | 0.197 | Old |
| 0.003055 | 0.863939 | 0.484 | 0.307 | 1 | 13 | Utp6 | 0.177 | Old |
| 0.003057 | 1.962214 | 0.115 | 0.018 | 1 | 13 | Procr | 0.097 | Old |
| 0.003106 | 0.470716 | 0.943 | 0.851 | 1 | 13 | Pld3 | 0.092 | Old |
| 0.003115 | 0.726833 | 0.623 | 0.43 | 1 | 13 | Ralgds | 0.193 | Old |
| 0.003121 | 0.50936 | 0.844 | 0.781 | 1 | 13 | Ifngr2 | 0.063 | Old |
| 0.003135 | 2.060827 | 0.131 | 0.026 | 1 | 13 | Myo10 | 0.105 | Old |
| 0.003145 | 1.974008 | 0.115 | 0.018 | 1 | 13 | Car6 | 0.097 | Old |
| 0.003169 | 0.699558 | 0.664 | 0.544 | 1 | 13 | Lpxn | 0.12 | Old |
| 0.003182 | 0.692371 | 0.361 | 0.167 | 1 | 13 | Ipo13 | 0.194 | Old |
| 0.00319 | 1.716418 | 0.115 | 0.018 | 1 | 13 | Creb3l1 | 0.097 | Old |
| 0.003209 | 2.11189 | 0.074 | 0 | 1 | 13 | Gm7967 | 0.074 | Old |
| 0.003209 | 2.007255 | 0.074 | 0 | 1 | 13 | Fbxl22 | 0.074 | Old |
| 0.003209 | 1.887202 | 0.074 | 0 | 1 | 13 | Hsd17b14 | 0.074 | Old |
| 0.003209 | 1.840888 | 0.074 | 0 | 1 | 13 | Gm10655 | 0.074 | Old |
| 0.003236 | 1.643365 | 0.115 | 0.018 | 1 | 13 | Gm12589 | 0.097 | Old |
| 0.003269 | 0.150356 | 1 | 1 | 1 | 13 | Actb | 0 | Old |
| 0.003311 | 0.445604 | 0.803 | 0.632 | 1 | 13 | Smarcd2 | 0.171 | Old |
| 0.0034 | 0.879534 | 0.311 | 0.14 | 1 | 13 | Edem3 | 0.171 | Old |
| 0.003415 | 0.671674 | 0.475 | 0.281 | 1 | 13 | Ttc5 | 0.194 | Old |
| 0.003453 | 1.307882 | 0.213 | 0.079 | 1 | 13 | Mccc1 | 0.134 | Old |
| 0.003521 | 1.644069 | 0.115 | 0.018 | 1 | 13 | Cpeb3 | 0.097 | Old |
| 0.003541 | 0.937086 | 0.566 | 0.395 | 1 | 13 | Hilpda | 0.171 | Old |
| 0.003573 | 0.932283 | 0.574 | 0.439 | 1 | 13 | Fam43a | 0.135 | Old |
| 0.003601 | 0.722797 | 0.672 | 0.509 | 1 | 13 | Rasa4 | 0.163 | Old |
| 0.003674 | 0.590848 | 0.803 | 0.693 | 1 | 13 | Igsf6 | 0.11 | Old |
| 0.003718 | 0.245275 | 1 | 0.991 | 1 | 13 | Eif3a | 0.009 | Old |
| 0.00374 | 0.790121 | 0.475 | 0.281 | 1 | 13 | Ccdc97 | 0.194 | Old |
| 0.00383 | 0.820819 | 0.459 | 0.281 | 1 | 13 | Bst1 | 0.178 | Old |
| 0.003841 | 1.053414 | 0.311 | 0.149 | 1 | 13 | Sort1 | 0.162 | Old |
| 0.003995 | 0.434139 | 0.877 | 0.816 | 1 | 13 | Ikzf1 | 0.061 | Old |
| 0.003998 | 0.302015 | 0.984 | 0.956 | 1 | 13 | P4hb | 0.028 | Old |
| 0.004118 | 0.732186 | 0.623 | 0.439 | 1 | 13 | Slc23a2 | 0.184 | Old |

| | | | | | | | | |
|---|---|---|---|---|---|---|---|---|
| 0.00412 | 1.328314 | 0.23 | 0.096 | 1 | 13 | Toe1 | 0.134 | Old |
| 0.004163 | 1.105727 | 0.148 | 0.035 | 1 | 13 | Src | 0.113 | Old |
| 0.004196 | 0.523097 | 0.852 | 0.623 | 1 | 13 | Rab20 | 0.229 | Old |
| 0.004287 | 0.738174 | 0.484 | 0.289 | 1 | 13 | Cpsf1 | 0.195 | Old |
| 0.00431 | 0.539365 | 0.73 | 0.579 | 1 | 13 | 9030624J0 | 0.151 | Old |
| 0.004358 | 0.904103 | 0.434 | 0.263 | 1 | 13 | Mmd | 0.171 | Old |
| 0.004383 | 0.661687 | 0.5 | 0.298 | 1 | 13 | Erlin2 | 0.202 | Old |
| 0.004389 | 0.504272 | 0.852 | 0.711 | 1 | 13 | AI314180 | 0.141 | Old |
| 0.004403 | 0.263433 | 1 | 1 | 1 | 13 | mt-Nd2 | 0 | Old |
| 0.004442 | 0.509102 | 0.828 | 0.684 | 1 | 13 | 2610001J0 | 0.144 | Old |
| 0.004589 | 0.448752 | 0.607 | 0.368 | 1 | 13 | Vps4a | 0.239 | Old |
| 0.00469 | 0.591207 | 0.73 | 0.623 | 1 | 13 | B4galt5 | 0.107 | Old |
| 0.004701 | 1.826734 | 0.09 | 0.009 | 1 | 13 | Pyroxd2 | 0.081 | Old |
| 0.004722 | 0.508848 | 0.795 | 0.623 | 1 | 13 | Foxn2 | 0.172 | Old |
| 0.004728 | 0.456394 | 0.885 | 0.789 | 1 | 13 | Gm21188 | 0.096 | Old |
| 0.004756 | 0.329108 | 0.967 | 0.93 | 1 | 13 | Tmem258 | 0.037 | Old |
| 0.004793 | 1.410791 | 0.221 | 0.088 | 1 | 13 | Ptgfrn | 0.133 | Old |
| 0.004896 | 0.501048 | 0.869 | 0.842 | 1 | 13 | Klf7 | 0.027 | Old |
| 0.0049 | 0.355456 | 0.951 | 0.956 | 1 | 13 | Vdac2 | -0.005 | Old |
| 0.004927 | 1.451219 | 0.09 | 0.009 | 1 | 13 | Gm6710 | 0.081 | Old |
| 0.00493 | 0.984722 | 0.205 | 0.07 | 1 | 13 | Pwp2 | 0.135 | Old |
| 0.004953 | 1.098447 | 0.213 | 0.079 | 1 | 13 | Fdxr | 0.134 | Old |
| 0.005005 | 0.862123 | 0.328 | 0.158 | 1 | 13 | Fbxl15 | 0.17 | Old |
| 0.005099 | 0.427871 | 0.82 | 0.693 | 1 | 13 | Tox4 | 0.127 | Old |
| 0.005115 | 0.523451 | 0.59 | 0.36 | 1 | 13 | Zswim8 | 0.23 | Old |
| 0.005236 | 0.694818 | 0.475 | 0.289 | 1 | 13 | Tom1 | 0.186 | Old |
| 0.005295 | 0.426014 | 0.918 | 0.86 | 1 | 13 | Mfsd1 | 0.058 | Old |
| 0.005322 | 0.340158 | 0.926 | 0.842 | 1 | 13 | Tuba1c | 0.084 | Old |
| 0.005358 | 1.117491 | 0.197 | 0.07 | 1 | 13 | Dnm1 | 0.127 | Old |
| 0.005417 | 0.754332 | 0.369 | 0.193 | 1 | 13 | Lsm14b | 0.176 | Old |
| 0.005419 | 0.647921 | 0.664 | 0.482 | 1 | 13 | Smu1 | 0.182 | Old |
| 0.005483 | 0.380754 | 0.869 | 0.807 | 1 | 13 | Rnpep | 0.062 | Old |
| 0.005508 | 0.407464 | 0.951 | 0.904 | 1 | 13 | Eno1 | 0.047 | Old |
| 0.005568 | 1.90782 | 0.066 | 0 | 1 | 13 | AC130815. | 0.066 | Old |
| 0.005568 | 1.707494 | 0.066 | 0 | 1 | 13 | Bvht | 0.066 | Old |
| 0.005623 | 0.884404 | 0.279 | 0.123 | 1 | 13 | Selenoo | 0.156 | Old |
| 0.005629 | 1.128142 | 0.197 | 0.07 | 1 | 13 | Ccdc191 | 0.127 | Old |
| 0.005662 | 0.806233 | 0.361 | 0.202 | 1 | 13 | Trappc2 | 0.159 | Old |
| 0.00569 | 1.244874 | 0.205 | 0.079 | 1 | 13 | Riox1 | 0.126 | Old |
| 0.005809 | 0.433718 | 0.885 | 0.825 | 1 | 13 | Vat1 | 0.06 | Old |
| 0.005815 | 0.29668 | 1 | 0.991 | 1 | 13 | Tpd52 | 0.009 | Old |
| 0.005815 | 0.663588 | 0.418 | 0.254 | 1 | 13 | Slc25a10 | 0.164 | Old |
| 0.005889 | 1.49066 | 0.164 | 0.053 | 1 | 13 | Wrb | 0.111 | Old |
| 0.005913 | 0.665287 | 0.443 | 0.254 | 1 | 13 | Ppid | 0.189 | Old |
| 0.005932 | 0.489843 | 0.844 | 0.746 | 1 | 13 | Tap1 | 0.098 | Old |

| | | | | | | | | |
|---|---|---|---|---|---|---|---|---|
| 0.006123 | 1.470363 | 0.139 | 0.035 | 1 | 13 | Kbtbd7 | 0.104 | Old |
| 0.006136 | 1.559654 | 0.123 | 0.026 | 1 | 13 | Mfsd13a | 0.097 | Old |
| 0.006163 | 0.249636 | 0.975 | 0.982 | 1 | 13 | Lyn | -0.007 | Old |
| 0.006364 | 0.82968 | 0.336 | 0.175 | 1 | 13 | Ccdc86 | 0.161 | Old |
| 0.0064 | 0.295669 | 0.992 | 0.974 | 1 | 13 | Atp5b | 0.018 | Old |
| 0.006417 | 0.512649 | 0.754 | 0.579 | 1 | 13 | Snd1 | 0.175 | Old |
| 0.00645 | 0.957796 | 0.713 | 0.596 | 1 | 13 | Tnfaip2 | 0.117 | Old |
| 0.006476 | 0.551197 | 0.73 | 0.535 | 1 | 13 | Slc35f6 | 0.195 | Old |
| 0.006506 | 0.259835 | 0.934 | 0.868 | 1 | 13 | Aamp | 0.066 | Old |
| 0.006534 | 1.612297 | 0.148 | 0.044 | 1 | 13 | Ahnak2 | 0.104 | Old |
| 0.006562 | 0.677173 | 0.779 | 0.658 | 1 | 13 | Cd300c2 | 0.121 | Old |
| 0.006627 | 0.552089 | 0.615 | 0.447 | 1 | 13 | Cyp51 | 0.168 | Old |
| 0.006673 | 1.027283 | 0.369 | 0.228 | 1 | 13 | Hebp1 | 0.141 | Old |
| 0.006762 | 0.412624 | 0.91 | 0.746 | 1 | 13 | Aebp2 | 0.164 | Old |
| 0.006787 | 0.370957 | 0.795 | 0.684 | 1 | 13 | Rreb1 | 0.111 | Old |
| 0.006826 | 0.323557 | 0.951 | 0.895 | 1 | 13 | Usmg5 | 0.056 | Old |
| 0.006885 | 0.327752 | 0.975 | 0.868 | 1 | 13 | Tpp1 | 0.107 | Old |
| 0.006955 | 0.812767 | 0.443 | 0.281 | 1 | 13 | Pogk | 0.162 | Old |
| 0.007009 | 0.856348 | 0.402 | 0.228 | 1 | 13 | Tpcn2 | 0.174 | Old |
| 0.007044 | 0.569999 | 0.672 | 0.518 | 1 | 13 | Bicd2 | 0.154 | Old |
| 0.007188 | 0.981065 | 0.221 | 0.088 | 1 | 13 | Rttn | 0.133 | Old |
| 0.007215 | 0.816118 | 0.32 | 0.167 | 1 | 13 | Tmem184c | 0.153 | Old |
| 0.007234 | 0.60367 | 0.5 | 0.333 | 1 | 13 | Mrpl38 | 0.167 | Old |
| 0.007241 | 0.408544 | 0.91 | 0.781 | 1 | 13 | Ttc7 | 0.129 | Old |
| 0.007324 | 0.240149 | 1 | 1 | 1 | 13 | Tm6sf1 | 0 | Old |
| 0.007382 | 0.783097 | 0.434 | 0.272 | 1 | 13 | Ddx41 | 0.162 | Old |
| 0.007415 | 1.032552 | 0.205 | 0.079 | 1 | 13 | Kptn | 0.126 | Old |
| 0.007644 | 0.219344 | 0.992 | 0.991 | 1 | 13 | Rpl27 | 0.001 | Old |
| 0.007715 | 0.605789 | 0.467 | 0.272 | 1 | 13 | Ceacam1 | 0.195 | Old |
| 0.007721 | 0.920958 | 0.32 | 0.167 | 1 | 13 | Mettl6 | 0.153 | Old |
| 0.007724 | 0.437811 | 0.943 | 0.816 | 1 | 13 | Apobec1 | 0.127 | Old |
| 0.007742 | 1.15965 | 0.238 | 0.105 | 1 | 13 | Wwox | 0.133 | Old |
| 0.007863 | 0.855906 | 0.27 | 0.123 | 1 | 13 | Gm9844 | 0.147 | Old |
| 0.007891 | 0.71429 | 0.459 | 0.272 | 1 | 13 | Hdac3 | 0.187 | Old |
| 0.007999 | 0.856001 | 0.295 | 0.149 | 1 | 13 | Slc12a4 | 0.146 | Old |
| 0.008021 | 0.241792 | 1 | 0.991 | 1 | 13 | Hnrnpab | 0.009 | Old |
| 0.008032 | 1.203084 | 0.189 | 0.07 | 1 | 13 | Tdp1 | 0.119 | Old |
| 0.008044 | 0.400098 | 0.861 | 0.737 | 1 | 13 | Ap3s1 | 0.124 | Old |
| 0.008093 | 0.701563 | 0.393 | 0.219 | 1 | 13 | Nfyb | 0.174 | Old |
| 0.008111 | 0.811819 | 0.418 | 0.246 | 1 | 13 | Sirpb1c | 0.172 | Old |
| 0.00812 | 0.592952 | 0.5 | 0.316 | 1 | 13 | Wdr20 | 0.184 | Old |
| 0.008122 | 0.891735 | 0.492 | 0.307 | 1 | 13 | Myo1e | 0.185 | Old |
| 0.008178 | 0.812548 | 0.402 | 0.237 | 1 | 13 | Odf2 | 0.165 | Old |
| 0.008178 | 0.476263 | 0.803 | 0.64 | 1 | 13 | Rnf213 | 0.163 | Old |
| 0.008179 | 0.551203 | 0.615 | 0.447 | 1 | 13 | Gps1 | 0.168 | Old |

| | | | | | | | | |
|---|---|---|---|---|---|---|---|---|
| 0.008233 | 0.274774 | 0.943 | 0.895 | 1 | 13 | Tcp1 | 0.048 | Old |
| 0.00825 | 0.166574 | 1 | 1 | 1 | 13 | Actr3 | 0 | Old |
| 0.008317 | 0.579148 | 0.582 | 0.386 | 1 | 13 | Tmem135 | 0.196 | Old |
| 0.008387 | 0.423016 | 0.902 | 0.807 | 1 | 13 | Edem1 | 0.095 | Old |
| 0.008405 | 1.502508 | 0.082 | 0.009 | 1 | 13 | Fn3krp | 0.073 | Old |
| 0.008438 | 0.256432 | 1 | 1 | 1 | 13 | Eef1g | 0 | Old |
| 0.008518 | 1.536778 | 0.156 | 0.053 | 1 | 13 | Ropn1l | 0.103 | Old |
| 0.008533 | 0.256329 | 0.984 | 0.991 | 1 | 13 | Pirb | -0.007 | Old |
| 0.008596 | 0.513593 | 0.787 | 0.596 | 1 | 13 | Kdelr2 | 0.191 | Old |
| 0.008607 | 0.490737 | 0.795 | 0.649 | 1 | 13 | Mgat4b | 0.146 | Old |
| 0.008776 | 0.237995 | 1 | 1 | 1 | 13 | Hspa5 | 0 | Old |
| 0.00889 | 0.940702 | 0.336 | 0.193 | 1 | 13 | Tmem38b | 0.143 | Old |
| 0.008897 | 0.349954 | 0.893 | 0.842 | 1 | 13 | Nadk | 0.051 | Old |
| 0.008968 | 0.781464 | 0.361 | 0.202 | 1 | 13 | Senp3 | 0.159 | Old |
| 0.009066 | 0.359752 | 0.926 | 0.947 | 1 | 13 | Jarid2 | -0.021 | Old |
| 0.009226 | 0.423479 | 0.828 | 0.632 | 1 | 13 | Cln8 | 0.196 | Old |
| 0.009396 | 1.21071 | 0.131 | 0.035 | 1 | 13 | Rab34 | 0.096 | Old |
| 0.009488 | 0.181891 | 1 | 1 | 1 | 13 | Hnrnpf | 0 | Old |
| 0.0095 | 1.404939 | 0.098 | 0.018 | 1 | 13 | Cttn | 0.08 | Old |
| 0.009541 | 0.225131 | 1 | 0.982 | 1 | 13 | Gas7 | 0.018 | Old |
| 0.00959 | 1.493384 | 0.115 | 0.026 | 1 | 13 | Shmt1 | 0.089 | Old |
| 0.00959 | 1.335615 | 0.115 | 0.026 | 1 | 13 | Arl6 | 0.089 | Old |
| 0.009676 | 1.799042 | 0.057 | 0 | 1 | 13 | Trmt5 | 0.057 | Old |
| 0.009676 | 1.691854 | 0.057 | 0 | 1 | 13 | Zscan18 | 0.057 | Old |
| 0.009676 | 1.67744 | 0.057 | 0 | 1 | 13 | Rxrg | 0.057 | Old |
| 0.009676 | 1.557333 | 0.057 | 0 | 1 | 13 | Fbxo36 | 0.057 | Old |
| 0.009676 | 1.472131 | 0.057 | 0 | 1 | 13 | Rpl13a-ps1 | 0.057 | Old |
| 0.009735 | 0.607924 | 0.508 | 0.342 | 1 | 13 | Zfp787 | 0.166 | Old |
| 0.009754 | 0.9961 | 0.254 | 0.123 | 1 | 13 | Gm16536 | 0.131 | Old |
| 0.009801 | 0.518198 | 0.59 | 0.412 | 1 | 13 | Vps37b | 0.178 | Old |
| 0.009809 | 1.008874 | 0.189 | 0.07 | 1 | 13 | D17H6S53I | 0.119 | Old |
| 0.00983 | 1.225203 | 0.115 | 0.026 | 1 | 13 | Park2 | 0.089 | Old |
| 0.009858 | 0.559781 | 0.549 | 0.333 | 1 | 13 | Ptpn9 | 0.216 | Old |
| 0.009917 | 0.209423 | 1 | 1 | 1 | 13 | Bri3 | 0 | Old |
| 0.009997 | 0.744472 | 0.385 | 0.228 | 1 | 13 | Atp8b2 | 0.157 | Old |
| 0.010023 | 0.661154 | 0.443 | 0.281 | 1 | 13 | Traf3 | 0.162 | Old |
| 0.010028 | 1.273214 | 0.098 | 0.018 | 1 | 13 | Capsl | 0.08 | Old |
| 0.010108 | 1.292665 | 0.279 | 0.149 | 1 | 13 | Bcar3 | 0.13 | Old |
| 0.010116 | 2.48871 | 0.27 | 0.14 | 1 | 13 | Acp5 | 0.13 | Old |
| 0.010297 | 0.712286 | 0.418 | 0.254 | 1 | 13 | Gpn1 | 0.164 | Old |
| 0.010307 | 0.212993 | 1 | 1 | 1 | 13 | Wdr1 | 0 | Old |
| 0.010364 | 0.188409 | 1 | 1 | 1 | 13 | Gnb1 | 0 | Old |
| 0.010386 | 0.424924 | 0.91 | 0.851 | 1 | 13 | Plec | 0.059 | Old |
| 0.01039 | 0.37014 | 0.541 | 0.36 | 1 | 13 | Rrm1 | 0.181 | Old |
| 0.010437 | 1.658973 | 0.139 | 0.044 | 1 | 13 | Ehf | 0.095 | Old |

| | | | | | | | | |
|---|---|---|---|---|---|---|---|---|
| 0.010438 | 0.664131 | 0.336 | 0.184 | 1 | 13 | Tmem97 | 0.152 | Old |
| 0.010458 | 0.726759 | 0.344 | 0.193 | 1 | 13 | Thoc5 | 0.151 | Old |
| 0.010503 | 1.059469 | 0.23 | 0.105 | 1 | 13 | Qpctl | 0.125 | Old |
| 0.010546 | 0.837688 | 0.418 | 0.272 | 1 | 13 | Prpf38a | 0.146 | Old |
| 0.010565 | 0.278178 | 0.992 | 0.974 | 1 | 13 | Metrnl | 0.018 | Old |
| 0.010593 | 0.489062 | 0.861 | 0.807 | 1 | 13 | Rab11fip5 | 0.054 | Old |
| 0.010664 | 0.884937 | 0.303 | 0.158 | 1 | 13 | Prmt7 | 0.145 | Old |
| 0.010733 | 0.59811 | 0.754 | 0.605 | 1 | 13 | Tmem51 | 0.149 | Old |
| 0.010851 | 0.585067 | 0.426 | 0.237 | 1 | 13 | Ggps1 | 0.189 | Old |
| 0.010909 | 0.516327 | 0.648 | 0.474 | 1 | 13 | Vars | 0.174 | Old |
| 0.011095 | 0.860534 | 0.27 | 0.132 | 1 | 13 | Inpp5a | 0.138 | Old |
| 0.011168 | 0.732885 | 0.385 | 0.219 | 1 | 13 | Abhd2 | 0.166 | Old |
| 0.011207 | 0.576062 | 0.557 | 0.377 | 1 | 13 | Cln3 | 0.18 | Old |
| 0.011248 | 0.90403 | 0.295 | 0.158 | 1 | 13 | Gzf1 | 0.137 | Old |
| 0.011319 | 0.843105 | 0.508 | 0.325 | 1 | 13 | Slc37a2 | 0.183 | Old |
| 0.011374 | 0.144661 | 1 | 1 | 1 | 13 | Gpx1 | 0 | Old |
| 0.011477 | 0.320952 | 0.951 | 0.886 | 1 | 13 | Tmed7 | 0.065 | Old |
| 0.011478 | 1.088248 | 0.172 | 0.061 | 1 | 13 | Oas3 | 0.111 | Old |
| 0.011536 | 0.477606 | 0.566 | 0.36 | 1 | 13 | Gsto1 | 0.206 | Old |
| 0.01163 | 1.258738 | 0.139 | 0.044 | 1 | 13 | Cyp2j6 | 0.095 | Old |
| 0.011685 | 0.342549 | 0.779 | 0.588 | 1 | 13 | Bag6 | 0.191 | Old |
| 0.011752 | 0.635133 | 0.516 | 0.333 | 1 | 13 | Pmpcb | 0.183 | Old |
| 0.011782 | 0.976026 | 0.246 | 0.114 | 1 | 13 | Pycrl | 0.132 | Old |
| 0.011972 | 0.309049 | 0.869 | 0.719 | 1 | 13 | Hadhb | 0.15 | Old |
| 0.012008 | 0.180876 | 1 | 0.991 | 1 | 13 | Grk2 | 0.009 | Old |
| 0.012164 | 1.515257 | 0.189 | 0.079 | 1 | 13 | Dcbld2 | 0.11 | Old |
| 0.012239 | 0.829224 | 0.344 | 0.202 | 1 | 13 | Fam78b | 0.142 | Old |
| 0.01226 | 1.816697 | 0.148 | 0.053 | 1 | 13 | Adgrl2 | 0.095 | Old |
| 0.012306 | 0.46877 | 0.656 | 0.482 | 1 | 13 | Tmem128 | 0.174 | Old |
| 0.01232 | 0.308811 | 0.934 | 0.921 | 1 | 13 | 1110008F1 | 0.013 | Old |
| 0.012332 | 0.253226 | 0.967 | 0.939 | 1 | 13 | Ewsr1 | 0.028 | Old |
| 0.012336 | 0.175891 | 1 | 1 | 1 | 13 | Cd47 | 0 | Old |
| 0.012348 | 0.73334 | 0.426 | 0.272 | 1 | 13 | Srprb | 0.154 | Old |
| 0.012521 | 0.254411 | 0.934 | 0.921 | 1 | 13 | Ubald2 | 0.013 | Old |
| 0.012537 | 0.487803 | 0.582 | 0.412 | 1 | 13 | Tfeb | 0.17 | Old |
| 0.012687 | 0.239958 | 0.943 | 0.904 | 1 | 13 | Ptpa | 0.039 | Old |
| 0.01295 | 0.248562 | 0.918 | 0.851 | 1 | 13 | Myo9b | 0.067 | Old |
| 0.013004 | 0.507567 | 0.705 | 0.544 | 1 | 13 | Hypk | 0.161 | Old |
| 0.013087 | 0.14552 | 1 | 1 | 1 | 13 | BC005537 | 0 | Old |
| 0.013355 | 0.58723 | 0.549 | 0.386 | 1 | 13 | Fam129b | 0.163 | Old |
| 0.013376 | 1.22572 | 0.164 | 0.061 | 1 | 13 | Hmgn3 | 0.103 | Old |
| 0.013417 | 0.695907 | 0.41 | 0.254 | 1 | 13 | Trim5 | 0.156 | Old |
| 0.013472 | 1.695924 | 0.074 | 0.009 | 1 | 13 | 4931414P1 | 0.065 | Old |
| 0.013489 | 0.484415 | 0.492 | 0.307 | 1 | 13 | Klhl28 | 0.185 | Old |
| 0.013581 | 0.248624 | 0.967 | 0.939 | 1 | 13 | Twf2 | 0.028 | Old |

| | | | | | | | | |
|---|---|---|---|---|---|---|---|---|
| 0.013592 | 0.458553 | 0.836 | 0.798 | 1 | 13 | Fgr | 0.038 | Old |
| 0.013707 | 0.688887 | 0.336 | 0.184 | 1 | 13 | Zbtb43 | 0.152 | Old |
| 0.013889 | 1.536369 | 0.074 | 0.009 | 1 | 13 | Lgi4 | 0.065 | Old |
| 0.013916 | 0.540146 | 0.656 | 0.5 | 1 | 13 | Ago3 | 0.156 | Old |
| 0.01396 | 0.714895 | 0.27 | 0.132 | 1 | 13 | Amd1 | 0.138 | Old |
| 0.013988 | 0.446689 | 0.664 | 0.509 | 1 | 13 | Phf20 | 0.155 | Old |
| 0.014041 | 1.552154 | 0.107 | 0.026 | 1 | 13 | Gpr162 | 0.081 | Old |
| 0.014289 | 0.174535 | 0.992 | 0.991 | 1 | 13 | Vcp | 0.001 | Old |
| 0.014301 | 0.623564 | 0.385 | 0.237 | 1 | 13 | Tbc1d8 | 0.148 | Old |
| 0.014318 | 0.610687 | 0.779 | 0.702 | 1 | 13 | Stat1 | 0.077 | Old |
| 0.014318 | 1.313462 | 0.074 | 0.009 | 1 | 13 | Tnfrsf14 | 0.065 | Old |
| 0.014511 | 0.670426 | 0.32 | 0.167 | 1 | 13 | Irf2bp1 | 0.153 | Old |
| 0.01462 | 0.872825 | 0.426 | 0.272 | 1 | 13 | Prkch | 0.154 | Old |
| 0.014627 | 0.268826 | 0.967 | 0.93 | 1 | 13 | Ube2n | 0.037 | Old |
| 0.014649 | 0.416597 | 0.754 | 0.649 | 1 | 13 | Btbd1 | 0.105 | Old |
| 0.014743 | 0.433048 | 0.623 | 0.447 | 1 | 13 | Rab43 | 0.176 | Old |
| 0.014788 | 0.386265 | 0.762 | 0.588 | 1 | 13 | Elf4 | 0.174 | Old |
| 0.014867 | 0.161727 | 1 | 1 | 1 | 13 | Arhgdia | 0 | Old |
| 0.014906 | 1.692782 | 0.09 | 0.018 | 1 | 13 | Plekha5 | 0.072 | Old |
| 0.014946 | 0.232637 | 1 | 0.991 | 1 | 13 | Ctsa | 0.009 | Old |
| 0.014994 | 0.533388 | 0.516 | 0.36 | 1 | 13 | Taok2 | 0.156 | Old |
| 0.015024 | 1.250491 | 0.148 | 0.053 | 1 | 13 | Nck2 | 0.095 | Old |
| 0.015127 | 0.399884 | 0.754 | 0.649 | 1 | 13 | Slamf8 | 0.105 | Old |
| 0.015183 | 0.71684 | 0.352 | 0.202 | 1 | 13 | Rnf19a | 0.15 | Old |
| 0.015184 | 0.402777 | 1 | 1 | 1 | 13 | Lgals3 | 0 | Old |
| 0.015258 | 0.667708 | 0.393 | 0.237 | 1 | 13 | Alg10b | 0.156 | Old |
| 0.015263 | 0.226002 | 0.975 | 0.965 | 1 | 13 | Xbp1 | 0.01 | Old |
| 0.015284 | 1.252244 | 0.107 | 0.026 | 1 | 13 | Rsad1 | 0.081 | Old |
| 0.015588 | 0.183249 | 0.992 | 0.982 | 1 | 13 | Arl6ip1 | 0.01 | Old |
| 0.015895 | 0.554919 | 0.672 | 0.535 | 1 | 13 | Gpr146 | 0.137 | Old |
| 0.015929 | 1.222468 | 0.09 | 0.018 | 1 | 13 | Kcng2 | 0.072 | Old |
| 0.015953 | 0.464554 | 0.836 | 0.702 | 1 | 13 | Man2a1 | 0.134 | Old |
| 0.015983 | 0.649272 | 0.492 | 0.342 | 1 | 13 | Nploc4 | 0.15 | Old |
| 0.016084 | 0.713786 | 0.369 | 0.228 | 1 | 13 | Tsc2 | 0.141 | Old |
| 0.016141 | 1.517498 | 0.09 | 0.018 | 1 | 13 | Ip6k3 | 0.072 | Old |
| 0.016158 | 0.370404 | 0.811 | 0.64 | 1 | 13 | Gatad1 | 0.171 | Old |
| 0.016274 | 0.956946 | 0.148 | 0.053 | 1 | 13 | Ssh3 | 0.095 | Old |
| 0.016461 | 0.917956 | 0.213 | 0.096 | 1 | 13 | C130026I2 | 0.117 | Old |
| 0.016518 | 3.358858 | 0.131 | 0.044 | 1 | 13 | Gpnmb | 0.087 | Old |
| 0.016626 | 0.880254 | 0.361 | 0.228 | 1 | 13 | Fnip2 | 0.133 | Old |
| 0.016722 | 0.871772 | 0.377 | 0.246 | 1 | 13 | Cryzl1 | 0.131 | Old |
| 0.01681 | 0.30689 | 0.91 | 0.851 | 1 | 13 | Arl8b | 0.059 | Old |
| 0.016864 | 2.044025 | 0.049 | 0 | 1 | 13 | Igf2bp2 | 0.049 | Old |
| 0.016864 | 2.035064 | 0.049 | 0 | 1 | 13 | Fzd8 | 0.049 | Old |
| 0.016864 | 1.585411 | 0.049 | 0 | 1 | 13 | Stab2 | 0.049 | Old |

| | | | | | | | | |
|---|---|---|---|---|---|---|---|---|
| 0.016864 | 1.528383 | 0.049 | 0 | 1 | 13 | Plp1 | 0.049 | Old |
| 0.016864 | 1.510581 | 0.049 | 0 | 1 | 13 | Cep170b | 0.049 | Old |
| 0.016864 | 1.432965 | 0.049 | 0 | 1 | 13 | Efna2 | 0.049 | Old |
| 0.016864 | 1.351111 | 0.049 | 0 | 1 | 13 | Mfsd9 | 0.049 | Old |
| 0.016864 | 1.302319 | 0.049 | 0 | 1 | 13 | Bub1 | 0.049 | Old |
| 0.016873 | 0.328739 | 0.877 | 0.789 | 1 | 13 | Gna12 | 0.088 | Old |
| 0.016887 | 0.565474 | 0.5 | 0.342 | 1 | 13 | Lrpprc | 0.158 | Old |
| 0.016929 | 0.5615 | 0.672 | 0.526 | 1 | 13 | Ckap4 | 0.146 | Old |
| 0.016945 | 0.129833 | 1 | 1 | 1 | 13 | H2-Aa | 0 | Old |
| 0.016998 | 0.817725 | 0.303 | 0.167 | 1 | 13 | Qser1 | 0.136 | Old |
| 0.017015 | 0.620234 | 0.434 | 0.281 | 1 | 13 | Ppa1 | 0.153 | Old |
| 0.017022 | 0.453998 | 0.795 | 0.649 | 1 | 13 | Arf2 | 0.146 | Old |
| 0.01706 | 0.451795 | 0.607 | 0.439 | 1 | 13 | Sar1b | 0.168 | Old |
| 0.017068 | 0.323492 | 0.828 | 0.754 | 1 | 13 | Psmc6 | 0.074 | Old |
| 0.017183 | 0.314295 | 0.852 | 0.711 | 1 | 13 | Nsf | 0.141 | Old |
| 0.017372 | 0.307867 | 0.795 | 0.64 | 1 | 13 | Atp6v1c1 | 0.155 | Old |
| 0.017449 | 0.64536 | 0.41 | 0.254 | 1 | 13 | Ap2a1 | 0.156 | Old |
| 0.017854 | 0.508662 | 0.713 | 0.553 | 1 | 13 | Noct | 0.16 | Old |
| 0.018004 | 0.416437 | 0.738 | 0.667 | 1 | 13 | Cope | 0.071 | Old |
| 0.018086 | 0.697227 | 0.369 | 0.211 | 1 | 13 | Casp9 | 0.158 | Old |
| 0.018168 | 0.798708 | 0.402 | 0.246 | 1 | 13 | Nuak1 | 0.156 | Old |
| 0.018219 | 0.602669 | 0.295 | 0.158 | 1 | 13 | B230307C2 | 0.137 | Old |
| 0.018335 | 0.270827 | 0.877 | 0.754 | 1 | 13 | Tm9sf2 | 0.123 | Old |
| 0.01834 | 0.273749 | 0.926 | 0.939 | 1 | 13 | Arap1 | -0.013 | Old |
| 0.018405 | 0.305985 | 0.984 | 0.982 | 1 | 13 | Dusp3 | 0.002 | Old |
| 0.018479 | 1.221991 | 0.189 | 0.088 | 1 | 13 | Card11 | 0.101 | Old |
| 0.018572 | 0.432588 | 0.82 | 0.746 | 1 | 13 | Pim3 | 0.074 | Old |
| 0.018578 | 0.857174 | 0.23 | 0.114 | 1 | 13 | Micu3 | 0.116 | Old |
| 0.018691 | 0.232206 | 1 | 0.991 | 1 | 13 | Tmsb10 | 0.009 | Old |
| 0.018704 | 0.444009 | 0.689 | 0.5 | 1 | 13 | Tbrg1 | 0.189 | Old |
| 0.018813 | 0.330454 | 0.951 | 0.833 | 1 | 13 | Nras | 0.118 | Old |
| 0.018843 | 1.491904 | 0.115 | 0.035 | 1 | 13 | Fbxl8 | 0.08 | Old |
| 0.018903 | 0.514656 | 0.68 | 0.509 | 1 | 13 | Mau2 | 0.171 | Old |
| 0.01892 | 1.032187 | 0.172 | 0.07 | 1 | 13 | Rbak | 0.102 | Old |
| 0.018923 | 1.243667 | 0.131 | 0.044 | 1 | 13 | Gm13431 | 0.087 | Old |
| 0.019028 | 0.351743 | 0.82 | 0.614 | 1 | 13 | Bub3 | 0.206 | Old |
| 0.01912 | 1.220369 | 0.131 | 0.044 | 1 | 13 | Fuz | 0.087 | Old |
| 0.019398 | 0.636242 | 0.295 | 0.158 | 1 | 13 | Cnot11 | 0.137 | Old |
| 0.019413 | 0.849162 | 0.311 | 0.184 | 1 | 13 | Rrp9 | 0.127 | Old |
| 0.019427 | 1.006231 | 0.148 | 0.053 | 1 | 13 | Nup35 | 0.095 | Old |
| 0.019441 | 0.585117 | 0.541 | 0.404 | 1 | 13 | Taf11 | 0.137 | Old |
| 0.019518 | 0.62458 | 0.164 | 0.061 | 1 | 13 | Bex6 | 0.103 | Old |
| 0.019532 | 0.800072 | 0.213 | 0.096 | 1 | 13 | Eefsec | 0.117 | Old |
| 0.019671 | 0.236443 | 1 | 0.991 | 1 | 13 | Eif4a1 | 0.009 | Old |
| 0.019772 | 0.154016 | 1 | 1 | 1 | 13 | Rrbp1 | 0 | Old |

| | | | | | | | | |
|---|---|---|---|---|---|---|---|---|
| 0.019833 | 0.69034 | 0.238 | 0.114 | 1 | 13 | 2010320M | 0.124 | Old |
| 0.020195 | 0.449903 | 0.689 | 0.518 | 1 | 13 | Ppp6c | 0.171 | Old |
| 0.020225 | 0.253339 | 0.975 | 0.939 | 1 | 13 | Larp4b | 0.036 | Old |
| 0.020302 | 0.354425 | 0.721 | 0.544 | 1 | 13 | Slc25a39 | 0.177 | Old |
| 0.020322 | 1.157051 | 0.156 | 0.061 | 1 | 13 | Qrsl1 | 0.095 | Old |
| 0.020341 | 1.15499 | 0.131 | 0.044 | 1 | 13 | Wdr92 | 0.087 | Old |
| 0.020356 | 0.423795 | 0.475 | 0.289 | 1 | 13 | Daxx | 0.186 | Old |
| 0.020556 | 0.493747 | 0.607 | 0.465 | 1 | 13 | Pdcd11 | 0.142 | Old |
| 0.020594 | 0.68785 | 0.385 | 0.246 | 1 | 13 | Bet1l | 0.139 | Old |
| 0.020656 | 0.624564 | 0.508 | 0.342 | 1 | 13 | Ap1g2 | 0.166 | Old |
| 0.020796 | 0.992349 | 0.238 | 0.123 | 1 | 13 | 2310040G2 | 0.115 | Old |
| 0.020823 | 0.602621 | 0.41 | 0.263 | 1 | 13 | Apeh | 0.147 | Old |
| 0.020919 | 0.419704 | 0.639 | 0.491 | 1 | 13 | Galnt10 | 0.148 | Old |
| 0.021022 | 0.63022 | 0.73 | 0.702 | 1 | 13 | Cd209g | 0.028 | Old |
| 0.021119 | 0.23598 | 1 | 0.982 | 1 | 13 | Mafb | 0.018 | Old |
| 0.021182 | 0.50073 | 0.631 | 0.474 | 1 | 13 | Ell2 | 0.157 | Old |
| 0.021311 | 0.857568 | 0.221 | 0.105 | 1 | 13 | Phykpl | 0.116 | Old |
| 0.021735 | 0.336488 | 0.893 | 0.816 | 1 | 13 | Gnpda1 | 0.077 | Old |
| 0.021876 | 0.214265 | 1 | 0.939 | 1 | 13 | Atp6ap2 | 0.061 | Old |
| 0.021935 | 1.589522 | 0.066 | 0.009 | 1 | 13 | Klhdc8b | 0.057 | Old |
| 0.021987 | 0.188622 | 1 | 0.965 | 1 | 13 | Clk1 | 0.035 | Old |
| 0.022025 | 0.584085 | 0.393 | 0.237 | 1 | 13 | Naa40 | 0.156 | Old |
| 0.022074 | 1.116071 | 0.131 | 0.044 | 1 | 13 | Btbd19 | 0.087 | Old |
| 0.02208 | 0.292786 | 0.877 | 0.737 | 1 | 13 | Mydgf | 0.14 | Old |
| 0.022153 | 0.213889 | 0.992 | 0.974 | 1 | 13 | Set | 0.018 | Old |
| 0.022224 | 0.569762 | 0.459 | 0.316 | 1 | 13 | Bckdha | 0.143 | Old |
| 0.02229 | 1.336429 | 0.098 | 0.026 | 1 | 13 | Tbc1d19 | 0.072 | Old |
| 0.022379 | 0.880037 | 0.287 | 0.158 | 1 | 13 | Mtor | 0.129 | Old |
| 0.022502 | 0.455524 | 0.811 | 0.658 | 1 | 13 | H2-M3 | 0.153 | Old |
| 0.022528 | 0.74994 | 0.205 | 0.096 | 1 | 13 | Gprc5b | 0.109 | Old |
| 0.022535 | 0.307982 | 0.697 | 0.465 | 1 | 13 | Cfdp1 | 0.232 | Old |
| 0.022938 | 0.669875 | 0.377 | 0.246 | 1 | 13 | Bloc1s1 | 0.131 | Old |
| 0.023001 | 0.198622 | 1 | 1 | 1 | 13 | Atp5f1 | 0 | Old |
| 0.023061 | 0.308029 | 0.918 | 0.895 | 1 | 13 | Ppig | 0.023 | Old |
| 0.023109 | 0.682608 | 0.303 | 0.167 | 1 | 13 | Sephs2 | 0.136 | Old |
| 0.023165 | 0.623276 | 0.418 | 0.263 | 1 | 13 | Exosc4 | 0.155 | Old |
| 0.023174 | 0.242053 | 0.975 | 0.991 | 1 | 13 | Itgam | -0.016 | Old |
| 0.023214 | 0.774291 | 0.197 | 0.088 | 1 | 13 | Alg11 | 0.109 | Old |
| 0.023231 | 0.186097 | 0.984 | 0.974 | 1 | 13 | Mtpn | 0.01 | Old |
| 0.023322 | 0.685961 | 0.148 | 0.053 | 1 | 13 | Cenpt | 0.095 | Old |
| 0.023389 | 0.857766 | 0.221 | 0.105 | 1 | 13 | Ano8 | 0.116 | Old |
| 0.023476 | 0.712796 | 0.287 | 0.158 | 1 | 13 | Ints8 | 0.129 | Old |
| 0.02375 | 0.501181 | 0.623 | 0.465 | 1 | 13 | Manba | 0.158 | Old |
| 0.023855 | 0.36954 | 0.844 | 0.754 | 1 | 13 | Atp6v0a1 | 0.09 | Old |
| 0.023881 | 0.686432 | 0.328 | 0.193 | 1 | 13 | Cdc42ep2 | 0.135 | Old |

| | | | | | | | | |
|---|---|---|---|---|---|---|---|---|
| 0.023928 | 0.475205 | 0.402 | 0.237 | 1 | 13 | Bcl2l13 | 0.165 | Old |
| 0.024009 | 1.317745 | 0.066 | 0.009 | 1 | 13 | Zfp773 | 0.057 | Old |
| 0.024009 | 1.292707 | 0.066 | 0.009 | 1 | 13 | 9130011E1 | 0.057 | Old |
| 0.024479 | 0.25735 | 0.91 | 0.851 | 1 | 13 | Pet100 | 0.059 | Old |
| 0.024737 | 1.304714 | 0.066 | 0.009 | 1 | 13 | Trim6 | 0.057 | Old |
| 0.024891 | 0.597221 | 0.393 | 0.254 | 1 | 13 | Abcf3 | 0.139 | Old |
| 0.024896 | 0.673289 | 0.393 | 0.263 | 1 | 13 | Mpv17 | 0.13 | Old |
| 0.024958 | 0.533301 | 0.262 | 0.132 | 1 | 13 | Rnf157 | 0.13 | Old |
| 0.025026 | 0.344272 | 0.984 | 0.965 | 1 | 13 | Rnf149 | 0.019 | Old |
| 0.025041 | 0.874994 | 0.262 | 0.149 | 1 | 13 | Mrpl1 | 0.113 | Old |
| 0.025083 | 0.328808 | 0.852 | 0.781 | 1 | 13 | Cdk11b | 0.071 | Old |
| 0.025223 | 0.850118 | 0.238 | 0.123 | 1 | 13 | Pvt1 | 0.115 | Old |
| 0.0253 | 0.322018 | 0.885 | 0.737 | 1 | 13 | Pcnp | 0.148 | Old |
| 0.025421 | 0.349965 | 0.697 | 0.544 | 1 | 13 | Chchd7 | 0.153 | Old |
| 0.025621 | 0.970592 | 0.213 | 0.105 | 1 | 13 | Gm7694 | 0.108 | Old |
| 0.025757 | 0.973016 | 0.197 | 0.096 | 1 | 13 | Tecpr2 | 0.101 | Old |
| 0.025791 | 0.40038 | 0.73 | 0.632 | 1 | 13 | Cyc1 | 0.098 | Old |
| 0.025844 | 0.580078 | 0.631 | 0.553 | 1 | 13 | Arrdc4 | 0.078 | Old |
| 0.025888 | 1.311601 | 0.082 | 0.018 | 1 | 13 | Dand5 | 0.064 | Old |
| 0.025888 | 1.043492 | 0.082 | 0.018 | 1 | 13 | Cdk18 | 0.064 | Old |
| 0.025978 | 1.073839 | 0.139 | 0.053 | 1 | 13 | 2900076A0 | 0.086 | Old |
| 0.026131 | 0.585042 | 0.779 | 0.737 | 1 | 13 | Cd209f | 0.042 | Old |
| 0.026143 | 0.301823 | 0.967 | 0.904 | 1 | 13 | Dpysl2 | 0.063 | Old |
| 0.026162 | 0.173906 | 1 | 0.991 | 1 | 13 | Dhrs3 | 0.009 | Old |
| 0.026194 | 0.785134 | 0.172 | 0.07 | 1 | 13 | Mkks | 0.102 | Old |
| 0.026224 | 1.233548 | 0.082 | 0.018 | 1 | 13 | Spib | 0.064 | Old |
| 0.026224 | 1.177745 | 0.082 | 0.018 | 1 | 13 | Kbtbd8 | 0.064 | Old |
| 0.026354 | 0.658976 | 0.238 | 0.123 | 1 | 13 | Msto1 | 0.115 | Old |
| 0.026378 | 1.236986 | 0.197 | 0.096 | 1 | 13 | Gm5150 | 0.101 | Old |
| 0.026619 | 0.462688 | 0.475 | 0.298 | 1 | 13 | Plcb3 | 0.177 | Old |
| 0.02674 | 0.838272 | 0.238 | 0.123 | 1 | 13 | Xrcc6 | 0.115 | Old |
| 0.026997 | 0.446819 | 0.836 | 0.763 | 1 | 13 | Parp14 | 0.073 | Old |
| 0.027186 | 0.557476 | 0.393 | 0.246 | 1 | 13 | Scd2 | 0.147 | Old |
| 0.027418 | 0.508677 | 0.615 | 0.456 | 1 | 13 | Fbxw11 | 0.159 | Old |
| 0.027424 | 0.327594 | 0.861 | 0.728 | 1 | 13 | Uqcrc1 | 0.133 | Old |
| 0.027424 | 0.881241 | 0.18 | 0.079 | 1 | 13 | P2ry14 | 0.101 | Old |
| 0.027454 | 1.064124 | 0.23 | 0.123 | 1 | 13 | Smyd5 | 0.107 | Old |
| 0.027466 | 0.667635 | 0.41 | 0.263 | 1 | 13 | Csf2rb2 | 0.147 | Old |
| 0.027528 | 0.764013 | 0.238 | 0.123 | 1 | 13 | Rai14 | 0.115 | Old |
| 0.027653 | 0.888406 | 0.18 | 0.079 | 1 | 13 | Enpep | 0.101 | Old |
| 0.027675 | 0.204823 | 0.992 | 0.974 | 1 | 13 | Pdcd6ip | 0.018 | Old |
| 0.027735 | 0.896141 | 0.254 | 0.14 | 1 | 13 | Vdr | 0.114 | Old |
| 0.027846 | 0.305019 | 0.705 | 0.561 | 1 | 13 | Znrd1 | 0.144 | Old |
| 0.027858 | 0.254842 | 0.951 | 0.921 | 1 | 13 | Ube3a | 0.03 | Old |
| 0.027864 | 0.262459 | 0.926 | 0.868 | 1 | 13 | Tnfrsf1b | 0.058 | Old |

| | | | | | | | | |
|---|---|---|---|---|---|---|---|---|
| 0.028097 | 0.43451 | 0.705 | 0.596 | 1 | 13 | Arhgap25 | 0.109 | Old |
| 0.02837 | 0.250842 | 0.902 | 0.772 | 1 | 13 | Ndufs2 | 0.13 | Old |
| 0.02842 | 0.345403 | 0.697 | 0.518 | 1 | 13 | Lrrc8c | 0.179 | Old |
| 0.028519 | 0.966259 | 0.254 | 0.14 | 1 | 13 | Pcid2 | 0.114 | Old |
| 0.028536 | 0.428985 | 0.697 | 0.544 | 1 | 13 | Snrpa | 0.153 | Old |
| 0.028702 | 0.19455 | 0.984 | 0.974 | 1 | 13 | Gusb | 0.01 | Old |
| 0.028779 | 0.455386 | 0.639 | 0.509 | 1 | 13 | Kcnab2 | 0.13 | Old |
| 0.028832 | 0.397808 | 0.738 | 0.605 | 1 | 13 | Plgrkt | 0.133 | Old |
| 0.028845 | 0.442742 | 0.852 | 0.763 | 1 | 13 | Lfng | 0.089 | Old |
| 0.028979 | 0.324616 | 0.779 | 0.667 | 1 | 13 | Acsl5 | 0.112 | Old |
| 0.029061 | 0.884588 | 0.287 | 0.175 | 1 | 13 | Dhx58 | 0.112 | Old |
| 0.029246 | 0.466764 | 0.746 | 0.544 | 1 | 13 | Ubxn6 | 0.202 | Old |
| 0.029261 | 0.525053 | 0.41 | 0.263 | 1 | 13 | Pik3r4 | 0.147 | Old |
| 0.029391 | 0.388528 | 0.721 | 0.57 | 1 | 13 | Tbc1d20 | 0.151 | Old |
| 0.029512 | 1.120755 | 0.123 | 0.044 | 1 | 13 | Inpp1 | 0.079 | Old |
| 0.029533 | 0.369966 | 0.516 | 0.351 | 1 | 13 | Pfdn4 | 0.165 | Old |
| 0.029543 | 1.714519 | 0.041 | 0 | 1 | 13 | Pcdhb22 | 0.041 | Old |
| 0.029543 | 1.520587 | 0.041 | 0 | 1 | 13 | Tmprss4 | 0.041 | Old |
| 0.029543 | 1.476412 | 0.041 | 0 | 1 | 13 | Gm10184 | 0.041 | Old |
| 0.029543 | 1.409874 | 0.041 | 0 | 1 | 13 | Dpyd | 0.041 | Old |
| 0.029543 | 1.400741 | 0.041 | 0 | 1 | 13 | Gm26542 | 0.041 | Old |
| 0.029543 | 1.296896 | 0.041 | 0 | 1 | 13 | Lyplal1 | 0.041 | Old |
| 0.029543 | 1.288179 | 0.041 | 0 | 1 | 13 | Bmp1 | 0.041 | Old |
| 0.029543 | 1.274628 | 0.041 | 0 | 1 | 13 | Cd300ld2 | 0.041 | Old |
| 0.029543 | 1.261537 | 0.041 | 0 | 1 | 13 | Jag2 | 0.041 | Old |
| 0.029543 | 1.251326 | 0.041 | 0 | 1 | 13 | Zfp507 | 0.041 | Old |
| 0.029543 | 1.242944 | 0.041 | 0 | 1 | 13 | Cyb5d1 | 0.041 | Old |
| 0.029543 | 1.198084 | 0.041 | 0 | 1 | 13 | Pidd1 | 0.041 | Old |
| 0.029547 | 0.84776 | 0.254 | 0.149 | 1 | 13 | Fam222b | 0.105 | Old |
| 0.029631 | 0.392061 | 0.697 | 0.57 | 1 | 13 | Pcna | 0.127 | Old |
| 0.029661 | 0.708231 | 0.508 | 0.395 | 1 | 13 | Mfsd12 | 0.113 | Old |
| 0.029682 | 0.420493 | 0.492 | 0.325 | 1 | 13 | Polr2b | 0.167 | Old |
| 0.029884 | 0.612783 | 0.303 | 0.175 | 1 | 13 | Tmem161b | 0.128 | Old |
| 0.02989 | 0.78441 | 0.205 | 0.096 | 1 | 13 | Ccnyl1 | 0.109 | Old |
| 0.029893 | 0.525963 | 0.361 | 0.219 | 1 | 13 | Cog8 | 0.142 | Old |
| 0.030097 | 0.460667 | 0.672 | 0.544 | 1 | 13 | Nupr1 | 0.128 | Old |
| 0.03027 | 0.508775 | 0.451 | 0.307 | 1 | 13 | Dhx16 | 0.144 | Old |
| 0.030277 | 0.26762 | 0.869 | 0.746 | 1 | 13 | Psmb2 | 0.123 | Old |
| 0.030375 | 0.377818 | 0.762 | 0.649 | 1 | 13 | Arf6 | 0.113 | Old |
| 0.030394 | 0.509955 | 0.467 | 0.316 | 1 | 13 | Mrpl55 | 0.151 | Old |
| 0.030408 | 1.154818 | 0.123 | 0.044 | 1 | 13 | Pdlim7 | 0.079 | Old |
| 0.030552 | 1.095918 | 0.516 | 0.404 | 1 | 13 | Mmp19 | 0.112 | Old |
| 0.030575 | 0.696717 | 0.164 | 0.07 | 1 | 13 | Cpne8 | 0.094 | Old |
| 0.030726 | 0.635191 | 0.328 | 0.211 | 1 | 13 | Ublcp1 | 0.117 | Old |
| 0.030792 | 0.329845 | 0.869 | 0.763 | 1 | 13 | Naa50 | 0.106 | Old |

| | | | | | | | | |
|---|---|---|---|---|---|---|---|---|
| 0.030797 | 0.761844 | 0.189 | 0.088 | 1 | 13 | Thop1 | 0.101 | Old |
| 0.030839 | 0.307652 | 0.746 | 0.588 | 1 | 13 | Arhgef12 | 0.158 | Old |
| 0.030938 | 0.403846 | 0.484 | 0.325 | 1 | 13 | Alg5 | 0.159 | Old |
| 0.031053 | 0.474265 | 0.697 | 0.561 | 1 | 13 | Hdac1 | 0.136 | Old |
| 0.031091 | 0.309113 | 0.861 | 0.684 | 1 | 13 | Lgals9 | 0.177 | Old |
| 0.031178 | 0.36501 | 0.77 | 0.632 | 1 | 13 | Stip1 | 0.138 | Old |
| 0.031453 | 0.170443 | 1 | 0.991 | 1 | 13 | Ywhah | 0.009 | Old |
| 0.031833 | 0.416498 | 0.5 | 0.316 | 1 | 13 | Evi2 | 0.184 | Old |
| 0.031958 | 0.217016 | 0.91 | 0.877 | 1 | 13 | Atp2a2 | 0.033 | Old |
| 0.031958 | 0.490861 | 0.615 | 0.474 | 1 | 13 | Tmem170k | 0.141 | Old |
| 0.031983 | 0.231541 | 0.975 | 0.991 | 1 | 13 | Itgb2 | -0.016 | Old |
| 0.03211 | 0.381782 | 0.516 | 0.325 | 1 | 13 | Zcchc2 | 0.191 | Old |
| 0.032208 | 0.324793 | 0.959 | 0.965 | 1 | 13 | Fyb | -0.006 | Old |
| 0.032238 | 0.695072 | 0.295 | 0.167 | 1 | 13 | Msantd4 | 0.128 | Old |
| 0.032271 | 1.03998 | 0.123 | 0.044 | 1 | 13 | Pus7l | 0.079 | Old |
| 0.032315 | 0.430862 | 0.574 | 0.404 | 1 | 13 | Kdm5c | 0.17 | Old |
| 0.03234 | 0.278356 | 0.91 | 0.798 | 1 | 13 | Lrrc25 | 0.112 | Old |
| 0.032366 | 0.238817 | 0.992 | 0.965 | 1 | 13 | Plxnb2 | 0.027 | Old |
| 0.03239 | 0.458394 | 0.664 | 0.544 | 1 | 13 | Acvrl1 | 0.12 | Old |
| 0.032419 | 0.413828 | 0.844 | 0.798 | 1 | 13 | Elmo2 | 0.046 | Old |
| 0.032438 | 0.464745 | 0.648 | 0.465 | 1 | 13 | Fbxo18 | 0.183 | Old |
| 0.032677 | 0.206428 | 0.992 | 1 | 1 | 13 | Tkt | -0.008 | Old |
| 0.032679 | 0.358224 | 0.713 | 0.518 | 1 | 13 | Nampt | 0.195 | Old |
| 0.032688 | 0.593994 | 0.32 | 0.193 | 1 | 13 | Arfgap3 | 0.127 | Old |
| 0.032822 | 0.762971 | 0.23 | 0.114 | 1 | 13 | Guf1 | 0.116 | Old |
| 0.033315 | 0.434412 | 0.393 | 0.237 | 1 | 13 | Dcaf1 | 0.156 | Old |
| 0.0337 | 0.445963 | 0.631 | 0.509 | 1 | 13 | Galc | 0.122 | Old |
| 0.033849 | 0.412111 | 0.623 | 0.474 | 1 | 13 | Itgb1bp1 | 0.149 | Old |
| 0.033966 | 0.867114 | 0.238 | 0.132 | 1 | 13 | Nol10 | 0.106 | Old |
| 0.034173 | 0.660095 | 0.434 | 0.289 | 1 | 13 | Cdc42ep3 | 0.145 | Old |
| 0.034349 | 0.538513 | 0.623 | 0.456 | 1 | 13 | Cops2 | 0.167 | Old |
| 0.034453 | 0.364452 | 0.779 | 0.64 | 1 | 13 | U2af1 | 0.139 | Old |
| 0.03484 | 0.634979 | 0.377 | 0.228 | 1 | 13 | Slc39a3 | 0.149 | Old |
| 0.034908 | 0.419962 | 0.664 | 0.579 | 1 | 13 | Prelid3b | 0.085 | Old |
| 0.03492 | 1.070527 | 0.148 | 0.061 | 1 | 13 | Zfp40 | 0.087 | Old |
| 0.034977 | 0.381442 | 0.803 | 0.737 | 1 | 13 | Pla2g15 | 0.066 | Old |
| 0.03504 | 0.24197 | 0.918 | 0.816 | 1 | 13 | Mapre2 | 0.102 | Old |
| 0.035062 | 0.408825 | 0.738 | 0.596 | 1 | 13 | Mff | 0.142 | Old |
| 0.035211 | 0.443169 | 0.607 | 0.474 | 1 | 13 | Idh3b | 0.133 | Old |
| 0.035435 | 0.33037 | 0.877 | 0.851 | 1 | 13 | Emilin2 | 0.026 | Old |
| 0.035623 | 0.309993 | 0.926 | 0.921 | 1 | 13 | Epn1 | 0.005 | Old |
| 0.035655 | 0.441897 | 0.73 | 0.605 | 1 | 13 | Mgat2 | 0.125 | Old |
| 0.035665 | 1.155011 | 0.107 | 0.035 | 1 | 13 | Arsg | 0.072 | Old |
| 0.035714 | 0.265874 | 0.959 | 0.895 | 1 | 13 | Stxbp3 | 0.064 | Old |
| 0.035741 | 0.613313 | 0.459 | 0.316 | 1 | 13 | C5ar2 | 0.143 | Old |

| | | | | | | | | |
|---|---|---|---|---|---|---|---|---|
| 0.03601 | 0.883236 | 0.172 | 0.079 | 1 | 13 | Plekha8 | 0.093 | Old |
| 0.036062 | 0.650731 | 0.32 | 0.193 | 1 | 13 | Cep295 | 0.127 | Old |
| 0.036156 | 0.524702 | 0.352 | 0.211 | 1 | 13 | Dusp10 | 0.141 | Old |
| 0.036303 | 0.988531 | 0.172 | 0.079 | 1 | 13 | Ftsj1 | 0.093 | Old |
| 0.036327 | 0.251582 | 0.934 | 0.86 | 1 | 13 | Morf4l2 | 0.074 | Old |
| 0.036376 | 0.274626 | 0.934 | 0.904 | 1 | 13 | Psmd2 | 0.03 | Old |
| 0.036425 | 0.292836 | 0.992 | 1 | 1 | 13 | Atp6v1b2 | -0.008 | Old |
| 0.036428 | 0.447343 | 0.5 | 0.342 | 1 | 13 | Mrpl2 | 0.158 | Old |
| 0.036555 | 0.25521 | 0.918 | 0.93 | 1 | 13 | Cct2 | -0.012 | Old |
| 0.03657 | 0.555392 | 0.467 | 0.325 | 1 | 13 | Dram1 | 0.142 | Old |
| 0.036784 | 0.381957 | 0.639 | 0.526 | 1 | 13 | Tbc1d13 | 0.113 | Old |
| 0.036808 | 0.888687 | 0.197 | 0.096 | 1 | 13 | Tfpi | 0.101 | Old |
| 0.036814 | 0.396681 | 0.82 | 0.693 | 1 | 13 | Slc20a1 | 0.127 | Old |
| 0.036891 | 0.52916 | 0.443 | 0.307 | 1 | 13 | Cfl2 | 0.136 | Old |
| 0.036936 | 0.224357 | 0.803 | 0.632 | 1 | 13 | Puf60 | 0.171 | Old |
| 0.037065 | 0.689252 | 0.328 | 0.211 | 1 | 13 | Nob1 | 0.117 | Old |
| 0.037144 | 0.30415 | 0.811 | 0.711 | 1 | 13 | Psmd14 | 0.1 | Old |
| 0.037461 | 0.165214 | 1 | 1 | 1 | 13 | Cox5a | 0 | Old |
| 0.037508 | 0.456348 | 0.492 | 0.325 | 1 | 13 | Usp50 | 0.167 | Old |
| 0.03751 | 1.021241 | 0.279 | 0.167 | 1 | 13 | Slc25a32 | 0.112 | Old |
| 0.037578 | 0.217436 | 0.967 | 0.86 | 1 | 13 | Eif4g1 | 0.107 | Old |
| 0.037812 | 0.14919 | 1 | 1 | 1 | 13 | Actr2 | 0 | Old |
| 0.037844 | 0.389558 | 0.672 | 0.535 | 1 | 13 | Pgrmc1 | 0.137 | Old |
| 0.037873 | 0.686692 | 0.23 | 0.123 | 1 | 13 | Gpatch1 | 0.107 | Old |
| 0.037906 | 0.685996 | 0.41 | 0.263 | 1 | 13 | Adpgk | 0.147 | Old |
| 0.037912 | 0.32847 | 0.566 | 0.395 | 1 | 13 | Tmem65 | 0.171 | Old |
| 0.037937 | 0.878156 | 0.238 | 0.132 | 1 | 13 | Gdpd1 | 0.106 | Old |
| 0.037955 | 1.636655 | 0.057 | 0.009 | 1 | 13 | Kif5c | 0.048 | Old |
| 0.037955 | 1.581244 | 0.057 | 0.009 | 1 | 13 | Scn3b | 0.048 | Old |
| 0.037981 | 0.589704 | 0.246 | 0.132 | 1 | 13 | Pola2 | 0.114 | Old |
| 0.038347 | 1.207267 | 0.131 | 0.053 | 1 | 13 | Oxld1 | 0.078 | Old |
| 0.038366 | 1.087397 | 0.107 | 0.035 | 1 | 13 | Cenpj | 0.072 | Old |
| 0.038453 | 0.855711 | 0.148 | 0.061 | 1 | 13 | Zc3h8 | 0.087 | Old |
| 0.038507 | 0.410629 | 0.533 | 0.386 | 1 | 13 | Lyst | 0.147 | Old |
| 0.038522 | 1.474875 | 0.057 | 0.009 | 1 | 13 | Rnf128 | 0.048 | Old |
| 0.038522 | 1.338044 | 0.057 | 0.009 | 1 | 13 | Lamb3 | 0.048 | Old |
| 0.038522 | 1.336549 | 0.057 | 0.009 | 1 | 13 | Kif11 | 0.048 | Old |
| 0.038554 | 0.388591 | 0.73 | 0.588 | 1 | 13 | 4933434E2 | 0.142 | Old |
| 0.038558 | 0.477809 | 0.344 | 0.202 | 1 | 13 | Uckl1 | 0.142 | Old |
| 0.038653 | 0.618007 | 0.402 | 0.281 | 1 | 13 | Mlx | 0.121 | Old |
| 0.038744 | 0.74713 | 0.221 | 0.114 | 1 | 13 | Zkscan6 | 0.107 | Old |
| 0.038857 | 0.220469 | 0.934 | 0.939 | 1 | 13 | Ighm | -0.005 | Old |
| 0.039062 | 0.106858 | 1 | 1 | 1 | 13 | Hnrnpa2b1 | 0 | Old |
| 0.039086 | 0.868282 | 0.18 | 0.088 | 1 | 13 | Evi5l | 0.092 | Old |
| 0.039305 | 0.531722 | 0.369 | 0.246 | 1 | 13 | Atg4d | 0.123 | Old |

| | | | | | | | | |
|---|---|---|---|---|---|---|---|---|
| 0.039318 | 0.872615 | 0.205 | 0.105 | 1 | 13 | Slc44a2 | 0.1 | Old |
| 0.039326 | 1.371162 | 0.074 | 0.018 | 1 | 13 | A530040E1 | 0.056 | Old |
| 0.039452 | 0.439481 | 0.648 | 0.518 | 1 | 13 | Prmt1 | 0.13 | Old |
| 0.039552 | 0.284081 | 0.779 | 0.684 | 1 | 13 | B4galnt1 | 0.095 | Old |
| 0.039639 | 0.401107 | 0.844 | 0.737 | 1 | 13 | Ak2 | 0.107 | Old |
| 0.039679 | 1.272842 | 0.057 | 0.009 | 1 | 13 | BC052040 | 0.048 | Old |
| 0.039679 | 0.363641 | 0.836 | 0.675 | 1 | 13 | Agtrap | 0.161 | Old |
| 0.039811 | 0.861369 | 0.148 | 0.061 | 1 | 13 | Nubpl | 0.087 | Old |
| 0.039827 | 1.227711 | 0.074 | 0.018 | 1 | 13 | Prkar1b | 0.056 | Old |
| 0.039962 | 0.49326 | 0.525 | 0.368 | 1 | 13 | Gyg | 0.157 | Old |
| 0.039964 | 0.275538 | 0.951 | 0.947 | 1 | 13 | Ccr5 | 0.004 | Old |
| 0.04005 | 0.825403 | 0.754 | 0.763 | 1 | 13 | Rgs1 | -0.009 | Old |
| 0.040157 | 0.793381 | 0.148 | 0.061 | 1 | 13 | Usp11 | 0.087 | Old |
| 0.040268 | 1.385238 | 0.057 | 0.009 | 1 | 13 | Klra3 | 0.048 | Old |
| 0.040334 | 1.273328 | 0.074 | 0.018 | 1 | 13 | Hspa12a | 0.056 | Old |
| 0.040649 | 0.539795 | 0.303 | 0.175 | 1 | 13 | Traf2 | 0.128 | Old |
| 0.040719 | 0.271569 | 0.975 | 0.965 | 1 | 13 | Gls | 0.01 | Old |
| 0.040811 | 0.385842 | 0.877 | 0.763 | 1 | 13 | Cpt1a | 0.114 | Old |
| 0.040846 | 4.037587 | 0.074 | 0.018 | 1 | 13 | Ctsk | 0.056 | Old |
| 0.040846 | 1.174844 | 0.074 | 0.018 | 1 | 13 | Pafah2 | 0.056 | Old |
| 0.04089 | 0.973959 | 0.131 | 0.053 | 1 | 13 | Nle1 | 0.078 | Old |
| 0.040959 | 0.306201 | 0.926 | 0.921 | 1 | 13 | Csf2rb | 0.005 | Old |
| 0.041003 | 0.217494 | 0.992 | 0.974 | 1 | 13 | Itm2c | 0.018 | Old |
| 0.041045 | 0.728429 | 0.393 | 0.289 | 1 | 13 | Il18 | 0.104 | Old |
| 0.041072 | 0.477289 | 0.361 | 0.228 | 1 | 13 | Rfxap | 0.133 | Old |
| 0.041174 | 0.850045 | 0.221 | 0.123 | 1 | 13 | Tnfsf13os | 0.098 | Old |
| 0.041183 | 0.44472 | 0.475 | 0.342 | 1 | 13 | Ruvbl1 | 0.133 | Old |
| 0.041219 | 0.48233 | 0.434 | 0.298 | 1 | 13 | Rab29 | 0.136 | Old |
| 0.041277 | 0.337778 | 0.623 | 0.465 | 1 | 13 | Pak1ip1 | 0.158 | Old |
| 0.041317 | 0.5643 | 0.27 | 0.149 | 1 | 13 | Ifi211 | 0.121 | Old |
| 0.041467 | 0.543985 | 0.475 | 0.333 | 1 | 13 | Adk | 0.142 | Old |
| 0.041468 | 1.241343 | 0.057 | 0.009 | 1 | 13 | Prr15 | 0.048 | Old |
| 0.041468 | 1.183022 | 0.057 | 0.009 | 1 | 13 | Fam120c | 0.048 | Old |
| 0.041468 | 1.020717 | 0.057 | 0.009 | 1 | 13 | Srd5a1 | 0.048 | Old |
| 0.041597 | 0.316205 | 0.77 | 0.728 | 1 | 13 | Kcnn4 | 0.042 | Old |
| 0.041887 | 1.175338 | 0.074 | 0.018 | 1 | 13 | Col16a1 | 0.056 | Old |
| 0.041887 | 1.117173 | 0.074 | 0.018 | 1 | 13 | CAAA0114 | 0.056 | Old |
| 0.041948 | 0.147248 | 0.992 | 0.921 | 1 | 13 | Atp5o.1 | 0.071 | Old |
| 0.042012 | 0.773122 | 0.238 | 0.132 | 1 | 13 | Nmral1 | 0.106 | Old |
| 0.042023 | 0.985796 | 0.131 | 0.053 | 1 | 13 | Tanc1 | 0.078 | Old |
| 0.042052 | 0.458913 | 0.566 | 0.412 | 1 | 13 | Tufm | 0.154 | Old |
| 0.042079 | 0.887925 | 0.057 | 0.009 | 1 | 13 | Eme2 | 0.048 | Old |
| 0.042103 | 0.295774 | 0.893 | 0.807 | 1 | 13 | Pnp | 0.086 | Old |
| 0.042161 | 0.959196 | 0.09 | 0.026 | 1 | 13 | 2610301B2 | 0.064 | Old |
| 0.042402 | 0.519075 | 0.402 | 0.281 | 1 | 13 | Gla | 0.121 | Old |

| | | | | | | | | |
|---|---|---|---|---|---|---|---|---|
| 0.042416 | 1.253504 | 0.074 | 0.018 | 1 | 13 | Pnpla1 | 0.056 | Old |
| 0.042586 | 0.942301 | 0.189 | 0.096 | 1 | 13 | Trmt11 | 0.093 | Old |
| 0.042649 | 0.834324 | 0.148 | 0.061 | 1 | 13 | Lrrc49 | 0.087 | Old |
| 0.042746 | 0.204698 | 0.918 | 0.895 | 1 | 13 | Rab31 | 0.023 | Old |
| 0.042748 | 0.716036 | 0.197 | 0.096 | 1 | 13 | Gm20707 | 0.101 | Old |
| 0.04281 | 1.17766 | 0.287 | 0.175 | 1 | 13 | Il10 | 0.112 | Old |
| 0.042892 | 0.244818 | 0.852 | 0.763 | 1 | 13 | Arfgef1 | 0.089 | Old |
| 0.042895 | 0.476716 | 0.475 | 0.316 | 1 | 13 | Uba3 | 0.159 | Old |
| 0.042931 | 0.582213 | 0.426 | 0.289 | 1 | 13 | Mettl1 | 0.137 | Old |
| 0.04295 | 0.95686 | 0.074 | 0.018 | 1 | 13 | Adipoq | 0.056 | Old |
| 0.04299 | 0.59609 | 0.475 | 0.333 | 1 | 13 | Cul2 | 0.142 | Old |
| 0.043182 | 0.881904 | 0.131 | 0.053 | 1 | 13 | Srfbp1 | 0.078 | Old |
| 0.043195 | 0.460603 | 0.672 | 0.518 | 1 | 13 | Cmip | 0.154 | Old |
| 0.043212 | 0.23677 | 0.975 | 0.93 | 1 | 13 | Cap1 | 0.045 | Old |
| 0.043526 | 0.348436 | 0.68 | 0.518 | 1 | 13 | Srf | 0.162 | Old |
| 0.043768 | 0.53689 | 0.467 | 0.342 | 1 | 13 | Ell | 0.125 | Old |
| 0.043839 | 0.412443 | 0.369 | 0.219 | 1 | 13 | Rrp15 | 0.15 | Old |
| 0.043941 | 0.265713 | 0.869 | 0.798 | 1 | 13 | Rab8a | 0.071 | Old |
| 0.044036 | 1.020126 | 0.074 | 0.018 | 1 | 13 | Mboat1 | 0.056 | Old |
| 0.044102 | 0.611949 | 0.336 | 0.211 | 1 | 13 | Caap1 | 0.125 | Old |
| 0.044244 | 0.499348 | 0.5 | 0.36 | 1 | 13 | Mak16 | 0.14 | Old |
| 0.04428 | 0.593978 | 0.221 | 0.114 | 1 | 13 | Ddx31 | 0.107 | Old |
| 0.044328 | 0.893647 | 0.156 | 0.07 | 1 | 13 | Ulbp1 | 0.086 | Old |
| 0.044452 | 0.418887 | 0.525 | 0.377 | 1 | 13 | Mrto4 | 0.148 | Old |
| 0.044577 | 1.01093 | 0.09 | 0.026 | 1 | 13 | Zfp946 | 0.064 | Old |
| 0.044595 | 0.651139 | 0.336 | 0.219 | 1 | 13 | Mmgt1 | 0.117 | Old |
| 0.044668 | 0.581892 | 0.32 | 0.184 | 1 | 13 | Atrn | 0.136 | Old |
| 0.044704 | 0.767837 | 0.213 | 0.114 | 1 | 13 | Fam198b | 0.099 | Old |
| 0.045165 | 0.519001 | 0.492 | 0.36 | 1 | 13 | Adap1 | 0.132 | Old |
| 0.045392 | 0.231502 | 0.934 | 0.912 | 1 | 13 | Psma6 | 0.022 | Old |
| 0.045422 | 0.319974 | 0.385 | 0.254 | 1 | 13 | Myg1 | 0.131 | Old |
| 0.045545 | 0.384439 | 0.385 | 0.237 | 1 | 13 | Gmppa | 0.148 | Old |
| 0.04564 | 0.187074 | 0.975 | 0.947 | 1 | 13 | Ywhag | 0.028 | Old |
| 0.045642 | 1.325984 | 0.098 | 0.035 | 1 | 13 | Rcn1 | 0.063 | Old |
| 0.04601 | 0.670673 | 0.197 | 0.096 | 1 | 13 | Atpaf1 | 0.101 | Old |
| 0.046091 | 0.460159 | 0.5 | 0.395 | 1 | 13 | Pin1 | 0.105 | Old |
| 0.046113 | 0.767461 | 0.352 | 0.237 | 1 | 13 | Gpx3 | 0.115 | Old |
| 0.04621 | 0.489469 | 0.746 | 0.763 | 1 | 13 | Pla2g2d | -0.017 | Old |
| 0.046406 | 0.768584 | 0.131 | 0.053 | 1 | 13 | Galnt6 | 0.078 | Old |
| 0.04655 | 0.54054 | 0.344 | 0.211 | 1 | 13 | Hacd3 | 0.133 | Old |
| 0.047079 | 0.265921 | 0.852 | 0.781 | 1 | 13 | Nfil3 | 0.071 | Old |
| 0.047138 | 0.504092 | 0.262 | 0.149 | 1 | 13 | Armc7 | 0.113 | Old |
| 0.047219 | 0.427635 | 0.5 | 0.36 | 1 | 13 | Hk3 | 0.14 | Old |
| 0.047295 | 0.256231 | 0.975 | 0.895 | 1 | 13 | Tubb2a | 0.08 | Old |
| 0.0473 | 1.232492 | 0.115 | 0.044 | 1 | 13 | Ly6c2 | 0.071 | Old |

| | | | | | | | | |
|---|---|---|---|---|---|---|---|---|
| 0.0473 | 1.061551 | 0.115 | 0.044 | 1 | 13 | Adamtsl4 | 0.071 | Old |
| 0.047418 | 0.966476 | 0.139 | 0.061 | 1 | 13 | Slc25a14 | 0.078 | Old |
| 0.047826 | 1.234723 | 0.139 | 0.061 | 1 | 13 | Gm42047 | 0.078 | Old |
| 0.047846 | 0.334656 | 0.787 | 0.772 | 1 | 13 | Rrp1 | 0.015 | Old |
| 0.047879 | 0.575482 | 0.27 | 0.158 | 1 | 13 | Dglucy | 0.112 | Old |
| 0.047924 | 0.696826 | 0.254 | 0.149 | 1 | 13 | Zbtb17 | 0.105 | Old |
| 0.047987 | 0.36483 | 0.77 | 0.693 | 1 | 13 | Napa | 0.077 | Old |
| 0.048017 | 0.711161 | 0.311 | 0.202 | 1 | 13 | Alg1 | 0.109 | Old |
| 0.048135 | 0.478109 | 0.393 | 0.263 | 1 | 13 | Nufip1 | 0.13 | Old |
| 0.04831 | 0.801725 | 0.189 | 0.096 | 1 | 13 | Abcb8 | 0.093 | Old |
| 0.048679 | 1.029658 | 0.172 | 0.088 | 1 | 13 | Ncapd2 | 0.084 | Old |
| 0.048712 | 0.261862 | 0.893 | 0.851 | 1 | 13 | Cmtm3 | 0.042 | Old |
| 0.04873 | 0.27948 | 0.926 | 0.877 | 1 | 13 | Hspd1 | 0.049 | Old |
| 0.048741 | 0.502103 | 0.443 | 0.307 | 1 | 13 | Tshz1 | 0.136 | Old |
| 0.048856 | 0.335319 | 0.779 | 0.693 | 1 | 13 | Blvra | 0.086 | Old |
| 0.049063 | 1.356753 | 0.098 | 0.035 | 1 | 13 | Clec1a | 0.063 | Old |
| 0.049143 | 0.338612 | 0.574 | 0.421 | 1 | 13 | Ptk2b | 0.153 | Old |
| 0.049179 | 0.457653 | 0.451 | 0.316 | 1 | 13 | Snhg12 | 0.135 | Old |
| 0.049306 | 0.327582 | 0.844 | 0.702 | 1 | 13 | Nus1 | 0.142 | Old |
| 0.049427 | 0.942658 | 0.172 | 0.088 | 1 | 13 | Ttbk2 | 0.084 | Old |
| 0.049435 | 0.521961 | 0.549 | 0.412 | 1 | 13 | Nacc1 | 0.137 | Old |
| 0.049544 | 0.531262 | 0.402 | 0.272 | 1 | 13 | Tvp23b | 0.13 | Old |
| 0.050137 | 0.284257 | 0.836 | 0.798 | 1 | 13 | Adssl1 | 0.038 | Old |
| 0.050337 | 0.925824 | 0.139 | 0.061 | 1 | 13 | Zbtb12 | 0.078 | Old |
| 0.050354 | 0.287654 | 0.811 | 0.693 | 1 | 13 | Ptprs | 0.118 | Old |
| 0.050567 | 0.259592 | 0.951 | 0.904 | 1 | 13 | Myof | 0.047 | Old |
| 0.050955 | 0.859194 | 0.287 | 0.184 | 1 | 13 | Map3k20 | 0.103 | Old |
| 0.050978 | 0.826514 | 0.23 | 0.132 | 1 | 13 | Zfand2a | 0.098 | Old |
| 0.051039 | 0.469281 | 0.418 | 0.272 | 1 | 13 | Chid1 | 0.146 | Old |
| 0.051113 | 1.233936 | 0.098 | 0.035 | 1 | 13 | Dtnb | 0.063 | Old |
| 0.05128 | 0.276948 | 0.861 | 0.737 | 1 | 13 | Srp72 | 0.124 | Old |
| 0.051353 | 0.36999 | 0.779 | 0.667 | 1 | 13 | Mrps18c | 0.112 | Old |
| 0.051414 | 0.325067 | 0.664 | 0.518 | 1 | 13 | Rcor1 | 0.146 | Old |
| 0.051614 | 0.609306 | 0.238 | 0.132 | 1 | 13 | Mrps2 | 0.106 | Old |
| 0.051717 | 0.56404 | 0.213 | 0.114 | 1 | 13 | Usp21 | 0.099 | Old |
| 0.051745 | 1.398511 | 0.123 | 0.053 | 1 | 13 | Uhrf1 | 0.07 | Old |
| 0.051915 | 0.242103 | 0.902 | 0.825 | 1 | 13 | Itch | 0.077 | Old |
| 0.05216 | 0.400235 | 0.533 | 0.377 | 1 | 13 | Mrpl40 | 0.156 | Old |
| 0.052164 | 1.032997 | 0.098 | 0.035 | 1 | 13 | Tldc1 | 0.063 | Old |
| 0.052208 | 3.476266 | 0.033 | 0 | 1 | 13 | Apol7c | 0.033 | Old |
| 0.052208 | 2.025188 | 0.033 | 0 | 1 | 13 | Dusp4 | 0.033 | Old |
| 0.052208 | 1.696699 | 0.033 | 0 | 1 | 13 | Gen1 | 0.033 | Old |
| 0.052208 | 1.541717 | 0.033 | 0 | 1 | 13 | Irak1bp1 | 0.033 | Old |
| 0.052208 | 1.396591 | 0.033 | 0 | 1 | 13 | Gm38336 | 0.033 | Old |
| 0.052208 | 1.381597 | 0.033 | 0 | 1 | 13 | AC168977. | 0.033 | Old |

| | | | | | | | | |
|---|---|---|---|---|---|---|---|---|
| 0.052208 | 1.358027 | 0.033 | 0 | 1 | 13 | Gm47815 | 0.033 | Old |
| 0.052208 | 1.316316 | 0.033 | 0 | 1 | 13 | Dclk1 | 0.033 | Old |
| 0.052208 | 1.315899 | 0.033 | 0 | 1 | 13 | Fhad1 | 0.033 | Old |
| 0.052208 | 1.305407 | 0.033 | 0 | 1 | 13 | Pnp2 | 0.033 | Old |
| 0.052208 | 1.294981 | 0.033 | 0 | 1 | 13 | Mirt1 | 0.033 | Old |
| 0.052208 | 1.294074 | 0.033 | 0 | 1 | 13 | Camk2n2 | 0.033 | Old |
| 0.052208 | 1.286432 | 0.033 | 0 | 1 | 13 | Fbxo48 | 0.033 | Old |
| 0.052208 | 1.266921 | 0.033 | 0 | 1 | 13 | Pde5a | 0.033 | Old |
| 0.052208 | 1.253328 | 0.033 | 0 | 1 | 13 | Pkia | 0.033 | Old |
| 0.052208 | 1.248287 | 0.033 | 0 | 1 | 13 | Sult1e1 | 0.033 | Old |
| 0.052208 | 1.238002 | 0.033 | 0 | 1 | 13 | Gm49322 | 0.033 | Old |
| 0.052208 | 1.234706 | 0.033 | 0 | 1 | 13 | Notch4 | 0.033 | Old |
| 0.052208 | 1.233215 | 0.033 | 0 | 1 | 13 | Gm15265 | 0.033 | Old |
| 0.052208 | 1.191919 | 0.033 | 0 | 1 | 13 | Tnfrsf9 | 0.033 | Old |
| 0.052208 | 1.184652 | 0.033 | 0 | 1 | 13 | Serpinc1 | 0.033 | Old |
| 0.052208 | 1.183413 | 0.033 | 0 | 1 | 13 | Gm21992 | 0.033 | Old |
| 0.052208 | 1.176766 | 0.033 | 0 | 1 | 13 | Zfp54 | 0.033 | Old |
| 0.052208 | 1.163496 | 0.033 | 0 | 1 | 13 | Zfp526 | 0.033 | Old |
| 0.052208 | 1.114923 | 0.033 | 0 | 1 | 13 | Gm32098 | 0.033 | Old |
| 0.052208 | 1.111992 | 0.033 | 0 | 1 | 13 | Zfp61 | 0.033 | Old |
| 0.052208 | 1.091259 | 0.033 | 0 | 1 | 13 | Gm43703 | 0.033 | Old |
| 0.052208 | 1.087177 | 0.033 | 0 | 1 | 13 | Brca1 | 0.033 | Old |
| 0.052208 | 1.014711 | 0.033 | 0 | 1 | 13 | Ceacam16 | 0.033 | Old |
| 0.052208 | 0.946339 | 0.033 | 0 | 1 | 13 | Elavl4 | 0.033 | Old |
| 0.052265 | 0.558064 | 0.336 | 0.219 | 1 | 13 | Rnf121 | 0.117 | Old |
| 0.052423 | 0.31085 | 0.811 | 0.693 | 1 | 13 | Piezo1 | 0.118 | Old |
| 0.05244 | 0.165545 | 1 | 1 | 1 | 13 | Slc6a6 | 0 | Old |
| 0.052497 | 0.316517 | 0.697 | 0.614 | 1 | 13 | Crybg1 | 0.083 | Old |
| 0.052514 | 0.842867 | 0.139 | 0.061 | 1 | 13 | Zfp523 | 0.078 | Old |
| 0.052534 | 0.642466 | 0.443 | 0.325 | 1 | 13 | Scn1b | 0.118 | Old |
| 0.052666 | 0.428254 | 0.426 | 0.298 | 1 | 13 | Tecpr1 | 0.128 | Old |
| 0.052801 | 0.475829 | 0.434 | 0.289 | 1 | 13 | Rpf1 | 0.145 | Old |
| 0.053011 | 0.306803 | 0.648 | 0.491 | 1 | 13 | Tars | 0.157 | Old |
| 0.053299 | 0.836887 | 0.32 | 0.211 | 1 | 13 | Vegfa | 0.109 | Old |
| 0.053397 | 0.56319 | 0.311 | 0.193 | 1 | 13 | Nos1ap | 0.118 | Old |
| 0.053505 | 0.354175 | 0.689 | 0.518 | 1 | 13 | Senp2 | 0.171 | Old |
| 0.053629 | 0.563979 | 0.41 | 0.281 | 1 | 13 | Fuom | 0.129 | Old |
| 0.053694 | 0.223072 | 0.893 | 0.807 | 1 | 13 | Ogdh | 0.086 | Old |
| 0.053706 | 0.579601 | 0.311 | 0.193 | 1 | 13 | Mtap | 0.118 | Old |
| 0.053965 | 0.231009 | 0.852 | 0.763 | 1 | 13 | Kpnb1 | 0.089 | Old |
| 0.054044 | 0.589312 | 0.254 | 0.149 | 1 | 13 | Zfp943 | 0.105 | Old |
| 0.054146 | 0.426864 | 0.451 | 0.316 | 1 | 13 | Ctso | 0.135 | Old |
| 0.054196 | 0.250975 | 0.91 | 0.833 | 1 | 13 | Cab39 | 0.077 | Old |
| 0.054344 | 0.19268 | 0.762 | 0.623 | 1 | 13 | Etnk1 | 0.139 | Old |
| 0.054483 | 0.864968 | 0.197 | 0.105 | 1 | 13 | Noa1 | 0.092 | Old |

| | | | | | | | | |
|---|---|---|---|---|---|---|---|---|
| 0.054538 | 0.215891 | 0.951 | 0.93 | 1 | 13 | Apbb1ip | 0.021 | Old |
| 0.05459 | 0.40122 | 0.492 | 0.342 | 1 | 13 | Phrf1 | 0.15 | Old |
| 0.054873 | 1.19489 | 0.098 | 0.035 | 1 | 13 | AU022793 | 0.063 | Old |
| 0.054873 | 0.959371 | 0.098 | 0.035 | 1 | 13 | Cdyl2 | 0.063 | Old |
| 0.055004 | 0.821744 | 0.164 | 0.079 | 1 | 13 | Rdh13 | 0.085 | Old |
| 0.055044 | 0.231525 | 0.992 | 0.965 | 1 | 13 | 9-Sep | 0.027 | Old |
| 0.055287 | 0.274424 | 1 | 0.965 | 1 | 13 | Lilr4b | 0.035 | Old |
| 0.055444 | 0.439995 | 0.459 | 0.316 | 1 | 13 | Katnbl1 | 0.143 | Old |
| 0.055503 | 0.425404 | 0.377 | 0.246 | 1 | 13 | Cks1b | 0.131 | Old |
| 0.055537 | 0.296798 | 0.795 | 0.711 | 1 | 13 | Ftl1-ps1 | 0.084 | Old |
| 0.055587 | 0.587835 | 0.549 | 0.43 | 1 | 13 | Man1b1 | 0.119 | Old |
| 0.055693 | 0.872477 | 0.139 | 0.061 | 1 | 13 | Cd320 | 0.078 | Old |
| 0.055887 | 0.368185 | 0.639 | 0.526 | 1 | 13 | Pcgf5 | 0.113 | Old |
| 0.055959 | 0.7088 | 0.205 | 0.114 | 1 | 13 | Prmt9 | 0.091 | Old |
| 0.056028 | 0.819362 | 0.115 | 0.044 | 1 | 13 | Nhej1 | 0.071 | Old |
| 0.056259 | 0.706003 | 0.189 | 0.096 | 1 | 13 | Hoxb4 | 0.093 | Old |
| 0.056343 | 0.593847 | 0.287 | 0.184 | 1 | 13 | Asap2 | 0.103 | Old |
| 0.056523 | 0.365819 | 0.852 | 0.746 | 1 | 13 | Rnh1 | 0.106 | Old |
| 0.056633 | 0.199529 | 0.975 | 0.974 | 1 | 13 | St13 | 0.001 | Old |
| 0.05695 | 0.561596 | 0.459 | 0.351 | 1 | 13 | Rnf139 | 0.108 | Old |
| 0.056991 | 0.934819 | 0.18 | 0.096 | 1 | 13 | Arrdc2 | 0.084 | Old |
| 0.057 | 0.15682 | 0.984 | 0.939 | 1 | 13 | Dnajc3 | 0.045 | Old |
| 0.05707 | 0.653914 | 0.189 | 0.096 | 1 | 13 | Fam160a2 | 0.093 | Old |
| 0.057257 | 0.126344 | 1 | 1 | 1 | 13 | Lcp1 | 0 | Old |
| 0.057512 | 0.267877 | 0.836 | 0.702 | 1 | 13 | Rdx | 0.134 | Old |
| 0.057597 | 0.340327 | 0.787 | 0.605 | 1 | 13 | Chd9 | 0.182 | Old |
| 0.05763 | 0.48954 | 0.467 | 0.36 | 1 | 13 | Hdac4 | 0.107 | Old |
| 0.057867 | 0.182234 | 0.959 | 0.939 | 1 | 13 | Fkbp1a | 0.02 | Old |
| 0.057889 | 0.924557 | 0.189 | 0.105 | 1 | 13 | Ppif | 0.084 | Old |
| 0.057892 | 0.35021 | 0.631 | 0.5 | 1 | 13 | Yipf5 | 0.131 | Old |
| 0.058221 | 0.32779 | 0.598 | 0.474 | 1 | 13 | Tubb4b | 0.124 | Old |
| 0.058278 | 0.350645 | 0.754 | 0.667 | 1 | 13 | Ctcf | 0.087 | Old |
| 0.058809 | 0.44372 | 0.361 | 0.237 | 1 | 13 | Icmt | 0.124 | Old |
| 0.058883 | 0.497761 | 0.287 | 0.175 | 1 | 13 | Bysl | 0.112 | Old |
| 0.058989 | 0.538634 | 0.492 | 0.368 | 1 | 13 | Pmvk | 0.124 | Old |
| 0.059139 | 0.145397 | 0.975 | 0.921 | 1 | 13 | Wipf1 | 0.054 | Old |
| 0.059497 | 1.307662 | 0.082 | 0.026 | 1 | 13 | Tbc1d30 | 0.056 | Old |
| 0.059497 | 1.171016 | 0.082 | 0.026 | 1 | 13 | Pou5f1 | 0.056 | Old |
| 0.059617 | 0.387675 | 0.672 | 0.623 | 1 | 13 | Cd226 | 0.049 | Old |
| 0.059878 | 0.476758 | 0.566 | 0.474 | 1 | 13 | Slfn8 | 0.092 | Old |
| 0.05988 | 0.465867 | 0.598 | 0.447 | 1 | 13 | Tnks | 0.151 | Old |
| 0.05991 | 0.318994 | 0.516 | 0.395 | 1 | 13 | Pvr | 0.121 | Old |
| 0.060032 | 0.327292 | 0.549 | 0.412 | 1 | 13 | Sptlc1 | 0.137 | Old |
| 0.060056 | 0.808442 | 0.172 | 0.088 | 1 | 13 | Mios | 0.084 | Old |
| 0.060254 | 0.302606 | 0.82 | 0.711 | 1 | 13 | Cat | 0.109 | Old |

| | | | | | | | | |
|---|---|---|---|---|---|---|---|---|
| 0.060332 | 0.314929 | 0.533 | 0.386 | 1 | 13 | Ak6 | 0.147 | Old |
| 0.060554 | 0.26738 | 0.951 | 0.93 | 1 | 13 | Serinc1 | 0.021 | Old |
| 0.060691 | 0.26548 | 0.68 | 0.588 | 1 | 13 | Pgap2 | 0.092 | Old |
| 0.060843 | 0.108296 | 0.984 | 0.991 | 1 | 13 | Prdx1 | -0.007 | Old |
| 0.061024 | 0.333806 | 0.762 | 0.658 | 1 | 13 | Ociad1 | 0.104 | Old |
| 0.061228 | 1.30249 | 0.123 | 0.053 | 1 | 13 | Gm26870 | 0.07 | Old |
| 0.061229 | 0.406878 | 0.475 | 0.351 | 1 | 13 | Snhg6 | 0.124 | Old |
| 0.061484 | 0.248196 | 0.869 | 0.781 | 1 | 13 | Cct7 | 0.088 | Old |
| 0.061514 | 0.682266 | 0.139 | 0.061 | 1 | 13 | Slc16a13 | 0.078 | Old |
| 0.061531 | 0.288612 | 0.844 | 0.737 | 1 | 13 | Klhl9 | 0.107 | Old |
| 0.061635 | 0.143042 | 1 | 1 | 1 | 13 | Tagln2 | 0 | Old |
| 0.06195 | 0.248965 | 0.885 | 0.877 | 1 | 13 | Vwa5a | 0.008 | Old |
| 0.062017 | 0.31289 | 0.828 | 0.825 | 1 | 13 | Ddx21 | 0.003 | Old |
| 0.062165 | 0.2451 | 0.984 | 0.956 | 1 | 13 | Ptprj | 0.028 | Old |
| 0.062168 | 0.119191 | 1 | 1 | 1 | 13 | Iqgap1 | 0 | Old |
| 0.062302 | 0.590408 | 0.451 | 0.333 | 1 | 13 | Pde4dip | 0.118 | Old |
| 0.062305 | 1.003611 | 0.123 | 0.053 | 1 | 13 | Scrn3 | 0.07 | Old |
| 0.062568 | 0.65601 | 0.148 | 0.07 | 1 | 13 | Prrg1 | 0.078 | Old |
| 0.062608 | 0.56966 | 0.164 | 0.079 | 1 | 13 | Mtg2 | 0.085 | Old |
| 0.062931 | 0.38558 | 0.443 | 0.316 | 1 | 13 | Ttc9c | 0.127 | Old |
| 0.063142 | 0.25077 | 0.869 | 0.746 | 1 | 13 | Hnrnpdl | 0.123 | Old |
| 0.063196 | 0.556055 | 0.344 | 0.228 | 1 | 13 | Smcr8 | 0.116 | Old |
| 0.063266 | 0.320034 | 0.795 | 0.772 | 1 | 13 | Slc11a1 | 0.023 | Old |
| 0.063398 | 0.808208 | 0.123 | 0.053 | 1 | 13 | Zfp874b | 0.07 | Old |
| 0.063471 | 0.577653 | 0.197 | 0.105 | 1 | 13 | Mad1l1 | 0.092 | Old |
| 0.063491 | 0.486337 | 0.377 | 0.254 | 1 | 13 | Psat1 | 0.123 | Old |
| 0.063521 | 0.35865 | 0.59 | 0.43 | 1 | 13 | Polr3c | 0.16 | Old |
| 0.063522 | 0.609153 | 0.41 | 0.298 | 1 | 13 | Nosip | 0.112 | Old |
| 0.063921 | 0.22235 | 0.992 | 0.939 | 1 | 13 | Cmpk1 | 0.053 | Old |
| 0.064063 | 0.05605 | 1 | 1 | 1 | 13 | Rps28 | 0 | Old |
| 0.064402 | 0.687391 | 0.262 | 0.158 | 1 | 13 | Hdac7 | 0.104 | Old |
| 0.064801 | 0.280361 | 0.73 | 0.596 | 1 | 13 | Usp47 | 0.134 | Old |
| 0.064907 | 0.922938 | 0.082 | 0.026 | 1 | 13 | Rom1 | 0.056 | Old |
| 0.064916 | 0.388626 | 0.525 | 0.421 | 1 | 13 | Snx13 | 0.104 | Old |
| 0.064935 | 0.595015 | 0.328 | 0.228 | 1 | 13 | Fam122a | 0.1 | Old |
| 0.065126 | 0.41892 | 0.762 | 0.684 | 1 | 13 | Stx12 | 0.078 | Old |
| 0.065245 | 0.299135 | 0.893 | 0.877 | 1 | 13 | Zfp703 | 0.016 | Old |
| 0.065278 | 0.270107 | 0.82 | 0.754 | 1 | 13 | Papd4 | 0.066 | Old |
| 0.065483 | 0.549405 | 0.369 | 0.246 | 1 | 13 | Cacna1d | 0.123 | Old |
| 0.065611 | 0.959433 | 0.082 | 0.026 | 1 | 13 | Uprt | 0.056 | Old |
| 0.065611 | 0.904638 | 0.082 | 0.026 | 1 | 13 | Arpin | 0.056 | Old |
| 0.065684 | 1.236181 | 0.049 | 0.009 | 1 | 13 | Gm10031 | 0.04 | Old |
| 0.065684 | 1.218138 | 0.049 | 0.009 | 1 | 13 | Angel1 | 0.04 | Old |
| 0.06571 | 0.345141 | 0.689 | 0.535 | 1 | 13 | Aqr | 0.154 | Old |
| 0.065724 | 0.426254 | 0.475 | 0.325 | 1 | 13 | Golt1b | 0.15 | Old |

| | | | | | | | | |
|---|---|---|---|---|---|---|---|---|
| 0.065755 | 0.480237 | 0.73 | 0.614 | 1 | 13 | Mdm2 | 0.116 | Old |
| 0.065824 | 1.361034 | 0.066 | 0.018 | 1 | 13 | Palb2 | 0.048 | Old |
| 0.066001 | 0.156058 | 0.984 | 0.947 | 1 | 13 | Rab1a | 0.037 | Old |
| 0.066177 | 0.890298 | 0.131 | 0.061 | 1 | 13 | Smg8 | 0.07 | Old |
| 0.066224 | 0.557788 | 0.328 | 0.202 | 1 | 13 | Zbtb34 | 0.126 | Old |
| 0.066225 | 0.292948 | 0.549 | 0.421 | 1 | 13 | Zmpste24 | 0.128 | Old |
| 0.066239 | 0.238259 | 0.238 | 0.132 | 1 | 13 | Rpgrip1 | 0.106 | Old |
| 0.066287 | 0.12688 | 0.984 | 1 | 1 | 13 | Atp5a1 | -0.016 | Old |
| 0.066334 | 0.549831 | 0.295 | 0.184 | 1 | 13 | Cd274 | 0.111 | Old |
| 0.066447 | 0.255085 | 0.697 | 0.588 | 1 | 13 | Ganab | 0.109 | Old |
| 0.066527 | 0.359029 | 0.361 | 0.228 | 1 | 13 | Pkd2 | 0.133 | Old |
| 0.066568 | 0.159624 | 1 | 0.947 | 1 | 13 | Capza1 | 0.053 | Old |
| 0.06657 | 0.092257 | 1 | 1 | 1 | 13 | Capzb | 0 | Old |
| 0.066642 | 1.232638 | 0.049 | 0.009 | 1 | 13 | Gm36839 | 0.04 | Old |
| 0.066642 | 1.118396 | 0.049 | 0.009 | 1 | 13 | Gm39459 | 0.04 | Old |
| 0.066642 | 1.033591 | 0.049 | 0.009 | 1 | 13 | Nap1l2 | 0.04 | Old |
| 0.066818 | 0.62464 | 0.238 | 0.14 | 1 | 13 | Ntmt1 | 0.098 | Old |
| 0.066859 | 0.341144 | 0.516 | 0.368 | 1 | 13 | Aspscr1 | 0.148 | Old |
| 0.066859 | 0.447626 | 0.328 | 0.193 | 1 | 13 | Med19 | 0.135 | Old |
| 0.066882 | 1.070536 | 0.107 | 0.044 | 1 | 13 | Tha1 | 0.063 | Old |
| 0.066882 | 1.055258 | 0.107 | 0.044 | 1 | 13 | Klhl8 | 0.063 | Old |
| 0.066989 | 0.531153 | 0.607 | 0.491 | 1 | 13 | Galnt7 | 0.116 | Old |
| 0.067313 | 0.559334 | 0.426 | 0.307 | 1 | 13 | A4galt | 0.119 | Old |
| 0.067485 | 0.521507 | 0.197 | 0.105 | 1 | 13 | Zbtb45 | 0.092 | Old |
| 0.067565 | 0.945522 | 0.156 | 0.079 | 1 | 13 | Helz2 | 0.077 | Old |
| 0.067675 | 0.134923 | 0.959 | 0.912 | 1 | 13 | Ctnnb1 | 0.047 | Old |
| 0.067678 | 0.981346 | 0.148 | 0.07 | 1 | 13 | Tnfsf14 | 0.078 | Old |
| 0.06771 | 0.170668 | 1 | 1 | 1 | 13 | Calm2 | 0 | Old |
| 0.067761 | 0.8876 | 0.082 | 0.026 | 1 | 13 | Harbi1 | 0.056 | Old |
| 0.067902 | 0.559735 | 0.377 | 0.281 | 1 | 13 | Zfp991 | 0.096 | Old |
| 0.068267 | 0.155622 | 0.951 | 0.947 | 1 | 13 | Prkar1a | 0.004 | Old |
| 0.06827 | 1.090166 | 0.066 | 0.018 | 1 | 13 | Zfp51 | 0.048 | Old |
| 0.068347 | 0.34798 | 0.623 | 0.491 | 1 | 13 | Dpp9 | 0.132 | Old |
| 0.068592 | 1.068022 | 0.049 | 0.009 | 1 | 13 | Tns2 | 0.04 | Old |
| 0.068649 | 0.442772 | 0.492 | 0.368 | 1 | 13 | Nop16 | 0.124 | Old |
| 0.068679 | 0.345232 | 0.738 | 0.649 | 1 | 13 | Cd200r1 | 0.089 | Old |
| 0.06898 | 0.439078 | 0.656 | 0.535 | 1 | 13 | 1810030O( | 0.121 | Old |
| 0.069081 | 0.299943 | 0.795 | 0.684 | 1 | 13 | Fam126a | 0.111 | Old |
| 0.069109 | 0.771306 | 0.156 | 0.079 | 1 | 13 | Vps33b | 0.077 | Old |
| 0.069134 | 0.274031 | 0.934 | 0.956 | 1 | 13 | Serpinb6a | -0.022 | Old |
| 0.069449 | 0.750122 | 0.254 | 0.158 | 1 | 13 | Foxred1 | 0.096 | Old |
| 0.069658 | 0.547204 | 0.32 | 0.202 | 1 | 13 | Rnf40 | 0.118 | Old |
| 0.069772 | 0.312516 | 0.615 | 0.447 | 1 | 13 | Rapgef2 | 0.168 | Old |
| 0.069804 | 0.744695 | 0.197 | 0.105 | 1 | 13 | Abca2 | 0.092 | Old |
| 0.069847 | 0.390599 | 0.639 | 0.526 | 1 | 13 | Slc1a5 | 0.113 | Old |

| | | | | | | | | |
|---|---|---|---|---|---|---|---|---|
| 0.069939 | 0.261847 | 0.934 | 0.886 | 1 | 13 | Trip12 | 0.048 | Old |
| 0.070033 | 0.208922 | 0.77 | 0.623 | 1 | 13 | Araf | 0.147 | Old |
| 0.070093 | 0.258178 | 0.861 | 0.798 | 1 | 13 | Glmp | 0.063 | Old |
| 0.07021 | 0.428046 | 0.77 | 0.649 | 1 | 13 | Gfpt1 | 0.121 | Old |
| 0.070252 | 0.452887 | 0.451 | 0.316 | 1 | 13 | Akr7a5 | 0.135 | Old |
| 0.070312 | 0.413901 | 0.5 | 0.368 | 1 | 13 | Gtf2h1 | 0.132 | Old |
| 0.070314 | 0.434927 | 0.295 | 0.175 | 1 | 13 | Ikbkap | 0.12 | Old |
| 0.070329 | 1.083354 | 0.205 | 0.123 | 1 | 13 | Iigp1 | 0.082 | Old |
| 0.070371 | 0.263463 | 0.869 | 0.763 | 1 | 13 | Ndufb2 | 0.106 | Old |
| 0.070446 | 0.448088 | 0.459 | 0.316 | 1 | 13 | Trp53inp2 | 0.143 | Old |
| 0.070495 | 0.37057 | 0.426 | 0.281 | 1 | 13 | Naa30 | 0.145 | Old |
| 0.070551 | 0.37925 | 0.664 | 0.482 | 1 | 13 | Clk3 | 0.182 | Old |
| 0.070569 | 0.466123 | 0.467 | 0.342 | 1 | 13 | Tk2 | 0.125 | Old |
| 0.070589 | 1.016637 | 0.049 | 0.009 | 1 | 13 | Rflnb | 0.04 | Old |
| 0.070589 | 0.982184 | 0.049 | 0.009 | 1 | 13 | 2010109A1 | 0.04 | Old |
| 0.070589 | 0.918654 | 0.049 | 0.009 | 1 | 13 | Gm11131 | 0.04 | Old |
| 0.070589 | 0.900199 | 0.049 | 0.009 | 1 | 13 | Ccsap | 0.04 | Old |
| 0.070763 | 0.350948 | 0.377 | 0.237 | 1 | 13 | Mtf1 | 0.14 | Old |
| 0.070791 | 1.044467 | 0.066 | 0.018 | 1 | 13 | Airn | 0.048 | Old |
| 0.070791 | 1.019229 | 0.066 | 0.018 | 1 | 13 | Gm9803 | 0.048 | Old |
| 0.070959 | 0.779693 | 0.246 | 0.158 | 1 | 13 | E2f3 | 0.088 | Old |
| 0.071054 | 0.24975 | 0.828 | 0.728 | 1 | 13 | Flii | 0.1 | Old |
| 0.07119 | 0.555667 | 0.23 | 0.132 | 1 | 13 | Fig4 | 0.098 | Old |
| 0.071472 | 0.796724 | 0.082 | 0.026 | 1 | 13 | Lysmd1 | 0.056 | Old |
| 0.07157 | 0.363028 | 0.41 | 0.281 | 1 | 13 | Appl2 | 0.129 | Old |
| 0.071591 | 0.635086 | 0.385 | 0.272 | 1 | 13 | Dtx2 | 0.113 | Old |
| 0.071605 | 0.880086 | 0.049 | 0.009 | 1 | 13 | 9130019O2 | 0.04 | Old |
| 0.071605 | 0.752768 | 0.049 | 0.009 | 1 | 13 | Dennd6b | 0.04 | Old |
| 0.071624 | 0.698377 | 0.279 | 0.184 | 1 | 13 | Alg13 | 0.095 | Old |
| 0.07167 | 0.527478 | 0.5 | 0.404 | 1 | 13 | Cdk17 | 0.096 | Old |
| 0.071706 | 0.474038 | 0.197 | 0.105 | 1 | 13 | Rcc1 | 0.092 | Old |
| 0.071803 | 0.466 | 0.475 | 0.368 | 1 | 13 | Rps19bp1 | 0.107 | Old |
| 0.07184 | 0.351923 | 0.779 | 0.675 | 1 | 13 | Smarca4 | 0.104 | Old |
| 0.072176 | 0.253803 | 0.656 | 0.474 | 1 | 13 | Ccdc22 | 0.182 | Old |
| 0.072228 | 0.428384 | 0.361 | 0.254 | 1 | 13 | Rexo4 | 0.107 | Old |
| 0.072259 | 0.57094 | 0.238 | 0.132 | 1 | 13 | D230025D1 | 0.106 | Old |
| 0.072295 | 0.322893 | 0.516 | 0.351 | 1 | 13 | Malsu1 | 0.165 | Old |
| 0.072327 | 0.323245 | 0.59 | 0.465 | 1 | 13 | Sash3 | 0.125 | Old |
| 0.0724 | 0.200596 | 0.934 | 0.947 | 1 | 13 | Atp6v1a | -0.013 | Old |
| 0.072513 | 1.014808 | 0.066 | 0.018 | 1 | 13 | Xrcc2 | 0.048 | Old |
| 0.072513 | 0.847069 | 0.066 | 0.018 | 1 | 13 | Pcdh7 | 0.048 | Old |
| 0.072513 | 0.807069 | 0.066 | 0.018 | 1 | 13 | Tmem144 | 0.048 | Old |
| 0.072615 | 0.488362 | 0.32 | 0.211 | 1 | 13 | Cog6 | 0.109 | Old |
| 0.072647 | 0.403664 | 0.475 | 0.351 | 1 | 13 | Shc1 | 0.124 | Old |
| 0.07287 | 0.343277 | 0.705 | 0.535 | 1 | 13 | Rhoq | 0.17 | Old |

| | | | | | | | | |
|---|---|---|---|---|---|---|---|---|
| 0.073256 | 0.277054 | 0.623 | 0.482 | 1 | 13 | Fnta | 0.141 | Old |
| 0.073318 | 0.328375 | 0.361 | 0.237 | 1 | 13 | Akr1e1 | 0.124 | Old |
| 0.07337 | 0.764944 | 0.156 | 0.079 | 1 | 13 | Ttyh2 | 0.077 | Old |
| 0.073387 | 0.858536 | 0.066 | 0.018 | 1 | 13 | Ppm1l | 0.048 | Old |
| 0.073433 | 0.413979 | 0.648 | 0.518 | 1 | 13 | Lin7c | 0.13 | Old |
| 0.073496 | 0.468883 | 0.5 | 0.377 | 1 | 13 | Agtpbp1 | 0.123 | Old |
| 0.073593 | 0.525091 | 0.213 | 0.114 | 1 | 13 | Sprtn | 0.099 | Old |
| 0.073731 | 0.317619 | 0.828 | 0.737 | 1 | 13 | Fam32a | 0.091 | Old |
| 0.073959 | 0.174443 | 1 | 0.965 | 1 | 13 | Txndc17 | 0.035 | Old |
| 0.073959 | 0.082074 | 1 | 1 | 1 | 13 | Cfl1 | 0 | Old |
| 0.073962 | 0.540874 | 0.123 | 0.053 | 1 | 13 | 2610021A( | 0.07 | Old |
| 0.074076 | 0.263027 | 0.369 | 0.228 | 1 | 13 | Utp14b | 0.141 | Old |
| 0.074131 | 0.330483 | 0.68 | 0.561 | 1 | 13 | Pisd | 0.119 | Old |
| 0.07421 | 0.958482 | 0.131 | 0.061 | 1 | 13 | Coq4 | 0.07 | Old |
| 0.074261 | 0.367642 | 0.557 | 0.43 | 1 | 13 | Akr1c13 | 0.127 | Old |
| 0.074355 | 0.431494 | 0.23 | 0.132 | 1 | 13 | Nek6 | 0.098 | Old |
| 0.074388 | 0.393977 | 0.336 | 0.219 | 1 | 13 | Gtpbp6 | 0.117 | Old |
| 0.074513 | 0.96977 | 0.197 | 0.114 | 1 | 13 | Ankmy2 | 0.083 | Old |
| 0.074588 | 0.441554 | 0.328 | 0.202 | 1 | 13 | Socs5 | 0.126 | Old |
| 0.074619 | 0.544923 | 0.238 | 0.14 | 1 | 13 | Gabpb1 | 0.098 | Old |
| 0.074668 | 0.487211 | 0.467 | 0.342 | 1 | 13 | Hax1 | 0.125 | Old |
| 0.07516 | 0.84766 | 0.066 | 0.018 | 1 | 13 | Sfxn5 | 0.048 | Old |
| 0.075197 | 0.749228 | 0.189 | 0.105 | 1 | 13 | Polm | 0.084 | Old |
| 0.07534 | 0.346923 | 0.328 | 0.193 | 1 | 13 | Wdyhv1 | 0.135 | Old |
| 0.075527 | 0.291243 | 0.623 | 0.482 | 1 | 13 | Rnf5 | 0.141 | Old |
| 0.075534 | 0.321735 | 0.803 | 0.702 | 1 | 13 | Pitpnc1 | 0.101 | Old |
| 0.075786 | 0.363865 | 0.541 | 0.404 | 1 | 13 | Tusc3 | 0.137 | Old |
| 0.075793 | 0.432878 | 0.32 | 0.211 | 1 | 13 | Fopnl | 0.109 | Old |
| 0.075902 | 0.219262 | 0.762 | 0.684 | 1 | 13 | Rela | 0.078 | Old |
| 0.075977 | 0.559442 | 0.516 | 0.404 | 1 | 13 | Sh3glb2 | 0.112 | Old |
| 0.076212 | 0.846458 | 0.172 | 0.096 | 1 | 13 | Hmga1 | 0.076 | Old |
| 0.076265 | 0.293951 | 0.492 | 0.351 | 1 | 13 | Rbm18 | 0.141 | Old |
| 0.076272 | 0.307062 | 0.574 | 0.412 | 1 | 13 | Spopl | 0.162 | Old |
| 0.076338 | 0.186431 | 0.885 | 0.825 | 1 | 13 | Copb2 | 0.06 | Old |
| 0.076545 | 0.300967 | 0.705 | 0.561 | 1 | 13 | Frrs1 | 0.144 | Old |
| 0.076936 | 0.786482 | 0.246 | 0.149 | 1 | 13 | Plac8 | 0.097 | Old |
| 0.077027 | 0.438734 | 0.5 | 0.351 | 1 | 13 | Ppip5k2 | 0.149 | Old |
| 0.077105 | 1.14995 | 0.115 | 0.053 | 1 | 13 | Cfap45 | 0.062 | Old |
| 0.077137 | 0.260244 | 0.738 | 0.596 | 1 | 13 | Zfp292 | 0.142 | Old |
| 0.077327 | 0.184924 | 0.475 | 0.333 | 1 | 13 | Uspl1 | 0.142 | Old |
| 0.077416 | 0.156299 | 1 | 1 | 1 | 13 | Unc93b1 | 0 | Old |
| 0.077635 | 0.539529 | 0.23 | 0.132 | 1 | 13 | Mad2l1bp | 0.098 | Old |
| 0.077888 | 0.245893 | 0.82 | 0.693 | 1 | 13 | Naa15 | 0.127 | Old |
| 0.077929 | 0.322506 | 0.82 | 0.754 | 1 | 13 | Pik3r5 | 0.066 | Old |
| 0.078035 | 0.303139 | 0.697 | 0.605 | 1 | 13 | Plxna1 | 0.092 | Old |

| | | | | | | | | |
|---|---|---|---|---|---|---|---|---|
| 0.078044 | 0.170884 | 0.811 | 0.728 | 1 | 13 | Ddx46 | 0.083 | Old |
| 0.078058 | 0.222538 | 0.984 | 0.982 | 1 | 13 | Cd68 | 0.002 | Old |
| 0.078086 | 0.534271 | 0.426 | 0.316 | 1 | 13 | Zfp219 | 0.11 | Old |
| 0.078242 | 0.387337 | 0.18 | 0.096 | 1 | 13 | Tmem35b | 0.084 | Old |
| 0.078338 | 0.22849 | 0.828 | 0.754 | 1 | 13 | Golph3 | 0.074 | Old |
| 0.078357 | 0.206415 | 0.975 | 0.912 | 1 | 13 | Surf4 | 0.063 | Old |
| 0.0787 | 0.35434 | 0.197 | 0.105 | 1 | 13 | Pex1 | 0.092 | Old |
| 0.078734 | 0.324841 | 0.672 | 0.561 | 1 | 13 | Shoc2 | 0.111 | Old |
| 0.078779 | 0.784945 | 0.18 | 0.096 | 1 | 13 | Dcun1d2 | 0.084 | Old |
| 0.078983 | 0.254756 | 0.525 | 0.412 | 1 | 13 | Ccdc34 | 0.113 | Old |
| 0.079016 | 0.19653 | 0.713 | 0.553 | 1 | 13 | Usp48 | 0.16 | Old |
| 0.079194 | 0.396457 | 0.639 | 0.544 | 1 | 13 | Sgta | 0.095 | Old |
| 0.079502 | 0.496208 | 0.746 | 0.667 | 1 | 13 | Nceh1 | 0.079 | Old |
| 0.079669 | 0.25832 | 0.959 | 0.947 | 1 | 13 | Spcs1 | 0.012 | Old |
| 0.079754 | 0.421609 | 0.311 | 0.193 | 1 | 13 | Xxylt1 | 0.118 | Old |
| 0.079849 | 0.300706 | 0.893 | 0.807 | 1 | 13 | Trim8 | 0.086 | Old |
| 0.079923 | 0.399651 | 0.574 | 0.421 | 1 | 13 | Lpcat3 | 0.153 | Old |
| 0.080089 | 0.310228 | 0.402 | 0.263 | 1 | 13 | Tmub2 | 0.139 | Old |
| 0.080163 | 0.185424 | 0.91 | 0.877 | 1 | 13 | Papola | 0.033 | Old |
| 0.08022 | 0.230686 | 0.902 | 0.807 | 1 | 13 | Snx10 | 0.095 | Old |
| 0.080342 | 0.162886 | 1 | 1 | 1 | 13 | Irf2bp2 | 0 | Old |
| 0.080342 | 0.121284 | 1 | 1 | 1 | 13 | Grn | 0 | Old |
| 0.080376 | 0.965776 | 0.139 | 0.07 | 1 | 13 | Prmt6 | 0.069 | Old |
| 0.080554 | 0.196779 | 0.861 | 0.728 | 1 | 13 | Vma21 | 0.133 | Old |
| 0.081115 | 0.638061 | 0.197 | 0.114 | 1 | 13 | Mov10 | 0.083 | Old |
| 0.081237 | 0.43676 | 0.393 | 0.263 | 1 | 13 | Taf2 | 0.13 | Old |
| 0.081325 | 0.327753 | 0.779 | 0.711 | 1 | 13 | Acaa2 | 0.068 | Old |
| 0.081337 | 0.57064 | 0.426 | 0.325 | 1 | 13 | Zswim4 | 0.101 | Old |
| 0.081341 | 0.708911 | 0.156 | 0.079 | 1 | 13 | Sumf2 | 0.077 | Old |
| 0.08151 | 0.523914 | 0.18 | 0.096 | 1 | 13 | Cdk20 | 0.084 | Old |
| 0.081553 | 0.518184 | 0.443 | 0.333 | 1 | 13 | Abhd17c | 0.11 | Old |
| 0.081596 | 0.27999 | 0.869 | 0.789 | 1 | 13 | Dock7 | 0.08 | Old |
| 0.081818 | 0.406799 | 0.689 | 0.588 | 1 | 13 | Arhgap10 | 0.101 | Old |
| 0.081845 | 0.968326 | 0.115 | 0.053 | 1 | 13 | Tnnt1 | 0.062 | Old |
| 0.082311 | 0.492231 | 0.254 | 0.158 | 1 | 13 | Mcm2 | 0.096 | Old |
| 0.082715 | 0.5318 | 0.418 | 0.307 | 1 | 13 | Zfp322a | 0.111 | Old |
| 0.082826 | 0.382499 | 0.689 | 0.544 | 1 | 13 | Gtf3c2 | 0.145 | Old |
| 0.082826 | 0.271933 | 0.705 | 0.605 | 1 | 13 | Dlgap4 | 0.1 | Old |
| 0.082844 | 0.607717 | 0.287 | 0.184 | 1 | 13 | Tfec | 0.103 | Old |
| 0.083132 | 0.638559 | 0.238 | 0.149 | 1 | 13 | Card9 | 0.089 | Old |
| 0.083279 | 0.4581 | 0.393 | 0.281 | 1 | 13 | Itpa | 0.112 | Old |
| 0.083296 | 0.511382 | 0.566 | 0.465 | 1 | 13 | Kcnq1ot1 | 0.101 | Old |
| 0.083324 | 0.305123 | 0.754 | 0.614 | 1 | 13 | Rara | 0.14 | Old |
| 0.083384 | 0.67446 | 0.221 | 0.132 | 1 | 13 | Sbk1 | 0.089 | Old |
| 0.083616 | 0.220609 | 0.852 | 0.868 | 1 | 13 | Cct3 | -0.016 | Old |

| | | | | | | | | |
|---|---|---|---|---|---|---|---|---|
| 0.083947 | 0.962622 | 0.115 | 0.053 | 1 | 13 | Snx33 | 0.062 | Old |
| 0.084037 | 0.400303 | 0.303 | 0.193 | 1 | 13 | Rbmx | 0.11 | Old |
| 0.084361 | 0.236223 | 0.967 | 0.93 | 1 | 13 | Eif1a | 0.037 | Old |
| 0.084889 | 0.268066 | 0.73 | 0.64 | 1 | 13 | Rps27rt | 0.09 | Old |
| 0.084916 | 0.298497 | 0.738 | 0.596 | 1 | 13 | Chmp1a | 0.142 | Old |
| 0.084926 | 0.642522 | 0.221 | 0.132 | 1 | 13 | Rhbdd2 | 0.089 | Old |
| 0.084957 | 0.697216 | 0.164 | 0.088 | 1 | 13 | Nxn | 0.076 | Old |
| 0.085018 | 0.637214 | 0.131 | 0.061 | 1 | 13 | Zfp747 | 0.07 | Old |
| 0.085018 | 0.430245 | 0.131 | 0.061 | 1 | 13 | Hells | 0.07 | Old |
| 0.085028 | 0.634243 | 0.811 | 0.737 | 1 | 13 | H2-Q7 | 0.074 | Old |
| 0.085184 | 0.930635 | 0.09 | 0.035 | 1 | 13 | Tmem132a | 0.055 | Old |
| 0.085338 | 0.305765 | 0.664 | 0.561 | 1 | 13 | Tmem165 | 0.103 | Old |
| 0.085445 | 0.721947 | 0.139 | 0.07 | 1 | 13 | Fbxo21 | 0.069 | Old |
| 0.085751 | 0.212463 | 0.852 | 0.728 | 1 | 13 | Ppp4r2 | 0.124 | Old |
| 0.085757 | 0.31528 | 0.902 | 0.833 | 1 | 13 | Spred1 | 0.069 | Old |
| 0.085759 | 0.590284 | 0.336 | 0.237 | 1 | 13 | Psmg1 | 0.099 | Old |
| 0.085762 | 0.108461 | 0.967 | 0.982 | 1 | 13 | Pum2 | -0.015 | Old |
| 0.085877 | 0.502151 | 0.566 | 0.456 | 1 | 13 | Uchl5 | 0.11 | Old |
| 0.085899 | 0.701925 | 0.23 | 0.14 | 1 | 13 | Alg6 | 0.09 | Old |
| 0.086078 | 0.389403 | 0.574 | 0.439 | 1 | 13 | Larp7 | 0.135 | Old |
| 0.086104 | 0.189218 | 0.984 | 0.939 | 1 | 13 | Atp6v0d1 | 0.045 | Old |
| 0.087371 | 0.839421 | 0.164 | 0.088 | 1 | 13 | Fahd1 | 0.076 | Old |
| 0.087548 | 0.514673 | 0.344 | 0.237 | 1 | 13 | Klra2 | 0.107 | Old |
| 0.087628 | 0.366574 | 0.434 | 0.307 | 1 | 13 | Tstd3 | 0.127 | Old |
| 0.087652 | 0.338834 | 0.566 | 0.395 | 1 | 13 | Fbxo3 | 0.171 | Old |
| 0.087668 | 0.860844 | 0.09 | 0.035 | 1 | 13 | Tmem263 | 0.055 | Old |
| 0.087668 | 0.8448 | 0.09 | 0.035 | 1 | 13 | A930007I1 | 0.055 | Old |
| 0.087677 | 0.178615 | 0.779 | 0.684 | 1 | 13 | Slc26a2 | 0.095 | Old |
| 0.088074 | 0.769014 | 0.139 | 0.07 | 1 | 13 | Zfp810 | 0.069 | Old |
| 0.088433 | 0.517494 | 0.287 | 0.184 | 1 | 13 | Sco2 | 0.103 | Old |
| 0.088584 | 0.471055 | 0.377 | 0.263 | 1 | 13 | Coq8b | 0.114 | Old |
| 0.088813 | 0.208391 | 0.934 | 0.877 | 1 | 13 | Vapa | 0.057 | Old |
| 0.089022 | 1.612916 | 0.115 | 0.053 | 1 | 13 | Hp | 0.062 | Old |
| 0.089093 | 0.27348 | 0.393 | 0.272 | 1 | 13 | Smurf2 | 0.121 | Old |
| 0.089176 | 0.475184 | 0.525 | 0.395 | 1 | 13 | Jmjd6 | 0.13 | Old |
| 0.089301 | 0.157755 | 1 | 1 | 1 | 13 | mt-Nd4l | 0 | Old |
| 0.089301 | 0.102662 | 1 | 1 | 1 | 13 | Npc2 | 0 | Old |
| 0.089355 | 0.752493 | 0.09 | 0.035 | 1 | 13 | Zfp251 | 0.055 | Old |
| 0.089417 | 0.490552 | 0.279 | 0.184 | 1 | 13 | Zbtb2 | 0.095 | Old |
| 0.089484 | 0.872742 | 0.18 | 0.105 | 1 | 13 | Kpna2 | 0.075 | Old |
| 0.089549 | 0.663072 | 0.23 | 0.14 | 1 | 13 | Gin1 | 0.09 | Old |
| 0.08969 | 0.527059 | 0.221 | 0.132 | 1 | 13 | Mill2 | 0.089 | Old |
| 0.089789 | 0.386049 | 0.238 | 0.14 | 1 | 13 | Nedd4l | 0.098 | Old |
| 0.089808 | 0.172886 | 0.902 | 0.877 | 1 | 13 | Ddx39b | 0.025 | Old |
| 0.090098 | 0.605596 | 0.246 | 0.149 | 1 | 13 | Gfm2 | 0.097 | Old |

| | | | | | | | | |
|---|---|---|---|---|---|---|---|---|
| 0.090209 | 0.799682 | 0.09 | 0.035 | 1 | 13 | A730063M | 0.055 | Old |
| 0.090317 | 0.616922 | 0.23 | 0.149 | 1 | 13 | Lrrc40 | 0.081 | Old |
| 0.090384 | 0.114532 | 0.984 | 0.982 | 1 | 13 | Romo1 | 0.002 | Old |
| 0.090516 | 0.864352 | 0.115 | 0.053 | 1 | 13 | Eid2b | 0.062 | Old |
| 0.090532 | 0.331355 | 0.336 | 0.219 | 1 | 13 | AC127341. | 0.117 | Old |
| 0.090602 | 0.411113 | 0.623 | 0.526 | 1 | 13 | Fchsd2 | 0.097 | Old |
| 0.090832 | 0.380544 | 0.508 | 0.404 | 1 | 13 | Dcaf13 | 0.104 | Old |
| 0.090883 | 0.677347 | 0.279 | 0.184 | 1 | 13 | Sirpb1b | 0.095 | Old |
| 0.090953 | 0.378299 | 0.434 | 0.316 | 1 | 13 | Isoc1 | 0.118 | Old |
| 0.091045 | 0.803066 | 0.148 | 0.079 | 1 | 13 | 2700038G2 | 0.069 | Old |
| 0.091069 | 0.761008 | 0.09 | 0.035 | 1 | 13 | E230032D2 | 0.055 | Old |
| 0.091151 | 0.589491 | 0.23 | 0.14 | 1 | 13 | Cutc | 0.09 | Old |
| 0.091203 | 1.14716 | 0.074 | 0.026 | 1 | 13 | Arg2 | 0.048 | Old |
| 0.091236 | 0.557019 | 0.131 | 0.061 | 1 | 13 | Mdm1 | 0.07 | Old |
| 0.091434 | 0.238323 | 0.738 | 0.675 | 1 | 13 | Hsd17b12 | 0.063 | Old |
| 0.091794 | 0.34325 | 0.434 | 0.316 | 1 | 13 | Rps6kc1 | 0.118 | Old |
| 0.091846 | 0.116883 | 1 | 1 | 1 | 13 | Hsp90b1 | 0 | Old |
| 0.092387 | 0.726142 | 0.205 | 0.123 | 1 | 13 | Macrod1 | 0.082 | Old |
| 0.092505 | 0.319834 | 0.475 | 0.342 | 1 | 13 | Nmd3 | 0.133 | Old |
| 0.092622 | 0.393173 | 0.484 | 0.351 | 1 | 13 | Rwdd4a | 0.133 | Old |
| 0.092855 | 0.564603 | 0.213 | 0.132 | 1 | 13 | Dolk | 0.081 | Old |
| 0.092917 | 0.583122 | 0.328 | 0.237 | 1 | 13 | Cdc14b | 0.091 | Old |
| 0.092931 | 0.585552 | 0.41 | 0.307 | 1 | 13 | Gstp1 | 0.103 | Old |
| 0.092954 | 0.167978 | 1 | 0.991 | 1 | 13 | Rnf130 | 0.009 | Old |
| 0.092961 | 0.454918 | 0.344 | 0.237 | 1 | 13 | Gnpat | 0.107 | Old |
| 0.093638 | 3.982663 | 0.025 | 0 | 1 | 13 | Wfdc21 | 0.025 | Old |
| 0.093638 | 2.087221 | 0.025 | 0 | 1 | 13 | Ctla4 | 0.025 | Old |
| 0.093638 | 1.834254 | 0.025 | 0 | 1 | 13 | Ksr2 | 0.025 | Old |
| 0.093638 | 1.491699 | 0.025 | 0 | 1 | 13 | Mgst2 | 0.025 | Old |
| 0.093638 | 1.331528 | 0.025 | 0 | 1 | 13 | Cndp1 | 0.025 | Old |
| 0.093638 | 1.265467 | 0.025 | 0 | 1 | 13 | Trnp1 | 0.025 | Old |
| 0.093638 | 1.208726 | 0.025 | 0 | 1 | 13 | Foxq1 | 0.025 | Old |
| 0.093638 | 1.186418 | 0.025 | 0 | 1 | 13 | C77080 | 0.025 | Old |
| 0.093638 | 1.183095 | 0.025 | 0 | 1 | 13 | Gimap9 | 0.025 | Old |
| 0.093638 | 1.167854 | 0.025 | 0 | 1 | 13 | Gm35339 | 0.025 | Old |
| 0.093638 | 1.160227 | 0.025 | 0 | 1 | 13 | Hmgb3 | 0.025 | Old |
| 0.093638 | 1.156514 | 0.025 | 0 | 1 | 13 | D030051J2 | 0.025 | Old |
| 0.093638 | 1.13938 | 0.025 | 0 | 1 | 13 | Vpreb1 | 0.025 | Old |
| 0.093638 | 1.138683 | 0.025 | 0 | 1 | 13 | Hid1 | 0.025 | Old |
| 0.093638 | 1.136773 | 0.025 | 0 | 1 | 13 | Gm16152 | 0.025 | Old |
| 0.093638 | 1.093278 | 0.025 | 0 | 1 | 13 | Gm16794 | 0.025 | Old |
| 0.093638 | 1.091396 | 0.025 | 0 | 1 | 13 | Rhov | 0.025 | Old |
| 0.093638 | 1.087116 | 0.025 | 0 | 1 | 13 | Gm10382 | 0.025 | Old |
| 0.093638 | 1.080603 | 0.025 | 0 | 1 | 13 | Gm17509 | 0.025 | Old |
| 0.093638 | 1.079177 | 0.025 | 0 | 1 | 13 | Slc23a1 | 0.025 | Old |

| | | | | | | | | |
|---|---|---|---|---|---|---|---|---|
| 0.093638 | 1.078533 | 0.025 | 0 | 1 | 13 | Efcc1 | 0.025 | Old |
| 0.093638 | 1.076442 | 0.025 | 0 | 1 | 13 | Nol3 | 0.025 | Old |
| 0.093638 | 1.07018 | 0.025 | 0 | 1 | 13 | Mkrn3 | 0.025 | Old |
| 0.093638 | 1.066883 | 0.025 | 0 | 1 | 13 | Gpr27 | 0.025 | Old |
| 0.093638 | 1.064126 | 0.025 | 0 | 1 | 13 | A130051J0 | 0.025 | Old |
| 0.093638 | 1.060326 | 0.025 | 0 | 1 | 13 | Gm26620 | 0.025 | Old |
| 0.093638 | 1.059224 | 0.025 | 0 | 1 | 13 | Gm26533 | 0.025 | Old |
| 0.093638 | 1.042252 | 0.025 | 0 | 1 | 13 | Tmem132c | 0.025 | Old |
| 0.093638 | 1.039101 | 0.025 | 0 | 1 | 13 | Zfand4 | 0.025 | Old |
| 0.093638 | 1.035109 | 0.025 | 0 | 1 | 13 | Gm35439 | 0.025 | Old |
| 0.093638 | 1.03147 | 0.025 | 0 | 1 | 13 | Rasl10a | 0.025 | Old |
| 0.093638 | 1.02668 | 0.025 | 0 | 1 | 13 | Gng7 | 0.025 | Old |
| 0.093638 | 1.026236 | 0.025 | 0 | 1 | 13 | Acoxl | 0.025 | Old |
| 0.093638 | 1.021334 | 0.025 | 0 | 1 | 13 | Slc9a5 | 0.025 | Old |
| 0.093638 | 1.012137 | 0.025 | 0 | 1 | 13 | Nxnl1 | 0.025 | Old |
| 0.093638 | 1.010069 | 0.025 | 0 | 1 | 13 | C230037L1 | 0.025 | Old |
| 0.093638 | 0.999115 | 0.025 | 0 | 1 | 13 | Hap1 | 0.025 | Old |
| 0.093638 | 0.994286 | 0.025 | 0 | 1 | 13 | Arhgap28 | 0.025 | Old |
| 0.093638 | 0.982238 | 0.025 | 0 | 1 | 13 | Slc5a11 | 0.025 | Old |
| 0.093638 | 0.977814 | 0.025 | 0 | 1 | 13 | Kdm8 | 0.025 | Old |
| 0.093638 | 0.975437 | 0.025 | 0 | 1 | 13 | Slc22a23 | 0.025 | Old |
| 0.093638 | 0.975393 | 0.025 | 0 | 1 | 13 | Traip | 0.025 | Old |
| 0.093638 | 0.969285 | 0.025 | 0 | 1 | 13 | Cdon | 0.025 | Old |
| 0.093638 | 0.961949 | 0.025 | 0 | 1 | 13 | Polq | 0.025 | Old |
| 0.093638 | 0.95989 | 0.025 | 0 | 1 | 13 | Bcar1 | 0.025 | Old |
| 0.093638 | 0.950403 | 0.025 | 0 | 1 | 13 | Gm29562 | 0.025 | Old |
| 0.093638 | 0.949169 | 0.025 | 0 | 1 | 13 | Gm14029 | 0.025 | Old |
| 0.093638 | 0.944595 | 0.025 | 0 | 1 | 13 | Pcdhga9 | 0.025 | Old |
| 0.093638 | 0.94336 | 0.025 | 0 | 1 | 13 | Vasn | 0.025 | Old |
| 0.093638 | 0.941477 | 0.025 | 0 | 1 | 13 | Plxnb1 | 0.025 | Old |
| 0.093638 | 0.930726 | 0.025 | 0 | 1 | 13 | 1700029I1 | 0.025 | Old |
| 0.093638 | 0.930556 | 0.025 | 0 | 1 | 13 | Gm44206 | 0.025 | Old |
| 0.093638 | 0.928744 | 0.025 | 0 | 1 | 13 | Pcdh17 | 0.025 | Old |
| 0.093638 | 0.924794 | 0.025 | 0 | 1 | 13 | AC113595. | 0.025 | Old |
| 0.093638 | 0.917761 | 0.025 | 0 | 1 | 13 | Slc1a4 | 0.025 | Old |
| 0.093638 | 0.915781 | 0.025 | 0 | 1 | 13 | Igha | 0.025 | Old |
| 0.093638 | 0.903204 | 0.025 | 0 | 1 | 13 | Nrg4 | 0.025 | Old |
| 0.093638 | 0.899592 | 0.025 | 0 | 1 | 13 | Gm31834 | 0.025 | Old |
| 0.093638 | 0.89676 | 0.025 | 0 | 1 | 13 | Gm3435 | 0.025 | Old |
| 0.093638 | 0.895115 | 0.025 | 0 | 1 | 13 | Zfp69 | 0.025 | Old |
| 0.093638 | 0.895027 | 0.025 | 0 | 1 | 13 | Gm26733 | 0.025 | Old |
| 0.093638 | 0.895007 | 0.025 | 0 | 1 | 13 | Gm4262 | 0.025 | Old |
| 0.093638 | 0.89011 | 0.025 | 0 | 1 | 13 | Rgs16 | 0.025 | Old |
| 0.093638 | 0.887927 | 0.025 | 0 | 1 | 13 | Nphp4 | 0.025 | Old |
| 0.093638 | 0.88453 | 0.025 | 0 | 1 | 13 | Dnajb3 | 0.025 | Old |

| | | | | | | | | |
|---|---|---|---|---|---|---|---|---|
| 0.093638 | 0.881871 | 0.025 | 0 | 1 | 13 | A430110L2 | 0.025 | Old |
| 0.093638 | 0.88169 | 0.025 | 0 | 1 | 13 | 1700040D1 | 0.025 | Old |
| 0.093638 | 0.880628 | 0.025 | 0 | 1 | 13 | Gm31410 | 0.025 | Old |
| 0.093638 | 0.877149 | 0.025 | 0 | 1 | 13 | Hunk | 0.025 | Old |
| 0.093638 | 0.872353 | 0.025 | 0 | 1 | 13 | Gm826 | 0.025 | Old |
| 0.093638 | 0.865702 | 0.025 | 0 | 1 | 13 | Mmp17 | 0.025 | Old |
| 0.093638 | 0.861939 | 0.025 | 0 | 1 | 13 | Nanos1 | 0.025 | Old |
| 0.093638 | 0.860214 | 0.025 | 0 | 1 | 13 | Dnhd1 | 0.025 | Old |
| 0.093638 | 0.855649 | 0.025 | 0 | 1 | 13 | Fstl1 | 0.025 | Old |
| 0.093638 | 0.851161 | 0.025 | 0 | 1 | 13 | 4833413G1 | 0.025 | Old |
| 0.093638 | 0.848322 | 0.025 | 0 | 1 | 13 | Ctxn1 | 0.025 | Old |
| 0.093638 | 0.846266 | 0.025 | 0 | 1 | 13 | Pars2 | 0.025 | Old |
| 0.093638 | 0.841284 | 0.025 | 0 | 1 | 13 | Zfp521 | 0.025 | Old |
| 0.093638 | 0.838463 | 0.025 | 0 | 1 | 13 | Gm28505 | 0.025 | Old |
| 0.093638 | 0.828757 | 0.025 | 0 | 1 | 13 | Gm9774 | 0.025 | Old |
| 0.093638 | 0.828555 | 0.025 | 0 | 1 | 13 | Esco2 | 0.025 | Old |
| 0.093638 | 0.82341 | 0.025 | 0 | 1 | 13 | Gm5127 | 0.025 | Old |
| 0.093638 | 0.797991 | 0.025 | 0 | 1 | 13 | 9130604C2 | 0.025 | Old |
| 0.093638 | 0.781628 | 0.025 | 0 | 1 | 13 | Gm26910 | 0.025 | Old |
| 0.093638 | 0.679836 | 0.025 | 0 | 1 | 13 | Zg16 | 0.025 | Old |
| 0.09369 | 0.758677 | 0.09 | 0.035 | 1 | 13 | Hgh1 | 0.055 | Old |
| 0.093828 | 0.473761 | 0.18 | 0.096 | 1 | 13 | Abhd18 | 0.084 | Old |
| 0.093997 | 0.280762 | 0.803 | 0.711 | 1 | 13 | Rap2b | 0.092 | Old |
| 0.094121 | 0.990478 | 0.074 | 0.026 | 1 | 13 | Agbl3 | 0.048 | Old |
| 0.094424 | 0.332377 | 0.475 | 0.325 | 1 | 13 | Ptbp2 | 0.15 | Old |
| 0.094546 | 0.478997 | 0.197 | 0.114 | 1 | 13 | Tinf2 | 0.083 | Old |
| 0.094666 | 0.42739 | 0.221 | 0.132 | 1 | 13 | Aspa | 0.089 | Old |
| 0.094836 | 0.22217 | 0.77 | 0.702 | 1 | 13 | Rnf4 | 0.068 | Old |
| 0.095091 | 0.262313 | 0.303 | 0.184 | 1 | 13 | Synj2 | 0.119 | Old |
| 0.09511 | 0.946311 | 0.074 | 0.026 | 1 | 13 | Gm26699 | 0.048 | Old |
| 0.09522 | 0.366009 | 0.607 | 0.474 | 1 | 13 | Fuca2 | 0.133 | Old |
| 0.095298 | 0.140093 | 0.352 | 0.219 | 1 | 13 | Hsdl2 | 0.133 | Old |
| 0.095421 | 0.271471 | 0.869 | 0.833 | 1 | 13 | Glrx3 | 0.036 | Old |
| 0.095477 | 0.557628 | 0.361 | 0.254 | 1 | 13 | Ppp2r3c | 0.107 | Old |
| 0.095736 | 0.545652 | 0.287 | 0.193 | 1 | 13 | Coq10a | 0.094 | Old |
| 0.095772 | 0.673736 | 0.262 | 0.167 | 1 | 13 | Pwp1 | 0.095 | Old |
| 0.095907 | 0.943619 | 0.115 | 0.053 | 1 | 13 | Ifi205 | 0.062 | Old |
| 0.096063 | 0.320622 | 0.541 | 0.404 | 1 | 13 | Gtf2f1 | 0.137 | Old |
| 0.09637 | 0.657963 | 0.09 | 0.035 | 1 | 13 | Cyp2s1 | 0.055 | Old |
| 0.096523 | 0.456621 | 0.426 | 0.307 | 1 | 13 | Nt5c2 | 0.119 | Old |
| 0.096684 | 0.313927 | 0.951 | 0.921 | 1 | 13 | Pla2g7 | 0.03 | Old |
| 0.096802 | 0.242544 | 0.656 | 0.482 | 1 | 13 | Dicer1 | 0.174 | Old |
| 0.097114 | 0.952821 | 0.074 | 0.026 | 1 | 13 | Zbtb48 | 0.048 | Old |
| 0.097265 | 0.537681 | 0.287 | 0.193 | 1 | 13 | Ampd2 | 0.094 | Old |
| 0.097565 | 0.248596 | 0.787 | 0.737 | 1 | 13 | Psmd7 | 0.05 | Old |

| | | | | | | | | |
|---|---|---|---|---|---|---|---|---|
| 0.097584 | 0.638361 | 0.164 | 0.088 | 1 | 13 | Rtn4ip1 | 0.076 | Old |
| 0.097714 | 0.46347 | 0.484 | 0.368 | 1 | 13 | Scarb1 | 0.116 | Old |
| 0.097941 | 0.469926 | 0.205 | 0.114 | 1 | 13 | Fars2 | 0.091 | Old |
| 0.097974 | 0.838424 | 0.23 | 0.149 | 1 | 13 | Angptl4 | 0.081 | Old |
| 0.098128 | 0.973683 | 0.074 | 0.026 | 1 | 13 | Intu | 0.048 | Old |
| 0.098128 | 0.885801 | 0.074 | 0.026 | 1 | 13 | 1110002L0 | 0.048 | Old |
| 0.098294 | 0.556081 | 0.287 | 0.193 | 1 | 13 | Mettl5 | 0.094 | Old |
| 0.098517 | 0.505574 | 0.443 | 0.342 | 1 | 13 | Ppp2cb | 0.101 | Old |
| 0.098521 | 0.577641 | 0.139 | 0.07 | 1 | 13 | Gfi1b | 0.069 | Old |
| 0.09894 | 0.333039 | 0.443 | 0.316 | 1 | 13 | Mtx2 | 0.127 | Old |
| 0.099044 | 0.987778 | 0.098 | 0.044 | 1 | 13 | Sag | 0.054 | Old |
| 0.099048 | 0.13888 | 1 | 1 | 1 | 13 | H2-Eb1 | 0 | Old |
| 0.099163 | 0.240742 | 0.844 | 0.763 | 1 | 13 | Cdipt | 0.081 | Old |
| 0.099224 | 0.35014 | 0.705 | 0.649 | 1 | 13 | Emc3 | 0.056 | Old |
| 0.099254 | 0.719687 | 0.148 | 0.079 | 1 | 13 | 1810011H1 | 0.069 | Old |
| 0.099353 | 0.423505 | 0.279 | 0.175 | 1 | 13 | Gpr89 | 0.104 | Old |
| 0.09942 | 0.378507 | 0.607 | 0.491 | 1 | 13 | Psmc2 | 0.116 | Old |
| 0.099637 | 0.411743 | 0.418 | 0.307 | 1 | 13 | Triap1 | 0.111 | Old |
| 0.09974 | 0.378237 | 0.492 | 0.395 | 1 | 13 | Polg | 0.097 | Old |
| 0.100064 | 0.278816 | 0.828 | 0.772 | 1 | 13 | Psmb7 | 0.056 | Old |
| 0.100181 | 0.904991 | 0.074 | 0.026 | 1 | 13 | Dpys | 0.048 | Old |
| 0.100181 | 0.857324 | 0.074 | 0.026 | 1 | 13 | C920006O1 | 0.048 | Old |
| 0.100222 | 0.412027 | 0.41 | 0.298 | 1 | 13 | Rab6b | 0.112 | Old |
| 0.100225 | 0.164485 | 1 | 1 | 1 | 13 | Gapdh | 0 | Old |
| 0.100288 | 0.363228 | 0.189 | 0.105 | 1 | 13 | Ccdc69 | 0.084 | Old |
| 0.100313 | 0.448753 | 0.484 | 0.368 | 1 | 13 | Usf1 | 0.116 | Old |
| 0.10033 | 0.414058 | 0.369 | 0.263 | 1 | 13 | Ttc13 | 0.106 | Old |
| 0.100401 | 0.573091 | 0.648 | 0.544 | 1 | 13 | Ints6 | 0.104 | Old |
| 0.100422 | 0.121618 | 0.975 | 0.991 | 1 | 13 | Sfpq | -0.016 | Old |
| 0.100506 | 0.517259 | 0.295 | 0.202 | 1 | 13 | Pgpep1 | 0.093 | Old |
| 0.10058 | 0.250196 | 0.82 | 0.754 | 1 | 13 | Txndc9 | 0.066 | Old |
| 0.100592 | 0.554305 | 0.344 | 0.237 | 1 | 13 | 1600002K0 | 0.107 | Old |
| 0.100638 | 0.850265 | 0.123 | 0.061 | 1 | 13 | Itfg2 | 0.062 | Old |
| 0.100665 | 0.603273 | 0.205 | 0.123 | 1 | 13 | Adck5 | 0.082 | Old |
| 0.100666 | 0.270663 | 0.828 | 0.719 | 1 | 13 | Ifi204 | 0.109 | Old |
| 0.100768 | 0.409127 | 0.393 | 0.272 | 1 | 13 | Taf4 | 0.121 | Old |
| 0.100882 | 0.376189 | 0.5 | 0.377 | 1 | 13 | Dck | 0.123 | Old |
| 0.100927 | 0.487133 | 0.189 | 0.105 | 1 | 13 | Atg4a | 0.084 | Old |
| 0.101221 | 0.916533 | 0.074 | 0.026 | 1 | 13 | Spc25 | 0.048 | Old |
| 0.101314 | 1.003341 | 0.197 | 0.123 | 1 | 13 | Cdt1 | 0.074 | Old |
| 0.101316 | 0.3954 | 0.525 | 0.377 | 1 | 13 | 9430038I0 | 0.148 | Old |
| 0.101552 | 0.691933 | 0.115 | 0.053 | 1 | 13 | Slc17a9 | 0.062 | Old |
| 0.102032 | 0.405737 | 0.32 | 0.193 | 1 | 13 | Kctd6 | 0.127 | Old |
| 0.102148 | 0.221676 | 0.959 | 0.895 | 1 | 13 | Prpf38b | 0.064 | Old |
| 0.102266 | 0.354076 | 0.574 | 0.43 | 1 | 13 | Hnrnph3 | 0.144 | Old |

| | | | | | | | | |
|---|---|---|---|---|---|---|---|---|
| 0.102269 | 0.904727 | 0.074 | 0.026 | 1 | 13 | Nynrin | 0.048 | Old |
| 0.102269 | 0.730899 | 0.074 | 0.026 | 1 | 13 | Prps1l3 | 0.048 | Old |
| 0.102297 | 0.282214 | 0.885 | 0.789 | 1 | 13 | Ist1 | 0.096 | Old |
| 0.102639 | 0.34947 | 0.5 | 0.377 | 1 | 13 | Zfp318 | 0.123 | Old |
| 0.102778 | 0.282833 | 0.869 | 0.763 | 1 | 13 | Rab3il1 | 0.106 | Old |
| 0.102832 | 0.239626 | 0.689 | 0.561 | 1 | 13 | Emc2 | 0.128 | Old |
| 0.102876 | 0.482899 | 0.426 | 0.307 | 1 | 13 | Trap1 | 0.119 | Old |
| 0.10298 | 0.157777 | 0.951 | 0.93 | 1 | 13 | Rtn3 | 0.021 | Old |
| 0.103135 | 0.503 | 0.205 | 0.123 | 1 | 13 | Diexf | 0.082 | Old |
| 0.103326 | 0.575757 | 0.074 | 0.026 | 1 | 13 | Bend3 | 0.048 | Old |
| 0.103761 | 0.402836 | 0.205 | 0.123 | 1 | 13 | Rad51b | 0.082 | Old |
| 0.103896 | 0.456394 | 0.566 | 0.456 | 1 | 13 | Nr2f6 | 0.11 | Old |
| 0.103915 | 0.482146 | 0.303 | 0.202 | 1 | 13 | Mief1 | 0.101 | Old |
| 0.10398 | 0.266102 | 0.557 | 0.395 | 1 | 13 | Plcl2 | 0.162 | Old |
| 0.104051 | 0.188524 | 0.115 | 0.053 | 1 | 13 | AW146154 | 0.062 | Old |
| 0.104221 | 0.215138 | 0.967 | 0.956 | 1 | 13 | Arf4 | 0.011 | Old |
| 0.104223 | 0.269186 | 0.828 | 0.684 | 1 | 13 | Clptm1 | 0.144 | Old |
| 0.104251 | 0.272678 | 0.5 | 0.368 | 1 | 13 | Uri1 | 0.132 | Old |
| 0.104265 | 0.310293 | 0.566 | 0.465 | 1 | 13 | Mindy3 | 0.101 | Old |
| 0.104459 | 0.715567 | 0.197 | 0.123 | 1 | 13 | Piga | 0.074 | Old |
| 0.105168 | 0.305643 | 0.467 | 0.351 | 1 | 13 | Prpf31 | 0.116 | Old |
| 0.10523 | 0.630993 | 0.221 | 0.14 | 1 | 13 | Aacs | 0.081 | Old |
| 0.105251 | 0.299454 | 0.566 | 0.447 | 1 | 13 | Zfp62 | 0.119 | Old |
| 0.105309 | 0.278643 | 0.828 | 0.772 | 1 | 13 | Zmat2 | 0.056 | Old |
| 0.10537 | 0.435689 | 0.393 | 0.289 | 1 | 13 | Stxbp5 | 0.104 | Old |
| 0.105382 | 0.349275 | 0.287 | 0.184 | 1 | 13 | Dxo | 0.103 | Old |
| 0.105463 | 0.209292 | 0.721 | 0.579 | 1 | 13 | Eif2a | 0.142 | Old |
| 0.105497 | 0.513203 | 0.189 | 0.105 | 1 | 13 | Dcp1b | 0.084 | Old |
| 0.105749 | 0.480137 | 0.287 | 0.193 | 1 | 13 | Zfp865 | 0.094 | Old |
| 0.105802 | 0.32212 | 0.525 | 0.404 | 1 | 13 | A430005L1 | 0.121 | Old |
| 0.106132 | 0.350545 | 0.492 | 0.36 | 1 | 13 | Stx11 | 0.132 | Old |
| 0.106189 | 0.323349 | 0.516 | 0.386 | 1 | 13 | Crkl | 0.13 | Old |
| 0.106206 | 0.256374 | 0.41 | 0.281 | 1 | 13 | Exoc7 | 0.129 | Old |
| 0.106485 | 0.091759 | 0.893 | 0.895 | 1 | 13 | Emc7 | -0.002 | Old |
| 0.106532 | 0.254896 | 0.451 | 0.307 | 1 | 13 | Gpr155 | 0.144 | Old |
| 0.106693 | 0.142248 | 1 | 1 | 1 | 13 | Sec61g | 0 | Old |
| 0.106958 | 0.259163 | 0.811 | 0.719 | 1 | 13 | Prdx6 | 0.092 | Old |
| 0.107413 | 0.330956 | 0.648 | 0.518 | 1 | 13 | Psmc4 | 0.13 | Old |
| 0.107471 | 0.089492 | 0.852 | 0.754 | 1 | 13 | Sbf2 | 0.098 | Old |
| 0.107624 | 0.269862 | 0.885 | 0.789 | 1 | 13 | Emc10 | 0.096 | Old |
| 0.10764 | 0.764356 | 0.074 | 0.026 | 1 | 13 | Pim2 | 0.048 | Old |
| 0.107987 | 0.824621 | 0.189 | 0.114 | 1 | 13 | Hsd17b7 | 0.075 | Old |
| 0.10812 | 0.31345 | 0.557 | 0.456 | 1 | 13 | Abhd17b | 0.101 | Old |
| 0.108307 | 0.415784 | 0.893 | 0.877 | 1 | 13 | Mif | 0.016 | Old |
| 0.108674 | 0.303406 | 0.844 | 0.833 | 1 | 13 | Sel1l | 0.011 | Old |

| | | | | | | | | |
|---|---|---|---|---|---|---|---|---|
| 0.109143 | 0.503503 | 0.139 | 0.07 | 1 | 13 | Med26 | 0.069 | Old |
| 0.109186 | 0.71921 | 0.238 | 0.158 | 1 | 13 | Ercc6l2 | 0.08 | Old |
| 0.109276 | 1.232713 | 0.057 | 0.018 | 1 | 13 | Gm15247 | 0.039 | Old |
| 0.109276 | 1.036872 | 0.057 | 0.018 | 1 | 13 | Plekhf1 | 0.039 | Old |
| 0.109414 | 0.319319 | 0.738 | 0.658 | 1 | 13 | Ythdf3 | 0.08 | Old |
| 0.109675 | 0.403446 | 0.59 | 0.509 | 1 | 13 | Slc27a1 | 0.081 | Old |
| 0.109723 | 0.4277 | 0.697 | 0.64 | 1 | 13 | Tns3 | 0.057 | Old |
| 0.109949 | 0.875371 | 0.098 | 0.044 | 1 | 13 | Hist1h2ap | 0.054 | Old |
| 0.110084 | 0.34507 | 0.779 | 0.737 | 1 | 13 | Coro1b | 0.042 | Old |
| 0.11032 | 0.463139 | 0.172 | 0.096 | 1 | 13 | Impa2 | 0.076 | Old |
| 0.110339 | 0.614221 | 0.23 | 0.14 | 1 | 13 | Tnfaip8l1 | 0.09 | Old |
| 0.110376 | 0.273764 | 0.828 | 0.737 | 1 | 13 | Otub1 | 0.091 | Old |
| 0.110408 | 0.323201 | 0.943 | 0.912 | 1 | 13 | Lst1 | 0.031 | Old |
| 0.110481 | 0.23493 | 0.828 | 0.728 | 1 | 13 | Vti1b | 0.1 | Old |
| 0.110555 | 0.895756 | 0.057 | 0.018 | 1 | 13 | Knl1 | 0.039 | Old |
| 0.110697 | 0.414357 | 0.459 | 0.342 | 1 | 13 | Fbxl17 | 0.117 | Old |
| 0.110767 | 0.352759 | 0.369 | 0.263 | 1 | 13 | Tmem39a | 0.106 | Old |
| 0.110809 | 0.398829 | 0.393 | 0.289 | 1 | 13 | Fbxo9 | 0.104 | Old |
| 0.110816 | 0.329404 | 0.59 | 0.447 | 1 | 13 | Atf2 | 0.143 | Old |
| 0.110818 | 0.405892 | 0.459 | 0.368 | 1 | 13 | Ddx39 | 0.091 | Old |
| 0.110942 | 0.273618 | 0.631 | 0.518 | 1 | 13 | Smarcb1 | 0.113 | Old |
| 0.110975 | 0.512748 | 0.213 | 0.132 | 1 | 13 | Fen1 | 0.081 | Old |
| 0.111252 | 1.786755 | 0.123 | 0.061 | 1 | 13 | Chil3 | 0.062 | Old |
| 0.111275 | 0.291452 | 0.525 | 0.404 | 1 | 13 | Stoml2 | 0.121 | Old |
| 0.111541 | 0.142995 | 0.975 | 0.974 | 1 | 13 | Ppp2ca | 0.001 | Old |
| 0.111573 | 0.285874 | 0.287 | 0.184 | 1 | 13 | Hps4 | 0.103 | Old |
| 0.11164 | 0.344024 | 0.566 | 0.421 | 1 | 13 | Slc25a28 | 0.145 | Old |
| 0.111845 | 1.096688 | 0.057 | 0.018 | 1 | 13 | Gca | 0.039 | Old |
| 0.111845 | 0.819829 | 0.057 | 0.018 | 1 | 13 | 1700020D( | 0.039 | Old |
| 0.111973 | 0.164121 | 0.992 | 0.991 | 1 | 13 | Eif5a | 0.001 | Old |
| 0.111984 | 0.387137 | 0.336 | 0.237 | 1 | 13 | Tsr1 | 0.099 | Old |
| 0.112001 | 0.34546 | 0.648 | 0.526 | 1 | 13 | Arid1b | 0.122 | Old |
| 0.112376 | 0.70397 | 0.156 | 0.088 | 1 | 13 | 5730480H( | 0.068 | Old |
| 0.112562 | 0.397412 | 0.541 | 0.447 | 1 | 13 | Srp68 | 0.094 | Old |
| 0.112626 | 0.388689 | 0.328 | 0.219 | 1 | 13 | Mlycd | 0.109 | Old |
| 0.112692 | 0.445288 | 0.689 | 0.561 | 1 | 13 | Pira2 | 0.128 | Old |
| 0.112694 | 0.245192 | 0.492 | 0.36 | 1 | 13 | Cmc2 | 0.132 | Old |
| 0.112709 | 0.452937 | 0.311 | 0.211 | 1 | 13 | Snx11 | 0.1 | Old |
| 0.112965 | 0.368467 | 0.451 | 0.333 | 1 | 13 | Tmem185b | 0.118 | Old |
| 0.112984 | 0.328948 | 0.672 | 0.553 | 1 | 13 | Birc2 | 0.119 | Old |
| 0.113019 | 0.269685 | 0.836 | 0.772 | 1 | 13 | Srgap2 | 0.064 | Old |
| 0.113049 | 0.152506 | 0.975 | 0.965 | 1 | 13 | Arpc4 | 0.01 | Old |
| 0.113053 | 0.201833 | 0.984 | 0.991 | 1 | 13 | Atf4 | -0.007 | Old |
| 0.113061 | 0.225206 | 0.738 | 0.614 | 1 | 13 | Gosr2 | 0.124 | Old |
| 0.113143 | 0.903297 | 0.131 | 0.07 | 1 | 13 | Taf9b | 0.061 | Old |

| | | | | | | | | |
|---|---|---|---|---|---|---|---|---|
| 0.113147 | 1.392624 | 0.057 | 0.018 | 1 | 13 | Cfb | 0.039 | Old |
| 0.113147 | 1.068061 | 0.057 | 0.018 | 1 | 13 | Fam83f | 0.039 | Old |
| 0.113147 | 1.015957 | 0.057 | 0.018 | 1 | 13 | Slc25a42 | 0.039 | Old |
| 0.113147 | 1.005047 | 0.057 | 0.018 | 1 | 13 | Gpd1 | 0.039 | Old |
| 0.113147 | 0.940321 | 0.057 | 0.018 | 1 | 13 | Slc9a7 | 0.039 | Old |
| 0.113147 | 0.906515 | 0.057 | 0.018 | 1 | 13 | Iqgap3 | 0.039 | Old |
| 0.113777 | 1.195464 | 0.041 | 0.009 | 1 | 13 | Mapk13 | 0.032 | Old |
| 0.113777 | 1.153664 | 0.041 | 0.009 | 1 | 13 | Mettl4 | 0.032 | Old |
| 0.113777 | 1.140472 | 0.041 | 0.009 | 1 | 13 | Ntng2 | 0.032 | Old |
| 0.114194 | 0.167294 | 0.934 | 0.868 | 1 | 13 | Pkn1 | 0.066 | Old |
| 0.114204 | 0.302306 | 0.713 | 0.658 | 1 | 13 | Mnt | 0.055 | Old |
| 0.114255 | 0.163187 | 0.598 | 0.456 | 1 | 13 | Dld | 0.142 | Old |
| 0.11428 | 0.293544 | 0.836 | 0.728 | 1 | 13 | Uba2 | 0.108 | Old |
| 0.114361 | 0.133928 | 1 | 0.991 | 1 | 13 | Ap2a2 | 0.009 | Old |
| 0.114426 | 0.384915 | 0.516 | 0.43 | 1 | 13 | Cask | 0.086 | Old |
| 0.114462 | 0.90495 | 0.057 | 0.018 | 1 | 13 | 0610040B1 | 0.039 | Old |
| 0.114462 | 0.860164 | 0.057 | 0.018 | 1 | 13 | Cdca5 | 0.039 | Old |
| 0.114556 | 0.316412 | 0.689 | 0.509 | 1 | 13 | Ubr3 | 0.18 | Old |
| 0.11469 | 0.617287 | 0.123 | 0.061 | 1 | 13 | Mocos | 0.062 | Old |
| 0.114995 | 0.23255 | 0.943 | 0.956 | 1 | 13 | Ninj1 | -0.013 | Old |
| 0.115049 | 0.716996 | 0.197 | 0.123 | 1 | 13 | AI597479 | 0.074 | Old |
| 0.115093 | 0.357739 | 0.451 | 0.351 | 1 | 13 | Kif13b | 0.1 | Old |
| 0.115104 | 0.468693 | 0.328 | 0.228 | 1 | 13 | Pdgfa | 0.1 | Old |
| 0.11514 | 0.246372 | 0.475 | 0.333 | 1 | 13 | AU019823 | 0.142 | Old |
| 0.115183 | 0.19155 | 0.885 | 0.833 | 1 | 13 | Csnk2b | 0.052 | Old |
| 0.11522 | 0.682257 | 0.107 | 0.053 | 1 | 13 | 5430416N0 | 0.054 | Old |
| 0.115347 | 0.358793 | 0.631 | 0.553 | 1 | 13 | Inppl1 | 0.078 | Old |
| 0.115366 | 0.319686 | 0.68 | 0.588 | 1 | 13 | Maea | 0.092 | Old |
| 0.115384 | 1.147906 | 0.041 | 0.009 | 1 | 13 | Lrig3 | 0.032 | Old |
| 0.115384 | 1.138755 | 0.041 | 0.009 | 1 | 13 | Gm4117 | 0.032 | Old |
| 0.115384 | 1.057882 | 0.041 | 0.009 | 1 | 13 | Pip5k1b | 0.032 | Old |
| 0.115595 | 0.385969 | 0.656 | 0.579 | 1 | 13 | Hmgcs1 | 0.077 | Old |
| 0.115675 | 0.175863 | 0.992 | 0.947 | 1 | 13 | Hsbp1 | 0.045 | Old |
| 0.115678 | 0.149725 | 1 | 0.956 | 1 | 13 | Myl12a | 0.044 | Old |
| 0.115753 | 0.285029 | 0.426 | 0.316 | 1 | 13 | Hat1 | 0.11 | Old |
| 0.115788 | 0.900569 | 0.057 | 0.018 | 1 | 13 | 1110035H1 | 0.039 | Old |
| 0.115788 | 0.858521 | 0.057 | 0.018 | 1 | 13 | 1700061G1 | 0.039 | Old |
| 0.115788 | 0.794803 | 0.057 | 0.018 | 1 | 13 | Pole | 0.039 | Old |
| 0.115869 | 0.274162 | 0.533 | 0.447 | 1 | 13 | Kmt2b | 0.086 | Old |
| 0.115924 | 0.492795 | 0.377 | 0.272 | 1 | 13 | Ric1 | 0.105 | Old |
| 0.116044 | 0.299462 | 0.361 | 0.263 | 1 | 13 | Rnf41 | 0.098 | Old |
| 0.116211 | 0.722519 | 0.156 | 0.088 | 1 | 13 | Gpr157 | 0.068 | Old |
| 0.11663 | 0.564958 | 0.148 | 0.079 | 1 | 13 | Galm | 0.069 | Old |
| 0.11677 | 0.331936 | 0.59 | 0.482 | 1 | 13 | Ergic2 | 0.108 | Old |
| 0.116854 | 0.362483 | 0.443 | 0.333 | 1 | 13 | Eif2b5 | 0.11 | Old |

| | | | | | | | | |
|---|---|---|---|---|---|---|---|---|
| 0.11701 | 1.393471 | 0.041 | 0.009 | 1 | 13 | Paqr9 | 0.032 | Old |
| 0.11701 | 1.234577 | 0.041 | 0.009 | 1 | 13 | Atp1b1 | 0.032 | Old |
| 0.11701 | 1.097732 | 0.041 | 0.009 | 1 | 13 | Eya1 | 0.032 | Old |
| 0.11701 | 0.954495 | 0.041 | 0.009 | 1 | 13 | Rpusd3 | 0.032 | Old |
| 0.11701 | 0.941994 | 0.041 | 0.009 | 1 | 13 | Nrap | 0.032 | Old |
| 0.117011 | 0.11694 | 1 | 1 | 1 | 13 | Hexa | 0 | Old |
| 0.117098 | 0.918586 | 0.107 | 0.053 | 1 | 13 | Etl4 | 0.054 | Old |
| 0.117881 | 0.102072 | 0.934 | 0.965 | 1 | 13 | Ap2s1 | -0.031 | Old |
| 0.117921 | 1.011743 | 0.082 | 0.035 | 1 | 13 | Rhebl1 | 0.047 | Old |
| 0.117956 | 0.319817 | 0.508 | 0.395 | 1 | 13 | Parp9 | 0.113 | Old |
| 0.118165 | 0.299055 | 0.73 | 0.702 | 1 | 13 | Nab1 | 0.028 | Old |
| 0.118177 | 0.164667 | 0.361 | 0.246 | 1 | 13 | Dhx29 | 0.115 | Old |
| 0.118489 | 0.496662 | 0.385 | 0.289 | 1 | 13 | Thumpd3 | 0.096 | Old |
| 0.118653 | 1.026954 | 0.041 | 0.009 | 1 | 13 | Itga2 | 0.032 | Old |
| 0.118653 | 0.536628 | 0.27 | 0.175 | 1 | 13 | Wdr7 | 0.095 | Old |
| 0.118752 | 0.248779 | 0.926 | 0.789 | 1 | 13 | Sdhc | 0.137 | Old |
| 0.118769 | 0.236017 | 0.689 | 0.632 | 1 | 13 | Etv6 | 0.057 | Old |
| 0.118878 | 0.237906 | 0.844 | 0.772 | 1 | 13 | Lamtor3 | 0.072 | Old |
| 0.118927 | 0.355852 | 0.566 | 0.456 | 1 | 13 | Kidins220 | 0.11 | Old |
| 0.119026 | 0.340591 | 0.697 | 0.57 | 1 | 13 | Skiv2l | 0.127 | Old |
| 0.11916 | 0.16755 | 0.934 | 0.904 | 1 | 13 | Llph | 0.03 | Old |
| 0.119167 | 0.446823 | 0.172 | 0.096 | 1 | 13 | Eif5a2 | 0.076 | Old |
| 0.119573 | 0.477208 | 0.492 | 0.36 | 1 | 13 | Rinl | 0.132 | Old |
| 0.119738 | 0.285083 | 0.541 | 0.43 | 1 | 13 | Psmd9 | 0.111 | Old |
| 0.119769 | 0.322884 | 0.311 | 0.202 | 1 | 13 | Tomm40l | 0.109 | Old |
| 0.119831 | 0.369366 | 0.336 | 0.228 | 1 | 13 | Cpped1 | 0.108 | Old |
| 0.119882 | 0.70004 | 0.148 | 0.079 | 1 | 13 | Cmtr2 | 0.069 | Old |
| 0.120065 | 0.256238 | 0.713 | 0.64 | 1 | 13 | Spty2d1 | 0.073 | Old |
| 0.120116 | 0.58852 | 0.254 | 0.167 | 1 | 13 | Rab40c | 0.087 | Old |
| 0.120145 | 0.190793 | 0.959 | 0.947 | 1 | 13 | Ube2l3 | 0.012 | Old |
| 0.120203 | 0.21528 | 0.443 | 0.316 | 1 | 13 | Eng | 0.127 | Old |
| 0.120315 | 1.000445 | 0.041 | 0.009 | 1 | 13 | Ect2 | 0.032 | Old |
| 0.120315 | 0.95473 | 0.041 | 0.009 | 1 | 13 | Tchh | 0.032 | Old |
| 0.120315 | 0.935891 | 0.041 | 0.009 | 1 | 13 | Gm8113 | 0.032 | Old |
| 0.120315 | 0.92306 | 0.041 | 0.009 | 1 | 13 | Sema7a | 0.032 | Old |
| 0.120315 | 0.91462 | 0.041 | 0.009 | 1 | 13 | Gm26526 | 0.032 | Old |
| 0.120315 | 0.884121 | 0.041 | 0.009 | 1 | 13 | Armcx6 | 0.032 | Old |
| 0.120315 | 0.847464 | 0.041 | 0.009 | 1 | 13 | Rnls | 0.032 | Old |
| 0.120403 | 0.429255 | 0.328 | 0.237 | 1 | 13 | Coq2 | 0.091 | Old |
| 0.12052 | 0.26018 | 0.836 | 0.763 | 1 | 13 | Calu | 0.073 | Old |
| 0.120786 | 0.709558 | 0.131 | 0.07 | 1 | 13 | Zmynd19 | 0.061 | Old |
| 0.12095 | 0.337403 | 0.467 | 0.36 | 1 | 13 | Ino80 | 0.107 | Old |
| 0.120968 | 0.187341 | 0.721 | 0.623 | 1 | 13 | Wwp2 | 0.098 | Old |
| 0.121075 | 0.402522 | 0.508 | 0.386 | 1 | 13 | Steap3 | 0.122 | Old |
| 0.121322 | 0.268493 | 0.623 | 0.509 | 1 | 13 | Fryl | 0.114 | Old |

| | | | | | | | | |
|---|---|---|---|---|---|---|---|---|
| 0.121435 | 0.419098 | 0.484 | 0.386 | 1 | 13 | Ints1 | 0.098 | Old |
| 0.121474 | 0.286161 | 0.352 | 0.246 | 1 | 13 | Elac2 | 0.106 | Old |
| 0.121558 | 0.40054 | 0.197 | 0.114 | 1 | 13 | Flad1 | 0.083 | Old |
| 0.121769 | 0.363995 | 0.27 | 0.175 | 1 | 13 | Dcun1d3 | 0.095 | Old |
| 0.121806 | 0.381512 | 0.393 | 0.289 | 1 | 13 | Snx24 | 0.104 | Old |
| 0.121945 | 0.215165 | 0.902 | 0.816 | 1 | 13 | Rcc2 | 0.086 | Old |
| 0.121994 | 0.919883 | 0.041 | 0.009 | 1 | 13 | 1-Mar | 0.032 | Old |
| 0.121994 | 0.836143 | 0.041 | 0.009 | 1 | 13 | Gm11175 | 0.032 | Old |
| 0.121994 | 0.817906 | 0.041 | 0.009 | 1 | 13 | Atp10a | 0.032 | Old |
| 0.121994 | 0.790386 | 0.041 | 0.009 | 1 | 13 | Tmem170 | 0.032 | Old |
| 0.121994 | 0.682118 | 0.041 | 0.009 | 1 | 13 | Htra1 | 0.032 | Old |
| 0.122132 | 0.379717 | 0.336 | 0.228 | 1 | 13 | 2410002F2 | 0.108 | Old |
| 0.122205 | 0.756753 | 0.164 | 0.096 | 1 | 13 | Abcg2 | 0.068 | Old |
| 0.122269 | 0.262871 | 0.762 | 0.675 | 1 | 13 | Arl1 | 0.087 | Old |
| 0.122344 | 0.134299 | 0.795 | 0.623 | 1 | 13 | Phf20l1 | 0.172 | Old |
| 0.122461 | 0.231488 | 0.525 | 0.439 | 1 | 13 | Pbdc1 | 0.086 | Old |
| 0.122507 | 0.67872 | 0.205 | 0.132 | 1 | 13 | Nepro | 0.073 | Old |
| 0.122826 | 0.318186 | 0.811 | 0.789 | 1 | 13 | Uhmk1 | 0.022 | Old |
| 0.122877 | 0.288104 | 0.861 | 0.702 | 1 | 13 | Exoc5 | 0.159 | Old |
| 0.122878 | 0.839667 | 0.107 | 0.053 | 1 | 13 | Cep41 | 0.054 | Old |
| 0.123168 | 0.389424 | 0.336 | 0.237 | 1 | 13 | Ubr7 | 0.099 | Old |
| 0.123219 | 0.326278 | 0.434 | 0.316 | 1 | 13 | Nhlrc2 | 0.118 | Old |
| 0.12325 | 0.158334 | 0.967 | 0.851 | 1 | 13 | Zfp91 | 0.116 | Old |
| 0.123767 | 0.318712 | 0.91 | 0.851 | 1 | 13 | Smpdl3a | 0.059 | Old |
| 0.123822 | 0.273452 | 0.754 | 0.64 | 1 | 13 | Rufy3 | 0.114 | Old |
| 0.124007 | 0.371492 | 0.418 | 0.307 | 1 | 13 | Yeats2 | 0.111 | Old |
| 0.124148 | 0.180722 | 0.574 | 0.439 | 1 | 13 | Smg7 | 0.135 | Old |
| 0.124288 | 0.326136 | 0.434 | 0.316 | 1 | 13 | Topbp1 | 0.118 | Old |
| 0.12431 | 0.729261 | 0.131 | 0.07 | 1 | 13 | Fzd1 | 0.061 | Old |
| 0.124449 | 0.248613 | 0.664 | 0.553 | 1 | 13 | Arnt | 0.111 | Old |
| 0.12449 | 0.397045 | 0.484 | 0.36 | 1 | 13 | Fzr1 | 0.124 | Old |
| 0.124637 | 0.937973 | 0.082 | 0.035 | 1 | 13 | Gbp8 | 0.047 | Old |
| 0.124638 | 0.537062 | 0.32 | 0.228 | 1 | 13 | 1190007I0 | 0.092 | Old |
| 0.124675 | 0.447754 | 0.205 | 0.123 | 1 | 13 | B4galt7 | 0.082 | Old |
| 0.124801 | 0.233785 | 0.525 | 0.386 | 1 | 13 | Scaf1 | 0.139 | Old |
| 0.124853 | 0.827324 | 0.107 | 0.053 | 1 | 13 | Lipt2 | 0.054 | Old |
| 0.125014 | 0.195432 | 0.959 | 0.991 | 1 | 13 | Hnrnpm | -0.032 | Old |
| 0.12557 | 0.143159 | 0.91 | 0.833 | 1 | 13 | Amfr | 0.077 | Old |
| 0.125573 | 0.628973 | 0.311 | 0.228 | 1 | 13 | Dzip3 | 0.083 | Old |
| 0.125677 | 0.242622 | 0.885 | 0.833 | 1 | 13 | Aco2 | 0.052 | Old |
| 0.125865 | 0.348259 | 0.59 | 0.474 | 1 | 13 | Akap8 | 0.116 | Old |
| 0.126065 | 0.570006 | 0.352 | 0.246 | 1 | 13 | Asb8 | 0.106 | Old |
| 0.126122 | 0.388508 | 0.623 | 0.535 | 1 | 13 | Xdh | 0.088 | Old |
| 0.126194 | 0.116593 | 1 | 1 | 1 | 13 | Anxa5 | 0 | Old |
| 0.126788 | 0.382601 | 0.41 | 0.307 | 1 | 13 | Cops7a | 0.103 | Old |

| | | | | | | | | |
|---|---|---|---|---|---|---|---|---|
| 0.126906 | 0.162124 | 0.582 | 0.43 | 1 | 13 | Stim1 | 0.152 | Old |
| 0.127097 | 0.389595 | 0.32 | 0.211 | 1 | 13 | Tradd | 0.109 | Old |
| 0.127113 | 0.318569 | 0.484 | 0.36 | 1 | 13 | Usp38 | 0.124 | Old |
| 0.127195 | 0.621597 | 0.238 | 0.158 | 1 | 13 | Gpr180 | 0.08 | Old |
| 0.127273 | 0.495258 | 0.385 | 0.289 | 1 | 13 | Hsf1 | 0.096 | Old |
| 0.127341 | 0.22268 | 0.836 | 0.746 | 1 | 13 | Ncf4 | 0.09 | Old |
| 0.127926 | 0.32703 | 0.377 | 0.281 | 1 | 13 | Zfp277 | 0.096 | Old |
| 0.127998 | 0.214746 | 0.926 | 0.912 | 1 | 13 | Ywhaq | 0.014 | Old |
| 0.128105 | 0.966912 | 0.082 | 0.035 | 1 | 13 | Gpc1 | 0.047 | Old |
| 0.128266 | 0.36339 | 0.451 | 0.342 | 1 | 13 | Xpnpep1 | 0.109 | Old |
| 0.128373 | 0.433503 | 0.557 | 0.509 | 1 | 13 | Ptdss1 | 0.048 | Old |
| 0.128617 | 0.200557 | 0.893 | 0.798 | 1 | 13 | Pdcd6 | 0.095 | Old |
| 0.128813 | 0.21996 | 0.992 | 0.956 | 1 | 13 | C5ar1 | 0.036 | Old |
| 0.128841 | 0.316516 | 0.213 | 0.132 | 1 | 13 | Gm45902 | 0.081 | Old |
| 0.129469 | 0.675802 | 0.23 | 0.158 | 1 | 13 | Slx4ip | 0.072 | Old |
| 0.129619 | 0.393903 | 0.492 | 0.386 | 1 | 13 | Dhcr24 | 0.106 | Old |
| 0.129737 | 0.61224 | 0.205 | 0.132 | 1 | 13 | Eif2ak4 | 0.073 | Old |
| 0.130027 | 0.485039 | 0.156 | 0.088 | 1 | 13 | Gins2 | 0.068 | Old |
| 0.130133 | 0.2467 | 0.5 | 0.36 | 1 | 13 | Dst | 0.14 | Old |
| 0.130206 | 0.58294 | 0.098 | 0.044 | 1 | 13 | Gm10053 | 0.054 | Old |
| 0.130384 | 0.204585 | 0.82 | 0.737 | 1 | 13 | Crat | 0.083 | Old |
| 0.130456 | 0.285515 | 0.336 | 0.228 | 1 | 13 | Galk1 | 0.108 | Old |
| 0.130459 | 0.801957 | 0.082 | 0.035 | 1 | 13 | Lig4 | 0.047 | Old |
| 0.130477 | 0.538991 | 0.205 | 0.132 | 1 | 13 | Rbm48 | 0.073 | Old |
| 0.130606 | 0.249863 | 0.689 | 0.614 | 1 | 13 | Cand1 | 0.075 | Old |
| 0.13063 | 0.342346 | 0.582 | 0.456 | 1 | 13 | Samd8 | 0.126 | Old |
| 0.130669 | 0.619223 | 0.131 | 0.07 | 1 | 13 | Lrrc28 | 0.061 | Old |
| 0.130711 | 0.526029 | 0.262 | 0.175 | 1 | 13 | Clp1 | 0.087 | Old |
| 0.130928 | 0.192124 | 0.82 | 0.781 | 1 | 13 | Dnajc13 | 0.039 | Old |
| 0.130952 | 0.224589 | 0.926 | 0.789 | 1 | 13 | Mdfic | 0.137 | Old |
| 0.131535 | 0.355421 | 0.787 | 0.719 | 1 | 13 | Abcc1 | 0.068 | Old |
| 0.131604 | 0.359654 | 0.369 | 0.254 | 1 | 13 | Pygo2 | 0.115 | Old |
| 0.13179 | 0.151769 | 0.943 | 0.86 | 1 | 13 | Bax | 0.083 | Old |
| 0.131843 | 0.361345 | 0.467 | 0.333 | 1 | 13 | Usp32 | 0.134 | Old |
| 0.131968 | 0.130008 | 0.984 | 0.991 | 1 | 13 | Colgalt1 | -0.007 | Old |
| 0.13204 | 0.365465 | 0.238 | 0.158 | 1 | 13 | Arl14ep | 0.08 | Old |
| 0.13212 | 0.320907 | 0.467 | 0.342 | 1 | 13 | Rap1gap2 | 0.125 | Old |
| 0.13237 | 0.266464 | 0.885 | 0.825 | 1 | 13 | Ghitm | 0.06 | Old |
| 0.13256 | 0.536288 | 0.32 | 0.228 | 1 | 13 | Cwc27 | 0.092 | Old |
| 0.13264 | 0.381277 | 0.172 | 0.096 | 1 | 13 | Mtrf1l | 0.076 | Old |
| 0.132846 | 0.769935 | 0.082 | 0.035 | 1 | 13 | C130013H( | 0.047 | Old |
| 0.132985 | 0.715385 | 0.139 | 0.079 | 1 | 13 | Lrmda | 0.06 | Old |
| 0.133026 | 0.230821 | 0.525 | 0.412 | 1 | 13 | Tcf20 | 0.113 | Old |
| 0.133443 | 0.124142 | 1 | 1 | 1 | 13 | H3f3b | 0 | Old |
| 0.13405 | 0.797615 | 0.107 | 0.053 | 1 | 13 | Cdc25b | 0.054 | Old |

| | | | | | | | | |
|---|---|---|---|---|---|---|---|---|
| 0.134052 | 1.115435 | 0.082 | 0.035 | 1 | 13 | Tpx2 | 0.047 | Old |
| 0.13407 | 0.271595 | 0.664 | 0.605 | 1 | 13 | Cpsf7 | 0.059 | Old |
| 0.134182 | 0.133606 | 0.992 | 0.956 | 1 | 13 | Bsg | 0.036 | Old |
| 0.134185 | 0.046051 | 0.992 | 0.991 | 1 | 13 | Shisa5 | 0.001 | Old |
| 0.134224 | 0.344124 | 0.393 | 0.289 | 1 | 13 | Fam53c | 0.104 | Old |
| 0.13426 | 0.415885 | 0.189 | 0.114 | 1 | 13 | Slc25a15 | 0.075 | Old |
| 0.13437 | 1.665749 | 0.115 | 0.061 | 1 | 13 | Tcim | 0.054 | Old |
| 0.13438 | 0.373489 | 0.664 | 0.588 | 1 | 13 | Kctd20 | 0.076 | Old |
| 0.134433 | 0.099653 | 1 | 1 | 1 | 13 | App | 0 | Old |
| 0.13447 | 0.250577 | 0.631 | 0.518 | 1 | 13 | Chfr | 0.113 | Old |
| 0.134799 | 0.428433 | 0.361 | 0.272 | 1 | 13 | Ascc2 | 0.089 | Old |
| 0.13493 | 0.159923 | 0.992 | 0.982 | 1 | 13 | Spi1 | 0.01 | Old |
| 0.134931 | 0.127433 | 0.992 | 1 | 1 | 13 | Celf2 | -0.008 | Old |
| 0.135177 | 0.966705 | 0.189 | 0.123 | 1 | 13 | Slc16a1 | 0.066 | Old |
| 0.135188 | 0.469573 | 0.287 | 0.184 | 1 | 13 | 3110062M | 0.103 | Old |
| 0.13525 | 0.465758 | 0.41 | 0.333 | 1 | 13 | Ugdh | 0.077 | Old |
| 0.135264 | 0.226005 | 0.779 | 0.728 | 1 | 13 | Rbm47 | 0.051 | Old |
| 0.13593 | 0.265138 | 0.68 | 0.535 | 1 | 13 | Egln2 | 0.145 | Old |
| 0.136073 | 0.437257 | 0.156 | 0.088 | 1 | 13 | Gatb | 0.068 | Old |
| 0.136219 | 0.504694 | 0.328 | 0.254 | 1 | 13 | Chst7 | 0.074 | Old |
| 0.136375 | 0.581117 | 0.197 | 0.123 | 1 | 13 | Coq9 | 0.074 | Old |
| 0.136489 | 0.764575 | 0.082 | 0.035 | 1 | 13 | Jmjd8 | 0.047 | Old |
| 0.136583 | 0.447866 | 0.393 | 0.289 | 1 | 13 | Med20 | 0.104 | Old |
| 0.136656 | 0.499601 | 0.189 | 0.114 | 1 | 13 | Scfd2 | 0.075 | Old |
| 0.136687 | 0.276552 | 0.205 | 0.123 | 1 | 13 | Pan2 | 0.082 | Old |
| 0.136754 | 0.436495 | 0.287 | 0.193 | 1 | 13 | Pef1 | 0.094 | Old |
| 0.137114 | 0.336285 | 0.434 | 0.316 | 1 | 13 | Ube2l6 | 0.118 | Old |
| 0.137145 | 0.273165 | 0.5 | 0.395 | 1 | 13 | Nabp2 | 0.105 | Old |
| 0.137149 | 0.451357 | 0.303 | 0.202 | 1 | 13 | Scnm1 | 0.101 | Old |
| 0.137229 | 0.67759 | 0.107 | 0.053 | 1 | 13 | Nicn1 | 0.054 | Old |
| 0.137406 | 0.189276 | 0.951 | 0.939 | 1 | 13 | Lbh | 0.012 | Old |
| 0.137412 | 1.060418 | 0.115 | 0.061 | 1 | 13 | Cep95 | 0.054 | Old |
| 0.137412 | 0.905316 | 0.115 | 0.061 | 1 | 13 | 1700066M | 0.054 | Old |
| 0.137579 | 0.217024 | 0.885 | 0.868 | 1 | 13 | Eif3b | 0.017 | Old |
| 0.137719 | 0.450601 | 0.336 | 0.237 | 1 | 13 | Ubxn7 | 0.099 | Old |
| 0.137721 | 0.646502 | 0.082 | 0.035 | 1 | 13 | Opn3 | 0.047 | Old |
| 0.137886 | 0.397982 | 0.434 | 0.333 | 1 | 13 | Nup93 | 0.101 | Old |
| 0.138463 | 0.319457 | 0.607 | 0.5 | 1 | 13 | Xpot | 0.107 | Old |
| 0.138468 | 0.162522 | 0.869 | 0.86 | 1 | 13 | Vdac3 | 0.009 | Old |
| 0.138778 | 0.218594 | 0.91 | 0.895 | 1 | 13 | Rbm8a | 0.015 | Old |
| 0.138819 | 0.222922 | 0.811 | 0.693 | 1 | 13 | Pa2g4 | 0.118 | Old |
| 0.138842 | 0.611331 | 0.221 | 0.149 | 1 | 13 | Naf1 | 0.072 | Old |
| 0.138899 | 0.269949 | 0.582 | 0.465 | 1 | 13 | Trnt1 | 0.117 | Old |
| 0.138909 | 0.316995 | 0.385 | 0.281 | 1 | 13 | Fem1a | 0.104 | Old |
| 0.138961 | 0.483837 | 0.082 | 0.035 | 1 | 13 | Fignl1 | 0.047 | Old |

| | | | | | | | | |
|---|---|---|---|---|---|---|---|---|
| 0.139078 | 0.26976 | 0.615 | 0.553 | 1 | 13 | Slain2 | 0.062 | Old |
| 0.139161 | 0.332567 | 0.197 | 0.114 | 1 | 13 | Coa7 | 0.083 | Old |
| 0.139171 | 0.310605 | 0.303 | 0.211 | 1 | 13 | Cog2 | 0.092 | Old |
| 0.139378 | 0.468417 | 0.213 | 0.14 | 1 | 13 | Gnpnat1 | 0.073 | Old |
| 0.139381 | 0.495895 | 0.107 | 0.053 | 1 | 13 | Gm31597 | 0.054 | Old |
| 0.13939 | 0.426385 | 0.344 | 0.254 | 1 | 13 | Pnkp | 0.09 | Old |
| 0.139471 | 0.06685 | 1 | 1 | 1 | 13 | Gm10076 | 0 | Old |
| 0.139685 | 0.371829 | 0.369 | 0.263 | 1 | 13 | Desi1 | 0.106 | Old |
| 0.139982 | 0.094752 | 1 | 0.974 | 1 | 13 | Rab5c | 0.026 | Old |
| 0.139982 | 0.069354 | 1 | 1 | 1 | 13 | Hnrnpk | 0 | Old |
| 0.140065 | 0.559686 | 0.27 | 0.184 | 1 | 13 | Polr3d | 0.086 | Old |
| 0.140114 | 0.405958 | 0.451 | 0.386 | 1 | 13 | Npl | 0.065 | Old |
| 0.140193 | 0.413232 | 0.475 | 0.351 | 1 | 13 | Sphk2 | 0.124 | Old |
| 0.140197 | 0.42197 | 0.697 | 0.64 | 1 | 13 | Ms4a6d | 0.057 | Old |
| 0.140212 | 0.424017 | 0.426 | 0.342 | 1 | 13 | Mrps5 | 0.084 | Old |
| 0.140275 | 0.491875 | 0.279 | 0.193 | 1 | 13 | Edc3 | 0.086 | Old |
| 0.140461 | 0.581304 | 0.164 | 0.096 | 1 | 13 | Fggy | 0.068 | Old |
| 0.140523 | 0.370089 | 0.156 | 0.088 | 1 | 13 | Trmt61a | 0.068 | Old |
| 0.140544 | 0.204353 | 0.852 | 0.711 | 1 | 13 | Ankrd13a | 0.141 | Old |
| 0.140682 | 0.164464 | 0.836 | 0.702 | 1 | 13 | Anapc5 | 0.134 | Old |
| 0.140861 | 0.173941 | 0.557 | 0.404 | 1 | 13 | Zyg11b | 0.153 | Old |
| 0.140876 | 0.33679 | 0.902 | 0.798 | 1 | 13 | Acp2 | 0.104 | Old |
| 0.140998 | 0.505558 | 0.32 | 0.219 | 1 | 13 | Mfn1 | 0.101 | Old |
| 0.141382 | 0.321448 | 0.549 | 0.447 | 1 | 13 | Glb1 | 0.102 | Old |
| 0.14148 | 0.396368 | 0.541 | 0.447 | 1 | 13 | Lap3 | 0.094 | Old |
| 0.141524 | 0.125777 | 0.975 | 0.982 | 1 | 13 | Taldo1 | -0.007 | Old |
| 0.141526 | 0.148559 | 0.992 | 1 | 1 | 13 | Ncl | -0.008 | Old |
| 0.141526 | 0.112509 | 0.992 | 1 | 1 | 13 | Nrros | -0.008 | Old |
| 0.141849 | 0.457111 | 0.861 | 0.798 | 1 | 13 | Eprs | 0.063 | Old |
| 0.142155 | 0.599273 | 0.131 | 0.07 | 1 | 13 | C1rl | 0.061 | Old |
| 0.142352 | 0.151103 | 0.918 | 0.877 | 1 | 13 | Itsn2 | 0.041 | Old |
| 0.142503 | 0.55418 | 0.23 | 0.158 | 1 | 13 | Pde6d | 0.072 | Old |
| 0.142562 | 0.105494 | 1 | 1 | 1 | 13 | Ywhaz | 0 | Old |
| 0.142589 | 0.318651 | 0.762 | 0.675 | 1 | 13 | Mcfd2 | 0.087 | Old |
| 0.142808 | 0.21839 | 0.762 | 0.632 | 1 | 13 | Il6st | 0.13 | Old |
| 0.142951 | 0.170663 | 0.893 | 0.807 | 1 | 13 | Gpsm3 | 0.086 | Old |
| 0.143082 | 0.113124 | 1 | 1 | 1 | 13 | Rhoa | 0 | Old |
| 0.143204 | 0.516418 | 0.189 | 0.114 | 1 | 13 | Telo2 | 0.075 | Old |
| 0.143603 | 0.130308 | 1 | 0.965 | 1 | 13 | Psmb1 | 0.035 | Old |
| 0.143995 | 0.532377 | 0.164 | 0.096 | 1 | 13 | Apba3 | 0.068 | Old |
| 0.144007 | 0.542855 | 0.082 | 0.035 | 1 | 13 | Jade3 | 0.047 | Old |
| 0.144126 | 0.394465 | 0.598 | 0.5 | 1 | 13 | Rnf44 | 0.098 | Old |
| 0.144148 | 1.108549 | 0.156 | 0.096 | 1 | 13 | Scarf1 | 0.06 | Old |
| 0.144281 | 0.936944 | 0.066 | 0.026 | 1 | 13 | Epas1 | 0.04 | Old |
| 0.144419 | 0.247118 | 0.689 | 0.596 | 1 | 13 | Slc25a51 | 0.093 | Old |

| | | | | | | | | |
|---|---|---|---|---|---|---|---|---|
| 0.144461 | 0.233411 | 0.877 | 0.781 | 1 | 13 | Srpr | 0.096 | Old |
| 0.144535 | 0.242185 | 0.836 | 0.754 | 1 | 13 | Kif1b | 0.082 | Old |
| 0.144574 | 0.422999 | 0.279 | 0.184 | 1 | 13 | Zfp954 | 0.095 | Old |
| 0.144719 | 0.212216 | 0.738 | 0.632 | 1 | 13 | Dctn2 | 0.106 | Old |
| 0.144877 | 0.200259 | 0.902 | 0.868 | 1 | 13 | Dnase2a | 0.034 | Old |
| 0.144889 | 0.521567 | 0.164 | 0.096 | 1 | 13 | Kremen1 | 0.068 | Old |
| 0.144897 | 0.165789 | 0.746 | 0.658 | 1 | 13 | Slc25a11 | 0.088 | Old |
| 0.145503 | 0.390986 | 0.221 | 0.14 | 1 | 13 | Naip6 | 0.081 | Old |
| 0.145937 | 0.558335 | 0.172 | 0.105 | 1 | 13 | Aldh6a1 | 0.067 | Old |
| 0.145967 | 0.165155 | 0.975 | 0.974 | 1 | 13 | Chmp4b | 0.001 | Old |
| 0.146235 | 0.110207 | 1 | 1 | 1 | 13 | Slc25a3 | 0 | Old |
| 0.146447 | 0.313676 | 0.361 | 0.237 | 1 | 13 | Gmnn | 0.124 | Old |
| 0.146493 | 0.683824 | 0.221 | 0.149 | 1 | 13 | Zfp280c | 0.072 | Old |
| 0.146494 | 0.214378 | 0.344 | 0.237 | 1 | 13 | Ormdl3 | 0.107 | Old |
| 0.146606 | 0.303441 | 0.689 | 0.57 | 1 | 13 | Tomm34 | 0.119 | Old |
| 0.14669 | 0.571958 | 0.164 | 0.096 | 1 | 13 | Mapkapk5 | 0.068 | Old |
| 0.146765 | 0.120192 | 1 | 1 | 1 | 13 | Rap1b | 0 | Old |
| 0.146874 | 0.606978 | 0.197 | 0.123 | 1 | 13 | Pdpr | 0.074 | Old |
| 0.147163 | 0.379052 | 0.131 | 0.07 | 1 | 13 | Rab38 | 0.061 | Old |
| 0.147174 | 1.070493 | 0.066 | 0.026 | 1 | 13 | Scrn2 | 0.04 | Old |
| 0.147174 | 0.948918 | 0.066 | 0.026 | 1 | 13 | Zfyve9 | 0.04 | Old |
| 0.147263 | 0.1528 | 0.959 | 0.93 | 1 | 13 | Drap1 | 0.029 | Old |
| 0.147277 | 0.953868 | 0.09 | 0.044 | 1 | 13 | 1810009A1 | 0.046 | Old |
| 0.147287 | 0.383487 | 0.295 | 0.202 | 1 | 13 | Lancl1 | 0.093 | Old |
| 0.147394 | 0.328082 | 0.369 | 0.272 | 1 | 13 | Nostrin | 0.097 | Old |
| 0.147492 | 0.264773 | 0.844 | 0.772 | 1 | 13 | Plaur | 0.072 | Old |
| 0.14757 | 0.398691 | 0.262 | 0.175 | 1 | 13 | Nup155 | 0.087 | Old |
| 0.147759 | 0.438985 | 0.344 | 0.254 | 1 | 13 | Tirap | 0.09 | Old |
| 0.147915 | 0.339299 | 0.418 | 0.298 | 1 | 13 | Arhgap21 | 0.12 | Old |
| 0.14797 | 0.467499 | 0.246 | 0.167 | 1 | 13 | Fktn | 0.079 | Old |
| 0.148061 | 0.614097 | 0.221 | 0.149 | 1 | 13 | Phospho2 | 0.072 | Old |
| 0.148365 | 0.081791 | 1 | 1 | 1 | 13 | Efhd2 | 0 | Old |
| 0.148437 | 0.223774 | 0.615 | 0.447 | 1 | 13 | Tor1a | 0.168 | Old |
| 0.148504 | 0.968154 | 0.09 | 0.044 | 1 | 13 | Snn | 0.046 | Old |
| 0.148504 | 0.947944 | 0.09 | 0.044 | 1 | 13 | Tk1 | 0.046 | Old |
| 0.148553 | 0.299503 | 0.697 | 0.596 | 1 | 13 | Pdha1 | 0.101 | Old |
| 0.148638 | 0.870274 | 0.066 | 0.026 | 1 | 13 | Gm43569 | 0.04 | Old |
| 0.148647 | 0.292463 | 0.516 | 0.412 | 1 | 13 | Atp7a | 0.104 | Old |
| 0.149412 | 0.255315 | 0.648 | 0.518 | 1 | 13 | Klhl6 | 0.13 | Old |
| 0.149438 | 0.160358 | 0.984 | 0.974 | 1 | 13 | Fermt3 | 0.01 | Old |
| 0.149523 | 0.347064 | 0.525 | 0.404 | 1 | 13 | Prpf19 | 0.121 | Old |
| 0.149978 | 0.100167 | 0.984 | 0.982 | 1 | 13 | Ap2m1 | 0.002 | Old |
| 0.150079 | 0.237881 | 0.852 | 0.711 | 1 | 13 | Evl | 0.141 | Old |
| 0.150099 | 0.563864 | 0.311 | 0.228 | 1 | 13 | Gna15 | 0.083 | Old |
| 0.150112 | 0.86848 | 0.066 | 0.026 | 1 | 13 | Tmem67 | 0.04 | Old |

| | | | | | | | | |
|---|---|---|---|---|---|---|---|---|
| 0.150186 | 0.871092 | 0.148 | 0.088 | 1 | 13 | Dhrs9 | 0.06 | Old |
| 0.150433 | 0.330661 | 0.541 | 0.474 | 1 | 13 | Mfsd10 | 0.067 | Old |
| 0.15052 | 0.15653 | 1 | 1 | 1 | 13 | mt-Nd5 | 0 | Old |
| 0.150846 | 0.320397 | 0.492 | 0.395 | 1 | 13 | Aggf1 | 0.097 | Old |
| 0.150853 | 0.326062 | 0.492 | 0.368 | 1 | 13 | Tmc6 | 0.124 | Old |
| 0.150914 | 0.676662 | 0.762 | 0.789 | 1 | 13 | Fgl2 | -0.027 | Old |
| 0.150936 | 0.26993 | 0.582 | 0.474 | 1 | 13 | Utp18 | 0.108 | Old |
| 0.151035 | 0.221238 | 0.459 | 0.342 | 1 | 13 | Prpsap1 | 0.117 | Old |
| 0.151597 | 0.910913 | 0.066 | 0.026 | 1 | 13 | Tnfsf15 | 0.04 | Old |
| 0.151597 | 0.868874 | 0.066 | 0.026 | 1 | 13 | Ace | 0.04 | Old |
| 0.151687 | 0.107148 | 0.902 | 0.833 | 1 | 13 | Psmd4 | 0.069 | Old |
| 0.151699 | 0.167106 | 0.869 | 0.789 | 1 | 13 | Erp44 | 0.08 | Old |
| 0.151726 | 0.407481 | 0.189 | 0.114 | 1 | 13 | Smyd2 | 0.075 | Old |
| 0.151837 | 0.899698 | 0.123 | 0.07 | 1 | 13 | Zrsr1 | 0.053 | Old |
| 0.152259 | 0.078688 | 0.926 | 0.886 | 1 | 13 | Midn | 0.04 | Old |
| 0.1524 | 0.250103 | 0.393 | 0.263 | 1 | 13 | Vamp7 | 0.13 | Old |
| 0.152579 | 0.494945 | 0.205 | 0.132 | 1 | 13 | Snip1 | 0.073 | Old |
| 0.152599 | 0.331691 | 0.189 | 0.114 | 1 | 13 | Slc35d2 | 0.075 | Old |
| 0.152692 | 0.464662 | 0.221 | 0.14 | 1 | 13 | Retsat | 0.081 | Old |
| 0.153094 | 0.861847 | 0.066 | 0.026 | 1 | 13 | Zfp607a | 0.04 | Old |
| 0.153292 | 0.287762 | 0.402 | 0.289 | 1 | 13 | Scaf4 | 0.113 | Old |
| 0.153305 | 0.178993 | 0.926 | 0.868 | 1 | 13 | Psmb10 | 0.058 | Old |
| 0.153422 | 0.684844 | 0.115 | 0.061 | 1 | 13 | Plekhh3 | 0.054 | Old |
| 0.153489 | 0.21537 | 0.893 | 0.877 | 1 | 13 | Ifnar1 | 0.016 | Old |
| 0.153529 | 0.420409 | 0.254 | 0.175 | 1 | 13 | Jak3 | 0.079 | Old |
| 0.153662 | 0.362561 | 0.287 | 0.202 | 1 | 13 | Rbbp5 | 0.085 | Old |
| 0.153839 | 0.367631 | 0.541 | 0.412 | 1 | 13 | Mbd1 | 0.129 | Old |
| 0.153947 | 0.180112 | 0.926 | 0.904 | 1 | 13 | Paip2 | 0.022 | Old |
| 0.154098 | 0.667026 | 0.139 | 0.079 | 1 | 13 | Smpd3 | 0.06 | Old |
| 0.154752 | 0.839623 | 0.09 | 0.044 | 1 | 13 | Oas1g | 0.046 | Old |
| 0.154874 | 0.106215 | 0.951 | 0.947 | 1 | 13 | Ndufs6 | 0.004 | Old |
| 0.155115 | 0.689642 | 0.139 | 0.079 | 1 | 13 | Dusp19 | 0.06 | Old |
| 0.155277 | 0.509533 | 0.221 | 0.149 | 1 | 13 | Cdkn2c | 0.072 | Old |
| 0.155374 | 0.138049 | 0.73 | 0.588 | 1 | 13 | Bbip1 | 0.142 | Old |
| 0.156025 | 0.970788 | 0.09 | 0.044 | 1 | 13 | Gemin8 | 0.046 | Old |
| 0.15612 | 0.805037 | 0.066 | 0.026 | 1 | 13 | Tmppe | 0.04 | Old |
| 0.15612 | 0.779557 | 0.066 | 0.026 | 1 | 13 | Dzip1 | 0.04 | Old |
| 0.156137 | 0.548933 | 0.139 | 0.079 | 1 | 13 | Phf11a | 0.06 | Old |
| 0.156676 | 0.325703 | 0.443 | 0.351 | 1 | 13 | Dnpep | 0.092 | Old |
| 0.156788 | 0.709963 | 0.115 | 0.061 | 1 | 13 | Pdss1 | 0.054 | Old |
| 0.157104 | 0.501194 | 0.197 | 0.123 | 1 | 13 | D2hgdh | 0.074 | Old |
| 0.157303 | 0.209363 | 0.27 | 0.175 | 1 | 13 | Adora2a | 0.095 | Old |
| 0.157306 | 0.970086 | 0.09 | 0.044 | 1 | 13 | Gm45871 | 0.046 | Old |
| 0.157331 | 0.63985 | 0.23 | 0.158 | 1 | 13 | Plekhg1 | 0.072 | Old |
| 0.157357 | 0.282684 | 0.533 | 0.43 | 1 | 13 | Lsm2 | 0.103 | Old |

| | | | | | | | | |
|---|---|---|---|---|---|---|---|---|
| 0.15765 | 0.776567 | 0.066 | 0.026 | 1 | 13 | Cfap126 | 0.04 | Old |
| 0.15765 | 0.764118 | 0.066 | 0.026 | 1 | 13 | B630019K0 | 0.04 | Old |
| 0.15765 | 0.695287 | 0.066 | 0.026 | 1 | 13 | Gm30074 | 0.04 | Old |
| 0.157672 | 0.038433 | 0.984 | 0.93 | 1 | 13 | Macf1 | 0.054 | Old |
| 0.158249 | 0.040085 | 1 | 1 | 1 | 13 | Rpl41 | 0 | Old |
| 0.158328 | 0.377596 | 0.27 | 0.184 | 1 | 13 | 5730455P1 | 0.086 | Old |
| 0.158571 | 0.583157 | 0.303 | 0.219 | 1 | 13 | Pelp1 | 0.084 | Old |
| 0.158595 | 0.711561 | 0.09 | 0.044 | 1 | 13 | Hist2h3c1 | 0.046 | Old |
| 0.158786 | 0.290516 | 0.328 | 0.246 | 1 | 13 | Tex10 | 0.082 | Old |
| 0.158792 | 0.281585 | 0.557 | 0.465 | 1 | 13 | Ammecr1l | 0.092 | Old |
| 0.159045 | 0.17755 | 0.779 | 0.711 | 1 | 13 | Fbxl3 | 0.068 | Old |
| 0.15906 | 0.233968 | 0.787 | 0.833 | 1 | 13 | Adam8 | -0.046 | Old |
| 0.159183 | 0.242342 | 0.492 | 0.351 | 1 | 13 | Fbrsl1 | 0.141 | Old |
| 0.159262 | 0.200438 | 0.885 | 0.816 | 1 | 13 | Sgk3 | 0.069 | Old |
| 0.15939 | 0.471564 | 0.205 | 0.132 | 1 | 13 | Hmgxb3 | 0.073 | Old |
| 0.159399 | 0.162057 | 0.73 | 0.596 | 1 | 13 | Kmt5a | 0.134 | Old |
| 0.159559 | 0.70229 | 0.189 | 0.123 | 1 | 13 | Dppa3 | 0.066 | Old |
| 0.159659 | 0.309412 | 0.361 | 0.254 | 1 | 13 | Mfap1b | 0.107 | Old |
| 0.15973 | 0.662107 | 0.205 | 0.14 | 1 | 13 | Shmt2 | 0.065 | Old |
| 0.159847 | 0.241475 | 0.336 | 0.237 | 1 | 13 | Heatr3 | 0.099 | Old |
| 0.159863 | 0.412437 | 0.27 | 0.184 | 1 | 13 | Mettl14 | 0.086 | Old |
| 0.159866 | 0.237986 | 0.902 | 0.956 | 1 | 13 | B4galt6 | -0.054 | Old |
| 0.160344 | 0.300732 | 0.525 | 0.404 | 1 | 13 | Prkaca | 0.121 | Old |
| 0.160491 | 0.147926 | 0.967 | 0.939 | 1 | 13 | Ptpra | 0.028 | Old |
| 0.160497 | 0.608815 | 0.172 | 0.105 | 1 | 13 | Fbxl19 | 0.067 | Old |
| 0.160744 | 0.145949 | 0.615 | 0.482 | 1 | 13 | Pip4k2b | 0.133 | Old |
| 0.161065 | 0.141046 | 0.967 | 0.947 | 1 | 13 | Kif5b | 0.02 | Old |
| 0.16107 | 0.19083 | 0.967 | 0.956 | 1 | 13 | Snx2 | 0.011 | Old |
| 0.16113 | 0.200521 | 0.902 | 0.825 | 1 | 13 | Aph1a | 0.077 | Old |
| 0.161239 | 0.143067 | 0.926 | 0.904 | 1 | 13 | Ppp1cb | 0.022 | Old |
| 0.161263 | 0.165704 | 0.902 | 0.851 | 1 | 13 | Nono | 0.051 | Old |
| 0.161398 | 0.238376 | 0.77 | 0.658 | 1 | 13 | Zmiz2 | 0.112 | Old |
| 0.161652 | 0.093037 | 1 | 1 | 1 | 13 | Emp3 | 0 | Old |
| 0.161987 | 0.195732 | 0.705 | 0.614 | 1 | 13 | Bfar | 0.091 | Old |
| 0.162105 | 0.211784 | 0.73 | 0.553 | 1 | 13 | Irak4 | 0.177 | Old |
| 0.162119 | 0.414805 | 0.213 | 0.14 | 1 | 13 | Zw10 | 0.073 | Old |
| 0.162309 | 0.336938 | 0.066 | 0.026 | 1 | 13 | Zfp820 | 0.04 | Old |
| 0.162635 | 0.134741 | 0.672 | 0.553 | 1 | 13 | Ube2g2 | 0.119 | Old |
| 0.162648 | 0.169629 | 0.631 | 0.509 | 1 | 13 | Creld2 | 0.122 | Old |
| 0.162663 | 0.289555 | 0.238 | 0.149 | 1 | 13 | Mrrf | 0.089 | Old |
| 0.162835 | 0.192865 | 0.689 | 0.623 | 1 | 13 | Pih1d1 | 0.066 | Old |
| 0.163381 | 0.159232 | 0.893 | 0.886 | 1 | 13 | Slc29a3 | 0.007 | Old |
| 0.163406 | 0.600229 | 0.541 | 0.465 | 1 | 13 | Anpep | 0.076 | Old |
| 0.163533 | 0.073047 | 0.557 | 0.456 | 1 | 13 | Hsph1 | 0.101 | Old |
| 0.163832 | 0.730341 | 0.09 | 0.044 | 1 | 13 | Zfp46 | 0.046 | Old |

| | | | | | | | | |
|---|---|---|---|---|---|---|---|---|
| 0.163885 | 0.797781 | 0.066 | 0.026 | 1 | 13 | Gbp4 | 0.04 | Old |
| 0.163885 | 0.62669 | 0.066 | 0.026 | 1 | 13 | Lins1 | 0.04 | Old |
| 0.163885 | 0.521787 | 0.066 | 0.026 | 1 | 13 | Rdh5 | 0.04 | Old |
| 0.163939 | 0.214854 | 0.77 | 0.667 | 1 | 13 | Ddx54 | 0.103 | Old |
| 0.164081 | 0.169881 | 0.836 | 0.825 | 1 | 13 | Grina | 0.011 | Old |
| 0.164413 | 0.186107 | 0.541 | 0.421 | 1 | 13 | Preb | 0.12 | Old |
| 0.164449 | 0.324765 | 0.459 | 0.325 | 1 | 13 | Gt(ROSA)2( | 0.134 | Old |
| 0.164603 | 0.198026 | 0.795 | 0.746 | 1 | 13 | Stom | 0.049 | Old |
| 0.164824 | 0.184597 | 0.484 | 0.377 | 1 | 13 | Golga3 | 0.107 | Old |
| 0.164831 | 0.304936 | 0.369 | 0.272 | 1 | 13 | Gpatch2 | 0.097 | Old |
| 0.164859 | 0.569604 | 0.115 | 0.061 | 1 | 13 | Rnf217 | 0.054 | Old |
| 0.164859 | 0.499056 | 0.115 | 0.061 | 1 | 13 | Pusl1 | 0.054 | Old |
| 0.164967 | 0.499013 | 0.434 | 0.368 | 1 | 13 | Ppp2r3d | 0.066 | Old |
| 0.16524 | 0.340039 | 0.262 | 0.175 | 1 | 13 | Rcc1l | 0.087 | Old |
| 0.165472 | 0.51751 | 0.066 | 0.026 | 1 | 13 | Rrm2 | 0.04 | Old |
| 0.16587 | 0.194437 | 0.779 | 0.719 | 1 | 13 | Prkx | 0.06 | Old |
| 0.165887 | 0.38089 | 0.328 | 0.254 | 1 | 13 | Ttf1 | 0.074 | Old |
| 0.165981 | 0.232398 | 0.279 | 0.193 | 1 | 13 | Aar2 | 0.086 | Old |
| 0.16616 | 0.213192 | 0.844 | 0.789 | 1 | 13 | Copb1 | 0.055 | Old |
| 0.166196 | 0.159614 | 0.967 | 0.886 | 1 | 13 | Trir | 0.081 | Old |
| 0.166231 | 0.55329 | 0.172 | 0.105 | 1 | 13 | Map4k3 | 0.067 | Old |
| 0.166294 | 0.520748 | 0.18 | 0.114 | 1 | 13 | Slc35e2 | 0.066 | Old |
| 0.166416 | 0.563294 | 0.238 | 0.167 | 1 | 13 | Med6 | 0.071 | Old |
| 0.166709 | 0.217876 | 0.533 | 0.421 | 1 | 13 | Dap3 | 0.112 | Old |
| 0.166765 | 0.440181 | 0.279 | 0.193 | 1 | 13 | Dhx34 | 0.086 | Old |
| 0.16686 | 0.105855 | 1 | 1 | 1 | 13 | H2-DMb1 | 0 | Old |
| 0.166923 | 0.241636 | 0.541 | 0.421 | 1 | 13 | Ado | 0.12 | Old |
| 0.167161 | 0.222929 | 0.467 | 0.342 | 1 | 13 | Pabpc4 | 0.125 | Old |
| 0.167222 | 0.283449 | 0.115 | 0.061 | 1 | 13 | 4933427D1 | 0.054 | Old |
| 0.167601 | 0.161115 | 0.516 | 0.404 | 1 | 13 | Tceal8 | 0.112 | Old |
| 0.168128 | 0.20232 | 0.787 | 0.737 | 1 | 13 | Srsf7 | 0.05 | Old |
| 0.168504 | 0.165594 | 0.574 | 0.439 | 1 | 13 | Dtx3l | 0.135 | Old |
| 0.169198 | 0.592878 | 0.09 | 0.044 | 1 | 13 | Gm15448 | 0.046 | Old |
| 0.169583 | 0.20451 | 0.951 | 0.842 | 1 | 13 | Atxn2l | 0.109 | Old |
| 0.169719 | 0.25642 | 0.393 | 0.272 | 1 | 13 | Strip1 | 0.121 | Old |
| 0.169807 | 0.137904 | 0.984 | 0.991 | 1 | 13 | Atp1b3 | -0.007 | Old |
| 0.169939 | 1.331129 | 0.049 | 0.018 | 1 | 13 | Mrnip | 0.031 | Old |
| 0.169939 | 1.310944 | 0.049 | 0.018 | 1 | 13 | Zfp455 | 0.031 | Old |
| 0.169939 | 0.921041 | 0.049 | 0.018 | 1 | 13 | Cdc45 | 0.031 | Old |
| 0.170077 | 0.073212 | 0.582 | 0.667 | 1 | 13 | Tnf | -0.085 | Old |
| 0.170215 | 0.274347 | 0.615 | 0.474 | 1 | 13 | Sp140 | 0.141 | Old |
| 0.170304 | 0.572968 | 0.066 | 0.026 | 1 | 13 | Zfp472 | 0.04 | Old |
| 0.17056 | 0.392133 | 0.09 | 0.044 | 1 | 13 | Rab11fip3 | 0.046 | Old |
| 0.170821 | 0.360165 | 0.508 | 0.412 | 1 | 13 | Tpst2 | 0.096 | Old |
| 0.170856 | 0.559491 | 0.361 | 0.281 | 1 | 13 | Lnpk | 0.08 | Old |

| | | | | | | | | |
|---|---|---|---|---|---|---|---|---|
| 0.170907 | 0.361622 | 0.582 | 0.447 | 1 | 13 | Trak1 | 0.135 | Old |
| 0.171062 | 0.309365 | 0.533 | 0.43 | 1 | 13 | Rabggtb | 0.103 | Old |
| 0.171155 | 0.426336 | 0.311 | 0.228 | 1 | 13 | Mbtps2 | 0.083 | Old |
| 0.171558 | 0.157582 | 0.959 | 0.93 | 1 | 13 | Evi2a | 0.029 | Old |
| 0.171586 | 0.125903 | 0.967 | 0.965 | 1 | 13 | Ndufa3 | 0.002 | Old |
| 0.171594 | 0.099674 | 1 | 1 | 1 | 13 | Canx | 0 | Old |
| 0.171594 | 0.084929 | 1 | 1 | 1 | 13 | Actg1 | 0 | Old |
| 0.17184 | 0.995606 | 0.049 | 0.018 | 1 | 13 | Plxnb3 | 0.031 | Old |
| 0.17184 | 0.929767 | 0.049 | 0.018 | 1 | 13 | Trp53i11 | 0.031 | Old |
| 0.172093 | 0.136578 | 0.852 | 0.781 | 1 | 13 | Stk4 | 0.071 | Old |
| 0.172319 | 0.223148 | 0.615 | 0.5 | 1 | 13 | Cntrl | 0.115 | Old |
| 0.172581 | 4.423098 | 0.016 | 0 | 1 | 13 | Lcn2 | 0.016 | Old |
| 0.172581 | 2.565003 | 0.016 | 0 | 1 | 13 | Gm15056 | 0.016 | Old |
| 0.172581 | 2.492056 | 0.016 | 0 | 1 | 13 | Il12b | 0.016 | Old |
| 0.172581 | 2.006282 | 0.016 | 0 | 1 | 13 | Hal | 0.016 | Old |
| 0.172581 | 1.456563 | 0.016 | 0 | 1 | 13 | Synpo2 | 0.016 | Old |
| 0.172581 | 1.241507 | 0.016 | 0 | 1 | 13 | Gm4841 | 0.016 | Old |
| 0.172581 | 1.23389 | 0.016 | 0 | 1 | 13 | B3galt2 | 0.016 | Old |
| 0.172581 | 1.125302 | 0.016 | 0 | 1 | 13 | Hoxb7 | 0.016 | Old |
| 0.172581 | 1.063705 | 0.016 | 0 | 1 | 13 | Hdgfl3 | 0.016 | Old |
| 0.172581 | 1.05968 | 0.016 | 0 | 1 | 13 | Adh1 | 0.016 | Old |
| 0.172581 | 1.059195 | 0.016 | 0 | 1 | 13 | Klra17 | 0.016 | Old |
| 0.172581 | 1.018044 | 0.016 | 0 | 1 | 13 | Olfml2b | 0.016 | Old |
| 0.172581 | 0.974155 | 0.016 | 0 | 1 | 13 | Rab26 | 0.016 | Old |
| 0.172581 | 0.935126 | 0.016 | 0 | 1 | 13 | Myo1d | 0.016 | Old |
| 0.172581 | 0.934032 | 0.016 | 0 | 1 | 13 | Zfhx4 | 0.016 | Old |
| 0.172581 | 0.928896 | 0.016 | 0 | 1 | 13 | Adgrl4 | 0.016 | Old |
| 0.172581 | 0.924784 | 0.016 | 0 | 1 | 13 | Gm26830 | 0.016 | Old |
| 0.172581 | 0.917344 | 0.016 | 0 | 1 | 13 | Fkbp11 | 0.016 | Old |
| 0.172581 | 0.914257 | 0.016 | 0 | 1 | 13 | Cd300e | 0.016 | Old |
| 0.172581 | 0.90862 | 0.016 | 0 | 1 | 13 | D730045B( | 0.016 | Old |
| 0.172581 | 0.905663 | 0.016 | 0 | 1 | 13 | Rell2 | 0.016 | Old |
| 0.172581 | 0.900393 | 0.016 | 0 | 1 | 13 | Fbxl21 | 0.016 | Old |
| 0.172581 | 0.899102 | 0.016 | 0 | 1 | 13 | Mtfp1 | 0.016 | Old |
| 0.172581 | 0.896662 | 0.016 | 0 | 1 | 13 | Gm44702 | 0.016 | Old |
| 0.172581 | 0.892407 | 0.016 | 0 | 1 | 13 | 2410018L1 | 0.016 | Old |
| 0.172581 | 0.886892 | 0.016 | 0 | 1 | 13 | Tmem231 | 0.016 | Old |
| 0.172581 | 0.886151 | 0.016 | 0 | 1 | 13 | Zfp970 | 0.016 | Old |
| 0.172581 | 0.880661 | 0.016 | 0 | 1 | 13 | Nid1 | 0.016 | Old |
| 0.172581 | 0.870868 | 0.016 | 0 | 1 | 13 | Catsper2 | 0.016 | Old |
| 0.172581 | 0.870512 | 0.016 | 0 | 1 | 13 | Sult5a1 | 0.016 | Old |
| 0.172581 | 0.869365 | 0.016 | 0 | 1 | 13 | Gm13620 | 0.016 | Old |
| 0.172581 | 0.866013 | 0.016 | 0 | 1 | 13 | Fermt2 | 0.016 | Old |
| 0.172581 | 0.858323 | 0.016 | 0 | 1 | 13 | A230020J2 | 0.016 | Old |
| 0.172581 | 0.856795 | 0.016 | 0 | 1 | 13 | Gm14154 | 0.016 | Old |

| | | | | | | | | |
|---|---|---|---|---|---|---|---|---|
| 0.172581 | 0.856526 | 0.016 | 0 | 1 | 13 | 0610039K1 | 0.016 | Old |
| 0.172581 | 0.850282 | 0.016 | 0 | 1 | 13 | Gm37035 | 0.016 | Old |
| 0.172581 | 0.844491 | 0.016 | 0 | 1 | 13 | Capns2 | 0.016 | Old |
| 0.172581 | 0.840676 | 0.016 | 0 | 1 | 13 | Gm14391 | 0.016 | Old |
| 0.172581 | 0.838261 | 0.016 | 0 | 1 | 13 | Gm11496 | 0.016 | Old |
| 0.172581 | 0.837069 | 0.016 | 0 | 1 | 13 | Gm26536 | 0.016 | Old |
| 0.172581 | 0.832108 | 0.016 | 0 | 1 | 13 | Zfp273 | 0.016 | Old |
| 0.172581 | 0.83034 | 0.016 | 0 | 1 | 13 | Mapk12 | 0.016 | Old |
| 0.172581 | 0.828046 | 0.016 | 0 | 1 | 13 | Pacrg | 0.016 | Old |
| 0.172581 | 0.821619 | 0.016 | 0 | 1 | 13 | Fam228b | 0.016 | Old |
| 0.172581 | 0.82106 | 0.016 | 0 | 1 | 13 | AC168977. | 0.016 | Old |
| 0.172581 | 0.819095 | 0.016 | 0 | 1 | 13 | Myrf | 0.016 | Old |
| 0.172581 | 0.815452 | 0.016 | 0 | 1 | 13 | Serinc4 | 0.016 | Old |
| 0.172581 | 0.812807 | 0.016 | 0 | 1 | 13 | Hspb7 | 0.016 | Old |
| 0.172581 | 0.807637 | 0.016 | 0 | 1 | 13 | Aqp3 | 0.016 | Old |
| 0.172581 | 0.807144 | 0.016 | 0 | 1 | 13 | Tgfa | 0.016 | Old |
| 0.172581 | 0.805523 | 0.016 | 0 | 1 | 13 | Ngfr | 0.016 | Old |
| 0.172581 | 0.804791 | 0.016 | 0 | 1 | 13 | A830052D1 | 0.016 | Old |
| 0.172581 | 0.804045 | 0.016 | 0 | 1 | 13 | Galnt14 | 0.016 | Old |
| 0.172581 | 0.797943 | 0.016 | 0 | 1 | 13 | Pik3r3 | 0.016 | Old |
| 0.172581 | 0.79658 | 0.016 | 0 | 1 | 13 | 3632451O( | 0.016 | Old |
| 0.172581 | 0.795079 | 0.016 | 0 | 1 | 13 | Cd109 | 0.016 | Old |
| 0.172581 | 0.792242 | 0.016 | 0 | 1 | 13 | Fosl1 | 0.016 | Old |
| 0.172581 | 0.790923 | 0.016 | 0 | 1 | 13 | Col9a3 | 0.016 | Old |
| 0.172581 | 0.789313 | 0.016 | 0 | 1 | 13 | Tusc5 | 0.016 | Old |
| 0.172581 | 0.788083 | 0.016 | 0 | 1 | 13 | Rufy4 | 0.016 | Old |
| 0.172581 | 0.787943 | 0.016 | 0 | 1 | 13 | Stra6l | 0.016 | Old |
| 0.172581 | 0.787073 | 0.016 | 0 | 1 | 13 | 9430078K2 | 0.016 | Old |
| 0.172581 | 0.784896 | 0.016 | 0 | 1 | 13 | Tek | 0.016 | Old |
| 0.172581 | 0.782815 | 0.016 | 0 | 1 | 13 | Acsbg1 | 0.016 | Old |
| 0.172581 | 0.781529 | 0.016 | 0 | 1 | 13 | Glra1 | 0.016 | Old |
| 0.172581 | 0.778944 | 0.016 | 0 | 1 | 13 | Ovgp1 | 0.016 | Old |
| 0.172581 | 0.775037 | 0.016 | 0 | 1 | 13 | Ube2cbp | 0.016 | Old |
| 0.172581 | 0.771282 | 0.016 | 0 | 1 | 13 | Peli3 | 0.016 | Old |
| 0.172581 | 0.769433 | 0.016 | 0 | 1 | 13 | Gm44672 | 0.016 | Old |
| 0.172581 | 0.768346 | 0.016 | 0 | 1 | 13 | Dtna | 0.016 | Old |
| 0.172581 | 0.767503 | 0.016 | 0 | 1 | 13 | Gm9873 | 0.016 | Old |
| 0.172581 | 0.76716 | 0.016 | 0 | 1 | 13 | Tpbg | 0.016 | Old |
| 0.172581 | 0.765491 | 0.016 | 0 | 1 | 13 | Gm3550 | 0.016 | Old |
| 0.172581 | 0.76511 | 0.016 | 0 | 1 | 13 | Gm5544 | 0.016 | Old |
| 0.172581 | 0.762796 | 0.016 | 0 | 1 | 13 | Serac1 | 0.016 | Old |
| 0.172581 | 0.761942 | 0.016 | 0 | 1 | 13 | Gm16299 | 0.016 | Old |
| 0.172581 | 0.749747 | 0.016 | 0 | 1 | 13 | Chrnb2 | 0.016 | Old |
| 0.172581 | 0.748093 | 0.016 | 0 | 1 | 13 | Gm128 | 0.016 | Old |
| 0.172581 | 0.747475 | 0.016 | 0 | 1 | 13 | Stil | 0.016 | Old |

| | | | | | | | | |
|---|---|---|---|---|---|---|---|---|
| 0.172581 | 0.745732 | 0.016 | 0 | 1 | 13 | Itgb3 | 0.016 | Old |
| 0.172581 | 0.741339 | 0.016 | 0 | 1 | 13 | Tjp1 | 0.016 | Old |
| 0.172581 | 0.741272 | 0.016 | 0 | 1 | 13 | Ldb2 | 0.016 | Old |
| 0.172581 | 0.740727 | 0.016 | 0 | 1 | 13 | Ccdc146 | 0.016 | Old |
| 0.172581 | 0.735816 | 0.016 | 0 | 1 | 13 | Ppic | 0.016 | Old |
| 0.172581 | 0.735106 | 0.016 | 0 | 1 | 13 | Cdk15 | 0.016 | Old |
| 0.172581 | 0.734111 | 0.016 | 0 | 1 | 13 | Jazf1 | 0.016 | Old |
| 0.172581 | 0.732558 | 0.016 | 0 | 1 | 13 | Tstd1 | 0.016 | Old |
| 0.172581 | 0.728055 | 0.016 | 0 | 1 | 13 | 2310034G( | 0.016 | Old |
| 0.172581 | 0.726828 | 0.016 | 0 | 1 | 13 | Lipt1 | 0.016 | Old |
| 0.172581 | 0.726194 | 0.016 | 0 | 1 | 13 | Rpl3l | 0.016 | Old |
| 0.172581 | 0.723884 | 0.016 | 0 | 1 | 13 | Ubd | 0.016 | Old |
| 0.172581 | 0.723641 | 0.016 | 0 | 1 | 13 | Cngb1 | 0.016 | Old |
| 0.172581 | 0.721623 | 0.016 | 0 | 1 | 13 | 2310047D( | 0.016 | Old |
| 0.172581 | 0.72135 | 0.016 | 0 | 1 | 13 | Gm46440 | 0.016 | Old |
| 0.172581 | 0.719544 | 0.016 | 0 | 1 | 13 | 4933404O1 | 0.016 | Old |
| 0.172581 | 0.719095 | 0.016 | 0 | 1 | 13 | Gm11127 | 0.016 | Old |
| 0.172581 | 0.717665 | 0.016 | 0 | 1 | 13 | 1810062G1 | 0.016 | Old |
| 0.172581 | 0.716857 | 0.016 | 0 | 1 | 13 | Itgb4 | 0.016 | Old |
| 0.172581 | 0.71502 | 0.016 | 0 | 1 | 13 | Col18a1 | 0.016 | Old |
| 0.172581 | 0.71243 | 0.016 | 0 | 1 | 13 | Gm10260 | 0.016 | Old |
| 0.172581 | 0.710325 | 0.016 | 0 | 1 | 13 | Zfp82 | 0.016 | Old |
| 0.172581 | 0.705277 | 0.016 | 0 | 1 | 13 | Tead1 | 0.016 | Old |
| 0.172581 | 0.702253 | 0.016 | 0 | 1 | 13 | Gm13889 | 0.016 | Old |
| 0.172581 | 0.698154 | 0.016 | 0 | 1 | 13 | Serpine2 | 0.016 | Old |
| 0.172581 | 0.697678 | 0.016 | 0 | 1 | 13 | Gm45495 | 0.016 | Old |
| 0.172581 | 0.697196 | 0.016 | 0 | 1 | 13 | Uchl4 | 0.016 | Old |
| 0.172581 | 0.694067 | 0.016 | 0 | 1 | 13 | Gm37800 | 0.016 | Old |
| 0.172581 | 0.694048 | 0.016 | 0 | 1 | 13 | Has1 | 0.016 | Old |
| 0.172581 | 0.693579 | 0.016 | 0 | 1 | 13 | Gm45605 | 0.016 | Old |
| 0.172581 | 0.69068 | 0.016 | 0 | 1 | 13 | Peg10 | 0.016 | Old |
| 0.172581 | 0.687564 | 0.016 | 0 | 1 | 13 | Emp2 | 0.016 | Old |
| 0.172581 | 0.68658 | 0.016 | 0 | 1 | 13 | Olfr77 | 0.016 | Old |
| 0.172581 | 0.685271 | 0.016 | 0 | 1 | 13 | A430057M | 0.016 | Old |
| 0.172581 | 0.685036 | 0.016 | 0 | 1 | 13 | Dnaaf2 | 0.016 | Old |
| 0.172581 | 0.683189 | 0.016 | 0 | 1 | 13 | Dpep1 | 0.016 | Old |
| 0.172581 | 0.680455 | 0.016 | 0 | 1 | 13 | Lat | 0.016 | Old |
| 0.172581 | 0.680207 | 0.016 | 0 | 1 | 13 | Npr2 | 0.016 | Old |
| 0.172581 | 0.67874 | 0.016 | 0 | 1 | 13 | Tm6sf2 | 0.016 | Old |
| 0.172581 | 0.677054 | 0.016 | 0 | 1 | 13 | Tmem25 | 0.016 | Old |
| 0.172581 | 0.672786 | 0.016 | 0 | 1 | 13 | Mreg | 0.016 | Old |
| 0.172581 | 0.672751 | 0.016 | 0 | 1 | 13 | Mok | 0.016 | Old |
| 0.172581 | 0.671276 | 0.016 | 0 | 1 | 13 | Pcdhgb6 | 0.016 | Old |
| 0.172581 | 0.668895 | 0.016 | 0 | 1 | 13 | Cldn23 | 0.016 | Old |
| 0.172581 | 0.668253 | 0.016 | 0 | 1 | 13 | Efcab11 | 0.016 | Old |

| | | | | | | | | |
|---|---|---|---|---|---|---|---|---|
| 0.172581 | 0.667466 | 0.016 | 0 | 1 | 13 | Hist1h3f | 0.016 | Old |
| 0.172581 | 0.666802 | 0.016 | 0 | 1 | 13 | 2700012I2( | 0.016 | Old |
| 0.172581 | 0.666585 | 0.016 | 0 | 1 | 13 | Gm20498 | 0.016 | Old |
| 0.172581 | 0.665961 | 0.016 | 0 | 1 | 13 | Necab3 | 0.016 | Old |
| 0.172581 | 0.665375 | 0.016 | 0 | 1 | 13 | Iqck | 0.016 | Old |
| 0.172581 | 0.664064 | 0.016 | 0 | 1 | 13 | Hoxb3os | 0.016 | Old |
| 0.172581 | 0.664015 | 0.016 | 0 | 1 | 13 | Gm16549 | 0.016 | Old |
| 0.172581 | 0.662292 | 0.016 | 0 | 1 | 13 | Kcnk5 | 0.016 | Old |
| 0.172581 | 0.659539 | 0.016 | 0 | 1 | 13 | Lox | 0.016 | Old |
| 0.172581 | 0.658803 | 0.016 | 0 | 1 | 13 | Srp54b | 0.016 | Old |
| 0.172581 | 0.655631 | 0.016 | 0 | 1 | 13 | Kctd4 | 0.016 | Old |
| 0.172581 | 0.655536 | 0.016 | 0 | 1 | 13 | Gm28727 | 0.016 | Old |
| 0.172581 | 0.64409 | 0.016 | 0 | 1 | 13 | Gm20457 | 0.016 | Old |
| 0.172581 | 0.643276 | 0.016 | 0 | 1 | 13 | Sox7 | 0.016 | Old |
| 0.172581 | 0.640783 | 0.016 | 0 | 1 | 13 | Gm15834 | 0.016 | Old |
| 0.172581 | 0.63888 | 0.016 | 0 | 1 | 13 | Itga2b | 0.016 | Old |
| 0.172581 | 0.638714 | 0.016 | 0 | 1 | 13 | Slc16a9 | 0.016 | Old |
| 0.172581 | 0.636014 | 0.016 | 0 | 1 | 13 | Smoc2 | 0.016 | Old |
| 0.172581 | 0.635059 | 0.016 | 0 | 1 | 13 | 2410004P0 | 0.016 | Old |
| 0.172581 | 0.633004 | 0.016 | 0 | 1 | 13 | Gm45237 | 0.016 | Old |
| 0.172581 | 0.630196 | 0.016 | 0 | 1 | 13 | Kifc1 | 0.016 | Old |
| 0.172581 | 0.626667 | 0.016 | 0 | 1 | 13 | AC150035. | 0.016 | Old |
| 0.172581 | 0.620767 | 0.016 | 0 | 1 | 13 | Hsd11b1 | 0.016 | Old |
| 0.172581 | 0.620767 | 0.016 | 0 | 1 | 13 | Crispld2 | 0.016 | Old |
| 0.172581 | 0.620142 | 0.016 | 0 | 1 | 13 | Postn | 0.016 | Old |
| 0.172581 | 0.620142 | 0.016 | 0 | 1 | 13 | Rec114 | 0.016 | Old |
| 0.172581 | 0.619039 | 0.016 | 0 | 1 | 13 | Thsd1 | 0.016 | Old |
| 0.172581 | 0.617328 | 0.016 | 0 | 1 | 13 | Apod | 0.016 | Old |
| 0.172581 | 0.616647 | 0.016 | 0 | 1 | 13 | Gm15133 | 0.016 | Old |
| 0.172581 | 0.615183 | 0.016 | 0 | 1 | 13 | Lrrtm2 | 0.016 | Old |
| 0.172581 | 0.614567 | 0.016 | 0 | 1 | 13 | 1700018L0 | 0.016 | Old |
| 0.172581 | 0.611969 | 0.016 | 0 | 1 | 13 | Tinagl1 | 0.016 | Old |
| 0.172581 | 0.611181 | 0.016 | 0 | 1 | 13 | Gm15655 | 0.016 | Old |
| 0.172581 | 0.606758 | 0.016 | 0 | 1 | 13 | Slc18a1 | 0.016 | Old |
| 0.172581 | 0.606467 | 0.016 | 0 | 1 | 13 | Ccdc63 | 0.016 | Old |
| 0.172581 | 0.604055 | 0.016 | 0 | 1 | 13 | Zfp78 | 0.016 | Old |
| 0.172581 | 0.603705 | 0.016 | 0 | 1 | 13 | Ccr3 | 0.016 | Old |
| 0.172581 | 0.602568 | 0.016 | 0 | 1 | 13 | Sntb1 | 0.016 | Old |
| 0.172581 | 0.600699 | 0.016 | 0 | 1 | 13 | Fxyd6 | 0.016 | Old |
| 0.172581 | 0.598029 | 0.016 | 0 | 1 | 13 | Ggn | 0.016 | Old |
| 0.172581 | 0.596931 | 0.016 | 0 | 1 | 13 | 1700013D2 | 0.016 | Old |
| 0.172581 | 0.596207 | 0.016 | 0 | 1 | 13 | Gm12708 | 0.016 | Old |
| 0.172581 | 0.594502 | 0.016 | 0 | 1 | 13 | AC152827. | 0.016 | Old |
| 0.172581 | 0.5895 | 0.016 | 0 | 1 | 13 | Gm45805 | 0.016 | Old |
| 0.172581 | 0.589428 | 0.016 | 0 | 1 | 13 | Ccdc141 | 0.016 | Old |

| | | | | | | | | |
|---|---|---|---|---|---|---|---|---|
| 0.172581 | 0.589097 | 0.016 | 0 | 1 | 13 | Adra1a | 0.016 | Old |
| 0.172581 | 0.584105 | 0.016 | 0 | 1 | 13 | Fscn3 | 0.016 | Old |
| 0.172581 | 0.583689 | 0.016 | 0 | 1 | 13 | Sgo1 | 0.016 | Old |
| 0.172581 | 0.579391 | 0.016 | 0 | 1 | 13 | F730311O2 | 0.016 | Old |
| 0.172581 | 0.579391 | 0.016 | 0 | 1 | 13 | Sapcd2 | 0.016 | Old |
| 0.172581 | 0.572737 | 0.016 | 0 | 1 | 13 | Tshz3 | 0.016 | Old |
| 0.172581 | 0.571955 | 0.016 | 0 | 1 | 13 | Lpar2 | 0.016 | Old |
| 0.172581 | 0.571789 | 0.016 | 0 | 1 | 13 | Lpar5 | 0.016 | Old |
| 0.172581 | 0.569755 | 0.016 | 0 | 1 | 13 | Sema3a | 0.016 | Old |
| 0.172581 | 0.5697 | 0.016 | 0 | 1 | 13 | Gm17455 | 0.016 | Old |
| 0.172581 | 0.569286 | 0.016 | 0 | 1 | 13 | Hook1 | 0.016 | Old |
| 0.172581 | 0.568912 | 0.016 | 0 | 1 | 13 | C8g | 0.016 | Old |
| 0.172581 | 0.564946 | 0.016 | 0 | 1 | 13 | Ikzf4 | 0.016 | Old |
| 0.172581 | 0.562502 | 0.016 | 0 | 1 | 13 | Man2c1os | 0.016 | Old |
| 0.172581 | 0.562294 | 0.016 | 0 | 1 | 13 | Gm5093 | 0.016 | Old |
| 0.172581 | 0.56028 | 0.016 | 0 | 1 | 13 | Ska3 | 0.016 | Old |
| 0.172581 | 0.559113 | 0.016 | 0 | 1 | 13 | 6030443J0 | 0.016 | Old |
| 0.172581 | 0.555546 | 0.016 | 0 | 1 | 13 | Gm9465 | 0.016 | Old |
| 0.172581 | 0.553256 | 0.016 | 0 | 1 | 13 | Htr7 | 0.016 | Old |
| 0.172581 | 0.551242 | 0.016 | 0 | 1 | 13 | Rnf225 | 0.016 | Old |
| 0.172581 | 0.54877 | 0.016 | 0 | 1 | 13 | Gm33370 | 0.016 | Old |
| 0.172581 | 0.543876 | 0.016 | 0 | 1 | 13 | Kcnab3 | 0.016 | Old |
| 0.172581 | 0.530954 | 0.016 | 0 | 1 | 13 | Gm28050 | 0.016 | Old |
| 0.172581 | 0.523991 | 0.016 | 0 | 1 | 13 | Nckap1 | 0.016 | Old |
| 0.172581 | 0.51603 | 0.016 | 0 | 1 | 13 | Gm15441 | 0.016 | Old |
| 0.172581 | 0.488157 | 0.016 | 0 | 1 | 13 | Rin1 | 0.016 | Old |
| 0.172581 | 0.47073 | 0.016 | 0 | 1 | 13 | Gm36423 | 0.016 | Old |
| 0.172581 | 0.415709 | 0.016 | 0 | 1 | 13 | Mpz | 0.016 | Old |
| 0.172793 | 0.135644 | 1 | 0.991 | 1 | 13 | Pdia3 | 0.009 | Old |
| 0.172818 | 0.124779 | 0.508 | 0.377 | 1 | 13 | Usp6nl | 0.131 | Old |
| 0.172871 | 0.220782 | 0.631 | 0.526 | 1 | 13 | Ap3m1 | 0.105 | Old |
| 0.173128 | 0.386695 | 0.41 | 0.316 | 1 | 13 | Agap1 | 0.094 | Old |
| 0.17324 | 0.425054 | 0.115 | 0.061 | 1 | 13 | Mms22l | 0.054 | Old |
| 0.173659 | 0.215836 | 0.68 | 0.632 | 1 | 13 | Psmd3 | 0.048 | Old |
| 0.173698 | 0.513936 | 0.205 | 0.132 | 1 | 13 | Ccdc28a | 0.073 | Old |
| 0.173698 | 0.260898 | 0.205 | 0.132 | 1 | 13 | Gpank1 | 0.073 | Old |
| 0.174062 | 0.207883 | 0.828 | 0.746 | 1 | 13 | Tecr | 0.082 | Old |
| 0.174364 | 0.081765 | 0.197 | 0.114 | 1 | 13 | Zbtb46 | 0.083 | Old |
| 0.174591 | 0.202994 | 0.967 | 0.956 | 1 | 13 | Sgpl1 | 0.011 | Old |
| 0.174863 | 0.580042 | 0.262 | 0.184 | 1 | 13 | Ints14 | 0.078 | Old |
| 0.174934 | 0.234118 | 0.721 | 0.623 | 1 | 13 | Tsen34 | 0.098 | Old |
| 0.175017 | 0.32264 | 0.508 | 0.421 | 1 | 13 | Dido1 | 0.087 | Old |
| 0.175112 | 0.221641 | 0.779 | 0.702 | 1 | 13 | Eif2s3x | 0.077 | Old |
| 0.175142 | 0.21453 | 0.361 | 0.263 | 1 | 13 | Inf2 | 0.098 | Old |
| 0.175691 | 0.908384 | 0.049 | 0.018 | 1 | 13 | Gm16845 | 0.031 | Old |

| | | | | | | | | |
|---|---|---|---|---|---|---|---|---|
| 0.175691 | 0.818112 | 0.049 | 0.018 | 1 | 13 | Gm43672 | 0.031 | Old |
| 0.175826 | 0.308053 | 0.484 | 0.36 | 1 | 13 | Mrpl45 | 0.124 | Old |
| 0.176083 | 0.554998 | 0.402 | 0.307 | 1 | 13 | Fdps | 0.095 | Old |
| 0.176491 | 0.488935 | 0.303 | 0.228 | 1 | 13 | Tma16 | 0.075 | Old |
| 0.176497 | 0.350917 | 0.443 | 0.351 | 1 | 13 | Gart | 0.092 | Old |
| 0.176579 | 0.640751 | 0.221 | 0.158 | 1 | 13 | Wbp1 | 0.063 | Old |
| 0.176725 | 0.169238 | 0.451 | 0.316 | 1 | 13 | Tmem68 | 0.135 | Old |
| 0.176782 | 0.285523 | 0.385 | 0.289 | 1 | 13 | Cdk2 | 0.096 | Old |
| 0.176809 | 0.55191 | 0.23 | 0.158 | 1 | 13 | Zfp64 | 0.072 | Old |
| 0.176927 | 0.493612 | 0.115 | 0.061 | 1 | 13 | Herc3 | 0.054 | Old |
| 0.176927 | 0.386508 | 0.115 | 0.061 | 1 | 13 | Sars2 | 0.054 | Old |
| 0.177185 | 0.266889 | 0.582 | 0.509 | 1 | 13 | Mitf | 0.073 | Old |
| 0.177185 | 1.579502 | 0.098 | 0.053 | 1 | 13 | Gm19951 | 0.045 | Old |
| 0.177219 | 0.22769 | 0.639 | 0.561 | 1 | 13 | Arhgap1 | 0.078 | Old |
| 0.17725 | 0.372672 | 0.189 | 0.123 | 1 | 13 | Hspbap1 | 0.066 | Old |
| 0.177455 | 0.344528 | 0.41 | 0.333 | 1 | 13 | Gngt2 | 0.077 | Old |
| 0.17763 | 0.119937 | 0.951 | 0.956 | 1 | 13 | Selenos | -0.005 | Old |
| 0.17764 | 1.066442 | 0.049 | 0.018 | 1 | 13 | Cavin3 | 0.031 | Old |
| 0.17764 | 1.059366 | 0.049 | 0.018 | 1 | 13 | Fads3 | 0.031 | Old |
| 0.17764 | 1.031489 | 0.049 | 0.018 | 1 | 13 | Hspa1l | 0.031 | Old |
| 0.177759 | 0.188793 | 0.459 | 0.342 | 1 | 13 | Sf3a2 | 0.117 | Old |
| 0.177801 | 0.328745 | 0.549 | 0.465 | 1 | 13 | Haus8 | 0.084 | Old |
| 0.177935 | 0.526711 | 0.148 | 0.088 | 1 | 13 | Msh3 | 0.06 | Old |
| 0.178025 | 0.259537 | 0.279 | 0.193 | 1 | 13 | Borcs5 | 0.086 | Old |
| 0.178094 | 0.540904 | 0.197 | 0.132 | 1 | 13 | Atf5 | 0.065 | Old |
| 0.178569 | 0.26533 | 0.336 | 0.246 | 1 | 13 | Cinp | 0.09 | Old |
| 0.178729 | 0.63826 | 0.139 | 0.088 | 1 | 13 | Gorab | 0.051 | Old |
| 0.178856 | 0.309223 | 0.311 | 0.219 | 1 | 13 | Wdr46 | 0.092 | Old |
| 0.178883 | 0.112801 | 1 | 1 | 1 | 13 | Gpi1 | 0 | Old |
| 0.179181 | 0.460255 | 0.172 | 0.105 | 1 | 13 | Ccar2 | 0.067 | Old |
| 0.179194 | 0.54005 | 0.189 | 0.123 | 1 | 13 | Nprl3 | 0.066 | Old |
| 0.17942 | 0.287537 | 0.533 | 0.439 | 1 | 13 | Sae1 | 0.094 | Old |
| 0.179605 | 1.733801 | 0.049 | 0.018 | 1 | 13 | H2-M2 | 0.031 | Old |
| 0.179605 | 0.916013 | 0.049 | 0.018 | 1 | 13 | Fut10 | 0.031 | Old |
| 0.179605 | 0.905826 | 0.049 | 0.018 | 1 | 13 | Tmem26 | 0.031 | Old |
| 0.180083 | 0.390538 | 0.27 | 0.193 | 1 | 13 | Urgcp | 0.077 | Old |
| 0.180092 | 0.485697 | 0.303 | 0.237 | 1 | 13 | Pex13 | 0.066 | Old |
| 0.180119 | 0.108763 | 1 | 0.991 | 1 | 13 | Tomm7 | 0.009 | Old |
| 0.180306 | 0.325338 | 0.697 | 0.57 | 1 | 13 | 8-Sep | 0.127 | Old |
| 0.180392 | 0.245624 | 0.492 | 0.614 | 1 | 13 | Cxcl1 | -0.122 | Old |
| 0.180434 | 0.151302 | 0.492 | 0.368 | 1 | 13 | Syap1 | 0.124 | Old |
| 0.180824 | 0.213457 | 0.705 | 0.596 | 1 | 13 | Ten1 | 0.109 | Old |
| 0.180924 | 0.348357 | 0.18 | 0.114 | 1 | 13 | Coq3 | 0.066 | Old |
| 0.180927 | 0.18561 | 0.787 | 0.675 | 1 | 13 | Cebpa | 0.112 | Old |
| 0.180988 | 0.188308 | 0.418 | 0.316 | 1 | 13 | Anapc10 | 0.102 | Old |

| | | | | | | | | |
|---|---|---|---|---|---|---|---|---|
| 0.181103 | 0.219413 | 0.926 | 0.86 | 1 | 13 | Pros1 | 0.066 | Old |
| 0.181136 | 0.361671 | 0.361 | 0.281 | 1 | 13 | Lmf2 | 0.08 | Old |
| 0.181235 | 0.512398 | 0.336 | 0.263 | 1 | 13 | Thbd | 0.073 | Old |
| 0.181466 | 0.280376 | 0.484 | 0.377 | 1 | 13 | Pmpca | 0.107 | Old |
| 0.181517 | 0.301464 | 0.484 | 0.386 | 1 | 13 | Camk2g | 0.098 | Old |
| 0.181587 | 1.100935 | 0.049 | 0.018 | 1 | 13 | Gm15564 | 0.031 | Old |
| 0.181587 | 0.916764 | 0.049 | 0.018 | 1 | 13 | Lonrf3 | 0.031 | Old |
| 0.181587 | 0.755675 | 0.049 | 0.018 | 1 | 13 | Cd46 | 0.031 | Old |
| 0.181587 | 0.724135 | 0.049 | 0.018 | 1 | 13 | Enkur | 0.031 | Old |
| 0.181587 | 0.646113 | 0.049 | 0.018 | 1 | 13 | Ppp1r3f | 0.031 | Old |
| 0.181598 | 0.379837 | 0.262 | 0.184 | 1 | 13 | Rbm10 | 0.078 | Old |
| 0.181611 | 0.415021 | 0.459 | 0.36 | 1 | 13 | Kri1 | 0.099 | Old |
| 0.181799 | 0.549879 | 0.156 | 0.096 | 1 | 13 | Meis3 | 0.06 | Old |
| 0.181799 | 0.508249 | 0.156 | 0.096 | 1 | 13 | Asb1 | 0.06 | Old |
| 0.181934 | 0.099566 | 0.91 | 0.851 | 1 | 13 | Map7d1 | 0.059 | Old |
| 0.181943 | 0.203512 | 0.434 | 0.342 | 1 | 13 | Atad2 | 0.092 | Old |
| 0.182438 | 0.276721 | 0.287 | 0.202 | 1 | 13 | Cdca4 | 0.085 | Old |
| 0.1828 | 0.188669 | 0.779 | 0.728 | 1 | 13 | Acbd3 | 0.051 | Old |
| 0.182841 | 0.22751 | 0.672 | 0.561 | 1 | 13 | Cpeb4 | 0.111 | Old |
| 0.183044 | 0.381442 | 0.459 | 0.386 | 1 | 13 | Phb | 0.073 | Old |
| 0.1832 | 0.372638 | 0.492 | 0.421 | 1 | 13 | Rars | 0.071 | Old |
| 0.183223 | 0.176964 | 0.967 | 0.947 | 1 | 13 | G3bp1 | 0.02 | Old |
| 0.183402 | 0.332193 | 0.598 | 0.465 | 1 | 13 | Nfatc1 | 0.133 | Old |
| 0.183468 | 0.297108 | 0.148 | 0.088 | 1 | 13 | Alyref2 | 0.06 | Old |
| 0.183476 | 0.259713 | 0.418 | 0.307 | 1 | 13 | Pigq | 0.111 | Old |
| 0.183585 | 0.801715 | 0.049 | 0.018 | 1 | 13 | Gm14286 | 0.031 | Old |
| 0.183585 | 0.782493 | 0.049 | 0.018 | 1 | 13 | Fam135a | 0.031 | Old |
| 0.183585 | 0.737045 | 0.049 | 0.018 | 1 | 13 | Gimap1 | 0.031 | Old |
| 0.183585 | 0.686788 | 0.049 | 0.018 | 1 | 13 | Spata2l | 0.031 | Old |
| 0.183665 | 0.216347 | 0.713 | 0.579 | 1 | 13 | Vps26a | 0.134 | Old |
| 0.183712 | 0.216044 | 0.779 | 0.711 | 1 | 13 | Skp1a | 0.068 | Old |
| 0.183732 | 0.192341 | 0.893 | 0.851 | 1 | 13 | Gna13 | 0.042 | Old |
| 0.184183 | 0.12097 | 0.992 | 0.991 | 1 | 13 | Srsf5 | 0.001 | Old |
| 0.18457 | 0.290962 | 0.344 | 0.246 | 1 | 13 | Slc9a6 | 0.098 | Old |
| 0.184588 | 0.251182 | 0.566 | 0.465 | 1 | 13 | Gas2l3 | 0.101 | Old |
| 0.184633 | 0.228841 | 0.09 | 0.044 | 1 | 13 | Cdc42bpa | 0.046 | Old |
| 0.184652 | 0.116251 | 0.508 | 0.386 | 1 | 13 | Gar1 | 0.122 | Old |
| 0.184965 | 0.340069 | 0.295 | 0.211 | 1 | 13 | Tfb2m | 0.084 | Old |
| 0.185099 | 0.241815 | 0.336 | 0.228 | 1 | 13 | Slc39a11 | 0.108 | Old |
| 0.185157 | 0.092965 | 0.77 | 0.684 | 1 | 13 | Kat2b | 0.086 | Old |
| 0.185216 | 0.23553 | 0.893 | 0.904 | 1 | 13 | Ap1b1 | -0.011 | Old |
| 0.185298 | 0.267856 | 0.369 | 0.272 | 1 | 13 | Slc17a5 | 0.097 | Old |
| 0.185341 | 0.227105 | 0.434 | 0.316 | 1 | 13 | Rtp4 | 0.118 | Old |
| 0.185439 | 0.251691 | 0.713 | 0.684 | 1 | 13 | Slc25a37 | 0.029 | Old |
| 0.185599 | 2.039895 | 0.049 | 0.018 | 1 | 13 | Pglyrp1 | 0.031 | Old |

| | | | | | | | | |
|---|---|---|---|---|---|---|---|---|
| 0.185599 | 0.838789 | 0.049 | 0.018 | 1 | 13 | Adap2os | 0.031 | Old |
| 0.18583 | 0.193614 | 0.607 | 0.544 | 1 | 13 | Gm34084 | 0.063 | Old |
| 0.185983 | 0.326268 | 0.541 | 0.421 | 1 | 13 | Zc3h4 | 0.12 | Old |
| 0.186087 | 0.471235 | 0.09 | 0.044 | 1 | 13 | Wdr59 | 0.046 | Old |
| 0.186451 | 0.300959 | 0.869 | 0.719 | 1 | 13 | Pkig | 0.15 | Old |
| 0.186753 | 0.195383 | 0.492 | 0.386 | 1 | 13 | Mtmr12 | 0.106 | Old |
| 0.186876 | 0.921585 | 0.074 | 0.035 | 1 | 13 | Trim13 | 0.039 | Old |
| 0.186882 | 0.437722 | 0.32 | 0.228 | 1 | 13 | Gm43848 | 0.092 | Old |
| 0.187036 | 0.302162 | 0.18 | 0.114 | 1 | 13 | Plpp6 | 0.066 | Old |
| 0.187129 | 0.407767 | 0.189 | 0.123 | 1 | 13 | Lyrm4 | 0.066 | Old |
| 0.187629 | 0.742032 | 0.049 | 0.018 | 1 | 13 | Gm26935 | 0.031 | Old |
| 0.187731 | 0.409093 | 0.311 | 0.228 | 1 | 13 | Lrch4 | 0.083 | Old |
| 0.188492 | 0.828013 | 0.074 | 0.035 | 1 | 13 | Naprt | 0.039 | Old |
| 0.188845 | 0.234384 | 0.262 | 0.175 | 1 | 13 | Adsl | 0.087 | Old |
| 0.18906 | 0.208372 | 0.541 | 0.439 | 1 | 13 | 8-Mar | 0.102 | Old |
| 0.189248 | 0.15429 | 0.975 | 0.921 | 1 | 13 | Nop53 | 0.054 | Old |
| 0.18937 | 0.394145 | 0.311 | 0.228 | 1 | 13 | Dvl3 | 0.083 | Old |
| 0.189466 | 0.720466 | 0.098 | 0.053 | 1 | 13 | Palm | 0.045 | Old |
| 0.189568 | 0.105687 | 0.238 | 0.158 | 1 | 13 | Galnt4 | 0.08 | Old |
| 0.189675 | 0.629813 | 0.049 | 0.018 | 1 | 13 | Gm16740 | 0.031 | Old |
| 0.189675 | 0.589238 | 0.049 | 0.018 | 1 | 13 | Aplnr | 0.031 | Old |
| 0.189675 | 0.537522 | 0.049 | 0.018 | 1 | 13 | Cit | 0.031 | Old |
| 0.189675 | 0.478225 | 0.049 | 0.018 | 1 | 13 | Ankrd55 | 0.031 | Old |
| 0.18993 | 0.333803 | 0.205 | 0.132 | 1 | 13 | Lrba | 0.073 | Old |
| 0.190009 | 0.469531 | 0.238 | 0.167 | 1 | 13 | Drg2 | 0.071 | Old |
| 0.190196 | 0.179412 | 0.885 | 0.877 | 1 | 13 | Psmd13 | 0.008 | Old |
| 0.190243 | 0.084823 | 1 | 1 | 1 | 13 | H2-Ab1 | 0 | Old |
| 0.190484 | 0.498657 | 0.131 | 0.079 | 1 | 13 | Cox18 | 0.052 | Old |
| 0.190645 | 0.163957 | 0.779 | 0.719 | 1 | 13 | Abcf1 | 0.06 | Old |
| 0.190682 | 0.136758 | 0.877 | 0.807 | 1 | 13 | Sidt2 | 0.07 | Old |
| 0.190713 | 0.042115 | 0.172 | 0.105 | 1 | 13 | Tacc3 | 0.067 | Old |
| 0.190853 | 0.154308 | 0.975 | 0.912 | 1 | 13 | Necap2 | 0.063 | Old |
| 0.190868 | 0.575039 | 0.098 | 0.053 | 1 | 13 | Large1 | 0.045 | Old |
| 0.190961 | 0.525853 | 0.385 | 0.289 | 1 | 13 | Dgat1 | 0.096 | Old |
| 0.191023 | 0.169633 | 0.533 | 0.395 | 1 | 13 | Pex7 | 0.138 | Old |
| 0.191026 | 0.385564 | 0.279 | 0.202 | 1 | 13 | Cpox | 0.077 | Old |
| 0.191111 | 0.197116 | 0.467 | 0.368 | 1 | 13 | Nol11 | 0.099 | Old |
| 0.191448 | 0.295395 | 0.451 | 0.36 | 1 | 13 | Strn | 0.091 | Old |
| 0.191537 | 0.240574 | 0.975 | 1 | 1 | 13 | Myadm | -0.025 | Old |
| 0.191556 | 0.248488 | 0.508 | 0.404 | 1 | 13 | Ube3c | 0.104 | Old |
| 0.191738 | 0.470386 | 0.049 | 0.018 | 1 | 13 | Helq | 0.031 | Old |
| 0.191798 | 0.359278 | 0.689 | 0.596 | 1 | 13 | Pcyt1a | 0.093 | Old |
| 0.19185 | 0.511793 | 0.238 | 0.167 | 1 | 13 | Slc25a22 | 0.071 | Old |
| 0.19218 | 0.414573 | 0.443 | 0.351 | 1 | 13 | Necap1 | 0.092 | Old |
| 0.192219 | 0.503752 | 0.221 | 0.158 | 1 | 13 | Akip1 | 0.063 | Old |

| | | | | | | | | |
|---|---|---|---|---|---|---|---|---|
| 0.192288 | 0.59276 | 0.18 | 0.123 | 1 | 13 | C130050O1 | 0.057 | Old |
| 0.192319 | 0.430615 | 0.361 | 0.272 | 1 | 13 | Dnajc30 | 0.089 | Old |
| 0.192592 | 0.141277 | 0.811 | 0.781 | 1 | 13 | Ifi27 | 0.03 | Old |
| 0.192756 | 0.201903 | 0.934 | 0.921 | 1 | 13 | Rhog | 0.013 | Old |
| 0.193139 | 0.219129 | 0.656 | 0.588 | 1 | 13 | Utp3 | 0.068 | Old |
| 0.193404 | 0.850987 | 0.074 | 0.035 | 1 | 13 | Jag1 | 0.039 | Old |
| 0.193613 | 0.206766 | 0.254 | 0.175 | 1 | 13 | Pinx1 | 0.079 | Old |
| 0.193627 | 0.375621 | 0.279 | 0.202 | 1 | 13 | Trmt2b | 0.077 | Old |
| 0.193818 | 0.533481 | 0.049 | 0.018 | 1 | 13 | Tedc1 | 0.031 | Old |
| 0.193819 | 0.142957 | 0.992 | 0.991 | 1 | 13 | Fam49b | 0.001 | Old |
| 0.193845 | 0.359187 | 0.262 | 0.184 | 1 | 13 | Slc25a13 | 0.078 | Old |
| 0.194134 | 0.110168 | 0.959 | 0.965 | 1 | 13 | Eif5b | -0.006 | Old |
| 0.194139 | 0.580297 | 0.131 | 0.079 | 1 | 13 | Kbtbd4 | 0.052 | Old |
| 0.194182 | 0.235662 | 0.574 | 0.421 | 1 | 13 | Mex3c | 0.153 | Old |
| 0.19435 | 0.214676 | 0.213 | 0.14 | 1 | 13 | Trpm2 | 0.073 | Old |
| 0.194529 | 0.336238 | 0.492 | 0.404 | 1 | 13 | Nolc1 | 0.088 | Old |
| 0.194747 | 0.375066 | 0.459 | 0.377 | 1 | 13 | Cramp1l | 0.082 | Old |
| 0.194827 | 0.345918 | 0.385 | 0.289 | 1 | 13 | Nubp2 | 0.096 | Old |
| 0.194919 | 0.155269 | 0.525 | 0.404 | 1 | 13 | Abl2 | 0.121 | Old |
| 0.194948 | 0.34363 | 0.393 | 0.316 | 1 | 13 | Slc25a17 | 0.077 | Old |
| 0.195005 | 0.121617 | 0.418 | 0.298 | 1 | 13 | Naglu | 0.12 | Old |
| 0.195062 | 0.816319 | 0.074 | 0.035 | 1 | 13 | Map7 | 0.039 | Old |
| 0.195121 | 0.552209 | 0.098 | 0.053 | 1 | 13 | Oas1c | 0.045 | Old |
| 0.195121 | 0.452583 | 0.098 | 0.053 | 1 | 13 | Fcho1 | 0.045 | Old |
| 0.19514 | 0.341722 | 0.549 | 0.474 | 1 | 13 | Htra2 | 0.075 | Old |
| 0.195168 | 0.40177 | 0.23 | 0.167 | 1 | 13 | Zkscan1 | 0.063 | Old |
| 0.195321 | 0.354548 | 0.189 | 0.123 | 1 | 13 | Ccdc163 | 0.066 | Old |
| 0.195421 | 0.332074 | 0.18 | 0.114 | 1 | 13 | Ippk | 0.066 | Old |
| 0.195434 | 0.108692 | 0.697 | 0.579 | 1 | 13 | Adrm1 | 0.118 | Old |
| 0.19546 | 0.066048 | 1 | 1 | 1 | 13 | Ptp4a2 | 0 | Old |
| 0.195578 | 0.684064 | 0.566 | 0.518 | 1 | 13 | Cp | 0.048 | Old |
| 0.195649 | 0.469493 | 0.467 | 0.421 | 1 | 13 | Tlr4 | 0.046 | Old |
| 0.195788 | 0.281962 | 0.639 | 0.518 | 1 | 13 | Tmem50b | 0.121 | Old |
| 0.195968 | 0.120182 | 0.385 | 0.272 | 1 | 13 | P2ry10b | 0.113 | Old |
| 0.196206 | 0.216865 | 0.68 | 0.535 | 1 | 13 | Fbxl5 | 0.145 | Old |
| 0.196221 | 0.181363 | 0.828 | 0.719 | 1 | 13 | Wbp2 | 0.109 | Old |
| 0.196324 | 0.269957 | 0.213 | 0.14 | 1 | 13 | Lrrc20 | 0.073 | Old |
| 0.196435 | 0.121534 | 0.549 | 0.395 | 1 | 13 | Oxsr1 | 0.154 | Old |
| 0.196441 | 0.125044 | 0.959 | 0.974 | 1 | 13 | Srsf2 | -0.015 | Old |
| 0.196554 | 0.575572 | 0.098 | 0.053 | 1 | 13 | Gale | 0.045 | Old |
| 0.19663 | 0.402504 | 0.443 | 0.351 | 1 | 13 | Ndufaf4 | 0.092 | Old |
| 0.19673 | 0.647051 | 0.074 | 0.035 | 1 | 13 | Abcc10 | 0.039 | Old |
| 0.19678 | 0.084517 | 1 | 1 | 1 | 13 | Cox7c | 0 | Old |
| 0.197007 | 0.515554 | 0.221 | 0.158 | 1 | 13 | Snx16 | 0.063 | Old |
| 0.197071 | 0.547518 | 0.23 | 0.158 | 1 | 13 | Slfn1 | 0.072 | Old |

| | | | | | | | | |
|---|---|---|---|---|---|---|---|---|
| 0.197076 | 0.570419 | 0.336 | 0.263 | 1 | 13 | Plp2 | 0.073 | Old |
| 0.197304 | 0.370178 | 0.607 | 0.544 | 1 | 13 | Drg1 | 0.063 | Old |
| 0.197413 | 0.164671 | 0.844 | 0.789 | 1 | 13 | Dhx9 | 0.055 | Old |
| 0.197444 | 1.400147 | 0.033 | 0.009 | 1 | 13 | 2200002D( | 0.024 | Old |
| 0.197444 | 1.016573 | 0.033 | 0.009 | 1 | 13 | Yes1 | 0.024 | Old |
| 0.197444 | 0.975363 | 0.033 | 0.009 | 1 | 13 | AC125149. | 0.024 | Old |
| 0.197444 | 0.970436 | 0.033 | 0.009 | 1 | 13 | Msantd1 | 0.024 | Old |
| 0.197444 | 0.888638 | 0.033 | 0.009 | 1 | 13 | Klri1 | 0.024 | Old |
| 0.197444 | 0.883213 | 0.033 | 0.009 | 1 | 13 | Slc39a12 | 0.024 | Old |
| 0.197444 | 0.797927 | 0.033 | 0.009 | 1 | 13 | Xrcc3 | 0.024 | Old |
| 0.197444 | 0.774464 | 0.033 | 0.009 | 1 | 13 | Chtf18 | 0.024 | Old |
| 0.197444 | 0.73176 | 0.033 | 0.009 | 1 | 13 | Msh5 | 0.024 | Old |
| 0.197844 | 0.638749 | 0.131 | 0.079 | 1 | 13 | Ankrd13b | 0.052 | Old |
| 0.197894 | 0.37142 | 0.344 | 0.254 | 1 | 13 | Slc11a2 | 0.09 | Old |
| 0.197907 | 0.135692 | 0.877 | 0.807 | 1 | 13 | Nucb1 | 0.07 | Old |
| 0.197924 | 0.547901 | 0.205 | 0.14 | 1 | 13 | Panx1 | 0.065 | Old |
| 0.197995 | 0.707899 | 0.098 | 0.053 | 1 | 13 | Flt1 | 0.045 | Old |
| 0.197995 | 0.538533 | 0.098 | 0.053 | 1 | 13 | Eml2 | 0.045 | Old |
| 0.198012 | 0.288031 | 0.77 | 0.702 | 1 | 13 | Noc2l | 0.068 | Old |
| 0.198018 | 0.561205 | 0.27 | 0.211 | 1 | 13 | Yars | 0.059 | Old |
| 0.198237 | 0.216006 | 0.533 | 0.43 | 1 | 13 | Rpl10-ps3 | 0.103 | Old |
| 0.198457 | 0.214718 | 0.754 | 0.658 | 1 | 13 | Nin | 0.096 | Old |
| 0.19846 | 0.240685 | 0.189 | 0.123 | 1 | 13 | Prune1 | 0.066 | Old |
| 0.198773 | 0.139603 | 1 | 1 | 1 | 13 | Hsp90aa1 | 0 | Old |
| 0.198773 | 0.08752 | 1 | 1 | 1 | 13 | Pfn1 | 0 | Old |
| 0.198876 | 0.590025 | 0.123 | 0.07 | 1 | 13 | Ccdc77 | 0.053 | Old |
| 0.198876 | 0.334521 | 0.123 | 0.07 | 1 | 13 | Prepl | 0.053 | Old |
| 0.198986 | 0.106698 | 0.689 | 0.561 | 1 | 13 | Snx30 | 0.128 | Old |
| 0.199022 | 0.147829 | 0.566 | 0.421 | 1 | 13 | Chsy1 | 0.145 | Old |
| 0.199218 | 0.395536 | 0.418 | 0.325 | 1 | 13 | Tnip1 | 0.093 | Old |
| 0.19944 | 0.054333 | 0.975 | 1 | 1 | 13 | Mcub | -0.025 | Old |
| 0.199659 | 0.225131 | 0.615 | 0.482 | 1 | 13 | Nagk | 0.133 | Old |
| 0.199945 | 0.607981 | 0.205 | 0.14 | 1 | 13 | Rora | 0.065 | Old |
| 0.200018 | 0.230152 | 0.861 | 0.798 | 1 | 13 | Rnasel | 0.063 | Old |
| 0.200117 | 1.288261 | 0.033 | 0.009 | 1 | 13 | Fam71a | 0.024 | Old |
| 0.200117 | 1.044742 | 0.033 | 0.009 | 1 | 13 | Gpr171 | 0.024 | Old |
| 0.200117 | 0.962638 | 0.033 | 0.009 | 1 | 13 | Actr3b | 0.024 | Old |
| 0.200117 | 0.87877 | 0.033 | 0.009 | 1 | 13 | Il17rc | 0.024 | Old |
| 0.200117 | 0.87478 | 0.033 | 0.009 | 1 | 13 | Neil3 | 0.024 | Old |
| 0.200117 | 0.856003 | 0.033 | 0.009 | 1 | 13 | Plppr2 | 0.024 | Old |
| 0.200117 | 0.813974 | 0.033 | 0.009 | 1 | 13 | Camkmt | 0.024 | Old |
| 0.200117 | 0.809148 | 0.033 | 0.009 | 1 | 13 | Lhfp | 0.024 | Old |
| 0.200146 | 0.25065 | 0.68 | 0.588 | 1 | 13 | Cept1 | 0.092 | Old |
| 0.200181 | 0.166774 | 0.852 | 0.789 | 1 | 13 | Psmd1 | 0.063 | Old |
| 0.200189 | 0.784025 | 0.172 | 0.114 | 1 | 13 | Cluap1 | 0.058 | Old |

| | | | | | | | | |
|---|---|---|---|---|---|---|---|---|
| 0.200296 | 0.305345 | 0.221 | 0.149 | 1 | 13 | Recql5 | 0.072 | Old |
| 0.200488 | 0.356013 | 0.721 | 0.693 | 1 | 13 | Stt3b | 0.028 | Old |
| 0.200692 | 0.42676 | 0.533 | 0.447 | 1 | 13 | Ipo11 | 0.086 | Old |
| 0.20088 | 0.166807 | 0.672 | 0.561 | 1 | 13 | Trpm7 | 0.111 | Old |
| 0.201333 | 0.274753 | 0.746 | 0.649 | 1 | 13 | Ap3b1 | 0.097 | Old |
| 0.201435 | 0.369108 | 0.361 | 0.281 | 1 | 13 | Fblim1 | 0.08 | Old |
| 0.201482 | 0.238817 | 0.598 | 0.491 | 1 | 13 | Sh3bp2 | 0.107 | Old |
| 0.201798 | 0.677503 | 0.074 | 0.035 | 1 | 13 | G430095P1 | 0.039 | Old |
| 0.20203 | 0.35269 | 0.148 | 0.088 | 1 | 13 | Cyb5d2 | 0.06 | Old |
| 0.202104 | 0.42261 | 0.434 | 0.351 | 1 | 13 | Dok3 | 0.083 | Old |
| 0.202142 | 0.2027 | 0.803 | 0.702 | 1 | 13 | Mcts1 | 0.101 | Old |
| 0.202251 | 0.205736 | 0.803 | 0.789 | 1 | 13 | Aldh3b1 | 0.014 | Old |
| 0.202266 | 0.347194 | 0.73 | 0.754 | 1 | 13 | Atp6v1d | -0.024 | Old |
| 0.202403 | 0.309626 | 0.197 | 0.123 | 1 | 13 | Clasrp | 0.074 | Old |
| 0.202816 | 1.593155 | 0.033 | 0.009 | 1 | 13 | Htra4 | 0.024 | Old |
| 0.202816 | 0.859673 | 0.033 | 0.009 | 1 | 13 | Fam89b | 0.024 | Old |
| 0.202816 | 0.843645 | 0.033 | 0.009 | 1 | 13 | Gabbr2 | 0.024 | Old |
| 0.202816 | 0.839683 | 0.033 | 0.009 | 1 | 13 | Gm527 | 0.024 | Old |
| 0.202816 | 0.815468 | 0.033 | 0.009 | 1 | 13 | Hspg2 | 0.024 | Old |
| 0.202816 | 0.791744 | 0.033 | 0.009 | 1 | 13 | Olfr1033 | 0.024 | Old |
| 0.202816 | 0.724015 | 0.033 | 0.009 | 1 | 13 | Mapk11 | 0.024 | Old |
| 0.202816 | 0.720111 | 0.033 | 0.009 | 1 | 13 | Ptpn3 | 0.024 | Old |
| 0.202816 | 0.661621 | 0.033 | 0.009 | 1 | 13 | Cystm1 | 0.024 | Old |
| 0.202816 | 0.645998 | 0.033 | 0.009 | 1 | 13 | Myo19 | 0.024 | Old |
| 0.20292 | 0.302194 | 0.27 | 0.184 | 1 | 13 | Cables2 | 0.086 | Old |
| 0.202943 | 0.14634 | 0.311 | 0.219 | 1 | 13 | Sun1 | 0.092 | Old |
| 0.20304 | 0.869952 | 0.115 | 0.07 | 1 | 13 | Zfp35 | 0.045 | Old |
| 0.203162 | 0.375375 | 0.156 | 0.096 | 1 | 13 | Twsg1 | 0.06 | Old |
| 0.203182 | 0.587916 | 0.246 | 0.175 | 1 | 13 | Ap3m2 | 0.071 | Old |
| 0.203266 | 0.289065 | 0.59 | 0.535 | 1 | 13 | Uba6 | 0.055 | Old |
| 0.203407 | 0.381394 | 0.41 | 0.342 | 1 | 13 | Dera | 0.068 | Old |
| 0.203508 | 0.595634 | 0.074 | 0.035 | 1 | 13 | Pkmyt1 | 0.039 | Old |
| 0.203959 | 0.275784 | 0.443 | 0.325 | 1 | 13 | E2f1 | 0.118 | Old |
| 0.204136 | 0.599266 | 0.131 | 0.079 | 1 | 13 | Nim1k | 0.052 | Old |
| 0.204289 | 0.392897 | 0.279 | 0.202 | 1 | 13 | Alg8 | 0.077 | Old |
| 0.204502 | 0.358782 | 0.631 | 0.588 | 1 | 13 | Lars2 | 0.043 | Old |
| 0.204781 | 0.179215 | 0.689 | 0.614 | 1 | 13 | Vrk2 | 0.075 | Old |
| 0.204887 | 0.21679 | 0.598 | 0.509 | 1 | 13 | Nr2c2 | 0.089 | Old |
| 0.205035 | 0.336354 | 0.172 | 0.105 | 1 | 13 | 1810032O( | 0.067 | Old |
| 0.205132 | 0.328205 | 0.508 | 0.456 | 1 | 13 | Esco1 | 0.052 | Old |
| 0.205229 | 0.584217 | 0.074 | 0.035 | 1 | 13 | Ttc30b | 0.039 | Old |
| 0.205229 | 0.465743 | 0.074 | 0.035 | 1 | 13 | Nudt15 | 0.039 | Old |
| 0.205314 | 0.472237 | 0.098 | 0.053 | 1 | 13 | Tmem143 | 0.045 | Old |
| 0.205443 | 0.308081 | 0.418 | 0.325 | 1 | 13 | Mpv17l2 | 0.093 | Old |
| 0.205497 | 0.327136 | 0.656 | 0.588 | 1 | 13 | Naga | 0.068 | Old |

| | | | | | | | | |
|---|---|---|---|---|---|---|---|---|
| 0.205533 | 0.183894 | 0.451 | 0.351 | 1 | 13 | Mnat1 | 0.1 | Old |
| 0.205541 | 0.924969 | 0.033 | 0.009 | 1 | 13 | Magi1 | 0.024 | Old |
| 0.205541 | 0.770934 | 0.033 | 0.009 | 1 | 13 | Gm12216 | 0.024 | Old |
| 0.205541 | 0.744489 | 0.033 | 0.009 | 1 | 13 | Scn7a | 0.024 | Old |
| 0.205541 | 0.730142 | 0.033 | 0.009 | 1 | 13 | Gm26808 | 0.024 | Old |
| 0.205541 | 0.702248 | 0.033 | 0.009 | 1 | 13 | Tpm2 | 0.024 | Old |
| 0.205541 | 0.661878 | 0.033 | 0.009 | 1 | 13 | Gm33103 | 0.024 | Old |
| 0.205541 | 0.654654 | 0.033 | 0.009 | 1 | 13 | Abca5 | 0.024 | Old |
| 0.205541 | 0.645524 | 0.033 | 0.009 | 1 | 13 | Gm43625 | 0.024 | Old |
| 0.205541 | 0.64205 | 0.033 | 0.009 | 1 | 13 | Mipol1 | 0.024 | Old |
| 0.205541 | 0.629963 | 0.033 | 0.009 | 1 | 13 | Plxna2 | 0.024 | Old |
| 0.205541 | 0.609822 | 0.033 | 0.009 | 1 | 13 | A230028O( | 0.024 | Old |
| 0.205541 | 0.572482 | 0.033 | 0.009 | 1 | 13 | AI463229 | 0.024 | Old |
| 0.205541 | 0.558557 | 0.033 | 0.009 | 1 | 13 | Serpinb12 | 0.024 | Old |
| 0.205541 | 0.53401 | 0.033 | 0.009 | 1 | 13 | Pbk | 0.024 | Old |
| 0.205541 | 0.480033 | 0.033 | 0.009 | 1 | 13 | Gm47507 | 0.024 | Old |
| 0.2056 | 0.507332 | 0.369 | 0.289 | 1 | 13 | Slc35a2 | 0.08 | Old |
| 0.205632 | 0.249905 | 0.295 | 0.219 | 1 | 13 | Mex3d | 0.076 | Old |
| 0.205753 | 0.37665 | 0.418 | 0.333 | 1 | 13 | Pycr2 | 0.085 | Old |
| 0.206014 | 0.201246 | 0.639 | 0.561 | 1 | 13 | Bet1 | 0.078 | Old |
| 0.206055 | 0.420928 | 0.352 | 0.281 | 1 | 13 | Lonp1 | 0.071 | Old |
| 0.206281 | 0.373474 | 0.221 | 0.149 | 1 | 13 | Papolg | 0.072 | Old |
| 0.206386 | 0.218416 | 0.869 | 0.781 | 1 | 13 | Rab1b | 0.088 | Old |
| 0.206428 | 0.298996 | 0.426 | 0.333 | 1 | 13 | Dohh | 0.093 | Old |
| 0.206791 | 0.263313 | 0.934 | 0.912 | 1 | 13 | AC149090. | 0.022 | Old |
| 0.206801 | 0.608184 | 0.098 | 0.053 | 1 | 13 | Nme6 | 0.045 | Old |
| 0.206926 | 0.242034 | 0.336 | 0.246 | 1 | 13 | Ppih | 0.09 | Old |
| 0.20696 | 0.483437 | 0.074 | 0.035 | 1 | 13 | Cdkl2 | 0.039 | Old |
| 0.207064 | 0.329695 | 0.352 | 0.254 | 1 | 13 | Herc6 | 0.098 | Old |
| 0.207214 | 0.414623 | 0.434 | 0.377 | 1 | 13 | Slc25a46 | 0.057 | Old |
| 0.207243 | 0.216026 | 0.844 | 0.868 | 1 | 13 | Smg1 | -0.024 | Old |
| 0.207324 | 0.176469 | 0.303 | 0.228 | 1 | 13 | Pparg | 0.075 | Old |
| 0.208267 | 0.097785 | 0.984 | 0.991 | 1 | 13 | Rpn2 | -0.007 | Old |
| 0.208293 | 0.674361 | 0.033 | 0.009 | 1 | 13 | Cenpi | 0.024 | Old |
| 0.208293 | 0.650348 | 0.033 | 0.009 | 1 | 13 | Carns1 | 0.024 | Old |
| 0.208293 | 0.641687 | 0.033 | 0.009 | 1 | 13 | Icam4 | 0.024 | Old |
| 0.208293 | 0.637198 | 0.033 | 0.009 | 1 | 13 | Cep97 | 0.024 | Old |
| 0.208293 | 0.607043 | 0.033 | 0.009 | 1 | 13 | Zfp947 | 0.024 | Old |
| 0.208293 | 0.599757 | 0.033 | 0.009 | 1 | 13 | Ankrd28 | 0.024 | Old |
| 0.208293 | 0.592004 | 0.033 | 0.009 | 1 | 13 | Neil2 | 0.024 | Old |
| 0.208293 | 0.536142 | 0.033 | 0.009 | 1 | 13 | A330069E1 | 0.024 | Old |
| 0.208293 | 0.518032 | 0.033 | 0.009 | 1 | 13 | Bcas1 | 0.024 | Old |
| 0.208293 | 0.504914 | 0.033 | 0.009 | 1 | 13 | Gpr150 | 0.024 | Old |
| 0.208293 | 0.237899 | 0.033 | 0.009 | 1 | 13 | Inhba | 0.024 | Old |
| 0.208293 | 0.224399 | 0.033 | 0.009 | 1 | 13 | Gm39469 | 0.024 | Old |

| | | | | | | | | |
|---|---|---|---|---|---|---|---|---|
| 0.208293 | 0.093326 | 0.033 | 0.009 | 1 | 13 | Iglc3 | 0.024 | Old |
| 0.208702 | 1.207001 | 0.074 | 0.035 | 1 | 13 | Cd69 | 0.039 | Old |
| 0.208727 | 0.300701 | 0.303 | 0.219 | 1 | 13 | Abhd13 | 0.084 | Old |
| 0.209029 | 0.373525 | 0.648 | 0.596 | 1 | 13 | Nsfl1c | 0.052 | Old |
| 0.209695 | 0.458659 | 0.23 | 0.167 | 1 | 13 | Lrrc61 | 0.063 | Old |
| 0.209792 | 0.201836 | 0.648 | 0.544 | 1 | 13 | Hivep1 | 0.104 | Old |
| 0.21015 | 0.210877 | 0.828 | 0.789 | 1 | 13 | Sdhd | 0.039 | Old |
| 0.210275 | 0.516518 | 0.205 | 0.14 | 1 | 13 | Smpd2 | 0.065 | Old |
| 0.210343 | 0.471737 | 0.221 | 0.149 | 1 | 13 | Nat10 | 0.072 | Old |
| 0.210416 | 0.098541 | 0.754 | 0.667 | 1 | 13 | Dynlt3 | 0.087 | Old |
| 0.210571 | 0.328048 | 0.131 | 0.079 | 1 | 13 | Gpr85 | 0.052 | Old |
| 0.210754 | 0.214667 | 0.803 | 0.684 | 1 | 13 | Psmd6 | 0.119 | Old |
| 0.210774 | 0.17616 | 0.672 | 0.535 | 1 | 13 | Mark3 | 0.137 | Old |
| 0.210988 | 0.173193 | 0.59 | 0.474 | 1 | 13 | Zdhhc5 | 0.116 | Old |
| 0.211547 | 0.306146 | 0.803 | 0.816 | 1 | 13 | Sulf2 | -0.013 | Old |
| 0.211737 | 0.101591 | 0.992 | 1 | 1 | 13 | Slc25a5 | -0.008 | Old |
| 0.21175 | 0.083529 | 0.844 | 0.789 | 1 | 13 | Ptpro | 0.055 | Old |
| 0.211894 | 0.2102 | 0.23 | 0.149 | 1 | 13 | Cela1 | 0.081 | Old |
| 0.212103 | 0.16168 | 0.893 | 0.772 | 1 | 13 | Lrp6 | 0.121 | Old |
| 0.212292 | 0.462701 | 0.557 | 0.535 | 1 | 13 | Rnd3 | 0.022 | Old |
| 0.212358 | 0.125806 | 0.557 | 0.447 | 1 | 13 | Knop1 | 0.11 | Old |
| 0.212425 | 0.390494 | 0.254 | 0.184 | 1 | 13 | Tmem199 | 0.07 | Old |
| 0.212425 | 0.302476 | 0.254 | 0.184 | 1 | 13 | 1190002N1 | 0.07 | Old |
| 0.212436 | 0.140915 | 0.975 | 1 | 1 | 13 | Arf1 | -0.025 | Old |
| 0.212644 | 0.714473 | 0.148 | 0.096 | 1 | 13 | Nod2 | 0.052 | Old |
| 0.212716 | 0.429097 | 0.385 | 0.307 | 1 | 13 | Ppp1r37 | 0.078 | Old |
| 0.212777 | 0.104368 | 0.975 | 0.956 | 1 | 13 | 7-Sep | 0.019 | Old |
| 0.212825 | 0.47569 | 0.213 | 0.149 | 1 | 13 | Kctd3 | 0.064 | Old |
| 0.212828 | 0.517747 | 0.098 | 0.053 | 1 | 13 | D11Wsu47 | 0.045 | Old |
| 0.212853 | 0.224035 | 0.467 | 0.36 | 1 | 13 | Cdk8 | 0.107 | Old |
| 0.212884 | 0.366616 | 0.467 | 0.412 | 1 | 13 | Wdr82 | 0.055 | Old |
| 0.212885 | 0.174557 | 0.697 | 0.632 | 1 | 13 | Lsm3 | 0.065 | Old |
| 0.213054 | 0.345364 | 0.148 | 0.088 | 1 | 13 | Abcb6 | 0.06 | Old |
| 0.213379 | 0.716864 | 0.107 | 0.061 | 1 | 13 | Cmtr1 | 0.046 | Old |
| 0.214231 | 0.16352 | 0.418 | 0.325 | 1 | 13 | Dnajc25 | 0.093 | Old |
| 0.214354 | 0.399336 | 0.098 | 0.053 | 1 | 13 | Gm12089 | 0.045 | Old |
| 0.21481 | 0.344538 | 0.197 | 0.132 | 1 | 13 | Gpn2 | 0.065 | Old |
| 0.214912 | 0.489693 | 0.516 | 0.482 | 1 | 13 | Psen2 | 0.034 | Old |
| 0.21511 | 0.551085 | 0.148 | 0.096 | 1 | 13 | Cep70 | 0.052 | Old |
| 0.215385 | 0.761985 | 0.115 | 0.07 | 1 | 13 | Pdia5 | 0.045 | Old |
| 0.215499 | 0.149077 | 0.672 | 0.57 | 1 | 13 | Suclg1 | 0.102 | Old |
| 0.216036 | 0.133099 | 0.918 | 0.798 | 1 | 13 | Pds5a | 0.12 | Old |
| 0.216052 | 0.22364 | 0.467 | 0.368 | 1 | 13 | Psmd10 | 0.099 | Old |
| 0.216193 | 0.048384 | 0.885 | 0.86 | 1 | 13 | Gatad2b | 0.025 | Old |
| 0.216224 | 0.333199 | 0.426 | 0.36 | 1 | 13 | Acox1 | 0.066 | Old |

| | | | | | | | | |
|---|---|---|---|---|---|---|---|---|
| 0.2163 | 0.648606 | 0.107 | 0.061 | 1 | 13 | Ifrd2 | 0.046 | Old |
| 0.216488 | 0.126195 | 0.516 | 0.386 | 1 | 13 | Ube4a | 0.13 | Old |
| 0.216495 | 0.338986 | 0.697 | 0.623 | 1 | 13 | Insig1 | 0.074 | Old |
| 0.21679 | 0.749517 | 0.115 | 0.07 | 1 | 13 | Zhx2 | 0.045 | Old |
| 0.216823 | 0.448223 | 0.311 | 0.237 | 1 | 13 | Plcb2 | 0.074 | Old |
| 0.216904 | 0.238835 | 0.287 | 0.202 | 1 | 13 | Fhod1 | 0.085 | Old |
| 0.217289 | 0.173717 | 0.934 | 0.921 | 1 | 13 | Tardbp | 0.013 | Old |
| 0.217431 | 0.493823 | 0.098 | 0.053 | 1 | 13 | Gm27003 | 0.045 | Old |
| 0.21747 | 0.18819 | 0.328 | 0.228 | 1 | 13 | Fbxw8 | 0.1 | Old |
| 0.217495 | 0.191729 | 0.779 | 0.711 | 1 | 13 | Swap70 | 0.068 | Old |
| 0.217564 | 0.310232 | 0.156 | 0.096 | 1 | 13 | Anln | 0.06 | Old |
| 0.217611 | 0.176462 | 0.828 | 0.719 | 1 | 13 | Txnl1 | 0.109 | Old |
| 0.217754 | 0.366235 | 0.525 | 0.43 | 1 | 13 | Fbxw7 | 0.095 | Old |
| 0.217771 | 0.718395 | 0.107 | 0.061 | 1 | 13 | Ift43 | 0.046 | Old |
| 0.217771 | 0.625819 | 0.107 | 0.061 | 1 | 13 | Phpt1 | 0.046 | Old |
| 0.217993 | 0.374279 | 0.434 | 0.36 | 1 | 13 | Reps1 | 0.074 | Old |
| 0.218057 | 0.23333 | 0.951 | 0.947 | 1 | 13 | P2rx4 | 0.004 | Old |
| 0.218087 | 0.096132 | 0.992 | 1 | 1 | 13 | Ostf1 | -0.008 | Old |
| 0.218133 | 0.237261 | 0.41 | 0.307 | 1 | 13 | Med17 | 0.103 | Old |
| 0.218382 | 0.381365 | 0.631 | 0.553 | 1 | 13 | Nfkbib | 0.078 | Old |
| 0.218475 | 0.120928 | 0.877 | 0.789 | 1 | 13 | Dlst | 0.088 | Old |
| 0.218736 | 0.296726 | 0.18 | 0.114 | 1 | 13 | Klhl36 | 0.066 | Old |
| 0.21925 | 0.562157 | 0.107 | 0.061 | 1 | 13 | Aste1 | 0.046 | Old |
| 0.219294 | 0.222635 | 0.254 | 0.175 | 1 | 13 | Fam114a1 | 0.079 | Old |
| 0.219377 | 0.44179 | 0.074 | 0.035 | 1 | 13 | Mfsd3 | 0.039 | Old |
| 0.219609 | 0.386684 | 0.328 | 0.263 | 1 | 13 | Tmem55a | 0.065 | Old |
| 0.219874 | 0.211094 | 0.975 | 0.991 | 1 | 13 | Egr1 | -0.016 | Old |
| 0.219913 | 0.253804 | 0.803 | 0.754 | 1 | 13 | Yeats4 | 0.049 | Old |
| 0.220034 | 0.371076 | 0.156 | 0.096 | 1 | 13 | Ndufaf5 | 0.06 | Old |
| 0.220233 | 0.134982 | 0.984 | 0.991 | 1 | 13 | Syngr2 | -0.007 | Old |
| 0.220561 | 0.755418 | 0.082 | 0.044 | 1 | 13 | Sirt5 | 0.038 | Old |
| 0.220901 | 0.123972 | 0.426 | 0.333 | 1 | 13 | Pygb | 0.093 | Old |
| 0.220953 | 0.124654 | 1 | 0.991 | 1 | 13 | Rpl13a | 0.009 | Old |
| 0.221058 | 0.455106 | 0.23 | 0.158 | 1 | 13 | Il1a | 0.072 | Old |
| 0.221905 | 0.194953 | 0.672 | 0.596 | 1 | 13 | Eri1 | 0.076 | Old |
| 0.222127 | 0.253927 | 0.623 | 0.509 | 1 | 13 | Hck | 0.114 | Old |
| 0.222143 | 0.231863 | 0.713 | 0.649 | 1 | 13 | Snx27 | 0.064 | Old |
| 0.222283 | 0.156194 | 0.91 | 0.833 | 1 | 13 | Notch2 | 0.077 | Old |
| 0.222436 | 0.180443 | 0.467 | 0.368 | 1 | 13 | Plagl2 | 0.099 | Old |
| 0.222628 | 0.743895 | 0.189 | 0.132 | 1 | 13 | Cd209e | 0.057 | Old |
| 0.222851 | 0.109644 | 0.525 | 0.412 | 1 | 13 | Pex6 | 0.113 | Old |
| 0.222866 | 0.193215 | 0.746 | 0.684 | 1 | 13 | Prpf8 | 0.062 | Old |
| 0.223011 | 0.453956 | 0.295 | 0.219 | 1 | 13 | Thoc1 | 0.076 | Old |
| 0.223662 | 0.376578 | 0.197 | 0.132 | 1 | 13 | Dimt1 | 0.065 | Old |
| 0.224733 | 0.542605 | 0.156 | 0.105 | 1 | 13 | Eps8l1 | 0.051 | Old |

| | | | | | | | | |
|---|---|---|---|---|---|---|---|---|
| 0.225031 | 0.302268 | 0.787 | 0.667 | 1 | 13 | Rasa1 | 0.12 | Old |
| 0.225243 | 0.475368 | 0.131 | 0.079 | 1 | 13 | Itga1 | 0.052 | Old |
| 0.225243 | 0.408763 | 0.131 | 0.079 | 1 | 13 | Ppp3cc | 0.052 | Old |
| 0.225243 | 0.351545 | 0.131 | 0.079 | 1 | 13 | Pigb | 0.052 | Old |
| 0.225302 | 0.066946 | 0.992 | 1 | 1 | 13 | Fos | -0.008 | Old |
| 0.22539 | 0.350963 | 0.336 | 0.263 | 1 | 13 | Neurl2 | 0.073 | Old |
| 0.225407 | 0.397723 | 0.172 | 0.114 | 1 | 13 | Plpp5 | 0.058 | Old |
| 0.225671 | 0.369434 | 0.377 | 0.298 | 1 | 13 | Exosc7 | 0.079 | Old |
| 0.225751 | 0.390467 | 0.18 | 0.114 | 1 | 13 | Gm16104 | 0.066 | Old |
| 0.226024 | 0.308516 | 0.434 | 0.351 | 1 | 13 | Cog7 | 0.083 | Old |
| 0.22609 | 0.561127 | 0.254 | 0.193 | 1 | 13 | Tmem242 | 0.061 | Old |
| 0.226554 | 0.15733 | 0.861 | 0.781 | 1 | 13 | Cacybp | 0.08 | Old |
| 0.226936 | 0.632795 | 0.279 | 0.211 | 1 | 13 | Tcf7l2 | 0.068 | Old |
| 0.227321 | 0.366806 | 0.369 | 0.298 | 1 | 13 | Maip1 | 0.071 | Old |
| 0.227361 | 0.391374 | 0.164 | 0.105 | 1 | 13 | Mtfr1 | 0.059 | Old |
| 0.227592 | 0.584761 | 0.139 | 0.088 | 1 | 13 | Ccdc126 | 0.051 | Old |
| 0.227645 | 0.28187 | 0.648 | 0.553 | 1 | 13 | Adam19 | 0.095 | Old |
| 0.227682 | 0.182955 | 0.5 | 0.395 | 1 | 13 | Atmin | 0.105 | Old |
| 0.227723 | 0.335171 | 0.352 | 0.272 | 1 | 13 | Tjp2 | 0.08 | Old |
| 0.22775 | 0.338587 | 0.861 | 0.789 | 1 | 13 | Aprt | 0.072 | Old |
| 0.22776 | 0.512786 | 0.213 | 0.149 | 1 | 13 | Irgm2 | 0.064 | Old |
| 0.227784 | 0.536238 | 0.23 | 0.175 | 1 | 13 | Spidr | 0.055 | Old |
| 0.228143 | 0.18405 | 0.238 | 0.158 | 1 | 13 | Nudt1 | 0.08 | Old |
| 0.228271 | 0.657779 | 0.107 | 0.061 | 1 | 13 | Cptp | 0.046 | Old |
| 0.2285 | 0.555738 | 0.197 | 0.14 | 1 | 13 | Rundc1 | 0.057 | Old |
| 0.228709 | 0.114588 | 0.541 | 0.412 | 1 | 13 | Sfxn1 | 0.129 | Old |
| 0.228759 | 0.234296 | 0.451 | 0.351 | 1 | 13 | Akap8l | 0.1 | Old |
| 0.228867 | 0.269143 | 0.689 | 0.553 | 1 | 13 | Glod4 | 0.136 | Old |
| 0.2289 | 0.113332 | 0.566 | 0.465 | 1 | 13 | Aldh16a1 | 0.101 | Old |
| 0.229014 | 0.355463 | 0.172 | 0.114 | 1 | 13 | Ddx20 | 0.058 | Old |
| 0.229108 | 0.111141 | 0.877 | 0.798 | 1 | 13 | Arglu1 | 0.079 | Old |
| 0.229342 | 0.083615 | 0.992 | 0.991 | 1 | 13 | Sh3bgrl | 0.001 | Old |
| 0.229855 | 0.28869 | 0.32 | 0.237 | 1 | 13 | Mterf3 | 0.083 | Old |
| 0.230122 | 0.318308 | 0.262 | 0.184 | 1 | 13 | 4930503L1 | 0.078 | Old |
| 0.230263 | 0.215446 | 0.139 | 0.088 | 1 | 13 | Tnfrsf18 | 0.051 | Old |
| 0.230366 | 0.141355 | 0.926 | 0.93 | 1 | 13 | C3ar1 | -0.004 | Old |
| 0.230435 | 0.098632 | 0.951 | 0.965 | 1 | 13 | 2900097C1 | -0.014 | Old |
| 0.230545 | 0.283057 | 0.525 | 0.447 | 1 | 13 | Fam204a | 0.078 | Old |
| 0.230545 | 0.213727 | 0.533 | 0.439 | 1 | 13 | Tbc1d15 | 0.094 | Old |
| 0.230885 | 0.113319 | 0.852 | 0.816 | 1 | 13 | Herc1 | 0.036 | Old |
| 0.230915 | 0.213776 | 0.631 | 0.5 | 1 | 13 | Hdac2 | 0.131 | Old |
| 0.230956 | 0.338634 | 0.205 | 0.14 | 1 | 13 | Parn | 0.065 | Old |
| 0.231619 | 0.305879 | 0.197 | 0.132 | 1 | 13 | Zmym1 | 0.065 | Old |
| 0.232013 | 0.298661 | 0.598 | 0.491 | 1 | 13 | Dpp3 | 0.107 | Old |
| 0.232083 | 0.356671 | 0.205 | 0.14 | 1 | 13 | Casp7 | 0.065 | Old |

| | | | | | | | | |
|---|---|---|---|---|---|---|---|---|
| 0.232184 | 0.35509 | 0.41 | 0.333 | 1 | 13 | Mphosph1 | 0.077 | Old |
| 0.232446 | 0.265597 | 0.615 | 0.553 | 1 | 13 | Ipo5 | 0.062 | Old |
| 0.232662 | 0.346662 | 0.172 | 0.114 | 1 | 13 | Ppm1f | 0.058 | Old |
| 0.232671 | 0.199988 | 0.344 | 0.254 | 1 | 13 | mt-Nd6 | 0.09 | Old |
| 0.232674 | 0.404468 | 0.32 | 0.237 | 1 | 13 | Fkrp | 0.083 | Old |
| 0.232682 | 0.111787 | 0.967 | 0.982 | 1 | 13 | Psma3 | -0.015 | Old |
| 0.232693 | 0.723619 | 0.082 | 0.044 | 1 | 13 | Rbm4 | 0.038 | Old |
| 0.232693 | 0.628376 | 0.082 | 0.044 | 1 | 13 | Mmab | 0.038 | Old |
| 0.233097 | 0.145676 | 0.475 | 0.36 | 1 | 13 | Vps54 | 0.115 | Old |
| 0.233217 | 0.153961 | 0.893 | 0.912 | 1 | 13 | Eif3m | -0.019 | Old |
| 0.233414 | 0.199686 | 0.82 | 0.807 | 1 | 13 | Scarb2 | 0.013 | Old |
| 0.233466 | 0.295384 | 0.23 | 0.167 | 1 | 13 | Zkscan14 | 0.063 | Old |
| 0.233727 | 0.218599 | 0.877 | 0.816 | 1 | 13 | Epb41l2 | 0.061 | Old |
| 0.23419 | 0.510414 | 0.197 | 0.14 | 1 | 13 | Lzic | 0.057 | Old |
| 0.234207 | 0.386906 | 0.41 | 0.316 | 1 | 13 | Gm46224 | 0.094 | Old |
| 0.234222 | 0.236443 | 0.664 | 0.579 | 1 | 13 | Mob1b | 0.085 | Old |
| 0.234364 | 0.212312 | 0.697 | 0.623 | 1 | 13 | Sik1 | 0.074 | Old |
| 0.234369 | 0.089073 | 0.91 | 0.904 | 1 | 13 | Smap2 | 0.006 | Old |
| 0.234384 | 0.18553 | 0.68 | 0.596 | 1 | 13 | Atp8b4 | 0.084 | Old |
| 0.23443 | 0.62575 | 0.107 | 0.061 | 1 | 13 | Ccdc130 | 0.046 | Old |
| 0.234656 | 0.203618 | 0.902 | 0.886 | 1 | 13 | Slc15a3 | 0.016 | Old |
| 0.234656 | 0.075356 | 0.238 | 0.167 | 1 | 13 | Slc30a4 | 0.071 | Old |
| 0.234794 | 0.328029 | 0.418 | 0.342 | 1 | 13 | Nup214 | 0.076 | Old |
| 0.234813 | 0.166317 | 0.852 | 0.763 | 1 | 13 | Mia3 | 0.089 | Old |
| 0.234854 | 0.335093 | 0.164 | 0.105 | 1 | 13 | Acp6 | 0.059 | Old |
| 0.234856 | 0.37589 | 0.18 | 0.123 | 1 | 13 | Nup107 | 0.057 | Old |
| 0.234931 | 0.101492 | 0.992 | 0.965 | 1 | 13 | Myl12b | 0.027 | Old |
| 0.235205 | 0.139433 | 0.885 | 0.86 | 1 | 13 | Esd | 0.025 | Old |
| 0.235297 | 0.20576 | 0.287 | 0.202 | 1 | 13 | Tbc1d24 | 0.085 | Old |
| 0.235298 | 0.087822 | 0.869 | 0.789 | 1 | 13 | Syf2 | 0.08 | Old |
| 0.235333 | 0.27393 | 0.631 | 0.57 | 1 | 13 | Ube2v2 | 0.061 | Old |
| 0.235495 | 0.701686 | 0.057 | 0.026 | 1 | 13 | Eri2 | 0.031 | Old |
| 0.235649 | 0.383143 | 0.115 | 0.07 | 1 | 13 | Ankrd33b | 0.045 | Old |
| 0.235673 | 0.493551 | 0.139 | 0.088 | 1 | 13 | Yars2 | 0.051 | Old |
| 0.235901 | 0.27161 | 0.664 | 0.605 | 1 | 13 | Dnase1l1 | 0.059 | Old |
| 0.236137 | 0.322017 | 0.434 | 0.351 | 1 | 13 | Ppp5c | 0.083 | Old |
| 0.236244 | 0.600043 | 0.082 | 0.044 | 1 | 13 | Tasp1 | 0.038 | Old |
| 0.236402 | 0.139675 | 0.951 | 0.939 | 1 | 13 | Sec62 | 0.012 | Old |
| 0.23658 | 0.36356 | 0.156 | 0.096 | 1 | 13 | Fkbp9 | 0.06 | Old |
| 0.236952 | 0.401214 | 0.41 | 0.351 | 1 | 13 | Lactb2 | 0.059 | Old |
| 0.236978 | 0.246179 | 0.705 | 0.632 | 1 | 13 | Hint2 | 0.073 | Old |
| 0.237044 | 0.206939 | 0.393 | 0.316 | 1 | 13 | Tubgcp4 | 0.077 | Old |
| 0.237105 | 0.883679 | 0.123 | 0.079 | 1 | 13 | Slc6a8 | 0.044 | Old |
| 0.237147 | 0.510242 | 0.115 | 0.07 | 1 | 13 | Hirip3 | 0.045 | Old |
| 0.237197 | 0.114038 | 1 | 0.991 | 1 | 13 | Serbp1 | 0.009 | Old |

| | | | | | | | | |
|---|---|---|---|---|---|---|---|---|
| 0.237554 | 0.305704 | 0.107 | 0.061 | 1 | 13 | Rida | 0.046 | Old |
| 0.23766 | 0.777753 | 0.057 | 0.026 | 1 | 13 | Nr3c2 | 0.031 | Old |
| 0.237968 | 0.469487 | 0.295 | 0.237 | 1 | 13 | Gnl2 | 0.058 | Old |
| 0.238377 | 0.103179 | 0.811 | 0.789 | 1 | 13 | Csnk1d | 0.022 | Old |
| 0.238407 | 0.369806 | 0.672 | 0.684 | 1 | 13 | Fam213b | -0.012 | Old |
| 0.238651 | 0.575849 | 0.115 | 0.07 | 1 | 13 | Dhx37 | 0.045 | Old |
| 0.238867 | 0.546971 | 0.189 | 0.132 | 1 | 13 | Ghdc | 0.057 | Old |
| 0.239209 | 0.320365 | 0.295 | 0.228 | 1 | 13 | Tpm3-rs7 | 0.067 | Old |
| 0.239279 | 0.175875 | 0.221 | 0.149 | 1 | 13 | Tysnd1 | 0.072 | Old |
| 0.23958 | 0.30054 | 0.525 | 0.465 | 1 | 13 | Mcrs1 | 0.06 | Old |
| 0.239605 | 0.194422 | 0.508 | 0.421 | 1 | 13 | Srebf1 | 0.087 | Old |
| 0.239828 | 0.123627 | 0.328 | 0.228 | 1 | 13 | Slc39a6 | 0.1 | Old |
| 0.239839 | 0.794777 | 0.057 | 0.026 | 1 | 13 | Ybey | 0.031 | Old |
| 0.239981 | 0.535845 | 0.156 | 0.105 | 1 | 13 | Rhbdd3 | 0.051 | Old |
| 0.240075 | 0.118605 | 0.705 | 0.632 | 1 | 13 | Ccnt2 | 0.073 | Old |
| 0.240707 | 0.612796 | 0.107 | 0.061 | 1 | 13 | Usp35 | 0.046 | Old |
| 0.240905 | 0.446082 | 0.303 | 0.228 | 1 | 13 | Rcan1 | 0.075 | Old |
| 0.241172 | 0.291363 | 0.139 | 0.088 | 1 | 13 | Slc36a3os | 0.051 | Old |
| 0.241257 | 0.211196 | 0.492 | 0.377 | 1 | 13 | Yif1a | 0.115 | Old |
| 0.24177 | 0.094468 | 1 | 0.991 | 1 | 13 | Flna | 0.009 | Old |
| 0.241838 | 0.200384 | 0.484 | 0.404 | 1 | 13 | Slc5a3 | 0.08 | Old |
| 0.242032 | 0.654226 | 0.057 | 0.026 | 1 | 13 | Ahrr | 0.031 | Old |
| 0.242032 | 0.6415 | 0.057 | 0.026 | 1 | 13 | Rnd2 | 0.031 | Old |
| 0.242032 | 0.638196 | 0.057 | 0.026 | 1 | 13 | Zfp28 | 0.031 | Old |
| 0.242107 | 0.323885 | 0.541 | 0.465 | 1 | 13 | Nasp | 0.076 | Old |
| 0.24217 | 0.194255 | 0.689 | 0.605 | 1 | 13 | Ncor2 | 0.084 | Old |
| 0.242434 | 0.24267 | 0.934 | 0.912 | 1 | 13 | Lrrc58 | 0.022 | Old |
| 0.242452 | 0.400705 | 0.189 | 0.132 | 1 | 13 | Gys1 | 0.057 | Old |
| 0.2425 | 0.394623 | 0.426 | 0.351 | 1 | 13 | Mb21d1 | 0.075 | Old |
| 0.242502 | 0.144333 | 0.721 | 0.596 | 1 | 13 | Ptpn2 | 0.125 | Old |
| 0.24275 | 0.155153 | 0.328 | 0.246 | 1 | 13 | Jade2 | 0.082 | Old |
| 0.24275 | 0.193753 | 0.713 | 0.632 | 1 | 13 | Ebp | 0.081 | Old |
| 0.242946 | 0.286874 | 0.221 | 0.158 | 1 | 13 | Nudt7 | 0.063 | Old |
| 0.243203 | 0.582247 | 0.115 | 0.07 | 1 | 13 | Prnp | 0.045 | Old |
| 0.243203 | 0.523634 | 0.115 | 0.07 | 1 | 13 | Armc6 | 0.045 | Old |
| 0.243286 | 0.283951 | 0.238 | 0.175 | 1 | 13 | Btg3 | 0.063 | Old |
| 0.2433 | 0.425601 | 0.369 | 0.281 | 1 | 13 | Mid1 | 0.088 | Old |
| 0.243427 | 0.344913 | 0.213 | 0.149 | 1 | 13 | Zswim7 | 0.064 | Old |
| 0.243461 | 0.473985 | 0.082 | 0.044 | 1 | 13 | Cep112 | 0.038 | Old |
| 0.243576 | 0.527616 | 0.254 | 0.193 | 1 | 13 | Bach2 | 0.061 | Old |
| 0.243584 | 0.749741 | 0.164 | 0.114 | 1 | 13 | Adcy3 | 0.05 | Old |
| 0.243603 | 0.499279 | 0.148 | 0.096 | 1 | 13 | Elovl6 | 0.052 | Old |
| 0.243643 | 0.119598 | 0.877 | 0.877 | 1 | 13 | Sptssa | 0 | Old |
| 0.243765 | 0.196369 | 0.918 | 0.86 | 1 | 13 | Gsk3b | 0.058 | Old |
| 0.243816 | 0.333618 | 0.164 | 0.105 | 1 | 13 | Grpel2 | 0.059 | Old |

| | | | | | | | | |
|---|---|---|---|---|---|---|---|---|
| 0.244151 | 0.13096 | 0.836 | 0.737 | 1 | 13 | Sumo3 | 0.099 | Old |
| 0.244636 | 0.272581 | 0.59 | 0.509 | 1 | 13 | Abcc5 | 0.081 | Old |
| 0.2449 | 0.269885 | 0.623 | 0.588 | 1 | 13 | Gripap1 | 0.035 | Old |
| 0.245491 | 0.507864 | 0.107 | 0.061 | 1 | 13 | Tchp | 0.046 | Old |
| 0.245759 | 0.256524 | 0.861 | 0.86 | 1 | 13 | Irf8 | 0.001 | Old |
| 0.246029 | 0.208116 | 0.73 | 0.675 | 1 | 13 | Tial1 | 0.055 | Old |
| 0.246175 | 0.20826 | 0.393 | 0.298 | 1 | 13 | Ahsa2 | 0.095 | Old |
| 0.246272 | 0.645514 | 0.115 | 0.07 | 1 | 13 | Mettl8 | 0.045 | Old |
| 0.246332 | 0.48902 | 0.221 | 0.158 | 1 | 13 | Krt10 | 0.063 | Old |
| 0.246461 | 0.670887 | 0.057 | 0.026 | 1 | 13 | Ptpdc1 | 0.031 | Old |
| 0.246461 | 0.608724 | 0.057 | 0.026 | 1 | 13 | Slc22a21 | 0.031 | Old |
| 0.246461 | 0.577269 | 0.057 | 0.026 | 1 | 13 | Mr1 | 0.031 | Old |
| 0.246914 | 0.285095 | 0.262 | 0.193 | 1 | 13 | Tmx2 | 0.069 | Old |
| 0.247015 | 0.263128 | 0.721 | 0.632 | 1 | 13 | Ythdf1 | 0.089 | Old |
| 0.24738 | 0.132768 | 0.844 | 0.763 | 1 | 13 | Ppm1a | 0.081 | Old |
| 0.24754 | 0.578557 | 0.336 | 0.272 | 1 | 13 | Hvcn1 | 0.064 | Old |
| 0.247633 | 0.39314 | 0.221 | 0.167 | 1 | 13 | Mgme1 | 0.054 | Old |
| 0.2478 | 0.332459 | 0.557 | 0.482 | 1 | 13 | Gadd45gip | 0.075 | Old |
| 0.247817 | 0.538097 | 0.115 | 0.07 | 1 | 13 | Tsen54 | 0.045 | Old |
| 0.247873 | 0.497751 | 0.156 | 0.105 | 1 | 13 | 1810024B0 | 0.051 | Old |
| 0.24789 | 0.103757 | 0.934 | 0.93 | 1 | 13 | Blnk | 0.004 | Old |
| 0.247941 | 0.19473 | 0.656 | 0.579 | 1 | 13 | Ebna1bp2 | 0.077 | Old |
| 0.247984 | 0.441807 | 0.279 | 0.211 | 1 | 13 | Ggta1 | 0.068 | Old |
| 0.248004 | 0.295987 | 0.336 | 0.246 | 1 | 13 | Prkrip1 | 0.09 | Old |
| 0.248381 | 0.282229 | 0.59 | 0.561 | 1 | 13 | Smarca2 | 0.029 | Old |
| 0.248398 | 0.212431 | 0.672 | 0.57 | 1 | 13 | Itpk1 | 0.102 | Old |
| 0.248697 | 0.563081 | 0.057 | 0.026 | 1 | 13 | Zbtb25 | 0.031 | Old |
| 0.248697 | 0.529704 | 0.057 | 0.026 | 1 | 13 | Frs3 | 0.031 | Old |
| 0.248697 | 0.399037 | 0.057 | 0.026 | 1 | 13 | Napepld | 0.031 | Old |
| 0.248825 | 0.372384 | 0.459 | 0.386 | 1 | 13 | Shprh | 0.073 | Old |
| 0.248902 | 0.246023 | 0.672 | 0.596 | 1 | 13 | Herpud2 | 0.076 | Old |
| 0.248975 | 0.584672 | 0.082 | 0.044 | 1 | 13 | Mblac1 | 0.038 | Old |
| 0.249368 | 0.5335 | 0.115 | 0.07 | 1 | 13 | Gm14326 | 0.045 | Old |
| 0.249368 | 0.245912 | 0.115 | 0.07 | 1 | 13 | 4930555A0 | 0.045 | Old |
| 0.249421 | 0.175693 | 0.639 | 0.5 | 1 | 13 | Bckdk | 0.139 | Old |
| 0.249589 | 0.424984 | 0.139 | 0.088 | 1 | 13 | C1ra | 0.051 | Old |
| 0.249665 | 0.392285 | 0.27 | 0.202 | 1 | 13 | Pdcd2 | 0.068 | Old |
| 0.24973 | 0.123672 | 0.926 | 0.886 | 1 | 13 | Rad23b | 0.04 | Old |
| 0.249943 | 0.284357 | 0.467 | 0.395 | 1 | 13 | Cabin1 | 0.072 | Old |
| 0.250009 | 0.304419 | 0.656 | 0.588 | 1 | 13 | 2310035C2 | 0.068 | Old |
| 0.250313 | 0.425576 | 0.254 | 0.184 | 1 | 13 | Dus3l | 0.07 | Old |
| 0.250315 | 0.355072 | 0.385 | 0.307 | 1 | 13 | Exosc9 | 0.078 | Old |
| 0.250331 | 0.268687 | 0.615 | 0.579 | 1 | 13 | Cers5 | 0.036 | Old |
| 0.25084 | 0.198518 | 0.385 | 0.298 | 1 | 13 | Ctnnbl1 | 0.087 | Old |
| 0.250873 | 0.173375 | 0.377 | 0.289 | 1 | 13 | Armc10 | 0.088 | Old |

| | | | | | | | | |
|---|---|---|---|---|---|---|---|---|
| 0.250878 | 0.318337 | 0.311 | 0.219 | 1 | 13 | 3110002H1 | 0.092 | Old |
| 0.250906 | 0.298035 | 0.344 | 0.263 | 1 | 13 | AY036118 | 0.081 | Old |
| 0.251091 | 0.385991 | 0.23 | 0.167 | 1 | 13 | Trp53bp1 | 0.063 | Old |
| 0.251276 | 0.611729 | 0.18 | 0.132 | 1 | 13 | Acvr2b | 0.048 | Old |
| 0.251604 | 0.149133 | 0.23 | 0.158 | 1 | 13 | Ppp1r9a | 0.072 | Old |
| 0.251633 | 0.433748 | 0.41 | 0.333 | 1 | 13 | Cluh | 0.077 | Old |
| 0.25188 | 0.268987 | 0.623 | 0.518 | 1 | 13 | Eif4e3 | 0.105 | Old |
| 0.251886 | 0.550551 | 0.156 | 0.105 | 1 | 13 | Zcchc4 | 0.051 | Old |
| 0.251975 | 0.190778 | 0.639 | 0.526 | 1 | 13 | Kif1c | 0.113 | Old |
| 0.252001 | 0.149966 | 0.82 | 0.754 | 1 | 13 | Ccnd3 | 0.066 | Old |
| 0.252681 | 0.149364 | 1 | 1 | 1 | 13 | Itm2b | 0 | Old |
| 0.252681 | 0.055937 | 1 | 1 | 1 | 13 | Rbms1 | 0 | Old |
| 0.252907 | 0.361721 | 0.303 | 0.228 | 1 | 13 | Scly | 0.075 | Old |
| 0.253192 | 0.471643 | 0.418 | 0.333 | 1 | 13 | Slc39a14 | 0.085 | Old |
| 0.253329 | 0.175046 | 0.352 | 0.254 | 1 | 13 | Helb | 0.098 | Old |
| 0.253358 | 0.315887 | 0.23 | 0.167 | 1 | 13 | Pnpla6 | 0.063 | Old |
| 0.253747 | 0.242583 | 0.541 | 0.447 | 1 | 13 | Smim20 | 0.094 | Old |
| 0.253772 | 0.354279 | 0.451 | 0.377 | 1 | 13 | Slc35a3 | 0.074 | Old |
| 0.25395 | 0.129123 | 0.902 | 0.877 | 1 | 13 | Hip1 | 0.025 | Old |
| 0.254062 | 0.416091 | 0.115 | 0.07 | 1 | 13 | Mad2l1 | 0.045 | Old |
| 0.254123 | 0.742134 | 0.172 | 0.123 | 1 | 13 | Plekhn1 | 0.049 | Old |
| 0.254269 | 0.112623 | 0.893 | 0.868 | 1 | 13 | Map2k2 | 0.025 | Old |
| 0.254382 | 0.207936 | 0.361 | 0.272 | 1 | 13 | Gbe1 | 0.089 | Old |
| 0.25453 | 0.414958 | 0.328 | 0.272 | 1 | 13 | Cdc42ep4 | 0.056 | Old |
| 0.254575 | 0.498519 | 0.082 | 0.044 | 1 | 13 | Cdk5rap1 | 0.038 | Old |
| 0.254875 | 0.255122 | 0.328 | 0.254 | 1 | 13 | Reep4 | 0.074 | Old |
| 0.254875 | 0.008086 | 0.631 | 0.693 | 1 | 13 | Ccl4 | -0.062 | Old |
| 0.255101 | 0.169773 | 0.795 | 0.868 | 1 | 13 | Pou2f2 | -0.073 | Old |
| 0.255294 | 0.355701 | 0.279 | 0.211 | 1 | 13 | Stoml1 | 0.068 | Old |
| 0.255317 | 0.277294 | 0.295 | 0.228 | 1 | 13 | Nr2c2ap | 0.067 | Old |
| 0.255361 | 0.134843 | 0.156 | 0.096 | 1 | 13 | Kat8 | 0.06 | Old |
| 0.255488 | 0.412625 | 0.057 | 0.026 | 1 | 13 | St5 | 0.031 | Old |
| 0.25554 | 0.138897 | 0.639 | 0.544 | 1 | 13 | Hnrnpll | 0.095 | Old |
| 0.255555 | 0.20448 | 0.811 | 0.737 | 1 | 13 | Sash1 | 0.074 | Old |
| 0.255987 | 0.275195 | 0.352 | 0.263 | 1 | 13 | Rgp1 | 0.089 | Old |
| 0.25639 | 0.702971 | 0.123 | 0.079 | 1 | 13 | Asb4 | 0.044 | Old |
| 0.256573 | 0.215834 | 0.311 | 0.228 | 1 | 13 | Rbm15b | 0.083 | Old |
| 0.256758 | 0.348436 | 0.139 | 0.088 | 1 | 13 | Zfp777 | 0.051 | Old |
| 0.257092 | 0.178685 | 0.492 | 0.386 | 1 | 13 | Cops3 | 0.106 | Old |
| 0.257125 | 0.319324 | 0.311 | 0.237 | 1 | 13 | Ring1 | 0.074 | Old |
| 0.257347 | 0.284509 | 0.492 | 0.395 | 1 | 13 | Ppp1r35 | 0.097 | Old |
| 0.257461 | 0.092012 | 1 | 1 | 1 | 13 | Ddx3x | 0 | Old |
| 0.257729 | 0.242874 | 0.672 | 0.649 | 1 | 13 | Rbm7 | 0.023 | Old |
| 0.257757 | 0.137076 | 0.943 | 0.904 | 1 | 13 | 2610507B1 | 0.039 | Old |
| 0.257857 | 0.404468 | 0.221 | 0.158 | 1 | 13 | Galnt9 | 0.063 | Old |

| | | | | | | | | |
|---|---|---|---|---|---|---|---|---|
| 0.257943 | 0.482171 | 0.164 | 0.114 | 1 | 13 | Arhgap22 | 0.05 | Old |
| 0.258224 | 0.124206 | 0.943 | 0.947 | 1 | 13 | Spcs2 | -0.004 | Old |
| 0.258472 | 0.509195 | 0.279 | 0.211 | 1 | 13 | Git1 | 0.068 | Old |
| 0.258675 | 0.451555 | 0.156 | 0.105 | 1 | 13 | Mvk | 0.051 | Old |
| 0.258778 | 0.265332 | 0.811 | 0.737 | 1 | 13 | Kmt2c | 0.074 | Old |
| 0.258818 | 0.452252 | 0.115 | 0.07 | 1 | 13 | Cxcr3 | 0.045 | Old |
| 0.259746 | 0.188587 | 0.549 | 0.439 | 1 | 13 | Lrrc47 | 0.11 | Old |
| 0.259912 | 0.321034 | 0.5 | 0.421 | 1 | 13 | Slc25a20 | 0.079 | Old |
| 0.25998 | 0.313015 | 0.238 | 0.175 | 1 | 13 | Rrs1 | 0.063 | Old |
| 0.260032 | 0.379668 | 0.385 | 0.316 | 1 | 13 | Paox | 0.069 | Old |
| 0.260536 | 0.266502 | 0.615 | 0.544 | 1 | 13 | Sirt7 | 0.071 | Old |
| 0.260759 | 0.699717 | 0.139 | 0.096 | 1 | 13 | Ankzf1 | 0.043 | Old |
| 0.261457 | 0.310935 | 0.336 | 0.263 | 1 | 13 | Spg20 | 0.073 | Old |
| 0.261482 | 0.09314 | 1 | 0.93 | 1 | 13 | Akr1b3 | 0.07 | Old |
| 0.261655 | 0.592052 | 0.197 | 0.14 | 1 | 13 | Atg2b | 0.057 | Old |
| 0.261685 | 0.252943 | 0.336 | 0.254 | 1 | 13 | Smarcd1 | 0.082 | Old |
| 0.261901 | 0.41939 | 0.451 | 0.368 | 1 | 13 | Oasl2 | 0.083 | Old |
| 0.26205 | 0.673329 | 0.131 | 0.088 | 1 | 13 | Zfp319 | 0.043 | Old |
| 0.262062 | 0.16447 | 0.721 | 0.623 | 1 | 13 | Parl | 0.098 | Old |
| 0.262407 | 0.345108 | 0.057 | 0.026 | 1 | 13 | Ffar4 | 0.031 | Old |
| 0.262407 | 0.170252 | 0.057 | 0.026 | 1 | 13 | Ddx11 | 0.031 | Old |
| 0.262537 | 0.366647 | 0.123 | 0.079 | 1 | 13 | Mpp7 | 0.044 | Old |
| 0.263372 | 0.280405 | 0.582 | 0.518 | 1 | 13 | Prpf39 | 0.064 | Old |
| 0.26354 | 0.167114 | 0.73 | 0.632 | 1 | 13 | Ubr4 | 0.098 | Old |
| 0.26376 | 0.373851 | 0.311 | 0.246 | 1 | 13 | Megf8 | 0.065 | Old |
| 0.263931 | 0.108787 | 1 | 1 | 1 | 13 | Ehd1 | 0 | Old |
| 0.263943 | 0.377073 | 0.426 | 0.333 | 1 | 13 | Ago1 | 0.093 | Old |
| 0.264007 | 0.263072 | 0.549 | 0.474 | 1 | 13 | Helz | 0.075 | Old |
| 0.264103 | 0.416873 | 0.082 | 0.044 | 1 | 13 | Pfkfb2 | 0.038 | Old |
| 0.264103 | 0.146036 | 0.852 | 0.719 | 1 | 13 | Tmem259 | 0.133 | Old |
| 0.264197 | 0.511859 | 0.156 | 0.105 | 1 | 13 | Dcaf6 | 0.051 | Old |
| 0.264264 | 0.256193 | 0.459 | 0.395 | 1 | 13 | Timm22 | 0.064 | Old |
| 0.264482 | 0.190788 | 0.656 | 0.509 | 1 | 13 | Ncoa6 | 0.147 | Old |
| 0.26449 | 0.368368 | 0.148 | 0.096 | 1 | 13 | Elp4 | 0.052 | Old |
| 0.264936 | 0.172826 | 0.836 | 0.798 | 1 | 13 | Dbnl | 0.038 | Old |
| 0.265048 | 0.513626 | 0.131 | 0.088 | 1 | 13 | Invs | 0.043 | Old |
| 0.265287 | 0.230993 | 0.762 | 0.649 | 1 | 13 | Ube2f | 0.113 | Old |
| 0.265319 | 0.373427 | 0.303 | 0.228 | 1 | 13 | Prkab1 | 0.075 | Old |
| 0.265319 | 0.210674 | 0.303 | 0.228 | 1 | 13 | AI606181 | 0.075 | Old |
| 0.265356 | 0.210144 | 0.311 | 0.237 | 1 | 13 | Yaf2 | 0.074 | Old |
| 0.265381 | 0.151742 | 0.738 | 0.605 | 1 | 13 | Sec24b | 0.133 | Old |
| 0.26542 | 0.15553 | 0.918 | 0.912 | 1 | 13 | Uqcrfs1 | 0.006 | Old |
| 0.265484 | 0.191529 | 0.877 | 0.868 | 1 | 13 | Fam96a | 0.009 | Old |
| 0.265548 | 0.367626 | 0.139 | 0.088 | 1 | 13 | Supv3l1 | 0.051 | Old |
| 0.26559 | 0.540834 | 0.156 | 0.105 | 1 | 13 | Acot11 | 0.051 | Old |

| | | | | | | | | |
|---|---|---|---|---|---|---|---|---|
| 0.265649 | 0.567481 | 0.123 | 0.079 | 1 | 13 | Wrap53 | 0.044 | Old |
| 0.265754 | 0.210761 | 0.475 | 0.368 | 1 | 13 | Elof1 | 0.107 | Old |
| 0.266033 | 0.223086 | 0.73 | 0.632 | 1 | 13 | Ei24 | 0.098 | Old |
| 0.26632 | 0.185741 | 0.467 | 0.368 | 1 | 13 | Arsa | 0.099 | Old |
| 0.266649 | 0.234588 | 0.197 | 0.14 | 1 | 13 | Slc37a4 | 0.057 | Old |
| 0.266943 | 0.269039 | 0.254 | 0.184 | 1 | 13 | Sugp2 | 0.07 | Old |
| 0.26695 | 0.081229 | 0.869 | 0.798 | 1 | 13 | Dnajc15 | 0.071 | Old |
| 0.267133 | 0.14511 | 0.934 | 0.93 | 1 | 13 | Ssr1 | 0.004 | Old |
| 0.267203 | 0.240667 | 0.369 | 0.281 | 1 | 13 | Incenp | 0.088 | Old |
| 0.267207 | 0.079574 | 1 | 1 | 1 | 13 | Gabarap | 0 | Old |
| 0.267366 | 0.376127 | 0.148 | 0.096 | 1 | 13 | Oxnad1 | 0.052 | Old |
| 0.267841 | 0.338979 | 0.377 | 0.316 | 1 | 13 | Ing3 | 0.061 | Old |
| 0.267846 | 0.160852 | 0.91 | 0.816 | 1 | 13 | Sdf2 | 0.094 | Old |
| 0.267982 | 0.402024 | 0.082 | 0.044 | 1 | 13 | Gpt | 0.038 | Old |
| 0.268202 | 0.279106 | 0.475 | 0.386 | 1 | 13 | Nln | 0.089 | Old |
| 0.268371 | 0.436945 | 0.41 | 0.333 | 1 | 13 | Lonp2 | 0.077 | Old |
| 0.269073 | 0.175861 | 0.697 | 0.596 | 1 | 13 | Cdc73 | 0.101 | Old |
| 0.269085 | 0.766496 | 0.066 | 0.035 | 1 | 13 | Taf1a | 0.031 | Old |
| 0.269105 | 0.313211 | 0.352 | 0.281 | 1 | 13 | Phkg2 | 0.071 | Old |
| 0.269317 | 0.43888 | 0.09 | 0.053 | 1 | 13 | Cd200r3 | 0.037 | Old |
| 0.269356 | 0.289049 | 0.32 | 0.246 | 1 | 13 | Prpf4 | 0.074 | Old |
| 0.269454 | 0.270389 | 0.721 | 0.693 | 1 | 13 | Iah1 | 0.028 | Old |
| 0.269661 | 0.441726 | 0.246 | 0.184 | 1 | 13 | Mpi | 0.062 | Old |
| 0.270018 | 0.518465 | 0.295 | 0.228 | 1 | 13 | Il27ra | 0.067 | Old |
| 0.270224 | 0.301665 | 0.525 | 0.447 | 1 | 13 | Magohb | 0.078 | Old |
| 0.270263 | 0.486214 | 0.148 | 0.096 | 1 | 13 | Fbxw9 | 0.052 | Old |
| 0.270488 | 0.145539 | 0.951 | 0.956 | 1 | 13 | Snrpg | -0.005 | Old |
| 0.270497 | 0.226366 | 0.59 | 0.474 | 1 | 13 | Gabpb2 | 0.116 | Old |
| 0.270554 | 0.233136 | 0.303 | 0.219 | 1 | 13 | Pcnx | 0.084 | Old |
| 0.270554 | 0.358142 | 0.336 | 0.263 | 1 | 13 | 2310001H1 | 0.073 | Old |
| 0.270697 | 0.424016 | 0.443 | 0.368 | 1 | 13 | Ikzf5 | 0.075 | Old |
| 0.271064 | 0.085479 | 0.59 | 0.491 | 1 | 13 | Pank2 | 0.099 | Old |
| 0.271167 | 0.547334 | 0.09 | 0.053 | 1 | 13 | Bora | 0.037 | Old |
| 0.271167 | 0.546574 | 0.09 | 0.053 | 1 | 13 | Vsig10 | 0.037 | Old |
| 0.271263 | 1.061328 | 0.066 | 0.035 | 1 | 13 | Plpp3 | 0.031 | Old |
| 0.271263 | 0.914286 | 0.066 | 0.035 | 1 | 13 | Stac2 | 0.031 | Old |
| 0.271296 | 0.233261 | 0.607 | 0.535 | 1 | 13 | Tbc1d23 | 0.072 | Old |
| 0.271663 | 0.156288 | 0.672 | 0.588 | 1 | 13 | Gars | 0.084 | Old |
| 0.271908 | 0.288156 | 0.492 | 0.404 | 1 | 13 | Plekhm1 | 0.088 | Old |
| 0.272025 | 0.103588 | 0.27 | 0.193 | 1 | 13 | Habp4 | 0.077 | Old |
| 0.272365 | 0.190839 | 0.484 | 0.412 | 1 | 13 | Washc1 | 0.072 | Old |
| 0.272373 | 0.528117 | 0.238 | 0.184 | 1 | 13 | Gm26782 | 0.054 | Old |
| 0.272517 | 0.193273 | 0.705 | 0.553 | 1 | 13 | Acp1 | 0.152 | Old |
| 0.272615 | 0.164103 | 0.484 | 0.404 | 1 | 13 | Qrich1 | 0.08 | Old |
| 0.27293 | 0.259338 | 0.525 | 0.465 | 1 | 13 | Naa10 | 0.06 | Old |

| | | | | | | | | |
|---|---|---|---|---|---|---|---|---|
| 0.272941 | 0.442474 | 0.344 | 0.272 | 1 | 13 | Limk1 | 0.072 | Old |
| 0.273025 | 0.482817 | 0.09 | 0.053 | 1 | 13 | Ift88 | 0.037 | Old |
| 0.27315 | 0.209317 | 0.27 | 0.193 | 1 | 13 | Dcp1a | 0.077 | Old |
| 0.273174 | 0.088016 | 0.926 | 0.868 | 1 | 13 | D17Wsu92 | 0.058 | Old |
| 0.273294 | 0.273549 | 0.623 | 0.561 | 1 | 13 | Setx | 0.062 | Old |
| 0.273779 | 0.25514 | 0.557 | 0.447 | 1 | 13 | Med30 | 0.11 | Old |
| 0.274237 | 0.08525 | 0.885 | 0.868 | 1 | 13 | Prkar2a | 0.017 | Old |
| 0.27438 | 0.46108 | 0.246 | 0.193 | 1 | 13 | Rdm1 | 0.053 | Old |
| 0.274892 | 0.616029 | 0.09 | 0.053 | 1 | 13 | Runx2 | 0.037 | Old |
| 0.274899 | 0.500631 | 0.295 | 0.237 | 1 | 13 | Usp39 | 0.058 | Old |
| 0.275146 | 0.36859 | 0.361 | 0.298 | 1 | 13 | Atp11a | 0.063 | Old |
| 0.275274 | 0.262721 | 0.574 | 0.465 | 1 | 13 | Cyp4f16 | 0.109 | Old |
| 0.275517 | 0.149887 | 0.984 | 0.965 | 1 | 13 | Glul | 0.019 | Old |
| 0.275556 | 0.720881 | 0.254 | 0.193 | 1 | 13 | Hes1 | 0.061 | Old |
| 0.275653 | 0.919347 | 0.066 | 0.035 | 1 | 13 | Myo1b | 0.031 | Old |
| 0.275728 | 0.267104 | 0.631 | 0.579 | 1 | 13 | Dpp8 | 0.052 | Old |
| 0.275966 | 0.113389 | 0.549 | 0.465 | 1 | 13 | Ccdc174 | 0.084 | Old |
| 0.276501 | 0.204401 | 0.475 | 0.386 | 1 | 13 | Ubap1 | 0.089 | Old |
| 0.276831 | 1.155018 | 0.041 | 0.018 | 1 | 13 | Mycl | 0.023 | Old |
| 0.276831 | 0.743293 | 0.041 | 0.018 | 1 | 13 | P2ry2 | 0.023 | Old |
| 0.276831 | 0.732106 | 0.041 | 0.018 | 1 | 13 | Sspn | 0.023 | Old |
| 0.276835 | 0.263468 | 0.197 | 0.14 | 1 | 13 | Kctd13 | 0.057 | Old |
| 0.27686 | 0.297768 | 0.443 | 0.36 | 1 | 13 | Mrpl10 | 0.083 | Old |
| 0.276884 | 0.143442 | 0.705 | 0.632 | 1 | 13 | Chp1 | 0.073 | Old |
| 0.277057 | 0.001768 | 0.189 | 0.123 | 1 | 13 | Zc3h3 | 0.066 | Old |
| 0.27706 | 0.307287 | 0.295 | 0.237 | 1 | 13 | Chkb | 0.058 | Old |
| 0.277365 | 0.208456 | 0.443 | 0.351 | 1 | 13 | Snx14 | 0.092 | Old |
| 0.277588 | 0.297756 | 0.139 | 0.088 | 1 | 13 | Ears2 | 0.051 | Old |
| 0.277599 | 0.283999 | 0.148 | 0.096 | 1 | 13 | Zfp53 | 0.052 | Old |
| 0.277793 | 0.329236 | 0.246 | 0.184 | 1 | 13 | Otud7b | 0.062 | Old |
| 0.278018 | 0.241954 | 0.951 | 0.947 | 1 | 13 | Rhob | 0.004 | Old |
| 0.278088 | 0.229478 | 0.451 | 0.404 | 1 | 13 | Tmem222 | 0.047 | Old |
| 0.278179 | 0.214886 | 0.639 | 0.553 | 1 | 13 | Ccz1 | 0.086 | Old |
| 0.278232 | 0.298513 | 0.361 | 0.281 | 1 | 13 | Akt1s1 | 0.08 | Old |
| 0.278407 | 0.235978 | 0.279 | 0.202 | 1 | 13 | 2410004B1 | 0.077 | Old |
| 0.278453 | 0.387504 | 0.164 | 0.114 | 1 | 13 | Nek4 | 0.05 | Old |
| 0.278454 | 0.221245 | 0.664 | 0.561 | 1 | 13 | Atg5 | 0.103 | Old |
| 0.278551 | 0.27242 | 0.664 | 0.596 | 1 | 13 | Derl2 | 0.068 | Old |
| 0.278832 | 0.480125 | 0.131 | 0.088 | 1 | 13 | Xndc1 | 0.043 | Old |
| 0.278895 | 0.127723 | 0.992 | 1 | 1 | 13 | Ccnl1 | -0.008 | Old |
| 0.279021 | 0.344524 | 0.254 | 0.193 | 1 | 13 | Gpatch4 | 0.061 | Old |
| 0.279179 | 0.123546 | 0.566 | 0.482 | 1 | 13 | Tdrd7 | 0.084 | Old |
| 0.279232 | 0.340453 | 0.295 | 0.237 | 1 | 13 | Itpkc | 0.058 | Old |
| 0.279313 | 0.129419 | 0.844 | 0.851 | 1 | 13 | Vps29 | -0.007 | Old |
| 0.279709 | 0.839262 | 0.041 | 0.018 | 1 | 13 | Il7 | 0.023 | Old |

| | | | | | | | | |
|---|---|---|---|---|---|---|---|---|
| 0.279713 | 0.238077 | 0.656 | 0.561 | 1 | 13 | Sec24c | 0.095 | Old |
| 0.27978 | 0.54056 | 0.262 | 0.211 | 1 | 13 | Rtca | 0.051 | Old |
| 0.280017 | 0.060414 | 0.557 | 0.456 | 1 | 13 | Cnot2 | 0.101 | Old |
| 0.280045 | 0.240282 | 0.623 | 0.544 | 1 | 13 | Kars | 0.079 | Old |
| 0.280308 | 0.155769 | 0.844 | 0.781 | 1 | 13 | Rap1gds1 | 0.063 | Old |
| 0.280593 | 0.065431 | 1 | 1 | 1 | 13 | Gnas | 0 | Old |
| 0.281238 | 0.111145 | 0.648 | 0.509 | 1 | 13 | Impad1 | 0.139 | Old |
| 0.281435 | 0.092293 | 0.967 | 0.965 | 1 | 13 | Pycard | 0.002 | Old |
| 0.281587 | 0.379876 | 0.123 | 0.079 | 1 | 13 | Rnf25 | 0.044 | Old |
| 0.281847 | 0.214804 | 0.664 | 0.544 | 1 | 13 | Arfgap2 | 0.12 | Old |
| 0.28223 | 0.241611 | 0.361 | 0.298 | 1 | 13 | Nudt13 | 0.063 | Old |
| 0.282242 | 0.320435 | 0.238 | 0.175 | 1 | 13 | Saal1 | 0.063 | Old |
| 0.282298 | 0.102448 | 1 | 1 | 1 | 13 | Rpl22l1 | 0 | Old |
| 0.28233 | 1.704959 | 0.066 | 0.035 | 1 | 13 | Acod1 | 0.031 | Old |
| 0.282358 | 0.060213 | 0.311 | 0.237 | 1 | 13 | Ppan | 0.074 | Old |
| 0.282447 | 0.994742 | 0.09 | 0.053 | 1 | 13 | Sell | 0.037 | Old |
| 0.282447 | 0.510511 | 0.09 | 0.053 | 1 | 13 | Slc7a5 | 0.037 | Old |
| 0.282447 | 0.493355 | 0.09 | 0.053 | 1 | 13 | Gtf2ird2 | 0.037 | Old |
| 0.282464 | 0.070292 | 0.172 | 0.114 | 1 | 13 | Zfp317 | 0.058 | Old |
| 0.282607 | 1.119962 | 0.041 | 0.018 | 1 | 13 | Hmmr | 0.023 | Old |
| 0.282607 | 0.818566 | 0.041 | 0.018 | 1 | 13 | Btnl9 | 0.023 | Old |
| 0.282607 | 0.777218 | 0.041 | 0.018 | 1 | 13 | Mterf1a | 0.023 | Old |
| 0.28283 | 0.15739 | 0.697 | 0.596 | 1 | 13 | Sf3b4 | 0.101 | Old |
| 0.282844 | 0.210238 | 0.525 | 0.43 | 1 | 13 | Swt1 | 0.095 | Old |
| 0.283035 | 0.09988 | 0.582 | 0.5 | 1 | 13 | Fam103a1 | 0.082 | Old |
| 0.283211 | 0.481095 | 0.23 | 0.175 | 1 | 13 | Btbd17 | 0.055 | Old |
| 0.28358 | 0.11445 | 0.975 | 0.991 | 1 | 13 | Nme1 | -0.016 | Old |
| 0.283894 | 0.233652 | 0.344 | 0.246 | 1 | 13 | Lias | 0.098 | Old |
| 0.28416 | 0.343162 | 0.156 | 0.105 | 1 | 13 | Dsel | 0.051 | Old |
| 0.284394 | 0.200724 | 0.361 | 0.281 | 1 | 13 | Yipf6 | 0.08 | Old |
| 0.28458 | 0.712423 | 0.066 | 0.035 | 1 | 13 | Gm16638 | 0.031 | Old |
| 0.28458 | 0.645862 | 0.066 | 0.035 | 1 | 13 | Fam120aos | 0.031 | Old |
| 0.28458 | 0.639087 | 0.066 | 0.035 | 1 | 13 | Armc9 | 0.031 | Old |
| 0.284674 | 0.414845 | 0.377 | 0.316 | 1 | 13 | Ruvbl2 | 0.061 | Old |
| 0.285193 | 0.283645 | 0.508 | 0.404 | 1 | 13 | Sil1 | 0.104 | Old |
| 0.285222 | 0.288299 | 0.115 | 0.07 | 1 | 13 | Slc7a6 | 0.045 | Old |
| 0.28528 | 0.167217 | 0.943 | 0.974 | 1 | 13 | Ptpn6 | -0.031 | Old |
| 0.28529 | 0.196717 | 0.574 | 0.509 | 1 | 13 | Prpf18 | 0.065 | Old |
| 0.285348 | 0.223392 | 0.762 | 0.667 | 1 | 13 | Mfsd5 | 0.095 | Old |
| 0.285526 | 0.8709 | 0.041 | 0.018 | 1 | 13 | Pomc | 0.023 | Old |
| 0.285526 | 0.81633 | 0.041 | 0.018 | 1 | 13 | Gm12184 | 0.023 | Old |
| 0.285526 | 0.728425 | 0.041 | 0.018 | 1 | 13 | Tmem201 | 0.023 | Old |
| 0.285526 | 0.668575 | 0.041 | 0.018 | 1 | 13 | Gm15326 | 0.023 | Old |
| 0.285816 | 0.253433 | 0.598 | 0.57 | 1 | 13 | Lyar | 0.028 | Old |
| 0.285857 | 0.263448 | 0.41 | 0.333 | 1 | 13 | Bcl2 | 0.077 | Old |

| | | | | | | | | |
|---|---|---|---|---|---|---|---|---|
| 0.285858 | 0.453572 | 0.205 | 0.158 | 1 | 13 | Fbxo42 | 0.047 | Old |
| 0.286117 | 0.119693 | 0.811 | 0.719 | 1 | 13 | Ypel5 | 0.092 | Old |
| 0.286266 | 0.287821 | 0.238 | 0.167 | 1 | 13 | Nt5m | 0.071 | Old |
| 0.286313 | 0.299443 | 0.566 | 0.456 | 1 | 13 | Flvcr1 | 0.11 | Old |
| 0.286347 | 0.198536 | 0.32 | 0.246 | 1 | 13 | Morc2a | 0.074 | Old |
| 0.286601 | 0.527358 | 0.098 | 0.061 | 1 | 13 | Ifit3 | 0.037 | Old |
| 0.286698 | 1.012946 | 0.131 | 0.088 | 1 | 13 | Il4i1 | 0.043 | Old |
| 0.286698 | 0.536013 | 0.131 | 0.088 | 1 | 13 | Cby1 | 0.043 | Old |
| 0.286988 | 0.003748 | 0.549 | 0.447 | 1 | 13 | Trabd | 0.102 | Old |
| 0.287088 | 0.338625 | 0.238 | 0.175 | 1 | 13 | Fam136a | 0.063 | Old |
| 0.287283 | 0.395544 | 0.197 | 0.14 | 1 | 13 | Fmn1 | 0.057 | Old |
| 0.28731 | 0.171457 | 0.795 | 0.754 | 1 | 13 | Bud31 | 0.041 | Old |
| 0.287351 | 0.20652 | 0.598 | 0.509 | 1 | 13 | Cbx4 | 0.089 | Old |
| 0.287481 | 0.187 | 0.623 | 0.535 | 1 | 13 | Snx12 | 0.088 | Old |
| 0.287829 | 0.190494 | 0.18 | 0.123 | 1 | 13 | Zfp426 | 0.057 | Old |
| 0.287831 | 0.31719 | 0.23 | 0.167 | 1 | 13 | Mto1 | 0.063 | Old |
| 0.287885 | 0.159779 | 0.992 | 0.974 | 1 | 13 | Ndufc2 | 0.018 | Old |
| 0.288204 | 0.360434 | 0.09 | 0.053 | 1 | 13 | 2700081O1 | 0.037 | Old |
| 0.288313 | 0.507139 | 0.279 | 0.211 | 1 | 13 | Styx | 0.068 | Old |
| 0.288316 | 0.354532 | 0.549 | 0.456 | 1 | 13 | Fem1b | 0.093 | Old |
| 0.288322 | 0.191141 | 0.762 | 0.649 | 1 | 13 | Med15 | 0.113 | Old |
| 0.28835 | 0.286577 | 0.893 | 0.868 | 1 | 13 | Icam1 | 0.025 | Old |
| 0.288434 | 0.591365 | 0.098 | 0.061 | 1 | 13 | Tmem14a | 0.037 | Old |
| 0.288453 | 0.26896 | 0.525 | 0.447 | 1 | 13 | Zufsp | 0.078 | Old |
| 0.288465 | 0.707051 | 0.041 | 0.018 | 1 | 13 | Smpd5 | 0.023 | Old |
| 0.288465 | 0.655191 | 0.041 | 0.018 | 1 | 13 | Car13 | 0.023 | Old |
| 0.288465 | 0.643321 | 0.041 | 0.018 | 1 | 13 | Ncapg | 0.023 | Old |
| 0.288465 | 0.635535 | 0.041 | 0.018 | 1 | 13 | Zbtb32 | 0.023 | Old |
| 0.289116 | 0.62809 | 0.066 | 0.035 | 1 | 13 | Taf4b | 0.031 | Old |
| 0.289116 | 0.600811 | 0.066 | 0.035 | 1 | 13 | Angptl6 | 0.031 | Old |
| 0.289146 | 0.117154 | 0.951 | 0.939 | 1 | 13 | Sh3kbp1 | 0.012 | Old |
| 0.289186 | 0.100021 | 1 | 0.991 | 1 | 13 | Ppp1ca | 0.009 | Old |
| 0.289214 | 0.177998 | 0.18 | 0.123 | 1 | 13 | Mon1a | 0.057 | Old |
| 0.289385 | 0.157331 | 0.533 | 0.439 | 1 | 13 | Fbrs | 0.094 | Old |
| 0.289994 | 0.115502 | 0.951 | 0.921 | 1 | 13 | Ptp4a1 | 0.03 | Old |
| 0.290053 | 0.084206 | 0.984 | 0.974 | 1 | 13 | Psmb4 | 0.01 | Old |
| 0.290278 | 0.271178 | 0.598 | 0.518 | 1 | 13 | Zdhhc3 | 0.08 | Old |
| 0.290309 | 0.196638 | 0.705 | 0.64 | 1 | 13 | Ncaph2 | 0.065 | Old |
| 0.290683 | 0.281847 | 0.344 | 0.281 | 1 | 13 | Dctn6 | 0.063 | Old |
| 0.290938 | 0.252751 | 0.32 | 0.237 | 1 | 13 | Fam53a | 0.083 | Old |
| 0.291011 | 0.140145 | 0.885 | 0.816 | 1 | 13 | Sirt2 | 0.069 | Old |
| 0.29113 | 0.321427 | 0.311 | 0.237 | 1 | 13 | Kat5 | 0.074 | Old |
| 0.29121 | 0.1567 | 0.869 | 0.763 | 1 | 13 | Smc6 | 0.106 | Old |
| 0.291402 | 0.57579 | 0.066 | 0.035 | 1 | 13 | Nr6a1 | 0.031 | Old |
| 0.291424 | 0.69732 | 0.041 | 0.018 | 1 | 13 | 9130230L2 | 0.023 | Old |

| | | | | | | | | |
|---|---|---|---|---|---|---|---|---|
| 0.291424 | 0.693307 | 0.041 | 0.018 | 1 | 13 | Btc | 0.023 | Old |
| 0.291424 | 0.626301 | 0.041 | 0.018 | 1 | 13 | Rexo5 | 0.023 | Old |
| 0.291455 | 0.421054 | 0.123 | 0.079 | 1 | 13 | Card6 | 0.044 | Old |
| 0.291767 | 0.246528 | 0.279 | 0.211 | 1 | 13 | 2410089E0 | 0.068 | Old |
| 0.291785 | 0.340719 | 0.549 | 0.719 | 1 | 13 | Clec4e | -0.17 | Old |
| 0.291789 | 0.18152 | 0.664 | 0.588 | 1 | 13 | Pqbp1 | 0.076 | Old |
| 0.291837 | 0.52579 | 0.295 | 0.246 | 1 | 13 | Mcm5 | 0.049 | Old |
| 0.292016 | 0.137165 | 0.762 | 0.64 | 1 | 13 | Krit1 | 0.122 | Old |
| 0.292087 | 0.231544 | 0.09 | 0.053 | 1 | 13 | Gpsm2 | 0.037 | Old |
| 0.292881 | 0.223639 | 0.484 | 0.386 | 1 | 13 | Cdkn2aipnl | 0.098 | Old |
| 0.293175 | 0.332811 | 0.254 | 0.193 | 1 | 13 | 3110040N1 | 0.061 | Old |
| 0.293514 | 0.325716 | 0.303 | 0.237 | 1 | 13 | Pcnx3 | 0.066 | Old |
| 0.293609 | 0.36142 | 0.361 | 0.298 | 1 | 13 | Klhdc2 | 0.063 | Old |
| 0.293686 | 0.05831 | 0.238 | 0.167 | 1 | 13 | Mul1 | 0.071 | Old |
| 0.2937 | 0.61375 | 0.066 | 0.035 | 1 | 13 | Gm13822 | 0.031 | Old |
| 0.2937 | 0.218382 | 0.066 | 0.035 | 1 | 13 | Top3a | 0.031 | Old |
| 0.293946 | 0.302087 | 0.197 | 0.14 | 1 | 13 | Dlg4 | 0.057 | Old |
| 0.29398 | 0.573032 | 0.098 | 0.061 | 1 | 13 | Cep19 | 0.037 | Old |
| 0.294213 | 0.323439 | 0.148 | 0.096 | 1 | 13 | Gm13373 | 0.052 | Old |
| 0.294403 | 0.747562 | 0.041 | 0.018 | 1 | 13 | Zfp993 | 0.023 | Old |
| 0.294403 | 0.674727 | 0.041 | 0.018 | 1 | 13 | Zfp968 | 0.023 | Old |
| 0.294403 | 0.623129 | 0.041 | 0.018 | 1 | 13 | Tbc1d2 | 0.023 | Old |
| 0.294403 | 0.015357 | 0.041 | 0.018 | 1 | 13 | Saa3 | 0.023 | Old |
| 0.294406 | 0.380914 | 0.352 | 0.272 | 1 | 13 | Dhdds | 0.08 | Old |
| 0.294498 | 0.184233 | 0.434 | 0.342 | 1 | 13 | Nop9 | 0.092 | Old |
| 0.294528 | 0.267558 | 0.557 | 0.474 | 1 | 13 | Bloc1s2 | 0.083 | Old |
| 0.294759 | 0.130554 | 0.861 | 0.877 | 1 | 13 | Tmem123 | -0.016 | Old |
| 0.294909 | 0.475859 | 0.205 | 0.149 | 1 | 13 | Prkdc | 0.056 | Old |
| 0.295102 | 0.340351 | 0.254 | 0.184 | 1 | 13 | Ogfod1 | 0.07 | Old |
| 0.295193 | 0.207326 | 0.664 | 0.64 | 1 | 13 | Mcmbp | 0.024 | Old |
| 0.295312 | 0.165282 | 0.664 | 0.553 | 1 | 13 | Stat5b | 0.111 | Old |
| 0.295316 | 0.27829 | 0.467 | 0.386 | 1 | 13 | Ralgapb | 0.081 | Old |
| 0.295348 | 0.188127 | 0.516 | 0.412 | 1 | 13 | Tnpo2 | 0.104 | Old |
| 0.295388 | 0.279168 | 0.672 | 0.57 | 1 | 13 | Gmip | 0.102 | Old |
| 0.295501 | 0.186658 | 0.836 | 0.789 | 1 | 13 | Thoc2 | 0.047 | Old |
| 0.295756 | 0.257433 | 0.148 | 0.096 | 1 | 13 | Urb2 | 0.052 | Old |
| 0.296004 | 0.341278 | 0.09 | 0.053 | 1 | 13 | Gm12689 | 0.037 | Old |
| 0.296004 | 0.309426 | 0.09 | 0.053 | 1 | 13 | Txnl4b | 0.037 | Old |
| 0.296066 | 0.190557 | 0.459 | 0.36 | 1 | 13 | Fam220a | 0.099 | Old |
| 0.296261 | 0.231049 | 0.549 | 0.491 | 1 | 13 | Rpl7l1 | 0.058 | Old |
| 0.296333 | 0.224222 | 0.262 | 0.202 | 1 | 13 | Sqle | 0.06 | Old |
| 0.296547 | 0.167362 | 0.352 | 0.272 | 1 | 13 | Traf6 | 0.08 | Old |
| 0.296763 | 0.310224 | 0.467 | 0.404 | 1 | 13 | Ftsj3 | 0.063 | Old |
| 0.297193 | 0.358263 | 0.434 | 0.377 | 1 | 13 | Thoc3 | 0.057 | Old |
| 0.297305 | 0.13038 | 0.803 | 0.64 | 1 | 13 | Stk24 | 0.163 | Old |

| | | | | | | | | |
|---|---|---|---|---|---|---|---|---|
| 0.297403 | 0.588953 | 0.041 | 0.018 | 1 | 13 | Rasl2-9 | 0.023 | Old |
| 0.297403 | 0.531197 | 0.041 | 0.018 | 1 | 13 | Stac3 | 0.023 | Old |
| 0.297403 | 0.496845 | 0.041 | 0.018 | 1 | 13 | Met | 0.023 | Old |
| 0.297403 | 0.490597 | 0.041 | 0.018 | 1 | 13 | Mlph | 0.023 | Old |
| 0.297479 | 0.322365 | 0.607 | 0.553 | 1 | 13 | Asxl2 | 0.054 | Old |
| 0.297479 | 0.213294 | 0.631 | 0.526 | 1 | 13 | Nsun2 | 0.105 | Old |
| 0.297716 | 0.752572 | 0.098 | 0.061 | 1 | 13 | Dnaaf5 | 0.037 | Old |
| 0.297716 | 0.474597 | 0.098 | 0.061 | 1 | 13 | Gm10521 | 0.037 | Old |
| 0.297766 | 0.26305 | 0.738 | 0.728 | 1 | 13 | Snw1 | 0.01 | Old |
| 0.297958 | 0.358154 | 0.131 | 0.088 | 1 | 13 | Gemin2 | 0.043 | Old |
| 0.297972 | 0.240062 | 0.615 | 0.518 | 1 | 13 | Acadvl | 0.097 | Old |
| 0.298333 | 0.467668 | 0.066 | 0.035 | 1 | 13 | Polr3g | 0.031 | Old |
| 0.298333 | 0.386312 | 0.066 | 0.035 | 1 | 13 | Kif22 | 0.031 | Old |
| 0.298352 | 0.242904 | 0.598 | 0.509 | 1 | 13 | Plekhm2 | 0.089 | Old |
| 0.298681 | 0.884171 | 0.074 | 0.044 | 1 | 13 | 1110008L1 | 0.03 | Old |
| 0.2989 | 0.156217 | 0.443 | 0.351 | 1 | 13 | Crebzf | 0.092 | Old |
| 0.299021 | 0.326173 | 0.361 | 0.289 | 1 | 13 | Trappc12 | 0.072 | Old |
| 0.299078 | 0.317533 | 0.59 | 0.491 | 1 | 13 | Echs1 | 0.099 | Old |
| 0.299488 | 0.296541 | 0.393 | 0.316 | 1 | 13 | Wdr11 | 0.077 | Old |
| 0.299545 | 0.226802 | 0.582 | 0.5 | 1 | 13 | Ifi209 | 0.082 | Old |
| 0.299631 | 0.108754 | 0.648 | 0.535 | 1 | 13 | Mrpl3 | 0.113 | Old |
| 0.299906 | 0.245393 | 0.631 | 0.596 | 1 | 13 | Dennd1a | 0.035 | Old |
| 0.299956 | 0.236102 | 0.09 | 0.053 | 1 | 13 | 4933417E1 | 0.037 | Old |
| 0.300069 | 0.238317 | 0.484 | 0.404 | 1 | 13 | Rfc2 | 0.08 | Old |
| 0.300069 | 0.19324 | 0.492 | 0.395 | 1 | 13 | Ddhd1 | 0.097 | Old |
| 0.300172 | 0.220735 | 0.393 | 0.325 | 1 | 13 | Heatr1 | 0.068 | Old |
| 0.30018 | 0.335503 | 0.27 | 0.202 | 1 | 13 | 1700037H( | 0.068 | Old |
| 0.300423 | 0.556568 | 0.041 | 0.018 | 1 | 13 | Gm49339 | 0.023 | Old |
| 0.300423 | 0.466585 | 0.041 | 0.018 | 1 | 13 | Cdkl5 | 0.023 | Old |
| 0.300445 | 0.256287 | 0.303 | 0.228 | 1 | 13 | Trpm4 | 0.075 | Old |
| 0.30049 | 0.399173 | 0.811 | 0.789 | 1 | 13 | Cebpd | 0.022 | Old |
| 0.30062 | 0.10759 | 1 | 1 | 1 | 13 | Malat1 | 0 | Old |
| 0.300668 | 0.462371 | 0.066 | 0.035 | 1 | 13 | Tnfrsf26 | 0.031 | Old |
| 0.300853 | 0.786817 | 0.074 | 0.044 | 1 | 13 | Ap5b1 | 0.03 | Old |
| 0.301016 | 0.087313 | 0.959 | 0.921 | 1 | 13 | Ptges3 | 0.038 | Old |
| 0.301085 | 0.326228 | 0.32 | 0.263 | 1 | 13 | Cgnl1 | 0.057 | Old |
| 0.301214 | 0.226538 | 0.475 | 0.377 | 1 | 13 | Pofut1 | 0.098 | Old |
| 0.301229 | 0.293383 | 0.131 | 0.088 | 1 | 13 | Phc1 | 0.043 | Old |
| 0.301512 | 0.062529 | 0.984 | 1 | 1 | 13 | Pim1 | -0.016 | Old |
| 0.30166 | 0.279923 | 0.541 | 0.439 | 1 | 13 | Itgav | 0.102 | Old |
| 0.30173 | 0.33998 | 0.68 | 0.588 | 1 | 13 | Mknk1 | 0.092 | Old |
| 0.301938 | 0.352147 | 0.246 | 0.184 | 1 | 13 | Zdhhc13 | 0.062 | Old |
| 0.301995 | 0.616598 | 0.295 | 0.246 | 1 | 13 | Flnb | 0.049 | Old |
| 0.302002 | 0.36645 | 0.213 | 0.158 | 1 | 13 | Utp4 | 0.055 | Old |
| 0.302206 | 0.329558 | 0.189 | 0.132 | 1 | 13 | Phlpp1 | 0.057 | Old |

| | | | | | | | | |
|---|---|---|---|---|---|---|---|---|
| 0.30228 | 0.116979 | 0.943 | 0.895 | 1 | 13 | Selenot | 0.048 | Old |
| 0.302334 | 0.419297 | 0.328 | 0.263 | 1 | 13 | Xaf1 | 0.065 | Old |
| 0.302344 | 0.146023 | 0.811 | 0.737 | 1 | 13 | Eif4e | 0.074 | Old |
| 0.302482 | 0.192715 | 0.762 | 0.693 | 1 | 13 | Cpsf2 | 0.069 | Old |
| 0.302881 | 0.727436 | 0.148 | 0.105 | 1 | 13 | P2ry1 | 0.043 | Old |
| 0.302981 | 0.266365 | 0.23 | 0.167 | 1 | 13 | Mettl21a | 0.063 | Old |
| 0.303015 | 0.466574 | 0.066 | 0.035 | 1 | 13 | Cavin2 | 0.031 | Old |
| 0.303015 | 0.443244 | 0.066 | 0.035 | 1 | 13 | Adgrl1 | 0.031 | Old |
| 0.303464 | 0.481086 | 0.041 | 0.018 | 1 | 13 | Ccdc114 | 0.023 | Old |
| 0.303464 | 0.378073 | 0.041 | 0.018 | 1 | 13 | Poc1a | 0.023 | Old |
| 0.303464 | 0.329325 | 0.041 | 0.018 | 1 | 13 | Guca1b | 0.023 | Old |
| 0.303518 | 0.181645 | 0.648 | 0.561 | 1 | 13 | Suds3 | 0.087 | Old |
| 0.303622 | 0.254424 | 0.418 | 0.333 | 1 | 13 | Bloc1s4 | 0.085 | Old |
| 0.303731 | 0.016099 | 0.869 | 0.772 | 1 | 13 | Ube2q1 | 0.097 | Old |
| 0.304439 | 0.150979 | 0.467 | 0.377 | 1 | 13 | Gle1 | 0.09 | Old |
| 0.304495 | 0.398753 | 0.164 | 0.114 | 1 | 13 | Trim30b | 0.05 | Old |
| 0.304649 | 0.197123 | 0.426 | 0.342 | 1 | 13 | Nmt2 | 0.084 | Old |
| 0.304673 | 0.152556 | 0.754 | 0.667 | 1 | 13 | Cd180 | 0.087 | Old |
| 0.304687 | 0.474037 | 0.221 | 0.167 | 1 | 13 | Abt1 | 0.054 | Old |
| 0.305097 | 0.13676 | 1 | 1 | 1 | 13 | C1qa | 0 | Old |
| 0.305177 | 0.333918 | 0.311 | 0.254 | 1 | 13 | Uxt | 0.057 | Old |
| 0.305395 | 0.093014 | 0.902 | 0.825 | 1 | 13 | Kdelr1 | 0.077 | Old |
| 0.30549 | 0.2461 | 0.213 | 0.149 | 1 | 13 | Rabif | 0.064 | Old |
| 0.305761 | 0.288222 | 0.238 | 0.175 | 1 | 13 | Stn1 | 0.063 | Old |
| 0.305869 | 0.148625 | 0.689 | 0.579 | 1 | 13 | Sppl3 | 0.11 | Old |
| 0.30645 | 0.300578 | 0.295 | 0.219 | 1 | 13 | Cyp4v3 | 0.076 | Old |
| 0.306525 | 0.453915 | 0.041 | 0.018 | 1 | 13 | Ogn | 0.023 | Old |
| 0.306525 | 0.326377 | 0.041 | 0.018 | 1 | 13 | Cysltr2 | 0.023 | Old |
| 0.306525 | 0.048616 | 0.041 | 0.018 | 1 | 13 | 1500002C1 | 0.023 | Old |
| 0.306793 | 0.2698 | 0.803 | 0.798 | 1 | 13 | Ms4a8a | 0.005 | Old |
| 0.306829 | 0.220687 | 0.5 | 0.412 | 1 | 13 | Cox16 | 0.088 | Old |
| 0.306929 | 0.117935 | 0.172 | 0.114 | 1 | 13 | Ncapg2 | 0.058 | Old |
| 0.307058 | 0.110934 | 0.574 | 0.491 | 1 | 13 | Sec24a | 0.083 | Old |
| 0.307161 | 0.318715 | 0.303 | 0.237 | 1 | 13 | Rapgef1 | 0.066 | Old |
| 0.307432 | 0.749104 | 0.074 | 0.044 | 1 | 13 | B3gnt2 | 0.03 | Old |
| 0.307517 | 0.143176 | 0.918 | 0.868 | 1 | 13 | Fam111a | 0.05 | Old |
| 0.307666 | 0.212216 | 0.221 | 0.158 | 1 | 13 | Scd1 | 0.063 | Old |
| 0.307684 | 0.223218 | 0.287 | 0.219 | 1 | 13 | Klhdc10 | 0.068 | Old |
| 0.308231 | 0.063175 | 0.393 | 0.307 | 1 | 13 | Rpa1 | 0.086 | Old |
| 0.308319 | 0.273027 | 0.369 | 0.298 | 1 | 13 | Nfyc | 0.071 | Old |
| 0.308612 | 0.290383 | 0.27 | 0.202 | 1 | 13 | Rrp12 | 0.068 | Old |
| 0.308708 | 0.079919 | 0.975 | 0.982 | 1 | 13 | Adam15 | -0.007 | Old |
| 0.3088 | 0.238335 | 0.41 | 0.342 | 1 | 13 | Zfp120 | 0.068 | Old |
| 0.308967 | 0.107356 | 0.197 | 0.14 | 1 | 13 | Gmppb | 0.057 | Old |
| 0.309 | 0.209925 | 0.607 | 0.544 | 1 | 13 | Supt6 | 0.063 | Old |

| | | | | | | | | |
|---|---|---|---|---|---|---|---|---|
| 0.309053 | 0.10671 | 0.787 | 0.693 | 1 | 13 | Ufm1 | 0.094 | Old |
| 0.309435 | 0.574833 | 0.156 | 0.114 | 1 | 13 | D930016D( | 0.042 | Old |
| 0.309647 | 1.129235 | 0.074 | 0.044 | 1 | 13 | Lhfpl2 | 0.03 | Old |
| 0.309877 | 0.135434 | 0.91 | 0.904 | 1 | 13 | Ppp1r12a | 0.006 | Old |
| 0.309989 | 0.230988 | 0.09 | 0.053 | 1 | 13 | Zfp408 | 0.037 | Old |
| 0.310128 | 0.374633 | 0.066 | 0.035 | 1 | 13 | Zfp235 | 0.031 | Old |
| 0.310128 | 0.030903 | 0.066 | 0.035 | 1 | 13 | Cdk1 | 0.031 | Old |
| 0.310196 | 0.273133 | 0.492 | 0.412 | 1 | 13 | Stat2 | 0.08 | Old |
| 0.310521 | 0.165939 | 0.385 | 0.325 | 1 | 13 | Stmn1 | 0.06 | Old |
| 0.310535 | 0.483103 | 0.607 | 0.553 | 1 | 13 | Slc16a3 | 0.054 | Old |
| 0.310767 | 0.235351 | 0.369 | 0.307 | 1 | 13 | Vkorc1l1 | 0.062 | Old |
| 0.311433 | 0.094521 | 0.975 | 0.965 | 1 | 13 | Hnrnph1 | 0.01 | Old |
| 0.311735 | 0.301855 | 0.451 | 0.386 | 1 | 13 | Adgre4 | 0.065 | Old |
| 0.311881 | 0.02488 | 0.123 | 0.079 | 1 | 13 | Brat1 | 0.044 | Old |
| 0.31195 | 0.202764 | 0.475 | 0.412 | 1 | 13 | Brd9 | 0.063 | Old |
| 0.31207 | 0.17639 | 0.721 | 0.737 | 1 | 13 | Smndc1 | -0.016 | Old |
| 0.312523 | 0.330819 | 0.066 | 0.035 | 1 | 13 | Cbwd1 | 0.031 | Old |
| 0.312523 | 0.021332 | 0.066 | 0.035 | 1 | 13 | Cdca3 | 0.031 | Old |
| 0.312818 | 0.118192 | 0.893 | 0.86 | 1 | 13 | Snrpd3 | 0.033 | Old |
| 0.312843 | 0.298792 | 0.607 | 0.518 | 1 | 13 | Hibadh | 0.089 | Old |
| 0.313185 | 0.280183 | 0.18 | 0.132 | 1 | 13 | Osbpl7 | 0.048 | Old |
| 0.31346 | 0.169509 | 0.705 | 0.605 | 1 | 13 | Atxn2 | 0.1 | Old |
| 0.313621 | 0.195006 | 0.689 | 0.605 | 1 | 13 | D1Ertd622( | 0.084 | Old |
| 0.313651 | 0.268922 | 0.615 | 0.57 | 1 | 13 | Trim12c | 0.045 | Old |
| 0.313712 | 0.179825 | 0.566 | 0.482 | 1 | 13 | Stx6 | 0.084 | Old |
| 0.31383 | 0.190138 | 0.828 | 0.825 | 1 | 13 | Ube2i | 0.003 | Old |
| 0.313877 | 0.365575 | 0.221 | 0.167 | 1 | 13 | Qtrt1 | 0.054 | Old |
| 0.314316 | 0.244795 | 0.32 | 0.246 | 1 | 13 | Aatf | 0.074 | Old |
| 0.31441 | 0.227513 | 0.156 | 0.105 | 1 | 13 | Cacna1f | 0.051 | Old |
| 0.31449 | 0.225694 | 0.803 | 0.763 | 1 | 13 | Baz1a | 0.04 | Old |
| 0.314634 | 0.302392 | 0.18 | 0.132 | 1 | 13 | Nr1d1 | 0.048 | Old |
| 0.314689 | 0.222613 | 0.32 | 0.237 | 1 | 13 | Clpp | 0.083 | Old |
| 0.314931 | 0.300531 | 0.066 | 0.035 | 1 | 13 | Gm15859 | 0.031 | Old |
| 0.314931 | 0.25331 | 0.066 | 0.035 | 1 | 13 | Cadm3 | 0.031 | Old |
| 0.315045 | 0.357475 | 0.164 | 0.114 | 1 | 13 | Rfx2 | 0.05 | Old |
| 0.315092 | 0.215101 | 0.172 | 0.123 | 1 | 13 | Pigh | 0.049 | Old |
| 0.315234 | 0.186008 | 0.59 | 0.491 | 1 | 13 | Ndufs1 | 0.099 | Old |
| 0.315418 | 0.126728 | 0.885 | 0.842 | 1 | 13 | Eif3l | 0.043 | Old |
| 0.315673 | 0.290919 | 0.475 | 0.395 | 1 | 13 | Mrpl9 | 0.08 | Old |
| 0.315725 | 0.186032 | 0.393 | 0.307 | 1 | 13 | Ralgapa1 | 0.086 | Old |
| 0.316037 | 0.292955 | 0.287 | 0.219 | 1 | 13 | Atr | 0.068 | Old |
| 0.31607 | 0.166212 | 0.705 | 0.623 | 1 | 13 | Klc1 | 0.082 | Old |
| 0.316352 | 0.581814 | 0.074 | 0.044 | 1 | 13 | Rtel1 | 0.03 | Old |
| 0.316399 | 0.112372 | 0.852 | 0.781 | 1 | 13 | Ssbp4 | 0.071 | Old |
| 0.31693 | 0.093245 | 0.975 | 0.947 | 1 | 13 | Coro1c | 0.028 | Old |

| | | | | | | | | |
|---|---|---|---|---|---|---|---|---|
| 0.317034 | 0.241267 | 0.115 | 0.07 | 1 | 13 | D3Ertd751 | 0.045 | Old |
| 0.317135 | 0.315722 | 0.402 | 0.325 | 1 | 13 | Clpx | 0.077 | Old |
| 0.317136 | 0.186874 | 0.352 | 0.289 | 1 | 13 | Mcm6 | 0.063 | Old |
| 0.31735 | 0.403541 | 0.066 | 0.035 | 1 | 13 | Clec3b | 0.031 | Old |
| 0.317423 | 0.163276 | 0.566 | 0.509 | 1 | 13 | Pum3 | 0.057 | Old |
| 0.317545 | 0.442965 | 0.18 | 0.132 | 1 | 13 | 2900089D1 | 0.048 | Old |
| 0.317562 | 0.281298 | 0.533 | 0.465 | 1 | 13 | Acadsb | 0.068 | Old |
| 0.317698 | 0.116752 | 0.861 | 0.807 | 1 | 13 | Supt16 | 0.054 | Old |
| 0.317832 | 0.187925 | 0.893 | 0.851 | 1 | 13 | Ddi2 | 0.042 | Old |
| 0.317849 | 0.453282 | 0.139 | 0.096 | 1 | 13 | Zfand1 | 0.043 | Old |
| 0.318075 | 0.493476 | 0.172 | 0.123 | 1 | 13 | Spice1 | 0.049 | Old |
| 0.318792 | 0.12837 | 1 | 0.991 | 1 | 13 | H2-T23 | 0.009 | Old |
| 0.319295 | 0.200384 | 0.418 | 0.342 | 1 | 13 | Cavin1 | 0.076 | Old |
| 0.319544 | 0.241274 | 0.262 | 0.202 | 1 | 13 | Mtmr9 | 0.06 | Old |
| 0.320232 | 0.236771 | 0.459 | 0.526 | 1 | 13 | Ccl3 | -0.067 | Old |
| 0.320292 | 0.184994 | 0.434 | 0.333 | 1 | 13 | Armc5 | 0.101 | Old |
| 0.320297 | 0.196345 | 0.697 | 0.588 | 1 | 13 | Mga | 0.109 | Old |
| 0.320612 | 0.326619 | 0.549 | 0.474 | 1 | 13 | Ubqln2 | 0.075 | Old |
| 0.320664 | 0.361546 | 0.123 | 0.079 | 1 | 13 | Timeless | 0.044 | Old |
| 0.320705 | 0.349635 | 0.156 | 0.105 | 1 | 13 | Mocs1 | 0.051 | Old |
| 0.320917 | 0.07047 | 0.893 | 0.825 | 1 | 13 | Sec11a | 0.068 | Old |
| 0.321507 | 0.382211 | 0.295 | 0.237 | 1 | 13 | Exosc2 | 0.058 | Old |
| 0.321579 | 0.088001 | 1 | 1 | 1 | 13 | Ndufa4 | 0 | Old |
| 0.321869 | 0.077162 | 0.779 | 0.816 | 1 | 13 | Gpbp1 | -0.037 | Old |
| 0.321961 | 0.463719 | 0.246 | 0.193 | 1 | 13 | AC165953. | 0.053 | Old |
| 0.322775 | 0.311327 | 0.098 | 0.061 | 1 | 13 | Naif1 | 0.037 | Old |
| 0.323055 | 0.239696 | 0.197 | 0.14 | 1 | 13 | Grwd1 | 0.057 | Old |
| 0.323152 | 0.618116 | 0.074 | 0.044 | 1 | 13 | Slc43a3 | 0.03 | Old |
| 0.323338 | 0.264341 | 0.377 | 0.325 | 1 | 13 | D6Wsu163 | 0.052 | Old |
| 0.323432 | 0.08654 | 0.992 | 0.93 | 1 | 13 | Cast | 0.062 | Old |
| 0.323497 | 0.072232 | 0.5 | 0.377 | 1 | 13 | Eef1akmt1 | 0.123 | Old |
| 0.323558 | 0.283224 | 0.295 | 0.228 | 1 | 13 | B3galnt2 | 0.067 | Old |
| 0.323591 | 0.218565 | 0.861 | 0.842 | 1 | 13 | Lima1 | 0.019 | Old |
| 0.323781 | 0.242001 | 0.566 | 0.491 | 1 | 13 | Hmgn2 | 0.075 | Old |
| 0.323983 | 0.124308 | 0.23 | 0.175 | 1 | 13 | Rmnd5b | 0.055 | Old |
| 0.324094 | 0.112995 | 0.91 | 0.877 | 1 | 13 | Rassf2 | 0.033 | Old |
| 0.324095 | 0.347852 | 0.172 | 0.123 | 1 | 13 | Cpt2 | 0.049 | Old |
| 0.324104 | 0.2201 | 0.459 | 0.377 | 1 | 13 | Btbd9 | 0.082 | Old |
| 0.324222 | 0.047722 | 0.123 | 0.079 | 1 | 13 | Efnb1 | 0.044 | Old |
| 0.324278 | 0.230996 | 0.164 | 0.114 | 1 | 13 | Snapc2 | 0.05 | Old |
| 0.324381 | 0.088877 | 0.984 | 1 | 1 | 13 | 1810037I1 | -0.016 | Old |
| 0.324537 | 0.272562 | 0.287 | 0.219 | 1 | 13 | Plrg1 | 0.068 | Old |
| 0.324758 | 0.429498 | 0.098 | 0.061 | 1 | 13 | Rnf122 | 0.037 | Old |
| 0.324793 | 0.329549 | 0.131 | 0.088 | 1 | 13 | Kdelc1 | 0.043 | Old |
| 0.324875 | 0.229516 | 0.77 | 0.711 | 1 | 13 | Paf1 | 0.059 | Old |

| | | | | | | | | |
|---|---|---|---|---|---|---|---|---|
| 0.324899 | 1.03293 | 0.18 | 0.132 | 1 | 13 | Mmp12 | 0.048 | Old |
| 0.325101 | 0.325608 | 0.598 | 0.509 | 1 | 13 | Mtmr10 | 0.089 | Old |
| 0.325165 | 0.217443 | 0.844 | 0.825 | 1 | 13 | Arhgef2 | 0.019 | Old |
| 0.325439 | 0.5792 | 0.074 | 0.044 | 1 | 13 | Rita1 | 0.03 | Old |
| 0.325439 | 0.576421 | 0.074 | 0.044 | 1 | 13 | Ss18l1 | 0.03 | Old |
| 0.325491 | 0.342026 | 0.549 | 0.5 | 1 | 13 | Mrps28 | 0.049 | Old |
| 0.326019 | 0.114931 | 0.852 | 0.807 | 1 | 13 | Eif3d | 0.045 | Old |
| 0.326168 | 0.185434 | 0.951 | 0.904 | 1 | 13 | Snx9 | 0.047 | Old |
| 0.326226 | 0.208656 | 0.779 | 0.719 | 1 | 13 | Psmb9 | 0.06 | Old |
| 0.326381 | 0.171885 | 0.713 | 0.623 | 1 | 13 | Phf23 | 0.09 | Old |
| 0.326477 | 0.573355 | 0.336 | 0.289 | 1 | 13 | F10 | 0.047 | Old |
| 0.326665 | 0.212826 | 0.189 | 0.14 | 1 | 13 | Skp2 | 0.049 | Old |
| 0.326681 | 0.558314 | 0.148 | 0.105 | 1 | 13 | Ppp1r12b | 0.043 | Old |
| 0.32677 | 0.298706 | 0.754 | 0.728 | 1 | 13 | Sec63 | 0.026 | Old |
| 0.326993 | 0.273331 | 0.287 | 0.219 | 1 | 13 | Crcp | 0.068 | Old |
| 0.32733 | 0.725992 | 0.164 | 0.123 | 1 | 13 | Tmem42 | 0.041 | Old |
| 0.327394 | 0.413803 | 0.205 | 0.158 | 1 | 13 | Pank1 | 0.047 | Old |
| 0.327514 | 0.277642 | 0.426 | 0.36 | 1 | 13 | Brwd1 | 0.066 | Old |
| 0.327524 | 0.111285 | 0.926 | 0.904 | 1 | 13 | Rilpl2 | 0.022 | Old |
| 0.327738 | 0.595292 | 0.074 | 0.044 | 1 | 13 | Bmpr1a | 0.03 | Old |
| 0.327826 | 0.425573 | 0.139 | 0.096 | 1 | 13 | Batf2 | 0.043 | Old |
| 0.327988 | 0.08166 | 0.844 | 0.781 | 1 | 13 | Hmox2 | 0.063 | Old |
| 0.328046 | 0.131817 | 0.254 | 0.193 | 1 | 13 | Zbtb16 | 0.061 | Old |
| 0.328133 | 0.100829 | 0.967 | 0.965 | 1 | 13 | Selplg | 0.002 | Old |
| 0.328142 | 0.057822 | 1 | 1 | 1 | 13 | Cox6b1 | 0 | Old |
| 0.329087 | 0.048394 | 1 | 1 | 1 | 13 | Rbpj | 0 | Old |
| 0.329453 | 0.166712 | 0.91 | 0.939 | 1 | 13 | Hnrnpl | -0.029 | Old |
| 0.330101 | 0.186293 | 0.467 | 0.404 | 1 | 13 | Sipa1l3 | 0.063 | Old |
| 0.330368 | 0.338823 | 0.262 | 0.211 | 1 | 13 | Mon1b | 0.051 | Old |
| 0.330646 | 0.232096 | 0.221 | 0.158 | 1 | 13 | Pnpt1 | 0.063 | Old |
| 0.330755 | 0.420427 | 0.098 | 0.061 | 1 | 13 | Ckap2l | 0.037 | Old |
| 0.330867 | 0.14911 | 0.68 | 0.605 | 1 | 13 | Mtfr1l | 0.075 | Old |
| 0.330981 | 0.020694 | 1 | 0.991 | 1 | 13 | Nfam1 | 0.009 | Old |
| 0.330986 | 0.264873 | 0.77 | 0.728 | 1 | 13 | Smarcc2 | 0.042 | Old |
| 0.331352 | 0.256938 | 0.705 | 0.579 | 1 | 13 | Parp1 | 0.126 | Old |
| 0.331559 | 0.281128 | 0.68 | 0.632 | 1 | 13 | Cyb5b | 0.048 | Old |
| 0.331584 | 0.11015 | 0.672 | 0.579 | 1 | 13 | Kpna1 | 0.093 | Old |
| 0.331586 | 0.169702 | 0.279 | 0.211 | 1 | 13 | Map4k2 | 0.068 | Old |
| 0.331635 | 0.218812 | 0.508 | 0.421 | 1 | 13 | Mrpl16 | 0.087 | Old |
| 0.331885 | 0.284227 | 0.238 | 0.175 | 1 | 13 | Rnf8 | 0.063 | Old |
| 0.331931 | 0.098685 | 1 | 1 | 1 | 13 | Pltp | 0 | Old |
| 0.332104 | 0.324132 | 0.279 | 0.219 | 1 | 13 | Kcnq1 | 0.06 | Old |
| 0.332109 | 0.465471 | 0.172 | 0.132 | 1 | 13 | Dtd2 | 0.04 | Old |
| 0.332183 | 0.123444 | 0.393 | 0.307 | 1 | 13 | Cdk10 | 0.086 | Old |
| 0.332274 | 0.297526 | 0.27 | 0.211 | 1 | 13 | Calcrl | 0.059 | Old |

| | | | | | | | | |
|---|---|---|---|---|---|---|---|---|
| 0.332366 | 0.712592 | 0.074 | 0.044 | 1 | 13 | Exog | 0.03 | Old |
| 0.332728 | 0.060676 | 0.721 | 0.57 | 1 | 13 | Tbc1d5 | 0.151 | Old |
| 0.333084 | 0.217039 | 0.648 | 0.596 | 1 | 13 | Tollip | 0.052 | Old |
| 0.333238 | 0.308252 | 0.566 | 0.509 | 1 | 13 | Copg1 | 0.057 | Old |
| 0.333388 | 0.235141 | 0.23 | 0.175 | 1 | 13 | Dusp12 | 0.055 | Old |
| 0.333685 | 0.288171 | 0.164 | 0.114 | 1 | 13 | 1500015A( | 0.05 | Old |
| 0.333694 | 0.320925 | 0.197 | 0.149 | 1 | 13 | Aasdh | 0.048 | Old |
| 0.333697 | 0.15143 | 0.5 | 0.421 | 1 | 13 | Parp2 | 0.079 | Old |
| 0.333702 | 0.19097 | 0.697 | 0.596 | 1 | 13 | Ahsa1 | 0.101 | Old |
| 0.334418 | 0.462385 | 0.393 | 0.307 | 1 | 13 | Srxn1 | 0.086 | Old |
| 0.334562 | 0.311525 | 0.41 | 0.325 | 1 | 13 | Lrp12 | 0.085 | Old |
| 0.334612 | 0.335256 | 0.279 | 0.219 | 1 | 13 | 5031425F1 | 0.06 | Old |
| 0.334623 | 0.131393 | 0.197 | 0.14 | 1 | 13 | Ctdspl | 0.057 | Old |
| 0.334695 | 0.628608 | 0.074 | 0.044 | 1 | 13 | Trim16 | 0.03 | Old |
| 0.334695 | 0.547235 | 0.074 | 0.044 | 1 | 13 | Trmt12 | 0.03 | Old |
| 0.334956 | 0.220015 | 0.443 | 0.368 | 1 | 13 | Tsfm | 0.075 | Old |
| 0.335046 | 0.339919 | 0.656 | 0.588 | 1 | 13 | Fnbp1l | 0.068 | Old |
| 0.33508 | 0.46555 | 0.107 | 0.07 | 1 | 13 | Tspyl3 | 0.037 | Old |
| 0.335088 | 0.212949 | 0.516 | 0.447 | 1 | 13 | Myo18a | 0.069 | Old |
| 0.335566 | 0.21282 | 0.361 | 0.289 | 1 | 13 | Nelfb | 0.072 | Old |
| 0.335674 | 0.122145 | 0.525 | 0.43 | 1 | 13 | Ddx10 | 0.095 | Old |
| 0.33612 | 0.181828 | 0.27 | 0.202 | 1 | 13 | Syne3 | 0.068 | Old |
| 0.336683 | 0.155112 | 0.59 | 0.526 | 1 | 13 | Zfx | 0.064 | Old |
| 0.336875 | 0.428717 | 0.336 | 0.289 | 1 | 13 | Ebag9 | 0.047 | Old |
| 0.337013 | 0.326607 | 0.131 | 0.088 | 1 | 13 | Sco1 | 0.043 | Old |
| 0.337021 | 0.422318 | 0.107 | 0.07 | 1 | 13 | Hus1 | 0.037 | Old |
| 0.337035 | 0.561186 | 0.074 | 0.044 | 1 | 13 | Ankrd66 | 0.03 | Old |
| 0.337035 | 0.488157 | 0.074 | 0.044 | 1 | 13 | 6330562C2 | 0.03 | Old |
| 0.337035 | 0.470103 | 0.074 | 0.044 | 1 | 13 | Ciart | 0.03 | Old |
| 0.337035 | 0.438687 | 0.074 | 0.044 | 1 | 13 | Ip6k2 | 0.03 | Old |
| 0.337061 | 0.314941 | 0.451 | 0.43 | 1 | 13 | Fads1 | 0.021 | Old |
| 0.337126 | 0.329358 | 0.377 | 0.316 | 1 | 13 | Pak4 | 0.061 | Old |
| 0.338338 | 0.28292 | 0.221 | 0.167 | 1 | 13 | Mlst8 | 0.054 | Old |
| 0.338344 | 0.243695 | 0.623 | 0.544 | 1 | 13 | Naa20 | 0.079 | Old |
| 0.338414 | 0.309362 | 0.18 | 0.132 | 1 | 13 | Abcc4 | 0.048 | Old |
| 0.338463 | 0.093995 | 0.41 | 0.316 | 1 | 13 | Akt3 | 0.094 | Old |
| 0.338528 | 0.227431 | 0.639 | 0.596 | 1 | 13 | Arhgef10l | 0.043 | Old |
| 0.33854 | 0.118746 | 0.303 | 0.228 | 1 | 13 | Fbxo8 | 0.075 | Old |
| 0.33863 | 0.080388 | 1 | 0.991 | 1 | 13 | Abracl | 0.009 | Old |
| 0.338735 | 0.157027 | 0.352 | 0.289 | 1 | 13 | Trim65 | 0.063 | Old |
| 0.338861 | 0.278237 | 0.098 | 0.061 | 1 | 13 | Rwdd2b | 0.037 | Old |
| 0.338969 | 0.299311 | 0.107 | 0.07 | 1 | 13 | Elp6 | 0.037 | Old |
| 0.339386 | 0.451251 | 0.074 | 0.044 | 1 | 13 | Fgfbp3 | 0.03 | Old |
| 0.339386 | 0.431517 | 0.074 | 0.044 | 1 | 13 | Gm12166 | 0.03 | Old |
| 0.33972 | 0.310146 | 0.139 | 0.096 | 1 | 13 | Rarb | 0.043 | Old |

| | | | | | | | | |
|---|---|---|---|---|---|---|---|---|
| 0.340141 | 0.298841 | 0.402 | 0.325 | 1 | 13 | Ttc32 | 0.077 | Old |
| 0.340332 | 0.216224 | 0.213 | 0.158 | 1 | 13 | Map4k5 | 0.055 | Old |
| 0.34052 | 0.104043 | 0.975 | 0.912 | 1 | 13 | Tmem234 | 0.063 | Old |
| 0.340528 | 0.109127 | 0.393 | 0.298 | 1 | 13 | Slc25a30 | 0.095 | Old |
| 0.340547 | 0.180353 | 0.959 | 0.965 | 1 | 13 | Ms4a6c | -0.006 | Old |
| 0.34074 | 0.418611 | 0.836 | 0.868 | 1 | 13 | Ccl7 | -0.032 | Old |
| 0.341265 | 0.081634 | 0.402 | 0.316 | 1 | 13 | Hjurp | 0.086 | Old |
| 0.341414 | 0.12749 | 0.844 | 0.781 | 1 | 13 | Anxa6 | 0.063 | Old |
| 0.341441 | 0.167651 | 0.139 | 0.096 | 1 | 13 | Nat8l | 0.043 | Old |
| 0.341576 | 1.377923 | 0.115 | 0.079 | 1 | 13 | Ly6a | 0.036 | Old |
| 0.341589 | 0.373892 | 0.205 | 0.158 | 1 | 13 | Ppil1 | 0.047 | Old |
| 0.342112 | 0.210028 | 0.549 | 0.482 | 1 | 13 | Galk2 | 0.067 | Old |
| 0.342258 | 0.398095 | 0.18 | 0.14 | 1 | 13 | Mamld1 | 0.04 | Old |
| 0.342529 | 0.066909 | 0.82 | 0.781 | 1 | 13 | Nudt3 | 0.039 | Old |
| 0.3426 | 0.332145 | 0.172 | 0.123 | 1 | 13 | Sec22c | 0.049 | Old |
| 0.342886 | 0.357629 | 0.107 | 0.07 | 1 | 13 | Elmod3 | 0.037 | Old |
| 0.342917 | 0.176971 | 0.566 | 0.491 | 1 | 13 | Atp11b | 0.075 | Old |
| 0.343401 | 0.186669 | 0.541 | 0.456 | 1 | 13 | Vps11 | 0.085 | Old |
| 0.343461 | 0.469049 | 0.115 | 0.079 | 1 | 13 | Tmem220 | 0.036 | Old |
| 0.343516 | 1.854392 | 0.025 | 0.009 | 1 | 13 | Upp1 | 0.016 | Old |
| 0.343516 | 0.949365 | 0.025 | 0.009 | 1 | 13 | Tsix | 0.016 | Old |
| 0.343516 | 0.829297 | 0.025 | 0.009 | 1 | 13 | Dctd | 0.016 | Old |
| 0.343516 | 0.775832 | 0.025 | 0.009 | 1 | 13 | Ripply3 | 0.016 | Old |
| 0.343516 | 0.760747 | 0.025 | 0.009 | 1 | 13 | AV099323 | 0.016 | Old |
| 0.343516 | 0.751252 | 0.025 | 0.009 | 1 | 13 | Gm37170 | 0.016 | Old |
| 0.343516 | 0.7329 | 0.025 | 0.009 | 1 | 13 | Alkbh4 | 0.016 | Old |
| 0.343516 | 0.711147 | 0.025 | 0.009 | 1 | 13 | Map1a | 0.016 | Old |
| 0.343516 | 0.688612 | 0.025 | 0.009 | 1 | 13 | Llgl2 | 0.016 | Old |
| 0.343516 | 0.675942 | 0.025 | 0.009 | 1 | 13 | Meig1 | 0.016 | Old |
| 0.343516 | 0.671038 | 0.025 | 0.009 | 1 | 13 | D130020L0 | 0.016 | Old |
| 0.343516 | 0.644767 | 0.025 | 0.009 | 1 | 13 | Zfp335os | 0.016 | Old |
| 0.343516 | 0.639318 | 0.025 | 0.009 | 1 | 13 | Parm1 | 0.016 | Old |
| 0.343516 | 0.609215 | 0.025 | 0.009 | 1 | 13 | Fap | 0.016 | Old |
| 0.343516 | 0.603479 | 0.025 | 0.009 | 1 | 13 | P2rx5 | 0.016 | Old |
| 0.343516 | 0.506564 | 0.025 | 0.009 | 1 | 13 | Mcmdc2 | 0.016 | Old |
| 0.343527 | 0.047433 | 0.639 | 0.561 | 1 | 13 | Zfp800 | 0.078 | Old |
| 0.343607 | 0.131502 | 0.713 | 0.64 | 1 | 13 | Nmt1 | 0.073 | Old |
| 0.343888 | 0.135279 | 0.27 | 0.202 | 1 | 13 | Phtf2 | 0.068 | Old |
| 0.344119 | 0.609981 | 0.074 | 0.044 | 1 | 13 | Gm9733 | 0.03 | Old |
| 0.344366 | 0.078041 | 0.918 | 0.947 | 1 | 13 | Ehbp1l1 | -0.029 | Old |
| 0.344597 | 0.142452 | 0.73 | 0.623 | 1 | 13 | Rnf6 | 0.107 | Old |
| 0.344642 | 0.283178 | 0.402 | 0.325 | 1 | 13 | Vps26b | 0.077 | Old |
| 0.344765 | 0.25984 | 0.279 | 0.219 | 1 | 13 | Kti12 | 0.06 | Old |
| 0.344796 | 0.246965 | 0.59 | 0.5 | 1 | 13 | Ric8a | 0.09 | Old |
| 0.344972 | 0.203304 | 0.582 | 0.518 | 1 | 13 | Sec13 | 0.064 | Old |

| | | | | | | | | |
|---|---|---|---|---|---|---|---|---|
| 0.345023 | 0.40099 | 0.098 | 0.061 | 1 | 13 | Hpn | 0.037 | Old |
| 0.345414 | 0.69046 | 0.082 | 0.053 | 1 | 13 | Trmu | 0.029 | Old |
| 0.345416 | 0.094834 | 1 | 1 | 1 | 13 | Ywhae | 0 | Old |
| 0.345605 | 0.092594 | 0.877 | 0.825 | 1 | 13 | Usp7 | 0.052 | Old |
| 0.345702 | 0.306996 | 0.344 | 0.281 | 1 | 13 | 4930453N2 | 0.063 | Old |
| 0.345736 | 0.232215 | 0.23 | 0.175 | 1 | 13 | Dcn | 0.055 | Old |
| 0.346048 | 0.143726 | 0.287 | 0.211 | 1 | 13 | Prdm4 | 0.076 | Old |
| 0.346305 | 0.168256 | 0.672 | 0.596 | 1 | 13 | Arl5a | 0.076 | Old |
| 0.34654 | 0.124278 | 0.713 | 0.579 | 1 | 13 | Dpy19l1 | 0.134 | Old |
| 0.346554 | 0.140887 | 0.639 | 0.579 | 1 | 13 | Nemf | 0.06 | Old |
| 0.347049 | 0.111383 | 0.549 | 0.447 | 1 | 13 | Ino80e | 0.102 | Old |
| 0.34713 | 1.162381 | 0.049 | 0.026 | 1 | 13 | Ccno | 0.023 | Old |
| 0.347485 | 0.146488 | 0.582 | 0.491 | 1 | 13 | Naxe | 0.091 | Old |
| 0.347684 | 0.143048 | 0.721 | 0.596 | 1 | 13 | Mrps17 | 0.125 | Old |
| 0.347881 | 1.440287 | 0.025 | 0.009 | 1 | 13 | Ereg | 0.016 | Old |
| 0.347881 | 0.961703 | 0.025 | 0.009 | 1 | 13 | Trp53cor1 | 0.016 | Old |
| 0.347881 | 0.918506 | 0.025 | 0.009 | 1 | 13 | Gm15228 | 0.016 | Old |
| 0.347881 | 0.768518 | 0.025 | 0.009 | 1 | 13 | Zfp937 | 0.016 | Old |
| 0.347881 | 0.76106 | 0.025 | 0.009 | 1 | 13 | Mfsd4b3 | 0.016 | Old |
| 0.347881 | 0.759904 | 0.025 | 0.009 | 1 | 13 | Ccnb1 | 0.016 | Old |
| 0.347881 | 0.729938 | 0.025 | 0.009 | 1 | 13 | Syde1 | 0.016 | Old |
| 0.347881 | 0.688001 | 0.025 | 0.009 | 1 | 13 | Zscan2 | 0.016 | Old |
| 0.347881 | 0.650729 | 0.025 | 0.009 | 1 | 13 | Gm35363 | 0.016 | Old |
| 0.347881 | 0.645298 | 0.025 | 0.009 | 1 | 13 | Tmem236 | 0.016 | Old |
| 0.347881 | 0.627228 | 0.025 | 0.009 | 1 | 13 | Ccdc80 | 0.016 | Old |
| 0.347881 | 0.592362 | 0.025 | 0.009 | 1 | 13 | B230303O1 | 0.016 | Old |
| 0.347881 | 0.569486 | 0.025 | 0.009 | 1 | 13 | Phyhd1 | 0.016 | Old |
| 0.347881 | 0.547803 | 0.025 | 0.009 | 1 | 13 | Dmtn | 0.016 | Old |
| 0.347881 | 0.541368 | 0.025 | 0.009 | 1 | 13 | Hoxa4 | 0.016 | Old |
| 0.347881 | 0.530701 | 0.025 | 0.009 | 1 | 13 | Gm26882 | 0.016 | Old |
| 0.347881 | 0.529352 | 0.025 | 0.009 | 1 | 13 | Bbs10 | 0.016 | Old |
| 0.347881 | 0.421577 | 0.025 | 0.009 | 1 | 13 | Tmem79 | 0.016 | Old |
| 0.348068 | 0.221615 | 0.393 | 0.325 | 1 | 13 | Wdr81 | 0.068 | Old |
| 0.348121 | 0.067949 | 0.164 | 0.114 | 1 | 13 | Gm45606 | 0.05 | Old |
| 0.348184 | 0.165344 | 0.697 | 0.623 | 1 | 13 | Ccdc115 | 0.074 | Old |
| 0.348191 | 0.265957 | 0.32 | 0.272 | 1 | 13 | Pex3 | 0.048 | Old |
| 0.34826 | 0.129841 | 0.918 | 0.939 | 1 | 13 | Cct8 | -0.021 | Old |
| 0.348382 | 0.278132 | 0.139 | 0.096 | 1 | 13 | Rbm45 | 0.043 | Old |
| 0.348519 | 0.089695 | 0.893 | 0.886 | 1 | 13 | Lman2 | 0.007 | Old |
| 0.348702 | 1.062009 | 0.189 | 0.149 | 1 | 13 | Fpr2 | 0.04 | Old |
| 0.348832 | 0.385028 | 0.205 | 0.158 | 1 | 13 | Calml4 | 0.047 | Old |
| 0.349539 | 0.114334 | 0.361 | 0.289 | 1 | 13 | Ctns | 0.072 | Old |
| 0.349641 | 0.094851 | 0.557 | 0.482 | 1 | 13 | Dscr3 | 0.075 | Old |
| 0.34979 | 0.200731 | 0.295 | 0.237 | 1 | 13 | Xrcc4 | 0.058 | Old |
| 0.349909 | 0.568139 | 0.082 | 0.053 | 1 | 13 | Pou6f1 | 0.029 | Old |

| | | | | | | | | |
|---|---|---|---|---|---|---|---|---|
| 0.350087 | 0.896192 | 0.049 | 0.026 | 1 | 13 | Rtl6 | 0.023 | Old |
| 0.350144 | 0.373485 | 0.27 | 0.219 | 1 | 13 | Mpp5 | 0.051 | Old |
| 0.350176 | 0.122743 | 0.32 | 0.246 | 1 | 13 | Slc35b4 | 0.074 | Old |
| 0.350381 | 0.250683 | 0.213 | 0.158 | 1 | 13 | Tlcd2 | 0.055 | Old |
| 0.350469 | 0.088446 | 0.344 | 0.281 | 1 | 13 | Lipo3 | 0.063 | Old |
| 0.350505 | 0.463806 | 0.172 | 0.123 | 1 | 13 | Sh3bp5l | 0.049 | Old |
| 0.350641 | 0.057632 | 0.893 | 0.825 | 1 | 13 | Mndal | 0.068 | Old |
| 0.350807 | 0.464979 | 0.107 | 0.07 | 1 | 13 | Hexdc | 0.037 | Old |
| 0.350807 | 0.427189 | 0.107 | 0.07 | 1 | 13 | Osgin1 | 0.037 | Old |
| 0.350898 | 0.123048 | 0.918 | 0.842 | 1 | 13 | Psmc3 | 0.076 | Old |
| 0.350917 | 0.1626 | 0.631 | 0.526 | 1 | 13 | Tfg | 0.105 | Old |
| 0.351069 | 0.463262 | 0.115 | 0.079 | 1 | 13 | Adam22 | 0.036 | Old |
| 0.351214 | 0.390351 | 0.295 | 0.246 | 1 | 13 | Lyrm9 | 0.049 | Old |
| 0.351297 | 0.27197 | 0.074 | 0.044 | 1 | 13 | Tgfbr3 | 0.03 | Old |
| 0.3513 | 0.092857 | 0.992 | 0.982 | 1 | 13 | Eif2s2 | 0.01 | Old |
| 0.351325 | 0.310609 | 0.303 | 0.246 | 1 | 13 | Bambi | 0.057 | Old |
| 0.351579 | 0.207757 | 0.434 | 0.351 | 1 | 13 | Ints9 | 0.083 | Old |
| 0.351651 | 0.262489 | 0.516 | 0.404 | 1 | 13 | Cpsf3 | 0.112 | Old |
| 0.35191 | 0.025938 | 0.566 | 0.447 | 1 | 13 | Casp8ap2 | 0.119 | Old |
| 0.352281 | 0.854677 | 0.025 | 0.009 | 1 | 13 | Cdkl4 | 0.016 | Old |
| 0.352281 | 0.8495 | 0.025 | 0.009 | 1 | 13 | Paqr5 | 0.016 | Old |
| 0.352281 | 0.610419 | 0.025 | 0.009 | 1 | 13 | Bdh1 | 0.016 | Old |
| 0.352281 | 0.567744 | 0.025 | 0.009 | 1 | 13 | Lca5 | 0.016 | Old |
| 0.352281 | 0.567272 | 0.025 | 0.009 | 1 | 13 | Hsf4 | 0.016 | Old |
| 0.352281 | 0.554986 | 0.025 | 0.009 | 1 | 13 | Thnsl2 | 0.016 | Old |
| 0.352281 | 0.532697 | 0.025 | 0.009 | 1 | 13 | Ccnf | 0.016 | Old |
| 0.352281 | 0.531501 | 0.025 | 0.009 | 1 | 13 | Ankdd1a | 0.016 | Old |
| 0.352281 | 0.505456 | 0.025 | 0.009 | 1 | 13 | Sugct | 0.016 | Old |
| 0.352281 | 0.502952 | 0.025 | 0.009 | 1 | 13 | Pacsin3 | 0.016 | Old |
| 0.352281 | 0.499212 | 0.025 | 0.009 | 1 | 13 | Neto2 | 0.016 | Old |
| 0.352281 | 0.496247 | 0.025 | 0.009 | 1 | 13 | Gm29994 | 0.016 | Old |
| 0.352281 | 0.465282 | 0.025 | 0.009 | 1 | 13 | Gm16701.1 | 0.016 | Old |
| 0.352281 | 0.456454 | 0.025 | 0.009 | 1 | 13 | Traf3ip2 | 0.016 | Old |
| 0.352456 | 0.147319 | 0.68 | 0.623 | 1 | 13 | Ubr5 | 0.057 | Old |
| 0.35265 | 0.191358 | 0.385 | 0.316 | 1 | 13 | Manea | 0.069 | Old |
| 0.352692 | 0.281668 | 0.607 | 0.491 | 1 | 13 | Rcn2 | 0.116 | Old |
| 0.352772 | 0.128779 | 0.975 | 0.956 | 1 | 13 | Ube2d2a | 0.019 | Old |
| 0.352806 | 0.343761 | 0.107 | 0.07 | 1 | 13 | Mccc2 | 0.037 | Old |
| 0.353002 | 0.034459 | 0.869 | 0.781 | 1 | 13 | 7-Mar | 0.088 | Old |
| 0.353017 | 0.243972 | 0.803 | 0.719 | 1 | 13 | Zcchc24 | 0.084 | Old |
| 0.35306 | 0.607823 | 0.049 | 0.026 | 1 | 13 | Tmsb15b1 | 0.023 | Old |
| 0.35371 | 0.227433 | 0.074 | 0.044 | 1 | 13 | Rpgrip1l | 0.03 | Old |
| 0.353794 | 0.391677 | 0.32 | 0.263 | 1 | 13 | Prmt3 | 0.057 | Old |
| 0.354321 | 0.321343 | 0.197 | 0.149 | 1 | 13 | Cep44 | 0.048 | Old |
| 0.354558 | 0.318597 | 0.254 | 0.202 | 1 | 13 | 3110082I1 | 0.052 | Old |

| | | | | | | | | |
|---|---|---|---|---|---|---|---|---|
| 0.354754 | 0.221894 | 0.336 | 0.263 | 1 | 13 | Mcoln1 | 0.073 | Old |
| 0.355388 | 0.28266 | 0.402 | 0.368 | 1 | 13 | Creb3 | 0.034 | Old |
| 0.355461 | 0.112055 | 0.82 | 0.789 | 1 | 13 | Gm16286 | 0.031 | Old |
| 0.355538 | 0.192275 | 0.639 | 0.614 | 1 | 13 | Gpr108 | 0.025 | Old |
| 0.355815 | 0.2559 | 0.525 | 0.482 | 1 | 13 | Mpdu1 | 0.043 | Old |
| 0.355822 | 0.080501 | 0.377 | 0.289 | 1 | 13 | Pctp | 0.088 | Old |
| 0.355859 | 0.216899 | 0.262 | 0.202 | 1 | 13 | Cpsf4 | 0.06 | Old |
| 0.356049 | 0.652906 | 0.049 | 0.026 | 1 | 13 | Pdk4 | 0.023 | Old |
| 0.356049 | 0.594827 | 0.049 | 0.026 | 1 | 13 | Hes2 | 0.023 | Old |
| 0.356134 | 0.317831 | 0.074 | 0.044 | 1 | 13 | A530013C2 | 0.03 | Old |
| 0.356152 | 0.166542 | 0.631 | 0.605 | 1 | 13 | Gpalpp1 | 0.026 | Old |
| 0.356702 | 0.171375 | 0.459 | 0.395 | 1 | 13 | Ift52 | 0.064 | Old |
| 0.356716 | 0.507381 | 0.025 | 0.009 | 1 | 13 | Ift122 | 0.016 | Old |
| 0.356716 | 0.504812 | 0.025 | 0.009 | 1 | 13 | Gm44949 | 0.016 | Old |
| 0.356716 | 0.492618 | 0.025 | 0.009 | 1 | 13 | Gm4349 | 0.016 | Old |
| 0.356716 | 0.491708 | 0.025 | 0.009 | 1 | 13 | Tmem72 | 0.016 | Old |
| 0.356716 | 0.460086 | 0.025 | 0.009 | 1 | 13 | Aqp7 | 0.016 | Old |
| 0.356716 | 0.449304 | 0.025 | 0.009 | 1 | 13 | Ehhadh | 0.016 | Old |
| 0.356716 | 0.401067 | 0.025 | 0.009 | 1 | 13 | Kif7 | 0.016 | Old |
| 0.356716 | 0.396251 | 0.025 | 0.009 | 1 | 13 | Gm46447 | 0.016 | Old |
| 0.356716 | 0.35792 | 0.025 | 0.009 | 1 | 13 | Gm30108 | 0.016 | Old |
| 0.356716 | 0.351928 | 0.025 | 0.009 | 1 | 13 | Cacna1b | 0.016 | Old |
| 0.356716 | 0.332761 | 0.025 | 0.009 | 1 | 13 | Dner | 0.016 | Old |
| 0.356716 | 0.305426 | 0.025 | 0.009 | 1 | 13 | Rwdd3 | 0.016 | Old |
| 0.356716 | 0.294481 | 0.025 | 0.009 | 1 | 13 | Slc16a12 | 0.016 | Old |
| 0.356716 | 0.289283 | 0.025 | 0.009 | 1 | 13 | Mylpf | 0.016 | Old |
| 0.356716 | 0.284467 | 0.025 | 0.009 | 1 | 13 | Adgb | 0.016 | Old |
| 0.356716 | 0.267128 | 0.025 | 0.009 | 1 | 13 | Rax | 0.016 | Old |
| 0.356716 | 0.247044 | 0.025 | 0.009 | 1 | 13 | Aldh1a3 | 0.016 | Old |
| 0.356716 | 0.244556 | 0.025 | 0.009 | 1 | 13 | Fam221a | 0.016 | Old |
| 0.356716 | 0.239232 | 0.025 | 0.009 | 1 | 13 | Tmem158 | 0.016 | Old |
| 0.356716 | 0.219927 | 0.025 | 0.009 | 1 | 13 | Gbp6 | 0.016 | Old |
| 0.356716 | 0.195437 | 0.025 | 0.009 | 1 | 13 | Gm14966 | 0.016 | Old |
| 0.356716 | 0.165496 | 0.025 | 0.009 | 1 | 13 | Cdc42ep5 | 0.016 | Old |
| 0.356716 | 0.164682 | 0.025 | 0.009 | 1 | 13 | Armcx2 | 0.016 | Old |
| 0.356845 | 0.478629 | 0.115 | 0.079 | 1 | 13 | Dhrs11 | 0.036 | Old |
| 0.356942 | 0.234703 | 0.607 | 0.535 | 1 | 13 | Ltbr | 0.072 | Old |
| 0.357025 | 0.081627 | 0.967 | 0.833 | 1 | 13 | Synj1 | 0.134 | Old |
| 0.357239 | 0.186319 | 0.648 | 0.57 | 1 | 13 | Ola1 | 0.078 | Old |
| 0.357261 | 0.324715 | 0.254 | 0.202 | 1 | 13 | Rpl36a-ps1 | 0.052 | Old |
| 0.357561 | 0.005137 | 0.098 | 0.061 | 1 | 13 | 9530036M | 0.037 | Old |
| 0.357627 | 0.485993 | 0.18 | 0.14 | 1 | 13 | Papss2 | 0.04 | Old |
| 0.357678 | 0.505371 | 0.123 | 0.088 | 1 | 13 | Wfdc18 | 0.035 | Old |
| 0.357885 | 0.167659 | 0.918 | 0.851 | 1 | 13 | Adam17 | 0.067 | Old |
| 0.357886 | 0.216328 | 0.615 | 0.518 | 1 | 13 | Dhrs1 | 0.097 | Old |

| | | | | | | | | |
|---|---|---|---|---|---|---|---|---|
| 0.35797 | 0.120976 | 0.73 | 0.605 | 1 | 13 | Maf1 | 0.125 | Old |
| 0.358245 | 0.21872 | 1 | 1 | 1 | 13 | Adgre1 | 0 | Old |
| 0.358451 | 0.338445 | 0.369 | 0.325 | 1 | 13 | Isca2 | 0.044 | Old |
| 0.358524 | 0.391401 | 0.148 | 0.105 | 1 | 13 | BC024978 | 0.043 | Old |
| 0.358569 | 0.359815 | 0.074 | 0.044 | 1 | 13 | Zfp442 | 0.03 | Old |
| 0.358569 | 0.335451 | 0.074 | 0.044 | 1 | 13 | Pkd1l2 | 0.03 | Old |
| 0.358791 | 0.142497 | 0.525 | 0.439 | 1 | 13 | Tspan5 | 0.086 | Old |
| 0.359053 | 0.552192 | 0.049 | 0.026 | 1 | 13 | Hspb6 | 0.023 | Old |
| 0.359053 | 0.542929 | 0.049 | 0.026 | 1 | 13 | Gm42722 | 0.023 | Old |
| 0.359053 | 0.53139 | 0.049 | 0.026 | 1 | 13 | Krtcap3 | 0.023 | Old |
| 0.359053 | 0.176988 | 0.049 | 0.026 | 1 | 13 | Gm17655 | 0.023 | Old |
| 0.359197 | 0.275274 | 0.336 | 0.281 | 1 | 13 | Cog3 | 0.055 | Old |
| 0.359313 | 0.459333 | 0.189 | 0.149 | 1 | 13 | Adnp2 | 0.04 | Old |
| 0.359383 | 0.056528 | 0.377 | 0.289 | 1 | 13 | Ap3s2 | 0.088 | Old |
| 0.35962 | 0.112337 | 0.164 | 0.114 | 1 | 13 | Knstrn | 0.05 | Old |
| 0.360029 | 0.353837 | 0.418 | 0.386 | 1 | 13 | Taf5l | 0.032 | Old |
| 0.360144 | 0.21152 | 0.172 | 0.123 | 1 | 13 | 5830432E0 | 0.049 | Old |
| 0.360145 | 0.584113 | 0.139 | 0.105 | 1 | 13 | Zbtb40 | 0.034 | Old |
| 0.360665 | 0.164451 | 0.59 | 0.518 | 1 | 13 | Vcl | 0.072 | Old |
| 0.360728 | 0.078313 | 0.959 | 0.956 | 1 | 13 | Slc3a2 | 0.003 | Old |
| 0.360761 | 0.210807 | 0.377 | 0.307 | 1 | 13 | Ddx19a | 0.07 | Old |
| 0.360837 | 0.084036 | 0.451 | 0.36 | 1 | 13 | Nip7 | 0.091 | Old |
| 0.360871 | 0.302593 | 0.107 | 0.07 | 1 | 13 | Fut4 | 0.037 | Old |
| 0.360893 | 0.365475 | 0.426 | 0.368 | 1 | 13 | Apba1 | 0.058 | Old |
| 0.361014 | 0.352291 | 0.074 | 0.044 | 1 | 13 | Znrd1as | 0.03 | Old |
| 0.361219 | 0.30008 | 0.23 | 0.175 | 1 | 13 | Rprd1a | 0.055 | Old |
| 0.361273 | 0.117135 | 0.393 | 0.333 | 1 | 13 | Nectin1 | 0.06 | Old |
| 0.361302 | 0.048165 | 0.705 | 0.789 | 1 | 13 | Dusp5 | -0.084 | Old |
| 0.361991 | 0.453204 | 0.213 | 0.167 | 1 | 13 | Gm6563 | 0.046 | Old |
| 0.362074 | 0.598631 | 0.049 | 0.026 | 1 | 13 | Rab23 | 0.023 | Old |
| 0.362074 | 0.514252 | 0.049 | 0.026 | 1 | 13 | Suv39h2 | 0.023 | Old |
| 0.362228 | 0.094409 | 0.951 | 0.956 | 1 | 13 | Caprin1 | -0.005 | Old |
| 0.362328 | 0.169859 | 0.131 | 0.088 | 1 | 13 | 4930522L1 | 0.043 | Old |
| 0.362534 | 0.203057 | 0.221 | 0.167 | 1 | 13 | Drosha | 0.054 | Old |
| 0.362973 | 0.165048 | 0.918 | 0.868 | 1 | 13 | Prrc2a | 0.05 | Old |
| 0.363173 | 0.151421 | 0.566 | 0.509 | 1 | 13 | Ndufv1 | 0.057 | Old |
| 0.363307 | 0.443444 | 0.123 | 0.088 | 1 | 13 | Mtif3 | 0.035 | Old |
| 0.363469 | 0.33824 | 0.074 | 0.044 | 1 | 13 | Tcaim | 0.03 | Old |
| 0.363469 | 0.295781 | 0.074 | 0.044 | 1 | 13 | Zfp607b | 0.03 | Old |
| 0.363545 | 0.186757 | 0.467 | 0.404 | 1 | 13 | Slc35e4 | 0.063 | Old |
| 0.363877 | 0.369328 | 0.238 | 0.184 | 1 | 13 | Nudt18 | 0.054 | Old |
| 0.364141 | 0.110155 | 0.951 | 0.886 | 1 | 13 | Sf1 | 0.065 | Old |
| 0.364638 | 0.420851 | 0.115 | 0.079 | 1 | 13 | Slc8a1 | 0.036 | Old |
| 0.364946 | 0.316636 | 0.107 | 0.07 | 1 | 13 | Capn10 | 0.037 | Old |
| 0.365111 | 1.337718 | 0.049 | 0.026 | 1 | 13 | Fabp7 | 0.023 | Old |

| | | | | | | | | |
|---|---|---|---|---|---|---|---|---|
| 0.365111 | 0.563962 | 0.049 | 0.026 | 1 | 13 | 4732471J0 | 0.023 | Old |
| 0.365111 | 0.523075 | 0.049 | 0.026 | 1 | 13 | Chst1 | 0.023 | Old |
| 0.365201 | 0.177855 | 0.951 | 0.912 | 1 | 13 | Pgk1 | 0.039 | Old |
| 0.36554 | 0.09715 | 0.68 | 0.526 | 1 | 13 | Cops4 | 0.154 | Old |
| 0.365637 | 0.289752 | 0.049 | 0.079 | 1 | 13 | Il18r1 | -0.03 | Old |
| 0.365739 | 0.112685 | 0.803 | 0.754 | 1 | 13 | Arl6ip5 | 0.049 | Old |
| 0.365948 | 0.214968 | 0.393 | 0.333 | 1 | 13 | Rufy1 | 0.06 | Old |
| 0.366186 | 0.115211 | 0.68 | 0.588 | 1 | 13 | Usp14 | 0.092 | Old |
| 0.366421 | 0.048207 | 0.689 | 0.553 | 1 | 13 | Actr10 | 0.136 | Old |
| 0.366649 | 0.192917 | 0.795 | 0.825 | 1 | 13 | Hfe | -0.03 | Old |
| 0.36682 | 0.233918 | 0.475 | 0.386 | 1 | 13 | BC003965 | 0.089 | Old |
| 0.366951 | 0.243673 | 0.705 | 0.667 | 1 | 13 | Suz12 | 0.038 | Old |
| 0.367136 | 0.267274 | 0.508 | 0.474 | 1 | 13 | Kansl3 | 0.034 | Old |
| 0.367193 | 0.283639 | 0.262 | 0.211 | 1 | 13 | Gon7 | 0.051 | Old |
| 0.36726 | 0.170647 | 0.377 | 0.298 | 1 | 13 | Fubp3 | 0.079 | Old |
| 0.3673 | 0.115101 | 0.992 | 1 | 1 | 13 | Ddx17 | -0.008 | Old |
| 0.367437 | 0.179148 | 0.402 | 0.342 | 1 | 13 | Prpsap2 | 0.06 | Old |
| 0.367828 | 0.130264 | 0.311 | 0.246 | 1 | 13 | Tacc2 | 0.065 | Old |
| 0.36792 | 0.124387 | 0.287 | 0.219 | 1 | 13 | Anapc7 | 0.068 | Old |
| 0.368163 | 0.562069 | 0.049 | 0.026 | 1 | 13 | Cenpm | 0.023 | Old |
| 0.368163 | 0.469477 | 0.049 | 0.026 | 1 | 13 | Ung | 0.023 | Old |
| 0.368163 | 0.40864 | 0.049 | 0.026 | 1 | 13 | Mageh1 | 0.023 | Old |
| 0.368163 | 0.405184 | 0.049 | 0.026 | 1 | 13 | Cenpk | 0.023 | Old |
| 0.368163 | 0.385046 | 0.049 | 0.026 | 1 | 13 | Gm14295 | 0.023 | Old |
| 0.368163 | 0.284948 | 0.049 | 0.026 | 1 | 13 | Ccdc17 | 0.023 | Old |
| 0.368163 | 0.283585 | 0.049 | 0.026 | 1 | 13 | Cchcr1 | 0.023 | Old |
| 0.369316 | 0.287495 | 0.525 | 0.456 | 1 | 13 | Csnk1g1 | 0.069 | Old |
| 0.369875 | 0.163056 | 0.459 | 0.377 | 1 | 13 | Ccdc25 | 0.082 | Old |
| 0.370897 | 0.494034 | 0.123 | 0.088 | 1 | 13 | Sirpb1a | 0.035 | Old |
| 0.371113 | 0.249469 | 0.107 | 0.07 | 1 | 13 | D130040H: | 0.037 | Old |
| 0.371115 | 0.001176 | 0.41 | 0.325 | 1 | 13 | Cebpe | 0.085 | Old |
| 0.371231 | 0.428414 | 0.049 | 0.026 | 1 | 13 | Bcl2a1c | 0.023 | Old |
| 0.371231 | 0.065779 | 0.049 | 0.026 | 1 | 13 | Rhou | 0.023 | Old |
| 0.371924 | 0.302189 | 0.205 | 0.149 | 1 | 13 | Tada2b | 0.056 | Old |
| 0.371948 | 0.109381 | 0.893 | 0.86 | 1 | 13 | Dnajc8 | 0.033 | Old |
| 0.372001 | 0.129399 | 0.762 | 0.667 | 1 | 13 | Atp6v1h | 0.095 | Old |
| 0.372304 | 0.255529 | 0.41 | 0.351 | 1 | 13 | Slc7a1 | 0.059 | Old |
| 0.372334 | 0.35583 | 0.303 | 0.254 | 1 | 13 | Wdsub1 | 0.049 | Old |
| 0.372392 | 0.119813 | 1 | 1 | 1 | 13 | Csf2ra | 0 | Old |
| 0.372411 | 0.149598 | 0.746 | 0.763 | 1 | 13 | mt-Atp8 | -0.017 | Old |
| 0.372537 | 0.281744 | 0.115 | 0.079 | 1 | 13 | Tyms | 0.036 | Old |
| 0.372805 | 0.349976 | 0.221 | 0.167 | 1 | 13 | Zbtb6 | 0.054 | Old |
| 0.372945 | 0.113128 | 0.18 | 0.132 | 1 | 13 | Stk39 | 0.048 | Old |
| 0.373156 | 0.407813 | 0.172 | 0.132 | 1 | 13 | Parp6 | 0.04 | Old |
| 0.373478 | 0.31323 | 0.352 | 0.289 | 1 | 13 | Dnajc11 | 0.063 | Old |

| | | | | | | | | |
|---|---|---|---|---|---|---|---|---|
| 0.373817 | 0.253763 | 0.648 | 0.588 | 1 | 13 | Efcab14 | 0.06 | Old |
| 0.373968 | 0.155626 | 0.631 | 0.57 | 1 | 13 | Phf10 | 0.061 | Old |
| 0.373976 | 0.321615 | 0.615 | 0.57 | 1 | 13 | Epb41l1 | 0.045 | Old |
| 0.374018 | 0.461428 | 0.131 | 0.096 | 1 | 13 | Primpol | 0.035 | Old |
| 0.374286 | 0.148243 | 0.221 | 0.167 | 1 | 13 | Stk38l | 0.054 | Old |
| 0.374315 | 0.399368 | 0.049 | 0.026 | 1 | 13 | Tmem240 | 0.023 | Old |
| 0.374315 | 0.333067 | 0.049 | 0.026 | 1 | 13 | Gm10320 | 0.023 | Old |
| 0.374729 | 0.311615 | 0.123 | 0.088 | 1 | 13 | Zfp609 | 0.035 | Old |
| 0.374945 | 0.069401 | 0.975 | 0.956 | 1 | 13 | Ndufb3 | 0.019 | Old |
| 0.374976 | 0.204921 | 0.525 | 0.491 | 1 | 13 | Tmem184b | 0.034 | Old |
| 0.375071 | 0.046709 | 0.672 | 0.596 | 1 | 13 | Slc30a5 | 0.076 | Old |
| 0.375606 | 0.195964 | 0.197 | 0.14 | 1 | 13 | Them4 | 0.057 | Old |
| 0.375942 | 0.144531 | 0.959 | 0.93 | 1 | 13 | Gsn | 0.029 | Old |
| 0.376019 | 0.422668 | 0.516 | 0.482 | 1 | 13 | Sestd1 | 0.034 | Old |
| 0.376417 | 0.0548 | 0.68 | 0.746 | 1 | 13 | Basp1 | -0.066 | Old |
| 0.376512 | 0.140598 | 0.861 | 0.693 | 1 | 13 | Usp4 | 0.168 | Old |
| 0.376901 | 0.038449 | 0.836 | 0.781 | 1 | 13 | Zc3h15 | 0.055 | Old |
| 0.377201 | 0.457875 | 0.09 | 0.061 | 1 | 13 | Natd1 | 0.029 | Old |
| 0.377415 | 0.211244 | 0.049 | 0.026 | 1 | 13 | Hist1h2be | 0.023 | Old |
| 0.377484 | 0.237779 | 0.574 | 0.491 | 1 | 13 | Ppm1h | 0.083 | Old |
| 0.377562 | 0.226911 | 0.369 | 0.325 | 1 | 13 | Ppfibp1 | 0.044 | Old |
| 0.37758 | 0.198262 | 0.32 | 0.368 | 1 | 13 | Cxcl10 | -0.048 | Old |
| 0.377702 | 0.302575 | 0.32 | 0.263 | 1 | 13 | Pign | 0.057 | Old |
| 0.377781 | 0.168614 | 0.869 | 0.833 | 1 | 13 | Soat1 | 0.036 | Old |
| 0.377816 | 0.128477 | 0.361 | 0.281 | 1 | 13 | Tmem167b | 0.08 | Old |
| 0.377833 | 0.06664 | 0.418 | 0.333 | 1 | 13 | Nsmce3 | 0.085 | Old |
| 0.377981 | 0.140123 | 0.639 | 0.605 | 1 | 13 | Bcl7b | 0.034 | Old |
| 0.378058 | 0.178898 | 0.32 | 0.246 | 1 | 13 | D430042O | 0.074 | Old |
| 0.378135 | 0.127858 | 0.475 | 0.386 | 1 | 13 | Fbxo22 | 0.089 | Old |
| 0.378339 | 0.287101 | 0.213 | 0.167 | 1 | 13 | Atpaf2 | 0.046 | Old |
| 0.378438 | 0.303392 | 0.352 | 0.289 | 1 | 13 | Slc35c1 | 0.063 | Old |
| 0.378775 | 0.168573 | 0.541 | 0.456 | 1 | 13 | Wdr41 | 0.085 | Old |
| 0.378805 | 0.284721 | 0.508 | 0.439 | 1 | 13 | Klhdc3 | 0.069 | Old |
| 0.379483 | 0.065282 | 0.311 | 0.246 | 1 | 13 | Prcc | 0.065 | Old |
| 0.379542 | 0.117284 | 0.451 | 0.351 | 1 | 13 | Klhdc4 | 0.1 | Old |
| 0.379999 | 0.332745 | 0.73 | 0.772 | 1 | 13 | Cxcr4 | -0.042 | Old |
| 0.380493 | 0.275647 | 0.426 | 0.368 | 1 | 13 | Tbck | 0.058 | Old |
| 0.380531 | 0.409515 | 0.049 | 0.026 | 1 | 13 | Trim62 | 0.023 | Old |
| 0.381616 | 0.185596 | 0.5 | 0.386 | 1 | 13 | Lipe | 0.114 | Old |
| 0.38224 | 0.04127 | 0.664 | 0.579 | 1 | 13 | Erlin1 | 0.085 | Old |
| 0.382304 | 0.165086 | 0.844 | 0.789 | 1 | 13 | Cox20 | 0.055 | Old |
| 0.382558 | 0.33537 | 0.115 | 0.079 | 1 | 13 | Rsl1 | 0.036 | Old |
| 0.382558 | 0.320255 | 0.115 | 0.079 | 1 | 13 | Slfn9 | 0.036 | Old |
| 0.382615 | 0.077684 | 0.492 | 0.421 | 1 | 13 | Otud6b | 0.071 | Old |
| 0.38304 | 0.337965 | 0.59 | 0.614 | 1 | 13 | Lpar1 | -0.024 | Old |

| | | | | | | | | |
|---|---|---|---|---|---|---|---|---|
| 0.384022 | 0.088298 | 0.713 | 0.702 | 1 | 13 | Apaf1 | 0.011 | Old |
| 0.384066 | 0.389313 | 0.262 | 0.219 | 1 | 13 | Ezh2 | 0.043 | Old |
| 0.384188 | 0.096382 | 0.803 | 0.693 | 1 | 13 | Wdr89 | 0.11 | Old |
| 0.384265 | 0.104745 | 0.361 | 0.298 | 1 | 13 | Ptgr1 | 0.063 | Old |
| 0.384507 | 0.149978 | 0.918 | 0.868 | 1 | 13 | Idh1 | 0.05 | Old |
| 0.384577 | 0.233463 | 0.393 | 0.325 | 1 | 13 | Eri3 | 0.068 | Old |
| 0.38475 | 0.115304 | 0.959 | 0.93 | 1 | 13 | Lims1 | 0.029 | Old |
| 0.384888 | 0.145216 | 0.508 | 0.404 | 1 | 13 | Bad | 0.104 | Old |
| 0.385027 | 0.196893 | 0.459 | 0.404 | 1 | 13 | Usp45 | 0.055 | Old |
| 0.385123 | 0.099805 | 0.869 | 0.842 | 1 | 13 | Camk1 | 0.027 | Old |
| 0.385753 | 0.185666 | 0.107 | 0.07 | 1 | 13 | Zfat | 0.037 | Old |
| 0.385918 | 0.282469 | 0.213 | 0.167 | 1 | 13 | Gtf2h4 | 0.046 | Old |
| 0.386117 | 0.168398 | 0.385 | 0.307 | 1 | 13 | Acot13 | 0.078 | Old |
| 0.386253 | 0.140677 | 0.295 | 0.246 | 1 | 13 | Fam8a1 | 0.049 | Old |
| 0.386271 | 0.15028 | 0.221 | 0.167 | 1 | 13 | Slc52a2 | 0.054 | Old |
| 0.386281 | 0.122478 | 0.139 | 0.096 | 1 | 13 | Polr3h | 0.043 | Old |
| 0.386631 | 0.053913 | 0.918 | 0.877 | 1 | 13 | Ddost | 0.041 | Old |
| 0.386732 | 0.24126 | 0.648 | 0.57 | 1 | 13 | Ogfr | 0.078 | Old |
| 0.386802 | 0.171611 | 0.148 | 0.105 | 1 | 13 | Zfp628 | 0.043 | Old |
| 0.386846 | 0.31507 | 0.23 | 0.184 | 1 | 13 | Tbrg4 | 0.046 | Old |
| 0.387318 | 0.264937 | 0.279 | 0.219 | 1 | 13 | Myef2 | 0.06 | Old |
| 0.387323 | 0.108876 | 0.705 | 0.623 | 1 | 13 | Ech1 | 0.082 | Old |
| 0.387604 | 0.085513 | 0.656 | 0.588 | 1 | 13 | Fbxw2 | 0.068 | Old |
| 0.388005 | 0.323105 | 0.18 | 0.14 | 1 | 13 | Tcta | 0.04 | Old |
| 0.388452 | 0.105184 | 0.992 | 0.991 | 1 | 13 | Ost4 | 0.001 | Old |
| 0.388459 | 0.024515 | 0.59 | 0.474 | 1 | 13 | Ccndbp1 | 0.116 | Old |
| 0.388854 | 0.130446 | 0.943 | 0.895 | 1 | 13 | Asap1 | 0.048 | Old |
| 0.389107 | 0.613693 | 0.877 | 0.886 | 1 | 13 | Ctsl | -0.009 | Old |
| 0.389252 | 0.179253 | 0.344 | 0.272 | 1 | 13 | Pld2 | 0.072 | Old |
| 0.389291 | 0.237558 | 0.525 | 0.456 | 1 | 13 | Dusp23 | 0.069 | Old |
| 0.389338 | 0.110859 | 0.18 | 0.132 | 1 | 13 | Kiz | 0.048 | Old |
| 0.389361 | 0.215467 | 0.533 | 0.465 | 1 | 13 | Brd8 | 0.068 | Old |
| 0.389452 | 0.213434 | 0.393 | 0.325 | 1 | 13 | Stx17 | 0.068 | Old |
| 0.389485 | 0.096661 | 0.975 | 0.939 | 1 | 13 | Ubap2l | 0.036 | Old |
| 0.389503 | 0.061178 | 0.992 | 0.991 | 1 | 13 | Dok2 | 0.001 | Old |
| 0.389514 | 0.239997 | 0.418 | 0.342 | 1 | 13 | Gapt | 0.076 | Old |
| 0.389726 | 0.139609 | 0.385 | 0.316 | 1 | 13 | Zfp672 | 0.069 | Old |
| 0.389735 | 0.164161 | 0.787 | 0.781 | 1 | 13 | Stat6 | 0.006 | Old |
| 0.389808 | 0.421493 | 0.197 | 0.158 | 1 | 13 | Phlpp2 | 0.039 | Old |
| 0.389885 | 0.254919 | 0.582 | 0.544 | 1 | 13 | Srd5a3 | 0.038 | Old |
| 0.390029 | 0.006076 | 1 | 0.982 | 1 | 13 | Prrc2c | 0.018 | Old |
| 0.390095 | 0.27499 | 0.615 | 0.579 | 1 | 13 | Cdk2ap1 | 0.036 | Old |
| 0.390146 | 0.348494 | 0.574 | 0.526 | 1 | 13 | Gclc | 0.048 | Old |
| 0.3903 | 0.365268 | 0.123 | 0.088 | 1 | 13 | Ccdc84 | 0.035 | Old |
| 0.39051 | 0.270478 | 0.213 | 0.167 | 1 | 13 | Usp42 | 0.046 | Old |

| | | | | | | | | |
|---|---|---|---|---|---|---|---|---|
| 0.390947 | 0.333524 | 0.131 | 0.096 | 1 | 13 | Zfp788 | 0.035 | Old |
| 0.390958 | 0.104708 | 0.951 | 0.886 | 1 | 13 | Syncrip | 0.065 | Old |
| 0.391058 | 0.174628 | 0.221 | 0.175 | 1 | 13 | Arid3b | 0.046 | Old |
| 0.391202 | 0.180491 | 0.615 | 0.57 | 1 | 13 | Dhx38 | 0.045 | Old |
| 0.391697 | 0.119087 | 0.918 | 0.833 | 1 | 13 | Ankrd17 | 0.085 | Old |
| 0.391782 | 0.154731 | 0.885 | 0.886 | 1 | 13 | Rsrc2 | -0.001 | Old |
| 0.391893 | 0.20322 | 0.139 | 0.096 | 1 | 13 | Mrps27 | 0.043 | Old |
| 0.392022 | 0.551633 | 0.082 | 0.053 | 1 | 13 | Nectin2 | 0.029 | Old |
| 0.392022 | 0.225413 | 0.082 | 0.053 | 1 | 13 | Sec16b | 0.029 | Old |
| 0.392913 | 0.029912 | 0.516 | 0.456 | 1 | 13 | Clcc1 | 0.06 | Old |
| 0.392971 | 0.361911 | 0.197 | 0.158 | 1 | 13 | Haus3 | 0.039 | Old |
| 0.393017 | 0.08064 | 0.459 | 0.377 | 1 | 13 | Ndrg3 | 0.082 | Old |
| 0.39307 | 0.116685 | 0.492 | 0.404 | 1 | 13 | Kif19a | 0.088 | Old |
| 0.39308 | 0.570198 | 0.09 | 0.061 | 1 | 13 | Mtrr | 0.029 | Old |
| 0.393339 | 0.106538 | 0.877 | 0.807 | 1 | 13 | Cd164 | 0.07 | Old |
| 0.393366 | 0.257683 | 0.68 | 0.667 | 1 | 13 | Coro2a | 0.013 | Old |
| 0.393377 | 0.207615 | 0.623 | 0.535 | 1 | 13 | Pes1 | 0.088 | Old |
| 0.39388 | 0.157513 | 0.221 | 0.167 | 1 | 13 | Tmco6 | 0.054 | Old |
| 0.393948 | 0.222846 | 0.41 | 0.325 | 1 | 13 | Slc30a1 | 0.085 | Old |
| 0.393995 | 0.082847 | 0.451 | 0.368 | 1 | 13 | Atg16l1 | 0.083 | Old |
| 0.394087 | 0.212031 | 0.221 | 0.175 | 1 | 13 | Micall1 | 0.046 | Old |
| 0.394122 | 0.124152 | 0.738 | 0.658 | 1 | 13 | Mrpl43 | 0.08 | Old |
| 0.39419 | 0.166104 | 0.951 | 0.921 | 1 | 13 | Mia2 | 0.03 | Old |
| 0.394243 | 0.128198 | 0.885 | 0.781 | 1 | 13 | Aldh9a1 | 0.104 | Old |
| 0.394259 | 0.401596 | 0.238 | 0.184 | 1 | 13 | Slc9a4 | 0.054 | Old |
| 0.394448 | 0.273272 | 0.082 | 0.053 | 1 | 13 | Lzts2 | 0.029 | Old |
| 0.39481 | 0.275527 | 0.172 | 0.132 | 1 | 13 | C1qtnf12 | 0.04 | Old |
| 0.39481 | 0.117897 | 0.172 | 0.132 | 1 | 13 | F3 | 0.04 | Old |
| 0.395271 | 0.266362 | 0.27 | 0.211 | 1 | 13 | Ppp2r1b | 0.059 | Old |
| 0.395314 | 0.151333 | 0.811 | 0.772 | 1 | 13 | Snf8 | 0.039 | Old |
| 0.395712 | 0.194193 | 0.443 | 0.342 | 1 | 13 | Decr1 | 0.101 | Old |
| 0.395713 | 0.091335 | 0.844 | 0.728 | 1 | 13 | Bcas2 | 0.116 | Old |
| 0.396631 | 0.245409 | 0.385 | 0.342 | 1 | 13 | Gm5547 | 0.043 | Old |
| 0.396901 | 0.087018 | 0.869 | 0.868 | 1 | 13 | Kras | 0.001 | Old |
| 0.396904 | 0.091632 | 0.992 | 0.991 | 1 | 13 | Psme1 | 0.001 | Old |
| 0.397015 | 0.406656 | 0.385 | 0.325 | 1 | 13 | Pprc1 | 0.06 | Old |
| 0.39723 | 0.455162 | 0.238 | 0.184 | 1 | 13 | Pdk1 | 0.054 | Old |
| 0.397245 | 0.069238 | 0.59 | 0.5 | 1 | 13 | Ranbp10 | 0.09 | Old |
| 0.397436 | 0.062385 | 1 | 1 | 1 | 13 | Tpm3 | 0 | Old |
| 0.397748 | 0.26169 | 0.148 | 0.105 | 1 | 13 | Esr1 | 0.043 | Old |
| 0.398023 | 0.183315 | 0.189 | 0.14 | 1 | 13 | Surf2 | 0.049 | Old |
| 0.398307 | 0.195653 | 0.418 | 0.333 | 1 | 13 | Tmem214 | 0.085 | Old |
| 0.398413 | 0.16988 | 0.918 | 0.939 | 1 | 13 | Wsb1 | -0.021 | Old |
| 0.39847 | 0.118693 | 0.803 | 0.667 | 1 | 13 | Tspan31 | 0.136 | Old |
| 0.398483 | 0.300647 | 0.393 | 0.333 | 1 | 13 | Hectd4 | 0.06 | Old |

| | | | | | | | | |
|---|---|---|---|---|---|---|---|---|
| 0.398679 | 0.246741 | 0.352 | 0.289 | 1 | 13 | Ptger2 | 0.063 | Old |
| 0.398875 | 1.078474 | 0.057 | 0.035 | 1 | 13 | Phactr1 | 0.022 | Old |
| 0.399018 | 0.708935 | 0.098 | 0.07 | 1 | 13 | Tarm1 | 0.028 | Old |
| 0.399034 | 0.111079 | 0.992 | 0.991 | 1 | 13 | Slfn2 | 0.001 | Old |
| 0.39908 | 0.302306 | 0.156 | 0.114 | 1 | 13 | Sh2b2 | 0.042 | Old |
| 0.399219 | 0.157593 | 0.59 | 0.526 | 1 | 13 | Naip2 | 0.064 | Old |
| 0.399582 | 0.40901 | 0.18 | 0.14 | 1 | 13 | Accs | 0.04 | Old |
| 0.399932 | 0.040444 | 0.852 | 0.86 | 1 | 13 | Luc7l3 | -0.008 | Old |
| 0.400036 | 0.259007 | 0.352 | 0.307 | 1 | 13 | Chchd4 | 0.045 | Old |
| 0.400058 | 0.100386 | 0.492 | 0.404 | 1 | 13 | Csad | 0.088 | Old |
| 0.400211 | 0.177905 | 0.516 | 0.439 | 1 | 13 | Rlf | 0.077 | Old |
| 0.400215 | 0.07236 | 0.238 | 0.184 | 1 | 13 | Lmbr1l | 0.054 | Old |
| 0.40056 | 0.182917 | 0.664 | 0.588 | 1 | 13 | Sun2 | 0.076 | Old |
| 0.400584 | 0.100974 | 0.893 | 0.833 | 1 | 13 | Rab21 | 0.06 | Old |
| 0.400641 | 0.120752 | 0.68 | 0.632 | 1 | 13 | Gnl1 | 0.048 | Old |
| 0.400695 | 0.195534 | 0.533 | 0.465 | 1 | 13 | Anapc1 | 0.068 | Old |
| 0.401109 | 0.117677 | 0.672 | 0.579 | 1 | 13 | Fbxo33 | 0.093 | Old |
| 0.401166 | 0.112835 | 0.197 | 0.149 | 1 | 13 | Ccdc91 | 0.048 | Old |
| 0.401729 | 0.171171 | 0.598 | 0.491 | 1 | 13 | Ube2j1 | 0.107 | Old |
| 0.401801 | 0.667439 | 0.057 | 0.035 | 1 | 13 | Simc1 | 0.022 | Old |
| 0.402064 | 0.151512 | 0.492 | 0.43 | 1 | 13 | AU040320 | 0.062 | Old |
| 0.402193 | 0.104595 | 0.943 | 0.939 | 1 | 13 | Srsf6 | 0.004 | Old |
| 0.402212 | 0.068372 | 0.328 | 0.263 | 1 | 13 | Kat2a | 0.065 | Old |
| 0.402232 | 0.378416 | 0.123 | 0.088 | 1 | 13 | Chaf1a | 0.035 | Old |
| 0.402295 | 0.056447 | 0.902 | 0.842 | 1 | 13 | Fam120a | 0.06 | Old |
| 0.402927 | 0.488278 | 0.18 | 0.14 | 1 | 13 | Socs2 | 0.04 | Old |
| 0.403245 | 0.060945 | 0.664 | 0.535 | 1 | 13 | Rnf20 | 0.129 | Old |
| 0.4035 | 0.166635 | 0.885 | 0.886 | 1 | 13 | Ldlr | -0.001 | Old |
| 0.403755 | 0.367344 | 0.189 | 0.149 | 1 | 13 | Gtf3c4 | 0.04 | Old |
| 0.404083 | 0.100352 | 0.295 | 0.228 | 1 | 13 | Lztr1 | 0.067 | Old |
| 0.404234 | 0.125067 | 0.664 | 0.553 | 1 | 13 | Pi4ka | 0.111 | Old |
| 0.404436 | 0.506898 | 0.246 | 0.211 | 1 | 13 | Gpr68 | 0.035 | Old |
| 0.404667 | 0.429754 | 0.09 | 0.061 | 1 | 13 | Nav2 | 0.029 | Old |
| 0.404667 | 0.126286 | 0.09 | 0.061 | 1 | 13 | Pclaf | 0.029 | Old |
| 0.404709 | 0.142552 | 0.541 | 0.456 | 1 | 13 | 0610030E2 | 0.085 | Old |
| 0.404927 | 0.05426 | 0.992 | 0.982 | 1 | 13 | Syk | 0.01 | Old |
| 0.404969 | 0.035087 | 0.475 | 0.377 | 1 | 13 | Golga2 | 0.098 | Old |
| 0.40497 | 0.122296 | 0.205 | 0.149 | 1 | 13 | Gmeb2 | 0.056 | Old |
| 0.405176 | 0.052629 | 0.795 | 0.737 | 1 | 13 | Pum1 | 0.058 | Old |
| 0.405465 | 0.112722 | 0.975 | 0.991 | 1 | 13 | Hif1a | -0.016 | Old |
| 0.405676 | 0.741009 | 0.098 | 0.07 | 1 | 13 | Chd3os | 0.028 | Old |
| 0.405769 | 0.254713 | 0.385 | 0.325 | 1 | 13 | Sec16a | 0.06 | Old |
| 0.406249 | 0.074647 | 0.779 | 0.702 | 1 | 13 | Morc3 | 0.077 | Old |
| 0.406356 | 0.16608 | 0.516 | 0.447 | 1 | 13 | Ilvbl | 0.069 | Old |
| 0.406586 | 0.029032 | 0.205 | 0.149 | 1 | 13 | Aven | 0.056 | Old |

| | | | | | | | | |
|---|---|---|---|---|---|---|---|---|
| 0.406802 | 0.33072 | 0.164 | 0.123 | 1 | 13 | Lmtk2 | 0.041 | Old |
| 0.407009 | 0.401306 | 0.09 | 0.061 | 1 | 13 | Ppp1r3e | 0.029 | Old |
| 0.407085 | 0.107417 | 0.762 | 0.667 | 1 | 13 | Nudcd3 | 0.095 | Old |
| 0.40714 | 0.151587 | 0.82 | 0.763 | 1 | 13 | Npepps | 0.057 | Old |
| 0.407275 | 0.161739 | 0.115 | 0.079 | 1 | 13 | Mettl25 | 0.036 | Old |
| 0.407681 | 0.149122 | 0.197 | 0.149 | 1 | 13 | Ap5z1 | 0.048 | Old |
| 0.407687 | 0.125439 | 0.451 | 0.36 | 1 | 13 | BC037034 | 0.091 | Old |
| 0.407693 | 0.609123 | 0.057 | 0.035 | 1 | 13 | Rad51ap1 | 0.022 | Old |
| 0.407697 | 0.179445 | 0.361 | 0.307 | 1 | 13 | Prps2 | 0.054 | Old |
| 0.407978 | 0.197689 | 0.189 | 0.14 | 1 | 13 | Sergef | 0.049 | Old |
| 0.408034 | 0.142106 | 0.68 | 0.588 | 1 | 13 | Tmem173 | 0.092 | Old |
| 0.408269 | 0.243212 | 0.557 | 0.518 | 1 | 13 | Mink1 | 0.039 | Old |
| 0.408278 | 0.048138 | 0.705 | 0.632 | 1 | 13 | Tceal9 | 0.073 | Old |
| 0.408322 | 0.28338 | 0.27 | 0.228 | 1 | 13 | Lig1 | 0.042 | Old |
| 0.408435 | 0.176358 | 0.77 | 0.763 | 1 | 13 | C1qbp | 0.007 | Old |
| 0.409135 | 0.15125 | 0.516 | 0.43 | 1 | 13 | Ap1ar | 0.086 | Old |
| 0.409956 | 0.462837 | 0.246 | 0.202 | 1 | 13 | Irf7 | 0.044 | Old |
| 0.410278 | 0.110438 | 0.5 | 0.421 | 1 | 13 | Thyn1 | 0.079 | Old |
| 0.410303 | 0.104961 | 0.352 | 0.281 | 1 | 13 | Fbxo6 | 0.071 | Old |
| 0.410345 | 0.034103 | 0.697 | 0.623 | 1 | 13 | Rabep1 | 0.074 | Old |
| 0.410346 | 0.219298 | 0.164 | 0.123 | 1 | 13 | Cad | 0.041 | Old |
| 0.410558 | 0.44798 | 0.205 | 0.167 | 1 | 13 | Zfp143 | 0.038 | Old |
| 0.410658 | 1.044875 | 0.057 | 0.035 | 1 | 13 | Spon1 | 0.022 | Old |
| 0.410956 | 0.277666 | 0.705 | 0.675 | 1 | 13 | Nacc2 | 0.03 | Old |
| 0.411416 | 0.056488 | 1 | 1 | 1 | 13 | Atp5l | 0 | Old |
| 0.411972 | 0.329095 | 0.172 | 0.132 | 1 | 13 | Tmem107 | 0.04 | Old |
| 0.412348 | 0.137239 | 0.877 | 0.833 | 1 | 13 | Phb2 | 0.044 | Old |
| 0.412401 | 0.552735 | 0.098 | 0.07 | 1 | 13 | Phf19 | 0.028 | Old |
| 0.41264 | 0.141077 | 0.385 | 0.298 | 1 | 13 | Gpatch2l | 0.087 | Old |
| 0.413017 | 0.125346 | 0.975 | 0.921 | 1 | 13 | Tubb6 | 0.054 | Old |
| 0.41315 | 0.313664 | 0.73 | 0.746 | 1 | 13 | R3hdm4 | -0.016 | Old |
| 0.413253 | 0.232072 | 0.68 | 0.658 | 1 | 13 | Rab32 | 0.022 | Old |
| 0.413267 | 0.123114 | 0.828 | 0.807 | 1 | 13 | Fxr1 | 0.021 | Old |
| 0.413313 | 0.181173 | 0.23 | 0.175 | 1 | 13 | Parp10 | 0.055 | Old |
| 0.413377 | 0.167081 | 0.467 | 0.386 | 1 | 13 | Usp33 | 0.081 | Old |
| 0.413577 | 0.173355 | 0.156 | 0.114 | 1 | 13 | F5 | 0.042 | Old |
| 0.413647 | 0.00129 | 0.557 | 0.447 | 1 | 13 | Uhrf1bp1l | 0.11 | Old |
| 0.41403 | 0.291249 | 0.279 | 0.228 | 1 | 13 | Vps8 | 0.051 | Old |
| 0.414032 | 0.282284 | 0.402 | 0.36 | 1 | 13 | Sumf1 | 0.042 | Old |
| 0.414082 | 0.340632 | 0.09 | 0.061 | 1 | 13 | Spryd4 | 0.029 | Old |
| 0.414381 | 0.346093 | 0.123 | 0.088 | 1 | 13 | Ggcx | 0.035 | Old |
| 0.414716 | 0.000966 | 0.697 | 0.57 | 1 | 13 | Ahcyl1 | 0.127 | Old |
| 0.414857 | 0.289676 | 0.344 | 0.281 | 1 | 13 | Plau | 0.063 | Old |
| 0.414864 | 0.171336 | 0.23 | 0.175 | 1 | 13 | Ints5.1 | 0.055 | Old |
| 0.415031 | 0.149341 | 0.828 | 0.675 | 1 | 13 | Dram2 | 0.153 | Old |

| | | | | | | | | |
|---|---|---|---|---|---|---|---|---|
| 0.415236 | 0.337209 | 0.139 | 0.105 | 1 | 13 | Otud1 | 0.034 | Old |
| 0.415352 | 0.195068 | 0.369 | 0.307 | 1 | 13 | Aifm1 | 0.062 | Old |
| 0.415607 | 0.179175 | 0.525 | 0.421 | 1 | 13 | Ddx47 | 0.104 | Old |
| 0.415715 | 0.089497 | 0.943 | 0.93 | 1 | 13 | Tor1aip1 | 0.013 | Old |
| 0.415768 | 0.22271 | 0.443 | 0.421 | 1 | 13 | Akap10 | 0.022 | Old |
| 0.415775 | 0.073152 | 1 | 1 | 1 | 13 | Dstn | 0 | Old |
| 0.416626 | 0.492035 | 0.057 | 0.035 | 1 | 13 | Tada2a | 0.022 | Old |
| 0.416748 | 0.091737 | 0.68 | 0.693 | 1 | 13 | Plcg2 | -0.013 | Old |
| 0.416869 | 0.043043 | 1 | 1 | 1 | 13 | Mob1a | 0 | Old |
| 0.416869 | 0.015292 | 1 | 1 | 1 | 13 | Tmem50a | 0 | Old |
| 0.416935 | 0.18218 | 0.393 | 0.325 | 1 | 13 | Serpinb1a | 0.068 | Old |
| 0.416935 | 0.145487 | 0.393 | 0.325 | 1 | 13 | Pggt1b | 0.068 | Old |
| 0.417016 | 0.150206 | 0.68 | 0.64 | 1 | 13 | Nfatc3 | 0.04 | Old |
| 0.417205 | 0.065913 | 0.852 | 0.851 | 1 | 13 | Eif1b | 0.001 | Old |
| 0.417382 | 0.19665 | 0.549 | 0.465 | 1 | 13 | Mrps23 | 0.084 | Old |
| 0.417487 | 0.127016 | 0.885 | 0.86 | 1 | 13 | Sbno1 | 0.025 | Old |
| 0.417844 | 0.124985 | 0.926 | 0.912 | 1 | 13 | Lasp1 | 0.014 | Old |
| 0.41796 | 0.422516 | 0.254 | 0.211 | 1 | 13 | Ms4a4c | 0.043 | Old |
| 0.418094 | 0.131063 | 0.738 | 0.693 | 1 | 13 | Fndc3b | 0.045 | Old |
| 0.418599 | 0.17726 | 0.361 | 0.316 | 1 | 13 | Aak1 | 0.045 | Old |
| 0.418614 | 0.06651 | 0.238 | 0.175 | 1 | 13 | Cdc23 | 0.063 | Old |
| 0.418711 | 0.386559 | 0.189 | 0.149 | 1 | 13 | Dip2c | 0.04 | Old |
| 0.41888 | 0.174276 | 0.664 | 0.544 | 1 | 13 | Ankib1 | 0.12 | Old |
| 0.419063 | 0.142553 | 0.992 | 1 | 1 | 13 | Fus | -0.008 | Old |
| 0.419284 | 0.196636 | 0.164 | 0.123 | 1 | 13 | Fbxl4 | 0.041 | Old |
| 0.419427 | 0.171047 | 0.77 | 0.711 | 1 | 13 | Hnrnpr | 0.059 | Old |
| 0.419441 | 0.390503 | 0.254 | 0.211 | 1 | 13 | Snupn | 0.043 | Old |
| 0.419489 | 0.334756 | 0.385 | 0.333 | 1 | 13 | Irgm1 | 0.052 | Old |
| 0.419612 | 0.108543 | 1 | 1 | 1 | 13 | Uqcr11 | 0 | Old |
| 0.419629 | 0.376857 | 0.057 | 0.035 | 1 | 13 | Atg9b | 0.022 | Old |
| 0.419795 | 0.100225 | 0.811 | 0.711 | 1 | 13 | BC017643 | 0.1 | Old |
| 0.420092 | 0.022047 | 0.713 | 0.658 | 1 | 13 | Dr1 | 0.055 | Old |
| 0.421429 | 0.27863 | 0.598 | 0.544 | 1 | 13 | Chka | 0.054 | Old |
| 0.421484 | 0.137176 | 0.238 | 0.184 | 1 | 13 | Nt5dc1 | 0.054 | Old |
| 0.421625 | 0.129685 | 0.516 | 0.447 | 1 | 13 | Iars2 | 0.069 | Old |
| 0.421961 | 0.277408 | 0.213 | 0.167 | 1 | 13 | Nol4l | 0.046 | Old |
| 0.42277 | 0.299698 | 0.549 | 0.447 | 1 | 13 | Dennd1c | 0.102 | Old |
| 0.422993 | 0.188635 | 0.885 | 0.86 | 1 | 13 | Rell1 | 0.025 | Old |
| 0.423185 | 0.071871 | 0.189 | 0.14 | 1 | 13 | Slf1 | 0.049 | Old |
| 0.423625 | 0.338861 | 0.09 | 0.061 | 1 | 13 | Zfp111 | 0.029 | Old |
| 0.4237 | 0.14947 | 0.656 | 0.579 | 1 | 13 | Xrn1 | 0.077 | Old |
| 0.423725 | 0.063377 | 0.68 | 0.588 | 1 | 13 | Slc25a36 | 0.092 | Old |
| 0.423835 | 0.025299 | 0.877 | 0.939 | 1 | 13 | Ccrl2 | -0.062 | Old |
| 0.423963 | 0.403244 | 0.279 | 0.237 | 1 | 13 | Dgcr8 | 0.042 | Old |
| 0.42401 | 0.233451 | 0.508 | 0.474 | 1 | 13 | Grk5 | 0.034 | Old |

| | | | | | | | | |
|---|---|---|---|---|---|---|---|---|
| 0.424157 | 0.100341 | 0.795 | 0.754 | 1 | 13 | Ppp1cc | 0.041 | Old |
| 0.424168 | 0.29519 | 0.131 | 0.096 | 1 | 13 | Gm5086 | 0.035 | Old |
| 0.424624 | 0.200717 | 0.377 | 0.342 | 1 | 13 | Emc4 | 0.035 | Old |
| 0.424755 | 0.1593 | 0.377 | 0.316 | 1 | 13 | Lcor | 0.061 | Old |
| 0.425673 | 0.529777 | 0.057 | 0.035 | 1 | 13 | Ascl2 | 0.022 | Old |
| 0.425673 | 0.378356 | 0.057 | 0.035 | 1 | 13 | Zfp27 | 0.022 | Old |
| 0.425843 | 0.132775 | 0.598 | 0.5 | 1 | 13 | Micu2 | 0.098 | Old |
| 0.426031 | 0.364407 | 0.09 | 0.061 | 1 | 13 | Rbks | 0.029 | Old |
| 0.426047 | 0.412631 | 0.098 | 0.07 | 1 | 13 | Zfp931 | 0.028 | Old |
| 0.426177 | 0.068693 | 0.582 | 0.456 | 1 | 13 | Asl | 0.126 | Old |
| 0.426605 | 0.117043 | 0.902 | 0.912 | 1 | 13 | Tmem189 | -0.01 | Old |
| 0.426823 | 0.572862 | 0.139 | 0.105 | 1 | 13 | Akr1b8 | 0.034 | Old |
| 0.427047 | 0.103838 | 0.41 | 0.342 | 1 | 13 | Hspa13 | 0.068 | Old |
| 0.427365 | 0.231371 | 0.377 | 0.316 | 1 | 13 | Tmem106c | 0.061 | Old |
| 0.42779 | 0.188874 | 0.566 | 0.5 | 1 | 13 | Clcn7 | 0.066 | Old |
| 0.427884 | 0.158902 | 0.91 | 0.816 | 1 | 13 | Arl8a | 0.094 | Old |
| 0.42809 | 0.370317 | 0.434 | 0.404 | 1 | 13 | Frmd4a | 0.03 | Old |
| 0.428123 | 0.153566 | 0.672 | 0.632 | 1 | 13 | Gsdmd | 0.04 | Old |
| 0.428125 | 0.123784 | 0.869 | 0.816 | 1 | 13 | Hprt | 0.053 | Old |
| 0.428337 | 0.219885 | 0.164 | 0.123 | 1 | 13 | Inafm2 | 0.041 | Old |
| 0.428347 | 0.532503 | 0.098 | 0.07 | 1 | 13 | A130010J1 | 0.028 | Old |
| 0.428347 | 0.230924 | 0.098 | 0.07 | 1 | 13 | Fut7 | 0.028 | Old |
| 0.428351 | 0.303162 | 0.205 | 0.167 | 1 | 13 | Fbxo25 | 0.038 | Old |
| 0.428444 | 0.370345 | 0.09 | 0.061 | 1 | 13 | Mtrf1 | 0.029 | Old |
| 0.428444 | 0.370127 | 0.09 | 0.061 | 1 | 13 | Ehbp1 | 0.029 | Old |
| 0.428663 | 0.532215 | 0.148 | 0.114 | 1 | 13 | Dclre1a | 0.034 | Old |
| 0.428714 | 0.355544 | 0.057 | 0.035 | 1 | 13 | A130077B1 | 0.022 | Old |
| 0.428719 | 0.119809 | 0.418 | 0.351 | 1 | 13 | Cyp4f13 | 0.067 | Old |
| 0.428959 | 0.056721 | 0.713 | 0.623 | 1 | 13 | Agfg1 | 0.09 | Old |
| 0.429569 | 0.26116 | 0.172 | 0.132 | 1 | 13 | Nprl2 | 0.04 | Old |
| 0.429734 | 0.001818 | 0.287 | 0.228 | 1 | 13 | Ccdc112 | 0.059 | Old |
| 0.429815 | 0.080183 | 0.516 | 0.412 | 1 | 13 | Trak2 | 0.104 | Old |
| 0.430028 | 0.154434 | 0.607 | 0.509 | 1 | 13 | Osbp | 0.098 | Old |
| 0.430046 | 0.36125 | 0.262 | 0.228 | 1 | 13 | Zfp358 | 0.034 | Old |
| 0.430215 | 0.162 | 0.893 | 0.851 | 1 | 13 | Tpi1 | 0.042 | Old |
| 0.430326 | 0.215102 | 0.68 | 0.649 | 1 | 13 | Twistnb | 0.031 | Old |
| 0.430799 | 0.221438 | 0.238 | 0.184 | 1 | 13 | Atm | 0.054 | Old |
| 0.430866 | 0.375111 | 0.09 | 0.061 | 1 | 13 | Rangrf | 0.029 | Old |
| 0.430866 | 0.146352 | 0.09 | 0.061 | 1 | 13 | C030006K1 | 0.029 | Old |
| 0.431208 | 0.103471 | 0.967 | 0.921 | 1 | 13 | Acer3 | 0.046 | Old |
| 0.431243 | 0.193165 | 0.361 | 0.307 | 1 | 13 | Mob2 | 0.054 | Old |
| 0.431688 | 0.088941 | 0.148 | 0.105 | 1 | 13 | Slc25a26 | 0.043 | Old |
| 0.431724 | 0.102825 | 0.779 | 0.702 | 1 | 13 | Map3k8 | 0.077 | Old |
| 0.431842 | 0.174947 | 0.852 | 0.886 | 1 | 13 | Malt1 | -0.034 | Old |
| 0.432058 | 0.266069 | 0.492 | 0.421 | 1 | 13 | 4833420G1 | 0.071 | Old |

| | | | | | | | | |
|---|---|---|---|---|---|---|---|---|
| 0.432164 | 0.358756 | 0.287 | 0.246 | 1 | 13 | Zc3h7b | 0.041 | Old |
| 0.433295 | 0.212047 | 0.09 | 0.061 | 1 | 13 | Fastkd1 | 0.029 | Old |
| 0.433325 | 0.132065 | 0.418 | 0.36 | 1 | 13 | Nudt5 | 0.058 | Old |
| 0.433538 | 0.220351 | 0.287 | 0.219 | 1 | 13 | Zbed4 | 0.068 | Old |
| 0.433602 | 0.213086 | 0.287 | 0.246 | 1 | 13 | Tmem161a | 0.041 | Old |
| 0.434026 | 0.066112 | 0.582 | 0.518 | 1 | 13 | Nup88 | 0.064 | Old |
| 0.434316 | 0.202588 | 0.451 | 0.386 | 1 | 13 | Zfp655 | 0.065 | Old |
| 0.434494 | 0.274306 | 0.508 | 0.447 | 1 | 13 | Ticam2 | 0.061 | Old |
| 0.435092 | 0.113911 | 0.246 | 0.184 | 1 | 13 | Cox10 | 0.062 | Old |
| 0.435103 | 0.205913 | 0.123 | 0.088 | 1 | 13 | Tnfsfm13 | 0.035 | Old |
| 0.435282 | 0.235018 | 0.492 | 0.439 | 1 | 13 | Hsdl1 | 0.053 | Old |
| 0.435501 | 0.329547 | 0.238 | 0.184 | 1 | 13 | Rnf144b | 0.054 | Old |
| 0.435573 | 0.1579 | 0.746 | 0.667 | 1 | 13 | Lrpap1 | 0.079 | Old |
| 0.435607 | 0.178781 | 0.664 | 0.579 | 1 | 13 | Cant1 | 0.085 | Old |
| 0.435647 | 0.102592 | 0.869 | 0.781 | 1 | 13 | Azin1 | 0.088 | Old |
| 0.435733 | 0.292342 | 0.09 | 0.061 | 1 | 13 | A430106G1 | 0.029 | Old |
| 0.43585 | 0.403359 | 0.23 | 0.184 | 1 | 13 | Naa16 | 0.046 | Old |
| 0.436067 | 0.141895 | 0.197 | 0.149 | 1 | 13 | Asb13 | 0.048 | Old |
| 0.436368 | 0.037695 | 0.32 | 0.246 | 1 | 13 | Hmbs | 0.074 | Old |
| 0.436494 | 0.030827 | 0.541 | 0.588 | 1 | 13 | Gmps | -0.047 | Old |
| 0.436577 | 0.215726 | 0.516 | 0.465 | 1 | 13 | Mrps18a | 0.051 | Old |
| 0.436585 | 0.201945 | 0.213 | 0.167 | 1 | 13 | Prpf3 | 0.046 | Old |
| 0.436846 | 0.173485 | 0.434 | 0.386 | 1 | 13 | Rrp7a | 0.048 | Old |
| 0.437072 | 0.089304 | 0.533 | 0.465 | 1 | 13 | Pnpla2 | 0.068 | Old |
| 0.437114 | 0.054185 | 0.516 | 0.412 | 1 | 13 | Dcun1d1 | 0.104 | Old |
| 0.437359 | 0.355806 | 0.27 | 0.228 | 1 | 13 | Prrg2 | 0.042 | Old |
| 0.437496 | 0.009391 | 0.492 | 0.395 | 1 | 13 | Wdr13 | 0.097 | Old |
| 0.437906 | 0.42088 | 0.418 | 0.368 | 1 | 13 | Kctd12b | 0.05 | Old |
| 0.437912 | 0.317775 | 0.057 | 0.035 | 1 | 13 | Gng10 | 0.022 | Old |
| 0.437912 | 0.2495 | 0.057 | 0.035 | 1 | 13 | Lgi3 | 0.022 | Old |
| 0.438049 | 0.106006 | 0.803 | 0.772 | 1 | 13 | Sec31a | 0.031 | Old |
| 0.43805 | 0.521843 | 0.107 | 0.079 | 1 | 13 | Tmtc4 | 0.028 | Old |
| 0.438178 | 0.328601 | 0.09 | 0.061 | 1 | 13 | Kcnd1 | 0.029 | Old |
| 0.438401 | 0.161137 | 0.328 | 0.272 | 1 | 13 | Sart3 | 0.056 | Old |
| 0.438603 | 0.604393 | 0.066 | 0.044 | 1 | 13 | Gvin1 | 0.022 | Old |
| 0.438607 | 0.290148 | 0.648 | 0.623 | 1 | 13 | H1f0 | 0.025 | Old |
| 0.438651 | 0.10512 | 0.238 | 0.184 | 1 | 13 | Ralgapa2 | 0.054 | Old |
| 0.439104 | 0.085973 | 0.967 | 0.991 | 1 | 13 | Hdgf | -0.024 | Old |
| 0.439569 | 0.201143 | 0.943 | 0.904 | 1 | 13 | Nrp2 | 0.039 | Old |
| 0.439655 | 0.045465 | 0.533 | 0.43 | 1 | 13 | Thoc6 | 0.103 | Old |
| 0.439789 | 0.282706 | 0.344 | 0.289 | 1 | 13 | Isyna1 | 0.055 | Old |
| 0.440176 | 0.143644 | 0.508 | 0.421 | 1 | 13 | Letm1 | 0.087 | Old |
| 0.440249 | 0.210735 | 0.393 | 0.325 | 1 | 13 | Nelfa | 0.068 | Old |
| 0.440961 | 0.113561 | 0.656 | 0.579 | 1 | 13 | Eftud2 | 0.077 | Old |
| 0.441004 | 0.164891 | 0.057 | 0.035 | 1 | 13 | Gm40304 | 0.022 | Old |

| | | | | | | | | |
|---|---|---|---|---|---|---|---|---|
| 0.441188 | 0.09526 | 0.582 | 0.535 | 1 | 13 | Slc12a7 | 0.047 | Old |
| 0.441435 | 0.226545 | 0.123 | 0.088 | 1 | 13 | Endov | 0.035 | Old |
| 0.441461 | 0.600928 | 0.066 | 0.044 | 1 | 13 | Tnfrsf22 | 0.022 | Old |
| 0.441815 | 0.357731 | 0.238 | 0.184 | 1 | 13 | Slc35a1 | 0.054 | Old |
| 0.44224 | 0.140494 | 0.77 | 0.675 | 1 | 13 | Secisbp2l | 0.095 | Old |
| 0.442404 | 0.218475 | 0.131 | 0.096 | 1 | 13 | Usp49 | 0.035 | Old |
| 0.442883 | 0.283946 | 0.279 | 0.237 | 1 | 13 | Mrps10 | 0.042 | Old |
| 0.443469 | 0.281092 | 0.303 | 0.263 | 1 | 13 | Pus1 | 0.04 | Old |
| 0.444107 | 0.22228 | 0.057 | 0.035 | 1 | 13 | Slc24a5 | 0.022 | Old |
| 0.444107 | 0.163951 | 0.057 | 0.035 | 1 | 13 | 2700046G( | 0.022 | Old |
| 0.444107 | 0.100091 | 0.057 | 0.035 | 1 | 13 | Mmachc | 0.022 | Old |
| 0.444173 | 0.066675 | 0.377 | 0.307 | 1 | 13 | Slc38a7 | 0.07 | Old |
| 0.44433 | 0.899132 | 0.066 | 0.044 | 1 | 13 | Vsig8 | 0.022 | Old |
| 0.444367 | 0.170266 | 0.18 | 0.14 | 1 | 13 | Ttc7b | 0.04 | Old |
| 0.444645 | 0.385508 | 0.098 | 0.07 | 1 | 13 | Gm10269 | 0.028 | Old |
| 0.444732 | 0.118682 | 0.951 | 0.939 | 1 | 13 | Herpud1 | 0.012 | Old |
| 0.444761 | 0.042952 | 0.557 | 0.57 | 1 | 13 | Errfi1 | -0.013 | Old |
| 0.445285 | 0.051427 | 0.893 | 0.825 | 1 | 13 | Camkk2 | 0.068 | Old |
| 0.445321 | 0.059817 | 0.959 | 0.956 | 1 | 13 | Tmem30a | 0.003 | Old |
| 0.445344 | 0.344847 | 0.27 | 0.237 | 1 | 13 | Orc3 | 0.033 | Old |
| 0.445381 | 0.093566 | 0.328 | 0.272 | 1 | 13 | Elp3 | 0.056 | Old |
| 0.445561 | 0.118979 | 0.09 | 0.061 | 1 | 13 | Fam84b | 0.029 | Old |
| 0.445907 | 0.074914 | 1 | 1 | 1 | 13 | Ube2d3 | 0 | Old |
| 0.445936 | 0.162626 | 0.615 | 0.526 | 1 | 13 | Pxk | 0.089 | Old |
| 0.446159 | 0.187587 | 0.639 | 0.632 | 1 | 13 | Glipr2 | 0.007 | Old |
| 0.44639 | 0.099131 | 0.516 | 0.43 | 1 | 13 | Eed | 0.086 | Old |
| 0.446425 | 0.219124 | 0.402 | 0.351 | 1 | 13 | Arhgap19 | 0.051 | Old |
| 0.446637 | 0.313575 | 0.156 | 0.123 | 1 | 13 | Tbl2 | 0.033 | Old |
| 0.446789 | 0.258642 | 0.336 | 0.281 | 1 | 13 | Fastk | 0.055 | Old |
| 0.446861 | 0.331092 | 0.369 | 0.316 | 1 | 13 | Rrp36 | 0.053 | Old |
| 0.447002 | 0.300096 | 0.098 | 0.07 | 1 | 13 | 1110002J0 | 0.028 | Old |
| 0.447209 | 0.916748 | 0.066 | 0.044 | 1 | 13 | Serpina3g | 0.022 | Old |
| 0.447223 | 0.188906 | 0.057 | 0.035 | 1 | 13 | Dph2 | 0.022 | Old |
| 0.447364 | 0.138173 | 0.254 | 0.202 | 1 | 13 | 6030458C1 | 0.052 | Old |
| 0.44741 | 0.376047 | 0.221 | 0.184 | 1 | 13 | Ppat | 0.037 | Old |
| 0.447713 | 0.93877 | 0.033 | 0.018 | 1 | 13 | Tlr12 | 0.015 | Old |
| 0.447713 | 0.762784 | 0.033 | 0.018 | 1 | 13 | 9430060I0 | 0.015 | Old |
| 0.447713 | 0.684698 | 0.033 | 0.018 | 1 | 13 | Timp3 | 0.015 | Old |
| 0.447713 | 0.591122 | 0.033 | 0.018 | 1 | 13 | Pole2 | 0.015 | Old |
| 0.448038 | 0.360687 | 0.09 | 0.061 | 1 | 13 | 9030025P2 | 0.029 | Old |
| 0.448131 | 0.140462 | 0.295 | 0.237 | 1 | 13 | Coprs | 0.058 | Old |
| 0.448179 | 0.083396 | 0.967 | 0.965 | 1 | 13 | Anp32a | 0.002 | Old |
| 0.44842 | 0.091958 | 0.779 | 0.684 | 1 | 13 | Coa3 | 0.095 | Old |
| 0.448704 | 0.165991 | 0.484 | 0.43 | 1 | 13 | Aldh3a2 | 0.054 | Old |
| 0.449232 | 0.078794 | 0.393 | 0.316 | 1 | 13 | Bicral | 0.077 | Old |

| | | | | | | | | |
|---|---|---|---|---|---|---|---|---|
| 0.449284 | 0.105326 | 0.852 | 0.798 | 1 | 13 | Strn3 | 0.054 | Old |
| 0.449284 | 0.228754 | 0.803 | 0.851 | 1 | 13 | Fxyd2 | -0.048 | Old |
| 0.449366 | 0.424791 | 0.098 | 0.07 | 1 | 13 | 4833445I0 | 0.028 | Old |
| 0.449782 | 0.213334 | 0.238 | 0.184 | 1 | 13 | Smim24 | 0.054 | Old |
| 0.449849 | 0.153935 | 0.402 | 0.36 | 1 | 13 | Dhx30 | 0.042 | Old |
| 0.450302 | 0.111839 | 0.467 | 0.404 | 1 | 13 | Syne2 | 0.063 | Old |
| 0.450474 | 0.089261 | 0.992 | 0.982 | 1 | 13 | Ptpn1 | 0.01 | Old |
| 0.450619 | 0.172666 | 0.549 | 0.544 | 1 | 13 | 2310011J0 | 0.005 | Old |
| 0.450733 | 0.120469 | 0.689 | 0.632 | 1 | 13 | Tmem9b | 0.057 | Old |
| 0.450747 | 0.119159 | 0.451 | 0.386 | 1 | 13 | Zfp984 | 0.065 | Old |
| 0.450916 | 0.058643 | 0.828 | 0.816 | 1 | 13 | Ppp1r15b | 0.012 | Old |
| 0.450976 | 0.151157 | 0.344 | 0.272 | 1 | 13 | Pias2 | 0.072 | Old |
| 0.450983 | 0.216268 | 0.557 | 0.465 | 1 | 13 | Ahctf1 | 0.092 | Old |
| 0.451563 | 0.129204 | 0.943 | 0.93 | 1 | 13 | Adgre5 | 0.013 | Old |
| 0.451574 | 0.012304 | 0.951 | 0.93 | 1 | 13 | Eif4h | 0.021 | Old |
| 0.451632 | 0.2227 | 0.205 | 0.167 | 1 | 13 | Blzf1 | 0.038 | Old |
| 0.451752 | 0.109027 | 0.27 | 0.219 | 1 | 13 | Pbx1 | 0.051 | Old |
| 0.451909 | 0.864157 | 0.033 | 0.018 | 1 | 13 | Cx3cl1 | 0.015 | Old |
| 0.451909 | 0.666548 | 0.033 | 0.018 | 1 | 13 | Mctp2 | 0.015 | Old |
| 0.451909 | 0.659682 | 0.033 | 0.018 | 1 | 13 | Gpr55 | 0.015 | Old |
| 0.451909 | 0.552647 | 0.033 | 0.018 | 1 | 13 | Prss30 | 0.015 | Old |
| 0.45215 | 0.110211 | 0.861 | 0.86 | 1 | 13 | Nsd1 | 0.001 | Old |
| 0.452177 | 0.078232 | 0.959 | 0.956 | 1 | 13 | Snx6 | 0.003 | Old |
| 0.453091 | 0.508891 | 0.189 | 0.149 | 1 | 13 | Fzd7 | 0.04 | Old |
| 0.453116 | 0.00594 | 0.615 | 0.509 | 1 | 13 | Zfp217 | 0.106 | Old |
| 0.453204 | 0.039082 | 0.721 | 0.667 | 1 | 13 | Cdk14 | 0.054 | Old |
| 0.453317 | 0.345308 | 0.475 | 0.43 | 1 | 13 | Sgk1 | 0.045 | Old |
| 0.453709 | 0.060911 | 0.902 | 0.904 | 1 | 13 | Akt1 | -0.002 | Old |
| 0.453759 | 0.033274 | 0.918 | 0.904 | 1 | 13 | Elavl1 | 0.014 | Old |
| 0.454049 | 0.150365 | 0.861 | 0.807 | 1 | 13 | Thrap3 | 0.054 | Old |
| 0.45432 | 0.265175 | 0.246 | 0.202 | 1 | 13 | Pag1 | 0.044 | Old |
| 0.454444 | 0.173799 | 0.557 | 0.491 | 1 | 13 | Atp13a1 | 0.066 | Old |
| 0.455031 | 0.074623 | 0.959 | 0.93 | 1 | 13 | Nav1 | 0.029 | Old |
| 0.455074 | 0.236856 | 0.689 | 0.623 | 1 | 13 | Ccng1 | 0.066 | Old |
| 0.455212 | 0.73784 | 0.074 | 0.053 | 1 | 13 | Fam109a | 0.021 | Old |
| 0.455212 | 0.693965 | 0.074 | 0.053 | 1 | 13 | Gm16973 | 0.021 | Old |
| 0.455251 | 0.120673 | 0.566 | 0.5 | 1 | 13 | Idi1 | 0.066 | Old |
| 0.455306 | 0.201488 | 0.68 | 0.623 | 1 | 13 | Rap2c | 0.057 | Old |
| 0.455515 | 0.098163 | 0.09 | 0.061 | 1 | 13 | Mast4 | 0.029 | Old |
| 0.456064 | 0.428086 | 0.164 | 0.132 | 1 | 13 | Fpgt | 0.032 | Old |
| 0.456072 | 0.130366 | 0.721 | 0.684 | 1 | 13 | Elk3 | 0.037 | Old |
| 0.456128 | 0.626597 | 0.033 | 0.018 | 1 | 13 | Tlr9 | 0.015 | Old |
| 0.456128 | 0.536727 | 0.033 | 0.018 | 1 | 13 | Usp27x | 0.015 | Old |
| 0.456128 | 0.515753 | 0.033 | 0.018 | 1 | 13 | Farp2 | 0.015 | Old |
| 0.456128 | 0.514731 | 0.033 | 0.018 | 1 | 13 | Zbtb49 | 0.015 | Old |

| | | | | | | | | |
|---|---|---|---|---|---|---|---|---|
| 0.456128 | 0.435786 | 0.033 | 0.018 | 1 | 13 | Dqx1 | 0.015 | Old |
| 0.456612 | 0.228514 | 0.369 | 0.342 | 1 | 13 | Ecd | 0.027 | Old |
| 0.456715 | 0.320213 | 0.59 | 0.518 | 1 | 13 | Dusp2 | 0.072 | Old |
| 0.456941 | 0.019317 | 0.475 | 0.377 | 1 | 13 | Mbd6 | 0.098 | Old |
| 0.457131 | 0.182635 | 0.68 | 0.596 | 1 | 13 | Ddx18 | 0.084 | Old |
| 0.457132 | 0.109886 | 0.623 | 0.544 | 1 | 13 | Plaa | 0.079 | Old |
| 0.457375 | 0.068957 | 1 | 1 | 1 | 13 | Hnrnpa3 | 0 | Old |
| 0.457441 | 0.262732 | 0.467 | 0.395 | 1 | 13 | Atrnl1 | 0.072 | Old |
| 0.458023 | 0.253972 | 0.09 | 0.061 | 1 | 13 | Oplah | 0.029 | Old |
| 0.458025 | 0.177767 | 0.23 | 0.175 | 1 | 13 | Thrb | 0.055 | Old |
| 0.458818 | 0.149484 | 0.557 | 0.5 | 1 | 13 | Atp9b | 0.057 | Old |
| 0.458831 | 0.444596 | 0.066 | 0.044 | 1 | 13 | M1ap | 0.022 | Old |
| 0.458833 | 0.16546 | 0.516 | 0.474 | 1 | 13 | Snapin | 0.042 | Old |
| 0.458893 | 0.28239 | 0.098 | 0.07 | 1 | 13 | Ipp | 0.028 | Old |
| 0.459196 | 0.169186 | 0.467 | 0.404 | 1 | 13 | Clasp1 | 0.063 | Old |
| 0.459262 | 0.178258 | 0.287 | 0.228 | 1 | 13 | Slc7a6os | 0.059 | Old |
| 0.459854 | 0.168531 | 0.516 | 0.465 | 1 | 13 | Smurf1 | 0.051 | Old |
| 0.45995 | 0.253466 | 0.295 | 0.237 | 1 | 13 | Caml | 0.058 | Old |
| 0.459989 | 0.359376 | 0.377 | 0.333 | 1 | 13 | Nde1 | 0.044 | Old |
| 0.460368 | 0.534412 | 0.033 | 0.018 | 1 | 13 | 1200007C1 | 0.015 | Old |
| 0.460368 | 0.498468 | 0.033 | 0.018 | 1 | 13 | Chek1 | 0.015 | Old |
| 0.460368 | 0.478146 | 0.033 | 0.018 | 1 | 13 | Mcm8 | 0.015 | Old |
| 0.460368 | 0.452385 | 0.033 | 0.018 | 1 | 13 | Cd22 | 0.015 | Old |
| 0.460368 | 0.402482 | 0.033 | 0.018 | 1 | 13 | B3gnt9 | 0.015 | Old |
| 0.460368 | 0.397577 | 0.033 | 0.018 | 1 | 13 | Penk | 0.015 | Old |
| 0.460368 | 0.373501 | 0.033 | 0.018 | 1 | 13 | Xkr6 | 0.015 | Old |
| 0.460368 | 0.358439 | 0.033 | 0.018 | 1 | 13 | Gpr18 | 0.015 | Old |
| 0.460368 | 0.266754 | 0.033 | 0.018 | 1 | 13 | Tm4sf1 | 0.015 | Old |
| 0.460368 | 0.134704 | 0.033 | 0.018 | 1 | 13 | Prodh | 0.015 | Old |
| 0.460368 | 0.101457 | 0.033 | 0.018 | 1 | 13 | Cenpn | 0.015 | Old |
| 0.460472 | 0.339155 | 0.434 | 0.395 | 1 | 13 | Ppfia1 | 0.039 | Old |
| 0.460831 | 0.002465 | 0.967 | 0.947 | 1 | 13 | Sppl2a | 0.02 | Old |
| 0.460901 | 0.119112 | 0.844 | 0.719 | 1 | 13 | Dctn1 | 0.125 | Old |
| 0.461031 | 0.212439 | 0.443 | 0.395 | 1 | 13 | Flot1 | 0.048 | Old |
| 0.4613 | 0.205766 | 0.623 | 0.596 | 1 | 13 | Mfsd11 | 0.027 | Old |
| 0.461378 | 0.341698 | 0.238 | 0.202 | 1 | 13 | Exoc6 | 0.036 | Old |
| 0.461594 | 0.458824 | 0.148 | 0.114 | 1 | 13 | Cald1 | 0.034 | Old |
| 0.461892 | 0.072168 | 0.27 | 0.211 | 1 | 13 | Aifm2 | 0.059 | Old |
| 0.461951 | 0.081678 | 0.951 | 0.921 | 1 | 13 | Etfb | 0.03 | Old |
| 0.462112 | 0.155102 | 0.525 | 0.447 | 1 | 13 | Smim19 | 0.078 | Old |
| 0.462238 | 0.2651 | 0.303 | 0.263 | 1 | 13 | Nt5dc3 | 0.04 | Old |
| 0.46249 | 0.184408 | 0.943 | 0.904 | 1 | 13 | Degs1 | 0.039 | Old |
| 0.462973 | 0.077573 | 0.639 | 0.544 | 1 | 13 | Osbpl11 | 0.095 | Old |
| 0.463171 | 0.150205 | 1 | 1 | 1 | 13 | Gm42418 | 0 | Old |
| 0.463401 | 0.100744 | 0.885 | 0.789 | 1 | 13 | Ctbp1 | 0.096 | Old |

| | | | | | | | | |
|---|---|---|---|---|---|---|---|---|
| 0.463832 | 0.19645 | 0.361 | 0.307 | 1 | 13 | Pitrm1 | 0.054 | Old |
| 0.463866 | 0.129236 | 0.656 | 0.614 | 1 | 13 | Pptc7 | 0.042 | Old |
| 0.464326 | 0.048926 | 0.787 | 0.711 | 1 | 13 | Dcun1d5 | 0.076 | Old |
| 0.464631 | 0.887374 | 0.033 | 0.018 | 1 | 13 | Ccr7 | 0.015 | Old |
| 0.464631 | 0.467141 | 0.033 | 0.018 | 1 | 13 | Satb1 | 0.015 | Old |
| 0.464631 | 0.410562 | 0.033 | 0.018 | 1 | 13 | Klrg2 | 0.015 | Old |
| 0.464631 | 0.380566 | 0.033 | 0.018 | 1 | 13 | Mss51 | 0.015 | Old |
| 0.464631 | 0.349681 | 0.033 | 0.018 | 1 | 13 | 2310068J1 | 0.015 | Old |
| 0.464631 | 0.199946 | 0.033 | 0.018 | 1 | 13 | Eda | 0.015 | Old |
| 0.464631 | 0.109849 | 0.033 | 0.018 | 1 | 13 | Serpinh1 | 0.015 | Old |
| 0.464659 | 0.143903 | 0.68 | 0.579 | 1 | 13 | Utrn | 0.101 | Old |
| 0.464755 | 0.139205 | 0.549 | 0.491 | 1 | 13 | Rhoh | 0.058 | Old |
| 0.465102 | 0.159126 | 0.443 | 0.377 | 1 | 13 | Apex1 | 0.066 | Old |
| 0.465502 | 0.102355 | 0.213 | 0.175 | 1 | 13 | Ahdc1 | 0.038 | Old |
| 0.465925 | 0.236097 | 0.295 | 0.237 | 1 | 13 | Psmf1 | 0.058 | Old |
| 0.466285 | 0.124574 | 0.23 | 0.175 | 1 | 13 | Mars | 0.055 | Old |
| 0.466645 | 0.112364 | 0.951 | 0.956 | 1 | 13 | Dnajb6 | -0.005 | Old |
| 0.467672 | 0.22973 | 0.369 | 0.307 | 1 | 13 | Brf1 | 0.062 | Old |
| 0.467733 | 0.281769 | 0.557 | 0.474 | 1 | 13 | Camk2d | 0.083 | Old |
| 0.467833 | 0.216173 | 0.426 | 0.377 | 1 | 13 | Med25 | 0.049 | Old |
| 0.468254 | 0.065316 | 0.672 | 0.588 | 1 | 13 | Polr2l | 0.084 | Old |
| 0.468562 | 0.172178 | 0.393 | 0.333 | 1 | 13 | Vopp1 | 0.06 | Old |
| 0.468691 | 0.227073 | 0.205 | 0.167 | 1 | 13 | Trmt13 | 0.038 | Old |
| 0.468915 | 0.65673 | 0.033 | 0.018 | 1 | 13 | Nrg1 | 0.015 | Old |
| 0.468915 | 0.366206 | 0.033 | 0.018 | 1 | 13 | Gbgt1 | 0.015 | Old |
| 0.468915 | 0.361501 | 0.033 | 0.018 | 1 | 13 | Gm49085 | 0.015 | Old |
| 0.468915 | 0.329248 | 0.033 | 0.018 | 1 | 13 | Mterf1b | 0.015 | Old |
| 0.468915 | 0.327339 | 0.033 | 0.018 | 1 | 13 | Gm26947 | 0.015 | Old |
| 0.468915 | 0.261381 | 0.033 | 0.018 | 1 | 13 | Kntc1 | 0.015 | Old |
| 0.468915 | 0.260204 | 0.033 | 0.018 | 1 | 13 | Mypop | 0.015 | Old |
| 0.468915 | 0.253203 | 0.033 | 0.018 | 1 | 13 | Gm12159 | 0.015 | Old |
| 0.468915 | 0.249051 | 0.033 | 0.018 | 1 | 13 | Zfp119a | 0.015 | Old |
| 0.468915 | 0.248098 | 0.033 | 0.018 | 1 | 13 | Mst1r | 0.015 | Old |
| 0.468915 | 0.204725 | 0.033 | 0.018 | 1 | 13 | A330084C1 | 0.015 | Old |
| 0.468915 | 0.186936 | 0.033 | 0.018 | 1 | 13 | Zfp37 | 0.015 | Old |
| 0.469278 | 0.115548 | 0.762 | 0.649 | 1 | 13 | Agps | 0.113 | Old |
| 0.469374 | 0.28061 | 0.336 | 0.281 | 1 | 13 | Adora2b | 0.055 | Old |
| 0.469722 | 0.163761 | 0.607 | 0.57 | 1 | 13 | Isy1 | 0.037 | Old |
| 0.469926 | 0.31825 | 0.197 | 0.158 | 1 | 13 | Bloc1s5 | 0.039 | Old |
| 0.470621 | 0.278267 | 0.066 | 0.044 | 1 | 13 | Eid2 | 0.022 | Old |
| 0.470859 | 0.13493 | 0.861 | 0.798 | 1 | 13 | Nufip2 | 0.063 | Old |
| 0.47097 | 0.373797 | 0.443 | 0.421 | 1 | 13 | 1700017B0 | 0.022 | Old |
| 0.471099 | 0.131831 | 0.32 | 0.254 | 1 | 13 | Mphosph6 | 0.066 | Old |
| 0.471456 | 0.203368 | 0.18 | 0.14 | 1 | 13 | Zbtb14 | 0.04 | Old |
| 0.471588 | 0.219952 | 0.156 | 0.123 | 1 | 13 | BC017158 | 0.033 | Old |

| | | | | | | | | |
|---|---|---|---|---|---|---|---|---|
| 0.471708 | 0.087979 | 0.926 | 0.868 | 1 | 13 | Cwc15 | 0.058 | Old |
| 0.471805 | 0.124207 | 0.852 | 0.851 | 1 | 13 | Mlf2 | 0.001 | Old |
| 0.471816 | 0.074056 | 0.721 | 0.623 | 1 | 13 | Rsbn1l | 0.098 | Old |
| 0.471876 | 0.214187 | 0.344 | 0.289 | 1 | 13 | Nfrkb | 0.055 | Old |
| 0.472122 | 0.139427 | 0.721 | 0.632 | 1 | 13 | Idnk | 0.089 | Old |
| 0.472163 | 0.234289 | 0.508 | 0.465 | 1 | 13 | Phf11b | 0.043 | Old |
| 0.472297 | 0.420772 | 0.107 | 0.079 | 1 | 13 | Ptar1 | 0.028 | Old |
| 0.472575 | 0.333752 | 0.311 | 0.272 | 1 | 13 | Aga | 0.039 | Old |
| 0.473023 | 0.268261 | 0.402 | 0.351 | 1 | 13 | Mut | 0.051 | Old |
| 0.473221 | 0.325437 | 0.033 | 0.018 | 1 | 13 | Dynlt1c | 0.015 | Old |
| 0.473221 | 0.2996 | 0.033 | 0.018 | 1 | 13 | 9130023H2 | 0.015 | Old |
| 0.473221 | 0.250503 | 0.033 | 0.018 | 1 | 13 | Dnajb13 | 0.015 | Old |
| 0.473221 | 0.247233 | 0.033 | 0.018 | 1 | 13 | Nup210l | 0.015 | Old |
| 0.473221 | 0.187001 | 0.033 | 0.018 | 1 | 13 | Tnfrsf8 | 0.015 | Old |
| 0.473221 | 0.161145 | 0.033 | 0.018 | 1 | 13 | Dtl | 0.015 | Old |
| 0.473221 | 0.065066 | 0.033 | 0.018 | 1 | 13 | Krba1 | 0.015 | Old |
| 0.473221 | 0.055572 | 0.033 | 0.018 | 1 | 13 | Akap5 | 0.015 | Old |
| 0.47344 | 0.213789 | 0.246 | 0.202 | 1 | 13 | Hars2 | 0.044 | Old |
| 0.473446 | 0.111384 | 0.574 | 0.465 | 1 | 13 | Naa60 | 0.109 | Old |
| 0.473539 | 0.217401 | 0.566 | 0.482 | 1 | 13 | Arl3 | 0.084 | Old |
| 0.47354 | 0.233875 | 0.172 | 0.132 | 1 | 13 | Def8 | 0.04 | Old |
| 0.473544 | 0.11616 | 0.844 | 0.789 | 1 | 13 | Dip2b | 0.055 | Old |
| 0.473594 | 0.353117 | 0.066 | 0.044 | 1 | 13 | Hist2h2ac | 0.022 | Old |
| 0.473594 | 0.345101 | 0.066 | 0.044 | 1 | 13 | Wdr73 | 0.022 | Old |
| 0.473635 | 0.137052 | 0.393 | 0.316 | 1 | 13 | Xpo4 | 0.077 | Old |
| 0.473683 | 0.087733 | 0.959 | 0.947 | 1 | 13 | Manf | 0.012 | Old |
| 0.473705 | 0.022331 | 1 | 1 | 1 | 13 | Mt1 | 0 | Old |
| 0.473735 | 0.289407 | 0.557 | 0.518 | 1 | 13 | Atic | 0.039 | Old |
| 0.473771 | 0.163855 | 0.131 | 0.096 | 1 | 13 | Ranbp6 | 0.035 | Old |
| 0.47383 | 0.118697 | 0.434 | 0.36 | 1 | 13 | Smc5 | 0.074 | Old |
| 0.474025 | 0.122209 | 0.508 | 0.421 | 1 | 13 | Exoc2 | 0.087 | Old |
| 0.474441 | 0.556748 | 0.189 | 0.158 | 1 | 13 | Cables1 | 0.031 | Old |
| 0.47468 | 0.051687 | 0.926 | 0.877 | 1 | 13 | Sumo1 | 0.049 | Old |
| 0.475046 | 0.52873 | 0.648 | 0.675 | 1 | 13 | Ccnd2 | -0.027 | Old |
| 0.475213 | 0.096443 | 0.893 | 0.789 | 1 | 13 | Usp34 | 0.104 | Old |
| 0.475465 | 0.244794 | 0.607 | 0.544 | 1 | 13 | Txndc11 | 0.063 | Old |
| 0.475494 | 0.319383 | 0.156 | 0.123 | 1 | 13 | Mcrip2 | 0.033 | Old |
| 0.475782 | 0.25905 | 0.418 | 0.386 | 1 | 13 | Bahd1 | 0.032 | Old |
| 0.476371 | 0.203726 | 0.738 | 0.719 | 1 | 13 | Golga7 | 0.019 | Old |
| 0.476578 | 0.368191 | 0.066 | 0.044 | 1 | 13 | Fastkd3 | 0.022 | Old |
| 0.476578 | 0.330767 | 0.066 | 0.044 | 1 | 13 | 1700037C1 | 0.022 | Old |
| 0.476578 | 0.320695 | 0.066 | 0.044 | 1 | 13 | Cox6a2 | 0.022 | Old |
| 0.476578 | 0.261658 | 0.066 | 0.044 | 1 | 13 | Mettl22 | 0.022 | Old |
| 0.477089 | 0.304336 | 0.139 | 0.105 | 1 | 13 | Acy1 | 0.034 | Old |
| 0.477163 | 0.115548 | 0.811 | 0.781 | 1 | 13 | Specc1l | 0.03 | Old |

| | | | | | | | | |
|---|---|---|---|---|---|---|---|---|
| 0.477518 | 0.094579 | 0.607 | 0.5 | 1 | 13 | Ccny | 0.107 | Old |
| 0.477549 | 0.297322 | 0.033 | 0.018 | 1 | 13 | 4933406I1ε | 0.015 | Old |
| 0.477549 | 0.27695 | 0.033 | 0.018 | 1 | 13 | Kyat3 | 0.015 | Old |
| 0.477549 | 0.246903 | 0.033 | 0.018 | 1 | 13 | Bbs5 | 0.015 | Old |
| 0.477549 | 0.190429 | 0.033 | 0.018 | 1 | 13 | Rbm8a2 | 0.015 | Old |
| 0.477549 | 0.171074 | 0.033 | 0.018 | 1 | 13 | Hist4h4 | 0.015 | Old |
| 0.478163 | 0.236442 | 0.779 | 0.675 | 1 | 13 | Wdfy4 | 0.104 | Old |
| 0.478226 | 0.076558 | 0.902 | 0.904 | 1 | 13 | Tbcb | -0.002 | Old |
| 0.478274 | 0.112223 | 0.918 | 0.904 | 1 | 13 | Cox17 | 0.014 | Old |
| 0.478295 | 0.128248 | 0.328 | 0.272 | 1 | 13 | Mrps11 | 0.056 | Old |
| 0.478506 | 0.143882 | 0.525 | 0.447 | 1 | 13 | Virma | 0.078 | Old |
| 0.478667 | 0.148293 | 0.648 | 0.579 | 1 | 13 | Mras | 0.069 | Old |
| 0.478997 | 0.228247 | 0.443 | 0.412 | 1 | 13 | Grsf1 | 0.031 | Old |
| 0.479045 | 0.181838 | 0.402 | 0.342 | 1 | 13 | Vti1a | 0.06 | Old |
| 0.479319 | 0.280237 | 0.107 | 0.079 | 1 | 13 | Ppp1r18os | 0.028 | Old |
| 0.479572 | 0.324234 | 0.066 | 0.044 | 1 | 13 | Zfp341 | 0.022 | Old |
| 0.479572 | 0.305934 | 0.066 | 0.044 | 1 | 13 | Wdr25 | 0.022 | Old |
| 0.479572 | 0.184513 | 0.066 | 0.044 | 1 | 13 | Xkr8 | 0.022 | Old |
| 0.479635 | 0.164137 | 0.811 | 0.711 | 1 | 13 | Plekha2 | 0.1 | Old |
| 0.479704 | 0.32527 | 0.426 | 0.386 | 1 | 13 | Brip1os | 0.04 | Old |
| 0.4801 | 0.151586 | 0.377 | 0.316 | 1 | 13 | Camta1 | 0.061 | Old |
| 0.480565 | 0.086507 | 0.156 | 0.114 | 1 | 13 | Slx4 | 0.042 | Old |
| 0.480649 | 0.087537 | 0.664 | 0.632 | 1 | 13 | Ormdl2 | 0.032 | Old |
| 0.4807 | 0.170795 | 0.467 | 0.421 | 1 | 13 | Usp5 | 0.046 | Old |
| 0.480764 | 0.132914 | 0.943 | 0.947 | 1 | 13 | Tomm20 | -0.004 | Old |
| 0.480806 | 0.097678 | 0.689 | 0.64 | 1 | 13 | Smim10l1 | 0.049 | Old |
| 0.48084 | 0.243964 | 0.205 | 0.167 | 1 | 13 | Numb | 0.038 | Old |
| 0.481083 | 0.472463 | 0.664 | 0.658 | 1 | 13 | Hk2 | 0.006 | Old |
| 0.481736 | 0.219687 | 0.492 | 0.439 | 1 | 13 | Zscan26 | 0.053 | Old |
| 0.481969 | 0.259667 | 0.295 | 0.246 | 1 | 13 | Agmo | 0.049 | Old |
| 0.481986 | 0.168935 | 0.451 | 0.386 | 1 | 13 | Abraxas2 | 0.065 | Old |
| 0.482512 | 0.194692 | 0.467 | 0.404 | 1 | 13 | Fam219b | 0.063 | Old |
| 0.482533 | 0.22383 | 0.336 | 0.272 | 1 | 13 | Pole3 | 0.064 | Old |
| 0.482549 | 0.132076 | 0.262 | 0.211 | 1 | 13 | Eif2b3 | 0.051 | Old |
| 0.482576 | 0.235126 | 0.066 | 0.044 | 1 | 13 | Zfp438 | 0.022 | Old |
| 0.482576 | 0.221779 | 0.066 | 0.044 | 1 | 13 | Mfsd6l | 0.022 | Old |
| 0.482576 | 0.21528 | 0.066 | 0.044 | 1 | 13 | Havcr2 | 0.022 | Old |
| 0.482576 | 0.004276 | 0.066 | 0.044 | 1 | 13 | Xlr3b | 0.022 | Old |
| 0.482909 | 0.677609 | 0.074 | 0.053 | 1 | 13 | Wdr62 | 0.021 | Old |
| 0.483046 | 0.132302 | 0.934 | 0.895 | 1 | 13 | 9530068E0 | 0.039 | Old |
| 0.483191 | 0.0385 | 0.689 | 0.553 | 1 | 13 | Chtf8 | 0.136 | Old |
| 0.483289 | 0.12454 | 0.549 | 0.474 | 1 | 13 | Tbk1 | 0.075 | Old |
| 0.483331 | 0.247714 | 0.139 | 0.105 | 1 | 13 | Efemp2 | 0.034 | Old |
| 0.483561 | 0.112177 | 0.664 | 0.614 | 1 | 13 | Rnf114 | 0.05 | Old |
| 0.485138 | 0.126591 | 0.803 | 0.807 | 1 | 13 | Ubn1 | -0.004 | Old |

| | | | | | | | | |
|---|---|---|---|---|---|---|---|---|
| 0.48535 | 0.213324 | 0.189 | 0.149 | 1 | 13 | Tmem185a | 0.04 | Old |
| 0.48559 | 0.238655 | 0.066 | 0.044 | 1 | 13 | Zgrf1 | 0.022 | Old |
| 0.48559 | 0.234705 | 0.066 | 0.044 | 1 | 13 | Klhl11 | 0.022 | Old |
| 0.485745 | 0.082494 | 0.262 | 0.211 | 1 | 13 | Ift22 | 0.051 | Old |
| 0.485825 | 0.187733 | 0.533 | 0.491 | 1 | 13 | Atg12 | 0.042 | Old |
| 0.486133 | 0.081271 | 0.443 | 0.386 | 1 | 13 | Cwf19l2 | 0.057 | Old |
| 0.486685 | 0.219603 | 0.41 | 0.36 | 1 | 13 | Iars | 0.05 | Old |
| 0.487016 | 0.071123 | 0.738 | 0.667 | 1 | 13 | Pole4 | 0.071 | Old |
| 0.487094 | 0.108863 | 0.328 | 0.272 | 1 | 13 | Zfp830 | 0.056 | Old |
| 0.487162 | 0.163631 | 0.697 | 0.623 | 1 | 13 | Ythdf2 | 0.074 | Old |
| 0.487162 | 0.142171 | 0.68 | 0.64 | 1 | 13 | Zfr | 0.04 | Old |
| 0.487228 | 0.133882 | 0.68 | 0.596 | 1 | 13 | Yipf4 | 0.084 | Old |
| 0.487353 | 0.136248 | 0.762 | 0.728 | 1 | 13 | Impact | 0.034 | Old |
| 0.487693 | 0.276453 | 0.197 | 0.158 | 1 | 13 | Usp31 | 0.039 | Old |
| 0.487693 | 0.242832 | 0.197 | 0.158 | 1 | 13 | Bag4 | 0.039 | Old |
| 0.487739 | 0.082042 | 0.82 | 0.807 | 1 | 13 | Osbpl9 | 0.013 | Old |
| 0.488066 | 0.243274 | 0.246 | 0.202 | 1 | 13 | Klc4 | 0.044 | Old |
| 0.48809 | 0.088979 | 0.516 | 0.43 | 1 | 13 | Phyh | 0.086 | Old |
| 0.48832 | 0.37959 | 0.115 | 0.088 | 1 | 13 | Zfp958 | 0.027 | Old |
| 0.48832 | 0.358535 | 0.115 | 0.088 | 1 | 13 | Piwil2 | 0.027 | Old |
| 0.488749 | 0.125342 | 0.582 | 0.491 | 1 | 13 | Sfswap | 0.091 | Old |
| 0.488768 | 0.019221 | 0.508 | 0.588 | 1 | 13 | Ncbp2 | -0.08 | Old |
| 0.488945 | 0.191404 | 0.721 | 0.711 | 1 | 13 | Alkbh5 | 0.01 | Old |
| 0.488951 | 0.06345 | 0.91 | 0.895 | 1 | 13 | Galnt1 | 0.015 | Old |
| 0.489009 | 0.215557 | 0.189 | 0.149 | 1 | 13 | Tars2 | 0.04 | Old |
| 0.489395 | 0.297989 | 0.336 | 0.289 | 1 | 13 | Sntb2 | 0.047 | Old |
| 0.489541 | 0.162763 | 0.779 | 0.763 | 1 | 13 | Tnpo3 | 0.016 | Old |
| 0.49001 | 0.092654 | 0.467 | 0.395 | 1 | 13 | Rab9 | 0.072 | Old |
| 0.490218 | 0.185703 | 0.615 | 0.588 | 1 | 13 | Mybbp1a | 0.027 | Old |
| 0.49045 | 0.021868 | 0.311 | 0.254 | 1 | 13 | Fam13b | 0.057 | Old |
| 0.49081 | 0.117105 | 0.533 | 0.482 | 1 | 13 | Chuk | 0.051 | Old |
| 0.491107 | 0.102561 | 0.902 | 0.842 | 1 | 13 | 2-Mar | 0.06 | Old |
| 0.491293 | 0.25658 | 0.156 | 0.123 | 1 | 13 | Cox15 | 0.033 | Old |
| 0.491548 | 0.097264 | 0.992 | 0.956 | 1 | 13 | Ndufc1 | 0.036 | Old |
| 0.491649 | 0.281231 | 0.066 | 0.044 | 1 | 13 | Gm26569 | 0.022 | Old |
| 0.491687 | 0.171209 | 0.32 | 0.272 | 1 | 13 | Dnajc16 | 0.048 | Old |
| 0.491786 | 0.51572 | 0.082 | 0.061 | 1 | 13 | Smco4 | 0.021 | Old |
| 0.491842 | 0.05183 | 0.607 | 0.509 | 1 | 13 | Bola3 | 0.098 | Old |
| 0.491919 | 0.053414 | 0.877 | 0.807 | 1 | 13 | Ccar1 | 0.07 | Old |
| 0.492047 | 0.010386 | 0.369 | 0.325 | 1 | 13 | Smpd1 | 0.044 | Old |
| 0.492465 | 0.069857 | 0.885 | 0.895 | 1 | 13 | Git2 | -0.01 | Old |
| 0.492479 | 0.330473 | 0.189 | 0.158 | 1 | 13 | Pold1 | 0.031 | Old |
| 0.492497 | 0.061642 | 0.492 | 0.43 | 1 | 13 | Nsrp1 | 0.062 | Old |
| 0.492995 | 0.486016 | 0.238 | 0.211 | 1 | 13 | Srbd1 | 0.027 | Old |
| 0.493533 | 0.214714 | 0.107 | 0.079 | 1 | 13 | Dhfr | 0.028 | Old |

| | | | | | | | | |
|---|---|---|---|---|---|---|---|---|
| 0.493615 | 0.111111 | 0.705 | 0.64 | 1 | 13 | Ssfa2 | 0.065 | Old |
| 0.493837 | 0.15544 | 0.697 | 0.605 | 1 | 13 | Bcat2 | 0.092 | Old |
| 0.494238 | 0.243319 | 0.074 | 0.053 | 1 | 13 | Zfp748 | 0.021 | Old |
| 0.494573 | 0.081474 | 0.361 | 0.298 | 1 | 13 | Mrps35 | 0.063 | Old |
| 0.494694 | 0.25686 | 0.066 | 0.044 | 1 | 13 | Gm42715 | 0.022 | Old |
| 0.494694 | 0.157193 | 0.066 | 0.044 | 1 | 13 | Cdc6 | 0.022 | Old |
| 0.494905 | 0.013377 | 0.426 | 0.342 | 1 | 13 | Rbm12 | 0.084 | Old |
| 0.495026 | 0.181917 | 0.23 | 0.193 | 1 | 13 | Mkrn2 | 0.037 | Old |
| 0.495164 | 0.038938 | 1 | 1 | 1 | 13 | Csde1 | 0 | Old |
| 0.495402 | 0.19741 | 0.418 | 0.377 | 1 | 13 | Sdad1 | 0.041 | Old |
| 0.495403 | 0.107531 | 0.803 | 0.693 | 1 | 13 | Ctdsp1 | 0.11 | Old |
| 0.495591 | 0.083868 | 0.377 | 0.316 | 1 | 13 | Ciapin1 | 0.061 | Old |
| 0.495608 | 0.217278 | 0.098 | 0.07 | 1 | 13 | Mettl17 | 0.028 | Old |
| 0.496052 | 0.091209 | 0.787 | 0.746 | 1 | 13 | Prkcb | 0.041 | Old |
| 0.497576 | 0.111736 | 0.967 | 0.982 | 1 | 13 | Atp6v1e1 | -0.015 | Old |
| 0.497749 | 0.152064 | 0.066 | 0.044 | 1 | 13 | Tmem41a | 0.022 | Old |
| 0.497877 | 0.209503 | 0.287 | 0.237 | 1 | 13 | Ccdc122 | 0.05 | Old |
| 0.498089 | 0.379481 | 0.123 | 0.096 | 1 | 13 | Unk | 0.027 | Old |
| 0.49832 | 0.191119 | 0.107 | 0.079 | 1 | 13 | AU041133 | 0.028 | Old |
| 0.498366 | 0.227257 | 0.402 | 0.342 | 1 | 13 | H2-Q6 | 0.06 | Old |
| 0.498472 | 0.238561 | 0.172 | 0.14 | 1 | 13 | Mus81 | 0.032 | Old |
| 0.498723 | 0.071091 | 0.951 | 0.912 | 1 | 13 | Cndp2 | 0.039 | Old |
| 0.498797 | 0.071815 | 0.828 | 0.763 | 1 | 13 | Vapb | 0.065 | Old |
| 0.49924 | 0.051973 | 0.885 | 0.807 | 1 | 13 | Sec61a1 | 0.078 | Old |
| 0.499916 | 0.135794 | 0.828 | 0.807 | 1 | 13 | Ube2k | 0.021 | Old |
| 0.499923 | 0.139422 | 0.779 | 0.702 | 1 | 13 | Cers6 | 0.077 | Old |
| 0.49995 | 0.063696 | 0.762 | 0.693 | 1 | 13 | Tomm70a | 0.069 | Old |
| 0.500002 | 0.035954 | 0.984 | 1 | 1 | 13 | Tma7 | -0.016 | Old |
| 0.50027 | 0.270196 | 0.631 | 0.623 | 1 | 13 | Amdhd2 | 0.008 | Old |
| 0.500365 | 0.066817 | 0.607 | 0.544 | 1 | 13 | Psen1 | 0.063 | Old |
| 0.500723 | 0.241105 | 0.107 | 0.079 | 1 | 13 | Zfp658 | 0.028 | Old |
| 0.500882 | 0.163427 | 0.361 | 0.316 | 1 | 13 | Avl9 | 0.045 | Old |
| 0.501049 | 0.173613 | 0.648 | 0.623 | 1 | 13 | Hist1h1c | 0.025 | Old |
| 0.501155 | 0.107637 | 0.418 | 0.342 | 1 | 13 | Cox19 | 0.076 | Old |
| 0.501683 | 0.008818 | 0.795 | 0.746 | 1 | 13 | Vamp3 | 0.049 | Old |
| 0.502047 | 0.266696 | 0.164 | 0.132 | 1 | 13 | Pcyox1l | 0.032 | Old |
| 0.502062 | 0.041625 | 0.205 | 0.167 | 1 | 13 | Mtg1 | 0.038 | Old |
| 0.502145 | 0.062078 | 0.877 | 0.798 | 1 | 13 | Ppp2r1a | 0.079 | Old |
| 0.5026 | 0.028306 | 0.262 | 0.307 | 1 | 13 | Tns4 | -0.045 | Old |
| 0.502859 | 0.123306 | 0.697 | 0.632 | 1 | 13 | Ano6 | 0.065 | Old |
| 0.503043 | 0.165613 | 0.369 | 0.307 | 1 | 13 | Timm9 | 0.062 | Old |
| 0.503328 | 0.149365 | 0.623 | 0.579 | 1 | 13 | Ebi3 | 0.044 | Old |
| 0.503389 | 0.11951 | 0.27 | 0.219 | 1 | 13 | Dhx33 | 0.051 | Old |
| 0.503451 | 0.072189 | 0.328 | 0.272 | 1 | 13 | Extl3 | 0.056 | Old |
| 0.503544 | 0.139435 | 0.221 | 0.175 | 1 | 13 | Pus10 | 0.046 | Old |

| | | | | | | | | |
|---|---|---|---|---|---|---|---|---|
| 0.503853 | 0.215664 | 0.434 | 0.386 | 1 | 13 | Mgst3 | 0.048 | Old |
| 0.503888 | 0.147414 | 0.066 | 0.044 | 1 | 13 | 1700113A1 | 0.022 | Old |
| 0.503996 | 0.067162 | 0.746 | 0.675 | 1 | 13 | Ppp3cb | 0.071 | Old |
| 0.504668 | 0.149696 | 0.902 | 0.904 | 1 | 13 | Ndufb6 | -0.002 | Old |
| 0.504839 | 0.14245 | 0.762 | 0.667 | 1 | 13 | Rmnd5a | 0.095 | Old |
| 0.50527 | 0.305778 | 0.566 | 0.526 | 1 | 13 | Slc28a2 | 0.04 | Old |
| 0.505291 | 0.172808 | 0.221 | 0.175 | 1 | 13 | Tom1l2 | 0.046 | Old |
| 0.505346 | 0.139693 | 0.156 | 0.123 | 1 | 13 | Btbd3 | 0.033 | Old |
| 0.505575 | 0.200021 | 0.254 | 0.211 | 1 | 13 | Bcas3 | 0.043 | Old |
| 0.505657 | 0.066353 | 0.098 | 0.07 | 1 | 13 | Taf1c | 0.028 | Old |
| 0.505708 | 0.400733 | 0.074 | 0.053 | 1 | 13 | Olfr1167 | 0.021 | Old |
| 0.505708 | 0.356898 | 0.074 | 0.053 | 1 | 13 | 4930524J0 | 0.021 | Old |
| 0.505752 | 0.25771 | 0.303 | 0.254 | 1 | 13 | Tusc2 | 0.049 | Old |
| 0.505808 | 0.05133 | 0.418 | 0.368 | 1 | 13 | E2f4 | 0.05 | Old |
| 0.505946 | 0.351567 | 0.246 | 0.211 | 1 | 13 | Slc39a13 | 0.035 | Old |
| 0.505948 | 0.169361 | 0.525 | 0.465 | 1 | 13 | Pmm2 | 0.06 | Old |
| 0.506584 | 0.146311 | 0.795 | 0.746 | 1 | 13 | Srp19 | 0.049 | Old |
| 0.506594 | 0.11747 | 0.139 | 0.105 | 1 | 13 | Qsox2 | 0.034 | Old |
| 0.506766 | 0.38236 | 0.115 | 0.088 | 1 | 13 | Tepsin | 0.027 | Old |
| 0.506895 | 0.199595 | 0.689 | 0.649 | 1 | 13 | Tnfrsf11a | 0.04 | Old |
| 0.507308 | 0.168889 | 0.738 | 0.693 | 1 | 13 | Cux1 | 0.045 | Old |
| 0.507364 | 0.111957 | 0.385 | 0.333 | 1 | 13 | Pik3c3 | 0.052 | Old |
| 0.507438 | 0.178272 | 0.607 | 0.535 | 1 | 13 | Smad2 | 0.072 | Old |
| 0.507446 | 0.095754 | 0.508 | 0.456 | 1 | 13 | Nsmce2 | 0.052 | Old |
| 0.508185 | 0.057536 | 0.098 | 0.07 | 1 | 13 | Vars2 | 0.028 | Old |
| 0.508378 | 0.133412 | 0.508 | 0.439 | 1 | 13 | Galnt11 | 0.069 | Old |
| 0.508597 | 0.363807 | 0.074 | 0.053 | 1 | 13 | Thumpd2 | 0.021 | Old |
| 0.508737 | 0.051741 | 0.139 | 0.105 | 1 | 13 | Pradc1 | 0.034 | Old |
| 0.509245 | 0.339279 | 0.205 | 0.167 | 1 | 13 | Pdp1 | 0.038 | Old |
| 0.509399 | 0.117797 | 0.156 | 0.123 | 1 | 13 | Aldh4a1 | 0.033 | Old |
| 0.509921 | 0.06593 | 0.5 | 0.43 | 1 | 13 | Hras | 0.07 | Old |
| 0.510035 | 0.055011 | 0.172 | 0.132 | 1 | 13 | Tmem237 | 0.04 | Old |
| 0.510035 | 0.049439 | 0.172 | 0.132 | 1 | 13 | Itgb3bp | 0.04 | Old |
| 0.510396 | 0.107802 | 0.107 | 0.079 | 1 | 13 | Pard6a | 0.028 | Old |
| 0.511008 | 0.09567 | 0.5 | 0.421 | 1 | 13 | Comtd1 | 0.079 | Old |
| 0.511024 | 0.13163 | 0.746 | 0.684 | 1 | 13 | Usp15 | 0.062 | Old |
| 0.511135 | 0.251847 | 0.344 | 0.289 | 1 | 13 | St7l | 0.055 | Old |
| 0.511275 | 0.132156 | 0.189 | 0.149 | 1 | 13 | Lrrc8b | 0.04 | Old |
| 0.511275 | 0.092202 | 0.189 | 0.149 | 1 | 13 | Fndc7 | 0.04 | Old |
| 0.511434 | 0.444305 | 0.115 | 0.088 | 1 | 13 | Osbpl1a | 0.027 | Old |
| 0.511434 | 0.230447 | 0.115 | 0.088 | 1 | 13 | 2700062C0 | 0.027 | Old |
| 0.511495 | 0.297631 | 0.074 | 0.053 | 1 | 13 | Zbtb26 | 0.021 | Old |
| 0.511685 | 0.135883 | 0.541 | 0.5 | 1 | 13 | Anks1 | 0.041 | Old |
| 0.511996 | 0.043437 | 0.172 | 0.132 | 1 | 13 | Ubxn2b | 0.04 | Old |
| 0.512062 | 0.450269 | 0.23 | 0.193 | 1 | 13 | Sema4c | 0.037 | Old |

| | | | | | | | | |
|---|---|---|---|---|---|---|---|---|
| 0.512127 | 0.172059 | 0.918 | 0.939 | 1 | 13 | Susd6 | -0.021 | Old |
| 0.512206 | 0.052751 | 1 | 1 | 1 | 13 | Cebpb | 0 | Old |
| 0.512316 | 0.138024 | 0.705 | 0.684 | 1 | 13 | Ep400 | 0.021 | Old |
| 0.512512 | 0.014439 | 0.689 | 0.632 | 1 | 13 | Kdm5a | 0.057 | Old |
| 0.512782 | 0.04887 | 0.18 | 0.14 | 1 | 13 | Zcchc10 | 0.04 | Old |
| 0.512805 | 0.089111 | 0.836 | 0.877 | 1 | 13 | Gfra2 | -0.041 | Old |
| 0.512856 | 0.121558 | 0.205 | 0.167 | 1 | 13 | Tmub1 | 0.038 | Old |
| 0.513096 | 0.143481 | 0.73 | 0.737 | 1 | 13 | Tsg101 | -0.007 | Old |
| 0.513212 | 0.160821 | 0.508 | 0.447 | 1 | 13 | Vta1 | 0.061 | Old |
| 0.513308 | 0.343015 | 0.418 | 0.377 | 1 | 13 | Sfxn2 | 0.041 | Old |
| 0.513566 | 0.255192 | 0.238 | 0.193 | 1 | 13 | Arel1 | 0.045 | Old |
| 0.513586 | 0.302815 | 0.295 | 0.254 | 1 | 13 | Tenm4 | 0.041 | Old |
| 0.514203 | 0.068393 | 0.484 | 0.412 | 1 | 13 | Gsap | 0.072 | Old |
| 0.514206 | 0.129134 | 0.689 | 0.579 | 1 | 13 | Maff | 0.11 | Old |
| 0.514314 | 0.213079 | 0.885 | 0.877 | 1 | 13 | Sp100 | 0.008 | Old |
| 0.514402 | 0.336843 | 0.074 | 0.053 | 1 | 13 | P3h1 | 0.021 | Old |
| 0.51454 | 0.027396 | 0.516 | 0.439 | 1 | 13 | Gan | 0.077 | Old |
| 0.514587 | 0.176216 | 0.902 | 0.956 | 1 | 13 | Rgs2 | -0.054 | Old |
| 0.514587 | 0.005021 | 0.926 | 0.93 | 1 | 13 | Parvb | -0.004 | Old |
| 0.514649 | 0.093867 | 0.984 | 0.93 | 1 | 13 | Tram1 | 0.054 | Old |
| 0.514665 | 0.076045 | 1 | 1 | 1 | 13 | Eif3e | 0 | Old |
| 0.514914 | 0.698314 | 0.041 | 0.026 | 1 | 13 | Asb10 | 0.015 | Old |
| 0.51492 | 0.175953 | 0.467 | 0.412 | 1 | 13 | Esrra | 0.055 | Old |
| 0.51514 | 0.140718 | 0.525 | 0.465 | 1 | 13 | Mrps7 | 0.06 | Old |
| 0.515259 | 0.208199 | 0.484 | 0.447 | 1 | 13 | Ece1 | 0.037 | Old |
| 0.515421 | 0.033075 | 0.885 | 0.851 | 1 | 13 | Rp2 | 0.034 | Old |
| 0.515696 | 0.128913 | 0.598 | 0.588 | 1 | 13 | Ktn1 | 0.01 | Old |
| 0.515802 | 0.130977 | 0.73 | 0.649 | 1 | 13 | Trappc6a | 0.081 | Old |
| 0.515983 | 0.105401 | 0.639 | 0.561 | 1 | 13 | C1galt1 | 0.078 | Old |
| 0.516788 | 0.172042 | 0.705 | 0.667 | 1 | 13 | Tec | 0.038 | Old |
| 0.516827 | 0.139808 | 0.754 | 0.711 | 1 | 13 | Stk11 | 0.043 | Old |
| 0.516864 | 0.190178 | 0.533 | 0.5 | 1 | 13 | Cds2 | 0.033 | Old |
| 0.517318 | 0.11179 | 0.893 | 0.851 | 1 | 13 | Dse | 0.042 | Old |
| 0.517338 | 0.246076 | 0.41 | 0.342 | 1 | 13 | Nsmf | 0.068 | Old |
| 0.517543 | 0.228352 | 0.238 | 0.202 | 1 | 13 | Rgmb | 0.036 | Old |
| 0.517673 | 0.244936 | 0.352 | 0.298 | 1 | 13 | Nelfcd | 0.054 | Old |
| 0.51779 | 0.776567 | 0.156 | 0.132 | 1 | 13 | Cadm1 | 0.024 | Old |
| 0.517955 | 0.080111 | 0.475 | 0.412 | 1 | 13 | Sec23ip | 0.063 | Old |
| 0.518124 | 0.225005 | 0.525 | 0.5 | 1 | 13 | Tmem192 | 0.025 | Old |
| 0.51834 | 0.359243 | 0.123 | 0.096 | 1 | 13 | Alms1 | 0.027 | Old |
| 0.518478 | 0.31925 | 0.115 | 0.088 | 1 | 13 | Wdr60 | 0.027 | Old |
| 0.518478 | 0.189742 | 0.115 | 0.088 | 1 | 13 | Akap1 | 0.027 | Old |
| 0.518653 | 0.161205 | 0.352 | 0.307 | 1 | 13 | Pacs1 | 0.045 | Old |
| 0.51885 | 0.697709 | 0.041 | 0.026 | 1 | 13 | Csf3 | 0.015 | Old |
| 0.51885 | 0.668616 | 0.041 | 0.026 | 1 | 13 | Gm29367 | 0.015 | Old |

| | | | | | | | | |
|---|---|---|---|---|---|---|---|---|
| 0.518979 | 0.075726 | 0.656 | 0.553 | 1 | 13 | Tmem230 | 0.103 | Old |
| 0.51941 | 0.04244 | 0.066 | 0.044 | 1 | 13 | Cep89 | 0.022 | Old |
| 0.519594 | 0.052454 | 0.967 | 0.965 | 1 | 13 | Timm13 | 0.002 | Old |
| 0.519688 | 0.034321 | 0.73 | 0.675 | 1 | 13 | Ptpn7 | 0.055 | Old |
| 0.52 | 0.381887 | 0.279 | 0.254 | 1 | 13 | Vegfb | 0.025 | Old |
| 0.520166 | 0.204713 | 0.107 | 0.079 | 1 | 13 | Katnb1 | 0.028 | Old |
| 0.520213 | 0.056851 | 0.975 | 0.956 | 1 | 13 | Ndufv3 | 0.019 | Old |
| 0.52024 | 0.118691 | 0.074 | 0.053 | 1 | 13 | Kif20b | 0.021 | Old |
| 0.52024 | 0.101436 | 0.074 | 0.053 | 1 | 13 | Nemp2 | 0.021 | Old |
| 0.520372 | 0.273668 | 0.434 | 0.386 | 1 | 13 | Pfkl | 0.048 | Old |
| 0.520837 | 0.133591 | 0.115 | 0.088 | 1 | 13 | Map2k3os | 0.027 | Old |
| 0.520898 | 0.149318 | 0.148 | 0.114 | 1 | 13 | Nbas | 0.034 | Old |
| 0.520987 | 0.218084 | 0.246 | 0.211 | 1 | 13 | Uqcc1 | 0.035 | Old |
| 0.521944 | 0.026439 | 0.557 | 0.447 | 1 | 13 | Lgals8 | 0.11 | Old |
| 0.522209 | 0.185029 | 0.434 | 0.395 | 1 | 13 | Nkap | 0.039 | Old |
| 0.522431 | 0.09985 | 0.18 | 0.14 | 1 | 13 | Klhl15 | 0.04 | Old |
| 0.522801 | 0.653093 | 0.041 | 0.026 | 1 | 13 | Txk | 0.015 | Old |
| 0.522801 | 0.5344 | 0.041 | 0.026 | 1 | 13 | 2210011C2 | 0.015 | Old |
| 0.522801 | 0.464754 | 0.041 | 0.026 | 1 | 13 | 1700003F1 | 0.015 | Old |
| 0.523321 | 0.044184 | 1 | 1 | 1 | 13 | Capza2 | 0 | Old |
| 0.52344 | 0.022315 | 0.631 | 0.649 | 1 | 13 | Dnttip2 | -0.018 | Old |
| 0.523596 | 0.108604 | 0.418 | 0.386 | 1 | 13 | Cstf2t | 0.032 | Old |
| 0.524058 | 0.063617 | 0.803 | 0.789 | 1 | 13 | Sdhb | 0.014 | Old |
| 0.524408 | 0.136959 | 0.23 | 0.184 | 1 | 13 | Bphl | 0.046 | Old |
| 0.526113 | 0.475242 | 0.074 | 0.053 | 1 | 13 | Rpp38 | 0.021 | Old |
| 0.526113 | 0.290594 | 0.074 | 0.053 | 1 | 13 | Zfp446 | 0.021 | Old |
| 0.526129 | 0.033847 | 0.475 | 0.412 | 1 | 13 | Borcs7 | 0.063 | Old |
| 0.526239 | 0.091281 | 0.418 | 0.36 | 1 | 13 | Map3k4 | 0.058 | Old |
| 0.526768 | 1.313048 | 0.041 | 0.026 | 1 | 13 | Plat | 0.015 | Old |
| 0.526768 | 0.444573 | 0.041 | 0.026 | 1 | 13 | Sgtb | 0.015 | Old |
| 0.527216 | 0.026922 | 0.844 | 0.807 | 1 | 13 | Srsf1 | 0.037 | Old |
| 0.527218 | 0.054437 | 0.279 | 0.228 | 1 | 13 | Arl2 | 0.051 | Old |
| 0.527346 | 0.07758 | 0.672 | 0.64 | 1 | 13 | Fip1l1 | 0.032 | Old |
| 0.527346 | 0.071015 | 0.697 | 0.614 | 1 | 13 | Snrnp27 | 0.083 | Old |
| 0.527369 | 0.105198 | 0.311 | 0.254 | 1 | 13 | Unc45a | 0.057 | Old |
| 0.527708 | 0.039218 | 0.762 | 0.684 | 1 | 13 | Pdcl3 | 0.078 | Old |
| 0.528184 | 0.053286 | 0.443 | 0.395 | 1 | 13 | Ctps2 | 0.048 | Old |
| 0.528378 | 0.223468 | 0.213 | 0.175 | 1 | 13 | Zfp938 | 0.038 | Old |
| 0.528526 | 0.032694 | 0.598 | 0.5 | 1 | 13 | Prdx3 | 0.098 | Old |
| 0.528728 | 0.1051 | 0.5 | 0.43 | 1 | 13 | Kctd2 | 0.07 | Old |
| 0.528743 | 0.082098 | 0.41 | 0.342 | 1 | 13 | Tmed4 | 0.068 | Old |
| 0.529062 | 0.265465 | 0.074 | 0.053 | 1 | 13 | Tnni3 | 0.021 | Old |
| 0.52913 | 0.049162 | 0.73 | 0.632 | 1 | 13 | Plbd2 | 0.098 | Old |
| 0.529274 | 0.115423 | 0.902 | 0.877 | 1 | 13 | Mfsd14b | 0.025 | Old |
| 0.529772 | 0.297747 | 0.123 | 0.096 | 1 | 13 | Hdac10 | 0.027 | Old |

| | | | | | | | | |
|---|---|---|---|---|---|---|---|---|
| 0.529772 | 0.251105 | 0.123 | 0.096 | 1 | 13 | Lancl2 | 0.027 | Old |
| 0.530066 | 0.226143 | 0.549 | 0.491 | 1 | 13 | Upf1 | 0.058 | Old |
| 0.530692 | 0.079372 | 0.943 | 0.895 | 1 | 13 | Dync1i2 | 0.048 | Old |
| 0.53075 | 0.577572 | 0.041 | 0.026 | 1 | 13 | Zfp532 | 0.015 | Old |
| 0.53075 | 0.490836 | 0.041 | 0.026 | 1 | 13 | Zfp503 | 0.015 | Old |
| 0.53075 | 0.396073 | 0.041 | 0.026 | 1 | 13 | L3hypdh | 0.015 | Old |
| 0.531079 | 0.084489 | 0.73 | 0.702 | 1 | 13 | Gnl3 | 0.028 | Old |
| 0.531153 | 0.120002 | 0.828 | 0.781 | 1 | 13 | Ap3d1 | 0.047 | Old |
| 0.531158 | 0.13984 | 0.27 | 0.219 | 1 | 13 | Pym1 | 0.051 | Old |
| 0.531552 | 0.280629 | 0.172 | 0.14 | 1 | 13 | Mrpl49 | 0.032 | Old |
| 0.531689 | 0.226853 | 0.361 | 0.325 | 1 | 13 | Arhgap23 | 0.036 | Old |
| 0.531921 | 0.377529 | 0.402 | 0.377 | 1 | 13 | Maoa | 0.025 | Old |
| 0.532142 | 0.07675 | 0.434 | 0.474 | 1 | 13 | Pdgfb | -0.04 | Old |
| 0.532371 | 4.546476 | 0.008 | 0.018 | 1 | 13 | Ltf | -0.01 | Old |
| 0.532371 | 0.538523 | 0.008 | 0.018 | 1 | 13 | Epha2 | -0.01 | Old |
| 0.532371 | 0.003363 | 0.008 | 0.018 | 1 | 13 | Gm19510 | -0.01 | Old |
| 0.532513 | 0.04511 | 0.107 | 0.079 | 1 | 13 | Lnx2 | 0.028 | Old |
| 0.532715 | 0.124817 | 0.115 | 0.088 | 1 | 13 | 6330418K0 | 0.027 | Old |
| 0.532761 | 0.108682 | 0.459 | 0.404 | 1 | 13 | Vps50 | 0.055 | Old |
| 0.533112 | 0.043408 | 0.451 | 0.404 | 1 | 13 | Topors | 0.047 | Old |
| 0.53406 | 0.053009 | 0.189 | 0.149 | 1 | 13 | Trim21 | 0.04 | Old |
| 0.534356 | 0.198263 | 0.648 | 0.649 | 1 | 13 | Prdx4 | -0.001 | Old |
| 0.534548 | 0.247882 | 0.246 | 0.211 | 1 | 13 | Inip | 0.035 | Old |
| 0.534639 | 0.258461 | 0.311 | 0.272 | 1 | 13 | Ints3 | 0.039 | Old |
| 0.534688 | 0.104364 | 0.689 | 0.614 | 1 | 13 | H2-Q4 | 0.075 | Old |
| 0.534747 | 0.354467 | 0.041 | 0.026 | 1 | 13 | E430024I0 | 0.015 | Old |
| 0.534747 | 0.33751 | 0.041 | 0.026 | 1 | 13 | Ak1 | 0.015 | Old |
| 0.534747 | 0.323657 | 0.041 | 0.026 | 1 | 13 | Gtf2ird1 | 0.015 | Old |
| 0.535202 | 0.49094 | 0.098 | 0.079 | 1 | 13 | Map3k15 | 0.019 | Old |
| 0.535352 | 0.118398 | 0.885 | 0.851 | 1 | 13 | Efr3a | 0.034 | Old |
| 0.535671 | 0.143004 | 0.877 | 0.947 | 1 | 13 | Cfh | -0.07 | Old |
| 0.535687 | 0.030107 | 0.77 | 0.693 | 1 | 13 | Rbm26 | 0.077 | Old |
| 0.535705 | 0.049911 | 0.639 | 0.57 | 1 | 13 | Trappc10 | 0.069 | Old |
| 0.536203 | 0.078285 | 0.418 | 0.36 | 1 | 13 | Ttc1 | 0.058 | Old |
| 0.536379 | 0.136714 | 0.828 | 0.798 | 1 | 13 | Sltm | 0.03 | Old |
| 0.53638 | 0.131279 | 0.926 | 0.939 | 1 | 13 | Mdh2 | -0.013 | Old |
| 0.536701 | 0.184812 | 0.344 | 0.289 | 1 | 13 | Smim8 | 0.055 | Old |
| 0.536757 | 0.024364 | 0.23 | 0.184 | 1 | 13 | Sft2d3 | 0.046 | Old |
| 0.536831 | 0.074682 | 0.893 | 0.895 | 1 | 13 | Stx4a | -0.002 | Old |
| 0.537504 | 0.231233 | 0.115 | 0.088 | 1 | 13 | Usp20 | 0.027 | Old |
| 0.537855 | 0.182566 | 0.27 | 0.228 | 1 | 13 | Memo1 | 0.042 | Old |
| 0.537959 | 0.269432 | 0.074 | 0.053 | 1 | 13 | Mgll | 0.021 | Old |
| 0.538319 | 0.054145 | 0.631 | 0.518 | 1 | 13 | Vps13d | 0.113 | Old |
| 0.538324 | 0.132809 | 1 | 0.93 | 1 | 13 | Usf2 | 0.07 | Old |
| 0.538331 | 0.038129 | 0.992 | 1 | 1 | 13 | Rap1a | -0.008 | Old |

| | | | | | | | | |
|---|---|---|---|---|---|---|---|---|
| 0.53876 | 0.57625 | 0.041 | 0.026 | 1 | 13 | Ccdc166 | 0.015 | Old |
| 0.53876 | 0.316489 | 0.041 | 0.026 | 1 | 13 | Trim45 | 0.015 | Old |
| 0.53876 | 0.311232 | 0.041 | 0.026 | 1 | 13 | 1700001L0 | 0.015 | Old |
| 0.53876 | 0.236159 | 0.041 | 0.026 | 1 | 13 | Ercc6l | 0.015 | Old |
| 0.538928 | 0.243774 | 0.443 | 0.439 | 1 | 13 | Bms1 | 0.004 | Old |
| 0.539565 | 0.053425 | 0.434 | 0.377 | 1 | 13 | Dlat | 0.057 | Old |
| 0.539703 | 0.358767 | 0.131 | 0.105 | 1 | 13 | Atrip | 0.026 | Old |
| 0.539732 | 0.141479 | 0.221 | 0.184 | 1 | 13 | Tlr8 | 0.037 | Old |
| 0.539902 | 0.253698 | 0.295 | 0.246 | 1 | 13 | Igtp | 0.049 | Old |
| 0.54022 | 0.134367 | 0.992 | 0.956 | 1 | 13 | Ran | 0.036 | Old |
| 0.540336 | 0.097733 | 0.279 | 0.228 | 1 | 13 | Calcoco1 | 0.051 | Old |
| 0.54036 | 0.003054 | 0.205 | 0.167 | 1 | 13 | Nup188 | 0.038 | Old |
| 0.540427 | 0.038157 | 0.648 | 0.526 | 1 | 13 | Rnf11 | 0.122 | Old |
| 0.54069 | 0.126885 | 0.918 | 0.895 | 1 | 13 | Ubash3b | 0.023 | Old |
| 0.540849 | 0.134907 | 0.967 | 0.974 | 1 | 13 | M6pr | -0.007 | Old |
| 0.541127 | 0.16235 | 0.615 | 0.588 | 1 | 13 | Pdhb | 0.027 | Old |
| 0.541132 | 0.046772 | 0.869 | 0.789 | 1 | 13 | Trappc2l | 0.08 | Old |
| 0.541488 | 0.201204 | 0.172 | 0.14 | 1 | 13 | Diras2 | 0.032 | Old |
| 0.541696 | 0.092651 | 0.582 | 0.518 | 1 | 13 | Rnft1 | 0.064 | Old |
| 0.541767 | 0.120254 | 0.189 | 0.149 | 1 | 13 | Zfp595 | 0.04 | Old |
| 0.542117 | 0.070719 | 0.992 | 1 | 1 | 13 | Vsir | -0.008 | Old |
| 0.542159 | 0.035369 | 0.828 | 0.807 | 1 | 13 | Saraf | 0.021 | Old |
| 0.542359 | 0.261263 | 0.557 | 0.544 | 1 | 13 | Txndc15 | 0.013 | Old |
| 0.542678 | 0.059005 | 0.492 | 0.421 | 1 | 13 | Tmem11 | 0.071 | Old |
| 0.5427 | 0.193009 | 0.213 | 0.184 | 1 | 13 | Slc9a3r2 | 0.029 | Old |
| 0.54277 | 0.439574 | 0.09 | 0.07 | 1 | 13 | Fchsd1 | 0.02 | Old |
| 0.542787 | 0.389664 | 0.041 | 0.026 | 1 | 13 | Zfp867 | 0.015 | Old |
| 0.542787 | 0.330284 | 0.041 | 0.026 | 1 | 13 | Nusap1 | 0.015 | Old |
| 0.542787 | 0.237076 | 0.041 | 0.026 | 1 | 13 | Gm9949 | 0.015 | Old |
| 0.542787 | 0.111108 | 0.041 | 0.026 | 1 | 13 | Tbc1d25 | 0.015 | Old |
| 0.54289 | 0.198152 | 0.443 | 0.412 | 1 | 13 | Zfp281 | 0.031 | Old |
| 0.543113 | 0.207516 | 0.246 | 0.211 | 1 | 13 | Miip | 0.035 | Old |
| 0.543377 | 0.061634 | 0.73 | 0.684 | 1 | 13 | Dnajb9 | 0.046 | Old |
| 0.543401 | 0.196161 | 0.238 | 0.202 | 1 | 13 | Etfrf1 | 0.036 | Old |
| 0.543627 | 0.183191 | 0.369 | 0.333 | 1 | 13 | Thumpd1 | 0.036 | Old |
| 0.543713 | 0.319879 | 0.287 | 0.263 | 1 | 13 | Atad3a | 0.024 | Old |
| 0.544834 | 0.164761 | 0.246 | 0.211 | 1 | 13 | Fer | 0.035 | Old |
| 0.54506 | 0.041678 | 0.828 | 0.781 | 1 | 13 | Psmc5 | 0.047 | Old |
| 0.545317 | 0.196139 | 0.689 | 0.623 | 1 | 13 | Abce1 | 0.066 | Old |
| 0.545897 | 0.012771 | 0.943 | 0.965 | 1 | 13 | Plekhb2 | -0.022 | Old |
| 0.545989 | 0.013042 | 0.123 | 0.096 | 1 | 13 | Ptrh1 | 0.027 | Old |
| 0.546532 | 0.044406 | 0.836 | 0.868 | 1 | 13 | Ppp1r12c | -0.032 | Old |
| 0.54683 | 0.185535 | 0.041 | 0.026 | 1 | 13 | Zfp287 | 0.015 | Old |
| 0.546897 | 0.143727 | 0.238 | 0.202 | 1 | 13 | Tspyl2 | 0.036 | Old |
| 0.547088 | 0.044202 | 0.484 | 0.412 | 1 | 13 | Smg5 | 0.072 | Old |

| | | | | | | | | |
|---|---|---|---|---|---|---|---|---|
| 0.547147 | 0.178224 | 0.115 | 0.088 | 1 | 13 | Tmem186 | 0.027 | Old |
| 0.547881 | 0.041405 | 0.5 | 0.43 | 1 | 13 | Traf7 | 0.07 | Old |
| 0.547928 | 0.067527 | 0.328 | 0.272 | 1 | 13 | Dis3l2 | 0.056 | Old |
| 0.548194 | 0.09935 | 0.607 | 0.544 | 1 | 13 | Dnajc14 | 0.063 | Old |
| 0.548459 | 0.09573 | 0.828 | 0.746 | 1 | 13 | Sys1 | 0.082 | Old |
| 0.548701 | 0.101357 | 0.484 | 0.421 | 1 | 13 | Prune2 | 0.063 | Old |
| 0.548738 | 0.227216 | 0.221 | 0.184 | 1 | 13 | Mcm7 | 0.037 | Old |
| 0.549093 | 0.067228 | 0.992 | 0.991 | 1 | 13 | Creg1 | 0.001 | Old |
| 0.549207 | 0.100455 | 0.697 | 0.614 | 1 | 13 | Nfya | 0.083 | Old |
| 0.549207 | 0.028713 | 0.705 | 0.605 | 1 | 13 | Por | 0.1 | Old |
| 0.549208 | 0.032464 | 0.631 | 0.561 | 1 | 13 | Rftn1 | 0.07 | Old |
| 0.549346 | 0.031185 | 0.754 | 0.64 | 1 | 13 | Sipa1 | 0.114 | Old |
| 0.549548 | 0.183395 | 0.738 | 0.719 | 1 | 13 | Hs6st1 | 0.019 | Old |
| 0.549566 | 0.173148 | 0.566 | 0.544 | 1 | 13 | Rbbp8 | 0.022 | Old |
| 0.550302 | 0.215937 | 0.352 | 0.325 | 1 | 13 | Kansl2 | 0.027 | Old |
| 0.550373 | 0.120928 | 0.164 | 0.132 | 1 | 13 | Zfp866 | 0.032 | Old |
| 0.550839 | 0.094951 | 0.672 | 0.614 | 1 | 13 | Rest | 0.058 | Old |
| 0.550887 | 0.267035 | 0.041 | 0.026 | 1 | 13 | Foxd2 | 0.015 | Old |
| 0.550887 | 0.239278 | 0.041 | 0.026 | 1 | 13 | Ubox5 | 0.015 | Old |
| 0.550887 | 0.22804 | 0.041 | 0.026 | 1 | 13 | Celsr3 | 0.015 | Old |
| 0.550887 | 0.212962 | 0.041 | 0.026 | 1 | 13 | Bbs12 | 0.015 | Old |
| 0.550887 | 0.207374 | 0.041 | 0.026 | 1 | 13 | Tgtp2 | 0.015 | Old |
| 0.550887 | 0.151451 | 0.041 | 0.026 | 1 | 13 | Gm16701 | 0.015 | Old |
| 0.550887 | 0.107686 | 0.041 | 0.026 | 1 | 13 | Prc1 | 0.015 | Old |
| 0.550952 | 0.065634 | 0.967 | 0.904 | 1 | 13 | Cdv3 | 0.063 | Old |
| 0.551003 | 0.048941 | 1 | 1 | 1 | 13 | Wnk1 | 0 | Old |
| 0.551003 | 0.030338 | 1 | 1 | 1 | 13 | Ywhab | 0 | Old |
| 0.551328 | 0.098852 | 0.598 | 0.561 | 1 | 13 | Ints6l | 0.037 | Old |
| 0.551639 | 0.017058 | 0.992 | 0.974 | 1 | 13 | Clec4a2 | 0.018 | Old |
| 0.55177 | 0.075527 | 0.811 | 0.772 | 1 | 13 | Hadha | 0.039 | Old |
| 0.55196 | 0.167906 | 0.672 | 0.684 | 1 | 13 | Dync1li1 | -0.012 | Old |
| 0.552278 | 0.014447 | 1 | 1 | 1 | 13 | Rac1 | 0 | Old |
| 0.552278 | 0.071459 | 1 | 1 | 1 | 13 | Hnrnpu | 0 | Old |
| 0.552338 | 0.02274 | 0.73 | 0.658 | 1 | 13 | Adprh | 0.072 | Old |
| 0.552401 | 0.000231 | 0.385 | 0.43 | 1 | 13 | Bcl2l1 | -0.045 | Old |
| 0.552439 | 0.209231 | 0.484 | 0.404 | 1 | 13 | Ccnh | 0.08 | Old |
| 0.552631 | 0.279333 | 0.082 | 0.061 | 1 | 13 | Tmem69 | 0.021 | Old |
| 0.55274 | 0.17865 | 0.721 | 0.658 | 1 | 13 | Dgkz | 0.063 | Old |
| 0.55288 | 0.121033 | 0.5 | 0.404 | 1 | 13 | Rexo1 | 0.096 | Old |
| 0.553555 | 0.021079 | 1 | 1 | 1 | 13 | Dazap2 | 0 | Old |
| 0.553562 | 0.194739 | 0.492 | 0.447 | 1 | 13 | Yrdc | 0.045 | Old |
| 0.55378 | 0.040908 | 0.68 | 0.614 | 1 | 13 | Tmem33 | 0.066 | Old |
| 0.553981 | 0.127761 | 0.615 | 0.544 | 1 | 13 | Tpcn1 | 0.071 | Old |
| 0.554043 | 0.036451 | 0.623 | 0.561 | 1 | 13 | Tbc1d2b | 0.062 | Old |
| 0.554435 | 0.191499 | 0.115 | 0.088 | 1 | 13 | Sec61a2 | 0.027 | Old |

| | | | | | | | | |
|---|---|---|---|---|---|---|---|---|
| 0.554959 | 0.177306 | 0.041 | 0.026 | 1 | 13 | Il20rb | 0.015 | Old |
| 0.554959 | 0.14589 | 0.041 | 0.026 | 1 | 13 | Cenpu | 0.015 | Old |
| 0.554959 | 0.071209 | 0.041 | 0.026 | 1 | 13 | Ajuba | 0.015 | Old |
| 0.55503 | 0.066055 | 0.303 | 0.246 | 1 | 13 | Bag5 | 0.057 | Old |
| 0.555245 | 0.215978 | 0.467 | 0.421 | 1 | 13 | Erg28 | 0.046 | Old |
| 0.555515 | 0.092386 | 0.311 | 0.246 | 1 | 13 | Snapc3 | 0.065 | Old |
| 0.555664 | 0.247421 | 0.607 | 0.535 | 1 | 13 | Tmem87b | 0.072 | Old |
| 0.555808 | 0.058997 | 0.639 | 0.553 | 1 | 13 | Gigyf2 | 0.086 | Old |
| 0.555995 | 0.190164 | 0.754 | 0.702 | 1 | 13 | Il1rn | 0.052 | Old |
| 0.556112 | 0.029167 | 1 | 1 | 1 | 13 | Arpc5 | 0 | Old |
| 0.556137 | 0.113394 | 0.295 | 0.246 | 1 | 13 | Gnptab | 0.049 | Old |
| 0.556373 | 0.173239 | 0.082 | 0.105 | 1 | 13 | Gcnt2 | -0.023 | Old |
| 0.557168 | 0.264642 | 0.303 | 0.272 | 1 | 13 | Trim11 | 0.031 | Old |
| 0.557354 | 0.010104 | 0.525 | 0.439 | 1 | 13 | Tcf3 | 0.086 | Old |
| 0.557993 | 0.00838 | 0.943 | 0.939 | 1 | 13 | Ncor1 | 0.004 | Old |
| 0.558123 | 0.108285 | 0.516 | 0.456 | 1 | 13 | Washc5 | 0.06 | Old |
| 0.558123 | 0.083363 | 0.525 | 0.447 | 1 | 13 | Sucla2 | 0.078 | Old |
| 0.558157 | 0.233455 | 0.582 | 0.57 | 1 | 13 | Cops5 | 0.012 | Old |
| 0.558236 | 0.074687 | 0.795 | 0.746 | 1 | 13 | Ncstn | 0.049 | Old |
| 0.558358 | 0.145261 | 0.828 | 0.833 | 1 | 13 | Uhrf2 | -0.005 | Old |
| 0.558643 | 0.138219 | 0.926 | 0.965 | 1 | 13 | Ccl2 | -0.039 | Old |
| 0.558661 | 0.063999 | 0.951 | 0.965 | 1 | 13 | Wdr83os | -0.014 | Old |
| 0.559007 | 0.114097 | 0.074 | 0.053 | 1 | 13 | Cntrob | 0.021 | Old |
| 0.559041 | 0.36216 | 0.156 | 0.132 | 1 | 13 | Ttf2 | 0.024 | Old |
| 0.559046 | 0.099661 | 0.041 | 0.026 | 1 | 13 | Bace1 | 0.015 | Old |
| 0.559046 | 0.071695 | 0.041 | 0.026 | 1 | 13 | Kctd1 | 0.015 | Old |
| 0.55967 | 0.156848 | 0.623 | 0.518 | 1 | 13 | Slc35c2 | 0.105 | Old |
| 0.559928 | 0.022668 | 0.672 | 0.579 | 1 | 13 | Usp22 | 0.093 | Old |
| 0.560082 | 0.203995 | 0.123 | 0.096 | 1 | 13 | Ppox | 0.027 | Old |
| 0.560483 | 0.048118 | 0.91 | 0.921 | 1 | 13 | Tmco1 | -0.011 | Old |
| 0.560937 | 0.412466 | 0.098 | 0.079 | 1 | 13 | Sh3rf1 | 0.019 | Old |
| 0.560997 | 0.179485 | 0.336 | 0.281 | 1 | 13 | Ankrd49 | 0.055 | Old |
| 0.561698 | 0.452608 | 0.09 | 0.07 | 1 | 13 | Zfp52 | 0.02 | Old |
| 0.561746 | 0.257117 | 0.139 | 0.114 | 1 | 13 | Paip2b | 0.025 | Old |
| 0.562065 | 0.12051 | 0.787 | 0.798 | 1 | 13 | Peli1 | -0.011 | Old |
| 0.562302 | 0.040399 | 0.689 | 0.588 | 1 | 13 | Zmym5 | 0.101 | Old |
| 0.562302 | 0.045836 | 0.68 | 0.596 | 1 | 13 | Pabpn1 | 0.084 | Old |
| 0.562416 | 0.177604 | 0.451 | 0.395 | 1 | 13 | Ergic1 | 0.056 | Old |
| 0.562462 | 0.112161 | 0.844 | 0.772 | 1 | 13 | Lpp | 0.072 | Old |
| 0.562603 | 0.04425 | 0.402 | 0.333 | 1 | 13 | Thap11 | 0.069 | Old |
| 0.563147 | 0.177188 | 0.041 | 0.026 | 1 | 13 | Armc2 | 0.015 | Old |
| 0.563147 | 0.121213 | 0.041 | 0.026 | 1 | 13 | Plpp1 | 0.015 | Old |
| 0.563203 | 0.031683 | 0.697 | 0.711 | 1 | 13 | Yme1l1 | -0.014 | Old |
| 0.563458 | 0.065834 | 0.852 | 0.904 | 1 | 13 | Casp8 | -0.052 | Old |
| 0.563633 | 0.078729 | 0.098 | 0.123 | 1 | 13 | Gprc5a | -0.025 | Old |

| | | | | | | | | |
|---|---|---|---|---|---|---|---|---|
| 0.563662 | 0.011216 | 0.779 | 0.781 | 1 | 13 | Inpp4a | -0.002 | Old |
| 0.563768 | 0.266283 | 0.525 | 0.5 | 1 | 13 | Rsbn1 | 0.025 | Old |
| 0.563819 | 0.058471 | 1 | 0.982 | 1 | 13 | Ifngr1 | 0.018 | Old |
| 0.564101 | 0.583469 | 0.049 | 0.035 | 1 | 13 | Hyls1 | 0.014 | Old |
| 0.564459 | 0.087155 | 0.525 | 0.474 | 1 | 13 | Vac14 | 0.051 | Old |
| 0.564565 | 0.075422 | 0.459 | 0.386 | 1 | 13 | Foxk2 | 0.073 | Old |
| 0.564595 | 0.150959 | 0.418 | 0.377 | 1 | 13 | Rptor | 0.041 | Old |
| 0.56469 | 0.177938 | 0.131 | 0.105 | 1 | 13 | Stau2 | 0.026 | Old |
| 0.564848 | 0.183968 | 0.582 | 0.57 | 1 | 13 | Sec23b | 0.012 | Old |
| 0.564887 | 0.129962 | 0.164 | 0.132 | 1 | 13 | Lclat1 | 0.032 | Old |
| 0.565088 | 0.005468 | 0.213 | 0.175 | 1 | 13 | Nkiras1 | 0.038 | Old |
| 0.565125 | 0.170512 | 0.533 | 0.456 | 1 | 13 | Relt | 0.077 | Old |
| 0.565439 | 0.225769 | 0.205 | 0.175 | 1 | 13 | Cipc | 0.03 | Old |
| 0.565614 | 0.019877 | 0.475 | 0.404 | 1 | 13 | Rbm14 | 0.071 | Old |
| 0.566389 | 0.070291 | 0.967 | 0.956 | 1 | 13 | Gpr132 | 0.011 | Old |
| 0.5664 | 0.03363 | 1 | 1 | 1 | 13 | Eif4g2 | 0 | Old |
| 0.566585 | 0.092632 | 0.648 | 0.623 | 1 | 13 | Dapp1 | 0.025 | Old |
| 0.566957 | 0.404511 | 0.082 | 0.061 | 1 | 13 | Ddit4 | 0.021 | Old |
| 0.566957 | 0.354173 | 0.082 | 0.061 | 1 | 13 | Loxl3 | 0.021 | Old |
| 0.566971 | 0.203692 | 0.221 | 0.184 | 1 | 13 | Gtpbp8 | 0.037 | Old |
| 0.567164 | 0.461117 | 0.09 | 0.07 | 1 | 13 | Ssx2ip | 0.02 | Old |
| 0.567164 | 0.225908 | 0.09 | 0.07 | 1 | 13 | Serpine1 | 0.02 | Old |
| 0.567345 | 0.081378 | 0.189 | 0.228 | 1 | 13 | Il7r | -0.039 | Old |
| 0.567397 | 0.143025 | 0.377 | 0.342 | 1 | 13 | Scyl1 | 0.035 | Old |
| 0.567427 | 0.061823 | 0.287 | 0.237 | 1 | 13 | Man2c1 | 0.05 | Old |
| 0.567537 | 0.178281 | 0.648 | 0.553 | 1 | 13 | Zzz3 | 0.095 | Old |
| 0.567581 | 0.24741 | 0.344 | 0.289 | 1 | 13 | Trim36 | 0.055 | Old |
| 0.567632 | 0.085014 | 0.311 | 0.263 | 1 | 13 | Daam1 | 0.048 | Old |
| 0.567874 | 0.295404 | 0.197 | 0.167 | 1 | 13 | Ydjc | 0.03 | Old |
| 0.568043 | 0.156697 | 0.779 | 0.728 | 1 | 13 | Zfp644 | 0.051 | Old |
| 0.568156 | 0.030792 | 0.451 | 0.395 | 1 | 13 | Nipsnap2 | 0.056 | Old |
| 0.568691 | 0.024072 | 0.795 | 0.868 | 1 | 13 | Slc38a1 | -0.073 | Old |
| 0.568757 | 0.099596 | 0.926 | 0.86 | 1 | 13 | Hectd1 | 0.066 | Old |
| 0.569152 | 0.044254 | 0.115 | 0.088 | 1 | 13 | Vill | 0.027 | Old |
| 0.569212 | 0.218129 | 0.697 | 0.667 | 1 | 13 | Mrpl48 | 0.03 | Old |
| 0.569272 | 0.241888 | 0.148 | 0.123 | 1 | 13 | Usp18 | 0.025 | Old |
| 0.569305 | 0.33965 | 0.344 | 0.307 | 1 | 13 | Tsga10 | 0.037 | Old |
| 0.569844 | 0.804942 | 0.082 | 0.061 | 1 | 13 | Chchd10 | 0.021 | Old |
| 0.569844 | 0.106105 | 0.082 | 0.061 | 1 | 13 | Tctn1 | 0.021 | Old |
| 0.569906 | 0.530025 | 0.09 | 0.07 | 1 | 13 | Sphk1 | 0.02 | Old |
| 0.569906 | 0.333128 | 0.09 | 0.07 | 1 | 13 | Sirt4 | 0.02 | Old |
| 0.569906 | 0.314898 | 0.09 | 0.07 | 1 | 13 | A930033H1 | 0.02 | Old |
| 0.569926 | 0.118062 | 0.738 | 0.702 | 1 | 13 | Clcn4 | 0.036 | Old |
| 0.569989 | 0.065546 | 0.844 | 0.816 | 1 | 13 | Mkln1 | 0.028 | Old |
| 0.569993 | 0.110608 | 0.902 | 0.868 | 1 | 13 | Stag2 | 0.034 | Old |

| | | | | | | | | |
|---|---|---|---|---|---|---|---|---|
| 0.570485 | 0.14315 | 0.607 | 0.561 | 1 | 13 | Brox | 0.046 | Old |
| 0.571051 | 0.065455 | 0.893 | 0.825 | 1 | 13 | Ago2 | 0.068 | Old |
| 0.571149 | 0.049807 | 0.844 | 0.737 | 1 | 13 | Rad21 | 0.107 | Old |
| 0.571583 | 0.171843 | 0.352 | 0.289 | 1 | 13 | Tmem181a | 0.063 | Old |
| 0.5716 | 0.159374 | 0.131 | 0.105 | 1 | 13 | Carf | 0.026 | Old |
| 0.5717 | 0.145293 | 0.5 | 0.439 | 1 | 13 | Hist3h2a | 0.061 | Old |
| 0.571991 | 0.024638 | 0.811 | 0.737 | 1 | 13 | Srsf10 | 0.074 | Old |
| 0.572079 | 0.281492 | 0.189 | 0.158 | 1 | 13 | Timd4 | 0.031 | Old |
| 0.572201 | 0.050181 | 0.943 | 0.956 | 1 | 13 | Shtn1 | -0.013 | Old |
| 0.572342 | 0.092661 | 0.705 | 0.675 | 1 | 13 | Pcmt1 | 0.03 | Old |
| 0.572737 | 0.189244 | 0.082 | 0.061 | 1 | 13 | Cep76 | 0.021 | Old |
| 0.57276 | 0.083362 | 0.787 | 0.728 | 1 | 13 | Csnk2a2 | 0.059 | Old |
| 0.573276 | 0.002676 | 0.516 | 0.439 | 1 | 13 | Papd5 | 0.077 | Old |
| 0.573625 | 0.13164 | 0.336 | 0.281 | 1 | 13 | Bzw2 | 0.055 | Old |
| 0.573812 | 0.104867 | 0.574 | 0.535 | 1 | 13 | Ankrd40 | 0.039 | Old |
| 0.573931 | 0.105947 | 0.344 | 0.298 | 1 | 13 | Pdzd11 | 0.046 | Old |
| 0.574007 | 0.178981 | 0.23 | 0.193 | 1 | 13 | Adprm | 0.037 | Old |
| 0.574026 | 0.416568 | 0.098 | 0.079 | 1 | 13 | Sccpdh | 0.019 | Old |
| 0.574175 | 0.088077 | 1 | 0.991 | 1 | 13 | Naaa | 0.009 | Old |
| 0.574454 | 0.20993 | 0.213 | 0.175 | 1 | 13 | Lig3 | 0.038 | Old |
| 0.574694 | 0.184288 | 0.295 | 0.254 | 1 | 13 | Fam60a | 0.041 | Old |
| 0.57489 | 0.232385 | 0.205 | 0.175 | 1 | 13 | Nsdhl | 0.03 | Old |
| 0.575498 | 0.270353 | 0.344 | 0.298 | 1 | 13 | Vps33a | 0.046 | Old |
| 0.575519 | 0.063098 | 0.803 | 0.719 | 1 | 13 | Ehmt2 | 0.084 | Old |
| 0.575706 | 0.012791 | 0.582 | 0.491 | 1 | 13 | Mthfd2 | 0.091 | Old |
| 0.576153 | 0.041422 | 0.18 | 0.149 | 1 | 13 | Cdpf1 | 0.031 | Old |
| 0.576167 | 0.063033 | 0.566 | 0.456 | 1 | 13 | Nudt9 | 0.11 | Old |
| 0.57666 | 0.318147 | 0.098 | 0.079 | 1 | 13 | Kazald1 | 0.019 | Old |
| 0.57676 | 0.334829 | 0.287 | 0.254 | 1 | 13 | Hmgn5 | 0.033 | Old |
| 0.57676 | 0.207034 | 0.377 | 0.325 | 1 | 13 | Apmap | 0.052 | Old |
| 0.576829 | 0.124609 | 0.582 | 0.509 | 1 | 13 | Parp8 | 0.073 | Old |
| 0.577052 | 0.200696 | 0.238 | 0.202 | 1 | 13 | Rfc3 | 0.036 | Old |
| 0.577052 | 0.137801 | 0.238 | 0.202 | 1 | 13 | Fhl3 | 0.036 | Old |
| 0.57741 | 0.333548 | 0.139 | 0.114 | 1 | 13 | Gtf3c5 | 0.025 | Old |
| 0.578041 | 0.151716 | 0.246 | 0.211 | 1 | 13 | Fam117a | 0.035 | Old |
| 0.57808 | 0.071683 | 0.992 | 0.965 | 1 | 13 | Birc3 | 0.027 | Old |
| 0.578081 | 0.073125 | 1 | 1 | 1 | 13 | Ifitm3 | 0 | Old |
| 0.578168 | 0.132661 | 0.18 | 0.149 | 1 | 13 | Gm47754 | 0.031 | Old |
| 0.578331 | 0.158518 | 0.279 | 0.246 | 1 | 13 | E4f1 | 0.033 | Old |
| 0.578552 | 0.040272 | 0.623 | 0.667 | 1 | 13 | Zrsr2 | -0.044 | Old |
| 0.57885 | 0.171633 | 0.238 | 0.202 | 1 | 13 | Papd7 | 0.036 | Old |
| 0.578968 | 0.465349 | 0.049 | 0.035 | 1 | 13 | Btbd11 | 0.014 | Old |
| 0.578968 | 0.308449 | 0.049 | 0.035 | 1 | 13 | Fam187b | 0.014 | Old |
| 0.57907 | 0.040701 | 0.918 | 0.842 | 1 | 13 | Ndufa5 | 0.076 | Old |
| 0.579269 | 0.124428 | 0.311 | 0.272 | 1 | 13 | Cfap36 | 0.039 | Old |

| | | | | | | | | |
|---|---|---|---|---|---|---|---|---|
| 0.579386 | 0.020863 | 1 | 0.991 | 1 | 13 | Ndufa1 | 0.009 | Old |
| 0.580114 | 0.092336 | 0.795 | 0.728 | 1 | 13 | Bcl3 | 0.067 | Old |
| 0.580389 | 0.048361 | 0.574 | 0.509 | 1 | 13 | Hcfc1 | 0.065 | Old |
| 0.580392 | 0.248703 | 0.508 | 0.465 | 1 | 13 | Gbp2 | 0.043 | Old |
| 0.580468 | 0.074431 | 0.91 | 0.877 | 1 | 13 | Esyt1 | 0.033 | Old |
| 0.58081 | 0.097369 | 0.705 | 0.605 | 1 | 13 | Sp1 | 0.1 | Old |
| 0.581171 | 0.178304 | 0.697 | 0.693 | 1 | 13 | Rras2 | 0.004 | Old |
| 0.581547 | 0.139146 | 0.123 | 0.096 | 1 | 13 | Gm30025 | 0.027 | Old |
| 0.581586 | 0.129856 | 0.328 | 0.289 | 1 | 13 | Agl | 0.039 | Old |
| 0.581694 | 0.053409 | 0.795 | 0.746 | 1 | 13 | Rbm6 | 0.049 | Old |
| 0.581698 | 0.003078 | 0.164 | 0.132 | 1 | 13 | Dnal4 | 0.032 | Old |
| 0.581717 | 0.090339 | 0.836 | 0.754 | 1 | 13 | Plin3 | 0.082 | Old |
| 0.581763 | 0.23815 | 0.221 | 0.184 | 1 | 13 | Acot2 | 0.037 | Old |
| 0.581834 | 0.121065 | 0.705 | 0.684 | 1 | 13 | Mrpl17 | 0.021 | Old |
| 0.581877 | 0.409924 | 0.123 | 0.105 | 1 | 13 | Mlkl | 0.018 | Old |
| 0.581896 | 0.151559 | 0.664 | 0.588 | 1 | 13 | Klhl5 | 0.076 | Old |
| 0.581947 | 0.258794 | 0.189 | 0.158 | 1 | 13 | Hsf2 | 0.031 | Old |
| 0.58238 | 0.173277 | 0.459 | 0.404 | 1 | 13 | Tmem120a | 0.055 | Old |
| 0.582713 | 0.508397 | 0.049 | 0.035 | 1 | 13 | Sbsn | 0.014 | Old |
| 0.582929 | 0.058959 | 0.82 | 0.763 | 1 | 13 | Mgea5 | 0.057 | Old |
| 0.584063 | 0.207805 | 0.385 | 0.333 | 1 | 13 | Zdhhc14 | 0.052 | Old |
| 0.584225 | 0.0025 | 0.525 | 0.579 | 1 | 13 | Cacul1 | -0.054 | Old |
| 0.584415 | 0.159331 | 0.205 | 0.175 | 1 | 13 | Copz2 | 0.03 | Old |
| 0.584442 | 0.01909 | 0.549 | 0.482 | 1 | 13 | Slc35b1 | 0.067 | Old |
| 0.584879 | 0.021599 | 0.623 | 0.561 | 1 | 13 | Rnps1 | 0.062 | Old |
| 0.585138 | 0.08769 | 0.557 | 0.474 | 1 | 13 | Sigmar1 | 0.083 | Old |
| 0.585921 | 0.018003 | 0.959 | 0.965 | 1 | 13 | Abi1 | -0.006 | Old |
| 0.585929 | 0.067741 | 0.984 | 0.965 | 1 | 13 | Scp2 | 0.019 | Old |
| 0.58647 | 0.447087 | 0.049 | 0.035 | 1 | 13 | Trim46 | 0.014 | Old |
| 0.58647 | 0.42756 | 0.049 | 0.035 | 1 | 13 | Nptxr | 0.014 | Old |
| 0.58647 | 0.402989 | 0.049 | 0.035 | 1 | 13 | Il15ra | 0.014 | Old |
| 0.58647 | 0.357286 | 0.049 | 0.035 | 1 | 13 | Rarres2 | 0.014 | Old |
| 0.58647 | 0.339031 | 0.049 | 0.035 | 1 | 13 | Dnajc28 | 0.014 | Old |
| 0.586857 | 0.119512 | 0.713 | 0.64 | 1 | 13 | Usp19 | 0.073 | Old |
| 0.586936 | 0.156729 | 0.246 | 0.211 | 1 | 13 | Ints10 | 0.035 | Old |
| 0.586977 | 0.199639 | 0.59 | 0.509 | 1 | 13 | Dusp16 | 0.081 | Old |
| 0.587758 | 0.178961 | 0.131 | 0.158 | 1 | 13 | Tspan33 | -0.027 | Old |
| 0.587908 | 0.041376 | 0.393 | 0.351 | 1 | 13 | Mrpl37 | 0.042 | Old |
| 0.588264 | 0.08185 | 0.77 | 0.728 | 1 | 13 | Gbp7 | 0.042 | Old |
| 0.5885 | 0.180499 | 0.418 | 0.386 | 1 | 13 | Hspbp1 | 0.032 | Old |
| 0.588557 | 0.110657 | 0.984 | 0.974 | 1 | 13 | Mapkapk2 | 0.01 | Old |
| 0.588723 | 0.216824 | 0.139 | 0.114 | 1 | 13 | Cenpl | 0.025 | Old |
| 0.588723 | 0.142627 | 0.139 | 0.114 | 1 | 13 | Armcx5 | 0.025 | Old |
| 0.589013 | 0.355563 | 0.148 | 0.123 | 1 | 13 | Socs1 | 0.025 | Old |
| 0.589226 | 0.078486 | 0.221 | 0.184 | 1 | 13 | Zfp687 | 0.037 | Old |

| | | | | | | | | |
|---|---|---|---|---|---|---|---|---|
| 0.589279 | 0.220074 | 0.09 | 0.07 | 1 | 13 | Trdmt1 | 0.02 | Old |
| 0.589279 | 0.211763 | 0.09 | 0.07 | 1 | 13 | Ficd | 0.02 | Old |
| 0.589953 | 0.15507 | 0.664 | 0.623 | 1 | 13 | Crtc3 | 0.041 | Old |
| 0.590071 | 0.15304 | 0.566 | 0.553 | 1 | 13 | Irf9 | 0.013 | Old |
| 0.590175 | 0.013891 | 0.656 | 0.553 | 1 | 13 | Spr | 0.103 | Old |
| 0.590217 | 0.00424 | 0.811 | 0.842 | 1 | 13 | Anxa3 | -0.031 | Old |
| 0.590237 | 4.506823 | 0.049 | 0.035 | 1 | 13 | S100a8 | 0.014 | Old |
| 0.590237 | 0.684966 | 0.049 | 0.035 | 1 | 13 | Zfp458 | 0.014 | Old |
| 0.590491 | 0.157428 | 0.23 | 0.193 | 1 | 13 | Flywch1 | 0.037 | Old |
| 0.590947 | 0.056593 | 0.861 | 0.798 | 1 | 13 | Huwe1 | 0.063 | Old |
| 0.591025 | 0.197259 | 0.361 | 0.316 | 1 | 13 | Ubap2 | 0.045 | Old |
| 0.592071 | 0.219728 | 0.09 | 0.07 | 1 | 13 | 3300005D( | 0.02 | Old |
| 0.592126 | 0.033419 | 0.27 | 0.228 | 1 | 13 | Rfng | 0.042 | Old |
| 0.592511 | 0.049852 | 1 | 1 | 1 | 13 | Junb | 0 | Old |
| 0.592511 | 0.028055 | 1 | 1 | 1 | 13 | Rpl38 | 0 | Old |
| 0.592589 | 0.380905 | 0.098 | 0.079 | 1 | 13 | 8030462N1 | 0.019 | Old |
| 0.592675 | 0.167406 | 0.623 | 0.526 | 1 | 13 | Tlk1 | 0.097 | Old |
| 0.592816 | 0.280627 | 0.32 | 0.298 | 1 | 13 | Las1l | 0.022 | Old |
| 0.593059 | 0.003803 | 0.885 | 0.789 | 1 | 13 | Pdcd10 | 0.096 | Old |
| 0.593131 | 0.096351 | 0.697 | 0.667 | 1 | 13 | Dennd4b | 0.03 | Old |
| 0.593287 | 0.330226 | 0.238 | 0.211 | 1 | 13 | Ap5s1 | 0.027 | Old |
| 0.593743 | 0.176308 | 0.107 | 0.132 | 1 | 13 | Nes | -0.025 | Old |
| 0.593831 | 0.03601 | 1 | 1 | 1 | 13 | Mpeg1 | 0 | Old |
| 0.593861 | 0.06735 | 0.27 | 0.228 | 1 | 13 | Polr3e | 0.042 | Old |
| 0.593994 | 0.058152 | 0.287 | 0.246 | 1 | 13 | Fam210a | 0.041 | Old |
| 0.594016 | 0.339307 | 0.049 | 0.035 | 1 | 13 | Gdpgp1 | 0.014 | Old |
| 0.594016 | 0.270008 | 0.049 | 0.035 | 1 | 13 | Chrna5 | 0.014 | Old |
| 0.594272 | 0.207425 | 0.721 | 0.684 | 1 | 13 | Rnf145 | 0.037 | Old |
| 0.594379 | 0.124931 | 0.82 | 0.728 | 1 | 13 | Lamtor5 | 0.092 | Old |
| 0.594499 | 0.067461 | 0.615 | 0.544 | 1 | 13 | Tia1 | 0.071 | Old |
| 0.59458 | 0.188814 | 0.164 | 0.14 | 1 | 13 | Ggct | 0.024 | Old |
| 0.594728 | 0.09602 | 0.541 | 0.509 | 1 | 13 | Asnsd1 | 0.032 | Old |
| 0.59487 | 0.187598 | 0.09 | 0.07 | 1 | 13 | Trip13 | 0.02 | Old |
| 0.595152 | 0.002824 | 1 | 1 | 1 | 13 | Ptafr | 0 | Old |
| 0.595266 | 0.150027 | 0.402 | 0.386 | 1 | 13 | Ptgir | 0.016 | Old |
| 0.595303 | 0.154993 | 0.385 | 0.342 | 1 | 13 | Mrps30 | 0.043 | Old |
| 0.59536 | 0.113984 | 0.836 | 0.798 | 1 | 13 | Commd8 | 0.038 | Old |
| 0.595544 | 0.006497 | 0.205 | 0.167 | 1 | 13 | Nup160 | 0.038 | Old |
| 0.595559 | 0.129716 | 0.139 | 0.114 | 1 | 13 | Adck2 | 0.025 | Old |
| 0.595777 | 0.018299 | 0.5 | 0.43 | 1 | 13 | Nudt16l1 | 0.07 | Old |
| 0.595813 | 0.11514 | 0.992 | 0.991 | 1 | 13 | Anp32b | 0.001 | Old |
| 0.596034 | 0.216252 | 0.23 | 0.193 | 1 | 13 | Heg1 | 0.037 | Old |
| 0.59619 | 0.180721 | 0.369 | 0.325 | 1 | 13 | Tmem154 | 0.044 | Old |
| 0.596378 | 0.192986 | 0.623 | 0.605 | 1 | 13 | Rnf166 | 0.018 | Old |
| 0.596474 | 0.054917 | 0.984 | 1 | 1 | 13 | Mat2a | -0.016 | Old |

| | | | | | | | | |
|---|---|---|---|---|---|---|---|---|
| 0.596775 | 0.283079 | 0.361 | 0.342 | 1 | 13 | Pkp4 | 0.019 | Old |
| 0.597055 | 0.149275 | 0.484 | 0.43 | 1 | 13 | Prr14 | 0.054 | Old |
| 0.597245 | 0.153374 | 0.525 | 0.482 | 1 | 13 | Irak3 | 0.043 | Old |
| 0.597522 | 0.121972 | 0.566 | 0.526 | 1 | 13 | Uba5 | 0.04 | Old |
| 0.597674 | 0.163011 | 0.09 | 0.07 | 1 | 13 | Fkbpl | 0.02 | Old |
| 0.597692 | 0.129174 | 0.246 | 0.211 | 1 | 13 | Cacfd1 | 0.035 | Old |
| 0.5979 | 0.175195 | 0.189 | 0.158 | 1 | 13 | Uck2 | 0.031 | Old |
| 0.598108 | 0.269861 | 0.541 | 0.518 | 1 | 13 | Tlr7 | 0.023 | Old |
| 0.598846 | 1.759545 | 0.016 | 0.009 | 1 | 13 | Mmp8 | 0.007 | Old |
| 0.598846 | 1.54837 | 0.016 | 0.009 | 1 | 13 | Ccl11 | 0.007 | Old |
| 0.598846 | 1.310835 | 0.016 | 0.009 | 1 | 13 | Rpl39l | 0.007 | Old |
| 0.598846 | 1.281785 | 0.016 | 0.009 | 1 | 13 | Ly6i | 0.007 | Old |
| 0.598846 | 0.768966 | 0.016 | 0.009 | 1 | 13 | Car2 | 0.007 | Old |
| 0.598846 | 0.757526 | 0.016 | 0.009 | 1 | 13 | D330045A2 | 0.007 | Old |
| 0.598846 | 0.703227 | 0.016 | 0.009 | 1 | 13 | Pimreg | 0.007 | Old |
| 0.598846 | 0.643528 | 0.016 | 0.009 | 1 | 13 | Mapk8ip1 | 0.007 | Old |
| 0.598846 | 0.558139 | 0.016 | 0.009 | 1 | 13 | G630030J0 | 0.007 | Old |
| 0.598846 | 0.555143 | 0.016 | 0.009 | 1 | 13 | Wt1 | 0.007 | Old |
| 0.598846 | 0.554138 | 0.016 | 0.009 | 1 | 13 | Gm20404 | 0.007 | Old |
| 0.598846 | 0.541404 | 0.016 | 0.009 | 1 | 13 | Gm14327 | 0.007 | Old |
| 0.598846 | 0.504338 | 0.016 | 0.009 | 1 | 13 | Zfp712 | 0.007 | Old |
| 0.598846 | 0.475479 | 0.016 | 0.009 | 1 | 13 | Ptpn13 | 0.007 | Old |
| 0.598846 | 0.459513 | 0.016 | 0.009 | 1 | 13 | Olfr373 | 0.007 | Old |
| 0.598846 | 0.456011 | 0.016 | 0.009 | 1 | 13 | Ager | 0.007 | Old |
| 0.598846 | 0.453337 | 0.016 | 0.009 | 1 | 13 | A430078I0 | 0.007 | Old |
| 0.598846 | 0.452873 | 0.016 | 0.009 | 1 | 13 | Slc7a10 | 0.007 | Old |
| 0.598846 | 0.446712 | 0.016 | 0.009 | 1 | 13 | Gspt2 | 0.007 | Old |
| 0.598846 | 0.440302 | 0.016 | 0.009 | 1 | 13 | Adgra2 | 0.007 | Old |
| 0.598846 | 0.435002 | 0.016 | 0.009 | 1 | 13 | Gm16230 | 0.007 | Old |
| 0.598846 | 0.429508 | 0.016 | 0.009 | 1 | 13 | 4933423P2 | 0.007 | Old |
| 0.598846 | 0.428903 | 0.016 | 0.009 | 1 | 13 | Rab19 | 0.007 | Old |
| 0.598846 | 0.420732 | 0.016 | 0.009 | 1 | 13 | Adam4 | 0.007 | Old |
| 0.598846 | 0.409648 | 0.016 | 0.009 | 1 | 13 | Sat2 | 0.007 | Old |
| 0.598846 | 0.400538 | 0.016 | 0.009 | 1 | 13 | Depdc1a | 0.007 | Old |
| 0.598846 | 0.393074 | 0.016 | 0.009 | 1 | 13 | Hyal3 | 0.007 | Old |
| 0.598846 | 0.386917 | 0.016 | 0.009 | 1 | 13 | Prelp | 0.007 | Old |
| 0.598846 | 0.384766 | 0.016 | 0.009 | 1 | 13 | Shisa8 | 0.007 | Old |
| 0.598846 | 0.373534 | 0.016 | 0.009 | 1 | 13 | Kcnj10 | 0.007 | Old |
| 0.598846 | 0.36876 | 0.016 | 0.009 | 1 | 13 | Zfp14 | 0.007 | Old |
| 0.598846 | 0.367656 | 0.016 | 0.009 | 1 | 13 | Angptl3 | 0.007 | Old |
| 0.598846 | 0.360827 | 0.016 | 0.009 | 1 | 13 | E130208F1 | 0.007 | Old |
| 0.598846 | 0.359764 | 0.016 | 0.009 | 1 | 13 | Tnnt3 | 0.007 | Old |
| 0.598846 | 0.340247 | 0.016 | 0.009 | 1 | 13 | Mxra8 | 0.007 | Old |
| 0.598846 | 0.335413 | 0.016 | 0.009 | 1 | 13 | Gm47079 | 0.007 | Old |
| 0.598846 | 0.335034 | 0.016 | 0.009 | 1 | 13 | Cpe | 0.007 | Old |

| | | | | | | | | |
|---|---|---|---|---|---|---|---|---|
| 0.598846 | 0.334568 | 0.016 | 0.009 | 1 | 13 | AC139347. | 0.007 | Old |
| 0.598846 | 0.323305 | 0.016 | 0.009 | 1 | 13 | 1700028J1 | 0.007 | Old |
| 0.598846 | 0.315372 | 0.016 | 0.009 | 1 | 13 | Ska1 | 0.007 | Old |
| 0.598846 | 0.314309 | 0.016 | 0.009 | 1 | 13 | Thap8 | 0.007 | Old |
| 0.598846 | 0.313125 | 0.016 | 0.009 | 1 | 13 | Cerkl | 0.007 | Old |
| 0.598846 | 0.264387 | 0.016 | 0.009 | 1 | 13 | Lrr1 | 0.007 | Old |
| 0.598846 | 0.251397 | 0.016 | 0.009 | 1 | 13 | Dlg5 | 0.007 | Old |
| 0.599087 | 0.056309 | 0.082 | 0.061 | 1 | 13 | Szt2 | 0.021 | Old |
| 0.599119 | 0.108556 | 0.984 | 0.956 | 1 | 13 | Cnbp | 0.028 | Old |
| 0.599232 | 0.175518 | 0.328 | 0.281 | 1 | 13 | Fbxo30 | 0.047 | Old |
| 0.599345 | 0.058524 | 0.836 | 0.798 | 1 | 13 | Bod1l | 0.038 | Old |
| 0.599626 | 0.239616 | 0.82 | 0.851 | 1 | 13 | Rab11fip1 | -0.031 | Old |
| 0.599882 | 0.090971 | 0.377 | 0.325 | 1 | 13 | Fgd3 | 0.052 | Old |
| 0.600021 | 0.036845 | 0.648 | 0.553 | 1 | 13 | Ubald1 | 0.095 | Old |
| 0.600104 | 0.234109 | 0.418 | 0.377 | 1 | 13 | Cav2 | 0.041 | Old |
| 0.600437 | 0.153757 | 0.967 | 0.947 | 1 | 13 | Lipa | 0.02 | Old |
| 0.600944 | 0.082248 | 0.123 | 0.096 | 1 | 13 | Shq1 | 0.027 | Old |
| 0.601562 | 0.091862 | 0.91 | 0.877 | 1 | 13 | Snrpb2 | 0.033 | Old |
| 0.601607 | 0.230132 | 0.049 | 0.035 | 1 | 13 | Znrf3 | 0.014 | Old |
| 0.601607 | 0.230044 | 0.049 | 0.035 | 1 | 13 | Gm31105 | 0.014 | Old |
| 0.601607 | 0.226739 | 0.049 | 0.035 | 1 | 13 | Tnfsf10 | 0.014 | Old |
| 0.601607 | 0.208011 | 0.049 | 0.035 | 1 | 13 | Gpx7 | 0.014 | Old |
| 0.601607 | 0.206001 | 0.049 | 0.035 | 1 | 13 | Sh2d3c | 0.014 | Old |
| 0.601729 | 0.088196 | 0.77 | 0.737 | 1 | 13 | Prkd3 | 0.033 | Old |
| 0.601841 | 0.217866 | 0.352 | 0.333 | 1 | 13 | Srm | 0.019 | Old |
| 0.601868 | 0.120104 | 0.754 | 0.763 | 1 | 13 | Larp4 | -0.009 | Old |
| 0.601999 | 0.1399 | 0.131 | 0.105 | 1 | 13 | Poll | 0.026 | Old |
| 0.602346 | 0.228465 | 0.148 | 0.123 | 1 | 13 | Dcaf4 | 0.025 | Old |
| 0.602532 | 0.071245 | 0.475 | 0.412 | 1 | 13 | Dnmt1 | 0.063 | Old |
| 0.602951 | 0.131334 | 0.197 | 0.167 | 1 | 13 | Sephs1 | 0.03 | Old |
| 0.603033 | 0.015111 | 0.164 | 0.132 | 1 | 13 | Taf5 | 0.032 | Old |
| 0.604049 | 0.175566 | 0.385 | 0.333 | 1 | 13 | Cdadc1 | 0.052 | Old |
| 0.604069 | 0.086719 | 0.32 | 0.281 | 1 | 13 | Zfp524 | 0.039 | Old |
| 0.6045 | 0.063068 | 0.525 | 0.447 | 1 | 13 | Cnot10 | 0.078 | Old |
| 0.604582 | 0.3029 | 0.148 | 0.123 | 1 | 13 | Hgf | 0.025 | Old |
| 0.604637 | 0.069539 | 0.59 | 0.553 | 1 | 13 | Fcf1 | 0.037 | Old |
| 0.60473 | 0.252267 | 0.139 | 0.114 | 1 | 13 | Gm46430 | 0.025 | Old |
| 0.604857 | 0.032928 | 0.352 | 0.289 | 1 | 13 | Ccdc85b | 0.063 | Old |
| 0.605024 | 0.035803 | 0.91 | 0.939 | 1 | 13 | Vasp | -0.029 | Old |
| 0.605418 | 0.247128 | 0.049 | 0.035 | 1 | 13 | Wdr5b | 0.014 | Old |
| 0.605418 | 0.225482 | 0.049 | 0.035 | 1 | 13 | Ifi214 | 0.014 | Old |
| 0.605418 | 0.217866 | 0.049 | 0.035 | 1 | 13 | Mief2 | 0.014 | Old |
| 0.605625 | 0.322475 | 0.205 | 0.175 | 1 | 13 | 5430427O1 | 0.03 | Old |
| 0.605695 | 2.163948 | 0.016 | 0.009 | 1 | 13 | Retnlg | 0.007 | Old |
| 0.605695 | 1.172842 | 0.016 | 0.009 | 1 | 13 | Ly6k | 0.007 | Old |

| | | | | | | | | |
|---|---|---|---|---|---|---|---|---|
| 0.605695 | 1.027252 | 0.016 | 0.009 | 1 | 13 | Gsdma | 0.007 | Old |
| 0.605695 | 0.698044 | 0.016 | 0.009 | 1 | 13 | Hexim2 | 0.007 | Old |
| 0.605695 | 0.671276 | 0.016 | 0.009 | 1 | 13 | 1700055D1 | 0.007 | Old |
| 0.605695 | 0.595408 | 0.016 | 0.009 | 1 | 13 | Chst2 | 0.007 | Old |
| 0.605695 | 0.562404 | 0.016 | 0.009 | 1 | 13 | Derl3 | 0.007 | Old |
| 0.605695 | 0.547121 | 0.016 | 0.009 | 1 | 13 | Mafa | 0.007 | Old |
| 0.605695 | 0.527742 | 0.016 | 0.009 | 1 | 13 | Akr1b7 | 0.007 | Old |
| 0.605695 | 0.468349 | 0.016 | 0.009 | 1 | 13 | Gm43597 | 0.007 | Old |
| 0.605695 | 0.465879 | 0.016 | 0.009 | 1 | 13 | Ccne2 | 0.007 | Old |
| 0.605695 | 0.44993 | 0.016 | 0.009 | 1 | 13 | H1fx | 0.007 | Old |
| 0.605695 | 0.449017 | 0.016 | 0.009 | 1 | 13 | Gm6598 | 0.007 | Old |
| 0.605695 | 0.447558 | 0.016 | 0.009 | 1 | 13 | Pdpn | 0.007 | Old |
| 0.605695 | 0.446176 | 0.016 | 0.009 | 1 | 13 | Cercam | 0.007 | Old |
| 0.605695 | 0.432981 | 0.016 | 0.009 | 1 | 13 | Loxl2 | 0.007 | Old |
| 0.605695 | 0.410285 | 0.016 | 0.009 | 1 | 13 | Jcad | 0.007 | Old |
| 0.605695 | 0.407496 | 0.016 | 0.009 | 1 | 13 | Ptcra | 0.007 | Old |
| 0.605695 | 0.404204 | 0.016 | 0.009 | 1 | 13 | Bach2os | 0.007 | Old |
| 0.605695 | 0.402401 | 0.016 | 0.009 | 1 | 13 | Cpt1b | 0.007 | Old |
| 0.605695 | 0.397247 | 0.016 | 0.009 | 1 | 13 | Mxd3 | 0.007 | Old |
| 0.605695 | 0.374497 | 0.016 | 0.009 | 1 | 13 | 1010001B2 | 0.007 | Old |
| 0.605695 | 0.354434 | 0.016 | 0.009 | 1 | 13 | 4732491K2 | 0.007 | Old |
| 0.605695 | 0.333748 | 0.016 | 0.009 | 1 | 13 | Esyt3 | 0.007 | Old |
| 0.605695 | 0.329791 | 0.016 | 0.009 | 1 | 13 | Hus1b | 0.007 | Old |
| 0.605695 | 0.320015 | 0.016 | 0.009 | 1 | 13 | Cldn12 | 0.007 | Old |
| 0.605695 | 0.312231 | 0.016 | 0.009 | 1 | 13 | Pcdhga7 | 0.007 | Old |
| 0.605695 | 0.301905 | 0.016 | 0.009 | 1 | 13 | Hba-a2 | 0.007 | Old |
| 0.605695 | 0.299722 | 0.016 | 0.009 | 1 | 13 | Calr3 | 0.007 | Old |
| 0.605695 | 0.297532 | 0.016 | 0.009 | 1 | 13 | Ablim2 | 0.007 | Old |
| 0.605695 | 0.288797 | 0.016 | 0.009 | 1 | 13 | Armcx1 | 0.007 | Old |
| 0.605695 | 0.286322 | 0.016 | 0.009 | 1 | 13 | Mindy4 | 0.007 | Old |
| 0.605695 | 0.286045 | 0.016 | 0.009 | 1 | 13 | Cdca2 | 0.007 | Old |
| 0.605695 | 0.281121 | 0.016 | 0.009 | 1 | 13 | Gm5113 | 0.007 | Old |
| 0.605695 | 0.272038 | 0.016 | 0.009 | 1 | 13 | Itga10 | 0.007 | Old |
| 0.605695 | 0.271441 | 0.016 | 0.009 | 1 | 13 | Gm43796 | 0.007 | Old |
| 0.605695 | 0.260889 | 0.016 | 0.009 | 1 | 13 | Gm16933 | 0.007 | Old |
| 0.605695 | 0.252782 | 0.016 | 0.009 | 1 | 13 | Ticrr | 0.007 | Old |
| 0.605695 | 0.23922 | 0.016 | 0.009 | 1 | 13 | Gm33280 | 0.007 | Old |
| 0.605695 | 0.231223 | 0.016 | 0.009 | 1 | 13 | Adcy4 | 0.007 | Old |
| 0.605695 | 0.230926 | 0.016 | 0.009 | 1 | 13 | 2010013B2 | 0.007 | Old |
| 0.605695 | 0.223249 | 0.016 | 0.009 | 1 | 13 | Raver1 | 0.007 | Old |
| 0.605695 | 0.223086 | 0.016 | 0.009 | 1 | 13 | Peg12 | 0.007 | Old |
| 0.605695 | 0.218663 | 0.016 | 0.009 | 1 | 13 | Cmya5 | 0.007 | Old |
| 0.605695 | 0.215067 | 0.016 | 0.009 | 1 | 13 | Esam | 0.007 | Old |
| 0.605695 | 0.214086 | 0.016 | 0.009 | 1 | 13 | Phf21b | 0.007 | Old |
| 0.605695 | 0.213334 | 0.016 | 0.009 | 1 | 13 | Mex3a | 0.007 | Old |

| | | | | | | | | |
|---|---|---|---|---|---|---|---|---|
| 0.605695 | 0.212714 | 0.016 | 0.009 | 1 | 13 | Gm45474 | 0.007 | Old |
| 0.605695 | 0.208546 | 0.016 | 0.009 | 1 | 13 | Kit | 0.007 | Old |
| 0.605695 | 0.206519 | 0.016 | 0.009 | 1 | 13 | Sel1l3 | 0.007 | Old |
| 0.605695 | 0.204109 | 0.016 | 0.009 | 1 | 13 | Fgfrl1 | 0.007 | Old |
| 0.605695 | 0.204106 | 0.016 | 0.009 | 1 | 13 | Tnfaip8l3 | 0.007 | Old |
| 0.605695 | 0.204094 | 0.016 | 0.009 | 1 | 13 | Srpx | 0.007 | Old |
| 0.605695 | 0.189977 | 0.016 | 0.009 | 1 | 13 | Gm26764 | 0.007 | Old |
| 0.605695 | 0.189726 | 0.016 | 0.009 | 1 | 13 | Tnfrsf4 | 0.007 | Old |
| 0.605695 | 0.182962 | 0.016 | 0.009 | 1 | 13 | Gm9484 | 0.007 | Old |
| 0.605695 | 0.131909 | 0.016 | 0.009 | 1 | 13 | Prrt1 | 0.007 | Old |
| 0.606014 | 0.206205 | 0.098 | 0.079 | 1 | 13 | Anxa9 | 0.019 | Old |
| 0.60615 | 0.09093 | 0.787 | 0.746 | 1 | 13 | Sf3b3 | 0.041 | Old |
| 0.606664 | 0.051561 | 0.508 | 0.456 | 1 | 13 | Tbc1d9b | 0.052 | Old |
| 0.606723 | 0.097634 | 0.246 | 0.211 | 1 | 13 | Nrde2 | 0.035 | Old |
| 0.606977 | 0.023874 | 0.926 | 0.895 | 1 | 13 | Sf3b5 | 0.031 | Old |
| 0.607033 | 0.291528 | 0.139 | 0.114 | 1 | 13 | Prkd2 | 0.025 | Old |
| 0.607172 | 0.161714 | 0.352 | 0.298 | 1 | 13 | Jagn1 | 0.054 | Old |
| 0.607196 | 0.044624 | 0.172 | 0.14 | 1 | 13 | Rnf135 | 0.032 | Old |
| 0.607293 | 0.109195 | 0.533 | 0.456 | 1 | 13 | Ylpm1 | 0.077 | Old |
| 0.607578 | 0.230483 | 0.107 | 0.088 | 1 | 13 | C330018D2 | 0.019 | Old |
| 0.607625 | 0.095881 | 0.926 | 0.886 | 1 | 13 | Mkrn1 | 0.04 | Old |
| 0.60783 | 0.0515 | 0.27 | 0.228 | 1 | 13 | Wdr12 | 0.042 | Old |
| 0.608016 | 0.329916 | 0.246 | 0.228 | 1 | 13 | Elac1 | 0.018 | Old |
| 0.608079 | 0.046973 | 0.549 | 0.5 | 1 | 13 | 3110043O2 | 0.049 | Old |
| 0.608546 | 0.013304 | 0.361 | 0.404 | 1 | 13 | Nagpa | -0.043 | Old |
| 0.608682 | 0.114776 | 0.451 | 0.377 | 1 | 13 | Kat7 | 0.074 | Old |
| 0.608833 | 0.021191 | 0.828 | 0.825 | 1 | 13 | Pacsin2 | 0.003 | Old |
| 0.609186 | 0.622695 | 0.057 | 0.044 | 1 | 13 | Clec2g | 0.013 | Old |
| 0.609241 | 0.344965 | 0.049 | 0.035 | 1 | 13 | Hist1h1b | 0.014 | Old |
| 0.609241 | 0.084929 | 0.049 | 0.035 | 1 | 13 | Dennd5b | 0.014 | Old |
| 0.609241 | 0.077693 | 0.049 | 0.035 | 1 | 13 | Shf | 0.014 | Old |
| 0.609241 | 0.038892 | 0.049 | 0.035 | 1 | 13 | Ccna2 | 0.014 | Old |
| 0.609431 | 0.024005 | 0.189 | 0.149 | 1 | 13 | Ttc27 | 0.04 | Old |
| 0.609608 | 0.096432 | 0.918 | 0.886 | 1 | 13 | Sdf4 | 0.032 | Old |
| 0.609877 | 0.107662 | 0.295 | 0.254 | 1 | 13 | Coq5 | 0.041 | Old |
| 0.609908 | 0.06152 | 0.893 | 0.816 | 1 | 13 | Sowahc | 0.077 | Old |
| 0.610205 | 0.098834 | 0.754 | 0.702 | 1 | 13 | Mob4 | 0.052 | Old |
| 0.610516 | 0.406784 | 0.139 | 0.167 | 1 | 13 | Gbp2b | -0.028 | Old |
| 0.610923 | 0.092431 | 0.443 | 0.386 | 1 | 13 | Phax | 0.057 | Old |
| 0.610988 | 0.027805 | 0.016 | 0.026 | 1 | 13 | Hoxa7 | -0.01 | Old |
| 0.61134 | 0.358473 | 0.279 | 0.263 | 1 | 13 | Mvd | 0.016 | Old |
| 0.611777 | 0.104902 | 0.295 | 0.263 | 1 | 13 | Tmem250- | 0.032 | Old |
| 0.611784 | 0.148556 | 0.09 | 0.07 | 1 | 13 | Fadd | 0.02 | Old |
| 0.611784 | 0.127097 | 0.09 | 0.07 | 1 | 13 | Setd4 | 0.02 | Old |
| 0.611844 | 0.008451 | 0.361 | 0.298 | 1 | 13 | Fam3a | 0.063 | Old |

| | | | | | | | | |
|---|---|---|---|---|---|---|---|---|
| 0.611897 | 0.147221 | 0.221 | 0.193 | 1 | 13 | Mier2 | 0.028 | Old |
| 0.612554 | 0.083428 | 0.82 | 0.798 | 1 | 13 | Eny2 | 0.022 | Old |
| 0.612579 | 0.280281 | 0.016 | 0.009 | 1 | 13 | Tgtp1 | 0.007 | Old |
| 0.612579 | 0.258797 | 0.016 | 0.009 | 1 | 13 | Acer2 | 0.007 | Old |
| 0.612579 | 0.249598 | 0.016 | 0.009 | 1 | 13 | Tex15 | 0.007 | Old |
| 0.612579 | 0.244095 | 0.016 | 0.009 | 1 | 13 | Tmeff1 | 0.007 | Old |
| 0.612579 | 0.236238 | 0.016 | 0.009 | 1 | 13 | Gm43403 | 0.007 | Old |
| 0.612579 | 0.232313 | 0.016 | 0.009 | 1 | 13 | Gm45799 | 0.007 | Old |
| 0.612579 | 0.230261 | 0.016 | 0.009 | 1 | 13 | Gm39121 | 0.007 | Old |
| 0.612579 | 0.228518 | 0.016 | 0.009 | 1 | 13 | Serpinb6b | 0.007 | Old |
| 0.612579 | 0.227107 | 0.016 | 0.009 | 1 | 13 | Gm26714 | 0.007 | Old |
| 0.612579 | 0.221367 | 0.016 | 0.009 | 1 | 13 | Rab4a | 0.007 | Old |
| 0.612579 | 0.220638 | 0.016 | 0.009 | 1 | 13 | Brdt | 0.007 | Old |
| 0.612579 | 0.21875 | 0.016 | 0.009 | 1 | 13 | Txlnb | 0.007 | Old |
| 0.612579 | 0.218637 | 0.016 | 0.009 | 1 | 13 | Als2cl | 0.007 | Old |
| 0.612579 | 0.212546 | 0.016 | 0.009 | 1 | 13 | Gm10605 | 0.007 | Old |
| 0.612579 | 0.210597 | 0.016 | 0.009 | 1 | 13 | Gm26797 | 0.007 | Old |
| 0.612579 | 0.204704 | 0.016 | 0.009 | 1 | 13 | Gm9888 | 0.007 | Old |
| 0.612579 | 0.20358 | 0.016 | 0.009 | 1 | 13 | Tmem82 | 0.007 | Old |
| 0.612579 | 0.199364 | 0.016 | 0.009 | 1 | 13 | B230369F2 | 0.007 | Old |
| 0.612579 | 0.196965 | 0.016 | 0.009 | 1 | 13 | Cstad | 0.007 | Old |
| 0.612579 | 0.193612 | 0.016 | 0.009 | 1 | 13 | Pcdhga6 | 0.007 | Old |
| 0.612579 | 0.193507 | 0.016 | 0.009 | 1 | 13 | Xcr1 | 0.007 | Old |
| 0.612579 | 0.190367 | 0.016 | 0.009 | 1 | 13 | Spns2 | 0.007 | Old |
| 0.612579 | 0.187449 | 0.016 | 0.009 | 1 | 13 | Slc4a8 | 0.007 | Old |
| 0.612579 | 0.187265 | 0.016 | 0.009 | 1 | 13 | Hist1h2bj | 0.007 | Old |
| 0.612579 | 0.175672 | 0.016 | 0.009 | 1 | 13 | 2010007H( | 0.007 | Old |
| 0.612579 | 0.173941 | 0.016 | 0.009 | 1 | 13 | Faah | 0.007 | Old |
| 0.612579 | 0.17305 | 0.016 | 0.009 | 1 | 13 | Grin2d | 0.007 | Old |
| 0.612579 | 0.172272 | 0.016 | 0.009 | 1 | 13 | 4930524O( | 0.007 | Old |
| 0.612579 | 0.16988 | 0.016 | 0.009 | 1 | 13 | Prss23 | 0.007 | Old |
| 0.612579 | 0.167535 | 0.016 | 0.009 | 1 | 13 | Egln3 | 0.007 | Old |
| 0.612579 | 0.166058 | 0.016 | 0.009 | 1 | 13 | Cux2 | 0.007 | Old |
| 0.612579 | 0.16366 | 0.016 | 0.009 | 1 | 13 | Pla1a | 0.007 | Old |
| 0.612579 | 0.163428 | 0.016 | 0.009 | 1 | 13 | Dync2li1 | 0.007 | Old |
| 0.612579 | 0.15769 | 0.016 | 0.009 | 1 | 13 | Igfbp5 | 0.007 | Old |
| 0.612579 | 0.155921 | 0.016 | 0.009 | 1 | 13 | Gm10037 | 0.007 | Old |
| 0.612579 | 0.154796 | 0.016 | 0.009 | 1 | 13 | Rmi2 | 0.007 | Old |
| 0.612579 | 0.15295 | 0.016 | 0.009 | 1 | 13 | Tgfbr3l | 0.007 | Old |
| 0.612579 | 0.149442 | 0.016 | 0.009 | 1 | 13 | Fgf1 | 0.007 | Old |
| 0.612579 | 0.147455 | 0.016 | 0.009 | 1 | 13 | Gm16982 | 0.007 | Old |
| 0.612579 | 0.134584 | 0.016 | 0.009 | 1 | 13 | Spsb1 | 0.007 | Old |
| 0.612579 | 0.1184 | 0.016 | 0.009 | 1 | 13 | Ccdc62 | 0.007 | Old |
| 0.612579 | 0.112404 | 0.016 | 0.009 | 1 | 13 | Zfp493 | 0.007 | Old |
| 0.612579 | 0.102148 | 0.016 | 0.009 | 1 | 13 | P2rx1 | 0.007 | Old |

| | | | | | | | | |
|---|---|---|---|---|---|---|---|---|
| 0.612579 | 0.101863 | 0.016 | 0.009 | 1 | 13 | Slc17a8 | 0.007 | Old |
| 0.612579 | 0.10174 | 0.016 | 0.009 | 1 | 13 | H3f3aos | 0.007 | Old |
| 0.612579 | 0.085223 | 0.016 | 0.009 | 1 | 13 | Fgf10 | 0.007 | Old |
| 0.612579 | 0.080027 | 0.016 | 0.009 | 1 | 13 | Kcna3 | 0.007 | Old |
| 0.612579 | 0.076626 | 0.016 | 0.009 | 1 | 13 | Tmem198 | 0.007 | Old |
| 0.612579 | 0.073326 | 0.016 | 0.009 | 1 | 13 | E2f7 | 0.007 | Old |
| 0.612579 | 0.055351 | 0.016 | 0.009 | 1 | 13 | D630033O: | 0.007 | Old |
| 0.612579 | 0.042612 | 0.016 | 0.009 | 1 | 13 | Vipr1 | 0.007 | Old |
| 0.612579 | 0.041684 | 0.016 | 0.009 | 1 | 13 | Nek2 | 0.007 | Old |
| 0.612579 | 0.03497 | 0.016 | 0.009 | 1 | 13 | Rab11fip4c | 0.007 | Old |
| 0.612579 | 0.022775 | 0.016 | 0.009 | 1 | 13 | Tacstd2 | 0.007 | Old |
| 0.612579 | 0.017133 | 0.016 | 0.009 | 1 | 13 | Tssk4 | 0.007 | Old |
| 0.612579 | 0.01585 | 0.016 | 0.009 | 1 | 13 | Grik5 | 0.007 | Old |
| 0.612579 | 0.014563 | 0.016 | 0.009 | 1 | 13 | Dact1 | 0.007 | Old |
| 0.612579 | 0.012746 | 0.016 | 0.009 | 1 | 13 | Glyctk | 0.007 | Old |
| 0.612579 | 0.008158 | 0.016 | 0.009 | 1 | 13 | Gm14418 | 0.007 | Old |
| 0.612644 | 0.17993 | 0.352 | 0.307 | 1 | 13 | Gphn | 0.045 | Old |
| 0.61302 | 0.022371 | 0.533 | 0.456 | 1 | 13 | Foxj2 | 0.077 | Old |
| 0.613074 | 0.312642 | 0.049 | 0.035 | 1 | 13 | Tmc8 | 0.014 | Old |
| 0.613074 | 0.148179 | 0.049 | 0.035 | 1 | 13 | Zfp428 | 0.014 | Old |
| 0.613074 | 0.075262 | 0.049 | 0.035 | 1 | 13 | AW011738 | 0.014 | Old |
| 0.61315 | 0.12701 | 0.484 | 0.456 | 1 | 13 | Nelfe | 0.028 | Old |
| 0.613392 | 0.138675 | 0.779 | 0.702 | 1 | 13 | Slc29a1 | 0.077 | Old |
| 0.613523 | 0.162301 | 0.893 | 0.877 | 1 | 13 | Cers2 | 0.016 | Old |
| 0.614049 | 0.133497 | 0.189 | 0.158 | 1 | 13 | Cdca8 | 0.031 | Old |
| 0.614388 | 0.057215 | 0.533 | 0.465 | 1 | 13 | Impa1 | 0.068 | Old |
| 0.614544 | 0.107226 | 0.664 | 0.553 | 1 | 13 | Rgl2 | 0.111 | Old |
| 0.614634 | 0.05755 | 0.459 | 0.404 | 1 | 13 | Med11 | 0.055 | Old |
| 0.615173 | 0.003531 | 0.844 | 0.851 | 1 | 13 | Eif3j1 | -0.007 | Old |
| 0.615379 | 0.585292 | 0.107 | 0.088 | 1 | 13 | Megf9 | 0.019 | Old |
| 0.615716 | 0.107782 | 0.434 | 0.377 | 1 | 13 | Plcb1 | 0.057 | Old |
| 0.615814 | 0.13553 | 0.148 | 0.123 | 1 | 13 | Phf7 | 0.025 | Old |
| 0.616087 | 0.082088 | 0.615 | 0.561 | 1 | 13 | Dars | 0.054 | Old |
| 0.616332 | 0.097233 | 0.336 | 0.289 | 1 | 13 | Kif1bp | 0.047 | Old |
| 0.61654 | 0.067166 | 0.525 | 0.474 | 1 | 13 | C330007P0 | 0.051 | Old |
| 0.616918 | 0.112795 | 0.049 | 0.035 | 1 | 13 | Abcb9 | 0.014 | Old |
| 0.616918 | 0.070885 | 0.049 | 0.035 | 1 | 13 | Dpf3 | 0.014 | Old |
| 0.617566 | 0.138529 | 0.648 | 0.632 | 1 | 13 | Tpm1 | 0.016 | Old |
| 0.61764 | 0.154299 | 0.623 | 0.588 | 1 | 13 | Ncoa1 | 0.035 | Old |
| 0.617686 | 0.205789 | 0.279 | 0.246 | 1 | 13 | Abhd5 | 0.033 | Old |
| 0.617989 | 0.343689 | 0.107 | 0.088 | 1 | 13 | Ctps | 0.019 | Old |
| 0.617989 | 0.231997 | 0.107 | 0.088 | 1 | 13 | 2310057M | 0.019 | Old |
| 0.618144 | 0.028069 | 0.557 | 0.482 | 1 | 13 | Guca1a | 0.075 | Old |
| 0.618151 | 0.165261 | 0.77 | 0.754 | 1 | 13 | Stt3a | 0.016 | Old |
| 0.618161 | 0.048774 | 0.779 | 0.667 | 1 | 13 | Guk1 | 0.112 | Old |

| | | | | | | | | |
|---|---|---|---|---|---|---|---|---|
| 0.618551 | 0.049206 | 0.393 | 0.351 | 1 | 13 | Calhm2 | 0.042 | Old |
| 0.618576 | 0.06999 | 0.59 | 0.553 | 1 | 13 | Sf3a3 | 0.037 | Old |
| 0.619497 | 0.21284 | 0.213 | 0.184 | 1 | 13 | Atat1 | 0.029 | Old |
| 0.619653 | 0.10691 | 0.852 | 0.807 | 1 | 13 | Fam104a | 0.045 | Old |
| 0.619754 | 0.290756 | 0.057 | 0.044 | 1 | 13 | 9130221H1 | 0.013 | Old |
| 0.619959 | 0.255155 | 0.541 | 0.553 | 1 | 13 | Katna1 | -0.012 | Old |
| 0.620057 | 0.045939 | 0.402 | 0.36 | 1 | 13 | Tmem86a | 0.042 | Old |
| 0.62032 | 0.249444 | 0.09 | 0.07 | 1 | 13 | Hist1h2ae | 0.02 | Old |
| 0.62032 | 0.14915 | 0.09 | 0.07 | 1 | 13 | Tmem62 | 0.02 | Old |
| 0.62038 | 0.118048 | 0.77 | 0.825 | 1 | 13 | Krcc1 | -0.055 | Old |
| 0.620507 | 0.171606 | 1 | 1 | 1 | 13 | Tmem176a | 0 | Old |
| 0.620628 | 0.133094 | 0.123 | 0.096 | 1 | 13 | Zfp668 | 0.027 | Old |
| 0.620772 | 0.119975 | 0.049 | 0.035 | 1 | 13 | Gm17586 | 0.014 | Old |
| 0.620931 | 0.114343 | 0.139 | 0.114 | 1 | 13 | Zkscan8 | 0.025 | Old |
| 0.621178 | 0.049342 | 0.975 | 0.982 | 1 | 13 | Rab8b | -0.007 | Old |
| 0.621261 | 0.083128 | 0.557 | 0.465 | 1 | 13 | Cdip1 | 0.092 | Old |
| 0.621261 | 0.056054 | 0.541 | 0.482 | 1 | 13 | Srp54a | 0.059 | Old |
| 0.621814 | 0.042068 | 0.344 | 0.298 | 1 | 13 | Ndufaf7 | 0.046 | Old |
| 0.621854 | 0.042914 | 1 | 1 | 1 | 13 | Cdc42 | 0 | Old |
| 0.622289 | 0.053848 | 0.664 | 0.632 | 1 | 13 | C1d | 0.032 | Old |
| 0.622302 | 0.181345 | 0.098 | 0.079 | 1 | 13 | Pdk2 | 0.019 | Old |
| 0.622302 | 0.170639 | 0.098 | 0.079 | 1 | 13 | Fpgs | 0.019 | Old |
| 0.622597 | 0.201352 | 0.148 | 0.123 | 1 | 13 | Nod1 | 0.025 | Old |
| 0.622851 | 0.064421 | 0.197 | 0.167 | 1 | 13 | 6720427I07 | 0.03 | Old |
| 0.622879 | 0.117881 | 0.369 | 0.298 | 1 | 13 | Cwc22 | 0.071 | Old |
| 0.622993 | 0.075187 | 0.82 | 0.763 | 1 | 13 | Asph | 0.057 | Old |
| 0.623062 | 0.043928 | 0.918 | 0.895 | 1 | 13 | Sqstm1 | 0.023 | Old |
| 0.623087 | 0.191838 | 0.803 | 0.789 | 1 | 13 | Zmynd8 | 0.014 | Old |
| 0.623177 | 0.001637 | 0.09 | 0.07 | 1 | 13 | Dnmt3b | 0.02 | Old |
| 0.623294 | 0.367028 | 0.057 | 0.044 | 1 | 13 | Usp54 | 0.013 | Old |
| 0.623294 | 0.345852 | 0.057 | 0.044 | 1 | 13 | Ska2 | 0.013 | Old |
| 0.623294 | 0.343769 | 0.057 | 0.044 | 1 | 13 | Numbl | 0.013 | Old |
| 0.623531 | 0.049967 | 0.525 | 0.491 | 1 | 13 | Slc25a1 | 0.034 | Old |
| 0.624637 | 0.173333 | 0.049 | 0.035 | 1 | 13 | Dnph1 | 0.014 | Old |
| 0.624637 | 0.023327 | 0.049 | 0.035 | 1 | 13 | Acsf2 | 0.014 | Old |
| 0.62476 | 0.206225 | 0.385 | 0.342 | 1 | 13 | Mrps34 | 0.043 | Old |
| 0.62495 | 0.147628 | 0.27 | 0.237 | 1 | 13 | Pithd1 | 0.033 | Old |
| 0.625119 | 0.100398 | 0.73 | 0.693 | 1 | 13 | Kat6a | 0.037 | Old |
| 0.625249 | 0.058931 | 0.689 | 0.658 | 1 | 13 | Rbm17 | 0.031 | Old |
| 0.625269 | 0.325734 | 0.443 | 0.439 | 1 | 13 | Dpep2 | 0.004 | Old |
| 0.625439 | 0.125145 | 0.336 | 0.281 | 1 | 13 | Prkaa1 | 0.055 | Old |
| 0.625908 | 0.086116 | 0.385 | 0.333 | 1 | 13 | Tada1 | 0.052 | Old |
| 0.626341 | 0.188974 | 0.648 | 0.588 | 1 | 13 | Kmt2d | 0.06 | Old |
| 0.626843 | 0.524749 | 0.057 | 0.044 | 1 | 13 | Xlr4a | 0.013 | Old |
| 0.627399 | 0.060778 | 0.77 | 0.737 | 1 | 13 | Tmem219 | 0.033 | Old |

| | | | | | | | | |
|---|---|---|---|---|---|---|---|---|
| 0.627482 | 0.095712 | 0.377 | 0.342 | 1 | 13 | Rnf170 | 0.035 | Old |
| 0.627482 | 0.079372 | 0.385 | 0.333 | 1 | 13 | Bmi1 | 0.052 | Old |
| 0.627752 | 0.096878 | 0.893 | 0.904 | 1 | 13 | Fkbp8 | -0.011 | Old |
| 0.627773 | 0.093324 | 0.098 | 0.079 | 1 | 13 | Gm47675 | 0.019 | Old |
| 0.628406 | 0.058279 | 0.918 | 0.868 | 1 | 13 | Mapk3 | 0.05 | Old |
| 0.628803 | 0.073929 | 0.164 | 0.14 | 1 | 13 | Btf3l4 | 0.024 | Old |
| 0.629106 | 0.05678 | 0.516 | 0.447 | 1 | 13 | Pam16 | 0.069 | Old |
| 0.629143 | 0.188505 | 0.336 | 0.307 | 1 | 13 | Stk16 | 0.029 | Old |
| 0.629143 | 0.025076 | 0.352 | 0.289 | 1 | 13 | Fam207a | 0.063 | Old |
| 0.629174 | 0.105609 | 0.246 | 0.219 | 1 | 13 | Sass6 | 0.027 | Old |
| 0.629428 | 0.081288 | 0.566 | 0.482 | 1 | 13 | Fam133b | 0.084 | Old |
| 0.629583 | 0.453441 | 0.172 | 0.158 | 1 | 13 | Uck1 | 0.014 | Old |
| 0.630097 | 0.186673 | 0.279 | 0.254 | 1 | 13 | Pde12 | 0.025 | Old |
| 0.630227 | 0.117687 | 0.238 | 0.202 | 1 | 13 | Rpa2 | 0.036 | Old |
| 0.6304 | 0.313888 | 0.057 | 0.044 | 1 | 13 | Fam234b | 0.013 | Old |
| 0.630485 | 0.087303 | 0.246 | 0.211 | 1 | 13 | Gramd1b | 0.035 | Old |
| 0.630516 | 0.158602 | 0.098 | 0.079 | 1 | 13 | Sgsm3 | 0.019 | Old |
| 0.630737 | 0.070444 | 0.811 | 0.737 | 1 | 13 | Immt | 0.074 | Old |
| 0.631326 | 0.071733 | 0.549 | 0.474 | 1 | 13 | Get4 | 0.075 | Old |
| 0.631549 | 0.141447 | 0.639 | 0.605 | 1 | 13 | B3gat3 | 0.034 | Old |
| 0.631772 | 0.266719 | 0.238 | 0.211 | 1 | 13 | Fdft1 | 0.027 | Old |
| 0.631782 | 0.036208 | 0.09 | 0.07 | 1 | 13 | Nkrf | 0.02 | Old |
| 0.632045 | 0.147645 | 0.467 | 0.421 | 1 | 13 | 9130401M | 0.046 | Old |
| 0.632055 | 0.066649 | 0.328 | 0.289 | 1 | 13 | Mapk9 | 0.039 | Old |
| 0.632411 | 0.19658 | 0.418 | 0.395 | 1 | 13 | Lrrc8a | 0.023 | Old |
| 0.632442 | 0.120072 | 0.189 | 0.158 | 1 | 13 | Zfp160 | 0.031 | Old |
| 0.632739 | 0.157749 | 0.557 | 0.561 | 1 | 13 | H2-T22 | -0.004 | Old |
| 0.63298 | 0.107422 | 0.631 | 0.579 | 1 | 13 | Nek9 | 0.052 | Old |
| 0.633017 | 0.161746 | 0.541 | 0.509 | 1 | 13 | Dag1 | 0.032 | Old |
| 0.63314 | 0.236489 | 0.164 | 0.14 | 1 | 13 | Smo | 0.024 | Old |
| 0.633615 | 0.061715 | 0.828 | 0.798 | 1 | 13 | Maz | 0.03 | Old |
| 0.633735 | 0.080599 | 0.393 | 0.412 | 1 | 13 | Trpv4 | -0.019 | Old |
| 0.63375 | 0.345602 | 0.107 | 0.088 | 1 | 13 | E330009J0 | 0.019 | Old |
| 0.63505 | 0.13658 | 0.418 | 0.377 | 1 | 13 | Vezt | 0.041 | Old |
| 0.635386 | 0.177019 | 0.598 | 0.588 | 1 | 13 | Erh | 0.01 | Old |
| 0.635579 | 0.186204 | 0.287 | 0.263 | 1 | 13 | Psmg2 | 0.024 | Old |
| 0.635699 | 0.115252 | 0.303 | 0.263 | 1 | 13 | Prr14l | 0.04 | Old |
| 0.635768 | 0.094052 | 0.385 | 0.342 | 1 | 13 | 4930402H2 | 0.043 | Old |
| 0.636024 | 0.120739 | 0.246 | 0.211 | 1 | 13 | Mtmr4 | 0.035 | Old |
| 0.636065 | 0.015343 | 0.959 | 0.956 | 1 | 13 | Cdc42se1 | 0.003 | Old |
| 0.636376 | 0.09847 | 0.869 | 0.868 | 1 | 13 | Socs6 | 0.001 | Old |
| 0.636413 | 0.080579 | 0.738 | 0.667 | 1 | 13 | Cyld | 0.071 | Old |
| 0.636715 | 0.081646 | 0.926 | 0.947 | 1 | 13 | Lcp2 | -0.021 | Old |
| 0.637078 | 0.114921 | 0.156 | 0.132 | 1 | 13 | Ddx51 | 0.024 | Old |
| 0.637351 | 0.553556 | 0.475 | 0.439 | 1 | 13 | Bmp2 | 0.036 | Old |

| | | | | | | | | |
|---|---|---|---|---|---|---|---|---|
| 0.637411 | 0.183582 | 0.295 | 0.272 | 1 | 13 | Ciao1 | 0.023 | Old |
| 0.637474 | 0.029266 | 0.77 | 0.684 | 1 | 13 | Htt | 0.086 | Old |
| 0.637541 | 0.281259 | 0.057 | 0.044 | 1 | 13 | A430033K( | 0.013 | Old |
| 0.637546 | 0.030318 | 0.09 | 0.07 | 1 | 13 | Plcg1 | 0.02 | Old |
| 0.637875 | 0.110046 | 0.246 | 0.211 | 1 | 13 | Strada | 0.035 | Old |
| 0.637972 | 0.17817 | 0.574 | 0.535 | 1 | 13 | Zfp451 | 0.039 | Old |
| 0.638051 | 0.130978 | 0.23 | 0.202 | 1 | 13 | Spred2 | 0.028 | Old |
| 0.638571 | 0.06749 | 0.73 | 0.649 | 1 | 13 | Dmxl1 | 0.081 | Old |
| 0.638972 | 0.182667 | 0.197 | 0.167 | 1 | 13 | Bcl9 | 0.03 | Old |
| 0.63945 | 0.262739 | 0.754 | 0.737 | 1 | 13 | Sarnp | 0.017 | Old |
| 0.639555 | 0.009434 | 0.287 | 0.237 | 1 | 13 | Dcaf10 | 0.05 | Old |
| 0.639916 | 0.08021 | 0.508 | 0.474 | 1 | 13 | Mgst1 | 0.034 | Old |
| 0.640092 | 0.021415 | 0.508 | 0.456 | 1 | 13 | BC029722 | 0.052 | Old |
| 0.64015 | 0.058825 | 0.82 | 0.86 | 1 | 13 | Ids | -0.04 | Old |
| 0.640425 | 0.165617 | 0.533 | 0.465 | 1 | 13 | Mapkapk3 | 0.068 | Old |
| 0.64057 | 0.019609 | 0.615 | 0.544 | 1 | 13 | Fbxo11 | 0.071 | Old |
| 0.641005 | 0.137887 | 0.492 | 0.447 | 1 | 13 | Ube4b | 0.045 | Old |
| 0.64154 | 0.047126 | 0.098 | 0.079 | 1 | 13 | Eef1akmt4 | 0.019 | Old |
| 0.642215 | 0.033921 | 1 | 0.991 | 1 | 13 | Picalm | 0.009 | Old |
| 0.642561 | 0.155345 | 0.467 | 0.447 | 1 | 13 | Rcbtb2 | 0.02 | Old |
| 0.642755 | 0.048031 | 0.533 | 0.561 | 1 | 13 | Smim27 | -0.028 | Old |
| 0.64276 | 0.029989 | 0.189 | 0.158 | 1 | 13 | Prkca | 0.031 | Old |
| 0.643029 | 0.076223 | 0.197 | 0.167 | 1 | 13 | Atg10 | 0.03 | Old |
| 0.643568 | 0.105699 | 0.18 | 0.158 | 1 | 13 | Zfp444 | 0.022 | Old |
| 0.643704 | 0.002152 | 0.336 | 0.281 | 1 | 13 | Wdtc1 | 0.055 | Old |
| 0.643875 | 0.069622 | 0.836 | 0.789 | 1 | 13 | Xrn2 | 0.047 | Old |
| 0.643998 | 0.103667 | 0.328 | 0.272 | 1 | 13 | Rhot2 | 0.056 | Old |
| 0.644081 | 0.05688 | 0.82 | 0.833 | 1 | 13 | Mapk1ip1l | -0.013 | Old |
| 0.644232 | 0.054101 | 0.926 | 0.956 | 1 | 13 | Itpkb | -0.03 | Old |
| 0.644531 | 0.210578 | 0.369 | 0.342 | 1 | 13 | Zfp608 | 0.027 | Old |
| 0.64461 | 0.041124 | 0.262 | 0.219 | 1 | 13 | Cryzl2 | 0.043 | Old |
| 0.644644 | 0.116012 | 0.893 | 0.86 | 1 | 13 | Mpp1 | 0.033 | Old |
| 0.644644 | 0.156718 | 0.402 | 0.351 | 1 | 13 | Stk25 | 0.051 | Old |
| 0.644831 | 0.138588 | 0.59 | 0.57 | 1 | 13 | Homer1 | 0.02 | Old |
| 0.644935 | 0.109492 | 0.508 | 0.482 | 1 | 13 | Pbx3 | 0.026 | Old |
| 0.64495 | 0.051652 | 1 | 0.974 | 1 | 13 | Pepd | 0.026 | Old |
| 0.644972 | 0.125927 | 0.344 | 0.289 | 1 | 13 | Tmcc2 | 0.055 | Old |
| 0.644972 | 0.213378 | 0.336 | 0.298 | 1 | 13 | Bloc1s6 | 0.038 | Old |
| 0.645313 | 0.109454 | 0.369 | 0.325 | 1 | 13 | Lztfl1 | 0.044 | Old |
| 0.645438 | 0.037792 | 0.148 | 0.123 | 1 | 13 | Fam206a | 0.025 | Old |
| 0.646124 | 0.203447 | 0.344 | 0.307 | 1 | 13 | Cd2bp2 | 0.037 | Old |
| 0.6462 | 0.443001 | 0.066 | 0.053 | 1 | 13 | Strip2 | 0.013 | Old |
| 0.646295 | 0.204799 | 0.475 | 0.421 | 1 | 13 | Gcsh | 0.054 | Old |
| 0.647034 | 0.06437 | 0.762 | 0.728 | 1 | 13 | Emd | 0.034 | Old |
| 0.647317 | 0.127533 | 0.664 | 0.632 | 1 | 13 | Cdk13 | 0.032 | Old |

| | | | | | | | | |
|---|---|---|---|---|---|---|---|---|
| 0.64736 | 0.320714 | 0.123 | 0.105 | 1 | 13 | Tmem268 | 0.018 | Old |
| 0.647404 | 0.037347 | 0.418 | 0.377 | 1 | 13 | Mfap1a | 0.041 | Old |
| 0.64745 | 0.03702 | 0.828 | 0.816 | 1 | 13 | Ndufv2 | 0.012 | Old |
| 0.647612 | 0.132471 | 0.287 | 0.246 | 1 | 13 | Tlr13 | 0.041 | Old |
| 0.647618 | 0.061194 | 0.918 | 0.93 | 1 | 13 | Tomm22 | -0.012 | Old |
| 0.647675 | 0.314716 | 0.295 | 0.281 | 1 | 13 | Vps45 | 0.014 | Old |
| 0.647883 | 0.112651 | 0.664 | 0.675 | 1 | 13 | Khsrp | -0.011 | Old |
| 0.648106 | 0.111075 | 0.369 | 0.316 | 1 | 13 | Fam219a | 0.053 | Old |
| 0.648316 | 0.183213 | 0.057 | 0.044 | 1 | 13 | Mfsd4b4 | 0.013 | Old |
| 0.648417 | 0.171831 | 0.164 | 0.14 | 1 | 13 | Gm4070 | 0.024 | Old |
| 0.64986 | 0.10916 | 0.098 | 0.079 | 1 | 13 | Tti1 | 0.019 | Old |
| 0.650351 | 0.0294 | 0.91 | 0.93 | 1 | 13 | Raly | -0.02 | Old |
| 0.650427 | 0.033333 | 0.967 | 0.956 | 1 | 13 | Lnpep | 0.011 | Old |
| 0.650682 | 0.139852 | 0.156 | 0.175 | 1 | 13 | Fbxo32 | -0.019 | Old |
| 0.650682 | 0.035748 | 0.18 | 0.149 | 1 | 13 | Sdhaf3 | 0.031 | Old |
| 0.650888 | 0.143465 | 0.41 | 0.368 | 1 | 13 | Lemd3 | 0.042 | Old |
| 0.651153 | 0.264115 | 0.189 | 0.167 | 1 | 13 | Shb | 0.022 | Old |
| 0.651581 | 0.170185 | 0.352 | 0.307 | 1 | 13 | Manbal | 0.045 | Old |
| 0.652385 | 0.027697 | 0.418 | 0.395 | 1 | 13 | Fbxl6 | 0.023 | Old |
| 0.653078 | 0.020209 | 0.795 | 0.728 | 1 | 13 | Emc6 | 0.067 | Old |
| 0.65317 | 0.04717 | 0.959 | 0.947 | 1 | 13 | Sdc4 | 0.012 | Old |
| 0.653219 | 0.113791 | 0.197 | 0.167 | 1 | 13 | Stradb | 0.03 | Old |
| 0.653513 | 0.182739 | 0.123 | 0.149 | 1 | 13 | 9530059O1 | -0.026 | Old |
| 0.654207 | 0.022543 | 0.557 | 0.482 | 1 | 13 | Rxrb | 0.075 | Old |
| 0.654508 | 0.285564 | 0.27 | 0.246 | 1 | 13 | Wdr75 | 0.024 | Old |
| 0.654601 | 0.209402 | 0.361 | 0.325 | 1 | 13 | Acot7 | 0.036 | Old |
| 0.654936 | 0.146666 | 0.18 | 0.149 | 1 | 13 | Frat2 | 0.031 | Old |
| 0.655348 | 0.197298 | 0.344 | 0.325 | 1 | 13 | Akr1b10 | 0.019 | Old |
| 0.655541 | 0.297475 | 0.057 | 0.044 | 1 | 13 | Zfp775 | 0.013 | Old |
| 0.655586 | 0.121855 | 0.254 | 0.228 | 1 | 13 | Dhx32 | 0.026 | Old |
| 0.655674 | 0.179117 | 0.23 | 0.211 | 1 | 13 | Zxdc | 0.019 | Old |
| 0.656271 | 0.344475 | 0.066 | 0.053 | 1 | 13 | Ddb2 | 0.013 | Old |
| 0.656444 | 0.079185 | 0.287 | 0.246 | 1 | 13 | Coq7 | 0.041 | Old |
| 0.656519 | 0.582307 | 0.025 | 0.035 | 1 | 13 | Sntg1 | -0.01 | Old |
| 0.657012 | 0.177267 | 0.754 | 0.711 | 1 | 13 | N4bp1 | 0.043 | Old |
| 0.657099 | 0.298693 | 0.23 | 0.202 | 1 | 13 | Crebl2 | 0.028 | Old |
| 0.657128 | 0.009984 | 0.541 | 0.5 | 1 | 13 | Cul5 | 0.041 | Old |
| 0.657287 | 0.046075 | 0.934 | 0.947 | 1 | 13 | Ezr | -0.013 | Old |
| 0.658035 | 0.286418 | 0.426 | 0.404 | 1 | 13 | Slc38a6 | 0.022 | Old |
| 0.65855 | 0.079274 | 0.754 | 0.719 | 1 | 13 | Dock11 | 0.035 | Old |
| 0.658694 | 0.088883 | 0.148 | 0.167 | 1 | 13 | Vwf | -0.019 | Old |
| 0.659334 | 0.372408 | 0.344 | 0.307 | 1 | 13 | Usp36 | 0.037 | Old |
| 0.659643 | 0.615907 | 0.066 | 0.053 | 1 | 13 | C1qtnf6 | 0.013 | Old |
| 0.659643 | 0.408092 | 0.066 | 0.053 | 1 | 13 | Gm26624 | 0.013 | Old |
| 0.660077 | 0.167294 | 0.566 | 0.526 | 1 | 13 | Xpo7 | 0.04 | Old |

| | | | | | | | | |
|---|---|---|---|---|---|---|---|---|
| 0.660082 | 0.089457 | 1 | 1 | 1 | 13 | Ubc | 0 | Old |
| 0.660886 | 0.224837 | 0.418 | 0.351 | 1 | 13 | Osgin2 | 0.067 | Old |
| 0.661122 | 0.070573 | 0.361 | 0.325 | 1 | 13 | Hscb | 0.036 | Old |
| 0.661428 | 0.004753 | 0.508 | 0.465 | 1 | 13 | Nudcd2 | 0.043 | Old |
| 0.661465 | 0.07856 | 1 | 1 | 1 | 13 | Selenop | 0 | Old |
| 0.661669 | 0.009894 | 0.639 | 0.544 | 1 | 13 | Ss18 | 0.095 | Old |
| 0.661807 | 0.033155 | 0.156 | 0.132 | 1 | 13 | Zbtb11os1 | 0.024 | Old |
| 0.66199 | 0.101293 | 0.877 | 0.921 | 1 | 13 | Ndufab1 | -0.044 | Old |
| 0.662108 | 0.207006 | 0.475 | 0.43 | 1 | 13 | Tbcel | 0.045 | Old |
| 0.662141 | 0.022191 | 0.246 | 0.211 | 1 | 13 | Riox2 | 0.035 | Old |
| 0.662372 | 0.105236 | 0.623 | 0.544 | 1 | 13 | Rangap1 | 0.079 | Old |
| 0.662756 | 0.067462 | 0.369 | 0.316 | 1 | 13 | Rpp25l | 0.053 | Old |
| 0.662797 | 0.145274 | 0.057 | 0.044 | 1 | 13 | Isoc2a | 0.013 | Old |
| 0.662916 | 0.061205 | 0.516 | 0.456 | 1 | 13 | Atg7 | 0.06 | Old |
| 0.663021 | 0.322161 | 0.066 | 0.053 | 1 | 13 | Ablim1 | 0.013 | Old |
| 0.66331 | 0.100605 | 0.172 | 0.149 | 1 | 13 | Dedd2 | 0.023 | Old |
| 0.664193 | 0.020364 | 0.918 | 0.956 | 1 | 13 | St8sia4 | -0.038 | Old |
| 0.664309 | 0.166499 | 0.23 | 0.193 | 1 | 13 | Foxp4 | 0.037 | Old |
| 0.664759 | 0.276856 | 0.311 | 0.272 | 1 | 13 | Dkc1 | 0.039 | Old |
| 0.664768 | 0.054011 | 0.369 | 0.325 | 1 | 13 | Rffl | 0.044 | Old |
| 0.66478 | 0.492934 | 0.623 | 0.605 | 1 | 13 | AW112010 | 0.018 | Old |
| 0.664815 | 0.049392 | 0.361 | 0.316 | 1 | 13 | Ercc3 | 0.045 | Old |
| 0.664968 | 0.096761 | 0.262 | 0.228 | 1 | 13 | Nr1d2 | 0.034 | Old |
| 0.665246 | 0.143248 | 0.287 | 0.263 | 1 | 13 | Ripor2 | 0.024 | Old |
| 0.665304 | 0.107364 | 0.762 | 0.658 | 1 | 13 | Ppt1 | 0.104 | Old |
| 0.665475 | 0.188353 | 0.172 | 0.149 | 1 | 13 | Zfp944 | 0.023 | Old |
| 0.665476 | 0.150868 | 0.172 | 0.149 | 1 | 13 | Thap7 | 0.023 | Old |
| 0.665619 | 0.033914 | 0.984 | 0.991 | 1 | 13 | Atp6v0e | -0.007 | Old |
| 0.66593 | 0.551281 | 0.025 | 0.035 | 1 | 13 | Cd177 | -0.01 | Old |
| 0.665946 | 0.073748 | 0.615 | 0.588 | 1 | 13 | Stard9 | 0.027 | Old |
| 0.666122 | 0.129142 | 0.82 | 0.825 | 1 | 13 | Fmnl1 | -0.005 | Old |
| 0.66613 | 0.10985 | 0.336 | 0.289 | 1 | 13 | Ttc17 | 0.047 | Old |
| 0.666148 | 0.096121 | 0.713 | 0.623 | 1 | 13 | Zfp263 | 0.09 | Old |
| 0.666186 | 0.031045 | 0.926 | 0.886 | 1 | 13 | Dhx15 | 0.04 | Old |
| 0.666406 | 0.285836 | 0.066 | 0.053 | 1 | 13 | Eml6 | 0.013 | Old |
| 0.666406 | 0.266395 | 0.066 | 0.053 | 1 | 13 | Cmbl | 0.013 | Old |
| 0.666438 | 0.045251 | 0.057 | 0.044 | 1 | 13 | Patj | 0.013 | Old |
| 0.667047 | 0.116417 | 0.844 | 0.833 | 1 | 13 | Aup1 | 0.011 | Old |
| 0.667566 | 0.16049 | 0.689 | 0.623 | 1 | 13 | Myd88 | 0.066 | Old |
| 0.667729 | 0.090611 | 0.189 | 0.167 | 1 | 13 | Tsr3 | 0.022 | Old |
| 0.66787 | 0.451237 | 0.074 | 0.061 | 1 | 13 | Layn | 0.013 | Old |
| 0.66817 | 0.053551 | 0.475 | 0.43 | 1 | 13 | Srpk1 | 0.045 | Old |
| 0.668363 | 0.054877 | 0.033 | 0.044 | 1 | 13 | Cryba4 | -0.011 | Old |
| 0.668397 | 0.033961 | 0.533 | 0.465 | 1 | 13 | Zfp330 | 0.068 | Old |
| 0.66864 | 0.008559 | 0.23 | 0.263 | 1 | 13 | 2510009E0 | -0.033 | Old |

| | | | | | | | | |
|---|---|---|---|---|---|---|---|---|
| 0.669379 | 0.010328 | 0.393 | 0.412 | 1 | 13 | Lyrm2 | -0.019 | Old |
| 0.669468 | 0.130086 | 0.664 | 0.553 | 1 | 13 | Cdk6 | 0.111 | Old |
| 0.669471 | 0.109479 | 0.484 | 0.447 | 1 | 13 | Parp4 | 0.037 | Old |
| 0.669733 | 0.157655 | 0.287 | 0.254 | 1 | 13 | Tpst1 | 0.033 | Old |
| 0.669845 | 0.199221 | 0.123 | 0.105 | 1 | 13 | Zc2hc1a | 0.018 | Old |
| 0.669941 | 0.013993 | 0.32 | 0.263 | 1 | 13 | Wdr77 | 0.057 | Old |
| 0.670214 | 0.066069 | 0.697 | 0.693 | 1 | 13 | Sp110 | 0.004 | Old |
| 0.670538 | 0.00964 | 0.295 | 0.254 | 1 | 13 | Snrnp35 | 0.041 | Old |
| 0.670789 | 0.071499 | 0.885 | 0.842 | 1 | 13 | Atp13a3 | 0.043 | Old |
| 0.671069 | 0.180431 | 0.475 | 0.447 | 1 | 13 | Ero1l | 0.028 | Old |
| 0.67108 | 0.061798 | 0.566 | 0.5 | 1 | 13 | Babam2 | 0.066 | Old |
| 0.671433 | 0.180337 | 0.139 | 0.123 | 1 | 13 | Coil | 0.016 | Old |
| 0.671605 | 0.031267 | 0.811 | 0.746 | 1 | 13 | Lsm5 | 0.065 | Old |
| 0.671906 | 0.102666 | 0.238 | 0.202 | 1 | 13 | Slc35b3 | 0.036 | Old |
| 0.67226 | 0.032563 | 0.098 | 0.079 | 1 | 13 | Tagap1 | 0.019 | Old |
| 0.672515 | 0.073692 | 0.426 | 0.377 | 1 | 13 | Nbr1 | 0.049 | Old |
| 0.672554 | 0.109956 | 0.033 | 0.044 | 1 | 13 | Bhlhe41 | -0.011 | Old |
| 0.673139 | 0.031529 | 0.926 | 0.886 | 1 | 13 | Epsti1 | 0.04 | Old |
| 0.673263 | 0.053685 | 0.992 | 0.991 | 1 | 13 | Cd36 | 0.001 | Old |
| 0.673362 | 0.003956 | 0.754 | 0.684 | 1 | 13 | Taf15 | 0.07 | Old |
| 0.673463 | 0.030678 | 0.246 | 0.211 | 1 | 13 | Slc2a1 | 0.035 | Old |
| 0.673742 | 0.049111 | 0.057 | 0.044 | 1 | 13 | Bbs2 | 0.013 | Old |
| 0.673742 | 0.013809 | 0.057 | 0.044 | 1 | 13 | Zfp316 | 0.013 | Old |
| 0.67396 | 0.023667 | 1 | 1 | 1 | 13 | Ptbp3 | 0 | Old |
| 0.674206 | 0.023164 | 0.811 | 0.728 | 1 | 13 | Dclre1c | 0.083 | Old |
| 0.674572 | 0.362847 | 0.197 | 0.175 | 1 | 13 | Fam26f | 0.022 | Old |
| 0.67495 | 0.09474 | 0.164 | 0.14 | 1 | 13 | Mapkbp1 | 0.024 | Old |
| 0.675098 | 0.029664 | 0.885 | 0.877 | 1 | 13 | Mafg | 0.008 | Old |
| 0.675354 | 0.065069 | 1 | 1 | 1 | 13 | Tmem176b | 0 | Old |
| 0.675611 | 0.282885 | 0.59 | 0.623 | 1 | 13 | Rasgef1b | -0.033 | Old |
| 0.676994 | 0.082718 | 0.705 | 0.64 | 1 | 13 | Arid4b | 0.065 | Old |
| 0.677319 | 0.193227 | 0.205 | 0.175 | 1 | 13 | Spaca9 | 0.03 | Old |
| 0.677351 | 0.180167 | 0.385 | 0.342 | 1 | 13 | Tlr1 | 0.043 | Old |
| 0.677407 | 0.442851 | 0.123 | 0.105 | 1 | 13 | Oasl1 | 0.018 | Old |
| 0.677865 | 0.059825 | 0.369 | 0.325 | 1 | 13 | Gm48065 | 0.044 | Old |
| 0.678277 | 0.10681 | 0.443 | 0.404 | 1 | 13 | Slc13a3 | 0.039 | Old |
| 0.678634 | 0.031662 | 0.287 | 0.254 | 1 | 13 | Ubac1 | 0.033 | Old |
| 0.67875 | 0.144871 | 0.533 | 0.57 | 1 | 13 | Plk2 | -0.037 | Old |
| 0.679413 | 0.023464 | 0.164 | 0.14 | 1 | 13 | Prdm10 | 0.024 | Old |
| 0.679754 | 0.209901 | 0.27 | 0.246 | 1 | 13 | Adamtsl5 | 0.024 | Old |
| 0.679755 | 0.007354 | 0.27 | 0.228 | 1 | 13 | Ddx19b | 0.042 | Old |
| 0.679838 | 0.091078 | 0.689 | 0.658 | 1 | 13 | Phc2 | 0.031 | Old |
| 0.679936 | 0.004229 | 0.754 | 0.658 | 1 | 13 | Thoc7 | 0.096 | Old |
| 0.680014 | 0.129079 | 0.066 | 0.053 | 1 | 13 | Top1mt | 0.013 | Old |
| 0.680014 | 5.81E-05 | 0.197 | 0.167 | 1 | 13 | Rab2b | 0.03 | Old |

| | | | | | | | | |
|---|---|---|---|---|---|---|---|---|
| 0.680029 | 0.12095 | 0.156 | 0.132 | 1 | 13 | Clip2 | 0.024 | Old |
| 0.680511 | 0.105874 | 0.508 | 0.482 | 1 | 13 | Rnmt | 0.026 | Old |
| 0.680592 | 0.077134 | 0.451 | 0.386 | 1 | 13 | Selenon | 0.065 | Old |
| 0.680711 | 0.031348 | 0.172 | 0.149 | 1 | 13 | Ncapd3 | 0.023 | Old |
| 0.680879 | 0.010709 | 0.328 | 0.289 | 1 | 13 | Nt5c3 | 0.039 | Old |
| 0.680941 | 0.044623 | 1 | 1 | 1 | 13 | Cltc | 0 | Old |
| 0.680966 | 0.038868 | 0.033 | 0.044 | 1 | 13 | 3010003L2 | -0.011 | Old |
| 0.681331 | 0.032026 | 0.877 | 0.868 | 1 | 13 | Serpinb8 | 0.009 | Old |
| 0.681333 | 0.13351 | 0.484 | 0.465 | 1 | 13 | Fry | 0.019 | Old |
| 0.682165 | 0.156376 | 0.434 | 0.412 | 1 | 13 | Zfp598 | 0.022 | Old |
| 0.682768 | 0.003589 | 0.82 | 0.807 | 1 | 13 | Prkcsh | 0.013 | Old |
| 0.682898 | 0.09909 | 0.172 | 0.149 | 1 | 13 | Gm26881 | 0.023 | Old |
| 0.683125 | 0.12915 | 0.246 | 0.219 | 1 | 13 | Zfp212 | 0.027 | Old |
| 0.683226 | 0.233428 | 0.279 | 0.254 | 1 | 13 | Efl1 | 0.025 | Old |
| 0.683255 | 0.085289 | 0.32 | 0.281 | 1 | 13 | Tbcd | 0.039 | Old |
| 0.683432 | 0.556275 | 0.066 | 0.053 | 1 | 13 | Il18rap | 0.013 | Old |
| 0.683432 | 0.220077 | 0.066 | 0.053 | 1 | 13 | Pkn3 | 0.013 | Old |
| 0.683586 | 0.03846 | 0.648 | 0.605 | 1 | 13 | Tcerg1 | 0.043 | Old |
| 0.684044 | 0.169665 | 0.393 | 0.386 | 1 | 13 | Zbtb22 | 0.007 | Old |
| 0.68436 | 0.339184 | 0.131 | 0.114 | 1 | 13 | Scn2b | 0.017 | Old |
| 0.684452 | 0.021126 | 0.525 | 0.456 | 1 | 13 | Braf | 0.069 | Old |
| 0.684614 | 0.071328 | 0.156 | 0.132 | 1 | 13 | Taf8 | 0.024 | Old |
| 0.685002 | 0.113344 | 0.123 | 0.105 | 1 | 13 | Zfp971 | 0.018 | Old |
| 0.685165 | 0.029974 | 0.115 | 0.096 | 1 | 13 | Gm3448 | 0.019 | Old |
| 0.685945 | 0.030334 | 0.721 | 0.64 | 1 | 13 | Mtch2 | 0.081 | Old |
| 0.686094 | 0.004207 | 0.557 | 0.5 | 1 | 13 | Hspa4l | 0.057 | Old |
| 0.686369 | 0.051687 | 0.893 | 0.895 | 1 | 13 | Kpna4 | -0.002 | Old |
| 0.686857 | 0.070852 | 0.066 | 0.053 | 1 | 13 | Gm15706 | 0.013 | Old |
| 0.686911 | 0.141921 | 0.156 | 0.132 | 1 | 13 | Zgpat | 0.024 | Old |
| 0.686934 | 0.148494 | 0.336 | 0.298 | 1 | 13 | Tsta3 | 0.038 | Old |
| 0.687093 | 0.47425 | 0.074 | 0.061 | 1 | 13 | AI480526 | 0.013 | Old |
| 0.687464 | 0.061863 | 0.107 | 0.088 | 1 | 13 | Lbx2 | 0.019 | Old |
| 0.687545 | 0.015475 | 0.426 | 0.386 | 1 | 13 | Timm44 | 0.04 | Old |
| 0.687845 | 0.025823 | 0.943 | 0.877 | 1 | 13 | Etf1 | 0.066 | Old |
| 0.688104 | 0.117214 | 0.139 | 0.123 | 1 | 13 | Zfp141 | 0.016 | Old |
| 0.688799 | 0.082815 | 0.697 | 0.667 | 1 | 13 | Lrrc59 | 0.03 | Old |
| 0.689251 | 0.058728 | 0.902 | 0.921 | 1 | 13 | Anxa7 | -0.019 | Old |
| 0.68933 | 0.04746 | 0.951 | 0.939 | 1 | 13 | Fgfr1op2 | 0.012 | Old |
| 0.68934 | 0.040849 | 0.959 | 0.947 | 1 | 13 | Nisch | 0.012 | Old |
| 0.689418 | 0.06425 | 0.033 | 0.044 | 1 | 13 | Otub2 | -0.011 | Old |
| 0.689423 | 0.000594 | 0.664 | 0.614 | 1 | 13 | Med1 | 0.05 | Old |
| 0.689476 | 0.136457 | 0.172 | 0.149 | 1 | 13 | Tmed1 | 0.023 | Old |
| 0.69113 | 0.07389 | 0.344 | 0.307 | 1 | 13 | Gcc1 | 0.037 | Old |
| 0.691467 | 0.091707 | 0.451 | 0.412 | 1 | 13 | Tm7sf3 | 0.039 | Old |
| 0.691495 | 0.016405 | 0.492 | 0.439 | 1 | 13 | Med29 | 0.053 | Old |

| | | | | | | | | |
|---|---|---|---|---|---|---|---|---|
| 0.691543 | 0.016119 | 0.328 | 0.281 | 1 | 13 | Zbtb21 | 0.047 | Old |
| 0.691669 | 0.409728 | 0.156 | 0.14 | 1 | 13 | Ndrg1 | 0.016 | Old |
| 0.691749 | 0.636738 | 0.131 | 0.114 | 1 | 13 | Serpinb9 | 0.017 | Old |
| 0.69183 | 0.125004 | 0.352 | 0.316 | 1 | 13 | Dcaf5 | 0.036 | Old |
| 0.691897 | 0.010162 | 0.852 | 0.904 | 1 | 13 | Dnajb14 | -0.052 | Old |
| 0.692148 | 0.058459 | 0.951 | 0.947 | 1 | 13 | Srsf9 | 0.004 | Old |
| 0.692725 | 0.13353 | 0.705 | 0.711 | 1 | 13 | Lpgat1 | -0.006 | Old |
| 0.692797 | 0.107131 | 0.68 | 0.614 | 1 | 13 | Ascc3 | 0.066 | Old |
| 0.692984 | 0.447517 | 0.057 | 0.07 | 1 | 13 | Cenpe | -0.013 | Old |
| 0.693546 | 0.064561 | 0.951 | 0.93 | 1 | 13 | Nop10 | 0.021 | Old |
| 0.693635 | 0.701563 | 0.025 | 0.018 | 1 | 13 | Gm38335 | 0.007 | Old |
| 0.693635 | 0.627464 | 0.025 | 0.018 | 1 | 13 | Tmem156 | 0.007 | Old |
| 0.693635 | 0.536538 | 0.025 | 0.018 | 1 | 13 | Gm17399 | 0.007 | Old |
| 0.693635 | 0.416269 | 0.025 | 0.018 | 1 | 13 | Hist1h3e | 0.007 | Old |
| 0.693727 | 0.13168 | 0.066 | 0.053 | 1 | 13 | Cacnb1 | 0.013 | Old |
| 0.693727 | 0.121569 | 0.066 | 0.053 | 1 | 13 | Ftx | 0.013 | Old |
| 0.693727 | 0.112655 | 0.066 | 0.053 | 1 | 13 | Osgepl1 | 0.013 | Old |
| 0.693746 | 0.01025 | 0.77 | 0.702 | 1 | 13 | Tmem164 | 0.068 | Old |
| 0.693994 | 0.048475 | 0.713 | 0.658 | 1 | 13 | Ankhd1 | 0.055 | Old |
| 0.694201 | 0.030749 | 0.574 | 0.526 | 1 | 13 | Ilf2 | 0.048 | Old |
| 0.694213 | 0.069924 | 0.279 | 0.246 | 1 | 13 | Nbn | 0.033 | Old |
| 0.694524 | 0.04561 | 0.434 | 0.439 | 1 | 13 | Scrib | -0.005 | Old |
| 0.694576 | 0.101988 | 0.18 | 0.158 | 1 | 13 | Metap1d | 0.022 | Old |
| 0.694588 | 0.079296 | 0.844 | 0.877 | 1 | 13 | Riok3 | -0.033 | Old |
| 0.694605 | 0.187894 | 0.557 | 0.553 | 1 | 13 | Pafah1b2 | 0.004 | Old |
| 0.694643 | 0.290546 | 0.295 | 0.281 | 1 | 13 | Lmf1 | 0.014 | Old |
| 0.694649 | 0.035998 | 0.5 | 0.465 | 1 | 13 | Tbc1d22a | 0.035 | Old |
| 0.694985 | 0.068817 | 1 | 1 | 1 | 13 | C1qc | 0 | Old |
| 0.695202 | 0.08121 | 0.295 | 0.263 | 1 | 13 | Tfdp1 | 0.032 | Old |
| 0.695579 | 0.021634 | 0.738 | 0.675 | 1 | 13 | Tap2 | 0.063 | Old |
| 0.696317 | 0.143776 | 0.057 | 0.07 | 1 | 13 | Plcxd2 | -0.013 | Old |
| 0.696331 | 0.038114 | 0.951 | 0.895 | 1 | 13 | Dynlrb1 | 0.056 | Old |
| 0.696407 | 0.167035 | 0.246 | 0.219 | 1 | 13 | Hdhd2 | 0.027 | Old |
| 0.696691 | 0.071499 | 0.131 | 0.114 | 1 | 13 | Ankrd39 | 0.017 | Old |
| 0.696696 | 0.163406 | 0.492 | 0.447 | 1 | 13 | Zbp1 | 0.045 | Old |
| 0.696734 | 0.11143 | 0.18 | 0.158 | 1 | 13 | Terf2ip | 0.022 | Old |
| 0.697171 | 0.068373 | 0.066 | 0.053 | 1 | 13 | Plxna3 | 0.013 | Old |
| 0.697803 | 0.156049 | 0.959 | 0.982 | 1 | 13 | Aldoa | -0.023 | Old |
| 0.697809 | 0.005678 | 0.426 | 0.377 | 1 | 13 | Hccs | 0.049 | Old |
| 0.698338 | 0.099564 | 0.115 | 0.096 | 1 | 13 | Gstk1 | 0.019 | Old |
| 0.698448 | 0.079761 | 0.41 | 0.368 | 1 | 13 | Bicra | 0.042 | Old |
| 0.698666 | 0.083895 | 0.861 | 0.825 | 1 | 13 | Emg1 | 0.036 | Old |
| 0.699118 | 0.0055 | 0.918 | 0.904 | 1 | 13 | Mapk14 | 0.014 | Old |
| 0.699291 | 1.023672 | 0.025 | 0.018 | 1 | 13 | Asns | 0.007 | Old |
| 0.699291 | 0.533172 | 0.025 | 0.018 | 1 | 13 | 9330175E1 | 0.007 | Old |

| | | | | | | | | |
|---|---|---|---|---|---|---|---|---|
| 0.699291 | 0.46758 | 0.025 | 0.018 | 1 | 13 | Xkrx | 0.007 | Old |
| 0.699291 | 0.451962 | 0.025 | 0.018 | 1 | 13 | Efr3b | 0.007 | Old |
| 0.699291 | 0.442164 | 0.025 | 0.018 | 1 | 13 | Gm33104 | 0.007 | Old |
| 0.699291 | 0.439292 | 0.025 | 0.018 | 1 | 13 | 5830454E0 | 0.007 | Old |
| 0.699291 | 0.415018 | 0.025 | 0.018 | 1 | 13 | Hist2h4 | 0.007 | Old |
| 0.699291 | 0.414339 | 0.025 | 0.018 | 1 | 13 | Xrcc5 | 0.007 | Old |
| 0.699291 | 0.353277 | 0.025 | 0.018 | 1 | 13 | Mpzl1 | 0.007 | Old |
| 0.69964 | 0.090661 | 0.852 | 0.763 | 1 | 13 | Ddb1 | 0.089 | Old |
| 0.700023 | 0.371338 | 0.074 | 0.061 | 1 | 13 | Stat4 | 0.013 | Old |
| 0.700281 | 0.03205 | 0.844 | 0.763 | 1 | 13 | Kcmf1 | 0.081 | Old |
| 0.700573 | 0.068483 | 0.467 | 0.421 | 1 | 13 | Fam53b | 0.046 | Old |
| 0.700576 | 0.023113 | 0.344 | 0.289 | 1 | 13 | Ampd3 | 0.055 | Old |
| 0.700579 | 0.115983 | 0.959 | 0.895 | 1 | 13 | Pik3cd | 0.064 | Old |
| 0.700621 | 0.10833 | 0.066 | 0.053 | 1 | 13 | Gatsl3 | 0.013 | Old |
| 0.700628 | 0.147871 | 0.148 | 0.132 | 1 | 13 | Spata5 | 0.016 | Old |
| 0.700829 | 0.086613 | 0.295 | 0.254 | 1 | 13 | Rtl8a | 0.041 | Old |
| 0.701324 | 0.240734 | 0.205 | 0.184 | 1 | 13 | Cc2d1a | 0.021 | Old |
| 0.701499 | 0.057197 | 0.852 | 0.833 | 1 | 13 | Adipor1 | 0.019 | Old |
| 0.701561 | 0.041097 | 0.525 | 0.518 | 1 | 13 | Myo7a | 0.007 | Old |
| 0.701894 | 0.029617 | 0.164 | 0.14 | 1 | 13 | Dnajc27 | 0.024 | Old |
| 0.702047 | 0.044821 | 1 | 1 | 1 | 13 | Ctsc | 0 | Old |
| 0.702127 | 0.087868 | 0.451 | 0.439 | 1 | 13 | Ss18l2 | 0.012 | Old |
| 0.702605 | 0.094408 | 0.877 | 0.921 | 1 | 13 | Man1a | -0.044 | Old |
| 0.702765 | 0.052259 | 0.59 | 0.535 | 1 | 13 | Irf2bpl | 0.055 | Old |
| 0.703185 | 0.013804 | 0.705 | 0.737 | 1 | 13 | Sp3 | -0.032 | Old |
| 0.703216 | 0.063584 | 0.066 | 0.079 | 1 | 13 | Slc25a53 | -0.013 | Old |
| 0.703269 | 0.20627 | 0.074 | 0.061 | 1 | 13 | Amotl2 | 0.013 | Old |
| 0.703371 | 0.036774 | 0.205 | 0.184 | 1 | 13 | Mettl3 | 0.021 | Old |
| 0.703633 | 0.14764 | 0.115 | 0.096 | 1 | 13 | Prr5 | 0.019 | Old |
| 0.703657 | 0.168346 | 0.566 | 0.5 | 1 | 13 | Mmadhc | 0.066 | Old |
| 0.703848 | 0.024642 | 0.77 | 0.711 | 1 | 13 | Lman1 | 0.059 | Old |
| 0.704078 | 0.068235 | 0.066 | 0.053 | 1 | 13 | Zfp518b | 0.013 | Old |
| 0.704243 | 0.035962 | 0.59 | 0.535 | 1 | 13 | Siah2 | 0.055 | Old |
| 0.704302 | 0.110433 | 0.656 | 0.588 | 1 | 13 | Ppil2 | 0.068 | Old |
| 0.704749 | 0.075039 | 0.902 | 0.904 | 1 | 13 | Ctdnep1 | -0.002 | Old |
| 0.704896 | 0.246235 | 0.041 | 0.053 | 1 | 13 | Il1r2 | -0.012 | Old |
| 0.704963 | 0.583711 | 0.025 | 0.018 | 1 | 13 | Nme3 | 0.007 | Old |
| 0.704963 | 0.360402 | 0.025 | 0.018 | 1 | 13 | Lzts3 | 0.007 | Old |
| 0.704963 | 0.359909 | 0.025 | 0.018 | 1 | 13 | Cmtm8 | 0.007 | Old |
| 0.704963 | 0.342702 | 0.025 | 0.018 | 1 | 13 | Gpat3 | 0.007 | Old |
| 0.704963 | 0.330247 | 0.025 | 0.018 | 1 | 13 | Gm48420 | 0.007 | Old |
| 0.704963 | 0.277544 | 0.025 | 0.018 | 1 | 13 | Gm20522 | 0.007 | Old |
| 0.705587 | 0.023849 | 0.992 | 0.991 | 1 | 13 | Ndufa2 | 0.001 | Old |
| 0.705736 | 0.21031 | 0.402 | 0.386 | 1 | 13 | 4931406P1 | 0.016 | Old |
| 0.706095 | 0.059405 | 0.607 | 0.605 | 1 | 13 | Gadd45g | 0.002 | Old |

| | | | | | | | | |
|---|---|---|---|---|---|---|---|---|
| 0.706282 | 0.030879 | 0.959 | 0.947 | 1 | 13 | Ssb | 0.012 | Old |
| 0.706286 | 0.475016 | 0.098 | 0.114 | 1 | 13 | Hspb1 | -0.016 | Old |
| 0.706286 | 0.114365 | 0.115 | 0.096 | 1 | 13 | Nat2 | 0.019 | Old |
| 0.70652 | 0.002669 | 0.074 | 0.061 | 1 | 13 | Prim1 | 0.013 | Old |
| 0.706616 | 0.076222 | 0.508 | 0.465 | 1 | 13 | Eef1e1 | 0.043 | Old |
| 0.706878 | 0.169547 | 0.705 | 0.632 | 1 | 13 | Herc2 | 0.073 | Old |
| 0.706907 | 0.148693 | 0.467 | 0.412 | 1 | 13 | Slc4a2 | 0.055 | Old |
| 0.707541 | 0.35841 | 0.066 | 0.053 | 1 | 13 | Ctif | 0.013 | Old |
| 0.707975 | 0.004123 | 0.123 | 0.105 | 1 | 13 | Amer1 | 0.018 | Old |
| 0.708512 | 0.136343 | 0.385 | 0.351 | 1 | 13 | Gpd2 | 0.034 | Old |
| 0.708687 | 0.079158 | 0.582 | 0.544 | 1 | 13 | Gtf2b | 0.038 | Old |
| 0.709324 | 0.129811 | 0.738 | 0.728 | 1 | 13 | Eif6 | 0.01 | Old |
| 0.709523 | 0.220686 | 0.205 | 0.184 | 1 | 13 | Mthfd1l | 0.021 | Old |
| 0.709777 | 0.192961 | 0.074 | 0.061 | 1 | 13 | Cep152 | 0.013 | Old |
| 0.709792 | 0.185161 | 0.197 | 0.175 | 1 | 13 | Taf1b | 0.022 | Old |
| 0.710029 | 0.12034 | 0.385 | 0.342 | 1 | 13 | Pygl | 0.043 | Old |
| 0.710397 | 0.355708 | 0.09 | 0.079 | 1 | 13 | Gm38832 | 0.011 | Old |
| 0.710458 | 0.072033 | 0.918 | 0.904 | 1 | 13 | Prex1 | 0.014 | Old |
| 0.710584 | 0.073983 | 0.418 | 0.386 | 1 | 13 | Eya3 | 0.032 | Old |
| 0.710652 | 4.559069 | 0.025 | 0.018 | 1 | 13 | S100a9 | 0.007 | Old |
| 0.710652 | 0.511018 | 0.025 | 0.018 | 1 | 13 | Gm37168 | 0.007 | Old |
| 0.710652 | 0.426737 | 0.025 | 0.018 | 1 | 13 | Myb | 0.007 | Old |
| 0.710652 | 0.294793 | 0.025 | 0.018 | 1 | 13 | Wwc1 | 0.007 | Old |
| 0.710652 | 0.233182 | 0.025 | 0.018 | 1 | 13 | Mnd1 | 0.007 | Old |
| 0.710652 | 0.226378 | 0.025 | 0.018 | 1 | 13 | Gm42726 | 0.007 | Old |
| 0.710652 | 0.221439 | 0.025 | 0.018 | 1 | 13 | Enkd1 | 0.007 | Old |
| 0.710652 | 0.172571 | 0.025 | 0.018 | 1 | 13 | Gm38251 | 0.007 | Old |
| 0.710652 | 0.171092 | 0.025 | 0.018 | 1 | 13 | Adamts5 | 0.007 | Old |
| 0.710652 | 0.16854 | 0.025 | 0.018 | 1 | 13 | Vma21-ps | 0.007 | Old |
| 0.710652 | 0.168235 | 0.025 | 0.018 | 1 | 13 | Tespa1 | 0.007 | Old |
| 0.710652 | 0.154288 | 0.025 | 0.018 | 1 | 13 | Tagln | 0.007 | Old |
| 0.710652 | 0.074636 | 0.025 | 0.018 | 1 | 13 | 9930022D1 | 0.007 | Old |
| 0.710884 | 0.186536 | 0.574 | 0.57 | 1 | 13 | Upf3a | 0.004 | Old |
| 0.711036 | 0.011274 | 0.746 | 0.649 | 1 | 13 | Cnih1 | 0.097 | Old |
| 0.711565 | 0.171715 | 0.582 | 0.579 | 1 | 13 | Rab12 | 0.003 | Old |
| 0.711594 | 0.25703 | 0.131 | 0.114 | 1 | 13 | Trim7 | 0.017 | Old |
| 0.711598 | 0.036736 | 0.467 | 0.412 | 1 | 13 | Stat5a | 0.055 | Old |
| 0.71165 | 0.085073 | 0.787 | 0.746 | 1 | 13 | Osgep | 0.041 | Old |
| 0.711678 | 0.093017 | 0.385 | 0.342 | 1 | 13 | Lsm1 | 0.043 | Old |
| 0.711864 | 0.240659 | 0.164 | 0.149 | 1 | 13 | Extl2 | 0.015 | Old |
| 0.711894 | 0.070819 | 0.607 | 0.561 | 1 | 13 | Ccdc88b | 0.046 | Old |
| 0.711925 | 0.058496 | 0.984 | 0.868 | 1 | 13 | Ppp1r9b | 0.116 | Old |
| 0.712221 | 0.04227 | 0.377 | 0.333 | 1 | 13 | Oxa1l | 0.044 | Old |
| 0.712737 | 0.065586 | 0.295 | 0.272 | 1 | 13 | Trim3 | 0.023 | Old |
| 0.712742 | 0.059556 | 0.541 | 0.456 | 1 | 13 | Wdr61 | 0.085 | Old |

| | | | | | | | | |
|---|---|---|---|---|---|---|---|---|
| 0.712835 | 0.074956 | 0.623 | 0.57 | 1 | 13 | Itpr1 | 0.053 | Old |
| 0.71359 | 0.177912 | 0.492 | 0.447 | 1 | 13 | Stk3 | 0.045 | Old |
| 0.71359 | 0.126346 | 0.484 | 0.456 | 1 | 13 | Map3k7 | 0.028 | Old |
| 0.713767 | 0.1174 | 0.279 | 0.263 | 1 | 13 | Rfx1 | 0.016 | Old |
| 0.714055 | 0.00459 | 0.623 | 0.658 | 1 | 13 | BC004004 | -0.035 | Old |
| 0.714263 | 0.00063 | 0.525 | 0.474 | 1 | 13 | Srpk2 | 0.051 | Old |
| 0.714309 | 0.053202 | 0.713 | 0.632 | 1 | 13 | Wtap | 0.081 | Old |
| 0.714484 | 0.214552 | 0.066 | 0.053 | 1 | 13 | Gm42670 | 0.013 | Old |
| 0.714526 | 0.007531 | 0.303 | 0.263 | 1 | 13 | Pip4k2c | 0.04 | Old |
| 0.714593 | 0.175252 | 0.311 | 0.289 | 1 | 13 | Racgap1 | 0.022 | Old |
| 0.714608 | 0.217857 | 0.139 | 0.123 | 1 | 13 | Thsd4 | 0.016 | Old |
| 0.715255 | 0.101855 | 0.844 | 0.904 | 1 | 13 | Mvp | -0.06 | Old |
| 0.715302 | 0.028926 | 0.836 | 0.781 | 1 | 13 | Sar1a | 0.055 | Old |
| 0.715423 | 0.075721 | 0.828 | 0.807 | 1 | 13 | Pdia4 | 0.021 | Old |
| 0.715535 | 0.019033 | 0.992 | 0.991 | 1 | 13 | Sumo2 | 0.001 | Old |
| 0.715594 | 0.020224 | 0.049 | 0.061 | 1 | 13 | Pcgf2 | -0.012 | Old |
| 0.715664 | 0.076877 | 0.811 | 0.763 | 1 | 13 | Ndufa12 | 0.048 | Old |
| 0.716274 | 0.077229 | 0.541 | 0.465 | 1 | 13 | Snrnp25 | 0.076 | Old |
| 0.716307 | 0.206997 | 0.074 | 0.061 | 1 | 13 | Gm11713 | 0.013 | Old |
| 0.716358 | 0.237141 | 0.025 | 0.018 | 1 | 13 | 2810405F1 | 0.007 | Old |
| 0.716358 | 0.21303 | 0.025 | 0.018 | 1 | 13 | Gm26514 | 0.007 | Old |
| 0.716358 | 0.174019 | 0.025 | 0.018 | 1 | 13 | Gm11491 | 0.007 | Old |
| 0.716358 | 0.171055 | 0.025 | 0.018 | 1 | 13 | Cib3 | 0.007 | Old |
| 0.716358 | 0.157985 | 0.025 | 0.018 | 1 | 13 | Fam189a2 | 0.007 | Old |
| 0.716358 | 0.156452 | 0.025 | 0.018 | 1 | 13 | Serpina3f | 0.007 | Old |
| 0.716358 | 0.155001 | 0.025 | 0.018 | 1 | 13 | Gm9725 | 0.007 | Old |
| 0.716358 | 0.137396 | 0.025 | 0.018 | 1 | 13 | 4930539J0 | 0.007 | Old |
| 0.716358 | 0.118212 | 0.025 | 0.018 | 1 | 13 | Gm9938 | 0.007 | Old |
| 0.716358 | 0.116517 | 0.025 | 0.018 | 1 | 13 | Dnaja4 | 0.007 | Old |
| 0.716358 | 0.109885 | 0.025 | 0.018 | 1 | 13 | Gm10384 | 0.007 | Old |
| 0.716358 | 0.068365 | 0.025 | 0.018 | 1 | 13 | 9230111E0 | 0.007 | Old |
| 0.716514 | 0.003228 | 0.385 | 0.333 | 1 | 13 | 1600012H0 | 0.052 | Old |
| 0.716586 | 0.152876 | 0.131 | 0.114 | 1 | 13 | Tab3 | 0.017 | Old |
| 0.716873 | 0.369002 | 0.156 | 0.14 | 1 | 13 | Sirt3 | 0.016 | Old |
| 0.716921 | 0.116765 | 0.631 | 0.596 | 1 | 13 | Vezf1 | 0.035 | Old |
| 0.717035 | 0.137577 | 0.139 | 0.123 | 1 | 13 | B3gnt8 | 0.016 | Old |
| 0.71712 | 0.161956 | 0.705 | 0.711 | 1 | 13 | Srrt | -0.006 | Old |
| 0.717716 | 0.162462 | 0.09 | 0.105 | 1 | 13 | Prr7 | -0.015 | Old |
| 0.717776 | 0.039892 | 0.393 | 0.421 | 1 | 13 | Acsl3 | -0.028 | Old |
| 0.717795 | 0.196126 | 0.525 | 0.482 | 1 | 13 | Tex264 | 0.043 | Old |
| 0.717907 | 0.074995 | 0.877 | 0.816 | 1 | 13 | Baz1b | 0.061 | Old |
| 0.718358 | 0.046182 | 0.943 | 0.939 | 1 | 13 | Hnrnpa1 | 0.004 | Old |
| 0.718654 | 0.064536 | 0.459 | 0.421 | 1 | 13 | Tspan4 | 0.038 | Old |
| 0.718817 | 0.006988 | 0.779 | 0.754 | 1 | 13 | Il4ra | 0.025 | Old |
| 0.719087 | 0.093067 | 0.131 | 0.114 | 1 | 13 | Cnst | 0.017 | Old |

| | | | | | | | | |
|---|---|---|---|---|---|---|---|---|
| 0.719099 | 0.002829 | 1 | 1 | 1 | 13 | Tspo | 0 | Old |
| 0.719465 | 0.162072 | 0.139 | 0.123 | 1 | 13 | Ulk3 | 0.016 | Old |
| 0.71958 | 0.121263 | 0.074 | 0.061 | 1 | 13 | Gne | 0.013 | Old |
| 0.719603 | 0.019345 | 0.115 | 0.096 | 1 | 13 | 3110009E1 | 0.019 | Old |
| 0.719907 | 0.084645 | 0.623 | 0.561 | 1 | 13 | Map3k2 | 0.062 | Old |
| 0.720228 | 0.028304 | 0.762 | 0.711 | 1 | 13 | Pan3 | 0.051 | Old |
| 0.720731 | 0.297195 | 0.098 | 0.088 | 1 | 13 | Spindoc | 0.01 | Old |
| 0.720825 | 0.080376 | 0.5 | 0.553 | 1 | 13 | Gm47283 | -0.053 | Old |
| 0.720856 | 0.144871 | 0.123 | 0.105 | 1 | 13 | Gemin6 | 0.018 | Old |
| 0.720916 | 0.067707 | 0.336 | 0.289 | 1 | 13 | Elp2 | 0.047 | Old |
| 0.721157 | 0.249911 | 0.18 | 0.167 | 1 | 13 | Wipi1 | 0.013 | Old |
| 0.721871 | 0.178848 | 0.393 | 0.377 | 1 | 13 | Arhgef7 | 0.016 | Old |
| 0.721921 | 0.062788 | 0.525 | 0.447 | 1 | 13 | Batf | 0.078 | Old |
| 0.721964 | 0.131436 | 0.287 | 0.254 | 1 | 13 | Trappc11 | 0.033 | Old |
| 0.722079 | 0.146358 | 0.025 | 0.018 | 1 | 13 | 1700086O( | 0.007 | Old |
| 0.722079 | 0.12559 | 0.025 | 0.018 | 1 | 13 | Marveld2 | 0.007 | Old |
| 0.722079 | 0.079915 | 0.025 | 0.018 | 1 | 13 | Neo1 | 0.007 | Old |
| 0.722079 | 0.075799 | 0.025 | 0.018 | 1 | 13 | Col3a1 | 0.007 | Old |
| 0.722079 | 0.021746 | 0.025 | 0.018 | 1 | 13 | Cdh5 | 0.007 | Old |
| 0.722101 | 0.031093 | 0.549 | 0.544 | 1 | 13 | Arid5b | 0.005 | Old |
| 0.722508 | 0.033709 | 0.738 | 0.746 | 1 | 13 | Hmgn1 | -0.008 | Old |
| 0.722562 | 0.068449 | 0.91 | 0.904 | 1 | 13 | Snu13 | 0.006 | Old |
| 0.722858 | 0.097492 | 0.074 | 0.061 | 1 | 13 | Per2 | 0.013 | Old |
| 0.722974 | 0.070971 | 0.451 | 0.404 | 1 | 13 | Aars | 0.047 | Old |
| 0.723131 | 0.176366 | 0.164 | 0.149 | 1 | 13 | Pias3 | 0.015 | Old |
| 0.723458 | 0.034754 | 0.525 | 0.447 | 1 | 13 | Otud4 | 0.078 | Old |
| 0.724044 | 0.017529 | 0.156 | 0.132 | 1 | 13 | Thap1 | 0.024 | Old |
| 0.724098 | 0.143635 | 0.131 | 0.114 | 1 | 13 | Dyrk1b | 0.017 | Old |
| 0.724449 | 0.064646 | 0.221 | 0.202 | 1 | 13 | Lin37 | 0.019 | Old |
| 0.724815 | 0.063449 | 1 | 1 | 1 | 13 | Grb2 | 0 | Old |
| 0.725134 | 0.235038 | 0.402 | 0.377 | 1 | 13 | Vrk3 | 0.025 | Old |
| 0.726141 | 0.005775 | 0.074 | 0.061 | 1 | 13 | Snap47 | 0.013 | Old |
| 0.726156 | 0.051893 | 0.303 | 0.281 | 1 | 13 | Banp | 0.022 | Old |
| 0.726156 | 0.036148 | 0.311 | 0.272 | 1 | 13 | Cog4 | 0.039 | Old |
| 0.726772 | 0.117087 | 0.139 | 0.123 | 1 | 13 | Gorasp1 | 0.016 | Old |
| 0.726788 | 0.06828 | 0.91 | 0.868 | 1 | 13 | Tex261 | 0.042 | Old |
| 0.726905 | 0.02957 | 0.926 | 0.921 | 1 | 13 | Runx1 | 0.005 | Old |
| 0.727417 | 0.289834 | 0.41 | 0.404 | 1 | 13 | Patl1 | 0.006 | Old |
| 0.727679 | 0.066753 | 0.992 | 1 | 1 | 13 | Txn1 | -0.008 | Old |
| 0.727816 | 0.083117 | 0.025 | 0.018 | 1 | 13 | Gm26609 | 0.007 | Old |
| 0.727816 | 0.05254 | 0.025 | 0.018 | 1 | 13 | Dnmt3aos | 0.007 | Old |
| 0.728879 | 0.238044 | 0.172 | 0.158 | 1 | 13 | Epn2 | 0.014 | Old |
| 0.729045 | 0.0466 | 0.951 | 0.886 | 1 | 13 | Magt1 | 0.065 | Old |
| 0.729209 | 0.080302 | 0.082 | 0.07 | 1 | 13 | Gm340 | 0.012 | Old |
| 0.729347 | 0.144325 | 0.295 | 0.263 | 1 | 13 | Pop4 | 0.032 | Old |

| | | | | | | | | |
|---|---|---|---|---|---|---|---|---|
| 0.729613 | 0.006889 | 0.861 | 0.904 | 1 | 13 | Mapk1 | -0.043 | Old |
| 0.730113 | 0.143083 | 0.508 | 0.474 | 1 | 13 | Trim30d | 0.034 | Old |
| 0.730166 | 0.152605 | 0.205 | 0.184 | 1 | 13 | Ddx28 | 0.021 | Old |
| 0.730592 | 0.106297 | 0.68 | 0.614 | 1 | 13 | Fam76a | 0.066 | Old |
| 0.730626 | 0.095967 | 0.5 | 0.439 | 1 | 13 | Frmd8 | 0.061 | Old |
| 0.730628 | 0.0896 | 0.451 | 0.421 | 1 | 13 | Cdc42bpb | 0.03 | Old |
| 0.731218 | 0.286708 | 0.123 | 0.105 | 1 | 13 | Gm10138 | 0.018 | Old |
| 0.731345 | 0.176215 | 0.525 | 0.509 | 1 | 13 | Dpagt1 | 0.016 | Old |
| 0.731581 | 0.075993 | 0.18 | 0.158 | 1 | 13 | Tlr5 | 0.022 | Old |
| 0.731603 | 0.002959 | 0.287 | 0.263 | 1 | 13 | Fra10ac1 | 0.024 | Old |
| 0.732477 | 0.039335 | 0.73 | 0.693 | 1 | 13 | Dcaf12 | 0.037 | Old |
| 0.732722 | 0.062571 | 0.074 | 0.061 | 1 | 13 | Cyfip2 | 0.013 | Old |
| 0.732764 | 0.184808 | 0.434 | 0.404 | 1 | 13 | Skiv2l2 | 0.03 | Old |
| 0.732764 | 0.055305 | 0.443 | 0.395 | 1 | 13 | Gtf3c1 | 0.048 | Old |
| 0.73302 | 0.071431 | 0.697 | 0.675 | 1 | 13 | Gatad2a | 0.022 | Old |
| 0.733314 | 0.093867 | 0.902 | 0.912 | 1 | 13 | Tiparp | -0.01 | Old |
| 0.733405 | 0.09199 | 0.959 | 0.947 | 1 | 13 | Pbrm1 | 0.012 | Old |
| 0.733673 | 0.038324 | 0.516 | 0.474 | 1 | 13 | 4833439L1 | 0.042 | Old |
| 0.733736 | 0.050163 | 0.672 | 0.632 | 1 | 13 | 4931406C0 | 0.04 | Old |
| 0.733842 | 0.053134 | 0.82 | 0.781 | 1 | 13 | Ncoa3 | 0.039 | Old |
| 0.734036 | 0.116809 | 0.795 | 0.746 | 1 | 13 | Cul1 | 0.049 | Old |
| 0.734324 | 0.143881 | 0.352 | 0.325 | 1 | 13 | Txndc12 | 0.027 | Old |
| 0.734861 | 0.137587 | 0.246 | 0.219 | 1 | 13 | Ripk3 | 0.027 | Old |
| 0.734896 | 0.046616 | 0.631 | 0.535 | 1 | 13 | Exosc5 | 0.096 | Old |
| 0.735491 | 0.216184 | 0.492 | 0.535 | 1 | 13 | Sdc1 | -0.043 | Old |
| 0.735573 | 0.137058 | 0.631 | 0.579 | 1 | 13 | Hmbox1 | 0.052 | Old |
| 0.735818 | 0.132837 | 0.533 | 0.482 | 1 | 13 | Atl3 | 0.051 | Old |
| 0.735993 | 0.059947 | 0.484 | 0.43 | 1 | 13 | Arhgap5 | 0.054 | Old |
| 0.73602 | 0.070265 | 0.074 | 0.061 | 1 | 13 | Ccdc138 | 0.013 | Old |
| 0.736802 | 0.034127 | 0.246 | 0.219 | 1 | 13 | Ormdl1 | 0.027 | Old |
| 0.737386 | 0.012742 | 0.869 | 0.851 | 1 | 13 | Cxcl16 | 0.018 | Old |
| 0.737925 | 0.085209 | 0.295 | 0.272 | 1 | 13 | Zfyve26 | 0.023 | Old |
| 0.738575 | 0.517304 | 0.082 | 0.07 | 1 | 13 | Coq8a | 0.012 | Old |
| 0.738575 | 0.088271 | 0.082 | 0.07 | 1 | 13 | A630072M | 0.012 | Old |
| 0.738646 | 0.046751 | 0.836 | 0.842 | 1 | 13 | Ccr1 | -0.006 | Old |
| 0.739171 | 0.041763 | 0.984 | 0.991 | 1 | 13 | Ptms | -0.007 | Old |
| 0.739201 | 0.028978 | 0.131 | 0.114 | 1 | 13 | Lhpp | 0.017 | Old |
| 0.739598 | 0.206818 | 0.213 | 0.193 | 1 | 13 | Alg14 | 0.02 | Old |
| 0.739905 | 0.033409 | 0.852 | 0.789 | 1 | 13 | Aurkaip1 | 0.063 | Old |
| 0.740194 | 0.030164 | 0.648 | 0.684 | 1 | 13 | Cdc26 | -0.036 | Old |
| 0.740612 | 0.076405 | 1 | 1 | 1 | 13 | Samhd1 | 0 | Old |
| 0.74069 | 0.084189 | 0.246 | 0.219 | 1 | 13 | Pigc | 0.027 | Old |
| 0.741461 | 0.100677 | 0.139 | 0.123 | 1 | 13 | Apex2 | 0.016 | Old |
| 0.741499 | 0.08306 | 0.508 | 0.456 | 1 | 13 | Nomo1 | 0.052 | Old |
| 0.741706 | 0.146205 | 0.082 | 0.07 | 1 | 13 | C2cd2 | 0.012 | Old |

| | | | | | | | | |
|---|---|---|---|---|---|---|---|---|
| 0.742054 | 0.025751 | 1 | 1 | 1 | 13 | Chchd2 | 0 | Old |
| 0.743047 | 0.163224 | 0.402 | 0.386 | 1 | 13 | Lats1 | 0.016 | Old |
| 0.743368 | 0.096819 | 0.77 | 0.763 | 1 | 13 | Slc35b2 | 0.007 | Old |
| 0.743417 | 0.095586 | 0.615 | 0.667 | 1 | 13 | Hist1h2bc | -0.052 | Old |
| 0.743491 | 0.048297 | 0.697 | 0.649 | 1 | 13 | Mdm4 | 0.048 | Old |
| 0.743493 | 0.023758 | 0.975 | 0.965 | 1 | 13 | Tra2b | 0.01 | Old |
| 0.743536 | 0.016594 | 0.738 | 0.693 | 1 | 13 | Pfdn1 | 0.045 | Old |
| 0.743918 | 0.11578 | 0.123 | 0.14 | 1 | 13 | Ube2e2 | -0.017 | Old |
| 0.744053 | 0.070636 | 0.91 | 0.868 | 1 | 13 | Copa | 0.042 | Old |
| 0.744147 | 0.13265 | 0.721 | 0.658 | 1 | 13 | Eif4ebp1 | 0.063 | Old |
| 0.744164 | 0.007866 | 0.943 | 0.904 | 1 | 13 | Rnf141 | 0.039 | Old |
| 0.744425 | 0.029574 | 0.836 | 0.842 | 1 | 13 | Cul3 | -0.006 | Old |
| 0.744638 | 0.049103 | 0.475 | 0.439 | 1 | 13 | Flcn | 0.036 | Old |
| 0.744866 | 0.145782 | 0.402 | 0.36 | 1 | 13 | Pon3 | 0.042 | Old |
| 0.745643 | 0.070402 | 0.197 | 0.175 | 1 | 13 | Tmem39b | 0.022 | Old |
| 0.746322 | 0.073349 | 0.418 | 0.368 | 1 | 13 | Zfp646 | 0.05 | Old |
| 0.74633 | 0.019091 | 0.918 | 0.93 | 1 | 13 | Tmem59 | -0.012 | Old |
| 0.746641 | 0.236161 | 0.5 | 0.474 | 1 | 13 | Scamp5 | 0.026 | Old |
| 0.746682 | 0.042785 | 0.172 | 0.158 | 1 | 13 | B4galt4 | 0.014 | Old |
| 0.746907 | 0.041349 | 0.5 | 0.509 | 1 | 13 | Zpr1 | -0.009 | Old |
| 0.747079 | 0.08618 | 0.705 | 0.675 | 1 | 13 | Snrnp200 | 0.03 | Old |
| 0.747114 | 0.071945 | 0.459 | 0.421 | 1 | 13 | Abcb7 | 0.038 | Old |
| 0.747696 | 0.166864 | 0.443 | 0.447 | 1 | 13 | Josd2 | -0.004 | Old |
| 0.74783 | 0.002618 | 1 | 1 | 1 | 13 | Ctsh | 0 | Old |
| 0.74793 | 0.303311 | 0.033 | 0.026 | 1 | 13 | Aspm | 0.007 | Old |
| 0.747981 | 0.261923 | 0.082 | 0.07 | 1 | 13 | 4632404H1 | 0.012 | Old |
| 0.748544 | 0.075353 | 0.943 | 0.974 | 1 | 13 | Arhgap30 | -0.031 | Old |
| 0.748659 | 0.05908 | 0.492 | 0.491 | 1 | 13 | Erap1 | 0.001 | Old |
| 0.748991 | 0.163097 | 0.09 | 0.079 | 1 | 13 | Chac2 | 0.011 | Old |
| 0.749114 | 0.080989 | 0.525 | 0.465 | 1 | 13 | Spast | 0.06 | Old |
| 0.74922 | 0.126023 | 0.18 | 0.158 | 1 | 13 | Ighmbp2 | 0.022 | Old |
| 0.74926 | 0.251708 | 0.074 | 0.061 | 1 | 13 | Tmem218 | 0.013 | Old |
| 0.74928 | 0.025061 | 0.582 | 0.57 | 1 | 13 | Chchd3 | 0.012 | Old |
| 0.749538 | 0.082373 | 0.582 | 0.535 | 1 | 13 | Mea1 | 0.047 | Old |
| 0.750277 | 0.023014 | 0.779 | 0.719 | 1 | 13 | Tmx3 | 0.06 | Old |
| 0.750328 | 0.148291 | 0.582 | 0.579 | 1 | 13 | D5Ertd579 | 0.003 | Old |
| 0.750543 | 0.072815 | 0.902 | 0.868 | 1 | 13 | Copz1 | 0.034 | Old |
| 0.750686 | 0.018438 | 0.951 | 0.912 | 1 | 13 | Comt | 0.039 | Old |
| 0.750931 | 0.100738 | 0.59 | 0.544 | 1 | 13 | Mlxip | 0.046 | Old |
| 0.750952 | 0.145815 | 0.279 | 0.254 | 1 | 13 | Prkci | 0.025 | Old |
| 0.751434 | 0.082349 | 0.18 | 0.158 | 1 | 13 | Zfp142 | 0.022 | Old |
| 0.751919 | 0.038852 | 0.5 | 0.456 | 1 | 13 | Dctn5 | 0.044 | Old |
| 0.751923 | 0.038213 | 0.811 | 0.702 | 1 | 13 | Ufsp2 | 0.109 | Old |
| 0.752171 | 0.034502 | 1 | 1 | 1 | 13 | Rtn4 | 0 | Old |
| 0.7527 | 0.138509 | 0.164 | 0.149 | 1 | 13 | Timmdc1 | 0.015 | Old |

| | | | | | | | | |
|---|---|---|---|---|---|---|---|---|
| 0.752854 | 0.506947 | 0.033 | 0.026 | 1 | 13 | Ifit3b | 0.007 | Old |
| 0.752854 | 0.469394 | 0.033 | 0.026 | 1 | 13 | Clec9a | 0.007 | Old |
| 0.752854 | 0.276569 | 0.033 | 0.026 | 1 | 13 | Rnf144a | 0.007 | Old |
| 0.752871 | 0.120536 | 0.148 | 0.132 | 1 | 13 | Clybl | 0.016 | Old |
| 0.753024 | 0.222087 | 0.459 | 0.439 | 1 | 13 | Adar | 0.02 | Old |
| 0.753319 | 0.108513 | 0.508 | 0.474 | 1 | 13 | Suco | 0.034 | Old |
| 0.753536 | 0.036828 | 0.91 | 0.912 | 1 | 13 | Nipa2 | -0.002 | Old |
| 0.753567 | 0.000425 | 0.934 | 0.912 | 1 | 13 | Sf3b6 | 0.022 | Old |
| 0.753776 | 0.056151 | 0.32 | 0.289 | 1 | 13 | Serhl | 0.031 | Old |
| 0.753966 | 0.307972 | 0.074 | 0.088 | 1 | 13 | Padi4 | -0.014 | Old |
| 0.754162 | 0.142711 | 0.197 | 0.175 | 1 | 13 | Dcakd | 0.022 | Old |
| 0.754515 | 0.083251 | 0.566 | 0.544 | 1 | 13 | Cir1 | 0.022 | Old |
| 0.754723 | 0.091739 | 0.402 | 0.368 | 1 | 13 | Man2b2 | 0.034 | Old |
| 0.755066 | 0.013694 | 0.959 | 0.982 | 1 | 13 | Nucks1 | -0.023 | Old |
| 0.755852 | 0.144233 | 0.73 | 0.693 | 1 | 13 | Pcmtd1 | 0.037 | Old |
| 0.755898 | 0.043367 | 0.738 | 0.64 | 1 | 13 | Anxa11 | 0.098 | Old |
| 0.75612 | 0.012905 | 0.844 | 0.728 | 1 | 13 | Appl1 | 0.116 | Old |
| 0.756479 | 0.027289 | 0.451 | 0.465 | 1 | 13 | Lypla1 | -0.014 | Old |
| 0.756518 | 0.0726 | 0.984 | 0.965 | 1 | 13 | Atp5c1 | 0.019 | Old |
| 0.75652 | 0.03676 | 1 | 1 | 1 | 13 | Timp2 | 0 | Old |
| 0.756957 | 0.033554 | 0.672 | 0.614 | 1 | 13 | Tbc1d10b | 0.058 | Old |
| 0.757296 | 0.350783 | 0.205 | 0.184 | 1 | 13 | Rcl1 | 0.021 | Old |
| 0.757693 | 0.035867 | 0.607 | 0.596 | 1 | 13 | Aco1 | 0.011 | Old |
| 0.757789 | 0.420425 | 0.033 | 0.026 | 1 | 13 | Gm20605 | 0.007 | Old |
| 0.757789 | 0.386954 | 0.033 | 0.026 | 1 | 13 | Dbndd2 | 0.007 | Old |
| 0.757789 | 0.341677 | 0.033 | 0.026 | 1 | 13 | 9330020H( | 0.007 | Old |
| 0.757789 | 0.313947 | 0.033 | 0.026 | 1 | 13 | Gm48960 | 0.007 | Old |
| 0.757789 | 0.288296 | 0.033 | 0.026 | 1 | 13 | Creg2 | 0.007 | Old |
| 0.758443 | 0.124484 | 0.656 | 0.632 | 1 | 13 | Fam105a | 0.024 | Old |
| 0.758492 | 0.203935 | 0.574 | 0.561 | 1 | 13 | Med8 | 0.013 | Old |
| 0.759122 | 0.034353 | 0.287 | 0.263 | 1 | 13 | Poc1b | 0.024 | Old |
| 0.75931 | 0.053357 | 0.467 | 0.404 | 1 | 13 | Tdg | 0.063 | Old |
| 0.759962 | 0.02614 | 0.123 | 0.105 | 1 | 13 | Gfod1 | 0.018 | Old |
| 0.759998 | 0.100315 | 0.77 | 0.684 | 1 | 13 | Strap | 0.086 | Old |
| 0.760174 | 0.113389 | 0.213 | 0.193 | 1 | 13 | Noc3l | 0.02 | Old |
| 0.760315 | 0.111414 | 0.73 | 0.754 | 1 | 13 | Taf1d | -0.024 | Old |
| 0.760433 | 0.078309 | 0.648 | 0.57 | 1 | 13 | Slc31a2 | 0.078 | Old |
| 0.76058 | 0.103911 | 0.082 | 0.07 | 1 | 13 | Btbd6 | 0.012 | Old |
| 0.760874 | 0.011072 | 0.975 | 0.965 | 1 | 13 | Hmox1 | 0.01 | Old |
| 0.760876 | 0.019964 | 1 | 1 | 1 | 13 | Son | 0 | Old |
| 0.760898 | 0.038047 | 0.32 | 0.289 | 1 | 13 | Ambra1 | 0.031 | Old |
| 0.760999 | 0.215676 | 0.09 | 0.079 | 1 | 13 | Smug1 | 0.011 | Old |
| 0.761818 | 0.214922 | 0.574 | 0.57 | 1 | 13 | Trim56 | 0.004 | Old |
| 0.761921 | 0.017381 | 0.869 | 0.825 | 1 | 13 | Zc3h7a | 0.044 | Old |
| 0.761938 | 0.153953 | 0.705 | 0.684 | 1 | 13 | Rassf5 | 0.021 | Old |

| | | | | | | | | |
|---|---|---|---|---|---|---|---|---|
| 0.762733 | 0.358632 | 0.033 | 0.026 | 1 | 13 | Gpr19 | 0.007 | Old |
| 0.762733 | 0.329823 | 0.033 | 0.026 | 1 | 13 | Igfbp7 | 0.007 | Old |
| 0.762733 | 0.237572 | 0.033 | 0.026 | 1 | 13 | Gm16253 | 0.007 | Old |
| 0.762937 | 0.075635 | 0.893 | 0.904 | 1 | 13 | Myo1g | -0.011 | Old |
| 0.763417 | 0.155495 | 0.279 | 0.263 | 1 | 13 | Pld1 | 0.016 | Old |
| 0.763418 | 0.188059 | 0.598 | 0.57 | 1 | 13 | Tfrc | 0.028 | Old |
| 0.763423 | 0.047589 | 0.566 | 0.518 | 1 | 13 | Gm49336 | 0.048 | Old |
| 0.763734 | 0.00618 | 0.934 | 0.912 | 1 | 13 | Pttg1ip | 0.022 | Old |
| 0.763784 | 0.055915 | 0.992 | 0.965 | 1 | 13 | Taok3 | 0.027 | Old |
| 0.76379 | 0.082732 | 0.508 | 0.465 | 1 | 13 | A630001G2 | 0.043 | Old |
| 0.764259 | 0.436237 | 0.221 | 0.219 | 1 | 13 | Myc | 0.002 | Old |
| 0.76468 | 0.057751 | 0.525 | 0.535 | 1 | 13 | Mrps9 | -0.01 | Old |
| 0.764914 | 0.219858 | 0.148 | 0.132 | 1 | 13 | Pigf | 0.016 | Old |
| 0.765337 | 0.008284 | 0.09 | 0.105 | 1 | 13 | Sv2b | -0.015 | Old |
| 0.765731 | 0.029512 | 0.779 | 0.728 | 1 | 13 | Rlim | 0.051 | Old |
| 0.765834 | 0.133182 | 0.156 | 0.14 | 1 | 13 | Exd2 | 0.016 | Old |
| 0.766254 | 0.105298 | 0.32 | 0.289 | 1 | 13 | Rbl1 | 0.031 | Old |
| 0.76644 | 0.00944 | 0.918 | 0.816 | 1 | 13 | Rassf1 | 0.102 | Old |
| 0.766448 | 0.093705 | 0.402 | 0.36 | 1 | 13 | Fbxo38 | 0.042 | Old |
| 0.7665 | 0.09821 | 0.098 | 0.088 | 1 | 13 | Pick1 | 0.01 | Old |
| 0.766842 | 0.004601 | 0.451 | 0.377 | 1 | 13 | Pop7 | 0.074 | Old |
| 0.766896 | 0.022373 | 0.369 | 0.342 | 1 | 13 | 0610037L1 | 0.027 | Old |
| 0.767397 | 0.074043 | 0.418 | 0.386 | 1 | 13 | Rnaseh2a | 0.032 | Old |
| 0.767562 | 0.013952 | 0.68 | 0.614 | 1 | 13 | Nr1h2 | 0.066 | Old |
| 0.767688 | 0.538491 | 0.033 | 0.026 | 1 | 13 | Dclk2 | 0.007 | Old |
| 0.767688 | 0.257531 | 0.033 | 0.026 | 1 | 13 | Camk2b | 0.007 | Old |
| 0.768036 | 0.028475 | 0.902 | 0.895 | 1 | 13 | Bcap31 | 0.007 | Old |
| 0.76819 | 0.076559 | 0.156 | 0.14 | 1 | 13 | Gm49359 | 0.016 | Old |
| 0.768967 | 0.114636 | 0.615 | 0.588 | 1 | 13 | Vav2 | 0.027 | Old |
| 0.769143 | 0.034024 | 0.197 | 0.175 | 1 | 13 | Nat6 | 0.022 | Old |
| 0.76928 | 0.179875 | 0.23 | 0.211 | 1 | 13 | Isca1 | 0.019 | Old |
| 0.769488 | 0.0527 | 0.77 | 0.746 | 1 | 13 | St3gal1 | 0.024 | Old |
| 0.770055 | 0.005134 | 0.484 | 0.491 | 1 | 13 | Rab3gap2 | -0.007 | Old |
| 0.770549 | 0.124291 | 0.156 | 0.14 | 1 | 13 | 1600014C1 | 0.016 | Old |
| 0.771082 | 0.115874 | 0.164 | 0.149 | 1 | 13 | Ccdc61 | 0.015 | Old |
| 0.771151 | 0.236833 | 0.197 | 0.184 | 1 | 13 | Mpst | 0.013 | Old |
| 0.771774 | 0.090786 | 0.762 | 0.763 | 1 | 13 | Mrpl51 | -0.001 | Old |
| 0.771862 | 0.158429 | 0.697 | 0.667 | 1 | 13 | Aida | 0.03 | Old |
| 0.77202 | 0.061253 | 0.246 | 0.219 | 1 | 13 | Arhgef18 | 0.027 | Old |
| 0.772119 | 0.148354 | 0.115 | 0.105 | 1 | 13 | Cep57l1 | 0.01 | Old |
| 0.772651 | 0.173647 | 0.033 | 0.026 | 1 | 13 | Plcd1 | 0.007 | Old |
| 0.772651 | 0.161775 | 0.033 | 0.026 | 1 | 13 | Chd5 | 0.007 | Old |
| 0.772651 | 0.15692 | 0.033 | 0.026 | 1 | 13 | Ccdc173 | 0.007 | Old |
| 0.772651 | 0.007644 | 0.033 | 0.026 | 1 | 13 | Cyp2j9 | 0.007 | Old |
| 0.772651 | 0.002975 | 0.033 | 0.026 | 1 | 13 | Ms4a4b | 0.007 | Old |

| | | | | | | | | |
|---|---|---|---|---|---|---|---|---|
| 0.772971 | 0.078107 | 0.59 | 0.623 | 1 | 13 | Mertk | -0.033 | Old |
| 0.773242 | 0.080276 | 0.082 | 0.07 | 1 | 13 | Grhpr | 0.012 | Old |
| 0.77328 | 0.13286 | 0.197 | 0.184 | 1 | 13 | Fam110b | 0.013 | Old |
| 0.773346 | 0.056646 | 0.639 | 0.649 | 1 | 13 | Tmpo | -0.01 | Old |
| 0.774654 | 0.097957 | 0.393 | 0.377 | 1 | 13 | Mtmr14 | 0.016 | Old |
| 0.775114 | 0.067294 | 0.697 | 0.684 | 1 | 13 | Nudc | 0.013 | Old |
| 0.775179 | 0.157756 | 0.098 | 0.088 | 1 | 13 | Rad1 | 0.01 | Old |
| 0.77545 | 0.121905 | 0.951 | 0.991 | 1 | 13 | Fn1 | -0.04 | Old |
| 0.775745 | 0.024235 | 0.803 | 0.754 | 1 | 13 | Dennd1b | 0.049 | Old |
| 0.775894 | 0.26963 | 0.377 | 0.368 | 1 | 13 | Smox | 0.009 | Old |
| 0.776418 | 0.042465 | 0.082 | 0.07 | 1 | 13 | Epop | 0.012 | Old |
| 0.776418 | 0.001453 | 0.082 | 0.07 | 1 | 13 | 1700008J0 | 0.012 | Old |
| 0.77646 | 0.038634 | 0.803 | 0.877 | 1 | 13 | Polr2k | -0.074 | Old |
| 0.776728 | 0.058563 | 0.467 | 0.412 | 1 | 13 | Smg9 | 0.055 | Old |
| 0.776896 | 0.17232 | 0.23 | 0.254 | 1 | 13 | Cd200r4 | -0.024 | Old |
| 0.777624 | 0.232358 | 0.033 | 0.026 | 1 | 13 | Clmn | 0.007 | Old |
| 0.777624 | 0.202825 | 0.033 | 0.026 | 1 | 13 | Zfp623 | 0.007 | Old |
| 0.777624 | 0.19785 | 0.033 | 0.026 | 1 | 13 | Rgs12 | 0.007 | Old |
| 0.777624 | 0.186579 | 0.033 | 0.026 | 1 | 13 | Col20a1 | 0.007 | Old |
| 0.777624 | 0.171388 | 0.033 | 0.026 | 1 | 13 | Pard3b | 0.007 | Old |
| 0.777624 | 0.142362 | 0.033 | 0.026 | 1 | 13 | Gnat3 | 0.007 | Old |
| 0.777938 | 0.044308 | 0.246 | 0.219 | 1 | 13 | Wiz | 0.027 | Old |
| 0.777975 | 0.045023 | 0.574 | 0.553 | 1 | 13 | Ppm1g | 0.021 | Old |
| 0.778177 | 0.013696 | 0.877 | 0.877 | 1 | 13 | Gnai3 | 0 | Old |
| 0.778936 | 0.08137 | 0.664 | 0.64 | 1 | 13 | Rps6kb1 | 0.024 | Old |
| 0.779022 | 0.023065 | 0.91 | 0.904 | 1 | 13 | Ythdc1 | 0.006 | Old |
| 0.779061 | 0.160491 | 0.492 | 0.474 | 1 | 13 | Taf13 | 0.018 | Old |
| 0.779119 | 0.178883 | 0.09 | 0.079 | 1 | 13 | Fam122b | 0.011 | Old |
| 0.779691 | 0.125842 | 0.361 | 0.351 | 1 | 13 | Kdm5b | 0.01 | Old |
| 0.779837 | 0.021903 | 0.967 | 0.965 | 1 | 13 | Prdx2 | 0.002 | Old |
| 0.779905 | 0.037224 | 0.434 | 0.395 | 1 | 13 | Grcc10 | 0.039 | Old |
| 0.780322 | 0.019574 | 0.164 | 0.149 | 1 | 13 | Edrf1 | 0.015 | Old |
| 0.780422 | 0.273643 | 0.172 | 0.158 | 1 | 13 | Spaca6 | 0.014 | Old |
| 0.780626 | 0.054605 | 0.361 | 0.342 | 1 | 13 | Gamt | 0.019 | Old |
| 0.780761 | 0.092289 | 0.541 | 0.535 | 1 | 13 | Flot2 | 0.006 | Old |
| 0.781567 | 0.078566 | 0.27 | 0.289 | 1 | 13 | Polr2d | -0.019 | Old |
| 0.781927 | 0.043372 | 0.877 | 0.921 | 1 | 13 | Adam10 | -0.044 | Old |
| 0.782128 | 0.106332 | 0.844 | 0.789 | 1 | 13 | Mtch1 | 0.055 | Old |
| 0.782153 | 0.028931 | 0.5 | 0.447 | 1 | 13 | Gab2 | 0.053 | Old |
| 0.78219 | 0.296085 | 0.107 | 0.096 | 1 | 13 | Colca2 | 0.011 | Old |
| 0.782375 | 0.034724 | 0.156 | 0.14 | 1 | 13 | Rilp | 0.016 | Old |
| 0.782408 | 0.019886 | 0.426 | 0.43 | 1 | 13 | Rprd1b | -0.004 | Old |
| 0.782553 | 0.028989 | 0.344 | 0.289 | 1 | 13 | Blcap | 0.055 | Old |
| 0.782606 | 0.05295 | 0.033 | 0.026 | 1 | 13 | Rgs5 | 0.007 | Old |
| 0.782606 | 0.016362 | 0.033 | 0.026 | 1 | 13 | Ube2v1 | 0.007 | Old |

| | | | | | | | | |
|---|---|---|---|---|---|---|---|---|
| 0.782771 | 0.091893 | 0.992 | 1 | 1 | 13 | Hspe1 | -0.008 | Old |
| 0.78347 | 0.035198 | 0.451 | 0.421 | 1 | 13 | Fbxo4 | 0.03 | Old |
| 0.783831 | 0.07317 | 0.811 | 0.877 | 1 | 13 | Ms4a7 | -0.066 | Old |
| 0.783966 | 0.029194 | 0.795 | 0.781 | 1 | 13 | Cs | 0.014 | Old |
| 0.784204 | 0.075829 | 0.951 | 0.912 | 1 | 13 | Tmed5 | 0.039 | Old |
| 0.784653 | 0.146532 | 0.098 | 0.114 | 1 | 13 | Slc30a6 | -0.016 | Old |
| 0.784954 | 0.136087 | 0.172 | 0.158 | 1 | 13 | Pmm1 | 0.014 | Old |
| 0.7853 | 0.071367 | 0.82 | 0.868 | 1 | 13 | Frmd4b | -0.048 | Old |
| 0.785504 | 0.033989 | 0.664 | 0.632 | 1 | 13 | Peak1 | 0.032 | Old |
| 0.785684 | 0.017388 | 0.475 | 0.465 | 1 | 13 | Tor3a | 0.01 | Old |
| 0.785888 | 0.023132 | 0.23 | 0.202 | 1 | 13 | Zfp729a | 0.028 | Old |
| 0.785898 | 0.315005 | 0.557 | 0.579 | 1 | 13 | Jdp2 | -0.022 | Old |
| 0.786284 | 0.067029 | 0.631 | 0.632 | 1 | 13 | Atad2b | -0.001 | Old |
| 0.787166 | 0.027262 | 0.967 | 0.956 | 1 | 13 | Eif4b | 0.011 | Old |
| 0.787598 | 0.030175 | 0.033 | 0.026 | 1 | 13 | 9530082P2 | 0.007 | Old |
| 0.787598 | 0.027641 | 0.033 | 0.026 | 1 | 13 | Gm34006 | 0.007 | Old |
| 0.787598 | 0.012444 | 0.033 | 0.026 | 1 | 13 | P4ha2 | 0.007 | Old |
| 0.78864 | 0.059078 | 1 | 1 | 1 | 13 | Dusp1 | 0 | Old |
| 0.788711 | 0.028612 | 0.385 | 0.351 | 1 | 13 | Tomm6 | 0.034 | Old |
| 0.78917 | 0.414696 | 0.041 | 0.035 | 1 | 13 | Gm15232 | 0.006 | Old |
| 0.789495 | 0.126617 | 0.156 | 0.14 | 1 | 13 | Zeb1 | 0.016 | Old |
| 0.7895 | 0.127285 | 0.402 | 0.377 | 1 | 13 | Ptpn23 | 0.025 | Old |
| 0.790003 | 0.004913 | 0.402 | 0.421 | 1 | 13 | Rnf181 | -0.019 | Old |
| 0.790109 | 0.010474 | 1 | 1 | 1 | 13 | Gnb2 | 0 | Old |
| 0.790109 | 0.009205 | 1 | 1 | 1 | 13 | Ucp2 | 0 | Old |
| 0.790127 | 0.102547 | 0.189 | 0.167 | 1 | 13 | Serinc5 | 0.022 | Old |
| 0.790304 | 0.027795 | 0.262 | 0.272 | 1 | 13 | Pck2 | -0.01 | Old |
| 0.790702 | 0.489008 | 0.131 | 0.123 | 1 | 13 | Dennd2a | 0.008 | Old |
| 0.790925 | 0.427126 | 0.09 | 0.105 | 1 | 13 | Gk5 | -0.015 | Old |
| 0.790945 | 0.033707 | 0.779 | 0.746 | 1 | 13 | Eif3g | 0.033 | Old |
| 0.790976 | 0.017184 | 0.303 | 0.263 | 1 | 13 | Phf11d | 0.04 | Old |
| 0.791094 | 0.01781 | 0.393 | 0.351 | 1 | 13 | Gipc1 | 0.042 | Old |
| 0.791189 | 0.063889 | 0.246 | 0.228 | 1 | 13 | Clpb | 0.018 | Old |
| 0.791267 | 0.107878 | 0.09 | 0.079 | 1 | 13 | Agrn | 0.011 | Old |
| 0.791353 | 0.012671 | 0.705 | 0.667 | 1 | 13 | Eif4e2 | 0.038 | Old |
| 0.791578 | 0.104431 | 0.984 | 0.982 | 1 | 13 | Lgals1 | 0.002 | Old |
| 0.791578 | 0.017108 | 1 | 1 | 1 | 13 | Ubl5 | 0 | Old |
| 0.791796 | 0.159924 | 0.238 | 0.228 | 1 | 13 | Gk | 0.01 | Old |
| 0.792488 | 0.059093 | 0.754 | 0.702 | 1 | 13 | Ube2m | 0.052 | Old |
| 0.792597 | 0.025804 | 0.033 | 0.026 | 1 | 13 | Epm2a | 0.007 | Old |
| 0.79262 | 0.053657 | 0.098 | 0.088 | 1 | 13 | Rbm41 | 0.01 | Old |
| 0.792671 | 0.07352 | 0.361 | 0.342 | 1 | 13 | C1galt1c1 | 0.019 | Old |
| 0.793024 | 0.006416 | 0.475 | 0.421 | 1 | 13 | Ddx23 | 0.054 | Old |
| 0.793057 | 0.127327 | 0.41 | 0.36 | 1 | 13 | Rbpms | 0.05 | Old |
| 0.793299 | 0.041955 | 0.434 | 0.386 | 1 | 13 | Spata2 | 0.048 | Old |

| | | | | | | | | |
|---|---|---|---|---|---|---|---|---|
| 0.793616 | 0.064108 | 0.926 | 0.833 | 1 | 13 | Ralb | 0.093 | Old |
| 0.79369 | 0.224405 | 0.115 | 0.105 | 1 | 13 | Zfp579 | 0.01 | Old |
| 0.793941 | 0.37013 | 0.189 | 0.184 | 1 | 13 | Ermard | 0.005 | Old |
| 0.793992 | 0.096943 | 0.869 | 0.781 | 1 | 13 | Sh3bp1 | 0.088 | Old |
| 0.794003 | 0.091504 | 0.689 | 0.579 | 1 | 13 | Stim2 | 0.11 | Old |
| 0.794669 | 0.193213 | 0.197 | 0.184 | 1 | 13 | Orc6 | 0.013 | Old |
| 0.795013 | 0.253857 | 0.32 | 0.298 | 1 | 13 | Mre11a | 0.022 | Old |
| 0.79524 | 0.060927 | 0.492 | 0.447 | 1 | 13 | Slc44a1 | 0.045 | Old |
| 0.795256 | 0.008496 | 1 | 0.991 | 1 | 13 | Itgb1 | 0.009 | Old |
| 0.795537 | 0.289414 | 0.098 | 0.088 | 1 | 13 | Vamp1 | 0.01 | Old |
| 0.795541 | 0.03881 | 0.885 | 0.781 | 1 | 13 | Zfand6 | 0.104 | Old |
| 0.796398 | 0.149188 | 0.115 | 0.105 | 1 | 13 | Sh3d19 | 0.01 | Old |
| 0.79661 | 0.035802 | 0.615 | 0.675 | 1 | 13 | Max | -0.06 | Old |
| 0.797631 | 0.110886 | 0.402 | 0.386 | 1 | 13 | Vps52 | 0.016 | Old |
| 0.797685 | 0.12415 | 0.27 | 0.254 | 1 | 13 | Mecr | 0.016 | Old |
| 0.79778 | 0.020107 | 0.533 | 0.544 | 1 | 13 | Mysm1 | -0.011 | Old |
| 0.798038 | 0.249991 | 0.041 | 0.035 | 1 | 13 | Il4 | 0.006 | Old |
| 0.798052 | 0.029271 | 0.82 | 0.763 | 1 | 13 | Fndc3a | 0.057 | Old |
| 0.798159 | 0.149691 | 0.287 | 0.263 | 1 | 13 | Mir142hg | 0.024 | Old |
| 0.798331 | 0.197511 | 0.131 | 0.123 | 1 | 13 | Mlh3 | 0.008 | Old |
| 0.798457 | 0.086667 | 0.098 | 0.088 | 1 | 13 | Zfp770 | 0.01 | Old |
| 0.798691 | 0.160482 | 0.254 | 0.237 | 1 | 13 | Zfp414 | 0.017 | Old |
| 0.799418 | 0.135914 | 0.697 | 0.719 | 1 | 13 | Wdfy2 | -0.022 | Old |
| 0.799577 | 0.03851 | 0.713 | 0.658 | 1 | 13 | Pigs | 0.055 | Old |
| 0.799632 | 0.09689 | 0.91 | 0.939 | 1 | 13 | Bclaf1 | -0.029 | Old |
| 0.799744 | 0.157018 | 0.238 | 0.228 | 1 | 13 | Rpp14 | 0.01 | Old |
| 0.799951 | 0.132133 | 0.377 | 0.351 | 1 | 13 | Tubgcp5 | 0.026 | Old |
| 0.80006 | 0.062656 | 0.352 | 0.316 | 1 | 13 | Kctd17 | 0.036 | Old |
| 0.800229 | 0.084193 | 0.295 | 0.272 | 1 | 13 | Siae | 0.023 | Old |
| 0.800411 | 0.107851 | 0.09 | 0.079 | 1 | 13 | Enpp4 | 0.011 | Old |
| 0.800742 | 0.109112 | 0.844 | 0.833 | 1 | 13 | Ripk1 | 0.011 | Old |
| 0.801304 | 0.038263 | 0.361 | 0.342 | 1 | 13 | Brwd3 | 0.019 | Old |
| 0.801401 | 0.214105 | 0.156 | 0.14 | 1 | 13 | Pigz | 0.016 | Old |
| 0.80153 | 0.006651 | 0.18 | 0.193 | 1 | 13 | Tmem110 | -0.013 | Old |
| 0.801599 | 0.011773 | 0.5 | 0.474 | 1 | 13 | Syvn1 | 0.026 | Old |
| 0.801851 | 0.033505 | 0.934 | 0.921 | 1 | 13 | Prelid1 | 0.013 | Old |
| 0.801992 | 0.047902 | 0.787 | 0.728 | 1 | 13 | Setd2 | 0.059 | Old |
| 0.802118 | 0.038561 | 0.41 | 0.395 | 1 | 13 | Nom1 | 0.015 | Old |
| 0.802481 | 0.302224 | 0.041 | 0.035 | 1 | 13 | Cd5l | 0.006 | Old |
| 0.802481 | 0.255147 | 0.041 | 0.035 | 1 | 13 | 1700109H( | 0.006 | Old |
| 0.802548 | 0.136276 | 0.82 | 0.772 | 1 | 13 | Cmklr1 | 0.048 | Old |
| 0.803134 | 0.081782 | 0.311 | 0.298 | 1 | 13 | Siah1a | 0.013 | Old |
| 0.803363 | 0.010637 | 1 | 0.991 | 1 | 13 | Zfp36l2 | 0.009 | Old |
| 0.803427 | 0.27856 | 0.131 | 0.123 | 1 | 13 | Zfp758 | 0.008 | Old |
| 0.803465 | 0.015549 | 0.09 | 0.079 | 1 | 13 | 2510046G1 | 0.011 | Old |

| | | | | | | | | |
|---|---|---|---|---|---|---|---|---|
| 0.803585 | 0.056097 | 0.459 | 0.456 | 1 | 13 | Itga9 | 0.003 | Old |
| 0.803822 | 0.138938 | 0.131 | 0.149 | 1 | 13 | Gga2 | -0.018 | Old |
| 0.803868 | 0.2344 | 0.287 | 0.272 | 1 | 13 | Usb1 | 0.015 | Old |
| 0.804051 | 0.239555 | 0.303 | 0.298 | 1 | 13 | Togaram1 | 0.005 | Old |
| 0.804173 | 0.082077 | 0.82 | 0.798 | 1 | 13 | Mfhas1 | 0.022 | Old |
| 0.804176 | 0.016786 | 0.623 | 0.561 | 1 | 13 | Inpp5k | 0.062 | Old |
| 0.804306 | 0.207963 | 0.098 | 0.088 | 1 | 13 | Slc22a18 | 0.01 | Old |
| 0.804536 | 0.07747 | 0.115 | 0.105 | 1 | 13 | Dus2 | 0.01 | Old |
| 0.804838 | 0.114307 | 0.984 | 0.965 | 1 | 13 | Rab7b | 0.019 | Old |
| 0.805317 | 0.003882 | 0.746 | 0.728 | 1 | 13 | Idh3g | 0.018 | Old |
| 0.805477 | 0.060869 | 0.91 | 0.886 | 1 | 13 | Nfe2l1 | 0.024 | Old |
| 0.805725 | 0.116821 | 0.418 | 0.377 | 1 | 13 | Tsc1 | 0.041 | Old |
| 0.806203 | 0.014793 | 0.877 | 0.912 | 1 | 13 | Cmtm7 | -0.035 | Old |
| 0.806311 | 0.06335 | 0.943 | 0.982 | 1 | 13 | Psme2 | -0.039 | Old |
| 0.806931 | 0.235826 | 0.041 | 0.035 | 1 | 13 | Bcl7a | 0.006 | Old |
| 0.807215 | 0.05921 | 0.27 | 0.254 | 1 | 13 | Zfp771 | 0.016 | Old |
| 0.807367 | 0.050177 | 0.844 | 0.825 | 1 | 13 | Zcrb1 | 0.019 | Old |
| 0.807581 | 0.007285 | 0.492 | 0.474 | 1 | 13 | Deptor | 0.018 | Old |
| 0.8077 | 0.011949 | 0.689 | 0.675 | 1 | 13 | Grk3 | 0.014 | Old |
| 0.808046 | 0.042384 | 0.426 | 0.421 | 1 | 13 | Zbtb18 | 0.005 | Old |
| 0.808051 | 0.196203 | 0.197 | 0.175 | 1 | 13 | Tsen15 | 0.022 | Old |
| 0.808419 | 0.097418 | 0.516 | 0.509 | 1 | 13 | Cisd1 | 0.007 | Old |
| 0.808481 | 0.1834 | 0.418 | 0.395 | 1 | 13 | Plekhg5 | 0.023 | Old |
| 0.809128 | 0.136772 | 0.762 | 0.719 | 1 | 13 | Mrpl12 | 0.043 | Old |
| 0.809469 | 0.114547 | 0.287 | 0.272 | 1 | 13 | Bid | 0.015 | Old |
| 0.809887 | 0.063352 | 0.877 | 0.904 | 1 | 13 | Pmp22 | -0.027 | Old |
| 0.81062 | 0.241692 | 0.73 | 0.754 | 1 | 13 | Emp1 | -0.024 | Old |
| 0.81074 | 0.072678 | 0.451 | 0.43 | 1 | 13 | Mafk | 0.021 | Old |
| 0.81092 | 0.081609 | 0.754 | 0.763 | 1 | 13 | Notch1 | -0.009 | Old |
| 0.810961 | 0.008168 | 0.23 | 0.246 | 1 | 13 | Mrpl46 | -0.016 | Old |
| 0.811284 | 0.048607 | 0.615 | 0.614 | 1 | 13 | Camta2 | 0.001 | Old |
| 0.811387 | 0.178408 | 0.041 | 0.035 | 1 | 13 | Cyp2r1 | 0.006 | Old |
| 0.811387 | 0.170505 | 0.041 | 0.035 | 1 | 13 | Tmlhe | 0.006 | Old |
| 0.811387 | 0.152792 | 0.041 | 0.035 | 1 | 13 | Plcb4 | 0.006 | Old |
| 0.811387 | 0.137812 | 0.041 | 0.035 | 1 | 13 | Gm4631 | 0.006 | Old |
| 0.811807 | 0.047173 | 0.418 | 0.395 | 1 | 13 | Stard3 | 0.023 | Old |
| 0.81213 | 0.333122 | 0.049 | 0.044 | 1 | 13 | Ndor1 | 0.005 | Old |
| 0.812169 | 0.029285 | 0.279 | 0.263 | 1 | 13 | Zfp26 | 0.016 | Old |
| 0.81236 | 0.001176 | 0.492 | 0.474 | 1 | 13 | Kbtbd11 | 0.018 | Old |
| 0.812942 | 0.036785 | 0.926 | 0.939 | 1 | 13 | Eif4a2 | -0.013 | Old |
| 0.813083 | 0.060152 | 0.107 | 0.096 | 1 | 13 | Aldh1b1 | 0.011 | Old |
| 0.813645 | 0.090561 | 0.131 | 0.123 | 1 | 13 | Setdb2 | 0.008 | Old |
| 0.813711 | 0.036181 | 1 | 1 | 1 | 13 | Pitpna | 0 | Old |
| 0.81427 | 0.035762 | 0.787 | 0.781 | 1 | 13 | Cep350 | 0.006 | Old |
| 0.814875 | 0.006916 | 0.426 | 0.491 | 1 | 13 | Bop1 | -0.065 | Old |

| | | | | | | | | |
|---|---|---|---|---|---|---|---|---|
| 0.815108 | 0.020331 | 0.926 | 0.877 | 1 | 13 | Zfp207 | 0.049 | Old |
| 0.81585 | 0.192145 | 0.041 | 0.035 | 1 | 13 | Tcaf1 | 0.006 | Old |
| 0.81585 | 0.162692 | 0.041 | 0.035 | 1 | 13 | Fhit | 0.006 | Old |
| 0.81585 | 0.141296 | 0.041 | 0.035 | 1 | 13 | Serping1 | 0.006 | Old |
| 0.816036 | 0.042857 | 0.098 | 0.088 | 1 | 13 | Tlr3 | 0.01 | Old |
| 0.816249 | 0.031885 | 0.508 | 0.474 | 1 | 13 | Babam1 | 0.034 | Old |
| 0.816452 | 0.019957 | 0.336 | 0.298 | 1 | 13 | Poglut1 | 0.038 | Old |
| 0.816644 | 0.038277 | 0.385 | 0.386 | 1 | 13 | Pno1 | -0.001 | Old |
| 0.817451 | 0.092474 | 0.426 | 0.412 | 1 | 13 | Rgs14 | 0.014 | Old |
| 0.817989 | 0.191191 | 0.213 | 0.202 | 1 | 13 | Acvr1b | 0.011 | Old |
| 0.818147 | 0.223807 | 0.115 | 0.105 | 1 | 13 | Tprn | 0.01 | Old |
| 0.818732 | 0.037949 | 0.107 | 0.096 | 1 | 13 | Il18bp | 0.011 | Old |
| 0.818886 | 0.053842 | 0.943 | 0.956 | 1 | 13 | Furin | -0.013 | Old |
| 0.818982 | 0.043433 | 0.549 | 0.588 | 1 | 13 | Mrpl32 | -0.039 | Old |
| 0.819296 | 0.016703 | 0.582 | 0.596 | 1 | 13 | Lpin1 | -0.014 | Old |
| 0.819482 | 0.054934 | 0.262 | 0.246 | 1 | 13 | Pgap1 | 0.016 | Old |
| 0.819639 | 0.036366 | 1 | 1 | 1 | 13 | Ptprc | 0 | Old |
| 0.8201 | 0.060827 | 0.566 | 0.526 | 1 | 13 | Dok1 | 0.04 | Old |
| 0.820243 | 0.273584 | 0.049 | 0.044 | 1 | 13 | Fsd1l | 0.005 | Old |
| 0.820243 | 0.137586 | 0.049 | 0.044 | 1 | 13 | BC030499 | 0.005 | Old |
| 0.820318 | 0.331309 | 0.041 | 0.035 | 1 | 13 | Fpr1 | 0.006 | Old |
| 0.820318 | 0.17527 | 0.041 | 0.035 | 1 | 13 | Zbtb39 | 0.006 | Old |
| 0.820318 | 0.170201 | 0.041 | 0.035 | 1 | 13 | 4931428F0 | 0.006 | Old |
| 0.820318 | 0.008789 | 0.041 | 0.035 | 1 | 13 | Cpeb1 | 0.006 | Old |
| 0.820342 | 0.043113 | 0.926 | 0.921 | 1 | 13 | Hivep2 | 0.005 | Old |
| 0.82113 | 0.1256 | 0.689 | 0.667 | 1 | 13 | Rsf1 | 0.022 | Old |
| 0.822322 | 0.025841 | 0.861 | 0.842 | 1 | 13 | Ddx24 | 0.019 | Old |
| 0.822519 | 0.108947 | 0.656 | 0.693 | 1 | 13 | 4932438A1 | -0.037 | Old |
| 0.822607 | 0.020506 | 1 | 1 | 1 | 13 | Tubb5 | 0 | Old |
| 0.822725 | 0.101639 | 0.713 | 0.789 | 1 | 13 | Rapgef6 | -0.076 | Old |
| 0.822765 | 0.052035 | 0.352 | 0.325 | 1 | 13 | Zfp945 | 0.027 | Old |
| 0.823073 | 0.083923 | 0.582 | 0.561 | 1 | 13 | Samd1 | 0.021 | Old |
| 0.823171 | 0.123627 | 0.877 | 0.868 | 1 | 13 | Hmgb2 | 0.009 | Old |
| 0.823301 | 0.014204 | 0.311 | 0.281 | 1 | 13 | Kank2 | 0.03 | Old |
| 0.823503 | 0.07647 | 0.607 | 0.553 | 1 | 13 | Cldnd1 | 0.054 | Old |
| 0.824452 | 0.007385 | 0.27 | 0.289 | 1 | 13 | Sec23a | -0.019 | Old |
| 0.824612 | 0.013908 | 0.459 | 0.421 | 1 | 13 | Cnot7 | 0.038 | Old |
| 0.824782 | 0.018807 | 0.926 | 0.904 | 1 | 13 | Lamtor1 | 0.022 | Old |
| 0.824791 | 0.078243 | 0.434 | 0.386 | 1 | 13 | Abhd4 | 0.048 | Old |
| 0.824792 | 0.130639 | 0.041 | 0.035 | 1 | 13 | Zfp280b | 0.006 | Old |
| 0.824792 | 0.067449 | 0.041 | 0.035 | 1 | 13 | Gm47232 | 0.006 | Old |
| 0.824792 | 0.053715 | 0.041 | 0.035 | 1 | 13 | Nnt | 0.006 | Old |
| 0.824828 | 0.021046 | 0.959 | 0.956 | 1 | 13 | Zmiz1 | 0.003 | Old |
| 0.824834 | 0.041951 | 0.975 | 0.991 | 1 | 13 | Aif1 | -0.016 | Old |
| 0.824908 | 0.106246 | 0.139 | 0.132 | 1 | 13 | Sap130 | 0.007 | Old |

| | | | | | | | | |
|---|---|---|---|---|---|---|---|---|
| 0.825406 | 0.119764 | 0.344 | 0.333 | 1 | 13 | Mettl16 | 0.011 | Old |
| 0.82546 | 0.018451 | 0.525 | 0.491 | 1 | 13 | Crtc2 | 0.034 | Old |
| 0.825839 | 0.011791 | 0.582 | 0.614 | 1 | 13 | Zc3h18 | -0.032 | Old |
| 0.826659 | 0.023078 | 0.221 | 0.202 | 1 | 13 | Sclt1 | 0.019 | Old |
| 0.826786 | 0.101414 | 0.869 | 0.833 | 1 | 13 | Gmfb | 0.036 | Old |
| 0.827098 | 0.050367 | 0.213 | 0.193 | 1 | 13 | Dnajb2 | 0.02 | Old |
| 0.827169 | 0.048494 | 0.328 | 0.351 | 1 | 13 | Trove2 | -0.023 | Old |
| 0.827397 | 0.023234 | 0.492 | 0.491 | 1 | 13 | Ncbp3 | 0.001 | Old |
| 0.827398 | 0.113939 | 0.672 | 0.702 | 1 | 13 | Ccl12 | -0.03 | Old |
| 0.827398 | 0.109758 | 0.705 | 0.667 | 1 | 13 | Fkbp3 | 0.038 | Old |
| 0.827412 | 0.042219 | 0.754 | 0.772 | 1 | 13 | AI467606 | -0.018 | Old |
| 0.828009 | 0.022756 | 0.393 | 0.342 | 1 | 13 | Rbm15 | 0.051 | Old |
| 0.82805 | 0.020298 | 0.352 | 0.325 | 1 | 13 | Eipr1 | 0.027 | Old |
| 0.828548 | 0.033614 | 0.975 | 0.965 | 1 | 13 | Bola2 | 0.01 | Old |
| 0.828918 | 0.111462 | 0.525 | 0.5 | 1 | 13 | Ap4s1 | 0.025 | Old |
| 0.829031 | 0.087227 | 0.328 | 0.307 | 1 | 13 | Trmt2a | 0.021 | Old |
| 0.829208 | 0.017806 | 0.189 | 0.193 | 1 | 13 | Gm37233 | -0.004 | Old |
| 0.829208 | 0.140752 | 0.197 | 0.184 | 1 | 13 | Susd1 | 0.013 | Old |
| 0.829272 | 0.098945 | 0.041 | 0.035 | 1 | 13 | Unc13b | 0.006 | Old |
| 0.829272 | 0.077948 | 0.041 | 0.035 | 1 | 13 | Magi3 | 0.006 | Old |
| 0.829272 | 0.027712 | 0.041 | 0.035 | 1 | 13 | Arv1 | 0.006 | Old |
| 0.829272 | 0.014768 | 0.041 | 0.035 | 1 | 13 | Pex10 | 0.006 | Old |
| 0.829917 | 0.012246 | 0.525 | 0.482 | 1 | 13 | Ppie | 0.043 | Old |
| 0.830007 | 0.036739 | 0.951 | 0.904 | 1 | 13 | Fubp1 | 0.047 | Old |
| 0.830082 | 0.031796 | 0.689 | 0.658 | 1 | 13 | Exoc4 | 0.031 | Old |
| 0.830732 | 0.021131 | 0.803 | 0.772 | 1 | 13 | Ccnt1 | 0.031 | Old |
| 0.831121 | 0.041829 | 0.852 | 0.807 | 1 | 13 | Ube2e1 | 0.045 | Old |
| 0.831483 | 0.017161 | 0.918 | 0.921 | 1 | 13 | Cct4 | -0.003 | Old |
| 0.831521 | 0.033553 | 0.943 | 0.982 | 1 | 13 | Bach1 | -0.039 | Old |
| 0.831556 | 0.063403 | 0.451 | 0.439 | 1 | 13 | Tspan17 | 0.012 | Old |
| 0.831567 | 0.034338 | 0.426 | 0.43 | 1 | 13 | Afg3l2 | -0.004 | Old |
| 0.831803 | 0.035066 | 0.861 | 0.781 | 1 | 13 | Aftph | 0.08 | Old |
| 0.831938 | 0.116435 | 0.574 | 0.553 | 1 | 13 | Usp12 | 0.021 | Old |
| 0.832145 | 0.027935 | 0.484 | 0.456 | 1 | 13 | Itfg1 | 0.028 | Old |
| 0.83222 | 0.056887 | 0.926 | 0.904 | 1 | 13 | Pde4b | 0.022 | Old |
| 0.832323 | 0.05104 | 0.467 | 0.439 | 1 | 13 | Sacm1l | 0.028 | Old |
| 0.83245 | 0.438695 | 0.049 | 0.044 | 1 | 13 | Pigl | 0.005 | Old |
| 0.833483 | 0.040486 | 0.23 | 0.219 | 1 | 13 | Taf6 | 0.011 | Old |
| 0.833595 | 0.081308 | 0.402 | 0.368 | 1 | 13 | Nmi | 0.034 | Old |
| 0.834039 | 0.065943 | 0.18 | 0.167 | 1 | 13 | 2810006K2 | 0.013 | Old |
| 0.834782 | 0.031719 | 0.393 | 0.368 | 1 | 13 | Senp5 | 0.025 | Old |
| 0.834831 | 5.15E-05 | 0.541 | 0.579 | 1 | 13 | Pbx2 | -0.038 | Old |
| 0.834972 | 0.000966 | 0.32 | 0.289 | 1 | 13 | Phf6 | 0.031 | Old |
| 0.834978 | 0.258653 | 0.197 | 0.193 | 1 | 13 | Ipcef1 | 0.004 | Old |
| 0.835046 | 0.133992 | 0.877 | 0.851 | 1 | 13 | Tle3 | 0.026 | Old |

| | | | | | | | | |
|---|---|---|---|---|---|---|---|---|
| 0.835266 | 0.089705 | 0.82 | 0.763 | 1 | 13 | Oat | 0.057 | Old |
| 0.835382 | 0.045612 | 0.492 | 0.491 | 1 | 13 | Parg | 0.001 | Old |
| 0.835531 | 0.021177 | 0.762 | 0.746 | 1 | 13 | Szrd1 | 0.016 | Old |
| 0.835652 | 0.067948 | 0.238 | 0.219 | 1 | 13 | Epm2aip1 | 0.019 | Old |
| 0.835737 | 0.048128 | 0.107 | 0.096 | 1 | 13 | E2f6 | 0.011 | Old |
| 0.835813 | 0.168513 | 0.123 | 0.114 | 1 | 13 | Gemin5 | 0.009 | Old |
| 0.836528 | 0.205506 | 0.049 | 0.044 | 1 | 13 | Eif3j2 | 0.005 | Old |
| 0.836569 | 0.004829 | 0.779 | 0.772 | 1 | 13 | Arhgap26 | 0.007 | Old |
| 0.836748 | 0.22374 | 0.131 | 0.123 | 1 | 13 | Tango6 | 0.008 | Old |
| 0.837106 | 0.04667 | 0.541 | 0.491 | 1 | 13 | Brpf1 | 0.05 | Old |
| 0.837294 | 0.167065 | 0.115 | 0.105 | 1 | 13 | Fmnl2 | 0.01 | Old |
| 0.837337 | 0.409564 | 0.066 | 0.061 | 1 | 13 | Rpusd1 | 0.005 | Old |
| 0.837448 | 0.017516 | 0.623 | 0.614 | 1 | 13 | Vps13a | 0.009 | Old |
| 0.837484 | 0.030258 | 1 | 0.991 | 1 | 13 | Ehd4 | 0.009 | Old |
| 0.837897 | 0.064211 | 0.533 | 0.5 | 1 | 13 | Fxr2 | 0.033 | Old |
| 0.838044 | 0.086819 | 0.328 | 0.307 | 1 | 13 | Il15 | 0.021 | Old |
| 0.838154 | 0.127872 | 0.164 | 0.158 | 1 | 13 | Gatc | 0.006 | Old |
| 0.838168 | 0.057764 | 0.787 | 0.781 | 1 | 13 | Mrps15 | 0.006 | Old |
| 0.839394 | 0.079144 | 0.32 | 0.333 | 1 | 13 | Mitd1 | -0.013 | Old |
| 0.83946 | 0.125367 | 0.541 | 0.518 | 1 | 13 | Stx8 | 0.023 | Old |
| 0.839973 | 0.132385 | 0.721 | 0.693 | 1 | 13 | Fcna | 0.028 | Old |
| 0.840288 | 0.040935 | 0.893 | 0.842 | 1 | 13 | Mknk2 | 0.051 | Old |
| 0.840374 | 0.08145 | 0.5 | 0.465 | 1 | 13 | Clns1a | 0.035 | Old |
| 0.84061 | 0.296371 | 0.049 | 0.044 | 1 | 13 | Klhl25 | 0.005 | Old |
| 0.840844 | 0.13281 | 0.246 | 0.228 | 1 | 13 | Cep290 | 0.018 | Old |
| 0.841046 | 0.119229 | 0.508 | 0.5 | 1 | 13 | Naxd | 0.008 | Old |
| 0.84108 | 0.026047 | 0.443 | 0.404 | 1 | 13 | Scaf8 | 0.039 | Old |
| 0.841095 | 0.163446 | 0.09 | 0.096 | 1 | 13 | Lmbr1 | -0.006 | Old |
| 0.841168 | 0.047304 | 0.475 | 0.447 | 1 | 13 | Chchd5 | 0.028 | Old |
| 0.841266 | 0.304649 | 0.057 | 0.053 | 1 | 13 | Ift81 | 0.004 | Old |
| 0.841485 | 0.037394 | 0.451 | 0.404 | 1 | 13 | Tnfaip1 | 0.047 | Old |
| 0.84193 | 0.020275 | 0.934 | 0.921 | 1 | 13 | Vps35 | 0.013 | Old |
| 0.842582 | 0.020633 | 0.41 | 0.368 | 1 | 13 | Tmem19 | 0.042 | Old |
| 0.842792 | 0.042943 | 0.484 | 0.439 | 1 | 13 | Fam160b1 | 0.045 | Old |
| 0.843236 | 0.090482 | 0.59 | 0.553 | 1 | 13 | Spg7 | 0.037 | Old |
| 0.843288 | 0.045969 | 0.713 | 0.658 | 1 | 13 | Elf2 | 0.055 | Old |
| 0.843538 | 0.123725 | 0.705 | 0.772 | 1 | 13 | Nbeal1 | -0.067 | Old |
| 0.843761 | 0.024527 | 0.238 | 0.219 | 1 | 13 | Galns | 0.019 | Old |
| 0.844167 | 0.03021 | 0.779 | 0.789 | 1 | 13 | Senp6 | -0.01 | Old |
| 0.844674 | 0.090737 | 0.73 | 0.702 | 1 | 13 | Polr2m | 0.028 | Old |
| 0.845042 | 0.163138 | 0.057 | 0.053 | 1 | 13 | Uhrf1bp1 | 0.004 | Old |
| 0.845112 | 0.093058 | 0.451 | 0.43 | 1 | 13 | Rpe | 0.021 | Old |
| 0.846246 | 0.008548 | 0.352 | 0.36 | 1 | 13 | Thap2 | -0.008 | Old |
| 0.846439 | 0.013622 | 0.992 | 1 | 1 | 13 | Fcgr3 | -0.008 | Old |
| 0.846484 | 0.023148 | 0.77 | 0.693 | 1 | 13 | Slc25a4 | 0.077 | Old |

| | | | | | | | | |
|---|---|---|---|---|---|---|---|---|
| 0.846569 | 0.314126 | 0.352 | 0.351 | 1 | 13 | Gdf15 | 0.001 | Old |
| 0.847477 | 0.014811 | 0.82 | 0.816 | 1 | 13 | Lbr | 0.004 | Old |
| 0.847735 | 0.019191 | 0.402 | 0.395 | 1 | 13 | Zfp280d | 0.007 | Old |
| 0.847764 | 0.12034 | 0.361 | 0.333 | 1 | 13 | Sos1 | 0.028 | Old |
| 0.847914 | 0.021933 | 0.951 | 0.921 | 1 | 13 | Swi5 | 0.03 | Old |
| 0.84792 | 0.060291 | 0.959 | 0.93 | 1 | 13 | Cerk | 0.029 | Old |
| 0.847965 | 0.106954 | 0.795 | 0.781 | 1 | 13 | Tsc22d1 | 0.014 | Old |
| 0.848081 | 0.02713 | 0.68 | 0.649 | 1 | 13 | Rarg | 0.031 | Old |
| 0.848279 | 0.017092 | 0.115 | 0.105 | 1 | 13 | Mblac2 | 0.01 | Old |
| 0.848446 | 0.046887 | 0.516 | 0.526 | 1 | 13 | Ak3 | -0.01 | Old |
| 0.848787 | 0.242266 | 0.049 | 0.044 | 1 | 13 | Arl4d | 0.005 | Old |
| 0.848787 | 0.105127 | 0.049 | 0.044 | 1 | 13 | Zfp975 | 0.005 | Old |
| 0.849534 | 0.275803 | 0.5 | 0.526 | 1 | 13 | Cd80 | -0.026 | Old |
| 0.849625 | 0.100195 | 0.811 | 0.798 | 1 | 13 | Cpsf6 | 0.013 | Old |
| 0.849769 | 0.039946 | 0.443 | 0.412 | 1 | 13 | Ccdc90b | 0.031 | Old |
| 0.849831 | 0.157661 | 0.246 | 0.237 | 1 | 13 | Ccdc167 | 0.009 | Old |
| 0.85074 | 0.218598 | 0.189 | 0.184 | 1 | 13 | Dip2a | 0.005 | Old |
| 0.850781 | 0.048968 | 0.615 | 0.553 | 1 | 13 | Raf1 | 0.062 | Old |
| 0.850829 | 0.085818 | 0.615 | 0.596 | 1 | 13 | Plekha1 | 0.019 | Old |
| 0.850921 | 0.072308 | 0.975 | 0.965 | 1 | 13 | Stat3 | 0.01 | Old |
| 0.850923 | 0.029203 | 0.984 | 0.982 | 1 | 13 | Srsf3 | 0.002 | Old |
| 0.851162 | 0.012337 | 0.082 | 0.088 | 1 | 13 | Gba2 | -0.006 | Old |
| 0.851542 | 0.057223 | 0.664 | 0.702 | 1 | 13 | Tsc22d2 | -0.038 | Old |
| 0.851721 | 0.02699 | 0.82 | 0.754 | 1 | 13 | Itsn1 | 0.066 | Old |
| 0.851765 | 0.007094 | 0.123 | 0.114 | 1 | 13 | 1700096K1 | 0.009 | Old |
| 0.851853 | 0.04209 | 0.541 | 0.509 | 1 | 13 | Lrrk1 | 0.032 | Old |
| 0.851944 | 0.044752 | 0.336 | 0.316 | 1 | 13 | Pcsk7 | 0.02 | Old |
| 0.852297 | 0.058386 | 0.885 | 0.877 | 1 | 13 | Rnf19b | 0.008 | Old |
| 0.852604 | 0.220169 | 0.057 | 0.053 | 1 | 13 | Klhl33 | 0.004 | Old |
| 0.852882 | 0.069046 | 0.049 | 0.044 | 1 | 13 | Aurka | 0.005 | Old |
| 0.852915 | 0.04655 | 0.713 | 0.711 | 1 | 13 | Chd1 | 0.002 | Old |
| 0.853582 | 0.089867 | 0.361 | 0.342 | 1 | 13 | Lxn | 0.019 | Old |
| 0.853703 | 0.057049 | 0.344 | 0.333 | 1 | 13 | Zdhhc18 | 0.011 | Old |
| 0.853902 | 0.059226 | 0.238 | 0.228 | 1 | 13 | Dnajc4 | 0.01 | Old |
| 0.853906 | 0.019211 | 0.967 | 0.93 | 1 | 13 | Cflar | 0.037 | Old |
| 0.854498 | 0.551978 | 0.164 | 0.158 | 1 | 13 | Chpf | 0.006 | Old |
| 0.854888 | 0.039478 | 0.426 | 0.395 | 1 | 13 | Pspc1 | 0.031 | Old |
| 0.855899 | 0.074533 | 0.525 | 0.5 | 1 | 13 | Ppp1r7 | 0.025 | Old |
| 0.856107 | 0.049639 | 0.574 | 0.518 | 1 | 13 | Dna2 | 0.056 | Old |
| 0.856391 | 0.381569 | 0.057 | 0.053 | 1 | 13 | Gm14325 | 0.004 | Old |
| 0.856435 | 0.330586 | 0.074 | 0.07 | 1 | 13 | Focad | 0.004 | Old |
| 0.856462 | 0.094295 | 0.721 | 0.675 | 1 | 13 | Ube2a | 0.046 | Old |
| 0.856679 | 0.017022 | 0.861 | 0.842 | 1 | 13 | Fam168b | 0.019 | Old |
| 0.8569 | 0.05601 | 0.811 | 0.746 | 1 | 13 | Cnot1 | 0.065 | Old |
| 0.856981 | 0.088521 | 0.049 | 0.044 | 1 | 13 | Paqr3 | 0.005 | Old |

| | | | | | | | | |
|---|---|---|---|---|---|---|---|---|
| 0.857096 | 0.055367 | 0.123 | 0.114 | 1 | 13 | Rnf123 | 0.009 | Old |
| 0.857411 | 0.07568 | 0.746 | 0.772 | 1 | 13 | Tmx1 | -0.026 | Old |
| 0.858304 | 0.030845 | 0.918 | 0.851 | 1 | 13 | Uba1 | 0.067 | Old |
| 0.858392 | 0.171689 | 0.287 | 0.272 | 1 | 13 | Siglec1 | 0.015 | Old |
| 0.85843 | 0.124428 | 0.115 | 0.123 | 1 | 13 | Wee1 | -0.008 | Old |
| 0.858544 | 0.537814 | 0.066 | 0.061 | 1 | 13 | Gm26756 | 0.005 | Old |
| 0.859338 | 0.201431 | 0.295 | 0.263 | 1 | 13 | Gbp5 | 0.032 | Old |
| 0.859902 | 0.042918 | 0.984 | 0.93 | 1 | 13 | Khdrbs1 | 0.054 | Old |
| 0.860913 | 0.015995 | 0.451 | 0.412 | 1 | 13 | Gm15987 | 0.039 | Old |
| 0.861084 | 0.202735 | 0.049 | 0.044 | 1 | 13 | Serpinb2 | 0.005 | Old |
| 0.861084 | 0.090555 | 0.049 | 0.044 | 1 | 13 | Zfp963 | 0.005 | Old |
| 0.861605 | 0.026841 | 0.189 | 0.175 | 1 | 13 | Tuft1 | 0.014 | Old |
| 0.861717 | 0.023842 | 0.393 | 0.36 | 1 | 13 | Hagh | 0.033 | Old |
| 0.86211 | 0.104384 | 0.557 | 0.596 | 1 | 13 | Afg3l1 | -0.039 | Old |
| 0.862361 | 0.285474 | 0.164 | 0.149 | 1 | 13 | Flrt2 | 0.015 | Old |
| 0.862367 | 0.439989 | 0.762 | 0.842 | 1 | 13 | Fabp4 | -0.08 | Old |
| 0.862434 | 0.174862 | 0.123 | 0.114 | 1 | 13 | Rtn4rl1 | 0.009 | Old |
| 0.862876 | 0.045976 | 0.918 | 0.93 | 1 | 13 | Daglb | -0.012 | Old |
| 0.863456 | 0.146234 | 0.516 | 0.465 | 1 | 13 | Mboat7 | 0.051 | Old |
| 0.863475 | 0.01163 | 0.352 | 0.325 | 1 | 13 | Wdr5 | 0.027 | Old |
| 0.863623 | 0.02041 | 0.344 | 0.351 | 1 | 13 | Ap5m1 | -0.007 | Old |
| 0.863706 | 0.047406 | 0.32 | 0.307 | 1 | 13 | Sirt1 | 0.013 | Old |
| 0.864402 | 0.036684 | 0.975 | 0.947 | 1 | 13 | Hnrnpc | 0.028 | Old |
| 0.864569 | 0.062064 | 0.549 | 0.535 | 1 | 13 | Ctbp2 | 0.014 | Old |
| 0.864812 | 0.180853 | 0.115 | 0.105 | 1 | 13 | Ppp2r3a | 0.01 | Old |
| 0.864919 | 0.034973 | 0.246 | 0.228 | 1 | 13 | Hic2 | 0.018 | Old |
| 0.864999 | 0.169755 | 0.516 | 0.491 | 1 | 13 | Mpnd | 0.025 | Old |
| 0.86519 | 0.033447 | 0.049 | 0.044 | 1 | 13 | Dgat2 | 0.005 | Old |
| 0.86519 | 0.031449 | 0.049 | 0.044 | 1 | 13 | Dnah2 | 0.005 | Old |
| 0.86519 | 0.016722 | 0.049 | 0.044 | 1 | 13 | Zfp324 | 0.005 | Old |
| 0.865636 | 0.21552 | 0.066 | 0.061 | 1 | 13 | Wdpcp | 0.005 | Old |
| 0.865882 | 0.124893 | 0.344 | 0.36 | 1 | 13 | Pja1 | -0.016 | Old |
| 0.866166 | 0.084536 | 0.5 | 0.465 | 1 | 13 | Arid2 | 0.035 | Old |
| 0.866451 | 0.001607 | 0.27 | 0.272 | 1 | 13 | Rgs3 | -0.002 | Old |
| 0.866473 | 0.033535 | 0.123 | 0.132 | 1 | 13 | BC002059 | -0.009 | Old |
| 0.866944 | 0.023816 | 0.713 | 0.667 | 1 | 13 | Trp53 | 0.046 | Old |
| 0.86705 | 0.17033 | 0.426 | 0.447 | 1 | 13 | Dolpp1 | -0.021 | Old |
| 0.867524 | 0.042267 | 0.615 | 0.684 | 1 | 13 | Capn1 | -0.069 | Old |
| 0.867743 | 0.021479 | 0.377 | 0.36 | 1 | 13 | Trappc9 | 0.017 | Old |
| 0.867769 | 0.129624 | 0.057 | 0.053 | 1 | 13 | Tubd1 | 0.004 | Old |
| 0.867769 | 0.03187 | 0.057 | 0.053 | 1 | 13 | Clspn | 0.004 | Old |
| 0.8693 | 0.061259 | 0.049 | 0.044 | 1 | 13 | Nek8 | 0.005 | Old |
| 0.8693 | 0.022044 | 0.049 | 0.044 | 1 | 13 | Prag1 | 0.005 | Old |
| 0.8693 | 0.006799 | 0.049 | 0.044 | 1 | 13 | Mocs3 | 0.005 | Old |
| 0.869328 | 0.200187 | 0.279 | 0.263 | 1 | 13 | Clec4d | 0.016 | Old |

| | | | | | | | | |
|---|---|---|---|---|---|---|---|---|
| 0.869573 | 0.114224 | 0.508 | 0.491 | 1 | 13 | Dlg1 | 0.017 | Old |
| 0.870376 | 0.009736 | 0.533 | 0.465 | 1 | 13 | 2810004N2 | 0.068 | Old |
| 0.871568 | 0.716689 | 0.057 | 0.053 | 1 | 13 | Dpt | 0.004 | Old |
| 0.871568 | 0.078014 | 0.057 | 0.053 | 1 | 13 | Gm26802 | 0.004 | Old |
| 0.871568 | 0.02949 | 0.057 | 0.053 | 1 | 13 | Pde4d | 0.004 | Old |
| 0.872195 | 0.045022 | 0.262 | 0.246 | 1 | 13 | 1110059E2 | 0.016 | Old |
| 0.872633 | 0.07785 | 0.467 | 0.421 | 1 | 13 | Polr3k | 0.046 | Old |
| 0.87274 | 0.224401 | 0.066 | 0.061 | 1 | 13 | Nme2 | 0.005 | Old |
| 0.872842 | 0.170919 | 0.205 | 0.193 | 1 | 13 | Entpd6 | 0.012 | Old |
| 0.87295 | 0.043983 | 0.254 | 0.263 | 1 | 13 | B3galnt1 | -0.009 | Old |
| 0.872953 | 0.037435 | 0.369 | 0.368 | 1 | 13 | Zfyve27 | 0.001 | Old |
| 0.873253 | 0.246531 | 0.164 | 0.158 | 1 | 13 | Tpk1 | 0.006 | Old |
| 0.873414 | 0.059826 | 0.049 | 0.044 | 1 | 13 | Ubtd2 | 0.005 | Old |
| 0.873872 | 0.03315 | 0.303 | 0.298 | 1 | 13 | Metap1 | 0.005 | Old |
| 0.874111 | 0.053756 | 0.402 | 0.377 | 1 | 13 | Arfip1 | 0.025 | Old |
| 0.874734 | 0.103367 | 0.885 | 0.825 | 1 | 13 | Tnpo1 | 0.06 | Old |
| 0.874737 | 0.053321 | 0.172 | 0.167 | 1 | 13 | Dmap1 | 0.005 | Old |
| 0.874818 | 0.032729 | 0.074 | 0.079 | 1 | 13 | Klk8 | -0.005 | Old |
| 0.874863 | 0.054198 | 0.721 | 0.693 | 1 | 13 | Pqlc1 | 0.028 | Old |
| 0.874917 | 0.030989 | 0.959 | 0.965 | 1 | 13 | Oxct1 | -0.006 | Old |
| 0.87492 | 0.002478 | 0.984 | 0.991 | 1 | 13 | Snrpb | -0.007 | Old |
| 0.874966 | 0.040455 | 0.41 | 0.368 | 1 | 13 | Tmem104 | 0.042 | Old |
| 0.874988 | 0.014834 | 0.246 | 0.228 | 1 | 13 | Nlrc5 | 0.018 | Old |
| 0.874998 | 0.055736 | 0.205 | 0.193 | 1 | 13 | Wdr55 | 0.012 | Old |
| 0.875007 | 0.047458 | 0.795 | 0.763 | 1 | 13 | Cenpb | 0.032 | Old |
| 0.875158 | 0.043569 | 0.139 | 0.132 | 1 | 13 | Paqr4 | 0.007 | Old |
| 0.87537 | 0.09967 | 0.057 | 0.053 | 1 | 13 | Zfp799 | 0.004 | Old |
| 0.87537 | 0.04938 | 0.057 | 0.053 | 1 | 13 | Mrm1 | 0.004 | Old |
| 0.876126 | 0.144338 | 0.262 | 0.246 | 1 | 13 | Cd79b | 0.016 | Old |
| 0.876273 | 0.339285 | 0.082 | 0.079 | 1 | 13 | Tmem86b | 0.003 | Old |
| 0.876527 | 0.223673 | 0.074 | 0.07 | 1 | 13 | Mrm2 | 0.004 | Old |
| 0.876921 | 0.00451 | 0.844 | 0.833 | 1 | 13 | Rbbp6 | 0.011 | Old |
| 0.877048 | 0.002065 | 0.484 | 0.439 | 1 | 13 | Lmbrd2 | 0.045 | Old |
| 0.877057 | 0.01124 | 0.533 | 0.509 | 1 | 13 | Dopey1 | 0.024 | Old |
| 0.877237 | 0.013745 | 0.811 | 0.737 | 1 | 13 | Uqcrc2 | 0.074 | Old |
| 0.877383 | 0.103972 | 0.779 | 0.807 | 1 | 13 | Lmbrd1 | -0.028 | Old |
| 0.877561 | 0.0106 | 0.844 | 0.789 | 1 | 13 | Zfp710 | 0.055 | Old |
| 0.877575 | 0.11116 | 0.713 | 0.763 | 1 | 13 | Ipo7 | -0.05 | Old |
| 0.878055 | 0.162964 | 0.443 | 0.412 | 1 | 13 | Rogdi | 0.031 | Old |
| 0.878055 | 0.051539 | 0.434 | 0.421 | 1 | 13 | Gcn1l1 | 0.013 | Old |
| 0.878709 | 0.303482 | 0.09 | 0.088 | 1 | 13 | Slc26a6 | 0.002 | Old |
| 0.879127 | 0.203419 | 0.213 | 0.202 | 1 | 13 | Yod1 | 0.011 | Old |
| 0.879175 | 0.297066 | 0.057 | 0.053 | 1 | 13 | Adra2a | 0.004 | Old |
| 0.879175 | 0.084017 | 0.057 | 0.053 | 1 | 13 | Htr1b | 0.004 | Old |
| 0.879175 | 0.040648 | 0.057 | 0.053 | 1 | 13 | Snx32 | 0.004 | Old |

| | | | | | | | | |
|---|---|---|---|---|---|---|---|---|
| 0.879217 | 0.016292 | 0.525 | 0.474 | 1 | 13 | Timm10 | 0.051 | Old |
| 0.879238 | 0.09478 | 0.869 | 0.833 | 1 | 13 | Sypl | 0.036 | Old |
| 0.879351 | 0.058819 | 0.615 | 0.588 | 1 | 13 | Spen | 0.027 | Old |
| 0.879405 | 0.039128 | 0.533 | 0.544 | 1 | 13 | Atxn7l3 | -0.011 | Old |
| 0.879433 | 0.089399 | 1 | 1 | 1 | 13 | Cybb | 0 | Old |
| 0.879459 | 0.073051 | 0.418 | 0.386 | 1 | 13 | Rab28 | 0.032 | Old |
| 0.880181 | 0.015695 | 0.959 | 0.956 | 1 | 13 | Laptm4a | 0.003 | Old |
| 0.881058 | 0.018508 | 0.779 | 0.781 | 1 | 13 | Cebpz | -0.002 | Old |
| 0.881202 | 0.044693 | 0.156 | 0.149 | 1 | 13 | Znhit3 | 0.007 | Old |
| 0.881845 | 0.05007 | 0.23 | 0.211 | 1 | 13 | Dpysl3 | 0.019 | Old |
| 0.882279 | 0.078273 | 0.623 | 0.623 | 1 | 13 | Atxn7l1 | 0 | Old |
| 0.882327 | 0.017535 | 0.795 | 0.711 | 1 | 13 | Sh3pxd2b | 0.084 | Old |
| 0.882505 | 0.055295 | 0.115 | 0.123 | 1 | 13 | Tut1 | -0.008 | Old |
| 0.882983 | 0.08988 | 0.057 | 0.053 | 1 | 13 | Adat1 | 0.004 | Old |
| 0.882983 | 0.033501 | 0.057 | 0.053 | 1 | 13 | Dagla | 0.004 | Old |
| 0.883243 | 0.177249 | 0.074 | 0.07 | 1 | 13 | Gm28707 | 0.004 | Old |
| 0.883413 | 0.305403 | 0.066 | 0.061 | 1 | 13 | Thnsl1 | 0.005 | Old |
| 0.883559 | 0.001771 | 0.574 | 0.553 | 1 | 13 | Sf3a1 | 0.021 | Old |
| 0.883933 | 0.364314 | 0.18 | 0.184 | 1 | 13 | Spic | -0.004 | Old |
| 0.884246 | 0.04888 | 0.68 | 0.596 | 1 | 13 | Xpo1 | 0.084 | Old |
| 0.88433 | 0.030207 | 0.795 | 0.833 | 1 | 13 | Txnrd1 | -0.038 | Old |
| 0.884455 | 0.178539 | 0.295 | 0.298 | 1 | 13 | Sik3 | -0.003 | Old |
| 0.88498 | 0.020897 | 0.27 | 0.237 | 1 | 13 | Tubgcp3 | 0.033 | Old |
| 0.885324 | 0.024952 | 0.598 | 0.588 | 1 | 13 | Slc50a1 | 0.01 | Old |
| 0.885355 | 0.058036 | 0.082 | 0.088 | 1 | 13 | Arhgap27 | -0.006 | Old |
| 0.885447 | 0.005134 | 0.951 | 0.939 | 1 | 13 | Nckap1l | 0.012 | Old |
| 0.885456 | 0.025678 | 0.975 | 1 | 1 | 13 | 5031439G( | -0.025 | Old |
| 0.88615 | 0.035086 | 0.893 | 0.904 | 1 | 13 | Kmt2e | -0.011 | Old |
| 0.886353 | 0.01466 | 0.623 | 0.579 | 1 | 13 | Carhsp1 | 0.044 | Old |
| 0.886639 | 0.010805 | 0.27 | 0.246 | 1 | 13 | 1110034G2 | 0.024 | Old |
| 0.887061 | 0.131894 | 0.475 | 0.518 | 1 | 13 | Med21 | -0.043 | Old |
| 0.887717 | 0.043283 | 1 | 0.991 | 1 | 13 | Ier5 | 0.009 | Old |
| 0.887808 | 0.096717 | 0.139 | 0.132 | 1 | 13 | Tatdn3 | 0.007 | Old |
| 0.887838 | 0.007088 | 0.762 | 0.789 | 1 | 13 | Cnppd1 | -0.027 | Old |
| 0.888028 | 0.023371 | 0.598 | 0.614 | 1 | 13 | Gm8186 | -0.016 | Old |
| 0.888321 | 0.043596 | 0.189 | 0.184 | 1 | 13 | Tmem87a | 0.005 | Old |
| 0.88847 | 0.041019 | 1 | 1 | 1 | 13 | Ftl1 | 0 | Old |
| 0.888477 | 0.010547 | 0.402 | 0.395 | 1 | 13 | Ddx56 | 0.007 | Old |
| 0.888686 | 0.077232 | 0.09 | 0.096 | 1 | 13 | Rfx3 | -0.006 | Old |
| 0.889306 | 0.130639 | 0.32 | 0.307 | 1 | 13 | Parp12 | 0.013 | Old |
| 0.889445 | 0.105574 | 0.5 | 0.456 | 1 | 13 | Igsf8 | 0.044 | Old |
| 0.889978 | 0.013309 | 0.992 | 1 | 1 | 13 | Cox5b | -0.008 | Old |
| 0.890541 | 0.100449 | 0.066 | 0.061 | 1 | 13 | 1-Sep | 0.005 | Old |
| 0.890606 | 0.000621 | 0.057 | 0.053 | 1 | 13 | Tmem177 | 0.004 | Old |
| 0.890617 | 0.339233 | 0.107 | 0.105 | 1 | 13 | Gm17745 | 0.002 | Old |

| | | | | | | | | |
|---|---|---|---|---|---|---|---|---|
| 0.890865 | 0.139376 | 0.172 | 0.167 | 1 | 13 | Gabbr1 | 0.005 | Old |
| 0.890895 | 0.029315 | 0.484 | 0.5 | 1 | 13 | Armcx3 | -0.016 | Old |
| 0.891429 | 0.109909 | 0.566 | 0.561 | 1 | 13 | Relb | 0.005 | Old |
| 0.891641 | 0.069749 | 0.705 | 0.711 | 1 | 13 | Phc3 | -0.006 | Old |
| 0.891766 | 0.027039 | 0.393 | 0.377 | 1 | 13 | Fam208b | 0.016 | Old |
| 0.891923 | 0.026908 | 0.295 | 0.298 | 1 | 13 | Dnajb12 | -0.003 | Old |
| 0.892107 | 0.046184 | 0.893 | 0.833 | 1 | 13 | Mef2d | 0.06 | Old |
| 0.89224 | 0.00851 | 0.975 | 0.991 | 1 | 13 | Ifrd1 | -0.016 | Old |
| 0.892452 | 0.113825 | 0.189 | 0.202 | 1 | 13 | Pam | -0.013 | Old |
| 0.892672 | 0.0231 | 0.82 | 0.816 | 1 | 13 | Ybx3 | 0.004 | Old |
| 0.892874 | 0.053259 | 0.869 | 0.868 | 1 | 13 | Adcy7 | 0.001 | Old |
| 0.892912 | 0.037126 | 0.344 | 0.333 | 1 | 13 | Ryk | 0.011 | Old |
| 0.89477 | 0.097794 | 0.426 | 0.43 | 1 | 13 | Taf3 | -0.004 | Old |
| 0.894835 | 0.007614 | 0.254 | 0.254 | 1 | 13 | Hnmt | 0 | Old |
| 0.895341 | 0.05695 | 0.434 | 0.386 | 1 | 13 | Tango2 | 0.048 | Old |
| 0.895632 | 0.00074 | 0.672 | 0.667 | 1 | 13 | Mtmr6 | 0.005 | Old |
| 0.895958 | 0.025746 | 0.492 | 0.465 | 1 | 13 | Cops8 | 0.027 | Old |
| 0.896012 | 0.052202 | 0.951 | 0.991 | 1 | 13 | Vmp1 | -0.04 | Old |
| 0.896013 | 0.007035 | 1 | 1 | 1 | 13 | Atp5e | 0 | Old |
| 0.896178 | 0.101666 | 0.057 | 0.061 | 1 | 13 | Cenpf | -0.004 | Old |
| 0.896266 | 0.347753 | 0.107 | 0.105 | 1 | 13 | Hdac6 | 0.002 | Old |
| 0.897213 | 0.045967 | 0.828 | 0.807 | 1 | 13 | Klf10 | 0.021 | Old |
| 0.897376 | 0.096436 | 0.189 | 0.175 | 1 | 13 | B230118H( | 0.014 | Old |
| 0.897487 | 0.070679 | 0.926 | 0.895 | 1 | 13 | Sin3b | 0.031 | Old |
| 0.897503 | 0.02242 | 0.926 | 0.93 | 1 | 13 | Arid1a | -0.004 | Old |
| 0.897561 | 0.01788 | 0.328 | 0.298 | 1 | 13 | AL590144.: | 0.03 | Old |
| 0.897678 | 0.110937 | 0.066 | 0.061 | 1 | 13 | Ube2t | 0.005 | Old |
| 0.897678 | 0.082003 | 0.066 | 0.061 | 1 | 13 | Lrrc32 | 0.005 | Old |
| 0.897678 | 0.040304 | 0.066 | 0.061 | 1 | 13 | Rhobtb2 | 0.005 | Old |
| 0.898768 | 0.05665 | 0.861 | 0.789 | 1 | 13 | Tank | 0.072 | Old |
| 0.898992 | 0.015769 | 0.5 | 0.482 | 1 | 13 | Zeb2os | 0.018 | Old |
| 0.899283 | 0.032217 | 0.705 | 0.711 | 1 | 13 | Polr2i | -0.006 | Old |
| 0.89931 | 0.059288 | 0.574 | 0.553 | 1 | 13 | Ilf3 | 0.021 | Old |
| 0.900487 | 0.027098 | 0.156 | 0.149 | 1 | 13 | Haus4 | 0.007 | Old |
| 0.900848 | 0.042364 | 0.484 | 0.474 | 1 | 13 | Fam20b | 0.01 | Old |
| 0.901145 | 0.018673 | 0.254 | 0.246 | 1 | 13 | Klhl21 | 0.008 | Old |
| 0.901197 | 0.067417 | 0.861 | 0.886 | 1 | 13 | Bhlhe40 | -0.025 | Old |
| 0.901283 | 0.006266 | 0.762 | 0.737 | 1 | 13 | Becn1 | 0.025 | Old |
| 0.901497 | 0.008542 | 0.787 | 0.763 | 1 | 13 | Poldip3 | 0.024 | Old |
| 0.901835 | 0.242809 | 0.082 | 0.079 | 1 | 13 | Nsun5 | 0.003 | Old |
| 0.902019 | 0.14241 | 0.91 | 0.912 | 1 | 13 | Id2 | -0.002 | Old |
| 0.90264 | 0.027548 | 0.426 | 0.412 | 1 | 13 | Zfp384 | 0.014 | Old |
| 0.902891 | 0.011695 | 0.262 | 0.263 | 1 | 13 | Slc25a44 | -0.001 | Old |
| 0.903547 | 0.017141 | 0.934 | 0.921 | 1 | 13 | Eloc | 0.013 | Old |
| 0.903565 | 0.007887 | 1 | 1 | 1 | 13 | Ddx5 | 0 | Old |

| | | | | | | | | |
|---|---|---|---|---|---|---|---|---|
| 0.903606 | 0.028463 | 0.615 | 0.596 | 1 | 13 | Ptpn22 | 0.019 | Old |
| 0.904551 | 0.121429 | 0.352 | 0.325 | 1 | 13 | Arntl | 0.027 | Old |
| 0.904823 | 1.035312 | 0.066 | 0.061 | 1 | 13 | Slc7a2 | 0.005 | Old |
| 0.904823 | 0.123406 | 0.066 | 0.061 | 1 | 13 | Gm29291 | 0.005 | Old |
| 0.904901 | 0.012712 | 0.164 | 0.167 | 1 | 13 | Slc12a2 | -0.003 | Old |
| 0.905068 | 0.023206 | 0.926 | 0.965 | 1 | 13 | Bag1 | -0.039 | Old |
| 0.905076 | 0.0042 | 0.992 | 0.982 | 1 | 13 | H2afy | 0.01 | Old |
| 0.905506 | 0.076934 | 0.705 | 0.728 | 1 | 13 | Trappc6b | -0.023 | Old |
| 0.905902 | 0.193233 | 0.311 | 0.298 | 1 | 13 | Mir17hg | 0.013 | Old |
| 0.906257 | 0.036489 | 0.443 | 0.404 | 1 | 13 | Ero1lb | 0.039 | Old |
| 0.906596 | 0.092346 | 0.68 | 0.684 | 1 | 13 | Plekhm3 | -0.004 | Old |
| 0.906811 | 0.12357 | 0.074 | 0.07 | 1 | 13 | Tfb1m | 0.004 | Old |
| 0.907859 | 0.054806 | 0.836 | 0.816 | 1 | 13 | Stk38 | 0.02 | Old |
| 0.908095 | 0.067084 | 0.959 | 0.947 | 1 | 13 | Rab11b | 0.012 | Old |
| 0.908218 | 0.571564 | 0.262 | 0.254 | 1 | 13 | Slc7a11 | 0.008 | Old |
| 0.908599 | 0.180974 | 0.041 | 0.044 | 1 | 13 | Poli | -0.003 | Old |
| 0.908599 | 0.095415 | 0.041 | 0.044 | 1 | 13 | Atxn7l2 | -0.003 | Old |
| 0.909015 | 0.068355 | 0.566 | 0.579 | 1 | 13 | Sart1 | -0.013 | Old |
| 0.909254 | 0.137125 | 0.361 | 0.368 | 1 | 13 | Naip5 | -0.007 | Old |
| 0.90961 | 0.112949 | 0.705 | 0.649 | 1 | 13 | Gm6377 | 0.056 | Old |
| 0.910185 | 0.663632 | 0.074 | 0.07 | 1 | 13 | Vcan | 0.004 | Old |
| 0.910185 | 0.143615 | 0.074 | 0.07 | 1 | 13 | Lag3 | 0.004 | Old |
| 0.910211 | 0.105778 | 0.049 | 0.053 | 1 | 13 | Sardh | -0.004 | Old |
| 0.910245 | 0.009542 | 0.197 | 0.184 | 1 | 13 | Tppp | 0.013 | Old |
| 0.911102 | 0.025824 | 0.926 | 0.912 | 1 | 13 | Ifi207 | 0.014 | Old |
| 0.911121 | 0.011125 | 0.623 | 0.561 | 1 | 13 | Pik3c2a | 0.062 | Old |
| 0.911449 | 0.310678 | 0.082 | 0.079 | 1 | 13 | B3gnt5 | 0.003 | Old |
| 0.91175 | 0.0132 | 0.877 | 0.833 | 1 | 13 | Rwdd1 | 0.044 | Old |
| 0.911952 | 0.00885 | 0.115 | 0.105 | 1 | 13 | Rubcnl | 0.01 | Old |
| 0.911976 | 0.019198 | 0.066 | 0.061 | 1 | 13 | Prpf40b | 0.005 | Old |
| 0.912409 | 0.020977 | 0.238 | 0.237 | 1 | 13 | Tmem70 | 0.001 | Old |
| 0.912453 | 0.110516 | 0.385 | 0.386 | 1 | 13 | Ifih1 | -0.001 | Old |
| 0.912934 | 0.064799 | 0.041 | 0.044 | 1 | 13 | Tmem267 | -0.003 | Old |
| 0.913076 | 0.099155 | 0.254 | 0.246 | 1 | 13 | Atg13 | 0.008 | Old |
| 0.913344 | 0.31372 | 0.107 | 0.114 | 1 | 13 | Slc7a13 | -0.007 | Old |
| 0.913845 | 0.013497 | 0.541 | 0.5 | 1 | 13 | D10Jhu81e | 0.041 | Old |
| 0.914013 | 0.023025 | 0.631 | 0.605 | 1 | 13 | Grk6 | 0.026 | Old |
| 0.914098 | 0.108575 | 0.402 | 0.395 | 1 | 13 | Gse1 | 0.007 | Old |
| 0.914152 | 0.009178 | 1 | 1 | 1 | 13 | Atp1a1 | 0 | Old |
| 0.914186 | 0.120776 | 0.049 | 0.053 | 1 | 13 | Zfp764 | -0.004 | Old |
| 0.914186 | 0.012133 | 0.049 | 0.053 | 1 | 13 | Sox12 | -0.004 | Old |
| 0.914498 | 0.141582 | 0.689 | 0.711 | 1 | 13 | Limd2 | -0.022 | Old |
| 0.914576 | 0.035515 | 0.066 | 0.07 | 1 | 13 | L2hgdh | -0.004 | Old |
| 0.914776 | 0.10795 | 0.385 | 0.377 | 1 | 13 | Ttc33 | 0.008 | Old |
| 0.914855 | 0.034027 | 0.746 | 0.719 | 1 | 13 | Pcbd2 | 0.027 | Old |

| | | | | | | | | |
|---|---|---|---|---|---|---|---|---|
| 0.915212 | 0.073416 | 0.574 | 0.561 | 1 | 13 | Bag3 | 0.013 | Old |
| 0.915677 | 0.040484 | 0.41 | 0.412 | 1 | 13 | Slc35e1 | -0.002 | Old |
| 0.915697 | 0.122679 | 0.164 | 0.158 | 1 | 13 | Csrp2 | 0.006 | Old |
| 0.916892 | 0.089257 | 0.803 | 0.816 | 1 | 13 | Sptbn1 | -0.013 | Old |
| 0.916901 | 0.037152 | 0.287 | 0.281 | 1 | 13 | Fbxo7 | 0.006 | Old |
| 0.916938 | 0.088828 | 0.074 | 0.07 | 1 | 13 | Bbs7 | 0.004 | Old |
| 0.917013 | 0.140563 | 0.336 | 0.316 | 1 | 13 | Rad54l2 | 0.02 | Old |
| 0.917164 | 0.03254 | 0.967 | 0.886 | 1 | 13 | Gnaq | 0.081 | Old |
| 0.917271 | 0.029783 | 0.041 | 0.044 | 1 | 13 | Fam149a | -0.003 | Old |
| 0.917271 | 0.007951 | 0.041 | 0.044 | 1 | 13 | Pcdhga12 | -0.003 | Old |
| 0.917468 | 0.064994 | 0.689 | 0.693 | 1 | 13 | Nhsl2 | -0.004 | Old |
| 0.918064 | 0.063883 | 0.164 | 0.158 | 1 | 13 | Mtfmt | 0.006 | Old |
| 0.918163 | 0.043744 | 0.049 | 0.053 | 1 | 13 | Gnb4 | -0.004 | Old |
| 0.918474 | 0.013737 | 0.598 | 0.544 | 1 | 13 | Sdccag3 | 0.054 | Old |
| 0.918498 | 0.014163 | 0.238 | 0.237 | 1 | 13 | Mthfsd | 0.001 | Old |
| 0.919504 | 0.01436 | 0.27 | 0.254 | 1 | 13 | Spsb3 | 0.016 | Old |
| 0.919576 | 0.002392 | 0.508 | 0.491 | 1 | 13 | Tmem183a | 0.017 | Old |
| 0.919903 | 0.186348 | 0.639 | 0.649 | 1 | 13 | Themis2 | -0.01 | Old |
| 0.919903 | 0.005618 | 0.639 | 0.649 | 1 | 13 | G6pdx | -0.01 | Old |
| 0.920028 | 0.00124 | 0.861 | 0.807 | 1 | 13 | Got1 | 0.054 | Old |
| 0.920317 | 0.103247 | 0.074 | 0.07 | 1 | 13 | Pex12 | 0.004 | Old |
| 0.920322 | 0.040814 | 0.41 | 0.395 | 1 | 13 | Edc4 | 0.015 | Old |
| 0.920685 | 0.061957 | 0.705 | 0.702 | 1 | 13 | Mrps36 | 0.003 | Old |
| 0.921213 | 0.084604 | 0.123 | 0.132 | 1 | 13 | Snhg9 | -0.009 | Old |
| 0.921622 | 0.145249 | 0.115 | 0.114 | 1 | 13 | Bloc1s3 | 0.001 | Old |
| 0.921716 | 0.09415 | 0.5 | 0.482 | 1 | 13 | Rnf146 | 0.018 | Old |
| 0.921819 | 0.027769 | 0.656 | 0.684 | 1 | 13 | 2210016F1 | -0.028 | Old |
| 0.9219 | 0.074247 | 0.549 | 0.553 | 1 | 13 | Iws1 | -0.004 | Old |
| 0.922142 | 0.101496 | 0.049 | 0.053 | 1 | 13 | Fsd2 | -0.004 | Old |
| 0.922262 | 0.035535 | 0.844 | 0.798 | 1 | 13 | Hsd17b11 | 0.046 | Old |
| 0.923238 | 0.023108 | 1 | 1 | 1 | 13 | Pomp | 0 | Old |
| 0.923306 | 0.027069 | 0.68 | 0.649 | 1 | 13 | Nop58 | 0.031 | Old |
| 0.923476 | 0.186349 | 0.533 | 0.509 | 1 | 13 | Trim12a | 0.024 | Old |
| 0.923624 | 0.263426 | 0.18 | 0.175 | 1 | 13 | Ube2o | 0.005 | Old |
| 0.923697 | 0.010483 | 0.074 | 0.07 | 1 | 13 | Zranb3 | 0.004 | Old |
| 0.923811 | 0.023125 | 0.828 | 0.833 | 1 | 13 | Sfxn3 | -0.005 | Old |
| 0.924067 | 0.008503 | 0.639 | 0.588 | 1 | 13 | Bsdc1 | 0.051 | Old |
| 0.924617 | 0.020527 | 0.672 | 0.623 | 1 | 13 | Atg2a | 0.049 | Old |
| 0.924754 | 0.031593 | 0.992 | 1 | 1 | 13 | Btg1 | -0.008 | Old |
| 0.924954 | 0.000217 | 0.746 | 0.763 | 1 | 13 | Nfkbie | -0.017 | Old |
| 0.92511 | 0.062934 | 0.672 | 0.702 | 1 | 13 | Zdhhc9 | -0.03 | Old |
| 0.925197 | 0.021643 | 0.828 | 0.763 | 1 | 13 | Rab18 | 0.065 | Old |
| 0.925245 | 0.081035 | 0.5 | 0.5 | 1 | 13 | Lpin2 | 0 | Old |
| 0.925739 | 0.094783 | 0.057 | 0.061 | 1 | 13 | Myom1 | -0.004 | Old |
| 0.925939 | 0.017152 | 0.779 | 0.807 | 1 | 13 | Dctn3 | -0.028 | Old |

| | | | | | | | | |
|---|---|---|---|---|---|---|---|---|
| 0.925953 | 0.141704 | 0.041 | 0.044 | 1 | 13 | Tmem150a | -0.003 | Old |
| 0.926093 | 0.192124 | 0.23 | 0.254 | 1 | 13 | Rhno1 | -0.024 | Old |
| 0.926123 | 0.147049 | 0.049 | 0.053 | 1 | 13 | Dnajc17 | -0.004 | Old |
| 0.926123 | 0.063109 | 0.049 | 0.053 | 1 | 13 | AC162302. | -0.004 | Old |
| 0.926365 | 0.003239 | 0.598 | 0.544 | 1 | 13 | Hdac5 | 0.054 | Old |
| 0.926506 | 0.029909 | 0.393 | 0.386 | 1 | 13 | Mroh1 | 0.007 | Old |
| 0.927001 | 0.134148 | 0.648 | 0.658 | 1 | 13 | Rhoc | -0.01 | Old |
| 0.927222 | 0.009279 | 0.697 | 0.64 | 1 | 13 | Azi2 | 0.057 | Old |
| 0.927271 | 0.010336 | 0.336 | 0.351 | 1 | 13 | Cdk16 | -0.015 | Old |
| 0.927545 | 0.160372 | 0.344 | 0.351 | 1 | 13 | Oma1 | -0.007 | Old |
| 0.92766 | 0.013819 | 0.09 | 0.096 | 1 | 13 | Pald1 | -0.006 | Old |
| 0.928543 | 0.004769 | 0.975 | 0.982 | 1 | 13 | Skap2 | -0.007 | Old |
| 0.92874 | 0.022157 | 0.516 | 0.5 | 1 | 13 | Foxo3 | 0.016 | Old |
| 0.929442 | 0.075891 | 0.057 | 0.061 | 1 | 13 | 1700052K1 | -0.004 | Old |
| 0.92968 | 0.507553 | 0.033 | 0.035 | 1 | 13 | Hdc | -0.002 | Old |
| 0.92968 | 0.150641 | 0.033 | 0.035 | 1 | 13 | Gm48604 | -0.002 | Old |
| 0.92968 | 0.059656 | 0.033 | 0.035 | 1 | 13 | Gnb1l | -0.002 | Old |
| 0.930106 | 0.084553 | 0.049 | 0.053 | 1 | 13 | Zfp951 | -0.004 | Old |
| 0.930297 | 0.141782 | 0.041 | 0.044 | 1 | 13 | Ppp1r3d | -0.003 | Old |
| 0.930297 | 0.055508 | 0.041 | 0.044 | 1 | 13 | Lysmd4 | -0.003 | Old |
| 0.930378 | 0.01626 | 0.779 | 0.754 | 1 | 13 | Wdr45b | 0.025 | Old |
| 0.930926 | 0.093591 | 0.574 | 0.553 | 1 | 13 | Ssbp3 | 0.021 | Old |
| 0.931355 | 0.023916 | 0.59 | 0.579 | 1 | 13 | Vbp1 | 0.011 | Old |
| 0.931437 | 0.049746 | 0.705 | 0.702 | 1 | 13 | Fundc2 | 0.003 | Old |
| 0.932819 | 0.092137 | 0.311 | 0.325 | 1 | 13 | Cep63 | -0.014 | Old |
| 0.933301 | 0.060941 | 0.639 | 0.57 | 1 | 13 | Pop5 | 0.069 | Old |
| 0.933741 | 0.027436 | 0.246 | 0.228 | 1 | 13 | Golim4 | 0.018 | Old |
| 0.933865 | 0.10626 | 0.623 | 0.667 | 1 | 13 | Appbp2 | -0.044 | Old |
| 0.934518 | 0.1898 | 0.033 | 0.035 | 1 | 13 | Tmem51os | -0.002 | Old |
| 0.934518 | 0.124733 | 0.033 | 0.035 | 1 | 13 | Abraxas1 | -0.002 | Old |
| 0.934518 | 0.050642 | 0.033 | 0.035 | 1 | 13 | Acot1 | -0.002 | Old |
| 0.934518 | 0.026946 | 0.033 | 0.035 | 1 | 13 | Jmjd4 | -0.002 | Old |
| 0.934518 | 0.016995 | 0.033 | 0.035 | 1 | 13 | Zfp784 | -0.002 | Old |
| 0.934518 | 0.014506 | 0.033 | 0.035 | 1 | 13 | Gm45894 | -0.002 | Old |
| 0.936275 | 0.005676 | 0.492 | 0.43 | 1 | 13 | Pgs1 | 0.062 | Old |
| 0.936796 | 0.005008 | 0.156 | 0.149 | 1 | 13 | Mgmt | 0.007 | Old |
| 0.936879 | 0.015666 | 0.943 | 0.93 | 1 | 13 | Xiap | 0.013 | Old |
| 0.936887 | 0.001794 | 0.992 | 0.965 | 1 | 13 | Rock2 | 0.027 | Old |
| 0.938402 | 0.003481 | 0.967 | 0.947 | 1 | 13 | Ccdc88a | 0.02 | Old |
| 0.938645 | 0.248384 | 0.025 | 0.026 | 1 | 13 | Tcea2 | -0.001 | Old |
| 0.938645 | 0.074161 | 0.025 | 0.026 | 1 | 13 | Stard10 | -0.001 | Old |
| 0.938645 | 0.039407 | 0.025 | 0.026 | 1 | 13 | Mb21d2 | -0.001 | Old |
| 0.938854 | 0.011899 | 0.713 | 0.632 | 1 | 13 | Myo9a | 0.081 | Old |
| 0.938986 | 0.011458 | 0.189 | 0.193 | 1 | 13 | Cry1 | -0.004 | Old |
| 0.938992 | 0.067505 | 0.041 | 0.044 | 1 | 13 | E2f8 | -0.003 | Old |

| | | | | | | | | |
|---|---|---|---|---|---|---|---|---|
| 0.939359 | 0.324194 | 0.033 | 0.035 | 1 | 13 | B130006D( | -0.002 | Old |
| 0.939359 | 0.147131 | 0.033 | 0.035 | 1 | 13 | Col27a1 | -0.002 | Old |
| 0.939359 | 0.021968 | 0.033 | 0.035 | 1 | 13 | Fam72a | -0.002 | Old |
| 0.939496 | 0.03479 | 0.172 | 0.167 | 1 | 13 | Dis3 | 0.005 | Old |
| 0.939546 | 0.006722 | 0.377 | 0.368 | 1 | 13 | Utp20 | 0.009 | Old |
| 0.940531 | 0.026128 | 0.648 | 0.588 | 1 | 13 | Dnajc21 | 0.06 | Old |
| 0.940617 | 0.144291 | 0.074 | 0.07 | 1 | 13 | Bub1b | 0.004 | Old |
| 0.940664 | 0.005425 | 0.943 | 0.886 | 1 | 13 | Atxn7l3b | 0.057 | Old |
| 0.940807 | 0.079394 | 0.385 | 0.377 | 1 | 13 | S100pbp | 0.008 | Old |
| 0.941149 | 0.071844 | 0.582 | 0.535 | 1 | 13 | Fyn | 0.047 | Old |
| 0.941309 | 0.114494 | 0.811 | 0.86 | 1 | 13 | Sod2 | -0.049 | Old |
| 0.941377 | 0.192425 | 0.279 | 0.281 | 1 | 13 | Crls1 | -0.002 | Old |
| 0.941648 | 0.140989 | 0.156 | 0.149 | 1 | 13 | 10-Sep | 0.007 | Old |
| 0.942254 | 0.010332 | 0.361 | 0.351 | 1 | 13 | Tmco3 | 0.01 | Old |
| 0.942509 | 0.050482 | 0.738 | 0.763 | 1 | 13 | Cdc5l | -0.025 | Old |
| 0.942884 | 0.072209 | 0.475 | 0.447 | 1 | 13 | Ppme1 | 0.028 | Old |
| 0.942904 | 0.038213 | 0.713 | 0.675 | 1 | 13 | Ppp4r3a | 0.038 | Old |
| 0.943833 | 0.031863 | 0.713 | 0.719 | 1 | 13 | Gtpbp4 | -0.006 | Old |
| 0.944213 | 0.252032 | 0.025 | 0.026 | 1 | 13 | D230017M | -0.001 | Old |
| 0.944213 | 0.105053 | 0.025 | 0.026 | 1 | 13 | Noxo1 | -0.001 | Old |
| 0.944213 | 0.02931 | 0.025 | 0.026 | 1 | 13 | Nutf2 | -0.001 | Old |
| 0.944213 | 0.007083 | 0.025 | 0.026 | 1 | 13 | Slc13a2 | -0.001 | Old |
| 0.944269 | 0.210613 | 0.057 | 0.061 | 1 | 13 | Zbtb10 | -0.004 | Old |
| 0.944478 | 0.080554 | 0.992 | 1 | 1 | 13 | S100a4 | -0.008 | Old |
| 0.944684 | 0.008351 | 0.73 | 0.728 | 1 | 13 | Tcf12 | 0.002 | Old |
| 0.94529 | 0.092235 | 0.148 | 0.158 | 1 | 13 | Pwwp2b | -0.01 | Old |
| 0.945631 | 0.060553 | 0.197 | 0.184 | 1 | 13 | Ptges2 | 0.013 | Old |
| 0.945941 | 0.051491 | 0.885 | 0.86 | 1 | 13 | Scpep1 | 0.025 | Old |
| 0.946053 | 0.247457 | 0.049 | 0.053 | 1 | 13 | Gm19325 | -0.004 | Old |
| 0.946317 | 0.021147 | 0.139 | 0.132 | 1 | 13 | Clstn1 | 0.007 | Old |
| 0.946345 | 0.327676 | 0.098 | 0.105 | 1 | 13 | Osbpl3 | -0.007 | Old |
| 0.946848 | 0.055701 | 0.705 | 0.711 | 1 | 13 | Fam50a | -0.006 | Old |
| 0.946939 | 0.062553 | 0.27 | 0.254 | 1 | 13 | Paxx | 0.016 | Old |
| 0.947289 | 0.06331 | 0.59 | 0.605 | 1 | 13 | Tmem205 | -0.015 | Old |
| 0.94738 | 0.030912 | 0.377 | 0.412 | 1 | 13 | Sema6d | -0.035 | Old |
| 0.947422 | 0.081226 | 0.205 | 0.211 | 1 | 13 | 1110032A( | -0.006 | Old |
| 0.947515 | 0.115148 | 0.967 | 0.991 | 1 | 13 | Nfkb1 | -0.024 | Old |
| 0.947757 | 0.062687 | 0.656 | 0.614 | 1 | 13 | Chmp2b | 0.042 | Old |
| 0.947843 | 0.135779 | 0.467 | 0.456 | 1 | 13 | Gm13212 | 0.011 | Old |
| 0.947897 | 0.102539 | 0.566 | 0.605 | 1 | 13 | Sdhaf4 | -0.039 | Old |
| 0.948432 | 0.159129 | 0.574 | 0.561 | 1 | 13 | Txlna | 0.013 | Old |
| 0.948783 | 0.036146 | 0.27 | 0.281 | 1 | 13 | Cnot9 | -0.011 | Old |
| 0.948861 | 0.071326 | 0.508 | 0.5 | 1 | 13 | Vps37a | 0.008 | Old |
| 0.948968 | 0.578621 | 0.016 | 0.018 | 1 | 13 | Cox4i2 | -0.002 | Old |
| 0.948968 | 0.179954 | 0.016 | 0.018 | 1 | 13 | Adig | -0.002 | Old |

| | | | | | | | | |
|---|---|---|---|---|---|---|---|---|
| 0.948968 | 0.078081 | 0.016 | 0.018 | 1 | 13 | C920021L1 | -0.002 | Old |
| 0.948968 | 0.072648 | 0.016 | 0.018 | 1 | 13 | Exoc3l4 | -0.002 | Old |
| 0.948968 | 0.072388 | 0.016 | 0.018 | 1 | 13 | Arc | -0.002 | Old |
| 0.948968 | 0.070766 | 0.016 | 0.018 | 1 | 13 | Ppef2 | -0.002 | Old |
| 0.948968 | 0.030102 | 0.016 | 0.018 | 1 | 13 | Cyyr1 | -0.002 | Old |
| 0.948968 | 0.020513 | 0.016 | 0.018 | 1 | 13 | 4930426I2 | -0.002 | Old |
| 0.948968 | 0.004918 | 0.016 | 0.018 | 1 | 13 | Tmem171 | -0.002 | Old |
| 0.948968 | 0.004837 | 0.016 | 0.018 | 1 | 13 | Mdk | -0.002 | Old |
| 0.948968 | 0.004823 | 0.016 | 0.018 | 1 | 13 | Adam23 | -0.002 | Old |
| 0.949047 | 0.172376 | 0.033 | 0.035 | 1 | 13 | Gprin3 | -0.002 | Old |
| 0.949047 | 0.084983 | 0.033 | 0.035 | 1 | 13 | Nadsyn1 | -0.002 | Old |
| 0.949784 | 0.426484 | 0.025 | 0.026 | 1 | 13 | K230015D( | -0.001 | Old |
| 0.949784 | 0.269539 | 0.025 | 0.026 | 1 | 13 | Olfml3 | -0.001 | Old |
| 0.949784 | 0.134225 | 0.025 | 0.026 | 1 | 13 | Tsku | -0.001 | Old |
| 0.949784 | 0.115145 | 0.025 | 0.026 | 1 | 13 | Spag1 | -0.001 | Old |
| 0.950019 | 0.06268 | 0.082 | 0.079 | 1 | 13 | Rabl3 | 0.003 | Old |
| 0.950045 | 0.025088 | 0.377 | 0.368 | 1 | 13 | Nvl | 0.009 | Old |
| 0.950466 | 0.029585 | 0.852 | 0.842 | 1 | 13 | Timm10b | 0.01 | Old |
| 0.951217 | 0.059012 | 0.098 | 0.096 | 1 | 13 | Abhd8 | 0.002 | Old |
| 0.95122 | 0.081796 | 0.664 | 0.684 | 1 | 13 | Mrps25 | -0.02 | Old |
| 0.951565 | 0.00184 | 0.262 | 0.254 | 1 | 13 | Trim34a | 0.008 | Old |
| 0.951768 | 0.03253 | 0.795 | 0.737 | 1 | 13 | Ccnl2 | 0.058 | Old |
| 0.952057 | 0.05327 | 0.918 | 0.904 | 1 | 13 | Ostc | 0.014 | Old |
| 0.952104 | 0.04319 | 0.508 | 0.5 | 1 | 13 | Usp1 | 0.008 | Old |
| 0.952828 | 0.063676 | 0.41 | 0.395 | 1 | 13 | Trmt1l | 0.015 | Old |
| 0.952828 | 0.015209 | 0.418 | 0.386 | 1 | 13 | Nlk | 0.032 | Old |
| 0.953082 | 0.017628 | 0.238 | 0.237 | 1 | 13 | Adamts10 | 0.001 | Old |
| 0.953097 | 0.17118 | 0.451 | 0.43 | 1 | 13 | Surf1 | 0.021 | Old |
| 0.953311 | 0.037245 | 0.533 | 0.518 | 1 | 13 | Zbtb7b | 0.015 | Old |
| 0.953539 | 0.131869 | 0.254 | 0.263 | 1 | 13 | Bend4 | -0.009 | Old |
| 0.954109 | 0.065072 | 0.5 | 0.465 | 1 | 13 | Sin3a | 0.035 | Old |
| 0.95417 | 0.125285 | 0.098 | 0.096 | 1 | 13 | Zfp760 | 0.002 | Old |
| 0.954349 | 0.214203 | 0.926 | 0.956 | 1 | 13 | Irf1 | -0.03 | Old |
| 0.954676 | 0.012112 | 0.361 | 0.351 | 1 | 13 | Cyb5r1 | 0.01 | Old |
| 0.95508 | 0.049758 | 0.885 | 0.886 | 1 | 13 | Acin1 | -0.001 | Old |
| 0.955113 | 0.056589 | 1 | 0.991 | 1 | 13 | Cct5 | 0.009 | Old |
| 0.955357 | 0.199164 | 0.025 | 0.026 | 1 | 13 | Rasgef1a | -0.001 | Old |
| 0.955357 | 0.19091 | 0.025 | 0.026 | 1 | 13 | Col6a2 | -0.001 | Old |
| 0.955357 | 0.165144 | 0.025 | 0.026 | 1 | 13 | AC146911. | -0.001 | Old |
| 0.955402 | 0.877183 | 0.057 | 0.061 | 1 | 13 | Sox4 | -0.004 | Old |
| 0.955506 | 0.015953 | 0.664 | 0.596 | 1 | 13 | Msl3 | 0.068 | Old |
| 0.955765 | 0.243658 | 0.016 | 0.018 | 1 | 13 | Abca8b | -0.002 | Old |
| 0.955765 | 0.212238 | 0.016 | 0.018 | 1 | 13 | Plk1 | -0.002 | Old |
| 0.955765 | 0.169469 | 0.016 | 0.018 | 1 | 13 | Rab4b | -0.002 | Old |
| 0.955765 | 0.167889 | 0.016 | 0.018 | 1 | 13 | Sptbn4 | -0.002 | Old |

| | | | | | | | | |
|---|---|---|---|---|---|---|---|---|
| 0.955765 | 0.131587 | 0.016 | 0.018 | 1 | 13 | Ptprb | -0.002 | Old |
| 0.955765 | 0.11725 | 0.016 | 0.018 | 1 | 13 | 1700099I0 | -0.002 | Old |
| 0.955765 | 0.117215 | 0.016 | 0.018 | 1 | 13 | Gm48682 | -0.002 | Old |
| 0.955765 | 0.10121 | 0.016 | 0.018 | 1 | 13 | Gm48079 | -0.002 | Old |
| 0.955765 | 0.089692 | 0.016 | 0.018 | 1 | 13 | Gm13710 | -0.002 | Old |
| 0.955765 | 0.079779 | 0.016 | 0.018 | 1 | 13 | Rpp40 | -0.002 | Old |
| 0.955765 | 0.077066 | 0.016 | 0.018 | 1 | 13 | Timp4 | -0.002 | Old |
| 0.955765 | 0.050862 | 0.016 | 0.018 | 1 | 13 | Gm16094 | -0.002 | Old |
| 0.955765 | 0.04733 | 0.016 | 0.018 | 1 | 13 | Ccr9 | -0.002 | Old |
| 0.955856 | 0.019388 | 0.926 | 0.886 | 1 | 13 | Tmed9 | 0.04 | Old |
| 0.956349 | 0.007928 | 0.336 | 0.325 | 1 | 13 | Gmeb1 | 0.011 | Old |
| 0.956479 | 0.072711 | 0.492 | 0.465 | 1 | 13 | Slu7 | 0.027 | Old |
| 0.956813 | 0.03854 | 0.5 | 0.491 | 1 | 13 | Pofut2 | 0.009 | Old |
| 0.957239 | 0.145291 | 0.27 | 0.272 | 1 | 13 | Map2k5 | -0.002 | Old |
| 0.957399 | 0.006217 | 0.541 | 0.518 | 1 | 13 | Imp4 | 0.023 | Old |
| 0.957461 | 0.175161 | 0.385 | 0.386 | 1 | 13 | Gsr | -0.001 | Old |
| 0.957667 | 0.02155 | 0.664 | 0.772 | 1 | 13 | Antxr2 | -0.108 | Old |
| 0.958088 | 5.62E-06 | 0.41 | 0.404 | 1 | 13 | Rnaset2a | 0.006 | Old |
| 0.959683 | 0.053835 | 0.41 | 0.395 | 1 | 13 | Foxk1 | 0.015 | Old |
| 0.959684 | 0.03918 | 0.082 | 0.079 | 1 | 13 | Syngr1 | 0.003 | Old |
| 0.960322 | 0.091709 | 0.156 | 0.158 | 1 | 13 | Rfc5 | -0.002 | Old |
| 0.960929 | 0.076812 | 0.631 | 0.623 | 1 | 13 | Dcp2 | 0.008 | Old |
| 0.960933 | 0.413865 | 0.025 | 0.026 | 1 | 13 | Chic1 | -0.001 | Old |
| 0.960933 | 0.186767 | 0.025 | 0.026 | 1 | 13 | Pcp4l1 | -0.001 | Old |
| 0.961159 | 0.070585 | 0.869 | 0.886 | 1 | 13 | Twf1 | -0.017 | Old |
| 0.961193 | 0.024342 | 0.984 | 0.939 | 1 | 13 | Capn2 | 0.045 | Old |
| 0.961259 | 0.108413 | 0.582 | 0.579 | 1 | 13 | Trim47 | 0.003 | Old |
| 0.961878 | 0.010096 | 0.656 | 0.64 | 1 | 13 | Trappc1 | 0.016 | Old |
| 0.962565 | 0.42989 | 0.016 | 0.018 | 1 | 13 | Gm26857 | -0.002 | Old |
| 0.962565 | 0.323985 | 0.016 | 0.018 | 1 | 13 | Syt7 | -0.002 | Old |
| 0.962565 | 0.238548 | 0.016 | 0.018 | 1 | 13 | Gm21917 | -0.002 | Old |
| 0.962565 | 0.178192 | 0.016 | 0.018 | 1 | 13 | Gm26716 | -0.002 | Old |
| 0.962565 | 0.161054 | 0.016 | 0.018 | 1 | 13 | Aarsd1 | -0.002 | Old |
| 0.962565 | 0.16021 | 0.016 | 0.018 | 1 | 13 | Lbhd1 | -0.002 | Old |
| 0.962565 | 0.15864 | 0.016 | 0.018 | 1 | 13 | Hoxaas3 | -0.002 | Old |
| 0.962565 | 0.154831 | 0.016 | 0.018 | 1 | 13 | Sox18 | -0.002 | Old |
| 0.962565 | 0.152477 | 0.016 | 0.018 | 1 | 13 | Tcf15 | -0.002 | Old |
| 0.962565 | 0.150133 | 0.016 | 0.018 | 1 | 13 | Lrrc63 | -0.002 | Old |
| 0.962565 | 0.146147 | 0.016 | 0.018 | 1 | 13 | Gm10399 | -0.002 | Old |
| 0.962565 | 0.138422 | 0.016 | 0.018 | 1 | 13 | Nudt17 | -0.002 | Old |
| 0.962565 | 0.131935 | 0.016 | 0.018 | 1 | 13 | Klc2 | -0.002 | Old |
| 0.962565 | 0.058802 | 0.016 | 0.018 | 1 | 13 | BC025920 | -0.002 | Old |
| 0.962624 | 0.148305 | 0.615 | 0.675 | 1 | 13 | Il3ra | -0.06 | Old |
| 0.962715 | 0.017493 | 1 | 1 | 1 | 13 | C1qb | 0 | Old |
| 0.96286 | 0.050422 | 0.336 | 0.333 | 1 | 13 | Zfpl1 | 0.003 | Old |

| | | | | | | | | |
|---|---|---|---|---|---|---|---|---|
| 0.963033 | 0.004574 | 0.098 | 0.096 | 1 | 13 | Nmnat1 | 0.002 | Old |
| 0.963327 | 0.101477 | 0.5 | 0.491 | 1 | 13 | Ssbp1 | 0.009 | Old |
| 0.964093 | 0.073866 | 0.574 | 0.526 | 1 | 13 | Slc36a1 | 0.048 | Old |
| 0.964225 | 0.174065 | 0.918 | 0.912 | 1 | 13 | Tuba1a | 0.006 | Old |
| 0.964444 | 0.022355 | 0.68 | 0.693 | 1 | 13 | Glrx | -0.013 | Old |
| 0.964814 | 0.128204 | 0.451 | 0.43 | 1 | 13 | Znrf1 | 0.021 | Old |
| 0.964829 | 0.137342 | 0.795 | 0.781 | 1 | 13 | Cycs | 0.014 | Old |
| 0.965204 | 0.027576 | 0.541 | 0.482 | 1 | 13 | Irs2 | 0.059 | Old |
| 0.965538 | 0.128932 | 0.779 | 0.754 | 1 | 13 | Nabp1 | 0.025 | Old |
| 0.965942 | 0.036197 | 0.156 | 0.149 | 1 | 13 | Tmem140 | 0.007 | Old |
| 0.965989 | 0.145223 | 0.098 | 0.096 | 1 | 13 | Casz1 | 0.002 | Old |
| 0.96613 | 0.002015 | 0.082 | 0.079 | 1 | 13 | Dync2h1 | 0.003 | Old |
| 0.966476 | 0.077522 | 0.861 | 0.868 | 1 | 13 | Smchd1 | -0.007 | Old |
| 0.966729 | 0.091404 | 0.418 | 0.404 | 1 | 13 | Slc30a9 | 0.014 | Old |
| 0.967131 | 0.013924 | 0.82 | 0.763 | 1 | 13 | Acat1 | 0.057 | Old |
| 0.967214 | 0.045569 | 0.18 | 0.184 | 1 | 13 | Tmtc3 | -0.004 | Old |
| 0.967399 | 0.08115 | 0.172 | 0.167 | 1 | 13 | Vwa8 | 0.005 | Old |
| 0.967554 | 0.019644 | 0.459 | 0.447 | 1 | 13 | Edem2 | 0.012 | Old |
| 0.967642 | 0.892068 | 0.09 | 0.088 | 1 | 13 | Fscn1 | 0.002 | Old |
| 0.968963 | 0.092287 | 0.557 | 0.561 | 1 | 13 | Atxn10 | -0.004 | Old |
| 0.968973 | 0.041271 | 0.246 | 0.254 | 1 | 13 | Ercc1 | -0.008 | Old |
| 0.969548 | 0.046112 | 0.639 | 0.623 | 1 | 13 | Psme2b | 0.016 | Old |
| 0.969841 | 0.087892 | 0.303 | 0.289 | 1 | 13 | Spryd7 | 0.014 | Old |
| 0.970029 | 0.123161 | 0.27 | 0.281 | 1 | 13 | Tmem2 | -0.011 | Old |
| 0.970267 | 0.054045 | 0.164 | 0.158 | 1 | 13 | Zfp780b | 0.006 | Old |
| 0.970287 | 0.049971 | 0.877 | 0.86 | 1 | 13 | Ppp3r1 | 0.017 | Old |
| 0.97032 | 0.076002 | 0.943 | 1 | 1 | 13 | Tnip3 | -0.057 | Old |
| 0.970876 | 0.063875 | 0.623 | 0.57 | 1 | 13 | Cd300lb | 0.053 | Old |
| 0.971078 | 0.051528 | 0.893 | 0.991 | 1 | 13 | Ear2 | -0.098 | Old |
| 0.971343 | 0.024823 | 0.352 | 0.342 | 1 | 13 | Ncoa5 | 0.01 | Old |
| 0.971603 | 0.003365 | 0.41 | 0.386 | 1 | 13 | Faap20 | 0.024 | Old |
| 0.971699 | 0.089617 | 0.787 | 0.781 | 1 | 13 | Dgkd | 0.006 | Old |
| 0.971998 | 0.073928 | 0.434 | 0.404 | 1 | 13 | Tlk2 | 0.03 | Old |
| 0.972594 | 0.113467 | 0.893 | 0.939 | 1 | 13 | Cd300ld | -0.046 | Old |
| 0.972666 | 0.049435 | 0.467 | 0.456 | 1 | 13 | Pfkfb3 | 0.011 | Old |
| 0.973803 | 0.040107 | 0.09 | 0.088 | 1 | 13 | Repin1 | 0.002 | Old |
| 0.973831 | 0.112893 | 0.426 | 0.43 | 1 | 13 | Sema4d | -0.004 | Old |
| 0.973836 | 0.20701 | 0.115 | 0.114 | 1 | 13 | Tgm1 | 0.001 | Old |
| 0.97423 | 0.206621 | 0.533 | 0.518 | 1 | 13 | Map3k14 | 0.015 | Old |
| 0.974891 | 0.005044 | 0.631 | 0.588 | 1 | 13 | Slc35a4 | 0.043 | Old |
| 0.974976 | 0.17582 | 0.246 | 0.254 | 1 | 13 | Maml3 | -0.008 | Old |
| 0.974976 | 0.169266 | 0.246 | 0.254 | 1 | 13 | Ppargc1b | -0.008 | Old |
| 0.97557 | 0.046399 | 0.639 | 0.544 | 1 | 13 | Mrpl13 | 0.095 | Old |
| 0.975686 | 0.006845 | 0.221 | 0.211 | 1 | 13 | Zer1 | 0.01 | Old |
| 0.975978 | 0.046699 | 0.467 | 0.456 | 1 | 13 | Faf1 | 0.011 | Old |

| | | | | | | | | |
|---|---|---|---|---|---|---|---|---|
| 0.976179 | 0.047916 | 0.738 | 0.728 | 1 | 13 | Atp13a2 | 0.01 | Old |
| 0.976272 | 0.020906 | 0.795 | 0.754 | 1 | 13 | Tpp2 | 0.041 | Old |
| 0.976349 | 0.101143 | 0.836 | 0.825 | 1 | 13 | Tppp3 | 0.011 | Old |
| 0.977866 | 0.051589 | 0.836 | 0.807 | 1 | 13 | P2ry6 | 0.029 | Old |
| 0.977903 | 0.037047 | 0.869 | 0.868 | 1 | 13 | Atf6 | 0.001 | Old |
| 0.977928 | 0.021291 | 1 | 1 | 1 | 13 | Sub1 | 0 | Old |
| 0.977958 | 0.138037 | 0.131 | 0.132 | 1 | 13 | Ifi206 | -0.001 | Old |
| 0.978147 | 0.025509 | 0.533 | 0.491 | 1 | 13 | Cc2d1b | 0.042 | Old |
| 0.978369 | 0.004894 | 0.27 | 0.254 | 1 | 13 | Mrs2 | 0.016 | Old |
| 0.978675 | 0.035945 | 0.893 | 0.886 | 1 | 13 | Wapl | 0.007 | Old |
| 0.979833 | 0.100947 | 0.525 | 0.518 | 1 | 13 | Zfp871 | 0.007 | Old |
| 0.980429 | 0.094214 | 0.426 | 0.404 | 1 | 13 | Wwc2 | 0.022 | Old |
| 0.982055 | 0.081056 | 0.639 | 0.632 | 1 | 13 | Dhrs7 | 0.007 | Old |
| 0.982429 | 0.074203 | 0.803 | 0.807 | 1 | 13 | Prpf4b | -0.004 | Old |
| 0.982437 | 0.017397 | 0.803 | 0.825 | 1 | 13 | Ppp4r3b | -0.022 | Old |
| 0.983184 | 0.037249 | 0.811 | 0.781 | 1 | 13 | Klhl24 | 0.03 | Old |
| 0.983672 | 0.055172 | 0.238 | 0.237 | 1 | 13 | Mynn | 0.001 | Old |
| 0.983683 | 0.07168 | 0.68 | 0.632 | 1 | 13 | Vamp4 | 0.048 | Old |
| 0.983788 | 0.019303 | 0.41 | 0.412 | 1 | 13 | Mta3 | -0.002 | Old |
| 0.984009 | 0.005082 | 0.902 | 0.904 | 1 | 13 | Ndufa11 | -0.002 | Old |
| 0.984598 | 0.063819 | 0.418 | 0.395 | 1 | 13 | Tctex1d2 | 0.023 | Old |
| 0.984754 | 0.039905 | 0.877 | 0.833 | 1 | 13 | Lat2 | 0.044 | Old |
| 0.984777 | 0.034574 | 0.975 | 0.974 | 1 | 13 | Asah1 | 0.001 | Old |
| 0.985318 | 0.139835 | 0.123 | 0.123 | 1 | 13 | Rnf152 | 0 | Old |
| 0.985537 | 0.03129 | 0.934 | 0.947 | 1 | 13 | Ier3ip1 | -0.013 | Old |
| 0.986034 | 0.016564 | 0.475 | 0.482 | 1 | 13 | Wdr33 | -0.007 | Old |
| 0.986688 | 0.025859 | 0.098 | 0.096 | 1 | 13 | Arvcf | 0.002 | Old |
| 0.986711 | 0.156631 | 0.59 | 0.667 | 1 | 13 | Ryr1 | -0.077 | Old |
| 0.986774 | 0.00877 | 0.467 | 0.465 | 1 | 13 | Zfp992 | 0.002 | Old |
| 0.986811 | 1.21E-05 | 0.656 | 0.675 | 1 | 13 | Slc25a24 | -0.019 | Old |
| 0.987002 | 0.083217 | 0.803 | 0.781 | 1 | 13 | Smim15 | 0.022 | Old |
| 0.987026 | 0.055897 | 0.811 | 0.842 | 1 | 13 | Pias1 | -0.031 | Old |
| 0.987031 | 0.103689 | 0.852 | 0.816 | 1 | 13 | Bak1 | 0.036 | Old |
| 0.987047 | 0.045753 | 0.869 | 0.877 | 1 | 13 | Nenf | -0.008 | Old |
| 0.987318 | 0.046194 | 0.172 | 0.175 | 1 | 13 | Pramef8 | -0.003 | Old |
| 0.987838 | 0.03952 | 0.508 | 0.509 | 1 | 13 | Fam3c | -0.001 | Old |
| 0.988229 | 0.100718 | 0.607 | 0.614 | 1 | 13 | Slc38a10 | -0.007 | Old |
| 0.988458 | 0.133959 | 0.27 | 0.289 | 1 | 13 | Actr8 | -0.019 | Old |
| 0.988483 | 0.067498 | 0.754 | 0.728 | 1 | 13 | Adipor2 | 0.026 | Old |
| 0.988501 | 0.062512 | 0.787 | 0.728 | 1 | 13 | Med28 | 0.059 | Old |
| 0.98855 | 0.035491 | 0.852 | 0.789 | 1 | 13 | Zfand3 | 0.063 | Old |
| 0.988582 | 0.026765 | 0.975 | 0.965 | 1 | 13 | Tpm4 | 0.01 | Old |
| 0.988583 | 0.017522 | 1 | 1 | 1 | 13 | Ahnak | 0 | Old |
| 0.988583 | 0.001594 | 1 | 1 | 1 | 13 | Ptma | 0 | Old |
| 0.98932 | 0.085521 | 0.475 | 0.482 | 1 | 13 | Mrps26 | -0.007 | Old |

| | | | | | | | | |
|---|---|---|---|---|---|---|---|---|
| 0.989546 | 0.118296 | 0.533 | 0.526 | 1 | 13 | Lilra6 | 0.007 | Old |
| 0.991171 | 0.080615 | 0.369 | 0.36 | 1 | 13 | Brap | 0.009 | Old |
| 0.991627 | 0.016824 | 0.984 | 0.982 | 1 | 13 | Rbm25 | 0.002 | Old |
| 0.992211 | 0.074653 | 0.664 | 0.623 | 1 | 13 | Hspa14 | 0.041 | Old |
| 0.99256 | 0.040591 | 0.459 | 0.474 | 1 | 13 | Ifi35 | -0.015 | Old |
| 0.993101 | 0.061347 | 0.779 | 0.737 | 1 | 13 | Stk40 | 0.042 | Old |
| 0.993146 | 0.06332 | 0.869 | 0.939 | 1 | 13 | Cbr2 | -0.07 | Old |
| 0.994353 | 0.177105 | 0.508 | 0.535 | 1 | 13 | Gcc2 | -0.027 | Old |
| 0.994569 | 0.013386 | 0.689 | 0.64 | 1 | 13 | Hbp1 | 0.049 | Old |
| 0.994672 | 0.043533 | 1 | 1 | 1 | 13 | Lyz2 | 0 | Old |
| 0.995373 | 0.064361 | 0.328 | 0.298 | 1 | 13 | Stx18 | 0.03 | Old |
| 0.99563 | 0.144771 | 0.385 | 0.368 | 1 | 13 | Csf3r | 0.017 | Old |
| 0.996143 | 0.108125 | 0.705 | 0.702 | 1 | 13 | Osbpl8 | 0.003 | Old |
| 0.996193 | 0.037168 | 0.902 | 0.921 | 1 | 13 | Ndufs7 | -0.019 | Old |
| 0.996213 | 0.306078 | 0.139 | 0.14 | 1 | 13 | Smad7 | -0.001 | Old |
| 0.996918 | 0.033377 | 0.23 | 0.237 | 1 | 13 | Mrps31 | -0.007 | Old |
| 0.997303 | 0.01661 | 0.344 | 0.342 | 1 | 13 | Pla2g4a | 0.002 | Old |
| 0.997394 | 0.013108 | 0.402 | 0.368 | 1 | 13 | Ctdspl2 | 0.034 | Old |
| 0.997572 | 0.24297 | 0.508 | 0.518 | 1 | 13 | Rnf38 | -0.01 | Old |
| 0.997697 | 0.022981 | 0.77 | 0.719 | 1 | 13 | Stx16 | 0.051 | Old |
| 0.99771 | 0.012761 | 0.82 | 0.825 | 1 | 13 | Npc1 | -0.005 | Old |
| 0.997716 | 0.014729 | 0.959 | 1 | 1 | 13 | Nktr | -0.041 | Old |
| 0.997716 | 0.036679 | 0.992 | 0.982 | 1 | 13 | Uqcr10 | 0.01 | Old |
| 0.997959 | 0.055856 | 0.238 | 0.237 | 1 | 13 | Asb7 | 0.001 | Old |
| 0.99818 | 0.050642 | 0.336 | 0.325 | 1 | 13 | Fam193b | 0.011 | Old |
| 0.998738 | 0.107645 | 0.139 | 0.14 | 1 | 13 | Snx7 | -0.001 | Old |
| 0.998769 | 0.120333 | 0.148 | 0.149 | 1 | 13 | Gpr141 | -0.001 | Old |
| 0.999229 | 0.127237 | 0.721 | 0.693 | 1 | 13 | Zc3h13 | 0.028 | Old |
| 0.999231 | 0.001127 | 0.73 | 0.737 | 1 | 13 | Ext1 | -0.007 | Old |
| 0.999237 | 0.016275 | 0.877 | 0.807 | 1 | 13 | Phf3 | 0.07 | Old |
| 3.48E-13 | 0.32908 | 1 | 1 | 1.08E-08 | 13 | Rps14 | 0 | Young |
| 4.59E-11 | 0.320835 | 1 | 1 | 1.43E-06 | 13 | Fau | 0 | Young |
| 1.50E-10 | 1.287156 | 0.851 | 0.598 | 4.67E-06 | 13 | Hpgd | 0.253 | Young |
| 2.04E-10 | 0.343592 | 1 | 1 | 6.33E-06 | 13 | Rpl18a | 0 | Young |
| 3.92E-10 | 0.331147 | 1 | 1 | 1.22E-05 | 13 | Rps4x | 0 | Young |
| 7.72E-10 | 0.296636 | 1 | 1 | 2.40E-05 | 13 | Rpl13 | 0 | Young |
| 2.60E-08 | 0.263527 | 1 | 1 | 0.000806 | 13 | Rpl17 | 0 | Young |
| 4.75E-08 | 1.287848 | 0.816 | 0.59 | 0.001477 | 13 | Ednrb | 0.226 | Young |
| 5.05E-08 | 2.237644 | 0.368 | 0.082 | 0.00157 | 13 | Cfd | 0.286 | Young |
| 6.16E-08 | 0.259881 | 1 | 1 | 0.001914 | 13 | Rps24 | 0 | Young |
| 6.78E-08 | 0.285677 | 1 | 1 | 0.002106 | 13 | Rpl18 | 0 | Young |
| 7.15E-08 | 0.222037 | 1 | 1 | 0.002221 | 13 | Rps3a1 | 0 | Young |
| 7.46E-08 | 0.348171 | 1 | 1 | 0.002317 | 13 | Cst3 | 0 | Young |
| 8.22E-08 | 0.931195 | 0.868 | 0.607 | 0.002553 | 13 | Gatm | 0.261 | Young |
| 8.38E-08 | 0.209045 | 1 | 1 | 0.002603 | 13 | Rps23 | 0 | Young |

| | | | | | | | | |
|---|---|---|---|---|---|---|---|---|
| 1.12E-07 | 0.671134 | 0.991 | 0.967 | 0.003493 | 13 | Apoe | 0.024 | Young |
| 1.32E-07 | 0.268183 | 1 | 1 | 0.004086 | 13 | Rpl26 | 0 | Young |
| 1.69E-07 | 0.264806 | 1 | 1 | 0.00524 | 13 | Rps16 | 0 | Young |
| 1.97E-07 | 0.285793 | 1 | 1 | 0.006115 | 13 | Rps13 | 0 | Young |
| 2.49E-07 | 1.205854 | 0.64 | 0.311 | 0.007743 | 13 | Plxdc2 | 0.329 | Young |
| 3.10E-07 | 0.942833 | 0.93 | 0.869 | 0.009632 | 13 | Ccl24 | 0.061 | Young |
| 4.00E-07 | 1.492181 | 0.491 | 0.18 | 0.012436 | 13 | Hepacam2 | 0.311 | Young |
| 4.51E-07 | 0.526006 | 0.982 | 0.951 | 0.014019 | 13 | Mef2c | 0.031 | Young |
| 8.10E-07 | 0.257003 | 0.965 | 0.779 | 0.025137 | 13 | Hspa1a | 0.186 | Young |
| 8.25E-07 | 0.221811 | 1 | 1 | 0.025616 | 13 | Rps9 | 0 | Young |
| 1.53E-06 | 0.675958 | 0.886 | 0.721 | 0.047528 | 13 | Nr4a3 | 0.165 | Young |
| 1.85E-06 | 0.254132 | 1 | 1 | 0.057461 | 13 | Rps10 | 0 | Young |
| 1.87E-06 | 0.236952 | 1 | 1 | 0.058008 | 13 | Rps5 | 0 | Young |
| 1.89E-06 | 0.252027 | 1 | 1 | 0.05856 | 13 | Rpl34 | 0 | Young |
| 2.32E-06 | 0.254126 | 1 | 1 | 0.072052 | 13 | Rpl19 | 0 | Young |
| 2.92E-06 | 1.232889 | 0.675 | 0.443 | 0.090665 | 13 | Cx3cr1 | 0.232 | Young |
| 3.16E-06 | 0.286612 | 1 | 1 | 0.098025 | 13 | Rpl11 | 0 | Young |
| 3.17E-06 | 0.226686 | 0.886 | 0.68 | 0.098396 | 13 | Hspa1b | 0.206 | Young |
| 3.81E-06 | 1.073228 | 0.377 | 0.115 | 0.118287 | 13 | Slc40a1 | 0.262 | Young |
| 3.90E-06 | 0.28456 | 1 | 1 | 0.121209 | 13 | Rps3 | 0 | Young |
| 4.52E-06 | 0.231792 | 1 | 1 | 0.140344 | 13 | Rpl30 | 0 | Young |
| 5.20E-06 | 0.708147 | 0.833 | 0.697 | 0.161618 | 13 | Rnase4 | 0.136 | Young |
| 5.69E-06 | 1.34508 | 0.482 | 0.23 | 0.176606 | 13 | Tslp | 0.252 | Young |
| 7.23E-06 | 0.231085 | 1 | 1 | 0.224599 | 13 | Rpl9 | 0 | Young |
| 7.83E-06 | 0.892474 | 0.895 | 0.762 | 0.243091 | 13 | Ltc4s | 0.133 | Young |
| 9.45E-06 | 0.236181 | 1 | 1 | 0.293328 | 13 | Rps11 | 0 | Young |
| 9.97E-06 | 0.741584 | 0.789 | 0.639 | 0.309692 | 13 | Cd33 | 0.15 | Young |
| 1.14E-05 | 0.202819 | 1 | 1 | 0.352939 | 13 | Rps20 | 0 | Young |
| 1.20E-05 | 0.591827 | 0.956 | 0.91 | 0.372463 | 13 | H2afv | 0.046 | Young |
| 1.50E-05 | 0.1835 | 1 | 1 | 0.4665 | 13 | Rpl7 | 0 | Young |
| 1.54E-05 | 0.251367 | 1 | 1 | 0.478773 | 13 | Rpl32 | 0 | Young |
| 1.69E-05 | 2.520814 | 0.298 | 0.09 | 0.526055 | 13 | Pmepa1 | 0.208 | Young |
| 1.80E-05 | 0.778581 | 0.684 | 0.426 | 0.557952 | 13 | Igf1r | 0.258 | Young |
| 1.98E-05 | 0.597803 | 0.956 | 0.803 | 0.614011 | 13 | Stab1 | 0.153 | Young |
| 2.10E-05 | 0.975533 | 0.675 | 0.525 | 0.652623 | 13 | Bmyc | 0.15 | Young |
| 2.39E-05 | 1.340434 | 0.474 | 0.221 | 0.742798 | 13 | Rasgrp3 | 0.253 | Young |
| 2.49E-05 | 0.839892 | 0.711 | 0.516 | 0.773361 | 13 | Cirbp | 0.195 | Young |
| 2.64E-05 | 0.619325 | 0.965 | 0.918 | 0.818437 | 13 | Dennd4a | 0.047 | Young |
| 2.78E-05 | 0.258674 | 1 | 1 | 0.862514 | 13 | Rps7 | 0 | Young |
| 2.92E-05 | 0.278749 | 1 | 1 | 0.907099 | 13 | Nr4a1 | 0 | Young |
| 3.15E-05 | 0.228926 | 1 | 1 | 0.978101 | 13 | Rpl28 | 0 | Young |
| 3.91E-05 | 0.260331 | 1 | 1 | 1 | 13 | Atp5h | 0 | Young |
| 4.44E-05 | 0.524755 | 0.921 | 0.852 | 1 | 13 | Higd2a | 0.069 | Young |
| 4.92E-05 | 0.19213 | 1 | 1 | 1 | 13 | Rpl23 | 0 | Young |
| 5.17E-05 | 0.260417 | 1 | 1 | 1 | 13 | Rsrp1 | 0 | Young |

| | | | | | | | | |
|---|---|---|---|---|---|---|---|---|
| 5.93E-05 | 0.559705 | 0.904 | 0.787 | 1 | 13 | Arl4c | 0.117 | Young |
| 7.75E-05 | 0.544921 | 0.877 | 0.738 | 1 | 13 | Cited2 | 0.139 | Young |
| 8.24E-05 | 0.270105 | 1 | 1 | 1 | 13 | Nfkbiz | 0 | Young |
| 8.31E-05 | 0.129632 | 1 | 1 | 1 | 13 | Tpt1 | 0 | Young |
| 8.58E-05 | 0.316006 | 1 | 1 | 1 | 13 | Pfdn5 | 0 | Young |
| 8.85E-05 | 0.378089 | 1 | 0.967 | 1 | 13 | Zfp36l1 | 0.033 | Young |
| 0.000103 | 0.145076 | 1 | 1 | 1 | 13 | Rplp1 | 0 | Young |
| 0.000117 | 0.273632 | 1 | 1 | 1 | 13 | Cd83 | 0 | Young |
| 0.000117 | 0.930555 | 0.553 | 0.344 | 1 | 13 | Lmo1 | 0.209 | Young |
| 0.000138 | 0.189414 | 1 | 1 | 1 | 13 | Rpl10a | 0 | Young |
| 0.00014 | 0.297481 | 1 | 0.984 | 1 | 13 | Gmfg | 0.016 | Young |
| 0.000149 | 0.782661 | 0.632 | 0.41 | 1 | 13 | Arhgap39 | 0.222 | Young |
| 0.000153 | 0.454591 | 0.991 | 0.984 | 1 | 13 | Txnip | 0.007 | Young |
| 0.000176 | 0.79377 | 0.579 | 0.328 | 1 | 13 | Serpinf1 | 0.251 | Young |
| 0.000211 | 0.775468 | 0.675 | 0.508 | 1 | 13 | Bri3bp | 0.167 | Young |
| 0.000219 | 0.387954 | 0.965 | 0.811 | 1 | 13 | Lyz1 | 0.154 | Young |
| 0.000223 | 2.046041 | 0.254 | 0.082 | 1 | 13 | Itga6 | 0.172 | Young |
| 0.000237 | 0.252609 | 1 | 1 | 1 | 13 | Rps18 | 0 | Young |
| 0.000252 | 0.82 | 0.632 | 0.451 | 1 | 13 | Mepce | 0.181 | Young |
| 0.000293 | 1.076206 | 0.544 | 0.328 | 1 | 13 | Ang | 0.216 | Young |
| 0.000298 | 0.216297 | 1 | 1 | 1 | 13 | Rpl21 | 0 | Young |
| 0.000305 | 0.19744 | 1 | 1 | 1 | 13 | Rpl12 | 0 | Young |
| 0.000321 | 0.191799 | 1 | 1 | 1 | 13 | Rps6 | 0 | Young |
| 0.000328 | 0.538857 | 0.842 | 0.639 | 1 | 13 | Tnfsf9 | 0.203 | Young |
| 0.000347 | 2.417145 | 0.184 | 0.041 | 1 | 13 | Col14a1 | 0.143 | Young |
| 0.000358 | 0.698328 | 0.719 | 0.541 | 1 | 13 | Bcl2a1a | 0.178 | Young |
| 0.000369 | 0.198401 | 1 | 1 | 1 | 13 | Rplp0 | 0 | Young |
| 0.000404 | 0.412294 | 0.93 | 0.91 | 1 | 13 | Rnf13 | 0.02 | Young |
| 0.000419 | 1.847243 | 0.184 | 0.041 | 1 | 13 | 3222401L1 | 0.143 | Young |
| 0.000464 | 0.472554 | 0.965 | 0.91 | 1 | 13 | Bcl2a1b | 0.055 | Young |
| 0.000499 | 0.155884 | 1 | 1 | 1 | 13 | Rpl35a | 0 | Young |
| 0.000499 | 0.494132 | 0.86 | 0.77 | 1 | 13 | Cebpg | 0.09 | Young |
| 0.000503 | 0.165803 | 1 | 1 | 1 | 13 | Rpl27a | 0 | Young |
| 0.000513 | 0.763448 | 0.544 | 0.328 | 1 | 13 | Gpr183 | 0.216 | Young |
| 0.000539 | 0.520831 | 0.816 | 0.689 | 1 | 13 | Slco2b1 | 0.127 | Young |
| 0.000564 | 0.497807 | 0.781 | 0.623 | 1 | 13 | Cd209a | 0.158 | Young |
| 0.000648 | 0.217406 | 1 | 1 | 1 | 13 | Rpl14 | 0 | Young |
| 0.000735 | 0.445364 | 0.956 | 0.893 | 1 | 13 | Tsc22d3 | 0.063 | Young |
| 0.00075 | 0.586145 | 0.877 | 0.795 | 1 | 13 | Aimp1 | 0.082 | Young |
| 0.000777 | 0.369202 | 0.991 | 0.951 | 1 | 13 | Maf | 0.04 | Young |
| 0.000777 | 0.519204 | 0.728 | 0.59 | 1 | 13 | Phactr2 | 0.138 | Young |
| 0.000785 | 1.051089 | 0.456 | 0.287 | 1 | 13 | Tmem238 | 0.169 | Young |
| 0.000852 | 0.448951 | 0.868 | 0.787 | 1 | 13 | Sft2d2 | 0.081 | Young |
| 0.000856 | 0.504294 | 0.912 | 0.836 | 1 | 13 | Ssh2 | 0.076 | Young |
| 0.000877 | 0.138328 | 0.061 | 0.213 | 1 | 13 | Igkc | -0.152 | Young |

| | | | | | | | | |
|---|---|---|---|---|---|---|---|---|
| 0.00095 | 0.387805 | 0.939 | 0.959 | 1 | 13 | Rtraf | -0.02 | Young |
| 0.001012 | 0.903277 | 0.632 | 0.492 | 1 | 13 | Skint3 | 0.14 | Young |
| 0.00102 | 0.348563 | 0.982 | 0.91 | 1 | 13 | Txn2 | 0.072 | Young |
| 0.001064 | 0.685961 | 0.684 | 0.549 | 1 | 13 | Nsmce4a | 0.135 | Young |
| 0.001157 | 0.294337 | 0.974 | 0.992 | 1 | 13 | Rel | -0.018 | Young |
| 0.001176 | 0.182393 | 1 | 1 | 1 | 13 | Rpl3 | 0 | Young |
| 0.001216 | 0.160672 | 1 | 1 | 1 | 13 | Tyrobp | 0 | Young |
| 0.001224 | 0.158559 | 1 | 1 | 1 | 13 | Rpsa | 0 | Young |
| 0.001242 | 1.776441 | 0.298 | 0.148 | 1 | 13 | Bckdhb | 0.15 | Young |
| 0.001242 | 0.496357 | 0.675 | 0.533 | 1 | 13 | Ncoa2 | 0.142 | Young |
| 0.001249 | 0.176686 | 1 | 1 | 1 | 13 | Eef1b2 | 0 | Young |
| 0.001329 | 1.052222 | 0.404 | 0.221 | 1 | 13 | Dusp8 | 0.183 | Young |
| 0.001362 | 0.231798 | 1 | 1 | 1 | 13 | Rpl15 | 0 | Young |
| 0.001389 | 0.265427 | 1 | 1 | 1 | 13 | Srgn | 0 | Young |
| 0.001426 | 0.30428 | 1 | 0.984 | 1 | 13 | Fnbp1 | 0.016 | Young |
| 0.001446 | 0.578002 | 0.746 | 0.598 | 1 | 13 | Fh1 | 0.148 | Young |
| 0.001474 | 0.290417 | 1 | 1 | 1 | 13 | Smdt1 | 0 | Young |
| 0.001564 | 0.195217 | 1 | 1 | 1 | 13 | Hint1 | 0 | Young |
| 0.001616 | 0.215772 | 1 | 1 | 1 | 13 | Ubb | 0 | Young |
| 0.001622 | 0.276653 | 0.956 | 0.885 | 1 | 13 | Ecm1 | 0.071 | Young |
| 0.001643 | 0.443442 | 0.904 | 0.697 | 1 | 13 | Ddx3y | 0.207 | Young |
| 0.001728 | 0.736717 | 0.64 | 0.492 | 1 | 13 | Arsb | 0.148 | Young |
| 0.001732 | 1.034818 | 0.404 | 0.23 | 1 | 13 | 6-Sep | 0.174 | Young |
| 0.001768 | 0.801506 | 0.596 | 0.475 | 1 | 13 | Ntpcr | 0.121 | Young |
| 0.001809 | 0.595839 | 0.675 | 0.549 | 1 | 13 | Rpp21 | 0.126 | Young |
| 0.001816 | 0.327092 | 0.982 | 0.951 | 1 | 13 | Batf3 | 0.031 | Young |
| 0.001961 | 0.40846 | 0.93 | 0.762 | 1 | 13 | Bcl2a1d | 0.168 | Young |
| 0.001988 | 0.269005 | 1 | 1 | 1 | 13 | Retnla | 0 | Young |
| 0.002008 | 0.300012 | 1 | 1 | 1 | 13 | F13a1 | 0 | Young |
| 0.002014 | 0.162451 | 1 | 1 | 1 | 13 | Kctd12 | 0 | Young |
| 0.002173 | 0.360228 | 0.982 | 0.902 | 1 | 13 | Use1 | 0.08 | Young |
| 0.002315 | 0.545091 | 0.667 | 0.508 | 1 | 13 | Vrk1 | 0.159 | Young |
| 0.002332 | 0.528176 | 0.763 | 0.656 | 1 | 13 | Klf9 | 0.107 | Young |
| 0.002372 | 0.370448 | 0.912 | 0.852 | 1 | 13 | Tlr2 | 0.06 | Young |
| 0.0024 | 1.008498 | 0.456 | 0.303 | 1 | 13 | Padi2 | 0.153 | Young |
| 0.002438 | 0.152547 | 1 | 1 | 1 | 13 | Rpl8 | 0 | Young |
| 0.002488 | 0.954246 | 0.368 | 0.213 | 1 | 13 | Metrn | 0.155 | Young |
| 0.002583 | 0.437752 | 0.825 | 0.639 | 1 | 13 | Elmsan1 | 0.186 | Young |
| 0.002603 | 0.775683 | 0.544 | 0.369 | 1 | 13 | Cxcl12 | 0.175 | Young |
| 0.00263 | 0.394006 | 1 | 0.975 | 1 | 13 | Nr3c1 | 0.025 | Young |
| 0.002697 | 0.127639 | 1 | 1 | 1 | 13 | Rps12 | 0 | Young |
| 0.002765 | 0.199644 | 1 | 1 | 1 | 13 | Alox5ap | 0 | Young |
| 0.002805 | 0.631958 | 0.377 | 0.213 | 1 | 13 | Siglech | 0.164 | Young |
| 0.002833 | 1.58805 | 0.193 | 0.066 | 1 | 13 | Klrd1 | 0.127 | Young |
| 0.002893 | 1.142376 | 0.333 | 0.189 | 1 | 13 | Tle1 | 0.144 | Young |

| | | | | | | | | |
|---|---|---|---|---|---|---|---|---|
| 0.003008 | 2.325039 | 0.07 | 0 | 1 | 13 | Cyr61 | 0.07 | Young |
| 0.003025 | 0.439167 | 0.781 | 0.639 | 1 | 13 | Rere | 0.142 | Young |
| 0.003047 | 0.67272 | 0.693 | 0.574 | 1 | 13 | Egr3 | 0.119 | Young |
| 0.003067 | 0.402556 | 0.789 | 0.754 | 1 | 13 | Msl2 | 0.035 | Young |
| 0.003099 | 0.501314 | 0.728 | 0.59 | 1 | 13 | Kdm4a | 0.138 | Young |
| 0.00317 | 0.129735 | 1 | 1 | 1 | 13 | Rps19 | 0 | Young |
| 0.003228 | 0.325486 | 0.974 | 0.934 | 1 | 13 | Abca9 | 0.04 | Young |
| 0.003351 | 0.131344 | 1 | 1 | 1 | 13 | Eif1 | 0 | Young |
| 0.003434 | 0.265036 | 1 | 0.975 | 1 | 13 | Fam46a | 0.025 | Young |
| 0.003455 | 0.227757 | 1 | 0.992 | 1 | 13 | Lamtor4 | 0.008 | Young |
| 0.003471 | 0.506735 | 0.649 | 0.5 | 1 | 13 | Gpr34 | 0.149 | Young |
| 0.003481 | 0.81823 | 0.404 | 0.246 | 1 | 13 | Map3k3 | 0.158 | Young |
| 0.003752 | 0.342845 | 0.991 | 0.959 | 1 | 13 | Pf4 | 0.032 | Young |
| 0.003777 | 0.374922 | 0.939 | 0.82 | 1 | 13 | Tgif1 | 0.119 | Young |
| 0.003942 | 0.544169 | 0.763 | 0.672 | 1 | 13 | Setd3 | 0.091 | Young |
| 0.003945 | 0.444007 | 0.886 | 0.738 | 1 | 13 | Cd163 | 0.148 | Young |
| 0.003946 | 1.742098 | 0.105 | 0.016 | 1 | 13 | Tmem138 | 0.089 | Young |
| 0.003982 | 0.978661 | 0.351 | 0.197 | 1 | 13 | Tmod1 | 0.154 | Young |
| 0.003985 | 0.836447 | 0.307 | 0.148 | 1 | 13 | Pkdcc | 0.159 | Young |
| 0.004002 | 0.982795 | 0.316 | 0.156 | 1 | 13 | Gm38843 | 0.16 | Young |
| 0.004009 | 0.31154 | 0.991 | 0.91 | 1 | 13 | Ssr4 | 0.081 | Young |
| 0.004053 | 0.753487 | 0.342 | 0.189 | 1 | 13 | Mcemp1 | 0.153 | Young |
| 0.004146 | 0.242033 | 1 | 0.984 | 1 | 13 | Gadd45b | 0.016 | Young |
| 0.004217 | 0.593286 | 0.64 | 0.533 | 1 | 13 | Ccm2 | 0.107 | Young |
| 0.004296 | 0.494265 | 0.939 | 0.836 | 1 | 13 | Unc119 | 0.103 | Young |
| 0.004377 | 0.273663 | 0.974 | 0.984 | 1 | 13 | Ogt | -0.01 | Young |
| 0.004483 | 0.261888 | 1 | 0.992 | 1 | 13 | Polr1d | 0.008 | Young |
| 0.0045 | 0.462173 | 0.754 | 0.689 | 1 | 13 | Dynll2 | 0.065 | Young |
| 0.004565 | 0.796342 | 0.412 | 0.246 | 1 | 13 | Arl10 | 0.166 | Young |
| 0.004618 | 0.656777 | 0.605 | 0.475 | 1 | 13 | Ccdc9 | 0.13 | Young |
| 0.004629 | 0.393889 | 0.798 | 0.705 | 1 | 13 | Cklf | 0.093 | Young |
| 0.00465 | 0.319983 | 0.912 | 0.877 | 1 | 13 | Arhgap45 | 0.035 | Young |
| 0.004731 | 0.120565 | 1 | 1 | 1 | 13 | Rps8 | 0 | Young |
| 0.004746 | 0.507155 | 0.798 | 0.721 | 1 | 13 | Krt80 | 0.077 | Young |
| 0.004837 | 0.508954 | 0.605 | 0.467 | 1 | 13 | Il13ra1 | 0.138 | Young |
| 0.004902 | 0.0921 | 1 | 1 | 1 | 13 | Rps29 | 0 | Young |
| 0.00502 | 0.162724 | 1 | 1 | 1 | 13 | Rpl10 | 0 | Young |
| 0.00514 | 0.225053 | 1 | 1 | 1 | 13 | H2afz | 0 | Young |
| 0.005272 | 0.734507 | 0.518 | 0.393 | 1 | 13 | Cib2 | 0.125 | Young |
| 0.005325 | 0.14469 | 1 | 1 | 1 | 13 | Rpl5 | 0 | Young |
| 0.005357 | 0.229077 | 1 | 1 | 1 | 13 | Tgfbi | 0 | Young |
| 0.005404 | 0.242438 | 0.991 | 0.959 | 1 | 13 | Lamtor2 | 0.032 | Young |
| 0.005491 | 0.705692 | 0.535 | 0.402 | 1 | 13 | Selenbp1 | 0.133 | Young |
| 0.00552 | 0.450961 | 0.711 | 0.557 | 1 | 13 | Tanc2 | 0.154 | Young |
| 0.005532 | 0.21858 | 0.991 | 0.975 | 1 | 13 | Slamf9 | 0.016 | Young |

| | | | | | | | | |
|---|---|---|---|---|---|---|---|---|
| 0.005541 | 1.405132 | 0.158 | 0.049 | 1 | 13 | 1810044D( | 0.109 | Young |
| 0.005549 | 0.501322 | 0.544 | 0.377 | 1 | 13 | Ppp4r1 | 0.167 | Young |
| 0.005623 | 1.998656 | 0.061 | 0 | 1 | 13 | Col1a2 | 0.061 | Young |
| 0.005623 | 1.841215 | 0.061 | 0 | 1 | 13 | Mybpc3 | 0.061 | Young |
| 0.005746 | 0.377058 | 0.851 | 0.811 | 1 | 13 | Smim14 | 0.04 | Young |
| 0.005747 | 0.212315 | 0.991 | 1 | 1 | 13 | Eif3k | -0.009 | Young |
| 0.005764 | 0.401522 | 0.851 | 0.738 | 1 | 13 | Clk4 | 0.113 | Young |
| 0.005776 | 0.328336 | 0.965 | 0.918 | 1 | 13 | Krtcap2 | 0.047 | Young |
| 0.005781 | 0.588466 | 0.491 | 0.303 | 1 | 13 | Dcxr | 0.188 | Young |
| 0.005787 | 0.357089 | 0.921 | 0.926 | 1 | 13 | Mpp6 | -0.005 | Young |
| 0.005844 | 1.209766 | 0.272 | 0.131 | 1 | 13 | AC163018. | 0.141 | Young |
| 0.005904 | 0.443854 | 0.667 | 0.525 | 1 | 13 | Smagp | 0.142 | Young |
| 0.00591 | 0.458281 | 0.693 | 0.557 | 1 | 13 | Nuak2 | 0.136 | Young |
| 0.005923 | 0.560973 | 0.667 | 0.533 | 1 | 13 | Nfxl1 | 0.134 | Young |
| 0.00602 | 0.742288 | 0.491 | 0.328 | 1 | 13 | Tagap | 0.163 | Young |
| 0.006067 | 0.52018 | 0.632 | 0.475 | 1 | 13 | Dpf2 | 0.157 | Young |
| 0.006262 | 0.566249 | 0.57 | 0.451 | 1 | 13 | Mdp1 | 0.119 | Young |
| 0.006297 | 1.335182 | 0.184 | 0.066 | 1 | 13 | Lix1 | 0.118 | Young |
| 0.006308 | 0.177528 | 1 | 1 | 1 | 13 | Srrm2 | 0 | Young |
| 0.006352 | 2.138023 | 0.096 | 0.016 | 1 | 13 | Siglecf | 0.08 | Young |
| 0.00637 | 0.384372 | 0.754 | 0.631 | 1 | 13 | Slc9a3r1 | 0.123 | Young |
| 0.006474 | 0.252311 | 0.974 | 0.992 | 1 | 13 | Rgs10 | -0.018 | Young |
| 0.006638 | 1.007707 | 0.281 | 0.139 | 1 | 13 | Nudt22 | 0.142 | Young |
| 0.00676 | 0.216915 | 1 | 0.975 | 1 | 13 | Kdm6b | 0.025 | Young |
| 0.006807 | 1.994309 | 0.079 | 0.008 | 1 | 13 | Prtn3 | 0.071 | Young |
| 0.00687 | 0.375918 | 0.842 | 0.828 | 1 | 13 | Zc3hav1 | 0.014 | Young |
| 0.00692 | 1.784447 | 0.079 | 0.008 | 1 | 13 | Slc46a1 | 0.071 | Young |
| 0.007024 | 0.791779 | 0.474 | 0.328 | 1 | 13 | Plxna4 | 0.146 | Young |
| 0.007136 | 0.358543 | 0.895 | 0.918 | 1 | 13 | Zfhx3 | -0.023 | Young |
| 0.007173 | 1.052075 | 0.237 | 0.107 | 1 | 13 | Polr3a | 0.13 | Young |
| 0.007208 | 0.38244 | 0.974 | 0.877 | 1 | 13 | Igfbp4 | 0.097 | Young |
| 0.007216 | 0.701101 | 0.342 | 0.189 | 1 | 13 | Lpcat2 | 0.153 | Young |
| 0.007303 | 1.121004 | 0.298 | 0.164 | 1 | 13 | Mtus1 | 0.134 | Young |
| 0.007496 | 0.361057 | 0.895 | 0.885 | 1 | 13 | Rex1bd | 0.01 | Young |
| 0.007596 | 0.256902 | 0.956 | 0.943 | 1 | 13 | Slk | 0.013 | Young |
| 0.007645 | 0.917284 | 0.412 | 0.279 | 1 | 13 | AI839979 | 0.133 | Young |
| 0.007675 | 0.401006 | 0.614 | 0.443 | 1 | 13 | Itga5 | 0.171 | Young |
| 0.007741 | 0.301369 | 0.93 | 0.926 | 1 | 13 | Cggbp1 | 0.004 | Young |
| 0.007763 | 0.855628 | 0.289 | 0.148 | 1 | 13 | Abcc3 | 0.141 | Young |
| 0.00778 | 1.633398 | 0.096 | 0.016 | 1 | 13 | Ch25h | 0.08 | Young |
| 0.007831 | 0.519817 | 0.658 | 0.533 | 1 | 13 | Dock1 | 0.125 | Young |
| 0.007975 | 0.138148 | 1 | 0.984 | 1 | 13 | Prdx5 | 0.016 | Young |
| 0.008045 | 0.329858 | 0.789 | 0.705 | 1 | 13 | Psd3 | 0.084 | Young |
| 0.008192 | 0.631132 | 0.64 | 0.516 | 1 | 13 | Adrb2 | 0.124 | Young |
| 0.008235 | 0.68836 | 0.553 | 0.467 | 1 | 13 | Fam96b | 0.086 | Young |

| | | | | | | | | |
|---|---|---|---|---|---|---|---|---|
| 0.00847 | 0.561949 | 0.544 | 0.393 | 1 | 13 | Senp1 | 0.151 | Young |
| 0.008589 | 0.330448 | 0.807 | 0.631 | 1 | 13 | Alox5 | 0.176 | Young |
| 0.008702 | 0.180793 | 0.991 | 0.992 | 1 | 13 | Man2b1 | -0.001 | Young |
| 0.008782 | 0.789276 | 0.43 | 0.303 | 1 | 13 | Gpaa1 | 0.127 | Young |
| 0.008939 | 0.842422 | 0.263 | 0.131 | 1 | 13 | Bcl11a | 0.132 | Young |
| 0.009024 | 0.246557 | 0.982 | 0.992 | 1 | 13 | Ndufb10 | -0.01 | Young |
| 0.009026 | 0.678661 | 0.623 | 0.484 | 1 | 13 | Upb1 | 0.139 | Young |
| 0.009134 | 0.902047 | 0.342 | 0.213 | 1 | 13 | Fam214a | 0.129 | Young |
| 0.009293 | 0.306219 | 0.842 | 0.779 | 1 | 13 | Ramp1 | 0.063 | Young |
| 0.009747 | 0.878508 | 0.184 | 0.066 | 1 | 13 | Colec12 | 0.118 | Young |
| 0.010251 | 0.237299 | 1 | 0.992 | 1 | 13 | S100a13 | 0.008 | Young |
| 0.010307 | 0.10144 | 1 | 1 | 1 | 13 | Fth1 | 0 | Young |
| 0.010314 | 1.591965 | 0.105 | 0.025 | 1 | 13 | Xpo5 | 0.08 | Young |
| 0.01035 | 0.413434 | 0.921 | 0.836 | 1 | 13 | Hexb | 0.085 | Young |
| 0.010537 | 0.428031 | 0.807 | 0.705 | 1 | 13 | Cd93 | 0.102 | Young |
| 0.01054 | 1.736919 | 0.053 | 0 | 1 | 13 | Tspan2 | 0.053 | Young |
| 0.01054 | 1.661863 | 0.053 | 0 | 1 | 13 | Amigo3 | 0.053 | Young |
| 0.01057 | 0.506102 | 0.746 | 0.689 | 1 | 13 | Rin3 | 0.057 | Young |
| 0.010595 | 1.523359 | 0.202 | 0.09 | 1 | 13 | Gm30978 | 0.112 | Young |
| 0.010653 | 0.225723 | 0.991 | 1 | 1 | 13 | Fcgr2b | -0.009 | Young |
| 0.010779 | 1.132088 | 0.246 | 0.123 | 1 | 13 | Ttll3 | 0.123 | Young |
| 0.011038 | 1.213028 | 0.158 | 0.057 | 1 | 13 | Stbd1 | 0.101 | Young |
| 0.01111 | 0.488706 | 0.754 | 0.68 | 1 | 13 | Stxbp6 | 0.074 | Young |
| 0.011147 | 1.989301 | 0.105 | 0.025 | 1 | 13 | F11r | 0.08 | Young |
| 0.011323 | 0.651539 | 0.509 | 0.385 | 1 | 13 | Dgcr2 | 0.124 | Young |
| 0.011498 | 0.19729 | 1 | 0.992 | 1 | 13 | H2afj | 0.008 | Young |
| 0.011513 | 0.558081 | 0.491 | 0.344 | 1 | 13 | Timm50 | 0.147 | Young |
| 0.011644 | 1.129976 | 0.123 | 0.033 | 1 | 13 | Atp2b4 | 0.09 | Young |
| 0.011783 | 1.708629 | 0.088 | 0.016 | 1 | 13 | 5830416I1! | 0.072 | Young |
| 0.011909 | 1.045597 | 0.254 | 0.131 | 1 | 13 | Slc27a3 | 0.123 | Young |
| 0.011943 | 0.188564 | 0.991 | 1 | 1 | 13 | Rbm3 | -0.009 | Young |
| 0.011943 | 0.122646 | 1 | 1 | 1 | 13 | Oaz1 | 0 | Young |
| 0.011951 | 1.710357 | 0.088 | 0.016 | 1 | 13 | Gdf9 | 0.072 | Young |
| 0.011967 | 0.581146 | 0.491 | 0.377 | 1 | 13 | Ate1 | 0.114 | Young |
| 0.011972 | 0.406078 | 0.842 | 0.754 | 1 | 13 | Rtf1 | 0.088 | Young |
| 0.012023 | 0.34601 | 0.816 | 0.762 | 1 | 13 | Ccni | 0.054 | Young |
| 0.012073 | 0.126219 | 1 | 1 | 1 | 13 | Rpl24 | 0 | Young |
| 0.012436 | 0.018401 | 0.079 | 0.205 | 1 | 13 | Rsad2 | -0.126 | Young |
| 0.01247 | 0.13932 | 1 | 1 | 1 | 13 | Fcer1g | 0 | Young |
| 0.012472 | 1.716105 | 0.07 | 0.008 | 1 | 13 | Lrrc1 | 0.062 | Young |
| 0.012583 | 0.45503 | 0.675 | 0.615 | 1 | 13 | Trappc5 | 0.06 | Young |
| 0.012721 | 1.190534 | 0.158 | 0.057 | 1 | 13 | Ctla2b | 0.101 | Young |
| 0.012828 | 0.528741 | 0.57 | 0.434 | 1 | 13 | Tmem243 | 0.136 | Young |
| 0.012862 | 0.248824 | 0.702 | 0.557 | 1 | 13 | Trem2 | 0.145 | Young |
| 0.012868 | 1.188058 | 0.167 | 0.066 | 1 | 13 | Fcor | 0.101 | Young |

| | | | | | | | | |
|---|---|---|---|---|---|---|---|---|
| 0.012925 | 0.374985 | 0.851 | 0.762 | 1 | 13 | Cnpy2 | 0.089 | Young |
| 0.013092 | 1.625813 | 0.07 | 0.008 | 1 | 13 | Nek3 | 0.062 | Young |
| 0.013092 | 1.617421 | 0.07 | 0.008 | 1 | 13 | P3h3 | 0.062 | Young |
| 0.013586 | 0.259366 | 0.982 | 0.984 | 1 | 13 | Wasf2 | -0.002 | Young |
| 0.013768 | 0.199212 | 0.991 | 0.992 | 1 | 13 | Jun | -0.001 | Young |
| 0.01379 | 0.619985 | 0.474 | 0.328 | 1 | 13 | Cyth2 | 0.146 | Young |
| 0.013913 | 0.269736 | 0.974 | 0.959 | 1 | 13 | Mycbp2 | 0.015 | Young |
| 0.013952 | 0.173862 | 1 | 0.992 | 1 | 13 | Fcrls | 0.008 | Young |
| 0.014027 | 0.132032 | 1 | 1 | 1 | 13 | Rps27a | 0 | Young |
| 0.014252 | 0.140562 | 1 | 1 | 1 | 13 | Cox4i1 | 0 | Young |
| 0.014272 | 0.6036 | 0.518 | 0.385 | 1 | 13 | Mcee | 0.133 | Young |
| 0.014321 | 0.308167 | 0.947 | 0.951 | 1 | 13 | Blvrb | -0.004 | Young |
| 0.014399 | 0.381779 | 0.781 | 0.73 | 1 | 13 | Tnfsf12 | 0.051 | Young |
| 0.014537 | 0.488456 | 0.009 | 0.074 | 1 | 13 | Jchain | -0.065 | Young |
| 0.014546 | 0.259185 | 0.965 | 0.918 | 1 | 13 | Spop | 0.047 | Young |
| 0.014975 | 0.318729 | 0.754 | 0.615 | 1 | 13 | H2-DMb2 | 0.139 | Young |
| 0.014993 | 0.264521 | 0.886 | 0.828 | 1 | 13 | Gpr65 | 0.058 | Young |
| 0.015025 | 0.220758 | 1 | 0.992 | 1 | 13 | Psma7 | 0.008 | Young |
| 0.015101 | 0.239531 | 0.991 | 0.934 | 1 | 13 | Glg1 | 0.057 | Young |
| 0.015207 | 1.026597 | 0.175 | 0.074 | 1 | 13 | Katnal1 | 0.101 | Young |
| 0.015264 | 0.088765 | 1 | 1 | 1 | 13 | Tmsb4x | 0 | Young |
| 0.015297 | 0.589969 | 0.553 | 0.451 | 1 | 13 | Dnajb4 | 0.102 | Young |
| 0.015343 | 0.223077 | 0.982 | 0.959 | 1 | 13 | Tmcc1 | 0.023 | Young |
| 0.015425 | 0.098829 | 1 | 1 | 1 | 13 | Rplp2 | 0 | Young |
| 0.015444 | 0.547921 | 0.325 | 0.189 | 1 | 13 | Dnaja3 | 0.136 | Young |
| 0.015777 | 1.108539 | 0.175 | 0.074 | 1 | 13 | 2900093K2 | 0.101 | Young |
| 0.015835 | 0.209161 | 1 | 1 | 1 | 13 | Srp9 | 0 | Young |
| 0.015844 | 0.506168 | 0.693 | 0.59 | 1 | 13 | 4930523C0 | 0.103 | Young |
| 0.016001 | 0.117007 | 1 | 1 | 1 | 13 | Rpl36 | 0 | Young |
| 0.016006 | 0.4194 | 0.789 | 0.697 | 1 | 13 | Il1rl1 | 0.092 | Young |
| 0.016085 | 0.219821 | 1 | 0.992 | 1 | 13 | Ndufa6 | 0.008 | Young |
| 0.016098 | 0.861098 | 0.386 | 0.246 | 1 | 13 | Dbt | 0.14 | Young |
| 0.016241 | 0.513845 | 0.579 | 0.443 | 1 | 13 | Rxra | 0.136 | Young |
| 0.016291 | 0.325886 | 0.877 | 0.852 | 1 | 13 | Top2b | 0.025 | Young |
| 0.016652 | 0.393682 | 0.851 | 0.77 | 1 | 13 | Ypel3 | 0.081 | Young |
| 0.016878 | 0.355076 | 0.772 | 0.672 | 1 | 13 | Snx8 | 0.1 | Young |
| 0.016931 | 1.257188 | 0.237 | 0.131 | 1 | 13 | Il1rl2 | 0.106 | Young |
| 0.017027 | 0.875445 | 0.298 | 0.172 | 1 | 13 | Dnase1l3 | 0.126 | Young |
| 0.017183 | 0.424325 | 0.816 | 0.754 | 1 | 13 | Arl5b | 0.062 | Young |
| 0.017252 | 0.384384 | 0.719 | 0.607 | 1 | 13 | Gabarapl1 | 0.112 | Young |
| 0.01727 | 0.206303 | 0.904 | 0.877 | 1 | 13 | Fcho2 | 0.027 | Young |
| 0.0173 | 0.199554 | 0.991 | 1 | 1 | 13 | Mrfap1 | -0.009 | Young |
| 0.017559 | 0.407818 | 0.702 | 0.664 | 1 | 13 | Mrps6 | 0.038 | Young |
| 0.017596 | 0.538888 | 0.447 | 0.32 | 1 | 13 | Rce1 | 0.127 | Young |
| 0.017753 | 0.183984 | 0.982 | 0.992 | 1 | 13 | Tacc1 | -0.01 | Young |

| | | | | | | | | |
|---|---|---|---|---|---|---|---|---|
| 0.01803 | 0.203601 | 1 | 0.967 | 1 | 13 | Il1b | 0.033 | Young |
| 0.018058 | 0.350262 | 0.868 | 0.828 | 1 | 13 | Arhgap31 | 0.04 | Young |
| 0.018094 | 0.558217 | 0.447 | 0.287 | 1 | 13 | Ivd | 0.16 | Young |
| 0.018217 | 0.114374 | 1 | 1 | 1 | 13 | Serf2 | 0 | Young |
| 0.018254 | 0.440035 | 0.789 | 0.656 | 1 | 13 | Ppp1r11 | 0.133 | Young |
| 0.018366 | 1.463324 | 0.096 | 0.025 | 1 | 13 | Fam13a | 0.071 | Young |
| 0.018583 | 0.353433 | 0.947 | 0.934 | 1 | 13 | Eps15 | 0.013 | Young |
| 0.018668 | 0.310085 | 0.833 | 0.82 | 1 | 13 | Smap1 | 0.013 | Young |
| 0.018684 | 0.661666 | 0.289 | 0.164 | 1 | 13 | Cdkn2aip | 0.125 | Young |
| 0.018944 | 0.647823 | 0.368 | 0.246 | 1 | 13 | Tspan13 | 0.122 | Young |
| 0.019318 | 0.627456 | 0.43 | 0.303 | 1 | 13 | Plpp2 | 0.127 | Young |
| 0.019333 | 1.189686 | 0.149 | 0.057 | 1 | 13 | Isg20 | 0.092 | Young |
| 0.019372 | 0.252285 | 1 | 0.984 | 1 | 13 | Cdkn1a | 0.016 | Young |
| 0.019372 | 0.202476 | 0.991 | 1 | 1 | 13 | Qk | -0.009 | Young |
| 0.019847 | 0.380797 | 0.895 | 0.852 | 1 | 13 | Anapc13 | 0.043 | Young |
| 0.019852 | 1.875494 | 0.044 | 0 | 1 | 13 | Fan1 | 0.044 | Young |
| 0.019852 | 1.777706 | 0.044 | 0 | 1 | 13 | Wdr19 | 0.044 | Young |
| 0.019852 | 1.560492 | 0.044 | 0 | 1 | 13 | 9830107B1 | 0.044 | Young |
| 0.019852 | 1.509535 | 0.044 | 0 | 1 | 13 | Treml4 | 0.044 | Young |
| 0.019852 | 1.459851 | 0.044 | 0 | 1 | 13 | Mutyh | 0.044 | Young |
| 0.019852 | 1.442213 | 0.044 | 0 | 1 | 13 | Ino80dos | 0.044 | Young |
| 0.019852 | 1.289736 | 0.044 | 0 | 1 | 13 | Gm17092 | 0.044 | Young |
| 0.019852 | 1.260562 | 0.044 | 0 | 1 | 13 | Fgf9 | 0.044 | Young |
| 0.01987 | 0.534112 | 0.719 | 0.689 | 1 | 13 | Gm26740 | 0.03 | Young |
| 0.0199 | 0.363665 | 0.772 | 0.631 | 1 | 13 | Ltb4r1 | 0.141 | Young |
| 0.019903 | 1.070513 | 0.228 | 0.115 | 1 | 13 | Slc25a33 | 0.113 | Young |
| 0.019904 | 0.345767 | 0.886 | 0.754 | 1 | 13 | Psmg4 | 0.132 | Young |
| 0.019974 | 0.215363 | 1 | 1 | 1 | 13 | Ptpn18 | 0 | Young |
| 0.020113 | 1.074635 | 0.193 | 0.09 | 1 | 13 | Uaca | 0.103 | Young |
| 0.020171 | 1.005483 | 0.211 | 0.107 | 1 | 13 | Asgr2 | 0.104 | Young |
| 0.020212 | 0.482073 | 0.64 | 0.508 | 1 | 13 | Rnase6 | 0.132 | Young |
| 0.020449 | 0.428056 | 0.737 | 0.656 | 1 | 13 | Cttnbp2nl | 0.081 | Young |
| 0.020488 | 0.109583 | 1 | 1 | 1 | 13 | Sh3bgrl3 | 0 | Young |
| 0.020498 | 0.553175 | 0.649 | 0.607 | 1 | 13 | Gdi1 | 0.042 | Young |
| 0.020573 | 0.758765 | 0.36 | 0.238 | 1 | 13 | Sh2b1 | 0.122 | Young |
| 0.020743 | 0.731507 | 0.377 | 0.246 | 1 | 13 | Rev1 | 0.131 | Young |
| 0.021007 | 0.29593 | 0.763 | 0.664 | 1 | 13 | Slc9a9 | 0.099 | Young |
| 0.021013 | 0.173729 | 0.974 | 1 | 1 | 13 | Bzw1 | -0.026 | Young |
| 0.021333 | 0.114512 | 1 | 1 | 1 | 13 | Rps25 | 0 | Young |
| 0.021494 | 0.236058 | 0.825 | 0.648 | 1 | 13 | Qpct | 0.177 | Young |
| 0.021543 | 0.272691 | 0.965 | 0.943 | 1 | 13 | Ets2 | 0.022 | Young |
| 0.021816 | 0.16724 | 0.93 | 0.877 | 1 | 13 | Zcchc11 | 0.053 | Young |
| 0.021906 | 0.443685 | 0.711 | 0.623 | 1 | 13 | Heca | 0.088 | Young |
| 0.021995 | 0.315665 | 0.772 | 0.705 | 1 | 13 | Id3 | 0.067 | Young |
| 0.022043 | 0.258958 | 0.895 | 0.836 | 1 | 13 | Ahcyl2 | 0.059 | Young |

| | | | | | | | | |
|---|---|---|---|---|---|---|---|---|
| 0.022411 | 0.951193 | 0.237 | 0.131 | 1 | 13 | Nop2 | 0.106 | Young |
| 0.022464 | 0.364666 | 0.904 | 0.844 | 1 | 13 | Arid4a | 0.06 | Young |
| 0.022634 | 0.472875 | 0.632 | 0.549 | 1 | 13 | Acss1 | 0.083 | Young |
| 0.022723 | 0.823281 | 0.325 | 0.213 | 1 | 13 | Pstk | 0.112 | Young |
| 0.022837 | 1.081566 | 0.202 | 0.098 | 1 | 13 | Afap1l1 | 0.104 | Young |
| 0.02304 | 0.415131 | 0.588 | 0.475 | 1 | 13 | Stx5a | 0.113 | Young |
| 0.023175 | 0.345598 | 0.798 | 0.656 | 1 | 13 | Lyve1 | 0.142 | Young |
| 0.023228 | 0.516479 | 0.561 | 0.475 | 1 | 13 | Tmem8 | 0.086 | Young |
| 0.023317 | 0.91379 | 0.307 | 0.18 | 1 | 13 | Mir155hg | 0.127 | Young |
| 0.023464 | 0.546625 | 0.404 | 0.27 | 1 | 13 | Phactr4 | 0.134 | Young |
| 0.023544 | 1.286735 | 0.079 | 0.016 | 1 | 13 | Rapsn | 0.063 | Young |
| 0.023546 | 0.272052 | 0.807 | 0.697 | 1 | 13 | Nlrp3 | 0.11 | Young |
| 0.023619 | 0.211519 | 0.939 | 0.926 | 1 | 13 | Hnrnpul2 | 0.013 | Young |
| 0.023889 | 0.868599 | 0.158 | 0.066 | 1 | 13 | Pxdc1 | 0.092 | Young |
| 0.024111 | 0.462325 | 0.535 | 0.443 | 1 | 13 | Nrf1 | 0.092 | Young |
| 0.024175 | 0.164282 | 1 | 0.967 | 1 | 13 | Mgl2 | 0.033 | Young |
| 0.024333 | 1.640825 | 0.061 | 0.008 | 1 | 13 | Ctla2a | 0.053 | Young |
| 0.024377 | 0.410994 | 0.807 | 0.754 | 1 | 13 | Glrx2 | 0.053 | Young |
| 0.024637 | 0.925443 | 0.289 | 0.18 | 1 | 13 | Rps6ka2 | 0.109 | Young |
| 0.024883 | 0.688373 | 0.342 | 0.238 | 1 | 13 | 1110065P2 | 0.104 | Young |
| 0.025156 | 1.054622 | 0.167 | 0.074 | 1 | 13 | Rwdd2a | 0.093 | Young |
| 0.025276 | 0.172883 | 0.991 | 0.984 | 1 | 13 | Rassf4 | 0.007 | Young |
| 0.025324 | 0.240314 | 0.93 | 0.869 | 1 | 13 | Zdhhc20 | 0.061 | Young |
| 0.025522 | 0.195836 | 0.982 | 0.943 | 1 | 13 | Ncf1 | 0.039 | Young |
| 0.025825 | 0.259634 | 0.904 | 0.77 | 1 | 13 | Sec14l1 | 0.134 | Young |
| 0.025915 | 1.153086 | 0.061 | 0.008 | 1 | 13 | Amot | 0.053 | Young |
| 0.026264 | 0.649327 | 0.342 | 0.23 | 1 | 13 | Timm29 | 0.112 | Young |
| 0.026376 | 0.278328 | 0.456 | 0.303 | 1 | 13 | Areg | 0.153 | Young |
| 0.026636 | 1.219613 | 0.254 | 0.156 | 1 | 13 | Mefv | 0.098 | Young |
| 0.026659 | 0.670134 | 0.36 | 0.238 | 1 | 13 | N4bp2 | 0.122 | Young |
| 0.026787 | 0.261285 | 0.93 | 0.934 | 1 | 13 | Aldh2 | -0.004 | Young |
| 0.026789 | 0.641104 | 0.447 | 0.32 | 1 | 13 | Grap | 0.127 | Young |
| 0.026992 | 0.183997 | 0.956 | 0.926 | 1 | 13 | Gng2 | 0.03 | Young |
| 0.027205 | 0.194906 | 0.965 | 0.975 | 1 | 13 | Srp14 | -0.01 | Young |
| 0.027608 | 0.162242 | 1 | 1 | 1 | 13 | H2-DMa | 0 | Young |
| 0.028014 | 0.078017 | 1 | 1 | 1 | 13 | Rps21 | 0 | Young |
| 0.028122 | 0.554563 | 0.412 | 0.295 | 1 | 13 | Chst12 | 0.117 | Young |
| 0.028205 | 0.236039 | 0.93 | 0.869 | 1 | 13 | Tmbim1 | 0.061 | Young |
| 0.028488 | 1.212971 | 0.158 | 0.074 | 1 | 13 | Ccdc170 | 0.084 | Young |
| 0.028624 | 0.386135 | 0.728 | 0.68 | 1 | 13 | Mrps12 | 0.048 | Young |
| 0.029125 | 0.593924 | 0.307 | 0.18 | 1 | 13 | Aff3 | 0.127 | Young |
| 0.029146 | 0.480624 | 0.544 | 0.434 | 1 | 13 | F630028O1 | 0.11 | Young |
| 0.029403 | 0.632606 | 0.325 | 0.197 | 1 | 13 | Eepd1 | 0.128 | Young |
| 0.029737 | 0.231449 | 0.465 | 0.328 | 1 | 13 | Slc15a2 | 0.137 | Young |
| 0.029749 | 0.669343 | 0.368 | 0.246 | 1 | 13 | D730003I1 | 0.122 | Young |

| | | | | | | | | |
|---|---|---|---|---|---|---|---|---|
| 0.029798 | 0.955466 | 0.237 | 0.131 | 1 | 13 | Peli2 | 0.106 | Young |
| 0.029965 | 0.218229 | 0.974 | 0.918 | 1 | 13 | Ctdsp2 | 0.056 | Young |
| 0.029975 | 0.512592 | 0.518 | 0.393 | 1 | 13 | Clec4n | 0.125 | Young |
| 0.029978 | 0.203919 | 0.991 | 0.984 | 1 | 13 | Elob | 0.007 | Young |
| 0.030065 | 1.480618 | 0.088 | 0.025 | 1 | 13 | Gins1 | 0.063 | Young |
| 0.030182 | 0.947748 | 0.175 | 0.082 | 1 | 13 | Rasgrf2 | 0.093 | Young |
| 0.030278 | 1.153824 | 0.149 | 0.066 | 1 | 13 | Fahd2a | 0.083 | Young |
| 0.030586 | 0.806162 | 0.351 | 0.254 | 1 | 13 | Rab33b | 0.097 | Young |
| 0.030806 | 0.54851 | 0.465 | 0.336 | 1 | 13 | Casd1 | 0.129 | Young |
| 0.030938 | 0.560628 | 0.43 | 0.303 | 1 | 13 | Tigd2 | 0.127 | Young |
| 0.031105 | 1.137905 | 0.149 | 0.066 | 1 | 13 | Zfp961 | 0.083 | Young |
| 0.031312 | 0.701513 | 0.412 | 0.295 | 1 | 13 | Unc119b | 0.117 | Young |
| 0.031406 | 0.734276 | 0.465 | 0.377 | 1 | 13 | Zfp704 | 0.088 | Young |
| 0.031412 | 0.969402 | 0.307 | 0.197 | 1 | 13 | Tbc1d10a | 0.11 | Young |
| 0.031425 | 0.66038 | 0.491 | 0.418 | 1 | 13 | Gm37494 | 0.073 | Young |
| 0.031561 | 1.446281 | 0.088 | 0.025 | 1 | 13 | Zfp12 | 0.063 | Young |
| 0.031886 | 0.387906 | 0.737 | 0.68 | 1 | 13 | Map2k1 | 0.057 | Young |
| 0.032064 | 0.565327 | 0.474 | 0.336 | 1 | 13 | Kank3 | 0.138 | Young |
| 0.032515 | 0.229825 | 0.982 | 0.934 | 1 | 13 | Nfkbid | 0.048 | Young |
| 0.032724 | 1.309268 | 0.088 | 0.025 | 1 | 13 | 1700030KC | 0.063 | Young |
| 0.032819 | 1.053334 | 0.149 | 0.066 | 1 | 13 | Sparc | 0.083 | Young |
| 0.03283 | 0.198266 | 0.974 | 0.967 | 1 | 13 | Ralbp1 | 0.007 | Young |
| 0.032971 | 0.247806 | 0.947 | 0.934 | 1 | 13 | Hacd4 | 0.013 | Young |
| 0.033028 | 0.683666 | 0.43 | 0.311 | 1 | 13 | Prkar2b | 0.119 | Young |
| 0.03312 | 1.298056 | 0.088 | 0.025 | 1 | 13 | Pcbp3 | 0.063 | Young |
| 0.033269 | 0.496492 | 0.544 | 0.402 | 1 | 13 | Cass4 | 0.142 | Young |
| 0.033466 | 0.667803 | 0.289 | 0.172 | 1 | 13 | Ifitm1 | 0.117 | Young |
| 0.033563 | 0.979618 | 0.219 | 0.123 | 1 | 13 | Fmnl3 | 0.096 | Young |
| 0.033595 | 0.29801 | 0.868 | 0.844 | 1 | 13 | B230219D2 | 0.024 | Young |
| 0.033675 | 0.530981 | 0.439 | 0.328 | 1 | 13 | Kmo | 0.111 | Young |
| 0.033677 | 0.578441 | 0.5 | 0.393 | 1 | 13 | Abcd2 | 0.107 | Young |
| 0.033688 | 0.493721 | 0.456 | 0.32 | 1 | 13 | Frmd6 | 0.136 | Young |
| 0.033944 | 0.378425 | 0.711 | 0.656 | 1 | 13 | Apc | 0.055 | Young |
| 0.034192 | 0.595865 | 0.447 | 0.361 | 1 | 13 | Rnf167 | 0.086 | Young |
| 0.034248 | 0.543186 | 0.386 | 0.27 | 1 | 13 | Kdm5d | 0.116 | Young |
| 0.034321 | 0.362181 | 0.772 | 0.68 | 1 | 13 | Npm3 | 0.092 | Young |
| 0.034417 | 0.273627 | 0.965 | 0.943 | 1 | 13 | Cmtm6 | 0.022 | Young |
| 0.034453 | 0.145624 | 0.939 | 0.893 | 1 | 13 | Cd86 | 0.046 | Young |
| 0.034697 | 0.778762 | 0.377 | 0.287 | 1 | 13 | Tgfbr1 | 0.09 | Young |
| 0.034737 | 1.218814 | 0.123 | 0.049 | 1 | 13 | Acad11 | 0.074 | Young |
| 0.035081 | 0.252949 | 1 | 0.967 | 1 | 13 | Fcgrt | 0.033 | Young |
| 0.035474 | 0.304699 | 0.623 | 0.467 | 1 | 13 | Itgax | 0.156 | Young |
| 0.035558 | 1.11343 | 0.114 | 0.041 | 1 | 13 | Gm26778 | 0.073 | Young |
| 0.03558 | 0.182039 | 1 | 1 | 1 | 13 | S100a10 | 0 | Young |
| 0.035746 | 0.423337 | 0.702 | 0.582 | 1 | 13 | Clcn5 | 0.12 | Young |

| | | | | | | | | |
|---|---|---|---|---|---|---|---|---|
| 0.035945 | 0.646869 | 0.307 | 0.205 | 1 | 13 | Capn5 | 0.102 | Young |
| 0.036095 | 0.227266 | 0.904 | 0.91 | 1 | 13 | Psmb3 | -0.006 | Young |
| 0.036248 | 0.698794 | 0.456 | 0.352 | 1 | 13 | Sh3bp5 | 0.104 | Young |
| 0.03627 | 0.31267 | 0.711 | 0.697 | 1 | 13 | Rnf10 | 0.014 | Young |
| 0.036278 | 0.721359 | 0.228 | 0.123 | 1 | 13 | Bnip1 | 0.105 | Young |
| 0.036685 | 0.52024 | 0.43 | 0.303 | 1 | 13 | Cracr2b | 0.127 | Young |
| 0.036784 | 1.31056 | 0.132 | 0.057 | 1 | 13 | Arl15 | 0.075 | Young |
| 0.037145 | 0.239945 | 0.886 | 0.885 | 1 | 13 | Mdh1 | 0.001 | Young |
| 0.037347 | 0.972492 | 0.228 | 0.131 | 1 | 13 | Map3k10 | 0.097 | Young |
| 0.037593 | 0.421812 | 0.596 | 0.467 | 1 | 13 | Trim33 | 0.129 | Young |
| 0.037712 | 1.395018 | 0.035 | 0 | 1 | 13 | Gm28086 | 0.035 | Young |
| 0.037713 | 1.453338 | 0.035 | 0 | 1 | 13 | Zfp566 | 0.035 | Young |
| 0.037713 | 1.451974 | 0.035 | 0 | 1 | 13 | Gm48529 | 0.035 | Young |
| 0.037713 | 1.383177 | 0.035 | 0 | 1 | 13 | Gm44777 | 0.035 | Young |
| 0.037713 | 1.381929 | 0.035 | 0 | 1 | 13 | Gstm2 | 0.035 | Young |
| 0.037713 | 1.343796 | 0.035 | 0 | 1 | 13 | AA543186 | 0.035 | Young |
| 0.037713 | 1.335816 | 0.035 | 0 | 1 | 13 | 5033406O( | 0.035 | Young |
| 0.037713 | 1.310461 | 0.035 | 0 | 1 | 13 | A530010L1 | 0.035 | Young |
| 0.037713 | 1.31001 | 0.035 | 0 | 1 | 13 | Nccrp1 | 0.035 | Young |
| 0.037713 | 1.294495 | 0.035 | 0 | 1 | 13 | Fancd2 | 0.035 | Young |
| 0.037713 | 1.281587 | 0.035 | 0 | 1 | 13 | Fmo1 | 0.035 | Young |
| 0.037713 | 1.258413 | 0.035 | 0 | 1 | 13 | Nnmt | 0.035 | Young |
| 0.037713 | 1.242063 | 0.035 | 0 | 1 | 13 | Xk | 0.035 | Young |
| 0.037713 | 1.221508 | 0.035 | 0 | 1 | 13 | Ccdc149 | 0.035 | Young |
| 0.037713 | 1.219437 | 0.035 | 0 | 1 | 13 | Gm16316 | 0.035 | Young |
| 0.037713 | 1.136613 | 0.035 | 0 | 1 | 13 | Atad3aos | 0.035 | Young |
| 0.038021 | 0.502071 | 0.412 | 0.279 | 1 | 13 | Ipo9 | 0.133 | Young |
| 0.038091 | 0.248687 | 0.807 | 0.68 | 1 | 13 | Napsa | 0.127 | Young |
| 0.038166 | 0.231074 | 0.991 | 1 | 1 | 13 | Atf3 | -0.009 | Young |
| 0.038195 | 1.128389 | 0.167 | 0.082 | 1 | 13 | Arap3 | 0.085 | Young |
| 0.038529 | 0.424659 | 0.64 | 0.557 | 1 | 13 | Gpr160 | 0.083 | Young |
| 0.038577 | 0.297192 | 0.64 | 0.574 | 1 | 13 | Gpat4 | 0.066 | Young |
| 0.039195 | 0.452279 | 0.421 | 0.303 | 1 | 13 | Zmym4 | 0.118 | Young |
| 0.039498 | 0.551016 | 0.421 | 0.32 | 1 | 13 | Arl4a | 0.101 | Young |
| 0.039859 | 0.256939 | 0.912 | 0.877 | 1 | 13 | Arpp19 | 0.035 | Young |
| 0.039977 | 0.097229 | 1 | 1 | 1 | 13 | H3f3a | 0 | Young |
| 0.040089 | 1.377725 | 0.07 | 0.016 | 1 | 13 | Pecam1 | 0.054 | Young |
| 0.040116 | 0.227682 | 0.86 | 0.82 | 1 | 13 | Nt5c | 0.04 | Young |
| 0.040179 | 0.732563 | 0.333 | 0.221 | 1 | 13 | Hira | 0.112 | Young |
| 0.040312 | 0.380637 | 0.693 | 0.68 | 1 | 13 | Smarce1 | 0.013 | Young |
| 0.040467 | 0.911148 | 0.158 | 0.074 | 1 | 13 | Izumo4 | 0.084 | Young |
| 0.040506 | 0.531101 | 0.632 | 0.541 | 1 | 13 | Gtf2i | 0.091 | Young |
| 0.040622 | 1.390535 | 0.07 | 0.016 | 1 | 13 | Cidec | 0.054 | Young |
| 0.040721 | 0.114861 | 1 | 1 | 1 | 13 | Eef2 | 0 | Young |
| 0.040938 | 0.869678 | 0.307 | 0.213 | 1 | 13 | Tfdp2 | 0.094 | Young |

| | | | | | | | | |
|---|---|---|---|---|---|---|---|---|
| 0.040949 | 0.59987 | 0.211 | 0.115 | 1 | 13 | Mettl13 | 0.096 | Young |
| 0.040992 | 0.698043 | 0.333 | 0.213 | 1 | 13 | Spn | 0.12 | Young |
| 0.041224 | 0.238508 | 0.86 | 0.869 | 1 | 13 | Mpc1 | -0.009 | Young |
| 0.041235 | 0.361992 | 0.754 | 0.73 | 1 | 13 | Sde2 | 0.024 | Young |
| 0.04144 | 0.30164 | 0.939 | 0.918 | 1 | 13 | Pmaip1 | 0.021 | Young |
| 0.041615 | 0.173887 | 0.947 | 0.893 | 1 | 13 | Tomm5 | 0.054 | Young |
| 0.041856 | 0.176813 | 0.965 | 0.967 | 1 | 13 | Clec4b1 | -0.002 | Young |
| 0.04186 | 0.147769 | 1 | 1 | 1 | 13 | Glud1 | 0 | Young |
| 0.042112 | 0.433916 | 0.386 | 0.27 | 1 | 13 | Mdc1 | 0.116 | Young |
| 0.042315 | 0.719263 | 0.439 | 0.344 | 1 | 13 | Thap3 | 0.095 | Young |
| 0.042437 | 0.187518 | 0.974 | 0.967 | 1 | 13 | Eid1 | 0.007 | Young |
| 0.042919 | 1.061039 | 0.123 | 0.049 | 1 | 13 | Gm36266 | 0.074 | Young |
| 0.043055 | 0.399916 | 0.114 | 0.041 | 1 | 13 | Gm26522 | 0.073 | Young |
| 0.043092 | 0.233718 | 0.93 | 0.934 | 1 | 13 | Mpc2 | -0.004 | Young |
| 0.043139 | 1.763996 | 0.053 | 0.008 | 1 | 13 | Ldlrad1 | 0.045 | Young |
| 0.043139 | 1.579338 | 0.053 | 0.008 | 1 | 13 | Mtcp1 | 0.045 | Young |
| 0.043139 | 1.437927 | 0.053 | 0.008 | 1 | 13 | Mlxipl | 0.045 | Young |
| 0.043572 | 0.209499 | 0.947 | 0.902 | 1 | 13 | Taf10 | 0.045 | Young |
| 0.043892 | 0.26693 | 0.921 | 0.869 | 1 | 13 | Crebbp | 0.052 | Young |
| 0.044152 | 1.020107 | 0.123 | 0.049 | 1 | 13 | Adnp | 0.074 | Young |
| 0.044568 | 1.048532 | 0.132 | 0.057 | 1 | 13 | Zfp81 | 0.075 | Young |
| 0.045132 | 0.906795 | 0.325 | 0.23 | 1 | 13 | Syce2 | 0.095 | Young |
| 0.04519 | 1.243178 | 0.053 | 0.008 | 1 | 13 | Lgalsl | 0.045 | Young |
| 0.045239 | 0.890344 | 0.333 | 0.23 | 1 | 13 | Fam118a | 0.103 | Young |
| 0.045485 | 0.273731 | 0.711 | 0.574 | 1 | 13 | Il6 | 0.137 | Young |
| 0.045545 | 0.269031 | 0.649 | 0.598 | 1 | 13 | Dnajb1 | 0.051 | Young |
| 0.045891 | 1.184257 | 0.053 | 0.008 | 1 | 13 | Garnl3 | 0.045 | Young |
| 0.046266 | 0.388348 | 0.623 | 0.533 | 1 | 13 | Hspb11 | 0.09 | Young |
| 0.046511 | 0.437213 | 0.526 | 0.393 | 1 | 13 | Selenom | 0.133 | Young |
| 0.046602 | 1.119524 | 0.053 | 0.008 | 1 | 13 | Pcx | 0.045 | Young |
| 0.046602 | 1.087729 | 0.053 | 0.008 | 1 | 13 | 8430408G2 | 0.045 | Young |
| 0.046783 | 0.711201 | 0.263 | 0.164 | 1 | 13 | Pianp | 0.099 | Young |
| 0.046811 | 1.111311 | 0.14 | 0.066 | 1 | 13 | Maged2 | 0.074 | Young |
| 0.046851 | 0.917761 | 0.175 | 0.09 | 1 | 13 | Ptpn4 | 0.085 | Young |
| 0.046956 | 0.30726 | 0.842 | 0.811 | 1 | 13 | Supt4a | 0.031 | Young |
| 0.046981 | 0.448705 | 0.588 | 0.484 | 1 | 13 | Ephx1 | 0.104 | Young |
| 0.047079 | 0.381265 | 0.798 | 0.836 | 1 | 13 | Ubxn4 | -0.038 | Young |
| 0.047092 | 0.497942 | 0.544 | 0.525 | 1 | 13 | Rabgef1 | 0.019 | Young |
| 0.047138 | 0.918136 | 0.254 | 0.164 | 1 | 13 | 2810402E2 | 0.09 | Young |
| 0.047246 | 0.265619 | 0.965 | 0.861 | 1 | 13 | Gas6 | 0.104 | Young |
| 0.047322 | 0.938487 | 0.053 | 0.008 | 1 | 13 | Epha4 | 0.045 | Young |
| 0.047328 | 0.653778 | 0.158 | 0.074 | 1 | 13 | Mical2 | 0.084 | Young |
| 0.047429 | 0.397868 | 0.579 | 0.5 | 1 | 13 | Crybg3 | 0.079 | Young |
| 0.047537 | 0.079108 | 1 | 1 | 1 | 13 | Rps27 | 0 | Young |
| 0.047633 | 0.534204 | 0.43 | 0.32 | 1 | 13 | Cd151 | 0.11 | Young |

| | | | | | | | | |
|---|---|---|---|---|---|---|---|---|
| 0.047785 | 0.232741 | 0.842 | 0.852 | 1 | 13 | Bmp2k | -0.01 | Young |
| 0.047795 | 0.838808 | 0.325 | 0.23 | 1 | 13 | Ppp1r8 | 0.095 | Young |
| 0.047966 | 0.157587 | 1 | 1 | 1 | 13 | Csf1r | 0 | Young |
| 0.04829 | 0.089919 | 1 | 1 | 1 | 13 | Rpl23a | 0 | Young |
| 0.04838 | 0.232004 | 0.939 | 0.951 | 1 | 13 | Tcf4 | -0.012 | Young |
| 0.048738 | 0.960005 | 0.132 | 0.057 | 1 | 13 | Trp53i13 | 0.075 | Young |
| 0.049273 | 0.169124 | 1 | 0.984 | 1 | 13 | Brk1 | 0.016 | Young |
| 0.049379 | 0.355567 | 0.561 | 0.443 | 1 | 13 | Axin1 | 0.118 | Young |
| 0.049634 | 0.371557 | 0.754 | 0.721 | 1 | 13 | N4bp2l2 | 0.033 | Young |
| 0.049699 | 1.276755 | 0.14 | 0.066 | 1 | 13 | Slc2a3 | 0.074 | Young |
| 0.049715 | 0.287888 | 0.64 | 0.533 | 1 | 13 | Washc3 | 0.107 | Young |
| 0.049753 | 0.517241 | 0.43 | 0.328 | 1 | 13 | Pafah1b3 | 0.102 | Young |
| 0.050251 | 0.607524 | 0.36 | 0.246 | 1 | 13 | Sesn3 | 0.114 | Young |
| 0.050279 | 0.218079 | 0.921 | 0.885 | 1 | 13 | Crlf2 | 0.036 | Young |
| 0.050383 | 0.149918 | 0.965 | 0.984 | 1 | 13 | Eif3c | -0.019 | Young |
| 0.050529 | 0.244364 | 0.921 | 0.902 | 1 | 13 | Nipbl | 0.019 | Young |
| 0.050938 | 0.327085 | 0.886 | 0.861 | 1 | 13 | Fam129a | 0.025 | Young |
| 0.051233 | 0.629885 | 0.395 | 0.311 | 1 | 13 | Phf1 | 0.084 | Young |
| 0.051269 | 0.504513 | 0.439 | 0.328 | 1 | 13 | Lysmd3 | 0.111 | Young |
| 0.051305 | 0.447442 | 0.526 | 0.418 | 1 | 13 | Ipmk | 0.108 | Young |
| 0.051349 | 0.774916 | 0.298 | 0.205 | 1 | 13 | Ctc1 | 0.093 | Young |
| 0.051517 | 0.441958 | 0.553 | 0.467 | 1 | 13 | Aim2 | 0.086 | Young |
| 0.052026 | 0.483875 | 0.553 | 0.434 | 1 | 13 | Pde3b | 0.119 | Young |
| 0.052284 | 0.230084 | 0.851 | 0.803 | 1 | 13 | Pkn2 | 0.048 | Young |
| 0.052336 | 0.676119 | 0.289 | 0.197 | 1 | 13 | Med24 | 0.092 | Young |
| 0.05247 | 0.809787 | 0.333 | 0.238 | 1 | 13 | Mettl26 | 0.095 | Young |
| 0.052493 | 0.557194 | 0.289 | 0.189 | 1 | 13 | Tatdn1 | 0.1 | Young |
| 0.052505 | 1.196776 | 0.105 | 0.041 | 1 | 13 | Zfp933 | 0.064 | Young |
| 0.052666 | 0.348591 | 0.43 | 0.311 | 1 | 13 | BC035044 | 0.119 | Young |
| 0.05303 | 1.449636 | 0.079 | 0.025 | 1 | 13 | Csf1 | 0.054 | Young |
| 0.053176 | 0.430674 | 0.518 | 0.402 | 1 | 13 | Icosl | 0.116 | Young |
| 0.053579 | 0.474486 | 0.579 | 0.467 | 1 | 13 | Snrk | 0.112 | Young |
| 0.053651 | 0.718551 | 0.36 | 0.279 | 1 | 13 | Tprkb | 0.081 | Young |
| 0.053769 | 0.942654 | 0.184 | 0.098 | 1 | 13 | Ccdc106 | 0.086 | Young |
| 0.05407 | 0.383852 | 0.491 | 0.377 | 1 | 13 | Fam120b | 0.114 | Young |
| 0.05408 | 1.071356 | 0.105 | 0.041 | 1 | 13 | Optn | 0.064 | Young |
| 0.054085 | 0.073224 | 1 | 1 | 1 | 13 | Rpl35 | 0 | Young |
| 0.054194 | 0.451052 | 0.535 | 0.418 | 1 | 13 | St3gal6 | 0.117 | Young |
| 0.054439 | 0.274644 | 0.816 | 0.77 | 1 | 13 | Kxd1 | 0.046 | Young |
| 0.055068 | 0.309439 | 0.675 | 0.582 | 1 | 13 | Chp2 | 0.093 | Young |
| 0.055144 | 0.618986 | 0.272 | 0.172 | 1 | 13 | Rac3 | 0.1 | Young |
| 0.055287 | 0.172067 | 0.982 | 0.992 | 1 | 13 | Tnfaip8 | -0.01 | Young |
| 0.05554 | 1.202419 | 0.079 | 0.025 | 1 | 13 | Carmil1 | 0.054 | Young |
| 0.055656 | 0.461095 | 0.667 | 0.648 | 1 | 13 | Marf1 | 0.019 | Young |
| 0.055693 | 1.046864 | 0.105 | 0.041 | 1 | 13 | Nedd4 | 0.064 | Young |

| | | | | | | | | |
|---|---|---|---|---|---|---|---|---|
| 0.055745 | 0.798639 | 0.272 | 0.18 | 1 | 13 | Ppm1d | 0.092 | Young |
| 0.055929 | 0.327399 | 0.86 | 0.861 | 1 | 13 | Foxn3 | -0.001 | Young |
| 0.055973 | 0.670073 | 0.351 | 0.262 | 1 | 13 | Gnl3l | 0.089 | Young |
| 0.055975 | 0.859397 | 0.228 | 0.148 | 1 | 13 | 0610009O2 | 0.08 | Young |
| 0.055981 | 0.100581 | 0.947 | 0.918 | 1 | 13 | Fuca1 | 0.029 | Young |
| 0.056119 | 0.45108 | 0.57 | 0.508 | 1 | 13 | Chd6 | 0.062 | Young |
| 0.056142 | 0.156829 | 0.991 | 0.992 | 1 | 13 | Selenow | -0.001 | Young |
| 0.056264 | 0.308633 | 0.675 | 0.598 | 1 | 13 | Pink1 | 0.077 | Young |
| 0.056472 | 0.380909 | 0.482 | 0.402 | 1 | 13 | Sympk | 0.08 | Young |
| 0.057 | 0.325319 | 0.667 | 0.541 | 1 | 13 | Susd3 | 0.126 | Young |
| 0.057108 | 0.813176 | 0.237 | 0.148 | 1 | 13 | Snhg20 | 0.089 | Young |
| 0.057255 | 0.128054 | 0.965 | 0.984 | 1 | 13 | Zcchc6 | -0.019 | Young |
| 0.057606 | 1.237969 | 0.114 | 0.049 | 1 | 13 | Gm15445 | 0.065 | Young |
| 0.057657 | 0.279937 | 0.912 | 0.902 | 1 | 13 | Cd2ap | 0.01 | Young |
| 0.057758 | 0.100484 | 1 | 1 | 1 | 13 | Rps15a | 0 | Young |
| 0.058014 | 1.021498 | 0.149 | 0.074 | 1 | 13 | Lin9 | 0.075 | Young |
| 0.058148 | 1.110457 | 0.079 | 0.025 | 1 | 13 | Acbd4 | 0.054 | Young |
| 0.058696 | 0.317744 | 0.596 | 0.5 | 1 | 13 | Map1s | 0.096 | Young |
| 0.058696 | 0.286372 | 0.868 | 0.779 | 1 | 13 | 1110004F1 | 0.089 | Young |
| 0.058875 | 0.44306 | 0.605 | 0.541 | 1 | 13 | Insr | 0.064 | Young |
| 0.05935 | 0.380037 | 0.64 | 0.59 | 1 | 13 | Dstyk | 0.05 | Young |
| 0.059412 | 0.469596 | 0.526 | 0.467 | 1 | 13 | Cstf3 | 0.059 | Young |
| 0.059482 | 0.284852 | 0.842 | 0.828 | 1 | 13 | Prkacb | 0.014 | Young |
| 0.059494 | 0.264744 | 0.904 | 0.951 | 1 | 13 | Pnisr | -0.047 | Young |
| 0.059759 | 0.724346 | 0.351 | 0.254 | 1 | 13 | Itgal | 0.097 | Young |
| 0.059822 | 0.175855 | 0.939 | 0.82 | 1 | 13 | Cyp4f18 | 0.119 | Young |
| 0.059943 | 0.603982 | 0.412 | 0.32 | 1 | 13 | Tuba4a | 0.092 | Young |
| 0.060029 | 0.21091 | 0.904 | 0.836 | 1 | 13 | Igf1 | 0.068 | Young |
| 0.060278 | 0.204227 | 0.886 | 0.893 | 1 | 13 | Ap1s2 | -0.007 | Young |
| 0.060375 | 0.786538 | 0.149 | 0.074 | 1 | 13 | Nipal3 | 0.075 | Young |
| 0.06061 | 0.271668 | 0.851 | 0.754 | 1 | 13 | Tet2 | 0.097 | Young |
| 0.061106 | 0.140414 | 1 | 1 | 1 | 13 | Npm1 | 0 | Young |
| 0.061243 | 0.709502 | 0.211 | 0.123 | 1 | 13 | Iqce | 0.088 | Young |
| 0.061261 | 0.952902 | 0.219 | 0.139 | 1 | 13 | Zbtb33 | 0.08 | Young |
| 0.06137 | 0.155692 | 0.982 | 0.984 | 1 | 13 | Nedd8 | -0.002 | Young |
| 0.061551 | 0.957316 | 0.079 | 0.025 | 1 | 13 | Ica1l | 0.054 | Young |
| 0.061577 | 0.97907 | 0.167 | 0.09 | 1 | 13 | Zfp182 | 0.077 | Young |
| 0.061675 | 0.667259 | 0.272 | 0.18 | 1 | 13 | Kansl1l | 0.092 | Young |
| 0.061901 | 0.145381 | 0.982 | 1 | 1 | 13 | Akap13 | -0.018 | Young |
| 0.062056 | 0.401227 | 0.526 | 0.402 | 1 | 13 | Mrpl35 | 0.124 | Young |
| 0.06209 | 0.45959 | 0.316 | 0.205 | 1 | 13 | C87436 | 0.111 | Young |
| 0.062628 | 1.416391 | 0.088 | 0.033 | 1 | 13 | Lrfn1 | 0.055 | Young |
| 0.063074 | 0.307283 | 0.596 | 0.475 | 1 | 13 | 5031425E2 | 0.121 | Young |
| 0.063089 | 0.35644 | 0.535 | 0.426 | 1 | 13 | Spin1 | 0.109 | Young |
| 0.063135 | 1.313006 | 0.114 | 0.049 | 1 | 13 | Tln2 | 0.065 | Young |

| | | | | | | | | |
|---|---|---|---|---|---|---|---|---|
| 0.063152 | 0.549716 | 0.307 | 0.205 | 1 | 13 | Arfgap1 | 0.102 | Young |
| 0.063289 | 0.286066 | 0.667 | 0.607 | 1 | 13 | Desi2 | 0.06 | Young |
| 0.063934 | 0.348563 | 0.596 | 0.525 | 1 | 13 | Cmah | 0.071 | Young |
| 0.063963 | 0.507856 | 0.526 | 0.467 | 1 | 13 | Rad50 | 0.059 | Young |
| 0.064019 | 0.23764 | 0.816 | 0.721 | 1 | 13 | Mbd2 | 0.095 | Young |
| 0.064214 | 0.156058 | 0.912 | 0.803 | 1 | 13 | Abca1 | 0.109 | Young |
| 0.064237 | 0.398727 | 0.711 | 0.68 | 1 | 13 | Arih2 | 0.031 | Young |
| 0.065499 | 0.462728 | 0.474 | 0.393 | 1 | 13 | Slc41a3 | 0.081 | Young |
| 0.06564 | 0.273457 | 0.886 | 0.877 | 1 | 13 | Orai1 | 0.009 | Young |
| 0.065666 | 0.331404 | 0.833 | 0.861 | 1 | 13 | Skil | -0.028 | Young |
| 0.065683 | 0.804632 | 0.281 | 0.197 | 1 | 13 | Jrkl | 0.084 | Young |
| 0.06575 | 0.877088 | 0.219 | 0.139 | 1 | 13 | Acap3 | 0.08 | Young |
| 0.066188 | 0.677701 | 0.246 | 0.156 | 1 | 13 | Mical1 | 0.09 | Young |
| 0.066615 | 0.253973 | 0.833 | 0.828 | 1 | 13 | 0610012G( | 0.005 | Young |
| 0.066657 | 0.919332 | 0.114 | 0.049 | 1 | 13 | Bcr | 0.065 | Young |
| 0.066861 | 0.753385 | 0.228 | 0.139 | 1 | 13 | Fam110a | 0.089 | Young |
| 0.067065 | 0.591115 | 0.333 | 0.23 | 1 | 13 | Rab11fip2 | 0.103 | Young |
| 0.067073 | 0.41697 | 0.553 | 0.475 | 1 | 13 | Ankle2 | 0.078 | Young |
| 0.067169 | 0.260437 | 0.895 | 0.861 | 1 | 13 | Park7 | 0.034 | Young |
| 0.067218 | 0.417386 | 0.386 | 0.279 | 1 | 13 | Nudt2 | 0.107 | Young |
| 0.06726 | 0.933053 | 0.114 | 0.049 | 1 | 13 | Btnl2 | 0.065 | Young |
| 0.067581 | 1.102665 | 0.184 | 0.107 | 1 | 13 | Gm27010 | 0.077 | Young |
| 0.067667 | 0.2359 | 0.93 | 0.934 | 1 | 13 | Scaf11 | -0.004 | Young |
| 0.067691 | 0.525177 | 0.474 | 0.393 | 1 | 13 | Arhgap11a | 0.081 | Young |
| 0.067742 | 0.58013 | 0.404 | 0.32 | 1 | 13 | Nat9 | 0.084 | Young |
| 0.068473 | 0.44613 | 0.614 | 0.525 | 1 | 13 | Dusp6 | 0.089 | Young |
| 0.068764 | 1.187129 | 0.061 | 0.016 | 1 | 13 | Gm16062 | 0.045 | Young |
| 0.069135 | 0.526623 | 0.404 | 0.303 | 1 | 13 | Atn1 | 0.101 | Young |
| 0.069153 | 0.194776 | 0.991 | 0.951 | 1 | 13 | Ier3 | 0.04 | Young |
| 0.06935 | 0.322621 | 0.719 | 0.623 | 1 | 13 | Tifa | 0.096 | Young |
| 0.069491 | 0.510903 | 0.456 | 0.393 | 1 | 13 | Kifap3 | 0.063 | Young |
| 0.069578 | 0.287861 | 0.614 | 0.574 | 1 | 13 | Cdc37l1 | 0.04 | Young |
| 0.069619 | 0.279719 | 0.895 | 0.82 | 1 | 13 | Pin4 | 0.075 | Young |
| 0.07007 | 0.97276 | 0.211 | 0.131 | 1 | 13 | Sema6b | 0.08 | Young |
| 0.070268 | 0.346261 | 0.456 | 0.352 | 1 | 13 | Hdac9 | 0.104 | Young |
| 0.070333 | 0.207546 | 1 | 0.984 | 1 | 13 | Myh9 | 0.016 | Young |
| 0.070534 | 1.956689 | 0.061 | 0.016 | 1 | 13 | Apold1 | 0.045 | Young |
| 0.070534 | 1.2276 | 0.061 | 0.016 | 1 | 13 | Homer2 | 0.045 | Young |
| 0.070631 | 0.323403 | 0.825 | 0.77 | 1 | 13 | Gabarapl2 | 0.055 | Young |
| 0.070927 | 0.173486 | 1 | 1 | 1 | 13 | Gm2a | 0 | Young |
| 0.071025 | 0.598851 | 0.342 | 0.254 | 1 | 13 | Zfp329 | 0.088 | Young |
| 0.071227 | 1.02116 | 0.193 | 0.115 | 1 | 13 | Gem | 0.078 | Young |
| 0.071433 | 1.153139 | 0.061 | 0.016 | 1 | 13 | Adgrg5 | 0.045 | Young |
| 0.071433 | 1.117279 | 0.061 | 0.016 | 1 | 13 | Tmem119 | 0.045 | Young |
| 0.071433 | 1.104463 | 0.061 | 0.016 | 1 | 13 | Gm28557 | 0.045 | Young |

| | | | | | | | | |
|---|---|---|---|---|---|---|---|---|
| 0.071433 | 1.09994 | 0.061 | 0.016 | 1 | 13 | Ccdc157 | 0.045 | Young |
| 0.071483 | 0.33643 | 0.605 | 0.549 | 1 | 13 | Actl6a | 0.056 | Young |
| 0.071969 | 0.358361 | 0.781 | 0.73 | 1 | 13 | Pdcd4 | 0.051 | Young |
| 0.071971 | 0.240505 | 0.746 | 0.607 | 1 | 13 | Cd209d | 0.139 | Young |
| 0.071983 | 0.424275 | 0.439 | 0.352 | 1 | 13 | Sord | 0.087 | Young |
| 0.072095 | 1.043849 | 0.14 | 0.074 | 1 | 13 | P2ry13 | 0.066 | Young |
| 0.072153 | 0.276792 | 0.351 | 0.23 | 1 | 13 | Sema4b | 0.121 | Young |
| 0.072327 | 0.390838 | 0.57 | 0.492 | 1 | 13 | Pdzd8 | 0.078 | Young |
| 0.07243 | 0.161169 | 0.991 | 1 | 1 | 13 | Tpr | -0.009 | Young |
| 0.072714 | 0.220394 | 0.895 | 0.844 | 1 | 13 | Ccnd1 | 0.051 | Young |
| 0.072732 | 1.889173 | 0.026 | 0 | 1 | 13 | Hbb-bs | 0.026 | Young |
| 0.072732 | 1.489384 | 0.026 | 0 | 1 | 13 | Scn4b | 0.026 | Young |
| 0.072732 | 1.48141 | 0.026 | 0 | 1 | 13 | AF529169 | 0.026 | Young |
| 0.072732 | 1.477964 | 0.026 | 0 | 1 | 13 | Slfn3 | 0.026 | Young |
| 0.072732 | 1.393004 | 0.026 | 0 | 1 | 13 | Zc3hav1l | 0.026 | Young |
| 0.072732 | 1.354743 | 0.026 | 0 | 1 | 13 | Gm26508 | 0.026 | Young |
| 0.072732 | 1.344026 | 0.026 | 0 | 1 | 13 | Zfp94 | 0.026 | Young |
| 0.072732 | 1.248735 | 0.026 | 0 | 1 | 13 | Gm20633 | 0.026 | Young |
| 0.072732 | 1.236123 | 0.026 | 0 | 1 | 13 | 9430037G( | 0.026 | Young |
| 0.072732 | 1.22605 | 0.026 | 0 | 1 | 13 | Gm5475 | 0.026 | Young |
| 0.072732 | 1.196317 | 0.026 | 0 | 1 | 13 | Podxl2 | 0.026 | Young |
| 0.072732 | 1.19521 | 0.026 | 0 | 1 | 13 | Sectm1a | 0.026 | Young |
| 0.072732 | 1.185496 | 0.026 | 0 | 1 | 13 | Mogat1 | 0.026 | Young |
| 0.072732 | 1.156518 | 0.026 | 0 | 1 | 13 | 9330104G( | 0.026 | Young |
| 0.072732 | 1.148304 | 0.026 | 0 | 1 | 13 | Gm11728 | 0.026 | Young |
| 0.072732 | 1.147999 | 0.026 | 0 | 1 | 13 | Gm8797 | 0.026 | Young |
| 0.072732 | 1.14019 | 0.026 | 0 | 1 | 13 | Gm47794 | 0.026 | Young |
| 0.072732 | 1.138416 | 0.026 | 0 | 1 | 13 | Dnal1 | 0.026 | Young |
| 0.072732 | 1.12732 | 0.026 | 0 | 1 | 13 | 3300002I0 | 0.026 | Young |
| 0.072732 | 1.104387 | 0.026 | 0 | 1 | 13 | Klkb1 | 0.026 | Young |
| 0.072732 | 1.09583 | 0.026 | 0 | 1 | 13 | Gm20449 | 0.026 | Young |
| 0.072732 | 1.095101 | 0.026 | 0 | 1 | 13 | Wdr35 | 0.026 | Young |
| 0.072732 | 1.093591 | 0.026 | 0 | 1 | 13 | Rbakdn | 0.026 | Young |
| 0.072732 | 1.080879 | 0.026 | 0 | 1 | 13 | Eml5 | 0.026 | Young |
| 0.072732 | 1.065889 | 0.026 | 0 | 1 | 13 | Mettl27 | 0.026 | Young |
| 0.072732 | 1.055003 | 0.026 | 0 | 1 | 13 | Proca1 | 0.026 | Young |
| 0.072732 | 1.054386 | 0.026 | 0 | 1 | 13 | Il1r1 | 0.026 | Young |
| 0.072732 | 1.050594 | 0.026 | 0 | 1 | 13 | Gm15551 | 0.026 | Young |
| 0.072732 | 1.039031 | 0.026 | 0 | 1 | 13 | Car4 | 0.026 | Young |
| 0.072732 | 1.032204 | 0.026 | 0 | 1 | 13 | Zfp973 | 0.026 | Young |
| 0.072732 | 1.026455 | 0.026 | 0 | 1 | 13 | Grin1 | 0.026 | Young |
| 0.072732 | 1.020812 | 0.026 | 0 | 1 | 13 | Zfp128 | 0.026 | Young |
| 0.072732 | 1.017905 | 0.026 | 0 | 1 | 13 | Setbp1 | 0.026 | Young |
| 0.072732 | 1.011895 | 0.026 | 0 | 1 | 13 | Hist1h2bh | 0.026 | Young |
| 0.072732 | 0.99352 | 0.026 | 0 | 1 | 13 | Serpinf2 | 0.026 | Young |

| | | | | | | | | |
|---|---|---|---|---|---|---|---|---|
| 0.072732 | 0.971682 | 0.026 | 0 | 1 | 13 | Slamf1 | 0.026 | Young |
| 0.072732 | 0.971612 | 0.026 | 0 | 1 | 13 | Plekhg6 | 0.026 | Young |
| 0.072732 | 0.971054 | 0.026 | 0 | 1 | 13 | Gm9889 | 0.026 | Young |
| 0.072732 | 0.952836 | 0.026 | 0 | 1 | 13 | Rasd1 | 0.026 | Young |
| 0.072732 | 0.946218 | 0.026 | 0 | 1 | 13 | Pdlim3 | 0.026 | Young |
| 0.072732 | 0.939178 | 0.026 | 0 | 1 | 13 | Nos3 | 0.026 | Young |
| 0.072732 | 0.93866 | 0.026 | 0 | 1 | 13 | Zfp248 | 0.026 | Young |
| 0.072732 | 0.909001 | 0.026 | 0 | 1 | 13 | Gm28809 | 0.026 | Young |
| 0.072732 | 0.849475 | 0.026 | 0 | 1 | 13 | Gm36028 | 0.026 | Young |
| 0.072732 | 0.831208 | 0.026 | 0 | 1 | 13 | Fam124a | 0.026 | Young |
| 0.072732 | 0.830674 | 0.026 | 0 | 1 | 13 | Gm45449 | 0.026 | Young |
| 0.072847 | 0.202972 | 0.886 | 0.893 | 1 | 13 | Cyb5a | -0.007 | Young |
| 0.072887 | 0.771368 | 0.114 | 0.049 | 1 | 13 | Dgke | 0.065 | Young |
| 0.073016 | 0.983279 | 0.132 | 0.066 | 1 | 13 | Tmtc2 | 0.066 | Young |
| 0.073138 | 0.50101 | 0.368 | 0.262 | 1 | 13 | Ssbp2 | 0.106 | Young |
| 0.073226 | 0.486095 | 0.535 | 0.451 | 1 | 13 | Stau1 | 0.084 | Young |
| 0.073335 | 0.900951 | 0.175 | 0.098 | 1 | 13 | 2610044O1 | 0.077 | Young |
| 0.07334 | 0.847201 | 0.123 | 0.057 | 1 | 13 | Uros | 0.066 | Young |
| 0.073344 | 0.127361 | 1 | 0.975 | 1 | 13 | Lsp1 | 0.025 | Young |
| 0.073668 | 0.413421 | 0.447 | 0.361 | 1 | 13 | Prkag2 | 0.086 | Young |
| 0.073809 | 0.444516 | 0.588 | 0.533 | 1 | 13 | Got2 | 0.055 | Young |
| 0.074322 | 0.194185 | 0.579 | 0.467 | 1 | 13 | Aoah | 0.112 | Young |
| 0.0746 | 0.380231 | 0.693 | 0.689 | 1 | 13 | Ube2e3 | 0.004 | Young |
| 0.074696 | 0.593231 | 0.465 | 0.385 | 1 | 13 | Stard4 | 0.08 | Young |
| 0.074728 | 0.197168 | 0.974 | 0.959 | 1 | 13 | Clic4 | 0.015 | Young |
| 0.074746 | 0.44994 | 0.447 | 0.352 | 1 | 13 | Txnrd2 | 0.095 | Young |
| 0.075117 | 0.325407 | 0.658 | 0.648 | 1 | 13 | Mettl23 | 0.01 | Young |
| 0.075215 | 0.255314 | 0.833 | 0.779 | 1 | 13 | Ppp6r3 | 0.054 | Young |
| 0.07522 | 1.007102 | 0.123 | 0.057 | 1 | 13 | Zfp580 | 0.066 | Young |
| 0.075424 | 0.280753 | 0.772 | 0.779 | 1 | 13 | St3gal4 | -0.007 | Young |
| 0.07553 | 0.644853 | 0.237 | 0.148 | 1 | 13 | Rnf185 | 0.089 | Young |
| 0.075565 | 0.71737 | 0.456 | 0.393 | 1 | 13 | Tmem71 | 0.063 | Young |
| 0.075583 | 0.474299 | 0.465 | 0.369 | 1 | 13 | Spire1 | 0.096 | Young |
| 0.075818 | 0.624352 | 0.351 | 0.262 | 1 | 13 | Ankrd46 | 0.089 | Young |
| 0.075829 | 0.180005 | 1 | 0.975 | 1 | 13 | Marcksl1 | 0.025 | Young |
| 0.075986 | 0.137935 | 0.991 | 0.992 | 1 | 13 | Dab2 | -0.001 | Young |
| 0.076069 | 1.444995 | 0.061 | 0.016 | 1 | 13 | Rnase2a | 0.045 | Young |
| 0.076069 | 0.965903 | 0.061 | 0.016 | 1 | 13 | Abcb4 | 0.045 | Young |
| 0.076161 | 0.3201 | 0.667 | 0.557 | 1 | 13 | Cdk12 | 0.11 | Young |
| 0.076283 | 0.210097 | 0.956 | 0.943 | 1 | 13 | Bnip2 | 0.013 | Young |
| 0.076357 | 0.235134 | 0.886 | 0.934 | 1 | 13 | Bptf | -0.048 | Young |
| 0.076958 | 0.965827 | 0.088 | 0.033 | 1 | 13 | Fndc4 | 0.055 | Young |
| 0.077247 | 0.17974 | 0.684 | 0.648 | 1 | 13 | Fcgr1 | 0.036 | Young |
| 0.07737 | 0.642567 | 0.36 | 0.287 | 1 | 13 | Ints12 | 0.073 | Young |
| 0.077502 | 0.727226 | 0.298 | 0.205 | 1 | 13 | Nsun6 | 0.093 | Young |

| | | | | | | | | |
|---|---|---|---|---|---|---|---|---|
| 0.077991 | 0.824376 | 0.061 | 0.016 | 1 | 13 | Spata7 | 0.045 | Young |
| 0.078166 | 0.587303 | 0.386 | 0.311 | 1 | 13 | Slc25a12 | 0.075 | Young |
| 0.078307 | 0.514735 | 0.404 | 0.328 | 1 | 13 | Gfer | 0.076 | Young |
| 0.078516 | 0.913787 | 0.132 | 0.066 | 1 | 13 | Ccdc73 | 0.066 | Young |
| 0.078633 | 0.858152 | 0.175 | 0.098 | 1 | 13 | Prickle3 | 0.077 | Young |
| 0.07882 | 0.214296 | 0.833 | 0.893 | 1 | 13 | Wac | -0.06 | Young |
| 0.079173 | 1.001278 | 0.149 | 0.082 | 1 | 13 | Exo5 | 0.067 | Young |
| 0.079446 | 0.422753 | 0.675 | 0.607 | 1 | 13 | Zfp395 | 0.068 | Young |
| 0.079787 | 0.264339 | 0.921 | 0.852 | 1 | 13 | Nars | 0.069 | Young |
| 0.079834 | 0.30914 | 0.746 | 0.68 | 1 | 13 | Nrip1 | 0.066 | Young |
| 0.080076 | 0.393628 | 0.544 | 0.5 | 1 | 13 | Celf4 | 0.044 | Young |
| 0.080225 | 1.375155 | 0.044 | 0.008 | 1 | 13 | Gpr137c | 0.036 | Young |
| 0.080225 | 1.369967 | 0.044 | 0.008 | 1 | 13 | Tal1 | 0.036 | Young |
| 0.080225 | 1.31906 | 0.044 | 0.008 | 1 | 13 | 1110046J0 | 0.036 | Young |
| 0.080225 | 1.172008 | 0.044 | 0.008 | 1 | 13 | Gm27168 | 0.036 | Young |
| 0.08043 | 0.798804 | 0.246 | 0.172 | 1 | 13 | Pgghg | 0.074 | Young |
| 0.080598 | 0.872097 | 0.211 | 0.131 | 1 | 13 | Egfr | 0.08 | Young |
| 0.080947 | 0.185575 | 0.947 | 0.91 | 1 | 13 | Clint1 | 0.037 | Young |
| 0.081047 | 0.244085 | 0.904 | 0.91 | 1 | 13 | Ankrd44 | -0.006 | Young |
| 0.081104 | 0.734608 | 0.404 | 0.344 | 1 | 13 | Foxo1 | 0.06 | Young |
| 0.081394 | 0.728232 | 0.272 | 0.189 | 1 | 13 | Txlng | 0.083 | Young |
| 0.081441 | 1.167676 | 0.044 | 0.008 | 1 | 13 | Lrp8 | 0.036 | Young |
| 0.081441 | 1.120162 | 0.044 | 0.008 | 1 | 13 | Zfp677 | 0.036 | Young |
| 0.081441 | 1.106918 | 0.044 | 0.008 | 1 | 13 | Rab42 | 0.036 | Young |
| 0.081715 | 0.988399 | 0.132 | 0.066 | 1 | 13 | Hist3h2ba | 0.066 | Young |
| 0.082105 | 0.262965 | 0.588 | 0.475 | 1 | 13 | Itgb5 | 0.113 | Young |
| 0.082509 | 0.492455 | 0.553 | 0.5 | 1 | 13 | Acox3 | 0.053 | Young |
| 0.082518 | 0.528519 | 0.421 | 0.352 | 1 | 13 | Foxj3 | 0.069 | Young |
| 0.082551 | 0.201981 | 0.851 | 0.713 | 1 | 13 | Cdkn1b | 0.138 | Young |
| 0.082672 | 3.851248 | 0.044 | 0.008 | 1 | 13 | Prg4 | 0.036 | Young |
| 0.082672 | 1.196997 | 0.044 | 0.008 | 1 | 13 | Zfp97 | 0.036 | Young |
| 0.082953 | 1.039265 | 0.096 | 0.041 | 1 | 13 | Rnf180 | 0.055 | Young |
| 0.083017 | 0.12992 | 1 | 0.992 | 1 | 13 | Tgfb1 | 0.008 | Young |
| 0.083019 | 0.203637 | 0.781 | 0.713 | 1 | 13 | Slc4a7 | 0.068 | Young |
| 0.083684 | 0.80491 | 0.132 | 0.066 | 1 | 13 | Dbp | 0.066 | Young |
| 0.083918 | 1.12393 | 0.044 | 0.008 | 1 | 13 | AI427809 | 0.036 | Young |
| 0.084039 | 0.101611 | 1 | 1 | 1 | 13 | Marcks | 0 | Young |
| 0.084238 | 0.224168 | 0.921 | 0.885 | 1 | 13 | 1810058I2 | 0.036 | Young |
| 0.085121 | 0.478565 | 0.491 | 0.41 | 1 | 13 | S1pr1 | 0.081 | Young |
| 0.085179 | 1.048579 | 0.044 | 0.008 | 1 | 13 | Hykk | 0.036 | Young |
| 0.085179 | 0.863684 | 0.044 | 0.008 | 1 | 13 | Pram1 | 0.036 | Young |
| 0.085199 | 0.588703 | 0.211 | 0.123 | 1 | 13 | Bod1 | 0.088 | Young |
| 0.085883 | 0.969339 | 0.149 | 0.082 | 1 | 13 | Trub1 | 0.067 | Young |
| 0.086456 | 1.055103 | 0.044 | 0.008 | 1 | 13 | Gm20939 | 0.036 | Young |
| 0.086769 | 0.522804 | 0.088 | 0.033 | 1 | 13 | Ms4a4a | 0.055 | Young |

| | | | | | | | | |
|---|---|---|---|---|---|---|---|---|
| 0.086916 | 0.278155 | 0.64 | 0.541 | 1 | 13 | AI413582 | 0.099 | Young |
| 0.08751 | 0.233706 | 0.737 | 0.762 | 1 | 13 | Elmo1 | -0.025 | Young |
| 0.087635 | 0.210125 | 0.912 | 0.861 | 1 | 13 | Plekhj1 | 0.051 | Young |
| 0.088017 | 0.235813 | 0.956 | 0.926 | 1 | 13 | Zbtb20 | 0.03 | Young |
| 0.088035 | 0.359781 | 0.746 | 0.721 | 1 | 13 | Phf21a | 0.025 | Young |
| 0.08805 | 0.083899 | 1 | 1 | 1 | 13 | Rpl39 | 0 | Young |
| 0.088159 | 0.903573 | 0.184 | 0.115 | 1 | 13 | Rnf26 | 0.069 | Young |
| 0.088202 | 0.27476 | 0.842 | 0.713 | 1 | 13 | Rgs18 | 0.129 | Young |
| 0.088902 | 0.550394 | 0.544 | 0.451 | 1 | 13 | Dcstamp | 0.093 | Young |
| 0.088973 | 0.226723 | 0.895 | 0.852 | 1 | 13 | Cbfa2t3 | 0.043 | Young |
| 0.089115 | 0.165495 | 0.991 | 0.943 | 1 | 13 | H13 | 0.048 | Young |
| 0.08956 | 1.273233 | 0.105 | 0.049 | 1 | 13 | Ptchd1 | 0.056 | Young |
| 0.089625 | 0.230234 | 0.737 | 0.738 | 1 | 13 | Pnn | -0.001 | Young |
| 0.089926 | 0.292451 | 0.781 | 0.779 | 1 | 13 | Erbin | 0.002 | Young |
| 0.090057 | 1.252228 | 0.07 | 0.025 | 1 | 13 | Kctd21 | 0.045 | Young |
| 0.090089 | 0.828363 | 0.14 | 0.074 | 1 | 13 | Sac3d1 | 0.066 | Young |
| 0.090098 | 0.902523 | 0.237 | 0.164 | 1 | 13 | Cul4b | 0.073 | Young |
| 0.090189 | 0.747493 | 0.096 | 0.041 | 1 | 13 | Mcf2l | 0.055 | Young |
| 0.090307 | 0.251795 | 0.868 | 0.852 | 1 | 13 | Hp1bp3 | 0.016 | Young |
| 0.090317 | 0.565161 | 0.237 | 0.148 | 1 | 13 | Gramd3 | 0.089 | Young |
| 0.090436 | 0.22678 | 0.939 | 0.877 | 1 | 13 | Bin1 | 0.062 | Young |
| 0.090574 | 0.446233 | 0.632 | 0.574 | 1 | 13 | Trim44 | 0.058 | Young |
| 0.09065 | 0.320525 | 0.544 | 0.459 | 1 | 13 | Nup153 | 0.085 | Young |
| 0.091137 | 1.060834 | 0.105 | 0.049 | 1 | 13 | Tlr11 | 0.056 | Young |
| 0.091642 | 0.435164 | 0.404 | 0.32 | 1 | 13 | Inafm1 | 0.084 | Young |
| 0.091743 | 0.310109 | 0.781 | 0.672 | 1 | 13 | Qdpr | 0.109 | Young |
| 0.092056 | 1.139131 | 0.07 | 0.025 | 1 | 13 | Nlgn2 | 0.045 | Young |
| 0.092132 | 0.553316 | 0.263 | 0.172 | 1 | 13 | Armt1 | 0.091 | Young |
| 0.092141 | 0.844676 | 0.14 | 0.074 | 1 | 13 | Dtx3 | 0.066 | Young |
| 0.092413 | 1.105658 | 0.114 | 0.057 | 1 | 13 | Hectd2 | 0.057 | Young |
| 0.092928 | 0.201375 | 0.737 | 0.68 | 1 | 13 | Pik3r6 | 0.057 | Young |
| 0.093069 | 1.121872 | 0.07 | 0.025 | 1 | 13 | Gm28187 | 0.045 | Young |
| 0.093069 | 1.081071 | 0.07 | 0.025 | 1 | 13 | Macrod2 | 0.045 | Young |
| 0.093437 | 0.844468 | 0.289 | 0.213 | 1 | 13 | Pot1a | 0.076 | Young |
| 0.093565 | 0.835265 | 0.096 | 0.041 | 1 | 13 | Fam161a | 0.055 | Young |
| 0.093588 | 0.310897 | 0.711 | 0.648 | 1 | 13 | Ivns1abp | 0.063 | Young |
| 0.093885 | 0.633748 | 0.325 | 0.254 | 1 | 13 | Endog | 0.071 | Young |
| 0.09409 | 1.165331 | 0.07 | 0.025 | 1 | 13 | Galnt12 | 0.045 | Young |
| 0.094508 | 0.190862 | 0.86 | 0.77 | 1 | 13 | Nfix | 0.09 | Young |
| 0.095574 | 0.392159 | 0.491 | 0.385 | 1 | 13 | Eci2 | 0.106 | Young |
| 0.095576 | 0.186973 | 0.693 | 0.615 | 1 | 13 | Pik3cg | 0.078 | Young |
| 0.095582 | 0.127261 | 1 | 1 | 1 | 13 | Cox7a2l | 0 | Young |
| 0.09575 | 0.567632 | 0.325 | 0.246 | 1 | 13 | Gga3 | 0.079 | Young |
| 0.095772 | 0.844057 | 0.254 | 0.18 | 1 | 13 | Kantr | 0.074 | Young |
| 0.095974 | 0.20051 | 0.868 | 0.803 | 1 | 13 | Lacc1 | 0.065 | Young |

| | | | | | | | | |
|---|---|---|---|---|---|---|---|---|
| 0.095989 | 0.341697 | 0.623 | 0.574 | 1 | 13 | Mbtps1 | 0.049 | Young |
| 0.096286 | 0.320227 | 0.737 | 0.713 | 1 | 13 | Luzp1 | 0.024 | Young |
| 0.097128 | 0.25599 | 0.816 | 0.828 | 1 | 13 | BC028528 | -0.012 | Young |
| 0.097687 | 0.515838 | 0.456 | 0.418 | 1 | 13 | Taf7 | 0.038 | Young |
| 0.09788 | 0.178948 | 0.982 | 0.975 | 1 | 13 | Aes | 0.007 | Young |
| 0.098027 | 0.449452 | 0.465 | 0.426 | 1 | 13 | Lrch3 | 0.039 | Young |
| 0.098447 | 0.357505 | 0.579 | 0.508 | 1 | 13 | Sesn1 | 0.071 | Young |
| 0.098463 | 0.165178 | 0.991 | 0.992 | 1 | 13 | Chd4 | -0.001 | Young |
| 0.09848 | 0.552345 | 0.36 | 0.295 | 1 | 13 | St3gal3 | 0.065 | Young |
| 0.098817 | 0.75757 | 0.096 | 0.041 | 1 | 13 | Cd101 | 0.055 | Young |
| 0.099301 | 0.180513 | 0.719 | 0.672 | 1 | 13 | Abhd17a | 0.047 | Young |
| 0.099634 | 0.162058 | 0.974 | 0.992 | 1 | 13 | Tcf25 | -0.018 | Young |
| 0.099662 | 0.274494 | 0.43 | 0.32 | 1 | 13 | Mxi1 | 0.11 | Young |
| 0.100225 | 0.072207 | 1 | 1 | 1 | 13 | Rpl6 | 0 | Young |
| 0.100294 | 0.4285 | 0.439 | 0.352 | 1 | 13 | Cherp | 0.087 | Young |
| 0.100482 | 0.185579 | 0.912 | 0.902 | 1 | 13 | Inpp5d | 0.01 | Young |
| 0.100591 | 0.254596 | 0.895 | 0.836 | 1 | 13 | Slc48a1 | 0.059 | Young |
| 0.100689 | 0.233299 | 0.746 | 0.639 | 1 | 13 | Zfp516 | 0.107 | Young |
| 0.100751 | 0.475621 | 0.465 | 0.393 | 1 | 13 | Zcchc9 | 0.072 | Young |
| 0.101003 | 0.489902 | 0.202 | 0.123 | 1 | 13 | Poc5 | 0.079 | Young |
| 0.101104 | 0.281998 | 0.728 | 0.672 | 1 | 13 | Rras | 0.056 | Young |
| 0.101131 | 0.32446 | 0.675 | 0.631 | 1 | 13 | N4bp2l1 | 0.044 | Young |
| 0.101497 | 0.334358 | 0.719 | 0.697 | 1 | 13 | Chd2 | 0.022 | Young |
| 0.101543 | 0.494176 | 0.509 | 0.443 | 1 | 13 | Dcaf11 | 0.066 | Young |
| 0.102175 | 0.272596 | 0.816 | 0.746 | 1 | 13 | Stxbp2 | 0.07 | Young |
| 0.102243 | 0.605165 | 0.184 | 0.107 | 1 | 13 | Gm16124 | 0.077 | Young |
| 0.102627 | 0.176183 | 0.5 | 0.41 | 1 | 13 | Mxd1 | 0.09 | Young |
| 0.102677 | 0.353697 | 0.737 | 0.697 | 1 | 13 | Baiap2 | 0.04 | Young |
| 0.102689 | 0.431227 | 0.167 | 0.09 | 1 | 13 | Zfp995 | 0.077 | Young |
| 0.103364 | 0.253128 | 0.605 | 0.541 | 1 | 13 | Setd5 | 0.064 | Young |
| 0.103384 | 0.436539 | 0.553 | 0.549 | 1 | 13 | Ip6k1 | 0.004 | Young |
| 0.103509 | 0.822579 | 0.175 | 0.107 | 1 | 13 | Enho | 0.068 | Young |
| 0.103614 | 0.498487 | 0.36 | 0.262 | 1 | 13 | Zbtb37 | 0.098 | Young |
| 0.103685 | 0.757805 | 0.105 | 0.049 | 1 | 13 | A430046D1 | 0.056 | Young |
| 0.103697 | 0.221911 | 0.789 | 0.779 | 1 | 13 | Rtf2 | 0.01 | Young |
| 0.103921 | 0.20753 | 0.737 | 0.738 | 1 | 13 | Znhit1 | -0.001 | Young |
| 0.104096 | 0.434345 | 0.465 | 0.385 | 1 | 13 | Irf3 | 0.08 | Young |
| 0.104174 | 0.171491 | 0.754 | 0.713 | 1 | 13 | Prrc2b | 0.041 | Young |
| 0.104214 | 0.610828 | 0.228 | 0.148 | 1 | 13 | Coro2b | 0.08 | Young |
| 0.104223 | 0.257363 | 0.798 | 0.721 | 1 | 13 | Taf6l | 0.077 | Young |
| 0.10423 | 0.092285 | 0.982 | 1 | 1 | 13 | Serp1 | -0.018 | Young |
| 0.104323 | 0.346941 | 0.693 | 0.672 | 1 | 13 | Lyl1 | 0.021 | Young |
| 0.10443 | 0.694993 | 0.237 | 0.164 | 1 | 13 | Gna11 | 0.073 | Young |
| 0.104473 | 0.277779 | 0.684 | 0.607 | 1 | 13 | Usp24 | 0.077 | Young |
| 0.104763 | 0.397786 | 0.57 | 0.5 | 1 | 13 | Rnf220 | 0.07 | Young |

| | | | | | | | | |
|---|---|---|---|---|---|---|---|---|
| 0.105356 | 0.499335 | 0.491 | 0.443 | 1 | 13 | Tesk1 | 0.048 | Young |
| 0.105417 | 0.565732 | 0.228 | 0.148 | 1 | 13 | Plk4 | 0.08 | Young |
| 0.105429 | 1.288888 | 0.079 | 0.033 | 1 | 13 | Tm9sf1 | 0.046 | Young |
| 0.105453 | 0.23586 | 0.886 | 0.811 | 1 | 13 | Arrb2 | 0.075 | Young |
| 0.105486 | 0.363204 | 0.588 | 0.557 | 1 | 13 | Oser1 | 0.031 | Young |
| 0.105494 | 0.309581 | 0.596 | 0.557 | 1 | 13 | Fam76b | 0.039 | Young |
| 0.105898 | 0.54528 | 0.219 | 0.139 | 1 | 13 | Cep128 | 0.08 | Young |
| 0.106061 | 0.669621 | 0.202 | 0.123 | 1 | 13 | Rpap2 | 0.079 | Young |
| 0.106247 | 0.656794 | 0.263 | 0.189 | 1 | 13 | Dclre1b | 0.074 | Young |
| 0.10628 | 0.173722 | 1 | 0.984 | 1 | 13 | Ndufb11 | 0.016 | Young |
| 0.106486 | 0.174655 | 0.991 | 0.975 | 1 | 13 | Klf3 | 0.016 | Young |
| 0.107108 | 0.1017 | 1 | 1 | 1 | 13 | Gltp | 0 | Young |
| 0.107108 | 0.084239 | 1 | 1 | 1 | 13 | Rps17 | 0 | Young |
| 0.107198 | 0.142954 | 0.86 | 0.73 | 1 | 13 | Tns1 | 0.13 | Young |
| 0.107254 | 0.790389 | 0.105 | 0.049 | 1 | 13 | Tnks1bp1 | 0.056 | Young |
| 0.107507 | 0.185846 | 0.974 | 0.934 | 1 | 13 | Mrpl23 | 0.04 | Young |
| 0.107643 | 0.362657 | 0.623 | 0.541 | 1 | 13 | Tcp11l2 | 0.082 | Young |
| 0.107733 | 0.188145 | 0.895 | 0.893 | 1 | 13 | Dnaja2 | 0.002 | Young |
| 0.107959 | 0.667923 | 0.281 | 0.205 | 1 | 13 | Ttc39c | 0.076 | Young |
| 0.107983 | 0.586026 | 0.5 | 0.459 | 1 | 13 | Sos2 | 0.041 | Young |
| 0.108037 | 0.334552 | 0.798 | 0.713 | 1 | 13 | Nfat5 | 0.085 | Young |
| 0.108282 | 0.399588 | 0.553 | 0.475 | 1 | 13 | Arrdc3 | 0.078 | Young |
| 0.108348 | 0.571141 | 0.5 | 0.443 | 1 | 13 | Itgb7 | 0.057 | Young |
| 0.108581 | 1.334093 | 0.079 | 0.033 | 1 | 13 | Barx2 | 0.046 | Young |
| 0.109036 | 0.817277 | 0.193 | 0.123 | 1 | 13 | Fbxo31 | 0.07 | Young |
| 0.109095 | 0.181446 | 0.895 | 0.82 | 1 | 13 | Derl1 | 0.075 | Young |
| 0.109537 | 0.279946 | 0.526 | 0.426 | 1 | 13 | Runx3 | 0.1 | Young |
| 0.110341 | 0.420838 | 0.456 | 0.377 | 1 | 13 | Wasl | 0.079 | Young |
| 0.110393 | 0.795292 | 0.175 | 0.107 | 1 | 13 | Rbm12b1 | 0.068 | Young |
| 0.110511 | 0.876785 | 0.114 | 0.057 | 1 | 13 | Fancc | 0.057 | Young |
| 0.110625 | 0.277624 | 0.711 | 0.639 | 1 | 13 | Sptlc2 | 0.072 | Young |
| 0.111767 | 0.319868 | 0.851 | 0.844 | 1 | 13 | Mapk6 | 0.007 | Young |
| 0.111806 | 1.006907 | 0.079 | 0.033 | 1 | 13 | AC129328. | 0.046 | Young |
| 0.112474 | 0.41257 | 0.43 | 0.361 | 1 | 13 | Ddit3 | 0.069 | Young |
| 0.112527 | 0.697054 | 0.175 | 0.107 | 1 | 13 | Rbm38 | 0.068 | Young |
| 0.112603 | 0.262437 | 0.974 | 0.934 | 1 | 13 | Sgms1 | 0.04 | Young |
| 0.112827 | 0.280329 | 0.833 | 0.828 | 1 | 13 | Mlec | 0.005 | Young |
| 0.112898 | 1.167659 | 0.079 | 0.033 | 1 | 13 | Senp8 | 0.046 | Young |
| 0.113065 | 0.655188 | 0.325 | 0.254 | 1 | 13 | Gm20696 | 0.071 | Young |
| 0.113271 | 0.521438 | 0.254 | 0.172 | 1 | 13 | Mthfd2l | 0.082 | Young |
| 0.114178 | 0.635173 | 0.202 | 0.131 | 1 | 13 | Kcne3 | 0.071 | Young |
| 0.114197 | 0.631242 | 0.228 | 0.156 | 1 | 13 | Lrp5 | 0.072 | Young |
| 0.114556 | 0.183661 | 0.956 | 0.943 | 1 | 13 | Cnn2 | 0.013 | Young |
| 0.114572 | 0.509567 | 0.395 | 0.328 | 1 | 13 | Zfp467 | 0.067 | Young |
| 0.114774 | 0.476015 | 0.281 | 0.197 | 1 | 13 | Tbkbp1 | 0.084 | Young |

| | | | | | | | | |
|---|---|---|---|---|---|---|---|---|
| 0.114915 | 0.173414 | 0.658 | 0.615 | 1 | 13 | Map4k1 | 0.043 | Young |
| 0.115333 | 0.445807 | 0.333 | 0.238 | 1 | 13 | Rnaseh2b | 0.095 | Young |
| 0.115585 | 0.170548 | 0.93 | 0.91 | 1 | 13 | Ilk | 0.02 | Young |
| 0.115679 | 0.161605 | 1 | 0.967 | 1 | 13 | Ptpre | 0.033 | Young |
| 0.115701 | 0.896253 | 0.149 | 0.09 | 1 | 13 | Ptpn21 | 0.059 | Young |
| 0.116224 | 0.965176 | 0.079 | 0.033 | 1 | 13 | Zfp3 | 0.046 | Young |
| 0.116489 | 0.83249 | 0.149 | 0.09 | 1 | 13 | AA467197 | 0.059 | Young |
| 0.116508 | 0.220441 | 0.781 | 0.705 | 1 | 13 | Vps36 | 0.076 | Young |
| 0.116677 | 0.320968 | 0.684 | 0.607 | 1 | 13 | Dnajc1 | 0.077 | Young |
| 0.116853 | 0.586496 | 0.474 | 0.41 | 1 | 13 | Plekhf2 | 0.064 | Young |
| 0.11687 | 0.170365 | 0.825 | 0.82 | 1 | 13 | Mrpl30 | 0.005 | Young |
| 0.116892 | 0.671688 | 0.175 | 0.107 | 1 | 13 | Klhl22 | 0.068 | Young |
| 0.117079 | 1.27445 | 0.053 | 0.016 | 1 | 13 | Slc39a8 | 0.037 | Young |
| 0.117079 | 1.147096 | 0.053 | 0.016 | 1 | 13 | Nipsnap1 | 0.037 | Young |
| 0.117083 | 0.274843 | 0.281 | 0.189 | 1 | 13 | Dpp4 | 0.092 | Young |
| 0.117193 | 0.392467 | 0.658 | 0.582 | 1 | 13 | Gch1 | 0.076 | Young |
| 0.117458 | 0.108829 | 1 | 1 | 1 | 13 | Pld4 | 0 | Young |
| 0.11758 | 0.259906 | 0.868 | 0.844 | 1 | 13 | Gm10073 | 0.024 | Young |
| 0.117673 | 0.567721 | 0.289 | 0.213 | 1 | 13 | Fiz1 | 0.076 | Young |
| 0.118117 | 0.690975 | 0.333 | 0.27 | 1 | 13 | Gpkow | 0.063 | Young |
| 0.118508 | 1.276816 | 0.053 | 0.016 | 1 | 13 | Parp16 | 0.037 | Young |
| 0.118508 | 1.163661 | 0.053 | 0.016 | 1 | 13 | Gm15545 | 0.037 | Young |
| 0.118709 | 0.263255 | 0.544 | 0.451 | 1 | 13 | Etv1 | 0.093 | Young |
| 0.119117 | 0.221625 | 0.711 | 0.639 | 1 | 13 | Lemd2 | 0.072 | Young |
| 0.119268 | 0.216672 | 0.561 | 0.533 | 1 | 13 | Atp11c | 0.028 | Young |
| 0.119485 | 0.892863 | 0.132 | 0.074 | 1 | 13 | Hist2h2be | 0.058 | Young |
| 0.119507 | 0.293769 | 0.684 | 0.648 | 1 | 13 | Psmd12 | 0.036 | Young |
| 0.119573 | 0.258392 | 0.825 | 0.82 | 1 | 13 | Sgpp1 | 0.005 | Young |
| 0.11993 | 0.337498 | 0.465 | 0.369 | 1 | 13 | Phf12 | 0.096 | Young |
| 0.120094 | 0.217281 | 0.728 | 0.648 | 1 | 13 | Sla | 0.08 | Young |
| 0.120164 | 0.095201 | 1 | 1 | 1 | 13 | Ybx1 | 0 | Young |
| 0.120777 | 1.06669 | 0.079 | 0.033 | 1 | 13 | Arhgap27o | 0.046 | Young |
| 0.120777 | 0.939513 | 0.079 | 0.033 | 1 | 13 | Slc5a6 | 0.046 | Young |
| 0.121222 | 0.323297 | 0.535 | 0.434 | 1 | 13 | Arid3a | 0.101 | Young |
| 0.121347 | 0.500299 | 0.219 | 0.148 | 1 | 13 | Stk11ip | 0.071 | Young |
| 0.121407 | 1.217639 | 0.053 | 0.016 | 1 | 13 | Sfn | 0.037 | Young |
| 0.121535 | 0.127142 | 1 | 1 | 1 | 13 | Tln1 | 0 | Young |
| 0.121719 | 0.379843 | 0.456 | 0.377 | 1 | 13 | Retreg1 | 0.079 | Young |
| 0.121995 | 0.171862 | 1 | 1 | 1 | 13 | Cd52 | 0 | Young |
| 0.121995 | 0.057129 | 1 | 1 | 1 | 13 | Rps2 | 0 | Young |
| 0.122203 | 0.394163 | 0.447 | 0.361 | 1 | 13 | Dcps | 0.086 | Young |
| 0.122456 | 0.679921 | 0.114 | 0.057 | 1 | 13 | Gm20513 | 0.057 | Young |
| 0.12271 | 0.339533 | 0.518 | 0.451 | 1 | 13 | Filip1l | 0.067 | Young |
| 0.122877 | 1.497889 | 0.053 | 0.016 | 1 | 13 | Creb3l3 | 0.037 | Young |
| 0.122877 | 1.058403 | 0.053 | 0.016 | 1 | 13 | Rhof | 0.037 | Young |

| | | | | | | | | |
|---|---|---|---|---|---|---|---|---|
| 0.122877 | 0.998427 | 0.053 | 0.016 | 1 | 13 | Erbb3 | 0.037 | Young |
| 0.122877 | 0.98329 | 0.053 | 0.016 | 1 | 13 | Zbtb3 | 0.037 | Young |
| 0.123378 | 0.150779 | 0.974 | 0.967 | 1 | 13 | Nsa2 | 0.007 | Young |
| 0.123527 | 0.511539 | 0.395 | 0.32 | 1 | 13 | Plxna4os1 | 0.075 | Young |
| 0.12383 | 0.179346 | 0.851 | 0.926 | 1 | 13 | Nxf1 | -0.075 | Young |
| 0.124266 | 0.146659 | 0.947 | 0.926 | 1 | 13 | Ifitm6 | 0.021 | Young |
| 0.124782 | 0.105788 | 1 | 1 | 1 | 13 | Rbm39 | 0 | Young |
| 0.124801 | 0.23259 | 0.825 | 0.836 | 1 | 13 | Mrps24 | -0.011 | Young |
| 0.12512 | 0.363361 | 0.474 | 0.393 | 1 | 13 | Stard7 | 0.081 | Young |
| 0.125241 | 0.621905 | 0.175 | 0.107 | 1 | 13 | Iqcb1 | 0.068 | Young |
| 0.125392 | 0.945272 | 0.175 | 0.115 | 1 | 13 | Tmem101 | 0.06 | Young |
| 0.12564 | 0.165836 | 0.825 | 0.779 | 1 | 13 | Csnk1e | 0.046 | Young |
| 0.125669 | 0.341861 | 0.518 | 0.467 | 1 | 13 | Zbtb11 | 0.051 | Young |
| 0.125859 | 1.019931 | 0.053 | 0.016 | 1 | 13 | Rbp1 | 0.037 | Young |
| 0.125859 | 0.902146 | 0.053 | 0.016 | 1 | 13 | Gm30329 | 0.037 | Young |
| 0.125859 | 0.854915 | 0.053 | 0.016 | 1 | 13 | Dchs1 | 0.037 | Young |
| 0.126141 | 0.192517 | 0.763 | 0.73 | 1 | 13 | Ppp1r10 | 0.033 | Young |
| 0.126494 | 0.254986 | 0.789 | 0.738 | 1 | 13 | Tnrc6c | 0.051 | Young |
| 0.12682 | 0.705659 | 0.211 | 0.148 | 1 | 13 | Tpbgl | 0.063 | Young |
| 0.126967 | 0.274081 | 0.456 | 0.361 | 1 | 13 | Atxn1 | 0.095 | Young |
| 0.127405 | 0.200814 | 0.746 | 0.762 | 1 | 13 | Ndufs3 | -0.016 | Young |
| 0.127781 | 0.336081 | 0.544 | 0.516 | 1 | 13 | Pom121 | 0.028 | Young |
| 0.1278 | 0.98376 | 0.088 | 0.041 | 1 | 13 | Gm32036 | 0.047 | Young |
| 0.127912 | 0.552669 | 0.36 | 0.287 | 1 | 13 | Isg20l2 | 0.073 | Young |
| 0.128044 | 0.318701 | 0.509 | 0.443 | 1 | 13 | Nsmaf | 0.066 | Young |
| 0.128097 | 0.06647 | 1 | 1 | 1 | 13 | Btg2 | 0 | Young |
| 0.128897 | 0.797661 | 0.053 | 0.016 | 1 | 13 | Zfp763 | 0.037 | Young |
| 0.129183 | 0.206328 | 0.895 | 0.861 | 1 | 13 | Ski | 0.034 | Young |
| 0.129294 | 0.691113 | 0.114 | 0.057 | 1 | 13 | Fzd4 | 0.057 | Young |
| 0.129634 | 0.332523 | 0.395 | 0.311 | 1 | 13 | Gpd1l | 0.084 | Young |
| 0.129862 | 0.187239 | 0.719 | 0.656 | 1 | 13 | Cnot4 | 0.063 | Young |
| 0.129958 | 0.207576 | 0.746 | 0.689 | 1 | 13 | Rbfa | 0.057 | Young |
| 0.130074 | 1.278641 | 0.088 | 0.041 | 1 | 13 | Cd28 | 0.047 | Young |
| 0.130303 | 0.473762 | 0.211 | 0.139 | 1 | 13 | Tbc1d32 | 0.072 | Young |
| 0.130362 | 0.278615 | 0.667 | 0.574 | 1 | 13 | Ccdc59 | 0.093 | Young |
| 0.130554 | 0.981318 | 0.149 | 0.09 | 1 | 13 | Platr17 | 0.059 | Young |
| 0.130554 | 0.270326 | 0.754 | 0.754 | 1 | 13 | Pikfyve | 0 | Young |
| 0.131222 | 0.950454 | 0.088 | 0.041 | 1 | 13 | Mks1 | 0.047 | Young |
| 0.131263 | 0.440013 | 0.526 | 0.492 | 1 | 13 | Ophn1 | 0.034 | Young |
| 0.131291 | 1.189689 | 0.096 | 0.049 | 1 | 13 | Bfsp1 | 0.047 | Young |
| 0.131506 | 0.266089 | 0.596 | 0.484 | 1 | 13 | Siglece | 0.112 | Young |
| 0.131677 | 0.479024 | 0.596 | 0.549 | 1 | 13 | Cks2 | 0.047 | Young |
| 0.131787 | 0.343549 | 0.491 | 0.418 | 1 | 13 | Nfia | 0.073 | Young |
| 0.131992 | 0.613181 | 0.053 | 0.016 | 1 | 13 | Rsph3a | 0.037 | Young |
| 0.132138 | 0.307771 | 0.658 | 0.623 | 1 | 13 | Micu1 | 0.035 | Young |

| | | | | | | | | |
|---|---|---|---|---|---|---|---|---|
| 0.132379 | 0.975312 | 0.088 | 0.041 | 1 | 13 | Abtb2 | 0.047 | Young |
| 0.132433 | 0.599201 | 0.184 | 0.115 | 1 | 13 | Zfhx2 | 0.069 | Young |
| 0.132458 | 0.099138 | 1 | 1 | 1 | 13 | Naca | 0 | Young |
| 0.132481 | 0.425335 | 0.289 | 0.213 | 1 | 13 | Sap30bp | 0.076 | Young |
| 0.13266 | 0.805225 | 0.123 | 0.066 | 1 | 13 | Inpp4b | 0.057 | Young |
| 0.132679 | 0.160294 | 0.956 | 0.943 | 1 | 13 | Pdcd5 | 0.013 | Young |
| 0.132947 | 0.119366 | 0.965 | 0.984 | 1 | 13 | Zfp706 | -0.019 | Young |
| 0.132974 | 0.2067 | 0.509 | 0.418 | 1 | 13 | Wnt11 | 0.091 | Young |
| 0.133036 | 0.200298 | 0.763 | 0.721 | 1 | 13 | Ift20 | 0.042 | Young |
| 0.133888 | 0.135847 | 0.939 | 0.934 | 1 | 13 | Mef2a | 0.005 | Young |
| 0.13402 | 0.614068 | 0.079 | 0.033 | 1 | 13 | D830025C( | 0.046 | Young |
| 0.134108 | 0.626738 | 0.342 | 0.27 | 1 | 13 | Slc14a1 | 0.072 | Young |
| 0.134433 | 0.163917 | 1 | 1 | 1 | 13 | Pnrc1 | 0 | Young |
| 0.135571 | 0.378657 | 0.491 | 0.418 | 1 | 13 | Ing5 | 0.073 | Young |
| 0.135659 | 0.379596 | 0.421 | 0.361 | 1 | 13 | AI837181 | 0.06 | Young |
| 0.135884 | 0.195628 | 0.816 | 0.787 | 1 | 13 | Fam173a | 0.029 | Young |
| 0.136272 | 0.326705 | 0.465 | 0.41 | 1 | 13 | Rnf169 | 0.055 | Young |
| 0.136491 | 0.541423 | 0.289 | 0.221 | 1 | 13 | Capn15 | 0.068 | Young |
| 0.13687 | 0.393111 | 0.684 | 0.672 | 1 | 13 | Zfp652 | 0.012 | Young |
| 0.137229 | 1.017645 | 0.105 | 0.057 | 1 | 13 | Pex11g | 0.048 | Young |
| 0.1373 | 0.588903 | 0.316 | 0.254 | 1 | 13 | Kdm4c | 0.062 | Young |
| 0.137368 | 0.725435 | 0.175 | 0.115 | 1 | 13 | Pik3ap1 | 0.06 | Young |
| 0.137609 | 0.294462 | 0.553 | 0.484 | 1 | 13 | 9930021J0 | 0.069 | Young |
| 0.137625 | 0.449121 | 0.404 | 0.32 | 1 | 13 | Chpt1 | 0.084 | Young |
| 0.137944 | 1.054091 | 0.096 | 0.049 | 1 | 13 | Arsi | 0.047 | Young |
| 0.13844 | 0.404947 | 0.482 | 0.451 | 1 | 13 | Alas1 | 0.031 | Young |
| 0.138488 | 0.111658 | 0.754 | 0.779 | 1 | 13 | Sdf2l1 | -0.025 | Young |
| 0.138772 | 0.522971 | 0.447 | 0.41 | 1 | 13 | Vgll4 | 0.037 | Young |
| 0.139033 | 0.531246 | 0.228 | 0.156 | 1 | 13 | Ptcd3 | 0.072 | Young |
| 0.139161 | 0.585319 | 0.193 | 0.123 | 1 | 13 | Zfp74 | 0.07 | Young |
| 0.139461 | 0.298277 | 0.623 | 0.566 | 1 | 13 | Lpcat1 | 0.057 | Young |
| 0.139778 | 0.491491 | 0.228 | 0.156 | 1 | 13 | Bpnt1 | 0.072 | Young |
| 0.139828 | 0.233549 | 0.544 | 0.451 | 1 | 13 | BC003331 | 0.093 | Young |
| 0.139995 | 0.469073 | 0.395 | 0.328 | 1 | 13 | BC005561 | 0.067 | Young |
| 0.140342 | 0.27253 | 0.649 | 0.607 | 1 | 13 | Pold4 | 0.042 | Young |
| 0.140347 | 0.770846 | 0.219 | 0.156 | 1 | 13 | Rab13 | 0.063 | Young |
| 0.140697 | 0.852904 | 0.088 | 0.041 | 1 | 13 | Apbb1 | 0.047 | Young |
| 0.140752 | 0.080274 | 1 | 1 | 1 | 13 | Rpl36a | 0 | Young |
| 0.141147 | 0.76089 | 0.193 | 0.131 | 1 | 13 | Flt3 | 0.062 | Young |
| 0.141327 | 0.29225 | 0.474 | 0.402 | 1 | 13 | Arl2bp | 0.072 | Young |
| 0.141743 | 0.474559 | 0.702 | 0.631 | 1 | 13 | Chd7 | 0.071 | Young |
| 0.141917 | 0.844138 | 0.088 | 0.041 | 1 | 13 | Nipa1 | 0.047 | Young |
| 0.14206 | 0.408536 | 0.43 | 0.361 | 1 | 13 | Tpgs1 | 0.069 | Young |
| 0.142282 | 0.395532 | 0.412 | 0.32 | 1 | 13 | Narf | 0.092 | Young |
| 0.142301 | 0.100946 | 0.991 | 0.975 | 1 | 13 | Tsc22d4 | 0.016 | Young |

| | | | | | | | | |
|---|---|---|---|---|---|---|---|---|
| 0.142302 | 0.112854 | 0.991 | 0.992 | 1 | 13 | Foxp1 | -0.001 | Young |
| 0.142312 | 0.333604 | 0.456 | 0.393 | 1 | 13 | Golga5 | 0.063 | Young |
| 0.142329 | 0.140789 | 0.807 | 0.779 | 1 | 13 | Sult1a1 | 0.028 | Young |
| 0.142505 | 0.30851 | 0.553 | 0.484 | 1 | 13 | Uimc1 | 0.069 | Young |
| 0.142562 | 0.082052 | 1 | 1 | 1 | 13 | Rpl22 | 0 | Young |
| 0.142781 | 0.899757 | 0.202 | 0.139 | 1 | 13 | Abhd14b | 0.063 | Young |
| 0.143149 | 0.399166 | 0.474 | 0.443 | 1 | 13 | Plk3 | 0.031 | Young |
| 0.143298 | 0.282942 | 0.693 | 0.631 | 1 | 13 | Commd4 | 0.062 | Young |
| 0.143871 | 0.681223 | 0.325 | 0.262 | 1 | 13 | Cep164 | 0.063 | Young |
| 0.144208 | 0.583043 | 0.36 | 0.287 | 1 | 13 | Jmy | 0.073 | Young |
| 0.144264 | 1.361465 | 0.018 | 0 | 1 | 13 | Dusp14 | 0.018 | Young |
| 0.144264 | 1.297607 | 0.018 | 0 | 1 | 13 | Fmr1nb | 0.018 | Young |
| 0.144264 | 1.098872 | 0.018 | 0 | 1 | 13 | Gm46404 | 0.018 | Young |
| 0.144264 | 1.076597 | 0.018 | 0 | 1 | 13 | A730011C1 | 0.018 | Young |
| 0.144264 | 1.066205 | 0.018 | 0 | 1 | 13 | Mybl2 | 0.018 | Young |
| 0.144264 | 1.046549 | 0.018 | 0 | 1 | 13 | Ly6g5b | 0.018 | Young |
| 0.144264 | 1.035307 | 0.018 | 0 | 1 | 13 | 9930004E1 | 0.018 | Young |
| 0.144264 | 1.013292 | 0.018 | 0 | 1 | 13 | Kif21a | 0.018 | Young |
| 0.144264 | 1.011384 | 0.018 | 0 | 1 | 13 | Ndufa4l2 | 0.018 | Young |
| 0.144264 | 1.010988 | 0.018 | 0 | 1 | 13 | Mfap5 | 0.018 | Young |
| 0.144264 | 1.005386 | 0.018 | 0 | 1 | 13 | Gm12002 | 0.018 | Young |
| 0.144264 | 0.999058 | 0.018 | 0 | 1 | 13 | Tmem91 | 0.018 | Young |
| 0.144264 | 0.992191 | 0.018 | 0 | 1 | 13 | Gm10603 | 0.018 | Young |
| 0.144264 | 0.981176 | 0.018 | 0 | 1 | 13 | 1700010I1 | 0.018 | Young |
| 0.144264 | 0.979139 | 0.018 | 0 | 1 | 13 | 1700064M | 0.018 | Young |
| 0.144264 | 0.971766 | 0.018 | 0 | 1 | 13 | Gm11361 | 0.018 | Young |
| 0.144264 | 0.967241 | 0.018 | 0 | 1 | 13 | Cfap99 | 0.018 | Young |
| 0.144264 | 0.954416 | 0.018 | 0 | 1 | 13 | 4930570N1 | 0.018 | Young |
| 0.144264 | 0.950411 | 0.018 | 0 | 1 | 13 | Gm44053 | 0.018 | Young |
| 0.144264 | 0.944366 | 0.018 | 0 | 1 | 13 | Atp2a1 | 0.018 | Young |
| 0.144264 | 0.943632 | 0.018 | 0 | 1 | 13 | Pcdhga2 | 0.018 | Young |
| 0.144264 | 0.941346 | 0.018 | 0 | 1 | 13 | Gm12905 | 0.018 | Young |
| 0.144264 | 0.934448 | 0.018 | 0 | 1 | 13 | Klk13 | 0.018 | Young |
| 0.144264 | 0.932013 | 0.018 | 0 | 1 | 13 | Adcyap1r1 | 0.018 | Young |
| 0.144264 | 0.93045 | 0.018 | 0 | 1 | 13 | Gm21859 | 0.018 | Young |
| 0.144264 | 0.924778 | 0.018 | 0 | 1 | 13 | Sema4g | 0.018 | Young |
| 0.144264 | 0.924649 | 0.018 | 0 | 1 | 13 | Gm10647 | 0.018 | Young |
| 0.144264 | 0.924493 | 0.018 | 0 | 1 | 13 | Majin | 0.018 | Young |
| 0.144264 | 0.923242 | 0.018 | 0 | 1 | 13 | Wdr54 | 0.018 | Young |
| 0.144264 | 0.92013 | 0.018 | 0 | 1 | 13 | Gm13713 | 0.018 | Young |
| 0.144264 | 0.909797 | 0.018 | 0 | 1 | 13 | Gm46218 | 0.018 | Young |
| 0.144264 | 0.902739 | 0.018 | 0 | 1 | 13 | Apoa2 | 0.018 | Young |
| 0.144264 | 0.901489 | 0.018 | 0 | 1 | 13 | Aldh1a1 | 0.018 | Young |
| 0.144264 | 0.900933 | 0.018 | 0 | 1 | 13 | Tnfaip6 | 0.018 | Young |
| 0.144264 | 0.899297 | 0.018 | 0 | 1 | 13 | Pcdh19 | 0.018 | Young |

| | | | | | | | | |
|---|---|---|---|---|---|---|---|---|
| 0.144264 | 0.898703 | 0.018 | 0 | 1 | 13 | Gm28192 | 0.018 | Young |
| 0.144264 | 0.898309 | 0.018 | 0 | 1 | 13 | Aoc3 | 0.018 | Young |
| 0.144264 | 0.89562 | 0.018 | 0 | 1 | 13 | Gm14167 | 0.018 | Young |
| 0.144264 | 0.894715 | 0.018 | 0 | 1 | 13 | Bicc1 | 0.018 | Young |
| 0.144264 | 0.892017 | 0.018 | 0 | 1 | 13 | Ccdc85a | 0.018 | Young |
| 0.144264 | 0.888814 | 0.018 | 0 | 1 | 13 | Fxyd1 | 0.018 | Young |
| 0.144264 | 0.883747 | 0.018 | 0 | 1 | 13 | Gm32401 | 0.018 | Young |
| 0.144264 | 0.871959 | 0.018 | 0 | 1 | 13 | Phgdh | 0.018 | Young |
| 0.144264 | 0.859483 | 0.018 | 0 | 1 | 13 | Spef2 | 0.018 | Young |
| 0.144264 | 0.85405 | 0.018 | 0 | 1 | 13 | Cdnf | 0.018 | Young |
| 0.144264 | 0.850132 | 0.018 | 0 | 1 | 13 | Col6a1 | 0.018 | Young |
| 0.144264 | 0.848246 | 0.018 | 0 | 1 | 13 | Klre1 | 0.018 | Young |
| 0.144264 | 0.845288 | 0.018 | 0 | 1 | 13 | Gm26724 | 0.018 | Young |
| 0.144264 | 0.84441 | 0.018 | 0 | 1 | 13 | Pdf | 0.018 | Young |
| 0.144264 | 0.844405 | 0.018 | 0 | 1 | 13 | Hcn4 | 0.018 | Young |
| 0.144264 | 0.843965 | 0.018 | 0 | 1 | 13 | Gm26785 | 0.018 | Young |
| 0.144264 | 0.839526 | 0.018 | 0 | 1 | 13 | Cd200r2 | 0.018 | Young |
| 0.144264 | 0.835252 | 0.018 | 0 | 1 | 13 | BC051537 | 0.018 | Young |
| 0.144264 | 0.834204 | 0.018 | 0 | 1 | 13 | Sdk1 | 0.018 | Young |
| 0.144264 | 0.829242 | 0.018 | 0 | 1 | 13 | 4930458B2 | 0.018 | Young |
| 0.144264 | 0.826693 | 0.018 | 0 | 1 | 13 | Gm13383 | 0.018 | Young |
| 0.144264 | 0.825984 | 0.018 | 0 | 1 | 13 | Mzf1 | 0.018 | Young |
| 0.144264 | 0.824773 | 0.018 | 0 | 1 | 13 | Ddit4l | 0.018 | Young |
| 0.144264 | 0.824197 | 0.018 | 0 | 1 | 13 | 4831440E1 | 0.018 | Young |
| 0.144264 | 0.822559 | 0.018 | 0 | 1 | 13 | Ninl | 0.018 | Young |
| 0.144264 | 0.821104 | 0.018 | 0 | 1 | 13 | Atcayos | 0.018 | Young |
| 0.144264 | 0.820292 | 0.018 | 0 | 1 | 13 | Nat14 | 0.018 | Young |
| 0.144264 | 0.818434 | 0.018 | 0 | 1 | 13 | Gm26582 | 0.018 | Young |
| 0.144264 | 0.813293 | 0.018 | 0 | 1 | 13 | Nwd1 | 0.018 | Young |
| 0.144264 | 0.807289 | 0.018 | 0 | 1 | 13 | 4930432J0 | 0.018 | Young |
| 0.144264 | 0.802789 | 0.018 | 0 | 1 | 13 | Zscan25 | 0.018 | Young |
| 0.144264 | 0.801504 | 0.018 | 0 | 1 | 13 | Gm12185 | 0.018 | Young |
| 0.144264 | 0.798846 | 0.018 | 0 | 1 | 13 | Rhbdl3 | 0.018 | Young |
| 0.144264 | 0.798411 | 0.018 | 0 | 1 | 13 | 6430550D2 | 0.018 | Young |
| 0.144264 | 0.793211 | 0.018 | 0 | 1 | 13 | Afmid | 0.018 | Young |
| 0.144264 | 0.792557 | 0.018 | 0 | 1 | 13 | Fbxl12os | 0.018 | Young |
| 0.144264 | 0.791827 | 0.018 | 0 | 1 | 13 | Zfp972 | 0.018 | Young |
| 0.144264 | 0.787527 | 0.018 | 0 | 1 | 13 | CT025521.1 | 0.018 | Young |
| 0.144264 | 0.786127 | 0.018 | 0 | 1 | 13 | Pdzd2 | 0.018 | Young |
| 0.144264 | 0.785649 | 0.018 | 0 | 1 | 13 | Ces1d | 0.018 | Young |
| 0.144264 | 0.774037 | 0.018 | 0 | 1 | 13 | Cyp2d22 | 0.018 | Young |
| 0.144264 | 0.773986 | 0.018 | 0 | 1 | 13 | Fstl3 | 0.018 | Young |
| 0.144264 | 0.771436 | 0.018 | 0 | 1 | 13 | Hdac11 | 0.018 | Young |
| 0.144264 | 0.766192 | 0.018 | 0 | 1 | 13 | Gm49004 | 0.018 | Young |
| 0.144264 | 0.765198 | 0.018 | 0 | 1 | 13 | Gfod2 | 0.018 | Young |

| | | | | | | | | |
|---|---|---|---|---|---|---|---|---|
| 0.144264 | 0.761068 | 0.018 | 0 | 1 | 13 | Smad6 | 0.018 | Young |
| 0.144264 | 0.759403 | 0.018 | 0 | 1 | 13 | 1700088E0 | 0.018 | Young |
| 0.144264 | 0.757723 | 0.018 | 0 | 1 | 13 | Zfp647 | 0.018 | Young |
| 0.144264 | 0.756715 | 0.018 | 0 | 1 | 13 | Unc45b | 0.018 | Young |
| 0.144264 | 0.756341 | 0.018 | 0 | 1 | 13 | Hist1h4n | 0.018 | Young |
| 0.144264 | 0.756058 | 0.018 | 0 | 1 | 13 | Aldh5a1 | 0.018 | Young |
| 0.144264 | 0.753799 | 0.018 | 0 | 1 | 13 | Smc2os | 0.018 | Young |
| 0.144264 | 0.746589 | 0.018 | 0 | 1 | 13 | Dab2ip | 0.018 | Young |
| 0.144264 | 0.743752 | 0.018 | 0 | 1 | 13 | C1qtnf7 | 0.018 | Young |
| 0.144264 | 0.740809 | 0.018 | 0 | 1 | 13 | Gm26637 | 0.018 | Young |
| 0.144264 | 0.738515 | 0.018 | 0 | 1 | 13 | Gm43573 | 0.018 | Young |
| 0.144264 | 0.735187 | 0.018 | 0 | 1 | 13 | Ackr3 | 0.018 | Young |
| 0.144264 | 0.733115 | 0.018 | 0 | 1 | 13 | Neu3 | 0.018 | Young |
| 0.144264 | 0.730145 | 0.018 | 0 | 1 | 13 | Zfr2 | 0.018 | Young |
| 0.144264 | 0.72086 | 0.018 | 0 | 1 | 13 | Frmd8os | 0.018 | Young |
| 0.144264 | 0.717998 | 0.018 | 0 | 1 | 13 | Chadl | 0.018 | Young |
| 0.144264 | 0.71541 | 0.018 | 0 | 1 | 13 | 2700016F2 | 0.018 | Young |
| 0.144264 | 0.714026 | 0.018 | 0 | 1 | 13 | Abat | 0.018 | Young |
| 0.144264 | 0.713172 | 0.018 | 0 | 1 | 13 | Epha10 | 0.018 | Young |
| 0.144264 | 0.710053 | 0.018 | 0 | 1 | 13 | Gm16894 | 0.018 | Young |
| 0.144264 | 0.708971 | 0.018 | 0 | 1 | 13 | Gm26676 | 0.018 | Young |
| 0.144264 | 0.708294 | 0.018 | 0 | 1 | 13 | Smc1b | 0.018 | Young |
| 0.144264 | 0.696029 | 0.018 | 0 | 1 | 13 | Limch1 | 0.018 | Young |
| 0.144264 | 0.695166 | 0.018 | 0 | 1 | 13 | Gm5426 | 0.018 | Young |
| 0.144264 | 0.691334 | 0.018 | 0 | 1 | 13 | S1pr4 | 0.018 | Young |
| 0.144264 | 0.686749 | 0.018 | 0 | 1 | 13 | B930094E0 | 0.018 | Young |
| 0.144264 | 0.683076 | 0.018 | 0 | 1 | 13 | Pygm | 0.018 | Young |
| 0.144264 | 0.682899 | 0.018 | 0 | 1 | 13 | Gnai1 | 0.018 | Young |
| 0.144264 | 0.680235 | 0.018 | 0 | 1 | 13 | Hmgcs2 | 0.018 | Young |
| 0.144264 | 0.677122 | 0.018 | 0 | 1 | 13 | 5033403F0 | 0.018 | Young |
| 0.144264 | 0.673801 | 0.018 | 0 | 1 | 13 | Gm10244 | 0.018 | Young |
| 0.144264 | 0.670083 | 0.018 | 0 | 1 | 13 | Fam57b | 0.018 | Young |
| 0.144264 | 0.668191 | 0.018 | 0 | 1 | 13 | Nnat | 0.018 | Young |
| 0.144264 | 0.661946 | 0.018 | 0 | 1 | 13 | Popdc2 | 0.018 | Young |
| 0.144264 | 0.658843 | 0.018 | 0 | 1 | 13 | Kif27 | 0.018 | Young |
| 0.144264 | 0.657939 | 0.018 | 0 | 1 | 13 | Lurap1 | 0.018 | Young |
| 0.144264 | 0.656714 | 0.018 | 0 | 1 | 13 | Dnase1l2 | 0.018 | Young |
| 0.144264 | 0.65383 | 0.018 | 0 | 1 | 13 | Ptgis | 0.018 | Young |
| 0.144264 | 0.651432 | 0.018 | 0 | 1 | 13 | Ttll7 | 0.018 | Young |
| 0.144264 | 0.649818 | 0.018 | 0 | 1 | 13 | 5430402O1 | 0.018 | Young |
| 0.144264 | 0.645527 | 0.018 | 0 | 1 | 13 | Gm8013 | 0.018 | Young |
| 0.144264 | 0.639556 | 0.018 | 0 | 1 | 13 | Cuedc1 | 0.018 | Young |
| 0.144264 | 0.639472 | 0.018 | 0 | 1 | 13 | Hpx | 0.018 | Young |
| 0.144264 | 0.631945 | 0.018 | 0 | 1 | 13 | Gm49077 | 0.018 | Young |
| 0.144264 | 0.628817 | 0.018 | 0 | 1 | 13 | Ramp2 | 0.018 | Young |

| | | | | | | | | |
|---|---|---|---|---|---|---|---|---|
| 0.144264 | 0.625956 | 0.018 | 0 | 1 | 13 | Mid2 | 0.018 | Young |
| 0.144264 | 0.576322 | 0.018 | 0 | 1 | 13 | Il2ra | 0.018 | Young |
| 0.144264 | 0.566655 | 0.018 | 0 | 1 | 13 | Gm15718 | 0.018 | Young |
| 0.144264 | 0.525949 | 0.018 | 0 | 1 | 13 | Impg1 | 0.018 | Young |
| 0.144382 | 0.745247 | 0.088 | 0.041 | 1 | 13 | Syp | 0.047 | Young |
| 0.144382 | 0.688296 | 0.088 | 0.041 | 1 | 13 | Nxpe3 | 0.047 | Young |
| 0.144524 | 0.1983 | 0.632 | 0.598 | 1 | 13 | Bmpr2 | 0.034 | Young |
| 0.14465 | 0.610939 | 0.263 | 0.197 | 1 | 13 | Cd1d1 | 0.066 | Young |
| 0.144854 | 0.860987 | 0.096 | 0.049 | 1 | 13 | Mpv17l | 0.047 | Young |
| 0.145174 | 0.425849 | 0.509 | 0.418 | 1 | 13 | Ddt | 0.091 | Young |
| 0.145307 | 0.304553 | 0.325 | 0.238 | 1 | 13 | Cars | 0.087 | Young |
| 0.145503 | 0.594096 | 0.219 | 0.148 | 1 | 13 | Gm44751 | 0.071 | Young |
| 0.145995 | 0.169532 | 0.921 | 0.869 | 1 | 13 | Dnajc19 | 0.052 | Young |
| 0.146866 | 0.297299 | 0.702 | 0.672 | 1 | 13 | Nck1 | 0.03 | Young |
| 0.147019 | 0.173699 | 0.605 | 0.582 | 1 | 13 | Arl5c | 0.023 | Young |
| 0.147203 | 0.615891 | 0.254 | 0.189 | 1 | 13 | Zfp994 | 0.065 | Young |
| 0.147244 | 0.27215 | 0.667 | 0.639 | 1 | 13 | Ccdc124 | 0.028 | Young |
| 0.147825 | 0.190228 | 0.947 | 0.992 | 1 | 13 | Top1 | -0.045 | Young |
| 0.147922 | 0.374024 | 0.553 | 0.492 | 1 | 13 | Vps53 | 0.061 | Young |
| 0.148358 | 0.151118 | 0.982 | 0.951 | 1 | 13 | Ppib | 0.031 | Young |
| 0.148367 | 0.096522 | 0.842 | 0.762 | 1 | 13 | Tiam1 | 0.08 | Young |
| 0.148445 | 0.784565 | 0.175 | 0.115 | 1 | 13 | Spout1 | 0.06 | Young |
| 0.148866 | 0.570422 | 0.211 | 0.139 | 1 | 13 | Ddah2 | 0.072 | Young |
| 0.148902 | 0.075315 | 1 | 1 | 1 | 13 | Atp2b1 | 0 | Young |
| 0.148945 | 1.038338 | 0.061 | 0.025 | 1 | 13 | Gm5141 | 0.036 | Young |
| 0.148945 | 1.001001 | 0.061 | 0.025 | 1 | 13 | Gm9796 | 0.036 | Young |
| 0.149094 | 0.231676 | 0.57 | 0.516 | 1 | 13 | Dpy30 | 0.054 | Young |
| 0.149303 | 0.317311 | 0.649 | 0.59 | 1 | 13 | Mctp1 | 0.059 | Young |
| 0.149345 | 1.275608 | 0.035 | 0.008 | 1 | 13 | Zfp58 | 0.027 | Young |
| 0.149345 | 1.147812 | 0.035 | 0.008 | 1 | 13 | Zfp354c | 0.027 | Young |
| 0.149345 | 1.107222 | 0.035 | 0.008 | 1 | 13 | Cdh24 | 0.027 | Young |
| 0.149345 | 1.085058 | 0.035 | 0.008 | 1 | 13 | Snhg18 | 0.027 | Young |
| 0.149345 | 1.036381 | 0.035 | 0.008 | 1 | 13 | Zkscan4 | 0.027 | Young |
| 0.149345 | 0.986637 | 0.035 | 0.008 | 1 | 13 | Palmd | 0.027 | Young |
| 0.149388 | 0.514482 | 0.289 | 0.23 | 1 | 13 | Fmo5 | 0.059 | Young |
| 0.149979 | 0.059763 | 1 | 1 | 1 | 13 | Cd74 | 0 | Young |
| 0.150196 | 0.443686 | 0.342 | 0.279 | 1 | 13 | Cxxc1 | 0.063 | Young |
| 0.150314 | 0.357262 | 0.421 | 0.336 | 1 | 13 | Hivep3 | 0.085 | Young |
| 0.150491 | 0.955087 | 0.061 | 0.025 | 1 | 13 | Mtbp | 0.036 | Young |
| 0.150504 | 0.23559 | 0.833 | 0.885 | 1 | 13 | Cnih4 | -0.052 | Young |
| 0.150509 | 0.073773 | 0.965 | 0.959 | 1 | 13 | Mtss1 | 0.006 | Young |
| 0.15052 | 0.07294 | 1 | 1 | 1 | 13 | Gpx4 | 0 | Young |
| 0.150712 | 0.143658 | 0.325 | 0.238 | 1 | 13 | Arl11 | 0.087 | Young |
| 0.150857 | 0.773113 | 0.184 | 0.123 | 1 | 13 | Rpap3 | 0.061 | Young |
| 0.15113 | 0.607032 | 0.325 | 0.246 | 1 | 13 | Abtb1 | 0.079 | Young |

| | | | | | | | | |
|---|---|---|---|---|---|---|---|---|
| 0.151362 | 0.418364 | 0.351 | 0.279 | 1 | 13 | Mpg | 0.072 | Young |
| 0.151515 | 1.140022 | 0.035 | 0.008 | 1 | 13 | Nt5e | 0.027 | Young |
| 0.151515 | 1.059901 | 0.035 | 0.008 | 1 | 13 | Dnajb7 | 0.027 | Young |
| 0.151515 | 1.004556 | 0.035 | 0.008 | 1 | 13 | 2210408I2: | 0.027 | Young |
| 0.151515 | 0.99747 | 0.035 | 0.008 | 1 | 13 | Gm19557 | 0.027 | Young |
| 0.151515 | 0.932044 | 0.035 | 0.008 | 1 | 13 | Fam229b | 0.027 | Young |
| 0.151515 | 0.926818 | 0.035 | 0.008 | 1 | 13 | Rasgrf1 | 0.027 | Young |
| 0.151515 | 0.8734 | 0.035 | 0.008 | 1 | 13 | Tmem246 | 0.027 | Young |
| 0.151852 | 0.113995 | 0.956 | 0.943 | 1 | 13 | Far1 | 0.013 | Young |
| 0.152048 | 1.031004 | 0.061 | 0.025 | 1 | 13 | Zcchc3 | 0.036 | Young |
| 0.15271 | 0.734779 | 0.167 | 0.107 | 1 | 13 | Zfp667 | 0.06 | Young |
| 0.152944 | 0.349427 | 0.447 | 0.385 | 1 | 13 | Igsf9 | 0.062 | Young |
| 0.152948 | 0.540552 | 0.158 | 0.098 | 1 | 13 | Ccdc180 | 0.06 | Young |
| 0.153124 | 0.72662 | 0.228 | 0.18 | 1 | 13 | F8a | 0.048 | Young |
| 0.153158 | 0.46189 | 0.36 | 0.279 | 1 | 13 | Msh2 | 0.081 | Young |
| 0.153235 | 0.134396 | 0.947 | 0.975 | 1 | 13 | Clec12a | -0.028 | Young |
| 0.153249 | 0.554235 | 0.395 | 0.344 | 1 | 13 | Borcs8 | 0.051 | Young |
| 0.153269 | 0.585828 | 0.088 | 0.041 | 1 | 13 | Fam167b | 0.047 | Young |
| 0.15342 | 0.215707 | 0.667 | 0.615 | 1 | 13 | Abcg1 | 0.052 | Young |
| 0.153709 | 2.535567 | 0.035 | 0.008 | 1 | 13 | Tnfsf18 | 0.027 | Young |
| 0.153709 | 1.035372 | 0.035 | 0.008 | 1 | 13 | Gm45844 | 0.027 | Young |
| 0.153709 | 0.930038 | 0.035 | 0.008 | 1 | 13 | Zfp772 | 0.027 | Young |
| 0.153709 | 0.847829 | 0.035 | 0.008 | 1 | 13 | Gm1720 | 0.027 | Young |
| 0.153709 | 0.839379 | 0.035 | 0.008 | 1 | 13 | Rtkn | 0.027 | Young |
| 0.153709 | 0.826631 | 0.035 | 0.008 | 1 | 13 | Msrb3 | 0.027 | Young |
| 0.153709 | 0.759516 | 0.035 | 0.008 | 1 | 13 | 2210008F0 | 0.027 | Young |
| 0.154348 | 0.074989 | 1 | 1 | 1 | 13 | Ubl3 | 0 | Young |
| 0.154497 | 0.700168 | 0.193 | 0.131 | 1 | 13 | Mbd4 | 0.062 | Young |
| 0.1552 | 0.891093 | 0.061 | 0.025 | 1 | 13 | Mansc1 | 0.036 | Young |
| 0.155348 | 0.265443 | 0.588 | 0.549 | 1 | 13 | Pex2 | 0.039 | Young |
| 0.155439 | 0.444444 | 0.57 | 0.533 | 1 | 13 | Rbm22 | 0.037 | Young |
| 0.155455 | 0.064849 | 0.991 | 1 | 1 | 13 | Cd14 | -0.009 | Young |
| 0.155906 | 0.515829 | 0.456 | 0.426 | 1 | 13 | Mosmo | 0.03 | Young |
| 0.155927 | 0.95917 | 0.035 | 0.008 | 1 | 13 | Scml4 | 0.027 | Young |
| 0.155927 | 0.913151 | 0.035 | 0.008 | 1 | 13 | Zfp955a | 0.027 | Young |
| 0.155927 | 0.690207 | 0.035 | 0.008 | 1 | 13 | Bicdl1 | 0.027 | Young |
| 0.156187 | 0.645755 | 0.211 | 0.148 | 1 | 13 | Pi4kb | 0.063 | Young |
| 0.156267 | 0.185684 | 0.842 | 0.738 | 1 | 13 | Ptpn11 | 0.104 | Young |
| 0.156594 | 0.435611 | 0.491 | 0.459 | 1 | 13 | 1700123O2 | 0.032 | Young |
| 0.156622 | 0.125625 | 0.333 | 0.467 | 1 | 13 | Isg15 | -0.134 | Young |
| 0.156788 | 1.079379 | 0.114 | 0.066 | 1 | 13 | Gm31243 | 0.048 | Young |
| 0.156795 | 0.973374 | 0.061 | 0.025 | 1 | 13 | Spata1 | 0.036 | Young |
| 0.156795 | 0.854382 | 0.061 | 0.025 | 1 | 13 | Map3k9 | 0.036 | Young |
| 0.156974 | 0.299212 | 0.395 | 0.32 | 1 | 13 | Cish | 0.075 | Young |
| 0.156974 | 0.289608 | 0.395 | 0.32 | 1 | 13 | Ints11 | 0.075 | Young |

| | | | | | | | | |
|---|---|---|---|---|---|---|---|---|
| 0.15701 | 0.232955 | 0.86 | 0.844 | 1 | 13 | Hcst | 0.016 | Young |
| 0.157774 | 0.472016 | 0.228 | 0.156 | 1 | 13 | Slco3a1 | 0.072 | Young |
| 0.157923 | 0.910776 | 0.114 | 0.066 | 1 | 13 | Thg1l | 0.048 | Young |
| 0.158118 | 0.443832 | 0.132 | 0.074 | 1 | 13 | Miga1 | 0.058 | Young |
| 0.15817 | 0.735208 | 0.035 | 0.008 | 1 | 13 | Kif18a | 0.027 | Young |
| 0.15817 | 0.45314 | 0.035 | 0.008 | 1 | 13 | Col1a1 | 0.027 | Young |
| 0.158208 | 0.682055 | 0.096 | 0.049 | 1 | 13 | Nudt6 | 0.047 | Young |
| 0.158217 | 0.303092 | 0.5 | 0.418 | 1 | 13 | Erf | 0.082 | Young |
| 0.158402 | 0.823292 | 0.061 | 0.025 | 1 | 13 | Cyp2e1 | 0.036 | Young |
| 0.158402 | 0.797377 | 0.061 | 0.025 | 1 | 13 | Zfp956 | 0.036 | Young |
| 0.158869 | 0.299143 | 0.465 | 0.385 | 1 | 13 | Plxnc1 | 0.08 | Young |
| 0.159685 | 0.269415 | 0.509 | 0.467 | 1 | 13 | 1110008P1 | 0.042 | Young |
| 0.159863 | 0.368511 | 0.272 | 0.189 | 1 | 13 | Dus1l | 0.083 | Young |
| 0.159922 | 0.458289 | 0.404 | 0.369 | 1 | 13 | Pdcd7 | 0.035 | Young |
| 0.15993 | 0.923214 | 0.105 | 0.057 | 1 | 13 | Dtwd1 | 0.048 | Young |
| 0.160021 | 0.859783 | 0.061 | 0.025 | 1 | 13 | Gm10645 | 0.036 | Young |
| 0.160021 | 0.791136 | 0.061 | 0.025 | 1 | 13 | Syt3 | 0.036 | Young |
| 0.160407 | 0.706263 | 0.211 | 0.148 | 1 | 13 | P2ry10 | 0.063 | Young |
| 0.16046 | 0.318039 | 0.64 | 0.664 | 1 | 13 | Mrpl21 | -0.024 | Young |
| 0.160801 | 0.110842 | 0.912 | 0.869 | 1 | 13 | Phlda1 | 0.043 | Young |
| 0.16096 | 0.275099 | 0.535 | 0.475 | 1 | 13 | Rcn3 | 0.06 | Young |
| 0.161372 | 0.138467 | 0.877 | 0.902 | 1 | 13 | Snrpd2 | -0.025 | Young |
| 0.161426 | 0.328699 | 0.526 | 0.475 | 1 | 13 | Bcap29 | 0.051 | Young |
| 0.161835 | 0.1587 | 0.868 | 0.885 | 1 | 13 | Sbds | -0.017 | Young |
| 0.162028 | 0.111441 | 0.754 | 0.705 | 1 | 13 | Csnk1g2 | 0.049 | Young |
| 0.162417 | 0.465277 | 0.377 | 0.328 | 1 | 13 | Hdgfl2 | 0.049 | Young |
| 0.162785 | 0.457131 | 0.544 | 0.525 | 1 | 13 | Wdr91 | 0.019 | Young |
| 0.162984 | 0.229176 | 0.737 | 0.713 | 1 | 13 | Cbx1 | 0.024 | Young |
| 0.163298 | 0.778792 | 0.061 | 0.025 | 1 | 13 | Tyw3 | 0.036 | Young |
| 0.163365 | 0.492564 | 0.202 | 0.131 | 1 | 13 | Ier5l | 0.071 | Young |
| 0.163387 | 0.373043 | 0.474 | 0.41 | 1 | 13 | Pou2f1 | 0.064 | Young |
| 0.163561 | 0.142681 | 0.842 | 0.91 | 1 | 13 | Vps4b | -0.068 | Young |
| 0.163688 | 0.885535 | 0.114 | 0.066 | 1 | 13 | Trp53rkb | 0.048 | Young |
| 0.163747 | 0.476415 | 0.316 | 0.238 | 1 | 13 | Frs2 | 0.078 | Young |
| 0.164059 | 0.690192 | 0.167 | 0.107 | 1 | 13 | Gimap6 | 0.06 | Young |
| 0.164426 | 0.364942 | 0.482 | 0.443 | 1 | 13 | Setd7 | 0.039 | Young |
| 0.164613 | 0.419365 | 0.518 | 0.459 | 1 | 13 | Kpna3 | 0.059 | Young |
| 0.164955 | 0.83157 | 0.061 | 0.025 | 1 | 13 | Tjp3 | 0.036 | Young |
| 0.165075 | 0.140798 | 0.965 | 0.926 | 1 | 13 | Mxd4 | 0.039 | Young |
| 0.165237 | 0.336187 | 0.395 | 0.336 | 1 | 13 | Smc2 | 0.059 | Young |
| 0.1654 | 0.104973 | 0.991 | 0.975 | 1 | 13 | Sec11c | 0.016 | Young |
| 0.165819 | 0.301039 | 0.675 | 0.697 | 1 | 13 | Wbp4 | -0.022 | Young |
| 0.165874 | 0.696976 | 0.096 | 0.049 | 1 | 13 | Zfp873 | 0.047 | Young |
| 0.166026 | 0.809417 | 0.105 | 0.057 | 1 | 13 | Tceal1 | 0.048 | Young |
| 0.166251 | 0.60036 | 0.36 | 0.303 | 1 | 13 | Nit1 | 0.057 | Young |

| | | | | | | | | |
|---|---|---|---|---|---|---|---|---|
| 0.166292 | 0.503163 | 0.368 | 0.328 | 1 | 13 | Cd82 | 0.04 | Young |
| 0.166549 | 0.221407 | 0.649 | 0.566 | 1 | 13 | Nucb2 | 0.083 | Young |
| 0.166735 | 0.441843 | 0.316 | 0.246 | 1 | 13 | Ermp1 | 0.07 | Young |
| 0.166785 | 0.054536 | 0.711 | 0.574 | 1 | 13 | Irf4 | 0.137 | Young |
| 0.166838 | 0.755214 | 0.184 | 0.131 | 1 | 13 | Usp30 | 0.053 | Young |
| 0.167137 | 0.196466 | 0.956 | 0.959 | 1 | 13 | Itga4 | -0.003 | Young |
| 0.167149 | 0.179963 | 0.974 | 0.975 | 1 | 13 | Sfr1 | -0.001 | Young |
| 0.167178 | 0.460318 | 0.096 | 0.049 | 1 | 13 | Tubb2b | 0.047 | Young |
| 0.167222 | 0.978044 | 0.114 | 0.066 | 1 | 13 | Chaf1b | 0.048 | Young |
| 0.167958 | 0.373891 | 0.474 | 0.385 | 1 | 13 | Dcaf15 | 0.089 | Young |
| 0.168282 | 0.138611 | 0.886 | 0.828 | 1 | 13 | Fgd4 | 0.058 | Young |
| 0.168586 | 0.345529 | 0.57 | 0.549 | 1 | 13 | Zbtb4 | 0.021 | Young |
| 0.169241 | 0.217869 | 0.649 | 0.582 | 1 | 13 | Rspry1 | 0.067 | Young |
| 0.169766 | 0.683161 | 0.105 | 0.057 | 1 | 13 | Gpr84 | 0.048 | Young |
| 0.16987 | 0.577836 | 0.149 | 0.09 | 1 | 13 | Urb1 | 0.059 | Young |
| 0.169967 | 0.589958 | 0.167 | 0.107 | 1 | 13 | Ms4a14 | 0.06 | Young |
| 0.170399 | 0.105963 | 0.982 | 0.975 | 1 | 13 | Mrpl52 | 0.007 | Young |
| 0.170531 | 0.446842 | 0.404 | 0.369 | 1 | 13 | Gm9493 | 0.035 | Young |
| 0.170674 | 1.218004 | 0.123 | 0.074 | 1 | 13 | Omp | 0.049 | Young |
| 0.170726 | 2.162835 | 0.132 | 0.082 | 1 | 13 | Ccl17 | 0.05 | Young |
| 0.170984 | 0.29684 | 0.728 | 0.689 | 1 | 13 | Nfu1 | 0.039 | Young |
| 0.170997 | 0.051313 | 1 | 1 | 1 | 13 | Pabpc1 | 0 | Young |
| 0.171096 | 0.388897 | 0.491 | 0.443 | 1 | 13 | Slc30a7 | 0.048 | Young |
| 0.171487 | 0.342635 | 0.439 | 0.377 | 1 | 13 | Wdr44 | 0.062 | Young |
| 0.171794 | 0.441495 | 0.474 | 0.41 | 1 | 13 | Dnlz | 0.064 | Young |
| 0.17213 | 3.164343 | 0.14 | 0.09 | 1 | 13 | Spp1 | 0.05 | Young |
| 0.172309 | 0.327916 | 0.535 | 0.516 | 1 | 13 | Wdr37 | 0.019 | Young |
| 0.172467 | 0.253615 | 0.518 | 0.443 | 1 | 13 | Idh3a | 0.075 | Young |
| 0.172474 | 0.506901 | 0.421 | 0.369 | 1 | 13 | Mthfd1 | 0.052 | Young |
| 0.172713 | 0.181679 | 0.965 | 0.893 | 1 | 13 | Bnip3l | 0.072 | Young |
| 0.173448 | 0.641762 | 0.193 | 0.131 | 1 | 13 | H2-Ob | 0.062 | Young |
| 0.173491 | 0.365945 | 0.316 | 0.246 | 1 | 13 | Tef | 0.07 | Young |
| 0.173836 | 0.212762 | 0.596 | 0.533 | 1 | 13 | Fam107b | 0.063 | Young |
| 0.174137 | 0.353764 | 0.149 | 0.09 | 1 | 13 | H2-Oa | 0.059 | Young |
| 0.174227 | 0.244762 | 0.596 | 0.557 | 1 | 13 | Gpr107 | 0.039 | Young |
| 0.17426 | 0.341137 | 0.316 | 0.238 | 1 | 13 | Sorl1 | 0.078 | Young |
| 0.175444 | 0.145367 | 0.667 | 0.631 | 1 | 13 | Rbbp7 | 0.036 | Young |
| 0.175522 | 0.461886 | 0.325 | 0.262 | 1 | 13 | Rpain | 0.063 | Young |
| 0.175552 | 0.504877 | 0.202 | 0.139 | 1 | 13 | Nsun3 | 0.063 | Young |
| 0.175708 | 0.58322 | 0.298 | 0.238 | 1 | 13 | 4921524J1 | 0.06 | Young |
| 0.175925 | 0.494422 | 0.377 | 0.311 | 1 | 13 | Ppm1k | 0.066 | Young |
| 0.176234 | 0.526928 | 0.289 | 0.23 | 1 | 13 | A530064D( | 0.059 | Young |
| 0.17633 | 0.124751 | 0.684 | 0.607 | 1 | 13 | Msr1 | 0.077 | Young |
| 0.177075 | 0.320029 | 0.57 | 0.582 | 1 | 13 | Lage3 | -0.012 | Young |
| 0.177423 | 0.390463 | 0.456 | 0.418 | 1 | 13 | Rsrc1 | 0.038 | Young |

| | | | | | | | | |
|---|---|---|---|---|---|---|---|---|
| 0.177616 | 0.140774 | 0.965 | 0.926 | 1 | 13 | Dek | 0.039 | Young |
| 0.178367 | 0.215105 | 0.544 | 0.5 | 1 | 13 | Aph1c | 0.044 | Young |
| 0.178382 | 0.912703 | 0.07 | 0.033 | 1 | 13 | Tle2 | 0.037 | Young |
| 0.178438 | 0.387996 | 0.272 | 0.205 | 1 | 13 | Dmac2 | 0.067 | Young |
| 0.178757 | 0.428471 | 0.263 | 0.197 | 1 | 13 | Tmem80 | 0.066 | Young |
| 0.179347 | 0.18213 | 0.895 | 0.926 | 1 | 13 | Cdc37 | -0.031 | Young |
| 0.179515 | 0.290535 | 0.658 | 0.598 | 1 | 13 | Tm9sf4 | 0.06 | Young |
| 0.179584 | 0.518671 | 0.386 | 0.361 | 1 | 13 | Prkce | 0.025 | Young |
| 0.179824 | 0.792166 | 0.132 | 0.082 | 1 | 13 | Gtpbp3 | 0.05 | Young |
| 0.17983 | 0.502673 | 0.281 | 0.221 | 1 | 13 | Tipin | 0.06 | Young |
| 0.180429 | 0.140683 | 0.991 | 0.975 | 1 | 13 | Reep3 | 0.016 | Young |
| 0.18074 | 0.100764 | 1 | 1 | 1 | 13 | Nfkbia | 0 | Young |
| 0.18135 | 0.151595 | 0.956 | 0.967 | 1 | 13 | Cox14 | -0.011 | Young |
| 0.181362 | 0.126426 | 1 | 0.992 | 1 | 13 | Ndufa13 | 0.008 | Young |
| 0.181638 | 1.022831 | 0.07 | 0.033 | 1 | 13 | Rapgef5 | 0.037 | Young |
| 0.181934 | 0.750938 | 0.114 | 0.066 | 1 | 13 | Gm20300 | 0.048 | Young |
| 0.182544 | 0.887955 | 0.123 | 0.074 | 1 | 13 | Ighd | 0.049 | Young |
| 0.182544 | 0.763955 | 0.123 | 0.074 | 1 | 13 | 4930512J1 | 0.049 | Young |
| 0.182612 | 0.107747 | 1 | 0.992 | 1 | 13 | S100a6 | 0.008 | Young |
| 0.182789 | 0.184299 | 0.702 | 0.639 | 1 | 13 | Rnf216 | 0.063 | Young |
| 0.183186 | 0.290206 | 0.237 | 0.164 | 1 | 13 | Klrb1b | 0.073 | Young |
| 0.184023 | 0.559853 | 0.105 | 0.057 | 1 | 13 | Ptges3l | 0.048 | Young |
| 0.184498 | 0.181086 | 0.991 | 1 | 1 | 13 | Il6ra | -0.009 | Young |
| 0.184695 | 0.492884 | 0.211 | 0.148 | 1 | 13 | Ofd1 | 0.063 | Young |
| 0.185476 | 0.125754 | 0.781 | 0.68 | 1 | 13 | Aldh1a2 | 0.101 | Young |
| 0.18568 | 0.181852 | 0.825 | 0.877 | 1 | 13 | Smim11 | -0.052 | Young |
| 0.18569 | 0.335373 | 0.132 | 0.082 | 1 | 13 | Ccl22 | 0.05 | Young |
| 0.186088 | 0.306625 | 0.614 | 0.566 | 1 | 13 | Trnau1ap | 0.048 | Young |
| 0.186264 | 0.377575 | 0.395 | 0.32 | 1 | 13 | Sbf1 | 0.075 | Young |
| 0.186399 | 0.096811 | 1 | 1 | 1 | 13 | Msn | 0 | Young |
| 0.186524 | 0.561644 | 0.368 | 0.336 | 1 | 13 | Carnmt1 | 0.032 | Young |
| 0.186604 | 0.776621 | 0.07 | 0.033 | 1 | 13 | 9430015G1 | 0.037 | Young |
| 0.187045 | 0.707296 | 0.114 | 0.066 | 1 | 13 | Fam102a | 0.048 | Young |
| 0.187108 | 0.20341 | 0.763 | 0.754 | 1 | 13 | Ugcg | 0.009 | Young |
| 0.187463 | 0.185806 | 0.693 | 0.59 | 1 | 13 | Mob3a | 0.103 | Young |
| 0.188404 | 0.192829 | 0.623 | 0.574 | 1 | 13 | 1700025G( | 0.049 | Young |
| 0.188645 | 0.604594 | 0.167 | 0.107 | 1 | 13 | Abhd6 | 0.06 | Young |
| 0.189091 | 0.235144 | 0.482 | 0.434 | 1 | 13 | Ice1 | 0.048 | Young |
| 0.189107 | 0.674924 | 0.175 | 0.123 | 1 | 13 | Alkbh2 | 0.052 | Young |
| 0.189515 | 0.245586 | 0.772 | 0.77 | 1 | 13 | Cnot6 | 0.002 | Young |
| 0.189812 | 0.461466 | 0.368 | 0.311 | 1 | 13 | Fdx1 | 0.057 | Young |
| 0.190011 | 0.158684 | 0.789 | 0.746 | 1 | 13 | Acadm | 0.043 | Young |
| 0.190143 | 0.828827 | 0.14 | 0.09 | 1 | 13 | Svip | 0.05 | Young |
| 0.190241 | 0.126344 | 0.974 | 0.984 | 1 | 13 | Tob2 | -0.01 | Young |
| 0.190533 | 0.156835 | 0.702 | 0.656 | 1 | 13 | Ocrl | 0.046 | Young |

| | | | | | | | | |
|---|---|---|---|---|---|---|---|---|
| 0.191193 | 0.65978 | 0.184 | 0.131 | 1 | 13 | Rpap1 | 0.053 | Young |
| 0.191422 | 0.626252 | 0.228 | 0.172 | 1 | 13 | Slc10a3 | 0.056 | Young |
| 0.191669 | 0.965389 | 0.07 | 0.033 | 1 | 13 | Card10 | 0.037 | Young |
| 0.192187 | 0.167553 | 1 | 0.992 | 1 | 13 | D8Ertd738 | 0.008 | Young |
| 0.192473 | 0.441958 | 0.123 | 0.074 | 1 | 13 | Dhodh | 0.049 | Young |
| 0.192554 | 0.183599 | 0.675 | 0.582 | 1 | 13 | Rbm33 | 0.093 | Young |
| 0.192984 | 0.701991 | 0.158 | 0.107 | 1 | 13 | Wdr24 | 0.051 | Young |
| 0.19338 | 0.797817 | 0.07 | 0.033 | 1 | 13 | Sh3bp4 | 0.037 | Young |
| 0.193476 | 0.237421 | 0.693 | 0.672 | 1 | 13 | Chn2 | 0.021 | Young |
| 0.193492 | 0.104189 | 1 | 1 | 1 | 13 | Clic1 | 0 | Young |
| 0.193508 | 0.244258 | 0.693 | 0.664 | 1 | 13 | Rbck1 | 0.029 | Young |
| 0.193834 | 0.34719 | 0.36 | 0.287 | 1 | 13 | Gnptg | 0.073 | Young |
| 0.194198 | 0.223606 | 0.833 | 0.844 | 1 | 13 | Ddrgk1 | -0.011 | Young |
| 0.194585 | 0.125521 | 0.789 | 0.713 | 1 | 13 | Neu1 | 0.076 | Young |
| 0.195151 | 0.365316 | 0.509 | 0.434 | 1 | 13 | Creb5 | 0.075 | Young |
| 0.195168 | 0.600122 | 0.228 | 0.172 | 1 | 13 | Aptx | 0.056 | Young |
| 0.195635 | 0.389539 | 0.386 | 0.32 | 1 | 13 | Maged1 | 0.066 | Young |
| 0.195697 | 0.256259 | 0.439 | 0.369 | 1 | 13 | Zbtb38 | 0.07 | Young |
| 0.195788 | 0.235185 | 0.588 | 0.574 | 1 | 13 | Kif2a | 0.014 | Young |
| 0.19584 | 0.461428 | 0.518 | 0.508 | 1 | 13 | Wipf2 | 0.01 | Young |
| 0.196324 | 0.481885 | 0.211 | 0.148 | 1 | 13 | 1810043G( | 0.063 | Young |
| 0.196329 | 0.184319 | 0.737 | 0.656 | 1 | 13 | Tgs1 | 0.081 | Young |
| 0.196406 | 0.269188 | 0.763 | 0.779 | 1 | 13 | Ubn2 | -0.016 | Young |
| 0.196507 | 0.489667 | 0.263 | 0.205 | 1 | 13 | Pcolce | 0.058 | Young |
| 0.196696 | 0.198245 | 0.816 | 0.746 | 1 | 13 | Nudt21 | 0.07 | Young |
| 0.196835 | 0.675479 | 0.07 | 0.033 | 1 | 13 | Zfp790 | 0.037 | Young |
| 0.196835 | 0.67279 | 0.07 | 0.033 | 1 | 13 | 4833418N( | 0.037 | Young |
| 0.197393 | 0.678972 | 0.254 | 0.189 | 1 | 13 | Pde8a | 0.065 | Young |
| 0.19755 | 0.56297 | 0.246 | 0.18 | 1 | 13 | Slc46a3 | 0.066 | Young |
| 0.19807 | 0.127785 | 0.982 | 0.91 | 1 | 13 | Dennd5a | 0.072 | Young |
| 0.198447 | 0.659163 | 0.14 | 0.09 | 1 | 13 | Rhbdf1 | 0.05 | Young |
| 0.198774 | 0.488748 | 0.342 | 0.279 | 1 | 13 | Mfn2 | 0.063 | Young |
| 0.198842 | 0.459407 | 0.43 | 0.385 | 1 | 13 | Car3 | 0.045 | Young |
| 0.198868 | 0.402357 | 0.404 | 0.336 | 1 | 13 | Foxred2 | 0.068 | Young |
| 0.199034 | 0.155035 | 0.789 | 0.77 | 1 | 13 | Mrpl42 | 0.019 | Young |
| 0.199981 | 0.910374 | 0.079 | 0.041 | 1 | 13 | Gm17025 | 0.038 | Young |
| 0.200444 | 0.165849 | 0.877 | 0.893 | 1 | 13 | Uqcc2 | -0.016 | Young |
| 0.200806 | 0.609138 | 0.289 | 0.238 | 1 | 13 | Rcor3 | 0.051 | Young |
| 0.201287 | 0.19664 | 0.5 | 0.418 | 1 | 13 | Zfp740 | 0.082 | Young |
| 0.201435 | 0.482909 | 0.351 | 0.295 | 1 | 13 | Orc4 | 0.056 | Young |
| 0.20179 | 0.083288 | 1 | 1 | 1 | 13 | Tnfaip3 | 0 | Young |
| 0.202125 | 0.117738 | 0.982 | 0.975 | 1 | 13 | Ckb | 0.007 | Young |
| 0.202227 | 0.643513 | 0.263 | 0.213 | 1 | 13 | Atg14 | 0.05 | Young |
| 0.202295 | 0.259078 | 0.693 | 0.664 | 1 | 13 | Avpi1 | 0.029 | Young |
| 0.202783 | 0.641276 | 0.167 | 0.115 | 1 | 13 | Trip6 | 0.052 | Young |

| | | | | | | | | |
|---|---|---|---|---|---|---|---|---|
| 0.203041 | 0.156418 | 0.956 | 0.893 | 1 | 13 | Hdlbp | 0.063 | Young |
| 0.203457 | 0.02101 | 0.974 | 0.934 | 1 | 13 | Slc38a2 | 0.04 | Young |
| 0.203501 | 0.52307 | 0.43 | 0.393 | 1 | 13 | Ccdc88c | 0.037 | Young |
| 0.20395 | 0.162705 | 0.921 | 0.885 | 1 | 13 | Hook3 | 0.036 | Young |
| 0.204145 | 0.35723 | 0.456 | 0.426 | 1 | 13 | H2-Ke6 | 0.03 | Young |
| 0.204352 | 1.249595 | 0.044 | 0.016 | 1 | 13 | Npr1 | 0.028 | Young |
| 0.204352 | 1.075841 | 0.044 | 0.016 | 1 | 13 | Ccdc15 | 0.028 | Young |
| 0.204352 | 1.021665 | 0.044 | 0.016 | 1 | 13 | Gm43254 | 0.028 | Young |
| 0.20472 | 0.197586 | 0.395 | 0.32 | 1 | 13 | Ginm1 | 0.075 | Young |
| 0.204922 | 0.123667 | 0.325 | 0.246 | 1 | 13 | Olr1 | 0.079 | Young |
| 0.204932 | 0.925553 | 0.079 | 0.041 | 1 | 13 | Fgf11 | 0.038 | Young |
| 0.20494 | 0.153221 | 0.667 | 0.598 | 1 | 13 | Mecp2 | 0.069 | Young |
| 0.205411 | 0.635608 | 0.132 | 0.082 | 1 | 13 | Depdc7 | 0.05 | Young |
| 0.205744 | 0.247242 | 0.675 | 0.607 | 1 | 13 | Aff1 | 0.068 | Young |
| 0.205803 | 0.38489 | 0.333 | 0.279 | 1 | 13 | Rfwd3 | 0.054 | Young |
| 0.20631 | 0.23985 | 0.921 | 0.877 | 1 | 13 | Eif3i | 0.044 | Young |
| 0.206368 | 0.297371 | 0.623 | 0.582 | 1 | 13 | Ppdpf | 0.041 | Young |
| 0.206676 | 1.156556 | 0.044 | 0.016 | 1 | 13 | Gm10643 | 0.028 | Young |
| 0.206676 | 0.970128 | 0.044 | 0.016 | 1 | 13 | Tonsl | 0.028 | Young |
| 0.206676 | 0.940972 | 0.044 | 0.016 | 1 | 13 | 4930477G( | 0.028 | Young |
| 0.207158 | 0.765587 | 0.114 | 0.066 | 1 | 13 | Zfyve28 | 0.048 | Young |
| 0.207428 | 0.546771 | 0.351 | 0.303 | 1 | 13 | Xylt2 | 0.048 | Young |
| 0.20759 | 1.004719 | 0.132 | 0.09 | 1 | 13 | Ogfod3 | 0.042 | Young |
| 0.208282 | 0.703507 | 0.079 | 0.041 | 1 | 13 | Tmem216 | 0.038 | Young |
| 0.208293 | 0.294842 | 0.009 | 0.033 | 1 | 13 | Iglc2 | -0.024 | Young |
| 0.208293 | 0.040541 | 0.009 | 0.033 | 1 | 13 | Gm36888 | -0.024 | Young |
| 0.208619 | 0.172256 | 0.658 | 0.639 | 1 | 13 | Impdh2 | 0.019 | Young |
| 0.208853 | 0.511796 | 0.219 | 0.164 | 1 | 13 | Nemp1 | 0.055 | Young |
| 0.20902 | 1.366223 | 0.044 | 0.016 | 1 | 13 | Hrh2 | 0.028 | Young |
| 0.209078 | 0.380397 | 0.43 | 0.361 | 1 | 13 | Cul4a | 0.069 | Young |
| 0.209663 | 0.108903 | 0.842 | 0.869 | 1 | 13 | Il10rb | -0.027 | Young |
| 0.209703 | 1.013124 | 0.088 | 0.049 | 1 | 13 | Negr1 | 0.039 | Young |
| 0.209768 | 0.955189 | 0.105 | 0.066 | 1 | 13 | Gm4285 | 0.039 | Young |
| 0.209955 | 0.118013 | 0.535 | 0.459 | 1 | 13 | Tmem57 | 0.076 | Young |
| 0.21016 | 0.152019 | 0.912 | 0.902 | 1 | 13 | Srsf11 | 0.01 | Young |
| 0.211113 | 0.160984 | 0.904 | 0.82 | 1 | 13 | Rsu1 | 0.084 | Young |
| 0.211298 | 0.963842 | 0.088 | 0.049 | 1 | 13 | Bbs9 | 0.039 | Young |
| 0.211382 | 1.051919 | 0.044 | 0.016 | 1 | 13 | Bard1 | 0.028 | Young |
| 0.211382 | 0.893383 | 0.044 | 0.016 | 1 | 13 | Enc1 | 0.028 | Young |
| 0.211725 | 0.24365 | 0.333 | 0.262 | 1 | 13 | Kdsr | 0.071 | Young |
| 0.212147 | 0.542908 | 0.123 | 0.074 | 1 | 13 | Crtc1 | 0.049 | Young |
| 0.212558 | 0.348818 | 0.202 | 0.139 | 1 | 13 | AC163354. | 0.063 | Young |
| 0.21264 | 0.144498 | 0.816 | 0.738 | 1 | 13 | Slamf7 | 0.078 | Young |
| 0.212902 | 0.816536 | 0.088 | 0.049 | 1 | 13 | 3-Mar | 0.039 | Young |
| 0.213137 | 0.101439 | 1 | 1 | 1 | 13 | Rpl29 | 0 | Young |

| | | | | | | | | |
|---|---|---|---|---|---|---|---|---|
| 0.213406 | 0.266325 | 0.781 | 0.844 | 1 | 13 | Jtb | -0.063 | Young |
| 0.213491 | 0.24535 | 0.807 | 0.754 | 1 | 13 | Usp16 | 0.053 | Young |
| 0.213764 | 1.101453 | 0.044 | 0.016 | 1 | 13 | Thrsp | 0.028 | Young |
| 0.213764 | 1.067129 | 0.044 | 0.016 | 1 | 13 | 6720489N1 | 0.028 | Young |
| 0.213764 | 0.801992 | 0.044 | 0.016 | 1 | 13 | Uggt2 | 0.028 | Young |
| 0.213764 | 0.760471 | 0.044 | 0.016 | 1 | 13 | Ppbp | 0.028 | Young |
| 0.213764 | 0.734555 | 0.044 | 0.016 | 1 | 13 | Gm6277 | 0.028 | Young |
| 0.214129 | 0.680937 | 0.307 | 0.246 | 1 | 13 | Ptcd2 | 0.061 | Young |
| 0.214231 | 0.28362 | 0.412 | 0.336 | 1 | 13 | Rtn1 | 0.076 | Young |
| 0.214933 | 0.361557 | 0.596 | 0.566 | 1 | 13 | Tbc1d9 | 0.03 | Young |
| 0.214959 | 0.757067 | 0.158 | 0.107 | 1 | 13 | Dynlt1a | 0.051 | Young |
| 0.215583 | 0.59668 | 0.202 | 0.148 | 1 | 13 | Fam217b | 0.054 | Young |
| 0.215598 | 0.130649 | 0.991 | 0.959 | 1 | 13 | Ppp1r14b | 0.032 | Young |
| 0.216136 | 1.041405 | 0.088 | 0.049 | 1 | 13 | Efnb2 | 0.039 | Young |
| 0.216164 | 0.652471 | 0.158 | 0.107 | 1 | 13 | Hebp2 | 0.051 | Young |
| 0.216166 | 1.019435 | 0.044 | 0.016 | 1 | 13 | G0s2 | 0.028 | Young |
| 0.216166 | 0.68071 | 0.044 | 0.016 | 1 | 13 | Selp | 0.028 | Young |
| 0.216166 | 0.559218 | 0.044 | 0.016 | 1 | 13 | Bhlhb9 | 0.028 | Young |
| 0.216292 | 0.099345 | 0.956 | 0.959 | 1 | 13 | Fkbp2 | -0.003 | Young |
| 0.216921 | 0.490482 | 0.228 | 0.164 | 1 | 13 | 1300002E1 | 0.064 | Young |
| 0.217102 | 0.105846 | 0.93 | 0.861 | 1 | 13 | Nr4a2 | 0.069 | Young |
| 0.217374 | 0.679983 | 0.158 | 0.107 | 1 | 13 | Gcdh | 0.051 | Young |
| 0.217395 | 0.36833 | 0.333 | 0.27 | 1 | 13 | Gtf3c3 | 0.063 | Young |
| 0.217424 | 0.105356 | 0.921 | 0.861 | 1 | 13 | Dot1l | 0.06 | Young |
| 0.217437 | 0.421799 | 0.281 | 0.221 | 1 | 13 | Nrm | 0.06 | Young |
| 0.21778 | 0.225174 | 0.895 | 0.885 | 1 | 13 | Dock8 | 0.01 | Young |
| 0.21795 | 0.429107 | 0.211 | 0.148 | 1 | 13 | Pdp2 | 0.063 | Young |
| 0.218227 | 0.265692 | 0.605 | 0.574 | 1 | 13 | Ubp1 | 0.031 | Young |
| 0.218494 | 0.233327 | 0.544 | 0.492 | 1 | 13 | Eif2s3y | 0.052 | Young |
| 0.218586 | 0.659512 | 0.044 | 0.016 | 1 | 13 | A930024E0 | 0.028 | Young |
| 0.218586 | 0.648621 | 0.044 | 0.016 | 1 | 13 | Fndc10 | 0.028 | Young |
| 0.21863 | 0.335174 | 0.649 | 0.631 | 1 | 13 | Usp3 | 0.018 | Young |
| 0.219711 | 0.149944 | 0.904 | 0.836 | 1 | 13 | Hcfc1r1 | 0.068 | Young |
| 0.219997 | 0.170809 | 0.649 | 0.574 | 1 | 13 | Atxn7 | 0.075 | Young |
| 0.220124 | 0.575527 | 0.167 | 0.115 | 1 | 13 | Nme4 | 0.052 | Young |
| 0.220461 | 0.467765 | 0.263 | 0.197 | 1 | 13 | Zfp597 | 0.066 | Young |
| 0.220856 | 0.237742 | 0.596 | 0.566 | 1 | 13 | St6galnac4 | 0.03 | Young |
| 0.22094 | 0.115295 | 0.965 | 0.959 | 1 | 13 | Wwp1 | 0.006 | Young |
| 0.220997 | 0.242924 | 0.763 | 0.795 | 1 | 13 | Rassf3 | -0.032 | Young |
| 0.221026 | 0.646043 | 0.044 | 0.016 | 1 | 13 | 4930430E1 | 0.028 | Young |
| 0.221026 | 0.618196 | 0.044 | 0.016 | 1 | 13 | Hist1h1d | 0.028 | Young |
| 0.221026 | 0.599833 | 0.044 | 0.016 | 1 | 13 | 4833407H1 | 0.028 | Young |
| 0.221313 | 0.063766 | 0.991 | 0.992 | 1 | 13 | Cxcl2 | -0.001 | Young |
| 0.221368 | 0.378177 | 0.368 | 0.311 | 1 | 13 | Cd72 | 0.057 | Young |
| 0.22154 | 0.350405 | 0.43 | 0.393 | 1 | 13 | Nudt19 | 0.037 | Young |

| | | | | | | | | |
|---|---|---|---|---|---|---|---|---|
| 0.221804 | 0.398033 | 0.298 | 0.238 | 1 | 13 | Mbd5 | 0.06 | Young |
| 0.222078 | 0.579485 | 0.079 | 0.041 | 1 | 13 | Arap2 | 0.038 | Young |
| 0.222236 | 0.559402 | 0.342 | 0.295 | 1 | 13 | Mier3 | 0.047 | Young |
| 0.222396 | 0.138595 | 1 | 0.992 | 1 | 13 | Trib1 | 0.008 | Young |
| 0.222446 | 0.2371 | 0.5 | 0.426 | 1 | 13 | Paip1 | 0.074 | Young |
| 0.222609 | 0.558076 | 0.184 | 0.131 | 1 | 13 | Cdk5rap2 | 0.053 | Young |
| 0.22289 | 0.814527 | 0.096 | 0.057 | 1 | 13 | Tmem53 | 0.039 | Young |
| 0.223086 | 0.361497 | 0.404 | 0.328 | 1 | 13 | Gaa | 0.076 | Young |
| 0.22312 | 0.094823 | 1 | 1 | 1 | 13 | Mrc1 | 0 | Young |
| 0.223453 | 0.126377 | 0.781 | 0.795 | 1 | 13 | Mrpl14 | -0.014 | Young |
| 0.224221 | 0.405303 | 0.351 | 0.295 | 1 | 13 | Dgcr6 | 0.056 | Young |
| 0.224267 | 0.284037 | 0.237 | 0.18 | 1 | 13 | Ccdc32 | 0.057 | Young |
| 0.224344 | 0.295903 | 0.114 | 0.066 | 1 | 13 | Gpsm1 | 0.048 | Young |
| 0.224385 | 0.159963 | 0.877 | 0.934 | 1 | 13 | Hspa4 | -0.057 | Young |
| 0.225179 | 0.542832 | 0.149 | 0.098 | 1 | 13 | Msl3l2 | 0.051 | Young |
| 0.225243 | 0.356656 | 0.132 | 0.082 | 1 | 13 | H2-T24 | 0.05 | Young |
| 0.225676 | 0.219368 | 0.509 | 0.459 | 1 | 13 | Tmem88 | 0.05 | Young |
| 0.226049 | 0.710529 | 0.088 | 0.049 | 1 | 13 | H2-Q10 | 0.039 | Young |
| 0.226327 | 0.086828 | 0.921 | 0.943 | 1 | 13 | Tet3 | -0.022 | Young |
| 0.227284 | 0.171467 | 0.912 | 0.893 | 1 | 13 | Stard8 | 0.019 | Young |
| 0.227647 | 0.845078 | 0.096 | 0.057 | 1 | 13 | Rnf24 | 0.039 | Young |
| 0.22801 | 0.215529 | 0.342 | 0.27 | 1 | 13 | Gm20186 | 0.072 | Young |
| 0.228083 | 0.455274 | 0.289 | 0.238 | 1 | 13 | Bud23 | 0.051 | Young |
| 0.228302 | 0.255291 | 0.544 | 0.533 | 1 | 13 | Brd3 | 0.011 | Young |
| 0.228399 | 0.474621 | 0.237 | 0.18 | 1 | 13 | Bcl9l | 0.057 | Young |
| 0.228599 | 0.171377 | 0.763 | 0.811 | 1 | 13 | Os9 | -0.048 | Young |
| 0.228995 | 0.349878 | 0.561 | 0.525 | 1 | 13 | Atp8a1 | 0.036 | Young |
| 0.229035 | 0.469536 | 0.289 | 0.238 | 1 | 13 | Bmf | 0.051 | Young |
| 0.229043 | 0.22507 | 0.64 | 0.557 | 1 | 13 | Ing4 | 0.083 | Young |
| 0.229283 | 0.337597 | 0.456 | 0.41 | 1 | 13 | Brix1 | 0.046 | Young |
| 0.229712 | 0.094193 | 1 | 1 | 1 | 13 | Hmgb1 | 0 | Young |
| 0.229715 | 0.178966 | 0.789 | 0.746 | 1 | 13 | Aff4 | 0.043 | Young |
| 0.229729 | 0.362243 | 0.219 | 0.164 | 1 | 13 | Rmnd1 | 0.055 | Young |
| 0.229737 | 0.321562 | 0.158 | 0.107 | 1 | 13 | Hspa2 | 0.051 | Young |
| 0.229783 | 0.480227 | 0.281 | 0.23 | 1 | 13 | Gde1 | 0.051 | Young |
| 0.230209 | 0.273277 | 0.702 | 0.607 | 1 | 13 | Pbxip1 | 0.095 | Young |
| 0.23047 | 0.436075 | 0.193 | 0.139 | 1 | 13 | Timm21 | 0.054 | Young |
| 0.231021 | 0.222712 | 0.5 | 0.434 | 1 | 13 | Zfp131 | 0.066 | Young |
| 0.231026 | 0.526431 | 0.079 | 0.041 | 1 | 13 | Lss | 0.038 | Young |
| 0.231126 | 0.809929 | 0.088 | 0.049 | 1 | 13 | Zbtb8a | 0.039 | Young |
| 0.231434 | 0.202107 | 0.456 | 0.385 | 1 | 13 | Ncbp1 | 0.071 | Young |
| 0.231502 | 0.262087 | 0.596 | 0.566 | 1 | 13 | Polr2j | 0.03 | Young |
| 0.231578 | 0.211268 | 0.605 | 0.607 | 1 | 13 | Cd300lg | -0.002 | Young |
| 0.231766 | 0.367301 | 0.491 | 0.459 | 1 | 13 | Nae1 | 0.032 | Young |
| 0.232171 | 0.483415 | 0.289 | 0.246 | 1 | 13 | Ttc4 | 0.043 | Young |

| | | | | | | | | |
|---|---|---|---|---|---|---|---|---|
| 0.232468 | 0.224458 | 0.588 | 0.549 | 1 | 13 | Mob3b | 0.039 | Young |
| 0.232685 | 0.076374 | 0.982 | 1 | 1 | 13 | Atp6v1g1 | -0.018 | Young |
| 0.232836 | 0.720439 | 0.088 | 0.049 | 1 | 13 | Lrsam1 | 0.039 | Young |
| 0.232846 | 0.612024 | 0.079 | 0.041 | 1 | 13 | 0610040J0 | 0.038 | Young |
| 0.233078 | 0.356505 | 0.412 | 0.352 | 1 | 13 | Gprc5c | 0.06 | Young |
| 0.233269 | 0.15386 | 0.904 | 0.918 | 1 | 13 | Ciita | -0.014 | Young |
| 0.233501 | 0.432779 | 0.36 | 0.311 | 1 | 13 | Brms1 | 0.049 | Young |
| 0.234077 | 0.638207 | 0.219 | 0.164 | 1 | 13 | Jaml | 0.055 | Young |
| 0.234263 | 0.387255 | 0.149 | 0.098 | 1 | 13 | Plscr1 | 0.051 | Young |
| 0.234554 | 0.147709 | 0.974 | 0.975 | 1 | 13 | Prr13 | -0.001 | Young |
| 0.234558 | 0.114162 | 0.991 | 0.992 | 1 | 13 | Dnaja1 | -0.001 | Young |
| 0.234928 | 0.120201 | 0.974 | 0.967 | 1 | 13 | Ccdc50 | 0.007 | Young |
| 0.235581 | 0.554809 | 0.149 | 0.098 | 1 | 13 | Rasal3 | 0.051 | Young |
| 0.235663 | 0.301189 | 0.57 | 0.516 | 1 | 13 | Pttg1 | 0.054 | Young |
| 0.235991 | 0.22954 | 0.596 | 0.598 | 1 | 13 | Oaz2 | -0.002 | Young |
| 0.236054 | 0.201831 | 0.825 | 0.836 | 1 | 13 | Ggnbp2 | -0.011 | Young |
| 0.236097 | 0.160474 | 0.719 | 0.639 | 1 | 13 | Cnpy3 | 0.08 | Young |
| 0.236631 | 0.528558 | 0.202 | 0.148 | 1 | 13 | Fbf1 | 0.054 | Young |
| 0.236777 | 0.205356 | 0.596 | 0.541 | 1 | 13 | Hmg20b | 0.055 | Young |
| 0.236886 | 0.39666 | 0.351 | 0.303 | 1 | 13 | Zfp260 | 0.048 | Young |
| 0.237197 | 0.052913 | 0.982 | 0.992 | 1 | 13 | Pak2 | -0.01 | Young |
| 0.237206 | 0.166166 | 0.553 | 0.492 | 1 | 13 | Clcn3 | 0.061 | Young |
| 0.237378 | 0.781265 | 0.096 | 0.057 | 1 | 13 | Zfp652os | 0.039 | Young |
| 0.237588 | 0.249621 | 0.702 | 0.697 | 1 | 13 | Dyrk2 | 0.005 | Young |
| 0.237732 | 0.633945 | 0.272 | 0.23 | 1 | 13 | Syt11 | 0.042 | Young |
| 0.237898 | 0.754605 | 0.114 | 0.074 | 1 | 13 | 4933433G1 | 0.04 | Young |
| 0.23804 | 0.239995 | 0.43 | 0.377 | 1 | 13 | Med12 | 0.053 | Young |
| 0.238255 | 0.162915 | 0.772 | 0.82 | 1 | 13 | Eif1ax | -0.048 | Young |
| 0.238273 | 0.314437 | 0.509 | 0.443 | 1 | 13 | Tgfbrap1 | 0.066 | Young |
| 0.238715 | 0.048006 | 1 | 1 | 1 | 13 | Hsp90ab1 | 0 | Young |
| 0.238746 | 0.263654 | 0.623 | 0.607 | 1 | 13 | Dhx36 | 0.016 | Young |
| 0.238875 | 0.525858 | 0.211 | 0.156 | 1 | 13 | Ccdc71 | 0.055 | Young |
| 0.238892 | 0.460233 | 0.36 | 0.311 | 1 | 13 | Gpatch11 | 0.049 | Young |
| 0.238897 | 0.0951 | 0.728 | 0.689 | 1 | 13 | Atf7ip | 0.039 | Young |
| 0.238985 | 0.356083 | 0.447 | 0.393 | 1 | 13 | Cebpzos | 0.054 | Young |
| 0.239025 | 0.621331 | 0.175 | 0.123 | 1 | 13 | Cdc25a | 0.052 | Young |
| 0.239498 | 0.291129 | 0.333 | 0.279 | 1 | 13 | Xylt1 | 0.054 | Young |
| 0.239766 | 0.657986 | 0.088 | 0.049 | 1 | 13 | E2f2 | 0.039 | Young |
| 0.239768 | 0.403805 | 0.167 | 0.115 | 1 | 13 | Trim39 | 0.052 | Young |
| 0.239836 | 0.221166 | 0.281 | 0.205 | 1 | 13 | Hcar2 | 0.076 | Young |
| 0.23994 | 0.477758 | 0.342 | 0.303 | 1 | 13 | Eif2ak3 | 0.039 | Young |
| 0.240239 | 0.04809 | 0.991 | 1 | 1 | 13 | Sat1 | -0.009 | Young |
| 0.240919 | 0.710614 | 0.114 | 0.074 | 1 | 13 | 1810034E1 | 0.04 | Young |
| 0.241004 | 0.081021 | 1 | 1 | 1 | 13 | Fxyd5 | 0 | Young |
| 0.241075 | 0.364039 | 0.439 | 0.41 | 1 | 13 | Klhl18 | 0.029 | Young |

| | | | | | | | | |
|---|---|---|---|---|---|---|---|---|
| 0.241174 | 0.156855 | 0.816 | 0.795 | 1 | 13 | Washc4 | 0.021 | Young |
| 0.24131 | 0.181008 | 0.421 | 0.344 | 1 | 13 | Secisbp2 | 0.077 | Young |
| 0.241768 | 0.069462 | 0.965 | 0.992 | 1 | 13 | Nfe2l2 | -0.027 | Young |
| 0.24177 | 0.128231 | 0.991 | 0.984 | 1 | 13 | Stk17b | 0.007 | Young |
| 0.243148 | 0.606875 | 0.219 | 0.172 | 1 | 13 | Scyl3 | 0.047 | Young |
| 0.243177 | 0.417155 | 0.325 | 0.262 | 1 | 13 | Zcchc8 | 0.063 | Young |
| 0.243681 | 0.271175 | 0.658 | 0.664 | 1 | 13 | Ncoa7 | -0.006 | Young |
| 0.243852 | 0.294591 | 0.746 | 0.738 | 1 | 13 | R3hdm1 | 0.008 | Young |
| 0.243904 | 0.505199 | 0.158 | 0.107 | 1 | 13 | Pigg | 0.051 | Young |
| 0.24419 | 0.212796 | 0.5 | 0.434 | 1 | 13 | Ssh1 | 0.066 | Young |
| 0.244204 | 0.393222 | 0.404 | 0.352 | 1 | 13 | Prep | 0.052 | Young |
| 0.244495 | 0.167587 | 0.807 | 0.746 | 1 | 13 | Cript | 0.061 | Young |
| 0.244696 | 0.047877 | 0.728 | 0.713 | 1 | 13 | Vav1 | 0.015 | Young |
| 0.244853 | 0.046933 | 1 | 1 | 1 | 13 | Gnai2 | 0 | Young |
| 0.244912 | 0.4886 | 0.263 | 0.205 | 1 | 13 | Dhx57 | 0.058 | Young |
| 0.244934 | 0.788963 | 0.237 | 0.205 | 1 | 13 | Cul9 | 0.032 | Young |
| 0.245435 | 0.267113 | 0.684 | 0.672 | 1 | 13 | Pigx | 0.012 | Young |
| 0.245437 | 0.119385 | 0.877 | 0.934 | 1 | 13 | Map4k4 | -0.057 | Young |
| 0.245502 | 0.952249 | 0.114 | 0.074 | 1 | 13 | Zfp825 | 0.04 | Young |
| 0.245713 | 0.405621 | 0.482 | 0.426 | 1 | 13 | Tmem64 | 0.056 | Young |
| 0.246345 | 0.357306 | 0.57 | 0.549 | 1 | 13 | Thap12 | 0.021 | Young |
| 0.246405 | 0.150409 | 0.991 | 0.984 | 1 | 13 | Tbca | 0.007 | Young |
| 0.246405 | 0.079257 | 1 | 1 | 1 | 13 | Cox8a | 0 | Young |
| 0.246432 | 0.284493 | 0.289 | 0.23 | 1 | 13 | Pou5f2 | 0.059 | Young |
| 0.24645 | 0.337178 | 0.482 | 0.459 | 1 | 13 | Dirc2 | 0.023 | Young |
| 0.246539 | 0.583192 | 0.246 | 0.197 | 1 | 13 | Foxo4 | 0.049 | Young |
| 0.246773 | 0.18471 | 0.772 | 0.705 | 1 | 13 | Entpd1 | 0.067 | Young |
| 0.246794 | 0.069399 | 0.991 | 0.992 | 1 | 13 | Cyth4 | -0.001 | Young |
| 0.246965 | 0.148063 | 0.886 | 0.918 | 1 | 13 | Cnot6l | -0.032 | Young |
| 0.247091 | 0.645649 | 0.175 | 0.131 | 1 | 13 | Tmem150b | 0.044 | Young |
| 0.247184 | 0.060155 | 1 | 1 | 1 | 13 | Rpl7a | 0 | Young |
| 0.247675 | 0.270753 | 0.623 | 0.639 | 1 | 13 | Rchy1 | -0.016 | Young |
| 0.247846 | 0.356917 | 0.544 | 0.492 | 1 | 13 | Ufl1 | 0.052 | Young |
| 0.248309 | 0.129434 | 0.535 | 0.484 | 1 | 13 | Ppm1b | 0.051 | Young |
| 0.24835 | 0.1126 | 0.772 | 0.738 | 1 | 13 | Btk | 0.034 | Young |
| 0.248459 | 0.490539 | 0.289 | 0.23 | 1 | 13 | Tbcc | 0.059 | Young |
| 0.248571 | 0.197752 | 0.526 | 0.451 | 1 | 13 | Tor1b | 0.075 | Young |
| 0.24869 | 0.677043 | 0.246 | 0.197 | 1 | 13 | Scai | 0.049 | Young |
| 0.248938 | 0.833554 | 0.053 | 0.025 | 1 | 13 | Dscc1 | 0.028 | Young |
| 0.249075 | 0.455505 | 0.447 | 0.418 | 1 | 13 | Nudt16 | 0.029 | Young |
| 0.249206 | 0.552666 | 0.158 | 0.107 | 1 | 13 | Ccdc152 | 0.051 | Young |
| 0.249538 | 0.483998 | 0.272 | 0.221 | 1 | 13 | Cog1 | 0.051 | Young |
| 0.249815 | 0.138944 | 0.605 | 0.615 | 1 | 13 | Ide | -0.01 | Young |
| 0.249857 | 0.152712 | 0.842 | 0.811 | 1 | 13 | Baz2b | 0.031 | Young |
| 0.250146 | 0.693199 | 0.114 | 0.074 | 1 | 13 | Gm6133 | 0.04 | Young |

| | | | | | | | | |
|---|---|---|---|---|---|---|---|---|
| 0.250221 | 0.132438 | 0.781 | 0.762 | 1 | 13 | Bcl2l11 | 0.019 | Young |
| 0.250617 | 0.161467 | 0.632 | 0.557 | 1 | 13 | Cltb | 0.075 | Young |
| 0.250651 | 0.50092 | 0.202 | 0.148 | 1 | 13 | Zkscan17 | 0.054 | Young |
| 0.250679 | 0.239255 | 0.807 | 0.787 | 1 | 13 | Mark2 | 0.02 | Young |
| 0.2507 | 0.186836 | 0.439 | 0.377 | 1 | 13 | Cnep1r1 | 0.062 | Young |
| 0.251521 | 0.598156 | 0.316 | 0.262 | 1 | 13 | Dym | 0.054 | Young |
| 0.251911 | 0.299484 | 0.465 | 0.434 | 1 | 13 | Zfp560 | 0.031 | Young |
| 0.25284 | 0.423758 | 0.175 | 0.123 | 1 | 13 | Rnf219 | 0.052 | Young |
| 0.253118 | 0.138671 | 0.904 | 0.869 | 1 | 13 | Rnf7 | 0.035 | Young |
| 0.253122 | 0.208812 | 0.798 | 0.828 | 1 | 13 | Ankrd12 | -0.03 | Young |
| 0.253409 | 0.227283 | 0.711 | 0.73 | 1 | 13 | Slbp | -0.019 | Young |
| 0.253551 | 0.352645 | 0.333 | 0.27 | 1 | 13 | Rubcn | 0.063 | Young |
| 0.25369 | 0.864645 | 0.053 | 0.025 | 1 | 13 | Brca2 | 0.028 | Young |
| 0.25369 | 0.859204 | 0.053 | 0.025 | 1 | 13 | Phldb3 | 0.028 | Young |
| 0.25369 | 0.765531 | 0.053 | 0.025 | 1 | 13 | Ildr2 | 0.028 | Young |
| 0.253852 | 0.041965 | 0.956 | 0.959 | 1 | 13 | 11-Sep | -0.003 | Young |
| 0.254449 | 0.255137 | 0.237 | 0.172 | 1 | 13 | Slc20a2 | 0.065 | Young |
| 0.25452 | 0.050138 | 0.781 | 0.721 | 1 | 13 | Tulp4 | 0.06 | Young |
| 0.254665 | 0.124711 | 0.991 | 0.992 | 1 | 13 | Snrnp70 | -0.001 | Young |
| 0.254839 | 0.381698 | 0.491 | 0.443 | 1 | 13 | Smim12 | 0.048 | Young |
| 0.255071 | 0.32859 | 0.211 | 0.156 | 1 | 13 | Rps6kb2 | 0.055 | Young |
| 0.255098 | 0.211054 | 0.377 | 0.303 | 1 | 13 | Tusc1 | 0.074 | Young |
| 0.25519 | 0.293434 | 0.789 | 0.787 | 1 | 13 | Ttc3 | 0.002 | Young |
| 0.25541 | 0.541697 | 0.175 | 0.123 | 1 | 13 | Mmp14 | 0.052 | Young |
| 0.255861 | 0.132914 | 1 | 1 | 1 | 13 | Lrp1 | 0 | Young |
| 0.255886 | 0.557461 | 0.088 | 0.049 | 1 | 13 | Lcmt2 | 0.039 | Young |
| 0.25609 | 0.877059 | 0.053 | 0.025 | 1 | 13 | Tspyl4 | 0.028 | Young |
| 0.256226 | 0.270477 | 0.719 | 0.721 | 1 | 13 | Smc4 | -0.002 | Young |
| 0.256324 | 0.264674 | 0.535 | 0.475 | 1 | 13 | Commd2 | 0.06 | Young |
| 0.256496 | 0.536019 | 0.325 | 0.279 | 1 | 13 | Ap4m1 | 0.046 | Young |
| 0.25681 | 0.16722 | 0.851 | 0.869 | 1 | 13 | Cbx3 | -0.018 | Young |
| 0.256817 | 0.207171 | 0.719 | 0.689 | 1 | 13 | St3gal5 | 0.03 | Young |
| 0.256849 | 0.278949 | 0.588 | 0.557 | 1 | 13 | Fibp | 0.031 | Young |
| 0.256955 | 0.091391 | 0.939 | 0.91 | 1 | 13 | Ndufs5 | 0.029 | Young |
| 0.257175 | 0.637179 | 0.14 | 0.098 | 1 | 13 | Fam131a | 0.042 | Young |
| 0.257216 | 0.463883 | 0.246 | 0.189 | 1 | 13 | Letmd1 | 0.057 | Young |
| 0.257307 | 0.347084 | 0.158 | 0.107 | 1 | 13 | Kbtbd3 | 0.051 | Young |
| 0.257419 | 0.242199 | 0.684 | 0.615 | 1 | 13 | Ing2 | 0.069 | Young |
| 0.257461 | 0.097635 | 1 | 1 | 1 | 13 | Serinc3 | 0 | Young |
| 0.257612 | 0.124494 | 0.886 | 0.91 | 1 | 13 | Kdm2a | -0.024 | Young |
| 0.257717 | 0.129748 | 0.798 | 0.762 | 1 | 13 | Ppp1r2 | 0.036 | Young |
| 0.257753 | 0.212358 | 0.728 | 0.697 | 1 | 13 | Ms4a6b | 0.031 | Young |
| 0.258034 | 0.286814 | 0.526 | 0.484 | 1 | 13 | Fam214b | 0.042 | Young |
| 0.258306 | 0.651719 | 0.202 | 0.156 | 1 | 13 | Ccdc58 | 0.046 | Young |
| 0.258505 | 0.734862 | 0.053 | 0.025 | 1 | 13 | Nphp1 | 0.028 | Young |

| | | | | | | | | |
|---|---|---|---|---|---|---|---|---|
| 0.258505 | 0.659661 | 0.053 | 0.025 | 1 | 13 | Gm28874 | 0.028 | Young |
| 0.258639 | 0.473469 | 0.211 | 0.156 | 1 | 13 | Nt5dc2 | 0.055 | Young |
| 0.259617 | 0.701735 | 0.114 | 0.074 | 1 | 13 | Tbc1d7 | 0.04 | Young |
| 0.259715 | 0.216548 | 0.57 | 0.525 | 1 | 13 | Oxr1 | 0.045 | Young |
| 0.260325 | 0.117694 | 0.886 | 0.885 | 1 | 13 | Pacs2 | 0.001 | Young |
| 0.260393 | 0.468437 | 0.254 | 0.205 | 1 | 13 | Dut | 0.049 | Young |
| 0.260536 | 0.259961 | 0.605 | 0.557 | 1 | 13 | Aaed1 | 0.048 | Young |
| 0.260615 | 0.499367 | 0.167 | 0.115 | 1 | 13 | Fancg | 0.052 | Young |
| 0.260936 | 0.683333 | 0.053 | 0.025 | 1 | 13 | Ttc21b | 0.028 | Young |
| 0.261055 | 0.149179 | 0.921 | 0.975 | 1 | 13 | Myo5a | -0.054 | Young |
| 0.261144 | 0.154417 | 0.57 | 0.516 | 1 | 13 | Qars | 0.054 | Young |
| 0.261219 | 0.649888 | 0.114 | 0.074 | 1 | 13 | Tgif2 | 0.04 | Young |
| 0.261362 | 0.222021 | 0.772 | 0.746 | 1 | 13 | Rc3h1 | 0.026 | Young |
| 0.261492 | 0.061319 | 1 | 1 | 1 | 13 | Pid1 | 0 | Young |
| 0.261671 | 0.350542 | 0.193 | 0.139 | 1 | 13 | Nek1 | 0.054 | Young |
| 0.261673 | 0.24532 | 0.474 | 0.451 | 1 | 13 | Rasgrp2 | 0.023 | Young |
| 0.261685 | 0.462901 | 0.325 | 0.27 | 1 | 13 | Smarcal1 | 0.055 | Young |
| 0.262195 | 0.087648 | 0.868 | 0.82 | 1 | 13 | Fam117b | 0.048 | Young |
| 0.262272 | 0.095348 | 0.947 | 0.951 | 1 | 13 | Arhgap18 | -0.004 | Young |
| 0.262347 | 0.338025 | 0.579 | 0.549 | 1 | 13 | Clasp2 | 0.03 | Young |
| 0.262359 | 0.191995 | 0.833 | 0.828 | 1 | 13 | U2surp | 0.005 | Young |
| 0.262675 | 0.184243 | 0.851 | 0.844 | 1 | 13 | Pdlim1 | 0.007 | Young |
| 0.262691 | 0.291472 | 0.325 | 0.27 | 1 | 13 | Hbegf | 0.055 | Young |
| 0.262759 | 0.568322 | 0.105 | 0.066 | 1 | 13 | Car9 | 0.039 | Young |
| 0.262801 | 0.349928 | 0.36 | 0.303 | 1 | 13 | Nup210 | 0.057 | Young |
| 0.263383 | 0.738103 | 0.053 | 0.025 | 1 | 13 | Zswim9 | 0.028 | Young |
| 0.263383 | 0.667894 | 0.053 | 0.025 | 1 | 13 | Lefty1 | 0.028 | Young |
| 0.264088 | 0.133377 | 0.904 | 0.893 | 1 | 13 | Cysltr1 | 0.011 | Young |
| 0.264444 | 0.70619 | 0.114 | 0.074 | 1 | 13 | Zfp953 | 0.04 | Young |
| 0.264662 | 0.389884 | 0.211 | 0.156 | 1 | 13 | Pex11b | 0.055 | Young |
| 0.264683 | 0.156124 | 0.64 | 0.582 | 1 | 13 | Grpel1 | 0.058 | Young |
| 0.264747 | 0.049677 | 1 | 1 | 1 | 13 | Arhgdib | 0 | Young |
| 0.264953 | 0.304838 | 0.518 | 0.492 | 1 | 13 | Nipsnap3b | 0.026 | Young |
| 0.265801 | 0.647721 | 0.132 | 0.09 | 1 | 13 | Nid2 | 0.042 | Young |
| 0.265826 | 0.311548 | 0.535 | 0.475 | 1 | 13 | Smim13 | 0.06 | Young |
| 0.266042 | 0.486644 | 0.184 | 0.139 | 1 | 13 | Armc3 | 0.045 | Young |
| 0.266053 | 0.390805 | 0.377 | 0.336 | 1 | 13 | B9d2 | 0.041 | Young |
| 0.266079 | 0.454079 | 0.228 | 0.172 | 1 | 13 | Umps | 0.056 | Young |
| 0.266519 | 0.156603 | 0.781 | 0.73 | 1 | 13 | Acsl4 | 0.051 | Young |
| 0.266747 | 0.052141 | 0.974 | 0.91 | 1 | 13 | Raph1 | 0.064 | Young |
| 0.267083 | 0.130297 | 0.93 | 0.91 | 1 | 13 | Matr3 | 0.02 | Young |
| 0.267101 | 0.114389 | 0.93 | 0.918 | 1 | 13 | Nol7 | 0.012 | Young |
| 0.267192 | 0.137624 | 0.842 | 0.779 | 1 | 13 | Tbl1xr1 | 0.063 | Young |
| 0.267446 | 0.217707 | 0.57 | 0.5 | 1 | 13 | Cd9 | 0.07 | Young |
| 0.267843 | 0.476195 | 0.105 | 0.066 | 1 | 13 | Tmem229a | 0.039 | Young |

| | | | | | | | | |
|---|---|---|---|---|---|---|---|---|
| 0.268209 | 0.174398 | 0.728 | 0.746 | 1 | 13 | Mtm1 | -0.018 | Young |
| 0.26832 | 0.543342 | 0.096 | 0.057 | 1 | 13 | Mgp | 0.039 | Young |
| 0.268324 | 0.607591 | 0.053 | 0.025 | 1 | 13 | Gm4707 | 0.028 | Young |
| 0.268324 | 0.598224 | 0.053 | 0.025 | 1 | 13 | Trmo | 0.028 | Young |
| 0.268324 | 0.560115 | 0.053 | 0.025 | 1 | 13 | Apoc1 | 0.028 | Young |
| 0.268324 | 0.551243 | 0.053 | 0.025 | 1 | 13 | Tcf19 | 0.028 | Young |
| 0.268324 | 0.543291 | 0.053 | 0.025 | 1 | 13 | Slc25a35 | 0.028 | Young |
| 0.268859 | 0.424628 | 0.228 | 0.18 | 1 | 13 | Mfge8 | 0.048 | Young |
| 0.269319 | 0.11705 | 0.807 | 0.787 | 1 | 13 | Leng8 | 0.02 | Young |
| 0.269804 | 0.27339 | 0.325 | 0.27 | 1 | 13 | Kdelc2 | 0.055 | Young |
| 0.269998 | 0.36144 | 0.465 | 0.451 | 1 | 13 | Hbs1l | 0.014 | Young |
| 0.270272 | 0.142663 | 0.746 | 0.713 | 1 | 13 | Hsd17b4 | 0.033 | Young |
| 0.270444 | 0.207454 | 0.763 | 0.754 | 1 | 13 | Mt2 | 0.009 | Young |
| 0.270819 | 0.529979 | 0.053 | 0.025 | 1 | 13 | Rasip1 | 0.028 | Young |
| 0.271338 | 0.101819 | 0.991 | 0.959 | 1 | 13 | Pgls | 0.032 | Young |
| 0.271685 | 0.249998 | 0.798 | 0.762 | 1 | 13 | Lilrb4a | 0.036 | Young |
| 0.271883 | 0.316471 | 0.614 | 0.566 | 1 | 13 | Dtymk | 0.048 | Young |
| 0.271982 | 0.208473 | 0.719 | 0.672 | 1 | 13 | Rps6ka1 | 0.047 | Young |
| 0.272591 | 0.097111 | 0.991 | 0.992 | 1 | 13 | Minos1 | -0.001 | Young |
| 0.273036 | 0.525981 | 0.412 | 0.402 | 1 | 13 | Ago4 | 0.01 | Young |
| 0.273083 | 0.097953 | 0.754 | 0.672 | 1 | 13 | Cd24a | 0.082 | Young |
| 0.273329 | 0.558677 | 0.053 | 0.025 | 1 | 13 | Hk1os | 0.028 | Young |
| 0.273708 | 0.685824 | 0.149 | 0.107 | 1 | 13 | Tmco4 | 0.042 | Young |
| 0.273716 | 0.150529 | 0.526 | 0.434 | 1 | 13 | Ddx1 | 0.092 | Young |
| 0.27422 | 0.709926 | 0.175 | 0.131 | 1 | 13 | Tnni2 | 0.044 | Young |
| 0.274283 | 0.037326 | 0.114 | 0.074 | 1 | 13 | Cxcl9 | 0.04 | Young |
| 0.274331 | 0.463727 | 0.325 | 0.287 | 1 | 13 | Bnip3 | 0.038 | Young |
| 0.274415 | 0.424655 | 0.193 | 0.139 | 1 | 13 | Wdr83 | 0.054 | Young |
| 0.274727 | 0.560959 | 0.105 | 0.066 | 1 | 13 | Thada | 0.039 | Young |
| 0.274781 | 0.210236 | 0.289 | 0.23 | 1 | 13 | Cd99l2 | 0.059 | Young |
| 0.274849 | 0.139922 | 0.904 | 0.893 | 1 | 13 | Uba52 | 0.011 | Young |
| 0.275077 | 0.289432 | 0.36 | 0.311 | 1 | 13 | Brcc3 | 0.049 | Young |
| 0.275101 | 0.200658 | 0.868 | 0.787 | 1 | 13 | Hopx | 0.081 | Young |
| 0.275155 | 0.879914 | 0.149 | 0.107 | 1 | 13 | Gtf2h2 | 0.042 | Young |
| 0.27534 | 0.155455 | 0.728 | 0.762 | 1 | 13 | Med13 | -0.034 | Young |
| 0.275431 | 0.107466 | 0.956 | 0.902 | 1 | 13 | Eif4ebp2 | 0.054 | Young |
| 0.275834 | 0.237761 | 0.456 | 0.418 | 1 | 13 | D16Ertd47 | 0.038 | Young |
| 0.276089 | 0.78306 | 0.061 | 0.033 | 1 | 13 | Pde2a | 0.028 | Young |
| 0.27656 | 0.388189 | 0.202 | 0.148 | 1 | 13 | 1110012L1 | 0.054 | Young |
| 0.276736 | 0.096245 | 0.789 | 0.795 | 1 | 13 | Ndufaf8 | -0.006 | Young |
| 0.276972 | 0.266541 | 0.395 | 0.328 | 1 | 13 | P2ry12 | 0.067 | Young |
| 0.277195 | 0.170897 | 0.5 | 0.443 | 1 | 13 | Lcorl | 0.057 | Young |
| 0.277543 | 0.503804 | 0.123 | 0.082 | 1 | 13 | R3hcc1 | 0.041 | Young |
| 0.277584 | 0.0641 | 0.798 | 0.762 | 1 | 13 | Wdfy3 | 0.036 | Young |
| 0.277618 | 0.556522 | 0.114 | 0.074 | 1 | 13 | Tsacc | 0.04 | Young |

| | | | | | | | | |
|---|---|---|---|---|---|---|---|---|
| 0.278314 | 0.133174 | 0.737 | 0.754 | 1 | 13 | Tmcc3 | -0.017 | Young |
| 0.278382 | 0.175359 | 0.763 | 0.77 | 1 | 13 | Tnfrsf21 | -0.007 | Young |
| 0.278462 | 1.660201 | 0.026 | 0.008 | 1 | 13 | Ifi44 | 0.018 | Young |
| 0.278462 | 1.019412 | 0.026 | 0.008 | 1 | 13 | 1700066B1 | 0.018 | Young |
| 0.278462 | 0.941055 | 0.026 | 0.008 | 1 | 13 | Ltbp4 | 0.018 | Young |
| 0.278462 | 0.936562 | 0.026 | 0.008 | 1 | 13 | Proser2 | 0.018 | Young |
| 0.278462 | 0.907534 | 0.026 | 0.008 | 1 | 13 | Polh | 0.018 | Young |
| 0.278462 | 0.901622 | 0.026 | 0.008 | 1 | 13 | U2af1l4 | 0.018 | Young |
| 0.278462 | 0.886609 | 0.026 | 0.008 | 1 | 13 | Zfp72 | 0.018 | Young |
| 0.278462 | 0.869488 | 0.026 | 0.008 | 1 | 13 | 5730422E0 | 0.018 | Young |
| 0.278462 | 0.863634 | 0.026 | 0.008 | 1 | 13 | Ttc23 | 0.018 | Young |
| 0.278462 | 0.861805 | 0.026 | 0.008 | 1 | 13 | Spns3 | 0.018 | Young |
| 0.278462 | 0.860574 | 0.026 | 0.008 | 1 | 13 | Gm15880 | 0.018 | Young |
| 0.278462 | 0.859 | 0.026 | 0.008 | 1 | 13 | Fmo2 | 0.018 | Young |
| 0.278462 | 0.851695 | 0.026 | 0.008 | 1 | 13 | Cracr2a | 0.018 | Young |
| 0.278462 | 0.848053 | 0.026 | 0.008 | 1 | 13 | Ptges | 0.018 | Young |
| 0.278462 | 0.808368 | 0.026 | 0.008 | 1 | 13 | Gm16174 | 0.018 | Young |
| 0.278462 | 0.797254 | 0.026 | 0.008 | 1 | 13 | Arhgap24 | 0.018 | Young |
| 0.278462 | 0.764162 | 0.026 | 0.008 | 1 | 13 | 1700030J2 | 0.018 | Young |
| 0.278462 | 0.757491 | 0.026 | 0.008 | 1 | 13 | Mustn1 | 0.018 | Young |
| 0.278462 | 0.749443 | 0.026 | 0.008 | 1 | 13 | 2810455O( | 0.018 | Young |
| 0.278462 | 0.742253 | 0.026 | 0.008 | 1 | 13 | E430024P1 | 0.018 | Young |
| 0.278462 | 0.72738 | 0.026 | 0.008 | 1 | 13 | Igsf10 | 0.018 | Young |
| 0.278462 | 0.717611 | 0.026 | 0.008 | 1 | 13 | Zfp955b | 0.018 | Young |
| 0.278462 | 0.697225 | 0.026 | 0.008 | 1 | 13 | Gm26563 | 0.018 | Young |
| 0.278462 | 0.670647 | 0.026 | 0.008 | 1 | 13 | Rdh12 | 0.018 | Young |
| 0.278462 | 0.662099 | 0.026 | 0.008 | 1 | 13 | Cdkl3 | 0.018 | Young |
| 0.278462 | 0.624828 | 0.026 | 0.008 | 1 | 13 | Foxm1 | 0.018 | Young |
| 0.279296 | 0.529047 | 0.114 | 0.074 | 1 | 13 | Cox11 | 0.04 | Young |
| 0.279347 | 0.188217 | 0.675 | 0.623 | 1 | 13 | Fam234a | 0.052 | Young |
| 0.280266 | 0.525139 | 0.333 | 0.303 | 1 | 13 | Zfp90 | 0.03 | Young |
| 0.281301 | 0.51278 | 0.333 | 0.311 | 1 | 13 | Pde4a | 0.022 | Young |
| 0.281415 | 0.274751 | 0.395 | 0.352 | 1 | 13 | Gm15417 | 0.043 | Young |
| 0.281866 | 0.316115 | 0.509 | 0.443 | 1 | 13 | Pilrb1 | 0.066 | Young |
| 0.281964 | 0.297808 | 0.333 | 0.27 | 1 | 13 | Zfp422 | 0.063 | Young |
| 0.282264 | 0.785588 | 0.026 | 0.008 | 1 | 13 | Akap12 | 0.018 | Young |
| 0.282264 | 0.773279 | 0.026 | 0.008 | 1 | 13 | Btla | 0.018 | Young |
| 0.282264 | 0.768189 | 0.026 | 0.008 | 1 | 13 | Kdr | 0.018 | Young |
| 0.282264 | 0.754201 | 0.026 | 0.008 | 1 | 13 | Fcrl1 | 0.018 | Young |
| 0.282264 | 0.749257 | 0.026 | 0.008 | 1 | 13 | D930028M | 0.018 | Young |
| 0.282264 | 0.747437 | 0.026 | 0.008 | 1 | 13 | Zfp870 | 0.018 | Young |
| 0.282264 | 0.747213 | 0.026 | 0.008 | 1 | 13 | Gm3604 | 0.018 | Young |
| 0.282264 | 0.725223 | 0.026 | 0.008 | 1 | 13 | Gm7324 | 0.018 | Young |
| 0.282264 | 0.721584 | 0.026 | 0.008 | 1 | 13 | AU019990 | 0.018 | Young |
| 0.282264 | 0.719147 | 0.026 | 0.008 | 1 | 13 | Gpr4 | 0.018 | Young |

| | | | | | | | | |
|---|---|---|---|---|---|---|---|---|
| 0.282264 | 0.713552 | 0.026 | 0.008 | 1 | 13 | Stxbp4 | 0.018 | Young |
| 0.282264 | 0.697298 | 0.026 | 0.008 | 1 | 13 | 2600006KC | 0.018 | Young |
| 0.282264 | 0.695046 | 0.026 | 0.008 | 1 | 13 | Gm1123 | 0.018 | Young |
| 0.282264 | 0.691227 | 0.026 | 0.008 | 1 | 13 | Gm12195 | 0.018 | Young |
| 0.282264 | 0.683573 | 0.026 | 0.008 | 1 | 13 | AC151836. | 0.018 | Young |
| 0.282264 | 0.679415 | 0.026 | 0.008 | 1 | 13 | Gm20517 | 0.018 | Young |
| 0.282264 | 0.671685 | 0.026 | 0.008 | 1 | 13 | Rasl11b | 0.018 | Young |
| 0.282264 | 0.658055 | 0.026 | 0.008 | 1 | 13 | Chrnb1 | 0.018 | Young |
| 0.282264 | 0.643325 | 0.026 | 0.008 | 1 | 13 | Arhgef39 | 0.018 | Young |
| 0.282264 | 0.615436 | 0.026 | 0.008 | 1 | 13 | Gm26674 | 0.018 | Young |
| 0.282264 | 0.587447 | 0.026 | 0.008 | 1 | 13 | Ccdc125 | 0.018 | Young |
| 0.282264 | 0.512845 | 0.026 | 0.008 | 1 | 13 | Amotl1 | 0.018 | Young |
| 0.28227 | 0.530872 | 0.254 | 0.213 | 1 | 13 | Tfip11 | 0.041 | Young |
| 0.282284 | 0.129751 | 0.982 | 0.943 | 1 | 13 | Smarca5 | 0.039 | Young |
| 0.28237 | 0.196218 | 0.772 | 0.68 | 1 | 13 | Bcl7c | 0.092 | Young |
| 0.2824 | 0.692327 | 0.123 | 0.082 | 1 | 13 | Etfbkmt | 0.041 | Young |
| 0.282464 | 0.488846 | 0.167 | 0.123 | 1 | 13 | Kdm4b | 0.044 | Young |
| 0.282517 | 0.216964 | 0.5 | 0.426 | 1 | 13 | Cyp27a1 | 0.074 | Young |
| 0.282672 | 0.464618 | 0.114 | 0.074 | 1 | 13 | Efcab2 | 0.04 | Young |
| 0.282698 | 0.497359 | 0.211 | 0.164 | 1 | 13 | Rhbdd1 | 0.047 | Young |
| 0.282709 | 0.151212 | 0.798 | 0.762 | 1 | 13 | Adss | 0.036 | Young |
| 0.283058 | 0.784601 | 0.061 | 0.033 | 1 | 13 | Gm26759 | 0.028 | Young |
| 0.283438 | 0.46013 | 0.368 | 0.336 | 1 | 13 | Pdgfc | 0.032 | Young |
| 0.283438 | 0.226563 | 0.377 | 0.328 | 1 | 13 | Chm | 0.049 | Young |
| 0.28346 | 0.200645 | 0.456 | 0.393 | 1 | 13 | Abcg3 | 0.063 | Young |
| 0.283565 | 1.006095 | 0.14 | 0.107 | 1 | 13 | Ndufaf1 | 0.033 | Young |
| 0.283779 | 0.222986 | 0.851 | 0.82 | 1 | 13 | Fam102b | 0.031 | Young |
| 0.2841 | 0.275835 | 0.728 | 0.721 | 1 | 13 | Col4a3bp | 0.007 | Young |
| 0.28437 | 0.53646 | 0.114 | 0.074 | 1 | 13 | BC147527 | 0.04 | Young |
| 0.285151 | 0.156438 | 0.921 | 0.91 | 1 | 13 | Cytip | 0.011 | Young |
| 0.285427 | 0.238774 | 0.728 | 0.738 | 1 | 13 | Me2 | -0.01 | Young |
| 0.285561 | 0.131318 | 0.675 | 0.697 | 1 | 13 | Rbm42 | -0.022 | Young |
| 0.28571 | 0.125164 | 0.982 | 0.934 | 1 | 13 | Fgfr1 | 0.048 | Young |
| 0.285719 | 0.137291 | 0.982 | 0.951 | 1 | 13 | Tmem160 | 0.031 | Young |
| 0.286101 | 1.009711 | 0.026 | 0.008 | 1 | 13 | Adamts4 | 0.018 | Young |
| 0.286101 | 0.663838 | 0.026 | 0.008 | 1 | 13 | Gm29394 | 0.018 | Young |
| 0.286101 | 0.662031 | 0.026 | 0.008 | 1 | 13 | Tspan18 | 0.018 | Young |
| 0.286101 | 0.618369 | 0.026 | 0.008 | 1 | 13 | Cep72 | 0.018 | Young |
| 0.286101 | 0.609595 | 0.026 | 0.008 | 1 | 13 | Ank1 | 0.018 | Young |
| 0.286101 | 0.582312 | 0.026 | 0.008 | 1 | 13 | Ispd | 0.018 | Young |
| 0.286101 | 0.558464 | 0.026 | 0.008 | 1 | 13 | Zfp850 | 0.018 | Young |
| 0.286101 | 0.53117 | 0.026 | 0.008 | 1 | 13 | Pdcd1lg2 | 0.018 | Young |
| 0.286101 | 0.486297 | 0.026 | 0.008 | 1 | 13 | Pfkfb1 | 0.018 | Young |
| 0.286101 | 0.484888 | 0.026 | 0.008 | 1 | 13 | Exo1 | 0.018 | Young |
| 0.286565 | 0.238982 | 0.43 | 0.361 | 1 | 13 | C1qtnf1 | 0.069 | Young |

| | | | | | | | | |
|---|---|---|---|---|---|---|---|---|
| 0.286733 | 0.108315 | 0.86 | 0.852 | 1 | 13 | Rala | 0.008 | Young |
| 0.286742 | 0.172373 | 0.798 | 0.77 | 1 | 13 | Lypla2 | 0.028 | Young |
| 0.287061 | 0.334966 | 0.105 | 0.066 | 1 | 13 | Ttll1 | 0.039 | Young |
| 0.287096 | 0.487804 | 0.184 | 0.139 | 1 | 13 | Zfp637 | 0.045 | Young |
| 0.288109 | 0.453957 | 0.228 | 0.18 | 1 | 13 | Trub2 | 0.048 | Young |
| 0.288404 | 0.310697 | 0.254 | 0.197 | 1 | 13 | Hddc2 | 0.057 | Young |
| 0.288537 | 0.450614 | 0.096 | 0.057 | 1 | 13 | Arhgap35 | 0.039 | Young |
| 0.288993 | 0.313112 | 0.579 | 0.566 | 1 | 13 | Gramd1a | 0.013 | Young |
| 0.289659 | 0.110188 | 0.868 | 0.787 | 1 | 13 | Rnf187 | 0.081 | Young |
| 0.289973 | 0.569254 | 0.026 | 0.008 | 1 | 13 | Ppp1r14a | 0.018 | Young |
| 0.289973 | 0.510837 | 0.026 | 0.008 | 1 | 13 | Gm26610 | 0.018 | Young |
| 0.289973 | 0.469684 | 0.026 | 0.008 | 1 | 13 | Best1 | 0.018 | Young |
| 0.289973 | 0.448154 | 0.026 | 0.008 | 1 | 13 | Rnf125 | 0.018 | Young |
| 0.289973 | 0.437862 | 0.026 | 0.008 | 1 | 13 | Efhd1 | 0.018 | Young |
| 0.289973 | 0.417613 | 0.026 | 0.008 | 1 | 13 | Rad9b | 0.018 | Young |
| 0.289973 | 0.410028 | 0.026 | 0.008 | 1 | 13 | 1700102P0 | 0.018 | Young |
| 0.289973 | 0.397165 | 0.026 | 0.008 | 1 | 13 | Gm43136 | 0.018 | Young |
| 0.289973 | 0.396755 | 0.026 | 0.008 | 1 | 13 | Gm34934 | 0.018 | Young |
| 0.289973 | 0.390731 | 0.026 | 0.008 | 1 | 13 | Itgb8 | 0.018 | Young |
| 0.289973 | 0.27832 | 0.026 | 0.008 | 1 | 13 | 2810457G0 | 0.018 | Young |
| 0.289973 | 0.269627 | 0.026 | 0.008 | 1 | 13 | Grtp1 | 0.018 | Young |
| 0.290055 | 0.075199 | 0.991 | 1 | 1 | 13 | Atpif1 | -0.009 | Young |
| 0.290144 | 0.691097 | 0.061 | 0.033 | 1 | 13 | Zfp39 | 0.028 | Young |
| 0.290144 | 0.657113 | 0.061 | 0.033 | 1 | 13 | Slc52a3 | 0.028 | Young |
| 0.290266 | 0.420005 | 0.289 | 0.246 | 1 | 13 | Nup133 | 0.043 | Young |
| 0.290416 | 0.175077 | 0.939 | 0.926 | 1 | 13 | Il10ra | 0.013 | Young |
| 0.290838 | 0.243629 | 0.649 | 0.582 | 1 | 13 | Slmap | 0.067 | Young |
| 0.290952 | 0.143974 | 0.561 | 0.516 | 1 | 13 | Itpr2 | 0.045 | Young |
| 0.290969 | 0.581553 | 0.202 | 0.156 | 1 | 13 | Rfc4 | 0.046 | Young |
| 0.293475 | 0.26881 | 0.579 | 0.541 | 1 | 13 | Zcchc17 | 0.038 | Young |
| 0.293684 | 0.605956 | 0.158 | 0.115 | 1 | 13 | Gm14548 | 0.043 | Young |
| 0.294104 | 0.332528 | 0.465 | 0.402 | 1 | 13 | 1110059G1 | 0.063 | Young |
| 0.294235 | 0.151733 | 0.719 | 0.689 | 1 | 13 | Madd | 0.03 | Young |
| 0.294275 | 0.381003 | 0.105 | 0.066 | 1 | 13 | Zdhhc1 | 0.039 | Young |
| 0.294275 | 0.31134 | 0.105 | 0.066 | 1 | 13 | Gm26853 | 0.039 | Young |
| 0.294935 | 0.699258 | 0.061 | 0.033 | 1 | 13 | Gm4610 | 0.028 | Young |
| 0.295273 | 0.5343 | 0.184 | 0.139 | 1 | 13 | I830077J02 | 0.045 | Young |
| 0.295843 | 0.116752 | 0.649 | 0.607 | 1 | 13 | Rab22a | 0.042 | Young |
| 0.296001 | 0.319125 | 0.404 | 0.361 | 1 | 13 | Gm12840 | 0.043 | Young |
| 0.296679 | 0.143566 | 0.588 | 0.582 | 1 | 13 | Dda1 | 0.006 | Young |
| 0.297024 | 0.172707 | 0.886 | 0.803 | 1 | 13 | Ash1l | 0.083 | Young |
| 0.297028 | 0.446698 | 0.447 | 0.418 | 1 | 13 | Pla2g16 | 0.029 | Young |
| 0.297317 | 0.423196 | 0.123 | 0.082 | 1 | 13 | Ifi213 | 0.041 | Young |
| 0.29735 | 0.609987 | 0.061 | 0.033 | 1 | 13 | Dlgap5 | 0.028 | Young |
| 0.29735 | 0.550427 | 0.061 | 0.033 | 1 | 13 | Samd10 | 0.028 | Young |

| | | | | | | | | |
|---|---|---|---|---|---|---|---|---|
| 0.29735 | 0.539416 | 0.061 | 0.033 | 1 | 13 | Col4a2 | 0.028 | Young |
| 0.297528 | 0.138153 | 0.728 | 0.672 | 1 | 13 | Abca7 | 0.056 | Young |
| 0.297799 | 0.139863 | 0.526 | 0.467 | 1 | 13 | Papss1 | 0.059 | Young |
| 0.29816 | 0.458866 | 0.333 | 0.295 | 1 | 13 | Med7 | 0.038 | Young |
| 0.298824 | 0.110671 | 0.947 | 0.967 | 1 | 13 | Psenen | -0.02 | Young |
| 0.298976 | 0.272652 | 0.614 | 0.557 | 1 | 13 | Apopt1 | 0.057 | Young |
| 0.298978 | 0.154427 | 0.728 | 0.787 | 1 | 13 | Trafd1 | -0.059 | Young |
| 0.299171 | 0.17246 | 0.632 | 0.557 | 1 | 13 | Dync1li2 | 0.075 | Young |
| 0.299698 | 0.3225 | 0.439 | 0.434 | 1 | 13 | Rabgap1 | 0.005 | Young |
| 0.29973 | 0.073477 | 1 | 0.984 | 1 | 13 | Cd81 | 0.016 | Young |
| 0.299778 | 0.779991 | 0.061 | 0.033 | 1 | 13 | Lrg1 | 0.028 | Young |
| 0.299778 | 0.562876 | 0.061 | 0.033 | 1 | 13 | Gnb5 | 0.028 | Young |
| 0.300383 | 0.194264 | 0.465 | 0.418 | 1 | 13 | Ttpal | 0.047 | Young |
| 0.30062 | 0.095746 | 1 | 0.984 | 1 | 13 | Selenok | 0.016 | Young |
| 0.30062 | 0.05752 | 1 | 1 | 1 | 13 | Uqcrh | 0 | Young |
| 0.300696 | 0.160183 | 0.605 | 0.598 | 1 | 13 | Srek1 | 0.007 | Young |
| 0.301597 | 0.195905 | 0.693 | 0.607 | 1 | 13 | Ddx42 | 0.086 | Young |
| 0.301661 | 0.437813 | 0.482 | 0.459 | 1 | 13 | Prpf6 | 0.023 | Young |
| 0.301699 | 0.593649 | 0.219 | 0.18 | 1 | 13 | Alkbh8 | 0.039 | Young |
| 0.302219 | 0.601561 | 0.061 | 0.033 | 1 | 13 | Tctn3 | 0.028 | Young |
| 0.302397 | 0.103514 | 0.982 | 0.951 | 1 | 13 | Tax1bp1 | 0.031 | Young |
| 0.302812 | 0.191279 | 0.588 | 0.516 | 1 | 13 | Pak1 | 0.072 | Young |
| 0.302903 | 0.031635 | 0.719 | 0.59 | 1 | 13 | Palld | 0.129 | Young |
| 0.302981 | 0.543553 | 0.219 | 0.18 | 1 | 13 | Otof | 0.039 | Young |
| 0.303164 | 0.180176 | 0.553 | 0.5 | 1 | 13 | Pgrmc2 | 0.053 | Young |
| 0.303462 | 0.200024 | 0.105 | 0.066 | 1 | 13 | Haus5 | 0.039 | Young |
| 0.303565 | 0.192651 | 0.64 | 0.631 | 1 | 13 | Rictor | 0.009 | Young |
| 0.303581 | 0.311984 | 0.395 | 0.352 | 1 | 13 | Gab1 | 0.043 | Young |
| 0.303622 | 0.3292 | 0.386 | 0.369 | 1 | 13 | Heatr6 | 0.017 | Young |
| 0.303911 | 0.084126 | 0.904 | 0.885 | 1 | 13 | Egr2 | 0.019 | Young |
| 0.304059 | 0.281407 | 0.14 | 0.098 | 1 | 13 | Smim5 | 0.042 | Young |
| 0.304113 | 0.456903 | 0.123 | 0.082 | 1 | 13 | Slc37a1 | 0.041 | Young |
| 0.304323 | 0.291725 | 0.553 | 0.574 | 1 | 13 | Znfx1 | -0.021 | Young |
| 0.304674 | 0.575392 | 0.061 | 0.033 | 1 | 13 | 4930579G2 | 0.028 | Young |
| 0.304674 | 0.547242 | 0.061 | 0.033 | 1 | 13 | Vmac | 0.028 | Young |
| 0.304674 | 0.360092 | 0.061 | 0.033 | 1 | 13 | Gm3336 | 0.028 | Young |
| 0.305119 | 0.260876 | 0.684 | 0.631 | 1 | 13 | Csgalnact2 | 0.053 | Young |
| 0.305382 | 0.301412 | 0.386 | 0.336 | 1 | 13 | Chpf2 | 0.05 | Young |
| 0.305424 | 0.250124 | 0.588 | 0.549 | 1 | 13 | Timm17a | 0.039 | Young |
| 0.305761 | 0.322475 | 0.228 | 0.189 | 1 | 13 | Brms1l | 0.039 | Young |
| 0.305891 | 0.068228 | 0.93 | 0.918 | 1 | 13 | Ppp2r5a | 0.012 | Young |
| 0.306525 | 0.012105 | 0.018 | 0.041 | 1 | 13 | Ildr1 | -0.023 | Young |
| 0.306542 | 0.318996 | 0.351 | 0.303 | 1 | 13 | Aqp9 | 0.048 | Young |
| 0.306894 | 0.070316 | 0.974 | 0.967 | 1 | 13 | Rac2 | 0.007 | Young |
| 0.307127 | 0.090155 | 0.605 | 0.59 | 1 | 13 | Rit1 | 0.015 | Young |

| | | | | | | | | |
|---|---|---|---|---|---|---|---|---|
| 0.307141 | 0.291051 | 0.061 | 0.033 | 1 | 13 | Dennd2d | 0.028 | Young |
| 0.307141 | 0.161463 | 0.061 | 0.033 | 1 | 13 | Cdkn2a | 0.028 | Young |
| 0.307151 | 0.415406 | 0.158 | 0.115 | 1 | 13 | Orc5 | 0.043 | Young |
| 0.307189 | 0.383916 | 0.105 | 0.066 | 1 | 13 | Bcorl1 | 0.039 | Young |
| 0.307234 | 0.233585 | 0.596 | 0.615 | 1 | 13 | Ino80d | -0.019 | Young |
| 0.307244 | 0.325546 | 0.404 | 0.361 | 1 | 13 | Ern1 | 0.043 | Young |
| 0.307271 | 0.198118 | 0.64 | 0.582 | 1 | 13 | Fez2 | 0.058 | Young |
| 0.30755 | 0.813643 | 0.123 | 0.082 | 1 | 13 | Cdkn1c | 0.041 | Young |
| 0.307754 | 0.069349 | 0.947 | 0.934 | 1 | 13 | Per1 | 0.013 | Young |
| 0.308057 | 0.222239 | 0.596 | 0.557 | 1 | 13 | Rab5b | 0.039 | Young |
| 0.308583 | 0.373388 | 0.307 | 0.262 | 1 | 13 | Zdhhc8 | 0.045 | Young |
| 0.308759 | 0.317191 | 0.272 | 0.213 | 1 | 13 | Glce | 0.059 | Young |
| 0.308798 | 0.227849 | 0.412 | 0.352 | 1 | 13 | 1600010M | 0.06 | Young |
| 0.30896 | 0.247204 | 0.667 | 0.631 | 1 | 13 | Rbbp4 | 0.036 | Young |
| 0.309063 | 0.462005 | 0.333 | 0.295 | 1 | 13 | Crot | 0.038 | Young |
| 0.309495 | 0.434576 | 0.298 | 0.262 | 1 | 13 | Iffo1 | 0.036 | Young |
| 0.309615 | 0.465383 | 0.342 | 0.287 | 1 | 13 | Uvssa | 0.055 | Young |
| 0.309622 | 0.407635 | 0.061 | 0.033 | 1 | 13 | Ttll5 | 0.028 | Young |
| 0.310402 | 0.149638 | 0.895 | 0.943 | 1 | 13 | Fli1 | -0.048 | Young |
| 0.310477 | 0.112273 | 0.965 | 0.918 | 1 | 13 | G3bp2 | 0.047 | Young |
| 0.310528 | 0.053138 | 1 | 1 | 1 | 13 | Mbnl1 | 0 | Young |
| 0.31085 | 0.289826 | 0.228 | 0.18 | 1 | 13 | Mfsd8 | 0.048 | Young |
| 0.311076 | 0.849357 | 0.079 | 0.049 | 1 | 13 | Bcs1l | 0.03 | Young |
| 0.311164 | 0.411276 | 0.281 | 0.238 | 1 | 13 | Ccdc28b | 0.043 | Young |
| 0.311399 | 0.100332 | 0.921 | 0.967 | 1 | 13 | Dock10 | -0.046 | Young |
| 0.311409 | 0.279048 | 0.491 | 0.443 | 1 | 13 | Wipi2 | 0.048 | Young |
| 0.311622 | 0.123626 | 0.579 | 0.5 | 1 | 13 | Rprd2 | 0.079 | Young |
| 0.311913 | 0.20199 | 0.623 | 0.648 | 1 | 13 | Ing1 | -0.025 | Young |
| 0.312116 | 0.798034 | 0.061 | 0.033 | 1 | 13 | Gadd45a | 0.028 | Young |
| 0.312116 | 0.524126 | 0.061 | 0.033 | 1 | 13 | F830016B0 | 0.028 | Young |
| 0.31223 | 0.139152 | 0.447 | 0.41 | 1 | 13 | Bcam | 0.037 | Young |
| 0.31235 | 0.195155 | 0.281 | 0.238 | 1 | 13 | Syne1 | 0.043 | Young |
| 0.313413 | 0.315742 | 0.544 | 0.5 | 1 | 13 | Cryl1 | 0.044 | Young |
| 0.313789 | 0.258908 | 0.64 | 0.631 | 1 | 13 | Trim35 | 0.009 | Young |
| 0.314064 | 0.958099 | 0.088 | 0.057 | 1 | 13 | Afg1l | 0.031 | Young |
| 0.31431 | 0.34336 | 0.246 | 0.197 | 1 | 13 | Rgcc | 0.049 | Young |
| 0.314321 | 0.658701 | 0.07 | 0.041 | 1 | 13 | Pter | 0.029 | Young |
| 0.314388 | 0.131142 | 0.904 | 0.893 | 1 | 13 | Fbl | 0.011 | Young |
| 0.314624 | 0.650055 | 0.061 | 0.033 | 1 | 13 | B3galt4 | 0.028 | Young |
| 0.314633 | 0.434773 | 0.149 | 0.107 | 1 | 13 | E130311K1 | 0.042 | Young |
| 0.314893 | 0.252616 | 0.693 | 0.656 | 1 | 13 | Sh2b3 | 0.037 | Young |
| 0.316648 | 0.822318 | 0.07 | 0.041 | 1 | 13 | Rhoj | 0.029 | Young |
| 0.316648 | 0.642062 | 0.07 | 0.041 | 1 | 13 | Gm21988 | 0.029 | Young |
| 0.316943 | 0.070188 | 0.991 | 1 | 1 | 13 | Dock2 | -0.009 | Young |
| 0.317027 | 0.390925 | 0.14 | 0.098 | 1 | 13 | Acss2 | 0.042 | Young |

| | | | | | | | | |
|---|---|---|---|---|---|---|---|---|
| 0.317135 | 0.487995 | 0.377 | 0.352 | 1 | 13 | Tbc1d4 | 0.025 | Young |
| 0.317527 | 0.260046 | 0.456 | 0.41 | 1 | 13 | Pcmtd2 | 0.046 | Young |
| 0.317586 | 0.734928 | 0.079 | 0.049 | 1 | 13 | Rmdn2 | 0.03 | Young |
| 0.317644 | 0.181815 | 0.57 | 0.557 | 1 | 13 | CAAA0111( | 0.013 | Young |
| 0.317867 | 0.072588 | 1 | 1 | 1 | 13 | Sf3b1 | 0 | Young |
| 0.31794 | 0.114581 | 0.596 | 0.59 | 1 | 13 | Keap1 | 0.006 | Young |
| 0.318075 | 1.006551 | 0.167 | 0.131 | 1 | 13 | Akr1c18 | 0.036 | Young |
| 0.318325 | 0.110296 | 0.789 | 0.738 | 1 | 13 | Dguok | 0.051 | Young |
| 0.318361 | 0.322877 | 0.246 | 0.189 | 1 | 13 | Zfp839 | 0.057 | Young |
| 0.318524 | 0.237332 | 0.737 | 0.689 | 1 | 13 | Kmt5b | 0.048 | Young |
| 0.319098 | 0.315426 | 0.333 | 0.295 | 1 | 13 | Rnf138 | 0.038 | Young |
| 0.319253 | 0.644085 | 0.132 | 0.098 | 1 | 13 | 2810454H( | 0.034 | Young |
| 0.319636 | 0.167234 | 0.772 | 0.73 | 1 | 13 | Trim28 | 0.042 | Young |
| 0.319678 | 0.398606 | 0.061 | 0.033 | 1 | 13 | Gm16867 | 0.028 | Young |
| 0.320041 | 0.256411 | 0.491 | 0.402 | 1 | 13 | Uchl3 | 0.089 | Young |
| 0.320104 | 0.295286 | 0.535 | 0.533 | 1 | 13 | Cnot8 | 0.002 | Young |
| 0.320648 | 0.039883 | 0.991 | 1 | 1 | 13 | Ly86 | -0.009 | Young |
| 0.320735 | 0.299797 | 0.289 | 0.23 | 1 | 13 | 0610009B2 | 0.059 | Young |
| 0.320827 | 0.242292 | 0.465 | 0.451 | 1 | 13 | Arfrp1 | 0.014 | Young |
| 0.321052 | 0.504973 | 0.211 | 0.172 | 1 | 13 | Slc25a38 | 0.039 | Young |
| 0.321105 | 0.173783 | 0.763 | 0.721 | 1 | 13 | Acbd5 | 0.042 | Young |
| 0.321334 | 0.195627 | 0.474 | 0.443 | 1 | 13 | Eif1ad | 0.031 | Young |
| 0.321335 | 0.7854 | 0.07 | 0.041 | 1 | 13 | Nars2 | 0.029 | Young |
| 0.321335 | 0.658445 | 0.07 | 0.041 | 1 | 13 | Hibch | 0.029 | Young |
| 0.321335 | 0.655789 | 0.07 | 0.041 | 1 | 13 | Gm11696 | 0.029 | Young |
| 0.321335 | 0.567725 | 0.07 | 0.041 | 1 | 13 | Smim10l2a | 0.029 | Young |
| 0.321343 | 0.005444 | 0.649 | 0.77 | 1 | 13 | Dnajb11 | -0.121 | Young |
| 0.321957 | 0.230395 | 0.36 | 0.295 | 1 | 13 | 2610524H( | 0.065 | Young |
| 0.322164 | 0.50178 | 0.211 | 0.164 | 1 | 13 | Ift172 | 0.047 | Young |
| 0.322189 | 0.329448 | 0.184 | 0.139 | 1 | 13 | Flrt3 | 0.045 | Young |
| 0.322731 | 0.352361 | 0.5 | 0.484 | 1 | 13 | Pcyox1 | 0.016 | Young |
| 0.323538 | 0.31975 | 0.211 | 0.164 | 1 | 13 | Ccl25 | 0.047 | Young |
| 0.323932 | 0.130768 | 0.851 | 0.902 | 1 | 13 | Tfe3 | -0.051 | Young |
| 0.324185 | 0.687878 | 0.079 | 0.049 | 1 | 13 | Ldlrad4 | 0.03 | Young |
| 0.324802 | 0.528128 | 0.289 | 0.254 | 1 | 13 | Wdr70 | 0.035 | Young |
| 0.325223 | 0.446477 | 0.228 | 0.189 | 1 | 13 | Zfp157 | 0.039 | Young |
| 0.325291 | 0.151333 | 0.851 | 0.893 | 1 | 13 | Traf1 | -0.042 | Young |
| 0.325319 | 0.062582 | 1 | 1 | 1 | 13 | Reep5 | 0 | Young |
| 0.325944 | 0.053333 | 0.798 | 0.713 | 1 | 13 | Lair1 | 0.085 | Young |
| 0.326067 | 0.5286 | 0.07 | 0.041 | 1 | 13 | Ubxn11 | 0.029 | Young |
| 0.32647 | 0.264559 | 0.43 | 0.402 | 1 | 13 | Tmx4 | 0.028 | Young |
| 0.327358 | 0.078249 | 0.061 | 0.033 | 1 | 13 | Tshz2 | 0.028 | Young |
| 0.327508 | 0.083876 | 0.772 | 0.705 | 1 | 13 | Spag7 | 0.067 | Young |
| 0.328633 | 0.713997 | 0.079 | 0.049 | 1 | 13 | Zfp930 | 0.03 | Young |
| 0.328633 | 0.652062 | 0.079 | 0.049 | 1 | 13 | Ankrd24 | 0.03 | Young |

| | | | | | | | | |
|---|---|---|---|---|---|---|---|---|
| 0.328633 | 0.297383 | 0.079 | 0.049 | 1 | 13 | Cryab | 0.03 | Young |
| 0.328712 | 0.139016 | 0.632 | 0.607 | 1 | 13 | Tyk2 | 0.025 | Young |
| 0.328796 | 0.446963 | 0.202 | 0.164 | 1 | 13 | Zscan21 | 0.038 | Young |
| 0.328951 | 0.292259 | 0.421 | 0.377 | 1 | 13 | Ngly1 | 0.044 | Young |
| 0.329087 | 0.053118 | 1 | 0.992 | 1 | 13 | Fosb | 0.008 | Young |
| 0.329365 | 0.149583 | 0.184 | 0.131 | 1 | 13 | Ttc38 | 0.053 | Young |
| 0.329694 | 0.506239 | 0.316 | 0.279 | 1 | 13 | Scmh1 | 0.037 | Young |
| 0.329733 | 0.385823 | 0.246 | 0.197 | 1 | 13 | Cops7b | 0.049 | Young |
| 0.330052 | 0.371929 | 0.219 | 0.172 | 1 | 13 | Tbc1d12 | 0.047 | Young |
| 0.330301 | 0.386439 | 0.491 | 0.492 | 1 | 13 | Cep120 | -0.001 | Young |
| 0.330439 | 0.369604 | 0.298 | 0.254 | 1 | 13 | Commd9 | 0.044 | Young |
| 0.330586 | 0.184784 | 0.772 | 0.754 | 1 | 13 | Mrpl34 | 0.018 | Young |
| 0.330872 | 0.664881 | 0.079 | 0.049 | 1 | 13 | Gstt1 | 0.03 | Young |
| 0.330872 | 0.64118 | 0.079 | 0.049 | 1 | 13 | Dffb | 0.03 | Young |
| 0.331388 | 0.227299 | 0.474 | 0.418 | 1 | 13 | Dhdh | 0.056 | Young |
| 0.331808 | 0.306079 | 0.412 | 0.377 | 1 | 13 | Mrpl19 | 0.035 | Young |
| 0.33187 | 0.076165 | 0.956 | 0.918 | 1 | 13 | Lifr | 0.038 | Young |
| 0.332503 | 0.290457 | 0.421 | 0.361 | 1 | 13 | Slc25a25 | 0.06 | Young |
| 0.332793 | 0.041749 | 0.912 | 0.943 | 1 | 13 | Neurl3 | -0.031 | Young |
| 0.332835 | 0.735851 | 0.088 | 0.057 | 1 | 13 | Egfl7 | 0.031 | Young |
| 0.33287 | 0.471313 | 0.175 | 0.131 | 1 | 13 | Acrbp | 0.044 | Young |
| 0.332887 | 0.469055 | 0.412 | 0.377 | 1 | 13 | Sipa1l1 | 0.035 | Young |
| 0.33325 | 0.505665 | 0.07 | 0.041 | 1 | 13 | Icos | 0.029 | Young |
| 0.33325 | 0.480781 | 0.07 | 0.041 | 1 | 13 | Ttc12 | 0.029 | Young |
| 0.334069 | 0.176025 | 0.816 | 0.803 | 1 | 13 | Wbp11 | 0.013 | Young |
| 0.334239 | 0.110593 | 0.904 | 0.959 | 1 | 13 | Tsn | -0.055 | Young |
| 0.335666 | 0.336394 | 0.07 | 0.041 | 1 | 13 | Wdr76 | 0.029 | Young |
| 0.335719 | 0.278832 | 0.439 | 0.402 | 1 | 13 | C2cd5 | 0.037 | Young |
| 0.335888 | 0.23893 | 0.167 | 0.123 | 1 | 13 | Prim2 | 0.044 | Young |
| 0.33609 | 0.346093 | 0.272 | 0.213 | 1 | 13 | Zfp180 | 0.059 | Young |
| 0.336181 | 0.097539 | 0.781 | 0.787 | 1 | 13 | Fes | -0.006 | Young |
| 0.336592 | 0.738582 | 0.193 | 0.156 | 1 | 13 | Supt3 | 0.037 | Young |
| 0.336706 | 0.144801 | 0.965 | 0.992 | 1 | 13 | Jak1 | -0.027 | Young |
| 0.336707 | 0.063194 | 1 | 0.975 | 1 | 13 | Luc7l2 | 0.025 | Young |
| 0.336906 | 0.151276 | 0.158 | 0.115 | 1 | 13 | Tgds | 0.043 | Young |
| 0.337021 | 0.756159 | 0.105 | 0.074 | 1 | 13 | Rab26os | 0.031 | Young |
| 0.337473 | 0.128747 | 0.912 | 0.902 | 1 | 13 | Lmo2 | 0.01 | Young |
| 0.338269 | 0.184669 | 0.895 | 0.877 | 1 | 13 | Pura | 0.018 | Young |
| 0.338453 | 0.498281 | 0.158 | 0.123 | 1 | 13 | Ankrd37 | 0.035 | Young |
| 0.339096 | 0.180484 | 0.825 | 0.811 | 1 | 13 | Siva1 | 0.014 | Young |
| 0.339127 | 0.1919 | 0.675 | 0.656 | 1 | 13 | Fam162a | 0.019 | Young |
| 0.339508 | 0.19053 | 0.553 | 0.533 | 1 | 13 | Yif1b | 0.02 | Young |
| 0.339545 | 0.085059 | 0.614 | 0.607 | 1 | 13 | Ykt6 | 0.007 | Young |
| 0.339801 | 0.4011 | 0.333 | 0.295 | 1 | 13 | Rcan3 | 0.038 | Young |
| 0.339993 | 0.19433 | 0.333 | 0.287 | 1 | 13 | Cdc16 | 0.046 | Young |

| | | | | | | | | |
|---|---|---|---|---|---|---|---|---|
| 0.340152 | 0.520779 | 0.202 | 0.164 | 1 | 13 | Dhx35 | 0.038 | Young |
| 0.340539 | 0.066963 | 0.175 | 0.131 | 1 | 13 | Xlr | 0.044 | Young |
| 0.340907 | 0.688959 | 0.096 | 0.066 | 1 | 13 | Tyw5 | 0.03 | Young |
| 0.340924 | 0.755977 | 0.105 | 0.074 | 1 | 13 | Wdhd1 | 0.031 | Young |
| 0.341729 | 0.602552 | 0.149 | 0.115 | 1 | 13 | Cenpv | 0.034 | Young |
| 0.342038 | 0.229654 | 0.482 | 0.426 | 1 | 13 | Rbl2 | 0.056 | Young |
| 0.342357 | 0.029932 | 0.263 | 0.352 | 1 | 13 | Paqr7 | -0.089 | Young |
| 0.342899 | 0.267099 | 0.43 | 0.418 | 1 | 13 | Smad1 | 0.012 | Young |
| 0.342967 | 0.359789 | 0.228 | 0.18 | 1 | 13 | Mrpl44 | 0.048 | Young |
| 0.343398 | 0.356095 | 0.158 | 0.115 | 1 | 13 | Nsl1 | 0.043 | Young |
| 0.344333 | 0.226734 | 0.456 | 0.385 | 1 | 13 | Arhgap6 | 0.071 | Young |
| 0.3446 | 0.168354 | 0.825 | 0.787 | 1 | 13 | Rheb | 0.038 | Young |
| 0.345047 | 0.168141 | 0.868 | 0.828 | 1 | 13 | Rab6a | 0.04 | Young |
| 0.345134 | 0.313243 | 0.474 | 0.475 | 1 | 13 | Mylip | -0.001 | Young |
| 0.345434 | 0.170307 | 0.658 | 0.648 | 1 | 13 | Zcchc7 | 0.01 | Young |
| 0.345486 | 0.28782 | 0.596 | 0.59 | 1 | 13 | Glo1 | 0.006 | Young |
| 0.34554 | 0.126339 | 0.596 | 0.607 | 1 | 13 | Casc3 | -0.011 | Young |
| 0.346583 | 0.13557 | 0.86 | 0.844 | 1 | 13 | Yy1 | 0.016 | Young |
| 0.346931 | 0.170244 | 0.649 | 0.656 | 1 | 13 | Nfatc2 | -0.007 | Young |
| 0.347093 | 0.804946 | 0.096 | 0.066 | 1 | 13 | Gm16287 | 0.03 | Young |
| 0.347802 | 0.074235 | 0.763 | 0.73 | 1 | 13 | Glyr1 | 0.033 | Young |
| 0.347925 | 0.560631 | 0.088 | 0.057 | 1 | 13 | C030034L1 | 0.031 | Young |
| 0.347995 | 0.262173 | 0.351 | 0.311 | 1 | 13 | Agfg2 | 0.04 | Young |
| 0.348191 | 0.191941 | 0.737 | 0.713 | 1 | 13 | 1810022KC | 0.024 | Young |
| 0.348279 | 0.201593 | 0.474 | 0.41 | 1 | 13 | Prorsd1 | 0.064 | Young |
| 0.34859 | 0.474861 | 0.167 | 0.123 | 1 | 13 | Ets1 | 0.044 | Young |
| 0.348599 | 0.082298 | 0.605 | 0.533 | 1 | 13 | Smim4 | 0.072 | Young |
| 0.34894 | 0.40402 | 0.298 | 0.27 | 1 | 13 | Cdyl | 0.028 | Young |
| 0.349891 | 0.343746 | 0.175 | 0.131 | 1 | 13 | Rasal2 | 0.044 | Young |
| 0.350109 | 0.448142 | 0.307 | 0.279 | 1 | 13 | Carm1 | 0.028 | Young |
| 0.350215 | 0.888505 | 0.035 | 0.016 | 1 | 13 | Mmp2 | 0.019 | Young |
| 0.350215 | 0.807113 | 0.035 | 0.016 | 1 | 13 | Gm15013 | 0.019 | Young |
| 0.350215 | 0.798687 | 0.035 | 0.016 | 1 | 13 | Bloc1s6os | 0.019 | Young |
| 0.350215 | 0.794117 | 0.035 | 0.016 | 1 | 13 | Kynu | 0.019 | Young |
| 0.350215 | 0.745581 | 0.035 | 0.016 | 1 | 13 | Arhgef4 | 0.019 | Young |
| 0.350448 | 0.381966 | 0.254 | 0.221 | 1 | 13 | Eef1akmt2 | 0.033 | Young |
| 0.350505 | 0.455836 | 0.167 | 0.131 | 1 | 13 | Irgq | 0.036 | Young |
| 0.350552 | 0.073264 | 0.667 | 0.574 | 1 | 13 | Sertad1 | 0.093 | Young |
| 0.350978 | 0.063538 | 0.877 | 0.902 | 1 | 13 | Ube2r2 | -0.025 | Young |
| 0.350992 | 0.092495 | 0.798 | 0.811 | 1 | 13 | Cbfb | -0.013 | Young |
| 0.351466 | 0.571246 | 0.079 | 0.049 | 1 | 13 | Slc12a5 | 0.03 | Young |
| 0.351466 | 0.458427 | 0.079 | 0.049 | 1 | 13 | Ptgs2os2 | 0.03 | Young |
| 0.351832 | 0.361708 | 0.211 | 0.164 | 1 | 13 | Gm35154 | 0.047 | Young |
| 0.352134 | 0.216772 | 0.614 | 0.598 | 1 | 13 | Angel2 | 0.016 | Young |
| 0.352283 | 0.084842 | 0.982 | 0.967 | 1 | 13 | Sod1 | 0.015 | Young |

| | | | | | | | | |
|---|---|---|---|---|---|---|---|---|
| 0.352766 | 0.460972 | 0.14 | 0.098 | 1 | 13 | Epor | 0.042 | Young |
| 0.353286 | 0.215424 | 0.158 | 0.115 | 1 | 13 | Hrh1 | 0.043 | Young |
| 0.353836 | 1.312602 | 0.035 | 0.016 | 1 | 13 | Nlrc3 | 0.019 | Young |
| 0.353836 | 1.282082 | 0.035 | 0.016 | 1 | 13 | Adm | 0.019 | Young |
| 0.353836 | 0.868415 | 0.035 | 0.016 | 1 | 13 | Kif15 | 0.019 | Young |
| 0.353836 | 0.682474 | 0.035 | 0.016 | 1 | 13 | Gm7467 | 0.019 | Young |
| 0.353836 | 0.664177 | 0.035 | 0.016 | 1 | 13 | Klhl3 | 0.019 | Young |
| 0.354252 | 0.088081 | 0.86 | 0.82 | 1 | 13 | Was | 0.04 | Young |
| 0.354529 | 0.178636 | 0.14 | 0.098 | 1 | 13 | Dcun1d4 | 0.042 | Young |
| 0.355087 | 0.049608 | 0.263 | 0.213 | 1 | 13 | Cep162 | 0.05 | Young |
| 0.355155 | 0.167886 | 0.868 | 0.861 | 1 | 13 | Ttc14 | 0.007 | Young |
| 0.355241 | 0.259211 | 0.342 | 0.303 | 1 | 13 | Rnf126 | 0.039 | Young |
| 0.355258 | 0.086063 | 1 | 1 | 1 | 13 | Gng5 | 0 | Young |
| 0.35615 | 0.425305 | 0.079 | 0.049 | 1 | 13 | Pecr | 0.03 | Young |
| 0.356716 | 0.547844 | 0.009 | 0.025 | 1 | 13 | Ly6d | -0.016 | Young |
| 0.356716 | 0.026151 | 0.009 | 0.025 | 1 | 13 | Cd79a | -0.016 | Young |
| 0.357192 | 0.142661 | 0.518 | 0.5 | 1 | 13 | Eif2ak2 | 0.018 | Young |
| 0.357481 | 0.855876 | 0.035 | 0.016 | 1 | 13 | Gm17146 | 0.019 | Young |
| 0.357481 | 0.834983 | 0.035 | 0.016 | 1 | 13 | Tnfrsf23 | 0.019 | Young |
| 0.357481 | 0.826215 | 0.035 | 0.016 | 1 | 13 | Acy3 | 0.019 | Young |
| 0.357481 | 0.659457 | 0.035 | 0.016 | 1 | 13 | Cobll1 | 0.019 | Young |
| 0.35754 | 0.445586 | 0.193 | 0.164 | 1 | 13 | Tnfrsf17 | 0.029 | Young |
| 0.357776 | 0.313791 | 0.43 | 0.377 | 1 | 13 | Snrnp40 | 0.053 | Young |
| 0.357999 | 0.585969 | 0.219 | 0.197 | 1 | 13 | Rpp30 | 0.022 | Young |
| 0.358009 | 0.392183 | 0.404 | 0.377 | 1 | 13 | Psip1 | 0.027 | Young |
| 0.358076 | 0.144702 | 0.904 | 0.918 | 1 | 13 | Tmem167 | -0.014 | Young |
| 0.358513 | 0.134452 | 0.781 | 0.77 | 1 | 13 | Api5 | 0.011 | Young |
| 0.358738 | 0.203495 | 0.404 | 0.344 | 1 | 13 | Sp2 | 0.06 | Young |
| 0.358844 | 0.727492 | 0.105 | 0.074 | 1 | 13 | Fads6 | 0.031 | Young |
| 0.358844 | 0.554073 | 0.105 | 0.074 | 1 | 13 | Tmem38a | 0.031 | Young |
| 0.358844 | 0.333484 | 0.105 | 0.074 | 1 | 13 | Limk2 | 0.031 | Young |
| 0.358966 | 0.55024 | 0.088 | 0.057 | 1 | 13 | Bahcc1 | 0.031 | Young |
| 0.359185 | 0.096998 | 0.816 | 0.811 | 1 | 13 | Hipk1 | 0.005 | Young |
| 0.359243 | 0.131251 | 1 | 0.967 | 1 | 13 | Chmp2a | 0.033 | Young |
| 0.359244 | 0.092952 | 0.991 | 0.984 | 1 | 13 | Atp6v1f | 0.007 | Young |
| 0.359579 | 0.179561 | 0.447 | 0.418 | 1 | 13 | Stk19 | 0.029 | Young |
| 0.359972 | 0.185517 | 0.693 | 0.73 | 1 | 13 | Rb1 | -0.037 | Young |
| 0.360143 | 0.550178 | 0.167 | 0.131 | 1 | 13 | 2810001G2 | 0.036 | Young |
| 0.360352 | 0.259352 | 0.456 | 0.443 | 1 | 13 | Arl6ip6 | 0.013 | Young |
| 0.360821 | 0.395426 | 0.36 | 0.344 | 1 | 13 | Mri1 | 0.016 | Young |
| 0.360871 | 0.592156 | 0.105 | 0.074 | 1 | 13 | Pced1b | 0.031 | Young |
| 0.360874 | 0.456305 | 0.079 | 0.049 | 1 | 13 | Slc22a17 | 0.03 | Young |
| 0.360874 | 0.435064 | 0.079 | 0.049 | 1 | 13 | Eno3 | 0.03 | Young |
| 0.360937 | 0.245359 | 0.544 | 0.508 | 1 | 13 | Fmr1 | 0.036 | Young |
| 0.361148 | 1.750747 | 0.035 | 0.016 | 1 | 13 | Cxcl14 | 0.019 | Young |

| | | | | | | | | |
|---|---|---|---|---|---|---|---|---|
| 0.361148 | 0.658767 | 0.035 | 0.016 | 1 | 13 | Bgn | 0.019 | Young |
| 0.361148 | 0.650752 | 0.035 | 0.016 | 1 | 13 | Gm38079 | 0.019 | Young |
| 0.361148 | 0.6463 | 0.035 | 0.016 | 1 | 13 | Gdpd5 | 0.019 | Young |
| 0.361148 | 0.576706 | 0.035 | 0.016 | 1 | 13 | Rad51 | 0.019 | Young |
| 0.36198 | 0.135892 | 0.439 | 0.402 | 1 | 13 | Itpripl1 | 0.037 | Young |
| 0.362157 | 0.205475 | 0.518 | 0.443 | 1 | 13 | Gskip | 0.075 | Young |
| 0.362246 | 0.05503 | 0.974 | 0.967 | 1 | 13 | Ndufb8 | 0.007 | Young |
| 0.362622 | 0.126914 | 0.404 | 0.352 | 1 | 13 | Agpat3 | 0.052 | Young |
| 0.362905 | 0.630919 | 0.105 | 0.074 | 1 | 13 | Zfp952 | 0.031 | Young |
| 0.362915 | 0.315864 | 0.386 | 0.352 | 1 | 13 | Stx3 | 0.034 | Young |
| 0.363866 | 0.232471 | 0.316 | 0.262 | 1 | 13 | Ctdp1 | 0.054 | Young |
| 0.363946 | 0.101871 | 0.912 | 0.869 | 1 | 13 | Hnrnpd | 0.043 | Young |
| 0.364141 | 0.109665 | 0.904 | 0.934 | 1 | 13 | Usp9x | -0.03 | Young |
| 0.364221 | 0.51082 | 0.175 | 0.131 | 1 | 13 | Sigirr | 0.044 | Young |
| 0.36426 | 0.054728 | 0.965 | 0.975 | 1 | 13 | Ndufb7 | -0.01 | Young |
| 0.364266 | 0.055292 | 0.991 | 1 | 1 | 13 | Zfand5 | -0.009 | Young |
| 0.364398 | 0.131737 | 0.658 | 0.582 | 1 | 13 | Tug1 | 0.076 | Young |
| 0.364493 | 0.187591 | 0.868 | 0.836 | 1 | 13 | Arcn1 | 0.032 | Young |
| 0.36473 | 0.198072 | 0.491 | 0.426 | 1 | 13 | Gon4l | 0.065 | Young |
| 0.36484 | 0.81764 | 0.035 | 0.016 | 1 | 13 | Kcnip3 | 0.019 | Young |
| 0.36484 | 0.686928 | 0.035 | 0.016 | 1 | 13 | Zscan22 | 0.019 | Young |
| 0.36484 | 0.590586 | 0.035 | 0.016 | 1 | 13 | Tcerg1l | 0.019 | Young |
| 0.36484 | 0.552236 | 0.035 | 0.016 | 1 | 13 | Rad51c | 0.019 | Young |
| 0.36484 | 0.506121 | 0.035 | 0.016 | 1 | 13 | Dock9 | 0.019 | Young |
| 0.36484 | 0.491046 | 0.035 | 0.016 | 1 | 13 | Cd200 | 0.019 | Young |
| 0.36484 | 0.490292 | 0.035 | 0.016 | 1 | 13 | Zfp296 | 0.019 | Young |
| 0.365141 | 0.22234 | 0.447 | 0.418 | 1 | 13 | Ccpg1 | 0.029 | Young |
| 0.365437 | 0.323668 | 0.605 | 0.631 | 1 | 13 | Rfk | -0.026 | Young |
| 0.365553 | 0.202868 | 0.588 | 0.607 | 1 | 13 | Vcpip1 | -0.019 | Young |
| 0.365901 | 0.072932 | 0.412 | 0.352 | 1 | 13 | Cd209b | 0.06 | Young |
| 0.366287 | 0.038472 | 1 | 1 | 1 | 13 | Arpc2 | 0 | Young |
| 0.366307 | 0.254688 | 0.535 | 0.5 | 1 | 13 | Armc1 | 0.035 | Young |
| 0.366523 | 0.13553 | 0.684 | 0.705 | 1 | 13 | Mgrn1 | -0.021 | Young |
| 0.366692 | 0.170288 | 0.377 | 0.336 | 1 | 13 | Ino80c | 0.041 | Young |
| 0.366709 | 0.202222 | 0.553 | 0.541 | 1 | 13 | Kif3b | 0.012 | Young |
| 0.36709 | 0.158225 | 0.684 | 0.721 | 1 | 13 | Fkbp4 | -0.037 | Young |
| 0.367213 | 0.565378 | 0.14 | 0.107 | 1 | 13 | Mvb12b | 0.033 | Young |
| 0.367284 | 0.173509 | 0.535 | 0.467 | 1 | 13 | Tmem41b | 0.068 | Young |
| 0.367498 | 0.239556 | 0.447 | 0.434 | 1 | 13 | Fam98b | 0.013 | Young |
| 0.367955 | 0.568786 | 0.088 | 0.057 | 1 | 13 | Zbed3 | 0.031 | Young |
| 0.368291 | 0.032615 | 0.184 | 0.254 | 1 | 13 | H2afx | -0.07 | Young |
| 0.368302 | 0.061648 | 0.974 | 0.951 | 1 | 13 | Eif3h | 0.023 | Young |
| 0.368555 | 0.578847 | 0.035 | 0.016 | 1 | 13 | Zkscan7 | 0.019 | Young |
| 0.368555 | 0.547187 | 0.035 | 0.016 | 1 | 13 | Dyrk3 | 0.019 | Young |
| 0.368555 | 0.537227 | 0.035 | 0.016 | 1 | 13 | Mmp25 | 0.019 | Young |

| | | | | | | | | |
|---|---|---|---|---|---|---|---|---|
| 0.368555 | 0.50121 | 0.035 | 0.016 | 1 | 13 | Ttc25 | 0.019 | Young |
| 0.368555 | 0.466542 | 0.035 | 0.016 | 1 | 13 | Gm14403 | 0.019 | Young |
| 0.368555 | 0.440592 | 0.035 | 0.016 | 1 | 13 | Erp27 | 0.019 | Young |
| 0.368555 | 0.409414 | 0.035 | 0.016 | 1 | 13 | Gm40773 | 0.019 | Young |
| 0.368555 | 0.318944 | 0.035 | 0.016 | 1 | 13 | Gramd1c | 0.019 | Young |
| 0.368912 | 0.282869 | 0.509 | 0.492 | 1 | 13 | Plscr3 | 0.017 | Young |
| 0.368979 | 0.171883 | 0.456 | 0.402 | 1 | 13 | Htatip2 | 0.054 | Young |
| 0.368991 | 0.558157 | 0.123 | 0.09 | 1 | 13 | Stk32c | 0.033 | Young |
| 0.369109 | 0.137763 | 0.377 | 0.328 | 1 | 13 | Klrk1 | 0.049 | Young |
| 0.369331 | 0.029532 | 0.991 | 0.984 | 1 | 13 | Ier2 | 0.007 | Young |
| 0.369612 | 0.203738 | 0.605 | 0.631 | 1 | 13 | Tmem248 | -0.026 | Young |
| 0.369619 | 0.39299 | 0.316 | 0.287 | 1 | 13 | Pogz | 0.029 | Young |
| 0.369625 | 0.11966 | 0.754 | 0.705 | 1 | 13 | Cuedc2 | 0.049 | Young |
| 0.370303 | 0.278536 | 0.368 | 0.336 | 1 | 13 | Dph5 | 0.032 | Young |
| 0.370303 | 0.171978 | 0.377 | 0.328 | 1 | 13 | Creb3l2 | 0.049 | Young |
| 0.37035 | 0.071924 | 0.974 | 1 | 1 | 13 | Ppp3ca | -0.026 | Young |
| 0.370522 | 0.365122 | 0.149 | 0.115 | 1 | 13 | Tapt1 | 0.034 | Young |
| 0.370527 | 0.13794 | 0.658 | 0.598 | 1 | 13 | Lsm8 | 0.06 | Young |
| 0.370844 | 0.240215 | 0.579 | 0.541 | 1 | 13 | Fundc1 | 0.038 | Young |
| 0.370897 | 0.479655 | 0.123 | 0.09 | 1 | 13 | B3galt6 | 0.033 | Young |
| 0.371189 | 0.187811 | 0.421 | 0.377 | 1 | 13 | Mzt1 | 0.044 | Young |
| 0.371229 | 0.313861 | 0.711 | 0.713 | 1 | 13 | Acadl | -0.002 | Young |
| 0.371293 | 0.074727 | 0.658 | 0.623 | 1 | 13 | Spata6 | 0.035 | Young |
| 0.371368 | 0.125804 | 0.982 | 0.975 | 1 | 13 | Eef1d | 0.007 | Young |
| 0.37137 | 0.098332 | 1 | 1 | 1 | 13 | Mrpl33 | 0 | Young |
| 0.371881 | 0.097994 | 0.991 | 0.992 | 1 | 13 | Cdk2ap2 | -0.001 | Young |
| 0.371904 | 0.18006 | 0.851 | 0.828 | 1 | 13 | Chic2 | 0.023 | Young |
| 0.372006 | 0.193277 | 0.675 | 0.68 | 1 | 13 | Mllt10 | -0.005 | Young |
| 0.372293 | 0.408948 | 0.035 | 0.016 | 1 | 13 | Elovl7 | 0.019 | Young |
| 0.372293 | 0.356076 | 0.035 | 0.016 | 1 | 13 | Gm33023 | 0.019 | Young |
| 0.372502 | 0.405171 | 0.088 | 0.057 | 1 | 13 | Tnk2 | 0.031 | Young |
| 0.372547 | 0.459524 | 0.096 | 0.066 | 1 | 13 | Pus3 | 0.03 | Young |
| 0.372587 | 0.257399 | 0.316 | 0.27 | 1 | 13 | Eif2b4 | 0.046 | Young |
| 0.372688 | 0.267556 | 0.482 | 0.467 | 1 | 13 | Zbtb1 | 0.015 | Young |
| 0.373458 | 0.134836 | 0.202 | 0.156 | 1 | 13 | Hist1h1e | 0.046 | Young |
| 0.374157 | 0.128315 | 0.886 | 0.902 | 1 | 13 | Anapc11 | -0.016 | Young |
| 0.374443 | 0.463037 | 0.246 | 0.213 | 1 | 13 | Pla2g12a | 0.033 | Young |
| 0.374719 | 0.551785 | 0.096 | 0.066 | 1 | 13 | 1110051M | 0.03 | Young |
| 0.374719 | 0.467171 | 0.096 | 0.066 | 1 | 13 | Wdr90 | 0.03 | Young |
| 0.374729 | 0.713365 | 0.123 | 0.09 | 1 | 13 | Slc2a9 | 0.033 | Young |
| 0.37495 | 0.060336 | 0.974 | 0.975 | 1 | 13 | Lsm4 | -0.001 | Young |
| 0.375372 | 0.453524 | 0.184 | 0.148 | 1 | 13 | Mbip | 0.036 | Young |
| 0.375404 | 0.179918 | 0.263 | 0.213 | 1 | 13 | Mthfr | 0.05 | Young |
| 0.375736 | 0.367458 | 0.193 | 0.156 | 1 | 13 | Tdrkh | 0.037 | Young |
| 0.375876 | 0.149362 | 0.132 | 0.098 | 1 | 13 | Slc22a15 | 0.034 | Young |

| | | | | | | | | |
|---|---|---|---|---|---|---|---|---|
| 0.376055 | 0.297959 | 0.035 | 0.016 | 1 | 13 | Gm2415 | 0.019 | Young |
| 0.376055 | 0.267189 | 0.035 | 0.016 | 1 | 13 | C2 | 0.019 | Young |
| 0.376055 | 0.189066 | 0.035 | 0.016 | 1 | 13 | Gpr21 | 0.019 | Young |
| 0.377004 | 0.284602 | 0.404 | 0.361 | 1 | 13 | Mtr | 0.043 | Young |
| 0.377415 | 0.241588 | 0.026 | 0.049 | 1 | 13 | Slc4a11 | -0.023 | Young |
| 0.377527 | 0.060669 | 0.991 | 1 | 1 | 13 | Atp5j | -0.009 | Young |
| 0.377985 | 0.269717 | 0.456 | 0.393 | 1 | 13 | Slc16a7 | 0.063 | Young |
| 0.37853 | 0.560711 | 0.114 | 0.082 | 1 | 13 | Pex11a | 0.032 | Young |
| 0.378559 | 0.071148 | 1 | 1 | 1 | 13 | Sh3glb1 | 0 | Young |
| 0.378859 | 0.357008 | 0.193 | 0.156 | 1 | 13 | 1600020E0 | 0.037 | Young |
| 0.37886 | 0.57967 | 0.193 | 0.164 | 1 | 13 | Ifitm5 | 0.029 | Young |
| 0.378998 | 0.258144 | 0.386 | 0.344 | 1 | 13 | Dhx8 | 0.042 | Young |
| 0.379269 | 0.619489 | 0.149 | 0.115 | 1 | 13 | Immp2l | 0.034 | Young |
| 0.379646 | 0.156067 | 0.667 | 0.574 | 1 | 13 | Zfp385a | 0.093 | Young |
| 0.37984 | 0.020072 | 0.035 | 0.016 | 1 | 13 | Mbnl3 | 0.019 | Young |
| 0.380424 | 0.255705 | 0.439 | 0.426 | 1 | 13 | Tmem43 | 0.013 | Young |
| 0.380577 | 0.112764 | 0.675 | 0.672 | 1 | 13 | Hps3 | 0.003 | Young |
| 0.380776 | 0.135576 | 0.649 | 0.631 | 1 | 13 | Pcm1 | 0.018 | Young |
| 0.381027 | 0.02354 | 0.333 | 0.41 | 1 | 13 | Plekhg2 | -0.077 | Young |
| 0.381087 | 0.465214 | 0.175 | 0.139 | 1 | 13 | Plscr4 | 0.036 | Young |
| 0.381191 | 0.115252 | 0.798 | 0.877 | 1 | 13 | Ptbp1 | -0.079 | Young |
| 0.381534 | 0.072027 | 0.07 | 0.107 | 1 | 13 | Adal | -0.037 | Young |
| 0.381625 | 0.243654 | 0.395 | 0.352 | 1 | 13 | Insig2 | 0.043 | Young |
| 0.381667 | 0.030205 | 1 | 1 | 1 | 13 | Ppia | 0 | Young |
| 0.381761 | 0.302304 | 0.184 | 0.148 | 1 | 13 | Gm49368 | 0.036 | Young |
| 0.38188 | 0.25842 | 0.421 | 0.385 | 1 | 13 | Dusp7 | 0.036 | Young |
| 0.382869 | 0.470915 | 0.246 | 0.213 | 1 | 13 | 2510002D2 | 0.033 | Young |
| 0.383746 | 0.062744 | 0.982 | 0.984 | 1 | 13 | Plekho1 | -0.002 | Young |
| 0.383899 | 0.457126 | 0.228 | 0.189 | 1 | 13 | Tnfsf13b | 0.039 | Young |
| 0.384395 | 0.375815 | 0.211 | 0.172 | 1 | 13 | Ctu2 | 0.039 | Young |
| 0.384611 | 0.258455 | 0.518 | 0.508 | 1 | 13 | Fam98c | 0.01 | Young |
| 0.384958 | 0.416273 | 0.167 | 0.131 | 1 | 13 | Atg16l2 | 0.036 | Young |
| 0.384968 | 0.205396 | 0.404 | 0.385 | 1 | 13 | B4galt3 | 0.019 | Young |
| 0.385067 | 0.106247 | 0.789 | 0.754 | 1 | 13 | Svil | 0.035 | Young |
| 0.385253 | 0.473982 | 0.132 | 0.098 | 1 | 13 | Cradd | 0.034 | Young |
| 0.385295 | 0.141998 | 0.912 | 0.828 | 1 | 13 | Gpcpd1 | 0.084 | Young |
| 0.385414 | 0.151638 | 0.333 | 0.287 | 1 | 13 | Bbc3 | 0.046 | Young |
| 0.385868 | 0.193235 | 0.561 | 0.516 | 1 | 13 | Hmgcr | 0.045 | Young |
| 0.386235 | 0.288097 | 0.386 | 0.344 | 1 | 13 | Prps1 | 0.042 | Young |
| 0.38635 | 0.465242 | 0.088 | 0.057 | 1 | 13 | Pex26 | 0.031 | Young |
| 0.386765 | 0.1342 | 0.649 | 0.623 | 1 | 13 | Nub1 | 0.026 | Young |
| 0.387145 | 0.621294 | 0.132 | 0.098 | 1 | 13 | Icam2 | 0.034 | Young |
| 0.387209 | 0.456171 | 0.298 | 0.27 | 1 | 13 | Atg4c | 0.028 | Young |
| 0.387353 | 0.118339 | 0.561 | 0.508 | 1 | 13 | Ranbp9 | 0.053 | Young |
| 0.387573 | 0.129744 | 0.886 | 0.885 | 1 | 13 | Samd4b | 0.001 | Young |

| | | | | | | | | |
|---|---|---|---|---|---|---|---|---|
| 0.387631 | 0.215748 | 0.289 | 0.246 | 1 | 13 | Rusc1 | 0.043 | Young |
| 0.387761 | 0.077946 | 0.895 | 0.926 | 1 | 13 | Psmb5 | -0.031 | Young |
| 0.387872 | 0.392917 | 0.105 | 0.074 | 1 | 13 | Cep78 | 0.031 | Young |
| 0.387894 | 0.156724 | 0.737 | 0.787 | 1 | 13 | Mfsd14a | -0.05 | Young |
| 0.387915 | 0.430244 | 0.096 | 0.066 | 1 | 13 | Uevld | 0.03 | Young |
| 0.387954 | 0.10805 | 0.719 | 0.721 | 1 | 13 | Lsm7 | -0.002 | Young |
| 0.388479 | 0.342045 | 0.579 | 0.607 | 1 | 13 | Nudt4 | -0.028 | Young |
| 0.388974 | 0.015469 | 0.982 | 0.967 | 1 | 13 | Kdm7a | 0.015 | Young |
| 0.389068 | 0.211062 | 0.719 | 0.762 | 1 | 13 | Tmem168 | -0.043 | Young |
| 0.389674 | 0.272425 | 0.456 | 0.426 | 1 | 13 | Snrnp48 | 0.03 | Young |
| 0.389972 | 0.089013 | 0.026 | 0.049 | 1 | 13 | Cldn5 | -0.023 | Young |
| 0.390444 | 0.201974 | 0.421 | 0.402 | 1 | 13 | Trappc3 | 0.019 | Young |
| 0.39082 | 0.882869 | 0.044 | 0.025 | 1 | 13 | Nrep | 0.019 | Young |
| 0.390841 | 0.212629 | 0.456 | 0.418 | 1 | 13 | Ggt5 | 0.038 | Young |
| 0.391465 | 0.113787 | 0.956 | 0.877 | 1 | 13 | 9930111J2 | 0.079 | Young |
| 0.391794 | 0.289944 | 0.482 | 0.467 | 1 | 13 | Mfap3 | 0.015 | Young |
| 0.392017 | 0.414686 | 0.158 | 0.123 | 1 | 13 | Ncs1 | 0.035 | Young |
| 0.392088 | 0.216382 | 0.614 | 0.615 | 1 | 13 | Slc15a4 | -0.001 | Young |
| 0.392133 | 0.420975 | 0.105 | 0.074 | 1 | 13 | Amigo1 | 0.031 | Young |
| 0.392274 | 0.56293 | 0.123 | 0.09 | 1 | 13 | Tsen2 | 0.033 | Young |
| 0.393017 | 0.195815 | 0.439 | 0.402 | 1 | 13 | Espl1 | 0.037 | Young |
| 0.393115 | 0.487006 | 0.211 | 0.18 | 1 | 13 | Eid3 | 0.031 | Young |
| 0.393132 | 0.102209 | 0.912 | 0.959 | 1 | 13 | Ankrd11 | -0.047 | Young |
| 0.393693 | 0.263882 | 0.351 | 0.32 | 1 | 13 | Mthfs | 0.031 | Young |
| 0.394085 | 0.111236 | 0.939 | 0.885 | 1 | 13 | Gapvd1 | 0.054 | Young |
| 0.394156 | 0.25327 | 0.491 | 0.516 | 1 | 13 | Prkag1 | -0.025 | Young |
| 0.394212 | 0.059087 | 0.939 | 0.951 | 1 | 13 | Tep1 | -0.012 | Young |
| 0.39422 | 0.126894 | 0.947 | 0.951 | 1 | 13 | Ccdc12 | -0.004 | Young |
| 0.394253 | 0.282282 | 0.123 | 0.09 | 1 | 13 | Fxn | 0.033 | Young |
| 0.395003 | 0.228272 | 0.518 | 0.484 | 1 | 13 | Usp37 | 0.034 | Young |
| 0.395014 | 0.173745 | 0.579 | 0.516 | 1 | 13 | Mospd3 | 0.063 | Young |
| 0.395292 | 0.201592 | 0.439 | 0.393 | 1 | 13 | Chml | 0.046 | Young |
| 0.395378 | 0.218018 | 0.447 | 0.393 | 1 | 13 | Slc12a9 | 0.054 | Young |
| 0.395744 | 0.45107 | 0.263 | 0.246 | 1 | 13 | Kif3a | 0.017 | Young |
| 0.395803 | 0.301745 | 0.518 | 0.508 | 1 | 13 | Phf8 | 0.01 | Young |
| 0.396141 | 0.013616 | 0.263 | 0.344 | 1 | 13 | Rnf168 | -0.081 | Young |
| 0.396235 | 0.223345 | 0.316 | 0.262 | 1 | 13 | Sh3pxd2a | 0.054 | Young |
| 0.39642 | 0.161887 | 0.518 | 0.492 | 1 | 13 | Apobr | 0.026 | Young |
| 0.396618 | 0.120389 | 0.667 | 0.631 | 1 | 13 | Gba | 0.036 | Young |
| 0.39673 | 0.437446 | 0.246 | 0.205 | 1 | 13 | Ift27 | 0.041 | Young |
| 0.397137 | 0.285053 | 0.465 | 0.434 | 1 | 13 | Kifc3 | 0.031 | Young |
| 0.397326 | 0.294786 | 0.693 | 0.738 | 1 | 13 | Ccdc47 | -0.045 | Young |
| 0.397576 | 0.709581 | 0.044 | 0.025 | 1 | 13 | Slc7a4 | 0.019 | Young |
| 0.397576 | 0.652174 | 0.044 | 0.025 | 1 | 13 | Zfp707 | 0.019 | Young |
| 0.397657 | 0.082231 | 0.412 | 0.361 | 1 | 13 | Nf1 | 0.051 | Young |

| | | | | | | | | |
|---|---|---|---|---|---|---|---|---|
| 0.398124 | 0.004087 | 0.36 | 0.426 | 1 | 13 | Hmg20a | -0.066 | Young |
| 0.39834 | 0.184217 | 0.798 | 0.697 | 1 | 13 | Mat2b | 0.101 | Young |
| 0.398543 | 0.366149 | 0.36 | 0.336 | 1 | 13 | Atg9a | 0.024 | Young |
| 0.398825 | 0.158064 | 0.649 | 0.672 | 1 | 13 | Mob3c | -0.023 | Young |
| 0.399316 | 0.21275 | 0.386 | 0.336 | 1 | 13 | Trmt6 | 0.05 | Young |
| 0.399658 | 0.099927 | 0.193 | 0.148 | 1 | 13 | Rft1 | 0.045 | Young |
| 0.399932 | 0.123515 | 0.842 | 0.869 | 1 | 13 | Eif4g3 | -0.027 | Young |
| 0.400328 | 0.130635 | 0.693 | 0.713 | 1 | 13 | Nans | -0.02 | Young |
| 0.400739 | 0.396492 | 0.105 | 0.074 | 1 | 13 | Snapc4 | 0.031 | Young |
| 0.400774 | 0.550381 | 0.149 | 0.115 | 1 | 13 | Zfp809 | 0.034 | Young |
| 0.40098 | 0.758432 | 0.044 | 0.025 | 1 | 13 | Napb | 0.019 | Young |
| 0.40139 | 0.329453 | 0.096 | 0.066 | 1 | 13 | Ccnb2 | 0.03 | Young |
| 0.401489 | 0.182773 | 0.877 | 0.926 | 1 | 13 | Tmod3 | -0.049 | Young |
| 0.402232 | 0.231263 | 0.088 | 0.123 | 1 | 13 | Ube2c | -0.035 | Young |
| 0.402464 | 0.1361 | 0.912 | 0.869 | 1 | 13 | Cd37 | 0.043 | Young |
| 0.402677 | 0.158424 | 0.702 | 0.664 | 1 | 13 | Polr2c | 0.038 | Young |
| 0.402697 | 0.140738 | 0.5 | 0.492 | 1 | 13 | Atad1 | 0.008 | Young |
| 0.402897 | 0.163751 | 0.088 | 0.057 | 1 | 13 | Ctnnd2 | 0.031 | Young |
| 0.403663 | 0.409391 | 0.096 | 0.066 | 1 | 13 | Wdr6 | 0.03 | Young |
| 0.404163 | 0.427544 | 0.193 | 0.164 | 1 | 13 | Msantd2 | 0.029 | Young |
| 0.404286 | 0.15739 | 0.904 | 0.943 | 1 | 13 | Mrpl20 | -0.039 | Young |
| 0.404401 | 0.681658 | 0.044 | 0.025 | 1 | 13 | Tesmin | 0.019 | Young |
| 0.404401 | 0.674431 | 0.044 | 0.025 | 1 | 13 | Dtwd2 | 0.019 | Young |
| 0.404526 | 0.25278 | 0.421 | 0.385 | 1 | 13 | Mis18a | 0.036 | Young |
| 0.405198 | 0.16496 | 0.693 | 0.73 | 1 | 13 | Cyb5r3 | -0.037 | Young |
| 0.405326 | 0.563823 | 0.316 | 0.295 | 1 | 13 | C2cd3 | 0.021 | Young |
| 0.405851 | 0.139068 | 0.518 | 0.467 | 1 | 13 | Htatsf1 | 0.051 | Young |
| 0.406153 | 0.188198 | 0.333 | 0.287 | 1 | 13 | Cdr2 | 0.046 | Young |
| 0.406918 | 0.118829 | 0.728 | 0.738 | 1 | 13 | Rtcb | -0.01 | Young |
| 0.406937 | 0.190554 | 0.509 | 0.459 | 1 | 13 | Il17ra | 0.05 | Young |
| 0.40718 | 0.118465 | 0.439 | 0.385 | 1 | 13 | Rnf103 | 0.054 | Young |
| 0.407275 | 0.388709 | 0.114 | 0.082 | 1 | 13 | Gm37305 | 0.032 | Young |
| 0.40784 | 0.731314 | 0.044 | 0.025 | 1 | 13 | Spc24 | 0.019 | Young |
| 0.40784 | 0.54509 | 0.044 | 0.025 | 1 | 13 | Adarb1 | 0.019 | Young |
| 0.40833 | 0.516765 | 0.272 | 0.246 | 1 | 13 | Lrrc42 | 0.026 | Young |
| 0.408392 | 0.084132 | 0.904 | 0.877 | 1 | 13 | Card19 | 0.027 | Young |
| 0.408392 | 0.277752 | 0.43 | 0.393 | 1 | 13 | Smad5 | 0.037 | Young |
| 0.40868 | 0.460074 | 0.289 | 0.254 | 1 | 13 | Pik3cb | 0.035 | Young |
| 0.408702 | 0.183826 | 0.807 | 0.811 | 1 | 13 | Adap2 | -0.004 | Young |
| 0.409038 | 0.406032 | 0.167 | 0.131 | 1 | 13 | Dgka | 0.036 | Young |
| 0.409384 | 0.312885 | 0.228 | 0.189 | 1 | 13 | Dexi | 0.039 | Young |
| 0.409458 | 0.13987 | 0.105 | 0.074 | 1 | 13 | Rad52 | 0.031 | Young |
| 0.409815 | 0.216683 | 0.149 | 0.107 | 1 | 13 | Klhl12 | 0.042 | Young |
| 0.41005 | 0.196807 | 0.395 | 0.344 | 1 | 13 | Mfsd6 | 0.051 | Young |
| 0.410291 | 0.099886 | 0.974 | 0.918 | 1 | 13 | Ubxn1 | 0.056 | Young |

| | | | | | | | | |
|---|---|---|---|---|---|---|---|---|
| 0.411259 | 0.094642 | 0.851 | 0.861 | 1 | 13 | Arhgap15 | -0.01 | Young |
| 0.411296 | 0.821453 | 0.044 | 0.025 | 1 | 13 | Mmp13 | 0.019 | Young |
| 0.411296 | 0.520568 | 0.044 | 0.025 | 1 | 13 | Magee1 | 0.019 | Young |
| 0.411296 | 0.505668 | 0.044 | 0.025 | 1 | 13 | Zcwpw1 | 0.019 | Young |
| 0.411296 | 0.264057 | 0.044 | 0.025 | 1 | 13 | Gm10791 | 0.019 | Young |
| 0.411416 | 0.046525 | 1 | 0.992 | 1 | 13 | Cox7a2 | 0.008 | Young |
| 0.411592 | 0.394976 | 0.377 | 0.352 | 1 | 13 | Sepsecs | 0.025 | Young |
| 0.411692 | 0.429704 | 0.307 | 0.27 | 1 | 13 | Abcb1b | 0.037 | Young |
| 0.411694 | 0.100437 | 0.675 | 0.68 | 1 | 13 | Ube2q2 | -0.005 | Young |
| 0.412549 | 0.293607 | 0.167 | 0.131 | 1 | 13 | 1110004E0 | 0.036 | Young |
| 0.413859 | 0.125006 | 0.07 | 0.107 | 1 | 13 | Pdss2 | -0.037 | Young |
| 0.414364 | 0.110218 | 0.675 | 0.664 | 1 | 13 | Tmf1 | 0.011 | Young |
| 0.414683 | 0.065827 | 1 | 0.984 | 1 | 13 | Plxnd1 | 0.016 | Young |
| 0.414733 | 0.112578 | 0.132 | 0.18 | 1 | 13 | Rabepk | -0.048 | Young |
| 0.41487 | 0.355466 | 0.211 | 0.18 | 1 | 13 | Oraov1 | 0.031 | Young |
| 0.415031 | 0.071949 | 0.763 | 0.746 | 1 | 13 | Ahr | 0.017 | Young |
| 0.41517 | 0.336393 | 0.158 | 0.123 | 1 | 13 | Zfp346 | 0.035 | Young |
| 0.415603 | 0.075114 | 0.772 | 0.795 | 1 | 13 | Trpc4ap | -0.023 | Young |
| 0.415687 | 0.358753 | 0.219 | 0.18 | 1 | 13 | Amd2 | 0.039 | Young |
| 0.415721 | 0.386331 | 0.114 | 0.082 | 1 | 13 | Leng9 | 0.032 | Young |
| 0.416132 | 0.251587 | 0.746 | 0.762 | 1 | 13 | Vav3 | -0.016 | Young |
| 0.416808 | 0.142744 | 0.702 | 0.631 | 1 | 13 | Magoh | 0.071 | Young |
| 0.416891 | 0.272475 | 0.5 | 0.467 | 1 | 13 | Pds5b | 0.033 | Young |
| 0.417276 | 0.105633 | 0.939 | 0.893 | 1 | 13 | Rps6ka3 | 0.046 | Young |
| 0.417561 | 0.203315 | 0.746 | 0.73 | 1 | 13 | Pilra | 0.016 | Young |
| 0.417849 | 0.558574 | 0.114 | 0.082 | 1 | 13 | Zfp101 | 0.032 | Young |
| 0.417965 | 0.093457 | 1 | 0.992 | 1 | 13 | Dynll1 | 0.008 | Young |
| 0.418261 | 0.466417 | 0.044 | 0.025 | 1 | 13 | Krt7 | 0.019 | Young |
| 0.418358 | 0.155328 | 0.675 | 0.68 | 1 | 13 | Vps41 | -0.005 | Young |
| 0.418752 | 0.153208 | 0.798 | 0.836 | 1 | 13 | Dnajc7 | -0.038 | Young |
| 0.419039 | 0.153777 | 0.974 | 0.934 | 1 | 13 | Map1lc3b | 0.04 | Young |
| 0.419063 | 0.056159 | 1 | 1 | 1 | 13 | Zfp36 | 0 | Young |
| 0.419078 | 0.380762 | 0.14 | 0.107 | 1 | 13 | Zfp276 | 0.033 | Young |
| 0.419243 | 0.470557 | 0.149 | 0.115 | 1 | 13 | Mapre3 | 0.034 | Young |
| 0.419489 | 0.076055 | 0.307 | 0.41 | 1 | 13 | Atl2 | -0.103 | Young |
| 0.419996 | 0.528069 | 0.36 | 0.344 | 1 | 13 | Arhgap4 | 0.016 | Young |
| 0.420504 | 0.093094 | 0.842 | 0.861 | 1 | 13 | Crlf3 | -0.019 | Young |
| 0.420987 | 0.110292 | 0.877 | 0.91 | 1 | 13 | Pten | -0.033 | Young |
| 0.421095 | 0.519169 | 0.272 | 0.246 | 1 | 13 | Adprhl2 | 0.026 | Young |
| 0.421233 | 0.391796 | 0.289 | 0.254 | 1 | 13 | Acpp | 0.035 | Young |
| 0.421277 | 0.129362 | 0.57 | 0.541 | 1 | 13 | Ezh1 | 0.029 | Young |
| 0.421382 | 0.169676 | 0.605 | 0.574 | 1 | 13 | Retreg3 | 0.031 | Young |
| 0.421769 | 0.387134 | 0.044 | 0.025 | 1 | 13 | Stk26 | 0.019 | Young |
| 0.421776 | 0.149858 | 0.447 | 0.426 | 1 | 13 | Cpd | 0.021 | Young |
| 0.421805 | 0.015756 | 0.956 | 0.975 | 1 | 13 | Brd2 | -0.019 | Young |

| | | | | | | | | |
|---|---|---|---|---|---|---|---|---|
| 0.421886 | 0.26426 | 0.219 | 0.189 | 1 | 13 | Srcap | 0.03 | Young |
| 0.422123 | 0.276286 | 0.114 | 0.082 | 1 | 13 | Tesk2 | 0.032 | Young |
| 0.422171 | 0.321695 | 0.342 | 0.311 | 1 | 13 | Rbsn | 0.031 | Young |
| 0.422225 | 0.165036 | 0.781 | 0.787 | 1 | 13 | Mrpl28 | -0.006 | Young |
| 0.422365 | 0.187022 | 0.474 | 0.443 | 1 | 13 | Ppm1m | 0.031 | Young |
| 0.422527 | 0.072611 | 0.281 | 0.238 | 1 | 13 | Gm45716 | 0.043 | Young |
| 0.42253 | 0.235937 | 0.561 | 0.566 | 1 | 13 | Rcsd1 | -0.005 | Young |
| 0.422599 | 0.358373 | 0.123 | 0.09 | 1 | 13 | Mfsd4a | 0.033 | Young |
| 0.422662 | 0.432176 | 0.254 | 0.23 | 1 | 13 | E2f5 | 0.024 | Young |
| 0.42304 | 0.010185 | 0.368 | 0.443 | 1 | 13 | Tmem203 | -0.075 | Young |
| 0.423187 | 0.644948 | 0.053 | 0.033 | 1 | 13 | Plppr3 | 0.02 | Young |
| 0.423411 | 0.00488 | 0.132 | 0.18 | 1 | 13 | Xpnpep3 | -0.048 | Young |
| 0.423608 | 0.263026 | 0.219 | 0.172 | 1 | 13 | Ercc4 | 0.047 | Young |
| 0.423873 | 0.122101 | 0.667 | 0.672 | 1 | 13 | Cops6 | -0.005 | Young |
| 0.424168 | 0.348647 | 0.132 | 0.098 | 1 | 13 | Enpp1 | 0.034 | Young |
| 0.424722 | 0.080102 | 0.667 | 0.639 | 1 | 13 | Bank1 | 0.028 | Young |
| 0.424871 | 0.239103 | 0.482 | 0.475 | 1 | 13 | Abl1 | 0.007 | Young |
| 0.424897 | 0.200933 | 0.333 | 0.303 | 1 | 13 | Zfp512 | 0.03 | Young |
| 0.425016 | 0.101459 | 0.281 | 0.254 | 1 | 13 | Lilra5 | 0.027 | Young |
| 0.425086 | 0.164782 | 0.64 | 0.713 | 1 | 13 | Diaph2 | -0.073 | Young |
| 0.425294 | 0.412309 | 0.044 | 0.025 | 1 | 13 | Abhd3 | 0.019 | Young |
| 0.425961 | 0.03999 | 0.254 | 0.205 | 1 | 13 | H6pd | 0.049 | Young |
| 0.426251 | 0.368864 | 0.272 | 0.238 | 1 | 13 | Zscan29 | 0.034 | Young |
| 0.426359 | 0.722715 | 0.053 | 0.033 | 1 | 13 | Tssk6 | 0.02 | Young |
| 0.426495 | 0.185728 | 0.728 | 0.77 | 1 | 13 | Ctnnd1 | -0.042 | Young |
| 0.426969 | 0.158621 | 0.649 | 0.631 | 1 | 13 | Casp3 | 0.018 | Young |
| 0.427009 | 0.163556 | 0.588 | 0.607 | 1 | 13 | Elp5 | -0.019 | Young |
| 0.427116 | 0.267256 | 0.588 | 0.59 | 1 | 13 | Ssna1 | -0.002 | Young |
| 0.42712 | 0.102314 | 0.728 | 0.672 | 1 | 13 | Tspyl1 | 0.056 | Young |
| 0.427551 | 0.277338 | 0.342 | 0.303 | 1 | 13 | Noc4l | 0.039 | Young |
| 0.427595 | 0.078046 | 0.895 | 0.885 | 1 | 13 | Ank | 0.01 | Young |
| 0.427616 | 0.076892 | 0.816 | 0.82 | 1 | 13 | Crk | -0.004 | Young |
| 0.427772 | 0.280956 | 0.482 | 0.459 | 1 | 13 | Seh1l | 0.023 | Young |
| 0.427903 | 0.05949 | 1 | 0.984 | 1 | 13 | Snrpe | 0.016 | Young |
| 0.428055 | 0.277412 | 0.351 | 0.311 | 1 | 13 | Gkap1 | 0.04 | Young |
| 0.428582 | 0.471939 | 0.114 | 0.082 | 1 | 13 | Adat2 | 0.032 | Young |
| 0.429016 | 0.046283 | 1 | 1 | 1 | 13 | Cox6c | 0 | Young |
| 0.429471 | 0.006547 | 0.956 | 0.893 | 1 | 13 | Gm26532 | 0.063 | Young |
| 0.4299 | 0.223126 | 0.254 | 0.213 | 1 | 13 | Rdh14 | 0.041 | Young |
| 0.430044 | 0.055521 | 0.939 | 0.918 | 1 | 13 | Cdc42se2 | 0.021 | Young |
| 0.430206 | 0.33709 | 0.605 | 0.615 | 1 | 13 | Crem | -0.01 | Young |
| 0.430532 | 0.094331 | 0.746 | 0.713 | 1 | 13 | Map3k11 | 0.033 | Young |
| 0.430785 | 0.140568 | 0.684 | 0.689 | 1 | 13 | Timm8b | -0.005 | Young |
| 0.430912 | 0.328793 | 0.123 | 0.09 | 1 | 13 | Trmt10b | 0.033 | Young |
| 0.430912 | 0.295322 | 0.123 | 0.09 | 1 | 13 | Slx1b | 0.033 | Young |

| | | | | | | | | |
|---|---|---|---|---|---|---|---|---|
| 0.431009 | 0.115971 | 0.772 | 0.787 | 1 | 13 | Slc12a6 | -0.015 | Young |
| 0.431246 | 0.06073 | 1 | 1 | 1 | 13 | Jpt1 | 0 | Young |
| 0.431428 | 0.122006 | 0.553 | 0.525 | 1 | 13 | Bcor | 0.028 | Young |
| 0.431526 | 0.144904 | 0.298 | 0.254 | 1 | 13 | Alg9 | 0.044 | Young |
| 0.431543 | 0.168286 | 0.412 | 0.377 | 1 | 13 | Cpne2 | 0.035 | Young |
| 0.431733 | 0.381967 | 0.351 | 0.328 | 1 | 13 | Mark4 | 0.023 | Young |
| 0.432396 | 0.246341 | 0.044 | 0.025 | 1 | 13 | Ttc30a1 | 0.019 | Young |
| 0.432579 | 0.325083 | 0.289 | 0.254 | 1 | 13 | Xrcc1 | 0.035 | Young |
| 0.433111 | 0.096226 | 0.798 | 0.754 | 1 | 13 | R3hdm2 | 0.044 | Young |
| 0.43389 | 0.129693 | 0.5 | 0.475 | 1 | 13 | Slc25a45 | 0.025 | Young |
| 0.434011 | 0.137373 | 0.114 | 0.156 | 1 | 13 | Nxpe4 | -0.042 | Young |
| 0.434154 | 0.142715 | 0.711 | 0.648 | 1 | 13 | Rybp | 0.063 | Young |
| 0.434244 | 0.245358 | 0.132 | 0.098 | 1 | 13 | Chchd6 | 0.034 | Young |
| 0.434316 | 0.246343 | 0.447 | 0.393 | 1 | 13 | Kdm3b | 0.054 | Young |
| 0.434479 | 0.019985 | 0.851 | 0.861 | 1 | 13 | Phip | -0.01 | Young |
| 0.4346 | 0.145485 | 0.412 | 0.369 | 1 | 13 | Glipr1 | 0.043 | Young |
| 0.434746 | 0.242365 | 0.219 | 0.18 | 1 | 13 | Trim30c | 0.039 | Young |
| 0.435516 | 0.078013 | 0.43 | 0.377 | 1 | 13 | Rnf14 | 0.053 | Young |
| 0.435958 | 0.62587 | 0.053 | 0.033 | 1 | 13 | Rab27a | 0.02 | Young |
| 0.435958 | 0.553215 | 0.053 | 0.033 | 1 | 13 | Asrgl1 | 0.02 | Young |
| 0.435972 | 0.18146 | 0.044 | 0.025 | 1 | 13 | Brip1 | 0.019 | Young |
| 0.43691 | 0.343093 | 0.307 | 0.287 | 1 | 13 | Rabgap1l | 0.02 | Young |
| 0.437225 | 0.065112 | 0.246 | 0.303 | 1 | 13 | Ptp4a3 | -0.057 | Young |
| 0.437231 | 0.97964 | 0.061 | 0.041 | 1 | 13 | Slc19a2 | 0.02 | Young |
| 0.437473 | 0.286839 | 0.474 | 0.467 | 1 | 13 | Ccnk | 0.007 | Young |
| 0.437501 | 0.166962 | 0.5 | 0.516 | 1 | 13 | Dnm1l | -0.016 | Young |
| 0.437513 | 0.151975 | 0.816 | 0.803 | 1 | 13 | Rb1cc1 | 0.013 | Young |
| 0.437984 | 0.152805 | 0.667 | 0.656 | 1 | 13 | Gtpbp2 | 0.011 | Young |
| 0.438044 | 0.095915 | 0.798 | 0.746 | 1 | 13 | Ptpn12 | 0.052 | Young |
| 0.438204 | 0.327464 | 0.149 | 0.115 | 1 | 13 | Cenpq | 0.034 | Young |
| 0.438261 | 0.217973 | 0.447 | 0.434 | 1 | 13 | Akap11 | 0.013 | Young |
| 0.438389 | 0.334944 | 0.351 | 0.328 | 1 | 13 | Casp6 | 0.023 | Young |
| 0.438681 | 0.0585 | 0.886 | 0.869 | 1 | 13 | Coq10b | 0.017 | Young |
| 0.438857 | 0.040677 | 0.833 | 0.861 | 1 | 13 | Brd1 | -0.028 | Young |
| 0.438882 | 0.325487 | 0.439 | 0.443 | 1 | 13 | Adcy9 | -0.004 | Young |
| 0.438951 | 0.322129 | 0.289 | 0.27 | 1 | 13 | Gmpr2 | 0.019 | Young |
| 0.439106 | 0.080128 | 1 | 1 | 1 | 13 | Jund | 0 | Young |
| 0.439106 | 0.066214 | 0.991 | 1 | 1 | 13 | Gdi2 | -0.009 | Young |
| 0.439185 | 0.660178 | 0.053 | 0.033 | 1 | 13 | Stox2 | 0.02 | Young |
| 0.439185 | 0.600282 | 0.053 | 0.033 | 1 | 13 | Cd276 | 0.02 | Young |
| 0.439185 | 0.582281 | 0.053 | 0.033 | 1 | 13 | Tspan9 | 0.02 | Young |
| 0.439345 | 0.209126 | 0.596 | 0.631 | 1 | 13 | Gpbp1l1 | -0.035 | Young |
| 0.439566 | 0.278167 | 0.044 | 0.025 | 1 | 13 | Ptprm | 0.019 | Young |
| 0.439934 | 0.160008 | 0.658 | 0.656 | 1 | 13 | Actr1a | 0.002 | Young |
| 0.440235 | 0.114052 | 0.991 | 0.984 | 1 | 13 | Tmed2 | 0.007 | Young |

| | | | | | | | | |
|---|---|---|---|---|---|---|---|---|
| 0.440533 | 0.922609 | 0.07 | 0.049 | 1 | 13 | Zdhhc24 | 0.021 | Young |
| 0.441749 | 0.213865 | 0.333 | 0.311 | 1 | 13 | Kif21b | 0.022 | Young |
| 0.441749 | 0.404699 | 0.404 | 0.385 | 1 | 13 | Pgm2l1 | 0.019 | Young |
| 0.442218 | 0.492333 | 0.219 | 0.197 | 1 | 13 | Mfng | 0.022 | Young |
| 0.442426 | 0.543874 | 0.053 | 0.033 | 1 | 13 | Gm30411 | 0.02 | Young |
| 0.442426 | 0.530477 | 0.053 | 0.033 | 1 | 13 | Afap1 | 0.02 | Young |
| 0.443035 | 0.344942 | 0.395 | 0.393 | 1 | 13 | Pcnt | 0.002 | Young |
| 0.443394 | 0.103785 | 0.912 | 0.885 | 1 | 13 | Ppp1r15a | 0.027 | Young |
| 0.443459 | 0.099714 | 0.772 | 0.787 | 1 | 13 | Tcn2 | -0.015 | Young |
| 0.443539 | 0.353758 | 0.263 | 0.221 | 1 | 13 | Ppp1r13b | 0.042 | Young |
| 0.443907 | 0.37615 | 0.368 | 0.344 | 1 | 13 | Sh2d1b1 | 0.024 | Young |
| 0.444559 | 0.107273 | 0.886 | 0.918 | 1 | 13 | Cuta | -0.032 | Young |
| 0.444596 | 0.53784 | 0.132 | 0.107 | 1 | 13 | BC051226 | 0.025 | Young |
| 0.444933 | 0.005181 | 0.456 | 0.541 | 1 | 13 | Mocs2 | -0.085 | Young |
| 0.445052 | 0.195921 | 0.263 | 0.221 | 1 | 13 | Iqsec2 | 0.042 | Young |
| 0.445078 | 0.007323 | 0.254 | 0.32 | 1 | 13 | Rad17 | -0.066 | Young |
| 0.445375 | 0.039468 | 0.772 | 0.713 | 1 | 13 | Agpat4 | 0.059 | Young |
| 0.445444 | 0.29832 | 0.105 | 0.074 | 1 | 13 | Gm42031 | 0.031 | Young |
| 0.445681 | 0.517923 | 0.053 | 0.033 | 1 | 13 | Ass1 | 0.02 | Young |
| 0.445715 | 0.115136 | 0.439 | 0.369 | 1 | 13 | Mdn1 | 0.07 | Young |
| 0.446483 | 0.113608 | 0.246 | 0.205 | 1 | 13 | Dmxl2 | 0.041 | Young |
| 0.446999 | 0.307348 | 0.412 | 0.393 | 1 | 13 | 2610037D( | 0.019 | Young |
| 0.447034 | 0.078701 | 0.947 | 0.975 | 1 | 13 | Zbtb7a | -0.028 | Young |
| 0.447046 | 0.095413 | 0.982 | 0.992 | 1 | 13 | Purb | -0.01 | Young |
| 0.447223 | 0.116553 | 0.035 | 0.057 | 1 | 13 | Zfp169 | -0.022 | Young |
| 0.447243 | 0.219099 | 0.614 | 0.631 | 1 | 13 | Anapc16 | -0.017 | Young |
| 0.447581 | 0.355023 | 0.342 | 0.32 | 1 | 13 | Mycbp | 0.022 | Young |
| 0.447725 | 0.14468 | 0.667 | 0.615 | 1 | 13 | Csnk1g3 | 0.052 | Young |
| 0.447762 | 0.210574 | 0.307 | 0.262 | 1 | 13 | Kat6b | 0.045 | Young |
| 0.448112 | 0.191863 | 0.632 | 0.656 | 1 | 13 | Ulk2 | -0.024 | Young |
| 0.448536 | 0.090643 | 0.482 | 0.459 | 1 | 13 | Paxbp1 | 0.023 | Young |
| 0.448849 | 0.296832 | 0.316 | 0.279 | 1 | 13 | Phkb | 0.037 | Young |
| 0.448949 | 0.431056 | 0.053 | 0.033 | 1 | 13 | Ckap2 | 0.02 | Young |
| 0.448949 | 0.407772 | 0.053 | 0.033 | 1 | 13 | Gstp2 | 0.02 | Young |
| 0.449826 | 0.085344 | 0.781 | 0.82 | 1 | 13 | Mrpl24 | -0.039 | Young |
| 0.450298 | 0.120603 | 0.754 | 0.697 | 1 | 13 | Rbm27 | 0.057 | Young |
| 0.450459 | 0.373808 | 0.211 | 0.18 | 1 | 13 | Ramp3 | 0.031 | Young |
| 0.450661 | 0.354333 | 0.219 | 0.189 | 1 | 13 | Entpd5 | 0.03 | Young |
| 0.451135 | 0.521709 | 0.254 | 0.23 | 1 | 13 | Rrp8 | 0.024 | Young |
| 0.451249 | 0.370674 | 0.246 | 0.213 | 1 | 13 | Idua | 0.033 | Young |
| 0.451683 | 0.296939 | 0.342 | 0.32 | 1 | 13 | Cbx7 | 0.022 | Young |
| 0.451891 | 0.258627 | 0.237 | 0.32 | 1 | 13 | Hif1an | -0.083 | Young |
| 0.452161 | 0.521882 | 0.061 | 0.041 | 1 | 13 | Tarsl2 | 0.02 | Young |
| 0.452424 | 0.382045 | 0.316 | 0.287 | 1 | 13 | Rrm2b | 0.029 | Young |
| 0.452619 | 0.262065 | 0.535 | 0.516 | 1 | 13 | Bcl6 | 0.019 | Young |

| | | | | | | | | |
|---|---|---|---|---|---|---|---|---|
| 0.452652 | 0.300747 | 0.219 | 0.18 | 1 | 13 | Dgcr14 | 0.039 | Young |
| 0.452677 | 0.137267 | 0.412 | 0.369 | 1 | 13 | Tmem37 | 0.043 | Young |
| 0.452768 | 0.064206 | 1 | 1 | 1 | 13 | Coro1a | 0 | Young |
| 0.453074 | 0.252177 | 0.518 | 0.492 | 1 | 13 | 5-Mar | 0.026 | Young |
| 0.453085 | 0.548946 | 0.149 | 0.123 | 1 | 13 | Zbtb41 | 0.026 | Young |
| 0.453488 | 0.462057 | 0.219 | 0.197 | 1 | 13 | 1500011BC | 0.022 | Young |
| 0.453701 | 0.133427 | 0.175 | 0.139 | 1 | 13 | Cox6b2 | 0.036 | Young |
| 0.453915 | 0.054186 | 0.991 | 0.967 | 1 | 13 | Cox7b | 0.024 | Young |
| 0.453917 | 0.028853 | 0.982 | 0.984 | 1 | 13 | Uqcrb | -0.002 | Young |
| 0.454507 | 0.034873 | 0.781 | 0.754 | 1 | 13 | Etfa | 0.027 | Young |
| 0.455181 | 0.632085 | 0.061 | 0.041 | 1 | 13 | Fam216a | 0.02 | Young |
| 0.455224 | 0.08944 | 0.825 | 0.754 | 1 | 13 | Mib1 | 0.071 | Young |
| 0.45552 | 0.106751 | 0.596 | 0.533 | 1 | 13 | Net1 | 0.063 | Young |
| 0.455525 | 0.462342 | 0.053 | 0.033 | 1 | 13 | Kif9 | 0.02 | Young |
| 0.455525 | 0.404172 | 0.053 | 0.033 | 1 | 13 | Ugt1a7c | 0.02 | Young |
| 0.455529 | 0.168041 | 0.667 | 0.631 | 1 | 13 | Dazap1 | 0.036 | Young |
| 0.455661 | 0.347388 | 0.43 | 0.426 | 1 | 13 | Pitpnm1 | 0.004 | Young |
| 0.455794 | 0.365533 | 0.289 | 0.254 | 1 | 13 | Clec5a | 0.035 | Young |
| 0.455952 | 0.34341 | 0.219 | 0.18 | 1 | 13 | Siglecg | 0.039 | Young |
| 0.45601 | 0.090917 | 0.816 | 0.82 | 1 | 13 | Chmp3 | -0.004 | Young |
| 0.456689 | 0.645731 | 0.079 | 0.057 | 1 | 13 | Slc10a6 | 0.022 | Young |
| 0.456712 | 0.335587 | 0.202 | 0.172 | 1 | 13 | Kif23 | 0.03 | Young |
| 0.456767 | 0.407628 | 0.167 | 0.139 | 1 | 13 | Trit1 | 0.028 | Young |
| 0.456955 | 0.524921 | 0.149 | 0.123 | 1 | 13 | Sgsm2 | 0.026 | Young |
| 0.457123 | 0.107004 | 0.623 | 0.598 | 1 | 13 | Gm2000 | 0.025 | Young |
| 0.457497 | 0.222701 | 0.509 | 0.459 | 1 | 13 | Cpeb2 | 0.05 | Young |
| 0.457696 | 0.663725 | 0.096 | 0.074 | 1 | 13 | Hyal1 | 0.022 | Young |
| 0.458031 | 0.614015 | 0.114 | 0.09 | 1 | 13 | Ccdc171 | 0.024 | Young |
| 0.458195 | 0.082813 | 0.86 | 0.91 | 1 | 13 | Arhgap9 | -0.05 | Young |
| 0.458505 | 0.300112 | 0.395 | 0.377 | 1 | 13 | Sdccag8 | 0.018 | Young |
| 0.45851 | 0.047403 | 0.561 | 0.541 | 1 | 13 | Epc1 | 0.02 | Young |
| 0.458574 | 0.067699 | 0.123 | 0.09 | 1 | 13 | Gm5431 | 0.033 | Young |
| 0.458639 | 0.342095 | 0.228 | 0.197 | 1 | 13 | St3gal2 | 0.031 | Young |
| 0.458727 | 0.328031 | 0.36 | 0.336 | 1 | 13 | Pkib | 0.024 | Young |
| 0.45876 | 0.117546 | 0.693 | 0.656 | 1 | 13 | Safb | 0.037 | Young |
| 0.458834 | 0.409997 | 0.053 | 0.033 | 1 | 13 | Gm29650 | 0.02 | Young |
| 0.458834 | 0.384093 | 0.053 | 0.033 | 1 | 13 | Zfp41 | 0.02 | Young |
| 0.458834 | 0.339837 | 0.053 | 0.033 | 1 | 13 | Tulp3 | 0.02 | Young |
| 0.459077 | 0.595191 | 0.246 | 0.213 | 1 | 13 | Lncpint | 0.033 | Young |
| 0.459284 | 0.429598 | 0.412 | 0.402 | 1 | 13 | Gclm | 0.01 | Young |
| 0.459292 | 0.166234 | 0.675 | 0.656 | 1 | 13 | Dcaf8 | 0.019 | Young |
| 0.45936 | 0.050746 | 0.386 | 0.475 | 1 | 13 | P3h2 | -0.089 | Young |
| 0.459689 | 0.033906 | 1 | 1 | 1 | 13 | Rpl4 | 0 | Young |
| 0.459724 | 0.058294 | 0.579 | 0.557 | 1 | 13 | Epb41 | 0.022 | Young |
| 0.459811 | 0.175764 | 0.035 | 0.057 | 1 | 13 | Gmpr | -0.022 | Young |

| | | | | | | | | |
|---|---|---|---|---|---|---|---|---|
| 0.459932 | 0.28852 | 0.184 | 0.148 | 1 | 13 | Ppp2r5b | 0.036 | Young |
| 0.461255 | 0.618779 | 0.061 | 0.041 | 1 | 13 | 2900060B1 | 0.02 | Young |
| 0.461579 | 0.23728 | 0.342 | 0.311 | 1 | 13 | Stambp | 0.031 | Young |
| 0.461851 | 0.109299 | 0.895 | 0.926 | 1 | 13 | Tnfrsf1a | -0.031 | Young |
| 0.462156 | 0.359641 | 0.053 | 0.033 | 1 | 13 | Ssc4d | 0.02 | Young |
| 0.462156 | 0.254523 | 0.053 | 0.033 | 1 | 13 | Ccpg1os | 0.02 | Young |
| 0.462447 | 0.531343 | 0.281 | 0.262 | 1 | 13 | Cnn3 | 0.019 | Young |
| 0.462631 | 0.1283 | 0.623 | 0.623 | 1 | 13 | Btbd7 | 0 | Young |
| 0.462941 | 0.09627 | 0.526 | 0.484 | 1 | 13 | Mapkap1 | 0.042 | Young |
| 0.463169 | 0.101467 | 1 | 0.959 | 1 | 13 | Ncf2 | 0.041 | Young |
| 0.46317 | 0.112399 | 0.728 | 0.746 | 1 | 13 | Smarcc1 | -0.018 | Young |
| 0.463795 | 0.069515 | 0.149 | 0.197 | 1 | 13 | Zfp942 | -0.048 | Young |
| 0.464261 | 0.371812 | 0.211 | 0.189 | 1 | 13 | Emc9 | 0.022 | Young |
| 0.464304 | 0.079651 | 0.728 | 0.721 | 1 | 13 | Lactb | 0.007 | Young |
| 0.464309 | 0.74473 | 0.061 | 0.041 | 1 | 13 | Tpmt | 0.02 | Young |
| 0.464309 | 0.520713 | 0.061 | 0.041 | 1 | 13 | Acad10 | 0.02 | Young |
| 0.464335 | 0.072626 | 1 | 0.984 | 1 | 13 | Ccr2 | 0.016 | Young |
| 0.464433 | 0.208418 | 0.605 | 0.607 | 1 | 13 | Hltf | -0.002 | Young |
| 0.464571 | 0.043264 | 0.579 | 0.557 | 1 | 13 | Bin3 | 0.022 | Young |
| 0.464609 | 0.256239 | 0.561 | 0.557 | 1 | 13 | Uxs1 | 0.004 | Young |
| 0.464812 | 0.357705 | 0.351 | 0.311 | 1 | 13 | Fam98a | 0.04 | Young |
| 0.465109 | 0.243908 | 0.728 | 0.73 | 1 | 13 | Map2k7 | -0.002 | Young |
| 0.465491 | 0.224188 | 0.053 | 0.033 | 1 | 13 | Csrnp2 | 0.02 | Young |
| 0.466364 | 0.032931 | 0.298 | 0.254 | 1 | 13 | Camsap2 | 0.044 | Young |
| 0.466534 | 0.347308 | 0.395 | 0.393 | 1 | 13 | Sssca1 | 0.002 | Young |
| 0.466585 | 0.081122 | 0.947 | 0.91 | 1 | 13 | Cd48 | 0.037 | Young |
| 0.466818 | 0.074879 | 0.281 | 0.238 | 1 | 13 | Gm1673 | 0.043 | Young |
| 0.467375 | 0.490588 | 0.061 | 0.041 | 1 | 13 | Gm17259 | 0.02 | Young |
| 0.467375 | 0.447951 | 0.061 | 0.041 | 1 | 13 | 6230400D1 | 0.02 | Young |
| 0.467658 | 0.127482 | 0.807 | 0.828 | 1 | 13 | Uqcc3 | -0.021 | Young |
| 0.468182 | 0.094724 | 0.772 | 0.811 | 1 | 13 | Zfp148 | -0.039 | Young |
| 0.468337 | 0.117188 | 0.851 | 0.795 | 1 | 13 | Axl | 0.056 | Young |
| 0.468338 | 0.323742 | 0.272 | 0.246 | 1 | 13 | Rmi1 | 0.026 | Young |
| 0.468367 | 0.212995 | 0.175 | 0.139 | 1 | 13 | Ints2 | 0.036 | Young |
| 0.468673 | 0.066484 | 0.579 | 0.582 | 1 | 13 | Selenoh | -0.003 | Young |
| 0.468768 | 0.204786 | 0.684 | 0.713 | 1 | 13 | Dcaf7 | -0.029 | Young |
| 0.46884 | 0.267057 | 0.053 | 0.033 | 1 | 13 | Zfp7 | 0.02 | Young |
| 0.46884 | 0.232401 | 0.053 | 0.033 | 1 | 13 | Dock6 | 0.02 | Young |
| 0.469125 | 0.602278 | 0.114 | 0.09 | 1 | 13 | Pcnx4 | 0.024 | Young |
| 0.469733 | 0.158157 | 0.325 | 0.393 | 1 | 13 | Scoc | -0.068 | Young |
| 0.469754 | 0.468741 | 0.351 | 0.328 | 1 | 13 | Lpar6 | 0.023 | Young |
| 0.469831 | 0.307899 | 0.605 | 0.639 | 1 | 13 | Zkscan3 | -0.034 | Young |
| 0.469859 | 0.017389 | 0.43 | 0.549 | 1 | 13 | Tmem131 | -0.119 | Young |
| 0.469926 | 0.366625 | 0.193 | 0.164 | 1 | 13 | Rtl8b | 0.029 | Young |
| 0.470451 | 0.451104 | 0.061 | 0.041 | 1 | 13 | Fuk | 0.02 | Young |

| | | | | | | | | |
|---|---|---|---|---|---|---|---|---|
| 0.471028 | 0.215446 | 0.316 | 0.287 | 1 | 13 | Psmd5 | 0.029 | Young |
| 0.47109 | 0.139552 | 0.904 | 0.885 | 1 | 13 | Limd1 | 0.019 | Young |
| 0.471099 | 0.183011 | 0.465 | 0.443 | 1 | 13 | Nrbf2 | 0.022 | Young |
| 0.471284 | 0.080528 | 0.93 | 0.934 | 1 | 13 | Snx20 | -0.004 | Young |
| 0.471507 | 0.445056 | 0.07 | 0.049 | 1 | 13 | Fam92a | 0.021 | Young |
| 0.47162 | 0.250111 | 0.447 | 0.418 | 1 | 13 | Ndst2 | 0.029 | Young |
| 0.472323 | 0.139395 | 0.912 | 0.893 | 1 | 13 | 1700020I14 | 0.019 | Young |
| 0.472659 | 0.354537 | 0.184 | 0.156 | 1 | 13 | Ano10 | 0.028 | Young |
| 0.472952 | 0.26182 | 0.14 | 0.107 | 1 | 13 | C030034I2 | 0.033 | Young |
| 0.473013 | 0.171914 | 0.693 | 0.656 | 1 | 13 | Wbp1l | 0.037 | Young |
| 0.473328 | 0.205848 | 0.377 | 0.352 | 1 | 13 | Dhrs4 | 0.025 | Young |
| 0.473539 | 0.405212 | 0.061 | 0.041 | 1 | 13 | Cdkn3 | 0.02 | Young |
| 0.473539 | 0.215282 | 0.061 | 0.041 | 1 | 13 | Traf3ip1 | 0.02 | Young |
| 0.473814 | 0.249078 | 0.289 | 0.262 | 1 | 13 | Zc3h10 | 0.027 | Young |
| 0.474041 | 0.374078 | 0.307 | 0.27 | 1 | 13 | Hhex | 0.037 | Young |
| 0.474064 | 0.35468 | 0.263 | 0.23 | 1 | 13 | Slc2a8 | 0.033 | Young |
| 0.474376 | 0.061418 | 0.404 | 0.352 | 1 | 13 | Nifk | 0.052 | Young |
| 0.474441 | 0.387214 | 0.184 | 0.164 | 1 | 13 | 2700097O0 | 0.02 | Young |
| 0.474597 | 0.171531 | 0.588 | 0.59 | 1 | 13 | Zfp950 | -0.002 | Young |
| 0.474861 | 0.131949 | 0.947 | 0.959 | 1 | 13 | Tra2a | -0.012 | Young |
| 0.474877 | 0.172858 | 0.711 | 0.713 | 1 | 13 | Gtf2a2 | -0.002 | Young |
| 0.474967 | 0.371139 | 0.281 | 0.254 | 1 | 13 | Prmt2 | 0.027 | Young |
| 0.475896 | 0.12271 | 0.202 | 0.164 | 1 | 13 | Dgkh | 0.038 | Young |
| 0.4762 | 0.055951 | 0.781 | 0.828 | 1 | 13 | Ergic3 | -0.047 | Young |
| 0.476585 | 0.424029 | 0.149 | 0.123 | 1 | 13 | Praf2 | 0.026 | Young |
| 0.476638 | 0.447028 | 0.061 | 0.041 | 1 | 13 | Crybb1 | 0.02 | Young |
| 0.476977 | 0.265716 | 0.307 | 0.279 | 1 | 13 | Cdc27 | 0.028 | Young |
| 0.477015 | 0.046305 | 0.693 | 0.607 | 1 | 13 | Reps2 | 0.086 | Young |
| 0.477076 | 0.808076 | 0.088 | 0.066 | 1 | 13 | Gm26982 | 0.022 | Young |
| 0.477076 | 0.584141 | 0.088 | 0.066 | 1 | 13 | E130308A1 | 0.022 | Young |
| 0.477178 | 0.351484 | 0.123 | 0.098 | 1 | 13 | 1110019D1 | 0.025 | Young |
| 0.477549 | 0.026302 | 0.018 | 0.033 | 1 | 13 | Gm568 | -0.015 | Young |
| 0.477884 | 0.193381 | 0.465 | 0.451 | 1 | 13 | Naa35 | 0.014 | Young |
| 0.478038 | 0.177648 | 0.605 | 0.59 | 1 | 13 | Orai3 | 0.015 | Young |
| 0.478298 | 0.233666 | 0.228 | 0.197 | 1 | 13 | Mettl7a1 | 0.031 | Young |
| 0.478766 | 0.119473 | 0.86 | 0.828 | 1 | 13 | Mien1 | 0.032 | Young |
| 0.478916 | 0.187749 | 0.623 | 0.648 | 1 | 13 | Cnot3 | -0.025 | Young |
| 0.478966 | 0.216409 | 0.053 | 0.033 | 1 | 13 | Gm26825 | 0.02 | Young |
| 0.478966 | 0.108246 | 0.053 | 0.033 | 1 | 13 | Il17rd | 0.02 | Young |
| 0.479649 | 0.532368 | 0.088 | 0.066 | 1 | 13 | Zfp229 | 0.022 | Young |
| 0.479748 | 0.410419 | 0.061 | 0.041 | 1 | 13 | Gm8369 | 0.02 | Young |
| 0.479765 | 0.141732 | 0.789 | 0.779 | 1 | 13 | Zranb2 | 0.01 | Young |
| 0.480313 | 0.011214 | 0.377 | 0.475 | 1 | 13 | Gopc | -0.098 | Young |
| 0.480662 | 0.275888 | 0.254 | 0.23 | 1 | 13 | Cep57 | 0.024 | Young |
| 0.481841 | 0.210895 | 0.544 | 0.574 | 1 | 13 | Smarcad1 | -0.03 | Young |

| | | | | | | | | |
|---|---|---|---|---|---|---|---|---|
| 0.482107 | 0.093077 | 0.675 | 0.615 | 1 | 13 | Ly96 | 0.06 | Young |
| 0.48237 | 0.262841 | 0.377 | 0.361 | 1 | 13 | Urod | 0.016 | Young |
| 0.482851 | 0.442921 | 0.105 | 0.082 | 1 | 13 | Tamm41 | 0.023 | Young |
| 0.482868 | 0.340277 | 0.061 | 0.041 | 1 | 13 | Grk4 | 0.02 | Young |
| 0.482868 | 0.327637 | 0.061 | 0.041 | 1 | 13 | Spa17 | 0.02 | Young |
| 0.48294 | 0.200598 | 0.377 | 0.352 | 1 | 13 | Fkbp1b | 0.025 | Young |
| 0.482949 | 0.169253 | 0.86 | 0.77 | 1 | 13 | C4b | 0.09 | Young |
| 0.482955 | 0.069885 | 0.737 | 0.746 | 1 | 13 | Tspan14 | -0.009 | Young |
| 0.483561 | 0.141837 | 0.719 | 0.738 | 1 | 13 | Baz2a | -0.019 | Young |
| 0.483742 | 0.096555 | 0.763 | 0.73 | 1 | 13 | Samm50 | 0.033 | Young |
| 0.483891 | 0.18926 | 0.596 | 0.615 | 1 | 13 | Cmas | -0.019 | Young |
| 0.484083 | 0.295145 | 0.237 | 0.205 | 1 | 13 | Cds1 | 0.032 | Young |
| 0.484314 | 0.074812 | 0.939 | 0.934 | 1 | 13 | Psmb8 | 0.005 | Young |
| 0.484566 | 0.042122 | 0.746 | 0.73 | 1 | 13 | Myo1c | 0.016 | Young |
| 0.485563 | 0.020957 | 0.982 | 1 | 1 | 13 | Pip4k2a | -0.018 | Young |
| 0.485694 | 0.111298 | 0.842 | 0.82 | 1 | 13 | Mrpl54 | 0.022 | Young |
| 0.485973 | 0.608361 | 0.07 | 0.049 | 1 | 13 | AC163032. | 0.021 | Young |
| 0.486 | 0.391538 | 0.061 | 0.041 | 1 | 13 | Fitm2 | 0.02 | Young |
| 0.486114 | 0.342458 | 0.132 | 0.107 | 1 | 13 | Hemk1 | 0.025 | Young |
| 0.486161 | 0.146183 | 0.886 | 0.836 | 1 | 13 | Snap23 | 0.05 | Young |
| 0.486336 | 0.467302 | 0.079 | 0.057 | 1 | 13 | Slc25a43 | 0.022 | Young |
| 0.48694 | 0.371449 | 0.158 | 0.131 | 1 | 13 | Ndst1 | 0.027 | Young |
| 0.487237 | 0.532365 | 0.175 | 0.156 | 1 | 13 | Tmed8 | 0.019 | Young |
| 0.487456 | 0.143357 | 0.535 | 0.533 | 1 | 13 | Ccng2 | 0.002 | Young |
| 0.487682 | 0.188575 | 0.623 | 0.639 | 1 | 13 | Mid1ip1 | -0.016 | Young |
| 0.487776 | 0.23126 | 0.272 | 0.23 | 1 | 13 | Tstd2 | 0.042 | Young |
| 0.487836 | 0.150368 | 0.482 | 0.459 | 1 | 13 | Eif2ak1 | 0.023 | Young |
| 0.487877 | 0.130156 | 0.772 | 0.779 | 1 | 13 | Kmt2a | -0.007 | Young |
| 0.487917 | 0.091652 | 0.939 | 0.951 | 1 | 13 | Ube2b | -0.012 | Young |
| 0.488282 | 0.159782 | 0.36 | 0.328 | 1 | 13 | Hs2st1 | 0.032 | Young |
| 0.488945 | 0.183907 | 0.711 | 0.721 | 1 | 13 | Rnf111 | -0.01 | Young |
| 0.488956 | 0.150219 | 0.412 | 0.41 | 1 | 13 | Rhot1 | 0.002 | Young |
| 0.489081 | 0.475293 | 0.079 | 0.057 | 1 | 13 | Lrp4 | 0.022 | Young |
| 0.489148 | 0.183017 | 0.342 | 0.328 | 1 | 13 | Msi2 | 0.014 | Young |
| 0.489894 | 0.012212 | 0.228 | 0.287 | 1 | 13 | Rnf214 | -0.059 | Young |
| 0.490329 | 0.351422 | 0.123 | 0.098 | 1 | 13 | Ap4b1 | 0.025 | Young |
| 0.490378 | 0.502375 | 0.132 | 0.107 | 1 | 13 | Klhl7 | 0.025 | Young |
| 0.490547 | 0.096169 | 0.421 | 0.385 | 1 | 13 | Fut8 | 0.036 | Young |
| 0.49068 | 0.334815 | 0.237 | 0.205 | 1 | 13 | Zfp398 | 0.032 | Young |
| 0.49071 | 0.297404 | 0.412 | 0.377 | 1 | 13 | Amn1 | 0.035 | Young |
| 0.490845 | 0.070585 | 0.149 | 0.189 | 1 | 13 | Srgap3 | -0.04 | Young |
| 0.491826 | 0.401075 | 0.07 | 0.049 | 1 | 13 | Ifi208 | 0.021 | Young |
| 0.491831 | 0.452965 | 0.14 | 0.115 | 1 | 13 | Zfp574 | 0.025 | Young |
| 0.492085 | 0.145007 | 0.939 | 0.926 | 1 | 13 | Alcam | 0.013 | Young |
| 0.492297 | 0.289529 | 0.061 | 0.041 | 1 | 13 | Sall2 | 0.02 | Young |

| | | | | | | | | |
|---|---|---|---|---|---|---|---|---|
| 0.492539 | 0.453173 | 0.123 | 0.098 | 1 | 13 | Arl16 | 0.025 | Young |
| 0.492626 | 0.392005 | 0.088 | 0.066 | 1 | 13 | Tmem191c | 0.022 | Young |
| 0.492658 | 0.054387 | 0.912 | 0.934 | 1 | 13 | Itpripl2 | -0.022 | Young |
| 0.492754 | 0.086728 | 1 | 0.984 | 1 | 13 | Calm3 | 0.016 | Young |
| 0.493094 | 0.299109 | 0.193 | 0.164 | 1 | 13 | Mipep | 0.029 | Young |
| 0.493164 | 0.028681 | 0.096 | 0.131 | 1 | 13 | Slamf6 | -0.035 | Young |
| 0.493439 | 0.395656 | 0.342 | 0.328 | 1 | 13 | Ppp2r5d | 0.014 | Young |
| 0.49353 | 0.274704 | 0.395 | 0.377 | 1 | 13 | Kin | 0.018 | Young |
| 0.493758 | 0.127903 | 0.877 | 0.926 | 1 | 13 | Rp9 | -0.049 | Young |
| 0.493761 | 0.184506 | 0.509 | 0.492 | 1 | 13 | Fmc1 | 0.017 | Young |
| 0.493917 | 0.251247 | 0.491 | 0.484 | 1 | 13 | Ubl4a | 0.007 | Young |
| 0.493996 | 0.287899 | 0.237 | 0.205 | 1 | 13 | Sp4 | 0.032 | Young |
| 0.494582 | 0.288635 | 0.272 | 0.246 | 1 | 13 | Sipa1l2 | 0.026 | Young |
| 0.494598 | 0.426674 | 0.079 | 0.057 | 1 | 13 | Klhl13 | 0.022 | Young |
| 0.494767 | 0.265146 | 0.07 | 0.049 | 1 | 13 | Mrm3 | 0.021 | Young |
| 0.494837 | 0.164071 | 0.772 | 0.762 | 1 | 13 | Ptger4 | 0.01 | Young |
| 0.495142 | 0.241292 | 0.351 | 0.328 | 1 | 13 | Aldh7a1 | 0.023 | Young |
| 0.495163 | 0.080034 | 0.982 | 0.984 | 1 | 13 | Rabac1 | -0.002 | Young |
| 0.495164 | 0.053167 | 1 | 0.992 | 1 | 13 | Snx5 | 0.008 | Young |
| 0.495244 | 0.502622 | 0.088 | 0.066 | 1 | 13 | Trim59 | 0.022 | Young |
| 0.495244 | 0.452853 | 0.088 | 0.066 | 1 | 13 | Pstpip2 | 0.022 | Young |
| 0.495462 | 0.095841 | 0.728 | 0.746 | 1 | 13 | Kdm2b | -0.018 | Young |
| 0.49613 | 0.153222 | 0.193 | 0.156 | 1 | 13 | Fzd5 | 0.037 | Young |
| 0.496226 | 0.153789 | 0.377 | 0.361 | 1 | 13 | Map3k5 | 0.016 | Young |
| 0.496295 | 0.163459 | 0.246 | 0.205 | 1 | 13 | Gtpbp1 | 0.041 | Young |
| 0.496761 | 0.137687 | 0.719 | 0.689 | 1 | 13 | Atraid | 0.03 | Young |
| 0.496808 | 0.203441 | 0.36 | 0.328 | 1 | 13 | Fam199x | 0.032 | Young |
| 0.496853 | 0.257224 | 0.377 | 0.352 | 1 | 13 | Rab10os | 0.025 | Young |
| 0.496974 | 0.052455 | 0.991 | 0.975 | 1 | 13 | Klf2 | 0.016 | Young |
| 0.4972 | 0.00014 | 0.158 | 0.205 | 1 | 13 | Rars2 | -0.047 | Young |
| 0.497369 | 0.438808 | 0.079 | 0.057 | 1 | 13 | 2610035D1 | 0.022 | Young |
| 0.49758 | 0.00078 | 1 | 1 | 1 | 13 | Hspa8 | 0 | Young |
| 0.497623 | 0.037509 | 0.5 | 0.467 | 1 | 13 | Supt20 | 0.033 | Young |
| 0.497717 | 0.372843 | 0.07 | 0.049 | 1 | 13 | Rab39 | 0.021 | Young |
| 0.497749 | 0.038617 | 0.044 | 0.066 | 1 | 13 | Gm11273 | -0.022 | Young |
| 0.497869 | 0.338181 | 0.088 | 0.066 | 1 | 13 | Nlrp10 | 0.022 | Young |
| 0.498183 | 0.034721 | 0.833 | 0.811 | 1 | 13 | Zc3h12a | 0.022 | Young |
| 0.498607 | 0.262929 | 0.36 | 0.344 | 1 | 13 | Gatsl2 | 0.016 | Young |
| 0.498686 | 0.257165 | 0.614 | 0.672 | 1 | 13 | Taf1 | -0.058 | Young |
| 0.498686 | 0.122934 | 0.649 | 0.639 | 1 | 13 | Hexim1 | 0.01 | Young |
| 0.49871 | 0.092018 | 0.904 | 0.951 | 1 | 13 | Klf13 | -0.047 | Young |
| 0.499368 | 0.124047 | 0.939 | 0.959 | 1 | 13 | Ndel1 | -0.02 | Young |
| 0.49992 | 0.122183 | 0.728 | 0.705 | 1 | 13 | Gga1 | 0.023 | Young |
| 0.500002 | 0.016594 | 1 | 1 | 1 | 13 | Btf3 | 0 | Young |
| 0.50006 | 0.174861 | 0.219 | 0.18 | 1 | 13 | Smim3 | 0.039 | Young |

| | | | | | | | | |
|---|---|---|---|---|---|---|---|---|
| 0.500148 | 0.484057 | 0.079 | 0.057 | 1 | 13 | Spata13 | 0.022 | Young |
| 0.500148 | 0.435047 | 0.079 | 0.057 | 1 | 13 | Lamb2 | 0.022 | Young |
| 0.500148 | 0.411929 | 0.079 | 0.057 | 1 | 13 | Map7d3 | 0.022 | Young |
| 0.500191 | 0.78199 | 0.132 | 0.115 | 1 | 13 | Tbc1d16 | 0.017 | Young |
| 0.500665 | 0.224392 | 0.237 | 0.205 | 1 | 13 | Phf13 | 0.032 | Young |
| 0.500676 | 0.364444 | 0.07 | 0.049 | 1 | 13 | Ank2 | 0.021 | Young |
| 0.500748 | 0.273466 | 0.158 | 0.131 | 1 | 13 | 3110001I2 | 0.027 | Young |
| 0.500814 | 0.050668 | 0.044 | 0.066 | 1 | 13 | Treml2 | -0.022 | Young |
| 0.501081 | 0.242784 | 0.342 | 0.32 | 1 | 13 | E130309D0 | 0.022 | Young |
| 0.501171 | 0.074426 | 0.93 | 0.951 | 1 | 13 | Dad1 | -0.021 | Young |
| 0.501517 | 0.075437 | 0.64 | 0.615 | 1 | 13 | Ube2d1 | 0.025 | Young |
| 0.501876 | 0.437028 | 0.096 | 0.074 | 1 | 13 | Lrrk2 | 0.022 | Young |
| 0.501934 | 0.17323 | 0.184 | 0.156 | 1 | 13 | Ttll4 | 0.028 | Young |
| 0.501996 | 0.096749 | 0.833 | 0.828 | 1 | 13 | AB124611 | 0.005 | Young |
| 0.50243 | 0.033885 | 1 | 1 | 1 | 13 | Mcl1 | 0 | Young |
| 0.502436 | 0.139521 | 0.781 | 0.77 | 1 | 13 | Tmem208 | 0.011 | Young |
| 0.502935 | 0.409682 | 0.079 | 0.057 | 1 | 13 | Nfe2 | 0.022 | Young |
| 0.502935 | 0.301431 | 0.079 | 0.057 | 1 | 13 | B3gnt3 | 0.022 | Young |
| 0.502971 | 0.288074 | 0.447 | 0.41 | 1 | 13 | Pex19 | 0.037 | Young |
| 0.50303 | 0.051977 | 0.974 | 0.959 | 1 | 13 | Atrx | 0.015 | Young |
| 0.503076 | 0.339641 | 0.254 | 0.23 | 1 | 13 | Tlr6 | 0.024 | Young |
| 0.50328 | 0.402411 | 0.114 | 0.09 | 1 | 13 | Ada | 0.024 | Young |
| 0.503363 | 0.299456 | 0.281 | 0.246 | 1 | 13 | Znhit6 | 0.035 | Young |
| 0.503582 | 0.314893 | 0.219 | 0.197 | 1 | 13 | Gss | 0.022 | Young |
| 0.503587 | 0.146059 | 0.509 | 0.5 | 1 | 13 | 0610010K1 | 0.009 | Young |
| 0.503645 | 0.209305 | 0.07 | 0.049 | 1 | 13 | E430018J2 | 0.021 | Young |
| 0.503853 | 0.005904 | 0.167 | 0.205 | 1 | 13 | Endod1 | -0.038 | Young |
| 0.504212 | 0.400458 | 0.289 | 0.27 | 1 | 13 | R3hcc1l | 0.019 | Young |
| 0.504256 | 0.275117 | 0.289 | 0.262 | 1 | 13 | Ppp6r2 | 0.027 | Young |
| 0.504726 | 0.230356 | 0.447 | 0.418 | 1 | 13 | Nubp1 | 0.029 | Young |
| 0.504825 | 0.32057 | 0.149 | 0.123 | 1 | 13 | Slc35g1 | 0.026 | Young |
| 0.504966 | 0.081448 | 0.833 | 0.82 | 1 | 13 | Irak1 | 0.013 | Young |
| 0.505613 | 0.152629 | 0.281 | 0.238 | 1 | 13 | Ngrn | 0.043 | Young |
| 0.505828 | 0.151875 | 0.833 | 0.787 | 1 | 13 | Dap | 0.046 | Young |
| 0.506513 | 0.215639 | 0.281 | 0.246 | 1 | 13 | Mxra7 | 0.035 | Young |
| 0.506889 | 0.070839 | 0.693 | 0.631 | 1 | 13 | Cd38 | 0.062 | Young |
| 0.50692 | 0.417395 | 0.096 | 0.074 | 1 | 13 | Rsph3b | 0.022 | Young |
| 0.507298 | 0.050984 | 0.965 | 0.967 | 1 | 13 | Pea15a | -0.002 | Young |
| 0.507669 | 0.089652 | 0.518 | 0.508 | 1 | 13 | Mrpl11 | 0.01 | Young |
| 0.507931 | 0.406638 | 0.114 | 0.09 | 1 | 13 | Ift57 | 0.024 | Young |
| 0.508283 | 0.271566 | 0.754 | 0.779 | 1 | 13 | Lmo4 | -0.025 | Young |
| 0.508402 | 0.045161 | 0.649 | 0.607 | 1 | 13 | Coro7 | 0.042 | Young |
| 0.508444 | 0.368599 | 0.088 | 0.066 | 1 | 13 | Sgce | 0.022 | Young |
| 0.508534 | 0.383967 | 0.079 | 0.057 | 1 | 13 | Ctu1 | 0.022 | Young |
| 0.50893 | 0.338128 | 0.149 | 0.123 | 1 | 13 | Dis3l | 0.026 | Young |

| | | | | | | | | |
|---|---|---|---|---|---|---|---|---|
| 0.50893 | 0.322328 | 0.149 | 0.123 | 1 | 13 | Haus6 | 0.026 | Young |
| 0.509254 | 0.424213 | 0.254 | 0.221 | 1 | 13 | Sugp1 | 0.033 | Young |
| 0.509939 | 0.198351 | 0.298 | 0.287 | 1 | 13 | Jpt2 | 0.011 | Young |
| 0.5109 | 0.10174 | 0.939 | 0.918 | 1 | 13 | Irf5 | 0.021 | Young |
| 0.510982 | 0.084936 | 0.316 | 0.287 | 1 | 13 | Slc9a8 | 0.029 | Young |
| 0.511106 | 0.376214 | 0.088 | 0.066 | 1 | 13 | Neil1 | 0.022 | Young |
| 0.511106 | 0.229335 | 0.088 | 0.066 | 1 | 13 | Nmrk1 | 0.022 | Young |
| 0.511135 | 0.27281 | 0.333 | 0.303 | 1 | 13 | Smpdl3b | 0.03 | Young |
| 0.511345 | 0.35905 | 0.079 | 0.057 | 1 | 13 | Tedc2 | 0.022 | Young |
| 0.51171 | 0.001972 | 0.325 | 0.41 | 1 | 13 | Pold3 | -0.085 | Young |
| 0.511788 | 0.183377 | 0.702 | 0.672 | 1 | 13 | Crebrf | 0.03 | Young |
| 0.511861 | 0.271271 | 0.421 | 0.41 | 1 | 13 | Rhbdf2 | 0.011 | Young |
| 0.511991 | 0.463007 | 0.096 | 0.074 | 1 | 13 | BC049352 | 0.022 | Young |
| 0.512543 | 0.078386 | 0.456 | 0.434 | 1 | 13 | Rassf8 | 0.022 | Young |
| 0.512607 | 0.290589 | 0.07 | 0.049 | 1 | 13 | Fignl2 | 0.021 | Young |
| 0.512835 | 0.438393 | 0.14 | 0.115 | 1 | 13 | Trmt10c | 0.025 | Young |
| 0.512945 | 0.207945 | 0.333 | 0.311 | 1 | 13 | Fbxw5 | 0.022 | Young |
| 0.512993 | 0.27705 | 0.307 | 0.279 | 1 | 13 | Zfp24 | 0.028 | Young |
| 0.513373 | 0.261039 | 0.325 | 0.295 | 1 | 13 | Triobp | 0.03 | Young |
| 0.513438 | 0.076371 | 0.202 | 0.254 | 1 | 13 | Trim37 | -0.052 | Young |
| 0.51366 | 0.30651 | 0.281 | 0.254 | 1 | 13 | Sfmbt1 | 0.027 | Young |
| 0.513725 | 0.303136 | 0.342 | 0.328 | 1 | 13 | Rem1 | 0.014 | Young |
| 0.513908 | 0.158702 | 0.728 | 0.746 | 1 | 13 | Ncoa4 | -0.018 | Young |
| 0.514035 | 0.135386 | 0.518 | 0.492 | 1 | 13 | Nhlrc3 | 0.026 | Young |
| 0.514524 | 0.411581 | 0.298 | 0.287 | 1 | 13 | Bcl2l12 | 0.011 | Young |
| 0.514675 | 0.184907 | 0.061 | 0.041 | 1 | 13 | 5430403G1 | 0.02 | Young |
| 0.514754 | 0.050978 | 0.754 | 0.68 | 1 | 13 | Pts | 0.074 | Young |
| 0.51496 | 0.203804 | 0.14 | 0.115 | 1 | 13 | Wdr53 | 0.025 | Young |
| 0.515836 | 0.213017 | 0.64 | 0.639 | 1 | 13 | 2810403A0 | 0.001 | Young |
| 0.515839 | 0.298363 | 0.211 | 0.189 | 1 | 13 | Tbp | 0.022 | Young |
| 0.515842 | 0.219945 | 0.254 | 0.221 | 1 | 13 | Zfp369 | 0.033 | Young |
| 0.515921 | 0.07382 | 0.395 | 0.361 | 1 | 13 | Rif1 | 0.034 | Young |
| 0.516192 | 0.226253 | 0.333 | 0.32 | 1 | 13 | Fech | 0.013 | Young |
| 0.516254 | 0.112878 | 0.842 | 0.836 | 1 | 13 | Stard3nl | 0.006 | Young |
| 0.51646 | 0.130722 | 0.518 | 0.475 | 1 | 13 | Espn | 0.043 | Young |
| 0.516491 | 0.370882 | 0.105 | 0.082 | 1 | 13 | Cep131 | 0.023 | Young |
| 0.516742 | 0.049693 | 0.877 | 0.877 | 1 | 13 | Ep300 | 0 | Young |
| 0.517301 | 0.546561 | 0.175 | 0.156 | 1 | 13 | Gab3 | 0.019 | Young |
| 0.517534 | 0.422376 | 0.202 | 0.18 | 1 | 13 | Zc3h6 | 0.022 | Young |
| 0.518127 | 0.269181 | 0.351 | 0.32 | 1 | 13 | Wdr18 | 0.031 | Young |
| 0.518235 | 0.1449 | 0.518 | 0.533 | 1 | 13 | Timm17b | -0.015 | Young |
| 0.51894 | 0.335285 | 0.105 | 0.082 | 1 | 13 | Tefm | 0.023 | Young |
| 0.519135 | 0.832164 | 0.088 | 0.066 | 1 | 13 | Clec4g | 0.022 | Young |
| 0.51921 | 0.163285 | 0.5 | 0.5 | 1 | 13 | Farsb | 0 | Young |
| 0.519539 | 1.223657 | 0.018 | 0.008 | 1 | 13 | Gm29554 | 0.01 | Young |

| | | | | | | | | |
|---|---|---|---|---|---|---|---|---|
| 0.519539 | 0.768397 | 0.018 | 0.008 | 1 | 13 | 4921516A( | 0.01 | Young |
| 0.519539 | 0.696965 | 0.018 | 0.008 | 1 | 13 | Rbp7 | 0.01 | Young |
| 0.519539 | 0.688207 | 0.018 | 0.008 | 1 | 13 | Glis3 | 0.01 | Young |
| 0.519539 | 0.668294 | 0.018 | 0.008 | 1 | 13 | Gm35315 | 0.01 | Young |
| 0.519539 | 0.661231 | 0.018 | 0.008 | 1 | 13 | Gm20324 | 0.01 | Young |
| 0.519539 | 0.656547 | 0.018 | 0.008 | 1 | 13 | Slit1 | 0.01 | Young |
| 0.519539 | 0.643907 | 0.018 | 0.008 | 1 | 13 | 2810021J2 | 0.01 | Young |
| 0.519539 | 0.636381 | 0.018 | 0.008 | 1 | 13 | Zfp661 | 0.01 | Young |
| 0.519539 | 0.632906 | 0.018 | 0.008 | 1 | 13 | AC122413. | 0.01 | Young |
| 0.519539 | 0.619471 | 0.018 | 0.008 | 1 | 13 | Lsmem1 | 0.01 | Young |
| 0.519539 | 0.613031 | 0.018 | 0.008 | 1 | 13 | Myo6 | 0.01 | Young |
| 0.519539 | 0.610896 | 0.018 | 0.008 | 1 | 13 | Gdf10 | 0.01 | Young |
| 0.519539 | 0.607505 | 0.018 | 0.008 | 1 | 13 | Il23a | 0.01 | Young |
| 0.519539 | 0.605765 | 0.018 | 0.008 | 1 | 13 | Gm34342 | 0.01 | Young |
| 0.519539 | 0.604062 | 0.018 | 0.008 | 1 | 13 | Sez6l2 | 0.01 | Young |
| 0.519539 | 0.60234 | 0.018 | 0.008 | 1 | 13 | Thbs3 | 0.01 | Young |
| 0.519539 | 0.599978 | 0.018 | 0.008 | 1 | 13 | Gm26909 | 0.01 | Young |
| 0.519539 | 0.597827 | 0.018 | 0.008 | 1 | 13 | Gimap5 | 0.01 | Young |
| 0.519539 | 0.595326 | 0.018 | 0.008 | 1 | 13 | Zfp2 | 0.01 | Young |
| 0.519539 | 0.593506 | 0.018 | 0.008 | 1 | 13 | Ric3 | 0.01 | Young |
| 0.519539 | 0.590871 | 0.018 | 0.008 | 1 | 13 | 4930589L2 | 0.01 | Young |
| 0.519539 | 0.589324 | 0.018 | 0.008 | 1 | 13 | Gm47469 | 0.01 | Young |
| 0.519539 | 0.583273 | 0.018 | 0.008 | 1 | 13 | Kctd14 | 0.01 | Young |
| 0.519539 | 0.581376 | 0.018 | 0.008 | 1 | 13 | Zfp651 | 0.01 | Young |
| 0.519539 | 0.564239 | 0.018 | 0.008 | 1 | 13 | Acap1 | 0.01 | Young |
| 0.519539 | 0.562332 | 0.018 | 0.008 | 1 | 13 | Kiss1r | 0.01 | Young |
| 0.519539 | 0.559837 | 0.018 | 0.008 | 1 | 13 | Unc5b | 0.01 | Young |
| 0.519539 | 0.55886 | 0.018 | 0.008 | 1 | 13 | Car7 | 0.01 | Young |
| 0.519539 | 0.556616 | 0.018 | 0.008 | 1 | 13 | AC127341. | 0.01 | Young |
| 0.519539 | 0.547905 | 0.018 | 0.008 | 1 | 13 | Kifc2 | 0.01 | Young |
| 0.519539 | 0.544599 | 0.018 | 0.008 | 1 | 13 | Gm10451 | 0.01 | Young |
| 0.519539 | 0.531971 | 0.018 | 0.008 | 1 | 13 | 2810408A1 | 0.01 | Young |
| 0.519539 | 0.522581 | 0.018 | 0.008 | 1 | 13 | Gm10282 | 0.01 | Young |
| 0.519539 | 0.519134 | 0.018 | 0.008 | 1 | 13 | Gm26541 | 0.01 | Young |
| 0.519539 | 0.517175 | 0.018 | 0.008 | 1 | 13 | Gm26664 | 0.01 | Young |
| 0.519539 | 0.516127 | 0.018 | 0.008 | 1 | 13 | 3110070M | 0.01 | Young |
| 0.519539 | 0.51316 | 0.018 | 0.008 | 1 | 13 | Prss53 | 0.01 | Young |
| 0.519539 | 0.508713 | 0.018 | 0.008 | 1 | 13 | 4930539E( | 0.01 | Young |
| 0.519539 | 0.507491 | 0.018 | 0.008 | 1 | 13 | Yjefn3 | 0.01 | Young |
| 0.519539 | 0.504494 | 0.018 | 0.008 | 1 | 13 | Zfp966 | 0.01 | Young |
| 0.519539 | 0.498662 | 0.018 | 0.008 | 1 | 13 | C130074G1 | 0.01 | Young |
| 0.519539 | 0.494919 | 0.018 | 0.008 | 1 | 13 | Pmel | 0.01 | Young |
| 0.519539 | 0.493045 | 0.018 | 0.008 | 1 | 13 | Pcna-ps2 | 0.01 | Young |
| 0.519539 | 0.483604 | 0.018 | 0.008 | 1 | 13 | Snhg10 | 0.01 | Young |
| 0.519539 | 0.477849 | 0.018 | 0.008 | 1 | 13 | Zfp759 | 0.01 | Young |

| | | | | | | | | |
|---|---|---|---|---|---|---|---|---|
| 0.519539 | 0.474542 | 0.018 | 0.008 | 1 | 13 | Syt17 | 0.01 | Young |
| 0.519539 | 0.472988 | 0.018 | 0.008 | 1 | 13 | Spats2l | 0.01 | Young |
| 0.519539 | 0.465642 | 0.018 | 0.008 | 1 | 13 | Gm44067 | 0.01 | Young |
| 0.519539 | 0.44563 | 0.018 | 0.008 | 1 | 13 | Fam178b | 0.01 | Young |
| 0.519539 | 0.435202 | 0.018 | 0.008 | 1 | 13 | 4930599N2 | 0.01 | Young |
| 0.519539 | 0.428061 | 0.018 | 0.008 | 1 | 13 | Etohd2 | 0.01 | Young |
| 0.519539 | 0.415401 | 0.018 | 0.008 | 1 | 13 | Cpt1c | 0.01 | Young |
| 0.519539 | 0.413707 | 0.018 | 0.008 | 1 | 13 | Zfp382 | 0.01 | Young |
| 0.519539 | 0.402129 | 0.018 | 0.008 | 1 | 13 | Zbed5 | 0.01 | Young |
| 0.519539 | 0.400746 | 0.018 | 0.008 | 1 | 13 | Ggact | 0.01 | Young |
| 0.519539 | 0.385026 | 0.018 | 0.008 | 1 | 13 | 4930537H2 | 0.01 | Young |
| 0.519539 | 0.384113 | 0.018 | 0.008 | 1 | 13 | Gm7879 | 0.01 | Young |
| 0.519539 | 0.35529 | 0.018 | 0.008 | 1 | 13 | Cacnb4 | 0.01 | Young |
| 0.519539 | 0.355171 | 0.018 | 0.008 | 1 | 13 | Gm5914 | 0.01 | Young |
| 0.519539 | 0.353501 | 0.018 | 0.008 | 1 | 13 | Cyb561 | 0.01 | Young |
| 0.519539 | 0.351043 | 0.018 | 0.008 | 1 | 13 | Hddc3 | 0.01 | Young |
| 0.519581 | 0.040464 | 0.956 | 0.951 | 1 | 13 | Taok1 | 0.005 | Young |
| 0.519647 | 0.344562 | 0.096 | 0.074 | 1 | 13 | Magi2 | 0.022 | Young |
| 0.519657 | 0.352799 | 0.114 | 0.09 | 1 | 13 | Pank4 | 0.024 | Young |
| 0.520673 | 0.117679 | 0.456 | 0.434 | 1 | 13 | Gatd1 | 0.022 | Young |
| 0.520986 | 0.190851 | 0.377 | 0.352 | 1 | 13 | Ubl7 | 0.025 | Young |
| 0.52101 | 0.210694 | 0.579 | 0.557 | 1 | 13 | Fyttd1 | 0.022 | Young |
| 0.521073 | 0.236019 | 0.342 | 0.328 | 1 | 13 | Sik2 | 0.014 | Young |
| 0.521392 | 0.223083 | 0.456 | 0.418 | 1 | 13 | Zfp592 | 0.038 | Young |
| 0.521661 | 0.192767 | 0.123 | 0.156 | 1 | 13 | Cmpk2 | -0.033 | Young |
| 0.521756 | 0.340723 | 0.123 | 0.098 | 1 | 13 | Zfp949 | 0.025 | Young |
| 0.521826 | 0.23214 | 0.088 | 0.066 | 1 | 13 | Prdm11 | 0.022 | Young |
| 0.521888 | 0.024547 | 0.956 | 0.852 | 1 | 13 | Vdac1 | 0.104 | Young |
| 0.522229 | 0.120592 | 0.816 | 0.836 | 1 | 13 | Rnf150 | -0.02 | Young |
| 0.522265 | 0.118723 | 0.596 | 0.582 | 1 | 13 | Dennd4c | 0.014 | Young |
| 0.522656 | 0.299756 | 0.219 | 0.197 | 1 | 13 | Zfp367 | 0.022 | Young |
| 0.522673 | 0.457257 | 0.079 | 0.057 | 1 | 13 | Gstm4 | 0.022 | Young |
| 0.522934 | 0.008212 | 0.175 | 0.221 | 1 | 13 | Ahi1 | -0.046 | Young |
| 0.523244 | 0.120866 | 0.947 | 0.91 | 1 | 13 | Arrb1 | 0.037 | Young |
| 0.523434 | 0.277312 | 0.149 | 0.123 | 1 | 13 | Psph | 0.026 | Young |
| 0.523509 | 0.292381 | 0.14 | 0.115 | 1 | 13 | 2610008E1 | 0.025 | Young |
| 0.523843 | 0.263145 | 0.167 | 0.139 | 1 | 13 | Trmt10a | 0.028 | Young |
| 0.524524 | 0.329049 | 0.088 | 0.066 | 1 | 13 | Nt5c3b | 0.022 | Young |
| 0.52478 | 0.299969 | 0.518 | 0.525 | 1 | 13 | Eif4enif1 | -0.007 | Young |
| 0.524783 | 0.356433 | 0.096 | 0.074 | 1 | 13 | Morn2 | 0.022 | Young |
| 0.524965 | 0.589571 | 0.175 | 0.156 | 1 | 13 | Fastkd2 | 0.019 | Young |
| 0.525178 | 0.057609 | 0.956 | 0.975 | 1 | 13 | Mrps21 | -0.019 | Young |
| 0.525338 | 0.131722 | 0.851 | 0.885 | 1 | 13 | Uvrag | -0.034 | Young |
| 0.525346 | 0.226047 | 0.737 | 0.672 | 1 | 13 | Commd7 | 0.065 | Young |
| 0.525662 | 0.044007 | 0.895 | 0.926 | 1 | 13 | Dnm2 | -0.031 | Young |

| | | | | | | | | |
|---|---|---|---|---|---|---|---|---|
| 0.525935 | 0.736664 | 0.018 | 0.008 | 1 | 13 | Etnk2 | 0.01 | Young |
| 0.525935 | 0.721884 | 0.018 | 0.008 | 1 | 13 | 1110020A2 | 0.01 | Young |
| 0.525935 | 0.545246 | 0.018 | 0.008 | 1 | 13 | Efcab7 | 0.01 | Young |
| 0.525935 | 0.52856 | 0.018 | 0.008 | 1 | 13 | Wwtr1 | 0.01 | Young |
| 0.525935 | 0.50096 | 0.018 | 0.008 | 1 | 13 | Rnf113a1 | 0.01 | Young |
| 0.525935 | 0.481588 | 0.018 | 0.008 | 1 | 13 | Rtkn2 | 0.01 | Young |
| 0.525935 | 0.481289 | 0.018 | 0.008 | 1 | 13 | Meox2 | 0.01 | Young |
| 0.525935 | 0.477285 | 0.018 | 0.008 | 1 | 13 | Kifc5b | 0.01 | Young |
| 0.525935 | 0.472445 | 0.018 | 0.008 | 1 | 13 | Pask | 0.01 | Young |
| 0.525935 | 0.446205 | 0.018 | 0.008 | 1 | 13 | Gm20655 | 0.01 | Young |
| 0.525935 | 0.442243 | 0.018 | 0.008 | 1 | 13 | Tctex1d4 | 0.01 | Young |
| 0.525935 | 0.442151 | 0.018 | 0.008 | 1 | 13 | BC030867 | 0.01 | Young |
| 0.525935 | 0.432071 | 0.018 | 0.008 | 1 | 13 | Gm49249 | 0.01 | Young |
| 0.525935 | 0.424256 | 0.018 | 0.008 | 1 | 13 | Gm10800 | 0.01 | Young |
| 0.525935 | 0.421904 | 0.018 | 0.008 | 1 | 13 | Gpr82 | 0.01 | Young |
| 0.525935 | 0.417995 | 0.018 | 0.008 | 1 | 13 | B9d1 | 0.01 | Young |
| 0.525935 | 0.414803 | 0.018 | 0.008 | 1 | 13 | Slc6a1 | 0.01 | Young |
| 0.525935 | 0.408686 | 0.018 | 0.008 | 1 | 13 | 6530409C1 | 0.01 | Young |
| 0.525935 | 0.404264 | 0.018 | 0.008 | 1 | 13 | Mcam | 0.01 | Young |
| 0.525935 | 0.399189 | 0.018 | 0.008 | 1 | 13 | Gm17552 | 0.01 | Young |
| 0.525935 | 0.396845 | 0.018 | 0.008 | 1 | 13 | Sorbs1 | 0.01 | Young |
| 0.525935 | 0.394221 | 0.018 | 0.008 | 1 | 13 | Pnma1 | 0.01 | Young |
| 0.525935 | 0.38646 | 0.018 | 0.008 | 1 | 13 | Ccdc120 | 0.01 | Young |
| 0.525935 | 0.369051 | 0.018 | 0.008 | 1 | 13 | AI661453 | 0.01 | Young |
| 0.525935 | 0.368485 | 0.018 | 0.008 | 1 | 13 | Rap1gap | 0.01 | Young |
| 0.525935 | 0.36132 | 0.018 | 0.008 | 1 | 13 | Hist1h2bb | 0.01 | Young |
| 0.525935 | 0.359337 | 0.018 | 0.008 | 1 | 13 | Flywch2 | 0.01 | Young |
| 0.525935 | 0.355256 | 0.018 | 0.008 | 1 | 13 | Abca14 | 0.01 | Young |
| 0.525935 | 0.354042 | 0.018 | 0.008 | 1 | 13 | Lbx1 | 0.01 | Young |
| 0.525935 | 0.353549 | 0.018 | 0.008 | 1 | 13 | Rep15 | 0.01 | Young |
| 0.525935 | 0.349196 | 0.018 | 0.008 | 1 | 13 | Fam212b | 0.01 | Young |
| 0.525935 | 0.344232 | 0.018 | 0.008 | 1 | 13 | Gm19412 | 0.01 | Young |
| 0.525935 | 0.341254 | 0.018 | 0.008 | 1 | 13 | Spag5 | 0.01 | Young |
| 0.525935 | 0.328615 | 0.018 | 0.008 | 1 | 13 | Gm47242 | 0.01 | Young |
| 0.525935 | 0.326649 | 0.018 | 0.008 | 1 | 13 | Nova2 | 0.01 | Young |
| 0.525935 | 0.324485 | 0.018 | 0.008 | 1 | 13 | Gm26576 | 0.01 | Young |
| 0.525935 | 0.309565 | 0.018 | 0.008 | 1 | 13 | Gm15283 | 0.01 | Young |
| 0.525935 | 0.299618 | 0.018 | 0.008 | 1 | 13 | Cenpp | 0.01 | Young |
| 0.525935 | 0.288057 | 0.018 | 0.008 | 1 | 13 | Pxylp1 | 0.01 | Young |
| 0.525935 | 0.282848 | 0.018 | 0.008 | 1 | 13 | Mycbpap | 0.01 | Young |
| 0.525935 | 0.282344 | 0.018 | 0.008 | 1 | 13 | Auts2 | 0.01 | Young |
| 0.525935 | 0.280276 | 0.018 | 0.008 | 1 | 13 | Fancb | 0.01 | Young |
| 0.527068 | 0.097026 | 0.719 | 0.705 | 1 | 13 | Ppp2r2a | 0.014 | Young |
| 0.527087 | 0.314179 | 0.281 | 0.262 | 1 | 13 | Zfyve21 | 0.019 | Young |
| 0.527915 | 0.169511 | 0.491 | 0.467 | 1 | 13 | Tob1 | 0.024 | Young |

| | | | | | | | | |
|---|---|---|---|---|---|---|---|---|
| 0.528132 | 0.091546 | 0.746 | 0.697 | 1 | 13 | Vps13c | 0.049 | Young |
| 0.528294 | 0.254727 | 0.667 | 0.639 | 1 | 13 | B4galt1 | 0.028 | Young |
| 0.528749 | 0.097452 | 0.518 | 0.533 | 1 | 13 | Trrap | -0.015 | Young |
| 0.529082 | 0.099334 | 0.868 | 0.869 | 1 | 13 | Ranbp2 | -0.001 | Young |
| 0.52924 | 0.230074 | 0.605 | 0.607 | 1 | 13 | Tlnrd1 | -0.002 | Young |
| 0.529369 | 0.138242 | 0.711 | 0.705 | 1 | 13 | Rap2a | 0.006 | Young |
| 0.529576 | 0.086741 | 0.789 | 0.754 | 1 | 13 | Igbp1 | 0.035 | Young |
| 0.529941 | 0.382564 | 0.088 | 0.066 | 1 | 13 | Wdr74 | 0.022 | Young |
| 0.529941 | 0.337731 | 0.088 | 0.066 | 1 | 13 | Ccl27a | 0.022 | Young |
| 0.530079 | 0.338571 | 0.211 | 0.189 | 1 | 13 | Rusc2 | 0.022 | Young |
| 0.530276 | 0.196428 | 0.281 | 0.262 | 1 | 13 | Gdap2 | 0.019 | Young |
| 0.530444 | 0.068452 | 0.272 | 0.336 | 1 | 13 | Fam45a | -0.064 | Young |
| 0.531035 | 0.205473 | 0.693 | 0.713 | 1 | 13 | Gramd4 | -0.02 | Young |
| 0.531172 | 0.112303 | 0.649 | 0.664 | 1 | 13 | Pstpip1 | -0.015 | Young |
| 0.531452 | 0.072144 | 0.36 | 0.311 | 1 | 13 | Smg6 | 0.049 | Young |
| 0.531462 | 0.125362 | 0.772 | 0.811 | 1 | 13 | Nek7 | -0.039 | Young |
| 0.531866 | 0.416639 | 0.316 | 0.311 | 1 | 13 | Bcl2l2 | 0.005 | Young |
| 0.532051 | 0.013923 | 1 | 1 | 1 | 13 | Dbi | 0 | Young |
| 0.532371 | 0.376164 | 0.018 | 0.008 | 1 | 13 | Zfp449 | 0.01 | Young |
| 0.532371 | 0.351404 | 0.018 | 0.008 | 1 | 13 | Tnfsf13 | 0.01 | Young |
| 0.532371 | 0.348406 | 0.018 | 0.008 | 1 | 13 | Dynlt1b | 0.01 | Young |
| 0.532371 | 0.344141 | 0.018 | 0.008 | 1 | 13 | Ghr | 0.01 | Young |
| 0.532371 | 0.30728 | 0.018 | 0.008 | 1 | 13 | Gsg1 | 0.01 | Young |
| 0.532371 | 0.303438 | 0.018 | 0.008 | 1 | 13 | Padi3 | 0.01 | Young |
| 0.532371 | 0.29729 | 0.018 | 0.008 | 1 | 13 | A930003A1 | 0.01 | Young |
| 0.532371 | 0.292034 | 0.018 | 0.008 | 1 | 13 | 1700008O( | 0.01 | Young |
| 0.532371 | 0.283359 | 0.018 | 0.008 | 1 | 13 | Zfp791 | 0.01 | Young |
| 0.532371 | 0.278176 | 0.018 | 0.008 | 1 | 13 | Kcnq3 | 0.01 | Young |
| 0.532371 | 0.264985 | 0.018 | 0.008 | 1 | 13 | Ermap | 0.01 | Young |
| 0.532371 | 0.260407 | 0.018 | 0.008 | 1 | 13 | Mcts2 | 0.01 | Young |
| 0.532371 | 0.258942 | 0.018 | 0.008 | 1 | 13 | 4930518I1! | 0.01 | Young |
| 0.532371 | 0.254791 | 0.018 | 0.008 | 1 | 13 | Plpp7 | 0.01 | Young |
| 0.532371 | 0.252497 | 0.018 | 0.008 | 1 | 13 | Mall | 0.01 | Young |
| 0.532371 | 0.248319 | 0.018 | 0.008 | 1 | 13 | Gm13012 | 0.01 | Young |
| 0.532371 | 0.245371 | 0.018 | 0.008 | 1 | 13 | Gm49087 | 0.01 | Young |
| 0.532371 | 0.242203 | 0.018 | 0.008 | 1 | 13 | Zpbp | 0.01 | Young |
| 0.532371 | 0.236543 | 0.018 | 0.008 | 1 | 13 | Apoc4 | 0.01 | Young |
| 0.532371 | 0.228668 | 0.018 | 0.008 | 1 | 13 | Peak1os | 0.01 | Young |
| 0.532371 | 0.217535 | 0.018 | 0.008 | 1 | 13 | Gm10273 | 0.01 | Young |
| 0.532371 | 0.216072 | 0.018 | 0.008 | 1 | 13 | Gm17354 | 0.01 | Young |
| 0.532371 | 0.215077 | 0.018 | 0.008 | 1 | 13 | Hist1h3a | 0.01 | Young |
| 0.532371 | 0.214166 | 0.018 | 0.008 | 1 | 13 | Lyrm7 | 0.01 | Young |
| 0.532371 | 0.207347 | 0.018 | 0.008 | 1 | 13 | Gm10851 | 0.01 | Young |
| 0.532371 | 0.182851 | 0.018 | 0.008 | 1 | 13 | Gm44851 | 0.01 | Young |
| 0.532371 | 0.181093 | 0.018 | 0.008 | 1 | 13 | Tox3 | 0.01 | Young |

| | | | | | | | | |
|---|---|---|---|---|---|---|---|---|
| 0.532371 | 0.166589 | 0.018 | 0.008 | 1 | 13 | Lrp11 | 0.01 | Young |
| 0.532371 | 0.133183 | 0.018 | 0.008 | 1 | 13 | Jam2 | 0.01 | Young |
| 0.532371 | 0.131815 | 0.018 | 0.008 | 1 | 13 | Casc1 | 0.01 | Young |
| 0.532371 | 0.123173 | 0.018 | 0.008 | 1 | 13 | Egfl8 | 0.01 | Young |
| 0.532371 | 0.114034 | 0.018 | 0.008 | 1 | 13 | Ctse | 0.01 | Young |
| 0.532371 | 0.109591 | 0.018 | 0.008 | 1 | 13 | Eno1b | 0.01 | Young |
| 0.532371 | 0.098181 | 0.018 | 0.008 | 1 | 13 | 4930480K2 | 0.01 | Young |
| 0.532371 | 0.059388 | 0.018 | 0.008 | 1 | 13 | Gm45620 | 0.01 | Young |
| 0.532371 | 0.016264 | 0.018 | 0.008 | 1 | 13 | Gm20234 | 0.01 | Young |
| 0.532661 | 0.286905 | 0.088 | 0.066 | 1 | 13 | Cd300lf | 0.022 | Young |
| 0.532937 | 0.157481 | 0.789 | 0.811 | 1 | 13 | Tm2d2 | -0.022 | Young |
| 0.533106 | 0.21771 | 0.298 | 0.262 | 1 | 13 | G2e3 | 0.036 | Young |
| 0.533169 | 0.269814 | 0.351 | 0.336 | 1 | 13 | Gpr137 | 0.015 | Young |
| 0.533226 | 0.518812 | 0.123 | 0.098 | 1 | 13 | Cd209c | 0.025 | Young |
| 0.533304 | 0.066753 | 1 | 0.984 | 1 | 13 | Rab14 | 0.016 | Young |
| 0.533343 | 0.151642 | 0.544 | 0.525 | 1 | 13 | Adh5 | 0.019 | Young |
| 0.534108 | 0.313347 | 0.333 | 0.32 | 1 | 13 | Tada3 | 0.013 | Young |
| 0.534355 | 0.347523 | 0.254 | 0.23 | 1 | 13 | Samd4 | 0.024 | Young |
| 0.535464 | 0.156676 | 0.447 | 0.434 | 1 | 13 | Ipo8 | 0.013 | Young |
| 0.535656 | 0.15059 | 0.807 | 0.811 | 1 | 13 | Cenpx | -0.004 | Young |
| 0.53615 | 0.201929 | 0.219 | 0.189 | 1 | 13 | Rad51d | 0.03 | Young |
| 0.536246 | 0.315754 | 0.105 | 0.082 | 1 | 13 | Coq6 | 0.023 | Young |
| 0.536379 | 0.091926 | 0.833 | 0.795 | 1 | 13 | Churc1 | 0.038 | Young |
| 0.537005 | 0.136763 | 0.746 | 0.787 | 1 | 13 | Cdc40 | -0.041 | Young |
| 0.53721 | 0.205091 | 0.667 | 0.672 | 1 | 13 | Vps13b | -0.005 | Young |
| 0.53785 | 0.310377 | 0.123 | 0.098 | 1 | 13 | 4930430F0 | 0.025 | Young |
| 0.537869 | 0.412953 | 0.281 | 0.279 | 1 | 13 | Alkbh7 | 0.002 | Young |
| 0.538504 | 0.270704 | 0.202 | 0.172 | 1 | 13 | Bclaf3 | 0.03 | Young |
| 0.53858 | 0.239162 | 0.132 | 0.107 | 1 | 13 | Gstcd | 0.025 | Young |
| 0.538642 | 0.285147 | 0.14 | 0.115 | 1 | 13 | Tmem241 | 0.025 | Young |
| 0.538943 | 0.156961 | 0.386 | 0.361 | 1 | 13 | Rae1 | 0.025 | Young |
| 0.539234 | 0.066237 | 0.465 | 0.451 | 1 | 13 | Milr1 | 0.014 | Young |
| 0.539592 | 0.061032 | 1 | 1 | 1 | 13 | Ifitm2 | 0 | Young |
| 0.539968 | 0.197786 | 0.386 | 0.385 | 1 | 13 | Sc5d | 0.001 | Young |
| 0.540274 | 0.257697 | 0.377 | 0.377 | 1 | 13 | Mkl1 | 0 | Young |
| 0.540826 | 0.249661 | 0.132 | 0.107 | 1 | 13 | Chst14 | 0.025 | Young |
| 0.540861 | 0.160461 | 0.088 | 0.066 | 1 | 13 | Gtf2h3 | 0.022 | Young |
| 0.541206 | 0.091041 | 0.158 | 0.197 | 1 | 13 | Zmym3 | -0.039 | Young |
| 0.541565 | 0.083914 | 0.886 | 0.836 | 1 | 13 | Camk1d | 0.05 | Young |
| 0.541807 | 0.233754 | 0.325 | 0.303 | 1 | 13 | Nckipsd | 0.022 | Young |
| 0.541855 | 0.162487 | 0.289 | 0.27 | 1 | 13 | Pnpo | 0.019 | Young |
| 0.541872 | 0.27474 | 0.167 | 0.139 | 1 | 13 | D930030I0 | 0.028 | Young |
| 0.541987 | 0.222492 | 0.263 | 0.246 | 1 | 13 | Cntln | 0.017 | Young |
| 0.542301 | 0.188845 | 0.395 | 0.369 | 1 | 13 | Suclg2 | 0.026 | Young |
| 0.542417 | 0.461376 | 0.175 | 0.156 | 1 | 13 | Sorbs3 | 0.019 | Young |

| | | | | | | | | |
|---|---|---|---|---|---|---|---|---|
| 0.542482 | 0.087858 | 0.789 | 0.77 | 1 | 13 | Rab5a | 0.019 | Young |
| 0.542493 | 0.323297 | 0.123 | 0.098 | 1 | 13 | Gas8 | 0.025 | Young |
| 0.542504 | 0.573944 | 0.114 | 0.098 | 1 | 13 | Tmem129 | 0.016 | Young |
| 0.542964 | 0.141435 | 0.096 | 0.074 | 1 | 13 | Zfp568 | 0.022 | Young |
| 0.543143 | 0.067922 | 0.921 | 0.869 | 1 | 13 | Mier1 | 0.052 | Young |
| 0.543183 | 0.250519 | 0.333 | 0.32 | 1 | 13 | Dnajc24 | 0.013 | Young |
| 0.543372 | 0.291998 | 0.518 | 0.5 | 1 | 13 | Rhobtb1 | 0.018 | Young |
| 0.544419 | 0.194866 | 0.491 | 0.484 | 1 | 13 | Ccdc186 | 0.007 | Young |
| 0.544649 | 0.020979 | 1 | 1 | 1 | 13 | Eef1a1 | 0 | Young |
| 0.544799 | 0.257212 | 0.307 | 0.295 | 1 | 13 | Kctd5 | 0.012 | Young |
| 0.545317 | 0.076545 | 0.667 | 0.648 | 1 | 13 | Dhx40 | 0.019 | Young |
| 0.545917 | 0.037924 | 1 | 1 | 1 | 13 | Uqcrq | 0 | Young |
| 0.545937 | 0.453553 | 0.114 | 0.09 | 1 | 13 | Zmym6 | 0.024 | Young |
| 0.546163 | 0.054493 | 0.07 | 0.049 | 1 | 13 | Vsig10l | 0.021 | Young |
| 0.546225 | 0.05551 | 0.298 | 0.352 | 1 | 13 | Ccnc | -0.054 | Young |
| 0.546335 | 0.313981 | 0.175 | 0.156 | 1 | 13 | Tab1 | 0.019 | Young |
| 0.546367 | 0.496695 | 0.281 | 0.262 | 1 | 13 | Rbm4b | 0.019 | Young |
| 0.546955 | 0.048401 | 0.228 | 0.279 | 1 | 13 | Faap100 | -0.051 | Young |
| 0.547191 | 0.343462 | 0.386 | 0.385 | 1 | 13 | Riok1 | 0.001 | Young |
| 0.547458 | 0.093629 | 0.184 | 0.23 | 1 | 13 | Gpt2 | -0.046 | Young |
| 0.547706 | 0.234826 | 0.395 | 0.385 | 1 | 13 | Mkl2 | 0.01 | Young |
| 0.547813 | 0.249864 | 0.254 | 0.23 | 1 | 13 | Paxip1 | 0.024 | Young |
| 0.547817 | 0.075815 | 0.202 | 0.172 | 1 | 13 | Ddr1 | 0.03 | Young |
| 0.54795 | 0.230448 | 0.167 | 0.139 | 1 | 13 | Abcb10 | 0.028 | Young |
| 0.547985 | 0.182363 | 0.561 | 0.566 | 1 | 13 | Luc7l | -0.005 | Young |
| 0.548418 | 0.183551 | 0.211 | 0.18 | 1 | 13 | Adi1 | 0.031 | Young |
| 0.548618 | 0.642155 | 0.193 | 0.189 | 1 | 13 | Mag | 0.004 | Young |
| 0.548782 | 0.162261 | 0.105 | 0.082 | 1 | 13 | A230050P2 | 0.023 | Young |
| 0.549507 | 0.262463 | 0.254 | 0.23 | 1 | 13 | Cmtm4 | 0.024 | Young |
| 0.549583 | 0.147359 | 0.36 | 0.328 | 1 | 13 | Tpra1 | 0.032 | Young |
| 0.550145 | 0.221523 | 0.526 | 0.5 | 1 | 13 | Il16 | 0.026 | Young |
| 0.550261 | 0.142443 | 0.719 | 0.77 | 1 | 13 | Slirp | -0.051 | Young |
| 0.550373 | 0.485558 | 0.158 | 0.139 | 1 | 13 | Prdm15 | 0.019 | Young |
| 0.550642 | 0.241867 | 0.491 | 0.492 | 1 | 13 | Zfp593 | -0.001 | Young |
| 0.550669 | 0.249738 | 0.237 | 0.213 | 1 | 13 | Depdc5 | 0.024 | Young |
| 0.550797 | 0.168616 | 0.482 | 0.459 | 1 | 13 | Ddhd2 | 0.023 | Young |
| 0.551048 | 0.312302 | 0.219 | 0.197 | 1 | 13 | Sec22a | 0.022 | Young |
| 0.551143 | 0.42589 | 0.289 | 0.279 | 1 | 13 | Cdc14a | 0.01 | Young |
| 0.551186 | 0.155752 | 0.263 | 0.238 | 1 | 13 | Lmnb1 | 0.025 | Young |
| 0.551633 | 0.059831 | 0.974 | 0.959 | 1 | 13 | Ppp1r18 | 0.015 | Young |
| 0.551672 | 0.214194 | 0.579 | 0.566 | 1 | 13 | Bdp1 | 0.013 | Young |
| 0.551684 | 0.285904 | 0.404 | 0.41 | 1 | 13 | Kmt5c | -0.006 | Young |
| 0.551927 | 0.336898 | 0.175 | 0.148 | 1 | 13 | Sacs | 0.027 | Young |
| 0.552277 | 0.040843 | 0.982 | 0.984 | 1 | 13 | Diaph1 | -0.002 | Young |
| 0.553269 | 0.072391 | 0.175 | 0.23 | 1 | 13 | Acad9 | -0.055 | Young |

| | | | | | | | | |
|---|---|---|---|---|---|---|---|---|
| 0.553371 | 0.051621 | 0.868 | 0.934 | 1 | 13 | Psmd8 | -0.066 | Young |
| 0.553866 | 0.032533 | 0.298 | 0.352 | 1 | 13 | Fbxl14 | -0.054 | Young |
| 0.554211 | 0.008259 | 0.281 | 0.32 | 1 | 13 | Mis12 | -0.039 | Young |
| 0.554466 | 0.05049 | 0.868 | 0.885 | 1 | 13 | Pdlim5 | -0.017 | Young |
| 0.554813 | 0.026643 | 0.93 | 0.975 | 1 | 13 | Psma2 | -0.045 | Young |
| 0.554959 | 0.007578 | 0.026 | 0.041 | 1 | 13 | Aqp1 | -0.015 | Young |
| 0.555221 | 0.499061 | 0.14 | 0.123 | 1 | 13 | Tubg1 | 0.017 | Young |
| 0.555273 | 0.293594 | 0.307 | 0.303 | 1 | 13 | Ttll12 | 0.004 | Young |
| 0.555565 | 0.220719 | 0.351 | 0.352 | 1 | 13 | Khk | -0.001 | Young |
| 0.555572 | 0.109603 | 0.763 | 0.738 | 1 | 13 | Naa38 | 0.025 | Young |
| 0.555592 | 0.16135 | 0.43 | 0.426 | 1 | 13 | Map2k4 | 0.004 | Young |
| 0.555662 | 0.189 | 0.412 | 0.41 | 1 | 13 | Lin54 | 0.002 | Young |
| 0.55605 | 0.379689 | 0.325 | 0.32 | 1 | 13 | Ndufaf2 | 0.005 | Young |
| 0.556061 | 0.19701 | 0.64 | 0.664 | 1 | 13 | Tifab | -0.024 | Young |
| 0.556063 | 0.405712 | 0.202 | 0.18 | 1 | 13 | Grap2 | 0.022 | Young |
| 0.556079 | 0.069689 | 0.93 | 0.959 | 1 | 13 | Stx7 | -0.029 | Young |
| 0.556459 | 0.112059 | 0.482 | 0.467 | 1 | 13 | Prkab2 | 0.015 | Young |
| 0.556567 | 0.355255 | 0.228 | 0.213 | 1 | 13 | Tbl3 | 0.015 | Young |
| 0.556789 | 0.103809 | 0.579 | 0.541 | 1 | 13 | Zfp326 | 0.038 | Young |
| 0.556871 | 0.198481 | 0.447 | 0.451 | 1 | 13 | Ppfia4 | -0.004 | Young |
| 0.557133 | 0.39594 | 0.281 | 0.279 | 1 | 13 | Ptk2 | 0.002 | Young |
| 0.557447 | 0.247816 | 0.088 | 0.066 | 1 | 13 | Ifit2 | 0.022 | Young |
| 0.557542 | 0.219224 | 0.377 | 0.352 | 1 | 13 | Wrn | 0.025 | Young |
| 0.557643 | 0.001036 | 0.079 | 0.107 | 1 | 13 | Arhgef5 | -0.028 | Young |
| 0.558186 | 0.139966 | 0.482 | 0.5 | 1 | 13 | Ube3b | -0.018 | Young |
| 0.558262 | 0.002235 | 0.184 | 0.23 | 1 | 13 | Mrpl39 | -0.046 | Young |
| 0.558535 | 0.133577 | 0.412 | 0.41 | 1 | 13 | Tmem251 | 0.002 | Young |
| 0.558676 | 0.033758 | 1 | 1 | 1 | 13 | Rps27l | 0 | Young |
| 0.558772 | 0.096267 | 0.307 | 0.27 | 1 | 13 | Aig1 | 0.037 | Young |
| 0.558977 | 0.03534 | 0.412 | 0.516 | 1 | 13 | Polr3gl | -0.104 | Young |
| 0.559071 | 0.21097 | 0.386 | 0.377 | 1 | 13 | Ltbp3 | 0.009 | Young |
| 0.559106 | 0.263739 | 0.272 | 0.254 | 1 | 13 | Hs1bp3 | 0.018 | Young |
| 0.559136 | 0.084747 | 0.772 | 0.787 | 1 | 13 | Wls | -0.015 | Young |
| 0.55931 | 0.031972 | 0.974 | 0.959 | 1 | 13 | Tbxas1 | 0.015 | Young |
| 0.559446 | 0.184997 | 0.535 | 0.533 | 1 | 13 | Gstm1 | 0.002 | Young |
| 0.559588 | 0.538451 | 0.14 | 0.123 | 1 | 13 | Ercc8 | 0.017 | Young |
| 0.559731 | 0.11894 | 0.728 | 0.697 | 1 | 13 | Ddx50 | 0.031 | Young |
| 0.559805 | 0.251088 | 0.202 | 0.18 | 1 | 13 | Ddx55 | 0.022 | Young |
| 0.560064 | 0.052422 | 0.719 | 0.746 | 1 | 13 | Setd1b | -0.027 | Young |
| 0.561245 | 0.037797 | 1 | 1 | 1 | 13 | Rpl37a | 0 | Young |
| 0.562339 | 0.012112 | 0.465 | 0.574 | 1 | 13 | Ncln | -0.109 | Young |
| 0.562738 | 0.061437 | 0.079 | 0.107 | 1 | 13 | Iqcc | -0.028 | Young |
| 0.562858 | 0.299342 | 0.14 | 0.115 | 1 | 13 | Tfap4 | 0.025 | Young |
| 0.563031 | 0.092801 | 0.088 | 0.066 | 1 | 13 | Grasp | 0.022 | Young |
| 0.563633 | 0.360247 | 0.123 | 0.098 | 1 | 13 | Ptch1 | 0.025 | Young |

| | | | | | | | | |
|---|---|---|---|---|---|---|---|---|
| 0.563633 | 0.343652 | 0.123 | 0.098 | 1 | 13 | Atp6v1g2 | 0.025 | Young |
| 0.563935 | 0.05308 | 0.43 | 0.385 | 1 | 13 | Txndc16 | 0.045 | Young |
| 0.564091 | 0.021365 | 0.237 | 0.287 | 1 | 13 | Dnajc9 | -0.05 | Young |
| 0.564358 | 0.093918 | 0.807 | 0.877 | 1 | 13 | Clptm1l | -0.07 | Young |
| 0.564868 | 0.298422 | 0.254 | 0.23 | 1 | 13 | Zcchc14 | 0.024 | Young |
| 0.565109 | 0.03581 | 1 | 1 | 1 | 13 | Prcp | 0 | Young |
| 0.566976 | 0.271129 | 0.158 | 0.139 | 1 | 13 | Pde1b | 0.019 | Young |
| 0.567031 | 0.045273 | 0.974 | 0.943 | 1 | 13 | Emb | 0.031 | Young |
| 0.567323 | 0.392512 | 0.202 | 0.18 | 1 | 13 | Alg2 | 0.022 | Young |
| 0.56766 | 0.107703 | 0.939 | 0.951 | 1 | 13 | Dpm3 | -0.012 | Young |
| 0.568361 | 0.614978 | 0.132 | 0.115 | 1 | 13 | Slc25a40 | 0.017 | Young |
| 0.568361 | 0.422123 | 0.132 | 0.115 | 1 | 13 | Nudcd1 | 0.017 | Young |
| 0.56868 | 0.015945 | 0.535 | 0.623 | 1 | 13 | Chmp1b | -0.088 | Young |
| 0.568986 | 0.049245 | 1 | 1 | 1 | 13 | Atp5g2 | 0 | Young |
| 0.568998 | 0.09546 | 0.719 | 0.738 | 1 | 13 | Rsl1d1 | -0.019 | Young |
| 0.569291 | 0.135859 | 0.728 | 0.664 | 1 | 13 | Mrps16 | 0.064 | Young |
| 0.569342 | 0.262707 | 0.351 | 0.311 | 1 | 13 | Gcnt1 | 0.04 | Young |
| 0.569372 | 0.105214 | 0.404 | 0.402 | 1 | 13 | Copg2 | 0.002 | Young |
| 0.569468 | 0.027942 | 0.096 | 0.074 | 1 | 13 | Cdc7 | 0.022 | Young |
| 0.569777 | 0.451636 | 0.254 | 0.246 | 1 | 13 | Alkbh3 | 0.008 | Young |
| 0.569847 | 0.0869 | 0.746 | 0.754 | 1 | 13 | Tnfrsf13b | -0.008 | Young |
| 0.570103 | 0.00052 | 0.939 | 0.869 | 1 | 13 | Stap1 | 0.07 | Young |
| 0.570392 | 0.14293 | 0.465 | 0.484 | 1 | 13 | Dapk3 | -0.019 | Young |
| 0.570766 | 0.296741 | 0.123 | 0.098 | 1 | 13 | Fdxacb1 | 0.025 | Young |
| 0.57095 | 0.063212 | 0.86 | 0.844 | 1 | 13 | Rragc | 0.016 | Young |
| 0.571368 | 0.232177 | 0.544 | 0.549 | 1 | 13 | Cse1l | -0.005 | Young |
| 0.571478 | 0.016026 | 0.912 | 0.926 | 1 | 13 | Bst2 | -0.014 | Young |
| 0.571709 | 0.003968 | 0.693 | 0.648 | 1 | 13 | Ccdc71l | 0.045 | Young |
| 0.57176 | 0.24535 | 0.254 | 0.23 | 1 | 13 | Ccdc93 | 0.024 | Young |
| 0.571845 | 0.365095 | 0.167 | 0.148 | 1 | 13 | Agk | 0.019 | Young |
| 0.571932 | 0.06975 | 0.421 | 0.393 | 1 | 13 | Proser1 | 0.028 | Young |
| 0.571991 | 0.143767 | 0.781 | 0.77 | 1 | 13 | Tnrc6a | 0.011 | Young |
| 0.572035 | 0.249841 | 0.175 | 0.148 | 1 | 13 | Ankrd54 | 0.027 | Young |
| 0.572133 | 0.368876 | 0.175 | 0.156 | 1 | 13 | Zfp429 | 0.019 | Young |
| 0.57264 | 0.140266 | 0.491 | 0.475 | 1 | 13 | Mrpl58 | 0.016 | Young |
| 0.572674 | 0.079082 | 0.895 | 0.902 | 1 | 13 | Pgd | -0.007 | Young |
| 0.572707 | 0.488084 | 0.246 | 0.23 | 1 | 13 | Llgl1 | 0.016 | Young |
| 0.572707 | 0.260721 | 0.211 | 0.262 | 1 | 13 | Abcd4 | -0.051 | Young |
| 0.57272 | 0.108572 | 0.93 | 0.885 | 1 | 13 | Brd4 | 0.045 | Young |
| 0.573488 | 0.251325 | 0.553 | 0.574 | 1 | 13 | Alkbh1 | -0.021 | Young |
| 0.573603 | 0.007451 | 0.526 | 0.623 | 1 | 13 | Cdc34 | -0.097 | Young |
| 0.573655 | 0.151921 | 0.588 | 0.533 | 1 | 13 | Tor1aip2 | 0.055 | Young |
| 0.573853 | 0.071572 | 0.588 | 0.582 | 1 | 13 | Zranb1 | 0.006 | Young |
| 0.573897 | 0.342597 | 0.167 | 0.148 | 1 | 13 | AW554918 | 0.019 | Young |
| 0.574126 | 0.326251 | 0.272 | 0.254 | 1 | 13 | Med31 | 0.018 | Young |

| | | | | | | | | |
|---|---|---|---|---|---|---|---|---|
| 0.574221 | 0.134249 | 0.553 | 0.533 | 1 | 13 | 6430548M | 0.02 | Young |
| 0.574661 | 0.144363 | 0.184 | 0.23 | 1 | 13 | Pskh1 | -0.046 | Young |
| 0.575001 | 0.015464 | 0.114 | 0.148 | 1 | 13 | Zfp146 | -0.034 | Young |
| 0.575346 | 0.298854 | 0.237 | 0.213 | 1 | 13 | Fasn | 0.024 | Young |
| 0.575346 | 0.267298 | 0.237 | 0.213 | 1 | 13 | Terf1 | 0.024 | Young |
| 0.575427 | 0.022966 | 0.93 | 0.943 | 1 | 13 | Arhgef6 | -0.013 | Young |
| 0.575468 | 0.235514 | 0.281 | 0.254 | 1 | 13 | Rnf34 | 0.027 | Young |
| 0.576153 | 0.445218 | 0.175 | 0.156 | 1 | 13 | Zfp626 | 0.019 | Young |
| 0.576379 | 0.718312 | 0.254 | 0.254 | 1 | 13 | Spink2 | 0 | Young |
| 0.576551 | 0.091343 | 0.64 | 0.648 | 1 | 13 | Man1a2 | -0.008 | Young |
| 0.576752 | 0.170759 | 0.167 | 0.139 | 1 | 13 | Fam210b | 0.028 | Young |
| 0.576778 | 0.139692 | 1 | 1 | 1 | 13 | Wfdc17 | 0 | Young |
| 0.576869 | 0.030475 | 0.719 | 0.738 | 1 | 13 | Zfp445 | -0.019 | Young |
| 0.577043 | 0.168964 | 0.158 | 0.131 | 1 | 13 | Cers4 | 0.027 | Young |
| 0.577412 | 0.127195 | 0.684 | 0.738 | 1 | 13 | Cyb5r4 | -0.054 | Young |
| 0.578168 | 0.326118 | 0.175 | 0.156 | 1 | 13 | Thap4 | 0.019 | Young |
| 0.578531 | 0.132171 | 0.605 | 0.59 | 1 | 13 | Pnrc2 | 0.015 | Young |
| 0.578987 | 0.353309 | 0.439 | 0.434 | 1 | 13 | Abi2 | 0.005 | Young |
| 0.579088 | 0.2815 | 0.377 | 0.361 | 1 | 13 | Nxpe5 | 0.016 | Young |
| 0.579199 | 0.010904 | 0.254 | 0.303 | 1 | 13 | Slc9a1 | -0.049 | Young |
| 0.579386 | 0.056471 | 0.991 | 0.984 | 1 | 13 | Lgmn | 0.007 | Young |
| 0.579448 | 0.192178 | 0.105 | 0.082 | 1 | 13 | Msrb2 | 0.023 | Young |
| 0.579512 | 0.573091 | 0.123 | 0.107 | 1 | 13 | 2700049A( | 0.016 | Young |
| 0.579584 | 0.478429 | 0.158 | 0.139 | 1 | 13 | St7 | 0.019 | Young |
| 0.580344 | 0.276493 | 0.123 | 0.098 | 1 | 13 | Acad12 | 0.025 | Young |
| 0.581459 | 0.748813 | 0.061 | 0.082 | 1 | 13 | Ccl5 | -0.021 | Young |
| 0.581774 | 0.268298 | 0.395 | 0.426 | 1 | 13 | Lsm10 | -0.031 | Young |
| 0.582 | 0.079126 | 1 | 1 | 1 | 13 | Eif3f | 0 | Young |
| 0.582322 | 0.058868 | 0.737 | 0.73 | 1 | 13 | Rnaseh2c | 0.007 | Young |
| 0.582929 | 0.225311 | 0.412 | 0.369 | 1 | 13 | Gm5617 | 0.043 | Young |
| 0.583015 | 0.262405 | 0.132 | 0.172 | 1 | 13 | Asb6 | -0.04 | Young |
| 0.583104 | 0.237273 | 0.447 | 0.418 | 1 | 13 | Slc2a6 | 0.029 | Young |
| 0.583205 | 0.138261 | 0.491 | 0.484 | 1 | 13 | Kbtbd2 | 0.007 | Young |
| 0.583285 | 0.02238 | 0.719 | 0.73 | 1 | 13 | Uso1 | -0.011 | Young |
| 0.583898 | 0.457505 | 0.14 | 0.123 | 1 | 13 | Gm29340 | 0.017 | Young |
| 0.583898 | 0.065331 | 0.114 | 0.148 | 1 | 13 | 5730409E( | -0.034 | Young |
| 0.58442 | 0.077817 | 0.447 | 0.492 | 1 | 13 | Wdr48 | -0.045 | Young |
| 0.584442 | 0.970973 | 0.026 | 0.016 | 1 | 13 | Marco | 0.01 | Young |
| 0.584442 | 0.796004 | 0.026 | 0.016 | 1 | 13 | Fabp3 | 0.01 | Young |
| 0.584442 | 0.782262 | 0.026 | 0.016 | 1 | 13 | Ebf1 | 0.01 | Young |
| 0.584442 | 0.780352 | 0.026 | 0.016 | 1 | 13 | Atp9a | 0.01 | Young |
| 0.584442 | 0.757381 | 0.026 | 0.016 | 1 | 13 | Pemt | 0.01 | Young |
| 0.584442 | 0.737616 | 0.026 | 0.016 | 1 | 13 | Rilpl1 | 0.01 | Young |
| 0.584442 | 0.704103 | 0.026 | 0.016 | 1 | 13 | Pde8b | 0.01 | Young |
| 0.584442 | 0.606551 | 0.026 | 0.016 | 1 | 13 | Gm49201 | 0.01 | Young |

| | | | | | | | | |
|---|---|---|---|---|---|---|---|---|
| 0.584442 | 0.604709 | 0.026 | 0.016 | 1 | 13 | Adgrf5 | 0.01 | Young |
| 0.584442 | 0.576175 | 0.026 | 0.016 | 1 | 13 | Slc22a4 | 0.01 | Young |
| 0.584442 | 0.532418 | 0.026 | 0.016 | 1 | 13 | Nbl1 | 0.01 | Young |
| 0.584442 | 0.524626 | 0.026 | 0.016 | 1 | 13 | Zfp11 | 0.01 | Young |
| 0.584442 | 0.519987 | 0.026 | 0.016 | 1 | 13 | 9930111J2 | 0.01 | Young |
| 0.584442 | 0.509354 | 0.026 | 0.016 | 1 | 13 | Tspan7 | 0.01 | Young |
| 0.584442 | 0.487575 | 0.026 | 0.016 | 1 | 13 | Phf11c | 0.01 | Young |
| 0.584442 | 0.476639 | 0.026 | 0.016 | 1 | 13 | Ppp1r13l | 0.01 | Young |
| 0.584442 | 0.467389 | 0.026 | 0.016 | 1 | 13 | Teddm2 | 0.01 | Young |
| 0.584442 | 0.454542 | 0.026 | 0.016 | 1 | 13 | S100a16 | 0.01 | Young |
| 0.584504 | 0.100405 | 0.544 | 0.484 | 1 | 13 | Epc2 | 0.06 | Young |
| 0.584566 | 0.066018 | 0.316 | 0.377 | 1 | 13 | Mfsd7a | -0.061 | Young |
| 0.585284 | 0.054195 | 0.439 | 0.402 | 1 | 13 | Upf2 | 0.037 | Young |
| 0.585291 | 0.521675 | 0.193 | 0.172 | 1 | 13 | Dgkg | 0.021 | Young |
| 0.585318 | 0.214912 | 0.14 | 0.115 | 1 | 13 | Lrrc45 | 0.025 | Young |
| 0.585386 | 0.111652 | 0.746 | 0.779 | 1 | 13 | Retreg2 | -0.033 | Young |
| 0.585428 | 0.253344 | 0.447 | 0.459 | 1 | 13 | Opa1 | -0.012 | Young |
| 0.585428 | 0.224505 | 0.456 | 0.451 | 1 | 13 | Tmem245 | 0.005 | Young |
| 0.585762 | 0.097958 | 0.57 | 0.557 | 1 | 13 | Mif4gd | 0.013 | Young |
| 0.586282 | 0.150688 | 0.316 | 0.295 | 1 | 13 | Tubgcp2 | 0.021 | Young |
| 0.586655 | 0.049223 | 0.342 | 0.393 | 1 | 13 | Usf3 | -0.051 | Young |
| 0.586973 | 0.012454 | 0.193 | 0.246 | 1 | 13 | Mib2 | -0.053 | Young |
| 0.587117 | 0.082684 | 0.658 | 0.664 | 1 | 13 | Sike1 | -0.006 | Young |
| 0.587659 | 0.102227 | 0.439 | 0.434 | 1 | 13 | Ciz1 | 0.005 | Young |
| 0.587902 | 0.027296 | 0.991 | 0.992 | 1 | 13 | Mtdh | -0.001 | Young |
| 0.588257 | 0.126651 | 0.693 | 0.689 | 1 | 13 | BC031181 | 0.004 | Young |
| 0.588291 | 0.433363 | 0.211 | 0.197 | 1 | 13 | Stk35 | 0.014 | Young |
| 0.588371 | 0.071979 | 0.114 | 0.148 | 1 | 13 | D030056L2 | -0.034 | Young |
| 0.58859 | 0.244754 | 0.675 | 0.77 | 1 | 13 | 1500011K1 | -0.095 | Young |
| 0.588661 | 0.030628 | 0.228 | 0.27 | 1 | 13 | Cbx6 | -0.042 | Young |
| 0.589248 | 0.102153 | 0.57 | 0.582 | 1 | 13 | Gbp3 | -0.012 | Young |
| 0.589523 | 0.434674 | 0.149 | 0.131 | 1 | 13 | Ercc2 | 0.018 | Young |
| 0.589523 | 0.374759 | 0.149 | 0.131 | 1 | 13 | Cnr2 | 0.018 | Young |
| 0.589655 | 0.356818 | 0.281 | 0.27 | 1 | 13 | Ercc5 | 0.011 | Young |
| 0.589708 | 0.673733 | 0.026 | 0.016 | 1 | 13 | Gm4950 | 0.01 | Young |
| 0.589708 | 0.579501 | 0.026 | 0.016 | 1 | 13 | 4930570G1 | 0.01 | Young |
| 0.589708 | 0.576023 | 0.026 | 0.016 | 1 | 13 | Igf2bp3 | 0.01 | Young |
| 0.589708 | 0.506289 | 0.026 | 0.016 | 1 | 13 | Rhox8 | 0.01 | Young |
| 0.589708 | 0.495818 | 0.026 | 0.016 | 1 | 13 | Agbl5 | 0.01 | Young |
| 0.589708 | 0.418877 | 0.026 | 0.016 | 1 | 13 | Tox2 | 0.01 | Young |
| 0.589708 | 0.362578 | 0.026 | 0.016 | 1 | 13 | Dok4 | 0.01 | Young |
| 0.589708 | 0.330546 | 0.026 | 0.016 | 1 | 13 | Mtus2 | 0.01 | Young |
| 0.58972 | 0.191911 | 0.272 | 0.246 | 1 | 13 | Rbm19 | 0.026 | Young |
| 0.59019 | 0.245341 | 0.158 | 0.139 | 1 | 13 | Zfp729b | 0.019 | Young |
| 0.590365 | 0.079226 | 0.561 | 0.574 | 1 | 13 | Atxn3 | -0.013 | Young |

| | | | | | | | | |
|---|---|---|---|---|---|---|---|---|
| 0.590845 | 0.10426 | 0.772 | 0.811 | 1 | 13 | Ptgs1 | -0.039 | Young |
| 0.591132 | 0.149318 | 0.395 | 0.402 | 1 | 13 | Ltn1 | -0.007 | Young |
| 0.59118 | 0.072051 | 0.798 | 0.762 | 1 | 13 | Polr2e | 0.036 | Young |
| 0.59157 | 0.093265 | 0.088 | 0.115 | 1 | 13 | Ccdc94 | -0.027 | Young |
| 0.592134 | 0.229816 | 0.298 | 0.279 | 1 | 13 | Camsap1 | 0.019 | Young |
| 0.592213 | 0.187131 | 0.561 | 0.582 | 1 | 13 | Nab2 | -0.021 | Young |
| 0.592333 | 0.185472 | 0.149 | 0.123 | 1 | 13 | Nsun4 | 0.026 | Young |
| 0.592658 | 0.226369 | 0.263 | 0.246 | 1 | 13 | Ints13 | 0.017 | Young |
| 0.592976 | 0.109846 | 0.886 | 0.91 | 1 | 13 | Msl1 | -0.024 | Young |
| 0.593199 | 0.375991 | 0.237 | 0.221 | 1 | 13 | Zmat3 | 0.016 | Young |
| 0.593575 | 0.591974 | 0.132 | 0.115 | 1 | 13 | Ppp1r16b | 0.017 | Young |
| 0.593807 | 0.040956 | 0.947 | 0.951 | 1 | 13 | Mrps33 | -0.004 | Young |
| 0.593863 | 0.024374 | 0.772 | 0.746 | 1 | 13 | Zfp638 | 0.026 | Young |
| 0.594468 | 0.328908 | 0.316 | 0.311 | 1 | 13 | Arhgef11 | 0.005 | Young |
| 0.594571 | 0.376562 | 0.193 | 0.18 | 1 | 13 | Nr2c1 | 0.013 | Young |
| 0.594996 | 0.662072 | 0.026 | 0.016 | 1 | 13 | Dntt | 0.01 | Young |
| 0.594996 | 0.621029 | 0.026 | 0.016 | 1 | 13 | Mir9-3hg | 0.01 | Young |
| 0.594996 | 0.598339 | 0.026 | 0.016 | 1 | 13 | Dcdc2b | 0.01 | Young |
| 0.594996 | 0.551431 | 0.026 | 0.016 | 1 | 13 | Cygb | 0.01 | Young |
| 0.594996 | 0.5124 | 0.026 | 0.016 | 1 | 13 | C1s1 | 0.01 | Young |
| 0.594996 | 0.496946 | 0.026 | 0.016 | 1 | 13 | Azin2 | 0.01 | Young |
| 0.594996 | 0.492807 | 0.026 | 0.016 | 1 | 13 | Rpl7a-ps5 | 0.01 | Young |
| 0.594996 | 0.476132 | 0.026 | 0.016 | 1 | 13 | 1110006O2 | 0.01 | Young |
| 0.594996 | 0.441742 | 0.026 | 0.016 | 1 | 13 | Satb2 | 0.01 | Young |
| 0.594996 | 0.435962 | 0.026 | 0.016 | 1 | 13 | Zfp976 | 0.01 | Young |
| 0.594996 | 0.430896 | 0.026 | 0.016 | 1 | 13 | AC163040. | 0.01 | Young |
| 0.594996 | 0.429878 | 0.026 | 0.016 | 1 | 13 | B230217C1 | 0.01 | Young |
| 0.594996 | 0.420599 | 0.026 | 0.016 | 1 | 13 | Gm10369 | 0.01 | Young |
| 0.594996 | 0.415917 | 0.026 | 0.016 | 1 | 13 | Egf | 0.01 | Young |
| 0.594996 | 0.378447 | 0.026 | 0.016 | 1 | 13 | Catsperg1 | 0.01 | Young |
| 0.594996 | 0.37679 | 0.026 | 0.016 | 1 | 13 | 4930426L0 | 0.01 | Young |
| 0.594996 | 0.366111 | 0.026 | 0.016 | 1 | 13 | Gm47204 | 0.01 | Young |
| 0.594996 | 0.359145 | 0.026 | 0.016 | 1 | 13 | Dus4l | 0.01 | Young |
| 0.594996 | 0.351904 | 0.026 | 0.016 | 1 | 13 | Xylb | 0.01 | Young |
| 0.594996 | 0.35129 | 0.026 | 0.016 | 1 | 13 | Il2 | 0.01 | Young |
| 0.594996 | 0.309082 | 0.026 | 0.016 | 1 | 13 | Tm4sf5 | 0.01 | Young |
| 0.594996 | 0.303573 | 0.026 | 0.016 | 1 | 13 | Lrrc3 | 0.01 | Young |
| 0.595262 | 0.234946 | 0.325 | 0.32 | 1 | 13 | Cgrrf1 | 0.005 | Young |
| 0.596461 | 0.046798 | 0.965 | 0.951 | 1 | 13 | Hcls1 | 0.014 | Young |
| 0.596698 | 0.320735 | 0.167 | 0.148 | 1 | 13 | Lsm11 | 0.019 | Young |
| 0.597886 | 0.364018 | 0.228 | 0.213 | 1 | 13 | Kctd18 | 0.015 | Young |
| 0.5979 | 0.3412 | 0.184 | 0.164 | 1 | 13 | Slc19a1 | 0.02 | Young |
| 0.597958 | 0.045125 | 0.254 | 0.23 | 1 | 13 | Tapbpl | 0.024 | Young |
| 0.598454 | 0.008748 | 0.991 | 0.943 | 1 | 13 | Etv3 | 0.048 | Young |
| 0.598521 | 0.024942 | 0.491 | 0.533 | 1 | 13 | Bbx | -0.042 | Young |

| | | | | | | | | |
|---|---|---|---|---|---|---|---|---|
| 0.598592 | 0.28221 | 0.237 | 0.221 | 1 | 13 | Mgat5 | 0.016 | Young |
| 0.598659 | 0.118958 | 0.5 | 0.508 | 1 | 13 | Ext2 | -0.008 | Young |
| 0.599099 | 0.09838 | 0.684 | 0.656 | 1 | 13 | Plpbp | 0.028 | Young |
| 0.599124 | 0.036639 | 1 | 1 | 1 | 13 | Clta | 0 | Young |
| 0.599252 | 0.229496 | 0.333 | 0.311 | 1 | 13 | Arsk | 0.022 | Young |
| 0.599495 | 0.181705 | 0.404 | 0.418 | 1 | 13 | Ptrhd1 | -0.014 | Young |
| 0.599648 | 0.159123 | 0.342 | 0.336 | 1 | 13 | Rfx7 | 0.006 | Young |
| 0.599855 | 0.158501 | 0.86 | 0.844 | 1 | 13 | Fam168a | 0.016 | Young |
| 0.600102 | 0.390066 | 0.202 | 0.189 | 1 | 13 | Zfp386 | 0.013 | Young |
| 0.600305 | 0.414506 | 0.026 | 0.016 | 1 | 13 | Traf4 | 0.01 | Young |
| 0.600305 | 0.386144 | 0.026 | 0.016 | 1 | 13 | Hist1h4c | 0.01 | Young |
| 0.600305 | 0.37565 | 0.026 | 0.016 | 1 | 13 | 0610038B2 | 0.01 | Young |
| 0.600305 | 0.344049 | 0.026 | 0.016 | 1 | 13 | Lgals4 | 0.01 | Young |
| 0.600305 | 0.318698 | 0.026 | 0.016 | 1 | 13 | Hs3st3b1 | 0.01 | Young |
| 0.600305 | 0.318239 | 0.026 | 0.016 | 1 | 13 | Rbpms2 | 0.01 | Young |
| 0.600305 | 0.296002 | 0.026 | 0.016 | 1 | 13 | Plagl1 | 0.01 | Young |
| 0.600305 | 0.289491 | 0.026 | 0.016 | 1 | 13 | Lum | 0.01 | Young |
| 0.600305 | 0.286869 | 0.026 | 0.016 | 1 | 13 | Pde7b | 0.01 | Young |
| 0.600305 | 0.283618 | 0.026 | 0.016 | 1 | 13 | Rgs7bp | 0.01 | Young |
| 0.600305 | 0.274795 | 0.026 | 0.016 | 1 | 13 | Med18 | 0.01 | Young |
| 0.600305 | 0.251899 | 0.026 | 0.016 | 1 | 13 | Gm15964 | 0.01 | Young |
| 0.600305 | 0.195671 | 0.026 | 0.016 | 1 | 13 | Arhgap29 | 0.01 | Young |
| 0.600547 | 0.252232 | 0.175 | 0.156 | 1 | 13 | Suv39h1 | 0.019 | Young |
| 0.600547 | 0.004457 | 0.149 | 0.18 | 1 | 13 | Deaf1 | -0.031 | Young |
| 0.601058 | 0.052488 | 0.921 | 0.943 | 1 | 13 | Tgoln1 | -0.022 | Young |
| 0.601073 | 0.168815 | 0.447 | 0.443 | 1 | 13 | Exosc10 | 0.004 | Young |
| 0.602237 | 0.144467 | 0.211 | 0.189 | 1 | 13 | Zc3h12c | 0.022 | Young |
| 0.602341 | 0.023617 | 0.675 | 0.639 | 1 | 13 | Rgs19 | 0.036 | Young |
| 0.603096 | 0.19751 | 0.535 | 0.557 | 1 | 13 | Napg | -0.022 | Young |
| 0.603933 | 0.254069 | 0.184 | 0.164 | 1 | 13 | Ppp1r16a | 0.02 | Young |
| 0.604148 | 0.299228 | 0.14 | 0.123 | 1 | 13 | Pls3 | 0.017 | Young |
| 0.604637 | 0.01718 | 0.5 | 0.639 | 1 | 13 | Sec22b | -0.139 | Young |
| 0.604781 | 0.150009 | 0.509 | 0.525 | 1 | 13 | Cnp | -0.016 | Young |
| 0.605467 | 0.100133 | 0.114 | 0.09 | 1 | 13 | Pms2 | 0.024 | Young |
| 0.605596 | 0.310421 | 0.246 | 0.221 | 1 | 13 | Mcm9 | 0.025 | Young |
| 0.605636 | 0.404481 | 0.026 | 0.016 | 1 | 13 | Acta2 | 0.01 | Young |
| 0.605636 | 0.355807 | 0.026 | 0.016 | 1 | 13 | Ifit1bl1 | 0.01 | Young |
| 0.605636 | 0.347855 | 0.026 | 0.016 | 1 | 13 | Gm26520 | 0.01 | Young |
| 0.605636 | 0.311765 | 0.026 | 0.016 | 1 | 13 | Zfp583 | 0.01 | Young |
| 0.605636 | 0.300481 | 0.026 | 0.016 | 1 | 13 | Gm40645 | 0.01 | Young |
| 0.605636 | 0.284456 | 0.026 | 0.016 | 1 | 13 | Gm16486 | 0.01 | Young |
| 0.605636 | 0.258156 | 0.026 | 0.016 | 1 | 13 | Lgr4 | 0.01 | Young |
| 0.605636 | 0.240866 | 0.026 | 0.016 | 1 | 13 | Psrc1 | 0.01 | Young |
| 0.605636 | 0.228913 | 0.026 | 0.016 | 1 | 13 | B230398E0 | 0.01 | Young |
| 0.605636 | 0.222286 | 0.026 | 0.016 | 1 | 13 | Gm15747 | 0.01 | Young |

| | | | | | | | | |
|---|---|---|---|---|---|---|---|---|
| 0.605636 | 0.204975 | 0.026 | 0.016 | 1 | 13 | 4930404N1 | 0.01 | Young |
| 0.605636 | 0.204119 | 0.026 | 0.016 | 1 | 13 | Id1 | 0.01 | Young |
| 0.605636 | 0.19151 | 0.026 | 0.016 | 1 | 13 | Lrrc75a | 0.01 | Young |
| 0.605636 | 0.188323 | 0.026 | 0.016 | 1 | 13 | Ly6g6d | 0.01 | Young |
| 0.605636 | 0.182054 | 0.026 | 0.016 | 1 | 13 | Neb | 0.01 | Young |
| 0.605636 | 0.119934 | 0.026 | 0.016 | 1 | 13 | Agap2 | 0.01 | Young |
| 0.605744 | 0.555773 | 0.114 | 0.098 | 1 | 13 | Cxxc5 | 0.016 | Young |
| 0.605805 | 0.02097 | 0.702 | 0.738 | 1 | 13 | Map4 | -0.036 | Young |
| 0.607267 | 0.176033 | 0.333 | 0.311 | 1 | 13 | Rnpc3 | 0.022 | Young |
| 0.607354 | 0.05483 | 0.272 | 0.328 | 1 | 13 | Rabggta | -0.056 | Young |
| 0.608123 | 0.221687 | 0.219 | 0.205 | 1 | 13 | Abhd10 | 0.014 | Young |
| 0.608123 | 0.099474 | 0.219 | 0.205 | 1 | 13 | Ttc37 | 0.014 | Young |
| 0.608183 | 0.265126 | 0.123 | 0.107 | 1 | 13 | Ubiad1 | 0.016 | Young |
| 0.608305 | 0.070431 | 0.561 | 0.549 | 1 | 13 | Cfap43 | 0.012 | Young |
| 0.608781 | 0.215902 | 0.175 | 0.156 | 1 | 13 | Rbm43 | 0.019 | Young |
| 0.609471 | 0.094361 | 0.193 | 0.164 | 1 | 13 | Vps37c | 0.029 | Young |
| 0.609471 | 0.076067 | 0.193 | 0.164 | 1 | 13 | Tor4a | 0.029 | Young |
| 0.609608 | 0.013646 | 0.904 | 0.902 | 1 | 13 | Mindy2 | 0.002 | Young |
| 0.609774 | 0.021697 | 1 | 1 | 1 | 13 | Pcbp2 | 0 | Young |
| 0.609882 | 0.345637 | 0.132 | 0.115 | 1 | 13 | Dtd1 | 0.017 | Young |
| 0.610022 | 0.1585 | 0.509 | 0.5 | 1 | 13 | Trim27 | 0.009 | Young |
| 0.61018 | 0.090851 | 0.789 | 0.746 | 1 | 13 | Rasgrp4 | 0.043 | Young |
| 0.610249 | 0.116517 | 0.465 | 0.475 | 1 | 13 | Pwwp2a | -0.01 | Young |
| 0.610298 | 0.670556 | 0.035 | 0.025 | 1 | 13 | Ly6c1 | 0.01 | Young |
| 0.610298 | 0.570868 | 0.035 | 0.025 | 1 | 13 | Timm23 | 0.01 | Young |
| 0.6106 | 0.396693 | 0.123 | 0.107 | 1 | 13 | Zfp709 | 0.016 | Young |
| 0.610648 | 0.181593 | 0.412 | 0.385 | 1 | 13 | Sap30 | 0.027 | Young |
| 0.610775 | 0.095748 | 0.632 | 0.615 | 1 | 13 | Taz | 0.017 | Young |
| 0.610988 | 0.235487 | 0.026 | 0.016 | 1 | 13 | Smarcd3 | 0.01 | Young |
| 0.610988 | 0.176901 | 0.026 | 0.016 | 1 | 13 | Bbs1 | 0.01 | Young |
| 0.610988 | 0.123481 | 0.026 | 0.016 | 1 | 13 | Cd59a | 0.01 | Young |
| 0.610988 | 0.077976 | 0.026 | 0.016 | 1 | 13 | Cd27 | 0.01 | Young |
| 0.611044 | 0.049324 | 0.307 | 0.352 | 1 | 13 | Plekha3 | -0.045 | Young |
| 0.611595 | 0.291236 | 0.377 | 0.369 | 1 | 13 | Crtap | 0.008 | Young |
| 0.612084 | 0.021953 | 0.895 | 0.852 | 1 | 13 | Ndufa8 | 0.043 | Young |
| 0.61225 | 0.119752 | 0.439 | 0.418 | 1 | 13 | Pnpla7 | 0.021 | Young |
| 0.612579 | 0.320003 | 0.009 | 0.016 | 1 | 13 | Iglc1 | -0.007 | Young |
| 0.612579 | 0.245833 | 0.009 | 0.016 | 1 | 13 | Npr3 | -0.007 | Young |
| 0.612579 | 0.110555 | 0.009 | 0.016 | 1 | 13 | Apol7d | -0.007 | Young |
| 0.612579 | 0.103979 | 0.009 | 0.016 | 1 | 13 | A630001O1 | -0.007 | Young |
| 0.612579 | 0.078058 | 0.009 | 0.016 | 1 | 13 | Gm32089 | -0.007 | Young |
| 0.612579 | 0.064804 | 0.009 | 0.016 | 1 | 13 | Tmem178b | -0.007 | Young |
| 0.612579 | 0.058742 | 0.009 | 0.016 | 1 | 13 | Kif20a | -0.007 | Young |
| 0.612579 | 0.056986 | 0.009 | 0.016 | 1 | 13 | C9 | -0.007 | Young |
| 0.612579 | 0.035781 | 0.009 | 0.016 | 1 | 13 | Pomgnt2 | -0.007 | Young |

| | | | | | | | | |
|---|---|---|---|---|---|---|---|---|
| 0.612689 | 0.065291 | 0.132 | 0.107 | 1 | 13 | Zfp512b | 0.025 | Young |
| 0.613518 | 0.385278 | 0.263 | 0.246 | 1 | 13 | Ypel2 | 0.017 | Young |
| 0.613782 | 0.062116 | 0.982 | 0.951 | 1 | 13 | Add3 | 0.031 | Young |
| 0.61396 | 0.238314 | 0.061 | 0.082 | 1 | 13 | 4930481A1 | -0.021 | Young |
| 0.614223 | 0.160958 | 0.886 | 0.893 | 1 | 13 | Anp32e | -0.007 | Young |
| 0.614509 | 0.034073 | 0.281 | 0.254 | 1 | 13 | Ankrd10 | 0.027 | Young |
| 0.614577 | 0.413035 | 0.132 | 0.115 | 1 | 13 | Maml2 | 0.017 | Young |
| 0.614589 | 0.028218 | 0.737 | 0.738 | 1 | 13 | Dennd6a | -0.001 | Young |
| 0.614624 | 0.67147 | 0.088 | 0.074 | 1 | 13 | Sirt6 | 0.014 | Young |
| 0.614626 | 0.035726 | 0.588 | 0.549 | 1 | 13 | Cenpa | 0.039 | Young |
| 0.614683 | 0.234751 | 0.526 | 0.549 | 1 | 13 | Ehmt1 | -0.023 | Young |
| 0.614855 | 0.755015 | 0.035 | 0.025 | 1 | 13 | C730034F0 | 0.01 | Young |
| 0.614855 | 0.678392 | 0.035 | 0.025 | 1 | 13 | Zfp974 | 0.01 | Young |
| 0.614962 | 0.065434 | 0.386 | 0.426 | 1 | 13 | Upf3b | -0.04 | Young |
| 0.615062 | 0.06003 | 0.833 | 0.844 | 1 | 13 | Psma1 | -0.011 | Young |
| 0.615379 | 0.426579 | 0.105 | 0.09 | 1 | 13 | Mtx3 | 0.015 | Young |
| 0.615716 | 0.176039 | 0.412 | 0.402 | 1 | 13 | Stx2 | 0.01 | Young |
| 0.615736 | 0.085187 | 0.447 | 0.443 | 1 | 13 | Mgat4a | 0.004 | Young |
| 0.616248 | 0.114164 | 0.737 | 0.754 | 1 | 13 | Cib1 | -0.017 | Young |
| 0.616361 | 0.159837 | 0.026 | 0.016 | 1 | 13 | Nupr1l | 0.01 | Young |
| 0.616974 | 0.093623 | 0.719 | 0.738 | 1 | 13 | Tmem109 | -0.019 | Young |
| 0.61706 | 0.037891 | 0.289 | 0.344 | 1 | 13 | Cep192 | -0.055 | Young |
| 0.617376 | 0.433288 | 0.263 | 0.254 | 1 | 13 | Wtip | 0.009 | Young |
| 0.617483 | 0.368173 | 0.202 | 0.189 | 1 | 13 | Ipo4 | 0.013 | Young |
| 0.617777 | 0.273024 | 0.167 | 0.148 | 1 | 13 | Ric8b | 0.019 | Young |
| 0.618783 | 0.310722 | 0.316 | 0.311 | 1 | 13 | Ikbip | 0.005 | Young |
| 0.618783 | 0.117261 | 0.228 | 0.279 | 1 | 13 | Dvl1 | -0.051 | Young |
| 0.6188 | 0.005912 | 0.86 | 0.885 | 1 | 13 | Samsn1 | -0.025 | Young |
| 0.618952 | 0.085366 | 0.877 | 0.91 | 1 | 13 | Myo1f | -0.033 | Young |
| 0.619427 | 0.612573 | 0.035 | 0.025 | 1 | 13 | Fbxo44 | 0.01 | Young |
| 0.619427 | 0.598592 | 0.035 | 0.025 | 1 | 13 | Ltb | 0.01 | Young |
| 0.619427 | 0.57059 | 0.035 | 0.025 | 1 | 13 | Ddias | 0.01 | Young |
| 0.619427 | 0.519129 | 0.035 | 0.025 | 1 | 13 | Cc2d2a | 0.01 | Young |
| 0.619427 | 0.414367 | 0.035 | 0.025 | 1 | 13 | Map1b | 0.01 | Young |
| 0.619954 | 0.533758 | 0.079 | 0.066 | 1 | 13 | Catip | 0.013 | Young |
| 0.62032 | 0.480052 | 0.088 | 0.074 | 1 | 13 | Zfp874a | 0.014 | Young |
| 0.620351 | 0.252593 | 0.158 | 0.139 | 1 | 13 | Dnmbp | 0.019 | Young |
| 0.620415 | 0.203801 | 0.561 | 0.59 | 1 | 13 | Rfc1 | -0.029 | Young |
| 0.620538 | 0.372872 | 0.184 | 0.172 | 1 | 13 | Klrb1a | 0.012 | Young |
| 0.620626 | 0.030186 | 0.684 | 0.648 | 1 | 13 | Rab24 | 0.036 | Young |
| 0.620793 | 0.463108 | 0.114 | 0.098 | 1 | 13 | Zfp868 | 0.016 | Young |
| 0.620845 | 0.625202 | 0.061 | 0.049 | 1 | 13 | Rccd1 | 0.012 | Young |
| 0.621007 | 0.351612 | 0.404 | 0.41 | 1 | 13 | Selenoi | -0.006 | Young |
| 0.621112 | 0.070026 | 0.702 | 0.697 | 1 | 13 | Dctn4 | 0.005 | Young |
| 0.621656 | 0.088609 | 0.342 | 0.311 | 1 | 13 | Rpusd4 | 0.031 | Young |

| | | | | | | | | |
|---|---|---|---|---|---|---|---|---|
| 0.621751 | 0.104428 | 0.184 | 0.156 | 1 | 13 | Top2a | 0.028 | Young |
| 0.621854 | 0.011873 | 0.991 | 0.984 | 1 | 13 | Cd84 | 0.007 | Young |
| 0.622007 | 0.121363 | 0.833 | 0.877 | 1 | 13 | Tprgl | -0.044 | Young |
| 0.622937 | 0.164962 | 0.439 | 0.434 | 1 | 13 | Ralgps2 | 0.005 | Young |
| 0.622967 | 0.595924 | 0.044 | 0.033 | 1 | 13 | Ccdc14 | 0.011 | Young |
| 0.623177 | 0.112109 | 0.07 | 0.09 | 1 | 13 | 1810055G( | -0.02 | Young |
| 0.624013 | 0.726792 | 0.035 | 0.025 | 1 | 13 | Htra3 | 0.01 | Young |
| 0.624013 | 0.434382 | 0.035 | 0.025 | 1 | 13 | Zfp691 | 0.01 | Young |
| 0.624163 | 0.02406 | 0.14 | 0.172 | 1 | 13 | Car5b | -0.032 | Young |
| 0.624191 | 0.131299 | 0.553 | 0.59 | 1 | 13 | Gigyf1 | -0.037 | Young |
| 0.624256 | 0.646065 | 0.061 | 0.049 | 1 | 13 | Homez | 0.012 | Young |
| 0.624359 | 0.080253 | 0.237 | 0.213 | 1 | 13 | Cep135 | 0.024 | Young |
| 0.624714 | 0.283312 | 0.158 | 0.139 | 1 | 13 | AU022252 | 0.019 | Young |
| 0.625699 | 0.011366 | 0.886 | 0.902 | 1 | 13 | Leprot | -0.016 | Young |
| 0.625973 | 0.525705 | 0.079 | 0.066 | 1 | 13 | Trmt44 | 0.013 | Young |
| 0.625987 | 0.064418 | 0.193 | 0.23 | 1 | 13 | Pdhx | -0.037 | Young |
| 0.62604 | 0.601707 | 0.088 | 0.074 | 1 | 13 | Fntb | 0.014 | Young |
| 0.626235 | 0.051772 | 0.395 | 0.369 | 1 | 13 | Slc35a5 | 0.026 | Young |
| 0.626864 | 0.313005 | 0.193 | 0.172 | 1 | 13 | Nlrp1b | 0.021 | Young |
| 0.627841 | 0.113301 | 0.211 | 0.262 | 1 | 13 | Pik3r2 | -0.051 | Young |
| 0.627931 | 0.055113 | 0.114 | 0.139 | 1 | 13 | Unc5a | -0.025 | Young |
| 0.627934 | 0.276789 | 0.333 | 0.328 | 1 | 13 | Cep68 | 0.005 | Young |
| 0.628061 | 0.290822 | 0.237 | 0.213 | 1 | 13 | Tyw1 | 0.024 | Young |
| 0.628149 | 0.416454 | 0.211 | 0.197 | 1 | 13 | Pola1 | 0.014 | Young |
| 0.628149 | 0.251794 | 0.211 | 0.197 | 1 | 13 | Cstf2 | 0.014 | Young |
| 0.628154 | 0.034148 | 0.535 | 0.508 | 1 | 13 | Hlcs | 0.027 | Young |
| 0.628276 | 0.134358 | 0.105 | 0.131 | 1 | 13 | Ick | -0.026 | Young |
| 0.628379 | 0.346093 | 0.114 | 0.098 | 1 | 13 | Slc37a3 | 0.016 | Young |
| 0.628478 | 0.397476 | 0.105 | 0.09 | 1 | 13 | Ralgps1 | 0.015 | Young |
| 0.628546 | 0.069099 | 0.412 | 0.434 | 1 | 13 | Ubtd1 | -0.022 | Young |
| 0.628614 | 4.976244 | 0.035 | 0.025 | 1 | 13 | Mucl1 | 0.01 | Young |
| 0.628614 | 0.399322 | 0.035 | 0.025 | 1 | 13 | 6430573P( | 0.01 | Young |
| 0.628614 | 0.384795 | 0.035 | 0.025 | 1 | 13 | Gpihbp1 | 0.01 | Young |
| 0.628614 | 0.383868 | 0.035 | 0.025 | 1 | 13 | Vps25 | 0.01 | Young |
| 0.628614 | 0.380127 | 0.035 | 0.025 | 1 | 13 | Lrrc29 | 0.01 | Young |
| 0.628614 | 0.368617 | 0.035 | 0.025 | 1 | 13 | Taco1 | 0.01 | Young |
| 0.628614 | 0.361908 | 0.035 | 0.025 | 1 | 13 | H2-Eb2 | 0.01 | Young |
| 0.628614 | 0.355398 | 0.035 | 0.025 | 1 | 13 | Prox2 | 0.01 | Young |
| 0.628614 | 0.242717 | 0.035 | 0.025 | 1 | 13 | Plag1 | 0.01 | Young |
| 0.628828 | 0.260932 | 0.219 | 0.205 | 1 | 13 | Wdcp | 0.014 | Young |
| 0.629725 | 0.049354 | 0.684 | 0.73 | 1 | 13 | Vps28 | -0.046 | Young |
| 0.629964 | 0.057905 | 1 | 0.975 | 1 | 13 | Scand1 | 0.025 | Young |
| 0.630495 | 0.056587 | 0.281 | 0.328 | 1 | 13 | Ccs | -0.047 | Young |
| 0.630516 | 0.450613 | 0.096 | 0.082 | 1 | 13 | Mcc | 0.014 | Young |
| 0.630574 | 0.219397 | 0.096 | 0.123 | 1 | 13 | Blm | -0.027 | Young |

| | | | | | | | | |
|---|---|---|---|---|---|---|---|---|
| 0.630863 | 0.015231 | 0.632 | 0.672 | 1 | 13 | Cdc123 | -0.04 | Young |
| 0.631112 | 0.361417 | 0.105 | 0.09 | 1 | 13 | Akap17b | 0.015 | Young |
| 0.631162 | 0.3701 | 0.228 | 0.213 | 1 | 13 | Alkbh6 | 0.015 | Young |
| 0.631189 | 0.014328 | 0.711 | 0.68 | 1 | 13 | Rabl6 | 0.031 | Young |
| 0.631394 | 0.028866 | 0.868 | 0.828 | 1 | 13 | Ppp1r21 | 0.04 | Young |
| 0.631475 | 0.220867 | 0.246 | 0.23 | 1 | 13 | Aktip | 0.016 | Young |
| 0.631782 | 0.459832 | 0.088 | 0.074 | 1 | 13 | Glb1l | 0.014 | Young |
| 0.631794 | 0.091266 | 0.325 | 0.385 | 1 | 13 | Med14 | -0.06 | Young |
| 0.632442 | 0.170192 | 0.184 | 0.164 | 1 | 13 | Coa4 | 0.02 | Young |
| 0.632821 | 0.230174 | 0.219 | 0.197 | 1 | 13 | Pex14 | 0.022 | Young |
| 0.633025 | 0.178511 | 0.333 | 0.32 | 1 | 13 | Krr1 | 0.013 | Young |
| 0.633069 | 0.176285 | 0.211 | 0.189 | 1 | 13 | Rnaseh1 | 0.022 | Young |
| 0.63323 | 0.468839 | 0.035 | 0.025 | 1 | 13 | Ehd2 | 0.01 | Young |
| 0.63323 | 0.377219 | 0.035 | 0.025 | 1 | 13 | Gm42595 | 0.01 | Young |
| 0.63323 | 0.370674 | 0.035 | 0.025 | 1 | 13 | Qprt | 0.01 | Young |
| 0.63323 | 0.348204 | 0.035 | 0.025 | 1 | 13 | Anks6 | 0.01 | Young |
| 0.63323 | 0.286124 | 0.035 | 0.025 | 1 | 13 | Dlg3 | 0.01 | Young |
| 0.633265 | 0.466098 | 0.096 | 0.082 | 1 | 13 | Aldoc | 0.014 | Young |
| 0.633303 | 0.054059 | 0.447 | 0.418 | 1 | 13 | Phka2 | 0.029 | Young |
| 0.633586 | 0.260266 | 0.447 | 0.451 | 1 | 13 | Zdhhc7 | -0.004 | Young |
| 0.63362 | 0.094329 | 0.456 | 0.451 | 1 | 13 | Dpm2 | 0.005 | Young |
| 0.634538 | 0.501721 | 0.061 | 0.049 | 1 | 13 | Nmb | 0.012 | Young |
| 0.634538 | 0.399558 | 0.061 | 0.049 | 1 | 13 | Zfp213 | 0.012 | Young |
| 0.634703 | 0.061713 | 0.421 | 0.393 | 1 | 13 | Ubqln4 | 0.028 | Young |
| 0.6351 | 0.151876 | 0.237 | 0.221 | 1 | 13 | Ift46 | 0.016 | Young |
| 0.635142 | 0.093831 | 0.667 | 0.689 | 1 | 13 | Dnajc2 | -0.022 | Young |
| 0.635216 | 0.723536 | 0.044 | 0.033 | 1 | 13 | Asb2 | 0.011 | Young |
| 0.635216 | 0.630137 | 0.044 | 0.033 | 1 | 13 | Zbtb7c | 0.011 | Young |
| 0.635216 | 0.55174 | 0.044 | 0.033 | 1 | 13 | Nckap5l | 0.011 | Young |
| 0.635216 | 0.441996 | 0.044 | 0.033 | 1 | 13 | Gm34961 | 0.011 | Young |
| 0.635216 | 0.36692 | 0.044 | 0.033 | 1 | 13 | Zswim3 | 0.011 | Young |
| 0.635714 | 0.167396 | 0.298 | 0.279 | 1 | 13 | Zfp407 | 0.019 | Young |
| 0.63574 | 0.092971 | 0.649 | 0.598 | 1 | 13 | Rab3d | 0.051 | Young |
| 0.635805 | 0.072745 | 0.842 | 0.803 | 1 | 13 | Akt2 | 0.039 | Young |
| 0.635882 | 0.086705 | 0.149 | 0.18 | 1 | 13 | Lrrc14 | -0.031 | Young |
| 0.635908 | 0.130497 | 0.596 | 0.557 | 1 | 13 | Tnfaip8l2 | 0.039 | Young |
| 0.636003 | 0.35719 | 0.114 | 0.098 | 1 | 13 | Cry2 | 0.016 | Young |
| 0.636398 | 0.244721 | 0.43 | 0.443 | 1 | 13 | Fam241a | -0.013 | Young |
| 0.636567 | 0.128978 | 0.447 | 0.451 | 1 | 13 | Tbpl1 | -0.004 | Young |
| 0.636758 | 0.037313 | 1 | 1 | 1 | 13 | Crip1 | 0 | Young |
| 0.637273 | 0.331014 | 0.07 | 0.057 | 1 | 13 | Itpr3 | 0.013 | Young |
| 0.637309 | 0.118753 | 0.289 | 0.262 | 1 | 13 | Hps1 | 0.027 | Young |
| 0.637355 | 0.300116 | 0.228 | 0.221 | 1 | 13 | Haus7 | 0.007 | Young |
| 0.637392 | 0.132667 | 0.386 | 0.385 | 1 | 13 | Sharpin | 0.001 | Young |
| 0.637468 | 0.369826 | 0.123 | 0.107 | 1 | 13 | Zfp932 | 0.016 | Young |

| | | | | | | | | |
|---|---|---|---|---|---|---|---|---|
| 0.637481 | 0.039731 | 0.439 | 0.402 | 1 | 13 | Gpr137b | 0.037 | Young |
| 0.637546 | 0.409666 | 0.088 | 0.074 | 1 | 13 | 9930104L0 | 0.014 | Young |
| 0.637546 | 0.265507 | 0.088 | 0.074 | 1 | 13 | Aaas | 0.014 | Young |
| 0.637725 | 0.180221 | 0.36 | 0.344 | 1 | 13 | Ccser2 | 0.016 | Young |
| 0.637859 | 0.317217 | 0.035 | 0.025 | 1 | 13 | Dlc1 | 0.01 | Young |
| 0.637859 | 0.309944 | 0.035 | 0.025 | 1 | 13 | Cenph | 0.01 | Young |
| 0.637859 | 0.280139 | 0.035 | 0.025 | 1 | 13 | Gm15873 | 0.01 | Young |
| 0.637882 | 0.100459 | 0.132 | 0.164 | 1 | 13 | B430306N( | -0.032 | Young |
| 0.638087 | 0.431196 | 0.079 | 0.066 | 1 | 13 | Tmem202 | 0.013 | Young |
| 0.638777 | 0.589304 | 0.096 | 0.082 | 1 | 13 | Pigo | 0.014 | Young |
| 0.639033 | 0.056455 | 0.693 | 0.811 | 1 | 13 | Herc4 | -0.118 | Young |
| 0.639321 | 0.620425 | 0.044 | 0.033 | 1 | 13 | Gm14296 | 0.011 | Young |
| 0.639321 | 0.507921 | 0.044 | 0.033 | 1 | 13 | Fam57a | 0.011 | Young |
| 0.639321 | 0.442795 | 0.044 | 0.033 | 1 | 13 | Mical3 | 0.011 | Young |
| 0.639321 | 0.389963 | 0.044 | 0.033 | 1 | 13 | Inmt | 0.011 | Young |
| 0.639321 | 0.353861 | 0.044 | 0.033 | 1 | 13 | Cep295nl | 0.011 | Young |
| 0.639321 | 0.331869 | 0.044 | 0.033 | 1 | 13 | Pms1 | 0.011 | Young |
| 0.639484 | 0.027463 | 1 | 0.984 | 1 | 13 | Csnk1a1 | 0.016 | Young |
| 0.639526 | 0.005353 | 0.447 | 0.492 | 1 | 13 | Asxl1 | -0.045 | Young |
| 0.640125 | 0.481045 | 0.053 | 0.041 | 1 | 13 | Gm26698 | 0.012 | Young |
| 0.640349 | 0.043903 | 0.842 | 0.869 | 1 | 13 | Snrpd1 | -0.027 | Young |
| 0.640437 | 0.316487 | 0.088 | 0.074 | 1 | 13 | Pigw | 0.014 | Young |
| 0.640504 | 0.226242 | 0.219 | 0.197 | 1 | 13 | Tfpt | 0.022 | Young |
| 0.640807 | 0.092451 | 0.789 | 0.803 | 1 | 13 | Tnrc6b | -0.014 | Young |
| 0.64084 | 0.061605 | 0.947 | 0.975 | 1 | 13 | Psma4 | -0.028 | Young |
| 0.641062 | 0.002961 | 0.421 | 0.5 | 1 | 13 | Matk | -0.079 | Young |
| 0.641131 | 0.576647 | 0.079 | 0.066 | 1 | 13 | Apoo | 0.013 | Young |
| 0.641432 | 0.537735 | 0.061 | 0.049 | 1 | 13 | Lekr1 | 0.012 | Young |
| 0.64197 | 0.186628 | 0.711 | 0.73 | 1 | 13 | Fam208a | -0.019 | Young |
| 0.642164 | 0.25541 | 0.561 | 0.574 | 1 | 13 | Dyrk1a | -0.013 | Young |
| 0.642212 | 0.178444 | 0.474 | 0.5 | 1 | 13 | Mphosph8 | -0.026 | Young |
| 0.6423 | 0.101115 | 0.667 | 0.697 | 1 | 13 | Hpf1 | -0.03 | Young |
| 0.642503 | 0.604478 | 0.035 | 0.025 | 1 | 13 | Dmkn | 0.01 | Young |
| 0.642503 | 0.440659 | 0.035 | 0.025 | 1 | 13 | Lif | 0.01 | Young |
| 0.642503 | 0.23827 | 0.035 | 0.025 | 1 | 13 | Nat8f1 | 0.01 | Young |
| 0.642947 | 0.253453 | 0.149 | 0.131 | 1 | 13 | Dennd2c | 0.018 | Young |
| 0.643073 | 0.209147 | 0.377 | 0.377 | 1 | 13 | Zfyve16 | 0 | Young |
| 0.643212 | 0.106426 | 0.728 | 0.648 | 1 | 13 | Eapp | 0.08 | Young |
| 0.643213 | 0.213396 | 0.5 | 0.508 | 1 | 13 | Lsr | -0.008 | Young |
| 0.643333 | 0.418824 | 0.088 | 0.074 | 1 | 13 | Pgap3 | 0.014 | Young |
| 0.643438 | 0.573595 | 0.044 | 0.033 | 1 | 13 | Apbb2 | 0.011 | Young |
| 0.643438 | 0.553643 | 0.044 | 0.033 | 1 | 13 | Gm28151 | 0.011 | Young |
| 0.643556 | 0.052678 | 0.351 | 0.393 | 1 | 13 | Bola1 | -0.042 | Young |
| 0.643609 | 0.221192 | 0.263 | 0.246 | 1 | 13 | Mrpl22 | 0.017 | Young |
| 0.643873 | 0.031168 | 0.702 | 0.73 | 1 | 13 | Kansl1 | -0.028 | Young |

| | | | | | | | | |
|---|---|---|---|---|---|---|---|---|
| 0.644498 | 0.047075 | 0.833 | 0.885 | 1 | 13 | Fem1c | -0.052 | Young |
| 0.644891 | 0.490051 | 0.061 | 0.049 | 1 | 13 | Tnfrsf13c | 0.012 | Young |
| 0.644891 | 0.448141 | 0.061 | 0.049 | 1 | 13 | Gm14305 | 0.012 | Young |
| 0.645185 | 0.006178 | 0.404 | 0.369 | 1 | 13 | Oard1 | 0.035 | Young |
| 0.645216 | 0.106643 | 0.149 | 0.131 | 1 | 13 | Mllt3 | 0.018 | Young |
| 0.645827 | 0.05275 | 0.842 | 0.869 | 1 | 13 | Ifi203 | -0.027 | Young |
| 0.645935 | 0.03542 | 0.816 | 0.811 | 1 | 13 | Sap18 | 0.005 | Young |
| 0.64623 | 0.304676 | 0.114 | 0.098 | 1 | 13 | Cdan1 | 0.016 | Young |
| 0.646234 | 0.309363 | 0.088 | 0.074 | 1 | 13 | Mex3b | 0.014 | Young |
| 0.647161 | 0.232081 | 0.035 | 0.025 | 1 | 13 | Tex14 | 0.01 | Young |
| 0.647161 | 0.076499 | 0.035 | 0.025 | 1 | 13 | Il31ra | 0.01 | Young |
| 0.647236 | 0.373062 | 0.079 | 0.066 | 1 | 13 | Rad18 | 0.013 | Young |
| 0.647565 | 0.289898 | 0.044 | 0.033 | 1 | 13 | Ccdc189 | 0.011 | Young |
| 0.647616 | 0.517329 | 0.053 | 0.041 | 1 | 13 | Cdca7l | 0.012 | Young |
| 0.647741 | 0.01121 | 0.123 | 0.148 | 1 | 13 | Syn1 | -0.025 | Young |
| 0.64829 | 0.180743 | 0.316 | 0.32 | 1 | 13 | Glt8d1 | -0.004 | Young |
| 0.648357 | 0.464169 | 0.061 | 0.049 | 1 | 13 | Adora3 | 0.012 | Young |
| 0.648489 | 0.013932 | 0.491 | 0.574 | 1 | 13 | Akna | -0.083 | Young |
| 0.648775 | 0.234625 | 0.377 | 0.361 | 1 | 13 | Nop14 | 0.016 | Young |
| 0.649057 | 0.040932 | 1 | 0.934 | 1 | 13 | Snrpf | 0.066 | Young |
| 0.650088 | 0.242147 | 0.254 | 0.23 | 1 | 13 | Adam33 | 0.024 | Young |
| 0.650213 | 0.475516 | 0.07 | 0.057 | 1 | 13 | Asah2 | 0.013 | Young |
| 0.650213 | 0.303109 | 0.07 | 0.057 | 1 | 13 | Rbfox2 | 0.013 | Young |
| 0.650298 | 0.518625 | 0.079 | 0.066 | 1 | 13 | Gm13479 | 0.013 | Young |
| 0.650401 | 0.165396 | 0.307 | 0.311 | 1 | 13 | Ctbs | -0.004 | Young |
| 0.650891 | 0.032731 | 0.842 | 0.803 | 1 | 13 | Ssrp1 | 0.039 | Young |
| 0.650901 | 0.053654 | 0.316 | 0.279 | 1 | 13 | Kat14 | 0.037 | Young |
| 0.65091 | 0.217521 | 0.149 | 0.189 | 1 | 13 | Dnajc12 | -0.04 | Young |
| 0.651375 | 0.273858 | 0.053 | 0.041 | 1 | 13 | Cldn10 | 0.012 | Young |
| 0.65145 | 0.025346 | 0.746 | 0.721 | 1 | 13 | 6-Mar | 0.025 | Young |
| 0.65158 | 0.027404 | 0.342 | 0.32 | 1 | 13 | Pex5 | 0.022 | Young |
| 0.651703 | 0.377843 | 0.044 | 0.033 | 1 | 13 | Hes7 | 0.011 | Young |
| 0.651703 | 0.317964 | 0.044 | 0.033 | 1 | 13 | Gm47230 | 0.011 | Young |
| 0.651831 | 0.608268 | 0.061 | 0.049 | 1 | 13 | Them6 | 0.012 | Young |
| 0.651833 | 0.041379 | 0.035 | 0.025 | 1 | 13 | Fam185a | 0.01 | Young |
| 0.652218 | 0.103502 | 0.605 | 0.648 | 1 | 13 | Tm2d1 | -0.043 | Young |
| 0.652715 | 0.12546 | 0.333 | 0.32 | 1 | 13 | Ppa2 | 0.013 | Young |
| 0.653148 | 0.061005 | 0.184 | 0.164 | 1 | 13 | Gstp3 | 0.02 | Young |
| 0.65328 | 0.147273 | 0.325 | 0.311 | 1 | 13 | Zfp511 | 0.014 | Young |
| 0.653281 | 0.005138 | 0.772 | 0.828 | 1 | 13 | Snrpc | -0.056 | Young |
| 0.654534 | 0.185309 | 0.447 | 0.426 | 1 | 13 | AW549877 | 0.021 | Young |
| 0.655112 | 0.223506 | 0.132 | 0.115 | 1 | 13 | Zmat1 | 0.017 | Young |
| 0.655143 | 0.430708 | 0.053 | 0.041 | 1 | 13 | Fam222a | 0.012 | Young |
| 0.655143 | 0.084366 | 0.053 | 0.041 | 1 | 13 | Sncg | 0.012 | Young |
| 0.655491 | 0.010948 | 0.482 | 0.492 | 1 | 13 | Polr2h | -0.01 | Young |

| | | | | | | | | |
|---|---|---|---|---|---|---|---|---|
| 0.655576 | 0.060107 | 0.614 | 0.598 | 1 | 13 | Tmem5 | 0.016 | Young |
| 0.655661 | 0.045167 | 0.693 | 0.705 | 1 | 13 | Commd3 | -0.012 | Young |
| 0.655852 | 0.323516 | 0.044 | 0.033 | 1 | 13 | Gm26606 | 0.011 | Young |
| 0.655852 | 0.284919 | 0.044 | 0.033 | 1 | 13 | Zfp105 | 0.011 | Young |
| 0.655852 | 0.187152 | 0.044 | 0.033 | 1 | 13 | Pla2g6 | 0.011 | Young |
| 0.655932 | 0.050803 | 0.982 | 0.943 | 1 | 13 | Sema4a | 0.039 | Young |
| 0.655941 | 0.025639 | 1 | 1 | 1 | 13 | Rack1 | 0 | Young |
| 0.656017 | 0.1788 | 0.254 | 0.246 | 1 | 13 | Tpgs2 | 0.008 | Young |
| 0.656157 | 0.187424 | 0.298 | 0.287 | 1 | 13 | Mum1 | 0.011 | Young |
| 0.656501 | 0.02629 | 0.904 | 0.91 | 1 | 13 | Tmed3 | -0.006 | Young |
| 0.656524 | 0.370976 | 0.114 | 0.098 | 1 | 13 | Mmgt2 | 0.016 | Young |
| 0.656524 | 0.209153 | 0.114 | 0.098 | 1 | 13 | Gstz1 | 0.016 | Young |
| 0.656636 | 0.135098 | 0.544 | 0.566 | 1 | 13 | Exosc3 | -0.022 | Young |
| 0.656957 | 0.030826 | 0.833 | 0.902 | 1 | 13 | Nup98 | -0.069 | Young |
| 0.657218 | 0.091744 | 0.667 | 0.68 | 1 | 13 | Egln1 | -0.013 | Young |
| 0.657576 | 0.145263 | 0.447 | 0.459 | 1 | 13 | Scyl2 | -0.012 | Young |
| 0.657803 | 0.102696 | 0.386 | 0.361 | 1 | 13 | Pde7a | 0.025 | Young |
| 0.657848 | 0.333879 | 0.158 | 0.139 | 1 | 13 | Pyroxd1 | 0.019 | Young |
| 0.658633 | 0.030666 | 0.36 | 0.344 | 1 | 13 | Parp3 | 0.016 | Young |
| 0.658642 | 0.031222 | 0.553 | 0.582 | 1 | 13 | Jkamp | -0.029 | Young |
| 0.658802 | 0.383335 | 0.061 | 0.049 | 1 | 13 | Gm26890 | 0.012 | Young |
| 0.659059 | 0.063561 | 0.702 | 0.664 | 1 | 13 | Kdm1a | 0.038 | Young |
| 0.659359 | 0.089912 | 0.404 | 0.41 | 1 | 13 | Soga1 | -0.006 | Young |
| 0.659519 | 0.329741 | 0.079 | 0.066 | 1 | 13 | Gm10642 | 0.013 | Young |
| 0.660012 | 0.182215 | 0.044 | 0.033 | 1 | 13 | Cdc20 | 0.011 | Young |
| 0.660078 | 0.001754 | 0.974 | 0.967 | 1 | 13 | Tab2 | 0.007 | Young |
| 0.660289 | 0.030305 | 0.272 | 0.32 | 1 | 13 | Prmt5 | -0.048 | Young |
| 0.66034 | 0.047652 | 0.535 | 0.549 | 1 | 13 | Tnip2 | -0.014 | Young |
| 0.660821 | 0.239868 | 0.088 | 0.074 | 1 | 13 | Parvg | 0.014 | Young |
| 0.661218 | 0.150968 | 0.035 | 0.025 | 1 | 13 | Gm16861 | 0.01 | Young |
| 0.661507 | 0.196443 | 0.184 | 0.164 | 1 | 13 | Tdrd3 | 0.02 | Young |
| 0.661874 | 0.16799 | 0.36 | 0.344 | 1 | 13 | Psd4 | 0.016 | Young |
| 0.662205 | 0.102297 | 0.816 | 0.779 | 1 | 13 | Nrbp1 | 0.037 | Young |
| 0.662216 | 0.133209 | 0.325 | 0.385 | 1 | 13 | Farsa | -0.06 | Young |
| 0.66229 | 0.068769 | 0.737 | 0.713 | 1 | 13 | Ankfy1 | 0.024 | Young |
| 0.662451 | 0.189918 | 0.237 | 0.23 | 1 | 13 | Wfs1 | 0.007 | Young |
| 0.662451 | 0.024259 | 0.219 | 0.246 | 1 | 13 | Zfp68 | -0.027 | Young |
| 0.662628 | 0.075343 | 0.5 | 0.492 | 1 | 13 | Nfx1 | 0.008 | Young |
| 0.662705 | 0.317934 | 0.053 | 0.041 | 1 | 13 | Zfp59 | 0.012 | Young |
| 0.662705 | 0.262907 | 0.053 | 0.041 | 1 | 13 | Caprin2 | 0.012 | Young |
| 0.662705 | 0.016939 | 0.053 | 0.041 | 1 | 13 | Cdk5r1 | 0.012 | Young |
| 0.662867 | 0.309206 | 0.202 | 0.189 | 1 | 13 | Gm20703 | 0.013 | Young |
| 0.663259 | 0.866207 | 0.07 | 0.057 | 1 | 13 | Vsig4 | 0.013 | Young |
| 0.663404 | 0.01166 | 0.816 | 0.836 | 1 | 13 | Atg3 | -0.02 | Young |
| 0.663825 | 0.035219 | 0.096 | 0.082 | 1 | 13 | Spef1 | 0.014 | Young |

| | | | | | | | | |
|---|---|---|---|---|---|---|---|---|
| 0.664182 | 0.27233 | 0.044 | 0.033 | 1 | 13 | Wdr27 | 0.011 | Young |
| 0.664182 | 0.237046 | 0.044 | 0.033 | 1 | 13 | Aplf | 0.011 | Young |
| 0.664182 | 0.209724 | 0.044 | 0.033 | 1 | 13 | Bco2 | 0.011 | Young |
| 0.664806 | 0.365097 | 0.132 | 0.115 | 1 | 13 | Ccdc66 | 0.017 | Young |
| 0.664806 | 0.185304 | 0.132 | 0.115 | 1 | 13 | Lonrf1 | 0.017 | Young |
| 0.66487 | 0.506324 | 0.202 | 0.189 | 1 | 13 | Ttc28 | 0.013 | Young |
| 0.665212 | 0.099422 | 0.263 | 0.246 | 1 | 13 | Neurl4 | 0.017 | Young |
| 0.665432 | 0.144507 | 0.307 | 0.287 | 1 | 13 | Ticam1 | 0.02 | Young |
| 0.665476 | 0.023788 | 0.149 | 0.172 | 1 | 13 | Rrnad1 | -0.023 | Young |
| 0.665802 | 0.366786 | 0.061 | 0.049 | 1 | 13 | Laptm4b | 0.012 | Young |
| 0.665802 | 0.362603 | 0.061 | 0.049 | 1 | 13 | Camk2n1 | 0.012 | Young |
| 0.665802 | 0.323533 | 0.061 | 0.049 | 1 | 13 | Bend6 | 0.012 | Young |
| 0.665802 | 0.218584 | 0.061 | 0.049 | 1 | 13 | Chek2 | 0.012 | Young |
| 0.666118 | 0.201852 | 0.254 | 0.254 | 1 | 13 | Scaper | 0 | Young |
| 0.666366 | 0.097836 | 0.482 | 0.459 | 1 | 13 | Hr | 0.023 | Young |
| 0.666438 | 3.056002 | 0.044 | 0.057 | 1 | 13 | Slpi | -0.013 | Young |
| 0.666499 | 0.287219 | 0.053 | 0.041 | 1 | 13 | Ugt1a6a | 0.012 | Young |
| 0.666499 | 0.285937 | 0.053 | 0.041 | 1 | 13 | Prelid3a | 0.012 | Young |
| 0.666537 | 0.47817 | 0.07 | 0.057 | 1 | 13 | Zfp948 | 0.013 | Young |
| 0.666692 | 0.315363 | 0.088 | 0.074 | 1 | 13 | D3Ertd254 | 0.014 | Young |
| 0.666693 | 0.29396 | 0.088 | 0.074 | 1 | 13 | Rab3a | 0.014 | Young |
| 0.666693 | 0.220392 | 0.07 | 0.09 | 1 | 13 | Zfp846 | -0.02 | Young |
| 0.666799 | 0.230001 | 0.289 | 0.279 | 1 | 13 | Mzt2 | 0.01 | Young |
| 0.666902 | 0.120533 | 0.421 | 0.377 | 1 | 13 | Crnkl1 | 0.044 | Young |
| 0.666929 | 0.062839 | 0.535 | 0.525 | 1 | 13 | Mast3 | 0.01 | Young |
| 0.667331 | 0.250424 | 0.123 | 0.107 | 1 | 13 | Cryz | 0.016 | Young |
| 0.668008 | 0.106486 | 0.237 | 0.287 | 1 | 13 | Hotairm1 | -0.05 | Young |
| 0.668008 | 0.105607 | 0.272 | 0.254 | 1 | 13 | Orc2 | 0.018 | Young |
| 0.668363 | 0.173334 | 0.044 | 0.033 | 1 | 13 | Plin4 | 0.011 | Young |
| 0.668363 | 0.155115 | 0.044 | 0.033 | 1 | 13 | Bag2 | 0.011 | Young |
| 0.669191 | 0.156426 | 0.202 | 0.18 | 1 | 13 | Dock4 | 0.022 | Young |
| 0.669306 | 0.079943 | 0.789 | 0.754 | 1 | 13 | Zmynd11 | 0.035 | Young |
| 0.669353 | 0.002219 | 0.5 | 0.574 | 1 | 13 | Bccip | -0.074 | Young |
| 0.669483 | 0.421092 | 0.114 | 0.098 | 1 | 13 | Dusp28 | 0.016 | Young |
| 0.669483 | 0.2676 | 0.114 | 0.098 | 1 | 13 | Tsr2 | 0.016 | Young |
| 0.670302 | 0.520215 | 0.053 | 0.041 | 1 | 13 | Nkapl | 0.012 | Young |
| 0.670302 | 0.267546 | 0.053 | 0.041 | 1 | 13 | Smkr-ps | 0.012 | Young |
| 0.670302 | 0.188395 | 0.053 | 0.041 | 1 | 13 | Disp1 | 0.012 | Young |
| 0.670676 | 0.191077 | 0.272 | 0.262 | 1 | 13 | Mrpl50 | 0.01 | Young |
| 0.67098 | 0.078347 | 0.298 | 0.352 | 1 | 13 | Engase | -0.054 | Young |
| 0.67111 | 0.029721 | 0.219 | 0.205 | 1 | 13 | Pomgnt1 | 0.014 | Young |
| 0.671168 | 0.029911 | 0.088 | 0.107 | 1 | 13 | Pcyt1b | -0.019 | Young |
| 0.671262 | 0.029005 | 0.807 | 0.787 | 1 | 13 | Ufc1 | 0.02 | Young |
| 0.671374 | 0.176893 | 0.36 | 0.385 | 1 | 13 | Ddx27 | -0.025 | Young |
| 0.671523 | 0.01478 | 0.623 | 0.615 | 1 | 13 | Akap9 | 0.008 | Young |

| | | | | | | | | |
|---|---|---|---|---|---|---|---|---|
| 0.671618 | 0.050642 | 0.219 | 0.197 | 1 | 13 | Zfp282 | 0.022 | Young |
| 0.671726 | 0.105101 | 0.596 | 0.59 | 1 | 13 | Ldb1 | 0.006 | Young |
| 0.671839 | 0.170683 | 0.167 | 0.148 | 1 | 13 | Zfyve1 | 0.019 | Young |
| 0.672023 | 0.143075 | 0.456 | 0.443 | 1 | 13 | Gm28875 | 0.013 | Young |
| 0.672192 | 0.246347 | 0.325 | 0.32 | 1 | 13 | Lsg1 | 0.005 | Young |
| 0.672585 | 0.23267 | 0.088 | 0.074 | 1 | 13 | Map2k6 | 0.014 | Young |
| 0.672811 | 0.073958 | 0.43 | 0.41 | 1 | 13 | Crim1 | 0.02 | Young |
| 0.672832 | 0.347771 | 0.061 | 0.049 | 1 | 13 | Ggnbp1 | 0.012 | Young |
| 0.672832 | 0.317379 | 0.061 | 0.049 | 1 | 13 | Eaf2 | 0.012 | Young |
| 0.672881 | 0.023887 | 0.447 | 0.418 | 1 | 13 | Prr5l | 0.029 | Young |
| 0.672906 | 0.219498 | 0.202 | 0.189 | 1 | 13 | Ppp1r3b | 0.013 | Young |
| 0.673111 | 0.371591 | 0.07 | 0.057 | 1 | 13 | Slc25a23 | 0.013 | Young |
| 0.673555 | 0.085778 | 0.404 | 0.369 | 1 | 13 | Zadh2 | 0.035 | Young |
| 0.673697 | 0.36628 | 0.237 | 0.23 | 1 | 13 | Wdr45 | 0.007 | Young |
| 0.674155 | 0.066455 | 0.667 | 0.68 | 1 | 13 | Eif4a3 | -0.013 | Young |
| 0.674172 | 0.184286 | 0.614 | 0.598 | 1 | 13 | P4ha1 | 0.016 | Young |
| 0.67426 | 0.044567 | 0.526 | 0.508 | 1 | 13 | Jak2 | 0.018 | Young |
| 0.674266 | 0.118953 | 0.474 | 0.492 | 1 | 13 | Trim26 | -0.018 | Young |
| 0.674693 | 0.120026 | 0.772 | 0.754 | 1 | 13 | Snx18 | 0.018 | Young |
| 0.675354 | 0.016892 | 1 | 1 | 1 | 13 | Atp5j2 | 0 | Young |
| 0.675538 | 0.477227 | 0.088 | 0.074 | 1 | 13 | Abi3 | 0.014 | Young |
| 0.675538 | 0.282625 | 0.088 | 0.074 | 1 | 13 | Bivm | 0.014 | Young |
| 0.676342 | 0.05025 | 0.404 | 0.393 | 1 | 13 | Aph1b | 0.011 | Young |
| 0.676358 | 0.228451 | 0.061 | 0.049 | 1 | 13 | Mtmr11 | 0.012 | Young |
| 0.676408 | 0.30311 | 0.07 | 0.057 | 1 | 13 | Zfp60 | 0.013 | Young |
| 0.676513 | 0.135209 | 0.64 | 0.639 | 1 | 13 | Erlec1 | 0.001 | Young |
| 0.676749 | 0.02702 | 1 | 1 | 1 | 13 | Arpc1b | 0 | Young |
| 0.676755 | 0.236665 | 0.044 | 0.033 | 1 | 13 | Gm47389 | 0.011 | Young |
| 0.677136 | 0.288361 | 0.465 | 0.459 | 1 | 13 | Slc16a6 | 0.006 | Young |
| 0.677261 | 0.114234 | 0.105 | 0.131 | 1 | 13 | Ift140 | -0.026 | Young |
| 0.677407 | 0.269843 | 0.123 | 0.107 | 1 | 13 | Tjap1 | 0.016 | Young |
| 0.677907 | 0.181881 | 0.096 | 0.082 | 1 | 13 | Lmln | 0.014 | Young |
| 0.677932 | 0.45787 | 0.053 | 0.041 | 1 | 13 | Gm26826 | 0.012 | Young |
| 0.678115 | 0.244707 | 0.079 | 0.066 | 1 | 13 | Acat2 | 0.013 | Young |
| 0.678141 | 0.03909 | 0.982 | 0.959 | 1 | 13 | Zyx | 0.023 | Young |
| 0.678805 | 0.16824 | 0.404 | 0.402 | 1 | 13 | Rpa3 | 0.002 | Young |
| 0.67929 | 0.046915 | 0.895 | 0.869 | 1 | 13 | Psmd11 | 0.026 | Young |
| 0.679304 | 0.116156 | 0.702 | 0.754 | 1 | 13 | Gxylt1 | -0.052 | Young |
| 0.679542 | 0.353982 | 0.175 | 0.164 | 1 | 13 | Wrap73 | 0.011 | Young |
| 0.679711 | 0.37112 | 0.07 | 0.057 | 1 | 13 | Phlda3 | 0.013 | Young |
| 0.679711 | 0.099646 | 0.07 | 0.057 | 1 | 13 | Zc3h11a | 0.013 | Young |
| 0.679754 | 0.204233 | 0.263 | 0.254 | 1 | 13 | Ccdc43 | 0.009 | Young |
| 0.679821 | 0.132572 | 0.465 | 0.484 | 1 | 13 | Kcnk13 | -0.019 | Young |
| 0.679831 | 0.218834 | 0.421 | 0.426 | 1 | 13 | Dopey2 | -0.005 | Young |
| 0.67989 | 0.224119 | 0.061 | 0.049 | 1 | 13 | Frat1 | 0.012 | Young |

| | | | | | | | | |
|---|---|---|---|---|---|---|---|---|
| 0.680024 | 0.004527 | 0.184 | 0.221 | 1 | 13 | Alad | -0.037 | Young |
| 0.680134 | 0.002221 | 0.588 | 0.648 | 1 | 13 | Txndc5 | -0.06 | Young |
| 0.680249 | 0.121029 | 0.614 | 0.607 | 1 | 13 | Tpd52l2 | 0.007 | Young |
| 0.680737 | 0.279726 | 0.096 | 0.082 | 1 | 13 | Ccdc102a | 0.014 | Young |
| 0.680737 | 0.080486 | 0.079 | 0.098 | 1 | 13 | Crocc | -0.019 | Young |
| 0.680966 | 0.019594 | 0.044 | 0.033 | 1 | 13 | Ldhb | 0.011 | Young |
| 0.681234 | 0.247476 | 0.079 | 0.066 | 1 | 13 | Gstt3 | 0.013 | Young |
| 0.68146 | 0.411891 | 0.088 | 0.074 | 1 | 13 | Retn | 0.014 | Young |
| 0.68146 | 0.24478 | 0.088 | 0.074 | 1 | 13 | Tkfc | 0.014 | Young |
| 0.681606 | 0.314497 | 0.482 | 0.525 | 1 | 13 | Ythdc2 | -0.043 | Young |
| 0.68176 | 0.223829 | 0.053 | 0.041 | 1 | 13 | Taco1os | 0.012 | Young |
| 0.68176 | 0.217162 | 0.053 | 0.041 | 1 | 13 | Gm43936 | 0.012 | Young |
| 0.68176 | 0.168694 | 0.053 | 0.041 | 1 | 13 | Klri2 | 0.012 | Young |
| 0.681977 | 0.140191 | 0.614 | 0.656 | 1 | 13 | Hmgcl | -0.042 | Young |
| 0.682288 | 0.134721 | 0.325 | 0.32 | 1 | 13 | Lrif1 | 0.005 | Young |
| 0.682466 | 0.128568 | 0.123 | 0.107 | 1 | 13 | Rdh10 | 0.016 | Young |
| 0.682529 | 0.051659 | 0.184 | 0.213 | 1 | 13 | Zfp746 | -0.029 | Young |
| 0.682541 | 0.264768 | 0.114 | 0.098 | 1 | 13 | Tbc1d8b | 0.016 | Young |
| 0.684024 | 0.005907 | 0.632 | 0.615 | 1 | 13 | 1810013L2 | 0.017 | Young |
| 0.684299 | 0.281809 | 0.307 | 0.311 | 1 | 13 | Setd1a | -0.004 | Young |
| 0.684338 | 0.120121 | 0.254 | 0.238 | 1 | 13 | Myh10 | 0.016 | Young |
| 0.684424 | 0.119548 | 0.228 | 0.213 | 1 | 13 | Clmp | 0.015 | Young |
| 0.684794 | 0.309127 | 0.219 | 0.205 | 1 | 13 | Utp23 | 0.014 | Young |
| 0.684842 | 0.09201 | 0.395 | 0.385 | 1 | 13 | Pml | 0.01 | Young |
| 0.685108 | 0.163023 | 0.447 | 0.443 | 1 | 13 | Timm8a1 | 0.004 | Young |
| 0.685596 | 0.142263 | 0.053 | 0.041 | 1 | 13 | A830010M | 0.012 | Young |
| 0.686012 | 0.149815 | 0.307 | 0.311 | 1 | 13 | Rps6ka5 | -0.004 | Young |
| 0.686458 | 0.084399 | 0.658 | 0.697 | 1 | 13 | Sms | -0.039 | Young |
| 0.687296 | 0.029498 | 0.5 | 0.566 | 1 | 13 | Pip5k1a | -0.066 | Young |
| 0.687402 | 0.157776 | 0.088 | 0.074 | 1 | 13 | Odf2l | 0.014 | Young |
| 0.687464 | 0.18609 | 0.105 | 0.09 | 1 | 13 | Ndufaf6 | 0.015 | Young |
| 0.687488 | 0.259457 | 0.079 | 0.066 | 1 | 13 | Gm4673 | 0.013 | Young |
| 0.687503 | 0.031491 | 0.719 | 0.721 | 1 | 13 | Cr1l | -0.002 | Young |
| 0.688039 | 0.105175 | 0.57 | 0.557 | 1 | 13 | Arpc5l | 0.013 | Young |
| 0.688132 | 0.075876 | 0.395 | 0.393 | 1 | 13 | Pigm | 0.002 | Young |
| 0.688186 | 0.024954 | 0.781 | 0.82 | 1 | 13 | Sugt1 | -0.039 | Young |
| 0.688674 | 0.051708 | 0.719 | 0.795 | 1 | 13 | 2510039O1 | -0.076 | Young |
| 0.688914 | 0.086265 | 0.86 | 0.852 | 1 | 13 | Lsm12 | 0.008 | Young |
| 0.689081 | 0.03787 | 0.86 | 0.893 | 1 | 13 | Trappc4 | -0.033 | Young |
| 0.689254 | 0.027041 | 0.096 | 0.082 | 1 | 13 | Txnrd3 | 0.014 | Young |
| 0.689262 | 0.08389 | 0.202 | 0.238 | 1 | 13 | Fam213a | -0.036 | Young |
| 0.689431 | 0.000135 | 0.579 | 0.615 | 1 | 13 | Atg101 | -0.036 | Young |
| 0.689615 | 0.133559 | 0.754 | 0.73 | 1 | 13 | Tbl1x | 0.024 | Young |
| 0.689656 | 0.260431 | 0.07 | 0.057 | 1 | 13 | AW209491 | 0.013 | Young |
| 0.689675 | 0.00711 | 0.632 | 0.656 | 1 | 13 | Gng12 | -0.024 | Young |

| | | | | | | | | |
|---|---|---|---|---|---|---|---|---|
| 0.690335 | 0.112668 | 0.325 | 0.311 | 1 | 13 | Msra | 0.014 | Young |
| 0.690428 | 0.188683 | 0.886 | 0.852 | 1 | 13 | Acly | 0.034 | Young |
| 0.690531 | 0.210392 | 0.061 | 0.049 | 1 | 13 | Wdr34 | 0.012 | Young |
| 0.690619 | 0.421825 | 0.158 | 0.148 | 1 | 13 | Kctd11 | 0.01 | Young |
| 0.690623 | 0.290491 | 0.079 | 0.066 | 1 | 13 | Hist1h4d | 0.013 | Young |
| 0.690623 | 0.188958 | 0.079 | 0.066 | 1 | 13 | Zfp553 | 0.013 | Young |
| 0.69068 | 0.188548 | 0.307 | 0.303 | 1 | 13 | Zfp110 | 0.004 | Young |
| 0.690752 | 0.02545 | 0.956 | 0.967 | 1 | 13 | Tmem256 | -0.011 | Young |
| 0.691027 | 0.019533 | 0.5 | 0.492 | 1 | 13 | Pqlc3 | 0.008 | Young |
| 0.69161 | 0.008408 | 0.482 | 0.541 | 1 | 13 | Mta1 | -0.059 | Young |
| 0.691731 | 0.11556 | 0.561 | 0.582 | 1 | 13 | Fto | -0.021 | Young |
| 0.691988 | 0.191043 | 0.491 | 0.492 | 1 | 13 | Nsd2 | -0.001 | Young |
| 0.69213 | 0.06174 | 0.93 | 0.943 | 1 | 13 | Sri | -0.013 | Young |
| 0.692245 | 0.005969 | 0.246 | 0.287 | 1 | 13 | Spg11 | -0.041 | Young |
| 0.692245 | 0.141442 | 0.439 | 0.426 | 1 | 13 | Dpy19l4 | 0.013 | Young |
| 0.692595 | 0.002346 | 0.184 | 0.213 | 1 | 13 | Nfatc2ip | -0.029 | Young |
| 0.692628 | 0.1988 | 0.123 | 0.107 | 1 | 13 | Polr1b | 0.016 | Young |
| 0.692968 | 0.291671 | 0.272 | 0.27 | 1 | 13 | Scimp | 0.002 | Young |
| 0.692984 | 0.176612 | 0.07 | 0.057 | 1 | 13 | Ddx59 | 0.013 | Young |
| 0.692984 | 0.110298 | 0.07 | 0.057 | 1 | 13 | Polr1e | 0.013 | Young |
| 0.693057 | 0.237521 | 0.114 | 0.098 | 1 | 13 | Cetn4 | 0.016 | Young |
| 0.693292 | 0.097081 | 0.053 | 0.041 | 1 | 13 | Tle6 | 0.012 | Young |
| 0.693421 | 0.111334 | 0.281 | 0.279 | 1 | 13 | Zc3hc1 | 0.002 | Young |
| 0.693576 | 0.009635 | 1 | 1 | 1 | 13 | Myl6 | 0 | Young |
| 0.693644 | 0.106532 | 0.684 | 0.705 | 1 | 13 | Tcof1 | -0.021 | Young |
| 0.693686 | 0.320181 | 0.202 | 0.18 | 1 | 13 | Sesn2 | 0.022 | Young |
| 0.694079 | 0.042516 | 0.518 | 0.525 | 1 | 13 | Dtnbp1 | -0.007 | Young |
| 0.694504 | 0.256319 | 0.237 | 0.23 | 1 | 13 | Dalrd3 | 0.007 | Young |
| 0.694815 | 0.074816 | 0.114 | 0.139 | 1 | 13 | Akr1c12 | -0.025 | Young |
| 0.695198 | 0.275148 | 0.202 | 0.189 | 1 | 13 | Rdh11 | 0.013 | Young |
| 0.695893 | 0.138687 | 0.316 | 0.311 | 1 | 13 | Osbpl2 | 0.005 | Young |
| 0.695972 | 0.028356 | 0.474 | 0.5 | 1 | 13 | Srsf4 | -0.026 | Young |
| 0.696202 | 0.077044 | 0.307 | 0.344 | 1 | 13 | Mtmr1 | -0.037 | Young |
| 0.696395 | 0.03104 | 0.982 | 1 | 1 | 13 | Ndufb9 | -0.018 | Young |
| 0.696909 | 0.128743 | 0.079 | 0.066 | 1 | 13 | Ppcs | 0.013 | Young |
| 0.69724 | 0.179579 | 0.281 | 0.27 | 1 | 13 | Zfp513 | 0.011 | Young |
| 0.697658 | 0.374917 | 0.061 | 0.049 | 1 | 13 | Nlrp1a | 0.012 | Young |
| 0.697796 | 0.052431 | 0.947 | 0.967 | 1 | 13 | Slc43a2 | -0.02 | Young |
| 0.697811 | 0.083955 | 0.096 | 0.082 | 1 | 13 | Gm26510 | 0.014 | Young |
| 0.697858 | 0.016262 | 0.246 | 0.279 | 1 | 13 | Zdhhc17 | -0.033 | Young |
| 0.69841 | 0.224575 | 0.105 | 0.09 | 1 | 13 | Mettl2 | 0.015 | Young |
| 0.698866 | 0.015126 | 0.289 | 0.344 | 1 | 13 | Mtpap | -0.055 | Young |
| 0.698894 | 0.312921 | 0.175 | 0.164 | 1 | 13 | Zfp335 | 0.011 | Young |
| 0.699007 | 0.047061 | 0.482 | 0.492 | 1 | 13 | AI504432 | -0.01 | Young |
| 0.699167 | 0.179195 | 0.132 | 0.115 | 1 | 13 | Rad9a | 0.017 | Young |

| | | | | | | | | |
|---|---|---|---|---|---|---|---|---|
| 0.699446 | 0.026791 | 0.57 | 0.566 | 1 | 13 | Gdap10 | 0.004 | Young |
| 0.699599 | 0.021167 | 0.307 | 0.279 | 1 | 13 | Eci1 | 0.028 | Young |
| 0.699656 | 0.162276 | 0.07 | 0.057 | 1 | 13 | Cdo1 | 0.013 | Young |
| 0.699839 | 0.025255 | 0.921 | 0.91 | 1 | 13 | Cep170 | 0.011 | Young |
| 0.699949 | 0.226086 | 0.228 | 0.213 | 1 | 13 | Gypc | 0.015 | Young |
| 0.700185 | 0.334911 | 0.307 | 0.32 | 1 | 13 | Saysd1 | -0.013 | Young |
| 0.700206 | 0.085488 | 0.877 | 0.836 | 1 | 13 | Tgfbr2 | 0.041 | Young |
| 0.700455 | 0.099125 | 0.588 | 0.59 | 1 | 13 | Gtf3c6 | -0.002 | Young |
| 0.70047 | 0.02548 | 0.623 | 0.639 | 1 | 13 | Lta4h | -0.016 | Young |
| 0.700559 | 0.189273 | 0.219 | 0.205 | 1 | 13 | Ust | 0.014 | Young |
| 0.700862 | 0.02413 | 0.57 | 0.533 | 1 | 13 | Fbxl20 | 0.037 | Young |
| 0.700976 | 0.0146 | 0.605 | 0.672 | 1 | 13 | Faim | -0.067 | Young |
| 0.701288 | 0.184838 | 0.325 | 0.328 | 1 | 13 | Nup50 | -0.003 | Young |
| 0.701603 | 0.327716 | 0.14 | 0.131 | 1 | 13 | Eral1 | 0.009 | Young |
| 0.701929 | 0.106579 | 0.316 | 0.295 | 1 | 13 | Med23 | 0.021 | Young |
| 0.702289 | 0.18771 | 0.316 | 0.32 | 1 | 13 | Kif16b | -0.004 | Young |
| 0.702341 | 0.042219 | 0.088 | 0.074 | 1 | 13 | Fam177a | 0.014 | Young |
| 0.702341 | 0.014908 | 0.816 | 0.787 | 1 | 13 | Ttyh3 | 0.029 | Young |
| 0.702561 | 0.067559 | 0.596 | 0.598 | 1 | 13 | Ndfip2 | -0.002 | Young |
| 0.702851 | 0.160133 | 0.658 | 0.664 | 1 | 13 | Rraga | -0.006 | Young |
| 0.703001 | 0.153811 | 0.07 | 0.057 | 1 | 13 | Otud3 | 0.013 | Young |
| 0.70375 | 0.046212 | 0.632 | 0.598 | 1 | 13 | Chrac1 | 0.034 | Young |
| 0.703815 | 0.048939 | 0.289 | 0.279 | 1 | 13 | Atp2a3 | 0.01 | Young |
| 0.704168 | 0.000648 | 0.956 | 0.992 | 1 | 13 | Eif5 | -0.036 | Young |
| 0.704336 | 0.018505 | 0.868 | 0.82 | 1 | 13 | Uggt1 | 0.048 | Young |
| 0.704813 | 0.040571 | 0.061 | 0.049 | 1 | 13 | Plppr4 | 0.012 | Young |
| 0.70528 | 0.008909 | 0.719 | 0.68 | 1 | 13 | Abcd1 | 0.039 | Young |
| 0.705799 | 0.14721 | 0.64 | 0.656 | 1 | 13 | Clock | -0.016 | Young |
| 0.706422 | 0.23968 | 0.158 | 0.148 | 1 | 13 | Dbf4 | 0.01 | Young |
| 0.706422 | 0.182272 | 0.158 | 0.148 | 1 | 13 | Clcn6 | 0.01 | Young |
| 0.706422 | 0.040805 | 0.14 | 0.164 | 1 | 13 | Rfesd | -0.024 | Young |
| 0.70683 | 0.106713 | 0.149 | 0.18 | 1 | 13 | Pdik1l | -0.031 | Young |
| 0.707009 | 0.131275 | 0.456 | 0.418 | 1 | 13 | Wdr3 | 0.038 | Young |
| 0.707219 | 0.19108 | 0.167 | 0.197 | 1 | 13 | Rpl9-ps6 | -0.03 | Young |
| 0.707344 | 0.193944 | 0.14 | 0.123 | 1 | 13 | Zfp653 | 0.017 | Young |
| 0.707541 | 0.018128 | 0.053 | 0.066 | 1 | 13 | Gm20632 | -0.013 | Young |
| 0.707559 | 0.207284 | 0.175 | 0.164 | 1 | 13 | Gm20342 | 0.011 | Young |
| 0.708231 | 0.080719 | 0.789 | 0.836 | 1 | 13 | Smc1a | -0.047 | Young |
| 0.708521 | 0.019686 | 0.667 | 0.68 | 1 | 13 | Xpr1 | -0.013 | Young |
| 0.708588 | 0.055023 | 0.746 | 0.77 | 1 | 13 | Creb1 | -0.024 | Young |
| 0.708691 | 0.081001 | 0.14 | 0.164 | 1 | 13 | 2-Sep | -0.024 | Young |
| 0.708779 | 0.170509 | 0.053 | 0.041 | 1 | 13 | Ccdc51 | 0.012 | Young |
| 0.708942 | 0.176144 | 0.114 | 0.098 | 1 | 13 | Strbp | 0.016 | Young |
| 0.70898 | 0.018031 | 0.535 | 0.549 | 1 | 13 | Atp6v0a2 | -0.014 | Young |
| 0.709083 | 0.066712 | 0.93 | 0.926 | 1 | 13 | Ctnna1 | 0.004 | Young |

| | | | | | | | | |
|---|---|---|---|---|---|---|---|---|
| 0.709133 | 0.042378 | 0.974 | 0.984 | 1 | 13 | Ddx6 | -0.01 | Young |
| 0.709435 | 0.026448 | 0.658 | 0.697 | 1 | 13 | Ubqln1 | -0.039 | Young |
| 0.709775 | 0.041773 | 0.404 | 0.41 | 1 | 13 | Acsl1 | -0.006 | Young |
| 0.709867 | 0.60169 | 0.123 | 0.115 | 1 | 13 | Tubgcp6 | 0.008 | Young |
| 0.710312 | 0.048836 | 0.482 | 0.467 | 1 | 13 | Wdr36 | 0.015 | Young |
| 0.710367 | 0.197229 | 0.184 | 0.164 | 1 | 13 | Fam35a | 0.02 | Young |
| 0.710508 | 0.165617 | 0.447 | 0.492 | 1 | 13 | Smim26 | -0.045 | Young |
| 0.71056 | 0.060266 | 0.456 | 0.443 | 1 | 13 | Cln5 | 0.013 | Young |
| 0.710652 | 0.118014 | 0.018 | 0.025 | 1 | 13 | Ptprcap | -0.007 | Young |
| 0.710777 | 0.125642 | 0.596 | 0.623 | 1 | 13 | Pigk | -0.027 | Young |
| 0.711266 | 0.009309 | 1 | 1 | 1 | 13 | Rps15 | 0 | Young |
| 0.711551 | 0.214816 | 0.246 | 0.238 | 1 | 13 | Vipas39 | 0.008 | Young |
| 0.711702 | 0.115832 | 0.237 | 0.23 | 1 | 13 | Cnrip1 | 0.007 | Young |
| 0.711711 | 0.094114 | 0.439 | 0.434 | 1 | 13 | Irak2 | 0.005 | Young |
| 0.712001 | 0.17109 | 0.184 | 0.172 | 1 | 13 | Trp53rka | 0.012 | Young |
| 0.712151 | 0.08541 | 0.395 | 0.385 | 1 | 13 | Josd1 | 0.01 | Young |
| 0.712869 | 0.216289 | 0.386 | 0.377 | 1 | 13 | Pkd1 | 0.009 | Young |
| 0.713235 | 0.275507 | 0.158 | 0.148 | 1 | 13 | Pus7 | 0.01 | Young |
| 0.713303 | 0.009468 | 0.877 | 0.943 | 1 | 13 | Pxn | -0.066 | Young |
| 0.713499 | 0.10853 | 0.667 | 0.648 | 1 | 13 | Slc8b1 | 0.019 | Young |
| 0.713499 | 0.094727 | 0.667 | 0.648 | 1 | 13 | Polb | 0.019 | Young |
| 0.714066 | 0.193868 | 0.272 | 0.262 | 1 | 13 | Gm38399 | 0.01 | Young |
| 0.714211 | 0.099202 | 0.754 | 0.754 | 1 | 13 | Ppp6r1 | 0 | Young |
| 0.714303 | 0.007794 | 0.412 | 0.492 | 1 | 13 | Trappc8 | -0.08 | Young |
| 0.714355 | 0.003624 | 0.404 | 0.443 | 1 | 13 | Cyth3 | -0.039 | Young |
| 0.714626 | 0.061961 | 0.509 | 0.615 | 1 | 13 | Mrpl15 | -0.106 | Young |
| 0.714676 | 0.082019 | 0.184 | 0.164 | 1 | 13 | Znhit2 | 0.02 | Young |
| 0.714815 | 0.016965 | 0.974 | 0.951 | 1 | 13 | Socs3 | 0.023 | Young |
| 0.714819 | 0.021892 | 0.956 | 0.984 | 1 | 13 | Klf4 | -0.028 | Young |
| 0.715087 | 0.013791 | 0.342 | 0.393 | 1 | 13 | Eef2k | -0.051 | Young |
| 0.715891 | 0.004993 | 0.079 | 0.066 | 1 | 13 | Fam189b | 0.013 | Young |
| 0.715918 | 0.323718 | 0.14 | 0.131 | 1 | 13 | Gm20604 | 0.009 | Young |
| 0.71696 | 0.04326 | 0.991 | 0.992 | 1 | 13 | Olfm1 | -0.001 | Young |
| 0.717064 | 0.174755 | 0.281 | 0.27 | 1 | 13 | Vps72 | 0.011 | Young |
| 0.717112 | 0.159387 | 0.149 | 0.131 | 1 | 13 | Zbtb24 | 0.018 | Young |
| 0.717616 | 0.37507 | 0.254 | 0.254 | 1 | 13 | Cdk5 | 0 | Young |
| 0.717789 | 0.027358 | 0.14 | 0.164 | 1 | 13 | Btrc | -0.024 | Young |
| 0.718106 | 0.217569 | 0.307 | 0.295 | 1 | 13 | Lrwd1 | 0.012 | Young |
| 0.7183 | 0.06934 | 0.561 | 0.582 | 1 | 13 | Inpp5f | -0.021 | Young |
| 0.718463 | 0.06509 | 0.377 | 0.369 | 1 | 13 | Klhl20 | 0.008 | Young |
| 0.718503 | 0.064123 | 0.281 | 0.279 | 1 | 13 | Fas | 0.002 | Young |
| 0.718529 | 0.028448 | 0.781 | 0.795 | 1 | 13 | Tbc1d14 | -0.014 | Young |
| 0.718548 | 0.223784 | 0.228 | 0.279 | 1 | 13 | Ccdc134 | -0.051 | Young |
| 0.718846 | 0.010776 | 0.281 | 0.336 | 1 | 13 | Rpia | -0.055 | Young |
| 0.718995 | 0.136177 | 0.702 | 0.738 | 1 | 13 | Ptov1 | -0.036 | Young |

| | | | | | | | | |
|---|---|---|---|---|---|---|---|---|
| 0.719393 | 0.012838 | 0.211 | 0.254 | 1 | 13 | Homer3 | -0.043 | Young |
| 0.719714 | 0.064761 | 0.807 | 0.828 | 1 | 13 | Psma5 | -0.021 | Young |
| 0.71981 | 0.071563 | 0.07 | 0.057 | 1 | 13 | Gm26669 | 0.013 | Young |
| 0.719823 | 0.00921 | 0.825 | 0.828 | 1 | 13 | Fgd2 | -0.003 | Young |
| 0.72001 | 0.026718 | 0.816 | 0.869 | 1 | 13 | Iqgap2 | -0.053 | Young |
| 0.720173 | 0.180082 | 0.439 | 0.484 | 1 | 13 | Mtx1 | -0.045 | Young |
| 0.720526 | 0.018871 | 0.982 | 0.984 | 1 | 13 | Ndufa7 | -0.002 | Young |
| 0.720527 | 0.00052 | 1 | 1 | 1 | 13 | Calm1 | 0 | Young |
| 0.720972 | 0.04867 | 0.518 | 0.598 | 1 | 13 | Aip | -0.08 | Young |
| 0.721452 | 0.914445 | 0.053 | 0.066 | 1 | 13 | Cxcl3 | -0.013 | Young |
| 0.721489 | 0.155756 | 0.158 | 0.139 | 1 | 13 | Cyp2ab1 | 0.019 | Young |
| 0.72158 | 0.167143 | 0.526 | 0.525 | 1 | 13 | Fam69a | 0.001 | Young |
| 0.721991 | 0.102111 | 0.307 | 0.303 | 1 | 13 | Abr | 0.004 | Young |
| 0.722075 | 0.121291 | 0.57 | 0.557 | 1 | 13 | 2310009B1 | 0.013 | Young |
| 0.722079 | 0.257222 | 0.018 | 0.025 | 1 | 13 | Kcnip4 | -0.007 | Young |
| 0.722079 | 0.022989 | 0.018 | 0.025 | 1 | 13 | Gm10093 | -0.007 | Young |
| 0.722481 | 0.006346 | 0.544 | 0.557 | 1 | 13 | Kctd10 | -0.013 | Young |
| 0.722529 | 0.000259 | 0.263 | 0.238 | 1 | 13 | Mamdc2 | 0.025 | Young |
| 0.722609 | 0.112321 | 0.351 | 0.336 | 1 | 13 | Rasgrp1 | 0.015 | Young |
| 0.722838 | 0.112592 | 0.605 | 0.648 | 1 | 13 | Elk4 | -0.043 | Young |
| 0.722929 | 0.352023 | 0.114 | 0.107 | 1 | 13 | Mturn | 0.007 | Young |
| 0.723442 | 0.106726 | 0.123 | 0.107 | 1 | 13 | Haus1 | 0.016 | Young |
| 0.723448 | 0.024219 | 0.088 | 0.074 | 1 | 13 | Hist2h3c2 | 0.014 | Young |
| 0.723479 | 0.277256 | 0.219 | 0.213 | 1 | 13 | Tbc1d31 | 0.006 | Young |
| 0.724044 | 0.367328 | 0.149 | 0.139 | 1 | 13 | Ttl | 0.01 | Young |
| 0.72446 | 0.16986 | 0.491 | 0.5 | 1 | 13 | Slc39a10 | -0.009 | Young |
| 0.724971 | 0.05492 | 0.316 | 0.361 | 1 | 13 | As3mt | -0.045 | Young |
| 0.725002 | 0.176095 | 0.175 | 0.164 | 1 | 13 | Gsdme | 0.011 | Young |
| 0.725085 | 0.4215 | 0.254 | 0.254 | 1 | 13 | Iqsec1 | 0 | Young |
| 0.725101 | 0.237272 | 0.123 | 0.115 | 1 | 13 | Ccp110 | 0.008 | Young |
| 0.725294 | 0.134934 | 0.289 | 0.279 | 1 | 13 | 2300009A0 | 0.01 | Young |
| 0.725473 | 0.07299 | 0.553 | 0.533 | 1 | 13 | Pdcd2l | 0.02 | Young |
| 0.725505 | 0.008436 | 0.719 | 0.77 | 1 | 13 | Lrrfip2 | -0.051 | Young |
| 0.725752 | 0.003471 | 0.658 | 0.623 | 1 | 13 | Otud5 | 0.035 | Young |
| 0.725779 | 0.114254 | 0.842 | 0.852 | 1 | 13 | Mcrip1 | -0.01 | Young |
| 0.725963 | 0.053833 | 0.535 | 0.631 | 1 | 13 | Opa3 | -0.096 | Young |
| 0.726116 | 0.164707 | 0.158 | 0.139 | 1 | 13 | Yipf2 | 0.019 | Young |
| 0.726368 | 0.04679 | 0.421 | 0.418 | 1 | 13 | Sap30l | 0.003 | Young |
| 0.726543 | 0.04252 | 0.421 | 0.402 | 1 | 13 | Cep250 | 0.019 | Young |
| 0.726575 | 0.064209 | 0.342 | 0.385 | 1 | 13 | Gtf2f2 | -0.043 | Young |
| 0.726607 | 0.101675 | 0.096 | 0.082 | 1 | 13 | Mmaa | 0.014 | Young |
| 0.727212 | 0.007922 | 0.491 | 0.549 | 1 | 13 | Kdm6a | -0.058 | Young |
| 0.727455 | 0.210144 | 0.219 | 0.213 | 1 | 13 | Hacd2 | 0.006 | Young |
| 0.727816 | 0.451355 | 0.018 | 0.025 | 1 | 13 | Frg2f1 | -0.007 | Young |
| 0.727816 | 0.320099 | 0.018 | 0.025 | 1 | 13 | Mttp | -0.007 | Young |

| | | | | | | | | |
|---|---|---|---|---|---|---|---|---|
| 0.727816 | 0.258213 | 0.018 | 0.025 | 1 | 13 | Art2b | -0.007 | Young |
| 0.727816 | 0.207596 | 0.018 | 0.025 | 1 | 13 | BC022687 | -0.007 | Young |
| 0.727816 | 0.081461 | 0.018 | 0.025 | 1 | 13 | Tube1 | -0.007 | Young |
| 0.727816 | 0.072578 | 0.018 | 0.025 | 1 | 13 | Scara5 | -0.007 | Young |
| 0.727816 | 0.056266 | 0.018 | 0.025 | 1 | 13 | Spry2 | -0.007 | Young |
| 0.727816 | 0.046747 | 0.018 | 0.025 | 1 | 13 | Cacna2d4 | -0.007 | Young |
| 0.727816 | 0.030313 | 0.018 | 0.025 | 1 | 13 | Trpt1 | -0.007 | Young |
| 0.727816 | 0.008561 | 0.018 | 0.025 | 1 | 13 | Ccnj | -0.007 | Young |
| 0.728011 | 0.084466 | 0.772 | 0.82 | 1 | 13 | Leprotl1 | -0.048 | Young |
| 0.728196 | 0.691889 | 0.114 | 0.107 | 1 | 13 | Slc26a11 | 0.007 | Young |
| 0.728871 | 0.083658 | 0.246 | 0.23 | 1 | 13 | Apool | 0.016 | Young |
| 0.729085 | 0.073202 | 0.588 | 0.607 | 1 | 13 | Synj2bp | -0.019 | Young |
| 0.729249 | 0.064999 | 0.711 | 0.738 | 1 | 13 | Fnip1 | -0.027 | Young |
| 0.729727 | 0.135638 | 0.447 | 0.467 | 1 | 13 | Prelid2 | -0.02 | Young |
| 0.729829 | 0.198056 | 0.158 | 0.189 | 1 | 13 | Cep104 | -0.031 | Young |
| 0.730336 | 0.022729 | 0.754 | 0.779 | 1 | 13 | Akirin2 | -0.025 | Young |
| 0.730547 | 0.007474 | 1 | 1 | 1 | 13 | Arf5 | 0 | Young |
| 0.730835 | 0.485006 | 0.114 | 0.107 | 1 | 13 | Cenpw | 0.007 | Young |
| 0.730984 | 0.16556 | 0.237 | 0.23 | 1 | 13 | Decr2 | 0.007 | Young |
| 0.731186 | 0.021982 | 0.904 | 0.926 | 1 | 13 | Arhgap17 | -0.022 | Young |
| 0.731275 | 0.170315 | 0.228 | 0.221 | 1 | 13 | Ankrd50 | 0.007 | Young |
| 0.731374 | 0.372291 | 0.228 | 0.213 | 1 | 13 | 2310061I0 | 0.015 | Young |
| 0.731577 | 0.320615 | 0.395 | 0.402 | 1 | 13 | Adgrg6 | -0.007 | Young |
| 0.732414 | 0.054589 | 0.096 | 0.082 | 1 | 13 | Fam109b | 0.014 | Young |
| 0.733351 | 0.098378 | 0.904 | 0.934 | 1 | 13 | Mbnl2 | -0.03 | Young |
| 0.73337 | 0.110473 | 0.921 | 0.934 | 1 | 13 | Celf1 | -0.013 | Young |
| 0.733636 | 0.18101 | 0.623 | 0.623 | 1 | 13 | Atf1 | 0 | Young |
| 0.733744 | 0.179643 | 0.272 | 0.254 | 1 | 13 | Slc27a4 | 0.018 | Young |
| 0.733816 | 0.258745 | 0.105 | 0.123 | 1 | 13 | Ifit1 | -0.018 | Young |
| 0.734627 | 0.093752 | 0.781 | 0.828 | 1 | 13 | Chtop | -0.047 | Young |
| 0.734855 | 0.028071 | 0.982 | 0.992 | 1 | 13 | Sf3b2 | -0.01 | Young |
| 0.735084 | 0.066164 | 0.298 | 0.287 | 1 | 13 | Ankrd16 | 0.011 | Young |
| 0.735302 | 0.133485 | 0.579 | 0.623 | 1 | 13 | Zhx1 | -0.044 | Young |
| 0.735385 | 0.096601 | 0.728 | 0.697 | 1 | 13 | Smad4 | 0.031 | Young |
| 0.736159 | 0.056455 | 0.684 | 0.664 | 1 | 13 | Fam91a1 | 0.02 | Young |
| 0.736659 | 0.260021 | 0.132 | 0.123 | 1 | 13 | Mrgbp | 0.009 | Young |
| 0.736778 | 0.076602 | 0.456 | 0.459 | 1 | 13 | Slc36a4 | -0.003 | Young |
| 0.736788 | 0.191831 | 0.579 | 0.623 | 1 | 13 | Ldah | -0.044 | Young |
| 0.737319 | 0.044801 | 0.833 | 0.869 | 1 | 13 | Rbm5 | -0.036 | Young |
| 0.737386 | 0.049362 | 0.86 | 0.861 | 1 | 13 | Birc6 | -0.001 | Young |
| 0.737443 | 0.045461 | 0.667 | 0.672 | 1 | 13 | Vamp2 | -0.005 | Young |
| 0.737445 | 0.040161 | 0.877 | 0.861 | 1 | 13 | Ik | 0.016 | Young |
| 0.737577 | 0.082142 | 0.623 | 0.607 | 1 | 13 | Ethe1 | 0.016 | Young |
| 0.737623 | 0.061198 | 0.623 | 0.721 | 1 | 13 | Cop1 | -0.098 | Young |
| 0.737672 | 0.072052 | 0.509 | 0.5 | 1 | 13 | Snrpa1 | 0.009 | Young |

| | | | | | | | | |
|---|---|---|---|---|---|---|---|---|
| 0.737893 | 0.187689 | 0.474 | 0.484 | 1 | 13 | Dhrs7b | -0.01 | Young |
| 0.737939 | 0.091358 | 0.632 | 0.664 | 1 | 13 | Snapc5 | -0.032 | Young |
| 0.738144 | 0.028624 | 0.395 | 0.393 | 1 | 13 | Tmem63b | 0.002 | Young |
| 0.738174 | 0.029027 | 0.798 | 0.803 | 1 | 13 | Usp8 | -0.005 | Young |
| 0.738201 | 0.184595 | 0.737 | 0.73 | 1 | 13 | Armc8 | 0.007 | Young |
| 0.738909 | 0.328027 | 0.272 | 0.279 | 1 | 13 | Lix1l | -0.007 | Young |
| 0.73897 | 0.066042 | 0.053 | 0.066 | 1 | 13 | BC048403 | -0.013 | Young |
| 0.739442 | 0.022915 | 0.342 | 0.336 | 1 | 13 | Eaf1 | 0.006 | Young |
| 0.739825 | 0.008803 | 0.596 | 0.566 | 1 | 13 | 4930455G( | 0.03 | Young |
| 0.740015 | 0.291984 | 0.333 | 0.328 | 1 | 13 | Snapc1 | 0.005 | Young |
| 0.74025 | 0.130623 | 0.263 | 0.262 | 1 | 13 | Erich1 | 0.001 | Young |
| 0.740296 | 0.062072 | 0.202 | 0.238 | 1 | 13 | 2010315B( | -0.036 | Young |
| 0.7406 | 0.018229 | 0.939 | 0.967 | 1 | 13 | Jmjd1c | -0.028 | Young |
| 0.740741 | 0.033767 | 0.333 | 0.393 | 1 | 13 | Enoph1 | -0.06 | Young |
| 0.741354 | 0.058488 | 0.447 | 0.443 | 1 | 13 | Agap3 | 0.004 | Young |
| 0.74194 | 0.061667 | 0.64 | 0.639 | 1 | 13 | Hipk3 | 0.001 | Young |
| 0.742029 | 0.081855 | 0.596 | 0.631 | 1 | 13 | Gpatch8 | -0.035 | Young |
| 0.74263 | 0.056105 | 0.061 | 0.074 | 1 | 13 | Polg2 | -0.013 | Young |
| 0.743058 | 0.103389 | 0.281 | 0.262 | 1 | 13 | Crbn | 0.019 | Young |
| 0.74315 | 0.081042 | 0.474 | 0.549 | 1 | 13 | Zwint | -0.075 | Young |
| 0.743561 | 0.109319 | 0.149 | 0.164 | 1 | 13 | Ankrd26 | -0.015 | Young |
| 0.743662 | 0.210626 | 0.246 | 0.246 | 1 | 13 | Abhd11 | 0 | Young |
| 0.744458 | 0.081077 | 0.605 | 0.656 | 1 | 13 | Mtmr3 | -0.051 | Young |
| 0.744617 | 0.028735 | 0.254 | 0.238 | 1 | 13 | Slc35d1 | 0.016 | Young |
| 0.745759 | 0.008337 | 0.561 | 0.557 | 1 | 13 | Sft2d1 | 0.004 | Young |
| 0.745785 | 0.008008 | 0.184 | 0.213 | 1 | 13 | Hace1 | -0.029 | Young |
| 0.746248 | 0.082122 | 0.64 | 0.664 | 1 | 13 | Sra1 | -0.024 | Young |
| 0.746298 | 0.061706 | 0.886 | 0.934 | 1 | 13 | Prpf40a | -0.048 | Young |
| 0.746385 | 0.014716 | 1 | 1 | 1 | 13 | Morf4l1 | 0 | Young |
| 0.746559 | 0.07932 | 0.36 | 0.393 | 1 | 13 | Klf16 | -0.033 | Young |
| 0.746674 | 0.096444 | 0.526 | 0.541 | 1 | 13 | Emsy | -0.015 | Young |
| 0.746914 | 0.056961 | 0.763 | 0.762 | 1 | 13 | Lrrfip1 | 0.001 | Young |
| 0.747007 | 0.106834 | 0.175 | 0.164 | 1 | 13 | Isoc2b | 0.011 | Young |
| 0.747038 | 0.197669 | 0.298 | 0.295 | 1 | 13 | Smad3 | 0.003 | Young |
| 0.747273 | 0.227209 | 0.14 | 0.131 | 1 | 13 | Hacd1 | 0.009 | Young |
| 0.7474 | 0.053833 | 0.333 | 0.377 | 1 | 13 | Commd10 | -0.044 | Young |
| 0.747654 | 0.235214 | 0.184 | 0.18 | 1 | 13 | Spryd3 | 0.004 | Young |
| 0.747708 | 0.140205 | 0.263 | 0.262 | 1 | 13 | Hectd3 | 0.001 | Young |
| 0.747961 | 0.202246 | 0.395 | 0.393 | 1 | 13 | 2310039H( | 0.002 | Young |
| 0.748206 | 0.230826 | 0.351 | 0.352 | 1 | 13 | Zfp397 | -0.001 | Young |
| 0.748835 | 0.276423 | 0.202 | 0.205 | 1 | 13 | Mcu | -0.003 | Young |
| 0.748856 | 0.195469 | 0.333 | 0.344 | 1 | 13 | Ttc19 | -0.011 | Young |
| 0.748878 | 0.280429 | 0.307 | 0.32 | 1 | 13 | Bap1 | -0.013 | Young |
| 0.749469 | 0.110904 | 0.105 | 0.123 | 1 | 13 | Fbxw17 | -0.018 | Young |
| 0.749684 | 0.382926 | 0.167 | 0.156 | 1 | 13 | Gtf2e1 | 0.011 | Young |

| | | | | | | | | |
|---|---|---|---|---|---|---|---|---|
| 0.74976 | 0.028732 | 0.193 | 0.221 | 1 | 13 | Ncdn | -0.028 | Young |
| 0.749999 | 0.073809 | 0.991 | 0.975 | 1 | 13 | Tm9sf3 | 0.016 | Young |
| 0.750258 | 0.076171 | 0.833 | 0.852 | 1 | 13 | Arhgef1 | -0.019 | Young |
| 0.750666 | 0.541685 | 0.096 | 0.09 | 1 | 13 | Gm20045 | 0.006 | Young |
| 0.751442 | 0.010958 | 0.956 | 0.975 | 1 | 13 | Irf2 | -0.019 | Young |
| 0.751658 | 0.024257 | 0.184 | 0.164 | 1 | 13 | Ecsit | 0.02 | Young |
| 0.752006 | 0.228867 | 0.202 | 0.197 | 1 | 13 | Tmem94 | 0.005 | Young |
| 0.752054 | 0.323777 | 0.114 | 0.107 | 1 | 13 | Ccdc181 | 0.007 | Young |
| 0.752171 | 0.006826 | 1 | 1 | 1 | 13 | Zeb2 | 0 | Young |
| 0.752208 | 0.165915 | 0.316 | 0.303 | 1 | 13 | Nfs1 | 0.013 | Young |
| 0.752208 | 0.181182 | 0.289 | 0.328 | 1 | 13 | Pmf1 | -0.039 | Young |
| 0.752291 | 0.109774 | 0.281 | 0.262 | 1 | 13 | Gosr1 | 0.019 | Young |
| 0.752292 | 0.116765 | 0.553 | 0.598 | 1 | 13 | Eps15l1 | -0.045 | Young |
| 0.752916 | 0.005049 | 0.167 | 0.189 | 1 | 13 | Elk1 | -0.022 | Young |
| 0.752988 | 0.195695 | 0.228 | 0.221 | 1 | 13 | Hook2 | 0.007 | Young |
| 0.75362 | 0.041431 | 1 | 1 | 1 | 13 | Erp29 | 0 | Young |
| 0.754347 | 0.000798 | 0.219 | 0.246 | 1 | 13 | Patz1 | -0.027 | Young |
| 0.754349 | 0.078999 | 0.798 | 0.836 | 1 | 13 | Polr2a | -0.038 | Young |
| 0.754515 | 0.00766 | 0.535 | 0.574 | 1 | 13 | Agpat5 | -0.039 | Young |
| 0.754737 | 0.086624 | 0.465 | 0.5 | 1 | 13 | Prdm1 | -0.035 | Young |
| 0.754737 | 0.064906 | 0.465 | 0.5 | 1 | 13 | Cog5 | -0.035 | Young |
| 0.754745 | 0.106745 | 0.781 | 0.934 | 1 | 13 | Nsd3 | -0.153 | Young |
| 0.755127 | 0.126615 | 0.579 | 0.59 | 1 | 13 | Abca3 | -0.011 | Young |
| 0.755245 | 0.049736 | 0.228 | 0.213 | 1 | 13 | Cisd3 | 0.015 | Young |
| 0.756444 | 0.020105 | 0.316 | 0.295 | 1 | 13 | Umad1 | 0.021 | Young |
| 0.756639 | 0.003153 | 0.333 | 0.352 | 1 | 13 | Pxmp4 | -0.019 | Young |
| 0.75667 | 0.105336 | 0.544 | 0.582 | 1 | 13 | Map1lc3a | -0.038 | Young |
| 0.756705 | 0.192225 | 0.36 | 0.352 | 1 | 13 | Gins4 | 0.008 | Young |
| 0.75692 | 0.152244 | 0.272 | 0.27 | 1 | 13 | Trip10 | 0.002 | Young |
| 0.757129 | 0.141239 | 0.298 | 0.303 | 1 | 13 | Cars2 | -0.005 | Young |
| 0.757245 | 0.132126 | 0.228 | 0.213 | 1 | 13 | Mast2 | 0.015 | Young |
| 0.75732 | 0.123725 | 0.693 | 0.73 | 1 | 13 | Chd8 | -0.037 | Young |
| 0.757371 | 0.067108 | 0.711 | 0.672 | 1 | 13 | Rc3h2 | 0.039 | Young |
| 0.757389 | 0.037137 | 0.623 | 0.623 | 1 | 13 | Tmem141 | 0 | Young |
| 0.757939 | 0.041378 | 0.219 | 0.197 | 1 | 13 | Cyb561d1 | 0.022 | Young |
| 0.757968 | 0.016111 | 0.974 | 0.967 | 1 | 13 | Atp5g1 | 0.007 | Young |
| 0.758246 | 0.061193 | 0.316 | 0.352 | 1 | 13 | Ankrd27 | -0.036 | Young |
| 0.758329 | 0.014258 | 0.43 | 0.475 | 1 | 13 | Rpl27-ps3 | -0.045 | Young |
| 0.758457 | 0.076473 | 0.851 | 0.893 | 1 | 13 | Denr | -0.042 | Young |
| 0.75873 | 0.148646 | 0.421 | 0.443 | 1 | 13 | Hps5 | -0.022 | Young |
| 0.759124 | 0.010165 | 0.658 | 0.697 | 1 | 13 | Exoc3 | -0.039 | Young |
| 0.75914 | 0.432865 | 0.132 | 0.123 | 1 | 13 | Abca6 | 0.009 | Young |
| 0.759239 | 0.240129 | 0.061 | 0.074 | 1 | 13 | Olfr920 | -0.013 | Young |
| 0.759665 | 0.067474 | 0.518 | 0.549 | 1 | 13 | Fam118b | -0.031 | Young |
| 0.759833 | 0.052834 | 0.816 | 0.77 | 1 | 13 | Iscu | 0.046 | Young |

| | | | | | | | | |
|---|---|---|---|---|---|---|---|---|
| 0.759918 | 0.058794 | 0.553 | 0.623 | 1 | 13 | Clip1 | -0.07 | Young |
| 0.760055 | 0.098383 | 0.096 | 0.115 | 1 | 13 | Gcat | -0.019 | Young |
| 0.760312 | 0.091571 | 0.175 | 0.164 | 1 | 13 | Npat | 0.011 | Young |
| 0.760358 | 0.031962 | 0.798 | 0.762 | 1 | 13 | Trip11 | 0.036 | Young |
| 0.761159 | 0.058871 | 0.754 | 0.811 | 1 | 13 | Ifnar2 | -0.057 | Young |
| 0.761251 | 0.029277 | 0.482 | 0.549 | 1 | 13 | Wdr43 | -0.067 | Young |
| 0.761534 | 0.214951 | 0.158 | 0.148 | 1 | 13 | Lmnb2 | 0.01 | Young |
| 0.761651 | 0.352931 | 0.132 | 0.123 | 1 | 13 | Pcgf1 | 0.009 | Young |
| 0.762038 | 0.165949 | 0.193 | 0.221 | 1 | 13 | Polr3b | -0.028 | Young |
| 0.762312 | 0.052469 | 0.412 | 0.402 | 1 | 13 | Fut11 | 0.01 | Young |
| 0.762371 | 0.23689 | 0.167 | 0.164 | 1 | 13 | Mrps22 | 0.003 | Young |
| 0.762559 | 0.154442 | 0.482 | 0.484 | 1 | 13 | Dctpp1 | -0.002 | Young |
| 0.762593 | 0.104727 | 0.105 | 0.123 | 1 | 13 | Cbr4 | -0.018 | Young |
| 0.762714 | 0.043432 | 0.851 | 0.861 | 1 | 13 | Hnrnpul1 | -0.01 | Young |
| 0.762924 | 0.243043 | 0.228 | 0.221 | 1 | 13 | Xpc | 0.007 | Young |
| 0.762975 | 0.029477 | 0.667 | 0.697 | 1 | 13 | Csnk2a1 | -0.03 | Young |
| 0.763125 | 0.204819 | 0.649 | 0.721 | 1 | 13 | Golgb1 | -0.072 | Young |
| 0.763362 | 0.210042 | 0.36 | 0.377 | 1 | 13 | Rbm34 | -0.017 | Young |
| 0.763594 | 0.072549 | 0.86 | 0.902 | 1 | 13 | Ssu72 | -0.042 | Young |
| 0.764062 | 0.047278 | 0.702 | 0.73 | 1 | 13 | 2-Mar | -0.028 | Young |
| 0.764413 | 0.009327 | 0.368 | 0.418 | 1 | 13 | Mapk8 | -0.05 | Young |
| 0.764772 | 0.081112 | 0.175 | 0.205 | 1 | 13 | Mcph1 | -0.03 | Young |
| 0.765226 | 0.042685 | 0.105 | 0.123 | 1 | 13 | Aurkb | -0.018 | Young |
| 0.765877 | 0.160815 | 0.281 | 0.279 | 1 | 13 | Nlrc4 | 0.002 | Young |
| 0.765914 | 0.259603 | 0.061 | 0.074 | 1 | 13 | Mdga1 | -0.013 | Young |
| 0.766697 | 0.015615 | 1 | 1 | 1 | 13 | Sec61b | 0 | Young |
| 0.768154 | 0.015105 | 1 | 1 | 1 | 13 | Hnrnpa0 | 0 | Young |
| 0.768338 | 0.153717 | 0.263 | 0.262 | 1 | 13 | Wdfy1 | 0.001 | Young |
| 0.768399 | 0.012925 | 0.298 | 0.336 | 1 | 13 | Mtmr2 | -0.038 | Young |
| 0.768551 | 0.123877 | 0.325 | 0.328 | 1 | 13 | Tmem159 | -0.003 | Young |
| 0.768872 | 0.104183 | 0.43 | 0.492 | 1 | 13 | Cyhr1 | -0.062 | Young |
| 0.769023 | 0.006444 | 0.175 | 0.205 | 1 | 13 | Cbfa2t2 | -0.03 | Young |
| 0.769551 | 0.029818 | 0.535 | 0.549 | 1 | 13 | Mesd | -0.014 | Young |
| 0.770071 | 0.112515 | 0.07 | 0.082 | 1 | 13 | Nmnat3 | -0.012 | Young |
| 0.770486 | 0.034723 | 0.211 | 0.246 | 1 | 13 | Srrd | -0.035 | Young |
| 0.770642 | 0.057367 | 0.833 | 0.852 | 1 | 13 | Higd1a | -0.019 | Young |
| 0.771733 | 0.215332 | 0.465 | 0.467 | 1 | 13 | Hyou1 | -0.002 | Young |
| 0.771929 | 0.04822 | 0.763 | 0.779 | 1 | 13 | Slc7a8 | -0.016 | Young |
| 0.771931 | 0.037395 | 0.298 | 0.336 | 1 | 13 | Usp40 | -0.038 | Young |
| 0.772168 | 0.04714 | 0.439 | 0.434 | 1 | 13 | Ulk1 | 0.005 | Young |
| 0.772399 | 0.224932 | 0.167 | 0.156 | 1 | 13 | Lppos | 0.011 | Young |
| 0.772453 | 0.003718 | 0.912 | 0.91 | 1 | 13 | Ensa | 0.002 | Young |
| 0.772559 | 0.226257 | 0.281 | 0.287 | 1 | 13 | Aasdhppt | -0.006 | Young |
| 0.772559 | 0.122447 | 0.263 | 0.303 | 1 | 13 | Hes6 | -0.04 | Young |
| 0.772673 | 0.017331 | 0.395 | 0.459 | 1 | 13 | Snx15 | -0.064 | Young |

| | | | | | | | | |
|---|---|---|---|---|---|---|---|---|
| 0.772736 | 0.159484 | 0.596 | 0.607 | 1 | 13 | Tes | -0.011 | Young |
| 0.772736 | 0.092782 | 0.596 | 0.607 | 1 | 13 | Kcnk6 | -0.011 | Young |
| 0.773855 | 0.001286 | 0.605 | 0.639 | 1 | 13 | Rnf115 | -0.034 | Young |
| 0.773973 | 0.070173 | 0.939 | 0.951 | 1 | 13 | Csk | -0.012 | Young |
| 0.774057 | 0.052219 | 0.737 | 0.795 | 1 | 13 | Nrd1 | -0.058 | Young |
| 0.774138 | 0.108668 | 0.14 | 0.131 | 1 | 13 | Map3k6 | 0.009 | Young |
| 0.774577 | 0.025745 | 0.43 | 0.508 | 1 | 13 | 3830406C1 | -0.078 | Young |
| 0.774606 | 0.135968 | 0.263 | 0.254 | 1 | 13 | Snx25 | 0.009 | Young |
| 0.774606 | 0.133341 | 0.263 | 0.254 | 1 | 13 | Qsox1 | 0.009 | Young |
| 0.774718 | 0.136658 | 0.702 | 0.754 | 1 | 13 | Smim7 | -0.052 | Young |
| 0.775811 | 0.043054 | 0.772 | 0.795 | 1 | 13 | Arid5a | -0.023 | Young |
| 0.776769 | 0.211989 | 0.132 | 0.123 | 1 | 13 | Zfp629 | 0.009 | Young |
| 0.776899 | 0.05751 | 0.211 | 0.238 | 1 | 13 | Rbms2 | -0.027 | Young |
| 0.777006 | 0.210772 | 0.43 | 0.443 | 1 | 13 | Cdk5rap3 | -0.013 | Young |
| 0.777006 | 0.065845 | 0.404 | 0.467 | 1 | 13 | Trpv2 | -0.063 | Young |
| 0.777112 | 0.07473 | 0.702 | 0.713 | 1 | 13 | Zzef1 | -0.011 | Young |
| 0.777624 | 0.021481 | 0.026 | 0.033 | 1 | 13 | Zfp202 | -0.007 | Young |
| 0.778161 | 0.056703 | 0.158 | 0.18 | 1 | 13 | Cmc4 | -0.022 | Young |
| 0.778199 | 0.027722 | 0.798 | 0.795 | 1 | 13 | Rab35 | 0.003 | Young |
| 0.778585 | 0.120743 | 0.123 | 0.139 | 1 | 13 | Gcfc2 | -0.016 | Young |
| 0.779149 | 0.183196 | 0.184 | 0.213 | 1 | 13 | Cactin | -0.029 | Young |
| 0.779297 | 0.104558 | 0.132 | 0.123 | 1 | 13 | Zfp959 | 0.009 | Young |
| 0.779491 | 0.193258 | 0.377 | 0.402 | 1 | 13 | Tm2d3 | -0.025 | Young |
| 0.779529 | 0.480239 | 0.096 | 0.09 | 1 | 13 | L3mbtl3 | 0.006 | Young |
| 0.779529 | 0.263604 | 0.096 | 0.09 | 1 | 13 | Neurl1b | 0.006 | Young |
| 0.779573 | 0.033696 | 0.596 | 0.582 | 1 | 13 | Plod1 | 0.014 | Young |
| 0.779597 | 0.029865 | 0.07 | 0.082 | 1 | 13 | Zc3h12d | -0.012 | Young |
| 0.779622 | 0.018209 | 0.711 | 0.787 | 1 | 13 | Ndufs4 | -0.076 | Young |
| 0.779837 | 0.062108 | 0.982 | 0.951 | 1 | 13 | Ikbkb | 0.031 | Young |
| 0.779911 | 0.244763 | 0.289 | 0.279 | 1 | 13 | Med4 | 0.01 | Young |
| 0.779913 | 0.116566 | 0.219 | 0.246 | 1 | 13 | Scap | -0.027 | Young |
| 0.780004 | 0.041391 | 0.465 | 0.459 | 1 | 13 | Faf2 | 0.006 | Young |
| 0.780193 | 0.030051 | 0.158 | 0.148 | 1 | 13 | Zfyve19 | 0.01 | Young |
| 0.780422 | 0.290944 | 0.167 | 0.164 | 1 | 13 | Actr5 | 0.003 | Young |
| 0.780517 | 0.033259 | 0.605 | 0.607 | 1 | 13 | Plekhg3 | -0.002 | Young |
| 0.781249 | 0.206072 | 0.202 | 0.197 | 1 | 13 | Cmss1 | 0.005 | Young |
| 0.781513 | 0.260568 | 0.14 | 0.131 | 1 | 13 | Pop1 | 0.009 | Young |
| 0.781855 | 0.167713 | 0.123 | 0.115 | 1 | 13 | Cox7a1 | 0.008 | Young |
| 0.782151 | 0.126207 | 0.079 | 0.09 | 1 | 13 | Steap4 | -0.011 | Young |
| 0.782394 | 0.065142 | 0.825 | 0.869 | 1 | 13 | Cdk9 | -0.044 | Young |
| 0.782433 | 0.323306 | 0.096 | 0.09 | 1 | 13 | Nphp3 | 0.006 | Young |
| 0.782506 | 0.089662 | 0.447 | 0.492 | 1 | 13 | Ndnf | -0.045 | Young |
| 0.782523 | 0.093599 | 0.114 | 0.131 | 1 | 13 | Spats2 | -0.017 | Young |
| 0.782606 | 0.10154 | 0.026 | 0.033 | 1 | 13 | Lepr | -0.007 | Young |
| 0.782606 | 0.010116 | 0.026 | 0.033 | 1 | 13 | Fam221b | -0.007 | Young |

| | | | | | | | | |
|---|---|---|---|---|---|---|---|---|
| 0.782633 | 0.080134 | 0.193 | 0.221 | 1 | 13 | Adck1 | -0.028 | Young |
| 0.782639 | 0.022835 | 0.351 | 0.393 | 1 | 13 | Jade1 | -0.042 | Young |
| 0.78335 | 0.147284 | 0.202 | 0.197 | 1 | 13 | Rnf113a2 | 0.005 | Young |
| 0.783527 | 0.035356 | 0.447 | 0.467 | 1 | 13 | Nup54 | -0.02 | Young |
| 0.783631 | 0.046138 | 0.605 | 0.557 | 1 | 13 | Snx29 | 0.048 | Young |
| 0.783738 | 0.09741 | 0.447 | 0.5 | 1 | 13 | Prrc1 | -0.053 | Young |
| 0.783886 | 0.038021 | 0.281 | 0.303 | 1 | 13 | Tfam | -0.022 | Young |
| 0.784102 | 0.003653 | 0.316 | 0.287 | 1 | 13 | Btbd10 | 0.029 | Young |
| 0.784116 | 0.061848 | 0.263 | 0.254 | 1 | 13 | Hinfp | 0.009 | Young |
| 0.784206 | 0.027407 | 0.491 | 0.484 | 1 | 13 | Rab3gap1 | 0.007 | Young |
| 0.784328 | 0.086584 | 0.368 | 0.377 | 1 | 13 | Ranbp3 | -0.009 | Young |
| 0.784465 | 0.241753 | 0.123 | 0.115 | 1 | 13 | Lin52 | 0.008 | Young |
| 0.784933 | 0.013252 | 0.553 | 0.533 | 1 | 13 | Rbmxl1 | 0.02 | Young |
| 0.785005 | 0.044325 | 0.351 | 0.344 | 1 | 13 | Taf12 | 0.007 | Young |
| 0.785066 | 0.006944 | 0.386 | 0.434 | 1 | 13 | Nol12 | -0.048 | Young |
| 0.785119 | 0.12976 | 0.421 | 0.426 | 1 | 13 | Bex3 | -0.005 | Young |
| 0.78551 | 0.008867 | 0.877 | 0.877 | 1 | 13 | Lrmp | 0 | Young |
| 0.785649 | 0.008672 | 0.904 | 0.934 | 1 | 13 | Rab11a | -0.03 | Young |
| 0.785774 | 0.07836 | 0.693 | 0.705 | 1 | 13 | Pfdn6 | -0.012 | Young |
| 0.785966 | 0.267535 | 0.07 | 0.082 | 1 | 13 | Map3k12 | -0.012 | Young |
| 0.785967 | 0.154598 | 0.509 | 0.541 | 1 | 13 | Slc31a1 | -0.032 | Young |
| 0.786306 | 0.026827 | 0.526 | 0.574 | 1 | 13 | Tmem131l | -0.048 | Young |
| 0.786456 | 0.053581 | 0.658 | 0.689 | 1 | 13 | St6gal1 | -0.031 | Young |
| 0.786849 | 0.15769 | 0.202 | 0.189 | 1 | 13 | Supt7l | 0.013 | Young |
| 0.786997 | 0.184776 | 0.316 | 0.328 | 1 | 13 | Zfp518a | -0.012 | Young |
| 0.787372 | 0.130269 | 0.386 | 0.377 | 1 | 13 | Med16 | 0.009 | Young |
| 0.787924 | 0.088971 | 0.167 | 0.189 | 1 | 13 | Exoc8 | -0.022 | Young |
| 0.78816 | 0.044124 | 0.149 | 0.139 | 1 | 13 | Pqlc2 | 0.01 | Young |
| 0.78825 | 0.239143 | 0.096 | 0.09 | 1 | 13 | Mcat | 0.006 | Young |
| 0.788634 | 0.037551 | 0.956 | 0.967 | 1 | 13 | Rbx1 | -0.011 | Young |
| 0.78884 | 0.052398 | 0.412 | 0.393 | 1 | 13 | Spns1 | 0.019 | Young |
| 0.789042 | 0.067727 | 0.105 | 0.123 | 1 | 13 | Det1 | -0.018 | Young |
| 0.789838 | 0.231735 | 0.237 | 0.279 | 1 | 13 | Mcm4 | -0.042 | Young |
| 0.790379 | 0.399362 | 0.193 | 0.189 | 1 | 13 | Fam46c | 0.004 | Young |
| 0.791094 | 0.173875 | 0.272 | 0.303 | 1 | 13 | Slc10a7 | -0.031 | Young |
| 0.791412 | 0.141553 | 0.439 | 0.459 | 1 | 13 | Dmtf1 | -0.02 | Young |
| 0.79151 | 0.107589 | 0.623 | 0.672 | 1 | 13 | Sav1 | -0.049 | Young |
| 0.79177 | 0.161734 | 0.202 | 0.197 | 1 | 13 | Ints4 | 0.005 | Young |
| 0.791933 | 0.027861 | 0.728 | 0.746 | 1 | 13 | Akirin1 | -0.018 | Young |
| 0.792009 | 0.031426 | 0.219 | 0.213 | 1 | 13 | Ints7 | 0.006 | Young |
| 0.792313 | 0.019363 | 0.974 | 0.992 | 1 | 13 | Selenof | -0.018 | Young |
| 0.792597 | 0.361164 | 0.026 | 0.033 | 1 | 13 | Haspin | -0.007 | Young |
| 0.792597 | 0.154143 | 0.026 | 0.033 | 1 | 13 | Zfpm1 | -0.007 | Young |
| 0.792821 | 0.053448 | 0.114 | 0.131 | 1 | 13 | Mterf4 | -0.017 | Young |
| 0.793043 | 0.019194 | 0.947 | 0.975 | 1 | 13 | Nptn | -0.028 | Young |

| | | | | | | | | |
|---|---|---|---|---|---|---|---|---|
| 0.793296 | 0.058703 | 0.579 | 0.549 | 1 | 13 | Hsd17b10 | 0.03 | Young |
| 0.793414 | 0.032461 | 0.5 | 0.541 | 1 | 13 | Auh | -0.041 | Young |
| 0.793429 | 0.161673 | 0.526 | 0.557 | 1 | 13 | Utp11 | -0.031 | Young |
| 0.793879 | 0.225801 | 0.202 | 0.197 | 1 | 13 | Flt3l | 0.005 | Young |
| 0.79406 | 0.070312 | 0.518 | 0.541 | 1 | 13 | Cd2 | -0.023 | Young |
| 0.794391 | 0.048725 | 0.623 | 0.631 | 1 | 13 | Phf5a | -0.008 | Young |
| 0.794773 | 0.182903 | 0.289 | 0.287 | 1 | 13 | Zfp362 | 0.002 | Young |
| 0.794969 | 0.13759 | 0.325 | 0.32 | 1 | 13 | Ocel1 | 0.005 | Young |
| 0.795064 | 0.135553 | 0.211 | 0.205 | 1 | 13 | Rbmx2 | 0.006 | Young |
| 0.795616 | 0.114703 | 0.465 | 0.467 | 1 | 13 | Socs7 | -0.002 | Young |
| 0.795742 | 0.027015 | 0.412 | 0.434 | 1 | 13 | Rtl5 | -0.022 | Young |
| 0.795782 | 0.35558 | 0.43 | 0.459 | 1 | 13 | Ubxn2a | -0.029 | Young |
| 0.796366 | 0.103102 | 0.553 | 0.598 | 1 | 13 | Anapc2 | -0.045 | Young |
| 0.796444 | 0.016025 | 0.228 | 0.254 | 1 | 13 | Dvl2 | -0.026 | Young |
| 0.796534 | 0.068126 | 0.518 | 0.525 | 1 | 13 | Dhps | -0.007 | Young |
| 0.796693 | 0.008264 | 0.316 | 0.336 | 1 | 13 | Terf2 | -0.02 | Young |
| 0.796702 | 0.15882 | 0.5 | 0.566 | 1 | 13 | Thra | -0.066 | Young |
| 0.796746 | 0.072862 | 0.184 | 0.172 | 1 | 13 | Synpo | 0.012 | Young |
| 0.797113 | 0.0448 | 0.842 | 0.852 | 1 | 13 | Elf1 | -0.01 | Young |
| 0.797146 | 0.474527 | 0.079 | 0.074 | 1 | 13 | Cenps | 0.005 | Young |
| 0.797753 | 0.33985 | 0.114 | 0.107 | 1 | 13 | Zfp1 | 0.007 | Young |
| 0.798388 | 0.215751 | 0.228 | 0.23 | 1 | 13 | Pgam5 | -0.002 | Young |
| 0.798429 | 0.045369 | 0.491 | 0.484 | 1 | 13 | Cdkn2d | 0.007 | Young |
| 0.798869 | 0.042551 | 0.895 | 0.926 | 1 | 13 | Tnrc18 | -0.031 | Young |
| 0.798956 | 0.043822 | 0.167 | 0.189 | 1 | 13 | Mllt1 | -0.022 | Young |
| 0.799631 | 0.14442 | 0.491 | 0.475 | 1 | 13 | Hlx | 0.016 | Young |
| 0.799974 | 0.072607 | 0.746 | 0.795 | 1 | 13 | Pip5k1c | -0.049 | Young |
| 0.800169 | 0.013731 | 0.877 | 0.852 | 1 | 13 | Map2k3 | 0.025 | Young |
| 0.800355 | 0.155921 | 0.281 | 0.303 | 1 | 13 | Vhl | -0.022 | Young |
| 0.80044 | 0.038369 | 0.386 | 0.443 | 1 | 13 | Tssc4 | -0.057 | Young |
| 0.800465 | 0.362663 | 0.114 | 0.107 | 1 | 13 | Acsf3 | 0.007 | Young |
| 0.800568 | 0.127376 | 0.114 | 0.131 | 1 | 13 | Oaf | -0.017 | Young |
| 0.800657 | 0.155728 | 0.412 | 0.434 | 1 | 13 | Pfkfb4 | -0.022 | Young |
| 0.800885 | 0.104299 | 0.518 | 0.557 | 1 | 13 | Il11ra1 | -0.039 | Young |
| 0.801242 | 0.074698 | 0.579 | 0.557 | 1 | 13 | Zfp622 | 0.022 | Young |
| 0.801275 | 0.21497 | 0.14 | 0.131 | 1 | 13 | Zhx3 | 0.009 | Young |
| 0.801298 | 0.028535 | 0.825 | 0.811 | 1 | 13 | Pfdn2 | 0.014 | Young |
| 0.8013 | 0.237161 | 0.211 | 0.205 | 1 | 13 | Anks3 | 0.006 | Young |
| 0.801549 | 0.088648 | 0.579 | 0.623 | 1 | 13 | Eml4 | -0.044 | Young |
| 0.801772 | 0.084755 | 0.886 | 0.902 | 1 | 13 | Dusp11 | -0.016 | Young |
| 0.801816 | 0.046718 | 0.325 | 0.344 | 1 | 13 | Kpna6 | -0.019 | Young |
| 0.801887 | 0.024244 | 1 | 1 | 1 | 13 | Cyba | 0 | Young |
| 0.802133 | 0.019996 | 0.307 | 0.328 | 1 | 13 | Coasy | -0.021 | Young |
| 0.802329 | 0.277186 | 0.202 | 0.197 | 1 | 13 | Traf3ip3 | 0.005 | Young |
| 0.802622 | 0.175451 | 0.026 | 0.033 | 1 | 13 | Gm38394 | -0.007 | Young |

| | | | | | | | | |
|---|---|---|---|---|---|---|---|---|
| 0.802622 | 0.086758 | 0.026 | 0.033 | 1 | 13 | Slc22a5 | -0.007 | Young |
| 0.802623 | 0.121246 | 0.149 | 0.139 | 1 | 13 | Rfx5 | 0.01 | Young |
| 0.802783 | 0.00606 | 0.509 | 0.533 | 1 | 13 | Acads | -0.024 | Young |
| 0.803027 | 0.058231 | 0.579 | 0.566 | 1 | 13 | Tiprl | 0.013 | Young |
| 0.803152 | 0.094262 | 0.167 | 0.164 | 1 | 13 | Cdc42bpg | 0.003 | Young |
| 0.803179 | 0.215184 | 0.114 | 0.107 | 1 | 13 | Chst15 | 0.007 | Young |
| 0.803465 | 0.19266 | 0.079 | 0.09 | 1 | 13 | Gm26518 | -0.011 | Young |
| 0.803465 | 0.067641 | 0.079 | 0.09 | 1 | 13 | Gm31718 | -0.011 | Young |
| 0.803477 | 0.068366 | 0.482 | 0.516 | 1 | 13 | Arhgap12 | -0.034 | Young |
| 0.803553 | 0.024711 | 0.491 | 0.492 | 1 | 13 | Cyp20a1 | -0.001 | Young |
| 0.803555 | 0.295901 | 0.158 | 0.156 | 1 | 13 | Kdm1b | 0.002 | Young |
| 0.803626 | 0.035815 | 0.474 | 0.475 | 1 | 13 | Xpa | -0.001 | Young |
| 0.803852 | 0.04056 | 0.202 | 0.221 | 1 | 13 | Polr3f | -0.019 | Young |
| 0.804176 | 0.129493 | 0.561 | 0.623 | 1 | 13 | Atf7 | -0.062 | Young |
| 0.804666 | 0.010632 | 0.781 | 0.795 | 1 | 13 | Banf1 | -0.014 | Young |
| 0.804702 | 0.02249 | 0.132 | 0.123 | 1 | 13 | Pced1a | 0.009 | Young |
| 0.804828 | 0.18185 | 0.377 | 0.393 | 1 | 13 | Senp7 | -0.016 | Young |
| 0.804836 | 0.205847 | 0.175 | 0.172 | 1 | 13 | Ptdss2 | 0.003 | Young |
| 0.804839 | 0.013901 | 1 | 1 | 1 | 13 | Rpl37 | 0 | Young |
| 0.805895 | 0.085277 | 0.114 | 0.107 | 1 | 13 | Cnnm2 | 0.007 | Young |
| 0.806014 | 0.054283 | 0.202 | 0.189 | 1 | 13 | Fam151b | 0.013 | Young |
| 0.806339 | 0.111488 | 0.202 | 0.23 | 1 | 13 | Zdhhc16 | -0.028 | Young |
| 0.806523 | 0.024346 | 0.754 | 0.77 | 1 | 13 | Pnpla8 | -0.016 | Young |
| 0.80679 | 0.149144 | 0.298 | 0.295 | 1 | 13 | Spsb2 | 0.003 | Young |
| 0.806822 | 0.102985 | 0.693 | 0.664 | 1 | 13 | Ankrd13c | 0.029 | Young |
| 0.806902 | 0.063625 | 0.544 | 0.574 | 1 | 13 | Unc50 | -0.03 | Young |
| 0.807048 | 0.010518 | 0.974 | 0.943 | 1 | 13 | S100a11 | 0.031 | Young |
| 0.807111 | 0.159981 | 0.386 | 0.418 | 1 | 13 | Gm17018 | -0.032 | Young |
| 0.80743 | 0.04562 | 0.825 | 0.861 | 1 | 13 | Washc2 | -0.036 | Young |
| 0.807621 | 0.013926 | 0.298 | 0.32 | 1 | 13 | Fgfr1op | -0.022 | Young |
| 0.807704 | 0.087839 | 0.272 | 0.311 | 1 | 13 | N6amt1 | -0.039 | Young |
| 0.807794 | 0.029388 | 1 | 0.992 | 1 | 13 | Atox1 | 0.008 | Young |
| 0.808418 | 0.30394 | 0.228 | 0.23 | 1 | 13 | Sppl2b | -0.002 | Young |
| 0.808722 | 0.15814 | 0.132 | 0.148 | 1 | 13 | F2rl2 | -0.016 | Young |
| 0.808722 | 0.050624 | 0.132 | 0.148 | 1 | 13 | Tti2 | -0.016 | Young |
| 0.808868 | 0.047981 | 0.412 | 0.434 | 1 | 13 | Actr1b | -0.022 | Young |
| 0.809536 | 0.2292 | 0.289 | 0.303 | 1 | 13 | Eef2kmt | -0.014 | Young |
| 0.809582 | 0.045639 | 0.079 | 0.09 | 1 | 13 | Dpy19l3 | -0.011 | Young |
| 0.809587 | 0.004124 | 0.333 | 0.344 | 1 | 13 | Mapk8ip3 | -0.011 | Young |
| 0.809955 | 0.090792 | 0.904 | 0.926 | 1 | 13 | Pigt | -0.022 | Young |
| 0.810227 | 0.061406 | 0.175 | 0.197 | 1 | 13 | Ston2 | -0.022 | Young |
| 0.810228 | 0.062432 | 0.211 | 0.197 | 1 | 13 | Gm4951 | 0.014 | Young |
| 0.810331 | 0.039001 | 0.816 | 0.852 | 1 | 13 | Eea1 | -0.036 | Young |
| 0.810428 | 0.263624 | 0.228 | 0.23 | 1 | 13 | Golga1 | -0.002 | Young |
| 0.810445 | 0.034903 | 0.395 | 0.41 | 1 | 13 | Psmg3 | -0.015 | Young |

| | | | | | | | | |
|---|---|---|---|---|---|---|---|---|
| 0.810446 | 0.193598 | 0.219 | 0.213 | 1 | 13 | Smpd4 | 0.006 | Young |
| 0.810462 | 0.127641 | 0.386 | 0.393 | 1 | 13 | Pigyl | -0.007 | Young |
| 0.810695 | 0.227819 | 0.123 | 0.115 | 1 | 13 | Rgl3 | 0.008 | Young |
| 0.810695 | 0.158527 | 0.184 | 0.18 | 1 | 13 | Whamm | 0.004 | Young |
| 0.810736 | 0.055959 | 0.921 | 0.967 | 1 | 13 | Rab2a | -0.046 | Young |
| 0.810751 | 0.044597 | 0.991 | 0.984 | 1 | 13 | Eps8 | 0.007 | Young |
| 0.810866 | 0.051502 | 0.325 | 0.328 | 1 | 13 | Asna1 | -0.003 | Young |
| 0.811025 | 0.132643 | 0.386 | 0.402 | 1 | 13 | Specc1 | -0.016 | Young |
| 0.811103 | 0.273547 | 0.421 | 0.451 | 1 | 13 | Exosc8 | -0.03 | Young |
| 0.811299 | 0.082083 | 0.342 | 0.377 | 1 | 13 | Acot8 | -0.035 | Young |
| 0.811403 | 0.492046 | 0.07 | 0.066 | 1 | 13 | Irgc1 | 0.004 | Young |
| 0.811484 | 0.039389 | 0.956 | 0.959 | 1 | 13 | Scamp2 | -0.003 | Young |
| 0.811491 | 0.013156 | 0.991 | 0.992 | 1 | 13 | Trf | -0.001 | Young |
| 0.811582 | 0.080264 | 0.07 | 0.082 | 1 | 13 | Zfp692 | -0.012 | Young |
| 0.811953 | 0.15033 | 0.237 | 0.238 | 1 | 13 | Brpf3 | -0.001 | Young |
| 0.812112 | 0.021261 | 0.719 | 0.762 | 1 | 13 | Hipk2 | -0.043 | Young |
| 0.812231 | 0.002708 | 1 | 0.984 | 1 | 13 | Snx3 | 0.016 | Young |
| 0.812332 | 0.069662 | 0.193 | 0.213 | 1 | 13 | Tex30 | -0.02 | Young |
| 0.812645 | 0.028573 | 0.079 | 0.09 | 1 | 13 | Ccdc137 | -0.011 | Young |
| 0.81304 | 0.067276 | 0.246 | 0.287 | 1 | 13 | Cnnm4 | -0.041 | Young |
| 0.813162 | 0.037579 | 0.316 | 0.287 | 1 | 13 | Fkbp5 | 0.029 | Young |
| 0.813504 | 0.103166 | 0.86 | 0.877 | 1 | 13 | Polr2f | -0.017 | Young |
| 0.813662 | 0.022208 | 0.351 | 0.377 | 1 | 13 | Ddx49 | -0.026 | Young |
| 0.813704 | 0.005964 | 0.219 | 0.246 | 1 | 13 | AU020206 | -0.027 | Young |
| 0.813732 | 0.022437 | 0.711 | 0.713 | 1 | 13 | Lsm14a | -0.002 | Young |
| 0.813829 | 0.124014 | 0.351 | 0.426 | 1 | 13 | Tmem63a | -0.075 | Young |
| 0.814178 | 0.428345 | 0.088 | 0.082 | 1 | 13 | Smyd4 | 0.006 | Young |
| 0.814443 | 0.135632 | 0.474 | 0.467 | 1 | 13 | Pgp | 0.007 | Young |
| 0.814558 | 0.242517 | 0.219 | 0.213 | 1 | 13 | Cep85 | 0.006 | Young |
| 0.814581 | 0.056383 | 0.158 | 0.172 | 1 | 13 | Tfcp2 | -0.014 | Young |
| 0.814602 | 0.002038 | 0.807 | 0.82 | 1 | 13 | Rin2 | -0.013 | Young |
| 0.8156 | 0.036364 | 0.711 | 0.746 | 1 | 13 | Snx4 | -0.035 | Young |
| 0.81585 | 0.018429 | 0.035 | 0.041 | 1 | 13 | Gm15050 | -0.006 | Young |
| 0.816036 | 0.133749 | 0.088 | 0.098 | 1 | 13 | Alg12 | -0.01 | Young |
| 0.816075 | 0.06177 | 0.351 | 0.41 | 1 | 13 | Pias4 | -0.059 | Young |
| 0.816085 | 0.137414 | 0.149 | 0.172 | 1 | 13 | Tmem9 | -0.023 | Young |
| 0.816408 | 0.313836 | 0.079 | 0.074 | 1 | 13 | Cbarp | 0.005 | Young |
| 0.81647 | 0.085211 | 0.158 | 0.18 | 1 | 13 | Mad2l2 | -0.022 | Young |
| 0.817303 | 0.083722 | 0.333 | 0.352 | 1 | 13 | Xab2 | -0.019 | Young |
| 0.817485 | 0.141107 | 0.132 | 0.123 | 1 | 13 | Nme7 | 0.009 | Young |
| 0.817492 | 0.136913 | 0.57 | 0.541 | 1 | 13 | Ptgs2 | 0.029 | Young |
| 0.817503 | 0.026096 | 0.553 | 0.533 | 1 | 13 | Nop56 | 0.02 | Young |
| 0.817598 | 0.22282 | 0.202 | 0.205 | 1 | 13 | Unc13d | -0.003 | Young |
| 0.817651 | 0.026364 | 0.825 | 0.82 | 1 | 13 | Msrb1 | 0.005 | Young |
| 0.817736 | 0.072413 | 0.605 | 0.639 | 1 | 13 | Gpr35 | -0.034 | Young |

| | | | | | | | | |
|---|---|---|---|---|---|---|---|---|
| 0.818146 | 0.057437 | 0.14 | 0.156 | 1 | 13 | Clcf1 | -0.016 | Young |
| 0.818758 | 0.065856 | 0.298 | 0.287 | 1 | 13 | Nup85 | 0.011 | Young |
| 0.81878 | 0.113913 | 0.079 | 0.09 | 1 | 13 | Cnbd2 | -0.011 | Young |
| 0.819104 | 0.120807 | 0.412 | 0.426 | 1 | 13 | Anapc15 | -0.014 | Young |
| 0.819482 | 0.006886 | 0.246 | 0.262 | 1 | 13 | Med9 | -0.016 | Young |
| 0.819582 | 0.097573 | 0.149 | 0.139 | 1 | 13 | Mapk1ip1 | 0.01 | Young |
| 0.819582 | 0.084199 | 0.149 | 0.139 | 1 | 13 | Acvr2a | 0.01 | Young |
| 0.819929 | 0.333828 | 0.193 | 0.197 | 1 | 13 | Usp46 | -0.004 | Young |
| 0.820081 | 0.135527 | 0.193 | 0.221 | 1 | 13 | Nudt14 | -0.028 | Young |
| 0.820288 | 0.03033 | 0.904 | 0.893 | 1 | 13 | Gtf2h5 | 0.011 | Young |
| 0.820316 | 0.060524 | 0.088 | 0.082 | 1 | 13 | Atad5 | 0.006 | Young |
| 0.820383 | 0.288273 | 0.342 | 0.361 | 1 | 13 | Aamdc | -0.019 | Young |
| 0.820445 | 0.151133 | 0.096 | 0.09 | 1 | 13 | Timp1 | 0.006 | Young |
| 0.820551 | 0.076925 | 0.807 | 0.82 | 1 | 13 | Eloa | -0.013 | Young |
| 0.820683 | 0.225657 | 0.281 | 0.287 | 1 | 13 | Uba7 | -0.006 | Young |
| 0.821103 | 0.526773 | 0.114 | 0.115 | 1 | 13 | Pcgf6 | -0.001 | Young |
| 0.821604 | 0.429175 | 0.07 | 0.066 | 1 | 13 | Gm10762 | 0.004 | Young |
| 0.822607 | 0.021628 | 0.991 | 1 | 1 | 13 | Rpl31 | -0.009 | Young |
| 0.822854 | 0.354422 | 0.079 | 0.074 | 1 | 13 | Zfp606 | 0.005 | Young |
| 0.822992 | 0.435602 | 0.061 | 0.057 | 1 | 13 | Hdhd5 | 0.004 | Young |
| 0.823073 | 0.011297 | 0.553 | 0.59 | 1 | 13 | Utp14a | -0.037 | Young |
| 0.823148 | 0.078339 | 0.184 | 0.205 | 1 | 13 | Narfl | -0.021 | Young |
| 0.823448 | 0.019961 | 0.789 | 0.82 | 1 | 13 | Cisd2 | -0.031 | Young |
| 0.823625 | 0.00034 | 0.658 | 0.713 | 1 | 13 | Paics | -0.055 | Young |
| 0.823744 | 0.112877 | 0.518 | 0.566 | 1 | 13 | Meaf6 | -0.048 | Young |
| 0.823804 | 0.123794 | 0.123 | 0.139 | 1 | 13 | Alg3 | -0.016 | Young |
| 0.823925 | 0.41114 | 0.123 | 0.123 | 1 | 13 | Nrg2 | 0 | Young |
| 0.824302 | 0.008422 | 0.596 | 0.598 | 1 | 13 | Ppp2r2d | -0.002 | Young |
| 0.824443 | 0.169625 | 0.149 | 0.139 | 1 | 13 | Zfp934 | 0.01 | Young |
| 0.824792 | 0.383164 | 0.035 | 0.041 | 1 | 13 | Trim68 | -0.006 | Young |
| 0.8248 | 0.026611 | 0.667 | 0.738 | 1 | 13 | Chd3 | -0.071 | Young |
| 0.825348 | 0.257054 | 0.149 | 0.148 | 1 | 13 | Actr6 | 0.001 | Young |
| 0.825401 | 0.029106 | 0.246 | 0.279 | 1 | 13 | Nedd1 | -0.033 | Young |
| 0.825432 | 0.011261 | 0.316 | 0.32 | 1 | 13 | Pcgf3 | -0.004 | Young |
| 0.825808 | 0.12074 | 0.105 | 0.098 | 1 | 13 | Mars2 | 0.007 | Young |
| 0.825988 | 0.039491 | 0.64 | 0.639 | 1 | 13 | Blmh | 0.001 | Young |
| 0.826333 | 0.284781 | 0.096 | 0.09 | 1 | 13 | Setd6 | 0.006 | Young |
| 0.82637 | 0.346102 | 0.193 | 0.197 | 1 | 13 | Eif2d | -0.004 | Young |
| 0.826377 | 0.08412 | 0.272 | 0.27 | 1 | 13 | Zdhhc4 | 0.002 | Young |
| 0.826924 | 0.06889 | 0.219 | 0.213 | 1 | 13 | Ndc1 | 0.006 | Young |
| 0.826964 | 0.058317 | 0.149 | 0.164 | 1 | 13 | Tgm2 | -0.015 | Young |
| 0.82698 | 0.122215 | 0.298 | 0.295 | 1 | 13 | Aimp2 | 0.003 | Young |
| 0.827321 | 0.177558 | 0.263 | 0.262 | 1 | 13 | Cnpy4 | 0.001 | Young |
| 0.827445 | 0.087439 | 0.202 | 0.189 | 1 | 13 | Snta1 | 0.013 | Young |
| 0.827708 | 0.109617 | 0.114 | 0.107 | 1 | 13 | Lamc1 | 0.007 | Young |

| | | | | | | | | |
|---|---|---|---|---|---|---|---|---|
| 0.828004 | 0.157164 | 0.456 | 0.5 | 1 | 13 | Ctnnbip1 | -0.044 | Young |
| 0.828642 | 0.353136 | 0.105 | 0.098 | 1 | 13 | Tmem206 | 0.007 | Young |
| 0.828861 | 0.077506 | 0.228 | 0.246 | 1 | 13 | Amz1 | -0.018 | Young |
| 0.829091 | 0.176598 | 0.351 | 0.352 | 1 | 13 | Haghl | -0.001 | Young |
| 0.829196 | 0.0275 | 0.289 | 0.336 | 1 | 13 | Mtif2 | -0.047 | Young |
| 0.829272 | 0.001706 | 0.035 | 0.041 | 1 | 13 | Tlcd1 | -0.006 | Young |
| 0.829312 | 0.361687 | 0.079 | 0.074 | 1 | 13 | Fastkd5 | 0.005 | Young |
| 0.829317 | 0.060537 | 0.64 | 0.656 | 1 | 13 | Psme3 | -0.016 | Young |
| 0.829582 | 0.010479 | 0.325 | 0.361 | 1 | 13 | Nup205 | -0.036 | Young |
| 0.82988 | 0.018035 | 0.193 | 0.18 | 1 | 13 | Creld1 | 0.013 | Young |
| 0.830007 | 0.064568 | 0.167 | 0.156 | 1 | 13 | Phtf1 | 0.011 | Young |
| 0.830017 | 0.2265 | 0.342 | 0.385 | 1 | 13 | Fam126b | -0.043 | Young |
| 0.830038 | 0.048576 | 1 | 1 | 1 | 13 | Arpc3 | 0 | Young |
| 0.830454 | 0.170526 | 0.307 | 0.344 | 1 | 13 | Sdhaf1 | -0.037 | Young |
| 0.830532 | 0.053532 | 0.465 | 0.475 | 1 | 13 | Gtf3a | -0.01 | Young |
| 0.830683 | 0.053888 | 0.763 | 0.803 | 1 | 13 | Fkbp15 | -0.04 | Young |
| 0.830699 | 0.091034 | 0.412 | 0.426 | 1 | 13 | Minpp1 | -0.014 | Young |
| 0.831379 | 0.019555 | 0.193 | 0.189 | 1 | 13 | Mki67 | 0.004 | Young |
| 0.8315 | 0.007549 | 0.947 | 0.918 | 1 | 13 | Fam174a | 0.029 | Young |
| 0.831665 | 0.168659 | 0.158 | 0.156 | 1 | 13 | Chd1l | 0.002 | Young |
| 0.831835 | 0.262008 | 0.123 | 0.115 | 1 | 13 | Letm2 | 0.008 | Young |
| 0.831835 | 0.092567 | 0.123 | 0.115 | 1 | 13 | Ogg1 | 0.008 | Young |
| 0.831836 | 0.401582 | 0.07 | 0.066 | 1 | 13 | Tm7sf2 | 0.004 | Young |
| 0.831836 | 0.347444 | 0.07 | 0.066 | 1 | 13 | Acyp2 | 0.004 | Young |
| 0.831976 | 0.037319 | 0.886 | 0.811 | 1 | 13 | Trim30a | 0.075 | Young |
| 0.83203 | 0.026118 | 0.272 | 0.279 | 1 | 13 | Ammecr1 | -0.007 | Young |
| 0.83207 | 0.11356 | 0.175 | 0.197 | 1 | 13 | Zfp84 | -0.022 | Young |
| 0.832083 | 0.151226 | 0.202 | 0.197 | 1 | 13 | Zfp617 | 0.005 | Young |
| 0.832634 | 0.045802 | 0.526 | 0.516 | 1 | 13 | Cd40 | 0.01 | Young |
| 0.832948 | 0.005932 | 0.895 | 0.918 | 1 | 13 | Hnrnph2 | -0.023 | Young |
| 0.833145 | 0.086547 | 0.289 | 0.32 | 1 | 13 | Leo1 | -0.031 | Young |
| 0.833445 | 0.032616 | 0.211 | 0.197 | 1 | 13 | B4gat1 | 0.014 | Young |
| 0.833707 | 0.111814 | 0.088 | 0.098 | 1 | 13 | Polrmt | -0.01 | Young |
| 0.833707 | 0.032301 | 0.088 | 0.098 | 1 | 13 | Gstm5 | -0.01 | Young |
| 0.833757 | 0.094976 | 0.035 | 0.041 | 1 | 13 | Tmem17 | -0.006 | Young |
| 0.833757 | 0.073582 | 0.035 | 0.041 | 1 | 13 | Smtn | -0.006 | Young |
| 0.833757 | 0.015469 | 0.035 | 0.041 | 1 | 13 | Fancf | -0.006 | Young |
| 0.834163 | 0.188755 | 0.07 | 0.082 | 1 | 13 | Gm26601 | -0.012 | Young |
| 0.834461 | 0.021122 | 0.912 | 0.926 | 1 | 13 | Trmt112 | -0.014 | Young |
| 0.835072 | 0.218932 | 0.588 | 0.607 | 1 | 13 | Fbxw4 | -0.019 | Young |
| 0.835072 | 0.03209 | 0.561 | 0.631 | 1 | 13 | Zdhhc21 | -0.07 | Young |
| 0.835844 | 0.032818 | 0.868 | 0.885 | 1 | 13 | Pon2 | -0.017 | Young |
| 0.836614 | 0.051463 | 0.342 | 0.393 | 1 | 13 | Poldip2 | -0.051 | Young |
| 0.836769 | 0.052392 | 0.667 | 0.73 | 1 | 13 | Stag1 | -0.063 | Young |
| 0.836806 | 0.049955 | 0.807 | 0.787 | 1 | 13 | Supt5 | 0.02 | Young |

| | | | | | | | | |
|---|---|---|---|---|---|---|---|---|
| 0.837459 | 0.021717 | 0.947 | 0.918 | 1 | 13 | Folr2 | 0.029 | Young |
| 0.837556 | 0.170691 | 0.167 | 0.164 | 1 | 13 | Pkp3 | 0.003 | Young |
| 0.837946 | 0.031367 | 0.833 | 0.861 | 1 | 13 | Cpne3 | -0.028 | Young |
| 0.838044 | 0.067668 | 0.316 | 0.32 | 1 | 13 | Nol8 | -0.004 | Young |
| 0.838138 | 0.177408 | 0.096 | 0.09 | 1 | 13 | Ift80 | 0.006 | Young |
| 0.838248 | 0.099801 | 0.035 | 0.041 | 1 | 13 | 4932422M | -0.006 | Young |
| 0.838248 | 0.030197 | 0.035 | 0.041 | 1 | 13 | Pdxp | -0.006 | Young |
| 0.838262 | 0.178164 | 0.316 | 0.344 | 1 | 13 | Ascc1 | -0.028 | Young |
| 0.838664 | 0.047473 | 0.14 | 0.131 | 1 | 13 | Rrp1b | 0.009 | Young |
| 0.838673 | 0.21431 | 0.07 | 0.066 | 1 | 13 | Glmn | 0.004 | Young |
| 0.838728 | 0.065644 | 0.149 | 0.164 | 1 | 13 | Trim14 | -0.015 | Young |
| 0.839099 | 0.099347 | 0.57 | 0.631 | 1 | 13 | Mogs | -0.061 | Young |
| 0.839217 | 0.151603 | 0.158 | 0.18 | 1 | 13 | Usp28 | -0.022 | Young |
| 0.839386 | 0.002791 | 0.184 | 0.18 | 1 | 13 | Zmynd15 | 0.004 | Young |
| 0.839617 | 0.022305 | 0.088 | 0.098 | 1 | 13 | Mphosph9 | -0.01 | Young |
| 0.839911 | 0.151392 | 0.237 | 0.262 | 1 | 13 | Cfap97 | -0.025 | Young |
| 0.840001 | 0.197621 | 0.105 | 0.098 | 1 | 13 | Gpatch3 | 0.007 | Young |
| 0.840337 | 0.318407 | 0.079 | 0.09 | 1 | 13 | Asf1b | -0.011 | Young |
| 0.840338 | 0.164072 | 0.64 | 0.631 | 1 | 13 | Epb41l4aos | 0.009 | Young |
| 0.840466 | 0.005841 | 0.982 | 0.984 | 1 | 13 | Capg | -0.002 | Young |
| 0.840508 | 0.037187 | 0.228 | 0.213 | 1 | 13 | Nxt2 | 0.015 | Young |
| 0.84064 | 0.13238 | 0.693 | 0.713 | 1 | 13 | Gtf2a1 | -0.02 | Young |
| 0.840718 | 0.18719 | 0.211 | 0.246 | 1 | 13 | Nfkbil1 | -0.035 | Young |
| 0.840813 | 0.23866 | 0.175 | 0.172 | 1 | 13 | Fance | 0.003 | Young |
| 0.840865 | 0.05975 | 0.482 | 0.475 | 1 | 13 | Cln6 | 0.007 | Young |
| 0.841353 | 0.07871 | 0.465 | 0.484 | 1 | 13 | Mllt6 | -0.019 | Young |
| 0.84141 | 0.126822 | 0.114 | 0.107 | 1 | 13 | Fkbp7 | 0.007 | Young |
| 0.841481 | 0.111642 | 0.579 | 0.557 | 1 | 13 | Dnajc10 | 0.022 | Young |
| 0.841882 | 0.289027 | 0.088 | 0.082 | 1 | 13 | Fancm | 0.006 | Young |
| 0.842097 | 0.323901 | 0.07 | 0.066 | 1 | 13 | Trim32 | 0.004 | Young |
| 0.842097 | 0.295868 | 0.07 | 0.066 | 1 | 13 | Ttc8 | 0.004 | Young |
| 0.842097 | 0.083311 | 0.07 | 0.066 | 1 | 13 | Zfp688 | 0.004 | Young |
| 0.842114 | 0.027616 | 0.526 | 0.516 | 1 | 13 | Ppp2r5e | 0.01 | Young |
| 0.842259 | 0.030238 | 0.535 | 0.598 | 1 | 13 | Hk1 | -0.063 | Young |
| 0.842324 | 0.125619 | 0.632 | 0.721 | 1 | 13 | Commd6 | -0.089 | Young |
| 0.842393 | 0.03403 | 0.64 | 0.664 | 1 | 13 | Ilkap | -0.024 | Young |
| 0.84245 | 0.073965 | 0.439 | 0.475 | 1 | 13 | Gps2 | -0.036 | Young |
| 0.842716 | 0.026241 | 0.263 | 0.262 | 1 | 13 | Gtdc1 | 0.001 | Young |
| 0.842744 | 0.263424 | 0.035 | 0.041 | 1 | 13 | Proser3 | -0.006 | Young |
| 0.842744 | 0.108657 | 0.035 | 0.041 | 1 | 13 | Pcbp4 | -0.006 | Young |
| 0.842744 | 0.106294 | 0.035 | 0.041 | 1 | 13 | Zscan20 | -0.006 | Young |
| 0.842744 | 0.051366 | 0.035 | 0.041 | 1 | 13 | Plxdc1 | -0.006 | Young |
| 0.842869 | 0.094011 | 0.228 | 0.262 | 1 | 13 | Cyb561d2 | -0.034 | Young |
| 0.843127 | 0.098546 | 0.456 | 0.475 | 1 | 13 | Setdb1 | -0.019 | Young |
| 0.843223 | 0.022758 | 0.307 | 0.336 | 1 | 13 | Zdhhc6 | -0.029 | Young |

| | | | | | | | | |
|---|---|---|---|---|---|---|---|---|
| 0.843427 | 0.302861 | 0.079 | 0.09 | 1 | 13 | Zfp113 | -0.011 | Young |
| 0.844269 | 0.043076 | 0.096 | 0.107 | 1 | 13 | Cenpo | -0.011 | Young |
| 0.844496 | 0.092035 | 0.596 | 0.607 | 1 | 13 | Tsnax | -0.011 | Young |
| 0.844768 | 0.058256 | 0.307 | 0.344 | 1 | 13 | Atg4b | -0.037 | Young |
| 0.84482 | 0.522513 | 0.123 | 0.123 | 1 | 13 | Rint1 | 0 | Young |
| 0.844854 | 0.032594 | 0.877 | 0.91 | 1 | 13 | Arih1 | -0.033 | Young |
| 0.84513 | 0.042635 | 0.377 | 0.402 | 1 | 13 | Thbs1 | -0.025 | Young |
| 0.845269 | 0.125081 | 0.316 | 0.32 | 1 | 13 | Parp11 | -0.004 | Young |
| 0.845335 | 0.124541 | 0.175 | 0.172 | 1 | 13 | Ankrd52 | 0.003 | Young |
| 0.845435 | 0.044851 | 0.281 | 0.311 | 1 | 13 | Leng1 | -0.03 | Young |
| 0.845507 | 0.233889 | 0.079 | 0.074 | 1 | 13 | L3mbtl2 | 0.005 | Young |
| 0.845836 | 0.031662 | 0.789 | 0.811 | 1 | 13 | Rad23a | -0.022 | Young |
| 0.846415 | 0.001863 | 0.298 | 0.328 | 1 | 13 | Arfgef2 | -0.03 | Young |
| 0.846575 | 0.031125 | 0.263 | 0.262 | 1 | 13 | Arhgef10 | 0.001 | Young |
| 0.846892 | 0.152835 | 0.281 | 0.287 | 1 | 13 | C2cd2l | -0.006 | Young |
| 0.847323 | 0.083833 | 0.307 | 0.32 | 1 | 13 | Dcaf17 | -0.013 | Young |
| 0.847669 | 0.300496 | 0.43 | 0.492 | 1 | 13 | Ube2w | -0.062 | Young |
| 0.847905 | 0.03117 | 0.895 | 0.959 | 1 | 13 | Arf3 | -0.064 | Young |
| 0.848135 | 0.032142 | 0.368 | 0.418 | 1 | 13 | Ptgr2 | -0.05 | Young |
| 0.849137 | 0.25943 | 0.281 | 0.311 | 1 | 13 | Tnfrsf12a | -0.03 | Young |
| 0.849459 | 0.091141 | 0.14 | 0.156 | 1 | 13 | Bud13 | -0.016 | Young |
| 0.850722 | 0.067307 | 0.728 | 0.705 | 1 | 13 | Evi5 | 0.023 | Young |
| 0.850923 | 0.0118 | 1 | 1 | 1 | 13 | Tmed10 | 0 | Young |
| 0.851213 | 0.060544 | 0.14 | 0.131 | 1 | 13 | Wdr47 | 0.009 | Young |
| 0.851223 | 0.115239 | 0.57 | 0.549 | 1 | 13 | Mvb12a | 0.021 | Young |
| 0.85131 | 0.410445 | 0.175 | 0.18 | 1 | 13 | Mx1 | -0.005 | Young |
| 0.851552 | 0.021328 | 0.447 | 0.492 | 1 | 13 | Ccdc82 | -0.045 | Young |
| 0.851751 | 0.3764 | 0.035 | 0.041 | 1 | 13 | C330011M | -0.006 | Young |
| 0.851751 | 0.262995 | 0.035 | 0.041 | 1 | 13 | Gm21781 | -0.006 | Young |
| 0.851751 | 0.045805 | 0.035 | 0.041 | 1 | 13 | Mppe1 | -0.006 | Young |
| 0.852003 | 0.262005 | 0.079 | 0.074 | 1 | 13 | Kif3c | 0.005 | Young |
| 0.852406 | 0.000133 | 0.114 | 0.107 | 1 | 13 | Ice2 | 0.007 | Young |
| 0.852751 | 0.194258 | 0.526 | 0.557 | 1 | 13 | Socs4 | -0.031 | Young |
| 0.852878 | 0.002674 | 0.36 | 0.361 | 1 | 13 | Rnf2 | -0.001 | Young |
| 0.853043 | 0.074407 | 0.754 | 0.787 | 1 | 13 | Imp3 | -0.033 | Young |
| 0.853088 | 0.039572 | 0.412 | 0.443 | 1 | 13 | Smn1 | -0.031 | Young |
| 0.853339 | 0.021683 | 0.684 | 0.705 | 1 | 13 | Ube2j2 | -0.021 | Young |
| 0.85346 | 0.184089 | 0.254 | 0.262 | 1 | 13 | Gm41307 | -0.008 | Young |
| 0.854498 | 0.404271 | 0.158 | 0.164 | 1 | 13 | Mllt11 | -0.006 | Young |
| 0.854871 | 0.058398 | 0.228 | 0.246 | 1 | 13 | Sgsh | -0.018 | Young |
| 0.855257 | 0.041728 | 0.149 | 0.164 | 1 | 13 | Kctd9 | -0.015 | Young |
| 0.855309 | 0.256082 | 0.123 | 0.123 | 1 | 13 | Dars2 | 0 | Young |
| 0.855346 | 0.164618 | 0.386 | 0.41 | 1 | 13 | Rasa2 | -0.024 | Young |
| 0.855376 | 0.037744 | 0.912 | 0.926 | 1 | 13 | Pebp1 | -0.014 | Young |
| 0.855391 | 0.012312 | 0.947 | 0.918 | 1 | 13 | Abhd12 | 0.029 | Young |

| | | | | | | | | |
|---|---|---|---|---|---|---|---|---|
| 0.855496 | 0.082597 | 0.456 | 0.459 | 1 | 13 | Tmem229b | -0.003 | Young |
| 0.855731 | 0.051852 | 0.237 | 0.254 | 1 | 13 | Dph6 | -0.017 | Young |
| 0.85577 | 0.118376 | 0.553 | 0.631 | 1 | 13 | Sptan1 | -0.078 | Young |
| 0.855775 | 0.298317 | 0.061 | 0.057 | 1 | 13 | Mis18bp1 | 0.004 | Young |
| 0.856237 | 0.114117 | 0.263 | 0.262 | 1 | 13 | Polk | 0.001 | Young |
| 0.856237 | 0.207319 | 0.263 | 0.262 | 1 | 13 | Pot1b | 0.001 | Young |
| 0.856262 | 0.114804 | 0.035 | 0.041 | 1 | 13 | Gm48551 | -0.006 | Young |
| 0.856312 | 0.062166 | 0.772 | 0.82 | 1 | 13 | Coa5 | -0.048 | Young |
| 0.856417 | 0.15015 | 0.307 | 0.32 | 1 | 13 | Mthfsl | -0.013 | Young |
| 0.85658 | 0.157387 | 0.193 | 0.197 | 1 | 13 | Trim23 | -0.004 | Young |
| 0.856703 | 0.066437 | 0.561 | 0.631 | 1 | 13 | Synrg | -0.07 | Young |
| 0.856839 | 0.261012 | 0.123 | 0.139 | 1 | 13 | Dbr1 | -0.016 | Young |
| 0.856951 | 0.05804 | 0.36 | 0.385 | 1 | 13 | Amz2 | -0.025 | Young |
| 0.856981 | 0.178501 | 0.044 | 0.049 | 1 | 13 | Rad54l | -0.005 | Young |
| 0.856981 | 0.143644 | 0.044 | 0.049 | 1 | 13 | Gm14005 | -0.005 | Young |
| 0.856981 | 0.051668 | 0.044 | 0.049 | 1 | 13 | Nfib | -0.005 | Young |
| 0.857298 | 0.022571 | 0.614 | 0.615 | 1 | 13 | Ogfrl1 | -0.001 | Young |
| 0.857335 | 0.140606 | 0.395 | 0.443 | 1 | 13 | 2210016L2 | -0.048 | Young |
| 0.857492 | 0.00163 | 0.518 | 0.508 | 1 | 13 | Zbtb8os | 0.01 | Young |
| 0.857557 | 0.048151 | 0.491 | 0.484 | 1 | 13 | Ubac2 | 0.007 | Young |
| 0.857557 | 0.048419 | 0.491 | 0.484 | 1 | 13 | Pdxdc1 | 0.007 | Young |
| 0.857606 | 0.05575 | 0.237 | 0.262 | 1 | 13 | 1700021F0 | -0.025 | Young |
| 0.857713 | 0.039305 | 0.237 | 0.254 | 1 | 13 | Fyco1 | -0.017 | Young |
| 0.857793 | 0.197956 | 0.412 | 0.418 | 1 | 13 | Cbx5 | -0.006 | Young |
| 0.858008 | 0.187687 | 0.167 | 0.156 | 1 | 13 | Per3 | 0.011 | Young |
| 0.858178 | 0.293791 | 0.281 | 0.287 | 1 | 13 | Slc25a19 | -0.006 | Young |
| 0.858778 | 0.414283 | 0.053 | 0.049 | 1 | 13 | Adamts6 | 0.004 | Young |
| 0.858778 | 0.331648 | 0.053 | 0.049 | 1 | 13 | Zswim1 | 0.004 | Young |
| 0.858884 | 0.110147 | 0.096 | 0.09 | 1 | 13 | Zfp768 | 0.006 | Young |
| 0.858932 | 0.049314 | 0.175 | 0.172 | 1 | 13 | Ubxn8 | 0.003 | Young |
| 0.859434 | 0.301628 | 0.061 | 0.057 | 1 | 13 | Col4a1 | 0.004 | Young |
| 0.859434 | 0.288501 | 0.061 | 0.057 | 1 | 13 | Gm13091 | 0.004 | Young |
| 0.859434 | 0.271615 | 0.061 | 0.057 | 1 | 13 | Klhl42 | 0.004 | Young |
| 0.859758 | 0.162072 | 0.298 | 0.336 | 1 | 13 | Fam160b2 | -0.038 | Young |
| 0.860659 | 0.070199 | 0.175 | 0.197 | 1 | 13 | Msh6 | -0.022 | Young |
| 0.860688 | 0.183661 | 0.246 | 0.246 | 1 | 13 | 9330160F1 | 0 | Young |
| 0.861233 | 0.107305 | 0.263 | 0.295 | 1 | 13 | Akap7 | -0.032 | Young |
| 0.861386 | 0.08038 | 0.096 | 0.107 | 1 | 13 | Clec16a | -0.011 | Young |
| 0.861727 | 0.024208 | 0.439 | 0.467 | 1 | 13 | Stam2 | -0.028 | Young |
| 0.862107 | 0.013378 | 0.667 | 0.689 | 1 | 13 | Fam49a | -0.022 | Young |
| 0.862118 | 0.057893 | 0.298 | 0.295 | 1 | 13 | Cblb | 0.003 | Young |
| 0.862155 | 0.075001 | 0.5 | 0.549 | 1 | 13 | Prdm2 | -0.049 | Young |
| 0.862716 | 0.423032 | 0.053 | 0.049 | 1 | 13 | A630089N0 | 0.004 | Young |
| 0.862816 | 0.079048 | 0.105 | 0.098 | 1 | 13 | E130307A1 | 0.007 | Young |
| 0.862825 | 0.035104 | 0.904 | 0.885 | 1 | 13 | Ranbp1 | 0.019 | Young |

| | | | | | | | | |
|---|---|---|---|---|---|---|---|---|
| 0.863077 | 0.130167 | 0.132 | 0.148 | 1 | 13 | Stambpl1 | -0.016 | Young |
| 0.863472 | 0.090662 | 0.456 | 0.5 | 1 | 13 | Hadh | -0.044 | Young |
| 0.863781 | 0.043813 | 0.526 | 0.557 | 1 | 13 | Pilrb2 | -0.031 | Young |
| 0.863938 | 0.032939 | 0.772 | 0.844 | 1 | 13 | Mgat1 | -0.072 | Young |
| 0.864081 | 0.023816 | 0.184 | 0.197 | 1 | 13 | Zfp410 | -0.013 | Young |
| 0.864103 | 0.100356 | 0.219 | 0.238 | 1 | 13 | Lrch1 | -0.019 | Young |
| 0.864186 | 0.069979 | 0.325 | 0.336 | 1 | 13 | Zfp639 | -0.011 | Young |
| 0.864207 | 0.066683 | 0.64 | 0.697 | 1 | 13 | Rasa3 | -0.057 | Young |
| 0.864216 | 0.019912 | 0.693 | 0.721 | 1 | 13 | Ube2z | -0.028 | Young |
| 0.864376 | 0.074779 | 0.421 | 0.426 | 1 | 13 | Abhd16a | -0.005 | Young |
| 0.86519 | 0.094559 | 0.044 | 0.049 | 1 | 13 | Eya4 | -0.005 | Young |
| 0.86519 | 0.047408 | 0.044 | 0.049 | 1 | 13 | Cldn34c1 | -0.005 | Young |
| 0.86519 | 0.04044 | 0.044 | 0.049 | 1 | 13 | B3gntl1 | -0.005 | Young |
| 0.865617 | 0.085259 | 0.5 | 0.475 | 1 | 13 | Mcm3 | 0.025 | Young |
| 0.865866 | 0.042769 | 0.947 | 0.885 | 1 | 13 | Psmb6 | 0.062 | Young |
| 0.865906 | 0.055092 | 0.982 | 0.984 | 1 | 13 | Nap1l1 | -0.002 | Young |
| 0.866139 | 0.183052 | 0.07 | 0.066 | 1 | 13 | Dph7 | 0.004 | Young |
| 0.866451 | 0.158015 | 0.377 | 0.385 | 1 | 13 | Pdcl | -0.008 | Young |
| 0.866466 | 0.046596 | 0.281 | 0.32 | 1 | 13 | Pibf1 | -0.039 | Young |
| 0.867129 | 0.040121 | 0.553 | 0.541 | 1 | 13 | Tbc1d1 | 0.012 | Young |
| 0.867407 | 0.176969 | 0.325 | 0.344 | 1 | 13 | Hint3 | -0.019 | Young |
| 0.867692 | 0.301326 | 0.281 | 0.311 | 1 | 13 | Nf2 | -0.03 | Young |
| 0.867806 | 0.049122 | 0.096 | 0.09 | 1 | 13 | Faap24 | 0.006 | Young |
| 0.868268 | 0.076014 | 0.298 | 0.328 | 1 | 13 | Fam173b | -0.03 | Young |
| 0.868288 | 0.069372 | 0.079 | 0.074 | 1 | 13 | Brf2 | 0.005 | Young |
| 0.86849 | 0.063546 | 0.202 | 0.221 | 1 | 13 | Unkl | -0.019 | Young |
| 0.868508 | 0.07123 | 0.184 | 0.205 | 1 | 13 | Hist1h4i | -0.021 | Young |
| 0.868524 | 0.207921 | 0.211 | 0.205 | 1 | 13 | Ppcdc | 0.006 | Young |
| 0.868534 | 0.090732 | 0.202 | 0.197 | 1 | 13 | Donson | 0.005 | Young |
| 0.868567 | 0.012392 | 0.807 | 0.844 | 1 | 13 | Man1c1 | -0.037 | Young |
| 0.868591 | 0.190356 | 0.474 | 0.492 | 1 | 13 | Atp1a3 | -0.018 | Young |
| 0.869047 | 0.072704 | 0.36 | 0.385 | 1 | 13 | Cspp1 | -0.025 | Young |
| 0.8694 | 0.04473 | 0.465 | 0.5 | 1 | 13 | Cpq | -0.035 | Young |
| 0.869496 | 0.031235 | 0.623 | 0.648 | 1 | 13 | Gemin7 | -0.025 | Young |
| 0.86966 | 0.015569 | 0.991 | 0.975 | 1 | 13 | Spag9 | 0.016 | Young |
| 0.86997 | 0.085354 | 0.096 | 0.107 | 1 | 13 | Qtrt2 | -0.011 | Young |
| 0.870254 | 0.124684 | 0.5 | 0.525 | 1 | 13 | Lrrc41 | -0.025 | Young |
| 0.870335 | 0.007737 | 0.851 | 0.934 | 1 | 13 | Rer1 | -0.083 | Young |
| 0.870359 | 0.001763 | 0.895 | 0.918 | 1 | 13 | Rexo2 | -0.023 | Young |
| 0.870384 | 0.025406 | 0.158 | 0.148 | 1 | 13 | Rmdn1 | 0.01 | Young |
| 0.870429 | 0.233226 | 0.061 | 0.057 | 1 | 13 | 4632415L0 | 0.004 | Young |
| 0.870602 | 0.250552 | 0.053 | 0.049 | 1 | 13 | Hacl1 | 0.004 | Young |
| 0.871608 | 0.040839 | 0.377 | 0.385 | 1 | 13 | Gtf2e2 | -0.008 | Young |
| 0.871643 | 0.116481 | 0.351 | 0.393 | 1 | 13 | Sp3os | -0.042 | Young |
| 0.87274 | 0.009014 | 0.772 | 0.787 | 1 | 13 | Mcur1 | -0.015 | Young |

| | | | | | | | | |
|---|---|---|---|---|---|---|---|---|
| 0.873032 | 0.03379 | 0.07 | 0.066 | 1 | 13 | Igfbp6 | 0.004 | Young |
| 0.873383 | 0.083039 | 0.351 | 0.361 | 1 | 13 | Ankra2 | -0.01 | Young |
| 0.873414 | 0.124257 | 0.044 | 0.049 | 1 | 13 | Zkscan5 | -0.005 | Young |
| 0.873414 | 0.116564 | 0.044 | 0.049 | 1 | 13 | 1700056N1 | -0.005 | Young |
| 0.873414 | 0.092715 | 0.044 | 0.049 | 1 | 13 | Gm32856 | -0.005 | Young |
| 0.873723 | 0.103235 | 0.684 | 0.779 | 1 | 13 | Psme4 | -0.095 | Young |
| 0.873774 | 0.061171 | 0.439 | 0.484 | 1 | 13 | Ppil4 | -0.045 | Young |
| 0.873804 | 0.153808 | 0.175 | 0.18 | 1 | 13 | Nlrx1 | -0.005 | Young |
| 0.873872 | 0.167961 | 0.298 | 0.303 | 1 | 13 | Ripk2 | -0.005 | Young |
| 0.87398 | 0.131437 | 0.237 | 0.238 | 1 | 13 | Btbd2 | -0.001 | Young |
| 0.874031 | 0.354917 | 0.044 | 0.041 | 1 | 13 | Zc4h2 | 0.003 | Young |
| 0.8741 | 0.295529 | 0.061 | 0.057 | 1 | 13 | Kcnj2 | 0.004 | Young |
| 0.874397 | 0.032664 | 0.404 | 0.443 | 1 | 13 | Scamp1 | -0.039 | Young |
| 0.87455 | 0.250428 | 0.053 | 0.049 | 1 | 13 | Cwf19l1 | 0.004 | Young |
| 0.874584 | 0.167785 | 0.219 | 0.221 | 1 | 13 | Nit2 | -0.002 | Young |
| 0.874818 | 0.0786 | 0.079 | 0.074 | 1 | 13 | Wars2 | 0.005 | Young |
| 0.875194 | 0.046031 | 0.544 | 0.557 | 1 | 13 | Hars | -0.013 | Young |
| 0.875559 | 0.014641 | 0.123 | 0.131 | 1 | 13 | Hyal2 | -0.008 | Young |
| 0.87559 | 0.048022 | 0.886 | 0.893 | 1 | 13 | Spcs3 | -0.007 | Young |
| 0.876067 | 0.152289 | 0.377 | 0.426 | 1 | 13 | 2310033P0 | -0.049 | Young |
| 0.876144 | 0.15245 | 0.14 | 0.156 | 1 | 13 | Galt | -0.016 | Young |
| 0.876243 | 0.064863 | 0.57 | 0.631 | 1 | 13 | Trmt1 | -0.061 | Young |
| 0.876423 | 0.013293 | 0.982 | 0.967 | 1 | 13 | Fosl2 | 0.015 | Young |
| 0.876596 | 0.079365 | 0.254 | 0.27 | 1 | 13 | Stam | -0.016 | Young |
| 0.876744 | 0.175976 | 0.096 | 0.09 | 1 | 13 | 1700006J1 | 0.006 | Young |
| 0.876778 | 0.258343 | 0.149 | 0.172 | 1 | 13 | Zfp87 | -0.023 | Young |
| 0.877135 | 0.079961 | 0.105 | 0.098 | 1 | 13 | Ankrd13d | 0.007 | Young |
| 0.877383 | 0.019556 | 0.781 | 0.803 | 1 | 13 | Chmp5 | -0.022 | Young |
| 0.877531 | 0.117325 | 0.044 | 0.049 | 1 | 13 | Prkcg | -0.005 | Young |
| 0.877976 | 0.063194 | 0.719 | 0.705 | 1 | 13 | Sertad2 | 0.014 | Young |
| 0.878066 | 0.05566 | 0.474 | 0.484 | 1 | 13 | Sh3tc1 | -0.01 | Young |
| 0.878086 | 0.261236 | 0.079 | 0.074 | 1 | 13 | Phka1 | 0.005 | Young |
| 0.878282 | 0.05738 | 0.614 | 0.68 | 1 | 13 | Mbd3 | -0.066 | Young |
| 0.878319 | 0.150447 | 0.307 | 0.32 | 1 | 13 | Traf5 | -0.013 | Young |
| 0.87834 | 0.313871 | 0.044 | 0.041 | 1 | 13 | Cdkn2b | 0.003 | Young |
| 0.87834 | 0.262291 | 0.044 | 0.041 | 1 | 13 | Hist2h2aa1 | 0.003 | Young |
| 0.878878 | 0.043863 | 0.219 | 0.213 | 1 | 13 | Cd3eap | 0.006 | Young |
| 0.878968 | 0.174353 | 0.368 | 0.385 | 1 | 13 | Agpat1 | -0.017 | Young |
| 0.878979 | 0.264852 | 0.158 | 0.156 | 1 | 13 | Echdc1 | 0.002 | Young |
| 0.879001 | 0.208971 | 0.123 | 0.123 | 1 | 13 | Hps6 | 0 | Young |
| 0.879074 | 0.028733 | 0.684 | 0.689 | 1 | 13 | Pcf11 | -0.005 | Young |
| 0.87912 | 0.035559 | 0.088 | 0.082 | 1 | 13 | 2410131K1 | 0.006 | Young |
| 0.879228 | 0.146654 | 0.202 | 0.205 | 1 | 13 | Nol6 | -0.003 | Young |
| 0.879404 | 0.018748 | 0.167 | 0.18 | 1 | 13 | Emilin1 | -0.013 | Young |
| 0.879404 | 0.011032 | 0.167 | 0.18 | 1 | 13 | Snx19 | -0.013 | Young |

| | | | | | | | | |
|---|---|---|---|---|---|---|---|---|
| 0.879574 | 0.210763 | 0.237 | 0.254 | 1 | 13 | Pgm2 | -0.017 | Young |
| 0.87973 | 0.075646 | 0.579 | 0.598 | 1 | 13 | Med10 | -0.019 | Young |
| 0.880811 | 0.039507 | 0.526 | 0.541 | 1 | 13 | Casp1 | -0.015 | Young |
| 0.880834 | 0.024477 | 0.281 | 0.287 | 1 | 13 | Gid4 | -0.006 | Young |
| 0.881219 | 0.115081 | 0.088 | 0.098 | 1 | 13 | Srr | -0.01 | Young |
| 0.881289 | 0.001574 | 0.307 | 0.295 | 1 | 13 | Agpat2 | 0.012 | Young |
| 0.881651 | 0.149957 | 0.044 | 0.049 | 1 | 13 | Trem1 | -0.005 | Young |
| 0.881651 | 0.013717 | 0.044 | 0.049 | 1 | 13 | Shpk | -0.005 | Young |
| 0.882137 | 0.043538 | 0.798 | 0.852 | 1 | 13 | Spg21 | -0.054 | Young |
| 0.882218 | 0.030683 | 0.825 | 0.861 | 1 | 13 | Pik3r1 | -0.036 | Young |
| 0.882237 | 0.038041 | 0.088 | 0.082 | 1 | 13 | Bpgm | 0.006 | Young |
| 0.882455 | 0.184447 | 0.053 | 0.049 | 1 | 13 | Gm9856 | 0.004 | Young |
| 0.882455 | 0.179381 | 0.053 | 0.049 | 1 | 13 | Ahcy | 0.004 | Young |
| 0.882573 | 0.175548 | 0.316 | 0.336 | 1 | 13 | D10Wsu10 | -0.02 | Young |
| 0.882653 | 0.327402 | 0.044 | 0.041 | 1 | 13 | Gm17435 | 0.003 | Young |
| 0.882653 | 0.281449 | 0.044 | 0.041 | 1 | 13 | Smim1 | 0.003 | Young |
| 0.882874 | 0.054182 | 0.105 | 0.098 | 1 | 13 | Ikzf2 | 0.007 | Young |
| 0.882894 | 0.088349 | 0.298 | 0.328 | 1 | 13 | 2610002M | -0.03 | Young |
| 0.88315 | 0.00289 | 0.904 | 0.91 | 1 | 13 | Tmem14c | -0.006 | Young |
| 0.883162 | 0.059168 | 0.605 | 0.648 | 1 | 13 | Pdpk1 | -0.043 | Young |
| 0.883185 | 0.097512 | 0.404 | 0.41 | 1 | 13 | Impdh1 | -0.006 | Young |
| 0.88339 | 0.026283 | 0.07 | 0.066 | 1 | 13 | Sgms2 | 0.004 | Young |
| 0.883573 | 0.036556 | 0.64 | 0.648 | 1 | 13 | Trio | -0.008 | Young |
| 0.883601 | 0.305619 | 0.412 | 0.434 | 1 | 13 | Osm | -0.022 | Young |
| 0.883727 | 0.016366 | 0.825 | 0.861 | 1 | 13 | Cyth1 | -0.036 | Young |
| 0.883737 | 0.215359 | 0.158 | 0.156 | 1 | 13 | Zfp821 | 0.002 | Young |
| 0.88395 | 0.024454 | 1 | 0.975 | 1 | 13 | Atp6ap1 | 0.025 | Young |
| 0.884174 | 0.075956 | 0.105 | 0.115 | 1 | 13 | Pfas | -0.01 | Young |
| 0.884205 | 0.190113 | 0.088 | 0.098 | 1 | 13 | Zfp275 | -0.01 | Young |
| 0.884754 | 0.098999 | 0.228 | 0.221 | 1 | 13 | Zdhhc12 | 0.007 | Young |
| 0.885128 | 0.197027 | 0.061 | 0.057 | 1 | 13 | Zfp605 | 0.004 | Young |
| 0.885128 | 0.081504 | 0.061 | 0.057 | 1 | 13 | Glis2 | 0.004 | Young |
| 0.885153 | 0.206629 | 0.211 | 0.221 | 1 | 13 | Wrnip1 | -0.01 | Young |
| 0.885226 | 0.047497 | 0.325 | 0.311 | 1 | 13 | Rnf31 | 0.014 | Young |
| 0.885698 | 0.122636 | 0.096 | 0.09 | 1 | 13 | Cd300ld3 | 0.006 | Young |
| 0.885775 | 0.252591 | 0.044 | 0.049 | 1 | 13 | A530032D1 | -0.005 | Young |
| 0.886233 | 0.287969 | 0.167 | 0.164 | 1 | 13 | Phldb1 | 0.003 | Young |
| 0.886273 | 0.31425 | 0.035 | 0.033 | 1 | 13 | 2210417A0 | 0.002 | Young |
| 0.886412 | 0.260892 | 0.053 | 0.049 | 1 | 13 | Gm33973 | 0.004 | Young |
| 0.886412 | 0.106849 | 0.053 | 0.049 | 1 | 13 | Tmem221 | 0.004 | Young |
| 0.886487 | 0.047228 | 0.351 | 0.377 | 1 | 13 | Ndufaf3 | -0.026 | Young |
| 0.886569 | 0.061886 | 0.36 | 0.352 | 1 | 13 | Pccb | 0.008 | Young |
| 0.886654 | 0.014866 | 0.342 | 0.336 | 1 | 13 | Prr12 | 0.006 | Young |
| 0.886724 | 0.197285 | 0.193 | 0.213 | 1 | 13 | Sgf29 | -0.02 | Young |
| 0.886847 | 0.0158 | 0.07 | 0.066 | 1 | 13 | Neurl1a | 0.004 | Young |

| | | | | | | | | |
|---|---|---|---|---|---|---|---|---|
| 0.886946 | 0.093113 | 0.105 | 0.115 | 1 | 13 | 1700056E2 | -0.01 | Young |
| 0.886969 | 0.536167 | 0.044 | 0.041 | 1 | 13 | Rnd1 | 0.003 | Young |
| 0.886992 | 0.051457 | 0.526 | 0.533 | 1 | 13 | Mplkip | -0.007 | Young |
| 0.887554 | 0.060284 | 0.325 | 0.336 | 1 | 13 | Vamp5 | -0.011 | Young |
| 0.887884 | 0.069837 | 0.675 | 0.746 | 1 | 13 | Trim41 | -0.071 | Young |
| 0.888016 | 0.17088 | 0.14 | 0.139 | 1 | 13 | A530017D2 | 0.001 | Young |
| 0.888109 | 0.095264 | 0.237 | 0.238 | 1 | 13 | Naa25 | -0.001 | Young |
| 0.8882 | 0.009244 | 0.825 | 0.836 | 1 | 13 | Gspt1 | -0.011 | Young |
| 0.888409 | 0.005636 | 0.281 | 0.287 | 1 | 13 | Ikbkg | -0.006 | Young |
| 0.88841 | 0.063522 | 0.175 | 0.189 | 1 | 13 | Rufy2 | -0.014 | Young |
| 0.88841 | 0.020265 | 0.184 | 0.18 | 1 | 13 | Tmem175 | 0.004 | Young |
| 0.888558 | 0.25021 | 0.167 | 0.172 | 1 | 13 | Ripor1 | -0.005 | Young |
| 0.888686 | 0.006941 | 0.096 | 0.09 | 1 | 13 | Slc18b1 | 0.006 | Young |
| 0.888704 | 0.071338 | 0.579 | 0.574 | 1 | 13 | Ap1g1 | 0.005 | Young |
| 0.888809 | 0.166968 | 0.061 | 0.057 | 1 | 13 | Hdac8 | 0.004 | Young |
| 0.88881 | 0.197089 | 0.342 | 0.361 | 1 | 13 | Lrrcc1 | -0.019 | Young |
| 0.888879 | 0.045531 | 0.474 | 0.525 | 1 | 13 | Bin2 | -0.051 | Young |
| 0.889241 | 0.227024 | 0.246 | 0.279 | 1 | 13 | Hmces | -0.033 | Young |
| 0.889389 | 0.081374 | 0.754 | 0.803 | 1 | 13 | Snx17 | -0.049 | Young |
| 0.889646 | 0.02248 | 0.825 | 0.811 | 1 | 13 | Chchd1 | 0.014 | Young |
| 0.889902 | 0.235536 | 0.044 | 0.049 | 1 | 13 | Map10 | -0.005 | Young |
| 0.890256 | 0.207152 | 0.307 | 0.336 | 1 | 13 | Tbce | -0.029 | Young |
| 0.890305 | 0.034029 | 0.412 | 0.434 | 1 | 13 | Tmem126a | -0.022 | Young |
| 0.890372 | 0.110202 | 0.053 | 0.049 | 1 | 13 | N4bp3 | 0.004 | Young |
| 0.890606 | 0.159671 | 0.053 | 0.057 | 1 | 13 | Lockd | -0.004 | Young |
| 0.891082 | 0.26819 | 0.035 | 0.033 | 1 | 13 | Spata24 | 0.002 | Young |
| 0.891289 | 0.291884 | 0.044 | 0.041 | 1 | 13 | Gm36445 | 0.003 | Young |
| 0.891597 | 0.038733 | 0.088 | 0.082 | 1 | 13 | Tigar | 0.006 | Young |
| 0.891657 | 0.149989 | 0.386 | 0.426 | 1 | 13 | Diablo | -0.04 | Young |
| 0.891707 | 0.046762 | 0.228 | 0.23 | 1 | 13 | Usp53 | -0.002 | Young |
| 0.892492 | 0.113926 | 0.061 | 0.057 | 1 | 13 | Tmem81 | 0.004 | Young |
| 0.892717 | 0.035086 | 0.544 | 0.566 | 1 | 13 | Tmem106b | -0.022 | Young |
| 0.892732 | 0.145542 | 0.219 | 0.238 | 1 | 13 | Golm1 | -0.019 | Young |
| 0.892862 | 0.033389 | 0.86 | 0.869 | 1 | 13 | Dnajc5 | -0.009 | Young |
| 0.892863 | 0.065436 | 0.254 | 0.254 | 1 | 13 | Ganc | 0 | Young |
| 0.892995 | 0.044364 | 0.982 | 0.992 | 1 | 13 | Pafah1b1 | -0.01 | Young |
| 0.892995 | 0.001692 | 1 | 1 | 1 | 13 | Laptm5 | 0 | Young |
| 0.893418 | 0.097045 | 0.298 | 0.32 | 1 | 13 | Txnl4a | -0.022 | Young |
| 0.894097 | 0.187766 | 0.272 | 0.295 | 1 | 13 | 2310022B0 | -0.023 | Young |
| 0.89417 | 0.032965 | 0.649 | 0.697 | 1 | 13 | Emc1 | -0.048 | Young |
| 0.894272 | 0.064968 | 0.491 | 0.533 | 1 | 13 | 1110038F1 | -0.042 | Young |
| 0.894335 | 0.127841 | 0.053 | 0.049 | 1 | 13 | Nthl1 | 0.004 | Young |
| 0.894335 | 0.081929 | 0.053 | 0.049 | 1 | 13 | Tarbp1 | 0.004 | Young |
| 0.894396 | 0.059723 | 0.86 | 0.885 | 1 | 13 | Nfic | -0.025 | Young |
| 0.894421 | 0.016608 | 0.053 | 0.057 | 1 | 13 | Bbs4 | -0.004 | Young |

| | | | | | | | | |
|---|---|---|---|---|---|---|---|---|
| 0.894421 | 0.003156 | 0.053 | 0.057 | 1 | 13 | A430078G: | -0.004 | Young |
| 0.894701 | 0.130784 | 0.316 | 0.361 | 1 | 13 | Zfp236 | -0.045 | Young |
| 0.8949 | 0.086698 | 0.263 | 0.287 | 1 | 13 | Tmem29 | -0.024 | Young |
| 0.89504 | 0.067301 | 0.833 | 0.844 | 1 | 13 | 2810474O: | -0.011 | Young |
| 0.895232 | 4.58E-05 | 0.596 | 0.697 | 1 | 13 | Xpo6 | -0.101 | Young |
| 0.895237 | 0.055922 | 0.754 | 0.754 | 1 | 13 | Borcs6 | 0 | Young |
| 0.895384 | 0.110695 | 0.175 | 0.172 | 1 | 13 | Zfp719 | 0.003 | Young |
| 0.895612 | 0.185531 | 0.044 | 0.041 | 1 | 13 | Pxmp2 | 0.003 | Young |
| 0.895894 | 0.474916 | 0.035 | 0.033 | 1 | 13 | Rabl2 | 0.002 | Young |
| 0.895894 | 0.384518 | 0.035 | 0.033 | 1 | 13 | Hist1h4a | 0.002 | Young |
| 0.896178 | 0.304241 | 0.061 | 0.057 | 1 | 13 | Tiam2 | 0.004 | Young |
| 0.896796 | 0.011966 | 0.158 | 0.164 | 1 | 13 | Crip2 | -0.006 | Young |
| 0.896817 | 0.065493 | 0.263 | 0.287 | 1 | 13 | Elmod2 | -0.024 | Young |
| 0.896833 | 0.020756 | 0.14 | 0.148 | 1 | 13 | Ercc6 | -0.008 | Young |
| 0.897458 | 0.135732 | 0.544 | 0.566 | 1 | 13 | Slc38a9 | -0.022 | Young |
| 0.897723 | 0.088012 | 0.211 | 0.221 | 1 | 13 | Acd | -0.01 | Young |
| 0.897779 | 0.028749 | 0.596 | 0.648 | 1 | 13 | Chordc1 | -0.052 | Young |
| 0.897838 | 0.171484 | 0.132 | 0.131 | 1 | 13 | Ppip5k1 | 0.001 | Young |
| 0.898024 | 0.274804 | 0.14 | 0.139 | 1 | 13 | Etaa1 | 0.001 | Young |
| 0.898036 | 0.048773 | 0.158 | 0.156 | 1 | 13 | Dnajc18 | 0.002 | Young |
| 0.898084 | 0.184771 | 0.149 | 0.148 | 1 | 13 | Gprasp1 | 0.001 | Young |
| 0.898108 | 0.172017 | 0.5 | 0.598 | 1 | 13 | Fam192a | -0.098 | Young |
| 0.898239 | 0.172958 | 0.053 | 0.057 | 1 | 13 | Ncaph | -0.004 | Young |
| 0.898239 | 0.089262 | 0.053 | 0.057 | 1 | 13 | Atp5s | -0.004 | Young |
| 0.898239 | 0.030259 | 0.053 | 0.057 | 1 | 13 | Sdr42e1 | -0.004 | Young |
| 0.898239 | 0.018279 | 0.053 | 0.057 | 1 | 13 | Prdm9 | -0.004 | Young |
| 0.898342 | 0.02002 | 0.675 | 0.713 | 1 | 13 | Numa1 | -0.038 | Young |
| 0.89844 | 0.082269 | 0.316 | 0.328 | 1 | 13 | Urm1 | -0.012 | Young |
| 0.898634 | 0.01977 | 0.807 | 0.795 | 1 | 13 | Usp25 | 0.012 | Young |
| 0.898654 | 0.107697 | 0.386 | 0.385 | 1 | 13 | Ddx60 | 0.001 | Young |
| 0.898871 | 0.123839 | 0.421 | 0.459 | 1 | 13 | Ddx52 | -0.038 | Young |
| 0.898886 | 0.035991 | 0.219 | 0.238 | 1 | 13 | Nol9 | -0.019 | Young |
| 0.899006 | 0.005947 | 0.939 | 0.902 | 1 | 13 | Znrf2 | 0.037 | Young |
| 0.899417 | 0.006735 | 0.474 | 0.492 | 1 | 13 | Enox2 | -0.018 | Young |
| 0.899596 | 0.008647 | 0.211 | 0.197 | 1 | 13 | Fancl | 0.014 | Young |
| 0.89983 | 0.085867 | 0.482 | 0.467 | 1 | 13 | Emc8 | 0.015 | Young |
| 0.899866 | 0.066283 | 0.061 | 0.057 | 1 | 13 | Tceanc | 0.004 | Young |
| 0.899961 | 0.063379 | 0.114 | 0.123 | 1 | 13 | Mospd1 | -0.009 | Young |
| 0.900283 | 0.024147 | 0.816 | 0.836 | 1 | 13 | Ggh | -0.02 | Young |
| 0.900419 | 0.061795 | 0.386 | 0.434 | 1 | 13 | Ctr9 | -0.048 | Young |
| 0.900697 | 0.014983 | 0.07 | 0.066 | 1 | 13 | Lyrm1 | 0.004 | Young |
| 0.900711 | 0.259126 | 0.035 | 0.033 | 1 | 13 | Zfp882 | 0.002 | Young |
| 0.900711 | 0.25038 | 0.035 | 0.033 | 1 | 13 | Glcci1 | 0.002 | Young |
| 0.901018 | 0.11316 | 0.079 | 0.074 | 1 | 13 | Adrb1 | 0.005 | Young |
| 0.902045 | 0.018281 | 0.939 | 0.951 | 1 | 13 | Metap2 | -0.012 | Young |

| | | | | | | | | |
|---|---|---|---|---|---|---|---|---|
| 0.90206 | 0.108722 | 0.053 | 0.057 | 1 | 13 | Ocstamp | -0.004 | Young |
| 0.90206 | 0.036927 | 0.053 | 0.057 | 1 | 13 | Ccne1 | -0.004 | Young |
| 0.902067 | 0.106647 | 0.509 | 0.59 | 1 | 13 | Nedd9 | -0.081 | Young |
| 0.902268 | 0.087225 | 0.053 | 0.049 | 1 | 13 | Klhdc1 | 0.004 | Young |
| 0.902299 | 0.298318 | 0.044 | 0.049 | 1 | 13 | Gm45051 | -0.005 | Young |
| 0.902299 | 0.26857 | 0.044 | 0.049 | 1 | 13 | Gm11772 | -0.005 | Young |
| 0.902498 | 0.093295 | 0.667 | 0.77 | 1 | 13 | Npepl1 | -0.103 | Young |
| 0.902527 | 0.14405 | 0.246 | 0.246 | 1 | 13 | Tex2 | 0 | Young |
| 0.902565 | 0.078798 | 0.158 | 0.172 | 1 | 13 | Tmem18 | -0.014 | Young |
| 0.902573 | 0.123775 | 0.272 | 0.279 | 1 | 13 | Tcp11l1 | -0.007 | Young |
| 0.902651 | 0.082921 | 0.114 | 0.123 | 1 | 13 | Atp23 | -0.009 | Young |
| 0.902801 | 0.251107 | 0.123 | 0.123 | 1 | 13 | Pold2 | 0 | Young |
| 0.902801 | 0.194525 | 0.123 | 0.123 | 1 | 13 | Surf6 | 0 | Young |
| 0.902985 | 0.240713 | 0.132 | 0.131 | 1 | 13 | E130102H2 | 0.001 | Young |
| 0.903213 | 0.033306 | 0.368 | 0.385 | 1 | 13 | Dpp7 | -0.017 | Young |
| 0.903556 | 0.098914 | 0.061 | 0.057 | 1 | 13 | Swsap1 | 0.004 | Young |
| 0.904088 | 0.032147 | 0.439 | 0.451 | 1 | 13 | Pdrg1 | -0.012 | Young |
| 0.904267 | 0.156671 | 0.044 | 0.041 | 1 | 13 | Gm4924 | 0.003 | Young |
| 0.904551 | 0.057787 | 0.272 | 0.311 | 1 | 13 | Aen | -0.039 | Young |
| 0.904567 | 0.013238 | 0.719 | 0.836 | 1 | 13 | Man2a2 | -0.117 | Young |
| 0.90479 | 0.007512 | 0.746 | 0.697 | 1 | 13 | Polr2g | 0.049 | Young |
| 0.904823 | 0.029276 | 0.061 | 0.066 | 1 | 13 | Klf8 | -0.005 | Young |
| 0.904823 | 0.018304 | 0.061 | 0.066 | 1 | 13 | Pomt1 | -0.005 | Young |
| 0.904958 | 0.015329 | 0.825 | 0.902 | 1 | 13 | Casp4 | -0.077 | Young |
| 0.905032 | 0.015153 | 0.912 | 0.893 | 1 | 13 | 2410015M | 0.019 | Young |
| 0.905066 | 0.00345 | 0.544 | 0.656 | 1 | 13 | Btaf1 | -0.112 | Young |
| 0.905119 | 0.128602 | 0.237 | 0.262 | 1 | 13 | Prr3 | -0.025 | Young |
| 0.905275 | 0.054755 | 0.342 | 0.352 | 1 | 13 | Rai1 | -0.01 | Young |
| 0.905531 | 0.24027 | 0.035 | 0.033 | 1 | 13 | 4732496C0 | 0.002 | Young |
| 0.905531 | 0.197159 | 0.035 | 0.033 | 1 | 13 | Psma8 | 0.002 | Young |
| 0.905531 | 0.138428 | 0.035 | 0.033 | 1 | 13 | Mpzl3 | 0.002 | Young |
| 0.905565 | 0.06069 | 0.132 | 0.139 | 1 | 13 | Zbtb5 | -0.007 | Young |
| 0.905831 | 0.017194 | 0.974 | 0.975 | 1 | 13 | Trps1 | -0.001 | Young |
| 0.905832 | 0.030536 | 0.991 | 0.975 | 1 | 13 | Cbl | 0.016 | Young |
| 0.905882 | 0.168015 | 0.053 | 0.057 | 1 | 13 | Zfp456 | -0.004 | Young |
| 0.905882 | 0.04967 | 0.053 | 0.057 | 1 | 13 | Spdl1 | -0.004 | Young |
| 0.90611 | 0.060422 | 0.465 | 0.508 | 1 | 13 | Nup62 | -0.043 | Young |
| 0.906117 | 0.034838 | 0.307 | 0.344 | 1 | 13 | Ppwd1 | -0.037 | Young |
| 0.906193 | 0.024485 | 0.64 | 0.672 | 1 | 13 | Arhgef3 | -0.032 | Young |
| 0.906193 | 0.005223 | 0.623 | 0.689 | 1 | 13 | Pja2 | -0.066 | Young |
| 0.906319 | 0.05668 | 0.728 | 0.713 | 1 | 13 | Brd7 | 0.015 | Young |
| 0.906433 | 0.033556 | 0.281 | 0.303 | 1 | 13 | Eva1b | -0.022 | Young |
| 0.906596 | 0.000871 | 0.667 | 0.697 | 1 | 13 | Sars | -0.03 | Young |
| 0.906717 | 0.184114 | 0.219 | 0.246 | 1 | 13 | Rabep2 | -0.027 | Young |
| 0.906909 | 0.094185 | 0.263 | 0.295 | 1 | 13 | Pphln1 | -0.032 | Young |

| | | | | | | | | |
|---|---|---|---|---|---|---|---|---|
| 0.907099 | 0.010934 | 0.465 | 0.516 | 1 | 13 | Nsmce1 | -0.051 | Young |
| 0.907216 | 0.070782 | 0.447 | 0.467 | 1 | 13 | Ash2l | -0.02 | Young |
| 0.907409 | 0.021632 | 0.421 | 0.467 | 1 | 13 | Ltv1 | -0.046 | Young |
| 0.907727 | 0.020536 | 0.518 | 0.541 | 1 | 13 | Rsl24d1 | -0.023 | Young |
| 0.908106 | 0.032183 | 0.132 | 0.139 | 1 | 13 | Ift74 | -0.007 | Young |
| 0.908308 | 0.087513 | 0.781 | 0.762 | 1 | 13 | Mon2 | 0.019 | Young |
| 0.908353 | 0.039763 | 0.237 | 0.238 | 1 | 13 | Sfi1 | -0.001 | Young |
| 0.908408 | 0.218126 | 0.526 | 0.574 | 1 | 13 | Arl6ip4 | -0.048 | Young |
| 0.908564 | 0.105858 | 0.579 | 0.656 | 1 | 13 | Strn4 | -0.077 | Young |
| 0.909161 | 0.017359 | 0.605 | 0.615 | 1 | 13 | Ibtk | -0.01 | Young |
| 0.909634 | 0.110421 | 0.211 | 0.213 | 1 | 13 | Champ1 | -0.002 | Young |
| 0.909707 | 0.265626 | 0.053 | 0.057 | 1 | 13 | Gm43660 | -0.004 | Young |
| 0.91 | 0.130797 | 0.465 | 0.5 | 1 | 13 | Srek1ip1 | -0.035 | Young |
| 0.910116 | 0.095675 | 0.316 | 0.344 | 1 | 13 | Tor2a | -0.028 | Young |
| 0.910211 | 0.047759 | 0.053 | 0.049 | 1 | 13 | Zwilch | 0.004 | Young |
| 0.910354 | 0.239028 | 0.035 | 0.033 | 1 | 13 | Cep55 | 0.002 | Young |
| 0.910354 | 0.237178 | 0.035 | 0.033 | 1 | 13 | Tbx6 | 0.002 | Young |
| 0.910557 | 0.129805 | 0.132 | 0.148 | 1 | 13 | Gbp9 | -0.016 | Young |
| 0.910848 | 0.137922 | 0.175 | 0.189 | 1 | 13 | Tbccd1 | -0.014 | Young |
| 0.910855 | 0.707639 | 0.026 | 0.025 | 1 | 13 | Gm45250 | 0.001 | Young |
| 0.910855 | 0.611999 | 0.026 | 0.025 | 1 | 13 | Gm45238 | 0.001 | Young |
| 0.910855 | 0.54733 | 0.026 | 0.025 | 1 | 13 | Gm13205 | 0.001 | Young |
| 0.910855 | 0.505442 | 0.026 | 0.025 | 1 | 13 | Zfp869 | 0.001 | Young |
| 0.910855 | 0.505087 | 0.026 | 0.025 | 1 | 13 | Hoxb3 | 0.001 | Young |
| 0.910855 | 0.484683 | 0.026 | 0.025 | 1 | 13 | Gm10602 | 0.001 | Young |
| 0.910855 | 0.368531 | 0.026 | 0.025 | 1 | 13 | Tmem8b | 0.001 | Young |
| 0.910855 | 0.337665 | 0.026 | 0.025 | 1 | 13 | Aatk | 0.001 | Young |
| 0.910855 | 0.293682 | 0.026 | 0.025 | 1 | 13 | Cd34 | 0.001 | Young |
| 0.910943 | 0.250433 | 0.061 | 0.057 | 1 | 13 | Shank1 | 0.004 | Young |
| 0.911121 | 0.153359 | 0.579 | 0.607 | 1 | 13 | Pcif1 | -0.028 | Young |
| 0.911407 | 0.026634 | 0.167 | 0.18 | 1 | 13 | Fbxl18 | -0.013 | Young |
| 0.911701 | 0.008982 | 0.263 | 0.279 | 1 | 13 | Def6 | -0.016 | Young |
| 0.911886 | 0.041969 | 0.36 | 0.369 | 1 | 13 | Slc4a1ap | -0.009 | Young |
| 0.91208 | 0.188648 | 0.289 | 0.295 | 1 | 13 | Chmp7 | -0.006 | Young |
| 0.912196 | 0.058622 | 0.763 | 0.803 | 1 | 13 | Glrx5 | -0.04 | Young |
| 0.912597 | 0.070846 | 0.421 | 0.475 | 1 | 13 | Exoc6b | -0.054 | Young |
| 0.912638 | 0.013611 | 1 | 1 | 1 | 13 | Atp6v0b | 0 | Young |
| 0.912934 | 0.119292 | 0.044 | 0.041 | 1 | 13 | Hhat | 0.003 | Young |
| 0.912934 | 0.102709 | 0.044 | 0.041 | 1 | 13 | Jmjd7 | 0.003 | Young |
| 0.912957 | 0.062941 | 0.325 | 0.361 | 1 | 13 | Klhl2 | -0.036 | Young |
| 0.913337 | 0.030761 | 0.886 | 0.893 | 1 | 13 | Srrm1 | -0.007 | Young |
| 0.913369 | 0.018519 | 0.412 | 0.418 | 1 | 13 | Ctsf | -0.006 | Young |
| 0.913386 | 0.012605 | 0.939 | 0.943 | 1 | 13 | Edf1 | -0.004 | Young |
| 0.913699 | 0.020551 | 0.175 | 0.172 | 1 | 13 | Sufu | 0.003 | Young |
| 0.913914 | 0.020752 | 0.632 | 0.68 | 1 | 13 | Mettl9 | -0.048 | Young |

| | | | | | | | | |
|---|---|---|---|---|---|---|---|---|
| 0.913986 | 0.028999 | 0.825 | 0.861 | 1 | 13 | Map3k1 | -0.036 | Young |
| 0.914072 | 0.003906 | 0.868 | 0.885 | 1 | 13 | Ube2s | -0.017 | Young |
| 0.914186 | 0.031667 | 0.053 | 0.049 | 1 | 13 | Fam71e1 | 0.004 | Young |
| 0.914255 | 0.258974 | 0.167 | 0.164 | 1 | 13 | Mapk7 | 0.003 | Young |
| 0.914255 | 0.154101 | 0.167 | 0.164 | 1 | 13 | Cd244 | 0.003 | Young |
| 0.914305 | 0.004382 | 0.184 | 0.205 | 1 | 13 | Orai2 | -0.021 | Young |
| 0.914382 | 0.128349 | 0.509 | 0.566 | 1 | 13 | Ppfibp2 | -0.057 | Young |
| 0.915153 | 0.069494 | 0.211 | 0.23 | 1 | 13 | Fbxo45 | -0.019 | Young |
| 0.915181 | 0.195285 | 0.035 | 0.033 | 1 | 13 | Gm29107 | 0.002 | Young |
| 0.915181 | 0.15778 | 0.035 | 0.033 | 1 | 13 | Gins3 | 0.002 | Young |
| 0.915181 | 0.135194 | 0.035 | 0.033 | 1 | 13 | Insl6 | 0.002 | Young |
| 0.915181 | 0.127845 | 0.035 | 0.033 | 1 | 13 | Adgra3 | 0.002 | Young |
| 0.91533 | 0.35092 | 0.202 | 0.213 | 1 | 13 | Pgm3 | -0.011 | Young |
| 0.91594 | 0.062458 | 0.412 | 0.451 | 1 | 13 | Zmym2 | -0.039 | Young |
| 0.916063 | 0.304254 | 0.123 | 0.123 | 1 | 13 | Rfxank | 0 | Young |
| 0.916405 | 0.364882 | 0.026 | 0.025 | 1 | 13 | Aldh1l1 | 0.001 | Young |
| 0.916405 | 0.339904 | 0.026 | 0.025 | 1 | 13 | Mettl18 | 0.001 | Young |
| 0.916405 | 0.32322 | 0.026 | 0.025 | 1 | 13 | Prdm16 | 0.001 | Young |
| 0.916405 | 0.29179 | 0.026 | 0.025 | 1 | 13 | E230001N( | 0.001 | Young |
| 0.916405 | 0.279022 | 0.026 | 0.025 | 1 | 13 | Mtfr2 | 0.001 | Young |
| 0.916405 | 0.238967 | 0.026 | 0.025 | 1 | 13 | Lrrc56 | 0.001 | Young |
| 0.916405 | 0.226211 | 0.026 | 0.025 | 1 | 13 | Casp12 | 0.001 | Young |
| 0.916896 | 0.058606 | 0.553 | 0.59 | 1 | 13 | Rbm28 | -0.037 | Young |
| 0.917194 | 0.017606 | 0.412 | 0.434 | 1 | 13 | Lman2l | -0.022 | Young |
| 0.917271 | 0.132639 | 0.044 | 0.041 | 1 | 13 | Tmem120b | 0.003 | Young |
| 0.917271 | 0.042805 | 0.044 | 0.041 | 1 | 13 | 4732440D( | 0.003 | Young |
| 0.917271 | 0.020484 | 0.044 | 0.041 | 1 | 13 | Ankrd9 | 0.003 | Young |
| 0.917322 | 0.239922 | 0.281 | 0.295 | 1 | 13 | Pyurf | -0.014 | Young |
| 0.917632 | 0.013414 | 0.816 | 0.795 | 1 | 13 | Mrpl57 | 0.021 | Young |
| 0.917761 | 0.019557 | 0.43 | 0.475 | 1 | 13 | Pdlim2 | -0.045 | Young |
| 0.917826 | 0.001404 | 0.614 | 0.615 | 1 | 13 | Rps6ka4 | -0.001 | Young |
| 0.917882 | 0.078593 | 0.851 | 0.926 | 1 | 13 | Jup | -0.075 | Young |
| 0.917945 | 0.017052 | 0.421 | 0.492 | 1 | 13 | Khnyn | -0.071 | Young |
| 0.918104 | 0.048232 | 0.509 | 0.5 | 1 | 13 | Acot9 | 0.009 | Young |
| 0.918382 | 0.041254 | 0.263 | 0.295 | 1 | 13 | Rnf215 | -0.032 | Young |
| 0.91851 | 0.005937 | 0.816 | 0.852 | 1 | 13 | Srebf2 | -0.036 | Young |
| 0.918694 | 0.037548 | 1 | 0.975 | 1 | 13 | Plek | 0.025 | Young |
| 0.918918 | 0.228641 | 0.544 | 0.516 | 1 | 13 | Hgsnat | 0.028 | Young |
| 0.919136 | 0.031767 | 0.061 | 0.066 | 1 | 13 | Tmc4 | -0.005 | Young |
| 0.919136 | 0.00626 | 0.061 | 0.066 | 1 | 13 | Hist1h4h | -0.005 | Young |
| 0.919403 | 0.060031 | 0.702 | 0.754 | 1 | 13 | Golga4 | -0.052 | Young |
| 0.91946 | 0.073836 | 0.465 | 0.484 | 1 | 13 | Cep83 | -0.019 | Young |
| 0.919576 | 0.132365 | 0.491 | 0.508 | 1 | 13 | Lcmt1 | -0.017 | Young |
| 0.920011 | 0.732599 | 0.035 | 0.033 | 1 | 13 | Tspoap1 | 0.002 | Young |
| 0.920011 | 0.113789 | 0.035 | 0.033 | 1 | 13 | Gm45774 | 0.002 | Young |

| | | | | | | | | |
|---|---|---|---|---|---|---|---|---|
| 0.920011 | 0.084846 | 0.035 | 0.033 | 1 | 13 | Zfp189 | 0.002 | Young |
| 0.920011 | 0.073169 | 0.035 | 0.033 | 1 | 13 | Xlr4c | 0.002 | Young |
| 0.920227 | 0.155914 | 0.211 | 0.213 | 1 | 13 | Cep85l | -0.002 | Young |
| 0.920519 | 0.081219 | 0.509 | 0.549 | 1 | 13 | Pnkd | -0.04 | Young |
| 0.920579 | 0.115029 | 0.175 | 0.172 | 1 | 13 | Mrvi1 | 0.003 | Young |
| 0.92073 | 0.056634 | 0.693 | 0.721 | 1 | 13 | Ube2h | -0.028 | Young |
| 0.920755 | 0.037472 | 0.298 | 0.311 | 1 | 13 | Hcfc2 | -0.013 | Young |
| 0.920933 | 0.077363 | 0.263 | 0.27 | 1 | 13 | Etfdh | -0.007 | Young |
| 0.920937 | 0.048142 | 0.5 | 0.533 | 1 | 13 | Ireb2 | -0.033 | Young |
| 0.921193 | 0.335633 | 0.053 | 0.057 | 1 | 13 | Cdca7 | -0.004 | Young |
| 0.921193 | 0.227076 | 0.053 | 0.057 | 1 | 13 | Cdiptos | -0.004 | Young |
| 0.921526 | 0.013021 | 0.07 | 0.066 | 1 | 13 | Usp2 | 0.004 | Young |
| 0.921591 | 0.050478 | 0.439 | 0.492 | 1 | 13 | Gid8 | -0.053 | Young |
| 0.921599 | 0.001846 | 0.14 | 0.148 | 1 | 13 | Inpp5b | -0.008 | Young |
| 0.921611 | 0.036978 | 0.044 | 0.041 | 1 | 13 | Dsn1 | 0.003 | Young |
| 0.921611 | 0.036501 | 0.044 | 0.041 | 1 | 13 | Gm29336 | 0.003 | Young |
| 0.921658 | 0.035354 | 0.842 | 0.918 | 1 | 13 | Zfp106 | -0.076 | Young |
| 0.921723 | 0.053605 | 1 | 0.984 | 1 | 13 | Wdr26 | 0.016 | Young |
| 0.92196 | 0.575283 | 0.026 | 0.025 | 1 | 13 | Dnd1 | 0.001 | Young |
| 0.92196 | 0.366665 | 0.026 | 0.025 | 1 | 13 | Mcm10 | 0.001 | Young |
| 0.92196 | 0.257551 | 0.026 | 0.025 | 1 | 13 | Lpcat4 | 0.001 | Young |
| 0.92196 | 0.219891 | 0.026 | 0.025 | 1 | 13 | Arhgef40 | 0.001 | Young |
| 0.92196 | 0.211542 | 0.026 | 0.025 | 1 | 13 | Il27 | 0.001 | Young |
| 0.922482 | 0.118486 | 0.333 | 0.369 | 1 | 13 | Chmp6 | -0.036 | Young |
| 0.922515 | 0.052347 | 0.57 | 0.582 | 1 | 13 | Esf1 | -0.012 | Young |
| 0.922545 | 0.05486 | 0.518 | 0.516 | 1 | 13 | Snap29 | 0.002 | Young |
| 0.922674 | 0.05589 | 0.491 | 0.5 | 1 | 13 | Ostm1 | -0.009 | Young |
| 0.922718 | 0.015782 | 0.061 | 0.066 | 1 | 13 | Zfp619 | -0.005 | Young |
| 0.923237 | 0.008346 | 0.982 | 0.959 | 1 | 13 | Csrnp1 | 0.023 | Young |
| 0.923263 | 0.018644 | 0.193 | 0.213 | 1 | 13 | Zfp654 | -0.02 | Young |
| 0.923383 | 0.057365 | 0.43 | 0.467 | 1 | 13 | Sh3gl1 | -0.037 | Young |
| 0.923697 | 0.031867 | 0.07 | 0.074 | 1 | 13 | Gm48066 | -0.004 | Young |
| 0.923729 | 0.094249 | 0.447 | 0.467 | 1 | 13 | Fnbp4 | -0.02 | Young |
| 0.924374 | 0.087308 | 0.404 | 0.426 | 1 | 13 | Slc7a7 | -0.022 | Young |
| 0.924558 | 0.034674 | 0.246 | 0.246 | 1 | 13 | Stxbp1 | 0 | Young |
| 0.924781 | 0.057043 | 0.386 | 0.393 | 1 | 13 | Vps16 | -0.007 | Young |
| 0.924844 | 0.266968 | 0.035 | 0.033 | 1 | 13 | Gm17089 | 0.002 | Young |
| 0.924844 | 0.049507 | 0.035 | 0.033 | 1 | 13 | Zfp119b | 0.002 | Young |
| 0.924844 | 0.032217 | 0.035 | 0.033 | 1 | 13 | Kctd7 | 0.002 | Young |
| 0.924844 | 0.026592 | 0.035 | 0.033 | 1 | 13 | Fbxo10 | 0.002 | Young |
| 0.925627 | 0.003169 | 0.737 | 0.746 | 1 | 13 | Kitl | -0.009 | Young |
| 0.92592 | 0.099568 | 0.465 | 0.525 | 1 | 13 | Cetn2 | -0.06 | Young |
| 0.92616 | 0.026302 | 0.842 | 0.869 | 1 | 13 | Tcea1 | -0.027 | Young |
| 0.926192 | 0.189564 | 0.132 | 0.131 | 1 | 13 | Haao | 0.001 | Young |
| 0.926302 | 0.159868 | 0.061 | 0.066 | 1 | 13 | Dact3 | -0.005 | Young |

| | | | | | | | | |
|---|---|---|---|---|---|---|---|---|
| 0.926302 | 0.079051 | 0.061 | 0.066 | 1 | 13 | Ovca2 | -0.005 | Young |
| 0.926311 | 0.100709 | 0.482 | 0.484 | 1 | 13 | Mrpl27 | -0.002 | Young |
| 0.926772 | 0.064907 | 0.254 | 0.279 | 1 | 13 | Haus2 | -0.025 | Young |
| 0.927021 | 0.054173 | 0.246 | 0.254 | 1 | 13 | Dtx4 | -0.008 | Young |
| 0.927068 | 0.131707 | 0.342 | 0.393 | 1 | 13 | Cwc25 | -0.051 | Young |
| 0.927079 | 0.04517 | 0.07 | 0.074 | 1 | 13 | E030030I0( | -0.004 | Young |
| 0.927518 | 0.366756 | 0.026 | 0.025 | 1 | 13 | Kif4 | 0.001 | Young |
| 0.927518 | 0.316618 | 0.026 | 0.025 | 1 | 13 | Ndc80 | 0.001 | Young |
| 0.927518 | 0.286864 | 0.026 | 0.025 | 1 | 13 | AC174678. | 0.001 | Young |
| 0.927518 | 0.186438 | 0.026 | 0.025 | 1 | 13 | Melk | 0.001 | Young |
| 0.927518 | 0.152098 | 0.026 | 0.025 | 1 | 13 | Sorcs2 | 0.001 | Young |
| 0.927518 | 0.145765 | 0.026 | 0.025 | 1 | 13 | Exd1 | 0.001 | Young |
| 0.927518 | 0.135026 | 0.026 | 0.025 | 1 | 13 | Gm2814 | 0.001 | Young |
| 0.927518 | 0.130401 | 0.026 | 0.025 | 1 | 13 | 1810010D( | 0.001 | Young |
| 0.927518 | 0.127377 | 0.026 | 0.025 | 1 | 13 | Psmc3ip | 0.001 | Young |
| 0.927518 | 0.099515 | 0.026 | 0.025 | 1 | 13 | Siah1b | 0.001 | Young |
| 0.927518 | 0.092635 | 0.026 | 0.025 | 1 | 13 | 2610020C( | 0.001 | Young |
| 0.927518 | 0.085259 | 0.026 | 0.025 | 1 | 13 | 2510017J1 | 0.001 | Young |
| 0.927518 | 0.042332 | 0.026 | 0.025 | 1 | 13 | Naip1 | 0.001 | Young |
| 0.927535 | 0.015698 | 0.816 | 0.803 | 1 | 13 | Safb2 | 0.013 | Young |
| 0.927635 | 0.056443 | 0.807 | 0.869 | 1 | 13 | Mrps14 | -0.062 | Young |
| 0.92792 | 0.261167 | 0.184 | 0.197 | 1 | 13 | Asf1a | -0.013 | Young |
| 0.928177 | 0.096631 | 0.509 | 0.525 | 1 | 13 | Cdk19 | -0.016 | Young |
| 0.928443 | 0.018976 | 0.719 | 0.705 | 1 | 13 | Ntan1 | 0.014 | Young |
| 0.928481 | 0.089649 | 0.693 | 0.738 | 1 | 13 | BC005624 | -0.045 | Young |
| 0.928652 | 0.162154 | 0.202 | 0.205 | 1 | 13 | Fam58b | -0.003 | Young |
| 0.928685 | 0.045878 | 0.719 | 0.762 | 1 | 13 | Tmem127 | -0.043 | Young |
| 0.92872 | 0.14493 | 0.263 | 0.27 | 1 | 13 | Gmcl1 | -0.007 | Young |
| 0.92886 | 0.31063 | 0.053 | 0.057 | 1 | 13 | Aars2 | -0.004 | Young |
| 0.928866 | 0.085581 | 0.386 | 0.418 | 1 | 13 | Anapc4 | -0.032 | Young |
| 0.929086 | 0.015678 | 0.632 | 0.639 | 1 | 13 | Yipf1 | -0.007 | Young |
| 0.929447 | 0.123477 | 0.246 | 0.262 | 1 | 13 | Utp15 | -0.016 | Young |
| 0.929487 | 0.064334 | 0.649 | 0.68 | 1 | 13 | Ubr2 | -0.031 | Young |
| 0.929487 | 0.202103 | 0.632 | 0.697 | 1 | 13 | Commd1 | -0.065 | Young |
| 0.929602 | 0.005858 | 0.623 | 0.615 | 1 | 13 | Gsk3a | 0.008 | Young |
| 0.92968 | 0.077991 | 0.035 | 0.033 | 1 | 13 | Mettl15 | 0.002 | Young |
| 0.92968 | 0.025551 | 0.035 | 0.033 | 1 | 13 | Zfp30 | 0.002 | Young |
| 0.92968 | 0.011176 | 0.035 | 0.033 | 1 | 13 | Iqcd | 0.002 | Young |
| 0.92985 | 0.180161 | 0.114 | 0.115 | 1 | 13 | Ap4e1 | -0.001 | Young |
| 0.929973 | 0.014529 | 0.868 | 0.861 | 1 | 13 | Cd302 | 0.007 | Young |
| 0.930236 | 0.032232 | 0.281 | 0.287 | 1 | 13 | Gfm1 | -0.006 | Young |
| 0.93038 | 0.166359 | 0.123 | 0.131 | 1 | 13 | Dnajb5 | -0.008 | Young |
| 0.931012 | 0.041943 | 0.132 | 0.139 | 1 | 13 | Bcdin3d | -0.007 | Young |
| 0.931695 | 0.206116 | 0.404 | 0.451 | 1 | 13 | Pknox1 | -0.047 | Young |
| 0.931715 | 0.001191 | 0.754 | 0.721 | 1 | 13 | Slc39a7 | 0.033 | Young |

| | | | | | | | | |
|---|---|---|---|---|---|---|---|---|
| 0.931845 | 0.01558 | 0.316 | 0.344 | 1 | 13 | Cnnm3 | -0.028 | Young |
| 0.931859 | 0.13026 | 0.342 | 0.377 | 1 | 13 | Fbxo34 | -0.035 | Young |
| 0.932259 | 0.004873 | 0.877 | 0.861 | 1 | 13 | Ubtf | 0.016 | Young |
| 0.932345 | 0.03118 | 0.184 | 0.197 | 1 | 13 | Cbr3 | -0.013 | Young |
| 0.93272 | 0.097375 | 0.175 | 0.18 | 1 | 13 | Lrrc57 | -0.005 | Young |
| 0.932774 | 0.067763 | 0.36 | 0.393 | 1 | 13 | Clec2i | -0.033 | Young |
| 0.933 | 0.311736 | 0.123 | 0.131 | 1 | 13 | Miga2 | -0.008 | Young |
| 0.93308 | 0.266663 | 0.026 | 0.025 | 1 | 13 | Rragd | 0.001 | Young |
| 0.93308 | 0.215412 | 0.026 | 0.025 | 1 | 13 | Nudt8.1 | 0.001 | Young |
| 0.93308 | 0.15572 | 0.026 | 0.025 | 1 | 13 | Gm48653 | 0.001 | Young |
| 0.93308 | 0.127055 | 0.026 | 0.025 | 1 | 13 | 6430531B1 | 0.001 | Young |
| 0.93308 | 0.114984 | 0.026 | 0.025 | 1 | 13 | Sparcl1 | 0.001 | Young |
| 0.93308 | 0.108792 | 0.026 | 0.025 | 1 | 13 | Gm15853 | 0.001 | Young |
| 0.93308 | 0.088996 | 0.026 | 0.025 | 1 | 13 | Meis1 | 0.001 | Young |
| 0.93308 | 0.06953 | 0.026 | 0.025 | 1 | 13 | Nr2f2 | 0.001 | Young |
| 0.93308 | 0.057272 | 0.026 | 0.025 | 1 | 13 | Rbp4 | 0.001 | Young |
| 0.93308 | 0.049026 | 0.026 | 0.025 | 1 | 13 | AL591952.3 | 0.001 | Young |
| 0.93308 | 0.037587 | 0.026 | 0.025 | 1 | 13 | Enah | 0.001 | Young |
| 0.93308 | 0.03652 | 0.026 | 0.025 | 1 | 13 | Rims3 | 0.001 | Young |
| 0.93308 | 0.022843 | 0.026 | 0.025 | 1 | 13 | AC139671. | 0.001 | Young |
| 0.933089 | 0.142887 | 0.605 | 0.68 | 1 | 13 | Nap1l4 | -0.075 | Young |
| 0.933147 | 0.004661 | 0.298 | 0.295 | 1 | 13 | Ppil3 | 0.003 | Young |
| 0.933357 | 0.018707 | 0.737 | 0.77 | 1 | 13 | Ssr2 | -0.033 | Young |
| 0.933438 | 0.052029 | 0.605 | 0.623 | 1 | 13 | Atp2c1 | -0.018 | Young |
| 0.933513 | 0.317717 | 0.096 | 0.098 | 1 | 13 | Spred3 | -0.002 | Young |
| 0.933871 | 0.04739 | 0.439 | 0.443 | 1 | 13 | Gabpa | -0.004 | Young |
| 0.93402 | 0.010178 | 0.202 | 0.221 | 1 | 13 | Zbtb42 | -0.019 | Young |
| 0.934604 | 0.043172 | 0.956 | 0.926 | 1 | 13 | Ndufb4 | 0.03 | Young |
| 0.934668 | 0.259204 | 0.123 | 0.123 | 1 | 13 | Dusp18 | 0 | Young |
| 0.934668 | 0.25473 | 0.123 | 0.123 | 1 | 13 | Ptcd1 | 0 | Young |
| 0.934709 | 0.115454 | 0.281 | 0.303 | 1 | 13 | Pdk3 | -0.022 | Young |
| 0.935 | 0.12757 | 0.596 | 0.631 | 1 | 13 | Ndufs8 | -0.035 | Young |
| 0.93516 | 0.068948 | 0.789 | 0.836 | 1 | 13 | 1-Mar | -0.047 | Young |
| 0.935386 | 2.502967 | 0.018 | 0.016 | 1 | 13 | Alox15 | 0.002 | Young |
| 0.935386 | 0.525733 | 0.018 | 0.016 | 1 | 13 | Me1 | 0.002 | Young |
| 0.935386 | 0.457846 | 0.018 | 0.016 | 1 | 13 | Hist1h4b | 0.002 | Young |
| 0.935386 | 0.447033 | 0.018 | 0.016 | 1 | 13 | Nme9 | 0.002 | Young |
| 0.935386 | 0.422771 | 0.018 | 0.016 | 1 | 13 | Hs3st3a1 | 0.002 | Young |
| 0.935386 | 0.406328 | 0.018 | 0.016 | 1 | 13 | Gm44175 | 0.002 | Young |
| 0.935386 | 0.40464 | 0.018 | 0.016 | 1 | 13 | Kcnc3 | 0.002 | Young |
| 0.935386 | 0.386063 | 0.018 | 0.016 | 1 | 13 | BC055324 | 0.002 | Young |
| 0.935386 | 0.37787 | 0.018 | 0.016 | 1 | 13 | Hecw2 | 0.002 | Young |
| 0.935386 | 0.373905 | 0.018 | 0.016 | 1 | 13 | Gnal | 0.002 | Young |
| 0.935386 | 0.368993 | 0.018 | 0.016 | 1 | 13 | Gm1070 | 0.002 | Young |
| 0.935386 | 0.355658 | 0.018 | 0.016 | 1 | 13 | Gm26690 | 0.002 | Young |

| | | | | | | | | |
|---|---|---|---|---|---|---|---|---|
| 0.935386 | 0.326502 | 0.018 | 0.016 | 1 | 13 | Tex9 | 0.002 | Young |
| 0.935386 | 0.319147 | 0.018 | 0.016 | 1 | 13 | Kif24 | 0.002 | Young |
| 0.935386 | 0.313822 | 0.018 | 0.016 | 1 | 13 | Nuf2 | 0.002 | Young |
| 0.935386 | 0.312947 | 0.018 | 0.016 | 1 | 13 | Slc25a27 | 0.002 | Young |
| 0.935386 | 0.311958 | 0.018 | 0.016 | 1 | 13 | 2900026A( | 0.002 | Young |
| 0.935386 | 0.305828 | 0.018 | 0.016 | 1 | 13 | Gm17275 | 0.002 | Young |
| 0.935386 | 0.301472 | 0.018 | 0.016 | 1 | 13 | Proscos | 0.002 | Young |
| 0.935386 | 0.282269 | 0.018 | 0.016 | 1 | 13 | 2810414N( | 0.002 | Young |
| 0.935386 | 0.281937 | 0.018 | 0.016 | 1 | 13 | Tgfb3 | 0.002 | Young |
| 0.935386 | 0.256375 | 0.018 | 0.016 | 1 | 13 | Ugt1a5 | 0.002 | Young |
| 0.935386 | 0.251821 | 0.018 | 0.016 | 1 | 13 | Rassf7 | 0.002 | Young |
| 0.935386 | 0.243704 | 0.018 | 0.016 | 1 | 13 | St18 | 0.002 | Young |
| 0.935386 | 0.242629 | 0.018 | 0.016 | 1 | 13 | Gm17767 | 0.002 | Young |
| 0.935386 | 0.237887 | 0.018 | 0.016 | 1 | 13 | Gm47428 | 0.002 | Young |
| 0.935386 | 0.216185 | 0.018 | 0.016 | 1 | 13 | Cped1 | 0.002 | Young |
| 0.935386 | 0.187108 | 0.018 | 0.016 | 1 | 13 | Gm30122 | 0.002 | Young |
| 0.935386 | 0.186636 | 0.018 | 0.016 | 1 | 13 | 4931440F1 | 0.002 | Young |
| 0.935386 | 0.173382 | 0.018 | 0.016 | 1 | 13 | Fkbp14 | 0.002 | Young |
| 0.935386 | 0.122402 | 0.018 | 0.016 | 1 | 13 | Phf24 | 0.002 | Young |
| 0.935391 | 0.007215 | 0.351 | 0.369 | 1 | 13 | Zfand2b | -0.018 | Young |
| 0.935934 | 0.118501 | 0.667 | 0.713 | 1 | 13 | Dph3 | -0.046 | Young |
| 0.936259 | 0.053288 | 0.368 | 0.385 | 1 | 13 | Ccdc107 | -0.017 | Young |
| 0.936334 | 0.10717 | 0.491 | 0.549 | 1 | 13 | Cab39l | -0.058 | Young |
| 0.936527 | 0.301928 | 0.132 | 0.131 | 1 | 13 | Rab3ip | 0.001 | Young |
| 0.936816 | 0.000596 | 0.851 | 0.885 | 1 | 13 | Med13l | -0.034 | Young |
| 0.937018 | 0.300874 | 0.158 | 0.164 | 1 | 13 | Epg5 | -0.006 | Young |
| 0.937063 | 0.259941 | 0.061 | 0.066 | 1 | 13 | Rpgr | -0.005 | Young |
| 0.937063 | 0.084402 | 0.061 | 0.066 | 1 | 13 | Platr25 | -0.005 | Young |
| 0.937208 | 0.110533 | 0.263 | 0.287 | 1 | 13 | Acaca | -0.024 | Young |
| 0.937231 | 0.044885 | 0.07 | 0.074 | 1 | 13 | Mypopos | -0.004 | Young |
| 0.937231 | 0.090983 | 0.07 | 0.074 | 1 | 13 | Als2 | -0.004 | Young |
| 0.937235 | 0.10064 | 0.368 | 0.426 | 1 | 13 | Tbc1d17 | -0.058 | Young |
| 0.93732 | 0.085226 | 0.193 | 0.197 | 1 | 13 | Rmdn3 | -0.004 | Young |
| 0.937508 | 0.076258 | 0.439 | 0.459 | 1 | 13 | Capn7 | -0.02 | Young |
| 0.937605 | 0.139102 | 0.237 | 0.254 | 1 | 13 | Rrn3 | -0.017 | Young |
| 0.937866 | 0.009152 | 0.588 | 0.607 | 1 | 13 | Mrpl4 | -0.019 | Young |
| 0.938471 | 0.06879 | 0.623 | 0.656 | 1 | 13 | Scamp3 | -0.033 | Young |
| 0.938589 | 0.050318 | 0.219 | 0.23 | 1 | 13 | Tdp2 | -0.011 | Young |
| 0.938645 | 0.126322 | 0.026 | 0.025 | 1 | 13 | Sgo2a | 0.001 | Young |
| 0.938645 | 0.082593 | 0.026 | 0.025 | 1 | 13 | Zfp85 | 0.001 | Young |
| 0.938645 | 0.021651 | 0.026 | 0.025 | 1 | 13 | Pcdhgb5 | 0.001 | Young |
| 0.938645 | 0.015482 | 0.026 | 0.025 | 1 | 13 | Stard13 | 0.001 | Young |
| 0.938937 | 0.042761 | 0.675 | 0.689 | 1 | 13 | Rnpepl1 | -0.014 | Young |
| 0.939076 | 0.111399 | 0.289 | 0.311 | 1 | 13 | Golph3l | -0.022 | Young |
| 0.939171 | 0.046855 | 0.439 | 0.459 | 1 | 13 | Nbdy | -0.02 | Young |

| | | | | | | | | |
|---|---|---|---|---|---|---|---|---|
| 0.939635 | 0.180435 | 0.298 | 0.32 | 1 | 13 | Ptrh2 | -0.022 | Young |
| 0.939809 | 0.050378 | 0.377 | 0.418 | 1 | 13 | Trim24 | -0.041 | Young |
| 0.940274 | 0.006 | 0.447 | 0.467 | 1 | 13 | Ap1m1 | -0.02 | Young |
| 0.940356 | 0.145872 | 0.649 | 0.689 | 1 | 13 | Fam172a | -0.04 | Young |
| 0.940362 | 0.362029 | 0.079 | 0.082 | 1 | 13 | Birc5 | -0.003 | Young |
| 0.940362 | 0.017369 | 0.079 | 0.082 | 1 | 13 | Atp6v0c | -0.003 | Young |
| 0.940416 | 0.228041 | 0.114 | 0.123 | 1 | 13 | B930036N1 | -0.009 | Young |
| 0.940482 | 0.054603 | 0.325 | 0.352 | 1 | 13 | S1pr2 | -0.027 | Young |
| 0.940543 | 0.061664 | 0.482 | 0.508 | 1 | 13 | Ckap5 | -0.026 | Young |
| 0.940653 | 0.090447 | 0.061 | 0.066 | 1 | 13 | Porcn | -0.005 | Young |
| 0.940722 | 0.117542 | 0.14 | 0.139 | 1 | 13 | Upp2 | 0.001 | Young |
| 0.940947 | 0.148427 | 0.289 | 0.311 | 1 | 13 | Pigu | -0.022 | Young |
| 0.940995 | 0.00687 | 0.202 | 0.213 | 1 | 13 | AI987944 | -0.011 | Young |
| 0.941253 | 0.06223 | 0.175 | 0.172 | 1 | 13 | Cbr1 | 0.003 | Young |
| 0.941304 | 0.06741 | 0.535 | 0.598 | 1 | 13 | Rps12-ps3 | -0.063 | Young |
| 0.942175 | 0.410458 | 0.018 | 0.016 | 1 | 13 | Hdhd3 | 0.002 | Young |
| 0.942175 | 0.25062 | 0.018 | 0.016 | 1 | 13 | Defb25 | 0.002 | Young |
| 0.942175 | 0.232535 | 0.018 | 0.016 | 1 | 13 | Cage1 | 0.002 | Young |
| 0.942175 | 0.223826 | 0.018 | 0.016 | 1 | 13 | Tnfrsf10b | 0.002 | Young |
| 0.942175 | 0.220038 | 0.018 | 0.016 | 1 | 13 | Npdc1 | 0.002 | Young |
| 0.942175 | 0.217426 | 0.018 | 0.016 | 1 | 13 | Srp54c | 0.002 | Young |
| 0.942175 | 0.182109 | 0.018 | 0.016 | 1 | 13 | Rarres1 | 0.002 | Young |
| 0.942175 | 0.18029 | 0.018 | 0.016 | 1 | 13 | D330050G2 | 0.002 | Young |
| 0.942175 | 0.174935 | 0.018 | 0.016 | 1 | 13 | Gm43971 | 0.002 | Young |
| 0.942175 | 0.172176 | 0.018 | 0.016 | 1 | 13 | Gm30246 | 0.002 | Young |
| 0.942175 | 0.1503 | 0.018 | 0.016 | 1 | 13 | B230322F0 | 0.002 | Young |
| 0.942175 | 0.148243 | 0.018 | 0.016 | 1 | 13 | D130017N0 | 0.002 | Young |
| 0.942175 | 0.147834 | 0.018 | 0.016 | 1 | 13 | Gm26660 | 0.002 | Young |
| 0.942175 | 0.11589 | 0.018 | 0.016 | 1 | 13 | Gm42067 | 0.002 | Young |
| 0.942175 | 0.115566 | 0.018 | 0.016 | 1 | 13 | B3gnt7 | 0.002 | Young |
| 0.942175 | 0.106666 | 0.018 | 0.016 | 1 | 13 | Ddr2 | 0.002 | Young |
| 0.942175 | 0.089383 | 0.018 | 0.016 | 1 | 13 | Lbp | 0.002 | Young |
| 0.942175 | 0.07196 | 0.018 | 0.016 | 1 | 13 | Hist1h2ac | 0.002 | Young |
| 0.942175 | 0.063125 | 0.018 | 0.016 | 1 | 13 | Itga8 | 0.002 | Young |
| 0.942175 | 0.062389 | 0.018 | 0.016 | 1 | 13 | Zfp940 | 0.002 | Young |
| 0.942265 | 0.479536 | 0.281 | 0.303 | 1 | 13 | Rrad | -0.022 | Young |
| 0.942364 | 0.044583 | 0.439 | 0.451 | 1 | 13 | Slf2 | -0.012 | Young |
| 0.942458 | 0.001562 | 0.342 | 0.377 | 1 | 13 | Apip | -0.035 | Young |
| 0.943148 | 0.058373 | 0.228 | 0.23 | 1 | 13 | Eif2b1 | -0.002 | Young |
| 0.94358 | 0.156243 | 0.211 | 0.213 | 1 | 13 | Acyp1 | -0.002 | Young |
| 0.943764 | 0.03492 | 0.132 | 0.139 | 1 | 13 | Dgkq | -0.007 | Young |
| 0.944005 | 0.064216 | 0.07 | 0.074 | 1 | 13 | Cav1 | -0.004 | Young |
| 0.944213 | 0.061653 | 0.026 | 0.025 | 1 | 13 | Fbxo5 | 0.001 | Young |
| 0.944213 | 0.032416 | 0.026 | 0.025 | 1 | 13 | Dhrs13 | 0.001 | Young |
| 0.944213 | 0.008223 | 0.026 | 0.025 | 1 | 13 | 9130008F2 | 0.001 | Young |

| | | | | | | | | |
|---|---|---|---|---|---|---|---|---|
| 0.944371 | 0.006958 | 0.833 | 0.852 | 1 | 13 | Acap2 | -0.019 | Young |
| 0.944509 | 0.343009 | 0.325 | 0.369 | 1 | 13 | Tceanc2 | -0.044 | Young |
| 0.944784 | 0.082461 | 0.605 | 0.689 | 1 | 13 | Heatr5a | -0.084 | Young |
| 0.944975 | 0.036388 | 0.36 | 0.393 | 1 | 13 | Abcf2 | -0.033 | Young |
| 0.945263 | 0.052878 | 0.544 | 0.59 | 1 | 13 | Slc45a4 | -0.046 | Young |
| 0.946118 | 0.087021 | 0.211 | 0.221 | 1 | 13 | Ino80b | -0.01 | Young |
| 0.946118 | 0.277458 | 0.211 | 0.221 | 1 | 13 | Zdhhc23 | -0.01 | Young |
| 0.946118 | 0.068936 | 0.211 | 0.221 | 1 | 13 | Casp2 | -0.01 | Young |
| 0.94628 | 0.147348 | 0.395 | 0.426 | 1 | 13 | Nqo2 | -0.031 | Young |
| 0.946317 | 0.166093 | 0.132 | 0.139 | 1 | 13 | Iba57 | -0.007 | Young |
| 0.94653 | 0.198696 | 0.456 | 0.492 | 1 | 13 | Larp1b | -0.036 | Young |
| 0.946677 | 0.156445 | 0.377 | 0.418 | 1 | 13 | 0610010F0 | -0.041 | Young |
| 0.946677 | 0.029507 | 0.395 | 0.402 | 1 | 13 | Exoc1 | -0.007 | Young |
| 0.946889 | 0.24822 | 0.175 | 0.189 | 1 | 13 | Heatr5b | -0.014 | Young |
| 0.947394 | 0.074184 | 0.07 | 0.074 | 1 | 13 | Gm7160 | -0.004 | Young |
| 0.947762 | 0.197249 | 0.351 | 0.369 | 1 | 13 | Dpm1 | -0.018 | Young |
| 0.947835 | 0.141978 | 0.061 | 0.066 | 1 | 13 | Cyb5rl | -0.005 | Young |
| 0.94798 | 0.049902 | 0.123 | 0.123 | 1 | 13 | Gtpbp10 | 0 | Young |
| 0.948014 | 0.061531 | 0.649 | 0.672 | 1 | 13 | Pik3ca | -0.023 | Young |
| 0.948085 | 0.057127 | 0.377 | 0.402 | 1 | 13 | Kdm3a | -0.025 | Young |
| 0.948861 | 0.133284 | 0.57 | 0.623 | 1 | 13 | Mprip | -0.053 | Young |
| 0.948968 | 0.502315 | 0.018 | 0.016 | 1 | 13 | Bcl6b | 0.002 | Young |
| 0.948968 | 0.225885 | 0.018 | 0.016 | 1 | 13 | Atp1a2 | 0.002 | Young |
| 0.948968 | 0.220206 | 0.018 | 0.016 | 1 | 13 | Ackr1 | 0.002 | Young |
| 0.948968 | 0.155624 | 0.018 | 0.016 | 1 | 13 | Sfrp1 | 0.002 | Young |
| 0.948968 | 0.150427 | 0.018 | 0.016 | 1 | 13 | BC043934 | 0.002 | Young |
| 0.948968 | 0.104472 | 0.018 | 0.016 | 1 | 13 | Gm21887 | 0.002 | Young |
| 0.948968 | 0.069294 | 0.018 | 0.016 | 1 | 13 | 4933406PC | 0.002 | Young |
| 0.948968 | 0.064418 | 0.018 | 0.016 | 1 | 13 | Fbxl2 | 0.002 | Young |
| 0.948968 | 0.059083 | 0.018 | 0.016 | 1 | 13 | Ulk4 | 0.002 | Young |
| 0.948968 | 0.056837 | 0.018 | 0.016 | 1 | 13 | Prdm8 | 0.002 | Young |
| 0.948968 | 0.053906 | 0.018 | 0.016 | 1 | 13 | Ccdc18 | 0.002 | Young |
| 0.948968 | 0.047042 | 0.018 | 0.016 | 1 | 13 | Cabp4 | 0.002 | Young |
| 0.948968 | 0.043407 | 0.018 | 0.016 | 1 | 13 | Slfn4 | 0.002 | Young |
| 0.948968 | 0.036212 | 0.018 | 0.016 | 1 | 13 | Adcy6 | 0.002 | Young |
| 0.948968 | 0.032164 | 0.018 | 0.016 | 1 | 13 | Phtf1os | 0.002 | Young |
| 0.948968 | 0.020364 | 0.018 | 0.016 | 1 | 13 | Clec1b | 0.002 | Young |
| 0.948968 | 0.015116 | 0.018 | 0.016 | 1 | 13 | Hist1h2ab | 0.002 | Young |
| 0.948968 | 0.013083 | 0.018 | 0.016 | 1 | 13 | Spaca1 | 0.002 | Young |
| 0.948968 | 0.000271 | 0.018 | 0.016 | 1 | 13 | Shcbp1 | 0.002 | Young |
| 0.949103 | 0.048527 | 0.386 | 0.402 | 1 | 13 | Ufd1 | -0.016 | Young |
| 0.949499 | 0.093743 | 0.377 | 0.385 | 1 | 13 | Svbp | -0.008 | Young |
| 0.949868 | 0.006822 | 0.675 | 0.721 | 1 | 13 | Nhp2 | -0.046 | Young |
| 0.950164 | 0.075657 | 0.737 | 0.762 | 1 | 13 | Phf14 | -0.025 | Young |
| 0.950538 | 0.206807 | 0.298 | 0.311 | 1 | 13 | Mavs | -0.013 | Young |

| | | | | | | | | |
|---|---|---|---|---|---|---|---|---|
| 0.950545 | 0.002792 | 0.93 | 0.926 | 1 | 13 | Hpcal1 | 0.004 | Young |
| 0.950731 | 0.12827 | 0.184 | 0.189 | 1 | 13 | Il1rap | -0.005 | Young |
| 0.950797 | 0.087415 | 0.14 | 0.139 | 1 | 13 | 1700019L1 | 0.001 | Young |
| 0.950915 | 0.029638 | 0.193 | 0.205 | 1 | 13 | Top3b | -0.012 | Young |
| 0.950977 | 0.102877 | 0.246 | 0.254 | 1 | 13 | Usp10 | -0.008 | Young |
| 0.951062 | 0.078996 | 0.219 | 0.23 | 1 | 13 | Mcm3ap | -0.011 | Young |
| 0.951115 | 0.246314 | 0.167 | 0.172 | 1 | 13 | L1cam | -0.005 | Young |
| 0.951563 | 0.02886 | 0.623 | 0.631 | 1 | 13 | Fam114a2 | -0.008 | Young |
| 0.951709 | 0.199032 | 0.202 | 0.213 | 1 | 13 | Fbxl12 | -0.011 | Young |
| 0.951825 | 0.112029 | 0.114 | 0.115 | 1 | 13 | Aldh18a1 | -0.001 | Young |
| 0.951845 | 0.150706 | 0.789 | 0.787 | 1 | 13 | AC160336. | 0.002 | Young |
| 0.952063 | 0.077912 | 0.228 | 0.246 | 1 | 13 | Med27 | -0.018 | Young |
| 0.952298 | 0.02479 | 0.605 | 0.639 | 1 | 13 | Zfc3h1 | -0.034 | Young |
| 0.952376 | 0.043172 | 0.228 | 0.254 | 1 | 13 | Slc35e3 | -0.026 | Young |
| 0.952399 | 0.022696 | 0.202 | 0.205 | 1 | 13 | Zfp738 | -0.003 | Young |
| 0.95283 | 0.140223 | 0.465 | 0.533 | 1 | 13 | Mms19 | -0.068 | Young |
| 0.952866 | 0.057662 | 0.518 | 0.574 | 1 | 13 | Fam193a | -0.056 | Young |
| 0.952998 | 0.021741 | 0.535 | 0.574 | 1 | 13 | Mrpl41 | -0.039 | Young |
| 0.953001 | 0.226574 | 0.105 | 0.107 | 1 | 13 | Disc1 | -0.002 | Young |
| 0.953139 | 0.107991 | 0.272 | 0.295 | 1 | 13 | Cbll1 | -0.023 | Young |
| 0.95324 | 0.148564 | 0.079 | 0.082 | 1 | 13 | Ldlrad3 | -0.003 | Young |
| 0.95344 | 0.106202 | 0.167 | 0.172 | 1 | 13 | Mrpl47 | -0.005 | Young |
| 0.95344 | 0.001791 | 0.167 | 0.172 | 1 | 13 | Fam20a | -0.005 | Young |
| 0.953822 | 0.082641 | 0.421 | 0.451 | 1 | 13 | Mtf2 | -0.03 | Young |
| 0.954053 | 0.032994 | 0.289 | 0.311 | 1 | 13 | Eif2b2 | -0.022 | Young |
| 0.954071 | 0.189878 | 0.263 | 0.27 | 1 | 13 | Sertad3 | -0.007 | Young |
| 0.954574 | 0.277052 | 0.263 | 0.287 | 1 | 13 | Rcbtb1 | -0.024 | Young |
| 0.95466 | 0.003581 | 0.719 | 0.738 | 1 | 13 | Ndufa10 | -0.019 | Young |
| 0.954876 | 0.177384 | 0.307 | 0.328 | 1 | 13 | Slc41a1 | -0.021 | Young |
| 0.955195 | 0.036416 | 0.535 | 0.549 | 1 | 13 | Acbd6 | -0.014 | Young |
| 0.955497 | 0.011479 | 0.175 | 0.18 | 1 | 13 | Fam149b | -0.005 | Young |
| 0.955765 | 0.024402 | 0.018 | 0.016 | 1 | 13 | Ngb | 0.002 | Young |
| 0.955765 | 0.017868 | 0.018 | 0.016 | 1 | 13 | Lrrc75aos2 | 0.002 | Young |
| 0.955765 | 0.003084 | 0.018 | 0.016 | 1 | 13 | 4930513N1 | 0.002 | Young |
| 0.955766 | 0.010253 | 0.167 | 0.172 | 1 | 13 | Wdr4 | -0.005 | Young |
| 0.955919 | 0.073999 | 0.175 | 0.189 | 1 | 13 | Ogfod2 | -0.014 | Young |
| 0.956255 | 0.060222 | 0.719 | 0.762 | 1 | 13 | Smc3 | -0.043 | Young |
| 0.956633 | 0.017186 | 0.974 | 1 | 1 | 13 | Mapre1 | -0.026 | Young |
| 0.957325 | 0.289643 | 0.114 | 0.115 | 1 | 13 | Tnfrsf25 | -0.001 | Young |
| 0.957721 | 0.04039 | 0.386 | 0.402 | 1 | 13 | Fbxo28 | -0.016 | Young |
| 0.957965 | 0.006192 | 0.789 | 0.795 | 1 | 13 | Ndfip1 | -0.006 | Young |
| 0.958025 | 0.031067 | 0.64 | 0.664 | 1 | 13 | Uap1 | -0.024 | Young |
| 0.958153 | 0.027761 | 0.982 | 0.959 | 1 | 13 | Ppp2r5c | 0.023 | Young |
| 0.958166 | 0.032154 | 0.325 | 0.336 | 1 | 13 | Gnpda2 | -0.011 | Young |
| 0.958177 | 0.278144 | 0.175 | 0.189 | 1 | 13 | Cnksr3 | -0.014 | Young |

| | | | | | | | | |
|---|---|---|---|---|---|---|---|---|
| 0.958177 | 0.17381 | 0.175 | 0.189 | 1 | 13 | Ak8 | -0.014 | Young |
| 0.958404 | 0.361103 | 0.088 | 0.09 | 1 | 13 | 6330403L0 | -0.002 | Young |
| 0.958637 | 0.052331 | 0.219 | 0.238 | 1 | 13 | Tmem126b | -0.019 | Young |
| 0.958637 | 0.016076 | 0.728 | 0.795 | 1 | 13 | Ugp2 | -0.067 | Young |
| 0.95864 | 0.119173 | 0.123 | 0.123 | 1 | 13 | Cul7 | 0 | Young |
| 0.958802 | 0.030957 | 0.649 | 0.656 | 1 | 13 | Tmem147 | -0.007 | Young |
| 0.958932 | 0.284058 | 0.184 | 0.197 | 1 | 13 | Nbeal2 | -0.013 | Young |
| 0.959067 | 0.029179 | 0.368 | 0.393 | 1 | 13 | Afdn | -0.025 | Young |
| 0.959222 | 0.12327 | 0.246 | 0.262 | 1 | 13 | Atxn1l | -0.016 | Young |
| 0.959284 | 0.019421 | 0.289 | 0.295 | 1 | 13 | Tmem260 | -0.006 | Young |
| 0.959354 | 0.19606 | 0.114 | 0.123 | 1 | 13 | Gm10036 | -0.009 | Young |
| 0.959386 | 0.071424 | 0.149 | 0.148 | 1 | 13 | Tram2 | 0.001 | Young |
| 0.959633 | 0.050547 | 0.193 | 0.205 | 1 | 13 | Vps51 | -0.012 | Young |
| 0.959951 | 0.019784 | 0.605 | 0.607 | 1 | 13 | Aagab | -0.002 | Young |
| 0.960152 | 0.100432 | 0.263 | 0.279 | 1 | 13 | Zfp266 | -0.016 | Young |
| 0.960475 | 0.031136 | 0.667 | 0.672 | 1 | 13 | Eif2s1 | -0.005 | Young |
| 0.961539 | 0.009991 | 0.272 | 0.287 | 1 | 13 | Acad8 | -0.015 | Young |
| 0.961627 | 0.157732 | 0.456 | 0.525 | 1 | 13 | Vps9d1 | -0.069 | Young |
| 0.961734 | 0.063789 | 0.237 | 0.254 | 1 | 13 | Slc25a16 | -0.017 | Young |
| 0.962062 | 0.364139 | 0.114 | 0.123 | 1 | 13 | Plcl1 | -0.009 | Young |
| 0.962097 | 0.061598 | 0.509 | 0.533 | 1 | 13 | Lrrc8d | -0.024 | Young |
| 0.962558 | 0.011279 | 0.772 | 0.82 | 1 | 13 | Arpc1a | -0.048 | Young |
| 0.962649 | 0.073428 | 0.825 | 0.869 | 1 | 13 | U2af2 | -0.044 | Young |
| 0.963033 | 0.034869 | 0.096 | 0.098 | 1 | 13 | Zfp65 | -0.002 | Young |
| 0.964071 | 0.227444 | 0.395 | 0.418 | 1 | 13 | Shkbp1 | -0.023 | Young |
| 0.964236 | 0.006156 | 0.982 | 0.959 | 1 | 13 | Fis1 | 0.023 | Young |
| 0.964236 | 0.012474 | 0.982 | 1 | 1 | 13 | Atp5d | -0.018 | Young |
| 0.964295 | 0.018352 | 0.64 | 0.68 | 1 | 13 | Ttc39b | -0.04 | Young |
| 0.964972 | 0.008986 | 0.246 | 0.254 | 1 | 13 | Clk2 | -0.008 | Young |
| 0.965043 | 0.0314 | 0.64 | 0.672 | 1 | 13 | Pigp | -0.032 | Young |
| 0.965437 | 0.118108 | 0.325 | 0.336 | 1 | 13 | Sec24d | -0.011 | Young |
| 0.965456 | 0.091452 | 0.342 | 0.377 | 1 | 13 | Cenpc1 | -0.035 | Young |
| 0.965594 | 0.23295 | 0.263 | 0.262 | 1 | 13 | Trp53inp1 | 0.001 | Young |
| 0.965706 | 0.009864 | 0.482 | 0.5 | 1 | 13 | Tatdn2 | -0.018 | Young |
| 0.965718 | 0.007343 | 0.868 | 0.869 | 1 | 13 | Add1 | -0.001 | Young |
| 0.965757 | 0.013932 | 1 | 1 | 1 | 13 | Sem1 | 0 | Young |
| 0.965833 | 0.129718 | 0.5 | 0.5 | 1 | 13 | Gm11808 | 0 | Young |
| 0.96613 | 0.127232 | 0.079 | 0.082 | 1 | 13 | Sap18b | -0.003 | Young |
| 0.966364 | 0.112832 | 0.596 | 0.648 | 1 | 13 | Frg1 | -0.052 | Young |
| 0.966688 | 0.118102 | 0.281 | 0.295 | 1 | 13 | Cfap20 | -0.014 | Young |
| 0.967128 | 0.076152 | 0.123 | 0.131 | 1 | 13 | Tmem115 | -0.008 | Young |
| 0.967179 | 0.0451 | 0.807 | 0.828 | 1 | 13 | Stub1 | -0.021 | Young |
| 0.967325 | 0.064392 | 0.298 | 0.311 | 1 | 13 | Tmem198b | -0.013 | Young |
| 0.9686 | 0.068322 | 0.746 | 0.787 | 1 | 13 | Mrpl36 | -0.041 | Young |
| 0.968621 | 0.021936 | 0.772 | 0.779 | 1 | 13 | Idh2 | -0.007 | Young |

| | | | | | | | | |
|---|---|---|---|---|---|---|---|---|
| 0.96869 | 0.042827 | 0.789 | 0.828 | 1 | 13 | Tmbim4 | -0.039 | Young |
| 0.968799 | 0.010566 | 1 | 1 | 1 | 13 | Vamp8 | 0 | Young |
| 0.968944 | 0.201735 | 0.096 | 0.098 | 1 | 13 | Vcpkmt | -0.002 | Young |
| 0.969161 | 0.068593 | 0.561 | 0.598 | 1 | 13 | Wars | -0.037 | Young |
| 0.969354 | 0.1498 | 0.079 | 0.082 | 1 | 13 | Gja1 | -0.003 | Young |
| 0.969354 | 0.024179 | 0.079 | 0.082 | 1 | 13 | Fbxo46 | -0.003 | Young |
| 0.969547 | 0.003217 | 0.904 | 0.91 | 1 | 13 | Dmac1 | -0.006 | Young |
| 0.969548 | 0.130289 | 0.456 | 0.492 | 1 | 13 | Maml1 | -0.036 | Young |
| 0.969792 | 0.001738 | 0.219 | 0.23 | 1 | 13 | Pigv | -0.011 | Young |
| 0.969843 | 0.079835 | 0.5 | 0.492 | 1 | 13 | Ppt2 | 0.008 | Young |
| 0.97017 | 0.022565 | 0.754 | 0.811 | 1 | 13 | Tmem55b | -0.057 | Young |
| 0.970187 | 0.251215 | 0.114 | 0.123 | 1 | 13 | Cbx8 | -0.009 | Young |
| 0.970211 | 0.029037 | 0.228 | 0.238 | 1 | 13 | Tmem209 | -0.01 | Young |
| 0.970258 | 0.100311 | 0.807 | 0.869 | 1 | 13 | Cic | -0.062 | Young |
| 0.970411 | 0.11816 | 0.228 | 0.246 | 1 | 13 | Cdk7 | -0.018 | Young |
| 0.970891 | 0.073648 | 0.325 | 0.336 | 1 | 13 | Pgm1 | -0.011 | Young |
| 0.971014 | 0.065922 | 0.202 | 0.213 | 1 | 13 | Zfp715 | -0.011 | Young |
| 0.971137 | 0.181744 | 0.07 | 0.074 | 1 | 13 | Zscan12 | -0.004 | Young |
| 0.971137 | 0.126857 | 0.07 | 0.074 | 1 | 13 | Pfkm | -0.004 | Young |
| 0.972388 | 0.104681 | 0.57 | 0.607 | 1 | 13 | Vkorc1 | -0.037 | Young |
| 0.972579 | 0.004395 | 0.079 | 0.082 | 1 | 13 | Nupl2 | -0.003 | Young |
| 0.972765 | 0.053921 | 0.649 | 0.648 | 1 | 13 | St6galnac6 | 0.001 | Young |
| 0.97332 | 0.023479 | 0.851 | 0.852 | 1 | 13 | S100a1 | -0.001 | Young |
| 0.973359 | 0.03645 | 0.947 | 0.918 | 1 | 13 | Bcl10 | 0.029 | Young |
| 0.973549 | 0.085849 | 0.5 | 0.574 | 1 | 13 | Esyt2 | -0.074 | Young |
| 0.973605 | 0.001231 | 0.64 | 0.68 | 1 | 13 | Elovl5 | -0.04 | Young |
| 0.973606 | 0.22576 | 0.254 | 0.279 | 1 | 13 | Dynlt1f | -0.025 | Young |
| 0.974026 | 0.092091 | 0.439 | 0.443 | 1 | 13 | Fgd6 | -0.004 | Young |
| 0.974536 | 0.1335 | 0.456 | 0.5 | 1 | 13 | Tax1bp3 | -0.044 | Young |
| 0.974551 | 0.081426 | 0.702 | 0.705 | 1 | 13 | Marveld1 | -0.003 | Young |
| 0.974732 | 0.020895 | 0.482 | 0.508 | 1 | 13 | Zmat5 | -0.026 | Young |
| 0.974858 | 0.173192 | 0.096 | 0.098 | 1 | 13 | Nup37 | -0.002 | Young |
| 0.974885 | 0.027922 | 1 | 1 | 1 | 13 | Klf6 | 0 | Young |
| 0.974894 | 0.05827 | 0.386 | 0.393 | 1 | 13 | Tmem60 | -0.007 | Young |
| 0.974923 | 0.02291 | 0.342 | 0.352 | 1 | 13 | Ccdc127 | -0.01 | Young |
| 0.975014 | 0.408808 | 0.123 | 0.131 | 1 | 13 | Rbbp9 | -0.008 | Young |
| 0.975761 | 0.014999 | 0.5 | 0.533 | 1 | 13 | Nkiras2 | -0.033 | Young |
| 0.97613 | 0.007071 | 0.474 | 0.475 | 1 | 13 | Ubfd1 | -0.001 | Young |
| 0.976168 | 0.070931 | 0.702 | 0.754 | 1 | 13 | Sqor | -0.052 | Young |
| 0.976313 | 0.001901 | 0.763 | 0.836 | 1 | 13 | Pdap1 | -0.073 | Young |
| 0.976321 | 0.102768 | 0.281 | 0.303 | 1 | 13 | Uty | -0.022 | Young |
| 0.976394 | 0.067838 | 0.895 | 0.902 | 1 | 13 | Nrp1 | -0.007 | Young |
| 0.976673 | 0.017295 | 0.623 | 0.68 | 1 | 13 | Bmt2 | -0.057 | Young |
| 0.976684 | 0.014436 | 0.237 | 0.246 | 1 | 13 | Exosc1 | -0.009 | Young |
| 0.976969 | 0.181063 | 0.307 | 0.328 | 1 | 13 | Dubr | -0.021 | Young |

| | | | | | | | | |
|---|---|---|---|---|---|---|---|---|
| 0.977265 | 0.111691 | 0.421 | 0.443 | 1 | 13 | Polr1c | -0.022 | Young |
| 0.977311 | 0.181189 | 0.123 | 0.123 | 1 | 13 | Ap1s3 | 0 | Young |
| 0.97749 | 0.13133 | 0.36 | 0.418 | 1 | 13 | Trip4 | -0.058 | Young |
| 0.977994 | 0.024485 | 0.281 | 0.287 | 1 | 13 | Alpk1 | -0.006 | Young |
| 0.978101 | 0.235303 | 0.149 | 0.156 | 1 | 13 | Arfip2 | -0.007 | Young |
| 0.978571 | 0.140513 | 0.228 | 0.246 | 1 | 13 | Acvr1 | -0.018 | Young |
| 0.978627 | 0.165255 | 0.105 | 0.107 | 1 | 13 | Twnk | -0.002 | Young |
| 0.978637 | 0.059719 | 0.798 | 0.861 | 1 | 13 | Tnks2 | -0.063 | Young |
| 0.979695 | 0.053579 | 0.482 | 0.525 | 1 | 13 | Zbtb44 | -0.043 | Young |
| 0.979779 | 0.054698 | 0.158 | 0.164 | 1 | 13 | Zxdb | -0.006 | Young |
| 0.980206 | 0.168367 | 0.219 | 0.23 | 1 | 13 | Pcyt2 | -0.011 | Young |
| 0.980568 | 0.012497 | 0.316 | 0.311 | 1 | 13 | Rngtt | 0.005 | Young |
| 0.980593 | 0.072874 | 0.175 | 0.18 | 1 | 13 | Xlr4b | -0.005 | Young |
| 0.980684 | 0.01565 | 0.675 | 0.705 | 1 | 13 | Cetn3 | -0.03 | Young |
| 0.980806 | 0.080733 | 0.719 | 0.762 | 1 | 13 | Dusp22 | -0.043 | Young |
| 0.98097 | 0.002547 | 0.947 | 0.934 | 1 | 13 | Ndufb5 | 0.013 | Young |
| 0.981066 | 0.130865 | 0.14 | 0.139 | 1 | 13 | Snx21 | 0.001 | Young |
| 0.981124 | 0.133765 | 0.184 | 0.197 | 1 | 13 | Cdkal1 | -0.013 | Young |
| 0.981533 | 0.067195 | 0.149 | 0.148 | 1 | 13 | Nif3l1 | 0.001 | Young |
| 0.982091 | 0.063542 | 0.254 | 0.254 | 1 | 13 | Rev3l | 0 | Young |
| 0.982223 | 0.034816 | 0.5 | 0.533 | 1 | 13 | Ngdn | -0.033 | Young |
| 0.982301 | 0.067671 | 0.254 | 0.27 | 1 | 13 | Asb3 | -0.016 | Young |
| 0.982316 | 0.015142 | 0.421 | 0.443 | 1 | 13 | Ccdc6 | -0.022 | Young |
| 0.982678 | 0.100911 | 0.447 | 0.492 | 1 | 13 | Arl13b | -0.045 | Young |
| 0.982903 | 0.278502 | 0.123 | 0.131 | 1 | 13 | B230354K1 | -0.008 | Young |
| 0.983047 | 1.512517 | 0.088 | 0.09 | 1 | 13 | Arg1 | -0.002 | Young |
| 0.983114 | 0.094887 | 0.491 | 0.566 | 1 | 13 | Vps18 | -0.075 | Young |
| 0.983142 | 0.043382 | 0.746 | 0.779 | 1 | 13 | Arrdc1 | -0.033 | Young |
| 0.983144 | 0.067304 | 0.132 | 0.131 | 1 | 13 | Nradd | 0.001 | Young |
| 0.98359 | 0.023098 | 0.14 | 0.139 | 1 | 13 | Eogt | 0.001 | Young |
| 0.9836 | 0.131834 | 0.623 | 0.639 | 1 | 13 | P2rx7 | -0.016 | Young |
| 0.983696 | 0.15234 | 0.167 | 0.172 | 1 | 13 | Serf1 | -0.005 | Young |
| 0.983959 | 0.042007 | 0.421 | 0.434 | 1 | 13 | Hikeshi | -0.013 | Young |
| 0.984745 | 0.156495 | 0.526 | 0.541 | 1 | 13 | Ebpl | -0.015 | Young |
| 0.984984 | 0.065624 | 0.246 | 0.254 | 1 | 13 | Gdpd3 | -0.008 | Young |
| 0.985346 | 0.028923 | 0.684 | 0.721 | 1 | 13 | Tle4 | -0.037 | Young |
| 0.985518 | 0.009824 | 0.842 | 0.877 | 1 | 13 | Mbtd1 | -0.035 | Young |
| 0.98606 | 0.008561 | 0.377 | 0.385 | 1 | 13 | Pi4k2b | -0.008 | Young |
| 0.986297 | 0.009271 | 0.325 | 0.328 | 1 | 13 | Tarbp2 | -0.003 | Young |
| 0.987031 | 0.065763 | 0.816 | 0.852 | 1 | 13 | Lsm6 | -0.036 | Young |
| 0.987336 | 0.173386 | 0.412 | 0.443 | 1 | 13 | Sdhaf2 | -0.031 | Young |
| 0.987407 | 0.088391 | 0.219 | 0.221 | 1 | 13 | Lrig2 | -0.002 | Young |
| 0.988172 | 0.090171 | 0.316 | 0.344 | 1 | 13 | Mrps18b | -0.028 | Young |
| 0.988312 | 0.053084 | 0.333 | 0.352 | 1 | 13 | Kif13a | -0.019 | Young |
| 0.988472 | 0.097593 | 0.728 | 0.738 | 1 | 13 | Mbp | -0.01 | Young |

| | | | | | | | | |
|---|---|---|---|---|---|---|---|---|
| 0.988707 | 0.229029 | 0.079 | 0.082 | 1 | 13 | Klhl26 | -0.003 | Young |
| 0.988707 | 0.176845 | 0.079 | 0.082 | 1 | 13 | Gm11423 | -0.003 | Young |
| 0.988778 | 0.054035 | 0.386 | 0.402 | 1 | 13 | Klf11 | -0.016 | Young |
| 0.988856 | 0.041856 | 0.553 | 0.566 | 1 | 13 | Atf6b | -0.013 | Young |
| 0.989211 | 0.294035 | 0.088 | 0.09 | 1 | 13 | Suox | -0.002 | Young |
| 0.989211 | 0.095672 | 0.088 | 0.09 | 1 | 13 | C330027C( | -0.002 | Young |
| 0.989245 | 0.073889 | 0.254 | 0.279 | 1 | 13 | Med22 | -0.025 | Young |
| 0.989409 | 0.36778 | 0.158 | 0.172 | 1 | 13 | Pcca | -0.014 | Young |
| 0.989528 | 0.047371 | 0.281 | 0.295 | 1 | 13 | Trp53bp2 | -0.014 | Young |
| 0.989765 | 0.030497 | 0.588 | 0.607 | 1 | 13 | Hpgds | -0.019 | Young |
| 0.989767 | 0.242122 | 0.132 | 0.139 | 1 | 13 | Igip | -0.007 | Young |
| 0.989847 | 0.03673 | 0.614 | 0.648 | 1 | 13 | Lats2 | -0.034 | Young |
| 0.989882 | 0.087415 | 0.623 | 0.672 | 1 | 13 | Samd9l | -0.049 | Young |
| 0.99005 | 0.314256 | 0.237 | 0.27 | 1 | 13 | Eml3 | -0.033 | Young |
| 0.99007 | 0.001571 | 0.825 | 0.795 | 1 | 13 | Stk10 | 0.03 | Young |
| 0.990167 | 0.037314 | 0.254 | 0.27 | 1 | 13 | Slc33a1 | -0.016 | Young |
| 0.990359 | 0.266106 | 0.114 | 0.115 | 1 | 13 | 3-Sep | -0.001 | Young |
| 0.990446 | 0.056557 | 0.368 | 0.402 | 1 | 13 | Zfp664 | -0.034 | Young |
| 0.990484 | 0.113977 | 0.211 | 0.221 | 1 | 13 | Ccdc117 | -0.01 | Young |
| 0.990811 | 0.018067 | 0.746 | 0.795 | 1 | 13 | Bscl2 | -0.049 | Young |
| 0.990999 | 0.091021 | 0.474 | 0.5 | 1 | 13 | Zc3h14 | -0.026 | Young |
| 0.99123 | 0.084448 | 0.36 | 0.385 | 1 | 13 | Coa6 | -0.025 | Young |
| 0.99123 | 0.067484 | 0.36 | 0.385 | 1 | 13 | Gucd1 | -0.025 | Young |
| 0.991275 | 0.081072 | 0.14 | 0.148 | 1 | 13 | Dffa | -0.008 | Young |
| 0.991621 | 0.063464 | 0.895 | 0.885 | 1 | 13 | Pfkp | 0.01 | Young |
| 0.991673 | 0.060145 | 0.158 | 0.164 | 1 | 13 | Tbc1d22b | -0.006 | Young |
| 0.991673 | 0.050658 | 0.158 | 0.164 | 1 | 13 | Dedd | -0.006 | Young |
| 0.992388 | 0.014701 | 0.991 | 0.992 | 1 | 13 | Rock1 | -0.001 | Young |
| 0.992459 | 0.075258 | 0.289 | 0.303 | 1 | 13 | Dnttip1 | -0.014 | Young |
| 0.992604 | 0.075967 | 0.096 | 0.098 | 1 | 13 | Nup43 | -0.002 | Young |
| 0.993046 | 0.019757 | 0.675 | 0.705 | 1 | 13 | Mta2 | -0.03 | Young |
| 0.993115 | 0.111453 | 0.763 | 0.803 | 1 | 13 | Lrp10 | -0.04 | Young |
| 0.993302 | 0.0935 | 0.272 | 0.295 | 1 | 13 | Fam78a | -0.023 | Young |
| 0.993578 | 0.0964 | 0.316 | 0.328 | 1 | 13 | Ubr1 | -0.012 | Young |
| 0.993861 | 0.038234 | 0.368 | 0.377 | 1 | 13 | Trappc13 | -0.009 | Young |
| 0.994134 | 0.20952 | 0.421 | 0.459 | 1 | 13 | Immp1l | -0.038 | Young |
| 0.994543 | 0.245713 | 0.193 | 0.205 | 1 | 13 | Men1 | -0.012 | Young |
| 0.994631 | 0.751029 | 0.202 | 0.213 | 1 | 13 | Yae1d1 | -0.011 | Young |
| 0.994672 | 0.020162 | 0.991 | 1 | 1 | 13 | Aplp2 | -0.009 | Young |
| 0.995111 | 0.154316 | 0.263 | 0.27 | 1 | 13 | Ppard | -0.007 | Young |
| 0.995562 | 0.22565 | 0.096 | 0.098 | 1 | 13 | Fam212a | -0.002 | Young |
| 0.995562 | 0.060008 | 0.096 | 0.098 | 1 | 13 | Pomt2 | -0.002 | Young |
| 0.995725 | 0.375344 | 0.105 | 0.107 | 1 | 13 | Tcea3 | -0.002 | Young |
| 0.995725 | 0.181351 | 0.105 | 0.107 | 1 | 13 | Btd | -0.002 | Young |
| 0.995868 | 0.063611 | 0.114 | 0.115 | 1 | 13 | Pomk | -0.001 | Young |

| | | | | | | | | |
|---|---|---|---|---|---|---|---|---|
| 0.995985 | 0.016728 | 0.526 | 0.541 | 1 | 13 | Pitpnb | -0.015 | Young |
| 0.996152 | 0.11852 | 0.272 | 0.287 | 1 | 13 | G6pc3 | -0.015 | Young |
| 0.996541 | 0.167263 | 0.167 | 0.18 | 1 | 13 | Smyd3 | -0.013 | Young |
| 0.996903 | 0.042931 | 0.658 | 0.697 | 1 | 13 | Mrpl18 | -0.039 | Young |
| 0.996955 | 0.018901 | 0.974 | 0.992 | 1 | 13 | Rab10 | -0.018 | Young |
| 0.997015 | 0.073426 | 0.246 | 0.262 | 1 | 13 | Gpn3 | -0.016 | Young |
| 0.997083 | 0.027575 | 0.263 | 0.279 | 1 | 13 | Vps39 | -0.016 | Young |
| 0.997454 | 0.062464 | 0.404 | 0.434 | 1 | 13 | Polr1a | -0.03 | Young |
| 0.997567 | 0.056105 | 0.491 | 0.525 | 1 | 13 | Phf2 | -0.034 | Young |
| 0.997576 | 0.009757 | 0.526 | 0.508 | 1 | 13 | Gbf1 | 0.018 | Young |
| 0.997648 | 0.08728 | 0.605 | 0.623 | 1 | 13 | Lars | -0.018 | Young |
| 0.998298 | 0.016199 | 0.412 | 0.418 | 1 | 13 | Ube2g1 | -0.006 | Young |
| 0.998459 | 0.186159 | 0.088 | 0.09 | 1 | 13 | Gm9530 | -0.002 | Young |
| 0.998703 | 0.21494 | 0.132 | 0.131 | 1 | 13 | Chst11 | 0.001 | Young |
| 0.998858 | 0.126742 | 0.175 | 0.18 | 1 | 13 | Gmds | -0.005 | Young |
| 0.998999 | 0.063411 | 0.246 | 0.254 | 1 | 13 | Iffo2 | -0.008 | Young |
| 0.999126 | 0.076586 | 0.368 | 0.385 | 1 | 13 | Ksr1 | -0.017 | Young |
| 0.999158 | 0.087209 | 0.404 | 0.459 | 1 | 13 | Scfd1 | -0.055 | Young |
| 1.99E-06 | 0.739721 | 0.978 | 0.88 | 0.06195 | 14 | Timp2 | 0.098 | Old |
| 3.54E-05 | 1.99264 | 0.519 | 0.2 | 1 | 14 | H2-T23 | 0.319 | Old |
| 4.96E-05 | 0.767332 | 0.885 | 0.62 | 1 | 14 | Celf2 | 0.265 | Old |
| 8.31E-05 | 0.812398 | 0.836 | 0.58 | 1 | 14 | Fbln2 | 0.256 | Old |
| 9.03E-05 | 0.67786 | 0.962 | 0.78 | 1 | 14 | Ahnak | 0.182 | Old |
| 0.000106 | 1.564572 | 0.645 | 0.4 | 1 | 14 | Ccl11 | 0.245 | Old |
| 0.000122 | 1.158227 | 0.825 | 0.52 | 1 | 14 | Aebp1 | 0.305 | Old |
| 0.000132 | 0.779925 | 0.852 | 0.74 | 1 | 14 | H2-K1 | 0.112 | Old |
| 0.000144 | 0.943322 | 0.77 | 0.5 | 1 | 14 | Ebf1 | 0.27 | Old |
| 0.000172 | 0.844925 | 0.874 | 0.62 | 1 | 14 | Neat1 | 0.254 | Old |
| 0.000219 | 0.860415 | 0.792 | 0.52 | 1 | 14 | Scara5 | 0.272 | Old |
| 0.00023 | 2.566737 | 0.437 | 0.18 | 1 | 14 | Vcam1 | 0.257 | Old |
| 0.000242 | 0.786502 | 0.852 | 0.68 | 1 | 14 | App | 0.172 | Old |
| 0.000278 | 0.935385 | 0.809 | 0.54 | 1 | 14 | Slc43a3 | 0.269 | Old |
| 0.00028 | 1.040964 | 0.77 | 0.56 | 1 | 14 | Slit3 | 0.21 | Old |
| 0.000289 | 0.914496 | 0.814 | 0.56 | 1 | 14 | Frmd6 | 0.254 | Old |
| 0.000373 | 0.924072 | 0.787 | 0.54 | 1 | 14 | Ifi204 | 0.247 | Old |
| 0.000383 | 0.710034 | 0.869 | 0.62 | 1 | 14 | Anxa1 | 0.249 | Old |
| 0.000409 | 0.445215 | 0.956 | 0.8 | 1 | 14 | C1s1 | 0.156 | Old |
| 0.000433 | 2.049238 | 0.415 | 0.16 | 1 | 14 | Pcsk6 | 0.255 | Old |
| 0.000468 | 1.948592 | 0.333 | 0.08 | 1 | 14 | Poldip3 | 0.253 | Old |
| 0.000505 | 0.748916 | 0.776 | 0.56 | 1 | 14 | Tnxb | 0.216 | Old |
| 0.000539 | 0.751664 | 0.847 | 0.66 | 1 | 14 | Dpep1 | 0.187 | Old |
| 0.000543 | 0.547097 | 0.754 | 0.5 | 1 | 14 | Ecm1 | 0.254 | Old |
| 0.000553 | 1.50329 | 0.519 | 0.28 | 1 | 14 | Scara3 | 0.239 | Old |
| 0.000627 | 0.755204 | 0.76 | 0.52 | 1 | 14 | Ddr2 | 0.24 | Old |
| 0.000638 | 0.675054 | 0.907 | 0.68 | 1 | 14 | C3 | 0.227 | Old |

| | | | | | | | | |
|---|---|---|---|---|---|---|---|---|
| 0.000961 | 0.682241 | 0.787 | 0.66 | 1 | 14 | Npc2 | 0.127 | Old |
| 0.001144 | 0.870752 | 0.645 | 0.4 | 1 | 14 | Amotl2 | 0.245 | Old |
| 0.001217 | 0.399144 | 0.47 | 0.22 | 1 | 14 | Tmod3 | 0.25 | Old |
| 0.001241 | 0.663251 | 0.699 | 0.46 | 1 | 14 | Cfh | 0.239 | Old |
| 0.001344 | 1.83033 | 0.443 | 0.24 | 1 | 14 | Rtraf | 0.203 | Old |
| 0.001345 | 0.963428 | 0.623 | 0.32 | 1 | 14 | Ifi207 | 0.303 | Old |
| 0.001416 | 0.520742 | 0.984 | 0.86 | 1 | 14 | Gsn | 0.124 | Old |
| 0.001421 | 0.533097 | 0.678 | 0.4 | 1 | 14 | Prnp | 0.278 | Old |
| 0.001777 | 0.897388 | 0.65 | 0.48 | 1 | 14 | Cyb5a | 0.17 | Old |
| 0.001801 | 1.152762 | 0.497 | 0.24 | 1 | 14 | Aldh1a3 | 0.257 | Old |
| 0.001893 | 0.404836 | 0.945 | 0.72 | 1 | 14 | Rnase4 | 0.225 | Old |
| 0.001935 | 0.595803 | 0.65 | 0.44 | 1 | 14 | Vldlr | 0.21 | Old |
| 0.002372 | 0.677983 | 0.552 | 0.28 | 1 | 14 | Lrrn4cl | 0.272 | Old |
| 0.002389 | 1.194886 | 0.574 | 0.36 | 1 | 14 | Pros1 | 0.214 | Old |
| 0.002497 | 1.642918 | 0.377 | 0.16 | 1 | 14 | Rtn1 | 0.217 | Old |
| 0.002566 | 0.704911 | 0.836 | 0.66 | 1 | 14 | Mbnl1 | 0.176 | Old |
| 0.002606 | 2.783832 | 0.251 | 0.06 | 1 | 14 | Irak3 | 0.191 | Old |
| 0.002609 | 0.500277 | 0.956 | 0.9 | 1 | 14 | Dcn | 0.056 | Old |
| 0.002981 | 0.911866 | 0.486 | 0.26 | 1 | 14 | Dact1 | 0.226 | Old |
| 0.003253 | 0.950404 | 0.689 | 0.44 | 1 | 14 | Nisch | 0.249 | Old |
| 0.003273 | 1.164713 | 0.448 | 0.24 | 1 | 14 | Nck1 | 0.208 | Old |
| 0.003275 | 0.619985 | 0.858 | 0.68 | 1 | 14 | Rsrp1 | 0.178 | Old |
| 0.00331 | 0.590082 | 0.372 | 0.18 | 1 | 14 | Ifi203 | 0.192 | Old |
| 0.003467 | 0.541344 | 0.902 | 0.78 | 1 | 14 | Nid1 | 0.122 | Old |
| 0.003495 | 0.458315 | 0.967 | 0.84 | 1 | 14 | Malat1 | 0.127 | Old |
| 0.003549 | 0.493488 | 0.874 | 0.82 | 1 | 14 | B2m | 0.054 | Old |
| 0.003716 | 0.646166 | 0.836 | 0.66 | 1 | 14 | Lamp1 | 0.176 | Old |
| 0.003758 | 0.616818 | 0.71 | 0.46 | 1 | 14 | Plat | 0.25 | Old |
| 0.003774 | 0.523663 | 0.792 | 0.62 | 1 | 14 | Gm10076 | 0.172 | Old |
| 0.003828 | 1.402745 | 0.404 | 0.18 | 1 | 14 | Dcbld2 | 0.224 | Old |
| 0.004219 | 0.453239 | 0.885 | 0.66 | 1 | 14 | Sptbn1 | 0.225 | Old |
| 0.004233 | 0.597041 | 0.667 | 0.52 | 1 | 14 | Rbms3 | 0.147 | Old |
| 0.004282 | 1.469944 | 0.492 | 0.28 | 1 | 14 | Cyp1b1 | 0.212 | Old |
| 0.004316 | 1.087955 | 0.667 | 0.44 | 1 | 14 | Svep1 | 0.227 | Old |
| 0.004493 | 0.789618 | 0.738 | 0.58 | 1 | 14 | Rtn4 | 0.158 | Old |
| 0.004622 | 0.860981 | 0.443 | 0.22 | 1 | 14 | Metrnl | 0.223 | Old |
| 0.004694 | 1.399339 | 0.415 | 0.22 | 1 | 14 | Fads1 | 0.195 | Old |
| 0.004718 | 0.915371 | 0.552 | 0.36 | 1 | 14 | Fbln5 | 0.192 | Old |
| 0.004909 | 5.083041 | 0.142 | 0 | 1 | 14 | Txlng | 0.142 | Old |
| 0.004915 | 0.64135 | 0.82 | 0.68 | 1 | 14 | Zfp36l2 | 0.14 | Old |
| 0.00502 | 0.799087 | 0.601 | 0.34 | 1 | 14 | Cdon | 0.261 | Old |
| 0.005222 | 0.644313 | 0.601 | 0.36 | 1 | 14 | Lima1 | 0.241 | Old |
| 0.005222 | 0.854488 | 0.448 | 0.22 | 1 | 14 | Smpd3 | 0.228 | Old |
| 0.005248 | 1.141921 | 0.781 | 0.62 | 1 | 14 | Gpx3 | 0.161 | Old |
| 0.005291 | 0.725477 | 0.82 | 0.68 | 1 | 14 | H2-D1 | 0.14 | Old |

| | | | | | | | | |
|---|---|---|---|---|---|---|---|---|
| 0.005322 | 0.447089 | 0.896 | 0.72 | 1 | 14 | Klf2 | 0.176 | Old |
| 0.005373 | 0.474397 | 0.891 | 0.78 | 1 | 14 | Serping1 | 0.111 | Old |
| 0.005597 | 0.653928 | 0.563 | 0.34 | 1 | 14 | Plau | 0.223 | Old |
| 0.005776 | 1.243703 | 0.41 | 0.2 | 1 | 14 | Arhgap24 | 0.21 | Old |
| 0.005788 | 0.796652 | 0.514 | 0.32 | 1 | 14 | Ctnna1 | 0.194 | Old |
| 0.00584 | 0.87002 | 0.689 | 0.56 | 1 | 14 | Fcgrt | 0.129 | Old |
| 0.005865 | 0.404284 | 0.869 | 0.74 | 1 | 14 | Lamp2 | 0.129 | Old |
| 0.00594 | 0.861568 | 0.623 | 0.42 | 1 | 14 | Sh3glb1 | 0.203 | Old |
| 0.006136 | 0.615278 | 0.77 | 0.56 | 1 | 14 | Ccdc80 | 0.21 | Old |
| 0.006169 | 1.057966 | 0.47 | 0.26 | 1 | 14 | Igf1r | 0.21 | Old |
| 0.00628 | 0.525274 | 0.508 | 0.3 | 1 | 14 | Cd302 | 0.208 | Old |
| 0.006529 | 1.105786 | 0.574 | 0.44 | 1 | 14 | Dst | 0.134 | Old |
| 0.006564 | 1.330539 | 0.366 | 0.16 | 1 | 14 | Pxn | 0.206 | Old |
| 0.006638 | 0.704935 | 0.913 | 0.74 | 1 | 14 | Selenop | 0.173 | Old |
| 0.006801 | 0.731089 | 0.852 | 0.6 | 1 | 14 | Mt1 | 0.252 | Old |
| 0.007007 | 0.721485 | 0.814 | 0.58 | 1 | 14 | Zeb2 | 0.234 | Old |
| 0.007136 | 5.547178 | 0.131 | 0 | 1 | 14 | Hhipl1 | 0.131 | Old |
| 0.007136 | 4.626819 | 0.131 | 0 | 1 | 14 | C130026I2 | 0.131 | Old |
| 0.007481 | 1.132107 | 0.415 | 0.22 | 1 | 14 | Tpp1 | 0.195 | Old |
| 0.007596 | 1.362501 | 0.454 | 0.26 | 1 | 14 | Sp3 | 0.194 | Old |
| 0.007627 | 0.398539 | 0.869 | 0.62 | 1 | 14 | Adamts5 | 0.249 | Old |
| 0.00774 | 0.519358 | 0.847 | 0.8 | 1 | 14 | Cd47 | 0.047 | Old |
| 0.007772 | 2.070036 | 0.317 | 0.14 | 1 | 14 | Tmem106a | 0.177 | Old |
| 0.007986 | 0.732576 | 0.366 | 0.14 | 1 | 14 | Jcad | 0.226 | Old |
| 0.008224 | 0.487946 | 0.852 | 0.62 | 1 | 14 | Ctsb | 0.232 | Old |
| 0.008312 | 0.902206 | 0.617 | 0.44 | 1 | 14 | Entpd2 | 0.177 | Old |
| 0.008371 | 0.572302 | 0.699 | 0.5 | 1 | 14 | C1ra | 0.199 | Old |
| 0.008409 | 0.437697 | 0.858 | 0.74 | 1 | 14 | Anxa5 | 0.118 | Old |
| 0.008492 | 0.540579 | 0.519 | 0.28 | 1 | 14 | Eif3k | 0.239 | Old |
| 0.008499 | 0.695747 | 0.607 | 0.38 | 1 | 14 | Mt2 | 0.227 | Old |
| 0.008593 | 4.877986 | 0.126 | 0 | 1 | 14 | Cgrrf1 | 0.126 | Old |
| 0.008793 | 0.750723 | 0.672 | 0.5 | 1 | 14 | Ltbp1 | 0.172 | Old |
| 0.009024 | 0.614033 | 0.792 | 0.56 | 1 | 14 | Klf6 | 0.232 | Old |
| 0.009076 | 1.262595 | 0.415 | 0.22 | 1 | 14 | Trio | 0.195 | Old |
| 0.009344 | 2.552201 | 0.191 | 0.04 | 1 | 14 | Gbp7 | 0.151 | Old |
| 0.009348 | 1.125 | 0.639 | 0.42 | 1 | 14 | Inmt | 0.219 | Old |
| 0.009602 | 0.932125 | 0.333 | 0.14 | 1 | 14 | Lpgat1 | 0.193 | Old |
| 0.009948 | 0.365494 | 0.716 | 0.54 | 1 | 14 | Tshz2 | 0.176 | Old |
| 0.010234 | 3.772543 | 0.153 | 0.02 | 1 | 14 | Tusc2 | 0.133 | Old |
| 0.010235 | 1.000533 | 0.568 | 0.38 | 1 | 14 | Anxa7 | 0.188 | Old |
| 0.010338 | 4.328758 | 0.12 | 0 | 1 | 14 | Ackr2 | 0.12 | Old |
| 0.01054 | 3.984207 | 0.18 | 0.04 | 1 | 14 | Pygo1 | 0.14 | Old |
| 0.010703 | 1.318144 | 0.35 | 0.16 | 1 | 14 | Tmeff2 | 0.19 | Old |
| 0.010719 | 0.485432 | 0.721 | 0.5 | 1 | 14 | Csde1 | 0.221 | Old |
| 0.011012 | 0.592116 | 0.683 | 0.46 | 1 | 14 | Abca8a | 0.223 | Old |

| | | | | | | | | |
|---|---|---|---|---|---|---|---|---|
| 0.011153 | 0.489919 | 0.995 | 0.96 | 1 | 14 | Jund | 0.035 | Old |
| 0.011209 | 1.193078 | 0.388 | 0.2 | 1 | 14 | Ahnak2 | 0.188 | Old |
| 0.011282 | 0.960157 | 0.459 | 0.26 | 1 | 14 | Wasf2 | 0.199 | Old |
| 0.011604 | 0.252512 | 0.754 | 0.6 | 1 | 14 | Bicc1 | 0.154 | Old |
| 0.011614 | 0.331354 | 0.989 | 0.92 | 1 | 14 | S100a6 | 0.069 | Old |
| 0.011969 | 0.130704 | 0.574 | 0.34 | 1 | 14 | Lhfp | 0.234 | Old |
| 0.012233 | 0.406923 | 0.792 | 0.62 | 1 | 14 | Xbp1 | 0.172 | Old |
| 0.0124 | 3.901798 | 0.148 | 0.02 | 1 | 14 | Large1 | 0.128 | Old |
| 0.01243 | 5.138595 | 0.115 | 0 | 1 | 14 | Gabre | 0.115 | Old |
| 0.01243 | 4.886127 | 0.115 | 0 | 1 | 14 | Kdm4b | 0.115 | Old |
| 0.01243 | 4.666805 | 0.115 | 0 | 1 | 14 | Sgk3 | 0.115 | Old |
| 0.01243 | 4.622848 | 0.115 | 0 | 1 | 14 | Commd4 | 0.115 | Old |
| 0.01243 | 4.368207 | 0.115 | 0 | 1 | 14 | Thap4 | 0.115 | Old |
| 0.012698 | 2.032399 | 0.213 | 0.06 | 1 | 14 | Lfng | 0.153 | Old |
| 0.012749 | 1.140302 | 0.426 | 0.24 | 1 | 14 | Sbds | 0.186 | Old |
| 0.012848 | 3.539858 | 0.148 | 0.02 | 1 | 14 | Trappc3 | 0.128 | Old |
| 0.013125 | 1.258368 | 0.366 | 0.2 | 1 | 14 | Cand1 | 0.166 | Old |
| 0.013255 | 1.436192 | 0.268 | 0.1 | 1 | 14 | Pamr1 | 0.168 | Old |
| 0.013309 | 3.524691 | 0.148 | 0.02 | 1 | 14 | Dpp7 | 0.128 | Old |
| 0.013544 | 1.135091 | 0.546 | 0.38 | 1 | 14 | Zyx | 0.166 | Old |
| 0.013625 | 3.065994 | 0.148 | 0.02 | 1 | 14 | Slc25a37 | 0.128 | Old |
| 0.013679 | 1.821621 | 0.273 | 0.12 | 1 | 14 | Abcc1 | 0.153 | Old |
| 0.013742 | 0.389146 | 0.825 | 0.7 | 1 | 14 | Efemp1 | 0.125 | Old |
| 0.013796 | 0.617065 | 0.607 | 0.46 | 1 | 14 | Irf2bpl | 0.147 | Old |
| 0.01401 | 0.42964 | 0.617 | 0.44 | 1 | 14 | Jpt1 | 0.177 | Old |
| 0.014073 | 0.650745 | 0.557 | 0.4 | 1 | 14 | Trip12 | 0.157 | Old |
| 0.014434 | 0.294693 | 0.721 | 0.48 | 1 | 14 | Adgrd1 | 0.241 | Old |
| 0.014935 | 4.161882 | 0.109 | 0 | 1 | 14 | Csprs | 0.109 | Old |
| 0.014935 | 4.155381 | 0.109 | 0 | 1 | 14 | Letmd1 | 0.109 | Old |
| 0.014949 | 1.393894 | 0.372 | 0.2 | 1 | 14 | Fmo2 | 0.172 | Old |
| 0.015083 | 1.949387 | 0.23 | 0.08 | 1 | 14 | Ogfr | 0.15 | Old |
| 0.015146 | 1.276635 | 0.383 | 0.22 | 1 | 14 | Zcchc7 | 0.163 | Old |
| 0.015204 | 1.64256 | 0.377 | 0.24 | 1 | 14 | Gpatch8 | 0.137 | Old |
| 0.015262 | 1.875182 | 0.339 | 0.18 | 1 | 14 | Hmgcs2 | 0.159 | Old |
| 0.015361 | 0.488448 | 0.607 | 0.44 | 1 | 14 | Mmp23 | 0.167 | Old |
| 0.01538 | 2.655672 | 0.175 | 0.04 | 1 | 14 | Iigp1 | 0.135 | Old |
| 0.0154 | 2.166709 | 0.377 | 0.22 | 1 | 14 | Adm | 0.157 | Old |
| 0.015691 | 0.972002 | 0.53 | 0.4 | 1 | 14 | Map4k4 | 0.13 | Old |
| 0.015822 | 1.091772 | 0.404 | 0.22 | 1 | 14 | Ncstn | 0.184 | Old |
| 0.015969 | 1.463024 | 0.301 | 0.14 | 1 | 14 | Eif2a | 0.161 | Old |
| 0.016843 | 1.512069 | 0.24 | 0.08 | 1 | 14 | Procr | 0.16 | Old |
| 0.016859 | 2.26652 | 0.224 | 0.08 | 1 | 14 | Kbtbd2 | 0.144 | Old |
| 0.017101 | 0.455513 | 0.634 | 0.44 | 1 | 14 | Ms4a4d | 0.194 | Old |
| 0.017129 | 2.825977 | 0.169 | 0.04 | 1 | 14 | 2610037D( | 0.129 | Old |
| 0.017136 | 1.390935 | 0.35 | 0.18 | 1 | 14 | Cltb | 0.17 | Old |

| | | | | | | | | |
|---|---|---|---|---|---|---|---|---|
| 0.017456 | 0.609968 | 0.601 | 0.42 | 1 | 14 | Sirpa | 0.181 | Old |
| 0.017682 | 0.288775 | 0.749 | 0.56 | 1 | 14 | Ace | 0.189 | Old |
| 0.017764 | 0.640625 | 0.033 | 0.12 | 1 | 14 | Dlg3 | -0.087 | Old |
| 0.017934 | 4.17704 | 0.104 | 0 | 1 | 14 | Caprin2 | 0.104 | Old |
| 0.017957 | 1.07573 | 0.35 | 0.18 | 1 | 14 | Pdlim4 | 0.17 | Old |
| 0.01797 | 0.428733 | 0.776 | 0.6 | 1 | 14 | Srsf3 | 0.176 | Old |
| 0.017983 | 1.376027 | 0.366 | 0.22 | 1 | 14 | Gpc6 | 0.146 | Old |
| 0.018156 | 0.644631 | 0.585 | 0.44 | 1 | 14 | Ifi27l2a | 0.145 | Old |
| 0.018176 | 0.887996 | 0.437 | 0.28 | 1 | 14 | Gdi1 | 0.157 | Old |
| 0.018206 | 0.804056 | 0.481 | 0.28 | 1 | 14 | Klhl9 | 0.201 | Old |
| 0.018288 | 2.388431 | 0.142 | 0.02 | 1 | 14 | 9930111J2 | 0.122 | Old |
| 0.018313 | 0.592726 | 0.776 | 0.62 | 1 | 14 | Nav1 | 0.156 | Old |
| 0.018578 | 1.644018 | 0.175 | 0.04 | 1 | 14 | Dpp9 | 0.135 | Old |
| 0.01858 | 3.509003 | 0.137 | 0.02 | 1 | 14 | Ddx56 | 0.117 | Old |
| 0.018671 | 2.169641 | 0.197 | 0.06 | 1 | 14 | Ranbp9 | 0.137 | Old |
| 0.018746 | 0.527401 | 0.339 | 0.18 | 1 | 14 | 2610001J0 | 0.159 | Old |
| 0.018797 | 3.558909 | 0.137 | 0.02 | 1 | 14 | Trim23 | 0.117 | Old |
| 0.019033 | 2.409212 | 0.169 | 0.04 | 1 | 14 | Ddhd1 | 0.129 | Old |
| 0.019058 | 0.347352 | 0.907 | 0.7 | 1 | 14 | Clec3b | 0.207 | Old |
| 0.019107 | 0.645269 | 0.628 | 0.42 | 1 | 14 | Creb5 | 0.208 | Old |
| 0.019381 | 0.791618 | 0.514 | 0.34 | 1 | 14 | Tnrc6a | 0.174 | Old |
| 0.019392 | 0.895253 | 0.47 | 0.32 | 1 | 14 | Crip2 | 0.15 | Old |
| 0.019435 | 2.461797 | 0.169 | 0.04 | 1 | 14 | Map2k1 | 0.129 | Old |
| 0.019481 | 0.48428 | 0.869 | 0.7 | 1 | 14 | Klf4 | 0.169 | Old |
| 0.019525 | 0.529492 | 0.71 | 0.52 | 1 | 14 | Man1a | 0.19 | Old |
| 0.019778 | 1.584118 | 0.257 | 0.1 | 1 | 14 | Ildr2 | 0.157 | Old |
| 0.020248 | 0.592049 | 0.754 | 0.58 | 1 | 14 | Mmp14 | 0.174 | Old |
| 0.020471 | 0.819056 | 0.557 | 0.38 | 1 | 14 | Sdc4 | 0.177 | Old |
| 0.020981 | 0.758088 | 0.607 | 0.42 | 1 | 14 | Sh3bgrl | 0.187 | Old |
| 0.020991 | 0.46074 | 0.88 | 0.64 | 1 | 14 | Maf | 0.24 | Old |
| 0.02102 | 2.55425 | 0.213 | 0.08 | 1 | 14 | Lacc1 | 0.133 | Old |
| 0.021098 | 1.909695 | 0.24 | 0.1 | 1 | 14 | B4galt5 | 0.14 | Old |
| 0.021456 | 1.998887 | 0.279 | 0.14 | 1 | 14 | Srgap1 | 0.139 | Old |
| 0.021525 | 4.765563 | 0.098 | 0 | 1 | 14 | Cd109 | 0.098 | Old |
| 0.021525 | 4.757167 | 0.098 | 0 | 1 | 14 | Med6 | 0.098 | Old |
| 0.021525 | 3.989953 | 0.098 | 0 | 1 | 14 | Vps45 | 0.098 | Old |
| 0.021536 | 0.958645 | 0.634 | 0.48 | 1 | 14 | Gm26917 | 0.154 | Old |
| 0.021547 | 2.268307 | 0.191 | 0.06 | 1 | 14 | Mocs2 | 0.131 | Old |
| 0.021666 | 0.386173 | 0.661 | 0.46 | 1 | 14 | Sntb2 | 0.201 | Old |
| 0.021696 | 0.323964 | 0.945 | 0.86 | 1 | 14 | Zbtb20 | 0.085 | Old |
| 0.021859 | 2.833131 | 0.164 | 0.04 | 1 | 14 | Zfp26 | 0.124 | Old |
| 0.021883 | 1.259247 | 0.383 | 0.22 | 1 | 14 | Sdhc | 0.163 | Old |
| 0.021916 | 0.866873 | 0.732 | 0.64 | 1 | 14 | Mat2a | 0.092 | Old |
| 0.021925 | 0.32595 | 0.831 | 0.64 | 1 | 14 | Pdgfra | 0.191 | Old |
| 0.022044 | 1.822337 | 0.24 | 0.1 | 1 | 14 | Pdlim2 | 0.14 | Old |

| | | | | | | | | |
|---|---|---|---|---|---|---|---|---|
| 0.022426 | 0.126195 | 0.727 | 0.48 | 1 | 14 | Mfap5 | 0.247 | Old |
| 0.022782 | 0.643186 | 0.311 | 0.16 | 1 | 14 | Pxdn | 0.151 | Old |
| 0.022924 | 0.398751 | 0.787 | 0.68 | 1 | 14 | Heg1 | 0.107 | Old |
| 0.023151 | 0.995045 | 0.366 | 0.2 | 1 | 14 | Gabra3 | 0.166 | Old |
| 0.023214 | 2.513095 | 0.186 | 0.06 | 1 | 14 | Morc3 | 0.126 | Old |
| 0.023235 | 3.318781 | 0.131 | 0.02 | 1 | 14 | Zfp330 | 0.111 | Old |
| 0.023265 | 1.1412 | 0.361 | 0.2 | 1 | 14 | Mcc | 0.161 | Old |
| 0.023299 | 0.98795 | 0.579 | 0.42 | 1 | 14 | Clec2d | 0.159 | Old |
| 0.023478 | 1.14493 | 0.328 | 0.16 | 1 | 14 | Cnot1 | 0.168 | Old |
| 0.023631 | 0.773338 | 0.716 | 0.56 | 1 | 14 | Id3 | 0.156 | Old |
| 0.023987 | 1.052503 | 0.361 | 0.18 | 1 | 14 | Chpt1 | 0.181 | Old |
| 0.024181 | 0.245497 | 0.137 | 0.02 | 1 | 14 | Mbd5 | 0.117 | Old |
| 0.024291 | 0.497995 | 0.617 | 0.46 | 1 | 14 | Antxr2 | 0.157 | Old |
| 0.024384 | 0.733301 | 0.404 | 0.22 | 1 | 14 | Hsd11b1 | 0.184 | Old |
| 0.024603 | 2.457306 | 0.186 | 0.06 | 1 | 14 | Psmc2 | 0.126 | Old |
| 0.024918 | 0.586864 | 0.628 | 0.42 | 1 | 14 | B4galt1 | 0.208 | Old |
| 0.025102 | 0.755807 | 0.35 | 0.18 | 1 | 14 | Osmr | 0.17 | Old |
| 0.025439 | 0.500949 | 0.765 | 0.62 | 1 | 14 | Slfn5 | 0.145 | Old |
| 0.025664 | 1.630878 | 0.279 | 0.14 | 1 | 14 | Med15 | 0.139 | Old |
| 0.025687 | 0.438239 | 0.716 | 0.54 | 1 | 14 | Rac1 | 0.176 | Old |
| 0.025826 | 4.322594 | 0.093 | 0 | 1 | 14 | Tmem87a | 0.093 | Old |
| 0.025938 | 0.616385 | 0.754 | 0.58 | 1 | 14 | Scn7a | 0.174 | Old |
| 0.026013 | 0.864186 | 0.612 | 0.44 | 1 | 14 | Abi3bp | 0.172 | Old |
| 0.026059 | 0.450794 | 0.645 | 0.5 | 1 | 14 | Aldh2 | 0.145 | Old |
| 0.026177 | 0.965611 | 0.279 | 0.12 | 1 | 14 | Naa15 | 0.159 | Old |
| 0.026295 | 1.185161 | 0.432 | 0.28 | 1 | 14 | Stk38 | 0.152 | Old |
| 0.026473 | 3.883474 | 0.126 | 0.02 | 1 | 14 | Tbc1d16 | 0.106 | Old |
| 0.026779 | 3.583816 | 0.126 | 0.02 | 1 | 14 | Rnf168 | 0.106 | Old |
| 0.026822 | 1.463278 | 0.246 | 0.1 | 1 | 14 | Magi3 | 0.146 | Old |
| 0.02708 | 2.505675 | 0.186 | 0.06 | 1 | 14 | Prkx | 0.126 | Old |
| 0.027089 | 3.549154 | 0.126 | 0.02 | 1 | 14 | Galnt17 | 0.106 | Old |
| 0.027092 | 1.908155 | 0.219 | 0.08 | 1 | 14 | Akap8 | 0.139 | Old |
| 0.027095 | 0.5177 | 0.705 | 0.6 | 1 | 14 | Rpl31 | 0.105 | Old |
| 0.027159 | 2.038686 | 0.235 | 0.1 | 1 | 14 | Siah1a | 0.135 | Old |
| 0.027349 | 1.317441 | 0.448 | 0.32 | 1 | 14 | Arid1a | 0.128 | Old |
| 0.02818 | 0.39341 | 0.546 | 0.34 | 1 | 14 | Pdpn | 0.206 | Old |
| 0.028185 | 3.03692 | 0.153 | 0.04 | 1 | 14 | Fam96b | 0.113 | Old |
| 0.02819 | 1.779967 | 0.131 | 0.02 | 1 | 14 | Arhgap28 | 0.111 | Old |
| 0.028347 | 1.271844 | 0.295 | 0.14 | 1 | 14 | Glrx5 | 0.155 | Old |
| 0.028606 | 0.794538 | 0.366 | 0.2 | 1 | 14 | Crebrf | 0.166 | Old |
| 0.028746 | 1.042868 | 0.546 | 0.4 | 1 | 14 | Gdf10 | 0.146 | Old |
| 0.028845 | 0.752792 | 0.372 | 0.2 | 1 | 14 | Spon2 | 0.172 | Old |
| 0.029076 | 1.531012 | 0.219 | 0.08 | 1 | 14 | Hmg20a | 0.139 | Old |
| 0.029265 | 1.652488 | 0.158 | 0.04 | 1 | 14 | Syap1 | 0.118 | Old |
| 0.029385 | 1.146152 | 0.372 | 0.22 | 1 | 14 | Atraid | 0.152 | Old |

| | | | | | | | | |
|---|---|---|---|---|---|---|---|---|
| 0.029533 | 0.807429 | 0.53 | 0.36 | 1 | 14 | Khdrbs1 | 0.17 | Old |
| 0.030009 | 1.486014 | 0.24 | 0.1 | 1 | 14 | Rspo1 | 0.14 | Old |
| 0.030019 | 2.801325 | 0.126 | 0.02 | 1 | 14 | Mex3b | 0.106 | Old |
| 0.030061 | 0.388001 | 0.525 | 0.3 | 1 | 14 | Emp1 | 0.225 | Old |
| 0.030106 | 1.353296 | 0.366 | 0.22 | 1 | 14 | Hexb | 0.146 | Old |
| 0.030221 | 0.512575 | 0.596 | 0.48 | 1 | 14 | Strn3 | 0.116 | Old |
| 0.030414 | 1.605432 | 0.328 | 0.2 | 1 | 14 | Ppp6r3 | 0.128 | Old |
| 0.030976 | 4.203441 | 0.087 | 0 | 1 | 14 | Rps6kc1 | 0.087 | Old |
| 0.031045 | 1.777894 | 0.317 | 0.2 | 1 | 14 | Eif1b | 0.117 | Old |
| 0.03121 | 0.092122 | 0.65 | 0.46 | 1 | 14 | Plxdc2 | 0.19 | Old |
| 0.031244 | 0.938324 | 0.437 | 0.28 | 1 | 14 | Tab2 | 0.157 | Old |
| 0.031417 | 0.554882 | 0.727 | 0.6 | 1 | 14 | Epb41l2 | 0.127 | Old |
| 0.031668 | 1.099689 | 0.47 | 0.34 | 1 | 14 | Manf | 0.13 | Old |
| 0.031816 | 1.161675 | 0.24 | 0.1 | 1 | 14 | Sgta | 0.14 | Old |
| 0.031974 | 1.887804 | 0.23 | 0.1 | 1 | 14 | Uhmk1 | 0.13 | Old |
| 0.032047 | 0.385542 | 0.667 | 0.5 | 1 | 14 | Chd4 | 0.167 | Old |
| 0.03277 | 0.337145 | 0.699 | 0.58 | 1 | 14 | Islr | 0.119 | Old |
| 0.032869 | 2.692764 | 0.153 | 0.04 | 1 | 14 | Tmem110 | 0.113 | Old |
| 0.032928 | 0.869428 | 0.503 | 0.32 | 1 | 14 | Ifi205 | 0.183 | Old |
| 0.032983 | 2.224549 | 0.18 | 0.06 | 1 | 14 | Atp13a1 | 0.12 | Old |
| 0.033044 | 3.164529 | 0.12 | 0.02 | 1 | 14 | Ptgir | 0.1 | Old |
| 0.033156 | 0.536001 | 0.574 | 0.38 | 1 | 14 | Emc10 | 0.194 | Old |
| 0.033324 | 3.209161 | 0.148 | 0.04 | 1 | 14 | Fpgt | 0.108 | Old |
| 0.033664 | 0.827944 | 0.546 | 0.4 | 1 | 14 | Copa | 0.146 | Old |
| 0.033782 | 1.003222 | 0.486 | 0.38 | 1 | 14 | Lrp6 | 0.106 | Old |
| 0.033792 | 0.16773 | 0.06 | 0.16 | 1 | 14 | Wtip | -0.1 | Old |
| 0.033843 | 0.866292 | 0.437 | 0.28 | 1 | 14 | Vasn | 0.157 | Old |
| 0.033903 | 1.692282 | 0.208 | 0.08 | 1 | 14 | Slc35a5 | 0.128 | Old |
| 0.034189 | 0.734375 | 0.393 | 0.22 | 1 | 14 | Ndufb5 | 0.173 | Old |
| 0.034362 | 1.018591 | 0.426 | 0.28 | 1 | 14 | Col6a6 | 0.146 | Old |
| 0.034548 | 1.598318 | 0.186 | 0.06 | 1 | 14 | Eya2 | 0.126 | Old |
| 0.034705 | 0.711484 | 0.656 | 0.52 | 1 | 14 | Ly6e | 0.136 | Old |
| 0.034717 | 3.188126 | 0.148 | 0.04 | 1 | 14 | Appl2 | 0.108 | Old |
| 0.034837 | 1.46841 | 0.393 | 0.3 | 1 | 14 | Sh3d19 | 0.093 | Old |
| 0.034901 | 1.42304 | 0.361 | 0.24 | 1 | 14 | Bola2 | 0.121 | Old |
| 0.034921 | 1.879994 | 0.153 | 0.04 | 1 | 14 | Map3k4 | 0.113 | Old |
| 0.035274 | 2.241226 | 0.153 | 0.04 | 1 | 14 | Snrnp40 | 0.113 | Old |
| 0.03539 | 0.486952 | 0.705 | 0.54 | 1 | 14 | Fgfr1 | 0.165 | Old |
| 0.035489 | 0.77678 | 0.448 | 0.3 | 1 | 14 | Dse | 0.148 | Old |
| 0.035515 | 1.693273 | 0.246 | 0.12 | 1 | 14 | Atrnl1 | 0.126 | Old |
| 0.035968 | 0.54562 | 0.645 | 0.5 | 1 | 14 | Rap1b | 0.145 | Old |
| 0.036002 | 1.339754 | 0.361 | 0.22 | 1 | 14 | Akr1b3 | 0.141 | Old |
| 0.036038 | 1.2044 | 0.208 | 0.08 | 1 | 14 | H2-M3 | 0.128 | Old |
| 0.036139 | 0.672604 | 0.421 | 0.28 | 1 | 14 | Slc4a4 | 0.141 | Old |
| 0.036504 | 0.874596 | 0.525 | 0.38 | 1 | 14 | Arpc5 | 0.145 | Old |

| | | | | | | | | |
|---|---|---|---|---|---|---|---|---|
| 0.037149 | 3.965752 | 0.082 | 0 | 1 | 14 | Sirt5 | 0.082 | Old |
| 0.037149 | 3.947067 | 0.082 | 0 | 1 | 14 | Tnip2 | 0.082 | Old |
| 0.037223 | 0.504707 | 0.738 | 0.58 | 1 | 14 | Ddx17 | 0.158 | Old |
| 0.037307 | 0.386696 | 0.672 | 0.54 | 1 | 14 | S100a16 | 0.132 | Old |
| 0.037633 | 0.984166 | 0.208 | 0.08 | 1 | 14 | Pfn2 | 0.128 | Old |
| 0.038034 | 2.509476 | 0.148 | 0.04 | 1 | 14 | Arhgef7 | 0.108 | Old |
| 0.038086 | 1.404845 | 0.251 | 0.12 | 1 | 14 | Scn1b | 0.131 | Old |
| 0.038107 | 0.675336 | 0.355 | 0.18 | 1 | 14 | Gm2a | 0.175 | Old |
| 0.038238 | 0.26354 | 0.918 | 0.72 | 1 | 14 | Serpinh1 | 0.198 | Old |
| 0.038302 | 0.853938 | 0.459 | 0.32 | 1 | 14 | Hspa9 | 0.139 | Old |
| 0.03834 | 0.429593 | 0.601 | 0.44 | 1 | 14 | Pura | 0.161 | Old |
| 0.038686 | 2.006927 | 0.12 | 0.02 | 1 | 14 | Zmat1 | 0.1 | Old |
| 0.038926 | 3.139257 | 0.115 | 0.02 | 1 | 14 | Abcd4 | 0.095 | Old |
| 0.03893 | 1.242785 | 0.372 | 0.24 | 1 | 14 | Tomm22 | 0.132 | Old |
| 0.039369 | 3.271456 | 0.115 | 0.02 | 1 | 14 | D3Ertd254 | 0.095 | Old |
| 0.039369 | 3.192339 | 0.115 | 0.02 | 1 | 14 | Ptcd3 | 0.095 | Old |
| 0.039497 | 0.912691 | 0.546 | 0.42 | 1 | 14 | Serpine2 | 0.126 | Old |
| 0.039557 | 0.497025 | 0.617 | 0.48 | 1 | 14 | Tmed2 | 0.137 | Old |
| 0.039915 | 1.788019 | 0.301 | 0.18 | 1 | 14 | Sav1 | 0.121 | Old |
| 0.040139 | 1.577035 | 0.153 | 0.04 | 1 | 14 | Mmp11 | 0.113 | Old |
| 0.040169 | 0.656774 | 0.486 | 0.34 | 1 | 14 | Pkd2 | 0.146 | Old |
| 0.040618 | 0.868716 | 0.322 | 0.18 | 1 | 14 | Mndal | 0.142 | Old |
| 0.040726 | 3.102501 | 0.115 | 0.02 | 1 | 14 | Olfml2a | 0.095 | Old |
| 0.040945 | 0.962734 | 0.306 | 0.16 | 1 | 14 | Ssr4 | 0.146 | Old |
| 0.04135 | 0.722564 | 0.574 | 0.48 | 1 | 14 | Tomm7 | 0.094 | Old |
| 0.041374 | 1.196111 | 0.372 | 0.24 | 1 | 14 | Plac8 | 0.132 | Old |
| 0.041653 | 3.104615 | 0.115 | 0.02 | 1 | 14 | Nfkb2 | 0.095 | Old |
| 0.041828 | 2.830527 | 0.142 | 0.04 | 1 | 14 | Adcyap1r1 | 0.102 | Old |
| 0.042096 | 0.461974 | 0.803 | 0.66 | 1 | 14 | Htra3 | 0.143 | Old |
| 0.042679 | 1.8245 | 0.197 | 0.08 | 1 | 14 | Slco2b1 | 0.117 | Old |
| 0.04294 | 0.270201 | 0.694 | 0.58 | 1 | 14 | Vat1 | 0.114 | Old |
| 0.043124 | 1.349242 | 0.29 | 0.16 | 1 | 14 | Mustn1 | 0.13 | Old |
| 0.043292 | 1.76992 | 0.317 | 0.2 | 1 | 14 | Nmb | 0.117 | Old |
| 0.043302 | 1.548359 | 0.202 | 0.08 | 1 | 14 | Mitf | 0.122 | Old |
| 0.043634 | 0.371179 | 0.76 | 0.56 | 1 | 14 | Serpinf1 | 0.2 | Old |
| 0.043772 | 1.72903 | 0.24 | 0.12 | 1 | 14 | Cpeb1 | 0.12 | Old |
| 0.043911 | 1.068294 | 0.393 | 0.26 | 1 | 14 | Kras | 0.133 | Old |
| 0.043967 | 0.335733 | 0.71 | 0.54 | 1 | 14 | Mxra8 | 0.17 | Old |
| 0.044011 | 0.841375 | 0.464 | 0.32 | 1 | 14 | Spcs2 | 0.144 | Old |
| 0.044139 | 0.150218 | 0.514 | 0.38 | 1 | 14 | Ppp1cc | 0.134 | Old |
| 0.044345 | 1.404662 | 0.246 | 0.12 | 1 | 14 | Gpnmb | 0.126 | Old |
| 0.044553 | 4.465661 | 0.077 | 0 | 1 | 14 | Spindoc | 0.077 | Old |
| 0.044553 | 3.997147 | 0.077 | 0 | 1 | 14 | Tmem126b | 0.077 | Old |
| 0.044553 | 3.960485 | 0.077 | 0 | 1 | 14 | Cby1 | 0.077 | Old |
| 0.044553 | 3.890102 | 0.077 | 0 | 1 | 14 | St7 | 0.077 | Old |

| | | | | | | | | |
|---|---|---|---|---|---|---|---|---|
| 0.044553 | 3.743902 | 0.077 | 0 | 1 | 14 | 1110051M | 0.077 | Old |
| 0.045295 | 0.897824 | 0.432 | 0.3 | 1 | 14 | Copb1 | 0.132 | Old |
| 0.04559 | 1.967552 | 0.169 | 0.06 | 1 | 14 | Mrpl18 | 0.109 | Old |
| 0.045867 | 0.932263 | 0.415 | 0.28 | 1 | 14 | Cdc42se1 | 0.135 | Old |
| 0.046356 | 3.428192 | 0.109 | 0.02 | 1 | 14 | Tbc1d8b | 0.089 | Old |
| 0.046499 | 0.287522 | 0.667 | 0.54 | 1 | 14 | Arpc1b | 0.127 | Old |
| 0.046794 | 0.225629 | 0.88 | 0.76 | 1 | 14 | Gstm1 | 0.12 | Old |
| 0.046882 | 3.163357 | 0.109 | 0.02 | 1 | 14 | Klhl26 | 0.089 | Old |
| 0.047153 | 2.491868 | 0.142 | 0.04 | 1 | 14 | Dnlz | 0.102 | Old |
| 0.047268 | 0.429439 | 0.639 | 0.52 | 1 | 14 | Actr2 | 0.119 | Old |
| 0.047412 | 3.241968 | 0.109 | 0.02 | 1 | 14 | Car8 | 0.089 | Old |
| 0.047568 | 0.486539 | 0.585 | 0.4 | 1 | 14 | Pcolce2 | 0.185 | Old |
| 0.047689 | 0.316058 | 0.76 | 0.64 | 1 | 14 | Tgfbr3 | 0.12 | Old |
| 0.047793 | 1.341786 | 0.35 | 0.24 | 1 | 14 | Prkcsh | 0.11 | Old |
| 0.047871 | 3.443441 | 0.137 | 0.04 | 1 | 14 | Ubr7 | 0.097 | Old |
| 0.048003 | 0.326019 | 0.628 | 0.44 | 1 | 14 | Adamts2 | 0.188 | Old |
| 0.04808 | 0.994089 | 0.322 | 0.18 | 1 | 14 | Leprot | 0.142 | Old |
| 0.048198 | 0.659799 | 0.661 | 0.5 | 1 | 14 | Nktr | 0.161 | Old |
| 0.048311 | 0.598495 | 0.525 | 0.4 | 1 | 14 | Dapk1 | 0.125 | Old |
| 0.048353 | 3.19406 | 0.137 | 0.04 | 1 | 14 | Hsf2 | 0.097 | Old |
| 0.048414 | 1.638316 | 0.295 | 0.18 | 1 | 14 | Frmd4b | 0.115 | Old |
| 0.04857 | 1.9414 | 0.142 | 0.04 | 1 | 14 | Sp3os | 0.102 | Old |
| 0.048855 | 1.255579 | 0.372 | 0.24 | 1 | 14 | C4b | 0.132 | Old |
| 0.049157 | 0.619915 | 0.443 | 0.26 | 1 | 14 | Gxylt2 | 0.183 | Old |
| 0.049446 | 0.324376 | 0.891 | 0.7 | 1 | 14 | S100a10 | 0.191 | Old |
| 0.049857 | 0.21466 | 0.612 | 0.46 | 1 | 14 | Bri3 | 0.152 | Old |
| 0.050569 | 0.335356 | 0.612 | 0.48 | 1 | 14 | Emilin2 | 0.132 | Old |
| 0.051244 | 0.815714 | 0.426 | 0.28 | 1 | 14 | Etfa | 0.146 | Old |
| 0.051293 | 1.042615 | 0.29 | 0.16 | 1 | 14 | Mllt6 | 0.13 | Old |
| 0.05166 | 1.169374 | 0.29 | 0.16 | 1 | 14 | Dennd6a | 0.13 | Old |
| 0.051711 | 0.263668 | 0.699 | 0.5 | 1 | 14 | Itih5 | 0.199 | Old |
| 0.051779 | 2.137183 | 0.186 | 0.08 | 1 | 14 | Eef2k | 0.106 | Old |
| 0.05214 | 2.116325 | 0.208 | 0.1 | 1 | 14 | Orai3 | 0.108 | Old |
| 0.0522 | 0.702969 | 0.508 | 0.38 | 1 | 14 | Lama4 | 0.128 | Old |
| 0.052415 | 1.987159 | 0.109 | 0.02 | 1 | 14 | Mta3 | 0.089 | Old |
| 0.052947 | 0.151173 | 0.989 | 1 | 1 | 14 | Rpl37a | -0.011 | Old |
| 0.053441 | 4.480444 | 0.071 | 0 | 1 | 14 | Trim65 | 0.071 | Old |
| 0.053441 | 4.474181 | 0.071 | 0 | 1 | 14 | Arvcf | 0.071 | Old |
| 0.053441 | 4.141324 | 0.071 | 0 | 1 | 14 | Msc | 0.071 | Old |
| 0.053441 | 4.117663 | 0.071 | 0 | 1 | 14 | Slc2a3 | 0.071 | Old |
| 0.053441 | 4.089534 | 0.071 | 0 | 1 | 14 | Bend6 | 0.071 | Old |
| 0.053441 | 4.07891 | 0.071 | 0 | 1 | 14 | 4930430F0 | 0.071 | Old |
| 0.053441 | 3.97077 | 0.071 | 0 | 1 | 14 | Fbxl8 | 0.071 | Old |
| 0.053441 | 3.841959 | 0.071 | 0 | 1 | 14 | Bicra | 0.071 | Old |
| 0.053992 | 0.435076 | 0.459 | 0.28 | 1 | 14 | Scpep1 | 0.179 | Old |

| | | | | | | | | |
|---|---|---|---|---|---|---|---|---|
| 0.054084 | 0.311181 | 0.033 | 0.1 | 1 | 14 | Nufip1 | -0.067 | Old |
| 0.054216 | 2.023613 | 0.164 | 0.06 | 1 | 14 | Slc7a1 | 0.104 | Old |
| 0.054327 | 1.496181 | 0.191 | 0.08 | 1 | 14 | Igip | 0.111 | Old |
| 0.054643 | 0.704518 | 0.344 | 0.2 | 1 | 14 | Gem | 0.144 | Old |
| 0.054871 | 1.468344 | 0.279 | 0.16 | 1 | 14 | Atp6v0a1 | 0.119 | Old |
| 0.055 | 2.333437 | 0.137 | 0.04 | 1 | 14 | Bmp6 | 0.097 | Old |
| 0.055272 | 0.692578 | 0.372 | 0.24 | 1 | 14 | Tmem43 | 0.132 | Old |
| 0.055664 | 1.369147 | 0.142 | 0.04 | 1 | 14 | Acer3 | 0.102 | Old |
| 0.055803 | 3.48837 | 0.104 | 0.02 | 1 | 14 | Hps5 | 0.084 | Old |
| 0.055873 | 1.589654 | 0.213 | 0.1 | 1 | 14 | Gpr153 | 0.113 | Old |
| 0.056087 | 2.181566 | 0.137 | 0.04 | 1 | 14 | Fibp | 0.097 | Old |
| 0.056089 | 1.08497 | 0.295 | 0.16 | 1 | 14 | Cetn3 | 0.135 | Old |
| 0.056199 | 1.967467 | 0.164 | 0.06 | 1 | 14 | Hs3st1 | 0.104 | Old |
| 0.056704 | 2.098588 | 0.164 | 0.06 | 1 | 14 | Cnpy3 | 0.104 | Old |
| 0.056764 | 0.342138 | 0.689 | 0.5 | 1 | 14 | Thbd | 0.189 | Old |
| 0.056823 | 1.258438 | 0.311 | 0.18 | 1 | 14 | Rnf145 | 0.131 | Old |
| 0.05722 | 0.698515 | 0.213 | 0.1 | 1 | 14 | Zfhx4 | 0.113 | Old |
| 0.057533 | 0.312848 | 0.885 | 0.74 | 1 | 14 | Ddx5 | 0.145 | Old |
| 0.057785 | 0.166488 | 0.005 | 0.04 | 1 | 14 | Map3k13 | -0.035 | Old |
| 0.057785 | 0.140626 | 0.005 | 0.04 | 1 | 14 | Grk4 | -0.035 | Old |
| 0.05821 | 1.555777 | 0.317 | 0.2 | 1 | 14 | Guk1 | 0.117 | Old |
| 0.05823 | 0.823733 | 0.383 | 0.28 | 1 | 14 | 2510009E0 | 0.103 | Old |
| 0.058315 | 1.977723 | 0.137 | 0.04 | 1 | 14 | Sema3e | 0.097 | Old |
| 0.058425 | 1.733495 | 0.235 | 0.12 | 1 | 14 | Ano6 | 0.115 | Old |
| 0.058527 | 0.482482 | 0.656 | 0.52 | 1 | 14 | Rps27l | 0.136 | Old |
| 0.058602 | 1.378394 | 0.317 | 0.2 | 1 | 14 | Sqor | 0.117 | Old |
| 0.058885 | 1.370749 | 0.262 | 0.14 | 1 | 14 | Acsl4 | 0.122 | Old |
| 0.059006 | 0.480481 | 0.044 | 0.12 | 1 | 14 | Acot6 | -0.076 | Old |
| 0.059091 | 0.047059 | 0.033 | 0.1 | 1 | 14 | Wdr4 | -0.067 | Old |
| 0.059333 | 1.08303 | 0.339 | 0.22 | 1 | 14 | Rictor | 0.119 | Old |
| 0.059789 | 0.925729 | 0.421 | 0.3 | 1 | 14 | Arl4a | 0.121 | Old |
| 0.060259 | 0.611452 | 0.322 | 0.2 | 1 | 14 | Gm16286 | 0.122 | Old |
| 0.060624 | 1.80492 | 0.224 | 0.12 | 1 | 14 | Hspa14 | 0.104 | Old |
| 0.060858 | 0.235619 | 0.486 | 0.3 | 1 | 14 | Gfpt2 | 0.186 | Old |
| 0.060987 | 2.509062 | 0.104 | 0.02 | 1 | 14 | Eya4 | 0.084 | Old |
| 0.061051 | 2.844767 | 0.131 | 0.04 | 1 | 14 | Ddx58 | 0.091 | Old |
| 0.061129 | 1.47433 | 0.235 | 0.12 | 1 | 14 | Coro2b | 0.115 | Old |
| 0.061271 | 0.497301 | 0.541 | 0.38 | 1 | 14 | Sult1e1 | 0.161 | Old |
| 0.061298 | 0.876748 | 0.459 | 0.34 | 1 | 14 | Bmp1 | 0.119 | Old |
| 0.061317 | 0.826163 | 0.432 | 0.3 | 1 | 14 | Mlec | 0.132 | Old |
| 0.061663 | 2.414071 | 0.104 | 0.02 | 1 | 14 | Akr1b10 | 0.084 | Old |
| 0.06171 | 0.242957 | 0.306 | 0.18 | 1 | 14 | Flnb | 0.126 | Old |
| 0.061721 | 0.93246 | 0.197 | 0.08 | 1 | 14 | Tex2 | 0.117 | Old |
| 0.061806 | 1.926951 | 0.186 | 0.08 | 1 | 14 | Ankra2 | 0.106 | Old |
| 0.061825 | 0.301197 | 0.142 | 0.04 | 1 | 14 | Abca7 | 0.102 | Old |

| | | | | | | | | |
|---|---|---|---|---|---|---|---|---|
| 0.062013 | 0.549009 | 0.661 | 0.5 | 1 | 14 | Pla1a | 0.161 | Old |
| 0.06252 | 1.374836 | 0.235 | 0.12 | 1 | 14 | Dram2 | 0.115 | Old |
| 0.06259 | 0.388332 | 0.568 | 0.44 | 1 | 14 | Cbx6 | 0.128 | Old |
| 0.062905 | 0.411507 | 0.355 | 0.22 | 1 | 14 | Slc50a1 | 0.135 | Old |
| 0.062915 | 0.992594 | 0.497 | 0.42 | 1 | 14 | Gpm6b | 0.077 | Old |
| 0.06292 | 0.178804 | 0.945 | 0.94 | 1 | 14 | Rpl39 | 0.005 | Old |
| 0.063104 | 0.679066 | 0.481 | 0.34 | 1 | 14 | Col6a5 | 0.141 | Old |
| 0.063309 | 0.888923 | 0.311 | 0.18 | 1 | 14 | Lmbrd1 | 0.131 | Old |
| 0.063798 | 0.275537 | 0.831 | 0.76 | 1 | 14 | Rpl10a | 0.071 | Old |
| 0.064023 | 0.651782 | 0.557 | 0.42 | 1 | 14 | Prelp | 0.137 | Old |
| 0.064129 | 4.125418 | 0.066 | 0 | 1 | 14 | Pkmyt1 | 0.066 | Old |
| 0.064129 | 3.958942 | 0.066 | 0 | 1 | 14 | Cyp4f13 | 0.066 | Old |
| 0.064129 | 3.954402 | 0.066 | 0 | 1 | 14 | AW146154 | 0.066 | Old |
| 0.064129 | 3.92584 | 0.066 | 0 | 1 | 14 | Rnf165 | 0.066 | Old |
| 0.064129 | 3.894864 | 0.066 | 0 | 1 | 14 | Zfp316 | 0.066 | Old |
| 0.064129 | 3.784556 | 0.066 | 0 | 1 | 14 | Trim16 | 0.066 | Old |
| 0.064129 | 3.685542 | 0.066 | 0 | 1 | 14 | Atmin | 0.066 | Old |
| 0.064182 | 1.193572 | 0.164 | 0.06 | 1 | 14 | Ipo9 | 0.104 | Old |
| 0.064608 | 0.797563 | 0.525 | 0.4 | 1 | 14 | Anxa3 | 0.125 | Old |
| 0.064644 | 1.087704 | 0.262 | 0.14 | 1 | 14 | Dpf2 | 0.122 | Old |
| 0.064796 | 0.351697 | 0.776 | 0.66 | 1 | 14 | Ywhae | 0.116 | Old |
| 0.064943 | 0.209699 | 0.355 | 0.22 | 1 | 14 | Nudt3 | 0.135 | Old |
| 0.065284 | 1.554523 | 0.246 | 0.14 | 1 | 14 | Man1c1 | 0.106 | Old |
| 0.065486 | 1.249969 | 0.186 | 0.08 | 1 | 14 | Glis2 | 0.106 | Old |
| 0.06597 | 1.840049 | 0.202 | 0.1 | 1 | 14 | Atg4b | 0.102 | Old |
| 0.066015 | 0.440398 | 0.415 | 0.3 | 1 | 14 | Dlgap4 | 0.115 | Old |
| 0.066163 | 0.858558 | 0.459 | 0.34 | 1 | 14 | Vwa5a | 0.119 | Old |
| 0.066209 | 0.155243 | 0.208 | 0.1 | 1 | 14 | Lrrc8c | 0.108 | Old |
| 0.06688 | 0.080777 | 0.071 | 0.16 | 1 | 14 | Qsox2 | -0.089 | Old |
| 0.067141 | 3.295299 | 0.098 | 0.02 | 1 | 14 | Zfp626 | 0.078 | Old |
| 0.067277 | 2.225301 | 0.131 | 0.04 | 1 | 14 | Sema6c | 0.091 | Old |
| 0.067316 | 0.088263 | 0.191 | 0.08 | 1 | 14 | Timm10b | 0.111 | Old |
| 0.067335 | 1.265537 | 0.137 | 0.04 | 1 | 14 | Blvra | 0.097 | Old |
| 0.067396 | 0.935268 | 0.322 | 0.2 | 1 | 14 | Xrn1 | 0.122 | Old |
| 0.067471 | 1.01439 | 0.426 | 0.32 | 1 | 14 | Psme1 | 0.106 | Old |
| 0.067524 | 0.506971 | 0.284 | 0.16 | 1 | 14 | C330007P0 | 0.124 | Old |
| 0.067835 | 0.564383 | 0.53 | 0.42 | 1 | 14 | Grina | 0.11 | Old |
| 0.067889 | 3.524821 | 0.098 | 0.02 | 1 | 14 | Tspyl4 | 0.078 | Old |
| 0.067928 | 1.86745 | 0.131 | 0.04 | 1 | 14 | Edem2 | 0.091 | Old |
| 0.067989 | 0.427442 | 0.88 | 0.72 | 1 | 14 | Igfbp4 | 0.16 | Old |
| 0.068457 | 1.447647 | 0.158 | 0.06 | 1 | 14 | Hmox2 | 0.098 | Old |
| 0.068662 | 1.441474 | 0.23 | 0.12 | 1 | 14 | Arhgap12 | 0.11 | Old |
| 0.06906 | 1.860916 | 0.158 | 0.06 | 1 | 14 | Slc30a7 | 0.098 | Old |
| 0.069089 | 2.934837 | 0.126 | 0.04 | 1 | 14 | Mcm3ap | 0.086 | Old |
| 0.069141 | 0.187736 | 0.645 | 0.48 | 1 | 14 | Sod1 | 0.165 | Old |

| | | | | | | | | |
|---|---|---|---|---|---|---|---|---|
| 0.06918 | 0.594448 | 0.224 | 0.1 | 1 | 14 | Hp | 0.124 | Old |
| 0.069393 | 1.310771 | 0.284 | 0.16 | 1 | 14 | Ccpg1 | 0.124 | Old |
| 0.069407 | 3.335456 | 0.098 | 0.02 | 1 | 14 | Dmkn | 0.078 | Old |
| 0.069667 | 0.238332 | 0.158 | 0.06 | 1 | 14 | Pip4k2c | 0.098 | Old |
| 0.06969 | 0.813343 | 0.464 | 0.36 | 1 | 14 | Rora | 0.104 | Old |
| 0.069742 | 0.480063 | 0.361 | 0.24 | 1 | 14 | Ltbp3 | 0.121 | Old |
| 0.070031 | 1.137424 | 0.399 | 0.28 | 1 | 14 | Lifr | 0.119 | Old |
| 0.070074 | 1.148838 | 0.279 | 0.16 | 1 | 14 | Dsel | 0.119 | Old |
| 0.070176 | 2.824227 | 0.098 | 0.02 | 1 | 14 | Bclaf3 | 0.078 | Old |
| 0.070224 | 0.884276 | 0.443 | 0.32 | 1 | 14 | Cyth3 | 0.123 | Old |
| 0.070253 | 0.175423 | 0.661 | 0.5 | 1 | 14 | Gnb1 | 0.161 | Old |
| 0.070254 | 1.287078 | 0.273 | 0.16 | 1 | 14 | Sgk1 | 0.113 | Old |
| 0.070447 | 3.627758 | 0.126 | 0.04 | 1 | 14 | Utp20 | 0.086 | Old |
| 0.070449 | 1.410476 | 0.208 | 0.1 | 1 | 14 | Vsig10 | 0.108 | Old |
| 0.071307 | 0.300451 | 0.738 | 0.66 | 1 | 14 | Tmbim6 | 0.078 | Old |
| 0.07131 | 0.447606 | 0.781 | 0.68 | 1 | 14 | Fn1 | 0.101 | Old |
| 0.071384 | 0.212976 | 0.355 | 0.22 | 1 | 14 | Psma4 | 0.135 | Old |
| 0.071576 | 0.414429 | 0.432 | 0.28 | 1 | 14 | Prdx6 | 0.152 | Old |
| 0.071585 | 0.959027 | 0.301 | 0.18 | 1 | 14 | Pde8a | 0.121 | Old |
| 0.072013 | 1.085978 | 0.279 | 0.16 | 1 | 14 | Med28 | 0.119 | Old |
| 0.07206 | 1.065362 | 0.301 | 0.18 | 1 | 14 | Synj2bp | 0.121 | Old |
| 0.072249 | 0.353642 | 0.607 | 0.48 | 1 | 14 | Antxr1 | 0.127 | Old |
| 0.072292 | 0.21553 | 0.749 | 0.6 | 1 | 14 | Medag | 0.149 | Old |
| 0.072508 | 0.854437 | 0.29 | 0.16 | 1 | 14 | Pcyt1a | 0.13 | Old |
| 0.072686 | 1.234073 | 0.257 | 0.14 | 1 | 14 | Ang | 0.117 | Old |
| 0.072789 | 1.152266 | 0.213 | 0.1 | 1 | 14 | Calhm2 | 0.113 | Old |
| 0.07326 | 0.830843 | 0.284 | 0.16 | 1 | 14 | Afap1 | 0.124 | Old |
| 0.073316 | 0.969284 | 0.503 | 0.4 | 1 | 14 | Tnfaip2 | 0.103 | Old |
| 0.073331 | 0.683814 | 0.53 | 0.42 | 1 | 14 | Fam120a | 0.11 | Old |
| 0.074017 | 1.562719 | 0.131 | 0.04 | 1 | 14 | Adora2b | 0.091 | Old |
| 0.074396 | 0.210897 | 0.787 | 0.6 | 1 | 14 | Serf2 | 0.187 | Old |
| 0.074655 | 0.615445 | 0.137 | 0.04 | 1 | 14 | 1600012H( | 0.097 | Old |
| 0.07558 | 1.21955 | 0.366 | 0.26 | 1 | 14 | Itch | 0.106 | Old |
| 0.075601 | 1.89464 | 0.153 | 0.06 | 1 | 14 | Gtf3a | 0.093 | Old |
| 0.075634 | 0.479303 | 0.311 | 0.18 | 1 | 14 | Ssc5d | 0.131 | Old |
| 0.076038 | 0.288624 | 0.918 | 0.82 | 1 | 14 | Rps27 | 0.098 | Old |
| 0.076475 | 1.768136 | 0.202 | 0.1 | 1 | 14 | Dock4 | 0.102 | Old |
| 0.076516 | 0.578558 | 0.448 | 0.32 | 1 | 14 | Hexa | 0.128 | Old |
| 0.077005 | 4.324839 | 0.06 | 0 | 1 | 14 | Ift122 | 0.06 | Old |
| 0.077005 | 4.113726 | 0.06 | 0 | 1 | 14 | Tspan17 | 0.06 | Old |
| 0.077005 | 4.022123 | 0.06 | 0 | 1 | 14 | Wdyhv1 | 0.06 | Old |
| 0.077005 | 4.014015 | 0.06 | 0 | 1 | 14 | Nmnat3 | 0.06 | Old |
| 0.077005 | 3.99544 | 0.06 | 0 | 1 | 14 | Tgfa | 0.06 | Old |
| 0.077005 | 3.957633 | 0.06 | 0 | 1 | 14 | Rmdn3 | 0.06 | Old |
| 0.077005 | 3.871364 | 0.06 | 0 | 1 | 14 | Cyp2u1 | 0.06 | Old |

| | | | | | | | | |
|---|---|---|---|---|---|---|---|---|
| 0.077005 | 3.85736 | 0.06 | 0 | 1 | 14 | Fam131a | 0.06 | Old |
| 0.077005 | 3.673185 | 0.06 | 0 | 1 | 14 | Cd300a | 0.06 | Old |
| 0.077005 | 3.591366 | 0.06 | 0 | 1 | 14 | Pfkfb4 | 0.06 | Old |
| 0.077005 | 3.456324 | 0.06 | 0 | 1 | 14 | Uevld | 0.06 | Old |
| 0.077005 | 3.347002 | 0.06 | 0 | 1 | 14 | 1110032A( | 0.06 | Old |
| 0.077005 | 3.332293 | 0.06 | 0 | 1 | 14 | Zfp40 | 0.06 | Old |
| 0.077005 | 3.052074 | 0.06 | 0 | 1 | 14 | Gm45187 | 0.06 | Old |
| 0.077156 | 0.280603 | 0.831 | 0.68 | 1 | 14 | Ly6a | 0.151 | Old |
| 0.077278 | 0.057739 | 0.383 | 0.26 | 1 | 14 | Cdc37 | 0.123 | Old |
| 0.077591 | 0.996543 | 0.295 | 0.18 | 1 | 14 | Smim15 | 0.115 | Old |
| 0.077651 | 0.623977 | 0.388 | 0.26 | 1 | 14 | Apbb1ip | 0.128 | Old |
| 0.077967 | 0.324595 | 0.858 | 0.7 | 1 | 14 | Hnrnpa2b1 | 0.158 | Old |
| 0.078129 | 0.461483 | 0.65 | 0.54 | 1 | 14 | Gpx4 | 0.11 | Old |
| 0.078257 | 1.960353 | 0.098 | 0.02 | 1 | 14 | Htra4 | 0.078 | Old |
| 0.07887 | 1.46628 | 0.186 | 0.08 | 1 | 14 | Eif4e3 | 0.106 | Old |
| 0.079255 | 0.290328 | 0.607 | 0.42 | 1 | 14 | Ggt5 | 0.187 | Old |
| 0.079411 | 0.791758 | 0.437 | 0.3 | 1 | 14 | Derl1 | 0.137 | Old |
| 0.079961 | 1.418792 | 0.098 | 0.02 | 1 | 14 | Amy1 | 0.078 | Old |
| 0.079993 | 0.243532 | 0.967 | 0.96 | 1 | 14 | Egr1 | 0.007 | Old |
| 0.080021 | 0.704861 | 0.333 | 0.2 | 1 | 14 | Ikbkb | 0.133 | Old |
| 0.08015 | 0.680055 | 0.475 | 0.34 | 1 | 14 | Nbeal1 | 0.135 | Old |
| 0.080367 | 0.042415 | 0.546 | 0.38 | 1 | 14 | Pea15a | 0.166 | Old |
| 0.080523 | 0.098163 | 0.333 | 0.2 | 1 | 14 | Opcml | 0.133 | Old |
| 0.0806 | 2.434593 | 0.148 | 0.06 | 1 | 14 | C1qtnf7 | 0.088 | Old |
| 0.080749 | 3.279852 | 0.093 | 0.02 | 1 | 14 | Ogfod2 | 0.073 | Old |
| 0.080749 | 3.141776 | 0.093 | 0.02 | 1 | 14 | Alms1 | 0.073 | Old |
| 0.080749 | 2.954539 | 0.093 | 0.02 | 1 | 14 | Hilpda | 0.073 | Old |
| 0.080795 | 0.073963 | 0.344 | 0.2 | 1 | 14 | Setd1b | 0.144 | Old |
| 0.080819 | 0.657523 | 0.53 | 0.4 | 1 | 14 | Zmynd11 | 0.13 | Old |
| 0.081359 | 0.193005 | 0.158 | 0.06 | 1 | 14 | Ncoa5 | 0.098 | Old |
| 0.081603 | 0.371493 | 0.792 | 0.72 | 1 | 14 | Hnrnpf | 0.072 | Old |
| 0.081641 | 2.98877 | 0.093 | 0.02 | 1 | 14 | Vangl2 | 0.073 | Old |
| 0.08168 | 0.95627 | 0.322 | 0.2 | 1 | 14 | Sbf2 | 0.122 | Old |
| 0.081695 | 0.638509 | 0.098 | 0.02 | 1 | 14 | Vwc2 | 0.078 | Old |
| 0.081908 | 1.535357 | 0.202 | 0.1 | 1 | 14 | Rabgap1l | 0.102 | Old |
| 0.081934 | 0.62273 | 0.333 | 0.22 | 1 | 14 | Cox17 | 0.113 | Old |
| 0.082021 | 2.674286 | 0.148 | 0.06 | 1 | 14 | Rb1 | 0.088 | Old |
| 0.082029 | 1.396281 | 0.208 | 0.1 | 1 | 14 | Cxcl2 | 0.108 | Old |
| 0.082056 | 1.158863 | 0.186 | 0.08 | 1 | 14 | Tnfsf13b | 0.106 | Old |
| 0.082541 | 3.206202 | 0.093 | 0.02 | 1 | 14 | Trmt10c | 0.073 | Old |
| 0.082541 | 3.066183 | 0.093 | 0.02 | 1 | 14 | Klhl42 | 0.073 | Old |
| 0.082544 | 0.397453 | 0.716 | 0.56 | 1 | 14 | Ccnl1 | 0.156 | Old |
| 0.082546 | 0.841799 | 0.355 | 0.24 | 1 | 14 | Ids | 0.115 | Old |
| 0.082649 | 0.875817 | 0.301 | 0.18 | 1 | 14 | Ranbp1 | 0.121 | Old |
| 0.082825 | 2.894823 | 0.12 | 0.04 | 1 | 14 | Sord | 0.08 | Old |

| | | | | | | | | |
|---|---|---|---|---|---|---|---|---|
| 0.083 | 1.262029 | 0.23 | 0.12 | 1 | 14 | Dnase2a | 0.11 | Old |
| 0.083 | 1.118643 | 0.23 | 0.12 | 1 | 14 | Cnppd1 | 0.11 | Old |
| 0.083054 | 0.498924 | 0.803 | 0.72 | 1 | 14 | Man2a1 | 0.083 | Old |
| 0.08324 | 0.860001 | 0.366 | 0.26 | 1 | 14 | Adam17 | 0.106 | Old |
| 0.083481 | 1.1684 | 0.268 | 0.16 | 1 | 14 | Nectin3 | 0.108 | Old |
| 0.083596 | 1.299603 | 0.23 | 0.12 | 1 | 14 | Jam2 | 0.11 | Old |
| 0.083905 | 1.184653 | 0.251 | 0.14 | 1 | 14 | Lrrk1 | 0.111 | Old |
| 0.084563 | 0.130623 | 0.956 | 0.9 | 1 | 14 | Rps27a | 0.056 | Old |
| 0.084817 | 0.476956 | 0.361 | 0.24 | 1 | 14 | Fis1 | 0.121 | Old |
| 0.085101 | 0.403286 | 0.432 | 0.3 | 1 | 14 | Foxn3 | 0.132 | Old |
| 0.085271 | 0.070594 | 0.126 | 0.04 | 1 | 14 | Aar2 | 0.086 | Old |
| 0.085817 | 0.581163 | 0.443 | 0.32 | 1 | 14 | Gja1 | 0.123 | Old |
| 0.086883 | 1.130531 | 0.126 | 0.04 | 1 | 14 | Parp10 | 0.086 | Old |
| 0.086906 | 0.607175 | 0.497 | 0.32 | 1 | 14 | Eif4g3 | 0.177 | Old |
| 0.087154 | 2.138608 | 0.148 | 0.06 | 1 | 14 | Lrpprc | 0.088 | Old |
| 0.08771 | 2.171515 | 0.12 | 0.04 | 1 | 14 | Bcl2l13 | 0.08 | Old |
| 0.087855 | 1.16232 | 0.339 | 0.24 | 1 | 14 | Lman2 | 0.099 | Old |
| 0.08787 | 0.948246 | 0.23 | 0.12 | 1 | 14 | Rbck1 | 0.11 | Old |
| 0.088199 | 0.930814 | 0.355 | 0.24 | 1 | 14 | Ccnl2 | 0.115 | Old |
| 0.088389 | 0.586656 | 0.628 | 0.5 | 1 | 14 | Pbrm1 | 0.128 | Old |
| 0.089019 | 1.14957 | 0.224 | 0.12 | 1 | 14 | Acss3 | 0.104 | Old |
| 0.089107 | 0.20184 | 0.049 | 0.12 | 1 | 14 | Erbb2 | -0.071 | Old |
| 0.089253 | 0.316677 | 0.557 | 0.42 | 1 | 14 | Twsg1 | 0.137 | Old |
| 0.089432 | 2.312953 | 0.148 | 0.06 | 1 | 14 | 6430548M | 0.088 | Old |
| 0.089609 | 0.553474 | 0.421 | 0.32 | 1 | 14 | Hcfc1r1 | 0.101 | Old |
| 0.089743 | 0.2278 | 0.891 | 0.7 | 1 | 14 | Ogn | 0.191 | Old |
| 0.089844 | 1.04857 | 0.251 | 0.14 | 1 | 14 | Gps1 | 0.111 | Old |
| 0.089987 | 0.166408 | 0.016 | 0.06 | 1 | 14 | Gtf2h4 | -0.044 | Old |
| 0.090034 | 2.506562 | 0.093 | 0.02 | 1 | 14 | Caap1 | 0.073 | Old |
| 0.090202 | 1.736313 | 0.148 | 0.06 | 1 | 14 | Zdhhc7 | 0.088 | Old |
| 0.090392 | 1.07809 | 0.284 | 0.18 | 1 | 14 | Golga7 | 0.104 | Old |
| 0.090529 | 2.494046 | 0.142 | 0.06 | 1 | 14 | Plekha8 | 0.082 | Old |
| 0.090714 | 0.794103 | 0.262 | 0.16 | 1 | 14 | Gkap1 | 0.102 | Old |
| 0.090996 | 0.560832 | 0.765 | 0.64 | 1 | 14 | Mcl1 | 0.125 | Old |
| 0.09102 | 1.134268 | 0.126 | 0.04 | 1 | 14 | Mzt2 | 0.086 | Old |
| 0.091042 | 0.271962 | 0.623 | 0.5 | 1 | 14 | Pnisr | 0.123 | Old |
| 0.091519 | 0.795064 | 0.481 | 0.38 | 1 | 14 | Prr13 | 0.101 | Old |
| 0.09163 | 2.320351 | 0.164 | 0.08 | 1 | 14 | Epb41l4aos | 0.084 | Old |
| 0.091634 | 0.392604 | 0.016 | 0.06 | 1 | 14 | Peg10 | -0.044 | Old |
| 0.091991 | 1.755905 | 0.093 | 0.02 | 1 | 14 | Trim37 | 0.073 | Old |
| 0.092075 | 1.203896 | 0.153 | 0.06 | 1 | 14 | Gpsm2 | 0.093 | Old |
| 0.092374 | 2.945045 | 0.164 | 0.08 | 1 | 14 | Lpin1 | 0.084 | Old |
| 0.092556 | 4.438845 | 0.055 | 0 | 1 | 14 | Fbln7 | 0.055 | Old |
| 0.092556 | 4.380307 | 0.055 | 0 | 1 | 14 | Zfp653 | 0.055 | Old |
| 0.092556 | 4.305429 | 0.055 | 0 | 1 | 14 | Rtn2 | 0.055 | Old |

| | | | | | | | | |
|---|---|---|---|---|---|---|---|---|
| 0.092556 | 4.2024 | 0.055 | 0 | 1 | 14 | Rsad2 | 0.055 | Old |
| 0.092556 | 3.731306 | 0.055 | 0 | 1 | 14 | Tldc1 | 0.055 | Old |
| 0.092556 | 3.726281 | 0.055 | 0 | 1 | 14 | Tmem201 | 0.055 | Old |
| 0.092556 | 3.543507 | 0.055 | 0 | 1 | 14 | Idua | 0.055 | Old |
| 0.092556 | 3.528522 | 0.055 | 0 | 1 | 14 | Trim39 | 0.055 | Old |
| 0.092556 | 3.527153 | 0.055 | 0 | 1 | 14 | Gca | 0.055 | Old |
| 0.092556 | 3.517318 | 0.055 | 0 | 1 | 14 | Sgtb | 0.055 | Old |
| 0.092556 | 3.464656 | 0.055 | 0 | 1 | 14 | Igkc | 0.055 | Old |
| 0.092556 | 3.369236 | 0.055 | 0 | 1 | 14 | Gtf2ird1 | 0.055 | Old |
| 0.092556 | 3.349049 | 0.055 | 0 | 1 | 14 | Lxn | 0.055 | Old |
| 0.092556 | 3.316945 | 0.055 | 0 | 1 | 14 | Hook2 | 0.055 | Old |
| 0.092556 | 3.313326 | 0.055 | 0 | 1 | 14 | Pola1 | 0.055 | Old |
| 0.092556 | 3.301909 | 0.055 | 0 | 1 | 14 | Nptxr | 0.055 | Old |
| 0.092556 | 3.034856 | 0.055 | 0 | 1 | 14 | Ercc6 | 0.055 | Old |
| 0.092556 | 2.996104 | 0.055 | 0 | 1 | 14 | Trim21 | 0.055 | Old |
| 0.092556 | 2.889013 | 0.055 | 0 | 1 | 14 | Map6 | 0.055 | Old |
| 0.092826 | 2.137845 | 0.12 | 0.04 | 1 | 14 | Srr | 0.08 | Old |
| 0.092916 | 0.780769 | 0.437 | 0.32 | 1 | 14 | S100a1 | 0.117 | Old |
| 0.092934 | 0.761052 | 0.197 | 0.1 | 1 | 14 | Siva1 | 0.097 | Old |
| 0.092938 | 0.359628 | 0.18 | 0.08 | 1 | 14 | Fam193b | 0.1 | Old |
| 0.092992 | 0.354057 | 0.683 | 0.54 | 1 | 14 | Sdcbp | 0.143 | Old |
| 0.093051 | 0.565725 | 0.257 | 0.14 | 1 | 14 | Aox3 | 0.117 | Old |
| 0.093336 | 1.853605 | 0.148 | 0.06 | 1 | 14 | Coq2 | 0.088 | Old |
| 0.093343 | 1.361874 | 0.284 | 0.18 | 1 | 14 | Prcp | 0.104 | Old |
| 0.093674 | 0.608812 | 0.388 | 0.26 | 1 | 14 | Cflar | 0.128 | Old |
| 0.093881 | 1.022737 | 0.399 | 0.3 | 1 | 14 | Igsf8 | 0.099 | Old |
| 0.093996 | 0.422168 | 0.689 | 0.58 | 1 | 14 | Tm9sf3 | 0.109 | Old |
| 0.094178 | 0.357864 | 0.656 | 0.54 | 1 | 14 | Atp5e | 0.116 | Old |
| 0.094389 | 1.018496 | 0.262 | 0.16 | 1 | 14 | Gstt1 | 0.102 | Old |
| 0.094405 | 1.114946 | 0.202 | 0.1 | 1 | 14 | Crebzf | 0.102 | Old |
| 0.094584 | 1.614686 | 0.12 | 0.04 | 1 | 14 | Vps16 | 0.08 | Old |
| 0.094854 | 1.009422 | 0.426 | 0.34 | 1 | 14 | Mindy2 | 0.086 | Old |
| 0.094935 | 1.716068 | 0.148 | 0.06 | 1 | 14 | Plekha6 | 0.088 | Old |
| 0.09531 | 0.670495 | 0.257 | 0.16 | 1 | 14 | Egln2 | 0.097 | Old |
| 0.095608 | 1.222225 | 0.235 | 0.12 | 1 | 14 | Usp33 | 0.115 | Old |
| 0.095743 | 1.264107 | 0.197 | 0.1 | 1 | 14 | Naga | 0.097 | Old |
| 0.096016 | 0.291247 | 0.399 | 0.28 | 1 | 14 | Pdlim5 | 0.119 | Old |
| 0.09625 | 0.174341 | 0.721 | 0.62 | 1 | 14 | Hnrnpu | 0.101 | Old |
| 0.096357 | 0.502386 | 0.508 | 0.36 | 1 | 14 | Cpxm1 | 0.148 | Old |
| 0.096407 | 1.707104 | 0.169 | 0.08 | 1 | 14 | Mtmr4 | 0.089 | Old |
| 0.096455 | 0.522464 | 0.366 | 0.26 | 1 | 14 | Lamtor4 | 0.106 | Old |
| 0.096511 | 1.03863 | 0.202 | 0.1 | 1 | 14 | Kctd20 | 0.102 | Old |
| 0.096555 | 1.88699 | 0.148 | 0.06 | 1 | 14 | Prdm8 | 0.088 | Old |
| 0.096716 | 0.722947 | 0.541 | 0.44 | 1 | 14 | Wt1 | 0.101 | Old |
| 0.097031 | 0.838189 | 0.093 | 0.02 | 1 | 14 | Wdr41 | 0.073 | Old |

| | | | | | | | | |
|---|---|---|---|---|---|---|---|---|
| 0.097086 | 3.082753 | 0.087 | 0.02 | 1 | 14 | 2510002D2 | 0.067 | Old |
| 0.097086 | 2.907297 | 0.087 | 0.02 | 1 | 14 | Pitpnm2 | 0.067 | Old |
| 0.09727 | 2.115133 | 0.12 | 0.04 | 1 | 14 | Mrpl58 | 0.08 | Old |
| 0.09727 | 1.611169 | 0.12 | 0.04 | 1 | 14 | Cyp39a1 | 0.08 | Old |
| 0.097292 | 3.116464 | 0.115 | 0.04 | 1 | 14 | Cyb5r1 | 0.075 | Old |
| 0.097308 | 0.680592 | 0.372 | 0.28 | 1 | 14 | Rc3h2 | 0.092 | Old |
| 0.097598 | 2.236819 | 0.186 | 0.1 | 1 | 14 | Akap8l | 0.086 | Old |
| 0.097694 | 0.446088 | 0.328 | 0.22 | 1 | 14 | Adam10 | 0.108 | Old |
| 0.097776 | 0.57369 | 0.437 | 0.34 | 1 | 14 | Use1 | 0.097 | Old |
| 0.097829 | 2.105847 | 0.208 | 0.12 | 1 | 14 | Cask | 0.088 | Old |
| 0.097832 | 2.415977 | 0.142 | 0.06 | 1 | 14 | Eogt | 0.082 | Old |
| 0.097832 | 2.315833 | 0.142 | 0.06 | 1 | 14 | Mrps23 | 0.082 | Old |
| 0.097896 | 0.68893 | 0.426 | 0.32 | 1 | 14 | Rnf130 | 0.106 | Old |
| 0.098073 | 1.297205 | 0.175 | 0.08 | 1 | 14 | Me1 | 0.095 | Old |
| 0.098148 | 3.048566 | 0.087 | 0.02 | 1 | 14 | Ttc30b | 0.067 | Old |
| 0.098414 | 0.326684 | 0.284 | 0.16 | 1 | 14 | Itfg1 | 0.124 | Old |
| 0.098501 | 2.278068 | 0.164 | 0.08 | 1 | 14 | Ripor1 | 0.084 | Old |
| 0.098655 | 0.629109 | 0.202 | 0.1 | 1 | 14 | Xpot | 0.102 | Old |
| 0.098671 | 2.134882 | 0.142 | 0.06 | 1 | 14 | Rdm1 | 0.082 | Old |
| 0.098847 | 0.001373 | 0.825 | 0.62 | 1 | 14 | Rhob | 0.205 | Old |
| 0.098865 | 0.357592 | 0.126 | 0.04 | 1 | 14 | Tnfrsf11b | 0.086 | Old |
| 0.09908 | 2.115036 | 0.186 | 0.1 | 1 | 14 | Rpl7l1 | 0.086 | Old |
| 0.09922 | 3.004184 | 0.087 | 0.02 | 1 | 14 | Scg3 | 0.067 | Old |
| 0.09922 | 2.941626 | 0.087 | 0.02 | 1 | 14 | Rrn3 | 0.067 | Old |
| 0.09922 | 2.879353 | 0.087 | 0.02 | 1 | 14 | Slc25a45 | 0.067 | Old |
| 0.09922 | 2.391947 | 0.087 | 0.02 | 1 | 14 | Nt5c3b | 0.067 | Old |
| 0.0998 | 0.305174 | 0.672 | 0.48 | 1 | 14 | Bgn | 0.192 | Old |
| 0.100051 | 0.232076 | 0.656 | 0.54 | 1 | 14 | Rtn3 | 0.116 | Old |
| 0.100155 | 0.854585 | 0.404 | 0.3 | 1 | 14 | Pdlim1 | 0.104 | Old |
| 0.100388 | 1.119128 | 0.175 | 0.08 | 1 | 14 | Casc4 | 0.095 | Old |
| 0.100552 | 0.594686 | 0.705 | 0.58 | 1 | 14 | Cebpb | 0.125 | Old |
| 0.100736 | 0.438359 | 0.601 | 0.5 | 1 | 14 | Mia3 | 0.101 | Old |
| 0.100949 | 0.979922 | 0.279 | 0.18 | 1 | 14 | Efhd1 | 0.099 | Old |
| 0.10104 | 2.547559 | 0.115 | 0.04 | 1 | 14 | Rbm34 | 0.075 | Old |
| 0.101042 | 1.214152 | 0.268 | 0.16 | 1 | 14 | Mindy1 | 0.108 | Old |
| 0.101391 | 2.844292 | 0.087 | 0.02 | 1 | 14 | Smpd4 | 0.067 | Old |
| 0.101691 | 1.162079 | 0.219 | 0.12 | 1 | 14 | Cpsf2 | 0.099 | Old |
| 0.101957 | 0.312613 | 0.563 | 0.42 | 1 | 14 | Hdgf | 0.143 | Old |
| 0.102396 | 0.059505 | 0.328 | 0.2 | 1 | 14 | Ptpn11 | 0.128 | Old |
| 0.102956 | 2.252386 | 0.142 | 0.06 | 1 | 14 | Nln | 0.082 | Old |
| 0.103072 | 0.22468 | 0.705 | 0.48 | 1 | 14 | Luc7l2 | 0.225 | Old |
| 0.103105 | 0.275839 | 0.328 | 0.22 | 1 | 14 | Gigyf1 | 0.108 | Old |
| 0.103209 | 0.896859 | 0.339 | 0.22 | 1 | 14 | Ccar1 | 0.119 | Old |
| 0.103367 | 0.373104 | 0.585 | 0.48 | 1 | 14 | Sf3b1 | 0.105 | Old |
| 0.103445 | 0.149846 | 0.814 | 0.64 | 1 | 14 | Lamc1 | 0.174 | Old |

| | | | | | | | | |
|---|---|---|---|---|---|---|---|---|
| 0.103615 | 0.160027 | 0.361 | 0.24 | 1 | 14 | Kmt2c | 0.121 | Old |
| 0.103831 | 1.967565 | 0.142 | 0.06 | 1 | 14 | Nt5c3 | 0.082 | Old |
| 0.103865 | 0.348977 | 0.749 | 0.62 | 1 | 14 | Axl | 0.129 | Old |
| 0.103996 | 0.427195 | 0.492 | 0.36 | 1 | 14 | Ackr3 | 0.132 | Old |
| 0.104125 | 1.28582 | 0.148 | 0.06 | 1 | 14 | Trabd | 0.088 | Old |
| 0.104145 | 0.67756 | 0.257 | 0.14 | 1 | 14 | Rbm42 | 0.117 | Old |
| 0.104711 | 2.095354 | 0.142 | 0.06 | 1 | 14 | Rhobtb3 | 0.082 | Old |
| 0.104719 | 2.296257 | 0.087 | 0.02 | 1 | 14 | Rabepk | 0.067 | Old |
| 0.104875 | 1.20731 | 0.23 | 0.14 | 1 | 14 | Gpbp1l1 | 0.09 | Old |
| 0.104903 | 2.842662 | 0.115 | 0.04 | 1 | 14 | Ttc21b | 0.075 | Old |
| 0.105811 | 0.270739 | 0.896 | 0.8 | 1 | 14 | Ctsl | 0.096 | Old |
| 0.105847 | 2.195918 | 0.087 | 0.02 | 1 | 14 | Pick1 | 0.067 | Old |
| 0.105879 | 0.795486 | 0.317 | 0.2 | 1 | 14 | Nampt | 0.117 | Old |
| 0.105887 | 2.643573 | 0.115 | 0.04 | 1 | 14 | Tppp | 0.075 | Old |
| 0.105973 | 1.787192 | 0.186 | 0.1 | 1 | 14 | Tmem106c | 0.086 | Old |
| 0.106666 | 0.29869 | 0.59 | 0.5 | 1 | 14 | Spag9 | 0.09 | Old |
| 0.106841 | 0.693879 | 0.202 | 0.1 | 1 | 14 | Odf2 | 0.102 | Old |
| 0.106868 | 0.537849 | 0.634 | 0.52 | 1 | 14 | Col5a3 | 0.114 | Old |
| 0.108292 | 0.602122 | 0.142 | 0.06 | 1 | 14 | Ptk2b | 0.082 | Old |
| 0.108438 | 1.702165 | 0.191 | 0.1 | 1 | 14 | Snx33 | 0.091 | Old |
| 0.108447 | 0.946462 | 0.197 | 0.1 | 1 | 14 | Phlpp1 | 0.097 | Old |
| 0.10863 | 1.259058 | 0.12 | 0.04 | 1 | 14 | Nip7 | 0.08 | Old |
| 0.109013 | 0.747494 | 0.219 | 0.12 | 1 | 14 | Trim25 | 0.099 | Old |
| 0.109281 | 0.283583 | 0.366 | 0.24 | 1 | 14 | Gprc5b | 0.126 | Old |
| 0.109319 | 0.205138 | 0.066 | 0.14 | 1 | 14 | Gnao1 | -0.074 | Old |
| 0.109592 | 0.494316 | 0.262 | 0.16 | 1 | 14 | Acadl | 0.102 | Old |
| 0.109963 | 1.792763 | 0.186 | 0.1 | 1 | 14 | Jak2 | 0.086 | Old |
| 0.110002 | 1.64503 | 0.164 | 0.08 | 1 | 14 | Cnnm3 | 0.084 | Old |
| 0.110422 | 0.095045 | 0.639 | 0.58 | 1 | 14 | Cox7a2l | 0.059 | Old |
| 0.110457 | 1.842758 | 0.087 | 0.02 | 1 | 14 | Cklf | 0.067 | Old |
| 0.110775 | 1.48579 | 0.186 | 0.1 | 1 | 14 | Clip3 | 0.086 | Old |
| 0.111249 | 0.720201 | 0.454 | 0.36 | 1 | 14 | Ppp2r1a | 0.094 | Old |
| 0.111401 | 4.468831 | 0.049 | 0 | 1 | 14 | Pik3r5 | 0.049 | Old |
| 0.111401 | 4.081722 | 0.049 | 0 | 1 | 14 | Slc12a7 | 0.049 | Old |
| 0.111401 | 4.020871 | 0.049 | 0 | 1 | 14 | Mterf4 | 0.049 | Old |
| 0.111401 | 3.88828 | 0.049 | 0 | 1 | 14 | Nkd1 | 0.049 | Old |
| 0.111401 | 3.871662 | 0.049 | 0 | 1 | 14 | E2f6 | 0.049 | Old |
| 0.111401 | 3.726589 | 0.049 | 0 | 1 | 14 | Zscan22 | 0.049 | Old |
| 0.111401 | 3.713151 | 0.049 | 0 | 1 | 14 | Rnf26 | 0.049 | Old |
| 0.111401 | 3.618238 | 0.049 | 0 | 1 | 14 | Saxo2 | 0.049 | Old |
| 0.111401 | 3.529646 | 0.049 | 0 | 1 | 14 | Tceanc | 0.049 | Old |
| 0.111401 | 3.51106 | 0.049 | 0 | 1 | 14 | Alad | 0.049 | Old |
| 0.111401 | 3.450194 | 0.049 | 0 | 1 | 14 | Omd | 0.049 | Old |
| 0.111401 | 3.426637 | 0.049 | 0 | 1 | 14 | Zfp747 | 0.049 | Old |
| 0.111401 | 3.410007 | 0.049 | 0 | 1 | 14 | Cdkn2c | 0.049 | Old |

| | | | | | | | | |
|---|---|---|---|---|---|---|---|---|
| 0.111401 | 3.399068 | 0.049 | 0 | 1 | 14 | Tuba4a | 0.049 | Old |
| 0.111401 | 3.395993 | 0.049 | 0 | 1 | 14 | Cox15 | 0.049 | Old |
| 0.111401 | 3.394283 | 0.049 | 0 | 1 | 14 | AW209491 | 0.049 | Old |
| 0.111401 | 3.362241 | 0.049 | 0 | 1 | 14 | Gemin5 | 0.049 | Old |
| 0.111401 | 3.344798 | 0.049 | 0 | 1 | 14 | Rangrf | 0.049 | Old |
| 0.111401 | 3.202715 | 0.049 | 0 | 1 | 14 | Fastkd2 | 0.049 | Old |
| 0.111401 | 3.170536 | 0.049 | 0 | 1 | 14 | Brat1 | 0.049 | Old |
| 0.111401 | 3.158307 | 0.049 | 0 | 1 | 14 | Wdr24 | 0.049 | Old |
| 0.111401 | 3.136266 | 0.049 | 0 | 1 | 14 | Gm9947 | 0.049 | Old |
| 0.111401 | 3.119286 | 0.049 | 0 | 1 | 14 | Zfp692 | 0.049 | Old |
| 0.111401 | 3.049435 | 0.049 | 0 | 1 | 14 | Cox10 | 0.049 | Old |
| 0.111401 | 2.73958 | 0.049 | 0 | 1 | 14 | Thoc5 | 0.049 | Old |
| 0.111401 | 2.522034 | 0.049 | 0 | 1 | 14 | Mrgprf | 0.049 | Old |
| 0.111473 | 0.945339 | 0.361 | 0.26 | 1 | 14 | Ubxn1 | 0.101 | Old |
| 0.111614 | 0.119664 | 0.956 | 0.94 | 1 | 14 | Rps4x | 0.016 | Old |
| 0.112188 | 1.280968 | 0.23 | 0.14 | 1 | 14 | Edem1 | 0.09 | Old |
| 0.112475 | 0.721684 | 0.481 | 0.4 | 1 | 14 | Ptp4a1 | 0.081 | Old |
| 0.112708 | 0.08979 | 0.47 | 0.36 | 1 | 14 | Ogdh | 0.11 | Old |
| 0.11282 | 1.277771 | 0.087 | 0.02 | 1 | 14 | Uchl5 | 0.067 | Old |
| 0.112999 | 0.270715 | 0.792 | 0.7 | 1 | 14 | Wnk1 | 0.092 | Old |
| 0.113058 | 1.320166 | 0.284 | 0.2 | 1 | 14 | Clasp1 | 0.084 | Old |
| 0.11314 | 0.24386 | 0.262 | 0.16 | 1 | 14 | Map3k2 | 0.102 | Old |
| 0.113249 | 0.445825 | 0.59 | 0.5 | 1 | 14 | Nr3c1 | 0.09 | Old |
| 0.113299 | 0.685494 | 0.47 | 0.36 | 1 | 14 | Mapk1 | 0.11 | Old |
| 0.113476 | 2.127858 | 0.164 | 0.08 | 1 | 14 | Tmem120a | 0.084 | Old |
| 0.113525 | 1.04881 | 0.169 | 0.08 | 1 | 14 | Slc30a9 | 0.089 | Old |
| 0.113772 | 0.73191 | 0.383 | 0.26 | 1 | 14 | Slc6a6 | 0.123 | Old |
| 0.113908 | 0.289465 | 0.672 | 0.54 | 1 | 14 | Fam102b | 0.132 | Old |
| 0.114804 | 0.201136 | 0.246 | 0.14 | 1 | 14 | Fam174a | 0.106 | Old |
| 0.115073 | 1.70619 | 0.115 | 0.04 | 1 | 14 | Rnf38 | 0.075 | Old |
| 0.115223 | 1.436649 | 0.23 | 0.14 | 1 | 14 | Bcl6 | 0.09 | Old |
| 0.115224 | 0.626175 | 0.087 | 0.02 | 1 | 14 | Dbr1 | 0.067 | Old |
| 0.115265 | 0.992662 | 0.322 | 0.22 | 1 | 14 | Abca9 | 0.102 | Old |
| 0.115312 | 0.964274 | 0.262 | 0.16 | 1 | 14 | Cpsf7 | 0.102 | Old |
| 0.116014 | 0.362588 | 0.432 | 0.28 | 1 | 14 | Ostc | 0.152 | Old |
| 0.116043 | 1.181621 | 0.262 | 0.16 | 1 | 14 | Ap3d1 | 0.102 | Old |
| 0.116125 | 1.084471 | 0.273 | 0.18 | 1 | 14 | Dnajc7 | 0.093 | Old |
| 0.116164 | 0.383415 | 0.421 | 0.3 | 1 | 14 | Mat2b | 0.121 | Old |
| 0.116292 | 0.464048 | 0.546 | 0.5 | 1 | 14 | Ywhaq | 0.046 | Old |
| 0.116343 | 2.591843 | 0.109 | 0.04 | 1 | 14 | Zfp74 | 0.069 | Old |
| 0.116371 | 0.339591 | 0.596 | 0.48 | 1 | 14 | Sulf2 | 0.116 | Old |
| 0.116455 | 0.639836 | 0.317 | 0.2 | 1 | 14 | Dpp4 | 0.117 | Old |
| 0.116494 | 0.152023 | 0.885 | 0.74 | 1 | 14 | Dpt | 0.145 | Old |
| 0.116522 | 0.78221 | 0.197 | 0.1 | 1 | 14 | Flt3l | 0.097 | Old |
| 0.116594 | 1.676727 | 0.186 | 0.1 | 1 | 14 | Agpat4 | 0.086 | Old |

| | | | | | | | | |
|---|---|---|---|---|---|---|---|---|
| 0.116703 | 1.376979 | 0.142 | 0.06 | 1 | 14 | Hace1 | 0.082 | Old |
| 0.116703 | 1.344566 | 0.142 | 0.06 | 1 | 14 | Plekha4 | 0.082 | Old |
| 0.117011 | 1.237518 | 0.251 | 0.16 | 1 | 14 | Sec23b | 0.091 | Old |
| 0.117198 | 1.68053 | 0.115 | 0.04 | 1 | 14 | Serpinb9 | 0.075 | Old |
| 0.117508 | 0.829694 | 0.448 | 0.34 | 1 | 14 | Gm26532 | 0.108 | Old |
| 0.117589 | 0.273514 | 0.847 | 0.78 | 1 | 14 | Nfia | 0.067 | Old |
| 0.11784 | 2.437354 | 0.158 | 0.08 | 1 | 14 | Engase | 0.078 | Old |
| 0.117932 | 0.739938 | 0.339 | 0.24 | 1 | 14 | Ensa | 0.099 | Old |
| 0.117969 | 3.115887 | 0.082 | 0.02 | 1 | 14 | Cfap43 | 0.062 | Old |
| 0.117969 | 2.785047 | 0.082 | 0.02 | 1 | 14 | Sdad1 | 0.062 | Old |
| 0.117969 | 2.561812 | 0.082 | 0.02 | 1 | 14 | Nphp3 | 0.062 | Old |
| 0.117985 | 0.796154 | 0.29 | 0.18 | 1 | 14 | Nipsnap2 | 0.11 | Old |
| 0.11799 | 1.114798 | 0.279 | 0.18 | 1 | 14 | Rab1b | 0.099 | Old |
| 0.118185 | 0.721545 | 0.344 | 0.24 | 1 | 14 | Gata6 | 0.104 | Old |
| 0.118196 | 1.345923 | 0.246 | 0.16 | 1 | 14 | Ice1 | 0.086 | Old |
| 0.118464 | 0.553956 | 0.601 | 0.54 | 1 | 14 | Selenok | 0.061 | Old |
| 0.118466 | 0.993811 | 0.235 | 0.14 | 1 | 14 | Efna5 | 0.095 | Old |
| 0.1188 | 0.535683 | 0.372 | 0.26 | 1 | 14 | Cuta | 0.112 | Old |
| 0.119103 | 1.918267 | 0.18 | 0.1 | 1 | 14 | Arfgap3 | 0.08 | Old |
| 0.119244 | 3.04427 | 0.082 | 0.02 | 1 | 14 | Rbbp5 | 0.062 | Old |
| 0.119244 | 2.889527 | 0.082 | 0.02 | 1 | 14 | Rffl | 0.062 | Old |
| 0.119352 | 1.027367 | 0.115 | 0.04 | 1 | 14 | Stat5a | 0.075 | Old |
| 0.119924 | 0.245508 | 0.809 | 0.74 | 1 | 14 | Gas1 | 0.069 | Old |
| 0.119952 | 0.635272 | 0.443 | 0.34 | 1 | 14 | 2310022B0 | 0.103 | Old |
| 0.119959 | 0.522298 | 0.596 | 0.46 | 1 | 14 | Oat | 0.136 | Old |
| 0.120413 | 0.211391 | 0.896 | 0.84 | 1 | 14 | Cavin1 | 0.056 | Old |
| 0.120732 | 0.443213 | 0.59 | 0.44 | 1 | 14 | Csf1 | 0.15 | Old |
| 0.120756 | 0.643771 | 0.328 | 0.2 | 1 | 14 | Emg1 | 0.128 | Old |
| 0.120817 | 1.318302 | 0.235 | 0.14 | 1 | 14 | Lyst | 0.095 | Old |
| 0.120907 | 2.174176 | 0.24 | 0.16 | 1 | 14 | Svbp | 0.08 | Old |
| 0.121393 | 1.980973 | 0.137 | 0.06 | 1 | 14 | Akap7 | 0.077 | Old |
| 0.121734 | 0.564232 | 0.585 | 0.5 | 1 | 14 | Tmem59 | 0.085 | Old |
| 0.121826 | 2.144705 | 0.082 | 0.02 | 1 | 14 | Gorab | 0.062 | Old |
| 0.12246 | 1.411337 | 0.208 | 0.12 | 1 | 14 | Rnf220 | 0.088 | Old |
| 0.122539 | 1.522666 | 0.164 | 0.08 | 1 | 14 | Samd14 | 0.084 | Old |
| 0.12265 | 1.743601 | 0.186 | 0.1 | 1 | 14 | Ankrd16 | 0.086 | Old |
| 0.122998 | 2.857155 | 0.109 | 0.04 | 1 | 14 | Ddx60 | 0.069 | Old |
| 0.123396 | 0.521361 | 0.508 | 0.38 | 1 | 14 | Hnrnpul1 | 0.128 | Old |
| 0.124889 | 0.327453 | 0.798 | 0.74 | 1 | 14 | Ctdsp2 | 0.058 | Old |
| 0.125229 | 0.991709 | 0.191 | 0.1 | 1 | 14 | Spred2 | 0.091 | Old |
| 0.125281 | 2.458702 | 0.109 | 0.04 | 1 | 14 | Satb1 | 0.069 | Old |
| 0.125781 | 2.290875 | 0.082 | 0.02 | 1 | 14 | Plscr2 | 0.062 | Old |
| 0.126004 | 1.336678 | 0.115 | 0.04 | 1 | 14 | Ptgr2 | 0.075 | Old |
| 0.12601 | 0.201039 | 0.306 | 0.18 | 1 | 14 | Clip1 | 0.126 | Old |
| 0.126022 | 0.535163 | 0.268 | 0.16 | 1 | 14 | Cant1 | 0.108 | Old |

| | | | | | | | | |
|---|---|---|---|---|---|---|---|---|
| 0.126073 | 0.487389 | 0.426 | 0.32 | 1 | 14 | Dap | 0.106 | Old |
| 0.12623 | 0.154977 | 0.546 | 0.44 | 1 | 14 | Ppp3ca | 0.106 | Old |
| 0.126319 | 1.235482 | 0.164 | 0.08 | 1 | 14 | Fnbp4 | 0.084 | Old |
| 0.126444 | 0.779296 | 0.235 | 0.14 | 1 | 14 | Apod | 0.095 | Old |
| 0.127265 | 0.523946 | 0.235 | 0.14 | 1 | 14 | Sik3 | 0.095 | Old |
| 0.127289 | 2.585267 | 0.131 | 0.06 | 1 | 14 | Ier5l | 0.071 | Old |
| 0.127558 | 1.915141 | 0.137 | 0.06 | 1 | 14 | Hsdl1 | 0.077 | Old |
| 0.127558 | 1.755439 | 0.137 | 0.06 | 1 | 14 | Myef2 | 0.077 | Old |
| 0.127702 | 0.379544 | 0.536 | 0.4 | 1 | 14 | Tsen34 | 0.136 | Old |
| 0.128072 | 1.757692 | 0.18 | 0.1 | 1 | 14 | Ubap1 | 0.08 | Old |
| 0.128247 | 0.250086 | 0.869 | 0.8 | 1 | 14 | Rpl36a | 0.069 | Old |
| 0.128272 | 1.746255 | 0.158 | 0.08 | 1 | 14 | Cebpa | 0.078 | Old |
| 0.128285 | 0.607592 | 0.421 | 0.32 | 1 | 14 | Rgl1 | 0.101 | Old |
| 0.128355 | 2.35999 | 0.131 | 0.06 | 1 | 14 | Eps15l1 | 0.071 | Old |
| 0.128419 | 0.385153 | 0.546 | 0.44 | 1 | 14 | Il11ra1 | 0.106 | Old |
| 0.128504 | 1.141307 | 0.355 | 0.28 | 1 | 14 | Smurf2 | 0.075 | Old |
| 0.128651 | 0.45128 | 0.464 | 0.34 | 1 | 14 | Enpp2 | 0.124 | Old |
| 0.128768 | 1.681684 | 0.109 | 0.04 | 1 | 14 | Terf1 | 0.069 | Old |
| 0.128997 | 1.610828 | 0.18 | 0.1 | 1 | 14 | 1110038F1 | 0.08 | Old |
| 0.129689 | 0.618489 | 0.454 | 0.34 | 1 | 14 | Adam15 | 0.114 | Old |
| 0.129748 | 1.225394 | 0.142 | 0.06 | 1 | 14 | Rbm15b | 0.082 | Old |
| 0.129752 | 0.217007 | 0.639 | 0.56 | 1 | 14 | Zcchc24 | 0.079 | Old |
| 0.129988 | 0.204294 | 0.557 | 0.42 | 1 | 14 | Arcn1 | 0.137 | Old |
| 0.130442 | 0.320389 | 0.617 | 0.52 | 1 | 14 | Ctnnb1 | 0.097 | Old |
| 0.130624 | 0.503766 | 0.546 | 0.46 | 1 | 14 | Slc44a1 | 0.086 | Old |
| 0.130731 | 1.589273 | 0.137 | 0.06 | 1 | 14 | Etfbkmt | 0.077 | Old |
| 0.130788 | 0.012036 | 0.645 | 0.58 | 1 | 14 | Nfe2l1 | 0.065 | Old |
| 0.130794 | 1.441343 | 0.186 | 0.1 | 1 | 14 | Atad2b | 0.086 | Old |
| 0.130863 | 1.549152 | 0.18 | 0.1 | 1 | 14 | Chmp7 | 0.08 | Old |
| 0.131079 | 1.392259 | 0.208 | 0.12 | 1 | 14 | Mturn | 0.088 | Old |
| 0.131093 | 0.902061 | 0.251 | 0.16 | 1 | 14 | Abhd17b | 0.091 | Old |
| 0.131209 | 1.38164 | 0.082 | 0.02 | 1 | 14 | Nckap5 | 0.062 | Old |
| 0.131776 | 0.686346 | 0.333 | 0.24 | 1 | 14 | Dnajc10 | 0.093 | Old |
| 0.132232 | 1.198423 | 0.23 | 0.14 | 1 | 14 | Snx21 | 0.09 | Old |
| 0.132329 | 1.938701 | 0.109 | 0.04 | 1 | 14 | Parp12 | 0.069 | Old |
| 0.132641 | 0.622327 | 0.475 | 0.38 | 1 | 14 | Actr3 | 0.095 | Old |
| 0.13266 | 1.151316 | 0.186 | 0.1 | 1 | 14 | Noc2l | 0.086 | Old |
| 0.132758 | 0.199098 | 0.995 | 0.9 | 1 | 14 | mt-Nd2 | 0.095 | Old |
| 0.13288 | 1.521486 | 0.137 | 0.06 | 1 | 14 | Larp6 | 0.077 | Old |
| 0.13288 | 1.412255 | 0.137 | 0.06 | 1 | 14 | Ago4 | 0.077 | Old |
| 0.13288 | 1.368379 | 0.137 | 0.06 | 1 | 14 | Atl2 | 0.077 | Old |
| 0.1336 | 1.352149 | 0.186 | 0.1 | 1 | 14 | Zfp62 | 0.086 | Old |
| 0.134186 | 0.198878 | 0.486 | 0.36 | 1 | 14 | Srsf6 | 0.126 | Old |
| 0.134338 | 3.839322 | 0.044 | 0 | 1 | 14 | Rab20 | 0.044 | Old |
| 0.134338 | 3.810474 | 0.044 | 0 | 1 | 14 | Igf2bp2 | 0.044 | Old |

| | | | | | | | | |
|---|---|---|---|---|---|---|---|---|
| 0.134338 | 3.712992 | 0.044 | 0 | 1 | 14 | Tmem240 | 0.044 | Old |
| 0.134338 | 3.619096 | 0.044 | 0 | 1 | 14 | Siglec1 | 0.044 | Old |
| 0.134338 | 3.561366 | 0.044 | 0 | 1 | 14 | Gm4951 | 0.044 | Old |
| 0.134338 | 3.54758 | 0.044 | 0 | 1 | 14 | Atoh8 | 0.044 | Old |
| 0.134338 | 3.463455 | 0.044 | 0 | 1 | 14 | Dffa | 0.044 | Old |
| 0.134338 | 3.444575 | 0.044 | 0 | 1 | 14 | Pianp | 0.044 | Old |
| 0.134338 | 3.330894 | 0.044 | 0 | 1 | 14 | Hes6 | 0.044 | Old |
| 0.134338 | 3.315451 | 0.044 | 0 | 1 | 14 | Bmp7 | 0.044 | Old |
| 0.134338 | 3.302004 | 0.044 | 0 | 1 | 14 | Nol3 | 0.044 | Old |
| 0.134338 | 3.273265 | 0.044 | 0 | 1 | 14 | Ggt7 | 0.044 | Old |
| 0.134338 | 3.248172 | 0.044 | 0 | 1 | 14 | Tfip11 | 0.044 | Old |
| 0.134338 | 3.201412 | 0.044 | 0 | 1 | 14 | Donson | 0.044 | Old |
| 0.134338 | 3.127475 | 0.044 | 0 | 1 | 14 | Aste1 | 0.044 | Old |
| 0.134338 | 3.087908 | 0.044 | 0 | 1 | 14 | Cd300c2 | 0.044 | Old |
| 0.134338 | 3.042941 | 0.044 | 0 | 1 | 14 | Fuk | 0.044 | Old |
| 0.134338 | 3.008964 | 0.044 | 0 | 1 | 14 | Zfp39 | 0.044 | Old |
| 0.134338 | 2.942757 | 0.044 | 0 | 1 | 14 | Smarca1 | 0.044 | Old |
| 0.134338 | 2.91988 | 0.044 | 0 | 1 | 14 | Gfer | 0.044 | Old |
| 0.134338 | 2.899668 | 0.044 | 0 | 1 | 14 | Gm20604 | 0.044 | Old |
| 0.134338 | 2.712543 | 0.044 | 0 | 1 | 14 | Amigo1 | 0.044 | Old |
| 0.134338 | 2.669641 | 0.044 | 0 | 1 | 14 | Gpr1 | 0.044 | Old |
| 0.134338 | 2.42223 | 0.044 | 0 | 1 | 14 | Slc35g1 | 0.044 | Old |
| 0.13455 | 1.170401 | 0.213 | 0.12 | 1 | 14 | Tmem219 | 0.093 | Old |
| 0.134655 | 1.33137 | 0.208 | 0.12 | 1 | 14 | Poglut1 | 0.088 | Old |
| 0.135397 | 0.425846 | 0.055 | 0.12 | 1 | 14 | Cluh | -0.065 | Old |
| 0.135465 | 0.309436 | 0.705 | 0.58 | 1 | 14 | Mrfap1 | 0.125 | Old |
| 0.135702 | 0.443968 | 0.388 | 0.28 | 1 | 14 | Gnai3 | 0.108 | Old |
| 0.135967 | 1.386946 | 0.109 | 0.04 | 1 | 14 | Lrp12 | 0.069 | Old |
| 0.136071 | 0.370568 | 0.421 | 0.3 | 1 | 14 | Ppp2r5c | 0.121 | Old |
| 0.136555 | 1.112551 | 0.23 | 0.14 | 1 | 14 | Ust | 0.09 | Old |
| 0.13676 | 1.016598 | 0.328 | 0.24 | 1 | 14 | Nrp2 | 0.088 | Old |
| 0.13676 | 0.527405 | 0.328 | 0.24 | 1 | 14 | Utrn | 0.088 | Old |
| 0.13726 | 1.083465 | 0.137 | 0.06 | 1 | 14 | Yae1d1 | 0.077 | Old |
| 0.137364 | 2.07125 | 0.153 | 0.08 | 1 | 14 | Plcb3 | 0.073 | Old |
| 0.137607 | 0.722155 | 0.35 | 0.24 | 1 | 14 | Nars | 0.11 | Old |
| 0.137945 | 0.546864 | 0.383 | 0.28 | 1 | 14 | Bst2 | 0.103 | Old |
| 0.138005 | 0.116636 | 0.339 | 0.24 | 1 | 14 | Epn1 | 0.099 | Old |
| 0.138264 | 2.231519 | 0.131 | 0.06 | 1 | 14 | Prdm5 | 0.071 | Old |
| 0.138602 | 2.022897 | 0.175 | 0.1 | 1 | 14 | Idnk | 0.075 | Old |
| 0.138782 | 0.347434 | 0.585 | 0.48 | 1 | 14 | Ugdh | 0.105 | Old |
| 0.139232 | 1.027605 | 0.164 | 0.08 | 1 | 14 | Prpf6 | 0.084 | Old |
| 0.139396 | 1.1276 | 0.284 | 0.2 | 1 | 14 | Ccdc6 | 0.084 | Old |
| 0.139681 | 0.74858 | 0.109 | 0.04 | 1 | 14 | Umps | 0.069 | Old |
| 0.140068 | 0.516973 | 0.191 | 0.1 | 1 | 14 | Psmb8 | 0.091 | Old |
| 0.140266 | 1.033742 | 0.164 | 0.08 | 1 | 14 | Rell1 | 0.084 | Old |

| | | | | | | | | |
|---|---|---|---|---|---|---|---|---|
| 0.140282 | 3.483936 | 0.077 | 0.02 | 1 | 14 | Mul1 | 0.057 | Old |
| 0.140282 | 3.477655 | 0.077 | 0.02 | 1 | 14 | Rbm19 | 0.057 | Old |
| 0.140282 | 2.960778 | 0.077 | 0.02 | 1 | 14 | Apba1 | 0.057 | Old |
| 0.140282 | 2.874099 | 0.077 | 0.02 | 1 | 14 | Duoxa1 | 0.057 | Old |
| 0.140596 | 2.034144 | 0.175 | 0.1 | 1 | 14 | Ppp1r21 | 0.075 | Old |
| 0.140745 | 0.573726 | 0.246 | 0.14 | 1 | 14 | Tfg | 0.106 | Old |
| 0.140936 | 1.014233 | 0.109 | 0.04 | 1 | 14 | Elk1 | 0.069 | Old |
| 0.140997 | 0.146082 | 0.169 | 0.08 | 1 | 14 | Lhfpl2 | 0.089 | Old |
| 0.141033 | 1.235822 | 0.284 | 0.2 | 1 | 14 | Plscr4 | 0.084 | Old |
| 0.141489 | 2.77416 | 0.104 | 0.04 | 1 | 14 | Arhgap15 | 0.064 | Old |
| 0.141492 | 0.197588 | 0.951 | 0.94 | 1 | 14 | Rpl38 | 0.011 | Old |
| 0.141784 | 2.549487 | 0.077 | 0.02 | 1 | 14 | Thrb | 0.057 | Old |
| 0.141784 | 2.511141 | 0.077 | 0.02 | 1 | 14 | Gtf2h3 | 0.057 | Old |
| 0.142179 | 1.714395 | 0.197 | 0.12 | 1 | 14 | Slc9a6 | 0.077 | Old |
| 0.1422 | 1.790261 | 0.109 | 0.04 | 1 | 14 | Timp1 | 0.069 | Old |
| 0.142828 | 0.292828 | 0.383 | 0.26 | 1 | 14 | Tcf21 | 0.123 | Old |
| 0.143115 | 0.115214 | 0.607 | 0.52 | 1 | 14 | Rabac1 | 0.087 | Old |
| 0.143294 | 0.266975 | 0.29 | 0.2 | 1 | 14 | Coa5 | 0.09 | Old |
| 0.1433 | 2.655933 | 0.077 | 0.02 | 1 | 14 | Metrn | 0.057 | Old |
| 0.144018 | 1.922397 | 0.131 | 0.06 | 1 | 14 | Metap1d | 0.071 | Old |
| 0.14451 | 0.86027 | 0.246 | 0.16 | 1 | 14 | Brd7 | 0.086 | Old |
| 0.144514 | 1.815288 | 0.18 | 0.1 | 1 | 14 | Flrt3 | 0.08 | Old |
| 0.14471 | 1.029814 | 0.355 | 0.26 | 1 | 14 | Zfp385a | 0.095 | Old |
| 0.144754 | 1.260747 | 0.109 | 0.04 | 1 | 14 | Pvr | 0.069 | Old |
| 0.144754 | 1.198557 | 0.109 | 0.04 | 1 | 14 | D630033O | 0.069 | Old |
| 0.144827 | 2.555131 | 0.077 | 0.02 | 1 | 14 | Ly96 | 0.057 | Old |
| 0.144827 | 2.386538 | 0.077 | 0.02 | 1 | 14 | Snn | 0.057 | Old |
| 0.144827 | 2.37895 | 0.077 | 0.02 | 1 | 14 | Ppp3cc | 0.057 | Old |
| 0.14488 | 1.066215 | 0.208 | 0.12 | 1 | 14 | Mapk14 | 0.088 | Old |
| 0.145013 | 0.19037 | 0.443 | 0.34 | 1 | 14 | Ggnbp2 | 0.103 | Old |
| 0.145177 | 0.150115 | 0.246 | 0.14 | 1 | 14 | Scrn1 | 0.106 | Old |
| 0.1454 | 2.630606 | 0.104 | 0.04 | 1 | 14 | Dnm3os | 0.064 | Old |
| 0.1454 | 2.1739 | 0.104 | 0.04 | 1 | 14 | Eif2s3y | 0.064 | Old |
| 0.145635 | 0.594525 | 0.557 | 0.5 | 1 | 14 | Amotl1 | 0.057 | Old |
| 0.145789 | 1.661466 | 0.158 | 0.08 | 1 | 14 | Chchd3 | 0.078 | Old |
| 0.145789 | 1.254558 | 0.158 | 0.08 | 1 | 14 | Zrsr2 | 0.078 | Old |
| 0.145939 | 1.789334 | 0.153 | 0.08 | 1 | 14 | Bnc2 | 0.073 | Old |
| 0.146218 | 0.358633 | 0.76 | 0.68 | 1 | 14 | Dnaja1 | 0.08 | Old |
| 0.146352 | 1.405308 | 0.137 | 0.06 | 1 | 14 | Hmcn2 | 0.077 | Old |
| 0.146367 | 2.240489 | 0.077 | 0.02 | 1 | 14 | Snhg9 | 0.057 | Old |
| 0.14642 | 0.357832 | 0.082 | 0.16 | 1 | 14 | H2-DMb1 | -0.078 | Old |
| 0.146858 | 0.384643 | 0.481 | 0.4 | 1 | 14 | Sec14l1 | 0.081 | Old |
| 0.147256 | 0.759802 | 0.279 | 0.18 | 1 | 14 | Ampd3 | 0.099 | Old |
| 0.147299 | 0.78052 | 0.317 | 0.22 | 1 | 14 | A4galt | 0.097 | Old |
| 0.147315 | 0.928195 | 0.186 | 0.1 | 1 | 14 | Il1rl2 | 0.086 | Old |

| | | | | | | | | |
|---|---|---|---|---|---|---|---|---|
| 0.147375 | 0.945723 | 0.23 | 0.14 | 1 | 14 | Rapgef6 | 0.09 | Old |
| 0.147576 | 1.452302 | 0.18 | 0.1 | 1 | 14 | Washc3 | 0.08 | Old |
| 0.147581 | 0.350069 | 0.596 | 0.52 | 1 | 14 | Snx18 | 0.076 | Old |
| 0.14792 | 2.541939 | 0.077 | 0.02 | 1 | 14 | Mapk12 | 0.057 | Old |
| 0.148292 | 1.761965 | 0.219 | 0.14 | 1 | 14 | Plod3 | 0.079 | Old |
| 0.148575 | 0.764566 | 0.328 | 0.24 | 1 | 14 | Ammecr1l | 0.088 | Old |
| 0.149021 | 1.897264 | 0.197 | 0.12 | 1 | 14 | Slc10a6 | 0.077 | Old |
| 0.149441 | 0.697131 | 0.432 | 0.34 | 1 | 14 | Paip2 | 0.092 | Old |
| 0.149485 | 1.92732 | 0.077 | 0.02 | 1 | 14 | Pmaip1 | 0.057 | Old |
| 0.149967 | 1.166269 | 0.109 | 0.04 | 1 | 14 | Zfp654 | 0.069 | Old |
| 0.15018 | 0.37481 | 0.23 | 0.14 | 1 | 14 | Dph3 | 0.09 | Old |
| 0.150197 | 1.716145 | 0.219 | 0.14 | 1 | 14 | Atp11a | 0.079 | Old |
| 0.150605 | 0.900556 | 0.262 | 0.18 | 1 | 14 | Psmb10 | 0.082 | Old |
| 0.151334 | 1.054588 | 0.317 | 0.24 | 1 | 14 | Fnip1 | 0.077 | Old |
| 0.151598 | 0.160701 | 0.596 | 0.46 | 1 | 14 | Steap3 | 0.136 | Old |
| 0.151603 | 0.746094 | 0.279 | 0.18 | 1 | 14 | Gpbp1 | 0.099 | Old |
| 0.151896 | 1.255837 | 0.202 | 0.12 | 1 | 14 | Ankrd44 | 0.082 | Old |
| 0.152653 | 2.101266 | 0.077 | 0.02 | 1 | 14 | Coro7 | 0.057 | Old |
| 0.152653 | 1.994021 | 0.077 | 0.02 | 1 | 14 | Madd | 0.057 | Old |
| 0.152756 | 2.409949 | 0.148 | 0.08 | 1 | 14 | Zfp780b | 0.068 | Old |
| 0.15277 | 0.162714 | 0.268 | 0.18 | 1 | 14 | Ube2v2 | 0.088 | Old |
| 0.153089 | 0.934401 | 0.224 | 0.14 | 1 | 14 | Nae1 | 0.084 | Old |
| 0.153139 | 0.872021 | 0.164 | 0.08 | 1 | 14 | Traf3 | 0.084 | Old |
| 0.153434 | 0.361018 | 0.623 | 0.52 | 1 | 14 | Bsg | 0.103 | Old |
| 0.153686 | 0.258873 | 0.383 | 0.26 | 1 | 14 | Wdr1 | 0.123 | Old |
| 0.154099 | 1.877491 | 0.175 | 0.1 | 1 | 14 | Zbtb37 | 0.075 | Old |
| 0.154813 | 0.764051 | 0.317 | 0.22 | 1 | 14 | Ddx24 | 0.097 | Old |
| 0.154888 | 0.50905 | 0.41 | 0.32 | 1 | 14 | Ghitm | 0.09 | Old |
| 0.154914 | 0.091817 | 0.525 | 0.4 | 1 | 14 | Dpysl3 | 0.125 | Old |
| 0.155029 | 0.248789 | 0.355 | 0.24 | 1 | 14 | Mib1 | 0.115 | Old |
| 0.155191 | 0.456351 | 0.492 | 0.4 | 1 | 14 | Ewsr1 | 0.092 | Old |
| 0.155322 | 0.943299 | 0.109 | 0.04 | 1 | 14 | Lyrm2 | 0.069 | Old |
| 0.155559 | 0.394995 | 0.24 | 0.14 | 1 | 14 | Fam126a | 0.1 | Old |
| 0.155775 | 1.130109 | 0.158 | 0.08 | 1 | 14 | Tbck | 0.078 | Old |
| 0.155776 | 0.324884 | 0.295 | 0.2 | 1 | 14 | Trim8 | 0.095 | Old |
| 0.155872 | 1.599203 | 0.077 | 0.02 | 1 | 14 | Gpatch1 | 0.057 | Old |
| 0.15606 | 0.092302 | 0.388 | 0.3 | 1 | 14 | Anxa4 | 0.088 | Old |
| 0.156076 | 0.207529 | 0.098 | 0.18 | 1 | 14 | Dgkh | -0.082 | Old |
| 0.156448 | 0.866712 | 0.393 | 0.32 | 1 | 14 | Lox | 0.073 | Old |
| 0.156545 | 0.950013 | 0.257 | 0.16 | 1 | 14 | Riok3 | 0.097 | Old |
| 0.156882 | 0.995533 | 0.23 | 0.14 | 1 | 14 | Iars2 | 0.09 | Old |
| 0.15696 | 1.265578 | 0.202 | 0.12 | 1 | 14 | Plekhh2 | 0.082 | Old |
| 0.156962 | 0.114829 | 0.448 | 0.34 | 1 | 14 | Sypl | 0.108 | Old |
| 0.157145 | 1.204241 | 0.197 | 0.12 | 1 | 14 | Prdx4 | 0.077 | Old |
| 0.157348 | 2.772181 | 0.126 | 0.06 | 1 | 14 | AI987944 | 0.066 | Old |

| | | | | | | | | |
|---|---|---|---|---|---|---|---|---|
| 0.157553 | 0.930429 | 0.306 | 0.22 | 1 | 14 | Pdcl3 | 0.086 | Old |
| 0.157632 | 1.815908 | 0.104 | 0.04 | 1 | 14 | Id1 | 0.064 | Old |
| 0.157632 | 1.580226 | 0.104 | 0.04 | 1 | 14 | Pex11b | 0.064 | Old |
| 0.157988 | 1.497571 | 0.202 | 0.12 | 1 | 14 | Lgals3 | 0.082 | Old |
| 0.158143 | 0.435427 | 0.18 | 0.1 | 1 | 14 | Snd1 | 0.08 | Old |
| 0.158143 | 0.303905 | 0.18 | 0.1 | 1 | 14 | Arsa | 0.08 | Old |
| 0.158225 | 0.976533 | 0.284 | 0.2 | 1 | 14 | Pla2r1 | 0.084 | Old |
| 0.159142 | 1.10764 | 0.077 | 0.02 | 1 | 14 | Hoxb2 | 0.057 | Old |
| 0.159409 | 0.303803 | 0.213 | 0.12 | 1 | 14 | Katnbl1 | 0.093 | Old |
| 0.16006 | 0.986476 | 0.268 | 0.18 | 1 | 14 | Cyp2d22 | 0.088 | Old |
| 0.1608 | 1.000372 | 0.208 | 0.12 | 1 | 14 | Cic | 0.088 | Old |
| 0.161045 | 1.503591 | 0.24 | 0.16 | 1 | 14 | Mrpl20 | 0.08 | Old |
| 0.161065 | 0.959401 | 0.306 | 0.22 | 1 | 14 | Otub1 | 0.086 | Old |
| 0.161117 | 1.138459 | 0.301 | 0.22 | 1 | 14 | Tusc3 | 0.081 | Old |
| 0.161168 | 1.264389 | 0.246 | 0.16 | 1 | 14 | Slc30a5 | 0.086 | Old |
| 0.161422 | 1.296347 | 0.18 | 0.1 | 1 | 14 | Cdc123 | 0.08 | Old |
| 0.161559 | 0.581311 | 0.541 | 0.44 | 1 | 14 | Txndc5 | 0.101 | Old |
| 0.161878 | 1.514035 | 0.104 | 0.04 | 1 | 14 | Eci1 | 0.064 | Old |
| 0.162219 | 0.378632 | 0.109 | 0.04 | 1 | 14 | H2-Q7 | 0.069 | Old |
| 0.16242 | 4.449806 | 0.038 | 0 | 1 | 14 | Krtdap | 0.038 | Old |
| 0.16242 | 4.047835 | 0.038 | 0 | 1 | 14 | Szt2 | 0.038 | Old |
| 0.16242 | 3.703261 | 0.038 | 0 | 1 | 14 | Pcdhgb4 | 0.038 | Old |
| 0.16242 | 3.571102 | 0.038 | 0 | 1 | 14 | Rbm41 | 0.038 | Old |
| 0.16242 | 3.469517 | 0.038 | 0 | 1 | 14 | Fam122b | 0.038 | Old |
| 0.16242 | 3.449088 | 0.038 | 0 | 1 | 14 | Whamm | 0.038 | Old |
| 0.16242 | 3.426647 | 0.038 | 0 | 1 | 14 | Tubgcp6 | 0.038 | Old |
| 0.16242 | 3.378459 | 0.038 | 0 | 1 | 14 | Hdac6 | 0.038 | Old |
| 0.16242 | 3.34436 | 0.038 | 0 | 1 | 14 | Fastkd3 | 0.038 | Old |
| 0.16242 | 3.33189 | 0.038 | 0 | 1 | 14 | Mthfsl | 0.038 | Old |
| 0.16242 | 3.315355 | 0.038 | 0 | 1 | 14 | Cdc25b | 0.038 | Old |
| 0.16242 | 3.291621 | 0.038 | 0 | 1 | 14 | Fchsd1 | 0.038 | Old |
| 0.16242 | 3.194486 | 0.038 | 0 | 1 | 14 | Yars2 | 0.038 | Old |
| 0.16242 | 3.175709 | 0.038 | 0 | 1 | 14 | Ugt1a6a | 0.038 | Old |
| 0.16242 | 3.157339 | 0.038 | 0 | 1 | 14 | Lonrf1 | 0.038 | Old |
| 0.16242 | 3.144304 | 0.038 | 0 | 1 | 14 | Lrfn3 | 0.038 | Old |
| 0.16242 | 3.132873 | 0.038 | 0 | 1 | 14 | Zbed4 | 0.038 | Old |
| 0.16242 | 3.122933 | 0.038 | 0 | 1 | 14 | Usp27x | 0.038 | Old |
| 0.16242 | 3.117441 | 0.038 | 0 | 1 | 14 | Gm45844 | 0.038 | Old |
| 0.16242 | 3.112748 | 0.038 | 0 | 1 | 14 | Lefty1 | 0.038 | Old |
| 0.16242 | 3.112008 | 0.038 | 0 | 1 | 14 | Gm45871 | 0.038 | Old |
| 0.16242 | 3.105561 | 0.038 | 0 | 1 | 14 | Ly9 | 0.038 | Old |
| 0.16242 | 3.100354 | 0.038 | 0 | 1 | 14 | Gstm6 | 0.038 | Old |
| 0.16242 | 3.100044 | 0.038 | 0 | 1 | 14 | Cdyl2 | 0.038 | Old |
| 0.16242 | 3.026142 | 0.038 | 0 | 1 | 14 | Celf4 | 0.038 | Old |
| 0.16242 | 3.01742 | 0.038 | 0 | 1 | 14 | Pcgf6 | 0.038 | Old |

| | | | | | | | | |
|---|---|---|---|---|---|---|---|---|
| 0.16242 | 3.010098 | 0.038 | 0 | 1 | 14 | Zfp867 | 0.038 | Old |
| 0.16242 | 2.950759 | 0.038 | 0 | 1 | 14 | Vmn1r90 | 0.038 | Old |
| 0.16242 | 2.864501 | 0.038 | 0 | 1 | 14 | Ttc26 | 0.038 | Old |
| 0.16242 | 2.824672 | 0.038 | 0 | 1 | 14 | Pcdhga6 | 0.038 | Old |
| 0.16242 | 2.765741 | 0.038 | 0 | 1 | 14 | Rtl5 | 0.038 | Old |
| 0.16242 | 2.737888 | 0.038 | 0 | 1 | 14 | Apobr | 0.038 | Old |
| 0.16242 | 2.722662 | 0.038 | 0 | 1 | 14 | Gm30025 | 0.038 | Old |
| 0.16242 | 2.695096 | 0.038 | 0 | 1 | 14 | Bbs1 | 0.038 | Old |
| 0.16242 | 2.67082 | 0.038 | 0 | 1 | 14 | Fam229b | 0.038 | Old |
| 0.16242 | 2.567338 | 0.038 | 0 | 1 | 14 | Camkk1 | 0.038 | Old |
| 0.16242 | 2.537755 | 0.038 | 0 | 1 | 14 | Cep95 | 0.038 | Old |
| 0.16242 | 2.503947 | 0.038 | 0 | 1 | 14 | Abcc10 | 0.038 | Old |
| 0.16242 | 2.498606 | 0.038 | 0 | 1 | 14 | Gm26935 | 0.038 | Old |
| 0.16242 | 2.430123 | 0.038 | 0 | 1 | 14 | Gcfc2 | 0.038 | Old |
| 0.162464 | 0.839307 | 0.077 | 0.02 | 1 | 14 | Usp31 | 0.057 | Old |
| 0.162679 | 1.384522 | 0.251 | 0.18 | 1 | 14 | Txnl1 | 0.071 | Old |
| 0.163056 | 0.62552 | 0.311 | 0.24 | 1 | 14 | Tmem165 | 0.071 | Old |
| 0.163293 | 0.657935 | 0.486 | 0.38 | 1 | 14 | Plec | 0.106 | Old |
| 0.16387 | 0.386368 | 0.033 | 0.08 | 1 | 14 | Klhl25 | -0.047 | Old |
| 0.164206 | 0.917433 | 0.361 | 0.28 | 1 | 14 | Vkorc1 | 0.081 | Old |
| 0.164432 | 1.073121 | 0.262 | 0.18 | 1 | 14 | Eif4enif1 | 0.082 | Old |
| 0.164633 | 0.682962 | 0.137 | 0.06 | 1 | 14 | Papss2 | 0.077 | Old |
| 0.164899 | 0.372306 | 0.568 | 0.44 | 1 | 14 | Rbm3 | 0.128 | Old |
| 0.165118 | 1.841872 | 0.126 | 0.06 | 1 | 14 | Trp53bp1 | 0.066 | Old |
| 0.165121 | 1.416304 | 0.213 | 0.14 | 1 | 14 | Camkk2 | 0.073 | Old |
| 0.16571 | 0.277979 | 0.126 | 0.22 | 1 | 14 | Mphosph8 | -0.094 | Old |
| 0.165726 | 1.173355 | 0.311 | 0.24 | 1 | 14 | Gmfb | 0.071 | Old |
| 0.166027 | 0.012616 | 0.913 | 0.74 | 1 | 14 | Nfib | 0.173 | Old |
| 0.16621 | 1.772916 | 0.104 | 0.04 | 1 | 14 | Ipmk | 0.064 | Old |
| 0.16621 | 1.572387 | 0.104 | 0.04 | 1 | 14 | Rnf41 | 0.064 | Old |
| 0.16621 | 1.554078 | 0.104 | 0.04 | 1 | 14 | Casp6 | 0.064 | Old |
| 0.166471 | 0.172052 | 0.011 | 0.04 | 1 | 14 | Arl5c | -0.029 | Old |
| 0.166723 | 1.337151 | 0.153 | 0.08 | 1 | 14 | Pex7 | 0.073 | Old |
| 0.166824 | 0.463024 | 0.355 | 0.28 | 1 | 14 | Mbd2 | 0.075 | Old |
| 0.166855 | 0.818199 | 0.35 | 0.28 | 1 | 14 | Itgb5 | 0.07 | Old |
| 0.166856 | 0.398819 | 0.415 | 0.28 | 1 | 14 | Tmem256 | 0.135 | Old |
| 0.166883 | 0.701411 | 0.317 | 0.24 | 1 | 14 | Psme2 | 0.077 | Old |
| 0.167453 | 0.632198 | 0.393 | 0.32 | 1 | 14 | Larp4b | 0.073 | Old |
| 0.167791 | 0.83371 | 0.268 | 0.16 | 1 | 14 | Prpf39 | 0.108 | Old |
| 0.16795 | 0.46831 | 0.301 | 0.2 | 1 | 14 | Tfpi | 0.101 | Old |
| 0.167991 | 0.391414 | 0.033 | 0.08 | 1 | 14 | Zfp882 | -0.047 | Old |
| 0.168523 | 0.135999 | 0.022 | 0.06 | 1 | 14 | Nav2 | -0.038 | Old |
| 0.168529 | 0.336138 | 0.344 | 0.24 | 1 | 14 | Vapb | 0.104 | Old |
| 0.168637 | 3.381388 | 0.071 | 0.02 | 1 | 14 | AI597479 | 0.051 | Old |
| 0.168637 | 3.364265 | 0.071 | 0.02 | 1 | 14 | Zbtb18 | 0.051 | Old |

| | | | | | | | | |
|---|---|---|---|---|---|---|---|---|
| 0.168637 | 3.335079 | 0.071 | 0.02 | 1 | 14 | Zfp991 | 0.051 | Old |
| 0.168637 | 3.328987 | 0.071 | 0.02 | 1 | 14 | Pinx1 | 0.051 | Old |
| 0.168637 | 2.918807 | 0.071 | 0.02 | 1 | 14 | Tpmt | 0.051 | Old |
| 0.168637 | 2.705724 | 0.071 | 0.02 | 1 | 14 | Snip1 | 0.051 | Old |
| 0.168648 | 2.444987 | 0.098 | 0.04 | 1 | 14 | Bnip1 | 0.058 | Old |
| 0.168971 | 0.886802 | 0.339 | 0.26 | 1 | 14 | Ccnt2 | 0.079 | Old |
| 0.169071 | 0.273613 | 0.831 | 0.68 | 1 | 14 | Lpl | 0.151 | Old |
| 0.16908 | 0.535418 | 0.672 | 0.6 | 1 | 14 | Mgst1 | 0.072 | Old |
| 0.169112 | 0.805573 | 0.415 | 0.36 | 1 | 14 | Ythdc1 | 0.055 | Old |
| 0.169232 | 0.331871 | 0.388 | 0.3 | 1 | 14 | Arsb | 0.088 | Old |
| 0.169255 | 0.740652 | 0.366 | 0.28 | 1 | 14 | Psip1 | 0.086 | Old |
| 0.16954 | 0.770332 | 0.268 | 0.18 | 1 | 14 | Gxylt1 | 0.088 | Old |
| 0.169718 | 0.631203 | 0.011 | 0.04 | 1 | 14 | Tpk1 | -0.029 | Old |
| 0.169718 | 0.414602 | 0.011 | 0.04 | 1 | 14 | Lhpp | -0.029 | Old |
| 0.169718 | 0.388892 | 0.011 | 0.04 | 1 | 14 | Itpk1 | -0.029 | Old |
| 0.169718 | 0.241927 | 0.011 | 0.04 | 1 | 14 | Fxyd7 | -0.029 | Old |
| 0.169718 | 0.229879 | 0.011 | 0.04 | 1 | 14 | Blm | -0.029 | Old |
| 0.169718 | 0.099908 | 0.011 | 0.04 | 1 | 14 | Trpc3 | -0.029 | Old |
| 0.169718 | 0.097116 | 0.011 | 0.04 | 1 | 14 | Sox18 | -0.029 | Old |
| 0.16972 | 1.210029 | 0.175 | 0.1 | 1 | 14 | AY036118 | 0.075 | Old |
| 0.16987 | 0.659012 | 0.186 | 0.1 | 1 | 14 | Stk11 | 0.086 | Old |
| 0.169976 | 1.061014 | 0.322 | 0.26 | 1 | 14 | Map4 | 0.062 | Old |
| 0.170296 | 0.092035 | 0.366 | 0.24 | 1 | 14 | Lysmd2 | 0.126 | Old |
| 0.17078 | 1.63593 | 0.148 | 0.08 | 1 | 14 | Mtf2 | 0.068 | Old |
| 0.170886 | 1.076093 | 0.311 | 0.22 | 1 | 14 | Cmklr1 | 0.091 | Old |
| 0.171012 | 0.321799 | 0.022 | 0.06 | 1 | 14 | Espn | -0.038 | Old |
| 0.171012 | 0.259035 | 0.022 | 0.06 | 1 | 14 | Msi1 | -0.038 | Old |
| 0.171105 | 0.729318 | 0.158 | 0.08 | 1 | 14 | Ccnk | 0.078 | Old |
| 0.17157 | 0.031776 | 0.995 | 0.96 | 1 | 14 | Rpl35a | 0.035 | Old |
| 0.171884 | 1.378482 | 0.235 | 0.16 | 1 | 14 | BC031181 | 0.075 | Old |
| 0.172222 | 2.910531 | 0.071 | 0.02 | 1 | 14 | Yipf2 | 0.051 | Old |
| 0.172222 | 2.836057 | 0.071 | 0.02 | 1 | 14 | Dusp23 | 0.051 | Old |
| 0.172854 | 0.276762 | 0.568 | 0.46 | 1 | 14 | Surf4 | 0.108 | Old |
| 0.172874 | 1.278588 | 0.153 | 0.08 | 1 | 14 | Snx13 | 0.073 | Old |
| 0.173174 | 1.956677 | 0.126 | 0.06 | 1 | 14 | Itpr3 | 0.066 | Old |
| 0.173174 | 0.754131 | 0.262 | 0.18 | 1 | 14 | Abcd3 | 0.082 | Old |
| 0.173521 | 0.728747 | 0.448 | 0.38 | 1 | 14 | Hivep2 | 0.068 | Old |
| 0.173529 | 0.242551 | 0.022 | 0.06 | 1 | 14 | Hspb2 | -0.038 | Old |
| 0.174035 | 2.102417 | 0.071 | 0.02 | 1 | 14 | Rusc1 | 0.051 | Old |
| 0.174073 | 0.706184 | 0.246 | 0.16 | 1 | 14 | Mrpl13 | 0.086 | Old |
| 0.174222 | 0.851163 | 0.131 | 0.06 | 1 | 14 | Scaper | 0.071 | Old |
| 0.174675 | 0.492226 | 0.585 | 0.5 | 1 | 14 | Dync1h1 | 0.085 | Old |
| 0.174759 | 2.167993 | 0.098 | 0.04 | 1 | 14 | Psmf1 | 0.058 | Old |
| 0.174759 | 1.828967 | 0.098 | 0.04 | 1 | 14 | Zmat5 | 0.058 | Old |
| 0.174798 | 1.173607 | 0.158 | 0.08 | 1 | 14 | Tank | 0.078 | Old |

| | | | | | | | | |
|---|---|---|---|---|---|---|---|---|
| 0.174817 | 0.590515 | 0.344 | 0.26 | 1 | 14 | Golph3 | 0.084 | Old |
| 0.175134 | 1.034295 | 0.104 | 0.04 | 1 | 14 | Pgm2l1 | 0.064 | Old |
| 0.175576 | 0.698044 | 0.131 | 0.06 | 1 | 14 | Dalrd3 | 0.071 | Old |
| 0.175923 | 1.632745 | 0.126 | 0.06 | 1 | 14 | Cwc25 | 0.066 | Old |
| 0.175956 | 0.342534 | 0.721 | 0.6 | 1 | 14 | Il6st | 0.121 | Old |
| 0.176072 | 0.04826 | 0.022 | 0.06 | 1 | 14 | F11r | -0.038 | Old |
| 0.176125 | 0.718949 | 0.246 | 0.16 | 1 | 14 | Timm8b | 0.086 | Old |
| 0.176645 | 1.367125 | 0.153 | 0.08 | 1 | 14 | Zcchc2 | 0.073 | Old |
| 0.176719 | 0.093088 | 0.372 | 0.26 | 1 | 14 | Ppm1a | 0.112 | Old |
| 0.176964 | 1.65754 | 0.191 | 0.12 | 1 | 14 | Ces1d | 0.071 | Old |
| 0.17731 | 1.706813 | 0.126 | 0.06 | 1 | 14 | Wrnip1 | 0.066 | Old |
| 0.177559 | 1.016492 | 0.224 | 0.14 | 1 | 14 | Boc | 0.084 | Old |
| 0.178004 | 1.085199 | 0.23 | 0.16 | 1 | 14 | Vps13a | 0.07 | Old |
| 0.178385 | 0.256366 | 0.383 | 0.28 | 1 | 14 | Adipor1 | 0.103 | Old |
| 0.178419 | 2.003414 | 0.148 | 0.08 | 1 | 14 | Lsm2 | 0.068 | Old |
| 0.178644 | 0.314473 | 0.022 | 0.06 | 1 | 14 | Slc16a6 | -0.038 | Old |
| 0.178919 | 0.131225 | 0.087 | 0.16 | 1 | 14 | Hoxa3 | -0.073 | Old |
| 0.179069 | 0.507702 | 0.235 | 0.14 | 1 | 14 | Dennd4c | 0.095 | Old |
| 0.179162 | 0.15735 | 0.694 | 0.6 | 1 | 14 | Pdia3 | 0.094 | Old |
| 0.179164 | 1.080489 | 0.175 | 0.1 | 1 | 14 | Smndc1 | 0.075 | Old |
| 0.17937 | 0.495864 | 0.344 | 0.24 | 1 | 14 | Sp1 | 0.104 | Old |
| 0.179561 | 1.957945 | 0.071 | 0.02 | 1 | 14 | Cdkn2aipnl | 0.051 | Old |
| 0.179685 | 1.234213 | 0.131 | 0.06 | 1 | 14 | Ndufv1 | 0.071 | Old |
| 0.179717 | 1.626534 | 0.148 | 0.08 | 1 | 14 | Stxbp5 | 0.068 | Old |
| 0.180111 | 1.881088 | 0.142 | 0.08 | 1 | 14 | Ctbs | 0.062 | Old |
| 0.181033 | 1.84667 | 0.098 | 0.04 | 1 | 14 | Med20 | 0.058 | Old |
| 0.181033 | 1.84479 | 0.098 | 0.04 | 1 | 14 | Gm17018 | 0.058 | Old |
| 0.181071 | 0.584679 | 0.131 | 0.06 | 1 | 14 | Nek1 | 0.071 | Old |
| 0.18189 | 0.550353 | 0.284 | 0.2 | 1 | 14 | Cttn | 0.084 | Old |
| 0.182393 | 0.515837 | 0.459 | 0.38 | 1 | 14 | Sp100 | 0.079 | Old |
| 0.182628 | 1.93965 | 0.098 | 0.04 | 1 | 14 | Ttll12 | 0.058 | Old |
| 0.182804 | 1.301181 | 0.175 | 0.1 | 1 | 14 | Nfu1 | 0.075 | Old |
| 0.183316 | 1.772851 | 0.071 | 0.02 | 1 | 14 | Smim8 | 0.051 | Old |
| 0.183501 | 0.312443 | 0.672 | 0.52 | 1 | 14 | Loxl1 | 0.152 | Old |
| 0.183569 | 0.730327 | 0.301 | 0.22 | 1 | 14 | Ppp1r9b | 0.081 | Old |
| 0.18372 | 0.054188 | 0.956 | 0.92 | 1 | 14 | Rpl30 | 0.036 | Old |
| 0.183999 | 0.223519 | 0.24 | 0.14 | 1 | 14 | Rnf139 | 0.1 | Old |
| 0.184233 | 1.936432 | 0.098 | 0.04 | 1 | 14 | 2410018L1 | 0.058 | Old |
| 0.184233 | 1.532099 | 0.098 | 0.04 | 1 | 14 | Dnal1 | 0.058 | Old |
| 0.184367 | 0.576596 | 0.126 | 0.06 | 1 | 14 | Gse1 | 0.066 | Old |
| 0.184513 | 0.250659 | 0.246 | 0.16 | 1 | 14 | Dlg1 | 0.086 | Old |
| 0.184556 | 0.942458 | 0.24 | 0.16 | 1 | 14 | Gcc2 | 0.08 | Old |
| 0.184683 | 2.132917 | 0.12 | 0.06 | 1 | 14 | Top3b | 0.06 | Old |
| 0.184755 | 0.162748 | 0.863 | 0.74 | 1 | 14 | Laptm4a | 0.123 | Old |
| 0.185108 | 1.010693 | 0.191 | 0.12 | 1 | 14 | Mrpl4 | 0.071 | Old |

| | | | | | | | | |
|---|---|---|---|---|---|---|---|---|
| 0.185305 | 0.161669 | 0.213 | 0.12 | 1 | 14 | Tbc1d9b | 0.093 | Old |
| 0.185848 | 1.901575 | 0.098 | 0.04 | 1 | 14 | Rhbdd2 | 0.058 | Old |
| 0.186065 | 0.242658 | 0.962 | 0.88 | 1 | 14 | Junb | 0.082 | Old |
| 0.186219 | 0.273678 | 0.672 | 0.58 | 1 | 14 | Cast | 0.092 | Old |
| 0.186499 | 1.318536 | 0.175 | 0.1 | 1 | 14 | Irf3 | 0.075 | Old |
| 0.186531 | 0.474157 | 0.301 | 0.22 | 1 | 14 | Ezh1 | 0.081 | Old |
| 0.186652 | 0.783628 | 0.322 | 0.24 | 1 | 14 | Ganab | 0.082 | Old |
| 0.187247 | 1.456669 | 0.126 | 0.06 | 1 | 14 | Trim5 | 0.066 | Old |
| 0.187458 | 0.786011 | 0.262 | 0.18 | 1 | 14 | Arntl | 0.082 | Old |
| 0.187618 | 1.821192 | 0.12 | 0.06 | 1 | 14 | Nceh1 | 0.06 | Old |
| 0.18762 | 0.321228 | 0.623 | 0.5 | 1 | 14 | Lasp1 | 0.123 | Old |
| 0.187626 | 0.370764 | 0.743 | 0.64 | 1 | 14 | Anxa2 | 0.103 | Old |
| 0.187743 | 1.011905 | 0.175 | 0.1 | 1 | 14 | Rpe | 0.075 | Old |
| 0.188892 | 1.048776 | 0.24 | 0.16 | 1 | 14 | Psen2 | 0.08 | Old |
| 0.188944 | 0.416024 | 0.361 | 0.26 | 1 | 14 | Eif3l | 0.101 | Old |
| 0.189058 | 0.878138 | 0.071 | 0.02 | 1 | 14 | Zfp229 | 0.051 | Old |
| 0.189059 | 0.089378 | 0.508 | 0.42 | 1 | 14 | Scarb2 | 0.088 | Old |
| 0.189098 | 1.964079 | 0.12 | 0.06 | 1 | 14 | Mccc1 | 0.06 | Old |
| 0.189259 | 0.93657 | 0.388 | 0.32 | 1 | 14 | Rbm5 | 0.068 | Old |
| 0.189328 | 1.202921 | 0.208 | 0.14 | 1 | 14 | Smim19 | 0.068 | Old |
| 0.190111 | 1.353677 | 0.251 | 0.18 | 1 | 14 | Otud7b | 0.071 | Old |
| 0.19016 | 1.127059 | 0.126 | 0.06 | 1 | 14 | Cxxc1 | 0.066 | Old |
| 0.190236 | 0.866791 | 0.339 | 0.26 | 1 | 14 | Crybg3 | 0.079 | Old |
| 0.190718 | 0.750396 | 0.295 | 0.22 | 1 | 14 | S1pr2 | 0.075 | Old |
| 0.190757 | 1.485368 | 0.098 | 0.04 | 1 | 14 | 1110059E2 | 0.058 | Old |
| 0.191083 | 0.840944 | 0.35 | 0.26 | 1 | 14 | Abca8b | 0.09 | Old |
| 0.191459 | 1.264104 | 0.268 | 0.2 | 1 | 14 | Kansl1 | 0.068 | Old |
| 0.191511 | 1.238076 | 0.175 | 0.1 | 1 | 14 | Gbp2 | 0.075 | Old |
| 0.191713 | 1.383637 | 0.148 | 0.08 | 1 | 14 | Usp2 | 0.068 | Old |
| 0.191981 | 0.486302 | 0.06 | 0.12 | 1 | 14 | Tmc6 | -0.06 | Old |
| 0.192084 | 2.166951 | 0.12 | 0.06 | 1 | 14 | Abcd2 | 0.06 | Old |
| 0.192233 | 0.278495 | 0.235 | 0.16 | 1 | 14 | Slc25a36 | 0.075 | Old |
| 0.192414 | 1.540028 | 0.098 | 0.04 | 1 | 14 | Fam118a | 0.058 | Old |
| 0.192414 | 1.236853 | 0.098 | 0.04 | 1 | 14 | Ppp1r7 | 0.058 | Old |
| 0.192414 | 0.310526 | 0.098 | 0.04 | 1 | 14 | 1810010H2 | 0.058 | Old |
| 0.192442 | 0.500652 | 0.131 | 0.06 | 1 | 14 | Sh2d4a | 0.071 | Old |
| 0.192694 | 0.228941 | 0.23 | 0.14 | 1 | 14 | Rnf20 | 0.09 | Old |
| 0.193107 | 1.222187 | 0.126 | 0.06 | 1 | 14 | Dexi | 0.066 | Old |
| 0.193658 | 0.285779 | 0.568 | 0.42 | 1 | 14 | Crtap | 0.148 | Old |
| 0.193832 | 0.691881 | 0.23 | 0.14 | 1 | 14 | Smurf1 | 0.09 | Old |
| 0.194131 | 0.211066 | 0.383 | 0.3 | 1 | 14 | Cd2ap | 0.083 | Old |
| 0.194182 | 0.327522 | 0.601 | 0.56 | 1 | 14 | Prrx1 | 0.041 | Old |
| 0.19423 | 1.248891 | 0.197 | 0.12 | 1 | 14 | Lamtor5 | 0.077 | Old |
| 0.194448 | 1.453455 | 0.251 | 0.18 | 1 | 14 | Ing1 | 0.071 | Old |
| 0.194776 | 0.519071 | 0.53 | 0.46 | 1 | 14 | Ryk | 0.07 | Old |

| | | | | | | | | |
|---|---|---|---|---|---|---|---|---|
| 0.195334 | 1.170065 | 0.175 | 0.1 | 1 | 14 | Apmap | 0.075 | Old |
| 0.195352 | 1.723269 | 0.202 | 0.14 | 1 | 14 | Trabd2b | 0.062 | Old |
| 0.195405 | 0.262951 | 0.541 | 0.46 | 1 | 14 | Nucb1 | 0.081 | Old |
| 0.195424 | 0.531139 | 0.41 | 0.34 | 1 | 14 | Elavl1 | 0.07 | Old |
| 0.195584 | 0.374361 | 0.454 | 0.36 | 1 | 14 | AC149090. | 0.094 | Old |
| 0.196029 | 0.246332 | 0.328 | 0.22 | 1 | 14 | Cox14 | 0.108 | Old |
| 0.196201 | 0.19241 | 0.617 | 0.48 | 1 | 14 | Lamb1 | 0.137 | Old |
| 0.196289 | 1.164163 | 0.273 | 0.2 | 1 | 14 | Adam33 | 0.073 | Old |
| 0.196659 | 0.659712 | 0.24 | 0.16 | 1 | 14 | Epb41l3 | 0.08 | Old |
| 0.19669 | 0.020705 | 0.366 | 0.28 | 1 | 14 | Hspb8 | 0.086 | Old |
| 0.196734 | 1.011234 | 0.235 | 0.16 | 1 | 14 | Ocrl | 0.075 | Old |
| 0.196748 | 1.010293 | 0.23 | 0.16 | 1 | 14 | Galnt16 | 0.07 | Old |
| 0.197036 | 0.12819 | 0.328 | 0.22 | 1 | 14 | Arid5b | 0.108 | Old |
| 0.197056 | 0.405765 | 0.426 | 0.34 | 1 | 14 | Psmd2 | 0.086 | Old |
| 0.197073 | 4.038176 | 0.033 | 0 | 1 | 14 | Wdr76 | 0.033 | Old |
| 0.197073 | 3.473969 | 0.033 | 0 | 1 | 14 | Epyc | 0.033 | Old |
| 0.197073 | 3.47162 | 0.033 | 0 | 1 | 14 | Ncf4 | 0.033 | Old |
| 0.197073 | 3.452463 | 0.033 | 0 | 1 | 14 | Mocs3 | 0.033 | Old |
| 0.197073 | 3.397394 | 0.033 | 0 | 1 | 14 | P2ry14 | 0.033 | Old |
| 0.197073 | 3.313849 | 0.033 | 0 | 1 | 14 | Lysmd1 | 0.033 | Old |
| 0.197073 | 3.188073 | 0.033 | 0 | 1 | 14 | 6330562C2 | 0.033 | Old |
| 0.197073 | 3.18222 | 0.033 | 0 | 1 | 14 | Card6 | 0.033 | Old |
| 0.197073 | 3.164178 | 0.033 | 0 | 1 | 14 | Tnfaip8l2 | 0.033 | Old |
| 0.197073 | 3.127402 | 0.033 | 0 | 1 | 14 | Bvht | 0.033 | Old |
| 0.197073 | 3.112233 | 0.033 | 0 | 1 | 14 | C920006O1 | 0.033 | Old |
| 0.197073 | 3.091905 | 0.033 | 0 | 1 | 14 | Slc6a4 | 0.033 | Old |
| 0.197073 | 3.063145 | 0.033 | 0 | 1 | 14 | Als2 | 0.033 | Old |
| 0.197073 | 3.021272 | 0.033 | 0 | 1 | 14 | Nxph3 | 0.033 | Old |
| 0.197073 | 3.01689 | 0.033 | 0 | 1 | 14 | 3110002H1 | 0.033 | Old |
| 0.197073 | 2.977859 | 0.033 | 0 | 1 | 14 | Abi3 | 0.033 | Old |
| 0.197073 | 2.938172 | 0.033 | 0 | 1 | 14 | Naip2 | 0.033 | Old |
| 0.197073 | 2.921167 | 0.033 | 0 | 1 | 14 | Gm16503 | 0.033 | Old |
| 0.197073 | 2.921141 | 0.033 | 0 | 1 | 14 | Foxf2 | 0.033 | Old |
| 0.197073 | 2.908898 | 0.033 | 0 | 1 | 14 | Fggy | 0.033 | Old |
| 0.197073 | 2.90704 | 0.033 | 0 | 1 | 14 | Fancf | 0.033 | Old |
| 0.197073 | 2.870267 | 0.033 | 0 | 1 | 14 | Rab3ip | 0.033 | Old |
| 0.197073 | 2.864868 | 0.033 | 0 | 1 | 14 | Nectin2 | 0.033 | Old |
| 0.197073 | 2.780872 | 0.033 | 0 | 1 | 14 | AU041133 | 0.033 | Old |
| 0.197073 | 2.765005 | 0.033 | 0 | 1 | 14 | Hps6 | 0.033 | Old |
| 0.197073 | 2.756244 | 0.033 | 0 | 1 | 14 | Stx1a | 0.033 | Old |
| 0.197073 | 2.753276 | 0.033 | 0 | 1 | 14 | Gm13212 | 0.033 | Old |
| 0.197073 | 2.736099 | 0.033 | 0 | 1 | 14 | E2f7 | 0.033 | Old |
| 0.197073 | 2.729626 | 0.033 | 0 | 1 | 14 | Dglucy | 0.033 | Old |
| 0.197073 | 2.70847 | 0.033 | 0 | 1 | 14 | AC130815. | 0.033 | Old |
| 0.197073 | 2.704529 | 0.033 | 0 | 1 | 14 | Baiap2l1 | 0.033 | Old |

| | | | | | | | | |
|---|---|---|---|---|---|---|---|---|
| 0.197073 | 2.637105 | 0.033 | 0 | 1 | 14 | Sla | 0.033 | Old |
| 0.197073 | 2.632264 | 0.033 | 0 | 1 | 14 | Gm16118 | 0.033 | Old |
| 0.197073 | 2.627394 | 0.033 | 0 | 1 | 14 | Gpr89 | 0.033 | Old |
| 0.197073 | 2.345635 | 0.033 | 0 | 1 | 14 | Sac3d1 | 0.033 | Old |
| 0.197073 | 2.327201 | 0.033 | 0 | 1 | 14 | Brinp3 | 0.033 | Old |
| 0.197073 | 2.288425 | 0.033 | 0 | 1 | 14 | Cenpl | 0.033 | Old |
| 0.197073 | 2.284397 | 0.033 | 0 | 1 | 14 | Abca2 | 0.033 | Old |
| 0.197073 | 2.105497 | 0.033 | 0 | 1 | 14 | Gstp2 | 0.033 | Old |
| 0.197227 | 2.435932 | 0.093 | 0.04 | 1 | 14 | Kri1 | 0.053 | Old |
| 0.197257 | 0.658509 | 0.202 | 0.12 | 1 | 14 | Gnl1 | 0.082 | Old |
| 0.197406 | 0.534245 | 0.372 | 0.28 | 1 | 14 | Trove2 | 0.092 | Old |
| 0.197589 | 0.813774 | 0.126 | 0.06 | 1 | 14 | Fer | 0.066 | Old |
| 0.1978 | 0.143443 | 0.464 | 0.36 | 1 | 14 | Numa1 | 0.104 | Old |
| 0.197914 | 0.814345 | 0.175 | 0.1 | 1 | 14 | Ccdc85b | 0.075 | Old |
| 0.198431 | 0.37423 | 0.23 | 0.14 | 1 | 14 | Maml1 | 0.09 | Old |
| 0.198764 | 0.270049 | 0.645 | 0.54 | 1 | 14 | Dnajc3 | 0.105 | Old |
| 0.198864 | 0.20323 | 0.792 | 0.7 | 1 | 14 | Rhoa | 0.092 | Old |
| 0.198966 | 2.303616 | 0.093 | 0.04 | 1 | 14 | Fam118b | 0.053 | Old |
| 0.199149 | 1.118919 | 0.098 | 0.04 | 1 | 14 | Fance | 0.058 | Old |
| 0.199331 | 1.94766 | 0.142 | 0.08 | 1 | 14 | Prmt3 | 0.062 | Old |
| 0.200475 | 0.276835 | 0.093 | 0.16 | 1 | 14 | Hoxd8 | -0.067 | Old |
| 0.200488 | 0.481953 | 0.344 | 0.26 | 1 | 14 | Smc1a | 0.084 | Old |
| 0.200759 | 2.534364 | 0.142 | 0.08 | 1 | 14 | Fam160b2 | 0.062 | Old |
| 0.200961 | 1.150816 | 0.191 | 0.12 | 1 | 14 | Dguok | 0.071 | Old |
| 0.201051 | 0.383313 | 0.574 | 0.48 | 1 | 14 | Kmt2e | 0.094 | Old |
| 0.201202 | 0.90649 | 0.24 | 0.16 | 1 | 14 | Hivep1 | 0.08 | Old |
| 0.20121 | 0.172218 | 0.623 | 0.54 | 1 | 14 | 2900097C1 | 0.083 | Old |
| 0.202165 | 0.707808 | 0.202 | 0.12 | 1 | 14 | Sobp | 0.082 | Old |
| 0.202273 | 0.426969 | 0.366 | 0.3 | 1 | 14 | Smdt1 | 0.066 | Old |
| 0.202389 | 1.398619 | 0.208 | 0.14 | 1 | 14 | Limch1 | 0.068 | Old |
| 0.202581 | 0.673004 | 0.098 | 0.04 | 1 | 14 | Gm15417 | 0.058 | Old |
| 0.202701 | 0.523525 | 0.268 | 0.18 | 1 | 14 | Ep400 | 0.088 | Old |
| 0.20278 | 3.133564 | 0.066 | 0.02 | 1 | 14 | Sntg1 | 0.046 | Old |
| 0.20278 | 2.911613 | 0.066 | 0.02 | 1 | 14 | Ptpn3 | 0.046 | Old |
| 0.20278 | 2.746029 | 0.066 | 0.02 | 1 | 14 | Eif2b1 | 0.046 | Old |
| 0.20278 | 2.734824 | 0.066 | 0.02 | 1 | 14 | Xrcc5 | 0.046 | Old |
| 0.20278 | 2.731829 | 0.066 | 0.02 | 1 | 14 | Gm21781 | 0.046 | Old |
| 0.20278 | 2.664303 | 0.066 | 0.02 | 1 | 14 | Nkain4 | 0.046 | Old |
| 0.20278 | 2.544115 | 0.066 | 0.02 | 1 | 14 | Gmeb1 | 0.046 | Old |
| 0.20278 | 2.409123 | 0.066 | 0.02 | 1 | 14 | Ldlrad3 | 0.046 | Old |
| 0.20278 | 2.270131 | 0.066 | 0.02 | 1 | 14 | Tfpt | 0.046 | Old |
| 0.20278 | 1.990287 | 0.066 | 0.02 | 1 | 14 | Mansc1 | 0.046 | Old |
| 0.202984 | 0.529077 | 0.404 | 0.34 | 1 | 14 | Acat1 | 0.064 | Old |
| 0.204249 | 2.212169 | 0.093 | 0.04 | 1 | 14 | 2010111I0 | 0.053 | Old |
| 0.204312 | 1.053068 | 0.098 | 0.04 | 1 | 14 | Mgme1 | 0.058 | Old |

| | | | | | | | | |
|---|---|---|---|---|---|---|---|---|
| 0.204376 | 1.218839 | 0.12 | 0.06 | 1 | 14 | Atxn7l1 | 0.06 | Old |
| 0.204651 | 0.713189 | 0.202 | 0.12 | 1 | 14 | Zfp397 | 0.082 | Old |
| 0.204796 | 1.004962 | 0.235 | 0.16 | 1 | 14 | Grk2 | 0.075 | Old |
| 0.204898 | 2.253539 | 0.066 | 0.02 | 1 | 14 | Stx11 | 0.046 | Old |
| 0.204898 | 2.080919 | 0.066 | 0.02 | 1 | 14 | Asna1 | 0.046 | Old |
| 0.204898 | 2.0302 | 0.066 | 0.02 | 1 | 14 | Tspyl3 | 0.046 | Old |
| 0.205307 | 1.511206 | 0.169 | 0.1 | 1 | 14 | Ndrg4 | 0.069 | Old |
| 0.205307 | 1.259052 | 0.169 | 0.1 | 1 | 14 | Fsd1l | 0.069 | Old |
| 0.205349 | 0.524726 | 0.432 | 0.34 | 1 | 14 | Golim4 | 0.092 | Old |
| 0.205683 | 0.088457 | 0.59 | 0.5 | 1 | 14 | Eef1d | 0.09 | Old |
| 0.20582 | 0.820926 | 0.24 | 0.16 | 1 | 14 | Pnrc2 | 0.08 | Old |
| 0.205952 | 1.334041 | 0.12 | 0.06 | 1 | 14 | Hoxa10 | 0.06 | Old |
| 0.206037 | 0.807888 | 0.268 | 0.18 | 1 | 14 | Klf10 | 0.088 | Old |
| 0.206055 | 0.502242 | 0.098 | 0.04 | 1 | 14 | Pex5 | 0.058 | Old |
| 0.206272 | 0.411637 | 0.536 | 0.44 | 1 | 14 | Tnrc6b | 0.096 | Old |
| 0.206545 | 0.225695 | 0.552 | 0.46 | 1 | 14 | Sec31a | 0.092 | Old |
| 0.206609 | 1.47303 | 0.186 | 0.12 | 1 | 14 | Eng | 0.066 | Old |
| 0.207033 | 2.706633 | 0.066 | 0.02 | 1 | 14 | 3110040N1 | 0.046 | Old |
| 0.207033 | 1.894635 | 0.066 | 0.02 | 1 | 14 | Zfp984 | 0.046 | Old |
| 0.207121 | 0.625832 | 0.448 | 0.38 | 1 | 14 | M6pr | 0.068 | Old |
| 0.207167 | 0.197263 | 0.317 | 0.22 | 1 | 14 | Snrpb2 | 0.097 | Old |
| 0.207177 | 0.719998 | 0.339 | 0.26 | 1 | 14 | Ubr5 | 0.079 | Old |
| 0.207254 | 0.632172 | 0.426 | 0.36 | 1 | 14 | Ube2d2a | 0.066 | Old |
| 0.208158 | 1.042159 | 0.24 | 0.16 | 1 | 14 | Rragc | 0.08 | Old |
| 0.208343 | 0.859606 | 0.126 | 0.06 | 1 | 14 | Ap3m2 | 0.066 | Old |
| 0.208609 | 1.620498 | 0.148 | 0.08 | 1 | 14 | Fbxo30 | 0.068 | Old |
| 0.209112 | 0.157195 | 0.628 | 0.54 | 1 | 14 | Capza2 | 0.088 | Old |
| 0.209183 | 2.553288 | 0.066 | 0.02 | 1 | 14 | Nfkbie | 0.046 | Old |
| 0.209183 | 2.319345 | 0.066 | 0.02 | 1 | 14 | Zfand2b | 0.046 | Old |
| 0.209492 | 0.38689 | 0.601 | 0.5 | 1 | 14 | Nfkbia | 0.101 | Old |
| 0.209572 | 0.818461 | 0.098 | 0.04 | 1 | 14 | Nadk2 | 0.058 | Old |
| 0.209634 | 2.079479 | 0.093 | 0.04 | 1 | 14 | Sox13 | 0.053 | Old |
| 0.209634 | 1.534187 | 0.093 | 0.04 | 1 | 14 | Dcps | 0.053 | Old |
| 0.209881 | 0.192383 | 0.858 | 0.76 | 1 | 14 | Ifitm3 | 0.098 | Old |
| 0.209913 | 0.737548 | 0.126 | 0.06 | 1 | 14 | Atxn1 | 0.066 | Old |
| 0.209919 | 0.621264 | 0.333 | 0.24 | 1 | 14 | Plekho2 | 0.093 | Old |
| 0.210293 | 0.608216 | 0.284 | 0.2 | 1 | 14 | Srp19 | 0.084 | Old |
| 0.210515 | 0.777698 | 0.273 | 0.2 | 1 | 14 | Tek | 0.073 | Old |
| 0.210515 | 0.673839 | 0.24 | 0.16 | 1 | 14 | Setx | 0.08 | Old |
| 0.210818 | 0.274986 | 0.104 | 0.18 | 1 | 14 | B4galt3 | -0.076 | Old |
| 0.211349 | 2.701979 | 0.066 | 0.02 | 1 | 14 | Alkbh6 | 0.046 | Old |
| 0.211349 | 2.582928 | 0.066 | 0.02 | 1 | 14 | Ptpdc1 | 0.046 | Old |
| 0.211349 | 2.029795 | 0.066 | 0.02 | 1 | 14 | Eef1e1 | 0.046 | Old |
| 0.211429 | 0.16516 | 0.503 | 0.4 | 1 | 14 | Eif3e | 0.103 | Old |
| 0.211451 | 1.885566 | 0.093 | 0.04 | 1 | 14 | Scnm1 | 0.053 | Old |

| | | | | | | | | |
|---|---|---|---|---|---|---|---|---|
| 0.211527 | 1.118517 | 0.148 | 0.08 | 1 | 14 | Nup133 | 0.068 | Old |
| 0.21153 | 0.319539 | 0.858 | 0.74 | 1 | 14 | Hspa5 | 0.118 | Old |
| 0.212179 | 0.587378 | 0.508 | 0.46 | 1 | 14 | Tor1aip1 | 0.048 | Old |
| 0.212352 | 0.158368 | 0.454 | 0.36 | 1 | 14 | Cct8 | 0.094 | Old |
| 0.212453 | 1.540926 | 0.142 | 0.08 | 1 | 14 | Rfx7 | 0.062 | Old |
| 0.21296 | 1.544905 | 0.164 | 0.1 | 1 | 14 | Lrrc8b | 0.064 | Old |
| 0.21328 | 1.852212 | 0.093 | 0.04 | 1 | 14 | B3gnt9 | 0.053 | Old |
| 0.213699 | 1.784497 | 0.158 | 0.1 | 1 | 14 | Akr1b8 | 0.058 | Old |
| 0.213963 | 1.892761 | 0.12 | 0.06 | 1 | 14 | Cxcl13 | 0.06 | Old |
| 0.213963 | 0.937245 | 0.12 | 0.06 | 1 | 14 | Ambra1 | 0.06 | Old |
| 0.213975 | 1.37222 | 0.262 | 0.2 | 1 | 14 | Nap1l4 | 0.062 | Old |
| 0.214653 | 1.865625 | 0.115 | 0.06 | 1 | 14 | Pnpla6 | 0.055 | Old |
| 0.214743 | 0.550844 | 0.388 | 0.3 | 1 | 14 | Rab5c | 0.088 | Old |
| 0.214793 | 0.710129 | 0.301 | 0.22 | 1 | 14 | Slbp | 0.081 | Old |
| 0.214823 | 1.159943 | 0.208 | 0.14 | 1 | 14 | Slc39a6 | 0.068 | Old |
| 0.215127 | 1.051015 | 0.262 | 0.2 | 1 | 14 | Cnih4 | 0.062 | Old |
| 0.215131 | 0.964055 | 0.191 | 0.12 | 1 | 14 | Aurkaip1 | 0.071 | Old |
| 0.215335 | 0.754456 | 0.372 | 0.28 | 1 | 14 | CT025619.1 | 0.092 | Old |
| 0.215452 | 1.463409 | 0.142 | 0.08 | 1 | 14 | Decr1 | 0.062 | Old |
| 0.215452 | 1.103138 | 0.142 | 0.08 | 1 | 14 | Tiam1 | 0.062 | Old |
| 0.215476 | 0.240169 | 0.049 | 0.1 | 1 | 14 | Sipa1l2 | -0.051 | Old |
| 0.215669 | 0.292251 | 0.503 | 0.38 | 1 | 14 | Olfml2b | 0.123 | Old |
| 0.215673 | 0.540675 | 0.197 | 0.12 | 1 | 14 | Trp53 | 0.077 | Old |
| 0.215731 | 1.914493 | 0.066 | 0.02 | 1 | 14 | Bloc1s2 | 0.046 | Old |
| 0.216309 | 0.378341 | 0.645 | 0.54 | 1 | 14 | Iqgap1 | 0.105 | Old |
| 0.216772 | 0.445525 | 0.388 | 0.3 | 1 | 14 | Hnrnpul2 | 0.088 | Old |
| 0.217425 | 0.833655 | 0.317 | 0.24 | 1 | 14 | Kat6a | 0.077 | Old |
| 0.217444 | 1.19824 | 0.262 | 0.2 | 1 | 14 | Rnf4 | 0.062 | Old |
| 0.217444 | 0.650867 | 0.268 | 0.18 | 1 | 14 | Imp3 | 0.088 | Old |
| 0.217451 | 0.960947 | 0.148 | 0.08 | 1 | 14 | Taf2 | 0.068 | Old |
| 0.217459 | 0.286564 | 0.53 | 0.44 | 1 | 14 | Asph | 0.09 | Old |
| 0.217613 | 0.824878 | 0.049 | 0.1 | 1 | 14 | Pus1 | -0.051 | Old |
| 0.217613 | 0.694848 | 0.049 | 0.1 | 1 | 14 | Serpinb8 | -0.051 | Old |
| 0.217777 | 0.644959 | 0.153 | 0.08 | 1 | 14 | Isoc1 | 0.073 | Old |
| 0.217946 | 2.396673 | 0.066 | 0.02 | 1 | 14 | Tada2a | 0.046 | Old |
| 0.217946 | 2.188247 | 0.066 | 0.02 | 1 | 14 | Pomt1 | 0.046 | Old |
| 0.217946 | 1.437591 | 0.066 | 0.02 | 1 | 14 | Ift140 | 0.046 | Old |
| 0.217988 | 1.798791 | 0.115 | 0.06 | 1 | 14 | Lcor | 0.055 | Old |
| 0.218004 | 0.538567 | 0.459 | 0.38 | 1 | 14 | Ece1 | 0.079 | Old |
| 0.218241 | 1.281182 | 0.246 | 0.18 | 1 | 14 | Ube2w | 0.066 | Old |
| 0.218373 | 0.244103 | 0.814 | 0.7 | 1 | 14 | Penk | 0.114 | Old |
| 0.218476 | 0.347259 | 0.273 | 0.2 | 1 | 14 | Itga9 | 0.073 | Old |
| 0.218585 | 0.297629 | 0.41 | 0.34 | 1 | 14 | Snx9 | 0.07 | Old |
| 0.218787 | 0.475618 | 0.366 | 0.28 | 1 | 14 | Dram1 | 0.086 | Old |
| 0.218835 | 1.661915 | 0.093 | 0.04 | 1 | 14 | Izumo4 | 0.053 | Old |

| | | | | | | | | |
|---|---|---|---|---|---|---|---|---|
| 0.218835 | 1.631854 | 0.093 | 0.04 | 1 | 14 | Alcam | 0.053 | Old |
| 0.218951 | 0.858188 | 0.148 | 0.08 | 1 | 14 | Clpp | 0.068 | Old |
| 0.21902 | 0.139535 | 0.661 | 0.5 | 1 | 14 | Egfr | 0.161 | Old |
| 0.219386 | 1.265302 | 0.301 | 0.24 | 1 | 14 | Rex1bd | 0.061 | Old |
| 0.21944 | 1.173038 | 0.246 | 0.18 | 1 | 14 | Cdc26 | 0.066 | Old |
| 0.219642 | 0.056356 | 0.432 | 0.32 | 1 | 14 | Ndufa3 | 0.112 | Old |
| 0.220007 | 1.245228 | 0.142 | 0.08 | 1 | 14 | Wdr11 | 0.062 | Old |
| 0.220177 | 1.413446 | 0.066 | 0.02 | 1 | 14 | Cutc | 0.046 | Old |
| 0.22071 | 1.705037 | 0.093 | 0.04 | 1 | 14 | Fbxo6 | 0.053 | Old |
| 0.22087 | 0.181689 | 0.628 | 0.58 | 1 | 14 | Glud1 | 0.048 | Old |
| 0.221219 | 0.598321 | 0.503 | 0.46 | 1 | 14 | Sfxn1 | 0.043 | Old |
| 0.221541 | 1.37235 | 0.142 | 0.08 | 1 | 14 | Ppp2cb | 0.062 | Old |
| 0.221917 | 0.344405 | 0.443 | 0.38 | 1 | 14 | Smad5 | 0.063 | Old |
| 0.222145 | 0.662624 | 0.262 | 0.18 | 1 | 14 | Emc2 | 0.082 | Old |
| 0.222179 | 0.228646 | 0.514 | 0.42 | 1 | 14 | Limd1 | 0.094 | Old |
| 0.222197 | 0.278258 | 0.12 | 0.06 | 1 | 14 | Filip1 | 0.06 | Old |
| 0.222425 | 1.412584 | 0.066 | 0.02 | 1 | 14 | Lekr1 | 0.046 | Old |
| 0.222425 | 1.215977 | 0.066 | 0.02 | 1 | 14 | A530020G: | 0.046 | Old |
| 0.222425 | 1.129569 | 0.066 | 0.02 | 1 | 14 | C2cd3 | 0.046 | Old |
| 0.222425 | 1.049706 | 0.066 | 0.02 | 1 | 14 | Tpcn2 | 0.046 | Old |
| 0.222495 | 1.94957 | 0.137 | 0.08 | 1 | 14 | Phf6 | 0.057 | Old |
| 0.222542 | 0.971167 | 0.208 | 0.14 | 1 | 14 | Arid5a | 0.068 | Old |
| 0.222596 | 1.477169 | 0.093 | 0.04 | 1 | 14 | Fgf11 | 0.053 | Old |
| 0.223157 | 1.170522 | 0.191 | 0.12 | 1 | 14 | Zfp9 | 0.071 | Old |
| 0.223532 | 0.150998 | 0.699 | 0.6 | 1 | 14 | Ltbp4 | 0.099 | Old |
| 0.223634 | 0.69252 | 0.197 | 0.12 | 1 | 14 | Paf1 | 0.077 | Old |
| 0.223832 | 0.505228 | 0.268 | 0.2 | 1 | 14 | Phactr2 | 0.068 | Old |
| 0.223898 | 0.47168 | 0.284 | 0.2 | 1 | 14 | Rab6b | 0.084 | Old |
| 0.224494 | 1.554508 | 0.093 | 0.04 | 1 | 14 | Ngrn | 0.053 | Old |
| 0.224494 | 1.391247 | 0.093 | 0.04 | 1 | 14 | Dcaf6 | 0.053 | Old |
| 0.224494 | 1.110394 | 0.093 | 0.04 | 1 | 14 | Cactin | 0.053 | Old |
| 0.224689 | 0.97376 | 0.066 | 0.02 | 1 | 14 | Cep97 | 0.046 | Old |
| 0.224689 | 0.868922 | 0.066 | 0.02 | 1 | 14 | Shroom3 | 0.046 | Old |
| 0.224689 | 0.507062 | 0.066 | 0.02 | 1 | 14 | Dyrk1b | 0.046 | Old |
| 0.224852 | 0.150563 | 0.109 | 0.18 | 1 | 14 | Afg3l1 | -0.071 | Old |
| 0.225064 | 0.705066 | 0.24 | 0.16 | 1 | 14 | Gadd45b | 0.08 | Old |
| 0.225322 | 1.034684 | 0.186 | 0.12 | 1 | 14 | Nfil3 | 0.066 | Old |
| 0.225731 | 0.341451 | 0.262 | 0.18 | 1 | 14 | Bin1 | 0.082 | Old |
| 0.225871 | 0.378202 | 0.525 | 0.48 | 1 | 14 | Ifngr1 | 0.045 | Old |
| 0.226188 | 1.349079 | 0.142 | 0.08 | 1 | 14 | Slc38a6 | 0.062 | Old |
| 0.226601 | 0.401693 | 0.219 | 0.14 | 1 | 14 | Zfyve9 | 0.079 | Old |
| 0.226703 | 1.163388 | 0.186 | 0.12 | 1 | 14 | Vps11 | 0.066 | Old |
| 0.226733 | 0.618236 | 0.475 | 0.4 | 1 | 14 | Trim35 | 0.075 | Old |
| 0.226926 | 0.149572 | 0.257 | 0.18 | 1 | 14 | Srp54a | 0.077 | Old |
| 0.22697 | 0.620474 | 0.066 | 0.02 | 1 | 14 | Abhd8 | 0.046 | Old |

| | | | | | | | | |
|---|---|---|---|---|---|---|---|---|
| 0.227229 | 0.54522 | 0.475 | 0.4 | 1 | 14 | Vegfd | 0.075 | Old |
| 0.227369 | 0.021525 | 0.628 | 0.58 | 1 | 14 | Eif3f | 0.048 | Old |
| 0.228021 | 0.084651 | 0.109 | 0.18 | 1 | 14 | Tspan14 | -0.071 | Old |
| 0.228411 | 0.276436 | 0.41 | 0.32 | 1 | 14 | Hsbp1 | 0.09 | Old |
| 0.228545 | 0.834431 | 0.213 | 0.14 | 1 | 14 | Dubr | 0.073 | Old |
| 0.228606 | 0.080003 | 0.656 | 0.58 | 1 | 14 | Arf4 | 0.076 | Old |
| 0.228819 | 2.039558 | 0.137 | 0.08 | 1 | 14 | Kdm3b | 0.057 | Old |
| 0.228819 | 1.524811 | 0.137 | 0.08 | 1 | 14 | Fgf7 | 0.057 | Old |
| 0.228995 | 0.908381 | 0.366 | 0.3 | 1 | 14 | Dpm3 | 0.066 | Old |
| 0.229267 | 0.185399 | 0.066 | 0.02 | 1 | 14 | Mdm1 | 0.046 | Old |
| 0.229581 | 0.002663 | 0.038 | 0.08 | 1 | 14 | Clec4a2 | -0.042 | Old |
| 0.229952 | 1.929211 | 0.115 | 0.06 | 1 | 14 | Kctd3 | 0.055 | Old |
| 0.230256 | 0.006679 | 0.093 | 0.04 | 1 | 14 | Lrrc32 | 0.053 | Old |
| 0.230451 | 1.355228 | 0.246 | 0.18 | 1 | 14 | Vps26a | 0.066 | Old |
| 0.230655 | 1.043158 | 0.12 | 0.06 | 1 | 14 | Taf4 | 0.06 | Old |
| 0.230703 | 0.874807 | 0.23 | 0.16 | 1 | 14 | Pi4ka | 0.07 | Old |
| 0.231477 | 0.185457 | 0.858 | 0.8 | 1 | 14 | Cd63 | 0.058 | Old |
| 0.232099 | 0.177086 | 0.623 | 0.58 | 1 | 14 | Fbn1 | 0.043 | Old |
| 0.232199 | 1.189078 | 0.093 | 0.04 | 1 | 14 | Sec22c | 0.053 | Old |
| 0.232212 | 0.393609 | 0.295 | 0.22 | 1 | 14 | Megf9 | 0.075 | Old |
| 0.232228 | 0.299874 | 0.492 | 0.42 | 1 | 14 | Ralbp1 | 0.072 | Old |
| 0.232318 | 0.259924 | 0.355 | 0.28 | 1 | 14 | Fhl1 | 0.075 | Old |
| 0.232492 | 1.37756 | 0.142 | 0.08 | 1 | 14 | Dip2b | 0.062 | Old |
| 0.232617 | 0.317812 | 0.825 | 0.76 | 1 | 14 | Col4a1 | 0.065 | Old |
| 0.232738 | 1.609039 | 0.158 | 0.1 | 1 | 14 | Gngt2 | 0.058 | Old |
| 0.232942 | 0.326179 | 0.268 | 0.18 | 1 | 14 | Uqcrfs1 | 0.088 | Old |
| 0.233033 | 0.065698 | 0.53 | 0.42 | 1 | 14 | Mapk3 | 0.11 | Old |
| 0.23314 | 1.023418 | 0.208 | 0.14 | 1 | 14 | Maml2 | 0.068 | Old |
| 0.233385 | 0.490244 | 0.295 | 0.22 | 1 | 14 | Itsn2 | 0.075 | Old |
| 0.234087 | 1.143798 | 0.142 | 0.08 | 1 | 14 | Aagab | 0.062 | Old |
| 0.234102 | 1.02793 | 0.12 | 0.06 | 1 | 14 | Gtf3c6 | 0.06 | Old |
| 0.234458 | 2.54873 | 0.087 | 0.04 | 1 | 14 | Rfc2 | 0.047 | Old |
| 0.235119 | 1.216932 | 0.186 | 0.12 | 1 | 14 | Maff | 0.066 | Old |
| 0.235132 | 0.21805 | 0.689 | 0.58 | 1 | 14 | Reep3 | 0.109 | Old |
| 0.236144 | 1.155927 | 0.224 | 0.16 | 1 | 14 | Deptor | 0.064 | Old |
| 0.236319 | 0.810346 | 0.426 | 0.36 | 1 | 14 | Sub1 | 0.066 | Old |
| 0.236477 | 2.761686 | 0.087 | 0.04 | 1 | 14 | Smg9 | 0.047 | Old |
| 0.236735 | 0.902769 | 0.246 | 0.18 | 1 | 14 | Epc2 | 0.066 | Old |
| 0.236815 | 0.05738 | 0.082 | 0.14 | 1 | 14 | Arl15 | -0.058 | Old |
| 0.236838 | 0.318624 | 0.235 | 0.16 | 1 | 14 | Aqr | 0.075 | Old |
| 0.236995 | 1.284548 | 0.115 | 0.06 | 1 | 14 | Hpse | 0.055 | Old |
| 0.237038 | 0.206277 | 0.754 | 0.64 | 1 | 14 | Tmem50a | 0.114 | Old |
| 0.237181 | 0.763902 | 0.23 | 0.16 | 1 | 14 | Pcbp4 | 0.07 | Old |
| 0.237214 | 0.198011 | 0.918 | 0.82 | 1 | 14 | Txnip | 0.098 | Old |
| 0.237714 | 0.749986 | 0.164 | 0.1 | 1 | 14 | Ap1ar | 0.064 | Old |

| | | | | | | | | |
|---|---|---|---|---|---|---|---|---|
| 0.238562 | 0.813476 | 0.29 | 0.22 | 1 | 14 | Mosmo | 0.07 | Old |
| 0.238705 | 0.632109 | 0.361 | 0.3 | 1 | 14 | Tle3 | 0.061 | Old |
| 0.238707 | 0.300455 | 0.273 | 0.2 | 1 | 14 | Mvp | 0.073 | Old |
| 0.23887 | 0.076762 | 0.393 | 0.3 | 1 | 14 | Rps6ka3 | 0.093 | Old |
| 0.239182 | 1.28788 | 0.202 | 0.14 | 1 | 14 | Hpcal1 | 0.062 | Old |
| 0.239182 | 0.289213 | 0.208 | 0.12 | 1 | 14 | Lgi4 | 0.088 | Old |
| 0.239846 | 0.548501 | 0.191 | 0.12 | 1 | 14 | Igbp1 | 0.071 | Old |
| 0.240078 | 0.431495 | 0.306 | 0.24 | 1 | 14 | D17Wsu92 | 0.066 | Old |
| 0.240091 | 1.49995 | 0.093 | 0.04 | 1 | 14 | Vopp1 | 0.053 | Old |
| 0.240091 | 1.469626 | 0.093 | 0.04 | 1 | 14 | Tmem70 | 0.053 | Old |
| 0.240179 | 0.217471 | 0.503 | 0.42 | 1 | 14 | Crebbp | 0.083 | Old |
| 0.240309 | 3.955291 | 0.027 | 0 | 1 | 14 | Dlx3 | 0.027 | Old |
| 0.240309 | 3.65724 | 0.027 | 0 | 1 | 14 | Riox1 | 0.027 | Old |
| 0.240309 | 3.531011 | 0.027 | 0 | 1 | 14 | Dock5 | 0.027 | Old |
| 0.240309 | 3.237901 | 0.027 | 0 | 1 | 14 | Cx3cr1 | 0.027 | Old |
| 0.240309 | 3.216022 | 0.027 | 0 | 1 | 14 | Meox1 | 0.027 | Old |
| 0.240309 | 3.214656 | 0.027 | 0 | 1 | 14 | Haus1 | 0.027 | Old |
| 0.240309 | 3.14068 | 0.027 | 0 | 1 | 14 | Osbp2 | 0.027 | Old |
| 0.240309 | 3.123232 | 0.027 | 0 | 1 | 14 | Atp1a3 | 0.027 | Old |
| 0.240309 | 3.121605 | 0.027 | 0 | 1 | 14 | Lca5l | 0.027 | Old |
| 0.240309 | 3.095047 | 0.027 | 0 | 1 | 14 | Nfatc2ip | 0.027 | Old |
| 0.240309 | 3.068971 | 0.027 | 0 | 1 | 14 | Slc12a8 | 0.027 | Old |
| 0.240309 | 3.055117 | 0.027 | 0 | 1 | 14 | Zfp87 | 0.027 | Old |
| 0.240309 | 3.039653 | 0.027 | 0 | 1 | 14 | A730020M | 0.027 | Old |
| 0.240309 | 3.037498 | 0.027 | 0 | 1 | 14 | Kif20b | 0.027 | Old |
| 0.240309 | 2.919433 | 0.027 | 0 | 1 | 14 | Gm9913 | 0.027 | Old |
| 0.240309 | 2.913124 | 0.027 | 0 | 1 | 14 | D130040H | 0.027 | Old |
| 0.240309 | 2.91 | 0.027 | 0 | 1 | 14 | Sync | 0.027 | Old |
| 0.240309 | 2.904213 | 0.027 | 0 | 1 | 14 | Taf5 | 0.027 | Old |
| 0.240309 | 2.904049 | 0.027 | 0 | 1 | 14 | Clcf1 | 0.027 | Old |
| 0.240309 | 2.897955 | 0.027 | 0 | 1 | 14 | Gm10710 | 0.027 | Old |
| 0.240309 | 2.888825 | 0.027 | 0 | 1 | 14 | Sdhaf3 | 0.027 | Old |
| 0.240309 | 2.88268 | 0.027 | 0 | 1 | 14 | Tmem229a | 0.027 | Old |
| 0.240309 | 2.857092 | 0.027 | 0 | 1 | 14 | Ttyh1 | 0.027 | Old |
| 0.240309 | 2.855198 | 0.027 | 0 | 1 | 14 | Surf6 | 0.027 | Old |
| 0.240309 | 2.854687 | 0.027 | 0 | 1 | 14 | Ric3 | 0.027 | Old |
| 0.240309 | 2.844776 | 0.027 | 0 | 1 | 14 | Enox1 | 0.027 | Old |
| 0.240309 | 2.842163 | 0.027 | 0 | 1 | 14 | Tmem98 | 0.027 | Old |
| 0.240309 | 2.824114 | 0.027 | 0 | 1 | 14 | Foxf1 | 0.027 | Old |
| 0.240309 | 2.79797 | 0.027 | 0 | 1 | 14 | Hist1h1e | 0.027 | Old |
| 0.240309 | 2.787874 | 0.027 | 0 | 1 | 14 | Pcdhb12 | 0.027 | Old |
| 0.240309 | 2.761368 | 0.027 | 0 | 1 | 14 | Gm42595 | 0.027 | Old |
| 0.240309 | 2.761199 | 0.027 | 0 | 1 | 14 | Gpr19 | 0.027 | Old |
| 0.240309 | 2.750274 | 0.027 | 0 | 1 | 14 | 2810021J2 | 0.027 | Old |
| 0.240309 | 2.683484 | 0.027 | 0 | 1 | 14 | Traf1 | 0.027 | Old |

| | | | | | | | | |
|---|---|---|---|---|---|---|---|---|
| 0.240309 | 2.679326 | 0.027 | 0 | 1 | 14 | Ifit3 | 0.027 | Old |
| 0.240309 | 2.673987 | 0.027 | 0 | 1 | 14 | Slc15a2 | 0.027 | Old |
| 0.240309 | 2.666406 | 0.027 | 0 | 1 | 14 | Zfp691 | 0.027 | Old |
| 0.240309 | 2.656866 | 0.027 | 0 | 1 | 14 | Ubtd1 | 0.027 | Old |
| 0.240309 | 2.654075 | 0.027 | 0 | 1 | 14 | Spata5 | 0.027 | Old |
| 0.240309 | 2.652365 | 0.027 | 0 | 1 | 14 | Mtrf1 | 0.027 | Old |
| 0.240309 | 2.652245 | 0.027 | 0 | 1 | 14 | Klhl11 | 0.027 | Old |
| 0.240309 | 2.651012 | 0.027 | 0 | 1 | 14 | Cyfip2 | 0.027 | Old |
| 0.240309 | 2.643947 | 0.027 | 0 | 1 | 14 | Syn3 | 0.027 | Old |
| 0.240309 | 2.626214 | 0.027 | 0 | 1 | 14 | 1600020E0 | 0.027 | Old |
| 0.240309 | 2.576787 | 0.027 | 0 | 1 | 14 | Mmaa | 0.027 | Old |
| 0.240309 | 2.563707 | 0.027 | 0 | 1 | 14 | Foxc2 | 0.027 | Old |
| 0.240309 | 2.557212 | 0.027 | 0 | 1 | 14 | Arid3a | 0.027 | Old |
| 0.240309 | 2.536511 | 0.027 | 0 | 1 | 14 | Astn2 | 0.027 | Old |
| 0.240309 | 2.532895 | 0.027 | 0 | 1 | 14 | Mis18a | 0.027 | Old |
| 0.240309 | 2.501602 | 0.027 | 0 | 1 | 14 | Zfp953 | 0.027 | Old |
| 0.240309 | 2.50022 | 0.027 | 0 | 1 | 14 | Hddc3 | 0.027 | Old |
| 0.240309 | 2.479471 | 0.027 | 0 | 1 | 14 | Acat2 | 0.027 | Old |
| 0.240309 | 2.446965 | 0.027 | 0 | 1 | 14 | Nnt | 0.027 | Old |
| 0.240309 | 2.43408 | 0.027 | 0 | 1 | 14 | Galnt12 | 0.027 | Old |
| 0.240309 | 2.428571 | 0.027 | 0 | 1 | 14 | 1700030K0 | 0.027 | Old |
| 0.240309 | 2.40706 | 0.027 | 0 | 1 | 14 | Pcdhga9 | 0.027 | Old |
| 0.240309 | 2.402845 | 0.027 | 0 | 1 | 14 | Lig4 | 0.027 | Old |
| 0.240309 | 2.395966 | 0.027 | 0 | 1 | 14 | Ccdc152 | 0.027 | Old |
| 0.240309 | 2.381919 | 0.027 | 0 | 1 | 14 | Fam35a | 0.027 | Old |
| 0.240309 | 2.370958 | 0.027 | 0 | 1 | 14 | Orai2 | 0.027 | Old |
| 0.240309 | 2.359025 | 0.027 | 0 | 1 | 14 | Lmnb2 | 0.027 | Old |
| 0.240309 | 2.334292 | 0.027 | 0 | 1 | 14 | Hlx | 0.027 | Old |
| 0.240309 | 2.327301 | 0.027 | 0 | 1 | 14 | Pira2 | 0.027 | Old |
| 0.240309 | 2.273851 | 0.027 | 0 | 1 | 14 | Zfp811 | 0.027 | Old |
| 0.240309 | 2.258741 | 0.027 | 0 | 1 | 14 | Mief2 | 0.027 | Old |
| 0.240309 | 2.256209 | 0.027 | 0 | 1 | 14 | Ttll5 | 0.027 | Old |
| 0.240309 | 2.250556 | 0.027 | 0 | 1 | 14 | Adck2 | 0.027 | Old |
| 0.240309 | 2.246426 | 0.027 | 0 | 1 | 14 | Haus5 | 0.027 | Old |
| 0.240309 | 2.226908 | 0.027 | 0 | 1 | 14 | BC022687 | 0.027 | Old |
| 0.240309 | 2.183906 | 0.027 | 0 | 1 | 14 | Ephx4 | 0.027 | Old |
| 0.240309 | 2.178912 | 0.027 | 0 | 1 | 14 | Zfp111 | 0.027 | Old |
| 0.240309 | 2.155181 | 0.027 | 0 | 1 | 14 | AC168977. | 0.027 | Old |
| 0.240309 | 2.14384 | 0.027 | 0 | 1 | 14 | Gm15706 | 0.027 | Old |
| 0.240309 | 2.143054 | 0.027 | 0 | 1 | 14 | Ptgfr | 0.027 | Old |
| 0.240309 | 2.072147 | 0.027 | 0 | 1 | 14 | Gm10330 | 0.027 | Old |
| 0.240309 | 2.036916 | 0.027 | 0 | 1 | 14 | Zfp873 | 0.027 | Old |
| 0.240309 | 2.028476 | 0.027 | 0 | 1 | 14 | Gm4425 | 0.027 | Old |
| 0.240309 | 1.981011 | 0.027 | 0 | 1 | 14 | Mrgpre | 0.027 | Old |
| 0.240309 | 1.942656 | 0.027 | 0 | 1 | 14 | C1rl | 0.027 | Old |

| | | | | | | | | |
|---|---|---|---|---|---|---|---|---|
| 0.240309 | 1.941381 | 0.027 | 0 | 1 | 14 | Zfp799 | 0.027 | Old |
| 0.240309 | 1.863804 | 0.027 | 0 | 1 | 14 | Sat2 | 0.027 | Old |
| 0.240309 | 1.767479 | 0.027 | 0 | 1 | 14 | Fam69b | 0.027 | Old |
| 0.240318 | 1.658734 | 0.18 | 0.12 | 1 | 14 | Adamtsl4 | 0.06 | Old |
| 0.240545 | 0.962739 | 0.142 | 0.08 | 1 | 14 | Zfp646 | 0.062 | Old |
| 0.240551 | 1.74133 | 0.087 | 0.04 | 1 | 14 | Gtpbp8 | 0.047 | Old |
| 0.240551 | 0.494668 | 0.066 | 0.12 | 1 | 14 | Zfp719 | -0.054 | Old |
| 0.24061 | 0.108688 | 0.699 | 0.58 | 1 | 14 | Dpysl2 | 0.119 | Old |
| 0.240904 | 0.725804 | 0.328 | 0.24 | 1 | 14 | Sox4 | 0.088 | Old |
| 0.241378 | 0.680051 | 0.35 | 0.26 | 1 | 14 | Vwa1 | 0.09 | Old |
| 0.241848 | 1.613902 | 0.137 | 0.08 | 1 | 14 | Trp53inp1 | 0.057 | Old |
| 0.242377 | 1.138792 | 0.115 | 0.06 | 1 | 14 | Oasl2 | 0.055 | Old |
| 0.242606 | 2.255715 | 0.087 | 0.04 | 1 | 14 | Abcb10 | 0.047 | Old |
| 0.242606 | 2.14002 | 0.087 | 0.04 | 1 | 14 | Eif2b4 | 0.047 | Old |
| 0.243455 | 0.11412 | 0.47 | 0.38 | 1 | 14 | Sdf4 | 0.09 | Old |
| 0.243513 | 1.395739 | 0.137 | 0.08 | 1 | 14 | Styx | 0.057 | Old |
| 0.243561 | 0.367942 | 0.579 | 0.44 | 1 | 14 | Smoc2 | 0.139 | Old |
| 0.243953 | 2.668699 | 0.06 | 0.02 | 1 | 14 | Hist3h2a | 0.04 | Old |
| 0.243953 | 2.647759 | 0.06 | 0.02 | 1 | 14 | Mrps18b | 0.04 | Old |
| 0.243953 | 2.410581 | 0.06 | 0.02 | 1 | 14 | Abhd3 | 0.04 | Old |
| 0.244169 | 0.318043 | 0.683 | 0.54 | 1 | 14 | Socs3 | 0.143 | Old |
| 0.244189 | 1.729591 | 0.115 | 0.06 | 1 | 14 | Clk3 | 0.055 | Old |
| 0.244564 | 0.567907 | 0.344 | 0.26 | 1 | 14 | Tmem100 | 0.084 | Old |
| 0.244674 | 2.295755 | 0.087 | 0.04 | 1 | 14 | Erc1 | 0.047 | Old |
| 0.244788 | 0.298762 | 0.328 | 0.26 | 1 | 14 | Csgalnact2 | 0.068 | Old |
| 0.244916 | 0.774443 | 0.038 | 0.08 | 1 | 14 | Pacsin3 | -0.042 | Old |
| 0.244916 | 0.273549 | 0.038 | 0.08 | 1 | 14 | Nat9 | -0.042 | Old |
| 0.244916 | 0.126979 | 0.038 | 0.08 | 1 | 14 | Amd1 | -0.042 | Old |
| 0.245471 | 1.073315 | 0.224 | 0.16 | 1 | 14 | Btbd1 | 0.064 | Old |
| 0.245885 | 0.541376 | 0.197 | 0.12 | 1 | 14 | Itga11 | 0.077 | Old |
| 0.246463 | 3.4649 | 0.06 | 0.02 | 1 | 14 | Adhfe1 | 0.04 | Old |
| 0.246463 | 2.768169 | 0.06 | 0.02 | 1 | 14 | Rmi1 | 0.04 | Old |
| 0.246463 | 2.225598 | 0.06 | 0.02 | 1 | 14 | Gsdme | 0.04 | Old |
| 0.246463 | 1.984 | 0.06 | 0.02 | 1 | 14 | Ogg1 | 0.04 | Old |
| 0.246656 | 0.944876 | 0.262 | 0.2 | 1 | 14 | Mex3c | 0.062 | Old |
| 0.246671 | 0.391026 | 0.448 | 0.36 | 1 | 14 | Eif4ebp2 | 0.088 | Old |
| 0.246681 | 0.925439 | 0.186 | 0.12 | 1 | 14 | Vps36 | 0.066 | Old |
| 0.247061 | 0.838906 | 0.268 | 0.2 | 1 | 14 | Rab31 | 0.068 | Old |
| 0.247173 | 0.60887 | 0.464 | 0.42 | 1 | 14 | Hdac7 | 0.044 | Old |
| 0.247173 | 0.322453 | 0.47 | 0.4 | 1 | 14 | Lamb2 | 0.07 | Old |
| 0.247375 | 0.260459 | 0.235 | 0.16 | 1 | 14 | Bdp1 | 0.075 | Old |
| 0.247411 | 0.806955 | 0.273 | 0.2 | 1 | 14 | Nf1 | 0.073 | Old |
| 0.247713 | 0.618273 | 0.617 | 0.58 | 1 | 14 | Ctsd | 0.037 | Old |
| 0.247773 | 0.317779 | 0.339 | 0.26 | 1 | 14 | Usp4 | 0.079 | Old |
| 0.247843 | 1.457027 | 0.115 | 0.06 | 1 | 14 | Thoc6 | 0.055 | Old |

| | | | | | | | | |
|---|---|---|---|---|---|---|---|---|
| 0.248097 | 0.121917 | 0.082 | 0.14 | 1 | 14 | Sdccag8 | -0.058 | Old |
| 0.248159 | 0.421943 | 0.492 | 0.4 | 1 | 14 | Klf7 | 0.092 | Old |
| 0.248181 | 1.348781 | 0.158 | 0.1 | 1 | 14 | Twist2 | 0.058 | Old |
| 0.248477 | 1.000455 | 0.164 | 0.1 | 1 | 14 | Umad1 | 0.064 | Old |
| 0.248992 | 2.351148 | 0.06 | 0.02 | 1 | 14 | Gbp5 | 0.04 | Old |
| 0.248992 | 2.323636 | 0.06 | 0.02 | 1 | 14 | D3Ertd751 | 0.04 | Old |
| 0.248992 | 2.156636 | 0.06 | 0.02 | 1 | 14 | Ncapg2 | 0.04 | Old |
| 0.248992 | 2.024345 | 0.06 | 0.02 | 1 | 14 | Zscan12 | 0.04 | Old |
| 0.248992 | 1.987869 | 0.06 | 0.02 | 1 | 14 | 9130008F2 | 0.04 | Old |
| 0.248992 | 1.751986 | 0.06 | 0.02 | 1 | 14 | Rfc3 | 0.04 | Old |
| 0.249544 | 1.297309 | 0.235 | 0.18 | 1 | 14 | Rchy1 | 0.055 | Old |
| 0.249633 | 0.634345 | 0.251 | 0.18 | 1 | 14 | Pdcd10 | 0.071 | Old |
| 0.249664 | 0.882348 | 0.246 | 0.18 | 1 | 14 | Cxcl1 | 0.066 | Old |
| 0.250188 | 0.188535 | 0.137 | 0.22 | 1 | 14 | Fam204a | -0.083 | Old |
| 0.250252 | 0.256646 | 0.137 | 0.08 | 1 | 14 | Gpr180 | 0.057 | Old |
| 0.250303 | 0.764699 | 0.213 | 0.14 | 1 | 14 | Med21 | 0.073 | Old |
| 0.250467 | 0.86956 | 0.142 | 0.08 | 1 | 14 | Eif2ak1 | 0.062 | Old |
| 0.250923 | 1.709286 | 0.109 | 0.06 | 1 | 14 | Ccdc91 | 0.049 | Old |
| 0.250951 | 1.553277 | 0.087 | 0.04 | 1 | 14 | Vrk3 | 0.047 | Old |
| 0.251117 | 1.102901 | 0.208 | 0.14 | 1 | 14 | Cul4a | 0.068 | Old |
| 0.25145 | 0.11434 | 0.191 | 0.12 | 1 | 14 | Dhx38 | 0.071 | Old |
| 0.251539 | 2.758354 | 0.06 | 0.02 | 1 | 14 | Pdik1l | 0.04 | Old |
| 0.251539 | 1.960367 | 0.06 | 0.02 | 1 | 14 | Gin1 | 0.04 | Old |
| 0.251539 | 1.946571 | 0.06 | 0.02 | 1 | 14 | Dzank1 | 0.04 | Old |
| 0.251539 | 1.933486 | 0.06 | 0.02 | 1 | 14 | Zbtb5 | 0.04 | Old |
| 0.251539 | 1.929906 | 0.06 | 0.02 | 1 | 14 | Hinfp | 0.04 | Old |
| 0.253077 | 0.457491 | 0.317 | 0.24 | 1 | 14 | Zdhhc3 | 0.077 | Old |
| 0.253151 | 0.602353 | 0.443 | 0.38 | 1 | 14 | Mafg | 0.063 | Old |
| 0.253396 | 1.143649 | 0.115 | 0.06 | 1 | 14 | Gm2000 | 0.055 | Old |
| 0.253836 | 0.593923 | 0.142 | 0.08 | 1 | 14 | Lrrk2 | 0.062 | Old |
| 0.253997 | 1.254921 | 0.197 | 0.14 | 1 | 14 | Zkscan1 | 0.057 | Old |
| 0.254103 | 1.856838 | 0.06 | 0.02 | 1 | 14 | Fmn1 | 0.04 | Old |
| 0.254237 | 1.041385 | 0.279 | 0.22 | 1 | 14 | Ube2q1 | 0.059 | Old |
| 0.254571 | 0.379877 | 0.333 | 0.26 | 1 | 14 | Rrp1 | 0.073 | Old |
| 0.255198 | 1.377784 | 0.087 | 0.04 | 1 | 14 | Taf9b | 0.047 | Old |
| 0.255698 | 0.404576 | 0.361 | 0.28 | 1 | 14 | Sgce | 0.081 | Old |
| 0.255961 | 0.349267 | 0.213 | 0.14 | 1 | 14 | Gtpbp2 | 0.073 | Old |
| 0.256296 | 0.198232 | 0.607 | 0.56 | 1 | 14 | Nptn | 0.047 | Old |
| 0.256445 | 0.217978 | 0.224 | 0.16 | 1 | 14 | Pcyox1 | 0.064 | Old |
| 0.256461 | 0.351559 | 0.295 | 0.22 | 1 | 14 | D5Ertd579 | 0.075 | Old |
| 0.256686 | 2.104877 | 0.06 | 0.02 | 1 | 14 | Fbxo46 | 0.04 | Old |
| 0.256686 | 1.930394 | 0.06 | 0.02 | 1 | 14 | Suox | 0.04 | Old |
| 0.256686 | 1.863413 | 0.06 | 0.02 | 1 | 14 | Zfp954 | 0.04 | Old |
| 0.256686 | 1.664155 | 0.06 | 0.02 | 1 | 14 | Atm | 0.04 | Old |
| 0.25712 | 0.845285 | 0.137 | 0.08 | 1 | 14 | Adar | 0.057 | Old |

| | | | | | | | | |
|---|---|---|---|---|---|---|---|---|
| 0.257395 | 0.408039 | 0.12 | 0.06 | 1 | 14 | Sp140 | 0.06 | Old |
| 0.257721 | 0.465506 | 0.393 | 0.32 | 1 | 14 | Atf5 | 0.073 | Old |
| 0.258265 | 1.175968 | 0.18 | 0.12 | 1 | 14 | Inppl1 | 0.06 | Old |
| 0.258554 | 1.900135 | 0.109 | 0.06 | 1 | 14 | Rgs10 | 0.049 | Old |
| 0.258751 | 0.850856 | 0.443 | 0.4 | 1 | 14 | Dusp3 | 0.043 | Old |
| 0.258766 | 0.163999 | 0.645 | 0.54 | 1 | 14 | Serinc1 | 0.105 | Old |
| 0.258857 | 1.234222 | 0.137 | 0.08 | 1 | 14 | Spata6 | 0.057 | Old |
| 0.259288 | 1.432096 | 0.06 | 0.02 | 1 | 14 | 2-Sep | 0.04 | Old |
| 0.259288 | 1.389529 | 0.06 | 0.02 | 1 | 14 | Per2 | 0.04 | Old |
| 0.259288 | 1.189261 | 0.06 | 0.02 | 1 | 14 | Lif | 0.04 | Old |
| 0.259569 | 0.736422 | 0.29 | 0.22 | 1 | 14 | Snrpd1 | 0.07 | Old |
| 0.259605 | 0.34135 | 0.268 | 0.18 | 1 | 14 | Hoxc4 | 0.088 | Old |
| 0.26016 | 0.088037 | 0.393 | 0.3 | 1 | 14 | Ecm2 | 0.093 | Old |
| 0.260487 | 1.87153 | 0.109 | 0.06 | 1 | 14 | Tpbgl | 0.049 | Old |
| 0.260602 | 1.100702 | 0.137 | 0.08 | 1 | 14 | Ndfip1 | 0.057 | Old |
| 0.260756 | 1.500863 | 0.153 | 0.1 | 1 | 14 | Dtx3l | 0.053 | Old |
| 0.26086 | 0.017635 | 0.448 | 0.36 | 1 | 14 | Ndufc1 | 0.088 | Old |
| 0.261661 | 1.056615 | 0.087 | 0.04 | 1 | 14 | Cyp2s1 | 0.047 | Old |
| 0.261907 | 1.714204 | 0.06 | 0.02 | 1 | 14 | Rsad1 | 0.04 | Old |
| 0.261907 | 1.335838 | 0.06 | 0.02 | 1 | 14 | Mrpl49 | 0.04 | Old |
| 0.261907 | 1.195267 | 0.06 | 0.02 | 1 | 14 | Zfp825 | 0.04 | Old |
| 0.262049 | 1.12583 | 0.202 | 0.14 | 1 | 14 | Fez2 | 0.062 | Old |
| 0.262409 | 0.481122 | 0.437 | 0.38 | 1 | 14 | Degs1 | 0.057 | Old |
| 0.262842 | 1.133047 | 0.197 | 0.14 | 1 | 14 | Pfdn1 | 0.057 | Old |
| 0.263206 | 1.114181 | 0.257 | 0.2 | 1 | 14 | Cwf19l2 | 0.057 | Old |
| 0.26332 | 0.000942 | 0.055 | 0.1 | 1 | 14 | Eif2ak4 | -0.045 | Old |
| 0.263716 | 0.052653 | 0.082 | 0.14 | 1 | 14 | Cln8 | -0.058 | Old |
| 0.263955 | 0.048715 | 0.377 | 0.3 | 1 | 14 | Notch2 | 0.077 | Old |
| 0.264117 | 1.467803 | 0.137 | 0.08 | 1 | 14 | Cpox | 0.057 | Old |
| 0.264257 | 0.305234 | 0.333 | 0.26 | 1 | 14 | Fundc2 | 0.073 | Old |
| 0.264312 | 0.141621 | 0.251 | 0.18 | 1 | 14 | Fbl | 0.071 | Old |
| 0.264383 | 1.874247 | 0.109 | 0.06 | 1 | 14 | Zxdc | 0.049 | Old |
| 0.264448 | 1.114458 | 0.18 | 0.12 | 1 | 14 | Heatr5a | 0.06 | Old |
| 0.264448 | 1.081988 | 0.18 | 0.12 | 1 | 14 | Srsf4 | 0.06 | Old |
| 0.264544 | 1.24158 | 0.06 | 0.02 | 1 | 14 | 3110082I1 | 0.04 | Old |
| 0.264614 | 0.568879 | 0.213 | 0.14 | 1 | 14 | Hoxa7 | 0.073 | Old |
| 0.265337 | 0.503015 | 0.448 | 0.36 | 1 | 14 | Mrc2 | 0.088 | Old |
| 0.265356 | 0.103231 | 0.224 | 0.14 | 1 | 14 | Csrp2 | 0.084 | Old |
| 0.26582 | 0.528893 | 0.246 | 0.18 | 1 | 14 | Ppp1r12c | 0.066 | Old |
| 0.265931 | 1.233865 | 0.235 | 0.18 | 1 | 14 | Myd88 | 0.055 | Old |
| 0.266032 | 1.359454 | 0.087 | 0.04 | 1 | 14 | Dhx57 | 0.047 | Old |
| 0.266053 | 0.435451 | 0.432 | 0.36 | 1 | 14 | Mast4 | 0.072 | Old |
| 0.266069 | 0.971355 | 0.164 | 0.1 | 1 | 14 | Mplkip | 0.064 | Old |
| 0.266318 | 0.724631 | 0.372 | 0.32 | 1 | 14 | Arpc1a | 0.052 | Old |
| 0.266345 | 1.72788 | 0.109 | 0.06 | 1 | 14 | Dip2a | 0.049 | Old |

| | | | | | | | | |
|---|---|---|---|---|---|---|---|---|
| 0.2672 | 0.586907 | 0.06 | 0.02 | 1 | 14 | Slc35e2 | 0.04 | Old |
| 0.267294 | 1.193082 | 0.24 | 0.18 | 1 | 14 | Mapk6 | 0.06 | Old |
| 0.26771 | 0.735256 | 0.164 | 0.1 | 1 | 14 | Ppp1r8 | 0.064 | Old |
| 0.267887 | 0.570856 | 0.284 | 0.22 | 1 | 14 | Ift20 | 0.064 | Old |
| 0.2681 | 0.217369 | 0.055 | 0.1 | 1 | 14 | Sssca1 | -0.045 | Old |
| 0.268416 | 0.064151 | 0.251 | 0.18 | 1 | 14 | Nsmaf | 0.071 | Old |
| 0.268454 | 0.247114 | 0.284 | 0.2 | 1 | 14 | Prelid1 | 0.084 | Old |
| 0.26925 | 0.231406 | 0.705 | 0.56 | 1 | 14 | Pttg1ip | 0.145 | Old |
| 0.269413 | 0.360064 | 0.366 | 0.28 | 1 | 14 | Rassf1 | 0.086 | Old |
| 0.269419 | 1.011151 | 0.186 | 0.12 | 1 | 14 | Ube4a | 0.066 | Old |
| 0.269874 | 0.777777 | 0.06 | 0.02 | 1 | 14 | Itfg2 | 0.04 | Old |
| 0.269874 | 0.707843 | 0.06 | 0.02 | 1 | 14 | D2hgdh | 0.04 | Old |
| 0.270337 | 0.318165 | 0.607 | 0.5 | 1 | 14 | Tagln2 | 0.107 | Old |
| 0.270366 | 0.394105 | 0.098 | 0.16 | 1 | 14 | Eef2kmt | -0.062 | Old |
| 0.270733 | 1.00338 | 0.18 | 0.12 | 1 | 14 | Timm29 | 0.06 | Old |
| 0.271097 | 0.250529 | 0.284 | 0.2 | 1 | 14 | Sgsm2 | 0.084 | Old |
| 0.271246 | 0.87373 | 0.137 | 0.08 | 1 | 14 | Lsm14b | 0.057 | Old |
| 0.271569 | 0.053984 | 0.027 | 0.06 | 1 | 14 | Ccdc181 | -0.033 | Old |
| 0.272294 | 1.455856 | 0.109 | 0.06 | 1 | 14 | Spire1 | 0.049 | Old |
| 0.272567 | 0.321973 | 0.06 | 0.02 | 1 | 14 | Zfp933 | 0.04 | Old |
| 0.272666 | 0.913925 | 0.158 | 0.1 | 1 | 14 | Mogs | 0.058 | Old |
| 0.272666 | 0.306258 | 0.158 | 0.1 | 1 | 14 | Atf6b | 0.058 | Old |
| 0.272683 | 0.670107 | 0.087 | 0.04 | 1 | 14 | Zfp407 | 0.047 | Old |
| 0.272683 | 0.580608 | 0.087 | 0.04 | 1 | 14 | Gstz1 | 0.047 | Old |
| 0.272933 | 0.209269 | 0.115 | 0.18 | 1 | 14 | Ldlrap1 | -0.065 | Old |
| 0.272985 | 1.51082 | 0.131 | 0.08 | 1 | 14 | Ranbp10 | 0.051 | Old |
| 0.273673 | 0.361877 | 0.328 | 0.24 | 1 | 14 | Kirrel | 0.088 | Old |
| 0.273757 | 0.029276 | 0.284 | 0.2 | 1 | 14 | Psmb5 | 0.084 | Old |
| 0.273914 | 0.959721 | 0.18 | 0.12 | 1 | 14 | Mum1 | 0.06 | Old |
| 0.273928 | 0.419073 | 0.426 | 0.36 | 1 | 14 | Abca1 | 0.066 | Old |
| 0.274148 | 0.274024 | 0.098 | 0.16 | 1 | 14 | Ttpal | -0.062 | Old |
| 0.274148 | 0.184198 | 0.098 | 0.16 | 1 | 14 | Ptger4 | -0.062 | Old |
| 0.274193 | 0.576994 | 0.333 | 0.26 | 1 | 14 | Npdc1 | 0.073 | Old |
| 0.274302 | 0.183229 | 0.563 | 0.48 | 1 | 14 | Nucks1 | 0.083 | Old |
| 0.274497 | 0.615014 | 0.404 | 0.34 | 1 | 14 | Ergic1 | 0.064 | Old |
| 0.274925 | 0.894467 | 0.087 | 0.04 | 1 | 14 | Zfp335 | 0.047 | Old |
| 0.274925 | 0.83589 | 0.087 | 0.04 | 1 | 14 | Slc15a4 | 0.047 | Old |
| 0.274942 | 0.509184 | 0.322 | 0.26 | 1 | 14 | Uqcrc2 | 0.062 | Old |
| 0.275068 | 0.109075 | 0.661 | 0.58 | 1 | 14 | Thra | 0.081 | Old |
| 0.275112 | 0.260683 | 0.503 | 0.4 | 1 | 14 | Hspd1 | 0.103 | Old |
| 0.275821 | 0.01382 | 0.164 | 0.24 | 1 | 14 | Sucla2 | -0.076 | Old |
| 0.276197 | 0.767998 | 0.301 | 0.24 | 1 | 14 | Chmp3 | 0.061 | Old |
| 0.276309 | 1.491418 | 0.109 | 0.06 | 1 | 14 | Mettl6 | 0.049 | Old |
| 0.276474 | 1.083911 | 0.284 | 0.24 | 1 | 14 | Dcaf7 | 0.044 | Old |
| 0.276564 | 0.554664 | 0.344 | 0.28 | 1 | 14 | Atxn2 | 0.064 | Old |

| | | | | | | | | |
|---|---|---|---|---|---|---|---|---|
| 0.27667 | 1.656692 | 0.131 | 0.08 | 1 | 14 | Polr1c | 0.051 | Old |
| 0.276677 | 0.754226 | 0.137 | 0.08 | 1 | 14 | Armc5 | 0.057 | Old |
| 0.277838 | 0.60152 | 0.055 | 0.1 | 1 | 14 | Kank3 | -0.045 | Old |
| 0.277866 | 0.158093 | 0.781 | 0.72 | 1 | 14 | Sem1 | 0.061 | Old |
| 0.27805 | 1.280791 | 0.197 | 0.14 | 1 | 14 | Eci2 | 0.057 | Old |
| 0.278167 | 1.317133 | 0.027 | 0.06 | 1 | 14 | Npy1r | -0.033 | Old |
| 0.278167 | 0.763886 | 0.027 | 0.06 | 1 | 14 | Stmn3 | -0.033 | Old |
| 0.278257 | 0.017403 | 0.891 | 0.8 | 1 | 14 | Rack1 | 0.091 | Old |
| 0.278266 | 2.743 | 0.082 | 0.04 | 1 | 14 | Ppm1f | 0.042 | Old |
| 0.278266 | 2.595899 | 0.082 | 0.04 | 1 | 14 | Bloc1s4 | 0.042 | Old |
| 0.278332 | 1.374508 | 0.109 | 0.06 | 1 | 14 | Cmc2 | 0.049 | Old |
| 0.279446 | 0.846739 | 0.087 | 0.04 | 1 | 14 | Tmem170b | 0.047 | Old |
| 0.280115 | 0.206771 | 0.115 | 0.18 | 1 | 14 | Slc38a9 | -0.065 | Old |
| 0.280193 | 0.171047 | 0.869 | 0.82 | 1 | 14 | Rpl26 | 0.049 | Old |
| 0.28021 | 0.724249 | 0.23 | 0.16 | 1 | 14 | Stx2 | 0.07 | Old |
| 0.280339 | 0.891265 | 0.137 | 0.08 | 1 | 14 | Tcirg1 | 0.057 | Old |
| 0.280528 | 0.123074 | 0.175 | 0.1 | 1 | 14 | Cenpx | 0.075 | Old |
| 0.2806 | 2.27475 | 0.082 | 0.04 | 1 | 14 | Copg2 | 0.042 | Old |
| 0.2806 | 2.177474 | 0.082 | 0.04 | 1 | 14 | Galt | 0.042 | Old |
| 0.281183 | 1.092512 | 0.158 | 0.1 | 1 | 14 | Atp9a | 0.058 | Old |
| 0.281183 | 0.880708 | 0.158 | 0.1 | 1 | 14 | Il17rc | 0.058 | Old |
| 0.281183 | 0.729069 | 0.158 | 0.1 | 1 | 14 | Ahi1 | 0.058 | Old |
| 0.281506 | 0.578858 | 0.027 | 0.06 | 1 | 14 | Mtfmt | -0.033 | Old |
| 0.281506 | 0.35502 | 0.027 | 0.06 | 1 | 14 | Cdc7 | -0.033 | Old |
| 0.281506 | 0.04267 | 0.027 | 0.06 | 1 | 14 | Cep41 | -0.033 | Old |
| 0.281634 | 0.534422 | 0.399 | 0.32 | 1 | 14 | Sptan1 | 0.079 | Old |
| 0.282147 | 2.045 | 0.126 | 0.08 | 1 | 14 | Kif1bp | 0.046 | Old |
| 0.282408 | 0.970056 | 0.109 | 0.06 | 1 | 14 | Ltv1 | 0.049 | Old |
| 0.282534 | 0.58131 | 0.284 | 0.22 | 1 | 14 | Dpy19l1 | 0.064 | Old |
| 0.282843 | 0.503815 | 0.519 | 0.46 | 1 | 14 | Slc29a1 | 0.059 | Old |
| 0.282893 | 0.793282 | 0.24 | 0.18 | 1 | 14 | Hibadh | 0.06 | Old |
| 0.282948 | 2.480132 | 0.082 | 0.04 | 1 | 14 | Zfp788 | 0.042 | Old |
| 0.282948 | 2.197867 | 0.082 | 0.04 | 1 | 14 | Slc25a44 | 0.042 | Old |
| 0.282948 | 2.140445 | 0.082 | 0.04 | 1 | 14 | Fuom | 0.042 | Old |
| 0.282948 | 1.898002 | 0.082 | 0.04 | 1 | 14 | Urm1 | 0.042 | Old |
| 0.283264 | 0.734439 | 0.372 | 0.32 | 1 | 14 | Rer1 | 0.052 | Old |
| 0.283555 | 0.149051 | 0.344 | 0.3 | 1 | 14 | Rad21 | 0.044 | Old |
| 0.283609 | 0.150297 | 0.18 | 0.12 | 1 | 14 | Kctd11 | 0.06 | Old |
| 0.283729 | 0.657039 | 0.377 | 0.32 | 1 | 14 | Mga | 0.057 | Old |
| 0.283763 | 0.364936 | 0.098 | 0.16 | 1 | 14 | Thoc1 | -0.062 | Old |
| 0.283823 | 0.228365 | 0.257 | 0.2 | 1 | 14 | Wtap | 0.057 | Old |
| 0.283859 | 0.498385 | 0.142 | 0.22 | 1 | 14 | Rnf141 | -0.078 | Old |
| 0.284034 | 0.767338 | 0.137 | 0.08 | 1 | 14 | Ccbe1 | 0.057 | Old |
| 0.284141 | 1.343826 | 0.131 | 0.08 | 1 | 14 | Gtf3c3 | 0.051 | Old |
| 0.284162 | 0.125147 | 0.377 | 0.34 | 1 | 14 | Rbbp6 | 0.037 | Old |

| | | | | | | | | |
|---|---|---|---|---|---|---|---|---|
| 0.284462 | 1.176476 | 0.109 | 0.06 | 1 | 14 | Vps53 | 0.049 | Old |
| 0.2845 | 0.735818 | 0.164 | 0.1 | 1 | 14 | Lemd3 | 0.064 | Old |
| 0.284647 | 0.899771 | 0.153 | 0.1 | 1 | 14 | Mms19 | 0.053 | Old |
| 0.285178 | 0.032433 | 0.393 | 0.3 | 1 | 14 | Fbxl3 | 0.093 | Old |
| 0.285308 | 2.0264 | 0.082 | 0.04 | 1 | 14 | Tmem192 | 0.042 | Old |
| 0.285308 | 1.934205 | 0.082 | 0.04 | 1 | 14 | Vps33a | 0.042 | Old |
| 0.285894 | 0.47996 | 0.137 | 0.08 | 1 | 14 | Casd1 | 0.057 | Old |
| 0.28603 | 1.106431 | 0.131 | 0.08 | 1 | 14 | Git1 | 0.051 | Old |
| 0.286597 | 1.022243 | 0.213 | 0.16 | 1 | 14 | Tmem9b | 0.053 | Old |
| 0.286653 | 0.924964 | 0.257 | 0.2 | 1 | 14 | Creb1 | 0.057 | Old |
| 0.287425 | 0.433701 | 0.115 | 0.18 | 1 | 14 | Egr2 | -0.065 | Old |
| 0.287425 | 0.125418 | 0.115 | 0.18 | 1 | 14 | Retreg3 | -0.065 | Old |
| 0.287682 | 1.911007 | 0.082 | 0.04 | 1 | 14 | Traf5 | 0.042 | Old |
| 0.288267 | 0.399647 | 0.027 | 0.06 | 1 | 14 | Xrcc4 | -0.033 | Old |
| 0.288267 | 0.257675 | 0.027 | 0.06 | 1 | 14 | Alg3 | -0.033 | Old |
| 0.288381 | 0.448649 | 0.372 | 0.3 | 1 | 14 | Tmod2 | 0.072 | Old |
| 0.28984 | 0.08014 | 0.104 | 0.16 | 1 | 14 | Cpt2 | -0.056 | Old |
| 0.289879 | 0.9074 | 0.158 | 0.1 | 1 | 14 | Klhl2 | 0.058 | Old |
| 0.289917 | 0.632578 | 0.246 | 0.18 | 1 | 14 | Becn1 | 0.066 | Old |
| 0.28993 | 0.616717 | 0.35 | 0.28 | 1 | 14 | Arhgap21 | 0.07 | Old |
| 0.290346 | 1.116364 | 0.055 | 0.1 | 1 | 14 | Fiz1 | -0.045 | Old |
| 0.290518 | 2.508383 | 0.104 | 0.06 | 1 | 14 | Zfp866 | 0.044 | Old |
| 0.291021 | 0.131214 | 0.415 | 0.32 | 1 | 14 | Nsd3 | 0.095 | Old |
| 0.291118 | 0.129632 | 0.344 | 0.26 | 1 | 14 | Snx4 | 0.084 | Old |
| 0.291312 | 0.842838 | 0.268 | 0.2 | 1 | 14 | Trim56 | 0.068 | Old |
| 0.291686 | 0.911154 | 0.191 | 0.14 | 1 | 14 | Ccdc174 | 0.051 | Old |
| 0.291714 | 1.019718 | 0.175 | 0.12 | 1 | 14 | Phf10 | 0.055 | Old |
| 0.291786 | 1.784188 | 0.126 | 0.08 | 1 | 14 | Sms | 0.046 | Old |
| 0.29247 | 1.811252 | 0.082 | 0.04 | 1 | 14 | Eid2 | 0.042 | Old |
| 0.2925 | 0.040283 | 0.219 | 0.3 | 1 | 14 | Sesn3 | -0.081 | Old |
| 0.292578 | 0.96162 | 0.257 | 0.22 | 1 | 14 | Pcf11 | 0.037 | Old |
| 0.292653 | 2.365201 | 0.104 | 0.06 | 1 | 14 | Slc41a2 | 0.044 | Old |
| 0.292742 | 0.16596 | 0.656 | 0.56 | 1 | 14 | Camk2n1 | 0.096 | Old |
| 0.292892 | 0.275425 | 0.055 | 0.1 | 1 | 14 | Vav2 | -0.045 | Old |
| 0.292949 | 0.328468 | 0.481 | 0.4 | 1 | 14 | Tspo | 0.081 | Old |
| 0.293278 | 0.407317 | 0.295 | 0.22 | 1 | 14 | Tcaf1 | 0.075 | Old |
| 0.293402 | 1.110815 | 0.175 | 0.12 | 1 | 14 | Fap | 0.055 | Old |
| 0.293415 | 0.690261 | 0.137 | 0.08 | 1 | 14 | 1700017B0 | 0.057 | Old |
| 0.293528 | 1.068365 | 0.153 | 0.1 | 1 | 14 | Tnip1 | 0.053 | Old |
| 0.293603 | 0.165581 | 0.098 | 0.16 | 1 | 14 | Nod1 | -0.062 | Old |
| 0.293671 | 1.432323 | 0.131 | 0.08 | 1 | 14 | Tbpl1 | 0.051 | Old |
| 0.293695 | 2.801745 | 0.055 | 0.02 | 1 | 14 | Rgs1 | 0.035 | Old |
| 0.293695 | 2.763696 | 0.055 | 0.02 | 1 | 14 | Mlkl | 0.035 | Old |
| 0.293695 | 2.709191 | 0.055 | 0.02 | 1 | 14 | Gmds | 0.035 | Old |
| 0.293695 | 2.655376 | 0.055 | 0.02 | 1 | 14 | Arhgap44 | 0.035 | Old |

| | | | | | | | | |
|---|---|---|---|---|---|---|---|---|
| 0.293695 | 2.598267 | 0.055 | 0.02 | 1 | 14 | Fam213b | 0.035 | Old |
| 0.293695 | 2.414513 | 0.055 | 0.02 | 1 | 14 | Mtcp1 | 0.035 | Old |
| 0.293695 | 2.337309 | 0.055 | 0.02 | 1 | 14 | Foxred1 | 0.035 | Old |
| 0.293695 | 2.312545 | 0.055 | 0.02 | 1 | 14 | Ift88 | 0.035 | Old |
| 0.293695 | 2.309284 | 0.055 | 0.02 | 1 | 14 | Kcnk2 | 0.035 | Old |
| 0.293695 | 2.259677 | 0.055 | 0.02 | 1 | 14 | Tet1 | 0.035 | Old |
| 0.293695 | 2.035228 | 0.055 | 0.02 | 1 | 14 | Ypel4 | 0.035 | Old |
| 0.294674 | 0.261428 | 0.705 | 0.6 | 1 | 14 | Lnpep | 0.105 | Old |
| 0.294883 | 1.570123 | 0.082 | 0.04 | 1 | 14 | Rdh11 | 0.042 | Old |
| 0.294903 | 0.403432 | 0.213 | 0.14 | 1 | 14 | Socs5 | 0.073 | Old |
| 0.295096 | 1.084688 | 0.175 | 0.12 | 1 | 14 | Plp2 | 0.055 | Old |
| 0.295131 | 3.82136 | 0.022 | 0 | 1 | 14 | Cldn10 | 0.022 | Old |
| 0.295131 | 3.619921 | 0.022 | 0 | 1 | 14 | Psd | 0.022 | Old |
| 0.295131 | 3.605265 | 0.022 | 0 | 1 | 14 | Hdx | 0.022 | Old |
| 0.295131 | 3.594956 | 0.022 | 0 | 1 | 14 | Serpinb6b | 0.022 | Old |
| 0.295131 | 3.487069 | 0.022 | 0 | 1 | 14 | Gm26771 | 0.022 | Old |
| 0.295131 | 3.435905 | 0.022 | 0 | 1 | 14 | 4632404H1 | 0.022 | Old |
| 0.295131 | 3.376724 | 0.022 | 0 | 1 | 14 | Rgs17 | 0.022 | Old |
| 0.295131 | 3.281347 | 0.022 | 0 | 1 | 14 | Tbrg4 | 0.022 | Old |
| 0.295131 | 3.163768 | 0.022 | 0 | 1 | 14 | D130020L0 | 0.022 | Old |
| 0.295131 | 3.102951 | 0.022 | 0 | 1 | 14 | Pcdhb4 | 0.022 | Old |
| 0.295131 | 3.065683 | 0.022 | 0 | 1 | 14 | Miga1 | 0.022 | Old |
| 0.295131 | 3.024073 | 0.022 | 0 | 1 | 14 | Itgal | 0.022 | Old |
| 0.295131 | 3.013155 | 0.022 | 0 | 1 | 14 | Scgb1a1 | 0.022 | Old |
| 0.295131 | 2.91496 | 0.022 | 0 | 1 | 14 | Slc45a3 | 0.022 | Old |
| 0.295131 | 2.886132 | 0.022 | 0 | 1 | 14 | Hoxa6 | 0.022 | Old |
| 0.295131 | 2.885769 | 0.022 | 0 | 1 | 14 | Chst10 | 0.022 | Old |
| 0.295131 | 2.851571 | 0.022 | 0 | 1 | 14 | Dlg2 | 0.022 | Old |
| 0.295131 | 2.842068 | 0.022 | 0 | 1 | 14 | Pard6a | 0.022 | Old |
| 0.295131 | 2.828966 | 0.022 | 0 | 1 | 14 | Lyl1 | 0.022 | Old |
| 0.295131 | 2.789655 | 0.022 | 0 | 1 | 14 | Trdmt1 | 0.022 | Old |
| 0.295131 | 2.778482 | 0.022 | 0 | 1 | 14 | Scn3b | 0.022 | Old |
| 0.295131 | 2.768953 | 0.022 | 0 | 1 | 14 | Cd72 | 0.022 | Old |
| 0.295131 | 2.759832 | 0.022 | 0 | 1 | 14 | Brms1 | 0.022 | Old |
| 0.295131 | 2.748393 | 0.022 | 0 | 1 | 14 | Ccdc173 | 0.022 | Old |
| 0.295131 | 2.729589 | 0.022 | 0 | 1 | 14 | Fkbp1b | 0.022 | Old |
| 0.295131 | 2.696962 | 0.022 | 0 | 1 | 14 | Adcy4 | 0.022 | Old |
| 0.295131 | 2.692412 | 0.022 | 0 | 1 | 14 | Tmem88b | 0.022 | Old |
| 0.295131 | 2.688429 | 0.022 | 0 | 1 | 14 | Gm20707 | 0.022 | Old |
| 0.295131 | 2.663237 | 0.022 | 0 | 1 | 14 | Mettl15 | 0.022 | Old |
| 0.295131 | 2.660277 | 0.022 | 0 | 1 | 14 | Fam110a | 0.022 | Old |
| 0.295131 | 2.658753 | 0.022 | 0 | 1 | 14 | Cptp | 0.022 | Old |
| 0.295131 | 2.64577 | 0.022 | 0 | 1 | 14 | Nos1 | 0.022 | Old |
| 0.295131 | 2.620464 | 0.022 | 0 | 1 | 14 | Dffb | 0.022 | Old |
| 0.295131 | 2.601464 | 0.022 | 0 | 1 | 14 | Lvrn | 0.022 | Old |

| | | | | | | | | |
|---|---|---|---|---|---|---|---|---|
| 0.295131 | 2.580777 | 0.022 | 0 | 1 | 14 | Bbs12 | 0.022 | Old |
| 0.295131 | 2.580558 | 0.022 | 0 | 1 | 14 | Fras1 | 0.022 | Old |
| 0.295131 | 2.573464 | 0.022 | 0 | 1 | 14 | Lonrf3 | 0.022 | Old |
| 0.295131 | 2.573313 | 0.022 | 0 | 1 | 14 | Tedc2 | 0.022 | Old |
| 0.295131 | 2.572815 | 0.022 | 0 | 1 | 14 | Best1 | 0.022 | Old |
| 0.295131 | 2.544791 | 0.022 | 0 | 1 | 14 | Zscan20 | 0.022 | Old |
| 0.295131 | 2.537556 | 0.022 | 0 | 1 | 14 | Mtif3 | 0.022 | Old |
| 0.295131 | 2.514512 | 0.022 | 0 | 1 | 14 | Clec5a | 0.022 | Old |
| 0.295131 | 2.513519 | 0.022 | 0 | 1 | 14 | Bend7 | 0.022 | Old |
| 0.295131 | 2.49756 | 0.022 | 0 | 1 | 14 | Cyb561d1 | 0.022 | Old |
| 0.295131 | 2.496598 | 0.022 | 0 | 1 | 14 | Mical2 | 0.022 | Old |
| 0.295131 | 2.494538 | 0.022 | 0 | 1 | 14 | Il20rb | 0.022 | Old |
| 0.295131 | 2.48677 | 0.022 | 0 | 1 | 14 | Rabggta | 0.022 | Old |
| 0.295131 | 2.478341 | 0.022 | 0 | 1 | 14 | Ccno | 0.022 | Old |
| 0.295131 | 2.461924 | 0.022 | 0 | 1 | 14 | Fen1 | 0.022 | Old |
| 0.295131 | 2.435107 | 0.022 | 0 | 1 | 14 | BC067074 | 0.022 | Old |
| 0.295131 | 2.421526 | 0.022 | 0 | 1 | 14 | Syt3 | 0.022 | Old |
| 0.295131 | 2.403166 | 0.022 | 0 | 1 | 14 | Dus4l | 0.022 | Old |
| 0.295131 | 2.397283 | 0.022 | 0 | 1 | 14 | Fbxo36 | 0.022 | Old |
| 0.295131 | 2.397021 | 0.022 | 0 | 1 | 14 | Zfp619 | 0.022 | Old |
| 0.295131 | 2.386585 | 0.022 | 0 | 1 | 14 | Nupl2 | 0.022 | Old |
| 0.295131 | 2.383062 | 0.022 | 0 | 1 | 14 | Frg2f1 | 0.022 | Old |
| 0.295131 | 2.358631 | 0.022 | 0 | 1 | 14 | Cldn3 | 0.022 | Old |
| 0.295131 | 2.358629 | 0.022 | 0 | 1 | 14 | Zranb3 | 0.022 | Old |
| 0.295131 | 2.3581 | 0.022 | 0 | 1 | 14 | Slc25a26 | 0.022 | Old |
| 0.295131 | 2.330722 | 0.022 | 0 | 1 | 14 | Pcdhgc5 | 0.022 | Old |
| 0.295131 | 2.320723 | 0.022 | 0 | 1 | 14 | Prrt4 | 0.022 | Old |
| 0.295131 | 2.310556 | 0.022 | 0 | 1 | 14 | Slc7a5 | 0.022 | Old |
| 0.295131 | 2.291769 | 0.022 | 0 | 1 | 14 | Dcdc2b | 0.022 | Old |
| 0.295131 | 2.289702 | 0.022 | 0 | 1 | 14 | Hyls1 | 0.022 | Old |
| 0.295131 | 2.282818 | 0.022 | 0 | 1 | 14 | Gm43936 | 0.022 | Old |
| 0.295131 | 2.278192 | 0.022 | 0 | 1 | 14 | Slc9a5 | 0.022 | Old |
| 0.295131 | 2.251217 | 0.022 | 0 | 1 | 14 | Wdr59 | 0.022 | Old |
| 0.295131 | 2.241958 | 0.022 | 0 | 1 | 14 | Gria4 | 0.022 | Old |
| 0.295131 | 2.223804 | 0.022 | 0 | 1 | 14 | Wdr31 | 0.022 | Old |
| 0.295131 | 2.223328 | 0.022 | 0 | 1 | 14 | Efna2 | 0.022 | Old |
| 0.295131 | 2.180302 | 0.022 | 0 | 1 | 14 | Rpusd1 | 0.022 | Old |
| 0.295131 | 2.172995 | 0.022 | 0 | 1 | 14 | Actn3 | 0.022 | Old |
| 0.295131 | 2.141257 | 0.022 | 0 | 1 | 14 | Kif1a | 0.022 | Old |
| 0.295131 | 2.119701 | 0.022 | 0 | 1 | 14 | Mypopos | 0.022 | Old |
| 0.295131 | 2.110551 | 0.022 | 0 | 1 | 14 | Tub | 0.022 | Old |
| 0.295131 | 2.101378 | 0.022 | 0 | 1 | 14 | Zfp61 | 0.022 | Old |
| 0.295131 | 2.092848 | 0.022 | 0 | 1 | 14 | Gm17334 | 0.022 | Old |
| 0.295131 | 2.09007 | 0.022 | 0 | 1 | 14 | Slc1a4 | 0.022 | Old |
| 0.295131 | 2.054444 | 0.022 | 0 | 1 | 14 | Snapc4 | 0.022 | Old |

| | | | | | | | | |
|---|---|---|---|---|---|---|---|---|
| 0.295131 | 2.050206 | 0.022 | 0 | 1 | 14 | Oas3 | 0.022 | Old |
| 0.295131 | 2.032393 | 0.022 | 0 | 1 | 14 | Gm45605 | 0.022 | Old |
| 0.295131 | 2.031643 | 0.022 | 0 | 1 | 14 | Kifc2 | 0.022 | Old |
| 0.295131 | 2.013777 | 0.022 | 0 | 1 | 14 | Gm6563 | 0.022 | Old |
| 0.295131 | 2.005311 | 0.022 | 0 | 1 | 14 | BC065403 | 0.022 | Old |
| 0.295131 | 1.996956 | 0.022 | 0 | 1 | 14 | Zbtb42 | 0.022 | Old |
| 0.295131 | 1.990015 | 0.022 | 0 | 1 | 14 | Vsig10l | 0.022 | Old |
| 0.295131 | 1.959622 | 0.022 | 0 | 1 | 14 | Gm42517 | 0.022 | Old |
| 0.295131 | 1.939192 | 0.022 | 0 | 1 | 14 | Miat | 0.022 | Old |
| 0.295131 | 1.926694 | 0.022 | 0 | 1 | 14 | Patj | 0.022 | Old |
| 0.295131 | 1.899489 | 0.022 | 0 | 1 | 14 | A630089N( | 0.022 | Old |
| 0.295131 | 1.887275 | 0.022 | 0 | 1 | 14 | Btk | 0.022 | Old |
| 0.295131 | 1.882021 | 0.022 | 0 | 1 | 14 | Reep2 | 0.022 | Old |
| 0.295131 | 1.877817 | 0.022 | 0 | 1 | 14 | Hist2h3c2 | 0.022 | Old |
| 0.295131 | 1.839504 | 0.022 | 0 | 1 | 14 | Map2k3os | 0.022 | Old |
| 0.295131 | 1.823758 | 0.022 | 0 | 1 | 14 | Cdc42bpg | 0.022 | Old |
| 0.295131 | 1.798597 | 0.022 | 0 | 1 | 14 | Polh | 0.022 | Old |
| 0.295131 | 1.790023 | 0.022 | 0 | 1 | 14 | Epha1 | 0.022 | Old |
| 0.295131 | 1.750657 | 0.022 | 0 | 1 | 14 | Lmx1a | 0.022 | Old |
| 0.295131 | 1.724382 | 0.022 | 0 | 1 | 14 | Cd200r1 | 0.022 | Old |
| 0.295131 | 1.709053 | 0.022 | 0 | 1 | 14 | Mettl25 | 0.022 | Old |
| 0.295131 | 1.704768 | 0.022 | 0 | 1 | 14 | Carmil3 | 0.022 | Old |
| 0.295131 | 1.70163 | 0.022 | 0 | 1 | 14 | Rbm47 | 0.022 | Old |
| 0.295131 | 1.656878 | 0.022 | 0 | 1 | 14 | Dusp28 | 0.022 | Old |
| 0.295131 | 1.596294 | 0.022 | 0 | 1 | 14 | Fam109a | 0.022 | Old |
| 0.295131 | 1.493253 | 0.022 | 0 | 1 | 14 | Il7 | 0.022 | Old |
| 0.296463 | 0.815162 | 0.251 | 0.2 | 1 | 14 | Dirc2 | 0.051 | Old |
| 0.296666 | 3.017316 | 0.055 | 0.02 | 1 | 14 | Snupn | 0.035 | Old |
| 0.296666 | 2.958514 | 0.055 | 0.02 | 1 | 14 | Nemp1 | 0.035 | Old |
| 0.296666 | 2.228243 | 0.055 | 0.02 | 1 | 14 | Slc17a7 | 0.035 | Old |
| 0.296797 | 1.380773 | 0.175 | 0.12 | 1 | 14 | Plbd2 | 0.055 | Old |
| 0.296881 | 0.694505 | 0.186 | 0.12 | 1 | 14 | Jup | 0.066 | Old |
| 0.296889 | 0.67713 | 0.262 | 0.2 | 1 | 14 | Cnot3 | 0.062 | Old |
| 0.297132 | 1.095045 | 0.153 | 0.1 | 1 | 14 | Capn7 | 0.053 | Old |
| 0.29731 | 1.626515 | 0.082 | 0.04 | 1 | 14 | Paox | 0.042 | Old |
| 0.297495 | 0.981473 | 0.202 | 0.14 | 1 | 14 | Ndufaf8 | 0.062 | Old |
| 0.297495 | 0.834477 | 0.202 | 0.14 | 1 | 14 | Sec16a | 0.062 | Old |
| 0.298117 | 0.009734 | 0.694 | 0.58 | 1 | 14 | Cygb | 0.114 | Old |
| 0.298172 | 0.095367 | 0.454 | 0.36 | 1 | 14 | Rbx1 | 0.094 | Old |
| 0.298755 | 1.141221 | 0.158 | 0.1 | 1 | 14 | Pla2g16 | 0.058 | Old |
| 0.299116 | 1.275655 | 0.109 | 0.06 | 1 | 14 | Sp110 | 0.049 | Old |
| 0.299116 | 1.036382 | 0.109 | 0.06 | 1 | 14 | Pld2 | 0.049 | Old |
| 0.299143 | 0.677901 | 0.137 | 0.08 | 1 | 14 | Pgghg | 0.057 | Old |
| 0.299375 | 1.304459 | 0.235 | 0.18 | 1 | 14 | Adipor2 | 0.055 | Old |
| 0.299637 | 0.365805 | 0.355 | 0.28 | 1 | 14 | Dyrk2 | 0.075 | Old |

| | | | | | | | | |
|---|---|---|---|---|---|---|---|---|
| 0.299658 | 2.533426 | 0.055 | 0.02 | 1 | 14 | 1600014C1 | 0.035 | Old |
| 0.299658 | 2.067533 | 0.055 | 0.02 | 1 | 14 | Map9 | 0.035 | Old |
| 0.299751 | 1.396366 | 0.082 | 0.04 | 1 | 14 | Zbtb8os | 0.042 | Old |
| 0.299751 | 1.304708 | 0.082 | 0.04 | 1 | 14 | Endov | 0.042 | Old |
| 0.300422 | 0.396603 | 0.525 | 0.46 | 1 | 14 | Fam198b | 0.065 | Old |
| 0.300621 | 0.239788 | 0.055 | 0.1 | 1 | 14 | Cd300ld | -0.045 | Old |
| 0.300765 | 1.070326 | 0.153 | 0.1 | 1 | 14 | Phlpp2 | 0.053 | Old |
| 0.30125 | 0.849868 | 0.109 | 0.06 | 1 | 14 | Zfp384 | 0.049 | Old |
| 0.301282 | 0.730644 | 0.23 | 0.16 | 1 | 14 | Pld3 | 0.07 | Old |
| 0.301479 | 0.57405 | 0.333 | 0.26 | 1 | 14 | Clock | 0.073 | Old |
| 0.301648 | 0.252691 | 0.519 | 0.44 | 1 | 14 | Erp29 | 0.079 | Old |
| 0.301967 | 0.876452 | 0.197 | 0.14 | 1 | 14 | Il3ra | 0.057 | Old |
| 0.302204 | 1.56607 | 0.082 | 0.04 | 1 | 14 | Nudt19 | 0.042 | Old |
| 0.302355 | 0.212706 | 0.158 | 0.1 | 1 | 14 | Cd320 | 0.058 | Old |
| 0.302539 | 0.061421 | 1 | 0.94 | 1 | 14 | mt-Co1 | 0.06 | Old |
| 0.30267 | 1.679479 | 0.055 | 0.02 | 1 | 14 | Retsat | 0.035 | Old |
| 0.302827 | 1.266421 | 0.224 | 0.18 | 1 | 14 | Ankib1 | 0.044 | Old |
| 0.302862 | 0.537949 | 0.464 | 0.4 | 1 | 14 | Pltp | 0.064 | Old |
| 0.302979 | 0.059912 | 0.568 | 0.44 | 1 | 14 | Igf1 | 0.128 | Old |
| 0.303394 | 0.906304 | 0.109 | 0.06 | 1 | 14 | Lyn | 0.049 | Old |
| 0.303486 | 2.225819 | 0.104 | 0.06 | 1 | 14 | Scly | 0.044 | Old |
| 0.303528 | 1.80766 | 0.169 | 0.12 | 1 | 14 | Pts | 0.049 | Old |
| 0.303688 | 0.315518 | 0.18 | 0.12 | 1 | 14 | Tor1aip2 | 0.06 | Old |
| 0.30455 | 0.340106 | 0.219 | 0.16 | 1 | 14 | Chd1 | 0.059 | Old |
| 0.304671 | 1.437176 | 0.082 | 0.04 | 1 | 14 | Casp3 | 0.042 | Old |
| 0.304671 | 1.229853 | 0.082 | 0.04 | 1 | 14 | Nfkbib | 0.042 | Old |
| 0.304979 | 0.656897 | 0.344 | 0.28 | 1 | 14 | Smchd1 | 0.064 | Old |
| 0.305386 | 1.077779 | 0.131 | 0.08 | 1 | 14 | Trim30d | 0.051 | Old |
| 0.305398 | 0.796952 | 0.175 | 0.12 | 1 | 14 | Nmt1 | 0.055 | Old |
| 0.305548 | 0.71633 | 0.109 | 0.06 | 1 | 14 | Fam149a | 0.049 | Old |
| 0.305631 | 0.683761 | 0.279 | 0.22 | 1 | 14 | Pip4k2b | 0.059 | Old |
| 0.305639 | 1.385798 | 0.257 | 0.22 | 1 | 14 | Insig2 | 0.037 | Old |
| 0.305702 | 2.23696 | 0.055 | 0.02 | 1 | 14 | Rps6ka2 | 0.035 | Old |
| 0.305702 | 1.855449 | 0.055 | 0.02 | 1 | 14 | Nol9 | 0.035 | Old |
| 0.305702 | 1.417261 | 0.055 | 0.02 | 1 | 14 | Helb | 0.035 | Old |
| 0.305932 | 0.102498 | 0.23 | 0.16 | 1 | 14 | Lrrc41 | 0.07 | Old |
| 0.305984 | 0.309214 | 0.131 | 0.2 | 1 | 14 | Tcf20 | -0.069 | Old |
| 0.305998 | 0.057163 | 0.601 | 0.5 | 1 | 14 | Mxd4 | 0.101 | Old |
| 0.30616 | 0.091459 | 0.016 | 0.04 | 1 | 14 | Mlf1 | -0.024 | Old |
| 0.306257 | 0.031678 | 0.115 | 0.18 | 1 | 14 | Ptprc | -0.065 | Old |
| 0.306631 | 0.725275 | 0.311 | 0.24 | 1 | 14 | Ythdf1 | 0.071 | Old |
| 0.307152 | 1.356806 | 0.082 | 0.04 | 1 | 14 | Ncf1 | 0.042 | Old |
| 0.307809 | 0.709046 | 0.158 | 0.1 | 1 | 14 | Card10 | 0.058 | Old |
| 0.308187 | 0.52027 | 0.29 | 0.22 | 1 | 14 | Wbp2 | 0.07 | Old |
| 0.308418 | 1.136571 | 0.301 | 0.26 | 1 | 14 | Mfsd14b | 0.041 | Old |

| | | | | | | | | |
|---|---|---|---|---|---|---|---|---|
| 0.30857 | 1.020936 | 0.197 | 0.14 | 1 | 14 | Echs1 | 0.057 | Old |
| 0.308754 | 2.109169 | 0.055 | 0.02 | 1 | 14 | 4930404H1 | 0.035 | Old |
| 0.308754 | 1.950162 | 0.055 | 0.02 | 1 | 14 | Pcdhgc3 | 0.035 | Old |
| 0.309338 | 0.99931 | 0.273 | 0.22 | 1 | 14 | Trim47 | 0.053 | Old |
| 0.309645 | 1.137503 | 0.082 | 0.04 | 1 | 14 | Rps19bp1 | 0.042 | Old |
| 0.309645 | 1.049561 | 0.082 | 0.04 | 1 | 14 | Rbm10 | 0.042 | Old |
| 0.309833 | 0.707091 | 0.191 | 0.14 | 1 | 14 | Eno1 | 0.051 | Old |
| 0.309886 | 0.583268 | 0.109 | 0.06 | 1 | 14 | Trappc11 | 0.049 | Old |
| 0.310177 | 0.249149 | 0.191 | 0.12 | 1 | 14 | Slc31a2 | 0.071 | Old |
| 0.310542 | 0.812819 | 0.202 | 0.14 | 1 | 14 | Apobec3 | 0.062 | Old |
| 0.311519 | 1.379744 | 0.191 | 0.14 | 1 | 14 | Fubp3 | 0.051 | Old |
| 0.311557 | 0.171067 | 0.88 | 0.84 | 1 | 14 | Rpl8 | 0.04 | Old |
| 0.311721 | 1.615567 | 0.126 | 0.08 | 1 | 14 | Arhgef28 | 0.046 | Old |
| 0.311826 | 1.626235 | 0.055 | 0.02 | 1 | 14 | Urb2 | 0.035 | Old |
| 0.311826 | 1.312517 | 0.055 | 0.02 | 1 | 14 | Zfp334 | 0.035 | Old |
| 0.311826 | 1.277487 | 0.055 | 0.02 | 1 | 14 | Tfeb | 0.035 | Old |
| 0.311826 | 1.253066 | 0.055 | 0.02 | 1 | 14 | Taf6 | 0.035 | Old |
| 0.312067 | 1.403895 | 0.148 | 0.1 | 1 | 14 | Alg14 | 0.048 | Old |
| 0.312071 | 0.402604 | 0.109 | 0.06 | 1 | 14 | Hspa12a | 0.049 | Old |
| 0.312153 | 1.128392 | 0.082 | 0.04 | 1 | 14 | Tmem185a | 0.042 | Old |
| 0.312153 | 0.832491 | 0.082 | 0.04 | 1 | 14 | Smim12 | 0.042 | Old |
| 0.312248 | 0.239751 | 0.328 | 0.26 | 1 | 14 | Vdac2 | 0.068 | Old |
| 0.312427 | 0.09294 | 0.071 | 0.12 | 1 | 14 | Nudt16 | -0.049 | Old |
| 0.312516 | 1.067873 | 0.208 | 0.16 | 1 | 14 | Phf2 | 0.048 | Old |
| 0.313708 | 0.640702 | 0.333 | 0.26 | 1 | 14 | Ddx3y | 0.073 | Old |
| 0.313709 | 0.393791 | 0.328 | 0.28 | 1 | 14 | Pcmtd2 | 0.048 | Old |
| 0.313763 | 1.64593 | 0.126 | 0.08 | 1 | 14 | 2310011J0 | 0.046 | Old |
| 0.314336 | 0.890312 | 0.279 | 0.22 | 1 | 14 | Mmp19 | 0.059 | Old |
| 0.314582 | 1.587264 | 0.104 | 0.06 | 1 | 14 | Klhl12 | 0.044 | Old |
| 0.314582 | 1.461399 | 0.104 | 0.06 | 1 | 14 | Frrs1 | 0.044 | Old |
| 0.314673 | 1.81081 | 0.082 | 0.04 | 1 | 14 | Stoml1 | 0.042 | Old |
| 0.314673 | 0.853149 | 0.082 | 0.04 | 1 | 14 | Zmym6 | 0.042 | Old |
| 0.314784 | 0.631214 | 0.224 | 0.16 | 1 | 14 | Zc3h14 | 0.064 | Old |
| 0.314918 | 2.056805 | 0.055 | 0.02 | 1 | 14 | Nphp1 | 0.035 | Old |
| 0.314918 | 1.189499 | 0.055 | 0.02 | 1 | 14 | Gpr135 | 0.035 | Old |
| 0.314918 | 1.167046 | 0.055 | 0.02 | 1 | 14 | Ddit4 | 0.035 | Old |
| 0.315048 | 0.233412 | 0.557 | 0.52 | 1 | 14 | Cxcl12 | 0.037 | Old |
| 0.315162 | 0.598337 | 0.016 | 0.04 | 1 | 14 | 2810405F1 | -0.024 | Old |
| 0.315162 | 0.385688 | 0.016 | 0.04 | 1 | 14 | Rragd | -0.024 | Old |
| 0.315162 | 0.312511 | 0.016 | 0.04 | 1 | 14 | Bend3 | -0.024 | Old |
| 0.315728 | 0.972961 | 0.219 | 0.16 | 1 | 14 | Tmem63a | 0.059 | Old |
| 0.315933 | 1.117239 | 0.169 | 0.12 | 1 | 14 | Snhg6 | 0.049 | Old |
| 0.315964 | 0.468556 | 0.596 | 0.56 | 1 | 14 | Col4a2 | 0.036 | Old |
| 0.316072 | 0.67236 | 0.268 | 0.2 | 1 | 14 | Ddi2 | 0.068 | Old |
| 0.316322 | 0.162126 | 0.208 | 0.28 | 1 | 14 | Htt | -0.072 | Old |

| | | | | | | | | |
|---|---|---|---|---|---|---|---|---|
| 0.316395 | 0.234373 | 0.41 | 0.36 | 1 | 14 | Ppp1r14b | 0.05 | Old |
| 0.316833 | 1.437985 | 0.104 | 0.06 | 1 | 14 | Mrto4 | 0.044 | Old |
| 0.317207 | 1.093652 | 0.082 | 0.04 | 1 | 14 | 2810006K2 | 0.042 | Old |
| 0.317207 | 0.74653 | 0.082 | 0.04 | 1 | 14 | Dhx33 | 0.042 | Old |
| 0.317207 | 0.348831 | 0.082 | 0.04 | 1 | 14 | Ints2 | 0.042 | Old |
| 0.31748 | 1.208369 | 0.153 | 0.1 | 1 | 14 | Map3k8 | 0.053 | Old |
| 0.317782 | 1.848738 | 0.148 | 0.1 | 1 | 14 | Leo1 | 0.048 | Old |
| 0.317872 | 1.351364 | 0.126 | 0.08 | 1 | 14 | Nf2 | 0.046 | Old |
| 0.31792 | 0.449455 | 0.421 | 0.36 | 1 | 14 | Tmed9 | 0.061 | Old |
| 0.318322 | 1.079857 | 0.191 | 0.14 | 1 | 14 | Uba3 | 0.051 | Old |
| 0.318471 | 0.426003 | 0.366 | 0.3 | 1 | 14 | Pfkp | 0.066 | Old |
| 0.319374 | 0.393951 | 0.153 | 0.1 | 1 | 14 | Ric1 | 0.053 | Old |
| 0.319728 | 1.640725 | 0.016 | 0.04 | 1 | 14 | Gm20186 | -0.024 | Old |
| 0.319754 | 0.109349 | 0.306 | 0.24 | 1 | 14 | Lgals3bp | 0.066 | Old |
| 0.319915 | 0.338271 | 0.115 | 0.18 | 1 | 14 | Napg | -0.065 | Old |
| 0.321162 | 1.34656 | 0.055 | 0.02 | 1 | 14 | Rnaseh1 | 0.035 | Old |
| 0.321162 | 0.887421 | 0.055 | 0.02 | 1 | 14 | Vav3 | 0.035 | Old |
| 0.321162 | 0.784836 | 0.055 | 0.02 | 1 | 14 | Slc16a10 | 0.035 | Old |
| 0.321162 | 0.63657 | 0.055 | 0.02 | 1 | 14 | Lpar3 | 0.035 | Old |
| 0.321162 | 0.1967 | 0.055 | 0.02 | 1 | 14 | Atl1 | 0.035 | Old |
| 0.321254 | 0.142255 | 0.732 | 0.6 | 1 | 14 | Ogt | 0.132 | Old |
| 0.321367 | 1.436001 | 0.104 | 0.06 | 1 | 14 | Urgcp | 0.044 | Old |
| 0.321367 | 1.052151 | 0.104 | 0.06 | 1 | 14 | Pot1a | 0.044 | Old |
| 0.321467 | 0.078645 | 0.656 | 0.56 | 1 | 14 | Hmgb1 | 0.096 | Old |
| 0.321578 | 0.399833 | 0.355 | 0.3 | 1 | 14 | Tmx3 | 0.055 | Old |
| 0.321601 | 0.269773 | 0.464 | 0.38 | 1 | 14 | Ptges3 | 0.084 | Old |
| 0.321631 | 1.115283 | 0.148 | 0.1 | 1 | 14 | Htra2 | 0.048 | Old |
| 0.321843 | 2.03465 | 0.142 | 0.1 | 1 | 14 | Rnf169 | 0.042 | Old |
| 0.322697 | 1.067148 | 0.284 | 0.24 | 1 | 14 | Ubn1 | 0.044 | Old |
| 0.322697 | 0.355197 | 0.29 | 0.22 | 1 | 14 | Birc2 | 0.07 | Old |
| 0.32317 | 1.053477 | 0.169 | 0.12 | 1 | 14 | Ltn1 | 0.049 | Old |
| 0.32317 | 0.295087 | 0.169 | 0.12 | 1 | 14 | Rassf5 | 0.049 | Old |
| 0.323385 | 0.379165 | 0.23 | 0.16 | 1 | 14 | Pikfyve | 0.07 | Old |
| 0.323489 | 0.980973 | 0.191 | 0.14 | 1 | 14 | Rpa3 | 0.051 | Old |
| 0.323508 | 0.06389 | 0.497 | 0.44 | 1 | 14 | Gpi1 | 0.057 | Old |
| 0.323772 | 0.576544 | 0.197 | 0.14 | 1 | 14 | Bicd1 | 0.057 | Old |
| 0.323797 | 0.258479 | 0.355 | 0.28 | 1 | 14 | Sec22b | 0.075 | Old |
| 0.32403 | 0.108604 | 0.383 | 0.32 | 1 | 14 | Nol7 | 0.063 | Old |
| 0.324314 | 0.521151 | 0.055 | 0.02 | 1 | 14 | Kat14 | 0.035 | Old |
| 0.324314 | 0.383913 | 0.055 | 0.02 | 1 | 14 | Tvp23a | 0.035 | Old |
| 0.324336 | 0.638747 | 0.016 | 0.04 | 1 | 14 | Mto1 | -0.024 | Old |
| 0.324336 | 0.30671 | 0.016 | 0.04 | 1 | 14 | Gm36529 | -0.024 | Old |
| 0.324336 | 0.099902 | 0.016 | 0.04 | 1 | 14 | Epha4 | -0.024 | Old |
| 0.324525 | 0.507227 | 0.071 | 0.12 | 1 | 14 | Snai1 | -0.049 | Old |
| 0.324818 | 0.139493 | 0.137 | 0.08 | 1 | 14 | Mfap1a | 0.057 | Old |

| | | | | | | | | |
|---|---|---|---|---|---|---|---|---|
| 0.324888 | 0.895893 | 0.082 | 0.04 | 1 | 14 | Aimp2 | 0.042 | Old |
| 0.324942 | 0.415969 | 0.295 | 0.24 | 1 | 14 | Rassf8 | 0.055 | Old |
| 0.325523 | 0.141014 | 0.273 | 0.36 | 1 | 14 | Dnajb4 | -0.087 | Old |
| 0.325592 | 0.750256 | 0.131 | 0.08 | 1 | 14 | Lynx1 | 0.051 | Old |
| 0.325665 | 0.628513 | 0.202 | 0.14 | 1 | 14 | Slc30a4 | 0.062 | Old |
| 0.326198 | 1.62627 | 0.126 | 0.08 | 1 | 14 | Washc1 | 0.046 | Old |
| 0.326505 | 0.804931 | 0.213 | 0.16 | 1 | 14 | Zmym2 | 0.053 | Old |
| 0.326658 | 0.325143 | 0.268 | 0.2 | 1 | 14 | Itga5 | 0.068 | Old |
| 0.326691 | 0.70446 | 0.284 | 0.22 | 1 | 14 | Iscu | 0.064 | Old |
| 0.326763 | 0.146018 | 0.781 | 0.7 | 1 | 14 | Fosb | 0.081 | Old |
| 0.326828 | 1.099846 | 0.169 | 0.12 | 1 | 14 | Fbxl7 | 0.049 | Old |
| 0.326832 | 1.661735 | 0.164 | 0.12 | 1 | 14 | Rreb1 | 0.044 | Old |
| 0.326947 | 0.432608 | 0.421 | 0.34 | 1 | 14 | Fgfr1op2 | 0.081 | Old |
| 0.32746 | 0.016014 | 0.126 | 0.18 | 1 | 14 | Clec10a | -0.054 | Old |
| 0.328186 | 0.820759 | 0.284 | 0.22 | 1 | 14 | Ccdc127 | 0.064 | Old |
| 0.328263 | 0.069123 | 0.257 | 0.2 | 1 | 14 | 9030624J0 | 0.057 | Old |
| 0.328301 | 1.523891 | 0.126 | 0.08 | 1 | 14 | Gpkow | 0.046 | Old |
| 0.328391 | 0.370376 | 0.426 | 0.36 | 1 | 14 | Rcn2 | 0.066 | Old |
| 0.328402 | 0.363286 | 0.432 | 0.4 | 1 | 14 | Aida | 0.032 | Old |
| 0.328575 | 1.836694 | 0.219 | 0.18 | 1 | 14 | Clk4 | 0.039 | Old |
| 0.328955 | 0.270639 | 0.153 | 0.1 | 1 | 14 | Nkap | 0.053 | Old |
| 0.329302 | 0.141454 | 0.519 | 0.46 | 1 | 14 | G3bp1 | 0.059 | Old |
| 0.329692 | 2.183957 | 0.077 | 0.04 | 1 | 14 | Zfp518b | 0.037 | Old |
| 0.329736 | 0.760319 | 0.131 | 0.08 | 1 | 14 | Def8 | 0.051 | Old |
| 0.329997 | 1.366705 | 0.202 | 0.16 | 1 | 14 | Dock11 | 0.042 | Old |
| 0.330808 | 1.291941 | 0.208 | 0.16 | 1 | 14 | Arsk | 0.048 | Old |
| 0.331815 | 1.297913 | 0.005 | 0.02 | 1 | 14 | 4932435O2 | -0.015 | Old |
| 0.331815 | 1.10682 | 0.005 | 0.02 | 1 | 14 | Hif3a | -0.015 | Old |
| 0.331815 | 0.798373 | 0.005 | 0.02 | 1 | 14 | Plvap | -0.015 | Old |
| 0.331815 | 0.744383 | 0.005 | 0.02 | 1 | 14 | Hoxaas3 | -0.015 | Old |
| 0.331815 | 0.682709 | 0.005 | 0.02 | 1 | 14 | Slco5a1 | -0.015 | Old |
| 0.331815 | 0.592658 | 0.005 | 0.02 | 1 | 14 | E430024I0 | -0.015 | Old |
| 0.331815 | 0.590906 | 0.005 | 0.02 | 1 | 14 | Rims4 | -0.015 | Old |
| 0.331815 | 0.590906 | 0.005 | 0.02 | 1 | 14 | Fat3 | -0.015 | Old |
| 0.331815 | 0.590906 | 0.005 | 0.02 | 1 | 14 | Asb2 | -0.015 | Old |
| 0.331815 | 0.528379 | 0.005 | 0.02 | 1 | 14 | Mtmr7 | -0.015 | Old |
| 0.331815 | 0.454775 | 0.005 | 0.02 | 1 | 14 | Arc | -0.015 | Old |
| 0.331815 | 0.379591 | 0.005 | 0.02 | 1 | 14 | Gm44647 | -0.015 | Old |
| 0.331815 | 0.376113 | 0.005 | 0.02 | 1 | 14 | C1ql2 | -0.015 | Old |
| 0.331815 | 0.325734 | 0.005 | 0.02 | 1 | 14 | Gm26779 | -0.015 | Old |
| 0.331815 | 0.274755 | 0.005 | 0.02 | 1 | 14 | Gm43703 | -0.015 | Old |
| 0.331815 | 0.267898 | 0.005 | 0.02 | 1 | 14 | 4930572G0 | -0.015 | Old |
| 0.331815 | 0.259694 | 0.005 | 0.02 | 1 | 14 | Ccdc14 | -0.015 | Old |
| 0.331815 | 0.217391 | 0.005 | 0.02 | 1 | 14 | Med12l | -0.015 | Old |
| 0.331815 | 0.217298 | 0.005 | 0.02 | 1 | 14 | Lmntd1 | -0.015 | Old |

| | | | | | | | | |
|---|---|---|---|---|---|---|---|---|
| 0.331815 | 0.196169 | 0.005 | 0.02 | 1 | 14 | Arhgap27o | -0.015 | Old |
| 0.331815 | 0.15994 | 0.005 | 0.02 | 1 | 14 | Tfec | -0.015 | Old |
| 0.331815 | 0.152527 | 0.005 | 0.02 | 1 | 14 | Slc6a17 | -0.015 | Old |
| 0.331815 | 0.118343 | 0.005 | 0.02 | 1 | 14 | Icos | -0.015 | Old |
| 0.331815 | 0.117919 | 0.005 | 0.02 | 1 | 14 | Ovgp1 | -0.015 | Old |
| 0.331815 | 0.09754 | 0.005 | 0.02 | 1 | 14 | Prkg2 | -0.015 | Old |
| 0.331815 | 0.088689 | 0.005 | 0.02 | 1 | 14 | Hyal3 | -0.015 | Old |
| 0.331815 | 0.08227 | 0.005 | 0.02 | 1 | 14 | Ramp3 | -0.015 | Old |
| 0.331815 | 0.058183 | 0.005 | 0.02 | 1 | 14 | Chn1os3 | -0.015 | Old |
| 0.331815 | 0.032476 | 0.005 | 0.02 | 1 | 14 | Slc29a2 | -0.015 | Old |
| 0.331815 | 0.025484 | 0.005 | 0.02 | 1 | 14 | Cgref1 | -0.015 | Old |
| 0.331815 | 0.006304 | 0.005 | 0.02 | 1 | 14 | Hus1b | -0.015 | Old |
| 0.331815 | 0.001812 | 0.005 | 0.02 | 1 | 14 | Neurl2 | -0.015 | Old |
| 0.33182 | 0.584195 | 0.131 | 0.08 | 1 | 14 | Cep120 | 0.051 | Old |
| 0.332062 | 0.122079 | 0.251 | 0.18 | 1 | 14 | Rufy3 | 0.071 | Old |
| 0.332379 | 1.947845 | 0.077 | 0.04 | 1 | 14 | Ttc23 | 0.037 | Old |
| 0.332534 | 0.019056 | 0.126 | 0.08 | 1 | 14 | Pcgf2 | 0.046 | Old |
| 0.333362 | 0.032202 | 0.044 | 0.08 | 1 | 14 | Cd300lg | -0.036 | Old |
| 0.334204 | 0.644538 | 0.268 | 0.22 | 1 | 14 | Ndufb2 | 0.048 | Old |
| 0.334476 | 0.103448 | 0.486 | 0.42 | 1 | 14 | Smc6 | 0.066 | Old |
| 0.334513 | 0.575689 | 0.257 | 0.2 | 1 | 14 | Dopey1 | 0.057 | Old |
| 0.334625 | 0.147541 | 0.153 | 0.22 | 1 | 14 | Ap3m1 | -0.067 | Old |
| 0.334758 | 0.557095 | 0.339 | 0.26 | 1 | 14 | Xdh | 0.079 | Old |
| 0.335079 | 1.848706 | 0.077 | 0.04 | 1 | 14 | Vps8 | 0.037 | Old |
| 0.33522 | 1.50522 | 0.104 | 0.06 | 1 | 14 | Xpc | 0.044 | Old |
| 0.335339 | 0.659435 | 0.148 | 0.1 | 1 | 14 | Msantd2 | 0.048 | Old |
| 0.335897 | 0.230207 | 0.464 | 0.36 | 1 | 14 | Foxp1 | 0.104 | Old |
| 0.336009 | 1.449756 | 0.197 | 0.16 | 1 | 14 | Samd9l | 0.037 | Old |
| 0.336316 | 0.151402 | 0.94 | 0.82 | 1 | 14 | mt-Nd3 | 0.12 | Old |
| 0.33646 | 0.093375 | 0.044 | 0.08 | 1 | 14 | Timmdc1 | -0.036 | Old |
| 0.337327 | 1.103919 | 0.148 | 0.1 | 1 | 14 | Zdhhc9 | 0.048 | Old |
| 0.337565 | 1.096844 | 0.104 | 0.06 | 1 | 14 | Ebag9 | 0.044 | Old |
| 0.337565 | 1.036999 | 0.104 | 0.06 | 1 | 14 | AU022252 | 0.044 | Old |
| 0.337956 | 0.081699 | 0.082 | 0.04 | 1 | 14 | Npat | 0.042 | Old |
| 0.337993 | 0.381068 | 0.295 | 0.22 | 1 | 14 | Rpl36al | 0.075 | Old |
| 0.338124 | 0.560187 | 0.131 | 0.08 | 1 | 14 | Gpc4 | 0.051 | Old |
| 0.338384 | 0.880734 | 0.311 | 0.28 | 1 | 14 | Zc3h7a | 0.031 | Old |
| 0.338443 | 0.555074 | 0.251 | 0.18 | 1 | 14 | Ube3c | 0.071 | Old |
| 0.3385 | 0.45516 | 0.399 | 0.34 | 1 | 14 | Smad4 | 0.059 | Old |
| 0.338721 | 0.895839 | 0.153 | 0.1 | 1 | 14 | Orc4 | 0.053 | Old |
| 0.338934 | 0.587249 | 0.415 | 0.36 | 1 | 14 | Ergic3 | 0.055 | Old |
| 0.338948 | 0.898786 | 0.126 | 0.08 | 1 | 14 | Psmb9 | 0.046 | Old |
| 0.339012 | 0.39985 | 0.383 | 0.32 | 1 | 14 | Acin1 | 0.063 | Old |
| 0.339034 | 0.42955 | 0.268 | 0.2 | 1 | 14 | Lmo2 | 0.068 | Old |
| 0.339576 | 0.620765 | 0.175 | 0.12 | 1 | 14 | Ifngr2 | 0.055 | Old |

| | | | | | | | | |
|---|---|---|---|---|---|---|---|---|
| 0.339921 | 0.086866 | 0.104 | 0.06 | 1 | 14 | Cntn1 | 0.044 | Old |
| 0.340004 | 0.856859 | 0.213 | 0.16 | 1 | 14 | Snx1 | 0.053 | Old |
| 0.340523 | 2.169451 | 0.077 | 0.04 | 1 | 14 | Tbc1d32 | 0.037 | Old |
| 0.340523 | 1.966801 | 0.077 | 0.04 | 1 | 14 | Zdhhc13 | 0.037 | Old |
| 0.341104 | 1.598838 | 0.126 | 0.08 | 1 | 14 | Gstm4 | 0.046 | Old |
| 0.341352 | 0.465545 | 0.322 | 0.28 | 1 | 14 | Rnf7 | 0.042 | Old |
| 0.34139 | 0.161482 | 0.109 | 0.06 | 1 | 14 | 1700021F0 | 0.049 | Old |
| 0.342062 | 0.534719 | 0.317 | 0.26 | 1 | 14 | Rwdd1 | 0.057 | Old |
| 0.342275 | 0.248227 | 0.219 | 0.16 | 1 | 14 | Gmps | 0.059 | Old |
| 0.342288 | 1.036128 | 0.104 | 0.06 | 1 | 14 | Pmvk | 0.044 | Old |
| 0.342603 | 0.361495 | 0.404 | 0.32 | 1 | 14 | Sf3b3 | 0.084 | Old |
| 0.343021 | 0.142027 | 0.202 | 0.14 | 1 | 14 | Aggf1 | 0.062 | Old |
| 0.343277 | 0.233687 | 0.082 | 0.04 | 1 | 14 | Ubxn8 | 0.042 | Old |
| 0.343277 | 0.000876 | 0.082 | 0.04 | 1 | 14 | Isyna1 | 0.042 | Old |
| 0.343303 | 0.217525 | 0.29 | 0.24 | 1 | 14 | Washc4 | 0.05 | Old |
| 0.3436 | 0.282036 | 0.383 | 0.3 | 1 | 14 | Lrp10 | 0.083 | Old |
| 0.343669 | 0.728828 | 0.366 | 0.32 | 1 | 14 | Slc39a7 | 0.046 | Old |
| 0.343981 | 0.565407 | 0.219 | 0.16 | 1 | 14 | Snap23 | 0.059 | Old |
| 0.34404 | 0.32736 | 0.574 | 0.48 | 1 | 14 | Kcnq1ot1 | 0.094 | Old |
| 0.344282 | 0.775413 | 0.322 | 0.28 | 1 | 14 | Cdk16 | 0.042 | Old |
| 0.344665 | 0.504998 | 0.104 | 0.06 | 1 | 14 | A230050P2 | 0.044 | Old |
| 0.344748 | 0.060778 | 0.525 | 0.46 | 1 | 14 | Mkln1 | 0.065 | Old |
| 0.345054 | 0.163949 | 0.825 | 0.76 | 1 | 14 | Rps6 | 0.065 | Old |
| 0.345109 | 0.482459 | 0.268 | 0.22 | 1 | 14 | Srrt | 0.048 | Old |
| 0.345136 | 0.140919 | 0.765 | 0.74 | 1 | 14 | Aplp2 | 0.025 | Old |
| 0.345353 | 2.286654 | 0.098 | 0.06 | 1 | 14 | Abhd17c | 0.038 | Old |
| 0.345353 | 2.104257 | 0.098 | 0.06 | 1 | 14 | Rint1 | 0.038 | Old |
| 0.346022 | 1.535073 | 0.077 | 0.04 | 1 | 14 | Rfx1 | 0.037 | Old |
| 0.346518 | 1.052183 | 0.224 | 0.18 | 1 | 14 | Arhgap18 | 0.044 | Old |
| 0.346665 | 0.321786 | 0.153 | 0.1 | 1 | 14 | Anks3 | 0.053 | Old |
| 0.347272 | 0.241572 | 0.579 | 0.54 | 1 | 14 | Prrc2c | 0.039 | Old |
| 0.347624 | 0.674968 | 0.126 | 0.08 | 1 | 14 | Zfp830 | 0.046 | Old |
| 0.347634 | 0.201528 | 0.279 | 0.22 | 1 | 14 | Mdfic | 0.059 | Old |
| 0.347794 | 1.793993 | 0.098 | 0.06 | 1 | 14 | Slc35b3 | 0.038 | Old |
| 0.348316 | 0.527888 | 0.191 | 0.14 | 1 | 14 | Ehmt1 | 0.051 | Old |
| 0.34862 | 0.94685 | 0.213 | 0.16 | 1 | 14 | Slc39a9 | 0.053 | Old |
| 0.348793 | 1.984483 | 0.077 | 0.04 | 1 | 14 | Rpa2 | 0.037 | Old |
| 0.348793 | 1.468523 | 0.077 | 0.04 | 1 | 14 | Pthlh | 0.037 | Old |
| 0.348852 | 0.025481 | 0.257 | 0.2 | 1 | 14 | Lamtor1 | 0.057 | Old |
| 0.349037 | 0.969453 | 0.044 | 0.08 | 1 | 14 | Galns | -0.036 | Old |
| 0.349037 | 0.245004 | 0.044 | 0.08 | 1 | 14 | Pno1 | -0.036 | Old |
| 0.349341 | 1.039545 | 0.169 | 0.12 | 1 | 14 | C1qtnf6 | 0.049 | Old |
| 0.349451 | 0.587237 | 0.104 | 0.06 | 1 | 14 | Sec61a2 | 0.044 | Old |
| 0.349494 | 0.430815 | 0.317 | 0.26 | 1 | 14 | Vma21 | 0.057 | Old |
| 0.349697 | 0.725793 | 0.24 | 0.18 | 1 | 14 | Stard3nl | 0.06 | Old |

| | | | | | | | | |
|---|---|---|---|---|---|---|---|---|
| 0.349815 | 0.205967 | 0.104 | 0.16 | 1 | 14 | Ddx55 | -0.056 | Old |
| 0.350247 | 1.900816 | 0.098 | 0.06 | 1 | 14 | P2ry1 | 0.038 | Old |
| 0.350792 | 0.816366 | 0.301 | 0.26 | 1 | 14 | Tmem123 | 0.041 | Old |
| 0.350907 | 0.783056 | 0.262 | 0.22 | 1 | 14 | Psmb7 | 0.042 | Old |
| 0.351458 | 0.930097 | 0.148 | 0.1 | 1 | 14 | Ccz1 | 0.048 | Old |
| 0.351578 | 1.563875 | 0.077 | 0.04 | 1 | 14 | Dcaf15 | 0.037 | Old |
| 0.351578 | 1.559121 | 0.077 | 0.04 | 1 | 14 | Ddx19b | 0.037 | Old |
| 0.35196 | 0.726578 | 0.191 | 0.14 | 1 | 14 | Asap2 | 0.051 | Old |
| 0.352067 | 0.033304 | 0.235 | 0.18 | 1 | 14 | Smg7 | 0.055 | Old |
| 0.352768 | 0.236018 | 0.377 | 0.32 | 1 | 14 | Galnt15 | 0.057 | Old |
| 0.353229 | 0.128913 | 0.661 | 0.54 | 1 | 14 | Sash1 | 0.121 | Old |
| 0.353856 | 0.731209 | 0.213 | 0.16 | 1 | 14 | Armcx2 | 0.053 | Old |
| 0.353928 | 0.1973 | 0.705 | 0.62 | 1 | 14 | Rpn2 | 0.085 | Old |
| 0.353931 | 3.471827 | 0.049 | 0.02 | 1 | 14 | Tmem255a | 0.029 | Old |
| 0.353931 | 3.03023 | 0.049 | 0.02 | 1 | 14 | Duox1 | 0.029 | Old |
| 0.353931 | 2.930455 | 0.049 | 0.02 | 1 | 14 | Dusp2 | 0.029 | Old |
| 0.353931 | 2.703588 | 0.049 | 0.02 | 1 | 14 | Urb1 | 0.029 | Old |
| 0.353931 | 2.31755 | 0.049 | 0.02 | 1 | 14 | Plpp7 | 0.029 | Old |
| 0.353931 | 2.233218 | 0.049 | 0.02 | 1 | 14 | Hacl1 | 0.029 | Old |
| 0.353931 | 2.093229 | 0.049 | 0.02 | 1 | 14 | Casp7 | 0.029 | Old |
| 0.353931 | 2.0433 | 0.049 | 0.02 | 1 | 14 | Camsap3 | 0.029 | Old |
| 0.353931 | 1.8922 | 0.049 | 0.02 | 1 | 14 | Ccdc58 | 0.029 | Old |
| 0.353931 | 1.866161 | 0.049 | 0.02 | 1 | 14 | Mgat5 | 0.029 | Old |
| 0.353938 | 0.782011 | 0.306 | 0.26 | 1 | 14 | Ap3s1 | 0.046 | Old |
| 0.354223 | 0.818991 | 0.126 | 0.08 | 1 | 14 | Foxo3 | 0.046 | Old |
| 0.354377 | 1.589121 | 0.077 | 0.04 | 1 | 14 | Enoph1 | 0.037 | Old |
| 0.354377 | 1.355967 | 0.077 | 0.04 | 1 | 14 | Tecpr2 | 0.037 | Old |
| 0.354674 | 0.017166 | 0.175 | 0.12 | 1 | 14 | Syne1 | 0.055 | Old |
| 0.354716 | 0.199498 | 0.224 | 0.16 | 1 | 14 | C1qtnf1 | 0.064 | Old |
| 0.355355 | 0.543169 | 0.372 | 0.32 | 1 | 14 | Wdr83os | 0.052 | Old |
| 0.355615 | 0.640714 | 0.197 | 0.14 | 1 | 14 | Tanc1 | 0.057 | Old |
| 0.356127 | 0.20639 | 0.53 | 0.48 | 1 | 14 | Cpq | 0.05 | Old |
| 0.356836 | 0.012723 | 0.574 | 0.48 | 1 | 14 | Mbnl2 | 0.094 | Old |
| 0.35719 | 2.225899 | 0.077 | 0.04 | 1 | 14 | Pafah2 | 0.037 | Old |
| 0.35719 | 0.111846 | 0.06 | 0.1 | 1 | 14 | Bcl2l12 | -0.04 | Old |
| 0.357374 | 0.753112 | 0.213 | 0.16 | 1 | 14 | Hspa13 | 0.053 | Old |
| 0.357404 | 1.628107 | 0.12 | 0.08 | 1 | 14 | Tmem184c | 0.04 | Old |
| 0.35742 | 1.27833 | 0.164 | 0.12 | 1 | 14 | Rbm4b | 0.044 | Old |
| 0.357442 | 3.181044 | 0.049 | 0.02 | 1 | 14 | Ifit1 | 0.029 | Old |
| 0.357442 | 2.474612 | 0.049 | 0.02 | 1 | 14 | Pdp2 | 0.029 | Old |
| 0.357442 | 2.294315 | 0.049 | 0.02 | 1 | 14 | Bbs7 | 0.029 | Old |
| 0.357442 | 2.13804 | 0.049 | 0.02 | 1 | 14 | Usp20 | 0.029 | Old |
| 0.357442 | 1.982477 | 0.049 | 0.02 | 1 | 14 | Wdr92 | 0.029 | Old |
| 0.357442 | 1.961557 | 0.049 | 0.02 | 1 | 14 | Gm26722 | 0.029 | Old |
| 0.357442 | 1.911406 | 0.049 | 0.02 | 1 | 14 | Fgf18 | 0.029 | Old |

| | | | | | | | | |
|---|---|---|---|---|---|---|---|---|
| 0.357442 | 1.885955 | 0.049 | 0.02 | 1 | 14 | Stradb | 0.029 | Old |
| 0.357442 | 1.790811 | 0.049 | 0.02 | 1 | 14 | Sergef | 0.029 | Old |
| 0.357442 | 1.687188 | 0.049 | 0.02 | 1 | 14 | Ankzf1 | 0.029 | Old |
| 0.357846 | 0.855671 | 0.268 | 0.22 | 1 | 14 | Tiprl | 0.048 | Old |
| 0.358051 | 0.413715 | 0.251 | 0.18 | 1 | 14 | Cd44 | 0.071 | Old |
| 0.358366 | 1.702899 | 0.142 | 0.1 | 1 | 14 | Rnf170 | 0.042 | Old |
| 0.358665 | 0.264321 | 0.372 | 0.3 | 1 | 14 | Zfp445 | 0.072 | Old |
| 0.358666 | 1.051416 | 0.126 | 0.08 | 1 | 14 | Pex16 | 0.046 | Old |
| 0.358666 | 0.807594 | 0.126 | 0.08 | 1 | 14 | E4f1 | 0.046 | Old |
| 0.358876 | 0.194613 | 0.153 | 0.22 | 1 | 14 | Glb1 | -0.067 | Old |
| 0.359459 | 0.720136 | 0.268 | 0.22 | 1 | 14 | Zfp704 | 0.048 | Old |
| 0.359671 | 1.739344 | 0.12 | 0.08 | 1 | 14 | Gnpda1 | 0.04 | Old |
| 0.359699 | 1.019949 | 0.148 | 0.1 | 1 | 14 | St7l | 0.048 | Old |
| 0.359699 | 0.791462 | 0.148 | 0.1 | 1 | 14 | Socs6 | 0.048 | Old |
| 0.360017 | 0.977929 | 0.077 | 0.04 | 1 | 14 | Shc2 | 0.037 | Old |
| 0.360072 | 0.551238 | 0.475 | 0.42 | 1 | 14 | Chmp4b | 0.055 | Old |
| 0.360362 | 0.549116 | 0.475 | 0.4 | 1 | 14 | Tbca | 0.075 | Old |
| 0.360439 | 0.064057 | 0.251 | 0.2 | 1 | 14 | Abcc5 | 0.051 | Old |
| 0.360489 | 0.054731 | 0.694 | 0.6 | 1 | 14 | Lpp | 0.094 | Old |
| 0.360975 | 2.025462 | 0.049 | 0.02 | 1 | 14 | Phf7 | 0.029 | Old |
| 0.361087 | 0.146649 | 0.552 | 0.46 | 1 | 14 | Ly6c1 | 0.092 | Old |
| 0.361104 | 0.184927 | 0.093 | 0.14 | 1 | 14 | Proser1 | -0.047 | Old |
| 0.361347 | 0.412165 | 0.514 | 0.46 | 1 | 14 | Spcs1 | 0.054 | Old |
| 0.361883 | 0.098331 | 0.929 | 0.84 | 1 | 14 | Eif1 | 0.089 | Old |
| 0.361892 | 0.406541 | 0.131 | 0.08 | 1 | 14 | Olfr1033 | 0.051 | Old |
| 0.361892 | 0.335807 | 0.131 | 0.08 | 1 | 14 | Terf2ip | 0.051 | Old |
| 0.362379 | 0.49661 | 0.175 | 0.12 | 1 | 14 | Plekhf1 | 0.055 | Old |
| 0.362579 | 1.430209 | 0.142 | 0.1 | 1 | 14 | Agap3 | 0.042 | Old |
| 0.362579 | 0.08821 | 0.12 | 0.18 | 1 | 14 | Ccm2 | -0.06 | Old |
| 0.362674 | 1.545251 | 0.098 | 0.06 | 1 | 14 | Endod1 | 0.038 | Old |
| 0.362859 | 1.074086 | 0.077 | 0.04 | 1 | 14 | Pank1 | 0.037 | Old |
| 0.362859 | 0.961642 | 0.077 | 0.04 | 1 | 14 | Ccdc22 | 0.037 | Old |
| 0.362859 | 0.814581 | 0.077 | 0.04 | 1 | 14 | Zfp113 | 0.037 | Old |
| 0.362917 | 0.72714 | 0.169 | 0.12 | 1 | 14 | Chst12 | 0.049 | Old |
| 0.362953 | 0.24522 | 0.328 | 0.26 | 1 | 14 | Dhx15 | 0.068 | Old |
| 0.36315 | 0.483722 | 0.24 | 0.18 | 1 | 14 | Pxk | 0.06 | Old |
| 0.364427 | 0.080135 | 0.311 | 0.4 | 1 | 14 | Dnajc1 | -0.089 | Old |
| 0.36453 | 1.575068 | 0.049 | 0.02 | 1 | 14 | St6gal1 | 0.029 | Old |
| 0.364677 | 0.212932 | 0.235 | 0.32 | 1 | 14 | Rbm33 | -0.085 | Old |
| 0.364736 | 0.515413 | 0.273 | 0.22 | 1 | 14 | Sh3kbp1 | 0.053 | Old |
| 0.3649 | 0.707807 | 0.191 | 0.14 | 1 | 14 | Kdm5c | 0.051 | Old |
| 0.364971 | 0.24762 | 0.153 | 0.1 | 1 | 14 | Gclm | 0.053 | Old |
| 0.365235 | 0.775342 | 0.257 | 0.2 | 1 | 14 | Abca6 | 0.057 | Old |
| 0.365415 | 0.900928 | 0.224 | 0.18 | 1 | 14 | Babam2 | 0.044 | Old |
| 0.365714 | 0.736084 | 0.06 | 0.1 | 1 | 14 | Rgcc | -0.04 | Old |

| | | | | | | | | |
|---|---|---|---|---|---|---|---|---|
| 0.365739 | 0.140149 | 0.656 | 0.58 | 1 | 14 | Ablim1 | 0.076 | Old |
| 0.365808 | 0.215959 | 0.475 | 0.4 | 1 | 14 | Mrpl33 | 0.075 | Old |
| 0.366271 | 0.33117 | 0.175 | 0.12 | 1 | 14 | Cbr3 | 0.055 | Old |
| 0.366419 | 4.73014 | 0.016 | 0 | 1 | 14 | Upk3b | 0.016 | Old |
| 0.366419 | 3.846112 | 0.016 | 0 | 1 | 14 | Ppp1r3c | 0.016 | Old |
| 0.366419 | 3.608767 | 0.016 | 0 | 1 | 14 | Trpt1 | 0.016 | Old |
| 0.366419 | 3.549405 | 0.016 | 0 | 1 | 14 | Tmem132b | 0.016 | Old |
| 0.366419 | 3.412151 | 0.016 | 0 | 1 | 14 | Dennd1c | 0.016 | Old |
| 0.366419 | 3.077663 | 0.016 | 0 | 1 | 14 | Kif9 | 0.016 | Old |
| 0.366419 | 3.071694 | 0.016 | 0 | 1 | 14 | 9330160F1 | 0.016 | Old |
| 0.366419 | 3.026897 | 0.016 | 0 | 1 | 14 | Iqck | 0.016 | Old |
| 0.366419 | 2.944622 | 0.016 | 0 | 1 | 14 | Tmem198 | 0.016 | Old |
| 0.366419 | 2.900192 | 0.016 | 0 | 1 | 14 | 4931440F1 | 0.016 | Old |
| 0.366419 | 2.864594 | 0.016 | 0 | 1 | 14 | Alpk1 | 0.016 | Old |
| 0.366419 | 2.858493 | 0.016 | 0 | 1 | 14 | Ddo | 0.016 | Old |
| 0.366419 | 2.825991 | 0.016 | 0 | 1 | 14 | Gm48604 | 0.016 | Old |
| 0.366419 | 2.791278 | 0.016 | 0 | 1 | 14 | Gm45716 | 0.016 | Old |
| 0.366419 | 2.788058 | 0.016 | 0 | 1 | 14 | Ackr4 | 0.016 | Old |
| 0.366419 | 2.787696 | 0.016 | 0 | 1 | 14 | Ddx28 | 0.016 | Old |
| 0.366419 | 2.774697 | 0.016 | 0 | 1 | 14 | Serpina3m | 0.016 | Old |
| 0.366419 | 2.742657 | 0.016 | 0 | 1 | 14 | Lmln | 0.016 | Old |
| 0.366419 | 2.708808 | 0.016 | 0 | 1 | 14 | Tmigd1 | 0.016 | Old |
| 0.366419 | 2.688047 | 0.016 | 0 | 1 | 14 | B230307C2 | 0.016 | Old |
| 0.366419 | 2.671087 | 0.016 | 0 | 1 | 14 | Gm48551 | 0.016 | Old |
| 0.366419 | 2.667275 | 0.016 | 0 | 1 | 14 | BC037039 | 0.016 | Old |
| 0.366419 | 2.666676 | 0.016 | 0 | 1 | 14 | Lcat | 0.016 | Old |
| 0.366419 | 2.662532 | 0.016 | 0 | 1 | 14 | Neto2 | 0.016 | Old |
| 0.366419 | 2.612656 | 0.016 | 0 | 1 | 14 | Ccdc166 | 0.016 | Old |
| 0.366419 | 2.575558 | 0.016 | 0 | 1 | 14 | Gm26901 | 0.016 | Old |
| 0.366419 | 2.548559 | 0.016 | 0 | 1 | 14 | Zfp709 | 0.016 | Old |
| 0.366419 | 2.541955 | 0.016 | 0 | 1 | 14 | 5033421B0 | 0.016 | Old |
| 0.366419 | 2.536438 | 0.016 | 0 | 1 | 14 | Spa17 | 0.016 | Old |
| 0.366419 | 2.520647 | 0.016 | 0 | 1 | 14 | Kcna1 | 0.016 | Old |
| 0.366419 | 2.517711 | 0.016 | 0 | 1 | 14 | Mkrn3 | 0.016 | Old |
| 0.366419 | 2.501455 | 0.016 | 0 | 1 | 14 | Gm11266 | 0.016 | Old |
| 0.366419 | 2.494906 | 0.016 | 0 | 1 | 14 | Sap30 | 0.016 | Old |
| 0.366419 | 2.489484 | 0.016 | 0 | 1 | 14 | Gm28707 | 0.016 | Old |
| 0.366419 | 2.482198 | 0.016 | 0 | 1 | 14 | Mok | 0.016 | Old |
| 0.366419 | 2.463674 | 0.016 | 0 | 1 | 14 | Zfp3 | 0.016 | Old |
| 0.366419 | 2.459843 | 0.016 | 0 | 1 | 14 | Stc2 | 0.016 | Old |
| 0.366419 | 2.441922 | 0.016 | 0 | 1 | 14 | 8030423F2 | 0.016 | Old |
| 0.366419 | 2.428641 | 0.016 | 0 | 1 | 14 | Osm | 0.016 | Old |
| 0.366419 | 2.424274 | 0.016 | 0 | 1 | 14 | Gm867 | 0.016 | Old |
| 0.366419 | 2.420577 | 0.016 | 0 | 1 | 14 | 4930579G2 | 0.016 | Old |
| 0.366419 | 2.414293 | 0.016 | 0 | 1 | 14 | Ydjc | 0.016 | Old |

| | | | | | | | | |
|---|---|---|---|---|---|---|---|---|
| 0.366419 | 2.413364 | 0.016 | 0 | 1 | 14 | Tor4a | 0.016 | Old |
| 0.366419 | 2.401339 | 0.016 | 0 | 1 | 14 | Armcx6 | 0.016 | Old |
| 0.366419 | 2.359316 | 0.016 | 0 | 1 | 14 | Gm5577 | 0.016 | Old |
| 0.366419 | 2.356249 | 0.016 | 0 | 1 | 14 | Thsd1 | 0.016 | Old |
| 0.366419 | 2.353367 | 0.016 | 0 | 1 | 14 | Neil2 | 0.016 | Old |
| 0.366419 | 2.344035 | 0.016 | 0 | 1 | 14 | Ccdc69 | 0.016 | Old |
| 0.366419 | 2.337584 | 0.016 | 0 | 1 | 14 | 1700088E0 | 0.016 | Old |
| 0.366419 | 2.320329 | 0.016 | 0 | 1 | 14 | Ticam2 | 0.016 | Old |
| 0.366419 | 2.314013 | 0.016 | 0 | 1 | 14 | Tonsl | 0.016 | Old |
| 0.366419 | 2.313488 | 0.016 | 0 | 1 | 14 | Tacc3 | 0.016 | Old |
| 0.366419 | 2.311616 | 0.016 | 0 | 1 | 14 | Nxpe3 | 0.016 | Old |
| 0.366419 | 2.292777 | 0.016 | 0 | 1 | 14 | Zfp781 | 0.016 | Old |
| 0.366419 | 2.274978 | 0.016 | 0 | 1 | 14 | Gm38251 | 0.016 | Old |
| 0.366419 | 2.272039 | 0.016 | 0 | 1 | 14 | Dnajc17 | 0.016 | Old |
| 0.366419 | 2.265778 | 0.016 | 0 | 1 | 14 | Ripor3 | 0.016 | Old |
| 0.366419 | 2.265501 | 0.016 | 0 | 1 | 14 | Gm36161 | 0.016 | Old |
| 0.366419 | 2.257521 | 0.016 | 0 | 1 | 14 | Trmo | 0.016 | Old |
| 0.366419 | 2.241764 | 0.016 | 0 | 1 | 14 | Tyro3 | 0.016 | Old |
| 0.366419 | 2.232734 | 0.016 | 0 | 1 | 14 | Gpr173 | 0.016 | Old |
| 0.366419 | 2.225457 | 0.016 | 0 | 1 | 14 | Gm20878 | 0.016 | Old |
| 0.366419 | 2.222196 | 0.016 | 0 | 1 | 14 | Gng8 | 0.016 | Old |
| 0.366419 | 2.221289 | 0.016 | 0 | 1 | 14 | Tc2n | 0.016 | Old |
| 0.366419 | 2.2165 | 0.016 | 0 | 1 | 14 | Ankrd1 | 0.016 | Old |
| 0.366419 | 2.214079 | 0.016 | 0 | 1 | 14 | Nlgn1 | 0.016 | Old |
| 0.366419 | 2.206486 | 0.016 | 0 | 1 | 14 | Prg4 | 0.016 | Old |
| 0.366419 | 2.171704 | 0.016 | 0 | 1 | 14 | Isoc2a | 0.016 | Old |
| 0.366419 | 2.169797 | 0.016 | 0 | 1 | 14 | Gm43660 | 0.016 | Old |
| 0.366419 | 2.167387 | 0.016 | 0 | 1 | 14 | Gm42726 | 0.016 | Old |
| 0.366419 | 2.163963 | 0.016 | 0 | 1 | 14 | Cdh24 | 0.016 | Old |
| 0.366419 | 2.160308 | 0.016 | 0 | 1 | 14 | Tnfsf10 | 0.016 | Old |
| 0.366419 | 2.156083 | 0.016 | 0 | 1 | 14 | Hexim2 | 0.016 | Old |
| 0.366419 | 2.152493 | 0.016 | 0 | 1 | 14 | Zfp27 | 0.016 | Old |
| 0.366419 | 2.146493 | 0.016 | 0 | 1 | 14 | Zfp354a | 0.016 | Old |
| 0.366419 | 2.142324 | 0.016 | 0 | 1 | 14 | Mlxipl | 0.016 | Old |
| 0.366419 | 2.13905 | 0.016 | 0 | 1 | 14 | Dbndd2 | 0.016 | Old |
| 0.366419 | 2.134786 | 0.016 | 0 | 1 | 14 | Lrrc36 | 0.016 | Old |
| 0.366419 | 2.131557 | 0.016 | 0 | 1 | 14 | Lrrc75a | 0.016 | Old |
| 0.366419 | 2.129586 | 0.016 | 0 | 1 | 14 | Hexdc | 0.016 | Old |
| 0.366419 | 2.126641 | 0.016 | 0 | 1 | 14 | Fgd6 | 0.016 | Old |
| 0.366419 | 2.120899 | 0.016 | 0 | 1 | 14 | Gm16141 | 0.016 | Old |
| 0.366419 | 2.118806 | 0.016 | 0 | 1 | 14 | Nphp4 | 0.016 | Old |
| 0.366419 | 2.110304 | 0.016 | 0 | 1 | 14 | Hist1h4i | 0.016 | Old |
| 0.366419 | 2.107201 | 0.016 | 0 | 1 | 14 | Gm10561 | 0.016 | Old |
| 0.366419 | 2.106569 | 0.016 | 0 | 1 | 14 | Rrm2 | 0.016 | Old |
| 0.366419 | 2.100113 | 0.016 | 0 | 1 | 14 | Clec4e | 0.016 | Old |

| | | | | | | | | |
|---|---|---|---|---|---|---|---|---|
| 0.366419 | 2.094308 | 0.016 | 0 | 1 | 14 | Plp1 | 0.016 | Old |
| 0.366419 | 2.093611 | 0.016 | 0 | 1 | 14 | Efr3b | 0.016 | Old |
| 0.366419 | 2.093517 | 0.016 | 0 | 1 | 14 | Kctd21 | 0.016 | Old |
| 0.366419 | 2.089789 | 0.016 | 0 | 1 | 14 | Sh2d6 | 0.016 | Old |
| 0.366419 | 2.088225 | 0.016 | 0 | 1 | 14 | 1600010M | 0.016 | Old |
| 0.366419 | 2.084599 | 0.016 | 0 | 1 | 14 | Celf5 | 0.016 | Old |
| 0.366419 | 2.054809 | 0.016 | 0 | 1 | 14 | Dsp | 0.016 | Old |
| 0.366419 | 2.051275 | 0.016 | 0 | 1 | 14 | Dnaaf3 | 0.016 | Old |
| 0.366419 | 2.049646 | 0.016 | 0 | 1 | 14 | Pigz | 0.016 | Old |
| 0.366419 | 2.047077 | 0.016 | 0 | 1 | 14 | 6330403L0 | 0.016 | Old |
| 0.366419 | 2.045599 | 0.016 | 0 | 1 | 14 | Carns1 | 0.016 | Old |
| 0.366419 | 2.043778 | 0.016 | 0 | 1 | 14 | Vpreb1 | 0.016 | Old |
| 0.366419 | 2.024573 | 0.016 | 0 | 1 | 14 | Ccdc94 | 0.016 | Old |
| 0.366419 | 2.024499 | 0.016 | 0 | 1 | 14 | Doc2g | 0.016 | Old |
| 0.366419 | 2.024166 | 0.016 | 0 | 1 | 14 | Cmtm8 | 0.016 | Old |
| 0.366419 | 2.016453 | 0.016 | 0 | 1 | 14 | Lcmt2 | 0.016 | Old |
| 0.366419 | 2.014149 | 0.016 | 0 | 1 | 14 | Gpr39 | 0.016 | Old |
| 0.366419 | 2.01237 | 0.016 | 0 | 1 | 14 | Ppargc1b | 0.016 | Old |
| 0.366419 | 2.012099 | 0.016 | 0 | 1 | 14 | Gm45495 | 0.016 | Old |
| 0.366419 | 1.992647 | 0.016 | 0 | 1 | 14 | Gm29650 | 0.016 | Old |
| 0.366419 | 1.990691 | 0.016 | 0 | 1 | 14 | Lsr | 0.016 | Old |
| 0.366419 | 1.980192 | 0.016 | 0 | 1 | 14 | Chek2 | 0.016 | Old |
| 0.366419 | 1.97293 | 0.016 | 0 | 1 | 14 | Zfp563 | 0.016 | Old |
| 0.366419 | 1.967167 | 0.016 | 0 | 1 | 14 | Sncg | 0.016 | Old |
| 0.366419 | 1.966567 | 0.016 | 0 | 1 | 14 | Fbxl13 | 0.016 | Old |
| 0.366419 | 1.959591 | 0.016 | 0 | 1 | 14 | C230035I1( | 0.016 | Old |
| 0.366419 | 1.955216 | 0.016 | 0 | 1 | 14 | Cks1brt | 0.016 | Old |
| 0.366419 | 1.948935 | 0.016 | 0 | 1 | 14 | Gnrh1 | 0.016 | Old |
| 0.366419 | 1.944571 | 0.016 | 0 | 1 | 14 | Trim66 | 0.016 | Old |
| 0.366419 | 1.928817 | 0.016 | 0 | 1 | 14 | Tubd1 | 0.016 | Old |
| 0.366419 | 1.901471 | 0.016 | 0 | 1 | 14 | Cenpq | 0.016 | Old |
| 0.366419 | 1.901182 | 0.016 | 0 | 1 | 14 | Arhgap4 | 0.016 | Old |
| 0.366419 | 1.897405 | 0.016 | 0 | 1 | 14 | Zfp551 | 0.016 | Old |
| 0.366419 | 1.894391 | 0.016 | 0 | 1 | 14 | Ube2t | 0.016 | Old |
| 0.366419 | 1.89176 | 0.016 | 0 | 1 | 14 | Crybg1 | 0.016 | Old |
| 0.366419 | 1.882469 | 0.016 | 0 | 1 | 14 | 9430037G( | 0.016 | Old |
| 0.366419 | 1.863492 | 0.016 | 0 | 1 | 14 | Gpat3 | 0.016 | Old |
| 0.366419 | 1.863233 | 0.016 | 0 | 1 | 14 | Ctla2a | 0.016 | Old |
| 0.366419 | 1.855293 | 0.016 | 0 | 1 | 14 | Zc3h11a | 0.016 | Old |
| 0.366419 | 1.855233 | 0.016 | 0 | 1 | 14 | Pcdhb19 | 0.016 | Old |
| 0.366419 | 1.850323 | 0.016 | 0 | 1 | 14 | Srl | 0.016 | Old |
| 0.366419 | 1.850048 | 0.016 | 0 | 1 | 14 | Arhgap25 | 0.016 | Old |
| 0.366419 | 1.845296 | 0.016 | 0 | 1 | 14 | Fam206a | 0.016 | Old |
| 0.366419 | 1.842851 | 0.016 | 0 | 1 | 14 | Myrf | 0.016 | Old |
| 0.366419 | 1.810501 | 0.016 | 0 | 1 | 14 | 5730507C( | 0.016 | Old |

| | | | | | | | | |
|---|---|---|---|---|---|---|---|---|
| 0.366419 | 1.801557 | 0.016 | 0 | 1 | 14 | Nupr1l | 0.016 | Old |
| 0.366419 | 1.797031 | 0.016 | 0 | 1 | 14 | A430046D1 | 0.016 | Old |
| 0.366419 | 1.785393 | 0.016 | 0 | 1 | 14 | Sars2 | 0.016 | Old |
| 0.366419 | 1.7696 | 0.016 | 0 | 1 | 14 | Ifi209 | 0.016 | Old |
| 0.366419 | 1.768141 | 0.016 | 0 | 1 | 14 | Efna4 | 0.016 | Old |
| 0.366419 | 1.763887 | 0.016 | 0 | 1 | 14 | Dcaf12l1 | 0.016 | Old |
| 0.366419 | 1.763454 | 0.016 | 0 | 1 | 14 | Sema4d | 0.016 | Old |
| 0.366419 | 1.755714 | 0.016 | 0 | 1 | 14 | Kif21a | 0.016 | Old |
| 0.366419 | 1.75524 | 0.016 | 0 | 1 | 14 | Glcci1 | 0.016 | Old |
| 0.366419 | 1.754468 | 0.016 | 0 | 1 | 14 | Gm10638 | 0.016 | Old |
| 0.366419 | 1.752758 | 0.016 | 0 | 1 | 14 | Fads2 | 0.016 | Old |
| 0.366419 | 1.75132 | 0.016 | 0 | 1 | 14 | Surf2 | 0.016 | Old |
| 0.366419 | 1.73145 | 0.016 | 0 | 1 | 14 | Plxna4os1 | 0.016 | Old |
| 0.366419 | 1.718841 | 0.016 | 0 | 1 | 14 | Noxo1 | 0.016 | Old |
| 0.366419 | 1.699466 | 0.016 | 0 | 1 | 14 | Gm15879 | 0.016 | Old |
| 0.366419 | 1.698402 | 0.016 | 0 | 1 | 14 | Gm43672 | 0.016 | Old |
| 0.366419 | 1.696555 | 0.016 | 0 | 1 | 14 | Rilp | 0.016 | Old |
| 0.366419 | 1.695955 | 0.016 | 0 | 1 | 14 | Klhl36 | 0.016 | Old |
| 0.366419 | 1.694725 | 0.016 | 0 | 1 | 14 | Nutf2 | 0.016 | Old |
| 0.366419 | 1.685843 | 0.016 | 0 | 1 | 14 | Ntf3 | 0.016 | Old |
| 0.366419 | 1.680703 | 0.016 | 0 | 1 | 14 | Olfr655 | 0.016 | Old |
| 0.366419 | 1.677214 | 0.016 | 0 | 1 | 14 | Zfp964 | 0.016 | Old |
| 0.366419 | 1.677019 | 0.016 | 0 | 1 | 14 | Sh3tc1 | 0.016 | Old |
| 0.366419 | 1.676658 | 0.016 | 0 | 1 | 14 | Gm14295 | 0.016 | Old |
| 0.366419 | 1.667078 | 0.016 | 0 | 1 | 14 | A930004D1 | 0.016 | Old |
| 0.366419 | 1.648729 | 0.016 | 0 | 1 | 14 | Cfap74 | 0.016 | Old |
| 0.366419 | 1.64815 | 0.016 | 0 | 1 | 14 | 6330418K0 | 0.016 | Old |
| 0.366419 | 1.642745 | 0.016 | 0 | 1 | 14 | Gm12689 | 0.016 | Old |
| 0.366419 | 1.624977 | 0.016 | 0 | 1 | 14 | Ccnj | 0.016 | Old |
| 0.366419 | 1.607091 | 0.016 | 0 | 1 | 14 | Primpol | 0.016 | Old |
| 0.366419 | 1.596589 | 0.016 | 0 | 1 | 14 | Thpo | 0.016 | Old |
| 0.366419 | 1.594157 | 0.016 | 0 | 1 | 14 | Syne3 | 0.016 | Old |
| 0.366419 | 1.592155 | 0.016 | 0 | 1 | 14 | Tnfrsf11a | 0.016 | Old |
| 0.366419 | 1.586638 | 0.016 | 0 | 1 | 14 | Cela1 | 0.016 | Old |
| 0.366419 | 1.586204 | 0.016 | 0 | 1 | 14 | Sytl3 | 0.016 | Old |
| 0.366419 | 1.585522 | 0.016 | 0 | 1 | 14 | AW551984 | 0.016 | Old |
| 0.366419 | 1.58229 | 0.016 | 0 | 1 | 14 | Sfn | 0.016 | Old |
| 0.366419 | 1.577015 | 0.016 | 0 | 1 | 14 | Hist1h4d | 0.016 | Old |
| 0.366419 | 1.571613 | 0.016 | 0 | 1 | 14 | Ahcy | 0.016 | Old |
| 0.366419 | 1.554595 | 0.016 | 0 | 1 | 14 | Zkscan4 | 0.016 | Old |
| 0.366419 | 1.546898 | 0.016 | 0 | 1 | 14 | Ppara | 0.016 | Old |
| 0.366419 | 1.54662 | 0.016 | 0 | 1 | 14 | Zfp456 | 0.016 | Old |
| 0.366419 | 1.543421 | 0.016 | 0 | 1 | 14 | Hoxa13 | 0.016 | Old |
| 0.366419 | 1.504098 | 0.016 | 0 | 1 | 14 | Gm26533 | 0.016 | Old |
| 0.366419 | 1.503303 | 0.016 | 0 | 1 | 14 | Bfsp1 | 0.016 | Old |

| | | | | | | | | |
|---|---|---|---|---|---|---|---|---|
| 0.366419 | 1.502684 | 0.016 | 0 | 1 | 14 | Zfp566 | 0.016 | Old |
| 0.366419 | 1.48049 | 0.016 | 0 | 1 | 14 | Fgf12 | 0.016 | Old |
| 0.366419 | 1.467893 | 0.016 | 0 | 1 | 14 | Gbp6 | 0.016 | Old |
| 0.366419 | 1.452979 | 0.016 | 0 | 1 | 14 | Tmem268 | 0.016 | Old |
| 0.366419 | 1.441746 | 0.016 | 0 | 1 | 14 | 5930412G1 | 0.016 | Old |
| 0.366419 | 1.437816 | 0.016 | 0 | 1 | 14 | Lrrn3 | 0.016 | Old |
| 0.366419 | 1.427178 | 0.016 | 0 | 1 | 14 | Gm29394 | 0.016 | Old |
| 0.366419 | 1.423098 | 0.016 | 0 | 1 | 14 | Ccdc136 | 0.016 | Old |
| 0.366419 | 1.420812 | 0.016 | 0 | 1 | 14 | Ndor1 | 0.016 | Old |
| 0.366419 | 1.390796 | 0.016 | 0 | 1 | 14 | Brd3os | 0.016 | Old |
| 0.366419 | 1.38907 | 0.016 | 0 | 1 | 14 | Gm42715 | 0.016 | Old |
| 0.366419 | 1.375422 | 0.016 | 0 | 1 | 14 | Gm43062 | 0.016 | Old |
| 0.366419 | 1.374004 | 0.016 | 0 | 1 | 14 | Milr1 | 0.016 | Old |
| 0.366419 | 1.35845 | 0.016 | 0 | 1 | 14 | Gm14322 | 0.016 | Old |
| 0.366419 | 1.355671 | 0.016 | 0 | 1 | 14 | AI467606 | 0.016 | Old |
| 0.366419 | 1.349365 | 0.016 | 0 | 1 | 14 | Fzd10 | 0.016 | Old |
| 0.366419 | 1.319934 | 0.016 | 0 | 1 | 14 | Zswim3 | 0.016 | Old |
| 0.366419 | 1.300203 | 0.016 | 0 | 1 | 14 | 5-Sep | 0.016 | Old |
| 0.366419 | 1.280429 | 0.016 | 0 | 1 | 14 | Snrpn | 0.016 | Old |
| 0.366419 | 1.275943 | 0.016 | 0 | 1 | 14 | Gm46447 | 0.016 | Old |
| 0.366419 | 1.249541 | 0.016 | 0 | 1 | 14 | BC051226 | 0.016 | Old |
| 0.366419 | 1.219096 | 0.016 | 0 | 1 | 14 | Cdk5rap1 | 0.016 | Old |
| 0.36656 | 0.408115 | 0.404 | 0.36 | 1 | 14 | Ranbp2 | 0.044 | Old |
| 0.366656 | 0.115264 | 0.847 | 0.8 | 1 | 14 | Rpl6 | 0.047 | Old |
| 0.366772 | 0.553417 | 0.191 | 0.14 | 1 | 14 | Tcof1 | 0.051 | Old |
| 0.366824 | 1.201024 | 0.142 | 0.1 | 1 | 14 | Tsc1 | 0.042 | Old |
| 0.366856 | 0.637117 | 0.169 | 0.12 | 1 | 14 | Utp3 | 0.049 | Old |
| 0.367722 | 1.349657 | 0.098 | 0.06 | 1 | 14 | Ddx19a | 0.038 | Old |
| 0.367926 | 0.060797 | 0.333 | 0.28 | 1 | 14 | Psmd4 | 0.053 | Old |
| 0.368061 | 0.536407 | 0.148 | 0.1 | 1 | 14 | Dda1 | 0.048 | Old |
| 0.368227 | 0.105503 | 0.093 | 0.14 | 1 | 14 | Ctdp1 | -0.047 | Old |
| 0.368515 | 0.908976 | 0.197 | 0.14 | 1 | 14 | Chpf2 | 0.057 | Old |
| 0.368583 | 0.97766 | 0.077 | 0.04 | 1 | 14 | Wdr18 | 0.037 | Old |
| 0.368583 | 0.910991 | 0.077 | 0.04 | 1 | 14 | Wdr73 | 0.037 | Old |
| 0.368583 | 0.837694 | 0.077 | 0.04 | 1 | 14 | AC130815. | 0.037 | Old |
| 0.368583 | 0.183655 | 0.06 | 0.1 | 1 | 14 | Tmem131l | -0.04 | Old |
| 0.368794 | 0.807933 | 0.251 | 0.2 | 1 | 14 | Rbm6 | 0.051 | Old |
| 0.368957 | 1.26645 | 0.142 | 0.1 | 1 | 14 | BC005561 | 0.042 | Old |
| 0.36912 | 0.432189 | 0.153 | 0.1 | 1 | 14 | Atp6v0a2 | 0.053 | Old |
| 0.369425 | 0.391247 | 0.208 | 0.16 | 1 | 14 | Pard3b | 0.048 | Old |
| 0.369926 | 0.801633 | 0.126 | 0.08 | 1 | 14 | Slc20a1 | 0.046 | Old |
| 0.370208 | 0.837954 | 0.328 | 0.3 | 1 | 14 | Ces2g | 0.028 | Old |
| 0.370262 | 1.262803 | 0.098 | 0.06 | 1 | 14 | Pitrm1 | 0.038 | Old |
| 0.370363 | 0.232165 | 0.372 | 0.3 | 1 | 14 | Rp9 | 0.072 | Old |
| 0.371238 | 0.098826 | 0.077 | 0.12 | 1 | 14 | Arl2 | -0.043 | Old |

| | | | | | | | | |
|---|---|---|---|---|---|---|---|---|
| 0.371577 | 0.691747 | 0.279 | 0.22 | 1 | 14 | Max | 0.059 | Old |
| 0.371664 | 0.524504 | 0.213 | 0.16 | 1 | 14 | Fbxo38 | 0.053 | Old |
| 0.37168 | 0.217654 | 0.23 | 0.18 | 1 | 14 | Hcfc1 | 0.05 | Old |
| 0.371706 | 1.551204 | 0.049 | 0.02 | 1 | 14 | Zfp579 | 0.029 | Old |
| 0.371706 | 1.355062 | 0.049 | 0.02 | 1 | 14 | Fstl3 | 0.029 | Old |
| 0.371706 | 0.874072 | 0.049 | 0.02 | 1 | 14 | Zfp11 | 0.029 | Old |
| 0.372132 | 0.299032 | 0.448 | 0.38 | 1 | 14 | Prdx5 | 0.068 | Old |
| 0.372157 | 0.393928 | 0.175 | 0.12 | 1 | 14 | Zbtb11 | 0.055 | Old |
| 0.372813 | 1.181689 | 0.098 | 0.06 | 1 | 14 | Htatip2 | 0.038 | Old |
| 0.373023 | 0.016146 | 0.093 | 0.14 | 1 | 14 | Cep290 | -0.047 | Old |
| 0.373165 | 0.11737 | 0.322 | 0.42 | 1 | 14 | Rnf11 | -0.098 | Old |
| 0.373369 | 0.914575 | 0.164 | 0.12 | 1 | 14 | Plpp5 | 0.044 | Old |
| 0.373687 | 0.235117 | 0.421 | 0.34 | 1 | 14 | Appbp2 | 0.081 | Old |
| 0.37401 | 0.42693 | 0.104 | 0.06 | 1 | 14 | Snrpa1 | 0.044 | Old |
| 0.374214 | 0.693199 | 0.224 | 0.18 | 1 | 14 | Fam160b1 | 0.044 | Old |
| 0.374363 | 0.958371 | 0.077 | 0.04 | 1 | 14 | Oplah | 0.037 | Old |
| 0.374363 | 0.842397 | 0.077 | 0.04 | 1 | 14 | Ralgps1 | 0.037 | Old |
| 0.374363 | 0.005115 | 0.06 | 0.1 | 1 | 14 | Cxcl14 | -0.04 | Old |
| 0.374783 | 0.290746 | 0.743 | 0.66 | 1 | 14 | Ptp4a2 | 0.083 | Old |
| 0.374997 | 0.437846 | 0.344 | 0.28 | 1 | 14 | Pde3a | 0.064 | Old |
| 0.375285 | 0.082599 | 0.973 | 0.92 | 1 | 14 | Ptma | 0.053 | Old |
| 0.375326 | 2.20234 | 0.049 | 0.02 | 1 | 14 | Col12a1 | 0.029 | Old |
| 0.375326 | 0.929708 | 0.049 | 0.02 | 1 | 14 | Fbxl15 | 0.029 | Old |
| 0.375326 | 0.907913 | 0.049 | 0.02 | 1 | 14 | Mtcl1 | 0.029 | Old |
| 0.375326 | 0.851636 | 0.049 | 0.02 | 1 | 14 | Anxa9 | 0.029 | Old |
| 0.375397 | 0.558295 | 0.153 | 0.1 | 1 | 14 | Nagk | 0.053 | Old |
| 0.376096 | 0.078078 | 0.235 | 0.18 | 1 | 14 | Mrps17 | 0.055 | Old |
| 0.376335 | 0.601553 | 0.328 | 0.3 | 1 | 14 | Fip1l1 | 0.028 | Old |
| 0.376985 | 0.864039 | 0.257 | 0.2 | 1 | 14 | Lats1 | 0.057 | Old |
| 0.377274 | 0.28462 | 0.077 | 0.04 | 1 | 14 | Apool | 0.037 | Old |
| 0.377274 | 0.230865 | 0.077 | 0.04 | 1 | 14 | Otud1 | 0.037 | Old |
| 0.377424 | 0.875386 | 0.164 | 0.12 | 1 | 14 | Ing3 | 0.044 | Old |
| 0.377842 | 0.527903 | 0.421 | 0.38 | 1 | 14 | Cct3 | 0.041 | Old |
| 0.378052 | 0.546845 | 0.301 | 0.24 | 1 | 14 | Ripk1 | 0.061 | Old |
| 0.378101 | 0.102136 | 0.175 | 0.12 | 1 | 14 | Ermp1 | 0.055 | Old |
| 0.378264 | 0.361177 | 0.153 | 0.22 | 1 | 14 | Bnip3 | -0.067 | Old |
| 0.378369 | 0.246888 | 0.284 | 0.22 | 1 | 14 | Xist | 0.064 | Old |
| 0.378681 | 0.840268 | 0.148 | 0.1 | 1 | 14 | Babam1 | 0.048 | Old |
| 0.378681 | 0.425349 | 0.148 | 0.1 | 1 | 14 | Fbxl14 | 0.048 | Old |
| 0.378968 | 1.191763 | 0.049 | 0.02 | 1 | 14 | Dhodh | 0.029 | Old |
| 0.378968 | 0.793275 | 0.049 | 0.02 | 1 | 14 | Tmem203 | 0.029 | Old |
| 0.37911 | 0.331069 | 0.077 | 0.12 | 1 | 14 | Trp53i13 | -0.043 | Old |
| 0.37911 | 0.08697 | 0.077 | 0.12 | 1 | 14 | Med29 | -0.043 | Old |
| 0.379221 | 0.803508 | 0.246 | 0.2 | 1 | 14 | Znhit1 | 0.046 | Old |
| 0.379619 | 0.252627 | 0.153 | 0.1 | 1 | 14 | Naxe | 0.053 | Old |

| | | | | | | | | |
|---|---|---|---|---|---|---|---|---|
| 0.379741 | 1.327518 | 0.142 | 0.1 | 1 | 14 | Cdc14b | 0.042 | Old |
| 0.379741 | 1.148545 | 0.142 | 0.1 | 1 | 14 | Vamp4 | 0.042 | Old |
| 0.380199 | 0.339141 | 0.077 | 0.04 | 1 | 14 | Ggct | 0.037 | Old |
| 0.380316 | 0.127815 | 0.765 | 0.74 | 1 | 14 | Srrm2 | 0.025 | Old |
| 0.380503 | 1.237746 | 0.23 | 0.18 | 1 | 14 | Agtrap | 0.05 | Old |
| 0.38192 | 1.283153 | 0.142 | 0.1 | 1 | 14 | Prr14 | 0.042 | Old |
| 0.382271 | 0.133026 | 0.596 | 0.54 | 1 | 14 | Bmpr2 | 0.056 | Old |
| 0.382632 | 0.668435 | 0.049 | 0.02 | 1 | 14 | Rnd1 | 0.029 | Old |
| 0.382632 | 0.661224 | 0.049 | 0.02 | 1 | 14 | Cep170b | 0.029 | Old |
| 0.382632 | 0.632556 | 0.049 | 0.02 | 1 | 14 | Isoc2b | 0.029 | Old |
| 0.382632 | 0.570806 | 0.049 | 0.02 | 1 | 14 | 6030419C1 | 0.029 | Old |
| 0.383124 | 1.009013 | 0.098 | 0.06 | 1 | 14 | Fam102a | 0.038 | Old |
| 0.383142 | 0.965677 | 0.224 | 0.18 | 1 | 14 | Abl2 | 0.044 | Old |
| 0.38321 | 0.154704 | 0.18 | 0.12 | 1 | 14 | Atg16l1 | 0.06 | Old |
| 0.384092 | 0.9643 | 0.186 | 0.14 | 1 | 14 | Dpy19l4 | 0.046 | Old |
| 0.384125 | 0.285665 | 0.104 | 0.06 | 1 | 14 | Hs1bp3 | 0.044 | Old |
| 0.384125 | 0.248455 | 0.104 | 0.06 | 1 | 14 | Hars2 | 0.044 | Old |
| 0.384404 | 0.673415 | 0.29 | 0.24 | 1 | 14 | Ubr2 | 0.05 | Old |
| 0.384414 | 0.342567 | 0.077 | 0.12 | 1 | 14 | Mapkbp1 | -0.043 | Old |
| 0.384455 | 0.093851 | 0.213 | 0.16 | 1 | 14 | Tmx1 | 0.053 | Old |
| 0.385143 | 0.55416 | 0.148 | 0.1 | 1 | 14 | Mapk8ip3 | 0.048 | Old |
| 0.385143 | 0.368959 | 0.148 | 0.1 | 1 | 14 | Slc35b4 | 0.048 | Old |
| 0.385383 | 0.687672 | 0.399 | 0.36 | 1 | 14 | Fuca1 | 0.039 | Old |
| 0.386053 | 0.385948 | 0.126 | 0.08 | 1 | 14 | Ostm1 | 0.046 | Old |
| 0.386196 | 0.186625 | 0.153 | 0.22 | 1 | 14 | Samm50 | -0.067 | Old |
| 0.386226 | 0.184913 | 0.525 | 0.48 | 1 | 14 | Pgrmc1 | 0.045 | Old |
| 0.386303 | 0.904415 | 0.142 | 0.1 | 1 | 14 | Steap2 | 0.042 | Old |
| 0.386317 | 0.248197 | 0.049 | 0.02 | 1 | 14 | Sgf29 | 0.029 | Old |
| 0.386679 | 0.242982 | 0.104 | 0.06 | 1 | 14 | Dusp7 | 0.044 | Old |
| 0.386696 | 0.213875 | 0.235 | 0.18 | 1 | 14 | Ilf3 | 0.055 | Old |
| 0.386749 | 0.81572 | 0.224 | 0.18 | 1 | 14 | Lin7c | 0.044 | Old |
| 0.386947 | 0.697083 | 0.169 | 0.12 | 1 | 14 | Cherp | 0.049 | Old |
| 0.387807 | 0.00375 | 0.65 | 0.56 | 1 | 14 | Glul | 0.09 | Old |
| 0.388076 | 0.091142 | 0.557 | 0.52 | 1 | 14 | Taok1 | 0.037 | Old |
| 0.388158 | 0.524035 | 0.213 | 0.16 | 1 | 14 | Tax1bp3 | 0.053 | Old |
| 0.388185 | 0.12856 | 0.388 | 0.32 | 1 | 14 | Ndrg1 | 0.068 | Old |
| 0.388345 | 1.352446 | 0.098 | 0.06 | 1 | 14 | Haghl | 0.038 | Old |
| 0.388505 | 1.26026 | 0.142 | 0.1 | 1 | 14 | Zfp217 | 0.042 | Old |
| 0.388561 | 0.906424 | 0.224 | 0.18 | 1 | 14 | Cc2d1b | 0.044 | Old |
| 0.388561 | 0.829793 | 0.224 | 0.18 | 1 | 14 | Ube2e2 | 0.044 | Old |
| 0.389488 | 0.753193 | 0.148 | 0.1 | 1 | 14 | Apopt1 | 0.048 | Old |
| 0.389488 | 0.164744 | 0.126 | 0.18 | 1 | 14 | Rbbp9 | -0.054 | Old |
| 0.389898 | 2.337934 | 0.071 | 0.04 | 1 | 14 | Plbd1 | 0.031 | Old |
| 0.389898 | 2.055928 | 0.071 | 0.04 | 1 | 14 | 6720427I0 | 0.031 | Old |
| 0.389898 | 1.84816 | 0.071 | 0.04 | 1 | 14 | Aifm1 | 0.031 | Old |

| | | | | | | | | |
|---|---|---|---|---|---|---|---|---|
| 0.389966 | 0.605114 | 0.186 | 0.14 | 1 | 14 | Mapk9 | 0.046 | Old |
| 0.390072 | 0.098956 | 0.552 | 0.48 | 1 | 14 | Emp3 | 0.072 | Old |
| 0.3902 | 0.490838 | 0.153 | 0.22 | 1 | 14 | Mn1 | -0.067 | Old |
| 0.390972 | 0.696358 | 0.098 | 0.06 | 1 | 14 | Lonp1 | 0.038 | Old |
| 0.391012 | 0.345139 | 0.24 | 0.18 | 1 | 14 | Flywch1 | 0.06 | Old |
| 0.391018 | 0.200519 | 0.421 | 0.36 | 1 | 14 | Cct5 | 0.061 | Old |
| 0.391911 | 0.504182 | 0.35 | 0.3 | 1 | 14 | Dstyk | 0.05 | Old |
| 0.392037 | 0.258542 | 0.077 | 0.04 | 1 | 14 | Cep19 | 0.037 | Old |
| 0.392095 | 0.077193 | 0.541 | 0.5 | 1 | 14 | Rab10 | 0.041 | Old |
| 0.392455 | 1.893436 | 0.093 | 0.06 | 1 | 14 | 2310009B1 | 0.033 | Old |
| 0.392934 | 1.151336 | 0.142 | 0.1 | 1 | 14 | Map1s | 0.042 | Old |
| 0.392973 | 2.324306 | 0.071 | 0.04 | 1 | 14 | Ccdc157 | 0.031 | Old |
| 0.393243 | 0.002824 | 0.792 | 0.7 | 1 | 14 | Jak1 | 0.092 | Old |
| 0.393609 | 1.210045 | 0.098 | 0.06 | 1 | 14 | Tceanc2 | 0.038 | Old |
| 0.393609 | 0.802506 | 0.098 | 0.06 | 1 | 14 | Cln6 | 0.038 | Old |
| 0.394119 | 0.13966 | 0.18 | 0.14 | 1 | 14 | Rps12-ps3 | 0.04 | Old |
| 0.394692 | 0.957419 | 0.197 | 0.16 | 1 | 14 | Nudcd2 | 0.037 | Old |
| 0.394754 | 1.877624 | 0.12 | 0.08 | 1 | 14 | Ing2 | 0.04 | Old |
| 0.39493 | 1.214311 | 0.219 | 0.18 | 1 | 14 | Tsg101 | 0.039 | Old |
| 0.394967 | 0.922418 | 0.383 | 0.34 | 1 | 14 | Nrp1 | 0.043 | Old |
| 0.395174 | 0.365563 | 0.197 | 0.14 | 1 | 14 | Sbsn | 0.057 | Old |
| 0.395401 | 0.175554 | 0.208 | 0.16 | 1 | 14 | Rcor1 | 0.048 | Old |
| 0.395446 | 0.047113 | 0.262 | 0.2 | 1 | 14 | Atg3 | 0.062 | Old |
| 0.395629 | 0.539347 | 0.213 | 0.16 | 1 | 14 | C77080 | 0.053 | Old |
| 0.395893 | 0.946597 | 0.186 | 0.14 | 1 | 14 | Trim33 | 0.046 | Old |
| 0.396061 | 0.007156 | 0.126 | 0.18 | 1 | 14 | Mrps5 | -0.054 | Old |
| 0.396063 | 1.842227 | 0.071 | 0.04 | 1 | 14 | 9430038I0 | 0.031 | Old |
| 0.396225 | 0.163518 | 0.579 | 0.5 | 1 | 14 | Lsp1 | 0.079 | Old |
| 0.396257 | 1.035901 | 0.098 | 0.06 | 1 | 14 | Zfp65 | 0.038 | Old |
| 0.396257 | 0.937085 | 0.098 | 0.06 | 1 | 14 | Tlr3 | 0.038 | Old |
| 0.396411 | 0.022378 | 0.437 | 0.36 | 1 | 14 | Angptl4 | 0.077 | Old |
| 0.39731 | 0.572881 | 0.393 | 0.36 | 1 | 14 | Ikbip | 0.033 | Old |
| 0.397501 | 0.810237 | 0.279 | 0.24 | 1 | 14 | Gna13 | 0.039 | Old |
| 0.397521 | 0.845474 | 0.262 | 0.22 | 1 | 14 | Zmat2 | 0.042 | Old |
| 0.397632 | 0.513235 | 0.29 | 0.24 | 1 | 14 | Slc25a24 | 0.05 | Old |
| 0.397803 | 0.06886 | 0.809 | 0.72 | 1 | 14 | Pcbp2 | 0.089 | Old |
| 0.397872 | 2.21653 | 0.093 | 0.06 | 1 | 14 | Plekhm3 | 0.033 | Old |
| 0.398034 | 0.587136 | 0.311 | 0.26 | 1 | 14 | Rc3h1 | 0.051 | Old |
| 0.398104 | 0.983407 | 0.164 | 0.12 | 1 | 14 | Ap1b1 | 0.044 | Old |
| 0.398104 | 0.741323 | 0.164 | 0.12 | 1 | 14 | Gdap10 | 0.044 | Old |
| 0.39813 | 1.166648 | 0.18 | 0.14 | 1 | 14 | Atp6v1c1 | 0.04 | Old |
| 0.398765 | 0.107798 | 0.033 | 0.06 | 1 | 14 | Zfp697 | -0.027 | Old |
| 0.398916 | 0.812537 | 0.098 | 0.06 | 1 | 14 | Stard5 | 0.038 | Old |
| 0.399168 | 1.782548 | 0.071 | 0.04 | 1 | 14 | Lmf1 | 0.031 | Old |
| 0.399318 | 0.720425 | 0.169 | 0.12 | 1 | 14 | Zfpl1 | 0.049 | Old |

| | | | | | | | | |
|---|---|---|---|---|---|---|---|---|
| 0.399583 | 0.99633 | 0.12 | 0.08 | 1 | 14 | Trpc1 | 0.04 | Old |
| 0.399633 | 1.413263 | 0.142 | 0.1 | 1 | 14 | Ap5m1 | 0.042 | Old |
| 0.39984 | 0.702713 | 0.306 | 0.26 | 1 | 14 | Tfe3 | 0.046 | Old |
| 0.400235 | 0.092314 | 0.459 | 0.38 | 1 | 14 | 1700020I14 | 0.079 | Old |
| 0.400597 | 0.441431 | 0.077 | 0.12 | 1 | 14 | Nbn | -0.043 | Old |
| 0.400597 | 0.319565 | 0.077 | 0.12 | 1 | 14 | Ap4m1 | -0.043 | Old |
| 0.400637 | 0.261571 | 0.459 | 0.4 | 1 | 14 | Scaf11 | 0.059 | Old |
| 0.40106 | 0.557206 | 0.06 | 0.1 | 1 | 14 | Rasa4 | -0.04 | Old |
| 0.401251 | 0.279163 | 0.268 | 0.22 | 1 | 14 | Uggt1 | 0.048 | Old |
| 0.401274 | 0.465312 | 0.454 | 0.38 | 1 | 14 | Osr1 | 0.074 | Old |
| 0.401496 | 0.135298 | 0.191 | 0.14 | 1 | 14 | Adamtsl3 | 0.051 | Old |
| 0.401586 | 0.726349 | 0.098 | 0.06 | 1 | 14 | Arhgef18 | 0.038 | Old |
| 0.401736 | 0.089025 | 0.284 | 0.22 | 1 | 14 | Ephx1 | 0.064 | Old |
| 0.402011 | 0.794975 | 0.12 | 0.08 | 1 | 14 | Surf1 | 0.04 | Old |
| 0.402198 | 0.331783 | 0.23 | 0.18 | 1 | 14 | Nov | 0.05 | Old |
| 0.402287 | 1.905909 | 0.071 | 0.04 | 1 | 14 | H1fx | 0.031 | Old |
| 0.402392 | 1.248306 | 0.24 | 0.2 | 1 | 14 | Alkal2 | 0.04 | Old |
| 0.40244 | 1.075308 | 0.197 | 0.16 | 1 | 14 | Xpo1 | 0.037 | Old |
| 0.402609 | 0.03534 | 0.093 | 0.14 | 1 | 14 | Atg2b | -0.047 | Old |
| 0.40274 | 0.449432 | 0.033 | 0.06 | 1 | 14 | Galm | -0.027 | Old |
| 0.402774 | 1.024638 | 0.202 | 0.16 | 1 | 14 | Nbea | 0.042 | Old |
| 0.402774 | 1.001872 | 0.202 | 0.16 | 1 | 14 | Vkorc1l1 | 0.042 | Old |
| 0.402786 | 0.007322 | 0.366 | 0.3 | 1 | 14 | Tacc2 | 0.066 | Old |
| 0.402981 | 0.050741 | 0.109 | 0.16 | 1 | 14 | Coro1a | -0.051 | Old |
| 0.403021 | 0.697771 | 0.208 | 0.16 | 1 | 14 | Tnfsf9 | 0.048 | Old |
| 0.403238 | 1.052171 | 0.224 | 0.18 | 1 | 14 | Thbs1 | 0.044 | Old |
| 0.404448 | 0.815659 | 0.12 | 0.08 | 1 | 14 | Tfdp1 | 0.04 | Old |
| 0.404876 | 0.061742 | 0.885 | 0.84 | 1 | 14 | Nfix | 0.045 | Old |
| 0.404929 | 0.606929 | 0.148 | 0.1 | 1 | 14 | Zfp318 | 0.048 | Old |
| 0.40494 | 0.887114 | 0.208 | 0.16 | 1 | 14 | Cxcl16 | 0.048 | Old |
| 0.405096 | 0.739706 | 0.224 | 0.18 | 1 | 14 | Arid2 | 0.044 | Old |
| 0.405193 | 0.126562 | 0.262 | 0.34 | 1 | 14 | Rab3gap1 | -0.078 | Old |
| 0.405287 | 0.981981 | 0.158 | 0.12 | 1 | 14 | Mrps18c | 0.038 | Old |
| 0.405397 | 0.844495 | 0.317 | 0.3 | 1 | 14 | Cab39 | 0.017 | Old |
| 0.405473 | 0.485788 | 0.191 | 0.14 | 1 | 14 | Gulp1 | 0.051 | Old |
| 0.405589 | 0.921689 | 0.251 | 0.22 | 1 | 14 | Synj1 | 0.031 | Old |
| 0.406144 | 1.757536 | 0.115 | 0.08 | 1 | 14 | Fam219a | 0.035 | Old |
| 0.406348 | 0.107055 | 0.202 | 0.14 | 1 | 14 | Crim1 | 0.062 | Old |
| 0.406739 | 0.557279 | 0.033 | 0.06 | 1 | 14 | Cysltr1 | -0.027 | Old |
| 0.406739 | 0.399758 | 0.033 | 0.06 | 1 | 14 | Zfp949 | -0.027 | Old |
| 0.406894 | 1.081806 | 0.12 | 0.08 | 1 | 14 | Mlh3 | 0.04 | Old |
| 0.406957 | 1.119716 | 0.098 | 0.06 | 1 | 14 | Map4k3 | 0.038 | Old |
| 0.406957 | 0.559714 | 0.098 | 0.06 | 1 | 14 | Wdr60 | 0.038 | Old |
| 0.407632 | 0.296456 | 0.317 | 0.26 | 1 | 14 | Zfpm2 | 0.057 | Old |
| 0.407885 | 0.087384 | 0.284 | 0.36 | 1 | 14 | P2rx4 | -0.076 | Old |

| | | | | | | | | |
|---|---|---|---|---|---|---|---|---|
| 0.40857 | 1.526013 | 0.071 | 0.04 | 1 | 14 | Utp18 | 0.031 | Old |
| 0.40857 | 1.41445 | 0.071 | 0.04 | 1 | 14 | Ttc38 | 0.031 | Old |
| 0.408695 | 0.769695 | 0.164 | 0.12 | 1 | 14 | Cyb561 | 0.044 | Old |
| 0.408695 | 0.116656 | 0.142 | 0.2 | 1 | 14 | Dcaf1 | -0.058 | Old |
| 0.40876 | 1.171273 | 0.246 | 0.22 | 1 | 14 | H2-T22 | 0.026 | Old |
| 0.40954 | 0.444053 | 0.311 | 0.26 | 1 | 14 | Heca | 0.051 | Old |
| 0.409868 | 0.140704 | 0.219 | 0.18 | 1 | 14 | Herpud2 | 0.039 | Old |
| 0.410219 | 1.281402 | 0.093 | 0.14 | 1 | 14 | Il6 | -0.047 | Old |
| 0.411239 | 0.220644 | 0.262 | 0.2 | 1 | 14 | Llph | 0.062 | Old |
| 0.411733 | 1.794993 | 0.071 | 0.04 | 1 | 14 | Eml2 | 0.031 | Old |
| 0.411733 | 1.786314 | 0.071 | 0.04 | 1 | 14 | Fgd1 | 0.031 | Old |
| 0.411733 | 1.573942 | 0.071 | 0.04 | 1 | 14 | Ermard | 0.031 | Old |
| 0.411733 | 1.450847 | 0.071 | 0.04 | 1 | 14 | Nprl2 | 0.031 | Old |
| 0.411733 | 1.361995 | 0.071 | 0.04 | 1 | 14 | Zdhhc16 | 0.031 | Old |
| 0.411733 | 1.216504 | 0.071 | 0.04 | 1 | 14 | Supt7l | 0.031 | Old |
| 0.411758 | 0.002333 | 0.29 | 0.36 | 1 | 14 | Peak1 | -0.07 | Old |
| 0.411845 | 0.709296 | 0.339 | 0.3 | 1 | 14 | Thbs3 | 0.039 | Old |
| 0.412578 | 0.624926 | 0.224 | 0.18 | 1 | 14 | Tet3 | 0.044 | Old |
| 0.41267 | 1.388716 | 0.175 | 0.14 | 1 | 14 | Brd9 | 0.035 | Old |
| 0.413106 | 0.89859 | 0.306 | 0.26 | 1 | 14 | Snx17 | 0.046 | Old |
| 0.413311 | 0.381009 | 0.301 | 0.26 | 1 | 14 | Ntn4 | 0.041 | Old |
| 0.414009 | 0.417774 | 0.432 | 0.4 | 1 | 14 | Ppp3cb | 0.032 | Old |
| 0.414157 | 0.414354 | 0.251 | 0.2 | 1 | 14 | Slain2 | 0.051 | Old |
| 0.414283 | 1.021283 | 0.12 | 0.08 | 1 | 14 | Apeh | 0.04 | Old |
| 0.414283 | 0.10911 | 0.12 | 0.08 | 1 | 14 | Dnajc9 | 0.04 | Old |
| 0.414806 | 1.427814 | 0.033 | 0.06 | 1 | 14 | Jph2 | -0.027 | Old |
| 0.414806 | 0.693635 | 0.033 | 0.06 | 1 | 14 | Myo7a | -0.027 | Old |
| 0.414911 | 1.248392 | 0.071 | 0.04 | 1 | 14 | Zc3hc1 | 0.031 | Old |
| 0.414997 | 0.11409 | 0.306 | 0.24 | 1 | 14 | Gstm5 | 0.066 | Old |
| 0.416021 | 0.068388 | 0.301 | 0.24 | 1 | 14 | Nr2f1 | 0.061 | Old |
| 0.416433 | 1.268269 | 0.202 | 0.16 | 1 | 14 | Polr2i | 0.042 | Old |
| 0.41662 | 0.407472 | 0.213 | 0.16 | 1 | 14 | Hsd17b4 | 0.053 | Old |
| 0.416764 | 0.630534 | 0.12 | 0.08 | 1 | 14 | Map3k5 | 0.04 | Old |
| 0.417241 | 0.45372 | 0.388 | 0.36 | 1 | 14 | Efemp2 | 0.028 | Old |
| 0.417576 | 0.403168 | 0.317 | 0.26 | 1 | 14 | Hnrnph3 | 0.057 | Old |
| 0.418304 | 0.85295 | 0.246 | 0.2 | 1 | 14 | Ubp1 | 0.046 | Old |
| 0.41849 | 0.270064 | 0.333 | 0.28 | 1 | 14 | Snx5 | 0.053 | Old |
| 0.418554 | 1.059647 | 0.18 | 0.14 | 1 | 14 | Apaf1 | 0.04 | Old |
| 0.418687 | 0.602578 | 0.366 | 0.32 | 1 | 14 | Sec24d | 0.046 | Old |
| 0.418729 | 0.832663 | 0.29 | 0.26 | 1 | 14 | Rab3d | 0.03 | Old |
| 0.419138 | 0.483869 | 0.344 | 0.28 | 1 | 14 | Arhgap5 | 0.064 | Old |
| 0.419836 | 0.675253 | 0.306 | 0.26 | 1 | 14 | Rnft1 | 0.046 | Old |
| 0.419978 | 0.440571 | 0.049 | 0.08 | 1 | 14 | Gys1 | -0.031 | Old |
| 0.420135 | 0.662557 | 0.246 | 0.2 | 1 | 14 | Fads6 | 0.046 | Old |
| 0.420143 | 0.826635 | 0.186 | 0.14 | 1 | 14 | Zfp281 | 0.046 | Old |

| | | | | | | | | |
|---|---|---|---|---|---|---|---|---|
| 0.420319 | 0.295598 | 0.557 | 0.48 | 1 | 14 | Purb | 0.077 | Old |
| 0.420412 | 0.488698 | 0.388 | 0.34 | 1 | 14 | Hspa4 | 0.048 | Old |
| 0.420457 | 0.149633 | 0.169 | 0.12 | 1 | 14 | Aox1 | 0.049 | Old |
| 0.420528 | 0.469946 | 0.268 | 0.22 | 1 | 14 | Il1r1 | 0.048 | Old |
| 0.421035 | 0.051384 | 0.552 | 0.44 | 1 | 14 | Set | 0.112 | Old |
| 0.421051 | 0.132118 | 0.552 | 0.48 | 1 | 14 | Tmem106b | 0.072 | Old |
| 0.421163 | 0.033586 | 0.131 | 0.18 | 1 | 14 | Creb3 | -0.049 | Old |
| 0.421309 | 1.646895 | 0.071 | 0.04 | 1 | 14 | Fabp5 | 0.031 | Old |
| 0.421309 | 0.936389 | 0.071 | 0.04 | 1 | 14 | Zfp532 | 0.031 | Old |
| 0.421461 | 0.124291 | 0.934 | 0.9 | 1 | 14 | Rpl13 | 0.034 | Old |
| 0.421547 | 0.026724 | 0.978 | 0.96 | 1 | 14 | Rps29 | 0.018 | Old |
| 0.421751 | 0.13852 | 0.12 | 0.08 | 1 | 14 | Anks1 | 0.04 | Old |
| 0.42176 | 0.744137 | 0.311 | 0.28 | 1 | 14 | Rab11fip2 | 0.031 | Old |
| 0.421856 | 0.082596 | 0.432 | 0.38 | 1 | 14 | Palm | 0.052 | Old |
| 0.421856 | 0.295218 | 0.426 | 0.4 | 1 | 14 | Xiap | 0.026 | Old |
| 0.421971 | 0.526832 | 0.246 | 0.2 | 1 | 14 | Bcl7b | 0.046 | Old |
| 0.421971 | 0.423416 | 0.246 | 0.2 | 1 | 14 | Cyhr1 | 0.046 | Old |
| 0.422205 | 1.043799 | 0.197 | 0.16 | 1 | 14 | Ola1 | 0.037 | Old |
| 0.422452 | 0.717531 | 0.208 | 0.16 | 1 | 14 | Unc50 | 0.048 | Old |
| 0.422456 | 0.518278 | 0.142 | 0.1 | 1 | 14 | Ebf3 | 0.042 | Old |
| 0.4228 | 1.693386 | 0.093 | 0.06 | 1 | 14 | Sema3b | 0.033 | Old |
| 0.4228 | 1.537387 | 0.093 | 0.06 | 1 | 14 | Tapt1 | 0.033 | Old |
| 0.422885 | 0.170294 | 0.743 | 0.66 | 1 | 14 | Rpl4 | 0.083 | Old |
| 0.42292 | 0.006884 | 0.995 | 1 | 1 | 14 | Tpt1 | -0.005 | Old |
| 0.423325 | 0.69387 | 0.098 | 0.06 | 1 | 14 | Dnajc16 | 0.038 | Old |
| 0.423325 | 0.55159 | 0.098 | 0.06 | 1 | 14 | Galnt10 | 0.038 | Old |
| 0.423325 | 0.400991 | 0.098 | 0.06 | 1 | 14 | Cxxc4 | 0.038 | Old |
| 0.423417 | 0.644644 | 0.049 | 0.08 | 1 | 14 | Neurl1b | -0.031 | Old |
| 0.423517 | 0.391027 | 0.492 | 0.44 | 1 | 14 | Pum1 | 0.052 | Old |
| 0.423798 | 1.471626 | 0.115 | 0.08 | 1 | 14 | Slc39a13 | 0.035 | Old |
| 0.423801 | 0.50297 | 0.164 | 0.12 | 1 | 14 | Ano10 | 0.044 | Old |
| 0.423801 | 0.124244 | 0.142 | 0.2 | 1 | 14 | Bod1 | -0.058 | Old |
| 0.424327 | 0.198836 | 0.191 | 0.26 | 1 | 14 | Dmpk | -0.069 | Old |
| 0.425463 | 0.297091 | 0.667 | 0.6 | 1 | 14 | Grn | 0.067 | Old |
| 0.425625 | 1.546731 | 0.093 | 0.06 | 1 | 14 | Tsen15 | 0.033 | Old |
| 0.425625 | 1.241191 | 0.093 | 0.06 | 1 | 14 | Nob1 | 0.033 | Old |
| 0.425705 | 0.173703 | 0.191 | 0.14 | 1 | 14 | Npc1 | 0.051 | Old |
| 0.425866 | 8.74E-05 | 0.202 | 0.26 | 1 | 14 | Mrc1 | -0.058 | Old |
| 0.425879 | 0.654183 | 0.268 | 0.22 | 1 | 14 | Ncor2 | 0.048 | Old |
| 0.425985 | 1.082668 | 0.164 | 0.12 | 1 | 14 | Atic | 0.044 | Old |
| 0.426088 | 0.07037 | 0.536 | 0.48 | 1 | 14 | Htra1 | 0.056 | Old |
| 0.426355 | 0.033633 | 0.18 | 0.24 | 1 | 14 | Mier3 | -0.06 | Old |
| 0.426872 | 0.222401 | 0.049 | 0.08 | 1 | 14 | Zfp764 | -0.031 | Old |
| 0.427084 | 2.877188 | 0.044 | 0.02 | 1 | 14 | N4bp2l1 | 0.024 | Old |
| 0.427084 | 2.438021 | 0.044 | 0.02 | 1 | 14 | 1110019D1 | 0.024 | Old |

| | | | | | | | | |
|---|---|---|---|---|---|---|---|---|
| 0.427084 | 2.317444 | 0.044 | 0.02 | 1 | 14 | Aptx | 0.024 | Old |
| 0.427084 | 2.305768 | 0.044 | 0.02 | 1 | 14 | Il15 | 0.024 | Old |
| 0.427084 | 2.295512 | 0.044 | 0.02 | 1 | 14 | Armc6 | 0.024 | Old |
| 0.427084 | 2.163691 | 0.044 | 0.02 | 1 | 14 | Trpm4 | 0.024 | Old |
| 0.427084 | 1.983445 | 0.044 | 0.02 | 1 | 14 | Ttf2 | 0.024 | Old |
| 0.427084 | 1.936054 | 0.044 | 0.02 | 1 | 14 | Pex26 | 0.024 | Old |
| 0.427084 | 1.821385 | 0.044 | 0.02 | 1 | 14 | Cep70 | 0.024 | Old |
| 0.427084 | 1.796082 | 0.044 | 0.02 | 1 | 14 | Tmco6 | 0.024 | Old |
| 0.427084 | 1.746378 | 0.044 | 0.02 | 1 | 14 | Napb | 0.024 | Old |
| 0.427084 | 1.541763 | 0.044 | 0.02 | 1 | 14 | Mapk1ip1 | 0.024 | Old |
| 0.427084 | 1.400018 | 0.044 | 0.02 | 1 | 14 | Rac3 | 0.024 | Old |
| 0.427765 | 0.904567 | 0.071 | 0.04 | 1 | 14 | 3110062M | 0.031 | Old |
| 0.427765 | 0.644125 | 0.071 | 0.04 | 1 | 14 | Mfsd8 | 0.031 | Old |
| 0.427882 | 0.051867 | 0.066 | 0.1 | 1 | 14 | Rpp30 | -0.034 | Old |
| 0.428153 | 0.113227 | 0.454 | 0.38 | 1 | 14 | Tex261 | 0.074 | Old |
| 0.428176 | 0.648867 | 0.164 | 0.12 | 1 | 14 | Vamp5 | 0.044 | Old |
| 0.428417 | 0.222513 | 0.186 | 0.14 | 1 | 14 | Paip1 | 0.046 | Old |
| 0.428461 | 1.850895 | 0.093 | 0.06 | 1 | 14 | Ccdc32 | 0.033 | Old |
| 0.428461 | 1.574032 | 0.093 | 0.06 | 1 | 14 | Invs | 0.033 | Old |
| 0.428896 | 0.05135 | 0.563 | 0.5 | 1 | 14 | Smim14 | 0.063 | Old |
| 0.428923 | 1.066454 | 0.115 | 0.08 | 1 | 14 | Nt5c | 0.035 | Old |
| 0.428993 | 0.675852 | 0.18 | 0.14 | 1 | 14 | Wdr37 | 0.04 | Old |
| 0.428993 | 0.072188 | 0.158 | 0.22 | 1 | 14 | Ppil2 | -0.062 | Old |
| 0.429387 | 0.769167 | 0.284 | 0.24 | 1 | 14 | Vps29 | 0.044 | Old |
| 0.430343 | 0.206869 | 0.049 | 0.08 | 1 | 14 | Kptn | -0.031 | Old |
| 0.430343 | 0.092584 | 0.049 | 0.08 | 1 | 14 | Acrbp | -0.031 | Old |
| 0.4305 | 0.144439 | 0.164 | 0.22 | 1 | 14 | Tmem55b | -0.056 | Old |
| 0.4305 | 0.024471 | 0.164 | 0.22 | 1 | 14 | Fryl | -0.056 | Old |
| 0.430717 | 0.128537 | 0.372 | 0.3 | 1 | 14 | Pdcd5 | 0.072 | Old |
| 0.430847 | 0.905804 | 0.251 | 0.22 | 1 | 14 | Clasp2 | 0.031 | Old |
| 0.430948 | 0.262678 | 0.066 | 0.1 | 1 | 14 | Fam161a | -0.034 | Old |
| 0.431014 | 1.130975 | 0.071 | 0.04 | 1 | 14 | Cdc42ep5 | 0.031 | Old |
| 0.431023 | 0.074462 | 0.47 | 0.4 | 1 | 14 | Abi1 | 0.07 | Old |
| 0.431223 | 2.635063 | 0.044 | 0.02 | 1 | 14 | Dcp1b | 0.024 | Old |
| 0.431223 | 2.548108 | 0.044 | 0.02 | 1 | 14 | Kcnd1 | 0.024 | Old |
| 0.431223 | 2.048915 | 0.044 | 0.02 | 1 | 14 | Dzip1l | 0.024 | Old |
| 0.431223 | 1.951341 | 0.044 | 0.02 | 1 | 14 | Neil1 | 0.024 | Old |
| 0.431223 | 1.943014 | 0.044 | 0.02 | 1 | 14 | Arhgap19 | 0.024 | Old |
| 0.431223 | 1.781276 | 0.044 | 0.02 | 1 | 14 | 3110009E1 | 0.024 | Old |
| 0.431223 | 1.730119 | 0.044 | 0.02 | 1 | 14 | Zc3h12b | 0.024 | Old |
| 0.431308 | 1.38331 | 0.093 | 0.06 | 1 | 14 | Tstd2 | 0.033 | Old |
| 0.431308 | 1.003324 | 0.093 | 0.06 | 1 | 14 | Tyk2 | 0.033 | Old |
| 0.431968 | 0.278151 | 0.454 | 0.38 | 1 | 14 | Tmed7 | 0.074 | Old |
| 0.432235 | 0.632888 | 0.126 | 0.18 | 1 | 14 | Tmem50b | -0.054 | Old |
| 0.433138 | 0.192182 | 0.311 | 0.26 | 1 | 14 | Khsrp | 0.051 | Old |

| | | | | | | | | |
|---|---|---|---|---|---|---|---|---|
| 0.433572 | 1.463159 | 0.137 | 0.1 | 1 | 14 | Kif3b | 0.037 | Old |
| 0.433831 | 0.497503 | 0.049 | 0.08 | 1 | 14 | Rnf126 | -0.031 | Old |
| 0.433831 | 0.280216 | 0.049 | 0.08 | 1 | 14 | Cpsf4 | -0.031 | Old |
| 0.434084 | 1.092736 | 0.115 | 0.08 | 1 | 14 | Senp3 | 0.035 | Old |
| 0.434166 | 1.560812 | 0.093 | 0.06 | 1 | 14 | Mre11a | 0.033 | Old |
| 0.434166 | 1.274721 | 0.093 | 0.06 | 1 | 14 | Zc3hav1l | 0.033 | Old |
| 0.434172 | 0.627811 | 0.23 | 0.18 | 1 | 14 | Dazap1 | 0.05 | Old |
| 0.434266 | 0.059085 | 0.448 | 0.4 | 1 | 14 | Rab7 | 0.048 | Old |
| 0.434278 | 0.660321 | 0.071 | 0.04 | 1 | 14 | Ercc2 | 0.031 | Old |
| 0.434278 | 0.617666 | 0.071 | 0.04 | 1 | 14 | Zfp938 | 0.031 | Old |
| 0.434278 | 0.47476 | 0.071 | 0.04 | 1 | 14 | Abcb7 | 0.031 | Old |
| 0.434318 | 0.603742 | 0.295 | 0.24 | 1 | 14 | Rraga | 0.055 | Old |
| 0.434428 | 0.349643 | 0.399 | 0.34 | 1 | 14 | Hadhb | 0.059 | Old |
| 0.434558 | 0.542335 | 0.148 | 0.1 | 1 | 14 | Dnajc2 | 0.048 | Old |
| 0.434558 | 0.37549 | 0.126 | 0.18 | 1 | 14 | Dgcr6 | -0.054 | Old |
| 0.434699 | 0.399206 | 0.257 | 0.22 | 1 | 14 | Tnfaip1 | 0.037 | Old |
| 0.435014 | 0.207077 | 0.421 | 0.36 | 1 | 14 | Ndufa5 | 0.061 | Old |
| 0.435161 | 0.03761 | 0.24 | 0.18 | 1 | 14 | Mia2 | 0.06 | Old |
| 0.435328 | 0.296225 | 0.18 | 0.14 | 1 | 14 | Cntln | 0.04 | Old |
| 0.435385 | 2.206896 | 0.044 | 0.02 | 1 | 14 | 3222401L1 | 0.024 | Old |
| 0.435385 | 2.023767 | 0.044 | 0.02 | 1 | 14 | Ubiad1 | 0.024 | Old |
| 0.435385 | 1.928516 | 0.044 | 0.02 | 1 | 14 | Mrm1 | 0.024 | Old |
| 0.435385 | 1.821878 | 0.044 | 0.02 | 1 | 14 | Col8a1 | 0.024 | Old |
| 0.435385 | 1.756347 | 0.044 | 0.02 | 1 | 14 | Slc28a2 | 0.024 | Old |
| 0.435385 | 1.668178 | 0.044 | 0.02 | 1 | 14 | Klhl33 | 0.024 | Old |
| 0.435385 | 1.583675 | 0.044 | 0.02 | 1 | 14 | Cbwd1 | 0.024 | Old |
| 0.435385 | 1.565835 | 0.044 | 0.02 | 1 | 14 | Gas8 | 0.024 | Old |
| 0.435385 | 1.481772 | 0.044 | 0.02 | 1 | 14 | Gm46323 | 0.024 | Old |
| 0.435385 | 1.401166 | 0.044 | 0.02 | 1 | 14 | Zfp446 | 0.024 | Old |
| 0.435385 | 1.280902 | 0.044 | 0.02 | 1 | 14 | Zdhhc24 | 0.024 | Old |
| 0.4354 | 0.50937 | 0.29 | 0.24 | 1 | 14 | Cers5 | 0.05 | Old |
| 0.436161 | 0.278084 | 0.251 | 0.2 | 1 | 14 | Dock1 | 0.051 | Old |
| 0.437035 | 1.395234 | 0.093 | 0.06 | 1 | 14 | Fut10 | 0.033 | Old |
| 0.437164 | 0.377037 | 0.29 | 0.24 | 1 | 14 | Slc12a6 | 0.05 | Old |
| 0.437335 | 0.809578 | 0.049 | 0.08 | 1 | 14 | Flvcr1 | -0.031 | Old |
| 0.437335 | 0.391903 | 0.049 | 0.08 | 1 | 14 | Ctif | -0.031 | Old |
| 0.437335 | 0.326096 | 0.049 | 0.08 | 1 | 14 | Mus81 | -0.031 | Old |
| 0.437335 | 0.114708 | 0.049 | 0.08 | 1 | 14 | Accs | -0.031 | Old |
| 0.438419 | 0.257864 | 0.175 | 0.24 | 1 | 14 | Arhgap32 | -0.065 | Old |
| 0.439149 | 0.142491 | 0.475 | 0.42 | 1 | 14 | H13 | 0.055 | Old |
| 0.439281 | 1.255648 | 0.115 | 0.08 | 1 | 14 | Acox3 | 0.035 | Old |
| 0.43932 | 0.076934 | 0.546 | 0.46 | 1 | 14 | Mapre1 | 0.086 | Old |
| 0.439571 | 1.838345 | 0.044 | 0.02 | 1 | 14 | Dis3l | 0.024 | Old |
| 0.439571 | 1.723858 | 0.044 | 0.02 | 1 | 14 | Npas2 | 0.024 | Old |
| 0.439571 | 1.470012 | 0.044 | 0.02 | 1 | 14 | Nek4 | 0.024 | Old |

| | | | | | | | | |
|---|---|---|---|---|---|---|---|---|
| 0.439571 | 1.050232 | 0.044 | 0.02 | 1 | 14 | Mrgprg | 0.024 | Old |
| 0.439581 | 0.862693 | 0.18 | 0.14 | 1 | 14 | Tlr4 | 0.04 | Old |
| 0.439915 | 1.366129 | 0.093 | 0.06 | 1 | 14 | Cmpk2 | 0.033 | Old |
| 0.439915 | 1.235661 | 0.093 | 0.06 | 1 | 14 | Dact2 | 0.033 | Old |
| 0.439915 | 0.827315 | 0.093 | 0.06 | 1 | 14 | Heph | 0.033 | Old |
| 0.440221 | 0.028846 | 0.066 | 0.1 | 1 | 14 | Apex1 | -0.034 | Old |
| 0.440464 | 0.625786 | 0.311 | 0.28 | 1 | 14 | Ifi211 | 0.031 | Old |
| 0.440586 | 0.007494 | 0.705 | 0.6 | 1 | 14 | Tppp3 | 0.105 | Old |
| 0.440848 | 0.848031 | 0.071 | 0.04 | 1 | 14 | Fyb | 0.031 | Old |
| 0.440848 | 0.139039 | 0.071 | 0.04 | 1 | 14 | B230118H( | 0.031 | Old |
| 0.440848 | 0.074894 | 0.071 | 0.04 | 1 | 14 | Ifi47 | 0.031 | Old |
| 0.440856 | 0.879861 | 0.049 | 0.08 | 1 | 14 | Zcchc4 | -0.031 | Old |
| 0.440856 | 0.711431 | 0.049 | 0.08 | 1 | 14 | Rbm48 | -0.031 | Old |
| 0.440856 | 0.353412 | 0.049 | 0.08 | 1 | 14 | Fbxw4 | -0.031 | Old |
| 0.440856 | 0.134246 | 0.049 | 0.08 | 1 | 14 | Ldlrad4 | -0.031 | Old |
| 0.441014 | 0.146755 | 0.415 | 0.34 | 1 | 14 | Pcmtd1 | 0.075 | Old |
| 0.441227 | 0.574002 | 0.322 | 0.28 | 1 | 14 | Asah1 | 0.042 | Old |
| 0.441769 | 1.342769 | 0.153 | 0.12 | 1 | 14 | Ppie | 0.033 | Old |
| 0.441892 | 1.329335 | 0.115 | 0.08 | 1 | 14 | Trappc12 | 0.035 | Old |
| 0.441892 | 1.215746 | 0.115 | 0.08 | 1 | 14 | Nfs1 | 0.035 | Old |
| 0.442048 | 0.457001 | 0.12 | 0.08 | 1 | 14 | Zfp518a | 0.04 | Old |
| 0.442406 | 0.151082 | 0.399 | 0.34 | 1 | 14 | Kif2a | 0.059 | Old |
| 0.4429 | 0.133633 | 0.098 | 0.06 | 1 | 14 | Dhcr24 | 0.038 | Old |
| 0.443156 | 0.474116 | 0.432 | 0.4 | 1 | 14 | Colec12 | 0.032 | Old |
| 0.443338 | 0.24787 | 0.066 | 0.1 | 1 | 14 | Isg15 | -0.034 | Old |
| 0.443402 | 0.068325 | 0.503 | 0.46 | 1 | 14 | Wdr26 | 0.043 | Old |
| 0.443643 | 0.477398 | 0.251 | 0.2 | 1 | 14 | Ncbp2 | 0.051 | Old |
| 0.443779 | 1.507464 | 0.044 | 0.02 | 1 | 14 | Slc39a8 | 0.024 | Old |
| 0.443779 | 1.448895 | 0.044 | 0.02 | 1 | 14 | Pik3c2b | 0.024 | Old |
| 0.443779 | 1.144445 | 0.044 | 0.02 | 1 | 14 | Vac14 | 0.024 | Old |
| 0.443779 | 0.935809 | 0.044 | 0.02 | 1 | 14 | Upp2 | 0.024 | Old |
| 0.443879 | 0.186196 | 0.279 | 0.22 | 1 | 14 | Mtch2 | 0.059 | Old |
| 0.443986 | 0.440579 | 0.262 | 0.22 | 1 | 14 | Ctcf | 0.042 | Old |
| 0.444189 | 0.039023 | 0.459 | 0.4 | 1 | 14 | Sel1l | 0.059 | Old |
| 0.444312 | 1.14637 | 0.23 | 0.2 | 1 | 14 | Scmh1 | 0.03 | Old |
| 0.444365 | 0.622872 | 0.246 | 0.2 | 1 | 14 | Cblb | 0.046 | Old |
| 0.444392 | 0.185693 | 0.049 | 0.08 | 1 | 14 | Fuz | -0.031 | Old |
| 0.444513 | 1.282721 | 0.115 | 0.08 | 1 | 14 | Bag4 | 0.035 | Old |
| 0.444895 | 0.39095 | 0.295 | 0.24 | 1 | 14 | Tcf7l1 | 0.055 | Old |
| 0.445223 | 0.31868 | 0.404 | 0.36 | 1 | 14 | Ypel3 | 0.044 | Old |
| 0.445707 | 1.187106 | 0.093 | 0.06 | 1 | 14 | Tpra1 | 0.033 | Old |
| 0.446467 | 0.165557 | 0.066 | 0.1 | 1 | 14 | Mid1 | -0.034 | Old |
| 0.447142 | 0.472221 | 0.115 | 0.08 | 1 | 14 | Mrps34 | 0.035 | Old |
| 0.447344 | 0.589708 | 0.158 | 0.12 | 1 | 14 | Dapk3 | 0.038 | Old |
| 0.447475 | 0.350201 | 0.071 | 0.04 | 1 | 14 | Dennd4b | 0.031 | Old |

| | | | | | | | | |
|---|---|---|---|---|---|---|---|---|
| 0.447527 | 0.25 | 0.65 | 0.62 | 1 | 14 | Pcbp1 | 0.03 | Old |
| 0.44801 | 1.032091 | 0.044 | 0.02 | 1 | 14 | Zc3h8 | 0.024 | Old |
| 0.44801 | 0.673114 | 0.044 | 0.02 | 1 | 14 | Nol10 | 0.024 | Old |
| 0.448187 | 0.663441 | 0.164 | 0.12 | 1 | 14 | Lnpk | 0.044 | Old |
| 0.44846 | 0.39171 | 0.098 | 0.14 | 1 | 14 | Brpf1 | -0.042 | Old |
| 0.44846 | 0.064923 | 0.098 | 0.14 | 1 | 14 | Btrc | -0.042 | Old |
| 0.448665 | 1.094492 | 0.153 | 0.12 | 1 | 14 | Fads3 | 0.033 | Old |
| 0.448898 | 0.068208 | 0.169 | 0.12 | 1 | 14 | Uxs1 | 0.049 | Old |
| 0.448932 | 0.135018 | 0.552 | 0.48 | 1 | 14 | Dync1i2 | 0.072 | Old |
| 0.449253 | 0.383181 | 0.109 | 0.16 | 1 | 14 | Klhl22 | -0.051 | Old |
| 0.449395 | 0.042438 | 0.956 | 0.94 | 1 | 14 | Rpsa | 0.016 | Old |
| 0.449609 | 0.295888 | 0.066 | 0.1 | 1 | 14 | Stard13 | -0.034 | Old |
| 0.449781 | 0.97718 | 0.115 | 0.08 | 1 | 14 | Rab40c | 0.035 | Old |
| 0.449781 | 0.442057 | 0.115 | 0.08 | 1 | 14 | Mak16 | 0.035 | Old |
| 0.449956 | 0.143553 | 0.158 | 0.2 | 1 | 14 | Loxl3 | -0.042 | Old |
| 0.449986 | 0.125999 | 0.59 | 0.54 | 1 | 14 | 8-Sep | 0.05 | Old |
| 0.450745 | 0.338429 | 0.191 | 0.14 | 1 | 14 | Tram2 | 0.051 | Old |
| 0.451049 | 1.229135 | 0.175 | 0.14 | 1 | 14 | Stard7 | 0.035 | Old |
| 0.451103 | 0.018607 | 0.098 | 0.14 | 1 | 14 | Kctd6 | -0.042 | Old |
| 0.45113 | 0.00252 | 0.148 | 0.2 | 1 | 14 | Man1b1 | -0.052 | Old |
| 0.451317 | 0.762852 | 0.251 | 0.22 | 1 | 14 | Hdac2 | 0.031 | Old |
| 0.451514 | 0.462263 | 0.049 | 0.08 | 1 | 14 | Abcb6 | -0.031 | Old |
| 0.451543 | 1.377357 | 0.093 | 0.06 | 1 | 14 | Jade2 | 0.033 | Old |
| 0.451543 | 0.889619 | 0.093 | 0.06 | 1 | 14 | Tpd52 | 0.033 | Old |
| 0.451651 | 0.460409 | 0.186 | 0.14 | 1 | 14 | Prex2 | 0.046 | Old |
| 0.452229 | 1.636454 | 0.109 | 0.08 | 1 | 14 | Scyl2 | 0.029 | Old |
| 0.452264 | 1.136909 | 0.044 | 0.02 | 1 | 14 | Fam217b | 0.024 | Old |
| 0.452264 | 0.660786 | 0.044 | 0.02 | 1 | 14 | Coprs | 0.024 | Old |
| 0.452264 | 0.540958 | 0.044 | 0.02 | 1 | 14 | Gemin2 | 0.024 | Old |
| 0.452264 | 0.304854 | 0.044 | 0.02 | 1 | 14 | Rfx2 | 0.024 | Old |
| 0.452599 | 0.333667 | 0.432 | 0.4 | 1 | 14 | Meis2 | 0.032 | Old |
| 0.452642 | 0.961287 | 0.219 | 0.18 | 1 | 14 | Rnd3 | 0.039 | Old |
| 0.452668 | 0.329634 | 0.503 | 0.5 | 1 | 14 | Kdelr1 | 0.003 | Old |
| 0.452879 | 0.164914 | 0.082 | 0.12 | 1 | 14 | Hoxb5 | -0.038 | Old |
| 0.452991 | 1.134805 | 0.273 | 0.24 | 1 | 14 | Nudt21 | 0.033 | Old |
| 0.453682 | 0.064512 | 0.115 | 0.16 | 1 | 14 | Khnyn | -0.045 | Old |
| 0.454136 | 0.136283 | 0.448 | 0.38 | 1 | 14 | Dlc1 | 0.068 | Old |
| 0.454158 | 0.42017 | 0.071 | 0.04 | 1 | 14 | Pcca | 0.031 | Old |
| 0.454384 | 0.159116 | 0.213 | 0.16 | 1 | 14 | Igsf10 | 0.053 | Old |
| 0.454476 | 0.132185 | 0.454 | 0.38 | 1 | 14 | Ehd2 | 0.074 | Old |
| 0.454476 | 0.779675 | 0.093 | 0.06 | 1 | 14 | H2afy2 | 0.033 | Old |
| 0.455083 | 1.166006 | 0.115 | 0.08 | 1 | 14 | Exosc9 | 0.035 | Old |
| 0.455083 | 0.877106 | 0.115 | 0.08 | 1 | 14 | Smarcal1 | 0.035 | Old |
| 0.455083 | 0.836449 | 0.115 | 0.08 | 1 | 14 | Zfp410 | 0.035 | Old |
| 0.455098 | 0.485839 | 0.049 | 0.08 | 1 | 14 | Cd48 | -0.031 | Old |

| | | | | | | | | |
|---|---|---|---|---|---|---|---|---|
| 0.455133 | 0.44594 | 0.262 | 0.22 | 1 | 14 | Usp22 | 0.042 | Old |
| 0.45551 | 0.350822 | 0.333 | 0.28 | 1 | 14 | Ctsa | 0.053 | Old |
| 0.455987 | 0.237495 | 0.372 | 0.36 | 1 | 14 | Cdkn1c | 0.012 | Old |
| 0.456209 | 0.339207 | 0.306 | 0.26 | 1 | 14 | Pafah1b2 | 0.046 | Old |
| 0.456541 | 0.58692 | 0.044 | 0.02 | 1 | 14 | Zfp992 | 0.024 | Old |
| 0.456541 | 0.473073 | 0.044 | 0.02 | 1 | 14 | Zkscan7 | 0.024 | Old |
| 0.456541 | 0.104174 | 0.044 | 0.02 | 1 | 14 | Zfp961 | 0.024 | Old |
| 0.456911 | 0.86665 | 0.246 | 0.22 | 1 | 14 | Atp2c1 | 0.026 | Old |
| 0.457148 | 0.941473 | 0.191 | 0.16 | 1 | 14 | Gpx7 | 0.031 | Old |
| 0.457421 | 1.180391 | 0.093 | 0.06 | 1 | 14 | Ccs | 0.033 | Old |
| 0.457646 | 1.47963 | 0.109 | 0.08 | 1 | 14 | Prrx2 | 0.029 | Old |
| 0.458964 | 0.398335 | 0.333 | 0.28 | 1 | 14 | Meis1 | 0.053 | Old |
| 0.460155 | 2.070838 | 0.066 | 0.04 | 1 | 14 | Nagpa | 0.026 | Old |
| 0.460155 | 1.969923 | 0.066 | 0.04 | 1 | 14 | Rorb | 0.026 | Old |
| 0.460155 | 1.580534 | 0.066 | 0.04 | 1 | 14 | Unc45b | 0.026 | Old |
| 0.460221 | 0.396875 | 0.306 | 0.24 | 1 | 14 | Ctso | 0.066 | Old |
| 0.460304 | 0.685629 | 0.137 | 0.1 | 1 | 14 | Sema6a | 0.037 | Old |
| 0.460486 | 0.182252 | 0.333 | 0.26 | 1 | 14 | Ddrgk1 | 0.073 | Old |
| 0.460518 | 0.299687 | 0.142 | 0.1 | 1 | 14 | Bcor | 0.042 | Old |
| 0.46084 | 0.112212 | 0.044 | 0.02 | 1 | 14 | Arl10 | 0.024 | Old |
| 0.460896 | 0.198299 | 0.071 | 0.04 | 1 | 14 | Gbp3 | 0.031 | Old |
| 0.460896 | 0.057408 | 0.071 | 0.04 | 1 | 14 | Ubqln4 | 0.031 | Old |
| 0.461124 | 1.039735 | 0.158 | 0.12 | 1 | 14 | Gsta3 | 0.038 | Old |
| 0.461124 | 0.717549 | 0.158 | 0.12 | 1 | 14 | Trappc5 | 0.038 | Old |
| 0.461179 | 0.2877 | 0.115 | 0.16 | 1 | 14 | Slc35b2 | -0.045 | Old |
| 0.462706 | 0.977977 | 0.257 | 0.22 | 1 | 14 | Gpsm3 | 0.037 | Old |
| 0.462779 | 0.77789 | 0.137 | 0.1 | 1 | 14 | Slc35e3 | 0.037 | Old |
| 0.463007 | 0.115247 | 0.984 | 0.92 | 1 | 14 | Itm2b | 0.064 | Old |
| 0.463312 | 3.545804 | 0.011 | 0 | 1 | 14 | Runx2 | 0.011 | Old |
| 0.463312 | 3.533982 | 0.011 | 0 | 1 | 14 | Slc2a9 | 0.011 | Old |
| 0.463312 | 3.530683 | 0.011 | 0 | 1 | 14 | Myl2 | 0.011 | Old |
| 0.463312 | 3.513632 | 0.011 | 0 | 1 | 14 | Enpp1 | 0.011 | Old |
| 0.463312 | 3.510456 | 0.011 | 0 | 1 | 14 | Gbp8 | 0.011 | Old |
| 0.463312 | 3.211482 | 0.011 | 0 | 1 | 14 | Incenp | 0.011 | Old |
| 0.463312 | 3.096948 | 0.011 | 0 | 1 | 14 | Lrrn4 | 0.011 | Old |
| 0.463312 | 3.079947 | 0.011 | 0 | 1 | 14 | Bdnf | 0.011 | Old |
| 0.463312 | 3.071063 | 0.011 | 0 | 1 | 14 | Gm4793 | 0.011 | Old |
| 0.463312 | 2.993847 | 0.011 | 0 | 1 | 14 | Tslrn1 | 0.011 | Old |
| 0.463312 | 2.989239 | 0.011 | 0 | 1 | 14 | Col28a1 | 0.011 | Old |
| 0.463312 | 2.97983 | 0.011 | 0 | 1 | 14 | Il18r1 | 0.011 | Old |
| 0.463312 | 2.968012 | 0.011 | 0 | 1 | 14 | Gm16124 | 0.011 | Old |
| 0.463312 | 2.95313 | 0.011 | 0 | 1 | 14 | Cxadr | 0.011 | Old |
| 0.463312 | 2.873475 | 0.011 | 0 | 1 | 14 | Lgals7 | 0.011 | Old |
| 0.463312 | 2.80607 | 0.011 | 0 | 1 | 14 | Shox2 | 0.011 | Old |
| 0.463312 | 2.755405 | 0.011 | 0 | 1 | 14 | Dnase1l3 | 0.011 | Old |

| | | | | | | | | |
|---|---|---|---|---|---|---|---|---|
| 0.463312 | 2.736828 | 0.011 | 0 | 1 | 14 | Zfp54 | 0.011 | Old |
| 0.463312 | 2.70697 | 0.011 | 0 | 1 | 14 | Gm26615 | 0.011 | Old |
| 0.463312 | 2.651586 | 0.011 | 0 | 1 | 14 | Hist4h4 | 0.011 | Old |
| 0.463312 | 2.609797 | 0.011 | 0 | 1 | 14 | Pkn3 | 0.011 | Old |
| 0.463312 | 2.575356 | 0.011 | 0 | 1 | 14 | Spic | 0.011 | Old |
| 0.463312 | 2.502587 | 0.011 | 0 | 1 | 14 | Gpr157 | 0.011 | Old |
| 0.463312 | 2.498864 | 0.011 | 0 | 1 | 14 | Ly6k | 0.011 | Old |
| 0.463312 | 2.472515 | 0.011 | 0 | 1 | 14 | Mgam | 0.011 | Old |
| 0.463312 | 2.456285 | 0.011 | 0 | 1 | 14 | Gm47972 | 0.011 | Old |
| 0.463312 | 2.413679 | 0.011 | 0 | 1 | 14 | Sntb1 | 0.011 | Old |
| 0.463312 | 2.368869 | 0.011 | 0 | 1 | 14 | Birc5 | 0.011 | Old |
| 0.463312 | 2.359947 | 0.011 | 0 | 1 | 14 | Gm2163 | 0.011 | Old |
| 0.463312 | 2.336151 | 0.011 | 0 | 1 | 14 | Prodh | 0.011 | Old |
| 0.463312 | 2.329889 | 0.011 | 0 | 1 | 14 | Cdkn2a | 0.011 | Old |
| 0.463312 | 2.326819 | 0.011 | 0 | 1 | 14 | Fam180a | 0.011 | Old |
| 0.463312 | 2.306369 | 0.011 | 0 | 1 | 14 | Sirpb1c | 0.011 | Old |
| 0.463312 | 2.296069 | 0.011 | 0 | 1 | 14 | Gm10399 | 0.011 | Old |
| 0.463312 | 2.282941 | 0.011 | 0 | 1 | 14 | Anln | 0.011 | Old |
| 0.463312 | 2.270942 | 0.011 | 0 | 1 | 14 | Stac3 | 0.011 | Old |
| 0.463312 | 2.270817 | 0.011 | 0 | 1 | 14 | Galnt2 | 0.011 | Old |
| 0.463312 | 2.264699 | 0.011 | 0 | 1 | 14 | Kif19a | 0.011 | Old |
| 0.463312 | 2.235777 | 0.011 | 0 | 1 | 14 | A330035P1 | 0.011 | Old |
| 0.463312 | 2.209392 | 0.011 | 0 | 1 | 14 | Gm7694 | 0.011 | Old |
| 0.463312 | 2.17662 | 0.011 | 0 | 1 | 14 | Ubxn11 | 0.011 | Old |
| 0.463312 | 2.168755 | 0.011 | 0 | 1 | 14 | Zbtb25 | 0.011 | Old |
| 0.463312 | 2.148763 | 0.011 | 0 | 1 | 14 | Sox6 | 0.011 | Old |
| 0.463312 | 2.140817 | 0.011 | 0 | 1 | 14 | Diras2 | 0.011 | Old |
| 0.463312 | 2.140757 | 0.011 | 0 | 1 | 14 | Tmem132c | 0.011 | Old |
| 0.463312 | 2.1135 | 0.011 | 0 | 1 | 14 | Cx3cl1 | 0.011 | Old |
| 0.463312 | 2.112038 | 0.011 | 0 | 1 | 14 | Acad10 | 0.011 | Old |
| 0.463312 | 2.097859 | 0.011 | 0 | 1 | 14 | Fhdc1 | 0.011 | Old |
| 0.463312 | 2.096772 | 0.011 | 0 | 1 | 14 | Agbl5 | 0.011 | Old |
| 0.463312 | 2.095609 | 0.011 | 0 | 1 | 14 | Prr16 | 0.011 | Old |
| 0.463312 | 2.093185 | 0.011 | 0 | 1 | 14 | Espl1 | 0.011 | Old |
| 0.463312 | 2.085997 | 0.011 | 0 | 1 | 14 | Egfros | 0.011 | Old |
| 0.463312 | 2.083891 | 0.011 | 0 | 1 | 14 | Gm4841 | 0.011 | Old |
| 0.463312 | 2.08373 | 0.011 | 0 | 1 | 14 | 2510046G1 | 0.011 | Old |
| 0.463312 | 2.071114 | 0.011 | 0 | 1 | 14 | Flvcr2 | 0.011 | Old |
| 0.463312 | 2.065415 | 0.011 | 0 | 1 | 14 | Slc47a1 | 0.011 | Old |
| 0.463312 | 2.063783 | 0.011 | 0 | 1 | 14 | AW011738 | 0.011 | Old |
| 0.463312 | 2.052297 | 0.011 | 0 | 1 | 14 | Nsl1 | 0.011 | Old |
| 0.463312 | 2.038177 | 0.011 | 0 | 1 | 14 | Wnk3 | 0.011 | Old |
| 0.463312 | 2.026992 | 0.011 | 0 | 1 | 14 | Igfbp2 | 0.011 | Old |
| 0.463312 | 2.023074 | 0.011 | 0 | 1 | 14 | C1qtnf4 | 0.011 | Old |
| 0.463312 | 2.0213 | 0.011 | 0 | 1 | 14 | Pcdhgb2 | 0.011 | Old |

| | | | | | | | | |
|---|---|---|---|---|---|---|---|---|
| 0.463312 | 2.017947 | 0.011 | 0 | 1 | 14 | Pcdhb5 | 0.011 | Old |
| 0.463312 | 2.010501 | 0.011 | 0 | 1 | 14 | Gm44386 | 0.011 | Old |
| 0.463312 | 2.003477 | 0.011 | 0 | 1 | 14 | Gprin2 | 0.011 | Old |
| 0.463312 | 2.000197 | 0.011 | 0 | 1 | 14 | Kcnq1 | 0.011 | Old |
| 0.463312 | 1.998291 | 0.011 | 0 | 1 | 14 | 5033428I2 | 0.011 | Old |
| 0.463312 | 1.997159 | 0.011 | 0 | 1 | 14 | Il2 | 0.011 | Old |
| 0.463312 | 1.982002 | 0.011 | 0 | 1 | 14 | Sox7 | 0.011 | Old |
| 0.463312 | 1.980824 | 0.011 | 0 | 1 | 14 | Cpne7 | 0.011 | Old |
| 0.463312 | 1.975117 | 0.011 | 0 | 1 | 14 | Gm6566 | 0.011 | Old |
| 0.463312 | 1.971339 | 0.011 | 0 | 1 | 14 | Rgs14 | 0.011 | Old |
| 0.463312 | 1.969506 | 0.011 | 0 | 1 | 14 | A330009N | 0.011 | Old |
| 0.463312 | 1.967044 | 0.011 | 0 | 1 | 14 | Lsamp | 0.011 | Old |
| 0.463312 | 1.965681 | 0.011 | 0 | 1 | 14 | Psrc1 | 0.011 | Old |
| 0.463312 | 1.964913 | 0.011 | 0 | 1 | 14 | Lpar2 | 0.011 | Old |
| 0.463312 | 1.962595 | 0.011 | 0 | 1 | 14 | Zfp955a | 0.011 | Old |
| 0.463312 | 1.95484 | 0.011 | 0 | 1 | 14 | Gm5141 | 0.011 | Old |
| 0.463312 | 1.945066 | 0.011 | 0 | 1 | 14 | Abcg3 | 0.011 | Old |
| 0.463312 | 1.940528 | 0.011 | 0 | 1 | 14 | Tlr13 | 0.011 | Old |
| 0.463312 | 1.935628 | 0.011 | 0 | 1 | 14 | Faxc | 0.011 | Old |
| 0.463312 | 1.927037 | 0.011 | 0 | 1 | 14 | Gm26707 | 0.011 | Old |
| 0.463312 | 1.926654 | 0.011 | 0 | 1 | 14 | Pla2g2e | 0.011 | Old |
| 0.463312 | 1.925724 | 0.011 | 0 | 1 | 14 | Sh3yl1 | 0.011 | Old |
| 0.463312 | 1.917831 | 0.011 | 0 | 1 | 14 | 1700010I1 | 0.011 | Old |
| 0.463312 | 1.905125 | 0.011 | 0 | 1 | 14 | Mefv | 0.011 | Old |
| 0.463312 | 1.898419 | 0.011 | 0 | 1 | 14 | Pim2 | 0.011 | Old |
| 0.463312 | 1.882483 | 0.011 | 0 | 1 | 14 | 4933406P0 | 0.011 | Old |
| 0.463312 | 1.882438 | 0.011 | 0 | 1 | 14 | Mettl22 | 0.011 | Old |
| 0.463312 | 1.87862 | 0.011 | 0 | 1 | 14 | Rnf182 | 0.011 | Old |
| 0.463312 | 1.874405 | 0.011 | 0 | 1 | 14 | BC049762 | 0.011 | Old |
| 0.463312 | 1.871375 | 0.011 | 0 | 1 | 14 | Gm7160 | 0.011 | Old |
| 0.463312 | 1.868265 | 0.011 | 0 | 1 | 14 | Gm31718 | 0.011 | Old |
| 0.463312 | 1.864407 | 0.011 | 0 | 1 | 14 | Dll1 | 0.011 | Old |
| 0.463312 | 1.857339 | 0.011 | 0 | 1 | 14 | Pde2a | 0.011 | Old |
| 0.463312 | 1.852359 | 0.011 | 0 | 1 | 14 | Ccdc160 | 0.011 | Old |
| 0.463312 | 1.852237 | 0.011 | 0 | 1 | 14 | Gm44899 | 0.011 | Old |
| 0.463312 | 1.850809 | 0.011 | 0 | 1 | 14 | Slpi | 0.011 | Old |
| 0.463312 | 1.848534 | 0.011 | 0 | 1 | 14 | Cox8b | 0.011 | Old |
| 0.463312 | 1.846885 | 0.011 | 0 | 1 | 14 | Napsa | 0.011 | Old |
| 0.463312 | 1.846579 | 0.011 | 0 | 1 | 14 | Krt7 | 0.011 | Old |
| 0.463312 | 1.846162 | 0.011 | 0 | 1 | 14 | Zbtb3 | 0.011 | Old |
| 0.463312 | 1.844309 | 0.011 | 0 | 1 | 14 | 5930422O | 0.011 | Old |
| 0.463312 | 1.835592 | 0.011 | 0 | 1 | 14 | Cerkl | 0.011 | Old |
| 0.463312 | 1.835592 | 0.011 | 0 | 1 | 14 | Pcdhb21 | 0.011 | Old |
| 0.463312 | 1.831292 | 0.011 | 0 | 1 | 14 | Mrln | 0.011 | Old |
| 0.463312 | 1.830457 | 0.011 | 0 | 1 | 14 | Tmem202 | 0.011 | Old |

| | | | | | | | | |
|---|---|---|---|---|---|---|---|---|
| 0.463312 | 1.820185 | 0.011 | 0 | 1 | 14 | Zfp493 | 0.011 | Old |
| 0.463312 | 1.81543 | 0.011 | 0 | 1 | 14 | Glrb | 0.011 | Old |
| 0.463312 | 1.810307 | 0.011 | 0 | 1 | 14 | Gm20075 | 0.011 | Old |
| 0.463312 | 1.804162 | 0.011 | 0 | 1 | 14 | Tnfsf13os | 0.011 | Old |
| 0.463312 | 1.803665 | 0.011 | 0 | 1 | 14 | Tcte2 | 0.011 | Old |
| 0.463312 | 1.796474 | 0.011 | 0 | 1 | 14 | Gm15867 | 0.011 | Old |
| 0.463312 | 1.79191 | 0.011 | 0 | 1 | 14 | Fam222a | 0.011 | Old |
| 0.463312 | 1.790034 | 0.011 | 0 | 1 | 14 | Sh3bgrl2 | 0.011 | Old |
| 0.463312 | 1.78023 | 0.011 | 0 | 1 | 14 | Trim14 | 0.011 | Old |
| 0.463312 | 1.777808 | 0.011 | 0 | 1 | 14 | Hist1h1d | 0.011 | Old |
| 0.463312 | 1.775293 | 0.011 | 0 | 1 | 14 | Gm26825 | 0.011 | Old |
| 0.463312 | 1.774886 | 0.011 | 0 | 1 | 14 | 1700056N1 | 0.011 | Old |
| 0.463312 | 1.773424 | 0.011 | 0 | 1 | 14 | Prkch | 0.011 | Old |
| 0.463312 | 1.772292 | 0.011 | 0 | 1 | 14 | Hoxa1 | 0.011 | Old |
| 0.463312 | 1.771817 | 0.011 | 0 | 1 | 14 | 1810062G1 | 0.011 | Old |
| 0.463312 | 1.771127 | 0.011 | 0 | 1 | 14 | Adamts3 | 0.011 | Old |
| 0.463312 | 1.768274 | 0.011 | 0 | 1 | 14 | Cenpe | 0.011 | Old |
| 0.463312 | 1.766731 | 0.011 | 0 | 1 | 14 | Gm14508 | 0.011 | Old |
| 0.463312 | 1.766351 | 0.011 | 0 | 1 | 14 | Fndc7 | 0.011 | Old |
| 0.463312 | 1.762657 | 0.011 | 0 | 1 | 14 | Cdkl1 | 0.011 | Old |
| 0.463312 | 1.761028 | 0.011 | 0 | 1 | 14 | K230015D0 | 0.011 | Old |
| 0.463312 | 1.76031 | 0.011 | 0 | 1 | 14 | Hrh1 | 0.011 | Old |
| 0.463312 | 1.756023 | 0.011 | 0 | 1 | 14 | Rbm38 | 0.011 | Old |
| 0.463312 | 1.755454 | 0.011 | 0 | 1 | 14 | Jakmip3 | 0.011 | Old |
| 0.463312 | 1.754257 | 0.011 | 0 | 1 | 14 | Cwf19l1 | 0.011 | Old |
| 0.463312 | 1.74626 | 0.011 | 0 | 1 | 14 | Gm29666 | 0.011 | Old |
| 0.463312 | 1.745496 | 0.011 | 0 | 1 | 14 | Nlgn3 | 0.011 | Old |
| 0.463312 | 1.742363 | 0.011 | 0 | 1 | 14 | 2310034G0 | 0.011 | Old |
| 0.463312 | 1.741138 | 0.011 | 0 | 1 | 14 | Il1rn | 0.011 | Old |
| 0.463312 | 1.739207 | 0.011 | 0 | 1 | 14 | C6 | 0.011 | Old |
| 0.463312 | 1.731885 | 0.011 | 0 | 1 | 14 | 2900060B1 | 0.011 | Old |
| 0.463312 | 1.729882 | 0.011 | 0 | 1 | 14 | Cntrob | 0.011 | Old |
| 0.463312 | 1.72805 | 0.011 | 0 | 1 | 14 | Gas2 | 0.011 | Old |
| 0.463312 | 1.727954 | 0.011 | 0 | 1 | 14 | Sarm1 | 0.011 | Old |
| 0.463312 | 1.72675 | 0.011 | 0 | 1 | 14 | A730017C2 | 0.011 | Old |
| 0.463312 | 1.721541 | 0.011 | 0 | 1 | 14 | 4933408B1 | 0.011 | Old |
| 0.463312 | 1.718938 | 0.011 | 0 | 1 | 14 | Nudt22 | 0.011 | Old |
| 0.463312 | 1.718242 | 0.011 | 0 | 1 | 14 | Dpf3 | 0.011 | Old |
| 0.463312 | 1.709351 | 0.011 | 0 | 1 | 14 | 2210406O1 | 0.011 | Old |
| 0.463312 | 1.707421 | 0.011 | 0 | 1 | 14 | Lpxn | 0.011 | Old |
| 0.463312 | 1.705055 | 0.011 | 0 | 1 | 14 | Fv1 | 0.011 | Old |
| 0.463312 | 1.70187 | 0.011 | 0 | 1 | 14 | Adora2a | 0.011 | Old |
| 0.463312 | 1.70187 | 0.011 | 0 | 1 | 14 | Mgat3 | 0.011 | Old |
| 0.463312 | 1.700298 | 0.011 | 0 | 1 | 14 | Nefm | 0.011 | Old |
| 0.463312 | 1.697112 | 0.011 | 0 | 1 | 14 | Arl11 | 0.011 | Old |

| | | | | | | | | |
|---|---|---|---|---|---|---|---|---|
| 0.463312 | 1.695213 | 0.011 | 0 | 1 | 14 | Parvg | 0.011 | Old |
| 0.463312 | 1.686518 | 0.011 | 0 | 1 | 14 | E430024P1 | 0.011 | Old |
| 0.463312 | 1.684413 | 0.011 | 0 | 1 | 14 | 9330179D1 | 0.011 | Old |
| 0.463312 | 1.68318 | 0.011 | 0 | 1 | 14 | Card14 | 0.011 | Old |
| 0.463312 | 1.67736 | 0.011 | 0 | 1 | 14 | Epm2a | 0.011 | Old |
| 0.463312 | 1.675618 | 0.011 | 0 | 1 | 14 | Slamf1 | 0.011 | Old |
| 0.463312 | 1.674405 | 0.011 | 0 | 1 | 14 | Paqr3 | 0.011 | Old |
| 0.463312 | 1.673844 | 0.011 | 0 | 1 | 14 | Wnt1 | 0.011 | Old |
| 0.463312 | 1.671197 | 0.011 | 0 | 1 | 14 | Svopl | 0.011 | Old |
| 0.463312 | 1.669175 | 0.011 | 0 | 1 | 14 | Kcns3 | 0.011 | Old |
| 0.463312 | 1.662913 | 0.011 | 0 | 1 | 14 | Disc1 | 0.011 | Old |
| 0.463312 | 1.66232 | 0.011 | 0 | 1 | 14 | Lockd | 0.011 | Old |
| 0.463312 | 1.661529 | 0.011 | 0 | 1 | 14 | Avil | 0.011 | Old |
| 0.463312 | 1.66093 | 0.011 | 0 | 1 | 14 | Adra2c | 0.011 | Old |
| 0.463312 | 1.657377 | 0.011 | 0 | 1 | 14 | Nrxn2 | 0.011 | Old |
| 0.463312 | 1.655122 | 0.011 | 0 | 1 | 14 | Gpatch3 | 0.011 | Old |
| 0.463312 | 1.653896 | 0.011 | 0 | 1 | 14 | Gm47754 | 0.011 | Old |
| 0.463312 | 1.647792 | 0.011 | 0 | 1 | 14 | Tm9sf1 | 0.011 | Old |
| 0.463312 | 1.644335 | 0.011 | 0 | 1 | 14 | Spns2 | 0.011 | Old |
| 0.463312 | 1.644335 | 0.011 | 0 | 1 | 14 | Ager | 0.011 | Old |
| 0.463312 | 1.637742 | 0.011 | 0 | 1 | 14 | Mapk11 | 0.011 | Old |
| 0.463312 | 1.636847 | 0.011 | 0 | 1 | 14 | 4732496C0 | 0.011 | Old |
| 0.463312 | 1.633383 | 0.011 | 0 | 1 | 14 | A430106G1 | 0.011 | Old |
| 0.463312 | 1.631912 | 0.011 | 0 | 1 | 14 | Steap1 | 0.011 | Old |
| 0.463312 | 1.625271 | 0.011 | 0 | 1 | 14 | Eme2 | 0.011 | Old |
| 0.463312 | 1.621637 | 0.011 | 0 | 1 | 14 | Zfp759 | 0.011 | Old |
| 0.463312 | 1.61959 | 0.011 | 0 | 1 | 14 | Gm9008 | 0.011 | Old |
| 0.463312 | 1.615076 | 0.011 | 0 | 1 | 14 | Scimp | 0.011 | Old |
| 0.463312 | 1.608782 | 0.011 | 0 | 1 | 14 | Col20a1 | 0.011 | Old |
| 0.463312 | 1.606017 | 0.011 | 0 | 1 | 14 | Fam72a | 0.011 | Old |
| 0.463312 | 1.603767 | 0.011 | 0 | 1 | 14 | G630016G0 | 0.011 | Old |
| 0.463312 | 1.599829 | 0.011 | 0 | 1 | 14 | Stk33 | 0.011 | Old |
| 0.463312 | 1.597067 | 0.011 | 0 | 1 | 14 | Spint2 | 0.011 | Old |
| 0.463312 | 1.594632 | 0.011 | 0 | 1 | 14 | Tigd5 | 0.011 | Old |
| 0.463312 | 1.5916 | 0.011 | 0 | 1 | 14 | Isl1 | 0.011 | Old |
| 0.463312 | 1.590441 | 0.011 | 0 | 1 | 14 | H19 | 0.011 | Old |
| 0.463312 | 1.584907 | 0.011 | 0 | 1 | 14 | Rxfp2 | 0.011 | Old |
| 0.463312 | 1.584774 | 0.011 | 0 | 1 | 14 | Smad9 | 0.011 | Old |
| 0.463312 | 1.579676 | 0.011 | 0 | 1 | 14 | Gm45606 | 0.011 | Old |
| 0.463312 | 1.578178 | 0.011 | 0 | 1 | 14 | Fcho1 | 0.011 | Old |
| 0.463312 | 1.576513 | 0.011 | 0 | 1 | 14 | Adgre4 | 0.011 | Old |
| 0.463312 | 1.57589 | 0.011 | 0 | 1 | 14 | Gm13373 | 0.011 | Old |
| 0.463312 | 1.572861 | 0.011 | 0 | 1 | 14 | 1810010D0 | 0.011 | Old |
| 0.463312 | 1.563615 | 0.011 | 0 | 1 | 14 | Slc6a12 | 0.011 | Old |
| 0.463312 | 1.562877 | 0.011 | 0 | 1 | 14 | Tpx2 | 0.011 | Old |

| | | | | | | | | |
|---|---|---|---|---|---|---|---|---|
| 0.463312 | 1.560345 | 0.011 | 0 | 1 | 14 | Rubcnl | 0.011 | Old |
| 0.463312 | 1.555622 | 0.011 | 0 | 1 | 14 | Gm15013 | 0.011 | Old |
| 0.463312 | 1.553774 | 0.011 | 0 | 1 | 14 | Grem2 | 0.011 | Old |
| 0.463312 | 1.552012 | 0.011 | 0 | 1 | 14 | Cyp2r1 | 0.011 | Old |
| 0.463312 | 1.544972 | 0.011 | 0 | 1 | 14 | 9030624G2 | 0.011 | Old |
| 0.463312 | 1.542454 | 0.011 | 0 | 1 | 14 | D830025C0 | 0.011 | Old |
| 0.463312 | 1.538385 | 0.011 | 0 | 1 | 14 | 9430015G1 | 0.011 | Old |
| 0.463312 | 1.532222 | 0.011 | 0 | 1 | 14 | Slc6a7 | 0.011 | Old |
| 0.463312 | 1.52852 | 0.011 | 0 | 1 | 14 | Gm7967 | 0.011 | Old |
| 0.463312 | 1.52852 | 0.011 | 0 | 1 | 14 | Slc7a13 | 0.011 | Old |
| 0.463312 | 1.52852 | 0.011 | 0 | 1 | 14 | Hhex | 0.011 | Old |
| 0.463312 | 1.526995 | 0.011 | 0 | 1 | 14 | Grtp1 | 0.011 | Old |
| 0.463312 | 1.522251 | 0.011 | 0 | 1 | 14 | Pcdhb7 | 0.011 | Old |
| 0.463312 | 1.510698 | 0.011 | 0 | 1 | 14 | Ebi3 | 0.011 | Old |
| 0.463312 | 1.50903 | 0.011 | 0 | 1 | 14 | Kif18a | 0.011 | Old |
| 0.463312 | 1.50136 | 0.011 | 0 | 1 | 14 | Rnf144b | 0.011 | Old |
| 0.463312 | 1.496732 | 0.011 | 0 | 1 | 14 | Gm38336 | 0.011 | Old |
| 0.463312 | 1.489591 | 0.011 | 0 | 1 | 14 | Vcpkmt | 0.011 | Old |
| 0.463312 | 1.488502 | 0.011 | 0 | 1 | 14 | Mfng | 0.011 | Old |
| 0.463312 | 1.4814 | 0.011 | 0 | 1 | 14 | Gm49085 | 0.011 | Old |
| 0.463312 | 1.471354 | 0.011 | 0 | 1 | 14 | Gm27010 | 0.011 | Old |
| 0.463312 | 1.461395 | 0.011 | 0 | 1 | 14 | Dhx58 | 0.011 | Old |
| 0.463312 | 1.447648 | 0.011 | 0 | 1 | 14 | Gm26534 | 0.011 | Old |
| 0.463312 | 1.436654 | 0.011 | 0 | 1 | 14 | E030030I0 | 0.011 | Old |
| 0.463312 | 1.433843 | 0.011 | 0 | 1 | 14 | Wdr62 | 0.011 | Old |
| 0.463312 | 1.430655 | 0.011 | 0 | 1 | 14 | Cntnap1 | 0.011 | Old |
| 0.463312 | 1.430445 | 0.011 | 0 | 1 | 14 | Ttn | 0.011 | Old |
| 0.463312 | 1.428715 | 0.011 | 0 | 1 | 14 | Lrp11 | 0.011 | Old |
| 0.463312 | 1.428112 | 0.011 | 0 | 1 | 14 | AC174678. | 0.011 | Old |
| 0.463312 | 1.419297 | 0.011 | 0 | 1 | 14 | Gm12355 | 0.011 | Old |
| 0.463312 | 1.415724 | 0.011 | 0 | 1 | 14 | Zfp712 | 0.011 | Old |
| 0.463312 | 1.412809 | 0.011 | 0 | 1 | 14 | Cdh9 | 0.011 | Old |
| 0.463312 | 1.410336 | 0.011 | 0 | 1 | 14 | Fpr2 | 0.011 | Old |
| 0.463312 | 1.409674 | 0.011 | 0 | 1 | 14 | Zfp454 | 0.011 | Old |
| 0.463312 | 1.394304 | 0.011 | 0 | 1 | 14 | Dnase1l2 | 0.011 | Old |
| 0.463312 | 1.393836 | 0.011 | 0 | 1 | 14 | Oas1c | 0.011 | Old |
| 0.463312 | 1.393799 | 0.011 | 0 | 1 | 14 | Gna15 | 0.011 | Old |
| 0.463312 | 1.39306 | 0.011 | 0 | 1 | 14 | Plxnb1 | 0.011 | Old |
| 0.463312 | 1.376953 | 0.011 | 0 | 1 | 14 | Clec4d | 0.011 | Old |
| 0.463312 | 1.375757 | 0.011 | 0 | 1 | 14 | Slc7a4 | 0.011 | Old |
| 0.463312 | 1.374983 | 0.011 | 0 | 1 | 14 | Pcdhb11 | 0.011 | Old |
| 0.463312 | 1.371791 | 0.011 | 0 | 1 | 14 | Rgs20 | 0.011 | Old |
| 0.463312 | 1.359406 | 0.011 | 0 | 1 | 14 | Tbx6 | 0.011 | Old |
| 0.463312 | 1.353391 | 0.011 | 0 | 1 | 14 | Oxld1 | 0.011 | Old |
| 0.463312 | 1.345458 | 0.011 | 0 | 1 | 14 | Amhr2 | 0.011 | Old |

| | | | | | | | | |
|---|---|---|---|---|---|---|---|---|
| 0.463312 | 1.34537 | 0.011 | 0 | 1 | 14 | Lgr5 | 0.011 | Old |
| 0.463312 | 1.331937 | 0.011 | 0 | 1 | 14 | Ropn1l | 0.011 | Old |
| 0.463312 | 1.329655 | 0.011 | 0 | 1 | 14 | Gramd1c | 0.011 | Old |
| 0.463312 | 1.317973 | 0.011 | 0 | 1 | 14 | 6-Sep | 0.011 | Old |
| 0.463312 | 1.316704 | 0.011 | 0 | 1 | 14 | Racgap1 | 0.011 | Old |
| 0.463312 | 1.316159 | 0.011 | 0 | 1 | 14 | Adam5 | 0.011 | Old |
| 0.463312 | 1.315233 | 0.011 | 0 | 1 | 14 | Nos1ap | 0.011 | Old |
| 0.463312 | 1.311741 | 0.011 | 0 | 1 | 14 | Lrrc9 | 0.011 | Old |
| 0.463312 | 1.306409 | 0.011 | 0 | 1 | 14 | Tnfrsf4 | 0.011 | Old |
| 0.463312 | 1.305827 | 0.011 | 0 | 1 | 14 | 2810414N( | 0.011 | Old |
| 0.463312 | 1.297874 | 0.011 | 0 | 1 | 14 | Tssk4 | 0.011 | Old |
| 0.463312 | 1.289942 | 0.011 | 0 | 1 | 14 | Slc25a47 | 0.011 | Old |
| 0.463312 | 1.283656 | 0.011 | 0 | 1 | 14 | Gm26829 | 0.011 | Old |
| 0.463312 | 1.281655 | 0.011 | 0 | 1 | 14 | Hpse2 | 0.011 | Old |
| 0.463312 | 1.27815 | 0.011 | 0 | 1 | 14 | Tnfrsf14 | 0.011 | Old |
| 0.463312 | 1.277638 | 0.011 | 0 | 1 | 14 | Zfp940 | 0.011 | Old |
| 0.463312 | 1.27539 | 0.011 | 0 | 1 | 14 | 2200002D( | 0.011 | Old |
| 0.463312 | 1.27539 | 0.011 | 0 | 1 | 14 | Tox3 | 0.011 | Old |
| 0.463312 | 1.272467 | 0.011 | 0 | 1 | 14 | Trim45 | 0.011 | Old |
| 0.463312 | 1.271078 | 0.011 | 0 | 1 | 14 | Tnfrsf10b | 0.011 | Old |
| 0.463312 | 1.269267 | 0.011 | 0 | 1 | 14 | Plpp2 | 0.011 | Old |
| 0.463312 | 1.268635 | 0.011 | 0 | 1 | 14 | Prkcz | 0.011 | Old |
| 0.463312 | 1.259417 | 0.011 | 0 | 1 | 14 | Plac9a | 0.011 | Old |
| 0.463312 | 1.258545 | 0.011 | 0 | 1 | 14 | Ppp1r3f | 0.011 | Old |
| 0.463312 | 1.254807 | 0.011 | 0 | 1 | 14 | Pgbd1 | 0.011 | Old |
| 0.463312 | 1.245895 | 0.011 | 0 | 1 | 14 | Gm3435 | 0.011 | Old |
| 0.463312 | 1.237294 | 0.011 | 0 | 1 | 14 | 0610040B1 | 0.011 | Old |
| 0.463312 | 1.234466 | 0.011 | 0 | 1 | 14 | Pcyt1b | 0.011 | Old |
| 0.463312 | 1.227896 | 0.011 | 0 | 1 | 14 | Zfp78 | 0.011 | Old |
| 0.463312 | 1.222748 | 0.011 | 0 | 1 | 14 | 4831440E1 | 0.011 | Old |
| 0.463312 | 1.218041 | 0.011 | 0 | 1 | 14 | Gm20324 | 0.011 | Old |
| 0.463312 | 1.216596 | 0.011 | 0 | 1 | 14 | D930048G: | 0.011 | Old |
| 0.463312 | 1.216466 | 0.011 | 0 | 1 | 14 | Nat8l | 0.011 | Old |
| 0.463312 | 1.215418 | 0.011 | 0 | 1 | 14 | Nudt8.1 | 0.011 | Old |
| 0.463312 | 1.213343 | 0.011 | 0 | 1 | 14 | Prph | 0.011 | Old |
| 0.463312 | 1.201315 | 0.011 | 0 | 1 | 14 | AC154507. | 0.011 | Old |
| 0.463312 | 1.195951 | 0.011 | 0 | 1 | 14 | Spata1 | 0.011 | Old |
| 0.463312 | 1.187023 | 0.011 | 0 | 1 | 14 | 2210039B( | 0.011 | Old |
| 0.463312 | 1.180991 | 0.011 | 0 | 1 | 14 | Trmt12 | 0.011 | Old |
| 0.463312 | 1.177685 | 0.011 | 0 | 1 | 14 | Gm6169 | 0.011 | Old |
| 0.463312 | 1.170142 | 0.011 | 0 | 1 | 14 | Chd3os | 0.011 | Old |
| 0.463312 | 1.159761 | 0.011 | 0 | 1 | 14 | Spag1 | 0.011 | Old |
| 0.463312 | 1.159102 | 0.011 | 0 | 1 | 14 | Cnga1 | 0.011 | Old |
| 0.463312 | 1.157357 | 0.011 | 0 | 1 | 14 | 0610043K1 | 0.011 | Old |
| 0.463312 | 1.149887 | 0.011 | 0 | 1 | 14 | Lhfpl3 | 0.011 | Old |

| | | | | | | | | |
|---|---|---|---|---|---|---|---|---|
| 0.463312 | 1.132965 | 0.011 | 0 | 1 | 14 | Lrriq3 | 0.011 | Old |
| 0.463312 | 1.130829 | 0.011 | 0 | 1 | 14 | Gm45250 | 0.011 | Old |
| 0.463312 | 1.125508 | 0.011 | 0 | 1 | 14 | CAAA0114[illegible] | 0.011 | Old |
| 0.463312 | 1.125071 | 0.011 | 0 | 1 | 14 | Plekhh1 | 0.011 | Old |
| 0.463312 | 1.117254 | 0.011 | 0 | 1 | 14 | Gale | 0.011 | Old |
| 0.463312 | 1.105139 | 0.011 | 0 | 1 | 14 | Gm26945 | 0.011 | Old |
| 0.463312 | 1.09926 | 0.011 | 0 | 1 | 14 | Slc5a11 | 0.011 | Old |
| 0.463312 | 1.087362 | 0.011 | 0 | 1 | 14 | Gdap1 | 0.011 | Old |
| 0.463312 | 1.085401 | 0.011 | 0 | 1 | 14 | Olr1 | 0.011 | Old |
| 0.463312 | 1.085401 | 0.011 | 0 | 1 | 14 | Glipr1 | 0.011 | Old |
| 0.463312 | 1.083029 | 0.011 | 0 | 1 | 14 | Gp1bb | 0.011 | Old |
| 0.463312 | 1.075712 | 0.011 | 0 | 1 | 14 | Gm10269 | 0.011 | Old |
| 0.463312 | 1.072034 | 0.011 | 0 | 1 | 14 | Tbc1d9 | 0.011 | Old |
| 0.463312 | 1.065896 | 0.011 | 0 | 1 | 14 | Fam221a | 0.011 | Old |
| 0.463312 | 1.065896 | 0.011 | 0 | 1 | 14 | Gm12359 | 0.011 | Old |
| 0.463312 | 1.057612 | 0.011 | 0 | 1 | 14 | Chchd10 | 0.011 | Old |
| 0.463312 | 1.056576 | 0.011 | 0 | 1 | 14 | BC030499 | 0.011 | Old |
| 0.463312 | 1.053639 | 0.011 | 0 | 1 | 14 | Gm44148 | 0.011 | Old |
| 0.463312 | 1.052559 | 0.011 | 0 | 1 | 14 | Fam120ao[illegible] | 0.011 | Old |
| 0.463312 | 1.051651 | 0.011 | 0 | 1 | 14 | Lrrc27 | 0.011 | Old |
| 0.463312 | 1.050875 | 0.011 | 0 | 1 | 14 | Cacng7 | 0.011 | Old |
| 0.463312 | 1.047055 | 0.011 | 0 | 1 | 14 | Snhg10 | 0.011 | Old |
| 0.463312 | 1.046341 | 0.011 | 0 | 1 | 14 | Wdr86 | 0.011 | Old |
| 0.463312 | 1.038691 | 0.011 | 0 | 1 | 14 | 3110001I2[illegible] | 0.011 | Old |
| 0.463312 | 1.031238 | 0.011 | 0 | 1 | 14 | A530032D[illegible] | 0.011 | Old |
| 0.463312 | 1.031238 | 0.011 | 0 | 1 | 14 | Olfr506 | 0.011 | Old |
| 0.463312 | 1.030649 | 0.011 | 0 | 1 | 14 | Gm16287 | 0.011 | Old |
| 0.463312 | 1.02996 | 0.011 | 0 | 1 | 14 | Osgin1 | 0.011 | Old |
| 0.463312 | 1.021589 | 0.011 | 0 | 1 | 14 | Gm26606 | 0.011 | Old |
| 0.463312 | 1.018631 | 0.011 | 0 | 1 | 14 | Proca1 | 0.011 | Old |
| 0.463312 | 1.009476 | 0.011 | 0 | 1 | 14 | Pcdhga5 | 0.011 | Old |
| 0.463312 | 1.00541 | 0.011 | 0 | 1 | 14 | Rtn4r | 0.011 | Old |
| 0.463312 | 1.003466 | 0.011 | 0 | 1 | 14 | Mppe1 | 0.011 | Old |
| 0.463312 | 1.001371 | 0.011 | 0 | 1 | 14 | Gm26691 | 0.011 | Old |
| 0.463312 | 1.000789 | 0.011 | 0 | 1 | 14 | Lyrm1 | 0.011 | Old |
| 0.463312 | 0.999638 | 0.011 | 0 | 1 | 14 | 2310068J1 | 0.011 | Old |
| 0.463312 | 0.997397 | 0.011 | 0 | 1 | 14 | Gm19276 | 0.011 | Old |
| 0.463312 | 0.996056 | 0.011 | 0 | 1 | 14 | Gm31410 | 0.011 | Old |
| 0.463312 | 0.986211 | 0.011 | 0 | 1 | 14 | Aass | 0.011 | Old |
| 0.463312 | 0.983902 | 0.011 | 0 | 1 | 14 | Catsper2 | 0.011 | Old |
| 0.463312 | 0.980165 | 0.011 | 0 | 1 | 14 | Matk | 0.011 | Old |
| 0.463312 | 0.9767 | 0.011 | 0 | 1 | 14 | Tlr5 | 0.011 | Old |
| 0.463312 | 0.976319 | 0.011 | 0 | 1 | 14 | Lrrc73 | 0.011 | Old |
| 0.463312 | 0.973432 | 0.011 | 0 | 1 | 14 | Hmga1 | 0.011 | Old |
| 0.463312 | 0.967736 | 0.011 | 0 | 1 | 14 | Dlx4 | 0.011 | Old |

| | | | | | | | | |
|---|---|---|---|---|---|---|---|---|
| 0.463312 | 0.948087 | 0.011 | 0 | 1 | 14 | Pacrg | 0.011 | Old |
| 0.463312 | 0.937065 | 0.011 | 0 | 1 | 14 | Ppef2 | 0.011 | Old |
| 0.463312 | 0.932307 | 0.011 | 0 | 1 | 14 | 1110006O2 | 0.011 | Old |
| 0.463312 | 0.927911 | 0.011 | 0 | 1 | 14 | Pars2 | 0.011 | Old |
| 0.463312 | 0.916362 | 0.011 | 0 | 1 | 14 | Gm31834 | 0.011 | Old |
| 0.463312 | 0.904746 | 0.011 | 0 | 1 | 14 | Amz1 | 0.011 | Old |
| 0.463312 | 0.904095 | 0.011 | 0 | 1 | 14 | Lrfn1 | 0.011 | Old |
| 0.463312 | 0.89399 | 0.011 | 0 | 1 | 14 | Eef1akmt4 | 0.011 | Old |
| 0.463312 | 0.893096 | 0.011 | 0 | 1 | 14 | Guca1b | 0.011 | Old |
| 0.463312 | 0.892398 | 0.011 | 0 | 1 | 14 | Aarsd1 | 0.011 | Old |
| 0.463312 | 0.88491 | 0.011 | 0 | 1 | 14 | Prss12 | 0.011 | Old |
| 0.463312 | 0.881309 | 0.011 | 0 | 1 | 14 | Gm10036 | 0.011 | Old |
| 0.463312 | 0.861303 | 0.011 | 0 | 1 | 14 | Gm11747 | 0.011 | Old |
| 0.463312 | 0.859164 | 0.011 | 0 | 1 | 14 | Gm13056 | 0.011 | Old |
| 0.463312 | 0.816125 | 0.011 | 0 | 1 | 14 | Zfp286 | 0.011 | Old |
| 0.463312 | 0.799999 | 0.011 | 0 | 1 | 14 | Ptgs2os2 | 0.011 | Old |
| 0.463342 | 0.732487 | 0.093 | 0.06 | 1 | 14 | Sft2d3 | 0.033 | Old |
| 0.463342 | 0.643585 | 0.093 | 0.06 | 1 | 14 | Gpatch2l | 0.033 | Old |
| 0.463654 | 2.358049 | 0.066 | 0.04 | 1 | 14 | Abhd11 | 0.026 | Old |
| 0.463654 | 2.33145 | 0.066 | 0.04 | 1 | 14 | Chmp6 | 0.026 | Old |
| 0.463654 | 1.934464 | 0.066 | 0.04 | 1 | 14 | Hemk1 | 0.026 | Old |
| 0.464319 | 0.247858 | 0.639 | 0.62 | 1 | 14 | Eif4a2 | 0.019 | Old |
| 0.46436 | 0.773186 | 0.24 | 0.2 | 1 | 14 | Bud31 | 0.04 | Old |
| 0.46436 | 0.714758 | 0.24 | 0.2 | 1 | 14 | Nin | 0.04 | Old |
| 0.464437 | 0.17974 | 0.082 | 0.12 | 1 | 14 | Tmco3 | -0.038 | Old |
| 0.464448 | 0.480591 | 0.098 | 0.14 | 1 | 14 | Ctnnbip1 | -0.042 | Old |
| 0.464815 | 0.133693 | 0.47 | 0.38 | 1 | 14 | Scp2 | 0.09 | Old |
| 0.465292 | 0.084457 | 0.18 | 0.24 | 1 | 14 | 2210016L2 | -0.06 | Old |
| 0.465503 | 0.269198 | 0.066 | 0.1 | 1 | 14 | Saal1 | -0.034 | Old |
| 0.465503 | 0.197763 | 0.066 | 0.1 | 1 | 14 | Cwc27 | -0.034 | Old |
| 0.465794 | 1.179254 | 0.115 | 0.08 | 1 | 14 | Rbsn | 0.035 | Old |
| 0.465948 | 0.625732 | 0.049 | 0.08 | 1 | 14 | AI480526 | -0.031 | Old |
| 0.465963 | 0.367257 | 0.481 | 0.46 | 1 | 14 | Ttc28 | 0.021 | Old |
| 0.466058 | 0.545436 | 0.443 | 0.4 | 1 | 14 | Ahcyl1 | 0.043 | Old |
| 0.466309 | 0.150773 | 0.268 | 0.22 | 1 | 14 | Bub3 | 0.048 | Old |
| 0.46631 | 0.576104 | 0.268 | 0.22 | 1 | 14 | Rnf216 | 0.048 | Old |
| 0.466319 | 1.270921 | 0.093 | 0.06 | 1 | 14 | Pxdc1 | 0.033 | Old |
| 0.46643 | 1.859989 | 0.087 | 0.06 | 1 | 14 | Rnaseh2a | 0.027 | Old |
| 0.467143 | 0.240351 | 0.098 | 0.14 | 1 | 14 | Nup205 | -0.042 | Old |
| 0.467167 | 1.620763 | 0.066 | 0.04 | 1 | 14 | Panx1 | 0.026 | Old |
| 0.467167 | 1.518392 | 0.066 | 0.04 | 1 | 14 | Casp9 | 0.026 | Old |
| 0.467395 | 0.453027 | 0.202 | 0.16 | 1 | 14 | Ttc19 | 0.042 | Old |
| 0.467803 | 0.833171 | 0.18 | 0.14 | 1 | 14 | BC017643 | 0.04 | Old |
| 0.468197 | 0.403436 | 0.268 | 0.22 | 1 | 14 | Stx12 | 0.048 | Old |
| 0.468493 | 0.841401 | 0.115 | 0.08 | 1 | 14 | Slc4a1ap | 0.035 | Old |

| | | | | | | | | |
|---|---|---|---|---|---|---|---|---|
| 0.468493 | 0.565237 | 0.115 | 0.08 | 1 | 14 | Traf6 | 0.035 | Old |
| 0.468589 | 1.759091 | 0.109 | 0.08 | 1 | 14 | Yrdc | 0.029 | Old |
| 0.468719 | 0.719219 | 0.066 | 0.1 | 1 | 14 | Evc2 | -0.034 | Old |
| 0.468719 | 0.473807 | 0.066 | 0.1 | 1 | 14 | Ganc | -0.034 | Old |
| 0.468987 | 0.603429 | 0.164 | 0.22 | 1 | 14 | Ets2 | -0.056 | Old |
| 0.468987 | 0.395283 | 0.186 | 0.14 | 1 | 14 | Fam114a2 | 0.046 | Old |
| 0.468987 | 0.341485 | 0.186 | 0.14 | 1 | 14 | Cpt1c | 0.046 | Old |
| 0.469059 | 0.974424 | 0.284 | 0.24 | 1 | 14 | Cops4 | 0.044 | Old |
| 0.469621 | 0.145388 | 0.29 | 0.24 | 1 | 14 | Reps2 | 0.05 | Old |
| 0.470014 | 1.69474 | 0.131 | 0.1 | 1 | 14 | Ctss | 0.031 | Old |
| 0.470014 | 1.272823 | 0.131 | 0.1 | 1 | 14 | Tmem55a | 0.031 | Old |
| 0.470088 | 0.456554 | 0.268 | 0.22 | 1 | 14 | Dtnbp1 | 0.048 | Old |
| 0.470249 | 0.211514 | 0.12 | 0.16 | 1 | 14 | Stk19 | -0.04 | Old |
| 0.470277 | 0.372956 | 0.082 | 0.12 | 1 | 14 | Tmem189 | -0.038 | Old |
| 0.470277 | 0.159133 | 0.082 | 0.12 | 1 | 14 | Pomk | -0.038 | Old |
| 0.470341 | 0.271784 | 0.257 | 0.22 | 1 | 14 | Mllt10 | 0.037 | Old |
| 0.470696 | 2.216849 | 0.066 | 0.04 | 1 | 14 | Irf7 | 0.026 | Old |
| 0.470696 | 1.716133 | 0.066 | 0.04 | 1 | 14 | Nckipsd | 0.026 | Old |
| 0.471201 | 0.394404 | 0.115 | 0.08 | 1 | 14 | Zfyve27 | 0.035 | Old |
| 0.471347 | 1.799374 | 0.109 | 0.08 | 1 | 14 | Angpt1 | 0.029 | Old |
| 0.471821 | 0.209317 | 0.443 | 0.36 | 1 | 14 | Ppp2ca | 0.083 | Old |
| 0.471947 | 0.311122 | 0.066 | 0.1 | 1 | 14 | Pdia5 | -0.034 | Old |
| 0.472233 | 0.703946 | 0.18 | 0.14 | 1 | 14 | Prps2 | 0.04 | Old |
| 0.472303 | 0.466197 | 0.093 | 0.06 | 1 | 14 | Clec4a3 | 0.033 | Old |
| 0.472325 | 0.125323 | 0.306 | 0.26 | 1 | 14 | Lamtor2 | 0.046 | Old |
| 0.472362 | 0.378329 | 0.492 | 0.44 | 1 | 14 | Phip | 0.052 | Old |
| 0.472558 | 1.948589 | 0.087 | 0.06 | 1 | 14 | D030056L2 | 0.027 | Old |
| 0.473034 | 0.377204 | 0.486 | 0.44 | 1 | 14 | Usp9x | 0.046 | Old |
| 0.473212 | 0.550477 | 0.082 | 0.12 | 1 | 14 | Eef1akmt2 | -0.038 | Old |
| 0.473346 | 0.589994 | 0.295 | 0.26 | 1 | 14 | Il4ra | 0.035 | Old |
| 0.473494 | 0.342081 | 0.421 | 0.36 | 1 | 14 | Hp1bp3 | 0.061 | Old |
| 0.474036 | 0.844867 | 0.197 | 0.16 | 1 | 14 | Epb41l1 | 0.037 | Old |
| 0.474191 | 0.263555 | 0.24 | 0.2 | 1 | 14 | Ube2e1 | 0.04 | Old |
| 0.474238 | 1.490207 | 0.066 | 0.04 | 1 | 14 | Diexf | 0.026 | Old |
| 0.474238 | 1.48365 | 0.066 | 0.04 | 1 | 14 | Stk10 | 0.026 | Old |
| 0.474238 | 1.385655 | 0.066 | 0.04 | 1 | 14 | Hagh | 0.026 | Old |
| 0.474264 | 1.066847 | 0.191 | 0.16 | 1 | 14 | Brpf3 | 0.031 | Old |
| 0.474456 | 0.776968 | 0.18 | 0.14 | 1 | 14 | Acox1 | 0.04 | Old |
| 0.474456 | 0.727042 | 0.18 | 0.14 | 1 | 14 | Cyp2j6 | 0.04 | Old |
| 0.474456 | 0.714159 | 0.18 | 0.14 | 1 | 14 | Nvl | 0.04 | Old |
| 0.474897 | 0.083746 | 0.82 | 0.7 | 1 | 14 | Fus | 0.12 | Old |
| 0.475041 | 1.229967 | 0.301 | 0.28 | 1 | 14 | Sned1 | 0.021 | Old |
| 0.475312 | 0.431217 | 0.093 | 0.06 | 1 | 14 | Jrkl | 0.033 | Old |
| 0.475463 | 0.833199 | 0.175 | 0.14 | 1 | 14 | Rabggtb | 0.035 | Old |
| 0.475638 | 1.535764 | 0.087 | 0.06 | 1 | 14 | Dhx29 | 0.027 | Old |

| | | | | | | | | |
|---|---|---|---|---|---|---|---|---|
| 0.475869 | 0.594778 | 0.23 | 0.2 | 1 | 14 | Cdc42bpb | 0.03 | Old |
| 0.475909 | 0.757167 | 0.279 | 0.24 | 1 | 14 | Csnk2b | 0.039 | Old |
| 0.476565 | 0.538205 | 0.022 | 0.04 | 1 | 14 | Svip | -0.018 | Old |
| 0.476643 | 0.753756 | 0.115 | 0.08 | 1 | 14 | Cars | 0.035 | Old |
| 0.47689 | 1.090935 | 0.109 | 0.08 | 1 | 14 | Trmt1 | 0.029 | Old |
| 0.477659 | 1.589101 | 0.131 | 0.1 | 1 | 14 | Inafm1 | 0.031 | Old |
| 0.477659 | 1.58724 | 0.131 | 0.1 | 1 | 14 | Hccs | 0.031 | Old |
| 0.477796 | 1.350504 | 0.066 | 0.04 | 1 | 14 | Borcs8 | 0.026 | Old |
| 0.477881 | 0.897193 | 0.23 | 0.2 | 1 | 14 | Aip | 0.03 | Old |
| 0.478092 | 0.28285 | 0.372 | 0.32 | 1 | 14 | Psmc6 | 0.052 | Old |
| 0.47833 | 0.575154 | 0.093 | 0.06 | 1 | 14 | Rcbtb1 | 0.033 | Old |
| 0.47833 | 0.223916 | 0.093 | 0.06 | 1 | 14 | Scamp5 | 0.033 | Old |
| 0.47844 | 0.652255 | 0.066 | 0.1 | 1 | 14 | Tmem38b | -0.034 | Old |
| 0.47873 | 1.989373 | 0.087 | 0.06 | 1 | 14 | Zcchc10 | 0.027 | Old |
| 0.478879 | 0.05224 | 0.104 | 0.14 | 1 | 14 | Apobec1 | -0.036 | Old |
| 0.478952 | 0.270914 | 0.514 | 0.5 | 1 | 14 | Sar1a | 0.014 | Old |
| 0.479112 | 0.680832 | 0.082 | 0.12 | 1 | 14 | 2700097O( | -0.038 | Old |
| 0.479112 | 0.606777 | 0.082 | 0.12 | 1 | 14 | Taf3 | -0.038 | Old |
| 0.479377 | 0.453068 | 0.115 | 0.08 | 1 | 14 | Wrb | 0.035 | Old |
| 0.479664 | 0.220279 | 0.279 | 0.24 | 1 | 14 | Abce1 | 0.039 | Old |
| 0.48022 | 0.050364 | 0.284 | 0.24 | 1 | 14 | Soat1 | 0.044 | Old |
| 0.480729 | 0.345175 | 0.29 | 0.24 | 1 | 14 | Pdpk1 | 0.05 | Old |
| 0.48116 | 0.52439 | 0.18 | 0.14 | 1 | 14 | Tbc1d5 | 0.04 | Old |
| 0.481368 | 1.656129 | 0.066 | 0.04 | 1 | 14 | Acy3 | 0.026 | Old |
| 0.481368 | 1.548773 | 0.066 | 0.04 | 1 | 14 | 2010315B( | 0.026 | Old |
| 0.481368 | 1.48026 | 0.066 | 0.04 | 1 | 14 | Slc25a19 | 0.026 | Old |
| 0.481368 | 1.267728 | 0.066 | 0.04 | 1 | 14 | Rfx3 | 0.026 | Old |
| 0.481368 | 1.090432 | 0.066 | 0.04 | 1 | 14 | Slx4ip | 0.026 | Old |
| 0.481974 | 0.241194 | 0.317 | 0.26 | 1 | 14 | Lrpap1 | 0.057 | Old |
| 0.482227 | 0.077322 | 0.186 | 0.14 | 1 | 14 | Ikzf5 | 0.046 | Old |
| 0.482469 | 1.627556 | 0.109 | 0.08 | 1 | 14 | Slf1 | 0.029 | Old |
| 0.482469 | 1.519507 | 0.109 | 0.08 | 1 | 14 | Mllt1 | 0.029 | Old |
| 0.482794 | 1.386263 | 0.131 | 0.1 | 1 | 14 | Jdp2 | 0.031 | Old |
| 0.48303 | 0.18506 | 0.257 | 0.2 | 1 | 14 | Pigk | 0.057 | Old |
| 0.483064 | 0.481121 | 0.235 | 0.2 | 1 | 14 | Mrpl30 | 0.035 | Old |
| 0.483196 | 0.327522 | 0.219 | 0.18 | 1 | 14 | Gstp1 | 0.039 | Old |
| 0.483444 | 0.841233 | 0.262 | 0.24 | 1 | 14 | Acsl5 | 0.022 | Old |
| 0.483493 | 0.382591 | 0.098 | 0.14 | 1 | 14 | Avpr1a | -0.042 | Old |
| 0.484048 | 0.975104 | 0.153 | 0.12 | 1 | 14 | Rbms2 | 0.033 | Old |
| 0.484694 | 0.373325 | 0.317 | 0.26 | 1 | 14 | Cebpg | 0.057 | Old |
| 0.484807 | 0.720928 | 0.197 | 0.16 | 1 | 14 | Zscan26 | 0.037 | Old |
| 0.484954 | 1.676228 | 0.066 | 0.04 | 1 | 14 | Nipal2 | 0.026 | Old |
| 0.4852 | 0.266647 | 0.448 | 0.42 | 1 | 14 | Angptl2 | 0.028 | Old |
| 0.485373 | 0.207431 | 0.131 | 0.1 | 1 | 14 | Ift57 | 0.031 | Old |
| 0.485386 | 0.814643 | 0.137 | 0.1 | 1 | 14 | Nup214 | 0.037 | Old |

| | | | | | | | | |
|---|---|---|---|---|---|---|---|---|
| 0.485796 | 0.638955 | 0.377 | 0.38 | 1 | 14 | Nova1 | -0.003 | Old |
| 0.486079 | 0.74329 | 0.246 | 0.22 | 1 | 14 | Dennd5b | 0.026 | Old |
| 0.486831 | 0.983871 | 0.224 | 0.2 | 1 | 14 | Eloa | 0.024 | Old |
| 0.487012 | 0.127426 | 0.131 | 0.18 | 1 | 14 | Rapgef1 | -0.049 | Old |
| 0.487318 | 1.125213 | 0.022 | 0.04 | 1 | 14 | Gon7 | -0.018 | Old |
| 0.487318 | 0.788859 | 0.022 | 0.04 | 1 | 14 | Btbd11 | -0.018 | Old |
| 0.487318 | 0.222888 | 0.022 | 0.04 | 1 | 14 | Adck5 | -0.018 | Old |
| 0.487318 | 0.150202 | 0.022 | 0.04 | 1 | 14 | Fastkd5 | -0.018 | Old |
| 0.487318 | 0.044786 | 0.022 | 0.04 | 1 | 14 | Dnajc4 | -0.018 | Old |
| 0.487398 | 0.055806 | 0.164 | 0.12 | 1 | 14 | Tmx2 | 0.044 | Old |
| 0.487934 | 0.866452 | 0.137 | 0.1 | 1 | 14 | Tada1 | 0.037 | Old |
| 0.48807 | 1.316201 | 0.087 | 0.06 | 1 | 14 | Micu3 | 0.027 | Old |
| 0.48807 | 1.301351 | 0.087 | 0.06 | 1 | 14 | Ppan | 0.027 | Old |
| 0.488555 | 1.822413 | 0.066 | 0.04 | 1 | 14 | B3galt1 | 0.026 | Old |
| 0.488555 | 1.749585 | 0.066 | 0.04 | 1 | 14 | Itgb3bp | 0.026 | Old |
| 0.488555 | 1.447306 | 0.066 | 0.04 | 1 | 14 | Tsr3 | 0.026 | Old |
| 0.488555 | 1.130108 | 0.066 | 0.04 | 1 | 14 | D730003I1 | 0.026 | Old |
| 0.48912 | 0.809684 | 0.235 | 0.2 | 1 | 14 | Raf1 | 0.035 | Old |
| 0.489121 | 0.504226 | 0.279 | 0.24 | 1 | 14 | Spsb1 | 0.039 | Old |
| 0.489121 | 0.128176 | 0.279 | 0.24 | 1 | 14 | Dnajb14 | 0.039 | Old |
| 0.489255 | 0.343752 | 0.115 | 0.16 | 1 | 14 | Map2k4 | -0.045 | Old |
| 0.489438 | 0.30718 | 0.219 | 0.18 | 1 | 14 | Maf1 | 0.039 | Old |
| 0.489462 | 0.338156 | 0.131 | 0.18 | 1 | 14 | Josd1 | -0.049 | Old |
| 0.489708 | 0.424691 | 0.224 | 0.18 | 1 | 14 | Stk25 | 0.044 | Old |
| 0.490009 | 0.215152 | 0.497 | 0.5 | 1 | 14 | Pkn2 | -0.003 | Old |
| 0.490052 | 0.045548 | 0.213 | 0.26 | 1 | 14 | Fgl2 | -0.047 | Old |
| 0.490248 | 0.184614 | 0.776 | 0.72 | 1 | 14 | Slc25a3 | 0.056 | Old |
| 0.490509 | 0.245679 | 0.093 | 0.06 | 1 | 14 | F2r | 0.033 | Old |
| 0.490509 | 0.237091 | 0.093 | 0.06 | 1 | 14 | Rars2 | 0.033 | Old |
| 0.490592 | 0.348029 | 0.306 | 0.26 | 1 | 14 | Trip10 | 0.046 | Old |
| 0.490615 | 0.438709 | 0.388 | 0.36 | 1 | 14 | Arf3 | 0.028 | Old |
| 0.490862 | 0.043409 | 0.672 | 0.64 | 1 | 14 | Sfpq | 0.032 | Old |
| 0.490896 | 0.584186 | 0.399 | 0.4 | 1 | 14 | Pim1 | -0.001 | Old |
| 0.491193 | 1.373458 | 0.164 | 0.14 | 1 | 14 | Mmgt1 | 0.024 | Old |
| 0.491302 | 1.010133 | 0.153 | 0.12 | 1 | 14 | Ophn1 | 0.033 | Old |
| 0.491571 | 0.660044 | 0.066 | 0.1 | 1 | 14 | Fzr1 | -0.034 | Old |
| 0.491571 | 0.568974 | 0.066 | 0.1 | 1 | 14 | D430042O( | -0.034 | Old |
| 0.491821 | 0.396169 | 0.295 | 0.26 | 1 | 14 | Hjurp | 0.035 | Old |
| 0.49217 | 1.029094 | 0.066 | 0.04 | 1 | 14 | Olfr920 | 0.026 | Old |
| 0.492477 | 0.071927 | 0.574 | 0.52 | 1 | 14 | Nfic | 0.054 | Old |
| 0.492492 | 1.405269 | 0.169 | 0.14 | 1 | 14 | Sap30l | 0.029 | Old |
| 0.492743 | 1.413177 | 0.022 | 0.04 | 1 | 14 | Haus6 | -0.018 | Old |
| 0.492743 | 0.916296 | 0.022 | 0.04 | 1 | 14 | Pycr1 | -0.018 | Old |
| 0.492743 | 0.69752 | 0.022 | 0.04 | 1 | 14 | Clcn2 | -0.018 | Old |
| 0.492743 | 0.677931 | 0.022 | 0.04 | 1 | 14 | Slc37a2 | -0.018 | Old |

| | | | | | | | | |
|---|---|---|---|---|---|---|---|---|
| 0.492743 | 0.49372 | 0.022 | 0.04 | 1 | 14 | Ptges3l | -0.018 | Old |
| 0.492743 | 0.39451 | 0.022 | 0.04 | 1 | 14 | Fam167a | -0.018 | Old |
| 0.492743 | 0.300518 | 0.022 | 0.04 | 1 | 14 | Plcg2 | -0.018 | Old |
| 0.492743 | 0.203286 | 0.022 | 0.04 | 1 | 14 | Mmp17 | -0.018 | Old |
| 0.492743 | 0.170522 | 0.022 | 0.04 | 1 | 14 | Napepld | -0.018 | Old |
| 0.492743 | 0.165982 | 0.022 | 0.04 | 1 | 14 | Dedd2 | -0.018 | Old |
| 0.492743 | 0.131901 | 0.022 | 0.04 | 1 | 14 | Exog | -0.018 | Old |
| 0.492743 | 0.116238 | 0.022 | 0.04 | 1 | 14 | Kctd7 | -0.018 | Old |
| 0.493053 | 0.293198 | 0.137 | 0.1 | 1 | 14 | Cep83 | 0.037 | Old |
| 0.493053 | 0.084249 | 0.12 | 0.16 | 1 | 14 | Ankrd49 | -0.04 | Old |
| 0.493153 | 1.191955 | 0.131 | 0.1 | 1 | 14 | Grsf1 | 0.031 | Old |
| 0.493153 | 0.349756 | 0.131 | 0.1 | 1 | 14 | Mthfd2l | 0.031 | Old |
| 0.493173 | 0.534895 | 0.115 | 0.08 | 1 | 14 | Hsf1 | 0.035 | Old |
| 0.493173 | 0.474226 | 0.115 | 0.08 | 1 | 14 | Pqlc3 | 0.035 | Old |
| 0.49318 | 0.184225 | 0.213 | 0.28 | 1 | 14 | Pip4k2a | -0.067 | Old |
| 0.493425 | 0.190823 | 0.404 | 0.38 | 1 | 14 | Setd5 | 0.024 | Old |
| 0.493598 | 0.133671 | 0.459 | 0.4 | 1 | 14 | Loxl2 | 0.059 | Old |
| 0.493665 | 0.550132 | 0.175 | 0.14 | 1 | 14 | Kxd1 | 0.035 | Old |
| 0.493731 | 1.432694 | 0.109 | 0.08 | 1 | 14 | Usp38 | 0.029 | Old |
| 0.493732 | 1.126351 | 0.153 | 0.12 | 1 | 14 | Drg1 | 0.033 | Old |
| 0.493732 | 1.041443 | 0.153 | 0.12 | 1 | 14 | Mettl16 | 0.033 | Old |
| 0.494351 | 1.356573 | 0.087 | 0.06 | 1 | 14 | Rexo4 | 0.027 | Old |
| 0.494351 | 0.426713 | 0.333 | 0.32 | 1 | 14 | Sppl3 | 0.013 | Old |
| 0.494668 | 0.325641 | 0.404 | 0.36 | 1 | 14 | Spred1 | 0.044 | Old |
| 0.494797 | 0.122959 | 0.628 | 0.6 | 1 | 14 | Sod3 | 0.028 | Old |
| 0.494891 | 0.531155 | 0.377 | 0.34 | 1 | 14 | Eps8 | 0.037 | Old |
| 0.495217 | 0.153646 | 0.213 | 0.28 | 1 | 14 | Oxr1 | -0.067 | Old |
| 0.495628 | 0.823313 | 0.257 | 0.22 | 1 | 14 | Phlda3 | 0.037 | Old |
| 0.495742 | 0.467509 | 0.388 | 0.36 | 1 | 14 | Wsb1 | 0.028 | Old |
| 0.495742 | 1.794097 | 0.126 | 0.1 | 1 | 14 | Slc25a32 | 0.026 | Old |
| 0.495762 | 1.106989 | 0.131 | 0.1 | 1 | 14 | Lpin2 | 0.031 | Old |
| 0.495762 | 1.031782 | 0.131 | 0.1 | 1 | 14 | Sh3glb2 | 0.031 | Old |
| 0.495762 | 0.746769 | 0.131 | 0.1 | 1 | 14 | Agtpbp1 | 0.031 | Old |
| 0.4958 | 0.789183 | 0.066 | 0.04 | 1 | 14 | Dph6 | 0.026 | Old |
| 0.4958 | 0.634899 | 0.066 | 0.04 | 1 | 14 | Ppat | 0.026 | Old |
| 0.495958 | 0.127463 | 0.115 | 0.08 | 1 | 14 | Gabbr1 | 0.035 | Old |
| 0.496151 | 0.448385 | 0.29 | 0.26 | 1 | 14 | Retreg2 | 0.03 | Old |
| 0.497507 | 1.722167 | 0.087 | 0.06 | 1 | 14 | Grik5 | 0.027 | Old |
| 0.497507 | 1.535945 | 0.087 | 0.06 | 1 | 14 | 2310061I0 | 0.027 | Old |
| 0.498135 | 0.450436 | 0.432 | 0.4 | 1 | 14 | Herpud1 | 0.032 | Old |
| 0.4982 | 1.510765 | 0.022 | 0.04 | 1 | 14 | Arhgef5 | -0.018 | Old |
| 0.4982 | 1.176596 | 0.022 | 0.04 | 1 | 14 | Cacna1c | -0.018 | Old |
| 0.4982 | 1.153602 | 0.022 | 0.04 | 1 | 14 | Gm37305 | -0.018 | Old |
| 0.4982 | 0.860517 | 0.022 | 0.04 | 1 | 14 | Slc25a29 | -0.018 | Old |
| 0.4982 | 0.851817 | 0.022 | 0.04 | 1 | 14 | Cdpf1 | -0.018 | Old |

| | | | | | | | | |
|---|---|---|---|---|---|---|---|---|
| 0.4982 | 0.630052 | 0.022 | 0.04 | 1 | 14 | 9130019O2 | -0.018 | Old |
| 0.4982 | 0.585904 | 0.022 | 0.04 | 1 | 14 | Ak5 | -0.018 | Old |
| 0.4982 | 0.547507 | 0.022 | 0.04 | 1 | 14 | Pstpip1 | -0.018 | Old |
| 0.4982 | 0.502007 | 0.022 | 0.04 | 1 | 14 | Cenpt | -0.018 | Old |
| 0.4982 | 0.390958 | 0.022 | 0.04 | 1 | 14 | Hdac11 | -0.018 | Old |
| 0.4982 | 0.351766 | 0.022 | 0.04 | 1 | 14 | Bcl2a1b | -0.018 | Old |
| 0.498377 | 0.842143 | 0.131 | 0.1 | 1 | 14 | Cdc23 | 0.031 | Old |
| 0.498751 | 0.378372 | 0.115 | 0.08 | 1 | 14 | Nudt16l1 | 0.035 | Old |
| 0.49895 | 0.53412 | 0.208 | 0.16 | 1 | 14 | Foxj2 | 0.048 | Old |
| 0.499444 | 1.124615 | 0.066 | 0.04 | 1 | 14 | Trmt10b | 0.026 | Old |
| 0.499444 | 0.854996 | 0.066 | 0.04 | 1 | 14 | Mettl26 | 0.026 | Old |
| 0.499444 | 0.843155 | 0.066 | 0.04 | 1 | 14 | Zfp90 | 0.026 | Old |
| 0.499444 | 0.822633 | 0.066 | 0.04 | 1 | 14 | Mettl3 | 0.026 | Old |
| 0.499444 | 0.776831 | 0.066 | 0.04 | 1 | 14 | Nop2 | 0.026 | Old |
| 0.499444 | 0.563098 | 0.066 | 0.04 | 1 | 14 | Thumpd2 | 0.026 | Old |
| 0.499559 | 0.380631 | 0.202 | 0.16 | 1 | 14 | Mcmbp | 0.042 | Old |
| 0.49973 | 0.154048 | 0.311 | 0.4 | 1 | 14 | Yme1l1 | -0.089 | Old |
| 0.500102 | 0.394432 | 0.273 | 0.24 | 1 | 14 | Safb | 0.033 | Old |
| 0.500675 | 1.775738 | 0.087 | 0.06 | 1 | 14 | Ccdc122 | 0.027 | Old |
| 0.500675 | 1.196468 | 0.087 | 0.06 | 1 | 14 | Mkl1 | 0.027 | Old |
| 0.501399 | 0.220596 | 0.776 | 0.7 | 1 | 14 | Ywhab | 0.076 | Old |
| 0.501648 | 0.198974 | 0.749 | 0.74 | 1 | 14 | Ier2 | 0.009 | Old |
| 0.501811 | 0.177123 | 0.213 | 0.18 | 1 | 14 | Zmpste24 | 0.033 | Old |
| 0.501819 | 0.977283 | 0.169 | 0.14 | 1 | 14 | Stat2 | 0.029 | Old |
| 0.502038 | 0.303766 | 0.251 | 0.2 | 1 | 14 | Prrg3 | 0.051 | Old |
| 0.503102 | 0.81654 | 0.066 | 0.04 | 1 | 14 | Pramef8 | 0.026 | Old |
| 0.503102 | 0.730554 | 0.066 | 0.04 | 1 | 14 | Actr6 | 0.026 | Old |
| 0.503102 | 0.664586 | 0.066 | 0.04 | 1 | 14 | Rad1 | 0.026 | Old |
| 0.503102 | 0.368097 | 0.066 | 0.04 | 1 | 14 | Arel1 | 0.026 | Old |
| 0.503102 | 0.169012 | 0.066 | 0.04 | 1 | 14 | Gnl2 | 0.026 | Old |
| 0.503853 | 1.525136 | 0.087 | 0.06 | 1 | 14 | Rcl1 | 0.027 | Old |
| 0.503853 | 1.339874 | 0.087 | 0.06 | 1 | 14 | Txndc16 | 0.027 | Old |
| 0.503853 | 0.793538 | 0.087 | 0.06 | 1 | 14 | Atat1 | 0.027 | Old |
| 0.503875 | 0.601319 | 0.18 | 0.14 | 1 | 14 | Has1 | 0.04 | Old |
| 0.504523 | 0.552418 | 0.197 | 0.16 | 1 | 14 | Pdcd11 | 0.037 | Old |
| 0.506302 | 0.297253 | 0.361 | 0.32 | 1 | 14 | Phldb1 | 0.041 | Old |
| 0.506437 | 1.050673 | 0.219 | 0.2 | 1 | 14 | Sars | 0.019 | Old |
| 0.506517 | 0.249342 | 0.208 | 0.26 | 1 | 14 | Dicer1 | -0.052 | Old |
| 0.506774 | 0.425411 | 0.066 | 0.04 | 1 | 14 | Paqr4 | 0.026 | Old |
| 0.507179 | 0.632895 | 0.115 | 0.08 | 1 | 14 | Btbd2 | 0.035 | Old |
| 0.507284 | 0.757358 | 0.191 | 0.16 | 1 | 14 | Nhsl1 | 0.031 | Old |
| 0.50736 | 0.245906 | 0.377 | 0.32 | 1 | 14 | Fxyd5 | 0.057 | Old |
| 0.507874 | 0.050843 | 0.273 | 0.24 | 1 | 14 | Srgap2 | 0.033 | Old |
| 0.508578 | 0.407112 | 0.137 | 0.1 | 1 | 14 | Hspa4l | 0.037 | Old |
| 0.508617 | 0.81989 | 0.23 | 0.2 | 1 | 14 | Cmtm6 | 0.03 | Old |

| | | | | | | | | |
|---|---|---|---|---|---|---|---|---|
| 0.508791 | 0.251826 | 0.322 | 0.28 | 1 | 14 | Mpdz | 0.042 | Old |
| 0.508883 | 0.241543 | 0.164 | 0.12 | 1 | 14 | Cept1 | 0.044 | Old |
| 0.508912 | 0.922501 | 0.131 | 0.1 | 1 | 14 | Sympk | 0.031 | Old |
| 0.509331 | 0.644449 | 0.268 | 0.24 | 1 | 14 | Cct7 | 0.028 | Old |
| 0.509893 | 0.57267 | 0.175 | 0.14 | 1 | 14 | Ralgds | 0.035 | Old |
| 0.51046 | 0.836347 | 0.066 | 0.04 | 1 | 14 | Zfp688 | 0.026 | Old |
| 0.51046 | 0.341195 | 0.066 | 0.04 | 1 | 14 | Lppos | 0.026 | Old |
| 0.510583 | 0.136344 | 0.377 | 0.34 | 1 | 14 | Slk | 0.037 | Old |
| 0.510798 | 0.271088 | 0.18 | 0.14 | 1 | 14 | Mmd | 0.04 | Old |
| 0.510899 | 0.232299 | 0.393 | 0.36 | 1 | 14 | Dynlrb1 | 0.033 | Old |
| 0.510974 | 0.104934 | 0.038 | 0.06 | 1 | 14 | Npr3 | -0.022 | Old |
| 0.511564 | 0.929699 | 0.131 | 0.1 | 1 | 14 | Ccdc124 | 0.031 | Old |
| 0.511737 | 0.23713 | 0.104 | 0.14 | 1 | 14 | Nup153 | -0.036 | Old |
| 0.511766 | 0.986411 | 0.224 | 0.2 | 1 | 14 | Tars | 0.024 | Old |
| 0.511811 | 0.527475 | 0.131 | 0.18 | 1 | 14 | Bckdha | -0.049 | Old |
| 0.512221 | 0.802993 | 0.306 | 0.28 | 1 | 14 | Bach1 | 0.026 | Old |
| 0.512435 | 0.417426 | 0.317 | 0.28 | 1 | 14 | Tmcc1 | 0.037 | Old |
| 0.512435 | 0.198816 | 0.317 | 0.28 | 1 | 14 | Bod1l | 0.037 | Old |
| 0.512615 | 0.238854 | 0.284 | 0.24 | 1 | 14 | Phldb2 | 0.044 | Old |
| 0.513429 | 0.104402 | 0.361 | 0.32 | 1 | 14 | Tead1 | 0.041 | Old |
| 0.513625 | 0.541599 | 0.158 | 0.12 | 1 | 14 | Rnf181 | 0.038 | Old |
| 0.514527 | 0.517217 | 0.104 | 0.14 | 1 | 14 | Fam149b | -0.036 | Old |
| 0.515463 | 0.375225 | 0.246 | 0.2 | 1 | 14 | Lrif1 | 0.046 | Old |
| 0.515983 | 0.874788 | 0.169 | 0.14 | 1 | 14 | Dusp22 | 0.029 | Old |
| 0.516231 | 3.394515 | 0.038 | 0.02 | 1 | 14 | Gm26870 | 0.018 | Old |
| 0.516231 | 2.727222 | 0.038 | 0.02 | 1 | 14 | Kat8 | 0.018 | Old |
| 0.516231 | 2.668467 | 0.038 | 0.02 | 1 | 14 | Dck | 0.018 | Old |
| 0.516231 | 2.54822 | 0.038 | 0.02 | 1 | 14 | Zfp821 | 0.018 | Old |
| 0.516231 | 2.423328 | 0.038 | 0.02 | 1 | 14 | Fn3krp | 0.018 | Old |
| 0.516231 | 2.404237 | 0.038 | 0.02 | 1 | 14 | Pced1b | 0.018 | Old |
| 0.516231 | 2.389586 | 0.038 | 0.02 | 1 | 14 | Capn10 | 0.018 | Old |
| 0.516231 | 2.354234 | 0.038 | 0.02 | 1 | 14 | Trnp1 | 0.018 | Old |
| 0.516231 | 2.041882 | 0.038 | 0.02 | 1 | 14 | Plxna4 | 0.018 | Old |
| 0.516231 | 2.039524 | 0.038 | 0.02 | 1 | 14 | Bivm | 0.018 | Old |
| 0.516231 | 2.030633 | 0.038 | 0.02 | 1 | 14 | Hid1 | 0.018 | Old |
| 0.516231 | 2.01435 | 0.038 | 0.02 | 1 | 14 | Ikzf1 | 0.018 | Old |
| 0.516231 | 1.980656 | 0.038 | 0.02 | 1 | 14 | H2-Q6 | 0.018 | Old |
| 0.516231 | 1.849523 | 0.038 | 0.02 | 1 | 14 | Wdr83 | 0.018 | Old |
| 0.516231 | 1.841627 | 0.038 | 0.02 | 1 | 14 | Scin | 0.018 | Old |
| 0.516231 | 1.83566 | 0.038 | 0.02 | 1 | 14 | Aff2 | 0.018 | Old |
| 0.516231 | 1.827391 | 0.038 | 0.02 | 1 | 14 | Rfc4 | 0.018 | Old |
| 0.516231 | 1.817437 | 0.038 | 0.02 | 1 | 14 | Slamf8 | 0.018 | Old |
| 0.516231 | 1.789011 | 0.038 | 0.02 | 1 | 14 | Alox5 | 0.018 | Old |
| 0.516231 | 1.629798 | 0.038 | 0.02 | 1 | 14 | Prdm10 | 0.018 | Old |
| 0.516231 | 1.623569 | 0.038 | 0.02 | 1 | 14 | Tmem42 | 0.018 | Old |

| | | | | | | | | |
|---|---|---|---|---|---|---|---|---|
| 0.516231 | 1.592138 | 0.038 | 0.02 | 1 | 14 | Pabpc4l | 0.018 | Old |
| 0.516231 | 1.510815 | 0.038 | 0.02 | 1 | 14 | Nup188 | 0.018 | Old |
| 0.516231 | 1.430957 | 0.038 | 0.02 | 1 | 14 | Lzic | 0.018 | Old |
| 0.516672 | 0.913255 | 0.109 | 0.08 | 1 | 14 | Zfp637 | 0.029 | Old |
| 0.516672 | 0.418582 | 0.109 | 0.08 | 1 | 14 | Fam13c | 0.029 | Old |
| 0.51689 | 0.706462 | 0.131 | 0.1 | 1 | 14 | Irgm1 | 0.031 | Old |
| 0.517678 | 0.037368 | 0.273 | 0.24 | 1 | 14 | St3gal1 | 0.033 | Old |
| 0.517874 | 0.484898 | 0.066 | 0.04 | 1 | 14 | Letm2 | 0.026 | Old |
| 0.517874 | 0.444369 | 0.066 | 0.04 | 1 | 14 | Blnk | 0.026 | Old |
| 0.517874 | 0.399398 | 0.066 | 0.04 | 1 | 14 | Fam57a | 0.026 | Old |
| 0.517874 | 0.144736 | 0.055 | 0.08 | 1 | 14 | Arhgap30 | -0.025 | Old |
| 0.517915 | 0.394115 | 0.235 | 0.2 | 1 | 14 | Fbxl5 | 0.035 | Old |
| 0.518226 | 0.209465 | 0.115 | 0.16 | 1 | 14 | Gopc | -0.045 | Old |
| 0.518533 | 1.237896 | 0.115 | 0.08 | 1 | 14 | Il1b | 0.035 | Old |
| 0.518976 | 0.059361 | 0.847 | 0.68 | 1 | 14 | Cebpd | 0.167 | Old |
| 0.519069 | 0.119714 | 0.137 | 0.1 | 1 | 14 | Dpm1 | 0.037 | Old |
| 0.519142 | 0.263799 | 0.186 | 0.16 | 1 | 14 | 4833439L1 | 0.026 | Old |
| 0.519179 | 0.126994 | 0.224 | 0.18 | 1 | 14 | Fam162a | 0.044 | Old |
| 0.519377 | 0.486 | 0.131 | 0.18 | 1 | 14 | Galnt11 | -0.049 | Old |
| 0.519563 | 0.890699 | 0.131 | 0.1 | 1 | 14 | Zfp512b | 0.031 | Old |
| 0.519563 | 0.449724 | 0.131 | 0.1 | 1 | 14 | Rab10os | 0.031 | Old |
| 0.519578 | 0.95421 | 0.109 | 0.08 | 1 | 14 | Pomt2 | 0.029 | Old |
| 0.519681 | 0.079352 | 0.038 | 0.06 | 1 | 14 | Ntpcr | -0.022 | Old |
| 0.519901 | 1.428882 | 0.087 | 0.06 | 1 | 14 | Gpatch11 | 0.027 | Old |
| 0.519901 | 1.262571 | 0.087 | 0.06 | 1 | 14 | E130307A1 | 0.027 | Old |
| 0.519901 | 0.58021 | 0.087 | 0.06 | 1 | 14 | Borcs6 | 0.027 | Old |
| 0.51998 | 0.52621 | 0.251 | 0.22 | 1 | 14 | Nipa2 | 0.031 | Old |
| 0.520146 | 0.571767 | 0.197 | 0.16 | 1 | 14 | Gm47283 | 0.037 | Old |
| 0.52061 | 0.91876 | 0.24 | 0.22 | 1 | 14 | Dctn2 | 0.02 | Old |
| 0.520709 | 0.309763 | 0.29 | 0.24 | 1 | 14 | Rtcb | 0.05 | Old |
| 0.521017 | 0.419354 | 0.158 | 0.12 | 1 | 14 | Mbd6 | 0.038 | Old |
| 0.521093 | 2.518607 | 0.038 | 0.02 | 1 | 14 | Mettl2 | 0.018 | Old |
| 0.521093 | 1.963049 | 0.038 | 0.02 | 1 | 14 | St8sia6 | 0.018 | Old |
| 0.521093 | 1.893281 | 0.038 | 0.02 | 1 | 14 | Gjc2 | 0.018 | Old |
| 0.521093 | 1.883668 | 0.038 | 0.02 | 1 | 14 | Dio3 | 0.018 | Old |
| 0.521093 | 1.847099 | 0.038 | 0.02 | 1 | 14 | Acot1 | 0.018 | Old |
| 0.521093 | 1.483262 | 0.038 | 0.02 | 1 | 14 | Gatb | 0.018 | Old |
| 0.521093 | 1.398035 | 0.038 | 0.02 | 1 | 14 | Mcm5 | 0.018 | Old |
| 0.521093 | 1.252387 | 0.038 | 0.02 | 1 | 14 | Smim24 | 0.018 | Old |
| 0.521093 | 1.229376 | 0.038 | 0.02 | 1 | 14 | Pfas | 0.018 | Old |
| 0.521602 | 0.138921 | 0.066 | 0.04 | 1 | 14 | Kantr | 0.026 | Old |
| 0.521648 | 0.016782 | 0.486 | 0.44 | 1 | 14 | Clic1 | 0.046 | Old |
| 0.521653 | 0.424539 | 0.175 | 0.14 | 1 | 14 | Bag6 | 0.035 | Old |
| 0.522446 | 0.193277 | 0.18 | 0.14 | 1 | 14 | Ncoa6 | 0.04 | Old |
| 0.522492 | 0.972079 | 0.109 | 0.08 | 1 | 14 | Zdhhc18 | 0.029 | Old |

| | | | | | | | | |
|---|---|---|---|---|---|---|---|---|
| 0.523142 | 1.055919 | 0.087 | 0.06 | 1 | 14 | Ccar2 | 0.027 | Old |
| 0.523142 | 0.292849 | 0.087 | 0.06 | 1 | 14 | Mtmr14 | 0.027 | Old |
| 0.523307 | 0.793711 | 0.23 | 0.2 | 1 | 14 | Usp48 | 0.03 | Old |
| 0.523413 | 2.429829 | 0.104 | 0.08 | 1 | 14 | Slco2a1 | 0.024 | Old |
| 0.524062 | 0.020834 | 0.038 | 0.06 | 1 | 14 | Cep76 | -0.022 | Old |
| 0.524983 | 0.32074 | 0.415 | 0.38 | 1 | 14 | Klc1 | 0.035 | Old |
| 0.525343 | 0.329484 | 0.066 | 0.04 | 1 | 14 | Xrcc1 | 0.026 | Old |
| 0.525343 | 0.043695 | 0.055 | 0.08 | 1 | 14 | Fign | -0.025 | Old |
| 0.525343 | 0.012002 | 0.055 | 0.08 | 1 | 14 | Bysl | -0.025 | Old |
| 0.525551 | 0.338834 | 0.322 | 0.28 | 1 | 14 | Plxnb2 | 0.042 | Old |
| 0.525726 | 1.009348 | 0.148 | 0.12 | 1 | 14 | Ash2l | 0.028 | Old |
| 0.525767 | 0.084215 | 0.104 | 0.14 | 1 | 14 | Gm16536 | -0.036 | Old |
| 0.525978 | 2.037665 | 0.038 | 0.02 | 1 | 14 | Gm4924 | 0.018 | Old |
| 0.525978 | 1.615989 | 0.038 | 0.02 | 1 | 14 | Hspbap1 | 0.018 | Old |
| 0.525978 | 1.268017 | 0.038 | 0.02 | 1 | 14 | Ptpn22 | 0.018 | Old |
| 0.525978 | 1.246428 | 0.038 | 0.02 | 1 | 14 | 2700049A( | 0.018 | Old |
| 0.525978 | 1.066528 | 0.038 | 0.02 | 1 | 14 | Alkbh3 | 0.018 | Old |
| 0.525978 | 1.047117 | 0.038 | 0.02 | 1 | 14 | Xkr8 | 0.018 | Old |
| 0.525978 | 0.97146 | 0.038 | 0.02 | 1 | 14 | Tubg1 | 0.018 | Old |
| 0.526289 | 0.217205 | 0.355 | 0.32 | 1 | 14 | Hnrnph2 | 0.035 | Old |
| 0.526393 | 0.969283 | 0.087 | 0.06 | 1 | 14 | Nckap5l | 0.027 | Old |
| 0.526393 | 0.742583 | 0.087 | 0.06 | 1 | 14 | Ppwd1 | 0.027 | Old |
| 0.526394 | 1.650967 | 0.104 | 0.08 | 1 | 14 | Mcm6 | 0.024 | Old |
| 0.526729 | 0.433962 | 0.262 | 0.24 | 1 | 14 | Tcn2 | 0.022 | Old |
| 0.527143 | 0.573583 | 0.18 | 0.14 | 1 | 14 | Psmd3 | 0.04 | Old |
| 0.527443 | 0.060062 | 0.322 | 0.26 | 1 | 14 | Ppp4r3b | 0.062 | Old |
| 0.527626 | 0.602733 | 0.131 | 0.1 | 1 | 14 | Ftsj3 | 0.031 | Old |
| 0.527713 | 0.041234 | 0.53 | 0.48 | 1 | 14 | Ssb | 0.05 | Old |
| 0.528346 | 0.624024 | 0.109 | 0.08 | 1 | 14 | Pcyt2 | 0.029 | Old |
| 0.528346 | 0.60351 | 0.109 | 0.08 | 1 | 14 | Gart | 0.029 | Old |
| 0.528373 | 0.04898 | 0.443 | 0.42 | 1 | 14 | Caprin1 | 0.023 | Old |
| 0.528407 | 0.231286 | 0.393 | 0.36 | 1 | 14 | Bnip2 | 0.033 | Old |
| 0.528463 | 0.230647 | 0.038 | 0.06 | 1 | 14 | Cep131 | -0.022 | Old |
| 0.528463 | 0.072051 | 0.038 | 0.06 | 1 | 14 | Nif3l1 | -0.022 | Old |
| 0.529098 | 0.259626 | 0.055 | 0.08 | 1 | 14 | Nrcam | -0.025 | Old |
| 0.529098 | 0.257913 | 0.055 | 0.08 | 1 | 14 | Trub1 | -0.025 | Old |
| 0.529098 | 0.069186 | 0.066 | 0.04 | 1 | 14 | Mtrr | 0.026 | Old |
| 0.529098 | 0.065689 | 0.066 | 0.04 | 1 | 14 | Asb13 | 0.026 | Old |
| 0.529655 | 0.870431 | 0.087 | 0.06 | 1 | 14 | Mbp | 0.027 | Old |
| 0.529655 | 0.680469 | 0.087 | 0.06 | 1 | 14 | 0610009O2 | 0.027 | Old |
| 0.529655 | 0.631482 | 0.087 | 0.06 | 1 | 14 | Cmc1 | 0.027 | Old |
| 0.529705 | 0.059008 | 0.208 | 0.16 | 1 | 14 | Preb | 0.048 | Old |
| 0.52995 | 0.067359 | 0.191 | 0.16 | 1 | 14 | Commd6 | 0.031 | Old |
| 0.530352 | 0.786947 | 0.169 | 0.14 | 1 | 14 | Pccb | 0.029 | Old |
| 0.530352 | 0.541576 | 0.169 | 0.14 | 1 | 14 | Zdhhc5 | 0.029 | Old |

| | | | | | | | | |
|---|---|---|---|---|---|---|---|---|
| 0.530488 | 0.308745 | 0.459 | 0.42 | 1 | 14 | Zfhx3 | 0.039 | Old |
| 0.530519 | 0.146414 | 0.235 | 0.2 | 1 | 14 | Lsm4 | 0.035 | Old |
| 0.530834 | 1.368046 | 0.148 | 0.12 | 1 | 14 | Tmem19 | 0.028 | Old |
| 0.530887 | 1.467756 | 0.038 | 0.02 | 1 | 14 | Nlrx1 | 0.018 | Old |
| 0.530887 | 1.463381 | 0.038 | 0.02 | 1 | 14 | Usp28 | 0.018 | Old |
| 0.530887 | 1.368639 | 0.038 | 0.02 | 1 | 14 | Ecscr | 0.018 | Old |
| 0.530887 | 1.243277 | 0.038 | 0.02 | 1 | 14 | Tyw3 | 0.018 | Old |
| 0.530887 | 1.14396 | 0.038 | 0.02 | 1 | 14 | Grpel2 | 0.018 | Old |
| 0.530887 | 1.133299 | 0.038 | 0.02 | 1 | 14 | Gm15446 | 0.018 | Old |
| 0.530887 | 0.94806 | 0.038 | 0.02 | 1 | 14 | Fam241a | 0.018 | Old |
| 0.530955 | 0.099347 | 0.137 | 0.18 | 1 | 14 | Ucp2 | -0.043 | Old |
| 0.530984 | 0.876106 | 0.24 | 0.22 | 1 | 14 | Fam104a | 0.02 | Old |
| 0.531028 | 0.172053 | 0.071 | 0.1 | 1 | 14 | Sdhaf1 | -0.029 | Old |
| 0.53116 | 0.361331 | 0.175 | 0.14 | 1 | 14 | Rab9 | 0.035 | Old |
| 0.531285 | 0.539533 | 0.109 | 0.08 | 1 | 14 | Helz2 | 0.029 | Old |
| 0.531285 | 0.406204 | 0.109 | 0.08 | 1 | 14 | Timm21 | 0.029 | Old |
| 0.531304 | 0.162807 | 0.404 | 0.38 | 1 | 14 | Ubn2 | 0.024 | Old |
| 0.531434 | 0.16346 | 0.104 | 0.14 | 1 | 14 | Usp37 | -0.036 | Old |
| 0.531565 | 0.206548 | 0.273 | 0.24 | 1 | 14 | Sipa1 | 0.033 | Old |
| 0.532882 | 0.480063 | 0.038 | 0.06 | 1 | 14 | Ndufaf5 | -0.022 | Old |
| 0.532926 | 0.267764 | 0.087 | 0.06 | 1 | 14 | Tgfb3 | 0.027 | Old |
| 0.53295 | 0.878484 | 0.295 | 0.28 | 1 | 14 | Spin1 | 0.015 | Old |
| 0.53329 | 0.169432 | 0.087 | 0.12 | 1 | 14 | Rrs1 | -0.033 | Old |
| 0.533422 | 0.468388 | 0.284 | 0.26 | 1 | 14 | Kank2 | 0.024 | Old |
| 0.533532 | 0.153412 | 0.612 | 0.56 | 1 | 14 | Cd248 | 0.052 | Old |
| 0.533603 | 1.245632 | 0.126 | 0.1 | 1 | 14 | Cib1 | 0.026 | Old |
| 0.533603 | 1.045343 | 0.126 | 0.1 | 1 | 14 | Mical1 | 0.026 | Old |
| 0.534325 | 0.277664 | 0.219 | 0.18 | 1 | 14 | Zfp326 | 0.039 | Old |
| 0.534332 | 0.599444 | 0.251 | 0.22 | 1 | 14 | AC160336. | 0.031 | Old |
| 0.534387 | 2.165092 | 0.082 | 0.06 | 1 | 14 | Tsr1 | 0.022 | Old |
| 0.534387 | 0.23867 | 0.071 | 0.1 | 1 | 14 | A830010M | -0.029 | Old |
| 0.535012 | 0.124053 | 0.12 | 0.16 | 1 | 14 | Abcf2 | -0.04 | Old |
| 0.535819 | 1.296581 | 0.038 | 0.02 | 1 | 14 | Lrch2 | 0.018 | Old |
| 0.535819 | 1.092439 | 0.038 | 0.02 | 1 | 14 | Fars2 | 0.018 | Old |
| 0.535819 | 1.023229 | 0.038 | 0.02 | 1 | 14 | Poll | 0.018 | Old |
| 0.535819 | 1.006859 | 0.038 | 0.02 | 1 | 14 | Zfp937 | 0.018 | Old |
| 0.535819 | 0.863037 | 0.038 | 0.02 | 1 | 14 | Clec4b1 | 0.018 | Old |
| 0.535946 | 0.161721 | 0.175 | 0.14 | 1 | 14 | Tmem140 | 0.035 | Old |
| 0.536014 | 0.070703 | 0.197 | 0.16 | 1 | 14 | 2410089E0 | 0.037 | Old |
| 0.536208 | 0.718859 | 0.087 | 0.06 | 1 | 14 | Emx2os | 0.027 | Old |
| 0.536208 | 0.275101 | 0.087 | 0.06 | 1 | 14 | Bbs9 | 0.027 | Old |
| 0.536439 | 0.097326 | 0.388 | 0.34 | 1 | 14 | Rab11a | 0.048 | Old |
| 0.536546 | 0.530767 | 0.213 | 0.18 | 1 | 14 | Mybbp1a | 0.033 | Old |
| 0.536567 | 0.021943 | 0.295 | 0.24 | 1 | 14 | Gan | 0.055 | Old |
| 0.53665 | 0.349865 | 0.055 | 0.08 | 1 | 14 | Shmt2 | -0.025 | Old |

| | | | | | | | | |
|---|---|---|---|---|---|---|---|---|
| 0.53665 | 0.122966 | 0.055 | 0.08 | 1 | 14 | Ccdc171 | -0.025 | Old |
| 0.537131 | 0.032421 | 0.104 | 0.14 | 1 | 14 | Rpp25l | -0.036 | Old |
| 0.53732 | 0.58243 | 0.038 | 0.06 | 1 | 14 | Frk | -0.022 | Old |
| 0.53732 | 0.520978 | 0.038 | 0.06 | 1 | 14 | Greb1l | -0.022 | Old |
| 0.53732 | 0.357076 | 0.038 | 0.06 | 1 | 14 | Mrps22 | -0.022 | Old |
| 0.537386 | 0.2186 | 0.202 | 0.16 | 1 | 14 | Gcn1l1 | 0.042 | Old |
| 0.537629 | 0.311336 | 0.213 | 0.16 | 1 | 14 | Has2 | 0.053 | Old |
| 0.537693 | 0.112043 | 0.12 | 0.16 | 1 | 14 | Kpna3 | -0.04 | Old |
| 0.537756 | 1.930594 | 0.082 | 0.06 | 1 | 14 | Atg4d | 0.022 | Old |
| 0.537756 | 1.572558 | 0.082 | 0.06 | 1 | 14 | Pde10a | 0.022 | Old |
| 0.538026 | 0.168976 | 0.344 | 0.3 | 1 | 14 | Smarcc2 | 0.044 | Old |
| 0.538212 | 0.611733 | 0.23 | 0.2 | 1 | 14 | Galk2 | 0.03 | Old |
| 0.538542 | 0.797412 | 0.148 | 0.12 | 1 | 14 | Zhx2 | 0.028 | Old |
| 0.539128 | 1.306665 | 0.126 | 0.1 | 1 | 14 | Chmp1b | 0.026 | Old |
| 0.539128 | 1.257592 | 0.126 | 0.1 | 1 | 14 | Eif2ak3 | 0.026 | Old |
| 0.539501 | 0.605019 | 0.087 | 0.06 | 1 | 14 | Pcid2 | 0.027 | Old |
| 0.539501 | 0.193899 | 0.087 | 0.06 | 1 | 14 | Ak6 | 0.027 | Old |
| 0.539501 | 0.095249 | 0.087 | 0.06 | 1 | 14 | Acot8 | 0.027 | Old |
| 0.539894 | 0.234444 | 0.437 | 0.38 | 1 | 14 | Yipf5 | 0.057 | Old |
| 0.539958 | 0.989741 | 0.186 | 0.16 | 1 | 14 | Syngr2 | 0.026 | Old |
| 0.540153 | 0.503089 | 0.109 | 0.08 | 1 | 14 | Asnsd1 | 0.029 | Old |
| 0.540153 | 0.22247 | 0.109 | 0.08 | 1 | 14 | Haus2 | 0.029 | Old |
| 0.540241 | 0.344168 | 0.519 | 0.5 | 1 | 14 | Ppp1r2 | 0.019 | Old |
| 0.540445 | 0.528548 | 0.055 | 0.08 | 1 | 14 | Rrad | -0.025 | Old |
| 0.540774 | 1.151356 | 0.038 | 0.02 | 1 | 14 | Chd1l | 0.018 | Old |
| 0.540774 | 0.819796 | 0.038 | 0.02 | 1 | 14 | Cep44 | 0.018 | Old |
| 0.540774 | 0.798919 | 0.038 | 0.02 | 1 | 14 | Olfr1034 | 0.018 | Old |
| 0.540774 | 0.470597 | 0.038 | 0.02 | 1 | 14 | Ovca2 | 0.018 | Old |
| 0.540774 | 0.395797 | 0.038 | 0.02 | 1 | 14 | Gm38394 | 0.018 | Old |
| 0.541124 | 0.853164 | 0.148 | 0.12 | 1 | 14 | Bri3bp | 0.028 | Old |
| 0.541561 | 0.204601 | 0.404 | 0.36 | 1 | 14 | Cers2 | 0.044 | Old |
| 0.541776 | 1.151018 | 0.038 | 0.06 | 1 | 14 | Sardh | -0.022 | Old |
| 0.541776 | 0.726152 | 0.038 | 0.06 | 1 | 14 | Lrrc28 | -0.022 | Old |
| 0.541776 | 0.585287 | 0.038 | 0.06 | 1 | 14 | Phtf2 | -0.022 | Old |
| 0.541776 | 0.574607 | 0.038 | 0.06 | 1 | 14 | Myo1d | -0.022 | Old |
| 0.541827 | 2.118842 | 0.06 | 0.04 | 1 | 14 | Noc3l | 0.02 | Old |
| 0.541827 | 1.895584 | 0.06 | 0.04 | 1 | 14 | Mars | 0.02 | Old |
| 0.541827 | 1.829698 | 0.06 | 0.04 | 1 | 14 | Slc25a40 | 0.02 | Old |
| 0.542435 | 0.883356 | 0.339 | 0.34 | 1 | 14 | Capn1 | -0.001 | Old |
| 0.542803 | 0.531059 | 0.087 | 0.06 | 1 | 14 | Nt5c2 | 0.027 | Old |
| 0.542859 | 0.450614 | 0.104 | 0.14 | 1 | 14 | Tm6sf1 | -0.036 | Old |
| 0.542859 | 0.132675 | 0.104 | 0.14 | 1 | 14 | Slc10a7 | -0.036 | Old |
| 0.543075 | 0.201191 | 0.12 | 0.16 | 1 | 14 | Hectd4 | -0.04 | Old |
| 0.543125 | 1.200059 | 0.109 | 0.08 | 1 | 14 | Zfp597 | 0.029 | Old |
| 0.543125 | 0.241031 | 0.109 | 0.08 | 1 | 14 | Rnf115 | 0.029 | Old |

| | | | | | | | | |
|---|---|---|---|---|---|---|---|---|
| 0.543215 | 0.790251 | 0.208 | 0.18 | 1 | 14 | Isy1 | 0.028 | Old |
| 0.543622 | 0.434618 | 0.339 | 0.32 | 1 | 14 | Ctsk | 0.019 | Old |
| 0.543793 | 0.645699 | 0.191 | 0.16 | 1 | 14 | Atp8b1 | 0.031 | Old |
| 0.544255 | 0.93866 | 0.055 | 0.08 | 1 | 14 | Slc35e4 | -0.025 | Old |
| 0.544255 | 0.405919 | 0.055 | 0.08 | 1 | 14 | Plcxd2 | -0.025 | Old |
| 0.544255 | 0.366502 | 0.055 | 0.08 | 1 | 14 | Wdsub1 | -0.025 | Old |
| 0.544255 | 0.354638 | 0.055 | 0.08 | 1 | 14 | Zfp523 | -0.025 | Old |
| 0.544255 | 0.222296 | 0.055 | 0.08 | 1 | 14 | 2610524H( | -0.025 | Old |
| 0.544255 | 0.044308 | 0.541 | 0.44 | 1 | 14 | Mrpl52 | 0.101 | Old |
| 0.544465 | 1.28409 | 0.104 | 0.08 | 1 | 14 | Dhdds | 0.024 | Old |
| 0.544526 | 1.822966 | 0.082 | 0.06 | 1 | 14 | Inf2 | 0.022 | Old |
| 0.544841 | 0.281718 | 0.454 | 0.4 | 1 | 14 | Crispld2 | 0.054 | Old |
| 0.544922 | 0.773247 | 0.169 | 0.14 | 1 | 14 | Phc2 | 0.029 | Old |
| 0.544922 | 0.702713 | 0.169 | 0.14 | 1 | 14 | Pcbd2 | 0.029 | Old |
| 0.544932 | 0.289153 | 0.426 | 0.4 | 1 | 14 | Twist1 | 0.026 | Old |
| 0.545189 | 0.306834 | 0.197 | 0.16 | 1 | 14 | Vps4a | 0.037 | Old |
| 0.545647 | 0.100185 | 0.284 | 0.34 | 1 | 14 | Zcchc11 | -0.056 | Old |
| 0.545752 | 0.726961 | 0.038 | 0.02 | 1 | 14 | Usp18 | 0.018 | Old |
| 0.545752 | 0.438325 | 0.038 | 0.02 | 1 | 14 | Gm30122 | 0.018 | Old |
| 0.545752 | 0.366323 | 0.038 | 0.02 | 1 | 14 | Pop7 | 0.018 | Old |
| 0.545776 | 0.151772 | 0.12 | 0.16 | 1 | 14 | Trmt1l | -0.04 | Old |
| 0.545777 | 1.495669 | 0.06 | 0.04 | 1 | 14 | Mrpl47 | 0.02 | Old |
| 0.546105 | 0.377309 | 0.109 | 0.08 | 1 | 14 | Pskh1 | 0.029 | Old |
| 0.546105 | 0.364536 | 0.109 | 0.08 | 1 | 14 | Guf1 | 0.029 | Old |
| 0.546115 | 0.298067 | 0.087 | 0.06 | 1 | 14 | Klf15 | 0.027 | Old |
| 0.546251 | 0.949508 | 0.038 | 0.06 | 1 | 14 | Evl | -0.022 | Old |
| 0.546251 | 0.633215 | 0.038 | 0.06 | 1 | 14 | Wee1 | -0.022 | Old |
| 0.546685 | 0.622659 | 0.131 | 0.1 | 1 | 14 | Klhl20 | 0.031 | Old |
| 0.546988 | 0.274059 | 0.169 | 0.22 | 1 | 14 | Fdxr | -0.051 | Old |
| 0.54747 | 0.97172 | 0.126 | 0.1 | 1 | 14 | Exd2 | 0.026 | Old |
| 0.547563 | 0.236682 | 0.235 | 0.2 | 1 | 14 | Syt11 | 0.035 | Old |
| 0.547705 | 0.021252 | 0.208 | 0.18 | 1 | 14 | Luc7l | 0.028 | Old |
| 0.547927 | 0.516141 | 0.071 | 0.1 | 1 | 14 | Dopey2 | -0.029 | Old |
| 0.547927 | 0.456173 | 0.071 | 0.1 | 1 | 14 | Necap1 | -0.029 | Old |
| 0.548077 | 0.069709 | 0.055 | 0.08 | 1 | 14 | Tdp1 | -0.025 | Old |
| 0.548617 | 0.291455 | 0.104 | 0.14 | 1 | 14 | Dpyd | -0.036 | Old |
| 0.548617 | 0.154948 | 0.104 | 0.14 | 1 | 14 | Tmem64 | -0.036 | Old |
| 0.548759 | 0.426079 | 0.126 | 0.16 | 1 | 14 | Bcr | -0.034 | Old |
| 0.548987 | 0.413387 | 0.23 | 0.2 | 1 | 14 | Lztfl1 | 0.03 | Old |
| 0.549094 | 0.2414 | 0.109 | 0.08 | 1 | 14 | Stxbp4 | 0.029 | Old |
| 0.549289 | 0.185652 | 0.23 | 0.18 | 1 | 14 | Filip1l | 0.05 | Old |
| 0.54942 | 0.695241 | 0.246 | 0.22 | 1 | 14 | Elf2 | 0.026 | Old |
| 0.549438 | 0.337036 | 0.087 | 0.06 | 1 | 14 | Hat1 | 0.027 | Old |
| 0.549736 | 0.311311 | 0.219 | 0.18 | 1 | 14 | Arhgap23 | 0.039 | Old |
| 0.549741 | 1.443243 | 0.06 | 0.04 | 1 | 14 | Fermt3 | 0.02 | Old |

| | | | | | | | | |
|---|---|---|---|---|---|---|---|---|
| 0.550125 | 0.678467 | 0.279 | 0.26 | 1 | 14 | Fat1 | 0.019 | Old |
| 0.550744 | 1.002835 | 0.038 | 0.06 | 1 | 14 | Rbm4 | -0.022 | Old |
| 0.550752 | 0.003453 | 0.038 | 0.02 | 1 | 14 | Gbp4 | 0.018 | Old |
| 0.551339 | 2.230267 | 0.082 | 0.06 | 1 | 14 | Mrpl40 | 0.022 | Old |
| 0.551339 | 1.526669 | 0.082 | 0.06 | 1 | 14 | Dgcr8 | 0.022 | Old |
| 0.551339 | 0.161081 | 0.071 | 0.1 | 1 | 14 | Ppa2 | -0.029 | Old |
| 0.551373 | 0.425323 | 0.153 | 0.12 | 1 | 14 | Arhgdib | 0.033 | Old |
| 0.551513 | 0.625457 | 0.148 | 0.12 | 1 | 14 | Selenoi | 0.028 | Old |
| 0.551513 | 0.590146 | 0.148 | 0.12 | 1 | 14 | Zdhhc8 | 0.028 | Old |
| 0.551643 | 0.232442 | 0.508 | 0.46 | 1 | 14 | Atp6v0b | 0.048 | Old |
| 0.5517 | 1.156227 | 0.186 | 0.16 | 1 | 14 | Ccdc90b | 0.026 | Old |
| 0.55209 | 0.805734 | 0.109 | 0.08 | 1 | 14 | Vhl | 0.029 | Old |
| 0.55209 | 0.436712 | 0.109 | 0.08 | 1 | 14 | Tsr2 | 0.029 | Old |
| 0.5523 | 0.575531 | 0.311 | 0.3 | 1 | 14 | Gtf2i | 0.011 | Old |
| 0.552396 | 0.361455 | 0.355 | 0.3 | 1 | 14 | Sspn | 0.055 | Old |
| 0.55256 | 0.919308 | 0.18 | 0.16 | 1 | 14 | Cdk19 | 0.02 | Old |
| 0.552737 | 0.08258 | 0.426 | 0.52 | 1 | 14 | Ankrd12 | -0.094 | Old |
| 0.553063 | 0.151889 | 0.393 | 0.34 | 1 | 14 | Krit1 | 0.053 | Old |
| 0.553066 | 1.022877 | 0.126 | 0.1 | 1 | 14 | Actr8 | 0.026 | Old |
| 0.553947 | 0.375737 | 0.224 | 0.18 | 1 | 14 | Vsir | 0.044 | Old |
| 0.553952 | 0.615697 | 0.153 | 0.12 | 1 | 14 | Herc4 | 0.033 | Old |
| 0.554096 | 0.208428 | 0.372 | 0.38 | 1 | 14 | Cct4 | -0.008 | Old |
| 0.554151 | 0.370833 | 0.284 | 0.24 | 1 | 14 | Cdk11b | 0.044 | Old |
| 0.554405 | 0.235556 | 0.104 | 0.14 | 1 | 14 | Timm9 | -0.036 | Old |
| 0.554476 | 0.128377 | 0.459 | 0.4 | 1 | 14 | Hif1a | 0.059 | Old |
| 0.55485 | 0.370711 | 0.35 | 0.34 | 1 | 14 | Prpf38b | 0.01 | Old |
| 0.555095 | 0.799087 | 0.109 | 0.08 | 1 | 14 | Asap3 | 0.029 | Old |
| 0.555562 | 0.442769 | 0.393 | 0.38 | 1 | 14 | Copz1 | 0.013 | Old |
| 0.55582 | 1.185974 | 0.158 | 0.14 | 1 | 14 | Pex2 | 0.018 | Old |
| 0.556537 | 1.037483 | 0.153 | 0.12 | 1 | 14 | Tmem167b | 0.033 | Old |
| 0.556537 | 0.446541 | 0.153 | 0.12 | 1 | 14 | Sec11c | 0.033 | Old |
| 0.556647 | 0.263786 | 0.12 | 0.16 | 1 | 14 | Rnf215 | -0.04 | Old |
| 0.556744 | 0.449207 | 0.148 | 0.12 | 1 | 14 | Gatad2a | 0.028 | Old |
| 0.557213 | 0.78034 | 0.169 | 0.14 | 1 | 14 | Sik1 | 0.029 | Old |
| 0.557711 | 2.06618 | 0.06 | 0.04 | 1 | 14 | BC037034 | 0.02 | Old |
| 0.557711 | 1.207339 | 0.06 | 0.04 | 1 | 14 | Map3k11 | 0.02 | Old |
| 0.558105 | 0.394287 | 0.087 | 0.12 | 1 | 14 | Med23 | -0.033 | Old |
| 0.558193 | 1.429012 | 0.082 | 0.06 | 1 | 14 | D10Wsu10 | 0.022 | Old |
| 0.558193 | 0.38622 | 0.071 | 0.1 | 1 | 14 | Exosc1 | -0.029 | Old |
| 0.558323 | 0.254563 | 0.536 | 0.48 | 1 | 14 | Selenos | 0.056 | Old |
| 0.558853 | 0.594246 | 0.213 | 0.18 | 1 | 14 | Zfp664 | 0.033 | Old |
| 0.559391 | 0.272312 | 0.251 | 0.22 | 1 | 14 | Ufm1 | 0.031 | Old |
| 0.559464 | 0.201169 | 0.087 | 0.06 | 1 | 14 | Sdhaf2 | 0.027 | Old |
| 0.559624 | 0.438572 | 0.055 | 0.08 | 1 | 14 | Pyroxd1 | -0.025 | Old |
| 0.559687 | 0.4093 | 0.169 | 0.14 | 1 | 14 | Uba52 | 0.029 | Old |

| | | | | | | | | |
|---|---|---|---|---|---|---|---|---|
| 0.560194 | 0.618216 | 0.311 | 0.28 | 1 | 14 | Uba2 | 0.031 | Old |
| 0.560216 | 0.410744 | 0.279 | 0.26 | 1 | 14 | Ntan1 | 0.019 | Old |
| 0.56068 | 0.286244 | 0.47 | 0.42 | 1 | 14 | Tkt | 0.05 | Old |
| 0.561246 | 0.81697 | 0.087 | 0.12 | 1 | 14 | Atg14 | -0.033 | Old |
| 0.561505 | 0.718402 | 0.251 | 0.22 | 1 | 14 | Ddx46 | 0.031 | Old |
| 0.561636 | 1.102222 | 0.082 | 0.06 | 1 | 14 | Mrpl19 | 0.022 | Old |
| 0.561679 | 0.094403 | 0.683 | 0.6 | 1 | 14 | Hnrnpa0 | 0.083 | Old |
| 0.561716 | 1.707124 | 0.06 | 0.04 | 1 | 14 | Fam151b | 0.02 | Old |
| 0.561716 | 1.676255 | 0.06 | 0.04 | 1 | 14 | Tbc1d24 | 0.02 | Old |
| 0.561716 | 1.436881 | 0.06 | 0.04 | 1 | 14 | Thap1 | 0.02 | Old |
| 0.561716 | 1.051384 | 0.06 | 0.04 | 1 | 14 | Ralgapa2 | 0.02 | Old |
| 0.561725 | 0.351627 | 0.153 | 0.12 | 1 | 14 | Slf2 | 0.033 | Old |
| 0.561725 | 0.03754 | 0.153 | 0.12 | 1 | 14 | AI837181 | 0.033 | Old |
| 0.561805 | 0.389167 | 0.383 | 0.36 | 1 | 14 | Csnk1d | 0.023 | Old |
| 0.562245 | 0.149175 | 0.284 | 0.24 | 1 | 14 | Marf1 | 0.044 | Old |
| 0.562839 | 1.092242 | 0.104 | 0.08 | 1 | 14 | Sh2b1 | 0.024 | Old |
| 0.563567 | 0.656026 | 0.186 | 0.16 | 1 | 14 | Ufsp2 | 0.026 | Old |
| 0.564342 | 0.731667 | 0.126 | 0.1 | 1 | 14 | AI413582 | 0.026 | Old |
| 0.564395 | 0.44693 | 0.087 | 0.12 | 1 | 14 | Alg1 | -0.033 | Old |
| 0.564395 | 0.097538 | 0.087 | 0.12 | 1 | 14 | Pmm1 | -0.033 | Old |
| 0.564402 | 0.332474 | 0.29 | 0.26 | 1 | 14 | Wdr45b | 0.03 | Old |
| 0.564553 | 0.877697 | 0.18 | 0.16 | 1 | 14 | Nabp2 | 0.02 | Old |
| 0.564834 | 1.487897 | 0.142 | 0.12 | 1 | 14 | Dcaf13 | 0.022 | Old |
| 0.564869 | 0.806404 | 0.12 | 0.16 | 1 | 14 | Trim3 | -0.04 | Old |
| 0.565089 | 1.248317 | 0.082 | 0.06 | 1 | 14 | Galnt4 | 0.022 | Old |
| 0.565105 | 0.372048 | 0.421 | 0.4 | 1 | 14 | Hnrnpdl | 0.021 | Old |
| 0.565274 | 1.531896 | 0.12 | 0.1 | 1 | 14 | Taf5l | 0.02 | Old |
| 0.565274 | 0.22873 | 0.109 | 0.14 | 1 | 14 | Nop56 | -0.031 | Old |
| 0.565736 | 1.586998 | 0.06 | 0.04 | 1 | 14 | Pnpla2 | 0.02 | Old |
| 0.565736 | 1.345136 | 0.06 | 0.04 | 1 | 14 | Naa40 | 0.02 | Old |
| 0.565736 | 1.001644 | 0.06 | 0.04 | 1 | 14 | Rnf113a2 | 0.02 | Old |
| 0.565736 | 0.882089 | 0.06 | 0.04 | 1 | 14 | Eya3 | 0.02 | Old |
| 0.565744 | 0.521208 | 0.251 | 0.22 | 1 | 14 | Diaph2 | 0.031 | Old |
| 0.56593 | 0.960116 | 0.104 | 0.08 | 1 | 14 | Rrp36 | 0.024 | Old |
| 0.56606 | 0.100888 | 0.257 | 0.3 | 1 | 14 | Cdkn1a | -0.043 | Old |
| 0.566071 | 0.5333 | 0.104 | 0.14 | 1 | 14 | Slc25a17 | -0.036 | Old |
| 0.566078 | 0.307642 | 0.131 | 0.1 | 1 | 14 | Mamld1 | 0.031 | Old |
| 0.567178 | 0.928797 | 0.126 | 0.1 | 1 | 14 | Pear1 | 0.026 | Old |
| 0.567253 | 0.334461 | 0.202 | 0.16 | 1 | 14 | Tspan5 | 0.042 | Old |
| 0.567386 | 0.829749 | 0.055 | 0.08 | 1 | 14 | Cd84 | -0.025 | Old |
| 0.567877 | 0.498486 | 0.317 | 0.28 | 1 | 14 | Emc1 | 0.037 | Old |
| 0.567946 | 0.526306 | 0.268 | 0.24 | 1 | 14 | Tgfbi | 0.028 | Old |
| 0.568442 | 0.298366 | 0.175 | 0.22 | 1 | 14 | Ugcg | -0.045 | Old |
| 0.568875 | 0.886875 | 0.131 | 0.1 | 1 | 14 | Slc33a1 | 0.031 | Old |
| 0.569029 | 1.145477 | 0.104 | 0.08 | 1 | 14 | Slc35a2 | 0.024 | Old |

| | | | | | | | | |
|---|---|---|---|---|---|---|---|---|
| 0.569475 | 0.035002 | 0.743 | 0.7 | 1 | 14 | Myadm | 0.043 | Old |
| 0.56977 | 0.948557 | 0.06 | 0.04 | 1 | 14 | Brdt | 0.02 | Old |
| 0.56977 | 0.867234 | 0.06 | 0.04 | 1 | 14 | Setdb2 | 0.02 | Old |
| 0.57011 | 0.699183 | 0.339 | 0.32 | 1 | 14 | Eif1ax | 0.019 | Old |
| 0.570236 | 0.200652 | 0.109 | 0.08 | 1 | 14 | Thumpd3 | 0.029 | Old |
| 0.570236 | 0.027812 | 0.109 | 0.08 | 1 | 14 | Lrba | 0.029 | Old |
| 0.570271 | 0.093546 | 0.377 | 0.34 | 1 | 14 | Trf | 0.037 | Old |
| 0.570424 | 0.625128 | 0.208 | 0.18 | 1 | 14 | Msl3 | 0.028 | Old |
| 0.570719 | 0.373409 | 0.087 | 0.12 | 1 | 14 | Mrps26 | -0.033 | Old |
| 0.571218 | 0.179224 | 0.393 | 0.48 | 1 | 14 | Hspb1 | -0.087 | Old |
| 0.571287 | 1.291619 | 0.055 | 0.08 | 1 | 14 | Zfp568 | -0.025 | Old |
| 0.571287 | 1.072997 | 0.055 | 0.08 | 1 | 14 | Rogdi | -0.025 | Old |
| 0.571656 | 1.059761 | 0.164 | 0.14 | 1 | 14 | Slc23a2 | 0.024 | Old |
| 0.571992 | 0.336147 | 0.311 | 0.28 | 1 | 14 | Msl2 | 0.031 | Old |
| 0.572007 | 0.394315 | 0.191 | 0.16 | 1 | 14 | Naa35 | 0.031 | Old |
| 0.572137 | 1.056886 | 0.104 | 0.08 | 1 | 14 | Manbal | 0.024 | Old |
| 0.572138 | 0.182168 | 0.169 | 0.14 | 1 | 14 | Meaf6 | 0.029 | Old |
| 0.57217 | 0.214482 | 0.137 | 0.18 | 1 | 14 | Yipf6 | -0.043 | Old |
| 0.57217 | 0.104041 | 0.153 | 0.12 | 1 | 14 | Fam3a | 0.033 | Old |
| 0.572273 | 0.190463 | 0.361 | 0.32 | 1 | 14 | Sbno2 | 0.041 | Old |
| 0.57258 | 0.69607 | 0.148 | 0.12 | 1 | 14 | Srm | 0.028 | Old |
| 0.572835 | 0.674695 | 0.246 | 0.22 | 1 | 14 | Ube2i | 0.026 | Old |
| 0.57296 | 0.050968 | 0.142 | 0.18 | 1 | 14 | Znrf2 | -0.038 | Old |
| 0.573013 | 0.040955 | 0.311 | 0.36 | 1 | 14 | Cisd2 | -0.049 | Old |
| 0.573079 | 0.030485 | 0.126 | 0.16 | 1 | 14 | Mrpl32 | -0.034 | Old |
| 0.573817 | 1.095718 | 0.06 | 0.04 | 1 | 14 | Orc6 | 0.02 | Old |
| 0.573817 | 0.948587 | 0.06 | 0.04 | 1 | 14 | Pirb | 0.02 | Old |
| 0.573817 | 0.683351 | 0.06 | 0.04 | 1 | 14 | Hoxa9 | 0.02 | Old |
| 0.573894 | 0.186969 | 0.087 | 0.12 | 1 | 14 | 9130401M | -0.033 | Old |
| 0.574011 | 0.215148 | 0.399 | 0.38 | 1 | 14 | Anapc5 | 0.019 | Old |
| 0.574389 | 0.429706 | 0.191 | 0.16 | 1 | 14 | Ncbp3 | 0.031 | Old |
| 0.574644 | 0.668612 | 0.169 | 0.14 | 1 | 14 | Nomo1 | 0.029 | Old |
| 0.574701 | 0.41574 | 0.383 | 0.36 | 1 | 14 | Atf7 | 0.023 | Old |
| 0.574731 | 0.450375 | 0.213 | 0.18 | 1 | 14 | Tex264 | 0.033 | Old |
| 0.574796 | 0.409392 | 0.153 | 0.12 | 1 | 14 | Parp6 | 0.033 | Old |
| 0.574896 | 0.590669 | 0.104 | 0.14 | 1 | 14 | Tanc2 | -0.036 | Old |
| 0.5752 | 0.824489 | 0.055 | 0.08 | 1 | 14 | Mtor | -0.025 | Old |
| 0.575508 | 1.51563 | 0.082 | 0.06 | 1 | 14 | Ccr2 | 0.022 | Old |
| 0.575508 | 0.617055 | 0.071 | 0.1 | 1 | 14 | Bcl2l11 | -0.029 | Old |
| 0.575728 | 0.12208 | 0.126 | 0.1 | 1 | 14 | Hspbp1 | 0.026 | Old |
| 0.575743 | 0.092066 | 0.317 | 0.28 | 1 | 14 | Eif1a | 0.037 | Old |
| 0.576369 | 0.300202 | 0.087 | 0.06 | 1 | 14 | Ormdl1 | 0.027 | Old |
| 0.576654 | 0.014406 | 0.219 | 0.18 | 1 | 14 | Rmnd5a | 0.039 | Old |
| 0.576727 | 0.936363 | 0.164 | 0.14 | 1 | 14 | Glt8d2 | 0.024 | Old |
| 0.576727 | 0.862992 | 0.164 | 0.14 | 1 | 14 | Aco1 | 0.024 | Old |

| | | | | | | | | |
|---|---|---|---|---|---|---|---|---|
| 0.576853 | 1.371979 | 0.12 | 0.1 | 1 | 14 | Mtmr12 | 0.02 | Old |
| 0.577729 | 0.09806 | 0.164 | 0.2 | 1 | 14 | Magoh | -0.036 | Old |
| 0.577853 | 0.45134 | 0.104 | 0.14 | 1 | 14 | Zdhhc6 | -0.036 | Old |
| 0.577853 | 0.280798 | 0.104 | 0.14 | 1 | 14 | Dimt1 | -0.036 | Old |
| 0.577878 | 0.96887 | 0.06 | 0.04 | 1 | 14 | Slc30a1 | 0.02 | Old |
| 0.577878 | 0.601153 | 0.06 | 0.04 | 1 | 14 | Ocel1 | 0.02 | Old |
| 0.577878 | 0.5636 | 0.06 | 0.04 | 1 | 14 | Tmem94 | 0.02 | Old |
| 0.577891 | 0.055182 | 0.94 | 0.86 | 1 | 14 | Zfp36 | 0.08 | Old |
| 0.577906 | 0.750673 | 0.148 | 0.12 | 1 | 14 | Ric8a | 0.028 | Old |
| 0.578212 | 0.577753 | 0.262 | 0.24 | 1 | 14 | Tnks1bp1 | 0.022 | Old |
| 0.57834 | 0.005358 | 0.503 | 0.46 | 1 | 14 | Hnrnpc | 0.043 | Old |
| 0.579001 | 1.259537 | 0.082 | 0.06 | 1 | 14 | Lactb | 0.022 | Old |
| 0.579001 | 1.018864 | 0.082 | 0.06 | 1 | 14 | Ern1 | 0.022 | Old |
| 0.579001 | 0.916205 | 0.082 | 0.06 | 1 | 14 | Trmt2b | 0.022 | Old |
| 0.579001 | 0.783354 | 0.082 | 0.06 | 1 | 14 | Zfp809 | 0.022 | Old |
| 0.579045 | 0.631562 | 0.437 | 0.44 | 1 | 14 | Itpripl2 | -0.003 | Old |
| 0.579696 | 0.001231 | 0.317 | 0.28 | 1 | 14 | Psenen | 0.037 | Old |
| 0.579765 | 1.215776 | 0.12 | 0.1 | 1 | 14 | Fem1a | 0.02 | Old |
| 0.580064 | 0.525291 | 0.137 | 0.18 | 1 | 14 | Sin3a | -0.043 | Old |
| 0.580233 | 0.433806 | 0.235 | 0.2 | 1 | 14 | Rhbdf1 | 0.035 | Old |
| 0.5807 | 0.483489 | 0.251 | 0.22 | 1 | 14 | Pdhb | 0.031 | Old |
| 0.580816 | 0.812366 | 0.104 | 0.14 | 1 | 14 | Crem | -0.036 | Old |
| 0.58123 | 0.467881 | 0.344 | 0.32 | 1 | 14 | Fam208a | 0.024 | Old |
| 0.581301 | 0.210903 | 0.202 | 0.16 | 1 | 14 | Taz | 0.042 | Old |
| 0.581564 | 0.553439 | 0.191 | 0.16 | 1 | 14 | Minpp1 | 0.031 | Old |
| 0.581952 | 0.704234 | 0.06 | 0.04 | 1 | 14 | Dus3l | 0.02 | Old |
| 0.581952 | 0.672926 | 0.06 | 0.04 | 1 | 14 | Nxt1 | 0.02 | Old |
| 0.581952 | 0.643428 | 0.06 | 0.04 | 1 | 14 | Krt80 | 0.02 | Old |
| 0.581952 | 0.570762 | 0.06 | 0.04 | 1 | 14 | Tmem199 | 0.02 | Old |
| 0.581952 | 0.537163 | 0.06 | 0.04 | 1 | 14 | 2310022A1 | 0.02 | Old |
| 0.581952 | 0.398406 | 0.06 | 0.04 | 1 | 14 | Mcm7 | 0.02 | Old |
| 0.582192 | 0.466155 | 0.169 | 0.14 | 1 | 14 | Mob3c | 0.029 | Old |
| 0.582238 | 0.366615 | 0.355 | 0.32 | 1 | 14 | Nras | 0.035 | Old |
| 0.582256 | 0.608854 | 0.257 | 0.24 | 1 | 14 | Sec23a | 0.017 | Old |
| 0.582504 | 0.926126 | 0.082 | 0.06 | 1 | 14 | Psmd10 | 0.022 | Old |
| 0.582959 | 0.220718 | 0.131 | 0.1 | 1 | 14 | Cdk10 | 0.031 | Old |
| 0.583255 | 0.627179 | 0.148 | 0.12 | 1 | 14 | Fbxo8 | 0.028 | Old |
| 0.583468 | 0.945363 | 0.087 | 0.12 | 1 | 14 | Gnb5 | -0.033 | Old |
| 0.584049 | 0.053558 | 0.126 | 0.16 | 1 | 14 | P3h4 | -0.034 | Old |
| 0.584276 | 0.436898 | 0.12 | 0.16 | 1 | 14 | Grpel1 | -0.04 | Old |
| 0.5847 | 0.481192 | 0.29 | 0.26 | 1 | 14 | Ptpn9 | 0.03 | Old |
| 0.585276 | 0.55764 | 0.219 | 0.2 | 1 | 14 | Rsl1d1 | 0.019 | Old |
| 0.585355 | 0.302695 | 0.153 | 0.12 | 1 | 14 | Upf2 | 0.033 | Old |
| 0.58561 | 0.267422 | 0.109 | 0.14 | 1 | 14 | 4-Sep | -0.031 | Old |
| 0.586017 | 0.921908 | 0.082 | 0.06 | 1 | 14 | Oxa1l | 0.022 | Old |

| | | | | | | | | |
|---|---|---|---|---|---|---|---|---|
| 0.586017 | 0.580389 | 0.082 | 0.06 | 1 | 14 | Alkbh1 | 0.022 | Old |
| 0.58604 | 0.920375 | 0.06 | 0.04 | 1 | 14 | Eif2b3 | 0.02 | Old |
| 0.58604 | 0.749798 | 0.06 | 0.04 | 1 | 14 | Car13 | 0.02 | Old |
| 0.58604 | 0.535758 | 0.06 | 0.04 | 1 | 14 | E130309D( | 0.02 | Old |
| 0.58604 | 0.387392 | 0.06 | 0.04 | 1 | 14 | Chrd | 0.02 | Old |
| 0.58604 | 0.014002 | 0.06 | 0.04 | 1 | 14 | Alg9 | 0.02 | Old |
| 0.58645 | 1.428605 | 0.18 | 0.16 | 1 | 14 | Serpine1 | 0.02 | Old |
| 0.586664 | 0.147687 | 0.262 | 0.24 | 1 | 14 | Fam8a1 | 0.022 | Old |
| 0.586676 | 1.68783 | 0.098 | 0.08 | 1 | 14 | Polr3f | 0.018 | Old |
| 0.586807 | 1.328675 | 0.137 | 0.12 | 1 | 14 | Gpd1l | 0.017 | Old |
| 0.587545 | 0.376216 | 0.219 | 0.2 | 1 | 14 | Tbc1d23 | 0.019 | Old |
| 0.587581 | 0.650675 | 0.262 | 0.22 | 1 | 14 | Sc5d | 0.042 | Old |
| 0.587661 | 0.154096 | 0.273 | 0.34 | 1 | 14 | Cbx1 | -0.067 | Old |
| 0.587661 | 0.004528 | 0.224 | 0.28 | 1 | 14 | Srebf1 | -0.056 | Old |
| 0.588614 | 0.521443 | 0.142 | 0.18 | 1 | 14 | Crot | -0.038 | Old |
| 0.588778 | 0.071652 | 0.863 | 0.82 | 1 | 14 | Rpl18 | 0.043 | Old |
| 0.589329 | 0.081989 | 0.251 | 0.22 | 1 | 14 | Atf6 | 0.031 | Old |
| 0.58954 | 0.533564 | 0.082 | 0.06 | 1 | 14 | Psme2b | 0.022 | Old |
| 0.589919 | 0.455301 | 0.284 | 0.26 | 1 | 14 | Acvrl1 | 0.024 | Old |
| 0.590141 | 0.355628 | 0.06 | 0.04 | 1 | 14 | Cdk20 | 0.02 | Old |
| 0.590668 | 0.420291 | 0.137 | 0.18 | 1 | 14 | Polb | -0.043 | Old |
| 0.590668 | 0.268584 | 0.153 | 0.12 | 1 | 14 | Fam49b | 0.033 | Old |
| 0.590787 | 0.574988 | 0.257 | 0.24 | 1 | 14 | Skiv2l | 0.017 | Old |
| 0.590994 | 0.457082 | 0.268 | 0.24 | 1 | 14 | Adam9 | 0.028 | Old |
| 0.592341 | 0.145952 | 0.126 | 0.16 | 1 | 14 | Kmt5c | -0.034 | Old |
| 0.592918 | 0.278032 | 0.415 | 0.38 | 1 | 14 | Adamts12 | 0.035 | Old |
| 0.593072 | 1.059956 | 0.082 | 0.06 | 1 | 14 | Slc12a9 | 0.022 | Old |
| 0.593072 | 0.660833 | 0.082 | 0.06 | 1 | 14 | Dvl2 | 0.022 | Old |
| 0.593116 | 0.599781 | 0.087 | 0.12 | 1 | 14 | Tmcc2 | -0.033 | Old |
| 0.593666 | 0.49848 | 0.251 | 0.22 | 1 | 14 | Smarcb1 | 0.031 | Old |
| 0.594022 | 0.658943 | 0.148 | 0.12 | 1 | 14 | Qars | 0.028 | Old |
| 0.594111 | 0.53212 | 0.104 | 0.08 | 1 | 14 | Ubac2 | 0.024 | Old |
| 0.594216 | 0.075668 | 0.645 | 0.6 | 1 | 14 | P4hb | 0.045 | Old |
| 0.594255 | 0.730667 | 0.06 | 0.04 | 1 | 14 | Il2ra | 0.02 | Old |
| 0.594255 | 0.145666 | 0.06 | 0.04 | 1 | 14 | Epsti1 | 0.02 | Old |
| 0.594603 | 0.106868 | 0.197 | 0.16 | 1 | 14 | Pi4k2b | 0.037 | Old |
| 0.59464 | 0.485141 | 0.164 | 0.14 | 1 | 14 | Erh | 0.024 | Old |
| 0.594875 | 0.273998 | 0.169 | 0.14 | 1 | 14 | Dyrk1a | 0.029 | Old |
| 0.594975 | 0.372657 | 0.29 | 0.26 | 1 | 14 | Lzts2 | 0.03 | Old |
| 0.595181 | 0.187179 | 0.262 | 0.32 | 1 | 14 | Sin3b | -0.058 | Old |
| 0.596148 | 0.346864 | 0.262 | 0.22 | 1 | 14 | Calml4 | 0.042 | Old |
| 0.596614 | 0.915021 | 0.082 | 0.06 | 1 | 14 | 9130221H1 | 0.022 | Old |
| 0.596614 | 0.765705 | 0.082 | 0.06 | 1 | 14 | Tmem67 | 0.022 | Old |
| 0.596614 | 0.541969 | 0.082 | 0.06 | 1 | 14 | Noc4l | 0.022 | Old |
| 0.596686 | 0.576882 | 0.295 | 0.28 | 1 | 14 | Akirin2 | 0.015 | Old |

| | | | | | | | | |
|---|---|---|---|---|---|---|---|---|
| 0.59722 | 0.735006 | 0.164 | 0.14 | 1 | 14 | Nynrin | 0.024 | Old |
| 0.597281 | 0.821118 | 0.104 | 0.08 | 1 | 14 | Pdcd2 | 0.024 | Old |
| 0.597281 | 0.252628 | 0.104 | 0.08 | 1 | 14 | Hacd1 | 0.024 | Old |
| 0.597362 | 1.323423 | 0.142 | 0.12 | 1 | 14 | Rcbtb2 | 0.022 | Old |
| 0.597381 | 0.208903 | 0.109 | 0.14 | 1 | 14 | Kmt2b | -0.031 | Old |
| 0.598307 | 0.259849 | 0.208 | 0.26 | 1 | 14 | Lsm8 | -0.052 | Old |
| 0.598382 | 0.453634 | 0.06 | 0.04 | 1 | 14 | Ppm1m | 0.02 | Old |
| 0.598382 | 0.240697 | 0.06 | 0.04 | 1 | 14 | Sik2 | 0.02 | Old |
| 0.598483 | 0.367707 | 0.525 | 0.46 | 1 | 14 | Klf13 | 0.065 | Old |
| 0.598744 | 0.39044 | 0.104 | 0.14 | 1 | 14 | Hacd2 | -0.036 | Old |
| 0.598958 | 0.023159 | 0.06 | 0.08 | 1 | 14 | Hpca | -0.02 | Old |
| 0.599435 | 0.146515 | 0.279 | 0.24 | 1 | 14 | Naa50 | 0.039 | Old |
| 0.599439 | 0.466189 | 0.148 | 0.12 | 1 | 14 | Ero1l | 0.028 | Old |
| 0.599439 | 0.448864 | 0.148 | 0.12 | 1 | 14 | Fam222b | 0.028 | Old |
| 0.599548 | 0.206856 | 0.355 | 0.32 | 1 | 14 | Papola | 0.035 | Old |
| 0.599766 | 1.149274 | 0.24 | 0.22 | 1 | 14 | Mbd1 | 0.02 | Old |
| 0.599984 | 0.38255 | 0.169 | 0.14 | 1 | 14 | Mospd2 | 0.029 | Old |
| 0.600111 | 1.007838 | 0.142 | 0.12 | 1 | 14 | Hmgn2 | 0.022 | Old |
| 0.600166 | 1.449395 | 0.082 | 0.06 | 1 | 14 | 1700037H[illegible] | 0.022 | Old |
| 0.600166 | 0.704471 | 0.082 | 0.06 | 1 | 14 | Focad | 0.022 | Old |
| 0.600166 | 0.336832 | 0.082 | 0.06 | 1 | 14 | Nol12 | 0.022 | Old |
| 0.600166 | 0.131248 | 0.082 | 0.06 | 1 | 14 | Phc1 | 0.022 | Old |
| 0.600198 | 0.306743 | 0.251 | 0.22 | 1 | 14 | Pdzd2 | 0.031 | Old |
| 0.600433 | 0.002336 | 0.077 | 0.1 | 1 | 14 | Fndc5 | -0.023 | Old |
| 0.600459 | 0.50281 | 0.104 | 0.08 | 1 | 14 | Prickle1 | 0.024 | Old |
| 0.600459 | 0.100858 | 0.093 | 0.12 | 1 | 14 | Tmem173 | -0.027 | Old |
| 0.600686 | 1.069446 | 0.137 | 0.12 | 1 | 14 | Camk1d | 0.017 | Old |
| 0.60129 | 0.487267 | 0.361 | 0.34 | 1 | 14 | Aco2 | 0.021 | Old |
| 0.601541 | 0.066152 | 0.279 | 0.24 | 1 | 14 | 2700081O[illegible] | 0.039 | Old |
| 0.601643 | 0.093684 | 0.956 | 0.94 | 1 | 14 | Rps15 | 0.016 | Old |
| 0.602151 | 0.288423 | 0.148 | 0.18 | 1 | 14 | Btbd3 | -0.032 | Old |
| 0.602508 | 0.195286 | 0.235 | 0.2 | 1 | 14 | Prdm2 | 0.035 | Old |
| 0.602522 | 0.061996 | 0.06 | 0.04 | 1 | 14 | Map3k14 | 0.02 | Old |
| 0.602865 | 0.75011 | 0.142 | 0.12 | 1 | 14 | Sos2 | 0.022 | Old |
| 0.603306 | 1.016741 | 0.12 | 0.1 | 1 | 14 | Wdr75 | 0.02 | Old |
| 0.603306 | 0.135602 | 0.109 | 0.14 | 1 | 14 | Snx7 | -0.031 | Old |
| 0.603443 | 0.701646 | 0.202 | 0.18 | 1 | 14 | Mapkap1 | 0.022 | Old |
| 0.603645 | 0.528453 | 0.104 | 0.08 | 1 | 14 | Slc25a20 | 0.024 | Old |
| 0.603645 | 0.184063 | 0.104 | 0.08 | 1 | 14 | Plk3 | 0.024 | Old |
| 0.603645 | 0.151272 | 0.093 | 0.12 | 1 | 14 | Pigyl | -0.027 | Old |
| 0.603727 | 0.468086 | 0.082 | 0.06 | 1 | 14 | Ephb3 | 0.022 | Old |
| 0.60388 | 0.653797 | 0.158 | 0.2 | 1 | 14 | Epc1 | -0.042 | Old |
| 0.604001 | 0.468742 | 0.224 | 0.2 | 1 | 14 | Il10rb | 0.024 | Old |
| 0.604406 | 0.230961 | 0.23 | 0.2 | 1 | 14 | Tnfrsf1b | 0.03 | Old |
| 0.604706 | 0.816889 | 0.273 | 0.26 | 1 | 14 | Esyt1 | 0.013 | Old |

| | | | | | | | | |
|---|---|---|---|---|---|---|---|---|
| 0.605112 | 0.341947 | 0.169 | 0.14 | 1 | 14 | Igsf3 | 0.029 | Old |
| 0.606374 | 0.106073 | 0.29 | 0.34 | 1 | 14 | Mgat2 | -0.05 | Old |
| 0.606674 | 0.946424 | 0.23 | 0.2 | 1 | 14 | Abhd5 | 0.03 | Old |
| 0.606838 | 0.502964 | 0.104 | 0.08 | 1 | 14 | Ttc8 | 0.024 | Old |
| 0.607076 | 0.200298 | 0.361 | 0.34 | 1 | 14 | Polr2a | 0.021 | Old |
| 0.607112 | 1.054276 | 0.208 | 0.2 | 1 | 14 | Man2b1 | 0.008 | Old |
| 0.607112 | 0.238511 | 0.213 | 0.18 | 1 | 14 | Rit1 | 0.033 | Old |
| 0.607123 | 0.09133 | 0.142 | 0.18 | 1 | 14 | Rnf166 | -0.038 | Old |
| 0.607298 | 0.760732 | 0.082 | 0.06 | 1 | 14 | Polr3e | 0.022 | Old |
| 0.607391 | 0.235715 | 0.077 | 0.1 | 1 | 14 | Acyp1 | -0.023 | Old |
| 0.607477 | 0.519583 | 0.328 | 0.3 | 1 | 14 | Usp19 | 0.028 | Old |
| 0.607589 | 0.780047 | 0.164 | 0.14 | 1 | 14 | Zmat3 | 0.024 | Old |
| 0.607683 | 0.664132 | 0.153 | 0.2 | 1 | 14 | Gtf2a2 | -0.047 | Old |
| 0.607999 | 0.363489 | 0.344 | 0.3 | 1 | 14 | AW112010 | 0.044 | Old |
| 0.608136 | 0.812367 | 0.23 | 0.22 | 1 | 14 | Casp4 | 0.01 | Old |
| 0.608271 | 0.058619 | 1 | 0.96 | 1 | 14 | mt-Co2 | 0.04 | Old |
| 0.608574 | 0.667994 | 0.224 | 0.2 | 1 | 14 | Gm10073 | 0.024 | Old |
| 0.60864 | 0.609523 | 0.24 | 0.22 | 1 | 14 | Nr2f6 | 0.02 | Old |
| 0.608727 | 0.54563 | 0.301 | 0.28 | 1 | 14 | Ctdsp1 | 0.021 | Old |
| 0.609502 | 0.397226 | 0.333 | 0.32 | 1 | 14 | Mut | 0.013 | Old |
| 0.609997 | 0.01826 | 0.279 | 0.24 | 1 | 14 | Bcl7c | 0.039 | Old |
| 0.610039 | 1.093003 | 0.158 | 0.14 | 1 | 14 | Qrich1 | 0.018 | Old |
| 0.610045 | 0.912097 | 0.208 | 0.18 | 1 | 14 | Cd83 | 0.028 | Old |
| 0.610243 | 0.506725 | 0.175 | 0.14 | 1 | 14 | Ttyh2 | 0.035 | Old |
| 0.610878 | 0.245733 | 0.082 | 0.06 | 1 | 14 | Rhot2 | 0.022 | Old |
| 0.610884 | 0.099287 | 0.077 | 0.1 | 1 | 14 | Egr3 | -0.023 | Old |
| 0.611058 | 0.281586 | 0.273 | 0.26 | 1 | 14 | Mgat4b | 0.013 | Old |
| 0.611584 | 0.624225 | 0.284 | 0.28 | 1 | 14 | Taf15 | 0.004 | Old |
| 0.611605 | 0.020078 | 0.131 | 0.1 | 1 | 14 | Bmp4 | 0.031 | Old |
| 0.612139 | 0.007138 | 0.153 | 0.12 | 1 | 14 | Susd2 | 0.033 | Old |
| 0.612243 | 0.892662 | 0.12 | 0.1 | 1 | 14 | Rnf40 | 0.02 | Old |
| 0.612243 | 0.283533 | 0.109 | 0.14 | 1 | 14 | Hsdl2 | -0.031 | Old |
| 0.612316 | 0.026939 | 0.202 | 0.24 | 1 | 14 | Sfswap | -0.038 | Old |
| 0.613223 | 0.156444 | 0.388 | 0.46 | 1 | 14 | Gabarapl1 | -0.072 | Old |
| 0.613939 | 0.090566 | 0.131 | 0.16 | 1 | 14 | Cfp | -0.029 | Old |
| 0.61395 | 0.426005 | 0.197 | 0.16 | 1 | 14 | Arl3 | 0.037 | Old |
| 0.61423 | 2.027237 | 0.077 | 0.06 | 1 | 14 | Btg3 | 0.017 | Old |
| 0.61436 | 0.458691 | 0.246 | 0.22 | 1 | 14 | Kitl | 0.026 | Old |
| 0.614385 | 0.226207 | 0.077 | 0.1 | 1 | 14 | Fbxo7 | -0.023 | Old |
| 0.614467 | 0.545086 | 0.082 | 0.06 | 1 | 14 | Slc2a8 | 0.022 | Old |
| 0.614467 | 0.12276 | 0.082 | 0.06 | 1 | 14 | Fbxw17 | 0.022 | Old |
| 0.61452 | 0.186259 | 0.257 | 0.22 | 1 | 14 | Sertad1 | 0.037 | Old |
| 0.614711 | 1.079455 | 0.137 | 0.12 | 1 | 14 | Sart1 | 0.017 | Old |
| 0.61502 | 0.326044 | 0.06 | 0.04 | 1 | 14 | Rbmx2 | 0.02 | Old |
| 0.615482 | 0.320846 | 0.328 | 0.3 | 1 | 14 | Kdm7a | 0.028 | Old |

| | | | | | | | | |
|---|---|---|---|---|---|---|---|---|
| 0.616055 | 0.143281 | 0.235 | 0.2 | 1 | 14 | Ncoa2 | 0.035 | Old |
| 0.616463 | 0.323044 | 0.104 | 0.08 | 1 | 14 | Bcat2 | 0.024 | Old |
| 0.616463 | 0.318962 | 0.104 | 0.08 | 1 | 14 | Oard1 | 0.024 | Old |
| 0.616617 | 0.290331 | 0.268 | 0.24 | 1 | 14 | Coro1c | 0.028 | Old |
| 0.616722 | 0.799095 | 0.142 | 0.12 | 1 | 14 | 3632451O( | 0.022 | Old |
| 0.617767 | 0.909956 | 0.202 | 0.18 | 1 | 14 | Ptpn12 | 0.022 | Old |
| 0.618066 | 0.386752 | 0.082 | 0.06 | 1 | 14 | Mark4 | 0.022 | Old |
| 0.618066 | 0.308388 | 0.082 | 0.06 | 1 | 14 | Mkl2 | 0.022 | Old |
| 0.618071 | 0.250958 | 0.23 | 0.2 | 1 | 14 | Lurap1l | 0.03 | Old |
| 0.618678 | 0.090559 | 0.612 | 0.66 | 1 | 14 | Klf3 | -0.048 | Old |
| 0.618755 | 0.00626 | 0.169 | 0.2 | 1 | 14 | Ube2j2 | -0.031 | Old |
| 0.619144 | 1.778878 | 0.011 | 0.02 | 1 | 14 | Gm28557 | -0.009 | Old |
| 0.619144 | 0.833009 | 0.011 | 0.02 | 1 | 14 | Hey2 | -0.009 | Old |
| 0.619144 | 0.754997 | 0.011 | 0.02 | 1 | 14 | Hspb6 | -0.009 | Old |
| 0.619144 | 0.618084 | 0.011 | 0.02 | 1 | 14 | AI182371 | -0.009 | Old |
| 0.619144 | 0.45469 | 0.011 | 0.02 | 1 | 14 | Gm34934 | -0.009 | Old |
| 0.619144 | 0.320775 | 0.011 | 0.02 | 1 | 14 | Tkfc | -0.009 | Old |
| 0.619144 | 0.245069 | 0.011 | 0.02 | 1 | 14 | Ltb4r1 | -0.009 | Old |
| 0.619144 | 0.139862 | 0.011 | 0.02 | 1 | 14 | Slc8a1 | -0.009 | Old |
| 0.619144 | 0.089736 | 0.011 | 0.02 | 1 | 14 | Aqp1 | -0.009 | Old |
| 0.619144 | 0.041658 | 0.011 | 0.02 | 1 | 14 | Mterf1a | -0.009 | Old |
| 0.619144 | 0.025814 | 0.011 | 0.02 | 1 | 14 | Dph2 | -0.009 | Old |
| 0.619144 | 0.001428 | 0.011 | 0.02 | 1 | 14 | Gm29107 | -0.009 | Old |
| 0.619194 | 1.1494 | 0.098 | 0.08 | 1 | 14 | Arfgef2 | 0.018 | Old |
| 0.61951 | 0.843325 | 0.142 | 0.12 | 1 | 14 | 5730455P1 | 0.022 | Old |
| 0.619542 | 0.571481 | 0.208 | 0.18 | 1 | 14 | Rsl24d1 | 0.028 | Old |
| 0.620221 | 0.06382 | 0.268 | 0.32 | 1 | 14 | Gatad1 | -0.052 | Old |
| 0.620238 | 0.156311 | 0.552 | 0.54 | 1 | 14 | Cyfip1 | 0.012 | Old |
| 0.620361 | 0.084187 | 0.213 | 0.26 | 1 | 14 | Vps41 | -0.047 | Old |
| 0.620365 | 0.271693 | 0.361 | 0.32 | 1 | 14 | Parva | 0.041 | Old |
| 0.620934 | 0.010522 | 0.268 | 0.24 | 1 | 14 | 2410015M | 0.028 | Old |
| 0.621036 | 0.359197 | 0.246 | 0.22 | 1 | 14 | Stard9 | 0.026 | Old |
| 0.621111 | 0.355496 | 0.257 | 0.22 | 1 | 14 | Lrrc8a | 0.037 | Old |
| 0.621235 | 0.21479 | 0.197 | 0.24 | 1 | 14 | Tmtc1 | -0.043 | Old |
| 0.621329 | 0.06862 | 0.148 | 0.12 | 1 | 14 | Axin2 | 0.028 | Old |
| 0.621419 | 0.086818 | 0.918 | 0.84 | 1 | 14 | Hsp90b1 | 0.078 | Old |
| 0.621628 | 1.297112 | 0.077 | 0.06 | 1 | 14 | Mrpl15 | 0.017 | Old |
| 0.621674 | 0.163164 | 0.082 | 0.06 | 1 | 14 | Dedd | 0.022 | Old |
| 0.622303 | 0.132362 | 0.131 | 0.16 | 1 | 14 | Crls1 | -0.029 | Old |
| 0.622321 | 0.077705 | 0.115 | 0.14 | 1 | 14 | Inpp5f | -0.025 | Old |
| 0.622488 | 1.164079 | 0.098 | 0.08 | 1 | 14 | Patl1 | 0.018 | Old |
| 0.623054 | 0.748516 | 0.235 | 0.22 | 1 | 14 | Atp6v1h | 0.015 | Old |
| 0.623213 | 0.765563 | 0.153 | 0.2 | 1 | 14 | Pofut2 | -0.047 | Old |
| 0.624315 | 0.538819 | 0.208 | 0.18 | 1 | 14 | Rnf122 | 0.028 | Old |
| 0.624388 | 0.43431 | 0.219 | 0.2 | 1 | 14 | Zc3h18 | 0.019 | Old |

| | | | | | | | | |
|---|---|---|---|---|---|---|---|---|
| 0.624506 | 0.103069 | 0.404 | 0.4 | 1 | 14 | Lgals9 | 0.004 | Old |
| 0.624543 | 0.004722 | 0.311 | 0.36 | 1 | 14 | Sdf2 | -0.049 | Old |
| 0.624925 | 0.049075 | 0.973 | 0.92 | 1 | 14 | Psap | 0.053 | Old |
| 0.624941 | 0.174149 | 0.077 | 0.1 | 1 | 14 | Slx1b | -0.023 | Old |
| 0.625063 | 0.097546 | 0.322 | 0.3 | 1 | 14 | Uqcc2 | 0.022 | Old |
| 0.625308 | 2.579524 | 0.033 | 0.02 | 1 | 14 | Zfp367 | 0.013 | Old |
| 0.625308 | 2.554492 | 0.033 | 0.02 | 1 | 14 | Helq | 0.013 | Old |
| 0.625308 | 2.405935 | 0.033 | 0.02 | 1 | 14 | Ntrk1 | 0.013 | Old |
| 0.625308 | 2.254569 | 0.033 | 0.02 | 1 | 14 | Camk2n2 | 0.013 | Old |
| 0.625308 | 2.226851 | 0.033 | 0.02 | 1 | 14 | Adrb2 | 0.013 | Old |
| 0.625308 | 2.166364 | 0.033 | 0.02 | 1 | 14 | Gm47204 | 0.013 | Old |
| 0.625308 | 2.078223 | 0.033 | 0.02 | 1 | 14 | Apex2 | 0.013 | Old |
| 0.625308 | 1.95634 | 0.033 | 0.02 | 1 | 14 | Ptpn4 | 0.013 | Old |
| 0.625308 | 1.881893 | 0.033 | 0.02 | 1 | 14 | Prrg2 | 0.013 | Old |
| 0.625308 | 1.880053 | 0.033 | 0.02 | 1 | 14 | Tmem267 | 0.013 | Old |
| 0.625308 | 1.868757 | 0.033 | 0.02 | 1 | 14 | Gm26660 | 0.013 | Old |
| 0.625308 | 1.818614 | 0.033 | 0.02 | 1 | 14 | Mb21d1 | 0.013 | Old |
| 0.625308 | 1.782134 | 0.033 | 0.02 | 1 | 14 | Rnf157 | 0.013 | Old |
| 0.625308 | 1.735793 | 0.033 | 0.02 | 1 | 14 | Cd46 | 0.013 | Old |
| 0.625308 | 1.7099 | 0.033 | 0.02 | 1 | 14 | Rnaset2a | 0.013 | Old |
| 0.625308 | 1.69929 | 0.033 | 0.02 | 1 | 14 | H2afx | 0.013 | Old |
| 0.625308 | 1.671695 | 0.033 | 0.02 | 1 | 14 | Nkd2 | 0.013 | Old |
| 0.625308 | 1.661774 | 0.033 | 0.02 | 1 | 14 | Rabl3 | 0.013 | Old |
| 0.625308 | 1.647274 | 0.033 | 0.02 | 1 | 14 | Ulk3 | 0.013 | Old |
| 0.625308 | 1.642913 | 0.033 | 0.02 | 1 | 14 | Shisa9 | 0.013 | Old |
| 0.625308 | 1.640954 | 0.033 | 0.02 | 1 | 14 | Phf11d | 0.013 | Old |
| 0.625308 | 1.605507 | 0.033 | 0.02 | 1 | 14 | Gm20696 | 0.013 | Old |
| 0.625308 | 1.585673 | 0.033 | 0.02 | 1 | 14 | Mettl13 | 0.013 | Old |
| 0.625308 | 1.583946 | 0.033 | 0.02 | 1 | 14 | Zfp958 | 0.013 | Old |
| 0.625308 | 1.576373 | 0.033 | 0.02 | 1 | 14 | Cxcr4 | 0.013 | Old |
| 0.625308 | 1.575831 | 0.033 | 0.02 | 1 | 14 | Gm12258 | 0.013 | Old |
| 0.625308 | 1.345468 | 0.033 | 0.02 | 1 | 14 | Fam227a | 0.013 | Old |
| 0.625308 | 1.337653 | 0.033 | 0.02 | 1 | 14 | Polr1b | 0.013 | Old |
| 0.625308 | 1.227423 | 0.033 | 0.02 | 1 | 14 | Cdkn2b | 0.013 | Old |
| 0.625342 | 1.510395 | 0.077 | 0.06 | 1 | 14 | Adprhl2 | 0.017 | Old |
| 0.625342 | 1.33842 | 0.077 | 0.06 | 1 | 14 | Tradd | 0.017 | Old |
| 0.625503 | 0.110047 | 0.41 | 0.4 | 1 | 14 | Rbfox2 | 0.01 | Old |
| 0.625925 | 0.254866 | 0.164 | 0.14 | 1 | 14 | Naa38 | 0.024 | Old |
| 0.625994 | 0.441526 | 0.273 | 0.26 | 1 | 14 | Rab11fip5 | 0.013 | Old |
| 0.626096 | 1.358037 | 0.115 | 0.1 | 1 | 14 | Gstt2 | 0.015 | Old |
| 0.626154 | 0.050368 | 0.104 | 0.08 | 1 | 14 | 2410002F2 | 0.024 | Old |
| 0.627719 | 1.958461 | 0.011 | 0.02 | 1 | 14 | Eef1akmt3 | -0.009 | Old |
| 0.627719 | 1.882406 | 0.011 | 0.02 | 1 | 14 | Fbxo5 | -0.009 | Old |
| 0.627719 | 1.739752 | 0.011 | 0.02 | 1 | 14 | Nthl1 | -0.009 | Old |
| 0.627719 | 1.347202 | 0.011 | 0.02 | 1 | 14 | Ston1 | -0.009 | Old |

| | | | | | | | | |
|---|---|---|---|---|---|---|---|---|
| 0.627719 | 1.343568 | 0.011 | 0.02 | 1 | 14 | Capn6 | -0.009 | Old |
| 0.627719 | 1.324129 | 0.011 | 0.02 | 1 | 14 | Gm15545 | -0.009 | Old |
| 0.627719 | 1.27905 | 0.011 | 0.02 | 1 | 14 | Rpl7a-ps5 | -0.009 | Old |
| 0.627719 | 1.210301 | 0.011 | 0.02 | 1 | 14 | Rpp25 | -0.009 | Old |
| 0.627719 | 1.201784 | 0.011 | 0.02 | 1 | 14 | Col13a1 | -0.009 | Old |
| 0.627719 | 0.965424 | 0.011 | 0.02 | 1 | 14 | Zfp773 | -0.009 | Old |
| 0.627719 | 0.931241 | 0.011 | 0.02 | 1 | 14 | Gprin3 | -0.009 | Old |
| 0.627719 | 0.910754 | 0.011 | 0.02 | 1 | 14 | Wrap53 | -0.009 | Old |
| 0.627719 | 0.788747 | 0.011 | 0.02 | 1 | 14 | Cyb5rl | -0.009 | Old |
| 0.627719 | 0.785856 | 0.011 | 0.02 | 1 | 14 | Zfp119b | -0.009 | Old |
| 0.627719 | 0.782893 | 0.011 | 0.02 | 1 | 14 | Zfp689 | -0.009 | Old |
| 0.627719 | 0.773374 | 0.011 | 0.02 | 1 | 14 | Bdkrb2 | -0.009 | Old |
| 0.627719 | 0.757522 | 0.011 | 0.02 | 1 | 14 | Insrr | -0.009 | Old |
| 0.627719 | 0.752287 | 0.011 | 0.02 | 1 | 14 | Coa4 | -0.009 | Old |
| 0.627719 | 0.667131 | 0.011 | 0.02 | 1 | 14 | Frmd7 | -0.009 | Old |
| 0.627719 | 0.651472 | 0.011 | 0.02 | 1 | 14 | Rassf9 | -0.009 | Old |
| 0.627719 | 0.643995 | 0.011 | 0.02 | 1 | 14 | 1700003F1 | -0.009 | Old |
| 0.627719 | 0.624688 | 0.011 | 0.02 | 1 | 14 | Lpcat4 | -0.009 | Old |
| 0.627719 | 0.610049 | 0.011 | 0.02 | 1 | 14 | Ddx59 | -0.009 | Old |
| 0.627719 | 0.604404 | 0.011 | 0.02 | 1 | 14 | Pik3cb | -0.009 | Old |
| 0.627719 | 0.603362 | 0.011 | 0.02 | 1 | 14 | Dgkb | -0.009 | Old |
| 0.627719 | 0.602183 | 0.011 | 0.02 | 1 | 14 | Ugt1a7c | -0.009 | Old |
| 0.627719 | 0.597768 | 0.011 | 0.02 | 1 | 14 | B3gnt2 | -0.009 | Old |
| 0.627719 | 0.585583 | 0.011 | 0.02 | 1 | 14 | Zfp553 | -0.009 | Old |
| 0.627719 | 0.580128 | 0.011 | 0.02 | 1 | 14 | 4833403J1 | -0.009 | Old |
| 0.627719 | 0.568908 | 0.011 | 0.02 | 1 | 14 | Cks2 | -0.009 | Old |
| 0.627719 | 0.562471 | 0.011 | 0.02 | 1 | 14 | Dusp14 | -0.009 | Old |
| 0.627719 | 0.506583 | 0.011 | 0.02 | 1 | 14 | 4932422M | -0.009 | Old |
| 0.627719 | 0.503708 | 0.011 | 0.02 | 1 | 14 | Hoxb8 | -0.009 | Old |
| 0.627719 | 0.499604 | 0.011 | 0.02 | 1 | 14 | St6galnac5 | -0.009 | Old |
| 0.627719 | 0.495796 | 0.011 | 0.02 | 1 | 14 | Pex10 | -0.009 | Old |
| 0.627719 | 0.479909 | 0.011 | 0.02 | 1 | 14 | Cd244 | -0.009 | Old |
| 0.627719 | 0.457576 | 0.011 | 0.02 | 1 | 14 | B3gnt3 | -0.009 | Old |
| 0.627719 | 0.455599 | 0.011 | 0.02 | 1 | 14 | Gm14812 | -0.009 | Old |
| 0.627719 | 0.446592 | 0.011 | 0.02 | 1 | 14 | Casz1 | -0.009 | Old |
| 0.627719 | 0.431775 | 0.011 | 0.02 | 1 | 14 | Phf24 | -0.009 | Old |
| 0.627719 | 0.425226 | 0.011 | 0.02 | 1 | 14 | 1010001N( | -0.009 | Old |
| 0.627719 | 0.416806 | 0.011 | 0.02 | 1 | 14 | Cep295nl | -0.009 | Old |
| 0.627719 | 0.410211 | 0.011 | 0.02 | 1 | 14 | Ada | -0.009 | Old |
| 0.627719 | 0.407696 | 0.011 | 0.02 | 1 | 14 | Uck2 | -0.009 | Old |
| 0.627719 | 0.397433 | 0.011 | 0.02 | 1 | 14 | Efcab2 | -0.009 | Old |
| 0.627719 | 0.373121 | 0.011 | 0.02 | 1 | 14 | Efhb | -0.009 | Old |
| 0.627719 | 0.28975 | 0.011 | 0.02 | 1 | 14 | Brca2 | -0.009 | Old |
| 0.627719 | 0.289736 | 0.011 | 0.02 | 1 | 14 | Fan1 | -0.009 | Old |
| 0.627719 | 0.273915 | 0.011 | 0.02 | 1 | 14 | Mdga1 | -0.009 | Old |

| | | | | | | | | |
|---|---|---|---|---|---|---|---|---|
| 0.627719 | 0.264367 | 0.011 | 0.02 | 1 | 14 | Vash1 | -0.009 | Old |
| 0.627719 | 0.255979 | 0.011 | 0.02 | 1 | 14 | Zbtb49 | -0.009 | Old |
| 0.627719 | 0.251762 | 0.011 | 0.02 | 1 | 14 | Cbx2 | -0.009 | Old |
| 0.627719 | 0.174849 | 0.011 | 0.02 | 1 | 14 | A430110L2 | -0.009 | Old |
| 0.627719 | 0.168948 | 0.011 | 0.02 | 1 | 14 | Ica1l | -0.009 | Old |
| 0.627719 | 0.160466 | 0.011 | 0.02 | 1 | 14 | Gper1 | -0.009 | Old |
| 0.627719 | 0.159673 | 0.011 | 0.02 | 1 | 14 | Shank2 | -0.009 | Old |
| 0.627719 | 0.157834 | 0.011 | 0.02 | 1 | 14 | Gm10687 | -0.009 | Old |
| 0.627719 | 0.150479 | 0.011 | 0.02 | 1 | 14 | Ctxn1 | -0.009 | Old |
| 0.627719 | 0.138328 | 0.011 | 0.02 | 1 | 14 | Gm17484 | -0.009 | Old |
| 0.627719 | 0.131637 | 0.011 | 0.02 | 1 | 14 | Jmjd4 | -0.009 | Old |
| 0.627719 | 0.116695 | 0.011 | 0.02 | 1 | 14 | Srrm4 | -0.009 | Old |
| 0.627719 | 0.115211 | 0.011 | 0.02 | 1 | 14 | Tube1 | -0.009 | Old |
| 0.627719 | 0.102602 | 0.011 | 0.02 | 1 | 14 | Vamp1 | -0.009 | Old |
| 0.627719 | 0.095112 | 0.011 | 0.02 | 1 | 14 | Nostrin | -0.009 | Old |
| 0.627719 | 0.091622 | 0.011 | 0.02 | 1 | 14 | C230037L1 | -0.009 | Old |
| 0.627719 | 0.090691 | 0.011 | 0.02 | 1 | 14 | Gm49322 | -0.009 | Old |
| 0.627719 | 0.045261 | 0.011 | 0.02 | 1 | 14 | Nim1k | -0.009 | Old |
| 0.627719 | 0.039677 | 0.011 | 0.02 | 1 | 14 | Mamstr | -0.009 | Old |
| 0.627719 | 0.033202 | 0.011 | 0.02 | 1 | 14 | Gm12796 | -0.009 | Old |
| 0.627719 | 0.019943 | 0.011 | 0.02 | 1 | 14 | Myo19 | -0.009 | Old |
| 0.627906 | 0.730335 | 0.142 | 0.12 | 1 | 14 | Aspscr1 | 0.022 | Old |
| 0.627932 | 0.048374 | 0.164 | 0.2 | 1 | 14 | Osbpl8 | -0.036 | Old |
| 0.627946 | 0.03291 | 0.191 | 0.16 | 1 | 14 | Sptlc1 | 0.031 | Old |
| 0.628808 | 0.126111 | 0.454 | 0.44 | 1 | 14 | Rb1cc1 | 0.014 | Old |
| 0.628878 | 0.327763 | 0.126 | 0.16 | 1 | 14 | Kdm3a | -0.034 | Old |
| 0.629065 | 1.645875 | 0.077 | 0.06 | 1 | 14 | Prrg1 | 0.017 | Old |
| 0.6291 | 1.057632 | 0.098 | 0.08 | 1 | 14 | Tnfaip8 | 0.018 | Old |
| 0.629399 | 0.708045 | 0.093 | 0.12 | 1 | 14 | Tspan7 | -0.027 | Old |
| 0.629399 | 0.082345 | 0.093 | 0.12 | 1 | 14 | Map3k3 | -0.027 | Old |
| 0.629567 | 0.581249 | 0.383 | 0.36 | 1 | 14 | Dennd5a | 0.023 | Old |
| 0.629959 | 0.471166 | 0.393 | 0.38 | 1 | 14 | Hipk1 | 0.013 | Old |
| 0.630248 | 0.103482 | 0.06 | 0.08 | 1 | 14 | Aldh18a1 | -0.02 | Old |
| 0.630491 | 0.241041 | 0.164 | 0.2 | 1 | 14 | Jag1 | -0.036 | Old |
| 0.630714 | 0.256939 | 0.372 | 0.32 | 1 | 14 | Mrps33 | 0.052 | Old |
| 0.630988 | 1.64525 | 0.033 | 0.02 | 1 | 14 | Slc5a6 | 0.013 | Old |
| 0.630988 | 1.475871 | 0.033 | 0.02 | 1 | 14 | Ccdc15 | 0.013 | Old |
| 0.630988 | 1.463018 | 0.033 | 0.02 | 1 | 14 | Pnpla7 | 0.013 | Old |
| 0.630988 | 1.260527 | 0.033 | 0.02 | 1 | 14 | Ppih | 0.013 | Old |
| 0.630988 | 1.104142 | 0.033 | 0.02 | 1 | 14 | Rasgrf1 | 0.013 | Old |
| 0.630988 | 1.056429 | 0.033 | 0.02 | 1 | 14 | Tubb3 | 0.013 | Old |
| 0.630988 | 0.893092 | 0.033 | 0.02 | 1 | 14 | Rasgrp4 | 0.013 | Old |
| 0.630988 | 0.836815 | 0.033 | 0.02 | 1 | 14 | Sco1 | 0.013 | Old |
| 0.630988 | 0.650563 | 0.033 | 0.02 | 1 | 14 | Dars2 | 0.013 | Old |
| 0.630988 | 0.623041 | 0.033 | 0.02 | 1 | 14 | Phpt1 | 0.013 | Old |

| | | | | | | | | |
|---|---|---|---|---|---|---|---|---|
| 0.631143 | 0.034578 | 0.432 | 0.4 | 1 | 14 | Uba1 | 0.032 | Old |
| 0.631294 | 0.307054 | 0.148 | 0.18 | 1 | 14 | Zmym4 | -0.032 | Old |
| 0.631728 | 0.861248 | 0.137 | 0.12 | 1 | 14 | Mfn1 | 0.017 | Old |
| 0.631918 | 0.314109 | 0.366 | 0.36 | 1 | 14 | Ube2n | 0.006 | Old |
| 0.632022 | 0.593278 | 0.077 | 0.1 | 1 | 14 | Mmp9 | -0.023 | Old |
| 0.632232 | 0.304796 | 0.23 | 0.28 | 1 | 14 | Skap2 | -0.05 | Old |
| 0.632552 | 0.144719 | 0.082 | 0.06 | 1 | 14 | Slc11a2 | 0.022 | Old |
| 0.63256 | 0.153943 | 0.563 | 0.5 | 1 | 14 | Dazap2 | 0.063 | Old |
| 0.632651 | 0.29794 | 0.104 | 0.08 | 1 | 14 | Sh2b3 | 0.024 | Old |
| 0.632724 | 0.202542 | 0.454 | 0.4 | 1 | 14 | Ccni | 0.054 | Old |
| 0.633531 | 0.705555 | 0.142 | 0.12 | 1 | 14 | Lsm3 | 0.022 | Old |
| 0.633531 | 0.594763 | 0.142 | 0.12 | 1 | 14 | Rpp21 | 0.022 | Old |
| 0.633568 | 0.181098 | 0.044 | 0.06 | 1 | 14 | Limd2 | -0.016 | Old |
| 0.63366 | 0.105613 | 0.169 | 0.14 | 1 | 14 | Dbnl | 0.029 | Old |
| 0.633973 | 0.715248 | 0.158 | 0.14 | 1 | 14 | Pgm2 | 0.018 | Old |
| 0.633992 | 0.606008 | 0.224 | 0.2 | 1 | 14 | Gtpbp4 | 0.024 | Old |
| 0.634006 | 0.436993 | 0.142 | 0.18 | 1 | 14 | Mfn2 | -0.038 | Old |
| 0.634181 | 0.220943 | 0.23 | 0.2 | 1 | 14 | Bcl2 | 0.03 | Old |
| 0.634207 | 0.038363 | 0.06 | 0.08 | 1 | 14 | Hic2 | -0.02 | Old |
| 0.63422 | 0.22932 | 0.126 | 0.1 | 1 | 14 | Plpp1 | 0.026 | Old |
| 0.634419 | 0.301339 | 0.235 | 0.22 | 1 | 14 | Icam1 | 0.015 | Old |
| 0.635184 | 0.000788 | 0.148 | 0.12 | 1 | 14 | Slc39a10 | 0.028 | Old |
| 0.63557 | 0.094318 | 0.251 | 0.22 | 1 | 14 | Rnasel | 0.031 | Old |
| 0.635575 | 0.195834 | 0.077 | 0.1 | 1 | 14 | Poldip2 | -0.023 | Old |
| 0.63591 | 0.231662 | 0.093 | 0.12 | 1 | 14 | Fam214b | -0.027 | Old |
| 0.636196 | 0.394063 | 0.082 | 0.06 | 1 | 14 | Rassf4 | 0.022 | Old |
| 0.636196 | 0.032818 | 0.082 | 0.06 | 1 | 14 | Adgrf5 | 0.022 | Old |
| 0.636309 | 2.051269 | 0.055 | 0.04 | 1 | 14 | Dcxr | 0.015 | Old |
| 0.636309 | 1.996851 | 0.055 | 0.04 | 1 | 14 | Sphk1 | 0.015 | Old |
| 0.636309 | 1.554886 | 0.055 | 0.04 | 1 | 14 | 2900076A( | 0.015 | Old |
| 0.6365 | 0.478331 | 0.29 | 0.28 | 1 | 14 | Psmc5 | 0.01 | Old |
| 0.63654 | 1.009475 | 0.077 | 0.06 | 1 | 14 | Pigu | 0.017 | Old |
| 0.636691 | 1.673879 | 0.033 | 0.02 | 1 | 14 | Dusp5 | 0.013 | Old |
| 0.636691 | 1.670776 | 0.033 | 0.02 | 1 | 14 | Cables1 | 0.013 | Old |
| 0.636691 | 1.660248 | 0.033 | 0.02 | 1 | 14 | Macrod1 | 0.013 | Old |
| 0.636691 | 1.004807 | 0.033 | 0.02 | 1 | 14 | Cep128 | 0.013 | Old |
| 0.636691 | 0.994864 | 0.033 | 0.02 | 1 | 14 | Mapkapk5 | 0.013 | Old |
| 0.636691 | 0.765294 | 0.033 | 0.02 | 1 | 14 | Hoxd11 | 0.013 | Old |
| 0.636734 | 0.206622 | 0.246 | 0.22 | 1 | 14 | Zbtb44 | 0.026 | Old |
| 0.637326 | 0.204275 | 0.913 | 0.8 | 1 | 14 | Lrp1 | 0.113 | Old |
| 0.637445 | 0.358802 | 0.126 | 0.16 | 1 | 14 | Ube2q2 | -0.034 | Old |
| 0.637445 | 0.220159 | 0.126 | 0.16 | 1 | 14 | Cops6 | -0.034 | Old |
| 0.637874 | 0.380927 | 0.24 | 0.22 | 1 | 14 | Sh3gl1 | 0.02 | Old |
| 0.638701 | 0.246824 | 0.257 | 0.24 | 1 | 14 | Mgea5 | 0.017 | Old |
| 0.638976 | 0.032846 | 0.732 | 0.68 | 1 | 14 | Eef2 | 0.052 | Old |

| | | | | | | | | |
|---|---|---|---|---|---|---|---|---|
| 0.639166 | 0.503102 | 0.164 | 0.14 | 1 | 14 | Trim26 | 0.024 | Old |
| 0.639434 | 0.110608 | 0.437 | 0.42 | 1 | 14 | Aamp | 0.017 | Old |
| 0.639536 | 0.124378 | 0.35 | 0.32 | 1 | 14 | Rcn1 | 0.03 | Old |
| 0.640312 | 1.34325 | 0.137 | 0.12 | 1 | 14 | Adpgk | 0.017 | Old |
| 0.640523 | 0.053773 | 0.268 | 0.24 | 1 | 14 | Anxa11 | 0.028 | Old |
| 0.640726 | 2.330558 | 0.055 | 0.04 | 1 | 14 | Nfkbil1 | 0.015 | Old |
| 0.640762 | 0.086224 | 0.148 | 0.12 | 1 | 14 | Tmub2 | 0.028 | Old |
| 0.641155 | 0.108913 | 0.628 | 0.6 | 1 | 14 | Rpn1 | 0.028 | Old |
| 0.641202 | 0.037133 | 0.541 | 0.5 | 1 | 14 | Ncl | 0.041 | Old |
| 0.642158 | 0.361522 | 0.06 | 0.08 | 1 | 14 | Hspb11 | -0.02 | Old |
| 0.642158 | 0.252252 | 0.06 | 0.08 | 1 | 14 | Cenpv | -0.02 | Old |
| 0.642158 | 0.204626 | 0.06 | 0.08 | 1 | 14 | Sema4b | -0.02 | Old |
| 0.642413 | 1.004935 | 0.098 | 0.08 | 1 | 14 | 2410004B1 | 0.018 | Old |
| 0.642413 | 0.748727 | 0.098 | 0.08 | 1 | 14 | Cpne8 | 0.018 | Old |
| 0.642415 | 1.719207 | 0.033 | 0.02 | 1 | 14 | Pus7l | 0.013 | Old |
| 0.642415 | 1.313126 | 0.033 | 0.02 | 1 | 14 | Zfp667 | 0.013 | Old |
| 0.642415 | 0.769989 | 0.033 | 0.02 | 1 | 14 | Zgpat | 0.013 | Old |
| 0.642415 | 0.55102 | 0.033 | 0.02 | 1 | 14 | Cmtr1 | 0.013 | Old |
| 0.642415 | 0.44265 | 0.033 | 0.02 | 1 | 14 | Matn4 | 0.013 | Old |
| 0.64245 | 0.577655 | 0.093 | 0.12 | 1 | 14 | Cldn15 | -0.027 | Old |
| 0.642457 | 0.253274 | 0.273 | 0.24 | 1 | 14 | Cwc15 | 0.033 | Old |
| 0.643184 | 1.032422 | 0.137 | 0.12 | 1 | 14 | Pgpep1 | 0.017 | Old |
| 0.643207 | 0.444946 | 0.126 | 0.1 | 1 | 14 | Fgf10 | 0.026 | Old |
| 0.643298 | 0.173279 | 0.24 | 0.28 | 1 | 14 | Nfatc4 | -0.04 | Old |
| 0.644121 | 0.011597 | 0.169 | 0.2 | 1 | 14 | Rras2 | -0.031 | Old |
| 0.644704 | 0.689887 | 0.24 | 0.22 | 1 | 14 | Nmt2 | 0.02 | Old |
| 0.644736 | 1.242752 | 0.158 | 0.14 | 1 | 14 | Ecd | 0.018 | Old |
| 0.644736 | 1.161042 | 0.158 | 0.14 | 1 | 14 | Dvl1 | 0.018 | Old |
| 0.644812 | 0.564991 | 0.186 | 0.16 | 1 | 14 | Mcu | 0.026 | Old |
| 0.644882 | 0.711371 | 0.322 | 0.32 | 1 | 14 | Vps4b | 0.002 | Old |
| 0.645156 | 1.87359 | 0.055 | 0.04 | 1 | 14 | Cd82 | 0.015 | Old |
| 0.645156 | 1.861256 | 0.055 | 0.04 | 1 | 14 | Tox | 0.015 | Old |
| 0.645156 | 1.768388 | 0.055 | 0.04 | 1 | 14 | Gm9844 | 0.015 | Old |
| 0.645177 | 0.048092 | 0.951 | 1 | 1 | 14 | Rps3a1 | -0.049 | Old |
| 0.645285 | 0.837395 | 0.027 | 0.04 | 1 | 14 | C130074G1 | -0.013 | Old |
| 0.645417 | 0.109678 | 0.415 | 0.38 | 1 | 14 | Srsf11 | 0.035 | Old |
| 0.645502 | 0.382829 | 0.109 | 0.14 | 1 | 14 | Gmcl1 | -0.031 | Old |
| 0.64573 | 0.425877 | 0.093 | 0.12 | 1 | 14 | Ms4a6c | -0.027 | Old |
| 0.64573 | 0.127595 | 0.093 | 0.12 | 1 | 14 | Slc45a4 | -0.027 | Old |
| 0.64576 | 1.019791 | 0.098 | 0.08 | 1 | 14 | Ddx39 | 0.018 | Old |
| 0.64576 | 0.887236 | 0.098 | 0.08 | 1 | 14 | Rfwd3 | 0.018 | Old |
| 0.645797 | 0.154751 | 0.224 | 0.22 | 1 | 14 | Cfdp1 | 0.004 | Old |
| 0.645935 | 0.122436 | 0.164 | 0.2 | 1 | 14 | Pik3r2 | -0.036 | Old |
| 0.646149 | 0.803585 | 0.06 | 0.08 | 1 | 14 | Nup155 | -0.02 | Old |
| 0.646149 | 0.444275 | 0.06 | 0.08 | 1 | 14 | Dis3l2 | -0.02 | Old |

| | | | | | | | | |
|---|---|---|---|---|---|---|---|---|
| 0.646149 | 0.102311 | 0.06 | 0.08 | 1 | 14 | Zfp729b | -0.02 | Old |
| 0.646149 | 0.04442 | 0.06 | 0.08 | 1 | 14 | Zfp846 | -0.02 | Old |
| 0.646285 | 0.365942 | 0.077 | 0.1 | 1 | 14 | Bcas3 | -0.023 | Old |
| 0.646642 | 0.075058 | 0.317 | 0.38 | 1 | 14 | Anapc11 | -0.063 | Old |
| 0.646931 | 0.389304 | 0.322 | 0.32 | 1 | 14 | Txn2 | 0.002 | Old |
| 0.647228 | 0.166806 | 0.29 | 0.36 | 1 | 14 | Dnm2 | -0.07 | Old |
| 0.647395 | 0.510518 | 0.044 | 0.06 | 1 | 14 | Fam43a | -0.016 | Old |
| 0.647395 | 0.29196 | 0.044 | 0.06 | 1 | 14 | F8a | -0.016 | Old |
| 0.647395 | 0.225542 | 0.044 | 0.06 | 1 | 14 | Adam22 | -0.016 | Old |
| 0.647725 | 0.068624 | 0.202 | 0.24 | 1 | 14 | Adamts4 | -0.038 | Old |
| 0.647821 | 1.325015 | 0.077 | 0.06 | 1 | 14 | Akap10 | 0.017 | Old |
| 0.647821 | 1.066844 | 0.077 | 0.06 | 1 | 14 | Zscan21 | 0.017 | Old |
| 0.647821 | 0.858321 | 0.077 | 0.06 | 1 | 14 | Vps72 | 0.017 | Old |
| 0.647909 | 0.651207 | 0.219 | 0.2 | 1 | 14 | Plekhj1 | 0.019 | Old |
| 0.648048 | 0.574523 | 0.246 | 0.22 | 1 | 14 | Ube2g2 | 0.026 | Old |
| 0.64813 | 0.257978 | 0.23 | 0.2 | 1 | 14 | Vps37a | 0.03 | Old |
| 0.64816 | 0.965821 | 0.033 | 0.02 | 1 | 14 | Msantd3 | 0.013 | Old |
| 0.64816 | 0.93463 | 0.033 | 0.02 | 1 | 14 | Clasrp | 0.013 | Old |
| 0.64816 | 0.879446 | 0.033 | 0.02 | 1 | 14 | Maneal | 0.013 | Old |
| 0.64816 | 0.578397 | 0.033 | 0.02 | 1 | 14 | Gm21188 | 0.013 | Old |
| 0.64816 | 0.546751 | 0.033 | 0.02 | 1 | 14 | Tmem254a | 0.013 | Old |
| 0.64816 | 0.414538 | 0.033 | 0.02 | 1 | 14 | Fam13a | 0.013 | Old |
| 0.64816 | 0.404048 | 0.033 | 0.02 | 1 | 14 | Mcoln2 | 0.013 | Old |
| 0.64816 | 0.278214 | 0.033 | 0.02 | 1 | 14 | Cd180 | 0.013 | Old |
| 0.64816 | 0.213909 | 0.033 | 0.02 | 1 | 14 | Rpgr | 0.013 | Old |
| 0.64816 | 0.194397 | 0.033 | 0.02 | 1 | 14 | Mvd | 0.013 | Old |
| 0.64816 | 0.185908 | 0.033 | 0.02 | 1 | 14 | Map7 | 0.013 | Old |
| 0.64816 | 0.069417 | 0.033 | 0.02 | 1 | 14 | Akr1c14 | 0.013 | Old |
| 0.64816 | 0.037999 | 0.033 | 0.02 | 1 | 14 | Clu | 0.013 | Old |
| 0.64816 | 0.034695 | 0.033 | 0.02 | 1 | 14 | Syngap1 | 0.013 | Old |
| 0.64816 | 0.020497 | 0.033 | 0.02 | 1 | 14 | Lca5 | 0.013 | Old |
| 0.648562 | 0.506641 | 0.109 | 0.14 | 1 | 14 | Dbt | -0.031 | Old |
| 0.649114 | 1.124893 | 0.098 | 0.08 | 1 | 14 | Saysd1 | 0.018 | Old |
| 0.649114 | 0.876389 | 0.098 | 0.08 | 1 | 14 | Tmem204 | 0.018 | Old |
| 0.649114 | 0.832476 | 0.098 | 0.08 | 1 | 14 | Mrpl2 | 0.018 | Old |
| 0.649228 | 0.522036 | 0.126 | 0.1 | 1 | 14 | Trim12a | 0.026 | Old |
| 0.649299 | 0.364184 | 0.202 | 0.18 | 1 | 14 | Uqcrc1 | 0.022 | Old |
| 0.649579 | 0.524312 | 0.262 | 0.24 | 1 | 14 | Ip6k1 | 0.022 | Old |
| 0.649598 | 1.658075 | 0.055 | 0.04 | 1 | 14 | Traf2 | 0.015 | Old |
| 0.649598 | 1.414347 | 0.055 | 0.04 | 1 | 14 | Ptges2 | 0.015 | Old |
| 0.649598 | 1.274985 | 0.055 | 0.04 | 1 | 14 | Ccdc170 | 0.015 | Old |
| 0.649598 | 1.24579 | 0.055 | 0.04 | 1 | 14 | Bloc1s6 | 0.015 | Old |
| 0.649892 | 0.064775 | 0.579 | 0.54 | 1 | 14 | Arf1 | 0.039 | Old |
| 0.65015 | 0.242143 | 0.06 | 0.08 | 1 | 14 | Arhgap9 | -0.02 | Old |
| 0.65015 | 0.241824 | 0.06 | 0.08 | 1 | 14 | Nr4a3 | -0.02 | Old |

| | | | | | | | | |
|---|---|---|---|---|---|---|---|---|
| 0.650203 | 0.091606 | 0.628 | 0.62 | 1 | 14 | Rab14 | 0.008 | Old |
| 0.65085 | 1.100799 | 0.115 | 0.1 | 1 | 14 | Eif2b5 | 0.015 | Old |
| 0.651041 | 0.588445 | 0.027 | 0.04 | 1 | 14 | Znrd1as | -0.013 | Old |
| 0.651041 | 0.572582 | 0.027 | 0.04 | 1 | 14 | Ftx | -0.013 | Old |
| 0.651041 | 0.279842 | 0.027 | 0.04 | 1 | 14 | 9330159F1 | -0.013 | Old |
| 0.651041 | 0.247496 | 0.027 | 0.04 | 1 | 14 | Angel1 | -0.013 | Old |
| 0.651041 | 0.240806 | 0.027 | 0.04 | 1 | 14 | Tspan32 | -0.013 | Old |
| 0.651041 | 0.058023 | 0.027 | 0.04 | 1 | 14 | Tmem71 | -0.013 | Old |
| 0.651041 | 0.030405 | 0.027 | 0.04 | 1 | 14 | Slc25a15 | -0.013 | Old |
| 0.6516 | 1.374167 | 0.077 | 0.06 | 1 | 14 | Gfm2 | 0.017 | Old |
| 0.6516 | 1.208978 | 0.077 | 0.06 | 1 | 14 | Ccp110 | 0.017 | Old |
| 0.6516 | 0.860984 | 0.077 | 0.06 | 1 | 14 | Zc3h12a | 0.017 | Old |
| 0.651979 | 0.033553 | 0.148 | 0.12 | 1 | 14 | Hbs1l | 0.028 | Old |
| 0.652006 | 0.124669 | 0.388 | 0.36 | 1 | 14 | Snrpf | 0.028 | Old |
| 0.652031 | 0.006357 | 0.044 | 0.06 | 1 | 14 | Zfp442 | -0.016 | Old |
| 0.652312 | 0.365555 | 0.213 | 0.18 | 1 | 14 | Hebp1 | 0.033 | Old |
| 0.652458 | 0.195885 | 0.448 | 0.4 | 1 | 14 | Ankrd17 | 0.048 | Old |
| 0.652502 | 0.329251 | 0.301 | 0.28 | 1 | 14 | Mrpl23 | 0.021 | Old |
| 0.65251 | 0.736747 | 0.213 | 0.2 | 1 | 14 | Wdfy2 | 0.013 | Old |
| 0.652657 | 0.538842 | 0.219 | 0.2 | 1 | 14 | Cdk12 | 0.019 | Old |
| 0.652783 | 0.008767 | 0.224 | 0.26 | 1 | 14 | Trrap | -0.036 | Old |
| 0.652858 | 0.971532 | 0.158 | 0.14 | 1 | 14 | Zfp598 | 0.018 | Old |
| 0.653098 | 0.260456 | 0.404 | 0.4 | 1 | 14 | Aldh1a1 | 0.004 | Old |
| 0.653723 | 0.058082 | 0.268 | 0.24 | 1 | 14 | Psme4 | 0.028 | Old |
| 0.653927 | 0.475828 | 0.033 | 0.02 | 1 | 14 | Myo1f | 0.013 | Old |
| 0.653927 | 0.384603 | 0.033 | 0.02 | 1 | 14 | Slc25a38 | 0.013 | Old |
| 0.653927 | 0.24007 | 0.033 | 0.02 | 1 | 14 | Hck | 0.013 | Old |
| 0.653927 | 0.219032 | 0.033 | 0.02 | 1 | 14 | Erbb3 | 0.013 | Old |
| 0.653927 | 0.164981 | 0.033 | 0.02 | 1 | 14 | A630001G2 | 0.013 | Old |
| 0.653927 | 0.032597 | 0.033 | 0.02 | 1 | 14 | Prcd | 0.013 | Old |
| 0.654162 | 0.669649 | 0.06 | 0.08 | 1 | 14 | Tctn1 | -0.02 | Old |
| 0.654162 | 0.419296 | 0.06 | 0.08 | 1 | 14 | Zeb2os | -0.02 | Old |
| 0.654162 | 0.3604 | 0.06 | 0.08 | 1 | 14 | Tsga10 | -0.02 | Old |
| 0.654162 | 0.226371 | 0.06 | 0.08 | 1 | 14 | Nup35 | -0.02 | Old |
| 0.654228 | 0.026694 | 0.35 | 0.38 | 1 | 14 | Mapk1ip1l | -0.03 | Old |
| 0.654514 | 0.236084 | 0.257 | 0.22 | 1 | 14 | Cnrip1 | 0.037 | Old |
| 0.654514 | 0.156108 | 0.24 | 0.28 | 1 | 14 | Fam32a | -0.04 | Old |
| 0.654701 | 0.548672 | 0.12 | 0.1 | 1 | 14 | Heatr6 | 0.02 | Old |
| 0.654701 | 0.423345 | 0.109 | 0.14 | 1 | 14 | Ciao1 | -0.031 | Old |
| 0.654728 | 0.176356 | 0.197 | 0.24 | 1 | 14 | Commd8 | -0.043 | Old |
| 0.655388 | 1.139946 | 0.077 | 0.06 | 1 | 14 | Nop9 | 0.017 | Old |
| 0.655842 | 1.105822 | 0.098 | 0.08 | 1 | 14 | Ank | 0.018 | Old |
| 0.65624 | 0.892225 | 0.142 | 0.12 | 1 | 14 | Golga1 | 0.022 | Old |
| 0.656681 | 0.283926 | 0.044 | 0.06 | 1 | 14 | Unk | -0.016 | Old |
| 0.656681 | 0.142995 | 0.044 | 0.06 | 1 | 14 | Rrp9 | -0.016 | Old |

| | | | | | | | | |
|---|---|---|---|---|---|---|---|---|
| 0.656767 | 0.288254 | 0.257 | 0.22 | 1 | 14 | Ccl2 | 0.037 | Old |
| 0.656818 | 0.857582 | 0.027 | 0.04 | 1 | 14 | Gch1 | -0.013 | Old |
| 0.656818 | 0.659522 | 0.027 | 0.04 | 1 | 14 | Zfp438 | -0.013 | Old |
| 0.656818 | 0.560142 | 0.027 | 0.04 | 1 | 14 | Gm20045 | -0.013 | Old |
| 0.656818 | 0.389431 | 0.027 | 0.04 | 1 | 14 | Chp2 | -0.013 | Old |
| 0.656818 | 0.136712 | 0.027 | 0.04 | 1 | 14 | Fgfbp3 | -0.013 | Old |
| 0.656964 | 0.761048 | 0.18 | 0.16 | 1 | 14 | Eif6 | 0.02 | Old |
| 0.656964 | 0.436828 | 0.18 | 0.16 | 1 | 14 | Plgrkt | 0.02 | Old |
| 0.657505 | 0.174659 | 0.317 | 0.3 | 1 | 14 | Tial1 | 0.017 | Old |
| 0.657623 | 0.321937 | 0.126 | 0.16 | 1 | 14 | Stk4 | -0.034 | Old |
| 0.657778 | 0.370885 | 0.109 | 0.14 | 1 | 14 | Slc25a30 | -0.031 | Old |
| 0.657784 | 0.261518 | 0.454 | 0.42 | 1 | 14 | Smarca5 | 0.034 | Old |
| 0.658183 | 0.117604 | 0.06 | 0.08 | 1 | 14 | Snapc5 | -0.02 | Old |
| 0.658296 | 0.130015 | 0.148 | 0.18 | 1 | 14 | Negr1 | -0.032 | Old |
| 0.658302 | 0.025507 | 0.126 | 0.1 | 1 | 14 | Serhl | 0.026 | Old |
| 0.658519 | 1.068051 | 0.055 | 0.04 | 1 | 14 | Tmc4 | 0.015 | Old |
| 0.658519 | 0.914583 | 0.055 | 0.04 | 1 | 14 | 1700066M | 0.015 | Old |
| 0.658702 | 0.021786 | 0.404 | 0.46 | 1 | 14 | Stag1 | -0.056 | Old |
| 0.65906 | 0.064076 | 0.47 | 0.42 | 1 | 14 | Nsa2 | 0.05 | Old |
| 0.659102 | 0.310606 | 0.142 | 0.12 | 1 | 14 | Mboat7 | 0.022 | Old |
| 0.659184 | 1.575817 | 0.077 | 0.06 | 1 | 14 | Mrps35 | 0.017 | Old |
| 0.659184 | 1.056265 | 0.077 | 0.06 | 1 | 14 | Cradd | 0.017 | Old |
| 0.659184 | 0.877888 | 0.077 | 0.06 | 1 | 14 | Apba3 | 0.017 | Old |
| 0.659217 | 0.886571 | 0.098 | 0.08 | 1 | 14 | Ttc5 | 0.018 | Old |
| 0.65938 | 0.313316 | 0.191 | 0.24 | 1 | 14 | Sun1 | -0.049 | Old |
| 0.659725 | 0.680911 | 0.235 | 0.22 | 1 | 14 | Eif4a3 | 0.015 | Old |
| 0.660526 | 0.883474 | 0.137 | 0.12 | 1 | 14 | Stx3 | 0.017 | Old |
| 0.660701 | 0.045944 | 0.251 | 0.28 | 1 | 14 | Gtf2a1 | -0.029 | Old |
| 0.661344 | 0.38273 | 0.044 | 0.06 | 1 | 14 | Atrip | -0.016 | Old |
| 0.661344 | 0.139169 | 0.044 | 0.06 | 1 | 14 | Smim3 | -0.016 | Old |
| 0.661344 | 0.124195 | 0.044 | 0.06 | 1 | 14 | Adamts14 | -0.016 | Old |
| 0.661534 | 0.232703 | 0.164 | 0.2 | 1 | 14 | Cpeb4 | -0.036 | Old |
| 0.661834 | 0.454779 | 0.175 | 0.22 | 1 | 14 | Arf6 | -0.045 | Old |
| 0.661918 | 0.129947 | 0.645 | 0.58 | 1 | 14 | Arhgdia | 0.065 | Old |
| 0.662045 | 0.720531 | 0.235 | 0.22 | 1 | 14 | Sec24c | 0.015 | Old |
| 0.662195 | 0.509002 | 0.219 | 0.2 | 1 | 14 | Tjp1 | 0.019 | Old |
| 0.662599 | 0.820716 | 0.098 | 0.08 | 1 | 14 | Zfp84 | 0.018 | Old |
| 0.662599 | 0.750717 | 0.098 | 0.08 | 1 | 14 | Plekhg2 | 0.018 | Old |
| 0.662615 | 1.69793 | 0.027 | 0.04 | 1 | 14 | Fam213a | -0.013 | Old |
| 0.662615 | 1.216833 | 0.027 | 0.04 | 1 | 14 | Tmem35a | -0.013 | Old |
| 0.662615 | 0.861619 | 0.027 | 0.04 | 1 | 14 | Trmt10a | -0.013 | Old |
| 0.662615 | 0.280182 | 0.027 | 0.04 | 1 | 14 | B4galnt1 | -0.013 | Old |
| 0.662615 | 0.279179 | 0.027 | 0.04 | 1 | 14 | Pusl1 | -0.013 | Old |
| 0.662615 | 0.165585 | 0.027 | 0.04 | 1 | 14 | Il1rl1 | -0.013 | Old |
| 0.662615 | 0.158516 | 0.027 | 0.04 | 1 | 14 | Zmynd15 | -0.013 | Old |

| | | | | | | | | |
|---|---|---|---|---|---|---|---|---|
| 0.662615 | 0.157506 | 0.027 | 0.04 | 1 | 14 | Pecr | -0.013 | Old |
| 0.662615 | 0.135101 | 0.027 | 0.04 | 1 | 14 | Atp23 | -0.013 | Old |
| 0.662806 | 0.040883 | 0.393 | 0.36 | 1 | 14 | Tmed5 | 0.033 | Old |
| 0.66299 | 0.677006 | 0.077 | 0.06 | 1 | 14 | Tmem101 | 0.017 | Old |
| 0.662998 | 1.604109 | 0.055 | 0.04 | 1 | 14 | Zfp81 | 0.015 | Old |
| 0.662998 | 1.193202 | 0.055 | 0.04 | 1 | 14 | Plcl1 | 0.015 | Old |
| 0.662998 | 0.659039 | 0.055 | 0.04 | 1 | 14 | B9d1 | 0.015 | Old |
| 0.663375 | 1.193617 | 0.115 | 0.1 | 1 | 14 | Ap4s1 | 0.015 | Old |
| 0.663487 | 0.255812 | 0.18 | 0.22 | 1 | 14 | Mrpl54 | -0.04 | Old |
| 0.663487 | 0.218087 | 0.18 | 0.22 | 1 | 14 | Slc4a7 | -0.04 | Old |
| 0.664059 | 0.483911 | 0.585 | 0.56 | 1 | 14 | Cp | 0.025 | Old |
| 0.664072 | 0.30272 | 0.202 | 0.18 | 1 | 14 | Wdr89 | 0.022 | Old |
| 0.664148 | 0.64229 | 0.175 | 0.16 | 1 | 14 | Vps13c | 0.015 | Old |
| 0.664379 | 0.038347 | 0.115 | 0.14 | 1 | 14 | Scaf4 | -0.025 | Old |
| 0.664485 | 0.122306 | 0.235 | 0.2 | 1 | 14 | Cacna2d1 | 0.035 | Old |
| 0.664546 | 0.202658 | 0.415 | 0.4 | 1 | 14 | Kdm2a | 0.015 | Old |
| 0.664719 | 0.101474 | 0.18 | 0.16 | 1 | 14 | Itgb1bp1 | 0.02 | Old |
| 0.664941 | 0.324883 | 0.251 | 0.22 | 1 | 14 | P4ha2 | 0.031 | Old |
| 0.665057 | 0.83263 | 0.197 | 0.18 | 1 | 14 | Bckdk | 0.017 | Old |
| 0.665988 | 1.016583 | 0.098 | 0.08 | 1 | 14 | Ints8 | 0.018 | Old |
| 0.665988 | 0.658187 | 0.098 | 0.08 | 1 | 14 | Msh2 | 0.018 | Old |
| 0.665988 | 0.421166 | 0.098 | 0.08 | 1 | 14 | Coa6 | 0.018 | Old |
| 0.666021 | 0.336493 | 0.044 | 0.06 | 1 | 14 | Brms1l | -0.016 | Old |
| 0.666022 | 0.191281 | 0.546 | 0.5 | 1 | 14 | 7-Sep | 0.046 | Old |
| 0.666255 | 0.774337 | 0.06 | 0.08 | 1 | 14 | Wdr46 | -0.02 | Old |
| 0.666255 | 0.537498 | 0.06 | 0.08 | 1 | 14 | Polr3h | -0.02 | Old |
| 0.666255 | 0.187471 | 0.06 | 0.08 | 1 | 14 | Phka2 | -0.02 | Old |
| 0.666516 | 0.221566 | 0.557 | 0.52 | 1 | 14 | Atxn7l3b | 0.037 | Old |
| 0.666547 | 0.550536 | 0.202 | 0.18 | 1 | 14 | 8-Mar | 0.022 | Old |
| 0.666732 | 0.398005 | 0.191 | 0.24 | 1 | 14 | Apc | -0.049 | Old |
| 0.666804 | 1.037827 | 0.077 | 0.06 | 1 | 14 | Mmgt2 | 0.017 | Old |
| 0.667047 | 0.395773 | 0.109 | 0.14 | 1 | 14 | Ankmy2 | -0.031 | Old |
| 0.667489 | 0.810085 | 0.055 | 0.04 | 1 | 14 | Cacnb4 | 0.015 | Old |
| 0.667716 | 0.505872 | 0.142 | 0.12 | 1 | 14 | Abat | 0.022 | Old |
| 0.667932 | 0.10008 | 0.262 | 0.32 | 1 | 14 | Rela | -0.058 | Old |
| 0.668117 | 0.101799 | 0.344 | 0.38 | 1 | 14 | Plce1 | -0.036 | Old |
| 0.668433 | 0.913935 | 0.027 | 0.04 | 1 | 14 | Klhdc1 | -0.013 | Old |
| 0.668433 | 0.875748 | 0.027 | 0.04 | 1 | 14 | Aldh5a1 | -0.013 | Old |
| 0.668433 | 0.640646 | 0.027 | 0.04 | 1 | 14 | Nme4 | -0.013 | Old |
| 0.668433 | 0.50862 | 0.027 | 0.04 | 1 | 14 | Gm26518 | -0.013 | Old |
| 0.668433 | 0.486403 | 0.027 | 0.04 | 1 | 14 | Acy1 | -0.013 | Old |
| 0.668433 | 0.419449 | 0.027 | 0.04 | 1 | 14 | Nmral1 | -0.013 | Old |
| 0.668433 | 0.293385 | 0.027 | 0.04 | 1 | 14 | Ms4a6d | -0.013 | Old |
| 0.668433 | 0.148533 | 0.027 | 0.04 | 1 | 14 | En1 | -0.013 | Old |
| 0.668433 | 0.101351 | 0.027 | 0.04 | 1 | 14 | Alpl | -0.013 | Old |

| | | | | | | | | |
|---|---|---|---|---|---|---|---|---|
| 0.668433 | 0.041982 | 0.027 | 0.04 | 1 | 14 | Gbp9 | -0.013 | Old |
| 0.668619 | 0.387604 | 0.23 | 0.28 | 1 | 14 | Unc93b1 | -0.05 | Old |
| 0.668882 | 0.430585 | 0.093 | 0.12 | 1 | 14 | Carnmt1 | -0.027 | Old |
| 0.668953 | 0.236793 | 0.148 | 0.12 | 1 | 14 | Kank1 | 0.028 | Old |
| 0.6693 | 0.426099 | 0.306 | 0.3 | 1 | 14 | Rabgap1 | 0.006 | Old |
| 0.669367 | 0.762451 | 0.213 | 0.2 | 1 | 14 | Birc3 | 0.013 | Old |
| 0.669389 | 0.638543 | 0.175 | 0.16 | 1 | 14 | Prr12 | 0.015 | Old |
| 0.669686 | 0.158516 | 0.301 | 0.34 | 1 | 14 | Gspt1 | -0.039 | Old |
| 0.670148 | 0.624479 | 0.12 | 0.1 | 1 | 14 | Cys1 | 0.02 | Old |
| 0.670306 | 0.939 | 0.06 | 0.08 | 1 | 14 | Spice1 | -0.02 | Old |
| 0.670306 | 0.291976 | 0.06 | 0.08 | 1 | 14 | Rcc1l | -0.02 | Old |
| 0.670348 | 0.413889 | 0.251 | 0.24 | 1 | 14 | Baiap2 | 0.011 | Old |
| 0.670381 | 0.212186 | 0.366 | 0.34 | 1 | 14 | Arl1 | 0.026 | Old |
| 0.670627 | 0.719525 | 0.077 | 0.06 | 1 | 14 | Zfp574 | 0.017 | Old |
| 0.670627 | 0.700078 | 0.077 | 0.06 | 1 | 14 | Spout1 | 0.017 | Old |
| 0.670627 | 0.637766 | 0.077 | 0.06 | 1 | 14 | Mkks | 0.017 | Old |
| 0.670627 | 0.338206 | 0.077 | 0.06 | 1 | 14 | Vwa8 | 0.017 | Old |
| 0.670627 | 0.334844 | 0.077 | 0.06 | 1 | 14 | Ppp4r1 | 0.017 | Old |
| 0.670627 | 0.252805 | 0.077 | 0.06 | 1 | 14 | Rab29 | 0.017 | Old |
| 0.67071 | 0.605576 | 0.044 | 0.06 | 1 | 14 | Cox11 | -0.016 | Old |
| 0.67071 | 0.517684 | 0.044 | 0.06 | 1 | 14 | Rtkn | -0.016 | Old |
| 0.671094 | 0.184998 | 0.301 | 0.36 | 1 | 14 | Dnajb9 | -0.059 | Old |
| 0.671412 | 0.037524 | 0.153 | 0.18 | 1 | 14 | Snx12 | -0.027 | Old |
| 0.671526 | 0.001107 | 0.235 | 0.2 | 1 | 14 | Mrpl28 | 0.035 | Old |
| 0.671558 | 0.572173 | 0.077 | 0.1 | 1 | 14 | Mast3 | -0.023 | Old |
| 0.671558 | 0.993432 | 0.077 | 0.1 | 1 | 14 | Cox16 | -0.023 | Old |
| 0.671558 | 0.58318 | 0.077 | 0.1 | 1 | 14 | Gpn1 | -0.023 | Old |
| 0.671992 | 0.987235 | 0.055 | 0.04 | 1 | 14 | B4galt2 | 0.015 | Old |
| 0.671992 | 0.734224 | 0.055 | 0.04 | 1 | 14 | Il27ra | 0.015 | Old |
| 0.671992 | 0.655452 | 0.055 | 0.04 | 1 | 14 | Zbtb34 | 0.015 | Old |
| 0.671992 | 0.387317 | 0.055 | 0.04 | 1 | 14 | Sgcd | 0.015 | Old |
| 0.671992 | 0.378914 | 0.055 | 0.04 | 1 | 14 | Wwp2 | 0.015 | Old |
| 0.67219 | 0.872795 | 0.137 | 0.12 | 1 | 14 | Siae | 0.017 | Old |
| 0.672216 | 1.035829 | 0.093 | 0.12 | 1 | 14 | Paxip1 | -0.027 | Old |
| 0.672939 | 0.244134 | 0.235 | 0.28 | 1 | 14 | Gsk3a | -0.045 | Old |
| 0.673063 | 0.014467 | 0.814 | 0.74 | 1 | 14 | Rrbp1 | 0.074 | Old |
| 0.673255 | 0.81049 | 0.109 | 0.14 | 1 | 14 | Rubcn | -0.031 | Old |
| 0.673337 | 0.069346 | 0.41 | 0.4 | 1 | 14 | Atp6v1b2 | 0.01 | Old |
| 0.67427 | 1.358442 | 0.027 | 0.04 | 1 | 14 | Asb6 | -0.013 | Old |
| 0.67427 | 1.056055 | 0.027 | 0.04 | 1 | 14 | Gm26881 | -0.013 | Old |
| 0.67427 | 1.007774 | 0.027 | 0.04 | 1 | 14 | Glb1l | -0.013 | Old |
| 0.67427 | 0.835094 | 0.027 | 0.04 | 1 | 14 | Pdss1 | -0.013 | Old |
| 0.67427 | 0.352268 | 0.027 | 0.04 | 1 | 14 | AB124611 | -0.013 | Old |
| 0.674367 | 0.539154 | 0.06 | 0.08 | 1 | 14 | Dxo | -0.02 | Old |
| 0.674458 | 0.950382 | 0.077 | 0.06 | 1 | 14 | Mis12 | 0.017 | Old |

| | | | | | | | | |
|---|---|---|---|---|---|---|---|---|
| 0.674458 | 0.824038 | 0.077 | 0.06 | 1 | 14 | Tchp | 0.017 | Old |
| 0.67453 | 0.581939 | 0.393 | 0.4 | 1 | 14 | Ppp1r15b | -0.007 | Old |
| 0.674568 | 0.989951 | 0.186 | 0.18 | 1 | 14 | Rnf19a | 0.006 | Old |
| 0.674847 | 0.089254 | 0.251 | 0.3 | 1 | 14 | Agpat3 | -0.049 | Old |
| 0.675119 | 0.869562 | 0.137 | 0.12 | 1 | 14 | Cenpc1 | 0.017 | Old |
| 0.675412 | 0.526464 | 0.044 | 0.06 | 1 | 14 | Rpgrip1l | -0.016 | Old |
| 0.675514 | 0.230905 | 0.186 | 0.16 | 1 | 14 | Wdr43 | 0.026 | Old |
| 0.67561 | 0.095581 | 0.29 | 0.26 | 1 | 14 | Ide | 0.03 | Old |
| 0.675932 | 0.541758 | 0.301 | 0.28 | 1 | 14 | Rbbp4 | 0.021 | Old |
| 0.676036 | 0.025396 | 0.224 | 0.26 | 1 | 14 | Acaa1a | -0.036 | Old |
| 0.676195 | 0.692402 | 0.098 | 0.08 | 1 | 14 | Rrp8 | 0.018 | Old |
| 0.676367 | 0.304395 | 0.12 | 0.1 | 1 | 14 | Magee1 | 0.02 | Old |
| 0.676507 | 0.805095 | 0.055 | 0.04 | 1 | 14 | Fbxo45 | 0.015 | Old |
| 0.676507 | 0.23492 | 0.055 | 0.04 | 1 | 14 | Apbb1 | 0.015 | Old |
| 0.676507 | 0.002172 | 0.055 | 0.04 | 1 | 14 | Eps8l1 | 0.015 | Old |
| 0.676638 | 0.877364 | 0.208 | 0.2 | 1 | 14 | Arl13b | 0.008 | Old |
| 0.676699 | 1.314736 | 0.131 | 0.12 | 1 | 14 | Pisd | 0.011 | Old |
| 0.677405 | 0.174405 | 0.399 | 0.4 | 1 | 14 | Cep170 | -0.001 | Old |
| 0.678028 | 1.023999 | 0.153 | 0.14 | 1 | 14 | Pkn1 | 0.013 | Old |
| 0.678053 | 0.989405 | 0.137 | 0.12 | 1 | 14 | Ruvbl1 | 0.017 | Old |
| 0.678053 | 0.988771 | 0.137 | 0.12 | 1 | 14 | Ino80c | 0.017 | Old |
| 0.678584 | 0.253534 | 0.459 | 0.4 | 1 | 14 | H2-Aa | 0.059 | Old |
| 0.678728 | 0.24004 | 0.295 | 0.28 | 1 | 14 | Git2 | 0.015 | Old |
| 0.67902 | 0.254895 | 0.437 | 0.44 | 1 | 14 | Gatad2b | -0.003 | Old |
| 0.679152 | 0.133615 | 0.295 | 0.3 | 1 | 14 | Ndufa8 | -0.005 | Old |
| 0.679271 | 0.237016 | 0.142 | 0.12 | 1 | 14 | Ap1s2 | 0.022 | Old |
| 0.679292 | 0.244371 | 0.24 | 0.22 | 1 | 14 | Trib1 | 0.02 | Old |
| 0.679611 | 0.842162 | 0.098 | 0.08 | 1 | 14 | Orai1 | 0.018 | Old |
| 0.679611 | 0.692639 | 0.098 | 0.08 | 1 | 14 | Gpr155 | 0.018 | Old |
| 0.679611 | 0.135441 | 0.098 | 0.08 | 1 | 14 | Rhbdd1 | 0.018 | Old |
| 0.679669 | 0.170251 | 0.115 | 0.14 | 1 | 14 | Renbp | -0.025 | Old |
| 0.679688 | 0.483088 | 0.175 | 0.22 | 1 | 14 | Rnf213 | -0.045 | Old |
| 0.679825 | 0.330005 | 0.246 | 0.24 | 1 | 14 | Arhgap10 | 0.006 | Old |
| 0.679964 | 0.11098 | 0.273 | 0.24 | 1 | 14 | Sox9 | 0.033 | Old |
| 0.680101 | 0.495728 | 0.197 | 0.18 | 1 | 14 | Hdac3 | 0.017 | Old |
| 0.680127 | 1.392119 | 0.044 | 0.06 | 1 | 14 | Net1 | -0.016 | Old |
| 0.680127 | 1.042624 | 0.044 | 0.06 | 1 | 14 | Ercc1 | -0.016 | Old |
| 0.680127 | 0.736534 | 0.044 | 0.06 | 1 | 14 | Acot7 | -0.016 | Old |
| 0.680253 | 0.345238 | 0.311 | 0.3 | 1 | 14 | Tbc1d2b | 0.011 | Old |
| 0.680336 | 0.668359 | 0.18 | 0.16 | 1 | 14 | Ifnar2 | 0.02 | Old |
| 0.681034 | 0.387946 | 0.055 | 0.04 | 1 | 14 | Zfp12 | 0.015 | Old |
| 0.681512 | 0.336646 | 0.202 | 0.24 | 1 | 14 | Mrps16 | -0.038 | Old |
| 0.682146 | 0.560078 | 0.077 | 0.06 | 1 | 14 | D930016D( | 0.017 | Old |
| 0.682146 | 0.44809 | 0.077 | 0.06 | 1 | 14 | Adprm | 0.017 | Old |
| 0.682146 | 0.080289 | 0.077 | 0.06 | 1 | 14 | Wbp1 | 0.017 | Old |

| | | | | | | | | |
|---|---|---|---|---|---|---|---|---|
| 0.682167 | 0.177636 | 0.579 | 0.54 | 1 | 14 | Ski | 0.039 | Old |
| 0.68234 | 0.86471 | 0.115 | 0.1 | 1 | 14 | Cdc27 | 0.015 | Old |
| 0.682516 | 0.805942 | 0.06 | 0.08 | 1 | 14 | Miga2 | -0.02 | Old |
| 0.682677 | 0.20763 | 0.317 | 0.3 | 1 | 14 | Ssh2 | 0.017 | Old |
| 0.682743 | 0.401416 | 0.115 | 0.14 | 1 | 14 | Ints13 | -0.025 | Old |
| 0.683033 | 0.438235 | 0.098 | 0.08 | 1 | 14 | Rae1 | 0.018 | Old |
| 0.683324 | 0.065094 | 0.235 | 0.2 | 1 | 14 | Cdk14 | 0.035 | Old |
| 0.683854 | 0.687994 | 0.219 | 0.2 | 1 | 14 | Zbtb16 | 0.019 | Old |
| 0.683934 | 0.239397 | 0.137 | 0.12 | 1 | 14 | Ftl1-ps1 | 0.017 | Old |
| 0.684535 | 0.581048 | 0.361 | 0.38 | 1 | 14 | Srsf7 | -0.019 | Old |
| 0.684854 | 1.181107 | 0.044 | 0.06 | 1 | 14 | Trp53rka | -0.016 | Old |
| 0.685077 | 0.174351 | 0.131 | 0.16 | 1 | 14 | Opa1 | -0.029 | Old |
| 0.68552 | 1.039016 | 0.115 | 0.1 | 1 | 14 | Serpina3n | 0.015 | Old |
| 0.685572 | 0.469954 | 0.055 | 0.04 | 1 | 14 | Rdh14 | 0.015 | Old |
| 0.685572 | 0.081131 | 0.055 | 0.04 | 1 | 14 | Zfp763 | 0.015 | Old |
| 0.685737 | 0.339599 | 0.12 | 0.1 | 1 | 14 | Tmem209 | 0.02 | Old |
| 0.685752 | 0.693711 | 0.148 | 0.18 | 1 | 14 | 4833420G1 | -0.032 | Old |
| 0.685752 | 0.022422 | 0.148 | 0.18 | 1 | 14 | Cfap20 | -0.032 | Old |
| 0.686002 | 0.487377 | 0.077 | 0.06 | 1 | 14 | Ogfod1 | 0.017 | Old |
| 0.686411 | 1.16491 | 0.153 | 0.14 | 1 | 14 | Erap1 | 0.013 | Old |
| 0.686882 | 0.455184 | 0.137 | 0.12 | 1 | 14 | Enpp4 | 0.017 | Old |
| 0.687157 | 1.626592 | 0.093 | 0.08 | 1 | 14 | Pelp1 | 0.013 | Old |
| 0.688197 | 0.232706 | 0.18 | 0.16 | 1 | 14 | Rbm17 | 0.02 | Old |
| 0.688242 | 0.318446 | 0.344 | 0.34 | 1 | 14 | Mcfd2 | 0.004 | Old |
| 0.688707 | 0.159396 | 0.115 | 0.1 | 1 | 14 | Lst1 | 0.015 | Old |
| 0.688871 | 0.591585 | 0.12 | 0.1 | 1 | 14 | Tlr2 | 0.02 | Old |
| 0.688981 | 0.724447 | 0.093 | 0.12 | 1 | 14 | Ints10 | -0.027 | Old |
| 0.689253 | 0.65577 | 0.23 | 0.22 | 1 | 14 | Lpcat1 | 0.01 | Old |
| 0.689442 | 0.376929 | 0.246 | 0.22 | 1 | 14 | Wsb2 | 0.026 | Old |
| 0.689594 | 0.807521 | 0.044 | 0.06 | 1 | 14 | Gpn2 | -0.016 | Old |
| 0.690122 | 0.764832 | 0.055 | 0.04 | 1 | 14 | Fam185a | 0.015 | Old |
| 0.690122 | 0.4087 | 0.055 | 0.04 | 1 | 14 | Wdr81 | 0.015 | Old |
| 0.690202 | 0.477304 | 0.197 | 0.18 | 1 | 14 | Impdh2 | 0.017 | Old |
| 0.690302 | 0.508403 | 0.35 | 0.32 | 1 | 14 | Tomm5 | 0.03 | Old |
| 0.69051 | 0.405504 | 0.153 | 0.18 | 1 | 14 | Nup50 | -0.027 | Old |
| 0.690825 | 0.586452 | 0.18 | 0.16 | 1 | 14 | Ssu72 | 0.02 | Old |
| 0.691294 | 0.525493 | 0.158 | 0.14 | 1 | 14 | Zdhhc4 | 0.018 | Old |
| 0.691392 | 0.121211 | 0.191 | 0.18 | 1 | 14 | Fam192a | 0.011 | Old |
| 0.691662 | 0.322455 | 0.311 | 0.28 | 1 | 14 | Armcx3 | 0.031 | Old |
| 0.69201 | 0.001721 | 0.12 | 0.1 | 1 | 14 | Cep112 | 0.02 | Old |
| 0.69221 | 0.023588 | 0.322 | 0.28 | 1 | 14 | Asap1 | 0.042 | Old |
| 0.692387 | 0.170042 | 0.169 | 0.14 | 1 | 14 | Smim13 | 0.029 | Old |
| 0.692501 | 0.323508 | 0.224 | 0.26 | 1 | 14 | Cbfb | -0.036 | Old |
| 0.692552 | 1.105483 | 0.186 | 0.18 | 1 | 14 | Cstf2t | 0.006 | Old |
| 0.692735 | 0.320484 | 0.197 | 0.18 | 1 | 14 | Polg | 0.017 | Old |

| | | | | | | | | |
|---|---|---|---|---|---|---|---|---|
| 0.693339 | 0.521483 | 0.098 | 0.08 | 1 | 14 | Slc35a3 | 0.018 | Old |
| 0.693821 | 0.121782 | 0.131 | 0.16 | 1 | 14 | Aldh9a1 | -0.029 | Old |
| 0.69395 | 0.313723 | 0.18 | 0.22 | 1 | 14 | Cops3 | -0.04 | Old |
| 0.694072 | 0.267 | 0.148 | 0.18 | 1 | 14 | Thap3 | -0.032 | Old |
| 0.694511 | 0.45397 | 0.262 | 0.24 | 1 | 14 | Ube2e3 | 0.022 | Old |
| 0.695092 | 0.276869 | 0.115 | 0.14 | 1 | 14 | Mtss1 | -0.025 | Old |
| 0.695096 | 1.070122 | 0.115 | 0.1 | 1 | 14 | Fam69a | 0.015 | Old |
| 0.695096 | 0.646503 | 0.115 | 0.1 | 1 | 14 | Bbip1 | 0.015 | Old |
| 0.695202 | 0.048921 | 1 | 0.98 | 1 | 14 | mt-Co3 | 0.02 | Old |
| 0.695364 | 0.052182 | 0.333 | 0.36 | 1 | 14 | Stt3a | -0.027 | Old |
| 0.695755 | 0.72669 | 0.137 | 0.12 | 1 | 14 | 2610008E1 | 0.017 | Old |
| 0.696004 | 0.229401 | 0.153 | 0.18 | 1 | 14 | Hgsnat | -0.027 | Old |
| 0.696004 | 0.040523 | 0.153 | 0.18 | 1 | 14 | Rbm22 | -0.027 | Old |
| 0.69629 | 0.086671 | 0.175 | 0.2 | 1 | 14 | Smim27 | -0.025 | Old |
| 0.696347 | 1.213707 | 0.208 | 0.2 | 1 | 14 | Thbs2 | 0.008 | Old |
| 0.696392 | 0.332209 | 0.219 | 0.26 | 1 | 14 | Gosr2 | -0.041 | Old |
| 0.696569 | 0.084247 | 0.514 | 0.48 | 1 | 14 | Add1 | 0.034 | Old |
| 0.696745 | 0.427784 | 0.142 | 0.12 | 1 | 14 | Bms1 | 0.022 | Old |
| 0.696745 | 0.131513 | 0.142 | 0.12 | 1 | 14 | Inpp5a | 0.022 | Old |
| 0.696854 | 0.592564 | 0.158 | 0.14 | 1 | 14 | Mien1 | 0.018 | Old |
| 0.697707 | 1.423554 | 0.093 | 0.08 | 1 | 14 | Six1 | 0.013 | Old |
| 0.697781 | 0.102406 | 0.464 | 0.44 | 1 | 14 | Uap1 | 0.024 | Old |
| 0.698587 | 0.020642 | 0.366 | 0.42 | 1 | 14 | Pnpla8 | -0.054 | Old |
| 0.698757 | 0.133661 | 0.153 | 0.18 | 1 | 14 | Fam129b | -0.027 | Old |
| 0.698855 | 0.013231 | 0.503 | 0.5 | 1 | 14 | Cavin3 | 0.003 | Old |
| 0.698904 | 0.210448 | 0.186 | 0.16 | 1 | 14 | Got2 | 0.026 | Old |
| 0.698904 | 0.195547 | 0.186 | 0.16 | 1 | 14 | Ube2g1 | 0.026 | Old |
| 0.698925 | 0.668974 | 0.06 | 0.08 | 1 | 14 | Rnf25 | -0.02 | Old |
| 0.699254 | 1.031975 | 0.055 | 0.04 | 1 | 14 | Chst11 | 0.015 | Old |
| 0.699254 | 0.248024 | 0.055 | 0.04 | 1 | 14 | Hdhd5 | 0.015 | Old |
| 0.699254 | 0.243158 | 0.055 | 0.04 | 1 | 14 | Fgd5 | 0.015 | Old |
| 0.699254 | 0.195457 | 0.055 | 0.04 | 1 | 14 | Ccnyl1 | 0.015 | Old |
| 0.699673 | 0.125937 | 0.142 | 0.12 | 1 | 14 | Manba | 0.022 | Old |
| 0.700141 | 0.176752 | 0.251 | 0.24 | 1 | 14 | Snx6 | 0.011 | Old |
| 0.700176 | 0.274254 | 0.437 | 0.44 | 1 | 14 | Psma3 | -0.003 | Old |
| 0.700241 | 0.767558 | 0.098 | 0.08 | 1 | 14 | Iffo2 | 0.018 | Old |
| 0.700241 | 0.231471 | 0.098 | 0.08 | 1 | 14 | Ifi35 | 0.018 | Old |
| 0.700433 | 0.018935 | 0.962 | 0.88 | 1 | 14 | Rpl7 | 0.082 | Old |
| 0.701197 | 1.215112 | 0.098 | 0.12 | 1 | 14 | Stx8 | -0.022 | Old |
| 0.701236 | 1.174381 | 0.093 | 0.08 | 1 | 14 | Slc25a25 | 0.013 | Old |
| 0.701509 | 0.173145 | 0.077 | 0.06 | 1 | 14 | Nipsnap3b | 0.017 | Old |
| 0.70208 | 0.311206 | 0.082 | 0.1 | 1 | 14 | Ikbkg | -0.018 | Old |
| 0.70208 | 0.141322 | 0.082 | 0.1 | 1 | 14 | AI606181 | -0.018 | Old |
| 0.702606 | 0.406363 | 0.142 | 0.12 | 1 | 14 | Nsmf | 0.022 | Old |
| 0.702631 | 0.428306 | 0.295 | 0.28 | 1 | 14 | Tbl1xr1 | 0.015 | Old |

| | | | | | | | | |
|---|---|---|---|---|---|---|---|---|
| 0.703328 | 0.334488 | 0.219 | 0.2 | 1 | 14 | Braf | 0.019 | Old |
| 0.703406 | 0.129302 | 0.137 | 0.16 | 1 | 14 | Polr2h | -0.023 | Old |
| 0.703702 | 0.300181 | 0.098 | 0.08 | 1 | 14 | Taf7 | 0.018 | Old |
| 0.703792 | 0.389833 | 0.197 | 0.24 | 1 | 14 | Map3k20 | -0.043 | Old |
| 0.70402 | 0.423452 | 0.169 | 0.2 | 1 | 14 | Idh3g | -0.031 | Old |
| 0.704132 | 0.252075 | 0.251 | 0.22 | 1 | 14 | Ccl9 | 0.031 | Old |
| 0.704141 | 0.147945 | 0.186 | 0.16 | 1 | 14 | Phf23 | 0.026 | Old |
| 0.704212 | 0.354849 | 0.257 | 0.24 | 1 | 14 | Fam171a1 | 0.017 | Old |
| 0.704407 | 0.73574 | 0.115 | 0.14 | 1 | 14 | Rnf150 | -0.025 | Old |
| 0.704407 | 0.200418 | 0.115 | 0.14 | 1 | 14 | Ankrd27 | -0.025 | Old |
| 0.70448 | 0.315959 | 0.503 | 0.5 | 1 | 14 | H2afz | 0.003 | Old |
| 0.70449 | 0.097245 | 0.596 | 0.56 | 1 | 14 | Akr1a1 | 0.036 | Old |
| 0.704621 | 0.88584 | 0.109 | 0.14 | 1 | 14 | Tango2 | -0.031 | Old |
| 0.704621 | 0.436578 | 0.12 | 0.1 | 1 | 14 | Ttc9c | 0.02 | Old |
| 0.704621 | 0.290051 | 0.12 | 0.1 | 1 | 14 | Fam117b | 0.02 | Old |
| 0.704997 | 0.476158 | 0.311 | 0.32 | 1 | 14 | Klhl24 | -0.009 | Old |
| 0.705394 | 0.233096 | 0.301 | 0.3 | 1 | 14 | Sgpp1 | 0.001 | Old |
| 0.705543 | 0.259976 | 0.142 | 0.12 | 1 | 14 | Ankle2 | 0.022 | Old |
| 0.705697 | 0.468833 | 0.082 | 0.1 | 1 | 14 | Klhdc4 | -0.018 | Old |
| 0.706118 | 0.056832 | 0.492 | 0.46 | 1 | 14 | Pja2 | 0.032 | Old |
| 0.706364 | 0.081288 | 0.667 | 0.56 | 1 | 14 | Serbp1 | 0.107 | Old |
| 0.70667 | 0.1676 | 0.169 | 0.2 | 1 | 14 | Rnf167 | -0.031 | Old |
| 0.707168 | 0.204291 | 0.098 | 0.08 | 1 | 14 | Fbxo25 | 0.018 | Old |
| 0.707428 | 1.514368 | 0.071 | 0.06 | 1 | 14 | Ctps2 | 0.011 | Old |
| 0.707522 | 0.143569 | 0.115 | 0.14 | 1 | 14 | Kcnt2 | -0.025 | Old |
| 0.707795 | 0.144956 | 0.279 | 0.32 | 1 | 14 | Sidt2 | -0.041 | Old |
| 0.707861 | 0.347934 | 0.098 | 0.12 | 1 | 14 | Nsmce1 | -0.022 | Old |
| 0.707939 | 0.723053 | 0.115 | 0.1 | 1 | 14 | Fbxo31 | 0.015 | Old |
| 0.708315 | 1.67223 | 0.093 | 0.08 | 1 | 14 | Eri1 | 0.013 | Old |
| 0.70843 | 0.007725 | 0.055 | 0.04 | 1 | 14 | Zbtb46 | 0.015 | Old |
| 0.708484 | 0.384712 | 0.142 | 0.12 | 1 | 14 | Frmd4a | 0.022 | Old |
| 0.708671 | 0.07553 | 0.377 | 0.38 | 1 | 14 | Eif3h | -0.003 | Old |
| 0.709373 | 0.816214 | 0.191 | 0.18 | 1 | 14 | Naxd | 0.011 | Old |
| 0.709807 | 0.279002 | 0.164 | 0.14 | 1 | 14 | C7 | 0.024 | Old |
| 0.709838 | 1.560492 | 0.148 | 0.14 | 1 | 14 | Dtymk | 0.008 | Old |
| 0.710635 | 0.603665 | 0.137 | 0.12 | 1 | 14 | Ube2l6 | 0.017 | Old |
| 0.710635 | 0.548441 | 0.137 | 0.12 | 1 | 14 | Ino80e | 0.017 | Old |
| 0.710642 | 0.668538 | 0.115 | 0.14 | 1 | 14 | Ubac1 | -0.025 | Old |
| 0.710642 | 0.354438 | 0.115 | 0.14 | 1 | 14 | Tarbp2 | -0.025 | Old |
| 0.710682 | 0.093087 | 0.208 | 0.24 | 1 | 14 | Polr2e | -0.032 | Old |
| 0.71104 | 0.113824 | 0.361 | 0.32 | 1 | 14 | Cul3 | 0.041 | Old |
| 0.711163 | 0.859912 | 0.115 | 0.1 | 1 | 14 | Spata2 | 0.015 | Old |
| 0.711163 | 0.746519 | 0.115 | 0.1 | 1 | 14 | Slc31a1 | 0.015 | Old |
| 0.711174 | 0.089145 | 0.563 | 0.54 | 1 | 14 | Srsf2 | 0.023 | Old |
| 0.711202 | 0.501053 | 0.098 | 0.12 | 1 | 14 | Smn1 | -0.022 | Old |

| | | | | | | | | |
|---|---|---|---|---|---|---|---|---|
| 0.71143 | 0.220537 | 0.131 | 0.16 | 1 | 14 | Ufd1 | -0.029 | Old |
| 0.712142 | 0.090486 | 0.268 | 0.24 | 1 | 14 | Pole4 | 0.028 | Old |
| 0.71238 | 0.787703 | 0.224 | 0.22 | 1 | 14 | Dcun1d5 | 0.004 | Old |
| 0.71258 | 0.02361 | 0.153 | 0.18 | 1 | 14 | Scfd1 | -0.027 | Old |
| 0.712951 | 0.461245 | 0.082 | 0.1 | 1 | 14 | Dnajc30 | -0.018 | Old |
| 0.713155 | 0.201984 | 0.219 | 0.26 | 1 | 14 | Cnot4 | -0.041 | Old |
| 0.713221 | 0.74326 | 0.077 | 0.06 | 1 | 14 | Sap30bp | 0.017 | Old |
| 0.713221 | 0.013988 | 0.077 | 0.06 | 1 | 14 | Zfp423 | 0.017 | Old |
| 0.713625 | 0.715115 | 0.137 | 0.12 | 1 | 14 | Zfp110 | 0.017 | Old |
| 0.713628 | 0.443024 | 0.148 | 0.18 | 1 | 14 | Swt1 | -0.032 | Old |
| 0.713721 | 0.106318 | 0.47 | 0.44 | 1 | 14 | Bptf | 0.03 | Old |
| 0.713817 | 0.032499 | 0.486 | 0.52 | 1 | 14 | Ube2b | -0.034 | Old |
| 0.71412 | 0.15209 | 0.098 | 0.08 | 1 | 14 | Cwc22 | 0.018 | Old |
| 0.71412 | 0.079399 | 0.098 | 0.08 | 1 | 14 | Myg1 | 0.018 | Old |
| 0.714392 | 0.603688 | 0.115 | 0.1 | 1 | 14 | Ubtd2 | 0.015 | Old |
| 0.714392 | 0.518479 | 0.115 | 0.1 | 1 | 14 | Lmtk2 | 0.015 | Old |
| 0.714541 | 0.340102 | 0.191 | 0.18 | 1 | 14 | Dido1 | 0.011 | Old |
| 0.714549 | 0.099899 | 0.098 | 0.12 | 1 | 14 | Kcnip3 | -0.022 | Old |
| 0.714877 | 0.161305 | 0.678 | 0.64 | 1 | 14 | Hspg2 | 0.038 | Old |
| 0.714894 | 0.040377 | 0.47 | 0.48 | 1 | 14 | H2-Ab1 | -0.01 | Old |
| 0.714895 | 0.057843 | 0.284 | 0.26 | 1 | 14 | Zfx | 0.024 | Old |
| 0.715032 | 0.11838 | 0.137 | 0.16 | 1 | 14 | Tmem150a | -0.023 | Old |
| 0.7156 | 0.515569 | 0.219 | 0.2 | 1 | 14 | Rasa1 | 0.019 | Old |
| 0.715693 | 0.604084 | 0.197 | 0.18 | 1 | 14 | Coro1b | 0.017 | Old |
| 0.716896 | 0.204218 | 0.115 | 0.14 | 1 | 14 | Hoxb4 | -0.025 | Old |
| 0.717141 | 0.183534 | 0.077 | 0.06 | 1 | 14 | Snx32 | 0.017 | Old |
| 0.717141 | 0.080106 | 0.077 | 0.06 | 1 | 14 | Mecr | 0.017 | Old |
| 0.717335 | 0.325266 | 0.131 | 0.16 | 1 | 14 | Tmem107 | -0.029 | Old |
| 0.717605 | 0.196741 | 0.098 | 0.08 | 1 | 14 | Aifm2 | 0.018 | Old |
| 0.7179 | 0.213624 | 0.098 | 0.12 | 1 | 14 | Dlat | -0.022 | Old |
| 0.718065 | 0.076495 | 0.246 | 0.28 | 1 | 14 | Hbp1 | -0.034 | Old |
| 0.71871 | 0.298767 | 0.24 | 0.28 | 1 | 14 | Ergic2 | -0.04 | Old |
| 0.718979 | 1.492673 | 0.093 | 0.08 | 1 | 14 | Taf12 | 0.013 | Old |
| 0.718979 | 0.721062 | 0.093 | 0.08 | 1 | 14 | Abcd1 | 0.013 | Old |
| 0.719 | 0.037917 | 0.268 | 0.24 | 1 | 14 | Mark3 | 0.028 | Old |
| 0.719251 | 0.62927 | 0.158 | 0.14 | 1 | 14 | Shprh | 0.018 | Old |
| 0.719511 | 1.403497 | 0.071 | 0.06 | 1 | 14 | Cox18 | 0.011 | Old |
| 0.719511 | 1.259627 | 0.071 | 0.06 | 1 | 14 | Capn5 | 0.011 | Old |
| 0.71957 | 0.260135 | 0.508 | 0.52 | 1 | 14 | Zfp207 | -0.012 | Old |
| 0.719702 | 0.622649 | 0.262 | 0.24 | 1 | 14 | Ppt1 | 0.022 | Old |
| 0.720294 | 0.616893 | 0.131 | 0.16 | 1 | 14 | Usp32 | -0.029 | Old |
| 0.720866 | 0.657492 | 0.115 | 0.1 | 1 | 14 | S100a4 | 0.015 | Old |
| 0.721068 | 0.062967 | 0.077 | 0.06 | 1 | 14 | Kiz | 0.017 | Old |
| 0.721273 | 0.613325 | 0.208 | 0.2 | 1 | 14 | Smim1 | 0.008 | Old |
| 0.721875 | 0.139652 | 0.191 | 0.22 | 1 | 14 | Psmc4 | -0.029 | Old |

| | | | | | | | | |
|---|---|---|---|---|---|---|---|---|
| 0.722069 | 0.35435 | 0.158 | 0.14 | 1 | 14 | Abhd17a | 0.018 | Old |
| 0.722069 | 0.226145 | 0.158 | 0.14 | 1 | 14 | D16Ertd47 | 0.018 | Old |
| 0.723257 | 0.37812 | 0.131 | 0.16 | 1 | 14 | Nup54 | -0.029 | Old |
| 0.723379 | 0.032712 | 0.24 | 0.28 | 1 | 14 | U2af1 | -0.04 | Old |
| 0.723555 | 1.28888 | 0.071 | 0.06 | 1 | 14 | Fam122a | 0.011 | Old |
| 0.72411 | 0.737115 | 0.115 | 0.1 | 1 | 14 | Dhps | 0.015 | Old |
| 0.72411 | 0.73339 | 0.115 | 0.1 | 1 | 14 | Dnase1l1 | 0.015 | Old |
| 0.72411 | 0.707783 | 0.115 | 0.1 | 1 | 14 | Nfya | 0.015 | Old |
| 0.72411 | 0.683355 | 0.115 | 0.1 | 1 | 14 | Prokr1 | 0.015 | Old |
| 0.72462 | 0.349416 | 0.098 | 0.12 | 1 | 14 | Hdac5 | -0.022 | Old |
| 0.725328 | 0.925334 | 0.169 | 0.16 | 1 | 14 | mt-Nd6 | 0.009 | Old |
| 0.725471 | 0.031231 | 0.219 | 0.2 | 1 | 14 | Cdc73 | 0.019 | Old |
| 0.725664 | 0.702986 | 0.328 | 0.34 | 1 | 14 | Araf | -0.012 | Old |
| 0.72612 | 0.734112 | 0.093 | 0.08 | 1 | 14 | Gpr137 | 0.013 | Old |
| 0.726983 | 0.130737 | 0.191 | 0.22 | 1 | 14 | Gtf3c1 | -0.029 | Old |
| 0.727537 | 0.358831 | 0.082 | 0.1 | 1 | 14 | Kctd5 | -0.018 | Old |
| 0.727606 | 1.266096 | 0.071 | 0.06 | 1 | 14 | Ttc7b | 0.011 | Old |
| 0.727606 | 1.126227 | 0.071 | 0.06 | 1 | 14 | Thyn1 | 0.011 | Old |
| 0.727606 | 0.214759 | 0.066 | 0.08 | 1 | 14 | Tpbg | -0.014 | Old |
| 0.727988 | 0.633886 | 0.098 | 0.12 | 1 | 14 | Ipo4 | -0.022 | Old |
| 0.728059 | 0.38654 | 0.24 | 0.28 | 1 | 14 | Nek9 | -0.04 | Old |
| 0.72806 | 0.048005 | 0.169 | 0.16 | 1 | 14 | Tmem9 | 0.009 | Old |
| 0.729699 | 0.408725 | 0.093 | 0.08 | 1 | 14 | Ttc7 | 0.013 | Old |
| 0.729925 | 0.093846 | 0.224 | 0.2 | 1 | 14 | Jtb | 0.024 | Old |
| 0.730537 | 0.053509 | 0.175 | 0.2 | 1 | 14 | Usp14 | -0.025 | Old |
| 0.730614 | 0.515537 | 0.115 | 0.1 | 1 | 14 | Dnajc18 | 0.015 | Old |
| 0.730674 | 0.459959 | 0.235 | 0.22 | 1 | 14 | Chordc1 | 0.015 | Old |
| 0.730738 | 0.066236 | 0.842 | 0.78 | 1 | 14 | Rplp0 | 0.062 | Old |
| 0.730978 | 0.274374 | 0.333 | 0.32 | 1 | 14 | Pnn | 0.013 | Old |
| 0.731249 | 0.070579 | 0.23 | 0.26 | 1 | 14 | Tollip | -0.03 | Old |
| 0.731665 | 1.072743 | 0.071 | 0.06 | 1 | 14 | Nup107 | 0.011 | Old |
| 0.731665 | 0.979979 | 0.071 | 0.06 | 1 | 14 | Sirt4 | 0.011 | Old |
| 0.731665 | 0.062475 | 0.066 | 0.08 | 1 | 14 | Fam207a | -0.014 | Old |
| 0.731665 | 0.002942 | 0.066 | 0.08 | 1 | 14 | Unc5a | -0.014 | Old |
| 0.731895 | 0.329369 | 0.35 | 0.34 | 1 | 14 | Eloc | 0.01 | Old |
| 0.731981 | 0.459968 | 0.268 | 0.26 | 1 | 14 | Zfp592 | 0.008 | Old |
| 0.732091 | 0.175217 | 0.153 | 0.18 | 1 | 14 | Bcl3 | -0.027 | Old |
| 0.732614 | 0.35052 | 0.115 | 0.14 | 1 | 14 | Med16 | -0.025 | Old |
| 0.733285 | 1.037317 | 0.093 | 0.08 | 1 | 14 | Cyth4 | 0.013 | Old |
| 0.733285 | 0.530167 | 0.093 | 0.08 | 1 | 14 | Pcnx | 0.013 | Old |
| 0.734361 | 1.036215 | 0.109 | 0.1 | 1 | 14 | Fcf1 | 0.009 | Old |
| 0.734578 | 0.427872 | 0.273 | 0.26 | 1 | 14 | Enpp5 | 0.013 | Old |
| 0.734668 | 0.089701 | 0.191 | 0.22 | 1 | 14 | Elmo2 | -0.029 | Old |
| 0.734867 | 0.221269 | 0.082 | 0.1 | 1 | 14 | Sgsh | -0.018 | Old |
| 0.734912 | 0.866184 | 0.23 | 0.22 | 1 | 14 | Sdhd | 0.01 | Old |

| | | | | | | | | |
|---|---|---|---|---|---|---|---|---|
| 0.734969 | 0.081948 | 0.12 | 0.14 | 1 | 14 | Osbpl1a | -0.02 | Old |
| 0.735342 | 0.908505 | 0.191 | 0.18 | 1 | 14 | Slc4a2 | 0.011 | Old |
| 0.735342 | 0.614967 | 0.191 | 0.18 | 1 | 14 | Nfx1 | 0.011 | Old |
| 0.735731 | 1.065383 | 0.071 | 0.06 | 1 | 14 | Galc | 0.011 | Old |
| 0.735731 | 0.043597 | 0.066 | 0.08 | 1 | 14 | Fcgr3 | -0.014 | Old |
| 0.735855 | 0.031666 | 0.175 | 0.2 | 1 | 14 | Dscr3 | -0.025 | Old |
| 0.735943 | 0.38482 | 0.421 | 0.4 | 1 | 14 | Gda | 0.021 | Old |
| 0.736046 | 0.246477 | 0.169 | 0.2 | 1 | 14 | Gypc | -0.031 | Old |
| 0.736188 | 0.353697 | 0.175 | 0.16 | 1 | 14 | Lmf2 | 0.015 | Old |
| 0.736211 | 0.333914 | 0.158 | 0.14 | 1 | 14 | Dnajc14 | 0.018 | Old |
| 0.736876 | 0.920489 | 0.093 | 0.08 | 1 | 14 | Dgat1 | 0.013 | Old |
| 0.736876 | 0.728946 | 0.093 | 0.08 | 1 | 14 | Xpnpep3 | 0.013 | Old |
| 0.737137 | 0.356596 | 0.115 | 0.1 | 1 | 14 | Fam20c | 0.015 | Old |
| 0.737164 | 0.770714 | 0.131 | 0.12 | 1 | 14 | Tmem181a | 0.011 | Old |
| 0.737164 | 0.734647 | 0.131 | 0.12 | 1 | 14 | Auh | 0.011 | Old |
| 0.73791 | 0.432987 | 0.235 | 0.22 | 1 | 14 | Thap12 | 0.015 | Old |
| 0.73791 | 0.24939 | 0.235 | 0.22 | 1 | 14 | Nbr1 | 0.015 | Old |
| 0.738123 | 0.331688 | 0.098 | 0.12 | 1 | 14 | Slc37a3 | -0.022 | Old |
| 0.738123 | 0.323856 | 0.098 | 0.12 | 1 | 14 | Agfg2 | -0.022 | Old |
| 0.738437 | 0.538589 | 0.23 | 0.26 | 1 | 14 | Arhgef1 | -0.03 | Old |
| 0.738555 | 0.467935 | 0.262 | 0.26 | 1 | 14 | Cdk4 | 0.002 | Old |
| 0.738736 | 0.043217 | 0.18 | 0.16 | 1 | 14 | Mkrn1 | 0.02 | Old |
| 0.738906 | 0.289702 | 0.175 | 0.16 | 1 | 14 | Fbxl17 | 0.015 | Old |
| 0.739805 | 1.537716 | 0.071 | 0.06 | 1 | 14 | Slc27a1 | 0.011 | Old |
| 0.739805 | 1.335181 | 0.071 | 0.06 | 1 | 14 | Tbc1d19 | 0.011 | Old |
| 0.739805 | 0.208933 | 0.066 | 0.08 | 1 | 14 | Mirg | -0.014 | Old |
| 0.739934 | 0.267235 | 0.421 | 0.4 | 1 | 14 | Trpm7 | 0.021 | Old |
| 0.740327 | 0.070985 | 0.224 | 0.26 | 1 | 14 | Pgrmc2 | -0.036 | Old |
| 0.740362 | 0.173499 | 0.208 | 0.24 | 1 | 14 | Zfp871 | -0.032 | Old |
| 0.740472 | 1.166278 | 0.093 | 0.08 | 1 | 14 | Vegfc | 0.013 | Old |
| 0.740472 | 0.566031 | 0.093 | 0.08 | 1 | 14 | Spg11 | 0.013 | Old |
| 0.740573 | 0.407208 | 0.18 | 0.22 | 1 | 14 | Scaf1 | -0.04 | Old |
| 0.740838 | 0.270863 | 0.388 | 0.38 | 1 | 14 | Eif5 | 0.008 | Old |
| 0.741358 | 0.114356 | 0.246 | 0.28 | 1 | 14 | Gna12 | -0.034 | Old |
| 0.741512 | 0.290443 | 0.098 | 0.12 | 1 | 14 | Arrb2 | -0.022 | Old |
| 0.741893 | 0.15971 | 0.148 | 0.18 | 1 | 14 | Mvb12a | -0.032 | Old |
| 0.742221 | 0.742037 | 0.082 | 0.1 | 1 | 14 | Nox4 | -0.018 | Old |
| 0.742221 | 0.590251 | 0.082 | 0.1 | 1 | 14 | Evi5l | -0.018 | Old |
| 0.742221 | 0.431572 | 0.082 | 0.1 | 1 | 14 | Nfrkb | -0.018 | Old |
| 0.743679 | 0.441291 | 0.115 | 0.1 | 1 | 14 | Zfp180 | 0.015 | Old |
| 0.743886 | 0.085725 | 0.066 | 0.08 | 1 | 14 | Akip1 | -0.014 | Old |
| 0.74432 | 0.379305 | 0.12 | 0.14 | 1 | 14 | Xpa | -0.02 | Old |
| 0.744341 | 1.437266 | 0.109 | 0.1 | 1 | 14 | Dhrs7b | 0.009 | Old |
| 0.744352 | 0.421062 | 0.175 | 0.16 | 1 | 14 | Msantd4 | 0.015 | Old |
| 0.744937 | 1.990378 | 0.049 | 0.04 | 1 | 14 | Zfp248 | 0.009 | Old |

| | | | | | | | | |
|---|---|---|---|---|---|---|---|---|
| 0.744937 | 1.849107 | 0.049 | 0.04 | 1 | 14 | Zbtb2 | 0.009 | Old |
| 0.744937 | 1.662083 | 0.049 | 0.04 | 1 | 14 | Epg5 | 0.009 | Old |
| 0.744937 | 1.618239 | 0.049 | 0.04 | 1 | 14 | Zscan29 | 0.009 | Old |
| 0.744937 | 1.595924 | 0.049 | 0.04 | 1 | 14 | Abhd10 | 0.009 | Old |
| 0.744937 | 1.471682 | 0.049 | 0.04 | 1 | 14 | Gm26853 | 0.009 | Old |
| 0.745648 | 0.296568 | 0.24 | 0.22 | 1 | 14 | C1d | 0.02 | Old |
| 0.745816 | 1.18198 | 0.191 | 0.18 | 1 | 14 | 1700025G( | 0.011 | Old |
| 0.746386 | 0.513449 | 0.131 | 0.12 | 1 | 14 | Chka | 0.011 | Old |
| 0.746704 | 0.409876 | 0.197 | 0.18 | 1 | 14 | Rnf111 | 0.017 | Old |
| 0.746704 | 0.280608 | 0.186 | 0.22 | 1 | 14 | Slc26a2 | -0.034 | Old |
| 0.747247 | 0.131575 | 0.432 | 0.4 | 1 | 14 | Ubl3 | 0.032 | Old |
| 0.747446 | 0.107416 | 0.12 | 0.14 | 1 | 14 | Faap100 | -0.02 | Old |
| 0.747518 | 0.589702 | 0.186 | 0.18 | 1 | 14 | Sdhb | 0.006 | Old |
| 0.747683 | 1.512016 | 0.093 | 0.08 | 1 | 14 | Coasy | 0.013 | Old |
| 0.747683 | 1.025808 | 0.093 | 0.08 | 1 | 14 | Exosc7 | 0.013 | Old |
| 0.747975 | 0.663702 | 0.071 | 0.06 | 1 | 14 | Bloc1s5 | 0.011 | Old |
| 0.748304 | 0.131673 | 0.098 | 0.12 | 1 | 14 | Wfs1 | -0.022 | Old |
| 0.748471 | 0.209013 | 0.399 | 0.38 | 1 | 14 | Tnpo1 | 0.019 | Old |
| 0.749285 | 0.665643 | 0.317 | 0.32 | 1 | 14 | Sltm | -0.003 | Old |
| 0.749599 | 1.432868 | 0.087 | 0.08 | 1 | 14 | Larp7 | 0.007 | Old |
| 0.749817 | 1.876668 | 0.049 | 0.04 | 1 | 14 | 5930430L0 | 0.009 | Old |
| 0.749817 | 1.763908 | 0.049 | 0.04 | 1 | 14 | Zxdb | 0.009 | Old |
| 0.749817 | 1.637062 | 0.049 | 0.04 | 1 | 14 | Map3k10 | 0.009 | Old |
| 0.749817 | 1.498618 | 0.049 | 0.04 | 1 | 14 | Spef1 | 0.009 | Old |
| 0.749817 | 1.476848 | 0.049 | 0.04 | 1 | 14 | Sirt3 | 0.009 | Old |
| 0.749831 | 0.34198 | 0.137 | 0.12 | 1 | 14 | Gnptg | 0.017 | Old |
| 0.750297 | 0.19892 | 0.137 | 0.16 | 1 | 14 | Tmem45a | -0.023 | Old |
| 0.750344 | 0.179879 | 0.366 | 0.34 | 1 | 14 | Pank3 | 0.026 | Old |
| 0.750506 | 0.361746 | 0.399 | 0.38 | 1 | 14 | Efcab14 | 0.019 | Old |
| 0.750575 | 0.228115 | 0.12 | 0.14 | 1 | 14 | 1110059G1 | -0.02 | Old |
| 0.750575 | 0.096685 | 0.12 | 0.14 | 1 | 14 | Pi4kb | -0.02 | Old |
| 0.750872 | 0.126161 | 0.208 | 0.18 | 1 | 14 | Cc2d2a | 0.028 | Old |
| 0.751297 | 0.661518 | 0.093 | 0.08 | 1 | 14 | Ror1 | 0.013 | Old |
| 0.751297 | 0.480302 | 0.093 | 0.08 | 1 | 14 | Garnl3 | 0.013 | Old |
| 0.751297 | 0.335905 | 0.093 | 0.08 | 1 | 14 | Ercc5 | 0.013 | Old |
| 0.751905 | 0.569594 | 0.153 | 0.14 | 1 | 14 | Gria3 | 0.013 | Old |
| 0.75207 | 0.957762 | 0.071 | 0.06 | 1 | 14 | Crat | 0.011 | Old |
| 0.75207 | 0.896728 | 0.071 | 0.06 | 1 | 14 | Tmem126a | 0.011 | Old |
| 0.75207 | 0.76359 | 0.066 | 0.08 | 1 | 14 | Cd163 | -0.014 | Old |
| 0.75207 | 0.219442 | 0.066 | 0.08 | 1 | 14 | Pank4 | -0.014 | Old |
| 0.752462 | 0.41039 | 0.262 | 0.26 | 1 | 14 | Rnf14 | 0.002 | Old |
| 0.752501 | 0.054777 | 0.514 | 0.58 | 1 | 14 | Btg1 | -0.066 | Old |
| 0.752808 | 0.601633 | 0.169 | 0.16 | 1 | 14 | Arhgap6 | 0.009 | Old |
| 0.753497 | 0.120857 | 0.213 | 0.2 | 1 | 14 | Cdc40 | 0.013 | Old |
| 0.753709 | 0.35078 | 0.12 | 0.14 | 1 | 14 | Kdelr3 | -0.02 | Old |

| | | | | | | | | |
|---|---|---|---|---|---|---|---|---|
| 0.753972 | 0.495676 | 0.251 | 0.24 | 1 | 14 | Vcpip1 | 0.011 | Old |
| 0.754432 | 0.704439 | 0.23 | 0.22 | 1 | 14 | Mdm2 | 0.01 | Old |
| 0.754644 | 0.107681 | 0.41 | 0.4 | 1 | 14 | Plin2 | 0.01 | Old |
| 0.754707 | 1.810361 | 0.049 | 0.04 | 1 | 14 | St8sia4 | 0.009 | Old |
| 0.754707 | 1.755303 | 0.049 | 0.04 | 1 | 14 | Rai2 | 0.009 | Old |
| 0.754707 | 1.713189 | 0.049 | 0.04 | 1 | 14 | Zbtb17 | 0.009 | Old |
| 0.754707 | 1.467132 | 0.049 | 0.04 | 1 | 14 | Fam173b | 0.009 | Old |
| 0.754707 | 1.426445 | 0.049 | 0.04 | 1 | 14 | Ogfod3 | 0.009 | Old |
| 0.754707 | 1.415693 | 0.049 | 0.04 | 1 | 14 | Eif5a2 | 0.009 | Old |
| 0.754707 | 1.155201 | 0.049 | 0.04 | 1 | 14 | Ptpro | 0.009 | Old |
| 0.754707 | 1.123436 | 0.049 | 0.04 | 1 | 14 | Sox12 | 0.009 | Old |
| 0.754781 | 0.065695 | 0.311 | 0.28 | 1 | 14 | Tmem5 | 0.031 | Old |
| 0.754916 | 0.428086 | 0.093 | 0.08 | 1 | 14 | Ormdl2 | 0.013 | Old |
| 0.755116 | 0.409524 | 0.098 | 0.12 | 1 | 14 | Kcnk6 | -0.022 | Old |
| 0.755259 | 0.227675 | 0.514 | 0.48 | 1 | 14 | Eif5a | 0.034 | Old |
| 0.755484 | 0.540201 | 0.186 | 0.18 | 1 | 14 | Rp2 | 0.006 | Old |
| 0.755771 | 0.203931 | 0.459 | 0.42 | 1 | 14 | 2-Mar | 0.039 | Old |
| 0.75582 | 0.281371 | 0.158 | 0.18 | 1 | 14 | Prkaca | -0.022 | Old |
| 0.756114 | 0.765292 | 0.131 | 0.16 | 1 | 14 | Lage3 | -0.029 | Old |
| 0.75616 | 0.626653 | 0.148 | 0.18 | 1 | 14 | Tmem230 | -0.032 | Old |
| 0.756173 | 0.804135 | 0.071 | 0.06 | 1 | 14 | Ankrd39 | 0.011 | Old |
| 0.756173 | 0.739623 | 0.071 | 0.06 | 1 | 14 | Fam210a | 0.011 | Old |
| 0.756173 | 0.673281 | 0.071 | 0.06 | 1 | 14 | Zkscan17 | 0.011 | Old |
| 0.756819 | 0.28648 | 0.115 | 0.1 | 1 | 14 | Pign | 0.015 | Old |
| 0.757001 | 0.582388 | 0.082 | 0.1 | 1 | 14 | Cdan1 | -0.018 | Old |
| 0.757001 | 0.277974 | 0.082 | 0.1 | 1 | 14 | Rmnd5b | -0.018 | Old |
| 0.757116 | 0.323293 | 0.262 | 0.26 | 1 | 14 | Rab8a | 0.002 | Old |
| 0.757219 | 0.292457 | 0.235 | 0.28 | 1 | 14 | Fam111a | -0.045 | Old |
| 0.757925 | 0.034728 | 0.328 | 0.3 | 1 | 14 | 2610507B1 | 0.028 | Old |
| 0.75854 | 0.619977 | 0.093 | 0.08 | 1 | 14 | Timm44 | 0.013 | Old |
| 0.758572 | 0.152542 | 0.197 | 0.22 | 1 | 14 | Ubald2 | -0.023 | Old |
| 0.758976 | 0.200141 | 0.191 | 0.18 | 1 | 14 | Rab23 | 0.011 | Old |
| 0.759203 | 0.75579 | 0.148 | 0.14 | 1 | 14 | Dcaf5 | 0.008 | Old |
| 0.759372 | 2.44183 | 0.027 | 0.02 | 1 | 14 | Dnaaf5 | 0.007 | Old |
| 0.759372 | 2.339354 | 0.027 | 0.02 | 1 | 14 | Bik | 0.007 | Old |
| 0.759372 | 2.214446 | 0.027 | 0.02 | 1 | 14 | Cdo1 | 0.007 | Old |
| 0.759372 | 2.14345 | 0.027 | 0.02 | 1 | 14 | Zfp146 | 0.007 | Old |
| 0.759372 | 2.141545 | 0.027 | 0.02 | 1 | 14 | Nudt7 | 0.007 | Old |
| 0.759372 | 2.11209 | 0.027 | 0.02 | 1 | 14 | Abtb2 | 0.007 | Old |
| 0.759372 | 2.088417 | 0.027 | 0.02 | 1 | 14 | BC024978 | 0.007 | Old |
| 0.759372 | 2.021735 | 0.027 | 0.02 | 1 | 14 | Tec | 0.007 | Old |
| 0.759372 | 1.975489 | 0.027 | 0.02 | 1 | 14 | Dmac2 | 0.007 | Old |
| 0.759372 | 1.935817 | 0.027 | 0.02 | 1 | 14 | Pigb | 0.007 | Old |
| 0.759372 | 1.910776 | 0.027 | 0.02 | 1 | 14 | Senp8 | 0.007 | Old |
| 0.759372 | 1.825978 | 0.027 | 0.02 | 1 | 14 | Setd4 | 0.007 | Old |

| | | | | | | | | |
|---|---|---|---|---|---|---|---|---|
| 0.759372 | 1.749519 | 0.027 | 0.02 | 1 | 14 | Gm48898 | 0.007 | Old |
| 0.759372 | 1.625817 | 0.027 | 0.02 | 1 | 14 | Lrp4 | 0.007 | Old |
| 0.759372 | 1.602921 | 0.027 | 0.02 | 1 | 14 | Eepd1 | 0.007 | Old |
| 0.759372 | 1.573331 | 0.027 | 0.02 | 1 | 14 | Lrmda | 0.007 | Old |
| 0.759372 | 1.543197 | 0.027 | 0.02 | 1 | 14 | Nat2 | 0.007 | Old |
| 0.759372 | 1.536289 | 0.027 | 0.02 | 1 | 14 | Mocos | 0.007 | Old |
| 0.759372 | 1.532854 | 0.027 | 0.02 | 1 | 14 | Ankrd24 | 0.007 | Old |
| 0.759372 | 1.496278 | 0.027 | 0.02 | 1 | 14 | Arhgap39 | 0.007 | Old |
| 0.759372 | 1.478267 | 0.027 | 0.02 | 1 | 14 | Dus2 | 0.007 | Old |
| 0.759372 | 1.465502 | 0.027 | 0.02 | 1 | 14 | Tti1 | 0.007 | Old |
| 0.759372 | 1.459713 | 0.027 | 0.02 | 1 | 14 | Txnrd2 | 0.007 | Old |
| 0.759372 | 1.434347 | 0.027 | 0.02 | 1 | 14 | Nat8f3 | 0.007 | Old |
| 0.759372 | 1.429962 | 0.027 | 0.02 | 1 | 14 | Ttc12 | 0.007 | Old |
| 0.759372 | 1.410209 | 0.027 | 0.02 | 1 | 14 | Gtdc1 | 0.007 | Old |
| 0.759372 | 1.385988 | 0.027 | 0.02 | 1 | 14 | Iglon5 | 0.007 | Old |
| 0.759372 | 1.376134 | 0.027 | 0.02 | 1 | 14 | Hsd11b2 | 0.007 | Old |
| 0.759372 | 1.37061 | 0.027 | 0.02 | 1 | 14 | Nubpl | 0.007 | Old |
| 0.759372 | 1.241389 | 0.027 | 0.02 | 1 | 14 | Gm14418 | 0.007 | Old |
| 0.759372 | 1.146813 | 0.027 | 0.02 | 1 | 14 | Mpv17l | 0.007 | Old |
| 0.759372 | 1.129221 | 0.027 | 0.02 | 1 | 14 | Trib3 | 0.007 | Old |
| 0.759372 | 1.127469 | 0.027 | 0.02 | 1 | 14 | Ms4a8a | 0.007 | Old |
| 0.759372 | 1.082033 | 0.027 | 0.02 | 1 | 14 | Itgb2 | 0.007 | Old |
| 0.759372 | 1.067895 | 0.027 | 0.02 | 1 | 14 | AI839979 | 0.007 | Old |
| 0.759372 | 1.01688 | 0.027 | 0.02 | 1 | 14 | Tmem69 | 0.007 | Old |
| 0.759372 | 1.0168 | 0.027 | 0.02 | 1 | 14 | Gtf2h2 | 0.007 | Old |
| 0.759372 | 0.994339 | 0.027 | 0.02 | 1 | 14 | Arhgap22 | 0.007 | Old |
| 0.759372 | 0.946422 | 0.027 | 0.02 | 1 | 14 | Pbx4 | 0.007 | Old |
| 0.759372 | 0.936434 | 0.027 | 0.02 | 1 | 14 | AC129328. | 0.007 | Old |
| 0.759372 | 0.581672 | 0.027 | 0.02 | 1 | 14 | Nlrc5 | 0.007 | Old |
| 0.759446 | 0.339739 | 0.262 | 0.26 | 1 | 14 | Rnf187 | 0.002 | Old |
| 0.759607 | 1.081435 | 0.049 | 0.04 | 1 | 14 | Chchd5 | 0.009 | Old |
| 0.759607 | 0.995663 | 0.049 | 0.04 | 1 | 14 | Psph | 0.009 | Old |
| 0.759607 | 0.859346 | 0.049 | 0.04 | 1 | 14 | Tmtc4 | 0.009 | Old |
| 0.759626 | 0.103742 | 0.049 | 0.06 | 1 | 14 | Adgrg6 | -0.011 | Old |
| 0.759751 | 0.283872 | 0.197 | 0.18 | 1 | 14 | Mcrs1 | 0.017 | Old |
| 0.760046 | 0.209771 | 0.404 | 0.4 | 1 | 14 | Tmed3 | 0.004 | Old |
| 0.760115 | 0.220907 | 0.115 | 0.1 | 1 | 14 | Ttc1 | 0.015 | Old |
| 0.760273 | 0.107667 | 0.284 | 0.32 | 1 | 14 | Os9 | -0.036 | Old |
| 0.760282 | 1.166277 | 0.066 | 0.08 | 1 | 14 | Dhx8 | -0.014 | Old |
| 0.760282 | 0.777753 | 0.071 | 0.06 | 1 | 14 | Dhx58os | 0.011 | Old |
| 0.760282 | 0.727562 | 0.071 | 0.06 | 1 | 14 | Soga1 | 0.011 | Old |
| 0.760282 | 0.641037 | 0.066 | 0.08 | 1 | 14 | Dok2 | -0.014 | Old |
| 0.760282 | 0.21417 | 0.066 | 0.08 | 1 | 14 | Ndufaf7 | -0.014 | Old |
| 0.760593 | 0.585421 | 0.153 | 0.14 | 1 | 14 | Knop1 | 0.013 | Old |
| 0.760709 | 0.519356 | 0.082 | 0.1 | 1 | 14 | Rgs12 | -0.018 | Old |

| | | | | | | | | |
|---|---|---|---|---|---|---|---|---|
| 0.760709 | 0.336791 | 0.082 | 0.1 | 1 | 14 | Zfp729a | -0.018 | Old |
| 0.761187 | 0.401969 | 0.115 | 0.14 | 1 | 14 | Ttc4 | -0.025 | Old |
| 0.761838 | 0.752668 | 0.131 | 0.12 | 1 | 14 | Tcta | 0.011 | Old |
| 0.761838 | 0.722776 | 0.131 | 0.12 | 1 | 14 | Pip5k1c | 0.011 | Old |
| 0.76217 | 0.420536 | 0.093 | 0.08 | 1 | 14 | Rbm12b1 | 0.013 | Old |
| 0.763415 | 0.494444 | 0.115 | 0.1 | 1 | 14 | Ssbp1 | 0.015 | Old |
| 0.763718 | 0.225466 | 0.208 | 0.18 | 1 | 14 | Ccndbp1 | 0.028 | Old |
| 0.763891 | 0.141978 | 0.169 | 0.16 | 1 | 14 | Xpnpep2 | 0.009 | Old |
| 0.764252 | 0.094531 | 0.23 | 0.22 | 1 | 14 | H2afj | 0.01 | Old |
| 0.764398 | 0.757501 | 0.071 | 0.06 | 1 | 14 | Stau2 | 0.011 | Old |
| 0.764398 | 0.368226 | 0.066 | 0.08 | 1 | 14 | Gm20342 | -0.014 | Old |
| 0.764398 | 0.25752 | 0.066 | 0.08 | 1 | 14 | Nuak2 | -0.014 | Old |
| 0.764424 | 1.462677 | 0.087 | 0.08 | 1 | 14 | Mtmr11 | 0.007 | Old |
| 0.764424 | 0.697626 | 0.082 | 0.1 | 1 | 14 | Ptprk | -0.018 | Old |
| 0.764517 | 2.072601 | 0.049 | 0.04 | 1 | 14 | Zfp141 | 0.009 | Old |
| 0.764517 | 1.404554 | 0.049 | 0.04 | 1 | 14 | Plcd3 | 0.009 | Old |
| 0.764517 | 1.244023 | 0.049 | 0.04 | 1 | 14 | Cops7b | 0.009 | Old |
| 0.764517 | 0.906598 | 0.049 | 0.04 | 1 | 14 | Taf1a | 0.009 | Old |
| 0.764517 | 0.855019 | 0.049 | 0.04 | 1 | 14 | 2900089D1 | 0.009 | Old |
| 0.76476 | 0.359827 | 0.158 | 0.14 | 1 | 14 | Fbxo22 | 0.018 | Old |
| 0.765151 | 0.301867 | 0.137 | 0.16 | 1 | 14 | Col27a1 | -0.023 | Old |
| 0.765369 | 0.649996 | 0.098 | 0.12 | 1 | 14 | Urod | -0.022 | Old |
| 0.765805 | 0.304391 | 0.093 | 0.08 | 1 | 14 | Mdc1 | 0.013 | Old |
| 0.765949 | 1.987092 | 0.027 | 0.02 | 1 | 14 | Dmap1 | 0.007 | Old |
| 0.765949 | 1.822236 | 0.027 | 0.02 | 1 | 14 | Suv39h1 | 0.007 | Old |
| 0.765949 | 1.39345 | 0.027 | 0.02 | 1 | 14 | Adnp | 0.007 | Old |
| 0.765949 | 1.337033 | 0.027 | 0.02 | 1 | 14 | Eda2r | 0.007 | Old |
| 0.765949 | 1.325674 | 0.027 | 0.02 | 1 | 14 | Jade3 | 0.007 | Old |
| 0.765949 | 1.297453 | 0.027 | 0.02 | 1 | 14 | Pop1 | 0.007 | Old |
| 0.765949 | 1.244921 | 0.027 | 0.02 | 1 | 14 | 1-Sep | 0.007 | Old |
| 0.765949 | 1.18965 | 0.027 | 0.02 | 1 | 14 | Ttll3 | 0.007 | Old |
| 0.765949 | 1.179567 | 0.027 | 0.02 | 1 | 14 | Sharpin | 0.007 | Old |
| 0.765949 | 1.175645 | 0.027 | 0.02 | 1 | 14 | Gab3 | 0.007 | Old |
| 0.765949 | 0.982785 | 0.027 | 0.02 | 1 | 14 | Coq3 | 0.007 | Old |
| 0.765949 | 0.946674 | 0.027 | 0.02 | 1 | 14 | Arhgef9 | 0.007 | Old |
| 0.765949 | 0.912117 | 0.027 | 0.02 | 1 | 14 | Cttnbp2 | 0.007 | Old |
| 0.765949 | 0.868282 | 0.027 | 0.02 | 1 | 14 | Dkk3 | 0.007 | Old |
| 0.765949 | 0.827583 | 0.027 | 0.02 | 1 | 14 | Ccdc130 | 0.007 | Old |
| 0.765949 | 0.741123 | 0.027 | 0.02 | 1 | 14 | Rsl1 | 0.007 | Old |
| 0.765949 | 0.540415 | 0.027 | 0.02 | 1 | 14 | Atp8b4 | 0.007 | Old |
| 0.766049 | 0.205655 | 0.519 | 0.52 | 1 | 14 | Atp6ap1 | -0.001 | Old |
| 0.766878 | 0.551043 | 0.208 | 0.2 | 1 | 14 | Ipo5 | 0.008 | Old |
| 0.767583 | 0.607522 | 0.115 | 0.14 | 1 | 14 | Agmo | -0.025 | Old |
| 0.767585 | 0.652246 | 0.202 | 0.2 | 1 | 14 | Cops8 | 0.002 | Old |
| 0.768024 | 0.404781 | 0.148 | 0.14 | 1 | 14 | Ints3 | 0.008 | Old |

| | | | | | | | | |
|---|---|---|---|---|---|---|---|---|
| 0.768046 | 0.64766 | 0.131 | 0.12 | 1 | 14 | 1700123O2 | 0.011 | Old |
| 0.768046 | 0.443758 | 0.131 | 0.12 | 1 | 14 | Mtpap | 0.011 | Old |
| 0.768521 | 0.921734 | 0.066 | 0.08 | 1 | 14 | Hcls1 | -0.014 | Old |
| 0.768521 | 0.5233 | 0.071 | 0.06 | 1 | 14 | Cln3 | 0.011 | Old |
| 0.768521 | 0.498666 | 0.071 | 0.06 | 1 | 14 | Tpgs2 | 0.011 | Old |
| 0.768521 | 0.405434 | 0.066 | 0.08 | 1 | 14 | Slc25a16 | -0.014 | Old |
| 0.768521 | 0.381326 | 0.071 | 0.06 | 1 | 14 | Sco2 | 0.011 | Old |
| 0.768521 | 0.37185 | 0.066 | 0.08 | 1 | 14 | Ticam1 | -0.014 | Old |
| 0.768521 | 0.354575 | 0.066 | 0.08 | 1 | 14 | Cdc42ep1 | -0.014 | Old |
| 0.768875 | 0.009187 | 0.705 | 0.7 | 1 | 14 | Prdx1 | 0.005 | Old |
| 0.769037 | 0.373082 | 0.273 | 0.26 | 1 | 14 | Pofut1 | 0.013 | Old |
| 0.769057 | 0.602901 | 0.049 | 0.06 | 1 | 14 | Pdss2 | -0.011 | Old |
| 0.769057 | 0.04713 | 0.049 | 0.06 | 1 | 14 | P2ry6 | -0.011 | Old |
| 0.769057 | 0.021899 | 0.049 | 0.06 | 1 | 14 | Fndc4 | -0.011 | Old |
| 0.769436 | 1.052455 | 0.049 | 0.04 | 1 | 14 | Lrrc25 | 0.009 | Old |
| 0.769436 | 0.839762 | 0.049 | 0.04 | 1 | 14 | Aph1c | 0.009 | Old |
| 0.769436 | 0.664364 | 0.049 | 0.04 | 1 | 14 | Chd7 | 0.009 | Old |
| 0.769843 | 0.105502 | 0.24 | 0.22 | 1 | 14 | Cir1 | 0.02 | Old |
| 0.770982 | 0.242847 | 0.262 | 0.24 | 1 | 14 | Blmh | 0.022 | Old |
| 0.771117 | 0.263288 | 0.137 | 0.16 | 1 | 14 | Prps1 | -0.023 | Old |
| 0.771163 | 0.869279 | 0.109 | 0.1 | 1 | 14 | Galnt7 | 0.009 | Old |
| 0.771215 | 0.008821 | 0.137 | 0.12 | 1 | 14 | C2cd2 | 0.017 | Old |
| 0.771869 | 0.829655 | 0.082 | 0.1 | 1 | 14 | Nudcd1 | -0.018 | Old |
| 0.772206 | 0.21442 | 0.191 | 0.18 | 1 | 14 | Cerk | 0.011 | Old |
| 0.772544 | 1.8327 | 0.027 | 0.02 | 1 | 14 | Gm13305 | 0.007 | Old |
| 0.772544 | 1.459417 | 0.027 | 0.02 | 1 | 14 | Akr1c18 | 0.007 | Old |
| 0.772544 | 1.188384 | 0.027 | 0.02 | 1 | 14 | Ak4 | 0.007 | Old |
| 0.772544 | 1.035569 | 0.027 | 0.02 | 1 | 14 | Tdo2 | 0.007 | Old |
| 0.772544 | 0.861876 | 0.027 | 0.02 | 1 | 14 | 1700109H( | 0.007 | Old |
| 0.772544 | 0.757933 | 0.027 | 0.02 | 1 | 14 | 1810024B( | 0.007 | Old |
| 0.772544 | 0.714709 | 0.027 | 0.02 | 1 | 14 | Raver2 | 0.007 | Old |
| 0.772544 | 0.701214 | 0.027 | 0.02 | 1 | 14 | Itgb7 | 0.007 | Old |
| 0.772544 | 0.599364 | 0.027 | 0.02 | 1 | 14 | Proser3 | 0.007 | Old |
| 0.772544 | 0.458445 | 0.027 | 0.02 | 1 | 14 | Mettl18 | 0.007 | Old |
| 0.772544 | 0.374769 | 0.027 | 0.02 | 1 | 14 | 1200007C1 | 0.007 | Old |
| 0.772544 | 0.357778 | 0.027 | 0.02 | 1 | 14 | Oas2 | 0.007 | Old |
| 0.772545 | 0.64097 | 0.24 | 0.24 | 1 | 14 | Lipa | 0 | Old |
| 0.77265 | 1.269071 | 0.066 | 0.08 | 1 | 14 | Phgdh | -0.014 | Old |
| 0.77265 | 0.473518 | 0.071 | 0.06 | 1 | 14 | Cetn4 | 0.011 | Old |
| 0.77265 | 0.343696 | 0.071 | 0.06 | 1 | 14 | Kctd18 | 0.011 | Old |
| 0.77265 | 0.213241 | 0.071 | 0.06 | 1 | 14 | Cluap1 | 0.011 | Old |
| 0.772663 | 0.851573 | 0.164 | 0.16 | 1 | 14 | Sod2 | 0.004 | Old |
| 0.77309 | 0.51602 | 0.093 | 0.08 | 1 | 14 | Bdh2 | 0.013 | Old |
| 0.77377 | 0.145712 | 0.213 | 0.2 | 1 | 14 | Utp11 | 0.013 | Old |
| 0.773785 | 0.193495 | 0.049 | 0.06 | 1 | 14 | Rpl36a-ps1 | -0.011 | Old |

| | | | | | | | | |
|---|---|---|---|---|---|---|---|---|
| 0.773785 | 0.034687 | 0.049 | 0.06 | 1 | 14 | Dusp18 | -0.011 | Old |
| 0.774365 | 1.108077 | 0.049 | 0.04 | 1 | 14 | Cd274 | 0.009 | Old |
| 0.774365 | 0.917733 | 0.049 | 0.04 | 1 | 14 | Etfdh | 0.009 | Old |
| 0.774365 | 0.730763 | 0.049 | 0.04 | 1 | 14 | Gnpat | 0.009 | Old |
| 0.774365 | 0.70082 | 0.049 | 0.04 | 1 | 14 | Tmem129 | 0.009 | Old |
| 0.774365 | 0.461668 | 0.049 | 0.04 | 1 | 14 | Ezh2 | 0.009 | Old |
| 0.774473 | 0.477394 | 0.153 | 0.18 | 1 | 14 | Polr1a | -0.027 | Old |
| 0.77486 | 0.713708 | 0.191 | 0.18 | 1 | 14 | Higd1a | 0.011 | Old |
| 0.77486 | 0.543828 | 0.191 | 0.18 | 1 | 14 | Ddx18 | 0.011 | Old |
| 0.774971 | 0.208881 | 0.372 | 0.36 | 1 | 14 | Srebf2 | 0.012 | Old |
| 0.774972 | 0.243676 | 0.372 | 0.36 | 1 | 14 | Stag2 | 0.012 | Old |
| 0.7756 | 1.25588 | 0.087 | 0.08 | 1 | 14 | Cryl1 | 0.007 | Old |
| 0.775756 | 0.258344 | 0.12 | 0.14 | 1 | 14 | Brap | -0.02 | Old |
| 0.776223 | 0.10407 | 0.104 | 0.12 | 1 | 14 | Cystm1 | -0.016 | Old |
| 0.776741 | 0.411519 | 0.093 | 0.08 | 1 | 14 | Zfp524 | 0.013 | Old |
| 0.776746 | 0.082242 | 0.41 | 0.46 | 1 | 14 | Mbtps1 | -0.05 | Old |
| 0.776786 | 0.979196 | 0.066 | 0.08 | 1 | 14 | C3ar1 | -0.014 | Old |
| 0.776786 | 0.638997 | 0.066 | 0.08 | 1 | 14 | Rnf219 | -0.014 | Old |
| 0.776786 | 0.359109 | 0.071 | 0.06 | 1 | 14 | Lancl2 | 0.011 | Old |
| 0.776786 | 0.295717 | 0.071 | 0.06 | 1 | 14 | Dock2 | 0.011 | Old |
| 0.777697 | 0.292913 | 0.322 | 0.3 | 1 | 14 | Zc3hav1 | 0.022 | Old |
| 0.777813 | 0.519967 | 0.169 | 0.16 | 1 | 14 | Micall1 | 0.009 | Old |
| 0.777912 | 1.542781 | 0.109 | 0.1 | 1 | 14 | Wrn | 0.009 | Old |
| 0.778521 | 0.508822 | 0.049 | 0.06 | 1 | 14 | Pola2 | -0.011 | Old |
| 0.778521 | 0.450001 | 0.049 | 0.06 | 1 | 14 | Apc2 | -0.011 | Old |
| 0.778521 | 0.299845 | 0.049 | 0.06 | 1 | 14 | Nsun5 | -0.011 | Old |
| 0.778521 | 0.20028 | 0.049 | 0.06 | 1 | 14 | Map3k6 | -0.011 | Old |
| 0.778521 | 0.17084 | 0.049 | 0.06 | 1 | 14 | Rnf214 | -0.011 | Old |
| 0.778521 | 0.024481 | 0.049 | 0.06 | 1 | 14 | Dok7 | -0.011 | Old |
| 0.779104 | 0.749601 | 0.098 | 0.12 | 1 | 14 | Rad17 | -0.022 | Old |
| 0.779155 | 1.004661 | 0.027 | 0.02 | 1 | 14 | Cacnb1 | 0.007 | Old |
| 0.779155 | 0.863486 | 0.027 | 0.02 | 1 | 14 | Ano3 | 0.007 | Old |
| 0.779155 | 0.841065 | 0.027 | 0.02 | 1 | 14 | Ighmbp2 | 0.007 | Old |
| 0.779155 | 0.795364 | 0.027 | 0.02 | 1 | 14 | Fmn2 | 0.007 | Old |
| 0.779155 | 0.480058 | 0.027 | 0.02 | 1 | 14 | Vars2 | 0.007 | Old |
| 0.779155 | 0.376335 | 0.027 | 0.02 | 1 | 14 | Zfhx2 | 0.007 | Old |
| 0.779155 | 0.177478 | 0.027 | 0.02 | 1 | 14 | Igsf9b | 0.007 | Old |
| 0.779155 | 0.175662 | 0.027 | 0.02 | 1 | 14 | E130308A1 | 0.007 | Old |
| 0.779155 | 0.098679 | 0.027 | 0.02 | 1 | 14 | AI661453 | 0.007 | Old |
| 0.779155 | 0.077576 | 0.027 | 0.02 | 1 | 14 | Gdpd3 | 0.007 | Old |
| 0.779155 | 0.062157 | 0.027 | 0.02 | 1 | 14 | Sorl1 | 0.007 | Old |
| 0.779158 | 0.875395 | 0.148 | 0.18 | 1 | 14 | Zfp329 | -0.032 | Old |
| 0.779302 | 1.186852 | 0.049 | 0.04 | 1 | 14 | Kif3c | 0.009 | Old |
| 0.779302 | 0.993047 | 0.049 | 0.04 | 1 | 14 | Bcar1 | 0.009 | Old |
| 0.779302 | 0.770254 | 0.049 | 0.04 | 1 | 14 | Akap17b | 0.009 | Old |

| | | | | | | | | |
|---|---|---|---|---|---|---|---|---|
| 0.779302 | 0.640747 | 0.049 | 0.04 | 1 | 14 | Prkd1 | 0.009 | Old |
| 0.779302 | 0.523353 | 0.049 | 0.04 | 1 | 14 | 4930481A1 | 0.009 | Old |
| 0.779302 | 0.503203 | 0.049 | 0.04 | 1 | 14 | L2hgdh | 0.009 | Old |
| 0.779475 | 0.13772 | 0.29 | 0.28 | 1 | 14 | Zfp451 | 0.01 | Old |
| 0.779535 | 0.760975 | 0.186 | 0.18 | 1 | 14 | Ddx54 | 0.006 | Old |
| 0.780179 | 0.550538 | 0.153 | 0.18 | 1 | 14 | Suds3 | -0.027 | Old |
| 0.780396 | 0.194663 | 0.093 | 0.08 | 1 | 14 | Prkg1 | 0.013 | Old |
| 0.780396 | 0.062246 | 0.093 | 0.08 | 1 | 14 | Nelfe | 0.013 | Old |
| 0.780928 | 0.895496 | 0.066 | 0.08 | 1 | 14 | Sugp1 | -0.014 | Old |
| 0.780928 | 0.619905 | 0.066 | 0.08 | 1 | 14 | Dnaja4 | -0.014 | Old |
| 0.780928 | 0.574204 | 0.071 | 0.06 | 1 | 14 | Pym1 | 0.011 | Old |
| 0.780928 | 0.376813 | 0.071 | 0.06 | 1 | 14 | Usf3 | 0.011 | Old |
| 0.780928 | 0.236473 | 0.071 | 0.06 | 1 | 14 | Slc18b1 | 0.011 | Old |
| 0.78227 | 0.265107 | 0.208 | 0.2 | 1 | 14 | Nrep | 0.008 | Old |
| 0.782549 | 0.71418 | 0.098 | 0.12 | 1 | 14 | Fbxo28 | -0.022 | Old |
| 0.782966 | 0.232739 | 0.273 | 0.28 | 1 | 14 | Sgpl1 | -0.007 | Old |
| 0.783076 | 0.803939 | 0.082 | 0.1 | 1 | 14 | Pdpr | -0.018 | Old |
| 0.783266 | 0.110774 | 0.049 | 0.06 | 1 | 14 | Slc25a22 | -0.011 | Old |
| 0.783266 | 0.107516 | 0.049 | 0.06 | 1 | 14 | Zfand2a | -0.011 | Old |
| 0.783266 | 0.029868 | 0.049 | 0.06 | 1 | 14 | 0610009B2 | -0.011 | Old |
| 0.783966 | 0.344181 | 0.213 | 0.2 | 1 | 14 | AW549877 | 0.013 | Old |
| 0.784041 | 0.059118 | 0.268 | 0.24 | 1 | 14 | Fkbp7 | 0.028 | Old |
| 0.784249 | 1.068211 | 0.049 | 0.04 | 1 | 14 | Itgb8 | 0.009 | Old |
| 0.784249 | 0.457868 | 0.049 | 0.04 | 1 | 14 | Kbtbd11 | 0.009 | Old |
| 0.784249 | 0.393714 | 0.049 | 0.04 | 1 | 14 | Rad18 | 0.009 | Old |
| 0.784249 | 0.112255 | 0.049 | 0.04 | 1 | 14 | 2310039H( | 0.009 | Old |
| 0.784679 | 0.694462 | 0.109 | 0.1 | 1 | 14 | Lrig1 | 0.009 | Old |
| 0.785077 | 1.104558 | 0.066 | 0.08 | 1 | 14 | Etaa1 | -0.014 | Old |
| 0.785077 | 0.157731 | 0.071 | 0.06 | 1 | 14 | Fra10ac1 | 0.011 | Old |
| 0.785077 | 0.090339 | 0.071 | 0.06 | 1 | 14 | Mtmr2 | 0.011 | Old |
| 0.785782 | 1.141545 | 0.027 | 0.02 | 1 | 14 | Atp5s | 0.007 | Old |
| 0.785782 | 0.559385 | 0.027 | 0.02 | 1 | 14 | Adat2 | 0.007 | Old |
| 0.785782 | 0.421929 | 0.027 | 0.02 | 1 | 14 | Ear2 | 0.007 | Old |
| 0.785782 | 0.405465 | 0.027 | 0.02 | 1 | 14 | Zfp790 | 0.007 | Old |
| 0.785782 | 0.202094 | 0.027 | 0.02 | 1 | 14 | Snrnp25 | 0.007 | Old |
| 0.785782 | 0.197921 | 0.027 | 0.02 | 1 | 14 | Dctd | 0.007 | Old |
| 0.785782 | 0.138318 | 0.027 | 0.02 | 1 | 14 | Ska2 | 0.007 | Old |
| 0.785782 | 0.082717 | 0.027 | 0.02 | 1 | 14 | Cd300lb | 0.007 | Old |
| 0.785782 | 0.061035 | 0.027 | 0.02 | 1 | 14 | Cd1d1 | 0.007 | Old |
| 0.786373 | 0.150588 | 0.104 | 0.12 | 1 | 14 | 1810055G( | -0.016 | Old |
| 0.786597 | 0.171186 | 0.137 | 0.12 | 1 | 14 | Dync2li1 | 0.017 | Old |
| 0.787601 | 0.161014 | 0.268 | 0.26 | 1 | 14 | Anapc1 | 0.008 | Old |
| 0.787684 | 0.539579 | 0.169 | 0.2 | 1 | 14 | Pde7a | -0.031 | Old |
| 0.788019 | 0.503841 | 0.049 | 0.06 | 1 | 14 | Traf3ip2 | -0.011 | Old |
| 0.788019 | 0.186717 | 0.049 | 0.06 | 1 | 14 | Cfb | -0.011 | Old |

| | | | | | | | | |
|---|---|---|---|---|---|---|---|---|
| 0.788019 | 0.186364 | 0.049 | 0.06 | 1 | 14 | P2rx7 | -0.011 | Old |
| 0.788019 | 0.137848 | 0.049 | 0.06 | 1 | 14 | Amot | -0.011 | Old |
| 0.788019 | 0.077413 | 0.049 | 0.06 | 1 | 14 | Fxyd2 | -0.011 | Old |
| 0.788019 | 0.018335 | 0.049 | 0.06 | 1 | 14 | Gm14964 | -0.011 | Old |
| 0.788019 | 0.003487 | 0.049 | 0.06 | 1 | 14 | Msto1 | -0.011 | Old |
| 0.788068 | 0.562567 | 0.109 | 0.1 | 1 | 14 | Efcc1 | 0.009 | Old |
| 0.788141 | 0.017518 | 0.219 | 0.2 | 1 | 14 | Ghr | 0.019 | Old |
| 0.788169 | 0.318747 | 0.191 | 0.18 | 1 | 14 | Map1a | 0.011 | Old |
| 0.788437 | 0.323964 | 0.12 | 0.14 | 1 | 14 | Mical3 | -0.02 | Old |
| 0.788442 | 0.142617 | 0.246 | 0.24 | 1 | 14 | Cspp1 | 0.006 | Old |
| 0.788897 | 0.258626 | 0.317 | 0.3 | 1 | 14 | Fam114a1 | 0.017 | Old |
| 0.789178 | 0.668242 | 0.235 | 0.22 | 1 | 14 | Mrps14 | 0.015 | Old |
| 0.789204 | 0.238088 | 0.049 | 0.04 | 1 | 14 | Gemin8 | 0.009 | Old |
| 0.789204 | 0.180627 | 0.049 | 0.04 | 1 | 14 | Zfp112 | 0.009 | Old |
| 0.789204 | 0.101112 | 0.049 | 0.04 | 1 | 14 | Srgap3 | 0.009 | Old |
| 0.789204 | 0.029293 | 0.049 | 0.04 | 1 | 14 | Mettl27 | 0.009 | Old |
| 0.789231 | 1.069961 | 0.066 | 0.08 | 1 | 14 | Kif16b | -0.014 | Old |
| 0.789234 | 0.393444 | 0.295 | 0.34 | 1 | 14 | Mrpl51 | -0.045 | Old |
| 0.789614 | 0.546816 | 0.164 | 0.16 | 1 | 14 | Zfp131 | 0.004 | Old |
| 0.789766 | 0.101033 | 0.153 | 0.14 | 1 | 14 | Snx14 | 0.013 | Old |
| 0.790404 | 0.020319 | 0.164 | 0.18 | 1 | 14 | Ppa1 | -0.016 | Old |
| 0.790431 | 0.088195 | 0.18 | 0.16 | 1 | 14 | Pgd | 0.02 | Old |
| 0.791018 | 0.228203 | 0.437 | 0.44 | 1 | 14 | Colgalt1 | -0.003 | Old |
| 0.791336 | 0.12924 | 0.197 | 0.18 | 1 | 14 | Idi1 | 0.017 | Old |
| 0.791644 | 0.135969 | 0.235 | 0.22 | 1 | 14 | Ets1 | 0.015 | Old |
| 0.791692 | 0.967954 | 0.148 | 0.14 | 1 | 14 | Wdr44 | 0.008 | Old |
| 0.79177 | 0.102965 | 0.295 | 0.3 | 1 | 14 | Ankrd13c | -0.005 | Old |
| 0.791812 | 0.073903 | 0.23 | 0.26 | 1 | 14 | Dap3 | -0.03 | Old |
| 0.792153 | 0.316635 | 0.399 | 0.4 | 1 | 14 | Map7d1 | -0.001 | Old |
| 0.792223 | 0.048083 | 0.732 | 0.66 | 1 | 14 | Ube2d3 | 0.072 | Old |
| 0.79229 | 0.354291 | 0.268 | 0.26 | 1 | 14 | Pcdh7 | 0.008 | Old |
| 0.792966 | 0.021847 | 0.333 | 0.36 | 1 | 14 | Ndufs2 | -0.027 | Old |
| 0.79316 | 0.37066 | 0.104 | 0.12 | 1 | 14 | Actl6a | -0.016 | Old |
| 0.79351 | 0.107321 | 0.191 | 0.18 | 1 | 14 | Sytl2 | 0.011 | Old |
| 0.794113 | 0.322108 | 0.224 | 0.26 | 1 | 14 | Dnajc19 | -0.036 | Old |
| 0.794168 | 0.878038 | 0.049 | 0.04 | 1 | 14 | Cd33 | 0.009 | Old |
| 0.794168 | 0.401983 | 0.049 | 0.04 | 1 | 14 | Tctn3 | 0.009 | Old |
| 0.794328 | 1.29503 | 0.087 | 0.08 | 1 | 14 | Slc38a7 | 0.007 | Old |
| 0.794339 | 0.0043 | 0.18 | 0.2 | 1 | 14 | Slc9a3r2 | -0.02 | Old |
| 0.794799 | 0.259233 | 0.12 | 0.14 | 1 | 14 | Rasal2 | -0.02 | Old |
| 0.794799 | 0.213678 | 0.12 | 0.14 | 1 | 14 | Ppp2r5d | -0.02 | Old |
| 0.794859 | 0.955319 | 0.109 | 0.1 | 1 | 14 | Relb | 0.009 | Old |
| 0.795064 | 0.119098 | 0.093 | 0.08 | 1 | 14 | Sh3bp2 | 0.013 | Old |
| 0.795627 | 0.178489 | 0.372 | 0.38 | 1 | 14 | Ndufb7 | -0.008 | Old |
| 0.7957 | 0.416538 | 0.033 | 0.04 | 1 | 14 | Tyms | -0.007 | Old |

| | | | | | | | | |
|---|---|---|---|---|---|---|---|---|
| 0.795931 | 0.423497 | 0.169 | 0.2 | 1 | 14 | Nsun2 | -0.031 | Old |
| 0.79616 | 0.166824 | 0.126 | 0.14 | 1 | 14 | Hmgcr | -0.014 | Old |
| 0.796391 | 0.935312 | 0.126 | 0.12 | 1 | 14 | Rpf1 | 0.006 | Old |
| 0.796526 | 0.130055 | 0.388 | 0.38 | 1 | 14 | Csnk1g2 | 0.008 | Old |
| 0.797548 | 0.687559 | 0.049 | 0.06 | 1 | 14 | Mblac2 | -0.011 | Old |
| 0.797548 | 0.577271 | 0.049 | 0.06 | 1 | 14 | Ccdc86 | -0.011 | Old |
| 0.797548 | 0.572721 | 0.049 | 0.06 | 1 | 14 | Ddx31 | -0.011 | Old |
| 0.797548 | 0.492913 | 0.049 | 0.06 | 1 | 14 | Shisa4 | -0.011 | Old |
| 0.797548 | 0.491312 | 0.049 | 0.06 | 1 | 14 | Pde4b | -0.011 | Old |
| 0.797548 | 0.453161 | 0.049 | 0.06 | 1 | 14 | Tbc1d31 | -0.011 | Old |
| 0.797748 | 0.392526 | 0.197 | 0.24 | 1 | 14 | Sra1 | -0.043 | Old |
| 0.797748 | 0.369448 | 0.208 | 0.2 | 1 | 14 | Fam193a | 0.008 | Old |
| 0.798089 | 0.760357 | 0.087 | 0.08 | 1 | 14 | Wdr12 | 0.007 | Old |
| 0.798571 | 0.904852 | 0.131 | 0.16 | 1 | 14 | Rpa1 | -0.029 | Old |
| 0.798743 | 0.289618 | 0.093 | 0.08 | 1 | 14 | Syk | 0.013 | Old |
| 0.798924 | 0.208619 | 0.404 | 0.4 | 1 | 14 | Arglu1 | 0.004 | Old |
| 0.79914 | 0.297889 | 0.049 | 0.04 | 1 | 14 | Xylt1 | 0.009 | Old |
| 0.79914 | 0.115082 | 0.049 | 0.04 | 1 | 14 | Ino80b | 0.009 | Old |
| 0.79914 | 0.045485 | 0.049 | 0.04 | 1 | 14 | Tmem231 | 0.009 | Old |
| 0.79914 | 0.0269 | 0.049 | 0.04 | 1 | 14 | Pcyox1l | 0.009 | Old |
| 0.799301 | 0.419019 | 0.131 | 0.12 | 1 | 14 | Gsr | 0.011 | Old |
| 0.799579 | 1.083493 | 0.126 | 0.12 | 1 | 14 | Magi2 | 0.006 | Old |
| 0.800239 | 0.643419 | 0.153 | 0.18 | 1 | 14 | Llgl1 | -0.027 | Old |
| 0.800618 | 0.56404 | 0.148 | 0.14 | 1 | 14 | Cetn2 | 0.008 | Old |
| 0.801441 | 0.541847 | 0.169 | 0.2 | 1 | 14 | Ube2f | -0.031 | Old |
| 0.801446 | 0.080586 | 0.404 | 0.38 | 1 | 14 | Trpc4ap | 0.024 | Old |
| 0.801496 | 0.094865 | 0.033 | 0.04 | 1 | 14 | Stn1 | -0.007 | Old |
| 0.801496 | 0.059781 | 0.033 | 0.04 | 1 | 14 | Tut1 | -0.007 | Old |
| 0.801665 | 0.577833 | 0.109 | 0.1 | 1 | 14 | Mapk8ip1 | 0.009 | Old |
| 0.80173 | 0.273055 | 0.071 | 0.06 | 1 | 14 | Cd209g | 0.011 | Old |
| 0.801854 | 1.667231 | 0.087 | 0.08 | 1 | 14 | Srbd1 | 0.007 | Old |
| 0.801854 | 1.022633 | 0.087 | 0.08 | 1 | 14 | Nudt13 | 0.007 | Old |
| 0.802299 | 0.230762 | 0.279 | 0.32 | 1 | 14 | Kmt5b | -0.041 | Old |
| 0.802324 | 2.034947 | 0.049 | 0.06 | 1 | 14 | Id4 | -0.011 | Old |
| 0.802324 | 0.368742 | 0.049 | 0.06 | 1 | 14 | Coq6 | -0.011 | Old |
| 0.802333 | 0.011754 | 0.213 | 0.24 | 1 | 14 | Ank2 | -0.027 | Old |
| 0.802925 | 0.142229 | 0.208 | 0.2 | 1 | 14 | Vangl1 | 0.008 | Old |
| 0.803094 | 0.214397 | 0.295 | 0.3 | 1 | 14 | Nadk | -0.005 | Old |
| 0.803369 | 0.501704 | 0.104 | 0.12 | 1 | 14 | Abhd18 | -0.016 | Old |
| 0.803405 | 0.516928 | 0.219 | 0.2 | 1 | 14 | Ccl6 | 0.019 | Old |
| 0.803599 | 0.576225 | 0.148 | 0.14 | 1 | 14 | Nacc1 | 0.008 | Old |
| 0.80412 | 0.690849 | 0.049 | 0.04 | 1 | 14 | Nr6a1 | 0.009 | Old |
| 0.804199 | 0.359661 | 0.169 | 0.2 | 1 | 14 | Rasa3 | -0.031 | Old |
| 0.804221 | 0.481333 | 0.191 | 0.18 | 1 | 14 | Cdip1 | 0.011 | Old |
| 0.80427 | 0.439563 | 0.087 | 0.1 | 1 | 14 | Edn1 | -0.013 | Old |

| | | | | | | | | |
|---|---|---|---|---|---|---|---|---|
| 0.80427 | 0.125079 | 0.087 | 0.1 | 1 | 14 | Smarcd1 | -0.013 | Old |
| 0.804561 | 0.293347 | 0.355 | 0.36 | 1 | 14 | Zfp950 | -0.005 | Old |
| 0.805074 | 0.61828 | 0.109 | 0.1 | 1 | 14 | Tnfrsf12a | 0.009 | Old |
| 0.805593 | 0.285716 | 0.131 | 0.12 | 1 | 14 | Zfp398 | 0.011 | Old |
| 0.805593 | 0.010515 | 0.131 | 0.12 | 1 | 14 | Hif1an | 0.011 | Old |
| 0.805624 | 1.613889 | 0.087 | 0.08 | 1 | 14 | Kat5 | 0.007 | Old |
| 0.805624 | 0.776323 | 0.087 | 0.08 | 1 | 14 | Tsc2 | 0.007 | Old |
| 0.805957 | 0.101369 | 0.213 | 0.22 | 1 | 14 | Hes1 | -0.007 | Old |
| 0.805965 | 0.684297 | 0.126 | 0.12 | 1 | 14 | Zpr1 | 0.006 | Old |
| 0.806545 | 0.028013 | 0.262 | 0.24 | 1 | 14 | Fnbp1 | 0.022 | Old |
| 0.806594 | 0.144108 | 0.24 | 0.24 | 1 | 14 | Tcf3 | 0 | Old |
| 0.80672 | 0.143168 | 0.344 | 0.34 | 1 | 14 | Usp7 | 0.004 | Old |
| 0.807108 | 1.097016 | 0.049 | 0.06 | 1 | 14 | Zfp946 | -0.011 | Old |
| 0.807108 | 0.889368 | 0.049 | 0.06 | 1 | 14 | Zfp41 | -0.011 | Old |
| 0.807108 | 0.665146 | 0.049 | 0.06 | 1 | 14 | Katnal1 | -0.011 | Old |
| 0.807108 | 0.659867 | 0.049 | 0.06 | 1 | 14 | Tmem39b | -0.011 | Old |
| 0.807108 | 0.637162 | 0.049 | 0.06 | 1 | 14 | Thnsl2 | -0.011 | Old |
| 0.807108 | 0.406993 | 0.049 | 0.06 | 1 | 14 | Smyd3 | -0.011 | Old |
| 0.807151 | 0.072745 | 0.443 | 0.42 | 1 | 14 | Uqcrq | 0.023 | Old |
| 0.807303 | 0.431741 | 0.033 | 0.04 | 1 | 14 | Kpna2 | -0.007 | Old |
| 0.807303 | 0.241347 | 0.033 | 0.04 | 1 | 14 | Utp14b | -0.007 | Old |
| 0.807635 | 0.000718 | 0.295 | 0.3 | 1 | 14 | Mon2 | -0.005 | Old |
| 0.808487 | 0.612674 | 0.109 | 0.1 | 1 | 14 | Asf1a | 0.009 | Old |
| 0.808687 | 0.159438 | 0.169 | 0.16 | 1 | 14 | St3gal6 | 0.009 | Old |
| 0.808743 | 0.100849 | 0.126 | 0.14 | 1 | 14 | Rpl10-ps3 | -0.014 | Old |
| 0.809109 | 0.364199 | 0.049 | 0.04 | 1 | 14 | Clybl | 0.009 | Old |
| 0.809398 | 1.025202 | 0.087 | 0.08 | 1 | 14 | Plekhm1 | 0.007 | Old |
| 0.809398 | 0.863563 | 0.087 | 0.08 | 1 | 14 | Pdk4 | 0.007 | Old |
| 0.810091 | 0.135724 | 0.071 | 0.06 | 1 | 14 | Pnp2 | 0.011 | Old |
| 0.810194 | 0.272319 | 0.104 | 0.12 | 1 | 14 | Dhx16 | -0.016 | Old |
| 0.810256 | 1.01471 | 0.104 | 0.1 | 1 | 14 | Ptdss2 | 0.004 | Old |
| 0.810708 | 0.326428 | 0.202 | 0.22 | 1 | 14 | Atl3 | -0.018 | Old |
| 0.81149 | 0.039046 | 0.235 | 0.26 | 1 | 14 | Ccdc59 | -0.025 | Old |
| 0.811898 | 0.535924 | 0.049 | 0.06 | 1 | 14 | Slamf9 | -0.011 | Old |
| 0.811903 | 0.382464 | 0.109 | 0.1 | 1 | 14 | Ttll4 | 0.009 | Old |
| 0.812363 | 0.713912 | 0.126 | 0.12 | 1 | 14 | Strn | 0.006 | Old |
| 0.812471 | 0.010422 | 0.224 | 0.2 | 1 | 14 | Prkcd | 0.024 | Old |
| 0.81256 | 0.603789 | 0.142 | 0.16 | 1 | 14 | Fbrsl1 | -0.018 | Old |
| 0.813121 | 0.116835 | 0.033 | 0.04 | 1 | 14 | Lss | -0.007 | Old |
| 0.813176 | 1.105193 | 0.087 | 0.08 | 1 | 14 | Dchs2 | 0.007 | Old |
| 0.813612 | 0.276322 | 0.104 | 0.12 | 1 | 14 | Wdr36 | -0.016 | Old |
| 0.813612 | 0.248147 | 0.104 | 0.12 | 1 | 14 | Cdk6 | -0.016 | Old |
| 0.813965 | 0.764585 | 0.12 | 0.14 | 1 | 14 | 9-Sep | -0.02 | Old |
| 0.81428 | 0.307413 | 0.071 | 0.06 | 1 | 14 | Fbxw9 | 0.011 | Old |
| 0.81428 | 0.153599 | 0.071 | 0.06 | 1 | 14 | Fibin | 0.011 | Old |

| | | | | | | | | |
|---|---|---|---|---|---|---|---|---|
| 0.815054 | 0.49912 | 0.131 | 0.12 | 1 | 14 | Kansl1l | 0.011 | Old |
| 0.815138 | 0.155876 | 0.279 | 0.3 | 1 | 14 | Sec24b | -0.021 | Old |
| 0.815192 | 1.265645 | 0.066 | 0.06 | 1 | 14 | 1810043G( | 0.006 | Old |
| 0.815323 | 0.695485 | 0.109 | 0.1 | 1 | 14 | Rdh10 | 0.009 | Old |
| 0.816219 | 0.224751 | 0.175 | 0.16 | 1 | 14 | Lrrc59 | 0.015 | Old |
| 0.816455 | 0.009144 | 0.399 | 0.38 | 1 | 14 | Rab18 | 0.019 | Old |
| 0.816696 | 1.069951 | 0.049 | 0.06 | 1 | 14 | Zfp608 | -0.011 | Old |
| 0.816696 | 0.984947 | 0.049 | 0.06 | 1 | 14 | Zfp618 | -0.011 | Old |
| 0.816959 | 0.626066 | 0.087 | 0.08 | 1 | 14 | Tmem242 | 0.007 | Old |
| 0.817452 | 0.048518 | 0.989 | 0.94 | 1 | 14 | Fos | 0.049 | Old |
| 0.817644 | 0.074275 | 0.497 | 0.44 | 1 | 14 | Fosl2 | 0.057 | Old |
| 0.817662 | 0.509981 | 0.191 | 0.18 | 1 | 14 | Kansl2 | 0.011 | Old |
| 0.817845 | 0.149742 | 0.306 | 0.34 | 1 | 14 | Bcl9l | -0.034 | Old |
| 0.818214 | 0.823724 | 0.131 | 0.12 | 1 | 14 | Alg11 | 0.011 | Old |
| 0.818474 | 1.42721 | 0.066 | 0.08 | 1 | 14 | Fam58b | -0.014 | Old |
| 0.818949 | 1.043656 | 0.033 | 0.04 | 1 | 14 | Zfp58 | -0.007 | Old |
| 0.818949 | 0.624182 | 0.033 | 0.04 | 1 | 14 | Tagap1 | -0.007 | Old |
| 0.818949 | 0.506622 | 0.033 | 0.04 | 1 | 14 | Zfp605 | -0.007 | Old |
| 0.818949 | 0.456269 | 0.033 | 0.04 | 1 | 14 | Bcs1l | -0.007 | Old |
| 0.818949 | 0.415131 | 0.033 | 0.04 | 1 | 14 | Klhl23 | -0.007 | Old |
| 0.818949 | 0.317556 | 0.033 | 0.04 | 1 | 14 | Zfp346 | -0.007 | Old |
| 0.818949 | 0.191212 | 0.033 | 0.04 | 1 | 14 | Nsun4 | -0.007 | Old |
| 0.818949 | 0.141565 | 0.033 | 0.04 | 1 | 14 | Cbx8 | -0.007 | Old |
| 0.818949 | 0.08768 | 0.033 | 0.04 | 1 | 14 | Dhfr | -0.007 | Old |
| 0.818949 | 0.02217 | 0.033 | 0.04 | 1 | 14 | Car5b | -0.007 | Old |
| 0.819043 | 0.5386 | 0.333 | 0.36 | 1 | 14 | Zfand6 | -0.027 | Old |
| 0.820133 | 0.422689 | 0.115 | 0.1 | 1 | 14 | Elac1 | 0.015 | Old |
| 0.820201 | 0.008068 | 0.94 | 0.98 | 1 | 14 | Rps24 | -0.04 | Old |
| 0.820459 | 0.791976 | 0.104 | 0.12 | 1 | 14 | Atxn7l3 | -0.016 | Old |
| 0.820459 | 0.459181 | 0.104 | 0.12 | 1 | 14 | Ephb4 | -0.016 | Old |
| 0.820595 | 0.470421 | 0.186 | 0.22 | 1 | 14 | Kctd2 | -0.034 | Old |
| 0.820739 | 0.330858 | 0.262 | 0.26 | 1 | 14 | Ppp4r3a | 0.002 | Old |
| 0.820747 | 0.697068 | 0.087 | 0.08 | 1 | 14 | Mavs | 0.007 | Old |
| 0.8208 | 1.046506 | 0.197 | 0.2 | 1 | 14 | Asxl2 | -0.003 | Old |
| 0.820945 | 0.727123 | 0.164 | 0.16 | 1 | 14 | Fam76b | 0.004 | Old |
| 0.821501 | 1.285653 | 0.049 | 0.06 | 1 | 14 | Dnajc24 | -0.011 | Old |
| 0.821501 | 1.177008 | 0.049 | 0.06 | 1 | 14 | Pus10 | -0.011 | Old |
| 0.821501 | 0.653391 | 0.049 | 0.06 | 1 | 14 | Tbc1d1 | -0.011 | Old |
| 0.821984 | 0.886774 | 0.126 | 0.12 | 1 | 14 | Xaf1 | 0.006 | Old |
| 0.821984 | 0.778392 | 0.126 | 0.12 | 1 | 14 | Gpn3 | 0.006 | Old |
| 0.821984 | 0.515199 | 0.126 | 0.12 | 1 | 14 | Banp | 0.006 | Old |
| 0.822344 | 0.377126 | 0.137 | 0.16 | 1 | 14 | Efs | -0.023 | Old |
| 0.822517 | 0.504381 | 0.175 | 0.2 | 1 | 14 | Zmiz2 | -0.025 | Old |
| 0.822804 | 0.464689 | 0.186 | 0.18 | 1 | 14 | Map4k5 | 0.006 | Old |
| 0.822804 | 0.370049 | 0.186 | 0.18 | 1 | 14 | Srp68 | 0.006 | Old |

| | | | | | | | | |
|---|---|---|---|---|---|---|---|---|
| 0.823928 | 0.006369 | 0.219 | 0.24 | 1 | 14 | Trafd1 | -0.021 | Old |
| 0.824201 | 1.144357 | 0.104 | 0.1 | 1 | 14 | Sirt1 | 0.004 | Old |
| 0.82424 | 0.253095 | 0.164 | 0.18 | 1 | 14 | C1qa | -0.016 | Old |
| 0.824538 | 0.958358 | 0.087 | 0.08 | 1 | 14 | Mpeg1 | 0.007 | Old |
| 0.824542 | 0.488722 | 0.131 | 0.12 | 1 | 14 | Mif4gd | 0.011 | Old |
| 0.824542 | 0.143782 | 0.126 | 0.14 | 1 | 14 | Papd7 | -0.014 | Old |
| 0.824787 | 1.83198 | 0.033 | 0.04 | 1 | 14 | Tmem141 | -0.007 | Old |
| 0.824787 | 0.753438 | 0.033 | 0.04 | 1 | 14 | Wnt9a | -0.007 | Old |
| 0.824787 | 0.56196 | 0.033 | 0.04 | 1 | 14 | R3hcc1 | -0.007 | Old |
| 0.824787 | 0.495979 | 0.033 | 0.04 | 1 | 14 | Zbtb40 | -0.007 | Old |
| 0.824787 | 0.481717 | 0.033 | 0.04 | 1 | 14 | Fam78b | -0.007 | Old |
| 0.824787 | 0.47726 | 0.033 | 0.04 | 1 | 14 | E330009J0 | -0.007 | Old |
| 0.824787 | 0.21781 | 0.033 | 0.04 | 1 | 14 | Hhat | -0.007 | Old |
| 0.824908 | 0.248468 | 0.339 | 0.32 | 1 | 14 | H2-Eb1 | 0.019 | Old |
| 0.825106 | 0.020517 | 0.262 | 0.3 | 1 | 14 | R3hdm1 | -0.038 | Old |
| 0.825321 | 0.265353 | 0.393 | 0.38 | 1 | 14 | Fbln1 | 0.013 | Old |
| 0.825603 | 0.745252 | 0.109 | 0.1 | 1 | 14 | Tfdp2 | 0.009 | Old |
| 0.826026 | 0.228554 | 0.355 | 0.36 | 1 | 14 | Ak3 | -0.005 | Old |
| 0.826229 | 0.114115 | 0.197 | 0.22 | 1 | 14 | Uimc1 | -0.023 | Old |
| 0.826312 | 1.291128 | 0.049 | 0.06 | 1 | 14 | Zfyve19 | -0.011 | Old |
| 0.826805 | 0.546596 | 0.12 | 0.14 | 1 | 14 | Etv6 | -0.02 | Old |
| 0.826805 | 0.092739 | 0.317 | 0.3 | 1 | 14 | Ccdc71l | 0.017 | Old |
| 0.827549 | 0.417589 | 0.148 | 0.14 | 1 | 14 | Tbc1d12 | 0.008 | Old |
| 0.82771 | 0.050715 | 0.126 | 0.14 | 1 | 14 | Pole3 | -0.014 | Old |
| 0.82813 | 0.056254 | 0.153 | 0.14 | 1 | 14 | Fbrs | 0.013 | Old |
| 0.828179 | 1.81548 | 0.066 | 0.06 | 1 | 14 | Rufy1 | 0.006 | Old |
| 0.828334 | 0.48814 | 0.087 | 0.08 | 1 | 14 | Aasdhppt | 0.007 | Old |
| 0.828457 | 0.365061 | 0.29 | 0.3 | 1 | 14 | Bmp2k | -0.01 | Old |
| 0.829037 | 0.479665 | 0.109 | 0.1 | 1 | 14 | Ikbke | 0.009 | Old |
| 0.829037 | 0.26818 | 0.109 | 0.1 | 1 | 14 | Gmip | 0.009 | Old |
| 0.829913 | 0.448043 | 0.164 | 0.18 | 1 | 14 | 10-Sep | -0.016 | Old |
| 0.830185 | 0.283653 | 0.087 | 0.1 | 1 | 14 | Bola1 | -0.013 | Old |
| 0.830435 | 0.374122 | 0.284 | 0.34 | 1 | 14 | Exoc5 | -0.056 | Old |
| 0.830554 | 0.636086 | 0.142 | 0.16 | 1 | 14 | Cdyl | -0.018 | Old |
| 0.830635 | 1.00916 | 0.033 | 0.04 | 1 | 14 | Lrrc61 | -0.007 | Old |
| 0.830635 | 0.692606 | 0.033 | 0.04 | 1 | 14 | Klf12 | -0.007 | Old |
| 0.830635 | 0.656705 | 0.033 | 0.04 | 1 | 14 | Prepl | -0.007 | Old |
| 0.830635 | 0.535945 | 0.033 | 0.04 | 1 | 14 | Leng9 | -0.007 | Old |
| 0.830635 | 0.394133 | 0.033 | 0.04 | 1 | 14 | Btbd6 | -0.007 | Old |
| 0.830635 | 0.251855 | 0.033 | 0.04 | 1 | 14 | Rmdn1 | -0.007 | Old |
| 0.831591 | 0.238667 | 0.372 | 0.38 | 1 | 14 | Kat2b | -0.008 | Old |
| 0.831632 | 0.6702 | 0.126 | 0.12 | 1 | 14 | Scyl1 | 0.006 | Old |
| 0.831659 | 0.153421 | 0.131 | 0.14 | 1 | 14 | Naf1 | -0.009 | Old |
| 0.832133 | 0.616595 | 0.087 | 0.08 | 1 | 14 | Cep192 | 0.007 | Old |
| 0.832133 | 0.076043 | 0.087 | 0.08 | 1 | 14 | Foxk2 | 0.007 | Old |

| | | | | | | | | |
|---|---|---|---|---|---|---|---|---|
| 0.832518 | 1.575638 | 0.066 | 0.06 | 1 | 14 | Cmtm4 | 0.006 | Old |
| 0.832518 | 1.326282 | 0.066 | 0.06 | 1 | 14 | Ctns | 0.006 | Old |
| 0.832518 | 1.188231 | 0.066 | 0.06 | 1 | 14 | Neurl3 | 0.006 | Old |
| 0.832758 | 0.279407 | 0.612 | 0.6 | 1 | 14 | Rhoj | 0.012 | Old |
| 0.832817 | 0.232734 | 0.279 | 0.28 | 1 | 14 | Ylpm1 | -0.001 | Old |
| 0.833922 | 0.209164 | 0.301 | 0.34 | 1 | 14 | Mrpl24 | -0.039 | Old |
| 0.834193 | 0.603987 | 0.104 | 0.12 | 1 | 14 | Tirap | -0.016 | Old |
| 0.834736 | 0.160751 | 0.361 | 0.38 | 1 | 14 | Vamp2 | -0.019 | Old |
| 0.834808 | 0.047313 | 0.268 | 0.26 | 1 | 14 | Taok2 | 0.008 | Old |
| 0.835077 | 0.288638 | 0.18 | 0.2 | 1 | 14 | Usp24 | -0.02 | Old |
| 0.835937 | 0.766261 | 0.087 | 0.08 | 1 | 14 | Stk39 | 0.007 | Old |
| 0.836444 | 0.677315 | 0.186 | 0.18 | 1 | 14 | Elk3 | 0.006 | Old |
| 0.836492 | 0.833818 | 0.033 | 0.04 | 1 | 14 | Tmem86a | -0.007 | Old |
| 0.836492 | 0.76494 | 0.033 | 0.04 | 1 | 14 | Slx4 | -0.007 | Old |
| 0.836541 | 0.243331 | 0.333 | 0.36 | 1 | 14 | Wwc2 | -0.027 | Old |
| 0.836863 | 1.473216 | 0.066 | 0.06 | 1 | 14 | Tmem178b | 0.006 | Old |
| 0.836863 | 1.081008 | 0.066 | 0.06 | 1 | 14 | Pdgfc | 0.006 | Old |
| 0.837486 | 0.557289 | 0.169 | 0.2 | 1 | 14 | Myo5a | -0.031 | Old |
| 0.837567 | 0.673256 | 0.137 | 0.16 | 1 | 14 | Dip2c | -0.023 | Old |
| 0.837625 | 0.715559 | 0.087 | 0.1 | 1 | 14 | Rexo1 | -0.013 | Old |
| 0.837625 | 0.524332 | 0.087 | 0.1 | 1 | 14 | Dclre1a | -0.013 | Old |
| 0.837625 | 0.348318 | 0.087 | 0.1 | 1 | 14 | G6pc3 | -0.013 | Old |
| 0.837625 | 0.283077 | 0.087 | 0.1 | 1 | 14 | Zfp386 | -0.013 | Old |
| 0.838006 | 1.119847 | 0.191 | 0.2 | 1 | 14 | Slc36a4 | -0.009 | Old |
| 0.838545 | 0.389166 | 0.24 | 0.24 | 1 | 14 | Prkab2 | 0 | Old |
| 0.839178 | 0.971543 | 0.186 | 0.18 | 1 | 14 | Cpped1 | 0.006 | Old |
| 0.839356 | 0.014283 | 0.109 | 0.1 | 1 | 14 | Ift43 | 0.009 | Old |
| 0.839397 | 0.074934 | 0.208 | 0.2 | 1 | 14 | Psmb2 | 0.008 | Old |
| 0.839805 | 0.196728 | 0.071 | 0.08 | 1 | 14 | Evi2a | -0.009 | Old |
| 0.839805 | 0.168955 | 0.071 | 0.08 | 1 | 14 | Rmnd1 | -0.009 | Old |
| 0.840022 | 0.246184 | 0.153 | 0.14 | 1 | 14 | Tbc1d14 | 0.013 | Old |
| 0.840395 | 0.372081 | 0.109 | 0.12 | 1 | 14 | Usp42 | -0.011 | Old |
| 0.840412 | 0.330117 | 0.126 | 0.14 | 1 | 14 | Cisd3 | -0.014 | Old |
| 0.840588 | 0.107929 | 0.251 | 0.24 | 1 | 14 | Plscr3 | 0.011 | Old |
| 0.841079 | 0.587293 | 0.104 | 0.12 | 1 | 14 | Snx30 | -0.016 | Old |
| 0.841079 | 0.580605 | 0.104 | 0.12 | 1 | 14 | Zfp251 | -0.016 | Old |
| 0.841213 | 1.460403 | 0.066 | 0.06 | 1 | 14 | Itpkc | 0.006 | Old |
| 0.841306 | 0.544654 | 0.126 | 0.12 | 1 | 14 | Pigm | 0.006 | Old |
| 0.84135 | 0.811545 | 0.087 | 0.1 | 1 | 14 | Lrrc20 | -0.013 | Old |
| 0.84135 | 0.414616 | 0.087 | 0.1 | 1 | 14 | Fzd8 | -0.013 | Old |
| 0.841376 | 0.074544 | 0.235 | 0.22 | 1 | 14 | Aimp1 | 0.015 | Old |
| 0.841511 | 0.033045 | 0.148 | 0.16 | 1 | 14 | Cnot7 | -0.012 | Old |
| 0.841542 | 0.123888 | 0.344 | 0.34 | 1 | 14 | Hexim1 | 0.004 | Old |
| 0.841669 | 1.00744 | 0.175 | 0.18 | 1 | 14 | Manea | -0.005 | Old |
| 0.841807 | 0.60511 | 0.18 | 0.18 | 1 | 14 | Rbm18 | 0 | Old |

| | | | | | | | | |
|---|---|---|---|---|---|---|---|---|
| 0.841807 | 0.052488 | 0.18 | 0.18 | 1 | 14 | Cul2 | 0 | Old |
| 0.842105 | 0.181021 | 0.317 | 0.32 | 1 | 14 | H6pd | -0.003 | Old |
| 0.842358 | 2.748583 | 0.033 | 0.04 | 1 | 14 | Rgs16 | -0.007 | Old |
| 0.842358 | 2.470346 | 0.033 | 0.04 | 1 | 14 | Nr1h4 | -0.007 | Old |
| 0.842358 | 1.729247 | 0.033 | 0.04 | 1 | 14 | Sap18b | -0.007 | Old |
| 0.842358 | 1.66796 | 0.033 | 0.04 | 1 | 14 | 9930104L0 | -0.007 | Old |
| 0.842358 | 1.467193 | 0.033 | 0.04 | 1 | 14 | Nt5dc1 | -0.007 | Old |
| 0.842358 | 1.255991 | 0.033 | 0.04 | 1 | 14 | Pctp | -0.007 | Old |
| 0.842358 | 1.227593 | 0.033 | 0.04 | 1 | 14 | Tnfrsf23 | -0.007 | Old |
| 0.842358 | 1.203329 | 0.033 | 0.04 | 1 | 14 | Unc5b | -0.007 | Old |
| 0.842358 | 1.122195 | 0.033 | 0.04 | 1 | 14 | Grap | -0.007 | Old |
| 0.842358 | 0.913852 | 0.033 | 0.04 | 1 | 14 | Stac2 | -0.007 | Old |
| 0.842358 | 0.879668 | 0.033 | 0.04 | 1 | 14 | Il18bp | -0.007 | Old |
| 0.842358 | 0.626601 | 0.033 | 0.04 | 1 | 14 | Ccdc191 | -0.007 | Old |
| 0.842358 | 0.622029 | 0.033 | 0.04 | 1 | 14 | Ptcd2 | -0.007 | Old |
| 0.842358 | 0.61875 | 0.033 | 0.04 | 1 | 14 | Snx11 | -0.007 | Old |
| 0.842358 | 0.452592 | 0.033 | 0.04 | 1 | 14 | Uhrf1bp1 | -0.007 | Old |
| 0.843485 | 0.209041 | 0.235 | 0.26 | 1 | 14 | Stat6 | -0.025 | Old |
| 0.843555 | 0.727693 | 0.087 | 0.08 | 1 | 14 | Cntrl | 0.007 | Old |
| 0.843555 | 0.299157 | 0.087 | 0.08 | 1 | 14 | Tex10 | 0.007 | Old |
| 0.843684 | 0.325691 | 0.268 | 0.3 | 1 | 14 | Ptpa | -0.032 | Old |
| 0.843912 | 0.019527 | 0.071 | 0.08 | 1 | 14 | Slc7a6os | -0.009 | Old |
| 0.844083 | 0.049802 | 0.333 | 0.32 | 1 | 14 | Ep300 | 0.013 | Old |
| 0.844637 | 0.342103 | 0.202 | 0.22 | 1 | 14 | Ppm1b | -0.018 | Old |
| 0.844702 | 0.233966 | 0.191 | 0.18 | 1 | 14 | Rab5b | 0.011 | Old |
| 0.845079 | 0.408373 | 0.087 | 0.1 | 1 | 14 | Tfcp2 | -0.013 | Old |
| 0.845079 | 0.026474 | 0.087 | 0.1 | 1 | 14 | Ruvbl2 | -0.013 | Old |
| 0.845567 | 0.902078 | 0.066 | 0.06 | 1 | 14 | Snx8 | 0.006 | Old |
| 0.845567 | 0.760672 | 0.066 | 0.06 | 1 | 14 | Cebpzos | 0.006 | Old |
| 0.845958 | 0.787233 | 0.24 | 0.24 | 1 | 14 | Ube2m | 0 | Old |
| 0.846006 | 0.251844 | 0.355 | 0.38 | 1 | 14 | R3hdm4 | -0.025 | Old |
| 0.846012 | 0.422202 | 0.202 | 0.2 | 1 | 14 | Prkag1 | 0.002 | Old |
| 0.846984 | 0.019498 | 0.164 | 0.18 | 1 | 14 | Mid1ip1 | -0.016 | Old |
| 0.847016 | 0.185624 | 0.546 | 0.62 | 1 | 14 | Phlda1 | -0.074 | Old |
| 0.847252 | 0.898044 | 0.175 | 0.18 | 1 | 14 | Agt | -0.005 | Old |
| 0.847474 | 0.004096 | 0.148 | 0.16 | 1 | 14 | Parp1 | -0.012 | Old |
| 0.848182 | 0.005112 | 0.607 | 0.64 | 1 | 14 | Atp2b1 | -0.033 | Old |
| 0.848358 | 0.156855 | 0.295 | 0.32 | 1 | 14 | Glyr1 | -0.025 | Old |
| 0.848515 | 0.382495 | 0.333 | 0.32 | 1 | 14 | Ppp1r18 | 0.013 | Old |
| 0.848638 | 0.464509 | 0.142 | 0.16 | 1 | 14 | Gtf2b | -0.018 | Old |
| 0.848658 | 0.205803 | 0.202 | 0.2 | 1 | 14 | Casc3 | 0.002 | Old |
| 0.848761 | 0.251185 | 0.18 | 0.2 | 1 | 14 | 1190002N1 | -0.02 | Old |
| 0.849124 | 0.182888 | 0.333 | 0.34 | 1 | 14 | Trim28 | -0.007 | Old |
| 0.849297 | 0.23215 | 0.224 | 0.22 | 1 | 14 | Elmsan1 | 0.004 | Old |
| 0.849485 | 0.022274 | 0.322 | 0.32 | 1 | 14 | Setd2 | 0.002 | Old |

| | | | | | | | | |
|---|---|---|---|---|---|---|---|---|
| 0.849642 | 0.088964 | 0.508 | 0.56 | 1 | 14 | Ash1l | -0.052 | Old |
| 0.849836 | 0.031659 | 0.164 | 0.18 | 1 | 14 | Trim30a | -0.016 | Old |
| 0.849926 | 1.446023 | 0.066 | 0.06 | 1 | 14 | Ptpn18 | 0.006 | Old |
| 0.849926 | 0.926456 | 0.066 | 0.06 | 1 | 14 | Cep135 | 0.006 | Old |
| 0.849926 | 0.68485 | 0.066 | 0.06 | 1 | 14 | Rpap1 | 0.006 | Old |
| 0.850047 | 0.732438 | 0.175 | 0.18 | 1 | 14 | Adh5 | -0.005 | Old |
| 0.850105 | 0.60028 | 0.18 | 0.18 | 1 | 14 | Insr | 0 | Old |
| 0.850133 | 0.343298 | 0.191 | 0.18 | 1 | 14 | Cdr2l | 0.011 | Old |
| 0.850229 | 0.106554 | 0.328 | 0.36 | 1 | 14 | D8Ertd738 | -0.032 | Old |
| 0.850733 | 0.271386 | 0.333 | 0.32 | 1 | 14 | Sulf1 | 0.013 | Old |
| 0.851003 | 0.645936 | 0.126 | 0.12 | 1 | 14 | Bcorl1 | 0.006 | Old |
| 0.851188 | 0.773672 | 0.087 | 0.08 | 1 | 14 | Slc9a9 | 0.007 | Old |
| 0.851188 | 0.327248 | 0.087 | 0.08 | 1 | 14 | Tfb2m | 0.007 | Old |
| 0.851188 | 0.213532 | 0.087 | 0.08 | 1 | 14 | Ranbp6 | 0.007 | Old |
| 0.852138 | 0.481622 | 0.071 | 0.08 | 1 | 14 | Sorbs3 | -0.009 | Old |
| 0.852138 | 0.206771 | 0.071 | 0.08 | 1 | 14 | AI464131 | -0.009 | Old |
| 0.852138 | 0.144304 | 0.071 | 0.08 | 1 | 14 | Mcm3 | -0.009 | Old |
| 0.852254 | 0.157888 | 0.104 | 0.1 | 1 | 14 | Gbe1 | 0.004 | Old |
| 0.8524 | 0.964757 | 0.153 | 0.18 | 1 | 14 | Dcp2 | -0.027 | Old |
| 0.852691 | 0.187865 | 0.164 | 0.18 | 1 | 14 | Rhot1 | -0.016 | Old |
| 0.852799 | 0.316457 | 0.24 | 0.28 | 1 | 14 | Ndst1 | -0.04 | Old |
| 0.852874 | 0.4133 | 0.18 | 0.18 | 1 | 14 | Xrn2 | 0 | Old |
| 0.853154 | 0.273578 | 0.126 | 0.14 | 1 | 14 | Letm1 | -0.014 | Old |
| 0.853154 | 0.047456 | 0.131 | 0.12 | 1 | 14 | Dusp16 | 0.011 | Old |
| 0.853556 | 0.01234 | 0.503 | 0.5 | 1 | 14 | Eln | 0.003 | Old |
| 0.854241 | 0.398345 | 0.126 | 0.12 | 1 | 14 | Mettl1 | 0.006 | Old |
| 0.85429 | 0.829167 | 0.066 | 0.06 | 1 | 14 | Zfp932 | 0.006 | Old |
| 0.85429 | 0.662301 | 0.066 | 0.06 | 1 | 14 | 1110034G2 | 0.006 | Old |
| 0.854604 | 0.304947 | 0.219 | 0.22 | 1 | 14 | Ankrd46 | -0.001 | Old |
| 0.85501 | 0.525532 | 0.087 | 0.08 | 1 | 14 | Faim | 0.007 | Old |
| 0.855407 | 0.207745 | 0.29 | 0.28 | 1 | 14 | Dnajc8 | 0.01 | Old |
| 0.855775 | 0.11504 | 0.104 | 0.1 | 1 | 14 | Ppp1r35 | 0.004 | Old |
| 0.855863 | 0.340021 | 0.24 | 0.24 | 1 | 14 | Sec24a | 0 | Old |
| 0.856257 | 1.181186 | 0.071 | 0.08 | 1 | 14 | Pth1r | -0.009 | Old |
| 0.856257 | 0.680247 | 0.071 | 0.08 | 1 | 14 | Plagl2 | -0.009 | Old |
| 0.856257 | 0.50127 | 0.071 | 0.08 | 1 | 14 | Gga3 | -0.009 | Old |
| 0.856257 | 0.34536 | 0.071 | 0.08 | 1 | 14 | Mlh1 | -0.009 | Old |
| 0.856257 | 0.24854 | 0.071 | 0.08 | 1 | 14 | Gmeb2 | -0.009 | Old |
| 0.856257 | 0.182464 | 0.071 | 0.08 | 1 | 14 | Cep162 | -0.009 | Old |
| 0.856258 | 0.053083 | 0.268 | 0.26 | 1 | 14 | Ccdc47 | 0.008 | Old |
| 0.856287 | 0.4883 | 0.087 | 0.1 | 1 | 14 | Mid2 | -0.013 | Old |
| 0.856287 | 0.403031 | 0.087 | 0.1 | 1 | 14 | Hmgn5 | -0.013 | Old |
| 0.856346 | 0.44781 | 0.131 | 0.12 | 1 | 14 | Zbtb22 | 0.011 | Old |
| 0.856433 | 0.028785 | 0.148 | 0.16 | 1 | 14 | Cyp51 | -0.012 | Old |
| 0.856832 | 0.13235 | 0.301 | 0.28 | 1 | 14 | Comt | 0.021 | Old |

| | | | | | | | | |
|---|---|---|---|---|---|---|---|---|
| 0.856976 | 0.353092 | 0.169 | 0.16 | 1 | 14 | Rars | 0.009 | Old |
| 0.857172 | 0.710688 | 0.219 | 0.22 | 1 | 14 | Arf2 | -0.001 | Old |
| 0.858343 | 0.080114 | 0.24 | 0.24 | 1 | 14 | Suco | 0 | Old |
| 0.858658 | 1.000424 | 0.066 | 0.06 | 1 | 14 | Pcdh1 | 0.006 | Old |
| 0.858658 | 0.83872 | 0.066 | 0.06 | 1 | 14 | Cnnm4 | 0.006 | Old |
| 0.858658 | 0.675414 | 0.066 | 0.06 | 1 | 14 | 4930503L1 | 0.006 | Old |
| 0.858658 | 0.59745 | 0.066 | 0.06 | 1 | 14 | Gm28875 | 0.006 | Old |
| 0.858658 | 0.583989 | 0.066 | 0.06 | 1 | 14 | D1Ertd622 | 0.006 | Old |
| 0.859298 | 0.825894 | 0.104 | 0.1 | 1 | 14 | Nr2c1 | 0.004 | Old |
| 0.859298 | 0.694783 | 0.104 | 0.1 | 1 | 14 | Ric8b | 0.004 | Old |
| 0.860029 | 0.655301 | 0.087 | 0.1 | 1 | 14 | Rgs19 | -0.013 | Old |
| 0.860399 | 0.040887 | 0.202 | 0.22 | 1 | 14 | Vamp3 | -0.018 | Old |
| 0.860514 | 0.365778 | 0.142 | 0.14 | 1 | 14 | BC029722 | 0.002 | Old |
| 0.860759 | 0.319939 | 0.29 | 0.3 | 1 | 14 | Stk40 | -0.01 | Old |
| 0.860954 | 0.270209 | 0.279 | 0.28 | 1 | 14 | Polr2f | -0.001 | Old |
| 0.861231 | 0.449679 | 0.158 | 0.18 | 1 | 14 | 5031425E2 | -0.022 | Old |
| 0.861421 | 0.021328 | 0.415 | 0.42 | 1 | 14 | Shisa5 | -0.005 | Old |
| 0.861758 | 0.120852 | 0.23 | 0.22 | 1 | 14 | Kremen1 | 0.01 | Old |
| 0.861803 | 0.199632 | 0.317 | 0.34 | 1 | 14 | Snx27 | -0.023 | Old |
| 0.861838 | 0.101051 | 0.322 | 0.36 | 1 | 14 | Mdm4 | -0.038 | Old |
| 0.861838 | 0.013978 | 0.322 | 0.36 | 1 | 14 | Fmo1 | -0.038 | Old |
| 0.862171 | 0.499181 | 0.016 | 0.02 | 1 | 14 | Ppp1r13b | -0.004 | Old |
| 0.862171 | 0.406272 | 0.016 | 0.02 | 1 | 14 | Ppp1r18os | -0.004 | Old |
| 0.862171 | 0.190777 | 0.016 | 0.02 | 1 | 14 | Piezo2 | -0.004 | Old |
| 0.862171 | 0.074327 | 0.016 | 0.02 | 1 | 14 | Il31ra | -0.004 | Old |
| 0.862171 | 0.043084 | 0.016 | 0.02 | 1 | 14 | Zfp85 | -0.004 | Old |
| 0.862491 | 0.170703 | 0.18 | 0.2 | 1 | 14 | Tmem131 | -0.02 | Old |
| 0.862664 | 0.215134 | 0.087 | 0.08 | 1 | 14 | Rab40b | 0.007 | Old |
| 0.862664 | 0.187649 | 0.087 | 0.08 | 1 | 14 | Clcn6 | 0.007 | Old |
| 0.862825 | 0.692959 | 0.104 | 0.1 | 1 | 14 | Dusp8 | 0.004 | Old |
| 0.863031 | 0.94115 | 0.066 | 0.06 | 1 | 14 | Scfd2 | 0.006 | Old |
| 0.863031 | 0.684881 | 0.066 | 0.06 | 1 | 14 | Ints14 | 0.006 | Old |
| 0.863031 | 0.519699 | 0.066 | 0.06 | 1 | 14 | Qtrt2 | 0.006 | Old |
| 0.863031 | 0.389651 | 0.066 | 0.06 | 1 | 14 | Smc2 | 0.006 | Old |
| 0.863031 | 0.022402 | 0.202 | 0.22 | 1 | 14 | Scaf8 | -0.018 | Old |
| 0.863076 | 0.266639 | 0.29 | 0.3 | 1 | 14 | Tnfsf12 | -0.01 | Old |
| 0.863076 | 0.120936 | 0.295 | 0.28 | 1 | 14 | Pgls | 0.015 | Old |
| 0.863913 | 0.367006 | 0.153 | 0.14 | 1 | 14 | Tmem164 | 0.013 | Old |
| 0.863913 | 0.166322 | 0.153 | 0.14 | 1 | 14 | Snrnp48 | 0.013 | Old |
| 0.864022 | 0.457051 | 0.279 | 0.3 | 1 | 14 | Plxna1 | -0.021 | Old |
| 0.864565 | 0.445845 | 0.202 | 0.2 | 1 | 14 | Mnt | 0.002 | Old |
| 0.864939 | 0.407914 | 0.273 | 0.28 | 1 | 14 | Fut11 | -0.007 | Old |
| 0.865274 | 0.704624 | 0.104 | 0.12 | 1 | 14 | B3gnt8 | -0.016 | Old |
| 0.865423 | 0.165456 | 0.24 | 0.26 | 1 | 14 | Pcmt1 | -0.02 | Old |
| 0.865893 | 0.317904 | 0.197 | 0.22 | 1 | 14 | Arl2bp | -0.023 | Old |

| | | | | | | | | |
|---|---|---|---|---|---|---|---|---|
| 0.866364 | 0.347989 | 0.284 | 0.28 | 1 | 14 | Stxbp3 | 0.004 | Old |
| 0.866496 | 0.385556 | 0.087 | 0.08 | 1 | 14 | Plscr1 | 0.007 | Old |
| 0.867007 | 0.159526 | 0.109 | 0.1 | 1 | 14 | Cpne2 | 0.009 | Old |
| 0.867408 | 0.889024 | 0.066 | 0.06 | 1 | 14 | Gm26669 | 0.006 | Old |
| 0.867408 | 0.582969 | 0.066 | 0.06 | 1 | 14 | B9d2 | 0.006 | Old |
| 0.867498 | 0.029889 | 0.251 | 0.24 | 1 | 14 | Maz | 0.011 | Old |
| 0.867523 | 0.477263 | 0.087 | 0.1 | 1 | 14 | Tmem175 | -0.013 | Old |
| 0.867971 | 0.054996 | 0.322 | 0.34 | 1 | 14 | Nek7 | -0.018 | Old |
| 0.868551 | 0.524998 | 0.202 | 0.2 | 1 | 14 | Arhgap31 | 0.002 | Old |
| 0.868784 | 0.078542 | 0.197 | 0.2 | 1 | 14 | Phf20 | -0.003 | Old |
| 0.868808 | 2.443968 | 0.044 | 0.04 | 1 | 14 | Ift172 | 0.004 | Old |
| 0.868808 | 2.441109 | 0.044 | 0.04 | 1 | 14 | Arpin | 0.004 | Old |
| 0.868808 | 1.697736 | 0.044 | 0.04 | 1 | 14 | Vps37c | 0.004 | Old |
| 0.868808 | 1.462841 | 0.044 | 0.04 | 1 | 14 | 2210408I2 | 0.004 | Old |
| 0.868808 | 1.428418 | 0.044 | 0.04 | 1 | 14 | Rap1gap2 | 0.004 | Old |
| 0.868808 | 1.419244 | 0.044 | 0.04 | 1 | 14 | Osbpl6 | 0.004 | Old |
| 0.868808 | 1.372034 | 0.044 | 0.04 | 1 | 14 | Gm16365 | 0.004 | Old |
| 0.868808 | 1.069628 | 0.044 | 0.04 | 1 | 14 | Mgst3 | 0.004 | Old |
| 0.868808 | 1.051511 | 0.044 | 0.04 | 1 | 14 | Sbk1 | 0.004 | Old |
| 0.868808 | 0.941073 | 0.044 | 0.04 | 1 | 14 | Stxbp2 | 0.004 | Old |
| 0.869015 | 0.18243 | 0.23 | 0.24 | 1 | 14 | Usp16 | -0.01 | Old |
| 0.869096 | 0.262517 | 0.317 | 0.32 | 1 | 14 | Snx2 | -0.003 | Old |
| 0.869137 | 0.168411 | 0.126 | 0.14 | 1 | 14 | Smap2 | -0.014 | Old |
| 0.869137 | 0.062309 | 0.131 | 0.12 | 1 | 14 | Wdtc1 | 0.011 | Old |
| 0.869903 | 0.530174 | 0.235 | 0.24 | 1 | 14 | Iah1 | -0.005 | Old |
| 0.870331 | 0.331487 | 0.087 | 0.08 | 1 | 14 | Wdr35 | 0.007 | Old |
| 0.870448 | 1.841561 | 0.016 | 0.02 | 1 | 14 | Colec11 | -0.004 | Old |
| 0.870448 | 0.921929 | 0.016 | 0.02 | 1 | 14 | Srd5a1 | -0.004 | Old |
| 0.870448 | 0.855695 | 0.016 | 0.02 | 1 | 14 | Tmem86b | -0.004 | Old |
| 0.870448 | 0.784433 | 0.016 | 0.02 | 1 | 14 | Clec2i | -0.004 | Old |
| 0.870448 | 0.784093 | 0.016 | 0.02 | 1 | 14 | Pus3 | -0.004 | Old |
| 0.870448 | 0.779063 | 0.016 | 0.02 | 1 | 14 | Mcrip2 | -0.004 | Old |
| 0.870448 | 0.762573 | 0.016 | 0.02 | 1 | 14 | Ccdc134 | -0.004 | Old |
| 0.870448 | 0.737128 | 0.016 | 0.02 | 1 | 14 | Irx2 | -0.004 | Old |
| 0.870448 | 0.667461 | 0.016 | 0.02 | 1 | 14 | Rpp38 | -0.004 | Old |
| 0.870448 | 0.640947 | 0.016 | 0.02 | 1 | 14 | Ddit4l | -0.004 | Old |
| 0.870448 | 0.570379 | 0.016 | 0.02 | 1 | 14 | Pilra | -0.004 | Old |
| 0.870448 | 0.566929 | 0.016 | 0.02 | 1 | 14 | Nudt6 | -0.004 | Old |
| 0.870448 | 0.472379 | 0.016 | 0.02 | 1 | 14 | Nepro | -0.004 | Old |
| 0.870448 | 0.445466 | 0.016 | 0.02 | 1 | 14 | Zfp119a | -0.004 | Old |
| 0.870448 | 0.440052 | 0.016 | 0.02 | 1 | 14 | Ngb | -0.004 | Old |
| 0.870448 | 0.434052 | 0.016 | 0.02 | 1 | 14 | Ulk4 | -0.004 | Old |
| 0.870448 | 0.387402 | 0.016 | 0.02 | 1 | 14 | Gsc | -0.004 | Old |
| 0.870448 | 0.320904 | 0.016 | 0.02 | 1 | 14 | 8030462N1 | -0.004 | Old |
| 0.870448 | 0.314064 | 0.016 | 0.02 | 1 | 14 | Zbtb11os1 | -0.004 | Old |

| | | | | | | | | |
|---|---|---|---|---|---|---|---|---|
| 0.870448 | 0.267355 | 0.016 | 0.02 | 1 | 14 | Ccdc84 | -0.004 | Old |
| 0.870448 | 0.247202 | 0.016 | 0.02 | 1 | 14 | Spata24 | -0.004 | Old |
| 0.870448 | 0.245642 | 0.016 | 0.02 | 1 | 14 | 1110008L1 | -0.004 | Old |
| 0.870448 | 0.217525 | 0.016 | 0.02 | 1 | 14 | Kifc1 | -0.004 | Old |
| 0.870448 | 0.210826 | 0.016 | 0.02 | 1 | 14 | Rhbdf2 | -0.004 | Old |
| 0.870448 | 0.195759 | 0.016 | 0.02 | 1 | 14 | Taf1c | -0.004 | Old |
| 0.870448 | 0.133548 | 0.016 | 0.02 | 1 | 14 | Pigl | -0.004 | Old |
| 0.870448 | 0.100565 | 0.016 | 0.02 | 1 | 14 | 2610020C0 | -0.004 | Old |
| 0.870448 | 0.087515 | 0.016 | 0.02 | 1 | 14 | Cyp4f18 | -0.004 | Old |
| 0.870448 | 0.080257 | 0.016 | 0.02 | 1 | 14 | Zfp971 | -0.004 | Old |
| 0.870448 | 0.027508 | 0.016 | 0.02 | 1 | 14 | Cnmd | -0.004 | Old |
| 0.870448 | 0.003689 | 0.016 | 0.02 | 1 | 14 | Ccpg1os | -0.004 | Old |
| 0.870475 | 0.025794 | 0.109 | 0.1 | 1 | 14 | Tns3 | 0.009 | Old |
| 0.870983 | 0.634488 | 0.109 | 0.12 | 1 | 14 | Ndufs8 | -0.011 | Old |
| 0.87121 | 1.516341 | 0.082 | 0.08 | 1 | 14 | Pigc | 0.002 | Old |
| 0.871789 | 1.161474 | 0.066 | 0.06 | 1 | 14 | 3110043O2 | 0.006 | Old |
| 0.871789 | 0.679021 | 0.066 | 0.06 | 1 | 14 | Orc3 | 0.006 | Old |
| 0.871789 | 0.4105 | 0.066 | 0.06 | 1 | 14 | Pdzd4 | 0.006 | Old |
| 0.871789 | 0.407346 | 0.066 | 0.06 | 1 | 14 | Ccdc61 | 0.006 | Old |
| 0.871789 | 0.316605 | 0.066 | 0.06 | 1 | 14 | Ptprf | 0.006 | Old |
| 0.871789 | 0.105628 | 0.066 | 0.06 | 1 | 14 | Dgkq | 0.006 | Old |
| 0.872802 | 0.706213 | 0.142 | 0.14 | 1 | 14 | Lonp2 | 0.002 | Old |
| 0.872812 | 0.444315 | 0.164 | 0.16 | 1 | 14 | Cog3 | 0.004 | Old |
| 0.872872 | 0.443025 | 0.142 | 0.16 | 1 | 14 | Yipf1 | -0.018 | Old |
| 0.87342 | 0.646644 | 0.104 | 0.1 | 1 | 14 | Arl6ip4 | 0.004 | Old |
| 0.87342 | 0.50985 | 0.104 | 0.1 | 1 | 14 | Nicn1 | 0.004 | Old |
| 0.873484 | 0.023707 | 0.333 | 0.34 | 1 | 14 | Bicd2 | -0.007 | Old |
| 0.874117 | 1.822604 | 0.044 | 0.04 | 1 | 14 | Rdh13 | 0.004 | Old |
| 0.874117 | 1.104983 | 0.044 | 0.04 | 1 | 14 | Ms4a6b | 0.004 | Old |
| 0.874169 | 0.058254 | 0.087 | 0.08 | 1 | 14 | Rnf19b | 0.007 | Old |
| 0.874395 | 0.665229 | 0.109 | 0.12 | 1 | 14 | Dpm2 | -0.011 | Old |
| 0.875029 | 0.983015 | 0.087 | 0.1 | 1 | 14 | Ipo13 | -0.013 | Old |
| 0.875091 | 0.206086 | 0.279 | 0.28 | 1 | 14 | Bcl10 | -0.001 | Old |
| 0.875093 | 0.513342 | 0.18 | 0.18 | 1 | 14 | Otud4 | 0 | Old |
| 0.875414 | 0.355565 | 0.257 | 0.3 | 1 | 14 | Slc16a2 | -0.043 | Old |
| 0.875592 | 0.436852 | 0.164 | 0.18 | 1 | 14 | Fbxl20 | -0.016 | Old |
| 0.875741 | 0.502159 | 0.24 | 0.24 | 1 | 14 | Abi2 | 0 | Old |
| 0.875872 | 1.081852 | 0.153 | 0.16 | 1 | 14 | Rbmx | -0.007 | Old |
| 0.876173 | 0.591054 | 0.066 | 0.06 | 1 | 14 | Mvb12b | 0.006 | Old |
| 0.876216 | 0.313033 | 0.202 | 0.22 | 1 | 14 | Dnmt3a | -0.018 | Old |
| 0.876676 | 0.409858 | 0.339 | 0.36 | 1 | 14 | Tmem234 | -0.021 | Old |
| 0.876909 | 0.46979 | 0.071 | 0.08 | 1 | 14 | Cstf2 | -0.009 | Old |
| 0.876909 | 0.467629 | 0.071 | 0.08 | 1 | 14 | Lrch1 | -0.009 | Old |
| 0.877022 | 0.088134 | 0.055 | 0.06 | 1 | 14 | Purg | -0.005 | Old |
| 0.877409 | 0.22607 | 0.148 | 0.16 | 1 | 14 | Cdk5rap3 | -0.012 | Old |

| | | | | | | | | |
|---|---|---|---|---|---|---|---|---|
| 0.877863 | 0.412515 | 0.202 | 0.2 | 1 | 14 | Csnk1g1 | 0.002 | Old |
| 0.877978 | 0.261277 | 0.366 | 0.42 | 1 | 14 | Tmem109 | -0.054 | Old |
| 0.878266 | 0.250668 | 0.525 | 0.5 | 1 | 14 | Samhd1 | 0.025 | Old |
| 0.878739 | 2.131792 | 0.016 | 0.02 | 1 | 14 | Tpm3-rs7 | -0.004 | Old |
| 0.878739 | 1.957188 | 0.016 | 0.02 | 1 | 14 | Cyp2f2 | -0.004 | Old |
| 0.878739 | 1.815728 | 0.016 | 0.02 | 1 | 14 | Stmn2 | -0.004 | Old |
| 0.878739 | 1.639915 | 0.016 | 0.02 | 1 | 14 | Slc25a53 | -0.004 | Old |
| 0.878739 | 1.589151 | 0.016 | 0.02 | 1 | 14 | Enpp6 | -0.004 | Old |
| 0.878739 | 1.500415 | 0.016 | 0.02 | 1 | 14 | Themis2 | -0.004 | Old |
| 0.878739 | 1.376268 | 0.016 | 0.02 | 1 | 14 | Cadm1 | -0.004 | Old |
| 0.878739 | 1.347834 | 0.016 | 0.02 | 1 | 14 | Tlr7 | -0.004 | Old |
| 0.878739 | 1.308524 | 0.016 | 0.02 | 1 | 14 | Egln3 | -0.004 | Old |
| 0.878739 | 1.282754 | 0.016 | 0.02 | 1 | 14 | Gm26682 | -0.004 | Old |
| 0.878739 | 1.266715 | 0.016 | 0.02 | 1 | 14 | Pcdhb16 | -0.004 | Old |
| 0.878739 | 1.24195 | 0.016 | 0.02 | 1 | 14 | Pot1b | -0.004 | Old |
| 0.878739 | 1.215621 | 0.016 | 0.02 | 1 | 14 | C920021L1 | -0.004 | Old |
| 0.878739 | 1.209728 | 0.016 | 0.02 | 1 | 14 | Gm42047 | -0.004 | Old |
| 0.878739 | 1.207038 | 0.016 | 0.02 | 1 | 14 | Cep126 | -0.004 | Old |
| 0.878739 | 1.147167 | 0.016 | 0.02 | 1 | 14 | Rtl4 | -0.004 | Old |
| 0.878739 | 1.032686 | 0.016 | 0.02 | 1 | 14 | Ccdc120 | -0.004 | Old |
| 0.878739 | 1.032286 | 0.016 | 0.02 | 1 | 14 | Sugct | -0.004 | Old |
| 0.878739 | 1.021458 | 0.016 | 0.02 | 1 | 14 | Nmnat1 | -0.004 | Old |
| 0.878739 | 0.99395 | 0.016 | 0.02 | 1 | 14 | Mad2l1 | -0.004 | Old |
| 0.878739 | 0.984254 | 0.016 | 0.02 | 1 | 14 | Slamf7 | -0.004 | Old |
| 0.878739 | 0.955517 | 0.016 | 0.02 | 1 | 14 | Dynlt1c | -0.004 | Old |
| 0.878739 | 0.929379 | 0.016 | 0.02 | 1 | 14 | Mmachc | -0.004 | Old |
| 0.878739 | 0.916234 | 0.016 | 0.02 | 1 | 14 | Gm10602 | -0.004 | Old |
| 0.878739 | 0.882025 | 0.016 | 0.02 | 1 | 14 | Uhrf1 | -0.004 | Old |
| 0.878739 | 0.869603 | 0.016 | 0.02 | 1 | 14 | Plppr3 | -0.004 | Old |
| 0.878739 | 0.866799 | 0.016 | 0.02 | 1 | 14 | Zfp35 | -0.004 | Old |
| 0.878739 | 0.842242 | 0.016 | 0.02 | 1 | 14 | Mpp2 | -0.004 | Old |
| 0.878739 | 0.83972 | 0.016 | 0.02 | 1 | 14 | Enho | -0.004 | Old |
| 0.878739 | 0.82936 | 0.016 | 0.02 | 1 | 14 | Gm4211 | -0.004 | Old |
| 0.878739 | 0.808288 | 0.016 | 0.02 | 1 | 14 | 4931414P1 | -0.004 | Old |
| 0.878739 | 0.799341 | 0.016 | 0.02 | 1 | 14 | Dtl | -0.004 | Old |
| 0.878739 | 0.798484 | 0.016 | 0.02 | 1 | 14 | Shq1 | -0.004 | Old |
| 0.878739 | 0.770212 | 0.016 | 0.02 | 1 | 14 | Gm36839 | -0.004 | Old |
| 0.878739 | 0.765872 | 0.016 | 0.02 | 1 | 14 | Neu3 | -0.004 | Old |
| 0.878739 | 0.746353 | 0.016 | 0.02 | 1 | 14 | Epb41l4b | -0.004 | Old |
| 0.878739 | 0.730335 | 0.016 | 0.02 | 1 | 14 | Gas2l3 | -0.004 | Old |
| 0.878739 | 0.723272 | 0.016 | 0.02 | 1 | 14 | Gm44777 | -0.004 | Old |
| 0.878739 | 0.707679 | 0.016 | 0.02 | 1 | 14 | Bcl2a1a | -0.004 | Old |
| 0.878739 | 0.685675 | 0.016 | 0.02 | 1 | 14 | Egfl7 | -0.004 | Old |
| 0.878739 | 0.654819 | 0.016 | 0.02 | 1 | 14 | Zfp429 | -0.004 | Old |
| 0.878739 | 0.648309 | 0.016 | 0.02 | 1 | 14 | Nme2 | -0.004 | Old |

| | | | | | | | | |
|---|---|---|---|---|---|---|---|---|
| 0.878739 | 0.630391 | 0.016 | 0.02 | 1 | 14 | Gm15050 | -0.004 | Old |
| 0.878739 | 0.628828 | 0.016 | 0.02 | 1 | 14 | Lpcat2 | -0.004 | Old |
| 0.878739 | 0.621373 | 0.016 | 0.02 | 1 | 14 | Hnmt | -0.004 | Old |
| 0.878739 | 0.581824 | 0.016 | 0.02 | 1 | 14 | Gm32856 | -0.004 | Old |
| 0.878739 | 0.540025 | 0.016 | 0.02 | 1 | 14 | Nudt14 | -0.004 | Old |
| 0.878739 | 0.528825 | 0.016 | 0.02 | 1 | 14 | Ybey | -0.004 | Old |
| 0.878739 | 0.520141 | 0.016 | 0.02 | 1 | 14 | Serp2 | -0.004 | Old |
| 0.878739 | 0.516654 | 0.016 | 0.02 | 1 | 14 | Gm9938 | -0.004 | Old |
| 0.878739 | 0.412422 | 0.016 | 0.02 | 1 | 14 | Fam60a | -0.004 | Old |
| 0.878739 | 0.401924 | 0.016 | 0.02 | 1 | 14 | Mks1 | -0.004 | Old |
| 0.878739 | 0.392417 | 0.016 | 0.02 | 1 | 14 | Gpr65 | -0.004 | Old |
| 0.878739 | 0.386226 | 0.016 | 0.02 | 1 | 14 | Mtfr1 | -0.004 | Old |
| 0.878739 | 0.376993 | 0.016 | 0.02 | 1 | 14 | Tm7sf2 | -0.004 | Old |
| 0.878739 | 0.35378 | 0.016 | 0.02 | 1 | 14 | Zfp934 | -0.004 | Old |
| 0.878739 | 0.311069 | 0.016 | 0.02 | 1 | 14 | Grin2d | -0.004 | Old |
| 0.878739 | 0.279547 | 0.016 | 0.02 | 1 | 14 | Gm32219 | -0.004 | Old |
| 0.878739 | 0.262266 | 0.016 | 0.02 | 1 | 14 | Smpdl3b | -0.004 | Old |
| 0.878739 | 0.255564 | 0.016 | 0.02 | 1 | 14 | Shank3 | -0.004 | Old |
| 0.878739 | 0.213691 | 0.016 | 0.02 | 1 | 14 | Gba2 | -0.004 | Old |
| 0.878739 | 0.181606 | 0.016 | 0.02 | 1 | 14 | Tekt2 | -0.004 | Old |
| 0.878839 | 0.070606 | 0.153 | 0.16 | 1 | 14 | Arpc5l | -0.007 | Old |
| 0.878862 | 0.040173 | 0.131 | 0.14 | 1 | 14 | Elovl6 | -0.009 | Old |
| 0.878951 | 0.323942 | 0.148 | 0.14 | 1 | 14 | Pkia | 0.008 | Old |
| 0.879196 | 0.003256 | 0.721 | 0.72 | 1 | 14 | Snrnp70 | 0.001 | Old |
| 0.879432 | 2.109251 | 0.044 | 0.04 | 1 | 14 | Mbd4 | 0.004 | Old |
| 0.879432 | 1.694602 | 0.044 | 0.04 | 1 | 14 | Polg2 | 0.004 | Old |
| 0.879432 | 1.550149 | 0.044 | 0.04 | 1 | 14 | Gcat | 0.004 | Old |
| 0.879432 | 1.118405 | 0.044 | 0.04 | 1 | 14 | Lrp5 | 0.004 | Old |
| 0.879432 | 1.114105 | 0.044 | 0.04 | 1 | 14 | Fam174b | 0.004 | Old |
| 0.879432 | 1.113829 | 0.044 | 0.04 | 1 | 14 | Tamm41 | 0.004 | Old |
| 0.879432 | 0.994385 | 0.044 | 0.04 | 1 | 14 | Parvb | 0.004 | Old |
| 0.879432 | 0.676041 | 0.044 | 0.04 | 1 | 14 | Pomgnt2 | 0.004 | Old |
| 0.879451 | 0.015641 | 0.093 | 0.1 | 1 | 14 | Mrpl16 | -0.007 | Old |
| 0.880205 | 0.086408 | 0.399 | 0.42 | 1 | 14 | Ldha | -0.021 | Old |
| 0.880207 | 0.597582 | 0.158 | 0.16 | 1 | 14 | Tubgcp5 | -0.002 | Old |
| 0.880376 | 0.14199 | 0.432 | 0.44 | 1 | 14 | Spry2 | -0.008 | Old |
| 0.880496 | 0.282033 | 0.104 | 0.1 | 1 | 14 | Fam171a2 | 0.004 | Old |
| 0.880527 | 0.353911 | 0.202 | 0.2 | 1 | 14 | Arfip1 | 0.002 | Old |
| 0.880562 | 0.375069 | 0.066 | 0.06 | 1 | 14 | Poc5 | 0.006 | Old |
| 0.880562 | 0.240868 | 0.066 | 0.06 | 1 | 14 | Cnst | 0.006 | Old |
| 0.880562 | 0.173852 | 0.066 | 0.06 | 1 | 14 | Hoxb6 | 0.006 | Old |
| 0.88105 | 0.767673 | 0.071 | 0.08 | 1 | 14 | Chchd6 | -0.009 | Old |
| 0.88105 | 0.669889 | 0.071 | 0.08 | 1 | 14 | Ctnnbl1 | -0.009 | Old |
| 0.881225 | 0.104592 | 0.109 | 0.12 | 1 | 14 | Sec22a | -0.011 | Old |
| 0.881992 | 0.228761 | 0.148 | 0.14 | 1 | 14 | 0610012G( | 0.008 | Old |

| | | | | | | | | |
|---|---|---|---|---|---|---|---|---|
| 0.882132 | 0.31328 | 0.311 | 0.32 | 1 | 14 | Arhgap29 | -0.009 | Old |
| 0.882839 | 0.143634 | 0.333 | 0.36 | 1 | 14 | Phf21a | -0.027 | Old |
| 0.883144 | 0.664239 | 0.158 | 0.16 | 1 | 14 | Cnot8 | -0.002 | Old |
| 0.883722 | 0.876548 | 0.175 | 0.18 | 1 | 14 | Rab35 | -0.005 | Old |
| 0.884038 | 0.475557 | 0.104 | 0.1 | 1 | 14 | Cry1 | 0.004 | Old |
| 0.884421 | 0.275949 | 0.164 | 0.16 | 1 | 14 | Myo6 | 0.004 | Old |
| 0.884704 | 0.045662 | 0.246 | 0.26 | 1 | 14 | Dmwd | -0.014 | Old |
| 0.884752 | 1.966359 | 0.044 | 0.04 | 1 | 14 | Lin52 | 0.004 | Old |
| 0.884752 | 1.382823 | 0.044 | 0.04 | 1 | 14 | Kbtbd3 | 0.004 | Old |
| 0.884752 | 1.201534 | 0.044 | 0.04 | 1 | 14 | Gstcd | 0.004 | Old |
| 0.884752 | 1.168449 | 0.044 | 0.04 | 1 | 14 | Gm26601 | 0.004 | Old |
| 0.884752 | 1.138666 | 0.044 | 0.04 | 1 | 14 | Lamc2 | 0.004 | Old |
| 0.884752 | 1.137818 | 0.044 | 0.04 | 1 | 14 | Ptpn7 | 0.004 | Old |
| 0.884752 | 1.084755 | 0.044 | 0.04 | 1 | 14 | Amdhd2 | 0.004 | Old |
| 0.884895 | 0.531283 | 0.197 | 0.2 | 1 | 14 | Sesn1 | -0.003 | Old |
| 0.884948 | 0.191756 | 0.235 | 0.24 | 1 | 14 | Qdpr | -0.005 | Old |
| 0.884954 | 0.545582 | 0.066 | 0.06 | 1 | 14 | Asb1 | 0.006 | Old |
| 0.884954 | 0.395401 | 0.066 | 0.06 | 1 | 14 | Vps9d1 | 0.006 | Old |
| 0.884954 | 0.280445 | 0.066 | 0.06 | 1 | 14 | Fes | 0.006 | Old |
| 0.884954 | 0.270843 | 0.066 | 0.06 | 1 | 14 | Zfp874b | 0.006 | Old |
| 0.885036 | 0.259793 | 0.142 | 0.16 | 1 | 14 | Ppp1r11 | -0.018 | Old |
| 0.885055 | 0.038194 | 0.333 | 0.36 | 1 | 14 | Rbm8a | -0.027 | Old |
| 0.885073 | 0.280126 | 0.224 | 0.22 | 1 | 14 | Slc25a51 | 0.004 | Old |
| 0.885173 | 0.217086 | 0.126 | 0.14 | 1 | 14 | Nsd2 | -0.014 | Old |
| 0.8857 | 0.184853 | 0.087 | 0.08 | 1 | 14 | Eif2s1 | 0.007 | Old |
| 0.885876 | 0.419814 | 0.448 | 0.48 | 1 | 14 | Bnip3l | -0.032 | Old |
| 0.885893 | 0.333471 | 0.317 | 0.36 | 1 | 14 | Fam168b | -0.043 | Old |
| 0.885926 | 0.365776 | 0.186 | 0.18 | 1 | 14 | Pik3ip1 | 0.006 | Old |
| 0.886293 | 0.243826 | 0.273 | 0.28 | 1 | 14 | Numbl | -0.007 | Old |
| 0.886415 | 0.204574 | 0.055 | 0.06 | 1 | 14 | 1110008P1 | -0.005 | Old |
| 0.886425 | 0.407011 | 0.148 | 0.16 | 1 | 14 | Mrps18a | -0.012 | Old |
| 0.886536 | 0.178119 | 0.317 | 0.34 | 1 | 14 | Crtc3 | -0.023 | Old |
| 0.886947 | 1.053183 | 0.082 | 0.08 | 1 | 14 | Tsen2 | 0.002 | Old |
| 0.887459 | 0.411725 | 0.235 | 0.24 | 1 | 14 | Map2k3 | -0.005 | Old |
| 0.887582 | 0.816159 | 0.104 | 0.1 | 1 | 14 | Washc5 | 0.004 | Old |
| 0.888318 | 0.730895 | 0.12 | 0.12 | 1 | 14 | Dnajc25 | 0 | Old |
| 0.888335 | 0.17183 | 0.186 | 0.2 | 1 | 14 | Dnttip2 | -0.014 | Old |
| 0.888386 | 0.418613 | 0.126 | 0.14 | 1 | 14 | Aup1 | -0.014 | Old |
| 0.888451 | 0.139846 | 0.306 | 0.32 | 1 | 14 | Ccnd3 | -0.014 | Old |
| 0.888522 | 0.001804 | 0.169 | 0.16 | 1 | 14 | Tmem248 | 0.009 | Old |
| 0.888671 | 0.314229 | 0.273 | 0.28 | 1 | 14 | Ndufa10 | -0.007 | Old |
| 0.889257 | 0.084707 | 0.415 | 0.4 | 1 | 14 | Mcrip1 | 0.015 | Old |
| 0.889631 | 0.026591 | 0.251 | 0.24 | 1 | 14 | 4931406P1 | 0.011 | Old |
| 0.88996 | 0.576382 | 0.164 | 0.18 | 1 | 14 | Arrb1 | -0.016 | Old |
| 0.890078 | 1.535456 | 0.044 | 0.04 | 1 | 14 | Gm43848 | 0.004 | Old |

| | | | | | | | | |
|---|---|---|---|---|---|---|---|---|
| 0.890078 | 1.245218 | 0.044 | 0.04 | 1 | 14 | Sp4 | 0.004 | Old |
| 0.890078 | 1.220289 | 0.044 | 0.04 | 1 | 14 | 1600002K0 | 0.004 | Old |
| 0.890078 | 1.21718 | 0.044 | 0.04 | 1 | 14 | Olfm2 | 0.004 | Old |
| 0.890078 | 0.894081 | 0.044 | 0.04 | 1 | 14 | Lin9 | 0.004 | Old |
| 0.890078 | 0.864467 | 0.044 | 0.04 | 1 | 14 | Pdk1 | 0.004 | Old |
| 0.890078 | 0.749643 | 0.044 | 0.04 | 1 | 14 | Ppp2r2b | 0.004 | Old |
| 0.890277 | 0.250661 | 0.197 | 0.2 | 1 | 14 | Mmadhc | -0.003 | Old |
| 0.890505 | 0.141856 | 0.093 | 0.1 | 1 | 14 | Mynn | -0.007 | Old |
| 0.890889 | 1.201004 | 0.082 | 0.08 | 1 | 14 | Samd1 | 0.002 | Old |
| 0.890889 | 0.847087 | 0.082 | 0.08 | 1 | 14 | Mov10 | 0.002 | Old |
| 0.891601 | 1.008955 | 0.126 | 0.14 | 1 | 14 | Ninj1 | -0.014 | Old |
| 0.891636 | 0.716674 | 0.12 | 0.14 | 1 | 14 | Rrm1 | -0.02 | Old |
| 0.891822 | 0.389096 | 0.18 | 0.18 | 1 | 14 | Stam2 | 0 | Old |
| 0.89297 | 0.409922 | 0.197 | 0.2 | 1 | 14 | Lims2 | -0.003 | Old |
| 0.893268 | 0.218472 | 0.126 | 0.12 | 1 | 14 | Axin1 | 0.006 | Old |
| 0.893407 | 0.517531 | 0.251 | 0.28 | 1 | 14 | Zranb1 | -0.029 | Old |
| 0.893492 | 0.536606 | 0.071 | 0.08 | 1 | 14 | Hspa12b | -0.009 | Old |
| 0.893573 | 0.079555 | 0.945 | 0.86 | 1 | 14 | Zfp36l1 | 0.085 | Old |
| 0.894177 | 0.52907 | 0.142 | 0.16 | 1 | 14 | Tbc1d17 | -0.018 | Old |
| 0.894203 | 0.11727 | 0.404 | 0.42 | 1 | 14 | Zfp148 | -0.016 | Old |
| 0.894378 | 0.519567 | 0.142 | 0.14 | 1 | 14 | Mrpl34 | 0.002 | Old |
| 0.89447 | 1.777579 | 0.098 | 0.1 | 1 | 14 | Mtr | -0.002 | Old |
| 0.894616 | 0.646074 | 0.18 | 0.18 | 1 | 14 | Bax | 0 | Old |
| 0.894676 | 0.71323 | 0.104 | 0.1 | 1 | 14 | Gsdmd | 0.004 | Old |
| 0.894676 | 0.169414 | 0.104 | 0.1 | 1 | 14 | E130102H2 | 0.004 | Old |
| 0.894732 | 0.236435 | 0.208 | 0.2 | 1 | 14 | Glo1 | 0.008 | Old |
| 0.894732 | 0.225086 | 0.202 | 0.22 | 1 | 14 | Xpo6 | -0.018 | Old |
| 0.894905 | 0.660737 | 0.158 | 0.16 | 1 | 14 | Trak1 | -0.002 | Old |
| 0.89491 | 0.425251 | 0.109 | 0.12 | 1 | 14 | Brwd1 | -0.011 | Old |
| 0.894946 | 0.865448 | 0.12 | 0.12 | 1 | 14 | Nrbf2 | 0 | Old |
| 0.895408 | 1.037458 | 0.044 | 0.04 | 1 | 14 | Glrx | 0.004 | Old |
| 0.895408 | 0.599949 | 0.044 | 0.04 | 1 | 14 | Zfp959 | 0.004 | Old |
| 0.895824 | 0.133632 | 0.055 | 0.06 | 1 | 14 | Pigv | -0.005 | Old |
| 0.896533 | 0.158186 | 0.126 | 0.12 | 1 | 14 | Arfgap2 | 0.006 | Old |
| 0.896998 | 0.001179 | 0.475 | 0.46 | 1 | 14 | Hectd1 | 0.015 | Old |
| 0.89755 | 0.149338 | 0.333 | 0.32 | 1 | 14 | Tnrc6c | 0.013 | Old |
| 0.897612 | 0.887163 | 0.087 | 0.1 | 1 | 14 | Grk5 | -0.013 | Old |
| 0.897645 | 1.259665 | 0.071 | 0.08 | 1 | 14 | Zbtb33 | -0.009 | Old |
| 0.897645 | 1.138415 | 0.071 | 0.08 | 1 | 14 | Armc9 | -0.009 | Old |
| 0.897704 | 0.161803 | 0.355 | 0.36 | 1 | 14 | Akt1 | -0.005 | Old |
| 0.897911 | 0.847373 | 0.219 | 0.24 | 1 | 14 | Nectin1 | -0.021 | Old |
| 0.898152 | 0.892679 | 0.066 | 0.06 | 1 | 14 | Fam26e | 0.006 | Old |
| 0.898152 | 0.332489 | 0.066 | 0.06 | 1 | 14 | Impdh1 | 0.006 | Old |
| 0.898152 | 0.136899 | 0.066 | 0.06 | 1 | 14 | Hlf | 0.006 | Old |
| 0.898262 | 0.913615 | 0.12 | 0.12 | 1 | 14 | Ctbp2 | 0 | Old |

| | | | | | | | | |
|---|---|---|---|---|---|---|---|---|
| 0.898641 | 0.4057 | 0.224 | 0.24 | 1 | 14 | Nrbp1 | -0.016 | Old |
| 0.89878 | 1.341488 | 0.082 | 0.08 | 1 | 14 | Mlycd | 0.002 | Old |
| 0.89878 | 1.242377 | 0.082 | 0.08 | 1 | 14 | Pdgfa | 0.002 | Old |
| 0.89878 | 1.065072 | 0.082 | 0.08 | 1 | 14 | Malsu1 | 0.002 | Old |
| 0.899301 | 0.184376 | 0.186 | 0.2 | 1 | 14 | Txlna | -0.014 | Old |
| 0.899301 | 0.142804 | 0.191 | 0.18 | 1 | 14 | Lbr | 0.011 | Old |
| 0.8998 | 0.062203 | 0.126 | 0.12 | 1 | 14 | Lman2l | 0.006 | Old |
| 0.900482 | 0.247473 | 0.224 | 0.22 | 1 | 14 | 2310035C2 | 0.004 | Old |
| 0.900534 | 0.309466 | 0.055 | 0.06 | 1 | 14 | Vstm2b | -0.005 | Old |
| 0.900534 | 0.147337 | 0.055 | 0.06 | 1 | 14 | Pfkfb3 | -0.005 | Old |
| 0.900534 | 0.01778 | 0.055 | 0.06 | 1 | 14 | Rnf144a | -0.005 | Old |
| 0.900558 | 0.326863 | 0.142 | 0.14 | 1 | 14 | Dhdh | 0.002 | Old |
| 0.900719 | 0.396149 | 0.251 | 0.28 | 1 | 14 | Herc2 | -0.029 | Old |
| 0.901049 | 0.261298 | 0.131 | 0.14 | 1 | 14 | Tdrd7 | -0.009 | Old |
| 0.901049 | 0.080741 | 0.131 | 0.14 | 1 | 14 | Pcdhga12 | -0.009 | Old |
| 0.901485 | 0.273199 | 0.148 | 0.16 | 1 | 14 | Utp6 | -0.012 | Old |
| 0.901581 | 0.726837 | 0.12 | 0.12 | 1 | 14 | Osbpl2 | 0 | Old |
| 0.901623 | 0.041414 | 0.328 | 0.32 | 1 | 14 | N4bp2l2 | 0.008 | Old |
| 0.901801 | 1.027005 | 0.071 | 0.08 | 1 | 14 | Cdca4 | -0.009 | Old |
| 0.901881 | 0.054913 | 0.164 | 0.16 | 1 | 14 | 1810013L2 | 0.004 | Old |
| 0.901885 | 0.236355 | 0.202 | 0.2 | 1 | 14 | Cops5 | 0.002 | Old |
| 0.901898 | 0.05105 | 0.852 | 0.9 | 1 | 14 | Rpl24 | -0.048 | Old |
| 0.902046 | 0.134332 | 0.186 | 0.2 | 1 | 14 | Acot9 | -0.014 | Old |
| 0.902143 | 0.02908 | 0.208 | 0.22 | 1 | 14 | Ahdc1 | -0.012 | Old |
| 0.902471 | 0.020767 | 0.23 | 0.22 | 1 | 14 | S100pbp | 0.01 | Old |
| 0.902529 | 0.432386 | 0.186 | 0.18 | 1 | 14 | Dnajb11 | 0.006 | Old |
| 0.902814 | 0.026548 | 0.333 | 0.36 | 1 | 14 | Pdcd6 | -0.027 | Old |
| 0.903753 | 0.587122 | 0.197 | 0.2 | 1 | 14 | Polr2c | -0.003 | Old |
| 0.90456 | 0.258659 | 0.202 | 0.2 | 1 | 14 | Chm | 0.002 | Old |
| 0.905071 | 0.225432 | 0.235 | 0.24 | 1 | 14 | Kidins220 | -0.005 | Old |
| 0.905196 | 0.045795 | 0.109 | 0.12 | 1 | 14 | Las1l | -0.011 | Old |
| 0.905248 | 0.322948 | 0.055 | 0.06 | 1 | 14 | Ttll1 | -0.005 | Old |
| 0.905248 | 0.141779 | 0.055 | 0.06 | 1 | 14 | Pald1 | -0.005 | Old |
| 0.905274 | 0.412642 | 0.366 | 0.38 | 1 | 14 | Psmd13 | -0.014 | Old |
| 0.905277 | 0.019858 | 0.093 | 0.1 | 1 | 14 | Zfp142 | -0.007 | Old |
| 0.905701 | 0.188139 | 0.24 | 0.24 | 1 | 14 | Myo1c | 0 | Old |
| 0.906083 | 1.269631 | 0.044 | 0.04 | 1 | 14 | Asrgl1 | 0.004 | Old |
| 0.906083 | 0.816328 | 0.044 | 0.04 | 1 | 14 | P4htm | 0.004 | Old |
| 0.906083 | 0.615005 | 0.044 | 0.04 | 1 | 14 | Tbxas1 | 0.004 | Old |
| 0.906083 | 0.465925 | 0.044 | 0.04 | 1 | 14 | Dgkz | 0.004 | Old |
| 0.906083 | 0.457844 | 0.044 | 0.04 | 1 | 14 | Thap7 | 0.004 | Old |
| 0.906083 | 0.288418 | 0.044 | 0.04 | 1 | 14 | Clec12a | 0.004 | Old |
| 0.906083 | 0.229297 | 0.044 | 0.04 | 1 | 14 | Afg1l | 0.004 | Old |
| 0.906083 | 0.115795 | 0.044 | 0.04 | 1 | 14 | Pfkl | 0.004 | Old |
| 0.906689 | 0.347479 | 0.158 | 0.16 | 1 | 14 | Usp40 | -0.002 | Old |

| | | | | | | | | |
|---|---|---|---|---|---|---|---|---|
| 0.906966 | 0.083124 | 0.066 | 0.06 | 1 | 14 | Rin3 | 0.006 | Old |
| 0.907403 | 0.20401 | 0.131 | 0.14 | 1 | 14 | Tor3a | -0.009 | Old |
| 0.90752 | 0.208653 | 0.148 | 0.16 | 1 | 14 | Phax | -0.012 | Old |
| 0.907539 | 0.116282 | 0.191 | 0.18 | 1 | 14 | Ppp2r5e | 0.011 | Old |
| 0.907712 | 0.159592 | 0.164 | 0.16 | 1 | 14 | Pigp | 0.004 | Old |
| 0.908223 | 0.842901 | 0.12 | 0.12 | 1 | 14 | Tubgcp3 | 0 | Old |
| 0.908223 | 0.618487 | 0.12 | 0.12 | 1 | 14 | Fkbp15 | 0 | Old |
| 0.908603 | 0.140087 | 0.153 | 0.16 | 1 | 14 | Cfap97 | -0.007 | Old |
| 0.90889 | 0.359896 | 0.104 | 0.1 | 1 | 14 | 4921524J1 | 0.004 | Old |
| 0.908936 | 1.143001 | 0.087 | 0.1 | 1 | 14 | Gm46430 | -0.013 | Old |
| 0.908957 | 1.081881 | 0.098 | 0.1 | 1 | 14 | Rnf31 | -0.002 | Old |
| 0.90946 | 0.203178 | 0.18 | 0.2 | 1 | 14 | Zufsp | -0.02 | Old |
| 0.909486 | 0.015928 | 0.333 | 0.36 | 1 | 14 | Apbb2 | -0.027 | Old |
| 0.909495 | 0.021516 | 0.23 | 0.24 | 1 | 14 | Sec23ip | -0.01 | Old |
| 0.909839 | 0.21227 | 0.142 | 0.14 | 1 | 14 | Sil1 | 0.002 | Old |
| 0.909965 | 0.258749 | 0.055 | 0.06 | 1 | 14 | Rbbp8 | -0.005 | Old |
| 0.910122 | 0.842226 | 0.071 | 0.08 | 1 | 14 | Cad | -0.009 | Old |
| 0.910583 | 0.081409 | 0.131 | 0.14 | 1 | 14 | Crtc2 | -0.009 | Old |
| 0.910629 | 0.231808 | 0.328 | 0.32 | 1 | 14 | Tril | 0.008 | Old |
| 0.910635 | 0.750262 | 0.082 | 0.08 | 1 | 14 | Zfp37 | 0.002 | Old |
| 0.910635 | 0.683771 | 0.082 | 0.08 | 1 | 14 | Nat10 | 0.002 | Old |
| 0.911427 | 0.986801 | 0.044 | 0.04 | 1 | 14 | Pla2g2d | 0.004 | Old |
| 0.912046 | 0.683828 | 0.213 | 0.22 | 1 | 14 | Map2k7 | -0.007 | Old |
| 0.912405 | 0.136959 | 0.29 | 0.32 | 1 | 14 | Tpd52l2 | -0.03 | Old |
| 0.912727 | 0.014594 | 0.508 | 0.52 | 1 | 14 | Emc7 | -0.012 | Old |
| 0.912885 | 0.019815 | 0.126 | 0.12 | 1 | 14 | Nup85 | 0.006 | Old |
| 0.912935 | 0.366978 | 0.142 | 0.14 | 1 | 14 | Bag5 | 0.002 | Old |
| 0.913037 | 0.071365 | 0.191 | 0.18 | 1 | 14 | Commd7 | 0.011 | Old |
| 0.913507 | 0.017147 | 0.115 | 0.12 | 1 | 14 | Tada3 | -0.005 | Old |
| 0.914591 | 0.67832 | 0.082 | 0.08 | 1 | 14 | Efl1 | 0.002 | Old |
| 0.914685 | 1.357749 | 0.055 | 0.06 | 1 | 14 | Kazald1 | -0.005 | Old |
| 0.914685 | 0.188357 | 0.055 | 0.06 | 1 | 14 | Actr5 | -0.005 | Old |
| 0.91526 | 0.014651 | 0.749 | 0.66 | 1 | 14 | Sparcl1 | 0.089 | Old |
| 0.915963 | 0.088982 | 0.208 | 0.2 | 1 | 14 | Copz2 | 0.008 | Old |
| 0.916082 | 0.136598 | 0.251 | 0.26 | 1 | 14 | Il17d | -0.009 | Old |
| 0.916377 | 0.392132 | 0.093 | 0.1 | 1 | 14 | Riok2 | -0.007 | Old |
| 0.916509 | 0.262148 | 0.077 | 0.08 | 1 | 14 | Plaur | -0.003 | Old |
| 0.916581 | 0.334238 | 0.148 | 0.16 | 1 | 14 | Rybp | -0.012 | Old |
| 0.916775 | 0.106454 | 0.044 | 0.04 | 1 | 14 | Haus7 | 0.004 | Old |
| 0.916946 | 0.254702 | 0.131 | 0.14 | 1 | 14 | Acbd6 | -0.009 | Old |
| 0.916946 | 0.239468 | 0.131 | 0.14 | 1 | 14 | Pogk | -0.009 | Old |
| 0.916946 | 0.005903 | 0.131 | 0.14 | 1 | 14 | Phrf1 | -0.009 | Old |
| 0.917811 | 0.157011 | 0.23 | 0.22 | 1 | 14 | Derl2 | 0.01 | Old |
| 0.918024 | 0.405731 | 0.262 | 0.3 | 1 | 14 | Acvr2a | -0.038 | Old |
| 0.918207 | 0.010607 | 0.169 | 0.18 | 1 | 14 | Hsph1 | -0.011 | Old |

| | | | | | | | | |
|---|---|---|---|---|---|---|---|---|
| 0.918549 | 0.932537 | 0.082 | 0.08 | 1 | 14 | Dera | 0.002 | Old |
| 0.918565 | 0.016791 | 0.366 | 0.4 | 1 | 14 | Timm13 | -0.034 | Old |
| 0.918617 | 0.121033 | 0.148 | 0.14 | 1 | 14 | Commd2 | 0.008 | Old |
| 0.918758 | 0.11369 | 0.246 | 0.24 | 1 | 14 | Pip5k1a | 0.006 | Old |
| 0.918989 | 0.010071 | 0.29 | 0.3 | 1 | 14 | BC005624 | -0.01 | Old |
| 0.919132 | 0.625887 | 0.142 | 0.14 | 1 | 14 | Ifih1 | 0.002 | Old |
| 0.919272 | 0.138539 | 0.246 | 0.26 | 1 | 14 | Ccl7 | -0.014 | Old |
| 0.919408 | 1.112074 | 0.055 | 0.06 | 1 | 14 | Tmem115 | -0.005 | Old |
| 0.919408 | 1.035532 | 0.055 | 0.06 | 1 | 14 | Inip | -0.005 | Old |
| 0.919408 | 0.884923 | 0.055 | 0.06 | 1 | 14 | Tubgcp2 | -0.005 | Old |
| 0.919408 | 0.884289 | 0.055 | 0.06 | 1 | 14 | Rab43 | -0.005 | Old |
| 0.919408 | 0.311248 | 0.055 | 0.06 | 1 | 14 | Ccnc | -0.005 | Old |
| 0.919959 | 0.279887 | 0.197 | 0.2 | 1 | 14 | Nup62 | -0.003 | Old |
| 0.920081 | 0.228242 | 0.093 | 0.1 | 1 | 14 | Pde12 | -0.007 | Old |
| 0.920129 | 0.014126 | 0.131 | 0.14 | 1 | 14 | Plekha3 | -0.009 | Old |
| 0.92044 | 0.380563 | 0.175 | 0.18 | 1 | 14 | Nol11 | -0.005 | Old |
| 0.920568 | 0.090353 | 0.077 | 0.08 | 1 | 14 | Zfp930 | -0.003 | Old |
| 0.921525 | 0.714491 | 0.12 | 0.12 | 1 | 14 | Tspyl2 | 0 | Old |
| 0.921688 | 0.028262 | 0.891 | 0.9 | 1 | 14 | Dusp1 | -0.009 | Old |
| 0.922372 | 0.340474 | 0.109 | 0.12 | 1 | 14 | Nit1 | -0.011 | Old |
| 0.92251 | 0.634064 | 0.082 | 0.08 | 1 | 14 | Mbtps2 | 0.002 | Old |
| 0.92251 | 0.484802 | 0.082 | 0.08 | 1 | 14 | Rftn2 | 0.002 | Old |
| 0.92262 | 1.321722 | 0.071 | 0.08 | 1 | 14 | Rilpl1 | -0.009 | Old |
| 0.922702 | 0.440328 | 0.251 | 0.28 | 1 | 14 | Ncoa7 | -0.029 | Old |
| 0.923839 | 1.203833 | 0.126 | 0.14 | 1 | 14 | Yars | -0.014 | Old |
| 0.923839 | 0.430838 | 0.126 | 0.14 | 1 | 14 | Irf2bp1 | -0.014 | Old |
| 0.924134 | 0.810743 | 0.055 | 0.06 | 1 | 14 | 1110065P2 | -0.005 | Old |
| 0.924134 | 0.685558 | 0.055 | 0.06 | 1 | 14 | Fuca2 | -0.005 | Old |
| 0.924134 | 0.456524 | 0.055 | 0.06 | 1 | 14 | Ccdc163 | -0.005 | Old |
| 0.924443 | 1.055238 | 0.06 | 0.06 | 1 | 14 | Dym | 0 | Old |
| 0.924629 | 0.019376 | 0.077 | 0.08 | 1 | 14 | Psmg3 | -0.003 | Old |
| 0.924854 | 1.139741 | 0.12 | 0.12 | 1 | 14 | Acap3 | 0 | Old |
| 0.924859 | 2.379528 | 0.022 | 0.02 | 1 | 14 | 2700038G2 | 0.002 | Old |
| 0.924859 | 2.308249 | 0.022 | 0.02 | 1 | 14 | Nusap1 | 0.002 | Old |
| 0.924859 | 2.181471 | 0.022 | 0.02 | 1 | 14 | Tdrp | 0.002 | Old |
| 0.924859 | 1.946779 | 0.022 | 0.02 | 1 | 14 | Nudt12 | 0.002 | Old |
| 0.924859 | 1.918513 | 0.022 | 0.02 | 1 | 14 | Nadsyn1 | 0.002 | Old |
| 0.924859 | 1.708228 | 0.022 | 0.02 | 1 | 14 | Ccdc39 | 0.002 | Old |
| 0.924859 | 1.683447 | 0.022 | 0.02 | 1 | 14 | Papln | 0.002 | Old |
| 0.924859 | 1.657754 | 0.022 | 0.02 | 1 | 14 | Lat2 | 0.002 | Old |
| 0.924859 | 1.544131 | 0.022 | 0.02 | 1 | 14 | Bend5 | 0.002 | Old |
| 0.924859 | 1.467929 | 0.022 | 0.02 | 1 | 14 | Mtfp1 | 0.002 | Old |
| 0.924859 | 1.451849 | 0.022 | 0.02 | 1 | 14 | Zfp810 | 0.002 | Old |
| 0.924859 | 1.426842 | 0.022 | 0.02 | 1 | 14 | Gins1 | 0.002 | Old |
| 0.924859 | 1.356785 | 0.022 | 0.02 | 1 | 14 | Ube2cbp | 0.002 | Old |

| | | | | | | | | |
|---|---|---|---|---|---|---|---|---|
| 0.924859 | 1.311178 | 0.022 | 0.02 | 1 | 14 | Tinf2 | 0.002 | Old |
| 0.924859 | 1.241038 | 0.022 | 0.02 | 1 | 14 | Gm6710 | 0.002 | Old |
| 0.924859 | 1.210597 | 0.022 | 0.02 | 1 | 14 | D11Wsu47 | 0.002 | Old |
| 0.924859 | 1.182328 | 0.022 | 0.02 | 1 | 14 | Kng2 | 0.002 | Old |
| 0.924859 | 1.154686 | 0.022 | 0.02 | 1 | 14 | Vav1 | 0.002 | Old |
| 0.924859 | 1.143682 | 0.022 | 0.02 | 1 | 14 | Siglech | 0.002 | Old |
| 0.924859 | 1.109087 | 0.022 | 0.02 | 1 | 14 | Nr3c2 | 0.002 | Old |
| 0.924859 | 1.089791 | 0.022 | 0.02 | 1 | 14 | Lrmp | 0.002 | Old |
| 0.924859 | 1.077248 | 0.022 | 0.02 | 1 | 14 | 5430416N( | 0.002 | Old |
| 0.924859 | 1.059629 | 0.022 | 0.02 | 1 | 14 | Ppcs | 0.002 | Old |
| 0.924859 | 1.05211 | 0.022 | 0.02 | 1 | 14 | 2510017J1 | 0.002 | Old |
| 0.924859 | 0.971704 | 0.022 | 0.02 | 1 | 14 | Ccl19 | 0.002 | Old |
| 0.924859 | 0.849338 | 0.022 | 0.02 | 1 | 14 | Mob3b | 0.002 | Old |
| 0.924859 | 0.843927 | 0.022 | 0.02 | 1 | 14 | Pclaf | 0.002 | Old |
| 0.924859 | 0.769478 | 0.022 | 0.02 | 1 | 14 | Tmem206 | 0.002 | Old |
| 0.924859 | 0.760881 | 0.022 | 0.02 | 1 | 14 | Bin2 | 0.002 | Old |
| 0.924859 | 0.744415 | 0.022 | 0.02 | 1 | 14 | Hoxd10 | 0.002 | Old |
| 0.924859 | 0.717623 | 0.022 | 0.02 | 1 | 14 | Ggnbp1 | 0.002 | Old |
| 0.925921 | 0.613481 | 0.137 | 0.14 | 1 | 14 | Srxn1 | -0.003 | Old |
| 0.925993 | 0.051759 | 0.126 | 0.12 | 1 | 14 | Abhd13 | 0.006 | Old |
| 0.926028 | 0.010249 | 0.169 | 0.16 | 1 | 14 | Ifnar1 | 0.009 | Old |
| 0.926472 | 0.666013 | 0.082 | 0.08 | 1 | 14 | Gdap2 | 0.002 | Old |
| 0.926686 | 0.18467 | 0.443 | 0.48 | 1 | 14 | Raph1 | -0.037 | Old |
| 0.926773 | 0.503384 | 0.224 | 0.24 | 1 | 14 | Med1 | -0.016 | Old |
| 0.927086 | 0.328845 | 0.383 | 0.42 | 1 | 14 | Chp1 | -0.037 | Old |
| 0.927339 | 0.147457 | 0.197 | 0.22 | 1 | 14 | Stx16 | -0.023 | Old |
| 0.927494 | 0.527209 | 0.093 | 0.1 | 1 | 14 | Hmcn1 | -0.007 | Old |
| 0.927869 | 0.449601 | 0.344 | 0.36 | 1 | 14 | Zadh2 | -0.016 | Old |
| 0.928226 | 0.22898 | 0.18 | 0.18 | 1 | 14 | Nsf | 0 | Old |
| 0.928863 | 1.284103 | 0.055 | 0.06 | 1 | 14 | Acd | -0.005 | Old |
| 0.928863 | 1.117121 | 0.055 | 0.06 | 1 | 14 | Nt5m | -0.005 | Old |
| 0.928863 | 0.771183 | 0.055 | 0.06 | 1 | 14 | Ints4 | -0.005 | Old |
| 0.92901 | 2.217954 | 0.06 | 0.06 | 1 | 14 | Plxna3 | 0 | Old |
| 0.92901 | 1.479619 | 0.06 | 0.06 | 1 | 14 | Eif2d | 0 | Old |
| 0.929597 | 0.12708 | 0.219 | 0.22 | 1 | 14 | Atp8b2 | -0.001 | Old |
| 0.929986 | 0.609166 | 0.279 | 0.3 | 1 | 14 | Zhx1 | -0.021 | Old |
| 0.930436 | 0.104906 | 0.082 | 0.08 | 1 | 14 | Dpp3 | 0.002 | Old |
| 0.93093 | 0.297567 | 0.377 | 0.4 | 1 | 14 | Srpx | -0.023 | Old |
| 0.931148 | 0.09944 | 0.208 | 0.22 | 1 | 14 | Rufy2 | -0.012 | Old |
| 0.931996 | 0.060166 | 0.279 | 0.28 | 1 | 14 | Ubr4 | -0.001 | Old |
| 0.932297 | 0.067314 | 0.344 | 0.36 | 1 | 14 | Psma2 | -0.016 | Old |
| 0.932354 | 0.29628 | 0.279 | 0.3 | 1 | 14 | Ggh | -0.021 | Old |
| 0.932354 | 1.787821 | 0.022 | 0.02 | 1 | 14 | Cxcl9 | 0.002 | Old |
| 0.932354 | 1.66542 | 0.022 | 0.02 | 1 | 14 | Rac2 | 0.002 | Old |
| 0.932354 | 1.423041 | 0.022 | 0.02 | 1 | 14 | Tom1l1 | 0.002 | Old |

| | | | | | | | | |
|---|---|---|---|---|---|---|---|---|
| 0.932354 | 1.358709 | 0.022 | 0.02 | 1 | 14 | Prr7 | 0.002 | Old |
| 0.932354 | 1.325329 | 0.022 | 0.02 | 1 | 14 | Ppfia2 | 0.002 | Old |
| 0.932354 | 1.319468 | 0.022 | 0.02 | 1 | 14 | Fam105a | 0.002 | Old |
| 0.932354 | 1.113009 | 0.022 | 0.02 | 1 | 14 | Ccdc114 | 0.002 | Old |
| 0.932354 | 1.060549 | 0.022 | 0.02 | 1 | 14 | Pnpo | 0.002 | Old |
| 0.932354 | 1.016508 | 0.022 | 0.02 | 1 | 14 | Gm10605 | 0.002 | Old |
| 0.932354 | 0.956435 | 0.022 | 0.02 | 1 | 14 | Zfp963 | 0.002 | Old |
| 0.932354 | 0.901634 | 0.022 | 0.02 | 1 | 14 | Aim2 | 0.002 | Old |
| 0.932354 | 0.848701 | 0.022 | 0.02 | 1 | 14 | Zfp59 | 0.002 | Old |
| 0.932354 | 0.841969 | 0.022 | 0.02 | 1 | 14 | Pik3r6 | 0.002 | Old |
| 0.932354 | 0.789326 | 0.022 | 0.02 | 1 | 14 | AL591952.: | 0.002 | Old |
| 0.932354 | 0.733159 | 0.022 | 0.02 | 1 | 14 | Mttp | 0.002 | Old |
| 0.932354 | 0.726079 | 0.022 | 0.02 | 1 | 14 | Bid | 0.002 | Old |
| 0.932354 | 0.719837 | 0.022 | 0.02 | 1 | 14 | Zfp13 | 0.002 | Old |
| 0.932354 | 0.710885 | 0.022 | 0.02 | 1 | 14 | Gemin6 | 0.002 | Old |
| 0.932354 | 0.694056 | 0.022 | 0.02 | 1 | 14 | Clec14a | 0.002 | Old |
| 0.932354 | 0.668787 | 0.022 | 0.02 | 1 | 14 | Samd10 | 0.002 | Old |
| 0.932354 | 0.632291 | 0.022 | 0.02 | 1 | 14 | Nmnat2 | 0.002 | Old |
| 0.932354 | 0.622381 | 0.022 | 0.02 | 1 | 14 | Jchain | 0.002 | Old |
| 0.932354 | 0.617199 | 0.022 | 0.02 | 1 | 14 | Alg8 | 0.002 | Old |
| 0.932354 | 0.594672 | 0.022 | 0.02 | 1 | 14 | Fcor | 0.002 | Old |
| 0.932354 | 0.582222 | 0.022 | 0.02 | 1 | 14 | Wdr74 | 0.002 | Old |
| 0.932354 | 0.557295 | 0.022 | 0.02 | 1 | 14 | Ppp4r4 | 0.002 | Old |
| 0.932354 | 0.465507 | 0.022 | 0.02 | 1 | 14 | AC154760. | 0.002 | Old |
| 0.932354 | 0.433478 | 0.022 | 0.02 | 1 | 14 | Zfp870 | 0.002 | Old |
| 0.932354 | 0.428882 | 0.022 | 0.02 | 1 | 14 | Zdhhc12 | 0.002 | Old |
| 0.932354 | 0.405897 | 0.022 | 0.02 | 1 | 14 | Ypel1 | 0.002 | Old |
| 0.932354 | 0.290899 | 0.022 | 0.02 | 1 | 14 | Eppk1 | 0.002 | Old |
| 0.932354 | 0.227575 | 0.022 | 0.02 | 1 | 14 | Asb4 | 0.002 | Old |
| 0.932452 | 0.215616 | 0.273 | 0.26 | 1 | 14 | Wwtr1 | 0.013 | Old |
| 0.932508 | 0.101038 | 0.153 | 0.16 | 1 | 14 | B3glct | -0.007 | Old |
| 0.932696 | 0.264835 | 0.109 | 0.12 | 1 | 14 | Parl | -0.011 | Old |
| 0.932756 | 0.276163 | 0.077 | 0.08 | 1 | 14 | Trpv4 | -0.003 | Old |
| 0.932875 | 0.088626 | 0.131 | 0.14 | 1 | 14 | Unc119b | -0.009 | Old |
| 0.933538 | 0.64598 | 0.126 | 0.14 | 1 | 14 | Tmem222 | -0.014 | Old |
| 0.933579 | 1.626186 | 0.06 | 0.06 | 1 | 14 | Med24 | 0 | Old |
| 0.933579 | 1.413443 | 0.06 | 0.06 | 1 | 14 | Ethe1 | 0 | Old |
| 0.933594 | 0.663608 | 0.055 | 0.06 | 1 | 14 | Plekhf2 | -0.005 | Old |
| 0.933832 | 0.249092 | 0.104 | 0.1 | 1 | 14 | Imp4 | 0.004 | Old |
| 0.934381 | 1.064153 | 0.098 | 0.1 | 1 | 14 | Tbk1 | -0.002 | Old |
| 0.934381 | 1.009157 | 0.098 | 0.1 | 1 | 14 | Ift46 | -0.002 | Old |
| 0.934381 | 0.823219 | 0.098 | 0.1 | 1 | 14 | Hmgcl | -0.002 | Old |
| 0.934402 | 0.431569 | 0.082 | 0.08 | 1 | 14 | Slc7a7 | 0.002 | Old |
| 0.93485 | 0.442336 | 0.12 | 0.12 | 1 | 14 | Adgra3 | 0 | Old |
| 0.934913 | 0.514752 | 0.093 | 0.1 | 1 | 14 | Mpdu1 | -0.007 | Old |

| | | | | | | | | |
|---|---|---|---|---|---|---|---|---|
| 0.934913 | 0.408053 | 0.093 | 0.1 | 1 | 14 | Wdr66 | -0.007 | Old |
| 0.934913 | 0.289283 | 0.093 | 0.1 | 1 | 14 | Tpgs1 | -0.007 | Old |
| 0.934913 | 0.199067 | 0.093 | 0.1 | 1 | 14 | Adgre5 | -0.007 | Old |
| 0.935385 | 0.114818 | 0.29 | 0.3 | 1 | 14 | Ndufab1 | -0.01 | Old |
| 0.935679 | 0.121492 | 0.366 | 0.38 | 1 | 14 | Slc38a10 | -0.014 | Old |
| 0.936065 | 0.227874 | 0.131 | 0.14 | 1 | 14 | Setbp1 | -0.009 | Old |
| 0.93614 | 0.661941 | 0.109 | 0.12 | 1 | 14 | 5-Mar | -0.011 | Old |
| 0.93614 | 0.652247 | 0.109 | 0.12 | 1 | 14 | Nab2 | -0.011 | Old |
| 0.936158 | 0.520979 | 0.164 | 0.18 | 1 | 14 | Snrpa | -0.016 | Old |
| 0.9362 | 0.509341 | 0.197 | 0.2 | 1 | 14 | Xpr1 | -0.003 | Old |
| 0.936433 | 0.125143 | 0.208 | 0.22 | 1 | 14 | Ech1 | -0.012 | Old |
| 0.936823 | 0.17477 | 0.077 | 0.08 | 1 | 14 | Rpap3 | -0.003 | Old |
| 0.937402 | 0.365069 | 0.104 | 0.1 | 1 | 14 | Bin3 | 0.004 | Old |
| 0.937402 | 0.205112 | 0.104 | 0.1 | 1 | 14 | Aktip | 0.004 | Old |
| 0.937762 | 0.249247 | 0.257 | 0.28 | 1 | 14 | Mta1 | -0.023 | Old |
| 0.93815 | 1.505225 | 0.06 | 0.06 | 1 | 14 | Arfip2 | 0 | Old |
| 0.93815 | 0.936543 | 0.06 | 0.06 | 1 | 14 | Nckap1l | 0 | Old |
| 0.938218 | 0.011535 | 0.77 | 0.7 | 1 | 14 | Rpl11 | 0.07 | Old |
| 0.938309 | 0.352697 | 0.235 | 0.26 | 1 | 14 | Osbpl5 | -0.025 | Old |
| 0.938327 | 0.867045 | 0.055 | 0.06 | 1 | 14 | Lin37 | -0.005 | Old |
| 0.938495 | 0.354212 | 0.153 | 0.16 | 1 | 14 | Necap2 | -0.007 | Old |
| 0.938624 | 0.472023 | 0.093 | 0.1 | 1 | 14 | Ppt2 | -0.007 | Old |
| 0.938624 | 0.30603 | 0.093 | 0.1 | 1 | 14 | Ubap2 | -0.007 | Old |
| 0.938829 | 0.038632 | 0.617 | 0.62 | 1 | 14 | Arhgef12 | -0.003 | Old |
| 0.939042 | 0.212877 | 0.388 | 0.42 | 1 | 14 | H2afy | -0.032 | Old |
| 0.939076 | 0.576874 | 0.208 | 0.22 | 1 | 14 | Bak1 | -0.012 | Old |
| 0.939139 | 0.137985 | 0.279 | 0.28 | 1 | 14 | Ibtk | -0.001 | Old |
| 0.939423 | 0.160393 | 0.202 | 0.2 | 1 | 14 | Erlin1 | 0.002 | Old |
| 0.939855 | 1.054728 | 0.022 | 0.02 | 1 | 14 | Myl4 | 0.002 | Old |
| 0.939855 | 0.544405 | 0.022 | 0.02 | 1 | 14 | Adat1 | 0.002 | Old |
| 0.939855 | 0.534448 | 0.022 | 0.02 | 1 | 14 | Entpd6 | 0.002 | Old |
| 0.939855 | 0.424834 | 0.022 | 0.02 | 1 | 14 | Tmem170 | 0.002 | Old |
| 0.939855 | 0.304976 | 0.022 | 0.02 | 1 | 14 | Tle6 | 0.002 | Old |
| 0.939855 | 0.280981 | 0.022 | 0.02 | 1 | 14 | Adam4 | 0.002 | Old |
| 0.939855 | 0.24091 | 0.022 | 0.02 | 1 | 14 | Abraxas1 | 0.002 | Old |
| 0.939855 | 0.125773 | 0.022 | 0.02 | 1 | 14 | Gm26740 | 0.002 | Old |
| 0.939855 | 0.107098 | 0.022 | 0.02 | 1 | 14 | Supt3 | 0.002 | Old |
| 0.939855 | 0.093049 | 0.022 | 0.02 | 1 | 14 | Tarbp1 | 0.002 | Old |
| 0.939855 | 0.053139 | 0.022 | 0.02 | 1 | 14 | Sfxn4 | 0.002 | Old |
| 0.939973 | 0.048031 | 0.322 | 0.34 | 1 | 14 | Fkbp8 | -0.018 | Old |
| 0.940006 | 0.304308 | 0.219 | 0.22 | 1 | 14 | Naa20 | -0.001 | Old |
| 0.940581 | 0.039275 | 0.115 | 0.12 | 1 | 14 | Lrp3 | -0.005 | Old |
| 0.940581 | 1.013003 | 0.186 | 0.2 | 1 | 14 | Cul7 | -0.014 | Old |
| 0.940648 | 0.179844 | 0.038 | 0.04 | 1 | 14 | Pnkp | -0.002 | Old |
| 0.940795 | 0.261198 | 0.148 | 0.16 | 1 | 14 | Slc35a1 | -0.012 | Old |

| | | | | | | | | |
|---|---|---|---|---|---|---|---|---|
| 0.941519 | 0.317967 | 0.12 | 0.12 | 1 | 14 | Zdhhc1 | 0 | Old |
| 0.941519 | 0.232742 | 0.12 | 0.12 | 1 | 14 | Baz1a | 0 | Old |
| 0.942337 | 0.708972 | 0.093 | 0.1 | 1 | 14 | Tomm40l | -0.007 | Old |
| 0.942337 | 0.278087 | 0.093 | 0.1 | 1 | 14 | Btbd9 | -0.007 | Old |
| 0.942338 | 0.539018 | 0.082 | 0.08 | 1 | 14 | Cpeb2 | 0.002 | Old |
| 0.942338 | 0.34766 | 0.082 | 0.08 | 1 | 14 | Mrs2 | 0.002 | Old |
| 0.942447 | 0.586228 | 0.131 | 0.14 | 1 | 14 | Mlxip | -0.009 | Old |
| 0.943031 | 0.513068 | 0.109 | 0.12 | 1 | 14 | Cinp | -0.011 | Old |
| 0.943063 | 1.132175 | 0.055 | 0.06 | 1 | 14 | Fyb2 | -0.005 | Old |
| 0.943063 | 1.074169 | 0.055 | 0.06 | 1 | 14 | Spi1 | -0.005 | Old |
| 0.943336 | 0.304954 | 0.235 | 0.26 | 1 | 14 | Cstf3 | -0.025 | Old |
| 0.943705 | 0.278017 | 0.246 | 0.24 | 1 | 14 | Chchd1 | 0.006 | Old |
| 0.943873 | 0.11312 | 0.295 | 0.3 | 1 | 14 | Psmb3 | -0.005 | Old |
| 0.943984 | 0.732686 | 0.169 | 0.18 | 1 | 14 | Laptm5 | -0.011 | Old |
| 0.94421 | 0.041362 | 0.383 | 0.42 | 1 | 14 | Zcchc6 | -0.037 | Old |
| 0.944543 | 0.162761 | 0.104 | 0.1 | 1 | 14 | Amn1 | 0.004 | Old |
| 0.944856 | 0.180096 | 0.12 | 0.12 | 1 | 14 | Wdfy1 | 0 | Old |
| 0.944961 | 0.411398 | 0.077 | 0.08 | 1 | 14 | Ccdc126 | -0.003 | Old |
| 0.944961 | 0.105456 | 0.077 | 0.08 | 1 | 14 | Plek | -0.003 | Old |
| 0.944961 | 0.086428 | 0.077 | 0.08 | 1 | 14 | Ror2 | -0.003 | Old |
| 0.945298 | 0.977356 | 0.098 | 0.1 | 1 | 14 | Cachd1 | -0.002 | Old |
| 0.945487 | 0.228329 | 0.235 | 0.24 | 1 | 14 | Ext2 | -0.005 | Old |
| 0.946052 | 0.430016 | 0.093 | 0.1 | 1 | 14 | Mrpl45 | -0.007 | Old |
| 0.946052 | 0.370503 | 0.093 | 0.1 | 1 | 14 | Patz1 | -0.007 | Old |
| 0.946292 | 0.719535 | 0.038 | 0.04 | 1 | 14 | Fbn2 | -0.002 | Old |
| 0.946292 | 0.042463 | 0.038 | 0.04 | 1 | 14 | Mcf2l | -0.002 | Old |
| 0.946292 | 0.03053 | 0.038 | 0.04 | 1 | 14 | Lrwd1 | -0.002 | Old |
| 0.946309 | 0.121003 | 0.082 | 0.08 | 1 | 14 | Cyp4f16 | 0.002 | Old |
| 0.946852 | 0.436657 | 0.169 | 0.18 | 1 | 14 | Hk2 | -0.011 | Old |
| 0.946858 | 0.12712 | 0.148 | 0.16 | 1 | 14 | Ebpl | -0.012 | Old |
| 0.947042 | 0.415724 | 0.197 | 0.2 | 1 | 14 | Cnot10 | -0.003 | Old |
| 0.947299 | 1.28186 | 0.06 | 0.06 | 1 | 14 | Adsl | 0 | Old |
| 0.947299 | 0.878531 | 0.06 | 0.06 | 1 | 14 | Gatsl3 | 0 | Old |
| 0.947361 | 0.419642 | 0.115 | 0.12 | 1 | 14 | Ppip5k2 | -0.005 | Old |
| 0.947362 | 1.170906 | 0.022 | 0.02 | 1 | 14 | B830012L1 | 0.002 | Old |
| 0.947362 | 0.822441 | 0.022 | 0.02 | 1 | 14 | Nfasc | 0.002 | Old |
| 0.947362 | 0.799657 | 0.022 | 0.02 | 1 | 14 | Taf4b | 0.002 | Old |
| 0.947362 | 0.796497 | 0.022 | 0.02 | 1 | 14 | Zfp324 | 0.002 | Old |
| 0.947362 | 0.302588 | 0.022 | 0.02 | 1 | 14 | Slc22a21 | 0.002 | Old |
| 0.947362 | 0.0627 | 0.022 | 0.02 | 1 | 14 | Mcm2 | 0.002 | Old |
| 0.947362 | 0.055042 | 0.022 | 0.02 | 1 | 14 | Tnfsfm13 | 0.002 | Old |
| 0.947362 | 0.019864 | 0.022 | 0.02 | 1 | 14 | B130006D( | 0.002 | Old |
| 0.947801 | 1.002473 | 0.055 | 0.06 | 1 | 14 | Pwp1 | -0.005 | Old |
| 0.948078 | 0.178342 | 0.158 | 0.16 | 1 | 14 | Zfp322a | -0.002 | Old |
| 0.94894 | 0.681971 | 0.098 | 0.1 | 1 | 14 | Fam199x | -0.002 | Old |

| | | | | | | | | |
|---|---|---|---|---|---|---|---|---|
| 0.949032 | 0.257741 | 0.077 | 0.08 | 1 | 14 | Ints12 | -0.003 | Old |
| 0.949122 | 0.692624 | 0.213 | 0.24 | 1 | 14 | Armc1 | -0.027 | Old |
| 0.949656 | 0.162256 | 0.208 | 0.22 | 1 | 14 | Fam208b | -0.012 | Old |
| 0.950215 | 0.002918 | 0.355 | 0.34 | 1 | 14 | Herc1 | 0.015 | Old |
| 0.950425 | 0.191056 | 0.219 | 0.22 | 1 | 14 | Seh1l | -0.001 | Old |
| 0.950479 | 0.246722 | 0.153 | 0.16 | 1 | 14 | Pmpca | -0.007 | Old |
| 0.950646 | 0.554256 | 0.164 | 0.18 | 1 | 14 | Arhgef10l | -0.016 | Old |
| 0.95138 | 0.047787 | 0.727 | 0.74 | 1 | 14 | Rps3 | -0.013 | Old |
| 0.951875 | 1.179662 | 0.06 | 0.06 | 1 | 14 | Zfp212 | 0 | Old |
| 0.951875 | 1.173638 | 0.06 | 0.06 | 1 | 14 | Nck2 | 0 | Old |
| 0.951875 | 1.076881 | 0.06 | 0.06 | 1 | 14 | Mettl21a | 0 | Old |
| 0.951875 | 0.936054 | 0.06 | 0.06 | 1 | 14 | Smad6 | 0 | Old |
| 0.951875 | 0.86628 | 0.06 | 0.06 | 1 | 14 | Apip | 0 | Old |
| 0.951938 | 0.032905 | 0.038 | 0.04 | 1 | 14 | Disp1 | -0.002 | Old |
| 0.952303 | 0.927978 | 0.208 | 0.22 | 1 | 14 | Srpk2 | -0.012 | Old |
| 0.95254 | 1.712515 | 0.055 | 0.06 | 1 | 14 | Platr25 | -0.005 | Old |
| 0.952583 | 0.965853 | 0.098 | 0.1 | 1 | 14 | Cdkn2d | -0.002 | Old |
| 0.952923 | 0.249583 | 0.148 | 0.16 | 1 | 14 | Card19 | -0.012 | Old |
| 0.953105 | 0.149901 | 0.077 | 0.08 | 1 | 14 | Entpd7 | -0.003 | Old |
| 0.953105 | 0.070103 | 0.077 | 0.08 | 1 | 14 | Vps18 | -0.003 | Old |
| 0.95396 | 0.665908 | 0.18 | 0.2 | 1 | 14 | Hltf | -0.02 | Old |
| 0.954145 | 0.179591 | 0.115 | 0.12 | 1 | 14 | Prkce | -0.005 | Old |
| 0.954382 | 0.1704 | 0.186 | 0.2 | 1 | 14 | Fam3c | -0.014 | Old |
| 0.954671 | 0.290525 | 0.219 | 0.24 | 1 | 14 | Ptgs1 | -0.021 | Old |
| 0.95487 | 0.354775 | 0.12 | 0.12 | 1 | 14 | Gt(ROSA)2( | 0 | Old |
| 0.955027 | 0.665553 | 0.306 | 0.32 | 1 | 14 | Pik3ca | -0.014 | Old |
| 0.955027 | 0.054283 | 0.306 | 0.32 | 1 | 14 | Yipf3 | -0.014 | Old |
| 0.955221 | 0.415229 | 0.131 | 0.14 | 1 | 14 | Mettl23 | -0.009 | Old |
| 0.955264 | 0.147541 | 0.104 | 0.1 | 1 | 14 | Cyp2j9 | 0.004 | Old |
| 0.955428 | 0.006739 | 0.377 | 0.36 | 1 | 14 | Gfpt1 | 0.017 | Old |
| 0.955618 | 0.295262 | 0.235 | 0.24 | 1 | 14 | Arl5b | -0.005 | Old |
| 0.956226 | 1.162217 | 0.098 | 0.1 | 1 | 14 | Mrpl55 | -0.002 | Old |
| 0.956226 | 0.476995 | 0.098 | 0.1 | 1 | 14 | Entpd5 | -0.002 | Old |
| 0.956454 | 1.209335 | 0.06 | 0.06 | 1 | 14 | Strada | 0 | Old |
| 0.956454 | 0.877723 | 0.06 | 0.06 | 1 | 14 | Asphd2 | 0 | Old |
| 0.956454 | 0.839487 | 0.06 | 0.06 | 1 | 14 | Cyp4v3 | 0 | Old |
| 0.956454 | 0.757303 | 0.06 | 0.06 | 1 | 14 | Zfyve26 | 0 | Old |
| 0.956475 | 0.264971 | 0.153 | 0.16 | 1 | 14 | Alg10b | -0.007 | Old |
| 0.95697 | 0.011766 | 0.158 | 0.16 | 1 | 14 | Cnot2 | -0.002 | Old |
| 0.95697 | 0.000691 | 0.158 | 0.16 | 1 | 14 | Smc5 | -0.002 | Old |
| 0.957178 | 1.200927 | 0.077 | 0.08 | 1 | 14 | Zfp157 | -0.003 | Old |
| 0.957178 | 0.041998 | 0.077 | 0.08 | 1 | 14 | Fbxl4 | -0.003 | Old |
| 0.957201 | 0.685666 | 0.093 | 0.1 | 1 | 14 | Fnip2 | -0.007 | Old |
| 0.957259 | 0.616675 | 0.219 | 0.24 | 1 | 14 | Usp15 | -0.021 | Old |
| 0.957259 | 0.26885 | 0.224 | 0.22 | 1 | 14 | Rfc1 | 0.004 | Old |

| | | | | | | | | |
|---|---|---|---|---|---|---|---|---|
| 0.957281 | 2.397033 | 0.055 | 0.06 | 1 | 14 | Ldb2 | -0.005 | Old |
| 0.957538 | 1.303997 | 0.115 | 0.12 | 1 | 14 | Kdm2b | -0.005 | Old |
| 0.957587 | 0.04386 | 0.038 | 0.04 | 1 | 14 | Cmss1 | -0.002 | Old |
| 0.957863 | 0.146488 | 0.23 | 0.24 | 1 | 14 | Trps1 | -0.01 | Old |
| 0.958416 | 0.235829 | 0.131 | 0.14 | 1 | 14 | Zfp521 | -0.009 | Old |
| 0.95884 | 0.105157 | 0.104 | 0.1 | 1 | 14 | Dhrs4 | 0.004 | Old |
| 0.958855 | 0.297964 | 0.29 | 0.3 | 1 | 14 | Peli1 | -0.01 | Old |
| 0.958977 | 0.048191 | 0.421 | 0.42 | 1 | 14 | Top2b | 0.001 | Old |
| 0.958992 | 0.395676 | 0.148 | 0.16 | 1 | 14 | Slc25a11 | -0.012 | Old |
| 0.95915 | 0.334358 | 0.18 | 0.18 | 1 | 14 | Mcur1 | 0 | Old |
| 0.95924 | 0.10213 | 0.213 | 0.22 | 1 | 14 | Pank2 | -0.007 | Old |
| 0.959465 | 0.119716 | 0.251 | 0.28 | 1 | 14 | Smu1 | -0.029 | Old |
| 0.959474 | 0.619449 | 0.153 | 0.16 | 1 | 14 | Mrps15 | -0.007 | Old |
| 0.959514 | 0.271087 | 0.142 | 0.14 | 1 | 14 | Pfdn6 | 0.002 | Old |
| 0.960013 | 0.097789 | 0.339 | 0.38 | 1 | 14 | Litaf | -0.041 | Old |
| 0.960555 | 0.560261 | 0.137 | 0.14 | 1 | 14 | Parp9 | -0.003 | Old |
| 0.960555 | 0.334744 | 0.137 | 0.14 | 1 | 14 | Med25 | -0.003 | Old |
| 0.96092 | 0.663735 | 0.093 | 0.1 | 1 | 14 | Kazn | -0.007 | Old |
| 0.961034 | 1.355903 | 0.06 | 0.06 | 1 | 14 | Ascc1 | 0 | Old |
| 0.961034 | 0.646247 | 0.06 | 0.06 | 1 | 14 | Tgif1 | 0 | Old |
| 0.961204 | 0.003246 | 0.29 | 0.3 | 1 | 14 | Clptm1l | -0.01 | Old |
| 0.961253 | 0.294319 | 0.077 | 0.08 | 1 | 14 | Taf1b | -0.003 | Old |
| 0.961253 | 0.163171 | 0.077 | 0.08 | 1 | 14 | Rbm12b2 | -0.003 | Old |
| 0.961613 | 0.467745 | 0.131 | 0.14 | 1 | 14 | Rnmt | -0.009 | Old |
| 0.962203 | 0.496509 | 0.082 | 0.08 | 1 | 14 | Prkaa2 | 0.002 | Old |
| 0.962203 | 0.29421 | 0.082 | 0.08 | 1 | 14 | Kdelc1 | 0.002 | Old |
| 0.962626 | 0.091334 | 0.142 | 0.14 | 1 | 14 | Ube2o | 0.002 | Old |
| 0.963237 | 0.846466 | 0.038 | 0.04 | 1 | 14 | Mrpl39 | -0.002 | Old |
| 0.963237 | 0.501071 | 0.038 | 0.04 | 1 | 14 | Rpusd4 | -0.002 | Old |
| 0.963237 | 0.191847 | 0.038 | 0.04 | 1 | 14 | Col4a6 | -0.002 | Old |
| 0.963237 | 0.038519 | 0.038 | 0.04 | 1 | 14 | Xndc1 | -0.002 | Old |
| 0.963872 | 0.044545 | 0.366 | 0.36 | 1 | 14 | Zmym5 | 0.006 | Old |
| 0.964685 | 0.27126 | 0.393 | 0.4 | 1 | 14 | Irf1 | -0.007 | Old |
| 0.964891 | 0.456805 | 0.12 | 0.12 | 1 | 14 | Phactr4 | 0 | Old |
| 0.965205 | 0.104291 | 0.372 | 0.38 | 1 | 14 | Sf1 | -0.008 | Old |
| 0.965329 | 0.133381 | 0.077 | 0.08 | 1 | 14 | Klf16 | -0.003 | Old |
| 0.965474 | 0.285058 | 0.153 | 0.16 | 1 | 14 | Slc6a8 | -0.007 | Old |
| 0.965615 | 1.565477 | 0.06 | 0.06 | 1 | 14 | Rnd2 | 0 | Old |
| 0.965615 | 0.901953 | 0.06 | 0.06 | 1 | 14 | Gfod1 | 0 | Old |
| 0.965615 | 0.629488 | 0.06 | 0.06 | 1 | 14 | Phykpl | 0 | Old |
| 0.965741 | 0.466958 | 0.191 | 0.2 | 1 | 14 | Esf1 | -0.009 | Old |
| 0.966179 | 0.4596 | 0.082 | 0.08 | 1 | 14 | Csgalnact1 | 0.002 | Old |
| 0.966403 | 0.134282 | 0.372 | 0.4 | 1 | 14 | Fzd4 | -0.028 | Old |
| 0.966551 | 0.382687 | 0.306 | 0.32 | 1 | 14 | Arl5a | -0.014 | Old |
| 0.966863 | 0.806128 | 0.137 | 0.14 | 1 | 14 | Srd5a3 | -0.003 | Old |

| | | | | | | | | |
|---|---|---|---|---|---|---|---|---|
| 0.966863 | 0.664931 | 0.137 | 0.14 | 1 | 14 | Nasp | -0.003 | Old |
| 0.967155 | 0.06502 | 0.361 | 0.38 | 1 | 14 | Pabpn1 | -0.019 | Old |
| 0.967597 | 0.05281 | 0.18 | 0.18 | 1 | 14 | C1qc | 0 | Old |
| 0.967723 | 1.01739 | 0.115 | 0.12 | 1 | 14 | Anapc2 | -0.005 | Old |
| 0.967723 | 0.611501 | 0.115 | 0.12 | 1 | 14 | Zkscan8 | -0.005 | Old |
| 0.967723 | 0.551975 | 0.115 | 0.12 | 1 | 14 | Iars | -0.005 | Old |
| 0.968022 | 0.170185 | 0.355 | 0.36 | 1 | 14 | Thoc2 | -0.005 | Old |
| 0.968191 | 0.06421 | 0.208 | 0.22 | 1 | 14 | Banf1 | -0.012 | Old |
| 0.968359 | 1.30279 | 0.093 | 0.1 | 1 | 14 | Nup160 | -0.007 | Old |
| 0.968851 | 0.184071 | 0.142 | 0.14 | 1 | 14 | Pknox1 | 0.002 | Old |
| 0.96889 | 0.463452 | 0.038 | 0.04 | 1 | 14 | Mad2l1bp | -0.002 | Old |
| 0.96889 | 0.439188 | 0.038 | 0.04 | 1 | 14 | Fbxl19 | -0.002 | Old |
| 0.969406 | 1.014175 | 0.077 | 0.08 | 1 | 14 | Exoc7 | -0.003 | Old |
| 0.969558 | 0.539764 | 0.23 | 0.26 | 1 | 14 | Spag7 | -0.03 | Old |
| 0.969753 | 0.022998 | 0.213 | 0.22 | 1 | 14 | Fkbp4 | -0.007 | Old |
| 0.970197 | 0.457102 | 0.06 | 0.06 | 1 | 14 | Gabpb1 | 0 | Old |
| 0.970302 | 0.277159 | 0.284 | 0.28 | 1 | 14 | Baz2b | 0.004 | Old |
| 0.970961 | 0.832381 | 0.186 | 0.2 | 1 | 14 | Sox5 | -0.014 | Old |
| 0.971119 | 0.285916 | 0.115 | 0.12 | 1 | 14 | Ip6k2 | -0.005 | Old |
| 0.97119 | 0.017327 | 0.399 | 0.4 | 1 | 14 | Jade1 | -0.001 | Old |
| 0.971286 | 0.5253 | 0.219 | 0.22 | 1 | 14 | Nr1h3 | -0.001 | Old |
| 0.971469 | 0.303022 | 0.197 | 0.2 | 1 | 14 | Aff1 | -0.003 | Old |
| 0.971544 | 0.662891 | 0.175 | 0.18 | 1 | 14 | Ankrd10 | -0.005 | Old |
| 0.971576 | 0.465313 | 0.12 | 0.12 | 1 | 14 | Rnf121 | 0 | Old |
| 0.972225 | 0.470263 | 0.344 | 0.36 | 1 | 14 | Capza1 | -0.016 | Old |
| 0.972441 | 0.098556 | 0.317 | 0.3 | 1 | 14 | Lyz2 | 0.017 | Old |
| 0.973172 | 0.57471 | 0.137 | 0.14 | 1 | 14 | Snapc3 | -0.003 | Old |
| 0.973484 | 0.444658 | 0.077 | 0.08 | 1 | 14 | L3mbtl3 | -0.003 | Old |
| 0.973539 | 0.044544 | 0.24 | 0.24 | 1 | 14 | Mtmr3 | 0 | Old |
| 0.974134 | 0.012679 | 0.082 | 0.08 | 1 | 14 | Slc35f6 | 0.002 | Old |
| 0.974477 | 0.136907 | 0.153 | 0.16 | 1 | 14 | Supt6 | -0.007 | Old |
| 0.974544 | 0.72078 | 0.038 | 0.04 | 1 | 14 | Scrn2 | -0.002 | Old |
| 0.974544 | 0.542907 | 0.038 | 0.04 | 1 | 14 | Asns | -0.002 | Old |
| 0.974544 | 0.454656 | 0.038 | 0.04 | 1 | 14 | Slc39a11 | -0.002 | Old |
| 0.974544 | 0.444961 | 0.038 | 0.04 | 1 | 14 | Bace2 | -0.002 | Old |
| 0.974544 | 0.431169 | 0.038 | 0.04 | 1 | 14 | Gpr34 | -0.002 | Old |
| 0.974544 | 0.413395 | 0.038 | 0.04 | 1 | 14 | Shf | -0.002 | Old |
| 0.974544 | 0.268968 | 0.038 | 0.04 | 1 | 14 | Dixdc1 | -0.002 | Old |
| 0.974767 | 0.281928 | 0.158 | 0.16 | 1 | 14 | Polr3gl | -0.002 | Old |
| 0.974919 | 0.10038 | 0.279 | 0.28 | 1 | 14 | Ralb | -0.001 | Old |
| 0.975277 | 0.146018 | 0.251 | 0.26 | 1 | 14 | Leprotl1 | -0.009 | Old |
| 0.9758 | 0.11252 | 0.421 | 0.42 | 1 | 14 | Hsd17b11 | 0.001 | Old |
| 0.975802 | 0.882336 | 0.093 | 0.1 | 1 | 14 | Aif1 | -0.007 | Old |
| 0.976047 | 0.181487 | 0.18 | 0.18 | 1 | 14 | Smad3 | 0 | Old |
| 0.976327 | 0.126561 | 0.137 | 0.14 | 1 | 14 | Kctd9 | -0.003 | Old |

| | | | | | | | | |
|---|---|---|---|---|---|---|---|---|
| 0.976902 | 0.01097 | 0.197 | 0.2 | 1 | 14 | Spaca6 | -0.003 | Old |
| 0.977084 | 0.139731 | 0.366 | 0.38 | 1 | 14 | Irak1 | -0.014 | Old |
| 0.977562 | 0.523561 | 0.077 | 0.08 | 1 | 14 | Utp15 | -0.003 | Old |
| 0.977562 | 0.464606 | 0.077 | 0.08 | 1 | 14 | Uba7 | -0.003 | Old |
| 0.977562 | 0.382398 | 0.077 | 0.08 | 1 | 14 | Aasdh | -0.003 | Old |
| 0.977656 | 0.304121 | 0.393 | 0.44 | 1 | 14 | Syne2 | -0.047 | Old |
| 0.977748 | 0.309936 | 0.251 | 0.26 | 1 | 14 | Rbm7 | -0.009 | Old |
| 0.978081 | 0.088101 | 0.306 | 0.32 | 1 | 14 | Zfr | -0.014 | Old |
| 0.97844 | 0.480161 | 0.169 | 0.18 | 1 | 14 | Slc25a1 | -0.011 | Old |
| 0.978864 | 0.080403 | 0.18 | 0.18 | 1 | 14 | Sema3d | 0 | Old |
| 0.979365 | 0.496198 | 0.06 | 0.06 | 1 | 14 | Ccl27a | 0 | Old |
| 0.979365 | 0.225734 | 0.06 | 0.06 | 1 | 14 | Cdkl2 | 0 | Old |
| 0.979523 | 0.532652 | 0.093 | 0.1 | 1 | 14 | Runx1 | -0.007 | Old |
| 0.9802 | 1.052152 | 0.038 | 0.04 | 1 | 14 | Gm527 | -0.002 | Old |
| 0.9802 | 0.921808 | 0.038 | 0.04 | 1 | 14 | Mcub | -0.002 | Old |
| 0.9802 | 0.819217 | 0.038 | 0.04 | 1 | 14 | Caskin2 | -0.002 | Old |
| 0.9802 | 0.759897 | 0.038 | 0.04 | 1 | 14 | Gpr137b | -0.002 | Old |
| 0.9802 | 0.665203 | 0.038 | 0.04 | 1 | 14 | BC017158 | -0.002 | Old |
| 0.9802 | 0.592498 | 0.038 | 0.04 | 1 | 14 | Dgka | -0.002 | Old |
| 0.9802 | 0.364457 | 0.038 | 0.04 | 1 | 14 | Polr1e | -0.002 | Old |
| 0.980271 | 0.118275 | 0.213 | 0.22 | 1 | 14 | Arhgef40 | -0.007 | Old |
| 0.980481 | 0.288644 | 0.153 | 0.16 | 1 | 14 | Rfk | -0.007 | Old |
| 0.980801 | 0.711415 | 0.131 | 0.14 | 1 | 14 | Med17 | -0.009 | Old |
| 0.980971 | 0.289985 | 0.235 | 0.24 | 1 | 14 | Pdcd4 | -0.005 | Old |
| 0.981641 | 0.518396 | 0.077 | 0.08 | 1 | 14 | Naa25 | -0.003 | Old |
| 0.981641 | 0.44018 | 0.077 | 0.08 | 1 | 14 | Ptk7 | -0.003 | Old |
| 0.981641 | 0.432117 | 0.077 | 0.08 | 1 | 14 | Trub2 | -0.003 | Old |
| 0.981678 | 0.697379 | 0.257 | 0.28 | 1 | 14 | Nab1 | -0.023 | Old |
| 0.981753 | 0.528596 | 0.098 | 0.1 | 1 | 14 | Naa30 | -0.002 | Old |
| 0.981753 | 0.30378 | 0.098 | 0.1 | 1 | 14 | Rnf217 | -0.002 | Old |
| 0.98203 | 0.015771 | 0.519 | 0.56 | 1 | 14 | Cyba | -0.041 | Old |
| 0.982336 | 0.22031 | 0.197 | 0.2 | 1 | 14 | Ints6 | -0.003 | Old |
| 0.982473 | 0.63351 | 0.24 | 0.26 | 1 | 14 | Ptk2 | -0.02 | Old |
| 0.982924 | 0.305689 | 0.175 | 0.18 | 1 | 14 | Zfp771 | -0.005 | Old |
| 0.98395 | 0.411535 | 0.06 | 0.06 | 1 | 14 | Irf4 | 0 | Old |
| 0.98395 | 0.26101 | 0.06 | 0.06 | 1 | 14 | Zw10 | 0 | Old |
| 0.984026 | 0.365776 | 0.251 | 0.28 | 1 | 14 | Sarnp | -0.029 | Old |
| 0.984189 | 0.428546 | 0.169 | 0.18 | 1 | 14 | Ctr9 | -0.011 | Old |
| 0.984336 | 0.051593 | 0.219 | 0.22 | 1 | 14 | Megf8 | -0.001 | Old |
| 0.984499 | 0.069169 | 0.18 | 0.18 | 1 | 14 | Rif1 | 0 | Old |
| 0.984563 | 0.004842 | 0.23 | 0.22 | 1 | 14 | Rtn4rl1 | 0.01 | Old |
| 0.984707 | 0.56299 | 0.115 | 0.12 | 1 | 14 | Fnbp1l | -0.005 | Old |
| 0.985301 | 0.432453 | 0.202 | 0.22 | 1 | 14 | Ppid | -0.018 | Old |
| 0.985402 | 0.433181 | 0.098 | 0.1 | 1 | 14 | Mterf3 | -0.002 | Old |
| 0.985402 | 0.427491 | 0.098 | 0.1 | 1 | 14 | Snx29 | -0.002 | Old |

| | | | | | | | | |
|---|---|---|---|---|---|---|---|---|
| 0.985641 | 0.602909 | 0.213 | 0.24 | 1 | 14 | Ndufv2 | -0.027 | Old |
| 0.985667 | 0.322059 | 0.273 | 0.3 | 1 | 14 | Lsm14a | -0.027 | Old |
| 0.98572 | 0.752789 | 0.077 | 0.08 | 1 | 14 | Smim4 | -0.003 | Old |
| 0.98577 | 0.331214 | 0.175 | 0.18 | 1 | 14 | Tmem176a | -0.005 | Old |
| 0.98577 | 0.276928 | 0.175 | 0.18 | 1 | 14 | Tnfaip3 | -0.005 | Old |
| 0.98577 | 0.221545 | 0.175 | 0.18 | 1 | 14 | Kmt2d | -0.005 | Old |
| 0.985856 | 1.661815 | 0.038 | 0.04 | 1 | 14 | Osbpl7 | -0.002 | Old |
| 0.985856 | 1.367646 | 0.038 | 0.04 | 1 | 14 | Edc3 | -0.002 | Old |
| 0.985856 | 1.155715 | 0.038 | 0.04 | 1 | 14 | Ninl | -0.002 | Old |
| 0.985856 | 1.099692 | 0.038 | 0.04 | 1 | 14 | Timm8a1 | -0.002 | Old |
| 0.985856 | 1.014356 | 0.038 | 0.04 | 1 | 14 | Smyd4 | -0.002 | Old |
| 0.985856 | 0.945769 | 0.038 | 0.04 | 1 | 14 | Flywch2 | -0.002 | Old |
| 0.985856 | 0.874661 | 0.038 | 0.04 | 1 | 14 | Pithd1 | -0.002 | Old |
| 0.985856 | 0.723629 | 0.038 | 0.04 | 1 | 14 | Shank1 | -0.002 | Old |
| 0.985856 | 0.573927 | 0.038 | 0.04 | 1 | 14 | Dph7 | -0.002 | Old |
| 0.986325 | 0.48245 | 0.148 | 0.16 | 1 | 14 | Med8 | -0.012 | Old |
| 0.986486 | 0.812113 | 0.153 | 0.16 | 1 | 14 | Foxn2 | -0.007 | Old |
| 0.986486 | 0.36845 | 0.153 | 0.16 | 1 | 14 | Gramd1a | -0.007 | Old |
| 0.986486 | 0.235847 | 0.153 | 0.16 | 1 | 14 | Trnt1 | -0.007 | Old |
| 0.98664 | 0.361813 | 0.158 | 0.16 | 1 | 14 | Lgals8 | -0.002 | Old |
| 0.986859 | 0.148069 | 0.361 | 0.38 | 1 | 14 | Sh3bgrl3 | -0.019 | Old |
| 0.987063 | 0.796458 | 0.169 | 0.18 | 1 | 14 | Cse1l | -0.011 | Old |
| 0.987193 | 0.145138 | 0.393 | 0.42 | 1 | 14 | Prpf4b | -0.027 | Old |
| 0.9872 | 0.666693 | 0.131 | 0.14 | 1 | 14 | Adarb1 | -0.009 | Old |
| 0.987317 | 0.251759 | 0.18 | 0.18 | 1 | 14 | Wipi2 | 0 | Old |
| 0.987552 | 0.408654 | 0.186 | 0.2 | 1 | 14 | Pwwp2a | -0.014 | Old |
| 0.987723 | 0.010264 | 1 | 0.94 | 1 | 14 | mt-Atp6 | 0.06 | Old |
| 0.98777 | 0.130346 | 0.197 | 0.2 | 1 | 14 | 5031439G( | -0.003 | Old |
| 0.987906 | 1.178012 | 0.109 | 0.12 | 1 | 14 | Cdh13 | -0.011 | Old |
| 0.988106 | 0.421997 | 0.115 | 0.12 | 1 | 14 | Ankrd28 | -0.005 | Old |
| 0.988294 | 0.126592 | 0.12 | 0.12 | 1 | 14 | Atg2a | 0 | Old |
| 0.988535 | 0.357314 | 0.06 | 0.06 | 1 | 14 | Snapc2 | 0 | Old |
| 0.988535 | 0.320515 | 0.06 | 0.06 | 1 | 14 | Agbl3 | 0 | Old |
| 0.988535 | 0.22468 | 0.06 | 0.06 | 1 | 14 | Qpctl | 0 | Old |
| 0.988804 | 0.232514 | 0.246 | 0.26 | 1 | 14 | Phf14 | -0.014 | Old |
| 0.989051 | 0.356995 | 0.098 | 0.1 | 1 | 14 | Cdk2 | -0.002 | Old |
| 0.989051 | 0.164785 | 0.098 | 0.1 | 1 | 14 | Nsmce2 | -0.002 | Old |
| 0.9898 | 1.134842 | 0.077 | 0.08 | 1 | 14 | Farp1 | -0.003 | Old |
| 0.9898 | 1.075165 | 0.077 | 0.08 | 1 | 14 | Mgl2 | -0.003 | Old |
| 0.989984 | 0.536703 | 0.24 | 0.26 | 1 | 14 | Pkig | -0.02 | Old |
| 0.991388 | 0.036818 | 0.716 | 0.66 | 1 | 14 | Csnk1a1 | 0.056 | Old |
| 0.991462 | 0.636208 | 0.175 | 0.18 | 1 | 14 | Osgep | -0.005 | Old |
| 0.991504 | 0.655167 | 0.115 | 0.12 | 1 | 14 | Cyth2 | -0.005 | Old |
| 0.991504 | 0.340711 | 0.115 | 0.12 | 1 | 14 | Srpk1 | -0.005 | Old |
| 0.991513 | 1.246953 | 0.038 | 0.04 | 1 | 14 | Anapc15 | -0.002 | Old |

| | | | | | | | | |
|---|---|---|---|---|---|---|---|---|
| 0.991513 | 1.116047 | 0.038 | 0.04 | 1 | 14 | Lrrc1 | -0.002 | Old |
| 0.991513 | 0.941241 | 0.038 | 0.04 | 1 | 14 | Mfsd4b4 | -0.002 | Old |
| 0.991848 | 0.251691 | 0.295 | 0.32 | 1 | 14 | Nsd1 | -0.025 | Old |
| 0.992108 | 0.34198 | 0.137 | 0.14 | 1 | 14 | Mtmr1 | -0.003 | Old |
| 0.992251 | 0.162551 | 0.35 | 0.36 | 1 | 14 | Cul1 | -0.01 | Old |
| 0.992577 | 0.176147 | 0.158 | 0.16 | 1 | 14 | Dnajc21 | -0.002 | Old |
| 0.992577 | 0.113865 | 0.158 | 0.16 | 1 | 14 | Psme3 | -0.002 | Old |
| 0.992701 | 0.14323 | 0.098 | 0.1 | 1 | 14 | Vrk2 | -0.002 | Old |
| 0.993121 | 0.170311 | 0.06 | 0.06 | 1 | 14 | Rps6kb2 | 0 | Old |
| 0.993121 | 0.072631 | 0.06 | 0.06 | 1 | 14 | Fbf1 | 0 | Old |
| 0.993146 | 0.332804 | 0.191 | 0.2 | 1 | 14 | Ccdc82 | -0.009 | Old |
| 0.993146 | 0.312853 | 0.191 | 0.2 | 1 | 14 | 2510039O1 | -0.009 | Old |
| 0.99388 | 0.925918 | 0.077 | 0.08 | 1 | 14 | Mfsd12 | -0.003 | Old |
| 0.99388 | 0.43924 | 0.077 | 0.08 | 1 | 14 | Slfn2 | -0.003 | Old |
| 0.99388 | 0.351577 | 0.077 | 0.08 | 1 | 14 | Wasf3 | -0.003 | Old |
| 0.994257 | 0.079871 | 0.311 | 0.32 | 1 | 14 | 2810474O1 | -0.009 | Old |
| 0.994257 | 0.040463 | 0.311 | 0.32 | 1 | 14 | Edf1 | -0.009 | Old |
| 0.994308 | 0.144901 | 0.175 | 0.18 | 1 | 14 | Gga2 | -0.005 | Old |
| 0.994415 | 0.782382 | 0.093 | 0.1 | 1 | 14 | Stx18 | -0.007 | Old |
| 0.995265 | 0.292096 | 0.137 | 0.14 | 1 | 14 | Sowahc | -0.003 | Old |
| 0.995619 | 0.170274 | 0.361 | 0.38 | 1 | 14 | Psmc3 | -0.019 | Old |
| 0.995688 | 0.522951 | 0.169 | 0.18 | 1 | 14 | Klhl28 | -0.011 | Old |
| 0.995958 | 0.056596 | 0.202 | 0.2 | 1 | 14 | Esr1 | 0.002 | Old |
| 0.996168 | 0.175551 | 0.23 | 0.24 | 1 | 14 | Exoc3 | -0.01 | Old |
| 0.99635 | 0.274542 | 0.098 | 0.1 | 1 | 14 | P3h1 | -0.002 | Old |
| 0.997171 | 1.521096 | 0.038 | 0.04 | 1 | 14 | Sfmbt1 | -0.002 | Old |
| 0.997171 | 1.432452 | 0.038 | 0.04 | 1 | 14 | Glis3 | -0.002 | Old |
| 0.997171 | 1.20642 | 0.038 | 0.04 | 1 | 14 | Ankrd29 | -0.002 | Old |
| 0.997171 | 1.105256 | 0.038 | 0.04 | 1 | 14 | Hoxa11 | -0.002 | Old |
| 0.997171 | 1.085071 | 0.038 | 0.04 | 1 | 14 | Cchcr1 | -0.002 | Old |
| 0.997171 | 1.009692 | 0.038 | 0.04 | 1 | 14 | Fam83f | -0.002 | Old |
| 0.997171 | 0.940297 | 0.038 | 0.04 | 1 | 14 | Tmem26 | -0.002 | Old |
| 0.997171 | 0.830925 | 0.038 | 0.04 | 1 | 14 | Wdr55 | -0.002 | Old |
| 0.997171 | 0.40843 | 0.038 | 0.04 | 1 | 14 | Rfc5 | -0.002 | Old |
| 0.997707 | 0.308317 | 0.06 | 0.06 | 1 | 14 | Kcnk5 | 0 | Old |
| 0.997707 | 0.287084 | 0.06 | 0.06 | 1 | 14 | Rxrb | 0 | Old |
| 0.997707 | 0.213643 | 0.06 | 0.06 | 1 | 14 | Irf5 | 0 | Old |
| 0.997707 | 0.070974 | 0.06 | 0.06 | 1 | 14 | Dhcr7 | 0 | Old |
| 0.99796 | 1.248069 | 0.077 | 0.08 | 1 | 14 | Lmo1 | -0.003 | Old |
| 0.99796 | 0.741335 | 0.077 | 0.08 | 1 | 14 | Lmnb1 | -0.003 | Old |
| 0.99796 | 0.503382 | 0.077 | 0.08 | 1 | 14 | Ctu2 | -0.003 | Old |
| 0.99796 | 0.433638 | 0.077 | 0.08 | 1 | 14 | Mpnd | -0.003 | Old |
| 0.998731 | 0.421032 | 0.235 | 0.24 | 1 | 14 | Pbx2 | -0.005 | Old |
| 0.998812 | 0.272488 | 0.279 | 0.3 | 1 | 14 | Brd1 | -0.021 | Old |
| 0.99883 | 0.49245 | 0.29 | 0.32 | 1 | 14 | S1pr3 | -0.03 | Old |

| | | | | | | | | |
|---|---|---|---|---|---|---|---|---|
| 4.42E-07 | 4.820427 | 0.2 | 0.016 | 0.013716 | 14 | Slc38a5 | 0.184 | Young |
| 4.70E-07 | 6.027374 | 0.16 | 0.005 | 0.014596 | 14 | Wfdc1 | 0.155 | Young |
| 7.92E-07 | 3.553631 | 0.36 | 0.087 | 0.024609 | 14 | Myl9 | 0.273 | Young |
| 2.22E-06 | 6.653414 | 0.12 | 0 | 0.068819 | 14 | Casq2 | 0.12 | Young |
| 3.99E-06 | 4.414786 | 0.14 | 0.005 | 0.124016 | 14 | Ptn | 0.135 | Young |
| 3.99E-06 | 4.387623 | 0.14 | 0.005 | 0.124016 | 14 | Hhip | 0.135 | Young |
| 4.14E-06 | 4.282512 | 0.14 | 0.005 | 0.128572 | 14 | Pdlim3 | 0.135 | Young |
| 4.19E-06 | 3.043806 | 0.18 | 0.016 | 0.129983 | 14 | Ednra | 0.164 | Young |
| 4.19E-06 | 2.946973 | 0.18 | 0.016 | 0.129983 | 14 | Col4a3 | 0.164 | Young |
| 4.93E-06 | 2.731065 | 0.16 | 0.011 | 0.15324 | 14 | Kcnd3 | 0.149 | Young |
| 8.84E-06 | 3.948525 | 0.2 | 0.027 | 0.274576 | 14 | Gng2 | 0.173 | Young |
| 1.63E-05 | 7.076578 | 0.1 | 0 | 0.506562 | 14 | Tnnt2 | 0.1 | Young |
| 1.63E-05 | 5.047348 | 0.1 | 0 | 0.506562 | 14 | Aatk | 0.1 | Young |
| 1.63E-05 | 4.212897 | 0.1 | 0 | 0.506562 | 14 | Myom1 | 0.1 | Young |
| 1.63E-05 | 4.183763 | 0.1 | 0 | 0.506562 | 14 | Cacnb2 | 0.1 | Young |
| 2.28E-05 | 3.450414 | 0.16 | 0.016 | 0.707719 | 14 | Cplx2 | 0.144 | Young |
| 2.45E-05 | 2.76502 | 0.3 | 0.077 | 0.761501 | 14 | Fam53b | 0.223 | Young |
| 2.98E-05 | 3.286954 | 0.12 | 0.005 | 0.925646 | 14 | Tspan13 | 0.115 | Young |
| 3.05E-05 | 3.565101 | 0.14 | 0.011 | 0.947926 | 14 | Poli | 0.129 | Young |
| 3.31E-05 | 2.451158 | 0.28 | 0.071 | 1 | 14 | Cpe | 0.209 | Young |
| 4.13E-05 | 0.754739 | 0.54 | 0.257 | 1 | 14 | Mgp | 0.283 | Young |
| 9.21E-05 | 3.422652 | 0.22 | 0.049 | 1 | 14 | Fbxo32 | 0.171 | Young |
| 0.000105 | 2.039385 | 0.16 | 0.022 | 1 | 14 | Gtpbp3 | 0.138 | Young |
| 0.00012 | 6.121438 | 0.08 | 0 | 1 | 14 | Car14 | 0.08 | Young |
| 0.00012 | 6.065042 | 0.08 | 0 | 1 | 14 | Cnn1 | 0.08 | Young |
| 0.00012 | 6.054244 | 0.08 | 0 | 1 | 14 | Wnt6 | 0.08 | Young |
| 0.00012 | 5.942795 | 0.08 | 0 | 1 | 14 | Npnt | 0.08 | Young |
| 0.00012 | 4.886039 | 0.08 | 0 | 1 | 14 | Chn1 | 0.08 | Young |
| 0.00012 | 4.717652 | 0.08 | 0 | 1 | 14 | Pgm5 | 0.08 | Young |
| 0.00012 | 3.599914 | 0.08 | 0 | 1 | 14 | Efna1 | 0.08 | Young |
| 0.000158 | 1.288194 | 0.5 | 0.24 | 1 | 14 | Chpf | 0.26 | Young |
| 0.000162 | 4.470325 | 0.12 | 0.011 | 1 | 14 | Dnmt3l | 0.109 | Young |
| 0.000167 | 4.123259 | 0.12 | 0.011 | 1 | 14 | Dtnb | 0.109 | Young |
| 0.000183 | 3.175814 | 0.12 | 0.011 | 1 | 14 | Pcbp3 | 0.109 | Young |
| 0.000184 | 4.268077 | 0.1 | 0.005 | 1 | 14 | Tenm2 | 0.095 | Young |
| 0.000197 | 4.370401 | 0.1 | 0.005 | 1 | 14 | Nrg2 | 0.095 | Young |
| 0.000197 | 3.568156 | 0.1 | 0.005 | 1 | 14 | Kcne4 | 0.095 | Young |
| 0.000211 | 2.401158 | 0.1 | 0.005 | 1 | 14 | Adgrg1 | 0.095 | Young |
| 0.000211 | 2.387649 | 0.1 | 0.005 | 1 | 14 | Unc119 | 0.095 | Young |
| 0.000223 | 2.202889 | 0.28 | 0.087 | 1 | 14 | Tpm2 | 0.193 | Young |
| 0.000225 | 1.443099 | 0.18 | 0.033 | 1 | 14 | Tagln | 0.147 | Young |
| 0.000234 | 2.904988 | 0.22 | 0.055 | 1 | 14 | Acta2 | 0.165 | Young |
| 0.000239 | 3.038756 | 0.24 | 0.066 | 1 | 14 | Itm2a | 0.174 | Young |
| 0.000315 | 1.581079 | 0.2 | 0.044 | 1 | 14 | Gnai1 | 0.156 | Young |
| 0.000338 | 3.496083 | 0.2 | 0.049 | 1 | 14 | Fry | 0.151 | Young |

| | | | | | | | | |
|---|---|---|---|---|---|---|---|---|
| 0.000478 | 2.120479 | 0.36 | 0.142 | 1 | 14 | Klhdc8b | 0.218 | Young |
| 0.000484 | 1.691611 | 0.2 | 0.049 | 1 | 14 | Rspo3 | 0.151 | Young |
| 0.000492 | 2.510888 | 0.14 | 0.022 | 1 | 14 | Gm26802 | 0.118 | Young |
| 0.000529 | 3.385945 | 0.14 | 0.022 | 1 | 14 | Narfl | 0.118 | Young |
| 0.000555 | 2.758783 | 0.14 | 0.022 | 1 | 14 | Cacna1d | 0.118 | Young |
| 0.000555 | 1.304319 | 0.14 | 0.022 | 1 | 14 | Nes | 0.118 | Young |
| 0.000576 | 2.135675 | 0.18 | 0.038 | 1 | 14 | Slc7a2 | 0.142 | Young |
| 0.000582 | 2.222899 | 0.14 | 0.022 | 1 | 14 | Tfap4 | 0.118 | Young |
| 0.000675 | 1.255177 | 0.62 | 0.399 | 1 | 14 | Hspa1a | 0.221 | Young |
| 0.000689 | 0.795377 | 0.18 | 0.038 | 1 | 14 | Mcam | 0.142 | Young |
| 0.000692 | 1.46563 | 0.4 | 0.186 | 1 | 14 | Cav1 | 0.214 | Young |
| 0.000709 | 3.407294 | 0.12 | 0.016 | 1 | 14 | Stk26 | 0.104 | Young |
| 0.000827 | 2.704876 | 0.22 | 0.066 | 1 | 14 | Myh11 | 0.154 | Young |
| 0.000889 | 1.822972 | 0.18 | 0.044 | 1 | 14 | Ank3 | 0.136 | Young |
| 0.000894 | 6.194052 | 0.06 | 0 | 1 | 14 | Ptpn5 | 0.06 | Young |
| 0.000894 | 5.774496 | 0.06 | 0 | 1 | 14 | Dlgap1 | 0.06 | Young |
| 0.000894 | 4.922402 | 0.06 | 0 | 1 | 14 | Itga8 | 0.06 | Young |
| 0.000894 | 4.786277 | 0.06 | 0 | 1 | 14 | Fbxl22 | 0.06 | Young |
| 0.000894 | 4.707133 | 0.06 | 0 | 1 | 14 | Pcp4l1 | 0.06 | Young |
| 0.000894 | 4.25536 | 0.06 | 0 | 1 | 14 | Tspan15 | 0.06 | Young |
| 0.000894 | 4.113329 | 0.06 | 0 | 1 | 14 | Amigo3 | 0.06 | Young |
| 0.000894 | 4.062059 | 0.06 | 0 | 1 | 14 | Stac | 0.06 | Young |
| 0.000894 | 4.014146 | 0.06 | 0 | 1 | 14 | Gja4 | 0.06 | Young |
| 0.000894 | 3.943253 | 0.06 | 0 | 1 | 14 | Chrnb1 | 0.06 | Young |
| 0.000894 | 3.885316 | 0.06 | 0 | 1 | 14 | Cap2 | 0.06 | Young |
| 0.000894 | 3.883751 | 0.06 | 0 | 1 | 14 | Abcc3 | 0.06 | Young |
| 0.000894 | 3.820249 | 0.06 | 0 | 1 | 14 | Gpr4 | 0.06 | Young |
| 0.000894 | 3.497144 | 0.06 | 0 | 1 | 14 | Pcdh11x | 0.06 | Young |
| 0.000894 | 3.333052 | 0.06 | 0 | 1 | 14 | Adgrl3 | 0.06 | Young |
| 0.000894 | 2.651368 | 0.06 | 0 | 1 | 14 | Clstn2 | 0.06 | Young |
| 0.000929 | 2.866699 | 0.2 | 0.055 | 1 | 14 | Crispld1 | 0.145 | Young |
| 0.000943 | 1.064535 | 0.52 | 0.251 | 1 | 14 | Exoc4 | 0.269 | Young |
| 0.001 | 1.362248 | 0.16 | 0.033 | 1 | 14 | Ccl12 | 0.127 | Young |
| 0.00103 | 4.406533 | 0.1 | 0.011 | 1 | 14 | Tnnt1 | 0.089 | Young |
| 0.00103 | 3.672555 | 0.1 | 0.011 | 1 | 14 | Rab36 | 0.089 | Young |
| 0.001059 | 4.002966 | 0.1 | 0.011 | 1 | 14 | Hbegf | 0.089 | Young |
| 0.001059 | 3.761048 | 0.1 | 0.011 | 1 | 14 | Prdm1 | 0.089 | Young |
| 0.001059 | 2.892444 | 0.1 | 0.011 | 1 | 14 | Zfp775 | 0.089 | Young |
| 0.00109 | 3.153014 | 0.1 | 0.011 | 1 | 14 | Mamdc2 | 0.089 | Young |
| 0.001117 | 1.62641 | 0.38 | 0.175 | 1 | 14 | Cul5 | 0.205 | Young |
| 0.001121 | 2.584449 | 0.1 | 0.011 | 1 | 14 | Rbpms2 | 0.089 | Young |
| 0.001153 | 2.496792 | 0.1 | 0.011 | 1 | 14 | Foxs1 | 0.089 | Young |
| 0.00122 | 2.917255 | 0.1 | 0.011 | 1 | 14 | 4933427D1 | 0.089 | Young |
| 0.001225 | 1.863661 | 0.2 | 0.055 | 1 | 14 | Mpzl1 | 0.145 | Young |
| 0.001278 | 3.618009 | 0.08 | 0.005 | 1 | 14 | Cldn11 | 0.075 | Young |

| | | | | | | | | |
|---|---|---|---|---|---|---|---|---|
| 0.001321 | 4.155502 | 0.08 | 0.005 | 1 | 14 | Lipg | 0.075 | Young |
| 0.001321 | 4.001201 | 0.08 | 0.005 | 1 | 14 | Emcn | 0.075 | Young |
| 0.001321 | 3.518863 | 0.08 | 0.005 | 1 | 14 | Arsg | 0.075 | Young |
| 0.001365 | 2.829487 | 0.08 | 0.005 | 1 | 14 | Nek8 | 0.075 | Young |
| 0.001365 | 2.828796 | 0.08 | 0.005 | 1 | 14 | Lmcd1 | 0.075 | Young |
| 0.001503 | 3.573362 | 0.14 | 0.027 | 1 | 14 | Qtrt1 | 0.113 | Young |
| 0.001566 | 3.024239 | 0.2 | 0.06 | 1 | 14 | Asl | 0.14 | Young |
| 0.001618 | 2.544781 | 0.16 | 0.038 | 1 | 14 | Mllt11 | 0.122 | Young |
| 0.001676 | 1.90187 | 0.34 | 0.158 | 1 | 14 | Srf | 0.182 | Young |
| 0.001732 | 1.975235 | 0.28 | 0.109 | 1 | 14 | Chsy1 | 0.171 | Young |
| 0.001766 | 2.409879 | 0.18 | 0.049 | 1 | 14 | Ano1 | 0.131 | Young |
| 0.001818 | 1.057921 | 0.14 | 0.027 | 1 | 14 | Ajuba | 0.113 | Young |
| 0.001896 | 0.890622 | 0.14 | 0.027 | 1 | 14 | Gla | 0.113 | Young |
| 0.002068 | 1.624623 | 0.16 | 0.038 | 1 | 14 | Abcc9 | 0.122 | Young |
| 0.002159 | 1.808161 | 0.22 | 0.071 | 1 | 14 | Zbtb21 | 0.149 | Young |
| 0.002183 | 2.647075 | 0.12 | 0.022 | 1 | 14 | Fgfr2 | 0.098 | Young |
| 0.002201 | 1.326347 | 0.34 | 0.142 | 1 | 14 | Ufl1 | 0.198 | Young |
| 0.002373 | 1.405936 | 0.24 | 0.082 | 1 | 14 | Vps50 | 0.158 | Young |
| 0.002663 | 1.624051 | 0.26 | 0.098 | 1 | 14 | Slc39a3 | 0.162 | Young |
| 0.002979 | 2.397568 | 0.14 | 0.033 | 1 | 14 | Ltc4s | 0.107 | Young |
| 0.003079 | 1.795222 | 0.36 | 0.18 | 1 | 14 | Lbh | 0.18 | Young |
| 0.003114 | 0.65695 | 0.12 | 0.022 | 1 | 14 | Gm13889 | 0.098 | Young |
| 0.003157 | 2.209269 | 0.14 | 0.033 | 1 | 14 | Vrk1 | 0.107 | Young |
| 0.003161 | 2.093212 | 0.2 | 0.066 | 1 | 14 | Polk | 0.134 | Young |
| 0.003161 | 1.79277 | 0.2 | 0.066 | 1 | 14 | Slc24a3 | 0.134 | Young |
| 0.003257 | 0.906183 | 0.42 | 0.224 | 1 | 14 | Kctd17 | 0.196 | Young |
| 0.003475 | 2.169038 | 0.14 | 0.033 | 1 | 14 | Carmn | 0.107 | Young |
| 0.003503 | 1.290425 | 0.24 | 0.087 | 1 | 14 | Cmas | 0.153 | Young |
| 0.003724 | 3.767548 | 0.1 | 0.016 | 1 | 14 | Gpld1 | 0.084 | Young |
| 0.003734 | 1.149842 | 0.26 | 0.098 | 1 | 14 | Neurl4 | 0.162 | Young |
| 0.003894 | 1.775664 | 0.14 | 0.033 | 1 | 14 | Pde5a | 0.107 | Young |
| 0.003894 | 1.243678 | 0.14 | 0.033 | 1 | 14 | Dapp1 | 0.107 | Young |
| 0.004097 | 2.176182 | 0.1 | 0.016 | 1 | 14 | Gsap | 0.084 | Young |
| 0.004102 | 2.281744 | 0.16 | 0.044 | 1 | 14 | Fxyd6 | 0.116 | Young |
| 0.004102 | 0.709875 | 0.16 | 0.044 | 1 | 14 | Exosc2 | 0.116 | Young |
| 0.004167 | 1.300238 | 0.24 | 0.087 | 1 | 14 | Usp21 | 0.153 | Young |
| 0.004184 | 1.987801 | 0.32 | 0.148 | 1 | 14 | Maged2 | 0.172 | Young |
| 0.004296 | 1.984824 | 0.1 | 0.016 | 1 | 14 | Rbl1 | 0.084 | Young |
| 0.004476 | 1.701014 | 0.4 | 0.208 | 1 | 14 | Cryab | 0.192 | Young |
| 0.004503 | 1.849739 | 0.1 | 0.016 | 1 | 14 | Atg10 | 0.084 | Young |
| 0.004536 | 2.729153 | 0.2 | 0.071 | 1 | 14 | Fblim1 | 0.129 | Young |
| 0.00463 | 1.144887 | 0.24 | 0.087 | 1 | 14 | Pigq | 0.153 | Young |
| 0.004767 | 0.872002 | 0.52 | 0.317 | 1 | 14 | Ncoa1 | 0.203 | Young |
| 0.004901 | 1.420453 | 0.24 | 0.093 | 1 | 14 | Emp2 | 0.147 | Young |
| 0.004935 | 1.099193 | 0.36 | 0.186 | 1 | 14 | Fam91a1 | 0.174 | Young |

| | | | | | | | | |
|---|---|---|---|---|---|---|---|---|
| 0.004953 | 1.186317 | 0.62 | 0.497 | 1 | 14 | Pfdn5 | 0.123 | Young |
| 0.005134 | 2.217349 | 0.54 | 0.344 | 1 | 14 | Tpm1 | 0.196 | Young |
| 0.005396 | 1.776598 | 0.3 | 0.142 | 1 | 14 | Sema5a | 0.158 | Young |
| 0.005838 | 3.838611 | 0.08 | 0.011 | 1 | 14 | Hopx | 0.069 | Young |
| 0.005838 | 3.669912 | 0.08 | 0.011 | 1 | 14 | Pygm | 0.069 | Young |
| 0.005985 | 1.453451 | 0.18 | 0.055 | 1 | 14 | BC034090 | 0.125 | Young |
| 0.005994 | 3.273787 | 0.08 | 0.011 | 1 | 14 | Tceal3 | 0.069 | Young |
| 0.005994 | 2.883131 | 0.08 | 0.011 | 1 | 14 | Tspan12 | 0.069 | Young |
| 0.006245 | 2.149975 | 0.12 | 0.027 | 1 | 14 | Lamc3 | 0.093 | Young |
| 0.006317 | 2.546288 | 0.08 | 0.011 | 1 | 14 | Jpt2 | 0.069 | Young |
| 0.006484 | 3.365168 | 0.08 | 0.011 | 1 | 14 | Angpt2 | 0.069 | Young |
| 0.006484 | 2.969646 | 0.08 | 0.011 | 1 | 14 | Reep1 | 0.069 | Young |
| 0.006484 | 2.6334 | 0.08 | 0.011 | 1 | 14 | Irak2 | 0.069 | Young |
| 0.006581 | 3.359219 | 0.24 | 0.104 | 1 | 14 | Ptch1 | 0.136 | Young |
| 0.006655 | 2.280265 | 0.08 | 0.011 | 1 | 14 | Mtss1l | 0.069 | Young |
| 0.006655 | 2.039547 | 0.08 | 0.011 | 1 | 14 | Aldh1l1 | 0.069 | Young |
| 0.006655 | 1.746795 | 0.08 | 0.011 | 1 | 14 | Itga7 | 0.069 | Young |
| 0.006725 | 2.520546 | 0.2 | 0.071 | 1 | 14 | Zfp651 | 0.129 | Young |
| 0.006831 | 1.556046 | 0.08 | 0.011 | 1 | 14 | Lair1 | 0.069 | Young |
| 0.006852 | 5.325355 | 0.04 | 0 | 1 | 14 | Nkapl | 0.04 | Young |
| 0.006852 | 5.13134 | 0.04 | 0 | 1 | 14 | Galr2 | 0.04 | Young |
| 0.006852 | 4.972091 | 0.04 | 0 | 1 | 14 | Gipr | 0.04 | Young |
| 0.006852 | 4.947752 | 0.04 | 0 | 1 | 14 | Tfcp2l1 | 0.04 | Young |
| 0.006852 | 4.867208 | 0.04 | 0 | 1 | 14 | 1700008J0 | 0.04 | Young |
| 0.006852 | 4.858104 | 0.04 | 0 | 1 | 14 | Frem1 | 0.04 | Young |
| 0.006852 | 4.813734 | 0.04 | 0 | 1 | 14 | Gm13861 | 0.04 | Young |
| 0.006852 | 4.813734 | 0.04 | 0 | 1 | 14 | Lin7a | 0.04 | Young |
| 0.006852 | 4.807481 | 0.04 | 0 | 1 | 14 | Pcsk9 | 0.04 | Young |
| 0.006852 | 4.807481 | 0.04 | 0 | 1 | 14 | Slc30a2 | 0.04 | Young |
| 0.006852 | 4.788558 | 0.04 | 0 | 1 | 14 | Jph1 | 0.04 | Young |
| 0.006852 | 4.480047 | 0.04 | 0 | 1 | 14 | Cacna1e | 0.04 | Young |
| 0.006852 | 4.451255 | 0.04 | 0 | 1 | 14 | Drd4 | 0.04 | Young |
| 0.006852 | 4.386787 | 0.04 | 0 | 1 | 14 | Gm48420 | 0.04 | Young |
| 0.006852 | 4.362993 | 0.04 | 0 | 1 | 14 | Ffar2 | 0.04 | Young |
| 0.006852 | 4.239028 | 0.04 | 0 | 1 | 14 | Musk | 0.04 | Young |
| 0.006852 | 4.106069 | 0.04 | 0 | 1 | 14 | Gm10827 | 0.04 | Young |
| 0.006852 | 4.096735 | 0.04 | 0 | 1 | 14 | Actg2 | 0.04 | Young |
| 0.006852 | 4.059597 | 0.04 | 0 | 1 | 14 | Kctd14 | 0.04 | Young |
| 0.006852 | 3.990121 | 0.04 | 0 | 1 | 14 | Rflna | 0.04 | Young |
| 0.006852 | 3.986046 | 0.04 | 0 | 1 | 14 | Gm12758 | 0.04 | Young |
| 0.006852 | 3.94448 | 0.04 | 0 | 1 | 14 | Dtx1 | 0.04 | Young |
| 0.006852 | 3.919169 | 0.04 | 0 | 1 | 14 | Tnni3 | 0.04 | Young |
| 0.006852 | 3.911021 | 0.04 | 0 | 1 | 14 | Tcf7 | 0.04 | Young |
| 0.006852 | 3.894348 | 0.04 | 0 | 1 | 14 | 2810454H( | 0.04 | Young |
| 0.006852 | 3.871112 | 0.04 | 0 | 1 | 14 | Homer2 | 0.04 | Young |

| | | | | | | | | |
|---|---|---|---|---|---|---|---|---|
| 0.006852 | 3.780841 | 0.04 | 0 | 1 | 14 | Ngef | 0.04 | Young |
| 0.006852 | 3.77605 | 0.04 | 0 | 1 | 14 | Susd1 | 0.04 | Young |
| 0.006852 | 3.773259 | 0.04 | 0 | 1 | 14 | Mmrn2 | 0.04 | Young |
| 0.006852 | 3.745728 | 0.04 | 0 | 1 | 14 | Myoz1 | 0.04 | Young |
| 0.006852 | 3.643463 | 0.04 | 0 | 1 | 14 | Adam11 | 0.04 | Young |
| 0.006852 | 3.602054 | 0.04 | 0 | 1 | 14 | Ankef1 | 0.04 | Young |
| 0.006852 | 3.521632 | 0.04 | 0 | 1 | 14 | Rpusd3 | 0.04 | Young |
| 0.006852 | 3.486224 | 0.04 | 0 | 1 | 14 | Mmp13 | 0.04 | Young |
| 0.006852 | 3.476192 | 0.04 | 0 | 1 | 14 | Synpo2l | 0.04 | Young |
| 0.006852 | 3.364339 | 0.04 | 0 | 1 | 14 | Fam46b | 0.04 | Young |
| 0.006852 | 3.244142 | 0.04 | 0 | 1 | 14 | Hcar1 | 0.04 | Young |
| 0.006852 | 3.244142 | 0.04 | 0 | 1 | 14 | Lgi1 | 0.04 | Young |
| 0.006852 | 3.236594 | 0.04 | 0 | 1 | 14 | Lilrb4a | 0.04 | Young |
| 0.006852 | 3.228421 | 0.04 | 0 | 1 | 14 | Prlr | 0.04 | Young |
| 0.006852 | 3.219449 | 0.04 | 0 | 1 | 14 | Glis1 | 0.04 | Young |
| 0.006852 | 3.219449 | 0.04 | 0 | 1 | 14 | Atp8a2 | 0.04 | Young |
| 0.006852 | 3.18503 | 0.04 | 0 | 1 | 14 | Acer2 | 0.04 | Young |
| 0.006852 | 3.122334 | 0.04 | 0 | 1 | 14 | Jazf1 | 0.04 | Young |
| 0.006852 | 3.108421 | 0.04 | 0 | 1 | 14 | Prelid3a | 0.04 | Young |
| 0.006852 | 3.08556 | 0.04 | 0 | 1 | 14 | Prx | 0.04 | Young |
| 0.006852 | 2.983875 | 0.04 | 0 | 1 | 14 | Peli3 | 0.04 | Young |
| 0.006852 | 2.913134 | 0.04 | 0 | 1 | 14 | Gm13830 | 0.04 | Young |
| 0.006852 | 2.886937 | 0.04 | 0 | 1 | 14 | Dusp15 | 0.04 | Young |
| 0.006852 | 2.866234 | 0.04 | 0 | 1 | 14 | Gm19705 | 0.04 | Young |
| 0.006852 | 2.821251 | 0.04 | 0 | 1 | 14 | Gm16740 | 0.04 | Young |
| 0.006852 | 2.759051 | 0.04 | 0 | 1 | 14 | P4ha3 | 0.04 | Young |
| 0.006852 | 2.728706 | 0.04 | 0 | 1 | 14 | Reep6 | 0.04 | Young |
| 0.006852 | 2.568598 | 0.04 | 0 | 1 | 14 | Hey1 | 0.04 | Young |
| 0.006852 | 2.568598 | 0.04 | 0 | 1 | 14 | Abcb1a | 0.04 | Young |
| 0.006852 | 2.568598 | 0.04 | 0 | 1 | 14 | Nrip2 | 0.04 | Young |
| 0.006852 | 2.559606 | 0.04 | 0 | 1 | 14 | Gm30524 | 0.04 | Young |
| 0.006852 | 2.551271 | 0.04 | 0 | 1 | 14 | Ldb3 | 0.04 | Young |
| 0.006852 | 2.442769 | 0.04 | 0 | 1 | 14 | Zwilch | 0.04 | Young |
| 0.006852 | 2.393493 | 0.04 | 0 | 1 | 14 | Hist1h4h | 0.04 | Young |
| 0.006852 | 2.363204 | 0.04 | 0 | 1 | 14 | Hoxb9 | 0.04 | Young |
| 0.006852 | 2.354188 | 0.04 | 0 | 1 | 14 | Aldh1l2 | 0.04 | Young |
| 0.006852 | 2.293807 | 0.04 | 0 | 1 | 14 | Tbx3os1 | 0.04 | Young |
| 0.006852 | 2.263288 | 0.04 | 0 | 1 | 14 | Tal1 | 0.04 | Young |
| 0.006852 | 2.248934 | 0.04 | 0 | 1 | 14 | Upb1 | 0.04 | Young |
| 0.006852 | 2.216587 | 0.04 | 0 | 1 | 14 | Rwdd2a | 0.04 | Young |
| 0.006852 | 2.195134 | 0.04 | 0 | 1 | 14 | Pnck | 0.04 | Young |
| 0.006852 | 2.116618 | 0.04 | 0 | 1 | 14 | Wdr27 | 0.04 | Young |
| 0.006852 | 2.020527 | 0.04 | 0 | 1 | 14 | Lyrm7 | 0.04 | Young |
| 0.006852 | 1.848713 | 0.04 | 0 | 1 | 14 | Ccdc159 | 0.04 | Young |
| 0.006852 | 1.581646 | 0.04 | 0 | 1 | 14 | Fbxo17 | 0.04 | Young |

| | | | | | | | | |
|---|---|---|---|---|---|---|---|---|
| 0.006852 | 1.581646 | 0.04 | 0 | 1 | 14 | Hunk | 0.04 | Young |
| 0.007085 | 2.726585 | 0.14 | 0.038 | 1 | 14 | Cgnl1 | 0.102 | Young |
| 0.007194 | 1.662024 | 0.08 | 0.011 | 1 | 14 | Pms2 | 0.069 | Young |
| 0.007194 | 1.475926 | 0.08 | 0.011 | 1 | 14 | Sh2d3c | 0.069 | Young |
| 0.007194 | 1.257195 | 0.08 | 0.011 | 1 | 14 | Slc2a4 | 0.069 | Young |
| 0.007194 | 1.173888 | 0.08 | 0.011 | 1 | 14 | Hpgds | 0.069 | Young |
| 0.007194 | 1.113546 | 0.08 | 0.011 | 1 | 14 | Klhl29 | 0.069 | Young |
| 0.007829 | 0.804962 | 0.58 | 0.388 | 1 | 14 | Wnt5a | 0.192 | Young |
| 0.007987 | 2.777707 | 0.14 | 0.038 | 1 | 14 | Usp35 | 0.102 | Young |
| 0.008096 | 1.169729 | 0.34 | 0.169 | 1 | 14 | Wipf2 | 0.171 | Young |
| 0.008193 | 1.690693 | 0.22 | 0.082 | 1 | 14 | Mtf1 | 0.138 | Young |
| 0.008203 | 2.048062 | 0.16 | 0.049 | 1 | 14 | Mvk | 0.111 | Young |
| 0.008227 | 1.321328 | 0.46 | 0.284 | 1 | 14 | Lbp | 0.176 | Young |
| 0.008395 | 4.456862 | 0.06 | 0.005 | 1 | 14 | Sh3bgr | 0.055 | Young |
| 0.008395 | 3.87831 | 0.06 | 0.005 | 1 | 14 | Bhlhe22 | 0.055 | Young |
| 0.008395 | 2.9671 | 0.06 | 0.005 | 1 | 14 | Gstm7 | 0.055 | Young |
| 0.008395 | 2.924945 | 0.06 | 0.005 | 1 | 14 | Slc2a6 | 0.055 | Young |
| 0.008593 | 1.066213 | 0.2 | 0.071 | 1 | 14 | Hmbs | 0.129 | Young |
| 0.008659 | 4.174318 | 0.06 | 0.005 | 1 | 14 | Ctnnal1 | 0.055 | Young |
| 0.008659 | 3.160344 | 0.06 | 0.005 | 1 | 14 | Chrna1os | 0.055 | Young |
| 0.008659 | 2.955813 | 0.06 | 0.005 | 1 | 14 | Zfp839 | 0.055 | Young |
| 0.008719 | 1.370654 | 0.54 | 0.393 | 1 | 14 | Ckb | 0.147 | Young |
| 0.008726 | 1.363517 | 0.16 | 0.049 | 1 | 14 | Ntmt1 | 0.111 | Young |
| 0.008931 | 2.704347 | 0.06 | 0.005 | 1 | 14 | Elmod1 | 0.055 | Young |
| 0.008931 | 2.668763 | 0.06 | 0.005 | 1 | 14 | Bcat1 | 0.055 | Young |
| 0.008931 | 2.657824 | 0.06 | 0.005 | 1 | 14 | Amigo2 | 0.055 | Young |
| 0.008931 | 2.167492 | 0.06 | 0.005 | 1 | 14 | Vps51 | 0.055 | Young |
| 0.008931 | 2.006439 | 0.06 | 0.005 | 1 | 14 | Lmo7 | 0.055 | Young |
| 0.008942 | 2.688427 | 0.18 | 0.06 | 1 | 14 | Tmem80 | 0.12 | Young |
| 0.009138 | 1.820113 | 0.16 | 0.049 | 1 | 14 | Mri1 | 0.111 | Young |
| 0.009199 | 0.840945 | 0.5 | 0.301 | 1 | 14 | Zfp703 | 0.199 | Young |
| 0.00921 | 2.514663 | 0.06 | 0.005 | 1 | 14 | Acsbg1 | 0.055 | Young |
| 0.00921 | 2.496272 | 0.06 | 0.005 | 1 | 14 | Cacna2d2 | 0.055 | Young |
| 0.00921 | 2.037023 | 0.06 | 0.005 | 1 | 14 | Tmem81 | 0.055 | Young |
| 0.00921 | 1.88298 | 0.06 | 0.005 | 1 | 14 | Dok4 | 0.055 | Young |
| 0.00921 | 1.736694 | 0.06 | 0.005 | 1 | 14 | Ino80dos | 0.055 | Young |
| 0.00921 | 1.486798 | 0.06 | 0.005 | 1 | 14 | 4931428F0 | 0.055 | Young |
| 0.00921 | 1.324438 | 0.06 | 0.005 | 1 | 14 | Tenm4 | 0.055 | Young |
| 0.00921 | 0.411804 | 0.06 | 0.005 | 1 | 14 | Cox4i2 | 0.055 | Young |
| 0.009279 | 1.098714 | 0.16 | 0.049 | 1 | 14 | Cav2 | 0.111 | Young |
| 0.009332 | 0.491537 | 0.18 | 0.06 | 1 | 14 | Snai2 | 0.12 | Young |
| 0.009469 | 0.659948 | 0.24 | 0.093 | 1 | 14 | Slc20a2 | 0.147 | Young |
| 0.009648 | 4.178855 | 0.1 | 0.022 | 1 | 14 | Gnaz | 0.078 | Young |
| 0.010052 | 3.362536 | 0.1 | 0.022 | 1 | 14 | Kcnn3 | 0.078 | Young |
| 0.010158 | 0.397174 | 0.22 | 0.082 | 1 | 14 | Usp39 | 0.138 | Young |

| | | | | | | | | |
|---|---|---|---|---|---|---|---|---|
| 0.010286 | 0.526047 | 0.22 | 0.082 | 1 | 14 | Ppp1r14a | 0.138 | Young |
| 0.010302 | 1.186829 | 0.4 | 0.235 | 1 | 14 | Adamtsl1 | 0.165 | Young |
| 0.010687 | 2.611493 | 0.1 | 0.022 | 1 | 14 | P3h2 | 0.078 | Young |
| 0.010714 | 1.365347 | 0.26 | 0.12 | 1 | 14 | Stk17b | 0.14 | Young |
| 0.010907 | 2.193399 | 0.1 | 0.022 | 1 | 14 | Rgs4 | 0.078 | Young |
| 0.01113 | 2.125964 | 0.1 | 0.022 | 1 | 14 | Vps37b | 0.078 | Young |
| 0.01113 | 1.012626 | 0.1 | 0.022 | 1 | 14 | Mnd1 | 0.078 | Young |
| 0.01127 | 0.688484 | 0.6 | 0.432 | 1 | 14 | Uqcrb | 0.168 | Young |
| 0.011358 | 2.259619 | 0.1 | 0.022 | 1 | 14 | Synpo | 0.078 | Young |
| 0.011358 | 1.991479 | 0.1 | 0.022 | 1 | 14 | Eif2b2 | 0.078 | Young |
| 0.011397 | 0.634973 | 0.42 | 0.235 | 1 | 14 | Kdm5b | 0.185 | Young |
| 0.01159 | 2.00126 | 0.1 | 0.022 | 1 | 14 | C5ar2 | 0.078 | Young |
| 0.011774 | 3.316515 | 0.12 | 0.033 | 1 | 14 | Tmco4 | 0.087 | Young |
| 0.011825 | 1.131494 | 0.22 | 0.087 | 1 | 14 | Prorsd1 | 0.133 | Young |
| 0.011826 | 1.677479 | 0.1 | 0.022 | 1 | 14 | Map7d2 | 0.078 | Young |
| 0.011842 | 1.589189 | 0.24 | 0.104 | 1 | 14 | Piezo1 | 0.136 | Young |
| 0.012066 | 0.237568 | 0.1 | 0.022 | 1 | 14 | Ndufa4l2 | 0.078 | Young |
| 0.012181 | 1.363508 | 0.18 | 0.066 | 1 | 14 | Pomgnt1 | 0.114 | Young |
| 0.012194 | 2.524004 | 0.12 | 0.033 | 1 | 14 | Specc1 | 0.087 | Young |
| 0.01231 | 1.99745 | 0.1 | 0.022 | 1 | 14 | Mrgbp | 0.078 | Young |
| 0.012347 | 2.100297 | 0.18 | 0.066 | 1 | 14 | Abr | 0.114 | Young |
| 0.012464 | 2.829276 | 0.14 | 0.044 | 1 | 14 | Rrnad1 | 0.096 | Young |
| 0.012559 | 0.791106 | 0.1 | 0.022 | 1 | 14 | Rab26os | 0.078 | Young |
| 0.013327 | 1.268302 | 0.28 | 0.131 | 1 | 14 | Uhrf1bp1l | 0.149 | Young |
| 0.013417 | 1.494163 | 0.16 | 0.055 | 1 | 14 | Mphosph1 | 0.105 | Young |
| 0.013478 | 2.01795 | 0.14 | 0.044 | 1 | 14 | Daam2 | 0.096 | Young |
| 0.01356 | 1.30502 | 0.18 | 0.066 | 1 | 14 | Farsa | 0.114 | Young |
| 0.013652 | 2.233235 | 0.18 | 0.071 | 1 | 14 | Bok | 0.109 | Young |
| 0.013785 | 1.465601 | 0.22 | 0.093 | 1 | 14 | Tom1 | 0.127 | Young |
| 0.013795 | 1.901295 | 0.14 | 0.044 | 1 | 14 | Sorbs1 | 0.096 | Young |
| 0.013834 | 1.33401 | 0.2 | 0.077 | 1 | 14 | Bzw2 | 0.123 | Young |
| 0.013903 | 1.776308 | 0.14 | 0.044 | 1 | 14 | Cbfa2t3 | 0.096 | Young |
| 0.014204 | 1.181821 | 0.16 | 0.055 | 1 | 14 | Cables2 | 0.105 | Young |
| 0.01449 | 1.212834 | 0.18 | 0.066 | 1 | 14 | Dnajc15 | 0.114 | Young |
| 0.014782 | 0.841577 | 0.22 | 0.093 | 1 | 14 | Ino80 | 0.127 | Young |
| 0.014789 | 2.120143 | 0.14 | 0.044 | 1 | 14 | Mccc2 | 0.096 | Young |
| 0.014907 | 0.819731 | 0.2 | 0.077 | 1 | 14 | Psmg2 | 0.123 | Young |
| 0.014955 | 1.461919 | 0.22 | 0.093 | 1 | 14 | Ubald1 | 0.127 | Young |
| 0.015246 | 1.22333 | 0.16 | 0.055 | 1 | 14 | Slc9a8 | 0.105 | Young |
| 0.015251 | 1.585922 | 0.14 | 0.044 | 1 | 14 | Cd209d | 0.096 | Young |
| 0.015486 | 1.021354 | 0.14 | 0.044 | 1 | 14 | Hivep3 | 0.096 | Young |
| 0.015582 | 1.131954 | 0.26 | 0.12 | 1 | 14 | Mrpl10 | 0.14 | Young |
| 0.015612 | 1.194368 | 0.3 | 0.153 | 1 | 14 | Mrpl12 | 0.147 | Young |
| 0.015674 | 0.413506 | 0.9 | 0.885 | 1 | 14 | Hspa8 | 0.015 | Young |
| 0.015725 | 2.278683 | 0.14 | 0.044 | 1 | 14 | Hspa2 | 0.096 | Young |

| | | | | | | | | |
|---|---|---|---|---|---|---|---|---|
| 0.015788 | 1.07332 | 0.12 | 0.033 | 1 | 14 | Pitpnm3 | 0.087 | Young |
| 0.015832 | 1.479077 | 0.2 | 0.082 | 1 | 14 | Ifitm1 | 0.118 | Young |
| 0.015902 | 0.97287 | 0.16 | 0.055 | 1 | 14 | Bicral | 0.105 | Young |
| 0.016358 | 1.052508 | 0.36 | 0.191 | 1 | 14 | Tomm70a | 0.169 | Young |
| 0.016587 | 1.475975 | 0.22 | 0.093 | 1 | 14 | Xylt2 | 0.127 | Young |
| 0.016649 | 1.452199 | 0.28 | 0.142 | 1 | 14 | Twistnb | 0.138 | Young |
| 0.016712 | 0.809225 | 0.14 | 0.044 | 1 | 14 | Arl4c | 0.096 | Young |
| 0.016813 | 1.657978 | 0.28 | 0.142 | 1 | 14 | 1810030O( | 0.138 | Young |
| 0.017004 | 1.295112 | 0.28 | 0.137 | 1 | 14 | Jam3 | 0.143 | Young |
| 0.017038 | 1.980908 | 0.22 | 0.104 | 1 | 14 | Col4a4 | 0.116 | Young |
| 0.01729 | 1.116315 | 0.26 | 0.12 | 1 | 14 | Rab3gap2 | 0.14 | Young |
| 0.017479 | 1.482031 | 0.38 | 0.224 | 1 | 14 | Smarce1 | 0.156 | Young |
| 0.017488 | 1.803672 | 0.14 | 0.044 | 1 | 14 | Mxd1 | 0.096 | Young |
| 0.017613 | 2.952207 | 0.08 | 0.016 | 1 | 14 | Btbd19 | 0.064 | Young |
| 0.017753 | 0.238459 | 0.14 | 0.044 | 1 | 14 | Fas | 0.096 | Young |
| 0.017777 | 0.816011 | 0.16 | 0.055 | 1 | 14 | Rptor | 0.105 | Young |
| 0.018391 | 3.457864 | 0.08 | 0.016 | 1 | 14 | Cyp4b1 | 0.064 | Young |
| 0.018471 | 1.024925 | 0.34 | 0.186 | 1 | 14 | Plekhg3 | 0.154 | Young |
| 0.018499 | 1.112205 | 0.2 | 0.082 | 1 | 14 | Ptafr | 0.118 | Young |
| 0.018791 | 2.16455 | 0.08 | 0.016 | 1 | 14 | Slc6a2 | 0.064 | Young |
| 0.018809 | 0.70172 | 0.34 | 0.18 | 1 | 14 | Tm9sf4 | 0.16 | Young |
| 0.018944 | 0.990304 | 0.58 | 0.432 | 1 | 14 | Snrpb | 0.148 | Young |
| 0.019199 | 2.501968 | 0.08 | 0.016 | 1 | 14 | Prmt6 | 0.064 | Young |
| 0.019199 | 2.299601 | 0.08 | 0.016 | 1 | 14 | Nme7 | 0.064 | Young |
| 0.019615 | 3.403554 | 0.08 | 0.016 | 1 | 14 | Gprasp2 | 0.064 | Young |
| 0.019615 | 2.428959 | 0.08 | 0.016 | 1 | 14 | Car3 | 0.064 | Young |
| 0.019733 | 0.35714 | 0.28 | 0.131 | 1 | 14 | Frs2 | 0.149 | Young |
| 0.019857 | 1.893715 | 0.2 | 0.082 | 1 | 14 | Maip1 | 0.118 | Young |
| 0.019857 | 1.275872 | 0.2 | 0.082 | 1 | 14 | Enah | 0.118 | Young |
| 0.019857 | 0.624559 | 0.2 | 0.082 | 1 | 14 | Plcg1 | 0.118 | Young |
| 0.020038 | 3.284473 | 0.08 | 0.016 | 1 | 14 | Zfp593 | 0.064 | Young |
| 0.020038 | 1.823408 | 0.08 | 0.016 | 1 | 14 | Timp4 | 0.064 | Young |
| 0.02047 | 1.970318 | 0.08 | 0.016 | 1 | 14 | Speg | 0.064 | Young |
| 0.02047 | 1.826217 | 0.08 | 0.016 | 1 | 14 | Nrm | 0.064 | Young |
| 0.02047 | 1.772905 | 0.08 | 0.016 | 1 | 14 | Cd59a | 0.064 | Young |
| 0.020652 | 1.443607 | 0.34 | 0.191 | 1 | 14 | Sf3a1 | 0.149 | Young |
| 0.020858 | 2.411717 | 0.2 | 0.087 | 1 | 14 | Oxsr1 | 0.113 | Young |
| 0.020903 | 1.268404 | 0.24 | 0.115 | 1 | 14 | Ap2a1 | 0.125 | Young |
| 0.020909 | 1.164426 | 0.08 | 0.016 | 1 | 14 | Higd1b | 0.064 | Young |
| 0.021328 | 2.251945 | 0.16 | 0.06 | 1 | 14 | Tma16 | 0.1 | Young |
| 0.021357 | 3.059403 | 0.08 | 0.016 | 1 | 14 | Fah | 0.064 | Young |
| 0.021357 | 2.433174 | 0.08 | 0.016 | 1 | 14 | Rasd1 | 0.064 | Young |
| 0.021357 | 1.192336 | 0.08 | 0.016 | 1 | 14 | Ap1g2 | 0.064 | Young |
| 0.021519 | 1.23222 | 0.6 | 0.508 | 1 | 14 | Picalm | 0.092 | Young |
| 0.021813 | 0.938097 | 0.08 | 0.016 | 1 | 14 | Rnaset2b | 0.064 | Young |

| | | | | | | | | |
|---|---|---|---|---|---|---|---|---|
| 0.022081 | 1.303874 | 0.18 | 0.071 | 1 | 14 | Gpr27 | 0.109 | Young |
| 0.022278 | 0.590239 | 0.08 | 0.016 | 1 | 14 | Mtg2 | 0.064 | Young |
| 0.022399 | 0.578029 | 0.86 | 0.76 | 1 | 14 | Rps11 | 0.1 | Young |
| 0.022751 | 1.022631 | 0.08 | 0.016 | 1 | 14 | Ncam1 | 0.064 | Young |
| 0.023293 | 2.256502 | 0.1 | 0.027 | 1 | 14 | Tigar | 0.073 | Young |
| 0.023293 | 2.095743 | 0.1 | 0.027 | 1 | 14 | Lyve1 | 0.073 | Young |
| 0.023491 | 0.778764 | 0.4 | 0.24 | 1 | 14 | Saraf | 0.16 | Young |
| 0.023913 | 1.058427 | 0.2 | 0.082 | 1 | 14 | Mylip | 0.118 | Young |
| 0.024126 | 2.532199 | 0.1 | 0.027 | 1 | 14 | Bcam | 0.073 | Young |
| 0.024126 | 2.341936 | 0.1 | 0.027 | 1 | 14 | Ehd3 | 0.073 | Young |
| 0.024126 | 2.116607 | 0.1 | 0.027 | 1 | 14 | Pycrl | 0.073 | Young |
| 0.024276 | 0.801443 | 0.26 | 0.126 | 1 | 14 | Wdr33 | 0.134 | Young |
| 0.024345 | 1.021909 | 0.18 | 0.071 | 1 | 14 | Irs1 | 0.109 | Young |
| 0.02437 | 1.117842 | 0.56 | 0.415 | 1 | 14 | Ttc3 | 0.145 | Young |
| 0.024552 | 1.773159 | 0.1 | 0.027 | 1 | 14 | Cd38 | 0.073 | Young |
| 0.024624 | 1.357243 | 0.16 | 0.06 | 1 | 14 | 2310057M | 0.1 | Young |
| 0.024716 | 0.989376 | 0.22 | 0.098 | 1 | 14 | Nbdy | 0.122 | Young |
| 0.024847 | 1.460616 | 0.14 | 0.049 | 1 | 14 | Sertad4 | 0.091 | Young |
| 0.024985 | 1.556538 | 0.1 | 0.027 | 1 | 14 | Gtf3c5 | 0.073 | Young |
| 0.025196 | 1.318323 | 0.14 | 0.049 | 1 | 14 | Aldh16a1 | 0.091 | Young |
| 0.025426 | 1.468796 | 0.12 | 0.038 | 1 | 14 | Msrb2 | 0.082 | Young |
| 0.025596 | 1.362627 | 0.16 | 0.06 | 1 | 14 | Lmod1 | 0.1 | Young |
| 0.025876 | 0.522398 | 0.22 | 0.093 | 1 | 14 | Nup93 | 0.127 | Young |
| 0.025902 | 0.602035 | 0.2 | 0.082 | 1 | 14 | Rara | 0.118 | Young |
| 0.026213 | 1.442401 | 0.18 | 0.077 | 1 | 14 | Uchl3 | 0.103 | Young |
| 0.026913 | 0.853549 | 0.22 | 0.098 | 1 | 14 | Phb | 0.122 | Young |
| 0.026943 | 0.520824 | 0.16 | 0.06 | 1 | 14 | Fli1 | 0.1 | Young |
| 0.027031 | 1.741188 | 0.12 | 0.038 | 1 | 14 | Kcna2 | 0.082 | Young |
| 0.027108 | 1.71567 | 0.38 | 0.257 | 1 | 14 | Mylk | 0.123 | Young |
| 0.027249 | 1.663109 | 0.1 | 0.027 | 1 | 14 | Mctp1 | 0.073 | Young |
| 0.027249 | 1.400344 | 0.1 | 0.027 | 1 | 14 | Cntfr | 0.073 | Young |
| 0.027386 | 1.137511 | 0.14 | 0.049 | 1 | 14 | Dusp12 | 0.091 | Young |
| 0.027386 | 0.996234 | 0.14 | 0.049 | 1 | 14 | Pop4 | 0.091 | Young |
| 0.027446 | 1.553939 | 0.12 | 0.038 | 1 | 14 | Zfp426 | 0.082 | Young |
| 0.027446 | 1.282988 | 0.12 | 0.038 | 1 | 14 | Zfp101 | 0.082 | Young |
| 0.027446 | 1.039926 | 0.12 | 0.038 | 1 | 14 | Mon1b | 0.082 | Young |
| 0.027446 | 0.558358 | 0.12 | 0.038 | 1 | 14 | Arhgap26 | 0.082 | Young |
| 0.027689 | 0.792957 | 0.5 | 0.366 | 1 | 14 | Cnpy2 | 0.134 | Young |
| 0.027723 | 1.767842 | 0.1 | 0.027 | 1 | 14 | Ptger2 | 0.073 | Young |
| 0.027766 | 0.910905 | 0.14 | 0.049 | 1 | 14 | Zc3h12c | 0.091 | Young |
| 0.027866 | 0.94357 | 0.12 | 0.038 | 1 | 14 | Inpp4b | 0.082 | Young |
| 0.028121 | 1.382881 | 0.18 | 0.077 | 1 | 14 | Nans | 0.103 | Young |
| 0.028135 | 1.560468 | 0.28 | 0.153 | 1 | 14 | Setdb1 | 0.127 | Young |
| 0.028151 | 1.136393 | 0.14 | 0.049 | 1 | 14 | Pparg | 0.091 | Young |
| 0.028292 | 1.441048 | 0.12 | 0.038 | 1 | 14 | Plpp6 | 0.082 | Young |

| | | | | | | | | |
|---|---|---|---|---|---|---|---|---|
| 0.028292 | 1.172129 | 0.12 | 0.038 | 1 | 14 | Cytip | 0.082 | Young |
| 0.028369 | 1.26393 | 0.22 | 0.098 | 1 | 14 | 2810004N2 | 0.122 | Young |
| 0.028451 | 2.021693 | 0.18 | 0.077 | 1 | 14 | Sccpdh | 0.103 | Young |
| 0.02868 | 1.921334 | 0.26 | 0.137 | 1 | 14 | Cabin1 | 0.123 | Young |
| 0.028714 | 0.888736 | 0.16 | 0.06 | 1 | 14 | Cd2bp2 | 0.1 | Young |
| 0.028783 | 1.439681 | 0.18 | 0.077 | 1 | 14 | Pus7 | 0.103 | Young |
| 0.028783 | 1.118661 | 0.18 | 0.077 | 1 | 14 | Uaca | 0.103 | Young |
| 0.029186 | 0.584065 | 0.1 | 0.027 | 1 | 14 | Pak1 | 0.073 | Young |
| 0.029215 | 1.775068 | 0.16 | 0.066 | 1 | 14 | Mink1 | 0.094 | Young |
| 0.029388 | 0.853612 | 0.38 | 0.235 | 1 | 14 | Mecp2 | 0.145 | Young |
| 0.02949 | 0.538523 | 0.26 | 0.131 | 1 | 14 | Ubxn2a | 0.129 | Young |
| 0.029603 | 0.412925 | 0.12 | 0.038 | 1 | 14 | Polr3a | 0.082 | Young |
| 0.029678 | 1.102593 | 0.22 | 0.104 | 1 | 14 | Trim27 | 0.116 | Young |
| 0.029689 | 0.310605 | 0.1 | 0.027 | 1 | 14 | Rasl12 | 0.073 | Young |
| 0.030187 | 1.138775 | 0.2 | 0.087 | 1 | 14 | Rilpl2 | 0.113 | Young |
| 0.030441 | 0.541656 | 0.58 | 0.415 | 1 | 14 | Pum2 | 0.165 | Young |
| 0.030506 | 1.160853 | 0.12 | 0.038 | 1 | 14 | Usp54 | 0.082 | Young |
| 0.030506 | 1.056015 | 0.12 | 0.038 | 1 | 14 | 1110012L1 | 0.082 | Young |
| 0.030518 | 0.847863 | 0.2 | 0.087 | 1 | 14 | Nelfcd | 0.113 | Young |
| 0.030969 | 0.561654 | 0.16 | 0.06 | 1 | 14 | Prkab1 | 0.1 | Young |
| 0.031129 | 0.897208 | 0.3 | 0.153 | 1 | 14 | Ascc3 | 0.147 | Young |
| 0.031407 | 0.774918 | 0.34 | 0.191 | 1 | 14 | Kifap3 | 0.149 | Young |
| 0.032082 | 0.969754 | 0.26 | 0.131 | 1 | 14 | Atrn | 0.129 | Young |
| 0.032083 | 0.221982 | 0.3 | 0.481 | 1 | 14 | Flrt2 | -0.181 | Young |
| 0.032984 | 1.277041 | 0.44 | 0.301 | 1 | 14 | Oaz2 | 0.139 | Young |
| 0.033047 | 3.133006 | 0.06 | 0.011 | 1 | 14 | Ajap1 | 0.049 | Young |
| 0.033047 | 2.331879 | 0.06 | 0.011 | 1 | 14 | Lysmd4 | 0.049 | Young |
| 0.033831 | 3.328545 | 0.06 | 0.011 | 1 | 14 | Dbp | 0.049 | Young |
| 0.033831 | 3.106973 | 0.06 | 0.011 | 1 | 14 | Rab39b | 0.049 | Young |
| 0.033831 | 2.823425 | 0.06 | 0.011 | 1 | 14 | Crocc | 0.049 | Young |
| 0.033831 | 2.783137 | 0.06 | 0.011 | 1 | 14 | Clspn | 0.049 | Young |
| 0.033831 | 2.697662 | 0.06 | 0.011 | 1 | 14 | Msh5 | 0.049 | Young |
| 0.033831 | 2.514259 | 0.06 | 0.011 | 1 | 14 | Zfp213 | 0.049 | Young |
| 0.033831 | 2.070354 | 0.06 | 0.011 | 1 | 14 | Was | 0.049 | Young |
| 0.033831 | 1.921108 | 0.06 | 0.011 | 1 | 14 | Swsap1 | 0.049 | Young |
| 0.03399 | 0.985738 | 0.56 | 0.459 | 1 | 14 | Tsc22d1 | 0.101 | Young |
| 0.034018 | 1.387696 | 0.48 | 0.328 | 1 | 14 | Tnpo3 | 0.152 | Young |
| 0.034042 | 0.570013 | 0.14 | 0.049 | 1 | 14 | Capn15 | 0.091 | Young |
| 0.034141 | 1.213179 | 0.24 | 0.115 | 1 | 14 | Eri3 | 0.125 | Young |
| 0.034493 | 0.920087 | 0.26 | 0.137 | 1 | 14 | Tspan18 | 0.123 | Young |
| 0.034579 | 1.211007 | 0.18 | 0.077 | 1 | 14 | Gata2 | 0.103 | Young |
| 0.034631 | 3.471417 | 0.06 | 0.011 | 1 | 14 | Zc3h10 | 0.049 | Young |
| 0.034631 | 2.653052 | 0.06 | 0.011 | 1 | 14 | Vps37d | 0.049 | Young |
| 0.034631 | 2.504206 | 0.06 | 0.011 | 1 | 14 | Col8a2 | 0.049 | Young |
| 0.034631 | 2.280517 | 0.06 | 0.011 | 1 | 14 | Mad2l2 | 0.049 | Young |

| | | | | | | | | |
|---|---|---|---|---|---|---|---|---|
| 0.034631 | 2.022449 | 0.06 | 0.011 | 1 | 14 | Slc14a1 | 0.049 | Young |
| 0.034631 | 1.939265 | 0.06 | 0.011 | 1 | 14 | Pecam1 | 0.049 | Young |
| 0.034631 | 1.643111 | 0.06 | 0.011 | 1 | 14 | Cdca7 | 0.049 | Young |
| 0.034631 | 1.61422 | 0.06 | 0.011 | 1 | 14 | Dnajc28 | 0.049 | Young |
| 0.034631 | 1.135253 | 0.06 | 0.011 | 1 | 14 | Ero1lb | 0.049 | Young |
| 0.034631 | 0.910993 | 0.06 | 0.011 | 1 | 14 | Pex11g | 0.049 | Young |
| 0.035447 | 2.909051 | 0.06 | 0.011 | 1 | 14 | Slc52a2 | 0.049 | Young |
| 0.035447 | 2.734082 | 0.06 | 0.011 | 1 | 14 | Gm14391 | 0.049 | Young |
| 0.035447 | 2.158136 | 0.06 | 0.011 | 1 | 14 | Tmem44 | 0.049 | Young |
| 0.035447 | 2.102697 | 0.06 | 0.011 | 1 | 14 | Zfp385b | 0.049 | Young |
| 0.035447 | 2.079913 | 0.06 | 0.011 | 1 | 14 | Map2k6 | 0.049 | Young |
| 0.035447 | 2.044459 | 0.06 | 0.011 | 1 | 14 | Csdc2 | 0.049 | Young |
| 0.035447 | 1.989595 | 0.06 | 0.011 | 1 | 14 | Dhtkd1 | 0.049 | Young |
| 0.035447 | 1.922192 | 0.06 | 0.011 | 1 | 14 | Rab11fip1 | 0.049 | Young |
| 0.035447 | 1.764864 | 0.06 | 0.011 | 1 | 14 | Tspan2 | 0.049 | Young |
| 0.035447 | 1.672421 | 0.06 | 0.011 | 1 | 14 | Fdxacb1 | 0.049 | Young |
| 0.035447 | 1.474927 | 0.06 | 0.011 | 1 | 14 | Nt5dc2 | 0.049 | Young |
| 0.035447 | 1.209013 | 0.06 | 0.011 | 1 | 14 | Trim6 | 0.049 | Young |
| 0.035478 | 1.922975 | 0.3 | 0.164 | 1 | 14 | Brox | 0.136 | Young |
| 0.035761 | 0.443186 | 0.5 | 0.333 | 1 | 14 | Larp1 | 0.167 | Young |
| 0.036165 | 2.376751 | 0.12 | 0.044 | 1 | 14 | Zfp1 | 0.076 | Young |
| 0.036279 | 1.63382 | 0.06 | 0.011 | 1 | 14 | Adssl1 | 0.049 | Young |
| 0.036279 | 1.451255 | 0.06 | 0.011 | 1 | 14 | Lrrtm3 | 0.049 | Young |
| 0.036279 | 1.39358 | 0.06 | 0.011 | 1 | 14 | Gm16973 | 0.049 | Young |
| 0.036279 | 1.382673 | 0.06 | 0.011 | 1 | 14 | Mmd2 | 0.049 | Young |
| 0.036279 | 1.354137 | 0.06 | 0.011 | 1 | 14 | Grhpr | 0.049 | Young |
| 0.036361 | 1.27644 | 0.16 | 0.066 | 1 | 14 | Sfxn2 | 0.094 | Young |
| 0.03712 | 1.481885 | 0.14 | 0.055 | 1 | 14 | Zfp609 | 0.085 | Young |
| 0.037128 | 1.671108 | 0.06 | 0.011 | 1 | 14 | Tmem182 | 0.049 | Young |
| 0.037128 | 1.559909 | 0.06 | 0.011 | 1 | 14 | Fgf9 | 0.049 | Young |
| 0.037128 | 1.486625 | 0.06 | 0.011 | 1 | 14 | Tmem88 | 0.049 | Young |
| 0.037128 | 1.28706 | 0.06 | 0.011 | 1 | 14 | Rasgef1b | 0.049 | Young |
| 0.037128 | 0.891839 | 0.06 | 0.011 | 1 | 14 | Dhx34 | 0.049 | Young |
| 0.037128 | 0.727093 | 0.06 | 0.011 | 1 | 14 | Trmu | 0.049 | Young |
| 0.037128 | 0.428908 | 0.06 | 0.011 | 1 | 14 | Slc30a6 | 0.049 | Young |
| 0.037269 | 1.00146 | 0.56 | 0.448 | 1 | 14 | Scd2 | 0.112 | Young |
| 0.037498 | 2.296488 | 0.16 | 0.071 | 1 | 14 | Nr4a2 | 0.089 | Young |
| 0.037512 | 0.724229 | 0.24 | 0.12 | 1 | 14 | Slc39a14 | 0.12 | Young |
| 0.037867 | 1.37975 | 0.24 | 0.12 | 1 | 14 | Rsrc1 | 0.12 | Young |
| 0.038138 | 1.065485 | 0.16 | 0.066 | 1 | 14 | Plekhg1 | 0.094 | Young |
| 0.038271 | 0.783057 | 0.32 | 0.18 | 1 | 14 | Gskip | 0.14 | Young |
| 0.038797 | 1.637529 | 0.12 | 0.044 | 1 | 14 | Prkcb | 0.076 | Young |
| 0.039452 | 0.960191 | 0.2 | 0.093 | 1 | 14 | Ahcyl2 | 0.107 | Young |
| 0.039555 | 0.697545 | 0.18 | 0.077 | 1 | 14 | Mast2 | 0.103 | Young |
| 0.039555 | 0.173939 | 0.18 | 0.077 | 1 | 14 | Tbx18 | 0.103 | Young |

| | | | | | | | | |
|---|---|---|---|---|---|---|---|---|
| 0.039572 | 1.320816 | 0.14 | 0.055 | 1 | 14 | Akna | 0.085 | Young |
| 0.039589 | 2.063525 | 0.18 | 0.087 | 1 | 14 | Ppm1g | 0.093 | Young |
| 0.040079 | 1.515442 | 0.14 | 0.055 | 1 | 14 | Sppl2b | 0.085 | Young |
| 0.040494 | 1.241699 | 0.18 | 0.082 | 1 | 14 | Dtx2 | 0.098 | Young |
| 0.040513 | 0.393769 | 0.32 | 0.169 | 1 | 14 | Osbpl11 | 0.151 | Young |
| 0.040813 | 0.733522 | 0.58 | 0.432 | 1 | 14 | Ubap2l | 0.148 | Young |
| 0.040849 | 2.772068 | 0.08 | 0.022 | 1 | 14 | Ccdc167 | 0.058 | Young |
| 0.040935 | 1.535353 | 0.18 | 0.082 | 1 | 14 | Atxn1l | 0.098 | Young |
| 0.041109 | 1.435328 | 0.14 | 0.055 | 1 | 14 | Mfap3l | 0.085 | Young |
| 0.041109 | 1.330648 | 0.14 | 0.055 | 1 | 14 | Radil | 0.085 | Young |
| 0.041163 | 0.846298 | 1 | 1 | 1 | 14 | Gm42418 | 0 | Young |
| 0.04183 | 0.821154 | 0.18 | 0.082 | 1 | 14 | Sufu | 0.098 | Young |
| 0.041963 | 1.378585 | 0.2 | 0.093 | 1 | 14 | Srek1ip1 | 0.107 | Young |
| 0.042169 | 1.968476 | 0.12 | 0.044 | 1 | 14 | Mrpl44 | 0.076 | Young |
| 0.042562 | 1.312948 | 0.26 | 0.142 | 1 | 14 | Jmy | 0.118 | Young |
| 0.042593 | 0.259828 | 0.98 | 0.989 | 1 | 14 | Actb | -0.009 | Young |
| 0.043094 | 2.308427 | 0.1 | 0.033 | 1 | 14 | Chrdl1 | 0.067 | Young |
| 0.04312 | 3.418667 | 0.08 | 0.022 | 1 | 14 | Col23a1 | 0.058 | Young |
| 0.04312 | 2.634509 | 0.08 | 0.022 | 1 | 14 | Sacs | 0.058 | Young |
| 0.04312 | 2.079087 | 0.08 | 0.022 | 1 | 14 | Gfod2 | 0.058 | Young |
| 0.043235 | 0.954597 | 0.14 | 0.055 | 1 | 14 | Tyw5 | 0.085 | Young |
| 0.043347 | 1.729707 | 0.12 | 0.044 | 1 | 14 | Dtwd1 | 0.076 | Young |
| 0.043519 | 0.834091 | 0.42 | 0.268 | 1 | 14 | Impact | 0.152 | Young |
| 0.043706 | 0.440975 | 0.38 | 0.235 | 1 | 14 | Ppp1r10 | 0.145 | Young |
| 0.043763 | 2.60277 | 0.1 | 0.033 | 1 | 14 | Ccl4 | 0.067 | Young |
| 0.043763 | 2.102874 | 0.1 | 0.033 | 1 | 14 | Ttc32 | 0.067 | Young |
| 0.043781 | 1.531794 | 0.14 | 0.055 | 1 | 14 | Nxn | 0.085 | Young |
| 0.0439 | 1.597697 | 0.08 | 0.022 | 1 | 14 | Optn | 0.058 | Young |
| 0.04444 | 2.118965 | 0.1 | 0.033 | 1 | 14 | Mfap4 | 0.067 | Young |
| 0.044692 | 1.818469 | 0.08 | 0.022 | 1 | 14 | Smtnl2 | 0.058 | Young |
| 0.044738 | 0.403081 | 0.92 | 0.907 | 1 | 14 | Hsp90ab1 | 0.013 | Young |
| 0.04483 | 0.925089 | 0.4 | 0.279 | 1 | 14 | Slc16a9 | 0.121 | Young |
| 0.044918 | 0.964016 | 0.22 | 0.109 | 1 | 14 | Tufm | 0.111 | Young |
| 0.045083 | 1.066319 | 0.36 | 0.213 | 1 | 14 | Dnpep | 0.147 | Young |
| 0.045166 | 2.361389 | 0.12 | 0.044 | 1 | 14 | Alg13 | 0.076 | Young |
| 0.045497 | 2.003309 | 0.08 | 0.022 | 1 | 14 | Esam | 0.058 | Young |
| 0.045633 | 0.607086 | 0.36 | 0.202 | 1 | 14 | Atf7ip | 0.158 | Young |
| 0.045786 | 3.075214 | 0.12 | 0.044 | 1 | 14 | Mbip | 0.076 | Young |
| 0.045786 | 1.475535 | 0.12 | 0.044 | 1 | 14 | Agpat2 | 0.076 | Young |
| 0.045786 | 0.812321 | 0.12 | 0.044 | 1 | 14 | Nkiras1 | 0.076 | Young |
| 0.045822 | 1.623313 | 0.1 | 0.033 | 1 | 14 | Tmem229b | 0.067 | Young |
| 0.046063 | 1.190723 | 0.18 | 0.082 | 1 | 14 | Atp11c | 0.098 | Young |
| 0.046259 | 0.85185 | 0.46 | 0.306 | 1 | 14 | Phf20l1 | 0.154 | Young |
| 0.046298 | 0.378047 | 0.72 | 0.639 | 1 | 14 | Cox6a1 | 0.081 | Young |
| 0.046313 | 2.216403 | 0.08 | 0.022 | 1 | 14 | Aaas | 0.058 | Young |

| | | | | | | | | |
|---|---|---|---|---|---|---|---|---|
| 0.046313 | 1.689915 | 0.08 | 0.022 | 1 | 14 | Kctd15 | 0.058 | Young |
| 0.046414 | 2.513053 | 0.12 | 0.044 | 1 | 14 | Tjap1 | 0.076 | Young |
| 0.046414 | 1.170261 | 0.12 | 0.044 | 1 | 14 | Prpf3 | 0.076 | Young |
| 0.046414 | 0.88409 | 0.12 | 0.044 | 1 | 14 | Pdzrn3 | 0.076 | Young |
| 0.046526 | 1.83787 | 0.1 | 0.033 | 1 | 14 | Hoxb3 | 0.067 | Young |
| 0.046861 | 1.647253 | 0.28 | 0.175 | 1 | 14 | Rnf10 | 0.105 | Young |
| 0.047049 | 2.605606 | 0.12 | 0.044 | 1 | 14 | Gtpbp6 | 0.076 | Young |
| 0.047049 | 2.124461 | 0.12 | 0.044 | 1 | 14 | Serf1 | 0.076 | Young |
| 0.047049 | 1.222036 | 0.12 | 0.044 | 1 | 14 | Stard3 | 0.076 | Young |
| 0.047089 | 1.251855 | 0.24 | 0.131 | 1 | 14 | Atg5 | 0.109 | Young |
| 0.047142 | 1.496295 | 0.08 | 0.022 | 1 | 14 | Tmod1 | 0.058 | Young |
| 0.047142 | 1.493112 | 0.08 | 0.022 | 1 | 14 | Tln2 | 0.058 | Young |
| 0.047142 | 1.456852 | 0.08 | 0.022 | 1 | 14 | Tepsin | 0.058 | Young |
| 0.047142 | 1.086849 | 0.08 | 0.022 | 1 | 14 | Dtna | 0.058 | Young |
| 0.04724 | 2.006002 | 0.1 | 0.033 | 1 | 14 | Cyp7b1 | 0.067 | Young |
| 0.047691 | 1.230245 | 0.12 | 0.044 | 1 | 14 | Pggt1b | 0.076 | Young |
| 0.047704 | 0.935987 | 0.26 | 0.148 | 1 | 14 | Mrpl21 | 0.112 | Young |
| 0.047768 | 0.520305 | 0.14 | 0.055 | 1 | 14 | Nop16 | 0.085 | Young |
| 0.047851 | 1.690902 | 0.18 | 0.087 | 1 | 14 | Lias | 0.093 | Young |
| 0.047855 | 1.015933 | 0.24 | 0.126 | 1 | 14 | Tob1 | 0.114 | Young |
| 0.047962 | 1.399801 | 0.1 | 0.033 | 1 | 14 | 2310033PC | 0.067 | Young |
| 0.047983 | 1.782888 | 0.08 | 0.022 | 1 | 14 | Acot11 | 0.058 | Young |
| 0.047983 | 1.636498 | 0.08 | 0.022 | 1 | 14 | Fzd5 | 0.058 | Young |
| 0.047983 | 1.263006 | 0.08 | 0.022 | 1 | 14 | Sirt6 | 0.058 | Young |
| 0.047983 | 1.111867 | 0.08 | 0.022 | 1 | 14 | Sorbs2 | 0.058 | Young |
| 0.048164 | 0.787298 | 0.16 | 0.066 | 1 | 14 | Echdc1 | 0.094 | Young |
| 0.04834 | 1.734124 | 0.12 | 0.044 | 1 | 14 | Exoc8 | 0.076 | Young |
| 0.048566 | 0.741905 | 0.18 | 0.082 | 1 | 14 | Trap1 | 0.098 | Young |
| 0.048582 | 1.54361 | 0.38 | 0.268 | 1 | 14 | Csrp1 | 0.112 | Young |
| 0.048694 | 2.1161 | 0.1 | 0.033 | 1 | 14 | Gcnt2 | 0.067 | Young |
| 0.048694 | 1.881242 | 0.1 | 0.033 | 1 | 14 | 9530077CC | 0.067 | Young |
| 0.048694 | 0.885895 | 0.1 | 0.033 | 1 | 14 | 4632415L0 | 0.067 | Young |
| 0.048837 | 0.971344 | 0.08 | 0.022 | 1 | 14 | Gm11627 | 0.058 | Young |
| 0.048837 | 0.794635 | 0.08 | 0.022 | 1 | 14 | Eid3 | 0.058 | Young |
| 0.048837 | 0.744654 | 0.08 | 0.022 | 1 | 14 | Kctd13 | 0.058 | Young |
| 0.048879 | 0.600192 | 0.48 | 0.333 | 1 | 14 | Ldb1 | 0.147 | Young |
| 0.04908 | 1.003273 | 0.18 | 0.082 | 1 | 14 | Pde1a | 0.098 | Young |
| 0.049435 | 1.364956 | 0.1 | 0.033 | 1 | 14 | Zfp617 | 0.067 | Young |
| 0.049435 | 1.264935 | 0.1 | 0.033 | 1 | 14 | Acvr1b | 0.067 | Young |
| 0.049435 | 0.838284 | 0.1 | 0.033 | 1 | 14 | Rtel1 | 0.067 | Young |
| 0.049563 | 0.812119 | 0.46 | 0.317 | 1 | 14 | Tspan31 | 0.143 | Young |
| 0.049703 | 1.623766 | 0.08 | 0.022 | 1 | 14 | Wdfy4 | 0.058 | Young |
| 0.049703 | 1.134364 | 0.08 | 0.022 | 1 | 14 | Tab1 | 0.058 | Young |
| 0.049703 | 0.745564 | 0.08 | 0.022 | 1 | 14 | Ccr1 | 0.058 | Young |
| 0.04996 | 1.799268 | 0.28 | 0.175 | 1 | 14 | Abca5 | 0.105 | Young |

| | | | | | | | | |
|---|---|---|---|---|---|---|---|---|
| 0.050186 | 1.603201 | 0.1 | 0.033 | 1 | 14 | Ndufaf1 | 0.067 | Young |
| 0.050186 | 1.133956 | 0.1 | 0.033 | 1 | 14 | Polrmt | 0.067 | Young |
| 0.050186 | 1.077523 | 0.1 | 0.033 | 1 | 14 | Angptl1 | 0.067 | Young |
| 0.050333 | 1.395045 | 0.12 | 0.044 | 1 | 14 | Mertk | 0.076 | Young |
| 0.050333 | 0.852209 | 0.12 | 0.044 | 1 | 14 | Tatdn1 | 0.076 | Young |
| 0.050348 | 0.375854 | 0.34 | 0.191 | 1 | 14 | Tsnax | 0.149 | Young |
| 0.050582 | 1.175268 | 0.08 | 0.022 | 1 | 14 | Csf2rb | 0.058 | Young |
| 0.050731 | 0.782671 | 0.32 | 0.197 | 1 | 14 | Alas1 | 0.123 | Young |
| 0.051183 | 0.865282 | 0.18 | 0.082 | 1 | 14 | Smc4 | 0.098 | Young |
| 0.051475 | 0.463373 | 0.08 | 0.022 | 1 | 14 | Thada | 0.058 | Young |
| 0.051475 | 0.296152 | 0.08 | 0.022 | 1 | 14 | Gm26890 | 0.058 | Young |
| 0.051475 | 0.19577 | 0.08 | 0.022 | 1 | 14 | Tbc1d7 | 0.058 | Young |
| 0.051475 | 0.194338 | 0.08 | 0.022 | 1 | 14 | Dgcr14 | 0.058 | Young |
| 0.0517 | 1.158474 | 0.12 | 0.044 | 1 | 14 | Ccdc28a | 0.076 | Young |
| 0.051716 | 1.041406 | 0.1 | 0.033 | 1 | 14 | Sdr39u1 | 0.067 | Young |
| 0.051744 | 1.221964 | 0.22 | 0.109 | 1 | 14 | Phf5a | 0.111 | Young |
| 0.052161 | 0.110575 | 0.16 | 0.066 | 1 | 14 | Rabgef1 | 0.094 | Young |
| 0.052262 | 1.050039 | 0.18 | 0.082 | 1 | 14 | Lncpint | 0.098 | Young |
| 0.052306 | 0.911213 | 0.26 | 0.142 | 1 | 14 | Pdcl | 0.118 | Young |
| 0.052358 | 0.816382 | 0.2 | 0.093 | 1 | 14 | Tmem161a | 0.107 | Young |
| 0.05238 | 1.013874 | 0.08 | 0.022 | 1 | 14 | Kifc3 | 0.058 | Young |
| 0.05238 | 0.560234 | 0.08 | 0.022 | 1 | 14 | Rpl9-ps6 | 0.058 | Young |
| 0.052495 | 0.838168 | 0.1 | 0.033 | 1 | 14 | Exoc3l4 | 0.067 | Young |
| 0.052495 | 0.708404 | 0.1 | 0.033 | 1 | 14 | Echdc2 | 0.067 | Young |
| 0.052618 | 0.912902 | 0.34 | 0.213 | 1 | 14 | Tmf1 | 0.127 | Young |
| 0.052754 | 0.495719 | 0.16 | 0.066 | 1 | 14 | Ss18l2 | 0.094 | Young |
| 0.052775 | 1.121914 | 0.26 | 0.148 | 1 | 14 | Cd36 | 0.112 | Young |
| 0.052811 | 1.40236 | 0.28 | 0.175 | 1 | 14 | Ralgps2 | 0.105 | Young |
| 0.052946 | 0.736692 | 0.82 | 0.738 | 1 | 14 | Ywhaz | 0.082 | Young |
| 0.053097 | 1.226625 | 0.12 | 0.044 | 1 | 14 | Tango6 | 0.076 | Young |
| 0.053098 | 1.717546 | 0.56 | 0.448 | 1 | 14 | Cbx5 | 0.112 | Young |
| 0.053284 | 0.7726 | 0.1 | 0.033 | 1 | 14 | Khk | 0.067 | Young |
| 0.053393 | 1.120839 | 0.34 | 0.213 | 1 | 14 | Zswim8 | 0.127 | Young |
| 0.053807 | 0.445748 | 0.12 | 0.044 | 1 | 14 | Ephb2 | 0.076 | Young |
| 0.054084 | 0.579286 | 0.1 | 0.033 | 1 | 14 | Park2 | 0.067 | Young |
| 0.054187 | 1.646586 | 0.14 | 0.06 | 1 | 14 | Eml3 | 0.08 | Young |
| 0.054659 | 4.231886 | 0.04 | 0.005 | 1 | 14 | Flrt1 | 0.035 | Young |
| 0.054659 | 3.871389 | 0.04 | 0.005 | 1 | 14 | Zfp469 | 0.035 | Young |
| 0.054659 | 3.838847 | 0.04 | 0.005 | 1 | 14 | Ablim2 | 0.035 | Young |
| 0.054659 | 3.838427 | 0.04 | 0.005 | 1 | 14 | Tnni2 | 0.035 | Young |
| 0.054659 | 3.648481 | 0.04 | 0.005 | 1 | 14 | Rab27a | 0.035 | Young |
| 0.054659 | 3.163313 | 0.04 | 0.005 | 1 | 14 | Dyx1c1 | 0.035 | Young |
| 0.054659 | 3.114761 | 0.04 | 0.005 | 1 | 14 | Ehhadh | 0.035 | Young |
| 0.054659 | 2.880112 | 0.04 | 0.005 | 1 | 14 | Prr5 | 0.035 | Young |
| 0.054659 | 2.836962 | 0.04 | 0.005 | 1 | 14 | Sucnr1 | 0.035 | Young |

| | | | | | | | | |
|---|---|---|---|---|---|---|---|---|
| 0.054659 | 2.611414 | 0.04 | 0.005 | 1 | 14 | Ddc | 0.035 | Young |
| 0.054659 | 2.568578 | 0.04 | 0.005 | 1 | 14 | Mark1 | 0.035 | Young |
| 0.054659 | 2.537209 | 0.04 | 0.005 | 1 | 14 | Klk8 | 0.035 | Young |
| 0.054659 | 2.496358 | 0.04 | 0.005 | 1 | 14 | Zfp174 | 0.035 | Young |
| 0.054659 | 2.482761 | 0.04 | 0.005 | 1 | 14 | Krtcap3 | 0.035 | Young |
| 0.054659 | 2.272682 | 0.04 | 0.005 | 1 | 14 | Sfmbt2 | 0.035 | Young |
| 0.054659 | 2.233428 | 0.04 | 0.005 | 1 | 14 | Polr3g | 0.035 | Young |
| 0.054659 | 1.778629 | 0.04 | 0.005 | 1 | 14 | Cd40 | 0.035 | Young |
| 0.054857 | 0.373858 | 0.82 | 0.765 | 1 | 14 | Cd81 | 0.055 | Young |
| 0.055712 | 1.035942 | 0.1 | 0.033 | 1 | 14 | Epb41l5 | 0.067 | Young |
| 0.05575 | 1.729413 | 0.22 | 0.115 | 1 | 14 | Wdr61 | 0.105 | Young |
| 0.056117 | 0.933211 | 0.38 | 0.246 | 1 | 14 | Aph1a | 0.134 | Young |
| 0.056202 | 0.916274 | 0.2 | 0.098 | 1 | 14 | Immp1l | 0.102 | Young |
| 0.056202 | 0.890247 | 0.2 | 0.098 | 1 | 14 | Zcchc9 | 0.102 | Young |
| 0.056204 | 3.965284 | 0.04 | 0.005 | 1 | 14 | Acp5 | 0.035 | Young |
| 0.056204 | 3.854177 | 0.04 | 0.005 | 1 | 14 | Tifab | 0.035 | Young |
| 0.056204 | 3.542982 | 0.04 | 0.005 | 1 | 14 | Serpini1 | 0.035 | Young |
| 0.056204 | 3.428839 | 0.04 | 0.005 | 1 | 14 | Rasgrp3 | 0.035 | Young |
| 0.056204 | 3.162769 | 0.04 | 0.005 | 1 | 14 | Adamts9 | 0.035 | Young |
| 0.056204 | 3.140487 | 0.04 | 0.005 | 1 | 14 | D630003M | 0.035 | Young |
| 0.056204 | 3.082087 | 0.04 | 0.005 | 1 | 14 | Rapgef4 | 0.035 | Young |
| 0.056204 | 3.054213 | 0.04 | 0.005 | 1 | 14 | Wipf3 | 0.035 | Young |
| 0.056204 | 2.962301 | 0.04 | 0.005 | 1 | 14 | Retreg1 | 0.035 | Young |
| 0.056204 | 2.856558 | 0.04 | 0.005 | 1 | 14 | Sapcd1 | 0.035 | Young |
| 0.056204 | 2.788249 | 0.04 | 0.005 | 1 | 14 | Igdcc3 | 0.035 | Young |
| 0.056204 | 2.67618 | 0.04 | 0.005 | 1 | 14 | Aldh1a7 | 0.035 | Young |
| 0.056204 | 2.556244 | 0.04 | 0.005 | 1 | 14 | Sorcs2 | 0.035 | Young |
| 0.056204 | 2.537765 | 0.04 | 0.005 | 1 | 14 | Rbp7 | 0.035 | Young |
| 0.056204 | 2.536647 | 0.04 | 0.005 | 1 | 14 | Cdk5r1 | 0.035 | Young |
| 0.056204 | 2.53431 | 0.04 | 0.005 | 1 | 14 | Peg12 | 0.035 | Young |
| 0.056204 | 2.51667 | 0.04 | 0.005 | 1 | 14 | Loxl4 | 0.035 | Young |
| 0.056204 | 2.497594 | 0.04 | 0.005 | 1 | 14 | Rgl3 | 0.035 | Young |
| 0.056204 | 2.4428 | 0.04 | 0.005 | 1 | 14 | Olfr558 | 0.035 | Young |
| 0.056204 | 2.404806 | 0.04 | 0.005 | 1 | 14 | 2600006K0 | 0.035 | Young |
| 0.056204 | 2.362982 | 0.04 | 0.005 | 1 | 14 | 1810011H1 | 0.035 | Young |
| 0.056204 | 2.307558 | 0.04 | 0.005 | 1 | 14 | Ltbp2 | 0.035 | Young |
| 0.056204 | 2.299926 | 0.04 | 0.005 | 1 | 14 | She | 0.035 | Young |
| 0.056204 | 2.234939 | 0.04 | 0.005 | 1 | 14 | Gtf2ird2 | 0.035 | Young |
| 0.056204 | 2.189075 | 0.04 | 0.005 | 1 | 14 | Cfap69 | 0.035 | Young |
| 0.056204 | 2.175114 | 0.04 | 0.005 | 1 | 14 | Sema3g | 0.035 | Young |
| 0.056204 | 2.114406 | 0.04 | 0.005 | 1 | 14 | Igsf9 | 0.035 | Young |
| 0.056204 | 1.949636 | 0.04 | 0.005 | 1 | 14 | Siglece | 0.035 | Young |
| 0.056204 | 1.880342 | 0.04 | 0.005 | 1 | 14 | Mapt | 0.035 | Young |
| 0.056204 | 1.865355 | 0.04 | 0.005 | 1 | 14 | Greb1 | 0.035 | Young |
| 0.056204 | 1.832306 | 0.04 | 0.005 | 1 | 14 | Dnaic2 | 0.035 | Young |

| | | | | | | | | |
|---|---|---|---|---|---|---|---|---|
| 0.056204 | 1.809873 | 0.04 | 0.005 | 1 | 14 | Hrct1 | 0.035 | Young |
| 0.056204 | 1.685532 | 0.04 | 0.005 | 1 | 14 | Oasl1 | 0.035 | Young |
| 0.056204 | 1.654337 | 0.04 | 0.005 | 1 | 14 | Slc23a1 | 0.035 | Young |
| 0.056204 | 1.630459 | 0.04 | 0.005 | 1 | 14 | Zfp93 | 0.035 | Young |
| 0.056204 | 1.560922 | 0.04 | 0.005 | 1 | 14 | Armc7 | 0.035 | Young |
| 0.056204 | 1.540922 | 0.04 | 0.005 | 1 | 14 | Ank1 | 0.035 | Young |
| 0.056541 | 0.066932 | 0.1 | 0.033 | 1 | 14 | Dclre1b | 0.067 | Young |
| 0.056915 | 0.830119 | 0.16 | 0.071 | 1 | 14 | Sft2d1 | 0.089 | Young |
| 0.057085 | 5.077818 | 0.02 | 0 | 1 | 14 | Edn3 | 0.02 | Young |
| 0.057085 | 4.844951 | 0.02 | 0 | 1 | 14 | Tmeff1 | 0.02 | Young |
| 0.057085 | 4.844951 | 0.02 | 0 | 1 | 14 | Sox30 | 0.02 | Young |
| 0.057085 | 4.710884 | 0.02 | 0 | 1 | 14 | Chrna4 | 0.02 | Young |
| 0.057085 | 4.710884 | 0.02 | 0 | 1 | 14 | Fhl5 | 0.02 | Young |
| 0.057085 | 4.710884 | 0.02 | 0 | 1 | 14 | Ppp1r1a | 0.02 | Young |
| 0.057085 | 4.502615 | 0.02 | 0 | 1 | 14 | Lmo3 | 0.02 | Young |
| 0.057085 | 4.502615 | 0.02 | 0 | 1 | 14 | Tnfrsf19 | 0.02 | Young |
| 0.057085 | 4.502615 | 0.02 | 0 | 1 | 14 | Wwc1 | 0.02 | Young |
| 0.057085 | 4.502615 | 0.02 | 0 | 1 | 14 | Itga3 | 0.02 | Young |
| 0.057085 | 4.502615 | 0.02 | 0 | 1 | 14 | Fam110c | 0.02 | Young |
| 0.057085 | 4.477457 | 0.02 | 0 | 1 | 14 | Brinp1 | 0.02 | Young |
| 0.057085 | 4.477457 | 0.02 | 0 | 1 | 14 | Hapln3 | 0.02 | Young |
| 0.057085 | 4.119914 | 0.02 | 0 | 1 | 14 | Lypd6 | 0.02 | Young |
| 0.057085 | 4.101663 | 0.02 | 0 | 1 | 14 | Gm39929 | 0.02 | Young |
| 0.057085 | 4.101663 | 0.02 | 0 | 1 | 14 | Gm15441 | 0.02 | Young |
| 0.057085 | 4.037632 | 0.02 | 0 | 1 | 14 | Tmem252 | 0.02 | Young |
| 0.057085 | 3.932308 | 0.02 | 0 | 1 | 14 | Pcdh10 | 0.02 | Young |
| 0.057085 | 3.827953 | 0.02 | 0 | 1 | 14 | Slc25a34 | 0.02 | Young |
| 0.057085 | 3.77424 | 0.02 | 0 | 1 | 14 | Mir670hg | 0.02 | Young |
| 0.057085 | 3.77424 | 0.02 | 0 | 1 | 14 | Frem2 | 0.02 | Young |
| 0.057085 | 3.77424 | 0.02 | 0 | 1 | 14 | Gria1 | 0.02 | Young |
| 0.057085 | 3.77424 | 0.02 | 0 | 1 | 14 | Zdhhc23 | 0.02 | Young |
| 0.057085 | 3.732137 | 0.02 | 0 | 1 | 14 | Rnf112 | 0.02 | Young |
| 0.057085 | 3.649133 | 0.02 | 0 | 1 | 14 | Pgr | 0.02 | Young |
| 0.057085 | 3.579651 | 0.02 | 0 | 1 | 14 | Spc25 | 0.02 | Young |
| 0.057085 | 3.579651 | 0.02 | 0 | 1 | 14 | Pip5k1b | 0.02 | Young |
| 0.057085 | 3.548549 | 0.02 | 0 | 1 | 14 | Mpz | 0.02 | Young |
| 0.057085 | 3.548549 | 0.02 | 0 | 1 | 14 | Sptlc3 | 0.02 | Young |
| 0.057085 | 3.548549 | 0.02 | 0 | 1 | 14 | Nudt11 | 0.02 | Young |
| 0.057085 | 3.548549 | 0.02 | 0 | 1 | 14 | 2010109A1 | 0.02 | Young |
| 0.057085 | 3.548549 | 0.02 | 0 | 1 | 14 | Gm42397 | 0.02 | Young |
| 0.057085 | 3.548549 | 0.02 | 0 | 1 | 14 | Ogdhl | 0.02 | Young |
| 0.057085 | 3.548549 | 0.02 | 0 | 1 | 14 | Fam198a | 0.02 | Young |
| 0.057085 | 3.548549 | 0.02 | 0 | 1 | 14 | Grhl1 | 0.02 | Young |
| 0.057085 | 3.548549 | 0.02 | 0 | 1 | 14 | Arhgap8 | 0.02 | Young |
| 0.057085 | 3.548549 | 0.02 | 0 | 1 | 14 | Krt18 | 0.02 | Young |

| | | | | | | | | |
|---|---|---|---|---|---|---|---|---|
| 0.057085 | 3.548549 | 0.02 | 0 | 1 | 14 | Aifm3 | 0.02 | Young |
| 0.057085 | 3.548549 | 0.02 | 0 | 1 | 14 | Slc22a8 | 0.02 | Young |
| 0.057085 | 3.538006 | 0.02 | 0 | 1 | 14 | Gm16070 | 0.02 | Young |
| 0.057085 | 3.538006 | 0.02 | 0 | 1 | 14 | Pappa2 | 0.02 | Young |
| 0.057085 | 3.538006 | 0.02 | 0 | 1 | 14 | Riiad1 | 0.02 | Young |
| 0.057085 | 3.538006 | 0.02 | 0 | 1 | 14 | 9530034E1 | 0.02 | Young |
| 0.057085 | 3.538006 | 0.02 | 0 | 1 | 14 | Sfxn5 | 0.02 | Young |
| 0.057085 | 3.538006 | 0.02 | 0 | 1 | 14 | Plekha7 | 0.02 | Young |
| 0.057085 | 3.538006 | 0.02 | 0 | 1 | 14 | Dleu7 | 0.02 | Young |
| 0.057085 | 3.538006 | 0.02 | 0 | 1 | 14 | Baalc | 0.02 | Young |
| 0.057085 | 3.452566 | 0.02 | 0 | 1 | 14 | Adamtsl2 | 0.02 | Young |
| 0.057085 | 3.320695 | 0.02 | 0 | 1 | 14 | C1qtnf9 | 0.02 | Young |
| 0.057085 | 3.215703 | 0.02 | 0 | 1 | 14 | Gm38037 | 0.02 | Young |
| 0.057085 | 3.215703 | 0.02 | 0 | 1 | 14 | Gpr150 | 0.02 | Young |
| 0.057085 | 3.215703 | 0.02 | 0 | 1 | 14 | Slc4a8 | 0.02 | Young |
| 0.057085 | 3.200593 | 0.02 | 0 | 1 | 14 | Bex4 | 0.02 | Young |
| 0.057085 | 3.200593 | 0.02 | 0 | 1 | 14 | Col11a1 | 0.02 | Young |
| 0.057085 | 3.200593 | 0.02 | 0 | 1 | 14 | Wscd2 | 0.02 | Young |
| 0.057085 | 3.200593 | 0.02 | 0 | 1 | 14 | D830026I1 | 0.02 | Young |
| 0.057085 | 3.200593 | 0.02 | 0 | 1 | 14 | Kcng4 | 0.02 | Young |
| 0.057085 | 3.200593 | 0.02 | 0 | 1 | 14 | Foxl2 | 0.02 | Young |
| 0.057085 | 3.200593 | 0.02 | 0 | 1 | 14 | 9030025P2 | 0.02 | Young |
| 0.057085 | 3.179713 | 0.02 | 0 | 1 | 14 | Gm9873 | 0.02 | Young |
| 0.057085 | 3.179713 | 0.02 | 0 | 1 | 14 | A930015D( | 0.02 | Young |
| 0.057085 | 3.127747 | 0.02 | 0 | 1 | 14 | Gja5 | 0.02 | Young |
| 0.057085 | 3.066963 | 0.02 | 0 | 1 | 14 | Cdca3 | 0.02 | Young |
| 0.057085 | 3.066963 | 0.02 | 0 | 1 | 14 | Gm31452 | 0.02 | Young |
| 0.057085 | 3.047441 | 0.02 | 0 | 1 | 14 | Angpt4 | 0.02 | Young |
| 0.057085 | 3.047441 | 0.02 | 0 | 1 | 14 | St6galnac3 | 0.02 | Young |
| 0.057085 | 3.047441 | 0.02 | 0 | 1 | 14 | Tarsl2 | 0.02 | Young |
| 0.057085 | 3.047441 | 0.02 | 0 | 1 | 14 | Gm38604 | 0.02 | Young |
| 0.057085 | 3.047441 | 0.02 | 0 | 1 | 14 | Trim9 | 0.02 | Young |
| 0.057085 | 3.047441 | 0.02 | 0 | 1 | 14 | Zfp641 | 0.02 | Young |
| 0.057085 | 3.047441 | 0.02 | 0 | 1 | 14 | Dscam | 0.02 | Young |
| 0.057085 | 3.018832 | 0.02 | 0 | 1 | 14 | Tnfaip8l3 | 0.02 | Young |
| 0.057085 | 2.926125 | 0.02 | 0 | 1 | 14 | Nap1l3 | 0.02 | Young |
| 0.057085 | 2.926125 | 0.02 | 0 | 1 | 14 | Aplp1 | 0.02 | Young |
| 0.057085 | 2.926125 | 0.02 | 0 | 1 | 14 | Scml4 | 0.02 | Young |
| 0.057085 | 2.926125 | 0.02 | 0 | 1 | 14 | Fam184a | 0.02 | Young |
| 0.057085 | 2.926125 | 0.02 | 0 | 1 | 14 | Gm3336 | 0.02 | Young |
| 0.057085 | 2.926125 | 0.02 | 0 | 1 | 14 | Vstm5 | 0.02 | Young |
| 0.057085 | 2.926125 | 0.02 | 0 | 1 | 14 | Sec14l4 | 0.02 | Young |
| 0.057085 | 2.926125 | 0.02 | 0 | 1 | 14 | Kcnmb1 | 0.02 | Young |
| 0.057085 | 2.926125 | 0.02 | 0 | 1 | 14 | Gm9796 | 0.02 | Young |
| 0.057085 | 2.926125 | 0.02 | 0 | 1 | 14 | Kcnj15 | 0.02 | Young |

| | | | | | | | | |
|---|---|---|---|---|---|---|---|---|
| 0.057085 | 2.926125 | 0.02 | 0 | 1 | 14 | Pcdhb15 | 0.02 | Young |
| 0.057085 | 2.91338 | 0.02 | 0 | 1 | 14 | Fam166b | 0.02 | Young |
| 0.057085 | 2.91338 | 0.02 | 0 | 1 | 14 | Gm15408 | 0.02 | Young |
| 0.057085 | 2.91338 | 0.02 | 0 | 1 | 14 | Gm15873 | 0.02 | Young |
| 0.057085 | 2.91338 | 0.02 | 0 | 1 | 14 | Cdhr5 | 0.02 | Young |
| 0.057085 | 2.91338 | 0.02 | 0 | 1 | 14 | Fam241b | 0.02 | Young |
| 0.057085 | 2.836811 | 0.02 | 0 | 1 | 14 | Prdm14 | 0.02 | Young |
| 0.057085 | 2.836811 | 0.02 | 0 | 1 | 14 | Lcn8 | 0.02 | Young |
| 0.057085 | 2.836811 | 0.02 | 0 | 1 | 14 | Adgrg5 | 0.02 | Young |
| 0.057085 | 2.836811 | 0.02 | 0 | 1 | 14 | Gm10353 | 0.02 | Young |
| 0.057085 | 2.806181 | 0.02 | 0 | 1 | 14 | Gm13594 | 0.02 | Young |
| 0.057085 | 2.806181 | 0.02 | 0 | 1 | 14 | Gm19935 | 0.02 | Young |
| 0.057085 | 2.769924 | 0.02 | 0 | 1 | 14 | 2610016A1 | 0.02 | Young |
| 0.057085 | 2.767419 | 0.02 | 0 | 1 | 14 | Afmid | 0.02 | Young |
| 0.057085 | 2.766704 | 0.02 | 0 | 1 | 14 | Tcf15 | 0.02 | Young |
| 0.057085 | 2.756748 | 0.02 | 0 | 1 | 14 | Pard3bos3 | 0.02 | Young |
| 0.057085 | 2.756748 | 0.02 | 0 | 1 | 14 | Gm10419 | 0.02 | Young |
| 0.057085 | 2.756748 | 0.02 | 0 | 1 | 14 | Gm17949 | 0.02 | Young |
| 0.057085 | 2.752068 | 0.02 | 0 | 1 | 14 | Dcstamp | 0.02 | Young |
| 0.057085 | 2.695538 | 0.02 | 0 | 1 | 14 | Ncaph | 0.02 | Young |
| 0.057085 | 2.695538 | 0.02 | 0 | 1 | 14 | A530013C2 | 0.02 | Young |
| 0.057085 | 2.695538 | 0.02 | 0 | 1 | 14 | Palm2 | 0.02 | Young |
| 0.057085 | 2.695538 | 0.02 | 0 | 1 | 14 | Aicda | 0.02 | Young |
| 0.057085 | 2.695538 | 0.02 | 0 | 1 | 14 | Gcgr | 0.02 | Young |
| 0.057085 | 2.695538 | 0.02 | 0 | 1 | 14 | Cabp4 | 0.02 | Young |
| 0.057085 | 2.686885 | 0.02 | 0 | 1 | 14 | Gm16263 | 0.02 | Young |
| 0.057085 | 2.618461 | 0.02 | 0 | 1 | 14 | Gm26768 | 0.02 | Young |
| 0.057085 | 2.618461 | 0.02 | 0 | 1 | 14 | Rbm24 | 0.02 | Young |
| 0.057085 | 2.59584 | 0.02 | 0 | 1 | 14 | Phex | 0.02 | Young |
| 0.057085 | 2.59584 | 0.02 | 0 | 1 | 14 | Nps | 0.02 | Young |
| 0.057085 | 2.59584 | 0.02 | 0 | 1 | 14 | Fitm1 | 0.02 | Young |
| 0.057085 | 2.575985 | 0.02 | 0 | 1 | 14 | Nyap1 | 0.02 | Young |
| 0.057085 | 2.575985 | 0.02 | 0 | 1 | 14 | Gm49282 | 0.02 | Young |
| 0.057085 | 2.575985 | 0.02 | 0 | 1 | 14 | Zfp983 | 0.02 | Young |
| 0.057085 | 2.575985 | 0.02 | 0 | 1 | 14 | Rasal3 | 0.02 | Young |
| 0.057085 | 2.524844 | 0.02 | 0 | 1 | 14 | Gm15834 | 0.02 | Young |
| 0.057085 | 2.524844 | 0.02 | 0 | 1 | 14 | Epor | 0.02 | Young |
| 0.057085 | 2.406705 | 0.02 | 0 | 1 | 14 | Atg9b | 0.02 | Young |
| 0.057085 | 2.363193 | 0.02 | 0 | 1 | 14 | Mreg | 0.02 | Young |
| 0.057085 | 2.363193 | 0.02 | 0 | 1 | 14 | Nr5a2 | 0.02 | Young |
| 0.057085 | 2.363193 | 0.02 | 0 | 1 | 14 | Bub1b | 0.02 | Young |
| 0.057085 | 2.363193 | 0.02 | 0 | 1 | 14 | Zfp536 | 0.02 | Young |
| 0.057085 | 2.363193 | 0.02 | 0 | 1 | 14 | Spn | 0.02 | Young |
| 0.057085 | 2.363193 | 0.02 | 0 | 1 | 14 | Pidd1 | 0.02 | Young |
| 0.057085 | 2.363193 | 0.02 | 0 | 1 | 14 | Tbc1d30 | 0.02 | Young |

| | | | | | | | | |
|---|---|---|---|---|---|---|---|---|
| 0.057085 | 2.363193 | 0.02 | 0 | 1 | 14 | Retn | 0.02 | Young |
| 0.057085 | 2.363193 | 0.02 | 0 | 1 | 14 | Sema5b | 0.02 | Young |
| 0.057085 | 2.303524 | 0.02 | 0 | 1 | 14 | 4930447C0 | 0.02 | Young |
| 0.057085 | 2.284024 | 0.02 | 0 | 1 | 14 | Atp2b2 | 0.02 | Young |
| 0.057085 | 2.284024 | 0.02 | 0 | 1 | 14 | Cck | 0.02 | Young |
| 0.057085 | 2.284024 | 0.02 | 0 | 1 | 14 | Icam2 | 0.02 | Young |
| 0.057085 | 2.284024 | 0.02 | 0 | 1 | 14 | Itgb4 | 0.02 | Young |
| 0.057085 | 2.284024 | 0.02 | 0 | 1 | 14 | Ablim3 | 0.02 | Young |
| 0.057085 | 2.251851 | 0.02 | 0 | 1 | 14 | Gm16638 | 0.02 | Young |
| 0.057085 | 2.251851 | 0.02 | 0 | 1 | 14 | Ascl1 | 0.02 | Young |
| 0.057085 | 2.251851 | 0.02 | 0 | 1 | 14 | Slc39a2 | 0.02 | Young |
| 0.057085 | 2.251851 | 0.02 | 0 | 1 | 14 | Fam212a | 0.02 | Young |
| 0.057085 | 2.24225 | 0.02 | 0 | 1 | 14 | Apoc1 | 0.02 | Young |
| 0.057085 | 2.24225 | 0.02 | 0 | 1 | 14 | Mapk13 | 0.02 | Young |
| 0.057085 | 2.186677 | 0.02 | 0 | 1 | 14 | Pclo | 0.02 | Young |
| 0.057085 | 2.146427 | 0.02 | 0 | 1 | 14 | Acvr1c | 0.02 | Young |
| 0.057085 | 2.146427 | 0.02 | 0 | 1 | 14 | Ntng1 | 0.02 | Young |
| 0.057085 | 2.146427 | 0.02 | 0 | 1 | 14 | Myocd | 0.02 | Young |
| 0.057085 | 2.146427 | 0.02 | 0 | 1 | 14 | Gmpr | 0.02 | Young |
| 0.057085 | 2.146427 | 0.02 | 0 | 1 | 14 | Popdc2 | 0.02 | Young |
| 0.057085 | 2.087077 | 0.02 | 0 | 1 | 14 | Misp | 0.02 | Young |
| 0.057085 | 2.087077 | 0.02 | 0 | 1 | 14 | Gm26520 | 0.02 | Young |
| 0.057085 | 2.087077 | 0.02 | 0 | 1 | 14 | Adrb1 | 0.02 | Young |
| 0.057085 | 1.998973 | 0.02 | 0 | 1 | 14 | Pabpc1l | 0.02 | Young |
| 0.057085 | 1.998973 | 0.02 | 0 | 1 | 14 | Oaz3 | 0.02 | Young |
| 0.057085 | 1.998973 | 0.02 | 0 | 1 | 14 | 2310047D0 | 0.02 | Young |
| 0.057085 | 1.998973 | 0.02 | 0 | 1 | 14 | Rinl | 0.02 | Young |
| 0.057085 | 1.998973 | 0.02 | 0 | 1 | 14 | Hrc | 0.02 | Young |
| 0.057085 | 1.998973 | 0.02 | 0 | 1 | 14 | Fam181b | 0.02 | Young |
| 0.057085 | 1.998973 | 0.02 | 0 | 1 | 14 | Aqp11 | 0.02 | Young |
| 0.057085 | 1.998973 | 0.02 | 0 | 1 | 14 | Lrrc3 | 0.02 | Young |
| 0.057085 | 1.998973 | 0.02 | 0 | 1 | 14 | 6430573F1 | 0.02 | Young |
| 0.057085 | 1.998973 | 0.02 | 0 | 1 | 14 | Ushbp1 | 0.02 | Young |
| 0.057085 | 1.998973 | 0.02 | 0 | 1 | 14 | Gm5464 | 0.02 | Young |
| 0.057085 | 1.998973 | 0.02 | 0 | 1 | 14 | Emid1 | 0.02 | Young |
| 0.057085 | 1.998973 | 0.02 | 0 | 1 | 14 | Arhgef15 | 0.02 | Young |
| 0.057085 | 1.998973 | 0.02 | 0 | 1 | 14 | Gmnn | 0.02 | Young |
| 0.057085 | 1.998973 | 0.02 | 0 | 1 | 14 | Pde8b | 0.02 | Young |
| 0.057085 | 1.998973 | 0.02 | 0 | 1 | 14 | Otub2 | 0.02 | Young |
| 0.057085 | 1.998973 | 0.02 | 0 | 1 | 14 | 1700001C1 | 0.02 | Young |
| 0.057085 | 1.995737 | 0.02 | 0 | 1 | 14 | Ipp | 0.02 | Young |
| 0.057085 | 1.995737 | 0.02 | 0 | 1 | 14 | Gabrd | 0.02 | Young |
| 0.057085 | 1.995737 | 0.02 | 0 | 1 | 14 | Lrrd1 | 0.02 | Young |
| 0.057085 | 1.995737 | 0.02 | 0 | 1 | 14 | Papolb | 0.02 | Young |
| 0.057085 | 1.995737 | 0.02 | 0 | 1 | 14 | Ccdc162 | 0.02 | Young |

| | | | | | | | | |
|---|---|---|---|---|---|---|---|---|
| 0.057085 | 1.995737 | 0.02 | 0 | 1 | 14 | Ctrl | 0.02 | Young |
| 0.057085 | 1.995737 | 0.02 | 0 | 1 | 14 | Wfdc18 | 0.02 | Young |
| 0.057085 | 1.995737 | 0.02 | 0 | 1 | 14 | Gm11940 | 0.02 | Young |
| 0.057085 | 1.995737 | 0.02 | 0 | 1 | 14 | Zfp455 | 0.02 | Young |
| 0.057085 | 1.962472 | 0.02 | 0 | 1 | 14 | Gm28874 | 0.02 | Young |
| 0.057085 | 1.962472 | 0.02 | 0 | 1 | 14 | Tac4 | 0.02 | Young |
| 0.057085 | 1.951741 | 0.02 | 0 | 1 | 14 | Gm44751 | 0.02 | Young |
| 0.057085 | 1.873584 | 0.02 | 0 | 1 | 14 | Siah3 | 0.02 | Young |
| 0.057085 | 1.854703 | 0.02 | 0 | 1 | 14 | B230398E0 | 0.02 | Young |
| 0.057085 | 1.854703 | 0.02 | 0 | 1 | 14 | Tnfrsf26 | 0.02 | Young |
| 0.057085 | 1.854703 | 0.02 | 0 | 1 | 14 | Gm42031 | 0.02 | Young |
| 0.057085 | 1.854703 | 0.02 | 0 | 1 | 14 | Adgrb1 | 0.02 | Young |
| 0.057085 | 1.854703 | 0.02 | 0 | 1 | 14 | 1700105P0 | 0.02 | Young |
| 0.057085 | 1.824541 | 0.02 | 0 | 1 | 14 | Sez6l | 0.02 | Young |
| 0.057085 | 1.824541 | 0.02 | 0 | 1 | 14 | Bora | 0.02 | Young |
| 0.057085 | 1.824541 | 0.02 | 0 | 1 | 14 | Gm12319 | 0.02 | Young |
| 0.057085 | 1.824541 | 0.02 | 0 | 1 | 14 | Gm26562 | 0.02 | Young |
| 0.057085 | 1.730796 | 0.02 | 0 | 1 | 14 | Cd28 | 0.02 | Young |
| 0.057085 | 1.730796 | 0.02 | 0 | 1 | 14 | Rufy4 | 0.02 | Young |
| 0.057085 | 1.730796 | 0.02 | 0 | 1 | 14 | Frmd5 | 0.02 | Young |
| 0.057085 | 1.730796 | 0.02 | 0 | 1 | 14 | Gm43569 | 0.02 | Young |
| 0.057085 | 1.730796 | 0.02 | 0 | 1 | 14 | Lilra5 | 0.02 | Young |
| 0.057085 | 1.730796 | 0.02 | 0 | 1 | 14 | Zscan2 | 0.02 | Young |
| 0.057085 | 1.730796 | 0.02 | 0 | 1 | 14 | Gm6793 | 0.02 | Young |
| 0.057085 | 1.730796 | 0.02 | 0 | 1 | 14 | Kif23 | 0.02 | Young |
| 0.057085 | 1.730796 | 0.02 | 0 | 1 | 14 | Hist3h2ba | 0.02 | Young |
| 0.057085 | 1.730796 | 0.02 | 0 | 1 | 14 | Slfn3 | 0.02 | Young |
| 0.057085 | 1.730796 | 0.02 | 0 | 1 | 14 | Brip1 | 0.02 | Young |
| 0.057085 | 1.730796 | 0.02 | 0 | 1 | 14 | Gnmt | 0.02 | Young |
| 0.057085 | 1.569642 | 0.02 | 0 | 1 | 14 | Pde1c | 0.02 | Young |
| 0.057085 | 1.569642 | 0.02 | 0 | 1 | 14 | Cox7a1 | 0.02 | Young |
| 0.057085 | 1.569642 | 0.02 | 0 | 1 | 14 | Ascl2 | 0.02 | Young |
| 0.057085 | 1.569642 | 0.02 | 0 | 1 | 14 | Ano4 | 0.02 | Young |
| 0.057085 | 1.569642 | 0.02 | 0 | 1 | 14 | Cngb1 | 0.02 | Young |
| 0.057085 | 1.569642 | 0.02 | 0 | 1 | 14 | Gm10371 | 0.02 | Young |
| 0.057085 | 1.569642 | 0.02 | 0 | 1 | 14 | Isl2 | 0.02 | Young |
| 0.057085 | 1.569642 | 0.02 | 0 | 1 | 14 | 9130213A2 | 0.02 | Young |
| 0.057085 | 1.569642 | 0.02 | 0 | 1 | 14 | Nrxn3 | 0.02 | Young |
| 0.057085 | 1.553445 | 0.02 | 0 | 1 | 14 | Thsd7b | 0.02 | Young |
| 0.057085 | 1.553445 | 0.02 | 0 | 1 | 14 | Cdk18 | 0.02 | Young |
| 0.057085 | 1.553445 | 0.02 | 0 | 1 | 14 | Map3k15 | 0.02 | Young |
| 0.057085 | 1.553445 | 0.02 | 0 | 1 | 14 | Chil5 | 0.02 | Young |
| 0.057085 | 1.553445 | 0.02 | 0 | 1 | 14 | Palmd | 0.02 | Young |
| 0.057085 | 1.553445 | 0.02 | 0 | 1 | 14 | Gm43614 | 0.02 | Young |
| 0.057085 | 1.553445 | 0.02 | 0 | 1 | 14 | Aqp7 | 0.02 | Young |

| | | | | | | | | |
|---|---|---|---|---|---|---|---|---|
| 0.057085 | 1.553445 | 0.02 | 0 | 1 | 14 | Tbc1d2 | 0.02 | Young |
| 0.057085 | 1.553445 | 0.02 | 0 | 1 | 14 | Zfp979 | 0.02 | Young |
| 0.057085 | 1.553445 | 0.02 | 0 | 1 | 14 | Fgfr3 | 0.02 | Young |
| 0.057085 | 1.553445 | 0.02 | 0 | 1 | 14 | Prom1 | 0.02 | Young |
| 0.057085 | 1.553445 | 0.02 | 0 | 1 | 14 | Pilrb1 | 0.02 | Young |
| 0.057085 | 1.553445 | 0.02 | 0 | 1 | 14 | Halr1 | 0.02 | Young |
| 0.057085 | 1.553445 | 0.02 | 0 | 1 | 14 | Gm45188 | 0.02 | Young |
| 0.057085 | 1.553445 | 0.02 | 0 | 1 | 14 | Trim30b | 0.02 | Young |
| 0.057085 | 1.553445 | 0.02 | 0 | 1 | 14 | Gm17745 | 0.02 | Young |
| 0.057085 | 1.553445 | 0.02 | 0 | 1 | 14 | Cpm | 0.02 | Young |
| 0.057085 | 1.553445 | 0.02 | 0 | 1 | 14 | Irx3 | 0.02 | Young |
| 0.057085 | 1.553445 | 0.02 | 0 | 1 | 14 | Adgrg3 | 0.02 | Young |
| 0.057085 | 1.553445 | 0.02 | 0 | 1 | 14 | Mmp15 | 0.02 | Young |
| 0.057085 | 1.553445 | 0.02 | 0 | 1 | 14 | Lrrc3b | 0.02 | Young |
| 0.057085 | 1.553445 | 0.02 | 0 | 1 | 14 | Esco2 | 0.02 | Young |
| 0.057085 | 1.553445 | 0.02 | 0 | 1 | 14 | Lrrc63 | 0.02 | Young |
| 0.057085 | 1.553445 | 0.02 | 0 | 1 | 14 | Sox21 | 0.02 | Young |
| 0.057085 | 1.553445 | 0.02 | 0 | 1 | 14 | Sema3f | 0.02 | Young |
| 0.057085 | 1.553445 | 0.02 | 0 | 1 | 14 | Ccdc51 | 0.02 | Young |
| 0.057085 | 1.553445 | 0.02 | 0 | 1 | 14 | Bcl6b | 0.02 | Young |
| 0.057085 | 1.553445 | 0.02 | 0 | 1 | 14 | Cdc6 | 0.02 | Young |
| 0.057085 | 1.553445 | 0.02 | 0 | 1 | 14 | Gm11713 | 0.02 | Young |
| 0.057085 | 1.553445 | 0.02 | 0 | 1 | 14 | Kcnj2 | 0.02 | Young |
| 0.057085 | 1.553445 | 0.02 | 0 | 1 | 14 | Rab37 | 0.02 | Young |
| 0.057085 | 1.553445 | 0.02 | 0 | 1 | 14 | Gm47675 | 0.02 | Young |
| 0.057085 | 1.553445 | 0.02 | 0 | 1 | 14 | Fzd6 | 0.02 | Young |
| 0.057085 | 1.553445 | 0.02 | 0 | 1 | 14 | Samd12 | 0.02 | Young |
| 0.057085 | 1.553445 | 0.02 | 0 | 1 | 14 | Baiap2l2 | 0.02 | Young |
| 0.057085 | 1.553445 | 0.02 | 0 | 1 | 14 | Prss27 | 0.02 | Young |
| 0.057085 | 1.553445 | 0.02 | 0 | 1 | 14 | Scube3 | 0.02 | Young |
| 0.057085 | 1.553445 | 0.02 | 0 | 1 | 14 | Col11a2 | 0.02 | Young |
| 0.057085 | 1.553445 | 0.02 | 0 | 1 | 14 | Ddx11 | 0.02 | Young |
| 0.057085 | 1.553445 | 0.02 | 0 | 1 | 14 | Gm26510 | 0.02 | Young |
| 0.057085 | 1.553445 | 0.02 | 0 | 1 | 14 | Tmem178 | 0.02 | Young |
| 0.057085 | 1.553445 | 0.02 | 0 | 1 | 14 | Trpm6 | 0.02 | Young |
| 0.057085 | 1.518873 | 0.02 | 0 | 1 | 14 | 9330175M | 0.02 | Young |
| 0.057085 | 1.518873 | 0.02 | 0 | 1 | 14 | Gm9530 | 0.02 | Young |
| 0.057085 | 1.518873 | 0.02 | 0 | 1 | 14 | Dmrtc1a | 0.02 | Young |
| 0.057085 | 1.518873 | 0.02 | 0 | 1 | 14 | C030034L1 | 0.02 | Young |
| 0.057085 | 1.518873 | 0.02 | 0 | 1 | 14 | Gm37035 | 0.02 | Young |
| 0.057085 | 1.518873 | 0.02 | 0 | 1 | 14 | Asb10 | 0.02 | Young |
| 0.057085 | 1.518873 | 0.02 | 0 | 1 | 14 | Exoc3l2 | 0.02 | Young |
| 0.057085 | 1.518873 | 0.02 | 0 | 1 | 14 | Fxyd3 | 0.02 | Young |
| 0.057085 | 1.518873 | 0.02 | 0 | 1 | 14 | Gldn | 0.02 | Young |
| 0.057085 | 1.518873 | 0.02 | 0 | 1 | 14 | Rdh12 | 0.02 | Young |

| | | | | | | | | |
|---|---|---|---|---|---|---|---|---|
| 0.057085 | 1.47681 | 0.02 | 0 | 1 | 14 | Slc7a3 | 0.02 | Young |
| 0.057085 | 1.47681 | 0.02 | 0 | 1 | 14 | Muc1 | 0.02 | Young |
| 0.057085 | 1.47681 | 0.02 | 0 | 1 | 14 | Zmynd12 | 0.02 | Young |
| 0.057085 | 1.47681 | 0.02 | 0 | 1 | 14 | Fabp3 | 0.02 | Young |
| 0.057085 | 1.47681 | 0.02 | 0 | 1 | 14 | Areg | 0.02 | Young |
| 0.057085 | 1.47681 | 0.02 | 0 | 1 | 14 | Hsd17b13 | 0.02 | Young |
| 0.057085 | 1.47681 | 0.02 | 0 | 1 | 14 | Gm43625 | 0.02 | Young |
| 0.057085 | 1.47681 | 0.02 | 0 | 1 | 14 | Ephb6 | 0.02 | Young |
| 0.057085 | 1.47681 | 0.02 | 0 | 1 | 14 | AI429214 | 0.02 | Young |
| 0.057085 | 1.47681 | 0.02 | 0 | 1 | 14 | Dand5 | 0.02 | Young |
| 0.057085 | 1.47681 | 0.02 | 0 | 1 | 14 | Mdga2 | 0.02 | Young |
| 0.057085 | 1.47681 | 0.02 | 0 | 1 | 14 | Ifit1bl2 | 0.02 | Young |
| 0.057085 | 1.473152 | 0.02 | 0 | 1 | 14 | Nrk | 0.02 | Young |
| 0.057085 | 1.473152 | 0.02 | 0 | 1 | 14 | AI115009 | 0.02 | Young |
| 0.057085 | 1.473152 | 0.02 | 0 | 1 | 14 | Ephb1 | 0.02 | Young |
| 0.057085 | 1.473152 | 0.02 | 0 | 1 | 14 | Sybu | 0.02 | Young |
| 0.057085 | 1.440234 | 0.02 | 0 | 1 | 14 | G730003C1 | 0.02 | Young |
| 0.057085 | 1.440234 | 0.02 | 0 | 1 | 14 | Brinp2 | 0.02 | Young |
| 0.057085 | 1.440234 | 0.02 | 0 | 1 | 14 | D430041D0 | 0.02 | Young |
| 0.057085 | 1.440234 | 0.02 | 0 | 1 | 14 | Scg5 | 0.02 | Young |
| 0.057085 | 1.440234 | 0.02 | 0 | 1 | 14 | Mcf2 | 0.02 | Young |
| 0.057085 | 1.440234 | 0.02 | 0 | 1 | 14 | 6530403H0 | 0.02 | Young |
| 0.057085 | 1.440234 | 0.02 | 0 | 1 | 14 | 4930570G1 | 0.02 | Young |
| 0.057085 | 1.440234 | 0.02 | 0 | 1 | 14 | Gm12454 | 0.02 | Young |
| 0.057085 | 1.440234 | 0.02 | 0 | 1 | 14 | Aldh1b1 | 0.02 | Young |
| 0.057085 | 1.440234 | 0.02 | 0 | 1 | 14 | Epha8 | 0.02 | Young |
| 0.057085 | 1.440234 | 0.02 | 0 | 1 | 14 | Gm13205 | 0.02 | Young |
| 0.057085 | 1.440234 | 0.02 | 0 | 1 | 14 | Tnfrsf25 | 0.02 | Young |
| 0.057085 | 1.440234 | 0.02 | 0 | 1 | 14 | Kcnh2 | 0.02 | Young |
| 0.057085 | 1.440234 | 0.02 | 0 | 1 | 14 | Gm15777 | 0.02 | Young |
| 0.057085 | 1.440234 | 0.02 | 0 | 1 | 14 | 5430427N1 | 0.02 | Young |
| 0.057085 | 1.440234 | 0.02 | 0 | 1 | 14 | Col26a1 | 0.02 | Young |
| 0.057085 | 1.440234 | 0.02 | 0 | 1 | 14 | Xntrpc | 0.02 | Young |
| 0.057085 | 1.440234 | 0.02 | 0 | 1 | 14 | Gm44623 | 0.02 | Young |
| 0.057085 | 1.440234 | 0.02 | 0 | 1 | 14 | Gm15677 | 0.02 | Young |
| 0.057085 | 1.440234 | 0.02 | 0 | 1 | 14 | Syce1 | 0.02 | Young |
| 0.057085 | 1.440234 | 0.02 | 0 | 1 | 14 | Ankrd37 | 0.02 | Young |
| 0.057085 | 1.440234 | 0.02 | 0 | 1 | 14 | Rasd2 | 0.02 | Young |
| 0.057085 | 1.440234 | 0.02 | 0 | 1 | 14 | Cdh3 | 0.02 | Young |
| 0.057085 | 1.440234 | 0.02 | 0 | 1 | 14 | 9930012K1 | 0.02 | Young |
| 0.057085 | 1.440234 | 0.02 | 0 | 1 | 14 | Jaml | 0.02 | Young |
| 0.057085 | 1.440234 | 0.02 | 0 | 1 | 14 | Scn2b | 0.02 | Young |
| 0.057085 | 1.440234 | 0.02 | 0 | 1 | 14 | Exph5 | 0.02 | Young |
| 0.057085 | 1.440234 | 0.02 | 0 | 1 | 14 | Asic2 | 0.02 | Young |
| 0.057085 | 1.440234 | 0.02 | 0 | 1 | 14 | Hoxb5os | 0.02 | Young |

| | | | | | | | | |
|---|---|---|---|---|---|---|---|---|
| 0.057085 | 1.440234 | 0.02 | 0 | 1 | 14 | Sdk2 | 0.02 | Young |
| 0.057085 | 1.440234 | 0.02 | 0 | 1 | 14 | Llgl2 | 0.02 | Young |
| 0.057085 | 1.440234 | 0.02 | 0 | 1 | 14 | Gm11725 | 0.02 | Young |
| 0.057085 | 1.440234 | 0.02 | 0 | 1 | 14 | Gm30655 | 0.02 | Young |
| 0.057085 | 1.440234 | 0.02 | 0 | 1 | 14 | Sptb | 0.02 | Young |
| 0.057085 | 1.440234 | 0.02 | 0 | 1 | 14 | Bbof1 | 0.02 | Young |
| 0.057085 | 1.440234 | 0.02 | 0 | 1 | 14 | AC163040. | 0.02 | Young |
| 0.057085 | 1.440234 | 0.02 | 0 | 1 | 14 | Gm10389 | 0.02 | Young |
| 0.057085 | 1.440234 | 0.02 | 0 | 1 | 14 | Cpne5 | 0.02 | Young |
| 0.057085 | 1.440234 | 0.02 | 0 | 1 | 14 | Vwce | 0.02 | Young |
| 0.057085 | 1.39128 | 0.02 | 0 | 1 | 14 | Ccdc34os | 0.02 | Young |
| 0.057085 | 1.39128 | 0.02 | 0 | 1 | 14 | Gm14419 | 0.02 | Young |
| 0.057085 | 1.39128 | 0.02 | 0 | 1 | 14 | Rnf113a1 | 0.02 | Young |
| 0.057085 | 1.39128 | 0.02 | 0 | 1 | 14 | Pxmp2 | 0.02 | Young |
| 0.057085 | 1.39128 | 0.02 | 0 | 1 | 14 | 9430002A1 | 0.02 | Young |
| 0.057085 | 1.39128 | 0.02 | 0 | 1 | 14 | Gm17233 | 0.02 | Young |
| 0.057085 | 1.39128 | 0.02 | 0 | 1 | 14 | Gm36107 | 0.02 | Young |
| 0.057085 | 1.39128 | 0.02 | 0 | 1 | 14 | 2010300F1 | 0.02 | Young |
| 0.057085 | 1.39128 | 0.02 | 0 | 1 | 14 | Rgs7bp | 0.02 | Young |
| 0.057085 | 1.39128 | 0.02 | 0 | 1 | 14 | Serac1 | 0.02 | Young |
| 0.057085 | 1.291984 | 0.02 | 0 | 1 | 14 | Tnnc2 | 0.02 | Young |
| 0.057085 | 1.291984 | 0.02 | 0 | 1 | 14 | Ugt2b36 | 0.02 | Young |
| 0.057085 | 1.291984 | 0.02 | 0 | 1 | 14 | Ufsp1 | 0.02 | Young |
| 0.057085 | 1.291984 | 0.02 | 0 | 1 | 14 | Prr15 | 0.02 | Young |
| 0.057085 | 1.291984 | 0.02 | 0 | 1 | 14 | Rep15 | 0.02 | Young |
| 0.057085 | 1.291984 | 0.02 | 0 | 1 | 14 | Rnase10 | 0.02 | Young |
| 0.057085 | 1.291984 | 0.02 | 0 | 1 | 14 | Krt222 | 0.02 | Young |
| 0.057085 | 1.291984 | 0.02 | 0 | 1 | 14 | Pcdhga8 | 0.02 | Young |
| 0.057085 | 1.283451 | 0.02 | 0 | 1 | 14 | Slc2a2 | 0.02 | Young |
| 0.057085 | 1.283451 | 0.02 | 0 | 1 | 14 | Tsacc | 0.02 | Young |
| 0.057085 | 1.283451 | 0.02 | 0 | 1 | 14 | 2310081O( | 0.02 | Young |
| 0.057085 | 1.283451 | 0.02 | 0 | 1 | 14 | Gm4876 | 0.02 | Young |
| 0.057085 | 1.283451 | 0.02 | 0 | 1 | 14 | Gm42372 | 0.02 | Young |
| 0.057085 | 1.283451 | 0.02 | 0 | 1 | 14 | Hsd3b7 | 0.02 | Young |
| 0.057085 | 1.283451 | 0.02 | 0 | 1 | 14 | Alx1 | 0.02 | Young |
| 0.057085 | 1.283451 | 0.02 | 0 | 1 | 14 | Tex15 | 0.02 | Young |
| 0.057085 | 1.283451 | 0.02 | 0 | 1 | 14 | 9330175E1 | 0.02 | Young |
| 0.057085 | 1.283451 | 0.02 | 0 | 1 | 14 | Ankdd1b | 0.02 | Young |
| 0.057085 | 1.283451 | 0.02 | 0 | 1 | 14 | Gm36495 | 0.02 | Young |
| 0.057085 | 1.283451 | 0.02 | 0 | 1 | 14 | A530016L2 | 0.02 | Young |
| 0.057085 | 1.283451 | 0.02 | 0 | 1 | 14 | B3gnt5 | 0.02 | Young |
| 0.057085 | 1.283451 | 0.02 | 0 | 1 | 14 | Bex6 | 0.02 | Young |
| 0.057085 | 1.283451 | 0.02 | 0 | 1 | 14 | Trpm3 | 0.02 | Young |
| 0.057085 | 1.219596 | 0.02 | 0 | 1 | 14 | Gm10134 | 0.02 | Young |
| 0.057085 | 1.219596 | 0.02 | 0 | 1 | 14 | Olfr73 | 0.02 | Young |

| | | | | | | | | |
|---|---|---|---|---|---|---|---|---|
| 0.057085 | 1.219596 | 0.02 | 0 | 1 | 14 | Oip5 | 0.02 | Young |
| 0.057085 | 1.219596 | 0.02 | 0 | 1 | 14 | Tceal5 | 0.02 | Young |
| 0.057085 | 1.219596 | 0.02 | 0 | 1 | 14 | B230334C0 | 0.02 | Young |
| 0.057085 | 1.219596 | 0.02 | 0 | 1 | 14 | Igsf21 | 0.02 | Young |
| 0.057085 | 1.219596 | 0.02 | 0 | 1 | 14 | Ttc34 | 0.02 | Young |
| 0.057085 | 1.219596 | 0.02 | 0 | 1 | 14 | 9430085M | 0.02 | Young |
| 0.057085 | 1.219596 | 0.02 | 0 | 1 | 14 | Gm9801 | 0.02 | Young |
| 0.057085 | 1.219596 | 0.02 | 0 | 1 | 14 | Tpte | 0.02 | Young |
| 0.057085 | 1.219596 | 0.02 | 0 | 1 | 14 | Ppp1r3b | 0.02 | Young |
| 0.057085 | 1.219596 | 0.02 | 0 | 1 | 14 | Car7 | 0.02 | Young |
| 0.057085 | 1.219596 | 0.02 | 0 | 1 | 14 | Gm26592 | 0.02 | Young |
| 0.057085 | 1.219596 | 0.02 | 0 | 1 | 14 | Ccdc92b | 0.02 | Young |
| 0.057085 | 1.219596 | 0.02 | 0 | 1 | 14 | Gm26836 | 0.02 | Young |
| 0.057085 | 1.219596 | 0.02 | 0 | 1 | 14 | Ccdc57 | 0.02 | Young |
| 0.057085 | 1.219596 | 0.02 | 0 | 1 | 14 | Slc17a3 | 0.02 | Young |
| 0.057085 | 1.219596 | 0.02 | 0 | 1 | 14 | Zfp457 | 0.02 | Young |
| 0.057085 | 1.219596 | 0.02 | 0 | 1 | 14 | Gm48633 | 0.02 | Young |
| 0.057085 | 1.219596 | 0.02 | 0 | 1 | 14 | Tmem117 | 0.02 | Young |
| 0.057085 | 1.073801 | 0.02 | 0 | 1 | 14 | Fam178b | 0.02 | Young |
| 0.057085 | 1.073801 | 0.02 | 0 | 1 | 14 | Rhov | 0.02 | Young |
| 0.057085 | 1.073801 | 0.02 | 0 | 1 | 14 | Arl9 | 0.02 | Young |
| 0.057085 | 1.073801 | 0.02 | 0 | 1 | 14 | Micall2 | 0.02 | Young |
| 0.057085 | 1.073801 | 0.02 | 0 | 1 | 14 | Gm38393 | 0.02 | Young |
| 0.057085 | 1.073801 | 0.02 | 0 | 1 | 14 | Gm48053 | 0.02 | Young |
| 0.057085 | 1.073801 | 0.02 | 0 | 1 | 14 | Kera | 0.02 | Young |
| 0.057085 | 1.073801 | 0.02 | 0 | 1 | 14 | Gm12153 | 0.02 | Young |
| 0.057085 | 1.073801 | 0.02 | 0 | 1 | 14 | Rnasek | 0.02 | Young |
| 0.057085 | 1.073801 | 0.02 | 0 | 1 | 14 | Serpinb9b | 0.02 | Young |
| 0.057085 | 1.073801 | 0.02 | 0 | 1 | 14 | Tert | 0.02 | Young |
| 0.057085 | 1.073801 | 0.02 | 0 | 1 | 14 | Prox2os | 0.02 | Young |
| 0.057085 | 1.073801 | 0.02 | 0 | 1 | 14 | Fosl1 | 0.02 | Young |
| 0.057085 | 0.916965 | 0.02 | 0 | 1 | 14 | Nppc | 0.02 | Young |
| 0.057085 | 0.916965 | 0.02 | 0 | 1 | 14 | Lgr6 | 0.02 | Young |
| 0.057085 | 0.916965 | 0.02 | 0 | 1 | 14 | B230369F2 | 0.02 | Young |
| 0.057085 | 0.916965 | 0.02 | 0 | 1 | 14 | Galnt5 | 0.02 | Young |
| 0.057085 | 0.916965 | 0.02 | 0 | 1 | 14 | Gm14703 | 0.02 | Young |
| 0.057085 | 0.916965 | 0.02 | 0 | 1 | 14 | Klhl4 | 0.02 | Young |
| 0.057085 | 0.916965 | 0.02 | 0 | 1 | 14 | Rragb | 0.02 | Young |
| 0.057085 | 0.916965 | 0.02 | 0 | 1 | 14 | Gm17146 | 0.02 | Young |
| 0.057085 | 0.916965 | 0.02 | 0 | 1 | 14 | Hist2h3c1 | 0.02 | Young |
| 0.057085 | 0.916965 | 0.02 | 0 | 1 | 14 | Col25a1 | 0.02 | Young |
| 0.057085 | 0.916965 | 0.02 | 0 | 1 | 14 | Spaca1 | 0.02 | Young |
| 0.057085 | 0.916965 | 0.02 | 0 | 1 | 14 | Hdhd3 | 0.02 | Young |
| 0.057085 | 0.916965 | 0.02 | 0 | 1 | 14 | 1500002C1 | 0.02 | Young |
| 0.057085 | 0.916965 | 0.02 | 0 | 1 | 14 | Sh2b2 | 0.02 | Young |

| | | | | | | | | |
|---|---|---|---|---|---|---|---|---|
| 0.057085 | 0.916965 | 0.02 | 0 | 1 | 14 | Gpr12 | 0.02 | Young |
| 0.057085 | 0.916965 | 0.02 | 0 | 1 | 14 | Gm38832 | 0.02 | Young |
| 0.057085 | 0.916965 | 0.02 | 0 | 1 | 14 | Cyb5r2 | 0.02 | Young |
| 0.057085 | 0.916965 | 0.02 | 0 | 1 | 14 | Gm5914 | 0.02 | Young |
| 0.057085 | 0.916965 | 0.02 | 0 | 1 | 14 | Chek1 | 0.02 | Young |
| 0.057085 | 0.916965 | 0.02 | 0 | 1 | 14 | Tmem266 | 0.02 | Young |
| 0.057085 | 0.916965 | 0.02 | 0 | 1 | 14 | Hbq1a | 0.02 | Young |
| 0.057085 | 0.916965 | 0.02 | 0 | 1 | 14 | Fam196b | 0.02 | Young |
| 0.057085 | 0.916965 | 0.02 | 0 | 1 | 14 | Ube2ql1 | 0.02 | Young |
| 0.057085 | 0.916965 | 0.02 | 0 | 1 | 14 | Gm20498 | 0.02 | Young |
| 0.057085 | 0.916965 | 0.02 | 0 | 1 | 14 | Tmem121 | 0.02 | Young |
| 0.057085 | 0.916965 | 0.02 | 0 | 1 | 14 | Phf21b | 0.02 | Young |
| 0.057085 | 0.916965 | 0.02 | 0 | 1 | 14 | Tomm6os | 0.02 | Young |
| 0.057538 | 1.416651 | 0.16 | 0.071 | 1 | 14 | Gtf2e1 | 0.089 | Young |
| 0.057566 | 0.443554 | 0.3 | 0.169 | 1 | 14 | Pi4k2a | 0.131 | Young |
| 0.057641 | 1.218323 | 0.28 | 0.169 | 1 | 14 | Adamts10 | 0.111 | Young |
| 0.057785 | 2.05543 | 0.04 | 0.005 | 1 | 14 | Slc16a12 | 0.035 | Young |
| 0.057785 | 1.920138 | 0.04 | 0.005 | 1 | 14 | Cyyr1 | 0.035 | Young |
| 0.057785 | 1.9104 | 0.04 | 0.005 | 1 | 14 | Lama5 | 0.035 | Young |
| 0.057785 | 1.723042 | 0.04 | 0.005 | 1 | 14 | Prps1l3 | 0.035 | Young |
| 0.057785 | 1.637145 | 0.04 | 0.005 | 1 | 14 | Dusp4 | 0.035 | Young |
| 0.057785 | 1.611359 | 0.04 | 0.005 | 1 | 14 | Arap3 | 0.035 | Young |
| 0.057785 | 1.576463 | 0.04 | 0.005 | 1 | 14 | Hoxa2 | 0.035 | Young |
| 0.057785 | 1.534264 | 0.04 | 0.005 | 1 | 14 | Batf2 | 0.035 | Young |
| 0.057785 | 1.506688 | 0.04 | 0.005 | 1 | 14 | Ssh3 | 0.035 | Young |
| 0.057785 | 1.494772 | 0.04 | 0.005 | 1 | 14 | Trpc4 | 0.035 | Young |
| 0.057785 | 1.467575 | 0.04 | 0.005 | 1 | 14 | Dna2 | 0.035 | Young |
| 0.057785 | 1.450406 | 0.04 | 0.005 | 1 | 14 | Lrrc56 | 0.035 | Young |
| 0.057785 | 1.394625 | 0.04 | 0.005 | 1 | 14 | Gpr85 | 0.035 | Young |
| 0.057785 | 1.29835 | 0.04 | 0.005 | 1 | 14 | Stmnd1 | 0.035 | Young |
| 0.057785 | 1.285337 | 0.04 | 0.005 | 1 | 14 | Hspb7 | 0.035 | Young |
| 0.057785 | 1.282693 | 0.04 | 0.005 | 1 | 14 | A230005M | 0.035 | Young |
| 0.057785 | 1.213237 | 0.04 | 0.005 | 1 | 14 | Mybl1 | 0.035 | Young |
| 0.057785 | 1.192225 | 0.04 | 0.005 | 1 | 14 | Gm16894 | 0.035 | Young |
| 0.057785 | 1.11951 | 0.04 | 0.005 | 1 | 14 | Folh1 | 0.035 | Young |
| 0.057785 | 1.07359 | 0.04 | 0.005 | 1 | 14 | Zfp947 | 0.035 | Young |
| 0.057785 | 0.766103 | 0.04 | 0.005 | 1 | 14 | Car11 | 0.035 | Young |
| 0.057785 | 0.680397 | 0.04 | 0.005 | 1 | 14 | Fancm | 0.035 | Young |
| 0.057785 | 0.470097 | 0.04 | 0.005 | 1 | 14 | Klhl6 | 0.035 | Young |
| 0.057785 | 0.412737 | 0.04 | 0.005 | 1 | 14 | Dennd3 | 0.035 | Young |
| 0.057785 | 0.259691 | 0.04 | 0.005 | 1 | 14 | Epha2 | 0.035 | Young |
| 0.057785 | 0.218824 | 0.04 | 0.005 | 1 | 14 | Zfp57 | 0.035 | Young |
| 0.057831 | 1.739906 | 0.16 | 0.077 | 1 | 14 | Etv5 | 0.083 | Young |
| 0.058138 | 1.249423 | 0.14 | 0.06 | 1 | 14 | Synpo2 | 0.08 | Young |
| 0.058231 | 0.303005 | 0.1 | 0.033 | 1 | 14 | Tcp11l1 | 0.067 | Young |

| | | | | | | | | |
|---|---|---|---|---|---|---|---|---|
| 0.05849 | 1.290981 | 0.36 | 0.224 | 1 | 14 | Arhgap1 | 0.136 | Young |
| 0.058846 | 2.500995 | 0.12 | 0.049 | 1 | 14 | Vstm4 | 0.071 | Young |
| 0.058923 | 1.02214 | 0.44 | 0.317 | 1 | 14 | Wasl | 0.123 | Young |
| 0.059014 | 1.26431 | 0.32 | 0.202 | 1 | 14 | Laptm4b | 0.118 | Young |
| 0.059058 | 1.020075 | 0.28 | 0.164 | 1 | 14 | Tmem135 | 0.116 | Young |
| 0.059127 | 0.917208 | 0.24 | 0.126 | 1 | 14 | Golt1b | 0.114 | Young |
| 0.059443 | 0.81608 | 0.16 | 0.071 | 1 | 14 | Atp13a2 | 0.089 | Young |
| 0.059522 | 1.321597 | 0.2 | 0.098 | 1 | 14 | Pibf1 | 0.102 | Young |
| 0.059696 | 0.745339 | 0.38 | 0.246 | 1 | 14 | Thoc7 | 0.134 | Young |
| 0.059789 | 0.648634 | 0.26 | 0.142 | 1 | 14 | Scai | 0.118 | Young |
| 0.060094 | 0.95981 | 0.24 | 0.131 | 1 | 14 | Fem1c | 0.109 | Young |
| 0.060239 | 0.768837 | 0.32 | 0.197 | 1 | 14 | Suclg1 | 0.123 | Young |
| 0.060247 | 0.360962 | 0.88 | 0.88 | 1 | 14 | Gnas | 0 | Young |
| 0.060326 | 1.42908 | 0.4 | 0.279 | 1 | 14 | Gng12 | 0.121 | Young |
| 0.060399 | 0.111457 | 0.4 | 0.251 | 1 | 14 | Adamts15 | 0.149 | Young |
| 0.060903 | 0.947905 | 0.14 | 0.06 | 1 | 14 | Zfp236 | 0.08 | Young |
| 0.060945 | 0.386384 | 0.96 | 0.951 | 1 | 14 | Sparc | 0.009 | Young |
| 0.06101 | 0.977337 | 0.22 | 0.115 | 1 | 14 | Klhl7 | 0.105 | Young |
| 0.061967 | 1.757249 | 0.22 | 0.12 | 1 | 14 | Golga5 | 0.1 | Young |
| 0.062175 | 0.842435 | 0.24 | 0.131 | 1 | 14 | Creld2 | 0.109 | Young |
| 0.063254 | 0.676851 | 0.76 | 0.749 | 1 | 14 | Igfbp7 | 0.011 | Young |
| 0.063308 | 0.544767 | 0.22 | 0.109 | 1 | 14 | Rad54l2 | 0.111 | Young |
| 0.06341 | 0.475881 | 0.16 | 0.071 | 1 | 14 | Zfp282 | 0.089 | Young |
| 0.063497 | 0.56106 | 0.18 | 0.082 | 1 | 14 | Mob2 | 0.098 | Young |
| 0.063603 | 0.540651 | 0.44 | 0.301 | 1 | 14 | Atp6v0d1 | 0.139 | Young |
| 0.063699 | 1.238758 | 0.3 | 0.186 | 1 | 14 | Ppfia1 | 0.114 | Young |
| 0.064066 | 0.715795 | 0.36 | 0.224 | 1 | 14 | Tmem159 | 0.136 | Young |
| 0.064644 | 0.606737 | 0.34 | 0.208 | 1 | 14 | Mcts1 | 0.132 | Young |
| 0.064888 | 1.295364 | 0.32 | 0.208 | 1 | 14 | Plcb4 | 0.112 | Young |
| 0.065286 | 1.398748 | 0.2 | 0.104 | 1 | 14 | Fmc1 | 0.096 | Young |
| 0.065509 | 0.627344 | 0.22 | 0.115 | 1 | 14 | Hdgfl2 | 0.105 | Young |
| 0.065555 | 0.515606 | 0.36 | 0.219 | 1 | 14 | Hs2st1 | 0.141 | Young |
| 0.065886 | 1.141642 | 0.22 | 0.12 | 1 | 14 | Aspn | 0.1 | Young |
| 0.066003 | 2.190381 | 0.14 | 0.06 | 1 | 14 | Gtpbp10 | 0.08 | Young |
| 0.066032 | 0.012814 | 0.2 | 0.098 | 1 | 14 | Mdn1 | 0.102 | Young |
| 0.066554 | 1.282606 | 0.34 | 0.23 | 1 | 14 | Scamp2 | 0.11 | Young |
| 0.066614 | 0.69089 | 0.5 | 0.333 | 1 | 14 | Cggbp1 | 0.167 | Young |
| 0.066619 | 0.912934 | 0.36 | 0.24 | 1 | 14 | Arid4b | 0.12 | Young |
| 0.06676 | 0.695587 | 0.14 | 0.06 | 1 | 14 | Fh1 | 0.08 | Young |
| 0.06688 | 0.326262 | 0.16 | 0.071 | 1 | 14 | Zc3h6 | 0.089 | Young |
| 0.067047 | 0.616798 | 0.48 | 0.372 | 1 | 14 | Anp32e | 0.108 | Young |
| 0.067531 | 1.619357 | 0.12 | 0.049 | 1 | 14 | Gucy1a1 | 0.071 | Young |
| 0.067599 | 0.995235 | 0.42 | 0.311 | 1 | 14 | Elk4 | 0.109 | Young |
| 0.068154 | 0.812623 | 0.3 | 0.18 | 1 | 14 | Scamp3 | 0.12 | Young |
| 0.068294 | 1.874043 | 0.14 | 0.06 | 1 | 14 | Spryd3 | 0.08 | Young |

| | | | | | | | | |
|---|---|---|---|---|---|---|---|---|
| 0.06837 | 2.35665 | 0.12 | 0.049 | 1 | 14 | Fhod1 | 0.071 | Young |
| 0.06866 | 1.282125 | 0.28 | 0.175 | 1 | 14 | Bcl2l1 | 0.105 | Young |
| 0.069218 | 3.153536 | 0.12 | 0.049 | 1 | 14 | Uqcc1 | 0.071 | Young |
| 0.069244 | 0.894393 | 0.18 | 0.093 | 1 | 14 | Actn1 | 0.087 | Young |
| 0.069406 | 1.043464 | 0.22 | 0.12 | 1 | 14 | Cers6 | 0.1 | Young |
| 0.070074 | 1.436706 | 0.12 | 0.049 | 1 | 14 | Rasl11a | 0.071 | Young |
| 0.070175 | 0.570916 | 0.36 | 0.251 | 1 | 14 | Rtf1 | 0.109 | Young |
| 0.070229 | 0.920597 | 0.28 | 0.175 | 1 | 14 | Hpgd | 0.105 | Young |
| 0.070439 | 0.427087 | 0.28 | 0.169 | 1 | 14 | Papss1 | 0.111 | Young |
| 0.070604 | 2.647803 | 0.1 | 0.038 | 1 | 14 | Rps6ka5 | 0.062 | Young |
| 0.070604 | 2.601147 | 0.1 | 0.038 | 1 | 14 | Prr5l | 0.062 | Young |
| 0.07065 | 0.549194 | 0.14 | 0.06 | 1 | 14 | Myo18a | 0.08 | Young |
| 0.07079 | 1.37789 | 0.18 | 0.087 | 1 | 14 | Hikeshi | 0.093 | Young |
| 0.070796 | 1.177713 | 0.34 | 0.23 | 1 | 14 | Lsm6 | 0.11 | Young |
| 0.070939 | 1.575104 | 0.12 | 0.049 | 1 | 14 | Smcr8 | 0.071 | Young |
| 0.071127 | 0.217452 | 0.2 | 0.098 | 1 | 14 | Cdc34 | 0.102 | Young |
| 0.071287 | 1.505683 | 0.16 | 0.077 | 1 | 14 | Cpeb3 | 0.083 | Young |
| 0.071509 | 0.556175 | 0.42 | 0.284 | 1 | 14 | Zmynd8 | 0.136 | Young |
| 0.071543 | 0.567451 | 0.2 | 0.104 | 1 | 14 | Nub1 | 0.096 | Young |
| 0.071813 | 0.880104 | 0.12 | 0.049 | 1 | 14 | Pnpt1 | 0.071 | Young |
| 0.071837 | 1.857108 | 0.22 | 0.12 | 1 | 14 | Armcx1 | 0.1 | Young |
| 0.072257 | 0.692463 | 0.14 | 0.06 | 1 | 14 | Cnot11 | 0.08 | Young |
| 0.072288 | 0.338388 | 0.24 | 0.131 | 1 | 14 | Glod4 | 0.109 | Young |
| 0.072359 | 0.427147 | 0.3 | 0.175 | 1 | 14 | Plekhm2 | 0.125 | Young |
| 0.072696 | 0.983349 | 0.12 | 0.049 | 1 | 14 | Aldh1a2 | 0.071 | Young |
| 0.073018 | 0.931904 | 0.38 | 0.246 | 1 | 14 | Atf2 | 0.134 | Young |
| 0.073237 | 1.081863 | 0.32 | 0.202 | 1 | 14 | Vegfa | 0.118 | Young |
| 0.073583 | 0.732193 | 0.18 | 0.087 | 1 | 14 | B4galt6 | 0.093 | Young |
| 0.073706 | 0.726154 | 0.22 | 0.12 | 1 | 14 | Gatsl2 | 0.1 | Young |
| 0.073792 | 0.180167 | 0.2 | 0.098 | 1 | 14 | Zfp263 | 0.102 | Young |
| 0.074098 | 0.785146 | 0.24 | 0.131 | 1 | 14 | Hist1h2bc | 0.109 | Young |
| 0.074481 | 2.01564 | 0.48 | 0.393 | 1 | 14 | Rbbp7 | 0.087 | Young |
| 0.074488 | 0.567449 | 0.12 | 0.049 | 1 | 14 | Sec16b | 0.071 | Young |
| 0.074539 | 1.658429 | 0.1 | 0.038 | 1 | 14 | Tinagl1 | 0.062 | Young |
| 0.074852 | 1.692529 | 0.2 | 0.104 | 1 | 14 | Bet1l | 0.096 | Young |
| 0.074983 | 1.158691 | 0.24 | 0.137 | 1 | 14 | Ppme1 | 0.103 | Young |
| 0.075025 | 1.090448 | 0.16 | 0.077 | 1 | 14 | Pef1 | 0.083 | Young |
| 0.075137 | 0.389394 | 0.78 | 0.787 | 1 | 14 | Hnrnpk | -0.007 | Young |
| 0.07516 | 0.967665 | 0.34 | 0.219 | 1 | 14 | Cd99l2 | 0.121 | Young |
| 0.075397 | 1.02191 | 0.12 | 0.049 | 1 | 14 | Rft1 | 0.071 | Young |
| 0.075398 | 0.443699 | 0.68 | 0.639 | 1 | 14 | Hspa1b | 0.041 | Young |
| 0.075479 | 1.334311 | 0.34 | 0.24 | 1 | 14 | Arl4d | 0.1 | Young |
| 0.075963 | 0.877633 | 0.26 | 0.148 | 1 | 14 | Twf2 | 0.112 | Young |
| 0.076143 | 1.023627 | 0.18 | 0.093 | 1 | 14 | Tk2 | 0.087 | Young |
| 0.076171 | 0.490603 | 0.28 | 0.164 | 1 | 14 | Psmd12 | 0.116 | Young |

| | | | | | | | | |
|---|---|---|---|---|---|---|---|---|
| 0.076274 | 1.377269 | 0.34 | 0.235 | 1 | 14 | Palld | 0.105 | Young |
| 0.076565 | 0.984425 | 0.16 | 0.077 | 1 | 14 | Ciapin1 | 0.083 | Young |
| 0.076573 | 1.689953 | 0.1 | 0.038 | 1 | 14 | Tusc5 | 0.062 | Young |
| 0.076573 | 1.686546 | 0.1 | 0.038 | 1 | 14 | Them4 | 0.062 | Young |
| 0.077161 | 0.796449 | 0.26 | 0.148 | 1 | 14 | Phf12 | 0.112 | Young |
| 0.077243 | 1.853921 | 0.12 | 0.049 | 1 | 14 | Ccdc117 | 0.071 | Young |
| 0.077243 | 0.665688 | 0.12 | 0.049 | 1 | 14 | Cd209a | 0.071 | Young |
| 0.077563 | 0.560033 | 0.22 | 0.12 | 1 | 14 | Phyh | 0.1 | Young |
| 0.077587 | 0.725546 | 0.2 | 0.104 | 1 | 14 | Elmod2 | 0.096 | Young |
| 0.077818 | 1.316315 | 0.2 | 0.109 | 1 | 14 | Rlf | 0.091 | Young |
| 0.077829 | 0.395789 | 0.24 | 0.131 | 1 | 14 | Slc7a8 | 0.109 | Young |
| 0.07818 | 1.661088 | 0.12 | 0.049 | 1 | 14 | Xrcc6 | 0.071 | Young |
| 0.078379 | 1.230076 | 0.62 | 0.492 | 1 | 14 | Eif3c | 0.128 | Young |
| 0.078506 | 1.288563 | 0.2 | 0.109 | 1 | 14 | Myo1b | 0.091 | Young |
| 0.078578 | 0.309558 | 0.94 | 0.984 | 1 | 14 | mt-Nd4 | -0.044 | Young |
| 0.078632 | 0.287472 | 0.22 | 0.115 | 1 | 14 | Isca1 | 0.105 | Young |
| 0.078822 | 0.382705 | 0.44 | 0.295 | 1 | 14 | Ino80d | 0.145 | Young |
| 0.079111 | 2.50404 | 0.06 | 0.016 | 1 | 14 | Magi1 | 0.044 | Young |
| 0.079126 | 0.899795 | 0.12 | 0.049 | 1 | 14 | Fcrls | 0.071 | Young |
| 0.079256 | 2.192146 | 0.08 | 0.027 | 1 | 14 | 0610010K1 | 0.053 | Young |
| 0.079552 | 0.696231 | 0.22 | 0.12 | 1 | 14 | Itpr1 | 0.1 | Young |
| 0.079604 | 0.581432 | 0.26 | 0.148 | 1 | 14 | Pdp1 | 0.112 | Young |
| 0.079692 | 0.63849 | 0.2 | 0.104 | 1 | 14 | Adam12 | 0.096 | Young |
| 0.079703 | 1.495524 | 0.22 | 0.126 | 1 | 14 | Mrps36 | 0.094 | Young |
| 0.080297 | 0.270595 | 0.48 | 0.306 | 1 | 14 | Aoc3 | 0.174 | Young |
| 0.080349 | 0.316249 | 0.8 | 0.749 | 1 | 14 | Chchd2 | 0.051 | Young |
| 0.080546 | 1.336179 | 0.18 | 0.093 | 1 | 14 | Mettl14 | 0.087 | Young |
| 0.080596 | 3.319686 | 0.06 | 0.016 | 1 | 14 | Hoxd3os1 | 0.044 | Young |
| 0.080597 | 0.721305 | 0.28 | 0.169 | 1 | 14 | Gak | 0.111 | Young |
| 0.080674 | 0.198572 | 0.28 | 0.153 | 1 | 14 | Gphn | 0.127 | Young |
| 0.080776 | 2.744716 | 0.1 | 0.038 | 1 | 14 | Pmf1 | 0.062 | Young |
| 0.080776 | 1.529594 | 0.1 | 0.038 | 1 | 14 | Znrf3 | 0.062 | Young |
| 0.080925 | 0.232902 | 0.14 | 0.268 | 1 | 14 | Cyld | -0.128 | Young |
| 0.080963 | 0.176272 | 0.18 | 0.087 | 1 | 14 | Rnf185 | 0.093 | Young |
| 0.081046 | 0.772899 | 0.12 | 0.049 | 1 | 14 | Fam136a | 0.071 | Young |
| 0.081271 | 0.818339 | 0.24 | 0.137 | 1 | 14 | Dnajb2 | 0.103 | Young |
| 0.081695 | 1.241137 | 0.14 | 0.066 | 1 | 14 | Ndufaf3 | 0.074 | Young |
| 0.081856 | 1.041069 | 0.1 | 0.038 | 1 | 14 | Hectd2 | 0.062 | Young |
| 0.082103 | 3.368927 | 0.06 | 0.016 | 1 | 14 | Dus1l | 0.044 | Young |
| 0.082103 | 2.138557 | 0.06 | 0.016 | 1 | 14 | Fam19a5 | 0.044 | Young |
| 0.082296 | 0.83909 | 0.42 | 0.306 | 1 | 14 | Ostf1 | 0.114 | Young |
| 0.08279 | 0.638482 | 0.5 | 0.355 | 1 | 14 | Lims1 | 0.145 | Young |
| 0.082947 | 1.978598 | 0.1 | 0.038 | 1 | 14 | Nol6 | 0.062 | Young |
| 0.082947 | 1.329097 | 0.1 | 0.038 | 1 | 14 | Slc25a14 | 0.062 | Young |
| 0.082947 | 1.192838 | 0.1 | 0.038 | 1 | 14 | Bpgm | 0.062 | Young |

| | | | | | | | | |
|---|---|---|---|---|---|---|---|---|
| 0.082947 | 0.461525 | 0.1 | 0.038 | 1 | 14 | Ypel2 | 0.062 | Young |
| 0.082959 | 0.658529 | 0.22 | 0.12 | 1 | 14 | Yes1 | 0.1 | Young |
| 0.082963 | 2.034312 | 0.08 | 0.027 | 1 | 14 | Pde3b | 0.053 | Young |
| 0.082963 | 1.320298 | 0.08 | 0.027 | 1 | 14 | Rcsd1 | 0.053 | Young |
| 0.082981 | 0.579818 | 0.16 | 0.077 | 1 | 14 | Pappa | 0.083 | Young |
| 0.083003 | 0.520952 | 0.12 | 0.049 | 1 | 14 | Usp30 | 0.071 | Young |
| 0.083097 | 2.525634 | 0.22 | 0.131 | 1 | 14 | Etv1 | 0.089 | Young |
| 0.083157 | 0.853775 | 0.26 | 0.158 | 1 | 14 | Vps28 | 0.102 | Young |
| 0.083343 | 0.76714 | 0.56 | 0.448 | 1 | 14 | Atp5k | 0.112 | Young |
| 0.083459 | 1.694515 | 0.14 | 0.066 | 1 | 14 | Smim26 | 0.074 | Young |
| 0.083459 | 1.247378 | 0.14 | 0.066 | 1 | 14 | Plk2 | 0.074 | Young |
| 0.083462 | 1.293312 | 0.2 | 0.109 | 1 | 14 | Papd5 | 0.091 | Young |
| 0.083634 | 3.533034 | 0.06 | 0.016 | 1 | 14 | Lgi3 | 0.044 | Young |
| 0.083634 | 2.48789 | 0.06 | 0.016 | 1 | 14 | Lyplal1 | 0.044 | Young |
| 0.083634 | 2.422207 | 0.06 | 0.016 | 1 | 14 | Endou | 0.044 | Young |
| 0.083634 | 2.308387 | 0.06 | 0.016 | 1 | 14 | Rnf152 | 0.044 | Young |
| 0.083634 | 1.86117 | 0.06 | 0.016 | 1 | 14 | Nup37 | 0.044 | Young |
| 0.083634 | 1.794748 | 0.06 | 0.016 | 1 | 14 | Tbc1d8 | 0.044 | Young |
| 0.083634 | 1.690668 | 0.06 | 0.016 | 1 | 14 | Coq4 | 0.044 | Young |
| 0.083634 | 1.529911 | 0.06 | 0.016 | 1 | 14 | Apold1 | 0.044 | Young |
| 0.083736 | 1.272556 | 0.3 | 0.191 | 1 | 14 | Tmed4 | 0.109 | Young |
| 0.083738 | 0.580303 | 0.48 | 0.372 | 1 | 14 | Slc3a2 | 0.108 | Young |
| 0.083804 | 0.851438 | 0.42 | 0.306 | 1 | 14 | Cdk2ap2 | 0.114 | Young |
| 0.083813 | 0.739685 | 0.16 | 0.077 | 1 | 14 | Cdk5 | 0.083 | Young |
| 0.083813 | 0.735639 | 0.16 | 0.077 | 1 | 14 | Acsl1 | 0.083 | Young |
| 0.084041 | 0.142671 | 0.2 | 0.104 | 1 | 14 | Gpcpd1 | 0.096 | Young |
| 0.08405 | 1.390613 | 0.1 | 0.038 | 1 | 14 | Fgfrl1 | 0.062 | Young |
| 0.08405 | 1.368304 | 0.1 | 0.038 | 1 | 14 | Plcxd3 | 0.062 | Young |
| 0.084078 | 0.432027 | 0.18 | 0.087 | 1 | 14 | Mzt1 | 0.093 | Young |
| 0.084229 | 2.178949 | 0.08 | 0.027 | 1 | 14 | Smyd5 | 0.053 | Young |
| 0.084229 | 1.917035 | 0.08 | 0.027 | 1 | 14 | Plekhg5 | 0.053 | Young |
| 0.084229 | 1.792689 | 0.08 | 0.027 | 1 | 14 | Ikzf4 | 0.053 | Young |
| 0.084452 | 1.256057 | 0.22 | 0.126 | 1 | 14 | Hprt | 0.094 | Young |
| 0.085187 | 2.45473 | 0.06 | 0.016 | 1 | 14 | Cspg4 | 0.044 | Young |
| 0.085187 | 1.907736 | 0.06 | 0.016 | 1 | 14 | A130010J1 | 0.044 | Young |
| 0.085187 | 1.905585 | 0.06 | 0.016 | 1 | 14 | Tbx2 | 0.044 | Young |
| 0.085187 | 1.79272 | 0.06 | 0.016 | 1 | 14 | AC127341. | 0.044 | Young |
| 0.085187 | 1.400704 | 0.06 | 0.016 | 1 | 14 | Rhpn2 | 0.044 | Young |
| 0.085187 | 1.319051 | 0.06 | 0.016 | 1 | 14 | Cep57l1 | 0.044 | Young |
| 0.085254 | 1.033185 | 0.14 | 0.066 | 1 | 14 | Fignl2 | 0.074 | Young |
| 0.085271 | 0.045261 | 0.04 | 0.126 | 1 | 14 | Wdr45 | -0.086 | Young |
| 0.085511 | 2.115953 | 0.08 | 0.027 | 1 | 14 | Spaar | 0.053 | Young |
| 0.085511 | 1.50148 | 0.08 | 0.027 | 1 | 14 | Dok3 | 0.053 | Young |
| 0.085511 | 1.490037 | 0.08 | 0.027 | 1 | 14 | Gdf11 | 0.053 | Young |
| 0.085511 | 1.376398 | 0.08 | 0.027 | 1 | 14 | Mfsd3 | 0.053 | Young |

| | | | | | | | | |
|---|---|---|---|---|---|---|---|---|
| 0.085511 | 1.37627 | 0.08 | 0.027 | 1 | 14 | Rfxank | 0.053 | Young |
| 0.086017 | 0.758418 | 0.3 | 0.18 | 1 | 14 | St6galnac6 | 0.12 | Young |
| 0.086405 | 0.935815 | 0.2 | 0.109 | 1 | 14 | Rbmxl1 | 0.091 | Young |
| 0.086556 | 1.646957 | 0.22 | 0.126 | 1 | 14 | Nelfb | 0.094 | Young |
| 0.086763 | 2.426418 | 0.06 | 0.016 | 1 | 14 | Gm14327 | 0.044 | Young |
| 0.086763 | 1.619358 | 0.06 | 0.016 | 1 | 14 | Stk11ip | 0.044 | Young |
| 0.086763 | 1.341228 | 0.06 | 0.016 | 1 | 14 | Acsf3 | 0.044 | Young |
| 0.086763 | 1.340298 | 0.06 | 0.016 | 1 | 14 | Mphosph9 | 0.044 | Young |
| 0.086763 | 1.252972 | 0.06 | 0.016 | 1 | 14 | Slc22a18 | 0.044 | Young |
| 0.086809 | 1.822625 | 0.08 | 0.027 | 1 | 14 | Commd9 | 0.053 | Young |
| 0.086809 | 1.8001 | 0.08 | 0.027 | 1 | 14 | Ngfr | 0.053 | Young |
| 0.086809 | 1.608056 | 0.08 | 0.027 | 1 | 14 | Znhit2 | 0.053 | Young |
| 0.08708 | 1.310433 | 0.14 | 0.066 | 1 | 14 | Esco1 | 0.074 | Young |
| 0.087395 | 0.60584 | 0.44 | 0.306 | 1 | 14 | Supt16 | 0.134 | Young |
| 0.08743 | 2.250616 | 0.1 | 0.038 | 1 | 14 | Kcnb1 | 0.062 | Young |
| 0.08743 | 0.987244 | 0.1 | 0.038 | 1 | 14 | Sphk2 | 0.062 | Young |
| 0.08743 | 0.90103 | 0.1 | 0.038 | 1 | 14 | Pcdh12 | 0.062 | Young |
| 0.08743 | 0.659422 | 0.1 | 0.038 | 1 | 14 | Zfp943 | 0.062 | Young |
| 0.087534 | 1.237836 | 0.18 | 0.093 | 1 | 14 | Osgin2 | 0.087 | Young |
| 0.087592 | 1.138783 | 0.28 | 0.175 | 1 | 14 | Arhgef17 | 0.105 | Young |
| 0.088123 | 1.559448 | 0.08 | 0.027 | 1 | 14 | Ercc4 | 0.053 | Young |
| 0.088123 | 1.510292 | 0.08 | 0.027 | 1 | 14 | Rtp4 | 0.053 | Young |
| 0.088123 | 1.409857 | 0.08 | 0.027 | 1 | 14 | Samsn1 | 0.053 | Young |
| 0.088339 | 0.47019 | 0.46 | 0.311 | 1 | 14 | Ube2k | 0.149 | Young |
| 0.088363 | 2.991771 | 0.06 | 0.016 | 1 | 14 | Gsto1 | 0.044 | Young |
| 0.088363 | 1.822908 | 0.06 | 0.016 | 1 | 14 | Art3 | 0.044 | Young |
| 0.088363 | 1.784545 | 0.06 | 0.016 | 1 | 14 | 1700052K1 | 0.044 | Young |
| 0.088363 | 1.567714 | 0.06 | 0.016 | 1 | 14 | Hoga1 | 0.044 | Young |
| 0.088363 | 1.301121 | 0.06 | 0.016 | 1 | 14 | Tle2 | 0.044 | Young |
| 0.088363 | 1.154869 | 0.06 | 0.016 | 1 | 14 | Mcat | 0.044 | Young |
| 0.088363 | 0.637543 | 0.06 | 0.016 | 1 | 14 | Pkp4 | 0.044 | Young |
| 0.088363 | 0.61545 | 0.06 | 0.016 | 1 | 14 | Arap2 | 0.044 | Young |
| 0.088462 | 0.876439 | 0.42 | 0.301 | 1 | 14 | Cyr61 | 0.119 | Young |
| 0.088964 | 0.66555 | 0.54 | 0.41 | 1 | 14 | Wipi1 | 0.13 | Young |
| 0.089356 | 0.974001 | 0.2 | 0.104 | 1 | 14 | Elovl5 | 0.096 | Young |
| 0.089356 | 0.382558 | 0.2 | 0.104 | 1 | 14 | Iqce | 0.096 | Young |
| 0.089452 | 2.032093 | 0.08 | 0.027 | 1 | 14 | Faap24 | 0.053 | Young |
| 0.089452 | 1.434287 | 0.08 | 0.027 | 1 | 14 | Pag1 | 0.053 | Young |
| 0.089452 | 0.94992 | 0.08 | 0.027 | 1 | 14 | Gpr88 | 0.053 | Young |
| 0.089878 | 0.961682 | 0.14 | 0.066 | 1 | 14 | Kif13a | 0.074 | Young |
| 0.089987 | 1.081849 | 0.06 | 0.016 | 1 | 14 | Itga6 | 0.044 | Young |
| 0.089987 | 0.998472 | 0.06 | 0.016 | 1 | 14 | Zfp189 | 0.044 | Young |
| 0.089987 | 0.959393 | 0.06 | 0.016 | 1 | 14 | Alkbh2 | 0.044 | Young |
| 0.089987 | 0.897265 | 0.06 | 0.016 | 1 | 14 | Dynlt1a | 0.044 | Young |
| 0.089987 | 0.876099 | 0.06 | 0.016 | 1 | 14 | Mpp3 | 0.044 | Young |

| | | | | | | | | |
|---|---|---|---|---|---|---|---|---|
| 0.089987 | 0.844521 | 0.06 | 0.016 | 1 | 14 | Gm17435 | 0.044 | Young |
| 0.089987 | 0.621522 | 0.06 | 0.016 | 1 | 14 | A230057D( | 0.044 | Young |
| 0.090155 | 1.285914 | 0.2 | 0.115 | 1 | 14 | Arl6ip6 | 0.085 | Young |
| 0.090422 | 1.692371 | 0.32 | 0.224 | 1 | 14 | Sh3bp5 | 0.096 | Young |
| 0.090798 | 2.468471 | 0.08 | 0.027 | 1 | 14 | Nradd | 0.053 | Young |
| 0.090798 | 1.510372 | 0.08 | 0.027 | 1 | 14 | Prickle3 | 0.053 | Young |
| 0.090798 | 0.793132 | 0.08 | 0.027 | 1 | 14 | Tsen54 | 0.053 | Young |
| 0.090826 | 0.706666 | 0.14 | 0.066 | 1 | 14 | Ccr5 | 0.074 | Young |
| 0.090904 | 0.59077 | 0.3 | 0.202 | 1 | 14 | Gpr107 | 0.098 | Young |
| 0.090919 | 0.624594 | 0.1 | 0.038 | 1 | 14 | Zfp868 | 0.062 | Young |
| 0.090978 | 1.185975 | 0.2 | 0.109 | 1 | 14 | Stx6 | 0.091 | Young |
| 0.091152 | 1.042657 | 0.44 | 0.322 | 1 | 14 | Tiparp | 0.118 | Young |
| 0.091608 | 0.894792 | 0.16 | 0.077 | 1 | 14 | Esrra | 0.083 | Young |
| 0.091623 | 0.933412 | 0.18 | 0.093 | 1 | 14 | Ptp4a3 | 0.087 | Young |
| 0.091634 | 0.871244 | 0.06 | 0.016 | 1 | 14 | Slc25a10 | 0.044 | Young |
| 0.09216 | 1.597996 | 0.08 | 0.027 | 1 | 14 | Adgrl2 | 0.053 | Young |
| 0.09216 | 1.260506 | 0.08 | 0.027 | 1 | 14 | 2410131K1 | 0.053 | Young |
| 0.09216 | 1.081715 | 0.08 | 0.027 | 1 | 14 | Triap1 | 0.053 | Young |
| 0.09216 | 1.07487 | 0.08 | 0.027 | 1 | 14 | Atg4a | 0.053 | Young |
| 0.092313 | 0.161032 | 0.14 | 0.251 | 1 | 14 | Rps27rt | -0.111 | Young |
| 0.092374 | 1.013292 | 0.22 | 0.12 | 1 | 14 | Cdc16 | 0.1 | Young |
| 0.092459 | 0.973691 | 0.18 | 0.093 | 1 | 14 | Lactb2 | 0.087 | Young |
| 0.092748 | 0.959513 | 0.14 | 0.066 | 1 | 14 | Bloc1s1 | 0.074 | Young |
| 0.093306 | 1.072505 | 0.06 | 0.016 | 1 | 14 | Tnfrsf22 | 0.044 | Young |
| 0.093306 | 1.068684 | 0.06 | 0.016 | 1 | 14 | Spp1 | 0.044 | Young |
| 0.093306 | 0.501144 | 0.06 | 0.016 | 1 | 14 | Hoxa4 | 0.044 | Young |
| 0.093306 | 0.145751 | 0.06 | 0.016 | 1 | 14 | Flad1 | 0.044 | Young |
| 0.093307 | 0.650399 | 0.1 | 0.038 | 1 | 14 | Rasgrp2 | 0.062 | Young |
| 0.093336 | 1.303144 | 0.2 | 0.109 | 1 | 14 | Gosr1 | 0.091 | Young |
| 0.094519 | 0.120873 | 0.1 | 0.038 | 1 | 14 | Odf2l | 0.062 | Young |
| 0.094651 | 1.724628 | 0.12 | 0.055 | 1 | 14 | Ccl24 | 0.065 | Young |
| 0.094933 | 1.201294 | 0.08 | 0.027 | 1 | 14 | Mmab | 0.053 | Young |
| 0.094933 | 1.10016 | 0.08 | 0.027 | 1 | 14 | Nxnl2 | 0.053 | Young |
| 0.094933 | 0.674087 | 0.08 | 0.027 | 1 | 14 | Fgd2 | 0.053 | Young |
| 0.094935 | 1.157238 | 0.2 | 0.109 | 1 | 14 | Aph1b | 0.091 | Young |
| 0.094935 | 0.843747 | 0.2 | 0.109 | 1 | 14 | Piga | 0.091 | Young |
| 0.094935 | 0.66878 | 0.2 | 0.109 | 1 | 14 | Ing5 | 0.091 | Young |
| 0.095004 | 1.16716 | 0.18 | 0.093 | 1 | 14 | Cacul1 | 0.087 | Young |
| 0.095107 | 1.326472 | 0.26 | 0.164 | 1 | 14 | Hsd17b12 | 0.096 | Young |
| 0.095572 | 0.717916 | 0.32 | 0.208 | 1 | 14 | Chd6 | 0.112 | Young |
| 0.095778 | 0.527349 | 0.46 | 0.317 | 1 | 14 | Tsc22d2 | 0.143 | Young |
| 0.09596 | 1.244863 | 0.28 | 0.169 | 1 | 14 | Pex19 | 0.111 | Young |
| 0.096167 | 1.239894 | 0.22 | 0.126 | 1 | 14 | Cdk2ap1 | 0.094 | Young |
| 0.096258 | 0.48776 | 0.52 | 0.41 | 1 | 14 | Cpd | 0.11 | Young |
| 0.096345 | 0.725171 | 0.08 | 0.027 | 1 | 14 | Smox | 0.053 | Young |

| | | | | | | | | |
|---|---|---|---|---|---|---|---|---|
| 0.096345 | 0.396141 | 0.08 | 0.027 | 1 | 14 | Plcb1 | 0.053 | Young |
| 0.096731 | 1.347538 | 0.18 | 0.093 | 1 | 14 | Tatdn2 | 0.087 | Young |
| 0.097079 | 2.479642 | 0.14 | 0.071 | 1 | 14 | Mme | 0.069 | Young |
| 0.097292 | 0.646978 | 0.16 | 0.082 | 1 | 14 | Hipk2 | 0.078 | Young |
| 0.097293 | 0.630277 | 0.36 | 0.235 | 1 | 14 | Nudc | 0.125 | Young |
| 0.09774 | 0.944692 | 0.32 | 0.202 | 1 | 14 | Pard3 | 0.118 | Young |
| 0.097773 | 0.785066 | 0.08 | 0.027 | 1 | 14 | C2cd2l | 0.053 | Young |
| 0.097773 | 0.653889 | 0.08 | 0.027 | 1 | 14 | Slc6a13 | 0.053 | Young |
| 0.097893 | 1.385191 | 0.12 | 0.055 | 1 | 14 | Zmym3 | 0.065 | Young |
| 0.098117 | 0.182568 | 0.08 | 0.18 | 1 | 14 | Nifk | -0.1 | Young |
| 0.098251 | 0.529767 | 0.2 | 0.104 | 1 | 14 | N6amt1 | 0.096 | Young |
| 0.098718 | 0.523153 | 0.44 | 0.317 | 1 | 14 | mt-Atp8 | 0.123 | Young |
| 0.099058 | 2.634236 | 0.14 | 0.071 | 1 | 14 | Cox19 | 0.069 | Young |
| 0.099101 | 0.936895 | 0.3 | 0.197 | 1 | 14 | Evi5 | 0.103 | Young |
| 0.099218 | 0.174116 | 0.08 | 0.027 | 1 | 14 | Zfp182 | 0.053 | Young |
| 0.099441 | 0.754704 | 0.44 | 0.322 | 1 | 14 | Zdhhc20 | 0.118 | Young |
| 0.099456 | 1.546512 | 0.32 | 0.208 | 1 | 14 | Uba5 | 0.112 | Young |
| 0.099731 | 0.746919 | 0.14 | 0.066 | 1 | 14 | Tbcd | 0.074 | Young |
| 0.099731 | 0.152984 | 0.14 | 0.066 | 1 | 14 | Gpaa1 | 0.074 | Young |
| 0.099864 | 1.087575 | 0.2 | 0.109 | 1 | 14 | Dab2ip | 0.091 | Young |
| 0.100102 | 1.544083 | 0.12 | 0.055 | 1 | 14 | Prr14l | 0.065 | Young |
| 0.100102 | 1.28001 | 0.12 | 0.055 | 1 | 14 | Med22 | 0.065 | Young |
| 0.100762 | 0.744292 | 0.14 | 0.066 | 1 | 14 | Pcgf1 | 0.074 | Young |
| 0.100762 | 0.716902 | 0.14 | 0.066 | 1 | 14 | Gcc1 | 0.074 | Young |
| 0.100981 | 1.514059 | 0.28 | 0.175 | 1 | 14 | Mrpl3 | 0.105 | Young |
| 0.101001 | 0.662714 | 0.4 | 0.257 | 1 | 14 | Nus1 | 0.143 | Young |
| 0.10104 | 2.135401 | 0.16 | 0.082 | 1 | 14 | Fam20a | 0.078 | Young |
| 0.10104 | 0.863712 | 0.16 | 0.082 | 1 | 14 | Fancc | 0.078 | Young |
| 0.101111 | 1.503418 | 0.4 | 0.301 | 1 | 14 | Matn2 | 0.099 | Young |
| 0.101169 | 0.002622 | 0.08 | 0.175 | 1 | 14 | Cdc42se2 | -0.095 | Young |
| 0.101613 | 1.25165 | 0.28 | 0.197 | 1 | 14 | Chuk | 0.083 | Young |
| 0.101752 | 1.237075 | 0.28 | 0.18 | 1 | 14 | Gna11 | 0.1 | Young |
| 0.102601 | 0.476201 | 0.42 | 0.284 | 1 | 14 | Cbl | 0.136 | Young |
| 0.102849 | 0.721041 | 0.14 | 0.066 | 1 | 14 | Ssna1 | 0.074 | Young |
| 0.102957 | 1.351267 | 0.16 | 0.082 | 1 | 14 | Slc41a3 | 0.078 | Young |
| 0.103011 | 1.020082 | 0.4 | 0.273 | 1 | 14 | Chd8 | 0.127 | Young |
| 0.103113 | 2.420307 | 0.14 | 0.071 | 1 | 14 | Lrig3 | 0.069 | Young |
| 0.103492 | 1.451832 | 0.12 | 0.055 | 1 | 14 | Atp1b2 | 0.065 | Young |
| 0.103492 | 1.380977 | 0.12 | 0.055 | 1 | 14 | Mthfd1 | 0.065 | Young |
| 0.104101 | 0.698957 | 0.44 | 0.322 | 1 | 14 | Postn | 0.118 | Young |
| 0.104147 | 2.218692 | 0.14 | 0.071 | 1 | 14 | Krt10 | 0.069 | Young |
| 0.104622 | 1.270463 | 0.18 | 0.098 | 1 | 14 | Akap1 | 0.082 | Young |
| 0.104642 | 1.486249 | 0.12 | 0.055 | 1 | 14 | Sugp2 | 0.065 | Young |
| 0.104642 | 1.037336 | 0.12 | 0.055 | 1 | 14 | Hibch | 0.065 | Young |
| 0.104683 | 0.910532 | 0.34 | 0.219 | 1 | 14 | Mettl9 | 0.121 | Young |

| | | | | | | | | |
|---|---|---|---|---|---|---|---|---|
| 0.104721 | 0.788906 | 0.2 | 0.115 | 1 | 14 | Pabpc4 | 0.085 | Young |
| 0.104874 | 0.4773 | 0.4 | 0.273 | 1 | 14 | Snw1 | 0.127 | Young |
| 0.104903 | 0.721309 | 0.16 | 0.082 | 1 | 14 | Usp11 | 0.078 | Young |
| 0.105163 | 0.924992 | 0.32 | 0.213 | 1 | 14 | Nsfl1c | 0.107 | Young |
| 0.105411 | 0.170365 | 0.46 | 0.596 | 1 | 14 | Rbp1 | -0.136 | Young |
| 0.10587 | 1.617129 | 0.2 | 0.109 | 1 | 14 | Fxr2 | 0.091 | Young |
| 0.106045 | 0.580869 | 0.14 | 0.066 | 1 | 14 | Pex14 | 0.074 | Young |
| 0.106076 | 0.801633 | 0.26 | 0.158 | 1 | 14 | Golga2 | 0.102 | Young |
| 0.106113 | 0.429328 | 0.26 | 0.153 | 1 | 14 | Tbc1d10b | 0.107 | Young |
| 0.106686 | 0.090082 | 0.18 | 0.093 | 1 | 14 | Btbd10 | 0.087 | Young |
| 0.106694 | 0.79996 | 0.3 | 0.197 | 1 | 14 | Ppp2r3a | 0.103 | Young |
| 0.106972 | 1.827437 | 0.12 | 0.055 | 1 | 14 | Mypop | 0.065 | Young |
| 0.107127 | 0.748572 | 0.14 | 0.066 | 1 | 14 | Plekho1 | 0.074 | Young |
| 0.107248 | 0.260172 | 0.16 | 0.273 | 1 | 14 | Ddx42 | -0.113 | Young |
| 0.107383 | 0.905909 | 0.18 | 0.098 | 1 | 14 | BC003965 | 0.082 | Young |
| 0.107876 | 0.800943 | 0.16 | 0.082 | 1 | 14 | Nudt5 | 0.078 | Young |
| 0.107929 | 0.525242 | 0.34 | 0.224 | 1 | 14 | Zfp260 | 0.116 | Young |
| 0.107988 | 1.743127 | 0.22 | 0.131 | 1 | 14 | Plac9b | 0.089 | Young |
| 0.107988 | 1.26925 | 0.22 | 0.131 | 1 | 14 | Spon1 | 0.089 | Young |
| 0.108262 | 0.827896 | 0.28 | 0.175 | 1 | 14 | Plaa | 0.105 | Young |
| 0.108945 | 0.641912 | 0.6 | 0.514 | 1 | 14 | Mprip | 0.086 | Young |
| 0.109273 | 0.499742 | 0.18 | 0.284 | 1 | 14 | Taf1 | -0.104 | Young |
| 0.109319 | 0.653408 | 0.14 | 0.066 | 1 | 14 | Pfkm | 0.074 | Young |
| 0.109429 | 1.35348 | 0.2 | 0.109 | 1 | 14 | Plrg1 | 0.091 | Young |
| 0.109429 | 1.214996 | 0.2 | 0.109 | 1 | 14 | Zbed3 | 0.091 | Young |
| 0.109767 | 0.644606 | 0.28 | 0.175 | 1 | 14 | Agpat5 | 0.105 | Young |
| 0.110169 | 0.784591 | 0.44 | 0.322 | 1 | 14 | Ntrk2 | 0.118 | Young |
| 0.110201 | 2.024816 | 0.18 | 0.098 | 1 | 14 | Slc25a23 | 0.082 | Young |
| 0.110295 | 0.933892 | 0.3 | 0.197 | 1 | 14 | Ubr3 | 0.103 | Young |
| 0.110543 | 0.961414 | 0.12 | 0.055 | 1 | 14 | B4galt4 | 0.065 | Young |
| 0.111729 | 1.013747 | 0.22 | 0.126 | 1 | 14 | Ppp6r2 | 0.094 | Young |
| 0.111944 | 1.029603 | 0.16 | 0.082 | 1 | 14 | Stx17 | 0.078 | Young |
| 0.111944 | 0.450101 | 0.16 | 0.082 | 1 | 14 | Wdr6 | 0.078 | Young |
| 0.112002 | 1.634613 | 0.24 | 0.148 | 1 | 14 | Gtf2h1 | 0.092 | Young |
| 0.112288 | 0.597403 | 0.24 | 0.142 | 1 | 14 | Adrm1 | 0.098 | Young |
| 0.112826 | 0.779276 | 0.28 | 0.175 | 1 | 14 | Atp6v1d | 0.105 | Young |
| 0.112977 | 1.751604 | 0.12 | 0.055 | 1 | 14 | Lyrm4 | 0.065 | Young |
| 0.113324 | 0.303204 | 0.42 | 0.284 | 1 | 14 | Zfp652 | 0.136 | Young |
| 0.113385 | 1.801339 | 0.1 | 0.044 | 1 | 14 | Zfp820 | 0.056 | Young |
| 0.113385 | 1.728436 | 0.1 | 0.044 | 1 | 14 | Kcnq4 | 0.056 | Young |
| 0.114023 | 1.270833 | 0.16 | 0.082 | 1 | 14 | Eed | 0.078 | Young |
| 0.114287 | 0.584349 | 0.62 | 0.508 | 1 | 14 | Nfkbiz | 0.112 | Young |
| 0.114691 | 0.191406 | 0.04 | 0.12 | 1 | 14 | Mea1 | -0.08 | Young |
| 0.114741 | 1.526206 | 0.3 | 0.197 | 1 | 14 | Abl1 | 0.103 | Young |
| 0.115027 | 0.500881 | 0.18 | 0.098 | 1 | 14 | Slc5a3 | 0.082 | Young |

| | | | | | | | | |
|---|---|---|---|---|---|---|---|---|
| 0.115224 | 0.092073 | 0.02 | 0.087 | 1 | 14 | Gm32618 | -0.067 | Young |
| 0.115452 | 0.812998 | 0.12 | 0.055 | 1 | 14 | Zfp595 | 0.065 | Young |
| 0.115564 | 0.453186 | 0.22 | 0.12 | 1 | 14 | Dot1l | 0.1 | Young |
| 0.115726 | 0.811419 | 0.04 | 0.12 | 1 | 14 | AC165953. | -0.08 | Young |
| 0.116123 | 2.359778 | 0.1 | 0.044 | 1 | 14 | Gtf2f2 | 0.056 | Young |
| 0.116413 | 0.186328 | 0.24 | 0.35 | 1 | 14 | Smarcd2 | -0.11 | Young |
| 0.116705 | 1.028651 | 0.12 | 0.055 | 1 | 14 | Irf8 | 0.065 | Young |
| 0.116705 | 1.012907 | 0.12 | 0.055 | 1 | 14 | Pigg | 0.065 | Young |
| 0.116705 | 0.777219 | 0.12 | 0.055 | 1 | 14 | Exosc4 | 0.065 | Young |
| 0.117108 | 0.687846 | 0.34 | 0.224 | 1 | 14 | Ccny | 0.116 | Young |
| 0.117208 | 1.040754 | 0.14 | 0.071 | 1 | 14 | Ccdc107 | 0.069 | Young |
| 0.117761 | 0.4995 | 0.14 | 0.257 | 1 | 14 | Ston2 | -0.117 | Young |
| 0.117969 | 0.439797 | 0.12 | 0.055 | 1 | 14 | Ispd | 0.065 | Young |
| 0.118089 | 0.359242 | 0.9 | 0.929 | 1 | 14 | Myl6 | -0.029 | Young |
| 0.118271 | 0.735604 | 0.16 | 0.082 | 1 | 14 | Nol8 | 0.078 | Young |
| 0.118877 | 1.60096 | 0.16 | 0.087 | 1 | 14 | Uggt2 | 0.073 | Young |
| 0.119006 | 0.621438 | 0.4 | 0.29 | 1 | 14 | Grb2 | 0.11 | Young |
| 0.119244 | 0.769594 | 0.12 | 0.055 | 1 | 14 | Ptpre | 0.065 | Young |
| 0.119244 | 0.498817 | 0.12 | 0.055 | 1 | 14 | Pik3cd | 0.065 | Young |
| 0.119382 | 0.503268 | 0.22 | 0.333 | 1 | 14 | Prpf40a | -0.113 | Young |
| 0.119977 | 0.744224 | 0.24 | 0.142 | 1 | 14 | Tomm40 | 0.098 | Young |
| 0.121084 | 0.049469 | 0.22 | 0.12 | 1 | 14 | Pih1d1 | 0.1 | Young |
| 0.121755 | 1.723435 | 0.1 | 0.044 | 1 | 14 | Triqk | 0.056 | Young |
| 0.122874 | 0.140042 | 0.22 | 0.328 | 1 | 14 | Tbcb | -0.108 | Young |
| 0.123093 | 0.619473 | 0.18 | 0.098 | 1 | 14 | Ralgapb | 0.082 | Young |
| 0.123097 | 1.232083 | 0.26 | 0.158 | 1 | 14 | Tmem41b | 0.102 | Young |
| 0.123182 | 1.028362 | 0.16 | 0.087 | 1 | 14 | Ddx23 | 0.073 | Young |
| 0.123232 | 1.442957 | 0.22 | 0.137 | 1 | 14 | Ndufa9 | 0.083 | Young |
| 0.123558 | 0.808255 | 0.32 | 0.219 | 1 | 14 | Ubr1 | 0.101 | Young |
| 0.123741 | 1.205293 | 0.42 | 0.322 | 1 | 14 | Cnih1 | 0.098 | Young |
| 0.123799 | 0.558437 | 0.52 | 0.415 | 1 | 14 | Zeb1 | 0.105 | Young |
| 0.123957 | 0.64745 | 0.26 | 0.158 | 1 | 14 | Tmem63b | 0.102 | Young |
| 0.124132 | 0.853741 | 0.18 | 0.098 | 1 | 14 | Heatr3 | 0.082 | Young |
| 0.124159 | 0.952718 | 0.42 | 0.328 | 1 | 14 | Plod2 | 0.092 | Young |
| 0.125125 | 0.560219 | 0.48 | 0.344 | 1 | 14 | Pkm | 0.136 | Young |
| 0.125177 | 0.820017 | 0.18 | 0.098 | 1 | 14 | Ssh1 | 0.082 | Young |
| 0.125377 | 0.625117 | 0.26 | 0.164 | 1 | 14 | Tbc1d10a | 0.096 | Young |
| 0.125401 | 1.530447 | 0.14 | 0.071 | 1 | 14 | Fam53a | 0.069 | Young |
| 0.125536 | 0.315918 | 0.3 | 0.186 | 1 | 14 | Gemin7 | 0.114 | Young |
| 0.125741 | 0.783626 | 0.28 | 0.175 | 1 | 14 | 4933434E2 | 0.105 | Young |
| 0.126055 | 1.809766 | 0.14 | 0.077 | 1 | 14 | Cdkn2aip | 0.063 | Young |
| 0.126117 | 1.46742 | 0.1 | 0.044 | 1 | 14 | Prdm15 | 0.056 | Young |
| 0.126117 | 1.221564 | 0.1 | 0.044 | 1 | 14 | Pkp2 | 0.056 | Young |
| 0.126117 | 1.125341 | 0.1 | 0.044 | 1 | 14 | C330018D2 | 0.056 | Young |
| 0.12649 | 1.680209 | 0.16 | 0.087 | 1 | 14 | Polr3d | 0.073 | Young |

| | | | | | | | | |
|---|---|---|---|---|---|---|---|---|
| 0.126494 | 1.318509 | 0.42 | 0.322 | 1 | 14 | Chd3 | 0.098 | Young |
| 0.126682 | 1.121559 | 0.22 | 0.131 | 1 | 14 | Kpna6 | 0.089 | Young |
| 0.127121 | 0.087119 | 0.12 | 0.055 | 1 | 14 | Mrpl22 | 0.065 | Young |
| 0.12714 | 0.498931 | 0.16 | 0.082 | 1 | 14 | Kdm5d | 0.078 | Young |
| 0.127198 | 1.146908 | 0.2 | 0.12 | 1 | 14 | Ifit2 | 0.08 | Young |
| 0.127598 | 1.448506 | 0.1 | 0.044 | 1 | 14 | Zmynd19 | 0.056 | Young |
| 0.128997 | 0.799453 | 0.42 | 0.35 | 1 | 14 | Efr3a | 0.07 | Young |
| 0.129092 | 1.074937 | 0.1 | 0.044 | 1 | 14 | Lnx2 | 0.056 | Young |
| 0.129092 | 0.873718 | 0.1 | 0.044 | 1 | 14 | Eef1akmt1 | 0.056 | Young |
| 0.129233 | 0.441867 | 0.34 | 0.224 | 1 | 14 | Napa | 0.116 | Young |
| 0.129319 | 1.01575 | 0.26 | 0.169 | 1 | 14 | Rap1gds1 | 0.091 | Young |
| 0.129428 | 0.535932 | 0.18 | 0.098 | 1 | 14 | Mdp1 | 0.082 | Young |
| 0.129867 | 1.155057 | 0.16 | 0.087 | 1 | 14 | Prep | 0.073 | Young |
| 0.129953 | 0.131671 | 0.16 | 0.268 | 1 | 14 | Scd1 | -0.108 | Young |
| 0.13028 | 1.51673 | 0.14 | 0.071 | 1 | 14 | Fkrp | 0.069 | Young |
| 0.13028 | 0.893527 | 0.14 | 0.071 | 1 | 14 | Ncf2 | 0.069 | Young |
| 0.1306 | 1.584611 | 0.1 | 0.044 | 1 | 14 | Traf4 | 0.056 | Young |
| 0.1306 | 1.030153 | 0.1 | 0.044 | 1 | 14 | Pbx3 | 0.056 | Young |
| 0.1306 | 0.908546 | 0.1 | 0.044 | 1 | 14 | Edrf1 | 0.056 | Young |
| 0.130729 | 0.779701 | 0.22 | 0.137 | 1 | 14 | Nqo2 | 0.083 | Young |
| 0.130731 | 1.575951 | 0.18 | 0.104 | 1 | 14 | Wdr3 | 0.076 | Young |
| 0.131015 | 0.36252 | 0.26 | 0.158 | 1 | 14 | Frg1 | 0.102 | Young |
| 0.132157 | 0.855002 | 0.16 | 0.087 | 1 | 14 | Acaca | 0.073 | Young |
| 0.132158 | 0.42703 | 0.22 | 0.126 | 1 | 14 | Cirbp | 0.094 | Young |
| 0.132773 | 0.969626 | 0.14 | 0.071 | 1 | 14 | Stk35 | 0.069 | Young |
| 0.132773 | 0.746108 | 0.14 | 0.071 | 1 | 14 | Sirt7 | 0.069 | Young |
| 0.133215 | 1.385155 | 0.32 | 0.23 | 1 | 14 | Fam20b | 0.09 | Young |
| 0.133432 | 0.980442 | 0.36 | 0.251 | 1 | 14 | Arhgap17 | 0.109 | Young |
| 0.133604 | 2.460101 | 0.08 | 0.033 | 1 | 14 | Gsta4 | 0.047 | Young |
| 0.133657 | 1.634659 | 0.1 | 0.044 | 1 | 14 | Pknox2 | 0.056 | Young |
| 0.133657 | 0.78163 | 0.1 | 0.044 | 1 | 14 | Gpr146 | 0.056 | Young |
| 0.133657 | 0.779634 | 0.1 | 0.044 | 1 | 14 | Rab4a | 0.056 | Young |
| 0.133871 | 1.407705 | 0.12 | 0.06 | 1 | 14 | Cdkal1 | 0.06 | Young |
| 0.133881 | 0.62075 | 0.42 | 0.301 | 1 | 14 | Uhrf2 | 0.119 | Young |
| 0.134034 | 1.014735 | 0.14 | 0.071 | 1 | 14 | Zfp777 | 0.069 | Young |
| 0.134034 | 0.805099 | 0.14 | 0.071 | 1 | 14 | Map2k5 | 0.069 | Young |
| 0.134034 | 0.537672 | 0.14 | 0.071 | 1 | 14 | St3gal5 | 0.069 | Young |
| 0.13412 | 0.618497 | 0.2 | 0.115 | 1 | 14 | Pim3 | 0.085 | Young |
| 0.134151 | 0.412375 | 0.42 | 0.311 | 1 | 14 | 7-Mar | 0.109 | Young |
| 0.134731 | 0.606688 | 0.28 | 0.186 | 1 | 14 | Mrpl42 | 0.094 | Young |
| 0.13504 | 0.590221 | 0.44 | 0.317 | 1 | 14 | Ube2l3 | 0.123 | Young |
| 0.135359 | 2.664211 | 0.08 | 0.033 | 1 | 14 | Scarb1 | 0.047 | Young |
| 0.13547 | 0.440997 | 0.6 | 0.508 | 1 | 14 | Drap1 | 0.092 | Young |
| 0.13565 | 0.92254 | 0.16 | 0.087 | 1 | 14 | Btaf1 | 0.073 | Young |
| 0.135937 | 0.220961 | 0.88 | 0.902 | 1 | 14 | Ubc | -0.022 | Young |

| | | | | | | | | |
|---|---|---|---|---|---|---|---|---|
| 0.136013 | 0.548248 | 0.18 | 0.098 | 1 | 14 | 5730409E0 | 0.082 | Young |
| 0.136154 | 0.878635 | 0.18 | 0.104 | 1 | 14 | Robo1 | 0.076 | Young |
| 0.136769 | 0.562994 | 0.1 | 0.044 | 1 | 14 | Zfp687 | 0.056 | Young |
| 0.136929 | 1.859603 | 0.16 | 0.093 | 1 | 14 | Cmc4 | 0.067 | Young |
| 0.136932 | 1.352824 | 0.24 | 0.148 | 1 | 14 | Dusp6 | 0.092 | Young |
| 0.136933 | 0.343348 | 0.24 | 0.148 | 1 | 14 | Pmpcb | 0.092 | Young |
| 0.137132 | 2.601444 | 0.08 | 0.033 | 1 | 14 | Adi1 | 0.047 | Young |
| 0.137132 | 2.342724 | 0.08 | 0.033 | 1 | 14 | Pced1a | 0.047 | Young |
| 0.137132 | 1.668271 | 0.08 | 0.033 | 1 | 14 | Rgs6 | 0.047 | Young |
| 0.137132 | 1.609771 | 0.08 | 0.033 | 1 | 14 | Zmym1 | 0.047 | Young |
| 0.137187 | 1.268108 | 0.24 | 0.153 | 1 | 14 | Ube3b | 0.087 | Young |
| 0.137227 | 0.43072 | 0.06 | 0.142 | 1 | 14 | Naa60 | -0.082 | Young |
| 0.137695 | 0.725048 | 0.28 | 0.18 | 1 | 14 | Tmem39a | 0.1 | Young |
| 0.138018 | 0.839615 | 0.16 | 0.087 | 1 | 14 | Cbll1 | 0.073 | Young |
| 0.13825 | 0.393958 | 0.3 | 0.197 | 1 | 14 | Trim2 | 0.103 | Young |
| 0.138757 | 0.772278 | 0.28 | 0.191 | 1 | 14 | Zfc3h1 | 0.089 | Young |
| 0.138922 | 1.521064 | 0.08 | 0.033 | 1 | 14 | St3gal3 | 0.047 | Young |
| 0.139169 | 1.668665 | 0.14 | 0.071 | 1 | 14 | Ccdc43 | 0.069 | Young |
| 0.139206 | 0.711422 | 0.34 | 0.459 | 1 | 14 | Anp32a | -0.119 | Young |
| 0.139214 | 1.186785 | 0.16 | 0.087 | 1 | 14 | Mst1r | 0.073 | Young |
| 0.139214 | 0.862532 | 0.16 | 0.087 | 1 | 14 | Pcnt | 0.073 | Young |
| 0.13946 | 1.147088 | 0.22 | 0.131 | 1 | 14 | Parp4 | 0.089 | Young |
| 0.139563 | 2.126417 | 0.14 | 0.077 | 1 | 14 | Zbtb10 | 0.063 | Young |
| 0.139693 | 0.752487 | 0.3 | 0.208 | 1 | 14 | Lamtor3 | 0.092 | Young |
| 0.139936 | 0.344392 | 0.1 | 0.044 | 1 | 14 | Spidr | 0.056 | Young |
| 0.139999 | 0.599131 | 0.3 | 0.197 | 1 | 14 | Clpx | 0.103 | Young |
| 0.140325 | 1.002861 | 0.3 | 0.213 | 1 | 14 | Iqsec1 | 0.087 | Young |
| 0.140352 | 1.03405 | 0.48 | 0.372 | 1 | 14 | Nckap1 | 0.108 | Young |
| 0.140418 | 0.993393 | 0.16 | 0.087 | 1 | 14 | Pld4 | 0.073 | Young |
| 0.140544 | 0.430638 | 0.18 | 0.098 | 1 | 14 | Secisbp2 | 0.082 | Young |
| 0.140731 | 1.335884 | 0.08 | 0.033 | 1 | 14 | Pde11a | 0.047 | Young |
| 0.140817 | 1.505113 | 0.12 | 0.06 | 1 | 14 | Ppil3 | 0.06 | Young |
| 0.142558 | 1.469917 | 0.08 | 0.033 | 1 | 14 | Zfp748 | 0.047 | Young |
| 0.142619 | 0.451893 | 0.26 | 0.164 | 1 | 14 | BC004004 | 0.096 | Young |
| 0.142968 | 0.048761 | 0.4 | 0.284 | 1 | 14 | Brd3 | 0.116 | Young |
| 0.143243 | 0.957619 | 0.28 | 0.191 | 1 | 14 | Rem1 | 0.089 | Young |
| 0.143434 | 0.87082 | 0.14 | 0.077 | 1 | 14 | Rab2b | 0.063 | Young |
| 0.143503 | 1.39256 | 0.46 | 0.393 | 1 | 14 | Ppp1r12a | 0.067 | Young |
| 0.144015 | 0.588379 | 0.34 | 0.224 | 1 | 14 | Vasp | 0.116 | Young |
| 0.144154 | 0.642533 | 0.28 | 0.191 | 1 | 14 | Rsf1 | 0.089 | Young |
| 0.144403 | 2.416922 | 0.08 | 0.033 | 1 | 14 | Bud13 | 0.047 | Young |
| 0.144403 | 1.367507 | 0.08 | 0.033 | 1 | 14 | Cmtr2 | 0.047 | Young |
| 0.144531 | 1.187087 | 0.48 | 0.421 | 1 | 14 | Nr2f2 | 0.059 | Young |
| 0.14471 | 1.820036 | 0.2 | 0.126 | 1 | 14 | Bcap29 | 0.074 | Young |
| 0.144956 | 0.793724 | 0.36 | 0.246 | 1 | 14 | Ndufb6 | 0.114 | Young |

| | | | | | | | | |
|---|---|---|---|---|---|---|---|---|
| 0.145115 | 1.209765 | 0.12 | 0.06 | 1 | 14 | Cdk5rap2 | 0.06 | Young |
| 0.145314 | 0.932958 | 0.16 | 0.087 | 1 | 14 | Ankrd50 | 0.073 | Young |
| 0.145314 | 0.884201 | 0.16 | 0.087 | 1 | 14 | Cisd1 | 0.073 | Young |
| 0.145314 | 0.510631 | 0.16 | 0.087 | 1 | 14 | Adcy9 | 0.073 | Young |
| 0.145517 | 0.537844 | 0.36 | 0.251 | 1 | 14 | Bcas2 | 0.109 | Young |
| 0.146266 | 1.666934 | 0.08 | 0.033 | 1 | 14 | Sephs2 | 0.047 | Young |
| 0.146266 | 1.458878 | 0.08 | 0.033 | 1 | 14 | Ptpru | 0.047 | Young |
| 0.146285 | 0.587076 | 0.66 | 0.574 | 1 | 14 | Gng5 | 0.086 | Young |
| 0.146352 | 0.827985 | 0.18 | 0.104 | 1 | 14 | Gfm1 | 0.076 | Young |
| 0.146594 | 0.187858 | 0.22 | 0.126 | 1 | 14 | Trnau1ap | 0.094 | Young |
| 0.147138 | 0.193184 | 0.42 | 0.295 | 1 | 14 | Lgmn | 0.125 | Young |
| 0.147828 | 1.337798 | 0.22 | 0.137 | 1 | 14 | Rab34 | 0.083 | Young |
| 0.148036 | 1.321319 | 0.12 | 0.06 | 1 | 14 | Gli2 | 0.06 | Young |
| 0.148036 | 1.186614 | 0.12 | 0.06 | 1 | 14 | Rgp1 | 0.06 | Young |
| 0.148147 | 2.10192 | 0.08 | 0.033 | 1 | 14 | Acss1 | 0.047 | Young |
| 0.148147 | 1.539486 | 0.08 | 0.033 | 1 | 14 | Slc35c1 | 0.047 | Young |
| 0.148696 | 1.139109 | 0.18 | 0.104 | 1 | 14 | Vars | 0.076 | Young |
| 0.148884 | 1.296802 | 0.22 | 0.137 | 1 | 14 | Polr2b | 0.083 | Young |
| 0.149071 | 0.619684 | 0.16 | 0.087 | 1 | 14 | Ramp1 | 0.073 | Young |
| 0.149153 | 0.901197 | 0.56 | 0.486 | 1 | 14 | Pten | 0.074 | Young |
| 0.149411 | 0.843283 | 0.26 | 0.164 | 1 | 14 | Tbcel | 0.096 | Young |
| 0.149513 | 1.099731 | 0.12 | 0.06 | 1 | 14 | Flcn | 0.06 | Young |
| 0.149855 | 0.150691 | 0.38 | 0.257 | 1 | 14 | Smg1 | 0.123 | Young |
| 0.149947 | 0.725197 | 0.22 | 0.137 | 1 | 14 | Nrf1 | 0.083 | Young |
| 0.150047 | 2.43673 | 0.08 | 0.033 | 1 | 14 | Gm5617 | 0.047 | Young |
| 0.150047 | 1.779534 | 0.08 | 0.033 | 1 | 14 | Hyal2 | 0.047 | Young |
| 0.150047 | 1.097186 | 0.08 | 0.033 | 1 | 14 | Ppif | 0.047 | Young |
| 0.150047 | 1.074745 | 0.08 | 0.033 | 1 | 14 | Shpk | 0.047 | Young |
| 0.150047 | 0.734436 | 0.08 | 0.033 | 1 | 14 | Sgsm3 | 0.047 | Young |
| 0.150339 | 1.412414 | 0.16 | 0.087 | 1 | 14 | Gcsh | 0.073 | Young |
| 0.150726 | 0.580302 | 0.34 | 0.235 | 1 | 14 | Hnrnpll | 0.105 | Young |
| 0.151001 | 0.893133 | 0.12 | 0.06 | 1 | 14 | Ppp2r5b | 0.06 | Young |
| 0.151503 | 0.963859 | 0.2 | 0.12 | 1 | 14 | Rnf138 | 0.08 | Young |
| 0.151551 | 2.197019 | 0.1 | 0.049 | 1 | 14 | Prickle2 | 0.051 | Young |
| 0.151835 | 0.336404 | 0.68 | 0.634 | 1 | 14 | Btg2 | 0.046 | Young |
| 0.151857 | 3.176096 | 0.06 | 0.022 | 1 | 14 | Chst2 | 0.038 | Young |
| 0.151965 | 1.703257 | 0.08 | 0.033 | 1 | 14 | Coil | 0.047 | Young |
| 0.151965 | 0.899677 | 0.08 | 0.033 | 1 | 14 | Cdh2 | 0.047 | Young |
| 0.151965 | 0.828016 | 0.08 | 0.033 | 1 | 14 | Gng7 | 0.047 | Young |
| 0.152059 | 0.61977 | 0.24 | 0.153 | 1 | 14 | Pex3 | 0.087 | Young |
| 0.152265 | 1.362059 | 0.18 | 0.104 | 1 | 14 | Mrpl43 | 0.076 | Young |
| 0.152501 | 1.511878 | 0.12 | 0.06 | 1 | 14 | Mief1 | 0.06 | Young |
| 0.152501 | 1.137556 | 0.12 | 0.06 | 1 | 14 | Camta1 | 0.06 | Young |
| 0.152682 | 0.521359 | 0.26 | 0.169 | 1 | 14 | Zc2hc1a | 0.091 | Young |
| 0.152682 | 0.397049 | 0.26 | 0.169 | 1 | 14 | Rap2b | 0.091 | Young |

| | | | | | | | | |
|---|---|---|---|---|---|---|---|---|
| 0.153061 | 0.637436 | 0.4 | 0.295 | 1 | 14 | Lars2 | 0.105 | Young |
| 0.154012 | 0.912395 | 0.12 | 0.06 | 1 | 14 | Tmed1 | 0.06 | Young |
| 0.154012 | 0.826449 | 0.12 | 0.06 | 1 | 14 | Prmt7 | 0.06 | Young |
| 0.154012 | 0.776718 | 0.12 | 0.06 | 1 | 14 | Prdm4 | 0.06 | Young |
| 0.154073 | 0.248061 | 0.14 | 0.071 | 1 | 14 | Rpp14 | 0.069 | Young |
| 0.154232 | 1.112981 | 0.3 | 0.219 | 1 | 14 | Ppp2r2d | 0.081 | Young |
| 0.154258 | 0.251561 | 0.34 | 0.235 | 1 | 14 | Tmcc3 | 0.105 | Young |
| 0.154636 | 0.371293 | 0.32 | 0.208 | 1 | 14 | Dnajc13 | 0.112 | Young |
| 0.154641 | 1.005872 | 0.2 | 0.126 | 1 | 14 | St3gal4 | 0.074 | Young |
| 0.15469 | 0.593874 | 0.38 | 0.279 | 1 | 14 | Qsox1 | 0.101 | Young |
| 0.154858 | 1.980483 | 0.1 | 0.049 | 1 | 14 | Mib2 | 0.051 | Young |
| 0.154919 | 1.492798 | 0.24 | 0.169 | 1 | 14 | Arhgap35 | 0.071 | Young |
| 0.155565 | 0.993777 | 0.3 | 0.213 | 1 | 14 | Nucb2 | 0.087 | Young |
| 0.155719 | 0.395467 | 0.82 | 0.77 | 1 | 14 | Rbm39 | 0.05 | Young |
| 0.155858 | 0.180342 | 0.08 | 0.033 | 1 | 14 | E2f3 | 0.047 | Young |
| 0.155899 | 0.701179 | 0.18 | 0.104 | 1 | 14 | Dynlt1f | 0.076 | Young |
| 0.156485 | 2.59241 | 0.06 | 0.022 | 1 | 14 | Mb21d2 | 0.038 | Young |
| 0.156485 | 1.755906 | 0.06 | 0.022 | 1 | 14 | Ddb2 | 0.038 | Young |
| 0.156485 | 1.574144 | 0.06 | 0.022 | 1 | 14 | Wasf1 | 0.038 | Young |
| 0.156821 | 0.35219 | 0.54 | 0.432 | 1 | 14 | Ubtf | 0.108 | Young |
| 0.156933 | 1.375601 | 0.14 | 0.077 | 1 | 14 | Itgbl1 | 0.063 | Young |
| 0.15701 | 3.849127 | 0.04 | 0.011 | 1 | 14 | Gm14443 | 0.029 | Young |
| 0.15701 | 3.815998 | 0.04 | 0.011 | 1 | 14 | Zbtb48 | 0.029 | Young |
| 0.15701 | 3.586539 | 0.04 | 0.011 | 1 | 14 | Zfp51 | 0.029 | Young |
| 0.15701 | 3.439258 | 0.04 | 0.011 | 1 | 14 | Ccl3 | 0.029 | Young |
| 0.15701 | 2.430208 | 0.04 | 0.011 | 1 | 14 | Gm14326 | 0.029 | Young |
| 0.15701 | 2.394263 | 0.04 | 0.011 | 1 | 14 | Nrtn | 0.029 | Young |
| 0.15701 | 2.382675 | 0.04 | 0.011 | 1 | 14 | Eral1 | 0.029 | Young |
| 0.15701 | 2.247416 | 0.04 | 0.011 | 1 | 14 | Fadd | 0.029 | Young |
| 0.15701 | 2.240437 | 0.04 | 0.011 | 1 | 14 | Prox2 | 0.029 | Young |
| 0.15701 | 2.135926 | 0.04 | 0.011 | 1 | 14 | Gm5113 | 0.029 | Young |
| 0.15701 | 2.133792 | 0.04 | 0.011 | 1 | 14 | Tmem30b | 0.029 | Young |
| 0.15701 | 1.867456 | 0.04 | 0.011 | 1 | 14 | Sag | 0.029 | Young |
| 0.15701 | 1.774363 | 0.04 | 0.011 | 1 | 14 | Gm46224 | 0.029 | Young |
| 0.157068 | 0.961846 | 0.12 | 0.06 | 1 | 14 | Gmppb | 0.06 | Young |
| 0.157068 | 0.625992 | 0.12 | 0.06 | 1 | 14 | Il33 | 0.06 | Young |
| 0.157073 | 0.351091 | 0.48 | 0.393 | 1 | 14 | Cd74 | 0.087 | Young |
| 0.157124 | 1.197896 | 0.18 | 0.104 | 1 | 14 | C1qtnf2 | 0.076 | Young |
| 0.157217 | 0.955598 | 0.24 | 0.148 | 1 | 14 | Usp1 | 0.092 | Young |
| 0.15737 | 0.609127 | 0.28 | 0.191 | 1 | 14 | Specc1l | 0.089 | Young |
| 0.157546 | 0.762801 | 0.22 | 0.137 | 1 | 14 | Nhlrc2 | 0.083 | Young |
| 0.157725 | 0.019332 | 0.52 | 0.361 | 1 | 14 | Gas6 | 0.159 | Young |
| 0.157833 | 1.191471 | 0.08 | 0.033 | 1 | 14 | Natd1 | 0.047 | Young |
| 0.157833 | 0.633933 | 0.08 | 0.033 | 1 | 14 | Riox2 | 0.047 | Young |
| 0.157833 | 0.347318 | 0.08 | 0.033 | 1 | 14 | Hyal1 | 0.047 | Young |

| | | | | | | | | |
|---|---|---|---|---|---|---|---|---|
| 0.158063 | 0.671518 | 0.2 | 0.126 | 1 | 14 | Dock10 | 0.074 | Young |
| 0.158334 | 1.262314 | 0.14 | 0.077 | 1 | 14 | Ncdn | 0.063 | Young |
| 0.15852 | 0.040005 | 0.42 | 0.552 | 1 | 14 | Vit | -0.132 | Young |
| 0.158839 | 1.876252 | 0.06 | 0.022 | 1 | 14 | Dcaf4 | 0.038 | Young |
| 0.158839 | 1.627958 | 0.06 | 0.022 | 1 | 14 | Rflnb | 0.038 | Young |
| 0.159432 | 0.713491 | 0.38 | 0.29 | 1 | 14 | Dusp11 | 0.09 | Young |
| 0.159548 | 0.617099 | 0.22 | 0.142 | 1 | 14 | Get4 | 0.078 | Young |
| 0.159602 | 0.297773 | 0.72 | 0.574 | 1 | 14 | Lrrc58 | 0.146 | Young |
| 0.159745 | 2.084358 | 0.14 | 0.077 | 1 | 14 | Psmd9 | 0.063 | Young |
| 0.159826 | 0.59353 | 0.08 | 0.033 | 1 | 14 | Adap2 | 0.047 | Young |
| 0.160117 | 3.538412 | 0.04 | 0.011 | 1 | 14 | Ankrd13b | 0.029 | Young |
| 0.160117 | 3.028398 | 0.04 | 0.011 | 1 | 14 | Mcm8 | 0.029 | Young |
| 0.160117 | 2.818979 | 0.04 | 0.011 | 1 | 14 | Il1r2 | 0.029 | Young |
| 0.160117 | 2.605108 | 0.04 | 0.011 | 1 | 14 | Rpp40 | 0.029 | Young |
| 0.160117 | 2.34593 | 0.04 | 0.011 | 1 | 14 | Coro2a | 0.029 | Young |
| 0.160117 | 2.202319 | 0.04 | 0.011 | 1 | 14 | Me3 | 0.029 | Young |
| 0.160117 | 2.030584 | 0.04 | 0.011 | 1 | 14 | Gli1 | 0.029 | Young |
| 0.160117 | 1.936007 | 0.04 | 0.011 | 1 | 14 | Map7d3 | 0.029 | Young |
| 0.160117 | 1.900759 | 0.04 | 0.011 | 1 | 14 | Gpihbp1 | 0.029 | Young |
| 0.160117 | 1.866654 | 0.04 | 0.011 | 1 | 14 | Pfkfb2 | 0.029 | Young |
| 0.160117 | 1.726494 | 0.04 | 0.011 | 1 | 14 | Gprc5a | 0.029 | Young |
| 0.160117 | 1.71454 | 0.04 | 0.011 | 1 | 14 | Mdfi | 0.029 | Young |
| 0.160117 | 1.653353 | 0.04 | 0.011 | 1 | 14 | Pik3r3 | 0.029 | Young |
| 0.160728 | 0.896066 | 0.4 | 0.301 | 1 | 14 | Igfbp5 | 0.099 | Young |
| 0.16099 | 0.372847 | 0.34 | 0.224 | 1 | 14 | Trim44 | 0.116 | Young |
| 0.161095 | 0.92469 | 0.44 | 0.344 | 1 | 14 | Mlf2 | 0.096 | Young |
| 0.161107 | 0.336842 | 0.06 | 0.131 | 1 | 14 | Prpf38a | -0.071 | Young |
| 0.16122 | 2.495766 | 0.06 | 0.022 | 1 | 14 | Sh3gl2 | 0.038 | Young |
| 0.16122 | 1.840853 | 0.06 | 0.022 | 1 | 14 | Zfp607a | 0.038 | Young |
| 0.16122 | 1.728384 | 0.06 | 0.022 | 1 | 14 | Sesn2 | 0.038 | Young |
| 0.16122 | 1.682807 | 0.06 | 0.022 | 1 | 14 | Rhbdl3 | 0.038 | Young |
| 0.16122 | 1.400516 | 0.06 | 0.022 | 1 | 14 | Fmnl3 | 0.038 | Young |
| 0.16122 | 1.285656 | 0.06 | 0.022 | 1 | 14 | Zfand1 | 0.038 | Young |
| 0.16122 | 1.285629 | 0.06 | 0.022 | 1 | 14 | Bank1 | 0.038 | Young |
| 0.16122 | 1.262057 | 0.06 | 0.022 | 1 | 14 | Gprc5c | 0.038 | Young |
| 0.161409 | 0.467798 | 0.86 | 0.885 | 1 | 14 | H3f3b | -0.025 | Young |
| 0.161415 | 0.849036 | 0.52 | 0.437 | 1 | 14 | Tbl1x | 0.083 | Young |
| 0.161637 | 2.430333 | 0.1 | 0.049 | 1 | 14 | Slc40a1 | 0.051 | Young |
| 0.161637 | 2.416714 | 0.1 | 0.049 | 1 | 14 | Hgf | 0.051 | Young |
| 0.161738 | 0.756737 | 0.12 | 0.06 | 1 | 14 | Utp14a | 0.06 | Young |
| 0.161738 | 0.577695 | 0.12 | 0.06 | 1 | 14 | Ppm1d | 0.06 | Young |
| 0.161839 | 0.401263 | 0.08 | 0.033 | 1 | 14 | Il10ra | 0.047 | Young |
| 0.161839 | 0.031553 | 0.08 | 0.033 | 1 | 14 | Slc35d2 | 0.047 | Young |
| 0.162167 | 0.342694 | 0.22 | 0.131 | 1 | 14 | Uap1l1 | 0.089 | Young |
| 0.162232 | 0.452242 | 0.44 | 0.333 | 1 | 14 | Psmb6 | 0.107 | Young |

| | | | | | | | | |
|---|---|---|---|---|---|---|---|---|
| 0.162716 | 1.24594 | 0.2 | 0.126 | 1 | 14 | Idh3a | 0.074 | Young |
| 0.16288 | 0.492065 | 0.26 | 0.169 | 1 | 14 | Phf1 | 0.091 | Young |
| 0.162903 | 1.125008 | 0.18 | 0.109 | 1 | 14 | Il15ra | 0.071 | Young |
| 0.163082 | 1.142211 | 0.16 | 0.093 | 1 | 14 | Mtx1 | 0.067 | Young |
| 0.163271 | 3.139462 | 0.04 | 0.011 | 1 | 14 | Snrnp35 | 0.029 | Young |
| 0.163271 | 2.330266 | 0.04 | 0.011 | 1 | 14 | Tmem177 | 0.029 | Young |
| 0.163271 | 2.207278 | 0.04 | 0.011 | 1 | 14 | Six4 | 0.029 | Young |
| 0.163271 | 2.123652 | 0.04 | 0.011 | 1 | 14 | Kiss1r | 0.029 | Young |
| 0.163271 | 1.985626 | 0.04 | 0.011 | 1 | 14 | Gm26645 | 0.029 | Young |
| 0.163271 | 1.918023 | 0.04 | 0.011 | 1 | 14 | Bloc1s3 | 0.029 | Young |
| 0.163271 | 1.874089 | 0.04 | 0.011 | 1 | 14 | Pcx | 0.029 | Young |
| 0.163271 | 1.861995 | 0.04 | 0.011 | 1 | 14 | Rnf123 | 0.029 | Young |
| 0.163271 | 1.846928 | 0.04 | 0.011 | 1 | 14 | Qprt | 0.029 | Young |
| 0.163271 | 1.846652 | 0.04 | 0.011 | 1 | 14 | Fhl3 | 0.029 | Young |
| 0.163271 | 1.837585 | 0.04 | 0.011 | 1 | 14 | Rccd1 | 0.029 | Young |
| 0.163271 | 1.788186 | 0.04 | 0.011 | 1 | 14 | Ptprb | 0.029 | Young |
| 0.163271 | 1.721139 | 0.04 | 0.011 | 1 | 14 | H2-DMb2 | 0.029 | Young |
| 0.163271 | 1.699369 | 0.04 | 0.011 | 1 | 14 | F630028O1 | 0.029 | Young |
| 0.163271 | 1.621315 | 0.04 | 0.011 | 1 | 14 | Wt1os | 0.029 | Young |
| 0.163271 | 1.61371 | 0.04 | 0.011 | 1 | 14 | Pde4d | 0.029 | Young |
| 0.163271 | 1.553019 | 0.04 | 0.011 | 1 | 14 | Cd24a | 0.029 | Young |
| 0.163271 | 1.525371 | 0.04 | 0.011 | 1 | 14 | Gins3 | 0.029 | Young |
| 0.163271 | 1.458138 | 0.04 | 0.011 | 1 | 14 | Susd3 | 0.029 | Young |
| 0.163271 | 1.45369 | 0.04 | 0.011 | 1 | 14 | Itga2 | 0.029 | Young |
| 0.163271 | 1.432421 | 0.04 | 0.011 | 1 | 14 | Samd5 | 0.029 | Young |
| 0.163271 | 1.387101 | 0.04 | 0.011 | 1 | 14 | Homez | 0.029 | Young |
| 0.163271 | 1.304779 | 0.04 | 0.011 | 1 | 14 | Ripor2 | 0.029 | Young |
| 0.163271 | 1.258552 | 0.04 | 0.011 | 1 | 14 | P2ry10b | 0.029 | Young |
| 0.163271 | 1.227658 | 0.04 | 0.011 | 1 | 14 | Zfp784 | 0.029 | Young |
| 0.163271 | 1.226189 | 0.04 | 0.011 | 1 | 14 | P2ry12 | 0.029 | Young |
| 0.163271 | 1.151336 | 0.04 | 0.011 | 1 | 14 | Tcaim | 0.029 | Young |
| 0.163271 | 1.121658 | 0.04 | 0.011 | 1 | 14 | Prelid2 | 0.029 | Young |
| 0.163271 | 1.011944 | 0.04 | 0.011 | 1 | 14 | Cdkl3 | 0.029 | Young |
| 0.163271 | 1.002591 | 0.04 | 0.011 | 1 | 14 | Myzap | 0.029 | Young |
| 0.163627 | 2.428406 | 0.06 | 0.022 | 1 | 14 | Tes | 0.038 | Young |
| 0.163627 | 1.419311 | 0.06 | 0.022 | 1 | 14 | Cd209b | 0.038 | Young |
| 0.163627 | 0.934271 | 0.06 | 0.022 | 1 | 14 | Msh3 | 0.038 | Young |
| 0.163627 | 0.85631 | 0.06 | 0.022 | 1 | 14 | Zbtb39 | 0.038 | Young |
| 0.163627 | 0.787596 | 0.06 | 0.022 | 1 | 14 | Bbc3 | 0.038 | Young |
| 0.163627 | 0.780846 | 0.06 | 0.022 | 1 | 14 | Colgalt2 | 0.038 | Young |
| 0.163921 | 0.752327 | 0.5 | 0.41 | 1 | 14 | Wipf1 | 0.09 | Young |
| 0.163977 | 0.204037 | 0.14 | 0.246 | 1 | 14 | Bhlhe40 | -0.106 | Young |
| 0.16429 | 0.918499 | 0.22 | 0.137 | 1 | 14 | Cul9 | 0.083 | Young |
| 0.164571 | 0.106301 | 0.28 | 0.18 | 1 | 14 | Cep350 | 0.1 | Young |
| 0.164633 | 0.483639 | 0.18 | 0.104 | 1 | 14 | Tmem47 | 0.076 | Young |

| | | | | | | | | |
|---|---|---|---|---|---|---|---|---|
| 0.164753 | 1.706945 | 0.22 | 0.148 | 1 | 14 | Cript | 0.072 | Young |
| 0.164909 | 1.269718 | 0.32 | 0.224 | 1 | 14 | Ctdspl | 0.096 | Young |
| 0.164909 | 0.543722 | 0.12 | 0.06 | 1 | 14 | Inafm2 | 0.06 | Young |
| 0.164909 | 0.438239 | 0.12 | 0.06 | 1 | 14 | Mthfr | 0.06 | Young |
| 0.165757 | 1.871265 | 0.16 | 0.093 | 1 | 14 | Acads | 0.067 | Young |
| 0.16591 | 0.412723 | 0.18 | 0.104 | 1 | 14 | Caml | 0.076 | Young |
| 0.165913 | 1.195433 | 0.24 | 0.153 | 1 | 14 | Fam53c | 0.087 | Young |
| 0.165921 | 0.000179 | 0.08 | 0.033 | 1 | 14 | Adam8 | 0.047 | Young |
| 0.166062 | 1.621159 | 0.06 | 0.022 | 1 | 14 | Ifitm6 | 0.038 | Young |
| 0.166062 | 1.335451 | 0.06 | 0.022 | 1 | 14 | Fastkd1 | 0.038 | Young |
| 0.166062 | 1.037129 | 0.06 | 0.022 | 1 | 14 | Zfp72 | 0.038 | Young |
| 0.166062 | 0.318231 | 0.06 | 0.022 | 1 | 14 | Gdpd1 | 0.038 | Young |
| 0.16621 | 0.363128 | 0.04 | 0.104 | 1 | 14 | Slfn8 | -0.064 | Young |
| 0.166211 | 0.228285 | 0.16 | 0.087 | 1 | 14 | Cybb | 0.073 | Young |
| 0.166471 | 1.258692 | 0.04 | 0.011 | 1 | 14 | Cldn5 | 0.029 | Young |
| 0.166471 | 1.246752 | 0.04 | 0.011 | 1 | 14 | Prune2 | 0.029 | Young |
| 0.166471 | 1.220419 | 0.04 | 0.011 | 1 | 14 | D630045J1 | 0.029 | Young |
| 0.166471 | 1.118879 | 0.04 | 0.011 | 1 | 14 | Cacna1g | 0.029 | Young |
| 0.166471 | 1.069493 | 0.04 | 0.011 | 1 | 14 | Gdpd2 | 0.029 | Young |
| 0.166471 | 1.064912 | 0.04 | 0.011 | 1 | 14 | Fam177a | 0.029 | Young |
| 0.166471 | 1.050049 | 0.04 | 0.011 | 1 | 14 | Atp2a3 | 0.029 | Young |
| 0.166471 | 1.033684 | 0.04 | 0.011 | 1 | 14 | Frs3 | 0.029 | Young |
| 0.166471 | 0.979701 | 0.04 | 0.011 | 1 | 14 | 1190005I0( | 0.029 | Young |
| 0.166471 | 0.969467 | 0.04 | 0.011 | 1 | 14 | Ceacam1 | 0.029 | Young |
| 0.166471 | 0.947426 | 0.04 | 0.011 | 1 | 14 | Paqr8 | 0.029 | Young |
| 0.166471 | 0.942338 | 0.04 | 0.011 | 1 | 14 | Rad9b | 0.029 | Young |
| 0.166471 | 0.936406 | 0.04 | 0.011 | 1 | 14 | Rtl3 | 0.029 | Young |
| 0.166471 | 0.875379 | 0.04 | 0.011 | 1 | 14 | 2310040G( | 0.029 | Young |
| 0.166471 | 0.652375 | 0.04 | 0.011 | 1 | 14 | Katnb1 | 0.029 | Young |
| 0.166471 | 0.369716 | 0.04 | 0.011 | 1 | 14 | Kcnj8 | 0.029 | Young |
| 0.166668 | 1.875234 | 0.18 | 0.109 | 1 | 14 | Mfap1b | 0.071 | Young |
| 0.167195 | 0.515682 | 0.18 | 0.104 | 1 | 14 | Senp5 | 0.076 | Young |
| 0.167195 | 0.140055 | 0.18 | 0.104 | 1 | 14 | Eif4ebp1 | 0.076 | Young |
| 0.167419 | 1.406559 | 0.22 | 0.142 | 1 | 14 | Gm8186 | 0.078 | Young |
| 0.16801 | 0.18129 | 0.32 | 0.213 | 1 | 14 | Ppp6c | 0.107 | Young |
| 0.168182 | 0.689445 | 0.46 | 0.372 | 1 | 14 | Mef2d | 0.088 | Young |
| 0.168408 | 0.760627 | 0.14 | 0.077 | 1 | 14 | Naa16 | 0.063 | Young |
| 0.168522 | 0.834112 | 0.3 | 0.208 | 1 | 14 | Npm3 | 0.092 | Young |
| 0.168523 | 1.486385 | 0.06 | 0.022 | 1 | 14 | Mthfs | 0.038 | Young |
| 0.168523 | 1.274082 | 0.06 | 0.022 | 1 | 14 | Begain | 0.038 | Young |
| 0.168523 | 1.072607 | 0.06 | 0.022 | 1 | 14 | 1810032O( | 0.038 | Young |
| 0.168523 | 0.817507 | 0.06 | 0.022 | 1 | 14 | Acp6 | 0.038 | Young |
| 0.168523 | 0.815877 | 0.06 | 0.022 | 1 | 14 | 2700062C( | 0.038 | Young |
| 0.168777 | 1.020932 | 0.38 | 0.295 | 1 | 14 | Bbx | 0.085 | Young |
| 0.168975 | 1.405375 | 0.16 | 0.087 | 1 | 14 | Pspc1 | 0.073 | Young |

| | | | | | | | | |
|---|---|---|---|---|---|---|---|---|
| 0.168975 | 1.402141 | 0.16 | 0.087 | 1 | 14 | Ubl4a | 0.073 | Young |
| 0.168975 | 0.784412 | 0.16 | 0.087 | 1 | 14 | Bambi | 0.073 | Young |
| 0.169271 | 0.659746 | 0.56 | 0.503 | 1 | 14 | Zmiz1 | 0.057 | Young |
| 0.169718 | 1.052266 | 0.04 | 0.011 | 1 | 14 | Ccdc62 | 0.029 | Young |
| 0.169718 | 0.915789 | 0.04 | 0.011 | 1 | 14 | Pkd1l2 | 0.029 | Young |
| 0.169718 | 0.915781 | 0.04 | 0.011 | 1 | 14 | Aoah | 0.029 | Young |
| 0.169718 | 0.900739 | 0.04 | 0.011 | 1 | 14 | Gm15551 | 0.029 | Young |
| 0.169718 | 0.789572 | 0.04 | 0.011 | 1 | 14 | Cenpa | 0.029 | Young |
| 0.169718 | 0.787357 | 0.04 | 0.011 | 1 | 14 | Map4k1 | 0.029 | Young |
| 0.169718 | 0.664356 | 0.04 | 0.011 | 1 | 14 | Chst3 | 0.029 | Young |
| 0.169718 | 0.623595 | 0.04 | 0.011 | 1 | 14 | Ears2 | 0.029 | Young |
| 0.169718 | 0.562214 | 0.04 | 0.011 | 1 | 14 | Drd1 | 0.029 | Young |
| 0.169718 | 0.47696 | 0.04 | 0.011 | 1 | 14 | Flt4 | 0.029 | Young |
| 0.169718 | 0.419686 | 0.04 | 0.011 | 1 | 14 | Cmbl | 0.029 | Young |
| 0.169718 | 0.358932 | 0.04 | 0.011 | 1 | 14 | Gm6598 | 0.029 | Young |
| 0.169718 | 0.349679 | 0.04 | 0.011 | 1 | 14 | Fgd3 | 0.029 | Young |
| 0.169718 | 0.326667 | 0.04 | 0.011 | 1 | 14 | Med18 | 0.029 | Young |
| 0.169718 | 0.161694 | 0.04 | 0.011 | 1 | 14 | Gm14399 | 0.029 | Young |
| 0.169718 | 0.002337 | 0.04 | 0.011 | 1 | 14 | Lrrc75b | 0.029 | Young |
| 0.169753 | 0.660117 | 0.12 | 0.06 | 1 | 14 | Ppp5c | 0.06 | Young |
| 0.169753 | 0.296451 | 0.12 | 0.06 | 1 | 14 | Commd10 | 0.06 | Young |
| 0.170326 | 0.369238 | 0.22 | 0.131 | 1 | 14 | Wbp11 | 0.089 | Young |
| 0.170422 | 2.446708 | 0.1 | 0.049 | 1 | 14 | Osgepl1 | 0.051 | Young |
| 0.171012 | 0.941076 | 0.06 | 0.022 | 1 | 14 | Oraov1 | 0.038 | Young |
| 0.171012 | 0.726724 | 0.06 | 0.022 | 1 | 14 | Nog | 0.038 | Young |
| 0.171012 | 0.700943 | 0.06 | 0.022 | 1 | 14 | Rab15 | 0.038 | Young |
| 0.171012 | 0.697181 | 0.06 | 0.022 | 1 | 14 | Mipol1 | 0.038 | Young |
| 0.171012 | 0.668948 | 0.06 | 0.022 | 1 | 14 | Map4k2 | 0.038 | Young |
| 0.171012 | 0.624852 | 0.06 | 0.022 | 1 | 14 | Cnnm2 | 0.038 | Young |
| 0.171012 | 0.5866 | 0.06 | 0.022 | 1 | 14 | Thop1 | 0.038 | Young |
| 0.171012 | 0.510755 | 0.06 | 0.022 | 1 | 14 | Rgs18 | 0.038 | Young |
| 0.171012 | 0.402991 | 0.06 | 0.022 | 1 | 14 | Aldh4a1 | 0.038 | Young |
| 0.171012 | 0.221757 | 0.06 | 0.022 | 1 | 14 | Zfp444 | 0.038 | Young |
| 0.171012 | 0.192436 | 0.06 | 0.022 | 1 | 14 | Epb41 | 0.038 | Young |
| 0.171012 | 0.133808 | 0.06 | 0.022 | 1 | 14 | Tmem143 | 0.038 | Young |
| 0.171012 | 0.121025 | 0.06 | 0.022 | 1 | 14 | E430018J2 | 0.038 | Young |
| 0.171206 | 1.432984 | 0.16 | 0.093 | 1 | 14 | Rnf114 | 0.067 | Young |
| 0.171373 | 1.187272 | 0.14 | 0.077 | 1 | 14 | St6galnac4 | 0.063 | Young |
| 0.171484 | 0.629765 | 0.28 | 0.191 | 1 | 14 | 6-Mar | 0.089 | Young |
| 0.171631 | 0.481006 | 0.2 | 0.12 | 1 | 14 | Sgcb | 0.08 | Young |
| 0.172019 | 0.500454 | 0.4 | 0.295 | 1 | 14 | Gpx8 | 0.105 | Young |
| 0.172045 | 0.101813 | 0.1 | 0.175 | 1 | 14 | Rtf2 | -0.075 | Young |
| 0.172099 | 0.736105 | 0.46 | 0.355 | 1 | 14 | Nme1 | 0.105 | Young |
| 0.172134 | 0.506029 | 0.5 | 0.404 | 1 | 14 | Sptssa | 0.096 | Young |
| 0.172409 | 0.788814 | 0.18 | 0.104 | 1 | 14 | Txndc12 | 0.076 | Young |

| | | | | | | | | |
|---|---|---|---|---|---|---|---|---|
| 0.172589 | 0.524726 | 0.16 | 0.093 | 1 | 14 | Slc27a4 | 0.067 | Young |
| 0.172619 | 0.362067 | 0.24 | 0.153 | 1 | 14 | B3galnt2 | 0.087 | Young |
| 0.172753 | 0.096228 | 0.14 | 0.235 | 1 | 14 | Ezr | -0.095 | Young |
| 0.17287 | 0.459074 | 0.14 | 0.077 | 1 | 14 | Exosc5 | 0.063 | Young |
| 0.173041 | 1.325972 | 0.12 | 0.06 | 1 | 14 | Dclre1c | 0.06 | Young |
| 0.173529 | 0.686447 | 0.06 | 0.022 | 1 | 14 | Olfr1167 | 0.038 | Young |
| 0.173529 | 0.294002 | 0.06 | 0.022 | 1 | 14 | Itga2b | 0.038 | Young |
| 0.174242 | 0.473011 | 0.74 | 0.743 | 1 | 14 | Gnai2 | -0.003 | Young |
| 0.174394 | 1.284375 | 0.18 | 0.109 | 1 | 14 | Msx1 | 0.071 | Young |
| 0.174605 | 0.192905 | 0.16 | 0.087 | 1 | 14 | Sprytd7 | 0.073 | Young |
| 0.175385 | 0.055873 | 0.28 | 0.366 | 1 | 14 | Arpp19 | -0.086 | Young |
| 0.175708 | 0.48517 | 0.38 | 0.262 | 1 | 14 | Gpat4 | 0.118 | Young |
| 0.175785 | 0.583679 | 0.26 | 0.169 | 1 | 14 | Baz2a | 0.091 | Young |
| 0.175863 | 2.179366 | 0.1 | 0.049 | 1 | 14 | BC028528 | 0.051 | Young |
| 0.175863 | 1.959169 | 0.1 | 0.049 | 1 | 14 | Man2c1 | 0.051 | Young |
| 0.175984 | 1.258203 | 0.34 | 0.246 | 1 | 14 | Bscl2 | 0.094 | Young |
| 0.176035 | 0.853823 | 0.16 | 0.087 | 1 | 14 | Pdrg1 | 0.073 | Young |
| 0.176072 | 0.399641 | 0.06 | 0.022 | 1 | 14 | Lix1 | 0.038 | Young |
| 0.176072 | 0.368894 | 0.06 | 0.022 | 1 | 14 | Fat4 | 0.038 | Young |
| 0.176072 | 0.3052 | 0.06 | 0.022 | 1 | 14 | Mir17hg | 0.038 | Young |
| 0.176376 | 0.409307 | 0.12 | 0.06 | 1 | 14 | Rundc1 | 0.06 | Young |
| 0.176398 | 0.949004 | 0.18 | 0.104 | 1 | 14 | Ppp6r1 | 0.076 | Young |
| 0.176484 | 0.629112 | 0.28 | 0.186 | 1 | 14 | Actr10 | 0.094 | Young |
| 0.176591 | 0.601256 | 0.46 | 0.366 | 1 | 14 | Cdc42bpa | 0.094 | Young |
| 0.176788 | 0.2751 | 0.16 | 0.093 | 1 | 14 | Hmgxb4 | 0.067 | Young |
| 0.176964 | 0.766459 | 0.24 | 0.158 | 1 | 14 | Hmox1 | 0.082 | Young |
| 0.17717 | 1.538107 | 0.3 | 0.23 | 1 | 14 | Htatsf1 | 0.07 | Young |
| 0.17742 | 0.991624 | 0.14 | 0.077 | 1 | 14 | Cog6 | 0.063 | Young |
| 0.17742 | 0.724613 | 0.14 | 0.077 | 1 | 14 | Snx16 | 0.063 | Young |
| 0.17742 | 0.496762 | 0.14 | 0.077 | 1 | 14 | Mrpl1 | 0.063 | Young |
| 0.177542 | 0.452157 | 0.24 | 0.35 | 1 | 14 | Cpt1a | -0.11 | Young |
| 0.17761 | 0.794408 | 0.38 | 0.284 | 1 | 14 | Kcmf1 | 0.096 | Young |
| 0.177704 | 2.039622 | 0.1 | 0.049 | 1 | 14 | Sh3rf1 | 0.051 | Young |
| 0.177704 | 1.336947 | 0.1 | 0.049 | 1 | 14 | Carf | 0.051 | Young |
| 0.177815 | 0.715449 | 0.28 | 0.191 | 1 | 14 | Gpalpp1 | 0.089 | Young |
| 0.17788 | 0.658677 | 0.32 | 0.219 | 1 | 14 | Spry1 | 0.101 | Young |
| 0.178361 | 0.513707 | 0.24 | 0.153 | 1 | 14 | D10Jhu81e | 0.087 | Young |
| 0.178593 | 0.819633 | 0.22 | 0.148 | 1 | 14 | Spry4 | 0.072 | Young |
| 0.178644 | 0.230139 | 0.06 | 0.022 | 1 | 14 | Cdc25a | 0.038 | Young |
| 0.178705 | 1.828954 | 0.16 | 0.098 | 1 | 14 | Tpst2 | 0.062 | Young |
| 0.178964 | 0.925496 | 0.46 | 0.41 | 1 | 14 | Kctd12 | 0.05 | Young |
| 0.179069 | 0.819346 | 0.28 | 0.197 | 1 | 14 | P3h3 | 0.083 | Young |
| 0.179096 | 0.629202 | 0.06 | 0.137 | 1 | 14 | Trappc1 | -0.077 | Young |
| 0.179127 | 0.847594 | 0.26 | 0.169 | 1 | 14 | Gclc | 0.091 | Young |
| 0.179276 | 0.458091 | 0.76 | 0.699 | 1 | 14 | Npm1 | 0.061 | Young |

| | | | | | | | | |
|---|---|---|---|---|---|---|---|---|
| 0.180064 | 0.759874 | 0.24 | 0.164 | 1 | 14 | Ppil4 | 0.076 | Young |
| 0.180127 | 0.998553 | 0.32 | 0.235 | 1 | 14 | Mrpl48 | 0.085 | Young |
| 0.180612 | 1.172897 | 0.3 | 0.219 | 1 | 14 | Cyp27a1 | 0.081 | Young |
| 0.181431 | 0.829779 | 0.1 | 0.049 | 1 | 14 | Prex1 | 0.051 | Young |
| 0.182746 | 0.323547 | 0.68 | 0.574 | 1 | 14 | Myl12a | 0.106 | Young |
| 0.183057 | 0.772395 | 0.6 | 0.481 | 1 | 14 | Apoe | 0.119 | Young |
| 0.183199 | 0.749211 | 0.18 | 0.104 | 1 | 14 | Carmil1 | 0.076 | Young |
| 0.183199 | 0.38069 | 0.18 | 0.104 | 1 | 14 | Igdcc4 | 0.076 | Young |
| 0.183625 | 0.666884 | 0.14 | 0.077 | 1 | 14 | Slc35d1 | 0.063 | Young |
| 0.184367 | 0.105707 | 0.06 | 0.126 | 1 | 14 | Parp11 | -0.066 | Young |
| 0.184798 | 0.368715 | 0.58 | 0.475 | 1 | 14 | Ncor1 | 0.105 | Young |
| 0.184898 | 0.522325 | 0.34 | 0.246 | 1 | 14 | Kif13b | 0.094 | Young |
| 0.184966 | 0.571524 | 0.4 | 0.295 | 1 | 14 | Med13l | 0.105 | Young |
| 0.185216 | 1.645092 | 0.1 | 0.049 | 1 | 14 | Myo10 | 0.051 | Young |
| 0.185216 | 0.922026 | 0.1 | 0.049 | 1 | 14 | Limk2 | 0.051 | Young |
| 0.185216 | 0.829453 | 0.1 | 0.049 | 1 | 14 | Sort1 | 0.051 | Young |
| 0.185261 | 0.504597 | 0.22 | 0.142 | 1 | 14 | Senp2 | 0.078 | Young |
| 0.185973 | 0.63396 | 0.18 | 0.104 | 1 | 14 | Ivns1abp | 0.076 | Young |
| 0.186198 | 1.768774 | 0.12 | 0.066 | 1 | 14 | Daxx | 0.054 | Young |
| 0.18648 | 1.460472 | 0.18 | 0.109 | 1 | 14 | Socs2 | 0.071 | Young |
| 0.186702 | 0.771261 | 0.5 | 0.426 | 1 | 14 | Skp1a | 0.074 | Young |
| 0.186715 | 0.538121 | 0.14 | 0.246 | 1 | 14 | Gm12840 | -0.106 | Young |
| 0.186786 | 0.468066 | 0.14 | 0.077 | 1 | 14 | Igfbp3 | 0.063 | Young |
| 0.186881 | 1.115952 | 0.16 | 0.093 | 1 | 14 | Uck1 | 0.067 | Young |
| 0.18713 | 0.916509 | 0.1 | 0.049 | 1 | 14 | Eda | 0.051 | Young |
| 0.18713 | 0.265986 | 0.1 | 0.049 | 1 | 14 | Jmjd8 | 0.051 | Young |
| 0.187316 | 0.378041 | 0.74 | 0.699 | 1 | 14 | Eef1g | 0.041 | Young |
| 0.18745 | 0.124172 | 0.28 | 0.186 | 1 | 14 | Stxbp1 | 0.094 | Young |
| 0.18786 | 0.679045 | 0.18 | 0.109 | 1 | 14 | Dcaf10 | 0.071 | Young |
| 0.188382 | 0.257483 | 0.14 | 0.077 | 1 | 14 | Ngf | 0.063 | Young |
| 0.188881 | 0.039721 | 0.12 | 0.202 | 1 | 14 | Gabpa | -0.082 | Young |
| 0.189058 | 0.923132 | 0.1 | 0.049 | 1 | 14 | Pan2 | 0.051 | Young |
| 0.189058 | 0.802693 | 0.1 | 0.049 | 1 | 14 | Foxo4 | 0.051 | Young |
| 0.189872 | 0.721055 | 0.28 | 0.191 | 1 | 14 | Dennd1b | 0.089 | Young |
| 0.189988 | 0.286349 | 0.14 | 0.077 | 1 | 14 | Cog5 | 0.063 | Young |
| 0.189994 | 0.871413 | 0.56 | 0.525 | 1 | 14 | Maged1 | 0.035 | Young |
| 0.19016 | 0.404938 | 0.06 | 0.126 | 1 | 14 | Arhgef11 | -0.066 | Young |
| 0.190643 | 0.596496 | 0.18 | 0.109 | 1 | 14 | Tbce | 0.071 | Young |
| 0.190792 | 0.518559 | 0.46 | 0.361 | 1 | 14 | Psma6 | 0.099 | Young |
| 0.190841 | 0.943296 | 0.42 | 0.317 | 1 | 14 | Scand1 | 0.103 | Young |
| 0.191001 | 1.377504 | 0.1 | 0.049 | 1 | 14 | Ccdc141 | 0.051 | Young |
| 0.191001 | 0.705828 | 0.1 | 0.049 | 1 | 14 | Ctps | 0.051 | Young |
| 0.191001 | 0.560841 | 0.1 | 0.049 | 1 | 14 | Phkg2 | 0.051 | Young |
| 0.191204 | 0.919389 | 0.38 | 0.306 | 1 | 14 | Golga4 | 0.074 | Young |
| 0.191604 | 0.251934 | 0.14 | 0.077 | 1 | 14 | Arl14ep | 0.063 | Young |

| | | | | | | | | |
|---|---|---|---|---|---|---|---|---|
| 0.191694 | 1.038171 | 0.3 | 0.23 | 1 | 14 | Adgra2 | 0.07 | Young |
| 0.192126 | 1.598201 | 0.14 | 0.082 | 1 | 14 | Slc12a2 | 0.058 | Young |
| 0.19216 | 0.357845 | 0.22 | 0.137 | 1 | 14 | Mpp5 | 0.083 | Young |
| 0.192214 | 0.523268 | 0.54 | 0.41 | 1 | 14 | Tcf25 | 0.13 | Young |
| 0.192306 | 0.691901 | 0.24 | 0.158 | 1 | 14 | Kansl3 | 0.082 | Young |
| 0.192343 | 0.269213 | 0.2 | 0.12 | 1 | 14 | Taf11 | 0.08 | Young |
| 0.192959 | 1.449184 | 0.1 | 0.049 | 1 | 14 | Ms4a7 | 0.051 | Young |
| 0.192959 | 0.210977 | 0.02 | 0.071 | 1 | 14 | Crcp | -0.051 | Young |
| 0.193102 | 1.360968 | 0.12 | 0.066 | 1 | 14 | Notch3 | 0.054 | Young |
| 0.19323 | 0.199334 | 0.14 | 0.077 | 1 | 14 | Akap2 | 0.063 | Young |
| 0.194402 | 0.483618 | 0.28 | 0.191 | 1 | 14 | Mob1b | 0.089 | Young |
| 0.194867 | 0.384064 | 0.14 | 0.077 | 1 | 14 | Pcsk7 | 0.063 | Young |
| 0.194932 | 0.970477 | 0.1 | 0.049 | 1 | 14 | Tead3 | 0.051 | Young |
| 0.194932 | 0.833466 | 0.1 | 0.049 | 1 | 14 | Plxna2 | 0.051 | Young |
| 0.194932 | 0.69762 | 0.1 | 0.049 | 1 | 14 | Ripk3 | 0.051 | Young |
| 0.194932 | 0.423098 | 0.1 | 0.049 | 1 | 14 | Lin54 | 0.051 | Young |
| 0.195091 | 2.220454 | 0.18 | 0.115 | 1 | 14 | Stim2 | 0.065 | Young |
| 0.195292 | 0.535903 | 0.78 | 0.776 | 1 | 14 | Ptms | 0.004 | Young |
| 0.195774 | 0.308616 | 0.7 | 0.699 | 1 | 14 | Rock2 | 0.001 | Young |
| 0.195865 | 0.482301 | 0.16 | 0.093 | 1 | 14 | Ttc33 | 0.067 | Young |
| 0.19615 | 0.887632 | 0.34 | 0.251 | 1 | 14 | Cope | 0.089 | Young |
| 0.196486 | 2.593345 | 0.08 | 0.038 | 1 | 14 | Tcf23 | 0.042 | Young |
| 0.196486 | 1.635158 | 0.08 | 0.038 | 1 | 14 | Cpxm2 | 0.042 | Young |
| 0.196919 | 1.247022 | 0.1 | 0.049 | 1 | 14 | Phka1 | 0.051 | Young |
| 0.196919 | 0.05849 | 0.1 | 0.049 | 1 | 14 | Ccdc66 | 0.051 | Young |
| 0.196941 | 1.730819 | 0.14 | 0.082 | 1 | 14 | Mr1 | 0.058 | Young |
| 0.197381 | 0.088046 | 0.18 | 0.104 | 1 | 14 | Trp53inp2 | 0.076 | Young |
| 0.197688 | 0.86914 | 0.24 | 0.153 | 1 | 14 | Cyc1 | 0.087 | Young |
| 0.197797 | 0.635372 | 0.2 | 0.12 | 1 | 14 | Psmd5 | 0.08 | Young |
| 0.198403 | 1.063337 | 0.12 | 0.066 | 1 | 14 | Elac2 | 0.054 | Young |
| 0.198403 | 0.760985 | 0.12 | 0.066 | 1 | 14 | Sepsecs | 0.054 | Young |
| 0.19846 | 0.665867 | 0.24 | 0.158 | 1 | 14 | Camta2 | 0.082 | Young |
| 0.198463 | 0.88083 | 0.34 | 0.273 | 1 | 14 | Srek1 | 0.067 | Young |
| 0.1991 | 1.600399 | 0.16 | 0.098 | 1 | 14 | Slc17a5 | 0.062 | Young |
| 0.199415 | 0.1808 | 0.32 | 0.213 | 1 | 14 | Klhdc2 | 0.107 | Young |
| 0.199708 | 1.143286 | 0.24 | 0.158 | 1 | 14 | Fam133b | 0.082 | Young |
| 0.199708 | 0.722279 | 0.24 | 0.158 | 1 | 14 | Anapc13 | 0.082 | Young |
| 0.200079 | 0.411878 | 0.38 | 0.284 | 1 | 14 | Dab2 | 0.096 | Young |
| 0.200367 | 0.206803 | 0.12 | 0.197 | 1 | 14 | Fez1 | -0.077 | Young |
| 0.200879 | 1.112712 | 0.26 | 0.18 | 1 | 14 | Rbl2 | 0.08 | Young |
| 0.200938 | 0.808165 | 0.1 | 0.049 | 1 | 14 | Nfam1 | 0.051 | Young |
| 0.200988 | 1.397096 | 0.08 | 0.038 | 1 | 14 | Nsun3 | 0.042 | Young |
| 0.201995 | 1.155989 | 0.12 | 0.066 | 1 | 14 | Ankrd54 | 0.054 | Young |
| 0.202029 | 0.468691 | 0.16 | 0.093 | 1 | 14 | Ints7 | 0.067 | Young |
| 0.202029 | 0.207904 | 0.16 | 0.093 | 1 | 14 | Ttbk2 | 0.067 | Young |

| | | | | | | | | |
|---|---|---|---|---|---|---|---|---|
| 0.202036 | 0.22079 | 0.62 | 0.781 | 1 | 14 | Hnrnph1 | -0.161 | Young |
| 0.202127 | 0.657082 | 0.38 | 0.284 | 1 | 14 | Egln1 | 0.096 | Young |
| 0.202142 | 0.005856 | 0.72 | 0.863 | 1 | 14 | Pabpc1 | -0.143 | Young |
| 0.202701 | 1.02354 | 0.32 | 0.23 | 1 | 14 | Dennd2a | 0.09 | Young |
| 0.20297 | 0.396534 | 0.1 | 0.049 | 1 | 14 | Gmfg | 0.051 | Young |
| 0.20297 | 0.353896 | 0.1 | 0.049 | 1 | 14 | Trim68 | 0.051 | Young |
| 0.2032 | 0.161692 | 0.14 | 0.077 | 1 | 14 | Arx | 0.063 | Young |
| 0.203273 | 0.038841 | 0.18 | 0.104 | 1 | 14 | Rbm43 | 0.076 | Young |
| 0.203591 | 0.767976 | 0.16 | 0.093 | 1 | 14 | Tigd2 | 0.067 | Young |
| 0.204473 | 0.539053 | 0.1 | 0.175 | 1 | 14 | Sh3bp4 | -0.075 | Young |
| 0.204501 | 1.334817 | 0.26 | 0.18 | 1 | 14 | Mgll | 0.08 | Young |
| 0.205014 | 0.442392 | 0.18 | 0.109 | 1 | 14 | Klhl5 | 0.071 | Young |
| 0.205088 | 0.991251 | 0.18 | 0.115 | 1 | 14 | Rprd1a | 0.065 | Young |
| 0.205093 | 0.691269 | 0.26 | 0.169 | 1 | 14 | Lars | 0.091 | Young |
| 0.205107 | 0.597012 | 0.2 | 0.126 | 1 | 14 | Supt4a | 0.074 | Young |
| 0.205216 | 0.169981 | 0.24 | 0.361 | 1 | 14 | Mesd | -0.121 | Young |
| 0.205373 | 0.016787 | 0.5 | 0.579 | 1 | 14 | Wls | -0.079 | Young |
| 0.205451 | 1.202893 | 0.32 | 0.246 | 1 | 14 | Zkscan3 | 0.074 | Young |
| 0.205564 | 1.308058 | 0.08 | 0.038 | 1 | 14 | Mgat4a | 0.042 | Young |
| 0.205564 | 1.267625 | 0.08 | 0.038 | 1 | 14 | Adck1 | 0.042 | Young |
| 0.205593 | 0.026288 | 0.54 | 0.628 | 1 | 14 | Calm3 | -0.088 | Young |
| 0.205635 | 1.677513 | 0.12 | 0.066 | 1 | 14 | Slc16a1 | 0.054 | Young |
| 0.205635 | 1.624031 | 0.12 | 0.066 | 1 | 14 | Rpf2 | 0.054 | Young |
| 0.20591 | 0.814648 | 0.44 | 0.333 | 1 | 14 | Ppp3r1 | 0.107 | Young |
| 0.206073 | 0.180144 | 0.18 | 0.273 | 1 | 14 | H2-Q4 | -0.093 | Young |
| 0.206434 | 0.937926 | 0.5 | 0.41 | 1 | 14 | Atp6v1f | 0.09 | Young |
| 0.206683 | 1.075388 | 0.2 | 0.131 | 1 | 14 | Nelfa | 0.069 | Young |
| 0.206846 | 0.410563 | 0.48 | 0.426 | 1 | 14 | Zfand5 | 0.054 | Young |
| 0.206931 | 1.11746 | 0.36 | 0.257 | 1 | 14 | Tpp2 | 0.103 | Young |
| 0.207186 | 1.124803 | 0.36 | 0.262 | 1 | 14 | Rbm28 | 0.098 | Young |
| 0.207545 | 0.546334 | 0.44 | 0.339 | 1 | 14 | Ubxn4 | 0.101 | Young |
| 0.20788 | 1.485743 | 0.08 | 0.038 | 1 | 14 | Shkbp1 | 0.042 | Young |
| 0.20788 | 1.166985 | 0.08 | 0.038 | 1 | 14 | Tmem41a | 0.042 | Young |
| 0.208011 | 0.878385 | 0.18 | 0.115 | 1 | 14 | Sipa1l3 | 0.065 | Young |
| 0.208013 | 0.152045 | 0.22 | 0.137 | 1 | 14 | Ick | 0.083 | Young |
| 0.20817 | 0.352823 | 0.26 | 0.18 | 1 | 14 | Ctsc | 0.08 | Young |
| 0.208332 | 0.351755 | 0.16 | 0.093 | 1 | 14 | Kat7 | 0.067 | Young |
| 0.208343 | 1.656152 | 0.16 | 0.098 | 1 | 14 | Iqsec2 | 0.062 | Young |
| 0.208483 | 0.969353 | 0.22 | 0.142 | 1 | 14 | Pom121 | 0.078 | Young |
| 0.209196 | 0.995106 | 0.24 | 0.164 | 1 | 14 | Eftud2 | 0.076 | Young |
| 0.209855 | 1.505923 | 0.38 | 0.301 | 1 | 14 | Efnb1 | 0.079 | Young |
| 0.209897 | 0.348082 | 0.24 | 0.158 | 1 | 14 | Sumf1 | 0.082 | Young |
| 0.209963 | 0.93705 | 0.5 | 0.415 | 1 | 14 | Fermt2 | 0.085 | Young |
| 0.210215 | 2.127309 | 0.08 | 0.038 | 1 | 14 | Ptpn14 | 0.042 | Young |
| 0.210293 | 0.292723 | 0.34 | 0.246 | 1 | 14 | Denr | 0.094 | Young |

| | | | | | | | | |
|---|---|---|---|---|---|---|---|---|
| 0.210554 | 0.341825 | 0.44 | 0.322 | 1 | 14 | Arih1 | 0.118 | Young |
| 0.211183 | 0.701997 | 0.12 | 0.066 | 1 | 14 | Lcorl | 0.054 | Young |
| 0.211183 | 0.020996 | 0.12 | 0.066 | 1 | 14 | Alg6 | 0.054 | Young |
| 0.211247 | 0.44643 | 0.1 | 0.049 | 1 | 14 | Acvr2b | 0.051 | Young |
| 0.211632 | 0.8004 | 0.3 | 0.23 | 1 | 14 | Vcl | 0.07 | Young |
| 0.211915 | 0.899638 | 0.14 | 0.082 | 1 | 14 | Strip1 | 0.058 | Young |
| 0.211926 | 0.082168 | 0.08 | 0.153 | 1 | 14 | Tmem128 | -0.073 | Young |
| 0.211946 | 0.631981 | 0.38 | 0.301 | 1 | 14 | Cdc37l1 | 0.079 | Young |
| 0.212025 | 0.034275 | 0.06 | 0.131 | 1 | 14 | Rpia | -0.071 | Young |
| 0.212488 | 0.482041 | 0.18 | 0.109 | 1 | 14 | Traf7 | 0.071 | Young |
| 0.212569 | 0.927761 | 0.08 | 0.038 | 1 | 14 | Gucy1b1 | 0.042 | Young |
| 0.21308 | 0.794378 | 0.16 | 0.098 | 1 | 14 | Rab13 | 0.062 | Young |
| 0.213551 | 0.349315 | 0.7 | 0.617 | 1 | 14 | Psmb4 | 0.083 | Young |
| 0.213844 | 1.240645 | 0.28 | 0.23 | 1 | 14 | Bmyc | 0.05 | Young |
| 0.214168 | 0.189768 | 0.52 | 0.448 | 1 | 14 | Tob2 | 0.072 | Young |
| 0.214497 | 0.080624 | 0.86 | 0.82 | 1 | 14 | Rpl21 | 0.04 | Young |
| 0.214942 | 1.392823 | 0.08 | 0.038 | 1 | 14 | Zswim1 | 0.042 | Young |
| 0.214942 | 1.077135 | 0.08 | 0.038 | 1 | 14 | Nxpe4 | 0.042 | Young |
| 0.215352 | 0.81982 | 0.14 | 0.082 | 1 | 14 | Il18 | 0.058 | Young |
| 0.215452 | 0.621412 | 0.18 | 0.115 | 1 | 14 | Mrps6 | 0.065 | Young |
| 0.216839 | 0.783809 | 0.12 | 0.066 | 1 | 14 | Rtca | 0.054 | Young |
| 0.216963 | 0.644505 | 0.18 | 0.115 | 1 | 14 | Gpd2 | 0.065 | Young |
| 0.217334 | 1.880922 | 0.08 | 0.038 | 1 | 14 | Met | 0.042 | Young |
| 0.217356 | 0.856302 | 0.3 | 0.224 | 1 | 14 | Cdipt | 0.076 | Young |
| 0.217898 | 1.471494 | 0.2 | 0.137 | 1 | 14 | Csf2ra | 0.063 | Young |
| 0.218828 | 1.345968 | 0.14 | 0.082 | 1 | 14 | Pycr2 | 0.058 | Young |
| 0.218828 | 0.785424 | 0.14 | 0.082 | 1 | 14 | Topbp1 | 0.058 | Young |
| 0.218839 | 0.40802 | 0.46 | 0.355 | 1 | 14 | Ppp2r2a | 0.105 | Young |
| 0.219724 | 0.100682 | 0.32 | 0.454 | 1 | 14 | Qpct | -0.134 | Young |
| 0.219745 | 1.448654 | 0.08 | 0.038 | 1 | 14 | Gm20300 | 0.042 | Young |
| 0.219745 | 1.129017 | 0.08 | 0.038 | 1 | 14 | Pwp2 | 0.042 | Young |
| 0.219745 | 0.776771 | 0.08 | 0.038 | 1 | 14 | Rhbdd3 | 0.042 | Young |
| 0.219779 | 0.855125 | 0.3 | 0.235 | 1 | 14 | Ppdpf | 0.065 | Young |
| 0.220312 | 0.575851 | 0.32 | 0.235 | 1 | 14 | Marveld1 | 0.085 | Young |
| 0.220458 | 0.429516 | 0.24 | 0.158 | 1 | 14 | Mfap3 | 0.082 | Young |
| 0.220581 | 0.797387 | 0.14 | 0.082 | 1 | 14 | Sema6d | 0.058 | Young |
| 0.2206 | 0.481589 | 0.46 | 0.377 | 1 | 14 | Atox1 | 0.083 | Young |
| 0.22067 | 0.826427 | 0.12 | 0.066 | 1 | 14 | Prkdc | 0.054 | Young |
| 0.22067 | 0.665064 | 0.12 | 0.066 | 1 | 14 | Mpv17l2 | 0.054 | Young |
| 0.22067 | 0.552106 | 0.12 | 0.066 | 1 | 14 | Nek6 | 0.054 | Young |
| 0.220708 | 0.647553 | 0.38 | 0.279 | 1 | 14 | Tceal8 | 0.101 | Young |
| 0.220746 | 0.119942 | 0.2 | 0.12 | 1 | 14 | Eif2ak2 | 0.08 | Young |
| 0.221367 | 0.439413 | 0.74 | 0.798 | 1 | 14 | mt-Nd5 | -0.058 | Young |
| 0.221481 | 0.512097 | 0.32 | 0.235 | 1 | 14 | Acp2 | 0.085 | Young |
| 0.221661 | 0.819927 | 0.52 | 0.432 | 1 | 14 | Ndufs6 | 0.088 | Young |

| | | | | | | | | |
|---|---|---|---|---|---|---|---|---|
| 0.221873 | 0.0012 | 0.4 | 0.503 | 1 | 14 | Galnt1 | -0.103 | Young |
| 0.222132 | 0.846128 | 0.32 | 0.23 | 1 | 14 | Gbf1 | 0.09 | Young |
| 0.222175 | 1.905881 | 0.08 | 0.038 | 1 | 14 | Palm3 | 0.042 | Young |
| 0.222175 | 0.867946 | 0.08 | 0.038 | 1 | 14 | Mmp3 | 0.042 | Young |
| 0.222293 | 1.16025 | 0.24 | 0.164 | 1 | 14 | Otud6b | 0.076 | Young |
| 0.222603 | 1.626105 | 0.12 | 0.066 | 1 | 14 | Tcea3 | 0.054 | Young |
| 0.222655 | 0.324237 | 0.72 | 0.842 | 1 | 14 | Dstn | -0.122 | Young |
| 0.222866 | 0.658705 | 0.56 | 0.454 | 1 | 14 | Mycbp2 | 0.106 | Young |
| 0.222939 | 0.019268 | 0.3 | 0.399 | 1 | 14 | Csnk2a1 | -0.099 | Young |
| 0.223806 | 1.47109 | 0.2 | 0.131 | 1 | 14 | Synm | 0.069 | Young |
| 0.223832 | 0.365329 | 0.32 | 0.235 | 1 | 14 | Cldnd1 | 0.085 | Young |
| 0.223847 | 0.430293 | 0.26 | 0.175 | 1 | 14 | Parp14 | 0.085 | Young |
| 0.223961 | 1.215078 | 0.22 | 0.153 | 1 | 14 | Atp7a | 0.067 | Young |
| 0.224017 | 0.310426 | 0.26 | 0.186 | 1 | 14 | Plxnd1 | 0.074 | Young |
| 0.224017 | 0.385976 | 0.48 | 0.377 | 1 | 14 | Atp6v0e | 0.103 | Young |
| 0.224118 | 0.359037 | 0.14 | 0.082 | 1 | 14 | Nt5e | 0.058 | Young |
| 0.224286 | 0.498919 | 0.3 | 0.213 | 1 | 14 | Zfp516 | 0.087 | Young |
| 0.224425 | 1.145829 | 0.3 | 0.219 | 1 | 14 | Ptprs | 0.081 | Young |
| 0.224548 | 0.926571 | 0.12 | 0.066 | 1 | 14 | Lig3 | 0.054 | Young |
| 0.224625 | 0.974125 | 0.08 | 0.038 | 1 | 14 | Tasp1 | 0.042 | Young |
| 0.224625 | 0.962917 | 0.08 | 0.038 | 1 | 14 | Zfp944 | 0.042 | Young |
| 0.224625 | 0.697511 | 0.08 | 0.038 | 1 | 14 | Dcun1d4 | 0.042 | Young |
| 0.224625 | 0.466403 | 0.08 | 0.038 | 1 | 14 | Pdhx | 0.042 | Young |
| 0.225094 | 0.951316 | 0.2 | 0.137 | 1 | 14 | Notch1 | 0.063 | Young |
| 0.225455 | 2.208454 | 0.14 | 0.087 | 1 | 14 | Pde6d | 0.053 | Young |
| 0.225641 | 0.158378 | 0.18 | 0.251 | 1 | 14 | Rnps1 | -0.071 | Young |
| 0.22579 | 1.925287 | 0.18 | 0.12 | 1 | 14 | Tgm2 | 0.06 | Young |
| 0.225902 | 1.057843 | 0.14 | 0.082 | 1 | 14 | Tfam | 0.058 | Young |
| 0.225923 | 1.18517 | 0.26 | 0.18 | 1 | 14 | Nup88 | 0.08 | Young |
| 0.225933 | 0.704928 | 0.66 | 0.617 | 1 | 14 | Flna | 0.043 | Young |
| 0.226101 | 0.474673 | 0.4 | 0.317 | 1 | 14 | Tmem167 | 0.083 | Young |
| 0.226398 | 0.15315 | 0.16 | 0.093 | 1 | 14 | Snx15 | 0.067 | Young |
| 0.226616 | 0.171159 | 0.26 | 0.344 | 1 | 14 | Rab21 | -0.084 | Young |
| 0.227093 | 0.697827 | 0.08 | 0.038 | 1 | 14 | Trmt61a | 0.042 | Young |
| 0.227142 | 0.110679 | 0.72 | 0.65 | 1 | 14 | Ddx3x | 0.07 | Young |
| 0.227193 | 1.215319 | 0.1 | 0.055 | 1 | 14 | Mrvi1 | 0.045 | Young |
| 0.227265 | 0.056649 | 0.46 | 0.585 | 1 | 14 | Hdlbp | -0.125 | Young |
| 0.227695 | 0.521561 | 0.14 | 0.082 | 1 | 14 | Nmd3 | 0.058 | Young |
| 0.228021 | 0.16689 | 0.18 | 0.109 | 1 | 14 | Hmgn3 | 0.071 | Young |
| 0.228092 | 0.031154 | 0.32 | 0.23 | 1 | 14 | Eif3b | 0.09 | Young |
| 0.228545 | 0.935672 | 0.26 | 0.18 | 1 | 14 | Fbxo33 | 0.08 | Young |
| 0.228762 | 0.396646 | 0.6 | 0.568 | 1 | 14 | Tra2a | 0.032 | Young |
| 0.229041 | 0.483787 | 0.3 | 0.208 | 1 | 14 | Gga1 | 0.092 | Young |
| 0.229423 | 0.349679 | 0.28 | 0.197 | 1 | 14 | Snx10 | 0.083 | Young |
| 0.229581 | 0.664579 | 0.08 | 0.038 | 1 | 14 | Zcchc3 | 0.042 | Young |

| | | | | | | | | |
|---|---|---|---|---|---|---|---|---|
| 0.229722 | 0.691723 | 0.52 | 0.443 | 1 | 14 | Tshz1 | 0.077 | Young |
| 0.229733 | 0.048046 | 0.2 | 0.131 | 1 | 14 | Kpna1 | 0.069 | Young |
| 0.230904 | 0.384861 | 0.18 | 0.115 | 1 | 14 | Rgs3 | 0.065 | Young |
| 0.231151 | 0.583111 | 0.78 | 0.721 | 1 | 14 | Sec62 | 0.059 | Young |
| 0.232177 | 1.061421 | 0.28 | 0.213 | 1 | 14 | Gnb4 | 0.067 | Young |
| 0.232199 | 0.950814 | 0.12 | 0.071 | 1 | 14 | Irak1bp1 | 0.049 | Young |
| 0.232387 | 0.819128 | 0.46 | 0.393 | 1 | 14 | Hoxc8 | 0.067 | Young |
| 0.232393 | 1.68667 | 0.22 | 0.153 | 1 | 14 | Chmp2b | 0.067 | Young |
| 0.232452 | 0.115516 | 0.12 | 0.066 | 1 | 14 | Hykk | 0.054 | Young |
| 0.232892 | 0.726885 | 0.2 | 0.126 | 1 | 14 | Uvssa | 0.074 | Young |
| 0.23295 | 0.722848 | 0.3 | 0.213 | 1 | 14 | Ndufs4 | 0.087 | Young |
| 0.232958 | 0.346407 | 0.4 | 0.311 | 1 | 14 | Svil | 0.089 | Young |
| 0.233137 | 0.985413 | 0.14 | 0.082 | 1 | 14 | Fbxw5 | 0.058 | Young |
| 0.233745 | 0.249514 | 0.22 | 0.142 | 1 | 14 | Rtl8b | 0.078 | Young |
| 0.234166 | 0.82167 | 0.3 | 0.235 | 1 | 14 | Fzd1 | 0.065 | Young |
| 0.234188 | 0.175528 | 0.24 | 0.158 | 1 | 14 | Lsm12 | 0.082 | Young |
| 0.234227 | 0.193275 | 0.3 | 0.443 | 1 | 14 | Zbtb4 | -0.143 | Young |
| 0.234503 | 0.550896 | 0.16 | 0.098 | 1 | 14 | Tyrobp | 0.062 | Young |
| 0.234574 | 0.920695 | 0.28 | 0.197 | 1 | 14 | Zfp945 | 0.083 | Young |
| 0.234615 | 0.504902 | 0.08 | 0.038 | 1 | 14 | Mrps27 | 0.042 | Young |
| 0.234615 | 0.290717 | 0.08 | 0.038 | 1 | 14 | Vipas39 | 0.042 | Young |
| 0.235153 | 0.436331 | 0.38 | 0.29 | 1 | 14 | Emsy | 0.09 | Young |
| 0.235285 | 0.334574 | 0.22 | 0.148 | 1 | 14 | Tvp23b | 0.072 | Young |
| 0.235875 | 1.049172 | 0.1 | 0.055 | 1 | 14 | Rgs5 | 0.045 | Young |
| 0.236122 | 0.942617 | 0.12 | 0.071 | 1 | 14 | Hlcs | 0.049 | Young |
| 0.236628 | 0.939573 | 0.52 | 0.475 | 1 | 14 | Mef2a | 0.045 | Young |
| 0.237161 | 0.302546 | 0.08 | 0.038 | 1 | 14 | S1pr1 | 0.042 | Young |
| 0.237441 | 0.258766 | 0.38 | 0.29 | 1 | 14 | Zbtb38 | 0.09 | Young |
| 0.237948 | 1.078746 | 0.14 | 0.087 | 1 | 14 | Zfp362 | 0.053 | Young |
| 0.238082 | 1.160346 | 0.1 | 0.055 | 1 | 14 | Dcun1d2 | 0.045 | Young |
| 0.238101 | 1.330063 | 0.12 | 0.071 | 1 | 14 | Sqle | 0.049 | Young |
| 0.238558 | 0.670171 | 0.54 | 0.475 | 1 | 14 | Sec61b | 0.065 | Young |
| 0.239009 | 0.796615 | 0.26 | 0.191 | 1 | 14 | Mbd3 | 0.069 | Young |
| 0.239427 | 0.223344 | 0.28 | 0.377 | 1 | 14 | Rnf6 | -0.097 | Young |
| 0.239669 | 0.814405 | 0.22 | 0.148 | 1 | 14 | Tulp3 | 0.072 | Young |
| 0.240303 | 0.814137 | 0.1 | 0.055 | 1 | 14 | Clp1 | 0.045 | Young |
| 0.240535 | 0.031707 | 0.14 | 0.082 | 1 | 14 | Wdr13 | 0.058 | Young |
| 0.240545 | 0.416551 | 0.18 | 0.115 | 1 | 14 | Tmem60 | 0.065 | Young |
| 0.240612 | 0.772691 | 0.26 | 0.18 | 1 | 14 | Ndufa12 | 0.08 | Young |
| 0.24074 | 0.303333 | 0.78 | 0.738 | 1 | 14 | Rpl10 | 0.042 | Young |
| 0.241118 | 1.591654 | 0.28 | 0.213 | 1 | 14 | Aldh7a1 | 0.067 | Young |
| 0.241281 | 0.260854 | 0.78 | 0.683 | 1 | 14 | Gnb2 | 0.097 | Young |
| 0.241887 | 0.414442 | 0.52 | 0.432 | 1 | 14 | Polr1d | 0.088 | Young |
| 0.242093 | 1.196565 | 0.12 | 0.071 | 1 | 14 | Paip2b | 0.049 | Young |
| 0.242312 | 0.319342 | 0.08 | 0.038 | 1 | 14 | Cep89 | 0.042 | Young |

| | | | | | | | | |
|---|---|---|---|---|---|---|---|---|
| 0.242312 | 0.317937 | 0.08 | 0.038 | 1 | 14 | Casp1 | 0.042 | Young |
| 0.242782 | 0.215569 | 0.44 | 0.344 | 1 | 14 | Dnajb1 | 0.096 | Young |
| 0.243027 | 0.534904 | 0.58 | 0.557 | 1 | 14 | Cox6b1 | 0.023 | Young |
| 0.243211 | 2.131171 | 0.06 | 0.027 | 1 | 14 | Simc1 | 0.033 | Young |
| 0.243389 | 0.297065 | 0.24 | 0.322 | 1 | 14 | Prrc1 | -0.082 | Young |
| 0.244039 | 1.669432 | 0.2 | 0.142 | 1 | 14 | Abhd14b | 0.058 | Young |
| 0.24405 | 0.277415 | 0.82 | 0.842 | 1 | 14 | Timp3 | -0.022 | Young |
| 0.244295 | 0.069799 | 0.14 | 0.082 | 1 | 14 | Unc45a | 0.058 | Young |
| 0.244388 | 0.059487 | 0.68 | 0.76 | 1 | 14 | Arf5 | -0.08 | Young |
| 0.244564 | 1.974824 | 0.18 | 0.131 | 1 | 14 | Exosc8 | 0.049 | Young |
| 0.244711 | 0.327663 | 0.28 | 0.202 | 1 | 14 | Dhx30 | 0.078 | Young |
| 0.244973 | 0.349423 | 0.52 | 0.426 | 1 | 14 | Syncrip | 0.094 | Young |
| 0.245037 | 0.389819 | 0.2 | 0.131 | 1 | 14 | Chrac1 | 0.069 | Young |
| 0.245214 | 1.320769 | 0.22 | 0.158 | 1 | 14 | Fam98b | 0.062 | Young |
| 0.245604 | 0.800832 | 0.22 | 0.148 | 1 | 14 | Slc48a1 | 0.072 | Young |
| 0.245604 | 0.44622 | 0.22 | 0.148 | 1 | 14 | Mfsd14a | 0.072 | Young |
| 0.245947 | 0.328105 | 0.68 | 0.689 | 1 | 14 | Cltc | -0.009 | Young |
| 0.246191 | 0.311985 | 0.14 | 0.082 | 1 | 14 | Ivd | 0.058 | Young |
| 0.246191 | 0.07686 | 0.14 | 0.082 | 1 | 14 | Atr | 0.058 | Young |
| 0.246452 | 0.313621 | 0.06 | 0.12 | 1 | 14 | Ccl8 | -0.06 | Young |
| 0.246463 | 0.921924 | 0.28 | 0.197 | 1 | 14 | Exoc6b | 0.083 | Young |
| 0.246641 | 0.513137 | 0.62 | 0.585 | 1 | 14 | Tacc1 | 0.035 | Young |
| 0.246661 | 0.509914 | 0.26 | 0.186 | 1 | 14 | Arhgef2 | 0.074 | Young |
| 0.246942 | 1.088608 | 0.22 | 0.153 | 1 | 14 | Hyou1 | 0.067 | Young |
| 0.24711 | 0.12033 | 0.6 | 0.732 | 1 | 14 | Ebf2 | -0.132 | Young |
| 0.247321 | 0.437981 | 0.6 | 0.503 | 1 | 14 | Ptpra | 0.097 | Young |
| 0.24754 | 0.185187 | 0.08 | 0.038 | 1 | 14 | 1190007I0 | 0.042 | Young |
| 0.24804 | 0.57802 | 0.18 | 0.12 | 1 | 14 | Socs7 | 0.06 | Young |
| 0.248097 | 0.378803 | 0.14 | 0.082 | 1 | 14 | Faf1 | 0.058 | Young |
| 0.24817 | 1.151485 | 0.12 | 0.071 | 1 | 14 | Taf13 | 0.049 | Young |
| 0.24817 | 0.937019 | 0.12 | 0.071 | 1 | 14 | Gm38399 | 0.049 | Young |
| 0.248181 | 0.346218 | 0.2 | 0.131 | 1 | 14 | Pnkd | 0.069 | Young |
| 0.248755 | 0.308799 | 0.86 | 0.858 | 1 | 14 | Ppia | 0.002 | Young |
| 0.24904 | 0.88867 | 0.14 | 0.087 | 1 | 14 | Mkrn2 | 0.053 | Young |
| 0.249325 | 2.554605 | 0.06 | 0.027 | 1 | 14 | C030014I2 | 0.033 | Young |
| 0.249325 | 1.658759 | 0.06 | 0.027 | 1 | 14 | Tbkbp1 | 0.033 | Young |
| 0.249325 | 1.595984 | 0.06 | 0.027 | 1 | 14 | Noa1 | 0.033 | Young |
| 0.249334 | 0.564472 | 0.1 | 0.055 | 1 | 14 | Ppfibp2 | 0.045 | Young |
| 0.249927 | 0.591285 | 0.22 | 0.153 | 1 | 14 | Mapk8 | 0.067 | Young |
| 0.250013 | 0.342643 | 0.14 | 0.082 | 1 | 14 | Rad50 | 0.058 | Young |
| 0.250026 | 0.910453 | 0.48 | 0.421 | 1 | 14 | Cstb | 0.059 | Young |
| 0.250122 | 0.404335 | 0.22 | 0.148 | 1 | 14 | Tmem205 | 0.072 | Young |
| 0.250923 | 1.040817 | 0.14 | 0.087 | 1 | 14 | Polr2d | 0.053 | Young |
| 0.25134 | 1.066903 | 0.18 | 0.12 | 1 | 14 | Slc12a4 | 0.06 | Young |
| 0.251514 | 0.03639 | 0.22 | 0.306 | 1 | 14 | Etf1 | -0.086 | Young |

| | | | | | | | | |
|---|---|---|---|---|---|---|---|---|
| 0.251535 | 0.207543 | 0.06 | 0.115 | 1 | 14 | Zfp358 | -0.055 | Young |
| 0.251628 | 0.771834 | 0.1 | 0.055 | 1 | 14 | Tmem104 | 0.045 | Young |
| 0.251682 | 0.363626 | 0.24 | 0.164 | 1 | 14 | Parp3 | 0.076 | Young |
| 0.252423 | 1.504041 | 0.06 | 0.027 | 1 | 14 | Zfp383 | 0.033 | Young |
| 0.252423 | 1.455946 | 0.06 | 0.027 | 1 | 14 | Ncapd2 | 0.033 | Young |
| 0.252423 | 1.313481 | 0.06 | 0.027 | 1 | 14 | Fam189a2 | 0.033 | Young |
| 0.25247 | 0.072062 | 0.32 | 0.399 | 1 | 14 | Usp25 | -0.079 | Young |
| 0.253209 | 0.400146 | 0.36 | 0.279 | 1 | 14 | Mgat1 | 0.081 | Young |
| 0.253274 | 0.195557 | 0.62 | 0.552 | 1 | 14 | Ube3a | 0.068 | Young |
| 0.253291 | 0.020305 | 0.1 | 0.175 | 1 | 14 | Kin | -0.075 | Young |
| 0.253368 | 0.292685 | 0.22 | 0.301 | 1 | 14 | Ptprg | -0.081 | Young |
| 0.253579 | 0.228742 | 0.3 | 0.219 | 1 | 14 | Fbxw11 | 0.081 | Young |
| 0.253616 | 1.793494 | 0.26 | 0.186 | 1 | 14 | Slirp | 0.074 | Young |
| 0.254719 | 0.835354 | 0.14 | 0.087 | 1 | 14 | Yif1b | 0.053 | Young |
| 0.25511 | 1.398759 | 0.34 | 0.301 | 1 | 14 | Des | 0.039 | Young |
| 0.255443 | 0.194187 | 0.4 | 0.301 | 1 | 14 | Por | 0.099 | Young |
| 0.255547 | 1.37596 | 0.06 | 0.027 | 1 | 14 | Tmem136 | 0.033 | Young |
| 0.255547 | 1.297264 | 0.06 | 0.027 | 1 | 14 | Herc3 | 0.033 | Young |
| 0.255547 | 1.148359 | 0.06 | 0.027 | 1 | 14 | Cbarp | 0.033 | Young |
| 0.255824 | 0.340249 | 0.14 | 0.082 | 1 | 14 | Orc5 | 0.058 | Young |
| 0.25626 | 0.664198 | 0.1 | 0.055 | 1 | 14 | Pex13 | 0.045 | Young |
| 0.25626 | 0.48155 | 0.1 | 0.055 | 1 | 14 | Kif21b | 0.045 | Young |
| 0.25626 | 0.379456 | 0.1 | 0.055 | 1 | 14 | Srcap | 0.045 | Young |
| 0.256316 | 1.085044 | 0.16 | 0.104 | 1 | 14 | Stoml2 | 0.056 | Young |
| 0.256347 | 1.787966 | 0.18 | 0.12 | 1 | 14 | Arhgef6 | 0.06 | Young |
| 0.256782 | 0.499661 | 0.32 | 0.251 | 1 | 14 | Golgb1 | 0.069 | Young |
| 0.25738 | 0.335967 | 0.28 | 0.197 | 1 | 14 | Spen | 0.083 | Young |
| 0.257654 | 0.215689 | 0.3 | 0.213 | 1 | 14 | Emilin1 | 0.087 | Young |
| 0.257718 | 0.499798 | 0.34 | 0.262 | 1 | 14 | Skil | 0.078 | Young |
| 0.257781 | 0.780183 | 0.22 | 0.148 | 1 | 14 | Lgalsl | 0.072 | Young |
| 0.257851 | 0.294924 | 0.64 | 0.59 | 1 | 14 | Glg1 | 0.05 | Young |
| 0.258147 | 1.155617 | 0.22 | 0.164 | 1 | 14 | Erf | 0.056 | Young |
| 0.258554 | 0.652802 | 0.14 | 0.087 | 1 | 14 | Ift52 | 0.053 | Young |
| 0.258599 | 2.192586 | 0.1 | 0.055 | 1 | 14 | Zfp976 | 0.045 | Young |
| 0.258599 | 0.834331 | 0.1 | 0.055 | 1 | 14 | Mios | 0.045 | Young |
| 0.258599 | 0.671775 | 0.1 | 0.055 | 1 | 14 | Ptrhd1 | 0.045 | Young |
| 0.258697 | 2.161813 | 0.06 | 0.027 | 1 | 14 | Dhx37 | 0.033 | Young |
| 0.258697 | 1.960552 | 0.06 | 0.027 | 1 | 14 | Gramd1b | 0.033 | Young |
| 0.258697 | 1.799935 | 0.06 | 0.027 | 1 | 14 | Cxcl10 | 0.033 | Young |
| 0.258697 | 1.328049 | 0.06 | 0.027 | 1 | 14 | Nhs | 0.033 | Young |
| 0.258697 | 1.188259 | 0.06 | 0.027 | 1 | 14 | Fahd2a | 0.033 | Young |
| 0.258697 | 1.021844 | 0.06 | 0.027 | 1 | 14 | Hoxb7 | 0.033 | Young |
| 0.258842 | 0.314157 | 0.34 | 0.251 | 1 | 14 | Olfm1 | 0.089 | Young |
| 0.258907 | 0.728249 | 0.3 | 0.219 | 1 | 14 | Dctn3 | 0.081 | Young |
| 0.259131 | 1.458624 | 0.24 | 0.169 | 1 | 14 | Pdzd8 | 0.071 | Young |

| | | | | | | | | |
|---|---|---|---|---|---|---|---|---|
| 0.259131 | 1.049792 | 0.24 | 0.169 | 1 | 14 | Pkdcc | 0.071 | Young |
| 0.259233 | 0.010307 | 0.16 | 0.098 | 1 | 14 | Nfxl1 | 0.062 | Young |
| 0.259291 | 0.312536 | 0.48 | 0.399 | 1 | 14 | Tsc22d3 | 0.081 | Young |
| 0.25976 | 0.496943 | 0.54 | 0.497 | 1 | 14 | Akap12 | 0.043 | Young |
| 0.259902 | 0.820999 | 0.16 | 0.104 | 1 | 14 | Dnajc11 | 0.056 | Young |
| 0.259902 | 0.781761 | 0.16 | 0.104 | 1 | 14 | Ccdc115 | 0.056 | Young |
| 0.260487 | 0.964635 | 0.14 | 0.087 | 1 | 14 | Usp36 | 0.053 | Young |
| 0.260642 | 0.898791 | 0.12 | 0.071 | 1 | 14 | Zfp143 | 0.049 | Young |
| 0.26089 | 0.1837 | 0.22 | 0.148 | 1 | 14 | Dcun1d1 | 0.072 | Young |
| 0.260952 | 1.256149 | 0.1 | 0.055 | 1 | 14 | Chchd4 | 0.045 | Young |
| 0.261117 | 0.410125 | 0.06 | 0.12 | 1 | 14 | Asb8 | -0.06 | Young |
| 0.261708 | 0.798049 | 0.26 | 0.18 | 1 | 14 | Gnpda2 | 0.08 | Young |
| 0.261754 | 0.607382 | 0.32 | 0.246 | 1 | 14 | Cnep1r1 | 0.074 | Young |
| 0.261875 | 1.935337 | 0.06 | 0.027 | 1 | 14 | Uros | 0.033 | Young |
| 0.261875 | 1.592506 | 0.06 | 0.027 | 1 | 14 | 2010320M | 0.033 | Young |
| 0.261875 | 1.265031 | 0.06 | 0.027 | 1 | 14 | Adal | 0.033 | Young |
| 0.261875 | 1.062811 | 0.06 | 0.027 | 1 | 14 | Pdgfrl | 0.033 | Young |
| 0.261875 | 1.03628 | 0.06 | 0.027 | 1 | 14 | Msr1 | 0.033 | Young |
| 0.261875 | 0.967448 | 0.06 | 0.027 | 1 | 14 | Adamts19 | 0.033 | Young |
| 0.261875 | 0.93679 | 0.06 | 0.027 | 1 | 14 | Plxnc1 | 0.033 | Young |
| 0.261875 | 0.885644 | 0.06 | 0.027 | 1 | 14 | Btc | 0.033 | Young |
| 0.261875 | 0.838431 | 0.06 | 0.027 | 1 | 14 | Wars2 | 0.033 | Young |
| 0.261875 | 0.71897 | 0.06 | 0.027 | 1 | 14 | Efnb2 | 0.033 | Young |
| 0.261907 | 4.63475 | 0.08 | 0.044 | 1 | 14 | Inhba | 0.036 | Young |
| 0.262393 | 0.128283 | 0.28 | 0.202 | 1 | 14 | Nop58 | 0.078 | Young |
| 0.262398 | 0.362754 | 0.18 | 0.115 | 1 | 14 | Nhp2 | 0.065 | Young |
| 0.262669 | 0.429054 | 0.4 | 0.301 | 1 | 14 | Park7 | 0.099 | Young |
| 0.262762 | 1.3 | 0.12 | 0.071 | 1 | 14 | Uspl1 | 0.049 | Young |
| 0.263192 | 0.703745 | 0.48 | 0.421 | 1 | 14 | Sema3c | 0.059 | Young |
| 0.26332 | 0.766895 | 0.1 | 0.055 | 1 | 14 | Rhobtb1 | 0.045 | Young |
| 0.263527 | 1.159667 | 0.24 | 0.175 | 1 | 14 | Odc1 | 0.065 | Young |
| 0.263588 | 0.716424 | 0.24 | 0.169 | 1 | 14 | Trappc8 | 0.071 | Young |
| 0.264134 | 0.188457 | 0.18 | 0.115 | 1 | 14 | Ubl7 | 0.065 | Young |
| 0.264327 | 0.254162 | 0.2 | 0.131 | 1 | 14 | Ddx27 | 0.069 | Young |
| 0.264351 | 0.263861 | 0.5 | 0.393 | 1 | 14 | Pak2 | 0.107 | Young |
| 0.264414 | 1.329227 | 0.2 | 0.142 | 1 | 14 | Faf2 | 0.058 | Young |
| 0.264523 | 0.254526 | 0.3 | 0.208 | 1 | 14 | Pcgf3 | 0.092 | Young |
| 0.264544 | 2.305464 | 0.08 | 0.044 | 1 | 14 | Hscb | 0.036 | Young |
| 0.265079 | 0.920951 | 0.06 | 0.027 | 1 | 14 | Hcst | 0.033 | Young |
| 0.265079 | 0.747995 | 0.06 | 0.027 | 1 | 14 | Tmem51 | 0.033 | Young |
| 0.265079 | 0.701247 | 0.06 | 0.027 | 1 | 14 | Elp6 | 0.033 | Young |
| 0.265079 | 0.669913 | 0.06 | 0.027 | 1 | 14 | Mier2 | 0.033 | Young |
| 0.265086 | 0.205933 | 0.24 | 0.169 | 1 | 14 | Syvn1 | 0.071 | Young |
| 0.265702 | 1.361667 | 0.1 | 0.055 | 1 | 14 | Ankrd26 | 0.045 | Young |
| 0.265702 | 0.71767 | 0.1 | 0.055 | 1 | 14 | Chkb | 0.045 | Young |

| | | | | | | | | |
|---|---|---|---|---|---|---|---|---|
| 0.26587 | 0.187053 | 0.1 | 0.175 | 1 | 14 | Rcan2 | -0.075 | Young |
| 0.266307 | 0.708266 | 0.22 | 0.158 | 1 | 14 | Idh3b | 0.062 | Young |
| 0.266888 | 0.42186 | 0.2 | 0.137 | 1 | 14 | Gars | 0.063 | Young |
| 0.267994 | 0.612388 | 0.24 | 0.175 | 1 | 14 | Sh3pxd2a | 0.065 | Young |
| 0.2681 | 0.392712 | 0.1 | 0.055 | 1 | 14 | Pla2g7 | 0.045 | Young |
| 0.268237 | 0.27287 | 0.04 | 0.087 | 1 | 14 | Pmm2 | -0.047 | Young |
| 0.268311 | 1.113074 | 0.06 | 0.027 | 1 | 14 | A630072M | 0.033 | Young |
| 0.268311 | 1.051006 | 0.06 | 0.027 | 1 | 14 | Rnf135 | 0.033 | Young |
| 0.268311 | 0.937579 | 0.06 | 0.027 | 1 | 14 | Traf3ip1 | 0.033 | Young |
| 0.268311 | 0.837404 | 0.06 | 0.027 | 1 | 14 | Eid2b | 0.033 | Young |
| 0.268311 | 0.577777 | 0.06 | 0.027 | 1 | 14 | Slc11a1 | 0.033 | Young |
| 0.268311 | 0.410443 | 0.06 | 0.027 | 1 | 14 | Gm49359 | 0.033 | Young |
| 0.268311 | 0.409062 | 0.06 | 0.027 | 1 | 14 | Rabep2 | 0.033 | Young |
| 0.268318 | 1.487056 | 0.14 | 0.087 | 1 | 14 | Chid1 | 0.053 | Young |
| 0.268533 | 0.557583 | 0.2 | 0.137 | 1 | 14 | Krr1 | 0.063 | Young |
| 0.268628 | 0.592778 | 0.28 | 0.197 | 1 | 14 | Iws1 | 0.083 | Young |
| 0.269019 | 0.707309 | 0.16 | 0.104 | 1 | 14 | Edem3 | 0.056 | Young |
| 0.269074 | 0.01975 | 0.22 | 0.306 | 1 | 14 | Pgap2 | -0.086 | Young |
| 0.269194 | 0.599938 | 0.12 | 0.071 | 1 | 14 | Cd209f | 0.049 | Young |
| 0.269333 | 1.238455 | 0.16 | 0.109 | 1 | 14 | Tbx3 | 0.051 | Young |
| 0.269495 | 0.948086 | 0.24 | 0.175 | 1 | 14 | Sf3a3 | 0.065 | Young |
| 0.269596 | 0.59443 | 0.76 | 0.694 | 1 | 14 | Rpl22l1 | 0.066 | Young |
| 0.269912 | 0.547142 | 0.26 | 0.197 | 1 | 14 | Atp11b | 0.063 | Young |
| 0.269942 | 0.547828 | 0.22 | 0.153 | 1 | 14 | Stau1 | 0.067 | Young |
| 0.270512 | 0.436349 | 0.1 | 0.055 | 1 | 14 | Scrn3 | 0.045 | Young |
| 0.270786 | 0.959737 | 0.24 | 0.18 | 1 | 14 | Kdm6a | 0.06 | Young |
| 0.270985 | 0.003806 | 0.12 | 0.186 | 1 | 14 | Tubb2b | -0.066 | Young |
| 0.271155 | 0.760839 | 0.16 | 0.109 | 1 | 14 | Ccdc102a | 0.051 | Young |
| 0.271569 | 1.837853 | 0.06 | 0.027 | 1 | 14 | Prim1 | 0.033 | Young |
| 0.271569 | 0.703718 | 0.06 | 0.027 | 1 | 14 | Tmem8 | 0.033 | Young |
| 0.271569 | 0.490494 | 0.06 | 0.027 | 1 | 14 | B230217C1 | 0.033 | Young |
| 0.272313 | 0.37507 | 0.3 | 0.224 | 1 | 14 | 4930402H2 | 0.076 | Young |
| 0.27261 | 0.72986 | 0.26 | 0.191 | 1 | 14 | Csrnp1 | 0.069 | Young |
| 0.272933 | 0.241135 | 0.18 | 0.115 | 1 | 14 | Dync2h1 | 0.065 | Young |
| 0.273247 | 0.516754 | 0.44 | 0.366 | 1 | 14 | P4ha1 | 0.074 | Young |
| 0.274148 | 0.08224 | 0.16 | 0.098 | 1 | 14 | Ngdn | 0.062 | Young |
| 0.274258 | 0.751379 | 0.52 | 0.459 | 1 | 14 | Srsf1 | 0.061 | Young |
| 0.274303 | 0.390244 | 0.6 | 0.514 | 1 | 14 | Atp5b | 0.086 | Young |
| 0.274855 | 0.345733 | 0.06 | 0.027 | 1 | 14 | Ifrd2 | 0.033 | Young |
| 0.275277 | 1.748589 | 0.08 | 0.044 | 1 | 14 | Nexn | 0.036 | Young |
| 0.275732 | 0.794242 | 0.12 | 0.071 | 1 | 14 | Mtm1 | 0.049 | Young |
| 0.276052 | 0.557094 | 0.1 | 0.158 | 1 | 14 | Stk16 | -0.058 | Young |
| 0.276052 | 0.377808 | 0.1 | 0.158 | 1 | 14 | Wnt11 | -0.058 | Young |
| 0.276216 | 0.922763 | 0.36 | 0.295 | 1 | 14 | Smtn | 0.065 | Young |
| 0.276547 | 0.487417 | 0.68 | 0.689 | 1 | 14 | Nenf | -0.009 | Young |

| | | | | | | | | |
|---|---|---|---|---|---|---|---|---|
| 0.276717 | 0.439494 | 0.38 | 0.301 | 1 | 14 | Ndufa1 | 0.079 | Young |
| 0.276821 | 0.70919 | 0.22 | 0.148 | 1 | 14 | Fut8 | 0.072 | Young |
| 0.277838 | 0.217693 | 0.1 | 0.055 | 1 | 14 | Wdr70 | 0.045 | Young |
| 0.277838 | 0.048143 | 0.1 | 0.055 | 1 | 14 | Tor2a | 0.045 | Young |
| 0.277936 | 0.746463 | 0.12 | 0.071 | 1 | 14 | Chst14 | 0.049 | Young |
| 0.277936 | 0.599238 | 0.12 | 0.071 | 1 | 14 | Zfp622 | 0.049 | Young |
| 0.278167 | 0.244936 | 0.06 | 0.027 | 1 | 14 | Pold1 | 0.033 | Young |
| 0.278352 | 1.21027 | 0.16 | 0.104 | 1 | 14 | Med14 | 0.056 | Young |
| 0.278352 | 0.554349 | 0.16 | 0.104 | 1 | 14 | Chml | 0.056 | Young |
| 0.278454 | 1.534439 | 0.14 | 0.093 | 1 | 14 | Adgrl1 | 0.047 | Young |
| 0.279477 | 1.023215 | 0.34 | 0.273 | 1 | 14 | Mpc1 | 0.067 | Young |
| 0.279741 | 0.564424 | 0.34 | 0.262 | 1 | 14 | Snrpc | 0.078 | Young |
| 0.280124 | 0.202225 | 0.32 | 0.235 | 1 | 14 | Plekha2 | 0.085 | Young |
| 0.280142 | 0.450358 | 0.28 | 0.208 | 1 | 14 | Dars | 0.072 | Young |
| 0.280151 | 0.5351 | 0.12 | 0.071 | 1 | 14 | Mycbp | 0.049 | Young |
| 0.280245 | 0.822752 | 0.16 | 0.104 | 1 | 14 | Tmem11 | 0.056 | Young |
| 0.280245 | 0.516687 | 0.16 | 0.104 | 1 | 14 | Fbxo21 | 0.056 | Young |
| 0.28031 | 1.549034 | 0.1 | 0.055 | 1 | 14 | Fbxo42 | 0.045 | Young |
| 0.280365 | 0.448362 | 0.14 | 0.087 | 1 | 14 | Neu1 | 0.053 | Young |
| 0.280497 | 0.301105 | 0.72 | 0.672 | 1 | 14 | BC005537 | 0.048 | Young |
| 0.280625 | 0.753786 | 0.18 | 0.12 | 1 | 14 | Mapre3 | 0.06 | Young |
| 0.280754 | 1.593984 | 0.08 | 0.044 | 1 | 14 | Trpv2 | 0.036 | Young |
| 0.280999 | 0.830141 | 0.4 | 0.339 | 1 | 14 | Ap2b1 | 0.061 | Young |
| 0.281446 | 0.651675 | 0.5 | 0.443 | 1 | 14 | Ccnd2 | 0.057 | Young |
| 0.281506 | 0.333612 | 0.06 | 0.027 | 1 | 14 | Rab3a | 0.033 | Young |
| 0.281506 | 0.115035 | 0.06 | 0.027 | 1 | 14 | Eif3j2 | 0.033 | Young |
| 0.281618 | 0.329095 | 0.68 | 0.596 | 1 | 14 | Pfn1 | 0.084 | Young |
| 0.281623 | 0.681324 | 0.26 | 0.197 | 1 | 14 | Ap1g1 | 0.063 | Young |
| 0.281937 | 0.915239 | 0.2 | 0.137 | 1 | 14 | Zfp560 | 0.063 | Young |
| 0.281937 | 0.370936 | 0.2 | 0.137 | 1 | 14 | Tle4 | 0.063 | Young |
| 0.282179 | 0.682851 | 0.62 | 0.563 | 1 | 14 | Rab2a | 0.057 | Young |
| 0.282266 | 0.042441 | 0.44 | 0.536 | 1 | 14 | Pnp | -0.096 | Young |
| 0.282379 | 0.396061 | 0.12 | 0.071 | 1 | 14 | Gorasp1 | 0.049 | Young |
| 0.282891 | 1.18331 | 0.46 | 0.404 | 1 | 14 | Sec63 | 0.056 | Young |
| 0.28352 | 1.549625 | 0.08 | 0.044 | 1 | 14 | Pyurf | 0.036 | Young |
| 0.283699 | 0.526022 | 0.26 | 0.175 | 1 | 14 | Eml4 | 0.085 | Young |
| 0.283757 | 0.02662 | 0.22 | 0.295 | 1 | 14 | Sugt1 | -0.075 | Young |
| 0.284057 | 1.370298 | 0.16 | 0.104 | 1 | 14 | Crlf3 | 0.056 | Young |
| 0.284237 | 0.817694 | 0.2 | 0.142 | 1 | 14 | Rrm2b | 0.058 | Young |
| 0.284462 | 1.180384 | 0.14 | 0.087 | 1 | 14 | Med7 | 0.053 | Young |
| 0.284618 | 0.703311 | 0.12 | 0.071 | 1 | 14 | Toe1 | 0.049 | Young |
| 0.284618 | 0.684508 | 0.12 | 0.071 | 1 | 14 | Nr1d2 | 0.049 | Young |
| 0.284618 | 0.608465 | 0.12 | 0.071 | 1 | 14 | Mipep | 0.049 | Young |
| 0.28479 | 0.885068 | 0.36 | 0.437 | 1 | 14 | St3gal2 | -0.077 | Young |
| 0.284873 | 0.142002 | 0.06 | 0.027 | 1 | 14 | Sprn | 0.033 | Young |

| | | | | | | | | |
|---|---|---|---|---|---|---|---|---|
| 0.284873 | 0.067005 | 0.06 | 0.027 | 1 | 14 | Gdf15 | 0.033 | Young |
| 0.285171 | 0.480743 | 0.32 | 0.421 | 1 | 14 | Btbd7 | -0.101 | Young |
| 0.285561 | 0.794332 | 0.3 | 0.213 | 1 | 14 | Mgrn1 | 0.087 | Young |
| 0.285673 | 0.283172 | 0.38 | 0.301 | 1 | 14 | Atp2b4 | 0.079 | Young |
| 0.28603 | 1.386245 | 0.16 | 0.109 | 1 | 14 | Hfe | 0.051 | Young |
| 0.286181 | 2.947171 | 0.1 | 0.06 | 1 | 14 | Rgs2 | 0.04 | Young |
| 0.286778 | 0.92999 | 0.38 | 0.295 | 1 | 14 | Scoc | 0.085 | Young |
| 0.286869 | 0.441198 | 0.12 | 0.071 | 1 | 14 | Tmem185b | 0.049 | Young |
| 0.287122 | 0.282181 | 0.26 | 0.18 | 1 | 14 | Bccip | 0.08 | Young |
| 0.287359 | 0.377944 | 0.48 | 0.372 | 1 | 14 | Atp5c1 | 0.108 | Young |
| 0.287362 | 0.50544 | 0.26 | 0.186 | 1 | 14 | Slc35b1 | 0.074 | Young |
| 0.287674 | 0.626303 | 0.68 | 0.716 | 1 | 14 | Col6a1 | -0.036 | Young |
| 0.287904 | 0.379146 | 0.16 | 0.104 | 1 | 14 | Prmt5 | 0.056 | Young |
| 0.287904 | 0.320493 | 0.16 | 0.104 | 1 | 14 | Commd1 | 0.056 | Young |
| 0.287948 | 0.776236 | 0.4 | 0.333 | 1 | 14 | Fyttd1 | 0.067 | Young |
| 0.288086 | 1.665892 | 0.18 | 0.126 | 1 | 14 | Hacd4 | 0.054 | Young |
| 0.288129 | 0.431879 | 0.36 | 0.268 | 1 | 14 | Dhx36 | 0.092 | Young |
| 0.288647 | 0.55288 | 0.38 | 0.322 | 1 | 14 | Clmp | 0.058 | Young |
| 0.289115 | 0.331917 | 0.28 | 0.197 | 1 | 14 | Slc35f5 | 0.083 | Young |
| 0.289133 | 1.312929 | 0.12 | 0.071 | 1 | 14 | Gm14296 | 0.049 | Young |
| 0.289133 | 0.582821 | 0.12 | 0.071 | 1 | 14 | Nbas | 0.049 | Young |
| 0.289133 | 0.522157 | 0.12 | 0.071 | 1 | 14 | Frmd8 | 0.049 | Young |
| 0.289133 | 0.13059 | 0.12 | 0.071 | 1 | 14 | Men1 | 0.049 | Young |
| 0.289663 | 0.740391 | 0.18 | 0.12 | 1 | 14 | Bola3 | 0.06 | Young |
| 0.290681 | 0.546634 | 0.14 | 0.087 | 1 | 14 | Casp8 | 0.053 | Young |
| 0.291118 | 0.831668 | 0.38 | 0.311 | 1 | 14 | Pcm1 | 0.069 | Young |
| 0.291276 | 0.170558 | 0.38 | 0.279 | 1 | 14 | Tspyl1 | 0.101 | Young |
| 0.29161 | 0.500476 | 0.48 | 0.388 | 1 | 14 | Ptbp3 | 0.092 | Young |
| 0.291928 | 3.218559 | 0.08 | 0.044 | 1 | 14 | 2810001G2 | 0.036 | Young |
| 0.291928 | 1.562058 | 0.08 | 0.044 | 1 | 14 | Lrsam1 | 0.036 | Young |
| 0.291928 | 1.362324 | 0.08 | 0.044 | 1 | 14 | Pitpnm1 | 0.036 | Young |
| 0.291928 | 1.010524 | 0.08 | 0.044 | 1 | 14 | Abt1 | 0.036 | Young |
| 0.292845 | 1.094305 | 0.28 | 0.219 | 1 | 14 | Stat5b | 0.061 | Young |
| 0.293505 | 0.77304 | 0.38 | 0.301 | 1 | 14 | Atp6v1a | 0.079 | Young |
| 0.293679 | 1.174516 | 0.12 | 0.077 | 1 | 14 | Ttc17 | 0.043 | Young |
| 0.29374 | 0.451203 | 0.16 | 0.104 | 1 | 14 | Obsl1 | 0.056 | Young |
| 0.294497 | 0.587764 | 0.6 | 0.601 | 1 | 14 | Ndufa4 | -0.001 | Young |
| 0.294522 | 0.354275 | 0.74 | 0.694 | 1 | 14 | Rpl7a | 0.046 | Young |
| 0.294653 | 0.313354 | 0.24 | 0.169 | 1 | 14 | Ing4 | 0.071 | Young |
| 0.294767 | 0.873262 | 0.08 | 0.044 | 1 | 14 | Spred3 | 0.036 | Young |
| 0.295163 | 0.588065 | 0.44 | 0.366 | 1 | 14 | Kdelr2 | 0.074 | Young |
| 0.295703 | 0.431133 | 0.16 | 0.104 | 1 | 14 | Elp5 | 0.056 | Young |
| 0.295925 | 1.865439 | 0.12 | 0.077 | 1 | 14 | H2-Ke6 | 0.043 | Young |
| 0.295925 | 1.184044 | 0.12 | 0.077 | 1 | 14 | Gtf2e2 | 0.043 | Young |
| 0.295995 | 0.167281 | 0.12 | 0.071 | 1 | 14 | Asb3 | 0.049 | Young |

| | | | | | | | | |
|---|---|---|---|---|---|---|---|---|
| 0.296148 | 0.283036 | 0.68 | 0.585 | 1 | 14 | Gdi2 | 0.095 | Young |
| 0.296852 | 1.257156 | 0.28 | 0.213 | 1 | 14 | Suclg2 | 0.067 | Young |
| 0.296881 | 0.780212 | 0.22 | 0.158 | 1 | 14 | Gle1 | 0.062 | Young |
| 0.296965 | 0.092791 | 0.1 | 0.158 | 1 | 14 | Timm10 | -0.058 | Young |
| 0.297032 | 1.005459 | 0.18 | 0.12 | 1 | 14 | Lyar | 0.06 | Young |
| 0.297331 | 2.886061 | 0.04 | 0.016 | 1 | 14 | Slc25a33 | 0.024 | Young |
| 0.297331 | 2.682658 | 0.04 | 0.016 | 1 | 14 | Klc2 | 0.024 | Young |
| 0.297331 | 1.470923 | 0.04 | 0.016 | 1 | 14 | Wdr78 | 0.024 | Young |
| 0.297331 | 1.417539 | 0.04 | 0.016 | 1 | 14 | Zfp607b | 0.024 | Young |
| 0.297438 | 0.181652 | 0.48 | 0.568 | 1 | 14 | Camk2d | -0.088 | Young |
| 0.297625 | 0.951699 | 0.08 | 0.044 | 1 | 14 | Zbtb43 | 0.036 | Young |
| 0.297625 | 0.943333 | 0.08 | 0.044 | 1 | 14 | Mad1l1 | 0.036 | Young |
| 0.298055 | 1.449179 | 0.24 | 0.186 | 1 | 14 | Bsdc1 | 0.054 | Young |
| 0.298311 | 1.568604 | 0.2 | 0.148 | 1 | 14 | Nav3 | 0.052 | Young |
| 0.298553 | 0.610811 | 0.22 | 0.158 | 1 | 14 | Dennd4a | 0.062 | Young |
| 0.298841 | 1.006971 | 0.14 | 0.093 | 1 | 14 | Ell2 | 0.047 | Young |
| 0.299116 | 0.289673 | 0.14 | 0.087 | 1 | 14 | Pigs | 0.053 | Young |
| 0.299375 | 0.728134 | 0.28 | 0.208 | 1 | 14 | Rnf146 | 0.072 | Young |
| 0.299827 | 0.630747 | 0.26 | 0.197 | 1 | 14 | Ccdc34 | 0.063 | Young |
| 0.299955 | 0.210603 | 0.32 | 0.24 | 1 | 14 | Brd8 | 0.08 | Young |
| 0.300417 | 0.574923 | 0.04 | 0.087 | 1 | 14 | Reep4 | -0.047 | Young |
| 0.300502 | 1.945245 | 0.08 | 0.044 | 1 | 14 | Hip1r | 0.036 | Young |
| 0.300502 | 0.942012 | 0.08 | 0.044 | 1 | 14 | Sgip1 | 0.036 | Young |
| 0.300814 | 0.347712 | 0.66 | 0.689 | 1 | 14 | Calm2 | -0.029 | Young |
| 0.300869 | 0.722317 | 0.3 | 0.224 | 1 | 14 | Fbxo18 | 0.076 | Young |
| 0.301073 | 2.389281 | 0.1 | 0.06 | 1 | 14 | Rpl27-ps3 | 0.04 | Young |
| 0.30125 | 0.332034 | 0.14 | 0.087 | 1 | 14 | Akt3 | 0.053 | Young |
| 0.30125 | 0.216734 | 0.14 | 0.087 | 1 | 14 | Ttc39c | 0.053 | Young |
| 0.301405 | 0.488965 | 0.26 | 0.191 | 1 | 14 | Mau2 | 0.069 | Young |
| 0.301486 | 0.415687 | 0.56 | 0.508 | 1 | 14 | Gsk3b | 0.052 | Young |
| 0.301505 | 0.640629 | 0.2 | 0.142 | 1 | 14 | Parg | 0.058 | Young |
| 0.301607 | 0.82017 | 0.34 | 0.262 | 1 | 14 | Ssrp1 | 0.078 | Young |
| 0.301644 | 0.593697 | 0.16 | 0.104 | 1 | 14 | Stk38l | 0.056 | Young |
| 0.301644 | 0.144599 | 0.16 | 0.104 | 1 | 14 | Stab1 | 0.056 | Young |
| 0.301724 | 2.512888 | 0.04 | 0.016 | 1 | 14 | Stmn1 | 0.024 | Young |
| 0.301724 | 1.73196 | 0.04 | 0.016 | 1 | 14 | Spryd4 | 0.024 | Young |
| 0.301724 | 1.534418 | 0.04 | 0.016 | 1 | 14 | Pik3cg | 0.024 | Young |
| 0.301724 | 1.343463 | 0.04 | 0.016 | 1 | 14 | Cfd | 0.024 | Young |
| 0.302346 | 0.645699 | 0.58 | 0.557 | 1 | 14 | Cox8a | 0.023 | Young |
| 0.302587 | 1.216601 | 0.22 | 0.164 | 1 | 14 | Akirin1 | 0.056 | Young |
| 0.302731 | 1.109776 | 0.12 | 0.077 | 1 | 14 | 4933431E2 | 0.043 | Young |
| 0.302731 | 0.778461 | 0.12 | 0.077 | 1 | 14 | Trim34a | 0.043 | Young |
| 0.303394 | 1.202628 | 0.06 | 0.109 | 1 | 14 | Sart3 | -0.049 | Young |
| 0.303394 | 0.70174 | 0.14 | 0.087 | 1 | 14 | Ptprd | 0.053 | Young |
| 0.303397 | 1.027412 | 0.08 | 0.044 | 1 | 14 | Cd93 | 0.036 | Young |

| | | | | | | | | |
|---|---|---|---|---|---|---|---|---|
| 0.303397 | 0.980105 | 0.08 | 0.044 | 1 | 14 | Tmem238 | 0.036 | Young |
| 0.303397 | 0.902346 | 0.08 | 0.044 | 1 | 14 | Tbc1d22a | 0.036 | Young |
| 0.303397 | 0.820761 | 0.08 | 0.044 | 1 | 14 | Dlg5 | 0.036 | Young |
| 0.303397 | 0.803871 | 0.08 | 0.044 | 1 | 14 | Bbs2 | 0.036 | Young |
| 0.303397 | 0.685945 | 0.08 | 0.044 | 1 | 14 | Zfp467 | 0.036 | Young |
| 0.303397 | 0.439903 | 0.08 | 0.044 | 1 | 14 | Stom | 0.036 | Young |
| 0.303606 | 0.47856 | 0.22 | 0.158 | 1 | 14 | Rcan1 | 0.062 | Young |
| 0.303642 | 0.798326 | 0.16 | 0.104 | 1 | 14 | Mrpl9 | 0.056 | Young |
| 0.303688 | 0.637251 | 0.12 | 0.18 | 1 | 14 | Zfp277 | -0.06 | Young |
| 0.303811 | 0.054832 | 0.32 | 0.426 | 1 | 14 | H2afv | -0.106 | Young |
| 0.304227 | 0.908611 | 0.48 | 0.432 | 1 | 14 | Oaf | 0.048 | Young |
| 0.304426 | 0.789725 | 0.24 | 0.186 | 1 | 14 | Sdc3 | 0.054 | Young |
| 0.304527 | 0.310935 | 0.26 | 0.186 | 1 | 14 | Ptdss1 | 0.074 | Young |
| 0.304673 | 0.242069 | 0.2 | 0.137 | 1 | 14 | Tmtc3 | 0.063 | Young |
| 0.305147 | 0.157425 | 0.06 | 0.115 | 1 | 14 | Clec4a1 | -0.055 | Young |
| 0.305216 | 0.58952 | 0.04 | 0.087 | 1 | 14 | Ikzf2 | -0.047 | Young |
| 0.30616 | 3.456679 | 0.04 | 0.016 | 1 | 14 | Gm15564 | 0.024 | Young |
| 0.30616 | 2.230514 | 0.04 | 0.016 | 1 | 14 | Cdkl5 | 0.024 | Young |
| 0.30616 | 1.631851 | 0.04 | 0.016 | 1 | 14 | Nrarp | 0.024 | Young |
| 0.30616 | 1.471045 | 0.04 | 0.016 | 1 | 14 | Pdgfb | 0.024 | Young |
| 0.30616 | 1.459903 | 0.04 | 0.016 | 1 | 14 | Bcl2a1d | 0.024 | Young |
| 0.30616 | 1.416443 | 0.04 | 0.016 | 1 | 14 | Nbeal2 | 0.024 | Young |
| 0.30616 | 1.345185 | 0.04 | 0.016 | 1 | 14 | Ppfia4 | 0.024 | Young |
| 0.30616 | 1.323641 | 0.04 | 0.016 | 1 | 14 | Pcdhb9 | 0.024 | Young |
| 0.30616 | 1.195851 | 0.04 | 0.016 | 1 | 14 | Adcy2 | 0.024 | Young |
| 0.30616 | 1.191305 | 0.04 | 0.016 | 1 | 14 | Harbi1 | 0.024 | Young |
| 0.30616 | 1.171532 | 0.04 | 0.016 | 1 | 14 | Slc16a3 | 0.024 | Young |
| 0.30616 | 1.149678 | 0.04 | 0.016 | 1 | 14 | Hr | 0.024 | Young |
| 0.30616 | 1.137657 | 0.04 | 0.016 | 1 | 14 | AC162302. | 0.024 | Young |
| 0.30616 | 1.041359 | 0.04 | 0.016 | 1 | 14 | Rbfox3 | 0.024 | Young |
| 0.30616 | 0.754084 | 0.04 | 0.016 | 1 | 14 | Rsph3b | 0.024 | Young |
| 0.30616 | 0.707342 | 0.04 | 0.016 | 1 | 14 | Gpc1 | 0.024 | Young |
| 0.30616 | 0.686891 | 0.04 | 0.016 | 1 | 14 | Rom1 | 0.024 | Young |
| 0.30631 | 0.767003 | 0.08 | 0.044 | 1 | 14 | Slc19a2 | 0.036 | Young |
| 0.30631 | 0.531655 | 0.08 | 0.044 | 1 | 14 | Ippk | 0.036 | Young |
| 0.306419 | 1.202512 | 0.18 | 0.126 | 1 | 14 | Mrps9 | 0.054 | Young |
| 0.307194 | 0.25814 | 0.32 | 0.24 | 1 | 14 | Psd3 | 0.08 | Young |
| 0.307246 | 0.019801 | 0.22 | 0.295 | 1 | 14 | Zc3h15 | -0.075 | Young |
| 0.307326 | 1.129785 | 0.12 | 0.077 | 1 | 14 | 2610044O1 | 0.043 | Young |
| 0.307639 | 0.221477 | 0.26 | 0.197 | 1 | 14 | Nsmce4a | 0.063 | Young |
| 0.307664 | 0.776286 | 0.16 | 0.104 | 1 | 14 | Nat6 | 0.056 | Young |
| 0.307664 | 0.580067 | 0.16 | 0.104 | 1 | 14 | Med31 | 0.056 | Young |
| 0.307712 | 0.264459 | 0.14 | 0.087 | 1 | 14 | Gpsm1 | 0.053 | Young |
| 0.307712 | 0.212619 | 0.14 | 0.087 | 1 | 14 | Ccdc9 | 0.053 | Young |
| 0.307716 | 0.577477 | 0.26 | 0.191 | 1 | 14 | Lemd2 | 0.069 | Young |

| | | | | | | | | |
|---|---|---|---|---|---|---|---|---|
| 0.307735 | 0.115668 | 0.3 | 0.23 | 1 | 14 | Stat1 | 0.07 | Young |
| 0.307964 | 0.513608 | 0.66 | 0.65 | 1 | 14 | Cnbp | 0.01 | Young |
| 0.308187 | 0.312127 | 0.34 | 0.257 | 1 | 14 | Zc3h13 | 0.083 | Young |
| 0.308279 | 0.32178 | 0.84 | 0.842 | 1 | 14 | Oaz1 | -0.002 | Young |
| 0.30861 | 0.534245 | 0.32 | 0.246 | 1 | 14 | C1qbp | 0.074 | Young |
| 0.308682 | 2.329253 | 0.14 | 0.098 | 1 | 14 | Thsd4 | 0.042 | Young |
| 0.308974 | 0.001746 | 0.24 | 0.317 | 1 | 14 | Ist1 | -0.077 | Young |
| 0.309039 | 0.621632 | 0.64 | 0.639 | 1 | 14 | Gpc3 | 0.001 | Young |
| 0.309058 | 0.830874 | 0.26 | 0.202 | 1 | 14 | Scrib | 0.058 | Young |
| 0.309391 | 0.21281 | 0.46 | 0.541 | 1 | 14 | Tspan3 | -0.081 | Young |
| 0.3094 | 0.771687 | 0.14 | 0.093 | 1 | 14 | Cd276 | 0.047 | Young |
| 0.309682 | 1.01091 | 0.38 | 0.311 | 1 | 14 | Fkbp2 | 0.069 | Young |
| 0.310466 | 0.000841 | 0.48 | 0.579 | 1 | 14 | Fyn | -0.099 | Young |
| 0.31064 | 2.345198 | 0.04 | 0.016 | 1 | 14 | Hgh1 | 0.024 | Young |
| 0.31064 | 2.259999 | 0.04 | 0.016 | 1 | 14 | Prkcg | 0.024 | Young |
| 0.31064 | 2.087156 | 0.04 | 0.016 | 1 | 14 | Col7a1 | 0.024 | Young |
| 0.31064 | 1.87129 | 0.04 | 0.016 | 1 | 14 | Cish | 0.024 | Young |
| 0.31064 | 1.838932 | 0.04 | 0.016 | 1 | 14 | Xkr6 | 0.024 | Young |
| 0.31064 | 1.772291 | 0.04 | 0.016 | 1 | 14 | Pwwp2b | 0.024 | Young |
| 0.31064 | 1.453052 | 0.04 | 0.016 | 1 | 14 | Zfp472 | 0.024 | Young |
| 0.31064 | 1.421504 | 0.04 | 0.016 | 1 | 14 | Gpr182 | 0.024 | Young |
| 0.31064 | 1.298316 | 0.04 | 0.016 | 1 | 14 | Nat14 | 0.024 | Young |
| 0.31064 | 1.119336 | 0.04 | 0.016 | 1 | 14 | Ubox5 | 0.024 | Young |
| 0.31064 | 1.118514 | 0.04 | 0.016 | 1 | 14 | Epha3 | 0.024 | Young |
| 0.31064 | 1.027213 | 0.04 | 0.016 | 1 | 14 | Cib2 | 0.024 | Young |
| 0.31064 | 1.015047 | 0.04 | 0.016 | 1 | 14 | Tmem2 | 0.024 | Young |
| 0.31064 | 0.988962 | 0.04 | 0.016 | 1 | 14 | Tlcd1 | 0.024 | Young |
| 0.31064 | 0.919227 | 0.04 | 0.016 | 1 | 14 | Dhrs11 | 0.024 | Young |
| 0.31064 | 0.913073 | 0.04 | 0.016 | 1 | 14 | Cdh5 | 0.024 | Young |
| 0.31064 | 0.908357 | 0.04 | 0.016 | 1 | 14 | Gm17501 | 0.024 | Young |
| 0.31064 | 0.869402 | 0.04 | 0.016 | 1 | 14 | Tnk1 | 0.024 | Young |
| 0.31064 | 0.869235 | 0.04 | 0.016 | 1 | 14 | Qrsl1 | 0.024 | Young |
| 0.31064 | 0.839532 | 0.04 | 0.016 | 1 | 14 | Zfp827 | 0.024 | Young |
| 0.31082 | 0.479331 | 0.08 | 0.137 | 1 | 14 | Dctn5 | -0.057 | Young |
| 0.311079 | 0.383819 | 0.42 | 0.35 | 1 | 14 | Epn2 | 0.07 | Young |
| 0.311282 | 0.995693 | 0.1 | 0.06 | 1 | 14 | Zfp160 | 0.04 | Young |
| 0.311529 | 0.767018 | 0.42 | 0.383 | 1 | 14 | Map1b | 0.037 | Young |
| 0.311541 | 0.721937 | 0.14 | 0.093 | 1 | 14 | Cbx4 | 0.047 | Young |
| 0.311907 | 0.803214 | 0.2 | 0.137 | 1 | 14 | Inpp5k | 0.063 | Young |
| 0.311967 | 1.738239 | 0.12 | 0.077 | 1 | 14 | Pphln1 | 0.043 | Young |
| 0.311967 | 1.164329 | 0.12 | 0.077 | 1 | 14 | Lipo3 | 0.043 | Young |
| 0.312192 | 0.933369 | 0.08 | 0.044 | 1 | 14 | Zswim9 | 0.036 | Young |
| 0.312192 | 0.840733 | 0.08 | 0.044 | 1 | 14 | Fdps | 0.036 | Young |
| 0.312192 | 0.833519 | 0.08 | 0.044 | 1 | 14 | Armcx4 | 0.036 | Young |
| 0.312192 | 0.676426 | 0.08 | 0.044 | 1 | 14 | Tbc1d22b | 0.036 | Young |

| | | | | | | | | |
|---|---|---|---|---|---|---|---|---|
| 0.312199 | 0.481417 | 0.24 | 0.175 | 1 | 14 | Mbtd1 | 0.065 | Young |
| 0.312365 | 1.165908 | 0.28 | 0.219 | 1 | 14 | Rev3l | 0.061 | Young |
| 0.312427 | 0.023893 | 0.12 | 0.071 | 1 | 14 | Lrrc14 | 0.049 | Young |
| 0.312507 | 0.260665 | 0.26 | 0.191 | 1 | 14 | Tet2 | 0.069 | Young |
| 0.31255 | 1.084258 | 0.26 | 0.186 | 1 | 14 | Mrps24 | 0.074 | Young |
| 0.313673 | 0.17581 | 0.4 | 0.317 | 1 | 14 | Cnot6l | 0.083 | Young |
| 0.313964 | 0.53405 | 0.18 | 0.126 | 1 | 14 | Gjc1 | 0.054 | Young |
| 0.314049 | 0.156119 | 0.14 | 0.208 | 1 | 14 | Fbxw2 | -0.068 | Young |
| 0.314305 | 1.910182 | 0.12 | 0.077 | 1 | 14 | Gramd3 | 0.043 | Young |
| 0.314649 | 0.455354 | 0.1 | 0.164 | 1 | 14 | Rcc2 | -0.064 | Young |
| 0.315161 | 0.728601 | 0.08 | 0.044 | 1 | 14 | Ankrd35 | 0.036 | Young |
| 0.315161 | 0.37123 | 0.08 | 0.044 | 1 | 14 | Tmem97 | 0.036 | Young |
| 0.315162 | 1.106306 | 0.04 | 0.016 | 1 | 14 | Podxl | 0.024 | Young |
| 0.315162 | 1.027845 | 0.04 | 0.016 | 1 | 14 | 6230400D1 | 0.024 | Young |
| 0.315162 | 1.008552 | 0.04 | 0.016 | 1 | 14 | Kdr | 0.024 | Young |
| 0.315162 | 0.918837 | 0.04 | 0.016 | 1 | 14 | Rsph9 | 0.024 | Young |
| 0.315162 | 0.878992 | 0.04 | 0.016 | 1 | 14 | Zfp668 | 0.024 | Young |
| 0.315162 | 0.723746 | 0.04 | 0.016 | 1 | 14 | Sh3d21 | 0.024 | Young |
| 0.315162 | 0.675146 | 0.04 | 0.016 | 1 | 14 | Gm3448 | 0.024 | Young |
| 0.315162 | 0.627676 | 0.04 | 0.016 | 1 | 14 | AC163032. | 0.024 | Young |
| 0.315162 | 0.596332 | 0.04 | 0.016 | 1 | 14 | Gm14286 | 0.024 | Young |
| 0.315162 | 0.578382 | 0.04 | 0.016 | 1 | 14 | Rttn | 0.024 | Young |
| 0.315162 | 0.565247 | 0.04 | 0.016 | 1 | 14 | Chst7 | 0.024 | Young |
| 0.315162 | 0.550648 | 0.04 | 0.016 | 1 | 14 | Rps6ka1 | 0.024 | Young |
| 0.315162 | 0.5497 | 0.04 | 0.016 | 1 | 14 | Nxpe5 | 0.024 | Young |
| 0.315162 | 0.544672 | 0.04 | 0.016 | 1 | 14 | Acsf2 | 0.024 | Young |
| 0.315162 | 0.523739 | 0.04 | 0.016 | 1 | 14 | C1qtnf12 | 0.024 | Young |
| 0.315162 | 0.51382 | 0.04 | 0.016 | 1 | 14 | Lyrm9 | 0.024 | Young |
| 0.315162 | 0.511298 | 0.04 | 0.016 | 1 | 14 | 9130023H2 | 0.024 | Young |
| 0.315162 | 0.491469 | 0.04 | 0.016 | 1 | 14 | Intu | 0.024 | Young |
| 0.315162 | 0.440061 | 0.04 | 0.016 | 1 | 14 | Tesk2 | 0.024 | Young |
| 0.315162 | 0.426029 | 0.04 | 0.016 | 1 | 14 | Mgmt | 0.024 | Young |
| 0.315162 | 0.326417 | 0.04 | 0.016 | 1 | 14 | Etl4 | 0.024 | Young |
| 0.315162 | 0.309953 | 0.04 | 0.016 | 1 | 14 | Cenpj | 0.024 | Young |
| 0.315162 | 0.152603 | 0.04 | 0.016 | 1 | 14 | Spata33 | 0.024 | Young |
| 0.315387 | 1.068786 | 0.32 | 0.262 | 1 | 14 | Acp1 | 0.058 | Young |
| 0.315714 | 0.033373 | 0.56 | 0.749 | 1 | 14 | Fndc3b | -0.189 | Young |
| 0.315803 | 0.819308 | 0.32 | 0.251 | 1 | 14 | Slmap | 0.069 | Young |
| 0.315852 | 1.45256 | 0.14 | 0.093 | 1 | 14 | Fbxl18 | 0.047 | Young |
| 0.315852 | 0.818577 | 0.14 | 0.093 | 1 | 14 | Usp45 | 0.047 | Young |
| 0.315852 | 0.594371 | 0.14 | 0.093 | 1 | 14 | As3mt | 0.047 | Young |
| 0.316472 | 0.313563 | 0.1 | 0.06 | 1 | 14 | Nmi | 0.04 | Young |
| 0.316653 | 0.049315 | 0.32 | 0.393 | 1 | 14 | Cmtm3 | -0.073 | Young |
| 0.316654 | 1.311428 | 0.12 | 0.077 | 1 | 14 | Wdr20 | 0.043 | Young |
| 0.316946 | 0.585458 | 0.54 | 0.53 | 1 | 14 | Msn | 0.01 | Young |

| | | | | | | | | |
|---|---|---|---|---|---|---|---|---|
| 0.316965 | 0.105076 | 0.28 | 0.197 | 1 | 14 | Smg6 | 0.083 | Young |
| 0.317346 | 0.360124 | 0.26 | 0.191 | 1 | 14 | Prdx3 | 0.069 | Young |
| 0.318022 | 0.760238 | 0.14 | 0.093 | 1 | 14 | Nlk | 0.047 | Young |
| 0.318148 | 1.049776 | 0.08 | 0.044 | 1 | 14 | Papolg | 0.036 | Young |
| 0.318148 | 0.570572 | 0.08 | 0.044 | 1 | 14 | 1500009L1 | 0.036 | Young |
| 0.31825 | 0.615468 | 0.28 | 0.213 | 1 | 14 | Hoxc6 | 0.067 | Young |
| 0.318321 | 0.535584 | 0.4 | 0.322 | 1 | 14 | Atp6v1e1 | 0.078 | Young |
| 0.318469 | 0.908706 | 0.22 | 0.153 | 1 | 14 | Golga3 | 0.067 | Young |
| 0.319088 | 0.731395 | 0.1 | 0.06 | 1 | 14 | Gdpd5 | 0.04 | Young |
| 0.319728 | 0.702155 | 0.04 | 0.016 | 1 | 14 | Slc25a35 | 0.024 | Young |
| 0.319728 | 0.595407 | 0.04 | 0.016 | 1 | 14 | 5430403G1 | 0.024 | Young |
| 0.319728 | 0.558285 | 0.04 | 0.016 | 1 | 14 | Mcph1 | 0.024 | Young |
| 0.319728 | 0.5556 | 0.04 | 0.016 | 1 | 14 | Sap130 | 0.024 | Young |
| 0.319728 | 0.428049 | 0.04 | 0.016 | 1 | 14 | AI504432 | 0.024 | Young |
| 0.319728 | 0.404829 | 0.04 | 0.016 | 1 | 14 | Pgf | 0.024 | Young |
| 0.319728 | 0.357782 | 0.04 | 0.016 | 1 | 14 | Exoc6 | 0.024 | Young |
| 0.319728 | 0.31397 | 0.04 | 0.016 | 1 | 14 | Krt36 | 0.024 | Young |
| 0.319728 | 0.278231 | 0.04 | 0.016 | 1 | 14 | Ttc27 | 0.024 | Young |
| 0.319728 | 0.276013 | 0.04 | 0.016 | 1 | 14 | Rtn4ip1 | 0.024 | Young |
| 0.319728 | 0.228358 | 0.04 | 0.016 | 1 | 14 | Tgif2 | 0.024 | Young |
| 0.319728 | 0.198809 | 0.04 | 0.016 | 1 | 14 | Bst1 | 0.024 | Young |
| 0.319728 | 0.167593 | 0.04 | 0.016 | 1 | 14 | 1700029I1 | 0.024 | Young |
| 0.319728 | 0.158155 | 0.04 | 0.016 | 1 | 14 | AC164424. | 0.024 | Young |
| 0.319728 | 0.025464 | 0.04 | 0.016 | 1 | 14 | Rtl6 | 0.024 | Young |
| 0.319728 | 0.01407 | 0.04 | 0.016 | 1 | 14 | Slc36a2 | 0.024 | Young |
| 0.321154 | 0.605497 | 0.08 | 0.044 | 1 | 14 | Casp2 | 0.036 | Young |
| 0.321292 | 0.395842 | 0.2 | 0.142 | 1 | 14 | Pitpnc1 | 0.058 | Young |
| 0.321387 | 1.471752 | 0.12 | 0.077 | 1 | 14 | Tceal1 | 0.043 | Young |
| 0.321387 | 0.450314 | 0.12 | 0.077 | 1 | 14 | Eno2 | 0.043 | Young |
| 0.321599 | 0.390845 | 0.4 | 0.344 | 1 | 14 | Hnrnpd | 0.056 | Young |
| 0.321718 | 0.777556 | 0.1 | 0.06 | 1 | 14 | Acbd4 | 0.04 | Young |
| 0.322616 | 1.025759 | 0.22 | 0.158 | 1 | 14 | Mospd3 | 0.062 | Young |
| 0.322697 | 0.60469 | 0.32 | 0.262 | 1 | 14 | Zcchc14 | 0.058 | Young |
| 0.323145 | 0.461562 | 0.14 | 0.087 | 1 | 14 | Dcaf11 | 0.053 | Young |
| 0.323534 | 0.500273 | 0.08 | 0.131 | 1 | 14 | Gtf3c4 | -0.051 | Young |
| 0.32377 | 0.620416 | 0.12 | 0.077 | 1 | 14 | Fcgr2b | 0.043 | Young |
| 0.323791 | 0.004493 | 0.42 | 0.497 | 1 | 14 | Spop | -0.077 | Young |
| 0.323961 | 0.259558 | 0.14 | 0.202 | 1 | 14 | Pum3 | -0.062 | Young |
| 0.324178 | 0.375627 | 0.08 | 0.044 | 1 | 14 | Meox2 | 0.036 | Young |
| 0.324178 | 0.025046 | 0.08 | 0.044 | 1 | 14 | Zbtb8a | 0.036 | Young |
| 0.324197 | 0.4332 | 0.18 | 0.251 | 1 | 14 | 1700019D0 | -0.071 | Young |
| 0.324314 | 1.447001 | 0.02 | 0.055 | 1 | 14 | Ccdc137 | -0.035 | Young |
| 0.324314 | 0.59366 | 0.02 | 0.055 | 1 | 14 | Rfx5 | -0.035 | Young |
| 0.324336 | 0.441623 | 0.04 | 0.016 | 1 | 14 | Alkbh4 | 0.024 | Young |
| 0.324336 | 0.343996 | 0.04 | 0.016 | 1 | 14 | Nkrf | 0.024 | Young |

| | | | | | | | | |
|---|---|---|---|---|---|---|---|---|
| 0.324336 | 0.318214 | 0.04 | 0.016 | 1 | 14 | Slc27a3 | 0.024 | Young |
| 0.324336 | 0.170907 | 0.04 | 0.016 | 1 | 14 | Mecom | 0.024 | Young |
| 0.324336 | 0.13747 | 0.04 | 0.016 | 1 | 14 | Recql5 | 0.024 | Young |
| 0.324336 | 0.123651 | 0.04 | 0.016 | 1 | 14 | Sh3rf3 | 0.024 | Young |
| 0.324336 | 0.077787 | 0.04 | 0.016 | 1 | 14 | Snx20 | 0.024 | Young |
| 0.324336 | 0.013789 | 0.04 | 0.016 | 1 | 14 | Nars2 | 0.024 | Young |
| 0.324362 | 1.414084 | 0.1 | 0.06 | 1 | 14 | Fgf1 | 0.04 | Young |
| 0.324382 | 1.016155 | 0.22 | 0.158 | 1 | 14 | Calcoco1 | 0.062 | Young |
| 0.324382 | 0.409662 | 0.22 | 0.158 | 1 | 14 | Tgs1 | 0.062 | Young |
| 0.324486 | 4.494599 | 0.02 | 0.005 | 1 | 14 | Cenps | 0.015 | Young |
| 0.324486 | 4.175089 | 0.02 | 0.005 | 1 | 14 | 1700023F0 | 0.015 | Young |
| 0.324486 | 3.840547 | 0.02 | 0.005 | 1 | 14 | Setmar | 0.015 | Young |
| 0.324486 | 3.632783 | 0.02 | 0.005 | 1 | 14 | Bnc1 | 0.015 | Young |
| 0.324486 | 3.632783 | 0.02 | 0.005 | 1 | 14 | Spock3 | 0.015 | Young |
| 0.324486 | 3.60796 | 0.02 | 0.005 | 1 | 14 | Mapk10 | 0.015 | Young |
| 0.324486 | 3.517921 | 0.02 | 0.005 | 1 | 14 | Egf | 0.015 | Young |
| 0.324486 | 3.440018 | 0.02 | 0.005 | 1 | 14 | 2700046G( | 0.015 | Young |
| 0.324486 | 3.230079 | 0.02 | 0.005 | 1 | 14 | 9130011E1 | 0.015 | Young |
| 0.324486 | 3.225846 | 0.02 | 0.005 | 1 | 14 | Ccdc187 | 0.015 | Young |
| 0.324486 | 3.154422 | 0.02 | 0.005 | 1 | 14 | 1500015A( | 0.015 | Young |
| 0.324486 | 3.121501 | 0.02 | 0.005 | 1 | 14 | Arnt2 | 0.015 | Young |
| 0.324486 | 3.105334 | 0.02 | 0.005 | 1 | 14 | 4930432K2 | 0.015 | Young |
| 0.324486 | 3.093958 | 0.02 | 0.005 | 1 | 14 | Tmem151a | 0.015 | Young |
| 0.324486 | 3.017536 | 0.02 | 0.005 | 1 | 14 | Wisp1 | 0.015 | Young |
| 0.324486 | 3.01352 | 0.02 | 0.005 | 1 | 14 | Apol10b | 0.015 | Young |
| 0.324486 | 2.95781 | 0.02 | 0.005 | 1 | 14 | Zfp97 | 0.015 | Young |
| 0.324486 | 2.924763 | 0.02 | 0.005 | 1 | 14 | Stum | 0.015 | Young |
| 0.324486 | 2.916624 | 0.02 | 0.005 | 1 | 14 | Sox11 | 0.015 | Young |
| 0.324486 | 2.913363 | 0.02 | 0.005 | 1 | 14 | AA986860 | 0.015 | Young |
| 0.324486 | 2.911639 | 0.02 | 0.005 | 1 | 14 | 4930518I1! | 0.015 | Young |
| 0.324486 | 2.904735 | 0.02 | 0.005 | 1 | 14 | D730045B( | 0.015 | Young |
| 0.324486 | 2.799198 | 0.02 | 0.005 | 1 | 14 | H2-Oa | 0.015 | Young |
| 0.324486 | 2.700243 | 0.02 | 0.005 | 1 | 14 | Cdh6 | 0.015 | Young |
| 0.324486 | 2.593252 | 0.02 | 0.005 | 1 | 14 | Gabrb2 | 0.015 | Young |
| 0.324486 | 2.564267 | 0.02 | 0.005 | 1 | 14 | Slc35g2 | 0.015 | Young |
| 0.324486 | 2.563683 | 0.02 | 0.005 | 1 | 14 | Gins2 | 0.015 | Young |
| 0.324486 | 2.54899 | 0.02 | 0.005 | 1 | 14 | Fbxo27 | 0.015 | Young |
| 0.324486 | 2.527447 | 0.02 | 0.005 | 1 | 14 | Gm26698 | 0.015 | Young |
| 0.324486 | 2.488656 | 0.02 | 0.005 | 1 | 14 | Slc17a9 | 0.015 | Young |
| 0.324486 | 2.479931 | 0.02 | 0.005 | 1 | 14 | Gm28151 | 0.015 | Young |
| 0.324486 | 2.471475 | 0.02 | 0.005 | 1 | 14 | Kcnab1 | 0.015 | Young |
| 0.324486 | 2.456466 | 0.02 | 0.005 | 1 | 14 | Gm10451 | 0.015 | Young |
| 0.324486 | 2.420307 | 0.02 | 0.005 | 1 | 14 | Dok5 | 0.015 | Young |
| 0.324486 | 2.321596 | 0.02 | 0.005 | 1 | 14 | Tbata | 0.015 | Young |
| 0.324486 | 2.306307 | 0.02 | 0.005 | 1 | 14 | Ifitm10 | 0.015 | Young |

| | | | | | | | | |
|---|---|---|---|---|---|---|---|---|
| 0.324486 | 2.300344 | 0.02 | 0.005 | 1 | 14 | Mybl2 | 0.015 | Young |
| 0.324486 | 2.299027 | 0.02 | 0.005 | 1 | 14 | 4732419C1 | 0.015 | Young |
| 0.324486 | 2.29277 | 0.02 | 0.005 | 1 | 14 | Lrch4 | 0.015 | Young |
| 0.324486 | 2.285829 | 0.02 | 0.005 | 1 | 14 | Dkk2 | 0.015 | Young |
| 0.324486 | 2.267788 | 0.02 | 0.005 | 1 | 14 | Rasip1 | 0.015 | Young |
| 0.324486 | 2.215587 | 0.02 | 0.005 | 1 | 14 | Sema7a | 0.015 | Young |
| 0.324486 | 2.130509 | 0.02 | 0.005 | 1 | 14 | Ccdc89 | 0.015 | Young |
| 0.324486 | 2.129893 | 0.02 | 0.005 | 1 | 14 | Zdhhc15 | 0.015 | Young |
| 0.324486 | 2.108339 | 0.02 | 0.005 | 1 | 14 | Mir142hg | 0.015 | Young |
| 0.324486 | 2.098005 | 0.02 | 0.005 | 1 | 14 | Nod2 | 0.015 | Young |
| 0.324486 | 2.074928 | 0.02 | 0.005 | 1 | 14 | Gm43126 | 0.015 | Young |
| 0.324486 | 2.069623 | 0.02 | 0.005 | 1 | 14 | Gimap9 | 0.015 | Young |
| 0.324486 | 2.061209 | 0.02 | 0.005 | 1 | 14 | Zfp661 | 0.015 | Young |
| 0.324486 | 2.061209 | 0.02 | 0.005 | 1 | 14 | Trim59 | 0.015 | Young |
| 0.324486 | 2.030825 | 0.02 | 0.005 | 1 | 14 | Ctu1 | 0.015 | Young |
| 0.324486 | 1.986656 | 0.02 | 0.005 | 1 | 14 | Gimap6 | 0.015 | Young |
| 0.324486 | 1.962392 | 0.02 | 0.005 | 1 | 14 | Etnk2 | 0.015 | Young |
| 0.324486 | 1.897587 | 0.02 | 0.005 | 1 | 14 | Gm44421 | 0.015 | Young |
| 0.324486 | 1.877331 | 0.02 | 0.005 | 1 | 14 | Clec4n | 0.015 | Young |
| 0.324486 | 1.867468 | 0.02 | 0.005 | 1 | 14 | Jak3 | 0.015 | Young |
| 0.324486 | 1.866277 | 0.02 | 0.005 | 1 | 14 | Kank4 | 0.015 | Young |
| 0.324486 | 1.85758 | 0.02 | 0.005 | 1 | 14 | 4933404O1 | 0.015 | Young |
| 0.324486 | 1.846496 | 0.02 | 0.005 | 1 | 14 | Lancl3 | 0.015 | Young |
| 0.324486 | 1.813705 | 0.02 | 0.005 | 1 | 14 | Fam71d | 0.015 | Young |
| 0.324486 | 1.806679 | 0.02 | 0.005 | 1 | 14 | Npas3 | 0.015 | Young |
| 0.324486 | 1.806347 | 0.02 | 0.005 | 1 | 14 | Slc2a12 | 0.015 | Young |
| 0.324486 | 1.796633 | 0.02 | 0.005 | 1 | 14 | Gm15512 | 0.015 | Young |
| 0.324486 | 1.776795 | 0.02 | 0.005 | 1 | 14 | Efcab7 | 0.015 | Young |
| 0.324486 | 1.756782 | 0.02 | 0.005 | 1 | 14 | Pde1b | 0.015 | Young |
| 0.324486 | 1.7431 | 0.02 | 0.005 | 1 | 14 | Zfp658 | 0.015 | Young |
| 0.324486 | 1.729922 | 0.02 | 0.005 | 1 | 14 | Ppp1r3d | 0.015 | Young |
| 0.324486 | 1.726846 | 0.02 | 0.005 | 1 | 14 | E230001N0 | 0.015 | Young |
| 0.324486 | 1.717351 | 0.02 | 0.005 | 1 | 14 | Tnnt3 | 0.015 | Young |
| 0.324486 | 1.715317 | 0.02 | 0.005 | 1 | 14 | Col6a4 | 0.015 | Young |
| 0.324486 | 1.679162 | 0.02 | 0.005 | 1 | 14 | Kif26a | 0.015 | Young |
| 0.324486 | 1.606246 | 0.02 | 0.005 | 1 | 14 | Olfr107 | 0.015 | Young |
| 0.324486 | 1.595659 | 0.02 | 0.005 | 1 | 14 | Gpam | 0.015 | Young |
| 0.324486 | 1.59156 | 0.02 | 0.005 | 1 | 14 | Gm19412 | 0.015 | Young |
| 0.324486 | 1.570527 | 0.02 | 0.005 | 1 | 14 | Zfp850 | 0.015 | Young |
| 0.324486 | 1.539914 | 0.02 | 0.005 | 1 | 14 | Mboat1 | 0.015 | Young |
| 0.324486 | 1.535237 | 0.02 | 0.005 | 1 | 14 | Sash3 | 0.015 | Young |
| 0.324486 | 1.534487 | 0.02 | 0.005 | 1 | 14 | Pih1d2 | 0.015 | Young |
| 0.324486 | 1.522839 | 0.02 | 0.005 | 1 | 14 | Mmp12 | 0.015 | Young |
| 0.324486 | 1.521389 | 0.02 | 0.005 | 1 | 14 | Gm4262 | 0.015 | Young |
| 0.324486 | 1.520683 | 0.02 | 0.005 | 1 | 14 | Myo1g | 0.015 | Young |

| | | | | | | | | |
|---|---|---|---|---|---|---|---|---|
| 0.324486 | 1.517467 | 0.02 | 0.005 | 1 | 14 | Xk | 0.015 | Young |
| 0.324486 | 1.47606 | 0.02 | 0.005 | 1 | 14 | Nnat | 0.015 | Young |
| 0.324486 | 1.463241 | 0.02 | 0.005 | 1 | 14 | Knl1 | 0.015 | Young |
| 0.324486 | 1.450517 | 0.02 | 0.005 | 1 | 14 | Ccdc85a | 0.015 | Young |
| 0.324486 | 1.442397 | 0.02 | 0.005 | 1 | 14 | Hpn | 0.015 | Young |
| 0.324486 | 1.437037 | 0.02 | 0.005 | 1 | 14 | Gm35248 | 0.015 | Young |
| 0.324486 | 1.411129 | 0.02 | 0.005 | 1 | 14 | Alx4 | 0.015 | Young |
| 0.324486 | 1.409832 | 0.02 | 0.005 | 1 | 14 | Zfp428 | 0.015 | Young |
| 0.324486 | 1.391892 | 0.02 | 0.005 | 1 | 14 | Tprn | 0.015 | Young |
| 0.324486 | 1.3554 | 0.02 | 0.005 | 1 | 14 | Prob1 | 0.015 | Young |
| 0.324486 | 1.3446 | 0.02 | 0.005 | 1 | 14 | Gm7324 | 0.015 | Young |
| 0.324486 | 1.336545 | 0.02 | 0.005 | 1 | 14 | Pemt | 0.015 | Young |
| 0.324486 | 1.335603 | 0.02 | 0.005 | 1 | 14 | Siah1b | 0.015 | Young |
| 0.324486 | 1.309997 | 0.02 | 0.005 | 1 | 14 | Gata3 | 0.015 | Young |
| 0.324486 | 1.292762 | 0.02 | 0.005 | 1 | 14 | Ifit3b | 0.015 | Young |
| 0.324486 | 1.251609 | 0.02 | 0.005 | 1 | 14 | Gm47040 | 0.015 | Young |
| 0.324486 | 1.243979 | 0.02 | 0.005 | 1 | 14 | Psd4 | 0.015 | Young |
| 0.324486 | 1.221893 | 0.02 | 0.005 | 1 | 14 | Opn3 | 0.015 | Young |
| 0.324486 | 1.212429 | 0.02 | 0.005 | 1 | 14 | Cd226 | 0.015 | Young |
| 0.324486 | 1.187631 | 0.02 | 0.005 | 1 | 14 | Arv1 | 0.015 | Young |
| 0.324486 | 1.162896 | 0.02 | 0.005 | 1 | 14 | Morn5 | 0.015 | Young |
| 0.324486 | 1.153137 | 0.02 | 0.005 | 1 | 14 | Kcnc4 | 0.015 | Young |
| 0.324486 | 1.151857 | 0.02 | 0.005 | 1 | 14 | Gm10762 | 0.015 | Young |
| 0.324486 | 1.13694 | 0.02 | 0.005 | 1 | 14 | Vwf | 0.015 | Young |
| 0.324486 | 1.12368 | 0.02 | 0.005 | 1 | 14 | 1700019L1 | 0.015 | Young |
| 0.324486 | 1.118796 | 0.02 | 0.005 | 1 | 14 | Gm973 | 0.015 | Young |
| 0.324486 | 1.118796 | 0.02 | 0.005 | 1 | 14 | 4930539J0 | 0.015 | Young |
| 0.324486 | 1.104881 | 0.02 | 0.005 | 1 | 14 | Pcdhga1 | 0.015 | Young |
| 0.324486 | 1.064083 | 0.02 | 0.005 | 1 | 14 | Kif7 | 0.015 | Young |
| 0.324486 | 1.058876 | 0.02 | 0.005 | 1 | 14 | Vsig4 | 0.015 | Young |
| 0.324486 | 0.983068 | 0.02 | 0.005 | 1 | 14 | Cspg5 | 0.015 | Young |
| 0.324486 | 0.866947 | 0.02 | 0.005 | 1 | 14 | Sigirr | 0.015 | Young |
| 0.324486 | 0.824403 | 0.02 | 0.005 | 1 | 14 | Gm46404 | 0.015 | Young |
| 0.324486 | 0.803092 | 0.02 | 0.005 | 1 | 14 | Gm34084 | 0.015 | Young |
| 0.325035 | 0.687958 | 0.3 | 0.246 | 1 | 14 | Mpv17 | 0.054 | Young |
| 0.325509 | 0.819621 | 0.18 | 0.126 | 1 | 14 | Ptpn2 | 0.054 | Young |
| 0.326166 | 1.037323 | 0.12 | 0.077 | 1 | 14 | Tfpi2 | 0.043 | Young |
| 0.326166 | 0.250028 | 0.12 | 0.077 | 1 | 14 | Usp6nl | 0.043 | Young |
| 0.326198 | 0.135002 | 0.16 | 0.104 | 1 | 14 | Prcc | 0.056 | Young |
| 0.326615 | 0.308349 | 0.4 | 0.339 | 1 | 14 | Emc3 | 0.061 | Young |
| 0.3268 | 1.676058 | 0.14 | 0.093 | 1 | 14 | Ggcx | 0.047 | Young |
| 0.32702 | 0.512982 | 0.1 | 0.06 | 1 | 14 | Arhgap45 | 0.04 | Young |
| 0.327221 | 1.306404 | 0.08 | 0.044 | 1 | 14 | Txnrd3 | 0.036 | Young |
| 0.327221 | 0.308896 | 0.08 | 0.044 | 1 | 14 | Shtn1 | 0.036 | Young |
| 0.327221 | 0.196165 | 0.08 | 0.044 | 1 | 14 | Wdr90 | 0.036 | Young |

| | | | | | | | | |
|---|---|---|---|---|---|---|---|---|
| 0.327221 | 0.013335 | 0.08 | 0.044 | 1 | 14 | Zfp449 | 0.036 | Young |
| 0.32737 | 0.754818 | 0.24 | 0.317 | 1 | 14 | Tuba1c | -0.077 | Young |
| 0.327374 | 1.236979 | 0.24 | 0.175 | 1 | 14 | Trip4 | 0.065 | Young |
| 0.328892 | 1.176258 | 0.16 | 0.115 | 1 | 14 | Ighm | 0.045 | Young |
| 0.328946 | 0.53034 | 0.46 | 0.426 | 1 | 14 | Eif4g1 | 0.034 | Young |
| 0.328999 | 0.393978 | 0.46 | 0.393 | 1 | 14 | Prkar2a | 0.067 | Young |
| 0.329399 | 0.665159 | 0.36 | 0.464 | 1 | 14 | Ssbp3 | -0.104 | Young |
| 0.329692 | 0.878677 | 0.1 | 0.06 | 1 | 14 | Atg4c | 0.04 | Young |
| 0.329692 | 0.688493 | 0.1 | 0.06 | 1 | 14 | Zbtb7c | 0.04 | Young |
| 0.329736 | 0.609389 | 0.16 | 0.109 | 1 | 14 | Grcc10 | 0.051 | Young |
| 0.329792 | 0.479824 | 0.7 | 0.678 | 1 | 14 | Cox7c | 0.022 | Young |
| 0.329997 | 0.652012 | 0.24 | 0.18 | 1 | 14 | Zbtb1 | 0.06 | Young |
| 0.330282 | 0.315382 | 0.08 | 0.044 | 1 | 14 | Pgam5 | 0.036 | Young |
| 0.330282 | 0.169654 | 0.08 | 0.044 | 1 | 14 | Abcb1b | 0.036 | Young |
| 0.330282 | 0.142649 | 0.08 | 0.044 | 1 | 14 | Zfp995 | 0.036 | Young |
| 0.330282 | 0.099695 | 0.08 | 0.044 | 1 | 14 | Polr3c | 0.036 | Young |
| 0.331247 | 1.39084 | 0.14 | 0.093 | 1 | 14 | Zfp422 | 0.047 | Young |
| 0.331384 | 0.163064 | 0.18 | 0.126 | 1 | 14 | Ptpn21 | 0.054 | Young |
| 0.331815 | 1.425669 | 0.02 | 0.005 | 1 | 14 | Cemip | 0.015 | Young |
| 0.331815 | 1.394581 | 0.02 | 0.005 | 1 | 14 | Rasgrf2 | 0.015 | Young |
| 0.331815 | 1.391027 | 0.02 | 0.005 | 1 | 14 | Card9 | 0.015 | Young |
| 0.331815 | 1.376075 | 0.02 | 0.005 | 1 | 14 | Wfikkn1 | 0.015 | Young |
| 0.331815 | 1.272348 | 0.02 | 0.005 | 1 | 14 | Spink2 | 0.015 | Young |
| 0.331815 | 1.250963 | 0.02 | 0.005 | 1 | 14 | Shisa6 | 0.015 | Young |
| 0.331815 | 1.174106 | 0.02 | 0.005 | 1 | 14 | Kcnk3 | 0.015 | Young |
| 0.331815 | 1.172688 | 0.02 | 0.005 | 1 | 14 | Foxd2 | 0.015 | Young |
| 0.331815 | 1.154998 | 0.02 | 0.005 | 1 | 14 | Fam167b | 0.015 | Young |
| 0.331815 | 1.154998 | 0.02 | 0.005 | 1 | 14 | Trim7 | 0.015 | Young |
| 0.331815 | 1.147173 | 0.02 | 0.005 | 1 | 14 | Cidec | 0.015 | Young |
| 0.331815 | 1.144737 | 0.02 | 0.005 | 1 | 14 | Slfn9 | 0.015 | Young |
| 0.331815 | 1.135476 | 0.02 | 0.005 | 1 | 14 | Mfsd4b3 | 0.015 | Young |
| 0.331815 | 1.129141 | 0.02 | 0.005 | 1 | 14 | Akr1c19 | 0.015 | Young |
| 0.331815 | 1.096167 | 0.02 | 0.005 | 1 | 14 | Mblac1 | 0.015 | Young |
| 0.331815 | 1.091783 | 0.02 | 0.005 | 1 | 14 | Skint3 | 0.015 | Young |
| 0.331815 | 1.06714 | 0.02 | 0.005 | 1 | 14 | A830035O: | 0.015 | Young |
| 0.331815 | 1.016301 | 0.02 | 0.005 | 1 | 14 | Ctf1 | 0.015 | Young |
| 0.331815 | 1.012929 | 0.02 | 0.005 | 1 | 14 | Kcng1 | 0.015 | Young |
| 0.331815 | 1.012578 | 0.02 | 0.005 | 1 | 14 | Asgr2 | 0.015 | Young |
| 0.331815 | 1.010951 | 0.02 | 0.005 | 1 | 14 | Zfp526 | 0.015 | Young |
| 0.331815 | 1.006376 | 0.02 | 0.005 | 1 | 14 | Mutyh | 0.015 | Young |
| 0.331815 | 0.999514 | 0.02 | 0.005 | 1 | 14 | 1700056E2 | 0.015 | Young |
| 0.331815 | 0.986669 | 0.02 | 0.005 | 1 | 14 | Gprin1 | 0.015 | Young |
| 0.331815 | 0.984871 | 0.02 | 0.005 | 1 | 14 | Eva1c | 0.015 | Young |
| 0.331815 | 0.918495 | 0.02 | 0.005 | 1 | 14 | Syt17 | 0.015 | Young |
| 0.331815 | 0.881525 | 0.02 | 0.005 | 1 | 14 | Gimap1 | 0.015 | Young |

| | | | | | | | | |
|---|---|---|---|---|---|---|---|---|
| 0.331815 | 0.862239 | 0.02 | 0.005 | 1 | 14 | Pcdhga2 | 0.015 | Young |
| 0.331815 | 0.853773 | 0.02 | 0.005 | 1 | 14 | Ccdc36 | 0.015 | Young |
| 0.331815 | 0.846954 | 0.02 | 0.005 | 1 | 14 | Timd4 | 0.015 | Young |
| 0.331815 | 0.837582 | 0.02 | 0.005 | 1 | 14 | Ttll7 | 0.015 | Young |
| 0.331815 | 0.831944 | 0.02 | 0.005 | 1 | 14 | 4933406C1 | 0.015 | Young |
| 0.331815 | 0.823399 | 0.02 | 0.005 | 1 | 14 | 4930525G2 | 0.015 | Young |
| 0.331815 | 0.81432 | 0.02 | 0.005 | 1 | 14 | Ccdc63 | 0.015 | Young |
| 0.331815 | 0.811507 | 0.02 | 0.005 | 1 | 14 | Gm30075 | 0.015 | Young |
| 0.331815 | 0.806937 | 0.02 | 0.005 | 1 | 14 | Rwdd2b | 0.015 | Young |
| 0.331815 | 0.80489 | 0.02 | 0.005 | 1 | 14 | Tuft1 | 0.015 | Young |
| 0.331815 | 0.792009 | 0.02 | 0.005 | 1 | 14 | Mpzl2 | 0.015 | Young |
| 0.331815 | 0.773521 | 0.02 | 0.005 | 1 | 14 | Ect2 | 0.015 | Young |
| 0.331815 | 0.773521 | 0.02 | 0.005 | 1 | 14 | 2700046A0 | 0.015 | Young |
| 0.331815 | 0.768314 | 0.02 | 0.005 | 1 | 14 | 9530026P0 | 0.015 | Young |
| 0.331815 | 0.768314 | 0.02 | 0.005 | 1 | 14 | Trank1 | 0.015 | Young |
| 0.331815 | 0.764351 | 0.02 | 0.005 | 1 | 14 | Thnsl1 | 0.015 | Young |
| 0.331815 | 0.752248 | 0.02 | 0.005 | 1 | 14 | Txk | 0.015 | Young |
| 0.331815 | 0.736558 | 0.02 | 0.005 | 1 | 14 | Eml6 | 0.015 | Young |
| 0.331815 | 0.736462 | 0.02 | 0.005 | 1 | 14 | Trpm2 | 0.015 | Young |
| 0.331815 | 0.736279 | 0.02 | 0.005 | 1 | 14 | Mir155hg | 0.015 | Young |
| 0.331815 | 0.734024 | 0.02 | 0.005 | 1 | 14 | Gm9856 | 0.015 | Young |
| 0.331815 | 0.732627 | 0.02 | 0.005 | 1 | 14 | Megf10 | 0.015 | Young |
| 0.331815 | 0.712228 | 0.02 | 0.005 | 1 | 14 | Etohd2 | 0.015 | Young |
| 0.331815 | 0.707642 | 0.02 | 0.005 | 1 | 14 | Dnah2 | 0.015 | Young |
| 0.331815 | 0.706497 | 0.02 | 0.005 | 1 | 14 | Tstd1 | 0.015 | Young |
| 0.331815 | 0.706404 | 0.02 | 0.005 | 1 | 14 | Trim13 | 0.015 | Young |
| 0.331815 | 0.69973 | 0.02 | 0.005 | 1 | 14 | Gm17586 | 0.015 | Young |
| 0.331815 | 0.686965 | 0.02 | 0.005 | 1 | 14 | Dclk2 | 0.015 | Young |
| 0.331815 | 0.686651 | 0.02 | 0.005 | 1 | 14 | Chac2 | 0.015 | Young |
| 0.331815 | 0.684804 | 0.02 | 0.005 | 1 | 14 | Ciart | 0.015 | Young |
| 0.331815 | 0.678237 | 0.02 | 0.005 | 1 | 14 | Dusp26 | 0.015 | Young |
| 0.331815 | 0.678237 | 0.02 | 0.005 | 1 | 14 | Egfl8 | 0.015 | Young |
| 0.331815 | 0.673724 | 0.02 | 0.005 | 1 | 14 | AC124739. | 0.015 | Young |
| 0.331815 | 0.672081 | 0.02 | 0.005 | 1 | 14 | B930082K0 | 0.015 | Young |
| 0.331815 | 0.671758 | 0.02 | 0.005 | 1 | 14 | Kcnab2 | 0.015 | Young |
| 0.331815 | 0.660594 | 0.02 | 0.005 | 1 | 14 | Tmem72 | 0.015 | Young |
| 0.331815 | 0.658915 | 0.02 | 0.005 | 1 | 14 | Slc46a1 | 0.015 | Young |
| 0.331815 | 0.657297 | 0.02 | 0.005 | 1 | 14 | Mfsd9 | 0.015 | Young |
| 0.331815 | 0.657297 | 0.02 | 0.005 | 1 | 14 | Gm4707 | 0.015 | Young |
| 0.331815 | 0.653493 | 0.02 | 0.005 | 1 | 14 | Klhl3 | 0.015 | Young |
| 0.331815 | 0.649041 | 0.02 | 0.005 | 1 | 14 | Hand2 | 0.015 | Young |
| 0.331815 | 0.648192 | 0.02 | 0.005 | 1 | 14 | Zfp184 | 0.015 | Young |
| 0.331815 | 0.643643 | 0.02 | 0.005 | 1 | 14 | Jpx | 0.015 | Young |
| 0.331815 | 0.640958 | 0.02 | 0.005 | 1 | 14 | Zfp955b | 0.015 | Young |
| 0.331815 | 0.634079 | 0.02 | 0.005 | 1 | 14 | Ryr1 | 0.015 | Young |

| | | | | | | | | |
|---|---|---|---|---|---|---|---|---|
| 0.331815 | 0.630339 | 0.02 | 0.005 | 1 | 14 | Gm9484 | 0.015 | Young |
| 0.331815 | 0.630016 | 0.02 | 0.005 | 1 | 14 | C78859 | 0.015 | Young |
| 0.331815 | 0.624663 | 0.02 | 0.005 | 1 | 14 | St6galnac2 | 0.015 | Young |
| 0.331815 | 0.624663 | 0.02 | 0.005 | 1 | 14 | Ankrd13d | 0.015 | Young |
| 0.331815 | 0.62316 | 0.02 | 0.005 | 1 | 14 | B3galt4 | 0.015 | Young |
| 0.331815 | 0.621367 | 0.02 | 0.005 | 1 | 14 | Acacb | 0.015 | Young |
| 0.331815 | 0.620844 | 0.02 | 0.005 | 1 | 14 | Hist2h2aa1 | 0.015 | Young |
| 0.331815 | 0.613185 | 0.02 | 0.005 | 1 | 14 | Jph4 | 0.015 | Young |
| 0.331815 | 0.611531 | 0.02 | 0.005 | 1 | 14 | Akr1c12 | 0.015 | Young |
| 0.331815 | 0.593682 | 0.02 | 0.005 | 1 | 14 | S100a3 | 0.015 | Young |
| 0.331815 | 0.579706 | 0.02 | 0.005 | 1 | 14 | S1pr4 | 0.015 | Young |
| 0.331815 | 0.579706 | 0.02 | 0.005 | 1 | 14 | Nek3 | 0.015 | Young |
| 0.331815 | 0.577241 | 0.02 | 0.005 | 1 | 14 | Tdrkh | 0.015 | Young |
| 0.331815 | 0.56944 | 0.02 | 0.005 | 1 | 14 | 3110021N2 | 0.015 | Young |
| 0.331815 | 0.567232 | 0.02 | 0.005 | 1 | 14 | Gm26809 | 0.015 | Young |
| 0.331815 | 0.560115 | 0.02 | 0.005 | 1 | 14 | Scarf1 | 0.015 | Young |
| 0.331815 | 0.54753 | 0.02 | 0.005 | 1 | 14 | Traip | 0.015 | Young |
| 0.331815 | 0.544258 | 0.02 | 0.005 | 1 | 14 | Prdm16 | 0.015 | Young |
| 0.331815 | 0.531206 | 0.02 | 0.005 | 1 | 14 | Zfp982 | 0.015 | Young |
| 0.331815 | 0.522683 | 0.02 | 0.005 | 1 | 14 | Gm26670 | 0.015 | Young |
| 0.331815 | 0.521822 | 0.02 | 0.005 | 1 | 14 | Fhit | 0.015 | Young |
| 0.331815 | 0.519366 | 0.02 | 0.005 | 1 | 14 | Pcdhga10 | 0.015 | Young |
| 0.331815 | 0.514905 | 0.02 | 0.005 | 1 | 14 | Prss53 | 0.015 | Young |
| 0.331815 | 0.514061 | 0.02 | 0.005 | 1 | 14 | 5430427O1 | 0.015 | Young |
| 0.331815 | 0.511834 | 0.02 | 0.005 | 1 | 14 | Gm37233 | 0.015 | Young |
| 0.331815 | 0.511452 | 0.02 | 0.005 | 1 | 14 | Gm10785 | 0.015 | Young |
| 0.331815 | 0.503758 | 0.02 | 0.005 | 1 | 14 | Gm33051 | 0.015 | Young |
| 0.331815 | 0.503758 | 0.02 | 0.005 | 1 | 14 | Apol9a | 0.015 | Young |
| 0.331815 | 0.493606 | 0.02 | 0.005 | 1 | 14 | Hist1h2be | 0.015 | Young |
| 0.331815 | 0.488895 | 0.02 | 0.005 | 1 | 14 | Smoc1 | 0.015 | Young |
| 0.331815 | 0.487423 | 0.02 | 0.005 | 1 | 14 | Siglecg | 0.015 | Young |
| 0.331815 | 0.458149 | 0.02 | 0.005 | 1 | 14 | Kif11 | 0.015 | Young |
| 0.331815 | 0.453984 | 0.02 | 0.005 | 1 | 14 | Nkain1 | 0.015 | Young |
| 0.331815 | 0.447638 | 0.02 | 0.005 | 1 | 14 | Slc22a3 | 0.015 | Young |
| 0.331815 | 0.441769 | 0.02 | 0.005 | 1 | 14 | Apol8 | 0.015 | Young |
| 0.331815 | 0.440531 | 0.02 | 0.005 | 1 | 14 | Rwdd3 | 0.015 | Young |
| 0.331815 | 0.418948 | 0.02 | 0.005 | 1 | 14 | Tfb1m | 0.015 | Young |
| 0.331815 | 0.41351 | 0.02 | 0.005 | 1 | 14 | Gm45767 | 0.015 | Young |
| 0.331815 | 0.407088 | 0.02 | 0.005 | 1 | 14 | Gm49207 | 0.015 | Young |
| 0.331815 | 0.403866 | 0.02 | 0.005 | 1 | 14 | Cd59b | 0.015 | Young |
| 0.331815 | 0.394128 | 0.02 | 0.005 | 1 | 14 | Zfp300 | 0.015 | Young |
| 0.331815 | 0.384178 | 0.02 | 0.005 | 1 | 14 | Ccdc149 | 0.015 | Young |
| 0.331815 | 0.365847 | 0.02 | 0.005 | 1 | 14 | Gm12184 | 0.015 | Young |
| 0.331815 | 0.349764 | 0.02 | 0.005 | 1 | 14 | Gm17092 | 0.015 | Young |
| 0.331815 | 0.332053 | 0.02 | 0.005 | 1 | 14 | Rbm46 | 0.015 | Young |

| | | | | | | | | |
|---|---|---|---|---|---|---|---|---|
| 0.331815 | 0.316347 | 0.02 | 0.005 | 1 | 14 | Nrg1 | 0.015 | Young |
| 0.331815 | 0.309713 | 0.02 | 0.005 | 1 | 14 | Dpep2 | 0.015 | Young |
| 0.331815 | 0.305878 | 0.02 | 0.005 | 1 | 14 | Cracr2a | 0.015 | Young |
| 0.331815 | 0.304157 | 0.02 | 0.005 | 1 | 14 | Ccdc148 | 0.015 | Young |
| 0.331815 | 0.299129 | 0.02 | 0.005 | 1 | 14 | Nat8f1 | 0.015 | Young |
| 0.331815 | 0.297148 | 0.02 | 0.005 | 1 | 14 | Mex3a | 0.015 | Young |
| 0.331815 | 0.296355 | 0.02 | 0.005 | 1 | 14 | Fignl1 | 0.015 | Young |
| 0.331815 | 0.295103 | 0.02 | 0.005 | 1 | 14 | Wdr25 | 0.015 | Young |
| 0.331815 | 0.292264 | 0.02 | 0.005 | 1 | 14 | Tnf | 0.015 | Young |
| 0.331815 | 0.281274 | 0.02 | 0.005 | 1 | 14 | 4930524J0 | 0.015 | Young |
| 0.331815 | 0.254358 | 0.02 | 0.005 | 1 | 14 | Gm10658 | 0.015 | Young |
| 0.331815 | 0.245045 | 0.02 | 0.005 | 1 | 14 | Zfp185 | 0.015 | Young |
| 0.331815 | 0.243287 | 0.02 | 0.005 | 1 | 14 | Zfp707 | 0.015 | Young |
| 0.331815 | 0.238984 | 0.02 | 0.005 | 1 | 14 | Pigw | 0.015 | Young |
| 0.331815 | 0.232773 | 0.02 | 0.005 | 1 | 14 | Zfp169 | 0.015 | Young |
| 0.331815 | 0.2217 | 0.02 | 0.005 | 1 | 14 | Phkg1 | 0.015 | Young |
| 0.331815 | 0.213515 | 0.02 | 0.005 | 1 | 14 | Pilrb2 | 0.015 | Young |
| 0.331815 | 0.206718 | 0.02 | 0.005 | 1 | 14 | Ttc9 | 0.015 | Young |
| 0.331815 | 0.205028 | 0.02 | 0.005 | 1 | 14 | Shmt1 | 0.015 | Young |
| 0.331815 | 0.203969 | 0.02 | 0.005 | 1 | 14 | Styk1 | 0.015 | Young |
| 0.331815 | 0.201654 | 0.02 | 0.005 | 1 | 14 | Kdm8 | 0.015 | Young |
| 0.331815 | 0.201654 | 0.02 | 0.005 | 1 | 14 | Zfp652os | 0.015 | Young |
| 0.331815 | 0.178867 | 0.02 | 0.005 | 1 | 14 | A730063M | 0.015 | Young |
| 0.331815 | 0.160426 | 0.02 | 0.005 | 1 | 14 | Aqp2 | 0.015 | Young |
| 0.331815 | 0.133898 | 0.02 | 0.005 | 1 | 14 | 2700033N1 | 0.015 | Young |
| 0.331815 | 0.13047 | 0.02 | 0.005 | 1 | 14 | Rhod | 0.015 | Young |
| 0.331815 | 0.127416 | 0.02 | 0.005 | 1 | 14 | Mag | 0.015 | Young |
| 0.331815 | 0.12712 | 0.02 | 0.005 | 1 | 14 | Acsl6 | 0.015 | Young |
| 0.331815 | 0.115286 | 0.02 | 0.005 | 1 | 14 | Atcayos | 0.015 | Young |
| 0.331815 | 0.100228 | 0.02 | 0.005 | 1 | 14 | Gm4285 | 0.015 | Young |
| 0.331815 | 0.09959 | 0.02 | 0.005 | 1 | 14 | Dennd2c | 0.015 | Young |
| 0.331815 | 0.065015 | 0.02 | 0.005 | 1 | 14 | Rad51c | 0.015 | Young |
| 0.331815 | 0.032014 | 0.02 | 0.005 | 1 | 14 | Akap6 | 0.015 | Young |
| 0.331815 | 0.002095 | 0.02 | 0.005 | 1 | 14 | Cecr2 | 0.015 | Young |
| 0.331815 | 0.000751 | 0.02 | 0.005 | 1 | 14 | Map2 | 0.015 | Young |
| 0.332366 | 0.647028 | 0.2 | 0.148 | 1 | 14 | Wars | 0.052 | Young |
| 0.332379 | 0.69366 | 0.1 | 0.06 | 1 | 14 | Gm4117 | 0.04 | Young |
| 0.332379 | 0.178902 | 0.1 | 0.06 | 1 | 14 | Abcc4 | 0.04 | Young |
| 0.332434 | 0.501495 | 0.2 | 0.142 | 1 | 14 | Spats2l | 0.058 | Young |
| 0.332538 | 0.224408 | 0.24 | 0.175 | 1 | 14 | Sf3b4 | 0.065 | Young |
| 0.332839 | 1.378811 | 0.18 | 0.131 | 1 | 14 | Smarca4 | 0.049 | Young |
| 0.333194 | 0.439812 | 0.52 | 0.459 | 1 | 14 | Lmna | 0.061 | Young |
| 0.333358 | 0.741702 | 0.18 | 0.126 | 1 | 14 | Aff3 | 0.054 | Young |
| 0.333362 | 0.180156 | 0.08 | 0.044 | 1 | 14 | Det1 | 0.036 | Young |
| 0.333362 | 0.135836 | 0.08 | 0.044 | 1 | 14 | Acss2 | 0.036 | Young |

| | | | | | | | | |
|---|---|---|---|---|---|---|---|---|
| 0.33342 | 1.011702 | 0.24 | 0.18 | 1 | 14 | Ccnh | 0.06 | Young |
| 0.33427 | 0.781247 | 0.24 | 0.175 | 1 | 14 | Pou2f1 | 0.065 | Young |
| 0.334428 | 0.187251 | 0.34 | 0.399 | 1 | 14 | Prkd3 | -0.059 | Young |
| 0.334663 | 0.835094 | 0.16 | 0.104 | 1 | 14 | Anapc10 | 0.056 | Young |
| 0.335079 | 0.815852 | 0.1 | 0.06 | 1 | 14 | Smug1 | 0.04 | Young |
| 0.335079 | 0.066429 | 0.1 | 0.06 | 1 | 14 | Dtd2 | 0.04 | Young |
| 0.335484 | 0.674659 | 0.22 | 0.164 | 1 | 14 | Ptpn13 | 0.056 | Young |
| 0.335854 | 0.030558 | 0.54 | 0.607 | 1 | 14 | Tgfbr2 | -0.067 | Young |
| 0.336014 | 1.100033 | 0.16 | 0.109 | 1 | 14 | Alg2 | 0.051 | Young |
| 0.336132 | 0.41557 | 0.34 | 0.257 | 1 | 14 | Flnc | 0.083 | Young |
| 0.336246 | 0.956351 | 0.36 | 0.295 | 1 | 14 | Dynll2 | 0.065 | Young |
| 0.33646 | 0.156641 | 0.08 | 0.044 | 1 | 14 | Ecsit | 0.036 | Young |
| 0.33684 | 1.256729 | 0.18 | 0.251 | 1 | 14 | Fmr1 | -0.071 | Young |
| 0.336868 | 0.537333 | 0.26 | 0.197 | 1 | 14 | Rxra | 0.063 | Young |
| 0.336919 | 0.753151 | 0.22 | 0.158 | 1 | 14 | G2e3 | 0.062 | Young |
| 0.337049 | 0.241747 | 0.34 | 0.273 | 1 | 14 | Snrpd2 | 0.067 | Young |
| 0.337794 | 1.003456 | 0.1 | 0.06 | 1 | 14 | Nabp1 | 0.04 | Young |
| 0.337992 | 1.218894 | 0.14 | 0.093 | 1 | 14 | Ago1 | 0.047 | Young |
| 0.338124 | 0.26037 | 0.08 | 0.131 | 1 | 14 | Mex3d | -0.051 | Young |
| 0.338542 | 0.952945 | 0.26 | 0.197 | 1 | 14 | Vps13b | 0.063 | Young |
| 0.33857 | 0.259081 | 0.24 | 0.169 | 1 | 14 | Crkl | 0.071 | Young |
| 0.338721 | 0.140798 | 0.1 | 0.153 | 1 | 14 | Med10 | -0.053 | Young |
| 0.338784 | 1.321592 | 0.26 | 0.213 | 1 | 14 | Ppp1r12b | 0.047 | Young |
| 0.33885 | 0.700393 | 0.32 | 0.24 | 1 | 14 | Szrd1 | 0.08 | Young |
| 0.339012 | 0.427169 | 0.42 | 0.355 | 1 | 14 | Bmper | 0.065 | Young |
| 0.340242 | 0.370333 | 0.16 | 0.109 | 1 | 14 | Abhd14a | 0.051 | Young |
| 0.340324 | 0.643222 | 0.44 | 0.399 | 1 | 14 | Uqcr10 | 0.041 | Young |
| 0.340324 | 0.512487 | 0.44 | 0.399 | 1 | 14 | Nufip2 | 0.041 | Young |
| 0.340396 | 0.718937 | 0.44 | 0.35 | 1 | 14 | Arl8a | 0.09 | Young |
| 0.340575 | 0.414293 | 0.26 | 0.191 | 1 | 14 | Tmem14c | 0.069 | Young |
| 0.340696 | 0.790239 | 0.18 | 0.131 | 1 | 14 | Farsb | 0.049 | Young |
| 0.340778 | 0.601049 | 0.12 | 0.077 | 1 | 14 | Ptpn23 | 0.043 | Young |
| 0.340975 | 0.274704 | 0.76 | 0.678 | 1 | 14 | Ubb | 0.082 | Young |
| 0.341198 | 0.451127 | 0.42 | 0.344 | 1 | 14 | Akt2 | 0.076 | Young |
| 0.341635 | 0.272293 | 0.34 | 0.421 | 1 | 14 | Tprgl | -0.081 | Young |
| 0.342019 | 0.553459 | 0.58 | 0.536 | 1 | 14 | Gls | 0.044 | Young |
| 0.342603 | 0.473077 | 0.44 | 0.372 | 1 | 14 | Pid1 | 0.068 | Young |
| 0.342934 | 0.141928 | 0.1 | 0.164 | 1 | 14 | Arfrp1 | -0.064 | Young |
| 0.343253 | 0.522428 | 0.12 | 0.077 | 1 | 14 | Smpd2 | 0.043 | Young |
| 0.343253 | 0.460842 | 0.12 | 0.077 | 1 | 14 | Cbfa2t2 | 0.043 | Young |
| 0.343253 | 0.457775 | 0.12 | 0.077 | 1 | 14 | Gatm | 0.043 | Young |
| 0.343253 | 0.389877 | 0.12 | 0.077 | 1 | 14 | Dock9 | 0.043 | Young |
| 0.343265 | 0.806627 | 0.1 | 0.06 | 1 | 14 | Bbs5 | 0.04 | Young |
| 0.343394 | 1.273031 | 0.14 | 0.098 | 1 | 14 | Itga1 | 0.042 | Young |
| 0.343495 | 0.432777 | 0.56 | 0.519 | 1 | 14 | Sqstm1 | 0.041 | Young |

| | | | | | | | | |
|---|---|---|---|---|---|---|---|---|
| 0.343537 | 0.40556 | 0.3 | 0.246 | 1 | 14 | Fkbp14 | 0.054 | Young |
| 0.343548 | 0.043416 | 0.38 | 0.295 | 1 | 14 | C1galt1 | 0.085 | Young |
| 0.343904 | 0.146231 | 0.2 | 0.137 | 1 | 14 | Fundc1 | 0.063 | Young |
| 0.344136 | 1.240877 | 0.12 | 0.082 | 1 | 14 | Ncald | 0.038 | Young |
| 0.344216 | 1.12249 | 0.46 | 0.377 | 1 | 14 | Mrps21 | 0.083 | Young |
| 0.344823 | 0.184007 | 0.14 | 0.093 | 1 | 14 | Adgre1 | 0.047 | Young |
| 0.344903 | 0.210352 | 0.6 | 0.667 | 1 | 14 | S100a11 | -0.067 | Young |
| 0.345537 | 0.134643 | 0.48 | 0.546 | 1 | 14 | Aes | -0.066 | Young |
| 0.345628 | 1.910748 | 0.06 | 0.033 | 1 | 14 | Gm4631 | 0.027 | Young |
| 0.345628 | 1.740022 | 0.06 | 0.033 | 1 | 14 | Kcnn1 | 0.027 | Young |
| 0.34574 | 0.993064 | 0.12 | 0.077 | 1 | 14 | Tbcc | 0.043 | Young |
| 0.34574 | 0.265196 | 0.12 | 0.077 | 1 | 14 | Mcoln1 | 0.043 | Young |
| 0.346022 | 0.342106 | 0.1 | 0.06 | 1 | 14 | Ap4e1 | 0.04 | Young |
| 0.346022 | 0.341957 | 0.1 | 0.06 | 1 | 14 | Agpat1 | 0.04 | Young |
| 0.346361 | 0.403264 | 0.24 | 0.29 | 1 | 14 | Lysmd3 | -0.05 | Young |
| 0.346405 | 0.102926 | 0.34 | 0.372 | 1 | 14 | Zfp638 | -0.032 | Young |
| 0.34712 | 0.452522 | 0.14 | 0.093 | 1 | 14 | Sclt1 | 0.047 | Young |
| 0.347592 | 0.827986 | 0.34 | 0.273 | 1 | 14 | Glipr2 | 0.067 | Young |
| 0.348316 | 0.708302 | 0.22 | 0.169 | 1 | 14 | Rnf44 | 0.051 | Young |
| 0.34867 | 1.069873 | 0.18 | 0.131 | 1 | 14 | Pogz | 0.049 | Young |
| 0.348793 | 1.286254 | 0.1 | 0.06 | 1 | 14 | R3hcc1l | 0.04 | Young |
| 0.348793 | 0.382383 | 0.1 | 0.06 | 1 | 14 | Fam46c | 0.04 | Young |
| 0.348793 | 0.362166 | 0.1 | 0.06 | 1 | 14 | Vta1 | 0.04 | Young |
| 0.348793 | 0.286008 | 0.1 | 0.06 | 1 | 14 | Ddx49 | 0.04 | Young |
| 0.348801 | 0.267288 | 0.16 | 0.109 | 1 | 14 | Senp1 | 0.051 | Young |
| 0.348866 | 0.359844 | 0.28 | 0.213 | 1 | 14 | Lypla2 | 0.067 | Young |
| 0.349267 | 1.996916 | 0.06 | 0.033 | 1 | 14 | Fkbp11 | 0.027 | Young |
| 0.349341 | 1.709045 | 0.2 | 0.148 | 1 | 14 | Vgll4 | 0.052 | Young |
| 0.350748 | 0.714482 | 0.12 | 0.077 | 1 | 14 | CAAA0111 | 0.043 | Young |
| 0.350961 | 0.520753 | 0.16 | 0.109 | 1 | 14 | Ercc6l2 | 0.051 | Young |
| 0.351358 | 0.357803 | 0.04 | 0.082 | 1 | 14 | Pcnx3 | -0.042 | Young |
| 0.351578 | 0.457752 | 0.1 | 0.06 | 1 | 14 | Ppox | 0.04 | Young |
| 0.351743 | 0.238946 | 0.14 | 0.093 | 1 | 14 | Nrbp2 | 0.047 | Young |
| 0.351886 | 1.210047 | 0.16 | 0.115 | 1 | 14 | Akr1e1 | 0.045 | Young |
| 0.35204 | 0.60255 | 0.38 | 0.311 | 1 | 14 | Ociad1 | 0.069 | Young |
| 0.352701 | 0.408196 | 0.18 | 0.131 | 1 | 14 | Pepd | 0.049 | Young |
| 0.353131 | 0.492294 | 0.16 | 0.109 | 1 | 14 | Afg3l2 | 0.051 | Young |
| 0.353131 | 0.04627 | 0.08 | 0.131 | 1 | 14 | Usp3 | -0.051 | Young |
| 0.35327 | 0.447415 | 0.12 | 0.077 | 1 | 14 | Gnptab | 0.043 | Young |
| 0.3537 | 0.490666 | 0.2 | 0.137 | 1 | 14 | Pla2g15 | 0.063 | Young |
| 0.353713 | 0.77282 | 0.24 | 0.175 | 1 | 14 | Tpcn1 | 0.065 | Young |
| 0.354026 | 0.90979 | 0.16 | 0.115 | 1 | 14 | Zfp462 | 0.045 | Young |
| 0.354068 | 0.46345 | 0.14 | 0.093 | 1 | 14 | Cog4 | 0.047 | Young |
| 0.354068 | 0.190041 | 0.14 | 0.093 | 1 | 14 | Grasp | 0.047 | Young |
| 0.354377 | 0.850456 | 0.1 | 0.06 | 1 | 14 | Rnf34 | 0.04 | Young |

| | | | | | | | | |
|---|---|---|---|---|---|---|---|---|
| 0.354377 | 0.231407 | 0.1 | 0.06 | 1 | 14 | Heatr1 | 0.04 | Young |
| 0.355119 | 0.852938 | 0.2 | 0.148 | 1 | 14 | Mrpl57 | 0.052 | Young |
| 0.356163 | 0.283399 | 0.34 | 0.257 | 1 | 14 | Ccser2 | 0.083 | Young |
| 0.356271 | 0.069852 | 0.18 | 0.12 | 1 | 14 | Phf13 | 0.06 | Young |
| 0.35719 | 0.137573 | 0.1 | 0.06 | 1 | 14 | Dnm3 | 0.04 | Young |
| 0.35742 | 0.230388 | 0.2 | 0.142 | 1 | 14 | Actr1b | 0.058 | Young |
| 0.357662 | 0.560722 | 0.36 | 0.301 | 1 | 14 | Fkbp5 | 0.059 | Young |
| 0.357732 | 0.426613 | 0.38 | 0.317 | 1 | 14 | Lix1l | 0.063 | Young |
| 0.358233 | 0.35496 | 0.54 | 0.519 | 1 | 14 | 2010107E0 | 0.021 | Young |
| 0.358346 | 0.057284 | 0.12 | 0.077 | 1 | 14 | Prkra | 0.043 | Young |
| 0.359052 | 0.36397 | 0.26 | 0.197 | 1 | 14 | Tmem176b | 0.063 | Young |
| 0.359119 | 0.063288 | 0.6 | 0.639 | 1 | 14 | Selenof | -0.039 | Young |
| 0.359318 | 0.668767 | 0.22 | 0.169 | 1 | 14 | Icmt | 0.051 | Young |
| 0.359561 | 0.337721 | 0.68 | 0.678 | 1 | 14 | Fkbp1a | 0.002 | Young |
| 0.359572 | 0.610081 | 0.26 | 0.191 | 1 | 14 | Dctn6 | 0.069 | Young |
| 0.360329 | 1.38653 | 0.06 | 0.033 | 1 | 14 | Zfp235 | 0.027 | Young |
| 0.360443 | 0.642161 | 0.2 | 0.153 | 1 | 14 | Cmtm7 | 0.047 | Young |
| 0.360472 | 0.394527 | 0.82 | 0.847 | 1 | 14 | Rps7 | -0.027 | Young |
| 0.360902 | 0.422915 | 0.12 | 0.077 | 1 | 14 | Tctex1d2 | 0.043 | Young |
| 0.360902 | 0.311432 | 0.12 | 0.077 | 1 | 14 | Zfp952 | 0.043 | Young |
| 0.360902 | 0.21351 | 0.12 | 0.077 | 1 | 14 | Wwox | 0.043 | Young |
| 0.36128 | 0.597614 | 0.4 | 0.328 | 1 | 14 | Wbp1l | 0.072 | Young |
| 0.361367 | 0.330291 | 0.2 | 0.142 | 1 | 14 | Wdr5 | 0.058 | Young |
| 0.361626 | 0.391394 | 0.46 | 0.383 | 1 | 14 | Fndc3a | 0.077 | Young |
| 0.361778 | 0.173836 | 0.1 | 0.148 | 1 | 14 | Ahsa2 | -0.048 | Young |
| 0.363025 | 0.597211 | 0.38 | 0.35 | 1 | 14 | Fcho2 | 0.03 | Young |
| 0.363389 | 0.616587 | 0.36 | 0.311 | 1 | 14 | Tbrg1 | 0.049 | Young |
| 0.363453 | 0.225886 | 0.42 | 0.333 | 1 | 14 | Pkd1 | 0.087 | Young |
| 0.363469 | 1.112271 | 0.12 | 0.077 | 1 | 14 | Klhdc10 | 0.043 | Young |
| 0.363469 | 0.240058 | 0.12 | 0.077 | 1 | 14 | B3galt6 | 0.043 | Young |
| 0.363865 | 0.36969 | 0.18 | 0.126 | 1 | 14 | Tprkb | 0.054 | Young |
| 0.363974 | 1.2131 | 0.14 | 0.098 | 1 | 14 | Nt5dc3 | 0.042 | Young |
| 0.364064 | 1.172442 | 0.06 | 0.033 | 1 | 14 | Comtd1 | 0.027 | Young |
| 0.364064 | 0.953785 | 0.06 | 0.033 | 1 | 14 | Nipsnap1 | 0.027 | Young |
| 0.364069 | 0.905827 | 0.08 | 0.049 | 1 | 14 | Nubp1 | 0.031 | Young |
| 0.364103 | 0.188868 | 0.16 | 0.109 | 1 | 14 | Fbxw7 | 0.051 | Young |
| 0.364299 | 0.83143 | 0.26 | 0.197 | 1 | 14 | Samd4 | 0.063 | Young |
| 0.364613 | 0.298868 | 0.24 | 0.175 | 1 | 14 | Myo9b | 0.065 | Young |
| 0.364883 | 0.891545 | 0.2 | 0.148 | 1 | 14 | Vps54 | 0.052 | Young |
| 0.364971 | 0.537595 | 0.1 | 0.153 | 1 | 14 | Ythdc2 | -0.053 | Young |
| 0.36507 | 0.408294 | 0.64 | 0.623 | 1 | 14 | Dynll1 | 0.017 | Young |
| 0.365675 | 0.491905 | 0.54 | 0.426 | 1 | 14 | Trir | 0.114 | Young |
| 0.365843 | 0.580876 | 0.14 | 0.093 | 1 | 14 | Ndc1 | 0.047 | Young |
| 0.365843 | 0.047009 | 0.14 | 0.093 | 1 | 14 | Slc35e1 | 0.047 | Young |
| 0.365959 | 0.438137 | 0.1 | 0.148 | 1 | 14 | Dzip1 | -0.048 | Young |

| | | | | | | | | |
|---|---|---|---|---|---|---|---|---|
| 0.366047 | 0.112032 | 0.12 | 0.077 | 1 | 14 | Pqlc1 | 0.043 | Young |
| 0.366307 | 0.966848 | 0.14 | 0.098 | 1 | 14 | Zcchc8 | 0.042 | Young |
| 0.366366 | 0.358246 | 0.54 | 0.53 | 1 | 14 | Fam129a | 0.01 | Young |
| 0.366515 | 0.279935 | 0.62 | 0.519 | 1 | 14 | Cops9 | 0.101 | Young |
| 0.367042 | 1.38033 | 0.18 | 0.131 | 1 | 14 | Clns1a | 0.049 | Young |
| 0.367364 | 0.736987 | 0.38 | 0.301 | 1 | 14 | Zfp106 | 0.079 | Young |
| 0.367824 | 1.092537 | 0.06 | 0.033 | 1 | 14 | Zfp513 | 0.027 | Young |
| 0.367824 | 0.828488 | 0.06 | 0.033 | 1 | 14 | Pak4 | 0.027 | Young |
| 0.368801 | 0.658721 | 0.32 | 0.246 | 1 | 14 | Taldo1 | 0.074 | Young |
| 0.369341 | 0.587455 | 0.2 | 0.142 | 1 | 14 | Sertad2 | 0.058 | Young |
| 0.369405 | 0.485603 | 0.3 | 0.23 | 1 | 14 | Ubqln2 | 0.07 | Young |
| 0.369616 | 0.5194 | 0.2 | 0.268 | 1 | 14 | Rab32 | -0.068 | Young |
| 0.370394 | 2.305466 | 0.08 | 0.049 | 1 | 14 | Hoxd9 | 0.031 | Young |
| 0.37062 | 0.326921 | 0.48 | 0.399 | 1 | 14 | Hnrnpl | 0.081 | Young |
| 0.37062 | 0.287093 | 0.14 | 0.093 | 1 | 14 | Slc43a2 | 0.047 | Young |
| 0.370679 | 0.309377 | 0.26 | 0.202 | 1 | 14 | Kat6b | 0.058 | Young |
| 0.370788 | 0.302588 | 0.16 | 0.109 | 1 | 14 | Timm17b | 0.051 | Young |
| 0.370788 | 0.094943 | 0.16 | 0.109 | 1 | 14 | Aars | 0.051 | Young |
| 0.371466 | 0.456086 | 0.1 | 0.06 | 1 | 14 | Tmem243 | 0.04 | Young |
| 0.371536 | 0.729128 | 0.12 | 0.082 | 1 | 14 | Snap47 | 0.038 | Young |
| 0.371608 | 1.472324 | 0.06 | 0.033 | 1 | 14 | Stap1 | 0.027 | Young |
| 0.371608 | 1.253599 | 0.06 | 0.033 | 1 | 14 | 1110002L0 | 0.027 | Young |
| 0.371608 | 1.17283 | 0.06 | 0.033 | 1 | 14 | Eno3 | 0.027 | Young |
| 0.371608 | 0.993521 | 0.06 | 0.033 | 1 | 14 | Recql | 0.027 | Young |
| 0.371608 | 0.828404 | 0.06 | 0.033 | 1 | 14 | Gmpr2 | 0.027 | Young |
| 0.371608 | 0.803618 | 0.06 | 0.033 | 1 | 14 | Cobll1 | 0.027 | Young |
| 0.371666 | 0.425506 | 0.38 | 0.333 | 1 | 14 | Acaa2 | 0.047 | Young |
| 0.37168 | 0.447663 | 0.26 | 0.208 | 1 | 14 | Ipo7 | 0.052 | Young |
| 0.372076 | 1.289028 | 0.18 | 0.137 | 1 | 14 | Tmem68 | 0.043 | Young |
| 0.372286 | 0.729724 | 0.18 | 0.126 | 1 | 14 | L3hypdh | 0.054 | Young |
| 0.372286 | 0.715163 | 0.18 | 0.126 | 1 | 14 | Cd200 | 0.054 | Young |
| 0.372504 | 1.300602 | 0.26 | 0.224 | 1 | 14 | Rest | 0.036 | Young |
| 0.372661 | 0.477136 | 0.2 | 0.29 | 1 | 14 | Hdac1 | -0.09 | Young |
| 0.373178 | 0.548434 | 0.36 | 0.29 | 1 | 14 | Tsc22d4 | 0.07 | Young |
| 0.373363 | 0.627805 | 0.14 | 0.098 | 1 | 14 | Fam234a | 0.042 | Young |
| 0.373582 | 1.087822 | 0.08 | 0.049 | 1 | 14 | Ttf1 | 0.031 | Young |
| 0.374092 | 1.044342 | 0.12 | 0.082 | 1 | 14 | Per3 | 0.038 | Young |
| 0.374363 | 0.098608 | 0.1 | 0.06 | 1 | 14 | Ppcdc | 0.04 | Young |
| 0.374389 | 0.238105 | 0.76 | 0.776 | 1 | 14 | Ifitm2 | -0.016 | Young |
| 0.374424 | 0.590683 | 0.22 | 0.164 | 1 | 14 | Fchsd2 | 0.056 | Young |
| 0.375211 | 0.707375 | 0.6 | 0.53 | 1 | 14 | Atpif1 | 0.07 | Young |
| 0.375416 | 2.430114 | 0.06 | 0.033 | 1 | 14 | Ephx2 | 0.027 | Young |
| 0.375416 | 1.922637 | 0.06 | 0.033 | 1 | 14 | Apln | 0.027 | Young |
| 0.375416 | 1.007445 | 0.06 | 0.033 | 1 | 14 | Kyat3 | 0.027 | Young |
| 0.375416 | 0.98063 | 0.06 | 0.033 | 1 | 14 | Amd2 | 0.027 | Young |

| | | | | | | | | |
|---|---|---|---|---|---|---|---|---|
| 0.375431 | 0.371165 | 0.3 | 0.24 | 1 | 14 | Ubfd1 | 0.06 | Young |
| 0.375947 | 0.258563 | 0.18 | 0.246 | 1 | 14 | Fgd4 | -0.066 | Young |
| 0.376659 | 1.051318 | 0.12 | 0.082 | 1 | 14 | Stard8 | 0.038 | Young |
| 0.376895 | 0.38748 | 0.52 | 0.481 | 1 | 14 | Cpsf6 | 0.039 | Young |
| 0.376928 | 0.571355 | 0.64 | 0.65 | 1 | 14 | Cox6c | -0.01 | Young |
| 0.377246 | 0.357408 | 0.6 | 0.536 | 1 | 14 | Hint1 | 0.064 | Young |
| 0.377408 | 0.259885 | 0.26 | 0.191 | 1 | 14 | Mark2 | 0.069 | Young |
| 0.37801 | 0.387141 | 0.48 | 0.432 | 1 | 14 | Ddost | 0.048 | Young |
| 0.378103 | 0.696897 | 0.16 | 0.115 | 1 | 14 | Emc6 | 0.045 | Young |
| 0.379221 | 0.71994 | 0.28 | 0.224 | 1 | 14 | Adcy6 | 0.056 | Young |
| 0.379248 | 1.606847 | 0.06 | 0.033 | 1 | 14 | Ociad2 | 0.027 | Young |
| 0.379248 | 1.54819 | 0.06 | 0.033 | 1 | 14 | Mlst8 | 0.027 | Young |
| 0.379248 | 1.426707 | 0.06 | 0.033 | 1 | 14 | Dmtn | 0.027 | Young |
| 0.379248 | 0.987499 | 0.06 | 0.033 | 1 | 14 | Zfp787 | 0.027 | Young |
| 0.379248 | 0.819417 | 0.06 | 0.033 | 1 | 14 | Gpank1 | 0.027 | Young |
| 0.379248 | 0.439452 | 0.06 | 0.033 | 1 | 14 | Azin2 | 0.027 | Young |
| 0.380084 | 0.353497 | 0.18 | 0.251 | 1 | 14 | Glt8d1 | -0.071 | Young |
| 0.380199 | 1.279308 | 0.04 | 0.077 | 1 | 14 | Trmt2a | -0.037 | Young |
| 0.38029 | 0.290199 | 0.14 | 0.093 | 1 | 14 | Zbtb7b | 0.047 | Young |
| 0.38085 | 0.223996 | 0.12 | 0.169 | 1 | 14 | Tmem260 | -0.049 | Young |
| 0.381938 | 0.417917 | 0.34 | 0.29 | 1 | 14 | Dlg4 | 0.05 | Young |
| 0.381968 | 1.567691 | 0.22 | 0.169 | 1 | 14 | Prkci | 0.051 | Young |
| 0.382098 | 0.316823 | 0.16 | 0.109 | 1 | 14 | Cep104 | 0.051 | Young |
| 0.382402 | 0.864957 | 0.44 | 0.388 | 1 | 14 | Eprs | 0.052 | Young |
| 0.382435 | 0.409603 | 0.3 | 0.421 | 1 | 14 | Tnfaip6 | -0.121 | Young |
| 0.382586 | 0.742052 | 0.16 | 0.115 | 1 | 14 | N4bp2 | 0.045 | Young |
| 0.382611 | 0.458773 | 0.24 | 0.191 | 1 | 14 | Srsf9 | 0.049 | Young |
| 0.382632 | 0.051953 | 0.02 | 0.049 | 1 | 14 | Zfp580 | -0.029 | Young |
| 0.382661 | 1.023496 | 0.36 | 0.322 | 1 | 14 | Pacs2 | 0.038 | Young |
| 0.382868 | 0.526748 | 0.52 | 0.475 | 1 | 14 | Tax1bp1 | 0.045 | Young |
| 0.382981 | 1.478584 | 0.1 | 0.148 | 1 | 14 | Rai14 | -0.048 | Young |
| 0.383035 | 0.750776 | 0.5 | 0.47 | 1 | 14 | Eif4b | 0.03 | Young |
| 0.383103 | 1.995421 | 0.06 | 0.033 | 1 | 14 | Oscp1 | 0.027 | Young |
| 0.383103 | 1.622365 | 0.06 | 0.033 | 1 | 14 | Ptch2 | 0.027 | Young |
| 0.383103 | 1.599671 | 0.06 | 0.033 | 1 | 14 | Eri2 | 0.027 | Young |
| 0.383103 | 1.550473 | 0.06 | 0.033 | 1 | 14 | 2610035D1 | 0.027 | Young |
| 0.383103 | 0.968706 | 0.06 | 0.033 | 1 | 14 | Elp4 | 0.027 | Young |
| 0.383103 | 0.910906 | 0.06 | 0.033 | 1 | 14 | Ccdc3 | 0.027 | Young |
| 0.383103 | 0.40622 | 0.06 | 0.033 | 1 | 14 | Alg12 | 0.027 | Young |
| 0.383103 | 0.275833 | 0.06 | 0.033 | 1 | 14 | Morn2 | 0.027 | Young |
| 0.383247 | 2.217914 | 0.08 | 0.049 | 1 | 14 | Mrps2 | 0.031 | Young |
| 0.383247 | 1.239767 | 0.08 | 0.049 | 1 | 14 | Eif1ad | 0.031 | Young |
| 0.384197 | 0.855049 | 0.2 | 0.164 | 1 | 14 | Dmd | 0.036 | Young |
| 0.384389 | 0.462232 | 0.5 | 0.41 | 1 | 14 | Stt3b | 0.09 | Young |
| 0.384426 | 0.692165 | 0.12 | 0.082 | 1 | 14 | Ttl | 0.038 | Young |

| | | | | | | | | |
|---|---|---|---|---|---|---|---|---|
| 0.384559 | 0.394349 | 0.4 | 0.344 | 1 | 14 | Thsd7a | 0.056 | Young |
| 0.384691 | 0.341575 | 0.66 | 0.694 | 1 | 14 | Tns1 | -0.034 | Young |
| 0.385183 | 0.178277 | 0.14 | 0.093 | 1 | 14 | Zfyve1 | 0.047 | Young |
| 0.385183 | 0.096997 | 0.14 | 0.093 | 1 | 14 | Rfxap | 0.047 | Young |
| 0.385582 | 0.064893 | 0.96 | 0.984 | 1 | 14 | Actg1 | -0.024 | Young |
| 0.385826 | 0.504275 | 0.22 | 0.169 | 1 | 14 | Zfp740 | 0.051 | Young |
| 0.386027 | 0.491597 | 0.48 | 0.41 | 1 | 14 | Atp1a2 | 0.07 | Young |
| 0.386044 | 0.417535 | 0.22 | 0.164 | 1 | 14 | Dpp8 | 0.056 | Young |
| 0.386095 | 0.018728 | 0.46 | 0.525 | 1 | 14 | Leng8 | -0.065 | Young |
| 0.38631 | 2.751215 | 0.1 | 0.066 | 1 | 14 | Dctpp1 | 0.034 | Young |
| 0.38631 | 0.895098 | 0.1 | 0.066 | 1 | 14 | Wdr91 | 0.034 | Young |
| 0.386317 | 1.228363 | 0.02 | 0.049 | 1 | 14 | Fam120c | -0.029 | Young |
| 0.386317 | 0.18204 | 0.02 | 0.049 | 1 | 14 | Nudt2 | -0.029 | Young |
| 0.386503 | 1.22257 | 0.08 | 0.049 | 1 | 14 | Prkd2 | 0.031 | Young |
| 0.386503 | 1.191344 | 0.08 | 0.049 | 1 | 14 | Aplf | 0.031 | Young |
| 0.386503 | 0.02226 | 0.08 | 0.049 | 1 | 14 | Dock6 | 0.031 | Young |
| 0.38668 | 0.108539 | 0.16 | 0.109 | 1 | 14 | Mrps28 | 0.051 | Young |
| 0.386983 | 2.177031 | 0.06 | 0.033 | 1 | 14 | Ptprm | 0.027 | Young |
| 0.386983 | 1.877039 | 0.06 | 0.033 | 1 | 14 | Ppp1r3e | 0.027 | Young |
| 0.386983 | 0.827559 | 0.06 | 0.033 | 1 | 14 | AI838599 | 0.027 | Young |
| 0.386983 | 0.537804 | 0.06 | 0.033 | 1 | 14 | Slc1a3 | 0.027 | Young |
| 0.386983 | 0.518717 | 0.06 | 0.033 | 1 | 14 | 4930522L1 | 0.027 | Young |
| 0.386983 | 0.503138 | 0.06 | 0.033 | 1 | 14 | Cadps | 0.027 | Young |
| 0.386983 | 0.277521 | 0.06 | 0.033 | 1 | 14 | Rrp12 | 0.027 | Young |
| 0.386983 | 0.212341 | 0.06 | 0.033 | 1 | 14 | Zfp458 | 0.027 | Young |
| 0.386983 | 0.169189 | 0.06 | 0.033 | 1 | 14 | Nubp2 | 0.027 | Young |
| 0.387036 | 1.144037 | 0.12 | 0.082 | 1 | 14 | Cryzl2 | 0.038 | Young |
| 0.387036 | 0.852292 | 0.12 | 0.082 | 1 | 14 | Cep295 | 0.038 | Young |
| 0.387072 | 0.376809 | 0.6 | 0.574 | 1 | 14 | Pebp1 | 0.026 | Young |
| 0.387782 | 0.460065 | 0.42 | 0.344 | 1 | 14 | Ppp4r2 | 0.076 | Young |
| 0.388345 | 0.558495 | 0.06 | 0.098 | 1 | 14 | Ccdc71 | -0.038 | Young |
| 0.388417 | 0.208956 | 0.32 | 0.262 | 1 | 14 | Psmd8 | 0.058 | Young |
| 0.388984 | 0.337877 | 0.16 | 0.109 | 1 | 14 | Pml | 0.051 | Young |
| 0.389336 | 0.027174 | 0.48 | 0.546 | 1 | 14 | Map1lc3b | -0.066 | Young |
| 0.389657 | 1.809253 | 0.12 | 0.082 | 1 | 14 | Bmi1 | 0.038 | Young |
| 0.390212 | 1.093674 | 0.24 | 0.197 | 1 | 14 | Ehbp1l1 | 0.043 | Young |
| 0.3903 | 0.640253 | 0.18 | 0.131 | 1 | 14 | Trappc10 | 0.049 | Young |
| 0.390312 | 0.052435 | 0.42 | 0.497 | 1 | 14 | Magt1 | -0.077 | Young |
| 0.390852 | 0.047764 | 0.24 | 0.175 | 1 | 14 | Tmem168 | 0.065 | Young |
| 0.390887 | 1.322266 | 0.06 | 0.033 | 1 | 14 | Armcx5 | 0.027 | Young |
| 0.390887 | 1.071834 | 0.06 | 0.033 | 1 | 14 | Wdr19 | 0.027 | Young |
| 0.390887 | 0.527691 | 0.06 | 0.033 | 1 | 14 | Nrn1 | 0.027 | Young |
| 0.390887 | 0.189388 | 0.06 | 0.033 | 1 | 14 | Hps1 | 0.027 | Young |
| 0.390887 | 0.162863 | 0.06 | 0.033 | 1 | 14 | Ddx20 | 0.027 | Young |
| 0.390887 | 0.004411 | 0.06 | 0.033 | 1 | 14 | Ptpn6 | 0.027 | Young |

| | | | | | | | | |
|---|---|---|---|---|---|---|---|---|
| 0.391095 | 0.468123 | 0.34 | 0.377 | 1 | 14 | Phf3 | -0.037 | Young |
| 0.391651 | 0.613225 | 0.16 | 0.115 | 1 | 14 | B3galnt1 | 0.045 | Young |
| 0.39183 | 0.238726 | 0.2 | 0.142 | 1 | 14 | Naa10 | 0.058 | Young |
| 0.392094 | 1.038617 | 0.1 | 0.066 | 1 | 14 | Mest | 0.034 | Young |
| 0.392289 | 0.890699 | 0.12 | 0.082 | 1 | 14 | Foxp2 | 0.038 | Young |
| 0.392594 | 0.051659 | 0.14 | 0.093 | 1 | 14 | Rhobtb2 | 0.047 | Young |
| 0.39277 | 0.290856 | 0.24 | 0.175 | 1 | 14 | 0610030E2 | 0.065 | Young |
| 0.393064 | 1.462818 | 0.08 | 0.049 | 1 | 14 | Nmrk1 | 0.031 | Young |
| 0.393064 | 1.214222 | 0.08 | 0.049 | 1 | 14 | Pcdhgc4 | 0.031 | Young |
| 0.393064 | 0.962574 | 0.08 | 0.049 | 1 | 14 | Zdhhc14 | 0.031 | Young |
| 0.393406 | 0.139853 | 0.36 | 0.273 | 1 | 14 | Gnpnat1 | 0.087 | Young |
| 0.393937 | 0.538212 | 0.16 | 0.115 | 1 | 14 | Rbm15 | 0.045 | Young |
| 0.394754 | 1.182764 | 0.14 | 0.104 | 1 | 14 | Kctd12b | 0.036 | Young |
| 0.394814 | 0.115695 | 0.06 | 0.033 | 1 | 14 | Rad9a | 0.027 | Young |
| 0.394814 | 0.105912 | 0.06 | 0.033 | 1 | 14 | Nol4l | 0.027 | Young |
| 0.394932 | 0.356592 | 0.12 | 0.082 | 1 | 14 | Ring1 | 0.038 | Young |
| 0.395005 | 1.438829 | 0.1 | 0.066 | 1 | 14 | Zfp414 | 0.034 | Young |
| 0.395031 | 0.565049 | 0.14 | 0.098 | 1 | 14 | Leng1 | 0.042 | Young |
| 0.39542 | 0.018854 | 0.26 | 0.333 | 1 | 14 | Pls3 | -0.073 | Young |
| 0.395806 | 0.029669 | 0.34 | 0.399 | 1 | 14 | Tulp4 | -0.059 | Young |
| 0.39637 | 2.306339 | 0.08 | 0.049 | 1 | 14 | Tgds | 0.031 | Young |
| 0.39637 | 0.92302 | 0.08 | 0.049 | 1 | 14 | Tnk2 | 0.031 | Young |
| 0.397485 | 0.931389 | 0.14 | 0.098 | 1 | 14 | Pdcd7 | 0.042 | Young |
| 0.397485 | 0.312865 | 0.14 | 0.098 | 1 | 14 | Dapk2 | 0.042 | Young |
| 0.398335 | 1.358361 | 0.2 | 0.153 | 1 | 14 | Fzd7 | 0.047 | Young |
| 0.39843 | 0.042215 | 0.34 | 0.257 | 1 | 14 | Tmem127 | 0.083 | Young |
| 0.398512 | 1.143165 | 0.16 | 0.126 | 1 | 14 | Socs1 | 0.034 | Young |
| 0.398765 | 0.132492 | 0.06 | 0.033 | 1 | 14 | Gm14305 | 0.027 | Young |
| 0.398765 | 0.098935 | 0.06 | 0.033 | 1 | 14 | Bphl | 0.027 | Young |
| 0.398765 | 0.02977 | 0.06 | 0.033 | 1 | 14 | Clec16a | 0.027 | Young |
| 0.398916 | 0.118664 | 0.06 | 0.098 | 1 | 14 | Repin1 | -0.038 | Young |
| 0.398974 | 0.373284 | 0.18 | 0.131 | 1 | 14 | Prkaa1 | 0.049 | Young |
| 0.399517 | 0.300326 | 0.14 | 0.191 | 1 | 14 | Nr1h2 | -0.051 | Young |
| 0.399693 | 1.505274 | 0.08 | 0.049 | 1 | 14 | Wnt5b | 0.031 | Young |
| 0.399947 | 1.754688 | 0.14 | 0.098 | 1 | 14 | Creld1 | 0.042 | Young |
| 0.400026 | 0.392293 | 0.46 | 0.388 | 1 | 14 | U2surp | 0.072 | Young |
| 0.400249 | 0.526151 | 0.12 | 0.082 | 1 | 14 | Fitm2 | 0.038 | Young |
| 0.400828 | 0.20744 | 0.38 | 0.29 | 1 | 14 | Pgk1 | 0.09 | Young |
| 0.400866 | 1.122285 | 0.1 | 0.066 | 1 | 14 | Atg16l2 | 0.034 | Young |
| 0.400882 | 1.054562 | 0.16 | 0.12 | 1 | 14 | Zdhhc17 | 0.04 | Young |
| 0.401161 | 0.117889 | 0.18 | 0.131 | 1 | 14 | Anapc4 | 0.049 | Young |
| 0.401184 | 0.006179 | 0.88 | 0.923 | 1 | 14 | Rpl23a | -0.043 | Young |
| 0.401854 | 0.089907 | 0.32 | 0.393 | 1 | 14 | U2af2 | -0.073 | Young |
| 0.402392 | 0.262384 | 0.28 | 0.219 | 1 | 14 | Itsn1 | 0.061 | Young |
| 0.40274 | 0.098361 | 0.06 | 0.033 | 1 | 14 | Brf2 | 0.027 | Young |

| | | | | | | | | |
|---|---|---|---|---|---|---|---|---|
| 0.40274 | 0.048443 | 0.06 | 0.033 | 1 | 14 | Mpi | 0.027 | Young |
| 0.402924 | 0.919698 | 0.12 | 0.082 | 1 | 14 | Ift81 | 0.038 | Young |
| 0.402935 | 0.0093 | 0.18 | 0.235 | 1 | 14 | Tnks | -0.055 | Young |
| 0.403032 | 0.707679 | 0.08 | 0.049 | 1 | 14 | Pradc1 | 0.031 | Young |
| 0.403032 | 0.54558 | 0.08 | 0.049 | 1 | 14 | Chrm3 | 0.031 | Young |
| 0.403166 | 0.58522 | 0.5 | 0.448 | 1 | 14 | Fst | 0.052 | Young |
| 0.403816 | 0.356761 | 0.1 | 0.066 | 1 | 14 | Gspt2 | 0.034 | Young |
| 0.403819 | 0.016395 | 0.3 | 0.23 | 1 | 14 | Ndfip2 | 0.07 | Young |
| 0.40429 | 1.559907 | 0.32 | 0.257 | 1 | 14 | Smarcc1 | 0.063 | Young |
| 0.40429 | 0.33097 | 0.32 | 0.257 | 1 | 14 | Csnk2a2 | 0.063 | Young |
| 0.405343 | 0.317793 | 0.08 | 0.131 | 1 | 14 | Mcee | -0.051 | Young |
| 0.405502 | 0.096376 | 0.36 | 0.432 | 1 | 14 | Rab12 | -0.072 | Young |
| 0.405556 | 0.395222 | 0.1 | 0.153 | 1 | 14 | Nfatc1 | -0.053 | Young |
| 0.405591 | 0.049067 | 0.12 | 0.169 | 1 | 14 | Mfhas1 | -0.049 | Young |
| 0.40561 | 0.448626 | 0.12 | 0.082 | 1 | 14 | 2610002M | 0.038 | Young |
| 0.405619 | 0.206129 | 0.28 | 0.224 | 1 | 14 | Hcfc2 | 0.056 | Young |
| 0.406102 | 0.162514 | 0.26 | 0.197 | 1 | 14 | Rgma | 0.063 | Young |
| 0.406388 | 1.884445 | 0.08 | 0.049 | 1 | 14 | Ano8 | 0.031 | Young |
| 0.406388 | 0.703649 | 0.08 | 0.049 | 1 | 14 | Ptgr1 | 0.031 | Young |
| 0.406388 | 0.646133 | 0.08 | 0.049 | 1 | 14 | Zfp931 | 0.031 | Young |
| 0.406388 | 0.492111 | 0.08 | 0.049 | 1 | 14 | Heyl | 0.031 | Young |
| 0.406388 | 0.469008 | 0.08 | 0.049 | 1 | 14 | Sumf2 | 0.031 | Young |
| 0.406564 | 0.132929 | 0.2 | 0.142 | 1 | 14 | Nsrp1 | 0.058 | Young |
| 0.407145 | 0.470965 | 0.04 | 0.077 | 1 | 14 | Emc9 | -0.037 | Young |
| 0.40739 | 0.258427 | 0.14 | 0.098 | 1 | 14 | Lrig2 | 0.042 | Young |
| 0.407443 | 0.3555 | 0.54 | 0.525 | 1 | 14 | Tma7 | 0.015 | Young |
| 0.407483 | 0.250777 | 0.4 | 0.339 | 1 | 14 | Rap2a | 0.061 | Young |
| 0.407764 | 0.089054 | 0.18 | 0.131 | 1 | 14 | F13a1 | 0.049 | Young |
| 0.407812 | 0.139756 | 0.18 | 0.246 | 1 | 14 | Mill2 | -0.066 | Young |
| 0.408268 | 0.341044 | 0.48 | 0.404 | 1 | 14 | Ndufb3 | 0.076 | Young |
| 0.408306 | 0.606795 | 0.12 | 0.082 | 1 | 14 | Isg20l2 | 0.038 | Young |
| 0.408325 | 0.324631 | 0.5 | 0.437 | 1 | 14 | Wac | 0.063 | Young |
| 0.408437 | 0.912132 | 0.26 | 0.213 | 1 | 14 | Ginm1 | 0.047 | Young |
| 0.408494 | 0.708831 | 0.3 | 0.257 | 1 | 14 | Mydgf | 0.043 | Young |
| 0.408912 | 1.056896 | 0.24 | 0.191 | 1 | 14 | Tlk2 | 0.049 | Young |
| 0.40893 | 1.011028 | 0.22 | 0.175 | 1 | 14 | B4gat1 | 0.045 | Young |
| 0.408984 | 0.416011 | 0.36 | 0.295 | 1 | 14 | Myo9a | 0.065 | Young |
| 0.409517 | 0.24963 | 0.44 | 0.355 | 1 | 14 | Ntn1 | 0.085 | Young |
| 0.409755 | 0.146456 | 0.1 | 0.066 | 1 | 14 | Ppp2r3c | 0.034 | Young |
| 0.409756 | 0.147804 | 0.76 | 0.863 | 1 | 14 | Tcf4 | -0.103 | Young |
| 0.409761 | 0.788478 | 0.08 | 0.049 | 1 | 14 | Ccdc138 | 0.031 | Young |
| 0.409761 | 0.719966 | 0.08 | 0.049 | 1 | 14 | Prpf40b | 0.031 | Young |
| 0.409761 | 0.514843 | 0.08 | 0.049 | 1 | 14 | Zfp768 | 0.031 | Young |
| 0.409761 | 0.373972 | 0.08 | 0.049 | 1 | 14 | Hdac10 | 0.031 | Young |
| 0.40989 | 0.514605 | 0.14 | 0.098 | 1 | 14 | Xab2 | 0.042 | Young |

| | | | | | | | | |
|---|---|---|---|---|---|---|---|---|
| 0.41001 | 0.203428 | 0.28 | 0.355 | 1 | 14 | Rcn3 | -0.075 | Young |
| 0.410109 | 0.258965 | 0.22 | 0.273 | 1 | 14 | Adss | -0.053 | Young |
| 0.410207 | 0.401185 | 0.04 | 0.077 | 1 | 14 | Setd6 | -0.037 | Young |
| 0.410699 | 0.787329 | 0.26 | 0.202 | 1 | 14 | Slu7 | 0.058 | Young |
| 0.410728 | 0.628738 | 0.24 | 0.186 | 1 | 14 | Pik3r4 | 0.054 | Young |
| 0.410761 | 0.218323 | 0.06 | 0.033 | 1 | 14 | Lpin3 | 0.027 | Young |
| 0.410918 | 0.080678 | 0.14 | 0.197 | 1 | 14 | Irf9 | -0.057 | Young |
| 0.411449 | 0.502109 | 0.28 | 0.219 | 1 | 14 | Clcn3 | 0.061 | Young |
| 0.411607 | 0.139217 | 0.4 | 0.322 | 1 | 14 | Arnt | 0.078 | Young |
| 0.411843 | 0.39135 | 0.42 | 0.361 | 1 | 14 | Chmp2a | 0.059 | Young |
| 0.412398 | 0.723751 | 0.14 | 0.098 | 1 | 14 | Bap1 | 0.042 | Young |
| 0.412398 | 0.579241 | 0.14 | 0.098 | 1 | 14 | 1300002E1 | 0.042 | Young |
| 0.412517 | 0.144078 | 0.16 | 0.115 | 1 | 14 | Hip1 | 0.045 | Young |
| 0.412744 | 0.53584 | 0.1 | 0.066 | 1 | 14 | Cacnb3 | 0.034 | Young |
| 0.412744 | 0.463923 | 0.1 | 0.066 | 1 | 14 | Tmem29 | 0.034 | Young |
| 0.41315 | 0.223551 | 0.08 | 0.049 | 1 | 14 | Hoxb3os | 0.031 | Young |
| 0.41315 | 0.060588 | 0.08 | 0.049 | 1 | 14 | Cul4b | 0.031 | Young |
| 0.413496 | 0.327717 | 0.22 | 0.169 | 1 | 14 | Trp53bp2 | 0.051 | Young |
| 0.41373 | 1.074049 | 0.12 | 0.082 | 1 | 14 | Cd14 | 0.038 | Young |
| 0.414024 | 0.162111 | 0.6 | 0.683 | 1 | 14 | Slc1a5 | -0.083 | Young |
| 0.414158 | 0.800368 | 0.38 | 0.366 | 1 | 14 | Pcnp | 0.014 | Young |
| 0.414275 | 0.148202 | 0.48 | 0.404 | 1 | 14 | Vezf1 | 0.076 | Young |
| 0.414431 | 0.837379 | 0.18 | 0.131 | 1 | 14 | Tmem198k | 0.049 | Young |
| 0.414656 | 0.230913 | 0.28 | 0.224 | 1 | 14 | Sult1a1 | 0.056 | Young |
| 0.414707 | 0.678757 | 0.16 | 0.12 | 1 | 14 | Cep85l | 0.04 | Young |
| 0.414916 | 0.47226 | 0.14 | 0.098 | 1 | 14 | Parn | 0.042 | Young |
| 0.41543 | 0.559529 | 0.68 | 0.71 | 1 | 14 | Serinc3 | -0.03 | Young |
| 0.415502 | 0.186076 | 0.28 | 0.213 | 1 | 14 | Ireb2 | 0.067 | Young |
| 0.416459 | 0.311401 | 0.12 | 0.082 | 1 | 14 | Alg5 | 0.038 | Young |
| 0.416459 | 0.28608 | 0.12 | 0.082 | 1 | 14 | Ccdc97 | 0.038 | Young |
| 0.416556 | 2.059194 | 0.08 | 0.049 | 1 | 14 | Fndc10 | 0.031 | Young |
| 0.416556 | 0.736368 | 0.08 | 0.049 | 1 | 14 | Parm1 | 0.031 | Young |
| 0.416668 | 0.064937 | 0.18 | 0.131 | 1 | 14 | Pygo2 | 0.049 | Young |
| 0.416916 | 0.799953 | 0.22 | 0.175 | 1 | 14 | Cers4 | 0.045 | Young |
| 0.417092 | 1.233201 | 0.2 | 0.153 | 1 | 14 | Lrrfip2 | 0.047 | Young |
| 0.417092 | 1.104393 | 0.2 | 0.153 | 1 | 14 | Rtl8a | 0.047 | Young |
| 0.417092 | 0.223551 | 0.2 | 0.153 | 1 | 14 | Arsi | 0.047 | Young |
| 0.418128 | 0.273929 | 0.5 | 0.454 | 1 | 14 | Tpr | 0.046 | Young |
| 0.418138 | 1.467003 | 0.12 | 0.087 | 1 | 14 | AU020206 | 0.033 | Young |
| 0.418183 | 0.108028 | 0.74 | 0.694 | 1 | 14 | Tapbp | 0.046 | Young |
| 0.418242 | 0.216971 | 0.26 | 0.202 | 1 | 14 | Cyp20a1 | 0.058 | Young |
| 0.418278 | 0.321629 | 0.18 | 0.24 | 1 | 14 | Dlst | -0.06 | Young |
| 0.418379 | 0.012339 | 0.24 | 0.306 | 1 | 14 | Tgfb1i1 | -0.066 | Young |
| 0.418522 | 0.329693 | 0.24 | 0.186 | 1 | 14 | Rnf149 | 0.054 | Young |
| 0.418761 | 0.977692 | 0.1 | 0.066 | 1 | 14 | Asah2 | 0.034 | Young |

| | | | | | | | | |
|---|---|---|---|---|---|---|---|---|
| 0.418761 | 0.684955 | 0.1 | 0.066 | 1 | 14 | Smg8 | 0.034 | Young |
| 0.418912 | 0.309875 | 0.18 | 0.131 | 1 | 14 | Spast | 0.049 | Young |
| 0.419782 | 0.206338 | 0.92 | 0.869 | 1 | 14 | Rpl18a | 0.051 | Young |
| 0.419978 | 0.889345 | 0.14 | 0.098 | 1 | 14 | Numb | 0.042 | Young |
| 0.419978 | 0.385593 | 0.08 | 0.049 | 1 | 14 | Tti2 | 0.031 | Young |
| 0.419978 | 0.374371 | 0.08 | 0.049 | 1 | 14 | Rabif | 0.031 | Young |
| 0.419978 | 0.321159 | 0.08 | 0.049 | 1 | 14 | Rpap2 | 0.031 | Young |
| 0.420015 | 0.179221 | 0.9 | 0.913 | 1 | 14 | Rps10 | -0.013 | Young |
| 0.420682 | 0.284204 | 0.24 | 0.311 | 1 | 14 | Alyref | -0.071 | Young |
| 0.42073 | 0.607174 | 0.1 | 0.148 | 1 | 14 | Wdr77 | -0.048 | Young |
| 0.421501 | 0.35671 | 0.92 | 0.945 | 1 | 14 | Col1a2 | -0.025 | Young |
| 0.421789 | 1.700539 | 0.1 | 0.066 | 1 | 14 | Dock8 | 0.034 | Young |
| 0.421789 | 0.349393 | 0.1 | 0.066 | 1 | 14 | Bop1 | 0.034 | Young |
| 0.422522 | 1.066936 | 0.14 | 0.098 | 1 | 14 | Zfp219 | 0.042 | Young |
| 0.422522 | 0.362678 | 0.14 | 0.098 | 1 | 14 | Fpgs | 0.042 | Young |
| 0.422522 | 0.148924 | 0.14 | 0.098 | 1 | 14 | Ints9 | 0.042 | Young |
| 0.422844 | 0.172483 | 0.42 | 0.481 | 1 | 14 | Ptgis | -0.061 | Young |
| 0.422966 | 0.142004 | 0.14 | 0.197 | 1 | 14 | Fam173a | -0.057 | Young |
| 0.423417 | 0.496778 | 0.08 | 0.049 | 1 | 14 | Ednrb | 0.031 | Young |
| 0.423417 | 0.20153 | 0.08 | 0.049 | 1 | 14 | Hirip3 | 0.031 | Young |
| 0.423417 | 0.144166 | 0.08 | 0.049 | 1 | 14 | Cep85 | 0.031 | Young |
| 0.423417 | 0.098595 | 0.08 | 0.049 | 1 | 14 | Zfp280b | 0.031 | Young |
| 0.423656 | 0.376456 | 0.22 | 0.169 | 1 | 14 | Cacna1a | 0.051 | Young |
| 0.423679 | 0.510006 | 0.18 | 0.137 | 1 | 14 | Clip2 | 0.043 | Young |
| 0.424258 | 1.643513 | 0.14 | 0.104 | 1 | 14 | Crbn | 0.036 | Young |
| 0.424706 | 0.168835 | 0.12 | 0.082 | 1 | 14 | Rel | 0.038 | Young |
| 0.424829 | 1.518973 | 0.1 | 0.066 | 1 | 14 | Prpf31 | 0.034 | Young |
| 0.424995 | 0.839247 | 0.22 | 0.175 | 1 | 14 | Maea | 0.045 | Young |
| 0.425076 | 0.015227 | 0.14 | 0.098 | 1 | 14 | Praf2 | 0.042 | Young |
| 0.426272 | 0.864814 | 0.24 | 0.197 | 1 | 14 | Pak1ip1 | 0.043 | Young |
| 0.42634 | 0.015346 | 0.22 | 0.164 | 1 | 14 | Fcer1g | 0.056 | Young |
| 0.42644 | 0.657597 | 0.16 | 0.12 | 1 | 14 | Ube2a | 0.04 | Young |
| 0.42652 | 0.114544 | 0.58 | 0.699 | 1 | 14 | Clic4 | -0.119 | Young |
| 0.426872 | 0.353834 | 0.08 | 0.049 | 1 | 14 | Lins1 | 0.031 | Young |
| 0.426872 | 0.2535 | 0.08 | 0.049 | 1 | 14 | Clec11a | 0.031 | Young |
| 0.426872 | 0.045522 | 0.08 | 0.049 | 1 | 14 | Lsm11 | 0.031 | Young |
| 0.426924 | 0.35896 | 0.2 | 0.148 | 1 | 14 | Kif3a | 0.052 | Young |
| 0.427785 | 0.205972 | 0.26 | 0.202 | 1 | 14 | Eif3g | 0.058 | Young |
| 0.427955 | 0.34929 | 0.4 | 0.328 | 1 | 14 | Tpi1 | 0.072 | Young |
| 0.428866 | 1.200339 | 0.06 | 0.098 | 1 | 14 | Foxk1 | -0.038 | Young |
| 0.428874 | 0.140385 | 0.38 | 0.443 | 1 | 14 | Zfand3 | -0.063 | Young |
| 0.429192 | 0.367024 | 0.16 | 0.115 | 1 | 14 | Klhl21 | 0.045 | Young |
| 0.429194 | 0.18255 | 0.56 | 0.617 | 1 | 14 | Ctsz | -0.057 | Young |
| 0.429298 | 0.55652 | 0.14 | 0.104 | 1 | 14 | AU040320 | 0.036 | Young |
| 0.429708 | 0.39736 | 0.26 | 0.202 | 1 | 14 | Cacybp | 0.058 | Young |

| | | | | | | | | |
|---|---|---|---|---|---|---|---|---|
| 0.43021 | 0.760875 | 0.14 | 0.098 | 1 | 14 | Gadd45a | 0.042 | Young |
| 0.43089 | 0.030974 | 0.18 | 0.235 | 1 | 14 | Cdc42ep2 | -0.055 | Young |
| 0.431113 | 0.917072 | 0.22 | 0.175 | 1 | 14 | Bace1 | 0.045 | Young |
| 0.431121 | 0.002139 | 0.32 | 0.41 | 1 | 14 | Clstn1 | -0.09 | Young |
| 0.431187 | 0.792011 | 0.16 | 0.12 | 1 | 14 | Map3k12 | 0.04 | Young |
| 0.431308 | 0.172516 | 0.5 | 0.448 | 1 | 14 | Reep5 | 0.052 | Young |
| 0.431606 | 1.091995 | 0.16 | 0.115 | 1 | 14 | Cdc42ep4 | 0.045 | Young |
| 0.431652 | 0.158136 | 0.06 | 0.098 | 1 | 14 | Vps52 | -0.038 | Young |
| 0.431888 | 0.296543 | 0.22 | 0.169 | 1 | 14 | Fam171b | 0.051 | Young |
| 0.432685 | 0.045847 | 0.3 | 0.23 | 1 | 14 | Emd | 0.07 | Young |
| 0.433049 | 0.202971 | 0.12 | 0.082 | 1 | 14 | Myo1e | 0.038 | Young |
| 0.433049 | 0.198239 | 0.12 | 0.082 | 1 | 14 | Fahd1 | 0.038 | Young |
| 0.433049 | 0.176743 | 0.12 | 0.082 | 1 | 14 | Zfp68 | 0.038 | Young |
| 0.433831 | 0.324935 | 0.08 | 0.049 | 1 | 14 | Doc2b | 0.031 | Young |
| 0.433909 | 0.472751 | 0.44 | 0.388 | 1 | 14 | Mdh1 | 0.052 | Young |
| 0.434026 | 0.341757 | 0.1 | 0.066 | 1 | 14 | Psmg1 | 0.034 | Young |
| 0.434266 | 0.027004 | 0.4 | 0.448 | 1 | 14 | Dbn1 | -0.048 | Young |
| 0.434683 | 0.089191 | 0.22 | 0.164 | 1 | 14 | Tox4 | 0.056 | Young |
| 0.435023 | 0.261499 | 0.38 | 0.443 | 1 | 14 | Myof | -0.063 | Young |
| 0.435056 | 0.092881 | 0.78 | 0.852 | 1 | 14 | Rbms1 | -0.072 | Young |
| 0.43538 | 0.186474 | 0.14 | 0.098 | 1 | 14 | Ulk1 | 0.042 | Young |
| 0.435688 | 0.542927 | 0.32 | 0.257 | 1 | 14 | Zfp292 | 0.063 | Young |
| 0.43585 | 0.346507 | 0.12 | 0.082 | 1 | 14 | Yaf2 | 0.038 | Young |
| 0.436373 | 0.383068 | 0.22 | 0.18 | 1 | 14 | Hsd17b10 | 0.04 | Young |
| 0.436508 | 0.400315 | 0.1 | 0.142 | 1 | 14 | Bad | -0.042 | Young |
| 0.436922 | 1.323352 | 0.08 | 0.12 | 1 | 14 | Atp6v0e2 | -0.04 | Young |
| 0.436962 | 0.640694 | 0.2 | 0.158 | 1 | 14 | Ebna1bp2 | 0.042 | Young |
| 0.437118 | 0.31131 | 0.1 | 0.066 | 1 | 14 | Lilr4b | 0.034 | Young |
| 0.437118 | 0.121722 | 0.1 | 0.066 | 1 | 14 | Ints5.1 | 0.034 | Young |
| 0.437978 | 0.3579 | 0.14 | 0.098 | 1 | 14 | Acyp2 | 0.042 | Young |
| 0.438052 | 0.828618 | 0.24 | 0.197 | 1 | 14 | Psen1 | 0.043 | Young |
| 0.438253 | 0.762451 | 0.24 | 0.191 | 1 | 14 | Ss18 | 0.049 | Young |
| 0.438663 | 0.428528 | 0.12 | 0.082 | 1 | 14 | Tubgcp4 | 0.038 | Young |
| 0.438704 | 0.342596 | 0.28 | 0.224 | 1 | 14 | Secisbp2l | 0.056 | Young |
| 0.438883 | 0.344382 | 0.48 | 0.426 | 1 | 14 | Msl1 | 0.054 | Young |
| 0.43963 | 0.062033 | 0.74 | 0.738 | 1 | 14 | Rarres2 | 0.002 | Young |
| 0.440073 | 0.512858 | 0.06 | 0.098 | 1 | 14 | D230025D: | -0.038 | Young |
| 0.440221 | 0.041424 | 0.1 | 0.066 | 1 | 14 | Mdk | 0.034 | Young |
| 0.440399 | 0.142366 | 0.24 | 0.186 | 1 | 14 | Msmo1 | 0.054 | Young |
| 0.440848 | 0.592882 | 0.04 | 0.071 | 1 | 14 | Parp2 | -0.031 | Young |
| 0.441014 | 0.974225 | 0.28 | 0.235 | 1 | 14 | Kdm1a | 0.045 | Young |
| 0.441075 | 0.162534 | 0.34 | 0.273 | 1 | 14 | Dctn4 | 0.067 | Young |
| 0.441251 | 0.305558 | 0.1 | 0.142 | 1 | 14 | Mpg | -0.042 | Young |
| 0.442019 | 0.932268 | 0.18 | 0.213 | 1 | 14 | Zwint | -0.033 | Young |
| 0.442141 | 0.680378 | 0.3 | 0.251 | 1 | 14 | Txndc11 | 0.049 | Young |

| | | | | | | | | |
|---|---|---|---|---|---|---|---|---|
| 0.443793 | 0.639215 | 0.16 | 0.115 | 1 | 14 | Pfdn4 | 0.045 | Young |
| 0.444059 | 0.638467 | 0.18 | 0.131 | 1 | 14 | Sh3bp1 | 0.049 | Young |
| 0.444059 | 0.012191 | 0.18 | 0.131 | 1 | 14 | Tbc1d13 | 0.049 | Young |
| 0.445585 | 0.767288 | 0.22 | 0.175 | 1 | 14 | Srgn | 0.045 | Young |
| 0.445707 | 1.68738 | 0.06 | 0.093 | 1 | 14 | Zfp710 | -0.033 | Young |
| 0.446248 | 1.708408 | 0.26 | 0.213 | 1 | 14 | Fbxw8 | 0.047 | Young |
| 0.446801 | 1.347353 | 0.48 | 0.459 | 1 | 14 | Atp1b3 | 0.021 | Young |
| 0.446969 | 0.509293 | 0.34 | 0.29 | 1 | 14 | Azin1 | 0.05 | Young |
| 0.447208 | 1.499301 | 0.14 | 0.104 | 1 | 14 | Rfng | 0.036 | Young |
| 0.447282 | 0.460886 | 0.2 | 0.153 | 1 | 14 | Rgl2 | 0.047 | Young |
| 0.447555 | 0.105328 | 0.3 | 0.23 | 1 | 14 | Dmxl1 | 0.07 | Young |
| 0.44846 | 1.225156 | 0.14 | 0.098 | 1 | 14 | Gpatch2 | 0.042 | Young |
| 0.448533 | 0.496469 | 0.46 | 0.437 | 1 | 14 | Dnajb6 | 0.023 | Young |
| 0.448722 | 1.758561 | 0.16 | 0.115 | 1 | 14 | Serpinb1a | 0.045 | Young |
| 0.448722 | 0.003932 | 0.16 | 0.115 | 1 | 14 | Oser1 | 0.045 | Young |
| 0.449771 | 1.020997 | 0.22 | 0.175 | 1 | 14 | Clcn7 | 0.045 | Young |
| 0.450015 | 0.597015 | 0.12 | 0.082 | 1 | 14 | Hs6st1 | 0.038 | Young |
| 0.450015 | 0.146795 | 0.12 | 0.082 | 1 | 14 | Mtap | 0.038 | Young |
| 0.450574 | 0.179511 | 0.38 | 0.311 | 1 | 14 | Gadd45g | 0.069 | Young |
| 0.45074 | 0.325433 | 0.58 | 0.552 | 1 | 14 | Mmp2 | 0.028 | Young |
| 0.450896 | 1.781917 | 0.14 | 0.109 | 1 | 14 | Dcakd | 0.031 | Young |
| 0.451119 | 0.670106 | 0.38 | 0.333 | 1 | 14 | Nop53 | 0.047 | Young |
| 0.451213 | 0.090321 | 0.38 | 0.306 | 1 | 14 | Ube2z | 0.074 | Young |
| 0.451366 | 0.794497 | 0.3 | 0.246 | 1 | 14 | Yipf4 | 0.054 | Young |
| 0.451446 | 1.330553 | 0.06 | 0.098 | 1 | 14 | Fam131b | -0.038 | Young |
| 0.451543 | 0.461936 | 0.06 | 0.093 | 1 | 14 | Polr3k | -0.033 | Young |
| 0.451609 | 0.592549 | 0.38 | 0.317 | 1 | 14 | Ndufb4 | 0.063 | Young |
| 0.452763 | 0.172096 | 0.1 | 0.066 | 1 | 14 | Ccl25 | 0.034 | Young |
| 0.452763 | 0.141457 | 0.1 | 0.066 | 1 | 14 | Tbp | 0.034 | Young |
| 0.452879 | 0.389021 | 0.12 | 0.082 | 1 | 14 | Slc43a1 | 0.038 | Young |
| 0.45309 | 0.020578 | 0.34 | 0.393 | 1 | 14 | Slc25a5 | -0.053 | Young |
| 0.453203 | 2.136974 | 0.08 | 0.055 | 1 | 14 | Acot2 | 0.025 | Young |
| 0.453685 | 0.451307 | 0.6 | 0.574 | 1 | 14 | Atp5g2 | 0.026 | Young |
| 0.453977 | 1.038547 | 0.34 | 0.29 | 1 | 14 | Slc25a39 | 0.05 | Young |
| 0.455455 | 3.303275 | 0.04 | 0.022 | 1 | 14 | Mss51 | 0.018 | Young |
| 0.455455 | 2.696281 | 0.04 | 0.022 | 1 | 14 | Nup43 | 0.018 | Young |
| 0.455455 | 2.57066 | 0.04 | 0.022 | 1 | 14 | Tmem191c | 0.018 | Young |
| 0.455455 | 1.839067 | 0.04 | 0.022 | 1 | 14 | Pard6g | 0.018 | Young |
| 0.455455 | 1.728856 | 0.04 | 0.022 | 1 | 14 | Pgap3 | 0.018 | Young |
| 0.455455 | 1.496148 | 0.04 | 0.022 | 1 | 14 | Zfp408 | 0.018 | Young |
| 0.455987 | 0.320362 | 0.42 | 0.355 | 1 | 14 | Idh1 | 0.065 | Young |
| 0.456067 | 0.563272 | 0.16 | 0.219 | 1 | 14 | Tle1 | -0.059 | Young |
| 0.45613 | 0.524143 | 0.2 | 0.153 | 1 | 14 | Snapc1 | 0.047 | Young |
| 0.45613 | 0.267304 | 0.2 | 0.153 | 1 | 14 | Tdp2 | 0.047 | Young |
| 0.456415 | 0.170178 | 0.14 | 0.098 | 1 | 14 | Usp50 | 0.042 | Young |

| | | | | | | | | |
|---|---|---|---|---|---|---|---|---|
| 0.456541 | 0.077035 | 0.02 | 0.044 | 1 | 14 | Il1rap | -0.024 | Young |
| 0.456566 | 0.559023 | 0.46 | 0.415 | 1 | 14 | Gnaq | 0.045 | Young |
| 0.4569 | 0.182415 | 0.48 | 0.393 | 1 | 14 | Zfp91 | 0.087 | Young |
| 0.457025 | 0.200319 | 0.3 | 0.235 | 1 | 14 | Mysm1 | 0.065 | Young |
| 0.457301 | 1.344885 | 0.12 | 0.093 | 1 | 14 | Lrrc57 | 0.027 | Young |
| 0.457944 | 0.852918 | 0.28 | 0.235 | 1 | 14 | Rad23a | 0.045 | Young |
| 0.458208 | 0.757919 | 0.22 | 0.175 | 1 | 14 | Olfml3 | 0.045 | Young |
| 0.458607 | 1.467558 | 0.14 | 0.109 | 1 | 14 | Trappc9 | 0.031 | Young |
| 0.458664 | 0.083264 | 0.18 | 0.219 | 1 | 14 | Gas2l1 | -0.039 | Young |
| 0.458777 | 0.418439 | 0.24 | 0.186 | 1 | 14 | Itpkb | 0.054 | Young |
| 0.459159 | 0.136166 | 0.26 | 0.202 | 1 | 14 | Sdccag3 | 0.058 | Young |
| 0.459268 | 0.320221 | 0.62 | 0.596 | 1 | 14 | Clk1 | 0.024 | Young |
| 0.459753 | 0.200145 | 0.48 | 0.372 | 1 | 14 | Med13 | 0.108 | Young |
| 0.460121 | 0.135076 | 0.12 | 0.169 | 1 | 14 | Osr2 | -0.049 | Young |
| 0.460249 | 0.32112 | 0.44 | 0.372 | 1 | 14 | Acadsb | 0.068 | Young |
| 0.460463 | 0.772086 | 0.4 | 0.377 | 1 | 14 | Cped1 | 0.023 | Young |
| 0.460683 | 1.30196 | 0.04 | 0.022 | 1 | 14 | 3-Mar | 0.018 | Young |
| 0.460683 | 1.250261 | 0.04 | 0.022 | 1 | 14 | Ap5s1 | 0.018 | Young |
| 0.4608 | 0.376242 | 0.5 | 0.421 | 1 | 14 | Rsrc2 | 0.079 | Young |
| 0.46084 | 1.941523 | 0.02 | 0.044 | 1 | 14 | Hcn4 | -0.024 | Young |
| 0.46084 | 1.542528 | 0.02 | 0.044 | 1 | 14 | Nudt18 | -0.024 | Young |
| 0.46084 | 0.955727 | 0.02 | 0.044 | 1 | 14 | Ncapd3 | -0.024 | Young |
| 0.46084 | 0.766316 | 0.02 | 0.044 | 1 | 14 | Zfp994 | -0.024 | Young |
| 0.46084 | 0.249076 | 0.02 | 0.044 | 1 | 14 | Bcdin3d | -0.024 | Young |
| 0.46084 | 0.127793 | 0.02 | 0.044 | 1 | 14 | Pcdhb20 | -0.024 | Young |
| 0.46084 | 0.006331 | 0.02 | 0.044 | 1 | 14 | Med11 | -0.024 | Young |
| 0.460874 | 0.203401 | 0.28 | 0.339 | 1 | 14 | Sf3b6 | -0.059 | Young |
| 0.460896 | 1.011664 | 0.04 | 0.071 | 1 | 14 | Lcp2 | -0.031 | Young |
| 0.461203 | 0.168051 | 0.14 | 0.18 | 1 | 14 | Trim12c | -0.04 | Young |
| 0.461572 | 0.106481 | 0.26 | 0.208 | 1 | 14 | Spty2d1 | 0.052 | Young |
| 0.461761 | 0.452277 | 0.14 | 0.098 | 1 | 14 | Parp16 | 0.042 | Young |
| 0.462163 | 0.653358 | 0.24 | 0.197 | 1 | 14 | Mfsd5 | 0.043 | Young |
| 0.462443 | 0.608843 | 0.22 | 0.164 | 1 | 14 | Dhrs1 | 0.056 | Young |
| 0.462831 | 0.239554 | 0.2 | 0.153 | 1 | 14 | Dmac1 | 0.047 | Young |
| 0.462918 | 0.319424 | 0.24 | 0.186 | 1 | 14 | Camsap1 | 0.054 | Young |
| 0.463054 | 1.533091 | 0.1 | 0.071 | 1 | 14 | Syde2 | 0.029 | Young |
| 0.463175 | 0.321822 | 0.62 | 0.574 | 1 | 14 | Slc25a4 | 0.046 | Young |
| 0.463893 | 0.064456 | 0.18 | 0.235 | 1 | 14 | Spopl | -0.055 | Young |
| 0.464228 | 0.511864 | 0.24 | 0.311 | 1 | 14 | Rbm26 | -0.071 | Young |
| 0.464448 | 0.05764 | 0.14 | 0.098 | 1 | 14 | Arfgap1 | 0.042 | Young |
| 0.464486 | 2.430117 | 0.06 | 0.038 | 1 | 14 | Gm6377 | 0.022 | Young |
| 0.465077 | 0.831693 | 0.2 | 0.153 | 1 | 14 | Ldah | 0.047 | Young |
| 0.46526 | 1.36586 | 0.18 | 0.131 | 1 | 14 | Prmt2 | 0.049 | Young |
| 0.46587 | 0.67073 | 0.46 | 0.437 | 1 | 14 | Myh9 | 0.023 | Young |
| 0.465944 | 1.75066 | 0.04 | 0.022 | 1 | 14 | Prdm11 | 0.018 | Young |

| | | | | | | | | |
|---|---|---|---|---|---|---|---|---|
| 0.465944 | 1.263169 | 0.04 | 0.022 | 1 | 14 | F830016B0 | 0.018 | Young |
| 0.465944 | 1.237753 | 0.04 | 0.022 | 1 | 14 | Cep78 | 0.018 | Young |
| 0.465944 | 1.237424 | 0.04 | 0.022 | 1 | 14 | Vdr | 0.018 | Young |
| 0.465944 | 1.132311 | 0.04 | 0.022 | 1 | 14 | 1-Mar | 0.018 | Young |
| 0.465944 | 1.029586 | 0.04 | 0.022 | 1 | 14 | Gm30648 | 0.018 | Young |
| 0.465944 | 0.933365 | 0.04 | 0.022 | 1 | 14 | Kcnk13 | 0.018 | Young |
| 0.465944 | 0.731739 | 0.04 | 0.022 | 1 | 14 | Colca2 | 0.018 | Young |
| 0.465944 | 0.549143 | 0.04 | 0.022 | 1 | 14 | Dhx35 | 0.018 | Young |
| 0.466296 | 0.1825 | 0.44 | 0.503 | 1 | 14 | Hspe1 | -0.063 | Young |
| 0.466317 | 0.160925 | 0.2 | 0.24 | 1 | 14 | Supt20 | -0.04 | Young |
| 0.466569 | 0.229195 | 0.94 | 0.934 | 1 | 14 | Rpl27a | 0.006 | Young |
| 0.46714 | 0.115682 | 0.3 | 0.372 | 1 | 14 | Adprh | -0.072 | Young |
| 0.467752 | 0.13783 | 0.16 | 0.12 | 1 | 14 | Slc7a6 | 0.04 | Young |
| 0.467866 | 0.670492 | 0.26 | 0.213 | 1 | 14 | Tm7sf3 | 0.047 | Young |
| 0.46793 | 0.153792 | 0.28 | 0.344 | 1 | 14 | Dnaja2 | -0.064 | Young |
| 0.468308 | 0.761963 | 0.32 | 0.273 | 1 | 14 | Lrch3 | 0.047 | Young |
| 0.468308 | 0.262238 | 0.32 | 0.273 | 1 | 14 | Cdkn1b | 0.047 | Young |
| 0.468493 | 0.553864 | 0.08 | 0.115 | 1 | 14 | Gstk1 | -0.035 | Young |
| 0.468612 | 2.08867 | 0.06 | 0.038 | 1 | 14 | Nedd1 | 0.022 | Young |
| 0.469092 | 0.64795 | 0.16 | 0.126 | 1 | 14 | Rab30 | 0.034 | Young |
| 0.469312 | 2.028572 | 0.1 | 0.071 | 1 | 14 | Tbl3 | 0.029 | Young |
| 0.469312 | 0.079176 | 0.1 | 0.071 | 1 | 14 | Decr2 | 0.029 | Young |
| 0.469431 | 0.53141 | 0.28 | 0.235 | 1 | 14 | Ier3 | 0.045 | Young |
| 0.469467 | 0.428839 | 0.4 | 0.355 | 1 | 14 | Brd2 | 0.045 | Young |
| 0.469716 | 0.218484 | 0.18 | 0.137 | 1 | 14 | Polr2l | 0.043 | Young |
| 0.469852 | 1.237216 | 0.2 | 0.158 | 1 | 14 | Ado | 0.042 | Young |
| 0.47006 | 0.525605 | 0.22 | 0.169 | 1 | 14 | Abhd2 | 0.051 | Young |
| 0.470238 | 0.185847 | 0.22 | 0.262 | 1 | 14 | Pptc7 | -0.042 | Young |
| 0.470249 | 0.452682 | 0.16 | 0.12 | 1 | 14 | Arhgef10 | 0.04 | Young |
| 0.470848 | 1.069717 | 0.14 | 0.104 | 1 | 14 | Phtf1 | 0.036 | Young |
| 0.470916 | 0.580157 | 0.12 | 0.087 | 1 | 14 | Stambp | 0.033 | Young |
| 0.471201 | 1.297648 | 0.08 | 0.115 | 1 | 14 | Spats2 | -0.035 | Young |
| 0.471238 | 1.501666 | 0.04 | 0.022 | 1 | 14 | Spata13 | 0.018 | Young |
| 0.471238 | 1.445018 | 0.04 | 0.022 | 1 | 14 | Apoo | 0.018 | Young |
| 0.471238 | 1.228392 | 0.04 | 0.022 | 1 | 14 | Rasl11b | 0.018 | Young |
| 0.471238 | 1.083307 | 0.04 | 0.022 | 1 | 14 | Tagap | 0.018 | Young |
| 0.471238 | 1.080347 | 0.04 | 0.022 | 1 | 14 | B3gntl1 | 0.018 | Young |
| 0.471238 | 1.06435 | 0.04 | 0.022 | 1 | 14 | Cnksr3 | 0.018 | Young |
| 0.47197 | 1.087821 | 0.32 | 0.273 | 1 | 14 | Raly | 0.047 | Young |
| 0.472165 | 0.251339 | 0.76 | 0.781 | 1 | 14 | Calu | -0.021 | Young |
| 0.472303 | 0.951758 | 0.06 | 0.093 | 1 | 14 | Cpsf1 | -0.033 | Young |
| 0.472343 | 0.14467 | 0.2 | 0.251 | 1 | 14 | Erlec1 | -0.051 | Young |
| 0.472458 | 0.8992 | 0.1 | 0.071 | 1 | 14 | Ascc2 | 0.029 | Young |
| 0.473351 | 0.806138 | 0.18 | 0.148 | 1 | 14 | Srpx2 | 0.032 | Young |
| 0.473792 | 0.875831 | 0.12 | 0.087 | 1 | 14 | Smg5 | 0.033 | Young |

| | | | | | | | | |
|---|---|---|---|---|---|---|---|---|
| 0.473793 | 0.004122 | 0.12 | 0.169 | 1 | 14 | Snx24 | -0.049 | Young |
| 0.474112 | 0.357212 | 0.42 | 0.508 | 1 | 14 | Dclk1 | -0.088 | Young |
| 0.474119 | 0.734396 | 0.2 | 0.153 | 1 | 14 | Sf3a2 | 0.047 | Young |
| 0.474222 | 0.615651 | 0.44 | 0.514 | 1 | 14 | Slc39a1 | -0.074 | Young |
| 0.474254 | 0.110641 | 0.6 | 0.585 | 1 | 14 | Tra2b | 0.015 | Young |
| 0.474338 | 0.40777 | 0.2 | 0.158 | 1 | 14 | Ddit3 | 0.042 | Young |
| 0.474467 | 0.340169 | 0.18 | 0.137 | 1 | 14 | Gba | 0.043 | Young |
| 0.474491 | 0.372925 | 0.24 | 0.197 | 1 | 14 | Msrb3 | 0.043 | Young |
| 0.47454 | 1.10511 | 0.04 | 0.071 | 1 | 14 | Adnp2 | -0.031 | Young |
| 0.474854 | 0.277987 | 0.44 | 0.377 | 1 | 14 | Ppig | 0.063 | Young |
| 0.475206 | 0.125402 | 0.38 | 0.454 | 1 | 14 | Rock1 | -0.074 | Young |
| 0.475317 | 0.21177 | 0.18 | 0.224 | 1 | 14 | Pqbp1 | -0.044 | Young |
| 0.47618 | 0.72488 | 0.22 | 0.18 | 1 | 14 | Ilkap | 0.04 | Young |
| 0.476251 | 0.200678 | 0.62 | 0.546 | 1 | 14 | Rab6a | 0.074 | Young |
| 0.476316 | 0.953637 | 0.18 | 0.142 | 1 | 14 | Gm49336 | 0.038 | Young |
| 0.476395 | 0.203343 | 0.2 | 0.153 | 1 | 14 | Mindy3 | 0.047 | Young |
| 0.476565 | 2.842198 | 0.04 | 0.022 | 1 | 14 | S100b | 0.018 | Young |
| 0.476565 | 2.833453 | 0.04 | 0.022 | 1 | 14 | Bicdl1 | 0.018 | Young |
| 0.476565 | 2.14357 | 0.04 | 0.022 | 1 | 14 | Zfp420 | 0.018 | Young |
| 0.476565 | 1.764354 | 0.04 | 0.022 | 1 | 14 | Zfp202 | 0.018 | Young |
| 0.476565 | 1.18107 | 0.04 | 0.022 | 1 | 14 | Cbr4 | 0.018 | Young |
| 0.476565 | 1.162772 | 0.04 | 0.022 | 1 | 14 | Med27 | 0.018 | Young |
| 0.476565 | 1.089331 | 0.04 | 0.022 | 1 | 14 | Rapgef5 | 0.018 | Young |
| 0.476565 | 1.068971 | 0.04 | 0.022 | 1 | 14 | Ptcd1 | 0.018 | Young |
| 0.476565 | 0.952758 | 0.04 | 0.022 | 1 | 14 | Slc19a1 | 0.018 | Young |
| 0.476565 | 0.851802 | 0.04 | 0.022 | 1 | 14 | Xrcc2 | 0.018 | Young |
| 0.476565 | 0.838858 | 0.04 | 0.022 | 1 | 14 | Oxtr | 0.018 | Young |
| 0.476565 | 0.825726 | 0.04 | 0.022 | 1 | 14 | Cacng4 | 0.018 | Young |
| 0.476565 | 0.754414 | 0.04 | 0.022 | 1 | 14 | P2ry2 | 0.018 | Young |
| 0.476565 | 0.702268 | 0.04 | 0.022 | 1 | 14 | Ercc8 | 0.018 | Young |
| 0.476565 | 0.698083 | 0.04 | 0.022 | 1 | 14 | Zfp647 | 0.018 | Young |
| 0.476565 | 0.676092 | 0.04 | 0.022 | 1 | 14 | Tmem220 | 0.018 | Young |
| 0.476565 | 0.655566 | 0.04 | 0.022 | 1 | 14 | Tspyl5 | 0.018 | Young |
| 0.476565 | 0.5916 | 0.04 | 0.022 | 1 | 14 | Dgke | 0.018 | Young |
| 0.476565 | 0.566052 | 0.04 | 0.022 | 1 | 14 | Gm34006 | 0.018 | Young |
| 0.476565 | 0.551926 | 0.04 | 0.022 | 1 | 14 | Notch4 | 0.018 | Young |
| 0.476565 | 0.512919 | 0.04 | 0.022 | 1 | 14 | Abcb9 | 0.018 | Young |
| 0.476565 | 0.384068 | 0.04 | 0.022 | 1 | 14 | Rnf125 | 0.018 | Young |
| 0.476677 | 0.664462 | 0.12 | 0.087 | 1 | 14 | Zfp760 | 0.033 | Young |
| 0.476926 | 0.985144 | 0.06 | 0.038 | 1 | 14 | Ccrl2 | 0.022 | Young |
| 0.476926 | 0.882011 | 0.06 | 0.038 | 1 | 14 | Tsix | 0.022 | Young |
| 0.477101 | 0.192826 | 0.24 | 0.191 | 1 | 14 | Psmd7 | 0.049 | Young |
| 0.477576 | 0.1911 | 0.16 | 0.208 | 1 | 14 | Cramp1l | -0.048 | Young |
| 0.477796 | 1.767627 | 0.08 | 0.055 | 1 | 14 | Tnik | 0.025 | Young |
| 0.477796 | 1.668432 | 0.08 | 0.055 | 1 | 14 | Drg2 | 0.025 | Young |

| | | | | | | | | |
|---|---|---|---|---|---|---|---|---|
| 0.478026 | 0.14436 | 0.46 | 0.399 | 1 | 14 | Bmpr1a | 0.061 | Young |
| 0.478149 | 0.080274 | 0.3 | 0.23 | 1 | 14 | Acbd3 | 0.07 | Young |
| 0.47832 | 0.471738 | 0.4 | 0.339 | 1 | 14 | Zfp503 | 0.061 | Young |
| 0.47833 | 0.70368 | 0.36 | 0.317 | 1 | 14 | Sfrp1 | 0.043 | Young |
| 0.47833 | 0.06486 | 0.06 | 0.093 | 1 | 14 | Slc4a3 | -0.033 | Young |
| 0.478879 | 0.506852 | 0.14 | 0.104 | 1 | 14 | Ddx47 | 0.036 | Young |
| 0.478999 | 0.686682 | 0.16 | 0.126 | 1 | 14 | Stard4 | 0.034 | Young |
| 0.479026 | 0.318335 | 0.82 | 0.858 | 1 | 14 | Eif4g2 | -0.038 | Young |
| 0.479768 | 0.627928 | 0.32 | 0.279 | 1 | 14 | Reck | 0.041 | Young |
| 0.480348 | 0.513519 | 0.5 | 0.481 | 1 | 14 | Psma7 | 0.019 | Young |
| 0.480723 | 0.438147 | 0.24 | 0.197 | 1 | 14 | Psmd6 | 0.043 | Young |
| 0.481113 | 2.319186 | 0.06 | 0.038 | 1 | 14 | Gss | 0.022 | Young |
| 0.481113 | 1.616039 | 0.06 | 0.038 | 1 | 14 | Zswim4 | 0.022 | Young |
| 0.481359 | 0.07718 | 0.06 | 0.093 | 1 | 14 | Myoc | -0.033 | Young |
| 0.481368 | 1.020936 | 0.08 | 0.055 | 1 | 14 | Nkiras2 | 0.025 | Young |
| 0.481572 | 0.446169 | 0.14 | 0.104 | 1 | 14 | Csf1r | 0.036 | Young |
| 0.48192 | 0.665349 | 0.26 | 0.213 | 1 | 14 | Pet100 | 0.047 | Young |
| 0.481925 | 0.70413 | 0.04 | 0.022 | 1 | 14 | Mtbp | 0.018 | Young |
| 0.481925 | 0.630403 | 0.04 | 0.022 | 1 | 14 | 2010007H( | 0.018 | Young |
| 0.481925 | 0.575108 | 0.04 | 0.022 | 1 | 14 | Fam234b | 0.018 | Young |
| 0.481925 | 0.551931 | 0.04 | 0.022 | 1 | 14 | Cadps2 | 0.018 | Young |
| 0.481925 | 0.45658 | 0.04 | 0.022 | 1 | 14 | Usp49 | 0.018 | Young |
| 0.481925 | 0.452492 | 0.04 | 0.022 | 1 | 14 | Relt | 0.018 | Young |
| 0.481925 | 0.433768 | 0.04 | 0.022 | 1 | 14 | Hoxc11 | 0.018 | Young |
| 0.481925 | 0.352164 | 0.04 | 0.022 | 1 | 14 | Flt1 | 0.018 | Young |
| 0.481925 | 0.344432 | 0.04 | 0.022 | 1 | 14 | Snx25 | 0.018 | Young |
| 0.481925 | 0.234041 | 0.04 | 0.022 | 1 | 14 | Tefm | 0.018 | Young |
| 0.481925 | 0.215677 | 0.04 | 0.022 | 1 | 14 | Naif1 | 0.018 | Young |
| 0.481925 | 0.017612 | 0.04 | 0.022 | 1 | 14 | Hdac9 | 0.018 | Young |
| 0.482477 | 0.594341 | 0.12 | 0.087 | 1 | 14 | Rhno1 | 0.033 | Young |
| 0.482477 | 0.095482 | 0.12 | 0.087 | 1 | 14 | Dact3 | 0.033 | Young |
| 0.482743 | 0.641922 | 0.1 | 0.142 | 1 | 14 | Elp3 | -0.042 | Young |
| 0.48281 | 0.354387 | 0.24 | 0.197 | 1 | 14 | Pan3 | 0.043 | Young |
| 0.482845 | 0.290262 | 0.16 | 0.12 | 1 | 14 | Scap | 0.04 | Young |
| 0.482976 | 0.410323 | 0.44 | 0.404 | 1 | 14 | Cox5a | 0.036 | Young |
| 0.483154 | 0.031556 | 0.28 | 0.224 | 1 | 14 | Gabpb2 | 0.056 | Young |
| 0.483186 | 1.052193 | 0.14 | 0.191 | 1 | 14 | Hpf1 | -0.051 | Young |
| 0.483381 | 0.688262 | 0.2 | 0.158 | 1 | 14 | Pygb | 0.042 | Young |
| 0.48346 | 0.312449 | 0.24 | 0.301 | 1 | 14 | Arpc4 | -0.061 | Young |
| 0.483493 | 1.340491 | 0.14 | 0.098 | 1 | 14 | Phkb | 0.042 | Young |
| 0.483801 | 0.025552 | 0.58 | 0.65 | 1 | 14 | Tubb5 | -0.07 | Young |
| 0.483802 | 0.100028 | 0.46 | 0.388 | 1 | 14 | Man2a2 | 0.072 | Young |
| 0.484465 | 0.056733 | 0.38 | 0.339 | 1 | 14 | Usp34 | 0.041 | Young |
| 0.484694 | 0.003161 | 0.28 | 0.311 | 1 | 14 | Trappc4 | -0.031 | Young |
| 0.484807 | 0.867445 | 0.22 | 0.18 | 1 | 14 | Snhg12 | 0.04 | Young |

| | | | | | | | | |
|---|---|---|---|---|---|---|---|---|
| 0.48489 | 0.11436 | 0.94 | 0.94 | 1 | 14 | Rps28 | 0 | Young |
| 0.484954 | 1.157429 | 0.08 | 0.055 | 1 | 14 | Abhd6 | 0.025 | Young |
| 0.484996 | 0.007641 | 0.72 | 0.803 | 1 | 14 | Morf4l1 | -0.083 | Young |
| 0.48509 | 0.099495 | 0.44 | 0.546 | 1 | 14 | Cd9 | -0.106 | Young |
| 0.485158 | 0.751491 | 0.1 | 0.071 | 1 | 14 | Spsb2 | 0.029 | Young |
| 0.48532 | 1.51299 | 0.06 | 0.038 | 1 | 14 | Slc2a13 | 0.022 | Young |
| 0.48532 | 1.121102 | 0.06 | 0.038 | 1 | 14 | Def6 | 0.022 | Young |
| 0.485392 | 1.241632 | 0.12 | 0.087 | 1 | 14 | Enox2 | 0.033 | Young |
| 0.486839 | 0.432856 | 0.4 | 0.344 | 1 | 14 | Cpne3 | 0.056 | Young |
| 0.487012 | 0.163299 | 0.18 | 0.131 | 1 | 14 | Sp2 | 0.049 | Young |
| 0.487318 | 0.425782 | 0.04 | 0.022 | 1 | 14 | Zkscan14 | 0.018 | Young |
| 0.487318 | 0.416316 | 0.04 | 0.022 | 1 | 14 | Ggact | 0.018 | Young |
| 0.487318 | 0.368438 | 0.04 | 0.022 | 1 | 14 | Zfp974 | 0.018 | Young |
| 0.487318 | 0.366024 | 0.04 | 0.022 | 1 | 14 | Stard10 | 0.018 | Young |
| 0.487318 | 0.224803 | 0.04 | 0.022 | 1 | 14 | Fam107a | 0.018 | Young |
| 0.487318 | 0.112007 | 0.04 | 0.022 | 1 | 14 | Lag3 | 0.018 | Young |
| 0.487318 | 0.030384 | 0.04 | 0.022 | 1 | 14 | Siah2 | 0.018 | Young |
| 0.487741 | 0.214593 | 0.24 | 0.284 | 1 | 14 | Bag1 | -0.044 | Young |
| 0.487934 | 0.285753 | 0.16 | 0.12 | 1 | 14 | Noct | 0.04 | Young |
| 0.488059 | 0.736492 | 0.28 | 0.23 | 1 | 14 | Rab3il1 | 0.05 | Young |
| 0.488152 | 0.412956 | 0.68 | 0.667 | 1 | 14 | Hnrnpa3 | 0.013 | Young |
| 0.488315 | 0.343473 | 0.12 | 0.087 | 1 | 14 | Clmn | 0.033 | Young |
| 0.488362 | 0.756551 | 0.1 | 0.071 | 1 | 14 | Stmn4 | 0.029 | Young |
| 0.488362 | 0.506102 | 0.1 | 0.071 | 1 | 14 | Src | 0.029 | Young |
| 0.488555 | 1.064802 | 0.08 | 0.055 | 1 | 14 | Ctgf | 0.025 | Young |
| 0.488555 | 0.958087 | 0.08 | 0.055 | 1 | 14 | Cox6b2 | 0.025 | Young |
| 0.488957 | 0.32732 | 0.44 | 0.607 | 1 | 14 | Prss23 | -0.167 | Young |
| 0.489156 | 0.232916 | 0.44 | 0.41 | 1 | 14 | Impad1 | 0.03 | Young |
| 0.489547 | 1.403803 | 0.06 | 0.038 | 1 | 14 | Pdk3 | 0.022 | Young |
| 0.489547 | 1.00183 | 0.06 | 0.038 | 1 | 14 | Mon1a | 0.022 | Young |
| 0.489631 | 0.061505 | 0.32 | 0.262 | 1 | 14 | Usp47 | 0.058 | Young |
| 0.49034 | 0.581089 | 0.24 | 0.186 | 1 | 14 | Ube2j1 | 0.054 | Young |
| 0.490777 | 0.168568 | 0.98 | 0.907 | 1 | 14 | Rpl17 | 0.073 | Young |
| 0.491168 | 0.339045 | 0.22 | 0.268 | 1 | 14 | Usp8 | -0.048 | Young |
| 0.491249 | 0.731977 | 0.12 | 0.087 | 1 | 14 | Cipc | 0.033 | Young |
| 0.491249 | 0.442247 | 0.12 | 0.087 | 1 | 14 | Crtc1 | 0.033 | Young |
| 0.491577 | 0.773785 | 0.1 | 0.071 | 1 | 14 | Mrps31 | 0.029 | Young |
| 0.491971 | 0.178478 | 0.26 | 0.208 | 1 | 14 | Vbp1 | 0.052 | Young |
| 0.49217 | 0.797982 | 0.08 | 0.055 | 1 | 14 | Nxt2 | 0.025 | Young |
| 0.492432 | 0.199256 | 0.66 | 0.71 | 1 | 14 | Atp5a1 | -0.05 | Young |
| 0.492517 | 0.500808 | 0.2 | 0.158 | 1 | 14 | Lsm5 | 0.042 | Young |
| 0.49255 | 0.47086 | 0.38 | 0.322 | 1 | 14 | Ndufa11 | 0.058 | Young |
| 0.492743 | 0.408941 | 0.04 | 0.022 | 1 | 14 | Rhoh | 0.018 | Young |
| 0.492858 | 0.550743 | 0.14 | 0.109 | 1 | 14 | Trappc6a | 0.031 | Young |
| 0.492979 | 0.879478 | 0.18 | 0.142 | 1 | 14 | Extl2 | 0.038 | Young |

| | | | | | | | | |
|---|---|---|---|---|---|---|---|---|
| 0.493053 | 0.227883 | 0.1 | 0.137 | 1 | 14 | Katna1 | -0.037 | Young |
| 0.493793 | 1.401834 | 0.06 | 0.038 | 1 | 14 | Lrrn1 | 0.022 | Young |
| 0.493793 | 0.828677 | 0.06 | 0.038 | 1 | 14 | A530017D2 | 0.022 | Young |
| 0.494291 | 0.083872 | 0.26 | 0.306 | 1 | 14 | Ythdf3 | -0.046 | Young |
| 0.494329 | 0.28921 | 0.38 | 0.322 | 1 | 14 | Snrpd3 | 0.058 | Young |
| 0.494803 | 0.718422 | 0.1 | 0.071 | 1 | 14 | Pin1 | 0.029 | Young |
| 0.494803 | 0.490334 | 0.1 | 0.071 | 1 | 14 | Irak4 | 0.029 | Young |
| 0.494815 | 0.105364 | 0.2 | 0.153 | 1 | 14 | Edc4 | 0.047 | Young |
| 0.494815 | 0.48169 | 0.2 | 0.158 | 1 | 14 | Fam126b | 0.042 | Young |
| 0.495537 | 0.550755 | 0.46 | 0.388 | 1 | 14 | Ppic | 0.072 | Young |
| 0.495548 | 0.947217 | 0.14 | 0.109 | 1 | 14 | Hras | 0.031 | Young |
| 0.495724 | 0.420545 | 0.5 | 0.596 | 1 | 14 | Rab1a | -0.096 | Young |
| 0.49723 | 0.265242 | 0.32 | 0.262 | 1 | 14 | Cdk9 | 0.058 | Young |
| 0.498041 | 0.763848 | 0.1 | 0.071 | 1 | 14 | Dph5 | 0.029 | Young |
| 0.498041 | 0.422424 | 0.1 | 0.071 | 1 | 14 | Gins4 | 0.029 | Young |
| 0.498059 | 1.893184 | 0.06 | 0.038 | 1 | 14 | Gfra4 | 0.022 | Young |
| 0.498059 | 1.447917 | 0.06 | 0.038 | 1 | 14 | Klhl8 | 0.022 | Young |
| 0.498059 | 1.158869 | 0.06 | 0.038 | 1 | 14 | Me2 | 0.022 | Young |
| 0.498059 | 1.037558 | 0.06 | 0.038 | 1 | 14 | Plppr2 | 0.022 | Young |
| 0.498059 | 0.976659 | 0.06 | 0.038 | 1 | 14 | Arsj | 0.022 | Young |
| 0.498059 | 0.901055 | 0.06 | 0.038 | 1 | 14 | Adap1 | 0.022 | Young |
| 0.498059 | 0.664183 | 0.06 | 0.038 | 1 | 14 | Kalrn | 0.022 | Young |
| 0.498059 | 0.599602 | 0.06 | 0.038 | 1 | 14 | Gm9493 | 0.022 | Young |
| 0.498059 | 0.597932 | 0.06 | 0.038 | 1 | 14 | Necab3 | 0.022 | Young |
| 0.498059 | 0.485827 | 0.06 | 0.038 | 1 | 14 | Lmbr1l | 0.022 | Young |
| 0.4982 | 0.123795 | 0.9 | 0.913 | 1 | 14 | Rpl37 | -0.013 | Young |
| 0.4982 | 0.020218 | 0.04 | 0.022 | 1 | 14 | Nlrp3 | 0.018 | Young |
| 0.498749 | 0.388542 | 0.22 | 0.169 | 1 | 14 | Adgrg2 | 0.051 | Young |
| 0.499151 | 0.102063 | 0.16 | 0.126 | 1 | 14 | Faap20 | 0.034 | Young |
| 0.499444 | 1.45761 | 0.08 | 0.055 | 1 | 14 | Rbm45 | 0.025 | Young |
| 0.499444 | 0.656978 | 0.08 | 0.055 | 1 | 14 | Tatdn3 | 0.025 | Young |
| 0.499444 | 0.647986 | 0.08 | 0.055 | 1 | 14 | Rpain | 0.025 | Young |
| 0.499444 | 0.643358 | 0.08 | 0.055 | 1 | 14 | Tenm3 | 0.025 | Young |
| 0.499444 | 0.557751 | 0.08 | 0.055 | 1 | 14 | Fam109b | 0.025 | Young |
| 0.499444 | 0.023706 | 0.08 | 0.055 | 1 | 14 | 1110046J0 | 0.025 | Young |
| 0.499681 | 0.005584 | 0.18 | 0.213 | 1 | 14 | Hk1 | -0.033 | Young |
| 0.500629 | 0.30741 | 0.26 | 0.35 | 1 | 14 | Auts2 | -0.09 | Young |
| 0.50129 | 1.750169 | 0.1 | 0.071 | 1 | 14 | Afap1l2 | 0.029 | Young |
| 0.50129 | 0.382569 | 0.1 | 0.071 | 1 | 14 | Tmem38a | 0.029 | Young |
| 0.501701 | 0.91539 | 0.16 | 0.126 | 1 | 14 | Wdr48 | 0.034 | Young |
| 0.501824 | 0.375881 | 0.22 | 0.251 | 1 | 14 | Zfp266 | -0.031 | Young |
| 0.502039 | 0.443598 | 0.3 | 0.257 | 1 | 14 | Eny2 | 0.043 | Young |
| 0.502345 | 0.684661 | 0.06 | 0.038 | 1 | 14 | 1700096K1 | 0.022 | Young |
| 0.502345 | 0.56763 | 0.06 | 0.038 | 1 | 14 | Rnaseh2b | 0.022 | Young |
| 0.502345 | 0.506599 | 0.06 | 0.038 | 1 | 14 | Kyat1 | 0.022 | Young |

| | | | | | | | | |
|---|---|---|---|---|---|---|---|---|
| 0.502528 | 0.160696 | 0.32 | 0.257 | 1 | 14 | Nfatc3 | 0.063 | Young |
| 0.502639 | 0.440701 | 0.18 | 0.142 | 1 | 14 | 1810022K0 | 0.038 | Young |
| 0.503102 | 2.476702 | 0.04 | 0.066 | 1 | 14 | Klc4 | -0.026 | Young |
| 0.503102 | 0.221072 | 0.08 | 0.055 | 1 | 14 | Emx2 | 0.025 | Young |
| 0.50352 | 0.014145 | 0.18 | 0.137 | 1 | 14 | 4930453N2 | 0.043 | Young |
| 0.50355 | 0.442535 | 0.58 | 0.574 | 1 | 14 | Nrip1 | 0.006 | Young |
| 0.504165 | 0.673533 | 0.2 | 0.153 | 1 | 14 | Extl3 | 0.047 | Young |
| 0.50455 | 0.621509 | 0.1 | 0.071 | 1 | 14 | Cmip | 0.029 | Young |
| 0.50455 | 0.255605 | 0.1 | 0.071 | 1 | 14 | Snx19 | 0.029 | Young |
| 0.504854 | 0.646186 | 0.38 | 0.333 | 1 | 14 | Dek | 0.047 | Young |
| 0.50488 | 0.241549 | 0.64 | 0.628 | 1 | 14 | Atf4 | 0.012 | Young |
| 0.504948 | 0.032757 | 0.16 | 0.115 | 1 | 14 | Ints6l | 0.045 | Young |
| 0.505015 | 0.015086 | 0.26 | 0.322 | 1 | 14 | Dnajc5 | -0.062 | Young |
| 0.506056 | 0.410796 | 0.12 | 0.087 | 1 | 14 | Nfkbid | 0.033 | Young |
| 0.506287 | 0.372857 | 0.12 | 0.158 | 1 | 14 | Bmt2 | -0.038 | Young |
| 0.506472 | 0.765149 | 0.18 | 0.148 | 1 | 14 | Prpsap1 | 0.032 | Young |
| 0.506482 | 0.139421 | 0.34 | 0.372 | 1 | 14 | Twf1 | -0.032 | Young |
| 0.506517 | 0.285194 | 0.2 | 0.153 | 1 | 14 | Upf3a | 0.047 | Young |
| 0.506642 | 0.566244 | 0.38 | 0.333 | 1 | 14 | Rin2 | 0.047 | Young |
| 0.50665 | 0.722704 | 0.06 | 0.038 | 1 | 14 | Tysnd1 | 0.022 | Young |
| 0.50665 | 0.679196 | 0.06 | 0.038 | 1 | 14 | Cd80 | 0.022 | Young |
| 0.506774 | 1.295753 | 0.08 | 0.055 | 1 | 14 | Amacr | 0.025 | Young |
| 0.506774 | 1.144936 | 0.08 | 0.055 | 1 | 14 | Gne | 0.025 | Young |
| 0.50768 | 0.650856 | 0.34 | 0.295 | 1 | 14 | Snrk | 0.045 | Young |
| 0.507821 | 1.040682 | 0.1 | 0.071 | 1 | 14 | Tada2b | 0.029 | Young |
| 0.507821 | 0.989025 | 0.1 | 0.071 | 1 | 14 | Acad8 | 0.029 | Young |
| 0.507821 | 0.185268 | 0.1 | 0.071 | 1 | 14 | Chic1 | 0.029 | Young |
| 0.507821 | 0.0923 | 0.1 | 0.071 | 1 | 14 | Dnttip1 | 0.029 | Young |
| 0.508635 | 0.402688 | 0.12 | 0.093 | 1 | 14 | Aen | 0.027 | Young |
| 0.508725 | 0.299164 | 0.2 | 0.158 | 1 | 14 | Sos1 | 0.042 | Young |
| 0.509183 | 0.048966 | 0.3 | 0.24 | 1 | 14 | Bfar | 0.06 | Young |
| 0.509305 | 0.222401 | 0.22 | 0.164 | 1 | 14 | Nr2c2 | 0.056 | Young |
| 0.509331 | 0.790395 | 0.28 | 0.257 | 1 | 14 | Pygl | 0.023 | Young |
| 0.50936 | 0.195091 | 0.86 | 0.891 | 1 | 14 | Rps13 | -0.031 | Young |
| 0.509489 | 0.632979 | 0.24 | 0.202 | 1 | 14 | Mef2c | 0.038 | Young |
| 0.51046 | 0.710873 | 0.04 | 0.066 | 1 | 14 | Vps33b | -0.026 | Young |
| 0.510635 | 0.260603 | 0.26 | 0.219 | 1 | 14 | Ndrg3 | 0.041 | Young |
| 0.510678 | 0.205535 | 0.32 | 0.262 | 1 | 14 | Gng11 | 0.058 | Young |
| 0.510911 | 0.019813 | 0.46 | 0.514 | 1 | 14 | Gas7 | -0.054 | Young |
| 0.510974 | 0.702116 | 0.06 | 0.038 | 1 | 14 | Naprt | 0.022 | Young |
| 0.510974 | 0.689546 | 0.06 | 0.038 | 1 | 14 | Anks6 | 0.022 | Young |
| 0.510974 | 0.474567 | 0.06 | 0.038 | 1 | 14 | Tmem251 | 0.022 | Young |
| 0.510974 | 0.434259 | 0.06 | 0.038 | 1 | 14 | Ppargc1a | 0.022 | Young |
| 0.510974 | 0.347779 | 0.06 | 0.038 | 1 | 14 | Zfp105 | 0.022 | Young |
| 0.510974 | 0.346035 | 0.06 | 0.038 | 1 | 14 | Fntb | 0.022 | Young |

| | | | | | | | | |
|---|---|---|---|---|---|---|---|---|
| 0.510974 | 0.31004 | 0.06 | 0.038 | 1 | 14 | Elmod3 | 0.022 | Young |
| 0.511063 | 0.265653 | 0.2 | 0.158 | 1 | 14 | Ttc39b | 0.042 | Young |
| 0.511104 | 0.258416 | 0.82 | 0.798 | 1 | 14 | Eef1b2 | 0.022 | Young |
| 0.51119 | 0.146809 | 0.16 | 0.12 | 1 | 14 | Selenoh | 0.04 | Young |
| 0.511766 | 0.235753 | 0.42 | 0.35 | 1 | 14 | Eea1 | 0.07 | Young |
| 0.511854 | 0.718122 | 0.14 | 0.109 | 1 | 14 | Vti1a | 0.031 | Young |
| 0.512345 | 0.551483 | 0.28 | 0.311 | 1 | 14 | Ktn1 | -0.031 | Young |
| 0.514237 | 0.032835 | 0.46 | 0.503 | 1 | 14 | Smarca2 | -0.043 | Young |
| 0.514397 | 0.738526 | 0.1 | 0.071 | 1 | 14 | Mrps30 | 0.029 | Young |
| 0.514397 | 0.413362 | 0.1 | 0.071 | 1 | 14 | Zfp120 | 0.029 | Young |
| 0.514555 | 0.566161 | 0.16 | 0.126 | 1 | 14 | Fmnl1 | 0.034 | Young |
| 0.516168 | 0.936149 | 0.28 | 0.246 | 1 | 14 | Ap3b1 | 0.034 | Young |
| 0.51686 | 0.143959 | 0.24 | 0.197 | 1 | 14 | Zfp280d | 0.043 | Young |
| 0.517361 | 0.76778 | 0.42 | 0.393 | 1 | 14 | Cdv3 | 0.027 | Young |
| 0.517678 | 0.41934 | 0.22 | 0.279 | 1 | 14 | Tug1 | -0.059 | Young |
| 0.517701 | 1.003375 | 0.1 | 0.071 | 1 | 14 | Aatf | 0.029 | Young |
| 0.517701 | 0.543005 | 0.1 | 0.071 | 1 | 14 | Mettl5 | 0.029 | Young |
| 0.517701 | 0.310293 | 0.1 | 0.071 | 1 | 14 | Mrpl37 | 0.029 | Young |
| 0.517874 | 1.839409 | 0.08 | 0.055 | 1 | 14 | Pyroxd2 | 0.025 | Young |
| 0.517874 | 0.811165 | 0.08 | 0.055 | 1 | 14 | Fam110b | 0.025 | Young |
| 0.517874 | 0.394384 | 0.08 | 0.055 | 1 | 14 | Gtpbp1 | 0.025 | Young |
| 0.517874 | 0.047958 | 0.08 | 0.055 | 1 | 14 | Kcp | 0.025 | Young |
| 0.518069 | 0.931068 | 0.12 | 0.087 | 1 | 14 | Usf1 | 0.033 | Young |
| 0.519028 | 0.220425 | 0.24 | 0.197 | 1 | 14 | 2810403A( | 0.043 | Young |
| 0.519681 | 0.957344 | 0.06 | 0.038 | 1 | 14 | 2300009A( | 0.022 | Young |
| 0.519681 | 0.777629 | 0.06 | 0.038 | 1 | 14 | Pex11a | 0.022 | Young |
| 0.519681 | 0.040541 | 0.06 | 0.038 | 1 | 14 | Atpaf2 | 0.022 | Young |
| 0.519744 | 1.457248 | 0.16 | 0.126 | 1 | 14 | Ciz1 | 0.034 | Young |
| 0.521017 | 0.140676 | 0.1 | 0.071 | 1 | 14 | Tmem218 | 0.029 | Young |
| 0.521095 | 0.223887 | 0.12 | 0.087 | 1 | 14 | Tars2 | 0.033 | Young |
| 0.521602 | 0.926446 | 0.08 | 0.055 | 1 | 14 | Fam13b | 0.025 | Young |
| 0.521602 | 0.181663 | 0.04 | 0.066 | 1 | 14 | Pou2f2 | -0.026 | Young |
| 0.521602 | 0.077351 | 0.08 | 0.055 | 1 | 14 | Nfatc2 | 0.025 | Young |
| 0.521893 | 0.012347 | 0.4 | 0.459 | 1 | 14 | Npepps | -0.059 | Young |
| 0.521959 | 0.503567 | 0.22 | 0.175 | 1 | 14 | Tssc4 | 0.045 | Young |
| 0.522019 | 0.409877 | 0.62 | 0.574 | 1 | 14 | 9530068E( | 0.046 | Young |
| 0.522349 | 0.773758 | 0.16 | 0.126 | 1 | 14 | Pigx | 0.034 | Young |
| 0.522407 | 0.245378 | 0.8 | 0.76 | 1 | 14 | Atp5l | 0.04 | Young |
| 0.522937 | 0.225281 | 0.26 | 0.224 | 1 | 14 | Sgms1 | 0.036 | Young |
| 0.523142 | 0.014427 | 0.2 | 0.153 | 1 | 14 | Gk5 | 0.047 | Young |
| 0.523148 | 1.018862 | 0.26 | 0.219 | 1 | 14 | Keap1 | 0.041 | Young |
| 0.523177 | 0.025792 | 0.44 | 0.519 | 1 | 14 | Pcdh19 | -0.079 | Young |
| 0.523398 | 0.107535 | 0.18 | 0.137 | 1 | 14 | Psmg4 | 0.043 | Young |
| 0.523413 | 0.989472 | 0.12 | 0.093 | 1 | 14 | Abcf3 | 0.027 | Young |
| 0.523413 | 0.125153 | 0.26 | 0.208 | 1 | 14 | Taf1d | 0.052 | Young |

| | | | | | | | | |
|---|---|---|---|---|---|---|---|---|
| 0.524062 | 0.216048 | 0.06 | 0.038 | 1 | 14 | Slc22a4 | 0.022 | Young |
| 0.524062 | 0.113962 | 0.06 | 0.038 | 1 | 14 | Gatc | 0.022 | Young |
| 0.52413 | 0.083879 | 0.12 | 0.087 | 1 | 14 | Atg101 | 0.033 | Young |
| 0.524343 | 0.400148 | 0.1 | 0.071 | 1 | 14 | Lsg1 | 0.029 | Young |
| 0.524343 | 0.242687 | 0.1 | 0.071 | 1 | 14 | Brwd3 | 0.029 | Young |
| 0.524343 | 0.170506 | 0.1 | 0.071 | 1 | 14 | Ikbkap | 0.029 | Young |
| 0.524343 | 0.168629 | 0.1 | 0.071 | 1 | 14 | Strbp | 0.029 | Young |
| 0.524842 | 0.660588 | 0.06 | 0.093 | 1 | 14 | Acad9 | -0.033 | Young |
| 0.524853 | 0.938058 | 0.32 | 0.273 | 1 | 14 | Tubb2a | 0.047 | Young |
| 0.525221 | 0.556185 | 0.3 | 0.24 | 1 | 14 | Sike1 | 0.06 | Young |
| 0.525343 | 0.074484 | 0.08 | 0.055 | 1 | 14 | Meis3 | 0.025 | Young |
| 0.525343 | 0.023555 | 0.08 | 0.055 | 1 | 14 | Il34 | 0.025 | Young |
| 0.525413 | 1.300595 | 0.16 | 0.131 | 1 | 14 | Gatd1 | 0.029 | Young |
| 0.52585 | 0.386272 | 0.36 | 0.421 | 1 | 14 | Tnks2 | -0.061 | Young |
| 0.525931 | 0.843112 | 0.18 | 0.148 | 1 | 14 | Rab24 | 0.032 | Young |
| 0.526158 | 0.106299 | 0.38 | 0.317 | 1 | 14 | Ptges | 0.063 | Young |
| 0.527143 | 0.849456 | 0.2 | 0.164 | 1 | 14 | Spock2 | 0.036 | Young |
| 0.527175 | 0.179753 | 0.12 | 0.087 | 1 | 14 | Wdr7 | 0.033 | Young |
| 0.527206 | 0.216795 | 0.72 | 0.787 | 1 | 14 | Cox4i1 | -0.067 | Young |
| 0.527427 | 0.249529 | 0.52 | 0.53 | 1 | 14 | Tceal9 | -0.01 | Young |
| 0.527477 | 0.793995 | 0.16 | 0.208 | 1 | 14 | Tmbim4 | -0.048 | Young |
| 0.527578 | 0.568202 | 0.16 | 0.126 | 1 | 14 | Tgfbrap1 | 0.034 | Young |
| 0.527578 | 0.371888 | 0.16 | 0.126 | 1 | 14 | Ranbp3 | 0.034 | Young |
| 0.527626 | 0.459839 | 0.1 | 0.131 | 1 | 14 | Mrpl41 | -0.031 | Young |
| 0.527648 | 0.156809 | 0.18 | 0.208 | 1 | 14 | Lsm7 | -0.028 | Young |
| 0.52768 | 0.029146 | 0.1 | 0.071 | 1 | 14 | Ccdc93 | 0.029 | Young |
| 0.528372 | 0.317129 | 0.34 | 0.295 | 1 | 14 | Luzp1 | 0.045 | Young |
| 0.528463 | 0.143221 | 0.06 | 0.038 | 1 | 14 | Tjp2 | 0.022 | Young |
| 0.528463 | 0.131836 | 0.06 | 0.038 | 1 | 14 | Wrap73 | 0.022 | Young |
| 0.528775 | 0.346944 | 0.14 | 0.175 | 1 | 14 | Sun2 | -0.035 | Young |
| 0.528805 | 0.542574 | 0.22 | 0.175 | 1 | 14 | Il13ra1 | 0.045 | Young |
| 0.529181 | 0.437644 | 0.36 | 0.311 | 1 | 14 | Polr2k | 0.049 | Young |
| 0.529183 | 0.344129 | 0.28 | 0.235 | 1 | 14 | Ythdf2 | 0.045 | Young |
| 0.529277 | 0.137573 | 0.26 | 0.311 | 1 | 14 | Fam76a | -0.051 | Young |
| 0.529385 | 1.580041 | 0.12 | 0.093 | 1 | 14 | Hax1 | 0.027 | Young |
| 0.5295 | 0.412299 | 0.2 | 0.164 | 1 | 14 | Stk24 | 0.036 | Young |
| 0.5295 | 0.106839 | 0.2 | 0.164 | 1 | 14 | Topors | 0.036 | Young |
| 0.529711 | 0.24828 | 0.18 | 0.142 | 1 | 14 | Cog7 | 0.038 | Young |
| 0.530228 | 0.284087 | 0.12 | 0.087 | 1 | 14 | Ap3s2 | 0.033 | Young |
| 0.530519 | 0.049625 | 0.2 | 0.235 | 1 | 14 | Actr1a | -0.035 | Young |
| 0.530587 | 0.929214 | 0.16 | 0.131 | 1 | 14 | Limk1 | 0.029 | Young |
| 0.530955 | 0.515594 | 0.18 | 0.137 | 1 | 14 | Ift27 | 0.043 | Young |
| 0.530964 | 0.138817 | 0.24 | 0.191 | 1 | 14 | Clptm1 | 0.049 | Young |
| 0.531784 | 0.311328 | 0.64 | 0.683 | 1 | 14 | Cd34 | -0.043 | Young |
| 0.532384 | 1.447696 | 0.12 | 0.093 | 1 | 14 | Nfyc | 0.027 | Young |

| | | | | | | | | |
|---|---|---|---|---|---|---|---|---|
| 0.532384 | 0.650992 | 0.12 | 0.093 | 1 | 14 | Zfp507 | 0.027 | Young |
| 0.532673 | 0.089641 | 0.9 | 0.934 | 1 | 14 | Vim | -0.034 | Young |
| 0.532797 | 0.498977 | 0.36 | 0.317 | 1 | 14 | Zfp706 | 0.043 | Young |
| 0.532867 | 0.456489 | 0.08 | 0.055 | 1 | 14 | Prune1 | 0.025 | Young |
| 0.532924 | 0.472211 | 0.36 | 0.311 | 1 | 14 | Itgav | 0.049 | Young |
| 0.533233 | 1.447782 | 0.24 | 0.202 | 1 | 14 | Ankrd13a | 0.038 | Young |
| 0.533394 | 0.079839 | 0.22 | 0.175 | 1 | 14 | Ykt6 | 0.045 | Young |
| 0.533732 | 0.35896 | 0.22 | 0.18 | 1 | 14 | Col18a1 | 0.04 | Young |
| 0.534022 | 1.216916 | 0.14 | 0.109 | 1 | 14 | Polr2j | 0.031 | Young |
| 0.535141 | 0.461649 | 0.38 | 0.41 | 1 | 14 | Adh1 | -0.03 | Young |
| 0.535472 | 0.731148 | 0.16 | 0.126 | 1 | 14 | Mrpl38 | 0.034 | Young |
| 0.535696 | 0.210711 | 0.22 | 0.175 | 1 | 14 | Rspry1 | 0.045 | Young |
| 0.535861 | 0.150362 | 1 | 0.962 | 1 | 14 | Rps8 | 0.038 | Young |
| 0.536001 | 0.633403 | 0.34 | 0.295 | 1 | 14 | Vamp8 | 0.045 | Young |
| 0.536243 | 0.477605 | 0.4 | 0.393 | 1 | 14 | Meg3 | 0.007 | Young |
| 0.536272 | 0.051475 | 0.26 | 0.208 | 1 | 14 | St5 | 0.052 | Young |
| 0.536361 | 0.104385 | 0.12 | 0.087 | 1 | 14 | Crlf2 | 0.033 | Young |
| 0.536601 | 0.22908 | 0.6 | 0.563 | 1 | 14 | Matr3 | 0.037 | Young |
| 0.53665 | 0.223584 | 0.08 | 0.055 | 1 | 14 | Kbtbd7 | 0.025 | Young |
| 0.53665 | 0.145767 | 0.08 | 0.055 | 1 | 14 | Trmt13 | 0.025 | Young |
| 0.537131 | 0.971883 | 0.14 | 0.104 | 1 | 14 | Josd2 | 0.036 | Young |
| 0.537574 | 0.267058 | 0.3 | 0.257 | 1 | 14 | Tgfb2 | 0.043 | Young |
| 0.537756 | 0.796364 | 0.1 | 0.071 | 1 | 14 | Dcp1a | 0.029 | Young |
| 0.537932 | 0.400632 | 0.56 | 0.525 | 1 | 14 | Ube2r2 | 0.035 | Young |
| 0.538713 | 0.76148 | 0.08 | 0.115 | 1 | 14 | Bahd1 | -0.035 | Young |
| 0.538713 | 0.028958 | 0.08 | 0.115 | 1 | 14 | Dut | -0.035 | Young |
| 0.538747 | 0.190765 | 0.64 | 0.639 | 1 | 14 | Atf3 | 0.001 | Young |
| 0.538844 | 0.326724 | 0.34 | 0.301 | 1 | 14 | Pcolce | 0.039 | Young |
| 0.539057 | 0.338486 | 0.42 | 0.377 | 1 | 14 | Psmd1 | 0.043 | Young |
| 0.539094 | 0.024906 | 0.32 | 0.355 | 1 | 14 | Npr1 | -0.035 | Young |
| 0.539449 | 0.027385 | 0.82 | 0.825 | 1 | 14 | Rpl28 | -0.005 | Young |
| 0.539638 | 1.064337 | 0.14 | 0.109 | 1 | 14 | Upf3b | 0.031 | Young |
| 0.539688 | 0.081726 | 0.4 | 0.426 | 1 | 14 | Ppfibp1 | -0.026 | Young |
| 0.540332 | 0.537067 | 0.32 | 0.268 | 1 | 14 | R3hdm2 | 0.052 | Young |
| 0.540332 | 0.070165 | 0.32 | 0.268 | 1 | 14 | Ankfy1 | 0.052 | Young |
| 0.540445 | 0.066026 | 0.08 | 0.055 | 1 | 14 | Ofd1 | 0.025 | Young |
| 0.542453 | 0.30455 | 0.4 | 0.339 | 1 | 14 | Amfr | 0.061 | Young |
| 0.542458 | 0.228568 | 0.14 | 0.109 | 1 | 14 | Eaf1 | 0.031 | Young |
| 0.54253 | 0.267976 | 0.12 | 0.087 | 1 | 14 | Nedd4l | 0.033 | Young |
| 0.543075 | 0.268241 | 0.1 | 0.137 | 1 | 14 | Fmo5 | -0.037 | Young |
| 0.543699 | 0.314986 | 0.34 | 0.284 | 1 | 14 | Akap9 | 0.056 | Young |
| 0.544255 | 0.002824 | 0.08 | 0.055 | 1 | 14 | Fgfr1op | 0.025 | Young |
| 0.544294 | 0.096412 | 0.4 | 0.339 | 1 | 14 | Atxn2l | 0.061 | Young |
| 0.544854 | 0.139199 | 0.3 | 0.372 | 1 | 14 | Epas1 | -0.072 | Young |
| 0.544991 | 0.720976 | 0.28 | 0.251 | 1 | 14 | Erp44 | 0.029 | Young |

| | | | | | | | | |
|---|---|---|---|---|---|---|---|---|
| 0.545189 | 0.220422 | 0.22 | 0.18 | 1 | 14 | Socs4 | 0.04 | Young |
| 0.545285 | 0.429093 | 0.14 | 0.109 | 1 | 14 | C1galt1c1 | 0.031 | Young |
| 0.545584 | 0.427296 | 0.14 | 0.175 | 1 | 14 | Cryzl1 | -0.035 | Young |
| 0.545734 | 1.875462 | 0.08 | 0.12 | 1 | 14 | Cnpy4 | -0.04 | Young |
| 0.545752 | 0.417854 | 0.02 | 0.038 | 1 | 14 | Trem2 | -0.018 | Young |
| 0.545752 | 0.137569 | 0.02 | 0.038 | 1 | 14 | Krba1 | -0.018 | Young |
| 0.545752 | 0.102938 | 0.02 | 0.038 | 1 | 14 | Fzd3 | -0.018 | Young |
| 0.545752 | 0.074424 | 0.02 | 0.038 | 1 | 14 | Lig1 | -0.018 | Young |
| 0.546115 | 1.763441 | 0.1 | 0.077 | 1 | 14 | Hebp2 | 0.023 | Young |
| 0.546115 | 0.517043 | 0.06 | 0.087 | 1 | 14 | Zbtb6 | -0.027 | Young |
| 0.546366 | 0.229004 | 0.9 | 0.858 | 1 | 14 | Fstl1 | 0.042 | Young |
| 0.546414 | 0.252061 | 0.36 | 0.301 | 1 | 14 | Cd151 | 0.059 | Young |
| 0.547315 | 1.139587 | 0.28 | 0.23 | 1 | 14 | Jarid2 | 0.05 | Young |
| 0.547495 | 0.07747 | 0.22 | 0.18 | 1 | 14 | Ddx10 | 0.04 | Young |
| 0.547503 | 0.029682 | 0.44 | 0.393 | 1 | 14 | Pds5a | 0.047 | Young |
| 0.547812 | 0.029277 | 0.5 | 0.579 | 1 | 14 | 1810037I1 | -0.079 | Young |
| 0.548013 | 0.37965 | 0.42 | 0.377 | 1 | 14 | Senp6 | 0.043 | Young |
| 0.548119 | 0.104831 | 0.14 | 0.109 | 1 | 14 | Bmp2 | 0.031 | Young |
| 0.548476 | 0.221499 | 0.4 | 0.464 | 1 | 14 | Acap2 | -0.064 | Young |
| 0.548663 | 1.108164 | 0.36 | 0.339 | 1 | 14 | Cox5b | 0.021 | Young |
| 0.549438 | 0.863759 | 0.1 | 0.077 | 1 | 14 | Ldhb | 0.023 | Young |
| 0.549438 | 0.800967 | 0.1 | 0.077 | 1 | 14 | Borcs7 | 0.023 | Young |
| 0.549438 | 0.768853 | 0.1 | 0.077 | 1 | 14 | Nipa1 | 0.023 | Young |
| 0.549676 | 1.162113 | 0.08 | 0.06 | 1 | 14 | Fhl2 | 0.02 | Young |
| 0.549927 | 0.141198 | 0.28 | 0.235 | 1 | 14 | Pja1 | 0.045 | Young |
| 0.550522 | 0.063841 | 0.26 | 0.301 | 1 | 14 | Puf60 | -0.041 | Young |
| 0.550752 | 1.022113 | 0.02 | 0.038 | 1 | 14 | 3930402G2 | -0.018 | Young |
| 0.550752 | 0.898737 | 0.02 | 0.038 | 1 | 14 | Grwd1 | -0.018 | Young |
| 0.550752 | 0.278564 | 0.02 | 0.038 | 1 | 14 | Tex9 | -0.018 | Young |
| 0.550752 | 0.245834 | 0.02 | 0.038 | 1 | 14 | Iqcc | -0.018 | Young |
| 0.550752 | 0.01034 | 0.02 | 0.038 | 1 | 14 | Alpk3 | -0.018 | Young |
| 0.550781 | 0.402098 | 0.18 | 0.148 | 1 | 14 | Chchd7 | 0.032 | Young |
| 0.55096 | 1.113531 | 0.14 | 0.109 | 1 | 14 | A430005L1 | 0.031 | Young |
| 0.551373 | 0.024797 | 0.12 | 0.153 | 1 | 14 | Reps1 | -0.033 | Young |
| 0.551587 | 1.14743 | 0.18 | 0.153 | 1 | 14 | Lrrc8d | 0.027 | Young |
| 0.552024 | 0.169739 | 0.28 | 0.235 | 1 | 14 | Fam172a | 0.045 | Young |
| 0.552425 | 0.258465 | 0.8 | 0.858 | 1 | 14 | Rps17 | -0.058 | Young |
| 0.55277 | 0.946046 | 0.1 | 0.077 | 1 | 14 | Sipa1l1 | 0.023 | Young |
| 0.55277 | 0.063326 | 0.06 | 0.087 | 1 | 14 | Gpr161 | -0.027 | Young |
| 0.55282 | 0.251457 | 0.64 | 0.568 | 1 | 14 | Sumo2 | 0.072 | Young |
| 0.553357 | 0.056696 | 0.2 | 0.164 | 1 | 14 | Syde1 | 0.036 | Young |
| 0.553745 | 0.480595 | 0.36 | 0.311 | 1 | 14 | Tubb6 | 0.049 | Young |
| 0.553809 | 0.615925 | 0.14 | 0.109 | 1 | 14 | Mrps7 | 0.031 | Young |
| 0.554382 | 0.046883 | 0.46 | 0.497 | 1 | 14 | Ndufa6 | -0.037 | Young |
| 0.554674 | 0.112375 | 0.7 | 0.661 | 1 | 14 | Eif3a | 0.039 | Young |

| | | | | | | | | |
|---|---|---|---|---|---|---|---|---|
| 0.554973 | 0.101858 | 0.12 | 0.087 | 1 | 14 | Coq5 | 0.033 | Young |
| 0.555104 | 0.258096 | 0.26 | 0.219 | 1 | 14 | Runx1t1 | 0.041 | Young |
| 0.555759 | 0.364746 | 0.28 | 0.23 | 1 | 14 | Tmem160 | 0.05 | Young |
| 0.556158 | 0.06925 | 0.56 | 0.656 | 1 | 14 | Qk | -0.096 | Young |
| 0.556288 | 0.207426 | 0.36 | 0.29 | 1 | 14 | Sirt2 | 0.07 | Young |
| 0.556664 | 0.387425 | 0.14 | 0.109 | 1 | 14 | Zfp738 | 0.031 | Young |
| 0.556825 | 0.533922 | 0.22 | 0.186 | 1 | 14 | Khdrbs3 | 0.034 | Young |
| 0.557127 | 2.335586 | 0.08 | 0.06 | 1 | 14 | Podxl2 | 0.02 | Young |
| 0.557523 | 0.477916 | 0.24 | 0.202 | 1 | 14 | Maob | 0.038 | Young |
| 0.558107 | 0.015394 | 0.08 | 0.109 | 1 | 14 | Kat2a | -0.029 | Young |
| 0.559464 | 0.335997 | 0.1 | 0.077 | 1 | 14 | Ly86 | 0.023 | Young |
| 0.559527 | 0.139283 | 0.14 | 0.109 | 1 | 14 | Tgfb1 | 0.031 | Young |
| 0.559558 | 0.405226 | 0.46 | 0.432 | 1 | 14 | Pomp | 0.028 | Young |
| 0.559662 | 0.427726 | 0.32 | 0.377 | 1 | 14 | Snrpg | -0.057 | Young |
| 0.559756 | 0.501657 | 0.12 | 0.093 | 1 | 14 | Pgs1 | 0.027 | Young |
| 0.559756 | 0.252445 | 0.12 | 0.093 | 1 | 14 | Dnmt1 | 0.027 | Young |
| 0.560616 | 0.762755 | 0.2 | 0.164 | 1 | 14 | Arhgap42 | 0.036 | Young |
| 0.560881 | 0.63792 | 0.18 | 0.148 | 1 | 14 | Acvr1 | 0.032 | Young |
| 0.561134 | 0.280602 | 0.4 | 0.41 | 1 | 14 | Ilk | -0.01 | Young |
| 0.561482 | 0.039498 | 0.22 | 0.268 | 1 | 14 | Pi15 | -0.048 | Young |
| 0.562207 | 0.425607 | 0.16 | 0.126 | 1 | 14 | Cyth1 | 0.034 | Young |
| 0.562264 | 0.084912 | 0.28 | 0.306 | 1 | 14 | Cyb5b | -0.026 | Young |
| 0.562397 | 0.307963 | 0.14 | 0.109 | 1 | 14 | Tipin | 0.031 | Young |
| 0.562468 | 0.208203 | 0.46 | 0.563 | 1 | 14 | Ptgfrn | -0.103 | Young |
| 0.562839 | 0.881276 | 0.12 | 0.093 | 1 | 14 | Ddr1 | 0.027 | Young |
| 0.562839 | 0.568711 | 0.12 | 0.093 | 1 | 14 | Mob3a | 0.027 | Young |
| 0.563045 | 0.083112 | 0.2 | 0.164 | 1 | 14 | Pir | 0.036 | Young |
| 0.563274 | 0.545147 | 0.26 | 0.224 | 1 | 14 | Erg28 | 0.036 | Young |
| 0.563367 | 0.122582 | 0.18 | 0.213 | 1 | 14 | Mtmr6 | -0.033 | Young |
| 0.563771 | 0.370951 | 0.16 | 0.197 | 1 | 14 | Enc1 | -0.037 | Young |
| 0.564927 | 0.271813 | 0.16 | 0.202 | 1 | 14 | Aldh6a1 | -0.042 | Young |
| 0.565089 | 0.788536 | 0.1 | 0.071 | 1 | 14 | Atg13 | 0.029 | Young |
| 0.566197 | 1.475482 | 0.1 | 0.077 | 1 | 14 | Anapc7 | 0.023 | Young |
| 0.566197 | 0.08211 | 0.1 | 0.077 | 1 | 14 | Aldh3b1 | 0.023 | Young |
| 0.566314 | 0.064647 | 0.32 | 0.262 | 1 | 14 | Tm2d2 | 0.058 | Young |
| 0.566446 | 0.991257 | 0.26 | 0.213 | 1 | 14 | Ube2d1 | 0.047 | Young |
| 0.567622 | 0.512509 | 0.1 | 0.137 | 1 | 14 | Elmo1 | -0.037 | Young |
| 0.567786 | 0.071944 | 0.18 | 0.142 | 1 | 14 | Fam98a | 0.038 | Young |
| 0.568159 | 0.766297 | 0.14 | 0.109 | 1 | 14 | Micu2 | 0.031 | Young |
| 0.568376 | 0.25034 | 0.32 | 0.284 | 1 | 14 | Col4a5 | 0.036 | Young |
| 0.568429 | 0.267069 | 0.46 | 0.475 | 1 | 14 | Stat3 | -0.015 | Young |
| 0.568464 | 0.360944 | 0.22 | 0.18 | 1 | 14 | Sap18 | 0.04 | Young |
| 0.568514 | 0.474933 | 0.18 | 0.148 | 1 | 14 | Ptbp2 | 0.032 | Young |
| 0.568535 | 0.160831 | 0.36 | 0.322 | 1 | 14 | Nudcd3 | 0.038 | Young |
| 0.568933 | 0.464855 | 0.28 | 0.235 | 1 | 14 | Uqcc3 | 0.045 | Young |

| | | | | | | | | |
|---|---|---|---|---|---|---|---|---|
| 0.569 | 0.236412 | 0.62 | 0.623 | 1 | 14 | Cd164 | -0.003 | Young |
| 0.569112 | 0.147747 | 1 | 0.989 | 1 | 14 | Eef1a1 | 0.011 | Young |
| 0.569578 | 0.528954 | 0.1 | 0.077 | 1 | 14 | Tesk1 | 0.023 | Young |
| 0.570759 | 0.691406 | 0.44 | 0.421 | 1 | 14 | Tns2 | 0.019 | Young |
| 0.570786 | 0.223505 | 0.58 | 0.563 | 1 | 14 | Fubp1 | 0.017 | Young |
| 0.571002 | 0.509058 | 0.4 | 0.372 | 1 | 14 | Eps15 | 0.028 | Young |
| 0.571069 | 1.34918 | 0.18 | 0.148 | 1 | 14 | Polr2g | 0.032 | Young |
| 0.571279 | 0.519979 | 0.8 | 0.831 | 1 | 14 | Col6a2 | -0.031 | Young |
| 0.571917 | 0.153052 | 0.16 | 0.202 | 1 | 14 | Aif1l | -0.042 | Young |
| 0.572137 | 1.163956 | 0.12 | 0.093 | 1 | 14 | Mlx | 0.027 | Young |
| 0.572137 | 0.903866 | 0.12 | 0.093 | 1 | 14 | Cbr1 | 0.027 | Young |
| 0.572137 | 0.556731 | 0.12 | 0.093 | 1 | 14 | Timm22 | 0.027 | Young |
| 0.572137 | 0.529691 | 0.12 | 0.093 | 1 | 14 | Borcs5 | 0.027 | Young |
| 0.572173 | 0.892943 | 0.08 | 0.06 | 1 | 14 | Hmces | 0.02 | Young |
| 0.572173 | 0.702818 | 0.08 | 0.06 | 1 | 14 | Ssx2ip | 0.02 | Young |
| 0.572596 | 0.39434 | 0.56 | 0.536 | 1 | 14 | Arl6ip1 | 0.024 | Young |
| 0.573148 | 0.098915 | 0.1 | 0.137 | 1 | 14 | Agtr1a | -0.037 | Young |
| 0.573287 | 0.705726 | 0.12 | 0.098 | 1 | 14 | Xpnpep1 | 0.022 | Young |
| 0.573836 | 0.09175 | 0.96 | 0.907 | 1 | 14 | Rps19 | 0.053 | Young |
| 0.573948 | 0.854771 | 0.14 | 0.109 | 1 | 14 | Tecpr1 | 0.031 | Young |
| 0.573948 | 0.404008 | 0.14 | 0.109 | 1 | 14 | Fam214a | 0.031 | Young |
| 0.574389 | 0.473642 | 0.16 | 0.191 | 1 | 14 | Micu1 | -0.031 | Young |
| 0.57478 | 0.219735 | 0.76 | 0.77 | 1 | 14 | Uqcrh | -0.01 | Young |
| 0.575406 | 0.049493 | 0.66 | 0.716 | 1 | 14 | Ar | -0.056 | Young |
| 0.575582 | 1.075189 | 0.16 | 0.131 | 1 | 14 | Ampd2 | 0.029 | Young |
| 0.575582 | 0.284089 | 0.16 | 0.131 | 1 | 14 | Hoxc10 | 0.029 | Young |
| 0.575728 | 0.735946 | 0.14 | 0.115 | 1 | 14 | Tmem237 | 0.025 | Young |
| 0.575865 | 0.211302 | 0.3 | 0.344 | 1 | 14 | Chmp5 | -0.044 | Young |
| 0.576406 | 0.042695 | 0.3 | 0.257 | 1 | 14 | Tmpo | 0.043 | Young |
| 0.576469 | 0.498397 | 0.38 | 0.415 | 1 | 14 | Naaa | -0.035 | Young |
| 0.576853 | 0.124461 | 0.14 | 0.109 | 1 | 14 | Pcna | 0.031 | Young |
| 0.576954 | 0.403715 | 0.22 | 0.262 | 1 | 14 | Nsg1 | -0.042 | Young |
| 0.577113 | 0.696486 | 0.62 | 0.53 | 1 | 14 | Macf1 | 0.09 | Young |
| 0.578767 | 1.393967 | 0.18 | 0.148 | 1 | 14 | Dhx32 | 0.032 | Young |
| 0.578767 | 0.263298 | 0.18 | 0.148 | 1 | 14 | Mxi1 | 0.032 | Young |
| 0.578767 | 0.111352 | 0.18 | 0.148 | 1 | 14 | Lcp1 | 0.032 | Young |
| 0.578829 | 0.180042 | 0.46 | 0.415 | 1 | 14 | Srpr | 0.045 | Young |
| 0.578925 | 0.510528 | 0.18 | 0.219 | 1 | 14 | Mrpl17 | -0.039 | Young |
| 0.580277 | 0.65447 | 0.16 | 0.197 | 1 | 14 | Kars | -0.037 | Young |
| 0.580569 | 0.430957 | 0.28 | 0.262 | 1 | 14 | Cs | 0.018 | Young |
| 0.580663 | 0.317694 | 0.32 | 0.262 | 1 | 14 | Naalad2 | 0.058 | Young |
| 0.582039 | 0.235271 | 0.28 | 0.24 | 1 | 14 | Arhgef25 | 0.04 | Young |
| 0.582192 | 0.57687 | 0.3 | 0.35 | 1 | 14 | Aebp2 | -0.05 | Young |
| 0.582416 | 0.175077 | 0.36 | 0.317 | 1 | 14 | Ramp2 | 0.043 | Young |
| 0.582479 | 0.629616 | 0.22 | 0.186 | 1 | 14 | Abhd16a | 0.034 | Young |

| | | | | | | | | |
|---|---|---|---|---|---|---|---|---|
| 0.582997 | 0.673637 | 0.62 | 0.634 | 1 | 14 | Pafah1b1 | -0.014 | Young |
| 0.583198 | 0.936452 | 0.1 | 0.077 | 1 | 14 | Ccnd1 | 0.023 | Young |
| 0.583582 | 2.275611 | 0.08 | 0.06 | 1 | 14 | Tomm6 | 0.02 | Young |
| 0.583582 | 1.967218 | 0.08 | 0.06 | 1 | 14 | Inpp5e | 0.02 | Young |
| 0.583582 | 1.371047 | 0.08 | 0.06 | 1 | 14 | Olfml1 | 0.02 | Young |
| 0.583582 | 0.993952 | 0.08 | 0.06 | 1 | 14 | Atp10a | 0.02 | Young |
| 0.583635 | 0.191395 | 0.92 | 0.962 | 1 | 14 | Rps14 | -0.042 | Young |
| 0.583661 | 0.208688 | 0.26 | 0.322 | 1 | 14 | Dhrs3 | -0.062 | Young |
| 0.583927 | 0.852792 | 0.18 | 0.148 | 1 | 14 | Tor1a | 0.032 | Young |
| 0.583985 | 1.660269 | 0.06 | 0.044 | 1 | 14 | Nsdhl | 0.016 | Young |
| 0.583985 | 1.400856 | 0.06 | 0.044 | 1 | 14 | Hps4 | 0.016 | Young |
| 0.584646 | 0.23327 | 0.12 | 0.093 | 1 | 14 | A830082K1 | 0.027 | Young |
| 0.584936 | 0.415131 | 0.74 | 0.792 | 1 | 14 | Cald1 | -0.052 | Young |
| 0.585034 | 0.424615 | 0.16 | 0.197 | 1 | 14 | Ctdspl2 | -0.037 | Young |
| 0.585034 | 0.073454 | 0.22 | 0.18 | 1 | 14 | Glce | 0.04 | Young |
| 0.585058 | 0.319782 | 0.34 | 0.311 | 1 | 14 | Syf2 | 0.029 | Young |
| 0.58528 | 0.028591 | 0.96 | 0.984 | 1 | 14 | mt-Nd1 | -0.024 | Young |
| 0.58604 | 0.526959 | 0.04 | 0.06 | 1 | 14 | B4galt7 | -0.02 | Young |
| 0.58604 | 0.314719 | 0.04 | 0.06 | 1 | 14 | Itga4 | -0.02 | Young |
| 0.586047 | 0.022559 | 0.38 | 0.437 | 1 | 14 | Psmd11 | -0.057 | Young |
| 0.586356 | 0.628132 | 0.34 | 0.306 | 1 | 14 | Swap70 | 0.034 | Young |
| 0.586626 | 1.13234 | 0.1 | 0.077 | 1 | 14 | Haus3 | 0.023 | Young |
| 0.586626 | 0.650827 | 0.1 | 0.077 | 1 | 14 | Prkrip1 | 0.023 | Young |
| 0.586626 | 0.255687 | 0.1 | 0.077 | 1 | 14 | Oxsm | 0.023 | Young |
| 0.586626 | 0.232731 | 0.1 | 0.077 | 1 | 14 | Prkag2 | 0.023 | Young |
| 0.586687 | 0.142833 | 0.36 | 0.311 | 1 | 14 | Sbno1 | 0.049 | Young |
| 0.586906 | 0.264747 | 0.24 | 0.202 | 1 | 14 | Ap1s1 | 0.038 | Young |
| 0.586932 | 0.522947 | 0.18 | 0.153 | 1 | 14 | Neo1 | 0.027 | Young |
| 0.58703 | 0.123506 | 0.28 | 0.328 | 1 | 14 | Gltp | -0.048 | Young |
| 0.587251 | 1.608518 | 0.18 | 0.158 | 1 | 14 | Ptgs2 | 0.022 | Young |
| 0.587793 | 0.908805 | 0.12 | 0.093 | 1 | 14 | Ttc13 | 0.027 | Young |
| 0.587793 | 0.352119 | 0.12 | 0.093 | 1 | 14 | Arhgef3 | 0.027 | Young |
| 0.588239 | 0.070779 | 0.5 | 0.546 | 1 | 14 | Aldoa | -0.046 | Young |
| 0.588637 | 0.076197 | 0.2 | 0.23 | 1 | 14 | Sae1 | -0.03 | Young |
| 0.589592 | 0.032254 | 0.48 | 0.404 | 1 | 14 | Dock7 | 0.076 | Young |
| 0.590064 | 1.160266 | 0.1 | 0.077 | 1 | 14 | Pold4 | 0.023 | Young |
| 0.590064 | 0.481416 | 0.1 | 0.077 | 1 | 14 | Ppp1r16a | 0.023 | Young |
| 0.590064 | 0.44252 | 0.1 | 0.077 | 1 | 14 | Ghdc | 0.023 | Young |
| 0.590102 | 0.373016 | 0.2 | 0.164 | 1 | 14 | Slc25a46 | 0.036 | Young |
| 0.590111 | 0.553522 | 0.14 | 0.115 | 1 | 14 | Pias3 | 0.025 | Young |
| 0.590141 | 0.438514 | 0.04 | 0.06 | 1 | 14 | 1500011B0 | -0.02 | Young |
| 0.590141 | 0.148994 | 0.04 | 0.06 | 1 | 14 | Zfp30 | -0.02 | Young |
| 0.590209 | 0.580728 | 0.26 | 0.219 | 1 | 14 | Timm17a | 0.041 | Young |
| 0.590209 | 0.565563 | 0.26 | 0.219 | 1 | 14 | Hmbox1 | 0.041 | Young |
| 0.590565 | 0.069988 | 0.26 | 0.301 | 1 | 14 | Foxo1 | -0.041 | Young |

| | | | | | | | | |
|---|---|---|---|---|---|---|---|---|
| 0.590878 | 0.256831 | 0.26 | 0.213 | 1 | 14 | Bcl2l2 | 0.047 | Young |
| 0.591322 | 1.14624 | 0.16 | 0.137 | 1 | 14 | Rasa2 | 0.023 | Young |
| 0.591758 | 0.680791 | 0.08 | 0.109 | 1 | 14 | Ptx3 | -0.029 | Young |
| 0.592292 | 0.375464 | 0.7 | 0.716 | 1 | 14 | Txn1 | -0.016 | Young |
| 0.592401 | 0.584622 | 0.24 | 0.311 | 1 | 14 | Anpep | -0.071 | Young |
| 0.592435 | 0.020391 | 0.6 | 0.689 | 1 | 14 | Fxyd1 | -0.089 | Young |
| 0.592865 | 1.32455 | 0.06 | 0.044 | 1 | 14 | Slc22a17 | 0.016 | Young |
| 0.592929 | 0.158416 | 0.24 | 0.257 | 1 | 14 | Rlim | -0.017 | Young |
| 0.593135 | 0.202152 | 0.18 | 0.213 | 1 | 14 | Fam92a | -0.033 | Young |
| 0.593211 | 0.025527 | 0.62 | 0.699 | 1 | 14 | Lum | -0.079 | Young |
| 0.593261 | 0.083288 | 0.58 | 0.678 | 1 | 14 | Atp2a2 | -0.098 | Young |
| 0.593293 | 0.555047 | 0.38 | 0.322 | 1 | 14 | Tnfrsf1a | 0.058 | Young |
| 0.593511 | 0.218495 | 0.1 | 0.077 | 1 | 14 | Hps3 | 0.023 | Young |
| 0.593511 | 0.200553 | 0.1 | 0.077 | 1 | 14 | Znhit6 | 0.023 | Young |
| 0.593639 | 1.818516 | 0.1 | 0.082 | 1 | 14 | Znrf1 | 0.018 | Young |
| 0.593978 | 0.856733 | 0.22 | 0.191 | 1 | 14 | Spr | 0.029 | Young |
| 0.594022 | 0.218644 | 0.12 | 0.148 | 1 | 14 | Carm1 | -0.028 | Young |
| 0.594111 | 0.504721 | 0.12 | 0.093 | 1 | 14 | Gm26523 | 0.027 | Young |
| 0.594428 | 0.337828 | 0.14 | 0.109 | 1 | 14 | Cdc42ep3 | 0.031 | Young |
| 0.594619 | 0.242608 | 0.16 | 0.131 | 1 | 14 | Cog8 | 0.029 | Young |
| 0.594947 | 0.252873 | 0.34 | 0.311 | 1 | 14 | Creb3l1 | 0.029 | Young |
| 0.595097 | 0.466204 | 0.08 | 0.06 | 1 | 14 | Dolpp1 | 0.02 | Young |
| 0.595116 | 0.294802 | 0.16 | 0.126 | 1 | 14 | Adcy7 | 0.034 | Young |
| 0.595551 | 0.083282 | 0.44 | 0.486 | 1 | 14 | Kmt2a | -0.046 | Young |
| 0.596661 | 0.831076 | 0.24 | 0.208 | 1 | 14 | Selenbp1 | 0.032 | Young |
| 0.596968 | 2.106635 | 0.1 | 0.077 | 1 | 14 | Acsl3 | 0.023 | Young |
| 0.596968 | 0.429865 | 0.1 | 0.077 | 1 | 14 | C5ar1 | 0.023 | Young |
| 0.596968 | 0.292587 | 0.1 | 0.077 | 1 | 14 | Lrrfip1 | 0.023 | Young |
| 0.597022 | 0.174103 | 0.74 | 0.749 | 1 | 14 | H3f3a | -0.009 | Young |
| 0.597022 | 0.02393 | 0.64 | 0.776 | 1 | 14 | Tuba1b | -0.136 | Young |
| 0.597328 | 1.307526 | 0.06 | 0.044 | 1 | 14 | Pcdh9 | 0.016 | Young |
| 0.597536 | 0.540557 | 0.34 | 0.317 | 1 | 14 | Lrrc17 | 0.023 | Young |
| 0.597547 | 0.088582 | 0.14 | 0.175 | 1 | 14 | Cilp | -0.035 | Young |
| 0.597975 | 1.598056 | 0.12 | 0.098 | 1 | 14 | Fam216a | 0.022 | Young |
| 0.598241 | 0.098326 | 0.48 | 0.525 | 1 | 14 | 11-Sep | -0.045 | Young |
| 0.598835 | 0.495282 | 0.46 | 0.437 | 1 | 14 | Tcp1 | 0.023 | Young |
| 0.600111 | 0.888184 | 0.16 | 0.131 | 1 | 14 | Tap1 | 0.029 | Young |
| 0.600174 | 0.12592 | 0.44 | 0.393 | 1 | 14 | Map1lc3a | 0.047 | Young |
| 0.600433 | 1.076541 | 0.1 | 0.077 | 1 | 14 | Ppm1k | 0.023 | Young |
| 0.600545 | 0.012046 | 0.3 | 0.251 | 1 | 14 | Spcs3 | 0.049 | Young |
| 0.601223 | 0.381518 | 0.44 | 0.399 | 1 | 14 | Prdx2 | 0.041 | Young |
| 0.601437 | 0.194756 | 0.92 | 0.885 | 1 | 14 | Nedd4 | 0.035 | Young |
| 0.601806 | 1.15895 | 0.06 | 0.044 | 1 | 14 | Itpripl1 | 0.016 | Young |
| 0.602151 | 0.119827 | 0.18 | 0.148 | 1 | 14 | Gripap1 | 0.032 | Young |
| 0.602156 | 0.525732 | 0.16 | 0.137 | 1 | 14 | Cacfd1 | 0.023 | Young |

| | | | | | | | | |
|---|---|---|---|---|---|---|---|---|
| 0.602437 | 0.322713 | 0.18 | 0.213 | 1 | 14 | Pdxk | -0.033 | Young |
| 0.602831 | 0.732758 | 0.08 | 0.06 | 1 | 14 | Lrrc45 | 0.02 | Young |
| 0.602913 | 0.609348 | 0.34 | 0.311 | 1 | 14 | Trmt112 | 0.029 | Young |
| 0.602913 | 0.30949 | 0.28 | 0.328 | 1 | 14 | Fgf2 | -0.048 | Young |
| 0.603264 | 0.083771 | 0.7 | 0.776 | 1 | 14 | Rpl14 | -0.076 | Young |
| 0.603332 | 0.448757 | 0.48 | 0.508 | 1 | 14 | Txndc17 | -0.028 | Young |
| 0.60357 | 0.378929 | 0.28 | 0.344 | 1 | 14 | Podn | -0.064 | Young |
| 0.603645 | 0.660356 | 0.08 | 0.104 | 1 | 14 | Usp53 | -0.024 | Young |
| 0.603645 | 0.131239 | 0.12 | 0.093 | 1 | 14 | Grk6 | 0.027 | Young |
| 0.603663 | 0.271512 | 0.76 | 0.743 | 1 | 14 | Rps18 | 0.017 | Young |
| 0.60387 | 0.729498 | 0.22 | 0.186 | 1 | 14 | Zzef1 | 0.034 | Young |
| 0.603908 | 0.366876 | 0.1 | 0.077 | 1 | 14 | Vezt | 0.023 | Young |
| 0.604222 | 0.620969 | 0.12 | 0.098 | 1 | 14 | Tapbpl | 0.022 | Young |
| 0.604235 | 0.100458 | 0.42 | 0.454 | 1 | 14 | Snu13 | -0.034 | Young |
| 0.604242 | 0.65848 | 0.22 | 0.18 | 1 | 14 | Zc3h4 | 0.04 | Young |
| 0.604298 | 0.458462 | 0.46 | 0.432 | 1 | 14 | Ext1 | 0.028 | Young |
| 0.60493 | 0.002734 | 0.36 | 0.421 | 1 | 14 | Col14a1 | -0.061 | Young |
| 0.605069 | 0.728312 | 0.16 | 0.202 | 1 | 14 | Rabl6 | -0.042 | Young |
| 0.605338 | 1.000744 | 0.24 | 0.202 | 1 | 14 | Smpd1 | 0.038 | Young |
| 0.606073 | 0.624264 | 0.42 | 0.421 | 1 | 14 | Ndufa13 | -0.001 | Young |
| 0.6063 | 1.456795 | 0.06 | 0.044 | 1 | 14 | Coa7 | 0.016 | Young |
| 0.6063 | 0.979051 | 0.06 | 0.044 | 1 | 14 | Wdcp | 0.016 | Young |
| 0.606452 | 0.081888 | 0.48 | 0.536 | 1 | 14 | Tmco1 | -0.056 | Young |
| 0.606714 | 1.269353 | 0.08 | 0.06 | 1 | 14 | Spata7 | 0.02 | Young |
| 0.606714 | 0.444016 | 0.08 | 0.06 | 1 | 14 | Slc26a11 | 0.02 | Young |
| 0.606714 | 0.359227 | 0.08 | 0.06 | 1 | 14 | Sdc1 | 0.02 | Young |
| 0.606714 | 0.340316 | 0.08 | 0.06 | 1 | 14 | Pxylp1 | 0.02 | Young |
| 0.606739 | 0.903664 | 0.12 | 0.153 | 1 | 14 | Amz2 | -0.033 | Young |
| 0.607056 | 0.325155 | 0.56 | 0.596 | 1 | 14 | Hook3 | -0.036 | Young |
| 0.607391 | 0.719587 | 0.1 | 0.077 | 1 | 14 | Ints1 | 0.023 | Young |
| 0.607404 | 0.920099 | 0.18 | 0.148 | 1 | 14 | Xxylt1 | 0.032 | Young |
| 0.607683 | 0.427584 | 0.18 | 0.158 | 1 | 14 | Ncln | 0.022 | Young |
| 0.607878 | 0.616033 | 0.28 | 0.268 | 1 | 14 | Ei24 | 0.012 | Young |
| 0.608391 | 0.019069 | 0.16 | 0.131 | 1 | 14 | Exoc2 | 0.029 | Young |
| 0.608535 | 0.356062 | 0.48 | 0.454 | 1 | 14 | Aff4 | 0.026 | Young |
| 0.608575 | 0.598979 | 0.42 | 0.503 | 1 | 14 | Cd55 | -0.083 | Young |
| 0.608663 | 0.19258 | 0.4 | 0.339 | 1 | 14 | Atp5o.1 | 0.061 | Young |
| 0.609763 | 0.580468 | 0.38 | 0.432 | 1 | 14 | Rhou | -0.052 | Young |
| 0.610609 | 0.402794 | 0.08 | 0.06 | 1 | 14 | Homer3 | 0.02 | Young |
| 0.610621 | 4.195048 | 0.02 | 0.011 | 1 | 14 | Upk1b | 0.009 | Young |
| 0.610621 | 3.432354 | 0.02 | 0.011 | 1 | 14 | Cldn1 | 0.009 | Young |
| 0.610621 | 3.400733 | 0.02 | 0.011 | 1 | 14 | Gm10031 | 0.009 | Young |
| 0.610621 | 3.344491 | 0.02 | 0.011 | 1 | 14 | 2010300C0 | 0.009 | Young |
| 0.610621 | 3.128851 | 0.02 | 0.011 | 1 | 14 | Fam155a | 0.009 | Young |
| 0.610621 | 2.764323 | 0.02 | 0.011 | 1 | 14 | Gm31105 | 0.009 | Young |

| | | | | | | | | |
|---|---|---|---|---|---|---|---|---|
| 0.610621 | 2.522986 | 0.02 | 0.011 | 1 | 14 | Krt19 | 0.009 | Young |
| 0.610621 | 2.495457 | 0.02 | 0.011 | 1 | 14 | Dpysl5 | 0.009 | Young |
| 0.610621 | 2.366307 | 0.02 | 0.011 | 1 | 14 | Bud23 | 0.009 | Young |
| 0.610621 | 2.297608 | 0.02 | 0.011 | 1 | 14 | Pqlc2 | 0.009 | Young |
| 0.610621 | 2.297119 | 0.02 | 0.011 | 1 | 14 | Top3a | 0.009 | Young |
| 0.610621 | 2.200127 | 0.02 | 0.011 | 1 | 14 | Gm34388 | 0.009 | Young |
| 0.610621 | 2.193266 | 0.02 | 0.011 | 1 | 14 | Nemp2 | 0.009 | Young |
| 0.610621 | 2.16215 | 0.02 | 0.011 | 1 | 14 | Ankrd55 | 0.009 | Young |
| 0.610621 | 2.004478 | 0.02 | 0.011 | 1 | 14 | Mgarp | 0.009 | Young |
| 0.610621 | 1.983969 | 0.02 | 0.011 | 1 | 14 | Gpm6a | 0.009 | Young |
| 0.610621 | 1.978517 | 0.02 | 0.011 | 1 | 14 | 1110035H1 | 0.009 | Young |
| 0.610621 | 1.876171 | 0.02 | 0.011 | 1 | 14 | Asxl3 | 0.009 | Young |
| 0.610621 | 1.772733 | 0.02 | 0.011 | 1 | 14 | Gm39469 | 0.009 | Young |
| 0.610621 | 1.765365 | 0.02 | 0.011 | 1 | 14 | Wfikkn2 | 0.009 | Young |
| 0.610621 | 1.755063 | 0.02 | 0.011 | 1 | 14 | Rbks | 0.009 | Young |
| 0.610621 | 1.726386 | 0.02 | 0.011 | 1 | 14 | Adamts7 | 0.009 | Young |
| 0.610621 | 1.673352 | 0.02 | 0.011 | 1 | 14 | Zcchc18 | 0.009 | Young |
| 0.610621 | 1.424814 | 0.02 | 0.011 | 1 | 14 | Fam26f | 0.009 | Young |
| 0.610621 | 1.416946 | 0.02 | 0.011 | 1 | 14 | Gm13199 | 0.009 | Young |
| 0.610621 | 1.357725 | 0.02 | 0.011 | 1 | 14 | Knstrn | 0.009 | Young |
| 0.610621 | 1.341423 | 0.02 | 0.011 | 1 | 14 | Rasgrp1 | 0.009 | Young |
| 0.610621 | 1.29161 | 0.02 | 0.011 | 1 | 14 | Wnk4 | 0.009 | Young |
| 0.610621 | 1.275586 | 0.02 | 0.011 | 1 | 14 | Mc2r | 0.009 | Young |
| 0.610621 | 1.257433 | 0.02 | 0.011 | 1 | 14 | Nova2 | 0.009 | Young |
| 0.610621 | 1.159367 | 0.02 | 0.011 | 1 | 14 | Trim29 | 0.009 | Young |
| 0.610621 | 1.152143 | 0.02 | 0.011 | 1 | 14 | Lrrc4 | 0.009 | Young |
| 0.610621 | 1.047145 | 0.02 | 0.011 | 1 | 14 | Ly6g6e | 0.009 | Young |
| 0.610621 | 1.029895 | 0.02 | 0.011 | 1 | 14 | Fjx1 | 0.009 | Young |
| 0.610621 | 1.022088 | 0.02 | 0.011 | 1 | 14 | Angptl8 | 0.009 | Young |
| 0.610621 | 0.954406 | 0.02 | 0.011 | 1 | 14 | Impa2 | 0.009 | Young |
| 0.610621 | 0.926877 | 0.02 | 0.011 | 1 | 14 | Exo5 | 0.009 | Young |
| 0.610621 | 0.913097 | 0.02 | 0.011 | 1 | 14 | Dsn1 | 0.009 | Young |
| 0.610621 | 0.904374 | 0.02 | 0.011 | 1 | 14 | Mastl | 0.009 | Young |
| 0.610621 | 0.89305 | 0.02 | 0.011 | 1 | 14 | Exd1 | 0.009 | Young |
| 0.610621 | 0.883728 | 0.02 | 0.011 | 1 | 14 | Kcnn4 | 0.009 | Young |
| 0.610621 | 0.839713 | 0.02 | 0.011 | 1 | 14 | Cenpo | 0.009 | Young |
| 0.610621 | 0.729813 | 0.02 | 0.011 | 1 | 14 | Bco1 | 0.009 | Young |
| 0.610621 | 0.653606 | 0.02 | 0.011 | 1 | 14 | Gm9780 | 0.009 | Young |
| 0.610621 | 0.547056 | 0.02 | 0.011 | 1 | 14 | Mterf2 | 0.009 | Young |
| 0.610808 | 1.11741 | 0.06 | 0.044 | 1 | 14 | Eipr1 | 0.016 | Young |
| 0.610808 | 0.697711 | 0.06 | 0.044 | 1 | 14 | Selplg | 0.016 | Young |
| 0.610884 | 0.34617 | 0.1 | 0.077 | 1 | 14 | Folr2 | 0.023 | Young |
| 0.610884 | 0.130918 | 0.1 | 0.077 | 1 | 14 | Tmem138 | 0.023 | Young |
| 0.61115 | 1.288254 | 0.22 | 0.208 | 1 | 14 | Stam | 0.012 | Young |
| 0.612035 | 0.163589 | 0.66 | 0.743 | 1 | 14 | Gabarap | -0.083 | Young |

| | | | | | | | | |
|---|---|---|---|---|---|---|---|---|
| 0.613247 | 1.180876 | 0.12 | 0.093 | 1 | 14 | Fbxl6 | 0.027 | Young |
| 0.613939 | 0.184298 | 0.16 | 0.131 | 1 | 14 | Gm12216 | 0.029 | Young |
| 0.613985 | 0.189073 | 0.36 | 0.317 | 1 | 14 | Stim1 | 0.043 | Young |
| 0.614463 | 0.195844 | 0.24 | 0.268 | 1 | 14 | Ube4b | -0.028 | Young |
| 0.614515 | 0.1014 | 0.08 | 0.06 | 1 | 14 | Pou5f2 | 0.02 | Young |
| 0.614515 | 0.077276 | 0.08 | 0.06 | 1 | 14 | Utp4 | 0.02 | Young |
| 0.614696 | 0.786275 | 0.16 | 0.202 | 1 | 14 | Stox2 | -0.042 | Young |
| 0.614847 | 0.171402 | 0.12 | 0.153 | 1 | 14 | Ublcp1 | -0.033 | Young |
| 0.614988 | 1.323721 | 0.14 | 0.186 | 1 | 14 | Abca3 | -0.046 | Young |
| 0.61502 | 1.823191 | 0.04 | 0.06 | 1 | 14 | Dnal4 | -0.02 | Young |
| 0.61502 | 1.056108 | 0.04 | 0.06 | 1 | 14 | Pold3 | -0.02 | Young |
| 0.615331 | 1.474909 | 0.06 | 0.044 | 1 | 14 | Ice2 | 0.016 | Young |
| 0.615331 | 1.159077 | 0.06 | 0.044 | 1 | 14 | Hvcn1 | 0.016 | Young |
| 0.615331 | 1.138304 | 0.06 | 0.044 | 1 | 14 | Emb | 0.016 | Young |
| 0.615363 | 0.145071 | 0.74 | 0.749 | 1 | 14 | Itm2c | -0.009 | Young |
| 0.615628 | 0.61359 | 0.22 | 0.262 | 1 | 14 | Sfrp2 | -0.042 | Young |
| 0.616119 | 0.540444 | 0.38 | 0.377 | 1 | 14 | Grb10 | 0.003 | Young |
| 0.616263 | 0.815402 | 0.46 | 0.475 | 1 | 14 | Mfge8 | -0.015 | Young |
| 0.61661 | 0.182057 | 0.34 | 0.29 | 1 | 14 | Hipk3 | 0.05 | Young |
| 0.616828 | 2.237559 | 0.04 | 0.027 | 1 | 14 | AC161607. | 0.013 | Young |
| 0.616828 | 1.546622 | 0.04 | 0.027 | 1 | 14 | Zfp772 | 0.013 | Young |
| 0.617071 | 0.366362 | 0.32 | 0.372 | 1 | 14 | Rab7b | -0.052 | Young |
| 0.617292 | 0.615571 | 0.32 | 0.284 | 1 | 14 | Ptbp1 | 0.036 | Young |
| 0.617408 | 0.230597 | 0.1 | 0.131 | 1 | 14 | Dmxl2 | -0.031 | Young |
| 0.617895 | 0.355052 | 0.1 | 0.077 | 1 | 14 | Zfp275 | 0.023 | Young |
| 0.617972 | 0.146481 | 0.18 | 0.148 | 1 | 14 | Gon4l | 0.032 | Young |
| 0.618066 | 1.279472 | 0.06 | 0.082 | 1 | 14 | Cdc14a | -0.022 | Young |
| 0.618416 | 0.430267 | 0.48 | 0.486 | 1 | 14 | Ssr1 | -0.006 | Young |
| 0.618432 | 0.38949 | 0.08 | 0.06 | 1 | 14 | Lsm10 | 0.02 | Young |
| 0.618432 | 0.386686 | 0.08 | 0.06 | 1 | 14 | Fbxo10 | 0.02 | Young |
| 0.618432 | 0.295524 | 0.08 | 0.06 | 1 | 14 | Tmem161b | 0.02 | Young |
| 0.618432 | 0.22929 | 0.08 | 0.06 | 1 | 14 | Tyw1 | 0.02 | Young |
| 0.619144 | 1.199652 | 0.02 | 0.011 | 1 | 14 | Spata2l | 0.009 | Young |
| 0.619144 | 1.031121 | 0.02 | 0.011 | 1 | 14 | Psmc3ip | 0.009 | Young |
| 0.619144 | 0.990985 | 0.02 | 0.011 | 1 | 14 | Slc25a21 | 0.009 | Young |
| 0.619144 | 0.917543 | 0.02 | 0.011 | 1 | 14 | A730056A( | 0.009 | Young |
| 0.619144 | 0.884007 | 0.02 | 0.011 | 1 | 14 | Tmem28 | 0.009 | Young |
| 0.619144 | 0.821547 | 0.02 | 0.011 | 1 | 14 | Vash2 | 0.009 | Young |
| 0.619144 | 0.762817 | 0.02 | 0.011 | 1 | 14 | BC035044 | 0.009 | Young |
| 0.619144 | 0.757282 | 0.02 | 0.011 | 1 | 14 | Lipt2 | 0.009 | Young |
| 0.619144 | 0.741654 | 0.02 | 0.011 | 1 | 14 | Ccnjl | 0.009 | Young |
| 0.619144 | 0.740198 | 0.02 | 0.011 | 1 | 14 | Oas1a | 0.009 | Young |
| 0.619144 | 0.739891 | 0.02 | 0.011 | 1 | 14 | Zfp94 | 0.009 | Young |
| 0.619144 | 0.73405 | 0.02 | 0.011 | 1 | 14 | Tro | 0.009 | Young |
| 0.619144 | 0.680358 | 0.02 | 0.011 | 1 | 14 | Itgax | 0.009 | Young |

| | | | | | | | | |
|---|---|---|---|---|---|---|---|---|
| 0.619144 | 0.646882 | 0.02 | 0.011 | 1 | 14 | Ccdc85c | 0.009 | Young |
| 0.619144 | 0.645395 | 0.02 | 0.011 | 1 | 14 | Kmo | 0.009 | Young |
| 0.619144 | 0.635311 | 0.02 | 0.011 | 1 | 14 | Gm29562 | 0.009 | Young |
| 0.619144 | 0.624037 | 0.02 | 0.011 | 1 | 14 | Galnt18 | 0.009 | Young |
| 0.619144 | 0.588023 | 0.02 | 0.011 | 1 | 14 | Gm37986 | 0.009 | Young |
| 0.619144 | 0.585648 | 0.02 | 0.011 | 1 | 14 | Hes7 | 0.009 | Young |
| 0.619144 | 0.572244 | 0.02 | 0.011 | 1 | 14 | Gm48529 | 0.009 | Young |
| 0.619144 | 0.571776 | 0.02 | 0.011 | 1 | 14 | Shroom4 | 0.009 | Young |
| 0.619144 | 0.541611 | 0.02 | 0.011 | 1 | 14 | Slc9a4 | 0.009 | Young |
| 0.619144 | 0.534361 | 0.02 | 0.011 | 1 | 14 | Slc38a1 | 0.009 | Young |
| 0.619144 | 0.528069 | 0.02 | 0.011 | 1 | 14 | Mars2 | 0.009 | Young |
| 0.619144 | 0.523685 | 0.02 | 0.011 | 1 | 14 | Amt | 0.009 | Young |
| 0.619144 | 0.507165 | 0.02 | 0.011 | 1 | 14 | Slc46a3 | 0.009 | Young |
| 0.619144 | 0.499368 | 0.02 | 0.011 | 1 | 14 | Agap2 | 0.009 | Young |
| 0.619144 | 0.491584 | 0.02 | 0.011 | 1 | 14 | Arhgap33 | 0.009 | Young |
| 0.619144 | 0.486617 | 0.02 | 0.011 | 1 | 14 | Gm9885 | 0.009 | Young |
| 0.619144 | 0.481813 | 0.02 | 0.011 | 1 | 14 | Cpz | 0.009 | Young |
| 0.619144 | 0.46712 | 0.02 | 0.011 | 1 | 14 | Gm36033 | 0.009 | Young |
| 0.619144 | 0.462247 | 0.02 | 0.011 | 1 | 14 | Zgrf1 | 0.009 | Young |
| 0.619144 | 0.423021 | 0.02 | 0.011 | 1 | 14 | Sema6b | 0.009 | Young |
| 0.619144 | 0.418646 | 0.02 | 0.011 | 1 | 14 | Gm10053 | 0.009 | Young |
| 0.619144 | 0.398929 | 0.02 | 0.011 | 1 | 14 | Pcdhga7 | 0.009 | Young |
| 0.619144 | 0.390223 | 0.02 | 0.011 | 1 | 14 | Ss18l1 | 0.009 | Young |
| 0.619144 | 0.359785 | 0.02 | 0.011 | 1 | 14 | Fut4 | 0.009 | Young |
| 0.619144 | 0.354963 | 0.02 | 0.011 | 1 | 14 | Tes3-ps | 0.009 | Young |
| 0.619144 | 0.351776 | 0.02 | 0.011 | 1 | 14 | Palb2 | 0.009 | Young |
| 0.619144 | 0.308046 | 0.02 | 0.011 | 1 | 14 | Foxred2 | 0.009 | Young |
| 0.619144 | 0.302345 | 0.02 | 0.011 | 1 | 14 | 9-Mar | 0.009 | Young |
| 0.619144 | 0.24476 | 0.02 | 0.011 | 1 | 14 | Gm5547 | 0.009 | Young |
| 0.619144 | 0.215824 | 0.02 | 0.011 | 1 | 14 | Gzmm | 0.009 | Young |
| 0.619144 | 0.086685 | 0.02 | 0.011 | 1 | 14 | Zfp2 | 0.009 | Young |
| 0.619144 | 0.040896 | 0.02 | 0.011 | 1 | 14 | Atad5 | 0.009 | Young |
| 0.619211 | 1.34481 | 0.04 | 0.06 | 1 | 14 | Crybb3 | -0.02 | Young |
| 0.619329 | 0.113294 | 0.24 | 0.202 | 1 | 14 | Gm37494 | 0.038 | Young |
| 0.619362 | 0.300188 | 0.14 | 0.115 | 1 | 14 | Rnf5 | 0.025 | Young |
| 0.61951 | 0.694099 | 0.16 | 0.131 | 1 | 14 | Nploc4 | 0.029 | Young |
| 0.61951 | 0.326603 | 0.16 | 0.131 | 1 | 14 | Hsd17b7 | 0.029 | Young |
| 0.619678 | 0.429716 | 0.2 | 0.175 | 1 | 14 | Uvrag | 0.025 | Young |
| 0.619813 | 0.168414 | 0.26 | 0.23 | 1 | 14 | Hmgb2 | 0.03 | Young |
| 0.619869 | 1.786002 | 0.06 | 0.044 | 1 | 14 | Grk3 | 0.016 | Young |
| 0.619869 | 1.654907 | 0.06 | 0.044 | 1 | 14 | Ppil1 | 0.016 | Young |
| 0.619869 | 1.277784 | 0.06 | 0.044 | 1 | 14 | Hoxc5 | 0.016 | Young |
| 0.619869 | 1.141412 | 0.06 | 0.044 | 1 | 14 | Homer1 | 0.016 | Young |
| 0.619869 | 0.679581 | 0.06 | 0.044 | 1 | 14 | Pvt1 | 0.016 | Young |
| 0.619869 | 0.588371 | 0.06 | 0.044 | 1 | 14 | Dusp10 | 0.016 | Young |

| | | | | | | | | |
|---|---|---|---|---|---|---|---|---|
| 0.620186 | 0.192139 | 0.5 | 0.454 | 1 | 14 | Nipbl | 0.046 | Young |
| 0.620891 | 1.236421 | 0.1 | 0.082 | 1 | 14 | Rcan3 | 0.018 | Young |
| 0.621414 | 0.988895 | 0.1 | 0.077 | 1 | 14 | Rad51d | 0.023 | Young |
| 0.621414 | 0.288724 | 0.1 | 0.077 | 1 | 14 | Fam135a | 0.023 | Young |
| 0.621414 | 0.286916 | 0.1 | 0.077 | 1 | 14 | Desi1 | 0.023 | Young |
| 0.621427 | 0.068762 | 0.46 | 0.525 | 1 | 14 | Minos1 | -0.065 | Young |
| 0.621674 | 0.196748 | 0.06 | 0.082 | 1 | 14 | Cldn12 | -0.022 | Young |
| 0.622056 | 0.518997 | 0.24 | 0.208 | 1 | 14 | N4bp1 | 0.032 | Young |
| 0.62236 | 0.76953 | 0.08 | 0.06 | 1 | 14 | Mras | 0.02 | Young |
| 0.622476 | 1.11594 | 0.04 | 0.027 | 1 | 14 | Pex12 | 0.013 | Young |
| 0.622828 | 0.240277 | 0.14 | 0.18 | 1 | 14 | Aga | -0.04 | Young |
| 0.622917 | 0.978357 | 0.12 | 0.093 | 1 | 14 | Gtf2f1 | 0.027 | Young |
| 0.623135 | 0.5696 | 0.08 | 0.109 | 1 | 14 | Ddt | -0.029 | Young |
| 0.623135 | 0.4142 | 0.12 | 0.098 | 1 | 14 | Poc1b | 0.022 | Young |
| 0.623279 | 0.21165 | 0.28 | 0.246 | 1 | 14 | Psmc1 | 0.034 | Young |
| 0.623414 | 0.229046 | 0.04 | 0.06 | 1 | 14 | Zfp287 | -0.02 | Young |
| 0.624421 | 0.432123 | 0.06 | 0.044 | 1 | 14 | Rida | 0.016 | Young |
| 0.624561 | 0.346057 | 0.42 | 0.404 | 1 | 14 | Ago2 | 0.016 | Young |
| 0.624984 | 0.679549 | 0.22 | 0.191 | 1 | 14 | Wbp4 | 0.029 | Young |
| 0.62501 | 0.227857 | 0.62 | 0.601 | 1 | 14 | Irf2bp2 | 0.019 | Young |
| 0.625102 | 0.028673 | 0.16 | 0.131 | 1 | 14 | 3830406C1 | 0.029 | Young |
| 0.625291 | 0.310884 | 0.06 | 0.082 | 1 | 14 | Ppip5k1 | -0.022 | Young |
| 0.625386 | 0.168882 | 0.28 | 0.251 | 1 | 14 | Tmem259 | 0.029 | Young |
| 0.625974 | 0.161301 | 0.18 | 0.213 | 1 | 14 | Dennd1a | -0.033 | Young |
| 0.626154 | 0.844492 | 0.08 | 0.104 | 1 | 14 | Vegfb | -0.024 | Young |
| 0.626154 | 0.284799 | 0.12 | 0.093 | 1 | 14 | Pprc1 | 0.027 | Young |
| 0.626243 | 0.663244 | 0.4 | 0.393 | 1 | 14 | Rad23b | 0.007 | Young |
| 0.626298 | 0.10892 | 0.08 | 0.06 | 1 | 14 | C87436 | 0.02 | Young |
| 0.626312 | 1.2167 | 0.12 | 0.098 | 1 | 14 | Endog | 0.022 | Young |
| 0.626723 | 0.477222 | 0.18 | 0.224 | 1 | 14 | Pdxdc1 | -0.044 | Young |
| 0.626834 | 0.182993 | 0.36 | 0.388 | 1 | 14 | Wapl | -0.028 | Young |
| 0.626855 | 0.164863 | 0.16 | 0.137 | 1 | 14 | Tmx4 | 0.023 | Young |
| 0.626961 | 0.429406 | 0.56 | 0.525 | 1 | 14 | Fkbp9 | 0.035 | Young |
| 0.627719 | 0.421528 | 0.02 | 0.011 | 1 | 14 | Fbxo2 | 0.009 | Young |
| 0.627719 | 0.410449 | 0.02 | 0.011 | 1 | 14 | B630019K0 | 0.009 | Young |
| 0.627719 | 0.392429 | 0.02 | 0.011 | 1 | 14 | Tlr1 | 0.009 | Young |
| 0.627719 | 0.391083 | 0.02 | 0.011 | 1 | 14 | BC052040 | 0.009 | Young |
| 0.627719 | 0.385908 | 0.02 | 0.011 | 1 | 14 | Lrrc51 | 0.009 | Young |
| 0.627719 | 0.355744 | 0.02 | 0.011 | 1 | 14 | Rab3b | 0.009 | Young |
| 0.627719 | 0.323657 | 0.02 | 0.011 | 1 | 14 | Arhgef19 | 0.009 | Young |
| 0.627719 | 0.305014 | 0.02 | 0.011 | 1 | 14 | Zfp956 | 0.009 | Young |
| 0.627719 | 0.259451 | 0.02 | 0.011 | 1 | 14 | Hsf4 | 0.009 | Young |
| 0.627719 | 0.214567 | 0.02 | 0.011 | 1 | 14 | AC153140. | 0.009 | Young |
| 0.627719 | 0.207987 | 0.02 | 0.011 | 1 | 14 | Epb41l4a | 0.009 | Young |
| 0.627719 | 0.204517 | 0.02 | 0.011 | 1 | 14 | Sema3a | 0.009 | Young |

| | | | | | | | | |
|---|---|---|---|---|---|---|---|---|
| 0.627719 | 0.195563 | 0.02 | 0.011 | 1 | 14 | Arhgap11a | 0.009 | Young |
| 0.627719 | 0.170437 | 0.02 | 0.011 | 1 | 14 | Csrnp2 | 0.009 | Young |
| 0.627719 | 0.151207 | 0.02 | 0.011 | 1 | 14 | Gm5784 | 0.009 | Young |
| 0.627719 | 0.141427 | 0.02 | 0.011 | 1 | 14 | Fgr | 0.009 | Young |
| 0.627719 | 0.125814 | 0.02 | 0.011 | 1 | 14 | Lcn2 | 0.009 | Young |
| 0.627719 | 0.11329 | 0.02 | 0.011 | 1 | 14 | Ripply3 | 0.009 | Young |
| 0.627719 | 0.080797 | 0.02 | 0.011 | 1 | 14 | Pigo | 0.009 | Young |
| 0.627719 | 0.068119 | 0.02 | 0.011 | 1 | 14 | Gpr183 | 0.009 | Young |
| 0.627719 | 0.067681 | 0.02 | 0.011 | 1 | 14 | Mthfd1l | 0.009 | Young |
| 0.627719 | 0.039256 | 0.02 | 0.011 | 1 | 14 | Gnb1l | 0.009 | Young |
| 0.627719 | 0.03683 | 0.02 | 0.011 | 1 | 14 | Arl16 | 0.009 | Young |
| 0.627719 | 0.036705 | 0.02 | 0.011 | 1 | 14 | Rnf208 | 0.009 | Young |
| 0.627719 | 0.001556 | 0.02 | 0.011 | 1 | 14 | Cand2 | 0.009 | Young |
| 0.627732 | 0.257868 | 0.28 | 0.24 | 1 | 14 | Ankrd40 | 0.04 | Young |
| 0.628145 | 1.505949 | 0.04 | 0.027 | 1 | 14 | Med26 | 0.013 | Young |
| 0.628145 | 1.316029 | 0.04 | 0.027 | 1 | 14 | Tbxa2r | 0.013 | Young |
| 0.628145 | 0.482619 | 0.04 | 0.027 | 1 | 14 | Ddx51 | 0.013 | Young |
| 0.628145 | 0.451392 | 0.04 | 0.027 | 1 | 14 | Tcf24 | 0.013 | Young |
| 0.628259 | 0.441346 | 0.14 | 0.115 | 1 | 14 | Yod1 | 0.025 | Young |
| 0.628396 | 0.393547 | 0.32 | 0.273 | 1 | 14 | Ndufb8 | 0.047 | Young |
| 0.628401 | 0.221161 | 0.72 | 0.738 | 1 | 14 | Calm1 | -0.018 | Young |
| 0.62862 | 0.026795 | 0.18 | 0.148 | 1 | 14 | Tmem208 | 0.032 | Young |
| 0.628641 | 0.413488 | 0.16 | 0.197 | 1 | 14 | Suz12 | -0.037 | Young |
| 0.629399 | 0.87733 | 0.08 | 0.104 | 1 | 14 | Cdk7 | -0.024 | Young |
| 0.629496 | 0.648744 | 0.12 | 0.098 | 1 | 14 | Fbxo9 | 0.022 | Young |
| 0.629598 | 0.3266 | 0.44 | 0.459 | 1 | 14 | Nfe2l2 | -0.019 | Young |
| 0.629681 | 0.381194 | 0.2 | 0.175 | 1 | 14 | Chst15 | 0.025 | Young |
| 0.629862 | 0.161343 | 0.2 | 0.24 | 1 | 14 | Abcf1 | -0.04 | Young |
| 0.629946 | 0.826443 | 0.22 | 0.257 | 1 | 14 | Fdx1 | -0.037 | Young |
| 0.630191 | 0.737379 | 0.36 | 0.35 | 1 | 14 | Eif3j1 | 0.01 | Young |
| 0.630248 | 0.057229 | 0.08 | 0.06 | 1 | 14 | Btf3l4 | 0.02 | Young |
| 0.630491 | 0.151079 | 0.2 | 0.164 | 1 | 14 | Slc35a4 | 0.036 | Young |
| 0.630716 | 0.05517 | 0.16 | 0.131 | 1 | 14 | Ttyh3 | 0.029 | Young |
| 0.631236 | 0.53446 | 0.1 | 0.126 | 1 | 14 | Pla2g12a | -0.026 | Young |
| 0.631384 | 1.331045 | 0.2 | 0.169 | 1 | 14 | Dr1 | 0.031 | Young |
| 0.631384 | 0.046125 | 0.2 | 0.169 | 1 | 14 | Plcd1 | 0.031 | Young |
| 0.631444 | 0.186181 | 0.88 | 0.896 | 1 | 14 | Rpl5 | -0.016 | Young |
| 0.631663 | 0.22749 | 0.24 | 0.213 | 1 | 14 | Ipo8 | 0.027 | Young |
| 0.631728 | 0.325062 | 0.16 | 0.126 | 1 | 14 | Yif1a | 0.034 | Young |
| 0.631857 | 1.199299 | 0.04 | 0.06 | 1 | 14 | Morc4 | -0.02 | Young |
| 0.631869 | 0.137054 | 0.26 | 0.213 | 1 | 14 | Shc1 | 0.047 | Young |
| 0.631874 | 0.505275 | 0.3 | 0.361 | 1 | 14 | Cebpz | -0.061 | Young |
| 0.631945 | 0.427656 | 0.28 | 0.251 | 1 | 14 | Srp72 | 0.029 | Young |
| 0.632068 | 0.568123 | 0.3 | 0.333 | 1 | 14 | H1f0 | -0.033 | Young |
| 0.632552 | 0.001981 | 0.06 | 0.082 | 1 | 14 | 1110004E0 | -0.022 | Young |

| | | | | | | | | |
|---|---|---|---|---|---|---|---|---|
| 0.632651 | 0.938413 | 0.12 | 0.093 | 1 | 14 | 0610010F0 | 0.027 | Young |
| 0.632687 | 0.585462 | 0.12 | 0.098 | 1 | 14 | Herc6 | 0.022 | Young |
| 0.632687 | 0.536284 | 0.12 | 0.098 | 1 | 14 | Cd53 | 0.022 | Young |
| 0.632919 | 0.227187 | 0.16 | 0.191 | 1 | 14 | Synrg | -0.031 | Young |
| 0.633252 | 0.091531 | 0.3 | 0.268 | 1 | 14 | Kctd10 | 0.032 | Young |
| 0.633321 | 0.502427 | 0.14 | 0.109 | 1 | 14 | Cep63 | 0.031 | Young |
| 0.633568 | 0.354697 | 0.06 | 0.044 | 1 | 14 | Bckdhb | 0.016 | Young |
| 0.633688 | 0.510506 | 0.62 | 0.601 | 1 | 14 | Sfr1 | 0.019 | Young |
| 0.633837 | 2.219356 | 0.04 | 0.027 | 1 | 14 | Inha | 0.013 | Young |
| 0.633837 | 1.846416 | 0.04 | 0.027 | 1 | 14 | Adamts6 | 0.013 | Young |
| 0.633837 | 1.774277 | 0.04 | 0.027 | 1 | 14 | Timeless | 0.013 | Young |
| 0.633837 | 1.577549 | 0.04 | 0.027 | 1 | 14 | Zfpm1 | 0.013 | Young |
| 0.633837 | 0.932382 | 0.04 | 0.027 | 1 | 14 | Echdc3 | 0.013 | Young |
| 0.633837 | 0.865369 | 0.04 | 0.027 | 1 | 14 | Cd86 | 0.013 | Young |
| 0.633837 | 0.798044 | 0.04 | 0.027 | 1 | 14 | Ap5z1 | 0.013 | Young |
| 0.633837 | 0.56681 | 0.04 | 0.027 | 1 | 14 | Rpgrip1 | 0.013 | Young |
| 0.633837 | 0.548481 | 0.04 | 0.027 | 1 | 14 | Necab1 | 0.013 | Young |
| 0.633837 | 0.305683 | 0.04 | 0.027 | 1 | 14 | Dnmt3b | 0.013 | Young |
| 0.633869 | 0.404145 | 0.42 | 0.388 | 1 | 14 | Dtx3 | 0.032 | Young |
| 0.633948 | 1.151209 | 0.16 | 0.142 | 1 | 14 | Camk2g | 0.018 | Young |
| 0.634181 | 0.202392 | 0.26 | 0.213 | 1 | 14 | Rcor3 | 0.047 | Young |
| 0.634207 | 0.277395 | 0.08 | 0.06 | 1 | 14 | Zfp365 | 0.02 | Young |
| 0.634417 | 0.20629 | 0.5 | 0.464 | 1 | 14 | Snx3 | 0.036 | Young |
| 0.635246 | 0.104393 | 0.28 | 0.328 | 1 | 14 | Nup98 | -0.048 | Young |
| 0.635375 | 0.066544 | 0.22 | 0.186 | 1 | 14 | Rnpep | 0.034 | Young |
| 0.635928 | 0.824883 | 0.3 | 0.257 | 1 | 14 | Rala | 0.043 | Young |
| 0.636099 | 0.630517 | 0.3 | 0.29 | 1 | 14 | Mif | 0.01 | Young |
| 0.63681 | 0.090657 | 0.4 | 0.426 | 1 | 14 | Plin3 | -0.026 | Young |
| 0.637093 | 0.734645 | 0.22 | 0.191 | 1 | 14 | Vps13d | 0.029 | Young |
| 0.63721 | 0.18764 | 0.14 | 0.115 | 1 | 14 | Akt1s1 | 0.025 | Young |
| 0.637227 | 0.02889 | 0.16 | 0.186 | 1 | 14 | Cmah | -0.026 | Young |
| 0.637307 | 0.265917 | 0.32 | 0.295 | 1 | 14 | Ilf2 | 0.025 | Young |
| 0.638163 | 0.551513 | 0.06 | 0.044 | 1 | 14 | Wdr53 | 0.016 | Young |
| 0.638163 | 0.376528 | 0.06 | 0.044 | 1 | 14 | Tstd3 | 0.016 | Young |
| 0.638163 | 0.255111 | 0.06 | 0.044 | 1 | 14 | Ap4b1 | 0.016 | Young |
| 0.638163 | 0.253309 | 0.06 | 0.044 | 1 | 14 | Synj2 | 0.016 | Young |
| 0.638178 | 0.068752 | 0.08 | 0.06 | 1 | 14 | Mtmr10 | 0.02 | Young |
| 0.638195 | 1.029685 | 0.1 | 0.082 | 1 | 14 | Coq7 | 0.018 | Young |
| 0.638678 | 0.025033 | 0.6 | 0.59 | 1 | 14 | Tgoln1 | 0.01 | Young |
| 0.638965 | 0.224964 | 0.68 | 0.798 | 1 | 14 | Ssr3 | -0.118 | Young |
| 0.639089 | 0.828959 | 0.12 | 0.098 | 1 | 14 | Hira | 0.022 | Young |
| 0.639166 | 0.534026 | 0.18 | 0.153 | 1 | 14 | Peli2 | 0.027 | Young |
| 0.639177 | 0.516555 | 0.12 | 0.093 | 1 | 14 | Ism1 | 0.027 | Young |
| 0.639399 | 0.427192 | 0.1 | 0.12 | 1 | 14 | Cttnbp2nl | -0.02 | Young |
| 0.63955 | 0.943109 | 0.04 | 0.027 | 1 | 14 | Trmt11 | 0.013 | Young |

| | | | | | | | | |
|---|---|---|---|---|---|---|---|---|
| 0.63955 | 0.800164 | 0.04 | 0.027 | 1 | 14 | Bmp5 | 0.013 | Young |
| 0.63955 | 0.618457 | 0.04 | 0.027 | 1 | 14 | Bag2 | 0.013 | Young |
| 0.63955 | 0.617288 | 0.04 | 0.027 | 1 | 14 | Atxn7l2 | 0.013 | Young |
| 0.63955 | 0.566184 | 0.04 | 0.027 | 1 | 14 | Gm4673 | 0.013 | Young |
| 0.63955 | 0.555563 | 0.04 | 0.027 | 1 | 14 | Zswim7 | 0.013 | Young |
| 0.63955 | 0.499687 | 0.04 | 0.027 | 1 | 14 | Gm45351 | 0.013 | Young |
| 0.63955 | 0.473742 | 0.04 | 0.027 | 1 | 14 | Cracr2b | 0.013 | Young |
| 0.63955 | 0.39432 | 0.04 | 0.027 | 1 | 14 | Mapkapk3 | 0.013 | Young |
| 0.63955 | 0.315027 | 0.04 | 0.027 | 1 | 14 | Bmp8a | 0.013 | Young |
| 0.640321 | 0.494001 | 0.38 | 0.35 | 1 | 14 | Ptov1 | 0.03 | Young |
| 0.640452 | 0.418093 | 0.32 | 0.306 | 1 | 14 | Rab22a | 0.014 | Young |
| 0.640762 | 0.475759 | 0.16 | 0.137 | 1 | 14 | AU019823 | 0.023 | Young |
| 0.640816 | 0.321416 | 0.46 | 0.448 | 1 | 14 | Snrpe | 0.012 | Young |
| 0.64168 | 1.039659 | 0.1 | 0.082 | 1 | 14 | Serpina3c | 0.018 | Young |
| 0.641829 | 0.175129 | 0.18 | 0.153 | 1 | 14 | Pbdc1 | 0.027 | Young |
| 0.642008 | 1.381604 | 0.16 | 0.131 | 1 | 14 | B230354K1 | 0.029 | Young |
| 0.642211 | 1.01896 | 0.16 | 0.142 | 1 | 14 | Il17rd | 0.018 | Young |
| 0.6423 | 1.026722 | 0.12 | 0.098 | 1 | 14 | Zbtb14 | 0.022 | Young |
| 0.642772 | 1.113964 | 0.06 | 0.044 | 1 | 14 | E330013P0 | 0.016 | Young |
| 0.642772 | 0.544664 | 0.06 | 0.044 | 1 | 14 | Slc16a7 | 0.016 | Young |
| 0.642772 | 0.246355 | 0.06 | 0.044 | 1 | 14 | BC002059 | 0.016 | Young |
| 0.642772 | 0.072895 | 0.06 | 0.044 | 1 | 14 | Gm14005 | 0.016 | Young |
| 0.643164 | 0.43595 | 0.6 | 0.579 | 1 | 14 | Ckap4 | 0.021 | Young |
| 0.643207 | 0.052021 | 0.14 | 0.115 | 1 | 14 | Atg12 | 0.025 | Young |
| 0.643266 | 0.632863 | 0.3 | 0.279 | 1 | 14 | Ptpn1 | 0.021 | Young |
| 0.643467 | 0.455753 | 0.24 | 0.219 | 1 | 14 | Uba6 | 0.021 | Young |
| 0.64348 | 0.249628 | 0.16 | 0.197 | 1 | 14 | Elovl1 | -0.037 | Young |
| 0.643508 | 0.12296 | 0.38 | 0.41 | 1 | 14 | Usmg5 | -0.03 | Young |
| 0.643559 | 0.666588 | 0.12 | 0.148 | 1 | 14 | Naaladl2 | -0.028 | Young |
| 0.643559 | 0.362577 | 0.12 | 0.148 | 1 | 14 | Pcgf5 | -0.028 | Young |
| 0.643798 | 0.440879 | 0.14 | 0.175 | 1 | 14 | Cdadc1 | -0.035 | Young |
| 0.643798 | 0.405494 | 0.14 | 0.175 | 1 | 14 | Marcksl1 | -0.035 | Young |
| 0.644405 | 0.040541 | 0.18 | 0.202 | 1 | 14 | Mtus1 | -0.022 | Young |
| 0.645285 | 1.441487 | 0.04 | 0.027 | 1 | 14 | Acad12 | 0.013 | Young |
| 0.645285 | 1.009984 | 0.04 | 0.027 | 1 | 14 | Ttc30a1 | 0.013 | Young |
| 0.645285 | 0.889878 | 0.04 | 0.027 | 1 | 14 | AL590144.1 | 0.013 | Young |
| 0.645285 | 0.675592 | 0.04 | 0.027 | 1 | 14 | Nalcn | 0.013 | Young |
| 0.645285 | 0.637702 | 0.04 | 0.027 | 1 | 14 | Mtg1 | 0.013 | Young |
| 0.645285 | 0.624052 | 0.04 | 0.027 | 1 | 14 | Cep152 | 0.013 | Young |
| 0.645285 | 0.474522 | 0.04 | 0.027 | 1 | 14 | Scml2 | 0.013 | Young |
| 0.645285 | 0.256574 | 0.04 | 0.027 | 1 | 14 | Il16 | 0.013 | Young |
| 0.646061 | 0.092623 | 0.16 | 0.126 | 1 | 14 | Lpar4 | 0.034 | Young |
| 0.646149 | 0.560096 | 0.08 | 0.06 | 1 | 14 | Prpsap2 | 0.02 | Young |
| 0.646285 | 0.587903 | 0.1 | 0.077 | 1 | 14 | Hectd3 | 0.023 | Young |
| 0.646682 | 0.372821 | 0.2 | 0.169 | 1 | 14 | Qser1 | 0.031 | Young |

| | | | | | | | | |
|---|---|---|---|---|---|---|---|---|
| 0.647128 | 0.318384 | 0.34 | 0.306 | 1 | 14 | Enpep | 0.034 | Young |
| 0.64718 | 0.138633 | 0.06 | 0.082 | 1 | 14 | Igtp | -0.022 | Young |
| 0.647693 | 0.210003 | 0.48 | 0.459 | 1 | 14 | Clta | 0.021 | Young |
| 0.64813 | 0.414656 | 0.24 | 0.219 | 1 | 14 | Mfap2 | 0.021 | Young |
| 0.648163 | 1.850577 | 0.08 | 0.066 | 1 | 14 | Plekhh3 | 0.014 | Young |
| 0.648175 | 0.036294 | 0.78 | 0.809 | 1 | 14 | Klf9 | -0.029 | Young |
| 0.648364 | 0.271674 | 0.54 | 0.503 | 1 | 14 | Ddx39b | 0.037 | Young |
| 0.648568 | 0.171294 | 0.74 | 0.792 | 1 | 14 | Canx | -0.052 | Young |
| 0.64877 | 0.250749 | 0.26 | 0.301 | 1 | 14 | Susd6 | -0.041 | Young |
| 0.648996 | 0.417053 | 0.36 | 0.328 | 1 | 14 | Mdh2 | 0.032 | Young |
| 0.649027 | 0.127436 | 0.14 | 0.175 | 1 | 14 | Rnf103 | -0.035 | Young |
| 0.649228 | 0.342829 | 0.14 | 0.115 | 1 | 14 | Mfsd10 | 0.025 | Young |
| 0.649317 | 0.042822 | 0.28 | 0.311 | 1 | 14 | Atn1 | -0.031 | Young |
| 0.64976 | 0.566807 | 0.44 | 0.432 | 1 | 14 | Ndufc2 | 0.008 | Young |
| 0.649843 | 1.298003 | 0.18 | 0.153 | 1 | 14 | Ddx52 | 0.027 | Young |
| 0.65015 | 0.195995 | 0.08 | 0.06 | 1 | 14 | E130311K1 | 0.02 | Young |
| 0.650305 | 0.1839 | 0.34 | 0.273 | 1 | 14 | Fam220a | 0.067 | Young |
| 0.650726 | 0.067531 | 0.4 | 0.361 | 1 | 14 | Sumo3 | 0.039 | Young |
| 0.651041 | 0.718857 | 0.04 | 0.027 | 1 | 14 | Tex30 | 0.013 | Young |
| 0.651041 | 0.544713 | 0.04 | 0.027 | 1 | 14 | Pla2g6 | 0.013 | Young |
| 0.651041 | 0.485536 | 0.04 | 0.027 | 1 | 14 | Tmub1 | 0.013 | Young |
| 0.651041 | 0.43038 | 0.04 | 0.027 | 1 | 14 | C030006K1 | 0.013 | Young |
| 0.651041 | 0.422329 | 0.04 | 0.027 | 1 | 14 | Gm45902 | 0.013 | Young |
| 0.651041 | 0.40859 | 0.04 | 0.027 | 1 | 14 | Gipc1 | 0.013 | Young |
| 0.651041 | 0.292093 | 0.04 | 0.027 | 1 | 14 | Gpr160 | 0.013 | Young |
| 0.651041 | 0.289878 | 0.04 | 0.027 | 1 | 14 | Gm2814 | 0.013 | Young |
| 0.651041 | 0.074397 | 0.04 | 0.027 | 1 | 14 | Fcgr1 | 0.013 | Young |
| 0.651176 | 0.369134 | 0.44 | 0.393 | 1 | 14 | Pigt | 0.047 | Young |
| 0.651752 | 0.371316 | 0.22 | 0.191 | 1 | 14 | Ggps1 | 0.029 | Young |
| 0.651771 | 0.27035 | 0.78 | 0.918 | 1 | 14 | Cst3 | -0.138 | Young |
| 0.651802 | 0.109736 | 0.42 | 0.464 | 1 | 14 | Mtpn | -0.044 | Young |
| 0.651953 | 0.186257 | 0.84 | 0.962 | 1 | 14 | Jun | -0.122 | Young |
| 0.652474 | 0.695492 | 0.08 | 0.098 | 1 | 14 | Atg9a | -0.018 | Young |
| 0.652523 | 0.139433 | 0.14 | 0.164 | 1 | 14 | Trak2 | -0.024 | Young |
| 0.652666 | 0.238642 | 0.3 | 0.268 | 1 | 14 | Mtfr1l | 0.032 | Young |
| 0.652978 | 0.078219 | 0.16 | 0.191 | 1 | 14 | Plod1 | -0.031 | Young |
| 0.653127 | 0.289871 | 0.6 | 0.596 | 1 | 14 | Top1 | 0.004 | Young |
| 0.653856 | 0.573585 | 0.26 | 0.23 | 1 | 14 | Dync1li2 | 0.03 | Young |
| 0.653927 | 0.139654 | 0.02 | 0.033 | 1 | 14 | Cyb561d2 | -0.013 | Young |
| 0.653927 | 0.136384 | 0.02 | 0.033 | 1 | 14 | Fam210b | -0.013 | Young |
| 0.653927 | 0.098682 | 0.02 | 0.033 | 1 | 14 | Ccdc77 | -0.013 | Young |
| 0.653927 | 0.018573 | 0.02 | 0.033 | 1 | 14 | Wdpcp | -0.013 | Young |
| 0.654208 | 0.160153 | 0.74 | 0.738 | 1 | 14 | Rpl35 | 0.002 | Young |
| 0.654387 | 0.043947 | 0.2 | 0.169 | 1 | 14 | Ksr1 | 0.031 | Young |
| 0.654796 | 0.139634 | 0.12 | 0.148 | 1 | 14 | Tgfbr1 | -0.028 | Young |

| | | | | | | | | |
|---|---|---|---|---|---|---|---|---|
| 0.654982 | 0.123822 | 0.16 | 0.186 | 1 | 14 | Tm2d3 | -0.026 | Young |
| 0.655152 | 0.733927 | 0.24 | 0.219 | 1 | 14 | Snrnp200 | 0.021 | Young |
| 0.655597 | 0.786501 | 0.3 | 0.257 | 1 | 14 | Arfgef1 | 0.043 | Young |
| 0.655698 | 1.135415 | 0.1 | 0.082 | 1 | 14 | Mllt3 | 0.018 | Young |
| 0.655699 | 0.744835 | 0.14 | 0.12 | 1 | 14 | Tspan6 | 0.02 | Young |
| 0.655699 | 0.412333 | 0.14 | 0.12 | 1 | 14 | Zfp655 | 0.02 | Young |
| 0.6557 | 0.453192 | 0.56 | 0.568 | 1 | 14 | Kif5b | -0.008 | Young |
| 0.656227 | 0.160663 | 0.44 | 0.437 | 1 | 14 | Dad1 | 0.003 | Young |
| 0.656325 | 0.517936 | 0.3 | 0.262 | 1 | 14 | Arrdc3 | 0.038 | Young |
| 0.656607 | 0.011884 | 0.38 | 0.437 | 1 | 14 | Srp9 | -0.057 | Young |
| 0.656681 | 0.34138 | 0.06 | 0.044 | 1 | 14 | Slc10a3 | 0.016 | Young |
| 0.656681 | 0.335881 | 0.06 | 0.044 | 1 | 14 | Cd37 | 0.016 | Young |
| 0.656681 | 0.084117 | 0.06 | 0.044 | 1 | 14 | Tifa | 0.016 | Young |
| 0.656818 | 0.277145 | 0.04 | 0.027 | 1 | 14 | Sytl4 | 0.013 | Young |
| 0.656818 | 0.175903 | 0.04 | 0.027 | 1 | 14 | Rbp4 | 0.013 | Young |
| 0.656838 | 0.06463 | 0.38 | 0.317 | 1 | 14 | Cr1l | 0.063 | Young |
| 0.656964 | 0.259171 | 0.2 | 0.169 | 1 | 14 | Idh2 | 0.031 | Young |
| 0.656964 | 0.023718 | 0.2 | 0.169 | 1 | 14 | Iqgap2 | 0.031 | Young |
| 0.657618 | 0.446245 | 0.16 | 0.137 | 1 | 14 | Adcy3 | 0.023 | Young |
| 0.657618 | 0.000665 | 0.12 | 0.148 | 1 | 14 | Lrrn2 | -0.028 | Young |
| 0.658302 | 0.295429 | 0.14 | 0.115 | 1 | 14 | Gzf1 | 0.025 | Young |
| 0.65847 | 0.274639 | 0.28 | 0.251 | 1 | 14 | Supt5 | 0.029 | Young |
| 0.658626 | 0.903207 | 0.38 | 0.366 | 1 | 14 | Aprt | 0.014 | Young |
| 0.658684 | 0.588902 | 0.14 | 0.12 | 1 | 14 | Tmem18 | 0.02 | Young |
| 0.659222 | 1.816347 | 0.1 | 0.082 | 1 | 14 | Pacs1 | 0.018 | Young |
| 0.65928 | 0.394659 | 0.28 | 0.333 | 1 | 14 | Tecr | -0.053 | Young |
| 0.659527 | 0.035315 | 0.44 | 0.475 | 1 | 14 | Mxra7 | -0.035 | Young |
| 0.659545 | 0.845725 | 0.2 | 0.169 | 1 | 14 | Snapin | 0.031 | Young |
| 0.659622 | 0.345971 | 0.46 | 0.519 | 1 | 14 | Eif5b | -0.059 | Young |
| 0.659714 | 2.332629 | 0.02 | 0.033 | 1 | 14 | Fxn | -0.013 | Young |
| 0.659714 | 1.976412 | 0.02 | 0.033 | 1 | 14 | Zc3h3 | -0.013 | Young |
| 0.659714 | 1.478629 | 0.02 | 0.033 | 1 | 14 | Dnajb5 | -0.013 | Young |
| 0.659714 | 1.273552 | 0.02 | 0.033 | 1 | 14 | Rhebl1 | -0.013 | Young |
| 0.659714 | 0.932784 | 0.02 | 0.033 | 1 | 14 | Sct | -0.013 | Young |
| 0.659714 | 0.92672 | 0.02 | 0.033 | 1 | 14 | Hist2h2be | -0.013 | Young |
| 0.659714 | 0.265593 | 0.02 | 0.033 | 1 | 14 | Cysltr2 | -0.013 | Young |
| 0.660291 | 0.066942 | 0.46 | 0.421 | 1 | 14 | Anp32b | 0.039 | Young |
| 0.660499 | 0.379389 | 0.38 | 0.361 | 1 | 14 | Smim11 | 0.019 | Young |
| 0.660592 | 0.10315 | 0.14 | 0.164 | 1 | 14 | Stx1b | -0.024 | Young |
| 0.660701 | 0.071956 | 0.24 | 0.262 | 1 | 14 | Blvrb | -0.022 | Young |
| 0.661207 | 0.766048 | 0.32 | 0.339 | 1 | 14 | Man1a2 | -0.019 | Young |
| 0.661338 | 1.141078 | 0.14 | 0.115 | 1 | 14 | Elof1 | 0.025 | Young |
| 0.662002 | 0.242198 | 0.3 | 0.273 | 1 | 14 | Tm4sf1 | 0.027 | Young |
| 0.662403 | 0.050649 | 0.76 | 0.716 | 1 | 14 | Ybx1 | 0.044 | Young |
| 0.662615 | 0.120028 | 0.04 | 0.027 | 1 | 14 | Tmem62 | 0.013 | Young |

| | | | | | | | | |
|---|---|---|---|---|---|---|---|---|
| 0.662716 | 0.235357 | 0.86 | 0.863 | 1 | 14 | Rpl19 | -0.003 | Young |
| 0.662754 | 0.271788 | 0.1 | 0.082 | 1 | 14 | Taf8 | 0.018 | Young |
| 0.662764 | 0.068132 | 0.18 | 0.158 | 1 | 14 | Armc10 | 0.022 | Young |
| 0.663291 | 0.909798 | 0.18 | 0.153 | 1 | 14 | Rprd1b | 0.027 | Young |
| 0.663656 | 0.263723 | 0.44 | 0.399 | 1 | 14 | Huwe1 | 0.041 | Young |
| 0.663682 | 0.7776 | 0.32 | 0.284 | 1 | 14 | Cndp2 | 0.036 | Young |
| 0.663752 | 0.348104 | 0.18 | 0.148 | 1 | 14 | Tmem119 | 0.032 | Young |
| 0.66379 | 0.036321 | 0.9 | 0.891 | 1 | 14 | Rpl22 | 0.009 | Young |
| 0.664018 | 0.571624 | 0.32 | 0.301 | 1 | 14 | Fkbp10 | 0.019 | Young |
| 0.66467 | 0.250741 | 0.14 | 0.12 | 1 | 14 | Mrpl35 | 0.02 | Young |
| 0.664766 | 0.209895 | 0.42 | 0.377 | 1 | 14 | Zranb2 | 0.043 | Young |
| 0.664963 | 1.217048 | 0.08 | 0.109 | 1 | 14 | Dpagt1 | -0.029 | Young |
| 0.664963 | 0.57765 | 0.12 | 0.098 | 1 | 14 | Ncbp1 | 0.022 | Young |
| 0.664963 | 0.248705 | 0.12 | 0.098 | 1 | 14 | Spns1 | 0.022 | Young |
| 0.664963 | 0.160124 | 0.12 | 0.098 | 1 | 14 | Psat1 | 0.022 | Young |
| 0.665657 | 0.137492 | 0.26 | 0.219 | 1 | 14 | Rnh1 | 0.041 | Young |
| 0.666112 | 0.092028 | 0.12 | 0.148 | 1 | 14 | Aspa | -0.028 | Young |
| 0.666227 | 0.191509 | 0.6 | 0.607 | 1 | 14 | Esyt2 | -0.007 | Young |
| 0.666293 | 0.772504 | 0.1 | 0.082 | 1 | 14 | Abraxas2 | 0.018 | Young |
| 0.666922 | 0.417903 | 0.2 | 0.23 | 1 | 14 | Dld | -0.03 | Young |
| 0.666999 | 0.209261 | 0.42 | 0.459 | 1 | 14 | Cbx3 | -0.039 | Young |
| 0.667426 | 0.830266 | 0.14 | 0.115 | 1 | 14 | Tubb4a | 0.025 | Young |
| 0.668002 | 0.445384 | 0.26 | 0.219 | 1 | 14 | Zzz3 | 0.041 | Young |
| 0.669176 | 0.054992 | 0.2 | 0.235 | 1 | 14 | Cadm3 | -0.035 | Young |
| 0.669262 | 0.044272 | 0.32 | 0.366 | 1 | 14 | 1110004F1 | -0.046 | Young |
| 0.669298 | 0.63924 | 0.46 | 0.437 | 1 | 14 | Ier3ip1 | 0.023 | Young |
| 0.669618 | 0.163011 | 0.26 | 0.295 | 1 | 14 | Ap2s1 | -0.035 | Young |
| 0.669623 | 0.23695 | 0.52 | 0.574 | 1 | 14 | Slc38a2 | -0.054 | Young |
| 0.669753 | 0.36034 | 0.78 | 0.814 | 1 | 14 | Rpl36 | -0.034 | Young |
| 0.670114 | 0.099014 | 0.32 | 0.284 | 1 | 14 | Abracl | 0.036 | Young |
| 0.670164 | 0.453521 | 0.5 | 0.486 | 1 | 14 | Lama2 | 0.014 | Young |
| 0.670627 | 1.016152 | 0.06 | 0.077 | 1 | 14 | 2610301B2 | -0.017 | Young |
| 0.670698 | 0.076004 | 0.32 | 0.279 | 1 | 14 | Larp4 | 0.041 | Young |
| 0.670793 | 0.508478 | 0.18 | 0.158 | 1 | 14 | Nom1 | 0.022 | Young |
| 0.671013 | 0.068582 | 0.34 | 0.29 | 1 | 14 | Ccdc88a | 0.05 | Young |
| 0.671418 | 0.112816 | 0.34 | 0.295 | 1 | 14 | Adk | 0.045 | Young |
| 0.671496 | 1.15537 | 0.12 | 0.098 | 1 | 14 | Fastk | 0.022 | Young |
| 0.672562 | 0.525683 | 0.22 | 0.186 | 1 | 14 | Gnl3 | 0.034 | Young |
| 0.672884 | 1.35181 | 0.22 | 0.197 | 1 | 14 | Aldh3a2 | 0.023 | Young |
| 0.672934 | 0.206459 | 0.2 | 0.175 | 1 | 14 | Ubqln1 | 0.025 | Young |
| 0.673194 | 0.213246 | 0.28 | 0.322 | 1 | 14 | Sec11a | -0.042 | Young |
| 0.673393 | 0.662507 | 0.1 | 0.082 | 1 | 14 | Parp8 | 0.018 | Young |
| 0.67369 | 0.774039 | 0.14 | 0.12 | 1 | 14 | Tef | 0.02 | Young |
| 0.673728 | 0.502489 | 0.28 | 0.257 | 1 | 14 | Sult5a1 | 0.023 | Young |
| 0.673893 | 0.109441 | 0.82 | 0.869 | 1 | 14 | Crip1 | -0.049 | Young |

| | | | | | | | | |
|---|---|---|---|---|---|---|---|---|
| 0.674401 | 0.201233 | 0.34 | 0.366 | 1 | 14 | Nxf1 | -0.026 | Young |
| 0.674739 | 0.320213 | 0.62 | 0.661 | 1 | 14 | Atp5h | -0.041 | Young |
| 0.675762 | 0.050921 | 0.38 | 0.328 | 1 | 14 | Hnrnpa1 | 0.052 | Young |
| 0.6766 | 0.003953 | 0.14 | 0.115 | 1 | 14 | Gar1 | 0.025 | Young |
| 0.676638 | 0.518553 | 0.22 | 0.202 | 1 | 14 | Trappc2l | 0.018 | Young |
| 0.676707 | 1.113363 | 0.14 | 0.12 | 1 | 14 | Golph3l | 0.02 | Young |
| 0.676803 | 0.088561 | 0.28 | 0.246 | 1 | 14 | Kpna4 | 0.034 | Young |
| 0.676954 | 1.128831 | 0.1 | 0.082 | 1 | 14 | Oxnad1 | 0.018 | Young |
| 0.676954 | 0.500316 | 0.1 | 0.082 | 1 | 14 | Nhlrc3 | 0.018 | Young |
| 0.67727 | 0.084812 | 0.3 | 0.273 | 1 | 14 | Arid1b | 0.027 | Young |
| 0.678054 | 0.182133 | 0.12 | 0.098 | 1 | 14 | Zfp369 | 0.022 | Young |
| 0.678054 | 0.15636 | 0.12 | 0.098 | 1 | 14 | Pycard | 0.022 | Young |
| 0.679208 | 0.591153 | 0.52 | 0.536 | 1 | 14 | Retnla | -0.016 | Young |
| 0.679669 | 0.424933 | 0.1 | 0.126 | 1 | 14 | Dhx40 | -0.026 | Young |
| 0.679678 | 0.277264 | 0.56 | 0.53 | 1 | 14 | Prrc2b | 0.03 | Young |
| 0.679729 | 0.853221 | 0.14 | 0.12 | 1 | 14 | Adamts16 | 0.02 | Young |
| 0.680307 | 1.730492 | 0.12 | 0.104 | 1 | 14 | Taf6l | 0.016 | Young |
| 0.680336 | 0.185196 | 0.2 | 0.169 | 1 | 14 | Kmt5a | 0.031 | Young |
| 0.681034 | 1.109897 | 0.04 | 0.055 | 1 | 14 | Nr2c2ap | -0.015 | Young |
| 0.681034 | 0.36242 | 0.04 | 0.055 | 1 | 14 | Meiob | -0.015 | Young |
| 0.681127 | 0.766844 | 0.28 | 0.273 | 1 | 14 | Mfsd1 | 0.007 | Young |
| 0.681194 | 0.657058 | 0.2 | 0.186 | 1 | 14 | Rnpepl1 | 0.014 | Young |
| 0.681472 | 0.834949 | 0.22 | 0.191 | 1 | 14 | Erlin2 | 0.029 | Young |
| 0.682141 | 0.103526 | 0.26 | 0.219 | 1 | 14 | Coq10b | 0.041 | Young |
| 0.682196 | 0.650601 | 0.3 | 0.257 | 1 | 14 | 4930523C0 | 0.043 | Young |
| 0.682374 | 0.006335 | 0.28 | 0.317 | 1 | 14 | Ldlr | -0.037 | Young |
| 0.682677 | 0.856371 | 0.34 | 0.306 | 1 | 14 | Yy1 | 0.034 | Young |
| 0.683229 | 0.583406 | 0.12 | 0.148 | 1 | 14 | Smo | -0.028 | Young |
| 0.683539 | 0.284173 | 0.24 | 0.219 | 1 | 14 | Dync1li1 | 0.021 | Young |
| 0.683617 | 0.561939 | 0.28 | 0.246 | 1 | 14 | Kif1c | 0.034 | Young |
| 0.684983 | 0.200823 | 0.54 | 0.623 | 1 | 14 | Tpm3 | -0.083 | Young |
| 0.685253 | 0.413199 | 0.22 | 0.197 | 1 | 14 | Pik3c2a | 0.023 | Young |
| 0.685426 | 0.116128 | 0.26 | 0.224 | 1 | 14 | Acadm | 0.036 | Young |
| 0.685882 | 0.110651 | 0.32 | 0.279 | 1 | 14 | Eif3i | 0.041 | Young |
| 0.685894 | 0.067209 | 0.28 | 0.246 | 1 | 14 | Csnk1g3 | 0.034 | Young |
| 0.68612 | 0.363852 | 0.24 | 0.213 | 1 | 14 | Kdsr | 0.027 | Young |
| 0.686295 | 0.325678 | 0.26 | 0.23 | 1 | 14 | Glmp | 0.03 | Young |
| 0.686497 | 0.20229 | 0.62 | 0.53 | 1 | 14 | Ttc14 | 0.09 | Young |
| 0.686964 | 0.764424 | 0.18 | 0.158 | 1 | 14 | Crebl2 | 0.022 | Young |
| 0.688471 | 0.558338 | 0.16 | 0.186 | 1 | 14 | Tmem57 | -0.026 | Young |
| 0.688635 | 0.054558 | 0.22 | 0.24 | 1 | 14 | Fam124a | -0.02 | Young |
| 0.689442 | 0.207462 | 0.26 | 0.235 | 1 | 14 | Higd2a | 0.025 | Young |
| 0.689555 | 0.098532 | 0.32 | 0.284 | 1 | 14 | Trip11 | 0.036 | Young |
| 0.689638 | 0.070836 | 0.48 | 0.519 | 1 | 14 | Ccdc50 | -0.039 | Young |
| 0.689846 | 0.577403 | 0.16 | 0.142 | 1 | 14 | Hypk | 0.018 | Young |

| | | | | | | | | |
|---|---|---|---|---|---|---|---|---|
| 0.689867 | 0.357013 | 0.06 | 0.077 | 1 | 14 | Atpaf1 | -0.017 | Young |
| 0.689897 | 0.147245 | 0.08 | 0.098 | 1 | 14 | Gyg | -0.018 | Young |
| 0.690024 | 0.754978 | 0.12 | 0.104 | 1 | 14 | Trim32 | 0.016 | Young |
| 0.690047 | 0.26948 | 0.36 | 0.328 | 1 | 14 | Ndel1 | 0.032 | Young |
| 0.690059 | 0.00071 | 0.66 | 0.765 | 1 | 14 | Calr | -0.105 | Young |
| 0.690122 | 1.37439 | 0.04 | 0.055 | 1 | 14 | Syp | -0.015 | Young |
| 0.69014 | 0.134708 | 0.26 | 0.224 | 1 | 14 | Ppp4c | 0.036 | Young |
| 0.690273 | 0.479414 | 0.32 | 0.295 | 1 | 14 | Ctdnep1 | 0.025 | Young |
| 0.690273 | 0.363656 | 0.26 | 0.311 | 1 | 14 | Basp1 | -0.051 | Young |
| 0.69051 | 0.233148 | 0.18 | 0.153 | 1 | 14 | Pde4dip | 0.027 | Young |
| 0.690939 | 0.578569 | 0.2 | 0.224 | 1 | 14 | Tomm34 | -0.024 | Young |
| 0.691074 | 0.640558 | 0.16 | 0.186 | 1 | 14 | Tmem147 | -0.026 | Young |
| 0.691074 | 0.270604 | 0.2 | 0.175 | 1 | 14 | Spg7 | 0.025 | Young |
| 0.69127 | 0.233557 | 0.1 | 0.082 | 1 | 14 | Nuak1 | 0.018 | Young |
| 0.691431 | 1.687541 | 0.08 | 0.066 | 1 | 14 | Mknk1 | 0.014 | Young |
| 0.691431 | 0.4304 | 0.08 | 0.066 | 1 | 14 | Slc22a15 | 0.014 | Young |
| 0.69185 | 0.410172 | 0.12 | 0.148 | 1 | 14 | Gm26782 | -0.028 | Young |
| 0.691997 | 0.856377 | 0.1 | 0.126 | 1 | 14 | Etfrf1 | -0.026 | Young |
| 0.69201 | 0.725135 | 0.1 | 0.12 | 1 | 14 | Arrdc1 | -0.02 | Young |
| 0.692518 | 0.487315 | 0.26 | 0.24 | 1 | 14 | 1810058I24 | 0.02 | Young |
| 0.692819 | 0.590827 | 0.24 | 0.224 | 1 | 14 | Mta2 | 0.016 | Young |
| 0.693457 | 0.140914 | 0.2 | 0.169 | 1 | 14 | Mnat1 | 0.031 | Young |
| 0.693594 | 0.138097 | 0.32 | 0.366 | 1 | 14 | Gorasp2 | -0.046 | Young |
| 0.693821 | 0.147691 | 0.12 | 0.142 | 1 | 14 | Pxmp4 | -0.022 | Young |
| 0.694072 | 0.309906 | 0.18 | 0.148 | 1 | 14 | Fnta | 0.032 | Young |
| 0.694555 | 0.714824 | 0.12 | 0.098 | 1 | 14 | Scn2a | 0.022 | Young |
| 0.694581 | 0.143519 | 0.28 | 0.306 | 1 | 14 | Cap1 | -0.026 | Young |
| 0.694682 | 1.322675 | 0.04 | 0.055 | 1 | 14 | Blzf1 | -0.015 | Young |
| 0.694682 | 0.13796 | 0.04 | 0.055 | 1 | 14 | Tab3 | -0.015 | Young |
| 0.694867 | 1.357863 | 0.1 | 0.082 | 1 | 14 | Actr3b | 0.018 | Young |
| 0.695092 | 0.303367 | 0.14 | 0.115 | 1 | 14 | Pde4a | 0.025 | Young |
| 0.695104 | 0.31564 | 0.18 | 0.158 | 1 | 14 | Ppp1r37 | 0.022 | Young |
| 0.695418 | 0.697471 | 0.08 | 0.066 | 1 | 14 | Dnajc27 | 0.014 | Young |
| 0.695418 | 0.553732 | 0.08 | 0.066 | 1 | 14 | Erich1 | 0.014 | Young |
| 0.695755 | 0.203651 | 0.12 | 0.137 | 1 | 14 | Yeats4 | -0.017 | Young |
| 0.69577 | 0.699315 | 0.24 | 0.213 | 1 | 14 | Commd3 | 0.027 | Young |
| 0.695989 | 0.15814 | 0.32 | 0.268 | 1 | 14 | Washc2 | 0.052 | Young |
| 0.696004 | 1.398359 | 0.14 | 0.164 | 1 | 14 | Smyd2 | -0.024 | Young |
| 0.69603 | 0.008863 | 0.94 | 0.929 | 1 | 14 | Rpl23 | 0.011 | Young |
| 0.696092 | 0.329034 | 0.16 | 0.18 | 1 | 14 | Nudt9 | -0.02 | Young |
| 0.696236 | 0.521666 | 0.58 | 0.656 | 1 | 14 | Morf4l2 | -0.076 | Young |
| 0.69653 | 1.825904 | 0.12 | 0.104 | 1 | 14 | Sgms2 | 0.016 | Young |
| 0.69653 | 0.529425 | 0.12 | 0.104 | 1 | 14 | Mitd1 | 0.016 | Young |
| 0.697155 | 0.59612 | 0.26 | 0.24 | 1 | 14 | Furin | 0.02 | Young |
| 0.698373 | 0.281531 | 0.36 | 0.328 | 1 | 14 | Taok3 | 0.032 | Young |

| | | | | | | | | |
|---|---|---|---|---|---|---|---|---|
| 0.69847 | 0.309167 | 0.1 | 0.082 | 1 | 14 | Fam189b | 0.018 | Young |
| 0.69847 | 0.193109 | 0.1 | 0.082 | 1 | 14 | Magohb | 0.018 | Young |
| 0.699067 | 1.381889 | 0.26 | 0.257 | 1 | 14 | Vti1b | 0.003 | Young |
| 0.699329 | 0.145741 | 0.54 | 0.552 | 1 | 14 | Tm9sf2 | -0.012 | Young |
| 0.699413 | 1.077958 | 0.08 | 0.066 | 1 | 14 | Cep250 | 0.014 | Young |
| 0.699603 | 0.076227 | 0.26 | 0.224 | 1 | 14 | Ncoa4 | 0.036 | Young |
| 0.700141 | 0.196464 | 0.22 | 0.257 | 1 | 14 | Api5 | -0.037 | Young |
| 0.70031 | 0.496982 | 0.24 | 0.219 | 1 | 14 | Nacc2 | 0.021 | Young |
| 0.70053 | 0.230482 | 0.72 | 0.787 | 1 | 14 | Col6a3 | -0.067 | Young |
| 0.700731 | 0.792237 | 0.32 | 0.361 | 1 | 14 | Creg1 | -0.041 | Young |
| 0.701217 | 0.159921 | 0.62 | 0.601 | 1 | 14 | Bclaf1 | 0.019 | Young |
| 0.701513 | 1.239822 | 0.18 | 0.153 | 1 | 14 | 1500011K1 | 0.027 | Young |
| 0.701854 | 0.68663 | 0.28 | 0.301 | 1 | 14 | Copg1 | -0.021 | Young |
| 0.70208 | 0.362711 | 0.1 | 0.082 | 1 | 14 | Cpsf3 | 0.018 | Young |
| 0.70208 | 0.222391 | 0.1 | 0.082 | 1 | 14 | Alkbh8 | 0.018 | Young |
| 0.702231 | 0.37088 | 0.46 | 0.464 | 1 | 14 | Kif1b | -0.004 | Young |
| 0.702314 | 0.664636 | 0.42 | 0.383 | 1 | 14 | Ubl5 | 0.037 | Young |
| 0.70241 | 0.364196 | 0.3 | 0.257 | 1 | 14 | Acadvl | 0.043 | Young |
| 0.702721 | 0.011144 | 0.66 | 0.683 | 1 | 14 | Pmp22 | -0.023 | Young |
| 0.702802 | 0.429411 | 0.38 | 0.35 | 1 | 14 | Smpdl3a | 0.03 | Young |
| 0.703058 | 0.869669 | 0.12 | 0.104 | 1 | 14 | Pop5 | 0.016 | Young |
| 0.703279 | 0.586886 | 0.14 | 0.169 | 1 | 14 | Slc44a2 | -0.029 | Young |
| 0.703416 | 0.713588 | 0.08 | 0.066 | 1 | 14 | Rrp15 | 0.014 | Young |
| 0.703416 | 0.594398 | 0.08 | 0.066 | 1 | 14 | Dohh | 0.014 | Young |
| 0.703416 | 0.444322 | 0.08 | 0.066 | 1 | 14 | Tnfrsf21 | 0.014 | Young |
| 0.703416 | 0.195878 | 0.08 | 0.066 | 1 | 14 | Osbpl3 | 0.014 | Young |
| 0.703485 | 0.347365 | 0.4 | 0.459 | 1 | 14 | Agap1 | -0.059 | Young |
| 0.703837 | 0.812449 | 0.04 | 0.055 | 1 | 14 | Amph | -0.015 | Young |
| 0.703837 | 0.311778 | 0.04 | 0.055 | 1 | 14 | Msh6 | -0.015 | Young |
| 0.704141 | 0.512286 | 0.2 | 0.175 | 1 | 14 | Map3k7 | 0.025 | Young |
| 0.704334 | 0.097252 | 0.5 | 0.568 | 1 | 14 | Ppp1cb | -0.068 | Young |
| 0.704631 | 0.248425 | 0.4 | 0.372 | 1 | 14 | Srrm1 | 0.028 | Young |
| 0.705099 | 0.03315 | 0.22 | 0.24 | 1 | 14 | Cmpk1 | -0.02 | Young |
| 0.705129 | 0.372574 | 0.24 | 0.219 | 1 | 14 | Ltbr | 0.021 | Young |
| 0.705697 | 0.64181 | 0.1 | 0.082 | 1 | 14 | Tbl2 | 0.018 | Young |
| 0.705697 | 0.026547 | 0.1 | 0.082 | 1 | 14 | Pck2 | 0.018 | Young |
| 0.705876 | 0.038252 | 0.5 | 0.552 | 1 | 14 | Tpm4 | -0.052 | Young |
| 0.706377 | 0.201134 | 0.54 | 0.536 | 1 | 14 | Rab11b | 0.004 | Young |
| 0.706722 | 0.117343 | 0.32 | 0.301 | 1 | 14 | Ube2s | 0.019 | Young |
| 0.707157 | 0.985047 | 0.14 | 0.12 | 1 | 14 | Rps6ka4 | 0.02 | Young |
| 0.707231 | 0.181127 | 0.4 | 0.443 | 1 | 14 | Csnk1e | -0.043 | Young |
| 0.707428 | 1.231809 | 0.08 | 0.066 | 1 | 14 | Tusc1 | 0.014 | Young |
| 0.707428 | 0.694133 | 0.08 | 0.066 | 1 | 14 | Ptrh2 | 0.014 | Young |
| 0.707902 | 0.290684 | 0.24 | 0.213 | 1 | 14 | Pla2g4a | 0.027 | Young |
| 0.70843 | 1.348387 | 0.04 | 0.055 | 1 | 14 | Eefsec | -0.015 | Young |

| | | | | | | | | |
|---|---|---|---|---|---|---|---|---|
| 0.70843 | 0.145263 | 0.04 | 0.055 | 1 | 14 | Ctc1 | -0.015 | Young |
| 0.708671 | 0.414522 | 0.32 | 0.393 | 1 | 14 | Rdx | -0.073 | Young |
| 0.708771 | 0.408081 | 0.22 | 0.202 | 1 | 14 | Ssr2 | 0.018 | Young |
| 0.708966 | 0.473675 | 0.34 | 0.35 | 1 | 14 | Immt | -0.01 | Young |
| 0.70931 | 0.021076 | 0.06 | 0.077 | 1 | 14 | Rdh5 | -0.017 | Young |
| 0.709393 | 0.030047 | 0.16 | 0.186 | 1 | 14 | Trip6 | -0.026 | Young |
| 0.709574 | 0.162997 | 0.4 | 0.377 | 1 | 14 | Elob | 0.023 | Young |
| 0.709726 | 1.529312 | 0.14 | 0.126 | 1 | 14 | Exoc1 | 0.014 | Young |
| 0.710243 | 0.60341 | 0.22 | 0.197 | 1 | 14 | Taf10 | 0.023 | Young |
| 0.710456 | 0.334405 | 0.38 | 0.388 | 1 | 14 | Tcea1 | -0.008 | Young |
| 0.710873 | 0.310717 | 0.34 | 0.377 | 1 | 14 | Bcap31 | -0.037 | Young |
| 0.71093 | 0.303819 | 0.36 | 0.328 | 1 | 14 | Hmgcs1 | 0.032 | Young |
| 0.711134 | 1.008011 | 0.22 | 0.251 | 1 | 14 | Tmem263 | -0.031 | Young |
| 0.711447 | 0.62067 | 0.08 | 0.066 | 1 | 14 | Aven | 0.014 | Young |
| 0.712024 | 0.180297 | 0.2 | 0.175 | 1 | 14 | Plekha1 | 0.025 | Young |
| 0.712661 | 0.315235 | 0.3 | 0.273 | 1 | 14 | Pdap1 | 0.027 | Young |
| 0.712719 | 0.271468 | 0.4 | 0.372 | 1 | 14 | Ppp1ca | 0.028 | Young |
| 0.713034 | 1.795657 | 0.06 | 0.049 | 1 | 14 | Paqr7 | 0.011 | Young |
| 0.713221 | 1.18473 | 0.06 | 0.077 | 1 | 14 | Prr3 | -0.017 | Young |
| 0.713221 | 0.121683 | 0.06 | 0.077 | 1 | 14 | Scyl3 | -0.017 | Young |
| 0.71364 | 0.999475 | 0.1 | 0.087 | 1 | 14 | Prrt1 | 0.013 | Young |
| 0.713766 | 0.498683 | 0.14 | 0.115 | 1 | 14 | Hddc2 | 0.025 | Young |
| 0.713939 | 0.226038 | 0.62 | 0.601 | 1 | 14 | Ddx6 | 0.019 | Young |
| 0.714132 | 0.068212 | 0.1 | 0.12 | 1 | 14 | Tdrd3 | -0.02 | Young |
| 0.714151 | 0.023429 | 0.3 | 0.262 | 1 | 14 | Tm2d1 | 0.038 | Young |
| 0.714296 | 0.227917 | 0.34 | 0.295 | 1 | 14 | Lmo4 | 0.045 | Young |
| 0.714358 | 0.543317 | 0.24 | 0.224 | 1 | 14 | Dcaf12 | 0.016 | Young |
| 0.714953 | 0.181753 | 0.42 | 0.355 | 1 | 14 | AI314180 | 0.065 | Young |
| 0.715217 | 0.328336 | 0.2 | 0.224 | 1 | 14 | Adamtsl5 | -0.024 | Young |
| 0.715404 | 0.53344 | 0.42 | 0.448 | 1 | 14 | Pgam1 | -0.028 | Young |
| 0.715587 | 0.202712 | 0.16 | 0.142 | 1 | 14 | Bex3 | 0.018 | Young |
| 0.715693 | 0.034592 | 0.18 | 0.197 | 1 | 14 | Sacm1l | -0.017 | Young |
| 0.716588 | 0.373596 | 0.1 | 0.082 | 1 | 14 | Zfp639 | 0.018 | Young |
| 0.716779 | 0.007711 | 0.18 | 0.202 | 1 | 14 | Epdr1 | -0.022 | Young |
| 0.71714 | 0.514228 | 0.28 | 0.301 | 1 | 14 | Osbpl9 | -0.021 | Young |
| 0.717238 | 0.184145 | 0.2 | 0.18 | 1 | 14 | Gramd4 | 0.02 | Young |
| 0.717313 | 0.089892 | 0.1 | 0.12 | 1 | 14 | Pstk | -0.02 | Young |
| 0.717649 | 1.311581 | 0.04 | 0.055 | 1 | 14 | Tnip3 | -0.015 | Young |
| 0.717649 | 1.301626 | 0.06 | 0.049 | 1 | 14 | Tmem186 | 0.011 | Young |
| 0.719251 | 0.373011 | 0.18 | 0.148 | 1 | 14 | Pias2 | 0.032 | Young |
| 0.719475 | 0.649489 | 0.12 | 0.104 | 1 | 14 | Avl9 | 0.016 | Young |
| 0.719511 | 0.755713 | 0.08 | 0.066 | 1 | 14 | Pdgfd | 0.014 | Young |
| 0.719702 | 0.096243 | 0.28 | 0.251 | 1 | 14 | Trappc6b | 0.029 | Young |
| 0.720231 | 0.461295 | 0.1 | 0.082 | 1 | 14 | Mtx3 | 0.018 | Young |
| 0.720498 | 0.747474 | 0.1 | 0.12 | 1 | 14 | Snap29 | -0.02 | Young |

| | | | | | | | | |
|---|---|---|---|---|---|---|---|---|
| 0.721068 | 0.260245 | 0.06 | 0.077 | 1 | 14 | Cep68 | -0.017 | Young |
| 0.721095 | 0.065784 | 0.08 | 0.098 | 1 | 14 | Naglu | -0.018 | Young |
| 0.721288 | 0.34154 | 0.3 | 0.268 | 1 | 14 | Eif2s3x | 0.032 | Young |
| 0.721568 | 0.572579 | 0.44 | 0.437 | 1 | 14 | Ndufb10 | 0.003 | Young |
| 0.721916 | 0.333797 | 0.46 | 0.53 | 1 | 14 | Romo1 | -0.07 | Young |
| 0.722007 | 0.615834 | 0.28 | 0.251 | 1 | 14 | Cds2 | 0.029 | Young |
| 0.722032 | 0.141699 | 0.44 | 0.41 | 1 | 14 | Sec61a1 | 0.03 | Young |
| 0.722274 | 1.180716 | 0.06 | 0.049 | 1 | 14 | Zfp53 | 0.011 | Young |
| 0.722563 | 1.241606 | 0.14 | 0.12 | 1 | 14 | Champ1 | 0.02 | Young |
| 0.722563 | 0.29016 | 0.14 | 0.12 | 1 | 14 | Adcy5 | 0.02 | Young |
| 0.722563 | 0.151602 | 0.14 | 0.12 | 1 | 14 | Iffo1 | 0.02 | Young |
| 0.722858 | 0.00699 | 0.18 | 0.208 | 1 | 14 | Xpo7 | -0.028 | Young |
| 0.723255 | 0.141499 | 0.34 | 0.306 | 1 | 14 | Dctn1 | 0.034 | Young |
| 0.723407 | 0.010826 | 0.24 | 0.202 | 1 | 14 | Eapp | 0.038 | Young |
| 0.723555 | 2.111736 | 0.08 | 0.066 | 1 | 14 | Inpp1 | 0.014 | Young |
| 0.723555 | 0.699905 | 0.08 | 0.066 | 1 | 14 | Fam98c | 0.014 | Young |
| 0.723555 | 0.359712 | 0.08 | 0.066 | 1 | 14 | Ift22 | 0.014 | Young |
| 0.723555 | 0.279104 | 0.08 | 0.066 | 1 | 14 | Opa3 | 0.014 | Young |
| 0.723555 | 0.123673 | 0.08 | 0.066 | 1 | 14 | Ftsj1 | 0.014 | Young |
| 0.723555 | 0.022259 | 0.08 | 0.066 | 1 | 14 | Cd52 | 0.014 | Young |
| 0.723688 | 0.676598 | 0.1 | 0.12 | 1 | 14 | Mafk | -0.02 | Young |
| 0.723795 | 0.823897 | 0.12 | 0.148 | 1 | 14 | Ebp | -0.028 | Young |
| 0.723795 | 0.165888 | 0.16 | 0.137 | 1 | 14 | Map3k1 | 0.023 | Young |
| 0.724085 | 0.070529 | 0.22 | 0.24 | 1 | 14 | Pttg1 | -0.02 | Young |
| 0.72489 | 0.384769 | 0.18 | 0.148 | 1 | 14 | Fktn | 0.032 | Young |
| 0.725231 | 0.171142 | 0.2 | 0.175 | 1 | 14 | Slc25a12 | 0.025 | Young |
| 0.725316 | 0.296679 | 0.16 | 0.18 | 1 | 14 | Rftn1 | -0.02 | Young |
| 0.725986 | 0.191802 | 0.9 | 0.923 | 1 | 14 | Fau | -0.023 | Young |
| 0.726039 | 0.135182 | 0.98 | 0.94 | 1 | 14 | Rps26 | 0.04 | Young |
| 0.726078 | 0.377618 | 0.12 | 0.104 | 1 | 14 | Tmem214 | 0.016 | Young |
| 0.726116 | 1.370135 | 0.4 | 0.437 | 1 | 14 | Fabp4 | -0.037 | Young |
| 0.726295 | 0.696007 | 0.22 | 0.202 | 1 | 14 | Angel2 | 0.018 | Young |
| 0.726909 | 1.134246 | 0.06 | 0.049 | 1 | 14 | Cyb5d2 | 0.011 | Young |
| 0.726909 | 1.075918 | 0.06 | 0.049 | 1 | 14 | Inpp5b | 0.011 | Young |
| 0.726909 | 0.910907 | 0.06 | 0.049 | 1 | 14 | Miip | 0.011 | Young |
| 0.726909 | 0.755821 | 0.06 | 0.049 | 1 | 14 | Iqcb1 | 0.011 | Young |
| 0.726983 | 0.328375 | 0.18 | 0.202 | 1 | 14 | Yeats2 | -0.022 | Young |
| 0.72726 | 0.34864 | 0.44 | 0.404 | 1 | 14 | Stx4a | 0.036 | Young |
| 0.727537 | 0.939875 | 0.1 | 0.082 | 1 | 14 | D6Wsu163 | 0.018 | Young |
| 0.727567 | 0.160749 | 0.34 | 0.361 | 1 | 14 | Tcf12 | -0.021 | Young |
| 0.727657 | 0.533534 | 0.3 | 0.322 | 1 | 14 | Arap1 | -0.022 | Young |
| 0.728009 | 0.299991 | 0.14 | 0.169 | 1 | 14 | Tmem245 | -0.029 | Young |
| 0.728054 | 0.412969 | 0.16 | 0.175 | 1 | 14 | Evc | -0.015 | Young |
| 0.728125 | 0.217681 | 0.66 | 0.721 | 1 | 14 | Sat1 | -0.061 | Young |
| 0.728224 | 0.455457 | 0.42 | 0.388 | 1 | 14 | Cnot6 | 0.032 | Young |

| | | | | | | | | |
|---|---|---|---|---|---|---|---|---|
| 0.728757 | 0.044804 | 0.14 | 0.12 | 1 | 14 | Pld1 | 0.02 | Young |
| 0.72881 | 0.105946 | 0.18 | 0.213 | 1 | 14 | Slit2 | -0.033 | Young |
| 0.728891 | 0.14139 | 0.52 | 0.585 | 1 | 14 | Sdc2 | -0.065 | Young |
| 0.729387 | 1.01412 | 0.12 | 0.104 | 1 | 14 | Pgm1 | 0.016 | Young |
| 0.729446 | 0.319767 | 0.4 | 0.388 | 1 | 14 | Ifrd1 | 0.012 | Young |
| 0.72968 | 0.426053 | 0.28 | 0.246 | 1 | 14 | Azi2 | 0.034 | Young |
| 0.731329 | 0.269558 | 0.22 | 0.202 | 1 | 14 | Ctsf | 0.018 | Young |
| 0.731554 | 0.806933 | 0.06 | 0.049 | 1 | 14 | Rab11fip3 | 0.011 | Young |
| 0.731554 | 0.545244 | 0.06 | 0.049 | 1 | 14 | Wnt2 | 0.011 | Young |
| 0.731637 | 0.167792 | 0.48 | 0.464 | 1 | 14 | Rpl29 | 0.016 | Young |
| 0.731665 | 0.296897 | 0.08 | 0.066 | 1 | 14 | Lrrc40 | 0.014 | Young |
| 0.731665 | 0.148445 | 0.08 | 0.066 | 1 | 14 | Zfp629 | 0.014 | Young |
| 0.731861 | 0.917155 | 0.14 | 0.12 | 1 | 14 | Irgm2 | 0.02 | Young |
| 0.731861 | 0.293299 | 0.14 | 0.12 | 1 | 14 | Pes1 | 0.02 | Young |
| 0.731861 | 0.075623 | 0.14 | 0.12 | 1 | 14 | Pias4 | 0.02 | Young |
| 0.732091 | 0.023032 | 0.18 | 0.153 | 1 | 14 | Entpd1 | 0.027 | Young |
| 0.73217 | 0.46289 | 0.12 | 0.142 | 1 | 14 | Stbd1 | -0.022 | Young |
| 0.732502 | 0.527395 | 0.5 | 0.546 | 1 | 14 | Lpar1 | -0.046 | Young |
| 0.732702 | 0.082724 | 0.12 | 0.104 | 1 | 14 | Itpa | 0.016 | Young |
| 0.732892 | 0.57127 | 0.06 | 0.077 | 1 | 14 | Fcna | -0.017 | Young |
| 0.733339 | 0.383804 | 0.4 | 0.432 | 1 | 14 | Fndc1 | -0.032 | Young |
| 0.733544 | 0.551854 | 0.18 | 0.158 | 1 | 14 | Glrx2 | 0.022 | Young |
| 0.734648 | 1.093157 | 0.26 | 0.24 | 1 | 14 | Sfxn3 | 0.02 | Young |
| 0.734867 | 0.480039 | 0.1 | 0.082 | 1 | 14 | Ift74 | 0.018 | Young |
| 0.735115 | 1.259356 | 0.08 | 0.098 | 1 | 14 | Pon3 | -0.018 | Young |
| 0.735115 | 0.871969 | 0.08 | 0.098 | 1 | 14 | Nfyb | -0.018 | Young |
| 0.735115 | 0.666688 | 0.08 | 0.098 | 1 | 14 | Med4 | -0.018 | Young |
| 0.73536 | 0.349305 | 0.46 | 0.459 | 1 | 14 | Rpl27 | 0.001 | Young |
| 0.735548 | 0.549758 | 0.22 | 0.197 | 1 | 14 | Scamp1 | 0.023 | Young |
| 0.735702 | 0.187761 | 0.28 | 0.295 | 1 | 14 | Prpf8 | -0.015 | Young |
| 0.735731 | 0.681144 | 0.08 | 0.066 | 1 | 14 | Nsun6 | 0.014 | Young |
| 0.735731 | 0.318216 | 0.08 | 0.066 | 1 | 14 | Amer1 | 0.014 | Young |
| 0.735855 | 0.38969 | 0.2 | 0.175 | 1 | 14 | Znrd1 | 0.025 | Young |
| 0.736021 | 0.274969 | 0.12 | 0.104 | 1 | 14 | Nosip | 0.016 | Young |
| 0.736169 | 0.146792 | 0.36 | 0.339 | 1 | 14 | Dcaf8 | 0.021 | Young |
| 0.736174 | 0.003579 | 0.28 | 0.317 | 1 | 14 | Pds5b | -0.037 | Young |
| 0.736209 | 0.148292 | 0.06 | 0.049 | 1 | 14 | Stambpl1 | 0.011 | Young |
| 0.736316 | 0.170846 | 0.18 | 0.158 | 1 | 14 | Ormdl3 | 0.022 | Young |
| 0.736567 | 0.169657 | 0.24 | 0.273 | 1 | 14 | Aak1 | -0.033 | Young |
| 0.737164 | 0.726692 | 0.14 | 0.126 | 1 | 14 | Cdh11 | 0.014 | Young |
| 0.737996 | 0.049511 | 0.36 | 0.388 | 1 | 14 | Selenon | -0.028 | Young |
| 0.738001 | 1.115245 | 0.24 | 0.279 | 1 | 14 | Msi2 | -0.039 | Young |
| 0.738217 | 0.306516 | 0.28 | 0.251 | 1 | 14 | Camk1 | 0.029 | Young |
| 0.738258 | 0.854676 | 0.2 | 0.18 | 1 | 14 | Nolc1 | 0.02 | Young |
| 0.738258 | 0.266591 | 0.2 | 0.18 | 1 | 14 | Casp8ap2 | 0.02 | Young |

| | | | | | | | | |
|---|---|---|---|---|---|---|---|---|
| 0.73852 | 0.13379 | 0.2 | 0.175 | 1 | 14 | Zdhhc21 | 0.025 | Young |
| 0.73852 | 0.125478 | 0.16 | 0.186 | 1 | 14 | Gm15675 | -0.026 | Young |
| 0.738673 | 0.937529 | 0.1 | 0.087 | 1 | 14 | Mapk7 | 0.013 | Young |
| 0.738927 | 0.195363 | 0.26 | 0.235 | 1 | 14 | Smad1 | 0.025 | Young |
| 0.739185 | 0.082903 | 0.24 | 0.219 | 1 | 14 | Churc1 | 0.021 | Young |
| 0.73966 | 0.003483 | 0.5 | 0.443 | 1 | 14 | Nfat5 | 0.057 | Young |
| 0.739805 | 0.975302 | 0.08 | 0.066 | 1 | 14 | Cep57 | 0.014 | Young |
| 0.739805 | 0.388085 | 0.08 | 0.066 | 1 | 14 | Spsb3 | 0.014 | Young |
| 0.739805 | 0.153792 | 0.08 | 0.066 | 1 | 14 | Pdcd2l | 0.014 | Young |
| 0.739805 | 0.087229 | 0.08 | 0.066 | 1 | 14 | Gm11808 | 0.014 | Young |
| 0.740874 | 2.265451 | 0.06 | 0.049 | 1 | 14 | Tmem246 | 0.011 | Young |
| 0.741199 | 0.04659 | 0.14 | 0.12 | 1 | 14 | Jagn1 | 0.02 | Young |
| 0.741428 | 0.515587 | 0.12 | 0.148 | 1 | 14 | Dgkd | -0.028 | Young |
| 0.741613 | 0.468777 | 0.42 | 0.464 | 1 | 14 | Ndufb9 | -0.044 | Young |
| 0.741628 | 0.254354 | 0.16 | 0.175 | 1 | 14 | Ndufs3 | -0.015 | Young |
| 0.741656 | 0.34646 | 0.16 | 0.142 | 1 | 14 | Rapgef2 | 0.018 | Young |
| 0.742157 | 0.592385 | 0.22 | 0.257 | 1 | 14 | Ndufs1 | -0.037 | Young |
| 0.742159 | 1.168524 | 0.08 | 0.098 | 1 | 14 | Ackr1 | -0.018 | Young |
| 0.742221 | 0.42137 | 0.1 | 0.082 | 1 | 14 | Deaf1 | 0.018 | Young |
| 0.742997 | 0.315084 | 0.48 | 0.443 | 1 | 14 | Ywhag | 0.037 | Young |
| 0.743198 | 0.492315 | 0.22 | 0.197 | 1 | 14 | Chmp1a | 0.023 | Young |
| 0.743857 | 0.07968 | 0.2 | 0.175 | 1 | 14 | Rhog | 0.025 | Young |
| 0.743964 | 0.465987 | 0.22 | 0.202 | 1 | 14 | Slc29a3 | 0.018 | Young |
| 0.744111 | 0.227511 | 0.12 | 0.142 | 1 | 14 | Pik3c3 | -0.022 | Young |
| 0.74432 | 0.413954 | 0.14 | 0.12 | 1 | 14 | Rwdd4a | 0.02 | Young |
| 0.74468 | 0.559037 | 0.22 | 0.208 | 1 | 14 | Prpf19 | 0.012 | Young |
| 0.744955 | 0.147076 | 0.18 | 0.202 | 1 | 14 | Clic6 | -0.022 | Young |
| 0.745548 | 0.943998 | 0.06 | 0.049 | 1 | 14 | Pigh | 0.011 | Young |
| 0.745548 | 0.496134 | 0.06 | 0.049 | 1 | 14 | Cstf1 | 0.011 | Young |
| 0.745695 | 0.062331 | 0.66 | 0.672 | 1 | 14 | Lgals1 | -0.012 | Young |
| 0.745816 | 1.073929 | 0.2 | 0.186 | 1 | 14 | Rsbn1 | 0.014 | Young |
| 0.745878 | 0.588812 | 0.1 | 0.087 | 1 | 14 | Maml3 | 0.013 | Young |
| 0.746414 | 0.454971 | 0.26 | 0.284 | 1 | 14 | Mettl7a1 | -0.024 | Young |
| 0.746827 | 0.680763 | 0.16 | 0.18 | 1 | 14 | Zcchc17 | -0.02 | Young |
| 0.747113 | 0.2817 | 0.32 | 0.317 | 1 | 14 | Ndufs5 | 0.003 | Young |
| 0.747337 | 0.565556 | 0.16 | 0.137 | 1 | 14 | Rangap1 | 0.023 | Young |
| 0.747446 | 0.298332 | 0.14 | 0.12 | 1 | 14 | Akr7a5 | 0.02 | Young |
| 0.747491 | 0.334648 | 0.16 | 0.142 | 1 | 14 | Hint2 | 0.018 | Young |
| 0.747876 | 0.221032 | 0.42 | 0.355 | 1 | 14 | Setd7 | 0.065 | Young |
| 0.747975 | 0.957292 | 0.08 | 0.066 | 1 | 14 | D17H6S53I | 0.014 | Young |
| 0.748572 | 0.19568 | 0.5 | 0.454 | 1 | 14 | Acly | 0.046 | Young |
| 0.748844 | 0.010853 | 0.2 | 0.18 | 1 | 14 | Gab2 | 0.02 | Young |
| 0.749173 | 0.097392 | 0.74 | 0.814 | 1 | 14 | Marcks | -0.074 | Young |
| 0.749348 | 0.001255 | 0.12 | 0.104 | 1 | 14 | Cbr2 | 0.016 | Young |
| 0.749489 | 0.478349 | 0.1 | 0.087 | 1 | 14 | Usp12 | 0.013 | Young |

| | | | | | | | | |
|---|---|---|---|---|---|---|---|---|
| 0.749868 | 0.016884 | 0.28 | 0.251 | 1 | 14 | Strn4 | 0.029 | Young |
| 0.750231 | 0.304155 | 0.18 | 0.158 | 1 | 14 | Hoxa5 | 0.022 | Young |
| 0.750232 | 0.548667 | 0.06 | 0.049 | 1 | 14 | Phospho2 | 0.011 | Young |
| 0.750232 | 0.37453 | 0.06 | 0.049 | 1 | 14 | Abtb1 | 0.011 | Young |
| 0.750232 | 0.367455 | 0.06 | 0.049 | 1 | 14 | Layn | 0.011 | Young |
| 0.75046 | 0.55388 | 0.16 | 0.148 | 1 | 14 | Otud5 | 0.012 | Young |
| 0.750468 | 0.327056 | 0.26 | 0.23 | 1 | 14 | Stk3 | 0.03 | Young |
| 0.751072 | 0.551032 | 0.2 | 0.186 | 1 | 14 | Clcn5 | 0.014 | Young |
| 0.751093 | 0.166554 | 0.38 | 0.328 | 1 | 14 | Kdm5a | 0.052 | Young |
| 0.751309 | 0.319979 | 0.26 | 0.301 | 1 | 14 | Rras | -0.041 | Young |
| 0.751426 | 0.675867 | 0.18 | 0.164 | 1 | 14 | Gprasp1 | 0.016 | Young |
| 0.751715 | 0.454033 | 0.54 | 0.514 | 1 | 14 | Eid1 | 0.026 | Young |
| 0.751777 | 0.441891 | 0.14 | 0.164 | 1 | 14 | Asb7 | -0.024 | Young |
| 0.752174 | 0.024102 | 0.24 | 0.213 | 1 | 14 | Ehbp1 | 0.027 | Young |
| 0.752435 | 0.328128 | 0.34 | 0.366 | 1 | 14 | Il6ra | -0.026 | Young |
| 0.752692 | 0.244184 | 0.12 | 0.104 | 1 | 14 | Mospd1 | 0.016 | Young |
| 0.752692 | 0.042339 | 0.12 | 0.104 | 1 | 14 | Coq8b | 0.016 | Young |
| 0.752701 | 0.321937 | 0.18 | 0.202 | 1 | 14 | Ddx1 | -0.022 | Young |
| 0.753105 | 0.336298 | 0.1 | 0.087 | 1 | 14 | Fam107b | 0.013 | Young |
| 0.753112 | 0.036435 | 0.86 | 0.814 | 1 | 14 | Rps25 | 0.046 | Young |
| 0.753299 | 0.059252 | 0.14 | 0.158 | 1 | 14 | Plpbp | -0.018 | Young |
| 0.753709 | 0.070822 | 0.14 | 0.12 | 1 | 14 | Lap3 | 0.02 | Young |
| 0.754645 | 0.205147 | 0.42 | 0.448 | 1 | 14 | Sppl2a | -0.028 | Young |
| 0.754913 | 0.352071 | 0.5 | 0.492 | 1 | 14 | Tnrc18 | 0.008 | Young |
| 0.754924 | 0.611509 | 0.06 | 0.049 | 1 | 14 | Bahcc1 | 0.011 | Young |
| 0.754924 | 0.592679 | 0.06 | 0.049 | 1 | 14 | Sema4a | 0.011 | Young |
| 0.754924 | 0.222366 | 0.06 | 0.049 | 1 | 14 | Mtrf1l | 0.011 | Young |
| 0.755216 | 0.178686 | 0.62 | 0.667 | 1 | 14 | Capzb | -0.047 | Young |
| 0.755679 | 0.126867 | 0.34 | 0.322 | 1 | 14 | Phc3 | 0.018 | Young |
| 0.75582 | 0.270972 | 0.18 | 0.158 | 1 | 14 | Airn | 0.022 | Young |
| 0.755922 | 0.321818 | 0.12 | 0.137 | 1 | 14 | Med30 | -0.017 | Young |
| 0.756114 | 0.667077 | 0.12 | 0.142 | 1 | 14 | Tmem183a | -0.022 | Young |
| 0.756173 | 0.089314 | 0.08 | 0.066 | 1 | 14 | Dnmbp | 0.014 | Young |
| 0.756412 | 0.024392 | 0.2 | 0.23 | 1 | 14 | Ehmt2 | -0.03 | Young |
| 0.756727 | 1.137206 | 0.1 | 0.087 | 1 | 14 | Mtif2 | 0.013 | Young |
| 0.757147 | 0.733524 | 0.24 | 0.213 | 1 | 14 | Zcrb1 | 0.027 | Young |
| 0.75786 | 0.279878 | 0.2 | 0.219 | 1 | 14 | Fbxo3 | -0.019 | Young |
| 0.757967 | 0.478623 | 0.3 | 0.284 | 1 | 14 | Dhx9 | 0.016 | Young |
| 0.758178 | 0.462191 | 0.42 | 0.454 | 1 | 14 | Snhg18 | -0.034 | Young |
| 0.758552 | 0.048688 | 0.5 | 0.514 | 1 | 14 | Sec61g | -0.014 | Young |
| 0.75862 | 0.318792 | 0.18 | 0.158 | 1 | 14 | Elp2 | 0.022 | Young |
| 0.75874 | 0.462414 | 0.12 | 0.131 | 1 | 14 | 0610037L1 | -0.011 | Young |
| 0.759023 | 0.425367 | 0.14 | 0.158 | 1 | 14 | Agps | -0.018 | Young |
| 0.759023 | 0.111261 | 0.14 | 0.158 | 1 | 14 | Gid4 | -0.018 | Young |
| 0.759163 | 0.119585 | 0.98 | 0.978 | 1 | 14 | Rpl41 | 0.002 | Young |

| | | | | | | | | |
|---|---|---|---|---|---|---|---|---|
| 0.759187 | 0.59734 | 0.32 | 0.29 | 1 | 14 | Wdr82 | 0.03 | Young |
| 0.759203 | 0.838899 | 0.16 | 0.142 | 1 | 14 | Usp10 | 0.018 | Young |
| 0.759348 | 0.007924 | 0.38 | 0.421 | 1 | 14 | Ddb1 | -0.041 | Young |
| 0.759626 | 1.562489 | 0.06 | 0.049 | 1 | 14 | Skp2 | 0.011 | Young |
| 0.759626 | 1.184956 | 0.06 | 0.049 | 1 | 14 | Kbtbd4 | 0.011 | Young |
| 0.759626 | 0.998282 | 0.06 | 0.049 | 1 | 14 | Tmem241 | 0.011 | Young |
| 0.759626 | 0.659463 | 0.06 | 0.049 | 1 | 14 | Ccdc88c | 0.011 | Young |
| 0.759626 | 0.277068 | 0.06 | 0.049 | 1 | 14 | Mfsd4a | 0.011 | Young |
| 0.759626 | 0.255749 | 0.06 | 0.049 | 1 | 14 | Hint3 | 0.011 | Young |
| 0.759892 | 0.236235 | 0.48 | 0.552 | 1 | 14 | Steap4 | -0.072 | Young |
| 0.760166 | 0.152742 | 0.24 | 0.257 | 1 | 14 | Col4a3bp | -0.017 | Young |
| 0.761304 | 0.332153 | 0.48 | 0.47 | 1 | 14 | Ran | 0.01 | Young |
| 0.762044 | 0.39849 | 0.16 | 0.148 | 1 | 14 | Xpo4 | 0.012 | Young |
| 0.762142 | 0.570212 | 0.2 | 0.18 | 1 | 14 | Tom1l2 | 0.02 | Young |
| 0.762194 | 0.556626 | 0.24 | 0.23 | 1 | 14 | Acbd5 | 0.01 | Young |
| 0.762521 | 0.110115 | 0.3 | 0.306 | 1 | 14 | Prkca | -0.006 | Young |
| 0.76275 | 0.349646 | 0.12 | 0.104 | 1 | 14 | Isca2 | 0.016 | Young |
| 0.763195 | 0.536646 | 0.26 | 0.24 | 1 | 14 | Cdc5l | 0.02 | Young |
| 0.763338 | 0.289169 | 0.56 | 0.525 | 1 | 14 | Ybx3 | 0.035 | Young |
| 0.763512 | 0.308202 | 0.18 | 0.169 | 1 | 14 | Ahctf1 | 0.011 | Young |
| 0.763557 | 0.553822 | 0.14 | 0.131 | 1 | 14 | Sbf1 | 0.009 | Young |
| 0.764337 | 1.623865 | 0.06 | 0.049 | 1 | 14 | Zfr2 | 0.011 | Young |
| 0.764337 | 0.028559 | 0.06 | 0.049 | 1 | 14 | Gcdh | 0.011 | Young |
| 0.76494 | 0.093967 | 0.14 | 0.126 | 1 | 14 | Rrp7a | 0.014 | Young |
| 0.765081 | 1.006954 | 0.16 | 0.142 | 1 | 14 | Calcrl | 0.018 | Young |
| 0.765896 | 0.026634 | 0.28 | 0.301 | 1 | 14 | Ndufs7 | -0.021 | Young |
| 0.766302 | 0.206987 | 0.32 | 0.366 | 1 | 14 | Alkbh5 | -0.046 | Young |
| 0.766547 | 0.099639 | 0.46 | 0.437 | 1 | 14 | Ndufb11 | 0.023 | Young |
| 0.766732 | 1.017515 | 0.08 | 0.071 | 1 | 14 | Atad2 | 0.009 | Young |
| 0.766823 | 0.0121 | 0.3 | 0.317 | 1 | 14 | Mpc2 | -0.017 | Young |
| 0.766906 | 0.681326 | 0.2 | 0.186 | 1 | 14 | Abhd4 | 0.014 | Young |
| 0.766906 | 0.29075 | 0.2 | 0.186 | 1 | 14 | Daglb | 0.014 | Young |
| 0.766979 | 0.515775 | 0.08 | 0.098 | 1 | 14 | C2cd5 | -0.018 | Young |
| 0.767037 | 0.122991 | 0.18 | 0.158 | 1 | 14 | Hars | 0.022 | Young |
| 0.767619 | 0.28126 | 0.24 | 0.268 | 1 | 14 | Ccdc12 | -0.028 | Young |
| 0.767857 | 0.787646 | 0.16 | 0.148 | 1 | 14 | Clip4 | 0.012 | Young |
| 0.768149 | 0.591461 | 0.12 | 0.137 | 1 | 14 | Timm50 | -0.017 | Young |
| 0.769057 | 0.264018 | 0.06 | 0.049 | 1 | 14 | Zbtb24 | 0.011 | Young |
| 0.769057 | 0.250141 | 0.06 | 0.049 | 1 | 14 | Malt1 | 0.011 | Young |
| 0.769057 | 0.091002 | 0.06 | 0.049 | 1 | 14 | Coq9 | 0.011 | Young |
| 0.769608 | 0.107364 | 0.32 | 0.333 | 1 | 14 | Cuedc1 | -0.013 | Young |
| 0.770128 | 0.134142 | 0.26 | 0.235 | 1 | 14 | Ufc1 | 0.025 | Young |
| 0.770945 | 0.076322 | 0.42 | 0.388 | 1 | 14 | Anxa6 | 0.032 | Young |
| 0.771524 | 0.71278 | 0.3 | 0.29 | 1 | 14 | Setd3 | 0.01 | Young |
| 0.772519 | 0.742624 | 0.42 | 0.448 | 1 | 14 | Ywhah | -0.028 | Young |

| | | | | | | | | |
|---|---|---|---|---|---|---|---|---|
| 0.77265 | 0.854116 | 0.08 | 0.066 | 1 | 14 | Ccdc112 | 0.014 | Young |
| 0.77265 | 0.425452 | 0.08 | 0.066 | 1 | 14 | Sephs1 | 0.014 | Young |
| 0.772848 | 0.136822 | 0.12 | 0.104 | 1 | 14 | Tlcd2 | 0.016 | Young |
| 0.773086 | 0.055452 | 0.48 | 0.448 | 1 | 14 | Atp6v1g1 | 0.032 | Young |
| 0.77309 | 0.369314 | 0.08 | 0.093 | 1 | 14 | Mtmr9 | -0.013 | Young |
| 0.773785 | 0.482511 | 0.06 | 0.049 | 1 | 14 | Inpp5d | 0.011 | Young |
| 0.773785 | 0.223933 | 0.06 | 0.049 | 1 | 14 | Uxt | 0.011 | Young |
| 0.773785 | 0.071926 | 0.06 | 0.049 | 1 | 14 | Gpatch4 | 0.011 | Young |
| 0.773785 | 0.028372 | 0.06 | 0.049 | 1 | 14 | Fancg | 0.011 | Young |
| 0.773922 | 0.704142 | 0.16 | 0.142 | 1 | 14 | Smim20 | 0.018 | Young |
| 0.773991 | 0.234201 | 0.66 | 0.667 | 1 | 14 | Prkar1a | -0.007 | Young |
| 0.774171 | 0.042758 | 0.18 | 0.186 | 1 | 14 | Trim24 | -0.006 | Young |
| 0.774207 | 0.071713 | 0.56 | 0.546 | 1 | 14 | Diaph1 | 0.014 | Young |
| 0.774269 | 0.53286 | 0.12 | 0.131 | 1 | 14 | Rnf8 | -0.011 | Young |
| 0.774526 | 0.373394 | 0.16 | 0.175 | 1 | 14 | Ccng1 | -0.015 | Young |
| 0.774827 | 0.655897 | 0.28 | 0.268 | 1 | 14 | Eif4e | 0.012 | Young |
| 0.774833 | 0.532601 | 0.34 | 0.333 | 1 | 14 | Fxr1 | 0.007 | Young |
| 0.774867 | 0.100772 | 0.28 | 0.301 | 1 | 14 | Slco3a1 | -0.021 | Young |
| 0.774963 | 0.005993 | 0.26 | 0.235 | 1 | 14 | Rheb | 0.025 | Young |
| 0.774974 | 0.016555 | 0.48 | 0.426 | 1 | 14 | Selenom | 0.054 | Young |
| 0.775066 | 0.721222 | 0.36 | 0.388 | 1 | 14 | Pi16 | -0.028 | Young |
| 0.77514 | 0.355155 | 0.14 | 0.153 | 1 | 14 | Ilvbl | -0.013 | Young |
| 0.775187 | 0.17724 | 0.26 | 0.24 | 1 | 14 | Wdfy3 | 0.02 | Young |
| 0.775246 | 0.039397 | 0.48 | 0.497 | 1 | 14 | Samd4b | -0.017 | Young |
| 0.775257 | 0.691786 | 0.32 | 0.355 | 1 | 14 | Cavin2 | -0.035 | Young |
| 0.776037 | 0.084276 | 0.26 | 0.279 | 1 | 14 | Arl6ip5 | -0.019 | Young |
| 0.776741 | 0.015162 | 0.24 | 0.257 | 1 | 14 | Pon2 | -0.017 | Young |
| 0.776991 | 0.380503 | 0.28 | 0.257 | 1 | 14 | Daam1 | 0.023 | Young |
| 0.777249 | 0.133434 | 0.98 | 0.978 | 1 | 14 | Rplp1 | 0.002 | Young |
| 0.777259 | 0.266399 | 0.12 | 0.142 | 1 | 14 | Smad7 | -0.022 | Young |
| 0.778059 | 0.106 | 0.28 | 0.251 | 1 | 14 | Pitpnb | 0.029 | Young |
| 0.778282 | 0.345858 | 0.6 | 0.672 | 1 | 14 | Pam | -0.072 | Young |
| 0.77838 | 1.000253 | 0.04 | 0.033 | 1 | 14 | Tst | 0.007 | Young |
| 0.779209 | 0.035664 | 0.18 | 0.164 | 1 | 14 | Terf2 | 0.016 | Young |
| 0.77952 | 0.609308 | 0.16 | 0.148 | 1 | 14 | Cfap36 | 0.012 | Young |
| 0.781015 | 0.068691 | 0.44 | 0.459 | 1 | 14 | Ndn | -0.019 | Young |
| 0.781085 | 0.092709 | 0.44 | 0.508 | 1 | 14 | Adamts1 | -0.068 | Young |
| 0.781311 | 0.164774 | 0.18 | 0.202 | 1 | 14 | Pex6 | -0.022 | Young |
| 0.781532 | 0.557629 | 0.3 | 0.339 | 1 | 14 | Pcsk5 | -0.039 | Young |
| 0.781737 | 0.042867 | 0.66 | 0.716 | 1 | 14 | Rpl15 | -0.056 | Young |
| 0.782369 | 0.406528 | 0.42 | 0.426 | 1 | 14 | B230219D2 | -0.006 | Young |
| 0.782787 | 0.030966 | 0.28 | 0.251 | 1 | 14 | Adam19 | 0.029 | Young |
| 0.782793 | 0.411505 | 0.16 | 0.142 | 1 | 14 | Mrps12 | 0.018 | Young |
| 0.782817 | 1.139085 | 0.08 | 0.071 | 1 | 14 | Coq10a | 0.009 | Young |
| 0.782837 | 0.354509 | 0.18 | 0.191 | 1 | 14 | 1110008F1 | -0.011 | Young |

| | | | | | | | | |
|---|---|---|---|---|---|---|---|---|
| 0.783918 | 0.183863 | 0.4 | 0.399 | 1 | 14 | Gapvd1 | 0.001 | Young |
| 0.784142 | 1.376639 | 0.04 | 0.033 | 1 | 14 | Supv3l1 | 0.007 | Young |
| 0.784142 | 1.180311 | 0.04 | 0.033 | 1 | 14 | Abcb8 | 0.007 | Young |
| 0.784142 | 0.99371 | 0.04 | 0.033 | 1 | 14 | A430105I1 | 0.007 | Young |
| 0.784142 | 0.928839 | 0.04 | 0.033 | 1 | 14 | Rita1 | 0.007 | Young |
| 0.784142 | 0.828347 | 0.04 | 0.033 | 1 | 14 | Ciita | 0.007 | Young |
| 0.78447 | 0.066169 | 0.22 | 0.24 | 1 | 14 | Cdk8 | -0.02 | Young |
| 0.784596 | 0.050189 | 0.34 | 0.355 | 1 | 14 | Ost4 | -0.015 | Young |
| 0.785077 | 0.098372 | 0.06 | 0.071 | 1 | 14 | Hmgxb3 | -0.011 | Young |
| 0.785141 | 0.353359 | 0.36 | 0.421 | 1 | 14 | Cox7a2 | -0.061 | Young |
| 0.78526 | 0.550846 | 0.24 | 0.273 | 1 | 14 | Chst1 | -0.033 | Young |
| 0.785586 | 0.358946 | 0.16 | 0.175 | 1 | 14 | Ubxn2b | -0.015 | Young |
| 0.785782 | 0.360417 | 0.02 | 0.027 | 1 | 14 | Zfp951 | -0.007 | Young |
| 0.785782 | 0.270814 | 0.02 | 0.027 | 1 | 14 | Zfp276 | -0.007 | Young |
| 0.785782 | 0.228268 | 0.02 | 0.027 | 1 | 14 | Nacad | -0.007 | Young |
| 0.785782 | 0.186825 | 0.02 | 0.027 | 1 | 14 | Ppm1h | -0.007 | Young |
| 0.785782 | 0.144887 | 0.02 | 0.027 | 1 | 14 | Pex1 | -0.007 | Young |
| 0.785782 | 0.071429 | 0.02 | 0.027 | 1 | 14 | 4732440D( | -0.007 | Young |
| 0.785782 | 0.040554 | 0.02 | 0.027 | 1 | 14 | Gm11131 | -0.007 | Young |
| 0.785782 | 0.010023 | 0.02 | 0.027 | 1 | 14 | Gpt2 | -0.007 | Young |
| 0.785888 | 0.304287 | 0.1 | 0.087 | 1 | 14 | H2-DMa | 0.013 | Young |
| 0.785888 | 0.075835 | 0.1 | 0.087 | 1 | 14 | Mfsd7a | 0.013 | Young |
| 0.786047 | 0.358996 | 0.18 | 0.197 | 1 | 14 | Stx5a | -0.017 | Young |
| 0.786259 | 0.244185 | 0.24 | 0.257 | 1 | 14 | Flot2 | -0.017 | Young |
| 0.786312 | 0.485862 | 0.18 | 0.175 | 1 | 14 | Etv3 | 0.005 | Young |
| 0.786373 | 0.729061 | 0.12 | 0.104 | 1 | 14 | Abcg2 | 0.016 | Young |
| 0.786546 | 0.745839 | 0.18 | 0.202 | 1 | 14 | Fkbp3 | -0.022 | Young |
| 0.786635 | 0.634504 | 0.12 | 0.109 | 1 | 14 | Scarf2 | 0.011 | Young |
| 0.786776 | 0.605875 | 0.36 | 0.399 | 1 | 14 | Rnf13 | -0.039 | Young |
| 0.786853 | 0.610041 | 0.08 | 0.071 | 1 | 14 | Slc2a1 | 0.009 | Young |
| 0.786983 | 0.960741 | 0.18 | 0.18 | 1 | 14 | Senp7 | 0 | Young |
| 0.787189 | 0.321752 | 0.24 | 0.213 | 1 | 14 | 9930021J0 | 0.027 | Young |
| 0.787295 | 0.116429 | 0.32 | 0.35 | 1 | 14 | Vdac1 | -0.03 | Young |
| 0.787412 | 0.100257 | 0.38 | 0.399 | 1 | 14 | Tsn | -0.019 | Young |
| 0.787601 | 0.063631 | 0.18 | 0.164 | 1 | 14 | Ppp2r3d | 0.016 | Young |
| 0.788169 | 0.565512 | 0.2 | 0.186 | 1 | 14 | Inhbb | 0.014 | Young |
| 0.788169 | 0.174493 | 0.2 | 0.186 | 1 | 14 | Upf1 | 0.014 | Young |
| 0.788434 | 0.865006 | 0.1 | 0.12 | 1 | 14 | Trmt6 | -0.02 | Young |
| 0.788446 | 1.581622 | 0.1 | 0.093 | 1 | 14 | Gstt3 | 0.007 | Young |
| 0.788446 | 1.319736 | 0.08 | 0.098 | 1 | 14 | Narf | -0.018 | Young |
| 0.788959 | 0.197852 | 0.36 | 0.383 | 1 | 14 | Rere | -0.023 | Young |
| 0.789204 | 0.574693 | 0.04 | 0.049 | 1 | 14 | Msra | -0.009 | Young |
| 0.789204 | 0.427902 | 0.04 | 0.049 | 1 | 14 | Six5 | -0.009 | Young |
| 0.789289 | 0.085099 | 0.8 | 0.814 | 1 | 14 | Tmsb10 | -0.014 | Young |
| 0.789431 | 0.061677 | 0.48 | 0.454 | 1 | 14 | Ugp2 | 0.026 | Young |

| | | | | | | | | |
|---|---|---|---|---|---|---|---|---|
| 0.789764 | 0.240021 | 0.12 | 0.104 | 1 | 14 | Bcl9 | 0.016 | Young |
| 0.789915 | 0.946141 | 0.04 | 0.033 | 1 | 14 | Abcg1 | 0.007 | Young |
| 0.789915 | 0.618814 | 0.04 | 0.033 | 1 | 14 | Trit1 | 0.007 | Young |
| 0.789915 | 0.462192 | 0.04 | 0.033 | 1 | 14 | Wdr34 | 0.007 | Young |
| 0.789968 | 0.796284 | 0.1 | 0.115 | 1 | 14 | Gps2 | -0.015 | Young |
| 0.79036 | 0.088315 | 0.38 | 0.355 | 1 | 14 | Nedd8 | 0.025 | Young |
| 0.790712 | 0.076963 | 0.54 | 0.557 | 1 | 14 | Sf3b2 | -0.017 | Young |
| 0.790732 | 0.896327 | 0.16 | 0.153 | 1 | 14 | Brip1os | 0.007 | Young |
| 0.790935 | 0.278442 | 0.42 | 0.421 | 1 | 14 | Far1 | -0.001 | Young |
| 0.791018 | 0.180791 | 0.44 | 0.437 | 1 | 14 | Arpc3 | 0.003 | Young |
| 0.791122 | 0.089963 | 0.62 | 0.661 | 1 | 14 | Atp5g3 | -0.041 | Young |
| 0.791229 | 0.851817 | 0.14 | 0.131 | 1 | 14 | Wnt10b | 0.009 | Young |
| 0.792424 | 2.309202 | 0.02 | 0.027 | 1 | 14 | Nipal3 | -0.007 | Young |
| 0.792424 | 2.237366 | 0.02 | 0.027 | 1 | 14 | Phf11b | -0.007 | Young |
| 0.792424 | 1.953225 | 0.02 | 0.027 | 1 | 14 | Trmt44 | -0.007 | Young |
| 0.792424 | 1.726428 | 0.02 | 0.027 | 1 | 14 | Iba57 | -0.007 | Young |
| 0.792424 | 1.712147 | 0.02 | 0.027 | 1 | 14 | Slc22a5 | -0.007 | Young |
| 0.792424 | 1.589286 | 0.02 | 0.027 | 1 | 14 | Tcf19 | -0.007 | Young |
| 0.792424 | 1.272864 | 0.02 | 0.027 | 1 | 14 | Hus1 | -0.007 | Young |
| 0.792424 | 1.271 | 0.02 | 0.027 | 1 | 14 | Il10 | -0.007 | Young |
| 0.792424 | 1.218864 | 0.02 | 0.027 | 1 | 14 | Bbs4 | -0.007 | Young |
| 0.792424 | 0.98353 | 0.02 | 0.027 | 1 | 14 | Mettl8 | -0.007 | Young |
| 0.792424 | 0.909524 | 0.02 | 0.027 | 1 | 14 | Slc25a27 | -0.007 | Young |
| 0.792424 | 0.88098 | 0.02 | 0.027 | 1 | 14 | Nr1d1 | -0.007 | Young |
| 0.792424 | 0.845222 | 0.02 | 0.027 | 1 | 14 | Tead2 | -0.007 | Young |
| 0.792424 | 0.823993 | 0.02 | 0.027 | 1 | 14 | Farp2 | -0.007 | Young |
| 0.792424 | 0.82225 | 0.02 | 0.027 | 1 | 14 | Fcgr4 | -0.007 | Young |
| 0.792424 | 0.676553 | 0.02 | 0.027 | 1 | 14 | Gucd1 | -0.007 | Young |
| 0.792424 | 0.596772 | 0.02 | 0.027 | 1 | 14 | Trmt5 | -0.007 | Young |
| 0.792424 | 0.42901 | 0.02 | 0.027 | 1 | 14 | Polr3b | -0.007 | Young |
| 0.792424 | 0.355095 | 0.02 | 0.027 | 1 | 14 | Vmac | -0.007 | Young |
| 0.792424 | 0.301826 | 0.02 | 0.027 | 1 | 14 | Arxes2 | -0.007 | Young |
| 0.792424 | 0.293627 | 0.02 | 0.027 | 1 | 14 | Wdr5b | -0.007 | Young |
| 0.792424 | 0.286115 | 0.02 | 0.027 | 1 | 14 | 2310001H1 | -0.007 | Young |
| 0.792424 | 0.227301 | 0.02 | 0.027 | 1 | 14 | Ndufaf6 | -0.007 | Young |
| 0.793038 | 0.017091 | 0.64 | 0.732 | 1 | 14 | Itgb1 | -0.092 | Young |
| 0.793227 | 1.014897 | 0.1 | 0.087 | 1 | 14 | Lmbrd2 | 0.013 | Young |
| 0.793227 | 0.384236 | 0.1 | 0.087 | 1 | 14 | Nprl3 | 0.013 | Young |
| 0.79351 | 0.241988 | 0.2 | 0.186 | 1 | 14 | Arid4a | 0.014 | Young |
| 0.793633 | 0.38729 | 0.14 | 0.158 | 1 | 14 | Ift80 | -0.018 | Young |
| 0.794168 | 0.651594 | 0.04 | 0.049 | 1 | 14 | Cc2d1a | -0.009 | Young |
| 0.794169 | 0.430145 | 0.16 | 0.148 | 1 | 14 | Maoa | 0.012 | Young |
| 0.794642 | 0.036689 | 0.26 | 0.257 | 1 | 14 | Sema4c | 0.003 | Young |
| 0.79528 | 0.628955 | 0.36 | 0.35 | 1 | 14 | Pmepa1 | 0.01 | Young |
| 0.79531 | 0.805071 | 0.2 | 0.191 | 1 | 14 | Tmem158 | 0.009 | Young |

| | | | | | | | | |
|---|---|---|---|---|---|---|---|---|
| 0.795638 | 0.565122 | 0.08 | 0.098 | 1 | 14 | Foxc1 | -0.018 | Young |
| 0.7957 | 1.610929 | 0.04 | 0.033 | 1 | 14 | Lyz1 | 0.007 | Young |
| 0.7957 | 0.92619 | 0.04 | 0.033 | 1 | 14 | Pcnx4 | 0.007 | Young |
| 0.7957 | 0.586789 | 0.04 | 0.033 | 1 | 14 | Cldn34c1 | 0.007 | Young |
| 0.795931 | 0.075062 | 0.2 | 0.169 | 1 | 14 | Epm2aip1 | 0.031 | Young |
| 0.796018 | 0.306047 | 0.18 | 0.164 | 1 | 14 | Ate1 | 0.016 | Young |
| 0.796018 | 0.294806 | 0.18 | 0.164 | 1 | 14 | Arrdc4 | 0.016 | Young |
| 0.796732 | 0.149502 | 0.34 | 0.366 | 1 | 14 | Ddx50 | -0.026 | Young |
| 0.79682 | 0.041638 | 0.52 | 0.59 | 1 | 14 | Rbpj | -0.07 | Young |
| 0.796903 | 0.73806 | 0.1 | 0.087 | 1 | 14 | Cdr2 | 0.013 | Young |
| 0.796903 | 0.016765 | 0.1 | 0.087 | 1 | 14 | Tmtc2 | 0.013 | Young |
| 0.797221 | 0.211613 | 0.56 | 0.514 | 1 | 14 | Nap1l1 | 0.046 | Young |
| 0.797432 | 0.875108 | 0.36 | 0.366 | 1 | 14 | Tmem258 | -0.006 | Young |
| 0.797558 | 1.481927 | 0.06 | 0.071 | 1 | 14 | Zfp606 | -0.011 | Young |
| 0.797558 | 0.415634 | 0.06 | 0.071 | 1 | 14 | Nedd9 | -0.011 | Young |
| 0.797667 | 1.016533 | 0.24 | 0.224 | 1 | 14 | Gabarapl2 | 0.016 | Young |
| 0.798109 | 0.230551 | 0.28 | 0.268 | 1 | 14 | Krcc1 | 0.012 | Young |
| 0.799301 | 0.683476 | 0.12 | 0.131 | 1 | 14 | Rbfa | -0.011 | Young |
| 0.799301 | 0.070033 | 0.12 | 0.131 | 1 | 14 | Tlnrd1 | -0.011 | Young |
| 0.799398 | 0.281095 | 0.26 | 0.257 | 1 | 14 | Pink1 | 0.003 | Young |
| 0.799472 | 0.128335 | 0.16 | 0.175 | 1 | 14 | Kdm4a | -0.015 | Young |
| 0.800074 | 0.12341 | 0.36 | 0.388 | 1 | 14 | Kdm6b | -0.028 | Young |
| 0.800188 | 0.187479 | 0.7 | 0.831 | 1 | 14 | Igfbp6 | -0.131 | Young |
| 0.800439 | 0.391769 | 0.3 | 0.306 | 1 | 14 | Ndufv3 | -0.006 | Young |
| 0.800616 | 0.308643 | 0.2 | 0.191 | 1 | 14 | Nemf | 0.009 | Young |
| 0.801404 | 0.227432 | 0.22 | 0.23 | 1 | 14 | Ulk2 | -0.01 | Young |
| 0.801487 | 0.148891 | 0.46 | 0.448 | 1 | 14 | Atp5j2 | 0.012 | Young |
| 0.801496 | 1.673356 | 0.04 | 0.033 | 1 | 14 | Mthfd2 | 0.007 | Young |
| 0.801496 | 0.571601 | 0.04 | 0.033 | 1 | 14 | Rad52 | 0.007 | Young |
| 0.801496 | 0.460461 | 0.04 | 0.033 | 1 | 14 | Atp6v1g2 | 0.007 | Young |
| 0.801496 | 0.425699 | 0.04 | 0.033 | 1 | 14 | Hdac8 | 0.007 | Young |
| 0.801496 | 0.370572 | 0.04 | 0.033 | 1 | 14 | Fancl | 0.007 | Young |
| 0.801496 | 0.305188 | 0.04 | 0.033 | 1 | 14 | Ahr | 0.007 | Young |
| 0.801496 | 0.263576 | 0.04 | 0.033 | 1 | 14 | Ptrh1 | 0.007 | Young |
| 0.801496 | 0.232834 | 0.04 | 0.033 | 1 | 14 | Avpi1 | 0.007 | Young |
| 0.801496 | 0.199304 | 0.04 | 0.033 | 1 | 14 | Tmlhe | 0.007 | Young |
| 0.801496 | 0.059648 | 0.04 | 0.033 | 1 | 14 | Lpar6 | 0.007 | Young |
| 0.801639 | 0.464904 | 0.54 | 0.546 | 1 | 14 | Celf1 | -0.006 | Young |
| 0.801642 | 0.454858 | 0.18 | 0.164 | 1 | 14 | Hmg20b | 0.016 | Young |
| 0.801712 | 0.410032 | 0.3 | 0.328 | 1 | 14 | Clcn4 | -0.028 | Young |
| 0.802063 | 0.680671 | 0.12 | 0.137 | 1 | 14 | Nde1 | -0.017 | Young |
| 0.802063 | 0.659757 | 0.12 | 0.137 | 1 | 14 | Tbc1d15 | -0.017 | Young |
| 0.802063 | 0.046181 | 0.12 | 0.137 | 1 | 14 | Alox5ap | -0.017 | Young |
| 0.802446 | 0.017212 | 0.14 | 0.126 | 1 | 14 | Nlgn2 | 0.014 | Young |
| 0.802446 | 0.646582 | 0.2 | 0.18 | 1 | 14 | Cuedc2 | 0.02 | Young |

| | | | | | | | | |
|---|---|---|---|---|---|---|---|---|
| 0.803053 | 0.96371 | 0.08 | 0.071 | 1 | 14 | Mrpl46 | 0.009 | Young |
| 0.803053 | 0.364253 | 0.08 | 0.071 | 1 | 14 | Hecw2 | 0.009 | Young |
| 0.803273 | 1.433542 | 0.2 | 0.191 | 1 | 14 | Anapc16 | 0.009 | Young |
| 0.803333 | 0.411687 | 0.18 | 0.18 | 1 | 14 | Dmtf1 | 0 | Young |
| 0.803337 | 0.300061 | 0.32 | 0.311 | 1 | 14 | Atp5g1 | 0.009 | Young |
| 0.803369 | 0.186368 | 0.12 | 0.104 | 1 | 14 | Mpp6 | 0.016 | Young |
| 0.803597 | 0.287046 | 0.3 | 0.279 | 1 | 14 | Flii | 0.021 | Young |
| 0.803599 | 0.129264 | 0.16 | 0.142 | 1 | 14 | Lpcat3 | 0.018 | Young |
| 0.803809 | 0.091274 | 0.32 | 0.301 | 1 | 14 | Etnk1 | 0.019 | Young |
| 0.80412 | 0.072447 | 0.04 | 0.049 | 1 | 14 | Lonrf2 | -0.009 | Young |
| 0.804871 | 0.685738 | 0.1 | 0.12 | 1 | 14 | Skiv2l2 | -0.02 | Young |
| 0.805248 | 0.457805 | 0.22 | 0.197 | 1 | 14 | Znfx1 | 0.023 | Young |
| 0.805593 | 0.189072 | 0.14 | 0.126 | 1 | 14 | Mageh1 | 0.014 | Young |
| 0.805593 | 0.105684 | 0.14 | 0.126 | 1 | 14 | Emc4 | 0.014 | Young |
| 0.805908 | 0.11457 | 0.06 | 0.071 | 1 | 14 | Cryz | -0.011 | Young |
| 0.806003 | 0.075284 | 0.32 | 0.361 | 1 | 14 | Fam46a | -0.041 | Young |
| 0.806691 | 1.001701 | 0.12 | 0.109 | 1 | 14 | Elf4 | 0.011 | Young |
| 0.806691 | 0.031298 | 0.1 | 0.115 | 1 | 14 | Usp46 | -0.015 | Young |
| 0.80678 | 0.131117 | 0.12 | 0.104 | 1 | 14 | Hotairm1 | 0.016 | Young |
| 0.807117 | 0.057513 | 0.08 | 0.071 | 1 | 14 | Spin4 | 0.009 | Young |
| 0.807262 | 0.229288 | 0.18 | 0.197 | 1 | 14 | Ankhd1 | -0.017 | Young |
| 0.807277 | 0.843534 | 0.18 | 0.164 | 1 | 14 | 2210016F1 | 0.016 | Young |
| 0.807303 | 1.483485 | 0.04 | 0.033 | 1 | 14 | Tmem17 | 0.007 | Young |
| 0.807303 | 1.258353 | 0.04 | 0.033 | 1 | 14 | Bcar3 | 0.007 | Young |
| 0.807303 | 0.735686 | 0.04 | 0.033 | 1 | 14 | Txnl4b | 0.007 | Young |
| 0.807303 | 0.572539 | 0.04 | 0.033 | 1 | 14 | Enkd1 | 0.007 | Young |
| 0.807303 | 0.570358 | 0.04 | 0.033 | 1 | 14 | Ankrd52 | 0.007 | Young |
| 0.807303 | 0.372157 | 0.04 | 0.033 | 1 | 14 | Glmn | 0.007 | Young |
| 0.807303 | 0.313238 | 0.04 | 0.033 | 1 | 14 | Zbtb26 | 0.007 | Young |
| 0.807303 | 0.25614 | 0.04 | 0.033 | 1 | 14 | Porcn | 0.007 | Young |
| 0.807862 | 0.657357 | 0.2 | 0.18 | 1 | 14 | Lztr1 | 0.02 | Young |
| 0.807863 | 0.171705 | 0.22 | 0.197 | 1 | 14 | Hdac4 | 0.023 | Young |
| 0.807875 | 0.025832 | 0.7 | 0.776 | 1 | 14 | Pbx1 | -0.076 | Young |
| 0.807959 | 0.12928 | 0.1 | 0.087 | 1 | 14 | Dis3 | 0.013 | Young |
| 0.808253 | 0.093114 | 0.92 | 0.951 | 1 | 14 | Rps21 | -0.031 | Young |
| 0.808338 | 0.284764 | 0.34 | 0.317 | 1 | 14 | Chtop | 0.023 | Young |
| 0.808487 | 0.031298 | 0.1 | 0.109 | 1 | 14 | Dzip3 | -0.009 | Young |
| 0.808743 | 0.774149 | 0.14 | 0.126 | 1 | 14 | Smarcad1 | 0.014 | Young |
| 0.808748 | 0.202735 | 0.5 | 0.481 | 1 | 14 | Smim10l1 | 0.019 | Young |
| 0.809109 | 0.811116 | 0.04 | 0.049 | 1 | 14 | Zfp628 | -0.009 | Young |
| 0.809109 | 0.215217 | 0.04 | 0.049 | 1 | 14 | Arntl2 | -0.009 | Young |
| 0.809239 | 0.440951 | 0.34 | 0.339 | 1 | 14 | Cotl1 | 0.001 | Young |
| 0.809454 | 0.150305 | 0.52 | 0.536 | 1 | 14 | Tardbp | -0.016 | Young |
| 0.809945 | 0.036627 | 0.66 | 0.623 | 1 | 14 | S100a13 | 0.037 | Young |
| 0.810041 | 0.255259 | 0.28 | 0.257 | 1 | 14 | Cxxc5 | 0.023 | Young |

| | | | | | | | | |
|---|---|---|---|---|---|---|---|---|
| 0.810091 | 0.538506 | 0.06 | 0.071 | 1 | 14 | Bhlhe41 | -0.011 | Young |
| 0.810091 | 0.366743 | 0.06 | 0.071 | 1 | 14 | Selenoo | -0.011 | Young |
| 0.811186 | 0.828208 | 0.08 | 0.071 | 1 | 14 | Egflam | 0.009 | Young |
| 0.811319 | 0.14877 | 0.24 | 0.262 | 1 | 14 | Dcun1d3 | -0.022 | Young |
| 0.811844 | 0.09401 | 0.16 | 0.148 | 1 | 14 | Inpp4a | 0.012 | Young |
| 0.811897 | 0.086053 | 0.14 | 0.126 | 1 | 14 | Slc9a1 | 0.014 | Young |
| 0.811903 | 0.116026 | 0.1 | 0.109 | 1 | 14 | Mrps11 | -0.009 | Young |
| 0.812279 | 0.629183 | 0.2 | 0.186 | 1 | 14 | 6030408B1 | 0.014 | Young |
| 0.812279 | 0.254692 | 0.2 | 0.186 | 1 | 14 | Rab28 | 0.014 | Young |
| 0.812369 | 0.159006 | 0.18 | 0.158 | 1 | 14 | Arih2 | 0.022 | Young |
| 0.813121 | 0.252565 | 0.04 | 0.033 | 1 | 14 | L3mbtl2 | 0.007 | Young |
| 0.813121 | 0.211893 | 0.04 | 0.033 | 1 | 14 | Pcdhgb5 | 0.007 | Young |
| 0.813121 | 0.16853 | 0.04 | 0.033 | 1 | 14 | Acad11 | 0.007 | Young |
| 0.813121 | 0.149875 | 0.04 | 0.033 | 1 | 14 | Chn2 | 0.007 | Young |
| 0.813288 | 0.09005 | 0.2 | 0.18 | 1 | 14 | Nnmt | 0.02 | Young |
| 0.813405 | 0.839631 | 0.12 | 0.109 | 1 | 14 | Zfp354c | 0.011 | Young |
| 0.813405 | 0.258462 | 0.12 | 0.109 | 1 | 14 | Zfp948 | 0.011 | Young |
| 0.813405 | 0.225794 | 0.12 | 0.109 | 1 | 14 | Btd | 0.011 | Young |
| 0.813422 | 0.662912 | 0.16 | 0.175 | 1 | 14 | Spg21 | -0.015 | Young |
| 0.813512 | 0.655025 | 0.56 | 0.568 | 1 | 14 | Rap1a | -0.008 | Young |
| 0.813898 | 0.322662 | 0.42 | 0.415 | 1 | 14 | Pitpna | 0.005 | Young |
| 0.813932 | 0.38696 | 0.24 | 0.23 | 1 | 14 | Tmem184b | 0.01 | Young |
| 0.814105 | 0.328844 | 0.04 | 0.049 | 1 | 14 | Dok1 | -0.009 | Young |
| 0.814105 | 0.172191 | 0.04 | 0.049 | 1 | 14 | Haus8 | -0.009 | Young |
| 0.814658 | 0.516728 | 0.72 | 0.798 | 1 | 14 | Col1a1 | -0.078 | Young |
| 0.815323 | 0.100247 | 0.1 | 0.109 | 1 | 14 | Mrps10 | -0.009 | Young |
| 0.815506 | 0.244104 | 0.18 | 0.202 | 1 | 14 | Snf8 | -0.022 | Young |
| 0.815928 | 0.324904 | 0.36 | 0.361 | 1 | 14 | Nupr1 | -0.001 | Young |
| 0.816377 | 0.110854 | 0.9 | 0.874 | 1 | 14 | Rpl9 | 0.026 | Young |
| 0.81659 | 0.685675 | 0.2 | 0.191 | 1 | 14 | Hacd3 | 0.009 | Young |
| 0.817202 | 0.245335 | 0.08 | 0.093 | 1 | 14 | Exosc3 | -0.013 | Young |
| 0.817236 | 0.095834 | 0.34 | 0.333 | 1 | 14 | Atp1a1 | 0.007 | Young |
| 0.817317 | 0.915128 | 0.1 | 0.093 | 1 | 14 | Zer1 | 0.007 | Young |
| 0.817994 | 0.124463 | 0.36 | 0.399 | 1 | 14 | Fto | -0.039 | Young |
| 0.818068 | 0.04292 | 0.38 | 0.355 | 1 | 14 | Smc3 | 0.025 | Young |
| 0.818634 | 0.333107 | 0.5 | 0.481 | 1 | 14 | Sdha | 0.019 | Young |
| 0.818875 | 0.65223 | 0.24 | 0.219 | 1 | 14 | Yap1 | 0.021 | Young |
| 0.818949 | 0.051655 | 0.04 | 0.033 | 1 | 14 | Lgals4 | 0.007 | Young |
| 0.819108 | 1.220123 | 0.04 | 0.049 | 1 | 14 | Rcc1 | -0.009 | Young |
| 0.819108 | 0.547358 | 0.04 | 0.049 | 1 | 14 | Zfp623 | -0.009 | Young |
| 0.819339 | 1.689789 | 0.08 | 0.071 | 1 | 14 | Tsku | 0.009 | Young |
| 0.819339 | 0.941883 | 0.08 | 0.071 | 1 | 14 | Ncs1 | 0.009 | Young |
| 0.819339 | 0.391983 | 0.08 | 0.071 | 1 | 14 | Rapgef3 | 0.009 | Young |
| 0.819739 | 1.721799 | 0.12 | 0.115 | 1 | 14 | Lrrc49 | 0.005 | Young |
| 0.821113 | 0.083477 | 0.2 | 0.208 | 1 | 14 | Ccdc186 | -0.008 | Young |

| | | | | | | | | |
|---|---|---|---|---|---|---|---|---|
| 0.821376 | 0.679354 | 0.12 | 0.131 | 1 | 14 | Thap11 | -0.011 | Young |
| 0.821872 | 0.049459 | 0.7 | 0.672 | 1 | 14 | Selenow | 0.028 | Young |
| 0.822517 | 0.218339 | 0.2 | 0.213 | 1 | 14 | B3gat3 | -0.013 | Young |
| 0.822673 | 0.148973 | 0.06 | 0.071 | 1 | 14 | Alkbh7 | -0.011 | Young |
| 0.823271 | 0.28232 | 0.5 | 0.492 | 1 | 14 | Tram1 | 0.008 | Young |
| 0.823311 | 0.300614 | 0.62 | 0.639 | 1 | 14 | Mtdh | -0.019 | Young |
| 0.823569 | 0.019683 | 0.4 | 0.361 | 1 | 14 | Creb3l2 | 0.039 | Young |
| 0.823887 | 0.406374 | 0.22 | 0.235 | 1 | 14 | Hdgfl3 | -0.015 | Young |
| 0.824119 | 1.603464 | 0.04 | 0.049 | 1 | 14 | Nptx2 | -0.009 | Young |
| 0.824576 | 1.909044 | 0.1 | 0.093 | 1 | 14 | Brcc3 | 0.007 | Young |
| 0.824721 | 0.044641 | 0.28 | 0.301 | 1 | 14 | Atp13a3 | -0.021 | Young |
| 0.824728 | 0.214841 | 0.38 | 0.377 | 1 | 14 | Igf2r | 0.003 | Young |
| 0.825313 | 0.47277 | 0.28 | 0.262 | 1 | 14 | Atxn10 | 0.018 | Young |
| 0.825603 | 0.679852 | 0.1 | 0.109 | 1 | 14 | Txnl4a | -0.009 | Young |
| 0.825657 | 0.246755 | 0.16 | 0.169 | 1 | 14 | Metap1 | -0.009 | Young |
| 0.825682 | 0.252265 | 0.26 | 0.257 | 1 | 14 | Armc8 | 0.003 | Young |
| 0.825704 | 1.260584 | 0.06 | 0.055 | 1 | 14 | Telo2 | 0.005 | Young |
| 0.826471 | 0.314366 | 0.1 | 0.087 | 1 | 14 | Gpr132 | 0.013 | Young |
| 0.826834 | 0.283759 | 0.46 | 0.437 | 1 | 14 | Ube2h | 0.023 | Young |
| 0.826875 | 0.096151 | 0.12 | 0.109 | 1 | 14 | Snta1 | 0.011 | Young |
| 0.826878 | 1.272558 | 0.06 | 0.071 | 1 | 14 | Atxn3 | -0.011 | Young |
| 0.826878 | 0.296222 | 0.06 | 0.071 | 1 | 14 | Srprb | -0.011 | Young |
| 0.826957 | 0.154013 | 0.58 | 0.563 | 1 | 14 | Dag1 | 0.017 | Young |
| 0.826969 | 0.354555 | 0.12 | 0.137 | 1 | 14 | Cdk17 | -0.017 | Young |
| 0.827075 | 0.279493 | 0.18 | 0.164 | 1 | 14 | Fasn | 0.016 | Young |
| 0.827409 | 0.327948 | 0.44 | 0.464 | 1 | 14 | Pbxip1 | -0.024 | Young |
| 0.827511 | 0.938762 | 0.08 | 0.071 | 1 | 14 | Zfp60 | 0.009 | Young |
| 0.827736 | 0.128623 | 0.2 | 0.224 | 1 | 14 | Sfrp4 | -0.024 | Young |
| 0.828494 | 0.024636 | 0.16 | 0.169 | 1 | 14 | Lmbr1 | -0.009 | Young |
| 0.828506 | 0.067017 | 0.96 | 0.995 | 1 | 14 | Tmsb4x | -0.035 | Young |
| 0.828987 | 0.368279 | 0.22 | 0.208 | 1 | 14 | Tshz3 | 0.012 | Young |
| 0.828988 | 0.439842 | 0.24 | 0.219 | 1 | 14 | Stip1 | 0.021 | Young |
| 0.829037 | 0.131295 | 0.1 | 0.109 | 1 | 14 | Snhg11 | -0.009 | Young |
| 0.82904 | 0.233737 | 0.48 | 0.448 | 1 | 14 | Sh3pxd2b | 0.032 | Young |
| 0.829137 | 0.603902 | 0.04 | 0.049 | 1 | 14 | Armt1 | -0.009 | Young |
| 0.829615 | 0.001048 | 0.16 | 0.148 | 1 | 14 | Pgp | 0.012 | Young |
| 0.830342 | 1.018984 | 0.06 | 0.055 | 1 | 14 | Tbccd1 | 0.005 | Young |
| 0.830554 | 0.105109 | 0.16 | 0.142 | 1 | 14 | Npepl1 | 0.018 | Young |
| 0.830847 | 0.412988 | 0.26 | 0.246 | 1 | 14 | Hadh | 0.014 | Young |
| 0.831005 | 0.011894 | 0.26 | 0.24 | 1 | 14 | Rsbn1l | 0.02 | Young |
| 0.831205 | 0.02584 | 0.9 | 0.907 | 1 | 14 | Rps16 | -0.007 | Young |
| 0.831605 | 0.401296 | 0.08 | 0.071 | 1 | 14 | Trim11 | 0.009 | Young |
| 0.831605 | 0.154752 | 0.08 | 0.071 | 1 | 14 | Aacs | 0.009 | Young |
| 0.831849 | 1.703579 | 0.1 | 0.093 | 1 | 14 | Smarcd3 | 0.007 | Young |
| 0.832473 | 0.433875 | 0.1 | 0.109 | 1 | 14 | Sdhaf4 | -0.009 | Young |

| | | | | | | | | |
|---|---|---|---|---|---|---|---|---|
| 0.832473 | 0.030526 | 0.1 | 0.109 | 1 | 14 | Slc8b1 | -0.009 | Young |
| 0.833455 | 0.058877 | 0.44 | 0.41 | 1 | 14 | Nop10 | 0.03 | Young |
| 0.833629 | 0.498081 | 0.12 | 0.109 | 1 | 14 | Setd1a | 0.011 | Young |
| 0.833629 | 0.041181 | 0.12 | 0.109 | 1 | 14 | Foxp4 | 0.011 | Young |
| 0.834072 | 0.393716 | 0.14 | 0.153 | 1 | 14 | Thap2 | -0.013 | Young |
| 0.8347 | 0.746389 | 0.16 | 0.18 | 1 | 14 | Coa3 | -0.02 | Young |
| 0.834789 | 0.079517 | 0.14 | 0.131 | 1 | 14 | Dpy30 | 0.009 | Young |
| 0.835224 | 0.33989 | 0.5 | 0.552 | 1 | 14 | Vcan | -0.052 | Young |
| 0.835301 | 0.526516 | 0.06 | 0.071 | 1 | 14 | Zfp942 | -0.011 | Young |
| 0.835558 | 1.547454 | 0.16 | 0.148 | 1 | 14 | Zfp24 | 0.012 | Young |
| 0.835578 | 0.071389 | 0.42 | 0.443 | 1 | 14 | Ehd1 | -0.023 | Young |
| 0.835718 | 0.054634 | 0.68 | 0.716 | 1 | 14 | Gpx1 | -0.036 | Young |
| 0.835818 | 0.575382 | 0.46 | 0.464 | 1 | 14 | Tpst1 | -0.004 | Young |
| 0.836093 | 0.895225 | 0.18 | 0.175 | 1 | 14 | Safb2 | 0.005 | Young |
| 0.836766 | 0.331132 | 0.22 | 0.208 | 1 | 14 | Psmd14 | 0.012 | Young |
| 0.836767 | 0.165735 | 0.22 | 0.197 | 1 | 14 | Cdk13 | 0.023 | Young |
| 0.83701 | 0.079045 | 0.12 | 0.109 | 1 | 14 | Dnajb12 | 0.011 | Young |
| 0.837016 | 0.42754 | 0.16 | 0.169 | 1 | 14 | Sft2d2 | -0.009 | Young |
| 0.837415 | 0.139547 | 0.46 | 0.415 | 1 | 14 | Bzw1 | 0.045 | Young |
| 0.837445 | 0.210608 | 0.36 | 0.383 | 1 | 14 | Arhgap20 | -0.023 | Young |
| 0.837638 | 0.668592 | 0.3 | 0.35 | 1 | 14 | G0s2 | -0.05 | Young |
| 0.837931 | 0.737478 | 0.26 | 0.251 | 1 | 14 | Aftph | 0.009 | Young |
| 0.838339 | 0.267758 | 0.28 | 0.284 | 1 | 14 | Sec13 | -0.004 | Young |
| 0.838534 | 1.726575 | 0.16 | 0.148 | 1 | 14 | Hadha | 0.012 | Young |
| 0.838644 | 0.069405 | 0.28 | 0.257 | 1 | 14 | Gusb | 0.023 | Young |
| 0.839356 | 0.912004 | 0.1 | 0.109 | 1 | 14 | Ell | -0.009 | Young |
| 0.839487 | 0.33853 | 0.08 | 0.093 | 1 | 14 | Ercc3 | -0.013 | Young |
| 0.839582 | 0.008707 | 0.54 | 0.552 | 1 | 14 | Ddah2 | -0.012 | Young |
| 0.839636 | 1.041611 | 0.06 | 0.055 | 1 | 14 | Peg3 | 0.005 | Young |
| 0.839862 | 0.855268 | 0.16 | 0.169 | 1 | 14 | Itpr2 | -0.009 | Young |
| 0.840092 | 0.186324 | 0.72 | 0.749 | 1 | 14 | Btf3 | -0.029 | Young |
| 0.840412 | 0.910646 | 0.14 | 0.126 | 1 | 14 | Lancl1 | 0.014 | Young |
| 0.840516 | 0.085246 | 0.94 | 0.973 | 1 | 14 | Fth1 | -0.033 | Young |
| 0.840692 | 0.436115 | 0.2 | 0.191 | 1 | 14 | Ptprj | 0.009 | Young |
| 0.840975 | 0.167993 | 0.42 | 0.393 | 1 | 14 | Pdia4 | 0.027 | Young |
| 0.841091 | 0.385063 | 0.5 | 0.492 | 1 | 14 | Eif2s2 | 0.008 | Young |
| 0.841542 | 0.208303 | 0.32 | 0.35 | 1 | 14 | Usf2 | -0.03 | Young |
| 0.84189 | 0.092751 | 0.3 | 0.295 | 1 | 14 | Psma5 | 0.005 | Young |
| 0.841945 | 0.135917 | 0.28 | 0.262 | 1 | 14 | Sys1 | 0.018 | Young |
| 0.842016 | 0.297495 | 0.22 | 0.202 | 1 | 14 | Rnf2 | 0.018 | Young |
| 0.842047 | 0.194977 | 0.22 | 0.197 | 1 | 14 | Pin4 | 0.023 | Young |
| 0.842803 | 0.939858 | 0.1 | 0.109 | 1 | 14 | Lrrcc1 | -0.009 | Young |
| 0.842803 | 0.816055 | 0.1 | 0.109 | 1 | 14 | Cercam | -0.009 | Young |
| 0.842995 | 2.137525 | 0.12 | 0.115 | 1 | 14 | Sdf2l1 | 0.005 | Young |
| 0.843165 | 0.016639 | 0.52 | 0.601 | 1 | 14 | Uqcr11 | -0.081 | Young |

| | | | | | | | | |
|---|---|---|---|---|---|---|---|---|
| 0.843593 | 0.725101 | 0.12 | 0.131 | 1 | 14 | Blcap | -0.011 | Young |
| 0.843593 | 0.165464 | 0.12 | 0.131 | 1 | 14 | Zfp46 | -0.011 | Young |
| 0.844491 | 0.253857 | 0.16 | 0.148 | 1 | 14 | Rarg | 0.012 | Young |
| 0.845884 | 0.491643 | 0.44 | 0.47 | 1 | 14 | Capn2 | -0.03 | Young |
| 0.846012 | 0.215246 | 0.2 | 0.202 | 1 | 14 | Dgcr2 | -0.002 | Young |
| 0.846536 | 0.118094 | 0.46 | 0.475 | 1 | 14 | Vgll3 | -0.015 | Young |
| 0.847135 | 1.261945 | 0.18 | 0.169 | 1 | 14 | Phf8 | 0.011 | Young |
| 0.847173 | 0.240597 | 0.12 | 0.109 | 1 | 14 | Cep164 | 0.011 | Young |
| 0.847173 | 0.109789 | 0.12 | 0.109 | 1 | 14 | Snhg20 | 0.011 | Young |
| 0.847387 | 1.045224 | 0.2 | 0.197 | 1 | 14 | Eml1 | 0.003 | Young |
| 0.847391 | 0.504493 | 0.18 | 0.186 | 1 | 14 | Mpp1 | -0.006 | Young |
| 0.847928 | 0.236144 | 0.3 | 0.306 | 1 | 14 | Agfg1 | -0.006 | Young |
| 0.847972 | 0.723849 | 0.06 | 0.071 | 1 | 14 | Fbxl12 | -0.011 | Young |
| 0.848022 | 0.148891 | 0.08 | 0.071 | 1 | 14 | Bmf | 0.009 | Young |
| 0.849246 | 0.441415 | 0.14 | 0.158 | 1 | 14 | Exosc10 | -0.018 | Young |
| 0.849679 | 0.085298 | 0.16 | 0.164 | 1 | 14 | Rev1 | -0.004 | Young |
| 0.849704 | 0.197372 | 0.1 | 0.109 | 1 | 14 | Fam160a2 | -0.009 | Young |
| 0.850095 | 0.385072 | 0.1 | 0.093 | 1 | 14 | E2f4 | 0.007 | Young |
| 0.850567 | 0.214009 | 0.1 | 0.115 | 1 | 14 | Mfsd6 | -0.015 | Young |
| 0.850834 | 0.147295 | 0.42 | 0.399 | 1 | 14 | Ankrd11 | 0.021 | Young |
| 0.851263 | 0.120675 | 0.16 | 0.169 | 1 | 14 | Med12 | -0.009 | Young |
| 0.851269 | 0.066521 | 0.78 | 0.842 | 1 | 14 | Srsf5 | -0.062 | Young |
| 0.851343 | 0.222749 | 0.32 | 0.301 | 1 | 14 | Eif3d | 0.019 | Young |
| 0.851843 | 0.320272 | 0.24 | 0.219 | 1 | 14 | Zhx3 | 0.021 | Young |
| 0.852138 | 0.657998 | 0.08 | 0.071 | 1 | 14 | Tslp | 0.009 | Young |
| 0.852229 | 0.147819 | 0.32 | 0.29 | 1 | 14 | Tor1b | 0.03 | Young |
| 0.85286 | 0.160396 | 0.44 | 0.421 | 1 | 14 | Brk1 | 0.019 | Young |
| 0.853497 | 0.176939 | 0.36 | 0.372 | 1 | 14 | Tubb4b | -0.012 | Young |
| 0.853618 | 1.706068 | 0.06 | 0.055 | 1 | 14 | Rbak | 0.005 | Young |
| 0.853622 | 0.92521 | 0.14 | 0.131 | 1 | 14 | Ccdc8 | 0.009 | Young |
| 0.853909 | 2.842447 | 0.02 | 0.016 | 1 | 14 | Ccdc189 | 0.004 | Young |
| 0.853909 | 2.402753 | 0.02 | 0.016 | 1 | 14 | Htr2a | 0.004 | Young |
| 0.853909 | 2.208354 | 0.02 | 0.016 | 1 | 14 | Ccdc68 | 0.004 | Young |
| 0.853909 | 2.16517 | 0.02 | 0.016 | 1 | 14 | Zfp52 | 0.004 | Young |
| 0.853909 | 2.143335 | 0.02 | 0.016 | 1 | 14 | Tmem132a | 0.004 | Young |
| 0.853909 | 2.121092 | 0.02 | 0.016 | 1 | 14 | Slc4a11 | 0.004 | Young |
| 0.853909 | 2.098416 | 0.02 | 0.016 | 1 | 14 | Arrdc2 | 0.004 | Young |
| 0.853909 | 2.090655 | 0.02 | 0.016 | 1 | 14 | Syce2 | 0.004 | Young |
| 0.853909 | 1.749624 | 0.02 | 0.016 | 1 | 14 | Ptger3 | 0.004 | Young |
| 0.853909 | 1.67085 | 0.02 | 0.016 | 1 | 14 | Cyp26b1 | 0.004 | Young |
| 0.853909 | 1.638639 | 0.02 | 0.016 | 1 | 14 | Gm26624 | 0.004 | Young |
| 0.853909 | 1.569615 | 0.02 | 0.016 | 1 | 14 | Efhc1 | 0.004 | Young |
| 0.853909 | 1.459013 | 0.02 | 0.016 | 1 | 14 | Hs3st3b1 | 0.004 | Young |
| 0.853909 | 1.403097 | 0.02 | 0.016 | 1 | 14 | B930036N1 | 0.004 | Young |
| 0.853909 | 1.382494 | 0.02 | 0.016 | 1 | 14 | Jmjd7 | 0.004 | Young |

| | | | | | | | | |
|---|---|---|---|---|---|---|---|---|
| 0.853909 | 1.375909 | 0.02 | 0.016 | 1 | 14 | Mmp16 | 0.004 | Young |
| 0.853909 | 1.365566 | 0.02 | 0.016 | 1 | 14 | Syn1 | 0.004 | Young |
| 0.853909 | 1.34233 | 0.02 | 0.016 | 1 | 14 | Lhfpl4 | 0.004 | Young |
| 0.853909 | 1.292046 | 0.02 | 0.016 | 1 | 14 | Il2rg | 0.004 | Young |
| 0.853909 | 1.273982 | 0.02 | 0.016 | 1 | 14 | Lrfn4 | 0.004 | Young |
| 0.853909 | 1.203175 | 0.02 | 0.016 | 1 | 14 | Ankrd42 | 0.004 | Young |
| 0.853909 | 1.186717 | 0.02 | 0.016 | 1 | 14 | Mrm2 | 0.004 | Young |
| 0.853909 | 1.117736 | 0.02 | 0.016 | 1 | 14 | Tmem216 | 0.004 | Young |
| 0.853909 | 1.045979 | 0.02 | 0.016 | 1 | 14 | Pawr | 0.004 | Young |
| 0.853909 | 1.044413 | 0.02 | 0.016 | 1 | 14 | Fig4 | 0.004 | Young |
| 0.853909 | 0.969269 | 0.02 | 0.016 | 1 | 14 | Tmem35b | 0.004 | Young |
| 0.853909 | 0.909131 | 0.02 | 0.016 | 1 | 14 | Enkur | 0.004 | Young |
| 0.853909 | 0.887971 | 0.02 | 0.016 | 1 | 14 | C030034I2 | 0.004 | Young |
| 0.853909 | 0.858919 | 0.02 | 0.016 | 1 | 14 | Pcdhb14 | 0.004 | Young |
| 0.853909 | 0.802948 | 0.02 | 0.016 | 1 | 14 | 4430402I1 | 0.004 | Young |
| 0.853909 | 0.757251 | 0.02 | 0.016 | 1 | 14 | Ppm1l | 0.004 | Young |
| 0.853909 | 0.740907 | 0.02 | 0.016 | 1 | 14 | Pnmal2 | 0.004 | Young |
| 0.853909 | 0.728983 | 0.02 | 0.016 | 1 | 14 | Whrn | 0.004 | Young |
| 0.853909 | 0.714353 | 0.02 | 0.016 | 1 | 14 | Gldc | 0.004 | Young |
| 0.853909 | 0.703154 | 0.02 | 0.016 | 1 | 14 | Tchh | 0.004 | Young |
| 0.853909 | 0.623119 | 0.02 | 0.016 | 1 | 14 | Bco2 | 0.004 | Young |
| 0.853909 | 0.618671 | 0.02 | 0.016 | 1 | 14 | Srfbp1 | 0.004 | Young |
| 0.853909 | 0.588989 | 0.02 | 0.016 | 1 | 14 | Ccdc17 | 0.004 | Young |
| 0.853909 | 0.587098 | 0.02 | 0.016 | 1 | 14 | 6720489N1 | 0.004 | Young |
| 0.853909 | 0.409698 | 0.02 | 0.016 | 1 | 14 | Ppp1r13l | 0.004 | Young |
| 0.853917 | 0.830048 | 0.24 | 0.23 | 1 | 14 | Atad1 | 0.01 | Young |
| 0.853963 | 1.182035 | 0.1 | 0.115 | 1 | 14 | Rassf3 | -0.015 | Young |
| 0.854329 | 0.787593 | 0.24 | 0.246 | 1 | 14 | Pa2g4 | -0.006 | Young |
| 0.854423 | 0.016272 | 0.94 | 0.973 | 1 | 14 | Rps23 | -0.033 | Young |
| 0.854542 | 0.070489 | 0.4 | 0.426 | 1 | 14 | Nfkb1 | -0.026 | Young |
| 0.855547 | 0.16929 | 0.18 | 0.164 | 1 | 14 | Diablo | 0.016 | Young |
| 0.856023 | 0.189916 | 0.46 | 0.448 | 1 | 14 | Mapre2 | 0.012 | Young |
| 0.856257 | 0.207916 | 0.08 | 0.071 | 1 | 14 | Kti12 | 0.009 | Young |
| 0.856258 | 0.111781 | 0.28 | 0.262 | 1 | 14 | Ddhd2 | 0.018 | Young |
| 0.85677 | 0.142997 | 0.14 | 0.131 | 1 | 14 | Afdn | 0.009 | Young |
| 0.857196 | 0.338385 | 0.38 | 0.372 | 1 | 14 | Vdac3 | 0.008 | Young |
| 0.857266 | 0.386725 | 0.42 | 0.415 | 1 | 14 | Nono | 0.005 | Young |
| 0.857416 | 0.065998 | 0.1 | 0.093 | 1 | 14 | Galk1 | 0.007 | Young |
| 0.85829 | 0.78757 | 0.06 | 0.055 | 1 | 14 | Tmem53 | 0.005 | Young |
| 0.85829 | 0.561372 | 0.06 | 0.055 | 1 | 14 | Gm10600 | 0.005 | Young |
| 0.858457 | 1.742751 | 0.2 | 0.224 | 1 | 14 | Myh10 | -0.024 | Young |
| 0.859207 | 0.320301 | 0.42 | 0.399 | 1 | 14 | Tcf7l2 | 0.021 | Young |
| 0.859415 | 0.371651 | 0.26 | 0.257 | 1 | 14 | Txndc9 | 0.003 | Young |
| 0.859424 | 0.613855 | 0.16 | 0.148 | 1 | 14 | Slc41a1 | 0.012 | Young |
| 0.859438 | 0.391124 | 0.28 | 0.268 | 1 | 14 | Wwp1 | 0.012 | Young |

| | | | | | | | | |
|---|---|---|---|---|---|---|---|---|
| 0.85956 | 0.307427 | 0.34 | 0.317 | 1 | 14 | Ppp2r5a | 0.023 | Young |
| 0.860381 | 0.42235 | 0.44 | 0.41 | 1 | 14 | Clint1 | 0.03 | Young |
| 0.861081 | 0.256951 | 0.1 | 0.093 | 1 | 14 | Nop14 | 0.007 | Young |
| 0.86148 | 0.133293 | 0.42 | 0.448 | 1 | 14 | Elf1 | -0.028 | Young |
| 0.862171 | 1.027417 | 0.02 | 0.016 | 1 | 14 | Zfat | 0.004 | Young |
| 0.862171 | 0.761545 | 0.02 | 0.016 | 1 | 14 | Top1mt | 0.004 | Young |
| 0.862171 | 0.737311 | 0.02 | 0.016 | 1 | 14 | Lepr | 0.004 | Young |
| 0.862171 | 0.729074 | 0.02 | 0.016 | 1 | 14 | 4930425PC | 0.004 | Young |
| 0.862171 | 0.709547 | 0.02 | 0.016 | 1 | 14 | 4930590J0 | 0.004 | Young |
| 0.862171 | 0.530515 | 0.02 | 0.016 | 1 | 14 | Rabl2 | 0.004 | Young |
| 0.862171 | 0.455351 | 0.02 | 0.016 | 1 | 14 | Ifitm5 | 0.004 | Young |
| 0.862171 | 0.409009 | 0.02 | 0.016 | 1 | 14 | Gng4 | 0.004 | Young |
| 0.862171 | 0.308277 | 0.02 | 0.016 | 1 | 14 | Cd3eap | 0.004 | Young |
| 0.862171 | 0.307437 | 0.02 | 0.016 | 1 | 14 | Npl | 0.004 | Young |
| 0.862171 | 0.295118 | 0.02 | 0.016 | 1 | 14 | Tmem8b | 0.004 | Young |
| 0.862171 | 0.249777 | 0.02 | 0.016 | 1 | 14 | Nle1 | 0.004 | Young |
| 0.862171 | 0.201767 | 0.02 | 0.016 | 1 | 14 | Plk4 | 0.004 | Young |
| 0.862171 | 0.162406 | 0.02 | 0.016 | 1 | 14 | Zfp14 | 0.004 | Young |
| 0.862171 | 0.139891 | 0.02 | 0.016 | 1 | 14 | Zfp7 | 0.004 | Young |
| 0.862171 | 0.023522 | 0.02 | 0.016 | 1 | 14 | Slc16a11 | 0.004 | Young |
| 0.86224 | 0.187383 | 0.5 | 0.497 | 1 | 14 | Selenot | 0.003 | Young |
| 0.862856 | 0.016291 | 0.42 | 0.437 | 1 | 14 | Vps35 | -0.017 | Young |
| 0.862966 | 1.463775 | 0.06 | 0.055 | 1 | 14 | Zfp280c | 0.005 | Young |
| 0.863468 | 0.387478 | 0.36 | 0.339 | 1 | 14 | Cyb5r3 | 0.021 | Young |
| 0.863541 | 0.257073 | 0.1 | 0.109 | 1 | 14 | Zfp746 | -0.009 | Young |
| 0.863687 | 0.04937 | 0.88 | 0.929 | 1 | 14 | Plpp3 | -0.049 | Young |
| 0.863867 | 0.075761 | 0.18 | 0.186 | 1 | 14 | Jmjd6 | -0.006 | Young |
| 0.863885 | 0.172786 | 0.4 | 0.443 | 1 | 14 | Atp5d | -0.043 | Young |
| 0.86397 | 0.834376 | 0.18 | 0.18 | 1 | 14 | Ndufaf4 | 0 | Young |
| 0.864102 | 0.286765 | 0.16 | 0.175 | 1 | 14 | Emc8 | -0.015 | Young |
| 0.864167 | 0.124125 | 0.12 | 0.109 | 1 | 14 | Rusc2 | 0.011 | Young |
| 0.864356 | 0.369911 | 0.44 | 0.475 | 1 | 14 | Psmb1 | -0.035 | Young |
| 0.864452 | 0.397875 | 0.58 | 0.623 | 1 | 14 | Dbi | -0.043 | Young |
| 0.86475 | 1.317556 | 0.1 | 0.093 | 1 | 14 | Asxl1 | 0.007 | Young |
| 0.86475 | 0.43082 | 0.1 | 0.093 | 1 | 14 | Ipo11 | 0.007 | Young |
| 0.86475 | 0.143188 | 0.1 | 0.093 | 1 | 14 | Bpnt1 | 0.007 | Young |
| 0.865443 | 0.198087 | 0.8 | 0.891 | 1 | 14 | Col3a1 | -0.091 | Young |
| 0.865651 | 0.534144 | 0.32 | 0.306 | 1 | 14 | Eif3m | 0.014 | Young |
| 0.865936 | 0.430446 | 0.12 | 0.131 | 1 | 14 | Ccdc25 | -0.011 | Young |
| 0.865936 | 0.068461 | 0.14 | 0.126 | 1 | 14 | Tspan9 | 0.014 | Young |
| 0.866388 | 0.24291 | 0.12 | 0.115 | 1 | 14 | Enpp3 | 0.005 | Young |
| 0.866391 | 0.026417 | 0.38 | 0.366 | 1 | 14 | Zfp395 | 0.014 | Young |
| 0.866599 | 0.238607 | 0.44 | 0.443 | 1 | 14 | Rpl13a | -0.003 | Young |
| 0.866766 | 0.013055 | 0.6 | 0.596 | 1 | 14 | Ndufa2 | 0.004 | Young |
| 0.866802 | 0.315521 | 0.14 | 0.148 | 1 | 14 | Atp1b1 | -0.008 | Young |

| | | | | | | | | |
|---|---|---|---|---|---|---|---|---|
| 0.866802 | 0.129887 | 0.14 | 0.148 | 1 | 14 | Chfr | -0.008 | Young |
| 0.866909 | 0.524677 | 0.14 | 0.153 | 1 | 14 | Crnkl1 | -0.013 | Young |
| 0.867093 | 0.287458 | 0.26 | 0.251 | 1 | 14 | Nrd1 | 0.009 | Young |
| 0.867222 | 1.097167 | 0.2 | 0.202 | 1 | 14 | Lats2 | -0.002 | Young |
| 0.867529 | 0.225798 | 0.46 | 0.454 | 1 | 14 | Rbm25 | 0.006 | Young |
| 0.867647 | 0.445599 | 0.06 | 0.055 | 1 | 14 | Oma1 | 0.005 | Young |
| 0.867647 | 0.225127 | 0.06 | 0.055 | 1 | 14 | Pcdhb17 | 0.005 | Young |
| 0.868378 | 0.173991 | 0.98 | 0.984 | 1 | 14 | Rplp2 | -0.004 | Young |
| 0.869381 | 0.242399 | 0.14 | 0.131 | 1 | 14 | 4931406CC | 0.009 | Young |
| 0.869381 | 0.121079 | 0.14 | 0.131 | 1 | 14 | Ttc37 | 0.009 | Young |
| 0.869662 | 1.407755 | 0.18 | 0.175 | 1 | 14 | Ypel5 | 0.005 | Young |
| 0.869941 | 0.181838 | 0.16 | 0.153 | 1 | 14 | Cks1b | 0.007 | Young |
| 0.87024 | 0.005047 | 0.96 | 0.967 | 1 | 14 | Rps12 | -0.007 | Young |
| 0.870331 | 1.313058 | 0.08 | 0.087 | 1 | 14 | Man2b2 | -0.007 | Young |
| 0.870331 | 0.018426 | 0.08 | 0.087 | 1 | 14 | Kdm1b | -0.007 | Young |
| 0.87035 | 0.01385 | 0.24 | 0.208 | 1 | 14 | Txndc15 | 0.032 | Young |
| 0.870448 | 0.30823 | 0.02 | 0.016 | 1 | 14 | Chadl | 0.004 | Young |
| 0.870448 | 0.233952 | 0.02 | 0.016 | 1 | 14 | Wnt4 | 0.004 | Young |
| 0.870448 | 0.153267 | 0.02 | 0.016 | 1 | 14 | Trp53i11 | 0.004 | Young |
| 0.870448 | 0.132753 | 0.02 | 0.016 | 1 | 14 | Zbtb12 | 0.004 | Young |
| 0.870448 | 0.080993 | 0.02 | 0.016 | 1 | 14 | Garem1 | 0.004 | Young |
| 0.870448 | 0.073878 | 0.02 | 0.016 | 1 | 14 | Cnbd2 | 0.004 | Young |
| 0.870448 | 0.030548 | 0.02 | 0.016 | 1 | 14 | Igsf6 | 0.004 | Young |
| 0.870464 | 0.306769 | 0.12 | 0.126 | 1 | 14 | Npr2 | -0.006 | Young |
| 0.870748 | 0.668115 | 0.3 | 0.306 | 1 | 14 | F3 | -0.006 | Young |
| 0.870938 | 0.134477 | 0.22 | 0.202 | 1 | 14 | Hoxc9 | 0.018 | Young |
| 0.870983 | 0.223188 | 0.12 | 0.109 | 1 | 14 | Agk | 0.011 | Young |
| 0.870985 | 0.134004 | 0.54 | 0.59 | 1 | 14 | Pdcd6ip | -0.05 | Young |
| 0.871077 | 0.230233 | 0.26 | 0.262 | 1 | 14 | Cited2 | -0.002 | Young |
| 0.871469 | 0.369857 | 0.44 | 0.421 | 1 | 14 | Baz1b | 0.019 | Young |
| 0.872095 | 0.790209 | 0.1 | 0.093 | 1 | 14 | Itgam | 0.007 | Young |
| 0.872332 | 1.698317 | 0.06 | 0.055 | 1 | 14 | Pdk2 | 0.005 | Young |
| 0.872332 | 1.406455 | 0.06 | 0.055 | 1 | 14 | Dchs1 | 0.005 | Young |
| 0.872332 | 1.112955 | 0.06 | 0.055 | 1 | 14 | Wdr47 | 0.005 | Young |
| 0.872332 | 0.308097 | 0.06 | 0.055 | 1 | 14 | Pdzd11 | 0.005 | Young |
| 0.872517 | 0.24758 | 0.4 | 0.415 | 1 | 14 | Ap2m1 | -0.015 | Young |
| 0.872537 | 0.005173 | 0.52 | 0.552 | 1 | 14 | Actn4 | -0.032 | Young |
| 0.872539 | 0.068624 | 0.14 | 0.131 | 1 | 14 | Slc35c2 | 0.009 | Young |
| 0.872604 | 0.029983 | 0.6 | 0.689 | 1 | 14 | Vcp | -0.089 | Young |
| 0.872609 | 0.430137 | 0.18 | 0.169 | 1 | 14 | Thumpd1 | 0.011 | Young |
| 0.872609 | 0.003999 | 0.18 | 0.169 | 1 | 14 | Ap1m1 | 0.011 | Young |
| 0.872812 | 0.318302 | 0.16 | 0.164 | 1 | 14 | Mphosph6 | -0.004 | Young |
| 0.872872 | 0.212945 | 0.14 | 0.148 | 1 | 14 | Dkc1 | -0.008 | Young |
| 0.873268 | 0.333898 | 0.22 | 0.208 | 1 | 14 | Cycs | 0.012 | Young |
| 0.873577 | 0.326405 | 0.6 | 0.639 | 1 | 14 | Eif4a1 | -0.039 | Young |

| | | | | | | | | |
|---|---|---|---|---|---|---|---|---|
| 0.874395 | 0.646201 | 0.12 | 0.109 | 1 | 14 | Dcaf17 | 0.011 | Young |
| 0.874695 | 0.012393 | 0.28 | 0.257 | 1 | 14 | Mff | 0.023 | Young |
| 0.875086 | 0.881734 | 0.12 | 0.12 | 1 | 14 | Lrrc47 | 0 | Young |
| 0.875201 | 1.263014 | 0.2 | 0.202 | 1 | 14 | Sestd1 | -0.002 | Young |
| 0.875201 | 0.141369 | 0.32 | 0.333 | 1 | 14 | Id2 | -0.013 | Young |
| 0.8757 | 0.148914 | 0.14 | 0.131 | 1 | 14 | G6pdx | 0.009 | Young |
| 0.8757 | 0.106312 | 0.14 | 0.131 | 1 | 14 | Cops7a | 0.009 | Young |
| 0.875765 | 0.03301 | 0.2 | 0.191 | 1 | 14 | Sar1b | 0.009 | Young |
| 0.876046 | 0.251047 | 0.34 | 0.35 | 1 | 14 | Appl1 | -0.01 | Young |
| 0.876057 | 1.092217 | 0.08 | 0.077 | 1 | 14 | Ten1 | 0.003 | Young |
| 0.876197 | 0.467513 | 0.36 | 0.333 | 1 | 14 | Glrx3 | 0.027 | Young |
| 0.87631 | 0.775068 | 0.36 | 0.361 | 1 | 14 | Cox7b | -0.001 | Young |
| 0.876456 | 0.046583 | 0.3 | 0.317 | 1 | 14 | Casp12 | -0.017 | Young |
| 0.877022 | 0.215968 | 0.06 | 0.055 | 1 | 14 | Mettl17 | 0.005 | Young |
| 0.877303 | 0.227522 | 0.34 | 0.339 | 1 | 14 | Cox20 | 0.001 | Young |
| 0.877409 | 0.684421 | 0.16 | 0.148 | 1 | 14 | Pcif1 | 0.012 | Young |
| 0.877409 | 0.452323 | 0.16 | 0.148 | 1 | 14 | Rnpc3 | 0.012 | Young |
| 0.877735 | 0.162056 | 0.46 | 0.497 | 1 | 14 | Gstm2 | -0.037 | Young |
| 0.878003 | 0.318551 | 0.4 | 0.437 | 1 | 14 | Midn | -0.037 | Young |
| 0.878614 | 0.214215 | 0.16 | 0.164 | 1 | 14 | Uchl1 | -0.004 | Young |
| 0.879743 | 0.003361 | 0.54 | 0.557 | 1 | 14 | Atp5f1 | -0.017 | Young |
| 0.879779 | 0.055894 | 0.34 | 0.317 | 1 | 14 | Ifi27 | 0.023 | Young |
| 0.880402 | 0.307255 | 0.18 | 0.186 | 1 | 14 | Nsmce3 | -0.006 | Young |
| 0.880444 | 0.828506 | 0.12 | 0.137 | 1 | 14 | Fmnl2 | -0.017 | Young |
| 0.88105 | 1.335077 | 0.08 | 0.071 | 1 | 14 | Shb | 0.009 | Young |
| 0.881494 | 0.287597 | 0.24 | 0.246 | 1 | 14 | Eif4e2 | -0.006 | Young |
| 0.881717 | 0.725567 | 0.06 | 0.055 | 1 | 14 | Pold2 | 0.005 | Young |
| 0.881809 | 0.674196 | 0.14 | 0.158 | 1 | 14 | Mepce | -0.018 | Young |
| 0.881853 | 0.275332 | 0.08 | 0.087 | 1 | 14 | Vps39 | -0.007 | Young |
| 0.882051 | 0.344868 | 0.14 | 0.137 | 1 | 14 | Rbm14 | 0.003 | Young |
| 0.88258 | 0.233907 | 0.64 | 0.71 | 1 | 14 | Col5a1 | -0.07 | Young |
| 0.882625 | 0.370006 | 0.58 | 0.579 | 1 | 14 | Nbl1 | 0.001 | Young |
| 0.882859 | 0.494287 | 0.2 | 0.186 | 1 | 14 | Osbp | 0.014 | Young |
| 0.883134 | 0.120314 | 0.1 | 0.093 | 1 | 14 | Fam45a | 0.007 | Young |
| 0.883216 | 0.373484 | 0.24 | 0.24 | 1 | 14 | Rabep1 | 0 | Young |
| 0.883751 | 0.06842 | 0.22 | 0.208 | 1 | 14 | Ppl | 0.012 | Young |
| 0.883751 | 0.035789 | 0.22 | 0.208 | 1 | 14 | Cab39l | 0.012 | Young |
| 0.883903 | 0.085424 | 0.8 | 0.825 | 1 | 14 | Errfi1 | -0.025 | Young |
| 0.884114 | 0.035425 | 0.62 | 0.634 | 1 | 14 | Col5a2 | -0.014 | Young |
| 0.884125 | 1.302077 | 0.08 | 0.077 | 1 | 14 | Tmem14a | 0.003 | Young |
| 0.885063 | 0.431923 | 0.2 | 0.213 | 1 | 14 | Fscn1 | -0.013 | Young |
| 0.885192 | 0.35746 | 0.14 | 0.131 | 1 | 14 | Thoc3 | 0.009 | Young |
| 0.8857 | 0.450453 | 0.08 | 0.087 | 1 | 14 | Mcm4 | -0.007 | Young |
| 0.88595 | 0.1758 | 0.22 | 0.213 | 1 | 14 | Plekhb2 | 0.007 | Young |
| 0.886415 | 0.672018 | 0.06 | 0.055 | 1 | 14 | Macrod2 | 0.005 | Young |

| | | | | | | | | |
|---|---|---|---|---|---|---|---|---|
| 0.887854 | 1.084373 | 0.1 | 0.109 | 1 | 14 | Rnaseh2c | -0.009 | Young |
| 0.888137 | 0.372451 | 0.32 | 0.328 | 1 | 14 | Esd | -0.008 | Young |
| 0.888441 | 0.247177 | 0.24 | 0.224 | 1 | 14 | Sptlc2 | 0.016 | Young |
| 0.888692 | 0.06592 | 0.38 | 0.393 | 1 | 14 | Tmem65 | -0.013 | Young |
| 0.890004 | 0.412292 | 0.12 | 0.126 | 1 | 14 | Hist1h1c | -0.006 | Young |
| 0.890505 | 0.266355 | 0.1 | 0.093 | 1 | 14 | 5730480H( | 0.007 | Young |
| 0.890939 | 0.524552 | 0.14 | 0.153 | 1 | 14 | Phactr1 | -0.013 | Young |
| 0.89099 | 0.33215 | 0.24 | 0.224 | 1 | 14 | Bet1 | 0.016 | Young |
| 0.891118 | 0.278228 | 0.06 | 0.055 | 1 | 14 | Larp1b | 0.005 | Young |
| 0.891118 | 0.008954 | 0.06 | 0.055 | 1 | 14 | Bhlhb9 | 0.005 | Young |
| 0.891486 | 0.919777 | 0.12 | 0.109 | 1 | 14 | Mrpl27 | 0.011 | Young |
| 0.892153 | 0.312521 | 0.46 | 0.481 | 1 | 14 | Pdgfrb | -0.021 | Young |
| 0.892205 | 0.625971 | 0.08 | 0.077 | 1 | 14 | Brf1 | 0.003 | Young |
| 0.892366 | 0.051386 | 0.8 | 0.831 | 1 | 14 | Rpl32 | -0.031 | Young |
| 0.892445 | 0.068338 | 0.16 | 0.148 | 1 | 14 | Eva1b | 0.012 | Young |
| 0.892842 | 0.153825 | 0.28 | 0.268 | 1 | 14 | Fam168a | 0.012 | Young |
| 0.893266 | 0.230446 | 0.12 | 0.115 | 1 | 14 | Itgb3 | 0.005 | Young |
| 0.893694 | 0.375128 | 0.58 | 0.596 | 1 | 14 | St13 | -0.016 | Young |
| 0.893749 | 0.418957 | 0.06 | 0.066 | 1 | 14 | Slc6a9 | -0.006 | Young |
| 0.893749 | 0.168013 | 0.06 | 0.066 | 1 | 14 | Ppp2r1b | -0.006 | Young |
| 0.893749 | 0.010907 | 0.06 | 0.066 | 1 | 14 | 6030458C1 | -0.006 | Young |
| 0.894177 | 0.151859 | 0.14 | 0.148 | 1 | 14 | Dnaja3 | -0.008 | Young |
| 0.894195 | 0.221042 | 0.1 | 0.093 | 1 | 14 | Gfra2 | 0.007 | Young |
| 0.89434 | 0.101375 | 0.34 | 0.344 | 1 | 14 | Etfb | -0.004 | Young |
| 0.894554 | 0.52698 | 0.14 | 0.137 | 1 | 14 | Vamp7 | 0.003 | Young |
| 0.894821 | 0.213459 | 0.1 | 0.109 | 1 | 14 | Wfdc17 | -0.009 | Young |
| 0.894947 | 0.180627 | 0.5 | 0.519 | 1 | 14 | Mob1a | -0.019 | Young |
| 0.895568 | 0.661812 | 0.2 | 0.213 | 1 | 14 | Klhl13 | -0.013 | Young |
| 0.895817 | 0.5452 | 0.4 | 0.383 | 1 | 14 | Ndufa7 | 0.017 | Young |
| 0.895824 | 0.100565 | 0.06 | 0.055 | 1 | 14 | Ints11 | 0.005 | Young |
| 0.896249 | 0.456694 | 0.08 | 0.077 | 1 | 14 | Zfp319 | 0.003 | Young |
| 0.896376 | 0.501764 | 0.22 | 0.213 | 1 | 14 | Cbx7 | 0.007 | Young |
| 0.896376 | 0.220656 | 0.22 | 0.213 | 1 | 14 | Trim41 | 0.007 | Young |
| 0.896455 | 0.155189 | 0.4 | 0.393 | 1 | 14 | Crk | 0.007 | Young |
| 0.89668 | 0.177822 | 0.16 | 0.153 | 1 | 14 | Virma | 0.007 | Young |
| 0.896815 | 0.375804 | 0.24 | 0.23 | 1 | 14 | Tbc1d20 | 0.01 | Young |
| 0.896989 | 0.826209 | 0.18 | 0.186 | 1 | 14 | Sdk1 | -0.006 | Young |
| 0.897028 | 0.05044 | 0.26 | 0.246 | 1 | 14 | Thy1 | 0.014 | Young |
| 0.89741 | 0.194635 | 0.18 | 0.18 | 1 | 14 | Ralgapa1 | 0 | Young |
| 0.897475 | 0.149183 | 0.2 | 0.191 | 1 | 14 | Ammecr1 | 0.009 | Young |
| 0.897874 | 0.019316 | 0.14 | 0.131 | 1 | 14 | Morc2a | 0.009 | Young |
| 0.897887 | 0.173582 | 0.1 | 0.093 | 1 | 14 | Uckl1 | 0.007 | Young |
| 0.898152 | 0.225978 | 0.06 | 0.066 | 1 | 14 | Med9 | -0.006 | Young |
| 0.898152 | 0.126702 | 0.06 | 0.066 | 1 | 14 | Pgap1 | -0.006 | Young |
| 0.898152 | 0.043858 | 0.06 | 0.066 | 1 | 14 | Plcl2 | -0.006 | Young |

| | | | | | | | | |
|---|---|---|---|---|---|---|---|---|
| 0.898359 | 0.405995 | 0.2 | 0.197 | 1 | 14 | Ogfrl1 | 0.003 | Young |
| 0.898985 | 0.118462 | 0.22 | 0.213 | 1 | 14 | Csk | 0.007 | Young |
| 0.899301 | 0.609496 | 0.18 | 0.191 | 1 | 14 | Tcerg1 | -0.011 | Young |
| 0.899892 | 0.155987 | 0.36 | 0.355 | 1 | 14 | Polr2m | 0.005 | Young |
| 0.900534 | 0.194197 | 0.06 | 0.055 | 1 | 14 | Lclat1 | 0.005 | Young |
| 0.900719 | 0.420242 | 0.26 | 0.257 | 1 | 14 | Akap11 | 0.003 | Young |
| 0.901702 | 0.041265 | 0.34 | 0.339 | 1 | 14 | Tmbim1 | 0.001 | Young |
| 0.901885 | 0.317136 | 0.2 | 0.202 | 1 | 14 | Desi2 | -0.002 | Young |
| 0.902046 | 0.263205 | 0.18 | 0.191 | 1 | 14 | Ak1 | -0.011 | Young |
| 0.902228 | 0.407647 | 0.38 | 0.388 | 1 | 14 | Kpnb1 | -0.008 | Young |
| 0.902557 | 0.626625 | 0.06 | 0.066 | 1 | 14 | Hdhd2 | -0.006 | Young |
| 0.902623 | 0.314594 | 0.38 | 0.415 | 1 | 14 | Hnrnpr | -0.035 | Young |
| 0.902827 | 0.27913 | 0.38 | 0.399 | 1 | 14 | Kdelc2 | -0.019 | Young |
| 0.903086 | 0.163139 | 0.46 | 0.475 | 1 | 14 | Ctsh | -0.015 | Young |
| 0.903467 | 0.300089 | 0.18 | 0.175 | 1 | 14 | Uri1 | 0.005 | Young |
| 0.903632 | 0.476929 | 0.22 | 0.219 | 1 | 14 | BC003331 | 0.001 | Young |
| 0.90427 | 0.013995 | 0.36 | 0.355 | 1 | 14 | Sf3b5 | 0.005 | Young |
| 0.904346 | 0.452852 | 0.08 | 0.077 | 1 | 14 | Cog1 | 0.003 | Young |
| 0.904788 | 0.257464 | 0.7 | 0.732 | 1 | 14 | Son | -0.032 | Young |
| 0.905025 | 0.667712 | 0.22 | 0.23 | 1 | 14 | Cux1 | -0.01 | Young |
| 0.905025 | 0.23874 | 0.22 | 0.23 | 1 | 14 | Tmem87b | -0.01 | Young |
| 0.905025 | 0.006272 | 0.24 | 0.257 | 1 | 14 | Psma1 | -0.017 | Young |
| 0.905248 | 1.546231 | 0.06 | 0.055 | 1 | 14 | Zfp317 | 0.005 | Young |
| 0.905277 | 0.027618 | 0.1 | 0.093 | 1 | 14 | Ddx41 | 0.007 | Young |
| 0.905287 | 0.015978 | 0.1 | 0.109 | 1 | 14 | Ppard | -0.009 | Young |
| 0.905598 | 0.388822 | 0.24 | 0.262 | 1 | 14 | Dnm1 | -0.022 | Young |
| 0.905627 | 0.092982 | 0.22 | 0.224 | 1 | 14 | Fem1b | -0.004 | Young |
| 0.905627 | 0.049352 | 0.22 | 0.224 | 1 | 14 | Mknk2 | -0.004 | Young |
| 0.906083 | 0.211312 | 0.04 | 0.044 | 1 | 14 | Tmem132c | -0.004 | Young |
| 0.906083 | 0.08419 | 0.04 | 0.044 | 1 | 14 | Utp23 | -0.004 | Young |
| 0.906083 | 0.009854 | 0.04 | 0.044 | 1 | 14 | Haus4 | -0.004 | Young |
| 0.906957 | 0.317602 | 0.22 | 0.235 | 1 | 14 | Tmem33 | -0.015 | Young |
| 0.906966 | 0.322504 | 0.06 | 0.066 | 1 | 14 | E2f1 | -0.006 | Young |
| 0.907539 | 1.131289 | 0.18 | 0.191 | 1 | 14 | Klhdc3 | -0.011 | Young |
| 0.907556 | 1.048137 | 0.22 | 0.251 | 1 | 14 | Fzd2 | -0.031 | Young |
| 0.907712 | 0.342886 | 0.16 | 0.164 | 1 | 14 | Togaram1 | -0.004 | Young |
| 0.90842 | 0.741035 | 0.26 | 0.273 | 1 | 14 | Cat | -0.013 | Young |
| 0.908779 | 0.179987 | 0.1 | 0.109 | 1 | 14 | Abhd12 | -0.009 | Young |
| 0.909932 | 0.289799 | 0.46 | 0.415 | 1 | 14 | Ap2a2 | 0.045 | Young |
| 0.909965 | 0.419328 | 0.06 | 0.055 | 1 | 14 | Cars2 | 0.005 | Young |
| 0.91022 | 0.083612 | 0.14 | 0.137 | 1 | 14 | Zfyve16 | 0.003 | Young |
| 0.910999 | 0.210258 | 0.32 | 0.333 | 1 | 14 | Ccng2 | -0.013 | Young |
| 0.911343 | 0.074747 | 0.3 | 0.311 | 1 | 14 | Srsf10 | -0.011 | Young |
| 0.911377 | 0.081162 | 0.06 | 0.066 | 1 | 14 | Bcl7a | -0.006 | Young |
| 0.911427 | 0.267726 | 0.04 | 0.044 | 1 | 14 | Mindy4 | -0.004 | Young |

| | | | | | | | | |
|---|---|---|---|---|---|---|---|---|
| 0.911427 | 0.210077 | 0.04 | 0.044 | 1 | 14 | Twnk | -0.004 | Young |
| 0.911427 | 0.099025 | 0.04 | 0.044 | 1 | 14 | Uty | -0.004 | Young |
| 0.911462 | 0.189343 | 0.54 | 0.53 | 1 | 14 | Rhoc | 0.01 | Young |
| 0.912273 | 1.184997 | 0.1 | 0.109 | 1 | 14 | Kdm4c | -0.009 | Young |
| 0.912306 | 0.046779 | 0.9 | 0.907 | 1 | 14 | Rps5 | -0.007 | Young |
| 0.912341 | 0.078965 | 0.96 | 0.967 | 1 | 14 | Rps9 | -0.007 | Young |
| 0.912452 | 0.444819 | 0.08 | 0.077 | 1 | 14 | Ccdc28b | 0.003 | Young |
| 0.912589 | 0.114721 | 0.16 | 0.158 | 1 | 14 | C1qb | 0.002 | Young |
| 0.912882 | 0.329578 | 0.3 | 0.333 | 1 | 14 | Ndrg2 | -0.033 | Young |
| 0.913036 | 0.463079 | 0.36 | 0.355 | 1 | 14 | Paics | 0.005 | Young |
| 0.913245 | 0.130878 | 0.48 | 0.47 | 1 | 14 | Rhoq | 0.01 | Young |
| 0.913306 | 0.052366 | 0.22 | 0.202 | 1 | 14 | Tnpo2 | 0.018 | Young |
| 0.913308 | 0.320635 | 0.56 | 0.612 | 1 | 14 | Mtch1 | -0.052 | Young |
| 0.913396 | 0.024041 | 0.62 | 0.667 | 1 | 14 | Tmem30a | -0.047 | Young |
| 0.913507 | 0.434219 | 0.12 | 0.115 | 1 | 14 | Ntrk3 | 0.005 | Young |
| 0.913507 | 0.023984 | 0.12 | 0.115 | 1 | 14 | Vps26b | 0.005 | Young |
| 0.914572 | 0.729225 | 0.16 | 0.153 | 1 | 14 | Fam96a | 0.007 | Young |
| 0.914572 | 0.715596 | 0.16 | 0.153 | 1 | 14 | Unkl | 0.007 | Young |
| 0.914778 | 0.306402 | 0.18 | 0.175 | 1 | 14 | Ubxn6 | 0.005 | Young |
| 0.915792 | 0.821807 | 0.06 | 0.066 | 1 | 14 | Mocs1 | -0.006 | Young |
| 0.915792 | 0.126119 | 0.06 | 0.066 | 1 | 14 | Mcm9 | -0.006 | Young |
| 0.916267 | 0.311077 | 0.24 | 0.246 | 1 | 14 | Jkamp | -0.006 | Young |
| 0.916496 | 0.029843 | 0.14 | 0.137 | 1 | 14 | Tlk1 | 0.003 | Young |
| 0.916509 | 0.451437 | 0.08 | 0.077 | 1 | 14 | Tmed8 | 0.003 | Young |
| 0.916509 | 0.448586 | 0.08 | 0.077 | 1 | 14 | Pter | 0.003 | Young |
| 0.916775 | 1.174717 | 0.04 | 0.044 | 1 | 14 | Akr1c13 | -0.004 | Young |
| 0.916775 | 0.043043 | 0.04 | 0.044 | 1 | 14 | Smco4 | -0.004 | Young |
| 0.916775 | 0.036881 | 0.04 | 0.044 | 1 | 14 | AW554918 | -0.004 | Young |
| 0.916946 | 0.170561 | 0.14 | 0.131 | 1 | 14 | Dvl3 | 0.009 | Young |
| 0.918004 | 0.107196 | 0.5 | 0.525 | 1 | 14 | Ppib | -0.025 | Young |
| 0.918092 | 0.504878 | 0.14 | 0.153 | 1 | 14 | Gpr108 | -0.013 | Young |
| 0.918106 | 1.667445 | 0.04 | 0.038 | 1 | 14 | Nit2 | 0.002 | Young |
| 0.91829 | 0.693983 | 0.38 | 0.366 | 1 | 14 | Rgmb | 0.014 | Young |
| 0.91829 | 0.286367 | 0.36 | 0.372 | 1 | 14 | Lman1 | -0.012 | Young |
| 0.919353 | 0.047982 | 0.4 | 0.383 | 1 | 14 | Rsu1 | 0.017 | Young |
| 0.92044 | 0.500734 | 0.18 | 0.175 | 1 | 14 | Fam120b | 0.005 | Young |
| 0.920568 | 0.591508 | 0.08 | 0.077 | 1 | 14 | Zfp874a | 0.003 | Young |
| 0.920568 | 0.097248 | 0.08 | 0.077 | 1 | 14 | Tcaf2 | 0.003 | Young |
| 0.921526 | 0.278669 | 0.36 | 0.35 | 1 | 14 | Aaed1 | 0.01 | Young |
| 0.921965 | 0.046136 | 0.6 | 0.667 | 1 | 14 | Tuba1a | -0.067 | Young |
| 0.922126 | 1.866549 | 0.04 | 0.044 | 1 | 14 | Rrp1b | -0.004 | Young |
| 0.922126 | 0.330825 | 0.04 | 0.044 | 1 | 14 | Ripk2 | -0.004 | Young |
| 0.922126 | 0.158048 | 0.04 | 0.044 | 1 | 14 | Batf3 | -0.004 | Young |
| 0.922126 | 0.023033 | 0.04 | 0.044 | 1 | 14 | Prkar1b | -0.004 | Young |
| 0.922204 | 0.012163 | 0.66 | 0.672 | 1 | 14 | Pdia6 | -0.012 | Young |

| | | | | | | | | |
|---|---|---|---|---|---|---|---|---|
| 0.922227 | 0.582109 | 0.32 | 0.333 | 1 | 14 | Ik | -0.013 | Young |
| 0.922232 | 0.593718 | 0.24 | 0.257 | 1 | 14 | Rab5a | -0.017 | Young |
| 0.922778 | 0.922581 | 0.14 | 0.137 | 1 | 14 | Ppp1r9a | 0.003 | Young |
| 0.923272 | 0.551935 | 0.18 | 0.175 | 1 | 14 | Tcp11l2 | 0.005 | Young |
| 0.923272 | 0.056906 | 0.18 | 0.175 | 1 | 14 | Pgm3 | 0.005 | Young |
| 0.923274 | 0.077335 | 0.42 | 0.393 | 1 | 14 | Metap2 | 0.027 | Young |
| 0.92373 | 0.124282 | 0.36 | 0.35 | 1 | 14 | Dhrs7 | 0.01 | Young |
| 0.92373 | 0.057 | 0.32 | 0.361 | 1 | 14 | Irs2 | -0.041 | Young |
| 0.923736 | 1.927424 | 0.04 | 0.038 | 1 | 14 | Fmod | 0.002 | Young |
| 0.923736 | 1.417232 | 0.04 | 0.038 | 1 | 14 | 2810402E2 | 0.002 | Young |
| 0.923736 | 0.858536 | 0.04 | 0.038 | 1 | 14 | Pigf | 0.002 | Young |
| 0.923736 | 0.728241 | 0.04 | 0.038 | 1 | 14 | Usb1 | 0.002 | Young |
| 0.923928 | 0.112676 | 0.16 | 0.175 | 1 | 14 | Ehd4 | -0.015 | Young |
| 0.924046 | 0.253437 | 0.32 | 0.29 | 1 | 14 | Sri | 0.03 | Young |
| 0.924628 | 1.640645 | 0.06 | 0.066 | 1 | 14 | Bach2 | -0.006 | Young |
| 0.924628 | 1.325378 | 0.06 | 0.066 | 1 | 14 | Zfp770 | -0.006 | Young |
| 0.925116 | 0.087404 | 0.3 | 0.311 | 1 | 14 | Sumo1 | -0.011 | Young |
| 0.925492 | 0.028692 | 0.22 | 0.23 | 1 | 14 | Phb2 | -0.01 | Young |
| 0.925867 | 0.190749 | 0.22 | 0.208 | 1 | 14 | Snrnp27 | 0.012 | Young |
| 0.925997 | 0.435016 | 0.2 | 0.202 | 1 | 14 | Adam23 | -0.002 | Young |
| 0.926472 | 0.134646 | 0.08 | 0.082 | 1 | 14 | Tsta3 | -0.002 | Young |
| 0.926807 | 0.227913 | 0.26 | 0.262 | 1 | 14 | Pcdh18 | -0.002 | Young |
| 0.927086 | 0.37391 | 0.38 | 0.393 | 1 | 14 | Ctbp1 | -0.013 | Young |
| 0.927482 | 0.515396 | 0.04 | 0.044 | 1 | 14 | Plekhn1 | -0.004 | Young |
| 0.927482 | 0.31564 | 0.04 | 0.044 | 1 | 14 | Zfp677 | -0.004 | Young |
| 0.927542 | 0.500672 | 0.3 | 0.295 | 1 | 14 | Uso1 | 0.005 | Young |
| 0.927682 | 0.356959 | 0.44 | 0.421 | 1 | 14 | Gapdh | 0.019 | Young |
| 0.927804 | 0.623236 | 0.14 | 0.148 | 1 | 14 | Serinc5 | -0.008 | Young |
| 0.927967 | 0.322909 | 0.54 | 0.563 | 1 | 14 | Prrc2a | -0.023 | Young |
| 0.928076 | 0.612977 | 0.2 | 0.197 | 1 | 14 | Cnn2 | 0.003 | Young |
| 0.928675 | 0.195557 | 0.32 | 0.328 | 1 | 14 | Vmp1 | -0.008 | Young |
| 0.928692 | 0.034296 | 0.08 | 0.077 | 1 | 14 | Klhl15 | 0.003 | Young |
| 0.928852 | 0.289021 | 0.5 | 0.519 | 1 | 14 | Hnrnpab | -0.019 | Young |
| 0.929049 | 0.450152 | 0.06 | 0.066 | 1 | 14 | Zbp1 | -0.006 | Young |
| 0.929049 | 0.156422 | 0.06 | 0.066 | 1 | 14 | Prmt9 | -0.006 | Young |
| 0.92937 | 0.731709 | 0.04 | 0.038 | 1 | 14 | Dpy19l3 | 0.002 | Young |
| 0.92937 | 0.66999 | 0.04 | 0.038 | 1 | 14 | Zfp715 | 0.002 | Young |
| 0.929506 | 0.190447 | 0.34 | 0.317 | 1 | 14 | Rps6kb1 | 0.023 | Young |
| 0.930306 | 0.04309 | 0.44 | 0.437 | 1 | 14 | Luc7l3 | 0.003 | Young |
| 0.930418 | 0.04568 | 0.12 | 0.115 | 1 | 14 | Gab1 | 0.005 | Young |
| 0.930436 | 0.681638 | 0.08 | 0.082 | 1 | 14 | A430033K0 | -0.002 | Young |
| 0.930934 | 0.055946 | 0.34 | 0.322 | 1 | 14 | Gli3 | 0.018 | Young |
| 0.93119 | 0.174402 | 0.34 | 0.35 | 1 | 14 | Carhsp1 | -0.01 | Young |
| 0.931213 | 0.340064 | 0.58 | 0.639 | 1 | 14 | Akap13 | -0.059 | Young |
| 0.931698 | 0.070504 | 0.34 | 0.361 | 1 | 14 | Oxct1 | -0.021 | Young |

| | | | | | | | | |
|---|---|---|---|---|---|---|---|---|
| 0.932209 | 0.167469 | 0.14 | 0.137 | 1 | 14 | Mfsd11 | 0.003 | Young |
| 0.932837 | 0.104847 | 0.24 | 0.235 | 1 | 14 | Shoc2 | 0.005 | Young |
| 0.93284 | 0.411223 | 0.04 | 0.044 | 1 | 14 | Msl3l2 | -0.004 | Young |
| 0.932993 | 0.322845 | 0.22 | 0.213 | 1 | 14 | Rap2c | 0.007 | Young |
| 0.933491 | 0.07368 | 0.2 | 0.197 | 1 | 14 | Ckap5 | 0.003 | Young |
| 0.93431 | 0.292872 | 0.3 | 0.311 | 1 | 14 | Gtf2h5 | -0.011 | Young |
| 0.934402 | 0.632935 | 0.08 | 0.082 | 1 | 14 | Clpb | -0.002 | Young |
| 0.934601 | 0.093907 | 0.78 | 0.792 | 1 | 14 | Rpl12 | -0.012 | Young |
| 0.9348 | 0.657689 | 0.22 | 0.219 | 1 | 14 | Atp9b | 0.001 | Young |
| 0.935007 | 3.03852 | 0.04 | 0.038 | 1 | 14 | C1qtnf3 | 0.002 | Young |
| 0.935007 | 1.901697 | 0.04 | 0.038 | 1 | 14 | Trib2 | 0.002 | Young |
| 0.935007 | 0.614109 | 0.04 | 0.038 | 1 | 14 | Prtg | 0.002 | Young |
| 0.935007 | 0.017111 | 0.04 | 0.038 | 1 | 14 | Gk | 0.002 | Young |
| 0.935501 | 0.275022 | 0.16 | 0.153 | 1 | 14 | Fopnl | 0.007 | Young |
| 0.935877 | 0.071971 | 0.42 | 0.421 | 1 | 14 | Bag3 | -0.001 | Young |
| 0.935918 | 1.109693 | 0.08 | 0.087 | 1 | 14 | Zfp865 | -0.007 | Young |
| 0.93815 | 1.605523 | 0.06 | 0.06 | 1 | 14 | Slc9a3r1 | 0 | Young |
| 0.93815 | 1.320009 | 0.06 | 0.06 | 1 | 14 | Agrn | 0 | Young |
| 0.93815 | 0.889538 | 0.06 | 0.06 | 1 | 14 | Nudt1 | 0 | Young |
| 0.938369 | 0.678325 | 0.08 | 0.082 | 1 | 14 | Cog2 | -0.002 | Young |
| 0.938469 | 0.214823 | 0.4 | 0.377 | 1 | 14 | Brd4 | 0.023 | Young |
| 0.93871 | 0.779886 | 0.24 | 0.246 | 1 | 14 | Strap | -0.006 | Young |
| 0.939255 | 0.719799 | 0.14 | 0.131 | 1 | 14 | Fbxo4 | 0.009 | Young |
| 0.939855 | 0.031573 | 0.02 | 0.022 | 1 | 14 | Prdm9 | -0.002 | Young |
| 0.939855 | 0.030755 | 0.02 | 0.022 | 1 | 14 | Trim46 | -0.002 | Young |
| 0.939855 | 0.011985 | 0.02 | 0.022 | 1 | 14 | Clic5 | -0.002 | Young |
| 0.939958 | 0.335912 | 0.26 | 0.24 | 1 | 14 | Cenpb | 0.02 | Young |
| 0.940648 | 0.831491 | 0.04 | 0.038 | 1 | 14 | Star | 0.002 | Young |
| 0.940648 | 0.33501 | 0.04 | 0.038 | 1 | 14 | Zfp758 | 0.002 | Young |
| 0.940882 | 0.303644 | 0.6 | 0.634 | 1 | 14 | Tmed10 | -0.034 | Young |
| 0.940891 | 0.610863 | 0.08 | 0.077 | 1 | 14 | Lrrc42 | 0.003 | Young |
| 0.940891 | 0.45949 | 0.08 | 0.077 | 1 | 14 | Heatr5b | 0.003 | Young |
| 0.941159 | 0.038947 | 0.36 | 0.344 | 1 | 14 | Thrap3 | 0.016 | Young |
| 0.94165 | 0.300233 | 0.14 | 0.137 | 1 | 14 | Vtn | 0.003 | Young |
| 0.942095 | 0.089849 | 0.3 | 0.306 | 1 | 14 | Pacsin2 | -0.006 | Young |
| 0.94213 | 0.079589 | 0.28 | 0.29 | 1 | 14 | Myc | -0.01 | Young |
| 0.942159 | 0.15734 | 0.24 | 0.224 | 1 | 14 | Chd2 | 0.016 | Young |
| 0.942338 | 0.111742 | 0.52 | 0.486 | 1 | 14 | Myl12b | 0.034 | Young |
| 0.942658 | 0.266794 | 0.46 | 0.47 | 1 | 14 | Copb2 | -0.01 | Young |
| 0.94338 | 0.034678 | 0.94 | 0.995 | 1 | 14 | mt-Cytb | -0.055 | Young |
| 0.943485 | 0.260184 | 0.22 | 0.213 | 1 | 14 | Per1 | 0.007 | Young |
| 0.943566 | 0.7745 | 0.04 | 0.044 | 1 | 14 | Ptar1 | -0.004 | Young |
| 0.943566 | 0.638326 | 0.04 | 0.044 | 1 | 14 | Rarb | -0.004 | Young |
| 0.943844 | 0.128348 | 0.2 | 0.191 | 1 | 14 | Spg20 | 0.009 | Young |
| 0.944159 | 0.151481 | 0.8 | 0.825 | 1 | 14 | Rpl34 | -0.025 | Young |

| | | | | | | | | |
|---|---|---|---|---|---|---|---|---|
| 0.944409 | 1.102499 | 0.24 | 0.257 | 1 | 14 | Pdlim7 | -0.017 | Young |
| 0.944543 | 0.149145 | 0.1 | 0.104 | 1 | 14 | Flot1 | -0.004 | Young |
| 0.944799 | 0.148741 | 0.14 | 0.137 | 1 | 14 | Gnl3l | 0.003 | Young |
| 0.944799 | 0.095493 | 0.14 | 0.137 | 1 | 14 | Gid8 | 0.003 | Young |
| 0.944856 | 0.709415 | 0.12 | 0.12 | 1 | 14 | Fdft1 | 0 | Young |
| 0.944961 | 0.398571 | 0.08 | 0.077 | 1 | 14 | Atad3a | 0.003 | Young |
| 0.945083 | 0.720709 | 0.18 | 0.18 | 1 | 14 | Slc25a28 | 0 | Young |
| 0.945251 | 0.678764 | 0.2 | 0.208 | 1 | 14 | Ssbp4 | -0.008 | Young |
| 0.945298 | 0.733077 | 0.1 | 0.098 | 1 | 14 | Gde1 | 0.002 | Young |
| 0.945389 | 0.074088 | 0.26 | 0.268 | 1 | 14 | Zc3h7b | -0.008 | Young |
| 0.945639 | 0.892677 | 0.14 | 0.131 | 1 | 14 | Klhl18 | 0.009 | Young |
| 0.945815 | 0.584768 | 0.28 | 0.317 | 1 | 14 | Rab8b | -0.037 | Young |
| 0.945851 | 0.250346 | 0.24 | 0.24 | 1 | 14 | Plekha5 | 0 | Young |
| 0.945997 | 0.200276 | 0.28 | 0.273 | 1 | 14 | Tbc1d4 | 0.007 | Young |
| 0.946078 | 0.435154 | 0.42 | 0.41 | 1 | 14 | Jmjd1c | 0.01 | Young |
| 0.946292 | 0.368693 | 0.04 | 0.038 | 1 | 14 | Pafah1b3 | 0.002 | Young |
| 0.946292 | 0.243187 | 0.04 | 0.038 | 1 | 14 | 2610021A0 | 0.002 | Young |
| 0.946292 | 0.164045 | 0.04 | 0.038 | 1 | 14 | Slc15a3 | 0.002 | Young |
| 0.946292 | 0.113558 | 0.04 | 0.038 | 1 | 14 | Dusp19 | 0.002 | Young |
| 0.946748 | 0.211783 | 0.46 | 0.47 | 1 | 14 | Swi5 | -0.01 | Young |
| 0.947114 | 0.31829 | 0.3 | 0.29 | 1 | 14 | Irf2 | 0.01 | Young |
| 0.947299 | 0.915777 | 0.06 | 0.06 | 1 | 14 | Zfp672 | 0 | Young |
| 0.947362 | 0.488386 | 0.02 | 0.022 | 1 | 14 | Smagp | -0.002 | Young |
| 0.947362 | 0.384587 | 0.02 | 0.022 | 1 | 14 | Zkscan5 | -0.002 | Young |
| 0.947362 | 0.332214 | 0.02 | 0.022 | 1 | 14 | Thg1l | -0.002 | Young |
| 0.947362 | 0.297795 | 0.02 | 0.022 | 1 | 14 | Slc25a42 | -0.002 | Young |
| 0.947362 | 0.250604 | 0.02 | 0.022 | 1 | 14 | Znhit3 | -0.002 | Young |
| 0.947362 | 0.215516 | 0.02 | 0.022 | 1 | 14 | Pcdhb22 | -0.002 | Young |
| 0.947362 | 0.168566 | 0.02 | 0.022 | 1 | 14 | Zkscan6 | -0.002 | Young |
| 0.947362 | 0.102268 | 0.02 | 0.022 | 1 | 14 | Naip5 | -0.002 | Young |
| 0.947482 | 0.384198 | 0.16 | 0.153 | 1 | 14 | Cln5 | 0.007 | Young |
| 0.947895 | 0.220451 | 0.18 | 0.18 | 1 | 14 | Ncoa3 | 0 | Young |
| 0.948078 | 0.13066 | 0.16 | 0.158 | 1 | 14 | Rbm12 | 0.002 | Young |
| 0.948333 | 0.13743 | 0.2 | 0.213 | 1 | 14 | Rai1 | -0.013 | Young |
| 0.948742 | 0.808574 | 0.38 | 0.443 | 1 | 14 | Pik3r1 | -0.063 | Young |
| 0.948933 | 0.721618 | 0.04 | 0.044 | 1 | 14 | Gamt | -0.004 | Young |
| 0.94898 | 0.080665 | 0.86 | 0.874 | 1 | 14 | Cdc42 | -0.014 | Young |
| 0.94902 | 0.220826 | 0.32 | 0.328 | 1 | 14 | Tap2 | -0.008 | Young |
| 0.949274 | 0.538801 | 0.14 | 0.148 | 1 | 14 | Fyco1 | -0.008 | Young |
| 0.94964 | 0.443568 | 0.32 | 0.344 | 1 | 14 | 4932438A1 | -0.024 | Young |
| 0.94978 | 0.188251 | 0.18 | 0.186 | 1 | 14 | Dtx4 | -0.006 | Young |
| 0.94978 | 0.103938 | 0.18 | 0.186 | 1 | 14 | Nid2 | -0.006 | Young |
| 0.950018 | 0.058382 | 0.4 | 0.404 | 1 | 14 | Smim7 | -0.004 | Young |
| 0.950028 | 0.173732 | 0.36 | 0.344 | 1 | 14 | Mob4 | 0.016 | Young |
| 0.950182 | 0.006536 | 0.14 | 0.142 | 1 | 14 | Ngly1 | -0.002 | Young |

| | | | | | | | | |
|---|---|---|---|---|---|---|---|---|
| 0.950438 | 0.288795 | 0.4 | 0.421 | 1 | 14 | Tln1 | -0.021 | Young |
| 0.950708 | 0.334399 | 0.18 | 0.18 | 1 | 14 | Tep1 | 0 | Young |
| 0.951099 | 0.650845 | 0.14 | 0.137 | 1 | 14 | Usp5 | 0.003 | Young |
| 0.951099 | 0.645581 | 0.14 | 0.137 | 1 | 14 | Got1 | 0.003 | Young |
| 0.951541 | 0.17634 | 0.42 | 0.464 | 1 | 14 | Serp1 | -0.044 | Young |
| 0.951621 | 0.523315 | 0.18 | 0.191 | 1 | 14 | Il17ra | -0.011 | Young |
| 0.95169 | 0.025852 | 0.1 | 0.104 | 1 | 14 | Ndufaf2 | -0.004 | Young |
| 0.951875 | 0.850352 | 0.06 | 0.06 | 1 | 14 | Mrrf | 0 | Young |
| 0.951938 | 0.030575 | 0.04 | 0.038 | 1 | 14 | Gucy1a2 | 0.002 | Young |
| 0.95217 | 0.160663 | 0.38 | 0.372 | 1 | 14 | Smap1 | 0.008 | Young |
| 0.952428 | 1.116515 | 0.24 | 0.224 | 1 | 14 | Ddx21 | 0.016 | Young |
| 0.953411 | 0.239019 | 0.36 | 0.383 | 1 | 14 | Vapa | -0.023 | Young |
| 0.95342 | 0.757329 | 0.18 | 0.197 | 1 | 14 | Impa1 | -0.017 | Young |
| 0.953441 | 0.60728 | 0.28 | 0.279 | 1 | 14 | Ssbp2 | 0.001 | Young |
| 0.953661 | 0.822636 | 0.3 | 0.306 | 1 | 14 | Zfp644 | -0.006 | Young |
| 0.9537 | 0.174295 | 0.24 | 0.246 | 1 | 14 | Nudt4 | -0.006 | Young |
| 0.954302 | 1.793903 | 0.04 | 0.044 | 1 | 14 | Polm | -0.004 | Young |
| 0.954485 | 0.316222 | 0.34 | 0.317 | 1 | 14 | Erbin | 0.023 | Young |
| 0.954753 | 0.136656 | 0.54 | 0.579 | 1 | 14 | Pnrc1 | -0.039 | Young |
| 0.954873 | 2.418079 | 0.02 | 0.022 | 1 | 14 | Cdt1 | -0.002 | Young |
| 0.954873 | 2.324613 | 0.02 | 0.022 | 1 | 14 | Them6 | -0.002 | Young |
| 0.954873 | 1.990603 | 0.02 | 0.022 | 1 | 14 | Mrm3 | -0.002 | Young |
| 0.954873 | 1.534761 | 0.02 | 0.022 | 1 | 14 | Prkar2b | -0.002 | Young |
| 0.954873 | 0.953387 | 0.02 | 0.022 | 1 | 14 | Gpr21 | -0.002 | Young |
| 0.954873 | 0.919058 | 0.02 | 0.022 | 1 | 14 | Gm28523 | -0.002 | Young |
| 0.954873 | 0.889187 | 0.02 | 0.022 | 1 | 14 | Fancd2 | -0.002 | Young |
| 0.954873 | 0.807631 | 0.02 | 0.022 | 1 | 14 | Trp53rkb | -0.002 | Young |
| 0.954873 | 0.790283 | 0.02 | 0.022 | 1 | 14 | Dagla | -0.002 | Young |
| 0.954873 | 0.733033 | 0.02 | 0.022 | 1 | 14 | Srrd | -0.002 | Young |
| 0.954873 | 0.65699 | 0.02 | 0.022 | 1 | 14 | Tnfrsf13b | -0.002 | Young |
| 0.954873 | 0.624055 | 0.02 | 0.022 | 1 | 14 | Gm17354 | -0.002 | Young |
| 0.954873 | 0.604784 | 0.02 | 0.022 | 1 | 14 | Dmrt2 | -0.002 | Young |
| 0.954873 | 0.57539 | 0.02 | 0.022 | 1 | 14 | Ccdc88b | -0.002 | Young |
| 0.954873 | 0.525388 | 0.02 | 0.022 | 1 | 14 | Gabrq | -0.002 | Young |
| 0.954873 | 0.46322 | 0.02 | 0.022 | 1 | 14 | Sass6 | -0.002 | Young |
| 0.954873 | 0.432462 | 0.02 | 0.022 | 1 | 14 | Gm29683 | -0.002 | Young |
| 0.955312 | 0.277038 | 0.24 | 0.23 | 1 | 14 | Zyg11b | 0.01 | Young |
| 0.95534 | 0.278379 | 0.28 | 0.268 | 1 | 14 | Gmppa | 0.012 | Young |
| 0.955622 | 0.332761 | 0.32 | 0.322 | 1 | 14 | Cfl2 | -0.002 | Young |
| 0.955859 | 0.511275 | 0.42 | 0.475 | 1 | 14 | Lgi2 | -0.055 | Young |
| 0.9562 | 0.385972 | 0.24 | 0.246 | 1 | 14 | Col16a1 | -0.006 | Young |
| 0.956226 | 0.57972 | 0.1 | 0.098 | 1 | 14 | Dtd1 | 0.002 | Young |
| 0.956879 | 0.235968 | 0.64 | 0.705 | 1 | 14 | Naca | -0.065 | Young |
| 0.957453 | 0.125848 | 0.84 | 0.858 | 1 | 14 | Rpl3 | -0.018 | Young |
| 0.957587 | 2.77828 | 0.04 | 0.038 | 1 | 14 | Ubash3b | 0.002 | Young |

| | | | | | | | | |
|---|---|---|---|---|---|---|---|---|
| 0.957587 | 1.61157 | 0.04 | 0.038 | 1 | 14 | Golm1 | 0.002 | Young |
| 0.957587 | 1.306261 | 0.04 | 0.038 | 1 | 14 | Rmdn2 | 0.002 | Young |
| 0.957587 | 1.210645 | 0.04 | 0.038 | 1 | 14 | Plagl1 | 0.002 | Young |
| 0.957587 | 1.171056 | 0.04 | 0.038 | 1 | 14 | Lzts3 | 0.002 | Young |
| 0.957587 | 0.541681 | 0.04 | 0.038 | 1 | 14 | Alyref2 | 0.002 | Young |
| 0.957587 | 0.204627 | 0.04 | 0.038 | 1 | 14 | Nrde2 | 0.002 | Young |
| 0.95809 | 0.367211 | 0.46 | 0.475 | 1 | 14 | G3bp2 | -0.015 | Young |
| 0.958151 | 0.30291 | 0.24 | 0.235 | 1 | 14 | Chtf8 | 0.005 | Young |
| 0.95884 | 1.045862 | 0.1 | 0.104 | 1 | 14 | Gm1673 | -0.004 | Young |
| 0.95884 | 0.144835 | 0.1 | 0.104 | 1 | 14 | Mpst | -0.004 | Young |
| 0.95915 | 0.877972 | 0.18 | 0.18 | 1 | 14 | Prmt1 | 0 | Young |
| 0.959353 | 0.069632 | 0.84 | 0.869 | 1 | 14 | Rps2 | -0.029 | Young |
| 0.959696 | 0.274884 | 0.42 | 0.404 | 1 | 14 | Mier1 | 0.016 | Young |
| 0.959871 | 0.531708 | 0.1 | 0.098 | 1 | 14 | E2f5 | 0.002 | Young |
| 0.960339 | 0.296264 | 0.42 | 0.437 | 1 | 14 | Gns | -0.017 | Young |
| 0.96093 | 0.174498 | 0.18 | 0.186 | 1 | 14 | Atxn7 | -0.006 | Young |
| 0.961204 | 0.360028 | 0.28 | 0.295 | 1 | 14 | Map2k2 | -0.015 | Young |
| 0.961549 | 0.12148 | 0.36 | 0.361 | 1 | 14 | Ppp1r15a | -0.001 | Young |
| 0.96155 | 0.957414 | 0.12 | 0.12 | 1 | 14 | Tsfm | 0 | Young |
| 0.96155 | 0.069336 | 0.12 | 0.12 | 1 | 14 | Ubxn7 | 0 | Young |
| 0.96155 | 0.026707 | 0.12 | 0.12 | 1 | 14 | Fam219b | 0 | Young |
| 0.961582 | 0.226574 | 0.3 | 0.301 | 1 | 14 | 2-Mar | -0.001 | Young |
| 0.962203 | 0.078177 | 0.08 | 0.082 | 1 | 14 | Rce1 | -0.002 | Young |
| 0.962416 | 0.783063 | 0.1 | 0.104 | 1 | 14 | Zbtb41 | -0.004 | Young |
| 0.9629 | 0.630708 | 0.16 | 0.158 | 1 | 14 | Trappc13 | 0.002 | Young |
| 0.96291 | 0.071142 | 0.34 | 0.333 | 1 | 14 | Birc6 | 0.007 | Young |
| 0.963709 | 0.265659 | 0.14 | 0.137 | 1 | 14 | Tmem250- | 0.003 | Young |
| 0.964496 | 0.39838 | 0.22 | 0.213 | 1 | 14 | Triobp | 0.007 | Young |
| 0.964989 | 0.130645 | 0.28 | 0.268 | 1 | 14 | Cry2 | 0.012 | Young |
| 0.966179 | 0.317062 | 0.08 | 0.082 | 1 | 14 | Med19 | -0.002 | Young |
| 0.966401 | 0.295088 | 0.46 | 0.486 | 1 | 14 | Atp5j | -0.026 | Young |
| 0.966824 | 0.600443 | 0.38 | 0.399 | 1 | 14 | Stub1 | -0.019 | Young |
| 0.966863 | 0.466523 | 0.14 | 0.137 | 1 | 14 | Ssfa2 | 0.003 | Young |
| 0.967162 | 0.691777 | 0.1 | 0.098 | 1 | 14 | Smad2 | 0.002 | Young |
| 0.967162 | 0.002692 | 0.1 | 0.098 | 1 | 14 | Orc2 | 0.002 | Young |
| 0.968137 | 0.413396 | 0.36 | 0.361 | 1 | 14 | Arl8b | -0.001 | Young |
| 0.969132 | 0.397875 | 0.22 | 0.23 | 1 | 14 | Fech | -0.01 | Young |
| 0.970017 | 0.057986 | 0.14 | 0.137 | 1 | 14 | Lgr4 | 0.003 | Young |
| 0.970156 | 0.893705 | 0.08 | 0.082 | 1 | 14 | Trappc2 | -0.002 | Young |
| 0.970156 | 0.730658 | 0.08 | 0.082 | 1 | 14 | Paxx | -0.002 | Young |
| 0.970197 | 1.147697 | 0.06 | 0.06 | 1 | 14 | Depdc5 | 0 | Young |
| 0.970334 | 0.198296 | 0.26 | 0.251 | 1 | 14 | Gigyf2 | 0.009 | Young |
| 0.970413 | 0.34028 | 0.18 | 0.18 | 1 | 14 | Ncaph2 | 0 | Young |
| 0.970781 | 0.142404 | 0.14 | 0.148 | 1 | 14 | Sde2 | -0.008 | Young |
| 0.970809 | 0.023447 | 0.1 | 0.098 | 1 | 14 | Clk2 | 0.002 | Young |

| | | | | | | | | |
|---|---|---|---|---|---|---|---|---|
| 0.97102 | 0.031857 | 0.24 | 0.24 | 1 | 14 | Msrb1 | 0 | Young |
| 0.971034 | 0.156123 | 0.62 | 0.656 | 1 | 14 | Nr4a1 | -0.036 | Young |
| 0.971119 | 0.245327 | 0.12 | 0.115 | 1 | 14 | Nrros | 0.005 | Young |
| 0.971703 | 0.267703 | 0.22 | 0.23 | 1 | 14 | Csad | -0.01 | Young |
| 0.9718 | 0.360002 | 0.16 | 0.158 | 1 | 14 | Lcmt1 | 0.002 | Young |
| 0.971885 | 0.226749 | 0.46 | 0.432 | 1 | 14 | Srp14 | 0.028 | Young |
| 0.972009 | 0.235268 | 0.36 | 0.333 | 1 | 14 | Pdha1 | 0.027 | Young |
| 0.97253 | 0.423763 | 0.36 | 0.361 | 1 | 14 | Efhd2 | -0.001 | Young |
| 0.972532 | 0.13539 | 0.28 | 0.279 | 1 | 14 | Pfdn2 | 0.001 | Young |
| 0.972991 | 0.241227 | 0.4 | 0.377 | 1 | 14 | Hic1 | 0.023 | Young |
| 0.97323 | 0.232534 | 0.18 | 0.18 | 1 | 14 | Klf11 | 0 | Young |
| 0.973401 | 0.150815 | 0.42 | 0.399 | 1 | 14 | Rexo2 | 0.021 | Young |
| 0.973819 | 0.336748 | 0.32 | 0.322 | 1 | 14 | Capg | -0.002 | Young |
| 0.973855 | 0.211235 | 0.14 | 0.148 | 1 | 14 | Foxj3 | -0.008 | Young |
| 0.974389 | 0.262818 | 0.18 | 0.175 | 1 | 14 | Mrpl36 | 0.005 | Young |
| 0.974643 | 0.197302 | 0.28 | 0.268 | 1 | 14 | Rbm27 | 0.012 | Young |
| 0.974781 | 1.442678 | 0.06 | 0.06 | 1 | 14 | Dolk | 0 | Young |
| 0.974781 | 1.108235 | 0.06 | 0.06 | 1 | 14 | Acot13 | 0 | Young |
| 0.974781 | 0.890767 | 0.06 | 0.06 | 1 | 14 | Mpp7 | 0 | Young |
| 0.974781 | 0.675734 | 0.06 | 0.06 | 1 | 14 | Arl6 | 0 | Young |
| 0.974781 | 0.297255 | 0.06 | 0.06 | 1 | 14 | Slc2a10 | 0 | Young |
| 0.974781 | 0.158438 | 0.06 | 0.06 | 1 | 14 | Isg20 | 0 | Young |
| 0.974918 | 0.482545 | 0.12 | 0.12 | 1 | 14 | Rab33b | 0 | Young |
| 0.974918 | 0.306963 | 0.12 | 0.12 | 1 | 14 | Gadd45gip | 0 | Young |
| 0.974918 | 0.155781 | 0.12 | 0.12 | 1 | 14 | Lsm1 | 0 | Young |
| 0.975079 | 0.99283 | 0.14 | 0.142 | 1 | 14 | Sigmar1 | -0.002 | Young |
| 0.975884 | 0.66438 | 0.32 | 0.311 | 1 | 14 | Cop1 | 0.009 | Young |
| 0.975898 | 0.582654 | 0.24 | 0.235 | 1 | 14 | C2 | 0.005 | Young |
| 0.976729 | 0.969417 | 0.1 | 0.104 | 1 | 14 | Wiz | -0.004 | Young |
| 0.977097 | 0.126399 | 0.2 | 0.202 | 1 | 14 | Mrpl14 | -0.002 | Young |
| 0.977234 | 1.445589 | 0.18 | 0.175 | 1 | 14 | Islr2 | 0.005 | Young |
| 0.977234 | 0.474484 | 0.18 | 0.175 | 1 | 14 | Papd4 | 0.005 | Young |
| 0.978014 | 0.398505 | 0.26 | 0.262 | 1 | 14 | Fam49a | -0.002 | Young |
| 0.978052 | 0.277538 | 0.68 | 0.738 | 1 | 14 | Cfl1 | -0.058 | Young |
| 0.978105 | 0.043416 | 0.1 | 0.098 | 1 | 14 | Ndst2 | 0.002 | Young |
| 0.978105 | 0.026307 | 0.1 | 0.098 | 1 | 14 | Sfi1 | 0.002 | Young |
| 0.978193 | 0.126422 | 0.14 | 0.142 | 1 | 14 | Zfyve21 | -0.002 | Young |
| 0.978864 | 0.362961 | 0.18 | 0.18 | 1 | 14 | Clcc1 | 0 | Young |
| 0.979365 | 1.757866 | 0.06 | 0.06 | 1 | 14 | Sh3bp5l | 0 | Young |
| 0.979365 | 0.484412 | 0.06 | 0.06 | 1 | 14 | Lipe | 0 | Young |
| 0.979436 | 0.394348 | 0.56 | 0.596 | 1 | 14 | Rbpms | -0.036 | Young |
| 0.979509 | 0.079986 | 0.34 | 0.317 | 1 | 14 | Ccnt1 | 0.023 | Young |
| 0.979681 | 0.323984 | 0.32 | 0.333 | 1 | 14 | Prkacb | -0.013 | Young |
| 0.980096 | 0.131109 | 0.5 | 0.546 | 1 | 14 | Hnrnpm | -0.046 | Young |
| 0.980203 | 0.440079 | 0.3 | 0.301 | 1 | 14 | Dynlt3 | -0.001 | Young |

| | | | | | | | | |
|---|---|---|---|---|---|---|---|---|
| 0.980271 | 0.512518 | 0.22 | 0.213 | 1 | 14 | Aig1 | 0.007 | Young |
| 0.980916 | 0.454238 | 0.16 | 0.164 | 1 | 14 | Samd8 | -0.004 | Young |
| 0.981991 | 0.061807 | 0.22 | 0.23 | 1 | 14 | Insig1 | -0.01 | Young |
| 0.982091 | 0.097282 | 0.08 | 0.082 | 1 | 14 | Fam117a | -0.002 | Young |
| 0.982485 | 0.484163 | 0.2 | 0.202 | 1 | 14 | 2900026A( | -0.002 | Young |
| 0.983277 | 0.110605 | 0.24 | 0.224 | 1 | 14 | Dnm1l | 0.016 | Young |
| 0.983724 | 0.070629 | 0.24 | 0.246 | 1 | 14 | Helz | -0.006 | Young |
| 0.98374 | 0.327112 | 0.36 | 0.383 | 1 | 14 | Chic2 | -0.023 | Young |
| 0.983888 | 1.155782 | 0.1 | 0.104 | 1 | 14 | Rngtt | -0.004 | Young |
| 0.983944 | 0.014777 | 0.4 | 0.393 | 1 | 14 | Krtcap2 | 0.007 | Young |
| 0.98395 | 1.502089 | 0.06 | 0.06 | 1 | 14 | Sall2 | 0 | Young |
| 0.98395 | 1.346317 | 0.06 | 0.06 | 1 | 14 | Zc4h2 | 0 | Young |
| 0.98395 | 0.985091 | 0.06 | 0.06 | 1 | 14 | Zbtb45 | 0 | Young |
| 0.98395 | 0.027599 | 0.06 | 0.06 | 1 | 14 | Drosha | 0 | Young |
| 0.984026 | 0.10235 | 0.26 | 0.257 | 1 | 14 | Txnrd1 | 0.003 | Young |
| 0.984336 | 0.037976 | 0.22 | 0.219 | 1 | 14 | Ak2 | 0.001 | Young |
| 0.984423 | 0.24613 | 0.14 | 0.142 | 1 | 14 | Zfp512 | -0.002 | Young |
| 0.985384 | 0.259775 | 0.34 | 0.333 | 1 | 14 | Tia1 | 0.007 | Young |
| 0.985609 | 0.465647 | 0.36 | 0.35 | 1 | 14 | Chd9 | 0.01 | Young |
| 0.985713 | 0.238749 | 0.38 | 0.355 | 1 | 14 | Atp6ap2 | 0.025 | Young |
| 0.985795 | 0.343225 | 0.14 | 0.137 | 1 | 14 | Mrpl11 | 0.003 | Young |
| 0.985833 | 0.128523 | 0.96 | 0.94 | 1 | 14 | Rps20 | 0.02 | Young |
| 0.98607 | 0.359335 | 0.08 | 0.082 | 1 | 14 | Zfp511 | -0.002 | Young |
| 0.98607 | 0.321886 | 0.08 | 0.082 | 1 | 14 | Memo1 | -0.002 | Young |
| 0.986946 | 0.268155 | 0.22 | 0.219 | 1 | 14 | Stx7 | 0.001 | Young |
| 0.987063 | 0.026429 | 0.18 | 0.169 | 1 | 14 | Tspan4 | 0.011 | Young |
| 0.987063 | 0.412244 | 0.18 | 0.169 | 1 | 14 | Riok1 | 0.011 | Young |
| 0.988294 | 0.273595 | 0.12 | 0.12 | 1 | 14 | Mrpl50 | 0 | Young |
| 0.988294 | 0.103397 | 0.12 | 0.12 | 1 | 14 | Agl | 0 | Young |
| 0.988361 | 0.257293 | 0.4 | 0.415 | 1 | 14 | Rassf2 | -0.015 | Young |
| 0.988863 | 0.361342 | 0.48 | 0.497 | 1 | 14 | Zbtb7a | -0.017 | Young |
| 0.989521 | 0.463316 | 0.68 | 0.76 | 1 | 14 | Serpinb6a | -0.08 | Young |
| 0.989723 | 0.104424 | 0.16 | 0.164 | 1 | 14 | Lypla1 | -0.004 | Young |
| 0.989938 | 0.710874 | 0.18 | 0.169 | 1 | 14 | Camsap2 | 0.011 | Young |
| 0.99005 | 1.04508 | 0.08 | 0.082 | 1 | 14 | Irgq | -0.002 | Young |
| 0.99005 | 0.305815 | 0.08 | 0.082 | 1 | 14 | Atp8a1 | -0.002 | Young |
| 0.990136 | 0.682935 | 0.18 | 0.18 | 1 | 14 | Stxbp6 | 0 | Young |
| 0.990207 | 0.266627 | 0.54 | 0.601 | 1 | 14 | Eif4h | -0.061 | Young |
| 0.990482 | 0.032203 | 0.7 | 0.765 | 1 | 14 | mt-Nd4l | -0.065 | Young |
| 0.990568 | 0.382256 | 0.2 | 0.202 | 1 | 14 | Ahsa1 | -0.002 | Young |
| 0.991239 | 0.805471 | 0.38 | 0.361 | 1 | 14 | Cct2 | 0.019 | Young |
| 0.991462 | 0.134875 | 0.18 | 0.175 | 1 | 14 | Cyb5r4 | 0.005 | Young |
| 0.991498 | 1.006782 | 0.26 | 0.268 | 1 | 14 | Gtf3c2 | -0.008 | Young |
| 0.991819 | 0.013483 | 0.46 | 0.475 | 1 | 14 | Col15a1 | -0.015 | Young |
| 0.992435 | 0.15281 | 0.78 | 0.885 | 1 | 14 | Ftl1 | -0.105 | Young |

| | | | | | | | | |
|---|---|---|---|---|---|---|---|---|
| 0.992507 | 0.076663 | 0.42 | 0.388 | 1 | 14 | Pias1 | 0.032 | Young |
| 0.992701 | 0.139716 | 0.1 | 0.098 | 1 | 14 | Otulin | 0.002 | Young |
| 0.993121 | 0.505406 | 0.06 | 0.06 | 1 | 14 | Cnot9 | 0 | Young |
| 0.993121 | 0.437178 | 0.06 | 0.06 | 1 | 14 | Mthfsd | 0 | Young |
| 0.993121 | 0.301504 | 0.06 | 0.06 | 1 | 14 | Cd68 | 0 | Young |
| 0.993121 | 0.228543 | 0.06 | 0.06 | 1 | 14 | Icosl | 0 | Young |
| 0.993121 | 0.011473 | 0.06 | 0.06 | 1 | 14 | 9530082P2 | 0 | Young |
| 0.993121 | 0.007347 | 0.06 | 0.06 | 1 | 14 | Ficd | 0 | Young |
| 0.993423 | 0.754318 | 0.22 | 0.213 | 1 | 14 | Cops2 | 0.007 | Young |
| 0.993769 | 0.983748 | 0.14 | 0.142 | 1 | 14 | Aamdc | -0.002 | Young |
| 0.99378 | 0.388327 | 0.24 | 0.251 | 1 | 14 | Ago3 | -0.011 | Young |
| 0.994163 | 0.102282 | 0.58 | 0.623 | 1 | 14 | Capns1 | -0.043 | Young |
| 0.99458 | 0.222907 | 0.38 | 0.377 | 1 | 14 | Ctnnd1 | 0.003 | Young |
| 0.994928 | 0.321506 | 0.46 | 0.497 | 1 | 14 | Cnn3 | -0.037 | Young |
| 0.994983 | 1.121807 | 0.12 | 0.12 | 1 | 14 | Mtx2 | 0 | Young |
| 0.99506 | 0.121597 | 0.5 | 0.574 | 1 | 14 | Atrx | -0.074 | Young |
| 0.995235 | 0.247573 | 0.68 | 0.754 | 1 | 14 | Hsp90aa1 | -0.074 | Young |
| 0.995265 | 0.537322 | 0.14 | 0.137 | 1 | 14 | Prpf4 | 0.003 | Young |
| 0.995546 | 0.067739 | 0.16 | 0.158 | 1 | 14 | Habp4 | 0.002 | Young |
| 0.995772 | 0.670842 | 0.18 | 0.18 | 1 | 14 | Nhsl2 | 0 | Young |
| 0.996194 | 0.023536 | 0.24 | 0.235 | 1 | 14 | Fbxo11 | 0.005 | Young |
| 0.99635 | 0.339851 | 0.1 | 0.098 | 1 | 14 | Atg7 | 0.002 | Young |
| 0.99635 | 0.110078 | 0.1 | 0.098 | 1 | 14 | Pam16 | 0.002 | Young |
| 0.9968 | 0.437266 | 0.14 | 0.131 | 1 | 14 | Lta4h | 0.009 | Young |
| 0.997126 | 0.01103 | 0.7 | 0.694 | 1 | 14 | Add3 | 0.006 | Young |
| 0.997707 | 0.134721 | 0.06 | 0.06 | 1 | 14 | Zfp975 | 0 | Young |
| 0.997707 | 0.104619 | 0.06 | 0.06 | 1 | 14 | Pou6f1 | 0 | Young |
| 0.997707 | 0.10316 | 0.06 | 0.06 | 1 | 14 | Tfrc | 0 | Young |
| 0.998301 | 0.280872 | 0.12 | 0.115 | 1 | 14 | Tdg | 0.005 | Young |
| 0.998328 | 0.161563 | 0.12 | 0.12 | 1 | 14 | Fam50a | 0 | Young |
| 0.998353 | 1.175161 | 0.12 | 0.126 | 1 | 14 | Adra1b | -0.006 | Young |
| 0.998532 | 0.446184 | 0.16 | 0.164 | 1 | 14 | Fam103a1 | -0.004 | Young |
| 0.998641 | 0.137079 | 0.2 | 0.197 | 1 | 14 | Mrps25 | 0.003 | Young |
| 0.998704 | 0.481944 | 0.22 | 0.224 | 1 | 14 | Atf1 | -0.004 | Young |
| 0.998905 | 0.368092 | 0.36 | 0.366 | 1 | 14 | Gaa | -0.006 | Young |
| 0.998945 | 0.247099 | 0.4 | 0.421 | 1 | 14 | Ier5 | -0.021 | Young |
| 0.999015 | 0.307671 | 0.58 | 0.563 | 1 | 14 | Hmgn1 | 0.017 | Young |
| 0.000103 | 1.655207 | 0.449 | 0.107 | 1 | 15 | Rps27rt | 0.342 | Old |
| 0.000163 | 4.324414 | 0.327 | 0.054 | 1 | 15 | Haus8 | 0.273 | Old |
| 0.000442 | 4.425221 | 0.245 | 0.018 | 1 | 15 | Trrap | 0.227 | Old |
| 0.001607 | 2.125778 | 0.286 | 0.054 | 1 | 15 | Usf1 | 0.232 | Old |
| 0.002211 | 2.270957 | 0.245 | 0.036 | 1 | 15 | Tbce | 0.209 | Old |
| 0.002269 | 2.193378 | 0.327 | 0.089 | 1 | 15 | Mysm1 | 0.238 | Old |
| 0.002382 | 1.566806 | 0.531 | 0.268 | 1 | 15 | Ddhd1 | 0.263 | Old |
| 0.002397 | 2.166609 | 0.367 | 0.125 | 1 | 15 | Rnf44 | 0.242 | Old |

| | | | | | | | | |
|---|---|---|---|---|---|---|---|---|
| 0.002487 | 2.391608 | 0.327 | 0.089 | 1 | 15 | Nr2c2 | 0.238 | Old |
| 0.002491 | 1.519179 | 0.49 | 0.214 | 1 | 15 | Psmd11 | 0.276 | Old |
| 0.002492 | 1.763933 | 0.408 | 0.161 | 1 | 15 | Cd200 | 0.247 | Old |
| 0.002846 | 2.54304 | 0.265 | 0.054 | 1 | 15 | Smarca2 | 0.211 | Old |
| 0.002877 | 3.187398 | 0.224 | 0.036 | 1 | 15 | Scaf1 | 0.188 | Old |
| 0.003684 | 7.208074 | 0.143 | 0 | 1 | 15 | Cxcl13 | 0.143 | Old |
| 0.003756 | 1.346285 | 0.531 | 0.286 | 1 | 15 | Irf2 | 0.245 | Old |
| 0.003885 | 0.505019 | 0.959 | 0.946 | 1 | 15 | H3f3b | 0.013 | Old |
| 0.004664 | 0.385461 | 1 | 1 | 1 | 15 | mt-Co3 | 0 | Old |
| 0.004952 | 0.381115 | 1 | 1 | 1 | 15 | H2-K1 | 0 | Old |
| 0.006916 | 1.672339 | 0.449 | 0.214 | 1 | 15 | Rab8a | 0.235 | Old |
| 0.007243 | 3.815084 | 0.163 | 0.018 | 1 | 15 | Pik3c3 | 0.145 | Old |
| 0.007359 | 1.21397 | 0.531 | 0.268 | 1 | 15 | Zfp106 | 0.263 | Old |
| 0.007485 | 4.94561 | 0.122 | 0 | 1 | 15 | Cep164 | 0.122 | Old |
| 0.007485 | 4.767756 | 0.122 | 0 | 1 | 15 | Ints11 | 0.122 | Old |
| 0.007485 | 4.568322 | 0.122 | 0 | 1 | 15 | Tbcc | 0.122 | Old |
| 0.007485 | 4.525667 | 0.122 | 0 | 1 | 15 | Cdca4 | 0.122 | Old |
| 0.007485 | 4.445766 | 0.122 | 0 | 1 | 15 | Dpp7 | 0.122 | Old |
| 0.007485 | 4.359038 | 0.122 | 0 | 1 | 15 | Zfp646 | 0.122 | Old |
| 0.007485 | 4.164738 | 0.122 | 0 | 1 | 15 | Ppil1 | 0.122 | Old |
| 0.007485 | 3.617512 | 0.122 | 0 | 1 | 15 | Mid1 | 0.122 | Old |
| 0.008166 | 1.861341 | 0.204 | 0.036 | 1 | 15 | Cdk19 | 0.168 | Old |
| 0.008268 | 1.272018 | 0.327 | 0.107 | 1 | 15 | Ivns1abp | 0.22 | Old |
| 0.008379 | 3.155395 | 0.224 | 0.054 | 1 | 15 | Babam2 | 0.17 | Old |
| 0.008641 | 0.779408 | 0.816 | 0.643 | 1 | 15 | Csde1 | 0.173 | Old |
| 0.00881 | 3.004577 | 0.163 | 0.018 | 1 | 15 | Bcl11a | 0.145 | Old |
| 0.008993 | 2.19481 | 0.286 | 0.089 | 1 | 15 | Eri1 | 0.197 | Old |
| 0.009142 | 1.570942 | 0.367 | 0.143 | 1 | 15 | Anapc11 | 0.224 | Old |
| 0.009158 | 2.628721 | 0.163 | 0.018 | 1 | 15 | Ebag9 | 0.145 | Old |
| 0.00922 | 2.719892 | 0.224 | 0.054 | 1 | 15 | Tdg | 0.17 | Old |
| 0.00951 | 2.830649 | 0.286 | 0.089 | 1 | 15 | Apc | 0.197 | Old |
| 0.009682 | 1.320537 | 0.204 | 0.036 | 1 | 15 | Rnf14 | 0.168 | Old |
| 0.011344 | 2.047153 | 0.245 | 0.071 | 1 | 15 | Aars | 0.174 | Old |
| 0.011482 | 2.317432 | 0.224 | 0.054 | 1 | 15 | Tmem11 | 0.17 | Old |
| 0.011528 | 1.161042 | 0.49 | 0.268 | 1 | 15 | Arhgap5 | 0.222 | Old |
| 0.012582 | 3.326858 | 0.184 | 0.036 | 1 | 15 | Lrpprc | 0.148 | Old |
| 0.012759 | 1.716184 | 0.388 | 0.179 | 1 | 15 | Elf4 | 0.209 | Old |
| 0.012975 | 1.749979 | 0.429 | 0.232 | 1 | 15 | Runx3 | 0.197 | Old |
| 0.013053 | 1.447792 | 0.469 | 0.268 | 1 | 15 | Hmgb2 | 0.201 | Old |
| 0.0135 | 2.571944 | 0.245 | 0.071 | 1 | 15 | Fhod1 | 0.174 | Old |
| 0.013523 | 0.296234 | 1 | 1 | 1 | 15 | mt-Co1 | 0 | Old |
| 0.013621 | 0.721928 | 0.347 | 0.125 | 1 | 15 | Exoc4 | 0.222 | Old |
| 0.013928 | 3.186773 | 0.184 | 0.036 | 1 | 15 | Gmcl1 | 0.148 | Old |
| 0.014666 | 2.001395 | 0.265 | 0.089 | 1 | 15 | Glrx2 | 0.176 | Old |
| 0.014894 | 2.190717 | 0.184 | 0.036 | 1 | 15 | Kmt5c | 0.148 | Old |

| | | | | | | | | |
|---|---|---|---|---|---|---|---|---|
| 0.015078 | 1.372393 | 0.286 | 0.089 | 1 | 15 | Uso1 | 0.197 | Old |
| 0.015175 | 4.845515 | 0.102 | 0 | 1 | 15 | Cd55 | 0.102 | Old |
| 0.015175 | 4.721618 | 0.102 | 0 | 1 | 15 | Otud7b | 0.102 | Old |
| 0.015175 | 4.493934 | 0.102 | 0 | 1 | 15 | Tmem97 | 0.102 | Old |
| 0.015175 | 4.426871 | 0.102 | 0 | 1 | 15 | Dnajb5 | 0.102 | Old |
| 0.015175 | 4.333112 | 0.102 | 0 | 1 | 15 | Zfp605 | 0.102 | Old |
| 0.015175 | 4.283865 | 0.102 | 0 | 1 | 15 | Mast2 | 0.102 | Old |
| 0.015175 | 4.240138 | 0.102 | 0 | 1 | 15 | Ints8 | 0.102 | Old |
| 0.015175 | 4.039978 | 0.102 | 0 | 1 | 15 | Ptpn23 | 0.102 | Old |
| 0.015175 | 3.882521 | 0.102 | 0 | 1 | 15 | Pls3 | 0.102 | Old |
| 0.015175 | 3.792543 | 0.102 | 0 | 1 | 15 | Ttll3 | 0.102 | Old |
| 0.015221 | 1.287278 | 0.469 | 0.25 | 1 | 15 | Tnrc6a | 0.219 | Old |
| 0.015387 | 1.643149 | 0.286 | 0.107 | 1 | 15 | Hsf2 | 0.179 | Old |
| 0.015398 | 2.095036 | 0.184 | 0.036 | 1 | 15 | 2010315B0 | 0.148 | Old |
| 0.015411 | 3.917398 | 0.143 | 0.018 | 1 | 15 | Pbx1 | 0.125 | Old |
| 0.015689 | 0.794243 | 0.755 | 0.571 | 1 | 15 | Srsf5 | 0.184 | Old |
| 0.016014 | 4.132974 | 0.143 | 0.018 | 1 | 15 | 2300009A0 | 0.125 | Old |
| 0.017044 | 1.160349 | 0.531 | 0.339 | 1 | 15 | Nfil3 | 0.192 | Old |
| 0.017188 | 0.046157 | 0.082 | 0.268 | 1 | 15 | Hspa1b | -0.186 | Old |
| 0.017947 | 2.669634 | 0.143 | 0.018 | 1 | 15 | Cog2 | 0.125 | Old |
| 0.018276 | 0.931204 | 0.388 | 0.179 | 1 | 15 | U2surp | 0.209 | Old |
| 0.018317 | 3.05629 | 0.224 | 0.071 | 1 | 15 | Rnf169 | 0.153 | Old |
| 0.018611 | 0.648602 | 0.531 | 0.321 | 1 | 15 | 1700020I14 | 0.21 | Old |
| 0.018628 | 0.974653 | 0.592 | 0.393 | 1 | 15 | Uvrag | 0.199 | Old |
| 0.018635 | 2.533662 | 0.143 | 0.018 | 1 | 15 | Xist | 0.125 | Old |
| 0.018635 | 2.188139 | 0.143 | 0.018 | 1 | 15 | Rngtt | 0.125 | Old |
| 0.018673 | 2.114288 | 0.306 | 0.125 | 1 | 15 | Szrd1 | 0.181 | Old |
| 0.018973 | 0.416071 | 0.551 | 0.304 | 1 | 15 | Vps4b | 0.247 | Old |
| 0.01934 | 0.762172 | 0.673 | 0.464 | 1 | 15 | Arl6ip1 | 0.209 | Old |
| 0.019541 | 2.248304 | 0.204 | 0.054 | 1 | 15 | Armc10 | 0.15 | Old |
| 0.020549 | 1.345872 | 0.673 | 0.518 | 1 | 15 | Gm26917 | 0.155 | Old |
| 0.020702 | 1.486969 | 0.265 | 0.089 | 1 | 15 | Ddx52 | 0.176 | Old |
| 0.021063 | 1.692381 | 0.306 | 0.125 | 1 | 15 | Rheb | 0.181 | Old |
| 0.022003 | 1.478289 | 0.367 | 0.179 | 1 | 15 | Arpc1a | 0.188 | Old |
| 0.022013 | 2.209708 | 0.204 | 0.054 | 1 | 15 | Man2c1 | 0.15 | Old |
| 0.023205 | 0.668107 | 0.898 | 0.839 | 1 | 15 | Ddx5 | 0.059 | Old |
| 0.023348 | 1.513918 | 0.204 | 0.054 | 1 | 15 | Znhit1 | 0.15 | Old |
| 0.023565 | 2.326649 | 0.224 | 0.071 | 1 | 15 | Rgl2 | 0.153 | Old |
| 0.023565 | 2.039604 | 0.224 | 0.071 | 1 | 15 | Ythdf1 | 0.153 | Old |
| 0.023586 | 3.006458 | 0.163 | 0.036 | 1 | 15 | Ddx49 | 0.127 | Old |
| 0.024408 | 1.206418 | 0.347 | 0.161 | 1 | 15 | Hist1h1c | 0.186 | Old |
| 0.024766 | 1.173128 | 0.286 | 0.107 | 1 | 15 | Nmt1 | 0.179 | Old |
| 0.024894 | 1.822142 | 0.224 | 0.071 | 1 | 15 | Faap20 | 0.153 | Old |
| 0.026016 | 2.13927 | 0.163 | 0.036 | 1 | 15 | Tmem184b | 0.127 | Old |
| 0.026288 | 1.984369 | 0.224 | 0.071 | 1 | 15 | Ppil2 | 0.153 | Old |

| | | | | | | | | |
|---|---|---|---|---|---|---|---|---|
| 0.026878 | 1.921194 | 0.245 | 0.089 | 1 | 15 | Slc25a20 | 0.156 | Old |
| 0.02701 | 1.514149 | 0.224 | 0.071 | 1 | 15 | Haghl | 0.153 | Old |
| 0.027165 | 1.138899 | 0.51 | 0.321 | 1 | 15 | Usp9x | 0.189 | Old |
| 0.027178 | 0.880354 | 0.429 | 0.25 | 1 | 15 | Strn3 | 0.179 | Old |
| 0.027754 | 1.5401 | 0.163 | 0.036 | 1 | 15 | Rtn1 | 0.127 | Old |
| 0.028659 | 2.426801 | 0.163 | 0.036 | 1 | 15 | Vamp7 | 0.127 | Old |
| 0.028659 | 2.068241 | 0.163 | 0.036 | 1 | 15 | Fbxl6 | 0.127 | Old |
| 0.029018 | 1.885926 | 0.245 | 0.089 | 1 | 15 | Zfp280d | 0.156 | Old |
| 0.029591 | 1.928193 | 0.163 | 0.036 | 1 | 15 | Dcaf13 | 0.127 | Old |
| 0.029591 | 1.902471 | 0.163 | 0.036 | 1 | 15 | Mrpl41 | 0.127 | Old |
| 0.029591 | 1.454992 | 0.163 | 0.036 | 1 | 15 | Lrba | 0.127 | Old |
| 0.030102 | 3.759145 | 0.122 | 0.018 | 1 | 15 | Foxj2 | 0.104 | Old |
| 0.030102 | 3.547685 | 0.122 | 0.018 | 1 | 15 | Trmt13 | 0.104 | Old |
| 0.030102 | 3.22458 | 0.122 | 0.018 | 1 | 15 | Cradd | 0.104 | Old |
| 0.030818 | 5.180994 | 0.082 | 0 | 1 | 15 | Tmem132a | 0.082 | Old |
| 0.030818 | 5.017968 | 0.082 | 0 | 1 | 15 | Sorl1 | 0.082 | Old |
| 0.030818 | 4.802291 | 0.082 | 0 | 1 | 15 | Tbc1d10c | 0.082 | Old |
| 0.030818 | 4.455651 | 0.082 | 0 | 1 | 15 | Ric8b | 0.082 | Old |
| 0.030818 | 4.450412 | 0.082 | 0 | 1 | 15 | Cdkal1 | 0.082 | Old |
| 0.030818 | 4.440937 | 0.082 | 0 | 1 | 15 | Frat2 | 0.082 | Old |
| 0.030818 | 4.394105 | 0.082 | 0 | 1 | 15 | Erc1 | 0.082 | Old |
| 0.030818 | 4.267573 | 0.082 | 0 | 1 | 15 | Hmgn5 | 0.082 | Old |
| 0.030818 | 4.253471 | 0.082 | 0 | 1 | 15 | Cdc42ep4 | 0.082 | Old |
| 0.030818 | 4.251342 | 0.082 | 0 | 1 | 15 | Kmo | 0.082 | Old |
| 0.030818 | 4.168073 | 0.082 | 0 | 1 | 15 | Nlrx1 | 0.082 | Old |
| 0.030818 | 4.11964 | 0.082 | 0 | 1 | 15 | Etaa1 | 0.082 | Old |
| 0.030818 | 4.076173 | 0.082 | 0 | 1 | 15 | Helb | 0.082 | Old |
| 0.030818 | 4.011792 | 0.082 | 0 | 1 | 15 | Ankrd35 | 0.082 | Old |
| 0.030818 | 4.011639 | 0.082 | 0 | 1 | 15 | Eef2kmt | 0.082 | Old |
| 0.030818 | 3.581139 | 0.082 | 0 | 1 | 15 | Cars | 0.082 | Old |
| 0.030818 | 3.543175 | 0.082 | 0 | 1 | 15 | Scmh1 | 0.082 | Old |
| 0.030818 | 3.460508 | 0.082 | 0 | 1 | 15 | Pgap1 | 0.082 | Old |
| 0.030818 | 3.40732 | 0.082 | 0 | 1 | 15 | Atg7 | 0.082 | Old |
| 0.030818 | 3.244685 | 0.082 | 0 | 1 | 15 | Zfp949 | 0.082 | Old |
| 0.031249 | 2.946676 | 0.122 | 0.018 | 1 | 15 | Aptx | 0.104 | Old |
| 0.031491 | 0.862321 | 0.51 | 0.304 | 1 | 15 | Flt3 | 0.206 | Old |
| 0.031786 | 1.365216 | 0.388 | 0.196 | 1 | 15 | Srp72 | 0.192 | Old |
| 0.032247 | 0.576974 | 0.612 | 0.357 | 1 | 15 | Pdcd6ip | 0.255 | Old |
| 0.032433 | 3.227332 | 0.122 | 0.018 | 1 | 15 | Mrpl55 | 0.104 | Old |
| 0.032433 | 2.668186 | 0.122 | 0.018 | 1 | 15 | Hinfp | 0.104 | Old |
| 0.033075 | 0.597712 | 0.714 | 0.536 | 1 | 15 | Prpf38b | 0.178 | Old |
| 0.033366 | 1.551529 | 0.367 | 0.196 | 1 | 15 | Uba2 | 0.171 | Old |
| 0.033655 | 2.44008 | 0.122 | 0.018 | 1 | 15 | Stam | 0.104 | Old |
| 0.033655 | 2.335318 | 0.122 | 0.018 | 1 | 15 | Zfp142 | 0.104 | Old |
| 0.034469 | 2.538323 | 0.204 | 0.071 | 1 | 15 | Knop1 | 0.133 | Old |

| | | | | | | | | |
|---|---|---|---|---|---|---|---|---|
| 0.034887 | 0.895684 | 0.531 | 0.321 | 1 | 15 | Snrpg | 0.21 | Old |
| 0.034917 | 2.309943 | 0.122 | 0.018 | 1 | 15 | Rfc3 | 0.104 | Old |
| 0.034917 | 1.977551 | 0.122 | 0.018 | 1 | 15 | Rrp7a | 0.104 | Old |
| 0.035509 | 1.01947 | 0.531 | 0.321 | 1 | 15 | Pten | 0.21 | Old |
| 0.036218 | 1.907613 | 0.122 | 0.018 | 1 | 15 | Pramef8 | 0.104 | Old |
| 0.036218 | 1.564233 | 0.122 | 0.018 | 1 | 15 | Asrgl1 | 0.104 | Old |
| 0.037218 | 1.185673 | 0.388 | 0.214 | 1 | 15 | Ankrd13a | 0.174 | Old |
| 0.037243 | 1.015055 | 0.245 | 0.089 | 1 | 15 | Kmt2d | 0.156 | Old |
| 0.037949 | 1.65252 | 0.184 | 0.054 | 1 | 15 | Wrn | 0.13 | Old |
| 0.038093 | 0.764215 | 0.531 | 0.321 | 1 | 15 | Srsf6 | 0.21 | Old |
| 0.039514 | 0.489244 | 0.959 | 0.929 | 1 | 15 | Pfn1 | 0.03 | Old |
| 0.039544 | 0.909567 | 0.714 | 0.536 | 1 | 15 | Jak2 | 0.178 | Old |
| 0.040155 | 2.185367 | 0.184 | 0.054 | 1 | 15 | Atp2c1 | 0.13 | Old |
| 0.040263 | 1.013963 | 0.531 | 0.339 | 1 | 15 | Coq10b | 0.192 | Old |
| 0.04033 | 1.589483 | 0.286 | 0.125 | 1 | 15 | Rpe | 0.161 | Old |
| 0.040655 | 1.142215 | 0.449 | 0.268 | 1 | 15 | H2-M3 | 0.181 | Old |
| 0.04105 | 1.169632 | 0.245 | 0.089 | 1 | 15 | Ankrd13c | 0.156 | Old |
| 0.041505 | 2.041067 | 0.204 | 0.071 | 1 | 15 | Samd8 | 0.133 | Old |
| 0.041547 | 1.002747 | 0.51 | 0.339 | 1 | 15 | Ccni | 0.171 | Old |
| 0.042013 | 1.41485 | 0.265 | 0.107 | 1 | 15 | Diablo | 0.158 | Old |
| 0.042469 | 1.72174 | 0.184 | 0.054 | 1 | 15 | Gbp9 | 0.13 | Old |
| 0.042604 | 1.716137 | 0.204 | 0.071 | 1 | 15 | Ipo7 | 0.133 | Old |
| 0.04347 | 1.125212 | 0.306 | 0.143 | 1 | 15 | Ly75 | 0.163 | Old |
| 0.043653 | 3.258013 | 0.143 | 0.036 | 1 | 15 | Hspa14 | 0.107 | Old |
| 0.043667 | 1.323255 | 0.184 | 0.054 | 1 | 15 | Pibf1 | 0.13 | Old |
| 0.044547 | 2.277534 | 0.286 | 0.143 | 1 | 15 | Stat4 | 0.143 | Old |
| 0.044874 | 1.786285 | 0.204 | 0.071 | 1 | 15 | Atr | 0.133 | Old |
| 0.045003 | 1.185642 | 0.469 | 0.321 | 1 | 15 | Aamp | 0.148 | Old |
| 0.045051 | 2.679923 | 0.143 | 0.036 | 1 | 15 | Tmub2 | 0.107 | Old |
| 0.045181 | 0.921226 | 0.245 | 0.089 | 1 | 15 | Ubac2 | 0.156 | Old |
| 0.046047 | 2.197714 | 0.204 | 0.071 | 1 | 15 | Zfp236 | 0.133 | Old |
| 0.046486 | 2.920247 | 0.143 | 0.036 | 1 | 15 | Cnot9 | 0.107 | Old |
| 0.046486 | 2.346694 | 0.143 | 0.036 | 1 | 15 | Cenpl | 0.107 | Old |
| 0.046528 | 2.015185 | 0.245 | 0.107 | 1 | 15 | Clptm1l | 0.138 | Old |
| 0.046748 | 0.257229 | 0.918 | 0.911 | 1 | 15 | Uqcrh | 0.007 | Old |
| 0.047084 | 0.947029 | 0.265 | 0.107 | 1 | 15 | Zrsr2 | 0.158 | Old |
| 0.047241 | 0.605176 | 0.714 | 0.571 | 1 | 15 | Pcbp1 | 0.143 | Old |
| 0.047435 | 0.912437 | 0.184 | 0.054 | 1 | 15 | Large1 | 0.13 | Old |
| 0.04796 | 3.08333 | 0.143 | 0.036 | 1 | 15 | Mut | 0.107 | Old |
| 0.04796 | 2.574922 | 0.143 | 0.036 | 1 | 15 | Malsu1 | 0.107 | Old |
| 0.048064 | 1.406479 | 0.286 | 0.125 | 1 | 15 | Trappc2l | 0.161 | Old |
| 0.048316 | 1.420389 | 0.388 | 0.232 | 1 | 15 | Hnrnpr | 0.156 | Old |
| 0.048471 | 2.239985 | 0.204 | 0.071 | 1 | 15 | Zfp260 | 0.133 | Old |
| 0.048918 | 1.980462 | 0.306 | 0.161 | 1 | 15 | Sart1 | 0.145 | Old |
| 0.048989 | 0.20848 | 1 | 1 | 1 | 15 | Rpl38 | 0 | Old |

| | | | | | | | | |
|---|---|---|---|---|---|---|---|---|
| 0.049184 | 1.645277 | 0.224 | 0.089 | 1 | 15 | Ankrd40 | 0.135 | Old |
| 0.049217 | 1.261276 | 0.408 | 0.25 | 1 | 15 | Prr14 | 0.158 | Old |
| 0.049473 | 2.593527 | 0.143 | 0.036 | 1 | 15 | Ddx56 | 0.107 | Old |
| 0.049473 | 2.269231 | 0.143 | 0.036 | 1 | 15 | Tmem106c | 0.107 | Old |
| 0.049723 | 1.816005 | 0.204 | 0.071 | 1 | 15 | Fam3a | 0.133 | Old |
| 0.051001 | 1.431442 | 0.204 | 0.071 | 1 | 15 | Slc7a1 | 0.133 | Old |
| 0.051026 | 3.189978 | 0.143 | 0.036 | 1 | 15 | Usp11 | 0.107 | Old |
| 0.051026 | 2.063205 | 0.143 | 0.036 | 1 | 15 | Zbtb11 | 0.107 | Old |
| 0.051026 | 1.882384 | 0.143 | 0.036 | 1 | 15 | Selenoh | 0.107 | Old |
| 0.051502 | 1.134222 | 0.265 | 0.107 | 1 | 15 | Ergic3 | 0.158 | Old |
| 0.051502 | 0.87989 | 0.265 | 0.107 | 1 | 15 | Crebl2 | 0.158 | Old |
| 0.051953 | 0.9728 | 0.408 | 0.25 | 1 | 15 | Mif4gd | 0.158 | Old |
| 0.052619 | 1.934686 | 0.143 | 0.036 | 1 | 15 | Prnp | 0.107 | Old |
| 0.052619 | 1.7184 | 0.143 | 0.036 | 1 | 15 | Cenpa | 0.107 | Old |
| 0.053939 | 0.58582 | 0.347 | 0.179 | 1 | 15 | Il4ra | 0.168 | Old |
| 0.054254 | 1.80837 | 0.143 | 0.036 | 1 | 15 | E130309D0 | 0.107 | Old |
| 0.054254 | 1.635163 | 0.143 | 0.036 | 1 | 15 | C3 | 0.107 | Old |
| 0.055357 | 1.12955 | 0.388 | 0.214 | 1 | 15 | Nfya | 0.174 | Old |
| 0.055412 | 1.397866 | 0.224 | 0.089 | 1 | 15 | Fbxo28 | 0.135 | Old |
| 0.055482 | 0.636171 | 0.469 | 0.286 | 1 | 15 | U2af2 | 0.183 | Old |
| 0.055835 | 1.924411 | 0.245 | 0.107 | 1 | 15 | Ufl1 | 0.138 | Old |
| 0.05593 | 1.589739 | 0.143 | 0.036 | 1 | 15 | Thoc6 | 0.107 | Old |
| 0.05593 | 1.573139 | 0.143 | 0.036 | 1 | 15 | Bahcc1 | 0.107 | Old |
| 0.056733 | 1.451527 | 0.224 | 0.089 | 1 | 15 | Asxl2 | 0.135 | Old |
| 0.057281 | 1.236425 | 0.429 | 0.268 | 1 | 15 | Ccnd1 | 0.161 | Old |
| 0.057281 | 1.107632 | 0.429 | 0.268 | 1 | 15 | Gtf2h5 | 0.161 | Old |
| 0.05765 | 1.896229 | 0.143 | 0.036 | 1 | 15 | Dhx35 | 0.107 | Old |
| 0.05765 | 1.2317 | 0.143 | 0.036 | 1 | 15 | Trpv2 | 0.107 | Old |
| 0.0578 | 2.167761 | 0.163 | 0.054 | 1 | 15 | Tnip1 | 0.109 | Old |
| 0.058081 | 1.284969 | 0.224 | 0.089 | 1 | 15 | Atad1 | 0.135 | Old |
| 0.058218 | 0.871362 | 0.286 | 0.125 | 1 | 15 | Stx8 | 0.161 | Old |
| 0.058566 | 0.920161 | 0.49 | 0.304 | 1 | 15 | Riok3 | 0.186 | Old |
| 0.059262 | 1.396702 | 0.204 | 0.071 | 1 | 15 | Stk38l | 0.133 | Old |
| 0.059413 | 2.272097 | 0.163 | 0.054 | 1 | 15 | Lats2 | 0.109 | Old |
| 0.059413 | 2.124827 | 0.163 | 0.054 | 1 | 15 | Orai2 | 0.109 | Old |
| 0.059839 | 1.437543 | 0.306 | 0.161 | 1 | 15 | Stx12 | 0.145 | Old |
| 0.060188 | 0.973238 | 0.551 | 0.393 | 1 | 15 | Strip2 | 0.158 | Old |
| 0.060642 | 4.060095 | 0.102 | 0.018 | 1 | 15 | Snapc1 | 0.084 | Old |
| 0.060642 | 3.599049 | 0.102 | 0.018 | 1 | 15 | Fastk | 0.084 | Old |
| 0.060642 | 3.403703 | 0.102 | 0.018 | 1 | 15 | Cc2d2a | 0.084 | Old |
| 0.060642 | 3.356923 | 0.102 | 0.018 | 1 | 15 | Nectin2 | 0.084 | Old |
| 0.060642 | 3.045092 | 0.102 | 0.018 | 1 | 15 | Atg9a | 0.084 | Old |
| 0.061062 | 2.408838 | 0.163 | 0.054 | 1 | 15 | Fbxo6 | 0.109 | Old |
| 0.061839 | 0.503265 | 0.857 | 0.696 | 1 | 15 | Ptbp3 | 0.161 | Old |
| 0.062786 | 1.560541 | 0.327 | 0.179 | 1 | 15 | Scarb2 | 0.148 | Old |

| | | | | | | | | |
|---|---|---|---|---|---|---|---|---|
| 0.062867 | 3.058006 | 0.102 | 0.018 | 1 | 15 | Cfap97 | 0.084 | Old |
| 0.06312 | 5.230489 | 0.061 | 0 | 1 | 15 | Mmaa | 0.061 | Old |
| 0.06312 | 5.217317 | 0.061 | 0 | 1 | 15 | Rrnad1 | 0.061 | Old |
| 0.06312 | 5.210729 | 0.061 | 0 | 1 | 15 | Armc7 | 0.061 | Old |
| 0.06312 | 5.124166 | 0.061 | 0 | 1 | 15 | Trpm2 | 0.061 | Old |
| 0.06312 | 4.999853 | 0.061 | 0 | 1 | 15 | Caprin2 | 0.061 | Old |
| 0.06312 | 4.711742 | 0.061 | 0 | 1 | 15 | Invs | 0.061 | Old |
| 0.06312 | 4.617032 | 0.061 | 0 | 1 | 15 | Abcb10 | 0.061 | Old |
| 0.06312 | 4.526175 | 0.061 | 0 | 1 | 15 | Platr25 | 0.061 | Old |
| 0.06312 | 4.430726 | 0.061 | 0 | 1 | 15 | Stard10 | 0.061 | Old |
| 0.06312 | 4.354477 | 0.061 | 0 | 1 | 15 | Mtor | 0.061 | Old |
| 0.06312 | 4.344747 | 0.061 | 0 | 1 | 15 | H2-Ob | 0.061 | Old |
| 0.06312 | 4.291034 | 0.061 | 0 | 1 | 15 | Polr2d | 0.061 | Old |
| 0.06312 | 4.274984 | 0.061 | 0 | 1 | 15 | Ppp3cc | 0.061 | Old |
| 0.06312 | 4.125309 | 0.061 | 0 | 1 | 15 | Nt5dc1 | 0.061 | Old |
| 0.06312 | 4.11504 | 0.061 | 0 | 1 | 15 | Lrrk2 | 0.061 | Old |
| 0.06312 | 4.091402 | 0.061 | 0 | 1 | 15 | Cstf1 | 0.061 | Old |
| 0.06312 | 4.089904 | 0.061 | 0 | 1 | 15 | Gcfc2 | 0.061 | Old |
| 0.06312 | 4.049322 | 0.061 | 0 | 1 | 15 | Tulp3 | 0.061 | Old |
| 0.06312 | 4.029669 | 0.061 | 0 | 1 | 15 | Mrm1 | 0.061 | Old |
| 0.06312 | 4.025549 | 0.061 | 0 | 1 | 15 | Orc5 | 0.061 | Old |
| 0.06312 | 4.005371 | 0.061 | 0 | 1 | 15 | Rbm43 | 0.061 | Old |
| 0.06312 | 4.005303 | 0.061 | 0 | 1 | 15 | Myom1 | 0.061 | Old |
| 0.06312 | 3.993429 | 0.061 | 0 | 1 | 15 | St7l | 0.061 | Old |
| 0.06312 | 3.94989 | 0.061 | 0 | 1 | 15 | Hdac10 | 0.061 | Old |
| 0.06312 | 3.881898 | 0.061 | 0 | 1 | 15 | Wiz | 0.061 | Old |
| 0.06312 | 3.864369 | 0.061 | 0 | 1 | 15 | Gm1673 | 0.061 | Old |
| 0.06312 | 3.859682 | 0.061 | 0 | 1 | 15 | Cacna1f | 0.061 | Old |
| 0.06312 | 3.832208 | 0.061 | 0 | 1 | 15 | Ddx20 | 0.061 | Old |
| 0.06312 | 3.825699 | 0.061 | 0 | 1 | 15 | Nfatc2ip | 0.061 | Old |
| 0.06312 | 3.788644 | 0.061 | 0 | 1 | 15 | Zfp553 | 0.061 | Old |
| 0.06312 | 3.787215 | 0.061 | 0 | 1 | 15 | Exoc6 | 0.061 | Old |
| 0.06312 | 3.773873 | 0.061 | 0 | 1 | 15 | Dock9 | 0.061 | Old |
| 0.06312 | 3.656266 | 0.061 | 0 | 1 | 15 | Samd10 | 0.061 | Old |
| 0.06312 | 3.63613 | 0.061 | 0 | 1 | 15 | Sirpb1c | 0.061 | Old |
| 0.06312 | 3.62174 | 0.061 | 0 | 1 | 15 | Gtpbp3 | 0.061 | Old |
| 0.06312 | 3.595421 | 0.061 | 0 | 1 | 15 | Cbwd1 | 0.061 | Old |
| 0.06312 | 3.589602 | 0.061 | 0 | 1 | 15 | Lpcat4 | 0.061 | Old |
| 0.06312 | 3.544932 | 0.061 | 0 | 1 | 15 | Bcl2l12 | 0.061 | Old |
| 0.06312 | 3.530865 | 0.061 | 0 | 1 | 15 | Tmem209 | 0.061 | Old |
| 0.06312 | 3.510959 | 0.061 | 0 | 1 | 15 | Nudt15 | 0.061 | Old |
| 0.06312 | 3.502383 | 0.061 | 0 | 1 | 15 | Ifi211 | 0.061 | Old |
| 0.06312 | 3.493451 | 0.061 | 0 | 1 | 15 | Hibch | 0.061 | Old |
| 0.06312 | 3.491124 | 0.061 | 0 | 1 | 15 | Plau | 0.061 | Old |
| 0.06312 | 3.482003 | 0.061 | 0 | 1 | 15 | Gm26542 | 0.061 | Old |

| | | | | | | | | |
|---|---|---|---|---|---|---|---|---|
| 0.06312 | 3.434972 | 0.061 | 0 | 1 | 15 | Snupn | 0.061 | Old |
| 0.06312 | 3.408272 | 0.061 | 0 | 1 | 15 | Fblim1 | 0.061 | Old |
| 0.06312 | 3.401475 | 0.061 | 0 | 1 | 15 | S1pr3 | 0.061 | Old |
| 0.06312 | 3.374675 | 0.061 | 0 | 1 | 15 | Slc38a7 | 0.061 | Old |
| 0.06312 | 3.371076 | 0.061 | 0 | 1 | 15 | Camta1 | 0.061 | Old |
| 0.06312 | 3.360624 | 0.061 | 0 | 1 | 15 | Pgm3 | 0.061 | Old |
| 0.06312 | 3.330099 | 0.061 | 0 | 1 | 15 | Acyp1 | 0.061 | Old |
| 0.06312 | 3.274675 | 0.061 | 0 | 1 | 15 | Cenpt | 0.061 | Old |
| 0.06312 | 3.227051 | 0.061 | 0 | 1 | 15 | Tcim | 0.061 | Old |
| 0.06312 | 3.219794 | 0.061 | 0 | 1 | 15 | Camk2a | 0.061 | Old |
| 0.06312 | 3.19984 | 0.061 | 0 | 1 | 15 | Ttc7b | 0.061 | Old |
| 0.06312 | 3.197802 | 0.061 | 0 | 1 | 15 | Trmt10a | 0.061 | Old |
| 0.06312 | 3.197802 | 0.061 | 0 | 1 | 15 | Clmn | 0.061 | Old |
| 0.06312 | 3.167782 | 0.061 | 0 | 1 | 15 | Fbf1 | 0.061 | Old |
| 0.06312 | 3.131005 | 0.061 | 0 | 1 | 15 | Ankzf1 | 0.061 | Old |
| 0.06312 | 3.123865 | 0.061 | 0 | 1 | 15 | Zfp41 | 0.061 | Old |
| 0.06312 | 3.107985 | 0.061 | 0 | 1 | 15 | B3glct | 0.061 | Old |
| 0.06312 | 3.082477 | 0.061 | 0 | 1 | 15 | Cnbd2 | 0.061 | Old |
| 0.06312 | 3.071941 | 0.061 | 0 | 1 | 15 | Thrb | 0.061 | Old |
| 0.06312 | 2.903538 | 0.061 | 0 | 1 | 15 | Gpatch3 | 0.061 | Old |
| 0.06312 | 2.86304 | 0.061 | 0 | 1 | 15 | Eif2b1 | 0.061 | Old |
| 0.06312 | 2.830352 | 0.061 | 0 | 1 | 15 | Rab11fip4 | 0.061 | Old |
| 0.06312 | 2.695586 | 0.061 | 0 | 1 | 15 | Zer1 | 0.061 | Old |
| 0.06312 | 2.561781 | 0.061 | 0 | 1 | 15 | Pfkfb2 | 0.061 | Old |
| 0.06312 | 2.434904 | 0.061 | 0 | 1 | 15 | Gmpr2 | 0.061 | Old |
| 0.063177 | 2.226457 | 0.184 | 0.071 | 1 | 15 | Nabp2 | 0.113 | Old |
| 0.063303 | 1.332765 | 0.347 | 0.232 | 1 | 15 | Rilpl2 | 0.115 | Old |
| 0.063303 | 1.168251 | 0.367 | 0.214 | 1 | 15 | Npepps | 0.153 | Old |
| 0.063571 | 0.490292 | 0.796 | 0.786 | 1 | 15 | Arpc5 | 0.01 | Old |
| 0.064121 | 1.368616 | 0.265 | 0.125 | 1 | 15 | Stt3a | 0.14 | Old |
| 0.064476 | 1.820329 | 0.163 | 0.054 | 1 | 15 | Arl3 | 0.109 | Old |
| 0.065016 | 0.573769 | 0.673 | 0.5 | 1 | 15 | Ndufa3 | 0.173 | Old |
| 0.065159 | 2.64754 | 0.102 | 0.018 | 1 | 15 | Ccpg1 | 0.084 | Old |
| 0.065159 | 2.530589 | 0.102 | 0.018 | 1 | 15 | Wrnip1 | 0.084 | Old |
| 0.065481 | 1.346149 | 0.265 | 0.125 | 1 | 15 | Ift52 | 0.14 | Old |
| 0.066433 | 2.049578 | 0.184 | 0.071 | 1 | 15 | Pam | 0.113 | Old |
| 0.06752 | 2.821669 | 0.102 | 0.018 | 1 | 15 | Tox2 | 0.084 | Old |
| 0.06752 | 2.597749 | 0.102 | 0.018 | 1 | 15 | Atad2 | 0.084 | Old |
| 0.06752 | 2.524031 | 0.102 | 0.018 | 1 | 15 | Nmnat3 | 0.084 | Old |
| 0.06752 | 2.393846 | 0.102 | 0.018 | 1 | 15 | Ggt5 | 0.084 | Old |
| 0.06752 | 2.188764 | 0.102 | 0.018 | 1 | 15 | Alyref2 | 0.084 | Old |
| 0.06752 | 2.101453 | 0.102 | 0.018 | 1 | 15 | Mpg | 0.084 | Old |
| 0.06752 | 1.96663 | 0.102 | 0.018 | 1 | 15 | Zfp809 | 0.084 | Old |
| 0.06752 | 1.77284 | 0.102 | 0.018 | 1 | 15 | Msto1 | 0.084 | Old |
| 0.067854 | 0.958743 | 0.592 | 0.464 | 1 | 15 | Glipr2 | 0.128 | Old |

| | | | | | | | | |
|---|---|---|---|---|---|---|---|---|
| 0.068275 | 1.069502 | 0.224 | 0.089 | 1 | 15 | CAAA0111! | 0.135 | Old |
| 0.068994 | 0.79119 | 0.429 | 0.25 | 1 | 15 | Pfdn2 | 0.179 | Old |
| 0.069379 | 0.392405 | 0.612 | 0.411 | 1 | 15 | Tab2 | 0.201 | Old |
| 0.0696 | 1.030627 | 0.245 | 0.107 | 1 | 15 | Fbxw7 | 0.138 | Old |
| 0.069844 | 0.732434 | 0.224 | 0.089 | 1 | 15 | Chchd3 | 0.135 | Old |
| 0.06989 | 1.658203 | 0.163 | 0.054 | 1 | 15 | Cnot10 | 0.109 | Old |
| 0.06995 | 2.47889 | 0.102 | 0.018 | 1 | 15 | Bcl9 | 0.084 | Old |
| 0.06995 | 2.255272 | 0.102 | 0.018 | 1 | 15 | Zfp787 | 0.084 | Old |
| 0.06995 | 2.16335 | 0.102 | 0.018 | 1 | 15 | Fam160b2 | 0.084 | Old |
| 0.06995 | 2.037553 | 0.102 | 0.018 | 1 | 15 | Zc3hc1 | 0.084 | Old |
| 0.06995 | 1.860275 | 0.102 | 0.018 | 1 | 15 | Tmem161t | 0.084 | Old |
| 0.06995 | 1.843448 | 0.102 | 0.018 | 1 | 15 | Acad8 | 0.084 | Old |
| 0.06995 | 1.82377 | 0.102 | 0.018 | 1 | 15 | 1810032O( | 0.084 | Old |
| 0.06995 | 1.188325 | 0.102 | 0.018 | 1 | 15 | Qsox2 | 0.084 | Old |
| 0.070254 | 1.181452 | 0.286 | 0.143 | 1 | 15 | Acly | 0.143 | Old |
| 0.071165 | 0.977959 | 0.408 | 0.25 | 1 | 15 | Safb2 | 0.158 | Old |
| 0.071612 | 0.707227 | 0.367 | 0.196 | 1 | 15 | mt-Atp8 | 0.171 | Old |
| 0.071776 | 1.610131 | 0.163 | 0.054 | 1 | 15 | Rnf139 | 0.109 | Old |
| 0.072452 | 1.531078 | 0.102 | 0.018 | 1 | 15 | Xylt1 | 0.084 | Old |
| 0.072452 | 0.887539 | 0.102 | 0.018 | 1 | 15 | Rai14 | 0.084 | Old |
| 0.073099 | 1.283586 | 0.286 | 0.143 | 1 | 15 | Osbpl9 | 0.143 | Old |
| 0.073099 | 0.970937 | 0.286 | 0.143 | 1 | 15 | Gpatch8 | 0.143 | Old |
| 0.073703 | 2.195225 | 0.163 | 0.054 | 1 | 15 | Cops7a | 0.109 | Old |
| 0.073886 | 1.057223 | 0.408 | 0.286 | 1 | 15 | Usp25 | 0.122 | Old |
| 0.074235 | 1.163762 | 0.245 | 0.107 | 1 | 15 | Nup85 | 0.138 | Old |
| 0.074385 | 0.888825 | 0.653 | 0.589 | 1 | 15 | Son | 0.064 | Old |
| 0.07443 | 1.397186 | 0.286 | 0.161 | 1 | 15 | Ube2g2 | 0.125 | Old |
| 0.074987 | 0.932526 | 0.449 | 0.286 | 1 | 15 | Aph1a | 0.163 | Old |
| 0.075092 | 0.433196 | 0.776 | 0.732 | 1 | 15 | Itm2c | 0.044 | Old |
| 0.075181 | 2.0085 | 0.184 | 0.071 | 1 | 15 | Rbl2 | 0.113 | Old |
| 0.075464 | 2.079545 | 0.224 | 0.107 | 1 | 15 | Ate1 | 0.117 | Old |
| 0.075672 | 1.633856 | 0.163 | 0.054 | 1 | 15 | Nmd3 | 0.109 | Old |
| 0.075675 | 0.872413 | 0.408 | 0.268 | 1 | 15 | Akt1 | 0.14 | Old |
| 0.076232 | 1.174371 | 0.51 | 0.339 | 1 | 15 | Nsd1 | 0.171 | Old |
| 0.076945 | 1.06956 | 0.408 | 0.268 | 1 | 15 | Alyref | 0.14 | Old |
| 0.077168 | 0.917269 | 0.388 | 0.25 | 1 | 15 | Ino80d | 0.138 | Old |
| 0.07853 | 0.518334 | 0.898 | 0.821 | 1 | 15 | Apobec3 | 0.077 | Old |
| 0.079072 | 1.4353 | 0.286 | 0.143 | 1 | 15 | Pnn | 0.143 | Old |
| 0.079072 | 0.807042 | 0.286 | 0.143 | 1 | 15 | Tcerg1 | 0.143 | Old |
| 0.079147 | 0.372402 | 0.408 | 0.232 | 1 | 15 | Fmnl1 | 0.176 | Old |
| 0.079739 | 1.040268 | 0.163 | 0.054 | 1 | 15 | Phkg2 | 0.109 | Old |
| 0.080805 | 0.677801 | 0.245 | 0.107 | 1 | 15 | Josd1 | 0.138 | Old |
| 0.080865 | 0.724361 | 0.571 | 0.393 | 1 | 15 | Fam60a | 0.178 | Old |
| 0.080867 | 2.577357 | 0.184 | 0.071 | 1 | 15 | Ube3c | 0.113 | Old |
| 0.080867 | 1.679803 | 0.184 | 0.071 | 1 | 15 | Fam50a | 0.113 | Old |

| | | | | | | | | |
|---|---|---|---|---|---|---|---|---|
| 0.080867 | 1.419329 | 0.184 | 0.071 | 1 | 15 | Sh3bp2 | 0.113 | Old |
| 0.080867 | 1.183677 | 0.184 | 0.071 | 1 | 15 | Ebna1bp2 | 0.113 | Old |
| 0.081155 | 0.359788 | 0.306 | 0.143 | 1 | 15 | Zfp524 | 0.163 | Old |
| 0.081589 | 1.564976 | 0.245 | 0.125 | 1 | 15 | Gps1 | 0.12 | Old |
| 0.082204 | 1.130114 | 0.286 | 0.143 | 1 | 15 | Slc35b2 | 0.143 | Old |
| 0.082382 | 0.381666 | 0.796 | 0.821 | 1 | 15 | Ier5 | -0.025 | Old |
| 0.083004 | 1.280434 | 0.306 | 0.161 | 1 | 15 | Cbx1 | 0.145 | Old |
| 0.083004 | 0.981582 | 0.306 | 0.161 | 1 | 15 | Slc4a8 | 0.145 | Old |
| 0.08492 | 0.731847 | 0.49 | 0.321 | 1 | 15 | Dnajc8 | 0.169 | Old |
| 0.085333 | 2.19151 | 0.204 | 0.089 | 1 | 15 | Emc1 | 0.115 | Old |
| 0.085333 | 1.773419 | 0.204 | 0.089 | 1 | 15 | Nsmaf | 0.115 | Old |
| 0.085333 | 1.668977 | 0.204 | 0.089 | 1 | 15 | Pfdn4 | 0.115 | Old |
| 0.086171 | 0.661876 | 0.163 | 0.054 | 1 | 15 | Manbal | 0.109 | Old |
| 0.08622 | 0.365939 | 0.02 | 0.107 | 1 | 15 | Gm6563 | -0.087 | Old |
| 0.08622 | 0.176518 | 0.02 | 0.107 | 1 | 15 | Ralgps2 | -0.087 | Old |
| 0.086802 | 1.139185 | 0.469 | 0.321 | 1 | 15 | Mkrn1 | 0.148 | Old |
| 0.086863 | 0.595635 | 0.857 | 0.75 | 1 | 15 | Arf5 | 0.107 | Old |
| 0.086897 | 2.125639 | 0.184 | 0.071 | 1 | 15 | Ppp2cb | 0.113 | Old |
| 0.087181 | 2.73543 | 0.122 | 0.036 | 1 | 15 | Med31 | 0.086 | Old |
| 0.087192 | 0.424619 | 0.837 | 0.804 | 1 | 15 | Slc25a3 | 0.033 | Old |
| 0.08729 | 0.740512 | 0.449 | 0.321 | 1 | 15 | Nipa2 | 0.128 | Old |
| 0.088321 | 1.10716 | 0.347 | 0.196 | 1 | 15 | Fbxo11 | 0.151 | Old |
| 0.089398 | 0.8593 | 0.306 | 0.161 | 1 | 15 | Cers2 | 0.145 | Old |
| 0.089792 | 2.349068 | 0.122 | 0.036 | 1 | 15 | Papd7 | 0.086 | Old |
| 0.089792 | 2.304596 | 0.122 | 0.036 | 1 | 15 | Sirt1 | 0.086 | Old |
| 0.090301 | 1.272478 | 0.469 | 0.339 | 1 | 15 | Ddi2 | 0.13 | Old |
| 0.090688 | 0.54056 | 0.163 | 0.054 | 1 | 15 | Akt3 | 0.109 | Old |
| 0.091101 | 0.583571 | 0.796 | 0.804 | 1 | 15 | Cnn2 | -0.008 | Old |
| 0.091115 | 1.158641 | 0.184 | 0.071 | 1 | 15 | Cebpzos | 0.113 | Old |
| 0.09157 | 1.6407 | 0.265 | 0.143 | 1 | 15 | Maea | 0.122 | Old |
| 0.092466 | 3.018254 | 0.122 | 0.036 | 1 | 15 | Pgam5 | 0.086 | Old |
| 0.092466 | 2.63361 | 0.122 | 0.036 | 1 | 15 | Rnf167 | 0.086 | Old |
| 0.092466 | 2.07172 | 0.122 | 0.036 | 1 | 15 | Zmym4 | 0.086 | Old |
| 0.092466 | 1.717986 | 0.122 | 0.036 | 1 | 15 | Cep295 | 0.086 | Old |
| 0.092756 | 0.831017 | 0.367 | 0.232 | 1 | 15 | Zc3h13 | 0.135 | Old |
| 0.093016 | 0.593634 | 0.163 | 0.054 | 1 | 15 | Tmem230 | 0.109 | Old |
| 0.093284 | 0.964053 | 0.184 | 0.071 | 1 | 15 | Rai1 | 0.113 | Old |
| 0.095203 | 2.129964 | 0.122 | 0.036 | 1 | 15 | Slc35b4 | 0.086 | Old |
| 0.095203 | 1.673344 | 0.122 | 0.036 | 1 | 15 | Scly | 0.086 | Old |
| 0.095205 | 0.774168 | 0.449 | 0.286 | 1 | 15 | Csnk1d | 0.163 | Old |
| 0.095362 | 1.172852 | 0.204 | 0.089 | 1 | 15 | Gps2 | 0.115 | Old |
| 0.095362 | 1.137116 | 0.204 | 0.089 | 1 | 15 | Nus1 | 0.115 | Old |
| 0.095555 | 0.17948 | 1 | 1 | 1 | 15 | Rpl41 | 0 | Old |
| 0.097427 | 0.918303 | 0.224 | 0.107 | 1 | 15 | 2210016F1 | 0.117 | Old |
| 0.097476 | 1.525587 | 0.204 | 0.089 | 1 | 15 | Usp37 | 0.115 | Old |

| | | | | | | | | |
|---|---|---|---|---|---|---|---|---|
| 0.097545 | 0.628485 | 0.612 | 0.464 | 1 | 15 | B4galnt1 | 0.148 | Old |
| 0.097748 | 0.67801 | 0.184 | 0.071 | 1 | 15 | Pik3cg | 0.113 | Old |
| 0.097748 | 0.330545 | 0.061 | 0.179 | 1 | 15 | Stard9 | -0.118 | Old |
| 0.098004 | 2.142652 | 0.122 | 0.036 | 1 | 15 | Dnajc4 | 0.086 | Old |
| 0.098004 | 1.778133 | 0.122 | 0.036 | 1 | 15 | Gm16685 | 0.086 | Old |
| 0.098004 | 1.764255 | 0.122 | 0.036 | 1 | 15 | Crbn | 0.086 | Old |
| 0.098004 | 1.762752 | 0.122 | 0.036 | 1 | 15 | Clcn7 | 0.086 | Old |
| 0.098004 | 1.660326 | 0.122 | 0.036 | 1 | 15 | Msr1 | 0.086 | Old |
| 0.098004 | 1.173721 | 0.122 | 0.036 | 1 | 15 | Cnnm4 | 0.086 | Old |
| 0.098773 | 1.594865 | 0.286 | 0.161 | 1 | 15 | Paf1 | 0.125 | Old |
| 0.098773 | 1.31795 | 0.286 | 0.161 | 1 | 15 | Ncaph2 | 0.125 | Old |
| 0.098862 | 1.472663 | 0.265 | 0.143 | 1 | 15 | Bbip1 | 0.122 | Old |
| 0.09897 | 0.602141 | 0.408 | 0.25 | 1 | 15 | Casc3 | 0.158 | Old |
| 0.099795 | 1.548012 | 0.245 | 0.125 | 1 | 15 | Elk4 | 0.12 | Old |
| 0.099947 | 0.619192 | 0.265 | 0.125 | 1 | 15 | Qdpr | 0.14 | Old |
| 0.100036 | 0.735715 | 0.449 | 0.304 | 1 | 15 | Bax | 0.145 | Old |
| 0.100044 | 0.776484 | 0.184 | 0.071 | 1 | 15 | Ap3m1 | 0.113 | Old |
| 0.100561 | 0.943849 | 0.408 | 0.25 | 1 | 15 | Ythdf3 | 0.158 | Old |
| 0.100871 | 2.352413 | 0.122 | 0.036 | 1 | 15 | Zc3h14 | 0.086 | Old |
| 0.100871 | 1.863982 | 0.122 | 0.036 | 1 | 15 | Gm19951 | 0.086 | Old |
| 0.100871 | 1.698456 | 0.122 | 0.036 | 1 | 15 | Zfp467 | 0.086 | Old |
| 0.100871 | 1.595466 | 0.122 | 0.036 | 1 | 15 | Tvp23b | 0.086 | Old |
| 0.100871 | 1.376664 | 0.122 | 0.036 | 1 | 15 | Med19 | 0.086 | Old |
| 0.101713 | 0.222639 | 0.122 | 0.268 | 1 | 15 | Susd6 | -0.146 | Old |
| 0.101818 | 1.491772 | 0.204 | 0.089 | 1 | 15 | Idh3b | 0.115 | Old |
| 0.101818 | 1.222704 | 0.204 | 0.089 | 1 | 15 | Trappc1 | 0.115 | Old |
| 0.102792 | 0.997487 | 0.367 | 0.214 | 1 | 15 | Ppm1b | 0.153 | Old |
| 0.103141 | 1.680851 | 0.306 | 0.179 | 1 | 15 | Chtop | 0.127 | Old |
| 0.103537 | 0.706145 | 0.429 | 0.268 | 1 | 15 | Mdm4 | 0.161 | Old |
| 0.103649 | 1.343335 | 0.224 | 0.107 | 1 | 15 | Eps15l1 | 0.117 | Old |
| 0.103649 | 1.057887 | 0.224 | 0.107 | 1 | 15 | Isy1 | 0.117 | Old |
| 0.103884 | 1.051151 | 0.347 | 0.214 | 1 | 15 | Kdm2b | 0.133 | Old |
| 0.104046 | 0.210333 | 0.204 | 0.089 | 1 | 15 | Fundc1 | 0.115 | Old |
| 0.104106 | 0.778697 | 0.367 | 0.232 | 1 | 15 | Nsf | 0.135 | Old |
| 0.104515 | 1.136234 | 0.327 | 0.179 | 1 | 15 | Arf6 | 0.148 | Old |
| 0.104629 | 1.11302 | 0.265 | 0.143 | 1 | 15 | Dr1 | 0.122 | Old |
| 0.105101 | 1.879923 | 0.143 | 0.054 | 1 | 15 | Dck | 0.089 | Old |
| 0.106313 | 0.660563 | 0.204 | 0.089 | 1 | 15 | Cetn2 | 0.115 | Old |
| 0.106641 | 0.498041 | 0.837 | 0.696 | 1 | 15 | Lcp1 | 0.141 | Old |
| 0.106806 | 1.413317 | 0.122 | 0.036 | 1 | 15 | Gpr146 | 0.086 | Old |
| 0.106806 | 0.929345 | 0.122 | 0.036 | 1 | 15 | Dcaf10 | 0.086 | Old |
| 0.107739 | 0.39735 | 0.755 | 0.625 | 1 | 15 | Myh9 | 0.13 | Old |
| 0.107835 | 2.339021 | 0.143 | 0.054 | 1 | 15 | Irak2 | 0.089 | Old |
| 0.107835 | 2.232139 | 0.143 | 0.054 | 1 | 15 | Mtx1 | 0.089 | Old |
| 0.108618 | 0.851881 | 0.204 | 0.089 | 1 | 15 | Mcm3 | 0.115 | Old |

| | | | | | | | | |
|---|---|---|---|---|---|---|---|---|
| 0.108828 | 0.322388 | 0.898 | 0.893 | 1 | 15 | Bhlhe40 | 0.005 | Old |
| 0.108941 | 1.048442 | 0.469 | 0.321 | 1 | 15 | Psmb6 | 0.148 | Old |
| 0.109496 | 0.749477 | 0.551 | 0.393 | 1 | 15 | Hnrnpdl | 0.158 | Old |
| 0.109875 | 0.835879 | 0.122 | 0.036 | 1 | 15 | Tspyl2 | 0.086 | Old |
| 0.109887 | 1.932389 | 0.163 | 0.071 | 1 | 15 | Mier3 | 0.092 | Old |
| 0.110137 | 1.062023 | 0.286 | 0.161 | 1 | 15 | Ccdc6 | 0.125 | Old |
| 0.110184 | 1.555533 | 0.224 | 0.107 | 1 | 15 | Cspp1 | 0.117 | Old |
| 0.110958 | 0.837968 | 0.429 | 0.304 | 1 | 15 | Fgfr1op2 | 0.125 | Old |
| 0.110963 | 0.598155 | 0.204 | 0.089 | 1 | 15 | Cc2d1b | 0.115 | Old |
| 0.112211 | 1.042971 | 0.367 | 0.25 | 1 | 15 | Cirbp | 0.117 | Old |
| 0.112433 | 1.280118 | 0.224 | 0.107 | 1 | 15 | Slf2 | 0.117 | Old |
| 0.113688 | 2.438391 | 0.184 | 0.089 | 1 | 15 | Rfwd3 | 0.095 | Old |
| 0.114032 | 1.25816 | 0.327 | 0.196 | 1 | 15 | Nsmce4a | 0.131 | Old |
| 0.114086 | 0.414101 | 0.388 | 0.25 | 1 | 15 | Pias1 | 0.138 | Old |
| 0.114223 | 0.049028 | 0.122 | 0.25 | 1 | 15 | Nemf | -0.128 | Old |
| 0.114704 | 1.3153 | 0.347 | 0.214 | 1 | 15 | Mreg | 0.133 | Old |
| 0.115629 | 0.715103 | 0.592 | 0.446 | 1 | 15 | Map3k1 | 0.146 | Old |
| 0.116222 | 0.931108 | 0.122 | 0.036 | 1 | 15 | Vps52 | 0.086 | Old |
| 0.116514 | 0.57766 | 0.755 | 0.679 | 1 | 15 | Clk1 | 0.076 | Old |
| 0.116578 | 1.692061 | 0.245 | 0.125 | 1 | 15 | Ubald1 | 0.12 | Old |
| 0.116578 | 0.644113 | 0.245 | 0.125 | 1 | 15 | Gngt2 | 0.12 | Old |
| 0.117021 | 0.471712 | 0.673 | 0.518 | 1 | 15 | Kdm6b | 0.155 | Old |
| 0.11827 | 0.593385 | 0.286 | 0.161 | 1 | 15 | Eapp | 0.125 | Old |
| 0.118288 | 0.613759 | 0.592 | 0.482 | 1 | 15 | Snx20 | 0.11 | Old |
| 0.118379 | 0.821486 | 0.429 | 0.286 | 1 | 15 | Cdk17 | 0.143 | Old |
| 0.118399 | 1.244089 | 0.367 | 0.232 | 1 | 15 | Zdhhc3 | 0.135 | Old |
| 0.118704 | 0.769587 | 0.51 | 0.393 | 1 | 15 | Scamp2 | 0.117 | Old |
| 0.119031 | 0.290629 | 0.306 | 0.161 | 1 | 15 | Yme1l1 | 0.145 | Old |
| 0.119053 | 0.406748 | 0.878 | 0.893 | 1 | 15 | Sub1 | -0.015 | Old |
| 0.119126 | 1.234419 | 0.265 | 0.143 | 1 | 15 | 2310035C2 | 0.122 | Old |
| 0.119339 | 1.697954 | 0.143 | 0.054 | 1 | 15 | Rab3gap1 | 0.089 | Old |
| 0.119339 | 1.493831 | 0.143 | 0.054 | 1 | 15 | Tmem18 | 0.089 | Old |
| 0.119517 | 1.319383 | 0.367 | 0.25 | 1 | 15 | Paip2 | 0.117 | Old |
| 0.119765 | 0.476426 | 0.347 | 0.196 | 1 | 15 | Ddx50 | 0.151 | Old |
| 0.120615 | 1.562124 | 0.163 | 0.071 | 1 | 15 | Trmt1l | 0.092 | Old |
| 0.121345 | 3.560626 | 0.082 | 0.018 | 1 | 15 | Arhgap21 | 0.064 | Old |
| 0.121345 | 3.166829 | 0.082 | 0.018 | 1 | 15 | Gpd1l | 0.064 | Old |
| 0.121345 | 3.002127 | 0.082 | 0.018 | 1 | 15 | Pgm1 | 0.064 | Old |
| 0.121345 | 2.940059 | 0.082 | 0.018 | 1 | 15 | Alg14 | 0.064 | Old |
| 0.121345 | 2.90929 | 0.082 | 0.018 | 1 | 15 | Fam129b | 0.064 | Old |
| 0.121345 | 2.778616 | 0.082 | 0.018 | 1 | 15 | Ppp1r16b | 0.064 | Old |
| 0.121345 | 2.744903 | 0.082 | 0.018 | 1 | 15 | Gnpnat1 | 0.064 | Old |
| 0.121345 | 2.53615 | 0.082 | 0.018 | 1 | 15 | Hace1 | 0.064 | Old |
| 0.121345 | 2.383662 | 0.082 | 0.018 | 1 | 15 | Uck1 | 0.064 | Old |
| 0.122361 | 1.808368 | 0.143 | 0.054 | 1 | 15 | Cabin1 | 0.089 | Old |

| | | | | | | | | |
|---|---|---|---|---|---|---|---|---|
| 0.122361 | 1.454816 | 0.143 | 0.054 | 1 | 15 | Phax | 0.089 | Old |
| 0.122509 | 1.027255 | 0.286 | 0.161 | 1 | 15 | Fermt3 | 0.125 | Old |
| 0.122635 | 0.970856 | 0.306 | 0.179 | 1 | 15 | Prmt1 | 0.127 | Old |
| 0.123182 | 0.339845 | 0.959 | 1 | 1 | 15 | Btg1 | -0.041 | Old |
| 0.123308 | 1.096861 | 0.367 | 0.25 | 1 | 15 | Prelid1 | 0.117 | Old |
| 0.123416 | 1.061353 | 0.245 | 0.125 | 1 | 15 | Kif1c | 0.12 | Old |
| 0.123422 | 1.89838 | 0.163 | 0.071 | 1 | 15 | Zfp397 | 0.092 | Old |
| 0.124744 | 0.365171 | 0.857 | 0.821 | 1 | 15 | Wnk1 | 0.036 | Old |
| 0.125238 | 0.959711 | 0.367 | 0.25 | 1 | 15 | Sun2 | 0.117 | Old |
| 0.125441 | 1.576922 | 0.143 | 0.054 | 1 | 15 | Zfp329 | 0.089 | Old |
| 0.125441 | 0.873175 | 0.143 | 0.054 | 1 | 15 | Abl2 | 0.089 | Old |
| 0.125585 | 3.638299 | 0.082 | 0.018 | 1 | 15 | Dnajc30 | 0.064 | Old |
| 0.125585 | 3.444136 | 0.082 | 0.018 | 1 | 15 | Nfyb | 0.064 | Old |
| 0.125585 | 3.29968 | 0.082 | 0.018 | 1 | 15 | Mlx | 0.064 | Old |
| 0.125585 | 2.886104 | 0.082 | 0.018 | 1 | 15 | Zbtb45 | 0.064 | Old |
| 0.125585 | 2.652716 | 0.082 | 0.018 | 1 | 15 | Nosip | 0.064 | Old |
| 0.125585 | 2.525512 | 0.082 | 0.018 | 1 | 15 | AU019823 | 0.064 | Old |
| 0.125585 | 2.521446 | 0.082 | 0.018 | 1 | 15 | Maoa | 0.064 | Old |
| 0.125585 | 2.481459 | 0.082 | 0.018 | 1 | 15 | Lxn | 0.064 | Old |
| 0.125585 | 2.410608 | 0.082 | 0.018 | 1 | 15 | Nrg1 | 0.064 | Old |
| 0.125585 | 2.348492 | 0.082 | 0.018 | 1 | 15 | Arhgef9 | 0.064 | Old |
| 0.125585 | 2.298951 | 0.082 | 0.018 | 1 | 15 | Crispld2 | 0.064 | Old |
| 0.125585 | 2.27783 | 0.082 | 0.018 | 1 | 15 | Rad51b | 0.064 | Old |
| 0.125585 | 2.263383 | 0.082 | 0.018 | 1 | 15 | Ccdc22 | 0.064 | Old |
| 0.125585 | 2.247168 | 0.082 | 0.018 | 1 | 15 | Ppa2 | 0.064 | Old |
| 0.125585 | 2.109933 | 0.082 | 0.018 | 1 | 15 | Kcnn1 | 0.064 | Old |
| 0.125585 | 2.092505 | 0.082 | 0.018 | 1 | 15 | Gtf3c5 | 0.064 | Old |
| 0.125881 | 0.325003 | 0.204 | 0.089 | 1 | 15 | Zfp277 | 0.115 | Old |
| 0.126279 | 2.264549 | 0.163 | 0.071 | 1 | 15 | Anapc1 | 0.092 | Old |
| 0.126279 | 1.698585 | 0.163 | 0.071 | 1 | 15 | Mettl9 | 0.092 | Old |
| 0.126859 | 0.895899 | 0.306 | 0.179 | 1 | 15 | Arrdc4 | 0.127 | Old |
| 0.126867 | 1.048262 | 0.286 | 0.161 | 1 | 15 | Esrra | 0.125 | Old |
| 0.128582 | 1.351974 | 0.143 | 0.054 | 1 | 15 | Kat6b | 0.089 | Old |
| 0.128582 | 1.273953 | 0.143 | 0.054 | 1 | 15 | Edrf1 | 0.089 | Old |
| 0.128582 | 1.231137 | 0.143 | 0.054 | 1 | 15 | Rexo1 | 0.089 | Old |
| 0.129187 | 1.845063 | 0.163 | 0.071 | 1 | 15 | Prelid3b | 0.092 | Old |
| 0.129187 | 1.8035 | 0.163 | 0.071 | 1 | 15 | Tmem222 | 0.092 | Old |
| 0.12994 | 2.838321 | 0.082 | 0.018 | 1 | 15 | Mlst8 | 0.064 | Old |
| 0.12994 | 2.683059 | 0.082 | 0.018 | 1 | 15 | Cdk16 | 0.064 | Old |
| 0.12994 | 2.545728 | 0.082 | 0.018 | 1 | 15 | Galnt10 | 0.064 | Old |
| 0.12994 | 2.529037 | 0.082 | 0.018 | 1 | 15 | Ikzf4 | 0.064 | Old |
| 0.12994 | 2.517574 | 0.082 | 0.018 | 1 | 15 | Unkl | 0.064 | Old |
| 0.12994 | 2.457696 | 0.082 | 0.018 | 1 | 15 | Ring1 | 0.064 | Old |
| 0.12994 | 2.319102 | 0.082 | 0.018 | 1 | 15 | Hist1h4i | 0.064 | Old |
| 0.12994 | 2.2131 | 0.082 | 0.018 | 1 | 15 | Lyrm9 | 0.064 | Old |

| | | | | | | | | |
|---|---|---|---|---|---|---|---|---|
| 0.12994 | 2.154092 | 0.082 | 0.018 | 1 | 15 | Ptcd3 | 0.064 | Old |
| 0.12994 | 2.141446 | 0.082 | 0.018 | 1 | 15 | Zfp655 | 0.064 | Old |
| 0.12994 | 2.098636 | 0.082 | 0.018 | 1 | 15 | Supt3 | 0.064 | Old |
| 0.12994 | 2.055367 | 0.082 | 0.018 | 1 | 15 | Cass4 | 0.064 | Old |
| 0.12994 | 2.05265 | 0.082 | 0.018 | 1 | 15 | Ppp2r3a | 0.064 | Old |
| 0.12994 | 1.954208 | 0.082 | 0.018 | 1 | 15 | Iqsec1 | 0.064 | Old |
| 0.12994 | 1.943395 | 0.082 | 0.018 | 1 | 15 | Coq2 | 0.064 | Old |
| 0.12994 | 1.934429 | 0.082 | 0.018 | 1 | 15 | Fuz | 0.064 | Old |
| 0.12994 | 1.909026 | 0.082 | 0.018 | 1 | 15 | Ercc4 | 0.064 | Old |
| 0.12994 | 1.902521 | 0.082 | 0.018 | 1 | 15 | Mfsd13a | 0.064 | Old |
| 0.12994 | 1.890192 | 0.082 | 0.018 | 1 | 15 | Mllt11 | 0.064 | Old |
| 0.12994 | 1.82414 | 0.082 | 0.018 | 1 | 15 | Kif13a | 0.064 | Old |
| 0.12994 | 1.810621 | 0.082 | 0.018 | 1 | 15 | Top3b | 0.064 | Old |
| 0.12994 | 1.674578 | 0.082 | 0.018 | 1 | 15 | 0610009B2 | 0.064 | Old |
| 0.130169 | 1.056488 | 0.327 | 0.196 | 1 | 15 | Maz | 0.131 | Old |
| 0.130335 | 0.560468 | 0.959 | 0.929 | 1 | 15 | Etv3 | 0.03 | Old |
| 0.131784 | 1.800256 | 0.143 | 0.054 | 1 | 15 | Mir17hg | 0.089 | Old |
| 0.131784 | 1.393411 | 0.143 | 0.054 | 1 | 15 | Fam46c | 0.089 | Old |
| 0.131784 | 0.738469 | 0.143 | 0.054 | 1 | 15 | Sertad3 | 0.089 | Old |
| 0.132147 | 1.710315 | 0.163 | 0.071 | 1 | 15 | Skiv2l2 | 0.092 | Old |
| 0.132184 | 5.431255 | 0.041 | 0 | 1 | 15 | Ldlrad3 | 0.041 | Old |
| 0.132184 | 5.266346 | 0.041 | 0 | 1 | 15 | Asb2 | 0.041 | Old |
| 0.132184 | 5.116008 | 0.041 | 0 | 1 | 15 | Arfip2 | 0.041 | Old |
| 0.132184 | 5.088097 | 0.041 | 0 | 1 | 15 | Mllt3 | 0.041 | Old |
| 0.132184 | 5.069747 | 0.041 | 0 | 1 | 15 | Upp2 | 0.041 | Old |
| 0.132184 | 4.966151 | 0.041 | 0 | 1 | 15 | Zfp839 | 0.041 | Old |
| 0.132184 | 4.939421 | 0.041 | 0 | 1 | 15 | Chil3 | 0.041 | Old |
| 0.132184 | 4.918916 | 0.041 | 0 | 1 | 15 | Fxn | 0.041 | Old |
| 0.132184 | 4.896316 | 0.041 | 0 | 1 | 15 | Neurl4 | 0.041 | Old |
| 0.132184 | 4.889796 | 0.041 | 0 | 1 | 15 | Synj2 | 0.041 | Old |
| 0.132184 | 4.830457 | 0.041 | 0 | 1 | 15 | Odf2l | 0.041 | Old |
| 0.132184 | 4.802978 | 0.041 | 0 | 1 | 15 | Ttc21b | 0.041 | Old |
| 0.132184 | 4.747616 | 0.041 | 0 | 1 | 15 | Wdfy1 | 0.041 | Old |
| 0.132184 | 4.68612 | 0.041 | 0 | 1 | 15 | Msantd2 | 0.041 | Old |
| 0.132184 | 4.568724 | 0.041 | 0 | 1 | 15 | Ppp1r3f | 0.041 | Old |
| 0.132184 | 4.469299 | 0.041 | 0 | 1 | 15 | Zc3h12d | 0.041 | Old |
| 0.132184 | 4.453127 | 0.041 | 0 | 1 | 15 | Casc1 | 0.041 | Old |
| 0.132184 | 4.451123 | 0.041 | 0 | 1 | 15 | Igkc | 0.041 | Old |
| 0.132184 | 4.42498 | 0.041 | 0 | 1 | 15 | Zfp580 | 0.041 | Old |
| 0.132184 | 4.42324 | 0.041 | 0 | 1 | 15 | Ugt1a7c | 0.041 | Old |
| 0.132184 | 4.303351 | 0.041 | 0 | 1 | 15 | 2900026A0 | 0.041 | Old |
| 0.132184 | 4.181687 | 0.041 | 0 | 1 | 15 | Ntn5 | 0.041 | Old |
| 0.132184 | 4.178741 | 0.041 | 0 | 1 | 15 | Cad | 0.041 | Old |
| 0.132184 | 4.170323 | 0.041 | 0 | 1 | 15 | Sntb1 | 0.041 | Old |
| 0.132184 | 4.149898 | 0.041 | 0 | 1 | 15 | Mtx3 | 0.041 | Old |

| | | | | | | | | |
|---|---|---|---|---|---|---|---|---|
| 0.132184 | 3.88025 | 0.041 | 0 | 1 | 15 | Gm11696 | 0.041 | Old |
| 0.132184 | 3.872305 | 0.041 | 0 | 1 | 15 | Zfp7 | 0.041 | Old |
| 0.132184 | 3.867146 | 0.041 | 0 | 1 | 15 | Lancl2 | 0.041 | Old |
| 0.132184 | 3.853854 | 0.041 | 0 | 1 | 15 | Tfb1m | 0.041 | Old |
| 0.132184 | 3.852878 | 0.041 | 0 | 1 | 15 | Nudt8.1 | 0.041 | Old |
| 0.132184 | 3.845075 | 0.041 | 0 | 1 | 15 | 6030468B1 | 0.041 | Old |
| 0.132184 | 3.836916 | 0.041 | 0 | 1 | 15 | Gm11131 | 0.041 | Old |
| 0.132184 | 3.821875 | 0.041 | 0 | 1 | 15 | S100a16 | 0.041 | Old |
| 0.132184 | 3.810712 | 0.041 | 0 | 1 | 15 | Sestd1 | 0.041 | Old |
| 0.132184 | 3.773071 | 0.041 | 0 | 1 | 15 | Gm15890 | 0.041 | Old |
| 0.132184 | 3.768918 | 0.041 | 0 | 1 | 15 | H2-T24 | 0.041 | Old |
| 0.132184 | 3.76848 | 0.041 | 0 | 1 | 15 | Fkbpl | 0.041 | Old |
| 0.132184 | 3.749653 | 0.041 | 0 | 1 | 15 | Ttn | 0.041 | Old |
| 0.132184 | 3.744631 | 0.041 | 0 | 1 | 15 | Mvb12b | 0.041 | Old |
| 0.132184 | 3.744086 | 0.041 | 0 | 1 | 15 | Gprc5b | 0.041 | Old |
| 0.132184 | 3.739994 | 0.041 | 0 | 1 | 15 | Tmem39b | 0.041 | Old |
| 0.132184 | 3.707436 | 0.041 | 0 | 1 | 15 | Atp1b1 | 0.041 | Old |
| 0.132184 | 3.706664 | 0.041 | 0 | 1 | 15 | Fam120aos | 0.041 | Old |
| 0.132184 | 3.686521 | 0.041 | 0 | 1 | 15 | Ypel1 | 0.041 | Old |
| 0.132184 | 3.669104 | 0.041 | 0 | 1 | 15 | Adamts10 | 0.041 | Old |
| 0.132184 | 3.659063 | 0.041 | 0 | 1 | 15 | Jcad | 0.041 | Old |
| 0.132184 | 3.63818 | 0.041 | 0 | 1 | 15 | A530017D2 | 0.041 | Old |
| 0.132184 | 3.637482 | 0.041 | 0 | 1 | 15 | Ffar2 | 0.041 | Old |
| 0.132184 | 3.583134 | 0.041 | 0 | 1 | 15 | Rab26os | 0.041 | Old |
| 0.132184 | 3.574504 | 0.041 | 0 | 1 | 15 | 5033406O0 | 0.041 | Old |
| 0.132184 | 3.568027 | 0.041 | 0 | 1 | 15 | Pdss2 | 0.041 | Old |
| 0.132184 | 3.560829 | 0.041 | 0 | 1 | 15 | Blzf1 | 0.041 | Old |
| 0.132184 | 3.548928 | 0.041 | 0 | 1 | 15 | Zfp456 | 0.041 | Old |
| 0.132184 | 3.530985 | 0.041 | 0 | 1 | 15 | 1700008J0 | 0.041 | Old |
| 0.132184 | 3.525877 | 0.041 | 0 | 1 | 15 | Cenph | 0.041 | Old |
| 0.132184 | 3.524189 | 0.041 | 0 | 1 | 15 | Glis2 | 0.041 | Old |
| 0.132184 | 3.517979 | 0.041 | 0 | 1 | 15 | Ncald | 0.041 | Old |
| 0.132184 | 3.498511 | 0.041 | 0 | 1 | 15 | Gm44067 | 0.041 | Old |
| 0.132184 | 3.481016 | 0.041 | 0 | 1 | 15 | Trmu | 0.041 | Old |
| 0.132184 | 3.468379 | 0.041 | 0 | 1 | 15 | Cep78 | 0.041 | Old |
| 0.132184 | 3.460195 | 0.041 | 0 | 1 | 15 | Gm43597 | 0.041 | Old |
| 0.132184 | 3.458912 | 0.041 | 0 | 1 | 15 | U2af1l4 | 0.041 | Old |
| 0.132184 | 3.449716 | 0.041 | 0 | 1 | 15 | Spata7 | 0.041 | Old |
| 0.132184 | 3.436279 | 0.041 | 0 | 1 | 15 | Tbl3 | 0.041 | Old |
| 0.132184 | 3.424155 | 0.041 | 0 | 1 | 15 | Pglyrp1 | 0.041 | Old |
| 0.132184 | 3.397297 | 0.041 | 0 | 1 | 15 | Ercc6 | 0.041 | Old |
| 0.132184 | 3.387268 | 0.041 | 0 | 1 | 15 | Zgpat | 0.041 | Old |
| 0.132184 | 3.386865 | 0.041 | 0 | 1 | 15 | Nup43 | 0.041 | Old |
| 0.132184 | 3.385918 | 0.041 | 0 | 1 | 15 | Gm42722 | 0.041 | Old |
| 0.132184 | 3.366456 | 0.041 | 0 | 1 | 15 | Tcta | 0.041 | Old |

| | | | | | | | | |
|---|---|---|---|---|---|---|---|---|
| 0.132184 | 3.363091 | 0.041 | 0 | 1 | 15 | Cmpk2 | 0.041 | Old |
| 0.132184 | 3.348721 | 0.041 | 0 | 1 | 15 | Epn2 | 0.041 | Old |
| 0.132184 | 3.339964 | 0.041 | 0 | 1 | 15 | Gm48505 | 0.041 | Old |
| 0.132184 | 3.33492 | 0.041 | 0 | 1 | 15 | Tgfb2 | 0.041 | Old |
| 0.132184 | 3.33235 | 0.041 | 0 | 1 | 15 | Cerkl | 0.041 | Old |
| 0.132184 | 3.329063 | 0.041 | 0 | 1 | 15 | Gpr55 | 0.041 | Old |
| 0.132184 | 3.309898 | 0.041 | 0 | 1 | 15 | Rubcnl | 0.041 | Old |
| 0.132184 | 3.297454 | 0.041 | 0 | 1 | 15 | Lzts2 | 0.041 | Old |
| 0.132184 | 3.294183 | 0.041 | 0 | 1 | 15 | Usp28 | 0.041 | Old |
| 0.132184 | 3.276947 | 0.041 | 0 | 1 | 15 | Mus81 | 0.041 | Old |
| 0.132184 | 3.272848 | 0.041 | 0 | 1 | 15 | Tjp3 | 0.041 | Old |
| 0.132184 | 3.258486 | 0.041 | 0 | 1 | 15 | Magi2 | 0.041 | Old |
| 0.132184 | 3.242688 | 0.041 | 0 | 1 | 15 | Vldlr | 0.041 | Old |
| 0.132184 | 3.22042 | 0.041 | 0 | 1 | 15 | Map4k2 | 0.041 | Old |
| 0.132184 | 3.216015 | 0.041 | 0 | 1 | 15 | Sag | 0.041 | Old |
| 0.132184 | 3.216015 | 0.041 | 0 | 1 | 15 | S1pr2 | 0.041 | Old |
| 0.132184 | 3.216015 | 0.041 | 0 | 1 | 15 | Abcd4 | 0.041 | Old |
| 0.132184 | 3.214937 | 0.041 | 0 | 1 | 15 | Gm7694 | 0.041 | Old |
| 0.132184 | 3.209162 | 0.041 | 0 | 1 | 15 | Tmem51os | 0.041 | Old |
| 0.132184 | 3.198369 | 0.041 | 0 | 1 | 15 | Ttc12 | 0.041 | Old |
| 0.132184 | 3.17902 | 0.041 | 0 | 1 | 15 | Il27ra | 0.041 | Old |
| 0.132184 | 3.17902 | 0.041 | 0 | 1 | 15 | Nudt7 | 0.041 | Old |
| 0.132184 | 3.17838 | 0.041 | 0 | 1 | 15 | Rragd | 0.041 | Old |
| 0.132184 | 3.177358 | 0.041 | 0 | 1 | 15 | Iqce | 0.041 | Old |
| 0.132184 | 3.17718 | 0.041 | 0 | 1 | 15 | Acad11 | 0.041 | Old |
| 0.132184 | 3.151543 | 0.041 | 0 | 1 | 15 | Ip6k2 | 0.041 | Old |
| 0.132184 | 3.147674 | 0.041 | 0 | 1 | 15 | Gstt3 | 0.041 | Old |
| 0.132184 | 3.133268 | 0.041 | 0 | 1 | 15 | Dus2 | 0.041 | Old |
| 0.132184 | 3.121948 | 0.041 | 0 | 1 | 15 | Efcab2 | 0.041 | Old |
| 0.132184 | 3.113653 | 0.041 | 0 | 1 | 15 | Ace | 0.041 | Old |
| 0.132184 | 3.110793 | 0.041 | 0 | 1 | 15 | Hmga1 | 0.041 | Old |
| 0.132184 | 3.098385 | 0.041 | 0 | 1 | 15 | Cpsf4 | 0.041 | Old |
| 0.132184 | 3.09289 | 0.041 | 0 | 1 | 15 | Atp6v0c | 0.041 | Old |
| 0.132184 | 3.089043 | 0.041 | 0 | 1 | 15 | Atad5 | 0.041 | Old |
| 0.132184 | 3.081477 | 0.041 | 0 | 1 | 15 | Dhodh | 0.041 | Old |
| 0.132184 | 3.079979 | 0.041 | 0 | 1 | 15 | Arhgap27 | 0.041 | Old |
| 0.132184 | 3.060894 | 0.041 | 0 | 1 | 15 | Thbd | 0.041 | Old |
| 0.132184 | 3.059064 | 0.041 | 0 | 1 | 15 | Rad54l | 0.041 | Old |
| 0.132184 | 3.053163 | 0.041 | 0 | 1 | 15 | Tm9sf1 | 0.041 | Old |
| 0.132184 | 3.046112 | 0.041 | 0 | 1 | 15 | Irak1bp1 | 0.041 | Old |
| 0.132184 | 3.045296 | 0.041 | 0 | 1 | 15 | Melk | 0.041 | Old |
| 0.132184 | 3.043173 | 0.041 | 0 | 1 | 15 | Flad1 | 0.041 | Old |
| 0.132184 | 3.043173 | 0.041 | 0 | 1 | 15 | Tmem246 | 0.041 | Old |
| 0.132184 | 3.042202 | 0.041 | 0 | 1 | 15 | Wwox | 0.041 | Old |
| 0.132184 | 3.03571 | 0.041 | 0 | 1 | 15 | Abca2 | 0.041 | Old |

| | | | | | | | | |
|---|---|---|---|---|---|---|---|---|
| 0.132184 | 3.03571 | 0.041 | 0 | 1 | 15 | Cep131 | 0.041 | Old |
| 0.132184 | 3.030191 | 0.041 | 0 | 1 | 15 | Neil1 | 0.041 | Old |
| 0.132184 | 3.029221 | 0.041 | 0 | 1 | 15 | Alkbh4 | 0.041 | Old |
| 0.132184 | 3.024653 | 0.041 | 0 | 1 | 15 | Zfp438 | 0.041 | Old |
| 0.132184 | 3.012188 | 0.041 | 0 | 1 | 15 | Plpp5 | 0.041 | Old |
| 0.132184 | 3.010249 | 0.041 | 0 | 1 | 15 | Gja1 | 0.041 | Old |
| 0.132184 | 3.000948 | 0.041 | 0 | 1 | 15 | Krt7 | 0.041 | Old |
| 0.132184 | 2.991337 | 0.041 | 0 | 1 | 15 | Lrrc49 | 0.041 | Old |
| 0.132184 | 2.988424 | 0.041 | 0 | 1 | 15 | Tmem268 | 0.041 | Old |
| 0.132184 | 2.97507 | 0.041 | 0 | 1 | 15 | Ccdc51 | 0.041 | Old |
| 0.132184 | 2.972754 | 0.041 | 0 | 1 | 15 | Hmgxb4 | 0.041 | Old |
| 0.132184 | 2.958579 | 0.041 | 0 | 1 | 15 | Shpk | 0.041 | Old |
| 0.132184 | 2.955624 | 0.041 | 0 | 1 | 15 | Primpol | 0.041 | Old |
| 0.132184 | 2.944312 | 0.041 | 0 | 1 | 15 | Ly6a | 0.041 | Old |
| 0.132184 | 2.933067 | 0.041 | 0 | 1 | 15 | Blm | 0.041 | Old |
| 0.132184 | 2.933067 | 0.041 | 0 | 1 | 15 | Pde8a | 0.041 | Old |
| 0.132184 | 2.931454 | 0.041 | 0 | 1 | 15 | Dcun1d3 | 0.041 | Old |
| 0.132184 | 2.910918 | 0.041 | 0 | 1 | 15 | Dhcr7 | 0.041 | Old |
| 0.132184 | 2.910918 | 0.041 | 0 | 1 | 15 | Znrd1as | 0.041 | Old |
| 0.132184 | 2.907476 | 0.041 | 0 | 1 | 15 | Gm10612 | 0.041 | Old |
| 0.132184 | 2.891502 | 0.041 | 0 | 1 | 15 | Esam | 0.041 | Old |
| 0.132184 | 2.885462 | 0.041 | 0 | 1 | 15 | Srr | 0.041 | Old |
| 0.132184 | 2.864613 | 0.041 | 0 | 1 | 15 | Rrn3 | 0.041 | Old |
| 0.132184 | 2.854682 | 0.041 | 0 | 1 | 15 | Ftx | 0.041 | Old |
| 0.132184 | 2.837858 | 0.041 | 0 | 1 | 15 | Rtn4ip1 | 0.041 | Old |
| 0.132184 | 2.837858 | 0.041 | 0 | 1 | 15 | Bbs9 | 0.041 | Old |
| 0.132184 | 2.837858 | 0.041 | 0 | 1 | 15 | B230217C1 | 0.041 | Old |
| 0.132184 | 2.837858 | 0.041 | 0 | 1 | 15 | Fasn | 0.041 | Old |
| 0.132184 | 2.83719 | 0.041 | 0 | 1 | 15 | Fam169b | 0.041 | Old |
| 0.132184 | 2.836749 | 0.041 | 0 | 1 | 15 | Qrsl1 | 0.041 | Old |
| 0.132184 | 2.829191 | 0.041 | 0 | 1 | 15 | Xpc | 0.041 | Old |
| 0.132184 | 2.822688 | 0.041 | 0 | 1 | 15 | 2810405F1 | 0.041 | Old |
| 0.132184 | 2.822688 | 0.041 | 0 | 1 | 15 | Htr7 | 0.041 | Old |
| 0.132184 | 2.818655 | 0.041 | 0 | 1 | 15 | Lysmd1 | 0.041 | Old |
| 0.132184 | 2.805374 | 0.041 | 0 | 1 | 15 | Golim4 | 0.041 | Old |
| 0.132184 | 2.799796 | 0.041 | 0 | 1 | 15 | Kctd1 | 0.041 | Old |
| 0.132184 | 2.793043 | 0.041 | 0 | 1 | 15 | Slc16a13 | 0.041 | Old |
| 0.132184 | 2.793043 | 0.041 | 0 | 1 | 15 | Trmt61a | 0.041 | Old |
| 0.132184 | 2.763392 | 0.041 | 0 | 1 | 15 | Als2 | 0.041 | Old |
| 0.132184 | 2.749269 | 0.041 | 0 | 1 | 15 | Zfp53 | 0.041 | Old |
| 0.132184 | 2.739001 | 0.041 | 0 | 1 | 15 | Icam2 | 0.041 | Old |
| 0.132184 | 2.732218 | 0.041 | 0 | 1 | 15 | Klhl8 | 0.041 | Old |
| 0.132184 | 2.732218 | 0.041 | 0 | 1 | 15 | Fadd | 0.041 | Old |
| 0.132184 | 2.725407 | 0.041 | 0 | 1 | 15 | Kat2a | 0.041 | Old |
| 0.132184 | 2.724473 | 0.041 | 0 | 1 | 15 | Aldoc | 0.041 | Old |

| | | | | | | | | |
|---|---|---|---|---|---|---|---|---|
| 0.132184 | 2.72177 | 0.041 | 0 | 1 | 15 | Zfp652os | 0.041 | Old |
| 0.132184 | 2.701299 | 0.041 | 0 | 1 | 15 | Gstz1 | 0.041 | Old |
| 0.132184 | 2.69713 | 0.041 | 0 | 1 | 15 | Tmem81 | 0.041 | Old |
| 0.132184 | 2.69713 | 0.041 | 0 | 1 | 15 | Fsd1l | 0.041 | Old |
| 0.132184 | 2.69713 | 0.041 | 0 | 1 | 15 | Kcnj2 | 0.041 | Old |
| 0.132184 | 2.671254 | 0.041 | 0 | 1 | 15 | Arhgef10 | 0.041 | Old |
| 0.132184 | 2.671254 | 0.041 | 0 | 1 | 15 | Kcnh3 | 0.041 | Old |
| 0.132184 | 2.658285 | 0.041 | 0 | 1 | 15 | Peg10 | 0.041 | Old |
| 0.132184 | 2.622763 | 0.041 | 0 | 1 | 15 | Zc3h8 | 0.041 | Old |
| 0.132184 | 2.598173 | 0.041 | 0 | 1 | 15 | Cd209e | 0.041 | Old |
| 0.132184 | 2.598173 | 0.041 | 0 | 1 | 15 | Gpt | 0.041 | Old |
| 0.132184 | 2.597962 | 0.041 | 0 | 1 | 15 | 4930570N1 | 0.041 | Old |
| 0.132184 | 2.556134 | 0.041 | 0 | 1 | 15 | Ccdc17 | 0.041 | Old |
| 0.132184 | 2.553509 | 0.041 | 0 | 1 | 15 | Rap1gap | 0.041 | Old |
| 0.132184 | 2.499868 | 0.041 | 0 | 1 | 15 | Sirpb1a | 0.041 | Old |
| 0.132184 | 2.499868 | 0.041 | 0 | 1 | 15 | Gm45606 | 0.041 | Old |
| 0.132184 | 2.491895 | 0.041 | 0 | 1 | 15 | Ptar1 | 0.041 | Old |
| 0.132184 | 2.487619 | 0.041 | 0 | 1 | 15 | Mterf4 | 0.041 | Old |
| 0.132184 | 2.483809 | 0.041 | 0 | 1 | 15 | Ifi206 | 0.041 | Old |
| 0.132184 | 2.483809 | 0.041 | 0 | 1 | 15 | Oas2 | 0.041 | Old |
| 0.132184 | 2.483809 | 0.041 | 0 | 1 | 15 | Sgf29 | 0.041 | Old |
| 0.132184 | 2.483522 | 0.041 | 0 | 1 | 15 | Colq | 0.041 | Old |
| 0.132184 | 2.483522 | 0.041 | 0 | 1 | 15 | Sh3bgrl2 | 0.041 | Old |
| 0.132184 | 2.470472 | 0.041 | 0 | 1 | 15 | Ccdc171 | 0.041 | Old |
| 0.132184 | 2.46351 | 0.041 | 0 | 1 | 15 | Edem3 | 0.041 | Old |
| 0.132184 | 2.46351 | 0.041 | 0 | 1 | 15 | Snx24 | 0.041 | Old |
| 0.132184 | 2.452583 | 0.041 | 0 | 1 | 15 | Bicd1 | 0.041 | Old |
| 0.132184 | 2.448522 | 0.041 | 0 | 1 | 15 | Cd4 | 0.041 | Old |
| 0.132184 | 2.448522 | 0.041 | 0 | 1 | 15 | Ifit1 | 0.041 | Old |
| 0.132184 | 2.422782 | 0.041 | 0 | 1 | 15 | Rfesd | 0.041 | Old |
| 0.132184 | 2.422782 | 0.041 | 0 | 1 | 15 | Twsg1 | 0.041 | Old |
| 0.132184 | 2.363076 | 0.041 | 0 | 1 | 15 | 2700049A0 | 0.041 | Old |
| 0.132184 | 2.348612 | 0.041 | 0 | 1 | 15 | Paqr3 | 0.041 | Old |
| 0.132184 | 2.339908 | 0.041 | 0 | 1 | 15 | Gtf2ird1 | 0.041 | Old |
| 0.132184 | 2.336631 | 0.041 | 0 | 1 | 15 | Gm9733 | 0.041 | Old |
| 0.132184 | 2.336631 | 0.041 | 0 | 1 | 15 | Limk2 | 0.041 | Old |
| 0.132184 | 2.32933 | 0.041 | 0 | 1 | 15 | Gm10184 | 0.041 | Old |
| 0.132184 | 2.296598 | 0.041 | 0 | 1 | 15 | Osbpl1a | 0.041 | Old |
| 0.132184 | 2.27213 | 0.041 | 0 | 1 | 15 | Wrap73 | 0.041 | Old |
| 0.132184 | 2.27213 | 0.041 | 0 | 1 | 15 | Nup107 | 0.041 | Old |
| 0.132184 | 2.258432 | 0.041 | 0 | 1 | 15 | Gxylt2 | 0.041 | Old |
| 0.132184 | 2.247237 | 0.041 | 0 | 1 | 15 | Ankmy2 | 0.041 | Old |
| 0.132184 | 2.202309 | 0.041 | 0 | 1 | 15 | Bcl2a1c | 0.041 | Old |
| 0.132184 | 2.186827 | 0.041 | 0 | 1 | 15 | Zbed4 | 0.041 | Old |
| 0.132184 | 2.176174 | 0.041 | 0 | 1 | 15 | Zfp874b | 0.041 | Old |

| | | | | | | | | |
|---|---|---|---|---|---|---|---|---|
| 0.132184 | 2.144241 | 0.041 | 0 | 1 | 15 | Zfp280c | 0.041 | Old |
| 0.132184 | 2.144241 | 0.041 | 0 | 1 | 15 | Lmf1 | 0.041 | Old |
| 0.132184 | 2.07218 | 0.041 | 0 | 1 | 15 | Hist2h2be | 0.041 | Old |
| 0.132184 | 2.061398 | 0.041 | 0 | 1 | 15 | Cep72 | 0.041 | Old |
| 0.132184 | 2.050539 | 0.041 | 0 | 1 | 15 | Slfn1 | 0.041 | Old |
| 0.132184 | 2.026354 | 0.041 | 0 | 1 | 15 | Hdhd5 | 0.041 | Old |
| 0.132184 | 1.910512 | 0.041 | 0 | 1 | 15 | Ift122 | 0.041 | Old |
| 0.132184 | 1.910512 | 0.041 | 0 | 1 | 15 | Gm31718 | 0.041 | Old |
| 0.132184 | 1.898445 | 0.041 | 0 | 1 | 15 | 9430060I0 | 0.041 | Old |
| 0.132184 | 1.886282 | 0.041 | 0 | 1 | 15 | Ppcs | 0.041 | Old |
| 0.132184 | 1.886282 | 0.041 | 0 | 1 | 15 | Spata2l | 0.041 | Old |
| 0.132184 | 1.792909 | 0.041 | 0 | 1 | 15 | Rnf113a1 | 0.041 | Old |
| 0.132184 | 1.792909 | 0.041 | 0 | 1 | 15 | E2f2 | 0.041 | Old |
| 0.132184 | 1.792909 | 0.041 | 0 | 1 | 15 | Prkd2 | 0.041 | Old |
| 0.132184 | 1.792909 | 0.041 | 0 | 1 | 15 | Mlycd | 0.041 | Old |
| 0.132184 | 1.792909 | 0.041 | 0 | 1 | 15 | Cenpo | 0.041 | Old |
| 0.132184 | 1.657322 | 0.041 | 0 | 1 | 15 | 1110002L0 | 0.041 | Old |
| 0.132482 | 1.006402 | 0.449 | 0.339 | 1 | 15 | Mapk1ip1l | 0.11 | Old |
| 0.13273 | 0.589639 | 0.571 | 0.429 | 1 | 15 | Dapp1 | 0.142 | Old |
| 0.13441 | 2.442477 | 0.082 | 0.018 | 1 | 15 | Pnpla7 | 0.064 | Old |
| 0.13441 | 2.234107 | 0.082 | 0.018 | 1 | 15 | Ormdl1 | 0.064 | Old |
| 0.13441 | 1.936088 | 0.082 | 0.018 | 1 | 15 | C330018D2 | 0.064 | Old |
| 0.13441 | 1.849353 | 0.082 | 0.018 | 1 | 15 | Fam20c | 0.064 | Old |
| 0.13441 | 1.713268 | 0.082 | 0.018 | 1 | 15 | Ern1 | 0.064 | Old |
| 0.13441 | 1.628454 | 0.082 | 0.018 | 1 | 15 | Zfp579 | 0.064 | Old |
| 0.13441 | 1.516312 | 0.082 | 0.018 | 1 | 15 | Zfp995 | 0.064 | Old |
| 0.13441 | 1.465803 | 0.082 | 0.018 | 1 | 15 | Pcyox1l | 0.064 | Old |
| 0.13441 | 1.452971 | 0.082 | 0.018 | 1 | 15 | Tfam | 0.064 | Old |
| 0.13441 | 1.428878 | 0.082 | 0.018 | 1 | 15 | Zfp429 | 0.064 | Old |
| 0.13441 | 1.410165 | 0.082 | 0.018 | 1 | 15 | Lrrc57 | 0.064 | Old |
| 0.13441 | 1.299487 | 0.082 | 0.018 | 1 | 15 | Mcm3ap | 0.064 | Old |
| 0.134523 | 0.316833 | 0.163 | 0.304 | 1 | 15 | Psme4 | -0.141 | Old |
| 0.135044 | 1.197626 | 0.347 | 0.232 | 1 | 15 | Rer1 | 0.115 | Old |
| 0.135044 | 1.151623 | 0.347 | 0.232 | 1 | 15 | Alcam | 0.115 | Old |
| 0.135047 | 1.049212 | 0.143 | 0.054 | 1 | 15 | Hdac3 | 0.089 | Old |
| 0.135047 | 1.01597 | 0.143 | 0.054 | 1 | 15 | Zfp384 | 0.089 | Old |
| 0.135151 | 1.162985 | 0.265 | 0.143 | 1 | 15 | Ppp1r11 | 0.122 | Old |
| 0.135151 | 0.3254 | 0.265 | 0.143 | 1 | 15 | Gnb4 | 0.122 | Old |
| 0.13516 | 1.571694 | 0.163 | 0.071 | 1 | 15 | Slc25a51 | 0.092 | Old |
| 0.135775 | 1.665273 | 0.224 | 0.125 | 1 | 15 | Gabpb2 | 0.099 | Old |
| 0.135775 | 1.289266 | 0.224 | 0.125 | 1 | 15 | Capn2 | 0.099 | Old |
| 0.137482 | 1.262208 | 0.265 | 0.161 | 1 | 15 | Nsmce3 | 0.104 | Old |
| 0.137812 | 1.544076 | 0.184 | 0.089 | 1 | 15 | Kdm5c | 0.095 | Old |
| 0.138876 | 0.601267 | 0.327 | 0.196 | 1 | 15 | Abca7 | 0.131 | Old |
| 0.138911 | 0.787768 | 0.388 | 0.268 | 1 | 15 | Col27a1 | 0.12 | Old |

| | | | | | | | | |
|---|---|---|---|---|---|---|---|---|
| 0.138999 | 1.603778 | 0.082 | 0.018 | 1 | 15 | Urgcp | 0.064 | Old |
| 0.138999 | 1.440271 | 0.082 | 0.018 | 1 | 15 | Dgkh | 0.064 | Old |
| 0.138999 | 1.227914 | 0.082 | 0.018 | 1 | 15 | C5ar2 | 0.064 | Old |
| 0.138999 | 0.834884 | 0.082 | 0.018 | 1 | 15 | Pde4d | 0.064 | Old |
| 0.138999 | 0.756417 | 0.082 | 0.018 | 1 | 15 | Hmces | 0.064 | Old |
| 0.138999 | 0.753341 | 0.082 | 0.018 | 1 | 15 | Fen1 | 0.064 | Old |
| 0.138999 | 0.673817 | 0.082 | 0.018 | 1 | 15 | Gm26853 | 0.064 | Old |
| 0.139889 | 1.472346 | 0.265 | 0.161 | 1 | 15 | Dnase1l1 | 0.104 | Old |
| 0.140339 | 0.819706 | 0.449 | 0.321 | 1 | 15 | Rpn2 | 0.128 | Old |
| 0.141343 | 1.225308 | 0.163 | 0.071 | 1 | 15 | Cdc14a | 0.092 | Old |
| 0.141343 | 1.192866 | 0.163 | 0.071 | 1 | 15 | Pcnt | 0.092 | Old |
| 0.141536 | 1.15356 | 0.367 | 0.25 | 1 | 15 | Ttc14 | 0.117 | Old |
| 0.14154 | 0.864454 | 0.408 | 0.268 | 1 | 15 | Cyth4 | 0.14 | Old |
| 0.141923 | 0.759881 | 0.51 | 0.393 | 1 | 15 | AC149090. | 0.117 | Old |
| 0.143183 | 0.759829 | 0.245 | 0.125 | 1 | 15 | Myof | 0.12 | Old |
| 0.143279 | 1.768271 | 0.204 | 0.107 | 1 | 15 | Dnaaf3 | 0.097 | Old |
| 0.143674 | 1.563168 | 0.184 | 0.089 | 1 | 15 | Parp1 | 0.095 | Old |
| 0.143674 | 1.335751 | 0.184 | 0.089 | 1 | 15 | Ncoa6 | 0.095 | Old |
| 0.143674 | 1.214279 | 0.184 | 0.089 | 1 | 15 | Tcof1 | 0.095 | Old |
| 0.143891 | 0.331573 | 0.694 | 0.607 | 1 | 15 | Lrrk1 | 0.087 | Old |
| 0.143933 | 0.292761 | 0.98 | 0.946 | 1 | 15 | H2afz | 0.034 | Old |
| 0.144026 | 0.272792 | 0.02 | 0.089 | 1 | 15 | Gpr137 | -0.069 | Old |
| 0.145211 | 0.644461 | 0.143 | 0.054 | 1 | 15 | Ulk2 | 0.089 | Old |
| 0.146676 | 1.571532 | 0.184 | 0.089 | 1 | 15 | Amdhd2 | 0.095 | Old |
| 0.147487 | 0.955894 | 0.327 | 0.214 | 1 | 15 | Impdh2 | 0.113 | Old |
| 0.147583 | 0.961685 | 0.265 | 0.143 | 1 | 15 | Pafah1b3 | 0.122 | Old |
| 0.148598 | 0.536455 | 0.673 | 0.607 | 1 | 15 | Tuba1b | 0.066 | Old |
| 0.148726 | 0.675109 | 0.143 | 0.054 | 1 | 15 | Scaper | 0.089 | Old |
| 0.148726 | 0.461821 | 0.143 | 0.054 | 1 | 15 | Dctn5 | 0.089 | Old |
| 0.149077 | 1.456831 | 0.224 | 0.125 | 1 | 15 | Zfp292 | 0.099 | Old |
| 0.149726 | 1.594804 | 0.184 | 0.089 | 1 | 15 | Slc38a10 | 0.095 | Old |
| 0.149917 | 0.468906 | 0.837 | 0.804 | 1 | 15 | Cox7a2l | 0.033 | Old |
| 0.14997 | 0.350203 | 0.388 | 0.25 | 1 | 15 | Tnpo3 | 0.138 | Old |
| 0.150136 | 0.930638 | 0.571 | 0.446 | 1 | 15 | Tpm1 | 0.125 | Old |
| 0.150203 | 0.731211 | 0.347 | 0.214 | 1 | 15 | Kdm6a | 0.133 | Old |
| 0.150403 | 0.848187 | 0.347 | 0.232 | 1 | 15 | Rwdd1 | 0.115 | Old |
| 0.150485 | 0.794664 | 0.286 | 0.161 | 1 | 15 | Acbd6 | 0.125 | Old |
| 0.150681 | 0.346817 | 0.224 | 0.107 | 1 | 15 | Tor1a | 0.117 | Old |
| 0.151023 | 1.68286 | 0.163 | 0.071 | 1 | 15 | Cd300c2 | 0.092 | Old |
| 0.151023 | 1.197807 | 0.163 | 0.071 | 1 | 15 | Map3k5 | 0.092 | Old |
| 0.151023 | 1.041807 | 0.163 | 0.071 | 1 | 15 | Fdps | 0.092 | Old |
| 0.151023 | 1.010806 | 0.163 | 0.071 | 1 | 15 | Ccdc28b | 0.092 | Old |
| 0.151283 | 1.069942 | 0.51 | 0.357 | 1 | 15 | Ankrd12 | 0.153 | Old |
| 0.151852 | 1.502234 | 0.224 | 0.125 | 1 | 15 | Slc48a1 | 0.099 | Old |
| 0.151852 | 1.328039 | 0.224 | 0.125 | 1 | 15 | Kpna1 | 0.099 | Old |

| | | | | | | | | |
|---|---|---|---|---|---|---|---|---|
| 0.151894 | 1.905458 | 0.204 | 0.107 | 1 | 15 | Osbpl11 | 0.097 | Old |
| 0.151894 | 0.519513 | 0.204 | 0.107 | 1 | 15 | Vti1a | 0.097 | Old |
| 0.151894 | 0.481862 | 0.204 | 0.107 | 1 | 15 | Shoc2 | 0.097 | Old |
| 0.152512 | 0.712328 | 0.469 | 0.357 | 1 | 15 | Chmp2a | 0.112 | Old |
| 0.152823 | 1.159016 | 0.184 | 0.089 | 1 | 15 | Braf | 0.095 | Old |
| 0.152857 | 1.295303 | 0.245 | 0.143 | 1 | 15 | H2-DMb2 | 0.102 | Old |
| 0.15302 | 0.925278 | 0.286 | 0.161 | 1 | 15 | Chuk | 0.125 | Old |
| 0.153333 | 1.256984 | 0.286 | 0.179 | 1 | 15 | Casp4 | 0.107 | Old |
| 0.15436 | 1.400641 | 0.163 | 0.071 | 1 | 15 | Dis3l2 | 0.092 | Old |
| 0.15436 | 1.204354 | 0.163 | 0.071 | 1 | 15 | Zbtb21 | 0.092 | Old |
| 0.15436 | 0.20397 | 0.163 | 0.071 | 1 | 15 | Zpr1 | 0.092 | Old |
| 0.154501 | 0.383284 | 0.592 | 0.518 | 1 | 15 | Picalm | 0.074 | Old |
| 0.155017 | 1.315369 | 0.265 | 0.161 | 1 | 15 | Got2 | 0.104 | Old |
| 0.15597 | 1.501869 | 0.184 | 0.089 | 1 | 15 | Xpo4 | 0.095 | Old |
| 0.15597 | 0.780148 | 0.184 | 0.089 | 1 | 15 | Sec23ip | 0.095 | Old |
| 0.156017 | 2.72617 | 0.102 | 0.036 | 1 | 15 | Trip4 | 0.066 | Old |
| 0.156387 | 0.400395 | 0.796 | 0.768 | 1 | 15 | Pik3r1 | 0.028 | Old |
| 0.156421 | 0.607917 | 0.49 | 0.375 | 1 | 15 | Ap2s1 | 0.115 | Old |
| 0.156586 | 0.486277 | 0.898 | 0.768 | 1 | 15 | Pmaip1 | 0.13 | Old |
| 0.158187 | 0.637953 | 0.286 | 0.161 | 1 | 15 | Washc1 | 0.125 | Old |
| 0.159538 | 0.873073 | 0.245 | 0.125 | 1 | 15 | Chfr | 0.12 | Old |
| 0.159683 | 0.770868 | 0.571 | 0.5 | 1 | 15 | Tmem131l | 0.071 | Old |
| 0.160396 | 3.270355 | 0.102 | 0.036 | 1 | 15 | Lmbr1l | 0.066 | Old |
| 0.160396 | 3.238453 | 0.102 | 0.036 | 1 | 15 | Nsun6 | 0.066 | Old |
| 0.160396 | 3.043966 | 0.102 | 0.036 | 1 | 15 | 1700096K1 | 0.066 | Old |
| 0.160396 | 2.689926 | 0.102 | 0.036 | 1 | 15 | Evi2 | 0.066 | Old |
| 0.160396 | 2.055654 | 0.102 | 0.036 | 1 | 15 | Rpia | 0.066 | Old |
| 0.160633 | 1.172431 | 0.306 | 0.196 | 1 | 15 | Mrpl38 | 0.11 | Old |
| 0.161017 | 1.342358 | 0.286 | 0.179 | 1 | 15 | Rabgef1 | 0.107 | Old |
| 0.161097 | 1.217176 | 0.245 | 0.143 | 1 | 15 | Ctcf | 0.102 | Old |
| 0.161574 | 0.705549 | 0.429 | 0.304 | 1 | 15 | Kif21b | 0.125 | Old |
| 0.162392 | 0.41414 | 0.224 | 0.107 | 1 | 15 | Puf60 | 0.117 | Old |
| 0.162409 | 1.229961 | 0.184 | 0.089 | 1 | 15 | Uhrf1bp1l | 0.095 | Old |
| 0.162409 | 0.863316 | 0.184 | 0.089 | 1 | 15 | Sdccag8 | 0.095 | Old |
| 0.16335 | 1.153871 | 0.224 | 0.125 | 1 | 15 | Zkscan3 | 0.099 | Old |
| 0.163917 | 1.450706 | 0.245 | 0.143 | 1 | 15 | Rnf181 | 0.102 | Old |
| 0.164319 | 0.894763 | 0.327 | 0.214 | 1 | 15 | Slc9a3r1 | 0.113 | Old |
| 0.164706 | 1.110848 | 0.163 | 0.071 | 1 | 15 | Tmed4 | 0.092 | Old |
| 0.164706 | 0.826104 | 0.163 | 0.071 | 1 | 15 | Upf3a | 0.092 | Old |
| 0.164867 | 2.224767 | 0.102 | 0.036 | 1 | 15 | Usb1 | 0.066 | Old |
| 0.164979 | 0.583396 | 0.429 | 0.286 | 1 | 15 | Usp34 | 0.143 | Old |
| 0.165703 | 0.692037 | 0.184 | 0.089 | 1 | 15 | Ccz1 | 0.095 | Old |
| 0.16575 | 1.28066 | 0.306 | 0.196 | 1 | 15 | Maff | 0.11 | Old |
| 0.166774 | 1.115615 | 0.245 | 0.143 | 1 | 15 | Ilf2 | 0.102 | Old |
| 0.166774 | 0.999559 | 0.245 | 0.143 | 1 | 15 | Rbm12 | 0.102 | Old |

| | | | | | | | | |
|---|---|---|---|---|---|---|---|---|
| 0.166837 | 0.845143 | 0.327 | 0.214 | 1 | 15 | Dcp2 | 0.113 | Old |
| 0.167121 | 1.685248 | 0.204 | 0.107 | 1 | 15 | Cx3cr1 | 0.097 | Old |
| 0.167239 | 0.666318 | 0.592 | 0.5 | 1 | 15 | Cox17 | 0.092 | Old |
| 0.168262 | 0.93244 | 0.51 | 0.446 | 1 | 15 | Marveld1 | 0.064 | Old |
| 0.168268 | 0.40024 | 0.163 | 0.071 | 1 | 15 | Ube4a | 0.092 | Old |
| 0.168353 | 0.816364 | 0.306 | 0.196 | 1 | 15 | Zfp91 | 0.11 | Old |
| 0.169047 | 1.390816 | 0.184 | 0.089 | 1 | 15 | Pak1ip1 | 0.095 | Old |
| 0.169047 | 0.458233 | 0.184 | 0.089 | 1 | 15 | 2610037D( | 0.095 | Old |
| 0.169432 | 2.024113 | 0.102 | 0.036 | 1 | 15 | Tbc1d17 | 0.066 | Old |
| 0.169432 | 1.745482 | 0.102 | 0.036 | 1 | 15 | Cish | 0.066 | Old |
| 0.169432 | 1.587373 | 0.102 | 0.036 | 1 | 15 | Map2k5 | 0.066 | Old |
| 0.169669 | 1.146729 | 0.245 | 0.143 | 1 | 15 | Casp8 | 0.102 | Old |
| 0.169773 | 1.032945 | 0.449 | 0.339 | 1 | 15 | Srsf11 | 0.11 | Old |
| 0.170197 | 0.9564 | 0.429 | 0.339 | 1 | 15 | Jmjd1c | 0.09 | Old |
| 0.171887 | 0.837085 | 0.163 | 0.071 | 1 | 15 | Ero1l | 0.092 | Old |
| 0.171887 | 0.547169 | 0.163 | 0.071 | 1 | 15 | Cxxc1 | 0.092 | Old |
| 0.172395 | 1.434929 | 0.224 | 0.125 | 1 | 15 | Ccdc71l | 0.099 | Old |
| 0.172395 | 0.764074 | 0.224 | 0.125 | 1 | 15 | Nudt16l1 | 0.099 | Old |
| 0.17349 | 0.674172 | 0.653 | 0.536 | 1 | 15 | Tank | 0.117 | Old |
| 0.173522 | 1.622535 | 0.204 | 0.107 | 1 | 15 | Ctnnd1 | 0.097 | Old |
| 0.173838 | 0.201013 | 0.776 | 0.839 | 1 | 15 | Aebp2 | -0.063 | Old |
| 0.17409 | 1.657537 | 0.102 | 0.036 | 1 | 15 | Gnl3l | 0.066 | Old |
| 0.17409 | 1.434493 | 0.102 | 0.036 | 1 | 15 | C1qtnf6 | 0.066 | Old |
| 0.17409 | 1.371894 | 0.102 | 0.036 | 1 | 15 | Isg15 | 0.066 | Old |
| 0.174262 | 0.793188 | 0.469 | 0.357 | 1 | 15 | Dpp4 | 0.112 | Old |
| 0.174466 | 0.966429 | 0.286 | 0.179 | 1 | 15 | Mgat4b | 0.107 | Old |
| 0.17557 | 0.420739 | 0.245 | 0.143 | 1 | 15 | Tlk1 | 0.102 | Old |
| 0.177047 | 0.859839 | 0.367 | 0.268 | 1 | 15 | Akap9 | 0.099 | Old |
| 0.177079 | 1.035934 | 0.265 | 0.161 | 1 | 15 | Zufsp | 0.104 | Old |
| 0.177253 | 0.489719 | 0.286 | 0.179 | 1 | 15 | Atp11b | 0.107 | Old |
| 0.178578 | 0.780496 | 0.245 | 0.143 | 1 | 15 | Stxbp6 | 0.102 | Old |
| 0.17863 | 0.92523 | 0.224 | 0.125 | 1 | 15 | Ube2j1 | 0.099 | Old |
| 0.178844 | 2.346541 | 0.102 | 0.036 | 1 | 15 | Zfp780b | 0.066 | Old |
| 0.178844 | 1.758304 | 0.102 | 0.036 | 1 | 15 | Dolpp1 | 0.066 | Old |
| 0.178844 | 1.742412 | 0.102 | 0.036 | 1 | 15 | D130040H: | 0.066 | Old |
| 0.178844 | 1.579843 | 0.102 | 0.036 | 1 | 15 | Sipa1l2 | 0.066 | Old |
| 0.178844 | 1.435179 | 0.102 | 0.036 | 1 | 15 | Mbip | 0.066 | Old |
| 0.178844 | 1.433319 | 0.102 | 0.036 | 1 | 15 | Pcca | 0.066 | Old |
| 0.178844 | 1.373191 | 0.102 | 0.036 | 1 | 15 | Batf | 0.066 | Old |
| 0.178844 | 1.194573 | 0.102 | 0.036 | 1 | 15 | Yif1a | 0.066 | Old |
| 0.178844 | 0.968823 | 0.102 | 0.036 | 1 | 15 | Mrpl58 | 0.066 | Old |
| 0.178844 | 0.767512 | 0.102 | 0.036 | 1 | 15 | Zfp101 | 0.066 | Old |
| 0.179383 | 0.85492 | 0.184 | 0.089 | 1 | 15 | Dync1li2 | 0.095 | Old |
| 0.179865 | 0.886786 | 0.327 | 0.214 | 1 | 15 | Gak | 0.113 | Old |
| 0.179885 | 0.51371 | 0.653 | 0.536 | 1 | 15 | Srp9 | 0.117 | Old |

| | | | | | | | | |
|---|---|---|---|---|---|---|---|---|
| 0.180106 | 0.920709 | 0.204 | 0.107 | 1 | 15 | Slu7 | 0.097 | Old |
| 0.180749 | 0.677845 | 0.367 | 0.232 | 1 | 15 | Dnajc2 | 0.135 | Old |
| 0.180832 | 1.059839 | 0.388 | 0.25 | 1 | 15 | Dusp22 | 0.138 | Old |
| 0.181495 | 0.83535 | 0.531 | 0.429 | 1 | 15 | Gpx4 | 0.102 | Old |
| 0.18151 | 1.698271 | 0.122 | 0.054 | 1 | 15 | Ubqln4 | 0.068 | Old |
| 0.18152 | 0.741121 | 0.388 | 0.286 | 1 | 15 | Zfp644 | 0.102 | Old |
| 0.181623 | 1.638884 | 0.245 | 0.143 | 1 | 15 | Naga | 0.102 | Old |
| 0.18181 | 0.953855 | 0.224 | 0.125 | 1 | 15 | Gxylt1 | 0.099 | Old |
| 0.181834 | 1.153554 | 0.306 | 0.196 | 1 | 15 | Atad2b | 0.11 | Old |
| 0.182931 | 0.749748 | 0.184 | 0.089 | 1 | 15 | Gtf2e2 | 0.095 | Old |
| 0.183693 | 1.686866 | 0.102 | 0.036 | 1 | 15 | Gmeb1 | 0.066 | Old |
| 0.183693 | 1.613717 | 0.102 | 0.036 | 1 | 15 | Ascc2 | 0.066 | Old |
| 0.183693 | 1.514405 | 0.102 | 0.036 | 1 | 15 | C2cd2l | 0.066 | Old |
| 0.183693 | 1.34784 | 0.102 | 0.036 | 1 | 15 | Plscr3 | 0.066 | Old |
| 0.183693 | 1.316576 | 0.102 | 0.036 | 1 | 15 | Fam173b | 0.066 | Old |
| 0.183693 | 1.188058 | 0.102 | 0.036 | 1 | 15 | Dnajb12 | 0.066 | Old |
| 0.183693 | 1.116038 | 0.102 | 0.036 | 1 | 15 | Zfp568 | 0.066 | Old |
| 0.183693 | 1.065325 | 0.102 | 0.036 | 1 | 15 | Pign | 0.066 | Old |
| 0.183693 | 0.365459 | 0.102 | 0.036 | 1 | 15 | Depdc5 | 0.066 | Old |
| 0.184707 | 1.146786 | 0.245 | 0.143 | 1 | 15 | Ergic1 | 0.102 | Old |
| 0.184716 | 0.150666 | 1 | 1 | 1 | 15 | Rpl37a | 0 | Old |
| 0.185815 | 0.963047 | 0.286 | 0.179 | 1 | 15 | Gsap | 0.107 | Old |
| 0.185872 | 1.694911 | 0.122 | 0.054 | 1 | 15 | Flot1 | 0.068 | Old |
| 0.18633 | 0.593216 | 0.51 | 0.429 | 1 | 15 | Fxr1 | 0.081 | Old |
| 0.186531 | 1.367503 | 0.184 | 0.089 | 1 | 15 | Mta3 | 0.095 | Old |
| 0.186531 | 0.60218 | 0.184 | 0.089 | 1 | 15 | Agpat1 | 0.095 | Old |
| 0.186531 | 0.520047 | 0.184 | 0.089 | 1 | 15 | Tmcc2 | 0.095 | Old |
| 0.186559 | 0.68861 | 0.571 | 0.464 | 1 | 15 | Map3k14 | 0.107 | Old |
| 0.186631 | 0.743915 | 0.327 | 0.232 | 1 | 15 | Syf2 | 0.095 | Old |
| 0.18783 | 0.786516 | 0.245 | 0.143 | 1 | 15 | Tmtc2 | 0.102 | Old |
| 0.187855 | 0.459576 | 0.694 | 0.607 | 1 | 15 | Ccdc50 | 0.087 | Old |
| 0.188295 | 0.721564 | 0.224 | 0.125 | 1 | 15 | Med16 | 0.099 | Old |
| 0.188295 | 0.521673 | 0.224 | 0.125 | 1 | 15 | Btaf1 | 0.099 | Old |
| 0.188509 | 0.821816 | 0.429 | 0.321 | 1 | 15 | Smarce1 | 0.108 | Old |
| 0.188639 | 1.364668 | 0.102 | 0.036 | 1 | 15 | Glb1l | 0.066 | Old |
| 0.188639 | 1.104512 | 0.102 | 0.036 | 1 | 15 | Nedd1 | 0.066 | Old |
| 0.188639 | 1.029479 | 0.102 | 0.036 | 1 | 15 | Agfg2 | 0.066 | Old |
| 0.188639 | 0.86069 | 0.102 | 0.036 | 1 | 15 | Pygb | 0.066 | Old |
| 0.188846 | 0.6095 | 0.551 | 0.482 | 1 | 15 | Ppp2ca | 0.069 | Old |
| 0.189005 | 0.216605 | 0.98 | 1 | 1 | 15 | Rpsa | -0.02 | Old |
| 0.189592 | 0.842941 | 0.449 | 0.375 | 1 | 15 | Polb | 0.074 | Old |
| 0.189814 | 0.056506 | 0.061 | 0.143 | 1 | 15 | Tshz1 | -0.082 | Old |
| 0.190247 | 0.853491 | 0.388 | 0.268 | 1 | 15 | Srsf1 | 0.12 | Old |
| 0.19031 | 2.021545 | 0.122 | 0.054 | 1 | 15 | Ston2 | 0.068 | Old |
| 0.19031 | 1.77605 | 0.122 | 0.054 | 1 | 15 | Fech | 0.068 | Old |

| | | | | | | | | |
|---|---|---|---|---|---|---|---|---|
| 0.190326 | 0.301914 | 0.204 | 0.107 | 1 | 15 | Ccl8 | 0.097 | Old |
| 0.192042 | 0.804055 | 0.367 | 0.25 | 1 | 15 | Tspan31 | 0.117 | Old |
| 0.193538 | 0.837846 | 0.429 | 0.321 | 1 | 15 | Arglu1 | 0.108 | Old |
| 0.193683 | 0.761623 | 0.102 | 0.036 | 1 | 15 | Tpm3-rs7 | 0.066 | Old |
| 0.193683 | 0.706912 | 0.102 | 0.036 | 1 | 15 | Rev1 | 0.066 | Old |
| 0.193826 | 0.130117 | 0.102 | 0.196 | 1 | 15 | Hsd17b12 | -0.094 | Old |
| 0.193886 | 0.058145 | 0.082 | 0.179 | 1 | 15 | Dpm2 | -0.097 | Old |
| 0.194386 | 0.148753 | 0.571 | 0.5 | 1 | 15 | Nedd8 | 0.071 | Old |
| 0.19463 | 0.186254 | 0.571 | 0.429 | 1 | 15 | Edem1 | 0.142 | Old |
| 0.194826 | 2.348335 | 0.122 | 0.054 | 1 | 15 | Hjurp | 0.068 | Old |
| 0.194826 | 2.184567 | 0.122 | 0.054 | 1 | 15 | Hmg20a | 0.068 | Old |
| 0.194826 | 1.408025 | 0.122 | 0.054 | 1 | 15 | Adam22 | 0.068 | Old |
| 0.195103 | 0.89569 | 0.265 | 0.161 | 1 | 15 | Mical3 | 0.104 | Old |
| 0.196698 | 0.914978 | 0.082 | 0.161 | 1 | 15 | Tspan32 | -0.079 | Old |
| 0.197431 | 1.152829 | 0.245 | 0.143 | 1 | 15 | Hsd17b4 | 0.102 | Old |
| 0.197431 | 1.065819 | 0.245 | 0.143 | 1 | 15 | Sike1 | 0.102 | Old |
| 0.19768 | 0.342855 | 0.776 | 0.786 | 1 | 15 | Plxnc1 | -0.01 | Old |
| 0.197978 | 0.192959 | 0.796 | 0.786 | 1 | 15 | BC005537 | 0.01 | Old |
| 0.198108 | 1.385967 | 0.143 | 0.071 | 1 | 15 | Fgd4 | 0.072 | Old |
| 0.198233 | 0.961514 | 0.265 | 0.161 | 1 | 15 | Dlg1 | 0.104 | Old |
| 0.198241 | 0.339107 | 0.592 | 0.5 | 1 | 15 | Ptprj | 0.092 | Old |
| 0.198781 | 0.856714 | 0.347 | 0.232 | 1 | 15 | Mecp2 | 0.115 | Old |
| 0.198825 | 0.625315 | 0.102 | 0.036 | 1 | 15 | Rnf34 | 0.066 | Old |
| 0.199421 | 2.385956 | 0.122 | 0.054 | 1 | 15 | Il18r1 | 0.068 | Old |
| 0.199421 | 1.899994 | 0.122 | 0.054 | 1 | 15 | Cep85l | 0.068 | Old |
| 0.199421 | 1.873462 | 0.122 | 0.054 | 1 | 15 | Pofut1 | 0.068 | Old |
| 0.199421 | 1.08172 | 0.122 | 0.054 | 1 | 15 | Pdlim2 | 0.068 | Old |
| 0.199611 | 0.318088 | 0.837 | 0.714 | 1 | 15 | Malt1 | 0.123 | Old |
| 0.200008 | 0.470803 | 0.816 | 0.696 | 1 | 15 | Fus | 0.12 | Old |
| 0.200659 | 1.190304 | 0.367 | 0.268 | 1 | 15 | Psmb5 | 0.099 | Old |
| 0.20071 | 0.754333 | 0.245 | 0.143 | 1 | 15 | Atg3 | 0.102 | Old |
| 0.20071 | 0.371914 | 0.245 | 0.143 | 1 | 15 | Ptbp2 | 0.102 | Old |
| 0.200967 | 1.305875 | 0.204 | 0.107 | 1 | 15 | Fbxw2 | 0.097 | Old |
| 0.201775 | 1.306441 | 0.224 | 0.125 | 1 | 15 | Cdk12 | 0.099 | Old |
| 0.202353 | 0.135017 | 0.061 | 0.143 | 1 | 15 | Rhot1 | -0.082 | Old |
| 0.203414 | 1.275729 | 0.204 | 0.125 | 1 | 15 | Nipsnap3b | 0.079 | Old |
| 0.203574 | 0.390093 | 0.51 | 0.393 | 1 | 15 | Rab10 | 0.117 | Old |
| 0.204066 | 0.38815 | 0.102 | 0.036 | 1 | 15 | Pcyt2 | 0.066 | Old |
| 0.204093 | 2.217174 | 0.122 | 0.054 | 1 | 15 | Atg4d | 0.068 | Old |
| 0.204093 | 1.674007 | 0.122 | 0.054 | 1 | 15 | Senp5 | 0.068 | Old |
| 0.204093 | 1.237885 | 0.122 | 0.054 | 1 | 15 | Cntln | 0.068 | Old |
| 0.204093 | 1.233351 | 0.122 | 0.054 | 1 | 15 | 1-Mar | 0.068 | Old |
| 0.204093 | 1.149973 | 0.122 | 0.054 | 1 | 15 | Bloc1s4 | 0.068 | Old |
| 0.204597 | 1.722679 | 0.163 | 0.089 | 1 | 15 | Acvr2a | 0.074 | Old |
| 0.204609 | 0.794783 | 0.204 | 0.107 | 1 | 15 | Pink1 | 0.097 | Old |

| | | | | | | | | |
|---|---|---|---|---|---|---|---|---|
| 0.205194 | 0.851797 | 0.327 | 0.214 | 1 | 15 | Rps12-ps3 | 0.113 | Old |
| 0.205238 | 1.4585 | 0.306 | 0.214 | 1 | 15 | Sys1 | 0.092 | Old |
| 0.206357 | 0.240063 | 0.041 | 0.107 | 1 | 15 | Coro7 | -0.066 | Old |
| 0.206663 | 1.320851 | 0.143 | 0.071 | 1 | 15 | Topors | 0.072 | Old |
| 0.207841 | 0.76719 | 0.388 | 0.286 | 1 | 15 | Ralbp1 | 0.102 | Old |
| 0.208361 | 0.693802 | 0.633 | 0.571 | 1 | 15 | Nop53 | 0.062 | Old |
| 0.208631 | 1.992317 | 0.163 | 0.089 | 1 | 15 | Tbpl1 | 0.074 | Old |
| 0.20882 | 0.589867 | 0.408 | 0.304 | 1 | 15 | Taf10 | 0.104 | Old |
| 0.208845 | 1.889925 | 0.122 | 0.054 | 1 | 15 | Nit1 | 0.068 | Old |
| 0.208845 | 1.70425 | 0.122 | 0.054 | 1 | 15 | Pkib | 0.068 | Old |
| 0.209293 | 0.749528 | 0.612 | 0.5 | 1 | 15 | Pfkfb3 | 0.112 | Old |
| 0.210436 | 0.320651 | 0.388 | 0.25 | 1 | 15 | Cd274 | 0.138 | Old |
| 0.210618 | 0.757737 | 0.388 | 0.286 | 1 | 15 | Cript | 0.102 | Old |
| 0.211139 | 1.063974 | 0.245 | 0.161 | 1 | 15 | Alkbh6 | 0.084 | Old |
| 0.211911 | 0.614493 | 0.796 | 0.679 | 1 | 15 | Birc2 | 0.117 | Old |
| 0.213677 | 1.467179 | 0.122 | 0.054 | 1 | 15 | Slc35b1 | 0.068 | Old |
| 0.213677 | 1.300183 | 0.122 | 0.054 | 1 | 15 | Med4 | 0.068 | Old |
| 0.213677 | 1.235148 | 0.122 | 0.054 | 1 | 15 | Chac2 | 0.068 | Old |
| 0.213677 | 1.056926 | 0.122 | 0.054 | 1 | 15 | Txlng | 0.068 | Old |
| 0.213677 | 1.007671 | 0.122 | 0.054 | 1 | 15 | Slc39a13 | 0.068 | Old |
| 0.213677 | 0.936785 | 0.122 | 0.054 | 1 | 15 | Zdhhc16 | 0.068 | Old |
| 0.214437 | 0.967306 | 0.265 | 0.161 | 1 | 15 | Mrps9 | 0.104 | Old |
| 0.214504 | 0.546703 | 0.633 | 0.554 | 1 | 15 | Spint2 | 0.079 | Old |
| 0.215484 | 2.423812 | 0.143 | 0.071 | 1 | 15 | Fam110a | 0.072 | Old |
| 0.21567 | 1.138118 | 0.265 | 0.179 | 1 | 15 | 1110004F1 | 0.086 | Old |
| 0.215822 | 0.35714 | 0.204 | 0.107 | 1 | 15 | Snrnp48 | 0.097 | Old |
| 0.216674 | 0.194055 | 0.898 | 0.804 | 1 | 15 | Map4k4 | 0.094 | Old |
| 0.216824 | 0.134654 | 0.347 | 0.214 | 1 | 15 | Tnfrsf9 | 0.133 | Old |
| 0.217248 | 0.993715 | 0.327 | 0.214 | 1 | 15 | Bsdc1 | 0.113 | Old |
| 0.217248 | 0.649165 | 0.327 | 0.214 | 1 | 15 | Tmem128 | 0.113 | Old |
| 0.217702 | 0.774847 | 0.245 | 0.143 | 1 | 15 | Trappc4 | 0.102 | Old |
| 0.217789 | 0.449235 | 0.265 | 0.161 | 1 | 15 | Mcrs1 | 0.104 | Old |
| 0.218589 | 1.305732 | 0.122 | 0.054 | 1 | 15 | Gm46430 | 0.068 | Old |
| 0.218589 | 1.281414 | 0.122 | 0.054 | 1 | 15 | Nf1 | 0.068 | Old |
| 0.218589 | 1.185349 | 0.122 | 0.054 | 1 | 15 | Slc36a1 | 0.068 | Old |
| 0.218589 | 1.029181 | 0.122 | 0.054 | 1 | 15 | Ncoa5 | 0.068 | Old |
| 0.218589 | 0.960575 | 0.122 | 0.054 | 1 | 15 | Adrb2 | 0.068 | Old |
| 0.218589 | 0.953254 | 0.122 | 0.054 | 1 | 15 | Gbp4 | 0.068 | Old |
| 0.218589 | 0.767268 | 0.122 | 0.054 | 1 | 15 | Kctd17 | 0.068 | Old |
| 0.218589 | 0.719031 | 0.122 | 0.054 | 1 | 15 | Cryzl1 | 0.068 | Old |
| 0.21868 | 0.346772 | 0.592 | 0.429 | 1 | 15 | Ptp4a1 | 0.163 | Old |
| 0.218741 | 0.998341 | 0.286 | 0.196 | 1 | 15 | Rhob | 0.09 | Old |
| 0.218804 | 2.223407 | 0.184 | 0.107 | 1 | 15 | Ptpn9 | 0.077 | Old |
| 0.219656 | 0.269897 | 0.204 | 0.107 | 1 | 15 | Akap8 | 0.097 | Old |
| 0.220554 | 0.89524 | 0.367 | 0.268 | 1 | 15 | Aimp1 | 0.099 | Old |

| | | | | | | | | |
|---|---|---|---|---|---|---|---|---|
| 0.22059 | 0.874965 | 0.306 | 0.214 | 1 | 15 | Wipf2 | 0.092 | Old |
| 0.221081 | 1.469661 | 0.163 | 0.089 | 1 | 15 | Evi5 | 0.074 | Old |
| 0.221081 | 0.208109 | 0.082 | 0.161 | 1 | 15 | Msmo1 | -0.079 | Old |
| 0.221102 | 0.208209 | 1 | 1 | 1 | 15 | mt-Atp6 | 0 | Old |
| 0.221355 | 1.189153 | 0.347 | 0.25 | 1 | 15 | Zfp638 | 0.097 | Old |
| 0.222934 | 0.365549 | 0.714 | 0.643 | 1 | 15 | 1810037I1 | 0.071 | Old |
| 0.223292 | 0.483553 | 0.224 | 0.125 | 1 | 15 | Dicer1 | 0.099 | Old |
| 0.223298 | 0.442133 | 0.49 | 0.375 | 1 | 15 | Gpcpd1 | 0.115 | Old |
| 0.223582 | 1.089571 | 0.122 | 0.054 | 1 | 15 | Gm17018 | 0.068 | Old |
| 0.223582 | 0.710383 | 0.122 | 0.054 | 1 | 15 | Ino80c | 0.068 | Old |
| 0.223582 | 0.529091 | 0.122 | 0.054 | 1 | 15 | Tceal8 | 0.068 | Old |
| 0.224563 | 0.772076 | 0.469 | 0.357 | 1 | 15 | Arih1 | 0.112 | Old |
| 0.224573 | 3.19201 | 0.143 | 0.071 | 1 | 15 | 4833420G1 | 0.072 | Old |
| 0.224573 | 1.781324 | 0.143 | 0.071 | 1 | 15 | Flnb | 0.072 | Old |
| 0.224573 | 1.051685 | 0.143 | 0.071 | 1 | 15 | Gramd3 | 0.072 | Old |
| 0.225347 | 0.785359 | 0.163 | 0.089 | 1 | 15 | Gga3 | 0.074 | Old |
| 0.22559 | 1.20565 | 0.204 | 0.125 | 1 | 15 | Ezh1 | 0.079 | Old |
| 0.22568 | 0.649866 | 0.49 | 0.411 | 1 | 15 | Ube2d2a | 0.079 | Old |
| 0.225989 | 0.197706 | 1 | 1 | 1 | 15 | Actb | 0 | Old |
| 0.2261 | 1.51451 | 0.224 | 0.143 | 1 | 15 | Ncbp3 | 0.081 | Old |
| 0.226513 | 0.609367 | 0.388 | 0.268 | 1 | 15 | Ube2n | 0.12 | Old |
| 0.227023 | 0.29329 | 0.939 | 0.875 | 1 | 15 | Btf3 | 0.064 | Old |
| 0.227057 | 0.711495 | 0.306 | 0.196 | 1 | 15 | Psmd6 | 0.11 | Old |
| 0.227784 | 0.195207 | 0.816 | 0.893 | 1 | 15 | Gabarap | -0.077 | Old |
| 0.229219 | 1.004994 | 0.143 | 0.071 | 1 | 15 | Tnfaip1 | 0.072 | Old |
| 0.229671 | 1.55538 | 0.163 | 0.089 | 1 | 15 | Ramp1 | 0.074 | Old |
| 0.229671 | 1.255387 | 0.163 | 0.089 | 1 | 15 | Unc50 | 0.074 | Old |
| 0.231574 | 0.709468 | 0.265 | 0.161 | 1 | 15 | Rb1 | 0.104 | Old |
| 0.23159 | 0.022055 | 0.735 | 0.857 | 1 | 15 | Naca | -0.122 | Old |
| 0.23221 | 0.493238 | 0.653 | 0.536 | 1 | 15 | Ywhah | 0.117 | Old |
| 0.233236 | 0.954513 | 0.286 | 0.179 | 1 | 15 | Mtmr4 | 0.107 | Old |
| 0.233353 | 2.288138 | 0.204 | 0.125 | 1 | 15 | Lat2 | 0.079 | Old |
| 0.233353 | 1.50952 | 0.204 | 0.125 | 1 | 15 | Gca | 0.079 | Old |
| 0.233353 | 1.429933 | 0.204 | 0.125 | 1 | 15 | Anapc16 | 0.079 | Old |
| 0.233525 | 1.132158 | 0.224 | 0.143 | 1 | 15 | Tial1 | 0.081 | Old |
| 0.233646 | 1.009292 | 0.306 | 0.196 | 1 | 15 | Srgap2 | 0.11 | Old |
| 0.23381 | 1.136126 | 0.122 | 0.054 | 1 | 15 | 6-Sep | 0.068 | Old |
| 0.23381 | 0.754093 | 0.122 | 0.054 | 1 | 15 | Elp2 | 0.068 | Old |
| 0.23381 | 0.60157 | 0.122 | 0.054 | 1 | 15 | Chst15 | 0.068 | Old |
| 0.233933 | 1.626058 | 0.143 | 0.071 | 1 | 15 | Sumf1 | 0.072 | Old |
| 0.233933 | 0.62638 | 0.143 | 0.071 | 1 | 15 | Iffo1 | 0.072 | Old |
| 0.234054 | 1.452837 | 0.163 | 0.089 | 1 | 15 | Snrnp200 | 0.074 | Old |
| 0.234946 | 0.318567 | 0.449 | 0.321 | 1 | 15 | Bsg | 0.128 | Old |
| 0.235115 | 0.407188 | 0.265 | 0.161 | 1 | 15 | Aqr | 0.104 | Old |
| 0.235481 | 0.230812 | 0.102 | 0.196 | 1 | 15 | Zdhhc9 | -0.094 | Old |

| | | | | | | | | |
|---|---|---|---|---|---|---|---|---|
| 0.236069 | 1.054606 | 0.265 | 0.179 | 1 | 15 | Hist1h2bc | 0.086 | Old |
| 0.236678 | 0.071252 | 0.163 | 0.286 | 1 | 15 | Cerk | -0.123 | Old |
| 0.236692 | 0.473397 | 0.429 | 0.304 | 1 | 15 | Ppig | 0.125 | Old |
| 0.23677 | 0.964448 | 0.347 | 0.25 | 1 | 15 | Herc1 | 0.097 | Old |
| 0.237381 | 0.918007 | 0.408 | 0.304 | 1 | 15 | Smg1 | 0.104 | Old |
| 0.237381 | 0.235551 | 0.408 | 0.304 | 1 | 15 | Vasp | 0.104 | Old |
| 0.23742 | 0.636485 | 0.592 | 0.518 | 1 | 15 | Lpp | 0.074 | Old |
| 0.238053 | 0.387752 | 0.041 | 0.107 | 1 | 15 | Larp1b | -0.066 | Old |
| 0.238496 | 1.763696 | 0.163 | 0.089 | 1 | 15 | Kdm4c | 0.074 | Old |
| 0.238496 | 1.710532 | 0.163 | 0.089 | 1 | 15 | Jade1 | 0.074 | Old |
| 0.238496 | 1.208895 | 0.163 | 0.089 | 1 | 15 | Ciz1 | 0.074 | Old |
| 0.238694 | 0.287573 | 0.265 | 0.161 | 1 | 15 | Chtf8 | 0.104 | Old |
| 0.238715 | 1.73598 | 0.143 | 0.071 | 1 | 15 | Ly9 | 0.072 | Old |
| 0.238715 | 1.691645 | 0.143 | 0.071 | 1 | 15 | Rfx5 | 0.072 | Old |
| 0.238715 | 1.446304 | 0.143 | 0.071 | 1 | 15 | Lsr | 0.072 | Old |
| 0.238715 | 1.00822 | 0.143 | 0.071 | 1 | 15 | Nfatc2 | 0.072 | Old |
| 0.239047 | 0.157628 | 0.122 | 0.054 | 1 | 15 | Klhdc4 | 0.068 | Old |
| 0.240027 | 1.009135 | 0.02 | 0.071 | 1 | 15 | Rora | -0.051 | Old |
| 0.240027 | 0.214334 | 0.02 | 0.071 | 1 | 15 | Dvl2 | -0.051 | Old |
| 0.240027 | 0.077095 | 0.02 | 0.071 | 1 | 15 | Uqcc1 | -0.051 | Old |
| 0.240155 | 0.432678 | 0.286 | 0.179 | 1 | 15 | Rab22a | 0.107 | Old |
| 0.241317 | 3.849146 | 0.061 | 0.018 | 1 | 15 | Polr3d | 0.043 | Old |
| 0.241317 | 3.139314 | 0.061 | 0.018 | 1 | 15 | Rbm48 | 0.043 | Old |
| 0.241317 | 2.947686 | 0.061 | 0.018 | 1 | 15 | Agap2 | 0.043 | Old |
| 0.241317 | 2.839062 | 0.061 | 0.018 | 1 | 15 | 2610528A1 | 0.043 | Old |
| 0.241317 | 2.722022 | 0.061 | 0.018 | 1 | 15 | Sft2d3 | 0.043 | Old |
| 0.241317 | 2.640343 | 0.061 | 0.018 | 1 | 15 | Tsr1 | 0.043 | Old |
| 0.241317 | 2.611272 | 0.061 | 0.018 | 1 | 15 | Gm15228 | 0.043 | Old |
| 0.241317 | 2.547248 | 0.061 | 0.018 | 1 | 15 | Eps8l1 | 0.043 | Old |
| 0.241317 | 2.504035 | 0.061 | 0.018 | 1 | 15 | Cmtr1 | 0.043 | Old |
| 0.241317 | 2.440646 | 0.061 | 0.018 | 1 | 15 | Tab3 | 0.043 | Old |
| 0.241317 | 2.412421 | 0.061 | 0.018 | 1 | 15 | Pithd1 | 0.043 | Old |
| 0.241317 | 2.400273 | 0.061 | 0.018 | 1 | 15 | Mtmr10 | 0.043 | Old |
| 0.241317 | 2.40008 | 0.061 | 0.018 | 1 | 15 | D130020L0 | 0.043 | Old |
| 0.241317 | 2.296177 | 0.061 | 0.018 | 1 | 15 | Cdcp1 | 0.043 | Old |
| 0.241317 | 2.280951 | 0.061 | 0.018 | 1 | 15 | Ttc39c | 0.043 | Old |
| 0.241317 | 2.264766 | 0.061 | 0.018 | 1 | 15 | Cadm1 | 0.043 | Old |
| 0.241317 | 2.26062 | 0.061 | 0.018 | 1 | 15 | Clec5a | 0.043 | Old |
| 0.241317 | 2.135118 | 0.061 | 0.018 | 1 | 15 | Vps72 | 0.043 | Old |
| 0.241317 | 2.017743 | 0.061 | 0.018 | 1 | 15 | Arhgap4 | 0.043 | Old |
| 0.241317 | 1.979161 | 0.061 | 0.018 | 1 | 15 | Cluap1 | 0.043 | Old |
| 0.241317 | 1.940582 | 0.061 | 0.018 | 1 | 15 | Cacna1d | 0.043 | Old |
| 0.241317 | 1.889254 | 0.061 | 0.018 | 1 | 15 | Gpx3 | 0.043 | Old |
| 0.24213 | 1.087907 | 0.286 | 0.196 | 1 | 15 | Atp13a3 | 0.09 | Old |
| 0.242595 | 0.707458 | 0.714 | 0.732 | 1 | 15 | Rbm39 | -0.018 | Old |

| | | | | | | | | |
|---|---|---|---|---|---|---|---|---|
| 0.242932 | 1.341186 | 0.245 | 0.161 | 1 | 15 | Mrps16 | 0.084 | Old |
| 0.242932 | 1.059528 | 0.245 | 0.161 | 1 | 15 | Baz2a | 0.084 | Old |
| 0.242997 | 1.272779 | 0.163 | 0.089 | 1 | 15 | Zfyve27 | 0.074 | Old |
| 0.242997 | 1.040049 | 0.163 | 0.089 | 1 | 15 | Cops3 | 0.074 | Old |
| 0.242997 | 1.039278 | 0.163 | 0.089 | 1 | 15 | Ddit4 | 0.074 | Old |
| 0.243566 | 1.413205 | 0.143 | 0.071 | 1 | 15 | Clns1a | 0.072 | Old |
| 0.243566 | 0.798498 | 0.143 | 0.071 | 1 | 15 | Tubgcp2 | 0.072 | Old |
| 0.243669 | 0.744115 | 0.286 | 0.179 | 1 | 15 | Hpf1 | 0.107 | Old |
| 0.243781 | 0.583102 | 0.306 | 0.196 | 1 | 15 | Nlrc5 | 0.11 | Old |
| 0.244366 | 1.558633 | 0.122 | 0.054 | 1 | 15 | Zfand2b | 0.068 | Old |
| 0.244366 | 0.575119 | 0.122 | 0.054 | 1 | 15 | Ahsa2 | 0.068 | Old |
| 0.244366 | 0.546802 | 0.122 | 0.054 | 1 | 15 | Pgs1 | 0.068 | Old |
| 0.24482 | 0.403236 | 0.388 | 0.268 | 1 | 15 | Ccar1 | 0.12 | Old |
| 0.245348 | 1.126954 | 0.204 | 0.125 | 1 | 15 | Acadsb | 0.079 | Old |
| 0.246403 | 0.412441 | 0.122 | 0.214 | 1 | 15 | Arhgap15 | -0.092 | Old |
| 0.246565 | 0.487497 | 0.51 | 0.393 | 1 | 15 | Got1 | 0.117 | Old |
| 0.246764 | 1.048106 | 0.265 | 0.179 | 1 | 15 | Ankfy1 | 0.086 | Old |
| 0.246764 | 0.774449 | 0.265 | 0.179 | 1 | 15 | Baz1b | 0.086 | Old |
| 0.247218 | 0.91609 | 0.286 | 0.179 | 1 | 15 | Cdc37l1 | 0.107 | Old |
| 0.247558 | 1.514903 | 0.163 | 0.089 | 1 | 15 | Hint3 | 0.074 | Old |
| 0.247558 | 1.420979 | 0.163 | 0.089 | 1 | 15 | Mrpl21 | 0.074 | Old |
| 0.247558 | 1.078227 | 0.163 | 0.089 | 1 | 15 | Pde7a | 0.074 | Old |
| 0.247558 | 0.911924 | 0.163 | 0.089 | 1 | 15 | Uckl1 | 0.074 | Old |
| 0.247752 | 0.812309 | 0.184 | 0.107 | 1 | 15 | Endod1 | 0.077 | Old |
| 0.248485 | 0.775556 | 0.143 | 0.071 | 1 | 15 | Trim30d | 0.072 | Old |
| 0.248485 | 0.66078 | 0.143 | 0.071 | 1 | 15 | Gpkow | 0.072 | Old |
| 0.24903 | 0.463898 | 0.551 | 0.5 | 1 | 15 | Eif3k | 0.051 | Old |
| 0.249184 | 2.805179 | 0.061 | 0.018 | 1 | 15 | 2310022B0 | 0.043 | Old |
| 0.249184 | 2.734347 | 0.061 | 0.018 | 1 | 15 | 3110082I1 | 0.043 | Old |
| 0.249184 | 2.688924 | 0.061 | 0.018 | 1 | 15 | Sssca1 | 0.043 | Old |
| 0.249184 | 2.513357 | 0.061 | 0.018 | 1 | 15 | Alad | 0.043 | Old |
| 0.249184 | 2.502485 | 0.061 | 0.018 | 1 | 15 | Med22 | 0.043 | Old |
| 0.249184 | 2.499956 | 0.061 | 0.018 | 1 | 15 | Phf11c | 0.043 | Old |
| 0.249184 | 2.482516 | 0.061 | 0.018 | 1 | 15 | Magee1 | 0.043 | Old |
| 0.249184 | 2.383726 | 0.061 | 0.018 | 1 | 15 | Nop9 | 0.043 | Old |
| 0.249184 | 2.382049 | 0.061 | 0.018 | 1 | 15 | Mettl7a1 | 0.043 | Old |
| 0.249184 | 2.372922 | 0.061 | 0.018 | 1 | 15 | Zhx2 | 0.043 | Old |
| 0.249184 | 2.35363 | 0.061 | 0.018 | 1 | 15 | Cnst | 0.043 | Old |
| 0.249184 | 2.341963 | 0.061 | 0.018 | 1 | 15 | Ubl4a | 0.043 | Old |
| 0.249184 | 2.332239 | 0.061 | 0.018 | 1 | 15 | Dscam | 0.043 | Old |
| 0.249184 | 2.272101 | 0.061 | 0.018 | 1 | 15 | Zfp846 | 0.043 | Old |
| 0.249184 | 2.261396 | 0.061 | 0.018 | 1 | 15 | Wfdc18 | 0.043 | Old |
| 0.249184 | 2.254159 | 0.061 | 0.018 | 1 | 15 | Alg13 | 0.043 | Old |
| 0.249184 | 2.157709 | 0.061 | 0.018 | 1 | 15 | Lilra6 | 0.043 | Old |
| 0.249184 | 2.119893 | 0.061 | 0.018 | 1 | 15 | Spred2 | 0.043 | Old |

| | | | | | | | | |
|---|---|---|---|---|---|---|---|---|
| 0.249184 | 2.045979 | 0.061 | 0.018 | 1 | 15 | Haus6 | 0.043 | Old |
| 0.249184 | 1.922712 | 0.061 | 0.018 | 1 | 15 | Ppp1r14a | 0.043 | Old |
| 0.249184 | 1.9225 | 0.061 | 0.018 | 1 | 15 | Tldc1 | 0.043 | Old |
| 0.249184 | 1.922063 | 0.061 | 0.018 | 1 | 15 | Nbas | 0.043 | Old |
| 0.249184 | 1.898898 | 0.061 | 0.018 | 1 | 15 | Rrp15 | 0.043 | Old |
| 0.249184 | 1.839934 | 0.061 | 0.018 | 1 | 15 | Sap130 | 0.043 | Old |
| 0.249184 | 1.82546 | 0.061 | 0.018 | 1 | 15 | Csprs | 0.043 | Old |
| 0.249184 | 1.810941 | 0.061 | 0.018 | 1 | 15 | 4930579C1 | 0.043 | Old |
| 0.249184 | 1.776127 | 0.061 | 0.018 | 1 | 15 | Ccdc28a | 0.043 | Old |
| 0.249184 | 1.766399 | 0.061 | 0.018 | 1 | 15 | Abi3 | 0.043 | Old |
| 0.249184 | 1.705822 | 0.061 | 0.018 | 1 | 15 | Mast3 | 0.043 | Old |
| 0.249184 | 1.693611 | 0.061 | 0.018 | 1 | 15 | Tti2 | 0.043 | Old |
| 0.249184 | 1.669543 | 0.061 | 0.018 | 1 | 15 | Aifm1 | 0.043 | Old |
| 0.249184 | 1.649144 | 0.061 | 0.018 | 1 | 15 | 2700081O1 | 0.043 | Old |
| 0.249184 | 1.645028 | 0.061 | 0.018 | 1 | 15 | Timm9 | 0.043 | Old |
| 0.249184 | 1.604092 | 0.061 | 0.018 | 1 | 15 | Ampd3 | 0.043 | Old |
| 0.249184 | 1.588883 | 0.061 | 0.018 | 1 | 15 | Ogfod1 | 0.043 | Old |
| 0.249184 | 1.533652 | 0.061 | 0.018 | 1 | 15 | Ano8 | 0.043 | Old |
| 0.249184 | 1.521065 | 0.061 | 0.018 | 1 | 15 | Gde1 | 0.043 | Old |
| 0.249184 | 1.422817 | 0.061 | 0.018 | 1 | 15 | Hoxb3 | 0.043 | Old |
| 0.249184 | 1.376974 | 0.061 | 0.018 | 1 | 15 | Lrig2 | 0.043 | Old |
| 0.249184 | 1.370035 | 0.061 | 0.018 | 1 | 15 | Tmco4 | 0.043 | Old |
| 0.249184 | 1.357645 | 0.061 | 0.018 | 1 | 15 | Leo1 | 0.043 | Old |
| 0.249638 | 0.713358 | 0.347 | 0.25 | 1 | 15 | Pkn2 | 0.097 | Old |
| 0.249767 | 0.047849 | 0.122 | 0.054 | 1 | 15 | Narf | 0.068 | Old |
| 0.250243 | 0.894622 | 0.449 | 0.357 | 1 | 15 | Tap1 | 0.092 | Old |
| 0.250709 | 0.624072 | 0.551 | 0.518 | 1 | 15 | Sf1 | 0.033 | Old |
| 0.25094 | 0.465228 | 0.347 | 0.232 | 1 | 15 | Gsn | 0.115 | Old |
| 0.252097 | 0.839017 | 0.184 | 0.107 | 1 | 15 | Dtx3l | 0.077 | Old |
| 0.252178 | 1.720226 | 0.163 | 0.089 | 1 | 15 | Ap1m1 | 0.074 | Old |
| 0.252178 | 1.333 | 0.163 | 0.089 | 1 | 15 | Parg | 0.074 | Old |
| 0.252178 | 1.10416 | 0.163 | 0.089 | 1 | 15 | Rab40c | 0.074 | Old |
| 0.252178 | 0.973166 | 0.163 | 0.089 | 1 | 15 | Cdc34 | 0.074 | Old |
| 0.252178 | 0.902003 | 0.163 | 0.089 | 1 | 15 | Gadd45gip | 0.074 | Old |
| 0.252345 | 1.291497 | 0.388 | 0.286 | 1 | 15 | Rras2 | 0.102 | Old |
| 0.252505 | 0.456689 | 0.633 | 0.5 | 1 | 15 | Use1 | 0.133 | Old |
| 0.252838 | 0.714967 | 0.224 | 0.143 | 1 | 15 | Stk40 | 0.081 | Old |
| 0.252902 | 0.201134 | 0.959 | 0.982 | 1 | 15 | Rps15 | -0.023 | Old |
| 0.253474 | 1.412579 | 0.143 | 0.071 | 1 | 15 | Dmxl1 | 0.072 | Old |
| 0.253474 | 0.884631 | 0.143 | 0.071 | 1 | 15 | Fkbp3 | 0.072 | Old |
| 0.253474 | 0.624999 | 0.143 | 0.071 | 1 | 15 | Ints12 | 0.072 | Old |
| 0.254242 | 1.502662 | 0.245 | 0.161 | 1 | 15 | Ago1 | 0.084 | Old |
| 0.254692 | 0.946855 | 0.347 | 0.268 | 1 | 15 | Rac2 | 0.079 | Old |
| 0.255252 | 0.011394 | 0.122 | 0.054 | 1 | 15 | Plcl1 | 0.068 | Old |
| 0.256128 | 0.595002 | 0.469 | 0.321 | 1 | 15 | Brd4 | 0.148 | Old |

| | | | | | | | | |
|---|---|---|---|---|---|---|---|---|
| 0.256494 | 0.189277 | 0.102 | 0.179 | 1 | 15 | Gemin7 | -0.077 | Old |
| 0.256494 | 0.122531 | 0.102 | 0.179 | 1 | 15 | Vac14 | -0.077 | Old |
| 0.25683 | 1.13393 | 0.224 | 0.143 | 1 | 15 | Snrk | 0.081 | Old |
| 0.256858 | 1.047085 | 0.163 | 0.089 | 1 | 15 | Ints3 | 0.074 | Old |
| 0.256858 | 0.811711 | 0.163 | 0.089 | 1 | 15 | Mrpl30 | 0.074 | Old |
| 0.257228 | 3.387303 | 0.061 | 0.018 | 1 | 15 | Stk35 | 0.043 | Old |
| 0.257228 | 2.519987 | 0.061 | 0.018 | 1 | 15 | B230118H( | 0.043 | Old |
| 0.257228 | 2.332931 | 0.061 | 0.018 | 1 | 15 | Tk2 | 0.043 | Old |
| 0.257228 | 2.030404 | 0.061 | 0.018 | 1 | 15 | Zc3h3 | 0.043 | Old |
| 0.257228 | 2.027984 | 0.061 | 0.018 | 1 | 15 | Ipo9 | 0.043 | Old |
| 0.257228 | 1.875126 | 0.061 | 0.018 | 1 | 15 | Actr6 | 0.043 | Old |
| 0.257228 | 1.799171 | 0.061 | 0.018 | 1 | 15 | Timd4 | 0.043 | Old |
| 0.257228 | 1.781618 | 0.061 | 0.018 | 1 | 15 | Zfp65 | 0.043 | Old |
| 0.257228 | 1.720927 | 0.061 | 0.018 | 1 | 15 | Fam45a | 0.043 | Old |
| 0.257228 | 1.716069 | 0.061 | 0.018 | 1 | 15 | Gm42047 | 0.043 | Old |
| 0.257228 | 1.682171 | 0.061 | 0.018 | 1 | 15 | Tor2a | 0.043 | Old |
| 0.257228 | 1.667185 | 0.061 | 0.018 | 1 | 15 | Cul4b | 0.043 | Old |
| 0.257228 | 1.539166 | 0.061 | 0.018 | 1 | 15 | Ppm1f | 0.043 | Old |
| 0.257228 | 1.457485 | 0.061 | 0.018 | 1 | 15 | Rrp12 | 0.043 | Old |
| 0.257228 | 1.409806 | 0.061 | 0.018 | 1 | 15 | Mief1 | 0.043 | Old |
| 0.257228 | 1.382392 | 0.061 | 0.018 | 1 | 15 | Pinx1 | 0.043 | Old |
| 0.257228 | 1.304116 | 0.061 | 0.018 | 1 | 15 | Zmym1 | 0.043 | Old |
| 0.257228 | 1.295816 | 0.061 | 0.018 | 1 | 15 | Rabepk | 0.043 | Old |
| 0.257228 | 1.249798 | 0.061 | 0.018 | 1 | 15 | Bcl2l13 | 0.043 | Old |
| 0.257228 | 1.220331 | 0.061 | 0.018 | 1 | 15 | Lin52 | 0.043 | Old |
| 0.257228 | 1.202331 | 0.061 | 0.018 | 1 | 15 | Med6 | 0.043 | Old |
| 0.257228 | 1.187863 | 0.061 | 0.018 | 1 | 15 | Mtfmt | 0.043 | Old |
| 0.257228 | 1.184519 | 0.061 | 0.018 | 1 | 15 | Nom1 | 0.043 | Old |
| 0.257228 | 1.17384 | 0.061 | 0.018 | 1 | 15 | Miga1 | 0.043 | Old |
| 0.257228 | 1.166537 | 0.061 | 0.018 | 1 | 15 | Slc25a45 | 0.043 | Old |
| 0.257228 | 1.151333 | 0.061 | 0.018 | 1 | 15 | Otud1 | 0.043 | Old |
| 0.257228 | 1.10987 | 0.061 | 0.018 | 1 | 15 | Zbtb24 | 0.043 | Old |
| 0.257228 | 1.064711 | 0.061 | 0.018 | 1 | 15 | Pcbp4 | 0.043 | Old |
| 0.257228 | 1.05341 | 0.061 | 0.018 | 1 | 15 | Zfp954 | 0.043 | Old |
| 0.257228 | 0.992465 | 0.061 | 0.018 | 1 | 15 | Nat10 | 0.043 | Old |
| 0.257228 | 0.972979 | 0.061 | 0.018 | 1 | 15 | Slc30a1 | 0.043 | Old |
| 0.257228 | 0.951863 | 0.061 | 0.018 | 1 | 15 | Zfp74 | 0.043 | Old |
| 0.257716 | 0.286227 | 0.878 | 0.857 | 1 | 15 | Gpx1 | 0.021 | Old |
| 0.258092 | 0.62171 | 0.245 | 0.161 | 1 | 15 | Tspyl1 | 0.084 | Old |
| 0.258532 | 0.950171 | 0.143 | 0.071 | 1 | 15 | Cog1 | 0.072 | Old |
| 0.258532 | 0.814096 | 0.143 | 0.071 | 1 | 15 | Ppwd1 | 0.072 | Old |
| 0.258532 | 0.754442 | 0.143 | 0.071 | 1 | 15 | Tmem260 | 0.072 | Old |
| 0.258532 | 0.6806 | 0.143 | 0.071 | 1 | 15 | Tmem70 | 0.072 | Old |
| 0.258532 | 0.680596 | 0.143 | 0.071 | 1 | 15 | Rnf122 | 0.072 | Old |
| 0.258532 | 0.571762 | 0.143 | 0.071 | 1 | 15 | Arhgap25 | 0.072 | Old |

| | | | | | | | | |
|---|---|---|---|---|---|---|---|---|
| 0.259321 | 0.593019 | 0.449 | 0.357 | 1 | 15 | Msl1 | 0.092 | Old |
| 0.259349 | 0.652228 | 0.531 | 0.429 | 1 | 15 | Fubp1 | 0.102 | Old |
| 0.259918 | 0.328256 | 0.327 | 0.214 | 1 | 15 | Ubash3b | 0.113 | Old |
| 0.260448 | 0.49751 | 0.653 | 0.625 | 1 | 15 | Arhgap30 | 0.028 | Old |
| 0.260866 | 1.276638 | 0.224 | 0.143 | 1 | 15 | Il3ra | 0.081 | Old |
| 0.260866 | 0.615348 | 0.224 | 0.143 | 1 | 15 | Ccdc82 | 0.081 | Old |
| 0.260945 | 0.831047 | 0.184 | 0.107 | 1 | 15 | Gpr68 | 0.077 | Old |
| 0.261195 | 0.91581 | 0.408 | 0.321 | 1 | 15 | Actn1 | 0.087 | Old |
| 0.261547 | 1.086196 | 0.265 | 0.179 | 1 | 15 | Pik3cb | 0.086 | Old |
| 0.261662 | 0.836568 | 0.469 | 0.411 | 1 | 15 | Kmt2e | 0.058 | Old |
| 0.261777 | 0.840159 | 0.286 | 0.179 | 1 | 15 | Relt | 0.107 | Old |
| 0.261982 | 1.005563 | 0.245 | 0.161 | 1 | 15 | Actr1a | 0.084 | Old |
| 0.262006 | 1.108477 | 0.204 | 0.125 | 1 | 15 | Get4 | 0.079 | Old |
| 0.262006 | 0.842154 | 0.204 | 0.125 | 1 | 15 | Lamtor5 | 0.079 | Old |
| 0.262064 | 1.21296 | 0.408 | 0.357 | 1 | 15 | Cxcr4 | 0.051 | Old |
| 0.263534 | 0.945143 | 0.286 | 0.196 | 1 | 15 | Smarca4 | 0.09 | Old |
| 0.263659 | 0.82787 | 0.143 | 0.071 | 1 | 15 | Donson | 0.072 | Old |
| 0.264946 | 0.76596 | 0.224 | 0.143 | 1 | 15 | Sde2 | 0.081 | Old |
| 0.265219 | 0.396364 | 0.612 | 0.5 | 1 | 15 | Hnrnpa1 | 0.112 | Old |
| 0.265336 | 0.813731 | 0.265 | 0.179 | 1 | 15 | Zfx | 0.086 | Old |
| 0.265449 | 0.854453 | 0.184 | 0.107 | 1 | 15 | Naa10 | 0.077 | Old |
| 0.265449 | 0.69358 | 0.184 | 0.107 | 1 | 15 | Hsd17b10 | 0.077 | Old |
| 0.265451 | 1.412958 | 0.061 | 0.018 | 1 | 15 | Timp3 | 0.043 | Old |
| 0.265451 | 1.081285 | 0.061 | 0.018 | 1 | 15 | Tbc1d31 | 0.043 | Old |
| 0.265451 | 1.060742 | 0.061 | 0.018 | 1 | 15 | Mypopos | 0.043 | Old |
| 0.265451 | 1.042581 | 0.061 | 0.018 | 1 | 15 | Agpat3 | 0.043 | Old |
| 0.265451 | 1.04122 | 0.061 | 0.018 | 1 | 15 | Zfp788 | 0.043 | Old |
| 0.265451 | 1.034963 | 0.061 | 0.018 | 1 | 15 | Btg3 | 0.043 | Old |
| 0.265451 | 1.003588 | 0.061 | 0.018 | 1 | 15 | Sort1 | 0.043 | Old |
| 0.265451 | 0.934809 | 0.061 | 0.018 | 1 | 15 | Zfp946 | 0.043 | Old |
| 0.265451 | 0.922474 | 0.061 | 0.018 | 1 | 15 | Arc | 0.043 | Old |
| 0.265451 | 0.91137 | 0.061 | 0.018 | 1 | 15 | Sugp1 | 0.043 | Old |
| 0.265451 | 0.904885 | 0.061 | 0.018 | 1 | 15 | Nup35 | 0.043 | Old |
| 0.265451 | 0.834409 | 0.061 | 0.018 | 1 | 15 | Nol9 | 0.043 | Old |
| 0.265451 | 0.82887 | 0.061 | 0.018 | 1 | 15 | Akap7 | 0.043 | Old |
| 0.265451 | 0.781775 | 0.061 | 0.018 | 1 | 15 | Rtl8b | 0.043 | Old |
| 0.265451 | 0.781725 | 0.061 | 0.018 | 1 | 15 | Dnaja3 | 0.043 | Old |
| 0.265451 | 0.781199 | 0.061 | 0.018 | 1 | 15 | Mphosph9 | 0.043 | Old |
| 0.265451 | 0.769441 | 0.061 | 0.018 | 1 | 15 | Gm26735 | 0.043 | Old |
| 0.265451 | 0.768905 | 0.061 | 0.018 | 1 | 15 | Gm26782 | 0.043 | Old |
| 0.265451 | 0.761997 | 0.061 | 0.018 | 1 | 15 | AC163040. | 0.043 | Old |
| 0.265451 | 0.711817 | 0.061 | 0.018 | 1 | 15 | Kit | 0.043 | Old |
| 0.265451 | 0.650998 | 0.061 | 0.018 | 1 | 15 | Tox3 | 0.043 | Old |
| 0.265451 | 0.603985 | 0.061 | 0.018 | 1 | 15 | Fance | 0.043 | Old |
| 0.265451 | 0.570852 | 0.061 | 0.018 | 1 | 15 | Arl14ep | 0.043 | Old |

| | | | | | | | | |
|---|---|---|---|---|---|---|---|---|
| 0.265451 | 0.533572 | 0.061 | 0.018 | 1 | 15 | Arfgef2 | 0.043 | Old |
| 0.265451 | 0.525289 | 0.061 | 0.018 | 1 | 15 | Zfp687 | 0.043 | Old |
| 0.265451 | 0.46917 | 0.061 | 0.018 | 1 | 15 | Gfod1 | 0.043 | Old |
| 0.265451 | 0.394989 | 0.061 | 0.018 | 1 | 15 | Akip1 | 0.043 | Old |
| 0.265451 | 0.33221 | 0.061 | 0.018 | 1 | 15 | Rhot2 | 0.043 | Old |
| 0.265451 | 0.309603 | 0.061 | 0.018 | 1 | 15 | Tex264 | 0.043 | Old |
| 0.265451 | 0.272066 | 0.061 | 0.018 | 1 | 15 | Ms4a6d | 0.043 | Old |
| 0.265451 | 0.201841 | 0.061 | 0.018 | 1 | 15 | Galt | 0.043 | Old |
| 0.265451 | 0.116093 | 0.061 | 0.018 | 1 | 15 | Tm7sf3 | 0.043 | Old |
| 0.265451 | 0.076169 | 0.061 | 0.018 | 1 | 15 | Prkca | 0.043 | Old |
| 0.265451 | 0.022233 | 0.061 | 0.018 | 1 | 15 | Capn15 | 0.043 | Old |
| 0.265498 | 0.101714 | 0.327 | 0.464 | 1 | 15 | Camkk2 | -0.137 | Old |
| 0.266199 | 0.376053 | 0.408 | 0.286 | 1 | 15 | Zc3hav1 | 0.122 | Old |
| 0.266289 | 0.712332 | 0.204 | 0.125 | 1 | 15 | Parp11 | 0.079 | Old |
| 0.266398 | 0.641868 | 0.163 | 0.089 | 1 | 15 | Stap2 | 0.074 | Old |
| 0.26645 | 0.256044 | 0.816 | 0.821 | 1 | 15 | Eef1g | -0.005 | Old |
| 0.267576 | 0.567105 | 0.327 | 0.232 | 1 | 15 | Gga1 | 0.095 | Old |
| 0.267905 | 1.19437 | 0.306 | 0.214 | 1 | 15 | Pabpn1 | 0.092 | Old |
| 0.268394 | 0.700657 | 0.49 | 0.357 | 1 | 15 | Fchsd2 | 0.133 | Old |
| 0.268857 | 0.055923 | 0.143 | 0.071 | 1 | 15 | Agap1 | 0.072 | Old |
| 0.26887 | 0.488282 | 0.735 | 0.732 | 1 | 15 | Grk3 | 0.003 | Old |
| 0.26907 | 0.643333 | 0.224 | 0.143 | 1 | 15 | D10Jhu81e | 0.081 | Old |
| 0.269163 | 0.799608 | 0.265 | 0.179 | 1 | 15 | Acer3 | 0.086 | Old |
| 0.269798 | 0.075591 | 0.061 | 0.125 | 1 | 15 | Cdk5r1 | -0.064 | Old |
| 0.270006 | 0.932327 | 0.184 | 0.107 | 1 | 15 | Sf3b4 | 0.077 | Old |
| 0.270649 | 0.249736 | 0.98 | 0.982 | 1 | 15 | Ccr7 | -0.002 | Old |
| 0.270711 | 0.251268 | 0.143 | 0.232 | 1 | 15 | Tbc1d20 | -0.089 | Old |
| 0.270821 | 0.595376 | 0.408 | 0.321 | 1 | 15 | Cop1 | 0.087 | Old |
| 0.270951 | 0.583752 | 0.286 | 0.196 | 1 | 15 | Zfp148 | 0.09 | Old |
| 0.271258 | 1.110716 | 0.163 | 0.089 | 1 | 15 | Dusp6 | 0.074 | Old |
| 0.271259 | 0.262146 | 0.531 | 0.429 | 1 | 15 | 7-Mar | 0.102 | Old |
| 0.273028 | 0.904576 | 0.265 | 0.179 | 1 | 15 | Tmem33 | 0.086 | Old |
| 0.273237 | 0.921778 | 0.224 | 0.143 | 1 | 15 | Hmgn1 | 0.081 | Old |
| 0.273237 | 0.816835 | 0.224 | 0.143 | 1 | 15 | Erp44 | 0.081 | Old |
| 0.273896 | 0.840232 | 0.245 | 0.161 | 1 | 15 | Egr2 | 0.084 | Old |
| 0.273896 | 0.691479 | 0.245 | 0.161 | 1 | 15 | Abcf1 | 0.084 | Old |
| 0.274186 | 0.974734 | 0.367 | 0.286 | 1 | 15 | Plekha2 | 0.081 | Old |
| 0.274616 | 0.729318 | 0.184 | 0.107 | 1 | 15 | Ppp1r35 | 0.077 | Old |
| 0.274616 | 0.445722 | 0.184 | 0.107 | 1 | 15 | Kcnq1ot1 | 0.077 | Old |
| 0.274951 | 0.922728 | 0.347 | 0.268 | 1 | 15 | Nbeal1 | 0.079 | Old |
| 0.276179 | 1.673059 | 0.163 | 0.089 | 1 | 15 | Ncor2 | 0.074 | Old |
| 0.276179 | 1.222253 | 0.163 | 0.089 | 1 | 15 | Gpr157 | 0.074 | Old |
| 0.276179 | 0.253757 | 0.163 | 0.089 | 1 | 15 | Traf7 | 0.074 | Old |
| 0.277448 | 0.589038 | 0.224 | 0.143 | 1 | 15 | Fut8 | 0.081 | Old |
| 0.277592 | 0.708984 | 0.388 | 0.268 | 1 | 15 | Stx11 | 0.12 | Old |

| | | | | | | | | |
|---|---|---|---|---|---|---|---|---|
| 0.277592 | 0.638416 | 0.367 | 0.286 | 1 | 15 | Tm9sf2 | 0.081 | Old |
| 0.278097 | 1.060018 | 0.367 | 0.268 | 1 | 15 | Wapl | 0.099 | Old |
| 0.278103 | 0.295518 | 0.898 | 0.786 | 1 | 15 | Srrm2 | 0.112 | Old |
| 0.278156 | 1.038327 | 0.429 | 0.357 | 1 | 15 | 2810474O1 | 0.072 | Old |
| 0.278431 | 0.782033 | 0.347 | 0.268 | 1 | 15 | Camta2 | 0.079 | Old |
| 0.27928 | 1.429554 | 0.184 | 0.107 | 1 | 15 | Gpd2 | 0.077 | Old |
| 0.27928 | 0.434482 | 0.184 | 0.107 | 1 | 15 | Erap1 | 0.077 | Old |
| 0.27928 | 0.347787 | 0.102 | 0.179 | 1 | 15 | Golga5 | -0.077 | Old |
| 0.279415 | 0.793405 | 0.408 | 0.339 | 1 | 15 | Ppp1r9b | 0.069 | Old |
| 0.280151 | 0.621482 | 0.449 | 0.429 | 1 | 15 | Ran | 0.02 | Old |
| 0.28116 | 0.661087 | 0.163 | 0.089 | 1 | 15 | Cul4a | 0.074 | Old |
| 0.28116 | 0.447695 | 0.163 | 0.089 | 1 | 15 | Gan | 0.074 | Old |
| 0.28116 | 0.422406 | 0.163 | 0.089 | 1 | 15 | Msantd4 | 0.074 | Old |
| 0.281569 | 1.046313 | 0.367 | 0.268 | 1 | 15 | Zfp516 | 0.099 | Old |
| 0.281703 | 1.033389 | 0.224 | 0.143 | 1 | 15 | Rbm6 | 0.081 | Old |
| 0.282041 | 0.703416 | 0.245 | 0.161 | 1 | 15 | AC160336. | 0.084 | Old |
| 0.282124 | 0.271596 | 0.347 | 0.232 | 1 | 15 | Ccdc88b | 0.115 | Old |
| 0.28392 | 0.595961 | 0.673 | 0.589 | 1 | 15 | Rbm25 | 0.084 | Old |
| 0.283997 | 0.690356 | 0.184 | 0.107 | 1 | 15 | Rnf38 | 0.077 | Old |
| 0.283997 | 0.613734 | 0.184 | 0.107 | 1 | 15 | Rfc2 | 0.077 | Old |
| 0.283997 | 0.609252 | 0.184 | 0.107 | 1 | 15 | Exosc5 | 0.077 | Old |
| 0.283997 | 0.227608 | 0.184 | 0.107 | 1 | 15 | Crybg3 | 0.077 | Old |
| 0.284798 | 0.114117 | 0.939 | 0.929 | 1 | 15 | Rel | 0.01 | Old |
| 0.284824 | 0.127967 | 0.959 | 0.929 | 1 | 15 | mt-Nd4 | 0.03 | Old |
| 0.284868 | 0.592288 | 0.143 | 0.071 | 1 | 15 | Dcn | 0.072 | Old |
| 0.285155 | 0.985413 | 0.306 | 0.232 | 1 | 15 | Sgk3 | 0.074 | Old |
| 0.286002 | 0.343825 | 0.224 | 0.143 | 1 | 15 | Abhd17b | 0.081 | Old |
| 0.286562 | 1.369908 | 0.061 | 0.125 | 1 | 15 | Rtl8a | -0.064 | Old |
| 0.286562 | 0.155542 | 0.061 | 0.125 | 1 | 15 | Nip7 | -0.064 | Old |
| 0.28739 | 1.340379 | 0.286 | 0.214 | 1 | 15 | Pole4 | 0.072 | Old |
| 0.28833 | 3.098773 | 0.082 | 0.036 | 1 | 15 | Hap1 | 0.046 | Old |
| 0.28833 | 2.927711 | 0.082 | 0.036 | 1 | 15 | Mindy3 | 0.046 | Old |
| 0.28833 | 2.231293 | 0.082 | 0.036 | 1 | 15 | Tmem80 | 0.046 | Old |
| 0.28833 | 2.20999 | 0.082 | 0.036 | 1 | 15 | Bet1l | 0.046 | Old |
| 0.28833 | 2.11552 | 0.082 | 0.036 | 1 | 15 | Dnajc16 | 0.046 | Old |
| 0.28833 | 2.027004 | 0.082 | 0.036 | 1 | 15 | Arhgap11a | 0.046 | Old |
| 0.288768 | 0.513794 | 0.184 | 0.107 | 1 | 15 | Cdk5rap3 | 0.077 | Old |
| 0.28894 | 0.790549 | 0.388 | 0.286 | 1 | 15 | Rassf1 | 0.102 | Old |
| 0.289186 | 0.190328 | 0.939 | 0.946 | 1 | 15 | Cfl1 | -0.007 | Old |
| 0.290346 | 0.709676 | 0.224 | 0.143 | 1 | 15 | Chmp1a | 0.081 | Old |
| 0.290346 | 0.633281 | 0.224 | 0.143 | 1 | 15 | Zmynd15 | 0.081 | Old |
| 0.290346 | 0.240506 | 0.143 | 0.071 | 1 | 15 | Egr3 | 0.072 | Old |
| 0.291139 | 0.511732 | 0.347 | 0.25 | 1 | 15 | Tia1 | 0.097 | Old |
| 0.291304 | 0.608049 | 0.163 | 0.089 | 1 | 15 | Tmem87b | 0.074 | Old |
| 0.291304 | 0.512371 | 0.082 | 0.161 | 1 | 15 | Gdap2 | -0.079 | Old |

| | | | | | | | | |
|---|---|---|---|---|---|---|---|---|
| 0.291304 | 0.510466 | 0.163 | 0.089 | 1 | 15 | Mthfsl | 0.074 | Old |
| 0.292919 | 0.97166 | 0.265 | 0.179 | 1 | 15 | Eif2a | 0.086 | Old |
| 0.292984 | 0.399771 | 0.327 | 0.232 | 1 | 15 | Zmynd8 | 0.095 | Old |
| 0.293739 | 5.85983 | 0.02 | 0 | 1 | 15 | Cd7 | 0.02 | Old |
| 0.293739 | 5.852861 | 0.02 | 0 | 1 | 15 | Cldn1 | 0.02 | Old |
| 0.293739 | 5.567879 | 0.02 | 0 | 1 | 15 | Aldh1a1 | 0.02 | Old |
| 0.293739 | 5.280326 | 0.02 | 0 | 1 | 15 | Senp8 | 0.02 | Old |
| 0.293739 | 5.070987 | 0.02 | 0 | 1 | 15 | Vegfa | 0.02 | Old |
| 0.293739 | 4.876963 | 0.02 | 0 | 1 | 15 | Prss23 | 0.02 | Old |
| 0.293739 | 4.700995 | 0.02 | 0 | 1 | 15 | Hspg2 | 0.02 | Old |
| 0.293739 | 4.700995 | 0.02 | 0 | 1 | 15 | 9130213A2 | 0.02 | Old |
| 0.293739 | 4.700995 | 0.02 | 0 | 1 | 15 | Gm20219 | 0.02 | Old |
| 0.293739 | 4.670186 | 0.02 | 0 | 1 | 15 | Aif1l | 0.02 | Old |
| 0.293739 | 4.670186 | 0.02 | 0 | 1 | 15 | Ccr6 | 0.02 | Old |
| 0.293739 | 4.597977 | 0.02 | 0 | 1 | 15 | Vill | 0.02 | Old |
| 0.293739 | 4.380132 | 0.02 | 0 | 1 | 15 | Kctd14 | 0.02 | Old |
| 0.293739 | 4.343301 | 0.02 | 0 | 1 | 15 | Gdap1 | 0.02 | Old |
| 0.293739 | 4.316979 | 0.02 | 0 | 1 | 15 | Gm37584 | 0.02 | Old |
| 0.293739 | 4.316979 | 0.02 | 0 | 1 | 15 | Bcl2l15 | 0.02 | Old |
| 0.293739 | 4.316979 | 0.02 | 0 | 1 | 15 | Cald1 | 0.02 | Old |
| 0.293739 | 4.316979 | 0.02 | 0 | 1 | 15 | Wdr62 | 0.02 | Old |
| 0.293739 | 4.316979 | 0.02 | 0 | 1 | 15 | Clec3b | 0.02 | Old |
| 0.293739 | 4.316979 | 0.02 | 0 | 1 | 15 | Gm12735 | 0.02 | Old |
| 0.293739 | 4.316979 | 0.02 | 0 | 1 | 15 | Abca13 | 0.02 | Old |
| 0.293739 | 4.316979 | 0.02 | 0 | 1 | 15 | Grhl1 | 0.02 | Old |
| 0.293739 | 4.316979 | 0.02 | 0 | 1 | 15 | Amn | 0.02 | Old |
| 0.293739 | 4.310638 | 0.02 | 0 | 1 | 15 | Ugt1a6a | 0.02 | Old |
| 0.293739 | 4.310638 | 0.02 | 0 | 1 | 15 | 5330413P1 | 0.02 | Old |
| 0.293739 | 4.310638 | 0.02 | 0 | 1 | 15 | Srpx | 0.02 | Old |
| 0.293739 | 4.310638 | 0.02 | 0 | 1 | 15 | Tmem240 | 0.02 | Old |
| 0.293739 | 4.310638 | 0.02 | 0 | 1 | 15 | 1110006O2 | 0.02 | Old |
| 0.293739 | 4.310638 | 0.02 | 0 | 1 | 15 | Car7 | 0.02 | Old |
| 0.293739 | 4.227341 | 0.02 | 0 | 1 | 15 | Rpl3l | 0.02 | Old |
| 0.293739 | 3.816656 | 0.02 | 0 | 1 | 15 | Xlr4a | 0.02 | Old |
| 0.293739 | 3.816656 | 0.02 | 0 | 1 | 15 | Xlr3b | 0.02 | Old |
| 0.293739 | 3.816656 | 0.02 | 0 | 1 | 15 | Ephb2 | 0.02 | Old |
| 0.293739 | 3.816656 | 0.02 | 0 | 1 | 15 | Mrpl22 | 0.02 | Old |
| 0.293739 | 3.816656 | 0.02 | 0 | 1 | 15 | Sac3d1 | 0.02 | Old |
| 0.293739 | 3.765834 | 0.02 | 0 | 1 | 15 | Cdhr1 | 0.02 | Old |
| 0.293739 | 3.629818 | 0.02 | 0 | 1 | 15 | Syde1 | 0.02 | Old |
| 0.293739 | 3.534888 | 0.02 | 0 | 1 | 15 | Adgrg5 | 0.02 | Old |
| 0.293739 | 3.447802 | 0.02 | 0 | 1 | 15 | Dph6 | 0.02 | Old |
| 0.293739 | 3.447802 | 0.02 | 0 | 1 | 15 | Grpr | 0.02 | Old |
| 0.293739 | 3.447802 | 0.02 | 0 | 1 | 15 | Hyal3 | 0.02 | Old |
| 0.293739 | 3.447802 | 0.02 | 0 | 1 | 15 | Trip13 | 0.02 | Old |

| | | | | | | | | |
|---|---|---|---|---|---|---|---|---|
| 0.293739 | 3.41268 | 0.02 | 0 | 1 | 15 | Lrsam1 | 0.02 | Old |
| 0.293739 | 3.41268 | 0.02 | 0 | 1 | 15 | Lekr1 | 0.02 | Old |
| 0.293739 | 3.41268 | 0.02 | 0 | 1 | 15 | Nxn | 0.02 | Old |
| 0.293739 | 3.41268 | 0.02 | 0 | 1 | 15 | Pdzd2 | 0.02 | Old |
| 0.293739 | 3.40713 | 0.02 | 0 | 1 | 15 | Igsf23 | 0.02 | Old |
| 0.293739 | 3.40713 | 0.02 | 0 | 1 | 15 | Ybey | 0.02 | Old |
| 0.293739 | 3.40713 | 0.02 | 0 | 1 | 15 | Gm32618 | 0.02 | Old |
| 0.293739 | 3.317683 | 0.02 | 0 | 1 | 15 | Psg16 | 0.02 | Old |
| 0.293739 | 3.317683 | 0.02 | 0 | 1 | 15 | 9130221H1 | 0.02 | Old |
| 0.293739 | 3.232949 | 0.02 | 0 | 1 | 15 | Gm12979 | 0.02 | Old |
| 0.293739 | 3.232949 | 0.02 | 0 | 1 | 15 | Mkrn3 | 0.02 | Old |
| 0.293739 | 3.205671 | 0.02 | 0 | 1 | 15 | Spef1 | 0.02 | Old |
| 0.293739 | 3.195862 | 0.02 | 0 | 1 | 15 | Dusp19 | 0.02 | Old |
| 0.293739 | 3.195862 | 0.02 | 0 | 1 | 15 | Extl2 | 0.02 | Old |
| 0.293739 | 3.195862 | 0.02 | 0 | 1 | 15 | Scube2 | 0.02 | Old |
| 0.293739 | 3.195862 | 0.02 | 0 | 1 | 15 | Sez6l2 | 0.02 | Old |
| 0.293739 | 3.195862 | 0.02 | 0 | 1 | 15 | B430219N1 | 0.02 | Old |
| 0.293739 | 3.195862 | 0.02 | 0 | 1 | 15 | Tnfrsf10b | 0.02 | Old |
| 0.293739 | 3.195862 | 0.02 | 0 | 1 | 15 | Zbed3 | 0.02 | Old |
| 0.293739 | 3.195862 | 0.02 | 0 | 1 | 15 | Gpd1 | 0.02 | Old |
| 0.293739 | 3.195862 | 0.02 | 0 | 1 | 15 | Chaf1a | 0.02 | Old |
| 0.293739 | 3.194321 | 0.02 | 0 | 1 | 15 | Npas2 | 0.02 | Old |
| 0.293739 | 3.194321 | 0.02 | 0 | 1 | 15 | Fcmr | 0.02 | Old |
| 0.293739 | 3.194321 | 0.02 | 0 | 1 | 15 | Usp20 | 0.02 | Old |
| 0.293739 | 3.194321 | 0.02 | 0 | 1 | 15 | Tasp1 | 0.02 | Old |
| 0.293739 | 3.194321 | 0.02 | 0 | 1 | 15 | Egfl6 | 0.02 | Old |
| 0.293739 | 3.194321 | 0.02 | 0 | 1 | 15 | Rorc | 0.02 | Old |
| 0.293739 | 3.194321 | 0.02 | 0 | 1 | 15 | Lingo4 | 0.02 | Old |
| 0.293739 | 3.194321 | 0.02 | 0 | 1 | 15 | I830077J02 | 0.02 | Old |
| 0.293739 | 3.194321 | 0.02 | 0 | 1 | 15 | Whrn | 0.02 | Old |
| 0.293739 | 3.194321 | 0.02 | 0 | 1 | 15 | Clstn1 | 0.02 | Old |
| 0.293739 | 3.194321 | 0.02 | 0 | 1 | 15 | Prdm16 | 0.02 | Old |
| 0.293739 | 3.194321 | 0.02 | 0 | 1 | 15 | Slc6a1 | 0.02 | Old |
| 0.293739 | 3.194321 | 0.02 | 0 | 1 | 15 | Klrg1 | 0.02 | Old |
| 0.293739 | 3.194321 | 0.02 | 0 | 1 | 15 | Chst8 | 0.02 | Old |
| 0.293739 | 3.194321 | 0.02 | 0 | 1 | 15 | Saal1 | 0.02 | Old |
| 0.293739 | 3.194321 | 0.02 | 0 | 1 | 15 | Pbld1 | 0.02 | Old |
| 0.293739 | 3.194321 | 0.02 | 0 | 1 | 15 | Rgcc | 0.02 | Old |
| 0.293739 | 3.194321 | 0.02 | 0 | 1 | 15 | Zfp599 | 0.02 | Old |
| 0.293739 | 3.194321 | 0.02 | 0 | 1 | 15 | N4bp3 | 0.02 | Old |
| 0.293739 | 3.194321 | 0.02 | 0 | 1 | 15 | D930048N1 | 0.02 | Old |
| 0.293739 | 3.194321 | 0.02 | 0 | 1 | 15 | Serpinb1c | 0.02 | Old |
| 0.293739 | 3.194321 | 0.02 | 0 | 1 | 15 | Smoc1 | 0.02 | Old |
| 0.293739 | 3.194321 | 0.02 | 0 | 1 | 15 | Il2rb | 0.02 | Old |
| 0.293739 | 3.194321 | 0.02 | 0 | 1 | 15 | Prepl | 0.02 | Old |

| | | | | | | | | |
|---|---|---|---|---|---|---|---|---|
| 0.293739 | 3.107801 | 0.02 | 0 | 1 | 15 | Atp6v0d2 | 0.02 | Old |
| 0.293739 | 3.088909 | 0.02 | 0 | 1 | 15 | Elp4 | 0.02 | Old |
| 0.293739 | 3.073093 | 0.02 | 0 | 1 | 15 | Eif2b3 | 0.02 | Old |
| 0.293739 | 3.073093 | 0.02 | 0 | 1 | 15 | Gm36445 | 0.02 | Old |
| 0.293739 | 3.073093 | 0.02 | 0 | 1 | 15 | 1110008L1 | 0.02 | Old |
| 0.293739 | 3.073093 | 0.02 | 0 | 1 | 15 | AC097366. | 0.02 | Old |
| 0.293739 | 3.065267 | 0.02 | 0 | 1 | 15 | Vsig4 | 0.02 | Old |
| 0.293739 | 3.065267 | 0.02 | 0 | 1 | 15 | AI427809 | 0.02 | Old |
| 0.293739 | 3.035386 | 0.02 | 0 | 1 | 15 | A530040E1 | 0.02 | Old |
| 0.293739 | 3.035386 | 0.02 | 0 | 1 | 15 | Atp1a2 | 0.02 | Old |
| 0.293739 | 3.035386 | 0.02 | 0 | 1 | 15 | Mecom | 0.02 | Old |
| 0.293739 | 3.035386 | 0.02 | 0 | 1 | 15 | Map9 | 0.02 | Old |
| 0.293739 | 3.035386 | 0.02 | 0 | 1 | 15 | Phf24 | 0.02 | Old |
| 0.293739 | 3.035386 | 0.02 | 0 | 1 | 15 | Cnr2 | 0.02 | Old |
| 0.293739 | 3.035386 | 0.02 | 0 | 1 | 15 | Cd27 | 0.02 | Old |
| 0.293739 | 3.035386 | 0.02 | 0 | 1 | 15 | Pclaf | 0.02 | Old |
| 0.293739 | 3.035386 | 0.02 | 0 | 1 | 15 | 4-Sep | 0.02 | Old |
| 0.293739 | 3.035386 | 0.02 | 0 | 1 | 15 | Esm1 | 0.02 | Old |
| 0.293739 | 3.035386 | 0.02 | 0 | 1 | 15 | 2700097O( | 0.02 | Old |
| 0.293739 | 3.035386 | 0.02 | 0 | 1 | 15 | Sytl3 | 0.02 | Old |
| 0.293739 | 3.035386 | 0.02 | 0 | 1 | 15 | Vars2 | 0.02 | Old |
| 0.293739 | 3.035386 | 0.02 | 0 | 1 | 15 | Vmac | 0.02 | Old |
| 0.293739 | 3.035386 | 0.02 | 0 | 1 | 15 | Iigp1 | 0.02 | Old |
| 0.293739 | 3.005405 | 0.02 | 0 | 1 | 15 | Prr15 | 0.02 | Old |
| 0.293739 | 3.005405 | 0.02 | 0 | 1 | 15 | Gm20506 | 0.02 | Old |
| 0.293739 | 3.005405 | 0.02 | 0 | 1 | 15 | Kif11 | 0.02 | Old |
| 0.293739 | 2.896473 | 0.02 | 0 | 1 | 15 | Gm37223 | 0.02 | Old |
| 0.293739 | 2.896473 | 0.02 | 0 | 1 | 15 | Exd1 | 0.02 | Old |
| 0.293739 | 2.896473 | 0.02 | 0 | 1 | 15 | Ctsk | 0.02 | Old |
| 0.293739 | 2.896473 | 0.02 | 0 | 1 | 15 | Crygn | 0.02 | Old |
| 0.293739 | 2.896473 | 0.02 | 0 | 1 | 15 | Zfp235 | 0.02 | Old |
| 0.293739 | 2.896473 | 0.02 | 0 | 1 | 15 | Rasip1 | 0.02 | Old |
| 0.293739 | 2.896473 | 0.02 | 0 | 1 | 15 | Proscos | 0.02 | Old |
| 0.293739 | 2.896473 | 0.02 | 0 | 1 | 15 | Gm26514 | 0.02 | Old |
| 0.293739 | 2.896473 | 0.02 | 0 | 1 | 15 | Fam227a | 0.02 | Old |
| 0.293739 | 2.896473 | 0.02 | 0 | 1 | 15 | Prdm9 | 0.02 | Old |
| 0.293739 | 2.888612 | 0.02 | 0 | 1 | 15 | Mybl1 | 0.02 | Old |
| 0.293739 | 2.888612 | 0.02 | 0 | 1 | 15 | Rbbp9 | 0.02 | Old |
| 0.293739 | 2.888612 | 0.02 | 0 | 1 | 15 | Gm15446 | 0.02 | Old |
| 0.293739 | 2.888612 | 0.02 | 0 | 1 | 15 | Fbln2 | 0.02 | Old |
| 0.293739 | 2.888612 | 0.02 | 0 | 1 | 15 | Zswim9 | 0.02 | Old |
| 0.293739 | 2.888612 | 0.02 | 0 | 1 | 15 | Wdr73 | 0.02 | Old |
| 0.293739 | 2.888612 | 0.02 | 0 | 1 | 15 | Gm48045 | 0.02 | Old |
| 0.293739 | 2.888612 | 0.02 | 0 | 1 | 15 | Gstt2 | 0.02 | Old |
| 0.293739 | 2.888612 | 0.02 | 0 | 1 | 15 | Apol8 | 0.02 | Old |

| | | | | | | | | |
|---|---|---|---|---|---|---|---|---|
| 0.293739 | 2.888612 | 0.02 | 0 | 1 | 15 | 4931406G( | 0.02 | Old |
| 0.293739 | 2.828042 | 0.02 | 0 | 1 | 15 | Gtf2ird2 | 0.02 | Old |
| 0.293739 | 2.828042 | 0.02 | 0 | 1 | 15 | Klra17 | 0.02 | Old |
| 0.293739 | 2.828042 | 0.02 | 0 | 1 | 15 | Gm12258 | 0.02 | Old |
| 0.293739 | 2.828042 | 0.02 | 0 | 1 | 15 | Unk | 0.02 | Old |
| 0.293739 | 2.828042 | 0.02 | 0 | 1 | 15 | Hist1h3f | 0.02 | Old |
| 0.293739 | 2.828042 | 0.02 | 0 | 1 | 15 | 4833418N( | 0.02 | Old |
| 0.293739 | 2.823235 | 0.02 | 0 | 1 | 15 | Siah1b | 0.02 | Old |
| 0.293739 | 2.798073 | 0.02 | 0 | 1 | 15 | BC034090 | 0.02 | Old |
| 0.293739 | 2.798073 | 0.02 | 0 | 1 | 15 | Ddr2 | 0.02 | Old |
| 0.293739 | 2.798073 | 0.02 | 0 | 1 | 15 | Hdac6 | 0.02 | Old |
| 0.293739 | 2.798073 | 0.02 | 0 | 1 | 15 | Egf | 0.02 | Old |
| 0.293739 | 2.798073 | 0.02 | 0 | 1 | 15 | Capn12 | 0.02 | Old |
| 0.293739 | 2.798073 | 0.02 | 0 | 1 | 15 | Tubb3 | 0.02 | Old |
| 0.293739 | 2.798073 | 0.02 | 0 | 1 | 15 | Ankrd36 | 0.02 | Old |
| 0.293739 | 2.798073 | 0.02 | 0 | 1 | 15 | Taf9 | 0.02 | Old |
| 0.293739 | 2.798073 | 0.02 | 0 | 1 | 15 | AI661453 | 0.02 | Old |
| 0.293739 | 2.798073 | 0.02 | 0 | 1 | 15 | Mrpl49 | 0.02 | Old |
| 0.293739 | 2.795553 | 0.02 | 0 | 1 | 15 | Rnf25 | 0.02 | Old |
| 0.293739 | 2.795553 | 0.02 | 0 | 1 | 15 | Gtf3c4 | 0.02 | Old |
| 0.293739 | 2.795553 | 0.02 | 0 | 1 | 15 | Gm21994 | 0.02 | Old |
| 0.293739 | 2.795553 | 0.02 | 0 | 1 | 15 | Trmt44 | 0.02 | Old |
| 0.293739 | 2.795553 | 0.02 | 0 | 1 | 15 | Rasl11a | 0.02 | Old |
| 0.293739 | 2.795553 | 0.02 | 0 | 1 | 15 | Gm26810 | 0.02 | Old |
| 0.293739 | 2.795553 | 0.02 | 0 | 1 | 15 | Taf1c | 0.02 | Old |
| 0.293739 | 2.795553 | 0.02 | 0 | 1 | 15 | Exog | 0.02 | Old |
| 0.293739 | 2.795553 | 0.02 | 0 | 1 | 15 | Immp2l | 0.02 | Old |
| 0.293739 | 2.741894 | 0.02 | 0 | 1 | 15 | Mcmdc2 | 0.02 | Old |
| 0.293739 | 2.741894 | 0.02 | 0 | 1 | 15 | Mageh1 | 0.02 | Old |
| 0.293739 | 2.741894 | 0.02 | 0 | 1 | 15 | Gm11413 | 0.02 | Old |
| 0.293739 | 2.741894 | 0.02 | 0 | 1 | 15 | Rasal1 | 0.02 | Old |
| 0.293739 | 2.741894 | 0.02 | 0 | 1 | 15 | 4930403D( | 0.02 | Old |
| 0.293739 | 2.741894 | 0.02 | 0 | 1 | 15 | Dnmt3aos | 0.02 | Old |
| 0.293739 | 2.741894 | 0.02 | 0 | 1 | 15 | AC163032. | 0.02 | Old |
| 0.293739 | 2.741894 | 0.02 | 0 | 1 | 15 | Npr3 | 0.02 | Old |
| 0.293739 | 2.741894 | 0.02 | 0 | 1 | 15 | Ccnf | 0.02 | Old |
| 0.293739 | 2.741894 | 0.02 | 0 | 1 | 15 | Vit | 0.02 | Old |
| 0.293739 | 2.708224 | 0.02 | 0 | 1 | 15 | Twist2 | 0.02 | Old |
| 0.293739 | 2.708224 | 0.02 | 0 | 1 | 15 | Gm14154 | 0.02 | Old |
| 0.293739 | 2.708224 | 0.02 | 0 | 1 | 15 | Syn1 | 0.02 | Old |
| 0.293739 | 2.708224 | 0.02 | 0 | 1 | 15 | Cysltr2 | 0.02 | Old |
| 0.293739 | 2.700464 | 0.02 | 0 | 1 | 15 | Prkcg | 0.02 | Old |
| 0.293739 | 2.700464 | 0.02 | 0 | 1 | 15 | Tescl | 0.02 | Old |
| 0.293739 | 2.700464 | 0.02 | 0 | 1 | 15 | Cdc6 | 0.02 | Old |
| 0.293739 | 2.684627 | 0.02 | 0 | 1 | 15 | Gm28706 | 0.02 | Old |

| | | | | | | | | |
|---|---|---|---|---|---|---|---|---|
| 0.293739 | 2.684627 | 0.02 | 0 | 1 | 15 | Map7d3 | 0.02 | Old |
| 0.293739 | 2.684627 | 0.02 | 0 | 1 | 15 | Gm14508 | 0.02 | Old |
| 0.293739 | 2.684627 | 0.02 | 0 | 1 | 15 | Zfand4 | 0.02 | Old |
| 0.293739 | 2.684627 | 0.02 | 0 | 1 | 15 | Gm15133 | 0.02 | Old |
| 0.293739 | 2.684627 | 0.02 | 0 | 1 | 15 | Bfsp2 | 0.02 | Old |
| 0.293739 | 2.684627 | 0.02 | 0 | 1 | 15 | Acsf2 | 0.02 | Old |
| 0.293739 | 2.684627 | 0.02 | 0 | 1 | 15 | Ptges3l | 0.02 | Old |
| 0.293739 | 2.684627 | 0.02 | 0 | 1 | 15 | Mrc2 | 0.02 | Old |
| 0.293739 | 2.684627 | 0.02 | 0 | 1 | 15 | Calm4 | 0.02 | Old |
| 0.293739 | 2.684627 | 0.02 | 0 | 1 | 15 | Gm21319 | 0.02 | Old |
| 0.293739 | 2.684627 | 0.02 | 0 | 1 | 15 | Rwdd2b | 0.02 | Old |
| 0.293739 | 2.684627 | 0.02 | 0 | 1 | 15 | Chaf1b | 0.02 | Old |
| 0.293739 | 2.684627 | 0.02 | 0 | 1 | 15 | Tedc2 | 0.02 | Old |
| 0.293739 | 2.684627 | 0.02 | 0 | 1 | 15 | Sema6a | 0.02 | Old |
| 0.293739 | 2.684627 | 0.02 | 0 | 1 | 15 | Syt7 | 0.02 | Old |
| 0.293739 | 2.65428 | 0.02 | 0 | 1 | 15 | Adamts13 | 0.02 | Old |
| 0.293739 | 2.65428 | 0.02 | 0 | 1 | 15 | Mrrf | 0.02 | Old |
| 0.293739 | 2.65428 | 0.02 | 0 | 1 | 15 | Gm13561 | 0.02 | Old |
| 0.293739 | 2.65428 | 0.02 | 0 | 1 | 15 | P3h1 | 0.02 | Old |
| 0.293739 | 2.65428 | 0.02 | 0 | 1 | 15 | Fndc5 | 0.02 | Old |
| 0.293739 | 2.65428 | 0.02 | 0 | 1 | 15 | Ctrc | 0.02 | Old |
| 0.293739 | 2.65428 | 0.02 | 0 | 1 | 15 | Gm16008 | 0.02 | Old |
| 0.293739 | 2.65428 | 0.02 | 0 | 1 | 15 | Clec4a4 | 0.02 | Old |
| 0.293739 | 2.65428 | 0.02 | 0 | 1 | 15 | Ccdc138 | 0.02 | Old |
| 0.293739 | 2.65428 | 0.02 | 0 | 1 | 15 | Hacl1 | 0.02 | Old |
| 0.293739 | 2.65428 | 0.02 | 0 | 1 | 15 | Mpzl2 | 0.02 | Old |
| 0.293739 | 2.65428 | 0.02 | 0 | 1 | 15 | Pou2af1 | 0.02 | Old |
| 0.293739 | 2.65428 | 0.02 | 0 | 1 | 15 | Gm9856 | 0.02 | Old |
| 0.293739 | 2.65428 | 0.02 | 0 | 1 | 15 | Rad51c | 0.02 | Old |
| 0.293739 | 2.65428 | 0.02 | 0 | 1 | 15 | Scrn2 | 0.02 | Old |
| 0.293739 | 2.65428 | 0.02 | 0 | 1 | 15 | Ahrr | 0.02 | Old |
| 0.293739 | 2.65428 | 0.02 | 0 | 1 | 15 | Enc1 | 0.02 | Old |
| 0.293739 | 2.65428 | 0.02 | 0 | 1 | 15 | Gm36287 | 0.02 | Old |
| 0.293739 | 2.65428 | 0.02 | 0 | 1 | 15 | Spag1 | 0.02 | Old |
| 0.293739 | 2.65428 | 0.02 | 0 | 1 | 15 | Stard6 | 0.02 | Old |
| 0.293739 | 2.65428 | 0.02 | 0 | 1 | 15 | Carns1 | 0.02 | Old |
| 0.293739 | 2.653021 | 0.02 | 0 | 1 | 15 | B430305J0 | 0.02 | Old |
| 0.293739 | 2.653021 | 0.02 | 0 | 1 | 15 | Pigo | 0.02 | Old |
| 0.293739 | 2.653021 | 0.02 | 0 | 1 | 15 | Ccsap | 0.02 | Old |
| 0.293739 | 2.653021 | 0.02 | 0 | 1 | 15 | Lrrc1 | 0.02 | Old |
| 0.293739 | 2.653021 | 0.02 | 0 | 1 | 15 | Map6d1 | 0.02 | Old |
| 0.293739 | 2.653021 | 0.02 | 0 | 1 | 15 | Mpp7 | 0.02 | Old |
| 0.293739 | 2.653021 | 0.02 | 0 | 1 | 15 | Poll | 0.02 | Old |
| 0.293739 | 2.632313 | 0.02 | 0 | 1 | 15 | Elobl | 0.02 | Old |
| 0.293739 | 2.623563 | 0.02 | 0 | 1 | 15 | Gm13564 | 0.02 | Old |

| | | | | | | | | |
|---|---|---|---|---|---|---|---|---|
| 0.293739 | 2.623563 | 0.02 | 0 | 1 | 15 | Afg1l | 0.02 | Old |
| 0.293739 | 2.623563 | 0.02 | 0 | 1 | 15 | Gm47392 | 0.02 | Old |
| 0.293739 | 2.623563 | 0.02 | 0 | 1 | 15 | Cygb | 0.02 | Old |
| 0.293739 | 2.623563 | 0.02 | 0 | 1 | 15 | Zfp623 | 0.02 | Old |
| 0.293739 | 2.623563 | 0.02 | 0 | 1 | 15 | Tmem241 | 0.02 | Old |
| 0.293739 | 2.539908 | 0.02 | 0 | 1 | 15 | Nkrf | 0.02 | Old |
| 0.293739 | 2.539908 | 0.02 | 0 | 1 | 15 | Ptger1 | 0.02 | Old |
| 0.293739 | 2.524799 | 0.02 | 0 | 1 | 15 | Mmgt1 | 0.02 | Old |
| 0.293739 | 2.524799 | 0.02 | 0 | 1 | 15 | Klhl4 | 0.02 | Old |
| 0.293739 | 2.524799 | 0.02 | 0 | 1 | 15 | 2610008E1 | 0.02 | Old |
| 0.293739 | 2.524799 | 0.02 | 0 | 1 | 15 | Lum | 0.02 | Old |
| 0.293739 | 2.524799 | 0.02 | 0 | 1 | 15 | Slc25a15 | 0.02 | Old |
| 0.293739 | 2.524799 | 0.02 | 0 | 1 | 15 | Brf2 | 0.02 | Old |
| 0.293739 | 2.524799 | 0.02 | 0 | 1 | 15 | Gm10131 | 0.02 | Old |
| 0.293739 | 2.524799 | 0.02 | 0 | 1 | 15 | Fhit | 0.02 | Old |
| 0.293739 | 2.524799 | 0.02 | 0 | 1 | 15 | Pomt2 | 0.02 | Old |
| 0.293739 | 2.524799 | 0.02 | 0 | 1 | 15 | Ctnnd2 | 0.02 | Old |
| 0.293739 | 2.524799 | 0.02 | 0 | 1 | 15 | Hectd2 | 0.02 | Old |
| 0.293739 | 2.493353 | 0.02 | 0 | 1 | 15 | Cd46 | 0.02 | Old |
| 0.293739 | 2.493353 | 0.02 | 0 | 1 | 15 | Ndor1 | 0.02 | Old |
| 0.293739 | 2.493353 | 0.02 | 0 | 1 | 15 | Lhfp | 0.02 | Old |
| 0.293739 | 2.493353 | 0.02 | 0 | 1 | 15 | Elavl4 | 0.02 | Old |
| 0.293739 | 2.493353 | 0.02 | 0 | 1 | 15 | Srm | 0.02 | Old |
| 0.293739 | 2.493353 | 0.02 | 0 | 1 | 15 | Zfp667 | 0.02 | Old |
| 0.293739 | 2.493353 | 0.02 | 0 | 1 | 15 | Fbxo17 | 0.02 | Old |
| 0.293739 | 2.493353 | 0.02 | 0 | 1 | 15 | Col6a2 | 0.02 | Old |
| 0.293739 | 2.493353 | 0.02 | 0 | 1 | 15 | Ddx28 | 0.02 | Old |
| 0.293739 | 2.493353 | 0.02 | 0 | 1 | 15 | Ift88 | 0.02 | Old |
| 0.293739 | 2.493353 | 0.02 | 0 | 1 | 15 | Adamts7 | 0.02 | Old |
| 0.293739 | 2.493353 | 0.02 | 0 | 1 | 15 | Gpr179 | 0.02 | Old |
| 0.293739 | 2.493353 | 0.02 | 0 | 1 | 15 | Adam11 | 0.02 | Old |
| 0.293739 | 2.493353 | 0.02 | 0 | 1 | 15 | Hist1h1d | 0.02 | Old |
| 0.293739 | 2.493353 | 0.02 | 0 | 1 | 15 | Tnk2 | 0.02 | Old |
| 0.293739 | 2.493353 | 0.02 | 0 | 1 | 15 | Urb1 | 0.02 | Old |
| 0.293739 | 2.493353 | 0.02 | 0 | 1 | 15 | Ccnj | 0.02 | Old |
| 0.293739 | 2.486511 | 0.02 | 0 | 1 | 15 | Gm2115 | 0.02 | Old |
| 0.293739 | 2.443333 | 0.02 | 0 | 1 | 15 | Gm12166 | 0.02 | Old |
| 0.293739 | 2.389869 | 0.02 | 0 | 1 | 15 | Dusp4 | 0.02 | Old |
| 0.293739 | 2.389869 | 0.02 | 0 | 1 | 15 | Csgalnact1 | 0.02 | Old |
| 0.293739 | 2.378772 | 0.02 | 0 | 1 | 15 | Gal3st2b | 0.02 | Old |
| 0.293739 | 2.378772 | 0.02 | 0 | 1 | 15 | Kcnb1 | 0.02 | Old |
| 0.293739 | 2.378772 | 0.02 | 0 | 1 | 15 | Gm15726 | 0.02 | Old |
| 0.293739 | 2.378772 | 0.02 | 0 | 1 | 15 | Rimkla | 0.02 | Old |
| 0.293739 | 2.378772 | 0.02 | 0 | 1 | 15 | Tssk3 | 0.02 | Old |
| 0.293739 | 2.378772 | 0.02 | 0 | 1 | 15 | Smkr-ps | 0.02 | Old |

| | | | | | | | | |
|---|---|---|---|---|---|---|---|---|
| 0.293739 | 2.378772 | 0.02 | 0 | 1 | 15 | Gm42688 | 0.02 | Old |
| 0.293739 | 2.378772 | 0.02 | 0 | 1 | 15 | Hpn | 0.02 | Old |
| 0.293739 | 2.378772 | 0.02 | 0 | 1 | 15 | Kcnc1 | 0.02 | Old |
| 0.293739 | 2.378772 | 0.02 | 0 | 1 | 15 | Whamm | 0.02 | Old |
| 0.293739 | 2.378772 | 0.02 | 0 | 1 | 15 | Chst3 | 0.02 | Old |
| 0.293739 | 2.378772 | 0.02 | 0 | 1 | 15 | 1500009L1 | 0.02 | Old |
| 0.293739 | 2.378772 | 0.02 | 0 | 1 | 15 | A530010L1 | 0.02 | Old |
| 0.293739 | 2.378772 | 0.02 | 0 | 1 | 15 | Carmil3 | 0.02 | Old |
| 0.293739 | 2.378772 | 0.02 | 0 | 1 | 15 | Gm26733 | 0.02 | Old |
| 0.293739 | 2.378772 | 0.02 | 0 | 1 | 15 | Nrgn | 0.02 | Old |
| 0.293739 | 2.378772 | 0.02 | 0 | 1 | 15 | Vipr1 | 0.02 | Old |
| 0.293739 | 2.378772 | 0.02 | 0 | 1 | 15 | Pigl | 0.02 | Old |
| 0.293739 | 2.378772 | 0.02 | 0 | 1 | 15 | Gm49233 | 0.02 | Old |
| 0.293739 | 2.378772 | 0.02 | 0 | 1 | 15 | Gm10791 | 0.02 | Old |
| 0.293739 | 2.378772 | 0.02 | 0 | 1 | 15 | Tcte1 | 0.02 | Old |
| 0.293739 | 2.378772 | 0.02 | 0 | 1 | 15 | Bvht | 0.02 | Old |
| 0.293739 | 2.378772 | 0.02 | 0 | 1 | 15 | Gm45871 | 0.02 | Old |
| 0.293739 | 2.378772 | 0.02 | 0 | 1 | 15 | Rad9a | 0.02 | Old |
| 0.293739 | 2.378772 | 0.02 | 0 | 1 | 15 | Taf5 | 0.02 | Old |
| 0.293739 | 2.354139 | 0.02 | 0 | 1 | 15 | Inpp1 | 0.02 | Old |
| 0.293739 | 2.354139 | 0.02 | 0 | 1 | 15 | Mars2 | 0.02 | Old |
| 0.293739 | 2.354139 | 0.02 | 0 | 1 | 15 | Atp2b4 | 0.02 | Old |
| 0.293739 | 2.354139 | 0.02 | 0 | 1 | 15 | Kif5c | 0.02 | Old |
| 0.293739 | 2.354139 | 0.02 | 0 | 1 | 15 | Gm29865 | 0.02 | Old |
| 0.293739 | 2.354139 | 0.02 | 0 | 1 | 15 | Gm8013 | 0.02 | Old |
| 0.293739 | 2.354139 | 0.02 | 0 | 1 | 15 | Tfip11 | 0.02 | Old |
| 0.293739 | 2.354139 | 0.02 | 0 | 1 | 15 | Zfp827 | 0.02 | Old |
| 0.293739 | 2.354139 | 0.02 | 0 | 1 | 15 | Pkd1l2 | 0.02 | Old |
| 0.293739 | 2.354139 | 0.02 | 0 | 1 | 15 | Gm10655 | 0.02 | Old |
| 0.293739 | 2.354139 | 0.02 | 0 | 1 | 15 | Ccnb2 | 0.02 | Old |
| 0.293739 | 2.354139 | 0.02 | 0 | 1 | 15 | Gm10277 | 0.02 | Old |
| 0.293739 | 2.354139 | 0.02 | 0 | 1 | 15 | 1700006E0 | 0.02 | Old |
| 0.293739 | 2.354139 | 0.02 | 0 | 1 | 15 | Fntb | 0.02 | Old |
| 0.293739 | 2.354139 | 0.02 | 0 | 1 | 15 | 3300005D0 | 0.02 | Old |
| 0.293739 | 2.289029 | 0.02 | 0 | 1 | 15 | Bard1 | 0.02 | Old |
| 0.293739 | 2.289029 | 0.02 | 0 | 1 | 15 | 2900009J0 | 0.02 | Old |
| 0.293739 | 2.289029 | 0.02 | 0 | 1 | 15 | Gm14325 | 0.02 | Old |
| 0.293739 | 2.289029 | 0.02 | 0 | 1 | 15 | Gpr171 | 0.02 | Old |
| 0.293739 | 2.289029 | 0.02 | 0 | 1 | 15 | Gm37696 | 0.02 | Old |
| 0.293739 | 2.289029 | 0.02 | 0 | 1 | 15 | Prkaa2 | 0.02 | Old |
| 0.293739 | 2.289029 | 0.02 | 0 | 1 | 15 | 8430408G2 | 0.02 | Old |
| 0.293739 | 2.289029 | 0.02 | 0 | 1 | 15 | Cbarp | 0.02 | Old |
| 0.293739 | 2.289029 | 0.02 | 0 | 1 | 15 | Foxc1 | 0.02 | Old |
| 0.293739 | 2.289029 | 0.02 | 0 | 1 | 15 | 5033430I1 | 0.02 | Old |
| 0.293739 | 2.289029 | 0.02 | 0 | 1 | 15 | Ccdc167 | 0.02 | Old |

| | | | | | | | | |
|---|---|---|---|---|---|---|---|---|
| 0.293739 | 2.288155 | 0.02 | 0 | 1 | 15 | Lrrc32 | 0.02 | Old |
| 0.293739 | 2.288155 | 0.02 | 0 | 1 | 15 | Ccdc84 | 0.02 | Old |
| 0.293739 | 2.254264 | 0.02 | 0 | 1 | 15 | Dtwd1 | 0.02 | Old |
| 0.293739 | 2.254264 | 0.02 | 0 | 1 | 15 | Gm14005 | 0.02 | Old |
| 0.293739 | 2.254264 | 0.02 | 0 | 1 | 15 | Spag4 | 0.02 | Old |
| 0.293739 | 2.254264 | 0.02 | 0 | 1 | 15 | Fam189b | 0.02 | Old |
| 0.293739 | 2.254264 | 0.02 | 0 | 1 | 15 | Arhgap29 | 0.02 | Old |
| 0.293739 | 2.254264 | 0.02 | 0 | 1 | 15 | 1700086P0 | 0.02 | Old |
| 0.293739 | 2.254264 | 0.02 | 0 | 1 | 15 | Hsd17b14 | 0.02 | Old |
| 0.293739 | 2.254264 | 0.02 | 0 | 1 | 15 | Grwd1 | 0.02 | Old |
| 0.293739 | 2.254264 | 0.02 | 0 | 1 | 15 | Cpeb1 | 0.02 | Old |
| 0.293739 | 2.254264 | 0.02 | 0 | 1 | 15 | Gm48066 | 0.02 | Old |
| 0.293739 | 2.254264 | 0.02 | 0 | 1 | 15 | Sap18b | 0.02 | Old |
| 0.293739 | 2.254264 | 0.02 | 0 | 1 | 15 | Cnrip1 | 0.02 | Old |
| 0.293739 | 2.254264 | 0.02 | 0 | 1 | 15 | Slfn9 | 0.02 | Old |
| 0.293739 | 2.254264 | 0.02 | 0 | 1 | 15 | Hus1b | 0.02 | Old |
| 0.293739 | 2.254264 | 0.02 | 0 | 1 | 15 | Dnajc27 | 0.02 | Old |
| 0.293739 | 2.254264 | 0.02 | 0 | 1 | 15 | Prdm15 | 0.02 | Old |
| 0.293739 | 2.249221 | 0.02 | 0 | 1 | 15 | Abcb6 | 0.02 | Old |
| 0.293739 | 2.249221 | 0.02 | 0 | 1 | 15 | Lrp11 | 0.02 | Old |
| 0.293739 | 2.249221 | 0.02 | 0 | 1 | 15 | Gm49087 | 0.02 | Old |
| 0.293739 | 2.249221 | 0.02 | 0 | 1 | 15 | Panx1 | 0.02 | Old |
| 0.293739 | 2.249221 | 0.02 | 0 | 1 | 15 | Elp6 | 0.02 | Old |
| 0.293739 | 2.249221 | 0.02 | 0 | 1 | 15 | Zpbp2 | 0.02 | Old |
| 0.293739 | 2.249221 | 0.02 | 0 | 1 | 15 | 4930426I24 | 0.02 | Old |
| 0.293739 | 2.249221 | 0.02 | 0 | 1 | 15 | Tfcp2 | 0.02 | Old |
| 0.293739 | 2.249221 | 0.02 | 0 | 1 | 15 | Gm6133 | 0.02 | Old |
| 0.293739 | 2.220737 | 0.02 | 0 | 1 | 15 | Zc3h11a | 0.02 | Old |
| 0.293739 | 2.220737 | 0.02 | 0 | 1 | 15 | Chrna1os | 0.02 | Old |
| 0.293739 | 2.220737 | 0.02 | 0 | 1 | 15 | Pex10 | 0.02 | Old |
| 0.293739 | 2.220737 | 0.02 | 0 | 1 | 15 | Gm42726 | 0.02 | Old |
| 0.293739 | 2.220737 | 0.02 | 0 | 1 | 15 | Cracr2a | 0.02 | Old |
| 0.293739 | 2.220737 | 0.02 | 0 | 1 | 15 | Kctd21 | 0.02 | Old |
| 0.293739 | 2.220737 | 0.02 | 0 | 1 | 15 | Slc12a3 | 0.02 | Old |
| 0.293739 | 2.220737 | 0.02 | 0 | 1 | 15 | Zfp54 | 0.02 | Old |
| 0.293739 | 2.220737 | 0.02 | 0 | 1 | 15 | Rab44 | 0.02 | Old |
| 0.293739 | 2.220737 | 0.02 | 0 | 1 | 15 | Zfp763 | 0.02 | Old |
| 0.293739 | 2.220737 | 0.02 | 0 | 1 | 15 | Xdh | 0.02 | Old |
| 0.293739 | 2.07784 | 0.02 | 0 | 1 | 15 | Rufy4 | 0.02 | Old |
| 0.293739 | 2.07784 | 0.02 | 0 | 1 | 15 | Hgf | 0.02 | Old |
| 0.293739 | 2.07784 | 0.02 | 0 | 1 | 15 | Gm26982 | 0.02 | Old |
| 0.293739 | 2.013904 | 0.02 | 0 | 1 | 15 | Zfp973 | 0.02 | Old |
| 0.293739 | 2.013904 | 0.02 | 0 | 1 | 15 | Tsix | 0.02 | Old |
| 0.293739 | 2.013904 | 0.02 | 0 | 1 | 15 | Jpx | 0.02 | Old |
| 0.293739 | 2.013904 | 0.02 | 0 | 1 | 15 | A430033K0 | 0.02 | Old |

| | | | | | | | | |
|---|---|---|---|---|---|---|---|---|
| 0.293739 | 2.013904 | 0.02 | 0 | 1 | 15 | Cidec | 0.02 | Old |
| 0.293739 | 2.013904 | 0.02 | 0 | 1 | 15 | Pak4 | 0.02 | Old |
| 0.293739 | 2.013904 | 0.02 | 0 | 1 | 15 | Lzts1 | 0.02 | Old |
| 0.293739 | 2.013904 | 0.02 | 0 | 1 | 15 | Timm8a2 | 0.02 | Old |
| 0.293739 | 2.013904 | 0.02 | 0 | 1 | 15 | 9930022D1 | 0.02 | Old |
| 0.293739 | 2.013904 | 0.02 | 0 | 1 | 15 | Gm48837 | 0.02 | Old |
| 0.293739 | 2.013904 | 0.02 | 0 | 1 | 15 | Dact1 | 0.02 | Old |
| 0.293739 | 2.013904 | 0.02 | 0 | 1 | 15 | Gm49077 | 0.02 | Old |
| 0.293739 | 2.013904 | 0.02 | 0 | 1 | 15 | Vwce | 0.02 | Old |
| 0.293739 | 1.90734 | 0.02 | 0 | 1 | 15 | Olfr1173 | 0.02 | Old |
| 0.293739 | 1.90734 | 0.02 | 0 | 1 | 15 | 1200007C1 | 0.02 | Old |
| 0.293739 | 1.90734 | 0.02 | 0 | 1 | 15 | Tcea2 | 0.02 | Old |
| 0.293739 | 1.90734 | 0.02 | 0 | 1 | 15 | Dlg3 | 0.02 | Old |
| 0.293739 | 1.90734 | 0.02 | 0 | 1 | 15 | Serpini1 | 0.02 | Old |
| 0.293739 | 1.90734 | 0.02 | 0 | 1 | 15 | Gjb3 | 0.02 | Old |
| 0.293739 | 1.90734 | 0.02 | 0 | 1 | 15 | Ficd | 0.02 | Old |
| 0.293739 | 1.90734 | 0.02 | 0 | 1 | 15 | Tmc4 | 0.02 | Old |
| 0.293739 | 1.90734 | 0.02 | 0 | 1 | 15 | Shcbp1 | 0.02 | Old |
| 0.293739 | 1.90734 | 0.02 | 0 | 1 | 15 | Sorbs2 | 0.02 | Old |
| 0.293739 | 1.90734 | 0.02 | 0 | 1 | 15 | Gm3235 | 0.02 | Old |
| 0.293739 | 1.90734 | 0.02 | 0 | 1 | 15 | Ankrd28 | 0.02 | Old |
| 0.293739 | 1.90734 | 0.02 | 0 | 1 | 15 | Gm48529 | 0.02 | Old |
| 0.293739 | 1.90734 | 0.02 | 0 | 1 | 15 | 4933427D1 | 0.02 | Old |
| 0.293739 | 1.90734 | 0.02 | 0 | 1 | 15 | Rnf43 | 0.02 | Old |
| 0.293739 | 1.90734 | 0.02 | 0 | 1 | 15 | Pcgf2 | 0.02 | Old |
| 0.293739 | 1.90734 | 0.02 | 0 | 1 | 15 | Gm28707 | 0.02 | Old |
| 0.293739 | 1.90734 | 0.02 | 0 | 1 | 15 | Adamts6 | 0.02 | Old |
| 0.293739 | 1.90734 | 0.02 | 0 | 1 | 15 | Bdkrb2 | 0.02 | Old |
| 0.293739 | 1.90734 | 0.02 | 0 | 1 | 15 | Tnfrsf13c | 0.02 | Old |
| 0.293739 | 1.90734 | 0.02 | 0 | 1 | 15 | Plcxd2 | 0.02 | Old |
| 0.293739 | 1.90734 | 0.02 | 0 | 1 | 15 | Mdga1 | 0.02 | Old |
| 0.293739 | 1.90734 | 0.02 | 0 | 1 | 15 | Tesmin | 0.02 | Old |
| 0.293739 | 1.90734 | 0.02 | 0 | 1 | 15 | Sptbn2 | 0.02 | Old |
| 0.293739 | 1.906912 | 0.02 | 0 | 1 | 15 | Gsta3 | 0.02 | Old |
| 0.293739 | 1.906912 | 0.02 | 0 | 1 | 15 | C1qtnf4 | 0.02 | Old |
| 0.293739 | 1.906912 | 0.02 | 0 | 1 | 15 | Mum1l1 | 0.02 | Old |
| 0.293739 | 1.906912 | 0.02 | 0 | 1 | 15 | Ccl27a | 0.02 | Old |
| 0.293739 | 1.906912 | 0.02 | 0 | 1 | 15 | Fggy | 0.02 | Old |
| 0.293739 | 1.906912 | 0.02 | 0 | 1 | 15 | Hook1 | 0.02 | Old |
| 0.293739 | 1.906912 | 0.02 | 0 | 1 | 15 | Kazn | 0.02 | Old |
| 0.293739 | 1.906912 | 0.02 | 0 | 1 | 15 | Gm26699 | 0.02 | Old |
| 0.293739 | 1.906912 | 0.02 | 0 | 1 | 15 | Gm43479 | 0.02 | Old |
| 0.293739 | 1.906912 | 0.02 | 0 | 1 | 15 | Dact3 | 0.02 | Old |
| 0.293739 | 1.906912 | 0.02 | 0 | 1 | 15 | Zfp60 | 0.02 | Old |
| 0.293739 | 1.906912 | 0.02 | 0 | 1 | 15 | Adamts14 | 0.02 | Old |

| | | | | | | | | |
|---|---|---|---|---|---|---|---|---|
| 0.293739 | 1.906912 | 0.02 | 0 | 1 | 15 | Zfr2 | 0.02 | Old |
| 0.293739 | 1.906912 | 0.02 | 0 | 1 | 15 | Tmtc4 | 0.02 | Old |
| 0.293739 | 1.906912 | 0.02 | 0 | 1 | 15 | Myo6 | 0.02 | Old |
| 0.293739 | 1.906912 | 0.02 | 0 | 1 | 15 | Lif | 0.02 | Old |
| 0.293739 | 1.906912 | 0.02 | 0 | 1 | 15 | Zfp2 | 0.02 | Old |
| 0.293739 | 1.906912 | 0.02 | 0 | 1 | 15 | Tbx21 | 0.02 | Old |
| 0.293739 | 1.906912 | 0.02 | 0 | 1 | 15 | Krt20 | 0.02 | Old |
| 0.293739 | 1.906912 | 0.02 | 0 | 1 | 15 | Arhgef28 | 0.02 | Old |
| 0.293739 | 1.906912 | 0.02 | 0 | 1 | 15 | Arg2 | 0.02 | Old |
| 0.293739 | 1.906912 | 0.02 | 0 | 1 | 15 | Ccdc152 | 0.02 | Old |
| 0.293739 | 1.906912 | 0.02 | 0 | 1 | 15 | Gm29019 | 0.02 | Old |
| 0.293739 | 1.906912 | 0.02 | 0 | 1 | 15 | Celsr1 | 0.02 | Old |
| 0.293739 | 1.906912 | 0.02 | 0 | 1 | 15 | Cpne8 | 0.02 | Old |
| 0.293739 | 1.906912 | 0.02 | 0 | 1 | 15 | Pus7l | 0.02 | Old |
| 0.293739 | 1.906912 | 0.02 | 0 | 1 | 15 | Ephb3 | 0.02 | Old |
| 0.293739 | 1.906912 | 0.02 | 0 | 1 | 15 | Riok2 | 0.02 | Old |
| 0.293739 | 1.906912 | 0.02 | 0 | 1 | 15 | Gm10269 | 0.02 | Old |
| 0.293739 | 1.906912 | 0.02 | 0 | 1 | 15 | Pstpip2 | 0.02 | Old |
| 0.293739 | 1.906912 | 0.02 | 0 | 1 | 15 | Ankrd13d | 0.02 | Old |
| 0.293739 | 1.906912 | 0.02 | 0 | 1 | 15 | Hspa12a | 0.02 | Old |
| 0.293739 | 1.848036 | 0.02 | 0 | 1 | 15 | Il27 | 0.02 | Old |
| 0.293739 | 1.848036 | 0.02 | 0 | 1 | 15 | Plscr2 | 0.02 | Old |
| 0.293739 | 1.723386 | 0.02 | 0 | 1 | 15 | 1110051M | 0.02 | Old |
| 0.293739 | 1.723386 | 0.02 | 0 | 1 | 15 | Rad51 | 0.02 | Old |
| 0.293739 | 1.723386 | 0.02 | 0 | 1 | 15 | Lcmt2 | 0.02 | Old |
| 0.293739 | 1.723386 | 0.02 | 0 | 1 | 15 | Cdk5rap1 | 0.02 | Old |
| 0.293739 | 1.723386 | 0.02 | 0 | 1 | 15 | Ahcy | 0.02 | Old |
| 0.293739 | 1.723386 | 0.02 | 0 | 1 | 15 | Gm11464 | 0.02 | Old |
| 0.293739 | 1.723386 | 0.02 | 0 | 1 | 15 | Gm14698 | 0.02 | Old |
| 0.293739 | 1.723386 | 0.02 | 0 | 1 | 15 | Acp6 | 0.02 | Old |
| 0.293739 | 1.723386 | 0.02 | 0 | 1 | 15 | Nccrp1 | 0.02 | Old |
| 0.293739 | 1.723386 | 0.02 | 0 | 1 | 15 | Zfp975 | 0.02 | Old |
| 0.293739 | 1.723386 | 0.02 | 0 | 1 | 15 | Gm47922 | 0.02 | Old |
| 0.293739 | 1.723386 | 0.02 | 0 | 1 | 15 | Tmem221 | 0.02 | Old |
| 0.293739 | 1.723386 | 0.02 | 0 | 1 | 15 | Fam89a | 0.02 | Old |
| 0.293739 | 1.723386 | 0.02 | 0 | 1 | 15 | Tgtp2 | 0.02 | Old |
| 0.293739 | 1.723386 | 0.02 | 0 | 1 | 15 | Gm15892 | 0.02 | Old |
| 0.293739 | 1.723386 | 0.02 | 0 | 1 | 15 | Gli3 | 0.02 | Old |
| 0.293739 | 1.723386 | 0.02 | 0 | 1 | 15 | Gen1 | 0.02 | Old |
| 0.293739 | 1.723386 | 0.02 | 0 | 1 | 15 | Them6 | 0.02 | Old |
| 0.293739 | 1.723386 | 0.02 | 0 | 1 | 15 | Ctsw | 0.02 | Old |
| 0.293739 | 1.723386 | 0.02 | 0 | 1 | 15 | Tmem252 | 0.02 | Old |
| 0.293739 | 1.723386 | 0.02 | 0 | 1 | 15 | Gsto2 | 0.02 | Old |
| 0.293739 | 1.68683 | 0.02 | 0 | 1 | 15 | Cacna1b | 0.02 | Old |
| 0.293739 | 1.68683 | 0.02 | 0 | 1 | 15 | Dhrs9 | 0.02 | Old |

| | | | | | | | | |
|---|---|---|---|---|---|---|---|---|
| 0.293739 | 1.68683 | 0.02 | 0 | 1 | 15 | Ubiad1 | 0.02 | Old |
| 0.293739 | 1.68683 | 0.02 | 0 | 1 | 15 | Zbtb49 | 0.02 | Old |
| 0.293739 | 1.68683 | 0.02 | 0 | 1 | 15 | Sspo | 0.02 | Old |
| 0.293739 | 1.68683 | 0.02 | 0 | 1 | 15 | Mfap5 | 0.02 | Old |
| 0.293739 | 1.68683 | 0.02 | 0 | 1 | 15 | Stx1b | 0.02 | Old |
| 0.293739 | 1.68683 | 0.02 | 0 | 1 | 15 | Zfp280b | 0.02 | Old |
| 0.293739 | 1.68683 | 0.02 | 0 | 1 | 15 | 4930404N1 | 0.02 | Old |
| 0.293739 | 1.68683 | 0.02 | 0 | 1 | 15 | Gm5427 | 0.02 | Old |
| 0.293739 | 1.68683 | 0.02 | 0 | 1 | 15 | Enkd1 | 0.02 | Old |
| 0.293739 | 1.68683 | 0.02 | 0 | 1 | 15 | Mrps22 | 0.02 | Old |
| 0.293739 | 1.68683 | 0.02 | 0 | 1 | 15 | Chrnb1 | 0.02 | Old |
| 0.293739 | 1.68683 | 0.02 | 0 | 1 | 15 | Trp53i13 | 0.02 | Old |
| 0.293739 | 1.68683 | 0.02 | 0 | 1 | 15 | Gm11496 | 0.02 | Old |
| 0.293739 | 1.68683 | 0.02 | 0 | 1 | 15 | Mpp3 | 0.02 | Old |
| 0.293739 | 1.68683 | 0.02 | 0 | 1 | 15 | Polg2 | 0.02 | Old |
| 0.293739 | 1.68683 | 0.02 | 0 | 1 | 15 | Caskin2 | 0.02 | Old |
| 0.293739 | 1.68683 | 0.02 | 0 | 1 | 15 | Nid1 | 0.02 | Old |
| 0.293739 | 1.68683 | 0.02 | 0 | 1 | 15 | Hist1h1b | 0.02 | Old |
| 0.293739 | 1.68683 | 0.02 | 0 | 1 | 15 | Hist1h3d | 0.02 | Old |
| 0.293739 | 1.68683 | 0.02 | 0 | 1 | 15 | Hist1h2ac | 0.02 | Old |
| 0.293739 | 1.68683 | 0.02 | 0 | 1 | 15 | Ntrk2 | 0.02 | Old |
| 0.293739 | 1.68683 | 0.02 | 0 | 1 | 15 | Kif3c | 0.02 | Old |
| 0.293739 | 1.68683 | 0.02 | 0 | 1 | 15 | Scin | 0.02 | Old |
| 0.293739 | 1.68683 | 0.02 | 0 | 1 | 15 | Ift27 | 0.02 | Old |
| 0.293739 | 1.68683 | 0.02 | 0 | 1 | 15 | Prrt1 | 0.02 | Old |
| 0.293739 | 1.68683 | 0.02 | 0 | 1 | 15 | Klc2 | 0.02 | Old |
| 0.293739 | 1.68683 | 0.02 | 0 | 1 | 15 | Plce1 | 0.02 | Old |
| 0.293739 | 1.68683 | 0.02 | 0 | 1 | 15 | AC125149. | 0.02 | Old |
| 0.293739 | 1.6418 | 0.02 | 0 | 1 | 15 | Pecr | 0.02 | Old |
| 0.293739 | 1.6418 | 0.02 | 0 | 1 | 15 | D2hgdh | 0.02 | Old |
| 0.293739 | 1.6418 | 0.02 | 0 | 1 | 15 | Vangl2 | 0.02 | Old |
| 0.293739 | 1.6418 | 0.02 | 0 | 1 | 15 | Fastkd1 | 0.02 | Old |
| 0.293739 | 1.6418 | 0.02 | 0 | 1 | 15 | Gm42893 | 0.02 | Old |
| 0.293739 | 1.6418 | 0.02 | 0 | 1 | 15 | Zswim5 | 0.02 | Old |
| 0.293739 | 1.6418 | 0.02 | 0 | 1 | 15 | Tlr12 | 0.02 | Old |
| 0.293739 | 1.6418 | 0.02 | 0 | 1 | 15 | Sumf2 | 0.02 | Old |
| 0.293739 | 1.6418 | 0.02 | 0 | 1 | 15 | Zbed5 | 0.02 | Old |
| 0.293739 | 1.6418 | 0.02 | 0 | 1 | 15 | BC024978 | 0.02 | Old |
| 0.293739 | 1.6418 | 0.02 | 0 | 1 | 15 | Fam187b | 0.02 | Old |
| 0.293739 | 1.6418 | 0.02 | 0 | 1 | 15 | Wdr83 | 0.02 | Old |
| 0.293739 | 1.6418 | 0.02 | 0 | 1 | 15 | Gm48935 | 0.02 | Old |
| 0.293739 | 1.6418 | 0.02 | 0 | 1 | 15 | Rpusd4 | 0.02 | Old |
| 0.293739 | 1.6418 | 0.02 | 0 | 1 | 15 | Inhba | 0.02 | Old |
| 0.293739 | 1.6418 | 0.02 | 0 | 1 | 15 | Plek2 | 0.02 | Old |
| 0.293739 | 1.6418 | 0.02 | 0 | 1 | 15 | Ttll5 | 0.02 | Old |

| | | | | | | | | |
|---|---|---|---|---|---|---|---|---|
| 0.293739 | 1.6418 | 0.02 | 0 | 1 | 15 | Myo10 | 0.02 | Old |
| 0.293739 | 1.6418 | 0.02 | 0 | 1 | 15 | Gm16386 | 0.02 | Old |
| 0.293739 | 1.6418 | 0.02 | 0 | 1 | 15 | Gm7334 | 0.02 | Old |
| 0.293739 | 1.6418 | 0.02 | 0 | 1 | 15 | Doc2g | 0.02 | Old |
| 0.293739 | 1.556873 | 0.02 | 0 | 1 | 15 | Dpt | 0.02 | Old |
| 0.293739 | 1.556873 | 0.02 | 0 | 1 | 15 | Ube4bos1 | 0.02 | Old |
| 0.293739 | 1.556873 | 0.02 | 0 | 1 | 15 | Usp30 | 0.02 | Old |
| 0.293739 | 1.556873 | 0.02 | 0 | 1 | 15 | Gsdme | 0.02 | Old |
| 0.293739 | 1.556873 | 0.02 | 0 | 1 | 15 | Zfp428 | 0.02 | Old |
| 0.293739 | 1.556873 | 0.02 | 0 | 1 | 15 | Fat1 | 0.02 | Old |
| 0.293739 | 1.556873 | 0.02 | 0 | 1 | 15 | Syce1l | 0.02 | Old |
| 0.293739 | 1.556873 | 0.02 | 0 | 1 | 15 | Mss51 | 0.02 | Old |
| 0.293739 | 1.556873 | 0.02 | 0 | 1 | 15 | Sncg | 0.02 | Old |
| 0.293739 | 1.556873 | 0.02 | 0 | 1 | 15 | Trav14-1 | 0.02 | Old |
| 0.293739 | 1.556873 | 0.02 | 0 | 1 | 15 | Tex14 | 0.02 | Old |
| 0.293739 | 1.556873 | 0.02 | 0 | 1 | 15 | Dimt1 | 0.02 | Old |
| 0.293739 | 1.556873 | 0.02 | 0 | 1 | 15 | Slc30a6 | 0.02 | Old |
| 0.293739 | 1.556873 | 0.02 | 0 | 1 | 15 | Gm17315 | 0.02 | Old |
| 0.293739 | 1.556873 | 0.02 | 0 | 1 | 15 | Colec12 | 0.02 | Old |
| 0.293739 | 1.556873 | 0.02 | 0 | 1 | 15 | Dtwd2 | 0.02 | Old |
| 0.293739 | 1.556873 | 0.02 | 0 | 1 | 15 | Pcx | 0.02 | Old |
| 0.293739 | 1.556873 | 0.02 | 0 | 1 | 15 | Trpt1 | 0.02 | Old |
| 0.293739 | 1.538622 | 0.02 | 0 | 1 | 15 | Serpinb12 | 0.02 | Old |
| 0.293739 | 1.538622 | 0.02 | 0 | 1 | 15 | Dnm3 | 0.02 | Old |
| 0.293739 | 1.538622 | 0.02 | 0 | 1 | 15 | Ccdc173 | 0.02 | Old |
| 0.293739 | 1.538622 | 0.02 | 0 | 1 | 15 | Gm4631 | 0.02 | Old |
| 0.293739 | 1.538622 | 0.02 | 0 | 1 | 15 | Creb3l4 | 0.02 | Old |
| 0.293739 | 1.538622 | 0.02 | 0 | 1 | 15 | Gm15234 | 0.02 | Old |
| 0.293739 | 1.538622 | 0.02 | 0 | 1 | 15 | Mcoln2 | 0.02 | Old |
| 0.293739 | 1.538622 | 0.02 | 0 | 1 | 15 | 2700016F2 | 0.02 | Old |
| 0.293739 | 1.538622 | 0.02 | 0 | 1 | 15 | Gnat3 | 0.02 | Old |
| 0.293739 | 1.538622 | 0.02 | 0 | 1 | 15 | Iqcd | 0.02 | Old |
| 0.293739 | 1.538622 | 0.02 | 0 | 1 | 15 | Acad10 | 0.02 | Old |
| 0.293739 | 1.538622 | 0.02 | 0 | 1 | 15 | Psph | 0.02 | Old |
| 0.293739 | 1.538622 | 0.02 | 0 | 1 | 15 | 4933404O1 | 0.02 | Old |
| 0.293739 | 1.538622 | 0.02 | 0 | 1 | 15 | Zkscan5 | 0.02 | Old |
| 0.293739 | 1.538622 | 0.02 | 0 | 1 | 15 | Numbl | 0.02 | Old |
| 0.293739 | 1.538622 | 0.02 | 0 | 1 | 15 | Dlg2 | 0.02 | Old |
| 0.293739 | 1.538622 | 0.02 | 0 | 1 | 15 | Tsku | 0.02 | Old |
| 0.293739 | 1.538622 | 0.02 | 0 | 1 | 15 | Cbr4 | 0.02 | Old |
| 0.293739 | 1.538622 | 0.02 | 0 | 1 | 15 | Mlkl | 0.02 | Old |
| 0.293739 | 1.538622 | 0.02 | 0 | 1 | 15 | Sdr42e1 | 0.02 | Old |
| 0.293739 | 1.538622 | 0.02 | 0 | 1 | 15 | Ccr3 | 0.02 | Old |
| 0.293739 | 1.538622 | 0.02 | 0 | 1 | 15 | Zkscan17 | 0.02 | Old |
| 0.293739 | 1.538622 | 0.02 | 0 | 1 | 15 | Atpaf2 | 0.02 | Old |

| | | | | | | | | |
|---|---|---|---|---|---|---|---|---|
| 0.293739 | 1.538622 | 0.02 | 0 | 1 | 15 | Scimp | 0.02 | Old |
| 0.293739 | 1.538622 | 0.02 | 0 | 1 | 15 | Mks1 | 0.02 | Old |
| 0.293739 | 1.538622 | 0.02 | 0 | 1 | 15 | Nt5c3b | 0.02 | Old |
| 0.293739 | 1.538622 | 0.02 | 0 | 1 | 15 | E430024I0 | 0.02 | Old |
| 0.293739 | 1.538622 | 0.02 | 0 | 1 | 15 | Prps1l3 | 0.02 | Old |
| 0.293739 | 1.538622 | 0.02 | 0 | 1 | 15 | Pop1 | 0.02 | Old |
| 0.293739 | 1.538622 | 0.02 | 0 | 1 | 15 | Gm4262 | 0.02 | Old |
| 0.293739 | 1.538622 | 0.02 | 0 | 1 | 15 | Tbc1d22b | 0.02 | Old |
| 0.293739 | 1.538622 | 0.02 | 0 | 1 | 15 | Slc37a1 | 0.02 | Old |
| 0.293739 | 1.538622 | 0.02 | 0 | 1 | 15 | Zbtb12 | 0.02 | Old |
| 0.293739 | 1.538622 | 0.02 | 0 | 1 | 15 | Pik3ap1 | 0.02 | Old |
| 0.293739 | 1.522988 | 0.02 | 0 | 1 | 15 | Pdcd1 | 0.02 | Old |
| 0.293739 | 1.522988 | 0.02 | 0 | 1 | 15 | Inafm2 | 0.02 | Old |
| 0.293739 | 1.522988 | 0.02 | 0 | 1 | 15 | Macrod2 | 0.02 | Old |
| 0.293739 | 1.522988 | 0.02 | 0 | 1 | 15 | Gm43403 | 0.02 | Old |
| 0.293739 | 1.522988 | 0.02 | 0 | 1 | 15 | Fam221b | 0.02 | Old |
| 0.293739 | 1.522988 | 0.02 | 0 | 1 | 15 | Pax5 | 0.02 | Old |
| 0.293739 | 1.522988 | 0.02 | 0 | 1 | 15 | Zfp982 | 0.02 | Old |
| 0.293739 | 1.522988 | 0.02 | 0 | 1 | 15 | 9530046B1 | 0.02 | Old |
| 0.293739 | 1.522988 | 0.02 | 0 | 1 | 15 | Gm26597 | 0.02 | Old |
| 0.293739 | 1.522988 | 0.02 | 0 | 1 | 15 | Zfp772 | 0.02 | Old |
| 0.293739 | 1.522988 | 0.02 | 0 | 1 | 15 | A730063M | 0.02 | Old |
| 0.293739 | 1.522988 | 0.02 | 0 | 1 | 15 | Cers4 | 0.02 | Old |
| 0.293739 | 1.522988 | 0.02 | 0 | 1 | 15 | Zfp963 | 0.02 | Old |
| 0.293739 | 1.522988 | 0.02 | 0 | 1 | 15 | Fuk | 0.02 | Old |
| 0.293739 | 1.522988 | 0.02 | 0 | 1 | 15 | Gm47957 | 0.02 | Old |
| 0.293739 | 1.522988 | 0.02 | 0 | 1 | 15 | Hist1h2ae | 0.02 | Old |
| 0.293739 | 1.522988 | 0.02 | 0 | 1 | 15 | Tc2n | 0.02 | Old |
| 0.293739 | 1.522988 | 0.02 | 0 | 1 | 15 | Vpreb1 | 0.02 | Old |
| 0.293739 | 1.522988 | 0.02 | 0 | 1 | 15 | Fam3b | 0.02 | Old |
| 0.293739 | 1.522988 | 0.02 | 0 | 1 | 15 | Tmem151b | 0.02 | Old |
| 0.293739 | 1.522988 | 0.02 | 0 | 1 | 15 | Mppe1 | 0.02 | Old |
| 0.293739 | 1.522988 | 0.02 | 0 | 1 | 15 | CAAA0114 | 0.02 | Old |
| 0.293739 | 1.507189 | 0.02 | 0 | 1 | 15 | Rabif | 0.02 | Old |
| 0.293739 | 1.507189 | 0.02 | 0 | 1 | 15 | Gm33973 | 0.02 | Old |
| 0.293739 | 1.507189 | 0.02 | 0 | 1 | 15 | A130010J1 | 0.02 | Old |
| 0.293739 | 1.507189 | 0.02 | 0 | 1 | 15 | Tnks1bp1 | 0.02 | Old |
| 0.293739 | 1.507189 | 0.02 | 0 | 1 | 15 | Kbtbd4 | 0.02 | Old |
| 0.293739 | 1.507189 | 0.02 | 0 | 1 | 15 | Wt1 | 0.02 | Old |
| 0.293739 | 1.507189 | 0.02 | 0 | 1 | 15 | Trp53rkb | 0.02 | Old |
| 0.293739 | 1.507189 | 0.02 | 0 | 1 | 15 | Gm42809 | 0.02 | Old |
| 0.293739 | 1.507189 | 0.02 | 0 | 1 | 15 | Car9 | 0.02 | Old |
| 0.293739 | 1.507189 | 0.02 | 0 | 1 | 15 | Tmem69 | 0.02 | Old |
| 0.293739 | 1.507189 | 0.02 | 0 | 1 | 15 | Rhbdl2 | 0.02 | Old |
| 0.293739 | 1.507189 | 0.02 | 0 | 1 | 15 | Pdik1l | 0.02 | Old |

| | | | | | | | | |
|---|---|---|---|---|---|---|---|---|
| 0.293739 | 1.507189 | 0.02 | 0 | 1 | 15 | Pqlc2 | 0.02 | Old |
| 0.293739 | 1.507189 | 0.02 | 0 | 1 | 15 | Zbtb48 | 0.02 | Old |
| 0.293739 | 1.507189 | 0.02 | 0 | 1 | 15 | Fzd1 | 0.02 | Old |
| 0.293739 | 1.507189 | 0.02 | 0 | 1 | 15 | Gm7467 | 0.02 | Old |
| 0.293739 | 1.507189 | 0.02 | 0 | 1 | 15 | Micall2 | 0.02 | Old |
| 0.293739 | 1.507189 | 0.02 | 0 | 1 | 15 | Elmod3 | 0.02 | Old |
| 0.293739 | 1.507189 | 0.02 | 0 | 1 | 15 | Lag3 | 0.02 | Old |
| 0.293739 | 1.507189 | 0.02 | 0 | 1 | 15 | Tspan9 | 0.02 | Old |
| 0.293739 | 1.507189 | 0.02 | 0 | 1 | 15 | Zfp420 | 0.02 | Old |
| 0.293739 | 1.507189 | 0.02 | 0 | 1 | 15 | Cd22 | 0.02 | Old |
| 0.293739 | 1.507189 | 0.02 | 0 | 1 | 15 | Tead1 | 0.02 | Old |
| 0.293739 | 1.507189 | 0.02 | 0 | 1 | 15 | B230208H1 | 0.02 | Old |
| 0.293739 | 1.507189 | 0.02 | 0 | 1 | 15 | Slc18a1 | 0.02 | Old |
| 0.293739 | 1.507189 | 0.02 | 0 | 1 | 15 | Rgl3 | 0.02 | Old |
| 0.293739 | 1.507189 | 0.02 | 0 | 1 | 15 | Kif23 | 0.02 | Old |
| 0.293739 | 1.507189 | 0.02 | 0 | 1 | 15 | Ctdspl | 0.02 | Old |
| 0.293739 | 1.507189 | 0.02 | 0 | 1 | 15 | Slc22a5 | 0.02 | Old |
| 0.293739 | 1.507189 | 0.02 | 0 | 1 | 15 | Nt5m | 0.02 | Old |
| 0.293739 | 1.507189 | 0.02 | 0 | 1 | 15 | Cyb5d2 | 0.02 | Old |
| 0.293739 | 1.507189 | 0.02 | 0 | 1 | 15 | Nme2 | 0.02 | Old |
| 0.293739 | 1.507189 | 0.02 | 0 | 1 | 15 | Gm12355 | 0.02 | Old |
| 0.293739 | 1.507189 | 0.02 | 0 | 1 | 15 | Hist1h4j | 0.02 | Old |
| 0.293739 | 1.507189 | 0.02 | 0 | 1 | 15 | Gm47460 | 0.02 | Old |
| 0.293739 | 1.507189 | 0.02 | 0 | 1 | 15 | 5430425K1 | 0.02 | Old |
| 0.293739 | 1.507189 | 0.02 | 0 | 1 | 15 | Lynx1 | 0.02 | Old |
| 0.293739 | 1.507189 | 0.02 | 0 | 1 | 15 | Fmnl3 | 0.02 | Old |
| 0.293739 | 1.507189 | 0.02 | 0 | 1 | 15 | AC126280. | 0.02 | Old |
| 0.293739 | 1.507189 | 0.02 | 0 | 1 | 15 | Cd200r4 | 0.02 | Old |
| 0.293739 | 1.507189 | 0.02 | 0 | 1 | 15 | Fahd1 | 0.02 | Old |
| 0.293739 | 1.507189 | 0.02 | 0 | 1 | 15 | Msh5 | 0.02 | Old |
| 0.293739 | 1.507189 | 0.02 | 0 | 1 | 15 | Psma8 | 0.02 | Old |
| 0.293739 | 1.507189 | 0.02 | 0 | 1 | 15 | Dagla | 0.02 | Old |
| 0.293739 | 1.507189 | 0.02 | 0 | 1 | 15 | Gm47242 | 0.02 | Old |
| 0.293739 | 1.4375 | 0.02 | 0 | 1 | 15 | Egfl7 | 0.02 | Old |
| 0.293739 | 1.4375 | 0.02 | 0 | 1 | 15 | Fahd2a | 0.02 | Old |
| 0.293739 | 1.4375 | 0.02 | 0 | 1 | 15 | Gm26881 | 0.02 | Old |
| 0.293739 | 1.4375 | 0.02 | 0 | 1 | 15 | Fam83f | 0.02 | Old |
| 0.293739 | 1.384533 | 0.02 | 0 | 1 | 15 | Sntg1 | 0.02 | Old |
| 0.293739 | 1.384533 | 0.02 | 0 | 1 | 15 | Snapc4 | 0.02 | Old |
| 0.293739 | 1.384533 | 0.02 | 0 | 1 | 15 | Ocstamp | 0.02 | Old |
| 0.293739 | 1.384533 | 0.02 | 0 | 1 | 15 | Mtg2 | 0.02 | Old |
| 0.293739 | 1.384533 | 0.02 | 0 | 1 | 15 | Car5b | 0.02 | Old |
| 0.293739 | 1.384533 | 0.02 | 0 | 1 | 15 | Cyp7b1 | 0.02 | Old |
| 0.293739 | 1.384533 | 0.02 | 0 | 1 | 15 | Dclk2 | 0.02 | Old |
| 0.293739 | 1.384533 | 0.02 | 0 | 1 | 15 | S100a3 | 0.02 | Old |

| | | | | | | | | |
|---|---|---|---|---|---|---|---|---|
| 0.293739 | 1.384533 | 0.02 | 0 | 1 | 15 | Gm12689 | 0.02 | Old |
| 0.293739 | 1.384533 | 0.02 | 0 | 1 | 15 | Afap1 | 0.02 | Old |
| 0.293739 | 1.384533 | 0.02 | 0 | 1 | 15 | Ppp1r9a | 0.02 | Old |
| 0.293739 | 1.384533 | 0.02 | 0 | 1 | 15 | Ceacam16 | 0.02 | Old |
| 0.293739 | 1.384533 | 0.02 | 0 | 1 | 15 | 4732496C0 | 0.02 | Old |
| 0.293739 | 1.384533 | 0.02 | 0 | 1 | 15 | Smpd2 | 0.02 | Old |
| 0.293739 | 1.384533 | 0.02 | 0 | 1 | 15 | Col4a2 | 0.02 | Old |
| 0.293739 | 1.384533 | 0.02 | 0 | 1 | 15 | Gm32050 | 0.02 | Old |
| 0.293739 | 1.384533 | 0.02 | 0 | 1 | 15 | 1810011H1 | 0.02 | Old |
| 0.293739 | 1.384533 | 0.02 | 0 | 1 | 15 | Gm17173 | 0.02 | Old |
| 0.293739 | 1.384533 | 0.02 | 0 | 1 | 15 | Thsd4 | 0.02 | Old |
| 0.293739 | 1.384533 | 0.02 | 0 | 1 | 15 | Fgf10 | 0.02 | Old |
| 0.293739 | 1.384533 | 0.02 | 0 | 1 | 15 | Dynlt1a | 0.02 | Old |
| 0.293739 | 1.384533 | 0.02 | 0 | 1 | 15 | Rsph3a | 0.02 | Old |
| 0.293739 | 1.384533 | 0.02 | 0 | 1 | 15 | Rsph9 | 0.02 | Old |
| 0.293739 | 1.384533 | 0.02 | 0 | 1 | 15 | Cnnm1 | 0.02 | Old |
| 0.293739 | 1.201664 | 0.02 | 0 | 1 | 15 | Tyw5 | 0.02 | Old |
| 0.293739 | 1.201664 | 0.02 | 0 | 1 | 15 | Kif1a | 0.02 | Old |
| 0.293739 | 1.201664 | 0.02 | 0 | 1 | 15 | Dars2 | 0.02 | Old |
| 0.293739 | 1.201664 | 0.02 | 0 | 1 | 15 | Hhat | 0.02 | Old |
| 0.293739 | 1.201664 | 0.02 | 0 | 1 | 15 | Cdca7 | 0.02 | Old |
| 0.293739 | 1.201664 | 0.02 | 0 | 1 | 15 | Gm614 | 0.02 | Old |
| 0.293739 | 1.201664 | 0.02 | 0 | 1 | 15 | 4632404H1 | 0.02 | Old |
| 0.293739 | 1.201664 | 0.02 | 0 | 1 | 15 | Adam30 | 0.02 | Old |
| 0.293739 | 1.201664 | 0.02 | 0 | 1 | 15 | Trmt10b | 0.02 | Old |
| 0.293739 | 1.201664 | 0.02 | 0 | 1 | 15 | Vma21-ps | 0.02 | Old |
| 0.293739 | 1.201664 | 0.02 | 0 | 1 | 15 | Tmem156 | 0.02 | Old |
| 0.293739 | 1.201664 | 0.02 | 0 | 1 | 15 | Auts2 | 0.02 | Old |
| 0.293739 | 1.201664 | 0.02 | 0 | 1 | 15 | Smo | 0.02 | Old |
| 0.293739 | 1.201664 | 0.02 | 0 | 1 | 15 | Rad51ap1 | 0.02 | Old |
| 0.293739 | 1.201664 | 0.02 | 0 | 1 | 15 | Nadsyn1 | 0.02 | Old |
| 0.293739 | 1.201664 | 0.02 | 0 | 1 | 15 | Cenpw | 0.02 | Old |
| 0.293739 | 1.201664 | 0.02 | 0 | 1 | 15 | Aifm2 | 0.02 | Old |
| 0.293739 | 1.201664 | 0.02 | 0 | 1 | 15 | Gm26541 | 0.02 | Old |
| 0.293739 | 1.201664 | 0.02 | 0 | 1 | 15 | Dpep2nb | 0.02 | Old |
| 0.293739 | 1.201664 | 0.02 | 0 | 1 | 15 | Rhou | 0.02 | Old |
| 0.293739 | 1.201664 | 0.02 | 0 | 1 | 15 | Ints14 | 0.02 | Old |
| 0.293739 | 1.201664 | 0.02 | 0 | 1 | 15 | Slc22a21 | 0.02 | Old |
| 0.293739 | 1.201664 | 0.02 | 0 | 1 | 15 | Pfas | 0.02 | Old |
| 0.293739 | 1.201664 | 0.02 | 0 | 1 | 15 | Dhrs13 | 0.02 | Old |
| 0.293739 | 1.201664 | 0.02 | 0 | 1 | 15 | Brca1 | 0.02 | Old |
| 0.293739 | 1.201664 | 0.02 | 0 | 1 | 15 | Mboat1 | 0.02 | Old |
| 0.293739 | 1.201664 | 0.02 | 0 | 1 | 15 | Nhlrc1 | 0.02 | Old |
| 0.293739 | 1.201664 | 0.02 | 0 | 1 | 15 | Zfp458 | 0.02 | Old |
| 0.293739 | 1.201664 | 0.02 | 0 | 1 | 15 | Zfp87 | 0.02 | Old |

| | | | | | | | | |
|---|---|---|---|---|---|---|---|---|
| 0.293739 | 1.201664 | 0.02 | 0 | 1 | 15 | 1700099I0! | 0.02 | Old |
| 0.293739 | 1.201664 | 0.02 | 0 | 1 | 15 | Slx4 | 0.02 | Old |
| 0.293739 | 1.201664 | 0.02 | 0 | 1 | 15 | 2610318N( | 0.02 | Old |
| 0.293739 | 1.201664 | 0.02 | 0 | 1 | 15 | Polq | 0.02 | Old |
| 0.293739 | 1.201664 | 0.02 | 0 | 1 | 15 | Col11a2 | 0.02 | Old |
| 0.293739 | 1.201664 | 0.02 | 0 | 1 | 15 | Trem3 | 0.02 | Old |
| 0.293739 | 1.201664 | 0.02 | 0 | 1 | 15 | Ms4a4b | 0.02 | Old |
| 0.293739 | 1.201664 | 0.02 | 0 | 1 | 15 | Sorbs1 | 0.02 | Old |
| 0.293739 | 0.89084 | 0.02 | 0 | 1 | 15 | Gm13391 | 0.02 | Old |
| 0.293739 | 0.89084 | 0.02 | 0 | 1 | 15 | Tti1 | 0.02 | Old |
| 0.293739 | 0.89084 | 0.02 | 0 | 1 | 15 | Lama5 | 0.02 | Old |
| 0.293739 | 0.89084 | 0.02 | 0 | 1 | 15 | Mbnl3 | 0.02 | Old |
| 0.293739 | 0.89084 | 0.02 | 0 | 1 | 15 | Zfp449 | 0.02 | Old |
| 0.293739 | 0.89084 | 0.02 | 0 | 1 | 15 | 4632415L0 | 0.02 | Old |
| 0.293739 | 0.89084 | 0.02 | 0 | 1 | 15 | P2ry13 | 0.02 | Old |
| 0.293739 | 0.89084 | 0.02 | 0 | 1 | 15 | Gm4349 | 0.02 | Old |
| 0.293739 | 0.89084 | 0.02 | 0 | 1 | 15 | Adamtsl4 | 0.02 | Old |
| 0.293739 | 0.89084 | 0.02 | 0 | 1 | 15 | Gm43569 | 0.02 | Old |
| 0.293739 | 0.89084 | 0.02 | 0 | 1 | 15 | Ddit4l | 0.02 | Old |
| 0.293739 | 0.89084 | 0.02 | 0 | 1 | 15 | Zfp618 | 0.02 | Old |
| 0.293739 | 0.89084 | 0.02 | 0 | 1 | 15 | Sfn | 0.02 | Old |
| 0.293739 | 0.89084 | 0.02 | 0 | 1 | 15 | Tnfrsf25 | 0.02 | Old |
| 0.293739 | 0.89084 | 0.02 | 0 | 1 | 15 | Ablim2 | 0.02 | Old |
| 0.293739 | 0.89084 | 0.02 | 0 | 1 | 15 | Gm40304 | 0.02 | Old |
| 0.293739 | 0.89084 | 0.02 | 0 | 1 | 15 | Gm42670 | 0.02 | Old |
| 0.293739 | 0.89084 | 0.02 | 0 | 1 | 15 | Brca2 | 0.02 | Old |
| 0.293739 | 0.89084 | 0.02 | 0 | 1 | 15 | Glcci1 | 0.02 | Old |
| 0.293739 | 0.89084 | 0.02 | 0 | 1 | 15 | Cep41 | 0.02 | Old |
| 0.293739 | 0.89084 | 0.02 | 0 | 1 | 15 | G630030J0 | 0.02 | Old |
| 0.293739 | 0.89084 | 0.02 | 0 | 1 | 15 | Serpinh1 | 0.02 | Old |
| 0.293739 | 0.89084 | 0.02 | 0 | 1 | 15 | Itgal | 0.02 | Old |
| 0.293739 | 0.89084 | 0.02 | 0 | 1 | 15 | Mapk1ip1 | 0.02 | Old |
| 0.293739 | 0.89084 | 0.02 | 0 | 1 | 15 | Slc18b1 | 0.02 | Old |
| 0.293739 | 0.89084 | 0.02 | 0 | 1 | 15 | Tet1 | 0.02 | Old |
| 0.293739 | 0.89084 | 0.02 | 0 | 1 | 15 | Bcr | 0.02 | Old |
| 0.293739 | 0.89084 | 0.02 | 0 | 1 | 15 | Creb3l3 | 0.02 | Old |
| 0.293739 | 0.89084 | 0.02 | 0 | 1 | 15 | Cib3 | 0.02 | Old |
| 0.293739 | 0.89084 | 0.02 | 0 | 1 | 15 | Gm31105 | 0.02 | Old |
| 0.293739 | 0.89084 | 0.02 | 0 | 1 | 15 | Cmtr2 | 0.02 | Old |
| 0.293739 | 0.89084 | 0.02 | 0 | 1 | 15 | Gins2 | 0.02 | Old |
| 0.293739 | 0.89084 | 0.02 | 0 | 1 | 15 | Zfp503 | 0.02 | Old |
| 0.293739 | 0.89084 | 0.02 | 0 | 1 | 15 | Emc9 | 0.02 | Old |
| 0.293739 | 0.89084 | 0.02 | 0 | 1 | 15 | Tipin | 0.02 | Old |
| 0.293739 | 0.89084 | 0.02 | 0 | 1 | 15 | Nme9 | 0.02 | Old |
| 0.293739 | 0.89084 | 0.02 | 0 | 1 | 15 | Il20rb | 0.02 | Old |

| | | | | | | | | |
|---|---|---|---|---|---|---|---|---|
| 0.293739 | 0.89084 | 0.02 | 0 | 1 | 15 | Mst1r | 0.02 | Old |
| 0.293739 | 0.89084 | 0.02 | 0 | 1 | 15 | Hist3h2ba | 0.02 | Old |
| 0.293739 | 0.89084 | 0.02 | 0 | 1 | 15 | Hoxb4 | 0.02 | Old |
| 0.293739 | 0.89084 | 0.02 | 0 | 1 | 15 | Abca6 | 0.02 | Old |
| 0.293739 | 0.89084 | 0.02 | 0 | 1 | 15 | Gm47675 | 0.02 | Old |
| 0.293739 | 0.89084 | 0.02 | 0 | 1 | 15 | Rnf182 | 0.02 | Old |
| 0.293739 | 0.89084 | 0.02 | 0 | 1 | 15 | Rgs14 | 0.02 | Old |
| 0.293739 | 0.89084 | 0.02 | 0 | 1 | 15 | Cdk20 | 0.02 | Old |
| 0.293739 | 0.89084 | 0.02 | 0 | 1 | 15 | Gpr150 | 0.02 | Old |
| 0.293739 | 0.89084 | 0.02 | 0 | 1 | 15 | Ift43 | 0.02 | Old |
| 0.293739 | 0.89084 | 0.02 | 0 | 1 | 15 | Adamts12 | 0.02 | Old |
| 0.293739 | 0.89084 | 0.02 | 0 | 1 | 15 | Nrbp2 | 0.02 | Old |
| 0.293739 | 0.89084 | 0.02 | 0 | 1 | 15 | Polr3h | 0.02 | Old |
| 0.293739 | 0.89084 | 0.02 | 0 | 1 | 15 | 1700037C1 | 0.02 | Old |
| 0.293739 | 0.89084 | 0.02 | 0 | 1 | 15 | Rabl3 | 0.02 | Old |
| 0.293739 | 0.89084 | 0.02 | 0 | 1 | 15 | Hunk | 0.02 | Old |
| 0.293739 | 0.89084 | 0.02 | 0 | 1 | 15 | Fpr2 | 0.02 | Old |
| 0.293739 | 0.89084 | 0.02 | 0 | 1 | 15 | Pacsin1 | 0.02 | Old |
| 0.293739 | 0.89084 | 0.02 | 0 | 1 | 15 | Ddx11 | 0.02 | Old |
| 0.293739 | 0.89084 | 0.02 | 0 | 1 | 15 | Mettl4 | 0.02 | Old |
| 0.293739 | 0.89084 | 0.02 | 0 | 1 | 15 | Mcc | 0.02 | Old |
| 0.293739 | 0.89084 | 0.02 | 0 | 1 | 15 | Eno1b | 0.02 | Old |
| 0.293739 | 0.89084 | 0.02 | 0 | 1 | 15 | Impa2 | 0.02 | Old |
| 0.293739 | 0.89084 | 0.02 | 0 | 1 | 15 | Tmem216 | 0.02 | Old |
| 0.293739 | 0.89084 | 0.02 | 0 | 1 | 15 | Stn1 | 0.02 | Old |
| 0.295642 | 2.669399 | 0.082 | 0.036 | 1 | 15 | Nmrk1 | 0.046 | Old |
| 0.295642 | 2.181727 | 0.082 | 0.036 | 1 | 15 | Rce1 | 0.046 | Old |
| 0.295642 | 2.160539 | 0.082 | 0.036 | 1 | 15 | Spryd3 | 0.046 | Old |
| 0.295642 | 2.058593 | 0.082 | 0.036 | 1 | 15 | Spidr | 0.046 | Old |
| 0.295642 | 1.751667 | 0.082 | 0.036 | 1 | 15 | Riox2 | 0.046 | Old |
| 0.295642 | 1.623939 | 0.082 | 0.036 | 1 | 15 | Mtpap | 0.046 | Old |
| 0.29629 | 0.503084 | 0.694 | 0.714 | 1 | 15 | Relb | -0.02 | Old |
| 0.296467 | 0.237139 | 0.163 | 0.089 | 1 | 15 | Ccdc9 | 0.074 | Old |
| 0.296467 | 0.22338 | 0.163 | 0.089 | 1 | 15 | Copg2 | 0.074 | Old |
| 0.296598 | 0.143425 | 0.98 | 1 | 1 | 15 | mt-Nd2 | -0.02 | Old |
| 0.296598 | 0.140723 | 1 | 1 | 1 | 15 | mt-Co2 | 0 | Old |
| 0.296742 | 0.484162 | 0.327 | 0.232 | 1 | 15 | Ddost | 0.095 | Old |
| 0.296794 | 0.224868 | 0.347 | 0.232 | 1 | 15 | Arl8a | 0.115 | Old |
| 0.297623 | 0.708696 | 0.204 | 0.125 | 1 | 15 | Kars | 0.079 | Old |
| 0.297623 | 0.181598 | 0.204 | 0.125 | 1 | 15 | Tmem64 | 0.079 | Old |
| 0.297872 | 0.958239 | 0.306 | 0.214 | 1 | 15 | Khsrp | 0.092 | Old |
| 0.300418 | 0.029082 | 0.429 | 0.554 | 1 | 15 | Mvp | -0.125 | Old |
| 0.30073 | 1.214021 | 0.265 | 0.196 | 1 | 15 | Ppp4r3a | 0.069 | Old |
| 0.301142 | 0.676687 | 0.265 | 0.179 | 1 | 15 | Vps28 | 0.086 | Old |
| 0.301142 | 0.552887 | 0.265 | 0.179 | 1 | 15 | Gapvd1 | 0.086 | Old |

| | | | | | | | | |
|---|---|---|---|---|---|---|---|---|
| 0.301691 | 0.469508 | 0.163 | 0.089 | 1 | 15 | Mrpl42 | 0.074 | Old |
| 0.301691 | 0.34156 | 0.163 | 0.089 | 1 | 15 | Specc1 | 0.074 | Old |
| 0.302077 | 0.283485 | 0.653 | 0.607 | 1 | 15 | Hnrnph1 | 0.046 | Old |
| 0.302153 | 0.168333 | 0.714 | 0.875 | 1 | 15 | Samhd1 | -0.161 | Old |
| 0.303076 | 2.309974 | 0.082 | 0.036 | 1 | 15 | Slc44a2 | 0.046 | Old |
| 0.303076 | 2.166053 | 0.082 | 0.036 | 1 | 15 | Gm48065 | 0.046 | Old |
| 0.303076 | 2.102825 | 0.082 | 0.036 | 1 | 15 | 2-Sep | 0.046 | Old |
| 0.303076 | 2.040321 | 0.082 | 0.036 | 1 | 15 | Edc3 | 0.046 | Old |
| 0.303076 | 1.949326 | 0.082 | 0.036 | 1 | 15 | Arhgef18 | 0.046 | Old |
| 0.303076 | 1.915813 | 0.082 | 0.036 | 1 | 15 | Tmem199 | 0.046 | Old |
| 0.303076 | 1.879688 | 0.082 | 0.036 | 1 | 15 | Parn | 0.046 | Old |
| 0.303076 | 1.861865 | 0.082 | 0.036 | 1 | 15 | Zfp182 | 0.046 | Old |
| 0.303076 | 1.774556 | 0.082 | 0.036 | 1 | 15 | Cdt1 | 0.046 | Old |
| 0.303076 | 1.750243 | 0.082 | 0.036 | 1 | 15 | Kctd11 | 0.046 | Old |
| 0.303076 | 1.743769 | 0.082 | 0.036 | 1 | 15 | Bank1 | 0.046 | Old |
| 0.303076 | 1.71468 | 0.082 | 0.036 | 1 | 15 | Tmem185b | 0.046 | Old |
| 0.303076 | 1.664907 | 0.082 | 0.036 | 1 | 15 | Zfp810 | 0.046 | Old |
| 0.303076 | 1.63267 | 0.082 | 0.036 | 1 | 15 | Clcc1 | 0.046 | Old |
| 0.303076 | 1.38477 | 0.082 | 0.036 | 1 | 15 | 1190007I0 | 0.046 | Old |
| 0.303076 | 1.267275 | 0.082 | 0.036 | 1 | 15 | Lcmt1 | 0.046 | Old |
| 0.304017 | 0.137216 | 0.061 | 0.125 | 1 | 15 | Trabd | -0.064 | Old |
| 0.304172 | 0.044945 | 0.776 | 0.857 | 1 | 15 | Ctsh | -0.081 | Old |
| 0.304647 | 0.345889 | 0.408 | 0.321 | 1 | 15 | Lsm4 | 0.087 | Old |
| 0.304929 | 0.04089 | 0.327 | 0.429 | 1 | 15 | Smarcd2 | -0.102 | Old |
| 0.30531 | 0.385043 | 0.265 | 0.179 | 1 | 15 | Znfx1 | 0.086 | Old |
| 0.3057 | 0.815739 | 0.306 | 0.214 | 1 | 15 | Olfm1 | 0.092 | Old |
| 0.3057 | 0.555078 | 0.306 | 0.214 | 1 | 15 | Mien1 | 0.092 | Old |
| 0.306086 | 0.833274 | 0.286 | 0.196 | 1 | 15 | Rnf187 | 0.09 | Old |
| 0.306692 | 1.759702 | 0.204 | 0.143 | 1 | 15 | Commd4 | 0.061 | Old |
| 0.307112 | 0.301091 | 0.939 | 0.982 | 1 | 15 | Ptma | -0.043 | Old |
| 0.307453 | 0.397477 | 0.245 | 0.161 | 1 | 15 | Mapk6 | 0.084 | Old |
| 0.307786 | 0.481026 | 0.347 | 0.464 | 1 | 15 | Ssb | -0.117 | Old |
| 0.308125 | 0.373261 | 0.347 | 0.232 | 1 | 15 | Glipr1 | 0.115 | Old |
| 0.308161 | 1.082962 | 0.224 | 0.143 | 1 | 15 | Mrpl9 | 0.081 | Old |
| 0.308207 | 0.820739 | 0.327 | 0.232 | 1 | 15 | Bcas2 | 0.095 | Old |
| 0.30839 | 0.841356 | 0.184 | 0.107 | 1 | 15 | Tatdn2 | 0.077 | Old |
| 0.30839 | 0.242135 | 0.102 | 0.179 | 1 | 15 | Atp7a | -0.077 | Old |
| 0.309754 | 0.7757 | 0.163 | 0.25 | 1 | 15 | Arhgap18 | -0.087 | Old |
| 0.309988 | 0.608942 | 0.061 | 0.125 | 1 | 15 | Fbxo8 | -0.064 | Old |
| 0.309988 | 0.439337 | 0.061 | 0.125 | 1 | 15 | Sgta | -0.064 | Old |
| 0.310168 | 1.072885 | 0.286 | 0.196 | 1 | 15 | Lmo2 | 0.09 | Old |
| 0.310402 | 0.402958 | 0.367 | 0.268 | 1 | 15 | Twf2 | 0.099 | Old |
| 0.310634 | 2.754403 | 0.082 | 0.036 | 1 | 15 | Mpi | 0.046 | Old |
| 0.310634 | 2.210328 | 0.082 | 0.036 | 1 | 15 | Ccdc163 | 0.046 | Old |
| 0.310634 | 1.994202 | 0.082 | 0.036 | 1 | 15 | Iqsec2 | 0.046 | Old |

| | | | | | | | | |
|---|---|---|---|---|---|---|---|---|
| 0.310634 | 1.923071 | 0.082 | 0.036 | 1 | 15 | Timm29 | 0.046 | Old |
| 0.310634 | 1.79352 | 0.082 | 0.036 | 1 | 15 | Trim11 | 0.046 | Old |
| 0.310634 | 1.370377 | 0.082 | 0.036 | 1 | 15 | Anapc4 | 0.046 | Old |
| 0.310634 | 1.346686 | 0.082 | 0.036 | 1 | 15 | 6030458C1 | 0.046 | Old |
| 0.310634 | 1.308787 | 0.082 | 0.036 | 1 | 15 | Rbsn | 0.046 | Old |
| 0.310634 | 1.263887 | 0.082 | 0.036 | 1 | 15 | Ell | 0.046 | Old |
| 0.310634 | 1.197269 | 0.082 | 0.036 | 1 | 15 | Sfmbt1 | 0.046 | Old |
| 0.311105 | 0.707907 | 0.347 | 0.268 | 1 | 15 | Ppp2r5a | 0.079 | Old |
| 0.311389 | 1.955584 | 0.102 | 0.054 | 1 | 15 | Pphln1 | 0.048 | Old |
| 0.311777 | 0.534787 | 0.204 | 0.125 | 1 | 15 | Mrpl57 | 0.079 | Old |
| 0.311777 | 0.428124 | 0.204 | 0.125 | 1 | 15 | Phf14 | 0.079 | Old |
| 0.311832 | 0.455082 | 0.245 | 0.161 | 1 | 15 | Ube2d1 | 0.084 | Old |
| 0.311999 | 0.756743 | 0.306 | 0.232 | 1 | 15 | Rbm42 | 0.074 | Old |
| 0.312726 | 0.524773 | 0.224 | 0.143 | 1 | 15 | Ube3b | 0.081 | Old |
| 0.313133 | 1.231584 | 0.265 | 0.196 | 1 | 15 | Emsy | 0.069 | Old |
| 0.314074 | 0.985202 | 0.245 | 0.179 | 1 | 15 | Inf2 | 0.066 | Old |
| 0.314885 | 0.062995 | 0.245 | 0.357 | 1 | 15 | Dmac1 | -0.112 | Old |
| 0.316251 | 0.239274 | 0.245 | 0.161 | 1 | 15 | Mark3 | 0.084 | Old |
| 0.316592 | 0.549328 | 0.204 | 0.125 | 1 | 15 | Hikeshi | 0.079 | Old |
| 0.317335 | 0.205933 | 0.224 | 0.143 | 1 | 15 | Strap | 0.081 | Old |
| 0.317642 | 0.883583 | 0.449 | 0.393 | 1 | 15 | Dnajb14 | 0.056 | Old |
| 0.31773 | 0.470114 | 0.163 | 0.089 | 1 | 15 | Mapkapk3 | 0.074 | Old |
| 0.31773 | 0.107717 | 0.163 | 0.089 | 1 | 15 | Cdkn2b | 0.074 | Old |
| 0.317905 | 0.291789 | 0.367 | 0.268 | 1 | 15 | Slc20a1 | 0.099 | Old |
| 0.31811 | 1.759881 | 0.102 | 0.054 | 1 | 15 | Itgb1bp1 | 0.048 | Old |
| 0.318155 | 0.30683 | 0.347 | 0.464 | 1 | 15 | Baz1a | -0.117 | Old |
| 0.318272 | 0.967125 | 0.367 | 0.321 | 1 | 15 | Ddx39b | 0.046 | Old |
| 0.318316 | 2.344403 | 0.082 | 0.036 | 1 | 15 | Tfdp2 | 0.046 | Old |
| 0.318316 | 2.044894 | 0.082 | 0.036 | 1 | 15 | Lysmd2 | 0.046 | Old |
| 0.318316 | 1.994046 | 0.082 | 0.036 | 1 | 15 | Fbxw17 | 0.046 | Old |
| 0.318316 | 1.71516 | 0.082 | 0.036 | 1 | 15 | Gm45902 | 0.046 | Old |
| 0.318316 | 1.708187 | 0.082 | 0.036 | 1 | 15 | Chpf2 | 0.046 | Old |
| 0.318316 | 1.52167 | 0.082 | 0.036 | 1 | 15 | Gtf2f1 | 0.046 | Old |
| 0.318316 | 1.371353 | 0.082 | 0.036 | 1 | 15 | Usp35 | 0.046 | Old |
| 0.318316 | 1.36344 | 0.082 | 0.036 | 1 | 15 | Ctr9 | 0.046 | Old |
| 0.318316 | 1.360963 | 0.082 | 0.036 | 1 | 15 | Coq7 | 0.046 | Old |
| 0.318316 | 1.19703 | 0.082 | 0.036 | 1 | 15 | H2afx | 0.046 | Old |
| 0.318316 | 1.089013 | 0.082 | 0.036 | 1 | 15 | Tfdp1 | 0.046 | Old |
| 0.318316 | 1.010373 | 0.082 | 0.036 | 1 | 15 | Manea | 0.046 | Old |
| 0.318316 | 0.948366 | 0.082 | 0.036 | 1 | 15 | Afg3l2 | 0.046 | Old |
| 0.318316 | 0.897277 | 0.082 | 0.036 | 1 | 15 | Narfl | 0.046 | Old |
| 0.318843 | 0.559216 | 0.429 | 0.357 | 1 | 15 | Gosr2 | 0.072 | Old |
| 0.31945 | 0.177075 | 0.98 | 0.982 | 1 | 15 | Rpl23a | -0.002 | Old |
| 0.319452 | 0.214262 | 0.98 | 0.982 | 1 | 15 | Rpl7 | -0.002 | Old |
| 0.320712 | 0.768164 | 0.245 | 0.161 | 1 | 15 | Hipk1 | 0.084 | Old |

| | | | | | | | | |
|---|---|---|---|---|---|---|---|---|
| 0.320712 | 0.306268 | 0.245 | 0.161 | 1 | 15 | Oat | 0.084 | Old |
| 0.32084 | 0.893373 | 0.408 | 0.339 | 1 | 15 | Trip12 | 0.069 | Old |
| 0.321456 | 0.655048 | 0.204 | 0.125 | 1 | 15 | Cx3cl1 | 0.079 | Old |
| 0.321456 | 0.442188 | 0.204 | 0.125 | 1 | 15 | D5Ertd579 | 0.079 | Old |
| 0.321456 | 0.441688 | 0.204 | 0.125 | 1 | 15 | Themis2 | 0.079 | Old |
| 0.3217 | 0.27933 | 0.367 | 0.268 | 1 | 15 | Napsa | 0.099 | Old |
| 0.32195 | 0.16995 | 0.224 | 0.321 | 1 | 15 | Sh3kbp1 | -0.097 | Old |
| 0.322163 | 1.259283 | 0.122 | 0.071 | 1 | 15 | Tmem243 | 0.051 | Old |
| 0.322226 | 0.419491 | 0.898 | 0.875 | 1 | 15 | mt-Nd3 | 0.023 | Old |
| 0.322364 | 0.561415 | 0.265 | 0.179 | 1 | 15 | Tmem63b | 0.086 | Old |
| 0.32257 | 0.575659 | 0.408 | 0.321 | 1 | 15 | Etf1 | 0.087 | Old |
| 0.322891 | 0.34191 | 0.673 | 0.625 | 1 | 15 | Hsp90b1 | 0.048 | Old |
| 0.323194 | 0.038216 | 0.367 | 0.5 | 1 | 15 | Rap1a | -0.133 | Old |
| 0.323673 | 0.830366 | 0.184 | 0.107 | 1 | 15 | Slc15a4 | 0.077 | Old |
| 0.323673 | 0.287192 | 0.102 | 0.179 | 1 | 15 | Fbxo18 | -0.077 | Old |
| 0.323945 | 0.778298 | 0.327 | 0.232 | 1 | 15 | Exoc5 | 0.095 | Old |
| 0.32463 | 0.59355 | 0.469 | 0.375 | 1 | 15 | Zfp217 | 0.094 | Old |
| 0.324926 | 1.638326 | 0.102 | 0.054 | 1 | 15 | Cib2 | 0.048 | Old |
| 0.324926 | 1.602142 | 0.102 | 0.054 | 1 | 15 | Dhcr24 | 0.048 | Old |
| 0.324926 | 1.183753 | 0.102 | 0.054 | 1 | 15 | Stx18 | 0.048 | Old |
| 0.326121 | 1.835316 | 0.082 | 0.036 | 1 | 15 | Prpf38a | 0.046 | Old |
| 0.326121 | 1.720242 | 0.082 | 0.036 | 1 | 15 | Zc3h6 | 0.046 | Old |
| 0.326121 | 1.552972 | 0.082 | 0.036 | 1 | 15 | Spdl1 | 0.046 | Old |
| 0.326121 | 0.9786 | 0.082 | 0.036 | 1 | 15 | Atg13 | 0.046 | Old |
| 0.326121 | 0.947414 | 0.082 | 0.036 | 1 | 15 | Ppp1r8 | 0.046 | Old |
| 0.326121 | 0.920151 | 0.082 | 0.036 | 1 | 15 | Zfp654 | 0.046 | Old |
| 0.326121 | 0.890289 | 0.082 | 0.036 | 1 | 15 | Galk1 | 0.046 | Old |
| 0.326121 | 0.848579 | 0.082 | 0.036 | 1 | 15 | Aph1b | 0.046 | Old |
| 0.326121 | 0.834833 | 0.082 | 0.036 | 1 | 15 | Plp2 | 0.046 | Old |
| 0.326121 | 0.465997 | 0.082 | 0.036 | 1 | 15 | Lap3 | 0.046 | Old |
| 0.326121 | 0.403856 | 0.082 | 0.036 | 1 | 15 | Strada | 0.046 | Old |
| 0.326121 | 0.335447 | 0.082 | 0.036 | 1 | 15 | Scfd2 | 0.046 | Old |
| 0.327215 | 0.261035 | 0.918 | 0.929 | 1 | 15 | Zfp36 | -0.011 | Old |
| 0.3273 | 1.525784 | 0.224 | 0.161 | 1 | 15 | Tob1 | 0.063 | Old |
| 0.3273 | 1.117031 | 0.224 | 0.161 | 1 | 15 | Ubr5 | 0.063 | Old |
| 0.3273 | 1.055901 | 0.224 | 0.161 | 1 | 15 | Ripk1 | 0.063 | Old |
| 0.328063 | 1.09933 | 0.306 | 0.232 | 1 | 15 | Sidt2 | 0.074 | Old |
| 0.328063 | 0.536279 | 0.306 | 0.232 | 1 | 15 | Acsl4 | 0.074 | Old |
| 0.328161 | 0.989906 | 0.449 | 0.393 | 1 | 15 | Ttyh3 | 0.056 | Old |
| 0.328366 | 1.753943 | 0.122 | 0.071 | 1 | 15 | Ralgapa2 | 0.051 | Old |
| 0.328366 | 1.492983 | 0.122 | 0.071 | 1 | 15 | Epb41l1 | 0.051 | Old |
| 0.328874 | 0.957332 | 0.184 | 0.107 | 1 | 15 | 0610037L1 | 0.077 | Old |
| 0.329755 | 0.49875 | 0.245 | 0.161 | 1 | 15 | Anapc13 | 0.084 | Old |
| 0.329755 | 0.328484 | 0.245 | 0.161 | 1 | 15 | Tnfrsf1a | 0.084 | Old |
| 0.329928 | 0.281645 | 0.408 | 0.321 | 1 | 15 | Map4 | 0.087 | Old |

| | | | | | | | | |
|---|---|---|---|---|---|---|---|---|
| 0.330187 | 0.605387 | 0.184 | 0.268 | 1 | 15 | Tcirg1 | -0.084 | Old |
| 0.330308 | 0.522685 | 0.347 | 0.268 | 1 | 15 | Pdcd5 | 0.079 | Old |
| 0.330545 | 1.364777 | 0.163 | 0.107 | 1 | 15 | Rrp36 | 0.056 | Old |
| 0.330545 | 0.981633 | 0.163 | 0.107 | 1 | 15 | Pin1 | 0.056 | Old |
| 0.330701 | 0.480768 | 0.571 | 0.464 | 1 | 15 | Mllt6 | 0.107 | Old |
| 0.331329 | 0.21548 | 0.204 | 0.125 | 1 | 15 | Nbdy | 0.079 | Old |
| 0.331475 | 0.468931 | 0.796 | 0.786 | 1 | 15 | Rassf2 | 0.01 | Old |
| 0.331624 | 1.849275 | 0.143 | 0.089 | 1 | 15 | Ccnh | 0.054 | Old |
| 0.331624 | 1.790324 | 0.143 | 0.089 | 1 | 15 | Dhrs1 | 0.054 | Old |
| 0.33172 | 1.841815 | 0.184 | 0.125 | 1 | 15 | Otud4 | 0.059 | Old |
| 0.331835 | 1.696484 | 0.102 | 0.054 | 1 | 15 | Zfp414 | 0.048 | Old |
| 0.331835 | 1.45837 | 0.102 | 0.054 | 1 | 15 | Sh3gl1 | 0.048 | Old |
| 0.331835 | 1.306236 | 0.102 | 0.054 | 1 | 15 | Acot8 | 0.048 | Old |
| 0.331835 | 1.064158 | 0.102 | 0.054 | 1 | 15 | Pias2 | 0.048 | Old |
| 0.332272 | 0.30664 | 0.816 | 0.821 | 1 | 15 | Prcp | -0.005 | Old |
| 0.333709 | 0.52849 | 0.469 | 0.411 | 1 | 15 | Mxd1 | 0.058 | Old |
| 0.334049 | 1.401037 | 0.082 | 0.036 | 1 | 15 | Gria3 | 0.046 | Old |
| 0.334049 | 0.861504 | 0.082 | 0.036 | 1 | 15 | Msl3l2 | 0.046 | Old |
| 0.334049 | 0.833337 | 0.082 | 0.036 | 1 | 15 | Pla2g4f | 0.046 | Old |
| 0.334049 | 0.786806 | 0.082 | 0.036 | 1 | 15 | Haus4 | 0.046 | Old |
| 0.334049 | 0.786362 | 0.082 | 0.036 | 1 | 15 | 2410002F2 | 0.046 | Old |
| 0.334049 | 0.774511 | 0.082 | 0.036 | 1 | 15 | Fam118a | 0.046 | Old |
| 0.334049 | 0.700459 | 0.082 | 0.036 | 1 | 15 | BC029722 | 0.046 | Old |
| 0.334049 | 0.636835 | 0.082 | 0.036 | 1 | 15 | Rfxank | 0.046 | Old |
| 0.334049 | 0.624845 | 0.082 | 0.036 | 1 | 15 | Zbtb14 | 0.046 | Old |
| 0.334049 | 0.413812 | 0.082 | 0.036 | 1 | 15 | Dcxr | 0.046 | Old |
| 0.334049 | 0.392944 | 0.082 | 0.036 | 1 | 15 | Acss2 | 0.046 | Old |
| 0.334049 | 0.366375 | 0.082 | 0.036 | 1 | 15 | Irgm2 | 0.046 | Old |
| 0.33413 | 0.38273 | 0.184 | 0.107 | 1 | 15 | Anxa11 | 0.077 | Old |
| 0.33413 | 0.305642 | 0.184 | 0.107 | 1 | 15 | Jmjd6 | 0.077 | Old |
| 0.334276 | 0.108952 | 0.041 | 0.089 | 1 | 15 | Ryr1 | -0.048 | Old |
| 0.334338 | 0.879379 | 0.245 | 0.161 | 1 | 15 | Ndrg1 | 0.084 | Old |
| 0.334338 | 0.362625 | 0.245 | 0.161 | 1 | 15 | Chd7 | 0.084 | Old |
| 0.334542 | 0.568929 | 0.265 | 0.196 | 1 | 15 | Klhl9 | 0.069 | Old |
| 0.334542 | 0.202606 | 0.184 | 0.268 | 1 | 15 | Lmo4 | -0.084 | Old |
| 0.33464 | 0.341756 | 0.653 | 0.589 | 1 | 15 | Sdc3 | 0.064 | Old |
| 0.33525 | 0.648211 | 0.51 | 0.411 | 1 | 15 | Fam168b | 0.099 | Old |
| 0.335976 | 1.187844 | 0.163 | 0.107 | 1 | 15 | Bicd2 | 0.056 | Old |
| 0.336213 | 0.433198 | 0.224 | 0.143 | 1 | 15 | Tbc1d5 | 0.081 | Old |
| 0.336268 | 0.707614 | 0.163 | 0.25 | 1 | 15 | Hbp1 | -0.087 | Old |
| 0.336871 | 0.133647 | 0.673 | 0.554 | 1 | 15 | Adam8 | 0.119 | Old |
| 0.337447 | 0.110651 | 0.082 | 0.143 | 1 | 15 | Nacc1 | -0.061 | Old |
| 0.337621 | 0.180221 | 0.837 | 0.732 | 1 | 15 | Kctd12 | 0.105 | Old |
| 0.338073 | 0.491567 | 0.49 | 0.411 | 1 | 15 | G3bp2 | 0.079 | Old |
| 0.33884 | 2.15231 | 0.102 | 0.054 | 1 | 15 | Desi1 | 0.048 | Old |

| | | | | | | | | |
|---|---|---|---|---|---|---|---|---|
| 0.33884 | 1.689818 | 0.102 | 0.054 | 1 | 15 | Akap8l | 0.048 | Old |
| 0.33884 | 1.52075 | 0.102 | 0.054 | 1 | 15 | Nfs1 | 0.048 | Old |
| 0.33884 | 1.1601 | 0.102 | 0.054 | 1 | 15 | Gm26535 | 0.048 | Old |
| 0.33884 | 1.150484 | 0.102 | 0.054 | 1 | 15 | Mphosph1 | 0.048 | Old |
| 0.33884 | 1.018572 | 0.102 | 0.054 | 1 | 15 | Lmnb1 | 0.048 | Old |
| 0.338962 | 0.726523 | 0.245 | 0.161 | 1 | 15 | Pank2 | 0.084 | Old |
| 0.339182 | 0.397964 | 0.408 | 0.304 | 1 | 15 | Tnfrsf11a | 0.104 | Old |
| 0.339724 | 0.614615 | 0.327 | 0.25 | 1 | 15 | Pgam1 | 0.077 | Old |
| 0.340167 | 0.145701 | 0.98 | 1 | 1 | 15 | Rack1 | -0.02 | Old |
| 0.340906 | 0.372102 | 0.429 | 0.339 | 1 | 15 | Gsto1 | 0.09 | Old |
| 0.341002 | 1.329608 | 0.122 | 0.071 | 1 | 15 | Hmgn2 | 0.051 | Old |
| 0.34112 | 0.638329 | 0.367 | 0.268 | 1 | 15 | Usf2 | 0.099 | Old |
| 0.341378 | 1.333059 | 0.224 | 0.161 | 1 | 15 | Frg1 | 0.063 | Old |
| 0.341396 | 0.316198 | 0.204 | 0.125 | 1 | 15 | Hadha | 0.079 | Old |
| 0.3421 | 0.602569 | 0.082 | 0.036 | 1 | 15 | Akap10 | 0.046 | Old |
| 0.3421 | 0.415052 | 0.082 | 0.036 | 1 | 15 | Gtdc1 | 0.046 | Old |
| 0.3421 | 0.374208 | 0.082 | 0.036 | 1 | 15 | Fbxo21 | 0.046 | Old |
| 0.3421 | 0.32274 | 0.082 | 0.036 | 1 | 15 | Dopey2 | 0.046 | Old |
| 0.3421 | 0.24228 | 0.082 | 0.036 | 1 | 15 | Prr14l | 0.046 | Old |
| 0.3421 | 0.178915 | 0.082 | 0.036 | 1 | 15 | Zfp688 | 0.046 | Old |
| 0.342534 | 0.597967 | 0.449 | 0.393 | 1 | 15 | 7-Sep | 0.056 | Old |
| 0.343626 | 0.456795 | 0.245 | 0.161 | 1 | 15 | Vps26a | 0.084 | Old |
| 0.343815 | 0.715581 | 0.327 | 0.25 | 1 | 15 | Ubxn2a | 0.077 | Old |
| 0.343815 | 0.690005 | 0.327 | 0.25 | 1 | 15 | Rgs19 | 0.077 | Old |
| 0.343819 | 0.057297 | 0.388 | 0.268 | 1 | 15 | Actn4 | 0.12 | Old |
| 0.344116 | 0.390441 | 0.286 | 0.196 | 1 | 15 | Emg1 | 0.09 | Old |
| 0.344116 | 0.376759 | 0.286 | 0.196 | 1 | 15 | Snx8 | 0.09 | Old |
| 0.344116 | 0.324784 | 0.286 | 0.196 | 1 | 15 | Slirp | 0.09 | Old |
| 0.344654 | 1.005132 | 0.306 | 0.232 | 1 | 15 | R3hdm2 | 0.074 | Old |
| 0.344654 | 0.675153 | 0.306 | 0.232 | 1 | 15 | Pdcd6 | 0.074 | Old |
| 0.344654 | 0.652367 | 0.306 | 0.232 | 1 | 15 | Ap3s1 | 0.074 | Old |
| 0.344735 | 1.719556 | 0.286 | 0.232 | 1 | 15 | Commd7 | 0.054 | Old |
| 0.345018 | 0.325066 | 0.714 | 0.643 | 1 | 15 | Cltc | 0.071 | Old |
| 0.345092 | 0.465675 | 0.367 | 0.268 | 1 | 15 | Selenos | 0.099 | Old |
| 0.345203 | 1.47033 | 0.204 | 0.143 | 1 | 15 | Irf3 | 0.061 | Old |
| 0.345203 | 1.195502 | 0.204 | 0.143 | 1 | 15 | Tmem109 | 0.061 | Old |
| 0.345918 | 0.081141 | 0.224 | 0.143 | 1 | 15 | Pogk | 0.081 | Old |
| 0.345938 | 1.776372 | 0.102 | 0.054 | 1 | 15 | 2310001H1 | 0.048 | Old |
| 0.345938 | 1.302453 | 0.102 | 0.054 | 1 | 15 | 1110038F1 | 0.048 | Old |
| 0.345938 | 0.934343 | 0.102 | 0.054 | 1 | 15 | Mgat5 | 0.048 | Old |
| 0.345938 | 0.809176 | 0.102 | 0.054 | 1 | 15 | Rbm14 | 0.048 | Old |
| 0.346072 | 0.303193 | 0.551 | 0.464 | 1 | 15 | Tra2b | 0.087 | Old |
| 0.346211 | 0.466532 | 0.347 | 0.268 | 1 | 15 | Mgea5 | 0.079 | Old |
| 0.347011 | 1.786626 | 0.163 | 0.107 | 1 | 15 | Phf12 | 0.056 | Old |
| 0.347011 | 0.029882 | 0.102 | 0.161 | 1 | 15 | Arhgef40 | -0.059 | Old |

| | | | | | | | | |
|---|---|---|---|---|---|---|---|---|
| 0.347435 | 2.237841 | 0.122 | 0.071 | 1 | 15 | Klri2 | 0.051 | Old |
| 0.347435 | 1.36836 | 0.122 | 0.071 | 1 | 15 | Fam53c | 0.051 | Old |
| 0.348415 | 0.052104 | 0.327 | 0.429 | 1 | 15 | Macf1 | -0.102 | Old |
| 0.349021 | 0.959517 | 0.286 | 0.232 | 1 | 15 | Cyp51 | 0.054 | Old |
| 0.349089 | 0.156626 | 0.265 | 0.179 | 1 | 15 | Guk1 | 0.086 | Old |
| 0.349161 | 0.324161 | 0.816 | 0.875 | 1 | 15 | Gpr132 | -0.059 | Old |
| 0.34929 | 0.906677 | 0.143 | 0.089 | 1 | 15 | Syvn1 | 0.054 | Old |
| 0.34929 | 0.68416 | 0.082 | 0.143 | 1 | 15 | Wdr91 | -0.061 | Old |
| 0.349326 | 0.207059 | 0.714 | 0.714 | 1 | 15 | Akr1a1 | 0 | Old |
| 0.349362 | 0.001515 | 0.041 | 0.089 | 1 | 15 | Pmf1 | -0.048 | Old |
| 0.350274 | 0.423073 | 0.082 | 0.036 | 1 | 15 | Ints5.1 | 0.046 | Old |
| 0.350274 | 0.146033 | 0.082 | 0.036 | 1 | 15 | Enpp4 | 0.046 | Old |
| 0.35056 | 0.260613 | 0.612 | 0.536 | 1 | 15 | Sirpa | 0.076 | Old |
| 0.352473 | 0.744717 | 0.449 | 0.446 | 1 | 15 | Ube2b | 0.003 | Old |
| 0.352614 | 1.723447 | 0.163 | 0.107 | 1 | 15 | Zfp362 | 0.056 | Old |
| 0.35313 | 1.938244 | 0.102 | 0.054 | 1 | 15 | Ddx39 | 0.048 | Old |
| 0.35313 | 1.694343 | 0.102 | 0.054 | 1 | 15 | Dgkz | 0.048 | Old |
| 0.35313 | 1.604876 | 0.102 | 0.054 | 1 | 15 | Mrpl2 | 0.048 | Old |
| 0.35313 | 1.528078 | 0.102 | 0.054 | 1 | 15 | Stau1 | 0.048 | Old |
| 0.35313 | 1.011492 | 0.102 | 0.054 | 1 | 15 | Tubb4b | 0.048 | Old |
| 0.35313 | 0.986477 | 0.102 | 0.054 | 1 | 15 | Mrpl50 | 0.048 | Old |
| 0.35313 | 0.82671 | 0.102 | 0.054 | 1 | 15 | Mki67 | 0.048 | Old |
| 0.35313 | 0.744811 | 0.102 | 0.054 | 1 | 15 | Preb | 0.048 | Old |
| 0.35313 | 0.657296 | 0.102 | 0.054 | 1 | 15 | Atp13a1 | 0.048 | Old |
| 0.354343 | 1.085349 | 0.327 | 0.286 | 1 | 15 | Cep350 | 0.041 | Old |
| 0.354357 | 0.335177 | 0.714 | 0.625 | 1 | 15 | Cops9 | 0.089 | Old |
| 0.354542 | 0.460939 | 0.449 | 0.411 | 1 | 15 | Mbp | 0.038 | Old |
| 0.354965 | 0.331833 | 0.571 | 0.5 | 1 | 15 | Atp5g3 | 0.071 | Old |
| 0.355309 | 1.785601 | 0.143 | 0.089 | 1 | 15 | Atp8a1 | 0.054 | Old |
| 0.355393 | 0.298287 | 0.49 | 0.393 | 1 | 15 | Cdc37 | 0.097 | Old |
| 0.355398 | 1.008315 | 0.327 | 0.268 | 1 | 15 | Cnih1 | 0.059 | Old |
| 0.355691 | 0.119319 | 0.102 | 0.179 | 1 | 15 | Fbxo3 | -0.077 | Old |
| 0.3558 | 0.337834 | 0.224 | 0.143 | 1 | 15 | Mrpl13 | 0.081 | Old |
| 0.3558 | 0.157558 | 0.224 | 0.143 | 1 | 15 | Pdgfb | 0.081 | Old |
| 0.355826 | 0.307389 | 0.878 | 0.821 | 1 | 15 | Cytip | 0.057 | Old |
| 0.355838 | 1.363977 | 0.224 | 0.161 | 1 | 15 | Tug1 | 0.063 | Old |
| 0.356028 | 0.017996 | 0.327 | 0.429 | 1 | 15 | Il13ra1 | -0.102 | Old |
| 0.356276 | 0.518094 | 0.327 | 0.25 | 1 | 15 | Icosl | 0.077 | Old |
| 0.357442 | 1.151564 | 0.306 | 0.232 | 1 | 15 | Akirin2 | 0.074 | Old |
| 0.357695 | 0.889594 | 0.286 | 0.232 | 1 | 15 | Rnf114 | 0.054 | Old |
| 0.358453 | 0.771158 | 0.347 | 0.268 | 1 | 15 | Zcrb1 | 0.079 | Old |
| 0.358453 | 0.701422 | 0.347 | 0.268 | 1 | 15 | Snap23 | 0.079 | Old |
| 0.359446 | 0.795328 | 0.245 | 0.179 | 1 | 15 | Vrk1 | 0.066 | Old |
| 0.360113 | 0.367721 | 0.633 | 0.554 | 1 | 15 | Zfand6 | 0.079 | Old |
| 0.360416 | 2.296285 | 0.102 | 0.054 | 1 | 15 | Rara | 0.048 | Old |

| | | | | | | | | |
|---|---|---|---|---|---|---|---|---|
| 0.360416 | 1.599974 | 0.102 | 0.054 | 1 | 15 | Nup155 | 0.048 | Old |
| 0.360416 | 1.513987 | 0.102 | 0.054 | 1 | 15 | Slc36a4 | 0.048 | Old |
| 0.360416 | 1.047939 | 0.102 | 0.054 | 1 | 15 | Rlf | 0.048 | Old |
| 0.360416 | 0.831035 | 0.102 | 0.054 | 1 | 15 | Cln3 | 0.048 | Old |
| 0.360416 | 0.81073 | 0.102 | 0.054 | 1 | 15 | Nrm | 0.048 | Old |
| 0.360416 | 0.615423 | 0.102 | 0.054 | 1 | 15 | Adk | 0.048 | Old |
| 0.360416 | 0.540554 | 0.102 | 0.054 | 1 | 15 | Rbm15 | 0.048 | Old |
| 0.360518 | 0.601254 | 0.347 | 0.268 | 1 | 15 | AI314180 | 0.079 | Old |
| 0.360532 | 1.40895 | 0.122 | 0.071 | 1 | 15 | Zfp597 | 0.051 | Old |
| 0.360532 | 1.218933 | 0.122 | 0.071 | 1 | 15 | Uspl1 | 0.051 | Old |
| 0.360532 | 0.713261 | 0.122 | 0.071 | 1 | 15 | Daxx | 0.051 | Old |
| 0.363991 | 1.084301 | 0.163 | 0.107 | 1 | 15 | Ppm1g | 0.056 | Old |
| 0.363991 | 0.889028 | 0.163 | 0.107 | 1 | 15 | Nipsnap2 | 0.056 | Old |
| 0.364199 | 0.024186 | 0.245 | 0.179 | 1 | 15 | Tpi1 | 0.066 | Old |
| 0.364288 | 0.272412 | 0.224 | 0.286 | 1 | 15 | Nup88 | -0.062 | Old |
| 0.364437 | 0.360297 | 0.388 | 0.304 | 1 | 15 | Sf3b6 | 0.084 | Old |
| 0.364873 | 0.354909 | 0.041 | 0.089 | 1 | 15 | Clk2 | -0.048 | Old |
| 0.365612 | 0.194429 | 0.837 | 0.821 | 1 | 15 | Tbc1d4 | 0.016 | Old |
| 0.36569 | 1.089397 | 0.224 | 0.161 | 1 | 15 | Commd6 | 0.063 | Old |
| 0.366063 | 0.528132 | 0.265 | 0.196 | 1 | 15 | Klrd1 | 0.069 | Old |
| 0.366267 | 0.899392 | 0.367 | 0.304 | 1 | 15 | Cdc42ep3 | 0.063 | Old |
| 0.366631 | 0.860751 | 0.286 | 0.214 | 1 | 15 | Ubxn4 | 0.072 | Old |
| 0.367195 | 1.948864 | 0.122 | 0.071 | 1 | 15 | Ap3s2 | 0.051 | Old |
| 0.367195 | 1.481816 | 0.122 | 0.071 | 1 | 15 | Exosc7 | 0.051 | Old |
| 0.367195 | 1.148358 | 0.122 | 0.071 | 1 | 15 | Uchl3 | 0.051 | Old |
| 0.367195 | 1.144066 | 0.122 | 0.071 | 1 | 15 | Gm9530 | 0.051 | Old |
| 0.367195 | 0.867074 | 0.122 | 0.071 | 1 | 15 | Ipo8 | 0.051 | Old |
| 0.367542 | 1.4345 | 0.143 | 0.089 | 1 | 15 | 8-Mar | 0.054 | Old |
| 0.367542 | 1.190078 | 0.143 | 0.089 | 1 | 15 | Gopc | 0.054 | Old |
| 0.36756 | 0.415453 | 0.163 | 0.232 | 1 | 15 | Csnk1g1 | -0.069 | Old |
| 0.367643 | 0.309984 | 0.429 | 0.339 | 1 | 15 | Crybg1 | 0.09 | Old |
| 0.367701 | 0.876627 | 0.306 | 0.25 | 1 | 15 | Zmynd11 | 0.056 | Old |
| 0.367796 | 0.865407 | 0.102 | 0.054 | 1 | 15 | Camk2d | 0.048 | Old |
| 0.367796 | 0.781378 | 0.102 | 0.054 | 1 | 15 | Zfp593 | 0.048 | Old |
| 0.367796 | 0.716412 | 0.102 | 0.054 | 1 | 15 | Zadh2 | 0.048 | Old |
| 0.367796 | 0.691106 | 0.102 | 0.054 | 1 | 15 | Mafk | 0.048 | Old |
| 0.367796 | 0.678566 | 0.102 | 0.054 | 1 | 15 | Med7 | 0.048 | Old |
| 0.367796 | 0.637103 | 0.102 | 0.054 | 1 | 15 | Ubl7 | 0.048 | Old |
| 0.36872 | 0.586392 | 0.51 | 0.446 | 1 | 15 | Ptprs | 0.064 | Old |
| 0.368992 | 0.544751 | 0.245 | 0.179 | 1 | 15 | Prdx6 | 0.066 | Old |
| 0.368992 | 0.210916 | 0.163 | 0.25 | 1 | 15 | Camk1d | -0.087 | Old |
| 0.369543 | 0.766262 | 0.449 | 0.411 | 1 | 15 | Frmd4a | 0.038 | Old |
| 0.370332 | 0.50927 | 0.49 | 0.393 | 1 | 15 | Lyst | 0.097 | Old |
| 0.371458 | 0.609793 | 0.449 | 0.375 | 1 | 15 | Pnisr | 0.074 | Old |
| 0.372008 | 0.801213 | 0.388 | 0.357 | 1 | 15 | Kdm2a | 0.031 | Old |

| | | | | | | | | |
|---|---|---|---|---|---|---|---|---|
| 0.372469 | 0.230515 | 0.163 | 0.232 | 1 | 15 | Wtap | -0.069 | Old |
| 0.37252 | 0.663901 | 0.388 | 0.304 | 1 | 15 | H2-Q4 | 0.084 | Old |
| 0.373327 | 0.171649 | 0.245 | 0.321 | 1 | 15 | Lamtor2 | -0.076 | Old |
| 0.373756 | 1.477816 | 0.143 | 0.089 | 1 | 15 | Rasa1 | 0.054 | Old |
| 0.373756 | 1.255854 | 0.143 | 0.089 | 1 | 15 | Nedd4l | 0.054 | Old |
| 0.373756 | 1.021881 | 0.143 | 0.089 | 1 | 15 | Rsbn1 | 0.054 | Old |
| 0.373756 | 0.67142 | 0.143 | 0.089 | 1 | 15 | Prkra | 0.054 | Old |
| 0.373756 | 0.392635 | 0.143 | 0.089 | 1 | 15 | Utp14a | 0.054 | Old |
| 0.373756 | 0.094209 | 0.082 | 0.143 | 1 | 15 | Adm | -0.061 | Old |
| 0.373934 | 1.676114 | 0.122 | 0.071 | 1 | 15 | Bpnt1 | 0.051 | Old |
| 0.373934 | 1.071905 | 0.122 | 0.071 | 1 | 15 | Trappc5 | 0.051 | Old |
| 0.373934 | 1.044813 | 0.122 | 0.071 | 1 | 15 | Ino80b | 0.051 | Old |
| 0.373934 | 0.946585 | 0.122 | 0.071 | 1 | 15 | Ifih1 | 0.051 | Old |
| 0.375261 | 0.543311 | 0.592 | 0.554 | 1 | 15 | Lnpep | 0.038 | Old |
| 0.37527 | 0.750743 | 0.102 | 0.054 | 1 | 15 | Cnnm3 | 0.048 | Old |
| 0.37527 | 0.70667 | 0.102 | 0.054 | 1 | 15 | Eif2b2 | 0.048 | Old |
| 0.37527 | 0.660843 | 0.102 | 0.054 | 1 | 15 | Atg4c | 0.048 | Old |
| 0.37527 | 0.614009 | 0.102 | 0.054 | 1 | 15 | Zdhhc4 | 0.048 | Old |
| 0.37527 | 0.581183 | 0.102 | 0.054 | 1 | 15 | Nfx1 | 0.048 | Old |
| 0.37527 | 0.4902 | 0.102 | 0.054 | 1 | 15 | Tdp2 | 0.048 | Old |
| 0.37527 | 0.356815 | 0.102 | 0.054 | 1 | 15 | Pfkl | 0.048 | Old |
| 0.375594 | 1.266958 | 0.163 | 0.107 | 1 | 15 | Nek9 | 0.056 | Old |
| 0.375594 | 1.086046 | 0.163 | 0.107 | 1 | 15 | Dpp3 | 0.056 | Old |
| 0.375594 | 1.066825 | 0.163 | 0.107 | 1 | 15 | Rbmxl1 | 0.056 | Old |
| 0.375594 | 0.928078 | 0.163 | 0.107 | 1 | 15 | Crkl | 0.056 | Old |
| 0.376025 | 0.846015 | 0.204 | 0.143 | 1 | 15 | Micall1 | 0.061 | Old |
| 0.376025 | 0.780752 | 0.204 | 0.143 | 1 | 15 | Vamp4 | 0.061 | Old |
| 0.376025 | 0.738667 | 0.204 | 0.143 | 1 | 15 | Cops4 | 0.061 | Old |
| 0.376025 | 0.259124 | 0.204 | 0.143 | 1 | 15 | Hspb11 | 0.061 | Old |
| 0.37609 | 0.409152 | 0.143 | 0.214 | 1 | 15 | Herc6 | -0.071 | Old |
| 0.378693 | 0.723664 | 0.245 | 0.179 | 1 | 15 | Fgr | 0.066 | Old |
| 0.379453 | 0.136584 | 0.347 | 0.268 | 1 | 15 | Il2ra | 0.079 | Old |
| 0.380034 | 1.321747 | 0.143 | 0.089 | 1 | 15 | Pde4a | 0.054 | Old |
| 0.380034 | 0.913814 | 0.143 | 0.089 | 1 | 15 | Ago4 | 0.054 | Old |
| 0.380034 | 0.850044 | 0.143 | 0.089 | 1 | 15 | Ercc3 | 0.054 | Old |
| 0.380034 | 0.7132 | 0.143 | 0.089 | 1 | 15 | Vav1 | 0.054 | Old |
| 0.380121 | 0.615615 | 0.265 | 0.196 | 1 | 15 | Atf7ip | 0.069 | Old |
| 0.380196 | 0.609972 | 0.449 | 0.393 | 1 | 15 | Arhgap45 | 0.056 | Old |
| 0.380749 | 0.870459 | 0.122 | 0.071 | 1 | 15 | Pla2g15 | 0.051 | Old |
| 0.38085 | 0.235447 | 0.224 | 0.321 | 1 | 15 | Ccnl2 | -0.097 | Old |
| 0.381322 | 1.329945 | 0.204 | 0.143 | 1 | 15 | Zfp263 | 0.061 | Old |
| 0.381466 | 0.28167 | 0.51 | 0.429 | 1 | 15 | Prrc2c | 0.081 | Old |
| 0.38148 | 1.842822 | 0.163 | 0.107 | 1 | 15 | Arpc5l | 0.056 | Old |
| 0.38148 | 0.820921 | 0.163 | 0.107 | 1 | 15 | Nlrp3 | 0.056 | Old |
| 0.38148 | 0.802273 | 0.163 | 0.107 | 1 | 15 | Cdk6 | 0.056 | Old |

| | | | | | | | | |
|---|---|---|---|---|---|---|---|---|
| 0.38148 | 0.740478 | 0.163 | 0.107 | 1 | 15 | Gfer | 0.056 | Old |
| 0.38148 | 0.012196 | 0.102 | 0.161 | 1 | 15 | Wdr48 | -0.059 | Old |
| 0.381993 | 0.56513 | 0.265 | 0.179 | 1 | 15 | Cpsf7 | 0.086 | Old |
| 0.382836 | 0.543419 | 0.102 | 0.054 | 1 | 15 | C1galt1c1 | 0.048 | Old |
| 0.382836 | 0.321954 | 0.102 | 0.054 | 1 | 15 | Ppip5k2 | 0.048 | Old |
| 0.382836 | 0.233771 | 0.102 | 0.054 | 1 | 15 | Zdhhc23 | 0.048 | Old |
| 0.383603 | 0.648792 | 0.245 | 0.179 | 1 | 15 | Mydgf | 0.066 | Old |
| 0.383603 | 0.571594 | 0.245 | 0.179 | 1 | 15 | Spag7 | 0.066 | Old |
| 0.383603 | 0.459808 | 0.245 | 0.179 | 1 | 15 | Gbp3 | 0.066 | Old |
| 0.383743 | 0.935856 | 0.347 | 0.268 | 1 | 15 | Rc3h2 | 0.079 | Old |
| 0.383869 | 0.356382 | 0.571 | 0.5 | 1 | 15 | Mrpl52 | 0.071 | Old |
| 0.383902 | 0.680525 | 0.306 | 0.232 | 1 | 15 | Cdk13 | 0.074 | Old |
| 0.385174 | 0.34928 | 0.327 | 0.268 | 1 | 15 | 2610001J0 | 0.059 | Old |
| 0.385427 | 0.709417 | 0.184 | 0.125 | 1 | 15 | Gcnt2 | 0.059 | Old |
| 0.385897 | 0.431723 | 0.224 | 0.161 | 1 | 15 | Mia3 | 0.063 | Old |
| 0.386377 | 1.173415 | 0.143 | 0.089 | 1 | 15 | Immp1l | 0.054 | Old |
| 0.386377 | 0.167939 | 0.082 | 0.143 | 1 | 15 | Ms4a8a | -0.061 | Old |
| 0.386663 | 0.393665 | 0.612 | 0.625 | 1 | 15 | Plxnd1 | -0.013 | Old |
| 0.386665 | 0.837298 | 0.204 | 0.143 | 1 | 15 | Cdc26 | 0.061 | Old |
| 0.386927 | 0.445489 | 0.367 | 0.304 | 1 | 15 | Bcl10 | 0.063 | Old |
| 0.387423 | 0.821395 | 0.163 | 0.107 | 1 | 15 | Lpgat1 | 0.056 | Old |
| 0.387423 | 0.535163 | 0.163 | 0.107 | 1 | 15 | Emc8 | 0.056 | Old |
| 0.38764 | 0.99563 | 0.122 | 0.071 | 1 | 15 | Banp | 0.051 | Old |
| 0.38764 | 0.765679 | 0.122 | 0.071 | 1 | 15 | Aak1 | 0.051 | Old |
| 0.38764 | 0.562449 | 0.122 | 0.071 | 1 | 15 | 2310033P0 | 0.051 | Old |
| 0.38764 | 0.558381 | 0.122 | 0.071 | 1 | 15 | Ostm1 | 0.051 | Old |
| 0.38764 | 0.520701 | 0.122 | 0.071 | 1 | 15 | Pip4k2c | 0.051 | Old |
| 0.38764 | 0.162882 | 0.122 | 0.071 | 1 | 15 | Exosc9 | 0.051 | Old |
| 0.388091 | 0.462352 | 0.429 | 0.321 | 1 | 15 | Uba1 | 0.108 | Old |
| 0.388425 | 0.017203 | 0.224 | 0.304 | 1 | 15 | Socs4 | -0.08 | Old |
| 0.388536 | 0.527702 | 0.408 | 0.321 | 1 | 15 | R3hdm4 | 0.087 | Old |
| 0.389086 | 0.181913 | 0.122 | 0.196 | 1 | 15 | S100pbp | -0.074 | Old |
| 0.390495 | 0.467962 | 0.102 | 0.054 | 1 | 15 | Med29 | 0.048 | Old |
| 0.390495 | 0.033895 | 0.102 | 0.054 | 1 | 15 | Hotairm1 | 0.048 | Old |
| 0.390495 | 0.018885 | 0.102 | 0.054 | 1 | 15 | Nt5c2 | 0.048 | Old |
| 0.390495 | 0.005275 | 0.102 | 0.054 | 1 | 15 | Zbtb42 | 0.048 | Old |
| 0.391414 | 0.118134 | 0.306 | 0.214 | 1 | 15 | Txndc9 | 0.092 | Old |
| 0.391768 | 0.079674 | 0.143 | 0.214 | 1 | 15 | Cab39l | -0.071 | Old |
| 0.392054 | 0.80698 | 0.204 | 0.143 | 1 | 15 | Btbd7 | 0.061 | Old |
| 0.392177 | 0.406896 | 0.429 | 0.321 | 1 | 15 | Psma4 | 0.108 | Old |
| 0.392784 | 0.772886 | 0.143 | 0.089 | 1 | 15 | Scfd1 | 0.054 | Old |
| 0.392784 | 0.596767 | 0.143 | 0.089 | 1 | 15 | Zbtb22 | 0.054 | Old |
| 0.392784 | 0.511791 | 0.143 | 0.089 | 1 | 15 | Hectd3 | 0.054 | Old |
| 0.393421 | 0.907925 | 0.163 | 0.107 | 1 | 15 | Commd2 | 0.056 | Old |
| 0.393421 | 0.598837 | 0.102 | 0.161 | 1 | 15 | Ppp4r1 | -0.059 | Old |

| | | | | | | | | |
|---|---|---|---|---|---|---|---|---|
| 0.393537 | 0.010234 | 0.163 | 0.25 | 1 | 15 | Slc35a3 | -0.087 | Old |
| 0.393729 | 1.247452 | 0.286 | 0.232 | 1 | 15 | Neurl3 | 0.054 | Old |
| 0.393729 | 0.023254 | 0.204 | 0.304 | 1 | 15 | Pdcd4 | -0.1 | Old |
| 0.394229 | 0.668235 | 0.408 | 0.339 | 1 | 15 | Plekho2 | 0.069 | Old |
| 0.394606 | 0.759208 | 0.122 | 0.071 | 1 | 15 | Ubap2 | 0.051 | Old |
| 0.394606 | 0.563612 | 0.122 | 0.071 | 1 | 15 | Thoc3 | 0.051 | Old |
| 0.395649 | 0.55171 | 0.408 | 0.357 | 1 | 15 | Washc2 | 0.051 | Old |
| 0.395994 | 0.114275 | 0.388 | 0.5 | 1 | 15 | Hnrnpc | -0.112 | Old |
| 0.396773 | 0.658424 | 0.184 | 0.125 | 1 | 15 | Snrnp25 | 0.059 | Old |
| 0.396803 | 0.260783 | 0.408 | 0.321 | 1 | 15 | Gm45716 | 0.087 | Old |
| 0.398246 | 0.260976 | 0.102 | 0.054 | 1 | 15 | Atf5 | 0.048 | Old |
| 0.398246 | 0.249122 | 0.102 | 0.054 | 1 | 15 | Coa3 | 0.048 | Old |
| 0.398246 | 0.21427 | 0.102 | 0.054 | 1 | 15 | Smad2 | 0.048 | Old |
| 0.398961 | 0.663468 | 0.429 | 0.357 | 1 | 15 | Snrpb2 | 0.072 | Old |
| 0.399135 | 0.917352 | 0.245 | 0.196 | 1 | 15 | Aldh9a1 | 0.049 | Old |
| 0.399255 | 1.793239 | 0.143 | 0.089 | 1 | 15 | Ankrd10 | 0.054 | Old |
| 0.399255 | 0.484704 | 0.143 | 0.089 | 1 | 15 | Ube2v2 | 0.054 | Old |
| 0.399476 | 0.839399 | 0.163 | 0.107 | 1 | 15 | Exoc3 | 0.056 | Old |
| 0.399529 | 1.281432 | 0.02 | 0.054 | 1 | 15 | Eml2 | -0.034 | Old |
| 0.399529 | 1.224784 | 0.02 | 0.054 | 1 | 15 | Mapkbp1 | -0.034 | Old |
| 0.399529 | 0.883803 | 0.02 | 0.054 | 1 | 15 | Areg | -0.034 | Old |
| 0.399529 | 0.51608 | 0.02 | 0.054 | 1 | 15 | Haus5 | -0.034 | Old |
| 0.399529 | 0.467495 | 0.02 | 0.054 | 1 | 15 | Gfm1 | -0.034 | Old |
| 0.399529 | 0.215209 | 0.02 | 0.054 | 1 | 15 | Tigd2 | -0.034 | Old |
| 0.400709 | 0.048537 | 0.367 | 0.482 | 1 | 15 | Cd53 | -0.115 | Old |
| 0.400723 | 0.535539 | 0.51 | 0.429 | 1 | 15 | Tpr | 0.081 | Old |
| 0.401492 | 1.080793 | 0.224 | 0.161 | 1 | 15 | D1Ertd622 | 0.063 | Old |
| 0.401648 | 0.628028 | 0.122 | 0.071 | 1 | 15 | Dopey1 | 0.051 | Old |
| 0.401648 | 0.427901 | 0.122 | 0.071 | 1 | 15 | Calcrl | 0.051 | Old |
| 0.40219 | 0.079846 | 0.306 | 0.232 | 1 | 15 | Ufm1 | 0.074 | Old |
| 0.403626 | 0.916583 | 0.245 | 0.179 | 1 | 15 | Rabep1 | 0.066 | Old |
| 0.403726 | 0.555499 | 0.286 | 0.214 | 1 | 15 | Herc4 | 0.072 | Old |
| 0.403726 | 0.449812 | 0.286 | 0.214 | 1 | 15 | Pim3 | 0.072 | Old |
| 0.404136 | 0.082706 | 0.286 | 0.196 | 1 | 15 | Hmgn3 | 0.09 | Old |
| 0.405436 | 0.091712 | 0.633 | 0.732 | 1 | 15 | Herpud1 | -0.099 | Old |
| 0.405457 | 0.205154 | 0.939 | 0.929 | 1 | 15 | Junb | 0.01 | Old |
| 0.405586 | 0.977063 | 0.163 | 0.107 | 1 | 15 | Zfp110 | 0.056 | Old |
| 0.405586 | 0.850684 | 0.163 | 0.107 | 1 | 15 | Ube2g1 | 0.056 | Old |
| 0.405586 | 0.621317 | 0.163 | 0.107 | 1 | 15 | Ilf3 | 0.056 | Old |
| 0.40563 | 0.658479 | 0.347 | 0.268 | 1 | 15 | Tmbim1 | 0.079 | Old |
| 0.40579 | 1.211194 | 0.143 | 0.089 | 1 | 15 | Ppp2r3c | 0.054 | Old |
| 0.40579 | 0.635946 | 0.143 | 0.089 | 1 | 15 | Rnf157 | 0.054 | Old |
| 0.40579 | 0.485592 | 0.143 | 0.089 | 1 | 15 | Alg9 | 0.054 | Old |
| 0.405947 | 0.40744 | 0.388 | 0.304 | 1 | 15 | Ggta1 | 0.084 | Old |
| 0.40609 | 0.254323 | 0.102 | 0.054 | 1 | 15 | Ptp4a3 | 0.048 | Old |

| | | | | | | | | |
|---|---|---|---|---|---|---|---|---|
| 0.406773 | 0.739579 | 0.224 | 0.161 | 1 | 15 | Arid1b | 0.063 | Old |
| 0.406773 | 0.501237 | 0.224 | 0.161 | 1 | 15 | Elmo2 | 0.063 | Old |
| 0.406903 | 0.119312 | 0.551 | 0.482 | 1 | 15 | P2ry10 | 0.069 | Old |
| 0.408319 | 0.422409 | 0.184 | 0.125 | 1 | 15 | Pigs | 0.059 | Old |
| 0.40849 | 1.062267 | 0.204 | 0.143 | 1 | 15 | Maml1 | 0.061 | Old |
| 0.40849 | 0.803142 | 0.204 | 0.143 | 1 | 15 | Bcor | 0.061 | Old |
| 0.40849 | 0.609094 | 0.204 | 0.143 | 1 | 15 | Cir1 | 0.061 | Old |
| 0.40849 | 0.558006 | 0.204 | 0.143 | 1 | 15 | Wdr82 | 0.061 | Old |
| 0.408728 | 0.627667 | 0.245 | 0.179 | 1 | 15 | Ola1 | 0.066 | Old |
| 0.408764 | 0.224831 | 0.122 | 0.071 | 1 | 15 | Syap1 | 0.051 | Old |
| 0.40909 | 0.2605 | 0.612 | 0.518 | 1 | 15 | Prkar1a | 0.094 | Old |
| 0.409161 | 0.099468 | 1 | 1 | 1 | 15 | B2m | 0 | Old |
| 0.409215 | 0.42598 | 0.265 | 0.196 | 1 | 15 | Slc15a3 | 0.069 | Old |
| 0.410897 | 0.479684 | 0.531 | 0.482 | 1 | 15 | Cyfip1 | 0.049 | Old |
| 0.411265 | 0.163361 | 0.367 | 0.446 | 1 | 15 | Ppp1r12a | -0.079 | Old |
| 0.411443 | 1.427267 | 0.265 | 0.214 | 1 | 15 | Uhrf2 | 0.051 | Old |
| 0.411527 | 0.469587 | 0.306 | 0.232 | 1 | 15 | Yeats2 | 0.074 | Old |
| 0.411752 | 0.458961 | 0.163 | 0.107 | 1 | 15 | Sar1b | 0.056 | Old |
| 0.412388 | 0.511274 | 0.143 | 0.089 | 1 | 15 | Btbd9 | 0.054 | Old |
| 0.412388 | 0.282719 | 0.082 | 0.143 | 1 | 15 | Lrmp | -0.061 | Old |
| 0.413404 | 0.507705 | 0.408 | 0.375 | 1 | 15 | Matr3 | 0.033 | Old |
| 0.413501 | 0.333499 | 0.327 | 0.25 | 1 | 15 | Esd | 0.077 | Old |
| 0.41366 | 0.56495 | 0.592 | 0.571 | 1 | 15 | Pirb | 0.021 | Old |
| 0.413868 | 0.889948 | 0.163 | 0.25 | 1 | 15 | Gtf2b | -0.087 | Old |
| 0.413868 | 0.446925 | 0.245 | 0.179 | 1 | 15 | Sppl3 | 0.066 | Old |
| 0.414059 | 1.122049 | 0.204 | 0.143 | 1 | 15 | Gm49336 | 0.061 | Old |
| 0.414167 | 0.710986 | 0.184 | 0.125 | 1 | 15 | Ndfip1 | 0.059 | Old |
| 0.414183 | 0.491232 | 0.388 | 0.321 | 1 | 15 | Cab39 | 0.067 | Old |
| 0.41419 | 0.885076 | 0.265 | 0.196 | 1 | 15 | Siglecg | 0.069 | Old |
| 0.414841 | 0.427789 | 0.347 | 0.286 | 1 | 15 | Atp6v0a1 | 0.061 | Old |
| 0.414915 | 0.767743 | 0.367 | 0.286 | 1 | 15 | Tbl1xr1 | 0.081 | Old |
| 0.415112 | 0.206606 | 0.408 | 0.339 | 1 | 15 | Tbc1d1 | 0.069 | Old |
| 0.415887 | 0.497552 | 0.061 | 0.107 | 1 | 15 | Idua | -0.046 | Old |
| 0.415887 | 0.005987 | 0.061 | 0.107 | 1 | 15 | St3gal5 | -0.046 | Old |
| 0.415955 | 0.439929 | 0.122 | 0.071 | 1 | 15 | Traf2 | 0.051 | Old |
| 0.415955 | 0.380032 | 0.122 | 0.071 | 1 | 15 | B3galnt2 | 0.051 | Old |
| 0.415955 | 0.359042 | 0.122 | 0.071 | 1 | 15 | Alas1 | 0.051 | Old |
| 0.415955 | 0.288009 | 0.122 | 0.071 | 1 | 15 | Scamp3 | 0.051 | Old |
| 0.415955 | 0.279054 | 0.122 | 0.071 | 1 | 15 | Ppih | 0.051 | Old |
| 0.415955 | 0.269999 | 0.122 | 0.071 | 1 | 15 | Nelfa | 0.051 | Old |
| 0.415955 | 0.116742 | 0.122 | 0.071 | 1 | 15 | Ap4m1 | 0.051 | Old |
| 0.417459 | 0.444629 | 0.224 | 0.161 | 1 | 15 | Ttc9c | 0.063 | Old |
| 0.417973 | 0.322259 | 0.163 | 0.107 | 1 | 15 | Tmem135 | 0.056 | Old |
| 0.419049 | 0.618249 | 0.143 | 0.089 | 1 | 15 | Mrps25 | 0.054 | Old |
| 0.4192 | 0.268125 | 0.184 | 0.268 | 1 | 15 | Actr10 | -0.084 | Old |

| | | | | | | | | |
|---|---|---|---|---|---|---|---|---|
| 0.419351 | 0.950391 | 0.347 | 0.304 | 1 | 15 | Nrbp1 | 0.043 | Old |
| 0.419351 | 0.176297 | 0.367 | 0.286 | 1 | 15 | Ago2 | 0.081 | Old |
| 0.420063 | 0.964731 | 0.184 | 0.125 | 1 | 15 | Rbm4b | 0.059 | Old |
| 0.421039 | 0.183382 | 0.694 | 0.607 | 1 | 15 | Hnrnpa3 | 0.087 | Old |
| 0.42157 | 0.343073 | 0.367 | 0.268 | 1 | 15 | Bcl2l11 | 0.099 | Old |
| 0.423087 | 0.320281 | 0.286 | 0.214 | 1 | 15 | Dpy19l1 | 0.072 | Old |
| 0.42322 | 0.23646 | 0.122 | 0.071 | 1 | 15 | Msh2 | 0.051 | Old |
| 0.42322 | 0.144188 | 0.122 | 0.071 | 1 | 15 | Slc12a7 | 0.051 | Old |
| 0.423317 | 0.48376 | 0.755 | 0.804 | 1 | 15 | Sod1 | -0.049 | Old |
| 0.423509 | 1.060286 | 0.061 | 0.107 | 1 | 15 | Cyp20a1 | -0.046 | Old |
| 0.423828 | 0.7661 | 0.347 | 0.286 | 1 | 15 | Hnrnpd | 0.061 | Old |
| 0.423898 | 0.133903 | 1 | 1 | 1 | 15 | Marcks | 0 | Old |
| 0.424246 | 0.781693 | 0.265 | 0.196 | 1 | 15 | Coro2a | 0.069 | Old |
| 0.424907 | 0.201933 | 0.408 | 0.357 | 1 | 15 | Ubap2l | 0.051 | Old |
| 0.425129 | 0.123732 | 0.082 | 0.125 | 1 | 15 | Mdc1 | -0.043 | Old |
| 0.42533 | 1.14917 | 0.204 | 0.143 | 1 | 15 | Mras | 0.061 | Old |
| 0.42533 | 0.818167 | 0.204 | 0.143 | 1 | 15 | Zmym5 | 0.061 | Old |
| 0.42533 | 0.588869 | 0.204 | 0.143 | 1 | 15 | Pde4dip | 0.061 | Old |
| 0.425622 | 0.251942 | 0.49 | 0.393 | 1 | 15 | Ctdnep1 | 0.097 | Old |
| 0.425774 | 0.308646 | 0.143 | 0.089 | 1 | 15 | Srek1ip1 | 0.054 | Old |
| 0.426009 | 0.422774 | 0.184 | 0.125 | 1 | 15 | Tubb2b | 0.059 | Old |
| 0.426221 | 0.442563 | 0.531 | 0.518 | 1 | 15 | Reep3 | 0.013 | Old |
| 0.426722 | 0.827387 | 0.306 | 0.25 | 1 | 15 | Spire1 | 0.056 | Old |
| 0.427193 | 0.191802 | 0.837 | 0.911 | 1 | 15 | Cd44 | -0.074 | Old |
| 0.42772 | 0.354724 | 0.367 | 0.321 | 1 | 15 | Pet100 | 0.046 | Old |
| 0.428308 | 0.805854 | 0.224 | 0.161 | 1 | 15 | Cux1 | 0.063 | Old |
| 0.428843 | 0.224952 | 0.122 | 0.196 | 1 | 15 | Derl2 | -0.074 | Old |
| 0.429327 | 1.27998 | 0.245 | 0.214 | 1 | 15 | Aurkaip1 | 0.031 | Old |
| 0.429517 | 0.679933 | 0.245 | 0.179 | 1 | 15 | Limd2 | 0.066 | Old |
| 0.429517 | 0.481977 | 0.245 | 0.179 | 1 | 15 | Ptpro | 0.066 | Old |
| 0.430156 | 0.498808 | 0.49 | 0.411 | 1 | 15 | Luc7l3 | 0.079 | Old |
| 0.43058 | 1.68376 | 0.163 | 0.107 | 1 | 15 | Tmem181a | 0.056 | Old |
| 0.43058 | 0.462741 | 0.163 | 0.107 | 1 | 15 | BC003965 | 0.056 | Old |
| 0.43058 | 0.055262 | 0.163 | 0.107 | 1 | 15 | Nr1h2 | 0.056 | Old |
| 0.430582 | 0.128155 | 0.224 | 0.304 | 1 | 15 | Yy1 | -0.08 | Old |
| 0.431212 | 0.470141 | 0.061 | 0.107 | 1 | 15 | Dcaf17 | -0.046 | Old |
| 0.431212 | 0.235181 | 0.061 | 0.107 | 1 | 15 | Tnfrsf12a | -0.046 | Old |
| 0.431212 | 0.103414 | 0.061 | 0.107 | 1 | 15 | P2ry12 | -0.046 | Old |
| 0.431315 | 0.014848 | 0.388 | 0.464 | 1 | 15 | Abi1 | -0.076 | Old |
| 0.431381 | 0.034817 | 0.98 | 1 | 1 | 15 | Rps21 | -0.02 | Old |
| 0.432003 | 1.616147 | 0.184 | 0.125 | 1 | 15 | Meaf6 | 0.059 | Old |
| 0.432003 | 0.263881 | 0.184 | 0.125 | 1 | 15 | Ddhd2 | 0.059 | Old |
| 0.43228 | 0.003366 | 0.327 | 0.411 | 1 | 15 | Sav1 | -0.084 | Old |
| 0.43256 | 0.65643 | 0.143 | 0.089 | 1 | 15 | Eaf1 | 0.054 | Old |
| 0.43256 | 0.222037 | 0.143 | 0.089 | 1 | 15 | Pkp3 | 0.054 | Old |

| | | | | | | | | |
|---|---|---|---|---|---|---|---|---|
| 0.432856 | 0.697755 | 0.327 | 0.286 | 1 | 15 | 9030624J0 | 0.041 | Old |
| 0.433794 | 0.19351 | 0.224 | 0.161 | 1 | 15 | Lrch1 | 0.063 | Old |
| 0.434444 | 0.488569 | 0.265 | 0.196 | 1 | 15 | Clk3 | 0.069 | Old |
| 0.434494 | 0.326188 | 0.163 | 0.232 | 1 | 15 | Emb | -0.069 | Old |
| 0.43474 | 2.274779 | 0.163 | 0.125 | 1 | 15 | Ccdc186 | 0.038 | Old |
| 0.434941 | 0.142895 | 0.245 | 0.339 | 1 | 15 | Spr | -0.094 | Old |
| 0.436966 | 0.434834 | 0.163 | 0.107 | 1 | 15 | Spg7 | 0.056 | Old |
| 0.436966 | 0.179408 | 0.163 | 0.107 | 1 | 15 | Ahsa1 | 0.056 | Old |
| 0.437288 | 0.21891 | 0.347 | 0.25 | 1 | 15 | Plekha1 | 0.097 | Old |
| 0.437288 | 0.144239 | 0.347 | 0.25 | 1 | 15 | Zc3h7b | 0.097 | Old |
| 0.437495 | 0.812292 | 0.327 | 0.286 | 1 | 15 | Smpdl3a | 0.041 | Old |
| 0.437972 | 0.418312 | 0.122 | 0.071 | 1 | 15 | Ebpl | 0.051 | Old |
| 0.438822 | 0.336282 | 0.429 | 0.321 | 1 | 15 | Rala | 0.108 | Old |
| 0.439259 | 0.393038 | 0.857 | 0.768 | 1 | 15 | Itgb1 | 0.089 | Old |
| 0.43933 | 0.623367 | 0.367 | 0.304 | 1 | 15 | Rdx | 0.063 | Old |
| 0.439642 | 0.476408 | 0.327 | 0.268 | 1 | 15 | Rnf6 | 0.059 | Old |
| 0.442313 | 0.308401 | 0.612 | 0.607 | 1 | 15 | Gltp | 0.005 | Old |
| 0.442568 | 0.570942 | 0.204 | 0.143 | 1 | 15 | Pdhb | 0.061 | Old |
| 0.442568 | 0.370278 | 0.204 | 0.143 | 1 | 15 | Smad1 | 0.061 | Old |
| 0.443405 | 0.31566 | 0.163 | 0.107 | 1 | 15 | Mthfs | 0.056 | Old |
| 0.443405 | 0.022592 | 0.163 | 0.107 | 1 | 15 | Ambra1 | 0.056 | Old |
| 0.444137 | 0.420797 | 0.122 | 0.179 | 1 | 15 | Fbxl12 | -0.057 | Old |
| 0.444137 | 0.173538 | 0.184 | 0.125 | 1 | 15 | Cebpd | 0.059 | Old |
| 0.444781 | 0.470869 | 0.184 | 0.268 | 1 | 15 | Pxn | -0.084 | Old |
| 0.445457 | 0.005291 | 0.122 | 0.071 | 1 | 15 | Ephx1 | 0.051 | Old |
| 0.446315 | 0.244872 | 0.898 | 0.893 | 1 | 15 | H2-Eb1 | 0.005 | Old |
| 0.44632 | 0.178096 | 0.143 | 0.089 | 1 | 15 | Arrdc3 | 0.054 | Old |
| 0.44632 | 0.113449 | 0.143 | 0.089 | 1 | 15 | Fah | 0.054 | Old |
| 0.446566 | 1.09072 | 0.265 | 0.214 | 1 | 15 | Psmd7 | 0.051 | Old |
| 0.446857 | 0.160164 | 0.347 | 0.268 | 1 | 15 | Sp100 | 0.079 | Old |
| 0.446859 | 0.146948 | 0.061 | 0.107 | 1 | 15 | Cadm3 | -0.046 | Old |
| 0.446859 | 0.024872 | 0.061 | 0.107 | 1 | 15 | Zfp318 | -0.046 | Old |
| 0.447068 | 0.503287 | 0.51 | 0.482 | 1 | 15 | Slc3a2 | 0.028 | Old |
| 0.448402 | 0.391956 | 0.204 | 0.143 | 1 | 15 | Mex3c | 0.061 | Old |
| 0.449254 | 0.879682 | 0.51 | 0.464 | 1 | 15 | Hnrnpab | 0.046 | Old |
| 0.449899 | 0.120942 | 0.163 | 0.107 | 1 | 15 | Snapc5 | 0.056 | Old |
| 0.450139 | 0.437621 | 0.306 | 0.232 | 1 | 15 | Pot1b | 0.074 | Old |
| 0.450139 | 0.322585 | 0.306 | 0.232 | 1 | 15 | Eps8 | 0.074 | Old |
| 0.450276 | 0.399517 | 0.184 | 0.125 | 1 | 15 | Jkamp | 0.059 | Old |
| 0.450276 | 0.344793 | 0.184 | 0.125 | 1 | 15 | Pnpla2 | 0.059 | Old |
| 0.450492 | 0.356664 | 0.224 | 0.161 | 1 | 15 | Rnf216 | 0.063 | Old |
| 0.451522 | 0.004322 | 0.265 | 0.321 | 1 | 15 | Parp8 | -0.056 | Old |
| 0.451729 | 0.452814 | 0.286 | 0.232 | 1 | 15 | Tmco1 | 0.054 | Old |
| 0.453036 | 0.528499 | 0.51 | 0.5 | 1 | 15 | Myl12a | 0.01 | Old |
| 0.453041 | 0.432064 | 0.367 | 0.304 | 1 | 15 | Akna | 0.063 | Old |

| | | | | | | | | |
|---|---|---|---|---|---|---|---|---|
| 0.453268 | 0.241963 | 0.327 | 0.232 | 1 | 15 | Gpr65 | 0.095 | Old |
| 0.453293 | 0.075487 | 0.143 | 0.089 | 1 | 15 | Rgmb | 0.054 | Old |
| 0.453649 | 0.976191 | 0.224 | 0.179 | 1 | 15 | Polr2l | 0.045 | Old |
| 0.45427 | 0.033492 | 0.51 | 0.607 | 1 | 15 | Rasa2 | -0.097 | Old |
| 0.454307 | 0.689614 | 0.408 | 0.339 | 1 | 15 | Kxd1 | 0.069 | Old |
| 0.454307 | 0.264647 | 0.408 | 0.339 | 1 | 15 | Tet2 | 0.069 | Old |
| 0.454724 | 0.602578 | 0.388 | 0.304 | 1 | 15 | Tbc1d9 | 0.084 | Old |
| 0.454802 | 0.414646 | 0.061 | 0.107 | 1 | 15 | Bmp2 | -0.046 | Old |
| 0.455258 | 0.209452 | 0.265 | 0.196 | 1 | 15 | Slc33a1 | 0.069 | Old |
| 0.455679 | 0.792327 | 0.286 | 0.214 | 1 | 15 | Mbtps1 | 0.072 | Old |
| 0.456446 | 0.127164 | 0.163 | 0.107 | 1 | 15 | Vps50 | 0.056 | Old |
| 0.456463 | 0.427767 | 0.122 | 0.179 | 1 | 15 | Ccdc174 | -0.057 | Old |
| 0.456947 | 0.428466 | 0.592 | 0.518 | 1 | 15 | Cnbp | 0.074 | Old |
| 0.457587 | 0.720805 | 0.388 | 0.339 | 1 | 15 | Plbd1 | 0.049 | Old |
| 0.457665 | 1.488676 | 0.367 | 0.304 | 1 | 15 | Chka | 0.063 | Old |
| 0.459316 | 0.196321 | 0.388 | 0.304 | 1 | 15 | Cap1 | 0.084 | Old |
| 0.459512 | 0.408237 | 0.204 | 0.268 | 1 | 15 | Rad23b | -0.064 | Old |
| 0.459585 | 0.057213 | 0.469 | 0.607 | 1 | 15 | Pbxip1 | -0.138 | Old |
| 0.460104 | 0.371007 | 0.306 | 0.232 | 1 | 15 | Notch2 | 0.074 | Old |
| 0.460199 | 0.624371 | 0.204 | 0.143 | 1 | 15 | Cept1 | 0.061 | Old |
| 0.460516 | 0.596557 | 0.347 | 0.304 | 1 | 15 | Sdf2 | 0.043 | Old |
| 0.460548 | 0.210835 | 0.265 | 0.196 | 1 | 15 | Kmt5a | 0.069 | Old |
| 0.460753 | 0.006029 | 0.633 | 0.679 | 1 | 15 | Serbp1 | -0.046 | Old |
| 0.462698 | 0.170568 | 0.184 | 0.125 | 1 | 15 | Cdip1 | 0.059 | Old |
| 0.462698 | 0.146714 | 0.184 | 0.125 | 1 | 15 | Arhgap12 | 0.059 | Old |
| 0.462903 | 0.47272 | 0.653 | 0.714 | 1 | 15 | Prex1 | -0.061 | Old |
| 0.463292 | 0.5013 | 0.408 | 0.339 | 1 | 15 | Irf5 | 0.069 | Old |
| 0.463581 | 0.501696 | 0.327 | 0.268 | 1 | 15 | Commd3 | 0.059 | Old |
| 0.463967 | 0.602589 | 0.469 | 0.446 | 1 | 15 | Fam32a | 0.023 | Old |
| 0.464649 | 0.787387 | 0.347 | 0.321 | 1 | 15 | Gusb | 0.026 | Old |
| 0.465134 | 0.206608 | 0.571 | 0.536 | 1 | 15 | Eif4a2 | 0.035 | Old |
| 0.465225 | 1.025925 | 0.347 | 0.304 | 1 | 15 | Snrpc | 0.043 | Old |
| 0.465339 | 0.380927 | 0.878 | 0.839 | 1 | 15 | Syngr2 | 0.039 | Old |
| 0.466889 | 0.19264 | 0.714 | 0.661 | 1 | 15 | Arf4 | 0.053 | Old |
| 0.467379 | 1.070054 | 0.265 | 0.214 | 1 | 15 | Osgep | 0.051 | Old |
| 0.468107 | 0.108911 | 0.449 | 0.518 | 1 | 15 | Snrpb | -0.069 | Old |
| 0.468823 | 0.466569 | 0.429 | 0.339 | 1 | 15 | Ncoa3 | 0.09 | Old |
| 0.46898 | 0.619759 | 0.184 | 0.125 | 1 | 15 | Mrpl32 | 0.059 | Old |
| 0.46898 | 0.150568 | 0.122 | 0.179 | 1 | 15 | Spin1 | -0.057 | Old |
| 0.469924 | 0.15939 | 1 | 1 | 1 | 15 | Actg1 | 0 | Old |
| 0.470434 | 0.001581 | 0.449 | 0.357 | 1 | 15 | Ldlr | 0.092 | Old |
| 0.470436 | 0.871657 | 0.347 | 0.286 | 1 | 15 | Gyg | 0.061 | Old |
| 0.470441 | 0.707278 | 0.224 | 0.179 | 1 | 15 | Fam193a | 0.045 | Old |
| 0.47123 | 0.068428 | 0.184 | 0.268 | 1 | 15 | Ypel5 | -0.084 | Old |
| 0.472123 | 0.138004 | 0.531 | 0.464 | 1 | 15 | Pum2 | 0.067 | Old |

| | | | | | | | | |
|---|---|---|---|---|---|---|---|---|
| 0.472507 | 0.345174 | 0.51 | 0.446 | 1 | 15 | Sfr1 | 0.064 | Old |
| 0.472563 | 0.014865 | 0.408 | 0.5 | 1 | 15 | Git2 | -0.092 | Old |
| 0.472888 | 0.089468 | 0.245 | 0.179 | 1 | 15 | Lad1 | 0.066 | Old |
| 0.473313 | 0.51905 | 0.224 | 0.161 | 1 | 15 | Sec24c | 0.063 | Old |
| 0.475929 | 1.971923 | 0.143 | 0.107 | 1 | 15 | Celf4 | 0.036 | Old |
| 0.476353 | 3.852025 | 0.041 | 0.018 | 1 | 15 | Dmwd | 0.023 | Old |
| 0.476353 | 3.656569 | 0.041 | 0.018 | 1 | 15 | Zfp12 | 0.023 | Old |
| 0.476353 | 3.567145 | 0.041 | 0.018 | 1 | 15 | Cd99l2 | 0.023 | Old |
| 0.476353 | 2.904909 | 0.041 | 0.018 | 1 | 15 | Cdkl5 | 0.023 | Old |
| 0.476353 | 2.759928 | 0.041 | 0.018 | 1 | 15 | Dync2h1 | 0.023 | Old |
| 0.476353 | 2.731498 | 0.041 | 0.018 | 1 | 15 | Med26 | 0.023 | Old |
| 0.476353 | 2.708187 | 0.041 | 0.018 | 1 | 15 | Kctd3 | 0.023 | Old |
| 0.476353 | 2.700488 | 0.041 | 0.018 | 1 | 15 | Plekha5 | 0.023 | Old |
| 0.476353 | 2.645155 | 0.041 | 0.018 | 1 | 15 | Vmn2r90 | 0.023 | Old |
| 0.476353 | 2.6205 | 0.041 | 0.018 | 1 | 15 | Zfp317 | 0.023 | Old |
| 0.476353 | 2.585482 | 0.041 | 0.018 | 1 | 15 | Dnajc25 | 0.023 | Old |
| 0.476353 | 2.5516 | 0.041 | 0.018 | 1 | 15 | Atrip | 0.023 | Old |
| 0.476353 | 2.500979 | 0.041 | 0.018 | 1 | 15 | Pde12 | 0.023 | Old |
| 0.476353 | 2.474026 | 0.041 | 0.018 | 1 | 15 | Zfp512b | 0.023 | Old |
| 0.476353 | 2.397642 | 0.041 | 0.018 | 1 | 15 | Top2a | 0.023 | Old |
| 0.476353 | 2.37909 | 0.041 | 0.018 | 1 | 15 | Gcdh | 0.023 | Old |
| 0.476353 | 2.336565 | 0.041 | 0.018 | 1 | 15 | Sars2 | 0.023 | Old |
| 0.476353 | 2.279686 | 0.041 | 0.018 | 1 | 15 | Cdadc1 | 0.023 | Old |
| 0.476353 | 2.262319 | 0.041 | 0.018 | 1 | 15 | Ccdc66 | 0.023 | Old |
| 0.476353 | 2.258883 | 0.041 | 0.018 | 1 | 15 | Slc52a2 | 0.023 | Old |
| 0.476353 | 2.22868 | 0.041 | 0.018 | 1 | 15 | Umps | 0.023 | Old |
| 0.476353 | 2.205559 | 0.041 | 0.018 | 1 | 15 | Bbc3 | 0.023 | Old |
| 0.476353 | 2.19857 | 0.041 | 0.018 | 1 | 15 | Rbm4 | 0.023 | Old |
| 0.476353 | 2.17765 | 0.041 | 0.018 | 1 | 15 | Tmx4 | 0.023 | Old |
| 0.476353 | 2.138245 | 0.041 | 0.018 | 1 | 15 | Zbtb43 | 0.023 | Old |
| 0.476353 | 2.135376 | 0.041 | 0.018 | 1 | 15 | Ears2 | 0.023 | Old |
| 0.476353 | 2.133587 | 0.041 | 0.018 | 1 | 15 | Ippk | 0.023 | Old |
| 0.476353 | 2.118102 | 0.041 | 0.018 | 1 | 15 | Pigh | 0.023 | Old |
| 0.476353 | 2.114778 | 0.041 | 0.018 | 1 | 15 | Cenpv | 0.023 | Old |
| 0.476353 | 2.089487 | 0.041 | 0.018 | 1 | 15 | Isg20 | 0.023 | Old |
| 0.476353 | 2.066341 | 0.041 | 0.018 | 1 | 15 | Timm21 | 0.023 | Old |
| 0.476353 | 2.059837 | 0.041 | 0.018 | 1 | 15 | Wdr24 | 0.023 | Old |
| 0.476353 | 2.028827 | 0.041 | 0.018 | 1 | 15 | Gm14548 | 0.023 | Old |
| 0.476353 | 2.028801 | 0.041 | 0.018 | 1 | 15 | Gpn2 | 0.023 | Old |
| 0.476353 | 2.016611 | 0.041 | 0.018 | 1 | 15 | Zmym6 | 0.023 | Old |
| 0.476353 | 1.999624 | 0.041 | 0.018 | 1 | 15 | Krt10 | 0.023 | Old |
| 0.476353 | 1.978791 | 0.041 | 0.018 | 1 | 15 | Tmem129 | 0.023 | Old |
| 0.476353 | 1.974849 | 0.041 | 0.018 | 1 | 15 | Mthfr | 0.023 | Old |
| 0.476353 | 1.97038 | 0.041 | 0.018 | 1 | 15 | Ankrd55 | 0.023 | Old |
| 0.476353 | 1.963638 | 0.041 | 0.018 | 1 | 15 | Plpp2 | 0.023 | Old |

| | | | | | | | | |
|---|---|---|---|---|---|---|---|---|
| 0.476353 | 1.944285 | 0.041 | 0.018 | 1 | 15 | C2cd3 | 0.023 | Old |
| 0.476353 | 1.885319 | 0.041 | 0.018 | 1 | 15 | Tatdn3 | 0.023 | Old |
| 0.476353 | 1.873227 | 0.041 | 0.018 | 1 | 15 | Evi5l | 0.023 | Old |
| 0.476353 | 1.862794 | 0.041 | 0.018 | 1 | 15 | Dnase1l2 | 0.023 | Old |
| 0.476353 | 1.852124 | 0.041 | 0.018 | 1 | 15 | Abtb1 | 0.023 | Old |
| 0.476353 | 1.83441 | 0.041 | 0.018 | 1 | 15 | Fmn1 | 0.023 | Old |
| 0.476353 | 1.832678 | 0.041 | 0.018 | 1 | 15 | Kat8 | 0.023 | Old |
| 0.476353 | 1.788915 | 0.041 | 0.018 | 1 | 15 | Cpeb3 | 0.023 | Old |
| 0.476353 | 1.764739 | 0.041 | 0.018 | 1 | 15 | Vps8 | 0.023 | Old |
| 0.476353 | 1.747789 | 0.041 | 0.018 | 1 | 15 | Snx7 | 0.023 | Old |
| 0.476353 | 1.699045 | 0.041 | 0.018 | 1 | 15 | Plcg1 | 0.023 | Old |
| 0.476353 | 1.694693 | 0.041 | 0.018 | 1 | 15 | Hivep3 | 0.023 | Old |
| 0.476353 | 1.676148 | 0.041 | 0.018 | 1 | 15 | Pdcd2 | 0.023 | Old |
| 0.476353 | 1.662326 | 0.041 | 0.018 | 1 | 15 | Hebp1 | 0.023 | Old |
| 0.476353 | 1.629665 | 0.041 | 0.018 | 1 | 15 | Ifitm5 | 0.023 | Old |
| 0.476353 | 1.626292 | 0.041 | 0.018 | 1 | 15 | Hist1h4d | 0.023 | Old |
| 0.476353 | 1.60807 | 0.041 | 0.018 | 1 | 15 | Zfp113 | 0.023 | Old |
| 0.476353 | 1.604757 | 0.041 | 0.018 | 1 | 15 | Rbm12b1 | 0.023 | Old |
| 0.476353 | 1.593122 | 0.041 | 0.018 | 1 | 15 | Vwf | 0.023 | Old |
| 0.476353 | 1.590924 | 0.041 | 0.018 | 1 | 15 | Pex12 | 0.023 | Old |
| 0.476353 | 1.564986 | 0.041 | 0.018 | 1 | 15 | Klhl33 | 0.023 | Old |
| 0.476353 | 1.563339 | 0.041 | 0.018 | 1 | 15 | Mms22l | 0.023 | Old |
| 0.476353 | 1.531625 | 0.041 | 0.018 | 1 | 15 | Echdc1 | 0.023 | Old |
| 0.476353 | 1.529616 | 0.041 | 0.018 | 1 | 15 | Cd79b | 0.023 | Old |
| 0.476353 | 1.529318 | 0.041 | 0.018 | 1 | 15 | Zfp455 | 0.023 | Old |
| 0.476353 | 1.500578 | 0.041 | 0.018 | 1 | 15 | Clec4d | 0.023 | Old |
| 0.476353 | 1.49562 | 0.041 | 0.018 | 1 | 15 | Gcc1 | 0.023 | Old |
| 0.476353 | 1.472873 | 0.041 | 0.018 | 1 | 15 | Tnr | 0.023 | Old |
| 0.476353 | 1.44745 | 0.041 | 0.018 | 1 | 15 | Dock4 | 0.023 | Old |
| 0.476353 | 1.434865 | 0.041 | 0.018 | 1 | 15 | Ctu2 | 0.023 | Old |
| 0.476353 | 1.375521 | 0.041 | 0.018 | 1 | 15 | Arhgap19 | 0.023 | Old |
| 0.476353 | 1.372239 | 0.041 | 0.018 | 1 | 15 | Slc25a44 | 0.023 | Old |
| 0.476353 | 1.335297 | 0.041 | 0.018 | 1 | 15 | Kif9 | 0.023 | Old |
| 0.476353 | 1.333076 | 0.041 | 0.018 | 1 | 15 | Dcaf6 | 0.023 | Old |
| 0.476353 | 1.321816 | 0.041 | 0.018 | 1 | 15 | Trp53rka | 0.023 | Old |
| 0.476353 | 1.242327 | 0.041 | 0.018 | 1 | 15 | Dtd2 | 0.023 | Old |
| 0.476353 | 1.225079 | 0.041 | 0.018 | 1 | 15 | Ccdc134 | 0.023 | Old |
| 0.476353 | 1.159059 | 0.041 | 0.018 | 1 | 15 | Chchd4 | 0.023 | Old |
| 0.476353 | 1.147854 | 0.041 | 0.018 | 1 | 15 | Sbk1 | 0.023 | Old |
| 0.476353 | 1.124653 | 0.041 | 0.018 | 1 | 15 | Lmnb2 | 0.023 | Old |
| 0.476353 | 1.090906 | 0.041 | 0.018 | 1 | 15 | Spout1 | 0.023 | Old |
| 0.476353 | 1.016645 | 0.041 | 0.018 | 1 | 15 | Slc35a1 | 0.023 | Old |
| 0.476353 | 0.928537 | 0.041 | 0.018 | 1 | 15 | Tnfaip8l1 | 0.023 | Old |
| 0.477682 | 0.748326 | 0.245 | 0.196 | 1 | 15 | Commd8 | 0.049 | Old |
| 0.478217 | 0.279712 | 0.204 | 0.143 | 1 | 15 | Snx1 | 0.061 | Old |

| | | | | | | | | |
|---|---|---|---|---|---|---|---|---|
| 0.478239 | 0.324102 | 0.571 | 0.518 | 1 | 15 | Rtn3 | 0.053 | Old |
| 0.4803 | 0.416587 | 0.531 | 0.482 | 1 | 15 | Stk17b | 0.049 | Old |
| 0.480431 | 0.373739 | 0.449 | 0.393 | 1 | 15 | Ptpra | 0.056 | Old |
| 0.480439 | 0.490407 | 0.184 | 0.25 | 1 | 15 | Appbp2 | -0.066 | Old |
| 0.4805 | 0.647182 | 0.367 | 0.357 | 1 | 15 | Lpcat1 | 0.01 | Old |
| 0.481033 | 0.837016 | 0.469 | 0.411 | 1 | 15 | Rlim | 0.058 | Old |
| 0.481107 | 0.031912 | 0.51 | 0.446 | 1 | 15 | Nono | 0.064 | Old |
| 0.481823 | 0.509962 | 0.224 | 0.179 | 1 | 15 | Brd3 | 0.045 | Old |
| 0.483258 | 0.307848 | 0.408 | 0.375 | 1 | 15 | Atxn7l3b | 0.033 | Old |
| 0.483336 | 0.670501 | 0.265 | 0.214 | 1 | 15 | Rbm22 | 0.051 | Old |
| 0.483377 | 0.131558 | 0.082 | 0.125 | 1 | 15 | Dscr3 | -0.043 | Old |
| 0.48392 | 0.423907 | 0.429 | 0.375 | 1 | 15 | Ube2i | 0.054 | Old |
| 0.484307 | 0.035418 | 0.204 | 0.143 | 1 | 15 | Qrich1 | 0.061 | Old |
| 0.484331 | 0.086233 | 0.286 | 0.214 | 1 | 15 | Acap2 | 0.072 | Old |
| 0.48757 | 0.65338 | 0.224 | 0.179 | 1 | 15 | Yipf3 | 0.045 | Old |
| 0.48757 | 0.626655 | 0.224 | 0.179 | 1 | 15 | Apopt1 | 0.045 | Old |
| 0.488105 | 0.22084 | 0.184 | 0.125 | 1 | 15 | Myo1f | 0.059 | Old |
| 0.488759 | 1.184137 | 0.245 | 0.196 | 1 | 15 | Skp1a | 0.049 | Old |
| 0.488759 | 0.978306 | 0.245 | 0.196 | 1 | 15 | Piezo1 | 0.049 | Old |
| 0.489753 | 0.255556 | 0.245 | 0.179 | 1 | 15 | Plxnb2 | 0.066 | Old |
| 0.490242 | 2.659468 | 0.041 | 0.018 | 1 | 15 | Nudt19 | 0.023 | Old |
| 0.490242 | 2.496708 | 0.041 | 0.018 | 1 | 15 | Dlg4 | 0.023 | Old |
| 0.490242 | 1.906255 | 0.041 | 0.018 | 1 | 15 | Ripor1 | 0.023 | Old |
| 0.490242 | 1.871762 | 0.041 | 0.018 | 1 | 15 | Gm10320 | 0.023 | Old |
| 0.490242 | 1.8157 | 0.041 | 0.018 | 1 | 15 | Snx21 | 0.023 | Old |
| 0.490242 | 1.80475 | 0.041 | 0.018 | 1 | 15 | Polr1e | 0.023 | Old |
| 0.490242 | 1.791599 | 0.041 | 0.018 | 1 | 15 | Mthfd2l | 0.023 | Old |
| 0.490242 | 1.765574 | 0.041 | 0.018 | 1 | 15 | Mief2 | 0.023 | Old |
| 0.490242 | 1.757124 | 0.041 | 0.018 | 1 | 15 | Nol6 | 0.023 | Old |
| 0.490242 | 1.740705 | 0.041 | 0.018 | 1 | 15 | Ttll1 | 0.023 | Old |
| 0.490242 | 1.680884 | 0.041 | 0.018 | 1 | 15 | Fbxo30 | 0.023 | Old |
| 0.490242 | 1.626924 | 0.041 | 0.018 | 1 | 15 | Rpap3 | 0.023 | Old |
| 0.490242 | 1.608308 | 0.041 | 0.018 | 1 | 15 | Ap4e1 | 0.023 | Old |
| 0.490242 | 1.593845 | 0.041 | 0.018 | 1 | 15 | Abhd6 | 0.023 | Old |
| 0.490242 | 1.576016 | 0.041 | 0.018 | 1 | 15 | Parp6 | 0.023 | Old |
| 0.490242 | 1.564385 | 0.041 | 0.018 | 1 | 15 | Smim24 | 0.023 | Old |
| 0.490242 | 1.56283 | 0.041 | 0.018 | 1 | 15 | Sec22c | 0.023 | Old |
| 0.490242 | 1.532922 | 0.041 | 0.018 | 1 | 15 | Tdrkh | 0.023 | Old |
| 0.490242 | 1.47449 | 0.041 | 0.018 | 1 | 15 | Rfx3 | 0.023 | Old |
| 0.490242 | 1.442563 | 0.041 | 0.018 | 1 | 15 | Cenpe | 0.023 | Old |
| 0.490242 | 1.435877 | 0.041 | 0.018 | 1 | 15 | Abtb2 | 0.023 | Old |
| 0.490242 | 1.4314 | 0.041 | 0.018 | 1 | 15 | Zkscan6 | 0.023 | Old |
| 0.490242 | 1.403891 | 0.041 | 0.018 | 1 | 15 | Car2 | 0.023 | Old |
| 0.490242 | 1.388233 | 0.041 | 0.018 | 1 | 15 | Ogfod2 | 0.023 | Old |
| 0.490242 | 1.384719 | 0.041 | 0.018 | 1 | 15 | Zfp943 | 0.023 | Old |

| | | | | | | | | |
|---|---|---|---|---|---|---|---|---|
| 0.490242 | 1.373509 | 0.041 | 0.018 | 1 | 15 | Akr1c12 | 0.023 | Old |
| 0.490242 | 1.347969 | 0.041 | 0.018 | 1 | 15 | AI480526 | 0.023 | Old |
| 0.490242 | 1.335825 | 0.041 | 0.018 | 1 | 15 | Arhgap10 | 0.023 | Old |
| 0.490242 | 1.30633 | 0.041 | 0.018 | 1 | 15 | Tsen54 | 0.023 | Old |
| 0.490242 | 1.304026 | 0.041 | 0.018 | 1 | 15 | Pop7 | 0.023 | Old |
| 0.490242 | 1.302979 | 0.041 | 0.018 | 1 | 15 | Cldn10 | 0.023 | Old |
| 0.490242 | 1.298787 | 0.041 | 0.018 | 1 | 15 | Crim1 | 0.023 | Old |
| 0.490242 | 1.293582 | 0.041 | 0.018 | 1 | 15 | Prickle3 | 0.023 | Old |
| 0.490242 | 1.289729 | 0.041 | 0.018 | 1 | 15 | Spic | 0.023 | Old |
| 0.490242 | 1.286888 | 0.041 | 0.018 | 1 | 15 | Tns2 | 0.023 | Old |
| 0.490242 | 1.273308 | 0.041 | 0.018 | 1 | 15 | Rab23 | 0.023 | Old |
| 0.490242 | 1.242598 | 0.041 | 0.018 | 1 | 15 | Zfp446 | 0.023 | Old |
| 0.490242 | 1.24104 | 0.041 | 0.018 | 1 | 15 | Gm32856 | 0.023 | Old |
| 0.490242 | 1.240597 | 0.041 | 0.018 | 1 | 15 | Extl3 | 0.023 | Old |
| 0.490242 | 1.219002 | 0.041 | 0.018 | 1 | 15 | Nsdhl | 0.023 | Old |
| 0.490242 | 1.217786 | 0.041 | 0.018 | 1 | 15 | Disc1 | 0.023 | Old |
| 0.490242 | 1.210252 | 0.041 | 0.018 | 1 | 15 | D3Ertd751 | 0.023 | Old |
| 0.490242 | 1.196857 | 0.041 | 0.018 | 1 | 15 | Prmt9 | 0.023 | Old |
| 0.490242 | 1.183932 | 0.041 | 0.018 | 1 | 15 | Asprv1 | 0.023 | Old |
| 0.490242 | 1.179791 | 0.041 | 0.018 | 1 | 15 | Cep85 | 0.023 | Old |
| 0.490242 | 1.166654 | 0.041 | 0.018 | 1 | 15 | Ighmbp2 | 0.023 | Old |
| 0.490242 | 1.141426 | 0.041 | 0.018 | 1 | 15 | Eri3 | 0.023 | Old |
| 0.490242 | 1.134492 | 0.041 | 0.018 | 1 | 15 | Efnb1 | 0.023 | Old |
| 0.490242 | 1.111569 | 0.041 | 0.018 | 1 | 15 | Cabp4 | 0.023 | Old |
| 0.490242 | 1.108791 | 0.041 | 0.018 | 1 | 15 | Rhobtb2 | 0.023 | Old |
| 0.490242 | 1.10049 | 0.041 | 0.018 | 1 | 15 | Mmp12 | 0.023 | Old |
| 0.490242 | 1.091422 | 0.041 | 0.018 | 1 | 15 | Spice1 | 0.023 | Old |
| 0.490242 | 1.089909 | 0.041 | 0.018 | 1 | 15 | Pole3 | 0.023 | Old |
| 0.490242 | 1.081973 | 0.041 | 0.018 | 1 | 15 | Hsf1 | 0.023 | Old |
| 0.490242 | 1.059529 | 0.041 | 0.018 | 1 | 15 | Guf1 | 0.023 | Old |
| 0.490242 | 1.059444 | 0.041 | 0.018 | 1 | 15 | Tnfsf4 | 0.023 | Old |
| 0.490242 | 1.038308 | 0.041 | 0.018 | 1 | 15 | Thumpd2 | 0.023 | Old |
| 0.490242 | 1.028569 | 0.041 | 0.018 | 1 | 15 | Etl4 | 0.023 | Old |
| 0.490242 | 1.028214 | 0.041 | 0.018 | 1 | 15 | Cpt2 | 0.023 | Old |
| 0.490242 | 1.020335 | 0.041 | 0.018 | 1 | 15 | Alg12 | 0.023 | Old |
| 0.490242 | 1.007383 | 0.041 | 0.018 | 1 | 15 | Coq3 | 0.023 | Old |
| 0.490242 | 1.002713 | 0.041 | 0.018 | 1 | 15 | Gtpbp10 | 0.023 | Old |
| 0.490242 | 0.98858 | 0.041 | 0.018 | 1 | 15 | Gstm4 | 0.023 | Old |
| 0.490242 | 0.974096 | 0.041 | 0.018 | 1 | 15 | Szt2 | 0.023 | Old |
| 0.490242 | 0.965304 | 0.041 | 0.018 | 1 | 15 | Ublcp1 | 0.023 | Old |
| 0.490242 | 0.906139 | 0.041 | 0.018 | 1 | 15 | Nhej1 | 0.023 | Old |
| 0.490242 | 0.896991 | 0.041 | 0.018 | 1 | 15 | Sparc | 0.023 | Old |
| 0.490242 | 0.864125 | 0.041 | 0.018 | 1 | 15 | Fkbp7 | 0.023 | Old |
| 0.490242 | 0.861175 | 0.041 | 0.018 | 1 | 15 | St5 | 0.023 | Old |
| 0.490242 | 0.859473 | 0.041 | 0.018 | 1 | 15 | Rdh10 | 0.023 | Old |

| | | | | | | | | |
|---|---|---|---|---|---|---|---|---|
| 0.490242 | 0.838632 | 0.041 | 0.018 | 1 | 15 | Ptpn3 | 0.023 | Old |
| 0.490242 | 0.837537 | 0.041 | 0.018 | 1 | 15 | Thop1 | 0.023 | Old |
| 0.490242 | 0.836644 | 0.041 | 0.018 | 1 | 15 | Klhl12 | 0.023 | Old |
| 0.490242 | 0.817746 | 0.041 | 0.018 | 1 | 15 | Ptprcap | 0.023 | Old |
| 0.490242 | 0.800699 | 0.041 | 0.018 | 1 | 15 | Cchcr1 | 0.023 | Old |
| 0.490242 | 0.790659 | 0.041 | 0.018 | 1 | 15 | Fam109a | 0.023 | Old |
| 0.490242 | 0.762538 | 0.041 | 0.018 | 1 | 15 | Ptprm | 0.023 | Old |
| 0.490242 | 0.761586 | 0.041 | 0.018 | 1 | 15 | Zfp952 | 0.023 | Old |
| 0.490242 | 0.743242 | 0.041 | 0.018 | 1 | 15 | Sorbs3 | 0.023 | Old |
| 0.490242 | 0.738899 | 0.041 | 0.018 | 1 | 15 | Plekha8 | 0.023 | Old |
| 0.490242 | 0.737202 | 0.041 | 0.018 | 1 | 15 | Tlr13 | 0.023 | Old |
| 0.490242 | 0.732249 | 0.041 | 0.018 | 1 | 15 | Gm20045 | 0.023 | Old |
| 0.490242 | 0.716328 | 0.041 | 0.018 | 1 | 15 | Pyroxd1 | 0.023 | Old |
| 0.490242 | 0.715578 | 0.041 | 0.018 | 1 | 15 | Nod2 | 0.023 | Old |
| 0.490242 | 0.681311 | 0.041 | 0.018 | 1 | 15 | Chst12 | 0.023 | Old |
| 0.490242 | 0.614786 | 0.041 | 0.018 | 1 | 15 | Nfe2l3 | 0.023 | Old |
| 0.490242 | 0.612456 | 0.041 | 0.018 | 1 | 15 | Sergef | 0.023 | Old |
| 0.490242 | 0.59164 | 0.041 | 0.018 | 1 | 15 | Naprt | 0.023 | Old |
| 0.490242 | 0.585587 | 0.041 | 0.018 | 1 | 15 | Hmbs | 0.023 | Old |
| 0.490242 | 0.583827 | 0.041 | 0.018 | 1 | 15 | Thg1l | 0.023 | Old |
| 0.490242 | 0.56136 | 0.041 | 0.018 | 1 | 15 | Apobr | 0.023 | Old |
| 0.490242 | 0.560163 | 0.041 | 0.018 | 1 | 15 | Pwp1 | 0.023 | Old |
| 0.490242 | 0.544144 | 0.041 | 0.018 | 1 | 15 | Eid2b | 0.023 | Old |
| 0.490242 | 0.485748 | 0.041 | 0.018 | 1 | 15 | Tacc3 | 0.023 | Old |
| 0.490242 | 0.422651 | 0.041 | 0.018 | 1 | 15 | Mipep | 0.023 | Old |
| 0.490455 | 0.738627 | 0.224 | 0.179 | 1 | 15 | Ap1s3 | 0.045 | Old |
| 0.490725 | 0.580556 | 0.306 | 0.232 | 1 | 15 | Pcf11 | 0.074 | Old |
| 0.490923 | 1.125352 | 0.306 | 0.268 | 1 | 15 | Tnrc6b | 0.038 | Old |
| 0.491065 | 0.433846 | 0.388 | 0.321 | 1 | 15 | Chp1 | 0.067 | Old |
| 0.492702 | 1.280888 | 0.184 | 0.143 | 1 | 15 | Fbxl20 | 0.041 | Old |
| 0.492702 | 1.243065 | 0.184 | 0.143 | 1 | 15 | Trio | 0.041 | Old |
| 0.492999 | 0.002922 | 0.265 | 0.196 | 1 | 15 | Sec23b | 0.069 | Old |
| 0.493242 | 0.552176 | 0.327 | 0.268 | 1 | 15 | Nae1 | 0.059 | Old |
| 0.493354 | 1.055711 | 0.224 | 0.179 | 1 | 15 | Ppp6r1 | 0.045 | Old |
| 0.493354 | 0.256488 | 0.224 | 0.179 | 1 | 15 | Heca | 0.045 | Old |
| 0.494036 | 0.49051 | 0.347 | 0.304 | 1 | 15 | Specc1l | 0.043 | Old |
| 0.494135 | 0.786549 | 0.265 | 0.214 | 1 | 15 | Pttg1ip | 0.051 | Old |
| 0.494349 | 0.525076 | 0.245 | 0.196 | 1 | 15 | Paics | 0.049 | Old |
| 0.494573 | 0.074454 | 0.122 | 0.179 | 1 | 15 | Cib1 | -0.057 | Old |
| 0.496007 | 0.222829 | 0.327 | 0.25 | 1 | 15 | Uhmk1 | 0.077 | Old |
| 0.49689 | 0.496314 | 0.204 | 0.161 | 1 | 15 | Stip1 | 0.043 | Old |
| 0.49702 | 1.599792 | 0.143 | 0.107 | 1 | 15 | Ccl12 | 0.036 | Old |
| 0.497156 | 0.023273 | 0.184 | 0.25 | 1 | 15 | Pld3 | -0.066 | Old |
| 0.498123 | 0.207865 | 0.673 | 0.661 | 1 | 15 | Morf4l1 | 0.012 | Old |
| 0.498414 | 0.153097 | 0.714 | 0.768 | 1 | 15 | Rassf3 | -0.054 | Old |

| | | | | | | | | |
|---|---|---|---|---|---|---|---|---|
| 0.498434 | 0.763778 | 0.449 | 0.393 | 1 | 15 | Gdi1 | 0.056 | Old |
| 0.499174 | 0.380306 | 0.224 | 0.179 | 1 | 15 | C5ar1 | 0.045 | Old |
| 0.499325 | 1.011466 | 0.163 | 0.125 | 1 | 15 | Pak1 | 0.038 | Old |
| 0.499325 | 0.973107 | 0.163 | 0.125 | 1 | 15 | Pxk | 0.038 | Old |
| 0.499583 | 0.694002 | 0.265 | 0.214 | 1 | 15 | Nuak1 | 0.051 | Old |
| 0.499972 | 0.821177 | 0.245 | 0.196 | 1 | 15 | Tmx1 | 0.049 | Old |
| 0.499972 | 0.479443 | 0.245 | 0.196 | 1 | 15 | Bdp1 | 0.049 | Old |
| 0.500361 | 0.131635 | 0.286 | 0.214 | 1 | 15 | Nsd3 | 0.072 | Old |
| 0.500779 | 0.132188 | 0.571 | 0.482 | 1 | 15 | Khdrbs1 | 0.089 | Old |
| 0.501086 | 0.093836 | 0.122 | 0.179 | 1 | 15 | Tm2d1 | -0.057 | Old |
| 0.501244 | 0.768499 | 0.306 | 0.25 | 1 | 15 | Ranbp2 | 0.056 | Old |
| 0.501795 | 0.852888 | 0.367 | 0.321 | 1 | 15 | Kmt2a | 0.046 | Old |
| 0.502098 | 0.175319 | 0.163 | 0.232 | 1 | 15 | Dyrk1a | -0.069 | Old |
| 0.502319 | 0.229767 | 0.204 | 0.268 | 1 | 15 | Sik1 | -0.064 | Old |
| 0.502711 | 0.642297 | 0.224 | 0.161 | 1 | 15 | Capn1 | 0.063 | Old |
| 0.502797 | 0.064388 | 0.184 | 0.25 | 1 | 15 | Acbd3 | -0.066 | Old |
| 0.502971 | 0.069274 | 0.204 | 0.161 | 1 | 15 | Ccl4 | 0.043 | Old |
| 0.502974 | 0.562203 | 0.51 | 0.446 | 1 | 15 | Tmem30a | 0.064 | Old |
| 0.503347 | 0.033293 | 0.408 | 0.464 | 1 | 15 | Hnrnpul2 | -0.056 | Old |
| 0.503351 | 0.840289 | 0.327 | 0.268 | 1 | 15 | Tmx3 | 0.059 | Old |
| 0.503546 | 0.559561 | 0.286 | 0.232 | 1 | 15 | Slc25a11 | 0.054 | Old |
| 0.503937 | 0.198188 | 0.776 | 0.786 | 1 | 15 | Tagln2 | -0.01 | Old |
| 0.504095 | 1.846077 | 0.102 | 0.071 | 1 | 15 | Myef2 | 0.031 | Old |
| 0.504159 | 0.871446 | 0.143 | 0.107 | 1 | 15 | Rgl1 | 0.036 | Old |
| 0.504345 | 0.875751 | 0.041 | 0.018 | 1 | 15 | Mtg1 | 0.023 | Old |
| 0.504345 | 0.737882 | 0.041 | 0.018 | 1 | 15 | Gm5547 | 0.023 | Old |
| 0.504345 | 0.70134 | 0.041 | 0.018 | 1 | 15 | Mex3d | 0.023 | Old |
| 0.504345 | 0.685917 | 0.041 | 0.018 | 1 | 15 | Gmppa | 0.023 | Old |
| 0.504345 | 0.653299 | 0.041 | 0.018 | 1 | 15 | Habp4 | 0.023 | Old |
| 0.504345 | 0.586858 | 0.041 | 0.018 | 1 | 15 | Pfkfb4 | 0.023 | Old |
| 0.504345 | 0.577108 | 0.041 | 0.018 | 1 | 15 | Zfp189 | 0.023 | Old |
| 0.504345 | 0.560859 | 0.041 | 0.018 | 1 | 15 | Slc43a3 | 0.023 | Old |
| 0.504345 | 0.534391 | 0.041 | 0.018 | 1 | 15 | Arhgap23 | 0.023 | Old |
| 0.504345 | 0.516428 | 0.041 | 0.018 | 1 | 15 | Dnaaf5 | 0.023 | Old |
| 0.504345 | 0.511556 | 0.041 | 0.018 | 1 | 15 | 6330562C2 | 0.023 | Old |
| 0.504345 | 0.496815 | 0.041 | 0.018 | 1 | 15 | Itfg2 | 0.023 | Old |
| 0.504345 | 0.479846 | 0.041 | 0.018 | 1 | 15 | Asb13 | 0.023 | Old |
| 0.504345 | 0.463272 | 0.041 | 0.018 | 1 | 15 | Pde5a | 0.023 | Old |
| 0.504345 | 0.42478 | 0.041 | 0.018 | 1 | 15 | Naa40 | 0.023 | Old |
| 0.504345 | 0.418899 | 0.041 | 0.018 | 1 | 15 | Pwwp2b | 0.023 | Old |
| 0.504345 | 0.4129 | 0.041 | 0.018 | 1 | 15 | Slc35e2 | 0.023 | Old |
| 0.504345 | 0.410681 | 0.041 | 0.018 | 1 | 15 | Sh2d4a | 0.023 | Old |
| 0.504345 | 0.406953 | 0.041 | 0.018 | 1 | 15 | Klre1 | 0.023 | Old |
| 0.504345 | 0.375424 | 0.041 | 0.018 | 1 | 15 | Rnf26 | 0.023 | Old |
| 0.504345 | 0.368302 | 0.041 | 0.018 | 1 | 15 | Tfb2m | 0.023 | Old |

| | | | | | | | | |
|---|---|---|---|---|---|---|---|---|
| 0.504345 | 0.328735 | 0.041 | 0.018 | 1 | 15 | Card10 | 0.023 | Old |
| 0.504345 | 0.312588 | 0.041 | 0.018 | 1 | 15 | Amacr | 0.023 | Old |
| 0.504345 | 0.310994 | 0.041 | 0.018 | 1 | 15 | Herc3 | 0.023 | Old |
| 0.504345 | 0.279107 | 0.041 | 0.018 | 1 | 15 | 1700021F0 | 0.023 | Old |
| 0.504345 | 0.275158 | 0.041 | 0.018 | 1 | 15 | Aurkb | 0.023 | Old |
| 0.504345 | 0.271852 | 0.041 | 0.018 | 1 | 15 | Creb3l2 | 0.023 | Old |
| 0.504345 | 0.222597 | 0.041 | 0.018 | 1 | 15 | Klhdc8b | 0.023 | Old |
| 0.504345 | 0.214015 | 0.041 | 0.018 | 1 | 15 | Dbf4 | 0.023 | Old |
| 0.504345 | 0.21023 | 0.041 | 0.018 | 1 | 15 | Smim5 | 0.023 | Old |
| 0.504345 | 0.208535 | 0.041 | 0.018 | 1 | 15 | B630019K0 | 0.023 | Old |
| 0.504345 | 0.158692 | 0.041 | 0.018 | 1 | 15 | Fbxw9 | 0.023 | Old |
| 0.504345 | 0.130501 | 0.041 | 0.018 | 1 | 15 | Usp21 | 0.023 | Old |
| 0.504345 | 0.109437 | 0.041 | 0.018 | 1 | 15 | Gm15445 | 0.023 | Old |
| 0.504345 | 0.10693 | 0.041 | 0.018 | 1 | 15 | Lhpp | 0.023 | Old |
| 0.504345 | 0.042407 | 0.041 | 0.018 | 1 | 15 | Thap1 | 0.023 | Old |
| 0.504345 | 0.005259 | 0.041 | 0.018 | 1 | 15 | Zfp275 | 0.023 | Old |
| 0.505419 | 0.945102 | 0.184 | 0.143 | 1 | 15 | Lpin1 | 0.041 | Old |
| 0.50563 | 0.963427 | 0.245 | 0.196 | 1 | 15 | Zfp871 | 0.049 | Old |
| 0.506183 | 0.116734 | 1 | 1 | 1 | 15 | Rplp2 | 0 | Old |
| 0.506288 | 1.839133 | 0.122 | 0.089 | 1 | 15 | Timm22 | 0.033 | Old |
| 0.506288 | 1.324657 | 0.122 | 0.089 | 1 | 15 | Fkbp1b | 0.033 | Old |
| 0.506288 | 0.167532 | 0.082 | 0.125 | 1 | 15 | Gm20186 | -0.043 | Old |
| 0.506446 | 0.432862 | 0.306 | 0.25 | 1 | 15 | Cs | 0.056 | Old |
| 0.506856 | 0.027836 | 0.367 | 0.429 | 1 | 15 | Mapkapk2 | -0.062 | Old |
| 0.507569 | 0.253981 | 0.449 | 0.393 | 1 | 15 | Lgals3 | 0.056 | Old |
| 0.507573 | 0.69443 | 0.041 | 0.071 | 1 | 15 | Nt5dc3 | -0.03 | Old |
| 0.507573 | 0.11211 | 0.041 | 0.071 | 1 | 15 | Grap | -0.03 | Old |
| 0.508895 | 0.297386 | 0.286 | 0.232 | 1 | 15 | Rnf111 | 0.054 | Old |
| 0.509084 | 0.198613 | 0.143 | 0.196 | 1 | 15 | Arhgap9 | -0.053 | Old |
| 0.509091 | 0.594786 | 0.204 | 0.161 | 1 | 15 | Pqlc3 | 0.043 | Old |
| 0.510825 | 0.181052 | 0.735 | 0.75 | 1 | 15 | Cbfa2t3 | -0.015 | Old |
| 0.51098 | 0.757926 | 0.592 | 0.571 | 1 | 15 | Arpc4 | 0.021 | Old |
| 0.511352 | 0.133404 | 0.102 | 0.143 | 1 | 15 | Naip2 | -0.041 | Old |
| 0.511866 | 0.342308 | 0.51 | 0.446 | 1 | 15 | Cct5 | 0.064 | Old |
| 0.512577 | 1.912018 | 0.102 | 0.071 | 1 | 15 | Rgs6 | 0.031 | Old |
| 0.512828 | 1.319946 | 0.163 | 0.125 | 1 | 15 | Rprd2 | 0.038 | Old |
| 0.513602 | 0.513522 | 0.388 | 0.339 | 1 | 15 | Foxn2 | 0.049 | Old |
| 0.513602 | 0.443859 | 0.388 | 0.339 | 1 | 15 | Ppp3cb | 0.049 | Old |
| 0.513928 | 3.184159 | 0.061 | 0.036 | 1 | 15 | Usp31 | 0.025 | Old |
| 0.513928 | 2.801992 | 0.061 | 0.036 | 1 | 15 | Jrkl | 0.025 | Old |
| 0.513928 | 2.299905 | 0.061 | 0.036 | 1 | 15 | Alpk1 | 0.025 | Old |
| 0.513928 | 2.072304 | 0.061 | 0.036 | 1 | 15 | Cpox | 0.025 | Old |
| 0.513928 | 2.058665 | 0.061 | 0.036 | 1 | 15 | Alg5 | 0.025 | Old |
| 0.513928 | 2.052555 | 0.061 | 0.036 | 1 | 15 | Insig2 | 0.025 | Old |
| 0.513928 | 1.639642 | 0.061 | 0.036 | 1 | 15 | Dhdds | 0.025 | Old |

| | | | | | | | | |
|---|---|---|---|---|---|---|---|---|
| 0.513928 | 1.589819 | 0.061 | 0.036 | 1 | 15 | Xrcc4 | 0.025 | Old |
| 0.513928 | 1.512288 | 0.061 | 0.036 | 1 | 15 | Acvr2b | 0.025 | Old |
| 0.513928 | 1.455831 | 0.061 | 0.036 | 1 | 15 | Vipas39 | 0.025 | Old |
| 0.513928 | 1.355111 | 0.061 | 0.036 | 1 | 15 | Chmp6 | 0.025 | Old |
| 0.513928 | 1.26308 | 0.061 | 0.036 | 1 | 15 | Dgcr6 | 0.025 | Old |
| 0.513928 | 1.180967 | 0.061 | 0.036 | 1 | 15 | Cep250 | 0.025 | Old |
| 0.513928 | 1.06211 | 0.061 | 0.036 | 1 | 15 | Stbd1 | 0.025 | Old |
| 0.51405 | 1.396158 | 0.122 | 0.089 | 1 | 15 | Chst7 | 0.033 | Old |
| 0.514696 | 0.299386 | 0.347 | 0.286 | 1 | 15 | Vdac3 | 0.061 | Old |
| 0.51525 | 0.795231 | 0.204 | 0.161 | 1 | 15 | Med1 | 0.043 | Old |
| 0.51679 | 0.261643 | 0.449 | 0.393 | 1 | 15 | Fcho2 | 0.056 | Old |
| 0.516852 | 0.962956 | 0.224 | 0.179 | 1 | 15 | Bfar | 0.045 | Old |
| 0.516956 | 0.126913 | 0.49 | 0.411 | 1 | 15 | Scpep1 | 0.079 | Old |
| 0.516993 | 0.175807 | 0.469 | 0.411 | 1 | 15 | Rock1 | 0.058 | Old |
| 0.517057 | 0.651786 | 0.286 | 0.25 | 1 | 15 | Mrps14 | 0.036 | Old |
| 0.517436 | 0.152793 | 0.612 | 0.554 | 1 | 15 | Eif4a1 | 0.058 | Old |
| 0.517487 | 0.562373 | 0.367 | 0.339 | 1 | 15 | Zmiz2 | 0.028 | Old |
| 0.517855 | 0.918268 | 0.041 | 0.071 | 1 | 15 | N6amt1 | -0.03 | Old |
| 0.518307 | 0.60638 | 0.184 | 0.143 | 1 | 15 | Pip5k1a | 0.041 | Old |
| 0.518323 | 0.356229 | 0.735 | 0.714 | 1 | 15 | Gnb1 | 0.021 | Old |
| 0.518874 | 0.488822 | 0.429 | 0.393 | 1 | 15 | Uqcrb | 0.036 | Old |
| 0.519236 | 0.353869 | 0.469 | 0.429 | 1 | 15 | Sbno2 | 0.04 | Old |
| 0.51965 | 0.875571 | 0.163 | 0.125 | 1 | 15 | Uimc1 | 0.038 | Old |
| 0.520802 | 0.106778 | 0.224 | 0.161 | 1 | 15 | Mx1 | 0.063 | Old |
| 0.520894 | 0.294655 | 0.122 | 0.179 | 1 | 15 | Herpud2 | -0.057 | Old |
| 0.52111 | 0.191917 | 0.592 | 0.679 | 1 | 15 | Nfkbid | -0.087 | Old |
| 0.521132 | 1.826437 | 0.102 | 0.071 | 1 | 15 | Cog3 | 0.031 | Old |
| 0.521132 | 1.211562 | 0.102 | 0.071 | 1 | 15 | Pcyox1 | 0.031 | Old |
| 0.521132 | 1.100626 | 0.102 | 0.071 | 1 | 15 | Thumpd3 | 0.031 | Old |
| 0.521321 | 0.447959 | 0.367 | 0.321 | 1 | 15 | Crebbp | 0.046 | Old |
| 0.521375 | 0.309073 | 0.51 | 0.411 | 1 | 15 | Fis1 | 0.099 | Old |
| 0.521689 | 0.853299 | 0.265 | 0.214 | 1 | 15 | Mon2 | 0.051 | Old |
| 0.521874 | 1.424213 | 0.122 | 0.089 | 1 | 15 | Gm43848 | 0.033 | Old |
| 0.521874 | 0.144928 | 0.082 | 0.125 | 1 | 15 | Copg1 | -0.043 | Old |
| 0.522163 | 0.113472 | 0.286 | 0.214 | 1 | 15 | Tsc22d1 | 0.072 | Old |
| 0.522409 | 0.272595 | 0.286 | 0.25 | 1 | 15 | P4hb | 0.036 | Old |
| 0.522802 | 1.057359 | 0.245 | 0.196 | 1 | 15 | Bmpr2 | 0.049 | Old |
| 0.522815 | 0.540414 | 0.224 | 0.179 | 1 | 15 | Dpysl2 | 0.045 | Old |
| 0.523813 | 0.309763 | 0.469 | 0.429 | 1 | 15 | Capza1 | 0.04 | Old |
| 0.524814 | 1.073168 | 0.184 | 0.143 | 1 | 15 | Cbx7 | 0.041 | Old |
| 0.524814 | 0.986511 | 0.184 | 0.143 | 1 | 15 | Ubp1 | 0.041 | Old |
| 0.524814 | 0.414679 | 0.184 | 0.143 | 1 | 15 | Cnot3 | 0.041 | Old |
| 0.524819 | 0.161828 | 0.306 | 0.393 | 1 | 15 | Rab1a | -0.087 | Old |
| 0.525215 | 2.689669 | 0.061 | 0.036 | 1 | 15 | Gm28875 | 0.025 | Old |
| 0.525215 | 2.086052 | 0.061 | 0.036 | 1 | 15 | Plcb2 | 0.025 | Old |

| | | | | | | | | |
|---|---|---|---|---|---|---|---|---|
| 0.525215 | 1.910488 | 0.061 | 0.036 | 1 | 15 | Jade2 | 0.025 | Old |
| 0.525215 | 1.81573 | 0.061 | 0.036 | 1 | 15 | Acvr1 | 0.025 | Old |
| 0.525215 | 1.80851 | 0.061 | 0.036 | 1 | 15 | Zfp821 | 0.025 | Old |
| 0.525215 | 1.706981 | 0.061 | 0.036 | 1 | 15 | Vamp1 | 0.025 | Old |
| 0.525215 | 1.576769 | 0.061 | 0.036 | 1 | 15 | Aig1 | 0.025 | Old |
| 0.525215 | 1.522964 | 0.061 | 0.036 | 1 | 15 | Slc35e3 | 0.025 | Old |
| 0.525215 | 1.497488 | 0.061 | 0.036 | 1 | 15 | Mri1 | 0.025 | Old |
| 0.525215 | 1.457019 | 0.061 | 0.036 | 1 | 15 | Zmat3 | 0.025 | Old |
| 0.525215 | 1.406299 | 0.061 | 0.036 | 1 | 15 | Ubac1 | 0.025 | Old |
| 0.525215 | 1.323832 | 0.061 | 0.036 | 1 | 15 | Fbxo25 | 0.025 | Old |
| 0.525215 | 1.138262 | 0.061 | 0.036 | 1 | 15 | Irf2bp1 | 0.025 | Old |
| 0.525897 | 1.437161 | 0.143 | 0.107 | 1 | 15 | Tmem158 | 0.036 | Old |
| 0.525897 | 0.896014 | 0.143 | 0.107 | 1 | 15 | Recql | 0.036 | Old |
| 0.525938 | 0.117728 | 0.429 | 0.482 | 1 | 15 | Nxf1 | -0.053 | Old |
| 0.526518 | 0.029316 | 0.122 | 0.161 | 1 | 15 | Fbxo45 | -0.039 | Old |
| 0.526923 | 0.117746 | 0.959 | 0.946 | 1 | 15 | Rpl37 | 0.013 | Old |
| 0.527096 | 0.018996 | 0.245 | 0.321 | 1 | 15 | Gns | -0.076 | Old |
| 0.527275 | 0.786388 | 0.367 | 0.339 | 1 | 15 | Xiap | 0.028 | Old |
| 0.527293 | 0.590932 | 0.265 | 0.214 | 1 | 15 | Aprt | 0.051 | Old |
| 0.527683 | 0.608452 | 0.204 | 0.161 | 1 | 15 | Khnyn | 0.043 | Old |
| 0.527777 | 0.07887 | 0.449 | 0.571 | 1 | 15 | Plek | -0.122 | Old |
| 0.528144 | 0.206656 | 0.306 | 0.411 | 1 | 15 | Zcchc11 | -0.105 | Old |
| 0.528244 | 0.397094 | 0.041 | 0.071 | 1 | 15 | Itgb5 | -0.03 | Old |
| 0.528244 | 0.346567 | 0.041 | 0.071 | 1 | 15 | Enoph1 | -0.03 | Old |
| 0.528244 | 0.344831 | 0.041 | 0.071 | 1 | 15 | Pex3 | -0.03 | Old |
| 0.528244 | 0.003454 | 0.041 | 0.071 | 1 | 15 | Ddit3 | -0.03 | Old |
| 0.528814 | 0.230182 | 0.224 | 0.179 | 1 | 15 | Sfxn3 | 0.045 | Old |
| 0.529022 | 0.009063 | 0.918 | 0.982 | 1 | 15 | Calm1 | -0.064 | Old |
| 0.529377 | 0.32487 | 0.633 | 0.554 | 1 | 15 | Rhog | 0.079 | Old |
| 0.529758 | 0.015483 | 0.082 | 0.125 | 1 | 15 | Cox16 | -0.043 | Old |
| 0.529758 | 0.008527 | 0.082 | 0.125 | 1 | 15 | Slc25a30 | -0.043 | Old |
| 0.529759 | 1.826411 | 0.102 | 0.071 | 1 | 15 | Fam43a | 0.031 | Old |
| 0.529759 | 1.283876 | 0.102 | 0.071 | 1 | 15 | Itga6 | 0.031 | Old |
| 0.529759 | 1.25688 | 0.102 | 0.071 | 1 | 15 | Rab3gap2 | 0.031 | Old |
| 0.529759 | 1.08278 | 0.102 | 0.071 | 1 | 15 | Pspc1 | 0.031 | Old |
| 0.529892 | 0.291108 | 0.51 | 0.446 | 1 | 15 | Adap1 | 0.064 | Old |
| 0.529975 | 0.454635 | 0.408 | 0.375 | 1 | 15 | Rftn1 | 0.033 | Old |
| 0.530331 | 0.865713 | 0.224 | 0.196 | 1 | 15 | Pycard | 0.028 | Old |
| 0.531362 | 0.498176 | 0.184 | 0.143 | 1 | 15 | Tcp11l1 | 0.041 | Old |
| 0.531731 | 0.211773 | 0.51 | 0.464 | 1 | 15 | Arrb2 | 0.046 | Old |
| 0.532355 | 0.765458 | 0.245 | 0.214 | 1 | 15 | Ext1 | 0.031 | Old |
| 0.532703 | 1.797223 | 0.082 | 0.054 | 1 | 15 | Yaf2 | 0.028 | Old |
| 0.532703 | 1.771805 | 0.082 | 0.054 | 1 | 15 | Cd151 | 0.028 | Old |
| 0.532703 | 1.616991 | 0.082 | 0.054 | 1 | 15 | Mrpl35 | 0.028 | Old |
| 0.532703 | 1.41729 | 0.082 | 0.054 | 1 | 15 | Letm1 | 0.028 | Old |

| | | | | | | | | |
|---|---|---|---|---|---|---|---|---|
| 0.532703 | 1.355166 | 0.082 | 0.054 | 1 | 15 | AC127341. | 0.028 | Old |
| 0.532703 | 1.29236 | 0.082 | 0.054 | 1 | 15 | Dhx8 | 0.028 | Old |
| 0.532703 | 1.214569 | 0.082 | 0.054 | 1 | 15 | Rfc4 | 0.028 | Old |
| 0.533433 | 0.679661 | 0.163 | 0.125 | 1 | 15 | Mrpl34 | 0.038 | Old |
| 0.533738 | 0.575753 | 0.408 | 0.339 | 1 | 15 | Zmat2 | 0.069 | Old |
| 0.534415 | 0.231558 | 0.245 | 0.196 | 1 | 15 | Zfp800 | 0.049 | Old |
| 0.534848 | 0.476083 | 0.224 | 0.179 | 1 | 15 | Denr | 0.045 | Old |
| 0.53504 | 0.39228 | 0.347 | 0.286 | 1 | 15 | Arcn1 | 0.061 | Old |
| 0.535369 | 0.002076 | 0.98 | 1 | 1 | 15 | Eif1 | -0.02 | Old |
| 0.535481 | 0.331532 | 0.469 | 0.411 | 1 | 15 | Sec11c | 0.058 | Old |
| 0.536264 | 0.095829 | 0.184 | 0.232 | 1 | 15 | Glod4 | -0.048 | Old |
| 0.536628 | 3.0947 | 0.061 | 0.036 | 1 | 15 | Kin | 0.025 | Old |
| 0.536628 | 3.036231 | 0.061 | 0.036 | 1 | 15 | Ltb | 0.025 | Old |
| 0.536628 | 2.249042 | 0.061 | 0.036 | 1 | 15 | Poc5 | 0.025 | Old |
| 0.536628 | 2.111905 | 0.061 | 0.036 | 1 | 15 | Tmx2 | 0.025 | Old |
| 0.536628 | 1.996563 | 0.061 | 0.036 | 1 | 15 | Mto1 | 0.025 | Old |
| 0.536628 | 1.625801 | 0.061 | 0.036 | 1 | 15 | Zmynd19 | 0.025 | Old |
| 0.536628 | 1.549319 | 0.061 | 0.036 | 1 | 15 | Dvl1 | 0.025 | Old |
| 0.536628 | 1.495046 | 0.061 | 0.036 | 1 | 15 | Fancl | 0.025 | Old |
| 0.536628 | 1.486264 | 0.061 | 0.036 | 1 | 15 | Tspan5 | 0.025 | Old |
| 0.536628 | 1.454862 | 0.061 | 0.036 | 1 | 15 | F11r | 0.025 | Old |
| 0.536628 | 1.448264 | 0.061 | 0.036 | 1 | 15 | Mgp | 0.025 | Old |
| 0.536628 | 1.440546 | 0.061 | 0.036 | 1 | 15 | Klf8 | 0.025 | Old |
| 0.536628 | 1.396033 | 0.061 | 0.036 | 1 | 15 | Tmem38a | 0.025 | Old |
| 0.536628 | 1.365829 | 0.061 | 0.036 | 1 | 15 | Gm15232 | 0.025 | Old |
| 0.536628 | 1.334662 | 0.061 | 0.036 | 1 | 15 | Syce2 | 0.025 | Old |
| 0.536628 | 1.311023 | 0.061 | 0.036 | 1 | 15 | Gdf15 | 0.025 | Old |
| 0.536628 | 1.310934 | 0.061 | 0.036 | 1 | 15 | Aen | 0.025 | Old |
| 0.536628 | 1.164663 | 0.061 | 0.036 | 1 | 15 | Kti12 | 0.025 | Old |
| 0.536628 | 1.122524 | 0.061 | 0.036 | 1 | 15 | Cdk5 | 0.025 | Old |
| 0.536628 | 1.117574 | 0.061 | 0.036 | 1 | 15 | Fam129c | 0.025 | Old |
| 0.536628 | 1.105292 | 0.061 | 0.036 | 1 | 15 | Rap1gap2 | 0.025 | Old |
| 0.536628 | 1.100476 | 0.061 | 0.036 | 1 | 15 | Ccdc43 | 0.025 | Old |
| 0.536628 | 1.095716 | 0.061 | 0.036 | 1 | 15 | Lclat1 | 0.025 | Old |
| 0.536628 | 1.082786 | 0.061 | 0.036 | 1 | 15 | Zfp426 | 0.025 | Old |
| 0.536628 | 1.069229 | 0.061 | 0.036 | 1 | 15 | Ikbkg | 0.025 | Old |
| 0.536628 | 1.062915 | 0.061 | 0.036 | 1 | 15 | Pola1 | 0.025 | Old |
| 0.536628 | 0.999601 | 0.061 | 0.036 | 1 | 15 | Pck2 | 0.025 | Old |
| 0.536628 | 0.98466 | 0.061 | 0.036 | 1 | 15 | Reep4 | 0.025 | Old |
| 0.536628 | 0.981826 | 0.061 | 0.036 | 1 | 15 | Sphk2 | 0.025 | Old |
| 0.536628 | 0.981164 | 0.061 | 0.036 | 1 | 15 | Decr1 | 0.025 | Old |
| 0.536628 | 0.975349 | 0.061 | 0.036 | 1 | 15 | Nr2c2ap | 0.025 | Old |
| 0.536628 | 0.949547 | 0.061 | 0.036 | 1 | 15 | Ddah2 | 0.025 | Old |
| 0.536628 | 0.926902 | 0.061 | 0.036 | 1 | 15 | Rnaset2b | 0.025 | Old |
| 0.536628 | 0.896161 | 0.061 | 0.036 | 1 | 15 | Dglucy | 0.025 | Old |

| | | | | | | | | |
|---|---|---|---|---|---|---|---|---|
| 0.536628 | 0.818618 | 0.061 | 0.036 | 1 | 15 | Bzw2 | 0.025 | Old |
| 0.536628 | 0.801873 | 0.061 | 0.036 | 1 | 15 | Lpcat3 | 0.025 | Old |
| 0.536628 | 0.789025 | 0.061 | 0.036 | 1 | 15 | Vps26b | 0.025 | Old |
| 0.536628 | 0.770167 | 0.061 | 0.036 | 1 | 15 | Ruvbl2 | 0.025 | Old |
| 0.536628 | 0.714403 | 0.061 | 0.036 | 1 | 15 | Tmem86a | 0.025 | Old |
| 0.536628 | 0.678011 | 0.061 | 0.036 | 1 | 15 | Ctps2 | 0.025 | Old |
| 0.536628 | 0.598148 | 0.061 | 0.036 | 1 | 15 | Brpf3 | 0.025 | Old |
| 0.537484 | 0.210087 | 0.612 | 0.75 | 1 | 15 | Ppp1r15a | -0.138 | Old |
| 0.537675 | 0.306094 | 0.306 | 0.268 | 1 | 15 | Fyb | 0.038 | Old |
| 0.537702 | 0.085796 | 0.082 | 0.125 | 1 | 15 | Bcl2l2 | -0.043 | Old |
| 0.537952 | 0.823443 | 0.184 | 0.143 | 1 | 15 | Emilin2 | 0.041 | Old |
| 0.538244 | 0.352145 | 0.224 | 0.321 | 1 | 15 | Tspan33 | -0.097 | Old |
| 0.538244 | 0.230751 | 0.306 | 0.25 | 1 | 15 | Phf20l1 | 0.056 | Old |
| 0.538459 | 2.033504 | 0.102 | 0.071 | 1 | 15 | Wdr53 | 0.031 | Old |
| 0.538459 | 1.030932 | 0.102 | 0.071 | 1 | 15 | Phospho2 | 0.031 | Old |
| 0.538739 | 0.31718 | 0.041 | 0.071 | 1 | 15 | Lnx2 | -0.03 | Old |
| 0.538739 | 0.296327 | 0.041 | 0.071 | 1 | 15 | Eid3 | -0.03 | Old |
| 0.539157 | 0.393993 | 0.592 | 0.536 | 1 | 15 | Synj1 | 0.056 | Old |
| 0.540267 | 0.956997 | 0.204 | 0.161 | 1 | 15 | Gtf3c2 | 0.043 | Old |
| 0.540393 | 0.664435 | 0.163 | 0.125 | 1 | 15 | 4833407H1 | 0.038 | Old |
| 0.540393 | 0.633054 | 0.163 | 0.125 | 1 | 15 | Xpnpep1 | 0.038 | Old |
| 0.540651 | 1.3521 | 0.143 | 0.107 | 1 | 15 | Fcf1 | 0.036 | Old |
| 0.540651 | 1.307043 | 0.143 | 0.107 | 1 | 15 | Rnf10 | 0.036 | Old |
| 0.540651 | 0.841097 | 0.143 | 0.107 | 1 | 15 | Slc2a6 | 0.036 | Old |
| 0.540651 | 0.687092 | 0.143 | 0.107 | 1 | 15 | Map3k7 | 0.036 | Old |
| 0.540651 | 0.57211 | 0.143 | 0.107 | 1 | 15 | Adrm1 | 0.036 | Old |
| 0.540916 | 0.966962 | 0.224 | 0.179 | 1 | 15 | Ndufa8 | 0.045 | Old |
| 0.540916 | 0.484978 | 0.224 | 0.179 | 1 | 15 | Vav3 | 0.045 | Old |
| 0.541268 | 0.182538 | 0.327 | 0.286 | 1 | 15 | Nadk | 0.041 | Old |
| 0.54221 | 0.134998 | 0.265 | 0.321 | 1 | 15 | Ik | -0.056 | Old |
| 0.542504 | 2.182535 | 0.082 | 0.054 | 1 | 15 | Gdpd3 | 0.028 | Old |
| 0.542504 | 2.005623 | 0.082 | 0.054 | 1 | 15 | Cwc22 | 0.028 | Old |
| 0.542504 | 1.189875 | 0.082 | 0.054 | 1 | 15 | Wdr77 | 0.028 | Old |
| 0.542504 | 1.151216 | 0.082 | 0.054 | 1 | 15 | Rasa3 | 0.028 | Old |
| 0.542504 | 1.043346 | 0.082 | 0.054 | 1 | 15 | Slc35f5 | 0.028 | Old |
| 0.542504 | 0.9163 | 0.082 | 0.054 | 1 | 15 | Mettl16 | 0.028 | Old |
| 0.542504 | 0.70284 | 0.082 | 0.054 | 1 | 15 | Abhd17c | 0.028 | Old |
| 0.543005 | 0.076591 | 0.245 | 0.321 | 1 | 15 | Ppp6r3 | -0.076 | Old |
| 0.543209 | 0.219127 | 0.184 | 0.25 | 1 | 15 | Gas7 | -0.066 | Old |
| 0.543857 | 0.27545 | 0.959 | 0.982 | 1 | 15 | Zfp36l1 | -0.023 | Old |
| 0.543885 | 0.186557 | 0.265 | 0.196 | 1 | 15 | Rab43 | 0.069 | Old |
| 0.545436 | 0.3377 | 0.163 | 0.214 | 1 | 15 | Luc7l | -0.051 | Old |
| 0.545706 | 0.852368 | 0.082 | 0.125 | 1 | 15 | Eipr1 | -0.043 | Old |
| 0.546156 | 0.468098 | 0.245 | 0.196 | 1 | 15 | Dctn4 | 0.049 | Old |
| 0.546615 | 0.32915 | 0.204 | 0.161 | 1 | 15 | Pdap1 | 0.043 | Old |

| | | | | | | | | |
|---|---|---|---|---|---|---|---|---|
| 0.547018 | 0.356068 | 0.224 | 0.179 | 1 | 15 | Pex16 | 0.045 | Old |
| 0.547147 | 0.021306 | 0.204 | 0.268 | 1 | 15 | Psen2 | -0.064 | Old |
| 0.54723 | 1.399513 | 0.102 | 0.071 | 1 | 15 | Efna2 | 0.031 | Old |
| 0.54723 | 1.069162 | 0.102 | 0.071 | 1 | 15 | Lix1l | 0.031 | Old |
| 0.54723 | 1.054167 | 0.102 | 0.071 | 1 | 15 | St6galnac4 | 0.031 | Old |
| 0.54723 | 1.016376 | 0.102 | 0.071 | 1 | 15 | Stoml2 | 0.031 | Old |
| 0.54723 | 0.722485 | 0.102 | 0.071 | 1 | 15 | Ccdc32 | 0.031 | Old |
| 0.547398 | 0.477369 | 0.163 | 0.125 | 1 | 15 | Mrpl36 | 0.038 | Old |
| 0.547853 | 0.411387 | 0.388 | 0.339 | 1 | 15 | Fbxl3 | 0.049 | Old |
| 0.548104 | 0.892055 | 0.143 | 0.107 | 1 | 15 | Akap11 | 0.036 | Old |
| 0.548104 | 0.16212 | 0.102 | 0.143 | 1 | 15 | Hypk | -0.041 | Old |
| 0.548164 | 1.712392 | 0.061 | 0.036 | 1 | 15 | Nudt14 | 0.025 | Old |
| 0.548164 | 1.312688 | 0.061 | 0.036 | 1 | 15 | Men1 | 0.025 | Old |
| 0.548164 | 1.229915 | 0.061 | 0.036 | 1 | 15 | Rpain | 0.025 | Old |
| 0.548164 | 1.135763 | 0.061 | 0.036 | 1 | 15 | Aasdh | 0.025 | Old |
| 0.548164 | 1.114419 | 0.061 | 0.036 | 1 | 15 | Spn | 0.025 | Old |
| 0.548164 | 1.107456 | 0.061 | 0.036 | 1 | 15 | Anapc7 | 0.025 | Old |
| 0.548164 | 1.019229 | 0.061 | 0.036 | 1 | 15 | Hirip3 | 0.025 | Old |
| 0.548164 | 0.966771 | 0.061 | 0.036 | 1 | 15 | Ccdc112 | 0.025 | Old |
| 0.548164 | 0.934498 | 0.061 | 0.036 | 1 | 15 | Hdac9 | 0.025 | Old |
| 0.548164 | 0.924509 | 0.061 | 0.036 | 1 | 15 | Bcl9l | 0.025 | Old |
| 0.548164 | 0.909415 | 0.061 | 0.036 | 1 | 15 | Nepro | 0.025 | Old |
| 0.548164 | 0.822968 | 0.061 | 0.036 | 1 | 15 | Flywch1 | 0.025 | Old |
| 0.548164 | 0.811064 | 0.061 | 0.036 | 1 | 15 | Fra10ac1 | 0.025 | Old |
| 0.548164 | 0.728526 | 0.061 | 0.036 | 1 | 15 | Paxip1 | 0.025 | Old |
| 0.548164 | 0.660176 | 0.061 | 0.036 | 1 | 15 | Terf2ip | 0.025 | Old |
| 0.548164 | 0.659124 | 0.061 | 0.036 | 1 | 15 | Stard5 | 0.025 | Old |
| 0.548164 | 0.636509 | 0.061 | 0.036 | 1 | 15 | Nup133 | 0.025 | Old |
| 0.548164 | 0.610103 | 0.061 | 0.036 | 1 | 15 | Mcm4 | 0.025 | Old |
| 0.548164 | 0.609294 | 0.061 | 0.036 | 1 | 15 | Ints7 | 0.025 | Old |
| 0.548164 | 0.6082 | 0.061 | 0.036 | 1 | 15 | Ppt2 | 0.025 | Old |
| 0.548164 | 0.534277 | 0.061 | 0.036 | 1 | 15 | 1500011B( | 0.025 | Old |
| 0.548164 | 0.519689 | 0.061 | 0.036 | 1 | 15 | Mgst3 | 0.025 | Old |
| 0.548164 | 0.465784 | 0.061 | 0.036 | 1 | 15 | Cactin | 0.025 | Old |
| 0.548164 | 0.291129 | 0.061 | 0.036 | 1 | 15 | Pgp | 0.025 | Old |
| 0.548164 | 0.212227 | 0.061 | 0.036 | 1 | 15 | Asna1 | 0.025 | Old |
| 0.548164 | 0.081713 | 0.061 | 0.036 | 1 | 15 | Jmjd8 | 0.025 | Old |
| 0.54818 | 0.01914 | 0.796 | 0.661 | 1 | 15 | Rgs1 | 0.135 | Old |
| 0.54823 | 0.197194 | 0.184 | 0.232 | 1 | 15 | Vapb | -0.048 | Old |
| 0.548924 | 0.36465 | 0.327 | 0.429 | 1 | 15 | Adam23 | -0.102 | Old |
| 0.549336 | 0.144288 | 0.041 | 0.071 | 1 | 15 | Prr3 | -0.03 | Old |
| 0.549377 | 0.177421 | 0.755 | 0.75 | 1 | 15 | Rpl31 | 0.005 | Old |
| 0.550023 | 0.470996 | 0.51 | 0.464 | 1 | 15 | Fam129a | 0.046 | Old |
| 0.551725 | 0.262721 | 0.531 | 0.464 | 1 | 15 | Csf2ra | 0.067 | Old |
| 0.551755 | 0.351249 | 0.327 | 0.286 | 1 | 15 | Metrnl | 0.041 | Old |

| | | | | | | | | |
|---|---|---|---|---|---|---|---|---|
| 0.552092 | 0.375599 | 0.347 | 0.464 | 1 | 15 | Acin1 | -0.117 | Old |
| 0.552394 | 2.811078 | 0.082 | 0.054 | 1 | 15 | Paxx | 0.028 | Old |
| 0.552394 | 1.521316 | 0.082 | 0.054 | 1 | 15 | Pex2 | 0.028 | Old |
| 0.552394 | 1.421844 | 0.082 | 0.054 | 1 | 15 | Zfp746 | 0.028 | Old |
| 0.552394 | 1.354014 | 0.082 | 0.054 | 1 | 15 | Alms1 | 0.028 | Old |
| 0.552394 | 1.103666 | 0.082 | 0.054 | 1 | 15 | Vta1 | 0.028 | Old |
| 0.552394 | 1.101391 | 0.082 | 0.054 | 1 | 15 | Fam13b | 0.028 | Old |
| 0.552394 | 0.997439 | 0.082 | 0.054 | 1 | 15 | G2e3 | 0.028 | Old |
| 0.552394 | 0.98865 | 0.082 | 0.054 | 1 | 15 | Tcf7 | 0.028 | Old |
| 0.552394 | 0.97892 | 0.082 | 0.054 | 1 | 15 | Eef1akmt1 | 0.028 | Old |
| 0.552394 | 0.891434 | 0.082 | 0.054 | 1 | 15 | Slc45a4 | 0.028 | Old |
| 0.552394 | 0.282469 | 0.082 | 0.054 | 1 | 15 | Gla | 0.028 | Old |
| 0.552415 | 0.04874 | 0.959 | 0.982 | 1 | 15 | Rpl6 | -0.023 | Old |
| 0.552741 | 0.373172 | 0.776 | 0.804 | 1 | 15 | Ddx6 | -0.028 | Old |
| 0.552999 | 0.437189 | 0.204 | 0.161 | 1 | 15 | Elovl5 | 0.043 | Old |
| 0.552999 | 0.124031 | 0.204 | 0.161 | 1 | 15 | Polr2c | 0.043 | Old |
| 0.553155 | 0.714889 | 0.224 | 0.179 | 1 | 15 | Utp3 | 0.045 | Old |
| 0.553167 | 0.09323 | 0.286 | 0.339 | 1 | 15 | Chd2 | -0.053 | Old |
| 0.553398 | 0.111918 | 0.327 | 0.411 | 1 | 15 | Il6ra | -0.084 | Old |
| 0.553768 | 1.188599 | 0.122 | 0.089 | 1 | 15 | Faf1 | 0.033 | Old |
| 0.553768 | 1.103897 | 0.122 | 0.089 | 1 | 15 | Ttc4 | 0.033 | Old |
| 0.553768 | 0.755012 | 0.122 | 0.089 | 1 | 15 | Blvra | 0.033 | Old |
| 0.553768 | 0.636779 | 0.122 | 0.089 | 1 | 15 | Foxo1 | 0.033 | Old |
| 0.553768 | 0.288258 | 0.082 | 0.125 | 1 | 15 | Edem2 | -0.043 | Old |
| 0.554448 | 0.880006 | 0.163 | 0.125 | 1 | 15 | Pla2g2d | 0.038 | Old |
| 0.554448 | 0.796977 | 0.163 | 0.125 | 1 | 15 | 1500011K1 | 0.038 | Old |
| 0.554789 | 0.545013 | 0.571 | 0.554 | 1 | 15 | 4930523C0 | 0.017 | Old |
| 0.555113 | 0.278903 | 0.286 | 0.25 | 1 | 15 | Ehmt2 | 0.036 | Old |
| 0.555113 | 0.023091 | 0.245 | 0.286 | 1 | 15 | Ube2e1 | -0.041 | Old |
| 0.555538 | 0.841759 | 0.245 | 0.214 | 1 | 15 | Tax1bp3 | 0.031 | Old |
| 0.555608 | 0.89578 | 0.102 | 0.143 | 1 | 15 | Prr5l | -0.041 | Old |
| 0.555608 | 0.666411 | 0.143 | 0.107 | 1 | 15 | Fam172a | 0.036 | Old |
| 0.555781 | 0.55314 | 0.347 | 0.286 | 1 | 15 | Ssfa2 | 0.061 | Old |
| 0.556072 | 1.065598 | 0.102 | 0.071 | 1 | 15 | Abcf2 | 0.031 | Old |
| 0.556072 | 1.064752 | 0.102 | 0.071 | 1 | 15 | Borcs8 | 0.031 | Old |
| 0.556072 | 0.987424 | 0.102 | 0.071 | 1 | 15 | Vps13c | 0.031 | Old |
| 0.556072 | 0.985593 | 0.102 | 0.071 | 1 | 15 | Tor3a | 0.031 | Old |
| 0.556072 | 0.946787 | 0.102 | 0.071 | 1 | 15 | Ttc19 | 0.031 | Old |
| 0.556072 | 0.850523 | 0.102 | 0.071 | 1 | 15 | Polr2b | 0.031 | Old |
| 0.556072 | 0.579542 | 0.102 | 0.071 | 1 | 15 | Dubr | 0.031 | Old |
| 0.556409 | 0.924227 | 0.347 | 0.339 | 1 | 15 | Sema7a | 0.008 | Old |
| 0.556409 | 0.049626 | 0.388 | 0.304 | 1 | 15 | Mcemp1 | 0.084 | Old |
| 0.556723 | 0.158032 | 0.959 | 1 | 1 | 15 | Tmem123 | -0.041 | Old |
| 0.557036 | 0.351018 | 0.327 | 0.286 | 1 | 15 | Il2rg | 0.041 | Old |
| 0.557192 | 0.436612 | 0.367 | 0.339 | 1 | 15 | Man1a | 0.028 | Old |

| | | | | | | | | |
|---|---|---|---|---|---|---|---|---|
| 0.558024 | 0.657965 | 0.245 | 0.196 | 1 | 15 | Osgin2 | 0.049 | Old |
| 0.558784 | 0.778422 | 0.449 | 0.446 | 1 | 15 | Tbcb | 0.003 | Old |
| 0.559068 | 0.413901 | 0.408 | 0.375 | 1 | 15 | Prrc2a | 0.033 | Old |
| 0.559821 | 2.384717 | 0.061 | 0.036 | 1 | 15 | Pank4 | 0.025 | Old |
| 0.559821 | 2.022285 | 0.061 | 0.036 | 1 | 15 | Atg4a | 0.025 | Old |
| 0.559821 | 0.972236 | 0.061 | 0.036 | 1 | 15 | Dusp7 | 0.025 | Old |
| 0.559821 | 0.959814 | 0.061 | 0.036 | 1 | 15 | Phb | 0.025 | Old |
| 0.559821 | 0.917723 | 0.061 | 0.036 | 1 | 15 | Gm45894 | 0.025 | Old |
| 0.559821 | 0.840737 | 0.061 | 0.036 | 1 | 15 | Entpd6 | 0.025 | Old |
| 0.559821 | 0.785157 | 0.061 | 0.036 | 1 | 15 | 1700017B0 | 0.025 | Old |
| 0.559821 | 0.740302 | 0.061 | 0.036 | 1 | 15 | Igsf3 | 0.025 | Old |
| 0.559821 | 0.675439 | 0.061 | 0.036 | 1 | 15 | Plrg1 | 0.025 | Old |
| 0.559821 | 0.544267 | 0.061 | 0.036 | 1 | 15 | Gprc5a | 0.025 | Old |
| 0.559821 | 0.502747 | 0.061 | 0.036 | 1 | 15 | Crls1 | 0.025 | Old |
| 0.559821 | 0.496527 | 0.061 | 0.036 | 1 | 15 | 1110032F0 | 0.025 | Old |
| 0.559821 | 0.486553 | 0.061 | 0.036 | 1 | 15 | Accs | 0.025 | Old |
| 0.559821 | 0.481842 | 0.061 | 0.036 | 1 | 15 | Klhl20 | 0.025 | Old |
| 0.559821 | 0.458627 | 0.061 | 0.036 | 1 | 15 | Mios | 0.025 | Old |
| 0.559821 | 0.418385 | 0.061 | 0.036 | 1 | 15 | Cdc42ep2 | 0.025 | Old |
| 0.559821 | 0.358927 | 0.061 | 0.036 | 1 | 15 | Nsun3 | 0.025 | Old |
| 0.559821 | 0.328499 | 0.061 | 0.036 | 1 | 15 | Ncapd3 | 0.025 | Old |
| 0.559821 | 0.325451 | 0.061 | 0.036 | 1 | 15 | Setdb2 | 0.025 | Old |
| 0.559821 | 0.296922 | 0.061 | 0.036 | 1 | 15 | Sgsh | 0.025 | Old |
| 0.559821 | 0.288218 | 0.061 | 0.036 | 1 | 15 | Rcan1 | 0.025 | Old |
| 0.559821 | 0.245013 | 0.061 | 0.036 | 1 | 15 | Adsl | 0.025 | Old |
| 0.559821 | 0.16897 | 0.061 | 0.036 | 1 | 15 | 5730480H0 | 0.025 | Old |
| 0.559821 | 0.166199 | 0.061 | 0.036 | 1 | 15 | Slc20a2 | 0.025 | Old |
| 0.559821 | 0.100215 | 0.061 | 0.036 | 1 | 15 | Cep104 | 0.025 | Old |
| 0.559821 | 0.08026 | 0.061 | 0.036 | 1 | 15 | Adnp2 | 0.025 | Old |
| 0.560213 | 0.205668 | 0.816 | 0.857 | 1 | 15 | Selplg | -0.041 | Old |
| 0.560865 | 0.141513 | 0.735 | 0.804 | 1 | 15 | Grb2 | -0.069 | Old |
| 0.561888 | 1.541397 | 0.122 | 0.089 | 1 | 15 | Tmco3 | 0.033 | Old |
| 0.561888 | 1.177069 | 0.122 | 0.089 | 1 | 15 | Apol10b | 0.033 | Old |
| 0.561888 | 0.817791 | 0.122 | 0.089 | 1 | 15 | Smim19 | 0.033 | Old |
| 0.561888 | 0.59311 | 0.122 | 0.089 | 1 | 15 | Gnl3 | 0.033 | Old |
| 0.561888 | 0.267348 | 0.122 | 0.089 | 1 | 15 | Washc5 | 0.033 | Old |
| 0.562227 | 0.161801 | 0.816 | 0.839 | 1 | 15 | Coro1a | -0.023 | Old |
| 0.562372 | 1.561105 | 0.082 | 0.054 | 1 | 15 | Taf2 | 0.028 | Old |
| 0.562372 | 1.475175 | 0.082 | 0.054 | 1 | 15 | AI606181 | 0.028 | Old |
| 0.562372 | 1.278956 | 0.082 | 0.054 | 1 | 15 | Bola1 | 0.028 | Old |
| 0.562372 | 0.976685 | 0.082 | 0.054 | 1 | 15 | Dtx4 | 0.028 | Old |
| 0.562372 | 0.968623 | 0.082 | 0.054 | 1 | 15 | Nifk | 0.028 | Old |
| 0.562372 | 0.879757 | 0.082 | 0.054 | 1 | 15 | Tmod2 | 0.028 | Old |
| 0.562372 | 0.778575 | 0.082 | 0.054 | 1 | 15 | Haao | 0.028 | Old |
| 0.562372 | 0.751931 | 0.082 | 0.054 | 1 | 15 | Cpped1 | 0.028 | Old |

| | | | | | | | | |
|---|---|---|---|---|---|---|---|---|
| 0.562372 | 0.715364 | 0.082 | 0.054 | 1 | 15 | Actr8 | 0.028 | Old |
| 0.563162 | 0.588222 | 0.143 | 0.107 | 1 | 15 | Aldh3a2 | 0.036 | Old |
| 0.563162 | 0.498802 | 0.102 | 0.143 | 1 | 15 | Eya3 | -0.041 | Old |
| 0.563162 | 0.231645 | 0.102 | 0.143 | 1 | 15 | Ccr1 | -0.041 | Old |
| 0.563174 | 0.180954 | 0.918 | 0.964 | 1 | 15 | Oaz1 | -0.046 | Old |
| 0.56353 | 0.194651 | 0.265 | 0.321 | 1 | 15 | Sipa1l3 | -0.056 | Old |
| 0.563819 | 0.321383 | 0.571 | 0.554 | 1 | 15 | Pak2 | 0.017 | Old |
| 0.564005 | 0.538886 | 0.245 | 0.196 | 1 | 15 | Gm15987 | 0.049 | Old |
| 0.564005 | 0.066686 | 0.245 | 0.196 | 1 | 15 | Vps29 | 0.049 | Old |
| 0.564146 | 0.339102 | 0.429 | 0.375 | 1 | 15 | Nuak2 | 0.054 | Old |
| 0.564582 | 0.021792 | 0.306 | 0.268 | 1 | 15 | Ppp1r2 | 0.038 | Old |
| 0.564714 | 0.749021 | 0.184 | 0.143 | 1 | 15 | Ccm2 | 0.041 | Old |
| 0.564982 | 1.76928 | 0.102 | 0.071 | 1 | 15 | Orc4 | 0.031 | Old |
| 0.564982 | 0.943772 | 0.102 | 0.071 | 1 | 15 | BC004004 | 0.031 | Old |
| 0.564982 | 0.856509 | 0.102 | 0.071 | 1 | 15 | Mrpl40 | 0.031 | Old |
| 0.564982 | 0.829704 | 0.102 | 0.071 | 1 | 15 | 1700037H( | 0.031 | Old |
| 0.564982 | 0.090991 | 0.102 | 0.071 | 1 | 15 | Mertk | 0.031 | Old |
| 0.566223 | 0.314882 | 0.327 | 0.268 | 1 | 15 | Bri3bp | 0.059 | Old |
| 0.566805 | 0.39303 | 0.224 | 0.286 | 1 | 15 | Ndufs7 | -0.062 | Old |
| 0.567167 | 0.410397 | 0.143 | 0.196 | 1 | 15 | Nudc | -0.053 | Old |
| 0.567188 | 0.888872 | 0.265 | 0.232 | 1 | 15 | Gpsm3 | 0.033 | Old |
| 0.56868 | 1.389555 | 0.163 | 0.125 | 1 | 15 | Crtc2 | 0.038 | Old |
| 0.56868 | 0.601892 | 0.163 | 0.125 | 1 | 15 | Ap1g2 | 0.038 | Old |
| 0.56868 | 0.337571 | 0.163 | 0.125 | 1 | 15 | Kctd20 | 0.038 | Old |
| 0.569015 | 0.597307 | 0.449 | 0.393 | 1 | 15 | Sepsecs | 0.056 | Old |
| 0.570016 | 0.68358 | 0.245 | 0.196 | 1 | 15 | Ccdc124 | 0.049 | Old |
| 0.570066 | 1.106539 | 0.122 | 0.089 | 1 | 15 | Fgfr1op | 0.033 | Old |
| 0.570066 | 0.902429 | 0.122 | 0.089 | 1 | 15 | Zc3h4 | 0.033 | Old |
| 0.570066 | 0.751601 | 0.122 | 0.089 | 1 | 15 | Cdk2 | 0.033 | Old |
| 0.570066 | 0.620622 | 0.122 | 0.089 | 1 | 15 | Wipi2 | 0.033 | Old |
| 0.570765 | 0.532643 | 0.143 | 0.107 | 1 | 15 | Naa38 | 0.036 | Old |
| 0.570765 | 0.519908 | 0.143 | 0.107 | 1 | 15 | Ilvbl | 0.036 | Old |
| 0.570765 | 0.409331 | 0.143 | 0.107 | 1 | 15 | Ube2m | 0.036 | Old |
| 0.571503 | 0.500864 | 0.184 | 0.143 | 1 | 15 | Rnf19a | 0.041 | Old |
| 0.571503 | 0.323054 | 0.184 | 0.143 | 1 | 15 | Npepl1 | 0.041 | Old |
| 0.571503 | 0.087267 | 0.184 | 0.143 | 1 | 15 | Serhl | 0.041 | Old |
| 0.571503 | 0.067375 | 0.143 | 0.179 | 1 | 15 | Setd7 | -0.036 | Old |
| 0.571596 | 0.34734 | 0.061 | 0.036 | 1 | 15 | 2810454H( | 0.025 | Old |
| 0.571596 | 0.344807 | 0.061 | 0.036 | 1 | 15 | Qpctl | 0.025 | Old |
| 0.571596 | 0.290748 | 0.061 | 0.036 | 1 | 15 | Ccdc191 | 0.025 | Old |
| 0.571596 | 0.175084 | 0.061 | 0.036 | 1 | 15 | Isg20l2 | 0.025 | Old |
| 0.571596 | 0.102883 | 0.061 | 0.036 | 1 | 15 | Coq8b | 0.025 | Old |
| 0.571596 | 0.061662 | 0.061 | 0.036 | 1 | 15 | Alkbh3 | 0.025 | Old |
| 0.571763 | 0.400273 | 0.224 | 0.179 | 1 | 15 | Dync1li1 | 0.045 | Old |
| 0.571791 | 0.33543 | 0.367 | 0.321 | 1 | 15 | Arid1a | 0.046 | Old |

| | | | | | | | | |
|---|---|---|---|---|---|---|---|---|
| 0.57237 | 0.892977 | 0.204 | 0.161 | 1 | 15 | Tmem173 | 0.043 | Old |
| 0.57237 | 0.84628 | 0.204 | 0.161 | 1 | 15 | Vcl | 0.043 | Old |
| 0.57237 | 0.54675 | 0.204 | 0.161 | 1 | 15 | Fam53b | 0.043 | Old |
| 0.572436 | 1.67265 | 0.082 | 0.054 | 1 | 15 | Zbtb37 | 0.028 | Old |
| 0.572436 | 1.616597 | 0.082 | 0.054 | 1 | 15 | Pde1b | 0.028 | Old |
| 0.572436 | 1.040689 | 0.082 | 0.054 | 1 | 15 | Fyco1 | 0.028 | Old |
| 0.572436 | 0.93969 | 0.082 | 0.054 | 1 | 15 | Slc39a3 | 0.028 | Old |
| 0.572436 | 0.775988 | 0.082 | 0.054 | 1 | 15 | Zfp46 | 0.028 | Old |
| 0.572436 | 0.757281 | 0.082 | 0.054 | 1 | 15 | Kynu | 0.028 | Old |
| 0.572436 | 0.704929 | 0.082 | 0.054 | 1 | 15 | Dera | 0.028 | Old |
| 0.572436 | 0.664209 | 0.082 | 0.054 | 1 | 15 | Mblac2 | 0.028 | Old |
| 0.572436 | 0.59373 | 0.082 | 0.054 | 1 | 15 | Txnrd2 | 0.028 | Old |
| 0.572436 | 0.588616 | 0.082 | 0.054 | 1 | 15 | Mtr | 0.028 | Old |
| 0.572436 | 0.570908 | 0.082 | 0.054 | 1 | 15 | Rnaset2a | 0.028 | Old |
| 0.572436 | 0.565362 | 0.082 | 0.054 | 1 | 15 | Rinl | 0.028 | Old |
| 0.572436 | 0.518876 | 0.082 | 0.054 | 1 | 15 | Ciao1 | 0.028 | Old |
| 0.572436 | 0.495335 | 0.082 | 0.054 | 1 | 15 | Slc35a5 | 0.028 | Old |
| 0.572436 | 0.449629 | 0.082 | 0.054 | 1 | 15 | Mycbp | 0.028 | Old |
| 0.572436 | 0.437605 | 0.082 | 0.054 | 1 | 15 | Pyurf | 0.028 | Old |
| 0.572436 | 0.348244 | 0.082 | 0.054 | 1 | 15 | Ercc5 | 0.028 | Old |
| 0.573204 | 0.683794 | 0.265 | 0.214 | 1 | 15 | Rnpepl1 | 0.051 | Old |
| 0.573204 | 0.143609 | 0.265 | 0.214 | 1 | 15 | Dnajc1 | 0.051 | Old |
| 0.573431 | 0.612678 | 0.449 | 0.429 | 1 | 15 | 9530068E0 | 0.02 | Old |
| 0.573431 | 0.025464 | 0.469 | 0.411 | 1 | 15 | Srrm1 | 0.058 | Old |
| 0.573515 | 0.384462 | 0.694 | 0.714 | 1 | 15 | H2-Q7 | -0.02 | Old |
| 0.573843 | 0.471324 | 0.612 | 0.571 | 1 | 15 | Ywhab | 0.041 | Old |
| 0.57396 | 0.980748 | 0.102 | 0.071 | 1 | 15 | Dcp1a | 0.031 | Old |
| 0.57396 | 0.852128 | 0.102 | 0.071 | 1 | 15 | Tprkb | 0.031 | Old |
| 0.57396 | 0.678601 | 0.102 | 0.071 | 1 | 15 | Mdn1 | 0.031 | Old |
| 0.57399 | 0.519182 | 0.408 | 0.393 | 1 | 15 | Ankrd11 | 0.015 | Old |
| 0.57399 | 0.509325 | 0.429 | 0.375 | 1 | 15 | Tnfsf9 | 0.054 | Old |
| 0.574339 | 0.049116 | 0.265 | 0.321 | 1 | 15 | Tmem59 | -0.056 | Old |
| 0.575861 | 0.726411 | 0.163 | 0.125 | 1 | 15 | Snrpa | 0.038 | Old |
| 0.575861 | 0.462421 | 0.163 | 0.125 | 1 | 15 | Fam193b | 0.038 | Old |
| 0.575861 | 0.458568 | 0.163 | 0.125 | 1 | 15 | Syne1 | 0.038 | Old |
| 0.575861 | 0.39615 | 0.163 | 0.125 | 1 | 15 | Izumo4 | 0.038 | Old |
| 0.577458 | 0.861741 | 0.286 | 0.25 | 1 | 15 | Atp2a2 | 0.036 | Old |
| 0.578299 | 1.095868 | 0.122 | 0.089 | 1 | 15 | Apip | 0.033 | Old |
| 0.578299 | 0.564236 | 0.122 | 0.089 | 1 | 15 | Xaf1 | 0.033 | Old |
| 0.578299 | 0.51154 | 0.122 | 0.089 | 1 | 15 | Atp8b2 | 0.033 | Old |
| 0.578299 | 0.372675 | 0.122 | 0.089 | 1 | 15 | Casp8ap2 | 0.033 | Old |
| 0.578299 | 0.126293 | 0.122 | 0.089 | 1 | 15 | Trim5 | 0.033 | Old |
| 0.578331 | 0.19736 | 0.184 | 0.143 | 1 | 15 | Zfp652 | 0.041 | Old |
| 0.578417 | 0.932141 | 0.143 | 0.107 | 1 | 15 | 2510009E0 | 0.036 | Old |
| 0.578417 | 0.863684 | 0.102 | 0.143 | 1 | 15 | Fibp | -0.041 | Old |

| | | | | | | | | |
|---|---|---|---|---|---|---|---|---|
| 0.578417 | 0.365879 | 0.143 | 0.107 | 1 | 15 | Ino80e | 0.036 | Old |
| 0.578417 | 0.239764 | 0.143 | 0.107 | 1 | 15 | Ubap1 | 0.036 | Old |
| 0.578417 | 0.040233 | 0.102 | 0.143 | 1 | 15 | Dcaf15 | -0.041 | Old |
| 0.57871 | 1.144534 | 0.224 | 0.196 | 1 | 15 | Ugp2 | 0.028 | Old |
| 0.578726 | 0.663001 | 0.408 | 0.357 | 1 | 15 | Anp32a | 0.051 | Old |
| 0.578775 | 0.049285 | 0.388 | 0.446 | 1 | 15 | Dusp2 | -0.058 | Old |
| 0.578862 | 0.304245 | 0.735 | 0.714 | 1 | 15 | Hnrnpa0 | 0.021 | Old |
| 0.578897 | 0.249696 | 0.204 | 0.161 | 1 | 15 | Mff | 0.043 | Old |
| 0.579652 | 0.487269 | 0.49 | 0.5 | 1 | 15 | Ddx3y | -0.01 | Old |
| 0.581073 | 0.151424 | 0.429 | 0.518 | 1 | 15 | Ywhaq | -0.089 | Old |
| 0.581267 | 0.554424 | 0.061 | 0.089 | 1 | 15 | Iars2 | -0.028 | Old |
| 0.582019 | 0.004685 | 0.204 | 0.268 | 1 | 15 | Tpp1 | -0.064 | Old |
| 0.582144 | 0.274675 | 0.306 | 0.25 | 1 | 15 | Sp110 | 0.056 | Old |
| 0.582157 | 0.410503 | 0.367 | 0.321 | 1 | 15 | Fbrsl1 | 0.046 | Old |
| 0.582585 | 1.009188 | 0.082 | 0.054 | 1 | 15 | Nol12 | 0.028 | Old |
| 0.582585 | 0.605354 | 0.082 | 0.054 | 1 | 15 | Fam210b | 0.028 | Old |
| 0.582585 | 0.511107 | 0.082 | 0.054 | 1 | 15 | Acd | 0.028 | Old |
| 0.582585 | 0.454509 | 0.082 | 0.054 | 1 | 15 | Thumpd1 | 0.028 | Old |
| 0.582585 | 0.379045 | 0.082 | 0.054 | 1 | 15 | Dcps | 0.028 | Old |
| 0.582762 | 0.377851 | 0.51 | 0.464 | 1 | 15 | Sri | 0.046 | Old |
| 0.583006 | 0.576473 | 0.102 | 0.071 | 1 | 15 | Proser1 | 0.031 | Old |
| 0.583006 | 0.520448 | 0.102 | 0.071 | 1 | 15 | Spib | 0.031 | Old |
| 0.583006 | 0.475435 | 0.102 | 0.071 | 1 | 15 | Zfp639 | 0.031 | Old |
| 0.583006 | 0.428676 | 0.102 | 0.071 | 1 | 15 | Taf4 | 0.031 | Old |
| 0.583006 | 0.411859 | 0.102 | 0.071 | 1 | 15 | Tmem161a | 0.031 | Old |
| 0.583072 | 0.575824 | 0.551 | 0.518 | 1 | 15 | Aftph | 0.033 | Old |
| 0.583084 | 0.899832 | 0.163 | 0.125 | 1 | 15 | Anapc2 | 0.038 | Old |
| 0.583084 | 0.602228 | 0.163 | 0.125 | 1 | 15 | Egln1 | 0.038 | Old |
| 0.583109 | 0.40263 | 0.286 | 0.25 | 1 | 15 | Adprh | 0.036 | Old |
| 0.583109 | 0.383885 | 0.245 | 0.286 | 1 | 15 | Golph3 | -0.041 | Old |
| 0.583109 | 0.174659 | 0.224 | 0.304 | 1 | 15 | Naa50 | -0.08 | Old |
| 0.583438 | 1.455362 | 0.204 | 0.179 | 1 | 15 | 5031425E2 | 0.025 | Old |
| 0.583438 | 0.07406 | 0.163 | 0.214 | 1 | 15 | Agfg1 | -0.051 | Old |
| 0.583489 | 0.245419 | 0.061 | 0.036 | 1 | 15 | Mrpl37 | 0.025 | Old |
| 0.583489 | 0.059017 | 0.061 | 0.036 | 1 | 15 | A930024E0 | 0.025 | Old |
| 0.583489 | 0.019416 | 0.061 | 0.036 | 1 | 15 | Pigu | 0.025 | Old |
| 0.583843 | 0.344149 | 0.449 | 0.375 | 1 | 15 | Nucks1 | 0.074 | Old |
| 0.584332 | 0.38755 | 0.224 | 0.179 | 1 | 15 | Fam76a | 0.045 | Old |
| 0.584332 | 0.36429 | 0.224 | 0.179 | 1 | 15 | Pkig | 0.045 | Old |
| 0.585192 | 0.477085 | 0.245 | 0.214 | 1 | 15 | Tpp2 | 0.031 | Old |
| 0.585197 | 0.796314 | 0.184 | 0.143 | 1 | 15 | Yipf4 | 0.041 | Old |
| 0.585197 | 0.666761 | 0.184 | 0.143 | 1 | 15 | Cdk2ap1 | 0.041 | Old |
| 0.58546 | 0.292836 | 0.204 | 0.161 | 1 | 15 | Rad50 | 0.043 | Old |
| 0.58546 | 0.270825 | 0.204 | 0.161 | 1 | 15 | Fam160b1 | 0.043 | Old |
| 0.58546 | 0.210257 | 0.204 | 0.161 | 1 | 15 | Trim24 | 0.043 | Old |

| | | | | | | | | |
|---|---|---|---|---|---|---|---|---|
| 0.58546 | 0.180257 | 0.204 | 0.161 | 1 | 15 | Ifnar2 | 0.043 | Old |
| 0.586117 | 0.980179 | 0.143 | 0.107 | 1 | 15 | 2410089E0 | 0.036 | Old |
| 0.586117 | 0.862408 | 0.143 | 0.107 | 1 | 15 | Fbrs | 0.036 | Old |
| 0.586117 | 0.489507 | 0.143 | 0.107 | 1 | 15 | Fnbp4 | 0.036 | Old |
| 0.586117 | 0.43445 | 0.143 | 0.107 | 1 | 15 | Cfl2 | 0.036 | Old |
| 0.586117 | 0.061726 | 0.102 | 0.143 | 1 | 15 | Exosc8 | -0.041 | Old |
| 0.586588 | 1.35324 | 0.122 | 0.089 | 1 | 15 | Rad54l2 | 0.033 | Old |
| 0.586588 | 0.537085 | 0.122 | 0.089 | 1 | 15 | Eif4e3 | 0.033 | Old |
| 0.586588 | 0.393328 | 0.122 | 0.089 | 1 | 15 | Map4k1 | 0.033 | Old |
| 0.586588 | 0.038098 | 0.122 | 0.089 | 1 | 15 | Usp33 | 0.033 | Old |
| 0.588332 | 0.230247 | 0.367 | 0.357 | 1 | 15 | Rab5c | 0.01 | Old |
| 0.588748 | 0.544272 | 0.551 | 0.5 | 1 | 15 | Ubxn1 | 0.051 | Old |
| 0.588786 | 0.343527 | 0.286 | 0.25 | 1 | 15 | Tpd52 | 0.036 | Old |
| 0.589712 | 1.938144 | 0.163 | 0.143 | 1 | 15 | Smg7 | 0.02 | Old |
| 0.590621 | 0.332331 | 0.469 | 0.482 | 1 | 15 | Bnip2 | -0.013 | Old |
| 0.591209 | 0.525603 | 0.245 | 0.214 | 1 | 15 | Eif1b | 0.031 | Old |
| 0.591677 | 0.183145 | 0.918 | 0.946 | 1 | 15 | Anxa2 | -0.028 | Old |
| 0.59173 | 0.035961 | 0.98 | 1 | 1 | 15 | Tpt1 | -0.02 | Old |
| 0.591884 | 0.235294 | 0.408 | 0.5 | 1 | 15 | Tcf25 | -0.092 | Old |
| 0.592057 | 0.178716 | 0.204 | 0.161 | 1 | 15 | Stard3nl | 0.043 | Old |
| 0.592057 | 0.09114 | 0.163 | 0.196 | 1 | 15 | Slbp | -0.033 | Old |
| 0.592101 | 0.685797 | 0.184 | 0.143 | 1 | 15 | Lrrc41 | 0.041 | Old |
| 0.592117 | 1.168915 | 0.102 | 0.071 | 1 | 15 | Arl10 | 0.031 | Old |
| 0.592117 | 0.493629 | 0.102 | 0.071 | 1 | 15 | 5730455P1 | 0.031 | Old |
| 0.592117 | 0.405577 | 0.102 | 0.071 | 1 | 15 | Zfp84 | 0.031 | Old |
| 0.592117 | 0.226291 | 0.102 | 0.071 | 1 | 15 | Camk2g | 0.031 | Old |
| 0.59212 | 0.775183 | 0.306 | 0.268 | 1 | 15 | Ccnd2 | 0.038 | Old |
| 0.592817 | 0.537365 | 0.082 | 0.054 | 1 | 15 | Emc4 | 0.028 | Old |
| 0.592817 | 0.352361 | 0.082 | 0.054 | 1 | 15 | Dnah2 | 0.028 | Old |
| 0.592817 | 0.304803 | 0.082 | 0.054 | 1 | 15 | Phtf1 | 0.028 | Old |
| 0.592817 | 0.266462 | 0.082 | 0.054 | 1 | 15 | Hps5 | 0.028 | Old |
| 0.592817 | 0.197628 | 0.082 | 0.054 | 1 | 15 | F630028O1 | 0.028 | Old |
| 0.592817 | 0.165271 | 0.082 | 0.054 | 1 | 15 | Gid8 | 0.028 | Old |
| 0.592817 | 0.082823 | 0.082 | 0.054 | 1 | 15 | Mknk1 | 0.028 | Old |
| 0.593841 | 0.277246 | 0.224 | 0.179 | 1 | 15 | Strbp | 0.045 | Old |
| 0.593863 | 0.810355 | 0.143 | 0.107 | 1 | 15 | Erlec1 | 0.036 | Old |
| 0.593863 | 0.162382 | 0.143 | 0.107 | 1 | 15 | Nrbf2 | 0.036 | Old |
| 0.594932 | 0.773469 | 0.122 | 0.089 | 1 | 15 | Ggps1 | 0.033 | Old |
| 0.594932 | 0.454881 | 0.122 | 0.089 | 1 | 15 | Nubp1 | 0.033 | Old |
| 0.594932 | 0.383202 | 0.122 | 0.089 | 1 | 15 | Nup214 | 0.033 | Old |
| 0.594932 | 0.338915 | 0.122 | 0.089 | 1 | 15 | Tlk2 | 0.033 | Old |
| 0.595449 | 0.286128 | 0.224 | 0.286 | 1 | 15 | Arl8b | -0.062 | Old |
| 0.595697 | 1.122905 | 0.327 | 0.304 | 1 | 15 | Mafg | 0.023 | Old |
| 0.595697 | 0.691907 | 0.327 | 0.304 | 1 | 15 | Dlst | 0.023 | Old |
| 0.596241 | 0.434372 | 0.265 | 0.232 | 1 | 15 | Glrx3 | 0.033 | Old |

| | | | | | | | | |
|---|---|---|---|---|---|---|---|---|
| 0.596489 | 0.159616 | 0.429 | 0.357 | 1 | 15 | Atp1b3 | 0.072 | Old |
| 0.596489 | 0.027221 | 0.429 | 0.357 | 1 | 15 | Snu13 | 0.072 | Old |
| 0.597028 | 0.562796 | 0.224 | 0.179 | 1 | 15 | Tceal9 | 0.045 | Old |
| 0.597066 | 0.074135 | 0.51 | 0.464 | 1 | 15 | Serp1 | 0.046 | Old |
| 0.59718 | 0.081095 | 0.51 | 0.589 | 1 | 15 | Rab11fip1 | -0.079 | Old |
| 0.597639 | 0.051804 | 0.633 | 0.679 | 1 | 15 | Ddx3x | -0.046 | Old |
| 0.597656 | 0.218131 | 0.122 | 0.161 | 1 | 15 | Eif4a3 | -0.039 | Old |
| 0.599041 | 0.263146 | 0.143 | 0.179 | 1 | 15 | Ahr | -0.036 | Old |
| 0.600217 | 0.509196 | 0.286 | 0.25 | 1 | 15 | Ap1b1 | 0.036 | Old |
| 0.600391 | 0.048163 | 0.694 | 0.714 | 1 | 15 | Psmb8 | -0.02 | Old |
| 0.600521 | 0.232783 | 0.204 | 0.232 | 1 | 15 | B4galt1 | -0.028 | Old |
| 0.60055 | 0.429685 | 0.061 | 0.089 | 1 | 15 | Slc9a1 | -0.028 | Old |
| 0.601293 | 1.25572 | 0.102 | 0.071 | 1 | 15 | Scap | 0.031 | Old |
| 0.601293 | 0.513439 | 0.102 | 0.071 | 1 | 15 | Fbxo42 | 0.031 | Old |
| 0.601293 | 0.500059 | 0.102 | 0.071 | 1 | 15 | Atg12 | 0.031 | Old |
| 0.601293 | 0.396515 | 0.102 | 0.071 | 1 | 15 | Hspa4l | 0.031 | Old |
| 0.601293 | 0.358999 | 0.102 | 0.071 | 1 | 15 | Rab11fip2 | 0.031 | Old |
| 0.601293 | 0.236019 | 0.102 | 0.071 | 1 | 15 | Traf5 | 0.031 | Old |
| 0.601293 | 0.227827 | 0.102 | 0.071 | 1 | 15 | Sra1 | 0.031 | Old |
| 0.601293 | 0.175791 | 0.102 | 0.071 | 1 | 15 | Dxo | 0.031 | Old |
| 0.601293 | 0.105074 | 0.102 | 0.071 | 1 | 15 | Scarb1 | 0.031 | Old |
| 0.601293 | 0.104381 | 0.102 | 0.071 | 1 | 15 | Adssl1 | 0.031 | Old |
| 0.601293 | 0.091101 | 0.102 | 0.071 | 1 | 15 | Ass1 | 0.031 | Old |
| 0.60154 | 0.186615 | 0.531 | 0.5 | 1 | 15 | Top1 | 0.031 | Old |
| 0.601657 | 0.178156 | 0.102 | 0.143 | 1 | 15 | Tuba4a | -0.041 | Old |
| 0.602705 | 0.190796 | 0.469 | 0.411 | 1 | 15 | Prdx2 | 0.058 | Old |
| 0.602864 | 0.056992 | 0.939 | 0.982 | 1 | 15 | mt-Cytb | -0.043 | Old |
| 0.60313 | 0.872401 | 0.082 | 0.054 | 1 | 15 | Tarbp2 | 0.028 | Old |
| 0.60313 | 0.821816 | 0.082 | 0.054 | 1 | 15 | Rnf217 | 0.028 | Old |
| 0.60313 | 0.24743 | 0.082 | 0.054 | 1 | 15 | A430005L1 | 0.028 | Old |
| 0.60313 | 0.239055 | 0.082 | 0.054 | 1 | 15 | Gar1 | 0.028 | Old |
| 0.60313 | 0.067677 | 0.082 | 0.054 | 1 | 15 | Zeb2os | 0.028 | Old |
| 0.60333 | 0.199362 | 0.122 | 0.089 | 1 | 15 | Slc4a1ap | 0.033 | Old |
| 0.60333 | 0.111464 | 0.122 | 0.089 | 1 | 15 | Asxl1 | 0.033 | Old |
| 0.60333 | 0.770841 | 0.245 | 0.214 | 1 | 15 | Cntrl | 0.031 | Old |
| 0.60333 | 0.670939 | 0.245 | 0.214 | 1 | 15 | Nck2 | 0.031 | Old |
| 0.60342 | 0.91335 | 0.51 | 0.482 | 1 | 15 | Nup98 | 0.028 | Old |
| 0.603423 | 0.303925 | 0.224 | 0.179 | 1 | 15 | Hint2 | 0.045 | Old |
| 0.603563 | 0.217102 | 0.367 | 0.339 | 1 | 15 | Magt1 | 0.028 | Old |
| 0.603653 | 0.226212 | 0.184 | 0.232 | 1 | 15 | 3830406C1 | -0.048 | Old |
| 0.603997 | 0.831148 | 0.408 | 0.393 | 1 | 15 | Gnai3 | 0.015 | Old |
| 0.60456 | 0.004695 | 0.327 | 0.268 | 1 | 15 | Psma1 | 0.059 | Old |
| 0.604704 | 0.0003 | 0.306 | 0.25 | 1 | 15 | Uba52 | 0.056 | Old |
| 0.605003 | 0.620036 | 0.122 | 0.161 | 1 | 15 | Kctd2 | -0.039 | Old |
| 0.605003 | 0.368578 | 0.163 | 0.125 | 1 | 15 | Pbx2 | 0.038 | Old |

| | | | | | | | | |
|---|---|---|---|---|---|---|---|---|
| 0.605003 | 0.215397 | 0.163 | 0.125 | 1 | 15 | Tbk1 | 0.038 | Old |
| 0.605003 | 0.097442 | 0.163 | 0.125 | 1 | 15 | Atrnl1 | 0.038 | Old |
| 0.605314 | 0.78108 | 0.571 | 0.554 | 1 | 15 | Nabp1 | 0.017 | Old |
| 0.605351 | 0.194227 | 0.204 | 0.161 | 1 | 15 | Rabl6 | 0.043 | Old |
| 0.60583 | 0.94355 | 0.245 | 0.232 | 1 | 15 | Slc38a1 | 0.013 | Old |
| 0.60583 | 0.493649 | 0.204 | 0.268 | 1 | 15 | Ubn1 | -0.064 | Old |
| 0.606019 | 0.487267 | 0.184 | 0.143 | 1 | 15 | Kif13b | 0.041 | Old |
| 0.606019 | 0.044528 | 0.143 | 0.179 | 1 | 15 | Phactr2 | -0.036 | Old |
| 0.606494 | 0.27324 | 0.469 | 0.446 | 1 | 15 | Tcp1 | 0.023 | Old |
| 0.606533 | 0.072519 | 0.714 | 0.768 | 1 | 15 | Ddx17 | -0.054 | Old |
| 0.607326 | 0.079928 | 0.347 | 0.304 | 1 | 15 | Aff4 | 0.043 | Old |
| 0.60805 | 0.846835 | 0.265 | 0.232 | 1 | 15 | Mtmr1 | 0.033 | Old |
| 0.608515 | 0.126582 | 0.449 | 0.393 | 1 | 15 | Usmg5 | 0.056 | Old |
| 0.608909 | 0.080257 | 0.49 | 0.571 | 1 | 15 | Ctsc | -0.081 | Old |
| 0.609432 | 0.246208 | 0.204 | 0.25 | 1 | 15 | Ncoa4 | -0.046 | Old |
| 0.609496 | 0.753114 | 0.143 | 0.107 | 1 | 15 | Hibadh | 0.036 | Old |
| 0.609496 | 0.600181 | 0.143 | 0.107 | 1 | 15 | Tmem65 | 0.036 | Old |
| 0.609496 | 0.320227 | 0.143 | 0.107 | 1 | 15 | Cnn3 | 0.036 | Old |
| 0.609496 | 0.253869 | 0.143 | 0.107 | 1 | 15 | Aagab | 0.036 | Old |
| 0.609496 | 0.214667 | 0.102 | 0.143 | 1 | 15 | Heatr5a | -0.041 | Old |
| 0.609848 | 0.117045 | 0.224 | 0.179 | 1 | 15 | Renbp | 0.045 | Old |
| 0.609848 | 0.029783 | 0.224 | 0.179 | 1 | 15 | Skiv2l | 0.045 | Old |
| 0.610015 | 0.200492 | 0.286 | 0.232 | 1 | 15 | Sptbn1 | 0.054 | Old |
| 0.610533 | 0.382233 | 0.102 | 0.071 | 1 | 15 | Hk3 | 0.031 | Old |
| 0.610533 | 0.367152 | 0.102 | 0.071 | 1 | 15 | Taf12 | 0.031 | Old |
| 0.611747 | 0.828226 | 0.286 | 0.25 | 1 | 15 | Irf9 | 0.036 | Old |
| 0.61178 | 0.454463 | 0.122 | 0.089 | 1 | 15 | Ralgds | 0.033 | Old |
| 0.61178 | 0.300447 | 0.122 | 0.089 | 1 | 15 | Golga2 | 0.033 | Old |
| 0.61178 | 0.085265 | 0.122 | 0.089 | 1 | 15 | Vdr | 0.033 | Old |
| 0.612239 | 0.058032 | 0.735 | 0.821 | 1 | 15 | mt-Nd5 | -0.086 | Old |
| 0.612391 | 1.554443 | 0.163 | 0.125 | 1 | 15 | Oxr1 | 0.038 | Old |
| 0.612927 | 0.012423 | 0.245 | 0.196 | 1 | 15 | Mau2 | 0.049 | Old |
| 0.613524 | 0.129127 | 0.082 | 0.054 | 1 | 15 | Anks3 | 0.028 | Old |
| 0.613524 | 0.122375 | 0.082 | 0.054 | 1 | 15 | Nf2 | 0.028 | Old |
| 0.613993 | 0.341916 | 0.265 | 0.232 | 1 | 15 | Smim15 | 0.033 | Old |
| 0.614098 | 0.030976 | 0.327 | 0.375 | 1 | 15 | Abhd12 | -0.048 | Old |
| 0.614154 | 0.191166 | 0.429 | 0.375 | 1 | 15 | Mrps21 | 0.054 | Old |
| 0.614585 | 0.163217 | 0.306 | 0.268 | 1 | 15 | Fam96a | 0.038 | Old |
| 0.614802 | 0.269323 | 0.429 | 0.518 | 1 | 15 | Ier2 | -0.089 | Old |
| 0.615724 | 0.116648 | 0.327 | 0.268 | 1 | 15 | Sf3b3 | 0.059 | Old |
| 0.616567 | 0.042575 | 0.653 | 0.679 | 1 | 15 | Shtn1 | -0.026 | Old |
| 0.616694 | 0.568022 | 0.327 | 0.286 | 1 | 15 | Zfp445 | 0.041 | Old |
| 0.617025 | 0.271708 | 0.388 | 0.357 | 1 | 15 | Mrpl24 | 0.031 | Old |
| 0.61759 | 0.091476 | 0.796 | 0.839 | 1 | 15 | Npm1 | -0.043 | Old |
| 0.617659 | 0.360777 | 0.673 | 0.696 | 1 | 15 | Canx | -0.023 | Old |

| | | | | | | | | |
|---|---|---|---|---|---|---|---|---|
| 0.618105 | 0.3909 | 0.551 | 0.589 | 1 | 15 | Fosl2 | -0.038 | Old |
| 0.618529 | 0.078712 | 0.265 | 0.321 | 1 | 15 | Dnajb9 | -0.056 | Old |
| 0.618778 | 0.955638 | 0.204 | 0.161 | 1 | 15 | Fmr1 | 0.043 | Old |
| 0.619172 | 0.422229 | 0.245 | 0.196 | 1 | 15 | Zfr | 0.049 | Old |
| 0.61926 | 0.644929 | 0.429 | 0.375 | 1 | 15 | Ewsr1 | 0.054 | Old |
| 0.619396 | 0.468419 | 0.367 | 0.339 | 1 | 15 | Snrpd2 | 0.028 | Old |
| 0.619835 | 0.091971 | 0.102 | 0.071 | 1 | 15 | Ctbp2 | 0.031 | Old |
| 0.619835 | 0.064393 | 0.102 | 0.071 | 1 | 15 | Ptk2 | 0.031 | Old |
| 0.620117 | 0.508015 | 0.061 | 0.089 | 1 | 15 | Tmem68 | -0.028 | Old |
| 0.620282 | 1.065272 | 0.122 | 0.089 | 1 | 15 | Tecpr1 | 0.033 | Old |
| 0.620282 | 0.524574 | 0.122 | 0.089 | 1 | 15 | Nsmce1 | 0.033 | Old |
| 0.620282 | 0.293658 | 0.122 | 0.089 | 1 | 15 | Tfrc | 0.033 | Old |
| 0.620507 | 0.443961 | 0.49 | 0.446 | 1 | 15 | Ppp1r18 | 0.044 | Old |
| 0.621263 | 0.064455 | 0.449 | 0.464 | 1 | 15 | Atxn7l1 | -0.015 | Old |
| 0.621345 | 0.08605 | 0.776 | 0.661 | 1 | 15 | Rap1b | 0.115 | Old |
| 0.622587 | 0.194917 | 0.408 | 0.464 | 1 | 15 | Leng8 | -0.056 | Old |
| 0.622629 | 0.805712 | 0.204 | 0.179 | 1 | 15 | Prdm1 | 0.025 | Old |
| 0.622671 | 0.024812 | 0.184 | 0.232 | 1 | 15 | Naaa | -0.048 | Old |
| 0.623147 | 0.246331 | 0.449 | 0.411 | 1 | 15 | Csrp1 | 0.038 | Old |
| 0.624448 | 0.256663 | 0.388 | 0.5 | 1 | 15 | Timm13 | -0.112 | Old |
| 0.624636 | 0.575276 | 0.408 | 0.375 | 1 | 15 | Snrpd3 | 0.033 | Old |
| 0.624714 | 0.253247 | 0.367 | 0.339 | 1 | 15 | Rab7 | 0.028 | Old |
| 0.6248 | 0.046702 | 0.347 | 0.411 | 1 | 15 | Myo5a | -0.064 | Old |
| 0.625308 | 0.510186 | 0.143 | 0.107 | 1 | 15 | Slc50a1 | 0.036 | Old |
| 0.625308 | 0.181735 | 0.143 | 0.107 | 1 | 15 | Tef | 0.036 | Old |
| 0.625437 | 0.133761 | 0.49 | 0.446 | 1 | 15 | Phf21a | 0.044 | Old |
| 0.62554 | 0.523712 | 0.204 | 0.161 | 1 | 15 | Suclg1 | 0.043 | Old |
| 0.625956 | 0.310446 | 0.306 | 0.268 | 1 | 15 | Ago3 | 0.038 | Old |
| 0.626221 | 0.149999 | 0.408 | 0.5 | 1 | 15 | Bmt2 | -0.092 | Old |
| 0.626602 | 0.201445 | 0.388 | 0.429 | 1 | 15 | Lsm6 | -0.041 | Old |
| 0.626826 | 0.452992 | 0.347 | 0.321 | 1 | 15 | Glyr1 | 0.026 | Old |
| 0.627166 | 0.800291 | 0.184 | 0.143 | 1 | 15 | Mrps17 | 0.041 | Old |
| 0.627166 | 0.002379 | 0.184 | 0.143 | 1 | 15 | Rnpep | 0.041 | Old |
| 0.627284 | 0.924514 | 0.163 | 0.125 | 1 | 15 | Ccdc122 | 0.038 | Old |
| 0.627284 | 0.439882 | 0.163 | 0.125 | 1 | 15 | Sema4d | 0.038 | Old |
| 0.627284 | 0.250107 | 0.163 | 0.125 | 1 | 15 | Snapc3 | 0.038 | Old |
| 0.627899 | 0.555693 | 0.245 | 0.214 | 1 | 15 | Tlr2 | 0.031 | Old |
| 0.628248 | 1.2204 | 0.184 | 0.161 | 1 | 15 | Mtch2 | 0.023 | Old |
| 0.628836 | 0.707068 | 0.122 | 0.089 | 1 | 15 | Gbp8 | 0.033 | Old |
| 0.628836 | 0.141687 | 0.122 | 0.089 | 1 | 15 | Antxr2 | 0.033 | Old |
| 0.628836 | 0.088387 | 0.122 | 0.089 | 1 | 15 | Metap1 | 0.033 | Old |
| 0.629199 | 0.36744 | 0.102 | 0.071 | 1 | 15 | Tmem167b | 0.031 | Old |
| 0.629199 | 0.192323 | 0.102 | 0.071 | 1 | 15 | Rubcn | 0.031 | Old |
| 0.629199 | 0.052507 | 0.102 | 0.071 | 1 | 15 | Ercc2 | 0.031 | Old |
| 0.629222 | 0.251919 | 0.286 | 0.25 | 1 | 15 | Ppp4r3b | 0.036 | Old |

| | | | | | | | | |
|---|---|---|---|---|---|---|---|---|
| 0.629271 | 0.726863 | 0.204 | 0.179 | 1 | 15 | Scyl2 | 0.025 | Old |
| 0.629271 | 0.020151 | 0.163 | 0.214 | 1 | 15 | Pigyl | -0.051 | Old |
| 0.629855 | 0.054544 | 0.306 | 0.375 | 1 | 15 | Sltm | -0.069 | Old |
| 0.629921 | 0.173617 | 0.939 | 0.911 | 1 | 15 | H2-Aa | 0.028 | Old |
| 0.630004 | 0.977153 | 0.061 | 0.089 | 1 | 15 | Kat5 | -0.028 | Old |
| 0.630004 | 0.814127 | 0.061 | 0.089 | 1 | 15 | Rhoq | -0.028 | Old |
| 0.630004 | 0.720345 | 0.061 | 0.089 | 1 | 15 | Fam234a | -0.028 | Old |
| 0.630004 | 0.365075 | 0.061 | 0.089 | 1 | 15 | Kif16b | -0.028 | Old |
| 0.630004 | 0.026462 | 0.061 | 0.089 | 1 | 15 | Ophn1 | -0.028 | Old |
| 0.630668 | 0.089994 | 0.224 | 0.286 | 1 | 15 | Cttnbp2nl | -0.062 | Old |
| 0.63197 | 0.459798 | 0.694 | 0.804 | 1 | 15 | Birc3 | -0.11 | Old |
| 0.631973 | 0.925782 | 0.265 | 0.232 | 1 | 15 | Smim7 | 0.033 | Old |
| 0.631973 | 0.518756 | 0.265 | 0.232 | 1 | 15 | Vps13a | 0.033 | Old |
| 0.632446 | 0.25616 | 0.673 | 0.732 | 1 | 15 | Akap13 | -0.059 | Old |
| 0.63328 | 0.818837 | 0.143 | 0.107 | 1 | 15 | Sfi1 | 0.036 | Old |
| 0.63328 | 0.218162 | 0.143 | 0.107 | 1 | 15 | Brpf1 | 0.036 | Old |
| 0.63328 | 0.098244 | 0.143 | 0.107 | 1 | 15 | Rbm18 | 0.036 | Old |
| 0.63328 | 0.082115 | 0.102 | 0.143 | 1 | 15 | Surf1 | -0.041 | Old |
| 0.634108 | 0.283003 | 0.245 | 0.214 | 1 | 15 | 2-Mar | 0.031 | Old |
| 0.634284 | 0.062644 | 0.143 | 0.179 | 1 | 15 | Cndp2 | -0.036 | Old |
| 0.634455 | 0.315 | 0.51 | 0.446 | 1 | 15 | Ifngr2 | 0.064 | Old |
| 0.634605 | 0.020148 | 0.367 | 0.286 | 1 | 15 | Cwc15 | 0.081 | Old |
| 0.634789 | 0.529873 | 0.122 | 0.161 | 1 | 15 | 1700025G( | -0.039 | Old |
| 0.635256 | 0.456135 | 0.347 | 0.339 | 1 | 15 | Prrc2b | 0.008 | Old |
| 0.635492 | 0.474218 | 0.224 | 0.196 | 1 | 15 | Erbin | 0.028 | Old |
| 0.635942 | 0.461035 | 0.163 | 0.214 | 1 | 15 | Ostc | -0.051 | Old |
| 0.637417 | 0.070944 | 0.327 | 0.25 | 1 | 15 | Ehmt1 | 0.077 | Old |
| 0.637439 | 0.034623 | 0.122 | 0.089 | 1 | 15 | Btbd10 | 0.033 | Old |
| 0.637617 | 0.342867 | 0.347 | 0.429 | 1 | 15 | Tprgl | -0.082 | Old |
| 0.638016 | 0.048627 | 0.224 | 0.268 | 1 | 15 | Ubr4 | -0.044 | Old |
| 0.638622 | 0.299127 | 0.102 | 0.071 | 1 | 15 | Rbmx2 | 0.031 | Old |
| 0.638622 | 0.279796 | 0.102 | 0.071 | 1 | 15 | Snx14 | 0.031 | Old |
| 0.638797 | 0.27073 | 0.551 | 0.536 | 1 | 15 | Ncl | 0.015 | Old |
| 0.638911 | 0.297642 | 0.592 | 0.589 | 1 | 15 | Rrbp1 | 0.003 | Old |
| 0.639158 | 0.152619 | 0.204 | 0.161 | 1 | 15 | 2200002D( | 0.043 | Old |
| 0.639158 | 0.084678 | 0.163 | 0.196 | 1 | 15 | Rnf220 | -0.033 | Old |
| 0.639958 | 0.743732 | 0.061 | 0.089 | 1 | 15 | Ppox | -0.028 | Old |
| 0.639958 | 0.737566 | 0.061 | 0.089 | 1 | 15 | Ube2e2 | -0.028 | Old |
| 0.639958 | 0.329164 | 0.061 | 0.089 | 1 | 15 | Isca2 | -0.028 | Old |
| 0.639958 | 0.176327 | 0.061 | 0.089 | 1 | 15 | Ints10 | -0.028 | Old |
| 0.639958 | 0.05302 | 0.061 | 0.089 | 1 | 15 | Tbc1d24 | -0.028 | Old |
| 0.640248 | 0.778094 | 0.163 | 0.143 | 1 | 15 | Fam208b | 0.02 | Old |
| 0.640343 | 0.04735 | 0.204 | 0.25 | 1 | 15 | Mvb12a | -0.046 | Old |
| 0.640546 | 0.108527 | 0.633 | 0.589 | 1 | 15 | Fam49b | 0.044 | Old |
| 0.640899 | 0.070554 | 0.714 | 0.768 | 1 | 15 | Sat1 | -0.054 | Old |

| | | | | | | | | |
|---|---|---|---|---|---|---|---|---|
| 0.64099 | 0.377057 | 0.286 | 0.25 | 1 | 15 | Dlgap4 | 0.036 | Old |
| 0.641212 | 0.204484 | 0.571 | 0.518 | 1 | 15 | Mbtd1 | 0.053 | Old |
| 0.641437 | 0.427723 | 0.143 | 0.179 | 1 | 15 | Pds5b | -0.036 | Old |
| 0.641944 | 0.435091 | 0.224 | 0.196 | 1 | 15 | Med10 | 0.028 | Old |
| 0.64242 | 0.119649 | 0.224 | 0.179 | 1 | 15 | Rab1b | 0.045 | Old |
| 0.642644 | 0.248382 | 0.163 | 0.214 | 1 | 15 | Pla2g7 | -0.051 | Old |
| 0.643344 | 0.440283 | 0.388 | 0.357 | 1 | 15 | Ptpn1 | 0.031 | Old |
| 0.644424 | 0.204776 | 0.204 | 0.232 | 1 | 15 | Pi4k2a | -0.028 | Old |
| 0.64528 | 0.370514 | 0.082 | 0.107 | 1 | 15 | Svbp | -0.025 | Old |
| 0.645981 | 0.120667 | 0.918 | 0.929 | 1 | 15 | Cd47 | -0.011 | Old |
| 0.646603 | 0.655971 | 0.245 | 0.214 | 1 | 15 | Parp14 | 0.031 | Old |
| 0.646603 | 0.355693 | 0.245 | 0.214 | 1 | 15 | Pip5k1c | 0.031 | Old |
| 0.648423 | 0.647841 | 0.224 | 0.196 | 1 | 15 | Golgb1 | 0.028 | Old |
| 0.648423 | 0.370166 | 0.224 | 0.196 | 1 | 15 | Tbrg1 | 0.028 | Old |
| 0.648966 | 0.724819 | 0.306 | 0.268 | 1 | 15 | Jarid2 | 0.038 | Old |
| 0.649178 | 0.041725 | 0.673 | 0.661 | 1 | 15 | Purb | 0.012 | Old |
| 0.649881 | 0.196968 | 0.469 | 0.554 | 1 | 15 | Nrip1 | -0.085 | Old |
| 0.649911 | 0.535025 | 0.163 | 0.125 | 1 | 15 | Pqbp1 | 0.038 | Old |
| 0.649911 | 0.341546 | 0.163 | 0.125 | 1 | 15 | Cnot7 | 0.038 | Old |
| 0.649911 | 0.05249 | 0.163 | 0.125 | 1 | 15 | Mbd1 | 0.038 | Old |
| 0.649976 | 1.386856 | 0.061 | 0.089 | 1 | 15 | Klhl6 | -0.028 | Old |
| 0.649976 | 1.106437 | 0.061 | 0.089 | 1 | 15 | Gcnt1 | -0.028 | Old |
| 0.649976 | 0.67295 | 0.061 | 0.089 | 1 | 15 | Pigc | -0.028 | Old |
| 0.649976 | 0.195432 | 0.061 | 0.089 | 1 | 15 | Ash2l | -0.028 | Old |
| 0.649976 | 0.020298 | 0.061 | 0.089 | 1 | 15 | Rbm15b | -0.028 | Old |
| 0.650371 | 0.443912 | 0.327 | 0.286 | 1 | 15 | Dock11 | 0.041 | Old |
| 0.650371 | 0.11575 | 0.327 | 0.286 | 1 | 15 | Arid4a | 0.041 | Old |
| 0.651083 | 0.102059 | 0.714 | 0.696 | 1 | 15 | Srsf3 | 0.018 | Old |
| 0.651217 | 0.438246 | 0.286 | 0.357 | 1 | 15 | Degs1 | -0.071 | Old |
| 0.651217 | 0.346754 | 0.347 | 0.304 | 1 | 15 | Cyld | 0.043 | Old |
| 0.652888 | 0.373909 | 0.245 | 0.214 | 1 | 15 | Ddx46 | 0.031 | Old |
| 0.653345 | 0.265847 | 0.551 | 0.536 | 1 | 15 | Tra2a | 0.015 | Old |
| 0.654357 | 0.346262 | 0.082 | 0.107 | 1 | 15 | Gm20465 | -0.025 | Old |
| 0.654772 | 0.113403 | 0.306 | 0.268 | 1 | 15 | N4bp2l2 | 0.038 | Old |
| 0.655978 | 0.32416 | 0.408 | 0.375 | 1 | 15 | Tomm22 | 0.033 | Old |
| 0.656133 | 0.56841 | 0.204 | 0.179 | 1 | 15 | Gba | 0.025 | Old |
| 0.656368 | 0.173585 | 0.408 | 0.357 | 1 | 15 | Il7r | 0.051 | Old |
| 0.657526 | 0.699683 | 0.122 | 0.161 | 1 | 15 | Zfp62 | -0.039 | Old |
| 0.657526 | 0.166833 | 0.122 | 0.161 | 1 | 15 | Sp3os | -0.039 | Old |
| 0.658368 | 0.153719 | 0.265 | 0.321 | 1 | 15 | C1qbp | -0.056 | Old |
| 0.658811 | 0.372314 | 0.245 | 0.286 | 1 | 15 | Mcur1 | -0.041 | Old |
| 0.659311 | 0.143916 | 0.347 | 0.286 | 1 | 15 | Kpna4 | 0.061 | Old |
| 0.659616 | 0.085828 | 0.571 | 0.643 | 1 | 15 | Pcgf5 | -0.072 | Old |
| 0.659717 | 1.741556 | 0.143 | 0.125 | 1 | 15 | Rcor1 | 0.018 | Old |
| 0.659814 | 0.26105 | 0.204 | 0.161 | 1 | 15 | Secisbp2l | 0.043 | Old |

| | | | | | | | | |
|---|---|---|---|---|---|---|---|---|
| 0.659814 | 0.067296 | 0.163 | 0.196 | 1 | 15 | Prkd3 | -0.033 | Old |
| 0.659903 | 0.093984 | 0.959 | 0.911 | 1 | 15 | Rps5 | 0.048 | Old |
| 0.660059 | 0.550517 | 0.061 | 0.089 | 1 | 15 | Dnajc12 | -0.028 | Old |
| 0.660599 | 0.225237 | 0.306 | 0.268 | 1 | 15 | Rybp | 0.038 | Old |
| 0.660676 | 0.651406 | 0.265 | 0.25 | 1 | 15 | Hipk2 | 0.015 | Old |
| 0.66093 | 1.231617 | 0.245 | 0.232 | 1 | 15 | Acadm | 0.013 | Old |
| 0.661692 | 0.089719 | 0.347 | 0.411 | 1 | 15 | Psmb10 | -0.064 | Old |
| 0.661903 | 0.143721 | 0.857 | 0.875 | 1 | 15 | Tmsb10 | -0.018 | Old |
| 0.662042 | 0.068441 | 0.857 | 0.911 | 1 | 15 | Ccdc88a | -0.054 | Old |
| 0.66255 | 0.422405 | 0.367 | 0.321 | 1 | 15 | N4bp1 | 0.046 | Old |
| 0.663094 | 0.061518 | 0.143 | 0.179 | 1 | 15 | 9930021J0 | -0.036 | Old |
| 0.663355 | 0.916548 | 0.184 | 0.161 | 1 | 15 | Fes | 0.023 | Old |
| 0.663486 | 0.354658 | 0.082 | 0.107 | 1 | 15 | Zfyve26 | -0.025 | Old |
| 0.66367 | 0.103026 | 0.592 | 0.607 | 1 | 15 | Csnk1a1 | -0.015 | Old |
| 0.663983 | 0.524688 | 0.408 | 0.446 | 1 | 15 | Tacc1 | -0.038 | Old |
| 0.6643 | 3.033905 | 0.02 | 0.036 | 1 | 15 | Top3a | -0.016 | Old |
| 0.6643 | 2.063286 | 0.02 | 0.036 | 1 | 15 | Taf8 | -0.016 | Old |
| 0.6643 | 1.995842 | 0.02 | 0.036 | 1 | 15 | Gatb | -0.016 | Old |
| 0.6643 | 1.95527 | 0.02 | 0.036 | 1 | 15 | AC163354. | -0.016 | Old |
| 0.6643 | 1.677724 | 0.02 | 0.036 | 1 | 15 | Il1rap | -0.016 | Old |
| 0.6643 | 1.41379 | 0.02 | 0.036 | 1 | 15 | Prcc | -0.016 | Old |
| 0.6643 | 1.397464 | 0.02 | 0.036 | 1 | 15 | AC163018. | -0.016 | Old |
| 0.6643 | 1.354545 | 0.02 | 0.036 | 1 | 15 | Zgrf1 | -0.016 | Old |
| 0.6643 | 1.270868 | 0.02 | 0.036 | 1 | 15 | Yars2 | -0.016 | Old |
| 0.6643 | 1.220378 | 0.02 | 0.036 | 1 | 15 | Gtpbp6 | -0.016 | Old |
| 0.6643 | 1.148869 | 0.02 | 0.036 | 1 | 15 | Mlh3 | -0.016 | Old |
| 0.6643 | 1.125465 | 0.02 | 0.036 | 1 | 15 | Rasgrp2 | -0.016 | Old |
| 0.6643 | 1.064725 | 0.02 | 0.036 | 1 | 15 | Srgap3 | -0.016 | Old |
| 0.6643 | 1.007329 | 0.02 | 0.036 | 1 | 15 | Ccnyl1 | -0.016 | Old |
| 0.6643 | 0.820828 | 0.02 | 0.036 | 1 | 15 | Tmem104 | -0.016 | Old |
| 0.6643 | 0.798803 | 0.02 | 0.036 | 1 | 15 | Ccdc126 | -0.016 | Old |
| 0.6643 | 0.70186 | 0.02 | 0.036 | 1 | 15 | 0610040B1 | -0.016 | Old |
| 0.6643 | 0.585175 | 0.02 | 0.036 | 1 | 15 | Cdh2 | -0.016 | Old |
| 0.6643 | 0.516855 | 0.02 | 0.036 | 1 | 15 | Ndufaf7 | -0.016 | Old |
| 0.6643 | 0.510171 | 0.02 | 0.036 | 1 | 15 | Zfp971 | -0.016 | Old |
| 0.6643 | 0.49061 | 0.02 | 0.036 | 1 | 15 | Mtmr9 | -0.016 | Old |
| 0.6643 | 0.480853 | 0.02 | 0.036 | 1 | 15 | Recql5 | -0.016 | Old |
| 0.6643 | 0.466538 | 0.02 | 0.036 | 1 | 15 | Zfp40 | -0.016 | Old |
| 0.6643 | 0.452454 | 0.02 | 0.036 | 1 | 15 | Fam222b | -0.016 | Old |
| 0.6643 | 0.435683 | 0.02 | 0.036 | 1 | 15 | Ercc6l | -0.016 | Old |
| 0.6643 | 0.36244 | 0.02 | 0.036 | 1 | 15 | Ipo4 | -0.016 | Old |
| 0.6643 | 0.341961 | 0.02 | 0.036 | 1 | 15 | C87436 | -0.016 | Old |
| 0.6643 | 0.336998 | 0.02 | 0.036 | 1 | 15 | 2410131K1 | -0.016 | Old |
| 0.6643 | 0.312338 | 0.02 | 0.036 | 1 | 15 | Ddx59 | -0.016 | Old |
| 0.6643 | 0.308375 | 0.02 | 0.036 | 1 | 15 | Tuft1 | -0.016 | Old |

| | | | | | | | | |
|---|---|---|---|---|---|---|---|---|
| 0.6643 | 0.283742 | 0.02 | 0.036 | 1 | 15 | Rpgrip1l | -0.016 | Old |
| 0.6643 | 0.243811 | 0.02 | 0.036 | 1 | 15 | Dock6 | -0.016 | Old |
| 0.6643 | 0.232838 | 0.02 | 0.036 | 1 | 15 | Zdhhc24 | -0.016 | Old |
| 0.6643 | 0.21249 | 0.02 | 0.036 | 1 | 15 | 2010111I0: | -0.016 | Old |
| 0.6643 | 0.194998 | 0.02 | 0.036 | 1 | 15 | Copz2 | -0.016 | Old |
| 0.6643 | 0.194998 | 0.02 | 0.036 | 1 | 15 | 1110004E0 | -0.016 | Old |
| 0.6643 | 0.14167 | 0.02 | 0.036 | 1 | 15 | Elovl6 | -0.016 | Old |
| 0.6643 | 0.122048 | 0.02 | 0.036 | 1 | 15 | Plpp6 | -0.016 | Old |
| 0.6643 | 0.112261 | 0.02 | 0.036 | 1 | 15 | Lonrf1 | -0.016 | Old |
| 0.6643 | 0.10435 | 0.02 | 0.036 | 1 | 15 | Enpep | -0.016 | Old |
| 0.6643 | 0.091905 | 0.02 | 0.036 | 1 | 15 | Gm12840 | -0.016 | Old |
| 0.6643 | 0.085871 | 0.02 | 0.036 | 1 | 15 | Eml3 | -0.016 | Old |
| 0.6643 | 0.067305 | 0.02 | 0.036 | 1 | 15 | D930016D( | -0.016 | Old |
| 0.6643 | 0.066138 | 0.02 | 0.036 | 1 | 15 | C920006O: | -0.016 | Old |
| 0.6643 | 0.017882 | 0.02 | 0.036 | 1 | 15 | Ndufaf2 | -0.016 | Old |
| 0.664728 | 0.417268 | 0.388 | 0.357 | 1 | 15 | Shisa5 | 0.031 | Old |
| 0.664867 | 0.382374 | 0.469 | 0.429 | 1 | 15 | Cnot6l | 0.04 | Old |
| 0.665533 | 0.129742 | 0.204 | 0.25 | 1 | 15 | Chd3 | -0.046 | Old |
| 0.665704 | 0.033262 | 0.245 | 0.304 | 1 | 15 | Klf7 | -0.059 | Old |
| 0.665948 | 0.164669 | 0.531 | 0.482 | 1 | 15 | Ifitm2 | 0.049 | Old |
| 0.666565 | 0.600068 | 0.408 | 0.375 | 1 | 15 | Apbb1ip | 0.033 | Old |
| 0.668004 | 0.217098 | 0.347 | 0.304 | 1 | 15 | Dctn1 | 0.043 | Old |
| 0.66802 | 0.392891 | 0.224 | 0.196 | 1 | 15 | Evl | 0.028 | Old |
| 0.668323 | 0.162113 | 0.449 | 0.429 | 1 | 15 | H2afv | 0.02 | Old |
| 0.668496 | 0.088265 | 0.592 | 0.607 | 1 | 15 | Ubl5 | -0.015 | Old |
| 0.669736 | 0.493538 | 0.204 | 0.179 | 1 | 15 | Acat1 | 0.025 | Old |
| 0.669736 | 0.382614 | 0.204 | 0.179 | 1 | 15 | Phc3 | 0.025 | Old |
| 0.669937 | 0.38777 | 0.306 | 0.286 | 1 | 15 | Pcyt1a | 0.02 | Old |
| 0.669939 | 0.361781 | 0.388 | 0.446 | 1 | 15 | Gpi1 | -0.058 | Old |
| 0.670203 | 0.883668 | 0.061 | 0.089 | 1 | 15 | Erich1 | -0.028 | Old |
| 0.670377 | 0.522179 | 0.143 | 0.179 | 1 | 15 | Zfp451 | -0.036 | Old |
| 0.670683 | 0.391357 | 0.388 | 0.321 | 1 | 15 | Bak1 | 0.067 | Old |
| 0.670876 | 0.471582 | 0.347 | 0.321 | 1 | 15 | Rcc2 | 0.026 | Old |
| 0.672315 | 0.540001 | 0.306 | 0.268 | 1 | 15 | Cycs | 0.038 | Old |
| 0.673486 | 0.095611 | 0.408 | 0.464 | 1 | 15 | Ndufb3 | -0.056 | Old |
| 0.673598 | 0.53663 | 0.367 | 0.321 | 1 | 15 | Ash1l | 0.046 | Old |
| 0.673673 | 0.268292 | 0.857 | 0.893 | 1 | 15 | Cxcl16 | -0.036 | Old |
| 0.674603 | 1.299517 | 0.224 | 0.196 | 1 | 15 | Dock7 | 0.028 | Old |
| 0.674614 | 0.005357 | 0.306 | 0.25 | 1 | 15 | Cul3 | 0.056 | Old |
| 0.674769 | 0.884646 | 0.265 | 0.232 | 1 | 15 | Phc2 | 0.033 | Old |
| 0.676579 | 0.236898 | 0.204 | 0.179 | 1 | 15 | Tbc1d23 | 0.025 | Old |
| 0.676601 | 0.350464 | 0.429 | 0.375 | 1 | 15 | Ap2b1 | 0.054 | Old |
| 0.677175 | 0.227085 | 0.653 | 0.589 | 1 | 15 | Ogfrl1 | 0.064 | Old |
| 0.677218 | 0.109142 | 0.367 | 0.411 | 1 | 15 | Srp14 | -0.044 | Old |
| 0.677594 | 0.124549 | 0.245 | 0.304 | 1 | 15 | Gucd1 | -0.059 | Old |

| | | | | | | | | |
|---|---|---|---|---|---|---|---|---|
| 0.677764 | 0.472189 | 0.388 | 0.375 | 1 | 15 | Cnot6 | 0.013 | Old |
| 0.678595 | 0.052556 | 0.918 | 0.929 | 1 | 15 | Rpl22 | -0.011 | Old |
| 0.679148 | 0.052939 | 0.306 | 0.375 | 1 | 15 | Rnf149 | -0.069 | Old |
| 0.679785 | 0.709447 | 0.102 | 0.125 | 1 | 15 | Gripap1 | -0.023 | Old |
| 0.680239 | 0.016665 | 0.592 | 0.661 | 1 | 15 | Spag9 | -0.069 | Old |
| 0.681586 | 0.414337 | 0.265 | 0.321 | 1 | 15 | Hsd17b11 | -0.056 | Old |
| 0.681892 | 0.900707 | 0.082 | 0.107 | 1 | 15 | Commd10 | -0.025 | Old |
| 0.682076 | 0.201909 | 0.306 | 0.357 | 1 | 15 | Coro1b | -0.051 | Old |
| 0.682476 | 0.065811 | 0.653 | 0.679 | 1 | 15 | Prr13 | -0.026 | Old |
| 0.682886 | 0.021122 | 0.224 | 0.25 | 1 | 15 | Ralb | -0.026 | Old |
| 0.68337 | 0.032307 | 0.959 | 1 | 1 | 15 | Eef1a1 | -0.041 | Old |
| 0.683502 | 0.50232 | 0.122 | 0.143 | 1 | 15 | Arhgef10l | -0.021 | Old |
| 0.684678 | 0.387451 | 0.245 | 0.214 | 1 | 15 | Kpna3 | 0.031 | Old |
| 0.684786 | 1.114403 | 0.184 | 0.161 | 1 | 15 | Pex6 | 0.023 | Old |
| 0.684935 | 0.05947 | 0.673 | 0.768 | 1 | 15 | Gfpt1 | -0.095 | Old |
| 0.684946 | 0.928232 | 0.163 | 0.143 | 1 | 15 | Tmem141 | 0.02 | Old |
| 0.684946 | 0.573085 | 0.163 | 0.143 | 1 | 15 | Nudcd2 | 0.02 | Old |
| 0.685036 | 0.211945 | 0.184 | 0.143 | 1 | 15 | Tbc1d10b | 0.041 | Old |
| 0.685036 | 0.007696 | 0.143 | 0.179 | 1 | 15 | Pold4 | -0.036 | Old |
| 0.686248 | 0.255604 | 0.653 | 0.643 | 1 | 15 | Skil | 0.01 | Old |
| 0.6872 | 0.302913 | 0.265 | 0.232 | 1 | 15 | Bud31 | 0.033 | Old |
| 0.687702 | 0.23398 | 0.347 | 0.321 | 1 | 15 | Nostrin | 0.026 | Old |
| 0.688428 | 0.026865 | 0.408 | 0.518 | 1 | 15 | Pafah1b1 | -0.11 | Old |
| 0.689602 | 0.020963 | 0.245 | 0.196 | 1 | 15 | Vamp3 | 0.049 | Old |
| 0.690285 | 0.182126 | 0.531 | 0.554 | 1 | 15 | Tmem50a | -0.023 | Old |
| 0.691074 | 0.159628 | 0.755 | 0.786 | 1 | 15 | Tpm3 | -0.031 | Old |
| 0.691106 | 0.196844 | 0.204 | 0.25 | 1 | 15 | Fmnl2 | -0.046 | Old |
| 0.691168 | 1.307051 | 0.082 | 0.107 | 1 | 15 | Sipa1l1 | -0.025 | Old |
| 0.691168 | 0.585626 | 0.082 | 0.107 | 1 | 15 | Papss1 | -0.025 | Old |
| 0.691168 | 0.323219 | 0.082 | 0.107 | 1 | 15 | Gm5617 | -0.025 | Old |
| 0.691502 | 0.819606 | 0.143 | 0.125 | 1 | 15 | Mbd6 | 0.018 | Old |
| 0.691502 | 0.683889 | 0.143 | 0.125 | 1 | 15 | Atg2a | 0.018 | Old |
| 0.691988 | 0.437718 | 0.184 | 0.161 | 1 | 15 | Chst11 | 0.023 | Old |
| 0.692108 | 0.049523 | 0.653 | 0.679 | 1 | 15 | Rpl27 | -0.026 | Old |
| 0.69241 | 0.274629 | 0.184 | 0.143 | 1 | 15 | Cope | 0.041 | Old |
| 0.692509 | 0.944735 | 0.163 | 0.143 | 1 | 15 | Phf10 | 0.02 | Old |
| 0.692702 | 0.315028 | 0.878 | 0.929 | 1 | 15 | Id2 | -0.051 | Old |
| 0.693447 | 0.002996 | 0.224 | 0.268 | 1 | 15 | Capg | -0.044 | Old |
| 0.693562 | 0.073049 | 0.469 | 0.536 | 1 | 15 | Ndufb9 | -0.067 | Old |
| 0.694503 | 0.647214 | 0.224 | 0.196 | 1 | 15 | Lsm3 | 0.028 | Old |
| 0.695281 | 0.349094 | 0.612 | 0.589 | 1 | 15 | Rpl29 | 0.023 | Old |
| 0.695973 | 0.019679 | 0.184 | 0.214 | 1 | 15 | Gmip | -0.03 | Old |
| 0.696127 | 0.315271 | 0.163 | 0.125 | 1 | 15 | Polr2j | 0.038 | Old |
| 0.696399 | 0.059614 | 0.327 | 0.286 | 1 | 15 | Hprt | 0.041 | Old |
| 0.696905 | 1.35599 | 0.122 | 0.107 | 1 | 15 | Eml4 | 0.015 | Old |

| | | | | | | | | |
|---|---|---|---|---|---|---|---|---|
| 0.697265 | 0.658067 | 0.204 | 0.179 | 1 | 15 | Prkcb | 0.025 | Old |
| 0.698737 | 0.535167 | 0.245 | 0.232 | 1 | 15 | Frmd4b | 0.013 | Old |
| 0.699201 | 0.839233 | 0.306 | 0.286 | 1 | 15 | Kmt2c | 0.02 | Old |
| 0.699537 | 1.427499 | 0.143 | 0.125 | 1 | 15 | Sertad2 | 0.018 | Old |
| 0.70049 | 0.438344 | 0.082 | 0.107 | 1 | 15 | Grpel1 | -0.025 | Old |
| 0.70049 | 0.192608 | 0.082 | 0.107 | 1 | 15 | Maged1 | -0.025 | Old |
| 0.700556 | 0.065676 | 0.449 | 0.411 | 1 | 15 | Ctnnb1 | 0.038 | Old |
| 0.700624 | 0.254889 | 0.388 | 0.339 | 1 | 15 | Marf1 | 0.049 | Old |
| 0.701184 | 0.066839 | 0.224 | 0.196 | 1 | 15 | Pik3cd | 0.028 | Old |
| 0.701193 | 0.095027 | 0.51 | 0.643 | 1 | 15 | Calr | -0.133 | Old |
| 0.702244 | 0.148864 | 0.673 | 0.679 | 1 | 15 | Eif3a | -0.006 | Old |
| 0.702553 | 0.035639 | 0.469 | 0.518 | 1 | 15 | Nol7 | -0.049 | Old |
| 0.703668 | 0.285851 | 0.531 | 0.625 | 1 | 15 | Gls | -0.094 | Old |
| 0.703896 | 0.063987 | 0.51 | 0.429 | 1 | 15 | Ogt | 0.081 | Old |
| 0.705523 | 1.862556 | 0.122 | 0.107 | 1 | 15 | Ccnc | 0.015 | Old |
| 0.705523 | 0.959195 | 0.102 | 0.125 | 1 | 15 | Cops6 | -0.023 | Old |
| 0.705523 | 0.455523 | 0.102 | 0.125 | 1 | 15 | Ppp2r3d | -0.023 | Old |
| 0.705523 | 0.246533 | 0.102 | 0.125 | 1 | 15 | Smim8 | -0.023 | Old |
| 0.705523 | 0.038378 | 0.102 | 0.125 | 1 | 15 | Smg5 | -0.023 | Old |
| 0.70602 | 0.746869 | 0.224 | 0.214 | 1 | 15 | Ms4a6c | 0.01 | Old |
| 0.706474 | 0.824715 | 0.184 | 0.161 | 1 | 15 | Stxbp1 | 0.023 | Old |
| 0.707087 | 0.066975 | 0.939 | 0.964 | 1 | 15 | Klf6 | -0.025 | Old |
| 0.707156 | 0.260804 | 0.327 | 0.357 | 1 | 15 | B230219D2 | -0.03 | Old |
| 0.707268 | 0.093663 | 0.286 | 0.25 | 1 | 15 | Lsm14a | 0.036 | Old |
| 0.707606 | 1.493592 | 0.143 | 0.125 | 1 | 15 | Spata6 | 0.018 | Old |
| 0.709009 | 0.010779 | 0.163 | 0.196 | 1 | 15 | AU040320 | -0.033 | Old |
| 0.709858 | 0.769845 | 0.082 | 0.107 | 1 | 15 | Atg4b | -0.025 | Old |
| 0.709858 | 0.380207 | 0.082 | 0.107 | 1 | 15 | Larp7 | -0.025 | Old |
| 0.709858 | 0.322163 | 0.082 | 0.107 | 1 | 15 | Dhx30 | -0.025 | Old |
| 0.709858 | 0.089143 | 0.082 | 0.107 | 1 | 15 | Psmd9 | -0.025 | Old |
| 0.709858 | 0.008322 | 0.082 | 0.107 | 1 | 15 | Nipal1 | -0.025 | Old |
| 0.710347 | 0.248483 | 0.347 | 0.411 | 1 | 15 | Bag1 | -0.064 | Old |
| 0.711183 | 0.888121 | 0.204 | 0.179 | 1 | 15 | Prpf8 | 0.025 | Old |
| 0.711183 | 0.511101 | 0.204 | 0.179 | 1 | 15 | Rnmt | 0.025 | Old |
| 0.711887 | 0.225044 | 0.531 | 0.536 | 1 | 15 | Uqcrq | -0.005 | Old |
| 0.713758 | 0.438463 | 0.184 | 0.161 | 1 | 15 | Tor1b | 0.023 | Old |
| 0.714933 | 0.339338 | 0.633 | 0.643 | 1 | 15 | Celf2 | -0.01 | Old |
| 0.715004 | 0.191057 | 0.571 | 0.625 | 1 | 15 | Taok3 | -0.054 | Old |
| 0.715707 | 0.111707 | 0.122 | 0.143 | 1 | 15 | Zscan26 | -0.021 | Old |
| 0.715943 | 0.158238 | 0.633 | 0.679 | 1 | 15 | Gdi2 | -0.046 | Old |
| 0.716675 | 0.110519 | 1 | 0.982 | 1 | 15 | Rps28 | 0.018 | Old |
| 0.71698 | 0.093234 | 0.306 | 0.286 | 1 | 15 | Thrap3 | 0.02 | Old |
| 0.717036 | 0.415855 | 0.245 | 0.214 | 1 | 15 | Cdc40 | 0.031 | Old |
| 0.717919 | 0.17142 | 0.327 | 0.375 | 1 | 15 | Atp6v0d1 | -0.048 | Old |
| 0.718167 | 0.186808 | 0.429 | 0.482 | 1 | 15 | Tes | -0.053 | Old |

| | | | | | | | | |
|---|---|---|---|---|---|---|---|---|
| 0.718179 | 0.330563 | 0.204 | 0.179 | 1 | 15 | Ei24 | 0.025 | Old |
| 0.718643 | 0.532428 | 0.265 | 0.232 | 1 | 15 | Zzef1 | 0.033 | Old |
| 0.71927 | 0.408812 | 0.082 | 0.107 | 1 | 15 | Nop56 | -0.025 | Old |
| 0.721068 | 0.716363 | 0.184 | 0.161 | 1 | 15 | Fzr1 | 0.023 | Old |
| 0.721068 | 0.25403 | 0.184 | 0.161 | 1 | 15 | Bckdha | 0.023 | Old |
| 0.721814 | 0.070786 | 0.306 | 0.357 | 1 | 15 | Psma5 | -0.051 | Old |
| 0.722873 | 1.220528 | 0.122 | 0.107 | 1 | 15 | Snrnp40 | 0.015 | Old |
| 0.722873 | 0.46637 | 0.102 | 0.125 | 1 | 15 | Klf11 | -0.023 | Old |
| 0.722873 | 0.278306 | 0.102 | 0.125 | 1 | 15 | Slc29a1 | -0.023 | Old |
| 0.722873 | 0.116889 | 0.102 | 0.125 | 1 | 15 | Acot9 | -0.023 | Old |
| 0.722878 | 0.309831 | 0.347 | 0.375 | 1 | 15 | Map1lc3b | -0.028 | Old |
| 0.722941 | 0.061334 | 0.306 | 0.286 | 1 | 15 | Lrp6 | 0.02 | Old |
| 0.723064 | 1.392736 | 0.163 | 0.143 | 1 | 15 | Hdac1 | 0.02 | Old |
| 0.723064 | 0.370092 | 0.163 | 0.143 | 1 | 15 | Pa2g4 | 0.02 | Old |
| 0.723413 | 0.033424 | 0.449 | 0.464 | 1 | 15 | Mat2a | -0.015 | Old |
| 0.723681 | 0.530915 | 0.551 | 0.679 | 1 | 15 | Fnbp1l | -0.128 | Old |
| 0.72469 | 0.297685 | 0.449 | 0.446 | 1 | 15 | Cct2 | 0.003 | Old |
| 0.725185 | 0.480881 | 0.531 | 0.5 | 1 | 15 | Cct4 | 0.031 | Old |
| 0.725199 | 0.526533 | 0.204 | 0.179 | 1 | 15 | Hnrnpll | 0.025 | Old |
| 0.725199 | 0.308076 | 0.204 | 0.179 | 1 | 15 | Setx | 0.025 | Old |
| 0.725199 | 0.303443 | 0.204 | 0.179 | 1 | 15 | Rnps1 | 0.025 | Old |
| 0.725904 | 0.965654 | 0.224 | 0.214 | 1 | 15 | Dstyk | 0.01 | Old |
| 0.725991 | 0.095541 | 0.306 | 0.268 | 1 | 15 | Arfgef1 | 0.038 | Old |
| 0.72627 | 0.066099 | 0.959 | 0.929 | 1 | 15 | Pim1 | 0.03 | Old |
| 0.728134 | 0.150116 | 0.224 | 0.196 | 1 | 15 | Ankhd1 | 0.028 | Old |
| 0.728589 | 0.57808 | 0.265 | 0.25 | 1 | 15 | Flii | 0.015 | Old |
| 0.728724 | 0.619107 | 0.082 | 0.107 | 1 | 15 | Nupr1 | -0.025 | Old |
| 0.728724 | 0.18794 | 0.082 | 0.107 | 1 | 15 | Brap | -0.025 | Old |
| 0.728919 | 0.196702 | 0.265 | 0.321 | 1 | 15 | Stk24 | -0.056 | Old |
| 0.729793 | 0.296315 | 0.429 | 0.429 | 1 | 15 | Add1 | 0 | Old |
| 0.730485 | 0.00603 | 0.163 | 0.196 | 1 | 15 | Mospd2 | -0.033 | Old |
| 0.730775 | 0.43958 | 0.143 | 0.161 | 1 | 15 | Parp9 | -0.018 | Old |
| 0.730775 | 0.35594 | 0.163 | 0.143 | 1 | 15 | Gm37494 | 0.02 | Old |
| 0.730829 | 0.950706 | 0.245 | 0.232 | 1 | 15 | Rnf213 | 0.013 | Old |
| 0.730934 | 0.082186 | 0.347 | 0.304 | 1 | 15 | Snd1 | 0.043 | Old |
| 0.730934 | 0.004666 | 0.306 | 0.339 | 1 | 15 | Eif4ebp2 | -0.033 | Old |
| 0.731603 | 0.953056 | 0.102 | 0.125 | 1 | 15 | Zcchc2 | -0.023 | Old |
| 0.731603 | 0.226596 | 0.102 | 0.125 | 1 | 15 | Gaa | -0.023 | Old |
| 0.732006 | 1.105511 | 0.143 | 0.125 | 1 | 15 | Rpa3 | 0.018 | Old |
| 0.732006 | 0.739518 | 0.143 | 0.125 | 1 | 15 | Osbp | 0.018 | Old |
| 0.732241 | 0.701013 | 0.204 | 0.179 | 1 | 15 | Sec24b | 0.025 | Old |
| 0.73324 | 0.078282 | 0.714 | 0.732 | 1 | 15 | Hnrnpf | -0.018 | Old |
| 0.733943 | 0.348703 | 0.469 | 0.446 | 1 | 15 | Htra2 | 0.023 | Old |
| 0.734086 | 1.484889 | 0.184 | 0.179 | 1 | 15 | Rbm47 | 0.005 | Old |
| 0.734094 | 0.028188 | 0.388 | 0.339 | 1 | 15 | Ppib | 0.049 | Old |

| | | | | | | | | |
|---|---|---|---|---|---|---|---|---|
| 0.735532 | 0.601683 | 0.122 | 0.161 | 1 | 15 | Nedd4 | -0.039 | Old |
| 0.735764 | 0.338685 | 0.184 | 0.161 | 1 | 15 | Mrpl45 | 0.023 | Old |
| 0.735882 | 0.086072 | 0.429 | 0.375 | 1 | 15 | Srsf2 | 0.054 | Old |
| 0.736053 | 0.145809 | 0.347 | 0.393 | 1 | 15 | Ythdc1 | -0.046 | Old |
| 0.736753 | 0.221477 | 0.306 | 0.339 | 1 | 15 | Leprot | -0.033 | Old |
| 0.737917 | 0.051359 | 0.286 | 0.304 | 1 | 15 | Uqcc2 | -0.018 | Old |
| 0.738115 | 0.278426 | 0.265 | 0.304 | 1 | 15 | Rab32 | -0.039 | Old |
| 0.738115 | 0.250371 | 0.265 | 0.304 | 1 | 15 | Ccrl2 | -0.039 | Old |
| 0.73822 | 1.473387 | 0.102 | 0.089 | 1 | 15 | Phf2 | 0.013 | Old |
| 0.73822 | 1.410211 | 0.102 | 0.089 | 1 | 15 | H2-Oa | 0.013 | Old |
| 0.738513 | 0.494339 | 0.163 | 0.143 | 1 | 15 | Rsl24d1 | 0.02 | Old |
| 0.739085 | 0.754546 | 0.347 | 0.321 | 1 | 15 | Csgalnact2 | 0.026 | Old |
| 0.740201 | 1.229836 | 0.143 | 0.125 | 1 | 15 | Fopnl | 0.018 | Old |
| 0.74054 | 0.412781 | 0.367 | 0.339 | 1 | 15 | Supt16 | 0.028 | Old |
| 0.740645 | 0.967094 | 0.204 | 0.196 | 1 | 15 | Pou2f2 | 0.008 | Old |
| 0.740725 | 0.095915 | 0.918 | 0.911 | 1 | 15 | Rpl36a | 0.007 | Old |
| 0.740767 | 0.011919 | 0.918 | 0.946 | 1 | 15 | Laptm5 | -0.028 | Old |
| 0.743148 | 0.046906 | 0.163 | 0.179 | 1 | 15 | Serpinb8 | -0.016 | Old |
| 0.743297 | 0.092038 | 0.245 | 0.214 | 1 | 15 | Kmt5b | 0.031 | Old |
| 0.743573 | 0.505111 | 0.306 | 0.304 | 1 | 15 | Ciita | 0.002 | Old |
| 0.744202 | 0.285919 | 0.306 | 0.268 | 1 | 15 | Dusp16 | 0.038 | Old |
| 0.744816 | 0.763755 | 0.143 | 0.179 | 1 | 15 | Mmp9 | -0.036 | Old |
| 0.744872 | 0.162294 | 0.306 | 0.357 | 1 | 15 | Glul | -0.051 | Old |
| 0.746227 | 0.047771 | 0.735 | 0.768 | 1 | 15 | Ahnak | -0.033 | Old |
| 0.746277 | 0.450876 | 0.163 | 0.143 | 1 | 15 | Atf6b | 0.02 | Old |
| 0.746394 | 0.201592 | 0.204 | 0.179 | 1 | 15 | Retreg2 | 0.025 | Old |
| 0.746844 | 0.190602 | 0.367 | 0.429 | 1 | 15 | Stard7 | -0.062 | Old |
| 0.74695 | 0.529091 | 0.306 | 0.286 | 1 | 15 | Rbck1 | 0.02 | Old |
| 0.747756 | 1.196717 | 0.102 | 0.089 | 1 | 15 | Gabpa | 0.013 | Old |
| 0.747756 | 0.557337 | 0.082 | 0.107 | 1 | 15 | Ddx47 | -0.025 | Old |
| 0.747964 | 0.225768 | 0.408 | 0.375 | 1 | 15 | Ncor1 | 0.033 | Old |
| 0.747967 | 0.398001 | 0.469 | 0.5 | 1 | 15 | Hdgf | -0.031 | Old |
| 0.74804 | 0.627876 | 0.592 | 0.625 | 1 | 15 | Sowahc | -0.033 | Old |
| 0.748426 | 0.791831 | 0.143 | 0.125 | 1 | 15 | Ankrd49 | 0.018 | Old |
| 0.748426 | 0.403275 | 0.143 | 0.125 | 1 | 15 | Irgm1 | 0.018 | Old |
| 0.748426 | 0.106895 | 0.122 | 0.143 | 1 | 15 | Pou2f1 | -0.021 | Old |
| 0.748573 | 0.566896 | 0.224 | 0.196 | 1 | 15 | Chd1 | 0.028 | Old |
| 0.749165 | 0.335059 | 0.122 | 0.107 | 1 | 15 | Ndufa12 | 0.015 | Old |
| 0.749335 | 0.25562 | 0.204 | 0.232 | 1 | 15 | Cops2 | -0.028 | Old |
| 0.75057 | 0.64641 | 0.224 | 0.268 | 1 | 15 | Ikzf1 | -0.044 | Old |
| 0.750684 | 0.40437 | 0.245 | 0.268 | 1 | 15 | Ubald2 | -0.023 | Old |
| 0.752137 | 0.055404 | 0.816 | 0.804 | 1 | 15 | Fabp5 | 0.012 | Old |
| 0.752174 | 0.231743 | 0.163 | 0.196 | 1 | 15 | Senp1 | -0.033 | Old |
| 0.753803 | 0.322288 | 0.245 | 0.286 | 1 | 15 | Amn1 | -0.041 | Old |
| 0.754702 | 0.09229 | 0.796 | 0.839 | 1 | 15 | Rogdi | -0.043 | Old |

| | | | | | | | | |
|---|---|---|---|---|---|---|---|---|
| 0.754902 | 0.046019 | 0.816 | 0.857 | 1 | 15 | Bmp2k | -0.041 | Old |
| 0.755527 | 0.127687 | 0.286 | 0.321 | 1 | 15 | Idh1 | -0.035 | Old |
| 0.756074 | 0.11274 | 0.204 | 0.232 | 1 | 15 | Bcl6 | -0.028 | Old |
| 0.756424 | 0.527805 | 0.286 | 0.286 | 1 | 15 | Anp32e | 0 | Old |
| 0.756424 | 0.516259 | 0.286 | 0.286 | 1 | 15 | Setd2 | 0 | Old |
| 0.756488 | 0.227468 | 0.347 | 0.321 | 1 | 15 | Mtch1 | 0.026 | Old |
| 0.756488 | 0.176954 | 0.306 | 0.357 | 1 | 15 | Ddb1 | -0.051 | Old |
| 0.75668 | 0.83661 | 0.143 | 0.125 | 1 | 15 | Erg28 | 0.018 | Old |
| 0.75668 | 0.235805 | 0.143 | 0.125 | 1 | 15 | Ufsp2 | 0.018 | Old |
| 0.756849 | 0.663722 | 0.245 | 0.232 | 1 | 15 | Gabarapl2 | 0.013 | Old |
| 0.757996 | 1.049754 | 0.122 | 0.107 | 1 | 15 | Rab3il1 | 0.015 | Old |
| 0.757996 | 0.805394 | 0.102 | 0.125 | 1 | 15 | Elof1 | -0.023 | Old |
| 0.757996 | 0.637157 | 0.122 | 0.107 | 1 | 15 | Bloc1s6 | 0.015 | Old |
| 0.757996 | 0.507074 | 0.122 | 0.107 | 1 | 15 | Rxrb | 0.015 | Old |
| 0.757996 | 0.474929 | 0.102 | 0.125 | 1 | 15 | Efnb2 | -0.023 | Old |
| 0.759428 | 0.268838 | 0.449 | 0.411 | 1 | 15 | Eif5 | 0.038 | Old |
| 0.760212 | 0.697664 | 0.367 | 0.393 | 1 | 15 | Trib1 | -0.026 | Old |
| 0.760212 | 0.425191 | 0.388 | 0.375 | 1 | 15 | Swi5 | 0.013 | Old |
| 0.760217 | 1.189572 | 0.265 | 0.25 | 1 | 15 | Rassf5 | 0.015 | Old |
| 0.760217 | 0.403731 | 0.265 | 0.25 | 1 | 15 | Cox20 | 0.015 | Old |
| 0.760633 | 0.955699 | 0.204 | 0.179 | 1 | 15 | Helz | 0.025 | Old |
| 0.76238 | 0.138202 | 0.388 | 0.339 | 1 | 15 | Eif5b | 0.049 | Old |
| 0.762558 | 0.31982 | 0.265 | 0.304 | 1 | 15 | Smndc1 | -0.039 | Old |
| 0.762941 | 0.414689 | 0.286 | 0.268 | 1 | 15 | Cd86 | 0.018 | Old |
| 0.764022 | 1.380168 | 0.082 | 0.071 | 1 | 15 | Scd2 | 0.011 | Old |
| 0.764342 | 0.50005 | 0.347 | 0.339 | 1 | 15 | Rspry1 | 0.008 | Old |
| 0.764342 | 0.252593 | 0.327 | 0.357 | 1 | 15 | Azin1 | -0.03 | Old |
| 0.764622 | 0.047535 | 0.367 | 0.411 | 1 | 15 | Ndufs6 | -0.044 | Old |
| 0.764961 | 1.378842 | 0.143 | 0.125 | 1 | 15 | St8sia1 | 0.018 | Old |
| 0.764961 | 1.161184 | 0.143 | 0.125 | 1 | 15 | Gpr107 | 0.018 | Old |
| 0.764961 | 0.657516 | 0.122 | 0.143 | 1 | 15 | Dpm1 | -0.021 | Old |
| 0.764961 | 0.541359 | 0.143 | 0.125 | 1 | 15 | Extl1 | 0.018 | Old |
| 0.764961 | 0.36595 | 0.143 | 0.125 | 1 | 15 | AY036118 | 0.018 | Old |
| 0.764961 | 0.261793 | 0.143 | 0.125 | 1 | 15 | Zc3h18 | 0.018 | Old |
| 0.764961 | 0.220212 | 0.143 | 0.125 | 1 | 15 | Cacul1 | 0.018 | Old |
| 0.764961 | 0.200062 | 0.143 | 0.125 | 1 | 15 | Faf2 | 0.018 | Old |
| 0.765442 | 0.666271 | 0.184 | 0.161 | 1 | 15 | Nfu1 | 0.023 | Old |
| 0.766071 | 0.265597 | 0.327 | 0.321 | 1 | 15 | Ankrd17 | 0.006 | Old |
| 0.766593 | 0.784832 | 0.224 | 0.286 | 1 | 15 | Cul1 | -0.062 | Old |
| 0.766593 | 0.104048 | 0.265 | 0.25 | 1 | 15 | Spcs1 | 0.015 | Old |
| 0.766858 | 0.711498 | 0.122 | 0.107 | 1 | 15 | Pam16 | 0.015 | Old |
| 0.766858 | 0.709315 | 0.122 | 0.107 | 1 | 15 | Esco1 | 0.015 | Old |
| 0.766858 | 0.438629 | 0.102 | 0.125 | 1 | 15 | Fam199x | -0.023 | Old |
| 0.766858 | 0.436071 | 0.122 | 0.107 | 1 | 15 | Pias3 | 0.015 | Old |
| 0.766858 | 0.111504 | 0.102 | 0.125 | 1 | 15 | Nacc2 | -0.023 | Old |

| | | | | | | | | |
|---|---|---|---|---|---|---|---|---|
| 0.766943 | 1.284666 | 0.102 | 0.089 | 1 | 15 | Mosmo | 0.013 | Old |
| 0.766943 | 0.712836 | 0.102 | 0.089 | 1 | 15 | Plekhg3 | 0.013 | Old |
| 0.767302 | 1.667875 | 0.041 | 0.054 | 1 | 15 | 9330175E1 | -0.013 | Old |
| 0.767302 | 0.7829 | 0.041 | 0.054 | 1 | 15 | Smim13 | -0.013 | Old |
| 0.767302 | 0.707611 | 0.041 | 0.054 | 1 | 15 | Cep68 | -0.013 | Old |
| 0.767302 | 0.652111 | 0.041 | 0.054 | 1 | 15 | Mib2 | -0.013 | Old |
| 0.767302 | 0.527564 | 0.041 | 0.054 | 1 | 15 | Setd1a | -0.013 | Old |
| 0.767302 | 0.480188 | 0.041 | 0.054 | 1 | 15 | Txndc12 | -0.013 | Old |
| 0.767302 | 0.228866 | 0.041 | 0.054 | 1 | 15 | Gabbr1 | -0.013 | Old |
| 0.767302 | 0.21797 | 0.041 | 0.054 | 1 | 15 | Ccp110 | -0.013 | Old |
| 0.767302 | 0.203692 | 0.041 | 0.054 | 1 | 15 | Pick1 | -0.013 | Old |
| 0.767302 | 0.146766 | 0.041 | 0.054 | 1 | 15 | Timm50 | -0.013 | Old |
| 0.767783 | 0.497916 | 0.204 | 0.179 | 1 | 15 | Fnip1 | 0.025 | Old |
| 0.767783 | 0.359014 | 0.204 | 0.179 | 1 | 15 | Ipmk | 0.025 | Old |
| 0.768077 | 0.172325 | 0.367 | 0.357 | 1 | 15 | Serinc1 | 0.01 | Old |
| 0.769189 | 1.129291 | 0.224 | 0.196 | 1 | 15 | Ncoa7 | 0.028 | Old |
| 0.769526 | 0.920653 | 0.265 | 0.268 | 1 | 15 | Arf2 | -0.003 | Old |
| 0.769526 | 0.816989 | 0.245 | 0.286 | 1 | 15 | Myo1c | -0.041 | Old |
| 0.769723 | 0.492997 | 0.163 | 0.143 | 1 | 15 | Zfp622 | 0.02 | Old |
| 0.770094 | 0.095372 | 1 | 1 | 1 | 15 | Rps26 | 0 | Old |
| 0.771722 | 0.372756 | 0.429 | 0.411 | 1 | 15 | Tomm20 | 0.018 | Old |
| 0.771977 | 0.671312 | 0.327 | 0.321 | 1 | 15 | Taf1d | 0.006 | Old |
| 0.772918 | 0.39622 | 0.163 | 0.179 | 1 | 15 | Wdtc1 | -0.016 | Old |
| 0.772918 | 0.091202 | 0.184 | 0.161 | 1 | 15 | Crat | 0.023 | Old |
| 0.772918 | 0.068989 | 0.184 | 0.161 | 1 | 15 | Rnase6 | 0.023 | Old |
| 0.773001 | 0.440177 | 0.224 | 0.214 | 1 | 15 | Cd164 | 0.01 | Old |
| 0.77327 | 0.248654 | 0.122 | 0.143 | 1 | 15 | Pcnx | -0.021 | Old |
| 0.77327 | 0.22473 | 0.143 | 0.125 | 1 | 15 | Azi2 | 0.018 | Old |
| 0.774257 | 0.003929 | 0.286 | 0.304 | 1 | 15 | Eif3d | -0.018 | Old |
| 0.774694 | 1.182576 | 0.082 | 0.071 | 1 | 15 | 3110062M | 0.011 | Old |
| 0.775401 | 0.220743 | 0.286 | 0.268 | 1 | 15 | Clec10a | 0.018 | Old |
| 0.775752 | 0.884972 | 0.122 | 0.107 | 1 | 15 | Mis18a | 0.015 | Old |
| 0.775752 | 0.701599 | 0.102 | 0.125 | 1 | 15 | Lilrb4a | -0.023 | Old |
| 0.775752 | 0.679894 | 0.122 | 0.107 | 1 | 15 | Zfp180 | 0.015 | Old |
| 0.775752 | 0.381005 | 0.102 | 0.125 | 1 | 15 | Atraid | -0.023 | Old |
| 0.775752 | 0.349367 | 0.122 | 0.107 | 1 | 15 | Zfp592 | 0.015 | Old |
| 0.7764 | 0.462687 | 0.204 | 0.232 | 1 | 15 | Wdr37 | -0.028 | Old |
| 0.7764 | 0.169595 | 0.204 | 0.232 | 1 | 15 | Kdm5a | -0.028 | Old |
| 0.776545 | 0.386754 | 0.245 | 0.214 | 1 | 15 | Sik3 | 0.031 | Old |
| 0.776591 | 1.734065 | 0.102 | 0.089 | 1 | 15 | Dhdh | 0.013 | Old |
| 0.776591 | 1.682039 | 0.102 | 0.089 | 1 | 15 | Vkorc1l1 | 0.013 | Old |
| 0.776591 | 0.821436 | 0.102 | 0.089 | 1 | 15 | Babam1 | 0.013 | Old |
| 0.776591 | 0.645525 | 0.102 | 0.089 | 1 | 15 | Tgfbrap1 | 0.013 | Old |
| 0.777586 | 1.139642 | 0.163 | 0.143 | 1 | 15 | Cnpy2 | 0.02 | Old |
| 0.777586 | 0.252107 | 0.163 | 0.143 | 1 | 15 | Vps37b | 0.02 | Old |

| | | | | | | | | |
|---|---|---|---|---|---|---|---|---|
| 0.777586 | 0.036255 | 0.143 | 0.161 | 1 | 15 | Setdb1 | -0.018 | Old |
| 0.777824 | 0.221322 | 0.592 | 0.607 | 1 | 15 | Cox6a1 | -0.015 | Old |
| 0.779148 | 0.041354 | 0.184 | 0.214 | 1 | 15 | Ift57 | -0.03 | Old |
| 0.779264 | 0.186136 | 0.551 | 0.536 | 1 | 15 | Fosb | 0.015 | Old |
| 0.779608 | 0.090427 | 0.327 | 0.286 | 1 | 15 | Hsbp1 | 0.041 | Old |
| 0.77968 | 0.259956 | 0.837 | 0.893 | 1 | 15 | Chchd2 | -0.056 | Old |
| 0.779683 | 0.065418 | 0.408 | 0.464 | 1 | 15 | Ssr3 | -0.056 | Old |
| 0.779717 | 0.216643 | 0.633 | 0.643 | 1 | 15 | Ubl3 | -0.01 | Old |
| 0.780415 | 0.109863 | 0.184 | 0.161 | 1 | 15 | Slc29a3 | 0.023 | Old |
| 0.780629 | 1.926831 | 0.041 | 0.054 | 1 | 15 | Ngrn | -0.013 | Old |
| 0.780629 | 1.021136 | 0.041 | 0.054 | 1 | 15 | Scai | -0.013 | Old |
| 0.780629 | 0.919035 | 0.041 | 0.054 | 1 | 15 | Eif5a2 | -0.013 | Old |
| 0.780629 | 0.86009 | 0.041 | 0.054 | 1 | 15 | Farsa | -0.013 | Old |
| 0.780629 | 0.824383 | 0.041 | 0.054 | 1 | 15 | Mrps18b | -0.013 | Old |
| 0.780629 | 0.616394 | 0.041 | 0.054 | 1 | 15 | Wdr6 | -0.013 | Old |
| 0.780629 | 0.525628 | 0.041 | 0.054 | 1 | 15 | Mapk8ip3 | -0.013 | Old |
| 0.780629 | 0.459945 | 0.041 | 0.054 | 1 | 15 | Cdc27 | -0.013 | Old |
| 0.780629 | 0.337755 | 0.041 | 0.054 | 1 | 15 | Cgrrf1 | -0.013 | Old |
| 0.780629 | 0.288226 | 0.041 | 0.054 | 1 | 15 | Cutc | -0.013 | Old |
| 0.780629 | 0.253092 | 0.041 | 0.054 | 1 | 15 | Cmtm4 | -0.013 | Old |
| 0.780629 | 0.237822 | 0.041 | 0.054 | 1 | 15 | Hopx | -0.013 | Old |
| 0.780629 | 0.19036 | 0.041 | 0.054 | 1 | 15 | Heatr1 | -0.013 | Old |
| 0.780629 | 0.173905 | 0.041 | 0.054 | 1 | 15 | Rbm34 | -0.013 | Old |
| 0.780629 | 0.092299 | 0.041 | 0.054 | 1 | 15 | Pnpla6 | -0.013 | Old |
| 0.780629 | 0.084434 | 0.041 | 0.054 | 1 | 15 | Fam8a1 | -0.013 | Old |
| 0.780629 | 0.065134 | 0.041 | 0.054 | 1 | 15 | Lamc1 | -0.013 | Old |
| 0.780629 | 0.043643 | 0.041 | 0.054 | 1 | 15 | Per3 | -0.013 | Old |
| 0.780629 | 0.026701 | 0.041 | 0.054 | 1 | 15 | Cul9 | -0.013 | Old |
| 0.780629 | 0.024727 | 0.041 | 0.054 | 1 | 15 | Tagap | -0.013 | Old |
| 0.780629 | 0.024565 | 0.041 | 0.054 | 1 | 15 | Smyd5 | -0.013 | Old |
| 0.780629 | 0.011171 | 0.041 | 0.054 | 1 | 15 | Cdr2 | -0.013 | Old |
| 0.781236 | 0.040855 | 0.571 | 0.571 | 1 | 15 | Emp3 | 0 | Old |
| 0.781605 | 0.243462 | 0.143 | 0.125 | 1 | 15 | Sod3 | 0.018 | Old |
| 0.781605 | 0.222415 | 0.143 | 0.125 | 1 | 15 | Zfp710 | 0.018 | Old |
| 0.782602 | 0.178995 | 0.245 | 0.268 | 1 | 15 | Fbxl5 | -0.023 | Old |
| 0.782713 | 0.232948 | 0.347 | 0.411 | 1 | 15 | H13 | -0.064 | Old |
| 0.783026 | 0.179843 | 0.224 | 0.196 | 1 | 15 | Srf | 0.028 | Old |
| 0.784195 | 0.137541 | 0.388 | 0.393 | 1 | 15 | Rp2 | -0.005 | Old |
| 0.784674 | 1.394887 | 0.122 | 0.107 | 1 | 15 | Cd82 | 0.015 | Old |
| 0.784674 | 1.101537 | 0.122 | 0.107 | 1 | 15 | Edc4 | 0.015 | Old |
| 0.784674 | 0.940206 | 0.122 | 0.107 | 1 | 15 | Hccs | 0.015 | Old |
| 0.784674 | 0.597751 | 0.122 | 0.107 | 1 | 15 | Blcap | 0.015 | Old |
| 0.784674 | 0.438802 | 0.122 | 0.107 | 1 | 15 | Dag1 | 0.015 | Old |
| 0.784674 | 0.102612 | 0.122 | 0.107 | 1 | 15 | Rab20 | 0.015 | Old |
| 0.785409 | 1.294106 | 0.082 | 0.071 | 1 | 15 | Bmi1 | 0.011 | Old |

| | | | | | | | | |
|---|---|---|---|---|---|---|---|---|
| 0.785409 | 1.117984 | 0.082 | 0.071 | 1 | 15 | Plk4 | 0.011 | Old |
| 0.785409 | 1.024372 | 0.082 | 0.071 | 1 | 15 | Sqor | 0.011 | Old |
| 0.785472 | 0.054525 | 0.143 | 0.161 | 1 | 15 | Echs1 | -0.018 | Old |
| 0.785603 | 0.387391 | 0.408 | 0.393 | 1 | 15 | Nisch | 0.015 | Old |
| 0.786274 | 1.865279 | 0.102 | 0.089 | 1 | 15 | Ncdn | 0.013 | Old |
| 0.786274 | 1.737446 | 0.102 | 0.089 | 1 | 15 | Rnf40 | 0.013 | Old |
| 0.786274 | 0.553417 | 0.102 | 0.089 | 1 | 15 | Ppme1 | 0.013 | Old |
| 0.786274 | 0.512274 | 0.102 | 0.089 | 1 | 15 | Tusc1 | 0.013 | Old |
| 0.786274 | 0.472064 | 0.102 | 0.089 | 1 | 15 | Ripor2 | 0.013 | Old |
| 0.786438 | 0.308535 | 0.612 | 0.625 | 1 | 15 | Dock2 | -0.013 | Old |
| 0.787309 | 0.240608 | 0.469 | 0.446 | 1 | 15 | Caprin1 | 0.023 | Old |
| 0.787602 | 0.490021 | 0.367 | 0.321 | 1 | 15 | Eif2s3x | 0.046 | Old |
| 0.787615 | 0.045216 | 0.694 | 0.75 | 1 | 15 | Tmbim6 | -0.056 | Old |
| 0.787919 | 0.34705 | 0.286 | 0.268 | 1 | 15 | Polr2g | 0.018 | Old |
| 0.788519 | 0.475748 | 0.245 | 0.286 | 1 | 15 | Cnot4 | -0.041 | Old |
| 0.788953 | 0.214383 | 0.408 | 0.482 | 1 | 15 | Mapk1 | -0.074 | Old |
| 0.78934 | 0.048854 | 0.347 | 0.357 | 1 | 15 | Pepd | -0.01 | Old |
| 0.78935 | 0.53739 | 0.204 | 0.179 | 1 | 15 | BC031181 | 0.025 | Old |
| 0.789421 | 0.233758 | 0.429 | 0.429 | 1 | 15 | Etnk1 | 0 | Old |
| 0.789963 | 0.890588 | 0.204 | 0.25 | 1 | 15 | Ckb | -0.046 | Old |
| 0.789963 | 0.00248 | 0.245 | 0.214 | 1 | 15 | Otulin | 0.031 | Old |
| 0.789964 | 0.813232 | 0.122 | 0.143 | 1 | 15 | Abhd4 | -0.021 | Old |
| 0.789964 | 0.209542 | 0.122 | 0.143 | 1 | 15 | Sec22b | -0.021 | Old |
| 0.789964 | 0.124071 | 0.143 | 0.125 | 1 | 15 | Slc38a9 | 0.018 | Old |
| 0.790333 | 0.072412 | 0.694 | 0.768 | 1 | 15 | Ptprc | -0.074 | Old |
| 0.790779 | 0.171632 | 0.327 | 0.304 | 1 | 15 | Emc10 | 0.023 | Old |
| 0.792164 | 0.139464 | 0.449 | 0.411 | 1 | 15 | Med13l | 0.038 | Old |
| 0.792238 | 0.318792 | 0.327 | 0.375 | 1 | 15 | Lrrc58 | -0.048 | Old |
| 0.792252 | 0.386355 | 0.265 | 0.25 | 1 | 15 | Ppm1a | 0.015 | Old |
| 0.79329 | 0.008444 | 0.184 | 0.214 | 1 | 15 | Eno1 | -0.03 | Old |
| 0.793381 | 0.410867 | 0.163 | 0.143 | 1 | 15 | Cd300a | 0.02 | Old |
| 0.793381 | 0.284562 | 0.163 | 0.143 | 1 | 15 | Phf8 | 0.02 | Old |
| 0.793444 | 0.110279 | 0.306 | 0.268 | 1 | 15 | Vamp2 | 0.038 | Old |
| 0.793625 | 1.009791 | 0.122 | 0.107 | 1 | 15 | Ltn1 | 0.015 | Old |
| 0.793625 | 0.820632 | 0.102 | 0.125 | 1 | 15 | Jaml | -0.023 | Old |
| 0.793625 | 0.042682 | 0.122 | 0.107 | 1 | 15 | Usp42 | 0.015 | Old |
| 0.79373 | 0.352717 | 0.388 | 0.446 | 1 | 15 | Zc3h7a | -0.058 | Old |
| 0.794021 | 1.715385 | 0.041 | 0.054 | 1 | 15 | Scnm1 | -0.013 | Old |
| 0.794021 | 0.8706 | 0.041 | 0.054 | 1 | 15 | Zfp830 | -0.013 | Old |
| 0.794021 | 0.767797 | 0.041 | 0.054 | 1 | 15 | Nln | -0.013 | Old |
| 0.794021 | 0.74869 | 0.041 | 0.054 | 1 | 15 | Tmem185a | -0.013 | Old |
| 0.794021 | 0.735369 | 0.041 | 0.054 | 1 | 15 | Sh3tc1 | -0.013 | Old |
| 0.794021 | 0.670856 | 0.041 | 0.054 | 1 | 15 | Ercc6l2 | -0.013 | Old |
| 0.794021 | 0.668737 | 0.041 | 0.054 | 1 | 15 | Polr2h | -0.013 | Old |
| 0.794021 | 0.606461 | 0.041 | 0.054 | 1 | 15 | Chid1 | -0.013 | Old |

| | | | | | | | | |
|---|---|---|---|---|---|---|---|---|
| 0.794021 | 0.493478 | 0.041 | 0.054 | 1 | 15 | Ackr1 | -0.013 | Old |
| 0.794021 | 0.409876 | 0.041 | 0.054 | 1 | 15 | Las1l | -0.013 | Old |
| 0.794021 | 0.373637 | 0.041 | 0.054 | 1 | 15 | D3Ertd254 | -0.013 | Old |
| 0.794021 | 0.333103 | 0.041 | 0.054 | 1 | 15 | Fam220a | -0.013 | Old |
| 0.794021 | 0.325037 | 0.041 | 0.054 | 1 | 15 | Paip2b | -0.013 | Old |
| 0.794021 | 0.317984 | 0.041 | 0.054 | 1 | 15 | Samd4 | -0.013 | Old |
| 0.794021 | 0.283137 | 0.041 | 0.054 | 1 | 15 | Kantr | -0.013 | Old |
| 0.794021 | 0.11542 | 0.041 | 0.054 | 1 | 15 | Bcas3 | -0.013 | Old |
| 0.794021 | 0.063147 | 0.041 | 0.054 | 1 | 15 | Klhdc2 | -0.013 | Old |
| 0.794021 | 0.06307 | 0.041 | 0.054 | 1 | 15 | Ppp2r1b | -0.013 | Old |
| 0.794612 | 0.124154 | 0.327 | 0.339 | 1 | 15 | Dnaja2 | -0.012 | Old |
| 0.794748 | 0.082093 | 0.939 | 0.964 | 1 | 15 | Rpl8 | -0.025 | Old |
| 0.794878 | 0.447677 | 0.265 | 0.268 | 1 | 15 | Stat5b | -0.003 | Old |
| 0.795469 | 0.756416 | 0.184 | 0.161 | 1 | 15 | Gpalpp1 | 0.023 | Old |
| 0.795983 | 0.585019 | 0.224 | 0.268 | 1 | 15 | Ramp3 | -0.044 | Old |
| 0.795983 | 0.115247 | 0.224 | 0.268 | 1 | 15 | Ttc3 | -0.044 | Old |
| 0.795989 | 1.229214 | 0.102 | 0.089 | 1 | 15 | Rwdd4a | 0.013 | Old |
| 0.795989 | 0.56186 | 0.102 | 0.089 | 1 | 15 | Hilpda | 0.013 | Old |
| 0.795989 | 0.49865 | 0.102 | 0.089 | 1 | 15 | Ctso | 0.013 | Old |
| 0.795989 | 0.430905 | 0.102 | 0.089 | 1 | 15 | Dhx16 | 0.013 | Old |
| 0.796164 | 1.765918 | 0.082 | 0.071 | 1 | 15 | Fam214a | 0.011 | Old |
| 0.796164 | 1.303406 | 0.082 | 0.071 | 1 | 15 | Mcm9 | 0.011 | Old |
| 0.796164 | 1.218781 | 0.082 | 0.071 | 1 | 15 | Mon1b | 0.011 | Old |
| 0.796164 | 0.877763 | 0.082 | 0.071 | 1 | 15 | Rcbtb2 | 0.011 | Old |
| 0.796164 | 0.789942 | 0.082 | 0.071 | 1 | 15 | Foxj3 | 0.011 | Old |
| 0.796164 | 0.589121 | 0.082 | 0.071 | 1 | 15 | Tex10 | 0.011 | Old |
| 0.796393 | 0.075169 | 0.245 | 0.232 | 1 | 15 | Spcs2 | 0.013 | Old |
| 0.796576 | 0.161708 | 0.204 | 0.179 | 1 | 15 | Daam1 | 0.025 | Old |
| 0.796796 | 0.234861 | 0.327 | 0.304 | 1 | 15 | Csnk2a2 | 0.023 | Old |
| 0.796836 | 0.381929 | 0.204 | 0.196 | 1 | 15 | Cant1 | 0.008 | Old |
| 0.79781 | 0.538601 | 0.265 | 0.339 | 1 | 15 | Lst1 | -0.074 | Old |
| 0.798348 | 0.201574 | 0.122 | 0.143 | 1 | 15 | Npc1 | -0.021 | Old |
| 0.798348 | 0.158917 | 0.122 | 0.143 | 1 | 15 | Plekhf2 | -0.021 | Old |
| 0.798348 | 0.079895 | 0.122 | 0.143 | 1 | 15 | Stx5a | -0.021 | Old |
| 0.798383 | 0.065969 | 0.735 | 0.786 | 1 | 15 | Arhgdia | -0.051 | Old |
| 0.798703 | 0.412287 | 0.265 | 0.25 | 1 | 15 | Ccnd3 | 0.015 | Old |
| 0.799304 | 0.032199 | 0.367 | 0.321 | 1 | 15 | Mdfic | 0.046 | Old |
| 0.799818 | 0.235024 | 0.449 | 0.446 | 1 | 15 | Sel1l | 0.003 | Old |
| 0.799823 | 0.918561 | 0.184 | 0.179 | 1 | 15 | B4galt3 | 0.005 | Old |
| 0.799823 | 0.488287 | 0.184 | 0.179 | 1 | 15 | Ilkap | 0.005 | Old |
| 0.800387 | 0.052705 | 0.184 | 0.214 | 1 | 15 | Itpk1 | -0.03 | Old |
| 0.800493 | 0.356755 | 0.286 | 0.268 | 1 | 15 | Ifi207 | 0.018 | Old |
| 0.800871 | 0.084384 | 0.469 | 0.5 | 1 | 15 | G3bp1 | -0.031 | Old |
| 0.801081 | 0.194245 | 0.388 | 0.393 | 1 | 15 | H2-DMb1 | -0.005 | Old |
| 0.80131 | 0.56012 | 0.143 | 0.161 | 1 | 15 | Tle4 | -0.018 | Old |

| | | | | | | | | |
|---|---|---|---|---|---|---|---|---|
| 0.801934 | 0.023735 | 0.245 | 0.268 | 1 | 15 | U2af1 | -0.023 | Old |
| 0.802604 | 0.286546 | 0.122 | 0.107 | 1 | 15 | Ccdc107 | 0.015 | Old |
| 0.802604 | 0.225839 | 0.122 | 0.107 | 1 | 15 | Zfp992 | 0.015 | Old |
| 0.803026 | 0.119917 | 0.184 | 0.161 | 1 | 15 | Lman2 | 0.023 | Old |
| 0.803444 | 0.098102 | 0.245 | 0.214 | 1 | 15 | Sptssa | 0.031 | Old |
| 0.803522 | 0.113944 | 0.163 | 0.196 | 1 | 15 | Clec4a2 | -0.033 | Old |
| 0.805737 | 0.59937 | 0.102 | 0.089 | 1 | 15 | Dnase1l3 | 0.013 | Old |
| 0.805737 | 0.546837 | 0.102 | 0.089 | 1 | 15 | Acadvl | 0.013 | Old |
| 0.805737 | 0.451926 | 0.102 | 0.089 | 1 | 15 | Tnfrsf4 | 0.013 | Old |
| 0.805737 | 0.320578 | 0.102 | 0.089 | 1 | 15 | Glb1 | 0.013 | Old |
| 0.806756 | 0.209503 | 0.122 | 0.143 | 1 | 15 | Tbcel | -0.021 | Old |
| 0.806756 | 0.057525 | 0.122 | 0.143 | 1 | 15 | Tbc1d2b | -0.021 | Old |
| 0.806756 | 0.023401 | 0.143 | 0.125 | 1 | 15 | Acaa2 | 0.018 | Old |
| 0.806959 | 1.984364 | 0.082 | 0.071 | 1 | 15 | Rab2b | 0.011 | Old |
| 0.806959 | 1.563378 | 0.082 | 0.071 | 1 | 15 | Scaf4 | 0.011 | Old |
| 0.806959 | 1.287228 | 0.082 | 0.071 | 1 | 15 | Tinf2 | 0.011 | Old |
| 0.806959 | 1.277736 | 0.082 | 0.071 | 1 | 15 | Ttc5 | 0.011 | Old |
| 0.806959 | 1.036101 | 0.082 | 0.071 | 1 | 15 | Creb5 | 0.011 | Old |
| 0.806959 | 0.913126 | 0.082 | 0.071 | 1 | 15 | Rcor3 | 0.011 | Old |
| 0.806959 | 0.740815 | 0.082 | 0.071 | 1 | 15 | Prmt3 | 0.011 | Old |
| 0.806959 | 0.708761 | 0.082 | 0.071 | 1 | 15 | Msrb3 | 0.011 | Old |
| 0.807225 | 0.346696 | 0.184 | 0.179 | 1 | 15 | Col4a3bp | 0.005 | Old |
| 0.807473 | 1.912155 | 0.041 | 0.054 | 1 | 15 | Agl | -0.013 | Old |
| 0.807473 | 1.312211 | 0.041 | 0.054 | 1 | 15 | Heatr5b | -0.013 | Old |
| 0.807473 | 1.287407 | 0.041 | 0.054 | 1 | 15 | Fam206a | -0.013 | Old |
| 0.807473 | 1.055049 | 0.041 | 0.054 | 1 | 15 | Usp36 | -0.013 | Old |
| 0.807473 | 0.816514 | 0.041 | 0.054 | 1 | 15 | Lrp5 | -0.013 | Old |
| 0.807473 | 0.805534 | 0.041 | 0.054 | 1 | 15 | Rcl1 | -0.013 | Old |
| 0.807473 | 0.800415 | 0.041 | 0.054 | 1 | 15 | Acox1 | -0.013 | Old |
| 0.807473 | 0.645433 | 0.041 | 0.054 | 1 | 15 | Iba57 | -0.013 | Old |
| 0.807473 | 0.472289 | 0.041 | 0.054 | 1 | 15 | Cd300lb | -0.013 | Old |
| 0.807473 | 0.460046 | 0.041 | 0.054 | 1 | 15 | Nadk2 | -0.013 | Old |
| 0.807473 | 0.436706 | 0.041 | 0.054 | 1 | 15 | Yes1 | -0.013 | Old |
| 0.807473 | 0.416435 | 0.041 | 0.054 | 1 | 15 | Hpgds | -0.013 | Old |
| 0.807473 | 0.124776 | 0.041 | 0.054 | 1 | 15 | Bet1 | -0.013 | Old |
| 0.807998 | 0.15142 | 0.367 | 0.429 | 1 | 15 | Pid1 | -0.062 | Old |
| 0.808674 | 0.332909 | 0.449 | 0.411 | 1 | 15 | Sbno1 | 0.038 | Old |
| 0.808865 | 0.091854 | 0.327 | 0.304 | 1 | 15 | Numa1 | 0.023 | Old |
| 0.808893 | 0.042414 | 0.49 | 0.482 | 1 | 15 | Tmed10 | 0.008 | Old |
| 0.809261 | 0.283488 | 0.163 | 0.143 | 1 | 15 | Plekha3 | 0.02 | Old |
| 0.809261 | 0.180159 | 0.143 | 0.161 | 1 | 15 | Dnajb11 | -0.018 | Old |
| 0.810147 | 0.463215 | 0.571 | 0.589 | 1 | 15 | Nr4a2 | -0.018 | Old |
| 0.810174 | 0.210679 | 0.633 | 0.643 | 1 | 15 | Stat3 | -0.01 | Old |
| 0.810453 | 0.361523 | 0.49 | 0.518 | 1 | 15 | Ehbp1l1 | -0.028 | Old |
| 0.810612 | 0.189155 | 0.204 | 0.232 | 1 | 15 | Setd5 | -0.028 | Old |

| | | | | | | | | |
|---|---|---|---|---|---|---|---|---|
| 0.810902 | 0.189917 | 0.224 | 0.196 | 1 | 15 | Kdm1a | 0.028 | Old |
| 0.810902 | 0.076905 | 0.204 | 0.214 | 1 | 15 | Rgs3 | -0.01 | Old |
| 0.811098 | 0.175229 | 0.286 | 0.304 | 1 | 15 | Bod1l | -0.018 | Old |
| 0.811608 | 0.67116 | 0.122 | 0.107 | 1 | 15 | Ifi204 | 0.015 | Old |
| 0.811608 | 0.347467 | 0.122 | 0.107 | 1 | 15 | Sgk1 | 0.015 | Old |
| 0.811608 | 0.232698 | 0.122 | 0.107 | 1 | 15 | Abcb7 | 0.015 | Old |
| 0.81311 | 0.252959 | 0.51 | 0.5 | 1 | 15 | Ppp1ca | 0.01 | Old |
| 0.813118 | 0.049845 | 0.286 | 0.268 | 1 | 15 | Psmc1 | 0.018 | Old |
| 0.813669 | 0.037893 | 0.306 | 0.339 | 1 | 15 | 2410015M | -0.033 | Old |
| 0.814601 | 0.02818 | 0.878 | 0.964 | 1 | 15 | Gm10076 | -0.086 | Old |
| 0.81463 | 0.389316 | 0.184 | 0.214 | 1 | 15 | Smarcc1 | -0.03 | Old |
| 0.814645 | 1.22529 | 0.184 | 0.179 | 1 | 15 | Cers5 | 0.005 | Old |
| 0.815186 | 0.586085 | 0.122 | 0.143 | 1 | 15 | Slc8b1 | -0.021 | Old |
| 0.815186 | 0.392009 | 0.122 | 0.143 | 1 | 15 | Trim36 | -0.021 | Old |
| 0.815186 | 0.310506 | 0.143 | 0.125 | 1 | 15 | Rae1 | 0.018 | Old |
| 0.815186 | 0.290049 | 0.122 | 0.143 | 1 | 15 | Ndufs8 | -0.021 | Old |
| 0.815186 | 0.279499 | 0.143 | 0.125 | 1 | 15 | E2f4 | 0.018 | Old |
| 0.815186 | 0.042622 | 0.122 | 0.143 | 1 | 15 | Imp4 | -0.021 | Old |
| 0.815515 | 1.056009 | 0.102 | 0.089 | 1 | 15 | Wdfy4 | 0.013 | Old |
| 0.815515 | 1.012203 | 0.102 | 0.089 | 1 | 15 | Gpn3 | 0.013 | Old |
| 0.815515 | 0.5165 | 0.102 | 0.089 | 1 | 15 | Vps54 | 0.013 | Old |
| 0.815757 | 0.112877 | 0.776 | 0.732 | 1 | 15 | Fnbp1 | 0.044 | Old |
| 0.815905 | 0.445372 | 0.408 | 0.429 | 1 | 15 | Slco5a1 | -0.021 | Old |
| 0.816912 | 2.629208 | 0.061 | 0.054 | 1 | 15 | Agk | 0.007 | Old |
| 0.816912 | 2.263129 | 0.061 | 0.054 | 1 | 15 | Tjp2 | 0.007 | Old |
| 0.816912 | 1.952355 | 0.061 | 0.054 | 1 | 15 | Vars | 0.007 | Old |
| 0.816912 | 1.479484 | 0.061 | 0.054 | 1 | 15 | Slc35a2 | 0.007 | Old |
| 0.816912 | 1.273064 | 0.061 | 0.054 | 1 | 15 | B4gat1 | 0.007 | Old |
| 0.816912 | 1.09315 | 0.061 | 0.054 | 1 | 15 | Lrwd1 | 0.007 | Old |
| 0.816912 | 0.835846 | 0.061 | 0.054 | 1 | 15 | Fubp3 | 0.007 | Old |
| 0.817231 | 0.472708 | 0.143 | 0.161 | 1 | 15 | Cmklr1 | -0.018 | Old |
| 0.817231 | 0.26002 | 0.163 | 0.143 | 1 | 15 | Sptan1 | 0.02 | Old |
| 0.817231 | 0.022354 | 0.143 | 0.161 | 1 | 15 | Erlin2 | -0.018 | Old |
| 0.817282 | 1.034525 | 0.286 | 0.304 | 1 | 15 | Pikfyve | -0.018 | Old |
| 0.817702 | 0.137374 | 0.49 | 0.5 | 1 | 15 | Creg1 | -0.01 | Old |
| 0.81779 | 1.696756 | 0.082 | 0.071 | 1 | 15 | Phlpp2 | 0.011 | Old |
| 0.81779 | 1.671435 | 0.082 | 0.071 | 1 | 15 | Nckipsd | 0.011 | Old |
| 0.81779 | 1.134955 | 0.082 | 0.071 | 1 | 15 | Cyfip2 | 0.011 | Old |
| 0.81779 | 1.107588 | 0.082 | 0.071 | 1 | 15 | Atp8b4 | 0.011 | Old |
| 0.81779 | 0.848728 | 0.082 | 0.071 | 1 | 15 | Exosc10 | 0.011 | Old |
| 0.81779 | 0.75692 | 0.082 | 0.071 | 1 | 15 | Zmym3 | 0.011 | Old |
| 0.81779 | 0.726433 | 0.082 | 0.071 | 1 | 15 | Hvcn1 | 0.011 | Old |
| 0.81779 | 0.473901 | 0.082 | 0.071 | 1 | 15 | Zbtb7b | 0.011 | Old |
| 0.817903 | 0.987638 | 0.061 | 0.071 | 1 | 15 | Mars | -0.01 | Old |
| 0.818193 | 0.297765 | 0.163 | 0.179 | 1 | 15 | Ip6k1 | -0.016 | Old |

| | | | | | | | | |
|---|---|---|---|---|---|---|---|---|
| 0.818392 | 0.519542 | 0.265 | 0.304 | 1 | 15 | Ndufs5 | -0.039 | Old |
| 0.81845 | 0.037715 | 0.347 | 0.357 | 1 | 15 | Ndufa11 | -0.01 | Old |
| 0.819449 | 0.121993 | 0.286 | 0.268 | 1 | 15 | BC005624 | 0.018 | Old |
| 0.819449 | 0.110561 | 0.286 | 0.268 | 1 | 15 | Lrrfip1 | 0.018 | Old |
| 0.819712 | 0.324191 | 0.224 | 0.25 | 1 | 15 | Ccl7 | -0.026 | Old |
| 0.820393 | 0.160611 | 0.388 | 0.429 | 1 | 15 | Mob3a | -0.041 | Old |
| 0.8209 | 0.055255 | 0.408 | 0.375 | 1 | 15 | Krcc1 | 0.033 | Old |
| 0.823057 | 0.183715 | 0.245 | 0.232 | 1 | 15 | Mib1 | 0.013 | Old |
| 0.823637 | 0.497812 | 0.143 | 0.125 | 1 | 15 | Mrpl18 | 0.018 | Old |
| 0.823637 | 0.276304 | 0.143 | 0.125 | 1 | 15 | 4833439L1 | 0.018 | Old |
| 0.823637 | 0.140917 | 0.122 | 0.143 | 1 | 15 | Lrrc8d | -0.021 | Old |
| 0.824641 | 0.267512 | 0.265 | 0.25 | 1 | 15 | 4932438A1 | 0.015 | Old |
| 0.825219 | 0.053119 | 0.163 | 0.143 | 1 | 15 | Ppm1k | 0.02 | Old |
| 0.825322 | 0.737761 | 0.102 | 0.089 | 1 | 15 | Smim3 | 0.013 | Old |
| 0.825322 | 0.467819 | 0.102 | 0.089 | 1 | 15 | Mocs2 | 0.013 | Old |
| 0.825322 | 0.319808 | 0.102 | 0.089 | 1 | 15 | Ddx10 | 0.013 | Old |
| 0.825322 | 0.291724 | 0.102 | 0.089 | 1 | 15 | Ifi47 | 0.013 | Old |
| 0.825322 | 0.228264 | 0.102 | 0.089 | 1 | 15 | Fam126b | 0.013 | Old |
| 0.825322 | 0.174324 | 0.102 | 0.089 | 1 | 15 | Ltbr | 0.013 | Old |
| 0.825353 | 0.433022 | 0.204 | 0.196 | 1 | 15 | Oxa1l | 0.008 | Old |
| 0.825802 | 0.494525 | 0.184 | 0.161 | 1 | 15 | Xpo6 | 0.023 | Old |
| 0.825805 | 0.577998 | 0.163 | 0.196 | 1 | 15 | Mgat1 | -0.033 | Old |
| 0.826405 | 0.267111 | 0.224 | 0.25 | 1 | 15 | Pdia4 | -0.026 | Old |
| 0.826405 | 0.059496 | 0.224 | 0.25 | 1 | 15 | Cpsf2 | -0.026 | Old |
| 0.826873 | 0.106111 | 0.245 | 0.286 | 1 | 15 | 4933434E2 | -0.041 | Old |
| 0.826927 | 0.111212 | 0.429 | 0.518 | 1 | 15 | Ak2 | -0.089 | Old |
| 0.826927 | 0.069818 | 0.469 | 0.482 | 1 | 15 | Zmiz1 | -0.013 | Old |
| 0.827049 | 0.022939 | 0.429 | 0.411 | 1 | 15 | Slc7a11 | 0.018 | Old |
| 0.827261 | 0.242639 | 0.347 | 0.321 | 1 | 15 | Taz | 0.026 | Old |
| 0.827832 | 0.112184 | 0.531 | 0.607 | 1 | 15 | Slc25a5 | -0.076 | Old |
| 0.827859 | 0.604824 | 0.224 | 0.214 | 1 | 15 | Xpr1 | 0.01 | Old |
| 0.828656 | 1.179376 | 0.082 | 0.071 | 1 | 15 | Actr5 | 0.011 | Old |
| 0.828656 | 1.097327 | 0.082 | 0.071 | 1 | 15 | Farsb | 0.011 | Old |
| 0.828656 | 0.957279 | 0.082 | 0.071 | 1 | 15 | Ppfia4 | 0.011 | Old |
| 0.828656 | 0.805655 | 0.082 | 0.071 | 1 | 15 | Mccc1 | 0.011 | Old |
| 0.828656 | 0.794266 | 0.082 | 0.071 | 1 | 15 | Carnmt1 | 0.011 | Old |
| 0.828656 | 0.571169 | 0.082 | 0.071 | 1 | 15 | Gatsl3 | 0.011 | Old |
| 0.828656 | 0.560683 | 0.082 | 0.071 | 1 | 15 | Pik3r4 | 0.011 | Old |
| 0.828656 | 0.476351 | 0.082 | 0.071 | 1 | 15 | Tbcd | 0.011 | Old |
| 0.828656 | 0.408336 | 0.082 | 0.071 | 1 | 15 | Nmt2 | 0.011 | Old |
| 0.828656 | 0.278362 | 0.082 | 0.071 | 1 | 15 | Dhx29 | 0.011 | Old |
| 0.828656 | 0.193047 | 0.082 | 0.071 | 1 | 15 | Crot | 0.011 | Old |
| 0.828656 | 0.160295 | 0.082 | 0.071 | 1 | 15 | Thbs1 | 0.011 | Old |
| 0.828934 | 0.089316 | 0.184 | 0.214 | 1 | 15 | Ear2 | -0.03 | Old |
| 0.829337 | 2.057905 | 0.061 | 0.054 | 1 | 15 | Mmp19 | 0.007 | Old |

| | | | | | | | | |
|---|---|---|---|---|---|---|---|---|
| 0.829337 | 1.972751 | 0.061 | 0.054 | 1 | 15 | Slc25a19 | 0.007 | Old |
| 0.829337 | 1.649015 | 0.061 | 0.054 | 1 | 15 | Ssna1 | 0.007 | Old |
| 0.829337 | 1.271936 | 0.061 | 0.054 | 1 | 15 | Rpp14 | 0.007 | Old |
| 0.829337 | 1.184582 | 0.061 | 0.054 | 1 | 15 | Asb7 | 0.007 | Old |
| 0.829337 | 1.170273 | 0.061 | 0.054 | 1 | 15 | Ppid | 0.007 | Old |
| 0.829337 | 0.850801 | 0.061 | 0.054 | 1 | 15 | Zfp608 | 0.007 | Old |
| 0.829337 | 0.81989 | 0.061 | 0.054 | 1 | 15 | Lig1 | 0.007 | Old |
| 0.829337 | 0.751907 | 0.061 | 0.054 | 1 | 15 | Fgd3 | 0.007 | Old |
| 0.829337 | 0.728012 | 0.061 | 0.054 | 1 | 15 | Endog | 0.007 | Old |
| 0.829691 | 0.539206 | 0.122 | 0.107 | 1 | 15 | Gm20342 | 0.015 | Old |
| 0.829691 | 0.344462 | 0.122 | 0.107 | 1 | 15 | Zmpste24 | 0.015 | Old |
| 0.829756 | 0.318145 | 0.245 | 0.232 | 1 | 15 | Cfh | 0.013 | Old |
| 0.829756 | 0.257635 | 0.245 | 0.232 | 1 | 15 | Fuca1 | 0.013 | Old |
| 0.829888 | 0.438522 | 0.367 | 0.375 | 1 | 15 | Atxn1 | -0.008 | Old |
| 0.830552 | 0.312426 | 0.367 | 0.429 | 1 | 15 | Llph | -0.062 | Old |
| 0.830552 | 0.131156 | 0.388 | 0.411 | 1 | 15 | Fkbp1a | -0.023 | Old |
| 0.831707 | 0.134657 | 0.306 | 0.339 | 1 | 15 | Ly6e | -0.033 | Old |
| 0.832108 | 0.253243 | 0.143 | 0.125 | 1 | 15 | Ahctf1 | 0.018 | Old |
| 0.832108 | 0.020211 | 0.143 | 0.125 | 1 | 15 | Zfp950 | 0.018 | Old |
| 0.832338 | 0.401532 | 0.469 | 0.482 | 1 | 15 | H2afj | -0.013 | Old |
| 0.832519 | 0.309544 | 0.204 | 0.196 | 1 | 15 | Pla2g16 | 0.008 | Old |
| 0.832519 | 0.285933 | 0.204 | 0.196 | 1 | 15 | B4galt6 | 0.008 | Old |
| 0.832956 | 0.276426 | 0.184 | 0.196 | 1 | 15 | Hacd4 | -0.012 | Old |
| 0.833226 | 0.543413 | 0.143 | 0.161 | 1 | 15 | Cep83 | -0.018 | Old |
| 0.833226 | 0.107019 | 0.163 | 0.143 | 1 | 15 | Rnf5 | 0.02 | Old |
| 0.833308 | 0.290543 | 0.265 | 0.268 | 1 | 15 | Larp1 | -0.003 | Old |
| 0.833428 | 0.517599 | 0.163 | 0.179 | 1 | 15 | Fem1b | -0.016 | Old |
| 0.833598 | 0.394755 | 0.429 | 0.429 | 1 | 15 | Scp2 | 0 | Old |
| 0.833983 | 0.155706 | 0.388 | 0.375 | 1 | 15 | Psmd4 | 0.013 | Old |
| 0.835156 | 1.046741 | 0.102 | 0.089 | 1 | 15 | Aco1 | 0.013 | Old |
| 0.835156 | 0.391618 | 0.102 | 0.089 | 1 | 15 | Phldb1 | 0.013 | Old |
| 0.835156 | 0.211746 | 0.102 | 0.089 | 1 | 15 | Mfsd12 | 0.013 | Old |
| 0.835156 | 0.152261 | 0.102 | 0.089 | 1 | 15 | Dexi | 0.013 | Old |
| 0.835156 | 0.133216 | 0.102 | 0.089 | 1 | 15 | Ncln | 0.013 | Old |
| 0.835156 | 0.069777 | 0.102 | 0.089 | 1 | 15 | Ndufs1 | 0.013 | Old |
| 0.835327 | 0.784498 | 0.265 | 0.286 | 1 | 15 | Atp6v1h | -0.021 | Old |
| 0.835474 | 0.636316 | 0.245 | 0.25 | 1 | 15 | Zbtb46 | -0.005 | Old |
| 0.836108 | 0.422437 | 0.184 | 0.214 | 1 | 15 | Fkbp2 | -0.03 | Old |
| 0.836221 | 0.102948 | 0.327 | 0.339 | 1 | 15 | Gsk3b | -0.012 | Old |
| 0.836378 | 0.057353 | 0.408 | 0.446 | 1 | 15 | Ppp3r1 | -0.038 | Old |
| 0.837001 | 0.667476 | 0.184 | 0.179 | 1 | 15 | Abhd2 | 0.005 | Old |
| 0.83728 | 0.590631 | 0.408 | 0.464 | 1 | 15 | Psmd8 | -0.056 | Old |
| 0.837785 | 0.140212 | 0.714 | 0.714 | 1 | 15 | Tspo | 0 | Old |
| 0.8383 | 0.240747 | 0.714 | 0.768 | 1 | 15 | Nsa2 | -0.054 | Old |
| 0.838338 | 0.399517 | 0.531 | 0.518 | 1 | 15 | Atf4 | 0.013 | Old |

| | | | | | | | | |
|---|---|---|---|---|---|---|---|---|
| 0.838348 | 0.003976 | 0.612 | 0.661 | 1 | 15 | Ptger4 | -0.049 | Old |
| 0.838511 | 0.385224 | 0.286 | 0.268 | 1 | 15 | Tomm70a | 0.018 | Old |
| 0.838767 | 0.639021 | 0.122 | 0.107 | 1 | 15 | Znrf1 | 0.015 | Old |
| 0.838767 | 0.433944 | 0.102 | 0.125 | 1 | 15 | Adam9 | -0.023 | Old |
| 0.838767 | 0.414782 | 0.122 | 0.107 | 1 | 15 | Cmas | 0.015 | Old |
| 0.838767 | 0.313126 | 0.122 | 0.107 | 1 | 15 | Faim | 0.015 | Old |
| 0.838767 | 0.036533 | 0.122 | 0.107 | 1 | 15 | Prpf6 | 0.015 | Old |
| 0.839555 | 0.936793 | 0.082 | 0.071 | 1 | 15 | 1600012H( | 0.011 | Old |
| 0.839555 | 0.922287 | 0.082 | 0.071 | 1 | 15 | Prrc1 | 0.011 | Old |
| 0.839555 | 0.807817 | 0.082 | 0.071 | 1 | 15 | Eif2ak3 | 0.011 | Old |
| 0.839555 | 0.365611 | 0.082 | 0.071 | 1 | 15 | Cd209b | 0.011 | Old |
| 0.839555 | 0.326469 | 0.082 | 0.071 | 1 | 15 | Olfr1167 | 0.011 | Old |
| 0.839555 | 0.232938 | 0.082 | 0.071 | 1 | 15 | Rhobtb1 | 0.011 | Old |
| 0.839555 | 0.000584 | 0.082 | 0.071 | 1 | 15 | Tbc1d9b | 0.011 | Old |
| 0.83966 | 0.076404 | 0.98 | 1 | 1 | 15 | Rps9 | -0.02 | Old |
| 0.839755 | 0.15355 | 0.286 | 0.25 | 1 | 15 | Efcab14 | 0.036 | Old |
| 0.8406 | 0.862676 | 0.122 | 0.143 | 1 | 15 | Kdm1b | -0.021 | Old |
| 0.8406 | 0.367395 | 0.122 | 0.143 | 1 | 15 | Mrps15 | -0.021 | Old |
| 0.8406 | 0.078603 | 0.143 | 0.125 | 1 | 15 | Kdelr2 | 0.018 | Old |
| 0.8406 | 0.038556 | 0.143 | 0.125 | 1 | 15 | Pkd1 | 0.018 | Old |
| 0.841063 | 0.762323 | 0.061 | 0.071 | 1 | 15 | Snrnp35 | -0.01 | Old |
| 0.841063 | 0.691112 | 0.061 | 0.071 | 1 | 15 | Tpra1 | -0.01 | Old |
| 0.841063 | 0.30537 | 0.061 | 0.071 | 1 | 15 | Ift46 | -0.01 | Old |
| 0.841063 | 0.254483 | 0.061 | 0.071 | 1 | 15 | Dbt | -0.01 | Old |
| 0.841063 | 0.174509 | 0.061 | 0.071 | 1 | 15 | Rufy2 | -0.01 | Old |
| 0.841063 | 0.116253 | 0.061 | 0.071 | 1 | 15 | 1600002K( | -0.01 | Old |
| 0.841173 | 0.001838 | 0.531 | 0.446 | 1 | 15 | H2-DMa | 0.085 | Old |
| 0.841248 | 0.571203 | 0.163 | 0.161 | 1 | 15 | Comt | 0.002 | Old |
| 0.841251 | 0.284436 | 0.143 | 0.161 | 1 | 15 | Ap3d1 | -0.018 | Old |
| 0.841805 | 1.486941 | 0.061 | 0.054 | 1 | 15 | Tmem63a | 0.007 | Old |
| 0.841805 | 1.47814 | 0.061 | 0.054 | 1 | 15 | Lipo3 | 0.007 | Old |
| 0.841805 | 1.257691 | 0.061 | 0.054 | 1 | 15 | Ing3 | 0.007 | Old |
| 0.841805 | 1.180607 | 0.061 | 0.054 | 1 | 15 | Armc1 | 0.007 | Old |
| 0.841805 | 1.143592 | 0.061 | 0.054 | 1 | 15 | Wdr90 | 0.007 | Old |
| 0.841805 | 1.14122 | 0.061 | 0.054 | 1 | 15 | Rnf215 | 0.007 | Old |
| 0.841805 | 0.865534 | 0.061 | 0.054 | 1 | 15 | Ankrd54 | 0.007 | Old |
| 0.841805 | 0.863318 | 0.061 | 0.054 | 1 | 15 | Trim3 | 0.007 | Old |
| 0.841805 | 0.804399 | 0.061 | 0.054 | 1 | 15 | Fam35a | 0.007 | Old |
| 0.841805 | 0.739626 | 0.061 | 0.054 | 1 | 15 | Wdr20 | 0.007 | Old |
| 0.841805 | 0.514911 | 0.061 | 0.054 | 1 | 15 | Fads1 | 0.007 | Old |
| 0.841805 | 0.397488 | 0.061 | 0.054 | 1 | 15 | Mical1 | 0.007 | Old |
| 0.841883 | 0.624022 | 0.367 | 0.429 | 1 | 15 | Tspan13 | -0.062 | Old |
| 0.843783 | 0.22994 | 0.327 | 0.321 | 1 | 15 | Ldha | 0.006 | Old |
| 0.843783 | 0.167439 | 0.347 | 0.304 | 1 | 15 | Cdv3 | 0.043 | Old |
| 0.844662 | 0.070555 | 0.918 | 0.946 | 1 | 15 | Rps7 | -0.028 | Old |

| | | | | | | | | |
|---|---|---|---|---|---|---|---|---|
| 0.845015 | 0.344875 | 0.102 | 0.089 | 1 | 15 | Dtx3 | 0.013 | Old |
| 0.845015 | 0.305608 | 0.102 | 0.089 | 1 | 15 | Ddx23 | 0.013 | Old |
| 0.845015 | 0.283248 | 0.102 | 0.089 | 1 | 15 | Zfyve16 | 0.013 | Old |
| 0.845015 | 0.158834 | 0.102 | 0.089 | 1 | 15 | Tlr11 | 0.013 | Old |
| 0.845015 | 0.127582 | 0.102 | 0.089 | 1 | 15 | BC003331 | 0.013 | Old |
| 0.845588 | 0.329295 | 0.449 | 0.429 | 1 | 15 | Cers6 | 0.02 | Old |
| 0.846543 | 0.040984 | 0.327 | 0.357 | 1 | 15 | Arrdc1 | -0.03 | Old |
| 0.846818 | 0.077855 | 0.816 | 0.839 | 1 | 15 | Capzb | -0.023 | Old |
| 0.846894 | 0.540372 | 0.204 | 0.196 | 1 | 15 | Clec4b1 | 0.008 | Old |
| 0.847179 | 0.204926 | 0.469 | 0.446 | 1 | 15 | Cbl | 0.023 | Old |
| 0.847496 | 0.201941 | 0.245 | 0.268 | 1 | 15 | Zranb1 | -0.023 | Old |
| 0.848363 | 0.244661 | 0.286 | 0.304 | 1 | 15 | Polr2m | -0.018 | Old |
| 0.848524 | 0.166345 | 0.429 | 0.393 | 1 | 15 | Pds5a | 0.036 | Old |
| 0.848727 | 0.028864 | 0.163 | 0.179 | 1 | 15 | Stk11 | -0.016 | Old |
| 0.849017 | 0.085726 | 0.367 | 0.357 | 1 | 15 | Slk | 0.01 | Old |
| 0.849109 | 0.187115 | 0.122 | 0.143 | 1 | 15 | Zdhhc6 | -0.021 | Old |
| 0.849163 | 0.098589 | 0.449 | 0.446 | 1 | 15 | Myo9b | 0.003 | Old |
| 0.849439 | 0.262738 | 0.408 | 0.429 | 1 | 15 | Trim25 | -0.021 | Old |
| 0.849711 | 0.007765 | 0.918 | 1 | 1 | 15 | Rpl39 | -0.082 | Old |
| 0.850485 | 1.015377 | 0.082 | 0.071 | 1 | 15 | Ptpn22 | 0.011 | Old |
| 0.850485 | 0.68115 | 0.082 | 0.071 | 1 | 15 | Plekhm3 | 0.011 | Old |
| 0.850485 | 0.585806 | 0.082 | 0.071 | 1 | 15 | Miip | 0.011 | Old |
| 0.850485 | 0.439833 | 0.082 | 0.071 | 1 | 15 | Gin1 | 0.011 | Old |
| 0.850485 | 0.347538 | 0.082 | 0.071 | 1 | 15 | Gstm1 | 0.011 | Old |
| 0.850485 | 0.324515 | 0.082 | 0.071 | 1 | 15 | Dhrs4 | 0.011 | Old |
| 0.850485 | 0.300885 | 0.082 | 0.071 | 1 | 15 | Zfp26 | 0.011 | Old |
| 0.850485 | 0.296785 | 0.082 | 0.071 | 1 | 15 | Sos1 | 0.011 | Old |
| 0.850485 | 0.221447 | 0.082 | 0.071 | 1 | 15 | Stx2 | 0.011 | Old |
| 0.850485 | 0.211688 | 0.082 | 0.071 | 1 | 15 | Kctd9 | 0.011 | Old |
| 0.850485 | 0.18702 | 0.082 | 0.071 | 1 | 15 | Exoc1 | 0.011 | Old |
| 0.850485 | 0.170592 | 0.082 | 0.071 | 1 | 15 | Rprd1b | 0.011 | Old |
| 0.850485 | 0.145733 | 0.082 | 0.071 | 1 | 15 | Nsun4 | 0.011 | Old |
| 0.850485 | 0.109789 | 0.082 | 0.071 | 1 | 15 | Dhps | 0.011 | Old |
| 0.850485 | 0.016552 | 0.082 | 0.071 | 1 | 15 | Atxn3 | 0.011 | Old |
| 0.850496 | 0.282965 | 0.184 | 0.214 | 1 | 15 | Plpbp | -0.03 | Old |
| 0.850649 | 0.250948 | 0.449 | 0.482 | 1 | 15 | Sulf2 | -0.033 | Old |
| 0.851064 | 0.054822 | 0.245 | 0.214 | 1 | 15 | Dnajc10 | 0.031 | Old |
| 0.851199 | 0.434574 | 0.347 | 0.321 | 1 | 15 | Cisd2 | 0.026 | Old |
| 0.852696 | 0.893289 | 0.061 | 0.071 | 1 | 15 | Selenom | -0.01 | Old |
| 0.852696 | 0.37905 | 0.061 | 0.071 | 1 | 15 | Gon7 | -0.01 | Old |
| 0.852696 | 0.350329 | 0.061 | 0.071 | 1 | 15 | Plekhg5 | -0.01 | Old |
| 0.852696 | 0.219991 | 0.061 | 0.071 | 1 | 15 | Ctc1 | -0.01 | Old |
| 0.852696 | 0.213653 | 0.061 | 0.071 | 1 | 15 | Gpr155 | -0.01 | Old |
| 0.852696 | 0.126653 | 0.061 | 0.071 | 1 | 15 | Gprc5c | -0.01 | Old |
| 0.852696 | 0.039638 | 0.061 | 0.071 | 1 | 15 | Tbck | -0.01 | Old |

| | | | | | | | | |
|---|---|---|---|---|---|---|---|---|
| 0.852696 | 0.008524 | 0.061 | 0.071 | 1 | 15 | Isca1 | -0.01 | Old |
| 0.853158 | 0.010683 | 0.204 | 0.214 | 1 | 15 | Mga | -0.01 | Old |
| 0.853247 | 0.181272 | 0.388 | 0.411 | 1 | 15 | Pdlim5 | -0.023 | Old |
| 0.854175 | 0.037199 | 0.735 | 0.857 | 1 | 15 | Hnrnpa2b1 | -0.122 | Old |
| 0.854202 | 0.244957 | 0.327 | 0.339 | 1 | 15 | Usp19 | -0.012 | Old |
| 0.854313 | 1.922443 | 0.061 | 0.054 | 1 | 15 | Fzd5 | 0.007 | Old |
| 0.854313 | 1.217222 | 0.061 | 0.054 | 1 | 15 | Tars2 | 0.007 | Old |
| 0.854313 | 1.041127 | 0.061 | 0.054 | 1 | 15 | Nkap | 0.007 | Old |
| 0.854313 | 1.038199 | 0.061 | 0.054 | 1 | 15 | Fam131a | 0.007 | Old |
| 0.854313 | 0.917067 | 0.061 | 0.054 | 1 | 15 | F8a | 0.007 | Old |
| 0.854313 | 0.611514 | 0.061 | 0.054 | 1 | 15 | Mocs1 | 0.007 | Old |
| 0.854313 | 0.597161 | 0.061 | 0.054 | 1 | 15 | Zfand2a | 0.007 | Old |
| 0.854313 | 0.570561 | 0.061 | 0.054 | 1 | 15 | Sdhaf4 | 0.007 | Old |
| 0.854313 | 0.521486 | 0.061 | 0.054 | 1 | 15 | D430042O( | 0.007 | Old |
| 0.854313 | 0.465171 | 0.061 | 0.054 | 1 | 15 | Dars | 0.007 | Old |
| 0.854313 | 0.45424 | 0.061 | 0.054 | 1 | 15 | Psmf1 | 0.007 | Old |
| 0.854313 | 0.444053 | 0.061 | 0.054 | 1 | 15 | Siae | 0.007 | Old |
| 0.854313 | 0.420621 | 0.061 | 0.054 | 1 | 15 | Pot1a | 0.007 | Old |
| 0.854313 | 0.41173 | 0.061 | 0.054 | 1 | 15 | Atg10 | 0.007 | Old |
| 0.854313 | 0.39536 | 0.061 | 0.054 | 1 | 15 | Strip1 | 0.007 | Old |
| 0.854313 | 0.04988 | 0.061 | 0.054 | 1 | 15 | Ctsf | 0.007 | Old |
| 0.854509 | 0.149068 | 0.306 | 0.321 | 1 | 15 | Dnm1l | -0.015 | Old |
| 0.8549 | 0.316716 | 0.102 | 0.089 | 1 | 15 | Arfgap1 | 0.013 | Old |
| 0.8549 | 0.165371 | 0.102 | 0.089 | 1 | 15 | Slc25a12 | 0.013 | Old |
| 0.8549 | 0.058354 | 0.102 | 0.089 | 1 | 15 | Zdhhc17 | 0.013 | Old |
| 0.854963 | 0.294037 | 0.184 | 0.196 | 1 | 15 | Etfa | -0.012 | Old |
| 0.856943 | 0.679287 | 0.163 | 0.161 | 1 | 15 | Anxa1 | 0.002 | Old |
| 0.856982 | 0.047703 | 0.122 | 0.107 | 1 | 15 | Arntl | 0.015 | Old |
| 0.857637 | 0.992595 | 0.122 | 0.143 | 1 | 15 | Dapk3 | -0.021 | Old |
| 0.857637 | 0.977174 | 0.122 | 0.143 | 1 | 15 | Nr1d2 | -0.021 | Old |
| 0.857637 | 0.841488 | 0.122 | 0.143 | 1 | 15 | Topbp1 | -0.021 | Old |
| 0.857637 | 0.315852 | 0.122 | 0.143 | 1 | 15 | Hps3 | -0.021 | Old |
| 0.857637 | 0.236715 | 0.122 | 0.143 | 1 | 15 | Avl9 | -0.021 | Old |
| 0.857669 | 0.458704 | 0.286 | 0.268 | 1 | 15 | Gtpbp2 | 0.018 | Old |
| 0.857669 | 0.185145 | 0.286 | 0.268 | 1 | 15 | 1110008F1 | 0.018 | Old |
| 0.857916 | 0.290107 | 0.224 | 0.232 | 1 | 15 | Fyn | -0.008 | Old |
| 0.859489 | 0.640745 | 0.184 | 0.179 | 1 | 15 | Imp3 | 0.005 | Old |
| 0.859774 | 0.065438 | 0.898 | 0.964 | 1 | 15 | Ctss | -0.066 | Old |
| 0.859833 | 0.156029 | 0.429 | 0.393 | 1 | 15 | Dynll1 | 0.036 | Old |
| 0.860049 | 0.383713 | 0.224 | 0.25 | 1 | 15 | Nup62 | -0.026 | Old |
| 0.860246 | 0.014664 | 0.224 | 0.196 | 1 | 15 | Cnppd1 | 0.028 | Old |
| 0.860675 | 0.330447 | 0.612 | 0.607 | 1 | 15 | Psmb1 | 0.005 | Old |
| 0.86132 | 0.260091 | 0.204 | 0.196 | 1 | 15 | Eif2s1 | 0.008 | Old |
| 0.861444 | 0.956702 | 0.082 | 0.071 | 1 | 15 | Serpina3g | 0.011 | Old |
| 0.861444 | 0.531519 | 0.082 | 0.071 | 1 | 15 | Ammecr1 | 0.011 | Old |

| | | | | | | | | |
|---|---|---|---|---|---|---|---|---|
| 0.861444 | 0.40101 | 0.082 | 0.071 | 1 | 15 | Por | 0.011 | Old |
| 0.861444 | 0.276929 | 0.082 | 0.071 | 1 | 15 | Kat7 | 0.011 | Old |
| 0.861444 | 0.228707 | 0.082 | 0.071 | 1 | 15 | Plod3 | 0.011 | Old |
| 0.861444 | 0.166766 | 0.082 | 0.071 | 1 | 15 | Armc5 | 0.011 | Old |
| 0.861444 | 0.071485 | 0.082 | 0.071 | 1 | 15 | Nudt5 | 0.011 | Old |
| 0.861444 | 0.046026 | 0.082 | 0.071 | 1 | 15 | Npl | 0.011 | Old |
| 0.861788 | 0.083332 | 0.388 | 0.357 | 1 | 15 | Nras | 0.031 | Old |
| 0.861816 | 0.23532 | 0.49 | 0.536 | 1 | 15 | Set | -0.046 | Old |
| 0.861816 | 0.006896 | 0.49 | 0.536 | 1 | 15 | Uqcr11 | -0.046 | Old |
| 0.862296 | 0.327397 | 0.633 | 0.696 | 1 | 15 | Csrnp1 | -0.063 | Old |
| 0.862324 | 0.213717 | 0.51 | 0.536 | 1 | 15 | Nktr | -0.026 | Old |
| 0.862596 | 0.417738 | 0.224 | 0.214 | 1 | 15 | Emc7 | 0.01 | Old |
| 0.862845 | 0.027785 | 0.49 | 0.482 | 1 | 15 | Ppp4r2 | 0.008 | Old |
| 0.863256 | 0.057906 | 0.714 | 0.714 | 1 | 15 | Rac1 | 0 | Old |
| 0.863424 | 0.046059 | 0.469 | 0.518 | 1 | 15 | N4bp2l1 | -0.049 | Old |
| 0.863428 | 0.077434 | 0.245 | 0.232 | 1 | 15 | Atl3 | 0.013 | Old |
| 0.863457 | 0.513156 | 0.082 | 0.089 | 1 | 15 | Klhl2 | -0.007 | Old |
| 0.863701 | 0.073509 | 0.551 | 0.554 | 1 | 15 | Sdcbp | -0.003 | Old |
| 0.863901 | 0.201159 | 0.265 | 0.25 | 1 | 15 | Nup210 | 0.015 | Old |
| 0.864074 | 0.375038 | 0.286 | 0.268 | 1 | 15 | Akt2 | 0.018 | Old |
| 0.864361 | 2.13198 | 0.061 | 0.071 | 1 | 15 | Gm10399 | -0.01 | Old |
| 0.864361 | 0.787753 | 0.061 | 0.071 | 1 | 15 | Chmp7 | -0.01 | Old |
| 0.864361 | 0.772942 | 0.061 | 0.071 | 1 | 15 | Bap1 | -0.01 | Old |
| 0.864361 | 0.498683 | 0.061 | 0.071 | 1 | 15 | Tmub1 | -0.01 | Old |
| 0.864361 | 0.376703 | 0.061 | 0.071 | 1 | 15 | Ivd | -0.01 | Old |
| 0.864361 | 0.124882 | 0.061 | 0.071 | 1 | 15 | Orc3 | -0.01 | Old |
| 0.864361 | 0.023317 | 0.061 | 0.071 | 1 | 15 | Mansc1 | -0.01 | Old |
| 0.864807 | 0.049931 | 0.102 | 0.089 | 1 | 15 | Pno1 | 0.013 | Old |
| 0.865419 | 0.077739 | 0.163 | 0.143 | 1 | 15 | Gas2l1 | 0.02 | Old |
| 0.865419 | 0.001932 | 0.143 | 0.161 | 1 | 15 | Shkbp1 | -0.018 | Old |
| 0.866119 | 0.387965 | 0.122 | 0.107 | 1 | 15 | Pus10 | 0.015 | Old |
| 0.866858 | 1.845373 | 0.061 | 0.054 | 1 | 15 | Dgke | 0.007 | Old |
| 0.866858 | 1.55585 | 0.061 | 0.054 | 1 | 15 | Tbc1d32 | 0.007 | Old |
| 0.866858 | 1.346472 | 0.061 | 0.054 | 1 | 15 | Il10 | 0.007 | Old |
| 0.866858 | 1.123666 | 0.061 | 0.054 | 1 | 15 | Tubgcp3 | 0.007 | Old |
| 0.866858 | 0.955307 | 0.061 | 0.054 | 1 | 15 | Klrb1b | 0.007 | Old |
| 0.866858 | 0.549242 | 0.061 | 0.054 | 1 | 15 | Atm | 0.007 | Old |
| 0.866858 | 0.467547 | 0.061 | 0.054 | 1 | 15 | Matk | 0.007 | Old |
| 0.866858 | 0.398313 | 0.061 | 0.054 | 1 | 15 | Gmeb2 | 0.007 | Old |
| 0.866858 | 0.373999 | 0.061 | 0.054 | 1 | 15 | Zw10 | 0.007 | Old |
| 0.866858 | 0.372608 | 0.061 | 0.054 | 1 | 15 | Arl4a | 0.007 | Old |
| 0.866858 | 0.319643 | 0.061 | 0.054 | 1 | 15 | Mark4 | 0.007 | Old |
| 0.866858 | 0.285737 | 0.061 | 0.054 | 1 | 15 | Zfp68 | 0.007 | Old |
| 0.866858 | 0.281688 | 0.061 | 0.054 | 1 | 15 | Nol8 | 0.007 | Old |
| 0.866858 | 0.257004 | 0.061 | 0.054 | 1 | 15 | 6430548M | 0.007 | Old |

| | | | | | | | | |
|---|---|---|---|---|---|---|---|---|
| 0.866858 | 0.13416 | 0.061 | 0.054 | 1 | 15 | Aktip | 0.007 | Old |
| 0.866858 | 0.0997 | 0.061 | 0.054 | 1 | 15 | Zfp617 | 0.007 | Old |
| 0.866858 | 0.076659 | 0.061 | 0.054 | 1 | 15 | Cdk5rap2 | 0.007 | Old |
| 0.866858 | 0.066485 | 0.061 | 0.054 | 1 | 15 | Rab27a | 0.007 | Old |
| 0.866858 | 0.003151 | 0.061 | 0.054 | 1 | 15 | Rarg | 0.007 | Old |
| 0.867344 | 0.572578 | 0.204 | 0.214 | 1 | 15 | Gnpda1 | -0.01 | Old |
| 0.868137 | 0.076086 | 0.531 | 0.536 | 1 | 15 | Dusp11 | -0.005 | Old |
| 0.868551 | 0.448618 | 0.204 | 0.196 | 1 | 15 | Tmem106b | 0.008 | Old |
| 0.868551 | 0.270918 | 0.204 | 0.196 | 1 | 15 | Psme2b | 0.008 | Old |
| 0.86919 | 0.055967 | 0.469 | 0.446 | 1 | 15 | Eif4h | 0.023 | Old |
| 0.869244 | 2.273366 | 0.041 | 0.036 | 1 | 15 | Scn1b | 0.005 | Old |
| 0.869244 | 2.206855 | 0.041 | 0.036 | 1 | 15 | Gm10036 | 0.005 | Old |
| 0.869244 | 2.097026 | 0.041 | 0.036 | 1 | 15 | Haus2 | 0.005 | Old |
| 0.869244 | 2.017885 | 0.041 | 0.036 | 1 | 15 | Pdk1 | 0.005 | Old |
| 0.869244 | 1.993045 | 0.041 | 0.036 | 1 | 15 | 2610002M | 0.005 | Old |
| 0.869244 | 1.812642 | 0.041 | 0.036 | 1 | 15 | Adck1 | 0.005 | Old |
| 0.869244 | 1.809515 | 0.041 | 0.036 | 1 | 15 | Btbd6 | 0.005 | Old |
| 0.869244 | 1.758807 | 0.041 | 0.036 | 1 | 15 | 9130008F2 | 0.005 | Old |
| 0.869244 | 1.703243 | 0.041 | 0.036 | 1 | 15 | L3mbtl3 | 0.005 | Old |
| 0.869244 | 1.642871 | 0.041 | 0.036 | 1 | 15 | Ndc1 | 0.005 | Old |
| 0.869244 | 1.565702 | 0.041 | 0.036 | 1 | 15 | Mettl26 | 0.005 | Old |
| 0.869244 | 1.483966 | 0.041 | 0.036 | 1 | 15 | Ppp1r12b | 0.005 | Old |
| 0.869244 | 1.3836 | 0.041 | 0.036 | 1 | 15 | Zfp770 | 0.005 | Old |
| 0.869244 | 1.299477 | 0.041 | 0.036 | 1 | 15 | Gm35154 | 0.005 | Old |
| 0.869244 | 1.299315 | 0.041 | 0.036 | 1 | 15 | Ddx19b | 0.005 | Old |
| 0.869244 | 1.274813 | 0.041 | 0.036 | 1 | 15 | Btbd2 | 0.005 | Old |
| 0.869244 | 1.187127 | 0.041 | 0.036 | 1 | 15 | Gm28043 | 0.005 | Old |
| 0.869244 | 1.136835 | 0.041 | 0.036 | 1 | 15 | Prmt5 | 0.005 | Old |
| 0.869244 | 1.098029 | 0.041 | 0.036 | 1 | 15 | Rps6kb2 | 0.005 | Old |
| 0.869244 | 1.0749 | 0.041 | 0.036 | 1 | 15 | Ntpcr | 0.005 | Old |
| 0.869244 | 1.06463 | 0.041 | 0.036 | 1 | 15 | Armcx5 | 0.005 | Old |
| 0.869244 | 1.055284 | 0.041 | 0.036 | 1 | 15 | Nos1ap | 0.005 | Old |
| 0.869244 | 0.980252 | 0.041 | 0.036 | 1 | 15 | Slc39a9 | 0.005 | Old |
| 0.869244 | 0.977735 | 0.041 | 0.036 | 1 | 15 | E2f1 | 0.005 | Old |
| 0.869244 | 0.957244 | 0.041 | 0.036 | 1 | 15 | Rcc1l | 0.005 | Old |
| 0.869244 | 0.931616 | 0.041 | 0.036 | 1 | 15 | Amz1 | 0.005 | Old |
| 0.869244 | 0.89877 | 0.041 | 0.036 | 1 | 15 | Fbxl18 | 0.005 | Old |
| 0.869244 | 0.846486 | 0.041 | 0.036 | 1 | 15 | Pola2 | 0.005 | Old |
| 0.869244 | 0.760144 | 0.041 | 0.036 | 1 | 15 | Ttll4 | 0.005 | Old |
| 0.869244 | 0.750988 | 0.041 | 0.036 | 1 | 15 | Tspan4 | 0.005 | Old |
| 0.869244 | 0.356342 | 0.041 | 0.036 | 1 | 15 | Hhip | 0.005 | Old |
| 0.869244 | 0.650954 | 0.347 | 0.321 | 1 | 15 | Rab12 | 0.026 | Old |
| 0.869577 | 0.523389 | 0.224 | 0.214 | 1 | 15 | Zdhhc20 | 0.01 | Old |
| 0.869623 | 0.223216 | 0.816 | 0.857 | 1 | 15 | Mob1a | -0.041 | Old |
| 0.870488 | 0.42154 | 0.286 | 0.268 | 1 | 15 | Baiap2 | 0.018 | Old |

| | | | | | | | | |
|---|---|---|---|---|---|---|---|---|
| 0.871 | 0.337142 | 0.469 | 0.5 | 1 | 15 | Hspa5 | -0.031 | Old |
| 0.87178 | 0.030077 | 0.184 | 0.161 | 1 | 15 | Mgat2 | 0.023 | Old |
| 0.87214 | 0.546071 | 0.265 | 0.268 | 1 | 15 | Clec4a3 | -0.003 | Old |
| 0.872429 | 0.593474 | 0.082 | 0.071 | 1 | 15 | Ethe1 | 0.011 | Old |
| 0.872429 | 0.258489 | 0.082 | 0.071 | 1 | 15 | Alox5 | 0.011 | Old |
| 0.872429 | 0.110101 | 0.082 | 0.071 | 1 | 15 | Fbxl17 | 0.011 | Old |
| 0.872694 | 0.718498 | 0.163 | 0.161 | 1 | 15 | Tgfbr2 | 0.002 | Old |
| 0.873401 | 0.185864 | 0.286 | 0.304 | 1 | 15 | Fam111a | -0.018 | Old |
| 0.873401 | 0.045026 | 0.286 | 0.304 | 1 | 15 | Suz12 | -0.018 | Old |
| 0.873505 | 0.064313 | 0.143 | 0.161 | 1 | 15 | Snrnp27 | -0.018 | Old |
| 0.873582 | 0.615401 | 0.224 | 0.25 | 1 | 15 | Tmem55b | -0.026 | Old |
| 0.873868 | 0.066607 | 0.082 | 0.089 | 1 | 15 | Cbll1 | -0.007 | Old |
| 0.873896 | 0.193052 | 0.551 | 0.536 | 1 | 15 | Peli1 | 0.015 | Old |
| 0.874112 | 0.131297 | 0.755 | 0.768 | 1 | 15 | Nap1l1 | -0.013 | Old |
| 0.874127 | 0.403667 | 0.327 | 0.321 | 1 | 15 | Nudt21 | 0.006 | Old |
| 0.874365 | 0.584441 | 0.388 | 0.375 | 1 | 15 | Eno3 | 0.013 | Old |
| 0.875163 | 0.175186 | 0.286 | 0.286 | 1 | 15 | Ube2h | 0 | Old |
| 0.875236 | 0.02632 | 0.388 | 0.393 | 1 | 15 | Mmd | -0.005 | Old |
| 0.875793 | 0.577074 | 0.204 | 0.196 | 1 | 15 | Tsg101 | 0.008 | Old |
| 0.876056 | 1.043638 | 0.061 | 0.071 | 1 | 15 | Syt11 | -0.01 | Old |
| 0.876056 | 0.77346 | 0.061 | 0.071 | 1 | 15 | Jmy | -0.01 | Old |
| 0.876056 | 0.517836 | 0.061 | 0.071 | 1 | 15 | Zfp160 | -0.01 | Old |
| 0.876056 | 0.517341 | 0.061 | 0.071 | 1 | 15 | Cln6 | -0.01 | Old |
| 0.876056 | 0.351734 | 0.061 | 0.071 | 1 | 15 | Ptgs1 | -0.01 | Old |
| 0.876056 | 0.310946 | 0.061 | 0.071 | 1 | 15 | Mcm6 | -0.01 | Old |
| 0.876056 | 0.272658 | 0.061 | 0.071 | 1 | 15 | Pgm2 | -0.01 | Old |
| 0.876056 | 0.143702 | 0.061 | 0.071 | 1 | 15 | Tnks | -0.01 | Old |
| 0.876056 | 0.122159 | 0.061 | 0.071 | 1 | 15 | 4930579G2 | -0.01 | Old |
| 0.87654 | 0.021156 | 0.306 | 0.286 | 1 | 15 | Acadl | 0.02 | Old |
| 0.876756 | 0.021338 | 0.796 | 0.75 | 1 | 15 | Snrpe | 0.046 | Old |
| 0.877389 | 0.057135 | 0.347 | 0.357 | 1 | 15 | Lamtor1 | -0.01 | Old |
| 0.878307 | 0.103569 | 0.429 | 0.429 | 1 | 15 | Myl12b | 0 | Old |
| 0.878355 | 0.90423 | 0.143 | 0.143 | 1 | 15 | Cnep1r1 | 0 | Old |
| 0.878387 | 0.051714 | 0.51 | 0.536 | 1 | 15 | Rpl13a | -0.026 | Old |
| 0.878645 | 0.338912 | 0.286 | 0.25 | 1 | 15 | Snx6 | 0.036 | Old |
| 0.879063 | 0.404704 | 0.306 | 0.321 | 1 | 15 | Smurf1 | -0.015 | Old |
| 0.879063 | 0.247812 | 0.306 | 0.321 | 1 | 15 | Clint1 | -0.015 | Old |
| 0.879436 | 1.283352 | 0.061 | 0.054 | 1 | 15 | Lias | 0.007 | Old |
| 0.879436 | 0.932119 | 0.061 | 0.054 | 1 | 15 | Sfxn2 | 0.007 | Old |
| 0.879436 | 0.753097 | 0.061 | 0.054 | 1 | 15 | Mllt1 | 0.007 | Old |
| 0.879436 | 0.691648 | 0.061 | 0.054 | 1 | 15 | Ints2 | 0.007 | Old |
| 0.879436 | 0.619543 | 0.061 | 0.054 | 1 | 15 | Lin37 | 0.007 | Old |
| 0.879436 | 0.203878 | 0.061 | 0.054 | 1 | 15 | Kdm4b | 0.007 | Old |
| 0.879436 | 0.19745 | 0.061 | 0.054 | 1 | 15 | Tmed8 | 0.007 | Old |
| 0.879436 | 0.182005 | 0.061 | 0.054 | 1 | 15 | Arsk | 0.007 | Old |

| | | | | | | | | |
|---|---|---|---|---|---|---|---|---|
| 0.879436 | 0.177917 | 0.061 | 0.054 | 1 | 15 | Nudt2 | 0.007 | Old |
| 0.879436 | 0.159066 | 0.061 | 0.054 | 1 | 15 | Ctnnbip1 | 0.007 | Old |
| 0.879436 | 0.136407 | 0.061 | 0.054 | 1 | 15 | Exosc4 | 0.007 | Old |
| 0.879436 | 0.098407 | 0.061 | 0.054 | 1 | 15 | Pdrg1 | 0.007 | Old |
| 0.879436 | 0.035624 | 0.061 | 0.054 | 1 | 15 | 2310039H( | 0.007 | Old |
| 0.87949 | 0.35508 | 0.163 | 0.179 | 1 | 15 | C330007P( | -0.016 | Old |
| 0.87949 | 0.145852 | 0.163 | 0.179 | 1 | 15 | Tcf3 | -0.016 | Old |
| 0.87949 | 0.043811 | 0.163 | 0.179 | 1 | 15 | Rictor | -0.016 | Old |
| 0.880362 | 1.05857 | 0.224 | 0.25 | 1 | 15 | Plec | -0.026 | Old |
| 0.880364 | 0.50573 | 0.245 | 0.268 | 1 | 15 | Furin | -0.023 | Old |
| 0.881604 | 0.218979 | 0.143 | 0.161 | 1 | 15 | Taf13 | -0.018 | Old |
| 0.883439 | 0.034757 | 0.082 | 0.071 | 1 | 15 | Clock | 0.011 | Old |
| 0.883756 | 0.091922 | 0.245 | 0.232 | 1 | 15 | Sp1 | 0.013 | Old |
| 0.884058 | 0.236466 | 0.286 | 0.321 | 1 | 15 | Prkar2a | -0.035 | Old |
| 0.884334 | 0.26539 | 0.327 | 0.339 | 1 | 15 | Sumo3 | -0.012 | Old |
| 0.884334 | 0.16069 | 0.327 | 0.339 | 1 | 15 | Rela | -0.012 | Old |
| 0.884478 | 0.307785 | 0.184 | 0.196 | 1 | 15 | Tnrc6c | -0.012 | Old |
| 0.884512 | 1.843159 | 0.041 | 0.036 | 1 | 15 | Pomgnt1 | 0.005 | Old |
| 0.884512 | 1.339892 | 0.041 | 0.036 | 1 | 15 | Amd1 | 0.005 | Old |
| 0.884512 | 1.331914 | 0.041 | 0.036 | 1 | 15 | Gm340 | 0.005 | Old |
| 0.884512 | 1.295684 | 0.041 | 0.036 | 1 | 15 | Fam78a | 0.005 | Old |
| 0.884512 | 1.288747 | 0.041 | 0.036 | 1 | 15 | B3gntl1 | 0.005 | Old |
| 0.884512 | 1.200073 | 0.041 | 0.036 | 1 | 15 | Slc16a1 | 0.005 | Old |
| 0.884512 | 1.194412 | 0.041 | 0.036 | 1 | 15 | 1110065P2 | 0.005 | Old |
| 0.884512 | 1.08565 | 0.041 | 0.036 | 1 | 15 | Cenpc1 | 0.005 | Old |
| 0.884512 | 1.047913 | 0.041 | 0.036 | 1 | 15 | Ttll12 | 0.005 | Old |
| 0.884512 | 0.921847 | 0.041 | 0.036 | 1 | 15 | Necap1 | 0.005 | Old |
| 0.884512 | 0.910396 | 0.041 | 0.036 | 1 | 15 | Cables1 | 0.005 | Old |
| 0.884512 | 0.908082 | 0.041 | 0.036 | 1 | 15 | Isoc1 | 0.005 | Old |
| 0.884512 | 0.897016 | 0.041 | 0.036 | 1 | 15 | Hbegf | 0.005 | Old |
| 0.884512 | 0.799928 | 0.041 | 0.036 | 1 | 15 | Pradc1 | 0.005 | Old |
| 0.884512 | 0.657428 | 0.041 | 0.036 | 1 | 15 | Zfp24 | 0.005 | Old |
| 0.884512 | 0.624078 | 0.041 | 0.036 | 1 | 15 | Ddx60 | 0.005 | Old |
| 0.884512 | 0.602904 | 0.041 | 0.036 | 1 | 15 | Ado | 0.005 | Old |
| 0.884512 | 0.564676 | 0.041 | 0.036 | 1 | 15 | Nxt1 | 0.005 | Old |
| 0.884512 | 0.563814 | 0.041 | 0.036 | 1 | 15 | Dbr1 | 0.005 | Old |
| 0.884512 | 0.498825 | 0.041 | 0.036 | 1 | 15 | Grap2 | 0.005 | Old |
| 0.884512 | 0.352086 | 0.041 | 0.036 | 1 | 15 | Zfyve21 | 0.005 | Old |
| 0.884512 | 0.283837 | 0.041 | 0.036 | 1 | 15 | Alkbh7 | 0.005 | Old |
| 0.884512 | 0.191352 | 0.041 | 0.036 | 1 | 15 | Sfxn1 | 0.005 | Old |
| 0.884549 | 0.290345 | 0.51 | 0.571 | 1 | 15 | Qk | -0.061 | Old |
| 0.885268 | 0.200088 | 0.551 | 0.571 | 1 | 15 | D17Wsu92 | -0.02 | Old |
| 0.886214 | 0.109738 | 0.592 | 0.589 | 1 | 15 | H2afy | 0.003 | Old |
| 0.886321 | 0.047478 | 0.469 | 0.464 | 1 | 15 | Rap2b | 0.005 | Old |
| 0.88656 | 0.164658 | 0.265 | 0.286 | 1 | 15 | Mier1 | -0.021 | Old |

| | | | | | | | | |
|---|---|---|---|---|---|---|---|---|
| 0.886674 | 0.136308 | 0.184 | 0.214 | 1 | 15 | Nr2f6 | -0.03 | Old |
| 0.886955 | 0.00884 | 0.776 | 0.768 | 1 | 15 | Rtn4 | 0.008 | Old |
| 0.887211 | 0.14316 | 0.163 | 0.179 | 1 | 15 | Rbm28 | -0.016 | Old |
| 0.887216 | 0.341424 | 0.531 | 0.554 | 1 | 15 | H2-Q6 | -0.023 | Old |
| 0.887429 | 0.041025 | 0.898 | 0.804 | 1 | 15 | Samsn1 | 0.094 | Old |
| 0.887778 | 1.305711 | 0.061 | 0.071 | 1 | 15 | Itga1 | -0.01 | Old |
| 0.887778 | 1.268328 | 0.061 | 0.071 | 1 | 15 | Phf11d | -0.01 | Old |
| 0.887778 | 1.194738 | 0.061 | 0.071 | 1 | 15 | Snx19 | -0.01 | Old |
| 0.887778 | 1.111442 | 0.061 | 0.071 | 1 | 15 | Tmem29 | -0.01 | Old |
| 0.887778 | 0.733342 | 0.061 | 0.071 | 1 | 15 | Slc28a2 | -0.01 | Old |
| 0.887778 | 0.681169 | 0.061 | 0.071 | 1 | 15 | Snhg20 | -0.01 | Old |
| 0.887778 | 0.50396 | 0.061 | 0.071 | 1 | 15 | Txndc16 | -0.01 | Old |
| 0.887778 | 0.491722 | 0.061 | 0.071 | 1 | 15 | Polg | -0.01 | Old |
| 0.887778 | 0.256109 | 0.061 | 0.071 | 1 | 15 | Zfp90 | -0.01 | Old |
| 0.888496 | 0.843354 | 0.163 | 0.161 | 1 | 15 | Mapk8 | 0.002 | Old |
| 0.888703 | 0.719399 | 0.204 | 0.214 | 1 | 15 | Tollip | -0.01 | Old |
| 0.888713 | 0.429168 | 0.51 | 0.518 | 1 | 15 | Gda | -0.008 | Old |
| 0.88912 | 0.207566 | 0.571 | 0.625 | 1 | 15 | Cacnb3 | -0.054 | Old |
| 0.889715 | 0.464597 | 0.143 | 0.161 | 1 | 15 | Gigyf1 | -0.018 | Old |
| 0.890307 | 0.904389 | 0.204 | 0.196 | 1 | 15 | Araf | 0.008 | Old |
| 0.890549 | 0.615957 | 0.245 | 0.232 | 1 | 15 | Cbfb | 0.013 | Old |
| 0.892044 | 0.700879 | 0.061 | 0.054 | 1 | 15 | 2900052N( | 0.007 | Old |
| 0.892044 | 0.319527 | 0.061 | 0.054 | 1 | 15 | Dym | 0.007 | Old |
| 0.892044 | 0.265045 | 0.061 | 0.054 | 1 | 15 | Acp5 | 0.007 | Old |
| 0.892044 | 0.123153 | 0.061 | 0.054 | 1 | 15 | Ranbp6 | 0.007 | Old |
| 0.892044 | 0.066831 | 0.061 | 0.054 | 1 | 15 | Tmod1 | 0.007 | Old |
| 0.892044 | 0.045413 | 0.061 | 0.054 | 1 | 15 | Pou5f2 | 0.007 | Old |
| 0.892325 | 0.172534 | 0.224 | 0.232 | 1 | 15 | Kctd10 | -0.008 | Old |
| 0.892426 | 0.191393 | 0.306 | 0.339 | 1 | 15 | Tmem19 | -0.033 | Old |
| 0.893915 | 0.198415 | 0.367 | 0.375 | 1 | 15 | Sbf2 | -0.008 | Old |
| 0.893941 | 0.292127 | 0.204 | 0.196 | 1 | 15 | Ntan1 | 0.008 | Old |
| 0.894472 | 0.646042 | 0.082 | 0.071 | 1 | 15 | Zfp410 | 0.011 | Old |
| 0.894649 | 0.047182 | 0.102 | 0.089 | 1 | 15 | Mboat7 | 0.013 | Old |
| 0.894756 | 0.485007 | 0.082 | 0.089 | 1 | 15 | Brf1 | -0.007 | Old |
| 0.894756 | 0.22287 | 0.082 | 0.089 | 1 | 15 | Gtf3c3 | -0.007 | Old |
| 0.894824 | 0.245648 | 0.796 | 0.804 | 1 | 15 | Dazap2 | -0.008 | Old |
| 0.895224 | 0.098987 | 0.551 | 0.536 | 1 | 15 | Dnajb6 | 0.015 | Old |
| 0.896179 | 0.356956 | 0.592 | 0.554 | 1 | 15 | Pip4k2a | 0.038 | Old |
| 0.896517 | 0.489275 | 0.286 | 0.321 | 1 | 15 | Emd | -0.035 | Old |
| 0.896846 | 0.554421 | 0.429 | 0.482 | 1 | 15 | Cox14 | -0.053 | Old |
| 0.897578 | 0.298521 | 0.204 | 0.196 | 1 | 15 | Nin | 0.008 | Old |
| 0.897824 | 0.293591 | 0.49 | 0.464 | 1 | 15 | Sgpl1 | 0.026 | Old |
| 0.897837 | 0.459088 | 0.143 | 0.161 | 1 | 15 | Mtmr14 | -0.018 | Old |
| 0.898084 | 0.271184 | 0.673 | 0.625 | 1 | 15 | Ccnl1 | 0.048 | Old |
| 0.898207 | 0.219028 | 0.265 | 0.268 | 1 | 15 | Slc41a1 | -0.003 | Old |

| | | | | | | | | |
|---|---|---|---|---|---|---|---|---|
| 0.899033 | 0.21469 | 0.102 | 0.107 | 1 | 15 | Spen | -0.005 | Old |
| 0.899524 | 0.866527 | 0.061 | 0.071 | 1 | 15 | Dnmt1 | -0.01 | Old |
| 0.899524 | 0.704149 | 0.061 | 0.071 | 1 | 15 | Hhex | -0.01 | Old |
| 0.899524 | 0.38891 | 0.061 | 0.071 | 1 | 15 | Hsd17b7 | -0.01 | Old |
| 0.899801 | 0.058489 | 0.755 | 0.804 | 1 | 15 | Brk1 | -0.049 | Old |
| 0.899823 | 1.869158 | 0.041 | 0.036 | 1 | 15 | Fam217b | 0.005 | Old |
| 0.899823 | 1.67086 | 0.041 | 0.036 | 1 | 15 | Nlrp10 | 0.005 | Old |
| 0.899823 | 1.239007 | 0.041 | 0.036 | 1 | 15 | Tmem41a | 0.005 | Old |
| 0.899823 | 1.238968 | 0.041 | 0.036 | 1 | 15 | Dennd1c | 0.005 | Old |
| 0.899823 | 1.117571 | 0.041 | 0.036 | 1 | 15 | Sun1 | 0.005 | Old |
| 0.899823 | 1.111695 | 0.041 | 0.036 | 1 | 15 | Zxdc | 0.005 | Old |
| 0.899823 | 1.107364 | 0.041 | 0.036 | 1 | 15 | Apex1 | 0.005 | Old |
| 0.899823 | 0.998399 | 0.041 | 0.036 | 1 | 15 | Rint1 | 0.005 | Old |
| 0.899823 | 0.991891 | 0.041 | 0.036 | 1 | 15 | Zfp942 | 0.005 | Old |
| 0.899823 | 0.830809 | 0.041 | 0.036 | 1 | 15 | Mettl5 | 0.005 | Old |
| 0.899823 | 0.811462 | 0.041 | 0.036 | 1 | 15 | Klhl15 | 0.005 | Old |
| 0.899823 | 0.802266 | 0.041 | 0.036 | 1 | 15 | Ubtd1 | 0.005 | Old |
| 0.899823 | 0.79089 | 0.041 | 0.036 | 1 | 15 | Slc14a1 | 0.005 | Old |
| 0.899823 | 0.764375 | 0.041 | 0.036 | 1 | 15 | Gm26870 | 0.005 | Old |
| 0.899823 | 0.752289 | 0.041 | 0.036 | 1 | 15 | Dtx2 | 0.005 | Old |
| 0.899823 | 0.737603 | 0.041 | 0.036 | 1 | 15 | Gm21781 | 0.005 | Old |
| 0.899823 | 0.733983 | 0.041 | 0.036 | 1 | 15 | Msh6 | 0.005 | Old |
| 0.899823 | 0.726528 | 0.041 | 0.036 | 1 | 15 | Wdr7 | 0.005 | Old |
| 0.899823 | 0.672185 | 0.041 | 0.036 | 1 | 15 | Spata5 | 0.005 | Old |
| 0.899823 | 0.637712 | 0.041 | 0.036 | 1 | 15 | Thoc1 | 0.005 | Old |
| 0.899823 | 0.626299 | 0.041 | 0.036 | 1 | 15 | Cog8 | 0.005 | Old |
| 0.899823 | 0.584677 | 0.041 | 0.036 | 1 | 15 | Cd244 | 0.005 | Old |
| 0.899823 | 0.526456 | 0.041 | 0.036 | 1 | 15 | Mrps27 | 0.005 | Old |
| 0.899823 | 0.52606 | 0.041 | 0.036 | 1 | 15 | Pex1 | 0.005 | Old |
| 0.899823 | 0.49387 | 0.041 | 0.036 | 1 | 15 | Ecsit | 0.005 | Old |
| 0.899823 | 0.492567 | 0.041 | 0.036 | 1 | 15 | Zdhhc13 | 0.005 | Old |
| 0.899823 | 0.483471 | 0.041 | 0.036 | 1 | 15 | Zfp771 | 0.005 | Old |
| 0.899823 | 0.481424 | 0.041 | 0.036 | 1 | 15 | 2410004B1 | 0.005 | Old |
| 0.899823 | 0.476421 | 0.041 | 0.036 | 1 | 15 | 2610301B2 | 0.005 | Old |
| 0.899823 | 0.451332 | 0.041 | 0.036 | 1 | 15 | Lsm11 | 0.005 | Old |
| 0.899823 | 0.409446 | 0.041 | 0.036 | 1 | 15 | AU022252 | 0.005 | Old |
| 0.899823 | 0.401056 | 0.041 | 0.036 | 1 | 15 | Brms1l | 0.005 | Old |
| 0.899823 | 0.396911 | 0.041 | 0.036 | 1 | 15 | Tle1 | 0.005 | Old |
| 0.899823 | 0.327483 | 0.041 | 0.036 | 1 | 15 | Nat9 | 0.005 | Old |
| 0.899823 | 0.323587 | 0.041 | 0.036 | 1 | 15 | Kif3a | 0.005 | Old |
| 0.899823 | 0.316944 | 0.041 | 0.036 | 1 | 15 | Ss18l1 | 0.005 | Old |
| 0.899823 | 0.311182 | 0.041 | 0.036 | 1 | 15 | Glmn | 0.005 | Old |
| 0.899823 | 0.301986 | 0.041 | 0.036 | 1 | 15 | Gm38394 | 0.005 | Old |
| 0.899823 | 0.297531 | 0.041 | 0.036 | 1 | 15 | Plekhg1 | 0.005 | Old |
| 0.899823 | 0.294597 | 0.041 | 0.036 | 1 | 15 | 1810024B0 | 0.005 | Old |

| | | | | | | | | |
|---|---|---|---|---|---|---|---|---|
| 0.899823 | 0.260985 | 0.041 | 0.036 | 1 | 15 | Polr1a | 0.005 | Old |
| 0.899823 | 0.23319 | 0.041 | 0.036 | 1 | 15 | Sned1 | 0.005 | Old |
| 0.899823 | 0.219781 | 0.041 | 0.036 | 1 | 15 | Gm46224 | 0.005 | Old |
| 0.899823 | 0.21067 | 0.041 | 0.036 | 1 | 15 | Dpep2 | 0.005 | Old |
| 0.899823 | 0.197098 | 0.041 | 0.036 | 1 | 15 | Rpusd1 | 0.005 | Old |
| 0.899823 | 0.188123 | 0.041 | 0.036 | 1 | 15 | Dock5 | 0.005 | Old |
| 0.899823 | 0.177619 | 0.041 | 0.036 | 1 | 15 | Ndufaf5 | 0.005 | Old |
| 0.899823 | 0.174821 | 0.041 | 0.036 | 1 | 15 | Lpcat2 | 0.005 | Old |
| 0.899823 | 0.163267 | 0.041 | 0.036 | 1 | 15 | Gm9844 | 0.005 | Old |
| 0.899823 | 0.162139 | 0.041 | 0.036 | 1 | 15 | Gm14286 | 0.005 | Old |
| 0.899823 | 0.15904 | 0.041 | 0.036 | 1 | 15 | Fbxl4 | 0.005 | Old |
| 0.899823 | 0.155504 | 0.041 | 0.036 | 1 | 15 | Zfp760 | 0.005 | Old |
| 0.899823 | 0.153156 | 0.041 | 0.036 | 1 | 15 | Nup160 | 0.005 | Old |
| 0.899823 | 0.145875 | 0.041 | 0.036 | 1 | 15 | Lman2l | 0.005 | Old |
| 0.899823 | 0.139317 | 0.041 | 0.036 | 1 | 15 | Fbxw8 | 0.005 | Old |
| 0.899823 | 0.133284 | 0.041 | 0.036 | 1 | 15 | Rufy1 | 0.005 | Old |
| 0.899823 | 0.123991 | 0.041 | 0.036 | 1 | 15 | Taf6 | 0.005 | Old |
| 0.899823 | 0.117257 | 0.041 | 0.036 | 1 | 15 | Alpk2 | 0.005 | Old |
| 0.899823 | 0.096252 | 0.041 | 0.036 | 1 | 15 | Arhgap39 | 0.005 | Old |
| 0.899823 | 0.079851 | 0.041 | 0.036 | 1 | 15 | Zfp799 | 0.005 | Old |
| 0.899823 | 0.058741 | 0.041 | 0.036 | 1 | 15 | Pmepa1 | 0.005 | Old |
| 0.899823 | 0.030336 | 0.041 | 0.036 | 1 | 15 | Bambi | 0.005 | Old |
| 0.899823 | 0.003067 | 0.041 | 0.036 | 1 | 15 | Lrp12 | 0.005 | Old |
| 0.900344 | 0.016407 | 0.347 | 0.339 | 1 | 15 | Pura | 0.008 | Old |
| 0.900754 | 0.606345 | 0.224 | 0.25 | 1 | 15 | Eif1a | -0.026 | Old |
| 0.900817 | 0.661357 | 0.429 | 0.429 | 1 | 15 | Krit1 | 0 | Old |
| 0.901198 | 0.435911 | 0.367 | 0.411 | 1 | 15 | Aldoa | -0.044 | Old |
| 0.902173 | 0.087226 | 0.245 | 0.25 | 1 | 15 | Armc8 | -0.005 | Old |
| 0.902678 | 0.189629 | 0.286 | 0.268 | 1 | 15 | Atp6v1d | 0.018 | Old |
| 0.902684 | 0.485531 | 0.163 | 0.179 | 1 | 15 | Bag6 | -0.016 | Old |
| 0.902698 | 0.040852 | 0.592 | 0.643 | 1 | 15 | Swap70 | -0.051 | Old |
| 0.902757 | 0.348256 | 0.286 | 0.321 | 1 | 15 | Rps6ka3 | -0.035 | Old |
| 0.902847 | 0.12566 | 0.898 | 0.946 | 1 | 15 | Rps17 | -0.048 | Old |
| 0.902987 | 0.006178 | 0.204 | 0.214 | 1 | 15 | Fas | -0.01 | Old |
| 0.903384 | 0.731872 | 0.143 | 0.143 | 1 | 15 | Gpr108 | 0 | Old |
| 0.903637 | 0.2429 | 0.449 | 0.411 | 1 | 15 | Ube2r2 | 0.038 | Old |
| 0.904658 | 0.028901 | 0.327 | 0.321 | 1 | 15 | Psenen | 0.006 | Old |
| 0.904791 | 2.177344 | 0.184 | 0.179 | 1 | 15 | Gclc | 0.005 | Old |
| 0.904791 | 0.314711 | 0.184 | 0.179 | 1 | 15 | Tnpo2 | 0.005 | Old |
| 0.904791 | 0.089425 | 0.184 | 0.179 | 1 | 15 | Gabarapl1 | 0.005 | Old |
| 0.904858 | 0.138054 | 0.204 | 0.196 | 1 | 15 | Stx16 | 0.008 | Old |
| 0.905228 | 0.987634 | 0.082 | 0.089 | 1 | 15 | Mospd3 | -0.007 | Old |
| 0.905228 | 0.048165 | 0.082 | 0.089 | 1 | 15 | Mcee | -0.007 | Old |
| 0.906723 | 0.064414 | 0.184 | 0.196 | 1 | 15 | Rabggtb | -0.012 | Old |
| 0.907002 | 0.578485 | 0.286 | 0.286 | 1 | 15 | Mrpl54 | 0 | Old |

| | | | | | | | | |
|---|---|---|---|---|---|---|---|---|
| 0.907033 | 0.015403 | 0.592 | 0.571 | 1 | 15 | Ccl9 | 0.021 | Old |
| 0.908032 | 0.003259 | 0.306 | 0.286 | 1 | 15 | Enox2 | 0.02 | Old |
| 0.908143 | 0.799044 | 0.204 | 0.232 | 1 | 15 | Uri1 | -0.028 | Old |
| 0.9085 | 0.800767 | 0.184 | 0.214 | 1 | 15 | Trp53inp2 | -0.03 | Old |
| 0.909896 | 0.203124 | 0.776 | 0.75 | 1 | 15 | Vcp | 0.026 | Old |
| 0.910142 | 0.590424 | 0.204 | 0.214 | 1 | 15 | Ppp1r21 | -0.01 | Old |
| 0.910435 | 0.637623 | 0.163 | 0.179 | 1 | 15 | Tcf7l2 | -0.016 | Old |
| 0.910477 | 0.024577 | 0.408 | 0.393 | 1 | 15 | Ifi27l2a | 0.015 | Old |
| 0.911283 | 0.517991 | 0.245 | 0.286 | 1 | 15 | Kat2b | -0.041 | Old |
| 0.911293 | 0.882264 | 0.061 | 0.071 | 1 | 15 | Mynn | -0.01 | Old |
| 0.911293 | 0.673862 | 0.061 | 0.071 | 1 | 15 | Hspbap1 | -0.01 | Old |
| 0.911749 | 1.423988 | 0.143 | 0.143 | 1 | 15 | Zcchc24 | 0 | Old |
| 0.912376 | 0.256903 | 0.184 | 0.179 | 1 | 15 | Arid5a | 0.005 | Old |
| 0.912385 | 0.330452 | 0.735 | 0.768 | 1 | 15 | Arl4c | -0.033 | Old |
| 0.913358 | 0.376496 | 0.592 | 0.661 | 1 | 15 | Spred1 | -0.069 | Old |
| 0.913441 | 0.266868 | 0.245 | 0.268 | 1 | 15 | Arl5b | -0.023 | Old |
| 0.913441 | 0.057352 | 0.245 | 0.268 | 1 | 15 | Il15ra | -0.023 | Old |
| 0.914928 | 0.076619 | 0.449 | 0.411 | 1 | 15 | Cdc42se1 | 0.038 | Old |
| 0.915077 | 0.048465 | 0.388 | 0.375 | 1 | 15 | Tle3 | 0.013 | Old |
| 0.915171 | 0.021103 | 0.041 | 0.036 | 1 | 15 | Nrg2 | 0.005 | Old |
| 0.915181 | 0.873624 | 0.204 | 0.232 | 1 | 15 | Ubqln2 | -0.028 | Old |
| 0.915254 | 0.619384 | 0.286 | 0.321 | 1 | 15 | Adam19 | -0.035 | Old |
| 0.915716 | 1.017657 | 0.082 | 0.089 | 1 | 15 | Grsf1 | -0.007 | Old |
| 0.915716 | 0.338827 | 0.082 | 0.089 | 1 | 15 | Dusp10 | -0.007 | Old |
| 0.915716 | 0.150969 | 0.082 | 0.089 | 1 | 15 | Rsad2 | -0.007 | Old |
| 0.917149 | 0.204727 | 0.551 | 0.571 | 1 | 15 | Atp6v1f | -0.02 | Old |
| 0.917799 | 0.60856 | 0.245 | 0.232 | 1 | 15 | Txn2 | 0.013 | Old |
| 0.918189 | 0.275696 | 0.102 | 0.107 | 1 | 15 | 1810030O( | -0.005 | Old |
| 0.918195 | 0.152749 | 0.163 | 0.179 | 1 | 15 | Dhx36 | -0.016 | Old |
| 0.919968 | 0.202234 | 0.184 | 0.179 | 1 | 15 | Bscl2 | 0.005 | Old |
| 0.919968 | 0.181592 | 0.184 | 0.179 | 1 | 15 | Mrpl48 | 0.005 | Old |
| 0.920003 | 0.107998 | 0.224 | 0.232 | 1 | 15 | Rbpms | -0.008 | Old |
| 0.920527 | 0.110445 | 0.49 | 0.5 | 1 | 15 | Cdkn1b | -0.01 | Old |
| 0.92058 | 0.242571 | 0.837 | 0.875 | 1 | 15 | H2-Ab1 | -0.038 | Old |
| 0.920712 | 0.225079 | 0.327 | 0.339 | 1 | 15 | Tnip3 | -0.012 | Old |
| 0.920914 | 0.038734 | 0.388 | 0.375 | 1 | 15 | Nipbl | 0.013 | Old |
| 0.921594 | 0.235049 | 0.184 | 0.196 | 1 | 15 | Ptpa | -0.012 | Old |
| 0.922062 | 0.511666 | 0.367 | 0.339 | 1 | 15 | Rpn1 | 0.028 | Old |
| 0.9223 | 0.12424 | 0.306 | 0.321 | 1 | 15 | Mknk2 | -0.015 | Old |
| 0.922341 | 0.150865 | 0.245 | 0.25 | 1 | 15 | Zc3h15 | -0.005 | Old |
| 0.923047 | 0.935733 | 0.306 | 0.339 | 1 | 15 | Lsm5 | -0.033 | Old |
| 0.923155 | 0.018371 | 0.816 | 0.911 | 1 | 15 | Cox7c | -0.095 | Old |
| 0.924359 | 3.359554 | 0.02 | 0.018 | 1 | 15 | Sell | 0.002 | Old |
| 0.924359 | 3.319417 | 0.02 | 0.018 | 1 | 15 | Cenpq | 0.002 | Old |
| 0.924359 | 3.278007 | 0.02 | 0.018 | 1 | 15 | Diras2 | 0.002 | Old |

| | | | | | | | | |
|---|---|---|---|---|---|---|---|---|
| 0.924359 | 3.143448 | 0.02 | 0.018 | 1 | 15 | Prpf40b | 0.002 | Old |
| 0.924359 | 3.134296 | 0.02 | 0.018 | 1 | 15 | 2900089D1 | 0.002 | Old |
| 0.924359 | 3.024282 | 0.02 | 0.018 | 1 | 15 | Cdca8 | 0.002 | Old |
| 0.924359 | 3.01571 | 0.02 | 0.018 | 1 | 15 | Slc27a3 | 0.002 | Old |
| 0.924359 | 2.781648 | 0.02 | 0.018 | 1 | 15 | Numb | 0.002 | Old |
| 0.924359 | 2.425645 | 0.02 | 0.018 | 1 | 15 | Mapkapk5 | 0.002 | Old |
| 0.924359 | 2.41644 | 0.02 | 0.018 | 1 | 15 | C130050O1 | 0.002 | Old |
| 0.924359 | 2.384973 | 0.02 | 0.018 | 1 | 15 | Pi16 | 0.002 | Old |
| 0.924359 | 2.289009 | 0.02 | 0.018 | 1 | 15 | Gdpd1 | 0.002 | Old |
| 0.924359 | 2.267038 | 0.02 | 0.018 | 1 | 15 | Scara3 | 0.002 | Old |
| 0.924359 | 2.168707 | 0.02 | 0.018 | 1 | 15 | Gm21992 | 0.002 | Old |
| 0.924359 | 2.155348 | 0.02 | 0.018 | 1 | 15 | Ubtd2 | 0.002 | Old |
| 0.924359 | 2.1187 | 0.02 | 0.018 | 1 | 15 | 1700016F1 | 0.002 | Old |
| 0.924359 | 2.034121 | 0.02 | 0.018 | 1 | 15 | Clmp | 0.002 | Old |
| 0.924359 | 2.034121 | 0.02 | 0.018 | 1 | 15 | Vcpkmt | 0.002 | Old |
| 0.924359 | 1.866455 | 0.02 | 0.018 | 1 | 15 | Acat2 | 0.002 | Old |
| 0.924359 | 1.857501 | 0.02 | 0.018 | 1 | 15 | Ccpg1os | 0.002 | Old |
| 0.924359 | 1.853511 | 0.02 | 0.018 | 1 | 15 | Zfp507 | 0.002 | Old |
| 0.924359 | 1.793385 | 0.02 | 0.018 | 1 | 15 | Lss | 0.002 | Old |
| 0.924359 | 1.759101 | 0.02 | 0.018 | 1 | 15 | Cdkn2c | 0.002 | Old |
| 0.924359 | 1.756581 | 0.02 | 0.018 | 1 | 15 | Telo2 | 0.002 | Old |
| 0.924359 | 1.753951 | 0.02 | 0.018 | 1 | 15 | Slc5a6 | 0.002 | Old |
| 0.924359 | 1.686067 | 0.02 | 0.018 | 1 | 15 | Spryd7 | 0.002 | Old |
| 0.924359 | 1.66247 | 0.02 | 0.018 | 1 | 15 | Dnal4 | 0.002 | Old |
| 0.924359 | 1.650189 | 0.02 | 0.018 | 1 | 15 | Satb2 | 0.002 | Old |
| 0.924359 | 1.645655 | 0.02 | 0.018 | 1 | 15 | Hmgxb3 | 0.002 | Old |
| 0.924359 | 1.630864 | 0.02 | 0.018 | 1 | 15 | Tmem191c | 0.002 | Old |
| 0.924359 | 1.601406 | 0.02 | 0.018 | 1 | 15 | Coq6 | 0.002 | Old |
| 0.924359 | 1.588326 | 0.02 | 0.018 | 1 | 15 | Disp1 | 0.002 | Old |
| 0.924359 | 1.584591 | 0.02 | 0.018 | 1 | 15 | Sh3d19 | 0.002 | Old |
| 0.924359 | 1.581315 | 0.02 | 0.018 | 1 | 15 | Plcb4 | 0.002 | Old |
| 0.924359 | 1.575679 | 0.02 | 0.018 | 1 | 15 | 6720489N1 | 0.002 | Old |
| 0.924359 | 1.51775 | 0.02 | 0.018 | 1 | 15 | Zfp719 | 0.002 | Old |
| 0.924359 | 1.513459 | 0.02 | 0.018 | 1 | 15 | Tmem138 | 0.002 | Old |
| 0.924359 | 1.471196 | 0.02 | 0.018 | 1 | 15 | F5 | 0.002 | Old |
| 0.924359 | 1.455346 | 0.02 | 0.018 | 1 | 15 | Havcr1 | 0.002 | Old |
| 0.924359 | 1.43171 | 0.02 | 0.018 | 1 | 15 | Amot | 0.002 | Old |
| 0.924359 | 1.404361 | 0.02 | 0.018 | 1 | 15 | D330050G2 | 0.002 | Old |
| 0.924359 | 1.382814 | 0.02 | 0.018 | 1 | 15 | Fastkd3 | 0.002 | Old |
| 0.924359 | 1.377175 | 0.02 | 0.018 | 1 | 15 | Inafm1 | 0.002 | Old |
| 0.924359 | 1.364361 | 0.02 | 0.018 | 1 | 15 | Wipi1 | 0.002 | Old |
| 0.924359 | 1.338927 | 0.02 | 0.018 | 1 | 15 | Klhl26 | 0.002 | Old |
| 0.924359 | 1.331982 | 0.02 | 0.018 | 1 | 15 | Zfyve28 | 0.002 | Old |
| 0.924359 | 1.331982 | 0.02 | 0.018 | 1 | 15 | Zfp934 | 0.002 | Old |
| 0.924359 | 1.315167 | 0.02 | 0.018 | 1 | 15 | 1700020D0 | 0.002 | Old |

| | | | | | | | | |
|---|---|---|---|---|---|---|---|---|
| 0.924359 | 1.31347 | 0.02 | 0.018 | 1 | 15 | Zfp958 | 0.002 | Old |
| 0.924359 | 1.301485 | 0.02 | 0.018 | 1 | 15 | Gm37759 | 0.002 | Old |
| 0.924359 | 1.300122 | 0.02 | 0.018 | 1 | 15 | Slco3a1 | 0.002 | Old |
| 0.924359 | 1.285308 | 0.02 | 0.018 | 1 | 15 | Spns3 | 0.002 | Old |
| 0.924359 | 1.282754 | 0.02 | 0.018 | 1 | 15 | Nat8l | 0.002 | Old |
| 0.924359 | 1.244543 | 0.02 | 0.018 | 1 | 15 | Tmem17 | 0.002 | Old |
| 0.924359 | 1.240527 | 0.02 | 0.018 | 1 | 15 | Xxylt1 | 0.002 | Old |
| 0.924359 | 1.232107 | 0.02 | 0.018 | 1 | 15 | Fbxo31 | 0.002 | Old |
| 0.924359 | 1.227064 | 0.02 | 0.018 | 1 | 15 | 2510046G1 | 0.002 | Old |
| 0.924359 | 1.225075 | 0.02 | 0.018 | 1 | 15 | Kpna2 | 0.002 | Old |
| 0.924359 | 1.210249 | 0.02 | 0.018 | 1 | 15 | Adat2 | 0.002 | Old |
| 0.924359 | 1.20491 | 0.02 | 0.018 | 1 | 15 | Rab13 | 0.002 | Old |
| 0.924359 | 1.19858 | 0.02 | 0.018 | 1 | 15 | Cbr3 | 0.002 | Old |
| 0.924359 | 1.184347 | 0.02 | 0.018 | 1 | 15 | Sco1 | 0.002 | Old |
| 0.924359 | 1.183989 | 0.02 | 0.018 | 1 | 15 | Zfp870 | 0.002 | Old |
| 0.924359 | 1.177806 | 0.02 | 0.018 | 1 | 15 | Capsl | 0.002 | Old |
| 0.924359 | 1.163099 | 0.02 | 0.018 | 1 | 15 | H1fx | 0.002 | Old |
| 0.924359 | 1.15755 | 0.02 | 0.018 | 1 | 15 | Kif15 | 0.002 | Old |
| 0.924359 | 1.115716 | 0.02 | 0.018 | 1 | 15 | Fam151b | 0.002 | Old |
| 0.924359 | 1.101185 | 0.02 | 0.018 | 1 | 15 | Epop | 0.002 | Old |
| 0.924359 | 1.076227 | 0.02 | 0.018 | 1 | 15 | Gm48604 | 0.002 | Old |
| 0.924359 | 1.068716 | 0.02 | 0.018 | 1 | 15 | Inmt | 0.002 | Old |
| 0.924359 | 1.066196 | 0.02 | 0.018 | 1 | 15 | Zfp146 | 0.002 | Old |
| 0.924359 | 1.061118 | 0.02 | 0.018 | 1 | 15 | A530099J1 | 0.002 | Old |
| 0.924359 | 1.056069 | 0.02 | 0.018 | 1 | 15 | St7 | 0.002 | Old |
| 0.924359 | 1.050606 | 0.02 | 0.018 | 1 | 15 | Tlcd2 | 0.002 | Old |
| 0.924359 | 1.048737 | 0.02 | 0.018 | 1 | 15 | Coq8a | 0.002 | Old |
| 0.924359 | 1.01333 | 0.02 | 0.018 | 1 | 15 | Gm44751 | 0.002 | Old |
| 0.924359 | 1.01333 | 0.02 | 0.018 | 1 | 15 | Trub1 | 0.002 | Old |
| 0.924359 | 1.012536 | 0.02 | 0.018 | 1 | 15 | Htr1b | 0.002 | Old |
| 0.924359 | 1.002991 | 0.02 | 0.018 | 1 | 15 | 3110001I2 | 0.002 | Old |
| 0.924359 | 1.001101 | 0.02 | 0.018 | 1 | 15 | Etfrf1 | 0.002 | Old |
| 0.924359 | 0.978867 | 0.02 | 0.018 | 1 | 15 | Dpep1 | 0.002 | Old |
| 0.924359 | 0.969332 | 0.02 | 0.018 | 1 | 15 | Bco2 | 0.002 | Old |
| 0.924359 | 0.967649 | 0.02 | 0.018 | 1 | 15 | Nnt | 0.002 | Old |
| 0.924359 | 0.967252 | 0.02 | 0.018 | 1 | 15 | Repin1 | 0.002 | Old |
| 0.924359 | 0.965138 | 0.02 | 0.018 | 1 | 15 | Zcchc3 | 0.002 | Old |
| 0.924359 | 0.965138 | 0.02 | 0.018 | 1 | 15 | Aldh18a1 | 0.002 | Old |
| 0.924359 | 0.935806 | 0.02 | 0.018 | 1 | 15 | Mettl25 | 0.002 | Old |
| 0.924359 | 0.93479 | 0.02 | 0.018 | 1 | 15 | Nemp2 | 0.002 | Old |
| 0.924359 | 0.926188 | 0.02 | 0.018 | 1 | 15 | Ulk3 | 0.002 | Old |
| 0.924359 | 0.925374 | 0.02 | 0.018 | 1 | 15 | Ppif | 0.002 | Old |
| 0.924359 | 0.924922 | 0.02 | 0.018 | 1 | 15 | Rab39b | 0.002 | Old |
| 0.924359 | 0.919659 | 0.02 | 0.018 | 1 | 15 | Zscan22 | 0.002 | Old |
| 0.924359 | 0.894206 | 0.02 | 0.018 | 1 | 15 | Asb4 | 0.002 | Old |

| | | | | | | | | |
|---|---|---|---|---|---|---|---|---|
| 0.924359 | 0.891895 | 0.02 | 0.018 | 1 | 15 | Smg8 | 0.002 | Old |
| 0.924359 | 0.885816 | 0.02 | 0.018 | 1 | 15 | Sdr39u1 | 0.002 | Old |
| 0.924359 | 0.885183 | 0.02 | 0.018 | 1 | 15 | Mrps2 | 0.002 | Old |
| 0.924359 | 0.885183 | 0.02 | 0.018 | 1 | 15 | Rffl | 0.002 | Old |
| 0.924359 | 0.879927 | 0.02 | 0.018 | 1 | 15 | Polr1b | 0.002 | Old |
| 0.924359 | 0.868368 | 0.02 | 0.018 | 1 | 15 | Mrvi1 | 0.002 | Old |
| 0.924359 | 0.868368 | 0.02 | 0.018 | 1 | 15 | Cdc42bpg | 0.002 | Old |
| 0.924359 | 0.86794 | 0.02 | 0.018 | 1 | 15 | Tor4a | 0.002 | Old |
| 0.924359 | 0.852539 | 0.02 | 0.018 | 1 | 15 | Ppp1r3e | 0.002 | Old |
| 0.924359 | 0.847695 | 0.02 | 0.018 | 1 | 15 | Gm17089 | 0.002 | Old |
| 0.924359 | 0.839417 | 0.02 | 0.018 | 1 | 15 | Enpp1 | 0.002 | Old |
| 0.924359 | 0.837951 | 0.02 | 0.018 | 1 | 15 | Psmg2 | 0.002 | Old |
| 0.924359 | 0.825879 | 0.02 | 0.018 | 1 | 15 | Cep290 | 0.002 | Old |
| 0.924359 | 0.81459 | 0.02 | 0.018 | 1 | 15 | BC002059 | 0.002 | Old |
| 0.924359 | 0.805078 | 0.02 | 0.018 | 1 | 15 | Anxa9 | 0.002 | Old |
| 0.924359 | 0.789312 | 0.02 | 0.018 | 1 | 15 | Znhit3 | 0.002 | Old |
| 0.924359 | 0.763996 | 0.02 | 0.018 | 1 | 15 | Gng11 | 0.002 | Old |
| 0.924359 | 0.751349 | 0.02 | 0.018 | 1 | 15 | Fzd7 | 0.002 | Old |
| 0.924359 | 0.737572 | 0.02 | 0.018 | 1 | 15 | Hps6 | 0.002 | Old |
| 0.924359 | 0.696718 | 0.02 | 0.018 | 1 | 15 | Adck2 | 0.002 | Old |
| 0.924359 | 0.696718 | 0.02 | 0.018 | 1 | 15 | Lhfpl2 | 0.002 | Old |
| 0.924359 | 0.692639 | 0.02 | 0.018 | 1 | 15 | Cd8a | 0.002 | Old |
| 0.924359 | 0.649415 | 0.02 | 0.018 | 1 | 15 | Tnfrsf17 | 0.002 | Old |
| 0.924359 | 0.647858 | 0.02 | 0.018 | 1 | 15 | Amer1 | 0.002 | Old |
| 0.924359 | 0.636768 | 0.02 | 0.018 | 1 | 15 | Cntrob | 0.002 | Old |
| 0.924359 | 0.634649 | 0.02 | 0.018 | 1 | 15 | Dip2c | 0.002 | Old |
| 0.924359 | 0.624782 | 0.02 | 0.018 | 1 | 15 | Tlr3 | 0.002 | Old |
| 0.924359 | 0.623435 | 0.02 | 0.018 | 1 | 15 | Mgmt | 0.002 | Old |
| 0.924359 | 0.619643 | 0.02 | 0.018 | 1 | 15 | Mpp2 | 0.002 | Old |
| 0.924359 | 0.617706 | 0.02 | 0.018 | 1 | 15 | Fancd2 | 0.002 | Old |
| 0.924359 | 0.616948 | 0.02 | 0.018 | 1 | 15 | Rbak | 0.002 | Old |
| 0.924359 | 0.614159 | 0.02 | 0.018 | 1 | 15 | Slc13a3 | 0.002 | Old |
| 0.924359 | 0.610337 | 0.02 | 0.018 | 1 | 15 | Dmpk | 0.002 | Old |
| 0.924359 | 0.60299 | 0.02 | 0.018 | 1 | 15 | Fdxr | 0.002 | Old |
| 0.924359 | 0.60299 | 0.02 | 0.018 | 1 | 15 | Pigf | 0.002 | Old |
| 0.924359 | 0.601085 | 0.02 | 0.018 | 1 | 15 | Ctu1 | 0.002 | Old |
| 0.924359 | 0.601085 | 0.02 | 0.018 | 1 | 15 | Gpt2 | 0.002 | Old |
| 0.924359 | 0.567557 | 0.02 | 0.018 | 1 | 15 | Nid2 | 0.002 | Old |
| 0.924359 | 0.566103 | 0.02 | 0.018 | 1 | 15 | Dhx58 | 0.002 | Old |
| 0.924359 | 0.562443 | 0.02 | 0.018 | 1 | 15 | Ror1 | 0.002 | Old |
| 0.924359 | 0.559671 | 0.02 | 0.018 | 1 | 15 | Lamb3 | 0.002 | Old |
| 0.924359 | 0.559671 | 0.02 | 0.018 | 1 | 15 | Ccdc14 | 0.002 | Old |
| 0.924359 | 0.546648 | 0.02 | 0.018 | 1 | 15 | Slc6a8 | 0.002 | Old |
| 0.924359 | 0.537677 | 0.02 | 0.018 | 1 | 15 | Adtrp | 0.002 | Old |
| 0.924359 | 0.534775 | 0.02 | 0.018 | 1 | 15 | Rrp1b | 0.002 | Old |

| | | | | | | | | |
|---|---|---|---|---|---|---|---|---|
| 0.924359 | 0.534716 | 0.02 | 0.018 | 1 | 15 | Nsl1 | 0.002 | Old |
| 0.924359 | 0.534716 | 0.02 | 0.018 | 1 | 15 | Gm26778 | 0.002 | Old |
| 0.924359 | 0.534716 | 0.02 | 0.018 | 1 | 15 | Lsmem1 | 0.002 | Old |
| 0.924359 | 0.529127 | 0.02 | 0.018 | 1 | 15 | Aldh1a3 | 0.002 | Old |
| 0.924359 | 0.524907 | 0.02 | 0.018 | 1 | 15 | Pwp2 | 0.002 | Old |
| 0.924359 | 0.516465 | 0.02 | 0.018 | 1 | 15 | Tmem218 | 0.002 | Old |
| 0.924359 | 0.507217 | 0.02 | 0.018 | 1 | 15 | Gm12981 | 0.002 | Old |
| 0.924359 | 0.505855 | 0.02 | 0.018 | 1 | 15 | Hdac8 | 0.002 | Old |
| 0.924359 | 0.504146 | 0.02 | 0.018 | 1 | 15 | Alg6 | 0.002 | Old |
| 0.924359 | 0.49965 | 0.02 | 0.018 | 1 | 15 | Spindoc | 0.002 | Old |
| 0.924359 | 0.491379 | 0.02 | 0.018 | 1 | 15 | Hexdc | 0.002 | Old |
| 0.924359 | 0.488726 | 0.02 | 0.018 | 1 | 15 | Gpatch4 | 0.002 | Old |
| 0.924359 | 0.48608 | 0.02 | 0.018 | 1 | 15 | Rtp4 | 0.002 | Old |
| 0.924359 | 0.485032 | 0.02 | 0.018 | 1 | 15 | Lipt2 | 0.002 | Old |
| 0.924359 | 0.485032 | 0.02 | 0.018 | 1 | 15 | 1700052K1 | 0.002 | Old |
| 0.924359 | 0.484015 | 0.02 | 0.018 | 1 | 15 | Zfp64 | 0.002 | Old |
| 0.924359 | 0.484015 | 0.02 | 0.018 | 1 | 15 | Tbkbp1 | 0.002 | Old |
| 0.924359 | 0.468217 | 0.02 | 0.018 | 1 | 15 | Dusp18 | 0.002 | Old |
| 0.924359 | 0.461409 | 0.02 | 0.018 | 1 | 15 | Endov | 0.002 | Old |
| 0.924359 | 0.457878 | 0.02 | 0.018 | 1 | 15 | Bend3 | 0.002 | Old |
| 0.924359 | 0.456358 | 0.02 | 0.018 | 1 | 15 | Gm29530 | 0.002 | Old |
| 0.924359 | 0.456358 | 0.02 | 0.018 | 1 | 15 | Selenoo | 0.002 | Old |
| 0.924359 | 0.453434 | 0.02 | 0.018 | 1 | 15 | Rassf7 | 0.002 | Old |
| 0.924359 | 0.448407 | 0.02 | 0.018 | 1 | 15 | Nemp1 | 0.002 | Old |
| 0.924359 | 0.443231 | 0.02 | 0.018 | 1 | 15 | Dsel | 0.002 | Old |
| 0.924359 | 0.437935 | 0.02 | 0.018 | 1 | 15 | Peak1os | 0.002 | Old |
| 0.924359 | 0.435047 | 0.02 | 0.018 | 1 | 15 | Plac8 | 0.002 | Old |
| 0.924359 | 0.415343 | 0.02 | 0.018 | 1 | 15 | Gins3 | 0.002 | Old |
| 0.924359 | 0.396157 | 0.02 | 0.018 | 1 | 15 | 9430078K2 | 0.002 | Old |
| 0.924359 | 0.395277 | 0.02 | 0.018 | 1 | 15 | Mill2 | 0.002 | Old |
| 0.924359 | 0.385333 | 0.02 | 0.018 | 1 | 15 | Ildr1 | 0.002 | Old |
| 0.924359 | 0.382577 | 0.02 | 0.018 | 1 | 15 | Tatdn1 | 0.002 | Old |
| 0.924359 | 0.355022 | 0.02 | 0.018 | 1 | 15 | Frmd5 | 0.002 | Old |
| 0.924359 | 0.349365 | 0.02 | 0.018 | 1 | 15 | Rhbdf1 | 0.002 | Old |
| 0.924359 | 0.346021 | 0.02 | 0.018 | 1 | 15 | Gm45187 | 0.002 | Old |
| 0.924359 | 0.345561 | 0.02 | 0.018 | 1 | 15 | Maml3 | 0.002 | Old |
| 0.924359 | 0.345561 | 0.02 | 0.018 | 1 | 15 | Abhd11 | 0.002 | Old |
| 0.924359 | 0.334565 | 0.02 | 0.018 | 1 | 15 | C1qtnf12 | 0.002 | Old |
| 0.924359 | 0.331928 | 0.02 | 0.018 | 1 | 15 | 1110002J0 | 0.002 | Old |
| 0.924359 | 0.331928 | 0.02 | 0.018 | 1 | 15 | Alg3 | 0.002 | Old |
| 0.924359 | 0.326538 | 0.02 | 0.018 | 1 | 15 | Tanc1 | 0.002 | Old |
| 0.924359 | 0.31845 | 0.02 | 0.018 | 1 | 15 | Zbtb2 | 0.002 | Old |
| 0.924359 | 0.318067 | 0.02 | 0.018 | 1 | 15 | Exo5 | 0.002 | Old |
| 0.924359 | 0.300991 | 0.02 | 0.018 | 1 | 15 | Cdc42bpa | 0.002 | Old |
| 0.924359 | 0.284565 | 0.02 | 0.018 | 1 | 15 | Map10 | 0.002 | Old |

| | | | | | | | | |
|---|---|---|---|---|---|---|---|---|
| 0.924359 | 0.268766 | 0.02 | 0.018 | 1 | 15 | Tnfsf13os | 0.002 | Old |
| 0.924359 | 0.258306 | 0.02 | 0.018 | 1 | 15 | Dgcr14 | 0.002 | Old |
| 0.924359 | 0.253733 | 0.02 | 0.018 | 1 | 15 | 2610021A( | 0.002 | Old |
| 0.924359 | 0.245146 | 0.02 | 0.018 | 1 | 15 | Mfsd4b4 | 0.002 | Old |
| 0.924359 | 0.187422 | 0.02 | 0.018 | 1 | 15 | Zcchc14 | 0.002 | Old |
| 0.924359 | 0.186909 | 0.02 | 0.018 | 1 | 15 | Zfp169 | 0.002 | Old |
| 0.924359 | 0.179507 | 0.02 | 0.018 | 1 | 15 | Megf9 | 0.002 | Old |
| 0.924359 | 0.179507 | 0.02 | 0.018 | 1 | 15 | Tepsin | 0.002 | Old |
| 0.924359 | 0.177982 | 0.02 | 0.018 | 1 | 15 | Toe1 | 0.002 | Old |
| 0.924359 | 0.177982 | 0.02 | 0.018 | 1 | 15 | Fam117a | 0.002 | Old |
| 0.924359 | 0.177555 | 0.02 | 0.018 | 1 | 15 | B3galnt1 | 0.002 | Old |
| 0.924359 | 0.177555 | 0.02 | 0.018 | 1 | 15 | Pon3 | 0.002 | Old |
| 0.924359 | 0.177555 | 0.02 | 0.018 | 1 | 15 | Cep112 | 0.002 | Old |
| 0.924359 | 0.16638 | 0.02 | 0.018 | 1 | 15 | Gk5 | 0.002 | Old |
| 0.924359 | 0.16638 | 0.02 | 0.018 | 1 | 15 | Fam20a | 0.002 | Old |
| 0.924359 | 0.162692 | 0.02 | 0.018 | 1 | 15 | Mterf1a | 0.002 | Old |
| 0.924359 | 0.149692 | 0.02 | 0.018 | 1 | 15 | 1700041G1 | 0.002 | Old |
| 0.924359 | 0.14611 | 0.02 | 0.018 | 1 | 15 | Eya4 | 0.002 | Old |
| 0.924359 | 0.14611 | 0.02 | 0.018 | 1 | 15 | Gm41307 | 0.002 | Old |
| 0.924359 | 0.135374 | 0.02 | 0.018 | 1 | 15 | AW146154 | 0.002 | Old |
| 0.924359 | 0.135374 | 0.02 | 0.018 | 1 | 15 | Hist1h2ap | 0.002 | Old |
| 0.924388 | 0.039692 | 0.673 | 0.661 | 1 | 15 | Plekhb2 | 0.012 | Old |
| 0.925051 | 0.315809 | 0.347 | 0.357 | 1 | 15 | Tap2 | -0.01 | Old |
| 0.925962 | 0.096041 | 0.163 | 0.179 | 1 | 15 | Stard8 | -0.016 | Old |
| 0.926219 | 0.789222 | 0.082 | 0.089 | 1 | 15 | Prkag2 | -0.007 | Old |
| 0.926219 | 0.478265 | 0.082 | 0.089 | 1 | 15 | Golph3l | -0.007 | Old |
| 0.926219 | 0.276212 | 0.082 | 0.089 | 1 | 15 | Slc11a1 | -0.007 | Old |
| 0.926292 | 0.176679 | 0.469 | 0.536 | 1 | 15 | Ktn1 | -0.067 | Old |
| 0.926742 | 0.1637 | 0.204 | 0.196 | 1 | 15 | Rbm26 | 0.008 | Old |
| 0.926939 | 0.160727 | 0.224 | 0.232 | 1 | 15 | Pgrmc1 | -0.008 | Old |
| 0.927771 | 0.373449 | 0.306 | 0.304 | 1 | 15 | Gpbp1l1 | 0.002 | Old |
| 0.927786 | 0.880549 | 0.102 | 0.107 | 1 | 15 | Snx13 | -0.005 | Old |
| 0.927786 | 0.695047 | 0.102 | 0.107 | 1 | 15 | Atxn10 | -0.005 | Old |
| 0.927786 | 0.32199 | 0.102 | 0.107 | 1 | 15 | Kctd18 | -0.005 | Old |
| 0.928181 | 0.14444 | 0.163 | 0.161 | 1 | 15 | 1810013L2 | 0.002 | Old |
| 0.928421 | 0.254198 | 0.449 | 0.518 | 1 | 15 | Zfp207 | -0.069 | Old |
| 0.928438 | 1.450404 | 0.122 | 0.125 | 1 | 15 | Ddx1 | -0.003 | Old |
| 0.928438 | 0.444905 | 0.122 | 0.125 | 1 | 15 | Phf1 | -0.003 | Old |
| 0.928508 | 1.076607 | 0.143 | 0.143 | 1 | 15 | Slc30a9 | 0 | Old |
| 0.928508 | 0.172261 | 0.143 | 0.143 | 1 | 15 | Ppp1r7 | 0 | Old |
| 0.928649 | 0.196297 | 0.408 | 0.429 | 1 | 15 | Mfhas1 | -0.021 | Old |
| 0.929041 | 0.230898 | 0.184 | 0.196 | 1 | 15 | Klhl28 | -0.012 | Old |
| 0.929041 | 0.214833 | 0.184 | 0.196 | 1 | 15 | Fam168a | -0.012 | Old |
| 0.929041 | 0.085915 | 0.184 | 0.196 | 1 | 15 | Rasal2 | -0.012 | Old |
| 0.929997 | 0.020912 | 0.714 | 0.696 | 1 | 15 | Colgalt1 | 0.018 | Old |

| | | | | | | | | |
|---|---|---|---|---|---|---|---|---|
| 0.930427 | 0.680676 | 0.143 | 0.161 | 1 | 15 | Cacybp | -0.018 | Old |
| 0.930427 | 0.561718 | 0.143 | 0.161 | 1 | 15 | Cnpy3 | -0.018 | Old |
| 0.93045 | 0.369609 | 0.327 | 0.357 | 1 | 15 | Laptm4a | -0.03 | Old |
| 0.930942 | 0.43519 | 0.265 | 0.268 | 1 | 15 | Metap2 | -0.003 | Old |
| 0.932584 | 0.138401 | 0.286 | 0.286 | 1 | 15 | Ighm | 0 | Old |
| 0.933391 | 0.046128 | 0.857 | 0.893 | 1 | 15 | Efhd2 | -0.036 | Old |
| 0.933517 | 0.053735 | 0.408 | 0.393 | 1 | 15 | Pebp1 | 0.015 | Old |
| 0.93388 | 0.30584 | 0.224 | 0.232 | 1 | 15 | Cct7 | -0.008 | Old |
| 0.934051 | 0.673363 | 0.204 | 0.196 | 1 | 15 | Micu1 | 0.008 | Old |
| 0.934792 | 0.153912 | 0.673 | 0.661 | 1 | 15 | Gnb2 | 0.012 | Old |
| 0.935814 | 0.171374 | 0.245 | 0.25 | 1 | 15 | Prkx | -0.005 | Old |
| 0.935814 | 0.094536 | 0.245 | 0.25 | 1 | 15 | Supt20 | -0.005 | Old |
| 0.935814 | 0.070993 | 0.245 | 0.25 | 1 | 15 | Thap2 | -0.005 | Old |
| 0.936494 | 0.682575 | 0.184 | 0.196 | 1 | 15 | Parp4 | -0.012 | Old |
| 0.936736 | 0.044102 | 0.082 | 0.089 | 1 | 15 | Lpar3 | -0.007 | Old |
| 0.9369 | 1.296952 | 0.143 | 0.143 | 1 | 15 | Pqlc1 | 0 | Old |
| 0.937364 | 0.779902 | 0.122 | 0.125 | 1 | 15 | Dennd1a | -0.003 | Old |
| 0.937364 | 0.232945 | 0.122 | 0.125 | 1 | 15 | Clcn5 | -0.003 | Old |
| 0.937364 | 0.03712 | 0.122 | 0.125 | 1 | 15 | Rbl1 | -0.003 | Old |
| 0.937393 | 0.595896 | 0.102 | 0.107 | 1 | 15 | Polr2i | -0.005 | Old |
| 0.937504 | 0.086624 | 0.265 | 0.268 | 1 | 15 | Usp4 | -0.003 | Old |
| 0.938449 | 0.424653 | 0.367 | 0.393 | 1 | 15 | Ly86 | -0.026 | Old |
| 0.938991 | 0.308391 | 0.286 | 0.286 | 1 | 15 | Nfkbie | 0 | Old |
| 0.940392 | 0.304937 | 0.429 | 0.429 | 1 | 15 | Pdcd10 | 0 | Old |
| 0.940881 | 0.036318 | 0.837 | 0.768 | 1 | 15 | Tapbp | 0.069 | Old |
| 0.941342 | 0.040022 | 0.449 | 0.464 | 1 | 15 | Dhx15 | -0.015 | Old |
| 0.941517 | 0.292717 | 0.163 | 0.179 | 1 | 15 | Eif4e | -0.016 | Old |
| 0.942838 | 0.47414 | 0.286 | 0.304 | 1 | 15 | Tiam1 | -0.018 | Old |
| 0.943952 | 0.194806 | 0.184 | 0.196 | 1 | 15 | Smarcc2 | -0.012 | Old |
| 0.944071 | 0.498466 | 0.265 | 0.268 | 1 | 15 | Klf13 | -0.003 | Old |
| 0.944071 | 0.181685 | 0.265 | 0.268 | 1 | 15 | Napa | -0.003 | Old |
| 0.944111 | 0.519049 | 0.163 | 0.161 | 1 | 15 | Ap1s1 | 0.002 | Old |
| 0.945299 | 0.727045 | 0.143 | 0.143 | 1 | 15 | Casp1 | 0 | Old |
| 0.945299 | 0.713516 | 0.143 | 0.143 | 1 | 15 | Naxd | 0 | Old |
| 0.945299 | 0.146362 | 0.143 | 0.143 | 1 | 15 | Abhd16a | 0 | Old |
| 0.945931 | 0.14818 | 0.02 | 0.018 | 1 | 15 | Ccdc58 | 0.002 | Old |
| 0.945931 | 0.129192 | 0.02 | 0.018 | 1 | 15 | Btla | 0.002 | Old |
| 0.945931 | 0.122314 | 0.02 | 0.018 | 1 | 15 | Gm29282 | 0.002 | Old |
| 0.945931 | 0.119706 | 0.02 | 0.018 | 1 | 15 | Treml4 | 0.002 | Old |
| 0.945931 | 0.116892 | 0.02 | 0.018 | 1 | 15 | Tlx2 | 0.002 | Old |
| 0.945931 | 0.105951 | 0.02 | 0.018 | 1 | 15 | Zbtb3 | 0.002 | Old |
| 0.945931 | 0.093192 | 0.02 | 0.018 | 1 | 15 | Svip | 0.002 | Old |
| 0.945931 | 0.093192 | 0.02 | 0.018 | 1 | 15 | Coil | 0.002 | Old |
| 0.945931 | 0.093192 | 0.02 | 0.018 | 1 | 15 | Clec16a | 0.002 | Old |
| 0.945931 | 0.090583 | 0.02 | 0.018 | 1 | 15 | Ccdc181 | 0.002 | Old |

| | | | | | | | | |
|---|---|---|---|---|---|---|---|---|
| 0.945931 | 0.080224 | 0.02 | 0.018 | 1 | 15 | Card11 | 0.002 | Old |
| 0.945931 | 0.080224 | 0.02 | 0.018 | 1 | 15 | Arhgap27o | 0.002 | Old |
| 0.945931 | 0.074941 | 0.02 | 0.018 | 1 | 15 | Zfp157 | 0.002 | Old |
| 0.945931 | 0.059396 | 0.02 | 0.018 | 1 | 15 | Abcb8 | 0.002 | Old |
| 0.945931 | 0.043724 | 0.02 | 0.018 | 1 | 15 | F10 | 0.002 | Old |
| 0.945931 | 0.043724 | 0.02 | 0.018 | 1 | 15 | Gorasp1 | 0.002 | Old |
| 0.945931 | 0.043508 | 0.02 | 0.018 | 1 | 15 | Spaca9 | 0.002 | Old |
| 0.945931 | 0.035214 | 0.02 | 0.018 | 1 | 15 | Ifi205 | 0.002 | Old |
| 0.945931 | 0.035066 | 0.02 | 0.018 | 1 | 15 | Simc1 | 0.002 | Old |
| 0.945931 | 0.028899 | 0.02 | 0.018 | 1 | 15 | Unc13d | 0.002 | Old |
| 0.945931 | 0.013141 | 0.02 | 0.018 | 1 | 15 | Cmss1 | 0.002 | Old |
| 0.945931 | 0.012713 | 0.02 | 0.018 | 1 | 15 | Tsacc | 0.002 | Old |
| 0.945931 | 0.005417 | 0.02 | 0.018 | 1 | 15 | Bmp1 | 0.002 | Old |
| 0.945931 | 0.002641 | 0.02 | 0.018 | 1 | 15 | Rapgef5 | 0.002 | Old |
| 0.946012 | 0.832056 | 0.204 | 0.214 | 1 | 15 | Hist3h2a | -0.01 | Old |
| 0.94658 | 0.360387 | 0.286 | 0.321 | 1 | 15 | Cyb5r4 | -0.035 | Old |
| 0.947009 | 1.090973 | 0.102 | 0.107 | 1 | 15 | Zfp330 | -0.005 | Old |
| 0.947009 | 0.345935 | 0.102 | 0.107 | 1 | 15 | Vgll4 | -0.005 | Old |
| 0.947009 | 0.09315 | 0.102 | 0.107 | 1 | 15 | Creld2 | -0.005 | Old |
| 0.947009 | 0.060534 | 0.102 | 0.107 | 1 | 15 | Rif1 | -0.005 | Old |
| 0.947263 | 1.418307 | 0.082 | 0.089 | 1 | 15 | Lyl1 | -0.007 | Old |
| 0.947263 | 1.339792 | 0.082 | 0.089 | 1 | 15 | Entpd7 | -0.007 | Old |
| 0.947263 | 0.565093 | 0.082 | 0.089 | 1 | 15 | Mbd5 | -0.007 | Old |
| 0.947425 | 0.051859 | 0.49 | 0.464 | 1 | 15 | Mrpl14 | 0.026 | Old |
| 0.947628 | 0.652714 | 0.306 | 0.339 | 1 | 15 | Mef2d | -0.033 | Old |
| 0.948564 | 0.135751 | 0.347 | 0.339 | 1 | 15 | Copz1 | 0.008 | Old |
| 0.948683 | 0.35383 | 0.204 | 0.196 | 1 | 15 | Inpp4a | 0.008 | Old |
| 0.948994 | 0.254618 | 0.347 | 0.357 | 1 | 15 | Srsf7 | -0.01 | Old |
| 0.949307 | 1.014937 | 0.245 | 0.25 | 1 | 15 | Per1 | -0.005 | Old |
| 0.949402 | 0.206714 | 0.347 | 0.375 | 1 | 15 | Srsf9 | -0.028 | Old |
| 0.950226 | 0.323205 | 0.306 | 0.321 | 1 | 15 | Arid4b | -0.015 | Old |
| 0.951991 | 0.265684 | 0.367 | 0.339 | 1 | 15 | Rcsd1 | 0.028 | Old |
| 0.952256 | 0.053031 | 0.469 | 0.429 | 1 | 15 | Ptges3 | 0.04 | Old |
| 0.953201 | 0.544931 | 0.204 | 0.214 | 1 | 15 | Rrad | -0.01 | Old |
| 0.953995 | 0.520939 | 0.224 | 0.214 | 1 | 15 | Med15 | 0.01 | Old |
| 0.953995 | 0.006792 | 0.224 | 0.214 | 1 | 15 | Slc25a10 | 0.01 | Old |
| 0.954067 | 0.1427 | 0.531 | 0.536 | 1 | 15 | Skap2 | -0.005 | Old |
| 0.954566 | 0.268943 | 0.408 | 0.446 | 1 | 15 | Rsrc2 | -0.038 | Old |
| 0.955105 | 0.03357 | 0.592 | 0.625 | 1 | 15 | Nfe2l1 | -0.033 | Old |
| 0.955237 | 0.655818 | 0.122 | 0.125 | 1 | 15 | Mrpl10 | -0.003 | Old |
| 0.95589 | 0.057952 | 0.673 | 0.732 | 1 | 15 | Rab8b | -0.059 | Old |
| 0.956633 | 0.345407 | 0.102 | 0.107 | 1 | 15 | Runx2 | -0.005 | Old |
| 0.956633 | 0.035848 | 0.102 | 0.107 | 1 | 15 | Inppl1 | -0.005 | Old |
| 0.957799 | 1.71592 | 0.082 | 0.089 | 1 | 15 | Slc31a2 | -0.007 | Old |
| 0.957799 | 1.563467 | 0.082 | 0.089 | 1 | 15 | Tcp11l2 | -0.007 | Old |

| | | | | | | | | |
|---|---|---|---|---|---|---|---|---|
| 0.957799 | 0.63144 | 0.082 | 0.089 | 1 | 15 | Axin1 | -0.007 | Old |
| 0.958028 | 0.089622 | 0.184 | 0.179 | 1 | 15 | Hmg20b | 0.005 | Old |
| 0.958083 | 0.053427 | 0.449 | 0.464 | 1 | 15 | Lfng | -0.015 | Old |
| 0.958882 | 0.541715 | 0.184 | 0.196 | 1 | 15 | Mtmr12 | -0.012 | Old |
| 0.959602 | 0.148524 | 0.551 | 0.536 | 1 | 15 | Adora2a | 0.015 | Old |
| 0.960063 | 0.126853 | 0.163 | 0.161 | 1 | 15 | Tmem147 | 0.002 | Old |
| 0.960395 | 0.724236 | 0.204 | 0.214 | 1 | 15 | Usp22 | -0.01 | Old |
| 0.960847 | 0.051727 | 0.653 | 0.679 | 1 | 15 | Tma7 | -0.026 | Old |
| 0.961075 | 0.075628 | 0.694 | 0.714 | 1 | 15 | Psma7 | -0.02 | Old |
| 0.961497 | 0.045111 | 0.816 | 0.857 | 1 | 15 | Dusp5 | -0.041 | Old |
| 0.961578 | 0.068214 | 0.959 | 0.982 | 1 | 15 | Myl6 | -0.023 | Old |
| 0.962115 | 0.741229 | 0.143 | 0.143 | 1 | 15 | Htatsf1 | 0 | Old |
| 0.962115 | 0.310914 | 0.143 | 0.143 | 1 | 15 | Cul5 | 0 | Old |
| 0.962813 | 0.384365 | 0.245 | 0.25 | 1 | 15 | Brd1 | -0.005 | Old |
| 0.962813 | 0.155834 | 0.245 | 0.25 | 1 | 15 | Klf9 | -0.005 | Old |
| 0.963794 | 0.164115 | 0.265 | 0.268 | 1 | 15 | Tiparp | -0.003 | Old |
| 0.964183 | 0.737426 | 0.122 | 0.125 | 1 | 15 | Dhrs7b | -0.003 | Old |
| 0.965204 | 0.258682 | 0.551 | 0.571 | 1 | 15 | Dync1h1 | -0.02 | Old |
| 0.965419 | 0.167121 | 0.306 | 0.304 | 1 | 15 | Ccnt1 | 0.002 | Old |
| 0.965654 | 0.385825 | 0.184 | 0.179 | 1 | 15 | Snx4 | 0.005 | Old |
| 0.965654 | 0.097624 | 0.184 | 0.179 | 1 | 15 | Crlf2 | 0.005 | Old |
| 0.965699 | 0.194671 | 0.245 | 0.232 | 1 | 15 | Traf3 | 0.013 | Old |
| 0.965699 | 0.024742 | 0.245 | 0.232 | 1 | 15 | Dennd1b | 0.013 | Old |
| 0.966118 | 0.082059 | 0.673 | 0.679 | 1 | 15 | Arhgdib | -0.006 | Old |
| 0.966264 | 0.18861 | 0.102 | 0.107 | 1 | 15 | Sephs1 | -0.005 | Old |
| 0.966264 | 0.093267 | 0.102 | 0.107 | 1 | 15 | Pogz | -0.005 | Old |
| 0.966264 | 0.05863 | 0.102 | 0.107 | 1 | 15 | Snapin | -0.005 | Old |
| 0.966658 | 0.080605 | 0.837 | 0.893 | 1 | 15 | Srgn | -0.056 | Old |
| 0.966693 | 0.031002 | 0.918 | 0.946 | 1 | 15 | Rpl28 | -0.028 | Old |
| 0.966704 | 0.063252 | 0.347 | 0.339 | 1 | 15 | Kras | 0.008 | Old |
| 0.968046 | 0.210687 | 0.163 | 0.161 | 1 | 15 | AI413582 | 0.002 | Old |
| 0.968046 | 0.199185 | 0.163 | 0.161 | 1 | 15 | Hdac2 | 0.002 | Old |
| 0.968046 | 0.16387 | 0.163 | 0.161 | 1 | 15 | Acaa1a | 0.002 | Old |
| 0.968652 | 0.374143 | 0.224 | 0.232 | 1 | 15 | Lin7c | -0.008 | Old |
| 0.968652 | 0.255804 | 0.224 | 0.232 | 1 | 15 | Clip1 | -0.008 | Old |
| 0.970373 | 0.111505 | 0.265 | 0.268 | 1 | 15 | Sec61a1 | -0.003 | Old |
| 0.970451 | 0.096678 | 0.367 | 0.375 | 1 | 15 | Slc26a2 | -0.008 | Old |
| 0.970464 | 0.096901 | 0.571 | 0.536 | 1 | 15 | Pitpna | 0.035 | Old |
| 0.970529 | 0.93726 | 0.143 | 0.143 | 1 | 15 | Mphosph8 | 0 | Old |
| 0.970529 | 0.374437 | 0.143 | 0.143 | 1 | 15 | Ap1ar | 0 | Old |
| 0.970662 | 0.205341 | 0.204 | 0.196 | 1 | 15 | Adgrg6 | 0.008 | Old |
| 0.970737 | 0.549278 | 0.265 | 0.286 | 1 | 15 | Ptpn2 | -0.021 | Old |
| 0.971819 | 0.02247 | 0.959 | 0.982 | 1 | 15 | Rpl13 | -0.023 | Old |
| 0.973133 | 1.419057 | 0.122 | 0.125 | 1 | 15 | Elmo1 | -0.003 | Old |
| 0.973133 | 1.130846 | 0.122 | 0.125 | 1 | 15 | Mmp25 | -0.003 | Old |

| | | | | | | | | |
|---|---|---|---|---|---|---|---|---|
| 0.973133 | 0.737735 | 0.122 | 0.125 | 1 | 15 | Mtf1 | -0.003 | Old |
| 0.973133 | 0.375829 | 0.122 | 0.125 | 1 | 15 | Eed | -0.003 | Old |
| 0.973133 | 0.349876 | 0.122 | 0.125 | 1 | 15 | Slc4a2 | -0.003 | Old |
| 0.973133 | 0.342864 | 0.122 | 0.125 | 1 | 15 | Igsf8 | -0.003 | Old |
| 0.973283 | 0.008692 | 0.184 | 0.179 | 1 | 15 | Itga5 | 0.005 | Old |
| 0.973405 | 0.042657 | 0.388 | 0.357 | 1 | 15 | Gdap10 | 0.031 | Old |
| 0.973751 | 0.20565 | 0.612 | 0.661 | 1 | 15 | Jpt1 | -0.049 | Old |
| 0.97479 | 0.190487 | 0.204 | 0.214 | 1 | 15 | Adipor2 | -0.01 | Old |
| 0.9754 | 0.220412 | 0.51 | 0.5 | 1 | 15 | Psmd2 | 0.01 | Old |
| 0.975899 | 0.48491 | 0.102 | 0.107 | 1 | 15 | Ralgapb | -0.005 | Old |
| 0.975899 | 0.43231 | 0.102 | 0.107 | 1 | 15 | Wdr44 | -0.005 | Old |
| 0.975899 | 0.278019 | 0.102 | 0.107 | 1 | 15 | Lrrcc1 | -0.005 | Old |
| 0.975899 | 0.241259 | 0.102 | 0.107 | 1 | 15 | Agpat5 | -0.005 | Old |
| 0.975899 | 0.124656 | 0.102 | 0.107 | 1 | 15 | Uba5 | -0.005 | Old |
| 0.976032 | 0.685108 | 0.163 | 0.161 | 1 | 15 | Dctn6 | 0.002 | Old |
| 0.976107 | 0.145452 | 0.633 | 0.554 | 1 | 15 | Polr1d | 0.079 | Old |
| 0.976331 | 0.094436 | 0.245 | 0.25 | 1 | 15 | Psmc5 | -0.005 | Old |
| 0.976359 | 0.101588 | 0.367 | 0.375 | 1 | 15 | Kdm7a | -0.008 | Old |
| 0.976955 | 0.304122 | 0.265 | 0.268 | 1 | 15 | Lcp2 | -0.003 | Old |
| 0.978212 | 0.376721 | 0.306 | 0.321 | 1 | 15 | Mdh1 | -0.015 | Old |
| 0.978892 | 1.328183 | 0.082 | 0.089 | 1 | 15 | Asb3 | -0.007 | Old |
| 0.978947 | 0.345213 | 0.143 | 0.143 | 1 | 15 | Desi2 | 0 | Old |
| 0.978947 | 0.188297 | 0.143 | 0.143 | 1 | 15 | Zbtb38 | 0 | Old |
| 0.978947 | 0.063233 | 0.143 | 0.143 | 1 | 15 | Klf16 | 0 | Old |
| 0.978947 | 0.038106 | 0.143 | 0.143 | 1 | 15 | Srpk1 | 0 | Old |
| 0.980225 | 0.342388 | 0.429 | 0.446 | 1 | 15 | Supt5 | -0.017 | Old |
| 0.980527 | 0.050795 | 0.469 | 0.464 | 1 | 15 | Tbc1d8 | 0.005 | Old |
| 0.980915 | 0.25933 | 0.184 | 0.179 | 1 | 15 | Acp1 | 0.005 | Old |
| 0.98094 | 0.191104 | 0.51 | 0.518 | 1 | 15 | Far1 | -0.008 | Old |
| 0.981674 | 0.138469 | 0.327 | 0.339 | 1 | 15 | Hnrnph2 | -0.012 | Old |
| 0.98191 | 0.061049 | 0.714 | 0.768 | 1 | 15 | Itga4 | -0.054 | Old |
| 0.981992 | 0.460397 | 0.204 | 0.214 | 1 | 15 | Cd209a | -0.01 | Old |
| 0.982087 | 0.32185 | 0.122 | 0.125 | 1 | 15 | Bahd1 | -0.003 | Old |
| 0.982087 | 0.191235 | 0.122 | 0.125 | 1 | 15 | Stim1 | -0.003 | Old |
| 0.982298 | 0.119286 | 0.224 | 0.214 | 1 | 15 | Myo9a | 0.01 | Old |
| 0.982298 | 0.105599 | 0.224 | 0.214 | 1 | 15 | Dpm3 | 0.01 | Old |
| 0.983092 | 0.185328 | 0.245 | 0.25 | 1 | 15 | Siva1 | -0.005 | Old |
| 0.983538 | 0.093015 | 0.265 | 0.268 | 1 | 15 | Sft2d2 | -0.003 | Old |
| 0.98402 | 0.017899 | 0.163 | 0.161 | 1 | 15 | Tmem41b | 0.002 | Old |
| 0.984278 | 0.135946 | 0.306 | 0.304 | 1 | 15 | Filip1l | 0.002 | Old |
| 0.984436 | 0.158142 | 0.306 | 0.321 | 1 | 15 | Adgre5 | -0.015 | Old |
| 0.985331 | 0.475577 | 0.367 | 0.393 | 1 | 15 | Phf3 | -0.026 | Old |
| 0.985331 | 0.443524 | 0.367 | 0.393 | 1 | 15 | Paxbp1 | -0.026 | Old |
| 0.985434 | 0.30868 | 0.388 | 0.393 | 1 | 15 | Creb1 | -0.005 | Old |
| 0.985538 | 0.803504 | 0.102 | 0.107 | 1 | 15 | Cuedc2 | -0.005 | Old |

| | | | | | | | | |
|---|---|---|---|---|---|---|---|---|
| 0.985538 | 0.385088 | 0.102 | 0.107 | 1 | 15 | Cavin1 | -0.005 | Old |
| 0.98595 | 0.105139 | 0.469 | 0.429 | 1 | 15 | Mkln1 | 0.04 | Old |
| 0.986022 | 0.818764 | 0.449 | 0.464 | 1 | 15 | Ap2m1 | -0.015 | Old |
| 0.986985 | 0.015968 | 0.673 | 0.696 | 1 | 15 | Rbpj | -0.023 | Old |
| 0.987367 | 0.244537 | 0.143 | 0.143 | 1 | 15 | Samd9l | 0 | Old |
| 0.987367 | 0.101725 | 0.143 | 0.143 | 1 | 15 | Ankrd27 | 0 | Old |
| 0.987782 | 0.172068 | 0.327 | 0.339 | 1 | 15 | 11-Sep | -0.012 | Old |
| 0.988499 | 0.225776 | 0.408 | 0.411 | 1 | 15 | Gna13 | -0.003 | Old |
| 0.988548 | 0.057687 | 0.184 | 0.179 | 1 | 15 | Atg5 | 0.005 | Old |
| 0.989194 | 0.32006 | 0.204 | 0.214 | 1 | 15 | Trim33 | -0.01 | Old |
| 0.989548 | 0.319554 | 0.224 | 0.232 | 1 | 15 | Fam117b | -0.008 | Old |
| 0.989548 | 0.181377 | 0.224 | 0.232 | 1 | 15 | Tmf1 | -0.008 | Old |
| 0.989548 | 0.115401 | 0.224 | 0.232 | 1 | 15 | Ctsl | -0.008 | Old |
| 0.989855 | 0.425595 | 0.245 | 0.25 | 1 | 15 | Chmp3 | -0.005 | Old |
| 0.989855 | 0.095479 | 0.245 | 0.25 | 1 | 15 | Stub1 | -0.005 | Old |
| 0.990122 | 0.258341 | 0.265 | 0.268 | 1 | 15 | Srpr | -0.003 | Old |
| 0.990244 | 0.166555 | 0.265 | 0.286 | 1 | 15 | Sox4 | -0.021 | Old |
| 0.990917 | 0.507837 | 0.327 | 0.357 | 1 | 15 | Plekhj1 | -0.03 | Old |
| 0.991043 | 0.650967 | 0.122 | 0.125 | 1 | 15 | Stxbp5 | -0.003 | Old |
| 0.991043 | 0.298538 | 0.122 | 0.125 | 1 | 15 | Plcl2 | -0.003 | Old |
| 0.991043 | 0.165067 | 0.122 | 0.125 | 1 | 15 | Eva1b | -0.003 | Old |
| 0.991043 | 0.156505 | 0.122 | 0.125 | 1 | 15 | Mrps7 | -0.003 | Old |
| 0.991043 | 0.114556 | 0.122 | 0.125 | 1 | 15 | Smurf2 | -0.003 | Old |
| 0.991065 | 0.002369 | 0.347 | 0.375 | 1 | 15 | Smchd1 | -0.028 | Old |
| 0.99126 | 0.21016 | 0.388 | 0.393 | 1 | 15 | Snx5 | -0.005 | Old |
| 0.992009 | 0.015736 | 0.163 | 0.161 | 1 | 15 | Ing1 | 0.002 | Old |
| 0.993138 | 0.128201 | 0.245 | 0.232 | 1 | 15 | Map7d1 | 0.013 | Old |
| 0.994611 | 0.082534 | 0.571 | 0.518 | 1 | 15 | Idi1 | 0.053 | Old |
| 0.995179 | 0.846824 | 0.102 | 0.107 | 1 | 15 | Gtf3c1 | -0.005 | Old |
| 0.995179 | 0.344059 | 0.102 | 0.107 | 1 | 15 | Nt5c3 | -0.005 | Old |
| 0.995789 | 0.564593 | 0.143 | 0.143 | 1 | 15 | Hyou1 | 0 | Old |
| 0.995789 | 0.150718 | 0.143 | 0.143 | 1 | 15 | Sdhc | 0 | Old |
| 0.99626 | 0.187262 | 0.184 | 0.196 | 1 | 15 | Soat1 | -0.012 | Old |
| 0.996398 | 0.454694 | 0.204 | 0.214 | 1 | 15 | Klc1 | -0.01 | Old |
| 0.996398 | 0.374379 | 0.204 | 0.214 | 1 | 15 | Traf6 | -0.01 | Old |
| 0.996459 | 0.113185 | 0.224 | 0.214 | 1 | 15 | Sec63 | 0.01 | Old |
| 0.996855 | 0.414509 | 0.306 | 0.304 | 1 | 15 | Yeats4 | 0.002 | Old |
| 0.996972 | 0.294868 | 0.347 | 0.339 | 1 | 15 | Pfkp | 0.008 | Old |
| 0.996998 | 0.82232 | 0.347 | 0.357 | 1 | 15 | Dnmt3a | -0.01 | Old |
| 0.996998 | 0.277372 | 0.347 | 0.357 | 1 | 15 | Gpr137b | -0.01 | Old |
| 0.997066 | 0.102724 | 0.388 | 0.375 | 1 | 15 | Ezr | 0.013 | Old |
| 0.997106 | 0.133476 | 0.388 | 0.411 | 1 | 15 | Pik3r5 | -0.023 | Old |
| 0.997142 | 0.059555 | 0.408 | 0.429 | 1 | 15 | H2-Eb2 | -0.021 | Old |
| 0.997255 | 0.024892 | 0.49 | 0.5 | 1 | 15 | Tmem168 | -0.01 | Old |
| 0.000627 | 0.605688 | 0.946 | 0.816 | 1 | 15 | Rpl35 | 0.13 | Young |

| | | | | | | | | |
|---|---|---|---|---|---|---|---|---|
| 0.00089 | 3.596614 | 0.304 | 0.061 | 1 | 15 | Fam162a | 0.243 | Young |
| 0.001233 | 1.290817 | 0.625 | 0.306 | 1 | 15 | Atp6v1b2 | 0.319 | Young |
| 0.001679 | 1.980492 | 0.339 | 0.082 | 1 | 15 | Flvcr1 | 0.257 | Young |
| 0.001811 | 1.391137 | 0.571 | 0.286 | 1 | 15 | Mpc1 | 0.285 | Young |
| 0.001899 | 0.613573 | 0.982 | 0.959 | 1 | 15 | Ftl1 | 0.023 | Young |
| 0.00191 | 2.743026 | 0.232 | 0.02 | 1 | 15 | Atp13a2 | 0.212 | Young |
| 0.002021 | 2.842143 | 0.339 | 0.102 | 1 | 15 | Vps36 | 0.237 | Young |
| 0.002551 | 2.164013 | 0.339 | 0.102 | 1 | 15 | Itgb2 | 0.237 | Young |
| 0.002576 | 4.3663 | 0.214 | 0.02 | 1 | 15 | Zfp560 | 0.194 | Young |
| 0.004221 | 2.742089 | 0.268 | 0.061 | 1 | 15 | Smim27 | 0.207 | Young |
| 0.004312 | 3.794557 | 0.196 | 0.02 | 1 | 15 | Fbxl14 | 0.176 | Young |
| 0.004351 | 1.77275 | 0.375 | 0.143 | 1 | 15 | Lipa | 0.232 | Young |
| 0.00468 | 0.563762 | 0.411 | 0.163 | 1 | 15 | Hspa1a | 0.248 | Young |
| 0.00495 | 0.468813 | 0.982 | 0.918 | 1 | 15 | Rps25 | 0.064 | Young |
| 0.005186 | 1.522432 | 0.536 | 0.306 | 1 | 15 | Casp3 | 0.23 | Young |
| 0.005502 | 2.56215 | 0.232 | 0.041 | 1 | 15 | Afdn | 0.191 | Young |
| 0.005515 | 1.056151 | 0.625 | 0.367 | 1 | 15 | Clic1 | 0.258 | Young |
| 0.005523 | 0.462247 | 0.893 | 0.837 | 1 | 15 | Rpl10 | 0.056 | Young |
| 0.0063 | 1.074936 | 0.446 | 0.184 | 1 | 15 | Rptor | 0.262 | Young |
| 0.006338 | 4.630651 | 0.143 | 0 | 1 | 15 | Hs1bp3 | 0.143 | Young |
| 0.006586 | 1.306094 | 0.607 | 0.388 | 1 | 15 | Tmem160 | 0.219 | Young |
| 0.00659 | 1.666226 | 0.357 | 0.122 | 1 | 15 | Ube2a | 0.235 | Young |
| 0.006665 | 2.367803 | 0.304 | 0.102 | 1 | 15 | Mettl23 | 0.202 | Young |
| 0.007282 | 1.471121 | 0.429 | 0.224 | 1 | 15 | Rex1bd | 0.205 | Young |
| 0.007412 | 0.517687 | 0.893 | 0.857 | 1 | 15 | Rpl10a | 0.036 | Young |
| 0.007709 | 4.522413 | 0.179 | 0.02 | 1 | 15 | Gle1 | 0.159 | Young |
| 0.007709 | 3.869504 | 0.179 | 0.02 | 1 | 15 | Utp18 | 0.159 | Young |
| 0.007756 | 0.441285 | 0.982 | 0.918 | 1 | 15 | Rps13 | 0.064 | Young |
| 0.009016 | 0.993952 | 0.625 | 0.408 | 1 | 15 | Ccser2 | 0.217 | Young |
| 0.009624 | 0.435233 | 0.929 | 0.776 | 1 | 15 | Eef2 | 0.153 | Young |
| 0.009818 | 0.396783 | 0.982 | 0.939 | 1 | 15 | Rpl23 | 0.043 | Young |
| 0.010286 | 0.593158 | 0.929 | 0.714 | 1 | 15 | Tubb5 | 0.215 | Young |
| 0.010367 | 0.486468 | 0.929 | 0.776 | 1 | 15 | Serf2 | 0.153 | Young |
| 0.010415 | 0.875214 | 0.429 | 0.204 | 1 | 15 | Mrpl23 | 0.225 | Young |
| 0.010594 | 1.171396 | 0.179 | 0.02 | 1 | 15 | Sin3a | 0.159 | Young |
| 0.010775 | 1.279256 | 0.393 | 0.163 | 1 | 15 | Slfn2 | 0.23 | Young |
| 0.011082 | 5.430457 | 0.125 | 0 | 1 | 15 | Dtymk | 0.125 | Young |
| 0.011082 | 5.376617 | 0.125 | 0 | 1 | 15 | 1700123O2 | 0.125 | Young |
| 0.011082 | 5.079354 | 0.125 | 0 | 1 | 15 | Tbc1d8b | 0.125 | Young |
| 0.011082 | 4.805916 | 0.125 | 0 | 1 | 15 | Plekhm1 | 0.125 | Young |
| 0.011082 | 4.763469 | 0.125 | 0 | 1 | 15 | Tmem192 | 0.125 | Young |
| 0.011082 | 4.389926 | 0.125 | 0 | 1 | 15 | Foxk2 | 0.125 | Young |
| 0.011082 | 4.001018 | 0.125 | 0 | 1 | 15 | Lpar1 | 0.125 | Young |
| 0.011495 | 0.45227 | 0.911 | 0.816 | 1 | 15 | Rpl15 | 0.095 | Young |
| 0.012045 | 1.181327 | 0.518 | 0.306 | 1 | 15 | Rb1cc1 | 0.212 | Young |

| | | | | | | | | |
|---|---|---|---|---|---|---|---|---|
| 0.012744 | 2.293923 | 0.232 | 0.061 | 1 | 15 | Mfap1b | 0.171 | Young |
| 0.012859 | 1.109918 | 0.589 | 0.367 | 1 | 15 | Nfe2l2 | 0.222 | Young |
| 0.013075 | 1.734259 | 0.393 | 0.184 | 1 | 15 | Pmvk | 0.209 | Young |
| 0.013299 | 1.155021 | 0.482 | 0.286 | 1 | 15 | Dok1 | 0.196 | Young |
| 0.013893 | 2.474123 | 0.232 | 0.061 | 1 | 15 | Slc5a3 | 0.171 | Young |
| 0.013893 | 1.96383 | 0.232 | 0.061 | 1 | 15 | Cenpx | 0.171 | Young |
| 0.01395 | 2.993817 | 0.196 | 0.041 | 1 | 15 | C4b | 0.155 | Young |
| 0.014523 | 0.808131 | 0.768 | 0.571 | 1 | 15 | Hspe1 | 0.197 | Young |
| 0.014611 | 1.21486 | 0.393 | 0.184 | 1 | 15 | Hmgcs1 | 0.209 | Young |
| 0.014657 | 2.056013 | 0.304 | 0.122 | 1 | 15 | Anapc5 | 0.182 | Young |
| 0.014672 | 3.241765 | 0.161 | 0.02 | 1 | 15 | Myo18a | 0.141 | Young |
| 0.014753 | 1.708183 | 0.446 | 0.245 | 1 | 15 | Ppp3ca | 0.201 | Young |
| 0.015019 | 2.014772 | 0.286 | 0.102 | 1 | 15 | Crtc3 | 0.184 | Young |
| 0.01524 | 0.754449 | 0.607 | 0.408 | 1 | 15 | Chd4 | 0.199 | Young |
| 0.015584 | 0.350138 | 0.964 | 0.918 | 1 | 15 | Rps11 | 0.046 | Young |
| 0.01573 | 1.271688 | 0.464 | 0.245 | 1 | 15 | Morf4l2 | 0.219 | Young |
| 0.01573 | 0.466646 | 0.875 | 0.714 | 1 | 15 | Rbm3 | 0.161 | Young |
| 0.016287 | 2.53009 | 0.196 | 0.041 | 1 | 15 | Vkorc1 | 0.155 | Young |
| 0.016474 | 0.575755 | 0.821 | 0.592 | 1 | 15 | Atpif1 | 0.229 | Young |
| 0.016793 | 0.778571 | 0.589 | 0.347 | 1 | 15 | Cox7a2 | 0.242 | Young |
| 0.01683 | 0.548463 | 0.714 | 0.531 | 1 | 15 | Hsp90aa1 | 0.183 | Young |
| 0.017077 | 1.240924 | 0.429 | 0.224 | 1 | 15 | Gm10073 | 0.205 | Young |
| 0.017439 | 1.951295 | 0.161 | 0.02 | 1 | 15 | Cpd | 0.141 | Young |
| 0.018214 | 0.401987 | 0.964 | 0.857 | 1 | 15 | Rps4x | 0.107 | Young |
| 0.018356 | 1.707157 | 0.286 | 0.102 | 1 | 15 | Atp11a | 0.184 | Young |
| 0.018804 | 0.48711 | 0.929 | 0.837 | 1 | 15 | Rpl32 | 0.092 | Young |
| 0.018923 | 1.681365 | 0.232 | 0.061 | 1 | 15 | Gfra2 | 0.171 | Young |
| 0.019006 | 1.589579 | 0.357 | 0.143 | 1 | 15 | Ptgs2 | 0.214 | Young |
| 0.019328 | 4.781826 | 0.107 | 0 | 1 | 15 | Gm29340 | 0.107 | Young |
| 0.019328 | 4.566912 | 0.107 | 0 | 1 | 15 | Cdan1 | 0.107 | Young |
| 0.019328 | 4.344801 | 0.107 | 0 | 1 | 15 | Dus3l | 0.107 | Young |
| 0.019328 | 4.197227 | 0.107 | 0 | 1 | 15 | Gm20703 | 0.107 | Young |
| 0.019328 | 4.182062 | 0.107 | 0 | 1 | 15 | Mthfd1 | 0.107 | Young |
| 0.019328 | 4.165136 | 0.107 | 0 | 1 | 15 | Prkci | 0.107 | Young |
| 0.019385 | 2.405543 | 0.214 | 0.061 | 1 | 15 | Fam98c | 0.153 | Young |
| 0.019406 | 0.947739 | 0.429 | 0.204 | 1 | 15 | Stat6 | 0.225 | Young |
| 0.019992 | 1.521214 | 0.232 | 0.061 | 1 | 15 | Iqgap2 | 0.171 | Young |
| 0.02164 | 0.624573 | 0.482 | 0.245 | 1 | 15 | Blnk | 0.237 | Young |
| 0.022951 | 1.924257 | 0.25 | 0.082 | 1 | 15 | Impad1 | 0.168 | Young |
| 0.023283 | 4.057431 | 0.143 | 0.02 | 1 | 15 | Ngly1 | 0.123 | Young |
| 0.023618 | 1.216894 | 0.464 | 0.265 | 1 | 15 | Rtcb | 0.199 | Young |
| 0.023876 | 1.983138 | 0.321 | 0.163 | 1 | 15 | Lrch3 | 0.158 | Young |
| 0.02416 | 0.292423 | 0.946 | 0.755 | 1 | 15 | Rps27l | 0.191 | Young |
| 0.024221 | 2.368478 | 0.214 | 0.061 | 1 | 15 | Mylip | 0.153 | Young |
| 0.024876 | 2.667817 | 0.232 | 0.082 | 1 | 15 | Thap3 | 0.15 | Young |

| | | | | | | | | |
|---|---|---|---|---|---|---|---|---|
| 0.024894 | 2.014678 | 0.214 | 0.061 | 1 | 15 | Usp14 | 0.153 | Young |
| 0.024942 | 3.705559 | 0.143 | 0.02 | 1 | 15 | Guca1a | 0.123 | Young |
| 0.024942 | 3.524869 | 0.143 | 0.02 | 1 | 15 | Plaur | 0.123 | Young |
| 0.024942 | 3.326946 | 0.143 | 0.02 | 1 | 15 | Zdhhc18 | 0.123 | Young |
| 0.02495 | 1.20366 | 0.339 | 0.143 | 1 | 15 | Snx18 | 0.196 | Young |
| 0.025062 | 1.472294 | 0.321 | 0.143 | 1 | 15 | Arrb1 | 0.178 | Young |
| 0.025209 | 2.744303 | 0.179 | 0.041 | 1 | 15 | Bola3 | 0.138 | Young |
| 0.025809 | 3.165044 | 0.143 | 0.02 | 1 | 15 | Nubp2 | 0.123 | Young |
| 0.02616 | 1.158748 | 0.411 | 0.204 | 1 | 15 | Uqcc3 | 0.207 | Young |
| 0.026245 | 1.721603 | 0.304 | 0.143 | 1 | 15 | Papss2 | 0.161 | Young |
| 0.026324 | 1.377535 | 0.357 | 0.184 | 1 | 15 | Hdlbp | 0.173 | Young |
| 0.026627 | 0.58573 | 0.786 | 0.653 | 1 | 15 | Arpc1b | 0.133 | Young |
| 0.026679 | 0.218867 | 0.982 | 0.898 | 1 | 15 | H3f3a | 0.084 | Young |
| 0.027006 | 0.904269 | 0.464 | 0.265 | 1 | 15 | Tram1 | 0.199 | Young |
| 0.02738 | 1.42702 | 0.321 | 0.143 | 1 | 15 | Ccng2 | 0.178 | Young |
| 0.027573 | 2.199367 | 0.179 | 0.041 | 1 | 15 | Fndc3a | 0.138 | Young |
| 0.027576 | 2.038194 | 0.232 | 0.082 | 1 | 15 | Dnajb1 | 0.15 | Young |
| 0.027749 | 2.165713 | 0.214 | 0.061 | 1 | 15 | Plk2 | 0.153 | Young |
| 0.028239 | 0.789588 | 0.679 | 0.551 | 1 | 15 | Gapdh | 0.128 | Young |
| 0.028403 | 2.024136 | 0.179 | 0.041 | 1 | 15 | Smad5 | 0.138 | Young |
| 0.028477 | 2.385263 | 0.196 | 0.061 | 1 | 15 | 8-Sep | 0.135 | Young |
| 0.028569 | 2.210468 | 0.143 | 0.02 | 1 | 15 | Etv1 | 0.123 | Young |
| 0.028612 | 1.551167 | 0.268 | 0.102 | 1 | 15 | Vav2 | 0.166 | Young |
| 0.028703 | 0.395361 | 0.929 | 0.918 | 1 | 15 | Rpl26 | 0.011 | Young |
| 0.028822 | 0.598356 | 0.75 | 0.612 | 1 | 15 | Gng5 | 0.138 | Young |
| 0.028922 | 0.816469 | 0.679 | 0.429 | 1 | 15 | Scand1 | 0.25 | Young |
| 0.029293 | 1.297666 | 0.393 | 0.204 | 1 | 15 | Gsdmd | 0.189 | Young |
| 0.029308 | 1.24602 | 0.375 | 0.184 | 1 | 15 | Fip1l1 | 0.191 | Young |
| 0.029308 | 1.171134 | 0.375 | 0.184 | 1 | 15 | Acsl5 | 0.191 | Young |
| 0.029532 | 0.543811 | 0.804 | 0.571 | 1 | 15 | Zeb2 | 0.233 | Young |
| 0.029532 | 0.479667 | 0.732 | 0.653 | 1 | 15 | Eif3e | 0.079 | Young |
| 0.029669 | 0.885002 | 0.661 | 0.449 | 1 | 15 | Serinc3 | 0.212 | Young |
| 0.029773 | 0.853189 | 0.5 | 0.306 | 1 | 15 | Adam17 | 0.194 | Young |
| 0.030127 | 2.001641 | 0.179 | 0.041 | 1 | 15 | Foxo3 | 0.138 | Young |
| 0.030721 | 1.940754 | 0.268 | 0.102 | 1 | 15 | Fbxo33 | 0.166 | Young |
| 0.031022 | 1.339691 | 0.179 | 0.041 | 1 | 15 | Pex19 | 0.138 | Young |
| 0.031388 | 1.02137 | 0.643 | 0.51 | 1 | 15 | Atp5j2 | 0.133 | Young |
| 0.031469 | 1.487108 | 0.375 | 0.204 | 1 | 15 | Ankib1 | 0.171 | Young |
| 0.031555 | 1.033085 | 0.357 | 0.163 | 1 | 15 | Wdr83os | 0.194 | Young |
| 0.031678 | 1.278081 | 0.446 | 0.286 | 1 | 15 | Cd81 | 0.16 | Young |
| 0.031812 | 0.518099 | 0.625 | 0.388 | 1 | 15 | Fgl2 | 0.237 | Young |
| 0.032269 | 0.491441 | 0.679 | 0.51 | 1 | 15 | Prdx1 | 0.169 | Young |
| 0.032689 | 0.322292 | 0.964 | 0.918 | 1 | 15 | Rpl34 | 0.046 | Young |
| 0.032747 | 0.987666 | 0.536 | 0.327 | 1 | 15 | Psmg4 | 0.209 | Young |
| 0.032962 | 1.37751 | 0.268 | 0.102 | 1 | 15 | Nfam1 | 0.166 | Young |

| | | | | | | | | |
|---|---|---|---|---|---|---|---|---|
| 0.032988 | 1.20188 | 0.393 | 0.204 | 1 | 15 | Cbx4 | 0.189 | Young |
| 0.033707 | 6.020094 | 0.089 | 0 | 1 | 15 | Hells | 0.089 | Young |
| 0.033707 | 4.940451 | 0.089 | 0 | 1 | 15 | Olr1 | 0.089 | Young |
| 0.033707 | 4.481844 | 0.089 | 0 | 1 | 15 | Spred3 | 0.089 | Young |
| 0.033707 | 4.441898 | 0.089 | 0 | 1 | 15 | Zbtb5 | 0.089 | Young |
| 0.033707 | 4.385199 | 0.089 | 0 | 1 | 15 | Yae1d1 | 0.089 | Young |
| 0.033707 | 4.282877 | 0.089 | 0 | 1 | 15 | Ddx41 | 0.089 | Young |
| 0.033707 | 4.200246 | 0.089 | 0 | 1 | 15 | Suclg2 | 0.089 | Young |
| 0.033707 | 4.124501 | 0.089 | 0 | 1 | 15 | Tmem115 | 0.089 | Young |
| 0.033707 | 4.123255 | 0.089 | 0 | 1 | 15 | Fry | 0.089 | Young |
| 0.033707 | 4.122364 | 0.089 | 0 | 1 | 15 | 4930455G( | 0.089 | Young |
| 0.033707 | 4.107695 | 0.089 | 0 | 1 | 15 | Tmem2 | 0.089 | Young |
| 0.033707 | 4.033445 | 0.089 | 0 | 1 | 15 | Diexf | 0.089 | Young |
| 0.033707 | 4.027778 | 0.089 | 0 | 1 | 15 | Clcf1 | 0.089 | Young |
| 0.033707 | 4.0175 | 0.089 | 0 | 1 | 15 | Tlr7 | 0.089 | Young |
| 0.033707 | 3.886274 | 0.089 | 0 | 1 | 15 | Naip6 | 0.089 | Young |
| 0.033707 | 3.833013 | 0.089 | 0 | 1 | 15 | Fgd6 | 0.089 | Young |
| 0.033707 | 3.773402 | 0.089 | 0 | 1 | 15 | Tmem71 | 0.089 | Young |
| 0.033707 | 3.748113 | 0.089 | 0 | 1 | 15 | Lgals8 | 0.089 | Young |
| 0.033707 | 3.61568 | 0.089 | 0 | 1 | 15 | D10Wsu10 | 0.089 | Young |
| 0.033707 | 3.548392 | 0.089 | 0 | 1 | 15 | Sema4b | 0.089 | Young |
| 0.033707 | 3.534094 | 0.089 | 0 | 1 | 15 | Pnpt1 | 0.089 | Young |
| 0.033707 | 3.532364 | 0.089 | 0 | 1 | 15 | Tmem237 | 0.089 | Young |
| 0.033707 | 3.488069 | 0.089 | 0 | 1 | 15 | Klc4 | 0.089 | Young |
| 0.033707 | 3.393948 | 0.089 | 0 | 1 | 15 | Pgghg | 0.089 | Young |
| 0.033707 | 3.17329 | 0.089 | 0 | 1 | 15 | Pcolce | 0.089 | Young |
| 0.034002 | 1.099767 | 0.321 | 0.143 | 1 | 15 | Phf5a | 0.178 | Young |
| 0.034393 | 0.879621 | 0.5 | 0.306 | 1 | 15 | Kif5b | 0.194 | Young |
| 0.03515 | 0.659517 | 0.571 | 0.367 | 1 | 15 | Zc3h12c | 0.204 | Young |
| 0.03516 | 0.71334 | 0.661 | 0.51 | 1 | 15 | Spi1 | 0.151 | Young |
| 0.035416 | 0.171111 | 1 | 1 | 1 | 15 | Rpl35a | 0 | Young |
| 0.036147 | 0.818464 | 0.214 | 0.061 | 1 | 15 | Ero1lb | 0.153 | Young |
| 0.036281 | 0.262115 | 0.518 | 0.327 | 1 | 15 | Xbp1 | 0.191 | Young |
| 0.036367 | 2.420375 | 0.196 | 0.061 | 1 | 15 | Upf1 | 0.135 | Young |
| 0.036885 | 0.311166 | 0.179 | 0.041 | 1 | 15 | Otud6b | 0.138 | Young |
| 0.037666 | 3.302324 | 0.161 | 0.041 | 1 | 15 | Mzt1 | 0.12 | Young |
| 0.037666 | 2.742005 | 0.161 | 0.041 | 1 | 15 | Atp11c | 0.12 | Young |
| 0.037815 | 1.08794 | 0.304 | 0.122 | 1 | 15 | Tcf12 | 0.182 | Young |
| 0.038134 | 0.374764 | 0.929 | 0.816 | 1 | 15 | Rpl11 | 0.113 | Young |
| 0.038165 | 1.682797 | 0.232 | 0.082 | 1 | 15 | Espl1 | 0.15 | Young |
| 0.038165 | 1.513899 | 0.232 | 0.082 | 1 | 15 | Ss18 | 0.15 | Young |
| 0.038353 | 2.207744 | 0.196 | 0.061 | 1 | 15 | Chd9 | 0.135 | Young |
| 0.038353 | 1.81262 | 0.196 | 0.061 | 1 | 15 | Snx12 | 0.135 | Young |
| 0.038642 | 0.813071 | 0.304 | 0.122 | 1 | 15 | Dok2 | 0.182 | Young |
| 0.038878 | 0.253218 | 0.929 | 0.918 | 1 | 15 | Rpl3 | 0.011 | Young |

| | | | | | | | | |
|---|---|---|---|---|---|---|---|---|
| 0.038995 | 0.824574 | 0.446 | 0.265 | 1 | 15 | Cebpg | 0.181 | Young |
| 0.039079 | 0.760521 | 0.643 | 0.449 | 1 | 15 | Net1 | 0.194 | Young |
| 0.03954 | 0.35073 | 0.714 | 0.612 | 1 | 15 | Atp6v0b | 0.102 | Young |
| 0.039988 | 0.355136 | 0.589 | 0.388 | 1 | 15 | Tmcc1 | 0.201 | Young |
| 0.040077 | 1.009671 | 0.429 | 0.245 | 1 | 15 | Kcnk6 | 0.184 | Young |
| 0.040135 | 0.775274 | 0.607 | 0.388 | 1 | 15 | Atp5b | 0.219 | Young |
| 0.040799 | 3.705967 | 0.125 | 0.02 | 1 | 15 | Tanc2 | 0.105 | Young |
| 0.040799 | 3.4431 | 0.125 | 0.02 | 1 | 15 | Fam219b | 0.105 | Young |
| 0.040799 | 3.288657 | 0.125 | 0.02 | 1 | 15 | Psmd5 | 0.105 | Young |
| 0.041116 | 1.208111 | 0.375 | 0.184 | 1 | 15 | Wdr89 | 0.191 | Young |
| 0.04113 | 2.363944 | 0.161 | 0.041 | 1 | 15 | Htatip2 | 0.12 | Young |
| 0.041266 | 0.453344 | 0.964 | 0.796 | 1 | 15 | Arpc2 | 0.168 | Young |
| 0.041547 | 0.855892 | 0.5 | 0.327 | 1 | 15 | Elavl1 | 0.173 | Young |
| 0.041867 | 0.654504 | 0.679 | 0.51 | 1 | 15 | Sqstm1 | 0.169 | Young |
| 0.042194 | 4.040722 | 0.125 | 0.02 | 1 | 15 | Cxcl1 | 0.105 | Young |
| 0.042194 | 3.330665 | 0.125 | 0.02 | 1 | 15 | Slc25a17 | 0.105 | Young |
| 0.042343 | 3.040344 | 0.161 | 0.041 | 1 | 15 | Slc1a5 | 0.12 | Young |
| 0.043242 | 1.405379 | 0.411 | 0.245 | 1 | 15 | Crlf3 | 0.166 | Young |
| 0.043441 | 0.908342 | 0.446 | 0.265 | 1 | 15 | Tor1aip1 | 0.181 | Young |
| 0.043586 | 2.457648 | 0.161 | 0.041 | 1 | 15 | Slc23a2 | 0.12 | Young |
| 0.043628 | 2.591113 | 0.125 | 0.02 | 1 | 15 | Zscan29 | 0.105 | Young |
| 0.043727 | 1.131359 | 0.196 | 0.061 | 1 | 15 | Cltb | 0.135 | Young |
| 0.043727 | 0.419681 | 0.196 | 0.061 | 1 | 15 | Dock1 | 0.135 | Young |
| 0.044331 | 2.044463 | 0.321 | 0.184 | 1 | 15 | Mpc2 | 0.137 | Young |
| 0.044874 | 1.387133 | 0.196 | 0.061 | 1 | 15 | Pmpca | 0.135 | Young |
| 0.045015 | 1.116942 | 0.429 | 0.286 | 1 | 15 | Pkm | 0.143 | Young |
| 0.045029 | 1.816213 | 0.268 | 0.122 | 1 | 15 | Dgkd | 0.146 | Young |
| 0.045104 | 2.77721 | 0.125 | 0.02 | 1 | 15 | Parp3 | 0.105 | Young |
| 0.045277 | 0.821052 | 0.375 | 0.184 | 1 | 15 | Fcgrt | 0.191 | Young |
| 0.045738 | 2.064044 | 0.214 | 0.082 | 1 | 15 | Umad1 | 0.132 | Young |
| 0.046091 | 0.358531 | 0.964 | 0.918 | 1 | 15 | Cst3 | 0.046 | Young |
| 0.046165 | 1.932202 | 0.161 | 0.041 | 1 | 15 | 9430038I0 | 0.12 | Young |
| 0.046515 | 0.472735 | 0.857 | 0.714 | 1 | 15 | Atp5h | 0.143 | Young |
| 0.047502 | 1.926339 | 0.161 | 0.041 | 1 | 15 | Prelid2 | 0.12 | Young |
| 0.047502 | 1.447267 | 0.161 | 0.041 | 1 | 15 | Plxna1 | 0.12 | Young |
| 0.047531 | 0.329079 | 1 | 0.959 | 1 | 15 | Rps14 | 0.041 | Young |
| 0.047773 | 1.318678 | 0.321 | 0.163 | 1 | 15 | Ccl24 | 0.158 | Young |
| 0.048183 | 2.119625 | 0.125 | 0.02 | 1 | 15 | Smarcal1 | 0.105 | Young |
| 0.048183 | 1.929931 | 0.125 | 0.02 | 1 | 15 | Akap2 | 0.105 | Young |
| 0.048221 | 1.292069 | 0.464 | 0.306 | 1 | 15 | Mir155hg | 0.158 | Young |
| 0.04862 | 0.171158 | 0.982 | 0.98 | 1 | 15 | Rps20 | 0.002 | Young |
| 0.048871 | 1.708989 | 0.161 | 0.041 | 1 | 15 | Wdr36 | 0.12 | Young |
| 0.049086 | 0.93265 | 0.375 | 0.204 | 1 | 15 | Usp7 | 0.171 | Young |
| 0.049184 | 1.644428 | 0.214 | 0.082 | 1 | 15 | Tssc4 | 0.132 | Young |
| 0.049777 | 0.889705 | 0.429 | 0.245 | 1 | 15 | Nck1 | 0.184 | Young |

| | | | | | | | | |
|---|---|---|---|---|---|---|---|---|
| 0.049787 | 1.760073 | 0.125 | 0.02 | 1 | 15 | Map3k3 | 0.105 | Young |
| 0.050832 | 0.766462 | 0.232 | 0.082 | 1 | 15 | Gab2 | 0.15 | Young |
| 0.050948 | 0.725041 | 0.643 | 0.51 | 1 | 15 | Mycbp2 | 0.133 | Young |
| 0.051004 | 2.050829 | 0.232 | 0.102 | 1 | 15 | Lyar | 0.13 | Young |
| 0.051436 | 1.553264 | 0.125 | 0.02 | 1 | 15 | Mitd1 | 0.105 | Young |
| 0.051436 | 1.483987 | 0.125 | 0.02 | 1 | 15 | Trmt6 | 0.105 | Young |
| 0.051436 | 0.84045 | 0.125 | 0.02 | 1 | 15 | Chml | 0.105 | Young |
| 0.051718 | 1.252763 | 0.321 | 0.163 | 1 | 15 | Cand1 | 0.158 | Young |
| 0.052804 | 2.716359 | 0.179 | 0.061 | 1 | 15 | Irak3 | 0.118 | Young |
| 0.052846 | 2.017598 | 0.214 | 0.082 | 1 | 15 | Sptlc1 | 0.132 | Young |
| 0.052846 | 1.784526 | 0.214 | 0.082 | 1 | 15 | Sec14l1 | 0.132 | Young |
| 0.054192 | 2.245999 | 0.179 | 0.061 | 1 | 15 | Oser1 | 0.118 | Young |
| 0.054363 | 0.583719 | 0.429 | 0.245 | 1 | 15 | Gpbp1 | 0.184 | Young |
| 0.054499 | 0.365852 | 0.232 | 0.082 | 1 | 15 | Slc25a25 | 0.15 | Young |
| 0.054684 | 1.356484 | 0.161 | 0.041 | 1 | 15 | Tnfsf12 | 0.12 | Young |
| 0.055003 | 0.937175 | 0.196 | 0.061 | 1 | 15 | Ebp | 0.135 | Young |
| 0.055412 | 1.678296 | 0.214 | 0.082 | 1 | 15 | Vps39 | 0.132 | Young |
| 0.055792 | 1.88858 | 0.286 | 0.143 | 1 | 15 | B3gat3 | 0.143 | Young |
| 0.055938 | 1.721165 | 0.375 | 0.224 | 1 | 15 | Csnk2b | 0.151 | Young |
| 0.056219 | 1.435312 | 0.304 | 0.163 | 1 | 15 | Cnih4 | 0.141 | Young |
| 0.056224 | 1.408941 | 0.161 | 0.041 | 1 | 15 | Snap29 | 0.12 | Young |
| 0.058914 | 2.480602 | 0.25 | 0.122 | 1 | 15 | C1d | 0.128 | Young |
| 0.058988 | 5.006017 | 0.071 | 0 | 1 | 15 | Itgb8 | 0.071 | Young |
| 0.058988 | 4.67734 | 0.071 | 0 | 1 | 15 | Kcna2 | 0.071 | Young |
| 0.058988 | 4.44907 | 0.071 | 0 | 1 | 15 | Ndufaf4 | 0.071 | Young |
| 0.058988 | 4.408708 | 0.071 | 0 | 1 | 15 | Slc2a3 | 0.071 | Young |
| 0.058988 | 4.401755 | 0.071 | 0 | 1 | 15 | Gins1 | 0.071 | Young |
| 0.058988 | 4.035213 | 0.071 | 0 | 1 | 15 | Hus1 | 0.071 | Young |
| 0.058988 | 4.02711 | 0.071 | 0 | 1 | 15 | Armt1 | 0.071 | Young |
| 0.058988 | 4.015618 | 0.071 | 0 | 1 | 15 | Pex26 | 0.071 | Young |
| 0.058988 | 4.011432 | 0.071 | 0 | 1 | 15 | Magohb | 0.071 | Young |
| 0.058988 | 3.971848 | 0.071 | 0 | 1 | 15 | Pard6a | 0.071 | Young |
| 0.058988 | 3.964398 | 0.071 | 0 | 1 | 15 | Man2b2 | 0.071 | Young |
| 0.058988 | 3.961376 | 0.071 | 0 | 1 | 15 | Tlr4 | 0.071 | Young |
| 0.058988 | 3.935738 | 0.071 | 0 | 1 | 15 | Aldh7a1 | 0.071 | Young |
| 0.058988 | 3.864438 | 0.071 | 0 | 1 | 15 | AW554918 | 0.071 | Young |
| 0.058988 | 3.801173 | 0.071 | 0 | 1 | 15 | Tm2d3 | 0.071 | Young |
| 0.058988 | 3.790891 | 0.071 | 0 | 1 | 15 | Fam118b | 0.071 | Young |
| 0.058988 | 3.788952 | 0.071 | 0 | 1 | 15 | Sass6 | 0.071 | Young |
| 0.058988 | 3.778938 | 0.071 | 0 | 1 | 15 | Unc45a | 0.071 | Young |
| 0.058988 | 3.75496 | 0.071 | 0 | 1 | 15 | Slc40a1 | 0.071 | Young |
| 0.058988 | 3.741255 | 0.071 | 0 | 1 | 15 | Mfsd7a | 0.071 | Young |
| 0.058988 | 3.723658 | 0.071 | 0 | 1 | 15 | Tlr1 | 0.071 | Young |
| 0.058988 | 3.691349 | 0.071 | 0 | 1 | 15 | Atp1a3 | 0.071 | Young |
| 0.058988 | 3.667116 | 0.071 | 0 | 1 | 15 | Ndst2 | 0.071 | Young |

| | | | | | | | | |
|---|---|---|---|---|---|---|---|---|
| 0.058988 | 3.631123 | 0.071 | 0 | 1 | 15 | Srrd | 0.071 | Young |
| 0.058988 | 3.582552 | 0.071 | 0 | 1 | 15 | Impdh1 | 0.071 | Young |
| 0.058988 | 3.541376 | 0.071 | 0 | 1 | 15 | Akr7a5 | 0.071 | Young |
| 0.058988 | 3.474667 | 0.071 | 0 | 1 | 15 | Fez2 | 0.071 | Young |
| 0.058988 | 3.436602 | 0.071 | 0 | 1 | 15 | Rem1 | 0.071 | Young |
| 0.058988 | 3.412681 | 0.071 | 0 | 1 | 15 | Dpysl3 | 0.071 | Young |
| 0.058988 | 3.409218 | 0.071 | 0 | 1 | 15 | Lgalsl | 0.071 | Young |
| 0.058988 | 3.386765 | 0.071 | 0 | 1 | 15 | Zkscan14 | 0.071 | Young |
| 0.058988 | 3.383652 | 0.071 | 0 | 1 | 15 | Zfp948 | 0.071 | Young |
| 0.058988 | 3.382215 | 0.071 | 0 | 1 | 15 | Sdhaf3 | 0.071 | Young |
| 0.058988 | 3.371431 | 0.071 | 0 | 1 | 15 | Kdelc2 | 0.071 | Young |
| 0.058988 | 3.173292 | 0.071 | 0 | 1 | 15 | Zfp653 | 0.071 | Young |
| 0.058988 | 3.166761 | 0.071 | 0 | 1 | 15 | Dbp | 0.071 | Young |
| 0.058988 | 3.139918 | 0.071 | 0 | 1 | 15 | Cd2 | 0.071 | Young |
| 0.058988 | 3.084008 | 0.071 | 0 | 1 | 15 | Sap30 | 0.071 | Young |
| 0.058988 | 3.083358 | 0.071 | 0 | 1 | 15 | Msra | 0.071 | Young |
| 0.058988 | 2.949332 | 0.071 | 0 | 1 | 15 | Ap5z1 | 0.071 | Young |
| 0.058988 | 2.90925 | 0.071 | 0 | 1 | 15 | Hs2st1 | 0.071 | Young |
| 0.059359 | 1.681915 | 0.321 | 0.184 | 1 | 15 | Igfbp4 | 0.137 | Young |
| 0.059454 | 1.480563 | 0.214 | 0.082 | 1 | 15 | Idh3g | 0.132 | Young |
| 0.059511 | 0.953254 | 0.393 | 0.224 | 1 | 15 | Cd36 | 0.169 | Young |
| 0.059619 | 0.484076 | 0.893 | 0.633 | 1 | 15 | Cox5a | 0.26 | Young |
| 0.05978 | 0.428058 | 0.464 | 0.327 | 1 | 15 | Clptm1 | 0.137 | Young |
| 0.059926 | 0.915219 | 0.339 | 0.163 | 1 | 15 | Nfix | 0.176 | Young |
| 0.060182 | 2.269255 | 0.25 | 0.122 | 1 | 15 | Mex3b | 0.128 | Young |
| 0.061045 | 0.584297 | 0.375 | 0.184 | 1 | 15 | Il6st | 0.191 | Young |
| 0.0616 | 1.919032 | 0.179 | 0.061 | 1 | 15 | Ide | 0.118 | Young |
| 0.061911 | 0.940588 | 0.446 | 0.286 | 1 | 15 | Ndufb8 | 0.16 | Young |
| 0.06205 | 0.345616 | 0.911 | 0.694 | 1 | 15 | Rpl4 | 0.217 | Young |
| 0.062235 | 0.888273 | 0.339 | 0.163 | 1 | 15 | Dcun1d5 | 0.176 | Young |
| 0.062509 | 1.183488 | 0.429 | 0.286 | 1 | 15 | Sar1a | 0.143 | Young |
| 0.062852 | 2.913399 | 0.143 | 0.041 | 1 | 15 | Cdyl2 | 0.102 | Young |
| 0.063192 | 0.383285 | 0.804 | 0.592 | 1 | 15 | Arl5c | 0.212 | Young |
| 0.063322 | 0.479873 | 0.643 | 0.531 | 1 | 15 | Eef1d | 0.112 | Young |
| 0.063507 | 0.648728 | 0.429 | 0.245 | 1 | 15 | Smarca5 | 0.184 | Young |
| 0.063918 | 0.538005 | 0.821 | 0.694 | 1 | 15 | Cdkn1a | 0.127 | Young |
| 0.064657 | 2.890513 | 0.143 | 0.041 | 1 | 15 | Usf3 | 0.102 | Young |
| 0.064716 | 2.546629 | 0.196 | 0.082 | 1 | 15 | Dnajc21 | 0.114 | Young |
| 0.064789 | 1.791875 | 0.179 | 0.061 | 1 | 15 | 1110008P1 | 0.118 | Young |
| 0.064806 | 0.534784 | 0.536 | 0.367 | 1 | 15 | 1810058I24 | 0.169 | Young |
| 0.065063 | 0.546559 | 0.786 | 0.612 | 1 | 15 | Asap1 | 0.174 | Young |
| 0.065205 | 1.453167 | 0.232 | 0.102 | 1 | 15 | Borcs6 | 0.13 | Young |
| 0.065526 | 1.183554 | 0.286 | 0.143 | 1 | 15 | Man2a1 | 0.143 | Young |
| 0.06638 | 0.621365 | 0.696 | 0.592 | 1 | 15 | Cox6b1 | 0.104 | Young |
| 0.066433 | 2.228617 | 0.179 | 0.061 | 1 | 15 | Trove2 | 0.118 | Young |

| | | | | | | | | |
|---|---|---|---|---|---|---|---|---|
| 0.066433 | 1.891208 | 0.179 | 0.061 | 1 | 15 | Cd33 | 0.118 | Young |
| 0.066456 | 0.738391 | 0.411 | 0.224 | 1 | 15 | Gbp2 | 0.187 | Young |
| 0.067297 | 1.042851 | 0.268 | 0.122 | 1 | 15 | Msrb1 | 0.146 | Young |
| 0.067608 | 1.078059 | 0.393 | 0.245 | 1 | 15 | Usp16 | 0.148 | Young |
| 0.067835 | 2.282098 | 0.196 | 0.082 | 1 | 15 | Orai3 | 0.114 | Young |
| 0.068109 | 2.070683 | 0.232 | 0.102 | 1 | 15 | Lyve1 | 0.13 | Young |
| 0.068112 | 1.742323 | 0.179 | 0.061 | 1 | 15 | Nelfb | 0.118 | Young |
| 0.068254 | 1.195103 | 0.304 | 0.163 | 1 | 15 | Ndufa5 | 0.141 | Young |
| 0.068394 | 2.473629 | 0.143 | 0.041 | 1 | 15 | Mak16 | 0.102 | Young |
| 0.068394 | 2.439908 | 0.143 | 0.041 | 1 | 15 | Tmem120b | 0.102 | Young |
| 0.068458 | 0.691385 | 0.393 | 0.224 | 1 | 15 | Alox5ap | 0.169 | Young |
| 0.068634 | 0.566562 | 0.571 | 0.388 | 1 | 15 | Zbtb20 | 0.183 | Young |
| 0.068684 | 0.828949 | 0.268 | 0.122 | 1 | 15 | Dirc2 | 0.146 | Young |
| 0.068779 | 0.87765 | 0.339 | 0.184 | 1 | 15 | Peak1 | 0.155 | Young |
| 0.0696 | 1.655766 | 0.232 | 0.102 | 1 | 15 | Usp47 | 0.13 | Young |
| 0.070171 | 1.196167 | 0.393 | 0.265 | 1 | 15 | Rbbp7 | 0.128 | Young |
| 0.070495 | 1.164613 | 0.357 | 0.204 | 1 | 15 | Mtss1 | 0.153 | Young |
| 0.071104 | 3.635675 | 0.107 | 0.02 | 1 | 15 | Dhx33 | 0.087 | Young |
| 0.071104 | 3.526616 | 0.107 | 0.02 | 1 | 15 | Nasp | 0.087 | Young |
| 0.071104 | 3.518927 | 0.107 | 0.02 | 1 | 15 | Lrrc8b | 0.087 | Young |
| 0.071104 | 3.487683 | 0.107 | 0.02 | 1 | 15 | Fuom | 0.087 | Young |
| 0.071104 | 3.476021 | 0.107 | 0.02 | 1 | 15 | Urod | 0.087 | Young |
| 0.071104 | 3.31237 | 0.107 | 0.02 | 1 | 15 | Abhd5 | 0.087 | Young |
| 0.071263 | 0.989371 | 0.446 | 0.327 | 1 | 15 | Zfp706 | 0.119 | Young |
| 0.071529 | 0.958541 | 0.268 | 0.122 | 1 | 15 | Stab1 | 0.146 | Young |
| 0.072308 | 2.120152 | 0.143 | 0.041 | 1 | 15 | Mfap1a | 0.102 | Young |
| 0.072308 | 1.873159 | 0.143 | 0.041 | 1 | 15 | Phka2 | 0.102 | Young |
| 0.072343 | 0.388845 | 0.929 | 0.816 | 1 | 15 | Rps3 | 0.113 | Young |
| 0.072744 | 2.295514 | 0.196 | 0.082 | 1 | 15 | Atf7 | 0.114 | Young |
| 0.072744 | 2.209422 | 0.196 | 0.082 | 1 | 15 | Hadhb | 0.114 | Young |
| 0.072997 | 0.868314 | 0.446 | 0.286 | 1 | 15 | Zbtb7a | 0.16 | Young |
| 0.073099 | 1.716663 | 0.268 | 0.143 | 1 | 15 | Ppm1h | 0.125 | Young |
| 0.073099 | 1.664235 | 0.268 | 0.143 | 1 | 15 | Cmip | 0.125 | Young |
| 0.073269 | 1.508864 | 0.411 | 0.286 | 1 | 15 | F13a1 | 0.125 | Young |
| 0.07336 | 1.166302 | 0.179 | 0.061 | 1 | 15 | Pros1 | 0.118 | Young |
| 0.07336 | 0.633364 | 0.179 | 0.061 | 1 | 15 | Pik3ca | 0.118 | Young |
| 0.073461 | 3.431414 | 0.107 | 0.02 | 1 | 15 | Bcam | 0.087 | Young |
| 0.073461 | 3.065298 | 0.107 | 0.02 | 1 | 15 | Ano6 | 0.087 | Young |
| 0.073461 | 2.719886 | 0.107 | 0.02 | 1 | 15 | Sap30bp | 0.087 | Young |
| 0.073461 | 2.571775 | 0.107 | 0.02 | 1 | 15 | Fuca2 | 0.087 | Young |
| 0.073461 | 2.193373 | 0.107 | 0.02 | 1 | 15 | Pdgfc | 0.087 | Young |
| 0.073484 | 0.463878 | 0.804 | 0.653 | 1 | 15 | Pnrc1 | 0.151 | Young |
| 0.07362 | 1.077675 | 0.393 | 0.245 | 1 | 15 | Timp2 | 0.148 | Young |
| 0.073711 | 1.146335 | 0.286 | 0.143 | 1 | 15 | Morc3 | 0.143 | Young |
| 0.073886 | 0.971719 | 0.411 | 0.265 | 1 | 15 | Sf3b5 | 0.146 | Young |

| | | | | | | | | |
|---|---|---|---|---|---|---|---|---|
| 0.074332 | 2.141976 | 0.143 | 0.041 | 1 | 15 | Carm1 | 0.102 | Young |
| 0.074332 | 1.914844 | 0.143 | 0.041 | 1 | 15 | Wbp1 | 0.102 | Young |
| 0.074332 | 1.850056 | 0.143 | 0.041 | 1 | 15 | Exoc7 | 0.102 | Young |
| 0.074442 | 1.212667 | 0.464 | 0.347 | 1 | 15 | Mrc1 | 0.117 | Young |
| 0.074484 | 1.139246 | 0.411 | 0.286 | 1 | 15 | Tsc22d4 | 0.125 | Young |
| 0.074767 | 0.603701 | 0.482 | 0.286 | 1 | 15 | C1qc | 0.196 | Young |
| 0.075142 | 0.14191 | 0.643 | 0.429 | 1 | 15 | Ccl6 | 0.214 | Young |
| 0.075729 | 0.611988 | 0.321 | 0.143 | 1 | 15 | Esyt2 | 0.178 | Young |
| 0.075881 | 3.058101 | 0.107 | 0.02 | 1 | 15 | Mfn2 | 0.087 | Young |
| 0.075881 | 2.502395 | 0.107 | 0.02 | 1 | 15 | Sdad1 | 0.087 | Young |
| 0.075881 | 2.379401 | 0.107 | 0.02 | 1 | 15 | Gtf2i | 0.087 | Young |
| 0.076175 | 1.819679 | 0.196 | 0.082 | 1 | 15 | Nudt16 | 0.114 | Young |
| 0.076403 | 2.424285 | 0.143 | 0.041 | 1 | 15 | Maip1 | 0.102 | Young |
| 0.076403 | 2.166312 | 0.143 | 0.041 | 1 | 15 | Adcy9 | 0.102 | Young |
| 0.076403 | 1.529242 | 0.143 | 0.041 | 1 | 15 | Gapt | 0.102 | Young |
| 0.076403 | 1.363883 | 0.143 | 0.041 | 1 | 15 | Kitl | 0.102 | Young |
| 0.077039 | 1.306961 | 0.179 | 0.061 | 1 | 15 | Cyp4f18 | 0.118 | Young |
| 0.077509 | 0.319243 | 0.268 | 0.122 | 1 | 15 | Aff1 | 0.146 | Young |
| 0.077616 | 1.062052 | 0.357 | 0.224 | 1 | 15 | Ppp1r14b | 0.133 | Young |
| 0.078344 | 0.379807 | 0.357 | 0.204 | 1 | 15 | Epc1 | 0.153 | Young |
| 0.078366 | 2.509783 | 0.107 | 0.02 | 1 | 15 | Rcn3 | 0.087 | Young |
| 0.078366 | 2.448122 | 0.107 | 0.02 | 1 | 15 | Pex7 | 0.087 | Young |
| 0.078366 | 2.202345 | 0.107 | 0.02 | 1 | 15 | Mfsd8 | 0.087 | Young |
| 0.078366 | 2.1202 | 0.107 | 0.02 | 1 | 15 | Xrcc1 | 0.087 | Young |
| 0.078391 | 1.235281 | 0.321 | 0.184 | 1 | 15 | Man2b1 | 0.137 | Young |
| 0.07852 | 2.305369 | 0.143 | 0.041 | 1 | 15 | Bcap29 | 0.102 | Young |
| 0.07852 | 1.807895 | 0.143 | 0.041 | 1 | 15 | Ube4b | 0.102 | Young |
| 0.07852 | 1.803556 | 0.143 | 0.041 | 1 | 15 | Scaf8 | 0.102 | Young |
| 0.07852 | 1.487798 | 0.143 | 0.041 | 1 | 15 | Gosr1 | 0.102 | Young |
| 0.078669 | 0.377039 | 0.714 | 0.49 | 1 | 15 | Arpp19 | 0.224 | Young |
| 0.078965 | 1.032971 | 0.357 | 0.224 | 1 | 15 | Nop10 | 0.133 | Young |
| 0.079066 | 0.889125 | 0.268 | 0.122 | 1 | 15 | Slc4a7 | 0.146 | Young |
| 0.079122 | 0.901527 | 0.446 | 0.286 | 1 | 15 | Vdac2 | 0.16 | Young |
| 0.079486 | 0.583952 | 0.625 | 0.469 | 1 | 15 | Mrfap1 | 0.156 | Young |
| 0.079537 | 0.456593 | 0.411 | 0.245 | 1 | 15 | Tmed7 | 0.166 | Young |
| 0.080045 | 0.550677 | 0.536 | 0.347 | 1 | 15 | Aif1 | 0.189 | Young |
| 0.080684 | 1.65813 | 0.143 | 0.041 | 1 | 15 | Lgals9 | 0.102 | Young |
| 0.080684 | 1.597776 | 0.143 | 0.041 | 1 | 15 | Cmah | 0.102 | Young |
| 0.080825 | 1.060461 | 0.304 | 0.163 | 1 | 15 | Prpf4b | 0.141 | Young |
| 0.080859 | 0.952927 | 0.411 | 0.245 | 1 | 15 | Rbm8a | 0.166 | Young |
| 0.081568 | 1.710748 | 0.196 | 0.082 | 1 | 15 | Tm9sf4 | 0.114 | Young |
| 0.082884 | 0.841797 | 0.411 | 0.265 | 1 | 15 | Gtpbp4 | 0.146 | Young |
| 0.082897 | 0.203615 | 0.143 | 0.041 | 1 | 15 | Sesn3 | 0.102 | Young |
| 0.083128 | 0.58168 | 0.357 | 0.224 | 1 | 15 | Scaf11 | 0.133 | Young |
| 0.083317 | 0.526529 | 0.768 | 0.571 | 1 | 15 | Bcl2a1d | 0.197 | Young |

| | | | | | | | | |
|---|---|---|---|---|---|---|---|---|
| 0.083534 | 1.936123 | 0.107 | 0.02 | 1 | 15 | Usp5 | 0.087 | Young |
| 0.085333 | 2.066048 | 0.196 | 0.082 | 1 | 15 | Tnfrsf18 | 0.114 | Young |
| 0.085333 | 1.513212 | 0.196 | 0.082 | 1 | 15 | Nhsl2 | 0.114 | Young |
| 0.085414 | 1.142689 | 0.25 | 0.122 | 1 | 15 | Mrpl12 | 0.128 | Young |
| 0.086036 | 1.028584 | 0.232 | 0.102 | 1 | 15 | Trappc10 | 0.13 | Young |
| 0.08622 | 1.370506 | 0.107 | 0.02 | 1 | 15 | Usp39 | 0.087 | Young |
| 0.08622 | 1.111607 | 0.107 | 0.02 | 1 | 15 | 2810004N2 | 0.087 | Young |
| 0.08622 | 1.026777 | 0.107 | 0.02 | 1 | 15 | Pisd | 0.087 | Young |
| 0.086328 | 0.559342 | 0.411 | 0.245 | 1 | 15 | Clec12a | 0.166 | Young |
| 0.08635 | 2.403535 | 0.161 | 0.061 | 1 | 15 | Golga3 | 0.1 | Young |
| 0.087066 | 0.604643 | 0.607 | 0.449 | 1 | 15 | Sf3b2 | 0.158 | Young |
| 0.087089 | 1.237906 | 0.268 | 0.143 | 1 | 15 | Smap1 | 0.125 | Young |
| 0.087131 | 1.310341 | 0.25 | 0.122 | 1 | 15 | Mif | 0.128 | Young |
| 0.087217 | 0.643032 | 0.5 | 0.367 | 1 | 15 | Cox5b | 0.133 | Young |
| 0.088388 | 0.807303 | 0.393 | 0.245 | 1 | 15 | Crebrf | 0.148 | Young |
| 0.088769 | 1.513963 | 0.268 | 0.143 | 1 | 15 | Me2 | 0.125 | Young |
| 0.089805 | 0.301 | 0.982 | 0.816 | 1 | 15 | Sem1 | 0.166 | Young |
| 0.090416 | 1.030194 | 0.321 | 0.184 | 1 | 15 | Cic | 0.137 | Young |
| 0.090467 | 0.564588 | 0.661 | 0.531 | 1 | 15 | Rbms1 | 0.13 | Young |
| 0.09086 | 0.590335 | 0.786 | 0.592 | 1 | 15 | Cstb | 0.194 | Young |
| 0.091242 | 1.381523 | 0.196 | 0.082 | 1 | 15 | Ndufb2 | 0.114 | Young |
| 0.091447 | 0.947549 | 0.339 | 0.184 | 1 | 15 | St6galnac6 | 0.155 | Young |
| 0.091449 | 0.801377 | 0.446 | 0.306 | 1 | 15 | Cast | 0.14 | Young |
| 0.09275 | 0.596373 | 0.411 | 0.265 | 1 | 15 | Ilk | 0.146 | Young |
| 0.092939 | 2.057038 | 0.161 | 0.061 | 1 | 15 | Snx29 | 0.1 | Young |
| 0.093284 | 1.201635 | 0.196 | 0.082 | 1 | 15 | Ints6 | 0.114 | Young |
| 0.093284 | 1.018187 | 0.179 | 0.061 | 1 | 15 | Rasgrp3 | 0.118 | Young |
| 0.093322 | 0.364608 | 0.821 | 0.612 | 1 | 15 | Gnas | 0.209 | Young |
| 0.093533 | 0.910887 | 0.357 | 0.224 | 1 | 15 | Rab5a | 0.133 | Young |
| 0.093644 | 0.790266 | 0.321 | 0.184 | 1 | 15 | Srsf10 | 0.137 | Young |
| 0.093926 | 1.217393 | 0.357 | 0.245 | 1 | 15 | Ssh1 | 0.112 | Young |
| 0.093963 | 0.271008 | 0.536 | 0.327 | 1 | 15 | Wfdc17 | 0.209 | Young |
| 0.094282 | 0.85544 | 0.25 | 0.122 | 1 | 15 | Rab18 | 0.128 | Young |
| 0.094661 | 0.648423 | 0.339 | 0.184 | 1 | 15 | Rbm27 | 0.155 | Young |
| 0.095224 | 2.368778 | 0.161 | 0.061 | 1 | 15 | Ibtk | 0.1 | Young |
| 0.095362 | 1.611023 | 0.196 | 0.082 | 1 | 15 | Abcd1 | 0.114 | Young |
| 0.095362 | 1.268962 | 0.196 | 0.082 | 1 | 15 | Lrrc47 | 0.114 | Young |
| 0.096182 | 1.341818 | 0.286 | 0.163 | 1 | 15 | Fcrls | 0.123 | Young |
| 0.096267 | 0.496087 | 0.536 | 0.388 | 1 | 15 | H2-M2 | 0.148 | Young |
| 0.096536 | 0.785289 | 0.536 | 0.429 | 1 | 15 | Eid1 | 0.107 | Young |
| 0.097227 | 0.135521 | 0.143 | 0.041 | 1 | 15 | Chn2 | 0.102 | Young |
| 0.097476 | 1.11735 | 0.196 | 0.082 | 1 | 15 | Rps6ka4 | 0.114 | Young |
| 0.09787 | 0.623367 | 0.286 | 0.143 | 1 | 15 | Chordc1 | 0.143 | Young |
| 0.098632 | 0.66827 | 0.554 | 0.408 | 1 | 15 | Rgs2 | 0.146 | Young |
| 0.098698 | 1.12508 | 0.464 | 0.367 | 1 | 15 | Jun | 0.097 | Young |

| | | | | | | | | |
|---|---|---|---|---|---|---|---|---|
| 0.098853 | 0.746819 | 0.446 | 0.327 | 1 | 15 | Pld4 | 0.119 | Young |
| 0.099628 | 1.36014 | 0.196 | 0.082 | 1 | 15 | Ldb1 | 0.114 | Young |
| 0.099628 | 0.955965 | 0.196 | 0.082 | 1 | 15 | Ing5 | 0.114 | Young |
| 0.099926 | 1.95504 | 0.161 | 0.061 | 1 | 15 | Sult1a1 | 0.1 | Young |
| 0.099926 | 1.936639 | 0.161 | 0.061 | 1 | 15 | Pabpc4 | 0.1 | Young |
| 0.099947 | 1.219829 | 0.25 | 0.122 | 1 | 15 | Jtb | 0.128 | Young |
| 0.100527 | 0.924055 | 0.536 | 0.388 | 1 | 15 | Bcl2a1a | 0.148 | Young |
| 0.101104 | 0.773895 | 0.357 | 0.204 | 1 | 15 | Tet3 | 0.153 | Young |
| 0.101361 | 1.170212 | 0.321 | 0.204 | 1 | 15 | Chic2 | 0.117 | Young |
| 0.101443 | 0.159311 | 0.982 | 0.98 | 1 | 15 | Rpl27a | 0.002 | Young |
| 0.101541 | 1.651775 | 0.214 | 0.102 | 1 | 15 | Man1a2 | 0.112 | Young |
| 0.101885 | 0.287287 | 0.804 | 0.673 | 1 | 15 | Sh3glb1 | 0.131 | Young |
| 0.101938 | 0.519877 | 0.411 | 0.265 | 1 | 15 | Dnajc7 | 0.146 | Young |
| 0.102115 | 0.230624 | 1 | 0.959 | 1 | 15 | Crip1 | 0.041 | Young |
| 0.102346 | 1.930614 | 0.161 | 0.061 | 1 | 15 | Bcl2l1 | 0.1 | Young |
| 0.102603 | 0.586044 | 0.518 | 0.347 | 1 | 15 | Ube2l3 | 0.171 | Young |
| 0.10347 | 0.112563 | 0.982 | 1 | 1 | 15 | H2-D1 | -0.018 | Young |
| 0.104235 | 5.049423 | 0.054 | 0 | 1 | 15 | Cks1b | 0.054 | Young |
| 0.104235 | 4.412134 | 0.054 | 0 | 1 | 15 | Cep44 | 0.054 | Young |
| 0.104235 | 4.278003 | 0.054 | 0 | 1 | 15 | Fcho1 | 0.054 | Young |
| 0.104235 | 4.271992 | 0.054 | 0 | 1 | 15 | Asf1b | 0.054 | Young |
| 0.104235 | 4.269878 | 0.054 | 0 | 1 | 15 | Tamm41 | 0.054 | Young |
| 0.104235 | 4.259222 | 0.054 | 0 | 1 | 15 | Alkbh8 | 0.054 | Young |
| 0.104235 | 4.209866 | 0.054 | 0 | 1 | 15 | Dcun1d4 | 0.054 | Young |
| 0.104235 | 4.209403 | 0.054 | 0 | 1 | 15 | Slc35d1 | 0.054 | Young |
| 0.104235 | 4.185299 | 0.054 | 0 | 1 | 15 | Fam114a1 | 0.054 | Young |
| 0.104235 | 4.165681 | 0.054 | 0 | 1 | 15 | Npat | 0.054 | Young |
| 0.104235 | 4.165383 | 0.054 | 0 | 1 | 15 | Cox10 | 0.054 | Young |
| 0.104235 | 4.058226 | 0.054 | 0 | 1 | 15 | Oas3 | 0.054 | Young |
| 0.104235 | 4.055175 | 0.054 | 0 | 1 | 15 | Nubpl | 0.054 | Young |
| 0.104235 | 4.03623 | 0.054 | 0 | 1 | 15 | Bcl2 | 0.054 | Young |
| 0.104235 | 4.031362 | 0.054 | 0 | 1 | 15 | Haus1 | 0.054 | Young |
| 0.104235 | 4.002654 | 0.054 | 0 | 1 | 15 | Deaf1 | 0.054 | Young |
| 0.104235 | 3.992214 | 0.054 | 0 | 1 | 15 | Il17re | 0.054 | Young |
| 0.104235 | 3.968942 | 0.054 | 0 | 1 | 15 | Rtl6 | 0.054 | Young |
| 0.104235 | 3.966733 | 0.054 | 0 | 1 | 15 | Gm26756 | 0.054 | Young |
| 0.104235 | 3.944723 | 0.054 | 0 | 1 | 15 | 1190002N1 | 0.054 | Young |
| 0.104235 | 3.902108 | 0.054 | 0 | 1 | 15 | Tmem42 | 0.054 | Young |
| 0.104235 | 3.878168 | 0.054 | 0 | 1 | 15 | Gtf2h4 | 0.054 | Young |
| 0.104235 | 3.868167 | 0.054 | 0 | 1 | 15 | Satb1 | 0.054 | Young |
| 0.104235 | 3.816095 | 0.054 | 0 | 1 | 15 | Dnajc9 | 0.054 | Young |
| 0.104235 | 3.751843 | 0.054 | 0 | 1 | 15 | Cry2 | 0.054 | Young |
| 0.104235 | 3.748246 | 0.054 | 0 | 1 | 15 | Tlr6 | 0.054 | Young |
| 0.104235 | 3.747995 | 0.054 | 0 | 1 | 15 | Slc17a9 | 0.054 | Young |
| 0.104235 | 3.74665 | 0.054 | 0 | 1 | 15 | Setbp1 | 0.054 | Young |

| | | | | | | | | |
|---|---|---|---|---|---|---|---|---|
| 0.104235 | 3.729161 | 0.054 | 0 | 1 | 15 | Frmd6 | 0.054 | Young |
| 0.104235 | 3.722707 | 0.054 | 0 | 1 | 15 | Tada2b | 0.054 | Young |
| 0.104235 | 3.710763 | 0.054 | 0 | 1 | 15 | Ispd | 0.054 | Young |
| 0.104235 | 3.706339 | 0.054 | 0 | 1 | 15 | Acot2 | 0.054 | Young |
| 0.104235 | 3.666834 | 0.054 | 0 | 1 | 15 | Saysd1 | 0.054 | Young |
| 0.104235 | 3.642159 | 0.054 | 0 | 1 | 15 | Gm31597 | 0.054 | Young |
| 0.104235 | 3.631558 | 0.054 | 0 | 1 | 15 | Atxn1l | 0.054 | Young |
| 0.104235 | 3.624072 | 0.054 | 0 | 1 | 15 | Ppp1r3b | 0.054 | Young |
| 0.104235 | 3.59892 | 0.054 | 0 | 1 | 15 | Heg1 | 0.054 | Young |
| 0.104235 | 3.589127 | 0.054 | 0 | 1 | 15 | Mtus1 | 0.054 | Young |
| 0.104235 | 3.571536 | 0.054 | 0 | 1 | 15 | Cinp | 0.054 | Young |
| 0.104235 | 3.567487 | 0.054 | 0 | 1 | 15 | Bicra | 0.054 | Young |
| 0.104235 | 3.533611 | 0.054 | 0 | 1 | 15 | Slc37a4 | 0.054 | Young |
| 0.104235 | 3.469851 | 0.054 | 0 | 1 | 15 | Pofut2 | 0.054 | Young |
| 0.104235 | 3.451722 | 0.054 | 0 | 1 | 15 | Sh3glb2 | 0.054 | Young |
| 0.104235 | 3.443906 | 0.054 | 0 | 1 | 15 | Cep95 | 0.054 | Young |
| 0.104235 | 3.435362 | 0.054 | 0 | 1 | 15 | Mfng | 0.054 | Young |
| 0.104235 | 3.427558 | 0.054 | 0 | 1 | 15 | Stambpl1 | 0.054 | Young |
| 0.104235 | 3.423828 | 0.054 | 0 | 1 | 15 | Fcrl1 | 0.054 | Young |
| 0.104235 | 3.422374 | 0.054 | 0 | 1 | 15 | Cela1 | 0.054 | Young |
| 0.104235 | 3.413429 | 0.054 | 0 | 1 | 15 | Fam136a | 0.054 | Young |
| 0.104235 | 3.407244 | 0.054 | 0 | 1 | 15 | Acsl3 | 0.054 | Young |
| 0.104235 | 3.397817 | 0.054 | 0 | 1 | 15 | Hr | 0.054 | Young |
| 0.104235 | 3.358171 | 0.054 | 0 | 1 | 15 | Kcnd1 | 0.054 | Young |
| 0.104235 | 3.350814 | 0.054 | 0 | 1 | 15 | Exoc3l4 | 0.054 | Young |
| 0.104235 | 3.332016 | 0.054 | 0 | 1 | 15 | B4galt4 | 0.054 | Young |
| 0.104235 | 3.327593 | 0.054 | 0 | 1 | 15 | Acot11 | 0.054 | Young |
| 0.104235 | 3.321669 | 0.054 | 0 | 1 | 15 | Arhgef4 | 0.054 | Young |
| 0.104235 | 3.27429 | 0.054 | 0 | 1 | 15 | Clec1a | 0.054 | Young |
| 0.104235 | 3.260474 | 0.054 | 0 | 1 | 15 | Gorab | 0.054 | Young |
| 0.104235 | 3.252343 | 0.054 | 0 | 1 | 15 | Tmem198b | 0.054 | Young |
| 0.104235 | 3.245981 | 0.054 | 0 | 1 | 15 | Gk | 0.054 | Young |
| 0.104235 | 3.222578 | 0.054 | 0 | 1 | 15 | AC162302. | 0.054 | Young |
| 0.104235 | 3.220785 | 0.054 | 0 | 1 | 15 | Rbpms2 | 0.054 | Young |
| 0.104235 | 3.194912 | 0.054 | 0 | 1 | 15 | Zfp933 | 0.054 | Young |
| 0.104235 | 3.194418 | 0.054 | 0 | 1 | 15 | Galnt4 | 0.054 | Young |
| 0.104235 | 3.193632 | 0.054 | 0 | 1 | 15 | Spata1 | 0.054 | Young |
| 0.104235 | 3.193499 | 0.054 | 0 | 1 | 15 | Aste1 | 0.054 | Young |
| 0.104235 | 3.165434 | 0.054 | 0 | 1 | 15 | Bcs1l | 0.054 | Young |
| 0.104235 | 3.134337 | 0.054 | 0 | 1 | 15 | Slc10a3 | 0.054 | Young |
| 0.104235 | 3.131179 | 0.054 | 0 | 1 | 15 | Katnal1 | 0.054 | Young |
| 0.104235 | 3.127159 | 0.054 | 0 | 1 | 15 | Ppp1r18os | 0.054 | Young |
| 0.104235 | 3.120145 | 0.054 | 0 | 1 | 15 | Bud13 | 0.054 | Young |
| 0.104235 | 3.112342 | 0.054 | 0 | 1 | 15 | Ldlrap1 | 0.054 | Young |
| 0.104235 | 3.0866 | 0.054 | 0 | 1 | 15 | Gpr89 | 0.054 | Young |

| | | | | | | | | |
|---|---|---|---|---|---|---|---|---|
| 0.104235 | 3.078968 | 0.054 | 0 | 1 | 15 | Isoc2b | 0.054 | Young |
| 0.104235 | 2.988394 | 0.054 | 0 | 1 | 15 | Adra2a | 0.054 | Young |
| 0.104235 | 2.923275 | 0.054 | 0 | 1 | 15 | Zfp944 | 0.054 | Young |
| 0.104235 | 2.90486 | 0.054 | 0 | 1 | 15 | Bphl | 0.054 | Young |
| 0.104235 | 2.885562 | 0.054 | 0 | 1 | 15 | Nectin4 | 0.054 | Young |
| 0.104235 | 2.863169 | 0.054 | 0 | 1 | 15 | Ikbip | 0.054 | Young |
| 0.104235 | 2.856404 | 0.054 | 0 | 1 | 15 | Wdr70 | 0.054 | Young |
| 0.104235 | 2.84674 | 0.054 | 0 | 1 | 15 | AW011738 | 0.054 | Young |
| 0.104235 | 2.846717 | 0.054 | 0 | 1 | 15 | Gemin6 | 0.054 | Young |
| 0.104235 | 2.798727 | 0.054 | 0 | 1 | 15 | Tenm4 | 0.054 | Young |
| 0.104235 | 2.791105 | 0.054 | 0 | 1 | 15 | Gm15448 | 0.054 | Young |
| 0.104235 | 2.777768 | 0.054 | 0 | 1 | 15 | Zswim7 | 0.054 | Young |
| 0.104235 | 2.774241 | 0.054 | 0 | 1 | 15 | Rcc1 | 0.054 | Young |
| 0.104235 | 2.696002 | 0.054 | 0 | 1 | 15 | Ptgr1 | 0.054 | Young |
| 0.104235 | 2.622432 | 0.054 | 0 | 1 | 15 | Lmo1 | 0.054 | Young |
| 0.104235 | 2.611063 | 0.054 | 0 | 1 | 15 | Cbfa2t2 | 0.054 | Young |
| 0.104235 | 2.599567 | 0.054 | 0 | 1 | 15 | Calml4 | 0.054 | Young |
| 0.104235 | 2.550183 | 0.054 | 0 | 1 | 15 | Btf3l4 | 0.054 | Young |
| 0.104235 | 2.525168 | 0.054 | 0 | 1 | 15 | Ppat | 0.054 | Young |
| 0.104235 | 2.518527 | 0.054 | 0 | 1 | 15 | 4933433G1 | 0.054 | Young |
| 0.104235 | 2.433365 | 0.054 | 0 | 1 | 15 | E2f3 | 0.054 | Young |
| 0.104235 | 2.339815 | 0.054 | 0 | 1 | 15 | Arl11 | 0.054 | Young |
| 0.104235 | 2.330342 | 0.054 | 0 | 1 | 15 | Mavs | 0.054 | Young |
| 0.104235 | 2.330342 | 0.054 | 0 | 1 | 15 | Agmo | 0.054 | Young |
| 0.104753 | 0.705061 | 0.339 | 0.204 | 1 | 15 | Dph5 | 0.135 | Young |
| 0.104813 | 1.591906 | 0.161 | 0.061 | 1 | 15 | Usp24 | 0.1 | Young |
| 0.105439 | 0.686407 | 0.607 | 0.49 | 1 | 15 | Apoe | 0.117 | Young |
| 0.105543 | 0.584489 | 0.589 | 0.449 | 1 | 15 | Spop | 0.14 | Young |
| 0.106313 | 1.183238 | 0.196 | 0.082 | 1 | 15 | Ubfd1 | 0.114 | Young |
| 0.106543 | 3.166463 | 0.125 | 0.041 | 1 | 15 | Lppos | 0.084 | Young |
| 0.107041 | 1.229905 | 0.268 | 0.143 | 1 | 15 | Nab1 | 0.125 | Young |
| 0.107326 | 1.718171 | 0.161 | 0.061 | 1 | 15 | Arl6ip6 | 0.1 | Young |
| 0.107326 | 1.483391 | 0.161 | 0.061 | 1 | 15 | Eno2 | 0.1 | Young |
| 0.107326 | 1.343505 | 0.161 | 0.061 | 1 | 15 | Ints6l | 0.1 | Young |
| 0.107619 | 0.210895 | 1 | 1 | 1 | 15 | Tmsb4x | 0 | Young |
| 0.108083 | 0.421964 | 0.554 | 0.388 | 1 | 15 | Gatsl2 | 0.166 | Young |
| 0.108289 | 0.693577 | 0.339 | 0.204 | 1 | 15 | Psmd1 | 0.135 | Young |
| 0.10862 | 1.974305 | 0.25 | 0.143 | 1 | 15 | Klf10 | 0.107 | Young |
| 0.108848 | 0.332883 | 0.786 | 0.694 | 1 | 15 | Cox6c | 0.092 | Young |
| 0.10901 | 0.962913 | 0.625 | 0.429 | 1 | 15 | Aldh1a2 | 0.196 | Young |
| 0.109296 | 0.474416 | 0.375 | 0.204 | 1 | 15 | Ccr2 | 0.171 | Young |
| 0.109887 | 1.475897 | 0.161 | 0.061 | 1 | 15 | Agtpbp1 | 0.1 | Young |
| 0.109989 | 0.670489 | 0.25 | 0.122 | 1 | 15 | Mdm2 | 0.128 | Young |
| 0.110062 | 1.043071 | 0.304 | 0.184 | 1 | 15 | Pdcl3 | 0.12 | Young |
| 0.110184 | 1.758028 | 0.214 | 0.102 | 1 | 15 | Slc25a39 | 0.112 | Young |

| | | | | | | | | |
|---|---|---|---|---|---|---|---|---|
| 0.11026 | 1.344187 | 0.321 | 0.204 | 1 | 15 | Nudt3 | 0.117 | Young |
| 0.110867 | 0.6423 | 0.5 | 0.408 | 1 | 15 | Diaph1 | 0.092 | Young |
| 0.110963 | 0.900879 | 0.196 | 0.082 | 1 | 15 | Pip4k2b | 0.114 | Young |
| 0.110984 | 1.232883 | 0.286 | 0.163 | 1 | 15 | Add3 | 0.123 | Young |
| 0.11103 | 1.27547 | 0.268 | 0.143 | 1 | 15 | St8sia6 | 0.125 | Young |
| 0.111585 | 0.899302 | 0.304 | 0.163 | 1 | 15 | Ddx54 | 0.141 | Young |
| 0.111585 | 0.742507 | 0.304 | 0.163 | 1 | 15 | Cuta | 0.141 | Young |
| 0.111864 | 0.524112 | 0.518 | 0.347 | 1 | 15 | Ltc4s | 0.171 | Young |
| 0.11199 | 0.348161 | 0.911 | 0.776 | 1 | 15 | Psme2 | 0.135 | Young |
| 0.11199 | 0.31922 | 0.911 | 0.776 | 1 | 15 | Rpl12 | 0.135 | Young |
| 0.112495 | 1.203796 | 0.161 | 0.061 | 1 | 15 | Urm1 | 0.1 | Young |
| 0.113347 | 0.746834 | 0.196 | 0.082 | 1 | 15 | Dnajb4 | 0.114 | Young |
| 0.113347 | 0.42675 | 0.196 | 0.082 | 1 | 15 | Ifitm6 | 0.114 | Young |
| 0.113574 | 0.268713 | 0.607 | 0.449 | 1 | 15 | Pnpla8 | 0.158 | Young |
| 0.113887 | 1.12823 | 0.304 | 0.184 | 1 | 15 | Efr3a | 0.12 | Young |
| 0.114165 | 0.471555 | 0.232 | 0.102 | 1 | 15 | Dnajc15 | 0.13 | Young |
| 0.114518 | 1.404016 | 0.286 | 0.184 | 1 | 15 | Idnk | 0.102 | Young |
| 0.114718 | 1.145767 | 0.214 | 0.102 | 1 | 15 | Agtrap | 0.112 | Young |
| 0.114718 | 0.954229 | 0.214 | 0.102 | 1 | 15 | Chrac1 | 0.112 | Young |
| 0.114952 | 0.50224 | 0.304 | 0.143 | 1 | 15 | Nhlrc2 | 0.161 | Young |
| 0.115153 | 1.27636 | 0.161 | 0.061 | 1 | 15 | Prmt2 | 0.1 | Young |
| 0.115153 | 0.938956 | 0.161 | 0.061 | 1 | 15 | Sc5d | 0.1 | Young |
| 0.115419 | 0.47476 | 0.429 | 0.306 | 1 | 15 | Rab2a | 0.123 | Young |
| 0.115432 | 2.134234 | 0.125 | 0.041 | 1 | 15 | Svil | 0.084 | Young |
| 0.115432 | 2.030701 | 0.125 | 0.041 | 1 | 15 | Tbrg4 | 0.084 | Young |
| 0.115432 | 2.002771 | 0.125 | 0.041 | 1 | 15 | Togaram1 | 0.084 | Young |
| 0.116102 | 0.197561 | 0.893 | 0.898 | 1 | 15 | Sh3bgrl3 | -0.005 | Young |
| 0.116182 | 0.380085 | 0.786 | 0.694 | 1 | 15 | Ubc | 0.092 | Young |
| 0.116192 | 2.205103 | 0.179 | 0.082 | 1 | 15 | Parvb | 0.097 | Young |
| 0.116578 | 1.488758 | 0.232 | 0.122 | 1 | 15 | Lta4h | 0.11 | Young |
| 0.116978 | 1.007572 | 0.286 | 0.163 | 1 | 15 | Saraf | 0.123 | Young |
| 0.116979 | 0.226467 | 0.946 | 0.898 | 1 | 15 | Rpl30 | 0.048 | Young |
| 0.117039 | 1.688756 | 0.214 | 0.102 | 1 | 15 | Ppp2r5e | 0.112 | Young |
| 0.117217 | 0.566463 | 0.589 | 0.429 | 1 | 15 | Tln1 | 0.16 | Young |
| 0.117657 | 0.737228 | 0.375 | 0.224 | 1 | 15 | Phb2 | 0.151 | Young |
| 0.117815 | 0.949869 | 0.339 | 0.184 | 1 | 15 | Clk4 | 0.155 | Young |
| 0.118521 | 2.471969 | 0.125 | 0.041 | 1 | 15 | Cnp | 0.084 | Young |
| 0.118521 | 2.174966 | 0.125 | 0.041 | 1 | 15 | Cdc14b | 0.084 | Young |
| 0.118739 | 1.857338 | 0.179 | 0.082 | 1 | 15 | Polr3gl | 0.097 | Young |
| 0.118739 | 1.620807 | 0.179 | 0.082 | 1 | 15 | Usp18 | 0.097 | Young |
| 0.118896 | 0.334764 | 0.321 | 0.184 | 1 | 15 | Gch1 | 0.137 | Young |
| 0.119006 | 1.017512 | 0.339 | 0.224 | 1 | 15 | Epn1 | 0.115 | Young |
| 0.119216 | 0.557631 | 0.5 | 0.327 | 1 | 15 | Rab11b | 0.173 | Young |
| 0.119765 | 0.53315 | 0.339 | 0.184 | 1 | 15 | Mgl2 | 0.155 | Young |
| 0.120476 | 0.874333 | 0.375 | 0.245 | 1 | 15 | Itpkb | 0.13 | Young |

| | | | | | | | | |
|---|---|---|---|---|---|---|---|---|
| 0.120615 | 1.170548 | 0.161 | 0.061 | 1 | 15 | Snx30 | 0.1 | Young |
| 0.120615 | 0.081942 | 0.161 | 0.061 | 1 | 15 | Tmem43 | 0.1 | Young |
| 0.120686 | 1.721191 | 0.786 | 0.796 | 1 | 15 | Retnla | -0.01 | Young |
| 0.120812 | 0.411929 | 0.25 | 0.122 | 1 | 15 | Tnf | 0.128 | Young |
| 0.12082 | 0.34082 | 0.554 | 0.388 | 1 | 15 | Zfc3h1 | 0.166 | Young |
| 0.121675 | 2.161974 | 0.125 | 0.041 | 1 | 15 | Man2a2 | 0.084 | Young |
| 0.121675 | 2.025652 | 0.125 | 0.041 | 1 | 15 | Tmem60 | 0.084 | Young |
| 0.121675 | 2.016728 | 0.125 | 0.041 | 1 | 15 | Hpgd | 0.084 | Young |
| 0.121675 | 1.960001 | 0.125 | 0.041 | 1 | 15 | Nedd9 | 0.084 | Young |
| 0.121675 | 1.807646 | 0.125 | 0.041 | 1 | 15 | Orc2 | 0.084 | Young |
| 0.123074 | 0.761738 | 0.25 | 0.122 | 1 | 15 | Dapk1 | 0.128 | Young |
| 0.123366 | 4.463404 | 0.089 | 0.02 | 1 | 15 | Lgi4 | 0.069 | Young |
| 0.123366 | 3.697042 | 0.089 | 0.02 | 1 | 15 | Nceh1 | 0.069 | Young |
| 0.123366 | 3.541001 | 0.089 | 0.02 | 1 | 15 | Gkap1 | 0.069 | Young |
| 0.123366 | 3.519902 | 0.089 | 0.02 | 1 | 15 | Zfp672 | 0.069 | Young |
| 0.123366 | 3.166861 | 0.089 | 0.02 | 1 | 15 | Gipc1 | 0.069 | Young |
| 0.123366 | 3.127434 | 0.089 | 0.02 | 1 | 15 | Mroh1 | 0.069 | Young |
| 0.123366 | 3.12018 | 0.089 | 0.02 | 1 | 15 | Eml5 | 0.069 | Young |
| 0.123366 | 3.070637 | 0.089 | 0.02 | 1 | 15 | Arpin | 0.069 | Young |
| 0.123366 | 3.065413 | 0.089 | 0.02 | 1 | 15 | Rnf219 | 0.069 | Young |
| 0.123366 | 2.951232 | 0.089 | 0.02 | 1 | 15 | Mob3b | 0.069 | Young |
| 0.123366 | 2.937648 | 0.089 | 0.02 | 1 | 15 | Pex11b | 0.069 | Young |
| 0.123366 | 2.937118 | 0.089 | 0.02 | 1 | 15 | Ccdc88c | 0.069 | Young |
| 0.123366 | 2.8431 | 0.089 | 0.02 | 1 | 15 | Mtap | 0.069 | Young |
| 0.123366 | 2.833762 | 0.089 | 0.02 | 1 | 15 | Slc25a16 | 0.069 | Young |
| 0.123366 | 2.797793 | 0.089 | 0.02 | 1 | 15 | Mthfd1l | 0.069 | Young |
| 0.123366 | 2.564454 | 0.089 | 0.02 | 1 | 15 | Mthfsd | 0.069 | Young |
| 0.123366 | 2.372426 | 0.089 | 0.02 | 1 | 15 | Cep135 | 0.069 | Young |
| 0.123545 | 1.281807 | 0.25 | 0.143 | 1 | 15 | Ndufs3 | 0.107 | Young |
| 0.123557 | 0.378327 | 0.804 | 0.755 | 1 | 15 | Ctsz | 0.049 | Young |
| 0.12464 | 2.138344 | 0.196 | 0.102 | 1 | 15 | Ednrb | 0.094 | Young |
| 0.124894 | 2.327763 | 0.125 | 0.041 | 1 | 15 | Cyp27a1 | 0.084 | Young |
| 0.124894 | 1.842753 | 0.125 | 0.041 | 1 | 15 | Cep162 | 0.084 | Young |
| 0.124947 | 0.311794 | 0.321 | 0.184 | 1 | 15 | Smc1a | 0.137 | Young |
| 0.125384 | 1.190802 | 0.339 | 0.204 | 1 | 15 | Ptbp1 | 0.135 | Young |
| 0.126378 | 0.517571 | 0.518 | 0.367 | 1 | 15 | Smim14 | 0.151 | Young |
| 0.126695 | 1.137961 | 0.214 | 0.102 | 1 | 15 | Fli1 | 0.112 | Young |
| 0.126769 | 0.528947 | 0.464 | 0.327 | 1 | 15 | Bst2 | 0.137 | Young |
| 0.12709 | 1.01717 | 0.357 | 0.224 | 1 | 15 | Mef2c | 0.133 | Young |
| 0.127302 | 2.821024 | 0.089 | 0.02 | 1 | 15 | Pilrb2 | 0.069 | Young |
| 0.127302 | 2.591963 | 0.089 | 0.02 | 1 | 15 | Tnni3 | 0.069 | Young |
| 0.127302 | 2.296838 | 0.089 | 0.02 | 1 | 15 | Thra | 0.069 | Young |
| 0.127302 | 2.268639 | 0.089 | 0.02 | 1 | 15 | Plxdc2 | 0.069 | Young |
| 0.128178 | 1.334692 | 0.125 | 0.041 | 1 | 15 | Anapc10 | 0.084 | Young |
| 0.128189 | 0.607356 | 0.554 | 0.449 | 1 | 15 | Ndufa4 | 0.105 | Young |

| | | | | | | | | |
|---|---|---|---|---|---|---|---|---|
| 0.128505 | 0.397575 | 0.696 | 0.51 | 1 | 15 | Stk4 | 0.186 | Young |
| 0.129367 | 1.679098 | 0.179 | 0.082 | 1 | 15 | Ppp2r2d | 0.097 | Young |
| 0.129367 | 1.565164 | 0.179 | 0.082 | 1 | 15 | Unc119 | 0.097 | Young |
| 0.129367 | 1.558167 | 0.179 | 0.082 | 1 | 15 | Abcg3 | 0.097 | Young |
| 0.129367 | 1.410637 | 0.179 | 0.082 | 1 | 15 | Nup54 | 0.097 | Young |
| 0.130178 | 0.515464 | 0.679 | 0.531 | 1 | 15 | S100a10 | 0.148 | Young |
| 0.130417 | 0.462839 | 0.482 | 0.306 | 1 | 15 | Znrf2 | 0.176 | Young |
| 0.13095 | 0.39779 | 0.679 | 0.51 | 1 | 15 | Tkt | 0.169 | Young |
| 0.130964 | 0.496473 | 0.714 | 0.551 | 1 | 15 | Pgap2 | 0.163 | Young |
| 0.131334 | 2.793216 | 0.089 | 0.02 | 1 | 15 | Lrrc20 | 0.069 | Young |
| 0.131334 | 2.499032 | 0.089 | 0.02 | 1 | 15 | Ppcdc | 0.069 | Young |
| 0.131334 | 2.467323 | 0.089 | 0.02 | 1 | 15 | Prune2 | 0.069 | Young |
| 0.131334 | 2.368429 | 0.089 | 0.02 | 1 | 15 | Cxcl12 | 0.069 | Young |
| 0.131334 | 2.259324 | 0.089 | 0.02 | 1 | 15 | Poc1b | 0.069 | Young |
| 0.131334 | 2.111823 | 0.089 | 0.02 | 1 | 15 | Slc41a3 | 0.069 | Young |
| 0.131334 | 2.033338 | 0.089 | 0.02 | 1 | 15 | Gm12089 | 0.069 | Young |
| 0.131334 | 2.004197 | 0.089 | 0.02 | 1 | 15 | Kifc3 | 0.069 | Young |
| 0.131334 | 1.914835 | 0.089 | 0.02 | 1 | 15 | C1qtnf1 | 0.069 | Young |
| 0.13153 | 1.533085 | 0.125 | 0.041 | 1 | 15 | Usp10 | 0.084 | Young |
| 0.13153 | 1.026474 | 0.125 | 0.041 | 1 | 15 | Tnfrsf13b | 0.084 | Young |
| 0.131869 | 0.570838 | 0.696 | 0.551 | 1 | 15 | Tbca | 0.145 | Young |
| 0.132136 | 1.362985 | 0.179 | 0.082 | 1 | 15 | Pdk3 | 0.097 | Young |
| 0.132136 | 1.324317 | 0.179 | 0.082 | 1 | 15 | Pggt1b | 0.097 | Young |
| 0.132147 | 0.647049 | 0.161 | 0.061 | 1 | 15 | Aspscr1 | 0.1 | Young |
| 0.132147 | 0.547131 | 0.161 | 0.061 | 1 | 15 | Itpa | 0.1 | Young |
| 0.132305 | 1.313818 | 0.304 | 0.204 | 1 | 15 | Ndufaf8 | 0.1 | Young |
| 0.132836 | 0.146375 | 0.982 | 0.98 | 1 | 15 | Rps27a | 0.002 | Young |
| 0.13352 | 0.434927 | 0.518 | 0.367 | 1 | 15 | Ndufa13 | 0.151 | Young |
| 0.133778 | 0.339073 | 0.857 | 0.735 | 1 | 15 | Ccl22 | 0.122 | Young |
| 0.134205 | 1.267759 | 0.286 | 0.163 | 1 | 15 | Abca1 | 0.123 | Young |
| 0.134951 | 1.592426 | 0.179 | 0.082 | 1 | 15 | Wdfy3 | 0.097 | Young |
| 0.13516 | 0.955008 | 0.161 | 0.061 | 1 | 15 | Tppp3 | 0.1 | Young |
| 0.135465 | 2.871489 | 0.089 | 0.02 | 1 | 15 | Scube3 | 0.069 | Young |
| 0.135465 | 2.603398 | 0.089 | 0.02 | 1 | 15 | Fam26f | 0.069 | Young |
| 0.135465 | 2.125508 | 0.089 | 0.02 | 1 | 15 | Ccdc91 | 0.069 | Young |
| 0.135465 | 1.945206 | 0.089 | 0.02 | 1 | 15 | Brwd1 | 0.069 | Young |
| 0.135465 | 1.87137 | 0.089 | 0.02 | 1 | 15 | Tdp1 | 0.069 | Young |
| 0.135465 | 1.694563 | 0.089 | 0.02 | 1 | 15 | Exoc8 | 0.069 | Young |
| 0.135503 | 1.074686 | 0.232 | 0.122 | 1 | 15 | Gtpbp1 | 0.11 | Young |
| 0.135602 | 0.502947 | 0.518 | 0.408 | 1 | 15 | Grn | 0.11 | Young |
| 0.136517 | 0.385059 | 0.75 | 0.633 | 1 | 15 | Nrp2 | 0.117 | Young |
| 0.136564 | 0.298665 | 0.893 | 0.837 | 1 | 15 | Tmem176k | 0.056 | Young |
| 0.137185 | 1.251797 | 0.393 | 0.306 | 1 | 15 | 9-Sep | 0.087 | Young |
| 0.137812 | 1.335769 | 0.179 | 0.082 | 1 | 15 | Atxn7l3 | 0.097 | Young |
| 0.138027 | 0.637721 | 0.232 | 0.122 | 1 | 15 | Stat2 | 0.11 | Young |

| | | | | | | | | |
|---|---|---|---|---|---|---|---|---|
| 0.138225 | 0.814374 | 0.161 | 0.061 | 1 | 15 | Cbx8 | 0.1 | Young |
| 0.138437 | 1.869716 | 0.125 | 0.041 | 1 | 15 | Sntb2 | 0.084 | Young |
| 0.138763 | 0.399471 | 0.732 | 0.673 | 1 | 15 | Atp2b1 | 0.059 | Young |
| 0.138999 | 0.005906 | 0.018 | 0.082 | 1 | 15 | Slc25a29 | -0.064 | Young |
| 0.139625 | 0.615486 | 0.214 | 0.102 | 1 | 15 | Pitpnb | 0.112 | Young |
| 0.139695 | 2.056327 | 0.089 | 0.02 | 1 | 15 | Socs1 | 0.069 | Young |
| 0.139695 | 1.50655 | 0.089 | 0.02 | 1 | 15 | Mpv17l2 | 0.069 | Young |
| 0.139695 | 1.439 | 0.089 | 0.02 | 1 | 15 | Isyna1 | 0.069 | Young |
| 0.139695 | 1.401988 | 0.089 | 0.02 | 1 | 15 | Mad2l1bp | 0.069 | Young |
| 0.139695 | 1.207344 | 0.089 | 0.02 | 1 | 15 | Zc3h10 | 0.069 | Young |
| 0.139695 | 1.118762 | 0.089 | 0.02 | 1 | 15 | Kat14 | 0.069 | Young |
| 0.139695 | 1.11398 | 0.089 | 0.02 | 1 | 15 | Ccs | 0.069 | Young |
| 0.140507 | 0.477589 | 0.75 | 0.653 | 1 | 15 | Fam49a | 0.097 | Young |
| 0.140587 | 1.076719 | 0.232 | 0.122 | 1 | 15 | Bcl3 | 0.11 | Young |
| 0.14072 | 1.431563 | 0.179 | 0.082 | 1 | 15 | Ecm1 | 0.097 | Young |
| 0.141235 | 0.266802 | 0.804 | 0.714 | 1 | 15 | Slc6a6 | 0.09 | Young |
| 0.141356 | 0.927071 | 0.304 | 0.184 | 1 | 15 | Ubxn6 | 0.12 | Young |
| 0.142043 | 2.705955 | 0.143 | 0.061 | 1 | 15 | Mms19 | 0.082 | Young |
| 0.142043 | 2.500829 | 0.143 | 0.061 | 1 | 15 | Mfap3 | 0.082 | Young |
| 0.142139 | 0.809011 | 0.446 | 0.327 | 1 | 15 | Pltp | 0.119 | Young |
| 0.142282 | 0.390648 | 0.339 | 0.204 | 1 | 15 | Rtf1 | 0.135 | Young |
| 0.142341 | 0.718633 | 0.589 | 0.469 | 1 | 15 | Icam1 | 0.12 | Young |
| 0.143525 | 0.887046 | 0.286 | 0.163 | 1 | 15 | Nhp2 | 0.123 | Young |
| 0.143534 | 0.340691 | 0.804 | 0.694 | 1 | 15 | Iscu | 0.11 | Young |
| 0.143674 | 0.559586 | 0.179 | 0.082 | 1 | 15 | Slc22a15 | 0.097 | Young |
| 0.143891 | 0.085668 | 0.732 | 0.551 | 1 | 15 | Arl5a | 0.181 | Young |
| 0.143987 | 0.300485 | 0.554 | 0.388 | 1 | 15 | Psmb3 | 0.166 | Young |
| 0.144026 | 1.298586 | 0.089 | 0.02 | 1 | 15 | Ankrd52 | 0.069 | Young |
| 0.144026 | 1.210471 | 0.089 | 0.02 | 1 | 15 | Arid5b | 0.069 | Young |
| 0.144026 | 0.887343 | 0.089 | 0.02 | 1 | 15 | Zfp991 | 0.069 | Young |
| 0.144026 | 0.881839 | 0.089 | 0.02 | 1 | 15 | Scd1 | 0.069 | Young |
| 0.144026 | 0.880948 | 0.089 | 0.02 | 1 | 15 | St3gal6 | 0.069 | Young |
| 0.144026 | 0.771905 | 0.089 | 0.02 | 1 | 15 | Cd180 | 0.069 | Young |
| 0.144026 | 0.700361 | 0.089 | 0.02 | 1 | 15 | Cbx6 | 0.069 | Young |
| 0.144026 | 0.690771 | 0.089 | 0.02 | 1 | 15 | Arhgap35 | 0.069 | Young |
| 0.144026 | 0.675237 | 0.089 | 0.02 | 1 | 15 | Rab33b | 0.069 | Young |
| 0.144026 | 0.562436 | 0.089 | 0.02 | 1 | 15 | Hepacam2 | 0.069 | Young |
| 0.144421 | 0.811693 | 0.411 | 0.306 | 1 | 15 | Plin2 | 0.105 | Young |
| 0.145354 | 2.427604 | 0.143 | 0.061 | 1 | 15 | Ofd1 | 0.082 | Young |
| 0.145619 | 0.841023 | 0.125 | 0.041 | 1 | 15 | Slc39a10 | 0.084 | Young |
| 0.145815 | 1.041815 | 0.232 | 0.122 | 1 | 15 | S100a1 | 0.11 | Young |
| 0.145815 | 0.978534 | 0.232 | 0.122 | 1 | 15 | Tns3 | 0.11 | Young |
| 0.145815 | 0.913805 | 0.232 | 0.122 | 1 | 15 | 2-Mar | 0.11 | Young |
| 0.145941 | 0.292121 | 0.75 | 0.592 | 1 | 15 | Flna | 0.158 | Young |
| 0.146857 | 0.354708 | 0.321 | 0.184 | 1 | 15 | Rmnd5a | 0.137 | Young |

| | | | | | | | | |
|---|---|---|---|---|---|---|---|---|
| 0.146886 | 0.919214 | 0.464 | 0.367 | 1 | 15 | Reep5 | 0.097 | Young |
| 0.147266 | 0.56061 | 0.464 | 0.388 | 1 | 15 | Tex261 | 0.076 | Young |
| 0.147434 | 0.389301 | 0.821 | 0.755 | 1 | 15 | Nfkbiz | 0.066 | Young |
| 0.147487 | 0.987755 | 0.321 | 0.204 | 1 | 15 | Tmed9 | 0.117 | Young |
| 0.147487 | 0.514822 | 0.321 | 0.204 | 1 | 15 | Pigt | 0.117 | Young |
| 0.147487 | 0.125391 | 0.321 | 0.204 | 1 | 15 | Gorasp2 | 0.117 | Young |
| 0.147518 | 0.921145 | 0.429 | 0.306 | 1 | 15 | Dab2 | 0.123 | Young |
| 0.147583 | 1.181424 | 0.25 | 0.143 | 1 | 15 | Pex13 | 0.107 | Young |
| 0.147583 | 1.088988 | 0.25 | 0.143 | 1 | 15 | Txndc11 | 0.107 | Young |
| 0.148485 | 1.16767 | 0.232 | 0.122 | 1 | 15 | Nucb1 | 0.11 | Young |
| 0.14898 | 1.527715 | 0.196 | 0.102 | 1 | 15 | Qars | 0.094 | Young |
| 0.150183 | 0.227708 | 0.25 | 0.122 | 1 | 15 | Ldah | 0.128 | Young |
| 0.150834 | 1.003507 | 0.286 | 0.163 | 1 | 15 | Elmsan1 | 0.123 | Young |
| 0.151361 | 0.525053 | 0.339 | 0.204 | 1 | 15 | Dnase2a | 0.135 | Young |
| 0.152156 | 0.722645 | 0.321 | 0.204 | 1 | 15 | Cbr2 | 0.117 | Young |
| 0.152702 | 0.971251 | 0.339 | 0.224 | 1 | 15 | Aco2 | 0.115 | Young |
| 0.152702 | 0.887439 | 0.339 | 0.224 | 1 | 15 | Ranbp1 | 0.115 | Young |
| 0.153084 | 0.513451 | 0.125 | 0.041 | 1 | 15 | Brwd3 | 0.084 | Young |
| 0.153184 | 0.729764 | 0.304 | 0.184 | 1 | 15 | Etfb | 0.12 | Young |
| 0.153323 | 0.455655 | 0.5 | 0.347 | 1 | 15 | Evi2a | 0.153 | Young |
| 0.153546 | 0.262996 | 0.214 | 0.102 | 1 | 15 | Zhx1 | 0.112 | Young |
| 0.154852 | 1.749425 | 0.196 | 0.102 | 1 | 15 | Lmbrd1 | 0.094 | Young |
| 0.154852 | 1.610495 | 0.196 | 0.102 | 1 | 15 | Sf3a3 | 0.094 | Young |
| 0.154852 | 1.303583 | 0.196 | 0.102 | 1 | 15 | Mrpl17 | 0.094 | Young |
| 0.15564 | 1.944113 | 0.143 | 0.061 | 1 | 15 | Nfia | 0.082 | Young |
| 0.15597 | 1.153013 | 0.179 | 0.082 | 1 | 15 | Tcn2 | 0.097 | Young |
| 0.15597 | 0.196157 | 0.179 | 0.082 | 1 | 15 | Lyz1 | 0.097 | Young |
| 0.156718 | 1.136695 | 0.232 | 0.122 | 1 | 15 | Gatm | 0.11 | Young |
| 0.156718 | 0.761364 | 0.232 | 0.122 | 1 | 15 | Ifi203 | 0.11 | Young |
| 0.157101 | 0.539334 | 0.696 | 0.571 | 1 | 15 | Calm3 | 0.125 | Young |
| 0.157128 | 1.449334 | 0.286 | 0.184 | 1 | 15 | Uqcrc2 | 0.102 | Young |
| 0.157654 | 0.511579 | 0.268 | 0.143 | 1 | 15 | Arhgap22 | 0.125 | Young |
| 0.158273 | 0.520808 | 0.464 | 0.327 | 1 | 15 | Fxyd5 | 0.137 | Young |
| 0.158437 | 0.345121 | 0.643 | 0.449 | 1 | 15 | Nfat5 | 0.194 | Young |
| 0.158539 | 0.200129 | 0.964 | 0.918 | 1 | 15 | Rpl9 | 0.046 | Young |
| 0.158554 | 0.312675 | 0.982 | 0.918 | 1 | 15 | Rps6 | 0.064 | Young |
| 0.159165 | 1.05417 | 0.179 | 0.082 | 1 | 15 | Ppt1 | 0.097 | Young |
| 0.159189 | 2.116829 | 0.143 | 0.061 | 1 | 15 | Plat | 0.082 | Young |
| 0.159189 | 1.731763 | 0.143 | 0.061 | 1 | 15 | Cstf2t | 0.082 | Young |
| 0.159328 | 0.157274 | 0.411 | 0.286 | 1 | 15 | Dync1i2 | 0.125 | Young |
| 0.159368 | 0.865466 | 0.321 | 0.204 | 1 | 15 | Socs3 | 0.117 | Young |
| 0.159538 | 0.695757 | 0.232 | 0.122 | 1 | 15 | Mndal | 0.11 | Young |
| 0.160021 | 0.742163 | 0.393 | 0.265 | 1 | 15 | Higd2a | 0.128 | Young |
| 0.1601 | 1.068649 | 0.304 | 0.204 | 1 | 15 | Serpinb6a | 0.1 | Young |
| 0.160324 | 0.546627 | 0.714 | 0.612 | 1 | 15 | Mapre1 | 0.102 | Young |

| | | | | | | | | |
|---|---|---|---|---|---|---|---|---|
| 0.160871 | 0.930362 | 0.25 | 0.143 | 1 | 15 | Lifr | 0.107 | Young |
| 0.160898 | 1.138595 | 0.196 | 0.102 | 1 | 15 | Ckap5 | 0.094 | Young |
| 0.16099 | 0.635153 | 0.482 | 0.408 | 1 | 15 | Hif1a | 0.074 | Young |
| 0.161348 | 0.414102 | 0.321 | 0.184 | 1 | 15 | Ifngr1 | 0.137 | Young |
| 0.161492 | 0.045264 | 0.786 | 0.735 | 1 | 15 | Hnrnpk | 0.051 | Young |
| 0.162087 | 0.508588 | 0.518 | 0.388 | 1 | 15 | Smdt1 | 0.13 | Young |
| 0.162799 | 1.641081 | 0.143 | 0.061 | 1 | 15 | Nvl | 0.082 | Young |
| 0.163063 | 0.733387 | 0.446 | 0.327 | 1 | 15 | Ppp1cb | 0.119 | Young |
| 0.163115 | 0.212708 | 0.679 | 0.571 | 1 | 15 | Glud1 | 0.108 | Young |
| 0.163176 | 0.518192 | 0.304 | 0.184 | 1 | 15 | Cdk9 | 0.12 | Young |
| 0.163822 | 0.574326 | 0.411 | 0.306 | 1 | 15 | Mia2 | 0.105 | Young |
| 0.164226 | 0.304586 | 0.839 | 0.612 | 1 | 15 | Pfdn5 | 0.227 | Young |
| 0.16575 | 0.661117 | 0.304 | 0.184 | 1 | 15 | Stag1 | 0.12 | Young |
| 0.16647 | 1.814148 | 0.143 | 0.061 | 1 | 15 | Dzip3 | 0.082 | Young |
| 0.16647 | 1.468312 | 0.143 | 0.061 | 1 | 15 | Fam69a | 0.082 | Young |
| 0.16647 | 1.145275 | 0.143 | 0.061 | 1 | 15 | Slco2b1 | 0.082 | Young |
| 0.167121 | 1.191369 | 0.196 | 0.102 | 1 | 15 | Pigk | 0.094 | Young |
| 0.167423 | 0.333038 | 0.589 | 0.449 | 1 | 15 | Psmb9 | 0.14 | Young |
| 0.168027 | 0.465637 | 0.536 | 0.429 | 1 | 15 | Ndufa7 | 0.107 | Young |
| 0.168256 | 0.208542 | 0.946 | 0.959 | 1 | 15 | Rps15a | -0.013 | Young |
| 0.168262 | 0.668208 | 0.518 | 0.429 | 1 | 15 | Dnaja1 | 0.089 | Young |
| 0.168975 | 0.034509 | 0.375 | 0.224 | 1 | 15 | Palld | 0.151 | Young |
| 0.169047 | 0.077621 | 0.179 | 0.082 | 1 | 15 | Cyth3 | 0.097 | Young |
| 0.16934 | 1.72198 | 0.214 | 0.122 | 1 | 15 | Alkbh1 | 0.092 | Young |
| 0.169384 | 1.053426 | 0.321 | 0.204 | 1 | 15 | Lsm12 | 0.117 | Young |
| 0.169384 | 0.802564 | 0.321 | 0.204 | 1 | 15 | Rp9 | 0.117 | Young |
| 0.169548 | 0.509271 | 0.339 | 0.224 | 1 | 15 | Dnm2 | 0.115 | Young |
| 0.169566 | 1.170231 | 0.321 | 0.224 | 1 | 15 | Zeb1 | 0.097 | Young |
| 0.170203 | 1.48382 | 0.143 | 0.061 | 1 | 15 | Diaph2 | 0.082 | Young |
| 0.170203 | 1.4226 | 0.143 | 0.061 | 1 | 15 | Icmt | 0.082 | Young |
| 0.170511 | 0.386661 | 0.446 | 0.286 | 1 | 15 | Smad4 | 0.16 | Young |
| 0.170812 | 1.98941 | 0.161 | 0.082 | 1 | 15 | Zfp131 | 0.079 | Young |
| 0.171711 | 0.39941 | 0.286 | 0.163 | 1 | 15 | Pin4 | 0.123 | Young |
| 0.172055 | 0.685532 | 0.446 | 0.327 | 1 | 15 | C1qa | 0.119 | Young |
| 0.172112 | 1.15926 | 0.321 | 0.224 | 1 | 15 | Tulp4 | 0.097 | Young |
| 0.172822 | 1.255526 | 0.286 | 0.184 | 1 | 15 | Ykt6 | 0.102 | Young |
| 0.173141 | 0.547817 | 0.411 | 0.286 | 1 | 15 | Tnrc18 | 0.125 | Young |
| 0.173268 | 0.358272 | 0.643 | 0.49 | 1 | 15 | Tax1bp1 | 0.153 | Young |
| 0.173999 | 1.21865 | 0.143 | 0.061 | 1 | 15 | BC035044 | 0.082 | Young |
| 0.173999 | 0.804529 | 0.143 | 0.061 | 1 | 15 | Aqp9 | 0.082 | Young |
| 0.174127 | 0.724204 | 0.393 | 0.265 | 1 | 15 | Rab6a | 0.128 | Young |
| 0.1742 | 1.417932 | 0.25 | 0.163 | 1 | 15 | Zfhx3 | 0.087 | Young |
| 0.174215 | 0.074537 | 0.232 | 0.122 | 1 | 15 | Gramd1a | 0.11 | Young |
| 0.174482 | 1.062273 | 0.268 | 0.163 | 1 | 15 | Cd84 | 0.105 | Young |
| 0.175318 | 0.195832 | 0.982 | 0.939 | 1 | 15 | Rps12 | 0.043 | Young |

| | | | | | | | | |
|---|---|---|---|---|---|---|---|---|
| 0.175506 | 0.331547 | 0.411 | 0.265 | 1 | 15 | Xrn2 | 0.146 | Young |
| 0.175547 | 0.958908 | 0.286 | 0.184 | 1 | 15 | Mrps6 | 0.102 | Young |
| 0.176349 | 0.740963 | 0.304 | 0.184 | 1 | 15 | Ssu72 | 0.12 | Young |
| 0.176791 | 1.16921 | 0.196 | 0.102 | 1 | 15 | Gm26740 | 0.094 | Young |
| 0.176791 | 0.481501 | 0.196 | 0.102 | 1 | 15 | Tlnrd1 | 0.094 | Young |
| 0.177291 | 0.94181 | 0.321 | 0.224 | 1 | 15 | Rapgef2 | 0.097 | Young |
| 0.177827 | 0.253705 | 0.554 | 0.408 | 1 | 15 | Atf3 | 0.146 | Young |
| 0.177844 | 2.602331 | 0.107 | 0.041 | 1 | 15 | Khk | 0.066 | Young |
| 0.177844 | 2.222538 | 0.107 | 0.041 | 1 | 15 | Fnta | 0.066 | Young |
| 0.177844 | 1.61083 | 0.107 | 0.041 | 1 | 15 | Samd1 | 0.066 | Young |
| 0.177857 | 1.293664 | 0.143 | 0.061 | 1 | 15 | Txlna | 0.082 | Young |
| 0.177857 | 1.280246 | 0.143 | 0.061 | 1 | 15 | Rnf138 | 0.082 | Young |
| 0.178128 | 0.05618 | 0.571 | 0.429 | 1 | 15 | Anp32b | 0.142 | Young |
| 0.178327 | 0.19618 | 0.554 | 0.367 | 1 | 15 | Lgmn | 0.187 | Young |
| 0.17863 | 1.225483 | 0.214 | 0.122 | 1 | 15 | Atxn2 | 0.092 | Young |
| 0.179455 | 0.150601 | 1 | 1 | 1 | 15 | Rps24 | 0 | Young |
| 0.179546 | 0.357368 | 0.446 | 0.306 | 1 | 15 | Tmem167 | 0.14 | Young |
| 0.179781 | 0.572855 | 0.339 | 0.224 | 1 | 15 | Vsir | 0.115 | Young |
| 0.180106 | 1.06543 | 0.196 | 0.102 | 1 | 15 | Mrps23 | 0.094 | Young |
| 0.180106 | 0.983109 | 0.196 | 0.102 | 1 | 15 | Ccdc115 | 0.094 | Young |
| 0.181147 | 0.180071 | 0.625 | 0.449 | 1 | 15 | Ccl17 | 0.176 | Young |
| 0.181779 | 1.059516 | 0.143 | 0.061 | 1 | 15 | Tpcn1 | 0.082 | Young |
| 0.181779 | 0.990659 | 0.143 | 0.061 | 1 | 15 | Yipf6 | 0.082 | Young |
| 0.181779 | 0.913032 | 0.143 | 0.061 | 1 | 15 | Fam120b | 0.082 | Young |
| 0.18205 | 0.540355 | 0.357 | 0.224 | 1 | 15 | Cct3 | 0.133 | Young |
| 0.182381 | 2.941445 | 0.107 | 0.041 | 1 | 15 | Cpeb2 | 0.066 | Young |
| 0.182381 | 2.503429 | 0.107 | 0.041 | 1 | 15 | Uty | 0.066 | Young |
| 0.182381 | 2.463804 | 0.107 | 0.041 | 1 | 15 | Fiz1 | 0.066 | Young |
| 0.182381 | 2.228229 | 0.107 | 0.041 | 1 | 15 | Nufip1 | 0.066 | Young |
| 0.182381 | 2.1687 | 0.107 | 0.041 | 1 | 15 | Ino80 | 0.066 | Young |
| 0.182942 | 0.626116 | 0.482 | 0.367 | 1 | 15 | Pea15a | 0.115 | Young |
| 0.182944 | 1.244919 | 0.25 | 0.163 | 1 | 15 | Sdhb | 0.087 | Young |
| 0.183565 | 0.485958 | 0.75 | 0.592 | 1 | 15 | Ywhae | 0.158 | Young |
| 0.183774 | 0.340669 | 0.554 | 0.388 | 1 | 15 | Poglut1 | 0.166 | Young |
| 0.184075 | 0.70846 | 0.393 | 0.306 | 1 | 15 | Uqcrfs1 | 0.087 | Young |
| 0.184176 | 0.440826 | 0.5 | 0.408 | 1 | 15 | Romo1 | 0.092 | Young |
| 0.185272 | 1.783256 | 0.161 | 0.082 | 1 | 15 | Psmd12 | 0.079 | Young |
| 0.185272 | 1.754992 | 0.161 | 0.082 | 1 | 15 | Cyth2 | 0.079 | Young |
| 0.185815 | 0.769246 | 0.286 | 0.163 | 1 | 15 | Ttc39b | 0.123 | Young |
| 0.18593 | 1.390138 | 0.25 | 0.163 | 1 | 15 | Mrps18c | 0.087 | Young |
| 0.18593 | 0.232046 | 0.268 | 0.143 | 1 | 15 | Vcam1 | 0.125 | Young |
| 0.185995 | 0.99015 | 0.339 | 0.245 | 1 | 15 | Ppp4c | 0.094 | Young |
| 0.186581 | 0.576999 | 0.482 | 0.388 | 1 | 15 | Ncf2 | 0.094 | Young |
| 0.186873 | 1.232998 | 0.196 | 0.102 | 1 | 15 | Gpat4 | 0.094 | Young |
| 0.186873 | 0.775588 | 0.107 | 0.204 | 1 | 15 | Ppfia1 | -0.097 | Young |

| | | | | | | | | |
|---|---|---|---|---|---|---|---|---|
| 0.187003 | 2.328312 | 0.107 | 0.041 | 1 | 15 | Nbn | 0.066 | Young |
| 0.187003 | 2.196681 | 0.107 | 0.041 | 1 | 15 | Slc35c2 | 0.066 | Young |
| 0.187003 | 2.119541 | 0.107 | 0.041 | 1 | 15 | Gm11772 | 0.066 | Young |
| 0.187003 | 1.979352 | 0.107 | 0.041 | 1 | 15 | Dpp9 | 0.066 | Young |
| 0.187003 | 1.896883 | 0.107 | 0.041 | 1 | 15 | Pik3r2 | 0.066 | Young |
| 0.187205 | 0.535835 | 0.696 | 0.633 | 1 | 15 | Tspan3 | 0.063 | Young |
| 0.187205 | 0.347837 | 0.714 | 0.612 | 1 | 15 | Aplp2 | 0.102 | Young |
| 0.187272 | 0.5492 | 0.375 | 0.245 | 1 | 15 | Irf4 | 0.13 | Young |
| 0.187551 | 0.30791 | 0.732 | 0.571 | 1 | 15 | Atp5k | 0.161 | Young |
| 0.188289 | 4.843392 | 0.036 | 0 | 1 | 15 | Gadd45a | 0.036 | Young |
| 0.188289 | 4.400041 | 0.036 | 0 | 1 | 15 | Mlh1 | 0.036 | Young |
| 0.188289 | 4.358078 | 0.036 | 0 | 1 | 15 | Bckdhb | 0.036 | Young |
| 0.188289 | 4.276855 | 0.036 | 0 | 1 | 15 | Gm39469 | 0.036 | Young |
| 0.188289 | 4.257858 | 0.036 | 0 | 1 | 15 | C530008M | 0.036 | Young |
| 0.188289 | 4.114293 | 0.036 | 0 | 1 | 15 | Gab3 | 0.036 | Young |
| 0.188289 | 3.963269 | 0.036 | 0 | 1 | 15 | Rmdn1 | 0.036 | Young |
| 0.188289 | 3.95719 | 0.036 | 0 | 1 | 15 | Rep15 | 0.036 | Young |
| 0.188289 | 3.927139 | 0.036 | 0 | 1 | 15 | Dao | 0.036 | Young |
| 0.188289 | 3.927139 | 0.036 | 0 | 1 | 15 | Cep70 | 0.036 | Young |
| 0.188289 | 3.862721 | 0.036 | 0 | 1 | 15 | Tmem140 | 0.036 | Young |
| 0.188289 | 3.75258 | 0.036 | 0 | 1 | 15 | 6330418K0 | 0.036 | Young |
| 0.188289 | 3.735357 | 0.036 | 0 | 1 | 15 | Fn3krp | 0.036 | Young |
| 0.188289 | 3.733774 | 0.036 | 0 | 1 | 15 | Ipcef1 | 0.036 | Young |
| 0.188289 | 3.698974 | 0.036 | 0 | 1 | 15 | Aldh4a1 | 0.036 | Young |
| 0.188289 | 3.693851 | 0.036 | 0 | 1 | 15 | Met | 0.036 | Young |
| 0.188289 | 3.682252 | 0.036 | 0 | 1 | 15 | E130102H2 | 0.036 | Young |
| 0.188289 | 3.66838 | 0.036 | 0 | 1 | 15 | Ggcx | 0.036 | Young |
| 0.188289 | 3.643739 | 0.036 | 0 | 1 | 15 | Ddx55 | 0.036 | Young |
| 0.188289 | 3.643545 | 0.036 | 0 | 1 | 15 | Sgsm2 | 0.036 | Young |
| 0.188289 | 3.613941 | 0.036 | 0 | 1 | 15 | Gm26890 | 0.036 | Young |
| 0.188289 | 3.600168 | 0.036 | 0 | 1 | 15 | Twist1 | 0.036 | Young |
| 0.188289 | 3.562903 | 0.036 | 0 | 1 | 15 | Flrt2 | 0.036 | Young |
| 0.188289 | 3.559508 | 0.036 | 0 | 1 | 15 | Shb | 0.036 | Young |
| 0.188289 | 3.53592 | 0.036 | 0 | 1 | 15 | Acvr1b | 0.036 | Young |
| 0.188289 | 3.515978 | 0.036 | 0 | 1 | 15 | Zfp58 | 0.036 | Young |
| 0.188289 | 3.50843 | 0.036 | 0 | 1 | 15 | Src | 0.036 | Young |
| 0.188289 | 3.504004 | 0.036 | 0 | 1 | 15 | Arap3 | 0.036 | Young |
| 0.188289 | 3.499545 | 0.036 | 0 | 1 | 15 | Gm29336 | 0.036 | Young |
| 0.188289 | 3.497199 | 0.036 | 0 | 1 | 15 | Pigw | 0.036 | Young |
| 0.188289 | 3.491324 | 0.036 | 0 | 1 | 15 | Mrpl46 | 0.036 | Young |
| 0.188289 | 3.48008 | 0.036 | 0 | 1 | 15 | Haus7 | 0.036 | Young |
| 0.188289 | 3.476699 | 0.036 | 0 | 1 | 15 | Gm20324 | 0.036 | Young |
| 0.188289 | 3.475496 | 0.036 | 0 | 1 | 15 | Ube2o | 0.036 | Young |
| 0.188289 | 3.471788 | 0.036 | 0 | 1 | 15 | Mylk | 0.036 | Young |
| 0.188289 | 3.468033 | 0.036 | 0 | 1 | 15 | Tmem242 | 0.036 | Young |

| | | | | | | | | |
|---|---|---|---|---|---|---|---|---|
| 0.188289 | 3.454675 | 0.036 | 0 | 1 | 15 | B130006D( | 0.036 | Young |
| 0.188289 | 3.453912 | 0.036 | 0 | 1 | 15 | C730034F( | 0.036 | Young |
| 0.188289 | 3.453668 | 0.036 | 0 | 1 | 15 | Agbl3 | 0.036 | Young |
| 0.188289 | 3.449059 | 0.036 | 0 | 1 | 15 | Fam102a | 0.036 | Young |
| 0.188289 | 3.432261 | 0.036 | 0 | 1 | 15 | Ccdc85a | 0.036 | Young |
| 0.188289 | 3.429208 | 0.036 | 0 | 1 | 15 | Rccd1 | 0.036 | Young |
| 0.188289 | 3.427294 | 0.036 | 0 | 1 | 15 | H2-Q10 | 0.036 | Young |
| 0.188289 | 3.416764 | 0.036 | 0 | 1 | 15 | Gstk1 | 0.036 | Young |
| 0.188289 | 3.406626 | 0.036 | 0 | 1 | 15 | Cep55 | 0.036 | Young |
| 0.188289 | 3.406227 | 0.036 | 0 | 1 | 15 | Ncs1 | 0.036 | Young |
| 0.188289 | 3.402112 | 0.036 | 0 | 1 | 15 | Utp23 | 0.036 | Young |
| 0.188289 | 3.400419 | 0.036 | 0 | 1 | 15 | Mad2l1 | 0.036 | Young |
| 0.188289 | 3.398729 | 0.036 | 0 | 1 | 15 | Cep19 | 0.036 | Young |
| 0.188289 | 3.393129 | 0.036 | 0 | 1 | 15 | Bcar3 | 0.036 | Young |
| 0.188289 | 3.38518 | 0.036 | 0 | 1 | 15 | Knl1 | 0.036 | Young |
| 0.188289 | 3.350996 | 0.036 | 0 | 1 | 15 | Poldip2 | 0.036 | Young |
| 0.188289 | 3.329818 | 0.036 | 0 | 1 | 15 | Thoc5 | 0.036 | Young |
| 0.188289 | 3.32574 | 0.036 | 0 | 1 | 15 | C2cd2 | 0.036 | Young |
| 0.188289 | 3.300548 | 0.036 | 0 | 1 | 15 | Cdc7 | 0.036 | Young |
| 0.188289 | 3.280711 | 0.036 | 0 | 1 | 15 | Hist2h2aa1 | 0.036 | Young |
| 0.188289 | 3.27713 | 0.036 | 0 | 1 | 15 | Carf | 0.036 | Young |
| 0.188289 | 3.271867 | 0.036 | 0 | 1 | 15 | Mfsd4a | 0.036 | Young |
| 0.188289 | 3.271867 | 0.036 | 0 | 1 | 15 | Zfp932 | 0.036 | Young |
| 0.188289 | 3.26623 | 0.036 | 0 | 1 | 15 | Arv1 | 0.036 | Young |
| 0.188289 | 3.26623 | 0.036 | 0 | 1 | 15 | 9130011E1 | 0.036 | Young |
| 0.188289 | 3.260624 | 0.036 | 0 | 1 | 15 | A930006I0 | 0.036 | Young |
| 0.188289 | 3.255839 | 0.036 | 0 | 1 | 15 | Gna11 | 0.036 | Young |
| 0.188289 | 3.253321 | 0.036 | 0 | 1 | 15 | Fstl1 | 0.036 | Young |
| 0.188289 | 3.249562 | 0.036 | 0 | 1 | 15 | Entpd5 | 0.036 | Young |
| 0.188289 | 3.248488 | 0.036 | 0 | 1 | 15 | Macc1 | 0.036 | Young |
| 0.188289 | 3.247256 | 0.036 | 0 | 1 | 15 | Brat1 | 0.036 | Young |
| 0.188289 | 3.239342 | 0.036 | 0 | 1 | 15 | Mum1 | 0.036 | Young |
| 0.188289 | 3.220725 | 0.036 | 0 | 1 | 15 | Chkb | 0.036 | Young |
| 0.188289 | 3.190128 | 0.036 | 0 | 1 | 15 | Oas1a | 0.036 | Young |
| 0.188289 | 3.184671 | 0.036 | 0 | 1 | 15 | Gm16536 | 0.036 | Young |
| 0.188289 | 3.18417 | 0.036 | 0 | 1 | 15 | Oaf | 0.036 | Young |
| 0.188289 | 3.174896 | 0.036 | 0 | 1 | 15 | Mr1 | 0.036 | Young |
| 0.188289 | 3.171056 | 0.036 | 0 | 1 | 15 | Vps33b | 0.036 | Young |
| 0.188289 | 3.169381 | 0.036 | 0 | 1 | 15 | Prpf3 | 0.036 | Young |
| 0.188289 | 3.169381 | 0.036 | 0 | 1 | 15 | Ifrd2 | 0.036 | Young |
| 0.188289 | 3.167421 | 0.036 | 0 | 1 | 15 | Fam110b | 0.036 | Young |
| 0.188289 | 3.162634 | 0.036 | 0 | 1 | 15 | Ccdc149 | 0.036 | Young |
| 0.188289 | 3.145416 | 0.036 | 0 | 1 | 15 | Trim32 | 0.036 | Young |
| 0.188289 | 3.139254 | 0.036 | 0 | 1 | 15 | Nckap1 | 0.036 | Young |
| 0.188289 | 3.130389 | 0.036 | 0 | 1 | 15 | Rbfox2 | 0.036 | Young |

| | | | | | | | | |
|---|---|---|---|---|---|---|---|---|
| 0.188289 | 3.12935 | 0.036 | 0 | 1 | 15 | Fer | 0.036 | Young |
| 0.188289 | 3.115182 | 0.036 | 0 | 1 | 15 | Gm9938 | 0.036 | Young |
| 0.188289 | 3.100905 | 0.036 | 0 | 1 | 15 | Xrcc6 | 0.036 | Young |
| 0.188289 | 3.096203 | 0.036 | 0 | 1 | 15 | Icam4 | 0.036 | Young |
| 0.188289 | 3.092709 | 0.036 | 0 | 1 | 15 | Ighd | 0.036 | Young |
| 0.188289 | 3.089137 | 0.036 | 0 | 1 | 15 | Inca1 | 0.036 | Young |
| 0.188289 | 3.077957 | 0.036 | 0 | 1 | 15 | Krtcap3 | 0.036 | Young |
| 0.188289 | 3.074887 | 0.036 | 0 | 1 | 15 | Mok | 0.036 | Young |
| 0.188289 | 3.061327 | 0.036 | 0 | 1 | 15 | Ebf1 | 0.036 | Young |
| 0.188289 | 3.047994 | 0.036 | 0 | 1 | 15 | H3f3aos | 0.036 | Young |
| 0.188289 | 3.046906 | 0.036 | 0 | 1 | 15 | Clic5 | 0.036 | Young |
| 0.188289 | 3.044046 | 0.036 | 0 | 1 | 15 | 0610010K1 | 0.036 | Young |
| 0.188289 | 3.023858 | 0.036 | 0 | 1 | 15 | Egfr | 0.036 | Young |
| 0.188289 | 3.020788 | 0.036 | 0 | 1 | 15 | Ttf2 | 0.036 | Young |
| 0.188289 | 3.014172 | 0.036 | 0 | 1 | 15 | Pgpep1 | 0.036 | Young |
| 0.188289 | 3.011556 | 0.036 | 0 | 1 | 15 | 4931440F1 | 0.036 | Young |
| 0.188289 | 2.987056 | 0.036 | 0 | 1 | 15 | Mettl1 | 0.036 | Young |
| 0.188289 | 2.986515 | 0.036 | 0 | 1 | 15 | Bmper | 0.036 | Young |
| 0.188289 | 2.984886 | 0.036 | 0 | 1 | 15 | Zcchc10 | 0.036 | Young |
| 0.188289 | 2.977948 | 0.036 | 0 | 1 | 15 | Siglec1 | 0.036 | Young |
| 0.188289 | 2.975315 | 0.036 | 0 | 1 | 15 | Trim39 | 0.036 | Young |
| 0.188289 | 2.96157 | 0.036 | 0 | 1 | 15 | Cox18 | 0.036 | Young |
| 0.188289 | 2.953874 | 0.036 | 0 | 1 | 15 | Shroom1 | 0.036 | Young |
| 0.188289 | 2.952383 | 0.036 | 0 | 1 | 15 | Gm1720 | 0.036 | Young |
| 0.188289 | 2.946114 | 0.036 | 0 | 1 | 15 | Jade3 | 0.036 | Young |
| 0.188289 | 2.944284 | 0.036 | 0 | 1 | 15 | Aatf | 0.036 | Young |
| 0.188289 | 2.935936 | 0.036 | 0 | 1 | 15 | Mcm7 | 0.036 | Young |
| 0.188289 | 2.927544 | 0.036 | 0 | 1 | 15 | Pms1 | 0.036 | Young |
| 0.188289 | 2.924367 | 0.036 | 0 | 1 | 15 | Zfp607a | 0.036 | Young |
| 0.188289 | 2.917145 | 0.036 | 0 | 1 | 15 | Bcas1 | 0.036 | Young |
| 0.188289 | 2.909514 | 0.036 | 0 | 1 | 15 | Bpgm | 0.036 | Young |
| 0.188289 | 2.901096 | 0.036 | 0 | 1 | 15 | Ssbp2 | 0.036 | Young |
| 0.188289 | 2.893614 | 0.036 | 0 | 1 | 15 | Mtmr11 | 0.036 | Young |
| 0.188289 | 2.888995 | 0.036 | 0 | 1 | 15 | Retn | 0.036 | Young |
| 0.188289 | 2.888672 | 0.036 | 0 | 1 | 15 | Bod1 | 0.036 | Young |
| 0.188289 | 2.883698 | 0.036 | 0 | 1 | 15 | Abca5 | 0.036 | Young |
| 0.188289 | 2.874451 | 0.036 | 0 | 1 | 15 | Ipo11 | 0.036 | Young |
| 0.188289 | 2.867649 | 0.036 | 0 | 1 | 15 | Tnfsf13b | 0.036 | Young |
| 0.188289 | 2.867127 | 0.036 | 0 | 1 | 15 | Fbxl19 | 0.036 | Young |
| 0.188289 | 2.862874 | 0.036 | 0 | 1 | 15 | Spp1 | 0.036 | Young |
| 0.188289 | 2.862874 | 0.036 | 0 | 1 | 15 | Gm4924 | 0.036 | Young |
| 0.188289 | 2.861395 | 0.036 | 0 | 1 | 15 | C030034L1 | 0.036 | Young |
| 0.188289 | 2.859131 | 0.036 | 0 | 1 | 15 | Smad3 | 0.036 | Young |
| 0.188289 | 2.857288 | 0.036 | 0 | 1 | 15 | Pcbp3 | 0.036 | Young |
| 0.188289 | 2.850113 | 0.036 | 0 | 1 | 15 | Yod1 | 0.036 | Young |

| | | | | | | | | |
|---|---|---|---|---|---|---|---|---|
| 0.188289 | 2.850021 | 0.036 | 0 | 1 | 15 | Zfp868 | 0.036 | Young |
| 0.188289 | 2.84625 | 0.036 | 0 | 1 | 15 | Gnptg | 0.036 | Young |
| 0.188289 | 2.844502 | 0.036 | 0 | 1 | 15 | Wdr4 | 0.036 | Young |
| 0.188289 | 2.844147 | 0.036 | 0 | 1 | 15 | Zhx3 | 0.036 | Young |
| 0.188289 | 2.840332 | 0.036 | 0 | 1 | 15 | Sgcb | 0.036 | Young |
| 0.188289 | 2.840326 | 0.036 | 0 | 1 | 15 | Cdo1 | 0.036 | Young |
| 0.188289 | 2.836529 | 0.036 | 0 | 1 | 15 | Pex14 | 0.036 | Young |
| 0.188289 | 2.83044 | 0.036 | 0 | 1 | 15 | Ddb2 | 0.036 | Young |
| 0.188289 | 2.825045 | 0.036 | 0 | 1 | 15 | Abhd15 | 0.036 | Young |
| 0.188289 | 2.82019 | 0.036 | 0 | 1 | 15 | Scyl3 | 0.036 | Young |
| 0.188289 | 2.82019 | 0.036 | 0 | 1 | 15 | Ier5l | 0.036 | Young |
| 0.188289 | 2.8175 | 0.036 | 0 | 1 | 15 | Gpr141 | 0.036 | Young |
| 0.188289 | 2.816147 | 0.036 | 0 | 1 | 15 | Bbs12 | 0.036 | Young |
| 0.188289 | 2.800117 | 0.036 | 0 | 1 | 15 | Lrrc14 | 0.036 | Young |
| 0.188289 | 2.774927 | 0.036 | 0 | 1 | 15 | Myh10 | 0.036 | Young |
| 0.188289 | 2.772728 | 0.036 | 0 | 1 | 15 | Trit1 | 0.036 | Young |
| 0.188289 | 2.772728 | 0.036 | 0 | 1 | 15 | Mturn | 0.036 | Young |
| 0.188289 | 2.772152 | 0.036 | 0 | 1 | 15 | Dcakd | 0.036 | Young |
| 0.188289 | 2.770702 | 0.036 | 0 | 1 | 15 | Htra1 | 0.036 | Young |
| 0.188289 | 2.769336 | 0.036 | 0 | 1 | 15 | Polr3g | 0.036 | Young |
| 0.188289 | 2.764764 | 0.036 | 0 | 1 | 15 | Tmem79 | 0.036 | Young |
| 0.188289 | 2.764288 | 0.036 | 0 | 1 | 15 | Trim62 | 0.036 | Young |
| 0.188289 | 2.763339 | 0.036 | 0 | 1 | 15 | Helq | 0.036 | Young |
| 0.188289 | 2.757714 | 0.036 | 0 | 1 | 15 | Nr1h3 | 0.036 | Young |
| 0.188289 | 2.757123 | 0.036 | 0 | 1 | 15 | Ndst1 | 0.036 | Young |
| 0.188289 | 2.736689 | 0.036 | 0 | 1 | 15 | Plscr4 | 0.036 | Young |
| 0.188289 | 2.717678 | 0.036 | 0 | 1 | 15 | Plcb1 | 0.036 | Young |
| 0.188289 | 2.71761 | 0.036 | 0 | 1 | 15 | Sccpdh | 0.036 | Young |
| 0.188289 | 2.711231 | 0.036 | 0 | 1 | 15 | Gm15564 | 0.036 | Young |
| 0.188289 | 2.701708 | 0.036 | 0 | 1 | 15 | Sirpb1b | 0.036 | Young |
| 0.188289 | 2.691199 | 0.036 | 0 | 1 | 15 | Tppp | 0.036 | Young |
| 0.188289 | 2.690511 | 0.036 | 0 | 1 | 15 | Abt1 | 0.036 | Young |
| 0.188289 | 2.690156 | 0.036 | 0 | 1 | 15 | Cd101 | 0.036 | Young |
| 0.188289 | 2.688593 | 0.036 | 0 | 1 | 15 | Zfp287 | 0.036 | Young |
| 0.188289 | 2.688465 | 0.036 | 0 | 1 | 15 | Tesk2 | 0.036 | Young |
| 0.188289 | 2.675408 | 0.036 | 0 | 1 | 15 | Hemk1 | 0.036 | Young |
| 0.188289 | 2.673934 | 0.036 | 0 | 1 | 15 | Serpine1 | 0.036 | Young |
| 0.188289 | 2.652688 | 0.036 | 0 | 1 | 15 | Trim45 | 0.036 | Young |
| 0.188289 | 2.645902 | 0.036 | 0 | 1 | 15 | 1600010M | 0.036 | Young |
| 0.188289 | 2.645585 | 0.036 | 0 | 1 | 15 | Hp | 0.036 | Young |
| 0.188289 | 2.645585 | 0.036 | 0 | 1 | 15 | Ccdc94 | 0.036 | Young |
| 0.188289 | 2.644801 | 0.036 | 0 | 1 | 15 | Zfp668 | 0.036 | Young |
| 0.188289 | 2.613778 | 0.036 | 0 | 1 | 15 | Naf1 | 0.036 | Young |
| 0.188289 | 2.612317 | 0.036 | 0 | 1 | 15 | Slc25a40 | 0.036 | Young |
| 0.188289 | 2.612317 | 0.036 | 0 | 1 | 15 | Snapc2 | 0.036 | Young |

| | | | | | | | | |
|---|---|---|---|---|---|---|---|---|
| 0.188289 | 2.597498 | 0.036 | 0 | 1 | 15 | Hes7 | 0.036 | Young |
| 0.188289 | 2.59454 | 0.036 | 0 | 1 | 15 | Gcat | 0.036 | Young |
| 0.188289 | 2.592733 | 0.036 | 0 | 1 | 15 | Bclaf3 | 0.036 | Young |
| 0.188289 | 2.551495 | 0.036 | 0 | 1 | 15 | Maml2 | 0.036 | Young |
| 0.188289 | 2.545385 | 0.036 | 0 | 1 | 15 | Apbb2 | 0.036 | Young |
| 0.188289 | 2.545385 | 0.036 | 0 | 1 | 15 | Wtip | 0.036 | Young |
| 0.188289 | 2.545385 | 0.036 | 0 | 1 | 15 | BC043934 | 0.036 | Young |
| 0.188289 | 2.539789 | 0.036 | 0 | 1 | 15 | Abhd14b | 0.036 | Young |
| 0.188289 | 2.537065 | 0.036 | 0 | 1 | 15 | Ctla2b | 0.036 | Young |
| 0.188289 | 2.515935 | 0.036 | 0 | 1 | 15 | Zfp931 | 0.036 | Young |
| 0.188289 | 2.515935 | 0.036 | 0 | 1 | 15 | Tmem53 | 0.036 | Young |
| 0.188289 | 2.509396 | 0.036 | 0 | 1 | 15 | Klhdc1 | 0.036 | Young |
| 0.188289 | 2.50261 | 0.036 | 0 | 1 | 15 | Gpm6b | 0.036 | Young |
| 0.188289 | 2.50261 | 0.036 | 0 | 1 | 15 | Ankrd26 | 0.036 | Young |
| 0.188289 | 2.50261 | 0.036 | 0 | 1 | 15 | Tecpr2 | 0.036 | Young |
| 0.188289 | 2.501005 | 0.036 | 0 | 1 | 15 | Ifi213 | 0.036 | Young |
| 0.188289 | 2.501005 | 0.036 | 0 | 1 | 15 | Elk1 | 0.036 | Young |
| 0.188289 | 2.496147 | 0.036 | 0 | 1 | 15 | Unc119b | 0.036 | Young |
| 0.188289 | 2.496147 | 0.036 | 0 | 1 | 15 | Gnal | 0.036 | Young |
| 0.188289 | 2.459282 | 0.036 | 0 | 1 | 15 | Tpk1 | 0.036 | Young |
| 0.188289 | 2.456873 | 0.036 | 0 | 1 | 15 | Urb2 | 0.036 | Young |
| 0.188289 | 2.450319 | 0.036 | 0 | 1 | 15 | Rsl1 | 0.036 | Young |
| 0.188289 | 2.443198 | 0.036 | 0 | 1 | 15 | Tbc1d7 | 0.036 | Young |
| 0.188289 | 2.436682 | 0.036 | 0 | 1 | 15 | Col4a1 | 0.036 | Young |
| 0.188289 | 2.422812 | 0.036 | 0 | 1 | 15 | Slc24a5 | 0.036 | Young |
| 0.188289 | 2.411875 | 0.036 | 0 | 1 | 15 | Dusp12 | 0.036 | Young |
| 0.188289 | 2.411875 | 0.036 | 0 | 1 | 15 | Pms2 | 0.036 | Young |
| 0.188289 | 2.403939 | 0.036 | 0 | 1 | 15 | Bivm | 0.036 | Young |
| 0.188289 | 2.403939 | 0.036 | 0 | 1 | 15 | Dennd2a | 0.036 | Young |
| 0.188289 | 2.403939 | 0.036 | 0 | 1 | 15 | Poli | 0.036 | Young |
| 0.188289 | 2.394879 | 0.036 | 0 | 1 | 15 | Mtrf1l | 0.036 | Young |
| 0.188289 | 2.390854 | 0.036 | 0 | 1 | 15 | Gm15545 | 0.036 | Young |
| 0.188289 | 2.390854 | 0.036 | 0 | 1 | 15 | Zfp764 | 0.036 | Young |
| 0.188289 | 2.390854 | 0.036 | 0 | 1 | 15 | Nt5dc2 | 0.036 | Young |
| 0.188289 | 2.38429 | 0.036 | 0 | 1 | 15 | Fastkd2 | 0.036 | Young |
| 0.188289 | 2.38429 | 0.036 | 0 | 1 | 15 | Aldh1b1 | 0.036 | Young |
| 0.188289 | 2.351376 | 0.036 | 0 | 1 | 15 | Gm14326 | 0.036 | Young |
| 0.188289 | 2.343099 | 0.036 | 0 | 1 | 15 | Il1a | 0.036 | Young |
| 0.188289 | 2.325525 | 0.036 | 0 | 1 | 15 | Lmbr1 | 0.036 | Young |
| 0.188289 | 2.298714 | 0.036 | 0 | 1 | 15 | Nek4 | 0.036 | Young |
| 0.188289 | 2.293774 | 0.036 | 0 | 1 | 15 | Dclre1b | 0.036 | Young |
| 0.188289 | 2.293774 | 0.036 | 0 | 1 | 15 | Asgr2 | 0.036 | Young |
| 0.188289 | 2.287249 | 0.036 | 0 | 1 | 15 | Ccl25 | 0.036 | Young |
| 0.188289 | 2.277608 | 0.036 | 0 | 1 | 15 | Anks6 | 0.036 | Young |
| 0.188289 | 2.277608 | 0.036 | 0 | 1 | 15 | Hic2 | 0.036 | Young |

| | | | | | | | | |
|---|---|---|---|---|---|---|---|---|
| 0.188289 | 2.269039 | 0.036 | 0 | 1 | 15 | Gm37422 | 0.036 | Young |
| 0.188289 | 2.254569 | 0.036 | 0 | 1 | 15 | Coro2b | 0.036 | Young |
| 0.188289 | 2.254569 | 0.036 | 0 | 1 | 15 | Rnd1 | 0.036 | Young |
| 0.188289 | 2.247353 | 0.036 | 0 | 1 | 15 | 1110034G2 | 0.036 | Young |
| 0.188289 | 2.243295 | 0.036 | 0 | 1 | 15 | Tln2 | 0.036 | Young |
| 0.188289 | 2.234011 | 0.036 | 0 | 1 | 15 | Als2cl | 0.036 | Young |
| 0.188289 | 2.225951 | 0.036 | 0 | 1 | 15 | Fars2 | 0.036 | Young |
| 0.188289 | 2.22503 | 0.036 | 0 | 1 | 15 | Gm10138 | 0.036 | Young |
| 0.188289 | 2.17142 | 0.036 | 0 | 1 | 15 | Fgf9 | 0.036 | Young |
| 0.188289 | 2.137425 | 0.036 | 0 | 1 | 15 | Polr3a | 0.036 | Young |
| 0.188289 | 2.129596 | 0.036 | 0 | 1 | 15 | Taf1a | 0.036 | Young |
| 0.188289 | 2.129596 | 0.036 | 0 | 1 | 15 | Klrb1a | 0.036 | Young |
| 0.188289 | 2.129596 | 0.036 | 0 | 1 | 15 | Zfp119b | 0.036 | Young |
| 0.188289 | 2.119939 | 0.036 | 0 | 1 | 15 | C130013H( | 0.036 | Young |
| 0.188289 | 2.119939 | 0.036 | 0 | 1 | 15 | Bdh1 | 0.036 | Young |
| 0.188289 | 2.112417 | 0.036 | 0 | 1 | 15 | Zfp335 | 0.036 | Young |
| 0.188289 | 2.105321 | 0.036 | 0 | 1 | 15 | Rpa2 | 0.036 | Young |
| 0.188289 | 2.101329 | 0.036 | 0 | 1 | 15 | Kbtbd8 | 0.036 | Young |
| 0.188289 | 2.070397 | 0.036 | 0 | 1 | 15 | Stradb | 0.036 | Young |
| 0.188289 | 2.062195 | 0.036 | 0 | 1 | 15 | Dolk | 0.036 | Young |
| 0.188289 | 2.062195 | 0.036 | 0 | 1 | 15 | Arsi | 0.036 | Young |
| 0.188289 | 2.043945 | 0.036 | 0 | 1 | 15 | Cnot11 | 0.036 | Young |
| 0.188289 | 2.041775 | 0.036 | 0 | 1 | 15 | Il1rl2 | 0.036 | Young |
| 0.188289 | 1.999384 | 0.036 | 0 | 1 | 15 | H6pd | 0.036 | Young |
| 0.188289 | 1.994902 | 0.036 | 0 | 1 | 15 | Slc9a4 | 0.036 | Young |
| 0.188289 | 1.994902 | 0.036 | 0 | 1 | 15 | Dedd | 0.036 | Young |
| 0.188289 | 1.986259 | 0.036 | 0 | 1 | 15 | Ada | 0.036 | Young |
| 0.188289 | 1.986259 | 0.036 | 0 | 1 | 15 | Fam57a | 0.036 | Young |
| 0.188289 | 1.960637 | 0.036 | 0 | 1 | 15 | Pkmyt1 | 0.036 | Young |
| 0.188289 | 1.930879 | 0.036 | 0 | 1 | 15 | Stac2 | 0.036 | Young |
| 0.188289 | 1.921842 | 0.036 | 0 | 1 | 15 | Thada | 0.036 | Young |
| 0.188289 | 1.843058 | 0.036 | 0 | 1 | 15 | Rusc2 | 0.036 | Young |
| 0.188289 | 1.833452 | 0.036 | 0 | 1 | 15 | Kif19a | 0.036 | Young |
| 0.188289 | 1.771534 | 0.036 | 0 | 1 | 15 | Trip6 | 0.036 | Young |
| 0.188289 | 1.761438 | 0.036 | 0 | 1 | 15 | Ptger2 | 0.036 | Young |
| 0.188295 | 1.650029 | 0.214 | 0.122 | 1 | 15 | Igf1r | 0.092 | Young |
| 0.189025 | 1.775007 | 0.161 | 0.082 | 1 | 15 | Opa3 | 0.079 | Young |
| 0.189592 | 0.204633 | 0.482 | 0.327 | 1 | 15 | Uap1 | 0.155 | Young |
| 0.189814 | 0.899035 | 0.143 | 0.061 | 1 | 15 | Rdm1 | 0.082 | Young |
| 0.189814 | 0.886517 | 0.143 | 0.061 | 1 | 15 | Zfp120 | 0.082 | Young |
| 0.189814 | 0.756533 | 0.143 | 0.061 | 1 | 15 | Socs6 | 0.082 | Young |
| 0.189814 | 0.298953 | 0.143 | 0.061 | 1 | 15 | Gtf2e1 | 0.082 | Young |
| 0.190298 | 0.632041 | 0.304 | 0.184 | 1 | 15 | Selenof | 0.12 | Young |
| 0.190298 | 0.436122 | 0.304 | 0.184 | 1 | 15 | Vwa5a | 0.12 | Young |
| 0.190326 | 1.135043 | 0.196 | 0.102 | 1 | 15 | Rnase4 | 0.094 | Young |

| | | | | | | | | |
|---|---|---|---|---|---|---|---|---|
| 0.191601 | 1.285988 | 0.214 | 0.122 | 1 | 15 | Slc12a6 | 0.092 | Young |
| 0.191646 | 0.230862 | 0.589 | 0.449 | 1 | 15 | Ghitm | 0.14 | Young |
| 0.191711 | 2.345883 | 0.107 | 0.041 | 1 | 15 | Flcn | 0.066 | Young |
| 0.191711 | 2.243457 | 0.107 | 0.041 | 1 | 15 | Uvssa | 0.066 | Young |
| 0.191711 | 1.996811 | 0.107 | 0.041 | 1 | 15 | Prkrip1 | 0.066 | Young |
| 0.191711 | 1.851494 | 0.107 | 0.041 | 1 | 15 | Eif2b5 | 0.066 | Young |
| 0.191711 | 1.595476 | 0.107 | 0.041 | 1 | 15 | Eepd1 | 0.066 | Young |
| 0.191999 | 1.363026 | 0.268 | 0.163 | 1 | 15 | Ndufs4 | 0.105 | Young |
| 0.191999 | 0.882956 | 0.268 | 0.163 | 1 | 15 | Sh2b3 | 0.105 | Young |
| 0.192579 | 0.723052 | 0.286 | 0.184 | 1 | 15 | Atmin | 0.102 | Young |
| 0.193271 | 0.284321 | 0.964 | 0.878 | 1 | 15 | Rpl36 | 0.086 | Young |
| 0.193826 | 0.805834 | 0.196 | 0.102 | 1 | 15 | Trim41 | 0.094 | Young |
| 0.193826 | 0.772518 | 0.196 | 0.102 | 1 | 15 | Nup93 | 0.094 | Young |
| 0.193928 | 0.837818 | 0.143 | 0.061 | 1 | 15 | Utp6 | 0.082 | Young |
| 0.193928 | 0.684808 | 0.143 | 0.061 | 1 | 15 | Ptrhd1 | 0.082 | Young |
| 0.194192 | 1.332423 | 0.232 | 0.143 | 1 | 15 | Sptlc2 | 0.089 | Young |
| 0.19587 | 0.344977 | 0.75 | 0.633 | 1 | 15 | Serpinb9 | 0.117 | Young |
| 0.196089 | 0.564398 | 0.429 | 0.306 | 1 | 15 | Lims1 | 0.123 | Young |
| 0.196454 | 0.178535 | 0.571 | 0.429 | 1 | 15 | Mthfd2 | 0.142 | Young |
| 0.196506 | 1.995633 | 0.107 | 0.041 | 1 | 15 | Igip | 0.066 | Young |
| 0.196506 | 1.799601 | 0.107 | 0.041 | 1 | 15 | Txnl4a | 0.066 | Young |
| 0.196506 | 1.71663 | 0.107 | 0.041 | 1 | 15 | Brip1os | 0.066 | Young |
| 0.196506 | 1.646666 | 0.107 | 0.041 | 1 | 15 | Ptgr2 | 0.066 | Young |
| 0.196506 | 1.609005 | 0.107 | 0.041 | 1 | 15 | Il1rl1 | 0.066 | Young |
| 0.196506 | 1.472 | 0.107 | 0.041 | 1 | 15 | Mrpl4 | 0.066 | Young |
| 0.196506 | 1.120858 | 0.107 | 0.041 | 1 | 15 | Kri1 | 0.066 | Young |
| 0.196698 | 1.47772 | 0.161 | 0.082 | 1 | 15 | Keap1 | 0.079 | Young |
| 0.197373 | 0.280267 | 0.196 | 0.102 | 1 | 15 | Fam98b | 0.094 | Young |
| 0.198108 | 0.888288 | 0.143 | 0.061 | 1 | 15 | 4931406P1 | 0.082 | Young |
| 0.198108 | 0.447908 | 0.143 | 0.061 | 1 | 15 | Oard1 | 0.082 | Young |
| 0.198493 | 0.266439 | 0.482 | 0.347 | 1 | 15 | Cpsf6 | 0.135 | Young |
| 0.198669 | 0.434682 | 0.5 | 0.367 | 1 | 15 | Cybb | 0.133 | Young |
| 0.198825 | 0.051466 | 0.036 | 0.102 | 1 | 15 | Siglech | -0.066 | Young |
| 0.199167 | 0.728326 | 0.321 | 0.224 | 1 | 15 | Ggnbp2 | 0.097 | Young |
| 0.19967 | 1.711279 | 0.179 | 0.102 | 1 | 15 | Lars | 0.077 | Young |
| 0.200619 | 2.166994 | 0.161 | 0.082 | 1 | 15 | Samm50 | 0.079 | Young |
| 0.200619 | 1.593666 | 0.161 | 0.082 | 1 | 15 | Mybbp1a | 0.079 | Young |
| 0.200619 | 1.353256 | 0.161 | 0.082 | 1 | 15 | Virma | 0.079 | Young |
| 0.20071 | 1.166278 | 0.232 | 0.143 | 1 | 15 | Ylpm1 | 0.089 | Young |
| 0.200967 | 0.897136 | 0.196 | 0.102 | 1 | 15 | Zfp366 | 0.094 | Young |
| 0.200967 | 0.792266 | 0.196 | 0.102 | 1 | 15 | Kdm3b | 0.094 | Young |
| 0.201388 | 2.091754 | 0.107 | 0.041 | 1 | 15 | Tfeb | 0.066 | Young |
| 0.201388 | 1.537656 | 0.107 | 0.041 | 1 | 15 | Eif4ebp1 | 0.066 | Young |
| 0.201388 | 1.487323 | 0.107 | 0.041 | 1 | 15 | Zfp358 | 0.066 | Young |
| 0.201388 | 1.477128 | 0.107 | 0.041 | 1 | 15 | Fbxo7 | 0.066 | Young |

| | | | | | | | | |
|---|---|---|---|---|---|---|---|---|
| 0.201388 | 1.406792 | 0.107 | 0.041 | 1 | 15 | Gpr160 | 0.066 | Young |
| 0.201388 | 1.28524 | 0.107 | 0.041 | 1 | 15 | Rpp21 | 0.066 | Young |
| 0.201388 | 1.080311 | 0.107 | 0.041 | 1 | 15 | Gem | 0.066 | Young |
| 0.201388 | 1.022549 | 0.107 | 0.041 | 1 | 15 | Eng | 0.066 | Young |
| 0.201531 | 0.320712 | 0.786 | 0.735 | 1 | 15 | Bcl2a1b | 0.051 | Young |
| 0.201775 | 1.100962 | 0.214 | 0.122 | 1 | 15 | Pnp | 0.092 | Young |
| 0.201775 | 0.991326 | 0.214 | 0.122 | 1 | 15 | Cd37 | 0.092 | Young |
| 0.201775 | 0.820734 | 0.214 | 0.122 | 1 | 15 | Slain2 | 0.092 | Young |
| 0.202353 | 0.435508 | 0.143 | 0.061 | 1 | 15 | Slc15a2 | 0.082 | Young |
| 0.202867 | 0.065109 | 0.25 | 0.143 | 1 | 15 | Ank | 0.107 | Young |
| 0.203419 | 0.174461 | 0.411 | 0.286 | 1 | 15 | Dbnl | 0.125 | Young |
| 0.204892 | 0.750055 | 0.554 | 0.449 | 1 | 15 | Irf1 | 0.105 | Young |
| 0.204892 | 0.59482 | 0.554 | 0.449 | 1 | 15 | Ehd1 | 0.105 | Young |
| 0.205243 | 0.35002 | 0.554 | 0.388 | 1 | 15 | Sdf4 | 0.166 | Young |
| 0.206032 | 0.388342 | 0.554 | 0.408 | 1 | 15 | Ndufb4 | 0.146 | Young |
| 0.206107 | 0.041009 | 0.411 | 0.286 | 1 | 15 | Runx1 | 0.125 | Young |
| 0.206357 | 1.577984 | 0.107 | 0.041 | 1 | 15 | Pom121 | 0.066 | Young |
| 0.206357 | 1.479492 | 0.107 | 0.041 | 1 | 15 | Lsm2 | 0.066 | Young |
| 0.206357 | 1.401815 | 0.107 | 0.041 | 1 | 15 | Nmi | 0.066 | Young |
| 0.206357 | 1.316679 | 0.107 | 0.041 | 1 | 15 | Slc31a1 | 0.066 | Young |
| 0.206357 | 1.281839 | 0.107 | 0.041 | 1 | 15 | Epm2aip1 | 0.066 | Young |
| 0.206357 | 0.84792 | 0.107 | 0.041 | 1 | 15 | Aamdc | 0.066 | Young |
| 0.206357 | 0.543282 | 0.107 | 0.041 | 1 | 15 | Pxmp4 | 0.066 | Young |
| 0.206357 | 0.207634 | 0.107 | 0.041 | 1 | 15 | Slx4ip | 0.066 | Young |
| 0.206357 | 0.0748 | 0.107 | 0.041 | 1 | 15 | Exosc3 | 0.066 | Young |
| 0.206663 | 0.104982 | 0.143 | 0.061 | 1 | 15 | Pygl | 0.082 | Young |
| 0.207171 | 1.710507 | 0.179 | 0.102 | 1 | 15 | Tbc1d22a | 0.077 | Young |
| 0.207674 | 1.093822 | 0.286 | 0.184 | 1 | 15 | Nans | 0.102 | Young |
| 0.207841 | 0.151494 | 0.393 | 0.265 | 1 | 15 | Fxyd2 | 0.128 | Young |
| 0.208631 | 1.683275 | 0.161 | 0.082 | 1 | 15 | Phf20 | 0.079 | Young |
| 0.208631 | 1.294405 | 0.161 | 0.082 | 1 | 15 | Gm6377 | 0.079 | Young |
| 0.208631 | 1.149248 | 0.161 | 0.082 | 1 | 15 | Irs2 | 0.079 | Young |
| 0.208845 | 0.151026 | 0.054 | 0.122 | 1 | 15 | Errfi1 | -0.068 | Young |
| 0.210134 | 1.154431 | 0.268 | 0.184 | 1 | 15 | Jup | 0.084 | Young |
| 0.210344 | 0.20728 | 0.982 | 0.898 | 1 | 15 | Rpl5 | 0.084 | Young |
| 0.210775 | 0.741342 | 0.268 | 0.163 | 1 | 15 | Folr2 | 0.105 | Young |
| 0.211001 | 0.882593 | 0.339 | 0.245 | 1 | 15 | Spcs3 | 0.094 | Young |
| 0.211416 | 1.383095 | 0.107 | 0.041 | 1 | 15 | mt-Nd6 | 0.066 | Young |
| 0.211416 | 1.355273 | 0.107 | 0.041 | 1 | 15 | Senp3 | 0.066 | Young |
| 0.211416 | 1.247016 | 0.107 | 0.041 | 1 | 15 | Dcaf1 | 0.066 | Young |
| 0.211416 | 1.234668 | 0.107 | 0.041 | 1 | 15 | Ptchd1 | 0.066 | Young |
| 0.211416 | 1.157479 | 0.107 | 0.041 | 1 | 15 | Ltv1 | 0.066 | Young |
| 0.211416 | 1.008374 | 0.107 | 0.041 | 1 | 15 | Rapgef1 | 0.066 | Young |
| 0.211416 | 0.835588 | 0.107 | 0.041 | 1 | 15 | Homer1 | 0.066 | Young |
| 0.211416 | 0.698213 | 0.107 | 0.041 | 1 | 15 | Commd1 | 0.066 | Young |

| | | | | | | | | |
|---|---|---|---|---|---|---|---|---|
| 0.211416 | 0.644613 | 0.107 | 0.041 | 1 | 15 | Zfp266 | 0.066 | Young |
| 0.212336 | 1.074156 | 0.214 | 0.122 | 1 | 15 | Glrx5 | 0.092 | Young |
| 0.212336 | 0.92217 | 0.214 | 0.122 | 1 | 15 | Syne2 | 0.092 | Young |
| 0.212336 | 0.904627 | 0.214 | 0.122 | 1 | 15 | Zbp1 | 0.092 | Young |
| 0.212724 | 1.263468 | 0.161 | 0.082 | 1 | 15 | Kdm5b | 0.079 | Young |
| 0.212724 | 1.147205 | 0.161 | 0.082 | 1 | 15 | 2210016L2 | 0.079 | Young |
| 0.212724 | 1.105173 | 0.161 | 0.082 | 1 | 15 | Uap1l1 | 0.079 | Young |
| 0.213292 | 0.78076 | 0.357 | 0.265 | 1 | 15 | Flrt3 | 0.092 | Young |
| 0.213299 | 3.677339 | 0.071 | 0.02 | 1 | 15 | Timm10 | 0.051 | Young |
| 0.213299 | 3.451437 | 0.071 | 0.02 | 1 | 15 | Siglece | 0.051 | Young |
| 0.213299 | 3.436648 | 0.071 | 0.02 | 1 | 15 | Clec4n | 0.051 | Young |
| 0.213299 | 3.42338 | 0.071 | 0.02 | 1 | 15 | Ecd | 0.051 | Young |
| 0.213299 | 3.298865 | 0.071 | 0.02 | 1 | 15 | Smim4 | 0.051 | Young |
| 0.213299 | 3.187382 | 0.071 | 0.02 | 1 | 15 | Il18bp | 0.051 | Young |
| 0.213299 | 3.137298 | 0.071 | 0.02 | 1 | 15 | Mpv17 | 0.051 | Young |
| 0.213299 | 2.842678 | 0.071 | 0.02 | 1 | 15 | Ccdc71 | 0.051 | Young |
| 0.213299 | 2.825256 | 0.071 | 0.02 | 1 | 15 | Gcsh | 0.051 | Young |
| 0.213299 | 2.770062 | 0.071 | 0.02 | 1 | 15 | Paox | 0.051 | Young |
| 0.213299 | 2.726975 | 0.071 | 0.02 | 1 | 15 | Zfp369 | 0.051 | Young |
| 0.213299 | 2.694108 | 0.071 | 0.02 | 1 | 15 | Cc2d1a | 0.051 | Young |
| 0.213299 | 2.665634 | 0.071 | 0.02 | 1 | 15 | Cpsf1 | 0.051 | Young |
| 0.213299 | 2.59002 | 0.071 | 0.02 | 1 | 15 | 2700038G2 | 0.051 | Young |
| 0.213299 | 2.588463 | 0.071 | 0.02 | 1 | 15 | Rcan3 | 0.051 | Young |
| 0.213299 | 2.531296 | 0.071 | 0.02 | 1 | 15 | Cdkn2a | 0.051 | Young |
| 0.213299 | 2.490568 | 0.071 | 0.02 | 1 | 15 | Itpripl1 | 0.051 | Young |
| 0.213299 | 2.480411 | 0.071 | 0.02 | 1 | 15 | Cep152 | 0.051 | Young |
| 0.213299 | 2.447282 | 0.071 | 0.02 | 1 | 15 | Sord | 0.051 | Young |
| 0.213299 | 2.423833 | 0.071 | 0.02 | 1 | 15 | Smpd1 | 0.051 | Young |
| 0.213299 | 2.377861 | 0.071 | 0.02 | 1 | 15 | Tom1 | 0.051 | Young |
| 0.213299 | 2.310062 | 0.071 | 0.02 | 1 | 15 | Abcb1b | 0.051 | Young |
| 0.213299 | 2.19322 | 0.071 | 0.02 | 1 | 15 | Psmg1 | 0.051 | Young |
| 0.213299 | 2.157658 | 0.071 | 0.02 | 1 | 15 | Prpf4 | 0.051 | Young |
| 0.213299 | 2.143817 | 0.071 | 0.02 | 1 | 15 | Plk3 | 0.051 | Young |
| 0.213299 | 1.891013 | 0.071 | 0.02 | 1 | 15 | Dennd6b | 0.051 | Young |
| 0.213477 | 0.407709 | 0.464 | 0.327 | 1 | 15 | Ndufv3 | 0.137 | Young |
| 0.213905 | 0.320558 | 0.714 | 0.694 | 1 | 15 | Cox8a | 0.02 | Young |
| 0.214421 | 0.580919 | 0.357 | 0.245 | 1 | 15 | Abcc1 | 0.112 | Young |
| 0.214834 | 0.569931 | 0.375 | 0.286 | 1 | 15 | Nr3c1 | 0.089 | Young |
| 0.214875 | 1.847777 | 0.179 | 0.102 | 1 | 15 | Ncbp2 | 0.077 | Young |
| 0.214875 | 1.816905 | 0.179 | 0.102 | 1 | 15 | Tsnax | 0.077 | Young |
| 0.214875 | 1.671718 | 0.179 | 0.102 | 1 | 15 | Tango2 | 0.077 | Young |
| 0.216292 | 0.149458 | 1 | 0.98 | 1 | 15 | Rplp1 | 0.02 | Young |
| 0.216563 | 1.675787 | 0.107 | 0.041 | 1 | 15 | Slc25a28 | 0.066 | Young |
| 0.216563 | 0.972114 | 0.107 | 0.041 | 1 | 15 | Pcgf3 | 0.066 | Young |
| 0.216563 | 0.902521 | 0.107 | 0.041 | 1 | 15 | Clpx | 0.066 | Young |

| | | | | | | | | |
|---|---|---|---|---|---|---|---|---|
| 0.216563 | 0.74945 | 0.107 | 0.041 | 1 | 15 | As3mt | 0.066 | Young |
| 0.216873 | 0.980254 | 0.161 | 0.082 | 1 | 15 | Ankle2 | 0.079 | Young |
| 0.216873 | 0.917977 | 0.161 | 0.082 | 1 | 15 | Ints13 | 0.079 | Young |
| 0.217134 | 0.536907 | 0.286 | 0.184 | 1 | 15 | Psmd14 | 0.102 | Young |
| 0.217678 | 1.019646 | 0.375 | 0.286 | 1 | 15 | Tbl1x | 0.089 | Young |
| 0.217678 | 0.754047 | 0.375 | 0.286 | 1 | 15 | Hspd1 | 0.089 | Young |
| 0.217702 | 1.135159 | 0.232 | 0.143 | 1 | 15 | Lacc1 | 0.089 | Young |
| 0.217789 | 1.262895 | 0.25 | 0.163 | 1 | 15 | Hat1 | 0.087 | Young |
| 0.218943 | 0.697754 | 0.714 | 0.592 | 1 | 15 | Epsti1 | 0.122 | Young |
| 0.219381 | 0.007742 | 0.625 | 0.49 | 1 | 15 | Celf1 | 0.135 | Young |
| 0.219771 | 3.044334 | 0.071 | 0.02 | 1 | 15 | Rnf8 | 0.051 | Young |
| 0.219771 | 2.881109 | 0.071 | 0.02 | 1 | 15 | Gzf1 | 0.051 | Young |
| 0.219771 | 2.774917 | 0.071 | 0.02 | 1 | 15 | Ubr7 | 0.051 | Young |
| 0.219771 | 2.698414 | 0.071 | 0.02 | 1 | 15 | Suv39h1 | 0.051 | Young |
| 0.219771 | 2.687625 | 0.071 | 0.02 | 1 | 15 | Grk5 | 0.051 | Young |
| 0.219771 | 2.486792 | 0.071 | 0.02 | 1 | 15 | Vps33a | 0.051 | Young |
| 0.219771 | 2.443048 | 0.071 | 0.02 | 1 | 15 | Fam122a | 0.051 | Young |
| 0.219771 | 2.337716 | 0.071 | 0.02 | 1 | 15 | Gm27010 | 0.051 | Young |
| 0.219771 | 2.175248 | 0.071 | 0.02 | 1 | 15 | Golga1 | 0.051 | Young |
| 0.219771 | 2.146164 | 0.071 | 0.02 | 1 | 15 | Pdpr | 0.051 | Young |
| 0.219771 | 2.110654 | 0.071 | 0.02 | 1 | 15 | Slc9a7 | 0.051 | Young |
| 0.219771 | 2.045641 | 0.071 | 0.02 | 1 | 15 | Adora2b | 0.051 | Young |
| 0.219771 | 2.014837 | 0.071 | 0.02 | 1 | 15 | Dus1l | 0.051 | Young |
| 0.219771 | 2.003714 | 0.071 | 0.02 | 1 | 15 | 1810043G( | 0.051 | Young |
| 0.219771 | 1.827076 | 0.071 | 0.02 | 1 | 15 | Mgst1 | 0.051 | Young |
| 0.219771 | 1.709419 | 0.071 | 0.02 | 1 | 15 | Aldh3b1 | 0.051 | Young |
| 0.220312 | 0.550214 | 0.393 | 0.286 | 1 | 15 | Dot1l | 0.107 | Young |
| 0.221081 | 1.111931 | 0.161 | 0.082 | 1 | 15 | Pcna | 0.079 | Young |
| 0.221081 | 0.247811 | 0.161 | 0.082 | 1 | 15 | Nbr1 | 0.079 | Young |
| 0.221221 | 0.966217 | 0.232 | 0.143 | 1 | 15 | Prdm2 | 0.089 | Young |
| 0.221779 | 1.962074 | 0.196 | 0.122 | 1 | 15 | Igf1 | 0.074 | Young |
| 0.2218 | 0.955305 | 0.107 | 0.041 | 1 | 15 | Slc25a33 | 0.066 | Young |
| 0.2218 | 0.948365 | 0.107 | 0.041 | 1 | 15 | Akr1e1 | 0.066 | Young |
| 0.2218 | 0.825557 | 0.107 | 0.041 | 1 | 15 | Krr1 | 0.066 | Young |
| 0.222783 | 1.646593 | 0.179 | 0.102 | 1 | 15 | Vps37a | 0.077 | Young |
| 0.222953 | 0.93333 | 0.321 | 0.224 | 1 | 15 | Plin3 | 0.097 | Young |
| 0.223554 | 0.415255 | 0.429 | 0.327 | 1 | 15 | Trmt112 | 0.102 | Young |
| 0.223851 | 0.316629 | 0.821 | 0.755 | 1 | 15 | Tyrobp | 0.066 | Young |
| 0.22401 | 0.787893 | 0.268 | 0.163 | 1 | 15 | Polr2k | 0.105 | Young |
| 0.224544 | 0.664551 | 0.482 | 0.388 | 1 | 15 | Klf3 | 0.094 | Young |
| 0.225347 | 1.124924 | 0.161 | 0.082 | 1 | 15 | Dram2 | 0.079 | Young |
| 0.225347 | 0.947491 | 0.161 | 0.082 | 1 | 15 | Dph3 | 0.079 | Young |
| 0.225347 | 0.258006 | 0.161 | 0.082 | 1 | 15 | Zfp664 | 0.079 | Young |
| 0.22629 | 0.427562 | 0.357 | 0.245 | 1 | 15 | Nub1 | 0.112 | Young |
| 0.226382 | 2.746724 | 0.071 | 0.02 | 1 | 15 | Manba | 0.051 | Young |

| | | | | | | | | |
|---|---|---|---|---|---|---|---|---|
| 0.226382 | 2.628564 | 0.071 | 0.02 | 1 | 15 | Rmnd5b | 0.051 | Young |
| 0.226382 | 2.314239 | 0.071 | 0.02 | 1 | 15 | Myb | 0.051 | Young |
| 0.226382 | 2.275168 | 0.071 | 0.02 | 1 | 15 | Trmt10c | 0.051 | Young |
| 0.226382 | 2.244202 | 0.071 | 0.02 | 1 | 15 | 4930453N2 | 0.051 | Young |
| 0.226382 | 2.194716 | 0.071 | 0.02 | 1 | 15 | Tirap | 0.051 | Young |
| 0.226382 | 2.113362 | 0.071 | 0.02 | 1 | 15 | Usp49 | 0.051 | Young |
| 0.226382 | 1.897506 | 0.071 | 0.02 | 1 | 15 | Ppil3 | 0.051 | Young |
| 0.226382 | 1.855693 | 0.071 | 0.02 | 1 | 15 | Plcb3 | 0.051 | Young |
| 0.226382 | 1.782847 | 0.071 | 0.02 | 1 | 15 | Bop1 | 0.051 | Young |
| 0.226382 | 1.674869 | 0.071 | 0.02 | 1 | 15 | Zscan21 | 0.051 | Young |
| 0.226382 | 1.614573 | 0.071 | 0.02 | 1 | 15 | Coq9 | 0.051 | Young |
| 0.226382 | 1.605629 | 0.071 | 0.02 | 1 | 15 | Dlat | 0.051 | Young |
| 0.226382 | 1.598127 | 0.071 | 0.02 | 1 | 15 | Ctnnbl1 | 0.051 | Young |
| 0.226382 | 1.433824 | 0.071 | 0.02 | 1 | 15 | Cdk7 | 0.051 | Young |
| 0.226382 | 1.296822 | 0.071 | 0.02 | 1 | 15 | Pkp4 | 0.051 | Young |
| 0.226382 | 1.259829 | 0.071 | 0.02 | 1 | 15 | Slc25a13 | 0.051 | Young |
| 0.226459 | 0.538409 | 0.268 | 0.184 | 1 | 15 | Washc4 | 0.084 | Young |
| 0.227032 | 0.718272 | 0.214 | 0.122 | 1 | 15 | Max | 0.092 | Young |
| 0.227127 | 0.378031 | 0.107 | 0.041 | 1 | 15 | Hspa13 | 0.066 | Young |
| 0.22718 | 0.111288 | 0.982 | 0.898 | 1 | 15 | Nfkbia | 0.084 | Young |
| 0.227409 | 0.157202 | 0.268 | 0.163 | 1 | 15 | Plekhm2 | 0.105 | Young |
| 0.227471 | 0.659974 | 0.196 | 0.102 | 1 | 15 | Hs6st1 | 0.094 | Young |
| 0.227936 | 0.304847 | 0.429 | 0.286 | 1 | 15 | Bach1 | 0.143 | Young |
| 0.227991 | 0.688043 | 0.357 | 0.265 | 1 | 15 | Samd4b | 0.092 | Young |
| 0.227991 | 0.188033 | 0.357 | 0.265 | 1 | 15 | Stat5a | 0.092 | Young |
| 0.228032 | 0.204836 | 0.804 | 0.571 | 1 | 15 | Bri3 | 0.233 | Young |
| 0.228426 | 1.997172 | 0.125 | 0.061 | 1 | 15 | Prkce | 0.064 | Young |
| 0.228638 | 0.481506 | 0.429 | 0.306 | 1 | 15 | Stk38 | 0.123 | Young |
| 0.229671 | 0.998915 | 0.161 | 0.082 | 1 | 15 | Synrg | 0.079 | Young |
| 0.229671 | 0.729836 | 0.161 | 0.082 | 1 | 15 | Taf6l | 0.079 | Young |
| 0.229804 | 0.982689 | 0.304 | 0.224 | 1 | 15 | Snx9 | 0.08 | Young |
| 0.23018 | 0.037573 | 0.482 | 0.347 | 1 | 15 | Prpf40a | 0.135 | Young |
| 0.230335 | 0.397194 | 0.304 | 0.184 | 1 | 15 | Zfand3 | 0.12 | Young |
| 0.230845 | 0.36595 | 0.268 | 0.163 | 1 | 15 | Krtcap2 | 0.105 | Young |
| 0.230898 | 1.240762 | 0.179 | 0.102 | 1 | 15 | Hmbox1 | 0.077 | Young |
| 0.231451 | 0.148703 | 0.196 | 0.102 | 1 | 15 | Atf1 | 0.094 | Young |
| 0.231922 | 0.364625 | 0.768 | 0.776 | 1 | 15 | Ybx1 | -0.008 | Young |
| 0.232022 | 0.868352 | 0.232 | 0.143 | 1 | 15 | Slc35a4 | 0.089 | Young |
| 0.232068 | 0.140849 | 0.339 | 0.245 | 1 | 15 | Rnf13 | 0.094 | Young |
| 0.232144 | 0.419983 | 0.518 | 0.388 | 1 | 15 | Rab11a | 0.13 | Young |
| 0.232377 | 0.569573 | 0.321 | 0.224 | 1 | 15 | Dctn3 | 0.097 | Young |
| 0.232544 | 0.645096 | 0.107 | 0.041 | 1 | 15 | Zfp444 | 0.066 | Young |
| 0.232544 | 0.10145 | 0.107 | 0.041 | 1 | 15 | 2810001G2 | 0.066 | Young |
| 0.233134 | 1.773865 | 0.071 | 0.02 | 1 | 15 | Gm47754 | 0.051 | Young |
| 0.233134 | 1.678435 | 0.071 | 0.02 | 1 | 15 | Cyb5r1 | 0.051 | Young |

| | | | | | | | | |
|---|---|---|---|---|---|---|---|---|
| 0.233134 | 1.662625 | 0.071 | 0.02 | 1 | 15 | Srcap | 0.051 | Young |
| 0.233134 | 1.57669 | 0.071 | 0.02 | 1 | 15 | Padi2 | 0.051 | Young |
| 0.233134 | 1.438945 | 0.071 | 0.02 | 1 | 15 | 4930503L1 | 0.051 | Young |
| 0.233134 | 1.435907 | 0.071 | 0.02 | 1 | 15 | Calcoco1 | 0.051 | Young |
| 0.233134 | 1.4333 | 0.071 | 0.02 | 1 | 15 | Med27 | 0.051 | Young |
| 0.233134 | 1.355372 | 0.071 | 0.02 | 1 | 15 | Insm1 | 0.051 | Young |
| 0.233134 | 1.312326 | 0.071 | 0.02 | 1 | 15 | Casp6 | 0.051 | Young |
| 0.233134 | 1.105659 | 0.071 | 0.02 | 1 | 15 | Heatr3 | 0.051 | Young |
| 0.233134 | 1.09562 | 0.071 | 0.02 | 1 | 15 | Wdr11 | 0.051 | Young |
| 0.233134 | 1.017617 | 0.071 | 0.02 | 1 | 15 | Ocel1 | 0.051 | Young |
| 0.233134 | 0.936651 | 0.071 | 0.02 | 1 | 15 | Dmac2 | 0.051 | Young |
| 0.233134 | 0.757261 | 0.071 | 0.02 | 1 | 15 | Ccdc85b | 0.051 | Young |
| 0.233353 | 1.459252 | 0.196 | 0.122 | 1 | 15 | Rnasel | 0.074 | Young |
| 0.233456 | 0.227988 | 0.982 | 0.959 | 1 | 15 | Pabpc1 | 0.023 | Young |
| 0.234054 | 1.098051 | 0.161 | 0.082 | 1 | 15 | Skint3 | 0.079 | Young |
| 0.234054 | 0.824974 | 0.161 | 0.082 | 1 | 15 | Atp6v0a2 | 0.079 | Young |
| 0.234318 | 0.343349 | 0.268 | 0.163 | 1 | 15 | Trip11 | 0.105 | Young |
| 0.234417 | 0.31444 | 0.429 | 0.306 | 1 | 15 | Psmb7 | 0.123 | Young |
| 0.234694 | 1.314528 | 0.321 | 0.245 | 1 | 15 | Cxcl2 | 0.076 | Young |
| 0.234694 | 0.922083 | 0.321 | 0.245 | 1 | 15 | Cask | 0.076 | Young |
| 0.235495 | 0.703788 | 0.339 | 0.265 | 1 | 15 | Ube2z | 0.074 | Young |
| 0.23561 | 0.541394 | 0.464 | 0.327 | 1 | 15 | Lima1 | 0.137 | Young |
| 0.235924 | 0.166581 | 0.946 | 0.918 | 1 | 15 | Rpl18a | 0.028 | Young |
| 0.236975 | 0.656238 | 0.286 | 0.184 | 1 | 15 | Ndnf | 0.102 | Young |
| 0.23715 | 0.806083 | 0.357 | 0.265 | 1 | 15 | Mrps33 | 0.092 | Young |
| 0.237657 | 0.259924 | 0.804 | 0.776 | 1 | 15 | Rnf19b | 0.028 | Young |
| 0.238319 | 2.127782 | 0.125 | 0.061 | 1 | 15 | Josd2 | 0.064 | Young |
| 0.238319 | 2.048494 | 0.125 | 0.061 | 1 | 15 | AI467606 | 0.064 | Young |
| 0.238319 | 1.671443 | 0.125 | 0.061 | 1 | 15 | Timm17a | 0.064 | Young |
| 0.238348 | 0.577356 | 0.482 | 0.388 | 1 | 15 | Cd9 | 0.094 | Young |
| 0.238496 | 0.744091 | 0.161 | 0.082 | 1 | 15 | Bag3 | 0.079 | Young |
| 0.238496 | 0.717678 | 0.161 | 0.082 | 1 | 15 | St6gal1 | 0.079 | Young |
| 0.23852 | 0.603005 | 0.214 | 0.122 | 1 | 15 | Adgre4 | 0.092 | Young |
| 0.238819 | 0.784815 | 0.411 | 0.327 | 1 | 15 | Mfsd14b | 0.084 | Young |
| 0.238876 | 0.349255 | 0.411 | 0.286 | 1 | 15 | Polr3c | 0.125 | Young |
| 0.23922 | 1.24264 | 0.179 | 0.102 | 1 | 15 | Lamtor3 | 0.077 | Young |
| 0.23922 | 1.187618 | 0.179 | 0.102 | 1 | 15 | Ppfibp2 | 0.077 | Young |
| 0.239428 | 0.506318 | 0.232 | 0.143 | 1 | 15 | Fundc2 | 0.089 | Young |
| 0.239663 | 0.239764 | 0.411 | 0.306 | 1 | 15 | Ctsa | 0.105 | Young |
| 0.240027 | 1.421183 | 0.071 | 0.02 | 1 | 15 | Brcc3 | 0.051 | Young |
| 0.240027 | 1.170304 | 0.071 | 0.02 | 1 | 15 | Mettl2 | 0.051 | Young |
| 0.240027 | 1.04517 | 0.071 | 0.02 | 1 | 15 | Slc38a6 | 0.051 | Young |
| 0.240027 | 0.918687 | 0.071 | 0.02 | 1 | 15 | Vps45 | 0.051 | Young |
| 0.240027 | 0.839659 | 0.071 | 0.02 | 1 | 15 | Mrps35 | 0.051 | Young |
| 0.240027 | 0.750251 | 0.071 | 0.02 | 1 | 15 | Sash3 | 0.051 | Young |

| | | | | | | | | |
|---|---|---|---|---|---|---|---|---|
| 0.240027 | 0.700316 | 0.071 | 0.02 | 1 | 15 | Ang | 0.051 | Young |
| 0.240027 | 0.680663 | 0.071 | 0.02 | 1 | 15 | Nprl3 | 0.051 | Young |
| 0.240027 | 0.486079 | 0.071 | 0.02 | 1 | 15 | Trmt2a | 0.051 | Young |
| 0.240027 | 0.383625 | 0.071 | 0.02 | 1 | 15 | Hcfc2 | 0.051 | Young |
| 0.240027 | 0.343913 | 0.071 | 0.02 | 1 | 15 | Tbc1d16 | 0.051 | Young |
| 0.240027 | 0.321643 | 0.071 | 0.02 | 1 | 15 | Trpv4 | 0.051 | Young |
| 0.240027 | 0.278044 | 0.071 | 0.02 | 1 | 15 | Fbxo9 | 0.051 | Young |
| 0.240027 | 0.261199 | 0.071 | 0.02 | 1 | 15 | Tmem250- | 0.051 | Young |
| 0.240027 | 0.156123 | 0.071 | 0.02 | 1 | 15 | Tslp | 0.051 | Young |
| 0.240027 | 0.155426 | 0.071 | 0.02 | 1 | 15 | Dusp8 | 0.051 | Young |
| 0.240027 | 0.093648 | 0.071 | 0.02 | 1 | 15 | Kctd6 | 0.051 | Young |
| 0.24026 | 1.145885 | 0.357 | 0.265 | 1 | 15 | Fn1 | 0.092 | Young |
| 0.24118 | 0.491333 | 0.482 | 0.388 | 1 | 15 | Ndufc2 | 0.094 | Young |
| 0.241303 | 0.842986 | 0.196 | 0.122 | 1 | 15 | Mlxip | 0.074 | Young |
| 0.241375 | 1.017327 | 0.268 | 0.163 | 1 | 15 | Dynlt3 | 0.105 | Young |
| 0.241538 | 0.296694 | 0.643 | 0.51 | 1 | 15 | Ctnna1 | 0.133 | Young |
| 0.242312 | 0.614496 | 0.25 | 0.163 | 1 | 15 | 2610507B1 | 0.087 | Young |
| 0.242439 | 0.00238 | 0.214 | 0.122 | 1 | 15 | Kansl3 | 0.092 | Young |
| 0.242997 | 0.870524 | 0.161 | 0.082 | 1 | 15 | Itgav | 0.079 | Young |
| 0.243377 | 2.118106 | 0.125 | 0.061 | 1 | 15 | Nup50 | 0.064 | Young |
| 0.243377 | 2.010006 | 0.125 | 0.061 | 1 | 15 | Wbp1l | 0.064 | Young |
| 0.243377 | 1.95264 | 0.125 | 0.061 | 1 | 15 | BC005561 | 0.064 | Young |
| 0.243377 | 1.737762 | 0.125 | 0.061 | 1 | 15 | Phlda1 | 0.064 | Young |
| 0.24346 | 1.575815 | 0.179 | 0.102 | 1 | 15 | Washc3 | 0.077 | Young |
| 0.244756 | 0.829364 | 0.339 | 0.245 | 1 | 15 | Ogdh | 0.094 | Young |
| 0.244756 | 0.56725 | 0.339 | 0.245 | 1 | 15 | Rnf7 | 0.094 | Young |
| 0.24482 | 0.799093 | 0.357 | 0.286 | 1 | 15 | Ptpn18 | 0.071 | Young |
| 0.244978 | 0.227416 | 0.375 | 0.245 | 1 | 15 | Lypla2 | 0.13 | Young |
| 0.245132 | 0.469327 | 0.482 | 0.408 | 1 | 15 | Bzw1 | 0.074 | Young |
| 0.245348 | 1.561427 | 0.196 | 0.122 | 1 | 15 | Gm38399 | 0.074 | Young |
| 0.245348 | 1.317705 | 0.196 | 0.122 | 1 | 15 | Gcn1l1 | 0.074 | Young |
| 0.245416 | 0.130884 | 0.75 | 0.653 | 1 | 15 | Tob2 | 0.097 | Young |
| 0.245593 | 0.532749 | 0.5 | 0.429 | 1 | 15 | Hexa | 0.071 | Young |
| 0.245616 | 0.30906 | 0.625 | 0.49 | 1 | 15 | Galnt1 | 0.135 | Young |
| 0.245859 | 0.188761 | 0.911 | 0.837 | 1 | 15 | Atp5l | 0.074 | Young |
| 0.245968 | 0.792406 | 0.25 | 0.163 | 1 | 15 | Gcc2 | 0.087 | Young |
| 0.246403 | 0.551182 | 0.214 | 0.122 | 1 | 15 | Trappc6a | 0.092 | Young |
| 0.247218 | 0.727838 | 0.268 | 0.184 | 1 | 15 | Emc2 | 0.084 | Young |
| 0.247513 | 0.19844 | 0.946 | 0.898 | 1 | 15 | Eef1b2 | 0.048 | Young |
| 0.247558 | 1.259708 | 0.161 | 0.082 | 1 | 15 | Slc7a7 | 0.079 | Young |
| 0.247752 | 1.131472 | 0.179 | 0.102 | 1 | 15 | Vcpip1 | 0.077 | Young |
| 0.247752 | 0.919521 | 0.179 | 0.102 | 1 | 15 | Nfic | 0.077 | Young |
| 0.247806 | 0.230782 | 0.554 | 0.388 | 1 | 15 | Tardbp | 0.166 | Young |
| 0.247925 | 0.354411 | 0.589 | 0.469 | 1 | 15 | Sema4a | 0.12 | Young |
| 0.24851 | 1.866197 | 0.125 | 0.061 | 1 | 15 | Gnl2 | 0.064 | Young |

| | | | | | | | | |
|---|---|---|---|---|---|---|---|---|
| 0.24851 | 1.646736 | 0.125 | 0.061 | 1 | 15 | C1galt1 | 0.064 | Young |
| 0.24851 | 1.634611 | 0.125 | 0.061 | 1 | 15 | Rnf170 | 0.064 | Young |
| 0.249441 | 2.049579 | 0.196 | 0.122 | 1 | 15 | Gm47283 | 0.074 | Young |
| 0.249441 | 1.265223 | 0.196 | 0.122 | 1 | 15 | Rbm33 | 0.074 | Young |
| 0.249591 | 0.468871 | 0.321 | 0.204 | 1 | 15 | Dnajc19 | 0.117 | Young |
| 0.250182 | 0.29875 | 0.821 | 0.653 | 1 | 15 | Ostf1 | 0.168 | Young |
| 0.250412 | 0.692025 | 0.214 | 0.122 | 1 | 15 | Uba6 | 0.092 | Young |
| 0.250412 | 0.389678 | 0.214 | 0.122 | 1 | 15 | Leprotl1 | 0.092 | Young |
| 0.250412 | 0.106714 | 0.214 | 0.122 | 1 | 15 | Crem | 0.092 | Young |
| 0.250804 | 0.790728 | 0.268 | 0.184 | 1 | 15 | Xpa | 0.084 | Young |
| 0.25084 | 0.832872 | 0.411 | 0.327 | 1 | 15 | Cd80 | 0.084 | Young |
| 0.251808 | 0.450567 | 0.411 | 0.306 | 1 | 15 | Cfp | 0.105 | Young |
| 0.252097 | 1.072515 | 0.179 | 0.102 | 1 | 15 | Man1b1 | 0.077 | Young |
| 0.252097 | 0.905438 | 0.179 | 0.102 | 1 | 15 | Mid1ip1 | 0.077 | Young |
| 0.252838 | 1.519471 | 0.214 | 0.143 | 1 | 15 | Rras | 0.071 | Young |
| 0.252986 | 0.009573 | 0.357 | 0.265 | 1 | 15 | Ctbp1 | 0.092 | Young |
| 0.253244 | 0.28875 | 0.464 | 0.327 | 1 | 15 | Pvr | 0.137 | Young |
| 0.25344 | 0.033698 | 0.75 | 0.653 | 1 | 15 | Dennd4a | 0.097 | Young |
| 0.253582 | 1.31946 | 0.196 | 0.122 | 1 | 15 | Nde1 | 0.074 | Young |
| 0.253718 | 1.614485 | 0.125 | 0.061 | 1 | 15 | Map3k20 | 0.064 | Young |
| 0.253718 | 1.51751 | 0.125 | 0.061 | 1 | 15 | Stoml1 | 0.064 | Young |
| 0.253718 | 1.490014 | 0.125 | 0.061 | 1 | 15 | Clip2 | 0.064 | Young |
| 0.253718 | 1.409112 | 0.125 | 0.061 | 1 | 15 | Rab10os | 0.064 | Young |
| 0.253819 | 0.170446 | 0.518 | 0.367 | 1 | 15 | Cct8 | 0.151 | Young |
| 0.25422 | 0.452277 | 0.286 | 0.204 | 1 | 15 | Sp140 | 0.082 | Young |
| 0.254425 | 0.705865 | 0.268 | 0.184 | 1 | 15 | Tomm5 | 0.084 | Young |
| 0.254466 | 0.642896 | 0.214 | 0.122 | 1 | 15 | Lrif1 | 0.092 | Young |
| 0.254574 | 0.256857 | 0.75 | 0.694 | 1 | 15 | Ptafr | 0.056 | Young |
| 0.254911 | 0.72443 | 0.411 | 0.306 | 1 | 15 | Psmd13 | 0.105 | Young |
| 0.256198 | 0.499096 | 0.464 | 0.367 | 1 | 15 | Tgif1 | 0.097 | Young |
| 0.256494 | 1.383518 | 0.179 | 0.102 | 1 | 15 | Synpo2 | 0.077 | Young |
| 0.256494 | 1.3095 | 0.179 | 0.102 | 1 | 15 | Cd209f | 0.077 | Young |
| 0.256653 | 0.07371 | 0.607 | 0.49 | 1 | 15 | Arhgef6 | 0.117 | Young |
| 0.257365 | 0.777047 | 0.393 | 0.306 | 1 | 15 | Dip2b | 0.087 | Young |
| 0.25777 | 1.021576 | 0.196 | 0.122 | 1 | 15 | Ddx27 | 0.074 | Young |
| 0.25879 | 0.529389 | 0.661 | 0.551 | 1 | 15 | Sec61g | 0.11 | Young |
| 0.258948 | 0.104113 | 0.429 | 0.327 | 1 | 15 | Elf1 | 0.102 | Young |
| 0.259002 | 1.885731 | 0.125 | 0.061 | 1 | 15 | Rnf166 | 0.064 | Young |
| 0.259002 | 1.522766 | 0.125 | 0.061 | 1 | 15 | Psmg3 | 0.064 | Young |
| 0.259002 | 0.329953 | 0.125 | 0.061 | 1 | 15 | Zswim4 | 0.064 | Young |
| 0.259504 | 0.027241 | 0.643 | 0.49 | 1 | 15 | Dock8 | 0.153 | Young |
| 0.260129 | 0.395759 | 0.411 | 0.327 | 1 | 15 | Ell2 | 0.084 | Young |
| 0.260866 | 1.17436 | 0.214 | 0.143 | 1 | 15 | Phyh | 0.071 | Young |
| 0.261547 | 1.329389 | 0.25 | 0.184 | 1 | 15 | Tec | 0.066 | Young |
| 0.261777 | 0.832629 | 0.268 | 0.184 | 1 | 15 | Tep1 | 0.084 | Young |

| | | | | | | | | |
|---|---|---|---|---|---|---|---|---|
| 0.262335 | 0.489129 | 0.321 | 0.224 | 1 | 15 | Tsn | 0.097 | Young |
| 0.262525 | 0.216143 | 0.304 | 0.204 | 1 | 15 | Gm21188 | 0.1 | Young |
| 0.263942 | 0.146513 | 0.25 | 0.143 | 1 | 15 | 6-Mar | 0.107 | Young |
| 0.264222 | 0.430716 | 0.5 | 0.388 | 1 | 15 | Ier3ip1 | 0.112 | Young |
| 0.264362 | 1.287445 | 0.125 | 0.061 | 1 | 15 | Hscb | 0.064 | Young |
| 0.264362 | 1.071722 | 0.125 | 0.061 | 1 | 15 | Pxdc1 | 0.064 | Young |
| 0.264827 | 0.838309 | 0.25 | 0.163 | 1 | 15 | Gm2000 | 0.087 | Young |
| 0.264827 | 0.54124 | 0.25 | 0.163 | 1 | 15 | Akirin1 | 0.087 | Young |
| 0.265083 | 0.083512 | 0.107 | 0.204 | 1 | 15 | Med12 | -0.097 | Young |
| 0.265192 | 0.193571 | 0.75 | 0.592 | 1 | 15 | Luc7l2 | 0.158 | Young |
| 0.265375 | 0.36816 | 0.286 | 0.184 | 1 | 15 | Vopp1 | 0.102 | Young |
| 0.265451 | 0.432451 | 0.018 | 0.061 | 1 | 15 | Tmem150b | -0.043 | Young |
| 0.265451 | 0.29258 | 0.018 | 0.061 | 1 | 15 | Gmnn | -0.043 | Young |
| 0.265451 | 0.131895 | 0.018 | 0.061 | 1 | 15 | 4921524J1 | -0.043 | Young |
| 0.265451 | 0.110808 | 0.018 | 0.061 | 1 | 15 | Mier2 | -0.043 | Young |
| 0.265451 | 0.065234 | 0.018 | 0.061 | 1 | 15 | Rbm10 | -0.043 | Young |
| 0.265451 | 0.022915 | 0.018 | 0.061 | 1 | 15 | Sirt5 | -0.043 | Young |
| 0.265821 | 0.622382 | 0.321 | 0.224 | 1 | 15 | Tm6sf1 | 0.097 | Young |
| 0.266289 | 0.897551 | 0.196 | 0.122 | 1 | 15 | Gatad1 | 0.074 | Young |
| 0.266289 | 0.707387 | 0.196 | 0.122 | 1 | 15 | Ptk2b | 0.074 | Young |
| 0.26645 | 0.396217 | 0.857 | 0.776 | 1 | 15 | Nr4a3 | 0.081 | Young |
| 0.266905 | 0.504166 | 0.375 | 0.286 | 1 | 15 | Impact | 0.089 | Young |
| 0.266966 | 0.896457 | 0.304 | 0.224 | 1 | 15 | Snx17 | 0.08 | Young |
| 0.26778 | 0.182714 | 0.821 | 0.714 | 1 | 15 | Fcer1g | 0.107 | Young |
| 0.267954 | 0.055948 | 0.339 | 0.224 | 1 | 15 | Nek7 | 0.115 | Young |
| 0.268596 | 0.073101 | 0.411 | 0.286 | 1 | 15 | Dusp3 | 0.125 | Young |
| 0.268624 | 0.461832 | 0.446 | 0.347 | 1 | 15 | Nme1 | 0.099 | Young |
| 0.269798 | 1.789046 | 0.125 | 0.061 | 1 | 15 | Pigx | 0.064 | Young |
| 0.269798 | 1.137184 | 0.125 | 0.061 | 1 | 15 | Hax1 | 0.064 | Young |
| 0.269884 | 0.978793 | 0.232 | 0.163 | 1 | 15 | Pmp22 | 0.069 | Young |
| 0.270006 | 0.851811 | 0.179 | 0.102 | 1 | 15 | Tifa | 0.077 | Young |
| 0.271504 | 0.669822 | 0.446 | 0.388 | 1 | 15 | Zcchc6 | 0.058 | Young |
| 0.271609 | 0.068333 | 0.786 | 0.735 | 1 | 15 | Mbnl1 | 0.051 | Young |
| 0.271776 | 0.63201 | 0.429 | 0.367 | 1 | 15 | Mrpl33 | 0.062 | Young |
| 0.27183 | 0.346841 | 0.482 | 0.408 | 1 | 15 | Ube2k | 0.074 | Young |
| 0.271879 | 1.624903 | 0.143 | 0.082 | 1 | 15 | Sirt7 | 0.061 | Young |
| 0.271946 | 0.203614 | 0.982 | 0.857 | 1 | 15 | Vim | 0.125 | Young |
| 0.272643 | 0.452637 | 0.25 | 0.163 | 1 | 15 | Tctex1d2 | 0.087 | Young |
| 0.272881 | 0.30795 | 0.411 | 0.327 | 1 | 15 | Tmem234 | 0.084 | Young |
| 0.273078 | 0.748602 | 0.268 | 0.184 | 1 | 15 | Tbc1d13 | 0.084 | Young |
| 0.273237 | 1.534914 | 0.214 | 0.143 | 1 | 15 | Dda1 | 0.071 | Young |
| 0.273237 | 1.189603 | 0.214 | 0.143 | 1 | 15 | Atf2 | 0.071 | Young |
| 0.274616 | 0.391843 | 0.179 | 0.102 | 1 | 15 | Cd1d1 | 0.077 | Young |
| 0.274654 | 1.237273 | 0.339 | 0.286 | 1 | 15 | Ids | 0.053 | Young |
| 0.274974 | 0.087944 | 0.268 | 0.163 | 1 | 15 | Rnf11 | 0.105 | Young |

| | | | | | | | | |
|---|---|---|---|---|---|---|---|---|
| 0.275001 | 0.996251 | 0.196 | 0.122 | 1 | 15 | Ugcg | 0.074 | Young |
| 0.275063 | 0.621704 | 0.339 | 0.245 | 1 | 15 | Gmfb | 0.094 | Young |
| 0.275309 | 1.054516 | 0.125 | 0.061 | 1 | 15 | Dennd4c | 0.064 | Young |
| 0.275309 | 0.988001 | 0.125 | 0.061 | 1 | 15 | Zyg11b | 0.064 | Young |
| 0.275309 | 0.956815 | 0.125 | 0.061 | 1 | 15 | Arid3a | 0.064 | Young |
| 0.276609 | 0.221112 | 0.25 | 0.163 | 1 | 15 | Srp54a | 0.087 | Young |
| 0.276918 | 0.300341 | 0.179 | 0.286 | 1 | 15 | Ndufab1 | -0.107 | Young |
| 0.276985 | 1.725946 | 0.143 | 0.082 | 1 | 15 | Zfp513 | 0.061 | Young |
| 0.277592 | 0.627158 | 0.357 | 0.286 | 1 | 15 | Fam46a | 0.071 | Young |
| 0.278023 | 0.347398 | 0.482 | 0.408 | 1 | 15 | Atp6ap1 | 0.074 | Young |
| 0.280795 | 0.896276 | 0.268 | 0.184 | 1 | 15 | Srrt | 0.084 | Young |
| 0.280898 | 1.36091 | 0.125 | 0.061 | 1 | 15 | Ocrl | 0.064 | Young |
| 0.280898 | 1.006717 | 0.125 | 0.061 | 1 | 15 | Plppr4 | 0.064 | Young |
| 0.280898 | 0.947406 | 0.125 | 0.061 | 1 | 15 | Reps1 | 0.064 | Young |
| 0.280898 | 0.814126 | 0.125 | 0.061 | 1 | 15 | Sharpin | 0.064 | Young |
| 0.280898 | 0.794883 | 0.125 | 0.061 | 1 | 15 | Mrps30 | 0.064 | Young |
| 0.280898 | 0.784492 | 0.125 | 0.061 | 1 | 15 | Iars | 0.064 | Young |
| 0.280898 | 0.52839 | 0.125 | 0.061 | 1 | 15 | Aven | 0.064 | Young |
| 0.280898 | 0.018104 | 0.125 | 0.061 | 1 | 15 | Med25 | 0.064 | Young |
| 0.281452 | 0.988225 | 0.304 | 0.224 | 1 | 15 | Igbp1 | 0.08 | Young |
| 0.281703 | 1.304559 | 0.214 | 0.143 | 1 | 15 | Luzp1 | 0.071 | Young |
| 0.282041 | 1.074597 | 0.232 | 0.163 | 1 | 15 | Tmem150c | 0.069 | Young |
| 0.282132 | 0.812377 | 0.339 | 0.245 | 1 | 15 | Lpl | 0.094 | Young |
| 0.282155 | 1.904864 | 0.143 | 0.082 | 1 | 15 | Zfp984 | 0.061 | Young |
| 0.282155 | 1.753425 | 0.143 | 0.082 | 1 | 15 | Prkdc | 0.061 | Young |
| 0.282155 | 1.471919 | 0.143 | 0.082 | 1 | 15 | Ppard | 0.061 | Young |
| 0.283316 | 0.538211 | 0.357 | 0.265 | 1 | 15 | Ccnt2 | 0.092 | Young |
| 0.283411 | 0.177732 | 0.982 | 0.939 | 1 | 15 | Rps10 | 0.043 | Young |
| 0.283524 | 0.375696 | 0.536 | 0.469 | 1 | 15 | Hint1 | 0.067 | Young |
| 0.283804 | 0.360778 | 0.482 | 0.367 | 1 | 15 | Uqcr10 | 0.115 | Young |
| 0.283905 | 0.893075 | 0.196 | 0.122 | 1 | 15 | Otub1 | 0.074 | Young |
| 0.284271 | 0.383569 | 0.286 | 0.184 | 1 | 15 | Tmem164 | 0.102 | Young |
| 0.284646 | 0.450812 | 0.518 | 0.469 | 1 | 15 | App | 0.049 | Young |
| 0.284659 | 0.743416 | 0.25 | 0.163 | 1 | 15 | Immt | 0.087 | Young |
| 0.284659 | 0.112279 | 0.25 | 0.163 | 1 | 15 | Herc2 | 0.087 | Young |
| 0.284709 | 0.193205 | 0.179 | 0.286 | 1 | 15 | Fam103a1 | -0.107 | Young |
| 0.284868 | 0.234339 | 0.071 | 0.143 | 1 | 15 | Ssbp1 | -0.072 | Young |
| 0.285155 | 0.640353 | 0.304 | 0.224 | 1 | 15 | Setd1b | 0.08 | Young |
| 0.285482 | 0.5831 | 0.339 | 0.265 | 1 | 15 | Tmem14c | 0.074 | Young |
| 0.285712 | 0.434692 | 0.339 | 0.245 | 1 | 15 | Trf | 0.094 | Young |
| 0.286562 | 1.018355 | 0.125 | 0.061 | 1 | 15 | Zfp395 | 0.064 | Young |
| 0.286562 | 0.852691 | 0.125 | 0.061 | 1 | 15 | Aggf1 | 0.064 | Young |
| 0.286562 | 0.844825 | 0.125 | 0.061 | 1 | 15 | Car3 | 0.064 | Young |
| 0.286562 | 0.807519 | 0.125 | 0.061 | 1 | 15 | Lztfl1 | 0.064 | Young |
| 0.286562 | 0.666358 | 0.125 | 0.061 | 1 | 15 | Trappc13 | 0.064 | Young |

| | | | | | | | | |
|---|---|---|---|---|---|---|---|---|
| 0.28739 | 1.589394 | 0.143 | 0.082 | 1 | 15 | Crebzf | 0.061 | Young |
| 0.28798 | 0.729126 | 0.429 | 0.347 | 1 | 15 | Adam10 | 0.082 | Young |
| 0.288429 | 0.862388 | 0.196 | 0.122 | 1 | 15 | Gm13546 | 0.074 | Young |
| 0.288768 | 0.532661 | 0.179 | 0.102 | 1 | 15 | Trpm7 | 0.077 | Young |
| 0.289396 | 0.669619 | 0.321 | 0.245 | 1 | 15 | Klhl5 | 0.076 | Young |
| 0.289504 | 0.25528 | 0.804 | 0.673 | 1 | 15 | Cd52 | 0.131 | Young |
| 0.290346 | 0.994368 | 0.214 | 0.143 | 1 | 15 | Cog5 | 0.071 | Young |
| 0.290348 | 1.172715 | 0.232 | 0.163 | 1 | 15 | Ing2 | 0.069 | Young |
| 0.290398 | 0.542166 | 0.518 | 0.449 | 1 | 15 | Rnf115 | 0.069 | Young |
| 0.291248 | 0.978838 | 0.286 | 0.204 | 1 | 15 | Il15 | 0.082 | Young |
| 0.29158 | 2.656274 | 0.089 | 0.041 | 1 | 15 | Sh2d5 | 0.048 | Young |
| 0.29158 | 2.543871 | 0.089 | 0.041 | 1 | 15 | Gm10851 | 0.048 | Young |
| 0.29158 | 2.482645 | 0.089 | 0.041 | 1 | 15 | Prpsap2 | 0.048 | Young |
| 0.29158 | 2.451399 | 0.089 | 0.041 | 1 | 15 | Pop4 | 0.048 | Young |
| 0.29158 | 2.089278 | 0.089 | 0.041 | 1 | 15 | Glrx | 0.048 | Young |
| 0.292304 | 0.860888 | 0.125 | 0.061 | 1 | 15 | Chp2 | 0.064 | Young |
| 0.292304 | 0.855086 | 0.125 | 0.061 | 1 | 15 | Il6 | 0.064 | Young |
| 0.292304 | 0.107537 | 0.125 | 0.061 | 1 | 15 | Cep192 | 0.064 | Young |
| 0.29266 | 0.830901 | 0.304 | 0.224 | 1 | 15 | Tmem51 | 0.08 | Young |
| 0.292691 | 1.438338 | 0.143 | 0.082 | 1 | 15 | Pdcd11 | 0.061 | Young |
| 0.292919 | 1.19722 | 0.25 | 0.184 | 1 | 15 | Ice1 | 0.066 | Young |
| 0.293002 | 0.641285 | 0.196 | 0.122 | 1 | 15 | Sipa1 | 0.074 | Young |
| 0.293002 | 0.526831 | 0.196 | 0.122 | 1 | 15 | Csnk1g3 | 0.074 | Young |
| 0.293002 | 0.496042 | 0.196 | 0.122 | 1 | 15 | Chchd7 | 0.074 | Young |
| 0.293079 | 0.851596 | 0.321 | 0.245 | 1 | 15 | Csf1r | 0.076 | Young |
| 0.293593 | 0.910783 | 0.179 | 0.102 | 1 | 15 | Mtm1 | 0.077 | Young |
| 0.293593 | 0.475999 | 0.179 | 0.102 | 1 | 15 | Gigyf2 | 0.077 | Young |
| 0.294008 | 1.016801 | 0.286 | 0.224 | 1 | 15 | Prdx5 | 0.062 | Young |
| 0.294563 | 1.140875 | 0.232 | 0.163 | 1 | 15 | Smad7 | 0.069 | Young |
| 0.294563 | 0.906311 | 0.232 | 0.163 | 1 | 15 | Ccdc59 | 0.069 | Young |
| 0.294733 | 0.704325 | 0.214 | 0.143 | 1 | 15 | Aaed1 | 0.071 | Young |
| 0.295141 | 0.853929 | 0.286 | 0.204 | 1 | 15 | Bcap31 | 0.082 | Young |
| 0.295938 | 0.263178 | 0.232 | 0.143 | 1 | 15 | Slc22a23 | 0.089 | Young |
| 0.296636 | 0.072464 | 0.339 | 0.245 | 1 | 15 | Dyrk2 | 0.094 | Young |
| 0.296669 | 0.306938 | 0.268 | 0.184 | 1 | 15 | Scn2b | 0.084 | Young |
| 0.298056 | 1.235741 | 0.143 | 0.082 | 1 | 15 | Zmat5 | 0.061 | Young |
| 0.298056 | 1.189678 | 0.143 | 0.082 | 1 | 15 | Taf3 | 0.061 | Young |
| 0.298122 | 0.834627 | 0.125 | 0.061 | 1 | 15 | Mkl2 | 0.064 | Young |
| 0.298122 | 0.625227 | 0.125 | 0.061 | 1 | 15 | Ncbp1 | 0.064 | Young |
| 0.298429 | 2.812896 | 0.089 | 0.041 | 1 | 15 | Smox | 0.048 | Young |
| 0.298429 | 2.545112 | 0.089 | 0.041 | 1 | 15 | Zfyve1 | 0.048 | Young |
| 0.298429 | 2.098411 | 0.089 | 0.041 | 1 | 15 | Nob1 | 0.048 | Young |
| 0.298429 | 2.01182 | 0.089 | 0.041 | 1 | 15 | Ampd2 | 0.048 | Young |
| 0.298429 | 1.991667 | 0.089 | 0.041 | 1 | 15 | Trappc12 | 0.048 | Young |
| 0.298429 | 1.957311 | 0.089 | 0.041 | 1 | 15 | Tmem245 | 0.048 | Young |

| | | | | | | | | |
|---|---|---|---|---|---|---|---|---|
| 0.298429 | 1.906385 | 0.089 | 0.041 | 1 | 15 | Lsm14b | 0.048 | Young |
| 0.298429 | 1.756996 | 0.089 | 0.041 | 1 | 15 | Ufd1 | 0.048 | Young |
| 0.298472 | 0.242104 | 0.179 | 0.102 | 1 | 15 | Itfg1 | 0.077 | Young |
| 0.298819 | 0.332809 | 0.232 | 0.163 | 1 | 15 | B4galt5 | 0.069 | Young |
| 0.29953 | 0.24878 | 0.732 | 0.592 | 1 | 15 | S100a11 | 0.14 | Young |
| 0.300974 | 0.215243 | 0.982 | 0.857 | 1 | 15 | Cox4i1 | 0.125 | Young |
| 0.30119 | 0.368708 | 0.357 | 0.265 | 1 | 15 | Arhgef1 | 0.092 | Young |
| 0.301278 | 0.496267 | 0.446 | 0.347 | 1 | 15 | Fam174a | 0.099 | Young |
| 0.303404 | 0.267449 | 0.179 | 0.102 | 1 | 15 | Pcm1 | 0.077 | Young |
| 0.303442 | 0.431895 | 0.464 | 0.388 | 1 | 15 | Larp4b | 0.076 | Young |
| 0.303487 | 1.506081 | 0.143 | 0.082 | 1 | 15 | Gpatch11 | 0.061 | Young |
| 0.303487 | 1.228174 | 0.143 | 0.082 | 1 | 15 | Vps9d1 | 0.061 | Young |
| 0.303487 | 0.946469 | 0.143 | 0.082 | 1 | 15 | Uxs1 | 0.061 | Young |
| 0.303634 | 0.631506 | 0.339 | 0.265 | 1 | 15 | Ccdc12 | 0.074 | Young |
| 0.303641 | 1.024598 | 0.214 | 0.143 | 1 | 15 | Fto | 0.071 | Young |
| 0.304578 | 0.609852 | 0.482 | 0.429 | 1 | 15 | Ndufb10 | 0.053 | Young |
| 0.305385 | 2.116342 | 0.089 | 0.041 | 1 | 15 | Aimp2 | 0.048 | Young |
| 0.305385 | 2.104582 | 0.089 | 0.041 | 1 | 15 | Pstpip1 | 0.048 | Young |
| 0.305385 | 2.026902 | 0.089 | 0.041 | 1 | 15 | Tns4 | 0.048 | Young |
| 0.305385 | 1.877109 | 0.089 | 0.041 | 1 | 15 | Ift140 | 0.048 | Young |
| 0.305385 | 1.751753 | 0.089 | 0.041 | 1 | 15 | Prrg2 | 0.048 | Young |
| 0.305385 | 1.535795 | 0.089 | 0.041 | 1 | 15 | Nop16 | 0.048 | Young |
| 0.307357 | 0.395731 | 0.393 | 0.286 | 1 | 15 | Cpeb4 | 0.107 | Young |
| 0.307845 | 0.210617 | 0.393 | 0.306 | 1 | 15 | Atp6v1a | 0.087 | Young |
| 0.308161 | 0.357942 | 0.214 | 0.143 | 1 | 15 | Pira2 | 0.071 | Young |
| 0.308161 | 0.321385 | 0.214 | 0.143 | 1 | 15 | Mcmbp | 0.071 | Young |
| 0.30839 | 0.804067 | 0.179 | 0.102 | 1 | 15 | BC017643 | 0.077 | Young |
| 0.308984 | 1.371201 | 0.143 | 0.082 | 1 | 15 | Map2k7 | 0.061 | Young |
| 0.308984 | 1.160443 | 0.143 | 0.082 | 1 | 15 | Pitrm1 | 0.061 | Young |
| 0.309754 | 0.3166 | 0.25 | 0.163 | 1 | 15 | Arih2 | 0.087 | Young |
| 0.309988 | 0.557337 | 0.125 | 0.061 | 1 | 15 | Rpl7l1 | 0.064 | Young |
| 0.309988 | 0.444973 | 0.125 | 0.061 | 1 | 15 | Bicral | 0.064 | Young |
| 0.309988 | 0.327029 | 0.125 | 0.061 | 1 | 15 | Zfp385a | 0.064 | Young |
| 0.310168 | 1.020121 | 0.268 | 0.204 | 1 | 15 | Ddrgk1 | 0.064 | Young |
| 0.311034 | 0.30844 | 0.589 | 0.49 | 1 | 15 | Dennd5a | 0.099 | Young |
| 0.31171 | 0.269651 | 1 | 1 | 1 | 15 | Gm42418 | 0 | Young |
| 0.311735 | 0.615169 | 0.393 | 0.327 | 1 | 15 | Hp1bp3 | 0.066 | Young |
| 0.311777 | 0.648687 | 0.196 | 0.122 | 1 | 15 | Impa1 | 0.074 | Young |
| 0.311832 | 1.447309 | 0.232 | 0.163 | 1 | 15 | Rsf1 | 0.069 | Young |
| 0.312448 | 2.234279 | 0.089 | 0.041 | 1 | 15 | Arhgap6 | 0.048 | Young |
| 0.312448 | 1.733501 | 0.089 | 0.041 | 1 | 15 | Dalrd3 | 0.048 | Young |
| 0.312448 | 1.571424 | 0.089 | 0.041 | 1 | 15 | Fam53a | 0.048 | Young |
| 0.312528 | 0.442713 | 0.5 | 0.449 | 1 | 15 | S100a13 | 0.051 | Young |
| 0.312726 | 0.835393 | 0.214 | 0.143 | 1 | 15 | Irak4 | 0.071 | Young |
| 0.313604 | 0.228361 | 0.232 | 0.143 | 1 | 15 | Ssbp3 | 0.089 | Young |

| | | | | | | | | |
|---|---|---|---|---|---|---|---|---|
| 0.313665 | 0.825783 | 0.286 | 0.224 | 1 | 15 | Pdxk | 0.062 | Young |
| 0.313761 | 0.211516 | 0.25 | 0.184 | 1 | 15 | Katna1 | 0.066 | Young |
| 0.314546 | 1.405637 | 0.143 | 0.082 | 1 | 15 | Tmem50b | 0.061 | Young |
| 0.314546 | 1.247228 | 0.143 | 0.082 | 1 | 15 | Pdzd8 | 0.061 | Young |
| 0.314546 | 1.209851 | 0.143 | 0.082 | 1 | 15 | Bckdk | 0.061 | Young |
| 0.314546 | 0.926958 | 0.143 | 0.082 | 1 | 15 | Idh2 | 0.061 | Young |
| 0.314588 | 0.213252 | 0.5 | 0.408 | 1 | 15 | Nckap1l | 0.092 | Young |
| 0.314885 | 0.107971 | 0.357 | 0.245 | 1 | 15 | Upb1 | 0.112 | Young |
| 0.316037 | 0.553057 | 0.125 | 0.061 | 1 | 15 | Hgsnat | 0.064 | Young |
| 0.316251 | 0.943967 | 0.232 | 0.163 | 1 | 15 | Psmb2 | 0.069 | Young |
| 0.316251 | 0.852831 | 0.232 | 0.163 | 1 | 15 | Map2k3 | 0.069 | Young |
| 0.316411 | 0.049408 | 0.446 | 0.367 | 1 | 15 | Crk | 0.079 | Young |
| 0.317308 | 0.149442 | 0.536 | 0.49 | 1 | 15 | Rock2 | 0.046 | Young |
| 0.318649 | 0.455192 | 0.518 | 0.408 | 1 | 15 | Vmp1 | 0.11 | Young |
| 0.319006 | 0.51882 | 0.411 | 0.347 | 1 | 15 | Sec62 | 0.064 | Young |
| 0.319312 | 0.36 | 0.339 | 0.245 | 1 | 15 | Bptf | 0.094 | Young |
| 0.319617 | 1.685919 | 0.089 | 0.041 | 1 | 15 | 1110032A0 | 0.048 | Young |
| 0.319617 | 1.554067 | 0.089 | 0.041 | 1 | 15 | Utp20 | 0.048 | Young |
| 0.319617 | 1.501543 | 0.089 | 0.041 | 1 | 15 | Gbe1 | 0.048 | Young |
| 0.319617 | 1.499472 | 0.089 | 0.041 | 1 | 15 | Arhgef11 | 0.048 | Young |
| 0.319617 | 1.188971 | 0.089 | 0.041 | 1 | 15 | Slamf8 | 0.048 | Young |
| 0.319617 | 1.091998 | 0.089 | 0.041 | 1 | 15 | Jpt2 | 0.048 | Young |
| 0.319617 | 1.056506 | 0.089 | 0.041 | 1 | 15 | Smim26 | 0.048 | Young |
| 0.319852 | 1.257121 | 0.161 | 0.102 | 1 | 15 | Dmtf1 | 0.059 | Young |
| 0.319962 | 1.033557 | 0.304 | 0.245 | 1 | 15 | Senp6 | 0.059 | Young |
| 0.320173 | 1.000348 | 0.143 | 0.082 | 1 | 15 | Rpl10-ps3 | 0.061 | Young |
| 0.320173 | 0.919563 | 0.143 | 0.082 | 1 | 15 | Secisbp2 | 0.061 | Young |
| 0.320173 | 0.91212 | 0.143 | 0.082 | 1 | 15 | Amz2 | 0.061 | Young |
| 0.320173 | 0.865895 | 0.143 | 0.082 | 1 | 15 | Cd302 | 0.061 | Young |
| 0.320173 | 0.356585 | 0.143 | 0.082 | 1 | 15 | Pafah1b2 | 0.061 | Young |
| 0.320704 | 0.255331 | 0.357 | 0.286 | 1 | 15 | Tmed5 | 0.071 | Young |
| 0.320712 | 0.885264 | 0.232 | 0.163 | 1 | 15 | Mgrn1 | 0.069 | Young |
| 0.322163 | 0.045743 | 0.125 | 0.061 | 1 | 15 | Znrd1 | 0.064 | Young |
| 0.322345 | 0.659627 | 0.393 | 0.306 | 1 | 15 | Coa5 | 0.087 | Young |
| 0.32263 | 0.815668 | 0.268 | 0.204 | 1 | 15 | Cd68 | 0.064 | Young |
| 0.322692 | 1.542039 | 0.214 | 0.163 | 1 | 15 | Card19 | 0.051 | Young |
| 0.322833 | 0.542143 | 0.25 | 0.163 | 1 | 15 | Pcmt1 | 0.087 | Young |
| 0.323194 | 0.147656 | 0.482 | 0.388 | 1 | 15 | Kdelr1 | 0.094 | Young |
| 0.323362 | 0.605183 | 0.286 | 0.204 | 1 | 15 | Ythdf2 | 0.082 | Young |
| 0.323673 | 0.381761 | 0.179 | 0.102 | 1 | 15 | Pcmtd2 | 0.077 | Young |
| 0.32463 | 0.206918 | 0.464 | 0.367 | 1 | 15 | Ssr4 | 0.097 | Young |
| 0.32517 | 1.242335 | 0.161 | 0.102 | 1 | 15 | Riok1 | 0.059 | Young |
| 0.32517 | 1.195668 | 0.161 | 0.102 | 1 | 15 | Hltf | 0.059 | Young |
| 0.325213 | 0.6895 | 0.232 | 0.163 | 1 | 15 | Tbxas1 | 0.069 | Young |
| 0.325866 | 0.871478 | 0.143 | 0.082 | 1 | 15 | Upf3b | 0.061 | Young |

| | | | | | | | | |
|---|---|---|---|---|---|---|---|---|
| 0.325866 | 0.699595 | 0.143 | 0.082 | 1 | 15 | Slc30a7 | 0.061 | Young |
| 0.326368 | 0.711928 | 0.196 | 0.122 | 1 | 15 | Vezf1 | 0.074 | Young |
| 0.326467 | 0.096348 | 0.893 | 0.796 | 1 | 15 | S100a6 | 0.097 | Young |
| 0.326893 | 1.990008 | 0.089 | 0.041 | 1 | 15 | Rnf126 | 0.048 | Young |
| 0.326893 | 1.772376 | 0.089 | 0.041 | 1 | 15 | Pias4 | 0.048 | Young |
| 0.326893 | 1.440185 | 0.089 | 0.041 | 1 | 15 | Xylt2 | 0.048 | Young |
| 0.326893 | 1.345271 | 0.089 | 0.041 | 1 | 15 | 4732440D( | 0.048 | Young |
| 0.326893 | 1.325164 | 0.089 | 0.041 | 1 | 15 | Snrpa1 | 0.048 | Young |
| 0.326893 | 1.237586 | 0.089 | 0.041 | 1 | 15 | Zfp518a | 0.048 | Young |
| 0.326893 | 1.053864 | 0.089 | 0.041 | 1 | 15 | Caml | 0.048 | Young |
| 0.326893 | 0.927482 | 0.089 | 0.041 | 1 | 15 | Srbd1 | 0.048 | Young |
| 0.326893 | 0.877594 | 0.089 | 0.041 | 1 | 15 | Pank1 | 0.048 | Young |
| 0.326893 | 0.864423 | 0.089 | 0.041 | 1 | 15 | Taf11 | 0.048 | Young |
| 0.326893 | 0.849296 | 0.089 | 0.041 | 1 | 15 | Tomm40 | 0.048 | Young |
| 0.326893 | 0.440086 | 0.089 | 0.041 | 1 | 15 | Cacfd1 | 0.048 | Young |
| 0.326893 | 0.000728 | 0.089 | 0.041 | 1 | 15 | Nfxl1 | 0.048 | Young |
| 0.327639 | 1.45828 | 0.321 | 0.245 | 1 | 15 | Ccl2 | 0.076 | Young |
| 0.328293 | 0.801362 | 0.357 | 0.286 | 1 | 15 | Med28 | 0.071 | Young |
| 0.328568 | 0.411337 | 0.625 | 0.51 | 1 | 15 | Pomp | 0.115 | Young |
| 0.328817 | 0.164565 | 0.571 | 0.429 | 1 | 15 | Mpeg1 | 0.142 | Young |
| 0.329755 | 0.966341 | 0.232 | 0.163 | 1 | 15 | Aldh2 | 0.069 | Young |
| 0.331427 | 0.857121 | 0.214 | 0.143 | 1 | 15 | Tbc1d14 | 0.071 | Young |
| 0.331624 | 0.004063 | 0.143 | 0.082 | 1 | 15 | Ranbp3 | 0.061 | Young |
| 0.332788 | 0.237616 | 0.768 | 0.714 | 1 | 15 | Ywhaz | 0.054 | Young |
| 0.334276 | 1.206881 | 0.089 | 0.041 | 1 | 15 | Fkbp5 | 0.048 | Young |
| 0.334276 | 1.020865 | 0.089 | 0.041 | 1 | 15 | Trim65 | 0.048 | Young |
| 0.334276 | 1.020642 | 0.089 | 0.041 | 1 | 15 | Ddx58 | 0.048 | Young |
| 0.334276 | 0.954345 | 0.089 | 0.041 | 1 | 15 | Fbxo46 | 0.048 | Young |
| 0.334276 | 0.923173 | 0.089 | 0.041 | 1 | 15 | Frrs1 | 0.048 | Young |
| 0.334276 | 0.890112 | 0.089 | 0.041 | 1 | 15 | Cks2 | 0.048 | Young |
| 0.334276 | 0.857545 | 0.089 | 0.041 | 1 | 15 | Dcstamp | 0.048 | Young |
| 0.334276 | 0.849248 | 0.089 | 0.041 | 1 | 15 | Creld1 | 0.048 | Young |
| 0.334276 | 0.790577 | 0.089 | 0.041 | 1 | 15 | Skp2 | 0.048 | Young |
| 0.334276 | 0.789399 | 0.089 | 0.041 | 1 | 15 | Atg14 | 0.048 | Young |
| 0.334276 | 0.783972 | 0.089 | 0.041 | 1 | 15 | Peli2 | 0.048 | Young |
| 0.334276 | 0.622335 | 0.089 | 0.041 | 1 | 15 | Dkk2 | 0.048 | Young |
| 0.334276 | 0.495283 | 0.089 | 0.041 | 1 | 15 | Comtd1 | 0.048 | Young |
| 0.335096 | 0.365142 | 0.714 | 0.633 | 1 | 15 | Txn1 | 0.081 | Young |
| 0.335298 | 1.15636 | 0.196 | 0.143 | 1 | 15 | Vat1 | 0.053 | Young |
| 0.3363 | 0.256813 | 0.5 | 0.388 | 1 | 15 | Atp5c1 | 0.112 | Young |
| 0.336338 | 0.046541 | 0.196 | 0.122 | 1 | 15 | Trappc2 | 0.074 | Young |
| 0.337272 | 0.03286 | 0.75 | 0.796 | 1 | 15 | Ccl5 | -0.046 | Young |
| 0.337447 | 1.017942 | 0.143 | 0.082 | 1 | 15 | BC028528 | 0.061 | Young |
| 0.337447 | 0.731508 | 0.143 | 0.082 | 1 | 15 | Ly96 | 0.061 | Young |
| 0.338525 | 0.453803 | 0.375 | 0.286 | 1 | 15 | Irak1 | 0.089 | Young |

| | | | | | | | | |
|---|---|---|---|---|---|---|---|---|
| 0.338935 | 0.789775 | 0.268 | 0.184 | 1 | 15 | Madd | 0.084 | Young |
| 0.33944 | 0.193219 | 0.107 | 0.184 | 1 | 15 | Insr | -0.077 | Young |
| 0.33944 | 0.029958 | 0.107 | 0.184 | 1 | 15 | Sars | -0.077 | Young |
| 0.341044 | 1.013735 | 0.214 | 0.143 | 1 | 15 | Insig1 | 0.071 | Young |
| 0.341465 | 1.120024 | 0.161 | 0.102 | 1 | 15 | Erh | 0.059 | Young |
| 0.341465 | 0.979794 | 0.161 | 0.102 | 1 | 15 | Ms4a7 | 0.059 | Young |
| 0.341766 | 1.037277 | 0.089 | 0.041 | 1 | 15 | Ttc1 | 0.048 | Young |
| 0.341766 | 0.933466 | 0.089 | 0.041 | 1 | 15 | Ankrd46 | 0.048 | Young |
| 0.341766 | 0.801196 | 0.089 | 0.041 | 1 | 15 | Sugp2 | 0.048 | Young |
| 0.341766 | 0.765318 | 0.089 | 0.041 | 1 | 15 | Ints1 | 0.048 | Young |
| 0.341766 | 0.717508 | 0.089 | 0.041 | 1 | 15 | Rad17 | 0.048 | Young |
| 0.341766 | 0.573742 | 0.089 | 0.041 | 1 | 15 | Procr | 0.048 | Young |
| 0.342534 | 0.711642 | 0.446 | 0.388 | 1 | 15 | Dad1 | 0.058 | Young |
| 0.342579 | 0.403095 | 0.768 | 0.714 | 1 | 15 | Socs2 | 0.054 | Young |
| 0.34286 | 0.132817 | 0.929 | 0.796 | 1 | 15 | Gadd45b | 0.133 | Young |
| 0.343336 | 0.863304 | 0.143 | 0.082 | 1 | 15 | Mtf2 | 0.061 | Young |
| 0.343336 | 0.675408 | 0.143 | 0.082 | 1 | 15 | Fam177a | 0.061 | Young |
| 0.343439 | 0.189434 | 0.821 | 0.694 | 1 | 15 | Cflar | 0.127 | Young |
| 0.343696 | 0.178836 | 0.607 | 0.531 | 1 | 15 | Dbi | 0.076 | Young |
| 0.343815 | 0.602487 | 0.321 | 0.245 | 1 | 15 | Adss | 0.076 | Young |
| 0.344116 | 0.949149 | 0.268 | 0.204 | 1 | 15 | Mat2b | 0.064 | Young |
| 0.344216 | 1.791743 | 0.107 | 0.061 | 1 | 15 | Slc25a38 | 0.046 | Young |
| 0.344353 | 0.943303 | 0.357 | 0.306 | 1 | 15 | C1qb | 0.051 | Young |
| 0.344539 | 1.122393 | 0.25 | 0.184 | 1 | 15 | Cd163 | 0.066 | Young |
| 0.344564 | 0.397036 | 0.286 | 0.204 | 1 | 15 | Pacsin2 | 0.082 | Young |
| 0.345918 | 0.435104 | 0.214 | 0.143 | 1 | 15 | Nfatc3 | 0.071 | Young |
| 0.346502 | 0.366465 | 0.196 | 0.122 | 1 | 15 | Zdhhc7 | 0.074 | Young |
| 0.34669 | 0.315309 | 0.732 | 0.612 | 1 | 15 | Ndufa6 | 0.12 | Young |
| 0.347299 | 1.121106 | 0.179 | 0.122 | 1 | 15 | Katnbl1 | 0.057 | Young |
| 0.347332 | 0.42293 | 0.518 | 0.449 | 1 | 15 | Eps15 | 0.069 | Young |
| 0.347831 | 0.854689 | 0.268 | 0.184 | 1 | 15 | Ehf | 0.084 | Young |
| 0.349107 | 0.02906 | 0.536 | 0.429 | 1 | 15 | Dhrs3 | 0.107 | Young |
| 0.34929 | 0.648983 | 0.143 | 0.082 | 1 | 15 | Cebpa | 0.061 | Young |
| 0.34929 | 0.586975 | 0.143 | 0.082 | 1 | 15 | Mre11a | 0.061 | Young |
| 0.34929 | 0.494975 | 0.143 | 0.082 | 1 | 15 | Tmem170b | 0.061 | Young |
| 0.34929 | 0.476701 | 0.143 | 0.082 | 1 | 15 | Bms1 | 0.061 | Young |
| 0.34929 | 0.366425 | 0.143 | 0.082 | 1 | 15 | Triap1 | 0.061 | Young |
| 0.34929 | 0.362481 | 0.143 | 0.082 | 1 | 15 | Was | 0.061 | Young |
| 0.34929 | 0.085451 | 0.143 | 0.082 | 1 | 15 | Ddx21 | 0.061 | Young |
| 0.349362 | 0.60145 | 0.089 | 0.041 | 1 | 15 | Serpinb9b | 0.048 | Young |
| 0.349362 | 0.585644 | 0.089 | 0.041 | 1 | 15 | Rpl36a-ps1 | 0.048 | Young |
| 0.349362 | 0.396431 | 0.089 | 0.041 | 1 | 15 | Tusc3 | 0.048 | Young |
| 0.349362 | 0.395638 | 0.089 | 0.041 | 1 | 15 | Pde6d | 0.048 | Young |
| 0.349362 | 0.109546 | 0.089 | 0.041 | 1 | 15 | Ercc1 | 0.048 | Young |
| 0.350408 | 0.120472 | 0.732 | 0.673 | 1 | 15 | Anxa5 | 0.059 | Young |

| | | | | | | | | |
|---|---|---|---|---|---|---|---|---|
| 0.350652 | 0.099071 | 0.5 | 0.388 | 1 | 15 | Abcg1 | 0.112 | Young |
| 0.350821 | 0.170332 | 0.875 | 0.816 | 1 | 15 | Clec2d | 0.059 | Young |
| 0.350837 | 0.606234 | 0.214 | 0.143 | 1 | 15 | Lbr | 0.071 | Young |
| 0.350837 | 0.216431 | 0.214 | 0.143 | 1 | 15 | Eif2s3y | 0.071 | Young |
| 0.350837 | 0.19678 | 0.214 | 0.143 | 1 | 15 | Mrpl51 | 0.071 | Young |
| 0.353073 | 0.196987 | 0.857 | 0.735 | 1 | 15 | Wdr1 | 0.122 | Young |
| 0.353078 | 0.664244 | 0.232 | 0.163 | 1 | 15 | Eif6 | 0.069 | Young |
| 0.353078 | 0.147657 | 0.232 | 0.163 | 1 | 15 | Emc3 | 0.069 | Young |
| 0.353124 | 0.168273 | 0.339 | 0.286 | 1 | 15 | Top2b | 0.053 | Young |
| 0.353287 | 0.31446 | 0.286 | 0.204 | 1 | 15 | Zmym2 | 0.082 | Young |
| 0.35437 | 0.40973 | 0.75 | 0.673 | 1 | 15 | Abracl | 0.077 | Young |
| 0.355309 | 0.001255 | 0.143 | 0.082 | 1 | 15 | Tmem87a | 0.061 | Young |
| 0.355514 | 1.249568 | 0.286 | 0.224 | 1 | 15 | Ube2q1 | 0.062 | Young |
| 0.355766 | 0.516598 | 0.357 | 0.286 | 1 | 15 | Ifnar1 | 0.071 | Young |
| 0.356465 | 0.338086 | 0.375 | 0.306 | 1 | 15 | Kdm4a | 0.069 | Young |
| 0.356465 | 0.247152 | 0.375 | 0.306 | 1 | 15 | Ttc7 | 0.069 | Young |
| 0.356572 | 0.387354 | 0.393 | 0.306 | 1 | 15 | Hnrnpul1 | 0.087 | Young |
| 0.356691 | 0.399565 | 0.839 | 0.776 | 1 | 15 | Cd63 | 0.063 | Young |
| 0.356859 | 0.177654 | 0.196 | 0.122 | 1 | 15 | Stap1 | 0.074 | Young |
| 0.357065 | 0.415412 | 0.089 | 0.041 | 1 | 15 | Grcc10 | 0.048 | Young |
| 0.357836 | 0.17993 | 0.643 | 0.551 | 1 | 15 | Rtraf | 0.092 | Young |
| 0.357865 | 0.232057 | 0.232 | 0.163 | 1 | 15 | Lipe | 0.069 | Young |
| 0.357882 | 1.756963 | 0.107 | 0.061 | 1 | 15 | Gfm2 | 0.046 | Young |
| 0.357882 | 1.750151 | 0.107 | 0.061 | 1 | 15 | Ube2q2 | 0.046 | Young |
| 0.357882 | 1.512846 | 0.107 | 0.061 | 1 | 15 | Gt(ROSA)2( | 0.046 | Young |
| 0.357882 | 1.501459 | 0.107 | 0.061 | 1 | 15 | Osbpl7 | 0.046 | Young |
| 0.358071 | 0.110079 | 0.554 | 0.408 | 1 | 15 | Mtpn | 0.146 | Young |
| 0.358274 | 1.021041 | 0.161 | 0.102 | 1 | 15 | Cdc73 | 0.059 | Young |
| 0.358274 | 0.589068 | 0.161 | 0.102 | 1 | 15 | Mrpl19 | 0.059 | Young |
| 0.358453 | 0.67105 | 0.339 | 0.265 | 1 | 15 | Becn1 | 0.074 | Young |
| 0.359497 | 4.852362 | 0.018 | 0 | 1 | 15 | Mucl1 | 0.018 | Young |
| 0.359497 | 4.710745 | 0.018 | 0 | 1 | 15 | Snx33 | 0.018 | Young |
| 0.359497 | 4.710745 | 0.018 | 0 | 1 | 15 | Cfb | 0.018 | Young |
| 0.359497 | 4.710745 | 0.018 | 0 | 1 | 15 | Plekhs1 | 0.018 | Young |
| 0.359497 | 4.437493 | 0.018 | 0 | 1 | 15 | Hba-a1 | 0.018 | Young |
| 0.359497 | 4.187714 | 0.018 | 0 | 1 | 15 | Hba-a2 | 0.018 | Young |
| 0.359497 | 4.053927 | 0.018 | 0 | 1 | 15 | Ppp1r14c | 0.018 | Young |
| 0.359497 | 4.053927 | 0.018 | 0 | 1 | 15 | 6230400D1 | 0.018 | Young |
| 0.359497 | 4.030441 | 0.018 | 0 | 1 | 15 | Gramd1c | 0.018 | Young |
| 0.359497 | 4.024926 | 0.018 | 0 | 1 | 15 | Ly6i | 0.018 | Young |
| 0.359497 | 3.599201 | 0.018 | 0 | 1 | 15 | Dnmt3b | 0.018 | Young |
| 0.359497 | 3.599201 | 0.018 | 0 | 1 | 15 | Mrpl47 | 0.018 | Young |
| 0.359497 | 3.599201 | 0.018 | 0 | 1 | 15 | Gm16845 | 0.018 | Young |
| 0.359497 | 3.599201 | 0.018 | 0 | 1 | 15 | Fam84b | 0.018 | Young |
| 0.359497 | 3.456288 | 0.018 | 0 | 1 | 15 | Id1 | 0.018 | Young |

| | | | | | | | | |
|---|---|---|---|---|---|---|---|---|
| 0.359497 | 3.442812 | 0.018 | 0 | 1 | 15 | Nmral1 | 0.018 | Young |
| 0.359497 | 3.398374 | 0.018 | 0 | 1 | 15 | F9 | 0.018 | Young |
| 0.359497 | 3.398374 | 0.018 | 0 | 1 | 15 | Mlf1 | 0.018 | Young |
| 0.359497 | 3.398374 | 0.018 | 0 | 1 | 15 | K230015D0 | 0.018 | Young |
| 0.359497 | 3.398374 | 0.018 | 0 | 1 | 15 | Ldlrad4 | 0.018 | Young |
| 0.359497 | 3.395933 | 0.018 | 0 | 1 | 15 | Prkg1 | 0.018 | Young |
| 0.359497 | 3.358565 | 0.018 | 0 | 1 | 15 | Mex3a | 0.018 | Young |
| 0.359497 | 3.358565 | 0.018 | 0 | 1 | 15 | Ndrg4 | 0.018 | Young |
| 0.359497 | 3.358565 | 0.018 | 0 | 1 | 15 | 2210408I2 | 0.018 | Young |
| 0.359497 | 3.358565 | 0.018 | 0 | 1 | 15 | Adipoq | 0.018 | Young |
| 0.359497 | 3.193221 | 0.018 | 0 | 1 | 15 | Metap1d | 0.018 | Young |
| 0.359497 | 3.193221 | 0.018 | 0 | 1 | 15 | Pcyt1b | 0.018 | Young |
| 0.359497 | 3.193221 | 0.018 | 0 | 1 | 15 | Nfib | 0.018 | Young |
| 0.359497 | 3.193221 | 0.018 | 0 | 1 | 15 | 4930512J1 | 0.018 | Young |
| 0.359497 | 3.193221 | 0.018 | 0 | 1 | 15 | Plekha4 | 0.018 | Young |
| 0.359497 | 3.193221 | 0.018 | 0 | 1 | 15 | Car11 | 0.018 | Young |
| 0.359497 | 3.193221 | 0.018 | 0 | 1 | 15 | Gm42715 | 0.018 | Young |
| 0.359497 | 3.193221 | 0.018 | 0 | 1 | 15 | Hebp2 | 0.018 | Young |
| 0.359497 | 3.193221 | 0.018 | 0 | 1 | 15 | A430057M | 0.018 | Young |
| 0.359497 | 3.193221 | 0.018 | 0 | 1 | 15 | Meis1 | 0.018 | Young |
| 0.359497 | 3.116131 | 0.018 | 0 | 1 | 15 | Elf3 | 0.018 | Young |
| 0.359497 | 3.116131 | 0.018 | 0 | 1 | 15 | Rbm45 | 0.018 | Young |
| 0.359497 | 3.116131 | 0.018 | 0 | 1 | 15 | Bhlhb9 | 0.018 | Young |
| 0.359497 | 3.116131 | 0.018 | 0 | 1 | 15 | Ptger3 | 0.018 | Young |
| 0.359497 | 3.116131 | 0.018 | 0 | 1 | 15 | Tpm2 | 0.018 | Young |
| 0.359497 | 3.116131 | 0.018 | 0 | 1 | 15 | Rad18 | 0.018 | Young |
| 0.359497 | 3.116131 | 0.018 | 0 | 1 | 15 | Cend1 | 0.018 | Young |
| 0.359497 | 3.116131 | 0.018 | 0 | 1 | 15 | Dhrs11 | 0.018 | Young |
| 0.359497 | 3.116131 | 0.018 | 0 | 1 | 15 | Gcnt4 | 0.018 | Young |
| 0.359497 | 3.116131 | 0.018 | 0 | 1 | 15 | 1700056N1 | 0.018 | Young |
| 0.359497 | 3.109026 | 0.018 | 0 | 1 | 15 | Cavin2 | 0.018 | Young |
| 0.359497 | 3.109026 | 0.018 | 0 | 1 | 15 | Pjvk | 0.018 | Young |
| 0.359497 | 3.109026 | 0.018 | 0 | 1 | 15 | 1700112D2 | 0.018 | Young |
| 0.359497 | 3.109026 | 0.018 | 0 | 1 | 15 | Ccne1 | 0.018 | Young |
| 0.359497 | 3.109026 | 0.018 | 0 | 1 | 15 | Trmt12 | 0.018 | Young |
| 0.359497 | 3.109026 | 0.018 | 0 | 1 | 15 | Rps6ka2 | 0.018 | Young |
| 0.359497 | 3.109026 | 0.018 | 0 | 1 | 15 | Ch25h | 0.018 | Young |
| 0.359497 | 3.072565 | 0.018 | 0 | 1 | 15 | 10-Sep | 0.018 | Young |
| 0.359497 | 3.072565 | 0.018 | 0 | 1 | 15 | Fam83h | 0.018 | Young |
| 0.359497 | 3.072565 | 0.018 | 0 | 1 | 15 | Amhr2 | 0.018 | Young |
| 0.359497 | 3.066513 | 0.018 | 0 | 1 | 15 | Pik3ip1 | 0.018 | Young |
| 0.359497 | 3.028069 | 0.018 | 0 | 1 | 15 | Plp1 | 0.018 | Young |
| 0.359497 | 3.028069 | 0.018 | 0 | 1 | 15 | Azin2 | 0.018 | Young |
| 0.359497 | 3.028069 | 0.018 | 0 | 1 | 15 | Tmem200b | 0.018 | Young |
| 0.359497 | 3.028069 | 0.018 | 0 | 1 | 15 | Gm48684 | 0.018 | Young |

| | | | | | | | | |
|---|---|---|---|---|---|---|---|---|
| 0.359497 | 3.028069 | 0.018 | 0 | 1 | 15 | Enpp5 | 0.018 | Young |
| 0.359497 | 3.028069 | 0.018 | 0 | 1 | 15 | Dnd1 | 0.018 | Young |
| 0.359497 | 3.028069 | 0.018 | 0 | 1 | 15 | Efemp2 | 0.018 | Young |
| 0.359497 | 3.008953 | 0.018 | 0 | 1 | 15 | Zbtb41 | 0.018 | Young |
| 0.359497 | 3.008953 | 0.018 | 0 | 1 | 15 | BC055324 | 0.018 | Young |
| 0.359497 | 3.008953 | 0.018 | 0 | 1 | 15 | Tyms | 0.018 | Young |
| 0.359497 | 3.008953 | 0.018 | 0 | 1 | 15 | Tead2 | 0.018 | Young |
| 0.359497 | 3.008953 | 0.018 | 0 | 1 | 15 | Gm15512 | 0.018 | Young |
| 0.359497 | 2.930002 | 0.018 | 0 | 1 | 15 | Gulp1 | 0.018 | Young |
| 0.359497 | 2.930002 | 0.018 | 0 | 1 | 15 | Slc2a1 | 0.018 | Young |
| 0.359497 | 2.930002 | 0.018 | 0 | 1 | 15 | Tagln | 0.018 | Young |
| 0.359497 | 2.930002 | 0.018 | 0 | 1 | 15 | Myo15 | 0.018 | Young |
| 0.359497 | 2.930002 | 0.018 | 0 | 1 | 15 | Appbp2os | 0.018 | Young |
| 0.359497 | 2.930002 | 0.018 | 0 | 1 | 15 | Abhd3 | 0.018 | Young |
| 0.359497 | 2.918444 | 0.018 | 0 | 1 | 15 | Vangl1 | 0.018 | Young |
| 0.359497 | 2.918444 | 0.018 | 0 | 1 | 15 | Zfp37 | 0.018 | Young |
| 0.359497 | 2.918444 | 0.018 | 0 | 1 | 15 | A730071L1 | 0.018 | Young |
| 0.359497 | 2.897488 | 0.018 | 0 | 1 | 15 | Ntng2 | 0.018 | Young |
| 0.359497 | 2.897488 | 0.018 | 0 | 1 | 15 | Ssx2ip | 0.018 | Young |
| 0.359497 | 2.897488 | 0.018 | 0 | 1 | 15 | Xrcc3 | 0.018 | Young |
| 0.359497 | 2.897488 | 0.018 | 0 | 1 | 15 | Kcng3 | 0.018 | Young |
| 0.359497 | 2.883587 | 0.018 | 0 | 1 | 15 | 1110019D1 | 0.018 | Young |
| 0.359497 | 2.868981 | 0.018 | 0 | 1 | 15 | 1700113A1 | 0.018 | Young |
| 0.359497 | 2.868981 | 0.018 | 0 | 1 | 15 | Sbk3 | 0.018 | Young |
| 0.359497 | 2.868981 | 0.018 | 0 | 1 | 15 | A630091E0 | 0.018 | Young |
| 0.359497 | 2.868981 | 0.018 | 0 | 1 | 15 | 9930111J2 | 0.018 | Young |
| 0.359497 | 2.851533 | 0.018 | 0 | 1 | 15 | 1110012L1 | 0.018 | Young |
| 0.359497 | 2.849371 | 0.018 | 0 | 1 | 15 | Ttc36 | 0.018 | Young |
| 0.359497 | 2.848508 | 0.018 | 0 | 1 | 15 | Gm7967 | 0.018 | Young |
| 0.359497 | 2.783598 | 0.018 | 0 | 1 | 15 | Gm6710 | 0.018 | Young |
| 0.359497 | 2.783598 | 0.018 | 0 | 1 | 15 | Syp | 0.018 | Young |
| 0.359497 | 2.783598 | 0.018 | 0 | 1 | 15 | Penk | 0.018 | Young |
| 0.359497 | 2.783598 | 0.018 | 0 | 1 | 15 | Etfbkmt | 0.018 | Young |
| 0.359497 | 2.783598 | 0.018 | 0 | 1 | 15 | Plagl1 | 0.018 | Young |
| 0.359497 | 2.783598 | 0.018 | 0 | 1 | 15 | Zfp882 | 0.018 | Young |
| 0.359497 | 2.783598 | 0.018 | 0 | 1 | 15 | Mylk4 | 0.018 | Young |
| 0.359497 | 2.696594 | 0.018 | 0 | 1 | 15 | Cercam | 0.018 | Young |
| 0.359497 | 2.696594 | 0.018 | 0 | 1 | 15 | 1700029I1 | 0.018 | Young |
| 0.359497 | 2.696594 | 0.018 | 0 | 1 | 15 | Phtf1os | 0.018 | Young |
| 0.359497 | 2.696594 | 0.018 | 0 | 1 | 15 | Xrcc2 | 0.018 | Young |
| 0.359497 | 2.696594 | 0.018 | 0 | 1 | 15 | Gm20696 | 0.018 | Young |
| 0.359497 | 2.696594 | 0.018 | 0 | 1 | 15 | Zscan2 | 0.018 | Young |
| 0.359497 | 2.696594 | 0.018 | 0 | 1 | 15 | Gm48689 | 0.018 | Young |
| 0.359497 | 2.696594 | 0.018 | 0 | 1 | 15 | Megf11 | 0.018 | Young |
| 0.359497 | 2.696594 | 0.018 | 0 | 1 | 15 | Ltbp2 | 0.018 | Young |

| | | | | | | | | |
|---|---|---|---|---|---|---|---|---|
| 0.359497 | 2.696594 | 0.018 | 0 | 1 | 15 | Ctif | 0.018 | Young |
| 0.359497 | 2.696594 | 0.018 | 0 | 1 | 15 | Casp7 | 0.018 | Young |
| 0.359497 | 2.59254 | 0.018 | 0 | 1 | 15 | Cdk15 | 0.018 | Young |
| 0.359497 | 2.59254 | 0.018 | 0 | 1 | 15 | Cdh22 | 0.018 | Young |
| 0.359497 | 2.59254 | 0.018 | 0 | 1 | 15 | Gspt2 | 0.018 | Young |
| 0.359497 | 2.59254 | 0.018 | 0 | 1 | 15 | Fam212b | 0.018 | Young |
| 0.359497 | 2.59254 | 0.018 | 0 | 1 | 15 | Cryz | 0.018 | Young |
| 0.359497 | 2.59254 | 0.018 | 0 | 1 | 15 | Rims3 | 0.018 | Young |
| 0.359497 | 2.59254 | 0.018 | 0 | 1 | 15 | Mad2l2 | 0.018 | Young |
| 0.359497 | 2.59254 | 0.018 | 0 | 1 | 15 | Zfp784 | 0.018 | Young |
| 0.359497 | 2.59254 | 0.018 | 0 | 1 | 15 | Dpy19l3 | 0.018 | Young |
| 0.359497 | 2.59254 | 0.018 | 0 | 1 | 15 | Xntrpc | 0.018 | Young |
| 0.359497 | 2.59254 | 0.018 | 0 | 1 | 15 | Mettl17 | 0.018 | Young |
| 0.359497 | 2.59254 | 0.018 | 0 | 1 | 15 | Tmem220 | 0.018 | Young |
| 0.359497 | 2.59254 | 0.018 | 0 | 1 | 15 | Hist1h4h | 0.018 | Young |
| 0.359497 | 2.582081 | 0.018 | 0 | 1 | 15 | Slc2a8 | 0.018 | Young |
| 0.359497 | 2.582081 | 0.018 | 0 | 1 | 15 | Aurka | 0.018 | Young |
| 0.359497 | 2.582081 | 0.018 | 0 | 1 | 15 | Ccdc77 | 0.018 | Young |
| 0.359497 | 2.582081 | 0.018 | 0 | 1 | 15 | Cyp2s1 | 0.018 | Young |
| 0.359497 | 2.582081 | 0.018 | 0 | 1 | 15 | Pald1 | 0.018 | Young |
| 0.359497 | 2.574056 | 0.018 | 0 | 1 | 15 | Tnnt3 | 0.018 | Young |
| 0.359497 | 2.569735 | 0.018 | 0 | 1 | 15 | Hist2h3c2 | 0.018 | Young |
| 0.359497 | 2.569735 | 0.018 | 0 | 1 | 15 | Abcg2 | 0.018 | Young |
| 0.359497 | 2.569735 | 0.018 | 0 | 1 | 15 | 4931428F0 | 0.018 | Young |
| 0.359497 | 2.569735 | 0.018 | 0 | 1 | 15 | Dis3 | 0.018 | Young |
| 0.359497 | 2.569735 | 0.018 | 0 | 1 | 15 | Mrm3 | 0.018 | Young |
| 0.359497 | 2.569735 | 0.018 | 0 | 1 | 15 | Dcaf4 | 0.018 | Young |
| 0.359497 | 2.569735 | 0.018 | 0 | 1 | 15 | Pnma1 | 0.018 | Young |
| 0.359497 | 2.569735 | 0.018 | 0 | 1 | 15 | Park2 | 0.018 | Young |
| 0.359497 | 2.569735 | 0.018 | 0 | 1 | 15 | Zfp35 | 0.018 | Young |
| 0.359497 | 2.54031 | 0.018 | 0 | 1 | 15 | A530032D1 | 0.018 | Young |
| 0.359497 | 2.54031 | 0.018 | 0 | 1 | 15 | Gm42957 | 0.018 | Young |
| 0.359497 | 2.54031 | 0.018 | 0 | 1 | 15 | Ampd1 | 0.018 | Young |
| 0.359497 | 2.54031 | 0.018 | 0 | 1 | 15 | Zfp974 | 0.018 | Young |
| 0.359497 | 2.54031 | 0.018 | 0 | 1 | 15 | Ctxn1 | 0.018 | Young |
| 0.359497 | 2.54031 | 0.018 | 0 | 1 | 15 | 2610044O1 | 0.018 | Young |
| 0.359497 | 2.54031 | 0.018 | 0 | 1 | 15 | Gm29562 | 0.018 | Young |
| 0.359497 | 2.54031 | 0.018 | 0 | 1 | 15 | Pgap3 | 0.018 | Young |
| 0.359497 | 2.54031 | 0.018 | 0 | 1 | 15 | Kif20a | 0.018 | Young |
| 0.359497 | 2.536129 | 0.018 | 0 | 1 | 15 | Plpp3 | 0.018 | Young |
| 0.359497 | 2.536129 | 0.018 | 0 | 1 | 15 | Sult1e1 | 0.018 | Young |
| 0.359497 | 2.536129 | 0.018 | 0 | 1 | 15 | Sparcl1 | 0.018 | Young |
| 0.359497 | 2.536129 | 0.018 | 0 | 1 | 15 | Lep | 0.018 | Young |
| 0.359497 | 2.536129 | 0.018 | 0 | 1 | 15 | Gm45155 | 0.018 | Young |
| 0.359497 | 2.536129 | 0.018 | 0 | 1 | 15 | Gfpt2 | 0.018 | Young |

| | | | | | | | | |
|---|---|---|---|---|---|---|---|---|
| 0.359497 | 2.536129 | 0.018 | 0 | 1 | 15 | Smpd5 | 0.018 | Young |
| 0.359497 | 2.536129 | 0.018 | 0 | 1 | 15 | Cby1 | 0.018 | Young |
| 0.359497 | 2.536129 | 0.018 | 0 | 1 | 15 | Vasn | 0.018 | Young |
| 0.359497 | 2.531965 | 0.018 | 0 | 1 | 15 | Gm38336 | 0.018 | Young |
| 0.359497 | 2.531965 | 0.018 | 0 | 1 | 15 | Cst7 | 0.018 | Young |
| 0.359497 | 2.531965 | 0.018 | 0 | 1 | 15 | Nxf2 | 0.018 | Young |
| 0.359497 | 2.531965 | 0.018 | 0 | 1 | 15 | Pdzk1ip1 | 0.018 | Young |
| 0.359497 | 2.526864 | 0.018 | 0 | 1 | 15 | C130074G1 | 0.018 | Young |
| 0.359497 | 2.526864 | 0.018 | 0 | 1 | 15 | Ipp | 0.018 | Young |
| 0.359497 | 2.526864 | 0.018 | 0 | 1 | 15 | Gm42846 | 0.018 | Young |
| 0.359497 | 2.526864 | 0.018 | 0 | 1 | 15 | Prc1 | 0.018 | Young |
| 0.359497 | 2.526864 | 0.018 | 0 | 1 | 15 | Gm30978 | 0.018 | Young |
| 0.359497 | 2.526864 | 0.018 | 0 | 1 | 15 | Gm8113 | 0.018 | Young |
| 0.359497 | 2.526864 | 0.018 | 0 | 1 | 15 | Carmil1 | 0.018 | Young |
| 0.359497 | 2.526864 | 0.018 | 0 | 1 | 15 | Etohd2 | 0.018 | Young |
| 0.359497 | 2.526864 | 0.018 | 0 | 1 | 15 | Clcn2 | 0.018 | Young |
| 0.359497 | 2.526864 | 0.018 | 0 | 1 | 15 | Zfp472 | 0.018 | Young |
| 0.359497 | 2.526864 | 0.018 | 0 | 1 | 15 | Lonp1 | 0.018 | Young |
| 0.359497 | 2.526864 | 0.018 | 0 | 1 | 15 | Mirt1 | 0.018 | Young |
| 0.359497 | 2.504841 | 0.018 | 0 | 1 | 15 | Gm17634 | 0.018 | Young |
| 0.359497 | 2.504841 | 0.018 | 0 | 1 | 15 | Klhdc8a | 0.018 | Young |
| 0.359497 | 2.504841 | 0.018 | 0 | 1 | 15 | Plxna3 | 0.018 | Young |
| 0.359497 | 2.504841 | 0.018 | 0 | 1 | 15 | 4930599N2 | 0.018 | Young |
| 0.359497 | 2.504841 | 0.018 | 0 | 1 | 15 | Apba3 | 0.018 | Young |
| 0.359497 | 2.504841 | 0.018 | 0 | 1 | 15 | Ano10 | 0.018 | Young |
| 0.359497 | 2.504841 | 0.018 | 0 | 1 | 15 | Adamts5 | 0.018 | Young |
| 0.359497 | 2.497709 | 0.018 | 0 | 1 | 15 | Ttc30b | 0.018 | Young |
| 0.359497 | 2.497709 | 0.018 | 0 | 1 | 15 | Lbp | 0.018 | Young |
| 0.359497 | 2.497709 | 0.018 | 0 | 1 | 15 | Gm9774 | 0.018 | Young |
| 0.359497 | 2.497709 | 0.018 | 0 | 1 | 15 | Ido1 | 0.018 | Young |
| 0.359497 | 2.497709 | 0.018 | 0 | 1 | 15 | Zfp276 | 0.018 | Young |
| 0.359497 | 2.497709 | 0.018 | 0 | 1 | 15 | Gm36839 | 0.018 | Young |
| 0.359497 | 2.497709 | 0.018 | 0 | 1 | 15 | Zfp346 | 0.018 | Young |
| 0.359497 | 2.497709 | 0.018 | 0 | 1 | 15 | Nme4 | 0.018 | Young |
| 0.359497 | 2.493545 | 0.018 | 0 | 1 | 15 | Zfp273 | 0.018 | Young |
| 0.359497 | 2.481919 | 0.018 | 0 | 1 | 15 | Zfp574 | 0.018 | Young |
| 0.359497 | 2.481919 | 0.018 | 0 | 1 | 15 | Mmgt2 | 0.018 | Young |
| 0.359497 | 2.481919 | 0.018 | 0 | 1 | 15 | Map3k9 | 0.018 | Young |
| 0.359497 | 2.439481 | 0.018 | 0 | 1 | 15 | Asb1 | 0.018 | Young |
| 0.359497 | 2.439481 | 0.018 | 0 | 1 | 15 | Gm26670 | 0.018 | Young |
| 0.359497 | 2.439481 | 0.018 | 0 | 1 | 15 | Ckmt1 | 0.018 | Young |
| 0.359497 | 2.439481 | 0.018 | 0 | 1 | 15 | Il11 | 0.018 | Young |
| 0.359497 | 2.439481 | 0.018 | 0 | 1 | 15 | Ltbp4 | 0.018 | Young |
| 0.359497 | 2.439481 | 0.018 | 0 | 1 | 15 | Gm47113 | 0.018 | Young |
| 0.359497 | 2.439481 | 0.018 | 0 | 1 | 15 | Hgd | 0.018 | Young |

| | | | | | | | | |
|---|---|---|---|---|---|---|---|---|
| 0.359497 | 2.439481 | 0.018 | 0 | 1 | 15 | A830052D1 | 0.018 | Young |
| 0.359497 | 2.439481 | 0.018 | 0 | 1 | 15 | 8030462N1 | 0.018 | Young |
| 0.359497 | 2.433957 | 0.018 | 0 | 1 | 15 | Cybrd1 | 0.018 | Young |
| 0.359497 | 2.433957 | 0.018 | 0 | 1 | 15 | Gm15952 | 0.018 | Young |
| 0.359497 | 2.433957 | 0.018 | 0 | 1 | 15 | Mrgpra2a | 0.018 | Young |
| 0.359497 | 2.433957 | 0.018 | 0 | 1 | 15 | Gm527 | 0.018 | Young |
| 0.359497 | 2.433957 | 0.018 | 0 | 1 | 15 | Ttc9 | 0.018 | Young |
| 0.359497 | 2.433957 | 0.018 | 0 | 1 | 15 | Gm26531 | 0.018 | Young |
| 0.359497 | 2.433957 | 0.018 | 0 | 1 | 15 | Cul7 | 0.018 | Young |
| 0.359497 | 2.414434 | 0.018 | 0 | 1 | 15 | Chst14 | 0.018 | Young |
| 0.359497 | 2.414434 | 0.018 | 0 | 1 | 15 | Stil | 0.018 | Young |
| 0.359497 | 2.414434 | 0.018 | 0 | 1 | 15 | 4632411P0 | 0.018 | Young |
| 0.359497 | 2.414434 | 0.018 | 0 | 1 | 15 | Gm17655 | 0.018 | Young |
| 0.359497 | 2.414434 | 0.018 | 0 | 1 | 15 | Gm45051 | 0.018 | Young |
| 0.359497 | 2.414434 | 0.018 | 0 | 1 | 15 | Nrip3 | 0.018 | Young |
| 0.359497 | 2.414434 | 0.018 | 0 | 1 | 15 | Gm33843 | 0.018 | Young |
| 0.359497 | 2.414434 | 0.018 | 0 | 1 | 15 | Tmie | 0.018 | Young |
| 0.359497 | 2.403409 | 0.018 | 0 | 1 | 15 | 3300002I0 | 0.018 | Young |
| 0.359497 | 2.403409 | 0.018 | 0 | 1 | 15 | Tnnt1 | 0.018 | Young |
| 0.359497 | 2.403409 | 0.018 | 0 | 1 | 15 | Trim43b | 0.018 | Young |
| 0.359497 | 2.346617 | 0.018 | 0 | 1 | 15 | Ildr2 | 0.018 | Young |
| 0.359497 | 2.346617 | 0.018 | 0 | 1 | 15 | Smyd3 | 0.018 | Young |
| 0.359497 | 2.346617 | 0.018 | 0 | 1 | 15 | Gm14327 | 0.018 | Young |
| 0.359497 | 2.346617 | 0.018 | 0 | 1 | 15 | Enho | 0.018 | Young |
| 0.359497 | 2.346617 | 0.018 | 0 | 1 | 15 | BC017158 | 0.018 | Young |
| 0.359497 | 2.346617 | 0.018 | 0 | 1 | 15 | AU041133 | 0.018 | Young |
| 0.359497 | 2.346617 | 0.018 | 0 | 1 | 15 | Parpbp | 0.018 | Young |
| 0.359497 | 2.346617 | 0.018 | 0 | 1 | 15 | Usp2 | 0.018 | Young |
| 0.359497 | 2.346617 | 0.018 | 0 | 1 | 15 | Cd300ld3 | 0.018 | Young |
| 0.359497 | 2.346617 | 0.018 | 0 | 1 | 15 | Ptpdc1 | 0.018 | Young |
| 0.359497 | 2.346617 | 0.018 | 0 | 1 | 15 | Vmn2r97 | 0.018 | Young |
| 0.359497 | 2.346617 | 0.018 | 0 | 1 | 15 | Pnpla1 | 0.018 | Young |
| 0.359497 | 2.342119 | 0.018 | 0 | 1 | 15 | Hsdl2 | 0.018 | Young |
| 0.359497 | 2.342119 | 0.018 | 0 | 1 | 15 | 0610043K1 | 0.018 | Young |
| 0.359497 | 2.342119 | 0.018 | 0 | 1 | 15 | Steap4 | 0.018 | Young |
| 0.359497 | 2.342119 | 0.018 | 0 | 1 | 15 | E030037K0 | 0.018 | Young |
| 0.359497 | 2.342119 | 0.018 | 0 | 1 | 15 | Has2 | 0.018 | Young |
| 0.359497 | 2.342119 | 0.018 | 0 | 1 | 15 | 1700025B1 | 0.018 | Young |
| 0.359497 | 2.308396 | 0.018 | 0 | 1 | 15 | Gm17435 | 0.018 | Young |
| 0.359497 | 2.292219 | 0.018 | 0 | 1 | 15 | Ptrh1 | 0.018 | Young |
| 0.359497 | 2.292219 | 0.018 | 0 | 1 | 15 | Gstm7 | 0.018 | Young |
| 0.359497 | 2.292219 | 0.018 | 0 | 1 | 15 | Henmt1 | 0.018 | Young |
| 0.359497 | 2.292219 | 0.018 | 0 | 1 | 15 | Ldhb | 0.018 | Young |
| 0.359497 | 2.292219 | 0.018 | 0 | 1 | 15 | Pex11g | 0.018 | Young |
| 0.359497 | 2.292219 | 0.018 | 0 | 1 | 15 | Gm8016 | 0.018 | Young |

| | | | | | | | | |
|---|---|---|---|---|---|---|---|---|
| 0.359497 | 2.292219 | 0.018 | 0 | 1 | 15 | Sdc2 | 0.018 | Young |
| 0.359497 | 2.292219 | 0.018 | 0 | 1 | 15 | Gm4950 | 0.018 | Young |
| 0.359497 | 2.288966 | 0.018 | 0 | 1 | 15 | Shq1 | 0.018 | Young |
| 0.359497 | 2.288966 | 0.018 | 0 | 1 | 15 | Top1mt | 0.018 | Young |
| 0.359497 | 2.281626 | 0.018 | 0 | 1 | 15 | Rarb | 0.018 | Young |
| 0.359497 | 2.268946 | 0.018 | 0 | 1 | 15 | Angptl2 | 0.018 | Young |
| 0.359497 | 2.268946 | 0.018 | 0 | 1 | 15 | Armcx2 | 0.018 | Young |
| 0.359497 | 2.268946 | 0.018 | 0 | 1 | 15 | Tmem38b | 0.018 | Young |
| 0.359497 | 2.268946 | 0.018 | 0 | 1 | 15 | Gm31363 | 0.018 | Young |
| 0.359497 | 2.268946 | 0.018 | 0 | 1 | 15 | Gm43936 | 0.018 | Young |
| 0.359497 | 2.268946 | 0.018 | 0 | 1 | 15 | Gstt1 | 0.018 | Young |
| 0.359497 | 2.268946 | 0.018 | 0 | 1 | 15 | Defb42 | 0.018 | Young |
| 0.359497 | 2.268946 | 0.018 | 0 | 1 | 15 | Mtrf1 | 0.018 | Young |
| 0.359497 | 2.268946 | 0.018 | 0 | 1 | 15 | Hrh2 | 0.018 | Young |
| 0.359497 | 2.268946 | 0.018 | 0 | 1 | 15 | Dok3 | 0.018 | Young |
| 0.359497 | 2.268946 | 0.018 | 0 | 1 | 15 | AL591952.: | 0.018 | Young |
| 0.359497 | 2.268946 | 0.018 | 0 | 1 | 15 | Nxpe3 | 0.018 | Young |
| 0.359497 | 2.24958 | 0.018 | 0 | 1 | 15 | Il1r2 | 0.018 | Young |
| 0.359497 | 2.24958 | 0.018 | 0 | 1 | 15 | Proser2 | 0.018 | Young |
| 0.359497 | 2.24958 | 0.018 | 0 | 1 | 15 | Gm15441 | 0.018 | Young |
| 0.359497 | 2.24958 | 0.018 | 0 | 1 | 15 | A030001D: | 0.018 | Young |
| 0.359497 | 2.24958 | 0.018 | 0 | 1 | 15 | Gm26882 | 0.018 | Young |
| 0.359497 | 2.24958 | 0.018 | 0 | 1 | 15 | Nprl2 | 0.018 | Young |
| 0.359497 | 2.24958 | 0.018 | 0 | 1 | 15 | Cmtm8 | 0.018 | Young |
| 0.359497 | 2.24958 | 0.018 | 0 | 1 | 15 | Ccr9 | 0.018 | Young |
| 0.359497 | 2.24958 | 0.018 | 0 | 1 | 15 | Meox1 | 0.018 | Young |
| 0.359497 | 2.24958 | 0.018 | 0 | 1 | 15 | Itgb3 | 0.018 | Young |
| 0.359497 | 2.24958 | 0.018 | 0 | 1 | 15 | BC147527 | 0.018 | Young |
| 0.359497 | 2.24958 | 0.018 | 0 | 1 | 15 | Ager | 0.018 | Young |
| 0.359497 | 2.229197 | 0.018 | 0 | 1 | 15 | Gm38079 | 0.018 | Young |
| 0.359497 | 2.229197 | 0.018 | 0 | 1 | 15 | Shf | 0.018 | Young |
| 0.359497 | 2.229197 | 0.018 | 0 | 1 | 15 | Gm9873 | 0.018 | Young |
| 0.359497 | 2.229197 | 0.018 | 0 | 1 | 15 | Gm17146 | 0.018 | Young |
| 0.359497 | 2.229197 | 0.018 | 0 | 1 | 15 | Acad12 | 0.018 | Young |
| 0.359497 | 2.229197 | 0.018 | 0 | 1 | 15 | Zfp976 | 0.018 | Young |
| 0.359497 | 2.229197 | 0.018 | 0 | 1 | 15 | Rexo5 | 0.018 | Young |
| 0.359497 | 2.229197 | 0.018 | 0 | 1 | 15 | Adarb1 | 0.018 | Young |
| 0.359497 | 2.229197 | 0.018 | 0 | 1 | 15 | Zfp781 | 0.018 | Young |
| 0.359497 | 2.229197 | 0.018 | 0 | 1 | 15 | Tssk4 | 0.018 | Young |
| 0.359497 | 2.229197 | 0.018 | 0 | 1 | 15 | Celsr3 | 0.018 | Young |
| 0.359497 | 2.229197 | 0.018 | 0 | 1 | 15 | Zfp3 | 0.018 | Young |
| 0.359497 | 2.229197 | 0.018 | 0 | 1 | 15 | Sarm1 | 0.018 | Young |
| 0.359497 | 2.229197 | 0.018 | 0 | 1 | 15 | Ctla2a | 0.018 | Young |
| 0.359497 | 2.229197 | 0.018 | 0 | 1 | 15 | Cep128 | 0.018 | Young |
| 0.359497 | 2.229197 | 0.018 | 0 | 1 | 15 | Spaca6 | 0.018 | Young |

| | | | | | | | | |
|---|---|---|---|---|---|---|---|---|
| 0.359497 | 2.229197 | 0.018 | 0 | 1 | 15 | Zfp820 | 0.018 | Young |
| 0.359497 | 2.229197 | 0.018 | 0 | 1 | 15 | A930029G2 | 0.018 | Young |
| 0.359497 | 2.229197 | 0.018 | 0 | 1 | 15 | Prelid3a | 0.018 | Young |
| 0.359497 | 2.192871 | 0.018 | 0 | 1 | 15 | Nudt6 | 0.018 | Young |
| 0.359497 | 2.192871 | 0.018 | 0 | 1 | 15 | Cd5l | 0.018 | Young |
| 0.359497 | 2.192871 | 0.018 | 0 | 1 | 15 | Zkscan1 | 0.018 | Young |
| 0.359497 | 2.192871 | 0.018 | 0 | 1 | 15 | Gm15889 | 0.018 | Young |
| 0.359497 | 2.192871 | 0.018 | 0 | 1 | 15 | Gnpat | 0.018 | Young |
| 0.359497 | 2.192871 | 0.018 | 0 | 1 | 15 | Me1 | 0.018 | Young |
| 0.359497 | 2.192871 | 0.018 | 0 | 1 | 15 | Shisa8 | 0.018 | Young |
| 0.359497 | 2.175117 | 0.018 | 0 | 1 | 15 | Arl4d | 0.018 | Young |
| 0.359497 | 2.152075 | 0.018 | 0 | 1 | 15 | Trim14 | 0.018 | Young |
| 0.359497 | 2.152075 | 0.018 | 0 | 1 | 15 | 9930104L0 | 0.018 | Young |
| 0.359497 | 2.152075 | 0.018 | 0 | 1 | 15 | Kif17 | 0.018 | Young |
| 0.359497 | 2.152075 | 0.018 | 0 | 1 | 15 | 4930524J0 | 0.018 | Young |
| 0.359497 | 2.152075 | 0.018 | 0 | 1 | 15 | Gm16251 | 0.018 | Young |
| 0.359497 | 2.152075 | 0.018 | 0 | 1 | 15 | Hbb-bs | 0.018 | Young |
| 0.359497 | 2.152075 | 0.018 | 0 | 1 | 15 | Armc6 | 0.018 | Young |
| 0.359497 | 2.152075 | 0.018 | 0 | 1 | 15 | Zfp1 | 0.018 | Young |
| 0.359497 | 2.152075 | 0.018 | 0 | 1 | 15 | Nkiras1 | 0.018 | Young |
| 0.359497 | 2.152075 | 0.018 | 0 | 1 | 15 | 2810457G0 | 0.018 | Young |
| 0.359497 | 2.152075 | 0.018 | 0 | 1 | 15 | Gm17259 | 0.018 | Young |
| 0.359497 | 2.152075 | 0.018 | 0 | 1 | 15 | Gpr135 | 0.018 | Young |
| 0.359497 | 2.152075 | 0.018 | 0 | 1 | 15 | Zfat | 0.018 | Young |
| 0.359497 | 2.152075 | 0.018 | 0 | 1 | 15 | AC113595. | 0.018 | Young |
| 0.359497 | 2.152075 | 0.018 | 0 | 1 | 15 | Gm17025 | 0.018 | Young |
| 0.359497 | 2.152075 | 0.018 | 0 | 1 | 15 | Cenpm | 0.018 | Young |
| 0.359497 | 2.152075 | 0.018 | 0 | 1 | 15 | Adrb1 | 0.018 | Young |
| 0.359497 | 2.066921 | 0.018 | 0 | 1 | 15 | Zfp212 | 0.018 | Young |
| 0.359497 | 2.066921 | 0.018 | 0 | 1 | 15 | A430078I0 | 0.018 | Young |
| 0.359497 | 2.066921 | 0.018 | 0 | 1 | 15 | Lrig1 | 0.018 | Young |
| 0.359497 | 2.066921 | 0.018 | 0 | 1 | 15 | Cep57l1 | 0.018 | Young |
| 0.359497 | 2.066921 | 0.018 | 0 | 1 | 15 | Stox2 | 0.018 | Young |
| 0.359497 | 2.066921 | 0.018 | 0 | 1 | 15 | Gm16486 | 0.018 | Young |
| 0.359497 | 2.066921 | 0.018 | 0 | 1 | 15 | Ttk | 0.018 | Young |
| 0.359497 | 2.066921 | 0.018 | 0 | 1 | 15 | Ankrd13b | 0.018 | Young |
| 0.359497 | 2.066921 | 0.018 | 0 | 1 | 15 | D11Wsu47 | 0.018 | Young |
| 0.359497 | 2.066921 | 0.018 | 0 | 1 | 15 | 1700001L0 | 0.018 | Young |
| 0.359497 | 2.066921 | 0.018 | 0 | 1 | 15 | Frmd8os | 0.018 | Young |
| 0.359497 | 2.038729 | 0.018 | 0 | 1 | 15 | Stkld1 | 0.018 | Young |
| 0.359497 | 2.038729 | 0.018 | 0 | 1 | 15 | Pkn3 | 0.018 | Young |
| 0.359497 | 2.038729 | 0.018 | 0 | 1 | 15 | Spc25 | 0.018 | Young |
| 0.359497 | 2.038729 | 0.018 | 0 | 1 | 15 | Them4 | 0.018 | Young |
| 0.359497 | 2.038729 | 0.018 | 0 | 1 | 15 | E130308A1 | 0.018 | Young |
| 0.359497 | 2.038729 | 0.018 | 0 | 1 | 15 | Dph2 | 0.018 | Young |

| | | | | | | | | |
|---|---|---|---|---|---|---|---|---|
| 0.359497 | 2.038729 | 0.018 | 0 | 1 | 15 | Adamts3 | 0.018 | Young |
| 0.359497 | 2.038729 | 0.018 | 0 | 1 | 15 | C1s1 | 0.018 | Young |
| 0.359497 | 2.038729 | 0.018 | 0 | 1 | 15 | Lins1 | 0.018 | Young |
| 0.359497 | 2.038729 | 0.018 | 0 | 1 | 15 | Hdgfl3 | 0.018 | Young |
| 0.359497 | 2.038729 | 0.018 | 0 | 1 | 15 | Capn5 | 0.018 | Young |
| 0.359497 | 2.038729 | 0.018 | 0 | 1 | 15 | Orc6 | 0.018 | Young |
| 0.359497 | 2.038729 | 0.018 | 0 | 1 | 15 | Lrmda | 0.018 | Young |
| 0.359497 | 2.038729 | 0.018 | 0 | 1 | 15 | Phf7 | 0.018 | Young |
| 0.359497 | 2.038729 | 0.018 | 0 | 1 | 15 | Fxyd6 | 0.018 | Young |
| 0.359497 | 2.038729 | 0.018 | 0 | 1 | 15 | Tmem202 | 0.018 | Young |
| 0.359497 | 2.038729 | 0.018 | 0 | 1 | 15 | Mafa | 0.018 | Young |
| 0.359497 | 2.038729 | 0.018 | 0 | 1 | 15 | Zfp119a | 0.018 | Young |
| 0.359497 | 2.036073 | 0.018 | 0 | 1 | 15 | Kdelc1 | 0.018 | Young |
| 0.359497 | 2.036073 | 0.018 | 0 | 1 | 15 | A430106G: | 0.018 | Young |
| 0.359497 | 2.036073 | 0.018 | 0 | 1 | 15 | Gfi1b | 0.018 | Young |
| 0.359497 | 2.036073 | 0.018 | 0 | 1 | 15 | Slc1a2 | 0.018 | Young |
| 0.359497 | 2.036073 | 0.018 | 0 | 1 | 15 | Rmnd1 | 0.018 | Young |
| 0.359497 | 2.036073 | 0.018 | 0 | 1 | 15 | Prag1 | 0.018 | Young |
| 0.359497 | 2.036073 | 0.018 | 0 | 1 | 15 | Gm47230 | 0.018 | Young |
| 0.359497 | 2.036073 | 0.018 | 0 | 1 | 15 | Gm16124 | 0.018 | Young |
| 0.359497 | 2.036073 | 0.018 | 0 | 1 | 15 | Gm11963 | 0.018 | Young |
| 0.359497 | 2.036073 | 0.018 | 0 | 1 | 15 | Vcan | 0.018 | Young |
| 0.359497 | 2.036073 | 0.018 | 0 | 1 | 15 | Myc | 0.018 | Young |
| 0.359497 | 2.036073 | 0.018 | 0 | 1 | 15 | Tab1 | 0.018 | Young |
| 0.359497 | 2.036073 | 0.018 | 0 | 1 | 15 | Ggnbp1 | 0.018 | Young |
| 0.359497 | 2.033109 | 0.018 | 0 | 1 | 15 | Tchh | 0.018 | Young |
| 0.359497 | 2.023563 | 0.018 | 0 | 1 | 15 | Hes1 | 0.018 | Young |
| 0.359497 | 1.894841 | 0.018 | 0 | 1 | 15 | Prim1 | 0.018 | Young |
| 0.359497 | 1.894199 | 0.018 | 0 | 1 | 15 | Cdk18 | 0.018 | Young |
| 0.359497 | 1.894199 | 0.018 | 0 | 1 | 15 | Catspere2 | 0.018 | Young |
| 0.359497 | 1.894199 | 0.018 | 0 | 1 | 15 | Gm13620 | 0.018 | Young |
| 0.359497 | 1.894199 | 0.018 | 0 | 1 | 15 | Zfyve19 | 0.018 | Young |
| 0.359497 | 1.894199 | 0.018 | 0 | 1 | 15 | Gm14410 | 0.018 | Young |
| 0.359497 | 1.894199 | 0.018 | 0 | 1 | 15 | Tbc1d25 | 0.018 | Young |
| 0.359497 | 1.894199 | 0.018 | 0 | 1 | 15 | Cxcl3 | 0.018 | Young |
| 0.359497 | 1.894199 | 0.018 | 0 | 1 | 15 | Thrsp | 0.018 | Young |
| 0.359497 | 1.894199 | 0.018 | 0 | 1 | 15 | Apbb1 | 0.018 | Young |
| 0.359497 | 1.894199 | 0.018 | 0 | 1 | 15 | Gm45449 | 0.018 | Young |
| 0.359497 | 1.894199 | 0.018 | 0 | 1 | 15 | C230057M | 0.018 | Young |
| 0.359497 | 1.894199 | 0.018 | 0 | 1 | 15 | Rft1 | 0.018 | Young |
| 0.359497 | 1.894199 | 0.018 | 0 | 1 | 15 | AC061963. | 0.018 | Young |
| 0.359497 | 1.894199 | 0.018 | 0 | 1 | 15 | Plxnb1 | 0.018 | Young |
| 0.359497 | 1.894199 | 0.018 | 0 | 1 | 15 | Gm12185 | 0.018 | Young |
| 0.359497 | 1.894199 | 0.018 | 0 | 1 | 15 | Gm11273 | 0.018 | Young |
| 0.359497 | 1.894199 | 0.018 | 0 | 1 | 15 | AC161165. | 0.018 | Young |

| | | | | | | | | |
|---|---|---|---|---|---|---|---|---|
| 0.359497 | 1.894199 | 0.018 | 0 | 1 | 15 | Gm19412 | 0.018 | Young |
| 0.359497 | 1.816329 | 0.018 | 0 | 1 | 15 | Suv39h2 | 0.018 | Young |
| 0.359497 | 1.816329 | 0.018 | 0 | 1 | 15 | Ubox5 | 0.018 | Young |
| 0.359497 | 1.816329 | 0.018 | 0 | 1 | 15 | Intu | 0.018 | Young |
| 0.359497 | 1.816329 | 0.018 | 0 | 1 | 15 | 4930522L1 | 0.018 | Young |
| 0.359497 | 1.816329 | 0.018 | 0 | 1 | 15 | Tsen2 | 0.018 | Young |
| 0.359497 | 1.816329 | 0.018 | 0 | 1 | 15 | Rad52 | 0.018 | Young |
| 0.359497 | 1.816329 | 0.018 | 0 | 1 | 15 | Nr2f2 | 0.018 | Young |
| 0.359497 | 1.816329 | 0.018 | 0 | 1 | 15 | A930033H1 | 0.018 | Young |
| 0.359497 | 1.816329 | 0.018 | 0 | 1 | 15 | 1700030K0 | 0.018 | Young |
| 0.359497 | 1.816329 | 0.018 | 0 | 1 | 15 | Nr3c2 | 0.018 | Young |
| 0.359497 | 1.816329 | 0.018 | 0 | 1 | 15 | B3gnt9 | 0.018 | Young |
| 0.359497 | 1.816329 | 0.018 | 0 | 1 | 15 | 4930432J0 | 0.018 | Young |
| 0.359497 | 1.816329 | 0.018 | 0 | 1 | 15 | Colca2 | 0.018 | Young |
| 0.359497 | 1.816329 | 0.018 | 0 | 1 | 15 | Gnb5 | 0.018 | Young |
| 0.359497 | 1.816329 | 0.018 | 0 | 1 | 15 | Mrnip | 0.018 | Young |
| 0.359497 | 1.816329 | 0.018 | 0 | 1 | 15 | Chd3os | 0.018 | Young |
| 0.359497 | 1.816329 | 0.018 | 0 | 1 | 15 | Card6 | 0.018 | Young |
| 0.359497 | 1.816329 | 0.018 | 0 | 1 | 15 | AC158554. | 0.018 | Young |
| 0.359497 | 1.816329 | 0.018 | 0 | 1 | 15 | Paqr4 | 0.018 | Young |
| 0.359497 | 1.816329 | 0.018 | 0 | 1 | 15 | Gpam | 0.018 | Young |
| 0.359497 | 1.742004 | 0.018 | 0 | 1 | 15 | Rftn2 | 0.018 | Young |
| 0.359497 | 1.742004 | 0.018 | 0 | 1 | 15 | Rcn1 | 0.018 | Young |
| 0.359497 | 1.742004 | 0.018 | 0 | 1 | 15 | Adnp | 0.018 | Young |
| 0.359497 | 1.742004 | 0.018 | 0 | 1 | 15 | 1700013H1 | 0.018 | Young |
| 0.359497 | 1.742004 | 0.018 | 0 | 1 | 15 | Rragb | 0.018 | Young |
| 0.359497 | 1.742004 | 0.018 | 0 | 1 | 15 | Ccdc180 | 0.018 | Young |
| 0.359497 | 1.742004 | 0.018 | 0 | 1 | 15 | Hes2 | 0.018 | Young |
| 0.359497 | 1.742004 | 0.018 | 0 | 1 | 15 | Tnfrsf14 | 0.018 | Young |
| 0.359497 | 1.742004 | 0.018 | 0 | 1 | 15 | Tespa1 | 0.018 | Young |
| 0.359497 | 1.742004 | 0.018 | 0 | 1 | 15 | Nlrp1b | 0.018 | Young |
| 0.359497 | 1.742004 | 0.018 | 0 | 1 | 15 | Tmem267 | 0.018 | Young |
| 0.359497 | 1.742004 | 0.018 | 0 | 1 | 15 | L3mbtl2 | 0.018 | Young |
| 0.359497 | 1.742004 | 0.018 | 0 | 1 | 15 | A630089N0 | 0.018 | Young |
| 0.359497 | 1.742004 | 0.018 | 0 | 1 | 15 | Uhrf1bp1 | 0.018 | Young |
| 0.359497 | 1.742004 | 0.018 | 0 | 1 | 15 | Slc8a1 | 0.018 | Young |
| 0.359497 | 1.729358 | 0.018 | 0 | 1 | 15 | Ptges2 | 0.018 | Young |
| 0.359497 | 1.729358 | 0.018 | 0 | 1 | 15 | Hdc | 0.018 | Young |
| 0.359497 | 1.729358 | 0.018 | 0 | 1 | 15 | Tspan7 | 0.018 | Young |
| 0.359497 | 1.729358 | 0.018 | 0 | 1 | 15 | Zfp691 | 0.018 | Young |
| 0.359497 | 1.729358 | 0.018 | 0 | 1 | 15 | Vps37d | 0.018 | Young |
| 0.359497 | 1.729358 | 0.018 | 0 | 1 | 15 | Pdk4 | 0.018 | Young |
| 0.359497 | 1.729358 | 0.018 | 0 | 1 | 15 | Lpar5 | 0.018 | Young |
| 0.359497 | 1.729358 | 0.018 | 0 | 1 | 15 | Tarm1 | 0.018 | Young |
| 0.359497 | 1.729358 | 0.018 | 0 | 1 | 15 | Tmem86b | 0.018 | Young |

| | | | | | | | | |
|---|---|---|---|---|---|---|---|---|
| 0.359497 | 1.729358 | 0.018 | 0 | 1 | 15 | Proser3 | 0.018 | Young |
| 0.359497 | 1.729358 | 0.018 | 0 | 1 | 15 | Zfp619 | 0.018 | Young |
| 0.359497 | 1.729358 | 0.018 | 0 | 1 | 15 | Gm4065 | 0.018 | Young |
| 0.359497 | 1.729358 | 0.018 | 0 | 1 | 15 | Lrrc36 | 0.018 | Young |
| 0.359497 | 1.729358 | 0.018 | 0 | 1 | 15 | Cuedc1 | 0.018 | Young |
| 0.359497 | 1.729358 | 0.018 | 0 | 1 | 15 | Rnd2 | 0.018 | Young |
| 0.359497 | 1.729358 | 0.018 | 0 | 1 | 15 | Zkscan8 | 0.018 | Young |
| 0.359497 | 1.729358 | 0.018 | 0 | 1 | 15 | Gm15908 | 0.018 | Young |
| 0.359497 | 1.729358 | 0.018 | 0 | 1 | 15 | Cxcl14 | 0.018 | Young |
| 0.359497 | 1.729358 | 0.018 | 0 | 1 | 15 | Tsr3 | 0.018 | Young |
| 0.359497 | 1.71949 | 0.018 | 0 | 1 | 15 | Armc3 | 0.018 | Young |
| 0.359497 | 1.71949 | 0.018 | 0 | 1 | 15 | Gemin8 | 0.018 | Young |
| 0.359497 | 1.71949 | 0.018 | 0 | 1 | 15 | Tmem55a | 0.018 | Young |
| 0.359497 | 1.71949 | 0.018 | 0 | 1 | 15 | Nupl2 | 0.018 | Young |
| 0.359497 | 1.71949 | 0.018 | 0 | 1 | 15 | Ptpn13 | 0.018 | Young |
| 0.359497 | 1.71949 | 0.018 | 0 | 1 | 15 | Cldn13 | 0.018 | Young |
| 0.359497 | 1.71949 | 0.018 | 0 | 1 | 15 | P3h3 | 0.018 | Young |
| 0.359497 | 1.71949 | 0.018 | 0 | 1 | 15 | Gdpgp1 | 0.018 | Young |
| 0.359497 | 1.71949 | 0.018 | 0 | 1 | 15 | Tacc2 | 0.018 | Young |
| 0.359497 | 1.71949 | 0.018 | 0 | 1 | 15 | Gm40645 | 0.018 | Young |
| 0.359497 | 1.71949 | 0.018 | 0 | 1 | 15 | Dlgap5 | 0.018 | Young |
| 0.359497 | 1.71949 | 0.018 | 0 | 1 | 15 | Sall2 | 0.018 | Young |
| 0.359497 | 1.71949 | 0.018 | 0 | 1 | 15 | Gm16867 | 0.018 | Young |
| 0.359497 | 1.71949 | 0.018 | 0 | 1 | 15 | Abr | 0.018 | Young |
| 0.359497 | 1.71949 | 0.018 | 0 | 1 | 15 | Nle1 | 0.018 | Young |
| 0.359497 | 1.71949 | 0.018 | 0 | 1 | 15 | Gm11491 | 0.018 | Young |
| 0.359497 | 1.71949 | 0.018 | 0 | 1 | 15 | Klhl11 | 0.018 | Young |
| 0.359497 | 1.71949 | 0.018 | 0 | 1 | 15 | Nfe2 | 0.018 | Young |
| 0.359497 | 1.71949 | 0.018 | 0 | 1 | 15 | Cbr1 | 0.018 | Young |
| 0.359497 | 1.71949 | 0.018 | 0 | 1 | 15 | Zfp955b | 0.018 | Young |
| 0.359497 | 1.71949 | 0.018 | 0 | 1 | 15 | Tbx10 | 0.018 | Young |
| 0.359497 | 1.653179 | 0.018 | 0 | 1 | 15 | Bag2 | 0.018 | Young |
| 0.359497 | 1.653179 | 0.018 | 0 | 1 | 15 | Gal3st2 | 0.018 | Young |
| 0.359497 | 1.653179 | 0.018 | 0 | 1 | 15 | Gm38251 | 0.018 | Young |
| 0.359497 | 1.653179 | 0.018 | 0 | 1 | 15 | Oip5 | 0.018 | Young |
| 0.359497 | 1.653179 | 0.018 | 0 | 1 | 15 | Tspyl3 | 0.018 | Young |
| 0.359497 | 1.653179 | 0.018 | 0 | 1 | 15 | Klhl13 | 0.018 | Young |
| 0.359497 | 1.653179 | 0.018 | 0 | 1 | 15 | Gm43825 | 0.018 | Young |
| 0.359497 | 1.653179 | 0.018 | 0 | 1 | 15 | Dlx5 | 0.018 | Young |
| 0.359497 | 1.653179 | 0.018 | 0 | 1 | 15 | Mmrn1 | 0.018 | Young |
| 0.359497 | 1.653179 | 0.018 | 0 | 1 | 15 | Rasgef1a | 0.018 | Young |
| 0.359497 | 1.653179 | 0.018 | 0 | 1 | 15 | Slc25a18 | 0.018 | Young |
| 0.359497 | 1.653179 | 0.018 | 0 | 1 | 15 | A930016O | 0.018 | Young |
| 0.359497 | 1.653179 | 0.018 | 0 | 1 | 15 | Zfp141 | 0.018 | Young |
| 0.359497 | 1.653179 | 0.018 | 0 | 1 | 15 | Coq10a | 0.018 | Young |

| | | | | | | | | |
|---|---|---|---|---|---|---|---|---|
| 0.359497 | 1.653179 | 0.018 | 0 | 1 | 15 | Zfp319 | 0.018 | Young |
| 0.359497 | 1.653179 | 0.018 | 0 | 1 | 15 | Gm5431 | 0.018 | Young |
| 0.359497 | 1.653179 | 0.018 | 0 | 1 | 15 | BC049762 | 0.018 | Young |
| 0.359497 | 1.653179 | 0.018 | 0 | 1 | 15 | Adap2os | 0.018 | Young |
| 0.359497 | 1.653179 | 0.018 | 0 | 1 | 15 | Rnf135 | 0.018 | Young |
| 0.359497 | 1.653179 | 0.018 | 0 | 1 | 15 | Ogfod3 | 0.018 | Young |
| 0.359497 | 1.653179 | 0.018 | 0 | 1 | 15 | Gm16907 | 0.018 | Young |
| 0.359497 | 1.653179 | 0.018 | 0 | 1 | 15 | 9330020H( | 0.018 | Young |
| 0.359497 | 1.653179 | 0.018 | 0 | 1 | 15 | Pou6f1 | 0.018 | Young |
| 0.359497 | 1.653179 | 0.018 | 0 | 1 | 15 | Map3k12 | 0.018 | Young |
| 0.359497 | 1.653179 | 0.018 | 0 | 1 | 15 | Dynlt1f | 0.018 | Young |
| 0.359497 | 1.653179 | 0.018 | 0 | 1 | 15 | Gm26682 | 0.018 | Young |
| 0.359497 | 1.653179 | 0.018 | 0 | 1 | 15 | Vegfb | 0.018 | Young |
| 0.359497 | 1.636695 | 0.018 | 0 | 1 | 15 | 2810414N( | 0.018 | Young |
| 0.359497 | 1.636695 | 0.018 | 0 | 1 | 15 | Rhoj | 0.018 | Young |
| 0.359497 | 1.617747 | 0.018 | 0 | 1 | 15 | Mybpc3 | 0.018 | Young |
| 0.359497 | 1.617747 | 0.018 | 0 | 1 | 15 | Zfp408 | 0.018 | Young |
| 0.359497 | 1.617747 | 0.018 | 0 | 1 | 15 | Ndp | 0.018 | Young |
| 0.359497 | 1.617747 | 0.018 | 0 | 1 | 15 | Slc25a14 | 0.018 | Young |
| 0.359497 | 1.617747 | 0.018 | 0 | 1 | 15 | Mttp | 0.018 | Young |
| 0.359497 | 1.617747 | 0.018 | 0 | 1 | 15 | Gm13067 | 0.018 | Young |
| 0.359497 | 1.617747 | 0.018 | 0 | 1 | 15 | Gm35082 | 0.018 | Young |
| 0.359497 | 1.617747 | 0.018 | 0 | 1 | 15 | Tgfbr3l | 0.018 | Young |
| 0.359497 | 1.617747 | 0.018 | 0 | 1 | 15 | Vegfc | 0.018 | Young |
| 0.359497 | 1.617747 | 0.018 | 0 | 1 | 15 | Slc1a4 | 0.018 | Young |
| 0.359497 | 1.617747 | 0.018 | 0 | 1 | 15 | Gm11423 | 0.018 | Young |
| 0.359497 | 1.617747 | 0.018 | 0 | 1 | 15 | Gm32184 | 0.018 | Young |
| 0.359497 | 1.617747 | 0.018 | 0 | 1 | 15 | Gm9465 | 0.018 | Young |
| 0.359497 | 1.617747 | 0.018 | 0 | 1 | 15 | Gm30025 | 0.018 | Young |
| 0.359497 | 1.617747 | 0.018 | 0 | 1 | 15 | Dcbld2 | 0.018 | Young |
| 0.359497 | 1.617747 | 0.018 | 0 | 1 | 15 | Rgs11 | 0.018 | Young |
| 0.359497 | 1.61443 | 0.018 | 0 | 1 | 15 | Sh2d1b1 | 0.018 | Young |
| 0.359497 | 1.61443 | 0.018 | 0 | 1 | 15 | Nes | 0.018 | Young |
| 0.359497 | 1.61443 | 0.018 | 0 | 1 | 15 | Faap24 | 0.018 | Young |
| 0.359497 | 1.61443 | 0.018 | 0 | 1 | 15 | Akr1c18 | 0.018 | Young |
| 0.359497 | 1.61443 | 0.018 | 0 | 1 | 15 | Acta2 | 0.018 | Young |
| 0.359497 | 1.573694 | 0.018 | 0 | 1 | 15 | Traf3ip1 | 0.018 | Young |
| 0.359497 | 1.573694 | 0.018 | 0 | 1 | 15 | Dusp28 | 0.018 | Young |
| 0.359497 | 1.573694 | 0.018 | 0 | 1 | 15 | Sardh | 0.018 | Young |
| 0.359497 | 1.573694 | 0.018 | 0 | 1 | 15 | Wdr34 | 0.018 | Young |
| 0.359497 | 1.573694 | 0.018 | 0 | 1 | 15 | Ncaph | 0.018 | Young |
| 0.359497 | 1.573694 | 0.018 | 0 | 1 | 15 | Gm14029 | 0.018 | Young |
| 0.359497 | 1.573694 | 0.018 | 0 | 1 | 15 | Fmo5 | 0.018 | Young |
| 0.359497 | 1.573694 | 0.018 | 0 | 1 | 15 | Slc7a13 | 0.018 | Young |
| 0.359497 | 1.573694 | 0.018 | 0 | 1 | 15 | Tbc1d19 | 0.018 | Young |

| | | | | | | | | |
|---|---|---|---|---|---|---|---|---|
| 0.359497 | 1.573694 | 0.018 | 0 | 1 | 15 | Rpap2 | 0.018 | Young |
| 0.359497 | 1.573694 | 0.018 | 0 | 1 | 15 | 5930430L0 | 0.018 | Young |
| 0.359497 | 1.573694 | 0.018 | 0 | 1 | 15 | Dyrk1b | 0.018 | Young |
| 0.359497 | 1.573694 | 0.018 | 0 | 1 | 15 | Ankrd37 | 0.018 | Young |
| 0.359497 | 1.573694 | 0.018 | 0 | 1 | 15 | Ccdc130 | 0.018 | Young |
| 0.359497 | 1.573694 | 0.018 | 0 | 1 | 15 | Txnl4b | 0.018 | Young |
| 0.359497 | 1.573694 | 0.018 | 0 | 1 | 15 | Gemin5 | 0.018 | Young |
| 0.359497 | 1.573694 | 0.018 | 0 | 1 | 15 | AW209491 | 0.018 | Young |
| 0.359497 | 1.573694 | 0.018 | 0 | 1 | 15 | Bloc1s5 | 0.018 | Young |
| 0.359497 | 1.573694 | 0.018 | 0 | 1 | 15 | Srd5a1 | 0.018 | Young |
| 0.359497 | 1.573694 | 0.018 | 0 | 1 | 15 | Akap5 | 0.018 | Young |
| 0.359497 | 1.573694 | 0.018 | 0 | 1 | 15 | AU022793 | 0.018 | Young |
| 0.359497 | 1.573694 | 0.018 | 0 | 1 | 15 | AC140186. | 0.018 | Young |
| 0.359497 | 1.573694 | 0.018 | 0 | 1 | 15 | Pacrg | 0.018 | Young |
| 0.359497 | 1.573694 | 0.018 | 0 | 1 | 15 | 9330136K2 | 0.018 | Young |
| 0.359497 | 1.573694 | 0.018 | 0 | 1 | 15 | Slc3a1 | 0.018 | Young |
| 0.359497 | 1.573694 | 0.018 | 0 | 1 | 15 | 0610009O2 | 0.018 | Young |
| 0.359497 | 1.573694 | 0.018 | 0 | 1 | 15 | Afap1l1 | 0.018 | Young |
| 0.359497 | 1.573694 | 0.018 | 0 | 1 | 15 | Tut1 | 0.018 | Young |
| 0.359497 | 1.557547 | 0.018 | 0 | 1 | 15 | St18 | 0.018 | Young |
| 0.359497 | 1.557547 | 0.018 | 0 | 1 | 15 | Opn3 | 0.018 | Young |
| 0.359497 | 1.557547 | 0.018 | 0 | 1 | 15 | Lin9 | 0.018 | Young |
| 0.359497 | 1.557547 | 0.018 | 0 | 1 | 15 | Surf2 | 0.018 | Young |
| 0.359497 | 1.557547 | 0.018 | 0 | 1 | 15 | Coq4 | 0.018 | Young |
| 0.359497 | 1.557547 | 0.018 | 0 | 1 | 15 | Nup188 | 0.018 | Young |
| 0.359497 | 1.557547 | 0.018 | 0 | 1 | 15 | Gm36266 | 0.018 | Young |
| 0.359497 | 1.557547 | 0.018 | 0 | 1 | 15 | Cnpy4 | 0.018 | Young |
| 0.359497 | 1.557547 | 0.018 | 0 | 1 | 15 | Cav1 | 0.018 | Young |
| 0.359497 | 1.557547 | 0.018 | 0 | 1 | 15 | Trim30c | 0.018 | Young |
| 0.359497 | 1.557547 | 0.018 | 0 | 1 | 15 | Slc22a18 | 0.018 | Young |
| 0.359497 | 1.557547 | 0.018 | 0 | 1 | 15 | Esr1 | 0.018 | Young |
| 0.359497 | 1.557547 | 0.018 | 0 | 1 | 15 | Mmp15 | 0.018 | Young |
| 0.359497 | 1.557547 | 0.018 | 0 | 1 | 15 | Tgm1 | 0.018 | Young |
| 0.359497 | 1.557547 | 0.018 | 0 | 1 | 15 | Parp16 | 0.018 | Young |
| 0.359497 | 1.557547 | 0.018 | 0 | 1 | 15 | Hyal1 | 0.018 | Young |
| 0.359497 | 1.557547 | 0.018 | 0 | 1 | 15 | Fam212a | 0.018 | Young |
| 0.359497 | 1.557547 | 0.018 | 0 | 1 | 15 | Fgf11 | 0.018 | Young |
| 0.359497 | 1.557547 | 0.018 | 0 | 1 | 15 | Gm21988 | 0.018 | Young |
| 0.359497 | 1.557547 | 0.018 | 0 | 1 | 15 | Gm9796 | 0.018 | Young |
| 0.359497 | 1.557547 | 0.018 | 0 | 1 | 15 | Dnal1 | 0.018 | Young |
| 0.359497 | 1.557547 | 0.018 | 0 | 1 | 15 | Kcnk13 | 0.018 | Young |
| 0.359497 | 1.557547 | 0.018 | 0 | 1 | 15 | Cdca5 | 0.018 | Young |
| 0.359497 | 1.557547 | 0.018 | 0 | 1 | 15 | Tctn3 | 0.018 | Young |
| 0.359497 | 1.557547 | 0.018 | 0 | 1 | 15 | Hoga1 | 0.018 | Young |
| 0.359497 | 1.551456 | 0.018 | 0 | 1 | 15 | Gm34342 | 0.018 | Young |

| | | | | | | | | |
|---|---|---|---|---|---|---|---|---|
| 0.359497 | 1.551456 | 0.018 | 0 | 1 | 15 | Pdzd4 | 0.018 | Young |
| 0.359497 | 1.551456 | 0.018 | 0 | 1 | 15 | Tceanc | 0.018 | Young |
| 0.359497 | 1.551456 | 0.018 | 0 | 1 | 15 | Sema6c | 0.018 | Young |
| 0.359497 | 1.551456 | 0.018 | 0 | 1 | 15 | St6galnac3 | 0.018 | Young |
| 0.359497 | 1.551456 | 0.018 | 0 | 1 | 15 | Fbxo44 | 0.018 | Young |
| 0.359497 | 1.551456 | 0.018 | 0 | 1 | 15 | Dffb | 0.018 | Young |
| 0.359497 | 1.551456 | 0.018 | 0 | 1 | 15 | 5430403G1 | 0.018 | Young |
| 0.359497 | 1.551456 | 0.018 | 0 | 1 | 15 | E030030I0 | 0.018 | Young |
| 0.359497 | 1.551456 | 0.018 | 0 | 1 | 15 | Chchd10 | 0.018 | Young |
| 0.359497 | 1.551456 | 0.018 | 0 | 1 | 15 | Tmem263 | 0.018 | Young |
| 0.359497 | 1.551456 | 0.018 | 0 | 1 | 15 | Fanca | 0.018 | Young |
| 0.359497 | 1.551456 | 0.018 | 0 | 1 | 15 | Piwil4 | 0.018 | Young |
| 0.359497 | 1.551456 | 0.018 | 0 | 1 | 15 | Jmjd4 | 0.018 | Young |
| 0.359497 | 1.551456 | 0.018 | 0 | 1 | 15 | Lrrc75a | 0.018 | Young |
| 0.359497 | 1.551456 | 0.018 | 0 | 1 | 15 | Erbb2 | 0.018 | Young |
| 0.359497 | 1.551456 | 0.018 | 0 | 1 | 15 | Pde8b | 0.018 | Young |
| 0.359497 | 1.551456 | 0.018 | 0 | 1 | 15 | Arl15 | 0.018 | Young |
| 0.359497 | 1.551456 | 0.018 | 0 | 1 | 15 | Ankrd9 | 0.018 | Young |
| 0.359497 | 1.551456 | 0.018 | 0 | 1 | 15 | Wrb | 0.018 | Young |
| 0.359497 | 1.551456 | 0.018 | 0 | 1 | 15 | Rsph1 | 0.018 | Young |
| 0.359497 | 1.551456 | 0.018 | 0 | 1 | 15 | A330032B1 | 0.018 | Young |
| 0.359497 | 1.463681 | 0.018 | 0 | 1 | 15 | Gm31812 | 0.018 | Young |
| 0.359497 | 1.463681 | 0.018 | 0 | 1 | 15 | Armc9 | 0.018 | Young |
| 0.359497 | 1.463681 | 0.018 | 0 | 1 | 15 | Gm10516 | 0.018 | Young |
| 0.359497 | 1.463681 | 0.018 | 0 | 1 | 15 | G0s2 | 0.018 | Young |
| 0.359497 | 1.463681 | 0.018 | 0 | 1 | 15 | Kyat1 | 0.018 | Young |
| 0.359497 | 1.463681 | 0.018 | 0 | 1 | 15 | Zfp335os | 0.018 | Young |
| 0.359497 | 1.463681 | 0.018 | 0 | 1 | 15 | Foxo4 | 0.018 | Young |
| 0.359497 | 1.463681 | 0.018 | 0 | 1 | 15 | Tmem154 | 0.018 | Young |
| 0.359497 | 1.463681 | 0.018 | 0 | 1 | 15 | Tgfbr3 | 0.018 | Young |
| 0.359497 | 1.463681 | 0.018 | 0 | 1 | 15 | Nupr1l | 0.018 | Young |
| 0.359497 | 1.463681 | 0.018 | 0 | 1 | 15 | Gpr85 | 0.018 | Young |
| 0.359497 | 1.463681 | 0.018 | 0 | 1 | 15 | Retsat | 0.018 | Young |
| 0.359497 | 1.463681 | 0.018 | 0 | 1 | 15 | Zfp607b | 0.018 | Young |
| 0.359497 | 1.463681 | 0.018 | 0 | 1 | 15 | Gm44851 | 0.018 | Young |
| 0.359497 | 1.463681 | 0.018 | 0 | 1 | 15 | Kcnq1 | 0.018 | Young |
| 0.359497 | 1.463681 | 0.018 | 0 | 1 | 15 | Tnfrsf22 | 0.018 | Young |
| 0.359497 | 1.463681 | 0.018 | 0 | 1 | 15 | Yjefn3 | 0.018 | Young |
| 0.359497 | 1.463681 | 0.018 | 0 | 1 | 15 | Adat1 | 0.018 | Young |
| 0.359497 | 1.463681 | 0.018 | 0 | 1 | 15 | Spsb4 | 0.018 | Young |
| 0.359497 | 1.463681 | 0.018 | 0 | 1 | 15 | Amotl2 | 0.018 | Young |
| 0.359497 | 1.463681 | 0.018 | 0 | 1 | 15 | Tmem107 | 0.018 | Young |
| 0.359497 | 1.463681 | 0.018 | 0 | 1 | 15 | Kif27 | 0.018 | Young |
| 0.359497 | 1.463681 | 0.018 | 0 | 1 | 15 | Mis18bp1 | 0.018 | Young |
| 0.359497 | 1.463681 | 0.018 | 0 | 1 | 15 | Gm26518 | 0.018 | Young |

| | | | | | | | | |
|---|---|---|---|---|---|---|---|---|
| 0.359497 | 1.463681 | 0.018 | 0 | 1 | 15 | Klhl22 | 0.018 | Young |
| 0.359497 | 1.463681 | 0.018 | 0 | 1 | 15 | 1810055G( | 0.018 | Young |
| 0.359497 | 1.340809 | 0.018 | 0 | 1 | 15 | Epha4 | 0.018 | Young |
| 0.359497 | 1.340809 | 0.018 | 0 | 1 | 15 | Per2 | 0.018 | Young |
| 0.359497 | 1.340809 | 0.018 | 0 | 1 | 15 | Serping1 | 0.018 | Young |
| 0.359497 | 1.340809 | 0.018 | 0 | 1 | 15 | Gm826 | 0.018 | Young |
| 0.359497 | 1.340809 | 0.018 | 0 | 1 | 15 | Fam209 | 0.018 | Young |
| 0.359497 | 1.340809 | 0.018 | 0 | 1 | 15 | Rnf128 | 0.018 | Young |
| 0.359497 | 1.340809 | 0.018 | 0 | 1 | 15 | Wdr78 | 0.018 | Young |
| 0.359497 | 1.340809 | 0.018 | 0 | 1 | 15 | Btbd19 | 0.018 | Young |
| 0.359497 | 1.340809 | 0.018 | 0 | 1 | 15 | Cacna2d4 | 0.018 | Young |
| 0.359497 | 1.340809 | 0.018 | 0 | 1 | 15 | Pianp | 0.018 | Young |
| 0.359497 | 1.340809 | 0.018 | 0 | 1 | 15 | Nop2 | 0.018 | Young |
| 0.359497 | 1.340809 | 0.018 | 0 | 1 | 15 | Cd3eap | 0.018 | Young |
| 0.359497 | 1.340809 | 0.018 | 0 | 1 | 15 | Sptbn4 | 0.018 | Young |
| 0.359497 | 1.340809 | 0.018 | 0 | 1 | 15 | 1700006J1 | 0.018 | Young |
| 0.359497 | 1.340809 | 0.018 | 0 | 1 | 15 | D830025C( | 0.018 | Young |
| 0.359497 | 1.340809 | 0.018 | 0 | 1 | 15 | Gm33103 | 0.018 | Young |
| 0.359497 | 1.340809 | 0.018 | 0 | 1 | 15 | Gas8 | 0.018 | Young |
| 0.359497 | 1.340809 | 0.018 | 0 | 1 | 15 | A230050P2 | 0.018 | Young |
| 0.359497 | 1.340809 | 0.018 | 0 | 1 | 15 | Rbp1 | 0.018 | Young |
| 0.359497 | 1.340809 | 0.018 | 0 | 1 | 15 | Tmppe | 0.018 | Young |
| 0.359497 | 1.340809 | 0.018 | 0 | 1 | 15 | Lpo | 0.018 | Young |
| 0.359497 | 1.340809 | 0.018 | 0 | 1 | 15 | Zfp825 | 0.018 | Young |
| 0.359497 | 1.340809 | 0.018 | 0 | 1 | 15 | Gm5086 | 0.018 | Young |
| 0.359497 | 1.340809 | 0.018 | 0 | 1 | 15 | Oplah | 0.018 | Young |
| 0.359497 | 1.340809 | 0.018 | 0 | 1 | 15 | Cdpf1 | 0.018 | Young |
| 0.359497 | 1.340809 | 0.018 | 0 | 1 | 15 | Rsph3b | 0.018 | Young |
| 0.359497 | 1.340809 | 0.018 | 0 | 1 | 15 | Dnajc18 | 0.018 | Young |
| 0.359497 | 1.340809 | 0.018 | 0 | 1 | 15 | Slc35g1 | 0.018 | Young |
| 0.359497 | 1.340809 | 0.018 | 0 | 1 | 15 | Neurl1a | 0.018 | Young |
| 0.359497 | 1.238423 | 0.018 | 0 | 1 | 15 | Zranb3 | 0.018 | Young |
| 0.359497 | 1.238423 | 0.018 | 0 | 1 | 15 | 1700056E2 | 0.018 | Young |
| 0.359497 | 1.238423 | 0.018 | 0 | 1 | 15 | Mettl8 | 0.018 | Young |
| 0.359497 | 1.238423 | 0.018 | 0 | 1 | 15 | Tcf15 | 0.018 | Young |
| 0.359497 | 1.238423 | 0.018 | 0 | 1 | 15 | 1700003F1 | 0.018 | Young |
| 0.359497 | 1.238423 | 0.018 | 0 | 1 | 15 | Ift80 | 0.018 | Young |
| 0.359497 | 1.238423 | 0.018 | 0 | 1 | 15 | Gstm2 | 0.018 | Young |
| 0.359497 | 1.238423 | 0.018 | 0 | 1 | 15 | Mmp17 | 0.018 | Young |
| 0.359497 | 1.238423 | 0.018 | 0 | 1 | 15 | Ssc4d | 0.018 | Young |
| 0.359497 | 1.238423 | 0.018 | 0 | 1 | 15 | Zfp777 | 0.018 | Young |
| 0.359497 | 1.238423 | 0.018 | 0 | 1 | 15 | Ggct | 0.018 | Young |
| 0.359497 | 1.238423 | 0.018 | 0 | 1 | 15 | Aplf | 0.018 | Young |
| 0.359497 | 1.238423 | 0.018 | 0 | 1 | 15 | 4930590J0 | 0.018 | Young |
| 0.359497 | 1.238423 | 0.018 | 0 | 1 | 15 | Gm4673 | 0.018 | Young |

| | | | | | | | | |
|---|---|---|---|---|---|---|---|---|
| 0.359497 | 1.238423 | 0.018 | 0 | 1 | 15 | Gm44763 | 0.018 | Young |
| 0.359497 | 1.238423 | 0.018 | 0 | 1 | 15 | Dctpp1 | 0.018 | Young |
| 0.359497 | 1.238423 | 0.018 | 0 | 1 | 15 | 9130019O2 | 0.018 | Young |
| 0.359497 | 1.238423 | 0.018 | 0 | 1 | 15 | 9130023H2 | 0.018 | Young |
| 0.359497 | 1.238423 | 0.018 | 0 | 1 | 15 | Dhx32 | 0.018 | Young |
| 0.359497 | 1.238423 | 0.018 | 0 | 1 | 15 | Mdm1 | 0.018 | Young |
| 0.359497 | 1.238423 | 0.018 | 0 | 1 | 15 | Rbms2 | 0.018 | Young |
| 0.359497 | 1.238423 | 0.018 | 0 | 1 | 15 | Timeless | 0.018 | Young |
| 0.359497 | 1.238423 | 0.018 | 0 | 1 | 15 | Slc7a2 | 0.018 | Young |
| 0.359497 | 1.238423 | 0.018 | 0 | 1 | 15 | Gm10643 | 0.018 | Young |
| 0.359497 | 1.238423 | 0.018 | 0 | 1 | 15 | 9930004E1 | 0.018 | Young |
| 0.359497 | 1.238423 | 0.018 | 0 | 1 | 15 | Fignl1 | 0.018 | Young |
| 0.359497 | 1.238423 | 0.018 | 0 | 1 | 15 | Csf3 | 0.018 | Young |
| 0.359497 | 1.238423 | 0.018 | 0 | 1 | 15 | Arsg | 0.018 | Young |
| 0.359497 | 1.238423 | 0.018 | 0 | 1 | 15 | Afmid | 0.018 | Young |
| 0.359497 | 1.238423 | 0.018 | 0 | 1 | 15 | Tpmt | 0.018 | Young |
| 0.359497 | 1.238423 | 0.018 | 0 | 1 | 15 | E2f6 | 0.018 | Young |
| 0.359497 | 1.238423 | 0.018 | 0 | 1 | 15 | Rida | 0.018 | Young |
| 0.359497 | 1.238423 | 0.018 | 0 | 1 | 15 | Gm48960 | 0.018 | Young |
| 0.359497 | 1.238423 | 0.018 | 0 | 1 | 15 | Gm26798 | 0.018 | Young |
| 0.359497 | 1.238423 | 0.018 | 0 | 1 | 15 | 9030025P2 | 0.018 | Young |
| 0.359497 | 1.238423 | 0.018 | 0 | 1 | 15 | BC051226 | 0.018 | Young |
| 0.359497 | 1.238423 | 0.018 | 0 | 1 | 15 | Gm17705 | 0.018 | Young |
| 0.359497 | 1.238423 | 0.018 | 0 | 1 | 15 | Cd70 | 0.018 | Young |
| 0.359497 | 1.038972 | 0.018 | 0 | 1 | 15 | Mfsd9 | 0.018 | Young |
| 0.359497 | 1.038972 | 0.018 | 0 | 1 | 15 | Sec16b | 0.018 | Young |
| 0.359497 | 1.038972 | 0.018 | 0 | 1 | 15 | Lrp4 | 0.018 | Young |
| 0.359497 | 1.038972 | 0.018 | 0 | 1 | 15 | Ccdc73 | 0.018 | Young |
| 0.359497 | 1.038972 | 0.018 | 0 | 1 | 15 | BC052040 | 0.018 | Young |
| 0.359497 | 1.038972 | 0.018 | 0 | 1 | 15 | Ttl | 0.018 | Young |
| 0.359497 | 1.038972 | 0.018 | 0 | 1 | 15 | A530013C2 | 0.018 | Young |
| 0.359497 | 1.038972 | 0.018 | 0 | 1 | 15 | Gm20633 | 0.018 | Young |
| 0.359497 | 1.038972 | 0.018 | 0 | 1 | 15 | Padi4 | 0.018 | Young |
| 0.359497 | 1.038972 | 0.018 | 0 | 1 | 15 | Pusl1 | 0.018 | Young |
| 0.359497 | 1.038972 | 0.018 | 0 | 1 | 15 | Hcar2 | 0.018 | Young |
| 0.359497 | 1.038972 | 0.018 | 0 | 1 | 15 | Aldh1l1 | 0.018 | Young |
| 0.359497 | 1.038972 | 0.018 | 0 | 1 | 15 | Slc25a26 | 0.018 | Young |
| 0.359497 | 1.038972 | 0.018 | 0 | 1 | 15 | Cdkn1c | 0.018 | Young |
| 0.359497 | 1.038972 | 0.018 | 0 | 1 | 15 | Slc19a1 | 0.018 | Young |
| 0.359497 | 1.038972 | 0.018 | 0 | 1 | 15 | Gm48551 | 0.018 | Young |
| 0.359497 | 1.038972 | 0.018 | 0 | 1 | 15 | Arhgef7 | 0.018 | Young |
| 0.359497 | 1.038972 | 0.018 | 0 | 1 | 15 | Cenpu | 0.018 | Young |
| 0.359497 | 1.038972 | 0.018 | 0 | 1 | 15 | Rab3a | 0.018 | Young |
| 0.359497 | 1.038972 | 0.018 | 0 | 1 | 15 | Nutf2 | 0.018 | Young |
| 0.359497 | 1.038972 | 0.018 | 0 | 1 | 15 | Nynrin | 0.018 | Young |

| | | | | | | | | |
|---|---|---|---|---|---|---|---|---|
| 0.359497 | 1.038972 | 0.018 | 0 | 1 | 15 | Pih1d2 | 0.018 | Young |
| 0.359497 | 1.038972 | 0.018 | 0 | 1 | 15 | Fam161a | 0.018 | Young |
| 0.359497 | 1.038972 | 0.018 | 0 | 1 | 15 | Mfsd6l | 0.018 | Young |
| 0.359497 | 1.038972 | 0.018 | 0 | 1 | 15 | Gmpr | 0.018 | Young |
| 0.359497 | 1.038972 | 0.018 | 0 | 1 | 15 | Zfp748 | 0.018 | Young |
| 0.359497 | 1.038972 | 0.018 | 0 | 1 | 15 | Gm32296 | 0.018 | Young |
| 0.359497 | 1.038972 | 0.018 | 0 | 1 | 15 | Gpihbp1 | 0.018 | Young |
| 0.359497 | 1.038972 | 0.018 | 0 | 1 | 15 | Pla2g6 | 0.018 | Young |
| 0.359497 | 1.038972 | 0.018 | 0 | 1 | 15 | Parvg | 0.018 | Young |
| 0.359497 | 1.038972 | 0.018 | 0 | 1 | 15 | AC123724. | 0.018 | Young |
| 0.359497 | 1.038972 | 0.018 | 0 | 1 | 15 | Slc7a4 | 0.018 | Young |
| 0.359497 | 1.038972 | 0.018 | 0 | 1 | 15 | Wdr5b | 0.018 | Young |
| 0.359497 | 1.038972 | 0.018 | 0 | 1 | 15 | Rbm11 | 0.018 | Young |
| 0.359497 | 1.038972 | 0.018 | 0 | 1 | 15 | Runx2os1 | 0.018 | Young |
| 0.359497 | 1.038972 | 0.018 | 0 | 1 | 15 | Eif3j2 | 0.018 | Young |
| 0.359497 | 1.038972 | 0.018 | 0 | 1 | 15 | Macrod1 | 0.018 | Young |
| 0.359497 | 1.022157 | 0.018 | 0 | 1 | 15 | Sulf1 | 0.018 | Young |
| 0.359497 | 1.022157 | 0.018 | 0 | 1 | 15 | Slc4a3 | 0.018 | Young |
| 0.359497 | 1.022157 | 0.018 | 0 | 1 | 15 | Prrx1 | 0.018 | Young |
| 0.359497 | 1.022157 | 0.018 | 0 | 1 | 15 | Creb3l1 | 0.018 | Young |
| 0.359497 | 1.022157 | 0.018 | 0 | 1 | 15 | Iqgap3 | 0.018 | Young |
| 0.359497 | 1.022157 | 0.018 | 0 | 1 | 15 | Ap4b1 | 0.018 | Young |
| 0.359497 | 1.022157 | 0.018 | 0 | 1 | 15 | Dmap1 | 0.018 | Young |
| 0.359497 | 1.022157 | 0.018 | 0 | 1 | 15 | Ubxn11 | 0.018 | Young |
| 0.359497 | 1.022157 | 0.018 | 0 | 1 | 15 | Crocc | 0.018 | Young |
| 0.359497 | 1.022157 | 0.018 | 0 | 1 | 15 | Fam185a | 0.018 | Young |
| 0.359497 | 1.022157 | 0.018 | 0 | 1 | 15 | Grk4 | 0.018 | Young |
| 0.359497 | 1.022157 | 0.018 | 0 | 1 | 15 | Jakmip1 | 0.018 | Young |
| 0.359497 | 1.022157 | 0.018 | 0 | 1 | 15 | Txk | 0.018 | Young |
| 0.359497 | 1.022157 | 0.018 | 0 | 1 | 15 | Pole | 0.018 | Young |
| 0.359497 | 1.022157 | 0.018 | 0 | 1 | 15 | Gm6598 | 0.018 | Young |
| 0.359497 | 1.022157 | 0.018 | 0 | 1 | 15 | Actg2 | 0.018 | Young |
| 0.359497 | 1.022157 | 0.018 | 0 | 1 | 15 | Eogt | 0.018 | Young |
| 0.359497 | 1.022157 | 0.018 | 0 | 1 | 15 | Ogg1 | 0.018 | Young |
| 0.359497 | 1.022157 | 0.018 | 0 | 1 | 15 | Zfp128 | 0.018 | Young |
| 0.359497 | 1.022157 | 0.018 | 0 | 1 | 15 | Zfp61 | 0.018 | Young |
| 0.359497 | 1.022157 | 0.018 | 0 | 1 | 15 | Cep89 | 0.018 | Young |
| 0.359497 | 1.022157 | 0.018 | 0 | 1 | 15 | Sv2b | 0.018 | Young |
| 0.359497 | 1.022157 | 0.018 | 0 | 1 | 15 | 1-Sep | 0.018 | Young |
| 0.359497 | 1.022157 | 0.018 | 0 | 1 | 15 | 9-Mar | 0.018 | Young |
| 0.359497 | 1.022157 | 0.018 | 0 | 1 | 15 | Zbtb39 | 0.018 | Young |
| 0.359497 | 1.022157 | 0.018 | 0 | 1 | 15 | Tssk6 | 0.018 | Young |
| 0.359497 | 1.022157 | 0.018 | 0 | 1 | 15 | Gm49004 | 0.018 | Young |
| 0.359497 | 1.022157 | 0.018 | 0 | 1 | 15 | Cebpe | 0.018 | Young |
| 0.359497 | 1.022157 | 0.018 | 0 | 1 | 15 | Lgi3 | 0.018 | Young |

| | | | | | | | | |
|---|---|---|---|---|---|---|---|---|
| 0.359497 | 1.022157 | 0.018 | 0 | 1 | 15 | Smco4 | 0.018 | Young |
| 0.359497 | 1.022157 | 0.018 | 0 | 1 | 15 | Hyls1 | 0.018 | Young |
| 0.359497 | 1.022157 | 0.018 | 0 | 1 | 15 | Arid3b | 0.018 | Young |
| 0.359497 | 1.022157 | 0.018 | 0 | 1 | 15 | Gm27211 | 0.018 | Young |
| 0.359497 | 1.022157 | 0.018 | 0 | 1 | 15 | P4htm | 0.018 | Young |
| 0.359497 | 1.022157 | 0.018 | 0 | 1 | 15 | Pold2 | 0.018 | Young |
| 0.359497 | 1.022157 | 0.018 | 0 | 1 | 15 | Wwc1 | 0.018 | Young |
| 0.359497 | 1.022157 | 0.018 | 0 | 1 | 15 | Wrap53 | 0.018 | Young |
| 0.359497 | 1.022157 | 0.018 | 0 | 1 | 15 | Slc46a1 | 0.018 | Young |
| 0.359497 | 1.022157 | 0.018 | 0 | 1 | 15 | Cep295nl | 0.018 | Young |
| 0.359497 | 1.022157 | 0.018 | 0 | 1 | 15 | Hist1h2bh | 0.018 | Young |
| 0.359497 | 1.022157 | 0.018 | 0 | 1 | 15 | Hist1h4c | 0.018 | Young |
| 0.359497 | 1.022157 | 0.018 | 0 | 1 | 15 | AI463229 | 0.018 | Young |
| 0.359497 | 1.022157 | 0.018 | 0 | 1 | 15 | Zfp712 | 0.018 | Young |
| 0.359497 | 1.022157 | 0.018 | 0 | 1 | 15 | Matn2 | 0.018 | Young |
| 0.359497 | 1.022157 | 0.018 | 0 | 1 | 15 | Gm10384 | 0.018 | Young |
| 0.359497 | 1.022157 | 0.018 | 0 | 1 | 15 | Hgh1 | 0.018 | Young |
| 0.359497 | 1.022157 | 0.018 | 0 | 1 | 15 | Timmdc1 | 0.018 | Young |
| 0.359497 | 1.022157 | 0.018 | 0 | 1 | 15 | Gm3448 | 0.018 | Young |
| 0.359497 | 1.022157 | 0.018 | 0 | 1 | 15 | D17H6S53I | 0.018 | Young |
| 0.359497 | 1.022157 | 0.018 | 0 | 1 | 15 | B230354K1 | 0.018 | Young |
| 0.359497 | 1.022157 | 0.018 | 0 | 1 | 15 | Abcc10 | 0.018 | Young |
| 0.359497 | 1.022157 | 0.018 | 0 | 1 | 15 | Ndc80 | 0.018 | Young |
| 0.359497 | 1.022157 | 0.018 | 0 | 1 | 15 | Gm1976 | 0.018 | Young |
| 0.359497 | 1.022157 | 0.018 | 0 | 1 | 15 | Ppargc1b | 0.018 | Young |
| 0.359497 | 1.022157 | 0.018 | 0 | 1 | 15 | Rnf165 | 0.018 | Young |
| 0.359751 | 0.041195 | 0.786 | 0.714 | 1 | 15 | Rsrp1 | 0.072 | Young |
| 0.359968 | 0.134384 | 0.946 | 0.918 | 1 | 15 | Psap | 0.028 | Young |
| 0.360122 | 0.247744 | 0.464 | 0.388 | 1 | 15 | Map2k2 | 0.076 | Young |
| 0.360356 | 0.39056 | 0.571 | 0.469 | 1 | 15 | Grk2 | 0.102 | Young |
| 0.361234 | 0.515034 | 0.304 | 0.245 | 1 | 15 | Ufc1 | 0.059 | Young |
| 0.361274 | 0.856445 | 0.339 | 0.286 | 1 | 15 | Cited2 | 0.053 | Young |
| 0.361393 | 0.435681 | 0.143 | 0.082 | 1 | 15 | Mindy1 | 0.061 | Young |
| 0.362459 | 0.203093 | 0.625 | 0.592 | 1 | 15 | Atp5j | 0.033 | Young |
| 0.363874 | 0.595202 | 0.357 | 0.286 | 1 | 15 | Slfn5 | 0.071 | Young |
| 0.363991 | 1.088841 | 0.161 | 0.102 | 1 | 15 | Itpr2 | 0.059 | Young |
| 0.363991 | 0.807709 | 0.161 | 0.102 | 1 | 15 | Slc39a7 | 0.059 | Young |
| 0.364609 | 0.210015 | 0.786 | 0.796 | 1 | 15 | Sec61b | -0.01 | Young |
| 0.36484 | 1.302784 | 0.107 | 0.061 | 1 | 15 | Ap5m1 | 0.046 | Young |
| 0.364873 | 0.245259 | 0.089 | 0.041 | 1 | 15 | Spns1 | 0.048 | Young |
| 0.364873 | 0.15131 | 0.089 | 0.041 | 1 | 15 | Lig3 | 0.048 | Young |
| 0.366764 | 0.526873 | 0.339 | 0.265 | 1 | 15 | Smc6 | 0.074 | Young |
| 0.367542 | 0.559159 | 0.143 | 0.082 | 1 | 15 | Casd1 | 0.061 | Young |
| 0.367542 | 0.552717 | 0.143 | 0.082 | 1 | 15 | 2310022A1 | 0.061 | Young |
| 0.36756 | 0.62717 | 0.232 | 0.163 | 1 | 15 | Trappc8 | 0.069 | Young |

| | | | | | | | | |
|---|---|---|---|---|---|---|---|---|
| 0.36756 | 0.18754 | 0.232 | 0.163 | 1 | 15 | Trim56 | 0.069 | Young |
| 0.367664 | 0.797556 | 0.25 | 0.184 | 1 | 15 | Egln2 | 0.066 | Young |
| 0.36782 | 3.208409 | 0.054 | 0.02 | 1 | 15 | Csf1 | 0.034 | Young |
| 0.36782 | 3.069971 | 0.054 | 0.02 | 1 | 15 | Tmem94 | 0.034 | Young |
| 0.36782 | 3.025694 | 0.054 | 0.02 | 1 | 15 | Selenoi | 0.034 | Young |
| 0.36782 | 2.995703 | 0.054 | 0.02 | 1 | 15 | Btbd3 | 0.034 | Young |
| 0.36782 | 2.917109 | 0.054 | 0.02 | 1 | 15 | Sec61a2 | 0.034 | Young |
| 0.36782 | 2.817053 | 0.054 | 0.02 | 1 | 15 | Rangrf | 0.034 | Young |
| 0.36782 | 2.778749 | 0.054 | 0.02 | 1 | 15 | Rab38 | 0.034 | Young |
| 0.36782 | 2.767557 | 0.054 | 0.02 | 1 | 15 | Ttc28 | 0.034 | Young |
| 0.36782 | 2.762812 | 0.054 | 0.02 | 1 | 15 | Mtfr1 | 0.034 | Young |
| 0.36782 | 2.696251 | 0.054 | 0.02 | 1 | 15 | Pdp1 | 0.034 | Young |
| 0.36782 | 2.689423 | 0.054 | 0.02 | 1 | 15 | Gm12216 | 0.034 | Young |
| 0.36782 | 2.659821 | 0.054 | 0.02 | 1 | 15 | Map4k3 | 0.034 | Young |
| 0.36782 | 2.645964 | 0.054 | 0.02 | 1 | 15 | Prim2 | 0.034 | Young |
| 0.36782 | 2.541201 | 0.054 | 0.02 | 1 | 15 | Fbxl8 | 0.034 | Young |
| 0.36782 | 2.526105 | 0.054 | 0.02 | 1 | 15 | Mterf3 | 0.034 | Young |
| 0.36782 | 2.525363 | 0.054 | 0.02 | 1 | 15 | Nat2 | 0.034 | Young |
| 0.36782 | 2.478868 | 0.054 | 0.02 | 1 | 15 | Elmod2 | 0.034 | Young |
| 0.36782 | 2.441851 | 0.054 | 0.02 | 1 | 15 | Etv5 | 0.034 | Young |
| 0.36782 | 2.356902 | 0.054 | 0.02 | 1 | 15 | Chpf | 0.034 | Young |
| 0.36782 | 2.353601 | 0.054 | 0.02 | 1 | 15 | Dcun1d2 | 0.034 | Young |
| 0.36782 | 2.343401 | 0.054 | 0.02 | 1 | 15 | Zscan20 | 0.034 | Young |
| 0.36782 | 2.322782 | 0.054 | 0.02 | 1 | 15 | Card9 | 0.034 | Young |
| 0.36782 | 2.318181 | 0.054 | 0.02 | 1 | 15 | Fam210a | 0.034 | Young |
| 0.36782 | 2.31465 | 0.054 | 0.02 | 1 | 15 | Mamdc2 | 0.034 | Young |
| 0.36782 | 2.303745 | 0.054 | 0.02 | 1 | 15 | Clpp | 0.034 | Young |
| 0.36782 | 2.257537 | 0.054 | 0.02 | 1 | 15 | Wdr45 | 0.034 | Young |
| 0.36782 | 2.23592 | 0.054 | 0.02 | 1 | 15 | Tusc2 | 0.034 | Young |
| 0.36782 | 2.187932 | 0.054 | 0.02 | 1 | 15 | Mbtps2 | 0.034 | Young |
| 0.36782 | 2.186526 | 0.054 | 0.02 | 1 | 15 | Rmi1 | 0.034 | Young |
| 0.36782 | 2.161705 | 0.054 | 0.02 | 1 | 15 | Polr3e | 0.034 | Young |
| 0.36782 | 2.108571 | 0.054 | 0.02 | 1 | 15 | Dedd2 | 0.034 | Young |
| 0.36782 | 2.097534 | 0.054 | 0.02 | 1 | 15 | AI597479 | 0.034 | Young |
| 0.36782 | 2.082761 | 0.054 | 0.02 | 1 | 15 | Iqcb1 | 0.034 | Young |
| 0.36782 | 2.073883 | 0.054 | 0.02 | 1 | 15 | Racgap1 | 0.034 | Young |
| 0.36782 | 2.046723 | 0.054 | 0.02 | 1 | 15 | Noxo1 | 0.034 | Young |
| 0.36782 | 2.027894 | 0.054 | 0.02 | 1 | 15 | Abhd8 | 0.034 | Young |
| 0.36782 | 2.01312 | 0.054 | 0.02 | 1 | 15 | Atp2a3 | 0.034 | Young |
| 0.36782 | 1.975936 | 0.054 | 0.02 | 1 | 15 | E130311K1 | 0.034 | Young |
| 0.36782 | 1.968755 | 0.054 | 0.02 | 1 | 15 | Acap3 | 0.034 | Young |
| 0.36782 | 1.940412 | 0.054 | 0.02 | 1 | 15 | Trp53bp1 | 0.034 | Young |
| 0.36782 | 1.937667 | 0.054 | 0.02 | 1 | 15 | Abhd13 | 0.034 | Young |
| 0.36782 | 1.937029 | 0.054 | 0.02 | 1 | 15 | Tstd2 | 0.034 | Young |
| 0.36782 | 1.936442 | 0.054 | 0.02 | 1 | 15 | Slc9a3r2 | 0.034 | Young |

| | | | | | | | | |
|---|---|---|---|---|---|---|---|---|
| 0.36782 | 1.805187 | 0.054 | 0.02 | 1 | 15 | Stk39 | 0.034 | Young |
| 0.36782 | 1.767812 | 0.054 | 0.02 | 1 | 15 | Scamp5 | 0.034 | Young |
| 0.36782 | 1.73736 | 0.054 | 0.02 | 1 | 15 | Tsga10 | 0.034 | Young |
| 0.36782 | 1.71905 | 0.054 | 0.02 | 1 | 15 | Cyb561d2 | 0.034 | Young |
| 0.36782 | 1.689975 | 0.054 | 0.02 | 1 | 15 | Prdm11 | 0.034 | Young |
| 0.36782 | 1.65715 | 0.054 | 0.02 | 1 | 15 | Serf1 | 0.034 | Young |
| 0.36782 | 1.6262 | 0.054 | 0.02 | 1 | 15 | Zfp51 | 0.034 | Young |
| 0.36782 | 1.574053 | 0.054 | 0.02 | 1 | 15 | Apba1 | 0.034 | Young |
| 0.36782 | 1.49587 | 0.054 | 0.02 | 1 | 15 | Mzt2 | 0.034 | Young |
| 0.36782 | 1.488656 | 0.054 | 0.02 | 1 | 15 | Rnf113a2 | 0.034 | Young |
| 0.36782 | 1.263404 | 0.054 | 0.02 | 1 | 15 | Fnip2 | 0.034 | Young |
| 0.36782 | 1.221491 | 0.054 | 0.02 | 1 | 15 | Ubr1 | 0.034 | Young |
| 0.368783 | 1.54202 | 0.179 | 0.122 | 1 | 15 | Tmem5 | 0.057 | Young |
| 0.368783 | 1.220448 | 0.179 | 0.122 | 1 | 15 | Rraga | 0.057 | Young |
| 0.368783 | 1.02752 | 0.179 | 0.122 | 1 | 15 | Myd88 | 0.057 | Young |
| 0.368783 | 1.024851 | 0.179 | 0.122 | 1 | 15 | Rab11fip5 | 0.057 | Young |
| 0.368783 | 0.996473 | 0.179 | 0.122 | 1 | 15 | Pitpnc1 | 0.057 | Young |
| 0.369018 | 0.56214 | 0.321 | 0.245 | 1 | 15 | Sec31a | 0.076 | Young |
| 0.369764 | 0.816471 | 0.161 | 0.102 | 1 | 15 | Odf2 | 0.059 | Young |
| 0.369764 | 0.688105 | 0.161 | 0.102 | 1 | 15 | Shprh | 0.059 | Young |
| 0.369764 | 0.604244 | 0.161 | 0.102 | 1 | 15 | Prkab2 | 0.059 | Young |
| 0.370952 | 0.463284 | 0.214 | 0.143 | 1 | 15 | Esyt1 | 0.071 | Young |
| 0.370952 | 0.277824 | 0.214 | 0.143 | 1 | 15 | Sf3a1 | 0.071 | Young |
| 0.371075 | 0.405393 | 0.268 | 0.204 | 1 | 15 | Il10rb | 0.064 | Young |
| 0.371883 | 1.948976 | 0.107 | 0.061 | 1 | 15 | Kbtbd3 | 0.046 | Young |
| 0.371883 | 1.744229 | 0.107 | 0.061 | 1 | 15 | Lnpk | 0.046 | Young |
| 0.371883 | 1.648307 | 0.107 | 0.061 | 1 | 15 | AI987944 | 0.046 | Young |
| 0.371883 | 1.596423 | 0.107 | 0.061 | 1 | 15 | Dohh | 0.046 | Young |
| 0.371883 | 1.45809 | 0.107 | 0.061 | 1 | 15 | Usp6nl | 0.046 | Young |
| 0.371883 | 1.381191 | 0.107 | 0.061 | 1 | 15 | Ak3 | 0.046 | Young |
| 0.371883 | 1.286548 | 0.107 | 0.061 | 1 | 15 | Plscr1 | 0.046 | Young |
| 0.371883 | 1.250615 | 0.107 | 0.061 | 1 | 15 | Serpinf1 | 0.046 | Young |
| 0.371883 | 1.028434 | 0.107 | 0.061 | 1 | 15 | Dnlz | 0.046 | Young |
| 0.372403 | 0.745146 | 0.25 | 0.184 | 1 | 15 | Plbd2 | 0.066 | Young |
| 0.372469 | 0.741377 | 0.232 | 0.163 | 1 | 15 | Auh | 0.069 | Young |
| 0.372469 | 0.544061 | 0.232 | 0.163 | 1 | 15 | Ndufv2 | 0.069 | Young |
| 0.373197 | 0.692128 | 0.286 | 0.224 | 1 | 15 | Cstf3 | 0.062 | Young |
| 0.373756 | 0.406161 | 0.143 | 0.082 | 1 | 15 | Dpy19l4 | 0.061 | Young |
| 0.374281 | 0.667649 | 0.179 | 0.122 | 1 | 15 | Fkbp15 | 0.057 | Young |
| 0.37527 | 0.123087 | 0.054 | 0.102 | 1 | 15 | Ppp1r37 | -0.048 | Young |
| 0.375394 | 0.157707 | 0.982 | 0.939 | 1 | 15 | Rpl21 | 0.043 | Young |
| 0.375594 | 0.918749 | 0.161 | 0.102 | 1 | 15 | Timm10b | 0.059 | Young |
| 0.375594 | 0.836599 | 0.161 | 0.102 | 1 | 15 | Rreb1 | 0.059 | Young |
| 0.375594 | 0.596204 | 0.161 | 0.102 | 1 | 15 | Nsd2 | 0.059 | Young |
| 0.375693 | 0.648329 | 0.268 | 0.204 | 1 | 15 | Lrp1 | 0.064 | Young |

| | | | | | | | | |
|---|---|---|---|---|---|---|---|---|
| 0.376087 | 0.018563 | 0.714 | 0.551 | 1 | 15 | Ankrd44 | 0.163 | Young |
| 0.376541 | 0.392225 | 0.393 | 0.306 | 1 | 15 | Tmem256 | 0.087 | Young |
| 0.376541 | 0.262659 | 0.393 | 0.306 | 1 | 15 | Dnajc3 | 0.087 | Young |
| 0.378152 | 0.184765 | 0.196 | 0.122 | 1 | 15 | Psmd3 | 0.074 | Young |
| 0.378211 | 3.044308 | 0.054 | 0.02 | 1 | 15 | Gpatch1 | 0.034 | Young |
| 0.378211 | 2.99656 | 0.054 | 0.02 | 1 | 15 | Nup205 | 0.034 | Young |
| 0.378211 | 2.787507 | 0.054 | 0.02 | 1 | 15 | P2ry1 | 0.034 | Young |
| 0.378211 | 2.385527 | 0.054 | 0.02 | 1 | 15 | Rpap1 | 0.034 | Young |
| 0.378211 | 2.213831 | 0.054 | 0.02 | 1 | 15 | Xlr4c | 0.034 | Young |
| 0.378211 | 2.188727 | 0.054 | 0.02 | 1 | 15 | Cdc16 | 0.034 | Young |
| 0.378211 | 2.161706 | 0.054 | 0.02 | 1 | 15 | Xlr4b | 0.034 | Young |
| 0.378211 | 2.088301 | 0.054 | 0.02 | 1 | 15 | Wdr59 | 0.034 | Young |
| 0.378211 | 2.056826 | 0.054 | 0.02 | 1 | 15 | Aar2 | 0.034 | Young |
| 0.378211 | 2.03409 | 0.054 | 0.02 | 1 | 15 | Pstk | 0.034 | Young |
| 0.378211 | 1.943926 | 0.054 | 0.02 | 1 | 15 | Plekhn1 | 0.034 | Young |
| 0.378211 | 1.899284 | 0.054 | 0.02 | 1 | 15 | Dennd2c | 0.034 | Young |
| 0.378211 | 1.859417 | 0.054 | 0.02 | 1 | 15 | Aldh6a1 | 0.034 | Young |
| 0.378211 | 1.753084 | 0.054 | 0.02 | 1 | 15 | Tomm40l | 0.034 | Young |
| 0.378211 | 1.6675 | 0.054 | 0.02 | 1 | 15 | Lrp8 | 0.034 | Young |
| 0.378211 | 1.663292 | 0.054 | 0.02 | 1 | 15 | A630072M | 0.034 | Young |
| 0.378211 | 1.659239 | 0.054 | 0.02 | 1 | 15 | Zfhx2 | 0.034 | Young |
| 0.378211 | 1.630442 | 0.054 | 0.02 | 1 | 15 | Mov10 | 0.034 | Young |
| 0.378211 | 1.590941 | 0.054 | 0.02 | 1 | 15 | Il16 | 0.034 | Young |
| 0.378211 | 1.575478 | 0.054 | 0.02 | 1 | 15 | Kcnn4 | 0.034 | Young |
| 0.378211 | 1.557205 | 0.054 | 0.02 | 1 | 15 | Oraov1 | 0.034 | Young |
| 0.378211 | 1.544539 | 0.054 | 0.02 | 1 | 15 | Ttc32 | 0.034 | Young |
| 0.378211 | 1.512341 | 0.054 | 0.02 | 1 | 15 | Elac1 | 0.034 | Young |
| 0.378211 | 1.498916 | 0.054 | 0.02 | 1 | 15 | Gm17767 | 0.034 | Young |
| 0.378211 | 1.497761 | 0.054 | 0.02 | 1 | 15 | Ttc27 | 0.034 | Young |
| 0.378211 | 1.495099 | 0.054 | 0.02 | 1 | 15 | Lrrc28 | 0.034 | Young |
| 0.378211 | 1.465483 | 0.054 | 0.02 | 1 | 15 | Bud23 | 0.034 | Young |
| 0.378211 | 1.454224 | 0.054 | 0.02 | 1 | 15 | Trmt2b | 0.034 | Young |
| 0.378211 | 1.357465 | 0.054 | 0.02 | 1 | 15 | Cmc4 | 0.034 | Young |
| 0.378211 | 1.353515 | 0.054 | 0.02 | 1 | 15 | Fkbp9 | 0.034 | Young |
| 0.378211 | 1.309996 | 0.054 | 0.02 | 1 | 15 | Mrgbp | 0.034 | Young |
| 0.378211 | 1.272825 | 0.054 | 0.02 | 1 | 15 | Hacd3 | 0.034 | Young |
| 0.378211 | 1.263475 | 0.054 | 0.02 | 1 | 15 | Zbtb26 | 0.034 | Young |
| 0.378211 | 1.259367 | 0.054 | 0.02 | 1 | 15 | Gstp1 | 0.034 | Young |
| 0.378211 | 1.2295 | 0.054 | 0.02 | 1 | 15 | Zfp953 | 0.034 | Young |
| 0.378211 | 1.219472 | 0.054 | 0.02 | 1 | 15 | Neurl1b | 0.034 | Young |
| 0.378211 | 1.199633 | 0.054 | 0.02 | 1 | 15 | Inpp5e | 0.034 | Young |
| 0.378211 | 1.190859 | 0.054 | 0.02 | 1 | 15 | C130026I2 | 0.034 | Young |
| 0.378211 | 1.119684 | 0.054 | 0.02 | 1 | 15 | R3hcc1l | 0.034 | Young |
| 0.37901 | 1.843255 | 0.107 | 0.061 | 1 | 15 | Cog7 | 0.046 | Young |
| 0.37901 | 1.78831 | 0.107 | 0.061 | 1 | 15 | Socs7 | 0.046 | Young |

| | | | | | | | | |
|---|---|---|---|---|---|---|---|---|
| 0.37901 | 1.60157 | 0.107 | 0.061 | 1 | 15 | Usp50 | 0.046 | Young |
| 0.37901 | 1.595132 | 0.107 | 0.061 | 1 | 15 | Terf1 | 0.046 | Young |
| 0.37901 | 1.554469 | 0.107 | 0.061 | 1 | 15 | Coa6 | 0.046 | Young |
| 0.37901 | 1.473863 | 0.107 | 0.061 | 1 | 15 | Srxn1 | 0.046 | Young |
| 0.37901 | 1.355632 | 0.107 | 0.061 | 1 | 15 | Aph1c | 0.046 | Young |
| 0.37901 | 1.346063 | 0.107 | 0.061 | 1 | 15 | Alg2 | 0.046 | Young |
| 0.37901 | 1.338343 | 0.107 | 0.061 | 1 | 15 | Mesd | 0.046 | Young |
| 0.379828 | 0.92644 | 0.179 | 0.122 | 1 | 15 | Capn7 | 0.057 | Young |
| 0.380034 | 0.346689 | 0.143 | 0.082 | 1 | 15 | Fbxw4 | 0.061 | Young |
| 0.380034 | 0.094269 | 0.143 | 0.082 | 1 | 15 | Pes1 | 0.061 | Young |
| 0.380333 | 0.426308 | 0.518 | 0.469 | 1 | 15 | Sh3bgrl | 0.049 | Young |
| 0.381272 | 0.380017 | 0.214 | 0.143 | 1 | 15 | Nploc4 | 0.071 | Young |
| 0.381272 | 0.294766 | 0.214 | 0.143 | 1 | 15 | Serinc5 | 0.071 | Young |
| 0.381322 | 1.153189 | 0.196 | 0.143 | 1 | 15 | Vps13d | 0.053 | Young |
| 0.38148 | 0.593432 | 0.161 | 0.102 | 1 | 15 | Trp53 | 0.059 | Young |
| 0.38148 | 0.325721 | 0.161 | 0.102 | 1 | 15 | Asb8 | 0.059 | Young |
| 0.38148 | 0.249551 | 0.161 | 0.102 | 1 | 15 | Slc41a2 | 0.059 | Young |
| 0.38206 | 0.468102 | 0.429 | 0.367 | 1 | 15 | Mapk3 | 0.062 | Young |
| 0.383743 | 0.601348 | 0.339 | 0.265 | 1 | 15 | Banf1 | 0.074 | Young |
| 0.383781 | 0.389362 | 0.643 | 0.571 | 1 | 15 | Ctsb | 0.072 | Young |
| 0.383807 | 0.164874 | 0.768 | 0.653 | 1 | 15 | Ndufa2 | 0.115 | Young |
| 0.385033 | 0.789514 | 0.268 | 0.204 | 1 | 15 | Msl2 | 0.064 | Young |
| 0.385033 | 0.001225 | 0.268 | 0.204 | 1 | 15 | Vti1b | 0.064 | Young |
| 0.38511 | 0.417238 | 0.429 | 0.388 | 1 | 15 | Cnot1 | 0.041 | Young |
| 0.385427 | 1.17899 | 0.179 | 0.122 | 1 | 15 | Ahcyl1 | 0.057 | Young |
| 0.385427 | 0.381214 | 0.179 | 0.122 | 1 | 15 | Sfswap | 0.057 | Young |
| 0.38622 | 1.970366 | 0.107 | 0.061 | 1 | 15 | Crtap | 0.046 | Young |
| 0.38622 | 1.8475 | 0.107 | 0.061 | 1 | 15 | Gid4 | 0.046 | Young |
| 0.38622 | 1.479896 | 0.107 | 0.061 | 1 | 15 | Arfgap2 | 0.046 | Young |
| 0.38622 | 1.261455 | 0.107 | 0.061 | 1 | 15 | Lrrc8a | 0.046 | Young |
| 0.38622 | 1.242635 | 0.107 | 0.061 | 1 | 15 | Map1s | 0.046 | Young |
| 0.38622 | 1.2312 | 0.107 | 0.061 | 1 | 15 | Nmnat1 | 0.046 | Young |
| 0.38622 | 1.119372 | 0.107 | 0.061 | 1 | 15 | Mvk | 0.046 | Young |
| 0.38622 | 0.892368 | 0.107 | 0.061 | 1 | 15 | Glt8d1 | 0.046 | Young |
| 0.38622 | 0.829485 | 0.107 | 0.061 | 1 | 15 | D230025D: | 0.046 | Young |
| 0.38622 | 0.743947 | 0.107 | 0.061 | 1 | 15 | Zfp626 | 0.046 | Young |
| 0.38622 | 0.725349 | 0.107 | 0.061 | 1 | 15 | Abcd2 | 0.046 | Young |
| 0.38622 | 0.219413 | 0.107 | 0.061 | 1 | 15 | Stx3 | 0.046 | Young |
| 0.386498 | 0.410778 | 0.214 | 0.143 | 1 | 15 | Rexo2 | 0.071 | Young |
| 0.386827 | 0.20688 | 0.429 | 0.347 | 1 | 15 | Baz2b | 0.082 | Young |
| 0.386844 | 0.520913 | 0.25 | 0.184 | 1 | 15 | Dclre1c | 0.066 | Young |
| 0.386884 | 0.38269 | 0.464 | 0.388 | 1 | 15 | Lamp2 | 0.076 | Young |
| 0.387086 | 0.226721 | 0.446 | 0.367 | 1 | 15 | Nampt | 0.079 | Young |
| 0.387423 | 0.933377 | 0.161 | 0.102 | 1 | 15 | Dguok | 0.059 | Young |
| 0.387437 | 0.002482 | 0.161 | 0.245 | 1 | 15 | Dido1 | -0.084 | Young |

| | | | | | | | | |
|---|---|---|---|---|---|---|---|---|
| 0.387531 | 0.379848 | 0.304 | 0.245 | 1 | 15 | Pon2 | 0.059 | Young |
| 0.388425 | 0.407145 | 0.304 | 0.224 | 1 | 15 | Copb2 | 0.08 | Young |
| 0.388781 | 1.983887 | 0.054 | 0.02 | 1 | 15 | Cfap36 | 0.034 | Young |
| 0.388781 | 1.711225 | 0.054 | 0.02 | 1 | 15 | Med17 | 0.034 | Young |
| 0.388781 | 1.527689 | 0.054 | 0.02 | 1 | 15 | Ttc39a | 0.034 | Young |
| 0.388781 | 1.441332 | 0.054 | 0.02 | 1 | 15 | Rab39 | 0.034 | Young |
| 0.388781 | 1.407452 | 0.054 | 0.02 | 1 | 15 | Trpm4 | 0.034 | Young |
| 0.388781 | 1.344769 | 0.054 | 0.02 | 1 | 15 | Pus7 | 0.034 | Young |
| 0.388781 | 1.285046 | 0.054 | 0.02 | 1 | 15 | Pef1 | 0.034 | Young |
| 0.388781 | 1.265743 | 0.054 | 0.02 | 1 | 15 | Klri1 | 0.034 | Young |
| 0.388781 | 1.252324 | 0.054 | 0.02 | 1 | 15 | Adamtsl5 | 0.034 | Young |
| 0.388781 | 1.212109 | 0.054 | 0.02 | 1 | 15 | Flt3l | 0.034 | Young |
| 0.388781 | 1.20882 | 0.054 | 0.02 | 1 | 15 | Hddc2 | 0.034 | Young |
| 0.388781 | 1.206138 | 0.054 | 0.02 | 1 | 15 | Ppa1 | 0.034 | Young |
| 0.388781 | 1.13976 | 0.054 | 0.02 | 1 | 15 | Mrpl1 | 0.034 | Young |
| 0.388781 | 1.123013 | 0.054 | 0.02 | 1 | 15 | Nlrc4 | 0.034 | Young |
| 0.388781 | 1.073502 | 0.054 | 0.02 | 1 | 15 | Slc25a32 | 0.034 | Young |
| 0.388781 | 1.06513 | 0.054 | 0.02 | 1 | 15 | Cxxc5 | 0.034 | Young |
| 0.388781 | 1.064545 | 0.054 | 0.02 | 1 | 15 | Clpb | 0.034 | Young |
| 0.388781 | 0.990405 | 0.054 | 0.02 | 1 | 15 | 1110059E2 | 0.034 | Young |
| 0.388781 | 0.982008 | 0.054 | 0.02 | 1 | 15 | Ncapd2 | 0.034 | Young |
| 0.388781 | 0.972748 | 0.054 | 0.02 | 1 | 15 | Tmem175 | 0.034 | Young |
| 0.388781 | 0.966535 | 0.054 | 0.02 | 1 | 15 | Eef2k | 0.034 | Young |
| 0.388781 | 0.958764 | 0.054 | 0.02 | 1 | 15 | Gm12159 | 0.034 | Young |
| 0.388781 | 0.884267 | 0.054 | 0.02 | 1 | 15 | Kiz | 0.034 | Young |
| 0.388781 | 0.848556 | 0.054 | 0.02 | 1 | 15 | Plxna4os1 | 0.034 | Young |
| 0.388781 | 0.733358 | 0.054 | 0.02 | 1 | 15 | Mtmr2 | 0.034 | Young |
| 0.388781 | 0.686394 | 0.054 | 0.02 | 1 | 15 | Pag1 | 0.034 | Young |
| 0.388781 | 0.678789 | 0.054 | 0.02 | 1 | 15 | 5031425F1 | 0.034 | Young |
| 0.388781 | 0.661159 | 0.054 | 0.02 | 1 | 15 | Atad3a | 0.034 | Young |
| 0.388781 | 0.547564 | 0.054 | 0.02 | 1 | 15 | Slc27a4 | 0.034 | Young |
| 0.389674 | 1.300371 | 0.25 | 0.204 | 1 | 15 | Ppp1r10 | 0.046 | Young |
| 0.390172 | 0.060303 | 0.839 | 0.837 | 1 | 15 | Iqgap1 | 0.002 | Young |
| 0.390311 | 0.40872 | 0.589 | 0.531 | 1 | 15 | Syk | 0.058 | Young |
| 0.390495 | 0.284489 | 0.054 | 0.102 | 1 | 15 | Prdm4 | -0.048 | Young |
| 0.390716 | 0.899843 | 0.339 | 0.286 | 1 | 15 | Itgax | 0.053 | Young |
| 0.391054 | 1.224145 | 0.214 | 0.163 | 1 | 15 | Smim11 | 0.051 | Young |
| 0.391075 | 0.010297 | 0.179 | 0.122 | 1 | 15 | Lsm7 | 0.057 | Young |
| 0.391414 | 0.680469 | 0.286 | 0.224 | 1 | 15 | Epb41l4ao: | 0.062 | Young |
| 0.391748 | 0.542043 | 0.321 | 0.265 | 1 | 15 | Slc25a4 | 0.056 | Young |
| 0.392054 | 1.369827 | 0.196 | 0.143 | 1 | 15 | Dap3 | 0.053 | Young |
| 0.392409 | 0.100862 | 0.339 | 0.265 | 1 | 15 | Polr2e | 0.074 | Young |
| 0.392507 | 0.613122 | 0.232 | 0.163 | 1 | 15 | Ptpn12 | 0.069 | Young |
| 0.392656 | 0.038363 | 0.411 | 0.306 | 1 | 15 | Supt4a | 0.105 | Young |
| 0.393421 | 1.877601 | 0.161 | 0.102 | 1 | 15 | Sesn1 | 0.059 | Young |

| | | | | | | | | |
|---|---|---|---|---|---|---|---|---|
| 0.393421 | 1.00433 | 0.161 | 0.102 | 1 | 15 | Tbc1d10a | 0.059 | Young |
| 0.393421 | 0.662802 | 0.161 | 0.102 | 1 | 15 | Epb41 | 0.059 | Young |
| 0.393459 | 0.273733 | 0.482 | 0.388 | 1 | 15 | Dnajc5 | 0.094 | Young |
| 0.393513 | 1.766209 | 0.107 | 0.061 | 1 | 15 | Zfp629 | 0.046 | Young |
| 0.393513 | 1.442232 | 0.107 | 0.061 | 1 | 15 | Pfdn6 | 0.046 | Young |
| 0.393513 | 1.304749 | 0.107 | 0.061 | 1 | 15 | Mob3c | 0.046 | Young |
| 0.393513 | 1.13296 | 0.107 | 0.061 | 1 | 15 | Akr1b10 | 0.046 | Young |
| 0.393513 | 0.586542 | 0.107 | 0.061 | 1 | 15 | Creb3 | 0.046 | Young |
| 0.394515 | 0.551504 | 0.268 | 0.204 | 1 | 15 | Slmap | 0.064 | Young |
| 0.394735 | 0.032412 | 0.982 | 1 | 1 | 15 | Rps3a1 | -0.018 | Young |
| 0.395443 | 0.009315 | 0.661 | 0.531 | 1 | 15 | Trps1 | 0.13 | Young |
| 0.396052 | 1.05808 | 0.286 | 0.224 | 1 | 15 | Blmh | 0.062 | Young |
| 0.396549 | 0.506719 | 0.304 | 0.245 | 1 | 15 | Birc6 | 0.059 | Young |
| 0.396773 | 0.964065 | 0.179 | 0.122 | 1 | 15 | Pts | 0.057 | Young |
| 0.396773 | 0.712029 | 0.179 | 0.122 | 1 | 15 | Yif1b | 0.057 | Young |
| 0.39708 | 0.287689 | 0.214 | 0.143 | 1 | 15 | Timm17b | 0.071 | Young |
| 0.397397 | 0.307268 | 0.464 | 0.408 | 1 | 15 | Ifitm3 | 0.056 | Young |
| 0.397449 | 0.295676 | 0.357 | 0.286 | 1 | 15 | Fkbp8 | 0.071 | Young |
| 0.397487 | 0.717599 | 0.196 | 0.143 | 1 | 15 | Ssbp4 | 0.053 | Young |
| 0.39757 | 0.268935 | 0.304 | 0.224 | 1 | 15 | Kcmf1 | 0.08 | Young |
| 0.398246 | 0.192198 | 0.054 | 0.102 | 1 | 15 | Decr2 | -0.048 | Young |
| 0.398847 | 0.147796 | 0.589 | 0.469 | 1 | 15 | Irf2bp2 | 0.12 | Young |
| 0.399476 | 0.654446 | 0.161 | 0.102 | 1 | 15 | Sos2 | 0.059 | Young |
| 0.399476 | 0.35404 | 0.161 | 0.102 | 1 | 15 | Ten1 | 0.059 | Young |
| 0.399476 | 0.165592 | 0.161 | 0.102 | 1 | 15 | Lcor | 0.059 | Young |
| 0.399529 | 0.889454 | 0.054 | 0.02 | 1 | 15 | Dcp1b | 0.034 | Young |
| 0.399529 | 0.83967 | 0.054 | 0.02 | 1 | 15 | Asf1a | 0.034 | Young |
| 0.399529 | 0.811809 | 0.054 | 0.02 | 1 | 15 | Incenp | 0.034 | Young |
| 0.399529 | 0.811368 | 0.054 | 0.02 | 1 | 15 | Ppp2r5d | 0.034 | Young |
| 0.399529 | 0.739076 | 0.054 | 0.02 | 1 | 15 | Ccdc114 | 0.034 | Young |
| 0.399529 | 0.738381 | 0.054 | 0.02 | 1 | 15 | Tmem35b | 0.034 | Young |
| 0.399529 | 0.737232 | 0.054 | 0.02 | 1 | 15 | Stk11ip | 0.034 | Young |
| 0.399529 | 0.704926 | 0.054 | 0.02 | 1 | 15 | Cnksr3 | 0.034 | Young |
| 0.399529 | 0.674089 | 0.054 | 0.02 | 1 | 15 | Gpank1 | 0.034 | Young |
| 0.399529 | 0.592069 | 0.054 | 0.02 | 1 | 15 | Uhrf1 | 0.034 | Young |
| 0.399529 | 0.521591 | 0.054 | 0.02 | 1 | 15 | 4930518I1 | 0.034 | Young |
| 0.399529 | 0.51733 | 0.054 | 0.02 | 1 | 15 | Atxn7l2 | 0.034 | Young |
| 0.399529 | 0.495878 | 0.054 | 0.02 | 1 | 15 | Irf7 | 0.034 | Young |
| 0.399529 | 0.437071 | 0.054 | 0.02 | 1 | 15 | Mctp1 | 0.034 | Young |
| 0.399529 | 0.421811 | 0.054 | 0.02 | 1 | 15 | Rdh11 | 0.034 | Young |
| 0.399529 | 0.382198 | 0.054 | 0.02 | 1 | 15 | Smug1 | 0.034 | Young |
| 0.399529 | 0.341095 | 0.054 | 0.02 | 1 | 15 | Rnf214 | 0.034 | Young |
| 0.399529 | 0.269136 | 0.054 | 0.02 | 1 | 15 | Qtrt1 | 0.034 | Young |
| 0.399529 | 0.179916 | 0.054 | 0.02 | 1 | 15 | Plod1 | 0.034 | Young |
| 0.399529 | 0.063868 | 0.054 | 0.02 | 1 | 15 | Pld1 | 0.034 | Young |

| | | | | | | | | |
|---|---|---|---|---|---|---|---|---|
| 0.399529 | 0.00969 | 0.054 | 0.02 | 1 | 15 | Vezt | 0.034 | Young |
| 0.400722 | 0.634914 | 0.286 | 0.224 | 1 | 15 | Gas6 | 0.062 | Young |
| 0.400888 | 1.419499 | 0.107 | 0.061 | 1 | 15 | Ppp5c | 0.046 | Young |
| 0.400888 | 1.228911 | 0.107 | 0.061 | 1 | 15 | Rexo4 | 0.046 | Young |
| 0.400888 | 0.975867 | 0.107 | 0.061 | 1 | 15 | Tmem251 | 0.046 | Young |
| 0.400888 | 0.965016 | 0.107 | 0.061 | 1 | 15 | Fmc1 | 0.046 | Young |
| 0.400888 | 0.787845 | 0.107 | 0.061 | 1 | 15 | Styx | 0.046 | Young |
| 0.400888 | 0.696873 | 0.107 | 0.061 | 1 | 15 | Tesk1 | 0.046 | Young |
| 0.400888 | 0.610805 | 0.107 | 0.061 | 1 | 15 | Wdr13 | 0.046 | Young |
| 0.400888 | 0.471152 | 0.107 | 0.061 | 1 | 15 | Fkbp4 | 0.046 | Young |
| 0.401194 | 0.355501 | 0.339 | 0.265 | 1 | 15 | Gnaq | 0.074 | Young |
| 0.401194 | 0.17557 | 0.339 | 0.265 | 1 | 15 | Rsu1 | 0.074 | Young |
| 0.40219 | 0.411139 | 0.304 | 0.224 | 1 | 15 | Tmem259 | 0.08 | Young |
| 0.402522 | 0.441835 | 0.179 | 0.122 | 1 | 15 | Pop5 | 0.057 | Young |
| 0.402766 | 0.042556 | 0.161 | 0.245 | 1 | 15 | Ssh2 | -0.084 | Young |
| 0.403955 | 0.227492 | 0.375 | 0.286 | 1 | 15 | Tmod3 | 0.089 | Young |
| 0.40439 | 1.582172 | 0.125 | 0.082 | 1 | 15 | Kbtbd11 | 0.043 | Young |
| 0.405112 | 0.149154 | 0.429 | 0.347 | 1 | 15 | Trir | 0.082 | Young |
| 0.405426 | 0.706863 | 0.286 | 0.224 | 1 | 15 | Nars | 0.062 | Young |
| 0.405501 | 0.040064 | 0.982 | 0.918 | 1 | 15 | Clic4 | 0.064 | Young |
| 0.405586 | 0.936 | 0.161 | 0.102 | 1 | 15 | Med21 | 0.059 | Young |
| 0.405586 | 0.221729 | 0.161 | 0.102 | 1 | 15 | Krt80 | 0.059 | Young |
| 0.40563 | 0.267506 | 0.339 | 0.265 | 1 | 15 | Hspa4 | 0.074 | Young |
| 0.40609 | 0.514656 | 0.054 | 0.102 | 1 | 15 | Eif2ak1 | -0.048 | Young |
| 0.40609 | 0.20026 | 0.054 | 0.102 | 1 | 15 | Trap1 | -0.048 | Young |
| 0.40609 | 0.137758 | 0.054 | 0.102 | 1 | 15 | Trp53inp1 | -0.048 | Young |
| 0.40609 | 0.058311 | 0.054 | 0.102 | 1 | 15 | Lsm10 | -0.048 | Young |
| 0.406126 | 0.53277 | 0.357 | 0.286 | 1 | 15 | Fcgr2b | 0.071 | Young |
| 0.406903 | 0.299776 | 0.554 | 0.469 | 1 | 15 | Necap2 | 0.085 | Young |
| 0.407177 | 0.238838 | 0.429 | 0.347 | 1 | 15 | Eif3m | 0.082 | Young |
| 0.407651 | 0.362994 | 0.571 | 0.531 | 1 | 15 | Ahcyl2 | 0.04 | Young |
| 0.408319 | 0.765817 | 0.179 | 0.122 | 1 | 15 | Tgs1 | 0.057 | Young |
| 0.408319 | 0.723478 | 0.179 | 0.122 | 1 | 15 | Seh1l | 0.057 | Young |
| 0.408347 | 1.209594 | 0.107 | 0.061 | 1 | 15 | Suox | 0.046 | Young |
| 0.408347 | 0.833519 | 0.107 | 0.061 | 1 | 15 | Rhno1 | 0.046 | Young |
| 0.408347 | 0.706109 | 0.107 | 0.061 | 1 | 15 | Rab29 | 0.046 | Young |
| 0.408347 | 0.398001 | 0.107 | 0.061 | 1 | 15 | Ust | 0.046 | Young |
| 0.408347 | 0.257875 | 0.107 | 0.061 | 1 | 15 | Igsf6 | 0.046 | Young |
| 0.40849 | 0.912313 | 0.196 | 0.143 | 1 | 15 | Cebpz | 0.053 | Young |
| 0.408728 | 0.999564 | 0.232 | 0.184 | 1 | 15 | C3ar1 | 0.048 | Young |
| 0.408764 | 0.063931 | 0.071 | 0.122 | 1 | 15 | Tmem203 | -0.051 | Young |
| 0.408951 | 0.28825 | 0.536 | 0.469 | 1 | 15 | Arf1 | 0.067 | Young |
| 0.409401 | 0.372034 | 0.393 | 0.327 | 1 | 15 | Eif4g1 | 0.066 | Young |
| 0.409401 | 0.348954 | 0.393 | 0.327 | 1 | 15 | Med13 | 0.066 | Young |
| 0.410544 | 0.030498 | 0.911 | 0.837 | 1 | 15 | Tmem176a | 0.074 | Young |

| | | | | | | | | |
|---|---|---|---|---|---|---|---|---|
| 0.410941 | 0.40131 | 0.518 | 0.469 | 1 | 15 | Trim35 | 0.049 | Young |
| 0.411235 | 1.835959 | 0.125 | 0.082 | 1 | 15 | Fdft1 | 0.043 | Young |
| 0.411235 | 1.193347 | 0.125 | 0.082 | 1 | 15 | Trip10 | 0.043 | Young |
| 0.411659 | 0.535554 | 0.25 | 0.184 | 1 | 15 | Fcgr3 | 0.066 | Young |
| 0.413184 | 0.321807 | 0.232 | 0.163 | 1 | 15 | Ybx3 | 0.069 | Young |
| 0.413184 | 0.011738 | 0.232 | 0.163 | 1 | 15 | Rgs12 | 0.069 | Young |
| 0.414059 | 1.025412 | 0.196 | 0.143 | 1 | 15 | Cep63 | 0.053 | Young |
| 0.414167 | 1.434407 | 0.179 | 0.122 | 1 | 15 | Hk2 | 0.057 | Young |
| 0.414167 | 0.565748 | 0.179 | 0.122 | 1 | 15 | Fam241a | 0.057 | Young |
| 0.414457 | 0.340654 | 0.429 | 0.367 | 1 | 15 | Pdia6 | 0.062 | Young |
| 0.415887 | 0.79883 | 0.107 | 0.061 | 1 | 15 | Phf23 | 0.046 | Young |
| 0.415887 | 0.719385 | 0.107 | 0.061 | 1 | 15 | Lats1 | 0.046 | Young |
| 0.415887 | 0.666627 | 0.107 | 0.061 | 1 | 15 | Tspan17 | 0.046 | Young |
| 0.415887 | 0.594415 | 0.107 | 0.061 | 1 | 15 | Loxl3 | 0.046 | Young |
| 0.415887 | 0.473557 | 0.107 | 0.061 | 1 | 15 | Abcf3 | 0.046 | Young |
| 0.415887 | 0.390604 | 0.107 | 0.061 | 1 | 15 | Dpf3 | 0.046 | Young |
| 0.415955 | 0.181383 | 0.071 | 0.122 | 1 | 15 | Sh3pxd2a | -0.051 | Young |
| 0.415955 | 0.029585 | 0.071 | 0.122 | 1 | 15 | Mrs2 | -0.051 | Young |
| 0.415955 | 0.022159 | 0.071 | 0.122 | 1 | 15 | Nek6 | -0.051 | Young |
| 0.417395 | 0.432638 | 0.821 | 0.714 | 1 | 15 | Dstn | 0.107 | Young |
| 0.417459 | 1.093618 | 0.214 | 0.163 | 1 | 15 | Tomm34 | 0.051 | Young |
| 0.417459 | 0.786298 | 0.214 | 0.163 | 1 | 15 | Ftl1-ps1 | 0.051 | Young |
| 0.418334 | 0.055281 | 0.982 | 0.98 | 1 | 15 | Cd83 | 0.002 | Young |
| 0.418545 | 0.029454 | 0.518 | 0.408 | 1 | 15 | Elob | 0.11 | Young |
| 0.418761 | 0.273252 | 0.214 | 0.143 | 1 | 15 | Gmps | 0.071 | Young |
| 0.418829 | 0.076571 | 0.268 | 0.204 | 1 | 15 | Snx2 | 0.064 | Young |
| 0.418829 | 0.01638 | 0.268 | 0.204 | 1 | 15 | Csk | 0.064 | Young |
| 0.418913 | 0.483304 | 0.464 | 0.388 | 1 | 15 | Egr1 | 0.076 | Young |
| 0.419113 | 0.160993 | 0.339 | 0.265 | 1 | 15 | Park7 | 0.074 | Young |
| 0.421961 | 0.298293 | 0.571 | 0.49 | 1 | 15 | Slamf7 | 0.081 | Young |
| 0.422162 | 1.303934 | 0.143 | 0.102 | 1 | 15 | Snhg6 | 0.041 | Young |
| 0.422278 | 0.231607 | 0.554 | 0.49 | 1 | 15 | Cmpk1 | 0.064 | Young |
| 0.422664 | 0.345572 | 0.696 | 0.633 | 1 | 15 | Ifrd1 | 0.063 | Young |
| 0.422863 | 0.77155 | 0.214 | 0.163 | 1 | 15 | Psen1 | 0.051 | Young |
| 0.423316 | 0.439964 | 0.321 | 0.265 | 1 | 15 | Rnf4 | 0.056 | Young |
| 0.423509 | 0.721696 | 0.107 | 0.061 | 1 | 15 | Pigv | 0.046 | Young |
| 0.423509 | 0.630975 | 0.107 | 0.061 | 1 | 15 | Slc39a6 | 0.046 | Young |
| 0.423509 | 0.528838 | 0.107 | 0.061 | 1 | 15 | Fam207a | 0.046 | Young |
| 0.423509 | 0.461625 | 0.107 | 0.061 | 1 | 15 | Slc10a7 | 0.046 | Young |
| 0.423509 | 0.280815 | 0.107 | 0.061 | 1 | 15 | Cyhr1 | 0.046 | Young |
| 0.423509 | 0.25306 | 0.107 | 0.061 | 1 | 15 | Sec23a | 0.046 | Young |
| 0.423795 | 0.348786 | 0.268 | 0.204 | 1 | 15 | Churc1 | 0.064 | Young |
| 0.423795 | 0.257437 | 0.268 | 0.204 | 1 | 15 | Larp4 | 0.064 | Young |
| 0.424566 | 0.597914 | 0.286 | 0.224 | 1 | 15 | Clasp1 | 0.062 | Young |
| 0.425021 | 0.858073 | 0.375 | 0.327 | 1 | 15 | Ube3a | 0.048 | Young |

| | | | | | | | | |
|---|---|---|---|---|---|---|---|---|
| 0.425129 | 1.382934 | 0.125 | 0.082 | 1 | 15 | Micu2 | 0.043 | Young |
| 0.425129 | 1.3114 | 0.125 | 0.082 | 1 | 15 | Ankrd16 | 0.043 | Young |
| 0.425129 | 1.258336 | 0.125 | 0.082 | 1 | 15 | Fam76b | 0.043 | Young |
| 0.425719 | 0.074813 | 0.982 | 0.898 | 1 | 15 | Hspa8 | 0.084 | Young |
| 0.425771 | 0.736137 | 0.286 | 0.245 | 1 | 15 | Mcub | 0.041 | Young |
| 0.425771 | 0.449511 | 0.304 | 0.224 | 1 | 15 | Rasa4 | 0.08 | Young |
| 0.425859 | 0.4393 | 0.643 | 0.592 | 1 | 15 | Pde4b | 0.051 | Young |
| 0.426009 | 0.631066 | 0.179 | 0.122 | 1 | 15 | AW549877 | 0.057 | Young |
| 0.426009 | 0.48312 | 0.179 | 0.122 | 1 | 15 | Ugdh | 0.057 | Young |
| 0.426092 | 0.701781 | 0.339 | 0.286 | 1 | 15 | Gmfg | 0.053 | Young |
| 0.426991 | 0.676732 | 0.25 | 0.184 | 1 | 15 | Cd48 | 0.066 | Young |
| 0.42772 | 0.633396 | 0.375 | 0.306 | 1 | 15 | Clu | 0.069 | Young |
| 0.428644 | 0.316179 | 0.5 | 0.429 | 1 | 15 | Hook3 | 0.071 | Young |
| 0.428796 | 0.56525 | 0.268 | 0.204 | 1 | 15 | Ap1s2 | 0.064 | Young |
| 0.429108 | 0.608819 | 0.232 | 0.163 | 1 | 15 | Wls | 0.069 | Young |
| 0.429327 | 0.708923 | 0.25 | 0.204 | 1 | 15 | Tsen34 | 0.046 | Young |
| 0.429517 | 0.853364 | 0.232 | 0.184 | 1 | 15 | Mink1 | 0.048 | Young |
| 0.430559 | 0.266729 | 0.071 | 0.122 | 1 | 15 | Fbxo34 | -0.051 | Young |
| 0.430559 | 0.187239 | 0.071 | 0.122 | 1 | 15 | Smarcd1 | -0.051 | Young |
| 0.430582 | 0.023461 | 0.304 | 0.224 | 1 | 15 | Mpp6 | 0.08 | Young |
| 0.431032 | 0.637225 | 0.196 | 0.143 | 1 | 15 | Magoh | 0.053 | Young |
| 0.431212 | 0.596169 | 0.107 | 0.061 | 1 | 15 | Smc5 | 0.046 | Young |
| 0.431212 | 0.209168 | 0.107 | 0.061 | 1 | 15 | Kifap3 | 0.046 | Young |
| 0.431212 | 0.027354 | 0.107 | 0.061 | 1 | 15 | Slc17a5 | 0.046 | Young |
| 0.432003 | 0.429018 | 0.179 | 0.122 | 1 | 15 | Nelfe | 0.057 | Young |
| 0.432174 | 0.463838 | 0.25 | 0.184 | 1 | 15 | Ece1 | 0.066 | Young |
| 0.432178 | 1.412715 | 0.125 | 0.082 | 1 | 15 | Psd3 | 0.043 | Young |
| 0.432178 | 1.10073 | 0.125 | 0.082 | 1 | 15 | 2310009B1 | 0.043 | Young |
| 0.432178 | 0.853562 | 0.125 | 0.082 | 1 | 15 | Gtf3a | 0.043 | Young |
| 0.432542 | 0.301328 | 0.643 | 0.571 | 1 | 15 | Csf2rb | 0.072 | Young |
| 0.433794 | 0.756303 | 0.214 | 0.163 | 1 | 15 | Xpo1 | 0.051 | Young |
| 0.433831 | 0.061981 | 0.268 | 0.204 | 1 | 15 | Lbh | 0.064 | Young |
| 0.434331 | 0.802853 | 0.286 | 0.224 | 1 | 15 | Med30 | 0.062 | Young |
| 0.434444 | 0.705017 | 0.25 | 0.204 | 1 | 15 | Elf2 | 0.046 | Young |
| 0.434454 | 0.372464 | 0.393 | 0.347 | 1 | 15 | Chmp5 | 0.046 | Young |
| 0.434494 | 0.296906 | 0.232 | 0.163 | 1 | 15 | Glmp | 0.069 | Young |
| 0.434494 | 0.162672 | 0.161 | 0.245 | 1 | 15 | Taf1 | -0.084 | Young |
| 0.435013 | 0.153534 | 0.768 | 0.714 | 1 | 15 | Atp5a1 | 0.054 | Young |
| 0.43522 | 0.281832 | 0.339 | 0.286 | 1 | 15 | Cdk11b | 0.053 | Young |
| 0.4353 | 0.16935 | 0.393 | 0.327 | 1 | 15 | Gpr183 | 0.066 | Young |
| 0.436778 | 1.182933 | 0.196 | 0.143 | 1 | 15 | Btk | 0.053 | Young |
| 0.436778 | 0.586935 | 0.196 | 0.143 | 1 | 15 | Cwf19l2 | 0.053 | Young |
| 0.437972 | 0.354156 | 0.071 | 0.122 | 1 | 15 | Mb21d1 | -0.051 | Young |
| 0.438046 | 0.654889 | 0.179 | 0.122 | 1 | 15 | Mtmr6 | 0.057 | Young |
| 0.438995 | 0.551268 | 0.107 | 0.061 | 1 | 15 | Cd93 | 0.046 | Young |

| | | | | | | | | |
|---|---|---|---|---|---|---|---|---|
| 0.439294 | 1.484133 | 0.125 | 0.082 | 1 | 15 | Dram1 | 0.043 | Young |
| 0.439294 | 1.32827 | 0.125 | 0.082 | 1 | 15 | Iws1 | 0.043 | Young |
| 0.439294 | 1.004373 | 0.125 | 0.082 | 1 | 15 | Mrpl27 | 0.043 | Young |
| 0.439294 | 0.913828 | 0.125 | 0.082 | 1 | 15 | Dhx57 | 0.043 | Young |
| 0.439409 | 0.482642 | 0.089 | 0.143 | 1 | 15 | Kctd12b | -0.054 | Young |
| 0.439409 | 0.183364 | 0.089 | 0.143 | 1 | 15 | Lrrc42 | -0.054 | Young |
| 0.439918 | 0.267185 | 0.232 | 0.163 | 1 | 15 | Wbp2 | 0.069 | Young |
| 0.441103 | 0.230257 | 0.375 | 0.306 | 1 | 15 | Txnrd1 | 0.069 | Young |
| 0.441895 | 1.543792 | 0.143 | 0.102 | 1 | 15 | Erf | 0.041 | Young |
| 0.442967 | 0.252362 | 0.536 | 0.449 | 1 | 15 | Wdr26 | 0.087 | Young |
| 0.443018 | 0.621809 | 0.482 | 0.449 | 1 | 15 | Sbds | 0.033 | Young |
| 0.443067 | 0.154673 | 0.518 | 0.449 | 1 | 15 | Klf2 | 0.069 | Young |
| 0.443405 | 0.120621 | 0.161 | 0.102 | 1 | 15 | Pigp | 0.059 | Young |
| 0.444504 | 0.187964 | 0.75 | 0.653 | 1 | 15 | Ikbkb | 0.097 | Young |
| 0.444781 | 0.899312 | 0.25 | 0.204 | 1 | 15 | Ube2e3 | 0.046 | Young |
| 0.444781 | 0.752616 | 0.25 | 0.204 | 1 | 15 | Ap1g1 | 0.046 | Young |
| 0.445457 | 0.28942 | 0.071 | 0.122 | 1 | 15 | Alg10b | -0.051 | Young |
| 0.44632 | 0.329234 | 0.089 | 0.143 | 1 | 15 | Galk2 | -0.054 | Young |
| 0.44632 | 0.005103 | 0.089 | 0.143 | 1 | 15 | Dclre1a | -0.054 | Young |
| 0.446477 | 1.858032 | 0.125 | 0.082 | 1 | 15 | Hif1an | 0.043 | Young |
| 0.446477 | 1.622985 | 0.125 | 0.082 | 1 | 15 | Fam214b | 0.043 | Young |
| 0.446477 | 1.30109 | 0.125 | 0.082 | 1 | 15 | Acvrl1 | 0.043 | Young |
| 0.446477 | 1.160594 | 0.125 | 0.082 | 1 | 15 | Abca3 | 0.043 | Young |
| 0.446477 | 0.871656 | 0.125 | 0.082 | 1 | 15 | Mphosph6 | 0.043 | Young |
| 0.446477 | 0.77528 | 0.125 | 0.082 | 1 | 15 | Pld2 | 0.043 | Young |
| 0.446859 | 0.515474 | 0.107 | 0.061 | 1 | 15 | Slc9a9 | 0.046 | Young |
| 0.446859 | 0.250067 | 0.107 | 0.061 | 1 | 15 | Fem1a | 0.046 | Young |
| 0.447188 | 0.552251 | 0.375 | 0.327 | 1 | 15 | Nufip2 | 0.048 | Young |
| 0.448402 | 0.631907 | 0.196 | 0.143 | 1 | 15 | Mrps12 | 0.053 | Young |
| 0.448588 | 0.541218 | 0.143 | 0.102 | 1 | 15 | Lrrc25 | 0.041 | Young |
| 0.44912 | 0.375713 | 0.339 | 0.286 | 1 | 15 | Hexim1 | 0.053 | Young |
| 0.449262 | 0.331996 | 0.5 | 0.429 | 1 | 15 | Eif3i | 0.071 | Young |
| 0.449459 | 0.223321 | 0.821 | 0.857 | 1 | 15 | Rps18 | -0.036 | Young |
| 0.449899 | 0.194277 | 0.161 | 0.102 | 1 | 15 | Ksr1 | 0.059 | Young |
| 0.4509 | 0.168852 | 0.768 | 0.653 | 1 | 15 | Btg2 | 0.115 | Young |
| 0.451327 | 0.198769 | 0.518 | 0.449 | 1 | 15 | Bola2 | 0.069 | Young |
| 0.45158 | 0.411698 | 0.339 | 0.265 | 1 | 15 | Coro1c | 0.074 | Young |
| 0.45158 | 0.344644 | 0.339 | 0.265 | 1 | 15 | Stx7 | 0.074 | Young |
| 0.453132 | 0.237912 | 0.625 | 0.571 | 1 | 15 | Clta | 0.054 | Young |
| 0.453726 | 1.123621 | 0.125 | 0.082 | 1 | 15 | Ganab | 0.043 | Young |
| 0.453726 | 1.077699 | 0.125 | 0.082 | 1 | 15 | Rrm2b | 0.043 | Young |
| 0.453726 | 0.879904 | 0.125 | 0.082 | 1 | 15 | Retreg3 | 0.043 | Young |
| 0.453726 | 0.871125 | 0.125 | 0.082 | 1 | 15 | Ankra2 | 0.043 | Young |
| 0.453726 | 0.827432 | 0.125 | 0.082 | 1 | 15 | Cfap43 | 0.043 | Young |
| 0.453726 | 0.817165 | 0.125 | 0.082 | 1 | 15 | Prkar2b | 0.043 | Young |

| | | | | | | | | |
|---|---|---|---|---|---|---|---|---|
| 0.454104 | 0.244859 | 0.446 | 0.388 | 1 | 15 | Zyx | 0.058 | Young |
| 0.454309 | 0.023226 | 0.268 | 0.204 | 1 | 15 | Ptpn6 | 0.064 | Young |
| 0.454802 | 0.054301 | 0.107 | 0.061 | 1 | 15 | Tm4sf5 | 0.046 | Young |
| 0.454988 | 0.223439 | 0.732 | 0.633 | 1 | 15 | Msn | 0.099 | Young |
| 0.455339 | 1.323893 | 0.143 | 0.102 | 1 | 15 | Wdfy2 | 0.041 | Young |
| 0.455458 | 0.579633 | 0.464 | 0.449 | 1 | 15 | Serpinb6b | 0.015 | Young |
| 0.455833 | 0.250491 | 0.357 | 0.286 | 1 | 15 | Phf11b | 0.071 | Young |
| 0.456138 | 0.819847 | 0.214 | 0.163 | 1 | 15 | Gspt1 | 0.051 | Young |
| 0.456446 | 0.039002 | 0.107 | 0.163 | 1 | 15 | Xpo7 | -0.056 | Young |
| 0.456463 | 0.189567 | 0.125 | 0.184 | 1 | 15 | Tmem57 | -0.059 | Young |
| 0.458193 | 0.394211 | 0.304 | 0.245 | 1 | 15 | Prpf39 | 0.059 | Young |
| 0.458336 | 0.313446 | 0.214 | 0.143 | 1 | 15 | 0610012G( | 0.071 | Young |
| 0.458434 | 0.689009 | 0.446 | 0.388 | 1 | 15 | Ywhag | 0.058 | Young |
| 0.459512 | 0.392573 | 0.268 | 0.204 | 1 | 15 | Raph1 | 0.064 | Young |
| 0.459778 | 0.036679 | 0.518 | 0.408 | 1 | 15 | Atp5d | 0.11 | Young |
| 0.460104 | 0.174935 | 0.232 | 0.306 | 1 | 15 | Ebi3 | -0.074 | Young |
| 0.460199 | 0.801744 | 0.196 | 0.143 | 1 | 15 | Ncf4 | 0.053 | Young |
| 0.460199 | 0.631367 | 0.196 | 0.143 | 1 | 15 | Atl2 | 0.053 | Young |
| 0.460199 | 0.567238 | 0.196 | 0.143 | 1 | 15 | Mapk14 | 0.053 | Young |
| 0.460199 | 0.349698 | 0.196 | 0.143 | 1 | 15 | Ssrp1 | 0.053 | Young |
| 0.460516 | 0.112277 | 0.357 | 0.286 | 1 | 15 | Atxn2l | 0.071 | Young |
| 0.460548 | 0.642193 | 0.25 | 0.204 | 1 | 15 | Epc2 | 0.046 | Young |
| 0.461041 | 0.932138 | 0.125 | 0.082 | 1 | 15 | Mvd | 0.043 | Young |
| 0.461041 | 0.920134 | 0.125 | 0.082 | 1 | 15 | Tmem120a | 0.043 | Young |
| 0.461041 | 0.854079 | 0.125 | 0.082 | 1 | 15 | Hps1 | 0.043 | Young |
| 0.461041 | 0.664789 | 0.125 | 0.082 | 1 | 15 | Slc16a3 | 0.043 | Young |
| 0.461041 | 0.65128 | 0.125 | 0.082 | 1 | 15 | Soga1 | 0.043 | Young |
| 0.461041 | 0.612378 | 0.125 | 0.082 | 1 | 15 | Acox3 | 0.043 | Young |
| 0.461041 | 0.213898 | 0.125 | 0.082 | 1 | 15 | Sec24a | 0.043 | Young |
| 0.461111 | 0.330502 | 0.339 | 0.265 | 1 | 15 | Acp2 | 0.074 | Young |
| 0.461824 | 0.692057 | 0.214 | 0.163 | 1 | 15 | Adh5 | 0.051 | Young |
| 0.461824 | 0.657686 | 0.214 | 0.163 | 1 | 15 | Ccng1 | 0.051 | Young |
| 0.461824 | 0.246944 | 0.214 | 0.163 | 1 | 15 | Brd8 | 0.051 | Young |
| 0.461824 | 0.903309 | 0.232 | 0.184 | 1 | 15 | Hmgcr | 0.048 | Young |
| 0.46212 | 0.40019 | 0.393 | 0.327 | 1 | 15 | Tgfb1 | 0.066 | Young |
| 0.462146 | 0.226745 | 0.143 | 0.102 | 1 | 15 | Fam133b | 0.041 | Young |
| 0.462698 | 0.386923 | 0.179 | 0.122 | 1 | 15 | Galnt12 | 0.057 | Young |
| 0.463694 | 0.267481 | 0.875 | 0.898 | 1 | 15 | Atp5e | -0.023 | Young |
| 0.464012 | 0.165429 | 0.982 | 0.939 | 1 | 15 | Rpl18 | 0.043 | Young |
| 0.464032 | 0.10015 | 0.179 | 0.265 | 1 | 15 | Cmtm7 | -0.086 | Young |
| 0.464864 | 0.145471 | 0.857 | 0.796 | 1 | 15 | 2010107E( | 0.061 | Young |
| 0.465132 | 0.347508 | 0.286 | 0.245 | 1 | 15 | Usp15 | 0.041 | Young |
| 0.465872 | 0.927672 | 0.25 | 0.204 | 1 | 15 | Mindy2 | 0.046 | Young |
| 0.466163 | 0.569592 | 0.196 | 0.143 | 1 | 15 | Pcif1 | 0.053 | Young |
| 0.466527 | 0.157955 | 0.786 | 0.714 | 1 | 15 | Gnai2 | 0.072 | Young |

| | | | | | | | | |
|---|---|---|---|---|---|---|---|---|
| 0.467539 | 2.32027 | 0.071 | 0.041 | 1 | 15 | Ptdss2 | 0.03 | Young |
| 0.467539 | 2.265273 | 0.071 | 0.041 | 1 | 15 | Map3k4 | 0.03 | Young |
| 0.467539 | 2.262688 | 0.071 | 0.041 | 1 | 15 | Cherp | 0.03 | Young |
| 0.467539 | 2.123903 | 0.071 | 0.041 | 1 | 15 | Ranbp10 | 0.03 | Young |
| 0.467539 | 1.910947 | 0.071 | 0.041 | 1 | 15 | Prdx4 | 0.03 | Young |
| 0.467539 | 1.897472 | 0.071 | 0.041 | 1 | 15 | Coq5 | 0.03 | Young |
| 0.467539 | 1.692254 | 0.071 | 0.041 | 1 | 15 | Pdhx | 0.03 | Young |
| 0.467539 | 1.605446 | 0.071 | 0.041 | 1 | 15 | Rfc5 | 0.03 | Young |
| 0.468422 | 1.189622 | 0.125 | 0.082 | 1 | 15 | Apeh | 0.043 | Young |
| 0.468422 | 1.013506 | 0.125 | 0.082 | 1 | 15 | 1110059G1 | 0.043 | Young |
| 0.468422 | 0.75471 | 0.125 | 0.082 | 1 | 15 | Lrrc61 | 0.043 | Young |
| 0.468422 | 0.544462 | 0.125 | 0.082 | 1 | 15 | Bend4 | 0.043 | Young |
| 0.468422 | 0.469258 | 0.125 | 0.082 | 1 | 15 | Tufm | 0.043 | Young |
| 0.468584 | 0.573111 | 0.268 | 0.224 | 1 | 15 | Supt6 | 0.044 | Young |
| 0.469009 | 1.106979 | 0.143 | 0.102 | 1 | 15 | Hcfc1 | 0.041 | Young |
| 0.469828 | 0.136612 | 0.946 | 0.898 | 1 | 15 | Cdc42 | 0.048 | Young |
| 0.469998 | 0.284996 | 0.375 | 0.327 | 1 | 15 | Akr1b3 | 0.048 | Young |
| 0.470754 | 0.547507 | 0.321 | 0.286 | 1 | 15 | Lamtor4 | 0.035 | Young |
| 0.470867 | 0.599429 | 0.304 | 0.265 | 1 | 15 | Bin1 | 0.039 | Young |
| 0.47123 | 0.652424 | 0.25 | 0.204 | 1 | 15 | Mcts1 | 0.046 | Young |
| 0.471712 | 0.069852 | 0.911 | 0.898 | 1 | 15 | Basp1 | 0.013 | Young |
| 0.472888 | 0.575442 | 0.232 | 0.184 | 1 | 15 | Msi2 | 0.048 | Young |
| 0.472888 | 0.338817 | 0.179 | 0.245 | 1 | 15 | L1cam | -0.066 | Young |
| 0.473273 | 0.17987 | 0.232 | 0.163 | 1 | 15 | Rusc1 | 0.069 | Young |
| 0.473313 | 0.632941 | 0.214 | 0.163 | 1 | 15 | Rab35 | 0.051 | Young |
| 0.473953 | 1.271793 | 0.179 | 0.143 | 1 | 15 | Csad | 0.036 | Young |
| 0.473953 | 0.600995 | 0.179 | 0.143 | 1 | 15 | Mlec | 0.036 | Young |
| 0.474665 | 0.108656 | 0.286 | 0.224 | 1 | 15 | Tox4 | 0.062 | Young |
| 0.475321 | 0.039692 | 0.268 | 0.204 | 1 | 15 | 1810022K0 | 0.064 | Young |
| 0.475503 | 0.112799 | 0.446 | 0.367 | 1 | 15 | Syncrip | 0.079 | Young |
| 0.475818 | 0.236443 | 0.321 | 0.265 | 1 | 15 | Cyb5a | 0.056 | Young |
| 0.475868 | 0.95701 | 0.125 | 0.082 | 1 | 15 | Tgif2 | 0.043 | Young |
| 0.475868 | 0.523083 | 0.125 | 0.082 | 1 | 15 | Wdr3 | 0.043 | Young |
| 0.475868 | 0.465823 | 0.125 | 0.082 | 1 | 15 | Mrps18a | 0.043 | Young |
| 0.475868 | 0.445376 | 0.125 | 0.082 | 1 | 15 | Ptpn7 | 0.043 | Young |
| 0.475929 | 1.214725 | 0.143 | 0.102 | 1 | 15 | Pih1d1 | 0.041 | Young |
| 0.475929 | 1.090309 | 0.143 | 0.102 | 1 | 15 | Gpr35 | 0.041 | Young |
| 0.476951 | 0.398041 | 0.393 | 0.347 | 1 | 15 | Cpne2 | 0.046 | Young |
| 0.47738 | 2.597699 | 0.071 | 0.041 | 1 | 15 | Mfsd10 | 0.03 | Young |
| 0.47738 | 2.273586 | 0.071 | 0.041 | 1 | 15 | Tceanc2 | 0.03 | Young |
| 0.47738 | 2.226084 | 0.071 | 0.041 | 1 | 15 | Coa4 | 0.03 | Young |
| 0.47738 | 2.120444 | 0.071 | 0.041 | 1 | 15 | Epg5 | 0.03 | Young |
| 0.47738 | 2.049565 | 0.071 | 0.041 | 1 | 15 | Tma16 | 0.03 | Young |
| 0.47738 | 2.030093 | 0.071 | 0.041 | 1 | 15 | G6pdx | 0.03 | Young |
| 0.47738 | 1.704969 | 0.071 | 0.041 | 1 | 15 | P2rx7 | 0.03 | Young |

| | | | | | | | | |
|---|---|---|---|---|---|---|---|---|
| 0.47738 | 1.517755 | 0.071 | 0.041 | 1 | 15 | Ptrh2 | 0.03 | Young |
| 0.47738 | 1.443974 | 0.071 | 0.041 | 1 | 15 | Swt1 | 0.03 | Young |
| 0.47738 | 1.382169 | 0.071 | 0.041 | 1 | 15 | 2810402E2 | 0.03 | Young |
| 0.478402 | 0.108386 | 0.696 | 0.551 | 1 | 15 | Selenok | 0.145 | Young |
| 0.478474 | 1.0637 | 0.232 | 0.184 | 1 | 15 | Dhx9 | 0.048 | Young |
| 0.479051 | 0.753596 | 0.268 | 0.224 | 1 | 15 | Tmed3 | 0.044 | Young |
| 0.479116 | 0.57398 | 0.214 | 0.163 | 1 | 15 | Arfrp1 | 0.051 | Young |
| 0.479401 | 0.186163 | 0.446 | 0.347 | 1 | 15 | 3110043O2 | 0.099 | Young |
| 0.480159 | 1.599118 | 0.179 | 0.143 | 1 | 15 | Dcun1d1 | 0.036 | Young |
| 0.480398 | 0.039808 | 0.304 | 0.224 | 1 | 15 | Poldip3 | 0.08 | Young |
| 0.480431 | 0.521 | 0.429 | 0.408 | 1 | 15 | Ndufa1 | 0.021 | Young |
| 0.4805 | 0.319534 | 0.393 | 0.327 | 1 | 15 | Cat | 0.066 | Young |
| 0.481033 | 0.207765 | 0.464 | 0.408 | 1 | 15 | Ndufb7 | 0.056 | Young |
| 0.481684 | 0.393064 | 0.179 | 0.122 | 1 | 15 | Trim7 | 0.057 | Young |
| 0.481684 | 0.216061 | 0.179 | 0.122 | 1 | 15 | Vps13b | 0.057 | Young |
| 0.482904 | 1.206508 | 0.143 | 0.102 | 1 | 15 | Msl3 | 0.041 | Young |
| 0.482904 | 0.873875 | 0.143 | 0.102 | 1 | 15 | Map1lc3a | 0.041 | Young |
| 0.483163 | 0.48651 | 0.107 | 0.163 | 1 | 15 | Dcaf11 | -0.056 | Young |
| 0.483163 | 0.240219 | 0.107 | 0.163 | 1 | 15 | Cln8 | -0.056 | Young |
| 0.483377 | 0.963121 | 0.125 | 0.082 | 1 | 15 | Glo1 | 0.043 | Young |
| 0.483377 | 0.659365 | 0.125 | 0.082 | 1 | 15 | Grk6 | 0.043 | Young |
| 0.483377 | 0.450721 | 0.125 | 0.082 | 1 | 15 | Gtf2f2 | 0.043 | Young |
| 0.483377 | 0.070708 | 0.125 | 0.082 | 1 | 15 | Itpr1 | 0.043 | Young |
| 0.483865 | 0.009033 | 0.393 | 0.347 | 1 | 15 | Surf4 | 0.046 | Young |
| 0.484095 | 0.711472 | 0.232 | 0.184 | 1 | 15 | Sypl | 0.048 | Young |
| 0.484307 | 0.245711 | 0.196 | 0.143 | 1 | 15 | Fam173a | 0.053 | Young |
| 0.484307 | 0.146302 | 0.196 | 0.143 | 1 | 15 | Zwint | 0.053 | Young |
| 0.484957 | 0.821783 | 0.214 | 0.163 | 1 | 15 | Mepce | 0.051 | Young |
| 0.485977 | 0.347368 | 0.25 | 0.184 | 1 | 15 | Papd4 | 0.066 | Young |
| 0.487334 | 1.959717 | 0.071 | 0.041 | 1 | 15 | Zcchc4 | 0.03 | Young |
| 0.487334 | 1.778711 | 0.071 | 0.041 | 1 | 15 | Wwc2 | 0.03 | Young |
| 0.487334 | 1.660735 | 0.071 | 0.041 | 1 | 15 | Xpnpep3 | 0.03 | Young |
| 0.487334 | 1.604605 | 0.071 | 0.041 | 1 | 15 | Apmap | 0.03 | Young |
| 0.487334 | 1.583367 | 0.071 | 0.041 | 1 | 15 | Leng1 | 0.03 | Young |
| 0.487334 | 1.576554 | 0.071 | 0.041 | 1 | 15 | Minpp1 | 0.03 | Young |
| 0.487334 | 1.48864 | 0.071 | 0.041 | 1 | 15 | Dcaf5 | 0.03 | Young |
| 0.487334 | 1.4392 | 0.071 | 0.041 | 1 | 15 | Med23 | 0.03 | Young |
| 0.487334 | 1.312804 | 0.071 | 0.041 | 1 | 15 | Nfrkb | 0.03 | Young |
| 0.487334 | 1.285914 | 0.071 | 0.041 | 1 | 15 | Adprm | 0.03 | Young |
| 0.487334 | 1.173112 | 0.071 | 0.041 | 1 | 15 | Llgl1 | 0.03 | Young |
| 0.487334 | 1.172939 | 0.071 | 0.041 | 1 | 15 | Nsmf | 0.03 | Young |
| 0.487334 | 1.146725 | 0.071 | 0.041 | 1 | 15 | Tsc2 | 0.03 | Young |
| 0.487334 | 1.025117 | 0.071 | 0.041 | 1 | 15 | Alg1 | 0.03 | Young |
| 0.487334 | 1.011177 | 0.071 | 0.041 | 1 | 15 | Nudt18 | 0.03 | Young |
| 0.487334 | 0.980759 | 0.071 | 0.041 | 1 | 15 | Agap3 | 0.03 | Young |

| | | | | | | | | |
|---|---|---|---|---|---|---|---|---|
| 0.488364 | 0.32575 | 0.304 | 0.245 | 1 | 15 | Tgfbi | 0.059 | Young |
| 0.488412 | 0.279308 | 0.536 | 0.49 | 1 | 15 | Plekho1 | 0.046 | Young |
| 0.49044 | 0.687612 | 0.196 | 0.143 | 1 | 15 | Xrn1 | 0.053 | Young |
| 0.490837 | 0.52964 | 0.214 | 0.163 | 1 | 15 | Stk3 | 0.051 | Young |
| 0.490837 | 0.188895 | 0.214 | 0.163 | 1 | 15 | Ski | 0.051 | Young |
| 0.490951 | 0.455116 | 0.125 | 0.082 | 1 | 15 | Oxsr1 | 0.043 | Young |
| 0.490951 | 0.380899 | 0.125 | 0.082 | 1 | 15 | Sbf1 | 0.043 | Young |
| 0.490951 | 0.26967 | 0.125 | 0.082 | 1 | 15 | Gclm | 0.043 | Young |
| 0.490951 | 0.127368 | 0.125 | 0.082 | 1 | 15 | Kpna6 | 0.043 | Young |
| 0.491423 | 0.216679 | 0.268 | 0.204 | 1 | 15 | Emc6 | 0.064 | Young |
| 0.493936 | 0.128861 | 1 | 0.98 | 1 | 15 | Rpl19 | 0.02 | Young |
| 0.494986 | 0.268491 | 0.268 | 0.224 | 1 | 15 | Ric1 | 0.044 | Young |
| 0.495506 | 0.061071 | 0.411 | 0.327 | 1 | 15 | Tsc22d3 | 0.084 | Young |
| 0.495759 | 0.449498 | 0.321 | 0.265 | 1 | 15 | Kansl1 | 0.056 | Young |
| 0.495759 | 0.321645 | 0.321 | 0.265 | 1 | 15 | Wnt11 | 0.056 | Young |
| 0.495933 | 0.591422 | 0.286 | 0.245 | 1 | 15 | Prkcsh | 0.041 | Young |
| 0.49626 | 0.242062 | 0.232 | 0.163 | 1 | 15 | Spopl | 0.069 | Young |
| 0.496613 | 0.148631 | 0.143 | 0.204 | 1 | 15 | Wasl | -0.061 | Young |
| 0.496855 | 0.244051 | 0.268 | 0.204 | 1 | 15 | Mnt | 0.064 | Young |
| 0.49689 | 1.028965 | 0.196 | 0.163 | 1 | 15 | Zbtb8os | 0.033 | Young |
| 0.49702 | 0.933703 | 0.143 | 0.102 | 1 | 15 | Noc2l | 0.041 | Young |
| 0.49702 | 0.705074 | 0.143 | 0.102 | 1 | 15 | Rnd3 | 0.041 | Young |
| 0.497027 | 0.171946 | 0.714 | 0.714 | 1 | 15 | Il4i1 | 0 | Young |
| 0.497398 | 1.923531 | 0.071 | 0.041 | 1 | 15 | Slc25a35 | 0.03 | Young |
| 0.497398 | 1.601362 | 0.071 | 0.041 | 1 | 15 | Mtx2 | 0.03 | Young |
| 0.497398 | 1.397293 | 0.071 | 0.041 | 1 | 15 | Ss18l2 | 0.03 | Young |
| 0.497398 | 1.287729 | 0.071 | 0.041 | 1 | 15 | Snx15 | 0.03 | Young |
| 0.497398 | 1.246551 | 0.071 | 0.041 | 1 | 15 | Ginm1 | 0.03 | Young |
| 0.497398 | 1.225901 | 0.071 | 0.041 | 1 | 15 | Traf3ip3 | 0.03 | Young |
| 0.497398 | 1.10768 | 0.071 | 0.041 | 1 | 15 | Mcph1 | 0.03 | Young |
| 0.497398 | 1.085049 | 0.071 | 0.041 | 1 | 15 | Ddr1 | 0.03 | Young |
| 0.497398 | 1.021427 | 0.071 | 0.041 | 1 | 15 | D6Wsu163 | 0.03 | Young |
| 0.497398 | 0.940831 | 0.071 | 0.041 | 1 | 15 | Rpl9-ps6 | 0.03 | Young |
| 0.497398 | 0.769428 | 0.071 | 0.041 | 1 | 15 | Cd226 | 0.03 | Young |
| 0.497398 | 0.699757 | 0.071 | 0.041 | 1 | 15 | Myg1 | 0.03 | Young |
| 0.497996 | 0.105484 | 0.982 | 0.959 | 1 | 15 | mt-Nd1 | 0.023 | Young |
| 0.498524 | 0.388914 | 0.25 | 0.204 | 1 | 15 | Trim26 | 0.046 | Young |
| 0.498588 | 0.337547 | 0.125 | 0.082 | 1 | 15 | Gphn | 0.043 | Young |
| 0.498654 | 0.067612 | 0.304 | 0.245 | 1 | 15 | Kif2a | 0.059 | Young |
| 0.499039 | 1.020881 | 0.179 | 0.143 | 1 | 15 | Yipf1 | 0.036 | Young |
| 0.499325 | 0.848708 | 0.161 | 0.122 | 1 | 15 | Neu1 | 0.039 | Young |
| 0.500361 | 0.269073 | 0.268 | 0.224 | 1 | 15 | Os9 | 0.044 | Young |
| 0.500949 | 0.072717 | 0.464 | 0.388 | 1 | 15 | Adcy6 | 0.076 | Young |
| 0.501174 | 0.562068 | 0.232 | 0.184 | 1 | 15 | Slc43a2 | 0.048 | Young |
| 0.502098 | 0.098641 | 0.232 | 0.163 | 1 | 15 | Sp3 | 0.069 | Young |

| | | | | | | | | |
|---|---|---|---|---|---|---|---|---|
| 0.502828 | 0.068682 | 0.196 | 0.143 | 1 | 15 | Polr1c | 0.053 | Young |
| 0.50299 | 0.669762 | 0.429 | 0.408 | 1 | 15 | Sin3b | 0.021 | Young |
| 0.50299 | 0.669282 | 0.429 | 0.408 | 1 | 15 | Txnip | 0.021 | Young |
| 0.50299 | 0.333795 | 0.446 | 0.388 | 1 | 15 | Selenop | 0.058 | Young |
| 0.504082 | 0.400982 | 0.25 | 0.204 | 1 | 15 | Wbp4 | 0.046 | Young |
| 0.504159 | 0.734546 | 0.143 | 0.102 | 1 | 15 | Rfx7 | 0.041 | Young |
| 0.504159 | 0.572536 | 0.143 | 0.102 | 1 | 15 | Carhsp1 | 0.041 | Young |
| 0.504345 | 1.361511 | 0.018 | 0.041 | 1 | 15 | Rps19bp1 | -0.023 | Young |
| 0.504345 | 1.29513 | 0.018 | 0.041 | 1 | 15 | Inpp5b | -0.023 | Young |
| 0.504345 | 1.225712 | 0.018 | 0.041 | 1 | 15 | H2-Ke6 | -0.023 | Young |
| 0.504345 | 0.986056 | 0.018 | 0.041 | 1 | 15 | Piga | -0.023 | Young |
| 0.504345 | 0.865897 | 0.018 | 0.041 | 1 | 15 | Ms4a4c | -0.023 | Young |
| 0.504345 | 0.761115 | 0.018 | 0.041 | 1 | 15 | Cacna1a | -0.023 | Young |
| 0.504345 | 0.711932 | 0.018 | 0.041 | 1 | 15 | Tfpt | -0.023 | Young |
| 0.504345 | 0.693891 | 0.018 | 0.041 | 1 | 15 | Catsperg1 | -0.023 | Young |
| 0.504345 | 0.641047 | 0.018 | 0.041 | 1 | 15 | Phlda3 | -0.023 | Young |
| 0.504345 | 0.552529 | 0.018 | 0.041 | 1 | 15 | Kdr | -0.023 | Young |
| 0.504345 | 0.545235 | 0.018 | 0.041 | 1 | 15 | 1110035H1 | -0.023 | Young |
| 0.504345 | 0.541324 | 0.018 | 0.041 | 1 | 15 | Omp | -0.023 | Young |
| 0.504345 | 0.501595 | 0.018 | 0.041 | 1 | 15 | Map3k10 | -0.023 | Young |
| 0.504345 | 0.43784 | 0.018 | 0.041 | 1 | 15 | Pla1a | -0.023 | Young |
| 0.504345 | 0.409035 | 0.018 | 0.041 | 1 | 15 | 2310061I0 | -0.023 | Young |
| 0.504345 | 0.353153 | 0.018 | 0.041 | 1 | 15 | Rrs1 | -0.023 | Young |
| 0.504345 | 0.335824 | 0.018 | 0.041 | 1 | 15 | Dis3l | -0.023 | Young |
| 0.504345 | 0.325305 | 0.018 | 0.041 | 1 | 15 | Lsg1 | -0.023 | Young |
| 0.504345 | 0.302662 | 0.018 | 0.041 | 1 | 15 | Trim2 | -0.023 | Young |
| 0.504345 | 0.274012 | 0.018 | 0.041 | 1 | 15 | Nol11 | -0.023 | Young |
| 0.504345 | 0.26897 | 0.018 | 0.041 | 1 | 15 | Smyd2 | -0.023 | Young |
| 0.504345 | 0.252692 | 0.018 | 0.041 | 1 | 15 | Angptl4 | -0.023 | Young |
| 0.504345 | 0.230016 | 0.018 | 0.041 | 1 | 15 | Trim30b | -0.023 | Young |
| 0.504345 | 0.195334 | 0.018 | 0.041 | 1 | 15 | Pou5f1 | -0.023 | Young |
| 0.504345 | 0.192261 | 0.018 | 0.041 | 1 | 15 | Acrbp | -0.023 | Young |
| 0.504345 | 0.177234 | 0.018 | 0.041 | 1 | 15 | 9430015G1 | -0.023 | Young |
| 0.504345 | 0.12069 | 0.018 | 0.041 | 1 | 15 | Ercc8 | -0.023 | Young |
| 0.504345 | 0.094792 | 0.018 | 0.041 | 1 | 15 | Rab26 | -0.023 | Young |
| 0.504345 | 0.086437 | 0.018 | 0.041 | 1 | 15 | Paqr7 | -0.023 | Young |
| 0.504345 | 0.038027 | 0.018 | 0.041 | 1 | 15 | Nxpe5 | -0.023 | Young |
| 0.504345 | 0.000957 | 0.018 | 0.041 | 1 | 15 | Tmem126b | -0.023 | Young |
| 0.505419 | 0.975376 | 0.179 | 0.143 | 1 | 15 | Sh3bp5 | 0.036 | Young |
| 0.506053 | 0.826773 | 0.161 | 0.122 | 1 | 15 | Ndufa9 | 0.039 | Young |
| 0.506053 | 0.322073 | 0.161 | 0.122 | 1 | 15 | Sdf2l1 | 0.039 | Young |
| 0.506288 | 0.414089 | 0.125 | 0.082 | 1 | 15 | Ttc17 | 0.043 | Young |
| 0.506288 | 0.359979 | 0.125 | 0.082 | 1 | 15 | Kdm3a | 0.043 | Young |
| 0.506288 | 0.285113 | 0.125 | 0.082 | 1 | 15 | Psd4 | 0.043 | Young |
| 0.507491 | 0.01137 | 0.911 | 0.776 | 1 | 15 | Rpl7a | 0.135 | Young |

| | | | | | | | | |
|---|---|---|---|---|---|---|---|---|
| 0.507573 | 1.331504 | 0.071 | 0.041 | 1 | 15 | Lonp2 | 0.03 | Young |
| 0.507573 | 1.325208 | 0.071 | 0.041 | 1 | 15 | Zfp865 | 0.03 | Young |
| 0.507573 | 1.091571 | 0.071 | 0.041 | 1 | 15 | Kdsr | 0.03 | Young |
| 0.507573 | 0.961636 | 0.071 | 0.041 | 1 | 15 | Nudt13 | 0.03 | Young |
| 0.507573 | 0.924455 | 0.071 | 0.041 | 1 | 15 | Pskh1 | 0.03 | Young |
| 0.507573 | 0.916946 | 0.071 | 0.041 | 1 | 15 | Ryk | 0.03 | Young |
| 0.507573 | 0.890118 | 0.071 | 0.041 | 1 | 15 | Smcr8 | 0.03 | Young |
| 0.507573 | 0.889265 | 0.071 | 0.041 | 1 | 15 | 3110040N1 | 0.03 | Young |
| 0.507573 | 0.800504 | 0.071 | 0.041 | 1 | 15 | Ruvbl1 | 0.03 | Young |
| 0.507573 | 0.797097 | 0.071 | 0.041 | 1 | 15 | Cetn4 | 0.03 | Young |
| 0.507573 | 0.749278 | 0.071 | 0.041 | 1 | 15 | E130307A1 | 0.03 | Young |
| 0.507573 | 0.745658 | 0.071 | 0.041 | 1 | 15 | Prep | 0.03 | Young |
| 0.507573 | 0.719408 | 0.071 | 0.041 | 1 | 15 | Rtca | 0.03 | Young |
| 0.507573 | 0.71552 | 0.071 | 0.041 | 1 | 15 | Vps51 | 0.03 | Young |
| 0.507573 | 0.686872 | 0.071 | 0.041 | 1 | 15 | Gpatch2 | 0.03 | Young |
| 0.507573 | 0.673335 | 0.071 | 0.041 | 1 | 15 | Trim47 | 0.03 | Young |
| 0.507573 | 0.614871 | 0.071 | 0.041 | 1 | 15 | Ccdc93 | 0.03 | Young |
| 0.507573 | 0.534809 | 0.071 | 0.041 | 1 | 15 | Ttc33 | 0.03 | Young |
| 0.507573 | 0.484504 | 0.071 | 0.041 | 1 | 15 | Sqle | 0.03 | Young |
| 0.507573 | 0.469647 | 0.071 | 0.041 | 1 | 15 | Fam3c | 0.03 | Young |
| 0.507573 | 0.455729 | 0.071 | 0.041 | 1 | 15 | Selenon | 0.03 | Young |
| 0.507573 | 0.449876 | 0.071 | 0.041 | 1 | 15 | Pctp | 0.03 | Young |
| 0.507573 | 0.356468 | 0.071 | 0.041 | 1 | 15 | Ift172 | 0.03 | Young |
| 0.507573 | 0.015621 | 0.071 | 0.041 | 1 | 15 | Selenbp1 | 0.03 | Young |
| 0.507644 | 0.257674 | 0.179 | 0.122 | 1 | 15 | Brox | 0.057 | Young |
| 0.508704 | 0.812061 | 0.214 | 0.163 | 1 | 15 | Psmd10 | 0.051 | Young |
| 0.508802 | 0.008016 | 0.393 | 0.327 | 1 | 15 | Mcrip1 | 0.066 | Young |
| 0.511003 | 0.305415 | 0.321 | 0.265 | 1 | 15 | Tyk2 | 0.056 | Young |
| 0.511352 | 0.785516 | 0.143 | 0.102 | 1 | 15 | Fcna | 0.041 | Young |
| 0.511352 | 0.469991 | 0.143 | 0.102 | 1 | 15 | Acsl1 | 0.041 | Young |
| 0.511352 | 0.194252 | 0.143 | 0.102 | 1 | 15 | Mea1 | 0.041 | Young |
| 0.511655 | 0.206565 | 0.571 | 0.469 | 1 | 15 | Rgs10 | 0.102 | Young |
| 0.511866 | 0.215797 | 0.5 | 0.449 | 1 | 15 | Il1b | 0.051 | Young |
| 0.512095 | 0.205491 | 0.607 | 0.571 | 1 | 15 | Rab14 | 0.036 | Young |
| 0.512598 | 0.101485 | 0.643 | 0.633 | 1 | 15 | Rabgap1l | 0.01 | Young |
| 0.512828 | 0.910922 | 0.161 | 0.122 | 1 | 15 | Actl6a | 0.039 | Young |
| 0.512828 | 0.729356 | 0.161 | 0.122 | 1 | 15 | Arhgef3 | 0.039 | Young |
| 0.51405 | 0.075864 | 0.125 | 0.082 | 1 | 15 | Tgfbr1 | 0.043 | Young |
| 0.51405 | 0.024297 | 0.125 | 0.082 | 1 | 15 | Nrde2 | 0.043 | Young |
| 0.514735 | 0.119642 | 0.161 | 0.224 | 1 | 15 | Polr2f | -0.063 | Young |
| 0.51525 | 0.805585 | 0.196 | 0.163 | 1 | 15 | Cnot8 | 0.033 | Young |
| 0.515296 | 0.579914 | 0.25 | 0.204 | 1 | 15 | Stom | 0.046 | Young |
| 0.515296 | 0.413434 | 0.25 | 0.204 | 1 | 15 | Eif3b | 0.046 | Young |
| 0.51538 | 0.378792 | 0.196 | 0.143 | 1 | 15 | Rrp1 | 0.053 | Young |
| 0.515549 | 0.195917 | 0.5 | 0.49 | 1 | 15 | Ppp2r5c | 0.01 | Young |

| | | | | | | | | |
|---|---|---|---|---|---|---|---|---|
| 0.516138 | 0.274819 | 0.321 | 0.265 | 1 | 15 | Klf4 | 0.056 | Young |
| 0.516138 | 0.245476 | 0.321 | 0.265 | 1 | 15 | Ppp2r1a | 0.056 | Young |
| 0.516623 | 0.375178 | 0.304 | 0.265 | 1 | 15 | Maf | 0.039 | Young |
| 0.516833 | 0.393342 | 0.482 | 0.429 | 1 | 15 | Lrrc8c | 0.053 | Young |
| 0.517855 | 0.721321 | 0.071 | 0.041 | 1 | 15 | Tubgcp5 | 0.03 | Young |
| 0.517855 | 0.549768 | 0.071 | 0.041 | 1 | 15 | Arfgap3 | 0.03 | Young |
| 0.517855 | 0.496036 | 0.071 | 0.041 | 1 | 15 | Cyb561d1 | 0.03 | Young |
| 0.517855 | 0.425703 | 0.071 | 0.041 | 1 | 15 | AI837181 | 0.03 | Young |
| 0.517855 | 0.424005 | 0.071 | 0.041 | 1 | 15 | Golm1 | 0.03 | Young |
| 0.517855 | 0.389788 | 0.071 | 0.041 | 1 | 15 | Exosc1 | 0.03 | Young |
| 0.517855 | 0.329435 | 0.071 | 0.041 | 1 | 15 | Mogs | 0.03 | Young |
| 0.517855 | 0.116857 | 0.071 | 0.041 | 1 | 15 | Arrdc2 | 0.03 | Young |
| 0.518307 | 0.975438 | 0.179 | 0.143 | 1 | 15 | Dpy30 | 0.036 | Young |
| 0.518307 | 0.791294 | 0.179 | 0.143 | 1 | 15 | Pitpnm1 | 0.036 | Young |
| 0.518598 | 0.644167 | 0.143 | 0.102 | 1 | 15 | Fam204a | 0.041 | Young |
| 0.518598 | 0.53962 | 0.143 | 0.102 | 1 | 15 | Raf1 | 0.041 | Young |
| 0.51965 | 1.080685 | 0.161 | 0.122 | 1 | 15 | Cbx5 | 0.039 | Young |
| 0.51965 | 0.793042 | 0.161 | 0.122 | 1 | 15 | Psmc2 | 0.039 | Young |
| 0.519744 | 0.122419 | 0.286 | 0.347 | 1 | 15 | Appl1 | -0.061 | Young |
| 0.520596 | 0.34484 | 0.321 | 0.286 | 1 | 15 | Ubtf | 0.035 | Young |
| 0.52095 | 0.354387 | 0.25 | 0.204 | 1 | 15 | Tsc22d2 | 0.046 | Young |
| 0.521433 | 0.308991 | 0.446 | 0.388 | 1 | 15 | Cd40 | 0.058 | Young |
| 0.521447 | 1.222398 | 0.196 | 0.163 | 1 | 15 | Cramp1l | 0.033 | Young |
| 0.521717 | 0.323867 | 0.196 | 0.143 | 1 | 15 | Golga7 | 0.053 | Young |
| 0.521874 | 0.427157 | 0.125 | 0.082 | 1 | 15 | Entpd1 | 0.043 | Young |
| 0.521874 | 0.216797 | 0.125 | 0.082 | 1 | 15 | Mmadhc | 0.043 | Young |
| 0.52258 | 0.159831 | 0.929 | 0.878 | 1 | 15 | Rps2 | 0.051 | Young |
| 0.522607 | 0.574812 | 0.357 | 0.306 | 1 | 15 | AA467197 | 0.051 | Young |
| 0.522815 | 0.838371 | 0.214 | 0.184 | 1 | 15 | Ap3b1 | 0.03 | Young |
| 0.524431 | 0.626593 | 0.232 | 0.184 | 1 | 15 | Rap1gds1 | 0.048 | Young |
| 0.524431 | 0.439651 | 0.232 | 0.184 | 1 | 15 | Pank3 | 0.048 | Young |
| 0.524431 | 0.369458 | 0.232 | 0.184 | 1 | 15 | 9930111J2 | 0.048 | Young |
| 0.525514 | 0.152505 | 0.786 | 0.837 | 1 | 15 | S100a4 | -0.051 | Young |
| 0.525897 | 1.159004 | 0.143 | 0.102 | 1 | 15 | Clec2i | 0.041 | Young |
| 0.525897 | 0.380052 | 0.143 | 0.102 | 1 | 15 | Naa35 | 0.041 | Young |
| 0.526336 | 0.174772 | 0.554 | 0.51 | 1 | 15 | Minos1 | 0.044 | Young |
| 0.526457 | 0.42527 | 0.393 | 0.347 | 1 | 15 | Sdhaf1 | 0.046 | Young |
| 0.526488 | 0.42987 | 0.321 | 0.265 | 1 | 15 | Ndufb6 | 0.056 | Young |
| 0.526488 | 0.146818 | 0.321 | 0.265 | 1 | 15 | Cfdp1 | 0.056 | Young |
| 0.526518 | 0.837938 | 0.161 | 0.122 | 1 | 15 | Snhg12 | 0.039 | Young |
| 0.526518 | 0.383256 | 0.161 | 0.122 | 1 | 15 | Ints9 | 0.039 | Young |
| 0.526637 | 0.449631 | 0.25 | 0.204 | 1 | 15 | Gm16286 | 0.046 | Young |
| 0.526907 | 0.296947 | 0.214 | 0.163 | 1 | 15 | Ptdss1 | 0.051 | Young |
| 0.527086 | 0.140602 | 0.554 | 0.49 | 1 | 15 | Zfand5 | 0.064 | Young |
| 0.527689 | 0.702687 | 0.268 | 0.224 | 1 | 15 | Mrps24 | 0.044 | Young |

| | | | | | | | | |
|---|---|---|---|---|---|---|---|---|
| 0.527848 | 0.158632 | 0.286 | 0.224 | 1 | 15 | Map3k8 | 0.062 | Young |
| 0.528093 | 0.390629 | 0.196 | 0.143 | 1 | 15 | Ppil4 | 0.053 | Young |
| 0.528244 | 0.672651 | 0.071 | 0.041 | 1 | 15 | Sacm1l | 0.03 | Young |
| 0.528244 | 0.521123 | 0.071 | 0.041 | 1 | 15 | Slamf1 | 0.03 | Young |
| 0.528244 | 0.393517 | 0.071 | 0.041 | 1 | 15 | Cyp4f13 | 0.03 | Young |
| 0.528244 | 0.364799 | 0.071 | 0.041 | 1 | 15 | 6430531B1 | 0.03 | Young |
| 0.528244 | 0.357353 | 0.071 | 0.041 | 1 | 15 | Cfd | 0.03 | Young |
| 0.528244 | 0.345414 | 0.071 | 0.041 | 1 | 15 | Kcnab2 | 0.03 | Young |
| 0.528244 | 0.341004 | 0.071 | 0.041 | 1 | 15 | S1pr1 | 0.03 | Young |
| 0.528244 | 0.338581 | 0.071 | 0.041 | 1 | 15 | Cpt1c | 0.03 | Young |
| 0.528244 | 0.159636 | 0.071 | 0.041 | 1 | 15 | Ndrg3 | 0.03 | Young |
| 0.528244 | 0.157701 | 0.071 | 0.041 | 1 | 15 | Gm36161 | 0.03 | Young |
| 0.528244 | 0.006554 | 0.071 | 0.041 | 1 | 15 | Lpar6 | 0.03 | Young |
| 0.528978 | 0.111608 | 0.839 | 0.714 | 1 | 15 | H2-T23 | 0.125 | Young |
| 0.529758 | 0.24285 | 0.125 | 0.082 | 1 | 15 | Acss1 | 0.043 | Young |
| 0.530202 | 0.447455 | 0.304 | 0.245 | 1 | 15 | Eif3j1 | 0.059 | Young |
| 0.530331 | 0.129977 | 0.179 | 0.245 | 1 | 15 | Nt5c | -0.066 | Young |
| 0.53035 | 0.113232 | 0.429 | 0.367 | 1 | 15 | Wac | 0.062 | Young |
| 0.532355 | 0.377569 | 0.25 | 0.204 | 1 | 15 | Ptpn11 | 0.046 | Young |
| 0.533037 | 0.729061 | 0.375 | 0.347 | 1 | 15 | Slc39a1 | 0.028 | Young |
| 0.533047 | 0.537159 | 0.161 | 0.224 | 1 | 15 | Ptov1 | -0.063 | Young |
| 0.533047 | 0.433202 | 0.214 | 0.163 | 1 | 15 | Mpp1 | 0.051 | Young |
| 0.533047 | 0.342482 | 0.214 | 0.163 | 1 | 15 | Cd300ld | 0.051 | Young |
| 0.5332 | 0.106419 | 0.232 | 0.306 | 1 | 15 | Gm26532 | -0.074 | Young |
| 0.533245 | 0.841817 | 0.268 | 0.224 | 1 | 15 | Ube2s | 0.044 | Young |
| 0.533245 | 0.641955 | 0.268 | 0.224 | 1 | 15 | Ralgapa1 | 0.044 | Young |
| 0.533248 | 0.444842 | 0.143 | 0.102 | 1 | 15 | Lancl1 | 0.041 | Young |
| 0.533248 | 0.442127 | 0.143 | 0.102 | 1 | 15 | Rps6ka5 | 0.041 | Young |
| 0.533256 | 0.031148 | 1 | 0.98 | 1 | 15 | Rps23 | 0.02 | Young |
| 0.533433 | 0.871038 | 0.161 | 0.122 | 1 | 15 | Adap2 | 0.039 | Young |
| 0.533433 | 0.818166 | 0.161 | 0.122 | 1 | 15 | Nup153 | 0.039 | Young |
| 0.533956 | 0.814299 | 0.196 | 0.163 | 1 | 15 | Ddt | 0.033 | Young |
| 0.53432 | 0.406674 | 0.125 | 0.184 | 1 | 15 | Ssr2 | -0.059 | Young |
| 0.534369 | 1.734833 | 0.089 | 0.061 | 1 | 15 | Cox19 | 0.028 | Young |
| 0.534369 | 1.664865 | 0.089 | 0.061 | 1 | 15 | Nelfcd | 0.028 | Young |
| 0.535481 | 0.491746 | 0.446 | 0.429 | 1 | 15 | Ndel1 | 0.017 | Young |
| 0.536176 | 0.247047 | 0.393 | 0.347 | 1 | 15 | Vdac1 | 0.046 | Young |
| 0.536264 | 0.249463 | 0.232 | 0.184 | 1 | 15 | Fam105a | 0.048 | Young |
| 0.537611 | 0.145823 | 0.339 | 0.286 | 1 | 15 | Rad23a | 0.053 | Young |
| 0.537675 | 0.548599 | 0.304 | 0.265 | 1 | 15 | Stag2 | 0.039 | Young |
| 0.537952 | 0.842055 | 0.179 | 0.143 | 1 | 15 | Twf1 | 0.036 | Young |
| 0.537952 | 0.712511 | 0.179 | 0.143 | 1 | 15 | Sh3bp4 | 0.036 | Young |
| 0.538828 | 0.196437 | 0.554 | 0.531 | 1 | 15 | Rabac1 | 0.023 | Young |
| 0.539224 | 0.028779 | 0.161 | 0.224 | 1 | 15 | Parl | -0.063 | Young |
| 0.539691 | 0.171605 | 0.571 | 0.49 | 1 | 15 | Selenow | 0.081 | Young |

| | | | | | | | | |
|---|---|---|---|---|---|---|---|---|
| 0.539747 | 0.476572 | 0.464 | 0.449 | 1 | 15 | Tubb2a | 0.015 | Young |
| 0.540267 | 0.554674 | 0.196 | 0.163 | 1 | 15 | Dcaf8 | 0.033 | Young |
| 0.54027 | 0.232846 | 0.196 | 0.245 | 1 | 15 | Chmp2b | -0.049 | Young |
| 0.54027 | 0.004074 | 0.196 | 0.245 | 1 | 15 | Trim28 | -0.049 | Young |
| 0.540393 | 0.867338 | 0.161 | 0.122 | 1 | 15 | Rasgrp4 | 0.039 | Young |
| 0.540393 | 0.676841 | 0.161 | 0.122 | 1 | 15 | Fbxo4 | 0.039 | Young |
| 0.540651 | 0.483907 | 0.143 | 0.102 | 1 | 15 | Mbd3 | 0.041 | Young |
| 0.540651 | 0.356032 | 0.143 | 0.102 | 1 | 15 | Pnkd | 0.041 | Young |
| 0.540916 | 0.273128 | 0.179 | 0.224 | 1 | 15 | Tor1aip2 | -0.045 | Young |
| 0.542231 | 0.284366 | 0.232 | 0.184 | 1 | 15 | Abca9 | 0.048 | Young |
| 0.542772 | 0.341605 | 0.339 | 0.286 | 1 | 15 | Mafb | 0.053 | Young |
| 0.543523 | 0.168724 | 0.339 | 0.408 | 1 | 15 | Psma6 | -0.069 | Young |
| 0.543594 | 1.684835 | 0.089 | 0.061 | 1 | 15 | Rars | 0.028 | Young |
| 0.543594 | 1.480512 | 0.089 | 0.061 | 1 | 15 | Timm8a1 | 0.028 | Young |
| 0.543594 | 1.138674 | 0.089 | 0.061 | 1 | 15 | Vps37c | 0.028 | Young |
| 0.544853 | 0.706972 | 0.304 | 0.286 | 1 | 15 | Zranb2 | 0.018 | Young |
| 0.546347 | 0.466268 | 0.304 | 0.245 | 1 | 15 | Taf15 | 0.059 | Young |
| 0.547398 | 0.99587 | 0.161 | 0.122 | 1 | 15 | Exoc6b | 0.039 | Young |
| 0.547398 | 0.630034 | 0.161 | 0.122 | 1 | 15 | Zfp281 | 0.039 | Young |
| 0.547398 | 0.618019 | 0.161 | 0.122 | 1 | 15 | 2510002D2 | 0.039 | Young |
| 0.547398 | 0.538482 | 0.161 | 0.122 | 1 | 15 | Arl1 | 0.039 | Young |
| 0.547502 | 0.257733 | 0.321 | 0.265 | 1 | 15 | Sec11a | 0.056 | Young |
| 0.548104 | 0.394598 | 0.143 | 0.102 | 1 | 15 | Brd9 | 0.041 | Young |
| 0.548104 | 0.36901 | 0.143 | 0.102 | 1 | 15 | Gbp7 | 0.041 | Young |
| 0.548104 | 0.258657 | 0.143 | 0.102 | 1 | 15 | Etfdh | 0.041 | Young |
| 0.548104 | 0.25183 | 0.143 | 0.102 | 1 | 15 | Ech1 | 0.041 | Young |
| 0.548104 | 0.15409 | 0.143 | 0.102 | 1 | 15 | Ammecr1l | 0.041 | Young |
| 0.548104 | 0.112357 | 0.143 | 0.102 | 1 | 15 | Sart3 | 0.041 | Young |
| 0.548164 | 0.04597 | 0.036 | 0.061 | 1 | 15 | Trub2 | -0.025 | Young |
| 0.548235 | 0.349271 | 0.5 | 0.449 | 1 | 15 | Snrpf | 0.051 | Young |
| 0.549336 | 0.132629 | 0.071 | 0.041 | 1 | 15 | Tdrd3 | 0.03 | Young |
| 0.549336 | 0.119188 | 0.071 | 0.041 | 1 | 15 | Fam149b | 0.03 | Young |
| 0.549336 | 0.025669 | 0.071 | 0.041 | 1 | 15 | Chchd6 | 0.03 | Young |
| 0.549594 | 0.406079 | 0.286 | 0.245 | 1 | 15 | Thoc7 | 0.041 | Young |
| 0.551253 | 0.779982 | 0.179 | 0.143 | 1 | 15 | Zfp326 | 0.036 | Young |
| 0.551326 | 0.526859 | 0.357 | 0.327 | 1 | 15 | Ctsd | 0.03 | Young |
| 0.552387 | 0.022115 | 0.643 | 0.551 | 1 | 15 | Hnrnpu | 0.092 | Young |
| 0.552897 | 1.339334 | 0.089 | 0.061 | 1 | 15 | Klhl7 | 0.028 | Young |
| 0.552897 | 1.264699 | 0.089 | 0.061 | 1 | 15 | Pold3 | 0.028 | Young |
| 0.552897 | 1.229356 | 0.089 | 0.061 | 1 | 15 | Gbf1 | 0.028 | Young |
| 0.552897 | 1.147725 | 0.089 | 0.061 | 1 | 15 | Abl1 | 0.028 | Young |
| 0.552897 | 1.093143 | 0.089 | 0.061 | 1 | 15 | Phf6 | 0.028 | Young |
| 0.552999 | 0.800729 | 0.196 | 0.163 | 1 | 15 | Csnk1e | 0.033 | Young |
| 0.553989 | 0.339926 | 0.196 | 0.143 | 1 | 15 | Cd209g | 0.053 | Young |
| 0.554165 | 0.423621 | 0.411 | 0.367 | 1 | 15 | Tmed2 | 0.044 | Young |

| | | | | | | | | |
|---|---|---|---|---|---|---|---|---|
| 0.554262 | 0.269855 | 0.232 | 0.184 | 1 | 15 | Nrp1 | 0.048 | Young |
| 0.554262 | 0.232767 | 0.232 | 0.184 | 1 | 15 | Plaa | 0.048 | Young |
| 0.554448 | 1.155975 | 0.161 | 0.122 | 1 | 15 | D16Ertd47 | 0.039 | Young |
| 0.554448 | 0.561075 | 0.161 | 0.122 | 1 | 15 | Sema6d | 0.039 | Young |
| 0.555538 | 0.598323 | 0.25 | 0.204 | 1 | 15 | Otud5 | 0.046 | Young |
| 0.555538 | 0.265392 | 0.25 | 0.204 | 1 | 15 | Hfe | 0.046 | Young |
| 0.555608 | 0.364531 | 0.143 | 0.102 | 1 | 15 | Stk25 | 0.041 | Young |
| 0.555608 | 0.354839 | 0.143 | 0.102 | 1 | 15 | Ms4a6b | 0.041 | Young |
| 0.555608 | 0.272368 | 0.143 | 0.102 | 1 | 15 | Zfp322a | 0.041 | Young |
| 0.555608 | 0.15276 | 0.143 | 0.102 | 1 | 15 | Pdpk1 | 0.041 | Young |
| 0.555608 | 0.123603 | 0.143 | 0.102 | 1 | 15 | Nop58 | 0.041 | Young |
| 0.556417 | 0.479394 | 0.464 | 0.429 | 1 | 15 | Tm9sf3 | 0.035 | Young |
| 0.557964 | 0.814803 | 0.179 | 0.143 | 1 | 15 | Gtf3c6 | 0.036 | Young |
| 0.557966 | 0.087106 | 0.214 | 0.163 | 1 | 15 | Atf6 | 0.051 | Young |
| 0.557966 | 0.055608 | 0.161 | 0.224 | 1 | 15 | Trappc6b | -0.063 | Young |
| 0.559051 | 0.059222 | 0.482 | 0.388 | 1 | 15 | Etv6 | 0.094 | Young |
| 0.559149 | 0.60125 | 0.304 | 0.265 | 1 | 15 | Rbm7 | 0.039 | Young |
| 0.55942 | 0.100898 | 0.161 | 0.204 | 1 | 15 | Rere | -0.043 | Young |
| 0.559776 | 0.179752 | 0.714 | 0.673 | 1 | 15 | Rab21 | 0.041 | Young |
| 0.559821 | 0.194365 | 0.036 | 0.061 | 1 | 15 | Tmem238 | -0.025 | Young |
| 0.559821 | 0.14512 | 0.036 | 0.061 | 1 | 15 | Akr1c13 | -0.025 | Young |
| 0.559821 | 0.137852 | 0.036 | 0.061 | 1 | 15 | Sufu | -0.025 | Young |
| 0.559821 | 0.110526 | 0.036 | 0.061 | 1 | 15 | Rad1 | -0.025 | Young |
| 0.559821 | 0.098139 | 0.036 | 0.061 | 1 | 15 | Rbm19 | -0.025 | Young |
| 0.559821 | 0.093764 | 0.036 | 0.061 | 1 | 15 | Acads | -0.025 | Young |
| 0.560659 | 0.563554 | 0.286 | 0.245 | 1 | 15 | Ddx24 | 0.041 | Young |
| 0.560958 | 0.045409 | 0.964 | 0.878 | 1 | 15 | Fth1 | 0.086 | Young |
| 0.561459 | 0.392931 | 0.268 | 0.224 | 1 | 15 | Txnl1 | 0.044 | Young |
| 0.561542 | 0.839527 | 0.161 | 0.122 | 1 | 15 | Vps18 | 0.039 | Young |
| 0.561542 | 0.748154 | 0.161 | 0.122 | 1 | 15 | Mfsd5 | 0.039 | Young |
| 0.561542 | 0.404992 | 0.161 | 0.122 | 1 | 15 | Rtl5 | 0.039 | Young |
| 0.562278 | 1.547661 | 0.089 | 0.061 | 1 | 15 | Mrps28 | 0.028 | Young |
| 0.562278 | 1.521181 | 0.089 | 0.061 | 1 | 15 | Siah2 | 0.028 | Young |
| 0.562278 | 1.349249 | 0.089 | 0.061 | 1 | 15 | Bnip3 | 0.028 | Young |
| 0.562278 | 1.189304 | 0.089 | 0.061 | 1 | 15 | Hpse | 0.028 | Young |
| 0.562278 | 1.166077 | 0.089 | 0.061 | 1 | 15 | Cp | 0.028 | Young |
| 0.562278 | 1.124265 | 0.089 | 0.061 | 1 | 15 | Pan2 | 0.028 | Young |
| 0.562278 | 0.990176 | 0.089 | 0.061 | 1 | 15 | Rbbp5 | 0.028 | Young |
| 0.562278 | 0.891513 | 0.089 | 0.061 | 1 | 15 | Trappc3 | 0.028 | Young |
| 0.562278 | 0.782057 | 0.089 | 0.061 | 1 | 15 | Brms1 | 0.028 | Young |
| 0.562278 | 0.54122 | 0.089 | 0.061 | 1 | 15 | Cfap20 | 0.028 | Young |
| 0.564582 | 0.072081 | 0.321 | 0.245 | 1 | 15 | Klhl24 | 0.076 | Young |
| 0.564714 | 0.358138 | 0.179 | 0.143 | 1 | 15 | Med11 | 0.036 | Young |
| 0.565527 | 0.38441 | 0.214 | 0.184 | 1 | 15 | Slc25a24 | 0.03 | Young |
| 0.565606 | 0.401172 | 0.5 | 0.49 | 1 | 15 | Hmgb1 | 0.01 | Young |

| | | | | | | | | |
|---|---|---|---|---|---|---|---|---|
| 0.566233 | 0.551582 | 0.286 | 0.245 | 1 | 15 | Spty2d1 | 0.041 | Young |
| 0.566233 | 0.23839 | 0.286 | 0.245 | 1 | 15 | Angel2 | 0.041 | Young |
| 0.567311 | 0.419131 | 0.25 | 0.204 | 1 | 15 | Wsb2 | 0.046 | Young |
| 0.567311 | 0.182993 | 0.25 | 0.204 | 1 | 15 | Hspa9 | 0.046 | Young |
| 0.567363 | 0.682953 | 0.25 | 0.224 | 1 | 15 | Chd8 | 0.026 | Young |
| 0.568632 | 0.131102 | 0.125 | 0.184 | 1 | 15 | Pnrc2 | -0.059 | Young |
| 0.56868 | 0.674572 | 0.161 | 0.122 | 1 | 15 | Usp12 | 0.039 | Young |
| 0.56868 | 0.526274 | 0.161 | 0.122 | 1 | 15 | Bmf | 0.039 | Young |
| 0.57004 | 0.307432 | 0.304 | 0.265 | 1 | 15 | Mbnl2 | 0.039 | Young |
| 0.570765 | 0.225892 | 0.143 | 0.102 | 1 | 15 | Rit1 | 0.041 | Young |
| 0.570765 | 0.069563 | 0.107 | 0.143 | 1 | 15 | Pum3 | -0.036 | Young |
| 0.571503 | 0.443798 | 0.179 | 0.143 | 1 | 15 | Zbtb1 | 0.036 | Young |
| 0.571596 | 0.604817 | 0.036 | 0.061 | 1 | 15 | Nol10 | -0.025 | Young |
| 0.571596 | 0.478568 | 0.036 | 0.061 | 1 | 15 | Bloc1s1 | -0.025 | Young |
| 0.571596 | 0.444338 | 0.036 | 0.061 | 1 | 15 | Slamf6 | -0.025 | Young |
| 0.571596 | 0.349417 | 0.036 | 0.061 | 1 | 15 | Med14 | -0.025 | Young |
| 0.571596 | 0.29262 | 0.036 | 0.061 | 1 | 15 | Cops7b | -0.025 | Young |
| 0.571596 | 0.237652 | 0.036 | 0.061 | 1 | 15 | E330009J0 | -0.025 | Young |
| 0.571596 | 0.197073 | 0.036 | 0.061 | 1 | 15 | Dffa | -0.025 | Young |
| 0.571596 | 0.135373 | 0.036 | 0.061 | 1 | 15 | Caap1 | -0.025 | Young |
| 0.571596 | 0.066109 | 0.036 | 0.061 | 1 | 15 | Ulbp1 | -0.025 | Young |
| 0.571596 | 0.047248 | 0.036 | 0.061 | 1 | 15 | Ptgir | -0.025 | Young |
| 0.571596 | 0.037688 | 0.036 | 0.061 | 1 | 15 | Stmn1 | -0.025 | Young |
| 0.571735 | 1.222291 | 0.089 | 0.061 | 1 | 15 | Lmbrd2 | 0.028 | Young |
| 0.571735 | 0.985436 | 0.089 | 0.061 | 1 | 15 | 1600020E0 | 0.028 | Young |
| 0.571735 | 0.954085 | 0.089 | 0.061 | 1 | 15 | Thyn1 | 0.028 | Young |
| 0.571735 | 0.941315 | 0.089 | 0.061 | 1 | 15 | Ppp1r16a | 0.028 | Young |
| 0.571735 | 0.801845 | 0.089 | 0.061 | 1 | 15 | Rab3d | 0.028 | Young |
| 0.571735 | 0.791035 | 0.089 | 0.061 | 1 | 15 | Drg1 | 0.028 | Young |
| 0.571735 | 0.768462 | 0.089 | 0.061 | 1 | 15 | Naglu | 0.028 | Young |
| 0.571735 | 0.737974 | 0.089 | 0.061 | 1 | 15 | Bad | 0.028 | Young |
| 0.571735 | 0.603853 | 0.089 | 0.061 | 1 | 15 | Camsap1 | 0.028 | Young |
| 0.571735 | 0.577705 | 0.089 | 0.061 | 1 | 15 | Fam219a | 0.028 | Young |
| 0.57237 | 1.264187 | 0.196 | 0.163 | 1 | 15 | Ggh | 0.033 | Young |
| 0.57237 | 0.224366 | 0.161 | 0.204 | 1 | 15 | Rsrc1 | -0.043 | Young |
| 0.572436 | 0.089128 | 0.054 | 0.082 | 1 | 15 | Sf3a2 | -0.028 | Young |
| 0.57285 | 0.087145 | 0.696 | 0.612 | 1 | 15 | Arpc3 | 0.084 | Young |
| 0.573544 | 0.253366 | 0.875 | 0.837 | 1 | 15 | Jund | 0.038 | Young |
| 0.57396 | 0.187713 | 0.071 | 0.102 | 1 | 15 | Sla | -0.031 | Young |
| 0.574952 | 0.514127 | 0.5 | 0.49 | 1 | 15 | Snx3 | 0.01 | Young |
| 0.575208 | 0.318525 | 0.571 | 0.551 | 1 | 15 | Rap2a | 0.02 | Young |
| 0.575861 | 0.667449 | 0.161 | 0.122 | 1 | 15 | Srp68 | 0.039 | Young |
| 0.575861 | 0.657695 | 0.161 | 0.122 | 1 | 15 | Atg16l1 | 0.039 | Young |
| 0.578331 | 0.753413 | 0.179 | 0.143 | 1 | 15 | Ctdspl2 | 0.036 | Young |
| 0.578331 | 0.700022 | 0.179 | 0.143 | 1 | 15 | Sae1 | 0.036 | Young |

| | | | | | | | | |
|---|---|---|---|---|---|---|---|---|
| 0.578402 | 0.125511 | 0.929 | 0.918 | 1 | 15 | Cyba | 0.011 | Young |
| 0.578417 | 0.37798 | 0.143 | 0.102 | 1 | 15 | Prpsap1 | 0.041 | Young |
| 0.578417 | 0.212875 | 0.143 | 0.102 | 1 | 15 | Abhd17a | 0.041 | Young |
| 0.578775 | 0.035637 | 0.429 | 0.408 | 1 | 15 | Cggbp1 | 0.021 | Young |
| 0.581073 | 0.057178 | 0.5 | 0.449 | 1 | 15 | Tnpo1 | 0.051 | Young |
| 0.581267 | 1.016399 | 0.089 | 0.061 | 1 | 15 | Naa30 | 0.028 | Young |
| 0.581267 | 0.963656 | 0.089 | 0.061 | 1 | 15 | Lysmd3 | 0.028 | Young |
| 0.581267 | 0.708585 | 0.089 | 0.061 | 1 | 15 | Zfp422 | 0.028 | Young |
| 0.581267 | 0.642055 | 0.089 | 0.061 | 1 | 15 | Mcu | 0.028 | Young |
| 0.581267 | 0.636564 | 0.089 | 0.061 | 1 | 15 | Tnfaip8l2 | 0.028 | Young |
| 0.581267 | 0.630182 | 0.089 | 0.061 | 1 | 15 | Letm2 | 0.028 | Young |
| 0.581267 | 0.570918 | 0.089 | 0.061 | 1 | 15 | Mrpl11 | 0.028 | Young |
| 0.581949 | 0.063344 | 0.607 | 0.551 | 1 | 15 | Tnfrsf1b | 0.056 | Young |
| 0.582129 | 0.091673 | 0.196 | 0.245 | 1 | 15 | Agpat4 | -0.049 | Young |
| 0.582585 | 0.356832 | 0.054 | 0.082 | 1 | 15 | Cdk8 | -0.028 | Young |
| 0.583006 | 0.590544 | 0.071 | 0.102 | 1 | 15 | Abraxas2 | -0.031 | Young |
| 0.583084 | 0.418671 | 0.161 | 0.122 | 1 | 15 | Rps6ka1 | 0.039 | Young |
| 0.583084 | 0.293765 | 0.161 | 0.122 | 1 | 15 | Napg | 0.039 | Young |
| 0.583272 | 1.452159 | 0.107 | 0.082 | 1 | 15 | Cd200r1 | 0.025 | Young |
| 0.583272 | 1.261773 | 0.107 | 0.082 | 1 | 15 | Pmpcb | 0.025 | Young |
| 0.58331 | 0.165193 | 0.75 | 0.694 | 1 | 15 | M6pr | 0.056 | Young |
| 0.583489 | 1.930237 | 0.036 | 0.061 | 1 | 15 | D030056L2 | -0.025 | Young |
| 0.583489 | 1.418091 | 0.036 | 0.061 | 1 | 15 | Casz1 | -0.025 | Young |
| 0.583489 | 1.250441 | 0.036 | 0.061 | 1 | 15 | Drg2 | -0.025 | Young |
| 0.583489 | 1.051022 | 0.036 | 0.061 | 1 | 15 | Tgds | -0.025 | Young |
| 0.583489 | 1.002519 | 0.036 | 0.061 | 1 | 15 | Deptor | -0.025 | Young |
| 0.583489 | 0.888489 | 0.036 | 0.061 | 1 | 15 | Usp45 | -0.025 | Young |
| 0.583489 | 0.792542 | 0.036 | 0.061 | 1 | 15 | Sp2 | -0.025 | Young |
| 0.583489 | 0.683758 | 0.036 | 0.061 | 1 | 15 | Borcs7 | -0.025 | Young |
| 0.583489 | 0.618733 | 0.036 | 0.061 | 1 | 15 | Zbtb40 | -0.025 | Young |
| 0.583489 | 0.472716 | 0.036 | 0.061 | 1 | 15 | 2610524H( | -0.025 | Young |
| 0.583489 | 0.363512 | 0.036 | 0.061 | 1 | 15 | Utp14b | -0.025 | Young |
| 0.583489 | 0.35126 | 0.036 | 0.061 | 1 | 15 | Zfp296 | -0.025 | Young |
| 0.583489 | 0.181969 | 0.036 | 0.061 | 1 | 15 | Fcgr1 | -0.025 | Young |
| 0.583489 | 0.176575 | 0.036 | 0.061 | 1 | 15 | Fancc | -0.025 | Young |
| 0.583489 | 0.137302 | 0.036 | 0.061 | 1 | 15 | Mtif2 | -0.025 | Young |
| 0.583489 | 0.136873 | 0.036 | 0.061 | 1 | 15 | Rps6kc1 | -0.025 | Young |
| 0.583489 | 0.132582 | 0.036 | 0.061 | 1 | 15 | Tmc6 | -0.025 | Young |
| 0.584068 | 0.321406 | 0.536 | 0.51 | 1 | 15 | Tmem39a | 0.026 | Young |
| 0.584332 | 0.902093 | 0.214 | 0.184 | 1 | 15 | Plgrkt | 0.03 | Young |
| 0.5849 | 0.261981 | 0.232 | 0.184 | 1 | 15 | Hnrnph3 | 0.048 | Young |
| 0.58507 | 0.113833 | 0.982 | 0.918 | 1 | 15 | Marcksl1 | 0.064 | Young |
| 0.585197 | 0.552182 | 0.179 | 0.143 | 1 | 15 | Bcl7b | 0.036 | Young |
| 0.585246 | 0.18889 | 0.321 | 0.265 | 1 | 15 | Stxbp2 | 0.056 | Young |
| 0.58546 | 0.423424 | 0.161 | 0.204 | 1 | 15 | Rnf20 | -0.043 | Young |

| | | | | | | | | |
|---|---|---|---|---|---|---|---|---|
| 0.586169 | 0.141321 | 0.661 | 0.531 | 1 | 15 | Myo1g | 0.13 | Young |
| 0.586387 | 0.121634 | 0.714 | 0.673 | 1 | 15 | Cotl1 | 0.041 | Young |
| 0.586406 | 0.106591 | 0.554 | 0.49 | 1 | 15 | Stat1 | 0.064 | Young |
| 0.587373 | 0.201922 | 0.375 | 0.306 | 1 | 15 | Nudt17 | 0.069 | Young |
| 0.58752 | 0.067659 | 0.607 | 0.571 | 1 | 15 | Litaf | 0.036 | Young |
| 0.587871 | 0.095938 | 0.589 | 0.51 | 1 | 15 | Rpl22l1 | 0.079 | Young |
| 0.587914 | 0.151366 | 0.375 | 0.327 | 1 | 15 | Arhgef2 | 0.048 | Young |
| 0.587987 | 0.024239 | 0.268 | 0.327 | 1 | 15 | Csnk2a1 | -0.059 | Young |
| 0.588332 | 0.491024 | 0.375 | 0.347 | 1 | 15 | Adipor1 | 0.028 | Young |
| 0.588338 | 0.2479 | 0.446 | 0.408 | 1 | 15 | Ankrd33b | 0.038 | Young |
| 0.588905 | 0.145598 | 0.411 | 0.388 | 1 | 15 | Il10ra | 0.023 | Young |
| 0.590068 | 0.080273 | 0.536 | 0.531 | 1 | 15 | Pgls | 0.005 | Young |
| 0.590102 | 0.167468 | 0.571 | 0.571 | 1 | 15 | 2900097C1 | 0 | Young |
| 0.5903 | 0.144473 | 0.339 | 0.286 | 1 | 15 | Rnf130 | 0.053 | Young |
| 0.590349 | 0.122136 | 0.161 | 0.122 | 1 | 15 | Pdlim1 | 0.039 | Young |
| 0.590872 | 1.061201 | 0.089 | 0.061 | 1 | 15 | Pcsk7 | 0.028 | Young |
| 0.590872 | 1.023932 | 0.089 | 0.061 | 1 | 15 | Letmd1 | 0.028 | Young |
| 0.590872 | 0.92737 | 0.089 | 0.061 | 1 | 15 | Lemd3 | 0.028 | Young |
| 0.590872 | 0.912209 | 0.089 | 0.061 | 1 | 15 | Ceacam1 | 0.028 | Young |
| 0.590872 | 0.696266 | 0.089 | 0.061 | 1 | 15 | Ubxn7 | 0.028 | Young |
| 0.590872 | 0.648103 | 0.089 | 0.061 | 1 | 15 | Rhof | 0.028 | Young |
| 0.590872 | 0.631528 | 0.089 | 0.061 | 1 | 15 | Smg9 | 0.028 | Young |
| 0.590872 | 0.56186 | 0.089 | 0.061 | 1 | 15 | Prorsd1 | 0.028 | Young |
| 0.590872 | 0.552069 | 0.089 | 0.061 | 1 | 15 | Cyp4v3 | 0.028 | Young |
| 0.590872 | 0.541415 | 0.089 | 0.061 | 1 | 15 | Rbm38 | 0.028 | Young |
| 0.590872 | 0.453661 | 0.089 | 0.061 | 1 | 15 | Ttf1 | 0.028 | Young |
| 0.590872 | 0.382442 | 0.089 | 0.061 | 1 | 15 | P2rx5 | 0.028 | Young |
| 0.591959 | 1.992203 | 0.107 | 0.082 | 1 | 15 | Tpgs1 | 0.025 | Young |
| 0.591959 | 1.312475 | 0.107 | 0.082 | 1 | 15 | Bcl2l14 | 0.025 | Young |
| 0.592057 | 0.267592 | 0.196 | 0.163 | 1 | 15 | Sigirr | 0.033 | Young |
| 0.592101 | 1.065334 | 0.179 | 0.143 | 1 | 15 | Ireb2 | 0.036 | Young |
| 0.592101 | 0.561542 | 0.179 | 0.143 | 1 | 15 | Bccip | 0.036 | Young |
| 0.592101 | 0.455277 | 0.179 | 0.143 | 1 | 15 | Rnaseh2b | 0.036 | Young |
| 0.592117 | 0.367851 | 0.071 | 0.102 | 1 | 15 | Arl6ip4 | -0.031 | Young |
| 0.592117 | 0.007313 | 0.071 | 0.102 | 1 | 15 | Prr12 | -0.031 | Young |
| 0.592817 | 0.394196 | 0.054 | 0.082 | 1 | 15 | A4galt | -0.028 | Young |
| 0.592817 | 0.130751 | 0.054 | 0.082 | 1 | 15 | Lsm1 | -0.028 | Young |
| 0.593863 | 0.018882 | 0.143 | 0.102 | 1 | 15 | Slc7a8 | 0.041 | Young |
| 0.594932 | 0.061063 | 0.089 | 0.122 | 1 | 15 | E2f5 | -0.033 | Young |
| 0.596247 | 0.168589 | 0.321 | 0.265 | 1 | 15 | Sgms1 | 0.056 | Young |
| 0.596788 | 1.470586 | 0.125 | 0.102 | 1 | 15 | Polr3k | 0.023 | Young |
| 0.597028 | 0.45115 | 0.214 | 0.184 | 1 | 15 | Ppp1cc | 0.03 | Young |
| 0.597372 | 0.217339 | 0.232 | 0.184 | 1 | 15 | Sap18 | 0.048 | Young |
| 0.597656 | 0.702265 | 0.161 | 0.122 | 1 | 15 | Zcchc17 | 0.039 | Young |
| 0.597701 | 0.330487 | 0.304 | 0.265 | 1 | 15 | Cd14 | 0.039 | Young |

| | | | | | | | | |
|---|---|---|---|---|---|---|---|---|
| 0.598663 | 0.100848 | 0.393 | 0.327 | 1 | 15 | Lmna | 0.066 | Young |
| 0.598687 | 0.359328 | 0.196 | 0.163 | 1 | 15 | Lemd2 | 0.033 | Young |
| 0.599024 | 0.260335 | 0.411 | 0.367 | 1 | 15 | Gatad2b | 0.044 | Young |
| 0.599041 | 0.658531 | 0.179 | 0.143 | 1 | 15 | Anpep | 0.036 | Young |
| 0.60055 | 0.92036 | 0.089 | 0.061 | 1 | 15 | Fbxw5 | 0.028 | Young |
| 0.60055 | 0.626651 | 0.089 | 0.061 | 1 | 15 | Ticam2 | 0.028 | Young |
| 0.60055 | 0.455087 | 0.089 | 0.061 | 1 | 15 | Aoah | 0.028 | Young |
| 0.60055 | 0.417858 | 0.089 | 0.061 | 1 | 15 | Hras | 0.028 | Young |
| 0.600585 | 0.125015 | 0.946 | 0.918 | 1 | 15 | Mcl1 | 0.028 | Young |
| 0.600704 | 0.81695 | 0.107 | 0.082 | 1 | 15 | Vps53 | 0.025 | Young |
| 0.601117 | 0.206718 | 0.339 | 0.286 | 1 | 15 | Cdc5l | 0.053 | Young |
| 0.60313 | 0.403576 | 0.054 | 0.082 | 1 | 15 | Ctns | -0.028 | Young |
| 0.60313 | 0.358353 | 0.054 | 0.082 | 1 | 15 | Akt1s1 | -0.028 | Young |
| 0.60313 | 0.000988 | 0.054 | 0.082 | 1 | 15 | Rnf185 | -0.028 | Young |
| 0.603148 | 0.181482 | 0.357 | 0.327 | 1 | 15 | Tubb6 | 0.03 | Young |
| 0.603653 | 0.640326 | 0.179 | 0.245 | 1 | 15 | Cd2bp2 | -0.066 | Young |
| 0.603938 | 0.357539 | 0.679 | 0.694 | 1 | 15 | Slc38a2 | -0.015 | Young |
| 0.604925 | 0.162926 | 0.732 | 0.673 | 1 | 15 | Ncf1 | 0.059 | Young |
| 0.604959 | 0.12673 | 0.929 | 0.878 | 1 | 15 | Atox1 | 0.051 | Young |
| 0.605003 | 0.168984 | 0.161 | 0.122 | 1 | 15 | Smc4 | 0.039 | Young |
| 0.605095 | 0.066654 | 0.946 | 0.959 | 1 | 15 | Dusp1 | -0.013 | Young |
| 0.605112 | 0.047788 | 0.982 | 0.98 | 1 | 15 | Rps19 | 0.002 | Young |
| 0.605113 | 0.098103 | 1 | 0.959 | 1 | 15 | Rplp0 | 0.041 | Young |
| 0.605351 | 1.062373 | 0.196 | 0.163 | 1 | 15 | Cnot2 | 0.033 | Young |
| 0.607342 | 0.094492 | 0.321 | 0.265 | 1 | 15 | Mt2 | 0.056 | Young |
| 0.607715 | 0.19529 | 0.839 | 0.714 | 1 | 15 | Ucp2 | 0.125 | Young |
| 0.609066 | 0.123493 | 0.375 | 0.429 | 1 | 15 | Rbm17 | -0.054 | Young |
| 0.609432 | 0.06943 | 0.25 | 0.204 | 1 | 15 | Trim27 | 0.046 | Young |
| 0.609508 | 1.293366 | 0.107 | 0.082 | 1 | 15 | Thap11 | 0.025 | Young |
| 0.609508 | 1.185804 | 0.107 | 0.082 | 1 | 15 | Hlx | 0.025 | Young |
| 0.609508 | 1.082044 | 0.107 | 0.082 | 1 | 15 | Wdr43 | 0.025 | Young |
| 0.609508 | 1.036429 | 0.107 | 0.082 | 1 | 15 | Cyc1 | 0.025 | Young |
| 0.609508 | 0.809554 | 0.107 | 0.082 | 1 | 15 | D730003I1 | 0.025 | Young |
| 0.609963 | 0.251992 | 0.232 | 0.184 | 1 | 15 | Rnft1 | 0.048 | Young |
| 0.610299 | 0.403214 | 0.089 | 0.061 | 1 | 15 | Tomm6 | 0.028 | Young |
| 0.610299 | 0.360294 | 0.089 | 0.061 | 1 | 15 | Cldnd1 | 0.028 | Young |
| 0.610299 | 0.341261 | 0.089 | 0.061 | 1 | 15 | Rhbdf2 | 0.028 | Young |
| 0.610299 | 0.267834 | 0.089 | 0.061 | 1 | 15 | Abi2 | 0.028 | Young |
| 0.610533 | 0.263719 | 0.071 | 0.102 | 1 | 15 | Dnajc11 | -0.031 | Young |
| 0.610533 | 0.075348 | 0.071 | 0.102 | 1 | 15 | Anapc15 | -0.031 | Young |
| 0.611357 | 0.117483 | 0.429 | 0.388 | 1 | 15 | Srebf2 | 0.041 | Young |
| 0.61178 | 1.092062 | 0.089 | 0.122 | 1 | 15 | Usp32 | -0.033 | Young |
| 0.612048 | 0.541962 | 0.196 | 0.163 | 1 | 15 | Pdlim4 | 0.033 | Young |
| 0.612048 | 0.521423 | 0.196 | 0.163 | 1 | 15 | Snx27 | 0.033 | Young |
| 0.612391 | 0.821409 | 0.161 | 0.122 | 1 | 15 | Osm | 0.039 | Young |

| | | | | | | | | |
|---|---|---|---|---|---|---|---|---|
| 0.612391 | 0.188793 | 0.161 | 0.122 | 1 | 15 | Phrf1 | 0.039 | Young |
| 0.612391 | 0.115097 | 0.161 | 0.122 | 1 | 15 | Lage3 | 0.039 | Young |
| 0.613017 | 1.500457 | 0.125 | 0.102 | 1 | 15 | Apool | 0.023 | Young |
| 0.613032 | 0.285858 | 0.179 | 0.143 | 1 | 15 | Gtf2a2 | 0.036 | Young |
| 0.613032 | 0.034902 | 0.179 | 0.143 | 1 | 15 | Adcy7 | 0.036 | Young |
| 0.613048 | 0.219204 | 0.518 | 0.469 | 1 | 15 | Anxa4 | 0.049 | Young |
| 0.613524 | 0.615117 | 0.054 | 0.082 | 1 | 15 | Aasdhppt | -0.028 | Young |
| 0.613524 | 0.360307 | 0.054 | 0.082 | 1 | 15 | B3gnt5 | -0.028 | Young |
| 0.613524 | 0.268744 | 0.054 | 0.082 | 1 | 15 | Retreg1 | -0.028 | Young |
| 0.613524 | 0.131846 | 0.054 | 0.082 | 1 | 15 | Arfip1 | -0.028 | Young |
| 0.613524 | 0.047855 | 0.054 | 0.082 | 1 | 15 | Gpn1 | -0.028 | Young |
| 0.613538 | 0.440922 | 0.429 | 0.408 | 1 | 15 | Edf1 | 0.021 | Young |
| 0.613993 | 0.342127 | 0.268 | 0.224 | 1 | 15 | Trim8 | 0.044 | Young |
| 0.61487 | 0.515322 | 0.25 | 0.224 | 1 | 15 | Ppp6c | 0.026 | Young |
| 0.616304 | 0.468082 | 0.214 | 0.184 | 1 | 15 | Fam102b | 0.03 | Young |
| 0.616336 | 0.100118 | 0.946 | 0.939 | 1 | 15 | Lamp1 | 0.007 | Young |
| 0.61738 | 0.17523 | 0.107 | 0.143 | 1 | 15 | Prdx3 | -0.036 | Young |
| 0.617548 | 0.166172 | 0.286 | 0.245 | 1 | 15 | Pptc7 | 0.041 | Young |
| 0.61759 | 0.356441 | 0.857 | 0.776 | 1 | 15 | Ptms | 0.081 | Young |
| 0.618368 | 1.117722 | 0.107 | 0.082 | 1 | 15 | Crnkl1 | 0.025 | Young |
| 0.618368 | 0.851484 | 0.107 | 0.082 | 1 | 15 | Trem2 | 0.025 | Young |
| 0.618778 | 0.481074 | 0.196 | 0.163 | 1 | 15 | Gatad2a | 0.033 | Young |
| 0.618778 | 0.423904 | 0.196 | 0.163 | 1 | 15 | Trim34a | 0.033 | Young |
| 0.619 | 0.133275 | 0.304 | 0.245 | 1 | 15 | Pacs2 | 0.059 | Young |
| 0.619111 | 0.607919 | 0.357 | 0.327 | 1 | 15 | Trpc4ap | 0.03 | Young |
| 0.619569 | 0.227854 | 0.375 | 0.347 | 1 | 15 | Wsb1 | 0.028 | Young |
| 0.619818 | 0.176652 | 0.161 | 0.122 | 1 | 15 | Eftud2 | 0.039 | Young |
| 0.619835 | 0.702201 | 0.071 | 0.102 | 1 | 15 | Rbfa | -0.031 | Young |
| 0.619835 | 0.223714 | 0.071 | 0.102 | 1 | 15 | Dtnbp1 | -0.031 | Young |
| 0.619835 | 0.119014 | 0.071 | 0.102 | 1 | 15 | Aacs | -0.031 | Young |
| 0.619835 | 0.02062 | 0.071 | 0.102 | 1 | 15 | Zfand1 | -0.031 | Young |
| 0.620081 | 0.308244 | 0.179 | 0.143 | 1 | 15 | Gm8186 | 0.036 | Young |
| 0.620117 | 0.506981 | 0.089 | 0.061 | 1 | 15 | Gins4 | 0.028 | Young |
| 0.620117 | 0.215577 | 0.089 | 0.061 | 1 | 15 | Rab28 | 0.028 | Young |
| 0.620117 | 0.189511 | 0.089 | 0.061 | 1 | 15 | Gab1 | 0.028 | Young |
| 0.620117 | 0.153692 | 0.089 | 0.061 | 1 | 15 | Camsap2 | 0.028 | Young |
| 0.620117 | 0.012399 | 0.089 | 0.061 | 1 | 15 | Bid | 0.028 | Young |
| 0.620282 | 0.216816 | 0.089 | 0.122 | 1 | 15 | AU020206 | -0.033 | Young |
| 0.622587 | 0.304877 | 0.446 | 0.429 | 1 | 15 | Trafd1 | 0.017 | Young |
| 0.622755 | 0.088158 | 0.911 | 0.816 | 1 | 15 | Traf1 | 0.095 | Young |
| 0.623996 | 0.435879 | 0.054 | 0.082 | 1 | 15 | Hacd2 | -0.028 | Young |
| 0.623996 | 0.042233 | 0.054 | 0.082 | 1 | 15 | Bin2 | -0.028 | Young |
| 0.624155 | 0.357835 | 0.321 | 0.265 | 1 | 15 | Eif3g | 0.056 | Young |
| 0.624385 | 0.291787 | 0.429 | 0.367 | 1 | 15 | Ier3 | 0.062 | Young |
| 0.62476 | 0.082326 | 0.786 | 0.694 | 1 | 15 | Tnfaip8 | 0.092 | Young |

| | | | | | | | | |
|---|---|---|---|---|---|---|---|---|
| 0.625308 | 0.603953 | 0.107 | 0.143 | 1 | 15 | Dgat1 | -0.036 | Young |
| 0.625308 | 0.186276 | 0.107 | 0.143 | 1 | 15 | Cln5 | -0.036 | Young |
| 0.62554 | 0.7803 | 0.196 | 0.163 | 1 | 15 | St3gal3 | 0.033 | Young |
| 0.627166 | 0.770638 | 0.179 | 0.143 | 1 | 15 | Slc30a5 | 0.036 | Young |
| 0.627166 | 0.180269 | 0.179 | 0.143 | 1 | 15 | Fut4 | 0.036 | Young |
| 0.627284 | 1.219932 | 0.107 | 0.082 | 1 | 15 | Gng12 | 0.025 | Young |
| 0.627284 | 0.778028 | 0.107 | 0.082 | 1 | 15 | Golt1b | 0.025 | Young |
| 0.627284 | 0.710197 | 0.107 | 0.082 | 1 | 15 | Pkd2 | 0.025 | Young |
| 0.627284 | 0.685139 | 0.107 | 0.082 | 1 | 15 | Gm47662 | 0.025 | Young |
| 0.627284 | 0.552285 | 0.107 | 0.082 | 1 | 15 | Pprc1 | 0.025 | Young |
| 0.627284 | 0.488332 | 0.107 | 0.082 | 1 | 15 | Itgb7 | 0.025 | Young |
| 0.627648 | 0.447373 | 0.286 | 0.265 | 1 | 15 | Amfr | 0.021 | Young |
| 0.627835 | 0.151179 | 0.321 | 0.286 | 1 | 15 | Wwp1 | 0.035 | Young |
| 0.628836 | 0.891481 | 0.089 | 0.122 | 1 | 15 | Inpp5f | -0.033 | Young |
| 0.628836 | 0.298825 | 0.089 | 0.122 | 1 | 15 | BC037034 | -0.033 | Young |
| 0.629067 | 0.40426 | 0.232 | 0.184 | 1 | 15 | Gnl1 | 0.048 | Young |
| 0.629199 | 0.2433 | 0.071 | 0.102 | 1 | 15 | Susd3 | -0.031 | Young |
| 0.629199 | 0.208088 | 0.071 | 0.102 | 1 | 15 | Zfp738 | -0.031 | Young |
| 0.629257 | 0.045639 | 0.661 | 0.633 | 1 | 15 | Tmcc3 | 0.028 | Young |
| 0.629964 | 0.061074 | 0.946 | 0.939 | 1 | 15 | Rps16 | 0.007 | Young |
| 0.630004 | 0.048678 | 0.089 | 0.061 | 1 | 15 | Afg3l1 | 0.028 | Young |
| 0.630016 | 0.369431 | 0.393 | 0.367 | 1 | 15 | Ctdsp2 | 0.026 | Young |
| 0.630278 | 0.066808 | 0.589 | 0.51 | 1 | 15 | Eif3c | 0.079 | Young |
| 0.631973 | 0.339059 | 0.268 | 0.224 | 1 | 15 | Igsf9 | 0.044 | Young |
| 0.632261 | 0.073925 | 0.964 | 0.939 | 1 | 15 | Ppia | 0.025 | Young |
| 0.632297 | 3.085514 | 0.036 | 0.02 | 1 | 15 | Ankrd50 | 0.016 | Young |
| 0.632297 | 2.732497 | 0.036 | 0.02 | 1 | 15 | Dnttip1 | 0.016 | Young |
| 0.632297 | 2.649035 | 0.036 | 0.02 | 1 | 15 | Mul1 | 0.016 | Young |
| 0.632297 | 2.606058 | 0.036 | 0.02 | 1 | 15 | Bex6 | 0.016 | Young |
| 0.632297 | 2.567833 | 0.036 | 0.02 | 1 | 15 | Nbeal2 | 0.016 | Young |
| 0.632297 | 2.565045 | 0.036 | 0.02 | 1 | 15 | Surf6 | 0.016 | Young |
| 0.632297 | 2.51067 | 0.036 | 0.02 | 1 | 15 | Slc25a22 | 0.016 | Young |
| 0.632297 | 2.485743 | 0.036 | 0.02 | 1 | 15 | Xlr | 0.016 | Young |
| 0.632297 | 2.42927 | 0.036 | 0.02 | 1 | 15 | Smpd4 | 0.016 | Young |
| 0.632297 | 2.395205 | 0.036 | 0.02 | 1 | 15 | Hsh2d | 0.016 | Young |
| 0.632297 | 2.365723 | 0.036 | 0.02 | 1 | 15 | Batf2 | 0.016 | Young |
| 0.632297 | 2.297556 | 0.036 | 0.02 | 1 | 15 | Coa7 | 0.016 | Young |
| 0.632297 | 2.284629 | 0.036 | 0.02 | 1 | 15 | Dennd3 | 0.016 | Young |
| 0.632297 | 2.240397 | 0.036 | 0.02 | 1 | 15 | Ndufaf1 | 0.016 | Young |
| 0.632297 | 2.220824 | 0.036 | 0.02 | 1 | 15 | Slc16a7 | 0.016 | Young |
| 0.632297 | 2.217512 | 0.036 | 0.02 | 1 | 15 | Foxred1 | 0.016 | Young |
| 0.632297 | 2.2167 | 0.036 | 0.02 | 1 | 15 | Phykpl | 0.016 | Young |
| 0.632297 | 2.202426 | 0.036 | 0.02 | 1 | 15 | Ndufaf3 | 0.016 | Young |
| 0.632297 | 2.183818 | 0.036 | 0.02 | 1 | 15 | Kptn | 0.016 | Young |
| 0.632297 | 2.174919 | 0.036 | 0.02 | 1 | 15 | Nqo2 | 0.016 | Young |

| | | | | | | | | |
|---|---|---|---|---|---|---|---|---|
| 0.632297 | 2.122942 | 0.036 | 0.02 | 1 | 15 | Prmt7 | 0.016 | Young |
| 0.632297 | 2.0804 | 0.036 | 0.02 | 1 | 15 | Pmm1 | 0.016 | Young |
| 0.632297 | 2.075648 | 0.036 | 0.02 | 1 | 15 | Mtss1l | 0.016 | Young |
| 0.632297 | 2.073113 | 0.036 | 0.02 | 1 | 15 | Nphp3 | 0.016 | Young |
| 0.632297 | 2.008789 | 0.036 | 0.02 | 1 | 15 | Pilrb1 | 0.016 | Young |
| 0.632297 | 1.96857 | 0.036 | 0.02 | 1 | 15 | Galns | 0.016 | Young |
| 0.632297 | 1.962598 | 0.036 | 0.02 | 1 | 15 | Tpcn2 | 0.016 | Young |
| 0.632297 | 1.955332 | 0.036 | 0.02 | 1 | 15 | 1700019L1 | 0.016 | Young |
| 0.632297 | 1.955035 | 0.036 | 0.02 | 1 | 15 | Cox15 | 0.016 | Young |
| 0.632297 | 1.937069 | 0.036 | 0.02 | 1 | 15 | Nit2 | 0.016 | Young |
| 0.632297 | 1.920103 | 0.036 | 0.02 | 1 | 15 | Rnf123 | 0.016 | Young |
| 0.632297 | 1.884029 | 0.036 | 0.02 | 1 | 15 | 1300002E1 | 0.016 | Young |
| 0.632297 | 1.865967 | 0.036 | 0.02 | 1 | 15 | Mad1l1 | 0.016 | Young |
| 0.632297 | 1.865611 | 0.036 | 0.02 | 1 | 15 | C230037L1 | 0.016 | Young |
| 0.632297 | 1.859902 | 0.036 | 0.02 | 1 | 15 | Tmtc3 | 0.016 | Young |
| 0.632297 | 1.852113 | 0.036 | 0.02 | 1 | 15 | Pus3 | 0.016 | Young |
| 0.632297 | 1.844907 | 0.036 | 0.02 | 1 | 15 | Zfp386 | 0.016 | Young |
| 0.632297 | 1.844033 | 0.036 | 0.02 | 1 | 15 | Nr1d1 | 0.016 | Young |
| 0.632297 | 1.832408 | 0.036 | 0.02 | 1 | 15 | Def6 | 0.016 | Young |
| 0.632297 | 1.829924 | 0.036 | 0.02 | 1 | 15 | Gm20707 | 0.016 | Young |
| 0.632297 | 1.818097 | 0.036 | 0.02 | 1 | 15 | Cdca7l | 0.016 | Young |
| 0.632297 | 1.812865 | 0.036 | 0.02 | 1 | 15 | 1600014C1 | 0.016 | Young |
| 0.632297 | 1.806545 | 0.036 | 0.02 | 1 | 15 | Mospd1 | 0.016 | Young |
| 0.632297 | 1.783824 | 0.036 | 0.02 | 1 | 15 | 2810006K2 | 0.016 | Young |
| 0.632297 | 1.760531 | 0.036 | 0.02 | 1 | 15 | Znhit2 | 0.016 | Young |
| 0.632297 | 1.75551 | 0.036 | 0.02 | 1 | 15 | Metrn | 0.016 | Young |
| 0.632297 | 1.753961 | 0.036 | 0.02 | 1 | 15 | Mcat | 0.016 | Young |
| 0.632297 | 1.742126 | 0.036 | 0.02 | 1 | 15 | Gpsm1 | 0.016 | Young |
| 0.632297 | 1.722769 | 0.036 | 0.02 | 1 | 15 | Phc1 | 0.016 | Young |
| 0.632297 | 1.69661 | 0.036 | 0.02 | 1 | 15 | Hrh1 | 0.016 | Young |
| 0.632297 | 1.687445 | 0.036 | 0.02 | 1 | 15 | Det1 | 0.016 | Young |
| 0.632297 | 1.668545 | 0.036 | 0.02 | 1 | 15 | Ccdc86 | 0.016 | Young |
| 0.632297 | 1.667583 | 0.036 | 0.02 | 1 | 15 | Ptgs2os2 | 0.016 | Young |
| 0.632297 | 1.664439 | 0.036 | 0.02 | 1 | 15 | Smg6 | 0.016 | Young |
| 0.632297 | 1.633402 | 0.036 | 0.02 | 1 | 15 | Trim23 | 0.016 | Young |
| 0.632297 | 1.616541 | 0.036 | 0.02 | 1 | 15 | Micu3 | 0.016 | Young |
| 0.632297 | 1.606624 | 0.036 | 0.02 | 1 | 15 | Cox7a1 | 0.016 | Young |
| 0.632297 | 1.59353 | 0.036 | 0.02 | 1 | 15 | Gpr180 | 0.016 | Young |
| 0.632297 | 1.591879 | 0.036 | 0.02 | 1 | 15 | Kank3 | 0.016 | Young |
| 0.632297 | 1.57812 | 0.036 | 0.02 | 1 | 15 | Ccdc61 | 0.016 | Young |
| 0.632297 | 1.572673 | 0.036 | 0.02 | 1 | 15 | Pla2g12a | 0.016 | Young |
| 0.632297 | 1.566607 | 0.036 | 0.02 | 1 | 15 | Katnb1 | 0.016 | Young |
| 0.632297 | 1.536857 | 0.036 | 0.02 | 1 | 15 | Cgnl1 | 0.016 | Young |
| 0.632297 | 1.517095 | 0.036 | 0.02 | 1 | 15 | Pkdcc | 0.016 | Young |
| 0.632297 | 1.512573 | 0.036 | 0.02 | 1 | 15 | Setd4 | 0.016 | Young |

| | | | | | | | | |
|---|---|---|---|---|---|---|---|---|
| 0.632297 | 1.49345 | 0.036 | 0.02 | 1 | 15 | Oxnad1 | 0.016 | Young |
| 0.632297 | 1.482175 | 0.036 | 0.02 | 1 | 15 | Wdyhv1 | 0.016 | Young |
| 0.632297 | 1.478867 | 0.036 | 0.02 | 1 | 15 | Scrn3 | 0.016 | Young |
| 0.632297 | 1.475423 | 0.036 | 0.02 | 1 | 15 | Mapk7 | 0.016 | Young |
| 0.632297 | 1.468443 | 0.036 | 0.02 | 1 | 15 | Pced1a | 0.016 | Young |
| 0.632297 | 1.459892 | 0.036 | 0.02 | 1 | 15 | Homer3 | 0.016 | Young |
| 0.632297 | 1.453011 | 0.036 | 0.02 | 1 | 15 | Mrps10 | 0.016 | Young |
| 0.632297 | 1.446288 | 0.036 | 0.02 | 1 | 15 | Pbdc1 | 0.016 | Young |
| 0.632297 | 1.44294 | 0.036 | 0.02 | 1 | 15 | Ciart | 0.016 | Young |
| 0.632297 | 1.428747 | 0.036 | 0.02 | 1 | 15 | Abhd14a | 0.016 | Young |
| 0.632297 | 1.427278 | 0.036 | 0.02 | 1 | 15 | Ctdp1 | 0.016 | Young |
| 0.632297 | 1.407092 | 0.036 | 0.02 | 1 | 15 | Snta1 | 0.016 | Young |
| 0.632297 | 1.402246 | 0.036 | 0.02 | 1 | 15 | Sirt6 | 0.016 | Young |
| 0.632297 | 1.381483 | 0.036 | 0.02 | 1 | 15 | Amd2 | 0.016 | Young |
| 0.632297 | 1.381483 | 0.036 | 0.02 | 1 | 15 | Abcc3 | 0.016 | Young |
| 0.632297 | 1.339408 | 0.036 | 0.02 | 1 | 15 | Atg16l2 | 0.016 | Young |
| 0.632297 | 1.335401 | 0.036 | 0.02 | 1 | 15 | Elac2 | 0.016 | Young |
| 0.632297 | 1.318666 | 0.036 | 0.02 | 1 | 15 | Ms4a4a | 0.016 | Young |
| 0.632297 | 1.270587 | 0.036 | 0.02 | 1 | 15 | Fndc10 | 0.016 | Young |
| 0.632297 | 1.247416 | 0.036 | 0.02 | 1 | 15 | Dgkg | 0.016 | Young |
| 0.632297 | 1.246992 | 0.036 | 0.02 | 1 | 15 | Wdhd1 | 0.016 | Young |
| 0.632297 | 1.230315 | 0.036 | 0.02 | 1 | 15 | Ppp2r5b | 0.016 | Young |
| 0.632297 | 1.224232 | 0.036 | 0.02 | 1 | 15 | Homez | 0.016 | Young |
| 0.632297 | 1.205741 | 0.036 | 0.02 | 1 | 15 | Exosc2 | 0.016 | Young |
| 0.632297 | 1.193251 | 0.036 | 0.02 | 1 | 15 | Wdr92 | 0.016 | Young |
| 0.632297 | 1.17032 | 0.036 | 0.02 | 1 | 15 | Mrpl44 | 0.016 | Young |
| 0.632297 | 1.143294 | 0.036 | 0.02 | 1 | 15 | Cpsf3 | 0.016 | Young |
| 0.632297 | 1.132201 | 0.036 | 0.02 | 1 | 15 | Spink2 | 0.016 | Young |
| 0.632297 | 1.10489 | 0.036 | 0.02 | 1 | 15 | Dhx37 | 0.016 | Young |
| 0.632297 | 1.098185 | 0.036 | 0.02 | 1 | 15 | Miga2 | 0.016 | Young |
| 0.632297 | 1.094851 | 0.036 | 0.02 | 1 | 15 | A830010M | 0.016 | Young |
| 0.632297 | 1.076259 | 0.036 | 0.02 | 1 | 15 | Stra6l | 0.016 | Young |
| 0.632297 | 1.04163 | 0.036 | 0.02 | 1 | 15 | Noc4l | 0.016 | Young |
| 0.632297 | 1.016801 | 0.036 | 0.02 | 1 | 15 | Cdc23 | 0.016 | Young |
| 0.632297 | 1.006941 | 0.036 | 0.02 | 1 | 15 | Zfp704 | 0.016 | Young |
| 0.632297 | 0.972864 | 0.036 | 0.02 | 1 | 15 | Fig4 | 0.016 | Young |
| 0.632297 | 0.969755 | 0.036 | 0.02 | 1 | 15 | Oasl2 | 0.016 | Young |
| 0.632297 | 0.884506 | 0.036 | 0.02 | 1 | 15 | Sdc1 | 0.016 | Young |
| 0.632297 | 0.773824 | 0.036 | 0.02 | 1 | 15 | Tex30 | 0.016 | Young |
| 0.632297 | 0.757679 | 0.036 | 0.02 | 1 | 15 | Limk1 | 0.016 | Young |
| 0.633266 | 0.224085 | 0.607 | 0.551 | 1 | 15 | Hnrnpm | 0.056 | Young |
| 0.63328 | 0.24485 | 0.107 | 0.143 | 1 | 15 | Eif2b4 | -0.036 | Young |
| 0.634284 | 0.741587 | 0.143 | 0.184 | 1 | 15 | Trim12a | -0.041 | Young |
| 0.634284 | 0.664704 | 0.143 | 0.184 | 1 | 15 | Pik3c2a | -0.041 | Young |
| 0.634284 | 0.486229 | 0.179 | 0.143 | 1 | 15 | Cklf | 0.036 | Young |

| | | | | | | | | |
|---|---|---|---|---|---|---|---|---|
| 0.634544 | 0.860474 | 0.054 | 0.082 | 1 | 15 | Hook2 | -0.028 | Young |
| 0.634544 | 0.781403 | 0.054 | 0.082 | 1 | 15 | Spg11 | -0.028 | Young |
| 0.634544 | 0.486542 | 0.054 | 0.082 | 1 | 15 | Nat6 | -0.028 | Young |
| 0.63525 | 0.021124 | 0.411 | 0.347 | 1 | 15 | Cebpb | 0.064 | Young |
| 0.635256 | 0.08831 | 0.375 | 0.306 | 1 | 15 | Eif3l | 0.069 | Young |
| 0.635492 | 0.09113 | 0.232 | 0.184 | 1 | 15 | Maf1 | 0.048 | Young |
| 0.635848 | 0.794074 | 0.214 | 0.184 | 1 | 15 | Dpp8 | 0.03 | Young |
| 0.635848 | 0.064736 | 0.214 | 0.184 | 1 | 15 | Ezh2 | 0.03 | Young |
| 0.636255 | 0.900354 | 0.107 | 0.082 | 1 | 15 | Arsb | 0.025 | Young |
| 0.636255 | 0.696769 | 0.107 | 0.082 | 1 | 15 | Ddx19a | 0.025 | Young |
| 0.636255 | 0.650531 | 0.107 | 0.082 | 1 | 15 | Tifab | 0.025 | Young |
| 0.636255 | 0.645403 | 0.107 | 0.082 | 1 | 15 | Scoc | 0.025 | Young |
| 0.636255 | 0.440562 | 0.107 | 0.082 | 1 | 15 | Trmt1 | 0.025 | Young |
| 0.637439 | 0.281009 | 0.089 | 0.122 | 1 | 15 | Rnf31 | -0.033 | Young |
| 0.638622 | 0.8624 | 0.071 | 0.102 | 1 | 15 | Trak2 | -0.031 | Young |
| 0.638622 | 0.41897 | 0.071 | 0.102 | 1 | 15 | Slfn8 | -0.031 | Young |
| 0.638622 | 0.168334 | 0.071 | 0.102 | 1 | 15 | Hdgfl2 | -0.031 | Young |
| 0.638622 | 0.090432 | 0.071 | 0.102 | 1 | 15 | Cars2 | -0.031 | Young |
| 0.63961 | 0.475453 | 0.268 | 0.245 | 1 | 15 | Bcat2 | 0.023 | Young |
| 0.639958 | 0.770183 | 0.089 | 0.061 | 1 | 15 | Sh3pxd2b | 0.028 | Young |
| 0.639958 | 0.0091 | 0.089 | 0.061 | 1 | 15 | Hspbp1 | 0.028 | Young |
| 0.640227 | 0.053254 | 0.411 | 0.367 | 1 | 15 | Bnip3l | 0.044 | Young |
| 0.64088 | 0.127865 | 0.464 | 0.388 | 1 | 15 | Nectin1 | 0.076 | Young |
| 0.641295 | 0.189689 | 0.107 | 0.143 | 1 | 15 | Ciapin1 | -0.036 | Young |
| 0.641437 | 0.506122 | 0.179 | 0.143 | 1 | 15 | Rin2 | 0.036 | Young |
| 0.642331 | 0.503664 | 0.125 | 0.163 | 1 | 15 | Gm13212 | -0.038 | Young |
| 0.642331 | 0.062987 | 0.161 | 0.122 | 1 | 15 | Rasgrp1 | 0.039 | Young |
| 0.642331 | 0.044471 | 0.161 | 0.122 | 1 | 15 | Ccdc47 | 0.039 | Young |
| 0.64528 | 0.768855 | 0.107 | 0.082 | 1 | 15 | Ubr3 | 0.025 | Young |
| 0.64528 | 0.717469 | 0.107 | 0.082 | 1 | 15 | Gnptab | 0.025 | Young |
| 0.64528 | 0.637235 | 0.107 | 0.082 | 1 | 15 | Rcn2 | 0.025 | Young |
| 0.64528 | 0.263293 | 0.107 | 0.082 | 1 | 15 | Slc16a6 | 0.025 | Young |
| 0.64528 | 0.256701 | 0.107 | 0.082 | 1 | 15 | Adpgk | 0.025 | Young |
| 0.64528 | 0.143108 | 0.107 | 0.082 | 1 | 15 | Hlcs | 0.025 | Young |
| 0.64546 | 0.036801 | 0.411 | 0.367 | 1 | 15 | Tcea1 | 0.044 | Young |
| 0.646014 | 0.410708 | 0.196 | 0.163 | 1 | 15 | Gsk3a | 0.033 | Young |
| 0.646014 | 0.205545 | 0.196 | 0.163 | 1 | 15 | Mrpl43 | 0.033 | Young |
| 0.646045 | 1.079414 | 0.125 | 0.102 | 1 | 15 | Strn | 0.023 | Young |
| 0.646049 | 0.145093 | 0.982 | 0.959 | 1 | 15 | Cd74 | 0.023 | Young |
| 0.646092 | 0.029628 | 0.089 | 0.122 | 1 | 15 | P4ha1 | -0.033 | Young |
| 0.646219 | 0.506486 | 0.446 | 0.469 | 1 | 15 | Nptn | -0.023 | Young |
| 0.646908 | 0.305827 | 0.286 | 0.245 | 1 | 15 | Kat6a | 0.041 | Young |
| 0.648104 | 1.914593 | 0.071 | 0.102 | 1 | 15 | Nop14 | -0.031 | Young |
| 0.648104 | 0.491946 | 0.071 | 0.102 | 1 | 15 | Rnf121 | -0.031 | Young |
| 0.648217 | 1.837771 | 0.036 | 0.02 | 1 | 15 | Zfp961 | 0.016 | Young |

| | | | | | | | | |
|---|---|---|---|---|---|---|---|---|
| 0.648217 | 1.647968 | 0.036 | 0.02 | 1 | 15 | Fancg | 0.016 | Young |
| 0.648217 | 1.609066 | 0.036 | 0.02 | 1 | 15 | B3gnt3 | 0.016 | Young |
| 0.648217 | 1.600354 | 0.036 | 0.02 | 1 | 15 | Zfp709 | 0.016 | Young |
| 0.648217 | 1.583206 | 0.036 | 0.02 | 1 | 15 | Dusp14 | 0.016 | Young |
| 0.648217 | 1.548545 | 0.036 | 0.02 | 1 | 15 | Hps4 | 0.016 | Young |
| 0.648217 | 1.534012 | 0.036 | 0.02 | 1 | 15 | Pfn2 | 0.016 | Young |
| 0.648217 | 1.3949 | 0.036 | 0.02 | 1 | 15 | Cox6b2 | 0.016 | Young |
| 0.648217 | 1.364955 | 0.036 | 0.02 | 1 | 15 | Zdhhc12 | 0.016 | Young |
| 0.648217 | 1.360512 | 0.036 | 0.02 | 1 | 15 | Zfp606 | 0.016 | Young |
| 0.648217 | 1.338619 | 0.036 | 0.02 | 1 | 15 | Gys1 | 0.016 | Young |
| 0.648217 | 1.302388 | 0.036 | 0.02 | 1 | 15 | Fbxl15 | 0.016 | Young |
| 0.648217 | 1.302057 | 0.036 | 0.02 | 1 | 15 | Spg20 | 0.016 | Young |
| 0.648217 | 1.242867 | 0.036 | 0.02 | 1 | 15 | Tmem62 | 0.016 | Young |
| 0.648217 | 1.234927 | 0.036 | 0.02 | 1 | 15 | Oxsm | 0.016 | Young |
| 0.648217 | 1.18794 | 0.036 | 0.02 | 1 | 15 | Fndc4 | 0.016 | Young |
| 0.648217 | 1.180989 | 0.036 | 0.02 | 1 | 15 | Rpgrip1 | 0.016 | Young |
| 0.648217 | 1.180162 | 0.036 | 0.02 | 1 | 15 | Il17rd | 0.016 | Young |
| 0.648217 | 1.156747 | 0.036 | 0.02 | 1 | 15 | Spats2 | 0.016 | Young |
| 0.648217 | 1.111228 | 0.036 | 0.02 | 1 | 15 | Trim59 | 0.016 | Young |
| 0.648217 | 1.107575 | 0.036 | 0.02 | 1 | 15 | Atg2b | 0.016 | Young |
| 0.648217 | 1.065845 | 0.036 | 0.02 | 1 | 15 | Rfx2 | 0.016 | Young |
| 0.648217 | 1.03638 | 0.036 | 0.02 | 1 | 15 | Mrpl39 | 0.016 | Young |
| 0.648217 | 1.035238 | 0.036 | 0.02 | 1 | 15 | Havcr2 | 0.016 | Young |
| 0.648217 | 1.027672 | 0.036 | 0.02 | 1 | 15 | Xpo5 | 0.016 | Young |
| 0.648217 | 1.003577 | 0.036 | 0.02 | 1 | 15 | Grpel2 | 0.016 | Young |
| 0.648217 | 0.985087 | 0.036 | 0.02 | 1 | 15 | Tex2 | 0.016 | Young |
| 0.648217 | 0.975053 | 0.036 | 0.02 | 1 | 15 | Eefsec | 0.016 | Young |
| 0.648217 | 0.970295 | 0.036 | 0.02 | 1 | 15 | Mapre3 | 0.016 | Young |
| 0.648217 | 0.954546 | 0.036 | 0.02 | 1 | 15 | Ankrd39 | 0.016 | Young |
| 0.648217 | 0.927932 | 0.036 | 0.02 | 1 | 15 | Sirt3 | 0.016 | Young |
| 0.648217 | 0.923835 | 0.036 | 0.02 | 1 | 15 | Rbm41 | 0.016 | Young |
| 0.648217 | 0.91285 | 0.036 | 0.02 | 1 | 15 | Jagn1 | 0.016 | Young |
| 0.648217 | 0.901515 | 0.036 | 0.02 | 1 | 15 | Fbln5 | 0.016 | Young |
| 0.648217 | 0.885152 | 0.036 | 0.02 | 1 | 15 | 6720427I0 | 0.016 | Young |
| 0.648217 | 0.854971 | 0.036 | 0.02 | 1 | 15 | Palm | 0.016 | Young |
| 0.648217 | 0.836676 | 0.036 | 0.02 | 1 | 15 | Tyw1 | 0.016 | Young |
| 0.648217 | 0.825645 | 0.036 | 0.02 | 1 | 15 | Zfp866 | 0.016 | Young |
| 0.648217 | 0.814484 | 0.036 | 0.02 | 1 | 15 | Gm29650 | 0.016 | Young |
| 0.648217 | 0.81335 | 0.036 | 0.02 | 1 | 15 | Gemin2 | 0.016 | Young |
| 0.648217 | 0.810698 | 0.036 | 0.02 | 1 | 15 | Coprs | 0.016 | Young |
| 0.648217 | 0.809355 | 0.036 | 0.02 | 1 | 15 | Ifi214 | 0.016 | Young |
| 0.648217 | 0.794466 | 0.036 | 0.02 | 1 | 15 | Utf1 | 0.016 | Young |
| 0.648217 | 0.775397 | 0.036 | 0.02 | 1 | 15 | Pml | 0.016 | Young |
| 0.648217 | 0.774576 | 0.036 | 0.02 | 1 | 15 | Snx25 | 0.016 | Young |
| 0.648217 | 0.748204 | 0.036 | 0.02 | 1 | 15 | Dgcr8 | 0.016 | Young |

| | | | | | | | | |
|---|---|---|---|---|---|---|---|---|
| 0.648217 | 0.745063 | 0.036 | 0.02 | 1 | 15 | Irgq | 0.016 | Young |
| 0.648217 | 0.735807 | 0.036 | 0.02 | 1 | 15 | Tlr8 | 0.016 | Young |
| 0.648217 | 0.729215 | 0.036 | 0.02 | 1 | 15 | Mrpl16 | 0.016 | Young |
| 0.648217 | 0.719452 | 0.036 | 0.02 | 1 | 15 | Prkch | 0.016 | Young |
| 0.648217 | 0.71819 | 0.036 | 0.02 | 1 | 15 | Smim20 | 0.016 | Young |
| 0.648217 | 0.643964 | 0.036 | 0.02 | 1 | 15 | Cdiptos | 0.016 | Young |
| 0.648217 | 0.643457 | 0.036 | 0.02 | 1 | 15 | Cfap45 | 0.016 | Young |
| 0.648217 | 0.633298 | 0.036 | 0.02 | 1 | 15 | Natd1 | 0.016 | Young |
| 0.648217 | 0.631541 | 0.036 | 0.02 | 1 | 15 | Birc5 | 0.016 | Young |
| 0.648217 | 0.625254 | 0.036 | 0.02 | 1 | 15 | Nek1 | 0.016 | Young |
| 0.648217 | 0.604832 | 0.036 | 0.02 | 1 | 15 | Cyp2ab1 | 0.016 | Young |
| 0.648217 | 0.602283 | 0.036 | 0.02 | 1 | 15 | Dph7 | 0.016 | Young |
| 0.648217 | 0.600415 | 0.036 | 0.02 | 1 | 15 | Nipa1 | 0.016 | Young |
| 0.648217 | 0.594236 | 0.036 | 0.02 | 1 | 15 | 5430427O1 | 0.016 | Young |
| 0.648217 | 0.593869 | 0.036 | 0.02 | 1 | 15 | Riox1 | 0.016 | Young |
| 0.648217 | 0.575824 | 0.036 | 0.02 | 1 | 15 | Nme7 | 0.016 | Young |
| 0.648217 | 0.56936 | 0.036 | 0.02 | 1 | 15 | Fpgt | 0.016 | Young |
| 0.648217 | 0.566258 | 0.036 | 0.02 | 1 | 15 | Rab3ip | 0.016 | Young |
| 0.648217 | 0.556104 | 0.036 | 0.02 | 1 | 15 | Cwc27 | 0.016 | Young |
| 0.648217 | 0.547482 | 0.036 | 0.02 | 1 | 15 | Ormdl3 | 0.016 | Young |
| 0.648217 | 0.534839 | 0.036 | 0.02 | 1 | 15 | Zfp715 | 0.016 | Young |
| 0.648217 | 0.528402 | 0.036 | 0.02 | 1 | 15 | B230307C2 | 0.016 | Young |
| 0.648217 | 0.528094 | 0.036 | 0.02 | 1 | 15 | Iqcc | 0.016 | Young |
| 0.648217 | 0.519971 | 0.036 | 0.02 | 1 | 15 | 5730409E0 | 0.016 | Young |
| 0.648217 | 0.506998 | 0.036 | 0.02 | 1 | 15 | Supt7l | 0.016 | Young |
| 0.648217 | 0.506748 | 0.036 | 0.02 | 1 | 15 | Snhg9 | 0.016 | Young |
| 0.648217 | 0.496478 | 0.036 | 0.02 | 1 | 15 | Sco2 | 0.016 | Young |
| 0.648217 | 0.487713 | 0.036 | 0.02 | 1 | 15 | Tubgcp6 | 0.016 | Young |
| 0.648217 | 0.469351 | 0.036 | 0.02 | 1 | 15 | Npr1 | 0.016 | Young |
| 0.648217 | 0.451868 | 0.036 | 0.02 | 1 | 15 | Eci1 | 0.016 | Young |
| 0.648217 | 0.445978 | 0.036 | 0.02 | 1 | 15 | Mbd4 | 0.016 | Young |
| 0.648217 | 0.42369 | 0.036 | 0.02 | 1 | 15 | Gm42031 | 0.016 | Young |
| 0.648217 | 0.388506 | 0.036 | 0.02 | 1 | 15 | Usp46 | 0.016 | Young |
| 0.648217 | 0.361738 | 0.036 | 0.02 | 1 | 15 | Zbtb33 | 0.016 | Young |
| 0.648217 | 0.345592 | 0.036 | 0.02 | 1 | 15 | Ticam1 | 0.016 | Young |
| 0.648217 | 0.332273 | 0.036 | 0.02 | 1 | 15 | Cog6 | 0.016 | Young |
| 0.648217 | 0.329112 | 0.036 | 0.02 | 1 | 15 | Zfp873 | 0.016 | Young |
| 0.648217 | 0.319317 | 0.036 | 0.02 | 1 | 15 | Zfp523 | 0.016 | Young |
| 0.648544 | 0.133194 | 0.446 | 0.408 | 1 | 15 | H1f0 | 0.038 | Young |
| 0.648623 | 0.145233 | 0.179 | 0.143 | 1 | 15 | Rsl1d1 | 0.036 | Young |
| 0.648883 | 0.578014 | 0.357 | 0.347 | 1 | 15 | Atp5o.1 | 0.01 | Young |
| 0.649352 | 0.937528 | 0.107 | 0.143 | 1 | 15 | Fbxo22 | -0.036 | Young |
| 0.649352 | 0.257518 | 0.107 | 0.143 | 1 | 15 | Zswim8 | -0.036 | Young |
| 0.649911 | 0.072278 | 0.125 | 0.163 | 1 | 15 | Clcn4 | -0.038 | Young |
| 0.650174 | 0.545765 | 0.25 | 0.245 | 1 | 15 | Cmc1 | 0.005 | Young |

| | | | | | | | | |
|---|---|---|---|---|---|---|---|---|
| 0.65071 | 0.109438 | 0.411 | 0.367 | 1 | 15 | Cep170 | 0.044 | Young |
| 0.650958 | 0.714186 | 0.286 | 0.265 | 1 | 15 | Psmc6 | 0.021 | Young |
| 0.651607 | 0.402972 | 0.25 | 0.224 | 1 | 15 | Fbxw11 | 0.026 | Young |
| 0.651863 | 1.048266 | 0.143 | 0.122 | 1 | 15 | Ociad1 | 0.021 | Young |
| 0.652849 | 0.244119 | 0.286 | 0.245 | 1 | 15 | Tfe3 | 0.041 | Young |
| 0.654347 | 0.098296 | 0.857 | 0.776 | 1 | 15 | Anxa3 | 0.081 | Young |
| 0.654357 | 0.631587 | 0.107 | 0.082 | 1 | 15 | Pus1 | 0.025 | Young |
| 0.654357 | 0.618049 | 0.107 | 0.082 | 1 | 15 | Ppie | 0.025 | Young |
| 0.654357 | 0.608694 | 0.107 | 0.082 | 1 | 15 | Pycr2 | 0.025 | Young |
| 0.654357 | 0.314783 | 0.107 | 0.082 | 1 | 15 | Parp2 | 0.025 | Young |
| 0.654357 | 0.219852 | 0.107 | 0.082 | 1 | 15 | Gars | 0.025 | Young |
| 0.654387 | 0.030945 | 0.589 | 0.51 | 1 | 15 | Eif2s2 | 0.079 | Young |
| 0.654415 | 1.002622 | 0.125 | 0.102 | 1 | 15 | Nsmce2 | 0.023 | Young |
| 0.654415 | 0.726374 | 0.125 | 0.102 | 1 | 15 | Usp38 | 0.023 | Young |
| 0.654772 | 0.19106 | 0.304 | 0.265 | 1 | 15 | Lilr4b | 0.039 | Young |
| 0.655648 | 0.408431 | 0.214 | 0.184 | 1 | 15 | Ipo5 | 0.03 | Young |
| 0.655842 | 0.296776 | 0.179 | 0.143 | 1 | 15 | Galnt11 | 0.036 | Young |
| 0.657526 | 0.190363 | 0.125 | 0.163 | 1 | 15 | Lman1 | -0.038 | Young |
| 0.659717 | 1.520655 | 0.143 | 0.122 | 1 | 15 | Rhoc | 0.021 | Young |
| 0.659747 | 0.171447 | 0.321 | 0.388 | 1 | 15 | Tnfaip2 | -0.067 | Young |
| 0.659814 | 0.280068 | 0.196 | 0.163 | 1 | 15 | Triobp | 0.033 | Young |
| 0.659814 | 0.123904 | 0.196 | 0.163 | 1 | 15 | Exoc2 | 0.033 | Young |
| 0.661263 | 0.344319 | 0.411 | 0.367 | 1 | 15 | P2rx4 | 0.044 | Young |
| 0.661462 | 0.180817 | 0.232 | 0.184 | 1 | 15 | Aip | 0.048 | Young |
| 0.662304 | 0.466468 | 0.179 | 0.224 | 1 | 15 | Ctdsp1 | -0.045 | Young |
| 0.662944 | 0.2019 | 0.839 | 0.714 | 1 | 15 | Atp5g2 | 0.125 | Young |
| 0.663486 | 0.730391 | 0.107 | 0.082 | 1 | 15 | Dgka | 0.025 | Young |
| 0.663486 | 0.600937 | 0.107 | 0.082 | 1 | 15 | Pdcl | 0.025 | Young |
| 0.663486 | 0.577695 | 0.107 | 0.082 | 1 | 15 | Mtmr3 | 0.025 | Young |
| 0.663638 | 0.622878 | 0.232 | 0.204 | 1 | 15 | Arap1 | 0.028 | Young |
| 0.663638 | 0.27391 | 0.232 | 0.204 | 1 | 15 | Ypel3 | 0.028 | Young |
| 0.663709 | 0.324361 | 0.268 | 0.245 | 1 | 15 | Csf2rb2 | 0.023 | Young |
| 0.664049 | 0.630603 | 0.25 | 0.224 | 1 | 15 | Sgpp1 | 0.026 | Young |
| 0.664049 | 0.381111 | 0.25 | 0.224 | 1 | 15 | Senp2 | 0.026 | Young |
| 0.664143 | 0.03929 | 0.321 | 0.265 | 1 | 15 | Cd24a | 0.056 | Young |
| 0.6643 | 0.499105 | 0.036 | 0.02 | 1 | 15 | R3hcc1 | 0.016 | Young |
| 0.6643 | 0.481754 | 0.036 | 0.02 | 1 | 15 | Tfec | 0.016 | Young |
| 0.6643 | 0.449773 | 0.036 | 0.02 | 1 | 15 | Ttc38 | 0.016 | Young |
| 0.6643 | 0.379157 | 0.036 | 0.02 | 1 | 15 | Nicn1 | 0.016 | Young |
| 0.6643 | 0.377771 | 0.036 | 0.02 | 1 | 15 | Ntmt1 | 0.016 | Young |
| 0.6643 | 0.374356 | 0.036 | 0.02 | 1 | 15 | Morc2a | 0.016 | Young |
| 0.6643 | 0.361035 | 0.036 | 0.02 | 1 | 15 | Tmem119 | 0.016 | Young |
| 0.6643 | 0.28024 | 0.036 | 0.02 | 1 | 15 | Praf2 | 0.016 | Young |
| 0.6643 | 0.268343 | 0.036 | 0.02 | 1 | 15 | Bcorl1 | 0.016 | Young |
| 0.6643 | 0.265703 | 0.036 | 0.02 | 1 | 15 | AL590144.: | 0.016 | Young |

| | | | | | | | | |
|---|---|---|---|---|---|---|---|---|
| 0.6643 | 0.254533 | 0.036 | 0.02 | 1 | 15 | Rrp9 | 0.016 | Young |
| 0.6643 | 0.245978 | 0.036 | 0.02 | 1 | 15 | Mettl6 | 0.016 | Young |
| 0.6643 | 0.230513 | 0.036 | 0.02 | 1 | 15 | Tmed1 | 0.016 | Young |
| 0.6643 | 0.222461 | 0.036 | 0.02 | 1 | 15 | Pi4kb | 0.016 | Young |
| 0.6643 | 0.207891 | 0.036 | 0.02 | 1 | 15 | Paqr8 | 0.016 | Young |
| 0.6643 | 0.189593 | 0.036 | 0.02 | 1 | 15 | Tmem186 | 0.016 | Young |
| 0.6643 | 0.188934 | 0.036 | 0.02 | 1 | 15 | Cobll1 | 0.016 | Young |
| 0.6643 | 0.163901 | 0.036 | 0.02 | 1 | 15 | Pelp1 | 0.016 | Young |
| 0.6643 | 0.155256 | 0.036 | 0.02 | 1 | 15 | Dnmbp | 0.016 | Young |
| 0.6643 | 0.134271 | 0.036 | 0.02 | 1 | 15 | Gm44777 | 0.016 | Young |
| 0.6643 | 0.128252 | 0.036 | 0.02 | 1 | 15 | 4930430F0 | 0.016 | Young |
| 0.6643 | 0.113915 | 0.036 | 0.02 | 1 | 15 | Nudcd1 | 0.016 | Young |
| 0.6643 | 0.112374 | 0.036 | 0.02 | 1 | 15 | Exd2 | 0.016 | Young |
| 0.6643 | 0.110031 | 0.036 | 0.02 | 1 | 15 | Tsen15 | 0.016 | Young |
| 0.6643 | 0.104145 | 0.036 | 0.02 | 1 | 15 | Pacs1 | 0.016 | Young |
| 0.6643 | 0.09927 | 0.036 | 0.02 | 1 | 15 | Haus3 | 0.016 | Young |
| 0.6643 | 0.05041 | 0.036 | 0.02 | 1 | 15 | Ppip5k1 | 0.016 | Young |
| 0.6643 | 0.044449 | 0.036 | 0.02 | 1 | 15 | Kank2 | 0.016 | Young |
| 0.6643 | 0.034527 | 0.036 | 0.02 | 1 | 15 | Car13 | 0.016 | Young |
| 0.6643 | 0.030151 | 0.036 | 0.02 | 1 | 15 | Adgrl1 | 0.016 | Young |
| 0.6643 | 0.02157 | 0.036 | 0.02 | 1 | 15 | Wdr25 | 0.016 | Young |
| 0.6643 | 0.007652 | 0.036 | 0.02 | 1 | 15 | Rars2 | 0.016 | Young |
| 0.665178 | 0.274943 | 0.125 | 0.163 | 1 | 15 | Sdccag3 | -0.038 | Young |
| 0.665178 | 0.151298 | 0.125 | 0.163 | 1 | 15 | Vps11 | -0.038 | Young |
| 0.665206 | 0.465971 | 0.357 | 0.347 | 1 | 15 | Polr2a | 0.01 | Young |
| 0.665206 | 0.281009 | 0.714 | 0.694 | 1 | 15 | Eif4b | 0.02 | Young |
| 0.665533 | 0.227744 | 0.25 | 0.204 | 1 | 15 | Rnh1 | 0.046 | Young |
| 0.666447 | 0.17758 | 0.304 | 0.265 | 1 | 15 | Mllt10 | 0.039 | Young |
| 0.66724 | 0.515622 | 0.071 | 0.102 | 1 | 15 | Lrch4 | -0.031 | Young |
| 0.668986 | 0.026103 | 0.214 | 0.184 | 1 | 15 | Cyb5b | 0.03 | Young |
| 0.66903 | 0.144152 | 0.571 | 0.531 | 1 | 15 | Calm2 | 0.04 | Young |
| 0.670377 | 0.127316 | 0.143 | 0.184 | 1 | 15 | Zdhhc21 | -0.041 | Young |
| 0.670377 | 0.077388 | 0.179 | 0.143 | 1 | 15 | Zfp703 | 0.036 | Young |
| 0.671287 | 1.109501 | 0.125 | 0.102 | 1 | 15 | Rnf146 | 0.023 | Young |
| 0.671287 | 0.825077 | 0.125 | 0.102 | 1 | 15 | Phf13 | 0.023 | Young |
| 0.671287 | 0.732322 | 0.125 | 0.102 | 1 | 15 | Ddx18 | 0.023 | Young |
| 0.671287 | 0.651104 | 0.125 | 0.102 | 1 | 15 | Zfpl1 | 0.023 | Young |
| 0.671287 | 0.527487 | 0.125 | 0.102 | 1 | 15 | Twistnb | 0.023 | Young |
| 0.671287 | 0.502941 | 0.125 | 0.102 | 1 | 15 | Senp7 | 0.023 | Young |
| 0.67158 | 0.331962 | 0.464 | 0.449 | 1 | 15 | Dek | 0.015 | Young |
| 0.672335 | 0.255566 | 0.089 | 0.122 | 1 | 15 | Ppm1m | -0.033 | Young |
| 0.672665 | 0.484543 | 0.107 | 0.082 | 1 | 15 | Fam20b | 0.025 | Young |
| 0.672665 | 0.309734 | 0.107 | 0.082 | 1 | 15 | Pla2g4a | 0.025 | Young |
| 0.672864 | 0.2803 | 0.125 | 0.163 | 1 | 15 | Asap2 | -0.038 | Young |
| 0.67338 | 0.118354 | 0.732 | 0.735 | 1 | 15 | Tomm7 | -0.003 | Young |

| | | | | | | | | |
|---|---|---|---|---|---|---|---|---|
| 0.673732 | 0.570174 | 0.161 | 0.204 | 1 | 15 | Daglb | -0.043 | Young |
| 0.675167 | 0.100513 | 0.446 | 0.388 | 1 | 15 | Brd2 | 0.058 | Young |
| 0.675537 | 1.025659 | 0.143 | 0.122 | 1 | 15 | Atic | 0.021 | Young |
| 0.675694 | 0.733723 | 0.179 | 0.224 | 1 | 15 | Mark2 | -0.045 | Young |
| 0.67689 | 0.726895 | 0.071 | 0.102 | 1 | 15 | Gas2l3 | -0.031 | Young |
| 0.678273 | 0.030494 | 0.25 | 0.204 | 1 | 15 | Hivep2 | 0.046 | Young |
| 0.679785 | 0.685287 | 0.125 | 0.102 | 1 | 15 | Naxe | 0.023 | Young |
| 0.680733 | 0.376735 | 0.196 | 0.163 | 1 | 15 | Rangap1 | 0.033 | Young |
| 0.680733 | 0.192845 | 0.196 | 0.163 | 1 | 15 | Psmc4 | 0.033 | Young |
| 0.681212 | 0.278661 | 0.232 | 0.184 | 1 | 15 | Higd1a | 0.048 | Young |
| 0.681892 | 0.950636 | 0.107 | 0.082 | 1 | 15 | Fdx1 | 0.025 | Young |
| 0.681892 | 0.234663 | 0.107 | 0.082 | 1 | 15 | Uchl5 | 0.025 | Young |
| 0.681892 | 0.217387 | 0.107 | 0.082 | 1 | 15 | Pde3b | 0.025 | Young |
| 0.681892 | 0.028315 | 0.107 | 0.082 | 1 | 15 | Mrto4 | 0.025 | Young |
| 0.682016 | 0.222857 | 0.268 | 0.245 | 1 | 15 | Arhgap1 | 0.023 | Young |
| 0.683075 | 0.669598 | 0.232 | 0.204 | 1 | 15 | Rell1 | 0.028 | Young |
| 0.683502 | 0.703031 | 0.143 | 0.122 | 1 | 15 | Slc27a1 | 0.021 | Young |
| 0.683502 | 0.699591 | 0.143 | 0.122 | 1 | 15 | Dnajc14 | 0.021 | Young |
| 0.685036 | 0.027526 | 0.143 | 0.184 | 1 | 15 | Elp5 | -0.041 | Young |
| 0.685457 | 0.066255 | 0.429 | 0.469 | 1 | 15 | Tuba1a | -0.04 | Young |
| 0.687244 | 0.343131 | 0.411 | 0.388 | 1 | 15 | Itpripl2 | 0.023 | Young |
| 0.688325 | 0.780882 | 0.125 | 0.102 | 1 | 15 | Cryl1 | 0.023 | Young |
| 0.688325 | 0.597388 | 0.125 | 0.102 | 1 | 15 | E4f1 | 0.023 | Young |
| 0.688325 | 0.443519 | 0.125 | 0.102 | 1 | 15 | Hars | 0.023 | Young |
| 0.689188 | 0.151614 | 0.214 | 0.184 | 1 | 15 | Sirt2 | 0.03 | Young |
| 0.68921 | 0.393905 | 0.25 | 0.224 | 1 | 15 | Hmox1 | 0.026 | Young |
| 0.689931 | 0.040352 | 0.643 | 0.612 | 1 | 15 | Psmb4 | 0.031 | Young |
| 0.691168 | 0.020897 | 0.107 | 0.082 | 1 | 15 | Fh1 | 0.025 | Young |
| 0.691502 | 0.561438 | 0.143 | 0.122 | 1 | 15 | Mpdu1 | 0.021 | Young |
| 0.69241 | 0.267335 | 0.143 | 0.184 | 1 | 15 | Utp11 | -0.041 | Young |
| 0.694326 | 0.194132 | 0.232 | 0.286 | 1 | 15 | Nudt9 | -0.054 | Young |
| 0.694693 | 0.03475 | 0.554 | 0.469 | 1 | 15 | Pbrm1 | 0.085 | Young |
| 0.695973 | 0.319133 | 0.214 | 0.184 | 1 | 15 | Mxd4 | 0.03 | Young |
| 0.695973 | 0.31735 | 0.214 | 0.184 | 1 | 15 | Ogfr | 0.03 | Young |
| 0.696147 | 0.405546 | 0.446 | 0.49 | 1 | 15 | Irf8 | -0.044 | Young |
| 0.696153 | 0.485958 | 0.232 | 0.204 | 1 | 15 | Mprip | 0.028 | Young |
| 0.696231 | 0.053777 | 0.393 | 0.449 | 1 | 15 | Psma2 | -0.056 | Young |
| 0.696399 | 0.117275 | 0.321 | 0.286 | 1 | 15 | Mbd2 | 0.035 | Young |
| 0.696793 | 0.159462 | 0.554 | 0.51 | 1 | 15 | Arhgap31 | 0.044 | Young |
| 0.696905 | 0.618094 | 0.125 | 0.102 | 1 | 15 | Usp48 | 0.023 | Young |
| 0.696905 | 0.42507 | 0.125 | 0.102 | 1 | 15 | Rnf103 | 0.023 | Young |
| 0.696905 | 0.038316 | 0.125 | 0.102 | 1 | 15 | Zfp945 | 0.023 | Young |
| 0.697318 | 0.151849 | 0.393 | 0.347 | 1 | 15 | Wasf2 | 0.046 | Young |
| 0.698984 | 0.970992 | 0.089 | 0.122 | 1 | 15 | Eif1ad | -0.033 | Young |
| 0.699201 | 0.006419 | 0.321 | 0.265 | 1 | 15 | Bub3 | 0.056 | Young |

| | | | | | | | | |
|---|---|---|---|---|---|---|---|---|
| 0.699537 | 0.642739 | 0.143 | 0.122 | 1 | 15 | Pdha1 | 0.021 | Young |
| 0.700103 | 0.758161 | 0.161 | 0.143 | 1 | 15 | Emp1 | 0.018 | Young |
| 0.70049 | 0.090711 | 0.107 | 0.082 | 1 | 15 | Tiprl | 0.025 | Young |
| 0.701901 | 0.139735 | 0.196 | 0.163 | 1 | 15 | Tns1 | 0.033 | Young |
| 0.701901 | 0.037158 | 0.196 | 0.163 | 1 | 15 | Tmpo | 0.033 | Young |
| 0.702726 | 0.108206 | 0.232 | 0.204 | 1 | 15 | Iah1 | 0.028 | Young |
| 0.70464 | 0.132639 | 0.286 | 0.265 | 1 | 15 | Vma21 | 0.021 | Young |
| 0.705215 | 0.011101 | 0.482 | 0.469 | 1 | 15 | Ndufb11 | 0.013 | Young |
| 0.705523 | 0.415809 | 0.125 | 0.102 | 1 | 15 | Oaz2 | 0.023 | Young |
| 0.706417 | 0.001594 | 0.625 | 0.612 | 1 | 15 | Sf3b1 | 0.013 | Young |
| 0.707156 | 0.069182 | 0.357 | 0.327 | 1 | 15 | Psmc3 | 0.03 | Young |
| 0.707268 | 0.081057 | 0.25 | 0.286 | 1 | 15 | Ube2j2 | -0.036 | Young |
| 0.707516 | 0.303089 | 0.714 | 0.653 | 1 | 15 | Tpm4 | 0.061 | Young |
| 0.707606 | 0.585039 | 0.143 | 0.122 | 1 | 15 | Cse1l | 0.021 | Young |
| 0.707606 | 0.548183 | 0.143 | 0.122 | 1 | 15 | Bmyc | 0.021 | Young |
| 0.707606 | 0.449903 | 0.143 | 0.122 | 1 | 15 | Hbs1l | 0.021 | Young |
| 0.707728 | 0.894656 | 0.161 | 0.143 | 1 | 15 | Ric8a | 0.018 | Young |
| 0.707728 | 0.593561 | 0.161 | 0.143 | 1 | 15 | M1ap | 0.018 | Young |
| 0.708311 | 0.337496 | 0.25 | 0.224 | 1 | 15 | Zcchc7 | 0.026 | Young |
| 0.709483 | 0.123846 | 0.982 | 0.939 | 1 | 15 | Lsp1 | 0.043 | Young |
| 0.710494 | 0.135066 | 0.393 | 0.429 | 1 | 15 | Lasp1 | -0.036 | Young |
| 0.711183 | 0.538073 | 0.196 | 0.184 | 1 | 15 | Rtf2 | 0.012 | Young |
| 0.711798 | 0.788856 | 0.125 | 0.163 | 1 | 15 | Ascc3 | -0.038 | Young |
| 0.712807 | 0.432266 | 0.339 | 0.347 | 1 | 15 | Cr1l | -0.008 | Young |
| 0.712943 | 0.247463 | 0.268 | 0.245 | 1 | 15 | Atp6v1c1 | 0.023 | Young |
| 0.713569 | 0.574774 | 0.321 | 0.327 | 1 | 15 | Snx10 | -0.006 | Young |
| 0.713758 | 0.503237 | 0.179 | 0.163 | 1 | 15 | Mcfd2 | 0.016 | Young |
| 0.71418 | 0.608213 | 0.125 | 0.102 | 1 | 15 | Dscaml1 | 0.023 | Young |
| 0.71418 | 0.470778 | 0.125 | 0.102 | 1 | 15 | Gskip | 0.023 | Young |
| 0.71418 | 0.334348 | 0.125 | 0.102 | 1 | 15 | Rgp1 | 0.023 | Young |
| 0.71418 | 0.269853 | 0.125 | 0.102 | 1 | 15 | Trappc11 | 0.023 | Young |
| 0.71418 | 0.077834 | 0.125 | 0.102 | 1 | 15 | Sema4c | 0.023 | Young |
| 0.71472 | 0.129477 | 0.214 | 0.265 | 1 | 15 | Usp3 | -0.051 | Young |
| 0.715707 | 0.553098 | 0.143 | 0.122 | 1 | 15 | Tmem205 | 0.021 | Young |
| 0.715707 | 0.517073 | 0.143 | 0.122 | 1 | 15 | Lncpint | 0.021 | Young |
| 0.715707 | 0.279498 | 0.143 | 0.122 | 1 | 15 | Tm2d2 | 0.021 | Young |
| 0.715938 | 0.522639 | 0.232 | 0.204 | 1 | 15 | Cmc2 | 0.028 | Young |
| 0.715938 | 0.095608 | 0.232 | 0.204 | 1 | 15 | Mapkap1 | 0.028 | Young |
| 0.716143 | 0.478991 | 0.161 | 0.204 | 1 | 15 | Fam96b | -0.043 | Young |
| 0.716143 | 0.159847 | 0.161 | 0.204 | 1 | 15 | Vps41 | -0.043 | Young |
| 0.716471 | 0.388317 | 0.214 | 0.184 | 1 | 15 | Il1rn | 0.03 | Young |
| 0.716784 | 0.190656 | 0.286 | 0.265 | 1 | 15 | Gramd4 | 0.021 | Young |
| 0.718179 | 0.00123 | 0.179 | 0.204 | 1 | 15 | Wwp2 | -0.025 | Young |
| 0.71927 | 0.960919 | 0.107 | 0.082 | 1 | 15 | Inip | 0.025 | Young |
| 0.719363 | 0.191858 | 0.393 | 0.347 | 1 | 15 | Apobec1 | 0.046 | Young |

| | | | | | | | | |
|---|---|---|---|---|---|---|---|---|
| 0.71968 | 0.546768 | 0.125 | 0.163 | 1 | 15 | Mfsd14a | -0.038 | Young |
| 0.71968 | 0.503124 | 0.125 | 0.163 | 1 | 15 | Eif2ak2 | -0.038 | Young |
| 0.719954 | 0.207669 | 0.304 | 0.265 | 1 | 15 | Ppp1r12c | 0.039 | Young |
| 0.721148 | 0.334993 | 0.25 | 0.224 | 1 | 15 | Thap12 | 0.026 | Young |
| 0.722576 | 0.239889 | 0.232 | 0.204 | 1 | 15 | Fbl | 0.028 | Young |
| 0.722873 | 0.549595 | 0.125 | 0.102 | 1 | 15 | Pttg1 | 0.023 | Young |
| 0.722873 | 0.474195 | 0.125 | 0.102 | 1 | 15 | Nrf1 | 0.023 | Young |
| 0.722873 | 0.17128 | 0.125 | 0.102 | 1 | 15 | Trim44 | 0.023 | Young |
| 0.722892 | 0.000604 | 0.446 | 0.388 | 1 | 15 | Capns1 | 0.058 | Young |
| 0.723841 | 0.759228 | 0.143 | 0.122 | 1 | 15 | 9530059O1 | 0.021 | Young |
| 0.723841 | 0.23462 | 0.143 | 0.122 | 1 | 15 | Taok2 | 0.021 | Young |
| 0.725444 | 0.433491 | 0.554 | 0.531 | 1 | 15 | Apol7c | 0.023 | Young |
| 0.725451 | 0.133672 | 0.268 | 0.245 | 1 | 15 | Gsr | 0.023 | Young |
| 0.727389 | 0.217702 | 0.714 | 0.673 | 1 | 15 | Stxbp3 | 0.041 | Young |
| 0.727597 | 0.700296 | 0.25 | 0.224 | 1 | 15 | Atxn7 | 0.026 | Young |
| 0.727747 | 2.828072 | 0.054 | 0.041 | 1 | 15 | Clec4e | 0.013 | Young |
| 0.727747 | 2.807809 | 0.054 | 0.041 | 1 | 15 | Calhm2 | 0.013 | Young |
| 0.727747 | 2.271765 | 0.054 | 0.041 | 1 | 15 | Thap4 | 0.013 | Young |
| 0.727747 | 2.042245 | 0.054 | 0.041 | 1 | 15 | Piwil2 | 0.013 | Young |
| 0.727747 | 1.884755 | 0.054 | 0.041 | 1 | 15 | Hdhd2 | 0.013 | Young |
| 0.727747 | 1.858414 | 0.054 | 0.041 | 1 | 15 | Gm49359 | 0.013 | Young |
| 0.727747 | 1.814801 | 0.054 | 0.041 | 1 | 15 | Gm9493 | 0.013 | Young |
| 0.727747 | 1.624267 | 0.054 | 0.041 | 1 | 15 | Nipsnap1 | 0.013 | Young |
| 0.727747 | 1.545348 | 0.054 | 0.041 | 1 | 15 | Socs5 | 0.013 | Young |
| 0.727747 | 1.512664 | 0.054 | 0.041 | 1 | 15 | Ppp1r13b | 0.013 | Young |
| 0.727747 | 1.453301 | 0.054 | 0.041 | 1 | 15 | Rhbdd3 | 0.013 | Young |
| 0.727747 | 1.326211 | 0.054 | 0.041 | 1 | 15 | Slc11a2 | 0.013 | Young |
| 0.727747 | 1.312466 | 0.054 | 0.041 | 1 | 15 | Ahi1 | 0.013 | Young |
| 0.727747 | 1.229454 | 0.054 | 0.041 | 1 | 15 | Osbpl3 | 0.013 | Young |
| 0.727747 | 1.159722 | 0.054 | 0.041 | 1 | 15 | Slc9a6 | 0.013 | Young |
| 0.727747 | 1.050984 | 0.054 | 0.041 | 1 | 15 | Sp4 | 0.013 | Young |
| 0.727747 | 0.925812 | 0.054 | 0.041 | 1 | 15 | Cryzl2 | 0.013 | Young |
| 0.727747 | 0.878548 | 0.054 | 0.041 | 1 | 15 | Cd300lg | 0.013 | Young |
| 0.727747 | 0.855304 | 0.054 | 0.041 | 1 | 15 | Insl6 | 0.013 | Young |
| 0.727747 | 0.751528 | 0.054 | 0.041 | 1 | 15 | Apex2 | 0.013 | Young |
| 0.728724 | 0.128876 | 0.107 | 0.082 | 1 | 15 | Lactb2 | 0.025 | Young |
| 0.728919 | 0.011249 | 0.321 | 0.265 | 1 | 15 | Adgre1 | 0.056 | Young |
| 0.729236 | 0.482277 | 0.232 | 0.204 | 1 | 15 | Mob4 | 0.028 | Young |
| 0.729978 | 0.182115 | 0.446 | 0.449 | 1 | 15 | H2-T22 | -0.003 | Young |
| 0.730252 | 0.214797 | 0.214 | 0.184 | 1 | 15 | Cmtm3 | 0.03 | Young |
| 0.730934 | 0.259788 | 0.321 | 0.327 | 1 | 15 | Fam120a | -0.006 | Young |
| 0.731603 | 0.254423 | 0.125 | 0.102 | 1 | 15 | Mrpl3 | 0.023 | Young |
| 0.731603 | 0.201532 | 0.125 | 0.102 | 1 | 15 | Rab5b | 0.023 | Young |
| 0.731603 | 0.199816 | 0.125 | 0.102 | 1 | 15 | Scrib | 0.023 | Young |
| 0.731603 | 0.143578 | 0.125 | 0.102 | 1 | 15 | Siah1a | 0.023 | Young |

| | | | | | | | | |
|---|---|---|---|---|---|---|---|---|
| 0.731603 | 0.137634 | 0.125 | 0.102 | 1 | 15 | Slc9a8 | 0.023 | Young |
| 0.731603 | 0.091599 | 0.125 | 0.102 | 1 | 15 | Pigm | 0.023 | Young |
| 0.731964 | 0.228239 | 0.286 | 0.245 | 1 | 15 | Drap1 | 0.041 | Young |
| 0.732006 | 0.372825 | 0.143 | 0.122 | 1 | 15 | Mpp5 | 0.021 | Young |
| 0.732006 | 0.357105 | 0.143 | 0.122 | 1 | 15 | Tfg | 0.021 | Young |
| 0.732568 | 0.105284 | 0.482 | 0.449 | 1 | 15 | Unc93b1 | 0.033 | Young |
| 0.733262 | 0.274091 | 0.375 | 0.367 | 1 | 15 | Mlf2 | 0.008 | Young |
| 0.734288 | 0.280787 | 0.321 | 0.306 | 1 | 15 | Manf | 0.015 | Young |
| 0.735764 | 0.857512 | 0.179 | 0.163 | 1 | 15 | Setd3 | 0.016 | Young |
| 0.736923 | 0.006804 | 0.375 | 0.408 | 1 | 15 | Dynlrb1 | -0.033 | Young |
| 0.7377 | 0.550663 | 0.357 | 0.347 | 1 | 15 | Fgfr1 | 0.01 | Young |
| 0.737912 | 0.160367 | 0.393 | 0.367 | 1 | 15 | Atp1a1 | 0.026 | Young |
| 0.737917 | 0.125933 | 0.304 | 0.286 | 1 | 15 | Eif1ax | 0.018 | Young |
| 0.738182 | 0.056291 | 0.286 | 0.245 | 1 | 15 | Atp5g1 | 0.041 | Young |
| 0.738513 | 0.562394 | 0.161 | 0.143 | 1 | 15 | Mta1 | 0.018 | Young |
| 0.738513 | 0.222271 | 0.161 | 0.143 | 1 | 15 | Ubqln1 | 0.018 | Young |
| 0.739307 | 0.567257 | 0.196 | 0.184 | 1 | 15 | Blvrb | 0.012 | Young |
| 0.739653 | 0.184163 | 0.714 | 0.673 | 1 | 15 | mt-Nd4l | 0.041 | Young |
| 0.740201 | 0.745669 | 0.143 | 0.122 | 1 | 15 | Med8 | 0.021 | Young |
| 0.740201 | 0.603863 | 0.143 | 0.122 | 1 | 15 | Aim2 | 0.021 | Young |
| 0.740201 | 0.426268 | 0.143 | 0.122 | 1 | 15 | Nudt4 | 0.021 | Young |
| 0.740367 | 0.200787 | 0.125 | 0.102 | 1 | 15 | Txndc15 | 0.023 | Young |
| 0.740858 | 2.166141 | 0.054 | 0.041 | 1 | 15 | Rnf152 | 0.013 | Young |
| 0.740858 | 2.064043 | 0.054 | 0.041 | 1 | 15 | Pdgfa | 0.013 | Young |
| 0.740858 | 1.987471 | 0.054 | 0.041 | 1 | 15 | Fam213a | 0.013 | Young |
| 0.740858 | 1.541596 | 0.054 | 0.041 | 1 | 15 | Zdhhc8 | 0.013 | Young |
| 0.740858 | 1.420394 | 0.054 | 0.041 | 1 | 15 | Lzic | 0.013 | Young |
| 0.740858 | 1.367703 | 0.054 | 0.041 | 1 | 15 | Wdcp | 0.013 | Young |
| 0.740858 | 1.300064 | 0.054 | 0.041 | 1 | 15 | Smagp | 0.013 | Young |
| 0.740858 | 1.251194 | 0.054 | 0.041 | 1 | 15 | Polr3b | 0.013 | Young |
| 0.740858 | 1.213904 | 0.054 | 0.041 | 1 | 15 | Qpct | 0.013 | Young |
| 0.740858 | 1.210933 | 0.054 | 0.041 | 1 | 15 | Srebf1 | 0.013 | Young |
| 0.740858 | 1.167567 | 0.054 | 0.041 | 1 | 15 | Clp1 | 0.013 | Young |
| 0.740858 | 1.076347 | 0.054 | 0.041 | 1 | 15 | Fbxo32 | 0.013 | Young |
| 0.740858 | 1.072582 | 0.054 | 0.041 | 1 | 15 | Rdh14 | 0.013 | Young |
| 0.740858 | 1.07061 | 0.054 | 0.041 | 1 | 15 | Rbm12b2 | 0.013 | Young |
| 0.740858 | 1.022036 | 0.054 | 0.041 | 1 | 15 | Cds1 | 0.013 | Young |
| 0.740858 | 1.001929 | 0.054 | 0.041 | 1 | 15 | Fndc7 | 0.013 | Young |
| 0.740858 | 0.971964 | 0.054 | 0.041 | 1 | 15 | Dst | 0.013 | Young |
| 0.740858 | 0.947639 | 0.054 | 0.041 | 1 | 15 | St3gal2 | 0.013 | Young |
| 0.740858 | 0.941824 | 0.054 | 0.041 | 1 | 15 | Sephs2 | 0.013 | Young |
| 0.740858 | 0.93496 | 0.054 | 0.041 | 1 | 15 | Wdr18 | 0.013 | Young |
| 0.740858 | 0.924481 | 0.054 | 0.041 | 1 | 15 | Uxt | 0.013 | Young |
| 0.740858 | 0.845339 | 0.054 | 0.041 | 1 | 15 | Tmem229b | 0.013 | Young |
| 0.740858 | 0.659362 | 0.054 | 0.041 | 1 | 15 | Ripk2 | 0.013 | Young |

| | | | | | | | | |
|---|---|---|---|---|---|---|---|---|
| 0.740858 | 0.562348 | 0.054 | 0.041 | 1 | 15 | Gart | 0.013 | Young |
| 0.742616 | 0.127244 | 0.232 | 0.204 | 1 | 15 | Gng2 | 0.028 | Young |
| 0.743148 | 0.271971 | 0.179 | 0.163 | 1 | 15 | Elovl1 | 0.016 | Young |
| 0.744202 | 0.545087 | 0.286 | 0.286 | 1 | 15 | Vapa | 0 | Young |
| 0.746277 | 0.51274 | 0.161 | 0.143 | 1 | 15 | Wars | 0.018 | Young |
| 0.746277 | 0.433358 | 0.161 | 0.143 | 1 | 15 | Rnaseh2c | 0.018 | Young |
| 0.746277 | 0.045615 | 0.143 | 0.163 | 1 | 15 | Dcaf7 | -0.02 | Young |
| 0.747452 | 0.336712 | 0.286 | 0.265 | 1 | 15 | Snn | 0.021 | Young |
| 0.747452 | 0.07484 | 0.286 | 0.265 | 1 | 15 | Apaf1 | 0.021 | Young |
| 0.748426 | 0.428784 | 0.143 | 0.122 | 1 | 15 | Slc25a1 | 0.021 | Young |
| 0.748808 | 0.407671 | 0.661 | 0.694 | 1 | 15 | Mt1 | -0.033 | Young |
| 0.749812 | 0.070242 | 0.857 | 0.776 | 1 | 15 | Eif4g2 | 0.081 | Young |
| 0.749911 | 0.089598 | 0.214 | 0.245 | 1 | 15 | Ppdpf | -0.031 | Young |
| 0.750671 | 0.139333 | 0.25 | 0.286 | 1 | 15 | Vsig10 | -0.036 | Young |
| 0.750733 | 0.402739 | 0.393 | 0.388 | 1 | 15 | Oxct1 | 0.005 | Young |
| 0.752303 | 0.226244 | 0.696 | 0.714 | 1 | 15 | Capza2 | -0.018 | Young |
| 0.754044 | 1.57568 | 0.054 | 0.041 | 1 | 15 | Noc3l | 0.013 | Young |
| 0.754044 | 1.353884 | 0.054 | 0.041 | 1 | 15 | Lmln | 0.013 | Young |
| 0.754044 | 1.305203 | 0.054 | 0.041 | 1 | 15 | Spsb2 | 0.013 | Young |
| 0.754044 | 1.248477 | 0.054 | 0.041 | 1 | 15 | Rhbdd1 | 0.013 | Young |
| 0.754044 | 1.24617 | 0.054 | 0.041 | 1 | 15 | Hagh | 0.013 | Young |
| 0.754044 | 1.212788 | 0.054 | 0.041 | 1 | 15 | Mcm2 | 0.013 | Young |
| 0.754044 | 1.170143 | 0.054 | 0.041 | 1 | 15 | Mecr | 0.013 | Young |
| 0.754044 | 1.126806 | 0.054 | 0.041 | 1 | 15 | Efl1 | 0.013 | Young |
| 0.754044 | 1.012515 | 0.054 | 0.041 | 1 | 15 | Igtp | 0.013 | Young |
| 0.754044 | 0.986018 | 0.054 | 0.041 | 1 | 15 | Naip5 | 0.013 | Young |
| 0.754044 | 0.95581 | 0.054 | 0.041 | 1 | 15 | Zmat1 | 0.013 | Young |
| 0.754044 | 0.927007 | 0.054 | 0.041 | 1 | 15 | Sh3bp5l | 0.013 | Young |
| 0.754044 | 0.922053 | 0.054 | 0.041 | 1 | 15 | Fhl3 | 0.013 | Young |
| 0.754044 | 0.912228 | 0.054 | 0.041 | 1 | 15 | Neurl2 | 0.013 | Young |
| 0.754044 | 0.858779 | 0.054 | 0.041 | 1 | 15 | Fam122b | 0.013 | Young |
| 0.754044 | 0.818331 | 0.054 | 0.041 | 1 | 15 | Pcnx3 | 0.013 | Young |
| 0.754044 | 0.806454 | 0.054 | 0.041 | 1 | 15 | Atat1 | 0.013 | Young |
| 0.754044 | 0.801051 | 0.054 | 0.041 | 1 | 15 | Mpst | 0.013 | Young |
| 0.754044 | 0.769031 | 0.054 | 0.041 | 1 | 15 | Ptpn4 | 0.013 | Young |
| 0.754044 | 0.757045 | 0.054 | 0.041 | 1 | 15 | Myo7a | 0.013 | Young |
| 0.754044 | 0.750973 | 0.054 | 0.041 | 1 | 15 | Def8 | 0.013 | Young |
| 0.754044 | 0.746933 | 0.054 | 0.041 | 1 | 15 | Dhx34 | 0.013 | Young |
| 0.754044 | 0.733656 | 0.054 | 0.041 | 1 | 15 | Pcgf1 | 0.013 | Young |
| 0.754044 | 0.719398 | 0.054 | 0.041 | 1 | 15 | Appl2 | 0.013 | Young |
| 0.754044 | 0.709619 | 0.054 | 0.041 | 1 | 15 | Sil1 | 0.013 | Young |
| 0.754044 | 0.691376 | 0.054 | 0.041 | 1 | 15 | Rassf8 | 0.013 | Young |
| 0.754044 | 0.684657 | 0.054 | 0.041 | 1 | 15 | Il23a | 0.013 | Young |
| 0.754044 | 0.652359 | 0.054 | 0.041 | 1 | 15 | Zfp729b | 0.013 | Young |
| 0.754044 | 0.646221 | 0.054 | 0.041 | 1 | 15 | Pcid2 | 0.013 | Young |

| | | | | | | | | |
|---|---|---|---|---|---|---|---|---|
| 0.754044 | 0.639307 | 0.054 | 0.041 | 1 | 15 | Mfn1 | 0.013 | Young |
| 0.754044 | 0.626664 | 0.054 | 0.041 | 1 | 15 | Armcx3 | 0.013 | Young |
| 0.754044 | 0.618151 | 0.054 | 0.041 | 1 | 15 | Lrrc45 | 0.013 | Young |
| 0.754044 | 0.607412 | 0.054 | 0.041 | 1 | 15 | AI839979 | 0.013 | Young |
| 0.754044 | 0.605833 | 0.054 | 0.041 | 1 | 15 | Stard3 | 0.013 | Young |
| 0.754044 | 0.57454 | 0.054 | 0.041 | 1 | 15 | Rmdn3 | 0.013 | Young |
| 0.754044 | 0.537971 | 0.054 | 0.041 | 1 | 15 | Hectd4 | 0.013 | Young |
| 0.754044 | 0.507521 | 0.054 | 0.041 | 1 | 15 | Rpf1 | 0.013 | Young |
| 0.754044 | 0.469272 | 0.054 | 0.041 | 1 | 15 | Elp3 | 0.013 | Young |
| 0.754044 | 0.457045 | 0.054 | 0.041 | 1 | 15 | Snip1 | 0.013 | Young |
| 0.754044 | 0.448648 | 0.054 | 0.041 | 1 | 15 | Zbtb6 | 0.013 | Young |
| 0.754044 | 0.421876 | 0.054 | 0.041 | 1 | 15 | Reps2 | 0.013 | Young |
| 0.754044 | 0.380371 | 0.054 | 0.041 | 1 | 15 | Zc2hc1a | 0.013 | Young |
| 0.754044 | 0.37701 | 0.054 | 0.041 | 1 | 15 | Sacs | 0.013 | Young |
| 0.754044 | 0.270771 | 0.054 | 0.041 | 1 | 15 | Ift22 | 0.013 | Young |
| 0.754068 | 0.647986 | 0.161 | 0.143 | 1 | 15 | Cops5 | 0.018 | Young |
| 0.754068 | 0.568647 | 0.161 | 0.143 | 1 | 15 | Rchy1 | 0.018 | Young |
| 0.755334 | 0.127487 | 0.429 | 0.408 | 1 | 15 | Zfp36l2 | 0.021 | Young |
| 0.755383 | 0.072695 | 0.964 | 0.918 | 1 | 15 | Rpl14 | 0.046 | Young |
| 0.756019 | 0.241279 | 0.304 | 0.286 | 1 | 15 | Smap2 | 0.018 | Young |
| 0.756118 | 0.012769 | 0.536 | 0.49 | 1 | 15 | Nrros | 0.046 | Young |
| 0.75668 | 0.335138 | 0.143 | 0.122 | 1 | 15 | Cisd1 | 0.021 | Young |
| 0.75668 | 0.302031 | 0.143 | 0.122 | 1 | 15 | Gse1 | 0.021 | Young |
| 0.757036 | 0.44514 | 0.268 | 0.245 | 1 | 15 | Fgd2 | 0.023 | Young |
| 0.757858 | 0.049096 | 0.375 | 0.429 | 1 | 15 | Dynll2 | -0.054 | Young |
| 0.757988 | 0.690629 | 0.179 | 0.163 | 1 | 15 | Ccnk | 0.016 | Young |
| 0.757988 | 0.501037 | 0.179 | 0.163 | 1 | 15 | Zfp740 | 0.016 | Young |
| 0.757988 | 0.433803 | 0.179 | 0.163 | 1 | 15 | Aida | 0.016 | Young |
| 0.757996 | 0.165924 | 0.125 | 0.102 | 1 | 15 | Hcst | 0.023 | Young |
| 0.757996 | 0.059759 | 0.125 | 0.102 | 1 | 15 | Faap100 | 0.023 | Young |
| 0.757996 | 0.049591 | 0.125 | 0.102 | 1 | 15 | Sdhaf2 | 0.023 | Young |
| 0.759461 | 0.243473 | 0.482 | 0.51 | 1 | 15 | Arf3 | -0.028 | Young |
| 0.759835 | 0.267153 | 0.286 | 0.265 | 1 | 15 | Alkbh5 | 0.021 | Young |
| 0.760122 | 0.192671 | 0.25 | 0.224 | 1 | 15 | Ing4 | 0.026 | Young |
| 0.760122 | 0.168178 | 0.214 | 0.265 | 1 | 15 | Fem1c | -0.051 | Young |
| 0.760122 | 0.046015 | 0.25 | 0.224 | 1 | 15 | Nsfl1c | 0.026 | Young |
| 0.760289 | 0.107115 | 1 | 0.939 | 1 | 15 | Rps8 | 0.061 | Young |
| 0.760292 | 0.072391 | 1 | 1 | 1 | 15 | Fau | 0 | Young |
| 0.761321 | 0.144082 | 0.375 | 0.367 | 1 | 15 | Cblb | 0.008 | Young |
| 0.761883 | 0.876501 | 0.161 | 0.143 | 1 | 15 | Gon4l | 0.018 | Young |
| 0.761883 | 0.262387 | 0.161 | 0.143 | 1 | 15 | Mpnd | 0.018 | Young |
| 0.76238 | 0.136224 | 0.339 | 0.388 | 1 | 15 | Fam107b | -0.049 | Young |
| 0.762558 | 0.561217 | 0.286 | 0.286 | 1 | 15 | Adam15 | 0 | Young |
| 0.762831 | 0.367857 | 0.232 | 0.204 | 1 | 15 | Orai1 | 0.028 | Young |
| 0.763431 | 0.058747 | 0.429 | 0.449 | 1 | 15 | Huwe1 | -0.02 | Young |

| | | | | | | | | |
|---|---|---|---|---|---|---|---|---|
| 0.763465 | 0.667968 | 0.25 | 0.245 | 1 | 15 | Cenpb | 0.005 | Young |
| 0.764961 | 0.507583 | 0.143 | 0.122 | 1 | 15 | 0610030E2 | 0.021 | Young |
| 0.766247 | 0.313051 | 0.411 | 0.429 | 1 | 15 | Tmem258 | -0.018 | Young |
| 0.767302 | 2.002786 | 0.054 | 0.041 | 1 | 15 | Inpp5a | 0.013 | Young |
| 0.767302 | 1.306587 | 0.054 | 0.041 | 1 | 15 | Pnpo | 0.013 | Young |
| 0.767302 | 1.018824 | 0.054 | 0.041 | 1 | 15 | Mrps5 | 0.013 | Young |
| 0.767302 | 0.921888 | 0.054 | 0.041 | 1 | 15 | Gamt | 0.013 | Young |
| 0.767302 | 0.871286 | 0.054 | 0.041 | 1 | 15 | Il18rap | 0.013 | Young |
| 0.767302 | 0.867248 | 0.054 | 0.041 | 1 | 15 | Mab21l3 | 0.013 | Young |
| 0.767302 | 0.687395 | 0.054 | 0.041 | 1 | 15 | Btrc | 0.013 | Young |
| 0.767302 | 0.658478 | 0.054 | 0.041 | 1 | 15 | Gpatch2l | 0.013 | Young |
| 0.767302 | 0.619386 | 0.054 | 0.041 | 1 | 15 | Nr2c1 | 0.013 | Young |
| 0.767302 | 0.617206 | 0.054 | 0.041 | 1 | 15 | Eif2ak4 | 0.013 | Young |
| 0.767302 | 0.53062 | 0.054 | 0.041 | 1 | 15 | Ssh3 | 0.013 | Young |
| 0.767302 | 0.510563 | 0.054 | 0.041 | 1 | 15 | Chm | 0.013 | Young |
| 0.767302 | 0.498129 | 0.054 | 0.041 | 1 | 15 | Trp53bp2 | 0.013 | Young |
| 0.767302 | 0.474639 | 0.054 | 0.041 | 1 | 15 | Cdc25a | 0.013 | Young |
| 0.767302 | 0.466682 | 0.054 | 0.041 | 1 | 15 | Nagpa | 0.013 | Young |
| 0.767302 | 0.443773 | 0.054 | 0.041 | 1 | 15 | Acot13 | 0.013 | Young |
| 0.767302 | 0.420796 | 0.054 | 0.041 | 1 | 15 | Zdhhc14 | 0.013 | Young |
| 0.767302 | 0.395409 | 0.054 | 0.041 | 1 | 15 | Zfp938 | 0.013 | Young |
| 0.767302 | 0.38448 | 0.054 | 0.041 | 1 | 15 | Hdac7 | 0.013 | Young |
| 0.767302 | 0.378956 | 0.054 | 0.041 | 1 | 15 | Ypel2 | 0.013 | Young |
| 0.767302 | 0.31244 | 0.054 | 0.041 | 1 | 15 | Wdr12 | 0.013 | Young |
| 0.767302 | 0.302299 | 0.054 | 0.041 | 1 | 15 | Ifitm1 | 0.013 | Young |
| 0.767302 | 0.265045 | 0.054 | 0.041 | 1 | 15 | Sft2d1 | 0.013 | Young |
| 0.767302 | 0.26238 | 0.054 | 0.041 | 1 | 15 | Eml6 | 0.013 | Young |
| 0.767302 | 0.259492 | 0.054 | 0.041 | 1 | 15 | Laptm4b | 0.013 | Young |
| 0.767302 | 0.175683 | 0.054 | 0.041 | 1 | 15 | Zfp595 | 0.013 | Young |
| 0.767302 | 0.160446 | 0.054 | 0.041 | 1 | 15 | Osbpl2 | 0.013 | Young |
| 0.767302 | 0.146978 | 0.054 | 0.041 | 1 | 15 | Gss | 0.013 | Young |
| 0.767783 | 0.717831 | 0.196 | 0.184 | 1 | 15 | R3hdm1 | 0.012 | Young |
| 0.767783 | 0.633243 | 0.196 | 0.184 | 1 | 15 | Rab7b | 0.012 | Young |
| 0.769526 | 0.351613 | 0.268 | 0.265 | 1 | 15 | Suco | 0.003 | Young |
| 0.769607 | 0.20292 | 0.232 | 0.204 | 1 | 15 | Dhx40 | 0.028 | Young |
| 0.769723 | 1.270026 | 0.143 | 0.163 | 1 | 15 | Flot2 | -0.02 | Young |
| 0.769821 | 0.215371 | 0.321 | 0.306 | 1 | 15 | Pcnp | 0.015 | Young |
| 0.76996 | 0.082389 | 0.5 | 0.49 | 1 | 15 | Epb41l2 | 0.01 | Young |
| 0.772076 | 1.195141 | 0.071 | 0.061 | 1 | 15 | Itpkc | 0.01 | Young |
| 0.772076 | 1.133027 | 0.071 | 0.061 | 1 | 15 | Sec22a | 0.01 | Young |
| 0.772076 | 1.023451 | 0.071 | 0.061 | 1 | 15 | Ap3m2 | 0.01 | Young |
| 0.772076 | 0.926358 | 0.071 | 0.061 | 1 | 15 | Msh3 | 0.01 | Young |
| 0.773255 | 0.003277 | 0.25 | 0.224 | 1 | 15 | Ccr5 | 0.026 | Young |
| 0.77327 | 0.107926 | 0.143 | 0.122 | 1 | 15 | Notch1 | 0.021 | Young |
| 0.77327 | 0.060706 | 0.125 | 0.143 | 1 | 15 | Pdxdc1 | -0.018 | Young |

| | | | | | | | | |
|---|---|---|---|---|---|---|---|---|
| 0.773567 | 0.271648 | 0.304 | 0.306 | 1 | 15 | Pi4k2b | -0.002 | Young |
| 0.773567 | 0.262389 | 0.286 | 0.327 | 1 | 15 | Mapre2 | -0.041 | Young |
| 0.77447 | 0.159177 | 0.375 | 0.429 | 1 | 15 | Smim10l1 | -0.054 | Young |
| 0.774953 | 0.163166 | 0.179 | 0.204 | 1 | 15 | Fam91a1 | -0.025 | Young |
| 0.775429 | 0.180597 | 0.464 | 0.429 | 1 | 15 | Eloc | 0.035 | Young |
| 0.775752 | 0.003618 | 0.125 | 0.102 | 1 | 15 | Srd5a3 | 0.023 | Young |
| 0.775793 | 0.080065 | 0.321 | 0.306 | 1 | 15 | Copa | 0.015 | Young |
| 0.775947 | 0.235418 | 0.357 | 0.327 | 1 | 15 | Snrpd1 | 0.03 | Young |
| 0.776426 | 0.479973 | 0.25 | 0.245 | 1 | 15 | Hcls1 | 0.005 | Young |
| 0.776469 | 0.047615 | 0.804 | 0.776 | 1 | 15 | Prkcd | 0.028 | Young |
| 0.777451 | 0.064289 | 0.946 | 0.939 | 1 | 15 | Rpl24 | 0.007 | Young |
| 0.777586 | 0.305088 | 0.143 | 0.163 | 1 | 15 | Phf11a | -0.02 | Young |
| 0.777586 | 0.281221 | 0.161 | 0.143 | 1 | 15 | Mrpl15 | 0.018 | Young |
| 0.777775 | 0.273048 | 0.357 | 0.347 | 1 | 15 | Tmem106a | 0.01 | Young |
| 0.780415 | 0.643408 | 0.179 | 0.163 | 1 | 15 | Calu | 0.016 | Young |
| 0.780415 | 0.362417 | 0.179 | 0.163 | 1 | 15 | Fxr2 | 0.016 | Young |
| 0.780415 | 0.273796 | 0.179 | 0.163 | 1 | 15 | Sarnp | 0.016 | Young |
| 0.780415 | 0.142352 | 0.179 | 0.163 | 1 | 15 | Gypc | 0.016 | Young |
| 0.780629 | 0.922854 | 0.054 | 0.041 | 1 | 15 | Mcrip2 | 0.013 | Young |
| 0.780629 | 0.560469 | 0.054 | 0.041 | 1 | 15 | Wars2 | 0.013 | Young |
| 0.780629 | 0.392264 | 0.054 | 0.041 | 1 | 15 | Cyp4f16 | 0.013 | Young |
| 0.780629 | 0.259913 | 0.054 | 0.041 | 1 | 15 | Phka1 | 0.013 | Young |
| 0.780629 | 0.242959 | 0.054 | 0.041 | 1 | 15 | Adam33 | 0.013 | Young |
| 0.780629 | 0.171953 | 0.054 | 0.041 | 1 | 15 | Dna2 | 0.013 | Young |
| 0.780629 | 0.128433 | 0.054 | 0.041 | 1 | 15 | Cdkn2d | 0.013 | Young |
| 0.780629 | 0.113941 | 0.054 | 0.041 | 1 | 15 | Cacnb1 | 0.013 | Young |
| 0.780629 | 0.09061 | 0.054 | 0.041 | 1 | 15 | 9330160F1 | 0.013 | Young |
| 0.780629 | 0.01492 | 0.054 | 0.041 | 1 | 15 | Rnf180 | 0.013 | Young |
| 0.780629 | 0.008087 | 0.054 | 0.041 | 1 | 15 | Mamld1 | 0.013 | Young |
| 0.781605 | 0.190078 | 0.143 | 0.122 | 1 | 15 | Trnt1 | 0.021 | Young |
| 0.781605 | 0.129816 | 0.125 | 0.143 | 1 | 15 | Cacna1s | -0.018 | Young |
| 0.782142 | 0.744474 | 0.196 | 0.184 | 1 | 15 | Il12b | 0.012 | Young |
| 0.782353 | 0.12883 | 0.946 | 0.898 | 1 | 15 | Hsp90ab1 | 0.048 | Young |
| 0.782602 | 0.299434 | 0.268 | 0.245 | 1 | 15 | Hexb | 0.023 | Young |
| 0.78293 | 0.148761 | 0.232 | 0.265 | 1 | 15 | Api5 | -0.033 | Young |
| 0.783026 | 0.732392 | 0.214 | 0.204 | 1 | 15 | Plagl2 | 0.01 | Young |
| 0.783246 | 0.766218 | 0.232 | 0.224 | 1 | 15 | Vhl | 0.008 | Young |
| 0.783246 | 0.327406 | 0.232 | 0.224 | 1 | 15 | Copb1 | 0.008 | Young |
| 0.783465 | 1.145409 | 0.071 | 0.061 | 1 | 15 | Tmem206 | 0.01 | Young |
| 0.784674 | 0.152743 | 0.107 | 0.122 | 1 | 15 | Rpl27-ps3 | -0.015 | Young |
| 0.785472 | 0.487662 | 0.161 | 0.143 | 1 | 15 | Dnajc13 | 0.018 | Young |
| 0.785472 | 0.24 | 0.143 | 0.163 | 1 | 15 | Cd209d | -0.02 | Young |
| 0.78621 | 0.506434 | 0.179 | 0.224 | 1 | 15 | Mir142hg | -0.045 | Young |
| 0.787003 | 0.707359 | 0.375 | 0.408 | 1 | 15 | Pf4 | -0.033 | Young |
| 0.78929 | 0.008538 | 0.5 | 0.449 | 1 | 15 | Rbx1 | 0.051 | Young |

| | | | | | | | | |
|---|---|---|---|---|---|---|---|---|
| 0.78935 | 0.427683 | 0.196 | 0.184 | 1 | 15 | Rap2c | 0.012 | Young |
| 0.78935 | 0.261941 | 0.196 | 0.184 | 1 | 15 | Nap1l4 | 0.012 | Young |
| 0.78935 | 0.013899 | 0.196 | 0.184 | 1 | 15 | Ubr2 | 0.012 | Young |
| 0.789964 | 0.052719 | 0.143 | 0.122 | 1 | 15 | Fam114a2 | 0.021 | Young |
| 0.78997 | 0.478625 | 0.214 | 0.204 | 1 | 15 | Npm3 | 0.01 | Young |
| 0.78997 | 0.046093 | 0.196 | 0.224 | 1 | 15 | Ifi209 | -0.028 | Young |
| 0.791058 | 0.142581 | 0.286 | 0.265 | 1 | 15 | Ndufa10 | 0.021 | Young |
| 0.791334 | 1.906053 | 0.089 | 0.082 | 1 | 15 | Scyl1 | 0.007 | Young |
| 0.791334 | 1.235054 | 0.089 | 0.082 | 1 | 15 | Slc35f6 | 0.007 | Young |
| 0.792252 | 0.797909 | 0.25 | 0.265 | 1 | 15 | Erlin1 | -0.015 | Young |
| 0.793381 | 0.409532 | 0.161 | 0.143 | 1 | 15 | Kdm5d | 0.018 | Young |
| 0.793381 | 0.388811 | 0.161 | 0.143 | 1 | 15 | Map3k2 | 0.018 | Young |
| 0.793381 | 0.086397 | 0.161 | 0.143 | 1 | 15 | Ccny | 0.018 | Young |
| 0.793381 | 0.061208 | 0.161 | 0.143 | 1 | 15 | A630001G2 | 0.018 | Young |
| 0.793625 | 0.46198 | 0.107 | 0.122 | 1 | 15 | Ap4s1 | -0.015 | Young |
| 0.794021 | 0.097647 | 0.054 | 0.041 | 1 | 15 | Dusp23 | 0.013 | Young |
| 0.794878 | 0.654787 | 0.268 | 0.265 | 1 | 15 | 5-Mar | 0.003 | Young |
| 0.794901 | 2.119346 | 0.071 | 0.061 | 1 | 15 | Zbtb17 | 0.01 | Young |
| 0.794901 | 1.332607 | 0.071 | 0.061 | 1 | 15 | Abcd3 | 0.01 | Young |
| 0.794901 | 1.280097 | 0.071 | 0.061 | 1 | 15 | Slc35e4 | 0.01 | Young |
| 0.794901 | 1.264843 | 0.071 | 0.061 | 1 | 15 | Mcoln1 | 0.01 | Young |
| 0.794901 | 0.884355 | 0.071 | 0.061 | 1 | 15 | Ppm1d | 0.01 | Young |
| 0.794901 | 0.703451 | 0.071 | 0.061 | 1 | 15 | Irf2bpl | 0.01 | Young |
| 0.794901 | 0.542953 | 0.071 | 0.061 | 1 | 15 | Gadd45g | 0.01 | Young |
| 0.795469 | 0.301658 | 0.179 | 0.163 | 1 | 15 | Phlpp1 | 0.016 | Young |
| 0.795983 | 0.550372 | 0.25 | 0.245 | 1 | 15 | Snw1 | 0.005 | Young |
| 0.795983 | 0.077121 | 0.232 | 0.265 | 1 | 15 | Nudcd3 | -0.033 | Young |
| 0.795989 | 0.141644 | 0.089 | 0.102 | 1 | 15 | Arap2 | -0.013 | Young |
| 0.795989 | 0.054961 | 0.089 | 0.102 | 1 | 15 | Tradd | -0.013 | Young |
| 0.796576 | 0.435675 | 0.196 | 0.184 | 1 | 15 | Lsm8 | 0.012 | Young |
| 0.796576 | 0.402228 | 0.179 | 0.204 | 1 | 15 | Ifi35 | -0.025 | Young |
| 0.797123 | 0.050637 | 0.911 | 0.878 | 1 | 15 | Actr3 | 0.033 | Young |
| 0.798348 | 0.009482 | 0.143 | 0.122 | 1 | 15 | Srsf4 | 0.021 | Young |
| 0.798767 | 0.379962 | 0.304 | 0.286 | 1 | 15 | Rnf141 | 0.018 | Young |
| 0.798811 | 0.006385 | 0.786 | 0.796 | 1 | 15 | Itm2b | -0.01 | Young |
| 0.80131 | 0.466514 | 0.161 | 0.143 | 1 | 15 | Rnf150 | 0.018 | Young |
| 0.80131 | 0.426492 | 0.143 | 0.163 | 1 | 15 | Cdk4 | -0.02 | Young |
| 0.80131 | 0.283324 | 0.143 | 0.163 | 1 | 15 | Cep120 | -0.02 | Young |
| 0.80131 | 0.130892 | 0.161 | 0.143 | 1 | 15 | Pdcd7 | 0.018 | Young |
| 0.80131 | 0.130629 | 0.161 | 0.143 | 1 | 15 | Ndfip2 | 0.018 | Young |
| 0.801711 | 0.03901 | 0.839 | 0.816 | 1 | 15 | St8sia4 | 0.023 | Young |
| 0.802532 | 0.202596 | 0.232 | 0.265 | 1 | 15 | Tmem189 | -0.033 | Young |
| 0.802604 | 0.710836 | 0.107 | 0.122 | 1 | 15 | AI504432 | -0.015 | Young |
| 0.803026 | 0.194466 | 0.179 | 0.163 | 1 | 15 | Dpf2 | 0.016 | Young |
| 0.803961 | 0.061478 | 0.536 | 0.51 | 1 | 15 | Txndc17 | 0.026 | Young |

| | | | | | | | | |
|---|---|---|---|---|---|---|---|---|
| 0.804674 | 0.126766 | 0.982 | 0.918 | 1 | 15 | Npc2 | 0.064 | Young |
| 0.804926 | 1.371472 | 0.125 | 0.122 | 1 | 15 | Anks1 | 0.003 | Young |
| 0.805917 | 0.297377 | 0.429 | 0.429 | 1 | 15 | Mdh2 | 0 | Young |
| 0.806381 | 1.201642 | 0.071 | 0.061 | 1 | 15 | Hars2 | 0.01 | Young |
| 0.806381 | 1.188203 | 0.071 | 0.061 | 1 | 15 | Mrps26 | 0.01 | Young |
| 0.806381 | 1.143897 | 0.071 | 0.061 | 1 | 15 | Pex5 | 0.01 | Young |
| 0.806381 | 1.143086 | 0.071 | 0.061 | 1 | 15 | Dnpep | 0.01 | Young |
| 0.806381 | 1.011656 | 0.071 | 0.061 | 1 | 15 | Naa16 | 0.01 | Young |
| 0.806381 | 0.998602 | 0.071 | 0.061 | 1 | 15 | Kctd5 | 0.01 | Young |
| 0.806381 | 0.916166 | 0.071 | 0.061 | 1 | 15 | Snx16 | 0.01 | Young |
| 0.806381 | 0.751696 | 0.071 | 0.061 | 1 | 15 | Polk | 0.01 | Young |
| 0.806381 | 0.696511 | 0.071 | 0.061 | 1 | 15 | Ikbkap | 0.01 | Young |
| 0.806381 | 0.645908 | 0.071 | 0.061 | 1 | 15 | Stk16 | 0.01 | Young |
| 0.806381 | 0.604442 | 0.071 | 0.061 | 1 | 15 | Vps4a | 0.01 | Young |
| 0.806381 | 0.56895 | 0.071 | 0.061 | 1 | 15 | Txndc5 | 0.01 | Young |
| 0.806381 | 0.519132 | 0.071 | 0.061 | 1 | 15 | Zxdb | 0.01 | Young |
| 0.806381 | 0.492834 | 0.071 | 0.061 | 1 | 15 | Naa20 | 0.01 | Young |
| 0.806381 | 0.459644 | 0.071 | 0.061 | 1 | 15 | Nfkbil1 | 0.01 | Young |
| 0.806799 | 0.052561 | 0.268 | 0.286 | 1 | 15 | Tnks2 | -0.018 | Young |
| 0.808059 | 0.103718 | 0.571 | 0.612 | 1 | 15 | Nav1 | -0.041 | Young |
| 0.808909 | 0.02361 | 0.804 | 0.796 | 1 | 15 | Psme1 | 0.008 | Young |
| 0.809094 | 0.275456 | 0.25 | 0.245 | 1 | 15 | Midn | 0.005 | Young |
| 0.809261 | 1.057284 | 0.161 | 0.143 | 1 | 15 | Rps6kb1 | 0.018 | Young |
| 0.809797 | 0.068231 | 0.625 | 0.633 | 1 | 15 | Atp5f1 | -0.008 | Young |
| 0.810722 | 0.02305 | 0.446 | 0.449 | 1 | 15 | Ensa | -0.003 | Young |
| 0.81108 | 0.010283 | 0.179 | 0.204 | 1 | 15 | Eloa | -0.025 | Young |
| 0.811788 | 1.242934 | 0.089 | 0.082 | 1 | 15 | Dhrs7 | 0.007 | Young |
| 0.811788 | 0.98188 | 0.089 | 0.082 | 1 | 15 | Ext2 | 0.007 | Young |
| 0.813711 | 1.785745 | 0.107 | 0.102 | 1 | 15 | Arhgap26 | 0.005 | Young |
| 0.814188 | 0.046812 | 0.286 | 0.306 | 1 | 15 | Sdhd | -0.02 | Young |
| 0.81489 | 0.219878 | 0.268 | 0.245 | 1 | 15 | Dennd6a | 0.023 | Young |
| 0.81489 | 0.060711 | 0.268 | 0.245 | 1 | 15 | Hcfc1r1 | 0.023 | Young |
| 0.815186 | 0.633082 | 0.125 | 0.143 | 1 | 15 | Abce1 | -0.018 | Young |
| 0.815186 | 0.205029 | 0.125 | 0.143 | 1 | 15 | Esf1 | -0.018 | Young |
| 0.815186 | 0.115417 | 0.143 | 0.122 | 1 | 15 | Lypla1 | 0.021 | Young |
| 0.815186 | 0.001994 | 0.125 | 0.143 | 1 | 15 | Ngdn | -0.018 | Young |
| 0.815477 | 0.033204 | 0.357 | 0.388 | 1 | 15 | Atp6ap2 | -0.031 | Young |
| 0.817001 | 0.006973 | 0.875 | 0.898 | 1 | 15 | Neat1 | -0.023 | Young |
| 0.817231 | 0.200016 | 0.161 | 0.143 | 1 | 15 | Atp9b | 0.018 | Young |
| 0.817231 | 0.091414 | 0.161 | 0.143 | 1 | 15 | Tnni2 | 0.018 | Young |
| 0.817702 | 0.119364 | 0.5 | 0.49 | 1 | 15 | Psma3 | 0.01 | Young |
| 0.81779 | 0.00999 | 0.071 | 0.082 | 1 | 15 | Slc16a10 | -0.011 | Young |
| 0.817903 | 0.846027 | 0.071 | 0.061 | 1 | 15 | Cluh | 0.01 | Young |
| 0.817903 | 0.814112 | 0.071 | 0.061 | 1 | 15 | Helz2 | 0.01 | Young |
| 0.817903 | 0.734153 | 0.071 | 0.061 | 1 | 15 | Slf1 | 0.01 | Young |

| | | | | | | | | |
|---|---|---|---|---|---|---|---|---|
| 0.817903 | 0.719427 | 0.071 | 0.061 | 1 | 15 | Mmp14 | 0.01 | Young |
| 0.817903 | 0.614938 | 0.071 | 0.061 | 1 | 15 | Depdc7 | 0.01 | Young |
| 0.817903 | 0.596703 | 0.071 | 0.061 | 1 | 15 | Pycrl | 0.01 | Young |
| 0.817903 | 0.592462 | 0.071 | 0.061 | 1 | 15 | Taf1b | 0.01 | Young |
| 0.817903 | 0.508591 | 0.071 | 0.061 | 1 | 15 | Mapk9 | 0.01 | Young |
| 0.817903 | 0.50827 | 0.071 | 0.061 | 1 | 15 | Taf7 | 0.01 | Young |
| 0.817903 | 0.488315 | 0.071 | 0.061 | 1 | 15 | Rhoh | 0.01 | Young |
| 0.817903 | 0.465006 | 0.071 | 0.061 | 1 | 15 | Sec16a | 0.01 | Young |
| 0.817903 | 0.414887 | 0.071 | 0.061 | 1 | 15 | Nolc1 | 0.01 | Young |
| 0.817903 | 0.287035 | 0.071 | 0.061 | 1 | 15 | Rxra | 0.01 | Young |
| 0.817903 | 0.037371 | 0.071 | 0.061 | 1 | 15 | Rpa1 | 0.01 | Young |
| 0.818354 | 0.717685 | 0.375 | 0.367 | 1 | 15 | Dap | 0.008 | Young |
| 0.818356 | 0.131942 | 0.179 | 0.204 | 1 | 15 | Noct | -0.025 | Young |
| 0.818356 | 0.047004 | 0.179 | 0.204 | 1 | 15 | Eif4g3 | -0.025 | Young |
| 0.818356 | 0.033853 | 0.179 | 0.204 | 1 | 15 | 2510039O1 | -0.025 | Young |
| 0.818564 | 0.08218 | 0.786 | 0.755 | 1 | 15 | Dock10 | 0.031 | Young |
| 0.820637 | 0.295712 | 0.107 | 0.122 | 1 | 15 | Kmt2b | -0.015 | Young |
| 0.820637 | 0.002578 | 0.107 | 0.122 | 1 | 15 | Gga2 | -0.015 | Young |
| 0.821095 | 0.243745 | 0.482 | 0.531 | 1 | 15 | Taldo1 | -0.049 | Young |
| 0.822349 | 1.104671 | 0.125 | 0.122 | 1 | 15 | Phtf2 | 0.003 | Young |
| 0.822843 | 0.01116 | 0.357 | 0.327 | 1 | 15 | Ehd4 | 0.03 | Young |
| 0.823774 | 0.414253 | 0.232 | 0.224 | 1 | 15 | Eif4e2 | 0.008 | Young |
| 0.823774 | 0.09393 | 0.214 | 0.245 | 1 | 15 | Rab31 | -0.031 | Young |
| 0.824933 | 0.500225 | 0.214 | 0.204 | 1 | 15 | Dennd4b | 0.01 | Young |
| 0.824933 | 0.381699 | 0.214 | 0.204 | 1 | 15 | Adar | 0.01 | Young |
| 0.824933 | 0.075428 | 0.196 | 0.224 | 1 | 15 | Fam192a | -0.028 | Young |
| 0.825076 | 0.032734 | 0.536 | 0.51 | 1 | 15 | Aes | 0.026 | Young |
| 0.825219 | 0.397313 | 0.161 | 0.143 | 1 | 15 | Stk10 | 0.018 | Young |
| 0.825219 | 0.143269 | 0.161 | 0.143 | 1 | 15 | Mgat4a | 0.018 | Young |
| 0.825684 | 0.429331 | 0.339 | 0.306 | 1 | 15 | Golga4 | 0.033 | Young |
| 0.826002 | 0.194968 | 0.429 | 0.388 | 1 | 15 | Hivep1 | 0.041 | Young |
| 0.826264 | 0.150076 | 0.571 | 0.51 | 1 | 15 | Atrx | 0.061 | Young |
| 0.827897 | 0.125733 | 0.268 | 0.245 | 1 | 15 | Nenf | 0.023 | Young |
| 0.828967 | 0.168193 | 0.286 | 0.265 | 1 | 15 | Hectd1 | 0.021 | Young |
| 0.829464 | 1.453786 | 0.071 | 0.061 | 1 | 15 | Cdkn2aip | 0.01 | Young |
| 0.829464 | 1.01818 | 0.071 | 0.061 | 1 | 15 | Nhlrc3 | 0.01 | Young |
| 0.829464 | 0.81792 | 0.071 | 0.061 | 1 | 15 | Memo1 | 0.01 | Young |
| 0.829464 | 0.659728 | 0.071 | 0.061 | 1 | 15 | Idh3a | 0.01 | Young |
| 0.829464 | 0.502323 | 0.071 | 0.061 | 1 | 15 | Ccdc90b | 0.01 | Young |
| 0.829464 | 0.492346 | 0.071 | 0.061 | 1 | 15 | Slc12a2 | 0.01 | Young |
| 0.829464 | 0.437949 | 0.071 | 0.061 | 1 | 15 | Lztr1 | 0.01 | Young |
| 0.829464 | 0.436741 | 0.071 | 0.061 | 1 | 15 | Sppl2b | 0.01 | Young |
| 0.829464 | 0.431875 | 0.071 | 0.061 | 1 | 15 | Pknox1 | 0.01 | Young |
| 0.829464 | 0.285951 | 0.071 | 0.061 | 1 | 15 | Dvl3 | 0.01 | Young |
| 0.829464 | 0.243745 | 0.071 | 0.061 | 1 | 15 | Gpaa1 | 0.01 | Young |

| | | | | | | | | |
|---|---|---|---|---|---|---|---|---|
| 0.829464 | 0.117145 | 0.071 | 0.061 | 1 | 15 | Pigq | 0.01 | Young |
| 0.829464 | 0.116556 | 0.071 | 0.061 | 1 | 15 | Qser1 | 0.01 | Young |
| 0.829464 | 0.071349 | 0.071 | 0.061 | 1 | 15 | Dip2a | 0.01 | Young |
| 0.829691 | 0.061814 | 0.107 | 0.122 | 1 | 15 | Stk19 | -0.015 | Young |
| 0.830578 | 0.186504 | 0.232 | 0.224 | 1 | 15 | Snf8 | 0.008 | Young |
| 0.831094 | 0.990476 | 0.125 | 0.122 | 1 | 15 | Nfyc | 0.003 | Young |
| 0.831094 | 0.640054 | 0.125 | 0.122 | 1 | 15 | Klhdc3 | 0.003 | Young |
| 0.831941 | 0.00176 | 0.875 | 0.857 | 1 | 15 | Actr2 | 0.018 | Young |
| 0.831969 | 0.016626 | 0.214 | 0.204 | 1 | 15 | Tecr | 0.01 | Young |
| 0.83237 | 1.122376 | 0.089 | 0.082 | 1 | 15 | Rnpc3 | 0.007 | Young |
| 0.83237 | 0.998143 | 0.089 | 0.082 | 1 | 15 | Mcm5 | 0.007 | Young |
| 0.83237 | 0.935245 | 0.089 | 0.082 | 1 | 15 | Ttc37 | 0.007 | Young |
| 0.83237 | 0.874982 | 0.089 | 0.082 | 1 | 15 | Eif2d | 0.007 | Young |
| 0.83237 | 0.59156 | 0.089 | 0.082 | 1 | 15 | P2ry6 | 0.007 | Young |
| 0.83237 | 0.519951 | 0.089 | 0.082 | 1 | 15 | Heatr6 | 0.007 | Young |
| 0.832518 | 1.183455 | 0.107 | 0.102 | 1 | 15 | Cdkn2aipnl | 0.005 | Young |
| 0.832518 | 1.158056 | 0.107 | 0.102 | 1 | 15 | Lgals3bp | 0.005 | Young |
| 0.832518 | 0.881148 | 0.107 | 0.102 | 1 | 15 | Smc2 | 0.005 | Young |
| 0.832956 | 0.169463 | 0.179 | 0.204 | 1 | 15 | Chd6 | -0.025 | Young |
| 0.832956 | 0.007189 | 0.196 | 0.184 | 1 | 15 | Fam126a | 0.012 | Young |
| 0.833428 | 0.38628 | 0.179 | 0.163 | 1 | 15 | 4931406C0 | 0.016 | Young |
| 0.833428 | 0.134581 | 0.179 | 0.163 | 1 | 15 | Arnt | 0.016 | Young |
| 0.833428 | 0.058868 | 0.161 | 0.184 | 1 | 15 | Gbp5 | -0.023 | Young |
| 0.833428 | 0.025487 | 0.161 | 0.184 | 1 | 15 | Usp8 | -0.023 | Young |
| 0.835156 | 0.032168 | 0.089 | 0.102 | 1 | 15 | Tars | -0.013 | Young |
| 0.836256 | 0.288706 | 0.536 | 0.531 | 1 | 15 | Limd1 | 0.005 | Young |
| 0.836908 | 0.000803 | 0.679 | 0.673 | 1 | 15 | Ptp4a2 | 0.006 | Young |
| 0.838767 | 0.191795 | 0.107 | 0.122 | 1 | 15 | Chpt1 | -0.015 | Young |
| 0.838767 | 0.084122 | 0.107 | 0.122 | 1 | 15 | Ccdc25 | -0.015 | Young |
| 0.83902 | 0.037439 | 0.196 | 0.224 | 1 | 15 | Rapgef6 | -0.028 | Young |
| 0.839659 | 0.140053 | 1 | 0.939 | 1 | 15 | Rps27 | 0.061 | Young |
| 0.839727 | 0.070473 | 0.375 | 0.388 | 1 | 15 | Selenot | -0.013 | Young |
| 0.839755 | 0.174728 | 0.268 | 0.265 | 1 | 15 | Inpp5k | 0.003 | Young |
| 0.840048 | 0.335442 | 0.446 | 0.429 | 1 | 15 | Pdia3 | 0.017 | Young |
| 0.840278 | 0.3736 | 0.196 | 0.184 | 1 | 15 | Srp19 | 0.012 | Young |
| 0.8406 | 0.175467 | 0.125 | 0.143 | 1 | 15 | Sik2 | -0.018 | Young |
| 0.8406 | 0.060394 | 0.125 | 0.143 | 1 | 15 | Mta2 | -0.018 | Young |
| 0.841063 | 1.007749 | 0.071 | 0.061 | 1 | 15 | Fkrp | 0.01 | Young |
| 0.841063 | 0.900557 | 0.071 | 0.061 | 1 | 15 | Kremen1 | 0.01 | Young |
| 0.841063 | 0.761726 | 0.071 | 0.061 | 1 | 15 | Megf8 | 0.01 | Young |
| 0.841063 | 0.57846 | 0.071 | 0.061 | 1 | 15 | P3h2 | 0.01 | Young |
| 0.841063 | 0.505997 | 0.071 | 0.061 | 1 | 15 | Atrn | 0.01 | Young |
| 0.841063 | 0.413174 | 0.071 | 0.061 | 1 | 15 | Zfp598 | 0.01 | Young |
| 0.841063 | 0.404884 | 0.071 | 0.061 | 1 | 15 | Hdac5 | 0.01 | Young |
| 0.841063 | 0.403438 | 0.071 | 0.061 | 1 | 15 | Ccdc127 | 0.01 | Young |

| | | | | | | | | |
|---|---|---|---|---|---|---|---|---|
| 0.841063 | 0.359617 | 0.071 | 0.061 | 1 | 15 | Fam213b | 0.01 | Young |
| 0.841063 | 0.276974 | 0.071 | 0.061 | 1 | 15 | Xab2 | 0.01 | Young |
| 0.841063 | 0.263396 | 0.071 | 0.061 | 1 | 15 | Ftsj3 | 0.01 | Young |
| 0.841063 | 0.165287 | 0.071 | 0.061 | 1 | 15 | Casp9 | 0.01 | Young |
| 0.841063 | 0.134936 | 0.071 | 0.061 | 1 | 15 | Rnf41 | 0.01 | Young |
| 0.841063 | 0.126527 | 0.071 | 0.061 | 1 | 15 | Stim2 | 0.01 | Young |
| 0.841063 | 0.106927 | 0.071 | 0.061 | 1 | 15 | G6pc3 | 0.01 | Young |
| 0.841063 | 0.103052 | 0.071 | 0.061 | 1 | 15 | 2700062CC | 0.01 | Young |
| 0.841063 | 0.059153 | 0.071 | 0.061 | 1 | 15 | Fktn | 0.01 | Young |
| 0.841063 | 0.045811 | 0.071 | 0.061 | 1 | 15 | Patl1 | 0.01 | Young |
| 0.841063 | 0.014022 | 0.071 | 0.061 | 1 | 15 | Rrm1 | 0.01 | Young |
| 0.84107 | 0.297596 | 0.161 | 0.184 | 1 | 15 | Uqcrc1 | -0.023 | Young |
| 0.84107 | 0.241257 | 0.179 | 0.163 | 1 | 15 | Ergic2 | 0.016 | Young |
| 0.84107 | 0.008249 | 0.161 | 0.184 | 1 | 15 | Tnip2 | -0.023 | Young |
| 0.841251 | 0.372532 | 0.143 | 0.163 | 1 | 15 | Trnau1ap | -0.02 | Young |
| 0.841251 | 0.000482 | 0.143 | 0.163 | 1 | 15 | Yipf5 | -0.02 | Young |
| 0.841717 | 0.002525 | 0.214 | 0.224 | 1 | 15 | Serpinb1a | -0.01 | Young |
| 0.842706 | 0.88495 | 0.089 | 0.082 | 1 | 15 | Hdac4 | 0.007 | Young |
| 0.842706 | 0.854966 | 0.089 | 0.082 | 1 | 15 | Ckap4 | 0.007 | Young |
| 0.842706 | 0.622424 | 0.089 | 0.082 | 1 | 15 | Nkiras2 | 0.007 | Young |
| 0.842706 | 0.165576 | 0.089 | 0.082 | 1 | 15 | Pi4ka | 0.007 | Young |
| 0.844867 | 0.048965 | 0.339 | 0.367 | 1 | 15 | Glg1 | -0.028 | Young |
| 0.845015 | 0.066231 | 0.089 | 0.102 | 1 | 15 | Prr7 | -0.013 | Young |
| 0.845176 | 0.755109 | 0.143 | 0.143 | 1 | 15 | Opa1 | 0 | Young |
| 0.84519 | 0.067567 | 0.232 | 0.204 | 1 | 15 | Mdp1 | 0.028 | Young |
| 0.846083 | 0.546922 | 0.214 | 0.204 | 1 | 15 | Prkacb | 0.01 | Young |
| 0.846855 | 0.25016 | 0.554 | 0.51 | 1 | 15 | Mef2a | 0.044 | Young |
| 0.847614 | 0.44554 | 0.196 | 0.184 | 1 | 15 | Dse | 0.012 | Young |
| 0.847813 | 0.226271 | 0.357 | 0.347 | 1 | 15 | Rad21 | 0.01 | Young |
| 0.847864 | 0.077439 | 0.107 | 0.122 | 1 | 15 | Sec13 | -0.015 | Young |
| 0.847864 | 0.048884 | 0.107 | 0.122 | 1 | 15 | Terf2 | -0.015 | Young |
| 0.847864 | 0.041845 | 0.107 | 0.122 | 1 | 15 | Eef1e1 | -0.015 | Young |
| 0.848363 | 0.252345 | 0.304 | 0.286 | 1 | 15 | Vrk2 | 0.018 | Young |
| 0.848646 | 1.481636 | 0.125 | 0.122 | 1 | 15 | Lrrfip2 | 0.003 | Young |
| 0.848646 | 0.767682 | 0.125 | 0.122 | 1 | 15 | Ormdl2 | 0.003 | Young |
| 0.849109 | 0.721948 | 0.125 | 0.143 | 1 | 15 | Gramd1b | -0.018 | Young |
| 0.849109 | 0.186314 | 0.125 | 0.143 | 1 | 15 | Gatd1 | -0.018 | Young |
| 0.849291 | 0.271279 | 0.161 | 0.143 | 1 | 15 | Cdc123 | 0.018 | Young |
| 0.849556 | 0.004507 | 0.607 | 0.571 | 1 | 15 | Foxp1 | 0.036 | Young |
| 0.850485 | 0.638167 | 0.071 | 0.082 | 1 | 15 | Usp40 | -0.011 | Young |
| 0.850485 | 0.33946 | 0.071 | 0.082 | 1 | 15 | Rnf168 | -0.011 | Young |
| 0.850485 | 0.338126 | 0.071 | 0.082 | 1 | 15 | Frs2 | -0.011 | Young |
| 0.850485 | 0.062516 | 0.071 | 0.082 | 1 | 15 | Pdcd2l | -0.011 | Young |
| 0.850762 | 0.419912 | 0.339 | 0.367 | 1 | 15 | Anxa7 | -0.028 | Young |
| 0.851064 | 0.378159 | 0.232 | 0.224 | 1 | 15 | Cdipt | 0.008 | Young |

| | | | | | | | | |
|---|---|---|---|---|---|---|---|---|
| 0.851064 | 0.055006 | 0.232 | 0.224 | 1 | 15 | Thoc2 | 0.008 | Young |
| 0.851422 | 0.554657 | 0.107 | 0.102 | 1 | 15 | Prkaca | 0.005 | Young |
| 0.851422 | 0.456626 | 0.107 | 0.102 | 1 | 15 | B9d2 | 0.005 | Young |
| 0.851778 | 0.269067 | 0.589 | 0.571 | 1 | 15 | Sdc4 | 0.018 | Young |
| 0.85227 | 0.037239 | 0.393 | 0.388 | 1 | 15 | Cdc42se2 | 0.005 | Young |
| 0.852696 | 0.545613 | 0.071 | 0.061 | 1 | 15 | Bag5 | 0.01 | Young |
| 0.852696 | 0.186262 | 0.071 | 0.061 | 1 | 15 | Usp53 | 0.01 | Young |
| 0.852696 | 0.174529 | 0.071 | 0.061 | 1 | 15 | Acsf3 | 0.01 | Young |
| 0.852696 | 0.010973 | 0.071 | 0.061 | 1 | 15 | Ccdc97 | 0.01 | Young |
| 0.852696 | 0.00582 | 0.071 | 0.061 | 1 | 15 | Borcs5 | 0.01 | Young |
| 0.853068 | 0.813816 | 0.089 | 0.082 | 1 | 15 | Parp10 | 0.007 | Young |
| 0.853068 | 0.729954 | 0.089 | 0.082 | 1 | 15 | Hsdl1 | 0.007 | Young |
| 0.853158 | 0.497744 | 0.214 | 0.204 | 1 | 15 | Nsun2 | 0.01 | Young |
| 0.853158 | 0.41095 | 0.214 | 0.204 | 1 | 15 | 2310011J0 | 0.01 | Young |
| 0.853158 | 0.410146 | 0.214 | 0.204 | 1 | 15 | Dcaf12 | 0.01 | Young |
| 0.853247 | 0.016913 | 0.393 | 0.408 | 1 | 15 | Cox7b | -0.015 | Young |
| 0.853448 | 0.76676 | 0.143 | 0.143 | 1 | 15 | Zbtb4 | 0 | Young |
| 0.8549 | 0.263946 | 0.089 | 0.102 | 1 | 15 | Cul2 | -0.013 | Young |
| 0.8549 | 0.206434 | 0.089 | 0.102 | 1 | 15 | Papd5 | -0.013 | Young |
| 0.8549 | 0.196186 | 0.089 | 0.102 | 1 | 15 | Rpp25l | -0.013 | Young |
| 0.854963 | 0.327007 | 0.196 | 0.184 | 1 | 15 | Chchd1 | 0.012 | Young |
| 0.854963 | 0.31391 | 0.196 | 0.184 | 1 | 15 | Pik3r6 | 0.012 | Young |
| 0.854963 | 0.038349 | 0.179 | 0.204 | 1 | 15 | Hmox2 | -0.025 | Young |
| 0.855895 | 0.205153 | 0.321 | 0.327 | 1 | 15 | Eprs | -0.006 | Young |
| 0.856398 | 0.201854 | 0.179 | 0.163 | 1 | 15 | Tmem9b | 0.016 | Young |
| 0.856783 | 0.078759 | 0.804 | 0.816 | 1 | 15 | Fscn1 | -0.012 | Young |
| 0.856943 | 0.822984 | 0.161 | 0.163 | 1 | 15 | Axl | -0.002 | Young |
| 0.856943 | 0.454562 | 0.161 | 0.163 | 1 | 15 | Tdrd7 | -0.002 | Young |
| 0.856982 | 0.26469 | 0.107 | 0.122 | 1 | 15 | Bin3 | -0.015 | Young |
| 0.856982 | 0.164804 | 0.107 | 0.122 | 1 | 15 | Prkag1 | -0.015 | Young |
| 0.856982 | 0.13779 | 0.107 | 0.122 | 1 | 15 | Lcorl | -0.015 | Young |
| 0.856982 | 0.111163 | 0.107 | 0.122 | 1 | 15 | Cdyl | -0.015 | Young |
| 0.856982 | 0.090511 | 0.107 | 0.122 | 1 | 15 | Pygo2 | -0.015 | Young |
| 0.857003 | 0.041168 | 0.411 | 0.408 | 1 | 15 | Tgoln1 | 0.003 | Young |
| 0.857012 | 0.008632 | 0.375 | 0.388 | 1 | 15 | Sumo1 | -0.013 | Young |
| 0.857206 | 0.170988 | 0.321 | 0.347 | 1 | 15 | Rbbp4 | -0.026 | Young |
| 0.857348 | 0.459965 | 0.143 | 0.163 | 1 | 15 | Man1c1 | -0.02 | Young |
| 0.857348 | 0.322299 | 0.143 | 0.163 | 1 | 15 | Rnf2 | -0.02 | Young |
| 0.857348 | 0.005164 | 0.161 | 0.143 | 1 | 15 | Slc25a37 | 0.018 | Young |
| 0.860246 | 0.073888 | 0.214 | 0.204 | 1 | 15 | Klrk1 | 0.01 | Young |
| 0.860635 | 0.034552 | 0.321 | 0.306 | 1 | 15 | Foxp4 | 0.015 | Young |
| 0.860733 | 0.28813 | 0.357 | 0.367 | 1 | 15 | Ube2f | -0.01 | Young |
| 0.860907 | 0.851638 | 0.107 | 0.102 | 1 | 15 | Zbtb44 | 0.005 | Young |
| 0.861444 | 0.565358 | 0.071 | 0.082 | 1 | 15 | Brix1 | -0.011 | Young |
| 0.861444 | 0.35359 | 0.071 | 0.082 | 1 | 15 | Hmgcl | -0.011 | Young |

| | | | | | | | | |
|---|---|---|---|---|---|---|---|---|
| 0.861444 | 0.043457 | 0.071 | 0.082 | 1 | 15 | Slc7a6 | -0.011 | Young |
| 0.861444 | 0.041863 | 0.071 | 0.082 | 1 | 15 | Patz1 | -0.011 | Young |
| 0.862324 | 0.136455 | 0.196 | 0.184 | 1 | 15 | Ncstn | 0.012 | Young |
| 0.863457 | 0.678874 | 0.089 | 0.082 | 1 | 15 | Stambp | 0.007 | Young |
| 0.863457 | 0.590143 | 0.089 | 0.082 | 1 | 15 | 0610010F0 | 0.007 | Young |
| 0.863457 | 0.427183 | 0.089 | 0.082 | 1 | 15 | Tada1 | 0.007 | Young |
| 0.863457 | 0.362419 | 0.089 | 0.082 | 1 | 15 | Tsta3 | 0.007 | Young |
| 0.863457 | 0.255392 | 0.089 | 0.082 | 1 | 15 | Tpst2 | 0.007 | Young |
| 0.864082 | 0.091347 | 0.179 | 0.163 | 1 | 15 | Sucla2 | 0.016 | Young |
| 0.864361 | 0.130977 | 0.071 | 0.061 | 1 | 15 | 4930402H2 | 0.01 | Young |
| 0.864361 | 0.021468 | 0.071 | 0.061 | 1 | 15 | Ttc13 | 0.01 | Young |
| 0.864807 | 0.377322 | 0.089 | 0.102 | 1 | 15 | Ccdc34 | -0.013 | Young |
| 0.864807 | 0.291252 | 0.089 | 0.102 | 1 | 15 | Nsrp1 | -0.013 | Young |
| 0.864807 | 0.245355 | 0.089 | 0.102 | 1 | 15 | Lin54 | -0.013 | Young |
| 0.864807 | 0.076513 | 0.089 | 0.102 | 1 | 15 | Hck | -0.013 | Young |
| 0.865419 | 0.251514 | 0.143 | 0.163 | 1 | 15 | Zbtb10 | -0.02 | Young |
| 0.865536 | 0.180329 | 0.357 | 0.347 | 1 | 15 | Vps35 | 0.01 | Young |
| 0.865644 | 0.049256 | 0.268 | 0.265 | 1 | 15 | Slc44a1 | 0.003 | Young |
| 0.866119 | 0.459377 | 0.107 | 0.122 | 1 | 15 | Rev3l | -0.015 | Young |
| 0.866119 | 0.153145 | 0.107 | 0.122 | 1 | 15 | Tsfm | -0.015 | Young |
| 0.866181 | 0.111453 | 0.125 | 0.143 | 1 | 15 | Pbx3 | -0.018 | Young |
| 0.866858 | 0.657162 | 0.054 | 0.061 | 1 | 15 | Ak6 | -0.007 | Young |
| 0.866858 | 0.159672 | 0.054 | 0.061 | 1 | 15 | Ermard | -0.007 | Young |
| 0.866858 | 0.079538 | 0.054 | 0.061 | 1 | 15 | Med24 | -0.007 | Young |
| 0.867344 | 0.263089 | 0.214 | 0.204 | 1 | 15 | Rsbn1l | 0.01 | Young |
| 0.867824 | 0.071329 | 0.464 | 0.408 | 1 | 15 | D8Ertd738 | 0.056 | Young |
| 0.868042 | 0.05416 | 0.321 | 0.327 | 1 | 15 | Ets2 | -0.006 | Young |
| 0.868706 | 0.070437 | 0.25 | 0.245 | 1 | 15 | Safb | 0.005 | Young |
| 0.869215 | 0.123675 | 0.786 | 0.776 | 1 | 15 | Tnfaip3 | 0.01 | Young |
| 0.869244 | 0.121915 | 0.321 | 0.347 | 1 | 15 | Kif1b | -0.026 | Young |
| 0.870225 | 0.051199 | 0.732 | 0.612 | 1 | 15 | Atp6v0e | 0.12 | Young |
| 0.870411 | 0.961059 | 0.107 | 0.102 | 1 | 15 | Bex3 | 0.005 | Young |
| 0.870411 | 0.747556 | 0.107 | 0.102 | 1 | 15 | Spast | 0.005 | Young |
| 0.871 | 0.344134 | 0.464 | 0.51 | 1 | 15 | Zbtb18 | -0.046 | Young |
| 0.871651 | 0.562256 | 0.232 | 0.224 | 1 | 15 | Fabp4 | 0.008 | Young |
| 0.87178 | 0.131841 | 0.179 | 0.163 | 1 | 15 | Tmem208 | 0.016 | Young |
| 0.872429 | 0.230592 | 0.071 | 0.082 | 1 | 15 | Naa60 | -0.011 | Young |
| 0.872429 | 0.164537 | 0.071 | 0.082 | 1 | 15 | Ctbs | -0.011 | Young |
| 0.872429 | 0.10478 | 0.071 | 0.082 | 1 | 15 | Fdxacb1 | -0.011 | Young |
| 0.872429 | 0.041688 | 0.071 | 0.082 | 1 | 15 | Smn1 | -0.011 | Young |
| 0.872558 | 0.116107 | 0.536 | 0.49 | 1 | 15 | Eif3h | 0.046 | Young |
| 0.873505 | 0.021233 | 0.143 | 0.163 | 1 | 15 | Eif4enif1 | -0.02 | Young |
| 0.873868 | 0.845472 | 0.089 | 0.082 | 1 | 15 | Yrdc | 0.007 | Young |
| 0.873868 | 0.844539 | 0.089 | 0.082 | 1 | 15 | Cog4 | 0.007 | Young |
| 0.873868 | 0.741109 | 0.089 | 0.082 | 1 | 15 | Mplkip | 0.007 | Young |

| | | | | | | | | |
|---|---|---|---|---|---|---|---|---|
| 0.873868 | 0.60012 | 0.089 | 0.082 | 1 | 15 | Cables2 | 0.007 | Young |
| 0.873868 | 0.571259 | 0.089 | 0.082 | 1 | 15 | Timm44 | 0.007 | Young |
| 0.873868 | 0.512942 | 0.089 | 0.082 | 1 | 15 | Tmem88 | 0.007 | Young |
| 0.873868 | 0.264973 | 0.089 | 0.082 | 1 | 15 | Prps2 | 0.007 | Young |
| 0.873868 | 0.26246 | 0.089 | 0.082 | 1 | 15 | Cwc25 | 0.007 | Young |
| 0.873868 | 0.224283 | 0.089 | 0.082 | 1 | 15 | Dld | 0.007 | Young |
| 0.873868 | 0.111376 | 0.089 | 0.082 | 1 | 15 | Kctd13 | 0.007 | Young |
| 0.874454 | 0.302682 | 0.196 | 0.224 | 1 | 15 | Rnf145 | -0.028 | Young |
| 0.874454 | 0.192773 | 0.214 | 0.204 | 1 | 15 | Slc25a36 | 0.01 | Young |
| 0.874735 | 0.254823 | 0.089 | 0.102 | 1 | 15 | Nomo1 | -0.013 | Young |
| 0.874741 | 0.521182 | 0.125 | 0.143 | 1 | 15 | Psip1 | -0.018 | Young |
| 0.874741 | 0.17884 | 0.125 | 0.143 | 1 | 15 | Sms | -0.018 | Young |
| 0.874964 | 0.168102 | 0.643 | 0.673 | 1 | 15 | Ubb | -0.03 | Young |
| 0.875113 | 1.240098 | 0.125 | 0.122 | 1 | 15 | Phkb | 0.003 | Young |
| 0.875113 | 1.198823 | 0.125 | 0.122 | 1 | 15 | Gpc1 | 0.003 | Young |
| 0.875274 | 0.361862 | 0.107 | 0.122 | 1 | 15 | Ffar4 | -0.015 | Young |
| 0.875274 | 0.351499 | 0.107 | 0.122 | 1 | 15 | Arsa | -0.015 | Young |
| 0.875274 | 0.348646 | 0.107 | 0.122 | 1 | 15 | Ythdc2 | -0.015 | Young |
| 0.875274 | 0.079525 | 0.107 | 0.122 | 1 | 15 | Sclt1 | -0.015 | Young |
| 0.876056 | 0.053738 | 0.071 | 0.061 | 1 | 15 | Prkaa1 | 0.01 | Young |
| 0.876056 | 0.026391 | 0.071 | 0.061 | 1 | 15 | Gm15417 | 0.01 | Young |
| 0.878355 | 0.71099 | 0.143 | 0.143 | 1 | 15 | Abcc5 | 0 | Young |
| 0.878814 | 0.395438 | 0.518 | 0.551 | 1 | 15 | Fos | -0.033 | Young |
| 0.879436 | 1.010943 | 0.054 | 0.061 | 1 | 15 | Klhl21 | -0.007 | Young |
| 0.879436 | 0.324077 | 0.054 | 0.061 | 1 | 15 | Ttpal | -0.007 | Young |
| 0.879436 | 0.275917 | 0.054 | 0.061 | 1 | 15 | Tmem126a | -0.007 | Young |
| 0.879436 | 0.241146 | 0.054 | 0.061 | 1 | 15 | Alg8 | -0.007 | Young |
| 0.879436 | 0.216741 | 0.054 | 0.061 | 1 | 15 | Rab24 | -0.007 | Young |
| 0.879436 | 0.140415 | 0.054 | 0.061 | 1 | 15 | Ltbp3 | -0.007 | Young |
| 0.879436 | 0.092495 | 0.054 | 0.061 | 1 | 15 | Dnajb2 | -0.007 | Young |
| 0.879436 | 0.070837 | 0.054 | 0.061 | 1 | 15 | Zfp398 | -0.007 | Young |
| 0.879436 | 0.064818 | 0.054 | 0.061 | 1 | 15 | Smim12 | -0.007 | Young |
| 0.879436 | 0.040361 | 0.054 | 0.061 | 1 | 15 | Adprhl2 | -0.007 | Young |
| 0.879436 | 0.034232 | 0.054 | 0.061 | 1 | 15 | Upf2 | -0.007 | Young |
| 0.879436 | 0.017656 | 0.054 | 0.061 | 1 | 15 | Chst2 | -0.007 | Young |
| 0.879436 | 0.015728 | 0.054 | 0.061 | 1 | 15 | Wdr61 | -0.007 | Young |
| 0.87949 | 0.034935 | 0.179 | 0.163 | 1 | 15 | Tmem165 | 0.016 | Young |
| 0.879935 | 1.079185 | 0.107 | 0.102 | 1 | 15 | Pja1 | 0.005 | Young |
| 0.879935 | 0.670647 | 0.107 | 0.102 | 1 | 15 | Arl13b | 0.005 | Young |
| 0.879935 | 0.3829 | 0.107 | 0.102 | 1 | 15 | Stam2 | 0.005 | Young |
| 0.881516 | 0.271874 | 0.286 | 0.286 | 1 | 15 | Eea1 | 0 | Young |
| 0.881604 | 0.463463 | 0.143 | 0.163 | 1 | 15 | Btbd1 | -0.02 | Young |
| 0.881604 | 0.438589 | 0.143 | 0.163 | 1 | 15 | Bbx | -0.02 | Young |
| 0.881604 | 0.348701 | 0.143 | 0.163 | 1 | 15 | Nab2 | -0.02 | Young |
| 0.881604 | 0.287395 | 0.161 | 0.143 | 1 | 15 | Cysltr1 | 0.018 | Young |

| | | | | | | | | |
|---|---|---|---|---|---|---|---|---|
| 0.882347 | 0.027529 | 0.339 | 0.347 | 1 | 15 | Pan3 | -0.008 | Young |
| 0.883315 | 0.3851 | 0.125 | 0.143 | 1 | 15 | Cdc42bpb | -0.018 | Young |
| 0.883439 | 1.260851 | 0.071 | 0.082 | 1 | 15 | Sigmar1 | -0.011 | Young |
| 0.883439 | 0.568683 | 0.071 | 0.082 | 1 | 15 | Utp15 | -0.011 | Young |
| 0.883439 | 0.211556 | 0.071 | 0.082 | 1 | 15 | Frmd8 | -0.011 | Young |
| 0.883439 | 0.14284 | 0.071 | 0.082 | 1 | 15 | Mfge8 | -0.011 | Young |
| 0.883439 | 0.137712 | 0.071 | 0.082 | 1 | 15 | Ascc1 | -0.011 | Young |
| 0.883968 | 0.600737 | 0.125 | 0.122 | 1 | 15 | Fbxo38 | 0.003 | Young |
| 0.883968 | 0.201279 | 0.125 | 0.122 | 1 | 15 | Smarcb1 | 0.003 | Young |
| 0.884058 | 0.387209 | 0.304 | 0.306 | 1 | 15 | Ncoa2 | -0.002 | Young |
| 0.884058 | 0.042797 | 0.304 | 0.306 | 1 | 15 | Papola | -0.002 | Young |
| 0.884302 | 0.718856 | 0.089 | 0.082 | 1 | 15 | Ganc | 0.007 | Young |
| 0.884302 | 0.59958 | 0.089 | 0.082 | 1 | 15 | Gnpda2 | 0.007 | Young |
| 0.884302 | 0.509411 | 0.089 | 0.082 | 1 | 15 | Cdk10 | 0.007 | Young |
| 0.884302 | 0.292519 | 0.089 | 0.082 | 1 | 15 | Scamp1 | 0.007 | Young |
| 0.884302 | 0.282576 | 0.089 | 0.082 | 1 | 15 | Il18 | 0.007 | Young |
| 0.884302 | 0.24509 | 0.089 | 0.082 | 1 | 15 | Sesn2 | 0.007 | Young |
| 0.884302 | 0.096754 | 0.089 | 0.082 | 1 | 15 | Zfp511 | 0.007 | Young |
| 0.884302 | 0.045112 | 0.089 | 0.082 | 1 | 15 | Shc1 | 0.007 | Young |
| 0.884302 | 0.030417 | 0.089 | 0.082 | 1 | 15 | Rcbtb1 | 0.007 | Young |
| 0.884445 | 0.541596 | 0.107 | 0.122 | 1 | 15 | Snx11 | -0.015 | Young |
| 0.884445 | 0.450544 | 0.107 | 0.122 | 1 | 15 | Myo1e | -0.015 | Young |
| 0.884445 | 0.229459 | 0.107 | 0.122 | 1 | 15 | Dpagt1 | -0.015 | Young |
| 0.884683 | 0.522895 | 0.089 | 0.102 | 1 | 15 | Parp12 | -0.013 | Young |
| 0.884683 | 0.462213 | 0.089 | 0.102 | 1 | 15 | Lair1 | -0.013 | Young |
| 0.884683 | 0.371748 | 0.089 | 0.102 | 1 | 15 | Smu1 | -0.013 | Young |
| 0.884683 | 0.339167 | 0.089 | 0.102 | 1 | 15 | Dgcr2 | -0.013 | Young |
| 0.88509 | 0.012761 | 1 | 1 | 1 | 15 | Rps29 | 0 | Young |
| 0.885425 | 0.738487 | 0.232 | 0.224 | 1 | 15 | Psme3 | 0.008 | Young |
| 0.88676 | 0.379125 | 0.393 | 0.388 | 1 | 15 | Cyth1 | 0.005 | Young |
| 0.887211 | 0.082416 | 0.179 | 0.163 | 1 | 15 | Csnk1g2 | 0.016 | Young |
| 0.887211 | 0.020301 | 0.179 | 0.163 | 1 | 15 | Arid2 | 0.016 | Young |
| 0.887512 | 0.130467 | 0.411 | 0.388 | 1 | 15 | Gm2a | 0.023 | Young |
| 0.887877 | 0.165686 | 0.268 | 0.306 | 1 | 15 | Tbc1d15 | -0.038 | Young |
| 0.887877 | 0.119127 | 0.286 | 0.286 | 1 | 15 | Ist1 | 0 | Young |
| 0.888703 | 0.07026 | 0.214 | 0.204 | 1 | 15 | Tmem127 | 0.01 | Young |
| 0.889476 | 0.478691 | 0.107 | 0.102 | 1 | 15 | Trappc9 | 0.005 | Young |
| 0.889476 | 0.402436 | 0.107 | 0.102 | 1 | 15 | Tmem183a | 0.005 | Young |
| 0.889476 | 0.298828 | 0.107 | 0.102 | 1 | 15 | Uba3 | 0.005 | Young |
| 0.889476 | 0.233547 | 0.107 | 0.102 | 1 | 15 | Nlk | 0.005 | Young |
| 0.889648 | 0.285415 | 0.179 | 0.184 | 1 | 15 | Mmp23 | -0.005 | Young |
| 0.889715 | 0.258495 | 0.143 | 0.163 | 1 | 15 | Gm34084 | -0.02 | Young |
| 0.889715 | 0.137603 | 0.143 | 0.163 | 1 | 15 | Papolg | -0.02 | Young |
| 0.891779 | 0.071809 | 0.375 | 0.388 | 1 | 15 | Itsn2 | -0.013 | Young |
| 0.891884 | 0.095509 | 0.196 | 0.184 | 1 | 15 | Rasgef1b | 0.012 | Young |

| | | | | | | | | |
|---|---|---|---|---|---|---|---|---|
| 0.892044 | 1.214874 | 0.054 | 0.061 | 1 | 15 | Wdr46 | -0.007 | Young |
| 0.892044 | 1.051743 | 0.054 | 0.061 | 1 | 15 | Hspa2 | -0.007 | Young |
| 0.892044 | 0.913077 | 0.054 | 0.061 | 1 | 15 | Tpgs2 | -0.007 | Young |
| 0.892044 | 0.516513 | 0.054 | 0.061 | 1 | 15 | Ccdc117 | -0.007 | Young |
| 0.892044 | 0.443253 | 0.054 | 0.061 | 1 | 15 | Qsox1 | -0.007 | Young |
| 0.892044 | 0.411503 | 0.054 | 0.061 | 1 | 15 | Gstcd | -0.007 | Young |
| 0.892044 | 0.372127 | 0.054 | 0.061 | 1 | 15 | Med20 | -0.007 | Young |
| 0.892044 | 0.297928 | 0.054 | 0.061 | 1 | 15 | Pgm2l1 | -0.007 | Young |
| 0.892044 | 0.29549 | 0.054 | 0.061 | 1 | 15 | Map4k5 | -0.007 | Young |
| 0.892044 | 0.277402 | 0.054 | 0.061 | 1 | 15 | Zfp959 | -0.007 | Young |
| 0.892044 | 0.226409 | 0.054 | 0.061 | 1 | 15 | Gm26669 | -0.007 | Young |
| 0.892044 | 0.14063 | 0.054 | 0.061 | 1 | 15 | Oma1 | -0.007 | Young |
| 0.892044 | 0.063616 | 0.054 | 0.061 | 1 | 15 | Mob2 | -0.007 | Young |
| 0.892044 | 0.037419 | 0.054 | 0.061 | 1 | 15 | Tmem214 | -0.007 | Young |
| 0.892044 | 0.016883 | 0.054 | 0.061 | 1 | 15 | Lrrc40 | -0.007 | Young |
| 0.892325 | 0.643651 | 0.232 | 0.224 | 1 | 15 | Nrd1 | 0.008 | Young |
| 0.892334 | 0.209432 | 0.464 | 0.49 | 1 | 15 | Atp6v1g1 | -0.026 | Young |
| 0.892837 | 0.97571 | 0.125 | 0.122 | 1 | 15 | Kbtbd2 | 0.003 | Young |
| 0.892837 | 0.968018 | 0.125 | 0.122 | 1 | 15 | Tmem131 | 0.003 | Young |
| 0.892837 | 0.819686 | 0.125 | 0.122 | 1 | 15 | Zcchc9 | 0.003 | Young |
| 0.892837 | 0.484497 | 0.125 | 0.122 | 1 | 15 | Slc25a46 | 0.003 | Young |
| 0.892837 | 0.425347 | 0.125 | 0.122 | 1 | 15 | Prpf18 | 0.003 | Young |
| 0.893573 | 0.196062 | 0.268 | 0.245 | 1 | 15 | St3gal4 | 0.023 | Young |
| 0.894471 | 0.105816 | 0.304 | 0.327 | 1 | 15 | Asah1 | -0.023 | Young |
| 0.894472 | 0.563166 | 0.071 | 0.082 | 1 | 15 | Hes6 | -0.011 | Young |
| 0.894472 | 0.554285 | 0.071 | 0.082 | 1 | 15 | Rad51d | -0.011 | Young |
| 0.894472 | 0.40743 | 0.071 | 0.082 | 1 | 15 | Wdr81 | -0.011 | Young |
| 0.894472 | 0.2941 | 0.071 | 0.082 | 1 | 15 | Slc35b3 | -0.011 | Young |
| 0.894472 | 0.262638 | 0.071 | 0.082 | 1 | 15 | Jag1 | -0.011 | Young |
| 0.894472 | 0.032976 | 0.071 | 0.082 | 1 | 15 | Aga | -0.011 | Young |
| 0.894756 | 0.871327 | 0.089 | 0.082 | 1 | 15 | Prpf19 | 0.007 | Young |
| 0.894756 | 0.247253 | 0.089 | 0.082 | 1 | 15 | 9130401M | 0.007 | Young |
| 0.894756 | 0.210114 | 0.089 | 0.082 | 1 | 15 | Zfp407 | 0.007 | Young |
| 0.894756 | 0.096451 | 0.089 | 0.082 | 1 | 15 | Srpk2 | 0.007 | Young |
| 0.895029 | 0.58122 | 0.143 | 0.143 | 1 | 15 | Pfdn1 | 0 | Young |
| 0.895841 | 0.031192 | 0.214 | 0.204 | 1 | 15 | Derl1 | 0.01 | Young |
| 0.897215 | 0.142448 | 0.179 | 0.184 | 1 | 15 | Grasp | -0.005 | Young |
| 0.897578 | 0.793148 | 0.196 | 0.204 | 1 | 15 | Uba7 | -0.008 | Young |
| 0.897595 | 0.257618 | 0.375 | 0.388 | 1 | 15 | Sdha | -0.013 | Young |
| 0.899033 | 0.743188 | 0.107 | 0.102 | 1 | 15 | Actr1b | 0.005 | Young |
| 0.899033 | 0.585704 | 0.107 | 0.102 | 1 | 15 | Map2k4 | 0.005 | Young |
| 0.899033 | 0.415471 | 0.107 | 0.102 | 1 | 15 | Mrps34 | 0.005 | Young |
| 0.899033 | 0.313976 | 0.107 | 0.102 | 1 | 15 | Lrpap1 | 0.005 | Young |
| 0.899033 | 0.292474 | 0.107 | 0.102 | 1 | 15 | Nagk | 0.005 | Young |
| 0.899451 | 0.105096 | 0.286 | 0.265 | 1 | 15 | Asph | 0.021 | Young |

| | | | | | | | | |
|---|---|---|---|---|---|---|---|---|
| 0.899823 | 1.106692 | 0.036 | 0.041 | 1 | 15 | Cisd3 | -0.005 | Young |
| 0.899823 | 1.105724 | 0.036 | 0.041 | 1 | 15 | Pym1 | -0.005 | Young |
| 0.899823 | 1.040665 | 0.036 | 0.041 | 1 | 15 | Prune1 | -0.005 | Young |
| 0.899823 | 0.98963 | 0.036 | 0.041 | 1 | 15 | Mkks | -0.005 | Young |
| 0.899823 | 0.75567 | 0.036 | 0.041 | 1 | 15 | Drosha | -0.005 | Young |
| 0.899823 | 0.700186 | 0.036 | 0.041 | 1 | 15 | Eci2 | -0.005 | Young |
| 0.899823 | 0.611578 | 0.036 | 0.041 | 1 | 15 | Slc12a4 | -0.005 | Young |
| 0.899823 | 0.607131 | 0.036 | 0.041 | 1 | 15 | Bnip1 | -0.005 | Young |
| 0.899823 | 0.55703 | 0.036 | 0.041 | 1 | 15 | Pparg | -0.005 | Young |
| 0.899823 | 0.460243 | 0.036 | 0.041 | 1 | 15 | Srprb | -0.005 | Young |
| 0.899823 | 0.38158 | 0.036 | 0.041 | 1 | 15 | Mocs3 | -0.005 | Young |
| 0.899823 | 0.381079 | 0.036 | 0.041 | 1 | 15 | Cox11 | -0.005 | Young |
| 0.899823 | 0.380842 | 0.036 | 0.041 | 1 | 15 | Trim37 | -0.005 | Young |
| 0.899823 | 0.28596 | 0.036 | 0.041 | 1 | 15 | Ap5s1 | -0.005 | Young |
| 0.899823 | 0.278794 | 0.036 | 0.041 | 1 | 15 | Klhdc10 | -0.005 | Young |
| 0.899823 | 0.247425 | 0.036 | 0.041 | 1 | 15 | Cenpj | -0.005 | Young |
| 0.899823 | 0.179804 | 0.036 | 0.041 | 1 | 15 | Glce | -0.005 | Young |
| 0.899823 | 0.135441 | 0.036 | 0.041 | 1 | 15 | Rabggta | -0.005 | Young |
| 0.899823 | 0.092663 | 0.036 | 0.041 | 1 | 15 | Ikbke | -0.005 | Young |
| 0.899823 | 0.0728 | 0.036 | 0.041 | 1 | 15 | Zfp994 | -0.005 | Young |
| 0.899823 | 0.022045 | 0.036 | 0.041 | 1 | 15 | Dut | -0.005 | Young |
| 0.900325 | 0.056831 | 0.982 | 1 | 1 | 15 | Malat1 | -0.018 | Young |
| 0.900817 | 0.002173 | 0.429 | 0.429 | 1 | 15 | Rpl36al | 0 | Young |
| 0.90172 | 0.363341 | 0.125 | 0.122 | 1 | 15 | Sash1 | 0.003 | Young |
| 0.90172 | 0.08151 | 0.125 | 0.122 | 1 | 15 | Rfc1 | 0.003 | Young |
| 0.902393 | 0.140298 | 0.464 | 0.449 | 1 | 15 | AW112010 | 0.015 | Young |
| 0.902684 | 0.026599 | 0.179 | 0.163 | 1 | 15 | Clec4a1 | 0.016 | Young |
| 0.902833 | 0.618629 | 0.107 | 0.122 | 1 | 15 | Hk1 | -0.015 | Young |
| 0.902833 | 0.380612 | 0.107 | 0.122 | 1 | 15 | Usp1 | -0.015 | Young |
| 0.903384 | 0.982546 | 0.143 | 0.143 | 1 | 15 | Rfk | 0 | Young |
| 0.903384 | 0.127246 | 0.143 | 0.143 | 1 | 15 | Lpin2 | 0 | Young |
| 0.904342 | 0.614753 | 0.161 | 0.163 | 1 | 15 | Mfsd6 | -0.002 | Young |
| 0.904627 | 0.502905 | 0.214 | 0.224 | 1 | 15 | Arl6ip5 | -0.01 | Young |
| 0.904627 | 0.063001 | 0.214 | 0.224 | 1 | 15 | Rabgap1 | -0.01 | Young |
| 0.904632 | 0.490534 | 0.089 | 0.102 | 1 | 15 | Cep57 | -0.013 | Young |
| 0.90468 | 1.377256 | 0.054 | 0.061 | 1 | 15 | Dkc1 | -0.007 | Young |
| 0.90468 | 1.307242 | 0.054 | 0.061 | 1 | 15 | Ubxn8 | -0.007 | Young |
| 0.90468 | 1.218432 | 0.054 | 0.061 | 1 | 15 | Thap7 | -0.007 | Young |
| 0.90468 | 1.011352 | 0.054 | 0.061 | 1 | 15 | Steap3 | -0.007 | Young |
| 0.90468 | 0.958029 | 0.054 | 0.061 | 1 | 15 | Spsb3 | -0.007 | Young |
| 0.90468 | 0.883613 | 0.054 | 0.061 | 1 | 15 | Med9 | -0.007 | Young |
| 0.90468 | 0.856345 | 0.054 | 0.061 | 1 | 15 | Mpzl3 | -0.007 | Young |
| 0.90468 | 0.64227 | 0.054 | 0.061 | 1 | 15 | Kansl1l | -0.007 | Young |
| 0.90468 | 0.61743 | 0.054 | 0.061 | 1 | 15 | Rdh13 | -0.007 | Young |
| 0.90468 | 0.579418 | 0.054 | 0.061 | 1 | 15 | Dgkq | -0.007 | Young |

| | | | | | | | | |
|---|---|---|---|---|---|---|---|---|
| 0.90468 | 0.37537 | 0.054 | 0.061 | 1 | 15 | Wee1 | -0.007 | Young |
| 0.90468 | 0.328378 | 0.054 | 0.061 | 1 | 15 | Tmem37 | -0.007 | Young |
| 0.90468 | 0.265603 | 0.054 | 0.061 | 1 | 15 | Tada3 | -0.007 | Young |
| 0.90468 | 0.101815 | 0.054 | 0.061 | 1 | 15 | Hsph1 | -0.007 | Young |
| 0.90468 | 0.069164 | 0.054 | 0.061 | 1 | 15 | Gabpb1 | -0.007 | Young |
| 0.90468 | 0.036967 | 0.054 | 0.061 | 1 | 15 | Zfp512 | -0.007 | Young |
| 0.904742 | 0.367902 | 0.25 | 0.286 | 1 | 15 | Rbm5 | -0.036 | Young |
| 0.904742 | 0.063014 | 0.268 | 0.265 | 1 | 15 | Ep300 | 0.003 | Young |
| 0.904875 | 0.1087 | 0.304 | 0.286 | 1 | 15 | Pum1 | 0.018 | Young |
| 0.905228 | 0.153421 | 0.089 | 0.082 | 1 | 15 | Nucb2 | 0.007 | Young |
| 0.905228 | 0.108065 | 0.089 | 0.082 | 1 | 15 | Ppp6r2 | 0.007 | Young |
| 0.905228 | 0.087271 | 0.089 | 0.082 | 1 | 15 | Git1 | 0.007 | Young |
| 0.905228 | 0.077285 | 0.089 | 0.082 | 1 | 15 | Cd72 | 0.007 | Young |
| 0.905525 | 0.439415 | 0.071 | 0.082 | 1 | 15 | Zfp219 | -0.011 | Young |
| 0.905525 | 0.300027 | 0.071 | 0.082 | 1 | 15 | Rfxap | -0.011 | Young |
| 0.907137 | 0.076749 | 0.357 | 0.347 | 1 | 15 | Hpcal1 | 0.01 | Young |
| 0.907474 | 0.015086 | 0.429 | 0.469 | 1 | 15 | Taok1 | -0.04 | Young |
| 0.907945 | 0.440163 | 0.464 | 0.449 | 1 | 15 | Bclaf1 | 0.015 | Young |
| 0.908032 | 0.434312 | 0.286 | 0.306 | 1 | 15 | Brd7 | -0.02 | Young |
| 0.908604 | 0.162322 | 0.107 | 0.102 | 1 | 15 | Mfsd11 | 0.005 | Young |
| 0.910125 | 0.153358 | 0.25 | 0.265 | 1 | 15 | Hipk3 | -0.015 | Young |
| 0.910142 | 0.184666 | 0.214 | 0.204 | 1 | 15 | Cyb5r3 | 0.01 | Young |
| 0.910615 | 1.054364 | 0.125 | 0.122 | 1 | 15 | Paip1 | 0.003 | Young |
| 0.910615 | 0.724214 | 0.125 | 0.122 | 1 | 15 | Id3 | 0.003 | Young |
| 0.910615 | 0.495759 | 0.125 | 0.122 | 1 | 15 | Rin3 | 0.003 | Young |
| 0.910615 | 0.493498 | 0.125 | 0.122 | 1 | 15 | Nfatc1 | 0.003 | Young |
| 0.910615 | 0.385545 | 0.125 | 0.122 | 1 | 15 | Stt3b | 0.003 | Young |
| 0.911599 | 0.187431 | 0.321 | 0.347 | 1 | 15 | St3gal1 | -0.026 | Young |
| 0.911749 | 0.495733 | 0.143 | 0.143 | 1 | 15 | Arhgap28 | 0 | Young |
| 0.912048 | 0.85402 | 0.107 | 0.122 | 1 | 15 | Smarcad1 | -0.015 | Young |
| 0.912048 | 0.66882 | 0.107 | 0.122 | 1 | 15 | Galc | -0.015 | Young |
| 0.912145 | 0.583384 | 0.196 | 0.204 | 1 | 15 | Cpne3 | -0.008 | Young |
| 0.912145 | 0.526952 | 0.196 | 0.204 | 1 | 15 | Arhgef12 | -0.008 | Young |
| 0.912145 | 0.215657 | 0.196 | 0.204 | 1 | 15 | Tcf20 | -0.008 | Young |
| 0.91228 | 0.78915 | 0.161 | 0.163 | 1 | 15 | Pja2 | -0.002 | Young |
| 0.913073 | 0.264956 | 0.232 | 0.224 | 1 | 15 | Ep400 | 0.008 | Young |
| 0.91463 | 1.142615 | 0.089 | 0.102 | 1 | 15 | Map3k11 | -0.013 | Young |
| 0.91463 | 0.543085 | 0.089 | 0.102 | 1 | 15 | Mefv | -0.013 | Young |
| 0.915171 | 1.246755 | 0.036 | 0.041 | 1 | 15 | Bach2 | -0.005 | Young |
| 0.915171 | 1.205775 | 0.036 | 0.041 | 1 | 15 | Diaph3 | -0.005 | Young |
| 0.915171 | 1.179503 | 0.036 | 0.041 | 1 | 15 | Ccar2 | -0.005 | Young |
| 0.915171 | 0.821197 | 0.036 | 0.041 | 1 | 15 | Mrps11 | -0.005 | Young |
| 0.915171 | 0.80856 | 0.036 | 0.041 | 1 | 15 | Sec24d | -0.005 | Young |
| 0.915171 | 0.749158 | 0.036 | 0.041 | 1 | 15 | Klra2 | -0.005 | Young |
| 0.915171 | 0.718985 | 0.036 | 0.041 | 1 | 15 | Wdsub1 | -0.005 | Young |

| | | | | | | | | |
|---|---|---|---|---|---|---|---|---|
| 0.915171 | 0.641168 | 0.036 | 0.041 | 1 | 15 | Zbtb16 | -0.005 | Young |
| 0.915171 | 0.623655 | 0.036 | 0.041 | 1 | 15 | Twnk | -0.005 | Young |
| 0.915171 | 0.62222 | 0.036 | 0.041 | 1 | 15 | Jak3 | -0.005 | Young |
| 0.915171 | 0.60606 | 0.036 | 0.041 | 1 | 15 | Yars | -0.005 | Young |
| 0.915171 | 0.560216 | 0.036 | 0.041 | 1 | 15 | Slc7a6os | -0.005 | Young |
| 0.915171 | 0.503018 | 0.036 | 0.041 | 1 | 15 | Tram2 | -0.005 | Young |
| 0.915171 | 0.498191 | 0.036 | 0.041 | 1 | 15 | Nol4l | -0.005 | Young |
| 0.915171 | 0.479981 | 0.036 | 0.041 | 1 | 15 | Wdr75 | -0.005 | Young |
| 0.915171 | 0.46262 | 0.036 | 0.041 | 1 | 15 | Tmem184c | -0.005 | Young |
| 0.915171 | 0.43257 | 0.036 | 0.041 | 1 | 15 | Naif1 | -0.005 | Young |
| 0.915171 | 0.416394 | 0.036 | 0.041 | 1 | 15 | Pigg | -0.005 | Young |
| 0.915171 | 0.413666 | 0.036 | 0.041 | 1 | 15 | Abhd10 | -0.005 | Young |
| 0.915171 | 0.410145 | 0.036 | 0.041 | 1 | 15 | Fam98a | -0.005 | Young |
| 0.915171 | 0.393406 | 0.036 | 0.041 | 1 | 15 | Mon1a | -0.005 | Young |
| 0.915171 | 0.365782 | 0.036 | 0.041 | 1 | 15 | Alg11 | -0.005 | Young |
| 0.915171 | 0.346462 | 0.036 | 0.041 | 1 | 15 | Tnfrsf21 | -0.005 | Young |
| 0.915171 | 0.339932 | 0.036 | 0.041 | 1 | 15 | Ccdc184 | -0.005 | Young |
| 0.915171 | 0.31515 | 0.036 | 0.041 | 1 | 15 | Nxt2 | -0.005 | Young |
| 0.915171 | 0.310403 | 0.036 | 0.041 | 1 | 15 | Tbp | -0.005 | Young |
| 0.915171 | 0.306445 | 0.036 | 0.041 | 1 | 15 | Ermp1 | -0.005 | Young |
| 0.915171 | 0.295709 | 0.036 | 0.041 | 1 | 15 | Clcn6 | -0.005 | Young |
| 0.915171 | 0.294727 | 0.036 | 0.041 | 1 | 15 | Ccdc102a | -0.005 | Young |
| 0.915171 | 0.267822 | 0.036 | 0.041 | 1 | 15 | Ptpn21 | -0.005 | Young |
| 0.915171 | 0.257397 | 0.036 | 0.041 | 1 | 15 | Cptp | -0.005 | Young |
| 0.915171 | 0.219539 | 0.036 | 0.041 | 1 | 15 | Mtif3 | -0.005 | Young |
| 0.915171 | 0.218326 | 0.036 | 0.041 | 1 | 15 | Zcchc8 | -0.005 | Young |
| 0.915171 | 0.154822 | 0.036 | 0.041 | 1 | 15 | Uck2 | -0.005 | Young |
| 0.915171 | 0.087814 | 0.036 | 0.041 | 1 | 15 | Ipo13 | -0.005 | Young |
| 0.915171 | 0.076863 | 0.036 | 0.041 | 1 | 15 | 4930481A1 | -0.005 | Young |
| 0.915171 | 0.073217 | 0.036 | 0.041 | 1 | 15 | Milr1 | -0.005 | Young |
| 0.915171 | 0.070968 | 0.036 | 0.041 | 1 | 15 | Gmppb | -0.005 | Young |
| 0.915171 | 0.061852 | 0.036 | 0.041 | 1 | 15 | Gm4707 | -0.005 | Young |
| 0.915171 | 0.025124 | 0.036 | 0.041 | 1 | 15 | Gtpbp8 | -0.005 | Young |
| 0.915171 | 0.021436 | 0.036 | 0.041 | 1 | 15 | Lmtk2 | -0.005 | Young |
| 0.915669 | 0.187138 | 0.393 | 0.388 | 1 | 15 | Hnrnpl | 0.005 | Young |
| 0.915716 | 0.007226 | 0.089 | 0.082 | 1 | 15 | Ulk1 | 0.007 | Young |
| 0.916596 | 0.940627 | 0.071 | 0.082 | 1 | 15 | Ppfibp1 | -0.011 | Young |
| 0.916596 | 0.748184 | 0.071 | 0.082 | 1 | 15 | Acaca | -0.011 | Young |
| 0.916758 | 0.51997 | 0.25 | 0.265 | 1 | 15 | Ndufs2 | -0.015 | Young |
| 0.917341 | 1.427313 | 0.054 | 0.061 | 1 | 15 | Ccl3 | -0.007 | Young |
| 0.917341 | 1.365123 | 0.054 | 0.061 | 1 | 15 | Cxcl9 | -0.007 | Young |
| 0.917341 | 1.000669 | 0.054 | 0.061 | 1 | 15 | N4bp2 | -0.007 | Young |
| 0.917341 | 0.899433 | 0.054 | 0.061 | 1 | 15 | Slc37a2 | -0.007 | Young |
| 0.917341 | 0.872861 | 0.054 | 0.061 | 1 | 15 | Itga9 | -0.007 | Young |
| 0.917341 | 0.560211 | 0.054 | 0.061 | 1 | 15 | Atpaf1 | -0.007 | Young |

| | | | | | | | | |
|---|---|---|---|---|---|---|---|---|
| 0.917341 | 0.440409 | 0.054 | 0.061 | 1 | 15 | AC133498. | -0.007 | Young |
| 0.917341 | 0.436263 | 0.054 | 0.061 | 1 | 15 | Mis12 | -0.007 | Young |
| 0.917341 | 0.367443 | 0.054 | 0.061 | 1 | 15 | Slc39a14 | -0.007 | Young |
| 0.917341 | 0.319213 | 0.054 | 0.061 | 1 | 15 | Pdcd1lg2 | -0.007 | Young |
| 0.917341 | 0.269323 | 0.054 | 0.061 | 1 | 15 | Crcp | -0.007 | Young |
| 0.918189 | 1.003346 | 0.107 | 0.102 | 1 | 15 | Lyrm2 | 0.005 | Young |
| 0.918189 | 0.244028 | 0.107 | 0.102 | 1 | 15 | Sympk | 0.005 | Young |
| 0.918703 | 0.151274 | 0.214 | 0.224 | 1 | 15 | Lars2 | -0.01 | Young |
| 0.918908 | 0.000699 | 0.661 | 0.633 | 1 | 15 | Sfpq | 0.028 | Young |
| 0.919075 | 0.050406 | 0.357 | 0.347 | 1 | 15 | Rbbp6 | 0.01 | Young |
| 0.91944 | 0.438448 | 0.196 | 0.204 | 1 | 15 | Acot7 | -0.008 | Young |
| 0.91944 | 0.214645 | 0.196 | 0.204 | 1 | 15 | Tmem248 | -0.008 | Young |
| 0.91944 | 0.009855 | 0.196 | 0.204 | 1 | 15 | Pwwp2a | -0.008 | Young |
| 0.919522 | 1.109239 | 0.125 | 0.122 | 1 | 15 | Mtfr1l | 0.003 | Young |
| 0.919522 | 0.591885 | 0.125 | 0.122 | 1 | 15 | Itsn1 | 0.003 | Young |
| 0.919522 | 0.496751 | 0.125 | 0.122 | 1 | 15 | Plcg2 | 0.003 | Young |
| 0.919782 | 0.105354 | 0.286 | 0.286 | 1 | 15 | Lyn | 0 | Young |
| 0.920124 | 0.646301 | 0.143 | 0.143 | 1 | 15 | 2810403A( | 0 | Young |
| 0.920226 | 0.283324 | 0.161 | 0.163 | 1 | 15 | Ptcd2 | -0.002 | Young |
| 0.92033 | 0.014427 | 0.821 | 0.755 | 1 | 15 | Vamp8 | 0.066 | Young |
| 0.920695 | 0.05517 | 1 | 0.98 | 1 | 15 | Rpl17 | 0.02 | Young |
| 0.921274 | 1.039663 | 0.107 | 0.122 | 1 | 15 | Tmem110 | -0.015 | Young |
| 0.921274 | 0.973378 | 0.107 | 0.122 | 1 | 15 | Pmm2 | -0.015 | Young |
| 0.921274 | 0.744972 | 0.107 | 0.122 | 1 | 15 | Prpf31 | -0.015 | Young |
| 0.92151 | 0.120861 | 0.304 | 0.306 | 1 | 15 | Ptpre | -0.002 | Young |
| 0.922341 | 0.157041 | 0.25 | 0.245 | 1 | 15 | Trim12c | 0.005 | Young |
| 0.922916 | 0.187206 | 0.571 | 0.592 | 1 | 15 | Nfkb1 | -0.021 | Young |
| 0.922931 | 0.159342 | 0.482 | 0.49 | 1 | 15 | Jak1 | -0.008 | Young |
| 0.924383 | 0.215587 | 0.268 | 0.265 | 1 | 15 | Cdk2ap2 | 0.003 | Young |
| 0.926181 | 0.190176 | 0.286 | 0.286 | 1 | 15 | Rufy3 | 0 | Young |
| 0.926217 | 0.029661 | 0.357 | 0.388 | 1 | 15 | Cd2ap | -0.031 | Young |
| 0.926742 | 0.553604 | 0.196 | 0.204 | 1 | 15 | Chchd5 | -0.008 | Young |
| 0.926939 | 0.241581 | 0.232 | 0.224 | 1 | 15 | Pcmtd1 | 0.008 | Young |
| 0.927568 | 0.713169 | 0.179 | 0.184 | 1 | 15 | Bloc1s2 | -0.005 | Young |
| 0.927684 | 1.059967 | 0.071 | 0.082 | 1 | 15 | Arel1 | -0.011 | Young |
| 0.927684 | 0.871794 | 0.071 | 0.082 | 1 | 15 | Prps1 | -0.011 | Young |
| 0.927786 | 1.177489 | 0.107 | 0.102 | 1 | 15 | Il11ra1 | 0.005 | Young |
| 0.927786 | 0.773629 | 0.107 | 0.102 | 1 | 15 | Pilra | 0.005 | Young |
| 0.927786 | 0.576785 | 0.107 | 0.102 | 1 | 15 | Plxna4 | 0.005 | Young |
| 0.927786 | 0.198072 | 0.107 | 0.102 | 1 | 15 | Wdr41 | 0.005 | Young |
| 0.927786 | 0.15339 | 0.107 | 0.102 | 1 | 15 | C2cd5 | 0.005 | Young |
| 0.927786 | 0.093096 | 0.107 | 0.102 | 1 | 15 | Slc12a9 | 0.005 | Young |
| 0.928438 | 0.103266 | 0.125 | 0.122 | 1 | 15 | Cds2 | 0.003 | Young |
| 0.928508 | 0.496458 | 0.143 | 0.143 | 1 | 15 | Hip1 | 0 | Young |
| 0.928508 | 0.495018 | 0.143 | 0.143 | 1 | 15 | Odc1 | 0 | Young |

| | | | | | | | | |
|---|---|---|---|---|---|---|---|---|
| 0.92916 | 0.093826 | 0.375 | 0.367 | 1 | 15 | Sugt1 | 0.008 | Young |
| 0.930022 | 1.446594 | 0.054 | 0.061 | 1 | 15 | Tbc1d12 | -0.007 | Young |
| 0.930022 | 1.279788 | 0.054 | 0.061 | 1 | 15 | Abhd18 | -0.007 | Young |
| 0.930022 | 1.239046 | 0.054 | 0.061 | 1 | 15 | Cxcl10 | -0.007 | Young |
| 0.930022 | 1.220982 | 0.054 | 0.061 | 1 | 15 | Gpr34 | -0.007 | Young |
| 0.930022 | 1.200293 | 0.054 | 0.061 | 1 | 15 | Kansl2 | -0.007 | Young |
| 0.930022 | 1.073536 | 0.054 | 0.061 | 1 | 15 | Dnajc24 | -0.007 | Young |
| 0.930022 | 1.007564 | 0.054 | 0.061 | 1 | 15 | Klhl18 | -0.007 | Young |
| 0.930022 | 0.959546 | 0.054 | 0.061 | 1 | 15 | Foxred2 | -0.007 | Young |
| 0.930022 | 0.924444 | 0.054 | 0.061 | 1 | 15 | Pold1 | -0.007 | Young |
| 0.930022 | 0.826535 | 0.054 | 0.061 | 1 | 15 | Klhl36 | -0.007 | Young |
| 0.930022 | 0.773537 | 0.054 | 0.061 | 1 | 15 | Stx17 | -0.007 | Young |
| 0.930022 | 0.679611 | 0.054 | 0.061 | 1 | 15 | Rfx1 | -0.007 | Young |
| 0.930022 | 0.308046 | 0.054 | 0.061 | 1 | 15 | Sh2b1 | -0.007 | Young |
| 0.93045 | 0.193973 | 0.339 | 0.347 | 1 | 15 | Rest | -0.008 | Young |
| 0.930551 | 1.971663 | 0.036 | 0.041 | 1 | 15 | Fgfbp3 | -0.005 | Young |
| 0.930551 | 1.827925 | 0.036 | 0.041 | 1 | 15 | Rprd1a | -0.005 | Young |
| 0.930551 | 1.71641 | 0.036 | 0.041 | 1 | 15 | Shmt2 | -0.005 | Young |
| 0.930551 | 1.648651 | 0.036 | 0.041 | 1 | 15 | Smpdl3b | -0.005 | Young |
| 0.930551 | 1.623683 | 0.036 | 0.041 | 1 | 15 | Pnkp | -0.005 | Young |
| 0.930551 | 1.360402 | 0.036 | 0.041 | 1 | 15 | 3110002H1 | -0.005 | Young |
| 0.930551 | 1.335882 | 0.036 | 0.041 | 1 | 15 | Gm37233 | -0.005 | Young |
| 0.930551 | 1.312091 | 0.036 | 0.041 | 1 | 15 | Cdon | -0.005 | Young |
| 0.930551 | 1.293792 | 0.036 | 0.041 | 1 | 15 | Naa25 | -0.005 | Young |
| 0.930551 | 1.256083 | 0.036 | 0.041 | 1 | 15 | Mettl14 | -0.005 | Young |
| 0.930551 | 1.201996 | 0.036 | 0.041 | 1 | 15 | Arl2 | -0.005 | Young |
| 0.930551 | 1.171792 | 0.036 | 0.041 | 1 | 15 | Atn1 | -0.005 | Young |
| 0.930551 | 1.14304 | 0.036 | 0.041 | 1 | 15 | Ttyh2 | -0.005 | Young |
| 0.930551 | 0.995747 | 0.036 | 0.041 | 1 | 15 | Nudt1 | -0.005 | Young |
| 0.930551 | 0.991825 | 0.036 | 0.041 | 1 | 15 | Tjap1 | -0.005 | Young |
| 0.930551 | 0.892704 | 0.036 | 0.041 | 1 | 15 | Aaas | -0.005 | Young |
| 0.930551 | 0.888555 | 0.036 | 0.041 | 1 | 15 | Jmjd7 | -0.005 | Young |
| 0.930551 | 0.886803 | 0.036 | 0.041 | 1 | 15 | Tm7sf2 | -0.005 | Young |
| 0.930551 | 0.84118 | 0.036 | 0.041 | 1 | 15 | D930030I0 | -0.005 | Young |
| 0.930551 | 0.789793 | 0.036 | 0.041 | 1 | 15 | 1810044D( | -0.005 | Young |
| 0.930551 | 0.753056 | 0.036 | 0.041 | 1 | 15 | Rhbdd2 | -0.005 | Young |
| 0.930551 | 0.730149 | 0.036 | 0.041 | 1 | 15 | Eef1akmt2 | -0.005 | Young |
| 0.930551 | 0.700956 | 0.036 | 0.041 | 1 | 15 | Prmt6 | -0.005 | Young |
| 0.930551 | 0.663651 | 0.036 | 0.041 | 1 | 15 | Rundc1 | -0.005 | Young |
| 0.930551 | 0.574583 | 0.036 | 0.041 | 1 | 15 | Cav2 | -0.005 | Young |
| 0.930551 | 0.533615 | 0.036 | 0.041 | 1 | 15 | Phactr4 | -0.005 | Young |
| 0.930551 | 0.462536 | 0.036 | 0.041 | 1 | 15 | Ncapg2 | -0.005 | Young |
| 0.930551 | 0.445368 | 0.036 | 0.041 | 1 | 15 | Chd1l | -0.005 | Young |
| 0.930551 | 0.436589 | 0.036 | 0.041 | 1 | 15 | Kif1bp | -0.005 | Young |
| 0.930551 | 0.402686 | 0.036 | 0.041 | 1 | 15 | Srfbp1 | -0.005 | Young |

| | | | | | | | | |
|---|---|---|---|---|---|---|---|---|
| 0.930551 | 0.252679 | 0.036 | 0.041 | 1 | 15 | Mrps31 | -0.005 | Young |
| 0.930551 | 0.238715 | 0.036 | 0.041 | 1 | 15 | Tsr2 | -0.005 | Young |
| 0.930551 | 0.190126 | 0.036 | 0.041 | 1 | 15 | Bst1 | -0.005 | Young |
| 0.930551 | 0.071426 | 0.036 | 0.041 | 1 | 15 | Crip2 | -0.005 | Young |
| 0.930859 | 0.024559 | 0.482 | 0.469 | 1 | 15 | Sh3bp1 | 0.013 | Young |
| 0.930942 | 0.054375 | 0.25 | 0.286 | 1 | 15 | Lrp10 | -0.036 | Young |
| 0.931582 | 0.275029 | 0.357 | 0.367 | 1 | 15 | Itch | -0.01 | Young |
| 0.931738 | 0.021465 | 0.5 | 0.51 | 1 | 15 | Osbpl8 | -0.01 | Young |
| 0.932494 | 0.105643 | 0.554 | 0.51 | 1 | 15 | Myadm | 0.044 | Young |
| 0.933737 | 0.370732 | 0.161 | 0.184 | 1 | 15 | Tubgcp4 | -0.023 | Young |
| 0.93388 | 0.107985 | 0.232 | 0.224 | 1 | 15 | Cetn3 | 0.008 | Young |
| 0.93388 | 0.08686 | 0.232 | 0.224 | 1 | 15 | Foxn3 | 0.008 | Young |
| 0.934037 | 0.146674 | 0.304 | 0.306 | 1 | 15 | Arhgap17 | -0.002 | Young |
| 0.934051 | 0.038572 | 0.196 | 0.204 | 1 | 15 | Cdk14 | -0.008 | Young |
| 0.935814 | 0.225031 | 0.25 | 0.245 | 1 | 15 | Grina | 0.005 | Young |
| 0.935846 | 0.020063 | 0.857 | 0.796 | 1 | 15 | Eif3f | 0.061 | Young |
| 0.936143 | 0.349748 | 0.161 | 0.163 | 1 | 15 | Naa15 | -0.002 | Young |
| 0.936143 | 0.118351 | 0.161 | 0.163 | 1 | 15 | Slc30a4 | -0.002 | Young |
| 0.9369 | 0.447554 | 0.143 | 0.143 | 1 | 15 | Aldh16a1 | 0 | Young |
| 0.937364 | 0.350934 | 0.125 | 0.122 | 1 | 15 | Srek1 | 0.003 | Young |
| 0.937364 | 0.155486 | 0.125 | 0.122 | 1 | 15 | Chmp1b | 0.003 | Young |
| 0.937364 | 0.099843 | 0.125 | 0.122 | 1 | 15 | AB124611 | 0.003 | Young |
| 0.937393 | 0.092437 | 0.107 | 0.102 | 1 | 15 | Ap2a1 | 0.005 | Young |
| 0.937393 | 0.082723 | 0.107 | 0.102 | 1 | 15 | Wdr33 | 0.005 | Young |
| 0.937393 | 0.015489 | 0.107 | 0.102 | 1 | 15 | Coasy | 0.005 | Young |
| 0.938301 | 0.190656 | 0.232 | 0.245 | 1 | 15 | Rragc | -0.013 | Young |
| 0.938595 | 0.536993 | 0.143 | 0.163 | 1 | 15 | Tmem219 | -0.02 | Young |
| 0.938786 | 0.901888 | 0.071 | 0.082 | 1 | 15 | Tmem159 | -0.011 | Young |
| 0.938786 | 0.85504 | 0.071 | 0.082 | 1 | 15 | Acad9 | -0.011 | Young |
| 0.938786 | 0.672005 | 0.071 | 0.082 | 1 | 15 | Cpq | -0.011 | Young |
| 0.939864 | 0.354497 | 0.214 | 0.224 | 1 | 15 | Asl | -0.01 | Young |
| 0.940009 | 0.141692 | 0.339 | 0.367 | 1 | 15 | Fam208a | -0.028 | Young |
| 0.940009 | 0.074588 | 0.339 | 0.367 | 1 | 15 | Agps | -0.028 | Young |
| 0.940042 | 0.047431 | 0.429 | 0.408 | 1 | 15 | Stx4a | 0.021 | Young |
| 0.940826 | 0.046907 | 0.232 | 0.224 | 1 | 15 | Tapbpl | 0.008 | Young |
| 0.941364 | 0.26232 | 0.196 | 0.204 | 1 | 15 | Foxk1 | -0.008 | Young |
| 0.941514 | 0.105018 | 0.268 | 0.286 | 1 | 15 | Kpnb1 | -0.018 | Young |
| 0.942788 | 0.286114 | 0.179 | 0.184 | 1 | 15 | Tpd52l2 | -0.005 | Young |
| 0.94334 | 0.275256 | 0.25 | 0.265 | 1 | 15 | Cbx3 | -0.015 | Young |
| 0.943459 | 0.356159 | 0.357 | 0.367 | 1 | 15 | Elk3 | -0.01 | Young |
| 0.943891 | 0.299309 | 0.357 | 0.388 | 1 | 15 | Raly | -0.031 | Young |
| 0.944111 | 0.500478 | 0.161 | 0.163 | 1 | 15 | Mrps36 | -0.002 | Young |
| 0.944111 | 0.35544 | 0.161 | 0.163 | 1 | 15 | Iffo2 | -0.002 | Young |
| 0.944301 | 0.119143 | 0.375 | 0.388 | 1 | 15 | Ost4 | -0.013 | Young |
| 0.945145 | 0.236447 | 0.232 | 0.245 | 1 | 15 | Itgam | -0.013 | Young |

| | | | | | | | | |
|---|---|---|---|---|---|---|---|---|
| 0.945299 | 0.385963 | 0.143 | 0.143 | 1 | 15 | Ikzf5 | 0 | Young |
| 0.945299 | 0.294052 | 0.143 | 0.143 | 1 | 15 | Stard4 | 0 | Young |
| 0.945299 | 0.203141 | 0.143 | 0.143 | 1 | 15 | Ndufv1 | 0 | Young |
| 0.945299 | 0.05596 | 0.143 | 0.143 | 1 | 15 | Rgs18 | 0 | Young |
| 0.945931 | 3.023915 | 0.018 | 0.02 | 1 | 15 | Prkab1 | -0.002 | Young |
| 0.945931 | 2.669394 | 0.018 | 0.02 | 1 | 15 | Tubd1 | -0.002 | Young |
| 0.945931 | 2.225291 | 0.018 | 0.02 | 1 | 15 | Tmem101 | -0.002 | Young |
| 0.945931 | 2.137229 | 0.018 | 0.02 | 1 | 15 | Osgepl1 | -0.002 | Young |
| 0.945931 | 2.039162 | 0.018 | 0.02 | 1 | 15 | Nsun5 | -0.002 | Young |
| 0.945931 | 2.039162 | 0.018 | 0.02 | 1 | 15 | Xndc1 | -0.002 | Young |
| 0.945931 | 1.974032 | 0.018 | 0.02 | 1 | 15 | Gm13012 | -0.002 | Young |
| 0.945931 | 1.914466 | 0.018 | 0.02 | 1 | 15 | Snap47 | -0.002 | Young |
| 0.945931 | 1.870901 | 0.018 | 0.02 | 1 | 15 | Commd9 | -0.002 | Young |
| 0.945931 | 1.826405 | 0.018 | 0.02 | 1 | 15 | Zfp637 | -0.002 | Young |
| 0.945931 | 1.692289 | 0.018 | 0.02 | 1 | 15 | Rab30 | -0.002 | Young |
| 0.945931 | 1.678895 | 0.018 | 0.02 | 1 | 15 | Tysnd1 | -0.002 | Young |
| 0.945931 | 1.671735 | 0.018 | 0.02 | 1 | 15 | Rom1 | -0.002 | Young |
| 0.945931 | 1.654599 | 0.018 | 0.02 | 1 | 15 | Gpnmb | -0.002 | Young |
| 0.945931 | 1.645289 | 0.018 | 0.02 | 1 | 15 | Gdf11 | -0.002 | Young |
| 0.945931 | 1.62442 | 0.018 | 0.02 | 1 | 15 | Epor | -0.002 | Young |
| 0.945931 | 1.608941 | 0.018 | 0.02 | 1 | 15 | Gm7160 | -0.002 | Young |
| 0.945931 | 1.606869 | 0.018 | 0.02 | 1 | 15 | Tpbg | -0.002 | Young |
| 0.945931 | 1.545469 | 0.018 | 0.02 | 1 | 15 | AC174678. | -0.002 | Young |
| 0.945931 | 1.52088 | 0.018 | 0.02 | 1 | 15 | Pdzd11 | -0.002 | Young |
| 0.945931 | 1.518599 | 0.018 | 0.02 | 1 | 15 | Tapt1 | -0.002 | Young |
| 0.945931 | 1.501764 | 0.018 | 0.02 | 1 | 15 | Prdm10 | -0.002 | Young |
| 0.945931 | 1.492502 | 0.018 | 0.02 | 1 | 15 | Ick | -0.002 | Young |
| 0.945931 | 1.455777 | 0.018 | 0.02 | 1 | 15 | Ralgps1 | -0.002 | Young |
| 0.945931 | 1.451279 | 0.018 | 0.02 | 1 | 15 | Bloc1s3 | -0.002 | Young |
| 0.945931 | 1.392171 | 0.018 | 0.02 | 1 | 15 | Htra3 | -0.002 | Young |
| 0.945931 | 1.374501 | 0.018 | 0.02 | 1 | 15 | Purg | -0.002 | Young |
| 0.945931 | 1.35874 | 0.018 | 0.02 | 1 | 15 | 2010013B2 | -0.002 | Young |
| 0.945931 | 1.353463 | 0.018 | 0.02 | 1 | 15 | Trdmt1 | -0.002 | Young |
| 0.945931 | 1.338357 | 0.018 | 0.02 | 1 | 15 | 2610035D1 | -0.002 | Young |
| 0.945931 | 1.334465 | 0.018 | 0.02 | 1 | 15 | Zbtb25 | -0.002 | Young |
| 0.945931 | 1.330301 | 0.018 | 0.02 | 1 | 15 | Tgm2 | -0.002 | Young |
| 0.945931 | 1.30203 | 0.018 | 0.02 | 1 | 15 | Tango6 | -0.002 | Young |
| 0.945931 | 1.296044 | 0.018 | 0.02 | 1 | 15 | Ccdc18 | -0.002 | Young |
| 0.945931 | 1.296044 | 0.018 | 0.02 | 1 | 15 | 1500015A0 | -0.002 | Young |
| 0.945931 | 1.286309 | 0.018 | 0.02 | 1 | 15 | Slc35c1 | -0.002 | Young |
| 0.945931 | 1.276409 | 0.018 | 0.02 | 1 | 15 | 5830432E0 | -0.002 | Young |
| 0.945931 | 1.261235 | 0.018 | 0.02 | 1 | 15 | Zbtb34 | -0.002 | Young |
| 0.945931 | 1.261235 | 0.018 | 0.02 | 1 | 15 | Sprtn | -0.002 | Young |
| 0.945931 | 1.261235 | 0.018 | 0.02 | 1 | 15 | Jrk | -0.002 | Young |
| 0.945931 | 1.232292 | 0.018 | 0.02 | 1 | 15 | Fbxl22 | -0.002 | Young |

| | | | | | | | | |
|---|---|---|---|---|---|---|---|---|
| 0.945931 | 1.232292 | 0.018 | 0.02 | 1 | 15 | Syne3 | -0.002 | Young |
| 0.945931 | 1.227181 | 0.018 | 0.02 | 1 | 15 | Ddx31 | -0.002 | Young |
| 0.945931 | 1.21277 | 0.018 | 0.02 | 1 | 15 | Mtbp | -0.002 | Young |
| 0.945931 | 1.179316 | 0.018 | 0.02 | 1 | 15 | Gm20300 | -0.002 | Young |
| 0.945931 | 1.147889 | 0.018 | 0.02 | 1 | 15 | Fzd4 | -0.002 | Young |
| 0.945931 | 1.137829 | 0.018 | 0.02 | 1 | 15 | 1700018L0 | -0.002 | Young |
| 0.945931 | 1.120729 | 0.018 | 0.02 | 1 | 15 | Zfp229 | -0.002 | Young |
| 0.945931 | 1.096769 | 0.018 | 0.02 | 1 | 15 | Spa17 | -0.002 | Young |
| 0.945931 | 1.066867 | 0.018 | 0.02 | 1 | 15 | Bag4 | -0.002 | Young |
| 0.945931 | 1.066867 | 0.018 | 0.02 | 1 | 15 | Gm48653 | -0.002 | Young |
| 0.945931 | 1.017323 | 0.018 | 0.02 | 1 | 15 | Oprd1 | -0.002 | Young |
| 0.945931 | 1.011104 | 0.018 | 0.02 | 1 | 15 | Pik3r3 | -0.002 | Young |
| 0.945931 | 1.004426 | 0.018 | 0.02 | 1 | 15 | Sfxn5 | -0.002 | Young |
| 0.945931 | 1.003359 | 0.018 | 0.02 | 1 | 15 | Gm29994 | -0.002 | Young |
| 0.945931 | 0.997508 | 0.018 | 0.02 | 1 | 15 | 4930532G1 | -0.002 | Young |
| 0.945931 | 0.997508 | 0.018 | 0.02 | 1 | 15 | Fap | -0.002 | Young |
| 0.945931 | 0.991206 | 0.018 | 0.02 | 1 | 15 | Ufsp1 | -0.002 | Young |
| 0.945931 | 0.991206 | 0.018 | 0.02 | 1 | 15 | Taf4b | -0.002 | Young |
| 0.945931 | 0.990519 | 0.018 | 0.02 | 1 | 15 | Samd14 | -0.002 | Young |
| 0.945931 | 0.983437 | 0.018 | 0.02 | 1 | 15 | Tpx2 | -0.002 | Young |
| 0.945931 | 0.979256 | 0.018 | 0.02 | 1 | 15 | Bicc1 | -0.002 | Young |
| 0.945931 | 0.979256 | 0.018 | 0.02 | 1 | 15 | Zfp729a | -0.002 | Young |
| 0.945931 | 0.962084 | 0.018 | 0.02 | 1 | 15 | L2hgdh | -0.002 | Young |
| 0.945931 | 0.959087 | 0.018 | 0.02 | 1 | 15 | Ahnak2 | -0.002 | Young |
| 0.945931 | 0.95041 | 0.018 | 0.02 | 1 | 15 | Abraxas1 | -0.002 | Young |
| 0.945931 | 0.95041 | 0.018 | 0.02 | 1 | 15 | Gm47232 | -0.002 | Young |
| 0.945931 | 0.940281 | 0.018 | 0.02 | 1 | 15 | B230398E0 | -0.002 | Young |
| 0.945931 | 0.925489 | 0.018 | 0.02 | 1 | 15 | Gm26522 | -0.002 | Young |
| 0.945931 | 0.925489 | 0.018 | 0.02 | 1 | 15 | Tbccd1 | -0.002 | Young |
| 0.945931 | 0.914574 | 0.018 | 0.02 | 1 | 15 | Scarf1 | -0.002 | Young |
| 0.945931 | 0.905021 | 0.018 | 0.02 | 1 | 15 | Zrsr1 | -0.002 | Young |
| 0.945931 | 0.898511 | 0.018 | 0.02 | 1 | 15 | Hcn3 | -0.002 | Young |
| 0.945931 | 0.895251 | 0.018 | 0.02 | 1 | 15 | Zfp775 | -0.002 | Young |
| 0.945931 | 0.869154 | 0.018 | 0.02 | 1 | 15 | Synpo | -0.002 | Young |
| 0.945931 | 0.851164 | 0.018 | 0.02 | 1 | 15 | Aebp1 | -0.002 | Young |
| 0.945931 | 0.85067 | 0.018 | 0.02 | 1 | 15 | Pgf | -0.002 | Young |
| 0.945931 | 0.837065 | 0.018 | 0.02 | 1 | 15 | Mettl13 | -0.002 | Young |
| 0.945931 | 0.83493 | 0.018 | 0.02 | 1 | 15 | Pim2 | -0.002 | Young |
| 0.945931 | 0.831446 | 0.018 | 0.02 | 1 | 15 | Dtnb | -0.002 | Young |
| 0.945931 | 0.82865 | 0.018 | 0.02 | 1 | 15 | Taf5l | -0.002 | Young |
| 0.945931 | 0.82363 | 0.018 | 0.02 | 1 | 15 | Mpv17l | -0.002 | Young |
| 0.945931 | 0.819131 | 0.018 | 0.02 | 1 | 15 | Rrp8 | -0.002 | Young |
| 0.945931 | 0.818301 | 0.018 | 0.02 | 1 | 15 | Tchp | -0.002 | Young |
| 0.945931 | 0.812081 | 0.018 | 0.02 | 1 | 15 | Agpat2 | -0.002 | Young |
| 0.945931 | 0.769231 | 0.018 | 0.02 | 1 | 15 | Thnsl1 | -0.002 | Young |

| | | | | | | | | |
|---|---|---|---|---|---|---|---|---|
| 0.945931 | 0.767542 | 0.018 | 0.02 | 1 | 15 | Cnnm2 | -0.002 | Young |
| 0.945931 | 0.762339 | 0.018 | 0.02 | 1 | 15 | Hyal2 | -0.002 | Young |
| 0.945931 | 0.762339 | 0.018 | 0.02 | 1 | 15 | Trmt5 | -0.002 | Young |
| 0.945931 | 0.752651 | 0.018 | 0.02 | 1 | 15 | Tom1l2 | -0.002 | Young |
| 0.945931 | 0.750344 | 0.018 | 0.02 | 1 | 15 | Tlr5 | -0.002 | Young |
| 0.945931 | 0.734045 | 0.018 | 0.02 | 1 | 15 | Bysl | -0.002 | Young |
| 0.945931 | 0.726907 | 0.018 | 0.02 | 1 | 15 | Ltb4r1 | -0.002 | Young |
| 0.945931 | 0.721449 | 0.018 | 0.02 | 1 | 15 | Taf9b | -0.002 | Young |
| 0.945931 | 0.712073 | 0.018 | 0.02 | 1 | 15 | Uevld | -0.002 | Young |
| 0.945931 | 0.710571 | 0.018 | 0.02 | 1 | 15 | Zfp367 | -0.002 | Young |
| 0.945931 | 0.704817 | 0.018 | 0.02 | 1 | 15 | Zfp442 | -0.002 | Young |
| 0.945931 | 0.699868 | 0.018 | 0.02 | 1 | 15 | Espn | -0.002 | Young |
| 0.945931 | 0.690576 | 0.018 | 0.02 | 1 | 15 | Mettl21a | -0.002 | Young |
| 0.945931 | 0.685628 | 0.018 | 0.02 | 1 | 15 | Vamp5 | -0.002 | Young |
| 0.945931 | 0.685201 | 0.018 | 0.02 | 1 | 15 | Mkrn2 | -0.002 | Young |
| 0.945931 | 0.669883 | 0.018 | 0.02 | 1 | 15 | 6530409C1 | -0.002 | Young |
| 0.945931 | 0.666707 | 0.018 | 0.02 | 1 | 15 | Gm29291 | -0.002 | Young |
| 0.945931 | 0.666707 | 0.018 | 0.02 | 1 | 15 | 1810034E1 | -0.002 | Young |
| 0.945931 | 0.662823 | 0.018 | 0.02 | 1 | 15 | Kif2c | -0.002 | Young |
| 0.945931 | 0.65154 | 0.018 | 0.02 | 1 | 15 | Rilp | -0.002 | Young |
| 0.945931 | 0.650419 | 0.018 | 0.02 | 1 | 15 | Slc29a2 | -0.002 | Young |
| 0.945931 | 0.630181 | 0.018 | 0.02 | 1 | 15 | Nlgn2 | -0.002 | Young |
| 0.945931 | 0.629217 | 0.018 | 0.02 | 1 | 15 | Mettl27 | -0.002 | Young |
| 0.945931 | 0.625053 | 0.018 | 0.02 | 1 | 15 | Mocos | -0.002 | Young |
| 0.945931 | 0.614716 | 0.018 | 0.02 | 1 | 15 | Mgme1 | -0.002 | Young |
| 0.945931 | 0.601229 | 0.018 | 0.02 | 1 | 15 | Tmem25 | -0.002 | Young |
| 0.945931 | 0.584227 | 0.018 | 0.02 | 1 | 15 | Slc25a53 | -0.002 | Young |
| 0.945931 | 0.55123 | 0.018 | 0.02 | 1 | 15 | Gm28874 | -0.002 | Young |
| 0.945931 | 0.551071 | 0.018 | 0.02 | 1 | 15 | Qtrt2 | -0.002 | Young |
| 0.945931 | 0.54034 | 0.018 | 0.02 | 1 | 15 | Flt1 | -0.002 | Young |
| 0.945931 | 0.534377 | 0.018 | 0.02 | 1 | 15 | Rpp38 | -0.002 | Young |
| 0.945931 | 0.528883 | 0.018 | 0.02 | 1 | 15 | Mtrr | -0.002 | Young |
| 0.945931 | 0.527693 | 0.018 | 0.02 | 1 | 15 | Tnfsfm13 | -0.002 | Young |
| 0.945931 | 0.527693 | 0.018 | 0.02 | 1 | 15 | B430306N( | -0.002 | Young |
| 0.945931 | 0.513085 | 0.018 | 0.02 | 1 | 15 | 2900093K2 | -0.002 | Young |
| 0.945931 | 0.510308 | 0.018 | 0.02 | 1 | 15 | Ank2 | -0.002 | Young |
| 0.945931 | 0.509666 | 0.018 | 0.02 | 1 | 15 | Vwa8 | -0.002 | Young |
| 0.945931 | 0.506041 | 0.018 | 0.02 | 1 | 15 | Ccnb1ip1 | -0.002 | Young |
| 0.945931 | 0.5019 | 0.018 | 0.02 | 1 | 15 | B3galt6 | -0.002 | Young |
| 0.945931 | 0.500107 | 0.018 | 0.02 | 1 | 15 | Harbi1 | -0.002 | Young |
| 0.945931 | 0.500107 | 0.018 | 0.02 | 1 | 15 | Igf2r | -0.002 | Young |
| 0.945931 | 0.496497 | 0.018 | 0.02 | 1 | 15 | C6 | -0.002 | Young |
| 0.945931 | 0.484997 | 0.018 | 0.02 | 1 | 15 | Rfng | -0.002 | Young |
| 0.945931 | 0.481856 | 0.018 | 0.02 | 1 | 15 | Cstf2 | -0.002 | Young |
| 0.945931 | 0.479199 | 0.018 | 0.02 | 1 | 15 | AC165953. | -0.002 | Young |

| | | | | | | | | |
|---|---|---|---|---|---|---|---|---|
| 0.945931 | 0.465245 | 0.018 | 0.02 | 1 | 15 | Hddc3 | -0.002 | Young |
| 0.945931 | 0.46311 | 0.018 | 0.02 | 1 | 15 | Mtcp1 | -0.002 | Young |
| 0.945931 | 0.46311 | 0.018 | 0.02 | 1 | 15 | Tkfc | -0.002 | Young |
| 0.945931 | 0.454125 | 0.018 | 0.02 | 1 | 15 | Hacd1 | -0.002 | Young |
| 0.945931 | 0.449969 | 0.018 | 0.02 | 1 | 15 | Noa1 | -0.002 | Young |
| 0.945931 | 0.439278 | 0.018 | 0.02 | 1 | 15 | Asb6 | -0.002 | Young |
| 0.945931 | 0.434779 | 0.018 | 0.02 | 1 | 15 | Rdh5 | -0.002 | Young |
| 0.945931 | 0.434779 | 0.018 | 0.02 | 1 | 15 | C430042M | -0.002 | Young |
| 0.945931 | 0.431504 | 0.018 | 0.02 | 1 | 15 | Mxra7 | -0.002 | Young |
| 0.945931 | 0.416083 | 0.018 | 0.02 | 1 | 15 | Gm8773 | -0.002 | Young |
| 0.945931 | 0.394273 | 0.018 | 0.02 | 1 | 15 | Gtf2h2 | -0.002 | Young |
| 0.945931 | 0.387009 | 0.018 | 0.02 | 1 | 15 | Uros | -0.002 | Young |
| 0.945931 | 0.385307 | 0.018 | 0.02 | 1 | 15 | Wdpcp | -0.002 | Young |
| 0.945931 | 0.37203 | 0.018 | 0.02 | 1 | 15 | BC048403 | -0.002 | Young |
| 0.945931 | 0.371211 | 0.018 | 0.02 | 1 | 15 | Rabep2 | -0.002 | Young |
| 0.945931 | 0.361606 | 0.018 | 0.02 | 1 | 15 | Amigo1 | -0.002 | Young |
| 0.945931 | 0.357472 | 0.018 | 0.02 | 1 | 15 | 2900076A( | -0.002 | Young |
| 0.945931 | 0.351899 | 0.018 | 0.02 | 1 | 15 | Gm15627 | -0.002 | Young |
| 0.945931 | 0.349792 | 0.018 | 0.02 | 1 | 15 | Tprn | -0.002 | Young |
| 0.945931 | 0.347583 | 0.018 | 0.02 | 1 | 15 | Gm10602 | -0.002 | Young |
| 0.945931 | 0.347583 | 0.018 | 0.02 | 1 | 15 | Map2k6 | -0.002 | Young |
| 0.945931 | 0.338276 | 0.018 | 0.02 | 1 | 15 | Smyd4 | -0.002 | Young |
| 0.945931 | 0.324835 | 0.018 | 0.02 | 1 | 15 | Slc35d2 | -0.002 | Young |
| 0.945931 | 0.322285 | 0.018 | 0.02 | 1 | 15 | L3mbtl1 | -0.002 | Young |
| 0.945931 | 0.313473 | 0.018 | 0.02 | 1 | 15 | Kctd7 | -0.002 | Young |
| 0.945931 | 0.303512 | 0.018 | 0.02 | 1 | 15 | 2410080I0 | -0.002 | Young |
| 0.945931 | 0.291858 | 0.018 | 0.02 | 1 | 15 | Engase | -0.002 | Young |
| 0.945931 | 0.291089 | 0.018 | 0.02 | 1 | 15 | Tspoap1 | -0.002 | Young |
| 0.945931 | 0.290245 | 0.018 | 0.02 | 1 | 15 | Fcgr4 | -0.002 | Young |
| 0.945931 | 0.285959 | 0.018 | 0.02 | 1 | 15 | Utp4 | -0.002 | Young |
| 0.945931 | 0.280984 | 0.018 | 0.02 | 1 | 15 | Fam58b | -0.002 | Young |
| 0.945931 | 0.277707 | 0.018 | 0.02 | 1 | 15 | Ets1 | -0.002 | Young |
| 0.945931 | 0.277701 | 0.018 | 0.02 | 1 | 15 | Ints4 | -0.002 | Young |
| 0.945931 | 0.268646 | 0.018 | 0.02 | 1 | 15 | Nav2 | -0.002 | Young |
| 0.945931 | 0.26505 | 0.018 | 0.02 | 1 | 15 | Myo1b | -0.002 | Young |
| 0.945931 | 0.262017 | 0.018 | 0.02 | 1 | 15 | Mccc2 | -0.002 | Young |
| 0.945931 | 0.245163 | 0.018 | 0.02 | 1 | 15 | Shmt1 | -0.002 | Young |
| 0.945931 | 0.243209 | 0.018 | 0.02 | 1 | 15 | Nme6 | -0.002 | Young |
| 0.945931 | 0.238009 | 0.018 | 0.02 | 1 | 15 | Prr5 | -0.002 | Young |
| 0.945931 | 0.2123 | 0.018 | 0.02 | 1 | 15 | Tubg1 | -0.002 | Young |
| 0.945931 | 0.203383 | 0.018 | 0.02 | 1 | 15 | Fam198b | -0.002 | Young |
| 0.945931 | 0.192212 | 0.018 | 0.02 | 1 | 15 | Gm42601 | -0.002 | Young |
| 0.945931 | 0.190736 | 0.018 | 0.02 | 1 | 15 | Wfs1 | -0.002 | Young |
| 0.945931 | 0.174529 | 0.018 | 0.02 | 1 | 15 | Zfp951 | -0.002 | Young |
| 0.945931 | 0.174529 | 0.018 | 0.02 | 1 | 15 | Wdr55 | -0.002 | Young |

| | | | | | | | | |
|---|---|---|---|---|---|---|---|---|
| 0.945931 | 0.173014 | 0.018 | 0.02 | 1 | 15 | Hadh | -0.002 | Young |
| 0.945931 | 0.173014 | 0.018 | 0.02 | 1 | 15 | Pccb | -0.002 | Young |
| 0.945931 | 0.172484 | 0.018 | 0.02 | 1 | 15 | Leng9 | -0.002 | Young |
| 0.945931 | 0.171455 | 0.018 | 0.02 | 1 | 15 | Wdr5 | -0.002 | Young |
| 0.945931 | 0.162617 | 0.018 | 0.02 | 1 | 15 | Gprasp1 | -0.002 | Young |
| 0.945931 | 0.156686 | 0.018 | 0.02 | 1 | 15 | Adck5 | -0.002 | Young |
| 0.945931 | 0.148132 | 0.018 | 0.02 | 1 | 15 | C1ra | -0.002 | Young |
| 0.945931 | 0.148132 | 0.018 | 0.02 | 1 | 15 | Ly6g6d | -0.002 | Young |
| 0.945931 | 0.139457 | 0.018 | 0.02 | 1 | 15 | Nbl1 | -0.002 | Young |
| 0.945931 | 0.139145 | 0.018 | 0.02 | 1 | 15 | Tigar | -0.002 | Young |
| 0.945931 | 0.139145 | 0.018 | 0.02 | 1 | 15 | Ripk3 | -0.002 | Young |
| 0.945931 | 0.131317 | 0.018 | 0.02 | 1 | 15 | Slx1b | -0.002 | Young |
| 0.945931 | 0.131317 | 0.018 | 0.02 | 1 | 15 | Ccdc69 | -0.002 | Young |
| 0.945931 | 0.126069 | 0.018 | 0.02 | 1 | 15 | Dmxl2 | -0.002 | Young |
| 0.945931 | 0.114557 | 0.018 | 0.02 | 1 | 15 | Fam222a | -0.002 | Young |
| 0.945931 | 0.114557 | 0.018 | 0.02 | 1 | 15 | Akr1b8 | -0.002 | Young |
| 0.945931 | 0.114557 | 0.018 | 0.02 | 1 | 15 | Cracr2b | -0.002 | Young |
| 0.945931 | 0.112479 | 0.018 | 0.02 | 1 | 15 | Zfp747 | -0.002 | Young |
| 0.945931 | 0.110558 | 0.018 | 0.02 | 1 | 15 | Zbtb7c | -0.002 | Young |
| 0.945931 | 0.100205 | 0.018 | 0.02 | 1 | 15 | Slc7a5 | -0.002 | Young |
| 0.945931 | 0.100205 | 0.018 | 0.02 | 1 | 15 | Sphk1 | -0.002 | Young |
| 0.945931 | 0.098971 | 0.018 | 0.02 | 1 | 15 | C030034I2 | -0.002 | Young |
| 0.945931 | 0.091443 | 0.018 | 0.02 | 1 | 15 | Mettl3 | -0.002 | Young |
| 0.945931 | 0.079818 | 0.018 | 0.02 | 1 | 15 | Gm14295 | -0.002 | Young |
| 0.945931 | 0.079148 | 0.018 | 0.02 | 1 | 15 | Rnf144b | -0.002 | Young |
| 0.945931 | 0.079148 | 0.018 | 0.02 | 1 | 15 | Zfp874a | -0.002 | Young |
| 0.945931 | 0.079125 | 0.018 | 0.02 | 1 | 15 | Chek2 | -0.002 | Young |
| 0.945931 | 0.066505 | 0.018 | 0.02 | 1 | 15 | Dnm1 | -0.002 | Young |
| 0.945931 | 0.055175 | 0.018 | 0.02 | 1 | 15 | Psat1 | -0.002 | Young |
| 0.945931 | 0.050707 | 0.018 | 0.02 | 1 | 15 | Foxc2 | -0.002 | Young |
| 0.945931 | 0.050357 | 0.018 | 0.02 | 1 | 15 | Sgms2 | -0.002 | Young |
| 0.945931 | 0.050357 | 0.018 | 0.02 | 1 | 15 | Trim21 | -0.002 | Young |
| 0.945931 | 0.050357 | 0.018 | 0.02 | 1 | 15 | Atcay | -0.002 | Young |
| 0.945931 | 0.050357 | 0.018 | 0.02 | 1 | 15 | Atp6v1g2 | -0.002 | Young |
| 0.945931 | 0.042174 | 0.018 | 0.02 | 1 | 15 | Atp5s | -0.002 | Young |
| 0.945931 | 0.037179 | 0.018 | 0.02 | 1 | 15 | Gtf2h3 | -0.002 | Young |
| 0.945931 | 0.036759 | 0.018 | 0.02 | 1 | 15 | Fam78b | -0.002 | Young |
| 0.945931 | 0.036759 | 0.018 | 0.02 | 1 | 15 | Gdf3 | -0.002 | Young |
| 0.945931 | 0.036759 | 0.018 | 0.02 | 1 | 15 | Zfp609 | -0.002 | Young |
| 0.945931 | 0.035072 | 0.018 | 0.02 | 1 | 15 | P2ry14 | -0.002 | Young |
| 0.945931 | 0.035072 | 0.018 | 0.02 | 1 | 15 | Plag1 | -0.002 | Young |
| 0.945931 | 0.029832 | 0.018 | 0.02 | 1 | 15 | Rasgrf2 | -0.002 | Young |
| 0.945931 | 0.024825 | 0.018 | 0.02 | 1 | 15 | Zfp282 | -0.002 | Young |
| 0.945931 | 0.018925 | 0.018 | 0.02 | 1 | 15 | Slc26a11 | -0.002 | Young |
| 0.945931 | 0.018618 | 0.018 | 0.02 | 1 | 15 | Ttbk2 | -0.002 | Young |

| | | | | | | | | |
|---|---|---|---|---|---|---|---|---|
| 0.945931 | 0.011379 | 0.018 | 0.02 | 1 | 15 | Fam160a2 | -0.002 | Young |
| 0.946009 | 0.603875 | 0.286 | 0.306 | 1 | 15 | Fam104a | -0.02 | Young |
| 0.946297 | 0.408498 | 0.125 | 0.122 | 1 | 15 | Dnttip2 | 0.003 | Young |
| 0.946297 | 0.137331 | 0.125 | 0.122 | 1 | 15 | Rbmx | 0.003 | Young |
| 0.947628 | 0.332855 | 0.321 | 0.327 | 1 | 15 | Zc3h12a | -0.006 | Young |
| 0.948109 | 0.183858 | 0.321 | 0.347 | 1 | 15 | Ssr1 | -0.026 | Young |
| 0.948683 | 0.591452 | 0.196 | 0.204 | 1 | 15 | Spsb1 | -0.008 | Young |
| 0.948683 | 0.461625 | 0.196 | 0.204 | 1 | 15 | Fndc3b | -0.008 | Young |
| 0.948683 | 0.282792 | 0.196 | 0.204 | 1 | 15 | Arl2bp | -0.008 | Young |
| 0.949307 | 0.496464 | 0.25 | 0.245 | 1 | 15 | Mxi1 | 0.005 | Young |
| 0.950405 | 0.322267 | 0.179 | 0.184 | 1 | 15 | Tspan14 | -0.005 | Young |
| 0.952085 | 0.46986 | 0.161 | 0.163 | 1 | 15 | Spata2 | -0.002 | Young |
| 0.952085 | 0.281704 | 0.161 | 0.163 | 1 | 15 | Htt | -0.002 | Young |
| 0.952085 | 0.211631 | 0.161 | 0.163 | 1 | 15 | Kidins220 | -0.002 | Young |
| 0.953333 | 0.273623 | 0.321 | 0.306 | 1 | 15 | Aup1 | 0.015 | Young |
| 0.953705 | 0.025288 | 0.143 | 0.143 | 1 | 15 | Rnaseh2a | 0 | Young |
| 0.953705 | 0.008953 | 0.143 | 0.143 | 1 | 15 | Ranbp9 | 0 | Young |
| 0.955237 | 0.356985 | 0.125 | 0.122 | 1 | 15 | Gtf2h1 | 0.003 | Young |
| 0.955237 | 0.219714 | 0.125 | 0.122 | 1 | 15 | Cd38 | 0.003 | Young |
| 0.955237 | 0.158422 | 0.125 | 0.122 | 1 | 15 | Lrrc59 | 0.003 | Young |
| 0.955237 | 0.148751 | 0.125 | 0.122 | 1 | 15 | Ncoa1 | 0.003 | Young |
| 0.955237 | 0.036059 | 0.125 | 0.122 | 1 | 15 | Itpr3 | 0.003 | Young |
| 0.956285 | 0.006981 | 0.821 | 0.776 | 1 | 15 | Pcbp2 | 0.045 | Young |
| 0.956633 | 0.212502 | 0.107 | 0.102 | 1 | 15 | Mitf | 0.005 | Young |
| 0.956633 | 0.121708 | 0.107 | 0.102 | 1 | 15 | Hira | 0.005 | Young |
| 0.956633 | 0.065362 | 0.107 | 0.102 | 1 | 15 | Il17ra | 0.005 | Young |
| 0.958013 | 0.108808 | 0.625 | 0.653 | 1 | 15 | Snrnp70 | -0.028 | Young |
| 0.959466 | 0.115448 | 0.571 | 0.49 | 1 | 15 | Rassf4 | 0.081 | Young |
| 0.959972 | 0.082293 | 0.589 | 0.551 | 1 | 15 | Mtdh | 0.038 | Young |
| 0.961067 | 0.079263 | 0.214 | 0.224 | 1 | 15 | Clasp2 | -0.01 | Young |
| 0.961297 | 0.230903 | 0.357 | 0.367 | 1 | 15 | Lactb | -0.01 | Young |
| 0.963333 | 0.377856 | 0.196 | 0.204 | 1 | 15 | Sod2 | -0.008 | Young |
| 0.963794 | 0.049591 | 0.268 | 0.265 | 1 | 15 | Suds3 | 0.003 | Young |
| 0.964656 | 0.216708 | 0.286 | 0.286 | 1 | 15 | 5031439G( | 0 | Young |
| 0.965654 | 0.417573 | 0.179 | 0.184 | 1 | 15 | Ccndbp1 | -0.005 | Young |
| 0.965654 | 0.223261 | 0.179 | 0.184 | 1 | 15 | Uggt1 | -0.005 | Young |
| 0.966217 | 0.004092 | 0.714 | 0.673 | 1 | 15 | Erp29 | 0.041 | Young |
| 0.966264 | 0.188574 | 0.107 | 0.102 | 1 | 15 | Sap30l | 0.005 | Young |
| 0.966264 | 0.040711 | 0.107 | 0.102 | 1 | 15 | Chsy1 | 0.005 | Young |
| 0.966264 | 0.039457 | 0.107 | 0.102 | 1 | 15 | Rab6b | 0.005 | Young |
| 0.96641 | 0.191437 | 0.321 | 0.347 | 1 | 15 | Pkn1 | -0.026 | Young |
| 0.967488 | 0.119269 | 0.268 | 0.286 | 1 | 15 | Fyttd1 | -0.018 | Young |
| 0.967591 | 0.054449 | 0.214 | 0.204 | 1 | 15 | Mob1b | 0.01 | Young |
| 0.967961 | 0.012165 | 0.393 | 0.388 | 1 | 15 | Il21r | 0.005 | Young |
| 0.968046 | 0.251768 | 0.161 | 0.163 | 1 | 15 | Cpt1a | -0.002 | Young |

| | | | | | | | | |
|---|---|---|---|---|---|---|---|---|
| 0.968046 | 0.057256 | 0.161 | 0.163 | 1 | 15 | Ifi27 | -0.002 | Young |
| 0.968062 | 0.187967 | 0.554 | 0.612 | 1 | 15 | Lyz2 | -0.058 | Young |
| 0.968141 | 0.021966 | 0.214 | 0.224 | 1 | 15 | Utrn | -0.01 | Young |
| 0.969404 | 0.17853 | 0.464 | 0.469 | 1 | 15 | Phip | -0.005 | Young |
| 0.969571 | 0.064596 | 0.25 | 0.245 | 1 | 15 | Spg21 | 0.005 | Young |
| 0.969983 | 0.708889 | 0.339 | 0.367 | 1 | 15 | Ube2l6 | -0.028 | Young |
| 0.969985 | 0.008473 | 0.25 | 0.265 | 1 | 15 | Tmem8 | -0.015 | Young |
| 0.970529 | 0.44701 | 0.143 | 0.143 | 1 | 15 | Mrpl28 | 0 | Young |
| 0.970529 | 0.262919 | 0.143 | 0.143 | 1 | 15 | Map2k1 | 0 | Young |
| 0.970529 | 0.250496 | 0.143 | 0.143 | 1 | 15 | Asnsd1 | 0 | Young |
| 0.970662 | 0.370044 | 0.196 | 0.204 | 1 | 15 | Zzz3 | -0.008 | Young |
| 0.971704 | 0.02503 | 0.304 | 0.306 | 1 | 15 | Lgals1 | -0.002 | Young |
| 0.972259 | 0.056097 | 0.304 | 0.347 | 1 | 15 | Ndufc1 | -0.043 | Young |
| 0.972556 | 0.167364 | 0.232 | 0.245 | 1 | 15 | Nfkbib | -0.013 | Young |
| 0.972755 | 0.212332 | 0.339 | 0.347 | 1 | 15 | Mkl1 | -0.008 | Young |
| 0.973231 | 0.093181 | 0.554 | 0.571 | 1 | 15 | Wipf1 | -0.017 | Young |
| 0.973283 | 0.339487 | 0.179 | 0.184 | 1 | 15 | Synj2bp | -0.005 | Young |
| 0.973283 | 0.055012 | 0.179 | 0.184 | 1 | 15 | Dhx38 | -0.005 | Young |
| 0.976331 | 0.485133 | 0.232 | 0.265 | 1 | 15 | Ppp1r15b | -0.033 | Young |
| 0.976639 | 0.03166 | 0.714 | 0.694 | 1 | 15 | Ube2d3 | 0.02 | Young |
| 0.976653 | 0.334177 | 0.25 | 0.265 | 1 | 15 | Acbd5 | -0.015 | Young |
| 0.977995 | 0.278857 | 0.196 | 0.204 | 1 | 15 | Ddx42 | -0.008 | Young |
| 0.978808 | 0.025058 | 0.339 | 0.347 | 1 | 15 | Tcf4 | -0.008 | Young |
| 0.978947 | 0.486341 | 0.143 | 0.143 | 1 | 15 | Anxa6 | 0 | Young |
| 0.978947 | 0.394166 | 0.143 | 0.143 | 1 | 15 | Jdp2 | 0 | Young |
| 0.978947 | 0.227232 | 0.143 | 0.143 | 1 | 15 | Zdhhc5 | 0 | Young |
| 0.978947 | 0.075484 | 0.143 | 0.143 | 1 | 15 | Rbbp8 | 0 | Young |
| 0.978947 | 0.015024 | 0.143 | 0.143 | 1 | 15 | Ift20 | 0 | Young |
| 0.979415 | 0.284952 | 0.232 | 0.245 | 1 | 15 | Wbp11 | -0.013 | Young |
| 0.980915 | 0.686413 | 0.179 | 0.184 | 1 | 15 | Ube2w | -0.005 | Young |
| 0.980915 | 0.173746 | 0.179 | 0.184 | 1 | 15 | Pgd | -0.005 | Young |
| 0.982087 | 0.699584 | 0.125 | 0.122 | 1 | 15 | Lmf2 | 0.003 | Young |
| 0.982087 | 0.474034 | 0.125 | 0.122 | 1 | 15 | Plekhg2 | 0.003 | Young |
| 0.982087 | 0.004884 | 0.125 | 0.122 | 1 | 15 | Lpxn | 0.003 | Young |
| 0.982298 | 0.393688 | 0.214 | 0.224 | 1 | 15 | Ppp2r2a | -0.01 | Young |
| 0.984109 | 0.392049 | 0.286 | 0.306 | 1 | 15 | Eny2 | -0.02 | Young |
| 0.984278 | 0.109158 | 0.304 | 0.306 | 1 | 15 | Vbp1 | -0.002 | Young |
| 0.984555 | 0.151861 | 0.839 | 0.735 | 1 | 15 | Rhoa | 0.104 | Young |
| 0.985331 | 0.00789 | 0.375 | 0.388 | 1 | 15 | Fryl | -0.013 | Young |
| 0.985795 | 0.110366 | 0.429 | 0.429 | 1 | 15 | Chmp4b | 0 | Young |
| 0.987367 | 0.105028 | 0.143 | 0.143 | 1 | 15 | Gm11808 | 0 | Young |
| 0.988548 | 0.013276 | 0.179 | 0.184 | 1 | 15 | Smc3 | -0.005 | Young |
| 0.989378 | 0.229988 | 0.214 | 0.224 | 1 | 15 | Pcbd2 | -0.01 | Young |
| 0.989572 | 0.091579 | 0.661 | 0.694 | 1 | 15 | Nr4a1 | -0.033 | Young |
| 0.989993 | 0.170196 | 0.25 | 0.265 | 1 | 15 | Wdr45b | -0.015 | Young |

| | | | | | | | | |
|---|---|---|---|---|---|---|---|---|
| 0.989993 | 0.147883 | 0.25 | 0.265 | 1 | 15 | Ubn2 | -0.015 | Young |
| 0.990122 | 0.182986 | 0.268 | 0.265 | 1 | 15 | Tmbim4 | 0.003 | Young |
| 0.990122 | 0.122876 | 0.268 | 0.265 | 1 | 15 | Rab9 | 0.003 | Young |
| 0.990465 | 0.211487 | 0.286 | 0.306 | 1 | 15 | Atp6v1e1 | -0.02 | Young |
| 0.991043 | 0.252163 | 0.125 | 0.122 | 1 | 15 | Sertad1 | 0.003 | Young |
| 0.991043 | 0.092804 | 0.125 | 0.122 | 1 | 15 | Atg101 | 0.003 | Young |
| 0.992009 | 0.340638 | 0.161 | 0.163 | 1 | 15 | Slc46a3 | -0.002 | Young |
| 0.992009 | 0.229666 | 0.161 | 0.163 | 1 | 15 | Slc35e1 | -0.002 | Young |
| 0.992009 | 0.061187 | 0.161 | 0.163 | 1 | 15 | Timm8b | -0.002 | Young |
| 0.992664 | 0.45352 | 0.196 | 0.204 | 1 | 15 | Clcn3 | -0.008 | Young |
| 0.995179 | 0.607589 | 0.107 | 0.102 | 1 | 15 | Mnat1 | 0.005 | Young |
| 0.995789 | 0.13807 | 0.143 | 0.143 | 1 | 15 | Trak1 | 0 | Young |
| 0.996618 | 0.017835 | 0.25 | 0.245 | 1 | 15 | Galnt7 | 0.005 | Young |
| 0.996822 | 0.567412 | 0.286 | 0.306 | 1 | 15 | Inpp5d | -0.02 | Young |
| 0.996822 | 0.180067 | 0.286 | 0.306 | 1 | 15 | Bcl7c | -0.02 | Young |
| 0.996972 | 0.049548 | 0.339 | 0.347 | 1 | 15 | Eif5a | -0.008 | Young |
| 0.997022 | 0.03235 | 0.357 | 0.367 | 1 | 15 | Ap2a2 | -0.01 | Young |
| 0.997255 | 0.105354 | 0.518 | 0.469 | 1 | 15 | Nfkb2 | 0.049 | Young |
| 0.997358 | 0.178692 | 0.607 | 0.612 | 1 | 15 | Sumo2 | -0.005 | Young |
| 2.83E-19 | 0.401507 | 0.969 | 0.93 | 8.79E-15 | 2 | B2m | 0.039 | Old |
| 8.83E-17 | 0.510706 | 0.815 | 0.715 | 2.74E-12 | 2 | Gda | 0.1 | Old |
| 7.32E-15 | 0.611758 | 0.858 | 0.765 | 2.27E-10 | 2 | Samhd1 | 0.093 | Old |
| 2.28E-11 | 0.263449 | 0.98 | 0.958 | 7.07E-07 | 2 | Gpx1 | 0.022 | Old |
| 6.08E-10 | 0.253342 | 0.934 | 0.906 | 1.89E-05 | 2 | H2-D1 | 0.028 | Old |
| 7.35E-10 | 0.465829 | 0.73 | 0.61 | 2.28E-05 | 2 | Srsf5 | 0.12 | Old |
| 7.52E-10 | 0.144872 | 1 | 0.999 | 2.34E-05 | 2 | mt-Co1 | 0.001 | Old |
| 1.46E-09 | 0.253685 | 0.989 | 0.99 | 4.53E-05 | 2 | Actg1 | -0.001 | Old |
| 1.62E-09 | 0.154722 | 0.998 | 0.999 | 5.05E-05 | 2 | Actb | -0.001 | Old |
| 1.88E-09 | 1.595976 | 0.079 | 0.017 | 5.84E-05 | 2 | Cxcl13 | 0.062 | Old |
| 6.74E-09 | 0.798669 | 0.3 | 0.184 | 0.000209 | 2 | Stat1 | 0.116 | Old |
| 1.35E-08 | 0.281732 | 0.942 | 0.93 | 0.00042 | 2 | Cebpb | 0.012 | Old |
| 1.37E-08 | 0.312795 | 0.837 | 0.766 | 0.000425 | 2 | Ywhaz | 0.071 | Old |
| 3.48E-08 | 0.835038 | 0.249 | 0.145 | 0.00108 | 2 | Ms4a4c | 0.104 | Old |
| 5.21E-08 | 0.79555 | 0.33 | 0.215 | 0.001619 | 2 | Plac8 | 0.115 | Old |
| 8.38E-08 | 0.165289 | 0.994 | 0.992 | 0.002603 | 2 | Psap | 0.002 | Old |
| 1.73E-07 | 0.762456 | 0.267 | 0.162 | 0.005358 | 2 | Rgs1 | 0.105 | Old |
| 2.90E-07 | 0.399462 | 0.711 | 0.613 | 0.009001 | 2 | Ms4a6c | 0.098 | Old |
| 3.08E-07 | 0.26122 | 0.882 | 0.824 | 0.009576 | 2 | Actr3 | 0.058 | Old |
| 3.79E-07 | 0.24881 | 0.941 | 0.91 | 0.011763 | 2 | H3f3b | 0.031 | Old |
| 4.03E-07 | 0.956026 | 0.219 | 0.129 | 0.012529 | 2 | Adgre4 | 0.09 | Old |
| 4.64E-07 | 0.508152 | 0.596 | 0.494 | 0.014421 | 2 | Klf2 | 0.102 | Old |
| 4.76E-07 | 0.32311 | 0.82 | 0.742 | 0.014773 | 2 | Cd44 | 0.078 | Old |
| 5.47E-07 | 0.238463 | 0.954 | 0.938 | 0.016987 | 2 | H2-K1 | 0.016 | Old |
| 5.81E-07 | 0.456496 | 0.435 | 0.321 | 0.018043 | 2 | Sp100 | 0.114 | Old |
| 6.11E-07 | 0.704482 | 0.4 | 0.29 | 0.018975 | 2 | Slfn5 | 0.11 | Old |

| | | | | | | | | |
|---|---|---|---|---|---|---|---|---|
| 8.84E-07 | 0.350747 | 0.681 | 0.606 | 0.027457 | 2 | Ifi27l2a | 0.075 | Old |
| 8.98E-07 | 0.389356 | 0.623 | 0.506 | 0.027899 | 2 | Xbp1 | 0.117 | Old |
| 9.46E-07 | 0.290835 | 0.889 | 0.867 | 0.029388 | 2 | BC005537 | 0.022 | Old |
| 1.20E-06 | 0.325919 | 0.858 | 0.849 | 0.037332 | 2 | Hsp90b1 | 0.009 | Old |
| 1.52E-06 | 0.447031 | 0.668 | 0.606 | 0.047079 | 2 | Ly6e | 0.062 | Old |
| 2.83E-06 | 0.171804 | 0.991 | 0.982 | 0.087893 | 2 | mt-Cytb | 0.009 | Old |
| 2.85E-06 | 0.435426 | 0.627 | 0.534 | 0.088431 | 2 | Clec4a3 | 0.093 | Old |
| 4.17E-06 | 0.234262 | 0.863 | 0.823 | 0.129351 | 2 | Capzb | 0.04 | Old |
| 6.26E-06 | 0.493951 | 0.444 | 0.333 | 0.194393 | 2 | Cd300ld | 0.111 | Old |
| 8.34E-06 | 0.434216 | 0.505 | 0.403 | 0.258875 | 2 | Prr13 | 0.102 | Old |
| 1.23E-05 | 0.651227 | 0.368 | 0.279 | 0.38277 | 2 | Ifitm6 | 0.089 | Old |
| 1.24E-05 | 0.266543 | 0.861 | 0.836 | 0.384487 | 2 | Neat1 | 0.025 | Old |
| 1.45E-05 | 0.714148 | 0.221 | 0.141 | 0.450968 | 2 | Ms4a6d | 0.08 | Old |
| 1.73E-05 | 0.443909 | 0.737 | 0.659 | 0.536047 | 2 | Msn | 0.078 | Old |
| 1.74E-05 | 1.003278 | 0.225 | 0.147 | 0.540243 | 2 | Chil3 | 0.078 | Old |
| 1.94E-05 | 0.79725 | 0.182 | 0.108 | 0.603065 | 2 | AW112010 | 0.074 | Old |
| 2.03E-05 | 0.356975 | 0.584 | 0.486 | 0.628974 | 2 | Atp5g3 | 0.098 | Old |
| 2.16E-05 | 0.382761 | 0.679 | 0.629 | 0.670669 | 2 | Naaa | 0.05 | Old |
| 2.39E-05 | 0.577559 | 0.415 | 0.322 | 0.742012 | 2 | Copz1 | 0.093 | Old |
| 2.66E-05 | 0.25663 | 0.897 | 0.855 | 0.824599 | 2 | mt-Nd3 | 0.042 | Old |
| 2.88E-05 | 0.513214 | 0.403 | 0.312 | 0.893717 | 2 | Ddx39b | 0.091 | Old |
| 2.94E-05 | 0.376238 | 0.47 | 0.373 | 0.911688 | 2 | Camkk2 | 0.097 | Old |
| 2.96E-05 | 0.237461 | 0.843 | 0.799 | 0.919693 | 2 | Rap1b | 0.044 | Old |
| 3.10E-05 | 2.526163 | 0.036 | 0.007 | 0.963052 | 2 | Pglyrp1 | 0.029 | Old |
| 3.14E-05 | 0.366651 | 0.745 | 0.704 | 0.975168 | 2 | Sdc3 | 0.041 | Old |
| 3.51E-05 | 0.219017 | 0.939 | 0.942 | 1 | 2 | Ctss | -0.003 | Old |
| 3.85E-05 | 0.117265 | 0.998 | 0.998 | 1 | 2 | mt-Co3 | 0 | Old |
| 3.88E-05 | 1.809496 | 0.047 | 0.013 | 1 | 2 | Igkc | 0.034 | Old |
| 4.43E-05 | 0.943911 | 0.086 | 0.038 | 1 | 2 | Usp50 | 0.048 | Old |
| 4.44E-05 | 0.251883 | 0.82 | 0.765 | 1 | 2 | Fam49b | 0.055 | Old |
| 5.37E-05 | 0.300403 | 0.647 | 0.563 | 1 | 2 | Metrnl | 0.084 | Old |
| 5.40E-05 | 0.196981 | 0.912 | 0.88 | 1 | 2 | Arpc2 | 0.032 | Old |
| 5.48E-05 | 0.456757 | 0.401 | 0.308 | 1 | 2 | Gm21188 | 0.093 | Old |
| 5.50E-05 | 0.422235 | 0.456 | 0.361 | 1 | 2 | Anxa1 | 0.095 | Old |
| 5.59E-05 | 1.059069 | 0.119 | 0.064 | 1 | 2 | Xist | 0.055 | Old |
| 5.71E-05 | 0.540411 | 0.33 | 0.241 | 1 | 2 | Gm26532 | 0.089 | Old |
| 6.38E-05 | 0.487056 | 0.551 | 0.471 | 1 | 2 | Cdk2ap2 | 0.08 | Old |
| 6.70E-05 | 0.815928 | 0.237 | 0.163 | 1 | 2 | Klra2 | 0.074 | Old |
| 6.96E-05 | 0.555138 | 0.249 | 0.171 | 1 | 2 | Plod3 | 0.078 | Old |
| 7.06E-05 | 0.618991 | 0.355 | 0.271 | 1 | 2 | Noct | 0.084 | Old |
| 7.36E-05 | 0.73846 | 0.245 | 0.171 | 1 | 2 | Agpat4 | 0.074 | Old |
| 7.87E-05 | 0.542374 | 0.379 | 0.3 | 1 | 2 | Kras | 0.079 | Old |
| 7.89E-05 | 0.993075 | 0.064 | 0.025 | 1 | 2 | Rgs3 | 0.039 | Old |
| 8.54E-05 | 0.780185 | 0.134 | 0.076 | 1 | 2 | Fyn | 0.058 | Old |
| 9.45E-05 | 0.715365 | 0.161 | 0.096 | 1 | 2 | Xdh | 0.065 | Old |

| | | | | | | | | |
|---|---|---|---|---|---|---|---|---|
| 0.000116 | 0.815917 | 0.146 | 0.087 | 1 | 2 | Ccl8 | 0.059 | Old |
| 0.000125 | 1.37469 | 0.064 | 0.026 | 1 | 2 | Tmem97 | 0.038 | Old |
| 0.000143 | 0.13351 | 0.98 | 0.983 | 1 | 2 | Laptm5 | -0.003 | Old |
| 0.000148 | 0.514088 | 0.198 | 0.128 | 1 | 2 | Dcp2 | 0.07 | Old |
| 0.000174 | 0.580508 | 0.253 | 0.182 | 1 | 2 | Prkar2a | 0.071 | Old |
| 0.000186 | 0.327894 | 0.31 | 0.227 | 1 | 2 | Tap1 | 0.083 | Old |
| 0.00019 | 0.241921 | 0.814 | 0.778 | 1 | 2 | Rhoa | 0.036 | Old |
| 0.000221 | 0.413217 | 0.332 | 0.252 | 1 | 2 | Elmo2 | 0.08 | Old |
| 0.000235 | 0.69746 | 0.206 | 0.141 | 1 | 2 | Scarb1 | 0.065 | Old |
| 0.000236 | 0.339026 | 0.608 | 0.524 | 1 | 2 | Arl6ip1 | 0.084 | Old |
| 0.000247 | 0.220961 | 0.862 | 0.818 | 1 | 2 | Dazap2 | 0.044 | Old |
| 0.000247 | 0.521053 | 0.308 | 0.229 | 1 | 2 | Gnpda1 | 0.079 | Old |
| 0.000262 | 0.573799 | 0.428 | 0.352 | 1 | 2 | Pou2f2 | 0.076 | Old |
| 0.000297 | 1.359074 | 0.06 | 0.025 | 1 | 2 | Dus1l | 0.035 | Old |
| 0.000318 | 0.332099 | 0.621 | 0.551 | 1 | 2 | Atp1a1 | 0.07 | Old |
| 0.00032 | 0.167049 | 0.97 | 0.973 | 1 | 2 | Junb | -0.003 | Old |
| 0.00035 | 2.935742 | 0.018 | 0.001 | 1 | 2 | Tgfbr3 | 0.017 | Old |
| 0.000353 | 1.13816 | 0.081 | 0.04 | 1 | 2 | Ldlrad3 | 0.041 | Old |
| 0.000354 | 0.818877 | 0.104 | 0.057 | 1 | 2 | Wdr20 | 0.047 | Old |
| 0.00036 | 0.299194 | 0.504 | 0.418 | 1 | 2 | Cap1 | 0.086 | Old |
| 0.000367 | 1.544897 | 0.024 | 0.004 | 1 | 2 | Tcta | 0.02 | Old |
| 0.000371 | 0.321655 | 0.673 | 0.619 | 1 | 2 | 2010107E0 | 0.054 | Old |
| 0.000406 | 0.390801 | 0.414 | 0.335 | 1 | 2 | Myl12a | 0.079 | Old |
| 0.000414 | 0.131497 | 0.964 | 0.96 | 1 | 2 | Crip1 | 0.004 | Old |
| 0.000423 | 0.490406 | 0.327 | 0.251 | 1 | 2 | B4galnt1 | 0.076 | Old |
| 0.000427 | 0.365149 | 0.292 | 0.215 | 1 | 2 | Aco2 | 0.077 | Old |
| 0.00043 | 0.745649 | 0.127 | 0.076 | 1 | 2 | P2ry10b | 0.051 | Old |
| 0.000435 | 0.898284 | 0.095 | 0.051 | 1 | 2 | Gm5150 | 0.044 | Old |
| 0.000448 | 0.47579 | 0.293 | 0.22 | 1 | 2 | Chmp5 | 0.073 | Old |
| 0.000449 | 0.392712 | 0.403 | 0.33 | 1 | 2 | 9-Sep | 0.073 | Old |
| 0.000463 | 0.328561 | 0.472 | 0.382 | 1 | 2 | Acer3 | 0.09 | Old |
| 0.000475 | 1.040158 | 0.051 | 0.02 | 1 | 2 | Gm26870 | 0.031 | Old |
| 0.000476 | 1.870743 | 0.02 | 0.002 | 1 | 2 | Phf11a | 0.018 | Old |
| 0.000499 | 0.501305 | 0.4 | 0.328 | 1 | 2 | Fgd4 | 0.072 | Old |
| 0.000514 | 0.308379 | 0.624 | 0.567 | 1 | 2 | Ccr5 | 0.057 | Old |
| 0.000522 | 0.171866 | 0.936 | 0.923 | 1 | 2 | Ccl6 | 0.013 | Old |
| 0.000527 | 0.53154 | 0.396 | 0.328 | 1 | 2 | Pla2g7 | 0.068 | Old |
| 0.000552 | 0.373062 | 0.377 | 0.298 | 1 | 2 | Ago2 | 0.079 | Old |
| 0.000556 | 0.485205 | 0.242 | 0.176 | 1 | 2 | Pld3 | 0.066 | Old |
| 0.000557 | 0.732744 | 0.225 | 0.163 | 1 | 2 | Ptk2b | 0.062 | Old |
| 0.000593 | 2.597193 | 0.014 | 0 | 1 | 2 | Tnfrsf14 | 0.014 | Old |
| 0.000597 | 0.80637 | 0.126 | 0.077 | 1 | 2 | Sf3a2 | 0.049 | Old |
| 0.000616 | 0.315631 | 0.503 | 0.425 | 1 | 2 | Myl12b | 0.078 | Old |
| 0.000617 | 0.239905 | 0.574 | 0.482 | 1 | 2 | Tnfrsf1b | 0.092 | Old |
| 0.000645 | 0.228282 | 0.787 | 0.724 | 1 | 2 | Hnrnpf | 0.063 | Old |

| | | | | | | | | |
|---|---|---|---|---|---|---|---|---|
| 0.000651 | 0.28756 | 0.589 | 0.523 | 1 | 2 | Arpc4 | 0.066 | Old |
| 0.000692 | 1.469102 | 0.054 | 0.023 | 1 | 2 | Tbp | 0.031 | Old |
| 0.000706 | 0.153548 | 0.805 | 0.742 | 1 | 2 | Cd53 | 0.063 | Old |
| 0.000732 | 0.455201 | 0.226 | 0.16 | 1 | 2 | Rnf44 | 0.066 | Old |
| 0.000746 | 0.519682 | 0.389 | 0.323 | 1 | 2 | Ifi204 | 0.066 | Old |
| 0.000765 | 1.832263 | 0.019 | 0.002 | 1 | 2 | Ahnak2 | 0.017 | Old |
| 0.000795 | 0.216662 | 0.741 | 0.672 | 1 | 2 | Tmem50a | 0.069 | Old |
| 0.000796 | 0.263015 | 0.669 | 0.596 | 1 | 2 | Ssr3 | 0.073 | Old |
| 0.000816 | 0.414212 | 0.281 | 0.209 | 1 | 2 | Ms4a6b | 0.072 | Old |
| 0.000826 | 0.616014 | 0.213 | 0.151 | 1 | 2 | Arnt | 0.062 | Old |
| 0.000836 | 0.230598 | 0.924 | 0.914 | 1 | 2 | Cybb | 0.01 | Old |
| 0.000852 | 0.549604 | 0.167 | 0.11 | 1 | 2 | Ccdc88c | 0.057 | Old |
| 0.000862 | 0.664551 | 0.11 | 0.065 | 1 | 2 | Rexo1 | 0.045 | Old |
| 0.000874 | 0.580482 | 0.191 | 0.131 | 1 | 2 | Araf | 0.06 | Old |
| 0.000894 | 0.650297 | 0.078 | 0.04 | 1 | 2 | Zdhhc6 | 0.038 | Old |
| 0.000916 | 0.713558 | 0.193 | 0.135 | 1 | 2 | Txndc9 | 0.058 | Old |
| 0.000944 | 0.370961 | 0.244 | 0.178 | 1 | 2 | Immt | 0.066 | Old |
| 0.000953 | 0.727755 | 0.106 | 0.062 | 1 | 2 | Pitpnm1 | 0.044 | Old |
| 0.000964 | 0.168178 | 0.909 | 0.899 | 1 | 2 | Ccl9 | 0.01 | Old |
| 0.00097 | 2.040304 | 0.013 | 0 | 1 | 2 | Colq | 0.013 | Old |
| 0.000971 | 0.274209 | 0.868 | 0.853 | 1 | 2 | Fos | 0.015 | Old |
| 0.000982 | 1.03761 | 0.076 | 0.039 | 1 | 2 | Sort1 | 0.037 | Old |
| 0.00099 | 2.270668 | 0.03 | 0.008 | 1 | 2 | Trem3 | 0.022 | Old |
| 0.000998 | 1.581122 | 0.024 | 0.005 | 1 | 2 | Gm3448 | 0.019 | Old |
| 0.001063 | 2.191401 | 0.054 | 0.024 | 1 | 2 | Ace | 0.03 | Old |
| 0.00108 | 0.65224 | 0.199 | 0.141 | 1 | 2 | AI467606 | 0.058 | Old |
| 0.001111 | 0.484197 | 0.275 | 0.211 | 1 | 2 | B4galt5 | 0.064 | Old |
| 0.001135 | 0.376365 | 0.415 | 0.336 | 1 | 2 | Ldlr | 0.079 | Old |
| 0.00119 | 0.244623 | 0.816 | 0.778 | 1 | 2 | Mafb | 0.038 | Old |
| 0.001214 | 0.670585 | 0.192 | 0.137 | 1 | 2 | Azi2 | 0.055 | Old |
| 0.00122 | 0.361684 | 0.519 | 0.443 | 1 | 2 | Klf13 | 0.076 | Old |
| 0.001229 | 0.56001 | 0.242 | 0.178 | 1 | 2 | Arf6 | 0.064 | Old |
| 0.001243 | 0.497716 | 0.373 | 0.306 | 1 | 2 | Msrb1 | 0.067 | Old |
| 0.001247 | 0.537714 | 0.194 | 0.135 | 1 | 2 | Smarce1 | 0.059 | Old |
| 0.001266 | 0.32823 | 0.58 | 0.518 | 1 | 2 | Jarid2 | 0.062 | Old |
| 0.001296 | 0.424656 | 0.199 | 0.14 | 1 | 2 | Crybg1 | 0.059 | Old |
| 0.001336 | 0.790075 | 0.075 | 0.039 | 1 | 2 | Cdc14b | 0.036 | Old |
| 0.001348 | 0.48168 | 0.397 | 0.333 | 1 | 2 | Smpdl3a | 0.064 | Old |
| 0.001381 | 0.650522 | 0.127 | 0.081 | 1 | 2 | Ndnf | 0.046 | Old |
| 0.001434 | 0.955066 | 0.041 | 0.015 | 1 | 2 | Igf2r | 0.026 | Old |
| 0.001451 | 0.656637 | 0.124 | 0.078 | 1 | 2 | Evi2 | 0.046 | Old |
| 0.001464 | 0.171135 | 0.892 | 0.874 | 1 | 2 | Vim | 0.018 | Old |
| 0.001477 | 0.625959 | 0.176 | 0.121 | 1 | 2 | Fcgr4 | 0.055 | Old |
| 0.001498 | 0.213744 | 0.808 | 0.777 | 1 | 2 | Ifitm3 | 0.031 | Old |
| 0.001501 | 0.746952 | 0.104 | 0.062 | 1 | 2 | Tmem222 | 0.042 | Old |

| | | | | | | | | |
|---|---|---|---|---|---|---|---|---|
| 0.001501 | 0.256624 | 0.708 | 0.663 | 1 | 2 | Flna | 0.045 | Old |
| 0.001512 | 0.843842 | 0.066 | 0.033 | 1 | 2 | Engase | 0.033 | Old |
| 0.001581 | 0.360382 | 0.225 | 0.163 | 1 | 2 | Lgals9 | 0.062 | Old |
| 0.00162 | 1.378384 | 0.025 | 0.006 | 1 | 2 | Cd55 | 0.019 | Old |
| 0.001685 | 0.419784 | 0.325 | 0.259 | 1 | 2 | Trim30a | 0.066 | Old |
| 0.001697 | 0.362091 | 0.353 | 0.283 | 1 | 2 | Tet2 | 0.07 | Old |
| 0.001725 | 0.330944 | 0.504 | 0.431 | 1 | 2 | Ppp2r5a | 0.073 | Old |
| 0.001743 | 0.111296 | 0.974 | 0.97 | 1 | 2 | mt-Nd2 | 0.004 | Old |
| 0.001806 | 0.605404 | 0.201 | 0.147 | 1 | 2 | Irf9 | 0.054 | Old |
| 0.001833 | 0.555537 | 0.233 | 0.178 | 1 | 2 | Gsr | 0.055 | Old |
| 0.001848 | 0.721312 | 0.064 | 0.032 | 1 | 2 | St3gal3 | 0.032 | Old |
| 0.001877 | 0.705224 | 0.113 | 0.07 | 1 | 2 | Tlr1 | 0.043 | Old |
| 0.001935 | 0.176582 | 0.835 | 0.809 | 1 | 2 | Capza2 | 0.026 | Old |
| 0.001959 | 0.688687 | 0.154 | 0.105 | 1 | 2 | Rap1gap2 | 0.049 | Old |
| 0.001979 | 0.286697 | 0.64 | 0.576 | 1 | 2 | Clec4a1 | 0.064 | Old |
| 0.002076 | 0.28771 | 0.526 | 0.448 | 1 | 2 | Myadm | 0.078 | Old |
| 0.002166 | 0.42045 | 0.202 | 0.145 | 1 | 2 | Lgals3bp | 0.057 | Old |
| 0.002247 | 1.790132 | 0.019 | 0.004 | 1 | 2 | Csprs | 0.015 | Old |
| 0.00225 | 0.29837 | 0.5 | 0.43 | 1 | 2 | Clec10a | 0.07 | Old |
| 0.002275 | 0.356193 | 0.473 | 0.405 | 1 | 2 | Nfam1 | 0.068 | Old |
| 0.002283 | 0.712384 | 0.197 | 0.144 | 1 | 2 | BC031181 | 0.053 | Old |
| 0.002303 | 0.224642 | 0.663 | 0.609 | 1 | 2 | Nrros | 0.054 | Old |
| 0.002316 | 0.196197 | 0.213 | 0.153 | 1 | 2 | Fgl2 | 0.06 | Old |
| 0.002376 | 0.222888 | 0.395 | 0.325 | 1 | 2 | Magt1 | 0.07 | Old |
| 0.002447 | 0.195952 | 0.897 | 0.887 | 1 | 2 | Ptprc | 0.01 | Old |
| 0.002546 | 0.492204 | 0.193 | 0.138 | 1 | 2 | Raf1 | 0.055 | Old |
| 0.002554 | 0.486282 | 0.275 | 0.214 | 1 | 2 | Ear2 | 0.061 | Old |
| 0.00258 | 0.530846 | 0.096 | 0.057 | 1 | 2 | Ttc13 | 0.039 | Old |
| 0.002615 | 2.643655 | 0.011 | 0 | 1 | 2 | Dmpk | 0.011 | Old |
| 0.002616 | 0.308151 | 0.732 | 0.709 | 1 | 2 | Hspa5 | 0.023 | Old |
| 0.002669 | 0.20497 | 0.588 | 0.507 | 1 | 2 | Plekho2 | 0.081 | Old |
| 0.002695 | 0.814325 | 0.046 | 0.02 | 1 | 2 | Trim30b | 0.026 | Old |
| 0.002699 | 0.640071 | 0.168 | 0.118 | 1 | 2 | Tyk2 | 0.05 | Old |
| 0.002702 | 0.624485 | 0.111 | 0.07 | 1 | 2 | Fpr2 | 0.041 | Old |
| 0.002809 | 0.118474 | 0.908 | 0.89 | 1 | 2 | Cdc42 | 0.018 | Old |
| 0.002858 | 0.244826 | 0.63 | 0.574 | 1 | 2 | Stk17b | 0.056 | Old |
| 0.002915 | 0.317598 | 0.357 | 0.292 | 1 | 2 | Lfng | 0.065 | Old |
| 0.002978 | 0.575796 | 0.091 | 0.053 | 1 | 2 | Nlrc5 | 0.038 | Old |
| 0.003 | 0.296924 | 0.262 | 0.198 | 1 | 2 | Esd | 0.064 | Old |
| 0.003017 | 0.634522 | 0.185 | 0.135 | 1 | 2 | Adap1 | 0.05 | Old |
| 0.003036 | 0.101528 | 0.996 | 0.995 | 1 | 2 | mt-Atp6 | 0.001 | Old |
| 0.003065 | 0.58439 | 0.244 | 0.185 | 1 | 2 | Gngt2 | 0.059 | Old |
| 0.003172 | 0.227148 | 0.562 | 0.486 | 1 | 2 | Rnf130 | 0.076 | Old |
| 0.003187 | 0.448085 | 0.182 | 0.129 | 1 | 2 | Chmp1a | 0.053 | Old |
| 0.003207 | 0.51334 | 0.103 | 0.063 | 1 | 2 | Irgm1 | 0.04 | Old |

| | | | | | | | | |
|---|---|---|---|---|---|---|---|---|
| 0.003235 | 0.17354 | 0.721 | 0.677 | 1 | 2 | Celf2 | 0.044 | Old |
| 0.003312 | 0.301295 | 0.275 | 0.21 | 1 | 2 | Napsa | 0.065 | Old |
| 0.003358 | 0.46914 | 0.11 | 0.07 | 1 | 2 | Sirpb1b | 0.04 | Old |
| 0.003369 | 1.307524 | 0.029 | 0.01 | 1 | 2 | Tbl2 | 0.019 | Old |
| 0.003385 | 0.31712 | 0.473 | 0.412 | 1 | 2 | Sppl2a | 0.061 | Old |
| 0.003556 | 0.750827 | 0.116 | 0.075 | 1 | 2 | Trim27 | 0.041 | Old |
| 0.003657 | 1.230941 | 0.048 | 0.023 | 1 | 2 | Rrp1b | 0.025 | Old |
| 0.003675 | 1.484971 | 0.025 | 0.007 | 1 | 2 | Ppm1f | 0.018 | Old |
| 0.003686 | 0.592196 | 0.11 | 0.07 | 1 | 2 | Rgs14 | 0.04 | Old |
| 0.003721 | 0.157494 | 0.796 | 0.779 | 1 | 2 | Alox5ap | 0.017 | Old |
| 0.003741 | 0.25965 | 0.212 | 0.154 | 1 | 2 | Wtap | 0.058 | Old |
| 0.003811 | 0.407503 | 0.382 | 0.323 | 1 | 2 | Usp25 | 0.059 | Old |
| 0.003818 | 0.283213 | 0.179 | 0.124 | 1 | 2 | Flvcr1 | 0.055 | Old |
| 0.003819 | 1.40558 | 0.076 | 0.044 | 1 | 2 | Tcf7l2 | 0.032 | Old |
| 0.003871 | 0.578534 | 0.209 | 0.156 | 1 | 2 | Fabp4 | 0.053 | Old |
| 0.003878 | 0.445023 | 0.252 | 0.194 | 1 | 2 | H2-Q7 | 0.058 | Old |
| 0.003896 | 0.434624 | 0.384 | 0.321 | 1 | 2 | Itch | 0.063 | Old |
| 0.00392 | 0.35124 | 0.345 | 0.278 | 1 | 2 | Sbno1 | 0.067 | Old |
| 0.003949 | 0.799239 | 0.045 | 0.02 | 1 | 2 | Fam78b | 0.025 | Old |
| 0.004052 | 0.683939 | 0.096 | 0.059 | 1 | 2 | Rtp4 | 0.037 | Old |
| 0.004095 | 0.826845 | 0.048 | 0.023 | 1 | 2 | 3110002H1 | 0.025 | Old |
| 0.004098 | 0.445059 | 0.211 | 0.158 | 1 | 2 | Suz12 | 0.053 | Old |
| 0.004109 | 0.84549 | 0.1 | 0.063 | 1 | 2 | Ap1g2 | 0.037 | Old |
| 0.004113 | 0.755699 | 0.086 | 0.051 | 1 | 2 | Slc35c1 | 0.035 | Old |
| 0.004174 | 0.863165 | 0.055 | 0.027 | 1 | 2 | Abi3 | 0.028 | Old |
| 0.004276 | 1.006012 | 0.037 | 0.015 | 1 | 2 | Treml4 | 0.022 | Old |
| 0.004311 | 2.2186 | 0.01 | 0 | 1 | 2 | Adap2os | 0.01 | Old |
| 0.004311 | 1.664258 | 0.01 | 0 | 1 | 2 | Fabp3 | 0.01 | Old |
| 0.004327 | 0.642729 | 0.133 | 0.089 | 1 | 2 | Gm36161 | 0.044 | Old |
| 0.004366 | 0.43174 | 0.169 | 0.12 | 1 | 2 | Tbc1d10b | 0.049 | Old |
| 0.004415 | 0.598467 | 0.177 | 0.128 | 1 | 2 | Vamp4 | 0.049 | Old |
| 0.004472 | 1.228351 | 0.041 | 0.018 | 1 | 2 | Kctd18 | 0.023 | Old |
| 0.00448 | 0.223823 | 0.2 | 0.146 | 1 | 2 | Hist1h1c | 0.054 | Old |
| 0.004619 | 0.155706 | 0.582 | 0.524 | 1 | 2 | Tapbp | 0.058 | Old |
| 0.004628 | 0.337424 | 0.335 | 0.276 | 1 | 2 | Cbfa2t3 | 0.059 | Old |
| 0.004672 | 0.254643 | 0.373 | 0.302 | 1 | 2 | Itsn2 | 0.071 | Old |
| 0.004705 | 0.601053 | 0.101 | 0.063 | 1 | 2 | Gdf3 | 0.038 | Old |
| 0.004736 | 0.398628 | 0.178 | 0.127 | 1 | 2 | Map2k1 | 0.051 | Old |
| 0.004874 | 0.462208 | 0.234 | 0.181 | 1 | 2 | Sik1 | 0.053 | Old |
| 0.005033 | 0.57598 | 0.175 | 0.127 | 1 | 2 | Nek9 | 0.048 | Old |
| 0.005036 | 1.507778 | 0.015 | 0.002 | 1 | 2 | Gm26510 | 0.013 | Old |
| 0.005049 | 0.361458 | 0.284 | 0.223 | 1 | 2 | Uhrf2 | 0.061 | Old |
| 0.005168 | 1.043086 | 0.06 | 0.032 | 1 | 2 | Dcn | 0.028 | Old |
| 0.005181 | 0.363092 | 0.396 | 0.324 | 1 | 2 | Limd1 | 0.072 | Old |
| 0.005259 | 0.228535 | 0.607 | 0.549 | 1 | 2 | Rnf149 | 0.058 | Old |

| | | | | | | | | |
|---|---|---|---|---|---|---|---|---|
| 0.005285 | 0.659455 | 0.062 | 0.033 | 1 | 2 | 6720427I0 | 0.029 | Old |
| 0.005329 | 0.41748 | 0.161 | 0.114 | 1 | 2 | Fbxw11 | 0.047 | Old |
| 0.005331 | 0.436877 | 0.109 | 0.07 | 1 | 2 | Tmem38b | 0.039 | Old |
| 0.00537 | 0.767596 | 0.152 | 0.108 | 1 | 2 | Cdc42ep3 | 0.044 | Old |
| 0.005413 | 2.004697 | 0.017 | 0.004 | 1 | 2 | Pnpla1 | 0.013 | Old |
| 0.005486 | 0.371737 | 0.156 | 0.109 | 1 | 2 | Rcn2 | 0.047 | Old |
| 0.005544 | 1.657269 | 0.017 | 0.004 | 1 | 2 | Dpt | 0.013 | Old |
| 0.005569 | 0.370029 | 0.29 | 0.23 | 1 | 2 | Polr2k | 0.06 | Old |
| 0.005581 | 0.29962 | 0.461 | 0.406 | 1 | 2 | Ptpn6 | 0.055 | Old |
| 0.005601 | 0.167743 | 0.9 | 0.881 | 1 | 2 | Pim1 | 0.019 | Old |
| 0.005624 | 0.420474 | 0.175 | 0.127 | 1 | 2 | Pira2 | 0.048 | Old |
| 0.005667 | 0.095409 | 0.987 | 0.974 | 1 | 2 | mt-Nd1 | 0.013 | Old |
| 0.005806 | 1.182967 | 0.021 | 0.006 | 1 | 2 | Ppp1r3e | 0.015 | Old |
| 0.005808 | 0.130025 | 0.878 | 0.876 | 1 | 2 | Calm1 | 0.002 | Old |
| 0.006 | 0.681583 | 0.054 | 0.027 | 1 | 2 | Dhx33 | 0.027 | Old |
| 0.006064 | 0.655528 | 0.081 | 0.049 | 1 | 2 | Agap1 | 0.032 | Old |
| 0.006073 | 1.386129 | 0.04 | 0.018 | 1 | 2 | Nudt1 | 0.022 | Old |
| 0.006103 | 0.781542 | 0.05 | 0.025 | 1 | 2 | Igtp | 0.025 | Old |
| 0.006142 | 0.273939 | 0.403 | 0.33 | 1 | 2 | Nfil3 | 0.073 | Old |
| 0.006154 | 0.779659 | 0.107 | 0.07 | 1 | 2 | Zbtb38 | 0.037 | Old |
| 0.006174 | 0.256124 | 0.636 | 0.584 | 1 | 2 | Cox6a1 | 0.052 | Old |
| 0.006174 | 0.173533 | 0.851 | 0.849 | 1 | 2 | Coro1a | 0.002 | Old |
| 0.006197 | 0.398515 | 0.299 | 0.24 | 1 | 2 | Man2a1 | 0.059 | Old |
| 0.006336 | 0.313283 | 0.348 | 0.284 | 1 | 2 | Tuba1c | 0.064 | Old |
| 0.006386 | 0.449625 | 0.153 | 0.108 | 1 | 2 | Gm9733 | 0.045 | Old |
| 0.006407 | 0.64197 | 0.067 | 0.038 | 1 | 2 | Rngtt | 0.029 | Old |
| 0.006695 | 0.202594 | 0.661 | 0.61 | 1 | 2 | Vmp1 | 0.051 | Old |
| 0.006751 | 0.375903 | 0.222 | 0.17 | 1 | 2 | Bub3 | 0.052 | Old |
| 0.006836 | 0.279594 | 0.333 | 0.266 | 1 | 2 | Actn4 | 0.067 | Old |
| 0.006965 | 0.764257 | 0.048 | 0.024 | 1 | 2 | Zbtb6 | 0.024 | Old |
| 0.006967 | 0.708435 | 0.069 | 0.039 | 1 | 2 | Nudt16 | 0.03 | Old |
| 0.00703 | 0.973898 | 0.133 | 0.094 | 1 | 2 | Rab29 | 0.039 | Old |
| 0.007062 | 0.427918 | 0.285 | 0.229 | 1 | 2 | Cdipt | 0.056 | Old |
| 0.007094 | 0.310878 | 0.33 | 0.268 | 1 | 2 | Naa50 | 0.062 | Old |
| 0.007101 | 0.505872 | 0.132 | 0.091 | 1 | 2 | Cops8 | 0.041 | Old |
| 0.007144 | 0.544346 | 0.172 | 0.127 | 1 | 2 | Srf | 0.045 | Old |
| 0.007166 | 0.496723 | 0.077 | 0.046 | 1 | 2 | Hist1h2bc | 0.031 | Old |
| 0.007221 | 0.253612 | 0.554 | 0.493 | 1 | 2 | Lyn | 0.061 | Old |
| 0.007286 | 0.233876 | 0.756 | 0.723 | 1 | 2 | Prkcd | 0.033 | Old |
| 0.007302 | 0.4241 | 0.221 | 0.169 | 1 | 2 | Ino80d | 0.052 | Old |
| 0.007397 | 0.462272 | 0.136 | 0.095 | 1 | 2 | Gpr146 | 0.041 | Old |
| 0.007424 | 0.46374 | 0.061 | 0.033 | 1 | 2 | Helz2 | 0.028 | Old |
| 0.007462 | 0.473642 | 0.163 | 0.118 | 1 | 2 | Ntan1 | 0.045 | Old |
| 0.007514 | 0.580993 | 0.11 | 0.074 | 1 | 2 | Ifi211 | 0.036 | Old |
| 0.007547 | 0.108888 | 0.91 | 0.872 | 1 | 2 | Btg1 | 0.038 | Old |

| | | | | | | | | |
|---|---|---|---|---|---|---|---|---|
| 0.0077 | 0.303524 | 0.262 | 0.207 | 1 | 2 | Stt3a | 0.055 | Old |
| 0.00773 | 0.377326 | 0.156 | 0.112 | 1 | 2 | Dnajc10 | 0.044 | Old |
| 0.007765 | 0.487968 | 0.244 | 0.191 | 1 | 2 | Ankfy1 | 0.053 | Old |
| 0.007777 | 0.287948 | 0.451 | 0.398 | 1 | 2 | Nipa2 | 0.053 | Old |
| 0.007841 | 0.225008 | 0.566 | 0.512 | 1 | 2 | Pip4k2a | 0.054 | Old |
| 0.007867 | 1.013069 | 0.025 | 0.008 | 1 | 2 | A530032D1 | 0.017 | Old |
| 0.007937 | 1.078548 | 0.032 | 0.013 | 1 | 2 | Svip | 0.019 | Old |
| 0.007991 | 1.959382 | 0.022 | 0.007 | 1 | 2 | Cpeb3 | 0.015 | Old |
| 0.008071 | 0.654578 | 0.121 | 0.083 | 1 | 2 | Abcb7 | 0.038 | Old |
| 0.008228 | 0.530067 | 0.077 | 0.046 | 1 | 2 | Vopp1 | 0.031 | Old |
| 0.008239 | 0.396446 | 0.241 | 0.189 | 1 | 2 | Arhgef10l | 0.052 | Old |
| 0.008293 | 0.367816 | 0.126 | 0.087 | 1 | 2 | Zbp1 | 0.039 | Old |
| 0.008329 | 0.192266 | 0.495 | 0.43 | 1 | 2 | Tmed5 | 0.065 | Old |
| 0.008334 | 0.451499 | 0.211 | 0.162 | 1 | 2 | Adprh | 0.049 | Old |
| 0.008374 | 0.718514 | 0.104 | 0.069 | 1 | 2 | Pilrb2 | 0.035 | Old |
| 0.008376 | 0.43097 | 0.137 | 0.096 | 1 | 2 | Tmem185b | 0.041 | Old |
| 0.008409 | 0.199854 | 0.7 | 0.667 | 1 | 2 | Lamp2 | 0.033 | Old |
| 0.008444 | 0.266553 | 0.079 | 0.048 | 1 | 2 | Tm2d3 | 0.031 | Old |
| 0.008468 | 0.218768 | 0.625 | 0.564 | 1 | 2 | Serbp1 | 0.061 | Old |
| 0.008593 | 0.191174 | 0.585 | 0.544 | 1 | 2 | H2-T23 | 0.041 | Old |
| 0.008672 | 1.564171 | 0.016 | 0.004 | 1 | 2 | Mag | 0.012 | Old |
| 0.008686 | 0.351087 | 0.201 | 0.151 | 1 | 2 | Chfr | 0.05 | Old |
| 0.008749 | 0.427818 | 0.081 | 0.05 | 1 | 2 | Llgl1 | 0.031 | Old |
| 0.008777 | 1.402809 | 0.045 | 0.023 | 1 | 2 | Ccdc163 | 0.022 | Old |
| 0.008876 | 0.282729 | 0.577 | 0.529 | 1 | 2 | Trim25 | 0.048 | Old |
| 0.008937 | 1.064608 | 0.018 | 0.005 | 1 | 2 | Cdc6 | 0.013 | Old |
| 0.008947 | 0.301 | 0.526 | 0.475 | 1 | 2 | Ptpn1 | 0.051 | Old |
| 0.009064 | 0.64953 | 0.101 | 0.067 | 1 | 2 | Sc5d | 0.034 | Old |
| 0.009115 | 0.231253 | 0.604 | 0.546 | 1 | 2 | Cyfip1 | 0.058 | Old |
| 0.009191 | 0.369055 | 0.31 | 0.255 | 1 | 2 | Eif3m | 0.055 | Old |
| 0.009318 | 0.344286 | 0.101 | 0.065 | 1 | 2 | Sdccag8 | 0.036 | Old |
| 0.009391 | 0.327267 | 0.243 | 0.189 | 1 | 2 | Srsf10 | 0.054 | Old |
| 0.009403 | 0.129594 | 0.818 | 0.779 | 1 | 2 | Arf5 | 0.039 | Old |
| 0.009404 | 0.379265 | 0.277 | 0.227 | 1 | 2 | Sdhd | 0.05 | Old |
| 0.009442 | 0.176674 | 0.792 | 0.777 | 1 | 2 | Actr2 | 0.015 | Old |
| 0.009459 | 0.570735 | 0.124 | 0.087 | 1 | 2 | Map3k14 | 0.037 | Old |
| 0.009493 | 0.353653 | 0.459 | 0.406 | 1 | 2 | Cd36 | 0.053 | Old |
| 0.009534 | 0.432084 | 0.158 | 0.115 | 1 | 2 | Aagab | 0.043 | Old |
| 0.00963 | 0.499806 | 0.091 | 0.058 | 1 | 2 | Nhlrc2 | 0.033 | Old |
| 0.009653 | 0.249143 | 0.501 | 0.442 | 1 | 2 | Mtpn | 0.059 | Old |
| 0.009775 | 0.414057 | 0.211 | 0.163 | 1 | 2 | Gak | 0.048 | Old |
| 0.009922 | 0.153424 | 0.664 | 0.614 | 1 | 2 | Atp5f1 | 0.05 | Old |
| 0.010154 | 0.523014 | 0.123 | 0.086 | 1 | 2 | Stx3 | 0.037 | Old |
| 0.0102 | 0.282884 | 0.564 | 0.512 | 1 | 2 | Zfand5 | 0.052 | Old |
| 0.010435 | 0.520474 | 0.269 | 0.221 | 1 | 2 | Slc16a10 | 0.048 | Old |

| | | | | | | | | |
|---|---|---|---|---|---|---|---|---|
| 0.010454 | 0.708145 | 0.089 | 0.057 | 1 | 2 | Bicra | 0.032 | Old |
| 0.010492 | 0.54911 | 0.091 | 0.058 | 1 | 2 | Haus8 | 0.033 | Old |
| 0.010534 | 0.340405 | 0.313 | 0.257 | 1 | 2 | Emc10 | 0.056 | Old |
| 0.010647 | 0.990187 | 0.031 | 0.013 | 1 | 2 | Rbm12b2 | 0.018 | Old |
| 0.01089 | 0.794971 | 0.201 | 0.158 | 1 | 2 | Dusp16 | 0.043 | Old |
| 0.010948 | 0.374722 | 0.355 | 0.304 | 1 | 2 | Fam174a | 0.051 | Old |
| 0.010967 | 0.946089 | 0.084 | 0.053 | 1 | 2 | Pparg | 0.031 | Old |
| 0.011258 | 0.347328 | 0.385 | 0.325 | 1 | 2 | Srsf6 | 0.06 | Old |
| 0.011279 | 0.639441 | 0.093 | 0.061 | 1 | 2 | Atp8b2 | 0.032 | Old |
| 0.011311 | 0.312269 | 0.151 | 0.108 | 1 | 2 | Rsl24d1 | 0.043 | Old |
| 0.011456 | 0.309151 | 0.333 | 0.278 | 1 | 2 | Spred1 | 0.055 | Old |
| 0.011813 | 0.273984 | 0.237 | 0.185 | 1 | 2 | Rnf213 | 0.052 | Old |
| 0.011936 | 0.552829 | 0.226 | 0.183 | 1 | 2 | Cept1 | 0.043 | Old |
| 0.012028 | 0.707619 | 0.107 | 0.074 | 1 | 2 | Gatd1 | 0.033 | Old |
| 0.012077 | 0.264687 | 0.429 | 0.372 | 1 | 2 | Alcam | 0.057 | Old |
| 0.012107 | 0.745941 | 0.075 | 0.046 | 1 | 2 | Cep85l | 0.029 | Old |
| 0.012136 | 0.603552 | 0.078 | 0.049 | 1 | 2 | Mkl2 | 0.029 | Old |
| 0.012188 | 1.012415 | 0.061 | 0.036 | 1 | 2 | Pi16 | 0.025 | Old |
| 0.012275 | 0.558799 | 0.049 | 0.026 | 1 | 2 | Scmh1 | 0.023 | Old |
| 0.012463 | 1.178285 | 0.058 | 0.033 | 1 | 2 | Naip6 | 0.025 | Old |
| 0.01259 | 1.852974 | 0.013 | 0.002 | 1 | 2 | Cenpe | 0.011 | Old |
| 0.012633 | 2.317973 | 0.013 | 0.002 | 1 | 2 | Adrb1 | 0.011 | Old |
| 0.012655 | 0.183137 | 0.342 | 0.279 | 1 | 2 | AB124611 | 0.063 | Old |
| 0.012661 | 0.346979 | 0.103 | 0.069 | 1 | 2 | Rhoh | 0.034 | Old |
| 0.012694 | 0.624275 | 0.092 | 0.061 | 1 | 2 | Gucd1 | 0.031 | Old |
| 0.012698 | 1.617553 | 0.013 | 0.002 | 1 | 2 | Mrm3 | 0.011 | Old |
| 0.012784 | 0.565189 | 0.021 | 0.007 | 1 | 2 | 2410131K1 | 0.014 | Old |
| 0.012906 | 1.378482 | 0.019 | 0.006 | 1 | 2 | Bmpr1a | 0.013 | Old |
| 0.012913 | 0.203502 | 0.446 | 0.382 | 1 | 2 | Surf4 | 0.064 | Old |
| 0.013036 | 0.861507 | 0.041 | 0.02 | 1 | 2 | Vps33b | 0.021 | Old |
| 0.013117 | 0.42259 | 0.268 | 0.219 | 1 | 2 | Prkx | 0.049 | Old |
| 0.013127 | 0.28516 | 0.232 | 0.182 | 1 | 2 | Dnajb11 | 0.05 | Old |
| 0.013131 | 0.269073 | 0.253 | 0.2 | 1 | 2 | Sidt2 | 0.053 | Old |
| 0.013336 | 0.426958 | 0.105 | 0.07 | 1 | 2 | Kmt2b | 0.035 | Old |
| 0.01338 | 0.228208 | 0.398 | 0.335 | 1 | 2 | Arhgap17 | 0.063 | Old |
| 0.013528 | 1.459146 | 0.015 | 0.004 | 1 | 2 | Nudt15 | 0.011 | Old |
| 0.013614 | 0.372578 | 0.181 | 0.137 | 1 | 2 | Lysmd3 | 0.044 | Old |
| 0.013706 | 0.467113 | 0.181 | 0.139 | 1 | 2 | Adam9 | 0.042 | Old |
| 0.013823 | 0.793584 | 0.047 | 0.025 | 1 | 2 | Rfx3 | 0.022 | Old |
| 0.014006 | 0.383523 | 0.213 | 0.166 | 1 | 2 | Piezo1 | 0.047 | Old |
| 0.014112 | 0.679495 | 0.081 | 0.052 | 1 | 2 | Slc35e3 | 0.029 | Old |
| 0.014166 | 0.195899 | 0.193 | 0.145 | 1 | 2 | Rnft1 | 0.048 | Old |
| 0.014264 | 0.310812 | 0.365 | 0.31 | 1 | 2 | Ppp1r15b | 0.055 | Old |
| 0.014314 | 0.817881 | 0.074 | 0.046 | 1 | 2 | Mitd1 | 0.028 | Old |
| 0.014366 | 0.41595 | 0.152 | 0.112 | 1 | 2 | Smg7 | 0.04 | Old |

| | | | | | | | | |
|---|---|---|---|---|---|---|---|---|
| 0.014397 | 0.360302 | 0.332 | 0.277 | 1 | 2 | Galnt1 | 0.055 | Old |
| 0.014404 | 0.158914 | 0.642 | 0.58 | 1 | 2 | Gnb1 | 0.062 | Old |
| 0.014432 | 0.542219 | 0.226 | 0.177 | 1 | 2 | Klc1 | 0.049 | Old |
| 0.014475 | 1.209788 | 0.039 | 0.019 | 1 | 2 | Zfp157 | 0.02 | Old |
| 0.014652 | 0.473765 | 0.154 | 0.115 | 1 | 2 | Sdccag3 | 0.039 | Old |
| 0.014892 | 0.321893 | 0.119 | 0.083 | 1 | 2 | F630028O1 | 0.036 | Old |
| 0.014912 | 0.556875 | 0.097 | 0.065 | 1 | 2 | Exoc6b | 0.032 | Old |
| 0.014928 | 0.40961 | 0.245 | 0.2 | 1 | 2 | Riok3 | 0.045 | Old |
| 0.0151 | 0.571151 | 0.106 | 0.072 | 1 | 2 | Mut | 0.034 | Old |
| 0.015133 | 0.962185 | 0.025 | 0.01 | 1 | 2 | Pde7b | 0.015 | Old |
| 0.015349 | 1.044894 | 0.067 | 0.042 | 1 | 2 | Zfyve9 | 0.025 | Old |
| 0.015356 | 0.921775 | 0.025 | 0.01 | 1 | 2 | Isg20 | 0.015 | Old |
| 0.015385 | 0.604142 | 0.079 | 0.05 | 1 | 2 | Dock5 | 0.029 | Old |
| 0.015429 | 0.175359 | 0.838 | 0.798 | 1 | 2 | Sirpa | 0.04 | Old |
| 0.015542 | 0.692499 | 0.066 | 0.04 | 1 | 2 | AL590144.1 | 0.026 | Old |
| 0.015804 | 0.991664 | 0.03 | 0.013 | 1 | 2 | Adsl | 0.017 | Old |
| 0.015877 | 0.239877 | 0.353 | 0.295 | 1 | 2 | Ppp1r12a | 0.058 | Old |
| 0.015907 | 0.8279 | 0.067 | 0.042 | 1 | 2 | Zfp954 | 0.025 | Old |
| 0.015939 | 0.252762 | 0.361 | 0.306 | 1 | 2 | Spcs1 | 0.055 | Old |
| 0.016013 | 0.774855 | 0.052 | 0.03 | 1 | 2 | Naa16 | 0.022 | Old |
| 0.016019 | 0.838956 | 0.06 | 0.036 | 1 | 2 | Elac1 | 0.024 | Old |
| 0.016148 | 0.220628 | 0.918 | 0.923 | 1 | 2 | Dusp1 | -0.005 | Old |
| 0.016154 | 1.188928 | 0.04 | 0.02 | 1 | 2 | Cyfip2 | 0.02 | Old |
| 0.016166 | 0.289348 | 0.355 | 0.298 | 1 | 2 | Stk38 | 0.057 | Old |
| 0.016257 | 0.787479 | 0.056 | 0.032 | 1 | 2 | Hmces | 0.024 | Old |
| 0.016489 | 0.476144 | 0.089 | 0.058 | 1 | 2 | Ptgr2 | 0.031 | Old |
| 0.016667 | 0.679555 | 0.104 | 0.072 | 1 | 2 | Taf11 | 0.032 | Old |
| 0.016711 | 1.09517 | 0.043 | 0.023 | 1 | 2 | Ccdc69 | 0.02 | Old |
| 0.016747 | 0.607467 | 0.101 | 0.069 | 1 | 2 | Ggta1 | 0.032 | Old |
| 0.016762 | 0.98839 | 0.033 | 0.015 | 1 | 2 | Recql5 | 0.018 | Old |
| 0.017004 | 0.346736 | 0.216 | 0.169 | 1 | 2 | Otulin | 0.047 | Old |
| 0.017053 | 0.262128 | 0.456 | 0.398 | 1 | 2 | Chmp4b | 0.058 | Old |
| 0.017113 | 0.560662 | 0.113 | 0.08 | 1 | 2 | Stat2 | 0.033 | Old |
| 0.017123 | 0.422694 | 0.155 | 0.114 | 1 | 2 | Gpr108 | 0.041 | Old |
| 0.017326 | 0.974048 | 0.057 | 0.033 | 1 | 2 | Tef | 0.024 | Old |
| 0.017327 | 0.664513 | 0.022 | 0.008 | 1 | 2 | Cox18 | 0.014 | Old |
| 0.017473 | 0.374703 | 0.224 | 0.179 | 1 | 2 | Rreb1 | 0.045 | Old |
| 0.017487 | 0.168287 | 0.714 | 0.675 | 1 | 2 | Plin2 | 0.039 | Old |
| 0.017607 | 1.520267 | 0.01 | 0.001 | 1 | 2 | Sparcl1 | 0.009 | Old |
| 0.017724 | 0.435189 | 0.127 | 0.091 | 1 | 2 | Glb1 | 0.036 | Old |
| 0.017726 | 0.603379 | 0.142 | 0.106 | 1 | 2 | Aqr | 0.036 | Old |
| 0.017756 | 1.101228 | 0.054 | 0.031 | 1 | 2 | Pon3 | 0.023 | Old |
| 0.017876 | 0.490501 | 0.254 | 0.205 | 1 | 2 | Nampt | 0.049 | Old |
| 0.017917 | 1.016039 | 0.065 | 0.04 | 1 | 2 | 1110059E2 | 0.025 | Old |
| 0.0181 | 0.341943 | 0.258 | 0.21 | 1 | 2 | Ckap4 | 0.048 | Old |

| | | | | | | | | |
|---|---|---|---|---|---|---|---|---|
| 0.018105 | 0.303358 | 0.415 | 0.362 | 1 | 2 | Mia2 | 0.053 | Old |
| 0.018149 | 1.240089 | 0.02 | 0.007 | 1 | 2 | Spef1 | 0.013 | Old |
| 0.01826 | 0.534007 | 0.18 | 0.141 | 1 | 2 | Fbxw2 | 0.039 | Old |
| 0.018286 | 0.767556 | 0.221 | 0.181 | 1 | 2 | Itgal | 0.04 | Old |
| 0.018326 | 0.148227 | 0.608 | 0.562 | 1 | 2 | Pirb | 0.046 | Old |
| 0.018539 | 0.277109 | 0.346 | 0.3 | 1 | 2 | Sypl | 0.046 | Old |
| 0.018801 | 0.292291 | 0.324 | 0.273 | 1 | 2 | Lypla2 | 0.051 | Old |
| 0.018953 | 0.825381 | 0.061 | 0.037 | 1 | 2 | Nob1 | 0.024 | Old |
| 0.019004 | 1.262327 | 0.026 | 0.011 | 1 | 2 | Dennd2a | 0.015 | Old |
| 0.019103 | 0.701335 | 0.064 | 0.039 | 1 | 2 | Sirpb1a | 0.025 | Old |
| 0.019143 | 0.558057 | 0.149 | 0.112 | 1 | 2 | Yrdc | 0.037 | Old |
| 0.019477 | 0.147236 | 0.696 | 0.635 | 1 | 2 | Rac1 | 0.061 | Old |
| 0.019647 | 0.412253 | 0.109 | 0.076 | 1 | 2 | Klhl2 | 0.033 | Old |
| 0.019911 | 1.374573 | 0.026 | 0.011 | 1 | 2 | Slc12a2 | 0.015 | Old |
| 0.019922 | 0.645877 | 0.097 | 0.067 | 1 | 2 | Pmm2 | 0.03 | Old |
| 0.020007 | 0.25636 | 0.642 | 0.591 | 1 | 2 | Klf4 | 0.051 | Old |
| 0.020016 | 0.20141 | 0.473 | 0.405 | 1 | 2 | Adgre5 | 0.068 | Old |
| 0.020037 | 1.735552 | 0.012 | 0.002 | 1 | 2 | 3110009E1 | 0.01 | Old |
| 0.020081 | 0.578503 | 0.103 | 0.071 | 1 | 2 | Mrpl10 | 0.032 | Old |
| 0.020085 | 0.188002 | 0.404 | 0.349 | 1 | 2 | Nadk | 0.055 | Old |
| 0.020098 | 0.424678 | 0.097 | 0.067 | 1 | 2 | Ddx58 | 0.03 | Old |
| 0.020108 | 0.47604 | 0.065 | 0.04 | 1 | 2 | Mrpl55 | 0.025 | Old |
| 0.020128 | 0.75142 | 0.037 | 0.019 | 1 | 2 | Rfx5 | 0.018 | Old |
| 0.02023 | 0.337694 | 0.278 | 0.229 | 1 | 2 | Esyt1 | 0.049 | Old |
| 0.020236 | 0.693203 | 0.041 | 0.021 | 1 | 2 | Tfip11 | 0.02 | Old |
| 0.020289 | 0.398402 | 0.031 | 0.014 | 1 | 2 | Gm9844 | 0.017 | Old |
| 0.020327 | 0.254511 | 0.31 | 0.259 | 1 | 2 | Anxa7 | 0.051 | Old |
| 0.020344 | 0.766282 | 0.047 | 0.026 | 1 | 2 | Rft1 | 0.021 | Old |
| 0.020375 | 1.187091 | 0.012 | 0.002 | 1 | 2 | Mettl22 | 0.01 | Old |
| 0.020376 | 0.366759 | 0.148 | 0.109 | 1 | 2 | Tmem127 | 0.039 | Old |
| 0.020428 | 0.507491 | 0.163 | 0.124 | 1 | 2 | Sdf2l1 | 0.039 | Old |
| 0.020486 | 0.445154 | 0.018 | 0.006 | 1 | 2 | Zfp688 | 0.012 | Old |
| 0.020614 | 0.816094 | 0.012 | 0.002 | 1 | 2 | D130020L0 | 0.01 | Old |
| 0.020643 | 0.451012 | 0.118 | 0.084 | 1 | 2 | Parp9 | 0.034 | Old |
| 0.020675 | 1.296627 | 0.016 | 0.005 | 1 | 2 | Acyp2 | 0.011 | Old |
| 0.020724 | 1.835476 | 0.014 | 0.004 | 1 | 2 | Nek8 | 0.01 | Old |
| 0.020731 | 0.363259 | 0.257 | 0.21 | 1 | 2 | Washc4 | 0.047 | Old |
| 0.020731 | 1.215207 | 0.029 | 0.013 | 1 | 2 | Ears2 | 0.016 | Old |
| 0.02087 | 0.68359 | 0.268 | 0.224 | 1 | 2 | Krt80 | 0.044 | Old |
| 0.021006 | 1.735193 | 0.014 | 0.004 | 1 | 2 | Mirt1 | 0.01 | Old |
| 0.021045 | 0.678154 | 0.059 | 0.036 | 1 | 2 | Nol6 | 0.023 | Old |
| 0.021344 | 0.124169 | 0.537 | 0.467 | 1 | 2 | Arf1 | 0.07 | Old |
| 0.021379 | 0.360762 | 0.16 | 0.12 | 1 | 2 | Atp8a1 | 0.04 | Old |
| 0.021411 | 0.884185 | 0.063 | 0.039 | 1 | 2 | 2010111I0 | 0.024 | Old |
| 0.021522 | 0.374661 | 0.193 | 0.153 | 1 | 2 | Cdk14 | 0.04 | Old |

| | | | | | | | | |
|---|---|---|---|---|---|---|---|---|
| 0.021538 | 0.363014 | 0.243 | 0.197 | 1 | 2 | Cyth1 | 0.046 | Old |
| 0.021581 | 0.281883 | 0.061 | 0.037 | 1 | 2 | Ctu2 | 0.024 | Old |
| 0.02166 | 0.75064 | 0.09 | 0.062 | 1 | 2 | 1110038F1 | 0.028 | Old |
| 0.021792 | 0.473486 | 0.161 | 0.122 | 1 | 2 | Derl2 | 0.039 | Old |
| 0.021806 | 0.242572 | 0.476 | 0.423 | 1 | 2 | Khdrbs1 | 0.053 | Old |
| 0.021987 | 0.310439 | 0.369 | 0.324 | 1 | 2 | Smchd1 | 0.045 | Old |
| 0.022079 | 0.11543 | 0.915 | 0.912 | 1 | 2 | Iqgap1 | 0.003 | Old |
| 0.022116 | 0.750425 | 0.066 | 0.042 | 1 | 2 | Sell | 0.024 | Old |
| 0.022134 | 0.474882 | 0.13 | 0.095 | 1 | 2 | Pla2g4a | 0.035 | Old |
| 0.022136 | 0.637168 | 0.066 | 0.042 | 1 | 2 | Srcap | 0.024 | Old |
| 0.022199 | 0.205819 | 0.283 | 0.229 | 1 | 2 | Vdac3 | 0.054 | Old |
| 0.022204 | 0.295904 | 0.236 | 0.188 | 1 | 2 | Senp2 | 0.048 | Old |
| 0.022376 | 0.453064 | 0.139 | 0.103 | 1 | 2 | Ltbr | 0.036 | Old |
| 0.022466 | 0.225394 | 0.253 | 0.202 | 1 | 2 | Cs | 0.051 | Old |
| 0.022766 | 0.19915 | 0.08 | 0.052 | 1 | 2 | Oas3 | 0.028 | Old |
| 0.022771 | 0.221554 | 0.224 | 0.176 | 1 | 2 | Map3k11 | 0.048 | Old |
| 0.023013 | 0.211686 | 0.453 | 0.397 | 1 | 2 | Atrx | 0.056 | Old |
| 0.023074 | 0.27531 | 0.255 | 0.205 | 1 | 2 | Smc6 | 0.05 | Old |
| 0.023244 | 0.21469 | 0.293 | 0.242 | 1 | 2 | Psmb3 | 0.051 | Old |
| 0.023266 | 0.244679 | 0.184 | 0.141 | 1 | 2 | Cope | 0.043 | Old |
| 0.023282 | 0.627887 | 0.051 | 0.03 | 1 | 2 | Dcun1d3 | 0.021 | Old |
| 0.023563 | 0.221726 | 0.428 | 0.375 | 1 | 2 | Selenos | 0.053 | Old |
| 0.023597 | 0.408462 | 0.142 | 0.106 | 1 | 2 | Ccz1 | 0.036 | Old |
| 0.023675 | 0.254556 | 0.522 | 0.473 | 1 | 2 | Pdcd6ip | 0.049 | Old |
| 0.023733 | 1.389645 | 0.027 | 0.012 | 1 | 2 | Gbp2b | 0.015 | Old |
| 0.023818 | 0.24022 | 0.279 | 0.233 | 1 | 2 | Thrap3 | 0.046 | Old |
| 0.023821 | 0.759088 | 0.06 | 0.037 | 1 | 2 | Isca1 | 0.023 | Old |
| 0.023908 | 0.426723 | 0.091 | 0.062 | 1 | 2 | Ppp2cb | 0.029 | Old |
| 0.023933 | 0.27478 | 0.115 | 0.081 | 1 | 2 | Stard3 | 0.034 | Old |
| 0.023997 | 1.397127 | 0.021 | 0.008 | 1 | 2 | Ifit1 | 0.013 | Old |
| 0.024058 | 0.309222 | 0.176 | 0.135 | 1 | 2 | Hyou1 | 0.041 | Old |
| 0.024244 | 1.29352 | 0.021 | 0.008 | 1 | 2 | Asb6 | 0.013 | Old |
| 0.024269 | 0.342822 | 0.164 | 0.125 | 1 | 2 | Idi1 | 0.039 | Old |
| 0.024561 | 0.27833 | 0.251 | 0.205 | 1 | 2 | Morc3 | 0.046 | Old |
| 0.024583 | 1.112643 | 0.046 | 0.026 | 1 | 2 | Snhg20 | 0.02 | Old |
| 0.024963 | 0.240839 | 0.543 | 0.487 | 1 | 2 | Cmpk1 | 0.056 | Old |
| 0.024992 | 0.347822 | 0.338 | 0.291 | 1 | 2 | Csk | 0.047 | Old |
| 0.025032 | 0.51949 | 0.08 | 0.053 | 1 | 2 | Slfn1 | 0.027 | Old |
| 0.025072 | 0.441841 | 0.13 | 0.095 | 1 | 2 | Ddhd1 | 0.035 | Old |
| 0.025133 | 0.247499 | 0.327 | 0.273 | 1 | 2 | Tm9sf2 | 0.054 | Old |
| 0.025162 | 0.286211 | 0.112 | 0.08 | 1 | 2 | Taf3 | 0.032 | Old |
| 0.025215 | 0.771766 | 0.088 | 0.061 | 1 | 2 | Crtap | 0.027 | Old |
| 0.025381 | 0.669533 | 0.153 | 0.116 | 1 | 2 | Ly6c2 | 0.037 | Old |
| 0.02543 | 0.474486 | 0.353 | 0.308 | 1 | 2 | Stk10 | 0.045 | Old |
| 0.025511 | 0.26938 | 0.287 | 0.236 | 1 | 2 | Polr2a | 0.051 | Old |

| | | | | | | | | |
|---|---|---|---|---|---|---|---|---|
| 0.02568 | 0.521795 | 0.119 | 0.087 | 1 | 2 | Crtc2 | 0.032 | Old |
| 0.02576 | 0.266738 | 0.045 | 0.025 | 1 | 2 | Tnfsf13os | 0.02 | Old |
| 0.025805 | 0.484962 | 0.107 | 0.076 | 1 | 2 | Sntb2 | 0.031 | Old |
| 0.025844 | 0.150613 | 0.081 | 0.053 | 1 | 2 | Prpf6 | 0.028 | Old |
| 0.025844 | 0.464267 | 0.125 | 0.091 | 1 | 2 | Mthfd2 | 0.034 | Old |
| 0.02597 | 0.48588 | 0.15 | 0.114 | 1 | 2 | Fam20b | 0.036 | Old |
| 0.026003 | 0.448861 | 0.175 | 0.137 | 1 | 2 | Sav1 | 0.038 | Old |
| 0.026027 | 0.612199 | 0.057 | 0.034 | 1 | 2 | Cog8 | 0.023 | Old |
| 0.026123 | 0.867643 | 0.03 | 0.014 | 1 | 2 | Sgcb | 0.016 | Old |
| 0.026188 | 0.379852 | 0.264 | 0.222 | 1 | 2 | Camk2d | 0.042 | Old |
| 0.026274 | 0.332474 | 0.353 | 0.304 | 1 | 2 | Azin1 | 0.049 | Old |
| 0.026302 | 0.366678 | 0.213 | 0.172 | 1 | 2 | Gdi1 | 0.041 | Old |
| 0.026338 | 0.111763 | 0.758 | 0.751 | 1 | 2 | Anxa2 | 0.007 | Old |
| 0.026356 | 0.400964 | 0.162 | 0.125 | 1 | 2 | Srd5a3 | 0.037 | Old |
| 0.026442 | 0.476469 | 0.079 | 0.052 | 1 | 2 | Stard5 | 0.027 | Old |
| 0.026474 | 0.488946 | 0.073 | 0.048 | 1 | 2 | Gpn3 | 0.025 | Old |
| 0.026492 | 0.858302 | 0.058 | 0.036 | 1 | 2 | Ddx60 | 0.022 | Old |
| 0.026575 | 0.179805 | 0.338 | 0.283 | 1 | 2 | Cpsf6 | 0.055 | Old |
| 0.026694 | 0.333553 | 0.063 | 0.039 | 1 | 2 | Fam26f | 0.024 | Old |
| 0.026724 | 0.919532 | 0.033 | 0.017 | 1 | 2 | Aacs | 0.016 | Old |
| 0.026794 | 0.23589 | 0.237 | 0.192 | 1 | 2 | Hopx | 0.045 | Old |
| 0.026804 | 0.116924 | 0.695 | 0.65 | 1 | 2 | Ubl3 | 0.045 | Old |
| 0.02711 | 0.388872 | 0.126 | 0.093 | 1 | 2 | Ifi47 | 0.033 | Old |
| 0.027152 | 0.460621 | 0.085 | 0.057 | 1 | 2 | Zfp524 | 0.028 | Old |
| 0.027321 | 0.276305 | 0.178 | 0.138 | 1 | 2 | Bsdc1 | 0.04 | Old |
| 0.027441 | 0.223848 | 0.076 | 0.05 | 1 | 2 | Irf7 | 0.026 | Old |
| 0.027489 | 0.44634 | 0.173 | 0.137 | 1 | 2 | Gigyf2 | 0.036 | Old |
| 0.02761 | 0.17394 | 0.056 | 0.033 | 1 | 2 | Dym | 0.023 | Old |
| 0.027697 | 0.157 | 0.335 | 0.279 | 1 | 2 | Fam96a | 0.056 | Old |
| 0.027724 | 0.911414 | 0.086 | 0.059 | 1 | 2 | Bcl2 | 0.027 | Old |
| 0.027974 | 0.414078 | 0.057 | 0.034 | 1 | 2 | Lias | 0.023 | Old |
| 0.028319 | 0.590685 | 0.04 | 0.021 | 1 | 2 | Pcca | 0.019 | Old |
| 0.028371 | 0.260444 | 0.485 | 0.452 | 1 | 2 | Tpm4 | 0.033 | Old |
| 0.028538 | 0.339039 | 0.346 | 0.298 | 1 | 2 | Nono | 0.048 | Old |
| 0.028578 | 0.613441 | 0.091 | 0.063 | 1 | 2 | Fbxo4 | 0.028 | Old |
| 0.028722 | 0.39839 | 0.158 | 0.121 | 1 | 2 | Rab43 | 0.037 | Old |
| 0.028801 | 0.154987 | 0.577 | 0.545 | 1 | 2 | Dock2 | 0.032 | Old |
| 0.02888 | 0.605041 | 0.05 | 0.03 | 1 | 2 | Espn | 0.02 | Old |
| 0.029099 | 0.273043 | 0.186 | 0.146 | 1 | 2 | Slc48a1 | 0.04 | Old |
| 0.029154 | 1.467424 | 0.017 | 0.006 | 1 | 2 | Gpnmb | 0.011 | Old |
| 0.029244 | 0.48127 | 0.135 | 0.101 | 1 | 2 | Mif4gd | 0.034 | Old |
| 0.029355 | 0.295073 | 0.233 | 0.19 | 1 | 2 | Sel1l | 0.043 | Old |
| 0.029372 | 0.296597 | 0.324 | 0.277 | 1 | 2 | Mdh2 | 0.047 | Old |
| 0.029417 | 1.361374 | 0.028 | 0.013 | 1 | 2 | Prr7 | 0.015 | Old |
| 0.029418 | 0.317356 | 0.117 | 0.084 | 1 | 2 | Atg7 | 0.033 | Old |

| | | | | | | | | |
|---|---|---|---|---|---|---|---|---|
| 0.029449 | 0.540619 | 0.14 | 0.107 | 1 | 2 | Fry | 0.033 | Old |
| 0.029663 | 0.549476 | 0.07 | 0.045 | 1 | 2 | Tpcn2 | 0.025 | Old |
| 0.02967 | 1.202966 | 0.017 | 0.006 | 1 | 2 | Dhrs9 | 0.011 | Old |
| 0.029873 | 0.266285 | 0.399 | 0.347 | 1 | 2 | Fgr | 0.052 | Old |
| 0.029899 | 0.288469 | 0.133 | 0.097 | 1 | 2 | Commd4 | 0.036 | Old |
| 0.03 | 0.379193 | 0.231 | 0.191 | 1 | 2 | Sp1 | 0.04 | Old |
| 0.030006 | 0.702161 | 0.017 | 0.006 | 1 | 2 | Cenpq | 0.011 | Old |
| 0.030024 | 0.245584 | 0.284 | 0.234 | 1 | 2 | Srgap2 | 0.05 | Old |
| 0.03011 | 0.459796 | 0.131 | 0.097 | 1 | 2 | Ptbp2 | 0.034 | Old |
| 0.030264 | 0.842584 | 0.028 | 0.013 | 1 | 2 | Gpatch3 | 0.015 | Old |
| 0.03053 | 0.757038 | 0.041 | 0.023 | 1 | 2 | Epg5 | 0.018 | Old |
| 0.030616 | 0.359049 | 0.148 | 0.112 | 1 | 2 | Bin2 | 0.036 | Old |
| 0.030754 | 0.513656 | 0.102 | 0.072 | 1 | 2 | Senp7 | 0.03 | Old |
| 0.030811 | 0.329349 | 0.028 | 0.013 | 1 | 2 | Cnst | 0.015 | Old |
| 0.030891 | 0.435995 | 0.202 | 0.164 | 1 | 2 | 6-Mar | 0.038 | Old |
| 0.031431 | 0.387012 | 0.157 | 0.121 | 1 | 2 | Pigk | 0.036 | Old |
| 0.031431 | 0.194839 | 0.337 | 0.284 | 1 | 2 | Psmb10 | 0.053 | Old |
| 0.031433 | 0.2594 | 0.125 | 0.091 | 1 | 2 | Pigm | 0.034 | Old |
| 0.03146 | 1.243817 | 0.015 | 0.005 | 1 | 2 | Znhit3 | 0.01 | Old |
| 0.031587 | 1.370342 | 0.015 | 0.005 | 1 | 2 | Tlr11 | 0.01 | Old |
| 0.031592 | 0.092425 | 0.425 | 0.366 | 1 | 2 | Rhog | 0.059 | Old |
| 0.031617 | 1.023728 | 0.052 | 0.032 | 1 | 2 | Mdm1 | 0.02 | Old |
| 0.031714 | 1.014247 | 0.015 | 0.005 | 1 | 2 | Gpsm2 | 0.01 | Old |
| 0.031884 | 1.004813 | 0.015 | 0.005 | 1 | 2 | P2ry14 | 0.01 | Old |
| 0.031914 | 0.339823 | 0.212 | 0.17 | 1 | 2 | Gfpt1 | 0.042 | Old |
| 0.032 | 0.108741 | 0.877 | 0.878 | 1 | 2 | Pfn1 | -0.001 | Old |
| 0.032064 | 0.148527 | 0.455 | 0.395 | 1 | 2 | Hdgf | 0.06 | Old |
| 0.032154 | 0.204966 | 0.216 | 0.173 | 1 | 2 | Zcchc7 | 0.043 | Old |
| 0.032215 | 2.275699 | 0.013 | 0.004 | 1 | 2 | Ctsk | 0.009 | Old |
| 0.032337 | 0.319492 | 0.106 | 0.076 | 1 | 2 | Zfp384 | 0.03 | Old |
| 0.032374 | 1.928146 | 0.011 | 0.002 | 1 | 2 | Hsd11b1 | 0.009 | Old |
| 0.032638 | 0.756317 | 0.011 | 0.002 | 1 | 2 | Fsd1l | 0.009 | Old |
| 0.032658 | 0.341714 | 0.128 | 0.096 | 1 | 2 | Zc3h18 | 0.032 | Old |
| 0.032789 | 0.37706 | 0.225 | 0.184 | 1 | 2 | D1Ertd622 | 0.041 | Old |
| 0.032928 | 1.280334 | 0.013 | 0.004 | 1 | 2 | Ptges3l | 0.009 | Old |
| 0.033031 | 0.792162 | 0.086 | 0.059 | 1 | 2 | Tmc6 | 0.027 | Old |
| 0.033368 | 0.609188 | 0.095 | 0.068 | 1 | 2 | Kitl | 0.027 | Old |
| 0.033423 | 0.535898 | 0.1 | 0.071 | 1 | 2 | Ccdc25 | 0.029 | Old |
| 0.033481 | 0.578194 | 0.086 | 0.059 | 1 | 2 | Sh3gl1 | 0.027 | Old |
| 0.033637 | 0.95576 | 0.054 | 0.033 | 1 | 2 | Cdc27 | 0.021 | Old |
| 0.033813 | 0.862115 | 0.049 | 0.03 | 1 | 2 | Golga1 | 0.019 | Old |
| 0.034023 | 0.507111 | 0.124 | 0.093 | 1 | 2 | Tbc1d2b | 0.031 | Old |
| 0.034043 | 0.143153 | 0.357 | 0.306 | 1 | 2 | Capn2 | 0.051 | Old |
| 0.034249 | 1.598147 | 0.02 | 0.008 | 1 | 2 | Limk2 | 0.012 | Old |
| 0.034331 | 0.42124 | 0.122 | 0.091 | 1 | 2 | Manba | 0.031 | Old |

| | | | | | | | | |
|---|---|---|---|---|---|---|---|---|
| 0.034399 | 0.365132 | 0.198 | 0.159 | 1 | 2 | Insig1 | 0.039 | Old |
| 0.034498 | 0.72568 | 0.026 | 0.012 | 1 | 2 | Primpol | 0.014 | Old |
| 0.034752 | 0.204496 | 0.572 | 0.527 | 1 | 2 | Clk1 | 0.045 | Old |
| 0.034859 | 1.22326 | 0.02 | 0.008 | 1 | 2 | Tnk2 | 0.012 | Old |
| 0.035141 | 0.222313 | 0.345 | 0.293 | 1 | 2 | Epsti1 | 0.052 | Old |
| 0.035156 | 0.421286 | 0.263 | 0.221 | 1 | 2 | Tmem51 | 0.042 | Old |
| 0.035334 | 0.400833 | 0.184 | 0.146 | 1 | 2 | Scamp3 | 0.038 | Old |
| 0.035505 | 0.255191 | 0.248 | 0.204 | 1 | 2 | Smarcd2 | 0.044 | Old |
| 0.035645 | 0.219851 | 0.383 | 0.334 | 1 | 2 | Atp13a3 | 0.049 | Old |
| 0.035717 | 0.318348 | 0.336 | 0.292 | 1 | 2 | Ikzf1 | 0.044 | Old |
| 0.035726 | 0.385085 | 0.091 | 0.064 | 1 | 2 | Cyth3 | 0.027 | Old |
| 0.03575 | 0.418152 | 0.214 | 0.178 | 1 | 2 | 2310035C2 | 0.036 | Old |
| 0.036147 | 0.382699 | 0.273 | 0.232 | 1 | 2 | Ptpn12 | 0.041 | Old |
| 0.036501 | 0.343878 | 0.263 | 0.217 | 1 | 2 | Lrrfip1 | 0.046 | Old |
| 0.036721 | 0.681603 | 0.032 | 0.017 | 1 | 2 | Iqsec2 | 0.015 | Old |
| 0.037162 | 0.768349 | 0.046 | 0.027 | 1 | 2 | Mapk7 | 0.019 | Old |
| 0.037259 | 0.466012 | 0.06 | 0.038 | 1 | 2 | Oas1a | 0.022 | Old |
| 0.037268 | 0.608631 | 0.107 | 0.078 | 1 | 2 | Naip5 | 0.029 | Old |
| 0.037277 | 0.241649 | 0.422 | 0.38 | 1 | 2 | Soat1 | 0.042 | Old |
| 0.037284 | 0.352962 | 0.346 | 0.303 | 1 | 2 | Lman2 | 0.043 | Old |
| 0.037539 | 0.229842 | 0.188 | 0.148 | 1 | 2 | Rnf6 | 0.04 | Old |
| 0.03759 | 0.388871 | 0.143 | 0.109 | 1 | 2 | Arhgap4 | 0.034 | Old |
| 0.037678 | 0.26198 | 0.216 | 0.176 | 1 | 2 | Med28 | 0.04 | Old |
| 0.037784 | 0.196277 | 0.178 | 0.14 | 1 | 2 | Guk1 | 0.038 | Old |
| 0.037811 | 0.418704 | 0.105 | 0.076 | 1 | 2 | Slc30a1 | 0.029 | Old |
| 0.038016 | 0.233901 | 0.405 | 0.371 | 1 | 2 | Atp5b | 0.034 | Old |
| 0.03815 | 0.219079 | 0.19 | 0.151 | 1 | 2 | Atp2c1 | 0.039 | Old |
| 0.038184 | 0.779704 | 0.106 | 0.078 | 1 | 2 | Secisbp2 | 0.028 | Old |
| 0.038363 | 0.389015 | 0.118 | 0.088 | 1 | 2 | Lmbrd2 | 0.03 | Old |
| 0.038411 | 0.335171 | 0.234 | 0.194 | 1 | 2 | Bcas2 | 0.04 | Old |
| 0.038536 | 0.232125 | 0.177 | 0.138 | 1 | 2 | Cmas | 0.039 | Old |
| 0.03866 | 0.525518 | 0.164 | 0.129 | 1 | 2 | Anxa11 | 0.035 | Old |
| 0.038789 | 0.255019 | 0.294 | 0.246 | 1 | 2 | Csnk2b | 0.048 | Old |
| 0.039184 | 0.573075 | 0.084 | 0.058 | 1 | 2 | Tubgcp3 | 0.026 | Old |
| 0.039249 | 0.149998 | 0.201 | 0.159 | 1 | 2 | Ufm1 | 0.042 | Old |
| 0.039281 | 0.209446 | 0.188 | 0.15 | 1 | 2 | Dennd1a | 0.038 | Old |
| 0.039333 | 0.081023 | 0.127 | 0.095 | 1 | 2 | Nudt16l1 | 0.032 | Old |
| 0.039367 | 0.36084 | 0.179 | 0.143 | 1 | 2 | Ing4 | 0.036 | Old |
| 0.039378 | 0.114162 | 0.809 | 0.806 | 1 | 2 | Arpc5 | 0.003 | Old |
| 0.039394 | 0.163254 | 0.846 | 0.834 | 1 | 2 | Lcp1 | 0.012 | Old |
| 0.039487 | 0.610868 | 0.024 | 0.011 | 1 | 2 | Cnbd2 | 0.013 | Old |
| 0.039521 | 0.146272 | 0.53 | 0.482 | 1 | 2 | Cd84 | 0.048 | Old |
| 0.039533 | 1.168818 | 0.018 | 0.007 | 1 | 2 | Rwdd2b | 0.011 | Old |
| 0.039533 | 1.136334 | 0.018 | 0.007 | 1 | 2 | Fam189b | 0.011 | Old |
| 0.039644 | 0.303668 | 0.287 | 0.243 | 1 | 2 | Spcs3 | 0.044 | Old |

| | | | | | | | | |
|---|---|---|---|---|---|---|---|---|
| 0.03976 | 0.660909 | 0.018 | 0.007 | 1 | 2 | Kdelc1 | 0.011 | Old |
| 0.039783 | 1.260923 | 0.027 | 0.013 | 1 | 2 | Cenpt | 0.014 | Old |
| 0.040037 | 0.192434 | 0.559 | 0.525 | 1 | 2 | Pdia3 | 0.034 | Old |
| 0.040141 | 0.392368 | 0.203 | 0.166 | 1 | 2 | Mgat2 | 0.037 | Old |
| 0.040162 | 0.386292 | 0.146 | 0.113 | 1 | 2 | Sestd1 | 0.033 | Old |
| 0.040267 | 0.111178 | 0.757 | 0.714 | 1 | 2 | Rbms1 | 0.043 | Old |
| 0.040326 | 0.633179 | 0.066 | 0.044 | 1 | 2 | Trp53inp1 | 0.022 | Old |
| 0.040396 | 0.447596 | 0.14 | 0.108 | 1 | 2 | Slc50a1 | 0.032 | Old |
| 0.04057 | 0.412407 | 0.061 | 0.039 | 1 | 2 | Zfp467 | 0.022 | Old |
| 0.040634 | 0.171945 | 0.018 | 0.007 | 1 | 2 | Pomt2 | 0.011 | Old |
| 0.040724 | 0.6988 | 0.09 | 0.064 | 1 | 2 | Mcoln1 | 0.026 | Old |
| 0.040808 | 0.42611 | 0.124 | 0.094 | 1 | 2 | Itfg1 | 0.03 | Old |
| 0.040945 | 0.772439 | 0.067 | 0.045 | 1 | 2 | C2cd3 | 0.022 | Old |
| 0.040986 | 0.515023 | 0.08 | 0.056 | 1 | 2 | Clpx | 0.024 | Old |
| 0.041051 | 0.26269 | 0.318 | 0.276 | 1 | 2 | Rnf19b | 0.042 | Old |
| 0.041148 | 1.045035 | 0.03 | 0.015 | 1 | 2 | L1cam | 0.015 | Old |
| 0.041179 | 0.139913 | 0.809 | 0.78 | 1 | 2 | Zfp36l2 | 0.029 | Old |
| 0.04118 | 0.393478 | 0.071 | 0.048 | 1 | 2 | Tmem209 | 0.023 | Old |
| 0.041307 | 0.890983 | 0.027 | 0.013 | 1 | 2 | Natd1 | 0.014 | Old |
| 0.041366 | 0.714504 | 0.036 | 0.02 | 1 | 2 | Wdr83 | 0.016 | Old |
| 0.041422 | 0.261582 | 0.178 | 0.14 | 1 | 2 | Cyp51 | 0.038 | Old |
| 0.041844 | 0.417292 | 0.091 | 0.064 | 1 | 2 | Hic2 | 0.027 | Old |
| 0.041945 | 0.655155 | 0.047 | 0.029 | 1 | 2 | Racgap1 | 0.018 | Old |
| 0.042044 | 0.207572 | 0.23 | 0.186 | 1 | 2 | Apaf1 | 0.044 | Old |
| 0.042112 | 0.238733 | 0.372 | 0.323 | 1 | 2 | Sgk3 | 0.049 | Old |
| 0.042155 | 0.502279 | 0.122 | 0.091 | 1 | 2 | Vcan | 0.031 | Old |
| 0.042192 | 0.424177 | 0.118 | 0.088 | 1 | 2 | Snx13 | 0.03 | Old |
| 0.042221 | 0.336155 | 0.079 | 0.055 | 1 | 2 | Gcc1 | 0.024 | Old |
| 0.042236 | 0.797471 | 0.036 | 0.02 | 1 | 2 | Ccar2 | 0.016 | Old |
| 0.042257 | 0.885015 | 0.063 | 0.042 | 1 | 2 | Il10 | 0.021 | Old |
| 0.042351 | 1.597159 | 0.016 | 0.006 | 1 | 2 | 4930522L1 | 0.01 | Old |
| 0.042359 | 0.107142 | 0.663 | 0.596 | 1 | 2 | Tkt | 0.067 | Old |
| 0.042367 | 0.286716 | 0.18 | 0.141 | 1 | 2 | Pdap1 | 0.039 | Old |
| 0.042473 | 0.706501 | 0.03 | 0.015 | 1 | 2 | Slc25a15 | 0.015 | Old |
| 0.042959 | 0.497291 | 0.027 | 0.013 | 1 | 2 | Nudt22 | 0.014 | Old |
| 0.042975 | 0.188292 | 0.195 | 0.157 | 1 | 2 | Rtcb | 0.038 | Old |
| 0.043021 | 0.553247 | 0.03 | 0.015 | 1 | 2 | Zfp932 | 0.015 | Old |
| 0.043182 | 0.411606 | 0.089 | 0.063 | 1 | 2 | Nsmce3 | 0.026 | Old |
| 0.043274 | 0.079479 | 0.949 | 0.943 | 1 | 2 | Ahnak | 0.006 | Old |
| 0.0433 | 0.104522 | 0.147 | 0.113 | 1 | 2 | Anks1 | 0.034 | Old |
| 0.043364 | 0.086456 | 0.746 | 0.716 | 1 | 2 | Ifitm2 | 0.03 | Old |
| 0.043414 | 0.454344 | 0.055 | 0.034 | 1 | 2 | Gpank1 | 0.021 | Old |
| 0.043445 | 1.208942 | 0.016 | 0.006 | 1 | 2 | Tada2a | 0.01 | Old |
| 0.043472 | 0.704627 | 0.067 | 0.045 | 1 | 2 | Tspan17 | 0.022 | Old |
| 0.043593 | 0.82807 | 0.056 | 0.036 | 1 | 2 | Trmt13 | 0.02 | Old |

| | | | | | | | | |
|---|---|---|---|---|---|---|---|---|
| 0.043595 | 0.821685 | 0.064 | 0.043 | 1 | 2 | Ngrn | 0.021 | Old |
| 0.043791 | 0.430274 | 0.093 | 0.067 | 1 | 2 | Tmem245 | 0.026 | Old |
| 0.044068 | 0.170486 | 0.578 | 0.532 | 1 | 2 | Ndufa4 | 0.046 | Old |
| 0.044085 | 0.267518 | 0.033 | 0.018 | 1 | 2 | Ccdc180 | 0.015 | Old |
| 0.044167 | 0.66698 | 0.056 | 0.036 | 1 | 2 | Gpatch11 | 0.02 | Old |
| 0.044504 | 0.269134 | 0.246 | 0.204 | 1 | 2 | Wbp2 | 0.042 | Old |
| 0.04454 | 0.726945 | 0.093 | 0.068 | 1 | 2 | Fam20c | 0.025 | Old |
| 0.044585 | 0.353484 | 0.03 | 0.015 | 1 | 2 | Jpx | 0.015 | Old |
| 0.044711 | 0.218025 | 0.182 | 0.144 | 1 | 2 | Rnf115 | 0.038 | Old |
| 0.045192 | 0.154544 | 0.363 | 0.309 | 1 | 2 | Hectd1 | 0.054 | Old |
| 0.045196 | 0.682224 | 0.021 | 0.01 | 1 | 2 | Zfp788 | 0.011 | Old |
| 0.045278 | 0.571532 | 0.067 | 0.045 | 1 | 2 | Terf1 | 0.022 | Old |
| 0.045557 | 0.170326 | 0.522 | 0.461 | 1 | 2 | Nptn | 0.061 | Old |
| 0.045596 | 0.047904 | 1 | 0.996 | 1 | 2 | mt-Co2 | 0.004 | Old |
| 0.045913 | 0.137429 | 0.574 | 0.53 | 1 | 2 | Atp1b3 | 0.044 | Old |
| 0.045987 | 0.705636 | 0.037 | 0.021 | 1 | 2 | Tmcc2 | 0.016 | Old |
| 0.046121 | 0.372742 | 0.231 | 0.191 | 1 | 2 | Gm6377 | 0.04 | Old |
| 0.046242 | 0.20279 | 0.558 | 0.533 | 1 | 2 | Shisa5 | 0.025 | Old |
| 0.046323 | 0.187917 | 0.036 | 0.02 | 1 | 2 | Dxo | 0.016 | Old |
| 0.046369 | 0.262112 | 0.245 | 0.203 | 1 | 2 | Acsl4 | 0.042 | Old |
| 0.046511 | 0.0993 | 0.919 | 0.907 | 1 | 2 | Myl6 | 0.012 | Old |
| 0.046711 | 0.26391 | 0.441 | 0.397 | 1 | 2 | Irf2 | 0.044 | Old |
| 0.046758 | 0.306417 | 0.109 | 0.081 | 1 | 2 | Pknox1 | 0.028 | Old |
| 0.046981 | 0.477266 | 0.142 | 0.11 | 1 | 2 | Ceacam1 | 0.032 | Old |
| 0.047205 | 0.137363 | 0.489 | 0.435 | 1 | 2 | Psme1 | 0.054 | Old |
| 0.047263 | 0.167521 | 0.274 | 0.228 | 1 | 2 | Anxa4 | 0.046 | Old |
| 0.04759 | 0.986227 | 0.031 | 0.017 | 1 | 2 | Ppif | 0.014 | Old |
| 0.047615 | 0.296974 | 0.108 | 0.08 | 1 | 2 | Yeats2 | 0.028 | Old |
| 0.047939 | 0.504951 | 0.037 | 0.021 | 1 | 2 | Leng9 | 0.016 | Old |
| 0.047949 | 0.705474 | 0.025 | 0.012 | 1 | 2 | Pinx1 | 0.013 | Old |
| 0.047982 | 0.311036 | 0.094 | 0.068 | 1 | 2 | Prpsap2 | 0.026 | Old |
| 0.047994 | 0.868464 | 0.025 | 0.012 | 1 | 2 | Gm26881 | 0.013 | Old |
| 0.04807 | 0.251392 | 0.277 | 0.234 | 1 | 2 | Mfsd14b | 0.043 | Old |
| 0.048079 | 0.491771 | 0.06 | 0.039 | 1 | 2 | Fdps | 0.021 | Old |
| 0.048158 | 0.056721 | 0.274 | 0.226 | 1 | 2 | Api5 | 0.048 | Old |
| 0.048186 | 0.545199 | 0.073 | 0.05 | 1 | 2 | Pdrg1 | 0.023 | Old |
| 0.048202 | 0.725749 | 0.014 | 0.005 | 1 | 2 | Mcat | 0.009 | Old |
| 0.048443 | 0.482401 | 0.025 | 0.012 | 1 | 2 | Kat8 | 0.013 | Old |
| 0.048503 | 0.352766 | 0.279 | 0.239 | 1 | 2 | Rhob | 0.04 | Old |
| 0.048552 | 0.749052 | 0.049 | 0.031 | 1 | 2 | D230025D: | 0.018 | Old |
| 0.048945 | 0.348268 | 0.165 | 0.131 | 1 | 2 | Rnps1 | 0.034 | Old |
| 0.049057 | 0.818543 | 0.028 | 0.014 | 1 | 2 | Fastkd2 | 0.014 | Old |
| 0.049415 | 0.461119 | 0.101 | 0.074 | 1 | 2 | Ncbp2 | 0.027 | Old |
| 0.049495 | 0.902968 | 0.019 | 0.008 | 1 | 2 | Aspa | 0.011 | Old |
| 0.049585 | 0.42717 | 0.182 | 0.15 | 1 | 2 | Tob1 | 0.032 | Old |

| | | | | | | | | |
|---|---|---|---|---|---|---|---|---|
| 0.049636 | 0.708763 | 0.031 | 0.017 | 1 | 2 | Usp46 | 0.014 | Old |
| 0.049733 | 0.534458 | 0.099 | 0.072 | 1 | 2 | Pgs1 | 0.027 | Old |
| 0.049778 | 0.245495 | 0.344 | 0.297 | 1 | 2 | Psma2 | 0.047 | Old |
| 0.050024 | 1.144507 | 0.019 | 0.008 | 1 | 2 | L3mbtl2 | 0.011 | Old |
| 0.050049 | 0.574697 | 0.104 | 0.077 | 1 | 2 | Nkiras2 | 0.027 | Old |
| 0.050286 | 0.467807 | 0.031 | 0.017 | 1 | 2 | Rnf123 | 0.014 | Old |
| 0.050341 | 1.468398 | 0.012 | 0.004 | 1 | 2 | Amacr | 0.008 | Old |
| 0.05044 | 0.396972 | 0.182 | 0.147 | 1 | 2 | Rnase6 | 0.035 | Old |
| 0.050573 | 0.41701 | 0.176 | 0.141 | 1 | 2 | Abhd17b | 0.035 | Old |
| 0.050619 | 0.276248 | 0.256 | 0.214 | 1 | 2 | Dram2 | 0.042 | Old |
| 0.050696 | 0.498813 | 0.054 | 0.034 | 1 | 2 | Alg8 | 0.02 | Old |
| 0.050786 | 0.213485 | 0.288 | 0.246 | 1 | 2 | Pnp | 0.042 | Old |
| 0.050797 | 0.132165 | 0.086 | 0.062 | 1 | 2 | Mmp9 | 0.024 | Old |
| 0.050974 | 1.194329 | 0.035 | 0.02 | 1 | 2 | Rab27a | 0.015 | Old |
| 0.051104 | 2.025681 | 0.01 | 0.002 | 1 | 2 | Cd300e | 0.008 | Old |
| 0.051185 | 1.433113 | 0.01 | 0.002 | 1 | 2 | Oasl1 | 0.008 | Old |
| 0.051318 | 0.206657 | 0.534 | 0.487 | 1 | 2 | Arhgap30 | 0.047 | Old |
| 0.051401 | 0.459765 | 0.106 | 0.078 | 1 | 2 | Sipa1l1 | 0.028 | Old |
| 0.051426 | 0.349706 | 0.155 | 0.122 | 1 | 2 | Rab22a | 0.033 | Old |
| 0.051593 | 1.194455 | 0.01 | 0.002 | 1 | 2 | Clec1a | 0.008 | Old |
| 0.051593 | 1.191747 | 0.01 | 0.002 | 1 | 2 | Rarres2 | 0.008 | Old |
| 0.051593 | 1.062418 | 0.01 | 0.002 | 1 | 2 | Chaf1a | 0.008 | Old |
| 0.051634 | 1.305512 | 0.01 | 0.002 | 1 | 2 | Spdl1 | 0.008 | Old |
| 0.051672 | 0.756887 | 0.046 | 0.029 | 1 | 2 | AI987944 | 0.017 | Old |
| 0.051844 | 0.449477 | 0.061 | 0.04 | 1 | 2 | Leo1 | 0.021 | Old |
| 0.051898 | 0.627876 | 0.071 | 0.049 | 1 | 2 | 2310033P0 | 0.022 | Old |
| 0.051921 | 1.173451 | 0.01 | 0.002 | 1 | 2 | Scnn1a | 0.008 | Old |
| 0.05201 | 0.736527 | 0.046 | 0.029 | 1 | 2 | Sbk1 | 0.017 | Old |
| 0.052042 | 0.211297 | 0.431 | 0.385 | 1 | 2 | Lims1 | 0.046 | Old |
| 0.052129 | 0.33329 | 0.218 | 0.181 | 1 | 2 | Khsrp | 0.037 | Old |
| 0.052246 | 0.242625 | 0.142 | 0.11 | 1 | 2 | Csnk1g1 | 0.032 | Old |
| 0.052338 | 0.426535 | 0.192 | 0.156 | 1 | 2 | Smim10l1 | 0.036 | Old |
| 0.052393 | 0.218671 | 0.329 | 0.284 | 1 | 2 | Spcs2 | 0.045 | Old |
| 0.052436 | 0.448514 | 0.249 | 0.215 | 1 | 2 | Dusp5 | 0.034 | Old |
| 0.05262 | 0.141336 | 0.575 | 0.54 | 1 | 2 | Rap1a | 0.035 | Old |
| 0.052667 | 0.259202 | 0.156 | 0.124 | 1 | 2 | Ints6 | 0.032 | Old |
| 0.052715 | 0.262654 | 0.183 | 0.147 | 1 | 2 | 2-Mar | 0.036 | Old |
| 0.05274 | 0.19094 | 0.156 | 0.122 | 1 | 2 | Eif2s1 | 0.034 | Old |
| 0.052754 | 0.612989 | 0.132 | 0.103 | 1 | 2 | Gm28875 | 0.029 | Old |
| 0.052937 | 0.384029 | 0.07 | 0.048 | 1 | 2 | Pex13 | 0.022 | Old |
| 0.053753 | 0.32254 | 0.268 | 0.229 | 1 | 2 | Dbnl | 0.039 | Old |
| 0.053917 | 0.368452 | 0.103 | 0.076 | 1 | 2 | Atmin | 0.027 | Old |
| 0.054188 | 0.597566 | 0.071 | 0.049 | 1 | 2 | Tmed4 | 0.022 | Old |
| 0.054225 | 0.464897 | 0.169 | 0.138 | 1 | 2 | Trim12c | 0.031 | Old |
| 0.05433 | 0.572291 | 0.039 | 0.023 | 1 | 2 | Trpm2 | 0.016 | Old |

| | | | | | | | | |
|---|---|---|---|---|---|---|---|---|
| 0.054548 | 0.478318 | 0.118 | 0.09 | 1 | 2 | Traf3 | 0.028 | Old |
| 0.054618 | 0.21937 | 0.18 | 0.144 | 1 | 2 | Gatad1 | 0.036 | Old |
| 0.054854 | 1.112586 | 0.026 | 0.013 | 1 | 2 | Wdr76 | 0.013 | Old |
| 0.054942 | 0.173828 | 0.762 | 0.742 | 1 | 2 | Ptbp3 | 0.02 | Old |
| 0.055115 | 0.313547 | 0.34 | 0.303 | 1 | 2 | Bst2 | 0.037 | Old |
| 0.055149 | 0.604036 | 0.118 | 0.091 | 1 | 2 | Thbd | 0.027 | Old |
| 0.05538 | 1.047097 | 0.043 | 0.026 | 1 | 2 | Heg1 | 0.017 | Old |
| 0.055391 | 0.830886 | 0.026 | 0.013 | 1 | 2 | Ugt1a7c | 0.013 | Old |
| 0.05561 | 0.336134 | 0.075 | 0.052 | 1 | 2 | Cpox | 0.023 | Old |
| 0.055637 | 0.774018 | 0.026 | 0.013 | 1 | 2 | Rnf144a | 0.013 | Old |
| 0.055868 | 0.893594 | 0.061 | 0.042 | 1 | 2 | Gosr1 | 0.019 | Old |
| 0.055868 | 0.330409 | 0.074 | 0.051 | 1 | 2 | Hdgfl2 | 0.023 | Old |
| 0.056091 | 0.098257 | 0.412 | 0.366 | 1 | 2 | Synj1 | 0.046 | Old |
| 0.056359 | 0.559123 | 0.133 | 0.105 | 1 | 2 | Nipsnap3b | 0.028 | Old |
| 0.056392 | 0.701064 | 0.029 | 0.015 | 1 | 2 | Smpd3 | 0.014 | Old |
| 0.056537 | 0.346366 | 0.049 | 0.031 | 1 | 2 | Gipc1 | 0.018 | Old |
| 0.056767 | 0.660793 | 0.043 | 0.026 | 1 | 2 | Golm1 | 0.017 | Old |
| 0.056925 | 0.300641 | 0.147 | 0.116 | 1 | 2 | Trnau1ap | 0.031 | Old |
| 0.057425 | 0.590574 | 0.122 | 0.095 | 1 | 2 | Tlr13 | 0.027 | Old |
| 0.057459 | 0.294848 | 0.23 | 0.192 | 1 | 2 | Lrmp | 0.038 | Old |
| 0.057771 | 0.287295 | 0.281 | 0.24 | 1 | 2 | Mtch1 | 0.041 | Old |
| 0.058519 | 0.260941 | 0.134 | 0.103 | 1 | 2 | Ndfip1 | 0.031 | Old |
| 0.058726 | 0.380577 | 0.262 | 0.226 | 1 | 2 | Adssl1 | 0.036 | Old |
| 0.058738 | 0.08196 | 0.225 | 0.183 | 1 | 2 | Cd300lb | 0.042 | Old |
| 0.058807 | 0.232509 | 0.287 | 0.247 | 1 | 2 | Nabp1 | 0.04 | Old |
| 0.058931 | 0.867699 | 0.036 | 0.021 | 1 | 2 | Gtf2h4 | 0.015 | Old |
| 0.05905 | 0.430375 | 0.047 | 0.03 | 1 | 2 | Cbfa2t2 | 0.017 | Old |
| 0.059108 | 0.187248 | 0.297 | 0.253 | 1 | 2 | Yy1 | 0.044 | Old |
| 0.059175 | 0.386728 | 0.643 | 0.624 | 1 | 2 | Lpl | 0.019 | Old |
| 0.059362 | 0.699058 | 0.056 | 0.037 | 1 | 2 | Tns4 | 0.019 | Old |
| 0.05946 | 0.535586 | 0.047 | 0.03 | 1 | 2 | Wdr12 | 0.017 | Old |
| 0.059698 | 0.670752 | 0.04 | 0.024 | 1 | 2 | Gm13373 | 0.016 | Old |
| 0.059783 | 0.366714 | 0.119 | 0.09 | 1 | 2 | Rars | 0.029 | Old |
| 0.059854 | 0.175841 | 0.306 | 0.264 | 1 | 2 | 2-Mar | 0.042 | Old |
| 0.059902 | 0.280993 | 0.221 | 0.184 | 1 | 2 | Pikfyve | 0.037 | Old |
| 0.060064 | 0.233267 | 0.084 | 0.059 | 1 | 2 | Psd4 | 0.025 | Old |
| 0.060126 | 0.320029 | 0.09 | 0.065 | 1 | 2 | Ctso | 0.025 | Old |
| 0.060314 | 0.335636 | 0.154 | 0.122 | 1 | 2 | Rpe | 0.032 | Old |
| 0.060338 | 0.276693 | 0.242 | 0.205 | 1 | 2 | Sertad1 | 0.037 | Old |
| 0.060575 | 0.27836 | 0.172 | 0.138 | 1 | 2 | Klhl5 | 0.034 | Old |
| 0.060578 | 0.178956 | 0.279 | 0.236 | 1 | 2 | Ostc | 0.043 | Old |
| 0.060636 | 0.171512 | 0.258 | 0.216 | 1 | 2 | Tpp2 | 0.042 | Old |
| 0.060707 | 0.2191 | 0.248 | 0.209 | 1 | 2 | Tle4 | 0.039 | Old |
| 0.060907 | 0.302355 | 0.148 | 0.116 | 1 | 2 | Slc35c2 | 0.032 | Old |
| 0.061526 | 0.33311 | 0.072 | 0.05 | 1 | 2 | Pigu | 0.022 | Old |

| | | | | | | | | |
|---|---|---|---|---|---|---|---|---|
| 0.06179 | 0.543168 | 0.084 | 0.061 | 1 | 2 | Cfl2 | 0.023 | Old |
| 0.061854 | 0.199452 | 0.232 | 0.192 | 1 | 2 | Oxct1 | 0.04 | Old |
| 0.062199 | 0.187791 | 0.452 | 0.406 | 1 | 2 | Mapk3 | 0.046 | Old |
| 0.062355 | 0.123023 | 0.378 | 0.331 | 1 | 2 | Snrpb2 | 0.047 | Old |
| 0.06238 | 0.397151 | 0.145 | 0.114 | 1 | 2 | Fryl | 0.031 | Old |
| 0.06249 | 1.081979 | 0.02 | 0.01 | 1 | 2 | Lss | 0.01 | Old |
| 0.062557 | 0.388038 | 0.196 | 0.16 | 1 | 2 | Snx27 | 0.036 | Old |
| 0.062632 | 0.319143 | 0.106 | 0.08 | 1 | 2 | Cdyl | 0.026 | Old |
| 0.062906 | 0.913394 | 0.062 | 0.043 | 1 | 2 | Manbal | 0.019 | Old |
| 0.062999 | 0.348126 | 0.123 | 0.095 | 1 | 2 | Sdad1 | 0.028 | Old |
| 0.063074 | 0.711696 | 0.105 | 0.081 | 1 | 2 | Pbx1 | 0.024 | Old |
| 0.063251 | 0.157668 | 0.166 | 0.132 | 1 | 2 | Polr2l | 0.034 | Old |
| 0.063282 | 0.286849 | 0.148 | 0.116 | 1 | 2 | Plagl2 | 0.032 | Old |
| 0.063357 | 0.78249 | 0.02 | 0.01 | 1 | 2 | C1qtnf6 | 0.01 | Old |
| 0.063531 | 0.379049 | 0.11 | 0.083 | 1 | 2 | Srebf1 | 0.027 | Old |
| 0.063799 | 0.816784 | 0.077 | 0.056 | 1 | 2 | Med16 | 0.021 | Old |
| 0.063903 | 0.429435 | 0.051 | 0.033 | 1 | 2 | Gamt | 0.018 | Old |
| 0.063937 | 0.434118 | 0.088 | 0.064 | 1 | 2 | Hcfc2 | 0.024 | Old |
| 0.063939 | 1.024974 | 0.015 | 0.006 | 1 | 2 | Satb1 | 0.009 | Old |
| 0.063981 | 0.244422 | 0.136 | 0.106 | 1 | 2 | Vps50 | 0.03 | Old |
| 0.063987 | 1.094781 | 0.024 | 0.012 | 1 | 2 | Mlkl | 0.012 | Old |
| 0.064139 | 0.311926 | 0.301 | 0.262 | 1 | 2 | Rassf5 | 0.039 | Old |
| 0.064238 | 1.036463 | 0.015 | 0.006 | 1 | 2 | Ttyh2 | 0.009 | Old |
| 0.064263 | 0.207114 | 0.253 | 0.21 | 1 | 2 | Sec31a | 0.043 | Old |
| 0.064291 | 0.431368 | 0.036 | 0.021 | 1 | 2 | Wwox | 0.015 | Old |
| 0.064298 | 0.266639 | 0.135 | 0.105 | 1 | 2 | Eif2ak2 | 0.03 | Old |
| 0.064312 | 1.02372 | 0.015 | 0.006 | 1 | 2 | Dhodh | 0.009 | Old |
| 0.064453 | 0.345831 | 0.293 | 0.257 | 1 | 2 | Sulf2 | 0.036 | Old |
| 0.064537 | 0.921515 | 0.024 | 0.012 | 1 | 2 | Iba57 | 0.012 | Old |
| 0.064746 | 0.328423 | 0.107 | 0.081 | 1 | 2 | Jkamp | 0.026 | Old |
| 0.064931 | 0.203326 | 0.081 | 0.058 | 1 | 2 | Xaf1 | 0.023 | Old |
| 0.064938 | 0.315991 | 0.227 | 0.19 | 1 | 2 | Tifab | 0.037 | Old |
| 0.065066 | 0.812309 | 0.015 | 0.006 | 1 | 2 | Tmem129 | 0.009 | Old |
| 0.065066 | 0.722791 | 0.015 | 0.006 | 1 | 2 | Acbd4 | 0.009 | Old |
| 0.06511 | 0.853251 | 0.027 | 0.014 | 1 | 2 | Trim30c | 0.013 | Old |
| 0.065119 | 0.490605 | 0.093 | 0.069 | 1 | 2 | Tspan5 | 0.024 | Old |
| 0.065385 | 0.997665 | 0.024 | 0.012 | 1 | 2 | Armcx5 | 0.012 | Old |
| 0.065538 | 0.352995 | 0.15 | 0.119 | 1 | 2 | Polr2c | 0.031 | Old |
| 0.065685 | 0.716339 | 0.108 | 0.083 | 1 | 2 | Fam160b1 | 0.025 | Old |
| 0.065837 | 0.078765 | 0.828 | 0.823 | 1 | 2 | Mbnl1 | 0.005 | Old |
| 0.065939 | 0.142041 | 0.758 | 0.749 | 1 | 2 | Fosl2 | 0.009 | Old |
| 0.065975 | 0.816022 | 0.024 | 0.012 | 1 | 2 | Med19 | 0.012 | Old |
| 0.066031 | 0.67318 | 0.052 | 0.034 | 1 | 2 | Gtf2e1 | 0.018 | Old |
| 0.066152 | 0.564357 | 0.024 | 0.012 | 1 | 2 | Agmo | 0.012 | Old |
| 0.066325 | 0.177566 | 0.273 | 0.229 | 1 | 2 | Rer1 | 0.044 | Old |

| | | | | | | | | |
|---|---|---|---|---|---|---|---|---|
| 0.06691 | 0.335443 | 0.192 | 0.159 | 1 | 2 | Wbp1l | 0.033 | Old |
| 0.066935 | 0.200442 | 0.051 | 0.033 | 1 | 2 | Limk1 | 0.018 | Old |
| 0.067103 | 0.318211 | 0.048 | 0.031 | 1 | 2 | Atp2a3 | 0.017 | Old |
| 0.067599 | 0.205942 | 0.449 | 0.409 | 1 | 2 | Cd300a | 0.04 | Old |
| 0.067625 | 0.196792 | 0.172 | 0.138 | 1 | 2 | Chuk | 0.034 | Old |
| 0.06771 | 0.204254 | 0.024 | 0.012 | 1 | 2 | Tbc1d12 | 0.012 | Old |
| 0.067721 | 0.477233 | 0.07 | 0.049 | 1 | 2 | Dhcr24 | 0.021 | Old |
| 0.067762 | 0.11786 | 0.355 | 0.309 | 1 | 2 | Arcn1 | 0.046 | Old |
| 0.067835 | 0.234265 | 0.264 | 0.221 | 1 | 2 | Trim8 | 0.043 | Old |
| 0.067887 | 0.234136 | 0.161 | 0.128 | 1 | 2 | Usp48 | 0.033 | Old |
| 0.068439 | 0.054139 | 0.527 | 0.475 | 1 | 2 | Syk | 0.052 | Old |
| 0.068675 | 0.15524 | 0.389 | 0.342 | 1 | 2 | Cox17 | 0.047 | Old |
| 0.068676 | 0.204134 | 0.296 | 0.249 | 1 | 2 | Arhgap31 | 0.047 | Old |
| 0.068784 | 0.318104 | 0.224 | 0.189 | 1 | 2 | Golga7 | 0.035 | Old |
| 0.06889 | 0.226971 | 0.489 | 0.448 | 1 | 2 | St8sia4 | 0.041 | Old |
| 0.068918 | 0.322215 | 0.246 | 0.21 | 1 | 2 | Stt3b | 0.036 | Old |
| 0.068937 | 0.709177 | 0.054 | 0.036 | 1 | 2 | Hsf2 | 0.018 | Old |
| 0.068959 | 0.06551 | 0.197 | 0.158 | 1 | 2 | Phc2 | 0.039 | Old |
| 0.069134 | 0.114676 | 0.316 | 0.273 | 1 | 2 | Plec | 0.043 | Old |
| 0.069255 | 0.247174 | 0.177 | 0.145 | 1 | 2 | Ss18 | 0.032 | Old |
| 0.069335 | 0.231114 | 0.173 | 0.141 | 1 | 2 | Gripap1 | 0.032 | Old |
| 0.069392 | 0.311451 | 0.074 | 0.052 | 1 | 2 | Ino80c | 0.022 | Old |
| 0.0697 | 0.483689 | 0.075 | 0.053 | 1 | 2 | Farsa | 0.022 | Old |
| 0.069918 | 0.158013 | 0.522 | 0.471 | 1 | 2 | Rab8b | 0.051 | Old |
| 0.070013 | 0.245426 | 0.135 | 0.106 | 1 | 2 | Ppdpf | 0.029 | Old |
| 0.070148 | 0.12952 | 0.33 | 0.287 | 1 | 2 | Shtn1 | 0.043 | Old |
| 0.070163 | 0.083831 | 0.526 | 0.482 | 1 | 2 | Eif5a | 0.044 | Old |
| 0.070206 | 0.153274 | 0.236 | 0.195 | 1 | 2 | Il2rg | 0.041 | Old |
| 0.070335 | 0.646533 | 0.037 | 0.023 | 1 | 2 | Pskh1 | 0.014 | Old |
| 0.070463 | 0.164232 | 0.684 | 0.644 | 1 | 2 | Ywhab | 0.04 | Old |
| 0.070549 | 0.430015 | 0.076 | 0.055 | 1 | 2 | Nlk | 0.021 | Old |
| 0.070565 | 0.275727 | 0.089 | 0.065 | 1 | 2 | Hhex | 0.024 | Old |
| 0.070637 | 0.215399 | 0.095 | 0.07 | 1 | 2 | Gpaa1 | 0.025 | Old |
| 0.070685 | 0.657657 | 0.034 | 0.02 | 1 | 2 | Bex6 | 0.014 | Old |
| 0.070703 | 0.235697 | 0.123 | 0.095 | 1 | 2 | Wars | 0.028 | Old |
| 0.070727 | 0.208512 | 0.354 | 0.309 | 1 | 2 | Gsk3b | 0.045 | Old |
| 0.070779 | 0.144989 | 0.171 | 0.138 | 1 | 2 | Mrps15 | 0.033 | Old |
| 0.070882 | 0.321577 | 0.116 | 0.089 | 1 | 2 | Aggf1 | 0.027 | Old |
| 0.07094 | 0.335665 | 0.09 | 0.067 | 1 | 2 | Steap3 | 0.023 | Old |
| 0.071018 | 1.327551 | 0.013 | 0.005 | 1 | 2 | Zhx2 | 0.008 | Old |
| 0.071018 | 0.164889 | 0.195 | 0.16 | 1 | 2 | Tmbim4 | 0.035 | Old |
| 0.071059 | 0.649598 | 0.094 | 0.071 | 1 | 2 | A430005L1 | 0.023 | Old |
| 0.071163 | 0.607241 | 0.12 | 0.094 | 1 | 2 | Kif2a | 0.026 | Old |
| 0.071174 | 1.032502 | 0.056 | 0.038 | 1 | 2 | Siglec1 | 0.018 | Old |
| 0.071275 | 0.203482 | 0.577 | 0.532 | 1 | 2 | Lst1 | 0.045 | Old |

| | | | | | | | | |
|---|---|---|---|---|---|---|---|---|
| 0.071709 | 1.027833 | 0.031 | 0.018 | 1 | 2 | Cnr2 | 0.013 | Old |
| 0.07173 | 0.900509 | 0.013 | 0.005 | 1 | 2 | Cdk5rap1 | 0.008 | Old |
| 0.071823 | 0.325097 | 0.052 | 0.034 | 1 | 2 | Spidr | 0.018 | Old |
| 0.071908 | 0.855819 | 0.013 | 0.005 | 1 | 2 | Enpp1 | 0.008 | Old |
| 0.071908 | 0.645601 | 0.013 | 0.005 | 1 | 2 | Gm17259 | 0.008 | Old |
| 0.071998 | 0.74227 | 0.013 | 0.005 | 1 | 2 | Zfp408 | 0.008 | Old |
| 0.072182 | 0.15886 | 0.58 | 0.538 | 1 | 2 | Csf2ra | 0.042 | Old |
| 0.07229 | 0.359183 | 0.037 | 0.023 | 1 | 2 | Ap4e1 | 0.014 | Old |
| 0.072346 | 0.496635 | 0.132 | 0.105 | 1 | 2 | Abcd2 | 0.027 | Old |
| 0.072357 | 0.400267 | 0.013 | 0.005 | 1 | 2 | Park2 | 0.008 | Old |
| 0.072427 | 0.726479 | 0.018 | 0.008 | 1 | 2 | Tspyl3 | 0.01 | Old |
| 0.072621 | 0.292643 | 0.138 | 0.108 | 1 | 2 | Zcchc2 | 0.03 | Old |
| 0.072993 | 0.370962 | 0.238 | 0.203 | 1 | 2 | Evl | 0.035 | Old |
| 0.073315 | 0.487081 | 0.018 | 0.008 | 1 | 2 | Tcaim | 0.01 | Old |
| 0.073617 | 0.138967 | 0.612 | 0.567 | 1 | 2 | Vamp8 | 0.045 | Old |
| 0.073627 | 0.204035 | 0.181 | 0.148 | 1 | 2 | Rgs19 | 0.033 | Old |
| 0.073737 | 0.398572 | 0.055 | 0.037 | 1 | 2 | Papd7 | 0.018 | Old |
| 0.073945 | 0.170655 | 0.059 | 0.04 | 1 | 2 | Mrpl40 | 0.019 | Old |
| 0.074463 | 0.158775 | 0.184 | 0.152 | 1 | 2 | Ly9 | 0.032 | Old |
| 0.07476 | 0.139454 | 0.427 | 0.379 | 1 | 2 | Slc39a1 | 0.048 | Old |
| 0.07479 | 0.439536 | 0.166 | 0.135 | 1 | 2 | Cops4 | 0.031 | Old |
| 0.07504 | 0.989224 | 0.025 | 0.013 | 1 | 2 | Slx4 | 0.012 | Old |
| 0.075167 | 1.091885 | 0.021 | 0.011 | 1 | 2 | Acod1 | 0.01 | Old |
| 0.0753 | 0.455356 | 0.076 | 0.055 | 1 | 2 | Mum1 | 0.021 | Old |
| 0.075342 | 0.714647 | 0.059 | 0.04 | 1 | 2 | Nup133 | 0.019 | Old |
| 0.075422 | 0.885809 | 0.028 | 0.015 | 1 | 2 | 1810011H1 | 0.013 | Old |
| 0.075615 | 0.240922 | 0.45 | 0.412 | 1 | 2 | Add3 | 0.038 | Old |
| 0.075662 | 0.446224 | 0.072 | 0.051 | 1 | 2 | Rptor | 0.021 | Old |
| 0.076466 | 0.701493 | 0.039 | 0.024 | 1 | 2 | Gabpb1 | 0.015 | Old |
| 0.076772 | 0.419878 | 0.021 | 0.011 | 1 | 2 | Zfp280b | 0.01 | Old |
| 0.076787 | 0.053825 | 0.106 | 0.081 | 1 | 2 | Gbp2 | 0.025 | Old |
| 0.076848 | 0.322529 | 0.184 | 0.152 | 1 | 2 | Slc25a28 | 0.032 | Old |
| 0.076933 | 0.166997 | 0.256 | 0.219 | 1 | 2 | Mob4 | 0.037 | Old |
| 0.077059 | 0.494849 | 0.046 | 0.03 | 1 | 2 | Sco2 | 0.016 | Old |
| 0.077066 | 0.306234 | 0.153 | 0.124 | 1 | 2 | Itpr1 | 0.029 | Old |
| 0.077123 | 0.269849 | 0.472 | 0.428 | 1 | 2 | Gsn | 0.044 | Old |
| 0.077171 | 0.619165 | 0.059 | 0.04 | 1 | 2 | Chmp6 | 0.019 | Old |
| 0.077183 | 0.41741 | 0.178 | 0.146 | 1 | 2 | Rnf114 | 0.032 | Old |
| 0.077184 | 0.899749 | 0.035 | 0.021 | 1 | 2 | Sft2d3 | 0.014 | Old |
| 0.077363 | 0.337382 | 0.223 | 0.19 | 1 | 2 | Stub1 | 0.033 | Old |
| 0.077776 | 0.886146 | 0.035 | 0.021 | 1 | 2 | Flad1 | 0.014 | Old |
| 0.077974 | 0.259478 | 0.2 | 0.166 | 1 | 2 | Anapc11 | 0.034 | Old |
| 0.078045 | 0.388224 | 0.076 | 0.055 | 1 | 2 | Mief1 | 0.021 | Old |
| 0.078071 | 0.21285 | 0.058 | 0.039 | 1 | 2 | Traf5 | 0.019 | Old |
| 0.078093 | 0.171778 | 0.406 | 0.357 | 1 | 2 | Hpcal1 | 0.049 | Old |

| | | | | | | | | |
|---|---|---|---|---|---|---|---|---|
| 0.07818 | 0.173883 | 0.621 | 0.583 | 1 | 2 | Ap2a2 | 0.038 | Old |
| 0.07838 | 0.834125 | 0.011 | 0.004 | 1 | 2 | Tnfrsf8 | 0.007 | Old |
| 0.078406 | 0.325085 | 0.226 | 0.195 | 1 | 2 | Golga4 | 0.031 | Old |
| 0.078486 | 0.743568 | 0.011 | 0.004 | 1 | 2 | Samd10 | 0.007 | Old |
| 0.078534 | 0.776446 | 0.035 | 0.021 | 1 | 2 | Cutc | 0.014 | Old |
| 0.078699 | 0.777012 | 0.011 | 0.004 | 1 | 2 | Pced1b | 0.007 | Old |
| 0.078913 | 0.955445 | 0.011 | 0.004 | 1 | 2 | Lif | 0.007 | Old |
| 0.078999 | 0.347966 | 0.194 | 0.164 | 1 | 2 | Pcna | 0.03 | Old |
| 0.079001 | 0.448881 | 0.067 | 0.048 | 1 | 2 | Mrps26 | 0.019 | Old |
| 0.079271 | 0.312928 | 0.182 | 0.151 | 1 | 2 | Slc30a5 | 0.031 | Old |
| 0.079641 | 0.239072 | 0.261 | 0.227 | 1 | 2 | Rsu1 | 0.034 | Old |
| 0.079667 | 0.021643 | 0.243 | 0.202 | 1 | 2 | Gm34084 | 0.041 | Old |
| 0.079667 | 0.435917 | 0.147 | 0.119 | 1 | 2 | Asxl2 | 0.028 | Old |
| 0.079731 | 0.186043 | 0.586 | 0.561 | 1 | 2 | Atp6v1b2 | 0.025 | Old |
| 0.079743 | 0.377951 | 0.082 | 0.061 | 1 | 2 | Bloc1s4 | 0.021 | Old |
| 0.079779 | 0.411272 | 0.105 | 0.081 | 1 | 2 | Wdr61 | 0.024 | Old |
| 0.080957 | 0.379357 | 0.12 | 0.094 | 1 | 2 | Pnpla2 | 0.026 | Old |
| 0.080981 | 0.258808 | 0.241 | 0.208 | 1 | 2 | Elk3 | 0.033 | Old |
| 0.081209 | 0.181262 | 0.42 | 0.385 | 1 | 2 | Asph | 0.035 | Old |
| 0.081239 | 0.163487 | 0.378 | 0.331 | 1 | 2 | Ogdh | 0.047 | Old |
| 0.0813 | 0.538576 | 0.086 | 0.064 | 1 | 2 | Smpdl3b | 0.022 | Old |
| 0.081364 | 0.501761 | 0.051 | 0.034 | 1 | 2 | Adnp2 | 0.017 | Old |
| 0.081613 | 1.167817 | 0.016 | 0.007 | 1 | 2 | Il15ra | 0.009 | Old |
| 0.081904 | 0.510648 | 0.056 | 0.038 | 1 | 2 | C130050O1 | 0.018 | Old |
| 0.082145 | 0.129586 | 0.14 | 0.11 | 1 | 2 | Kif1c | 0.03 | Old |
| 0.082314 | 0.384166 | 0.119 | 0.093 | 1 | 2 | Tor3a | 0.026 | Old |
| 0.082404 | 0.531942 | 0.061 | 0.043 | 1 | 2 | Pja1 | 0.018 | Old |
| 0.082438 | 0.283456 | 0.207 | 0.172 | 1 | 2 | Ppp4r3a | 0.035 | Old |
| 0.082622 | 0.386809 | 0.112 | 0.087 | 1 | 2 | Washc3 | 0.025 | Old |
| 0.082709 | 0.39653 | 0.043 | 0.027 | 1 | 2 | Phf11d | 0.016 | Old |
| 0.082747 | 0.769073 | 0.016 | 0.007 | 1 | 2 | Slc25a14 | 0.009 | Old |
| 0.082755 | 0.168797 | 0.247 | 0.213 | 1 | 2 | Cln8 | 0.034 | Old |
| 0.082896 | 0.382443 | 0.074 | 0.053 | 1 | 2 | Il15 | 0.021 | Old |
| 0.08323 | 0.6061 | 0.052 | 0.036 | 1 | 2 | Rrp12 | 0.016 | Old |
| 0.083299 | 0.165877 | 0.535 | 0.492 | 1 | 2 | Tacc1 | 0.043 | Old |
| 0.083394 | 0.181394 | 0.184 | 0.151 | 1 | 2 | Unc119 | 0.033 | Old |
| 0.083489 | 0.138258 | 0.179 | 0.145 | 1 | 2 | Csf3r | 0.034 | Old |
| 0.083555 | 0.557796 | 0.056 | 0.038 | 1 | 2 | Zfp738 | 0.018 | Old |
| 0.083558 | 0.284837 | 0.043 | 0.027 | 1 | 2 | Gnpnat1 | 0.016 | Old |
| 0.083593 | 0.158184 | 0.269 | 0.23 | 1 | 2 | Cct7 | 0.039 | Old |
| 0.083796 | 0.000923 | 0.217 | 0.179 | 1 | 2 | Pcgf5 | 0.038 | Old |
| 0.083804 | 0.718058 | 0.016 | 0.007 | 1 | 2 | 2700062C0 | 0.009 | Old |
| 0.084136 | 0.31647 | 0.294 | 0.26 | 1 | 2 | Mndal | 0.034 | Old |
| 0.084168 | 0.240861 | 0.14 | 0.112 | 1 | 2 | Ppm1h | 0.028 | Old |
| 0.084269 | 0.457817 | 0.157 | 0.129 | 1 | 2 | Pik3r6 | 0.028 | Old |

| | | | | | | | | |
|---|---|---|---|---|---|---|---|---|
| 0.084287 | 0.5345 | 0.044 | 0.029 | 1 | 2 | Eef1akmt2 | 0.015 | Old |
| 0.084832 | 0.182316 | 0.379 | 0.337 | 1 | 2 | Psmd13 | 0.042 | Old |
| 0.084863 | 0.344689 | 0.032 | 0.019 | 1 | 2 | Acap3 | 0.013 | Old |
| 0.084923 | 0.232889 | 0.12 | 0.093 | 1 | 2 | Fam69a | 0.027 | Old |
| 0.084969 | 0.34536 | 0.193 | 0.163 | 1 | 2 | Itgav | 0.03 | Old |
| 0.085018 | 0.220101 | 0.443 | 0.407 | 1 | 2 | Mapre1 | 0.036 | Old |
| 0.085058 | 0.76945 | 0.044 | 0.029 | 1 | 2 | Cavin1 | 0.015 | Old |
| 0.085255 | 0.309824 | 0.224 | 0.195 | 1 | 2 | Hps3 | 0.029 | Old |
| 0.085289 | 0.301556 | 0.175 | 0.144 | 1 | 2 | Csnk1g2 | 0.031 | Old |
| 0.085296 | 0.490602 | 0.036 | 0.023 | 1 | 2 | Dedd2 | 0.013 | Old |
| 0.085671 | 1.463297 | 0.019 | 0.01 | 1 | 2 | B230118H( | 0.009 | Old |
| 0.085753 | 0.979654 | 0.019 | 0.01 | 1 | 2 | Gm42715 | 0.009 | Old |
| 0.08582 | 0.636362 | 0.029 | 0.017 | 1 | 2 | Zmynd19 | 0.012 | Old |
| 0.086078 | 0.33322 | 0.094 | 0.071 | 1 | 2 | Dopey1 | 0.023 | Old |
| 0.086222 | 1.010137 | 0.026 | 0.014 | 1 | 2 | Tmem51os | 0.012 | Old |
| 0.086256 | 0.263602 | 0.149 | 0.12 | 1 | 2 | Fkbp4 | 0.029 | Old |
| 0.086279 | 0.604115 | 0.072 | 0.052 | 1 | 2 | Adgrg6 | 0.02 | Old |
| 0.086347 | 0.373872 | 0.115 | 0.09 | 1 | 2 | Hypk | 0.025 | Old |
| 0.086549 | 0.381557 | 0.105 | 0.081 | 1 | 2 | Plk2 | 0.024 | Old |
| 0.086615 | 0.183213 | 0.466 | 0.417 | 1 | 2 | Tor1aip1 | 0.049 | Old |
| 0.086809 | 0.106112 | 0.852 | 0.836 | 1 | 2 | Mpeg1 | 0.016 | Old |
| 0.08699 | 0.752559 | 0.026 | 0.014 | 1 | 2 | Zfp53 | 0.012 | Old |
| 0.08701 | 0.534561 | 0.125 | 0.1 | 1 | 2 | Vcl | 0.025 | Old |
| 0.087117 | 0.746291 | 0.058 | 0.04 | 1 | 2 | Fam78a | 0.018 | Old |
| 0.087142 | 0.949792 | 0.019 | 0.01 | 1 | 2 | Lysmd1 | 0.009 | Old |
| 0.08716 | 0.263651 | 0.147 | 0.118 | 1 | 2 | Med8 | 0.029 | Old |
| 0.087224 | 0.750117 | 0.019 | 0.01 | 1 | 2 | D3Ertd751( | 0.009 | Old |
| 0.087445 | 0.156675 | 0.128 | 0.101 | 1 | 2 | Dmtf1 | 0.027 | Old |
| 0.087975 | 0.458479 | 0.026 | 0.014 | 1 | 2 | Il18bp | 0.012 | Old |
| 0.088193 | 0.258317 | 0.198 | 0.166 | 1 | 2 | Themis2 | 0.032 | Old |
| 0.088213 | 1.111128 | 0.022 | 0.012 | 1 | 2 | Smyd3 | 0.01 | Old |
| 0.088468 | 0.167496 | 0.36 | 0.318 | 1 | 2 | Tcirg1 | 0.042 | Old |
| 0.088494 | 0.173089 | 0.234 | 0.198 | 1 | 2 | Ppp1r21 | 0.036 | Old |
| 0.088653 | 0.215087 | 0.128 | 0.102 | 1 | 2 | Hp | 0.026 | Old |
| 0.088748 | 0.881362 | 0.022 | 0.012 | 1 | 2 | Maml2 | 0.01 | Old |
| 0.088755 | 0.681668 | 0.026 | 0.014 | 1 | 2 | C130026I2 | 0.012 | Old |
| 0.0888 | 0.791977 | 0.019 | 0.01 | 1 | 2 | Gm26756 | 0.009 | Old |
| 0.089062 | 0.249733 | 0.04 | 0.025 | 1 | 2 | Gmnn | 0.015 | Old |
| 0.08909 | 0.196037 | 0.55 | 0.517 | 1 | 2 | Emilin2 | 0.033 | Old |
| 0.089121 | 0.311325 | 0.141 | 0.114 | 1 | 2 | Hars | 0.027 | Old |
| 0.089254 | 0.446344 | 0.026 | 0.014 | 1 | 2 | Adi1 | 0.012 | Old |
| 0.089362 | 0.155986 | 0.151 | 0.121 | 1 | 2 | Phf5a | 0.03 | Old |
| 0.089443 | 0.145383 | 0.253 | 0.214 | 1 | 2 | Brd7 | 0.039 | Old |
| 0.089573 | 0.424683 | 0.048 | 0.032 | 1 | 2 | Tradd | 0.016 | Old |
| 0.089735 | 0.230445 | 0.246 | 0.209 | 1 | 2 | H1f0 | 0.037 | Old |

| | | | | | | | | |
|---|---|---|---|---|---|---|---|---|
| 0.089771 | 0.11584 | 0.704 | 0.677 | 1 | 2 | Cd47 | 0.027 | Old |
| 0.089781 | 0.216934 | 0.175 | 0.143 | 1 | 2 | Chsy1 | 0.032 | Old |
| 0.089865 | 0.331586 | 0.121 | 0.095 | 1 | 2 | Ldlrap1 | 0.026 | Old |
| 0.090041 | 0.512365 | 0.136 | 0.112 | 1 | 2 | 6430548M | 0.024 | Old |
| 0.090382 | 0.171606 | 0.163 | 0.133 | 1 | 2 | Inppl1 | 0.03 | Old |
| 0.09046 | 0.297369 | 0.078 | 0.057 | 1 | 2 | Ndufv1 | 0.021 | Old |
| 0.090515 | 0.196457 | 0.102 | 0.077 | 1 | 2 | Galnt10 | 0.025 | Old |
| 0.09079 | 0.247328 | 0.209 | 0.173 | 1 | 2 | Zmynd11 | 0.036 | Old |
| 0.090916 | 0.792283 | 0.059 | 0.042 | 1 | 2 | Rhbdd1 | 0.017 | Old |
| 0.091091 | 0.280009 | 0.192 | 0.159 | 1 | 2 | Mdm4 | 0.033 | Old |
| 0.091302 | 0.266346 | 0.283 | 0.245 | 1 | 2 | Ppp3r1 | 0.038 | Old |
| 0.09139 | 0.187277 | 0.271 | 0.233 | 1 | 2 | Siva1 | 0.038 | Old |
| 0.091505 | 0.51549 | 0.081 | 0.061 | 1 | 2 | Cdc14a | 0.02 | Old |
| 0.091575 | 0.372737 | 0.066 | 0.048 | 1 | 2 | Dcaf10 | 0.018 | Old |
| 0.091583 | 0.116991 | 0.244 | 0.205 | 1 | 2 | Sec11a | 0.039 | Old |
| 0.091874 | 0.532292 | 0.088 | 0.067 | 1 | 2 | 2310001H1 | 0.021 | Old |
| 0.092064 | 0.347911 | 0.257 | 0.223 | 1 | 2 | Cop1 | 0.034 | Old |
| 0.092205 | 0.419108 | 0.136 | 0.11 | 1 | 2 | Nae1 | 0.026 | Old |
| 0.092382 | 0.172695 | 0.272 | 0.234 | 1 | 2 | Arrdc1 | 0.038 | Old |
| 0.092453 | 0.389401 | 0.088 | 0.067 | 1 | 2 | Dhx8 | 0.021 | Old |
| 0.092571 | 0.091366 | 0.204 | 0.17 | 1 | 2 | Trappc2l | 0.034 | Old |
| 0.092816 | 0.659161 | 0.033 | 0.02 | 1 | 2 | Fam149b | 0.013 | Old |
| 0.09288 | 0.770487 | 0.033 | 0.02 | 1 | 2 | Acp5 | 0.013 | Old |
| 0.092885 | 1.064557 | 0.014 | 0.006 | 1 | 2 | Mkks | 0.008 | Old |
| 0.092985 | 0.139563 | 0.319 | 0.278 | 1 | 2 | Aph1a | 0.041 | Old |
| 0.093178 | 0.435406 | 0.077 | 0.057 | 1 | 2 | Minpp1 | 0.02 | Old |
| 0.093301 | 1.052033 | 0.014 | 0.006 | 1 | 2 | Krt10 | 0.008 | Old |
| 0.093381 | 0.463775 | 0.15 | 0.125 | 1 | 2 | Rasa1 | 0.025 | Old |
| 0.093631 | 0.219695 | 0.105 | 0.081 | 1 | 2 | Cep83 | 0.024 | Old |
| 0.093667 | 0.442103 | 0.045 | 0.03 | 1 | 2 | Manea | 0.015 | Old |
| 0.093814 | 0.957167 | 0.03 | 0.018 | 1 | 2 | Cbr3 | 0.012 | Old |
| 0.093824 | 0.526785 | 0.014 | 0.006 | 1 | 2 | Mcf2l | 0.008 | Old |
| 0.093892 | 0.159934 | 0.741 | 0.733 | 1 | 2 | S100a4 | 0.008 | Old |
| 0.093903 | 0.581128 | 0.072 | 0.052 | 1 | 2 | Gm5547 | 0.02 | Old |
| 0.093996 | 0.226302 | 0.074 | 0.053 | 1 | 2 | Alg10b | 0.021 | Old |
| 0.094175 | 0.574211 | 0.041 | 0.026 | 1 | 2 | Gmds | 0.015 | Old |
| 0.094221 | 0.31119 | 0.057 | 0.039 | 1 | 2 | Palm | 0.018 | Old |
| 0.094257 | 0.402228 | 0.123 | 0.1 | 1 | 2 | Atxn10 | 0.023 | Old |
| 0.094266 | 0.743807 | 0.046 | 0.031 | 1 | 2 | Pnpt1 | 0.015 | Old |
| 0.09436 | 0.198259 | 0.228 | 0.194 | 1 | 2 | Dr1 | 0.034 | Old |
| 0.094362 | 0.246238 | 0.301 | 0.266 | 1 | 2 | Cnih4 | 0.035 | Old |
| 0.094439 | 0.21822 | 0.048 | 0.032 | 1 | 2 | B230307C2 | 0.016 | Old |
| 0.094559 | 0.746887 | 0.014 | 0.006 | 1 | 2 | Tha1 | 0.008 | Old |
| 0.094749 | 0.49375 | 0.101 | 0.078 | 1 | 2 | G6pdx | 0.023 | Old |
| 0.095052 | 0.395632 | 0.073 | 0.053 | 1 | 2 | Zdhhc13 | 0.02 | Old |

| | | | | | | | | |
|---|---|---|---|---|---|---|---|---|
| 0.095319 | 0.576715 | 0.065 | 0.048 | 1 | 2 | Ptger2 | 0.017 | Old |
| 0.095545 | 0.362754 | 0.096 | 0.074 | 1 | 2 | Slc12a7 | 0.022 | Old |
| 0.095638 | 0.423237 | 0.033 | 0.02 | 1 | 2 | Lrch4 | 0.013 | Old |
| 0.095666 | 0.306665 | 0.208 | 0.175 | 1 | 2 | Npm3 | 0.033 | Old |
| 0.095712 | 0.126946 | 0.049 | 0.033 | 1 | 2 | Parn | 0.016 | Old |
| 0.09628 | 0.191328 | 0.433 | 0.392 | 1 | 2 | Atp6ap2 | 0.041 | Old |
| 0.096316 | 0.777486 | 0.092 | 0.071 | 1 | 2 | Nxpe4 | 0.021 | Old |
| 0.096351 | 0.323737 | 0.134 | 0.108 | 1 | 2 | Rpl7l1 | 0.026 | Old |
| 0.096397 | 0.15599 | 0.121 | 0.095 | 1 | 2 | Dlgap4 | 0.026 | Old |
| 0.096481 | 0.143315 | 0.097 | 0.074 | 1 | 2 | Bop1 | 0.023 | Old |
| 0.096506 | 0.137163 | 0.103 | 0.08 | 1 | 2 | Zmym2 | 0.023 | Old |
| 0.096522 | 0.586886 | 0.063 | 0.045 | 1 | 2 | Pml | 0.018 | Old |
| 0.096547 | 0.287941 | 0.353 | 0.315 | 1 | 2 | Dse | 0.038 | Old |
| 0.096871 | 0.219374 | 0.204 | 0.172 | 1 | 2 | Ube2j2 | 0.032 | Old |
| 0.097102 | 0.173698 | 0.591 | 0.563 | 1 | 2 | H2afy | 0.028 | Old |
| 0.097241 | 0.300182 | 0.304 | 0.271 | 1 | 2 | D17Wsu92 | 0.033 | Old |
| 0.09747 | 0.437686 | 0.034 | 0.021 | 1 | 2 | Pank4 | 0.013 | Old |
| 0.09759 | 0.096933 | 0.338 | 0.293 | 1 | 2 | Cdc42se2 | 0.045 | Old |
| 0.097621 | 0.470163 | 0.203 | 0.177 | 1 | 2 | Csgalnact2 | 0.026 | Old |
| 0.097661 | 0.438439 | 0.079 | 0.059 | 1 | 2 | Sfi1 | 0.02 | Old |
| 0.097679 | 0.228452 | 0.126 | 0.101 | 1 | 2 | Suco | 0.025 | Old |
| 0.097702 | 0.169411 | 0.171 | 0.141 | 1 | 2 | Hmgcr | 0.03 | Old |
| 0.097707 | 0.235613 | 0.368 | 0.33 | 1 | 2 | Vwa5a | 0.038 | Old |
| 0.098 | 0.212842 | 0.335 | 0.298 | 1 | 2 | Ranbp2 | 0.037 | Old |
| 0.09804 | 0.115688 | 0.096 | 0.074 | 1 | 2 | Gga3 | 0.022 | Old |
| 0.09887 | 0.250526 | 0.13 | 0.103 | 1 | 2 | Ebna1bp2 | 0.027 | Old |
| 0.098911 | 0.205399 | 0.17 | 0.141 | 1 | 2 | Ubr4 | 0.029 | Old |
| 0.099005 | 0.8073 | 0.047 | 0.032 | 1 | 2 | Mrpl22 | 0.015 | Old |
| 0.099176 | 0.524149 | 0.142 | 0.118 | 1 | 2 | Gsap | 0.024 | Old |
| 0.099339 | 0.386443 | 0.063 | 0.045 | 1 | 2 | Lmbr1l | 0.018 | Old |
| 0.09939 | 1.103874 | 0.024 | 0.013 | 1 | 2 | F7 | 0.011 | Old |
| 0.099732 | 0.570924 | 0.126 | 0.103 | 1 | 2 | Bin3 | 0.023 | Old |
| 0.099846 | 0.133197 | 0.301 | 0.265 | 1 | 2 | Sin3b | 0.036 | Old |
| 0.099856 | 0.33221 | 0.055 | 0.038 | 1 | 2 | Taf1b | 0.017 | Old |
| 0.100326 | 0.175528 | 0.325 | 0.289 | 1 | 2 | Nsd3 | 0.036 | Old |
| 0.100333 | 0.109408 | 0.105 | 0.081 | 1 | 2 | Pgrmc2 | 0.024 | Old |
| 0.100356 | 0.587 | 0.027 | 0.015 | 1 | 2 | BC051226 | 0.012 | Old |
| 0.100439 | 0.166072 | 0.321 | 0.281 | 1 | 2 | Tomm22 | 0.04 | Old |
| 0.100531 | 0.113109 | 0.196 | 0.164 | 1 | 2 | Mogs | 0.032 | Old |
| 0.100671 | 0.175645 | 0.283 | 0.247 | 1 | 2 | Acly | 0.036 | Old |
| 0.100751 | 0.21274 | 0.142 | 0.115 | 1 | 2 | Pak1ip1 | 0.027 | Old |
| 0.100872 | 0.183821 | 0.245 | 0.208 | 1 | 2 | Arhgap26 | 0.037 | Old |
| 0.100971 | 0.299185 | 0.232 | 0.203 | 1 | 2 | Trappc4 | 0.029 | Old |
| 0.101161 | 0.682892 | 0.027 | 0.015 | 1 | 2 | Slc37a3 | 0.012 | Old |
| 0.101244 | 1.214898 | 0.02 | 0.011 | 1 | 2 | Tfap4 | 0.009 | Old |

| | | | | | | | | |
|---|---|---|---|---|---|---|---|---|
| 0.101254 | 0.163481 | 0.404 | 0.362 | 1 | 2 | Eif4a2 | 0.042 | Old |
| 0.101303 | 0.343773 | 0.037 | 0.024 | 1 | 2 | Nod2 | 0.013 | Old |
| 0.101349 | 0.415733 | 0.147 | 0.121 | 1 | 2 | Cdk17 | 0.026 | Old |
| 0.101357 | 1.049551 | 0.024 | 0.013 | 1 | 2 | Gatb | 0.011 | Old |
| 0.101365 | 0.144038 | 0.319 | 0.28 | 1 | 2 | Abhd12 | 0.039 | Old |
| 0.101517 | 0.112968 | 0.057 | 0.039 | 1 | 2 | Pdp2 | 0.018 | Old |
| 0.101547 | 0.58097 | 0.027 | 0.015 | 1 | 2 | Rab13 | 0.012 | Old |
| 0.101716 | 0.325537 | 0.077 | 0.057 | 1 | 2 | Tnip1 | 0.02 | Old |
| 0.101761 | 0.418513 | 0.069 | 0.05 | 1 | 2 | Armc10 | 0.019 | Old |
| 0.101904 | 0.395464 | 0.042 | 0.027 | 1 | 2 | Zfp142 | 0.015 | Old |
| 0.101987 | 0.277533 | 0.145 | 0.116 | 1 | 2 | Abcc5 | 0.029 | Old |
| 0.102178 | 0.530655 | 0.042 | 0.027 | 1 | 2 | Trp53bp1 | 0.015 | Old |
| 0.102185 | 0.725408 | 0.024 | 0.013 | 1 | 2 | Zfp35 | 0.011 | Old |
| 0.102332 | 0.701715 | 0.105 | 0.084 | 1 | 2 | Usp37 | 0.021 | Old |
| 0.102526 | 0.789162 | 0.017 | 0.008 | 1 | 2 | Rasal2 | 0.009 | Old |
| 0.102641 | 0.229001 | 0.096 | 0.074 | 1 | 2 | Brf1 | 0.022 | Old |
| 0.102725 | 0.537751 | 0.017 | 0.008 | 1 | 2 | Il27 | 0.009 | Old |
| 0.102757 | 0.281994 | 0.069 | 0.05 | 1 | 2 | Pi4k2b | 0.019 | Old |
| 0.102798 | 0.237874 | 0.227 | 0.194 | 1 | 2 | St3gal4 | 0.033 | Old |
| 0.102825 | 0.578418 | 0.017 | 0.008 | 1 | 2 | Magi3 | 0.009 | Old |
| 0.102845 | 0.246299 | 0.109 | 0.086 | 1 | 2 | Slu7 | 0.023 | Old |
| 0.103054 | 0.095289 | 0.317 | 0.277 | 1 | 2 | 2610507B1 | 0.04 | Old |
| 0.103099 | 0.255557 | 0.212 | 0.179 | 1 | 2 | Dctn4 | 0.033 | Old |
| 0.103122 | 0.398146 | 0.073 | 0.053 | 1 | 2 | Phkg2 | 0.02 | Old |
| 0.103178 | 0.348195 | 0.027 | 0.015 | 1 | 2 | Smo | 0.012 | Old |
| 0.103179 | 0.594619 | 0.047 | 0.032 | 1 | 2 | Dnaja3 | 0.015 | Old |
| 0.103185 | 0.108556 | 0.273 | 0.23 | 1 | 2 | Pdcd10 | 0.043 | Old |
| 0.103469 | 0.55611 | 0.042 | 0.027 | 1 | 2 | Ly6a | 0.015 | Old |
| 0.10354 | 0.217578 | 0.281 | 0.243 | 1 | 2 | Mbp | 0.038 | Old |
| 0.103554 | 0.107479 | 0.458 | 0.41 | 1 | 2 | Gusb | 0.048 | Old |
| 0.103699 | 0.475836 | 0.097 | 0.076 | 1 | 2 | Tmem229b | 0.021 | Old |
| 0.103708 | 0.442611 | 0.24 | 0.214 | 1 | 2 | Fam49a | 0.026 | Old |
| 0.103784 | 0.379147 | 0.05 | 0.034 | 1 | 2 | Bloc1s6 | 0.016 | Old |
| 0.103804 | 0.172584 | 0.157 | 0.129 | 1 | 2 | C1d | 0.028 | Old |
| 0.104065 | 0.523295 | 0.091 | 0.07 | 1 | 2 | Parp12 | 0.021 | Old |
| 0.104137 | 0.156062 | 0.429 | 0.387 | 1 | 2 | Ifi207 | 0.042 | Old |
| 0.104418 | 0.56606 | 0.051 | 0.036 | 1 | 2 | Glce | 0.015 | Old |
| 0.104487 | 0.242676 | 0.298 | 0.261 | 1 | 2 | Leprot | 0.037 | Old |
| 0.104666 | 0.547728 | 0.034 | 0.021 | 1 | 2 | Cryzl2 | 0.013 | Old |
| 0.104785 | 0.141539 | 0.848 | 0.825 | 1 | 2 | Nr4a1 | 0.023 | Old |
| 0.105157 | 0.504822 | 0.06 | 0.043 | 1 | 2 | Zfp511 | 0.017 | Old |
| 0.105266 | 0.138709 | 0.531 | 0.494 | 1 | 2 | 5031439G0 | 0.037 | Old |
| 0.105303 | 1.355796 | 0.012 | 0.005 | 1 | 2 | Arl4d | 0.007 | Old |
| 0.105337 | 0.175458 | 0.43 | 0.393 | 1 | 2 | Edem1 | 0.037 | Old |
| 0.105428 | 1.639883 | 0.012 | 0.005 | 1 | 2 | S1pr5 | 0.007 | Old |

| | | | | | | | | |
|---|---|---|---|---|---|---|---|---|
| 0.105466 | 0.334016 | 0.192 | 0.162 | 1 | 2 | Elf2 | 0.03 | Old |
| 0.105492 | 0.417058 | 0.142 | 0.118 | 1 | 2 | Gas2l3 | 0.024 | Old |
| 0.105569 | 0.251413 | 0.233 | 0.2 | 1 | 2 | Bak1 | 0.033 | Old |
| 0.105832 | 1.274282 | 0.028 | 0.017 | 1 | 2 | Dnaaf3 | 0.011 | Old |
| 0.105855 | 0.261043 | 0.193 | 0.162 | 1 | 2 | U2af1 | 0.031 | Old |
| 0.106117 | 0.179082 | 0.282 | 0.249 | 1 | 2 | Swi5 | 0.033 | Old |
| 0.106185 | 0.937791 | 0.012 | 0.005 | 1 | 2 | Bfsp1 | 0.007 | Old |
| 0.106187 | 0.466207 | 0.047 | 0.032 | 1 | 2 | Zc3h7b | 0.015 | Old |
| 0.106247 | 0.134986 | 0.162 | 0.134 | 1 | 2 | Hmgcs1 | 0.028 | Old |
| 0.106248 | 0.448359 | 0.118 | 0.095 | 1 | 2 | Slc25a1 | 0.023 | Old |
| 0.106279 | 0.313499 | 0.07 | 0.051 | 1 | 2 | Fam3a | 0.019 | Old |
| 0.106438 | 0.839887 | 0.012 | 0.005 | 1 | 2 | Sgsm3 | 0.007 | Old |
| 0.10644 | 0.091606 | 0.232 | 0.197 | 1 | 2 | Sys1 | 0.035 | Old |
| 0.1066 | 0.230999 | 0.074 | 0.055 | 1 | 2 | Slc35a2 | 0.019 | Old |
| 0.106692 | 0.756956 | 0.012 | 0.005 | 1 | 2 | Tnfrsf18 | 0.007 | Old |
| 0.106716 | 0.468417 | 0.061 | 0.044 | 1 | 2 | Prpf4 | 0.017 | Old |
| 0.106755 | 0.287366 | 0.077 | 0.057 | 1 | 2 | Heatr6 | 0.02 | Old |
| 0.10686 | 0.871733 | 0.044 | 0.03 | 1 | 2 | Rnf144b | 0.014 | Old |
| 0.106903 | 0.024023 | 0.011 | 0.02 | 1 | 2 | Dnaaf5 | -0.009 | Old |
| 0.107246 | 0.196732 | 0.467 | 0.439 | 1 | 2 | Itgb2 | 0.028 | Old |
| 0.107367 | 0.424889 | 0.13 | 0.106 | 1 | 2 | Cdip1 | 0.024 | Old |
| 0.107839 | 0.722414 | 0.012 | 0.005 | 1 | 2 | Xlr4a | 0.007 | Old |
| 0.107904 | 0.352385 | 0.094 | 0.072 | 1 | 2 | Slc41a3 | 0.022 | Old |
| 0.107945 | 0.335209 | 0.115 | 0.09 | 1 | 2 | Lilra6 | 0.025 | Old |
| 0.10802 | 0.37763 | 0.091 | 0.07 | 1 | 2 | Mrpl41 | 0.021 | Old |
| 0.108045 | 0.241891 | 0.304 | 0.262 | 1 | 2 | Usp7 | 0.042 | Old |
| 0.108058 | 0.02833 | 0.362 | 0.329 | 1 | 2 | Dock8 | 0.033 | Old |
| 0.108059 | 0.577123 | 0.043 | 0.029 | 1 | 2 | Nol10 | 0.014 | Old |
| 0.108261 | 0.175101 | 0.12 | 0.096 | 1 | 2 | C5ar2 | 0.024 | Old |
| 0.108281 | 0.415813 | 0.056 | 0.039 | 1 | 2 | Mfn1 | 0.017 | Old |
| 0.108353 | 0.171805 | 0.012 | 0.005 | 1 | 2 | Klhdc1 | 0.007 | Old |
| 0.108591 | 0.108915 | 0.219 | 0.186 | 1 | 2 | Rbm27 | 0.033 | Old |
| 0.108616 | 0.210667 | 0.138 | 0.112 | 1 | 2 | Mrpl17 | 0.026 | Old |
| 0.108905 | 0.172089 | 0.078 | 0.058 | 1 | 2 | Sec23a | 0.02 | Old |
| 0.109009 | 0.357918 | 0.121 | 0.097 | 1 | 2 | Zmpste24 | 0.024 | Old |
| 0.109029 | 0.360069 | 0.097 | 0.076 | 1 | 2 | Ankra2 | 0.021 | Old |
| 0.109039 | 0.388264 | 0.122 | 0.097 | 1 | 2 | Creld2 | 0.025 | Old |
| 0.10916 | 0.248766 | 0.239 | 0.209 | 1 | 2 | Tspan14 | 0.03 | Old |
| 0.109471 | 0.073955 | 0.192 | 0.159 | 1 | 2 | Cd180 | 0.033 | Old |
| 0.109574 | 0.002232 | 0.111 | 0.087 | 1 | 2 | Ufl1 | 0.024 | Old |
| 0.109599 | 0.177424 | 0.151 | 0.124 | 1 | 2 | Kcnk6 | 0.027 | Old |
| 0.109603 | 0.473004 | 0.075 | 0.056 | 1 | 2 | Slc9a8 | 0.019 | Old |
| 0.109605 | 0.189681 | 0.081 | 0.061 | 1 | 2 | Sp2 | 0.02 | Old |
| 0.109606 | 0.976343 | 0.04 | 0.026 | 1 | 2 | Zfp595 | 0.014 | Old |
| 0.109779 | 0.935179 | 0.028 | 0.017 | 1 | 2 | Jag1 | 0.011 | Old |

| | | | | | | | | |
|---|---|---|---|---|---|---|---|---|
| 0.109927 | 0.075541 | 0.072 | 0.052 | 1 | 2 | Cdc42bpb | 0.02 | Old |
| 0.110082 | 0.309486 | 0.075 | 0.056 | 1 | 2 | Kdm4c | 0.019 | Old |
| 0.110178 | 0.362205 | 0.178 | 0.148 | 1 | 2 | Gpr141 | 0.03 | Old |
| 0.110187 | 0.43732 | 0.031 | 0.019 | 1 | 2 | Rhbdd3 | 0.012 | Old |
| 0.11022 | 0.954292 | 0.028 | 0.017 | 1 | 2 | Abcb10 | 0.011 | Old |
| 0.110511 | 0.200722 | 0.438 | 0.398 | 1 | 2 | Far1 | 0.04 | Old |
| 0.110599 | 0.231141 | 0.338 | 0.303 | 1 | 2 | Idh1 | 0.035 | Old |
| 0.110962 | 0.390188 | 0.162 | 0.135 | 1 | 2 | H2-T22 | 0.027 | Old |
| 0.11097 | 0.400335 | 0.052 | 0.037 | 1 | 2 | Lrig2 | 0.015 | Old |
| 0.110995 | 0.11556 | 0.216 | 0.184 | 1 | 2 | Rbbp4 | 0.032 | Old |
| 0.111123 | 0.242627 | 0.335 | 0.3 | 1 | 2 | Stx4a | 0.035 | Old |
| 0.111209 | 0.26623 | 0.147 | 0.12 | 1 | 2 | Acadl | 0.027 | Old |
| 0.111252 | 0.082063 | 0.031 | 0.019 | 1 | 2 | Ccdc167 | 0.012 | Old |
| 0.111269 | 0.292069 | 0.106 | 0.083 | 1 | 2 | Nsrp1 | 0.023 | Old |
| 0.111356 | 0.072306 | 0.298 | 0.259 | 1 | 2 | Tbcb | 0.039 | Old |
| 0.11139 | 1.027753 | 0.028 | 0.017 | 1 | 2 | Cradd | 0.011 | Old |
| 0.111412 | 0.225801 | 0.106 | 0.083 | 1 | 2 | Fbxo38 | 0.023 | Old |
| 0.111773 | 0.117142 | 0.046 | 0.031 | 1 | 2 | Rai14 | 0.015 | Old |
| 0.111841 | 0.601341 | 0.035 | 0.023 | 1 | 2 | Nfe2 | 0.012 | Old |
| 0.111883 | 0.343168 | 0.173 | 0.145 | 1 | 2 | Gpat4 | 0.028 | Old |
| 0.112305 | 0.297812 | 0.165 | 0.138 | 1 | 2 | Ireb2 | 0.027 | Old |
| 0.112571 | 0.275279 | 0.299 | 0.266 | 1 | 2 | Aftph | 0.033 | Old |
| 0.112772 | 0.495285 | 0.052 | 0.037 | 1 | 2 | Ivd | 0.015 | Old |
| 0.113019 | 0.680973 | 0.028 | 0.017 | 1 | 2 | Zfyve21 | 0.011 | Old |
| 0.11313 | 0.515504 | 0.067 | 0.05 | 1 | 2 | Pnkp | 0.017 | Old |
| 0.113218 | 0.164551 | 0.364 | 0.329 | 1 | 2 | Ctsl | 0.035 | Old |
| 0.113307 | 0.444064 | 0.088 | 0.068 | 1 | 2 | Cryzl1 | 0.02 | Old |
| 0.113671 | 0.910622 | 0.055 | 0.039 | 1 | 2 | Micall1 | 0.016 | Old |
| 0.114774 | 0.302384 | 0.146 | 0.12 | 1 | 2 | Dnajc15 | 0.026 | Old |
| 0.114862 | 0.386548 | 0.066 | 0.049 | 1 | 2 | Naxe | 0.017 | Old |
| 0.114952 | 0.454301 | 0.072 | 0.053 | 1 | 2 | Myo7a | 0.019 | Old |
| 0.115242 | 0.386207 | 0.035 | 0.023 | 1 | 2 | Arid3b | 0.012 | Old |
| 0.115677 | 0.361226 | 0.048 | 0.033 | 1 | 2 | Palld | 0.015 | Old |
| 0.115816 | 0.16593 | 0.3 | 0.265 | 1 | 2 | Tmem123 | 0.035 | Old |
| 0.115866 | 0.38496 | 0.066 | 0.049 | 1 | 2 | Adk | 0.017 | Old |
| 0.116075 | 0.576241 | 0.015 | 0.007 | 1 | 2 | A930007I1 | 0.008 | Old |
| 0.116198 | 0.679134 | 0.025 | 0.014 | 1 | 2 | Rnf152 | 0.011 | Old |
| 0.116225 | 0.123161 | 0.043 | 0.029 | 1 | 2 | H2afx | 0.014 | Old |
| 0.116239 | 0.145196 | 0.035 | 0.051 | 1 | 2 | Map4k5 | -0.016 | Old |
| 0.11631 | 1.116153 | 0.015 | 0.007 | 1 | 2 | Mis18bp1 | 0.008 | Old |
| 0.116484 | 0.218487 | 0.047 | 0.032 | 1 | 2 | Dnajc24 | 0.015 | Old |
| 0.117063 | 0.329443 | 0.127 | 0.103 | 1 | 2 | Fabp5 | 0.024 | Old |
| 0.117138 | 1.02855 | 0.015 | 0.007 | 1 | 2 | Gm31597 | 0.008 | Old |
| 0.117373 | 0.400193 | 0.115 | 0.093 | 1 | 2 | Htatsf1 | 0.022 | Old |
| 0.117513 | 0.355657 | 0.248 | 0.217 | 1 | 2 | Herc4 | 0.031 | Old |

| | | | | | | | | |
|---|---|---|---|---|---|---|---|---|
| 0.117552 | 1.365627 | 0.018 | 0.01 | 1 | 2 | Plekha8 | 0.008 | Old |
| 0.117569 | 0.069531 | 0.097 | 0.075 | 1 | 2 | Rit1 | 0.022 | Old |
| 0.117796 | 1.305954 | 0.01 | 0.004 | 1 | 2 | Wdr19 | 0.006 | Old |
| 0.117993 | 0.322888 | 0.142 | 0.118 | 1 | 2 | Oasl2 | 0.024 | Old |
| 0.118001 | 0.366329 | 0.062 | 0.045 | 1 | 2 | Coq2 | 0.017 | Old |
| 0.118056 | 0.407419 | 0.066 | 0.049 | 1 | 2 | Senp3 | 0.017 | Old |
| 0.118063 | 0.366071 | 0.093 | 0.072 | 1 | 2 | Inf2 | 0.021 | Old |
| 0.118257 | 1.348634 | 0.01 | 0.004 | 1 | 2 | Gimap6 | 0.006 | Old |
| 0.118411 | 1.527985 | 0.01 | 0.004 | 1 | 2 | C920006O1 | 0.006 | Old |
| 0.118411 | 1.344347 | 0.01 | 0.004 | 1 | 2 | Slc12a5 | 0.006 | Old |
| 0.118411 | 1.235775 | 0.01 | 0.004 | 1 | 2 | Isoc2a | 0.006 | Old |
| 0.118523 | 0.142061 | 0.139 | 0.112 | 1 | 2 | Mrps25 | 0.027 | Old |
| 0.118567 | 0.672028 | 0.015 | 0.007 | 1 | 2 | Galm | 0.008 | Old |
| 0.118728 | 1.234425 | 0.018 | 0.01 | 1 | 2 | Ss18l1 | 0.008 | Old |
| 0.118795 | 0.467597 | 0.077 | 0.058 | 1 | 2 | Gm43848 | 0.019 | Old |
| 0.118969 | 0.427448 | 0.067 | 0.05 | 1 | 2 | Hmbs | 0.017 | Old |
| 0.119338 | 1.056681 | 0.01 | 0.004 | 1 | 2 | Hist1h4h | 0.006 | Old |
| 0.119782 | 0.13186 | 0.227 | 0.19 | 1 | 2 | Aldh9a1 | 0.037 | Old |
| 0.120021 | 0.824114 | 0.018 | 0.01 | 1 | 2 | Armc3 | 0.008 | Old |
| 0.120087 | 0.318143 | 0.152 | 0.126 | 1 | 2 | Rest | 0.026 | Old |
| 0.120089 | 0.255371 | 0.188 | 0.159 | 1 | 2 | Nedd9 | 0.029 | Old |
| 0.120184 | 0.189631 | 0.255 | 0.221 | 1 | 2 | Setd2 | 0.034 | Old |
| 0.120238 | 0.860619 | 0.018 | 0.01 | 1 | 2 | Lpcat4 | 0.008 | Old |
| 0.120246 | 0.161969 | 0.212 | 0.179 | 1 | 2 | Mtmr6 | 0.033 | Old |
| 0.12027 | 0.414805 | 0.01 | 0.004 | 1 | 2 | Selp | 0.006 | Old |
| 0.12027 | 0.370324 | 0.01 | 0.004 | 1 | 2 | Magi2 | 0.006 | Old |
| 0.12027 | 0.327294 | 0.01 | 0.004 | 1 | 2 | Klrb1a | 0.006 | Old |
| 0.120398 | 0.340881 | 0.052 | 0.037 | 1 | 2 | Nsun6 | 0.015 | Old |
| 0.120418 | 0.396952 | 0.112 | 0.09 | 1 | 2 | H2-Q4 | 0.022 | Old |
| 0.120563 | 1.397951 | 0.018 | 0.01 | 1 | 2 | Aaas | 0.008 | Old |
| 0.120689 | 0.56189 | 0.032 | 0.02 | 1 | 2 | Ptdss2 | 0.012 | Old |
| 0.120749 | 0.66511 | 0.036 | 0.024 | 1 | 2 | Prune1 | 0.012 | Old |
| 0.120771 | 0.165929 | 0.066 | 0.049 | 1 | 2 | Tomm6 | 0.017 | Old |
| 0.120774 | 0.497551 | 0.044 | 0.03 | 1 | 2 | Rars2 | 0.014 | Old |
| 0.121032 | 0.244744 | 0.081 | 0.062 | 1 | 2 | Lnpk | 0.019 | Old |
| 0.121158 | 0.613577 | 0.021 | 0.012 | 1 | 2 | Hibch | 0.009 | Old |
| 0.121168 | 0.260775 | 0.135 | 0.109 | 1 | 2 | Scd2 | 0.026 | Old |
| 0.121176 | 0.370054 | 0.153 | 0.127 | 1 | 2 | Nsfl1c | 0.026 | Old |
| 0.121476 | 0.447616 | 0.072 | 0.053 | 1 | 2 | Socs5 | 0.019 | Old |
| 0.121758 | 0.149446 | 0.108 | 0.084 | 1 | 2 | S100pbp | 0.024 | Old |
| 0.122091 | 0.477725 | 0.018 | 0.01 | 1 | 2 | Fam177a | 0.008 | Old |
| 0.122132 | 0.244875 | 0.182 | 0.156 | 1 | 2 | Dcaf7 | 0.026 | Old |
| 0.122158 | 0.06694 | 0.131 | 0.106 | 1 | 2 | Babam1 | 0.025 | Old |
| 0.122224 | 0.224729 | 0.167 | 0.139 | 1 | 2 | Pik3cg | 0.028 | Old |
| 0.122347 | 0.175332 | 0.173 | 0.144 | 1 | 2 | Usp22 | 0.029 | Old |

| | | | | | | | | |
|---|---|---|---|---|---|---|---|---|
| 0.122408 | 0.019344 | 0.044 | 0.03 | 1 | 2 | Phlpp1 | 0.014 | Old |
| 0.122453 | 0.12144 | 0.638 | 0.606 | 1 | 2 | Grk2 | 0.032 | Old |
| 0.122995 | 0.321451 | 0.122 | 0.099 | 1 | 2 | Esco1 | 0.023 | Old |
| 0.123039 | 0.227134 | 0.082 | 0.063 | 1 | 2 | Tceal8 | 0.019 | Old |
| 0.123123 | 0.655537 | 0.065 | 0.049 | 1 | 2 | B3glct | 0.016 | Old |
| 0.123382 | 0.132061 | 0.596 | 0.557 | 1 | 2 | Diaph1 | 0.039 | Old |
| 0.123603 | 0.13586 | 0.34 | 0.304 | 1 | 2 | Ube2n | 0.036 | Old |
| 0.123609 | 0.456829 | 0.036 | 0.024 | 1 | 2 | Atpaf2 | 0.012 | Old |
| 0.123741 | 0.87009 | 0.056 | 0.04 | 1 | 2 | Hgf | 0.016 | Old |
| 0.123835 | 0.264958 | 0.054 | 0.038 | 1 | 2 | Dnajc30 | 0.016 | Old |
| 0.124069 | 0.401296 | 0.045 | 0.031 | 1 | 2 | Ermard | 0.014 | Old |
| 0.124256 | 0.441698 | 0.177 | 0.15 | 1 | 2 | Pts | 0.027 | Old |
| 0.124483 | 0.151665 | 0.061 | 0.044 | 1 | 2 | Snapc3 | 0.017 | Old |
| 0.12456 | 0.509464 | 0.095 | 0.076 | 1 | 2 | Dhrs7b | 0.019 | Old |
| 0.124788 | 0.132826 | 0.021 | 0.012 | 1 | 2 | Thap7 | 0.009 | Old |
| 0.124788 | 0.095231 | 0.021 | 0.012 | 1 | 2 | Abcb8 | 0.009 | Old |
| 0.124868 | 0.57351 | 0.059 | 0.043 | 1 | 2 | Pot1a | 0.016 | Old |
| 0.124946 | 0.293451 | 0.118 | 0.094 | 1 | 2 | Usp6nl | 0.024 | Old |
| 0.124972 | 0.232214 | 0.209 | 0.177 | 1 | 2 | Bscl2 | 0.032 | Old |
| 0.125211 | 0.119016 | 0.467 | 0.429 | 1 | 2 | Apbb1ip | 0.038 | Old |
| 0.125607 | 0.640664 | 0.06 | 0.044 | 1 | 2 | Nup93 | 0.016 | Old |
| 0.125965 | 0.910288 | 0.037 | 0.025 | 1 | 2 | Pan2 | 0.012 | Old |
| 0.126044 | 0.208496 | 0.279 | 0.246 | 1 | 2 | Pros1 | 0.033 | Old |
| 0.126097 | 0.603839 | 0.029 | 0.018 | 1 | 2 | BC024978 | 0.011 | Old |
| 0.126285 | 0.814058 | 0.033 | 0.021 | 1 | 2 | Nod1 | 0.012 | Old |
| 0.126587 | 0.046622 | 0.413 | 0.369 | 1 | 2 | Wsb1 | 0.044 | Old |
| 0.126736 | 0.134429 | 0.907 | 0.897 | 1 | 2 | Zfp36 | 0.01 | Old |
| 0.126796 | 0.242611 | 0.029 | 0.018 | 1 | 2 | Pola2 | 0.011 | Old |
| 0.126875 | 0.304863 | 0.175 | 0.147 | 1 | 2 | Snx4 | 0.028 | Old |
| 0.126981 | 0.380431 | 0.079 | 0.061 | 1 | 2 | Fam220a | 0.018 | Old |
| 0.127202 | 0.10518 | 0.525 | 0.488 | 1 | 2 | Nme1 | 0.037 | Old |
| 0.127367 | 0.25595 | 0.128 | 0.105 | 1 | 2 | Sft2d1 | 0.023 | Old |
| 0.127394 | 0.129754 | 0.224 | 0.19 | 1 | 2 | Zfp263 | 0.034 | Old |
| 0.127452 | 0.15824 | 0.177 | 0.148 | 1 | 2 | Usp47 | 0.029 | Old |
| 0.1276 | 0.08986 | 0.296 | 0.253 | 1 | 2 | Ythdf3 | 0.043 | Old |
| 0.128492 | 0.170415 | 0.245 | 0.211 | 1 | 2 | Cbfb | 0.034 | Old |
| 0.12856 | 0.499098 | 0.055 | 0.039 | 1 | 2 | Rsad2 | 0.016 | Old |
| 0.128636 | 0.303375 | 0.201 | 0.175 | 1 | 2 | Cd37 | 0.026 | Old |
| 0.128701 | 0.192358 | 0.045 | 0.031 | 1 | 2 | Xrcc4 | 0.014 | Old |
| 0.128754 | 0.354004 | 0.063 | 0.046 | 1 | 2 | Dnlz | 0.017 | Old |
| 0.128786 | 0.217472 | 0.162 | 0.135 | 1 | 2 | Bst1 | 0.027 | Old |
| 0.128941 | 0.876156 | 0.086 | 0.067 | 1 | 2 | Spn | 0.019 | Old |
| 0.129002 | 0.230751 | 0.208 | 0.178 | 1 | 2 | Sp110 | 0.03 | Old |
| 0.129377 | 0.104949 | 0.032 | 0.02 | 1 | 2 | Apba3 | 0.012 | Old |
| 0.12958 | 0.144161 | 0.255 | 0.219 | 1 | 2 | Ppp1r2 | 0.036 | Old |

| | | | | | | | | |
|---|---|---|---|---|---|---|---|---|
| 0.12959 | 0.289132 | 0.262 | 0.23 | 1 | 2 | Hip1 | 0.032 | Old |
| 0.129624 | 0.275136 | 0.049 | 0.034 | 1 | 2 | Cstf1 | 0.015 | Old |
| 0.129729 | 0.303215 | 0.063 | 0.046 | 1 | 2 | Dtx2 | 0.017 | Old |
| 0.129744 | 0.321692 | 0.105 | 0.083 | 1 | 2 | Thoc6 | 0.022 | Old |
| 0.129756 | 0.132412 | 0.525 | 0.487 | 1 | 2 | Atp6ap1 | 0.038 | Old |
| 0.130402 | 0.1075 | 0.261 | 0.224 | 1 | 2 | Cux1 | 0.037 | Old |
| 0.131017 | 0.1174 | 0.704 | 0.678 | 1 | 2 | H2-DMb1 | 0.026 | Old |
| 0.131337 | 0.552644 | 0.06 | 0.044 | 1 | 2 | I830077J02 | 0.016 | Old |
| 0.131364 | 0.340611 | 0.102 | 0.081 | 1 | 2 | C2cd5 | 0.021 | Old |
| 0.13137 | 0.408026 | 0.077 | 0.058 | 1 | 2 | Tex10 | 0.019 | Old |
| 0.131433 | 0.267926 | 0.109 | 0.088 | 1 | 2 | Apobr | 0.021 | Old |
| 0.131823 | 0.512383 | 0.064 | 0.048 | 1 | 2 | Sipa1l2 | 0.016 | Old |
| 0.131974 | 0.498742 | 0.11 | 0.089 | 1 | 2 | Dnm1l | 0.021 | Old |
| 0.132082 | 0.211689 | 0.071 | 0.052 | 1 | 2 | Uba7 | 0.019 | Old |
| 0.132112 | 0.157634 | 0.218 | 0.189 | 1 | 2 | Tap2 | 0.029 | Old |
| 0.132368 | 0.184798 | 0.125 | 0.101 | 1 | 2 | Dennd1c | 0.024 | Old |
| 0.132731 | 0.424677 | 0.091 | 0.071 | 1 | 2 | Ddx56 | 0.02 | Old |
| 0.132763 | 0.269731 | 0.103 | 0.082 | 1 | 2 | Keap1 | 0.021 | Old |
| 0.132804 | 0.540325 | 0.056 | 0.04 | 1 | 2 | Cdca4 | 0.016 | Old |
| 0.132827 | 0.512447 | 0.046 | 0.032 | 1 | 2 | Cfap97 | 0.014 | Old |
| 0.133067 | 0.122928 | 0.418 | 0.372 | 1 | 2 | Anp32a | 0.046 | Old |
| 0.133168 | 0.093127 | 0.258 | 0.223 | 1 | 2 | Snrpd2 | 0.035 | Old |
| 0.133317 | 0.14507 | 0.456 | 0.411 | 1 | 2 | Kif5b | 0.045 | Old |
| 0.133322 | 0.047605 | 0.088 | 0.068 | 1 | 2 | Mtf1 | 0.02 | Old |
| 0.133323 | 0.542927 | 0.037 | 0.025 | 1 | 2 | Stambpl1 | 0.012 | Old |
| 0.133329 | 0.103299 | 0.177 | 0.147 | 1 | 2 | Pdha1 | 0.03 | Old |
| 0.133367 | 0.518084 | 0.089 | 0.07 | 1 | 2 | Ddx19a | 0.019 | Old |
| 0.133388 | 0.219535 | 0.097 | 0.077 | 1 | 2 | Gpr137b | 0.02 | Old |
| 0.133411 | 0.291013 | 0.091 | 0.071 | 1 | 2 | Tmub2 | 0.02 | Old |
| 0.133432 | 0.430569 | 0.075 | 0.057 | 1 | 2 | Ccdc127 | 0.018 | Old |
| 0.133493 | 0.266352 | 0.176 | 0.15 | 1 | 2 | Fam173a | 0.026 | Old |
| 0.133744 | 0.152851 | 0.466 | 0.418 | 1 | 2 | P4hb | 0.048 | Old |
| 0.134261 | 0.285896 | 0.055 | 0.039 | 1 | 2 | Klhdc10 | 0.016 | Old |
| 0.13432 | 0.179498 | 0.48 | 0.438 | 1 | 2 | Tra2b | 0.042 | Old |
| 0.134327 | 0.465603 | 0.071 | 0.053 | 1 | 2 | Abhd4 | 0.018 | Old |
| 0.134344 | 0.385953 | 0.118 | 0.096 | 1 | 2 | 4833420G1 | 0.022 | Old |
| 0.134416 | 0.054962 | 0.026 | 0.015 | 1 | 2 | Yars2 | 0.011 | Old |
| 0.134588 | 0.427932 | 0.08 | 0.062 | 1 | 2 | Sbf1 | 0.018 | Old |
| 0.134612 | 0.421563 | 0.056 | 0.04 | 1 | 2 | Slc39a3 | 0.016 | Old |
| 0.134679 | 0.930639 | 0.013 | 0.006 | 1 | 2 | Ankrd66 | 0.007 | Old |
| 0.134752 | 0.117462 | 0.58 | 0.54 | 1 | 2 | Hnrnph1 | 0.04 | Old |
| 0.134822 | 0.862033 | 0.013 | 0.006 | 1 | 2 | Anln | 0.007 | Old |
| 0.134965 | 1.06899 | 0.013 | 0.006 | 1 | 2 | Homez | 0.007 | Old |
| 0.135253 | 0.651509 | 0.013 | 0.006 | 1 | 2 | Zfp438 | 0.007 | Old |
| 0.135407 | 0.771304 | 0.03 | 0.019 | 1 | 2 | Telo2 | 0.011 | Old |

| | | | | | | | | |
|---|---|---|---|---|---|---|---|---|
| 0.135553 | 0.158055 | 0.05 | 0.036 | 1 | 2 | Ikbip | 0.014 | Old |
| 0.135558 | 0.300539 | 0.107 | 0.086 | 1 | 2 | Ddx47 | 0.021 | Old |
| 0.135575 | 0.024907 | 0.022 | 0.034 | 1 | 2 | Ahi1 | -0.012 | Old |
| 0.135589 | 0.371402 | 0.026 | 0.015 | 1 | 2 | Nepro | 0.011 | Old |
| 0.135684 | 0.757489 | 0.013 | 0.006 | 1 | 2 | 9330020H( | 0.007 | Old |
| 0.135909 | 0.277262 | 0.138 | 0.114 | 1 | 2 | Echs1 | 0.024 | Old |
| 0.135972 | 0.554884 | 0.071 | 0.053 | 1 | 2 | Grcc10 | 0.018 | Old |
| 0.136079 | 0.168241 | 0.197 | 0.167 | 1 | 2 | Gtf2a1 | 0.03 | Old |
| 0.136289 | 0.006348 | 0.069 | 0.088 | 1 | 2 | Gm37494 | -0.019 | Old |
| 0.13655 | 0.713188 | 0.013 | 0.006 | 1 | 2 | Dusp19 | 0.007 | Old |
| 0.136582 | 0.54893 | 0.03 | 0.019 | 1 | 2 | Hoxb4 | 0.011 | Old |
| 0.136981 | 0.178223 | 0.358 | 0.323 | 1 | 2 | Hnrnpd | 0.035 | Old |
| 0.13713 | 0.468688 | 0.013 | 0.006 | 1 | 2 | Rad52 | 0.007 | Old |
| 0.137134 | 0.404608 | 0.135 | 0.113 | 1 | 2 | Cxcl1 | 0.022 | Old |
| 0.137365 | 0.128911 | 0.026 | 0.015 | 1 | 2 | Foxp4 | 0.011 | Old |
| 0.137366 | 0.290672 | 0.101 | 0.08 | 1 | 2 | Gyg | 0.021 | Old |
| 0.137392 | 0.401236 | 0.056 | 0.04 | 1 | 2 | Fam53a | 0.016 | Old |
| 0.137553 | 0.448204 | 0.06 | 0.044 | 1 | 2 | Sap30bp | 0.016 | Old |
| 0.137715 | 0.191117 | 0.186 | 0.158 | 1 | 2 | Slc9a3r1 | 0.028 | Old |
| 0.137803 | 0.099072 | 0.084 | 0.064 | 1 | 2 | Chid1 | 0.02 | Old |
| 0.138002 | 0.140793 | 0.815 | 0.802 | 1 | 2 | Hnrnpa2b1 | 0.013 | Old |
| 0.138093 | 0.115916 | 0.2 | 0.17 | 1 | 2 | Mrpl14 | 0.03 | Old |
| 0.138224 | 0.313762 | 0.125 | 0.102 | 1 | 2 | Rangap1 | 0.023 | Old |
| 0.138636 | 0.370865 | 0.175 | 0.151 | 1 | 2 | Sqor | 0.024 | Old |
| 0.138653 | 0.776141 | 0.039 | 0.026 | 1 | 2 | Plscr1 | 0.013 | Old |
| 0.139057 | 0.740795 | 0.034 | 0.023 | 1 | 2 | Atat1 | 0.011 | Old |
| 0.139396 | 0.688818 | 0.019 | 0.011 | 1 | 2 | Myo10 | 0.008 | Old |
| 0.139546 | 0.094243 | 0.784 | 0.772 | 1 | 2 | Slc25a3 | 0.012 | Old |
| 0.139717 | 0.439444 | 0.13 | 0.107 | 1 | 2 | Dph3 | 0.023 | Old |
| 0.139753 | 0.158862 | 0.36 | 0.321 | 1 | 2 | Ddi2 | 0.039 | Old |
| 0.139771 | 0.347995 | 0.101 | 0.08 | 1 | 2 | Prkch | 0.021 | Old |
| 0.139782 | 0.302776 | 0.12 | 0.097 | 1 | 2 | Rasgrp2 | 0.023 | Old |
| 0.139992 | 0.121461 | 0.5 | 0.462 | 1 | 2 | Macf1 | 0.038 | Old |
| 0.140122 | 0.099229 | 0.055 | 0.039 | 1 | 2 | Pphln1 | 0.016 | Old |
| 0.140335 | 0.537344 | 0.019 | 0.011 | 1 | 2 | C1ra | 0.008 | Old |
| 0.140453 | 0.48813 | 0.019 | 0.011 | 1 | 2 | Slc2a3 | 0.008 | Old |
| 0.140607 | 0.145179 | 0.329 | 0.29 | 1 | 2 | Copa | 0.039 | Old |
| 0.140912 | 0.135599 | 0.022 | 0.013 | 1 | 2 | Impa2 | 0.009 | Old |
| 0.140923 | 0.256881 | 0.06 | 0.044 | 1 | 2 | Tmub1 | 0.016 | Old |
| 0.141164 | 0.316349 | 0.211 | 0.183 | 1 | 2 | Lsm14a | 0.028 | Old |
| 0.14136 | 0.57178 | 0.052 | 0.038 | 1 | 2 | Cstf2 | 0.014 | Old |
| 0.141453 | 0.083659 | 0.03 | 0.019 | 1 | 2 | 6330418K( | 0.011 | Old |
| 0.141753 | 0.19979 | 0.566 | 0.54 | 1 | 2 | Cytip | 0.026 | Old |
| 0.1418 | 0.346052 | 0.126 | 0.103 | 1 | 2 | Inpp5f | 0.023 | Old |
| 0.141846 | 0.588229 | 0.016 | 0.008 | 1 | 2 | Ccdc137 | 0.008 | Old |

| | | | | | | | | |
|---|---|---|---|---|---|---|---|---|
| 0.141846 | 0.391698 | 0.016 | 0.008 | 1 | 2 | Oas1g | 0.008 | Old |
| 0.141849 | 0.356269 | 0.115 | 0.094 | 1 | 2 | Cdkn2d | 0.021 | Old |
| 0.141967 | 0.721193 | 0.08 | 0.063 | 1 | 2 | Gcnt2 | 0.017 | Old |
| 0.14199 | 0.224705 | 0.051 | 0.037 | 1 | 2 | Cdadc1 | 0.014 | Old |
| 0.14202 | 0.161701 | 0.241 | 0.207 | 1 | 2 | Sbds | 0.034 | Old |
| 0.142031 | 0.149678 | 0.042 | 0.029 | 1 | 2 | Tipin | 0.013 | Old |
| 0.142169 | 0.100676 | 0.037 | 0.025 | 1 | 2 | Rnf217 | 0.012 | Old |
| 0.14224 | 0.368859 | 0.016 | 0.008 | 1 | 2 | Mmaa | 0.008 | Old |
| 0.142291 | 0.143626 | 0.084 | 0.064 | 1 | 2 | Tmem181a | 0.02 | Old |
| 0.142301 | 0.57404 | 0.048 | 0.034 | 1 | 2 | Cry1 | 0.014 | Old |
| 0.142554 | 0.120864 | 0.248 | 0.216 | 1 | 2 | Psmb5 | 0.032 | Old |
| 0.142584 | 0.24323 | 0.019 | 0.011 | 1 | 2 | Smug1 | 0.008 | Old |
| 0.142985 | 0.099816 | 0.022 | 0.013 | 1 | 2 | Prepl | 0.009 | Old |
| 0.143069 | 0.206589 | 0.375 | 0.342 | 1 | 2 | Morf4l2 | 0.033 | Old |
| 0.143175 | 0.298594 | 0.077 | 0.059 | 1 | 2 | Eipr1 | 0.018 | Old |
| 0.143183 | 0.155642 | 0.225 | 0.192 | 1 | 2 | Ncstn | 0.033 | Old |
| 0.143268 | 0.360921 | 0.134 | 0.112 | 1 | 2 | Fnbp1l | 0.022 | Old |
| 0.14341 | 0.322981 | 0.047 | 0.033 | 1 | 2 | Srm | 0.014 | Old |
| 0.143539 | 0.12015 | 0.055 | 0.039 | 1 | 2 | Zfp729a | 0.016 | Old |
| 0.143656 | 0.448011 | 0.084 | 0.065 | 1 | 2 | Tsfm | 0.019 | Old |
| 0.143789 | 0.090749 | 0.118 | 0.095 | 1 | 2 | Slc37a2 | 0.023 | Old |
| 0.143954 | 0.210569 | 0.481 | 0.464 | 1 | 2 | Srsf2 | 0.017 | Old |
| 0.144194 | 0.392433 | 0.061 | 0.045 | 1 | 2 | Nr1h3 | 0.016 | Old |
| 0.144488 | 0.496341 | 0.072 | 0.055 | 1 | 2 | Trappc12 | 0.017 | Old |
| 0.144566 | 0.193988 | 0.376 | 0.334 | 1 | 2 | Sf3b6 | 0.042 | Old |
| 0.144638 | 0.265379 | 0.047 | 0.033 | 1 | 2 | Plekhg1 | 0.014 | Old |
| 0.144678 | 0.660411 | 0.106 | 0.087 | 1 | 2 | Cwf19l2 | 0.019 | Old |
| 0.144882 | 0.526626 | 0.034 | 0.023 | 1 | 2 | Arl14ep | 0.011 | Old |
| 0.144973 | 0.051626 | 0.262 | 0.228 | 1 | 2 | Supt5 | 0.034 | Old |
| 0.145124 | 0.138961 | 0.283 | 0.246 | 1 | 2 | Acap2 | 0.037 | Old |
| 0.145352 | 0.384776 | 0.077 | 0.059 | 1 | 2 | Lyrm2 | 0.018 | Old |
| 0.145399 | 0.378782 | 0.126 | 0.105 | 1 | 2 | Trappc8 | 0.021 | Old |
| 0.145419 | 0.968769 | 0.031 | 0.02 | 1 | 2 | Rcl1 | 0.011 | Old |
| 0.145509 | 0.096618 | 0.536 | 0.488 | 1 | 2 | Pak2 | 0.048 | Old |
| 0.145685 | 0.619086 | 0.058 | 0.043 | 1 | 2 | Abcc4 | 0.015 | Old |
| 0.14596 | 0.802178 | 0.04 | 0.027 | 1 | 2 | Pafah1b3 | 0.013 | Old |
| 0.146211 | 0.623777 | 0.057 | 0.042 | 1 | 2 | Prkab1 | 0.015 | Old |
| 0.146237 | 0.118601 | 0.147 | 0.122 | 1 | 2 | Fyttd1 | 0.025 | Old |
| 0.146364 | 0.167293 | 0.236 | 0.207 | 1 | 2 | Mdm2 | 0.029 | Old |
| 0.14661 | 0.150524 | 0.335 | 0.293 | 1 | 2 | Tmod3 | 0.042 | Old |
| 0.146667 | 0.125788 | 0.97 | 0.957 | 1 | 2 | Jund | 0.013 | Old |
| 0.146799 | 0.230109 | 0.12 | 0.097 | 1 | 2 | Map1lc3a | 0.023 | Old |
| 0.14692 | 0.237119 | 0.151 | 0.126 | 1 | 2 | Auh | 0.025 | Old |
| 0.147591 | 0.221225 | 0.255 | 0.223 | 1 | 2 | Higd2a | 0.032 | Old |
| 0.147726 | 0.189285 | 0.238 | 0.208 | 1 | 2 | Erp44 | 0.03 | Old |

| | | | | | | | | |
|---|---|---|---|---|---|---|---|---|
| 0.147729 | 0.305353 | 0.078 | 0.061 | 1 | 2 | Erh | 0.017 | Old |
| 0.14803 | 0.582638 | 0.054 | 0.039 | 1 | 2 | Actr8 | 0.015 | Old |
| 0.148076 | 0.454728 | 0.102 | 0.082 | 1 | 2 | Arfip1 | 0.02 | Old |
| 0.148165 | 0.019519 | 0.043 | 0.03 | 1 | 2 | Irgm2 | 0.013 | Old |
| 0.14835 | 0.164357 | 0.882 | 0.881 | 1 | 2 | Ctsb | 0.001 | Old |
| 0.148717 | 0.30011 | 0.099 | 0.078 | 1 | 2 | Dhx30 | 0.021 | Old |
| 0.148761 | 0.164759 | 0.243 | 0.21 | 1 | 2 | Rell1 | 0.033 | Old |
| 0.148961 | 0.077197 | 0.076 | 0.058 | 1 | 2 | Dnajc11 | 0.018 | Old |
| 0.148962 | 0.126072 | 0.227 | 0.196 | 1 | 2 | Mat2b | 0.031 | Old |
| 0.149392 | 0.326495 | 0.079 | 0.062 | 1 | 2 | Washc5 | 0.017 | Old |
| 0.149462 | 0.660492 | 0.089 | 0.071 | 1 | 2 | Pcgf3 | 0.018 | Old |
| 0.149536 | 0.170245 | 0.303 | 0.268 | 1 | 2 | Luc7l3 | 0.035 | Old |
| 0.14963 | 0.182664 | 0.281 | 0.251 | 1 | 2 | Dnmt3a | 0.03 | Old |
| 0.150055 | 0.182659 | 0.289 | 0.254 | 1 | 2 | Ccnt1 | 0.035 | Old |
| 0.150089 | 0.19404 | 0.317 | 0.283 | 1 | 2 | Leng8 | 0.034 | Old |
| 0.150145 | 0.0356 | 0.199 | 0.166 | 1 | 2 | BC005624 | 0.033 | Old |
| 0.150294 | 0.35262 | 0.065 | 0.049 | 1 | 2 | Anapc15 | 0.016 | Old |
| 0.15031 | 0.638324 | 0.035 | 0.024 | 1 | 2 | AI597479 | 0.011 | Old |
| 0.150348 | 0.591996 | 0.044 | 0.031 | 1 | 2 | Cactin | 0.013 | Old |
| 0.150355 | 0.206962 | 0.062 | 0.046 | 1 | 2 | Plp2 | 0.016 | Old |
| 0.150405 | 0.423775 | 0.048 | 0.034 | 1 | 2 | Tyw1 | 0.014 | Old |
| 0.15057 | 0.210593 | 0.109 | 0.088 | 1 | 2 | Ece1 | 0.021 | Old |
| 0.150817 | 0.294327 | 0.163 | 0.138 | 1 | 2 | Eapp | 0.025 | Old |
| 0.150977 | 0.049388 | 0.047 | 0.033 | 1 | 2 | 1600002K0 | 0.014 | Old |
| 0.151105 | 0.121058 | 0.204 | 0.175 | 1 | 2 | Kmt5b | 0.029 | Old |
| 0.151203 | 0.548462 | 0.035 | 0.024 | 1 | 2 | Men1 | 0.011 | Old |
| 0.151631 | 0.269088 | 0.127 | 0.106 | 1 | 2 | Sap30 | 0.021 | Old |
| 0.151668 | 0.544734 | 0.027 | 0.017 | 1 | 2 | Fut7 | 0.01 | Old |
| 0.152057 | 0.17613 | 0.304 | 0.273 | 1 | 2 | Arglu1 | 0.031 | Old |
| 0.152236 | 0.713065 | 0.035 | 0.024 | 1 | 2 | Polm | 0.011 | Old |
| 0.152384 | 0.193859 | 0.399 | 0.367 | 1 | 2 | Pik3cd | 0.032 | Old |
| 0.152464 | 0.497156 | 0.094 | 0.076 | 1 | 2 | Prkaca | 0.018 | Old |
| 0.152549 | 0.425563 | 0.143 | 0.121 | 1 | 2 | Cln3 | 0.022 | Old |
| 0.152662 | 0.01201 | 0.173 | 0.146 | 1 | 2 | Sde2 | 0.027 | Old |
| 0.153381 | 0.199382 | 0.181 | 0.153 | 1 | 2 | Twf1 | 0.028 | Old |
| 0.153413 | 0.401768 | 0.074 | 0.057 | 1 | 2 | Gpatch2l | 0.017 | Old |
| 0.153469 | 0.138222 | 0.113 | 0.091 | 1 | 2 | Pdcd11 | 0.022 | Old |
| 0.153524 | 0.199656 | 0.476 | 0.444 | 1 | 2 | G3bp2 | 0.032 | Old |
| 0.153774 | 0.398196 | 0.048 | 0.034 | 1 | 2 | Zdhhc8 | 0.014 | Old |
| 0.154417 | 0.232059 | 0.223 | 0.194 | 1 | 2 | Nucb1 | 0.029 | Old |
| 0.154511 | 0.191368 | 0.215 | 0.186 | 1 | 2 | Tmpo | 0.029 | Old |
| 0.154895 | 0.21162 | 0.066 | 0.05 | 1 | 2 | Gdap2 | 0.016 | Old |
| 0.154929 | 0.385944 | 0.075 | 0.058 | 1 | 2 | Taf4 | 0.017 | Old |
| 0.154981 | 0.332094 | 0.125 | 0.103 | 1 | 2 | Rabggtb | 0.022 | Old |
| 0.155037 | 0.671681 | 0.011 | 0.005 | 1 | 2 | Gm49339 | 0.006 | Old |

| | | | | | | | | |
|---|---|---|---|---|---|---|---|---|
| 0.155163 | 0.437879 | 0.052 | 0.038 | 1 | 2 | Pgam5 | 0.014 | Old |
| 0.155254 | 0.409226 | 0.031 | 0.02 | 1 | 2 | Mllt3 | 0.011 | Old |
| 0.155388 | 1.136185 | 0.011 | 0.005 | 1 | 2 | Fam185a | 0.006 | Old |
| 0.155564 | 1.166006 | 0.011 | 0.005 | 1 | 2 | Hes7 | 0.006 | Old |
| 0.155609 | 0.408523 | 0.049 | 0.036 | 1 | 2 | Recql | 0.013 | Old |
| 0.155741 | 1.238486 | 0.011 | 0.005 | 1 | 2 | Fancm | 0.006 | Old |
| 0.155842 | 0.123891 | 0.031 | 0.02 | 1 | 2 | Ifi205 | 0.011 | Old |
| 0.155917 | 1.094555 | 0.011 | 0.005 | 1 | 2 | Nr6a1 | 0.006 | Old |
| 0.156093 | 1.236283 | 0.011 | 0.005 | 1 | 2 | Tarbp1 | 0.006 | Old |
| 0.156725 | 0.018326 | 0.024 | 0.014 | 1 | 2 | Hps6 | 0.01 | Old |
| 0.156886 | 0.364628 | 0.045 | 0.032 | 1 | 2 | Dcaf1 | 0.013 | Old |
| 0.157099 | 0.135863 | 0.533 | 0.5 | 1 | 2 | Olfm1 | 0.033 | Old |
| 0.157288 | 0.137814 | 0.065 | 0.049 | 1 | 2 | 2210016L2 | 0.016 | Old |
| 0.15751 | 0.665452 | 0.011 | 0.005 | 1 | 2 | Gspt2 | 0.006 | Old |
| 0.15751 | 0.409734 | 0.011 | 0.005 | 1 | 2 | Asah2 | 0.006 | Old |
| 0.157519 | 0.859242 | 0.02 | 0.012 | 1 | 2 | Gm14286 | 0.008 | Old |
| 0.157737 | 0.371582 | 0.059 | 0.044 | 1 | 2 | Srprb | 0.015 | Old |
| 0.157805 | 0.16894 | 0.228 | 0.2 | 1 | 2 | Rara | 0.028 | Old |
| 0.15783 | 0.392793 | 0.048 | 0.034 | 1 | 2 | Utp14b | 0.014 | Old |
| 0.157866 | 0.222441 | 0.011 | 0.005 | 1 | 2 | Tsix | 0.006 | Old |
| 0.157932 | 0.225963 | 0.278 | 0.252 | 1 | 2 | Smg1 | 0.026 | Old |
| 0.158011 | 0.262544 | 0.063 | 0.048 | 1 | 2 | Naa25 | 0.015 | Old |
| 0.158159 | 0.26962 | 0.118 | 0.096 | 1 | 2 | Nsd2 | 0.022 | Old |
| 0.158338 | 0.925551 | 0.036 | 0.025 | 1 | 2 | Lrba | 0.011 | Old |
| 0.158511 | 0.740635 | 0.028 | 0.018 | 1 | 2 | Tmem156 | 0.01 | Old |
| 0.15859 | 0.306303 | 0.142 | 0.12 | 1 | 2 | Cse1l | 0.022 | Old |
| 0.158706 | 0.152925 | 0.152 | 0.127 | 1 | 2 | Mrpl32 | 0.025 | Old |
| 0.159356 | 0.136324 | 0.573 | 0.539 | 1 | 2 | Nap1l1 | 0.034 | Old |
| 0.159488 | 0.55342 | 0.093 | 0.075 | 1 | 2 | Med21 | 0.018 | Old |
| 0.159678 | 0.165574 | 0.088 | 0.069 | 1 | 2 | Zfp593 | 0.019 | Old |
| 0.159881 | 0.267799 | 0.519 | 0.496 | 1 | 2 | Ptpre | 0.023 | Old |
| 0.16002 | 0.284334 | 0.089 | 0.07 | 1 | 2 | Copg2 | 0.019 | Old |
| 0.160398 | 1.057788 | 0.017 | 0.01 | 1 | 2 | Chst15 | 0.007 | Old |
| 0.160517 | 0.638374 | 0.02 | 0.012 | 1 | 2 | Trit1 | 0.008 | Old |
| 0.160724 | 0.077679 | 0.076 | 0.058 | 1 | 2 | Foxk2 | 0.018 | Old |
| 0.161526 | 0.61118 | 0.02 | 0.012 | 1 | 2 | B4galt4 | 0.008 | Old |
| 0.161544 | 0.494143 | 0.089 | 0.071 | 1 | 2 | Mroh1 | 0.018 | Old |
| 0.161906 | 0.483775 | 0.05 | 0.037 | 1 | 2 | Galk1 | 0.013 | Old |
| 0.162159 | 0.536673 | 0.02 | 0.012 | 1 | 2 | Dhx37 | 0.008 | Old |
| 0.162354 | 0.809864 | 0.032 | 0.021 | 1 | 2 | Ttc7b | 0.011 | Old |
| 0.162708 | 0.939622 | 0.014 | 0.007 | 1 | 2 | Srr | 0.007 | Old |
| 0.16295 | 0.200149 | 0.334 | 0.3 | 1 | 2 | Zfp703 | 0.034 | Old |
| 0.163049 | 0.277468 | 0.02 | 0.012 | 1 | 2 | Tchp | 0.008 | Old |
| 0.163198 | 0.224823 | 0.38 | 0.349 | 1 | 2 | Ube2l3 | 0.031 | Old |
| 0.163284 | 0.361173 | 0.093 | 0.075 | 1 | 2 | Nectin1 | 0.018 | Old |

| | | | | | | | | |
|---|---|---|---|---|---|---|---|---|
| 0.16347 | 0.295763 | 0.086 | 0.068 | 1 | 2 | Scyl1 | 0.018 | Old |
| 0.163473 | 0.04895 | 0.256 | 0.226 | 1 | 2 | Krit1 | 0.03 | Old |
| 0.16428 | 0.896703 | 0.014 | 0.007 | 1 | 2 | Elp6 | 0.007 | Old |
| 0.164307 | 0.267352 | 0.074 | 0.057 | 1 | 2 | Mtif2 | 0.017 | Old |
| 0.164336 | 0.096592 | 0.188 | 0.16 | 1 | 2 | Atp11b | 0.028 | Old |
| 0.164437 | 0.769674 | 0.014 | 0.007 | 1 | 2 | Gm15964 | 0.007 | Old |
| 0.164753 | 0.738239 | 0.014 | 0.007 | 1 | 2 | Dnm1 | 0.007 | Old |
| 0.164861 | 0.69664 | 0.017 | 0.01 | 1 | 2 | Fbxw9 | 0.007 | Old |
| 0.164912 | 1.152638 | 0.014 | 0.007 | 1 | 2 | Zfp606 | 0.007 | Old |
| 0.16501 | 0.139218 | 0.049 | 0.036 | 1 | 2 | Utp15 | 0.013 | Old |
| 0.165095 | 0.17049 | 0.02 | 0.012 | 1 | 2 | Eral1 | 0.008 | Old |
| 0.165103 | 0.223642 | 0.059 | 0.044 | 1 | 2 | Slc25a19 | 0.015 | Old |
| 0.165131 | 0.235274 | 0.237 | 0.209 | 1 | 2 | Ddx24 | 0.028 | Old |
| 0.165252 | 0.473862 | 0.195 | 0.172 | 1 | 2 | Dck | 0.023 | Old |
| 0.1653 | 0.141484 | 0.064 | 0.049 | 1 | 2 | Yif1b | 0.015 | Old |
| 0.165658 | 0.695969 | 0.036 | 0.025 | 1 | 2 | Lrrc57 | 0.011 | Old |
| 0.165704 | 0.062975 | 0.014 | 0.007 | 1 | 2 | Tnni3 | 0.007 | Old |
| 0.165993 | 0.096515 | 0.573 | 0.544 | 1 | 2 | Tuba1b | 0.029 | Old |
| 0.166057 | 0.220844 | 0.382 | 0.354 | 1 | 2 | Snx2 | 0.028 | Old |
| 0.166091 | 0.427499 | 0.046 | 0.033 | 1 | 2 | Tomm40l | 0.013 | Old |
| 0.166175 | 0.462613 | 0.075 | 0.058 | 1 | 2 | Kdm5b | 0.017 | Old |
| 0.166194 | 0.049914 | 0.224 | 0.191 | 1 | 2 | H2-M3 | 0.033 | Old |
| 0.166379 | 0.212891 | 0.303 | 0.274 | 1 | 2 | Fmnl1 | 0.029 | Old |
| 0.166701 | 0.358033 | 0.017 | 0.01 | 1 | 2 | Haus1 | 0.007 | Old |
| 0.167128 | 0.726364 | 0.017 | 0.01 | 1 | 2 | 4632404H1 | 0.007 | Old |
| 0.167267 | 0.052628 | 0.155 | 0.129 | 1 | 2 | Dennd4c | 0.026 | Old |
| 0.167273 | 0.223741 | 0.121 | 0.1 | 1 | 2 | Cox16 | 0.021 | Old |
| 0.167457 | 0.603744 | 0.014 | 0.007 | 1 | 2 | Gm8369 | 0.007 | Old |
| 0.167841 | 0.37181 | 0.017 | 0.01 | 1 | 2 | Ptar1 | 0.007 | Old |
| 0.167952 | 0.029646 | 0.063 | 0.048 | 1 | 2 | Sec24d | 0.015 | Old |
| 0.168098 | 0.30843 | 0.014 | 0.007 | 1 | 2 | Fam234b | 0.007 | Old |
| 0.168484 | 0.729605 | 0.046 | 0.033 | 1 | 2 | Gnptg | 0.013 | Old |
| 0.16854 | 0.106938 | 0.357 | 0.321 | 1 | 2 | Tmco1 | 0.036 | Old |
| 0.16858 | 0.436394 | 0.014 | 0.007 | 1 | 2 | 2810405F1 | 0.007 | Old |
| 0.168593 | 0.111329 | 0.468 | 0.429 | 1 | 2 | Rpn2 | 0.039 | Old |
| 0.169409 | 0.361154 | 0.032 | 0.021 | 1 | 2 | Ankmy2 | 0.011 | Old |
| 0.169446 | 0.379819 | 0.133 | 0.112 | 1 | 2 | Necap1 | 0.021 | Old |
| 0.169459 | 0.84006 | 0.042 | 0.03 | 1 | 2 | Cog1 | 0.012 | Old |
| 0.169967 | 0.630697 | 0.025 | 0.015 | 1 | 2 | Fmn1 | 0.01 | Old |
| 0.169989 | 0.607193 | 0.037 | 0.026 | 1 | 2 | Clcn6 | 0.011 | Old |
| 0.170143 | 0.23332 | 0.146 | 0.122 | 1 | 2 | Ccndbp1 | 0.024 | Old |
| 0.170301 | 0.2578 | 0.11 | 0.09 | 1 | 2 | Cramp1l | 0.02 | Old |
| 0.170696 | 0.178483 | 0.359 | 0.328 | 1 | 2 | Aamp | 0.031 | Old |
| 0.170853 | 0.273934 | 0.185 | 0.162 | 1 | 2 | Hfe | 0.023 | Old |
| 0.170881 | 0.450342 | 0.051 | 0.038 | 1 | 2 | Amd1 | 0.013 | Old |

| | | | | | | | | |
|---|---|---|---|---|---|---|---|---|
| 0.170933 | 0.442428 | 0.087 | 0.069 | 1 | 2 | Exoc1 | 0.018 | Old |
| 0.171025 | 0.235704 | 0.06 | 0.045 | 1 | 2 | Cdc42ep2 | 0.015 | Old |
| 0.17139 | 0.096238 | 0.336 | 0.297 | 1 | 2 | Nsf | 0.039 | Old |
| 0.171628 | 0.31104 | 0.258 | 0.233 | 1 | 2 | Casp8 | 0.025 | Old |
| 0.1721 | 0.467054 | 0.139 | 0.119 | 1 | 2 | Lyl1 | 0.02 | Old |
| 0.172462 | 0.175988 | 0.025 | 0.015 | 1 | 2 | Elmod3 | 0.01 | Old |
| 0.172622 | 0.271335 | 0.084 | 0.065 | 1 | 2 | Hpse | 0.019 | Old |
| 0.172901 | 0.19089 | 0.254 | 0.227 | 1 | 2 | Klf7 | 0.027 | Old |
| 0.172986 | 0.275755 | 0.05 | 0.037 | 1 | 2 | Pex14 | 0.013 | Old |
| 0.173732 | 1.622741 | 0.026 | 0.017 | 1 | 2 | Eno3 | 0.009 | Old |
| 0.173785 | 0.069597 | 0.179 | 0.151 | 1 | 2 | Ctcf | 0.028 | Old |
| 0.173898 | 0.003321 | 0.374 | 0.338 | 1 | 2 | Mbnl2 | 0.036 | Old |
| 0.173974 | 0.670116 | 0.058 | 0.044 | 1 | 2 | Aasdhppt | 0.014 | Old |
| 0.17413 | 0.215149 | 0.163 | 0.138 | 1 | 2 | Hs6st1 | 0.025 | Old |
| 0.174401 | 0.191343 | 0.076 | 0.059 | 1 | 2 | Brip1os | 0.017 | Old |
| 0.174905 | 0.727333 | 0.037 | 0.026 | 1 | 2 | Ublcp1 | 0.011 | Old |
| 0.174973 | 0.277699 | 0.056 | 0.042 | 1 | 2 | Donson | 0.014 | Old |
| 0.174983 | 0.525751 | 0.025 | 0.015 | 1 | 2 | Inpp5e | 0.01 | Old |
| 0.17516 | 0.403598 | 0.062 | 0.048 | 1 | 2 | Crbn | 0.014 | Old |
| 0.175193 | 0.788005 | 0.082 | 0.067 | 1 | 2 | Tusc1 | 0.015 | Old |
| 0.175467 | 0.460946 | 0.139 | 0.119 | 1 | 2 | Aip | 0.02 | Old |
| 0.17561 | 0.145058 | 0.148 | 0.124 | 1 | 2 | Ikbke | 0.024 | Old |
| 0.175846 | 0.322546 | 0.096 | 0.078 | 1 | 2 | Vps53 | 0.018 | Old |
| 0.175949 | 0.451633 | 0.029 | 0.019 | 1 | 2 | Dgkq | 0.01 | Old |
| 0.176277 | 0.225021 | 0.091 | 0.072 | 1 | 2 | Btbd9 | 0.019 | Old |
| 0.17639 | 0.218046 | 0.206 | 0.179 | 1 | 2 | Vps26a | 0.027 | Old |
| 0.176417 | 0.249212 | 0.071 | 0.055 | 1 | 2 | Ammecr1 | 0.016 | Old |
| 0.176427 | 0.26678 | 0.046 | 0.033 | 1 | 2 | Fdxr | 0.013 | Old |
| 0.176575 | 0.287521 | 0.268 | 0.242 | 1 | 2 | Atf7ip | 0.026 | Old |
| 0.176722 | 0.130898 | 0.407 | 0.365 | 1 | 2 | Arap1 | 0.042 | Old |
| 0.176821 | 0.341318 | 0.046 | 0.033 | 1 | 2 | Ptcd2 | 0.013 | Old |
| 0.176884 | 0.612904 | 0.042 | 0.03 | 1 | 2 | Rnaset2b | 0.012 | Old |
| 0.176962 | 0.481996 | 0.051 | 0.038 | 1 | 2 | Msantd4 | 0.013 | Old |
| 0.177066 | 0.281314 | 0.124 | 0.103 | 1 | 2 | Slc25a45 | 0.021 | Old |
| 0.17709 | 0.03161 | 0.178 | 0.15 | 1 | 2 | Mrps24 | 0.028 | Old |
| 0.177269 | 0.616536 | 0.033 | 0.023 | 1 | 2 | Tcea3 | 0.01 | Old |
| 0.177588 | 0.256416 | 0.131 | 0.11 | 1 | 2 | Cenpa | 0.021 | Old |
| 0.177938 | 0.299001 | 0.268 | 0.241 | 1 | 2 | Gpr35 | 0.027 | Old |
| 0.177946 | 0.336392 | 0.095 | 0.077 | 1 | 2 | Selenoi | 0.018 | Old |
| 0.178355 | 0.547496 | 0.05 | 0.037 | 1 | 2 | Ipcef1 | 0.013 | Old |
| 0.178425 | 0.0752 | 0.122 | 0.1 | 1 | 2 | Slc35b1 | 0.022 | Old |
| 0.178482 | 0.029793 | 0.064 | 0.049 | 1 | 2 | Smcr8 | 0.015 | Old |
| 0.1786 | 0.420676 | 0.149 | 0.127 | 1 | 2 | Gramd4 | 0.022 | Old |
| 0.178785 | 0.045738 | 0.145 | 0.12 | 1 | 2 | Cyb5r3 | 0.025 | Old |
| 0.178937 | 0.118886 | 0.033 | 0.023 | 1 | 2 | Usp18 | 0.01 | Old |

| | | | | | | | | |
|---|---|---|---|---|---|---|---|---|
| 0.179137 | 0.285739 | 0.061 | 0.046 | 1 | 2 | Spg20 | 0.015 | Old |
| 0.179366 | 0.466413 | 0.096 | 0.078 | 1 | 2 | Kpna3 | 0.018 | Old |
| 0.179548 | 0.611917 | 0.021 | 0.013 | 1 | 2 | Fam131a | 0.008 | Old |
| 0.179713 | 0.061967 | 0.221 | 0.19 | 1 | 2 | Ptpn11 | 0.031 | Old |
| 0.179788 | 0.103076 | 0.79 | 0.754 | 1 | 2 | Anxa5 | 0.036 | Old |
| 0.180515 | 0.188858 | 0.131 | 0.109 | 1 | 2 | Bccip | 0.022 | Old |
| 0.18054 | 0.234408 | 0.072 | 0.056 | 1 | 2 | Smad3 | 0.016 | Old |
| 0.180629 | 0.284111 | 0.089 | 0.071 | 1 | 2 | Otud6b | 0.018 | Old |
| 0.180929 | 0.042111 | 0.215 | 0.185 | 1 | 2 | Crlf2 | 0.03 | Old |
| 0.181071 | 0.064552 | 0.08 | 0.063 | 1 | 2 | Nom1 | 0.017 | Old |
| 0.18126 | 0.261808 | 0.115 | 0.095 | 1 | 2 | Tnpo2 | 0.02 | Old |
| 0.181264 | 0.211844 | 0.106 | 0.087 | 1 | 2 | Spin1 | 0.019 | Old |
| 0.181543 | 0.521262 | 0.021 | 0.013 | 1 | 2 | Tamm41 | 0.008 | Old |
| 0.181644 | 0.230354 | 0.116 | 0.095 | 1 | 2 | Exoc2 | 0.021 | Old |
| 0.181811 | 0.284343 | 0.427 | 0.399 | 1 | 2 | Lbh | 0.028 | Old |
| 0.181857 | 0.05951 | 0.042 | 0.03 | 1 | 2 | Mettl5 | 0.012 | Old |
| 0.182154 | 0.083447 | 0.253 | 0.221 | 1 | 2 | Agps | 0.032 | Old |
| 0.182179 | 0.095317 | 0.411 | 0.385 | 1 | 2 | Ankrd11 | 0.026 | Old |
| 0.182343 | 0.289506 | 0.117 | 0.097 | 1 | 2 | Tbk1 | 0.02 | Old |
| 0.182613 | 0.484472 | 0.021 | 0.013 | 1 | 2 | Smpd2 | 0.008 | Old |
| 0.182634 | 0.142357 | 0.455 | 0.419 | 1 | 2 | Dync1h1 | 0.036 | Old |
| 0.182881 | 0.178061 | 0.281 | 0.248 | 1 | 2 | Pfkp | 0.033 | Old |
| 0.182965 | 0.132084 | 0.076 | 0.059 | 1 | 2 | Pctp | 0.017 | Old |
| 0.182967 | 0.143532 | 0.077 | 0.061 | 1 | 2 | Isg15 | 0.016 | Old |
| 0.183126 | 0.416453 | 0.124 | 0.105 | 1 | 2 | Dap3 | 0.019 | Old |
| 0.18315 | 0.432896 | 0.021 | 0.013 | 1 | 2 | Gm20300 | 0.008 | Old |
| 0.183426 | 0.097386 | 0.611 | 0.594 | 1 | 2 | Rassf4 | 0.017 | Old |
| 0.183444 | 0.019776 | 0.16 | 0.133 | 1 | 2 | Ube2j1 | 0.027 | Old |
| 0.183446 | 0.099766 | 0.19 | 0.162 | 1 | 2 | Etv6 | 0.028 | Old |
| 0.183503 | 0.23314 | 0.105 | 0.086 | 1 | 2 | Sash3 | 0.019 | Old |
| 0.183523 | 0.450103 | 0.043 | 0.031 | 1 | 2 | Stn1 | 0.012 | Old |
| 0.183759 | 0.121901 | 0.239 | 0.211 | 1 | 2 | Zfp638 | 0.028 | Old |
| 0.183773 | 0.22794 | 0.061 | 0.046 | 1 | 2 | Lonp1 | 0.015 | Old |
| 0.18414 | 0.232125 | 0.089 | 0.071 | 1 | 2 | Bad | 0.018 | Old |
| 0.184205 | 0.473676 | 0.08 | 0.064 | 1 | 2 | Actr1b | 0.016 | Old |
| 0.184241 | 0.238882 | 0.061 | 0.046 | 1 | 2 | Abcf3 | 0.015 | Old |
| 0.18446 | 0.025677 | 0.277 | 0.243 | 1 | 2 | Ccnt2 | 0.034 | Old |
| 0.185057 | 0.442241 | 0.08 | 0.064 | 1 | 2 | Apba1 | 0.016 | Old |
| 0.185269 | 0.960535 | 0.018 | 0.011 | 1 | 2 | Gm26982 | 0.007 | Old |
| 0.185354 | 0.886987 | 0.026 | 0.017 | 1 | 2 | Parvg | 0.009 | Old |
| 0.185407 | 0.087088 | 0.216 | 0.189 | 1 | 2 | Rrp1 | 0.027 | Old |
| 0.185416 | 0.301544 | 0.046 | 0.033 | 1 | 2 | Wrap73 | 0.013 | Old |
| 0.185469 | 0.195481 | 0.107 | 0.088 | 1 | 2 | Pds5b | 0.019 | Old |
| 0.185642 | 0.20135 | 0.182 | 0.158 | 1 | 2 | Mon2 | 0.024 | Old |
| 0.185683 | 0.40204 | 0.105 | 0.087 | 1 | 2 | Hccs | 0.018 | Old |

| | | | | | | | | |
|---|---|---|---|---|---|---|---|---|
| 0.185764 | 0.475754 | 0.104 | 0.086 | 1 | 2 | Il11ra1 | 0.018 | Old |
| 0.185837 | 0.149939 | 0.426 | 0.394 | 1 | 2 | Zyx | 0.032 | Old |
| 0.18594 | 0.624773 | 0.039 | 0.027 | 1 | 2 | Ric8b | 0.012 | Old |
| 0.186027 | 0.306807 | 0.063 | 0.049 | 1 | 2 | Sirt1 | 0.014 | Old |
| 0.186172 | 0.286383 | 0.073 | 0.057 | 1 | 2 | Pkd1 | 0.016 | Old |
| 0.186348 | 0.239027 | 0.361 | 0.336 | 1 | 2 | Notch2 | 0.025 | Old |
| 0.187657 | 0.613924 | 0.03 | 0.02 | 1 | 2 | Lrrc14 | 0.01 | Old |
| 0.187705 | 0.26685 | 0.156 | 0.133 | 1 | 2 | Yipf4 | 0.023 | Old |
| 0.187731 | 0.13901 | 0.169 | 0.145 | 1 | 2 | Rnpepl1 | 0.024 | Old |
| 0.187794 | 0.867297 | 0.018 | 0.011 | 1 | 2 | Cyb5d2 | 0.007 | Old |
| 0.188021 | 0.434457 | 0.113 | 0.095 | 1 | 2 | Smc5 | 0.018 | Old |
| 0.188494 | 0.187239 | 0.325 | 0.296 | 1 | 2 | Map3k1 | 0.029 | Old |
| 0.188678 | 0.112547 | 0.282 | 0.252 | 1 | 2 | Ifi203 | 0.03 | Old |
| 0.188766 | 0.322628 | 0.018 | 0.011 | 1 | 2 | Sigirr | 0.007 | Old |
| 0.188931 | 0.201649 | 0.047 | 0.034 | 1 | 2 | Ppp1r13b | 0.013 | Old |
| 0.188959 | 0.113545 | 0.037 | 0.026 | 1 | 2 | Scarf1 | 0.011 | Old |
| 0.189052 | 0.198948 | 0.411 | 0.385 | 1 | 2 | Ankrd44 | 0.026 | Old |
| 0.189322 | 0.196269 | 0.103 | 0.084 | 1 | 2 | Ltn1 | 0.019 | Old |
| 0.189457 | 0.111228 | 0.097 | 0.078 | 1 | 2 | Capn7 | 0.019 | Old |
| 0.189471 | 0.098426 | 0.082 | 0.065 | 1 | 2 | Nelfcd | 0.017 | Old |
| 0.189859 | 0.235211 | 0.175 | 0.152 | 1 | 2 | Tbc1d23 | 0.023 | Old |
| 0.190088 | 1.622801 | 0.015 | 0.008 | 1 | 2 | 4833407H1 | 0.007 | Old |
| 0.190309 | 0.174634 | 0.093 | 0.075 | 1 | 2 | Kri1 | 0.018 | Old |
| 0.19059 | 0.974412 | 0.015 | 0.008 | 1 | 2 | Gdpd1 | 0.007 | Old |
| 0.190746 | 0.058002 | 0.23 | 0.201 | 1 | 2 | Cyb5r4 | 0.029 | Old |
| 0.190811 | 0.266765 | 0.026 | 0.017 | 1 | 2 | Tbc1d22b | 0.009 | Old |
| 0.190835 | 0.27069 | 0.052 | 0.039 | 1 | 2 | B3gnt8 | 0.013 | Old |
| 0.191158 | 0.156114 | 0.219 | 0.191 | 1 | 2 | Smim15 | 0.028 | Old |
| 0.191402 | 0.017097 | 0.018 | 0.011 | 1 | 2 | Tepsin | 0.007 | Old |
| 0.191626 | 0.25687 | 0.094 | 0.076 | 1 | 2 | Ttpal | 0.018 | Old |
| 0.191848 | 0.132239 | 0.195 | 0.169 | 1 | 2 | Arid5a | 0.026 | Old |
| 0.191857 | 0.215883 | 0.018 | 0.011 | 1 | 2 | Bcdin3d | 0.007 | Old |
| 0.191933 | 0.188208 | 0.034 | 0.024 | 1 | 2 | Gadd45a | 0.01 | Old |
| 0.192183 | 0.143095 | 0.175 | 0.15 | 1 | 2 | Rbm42 | 0.025 | Old |
| 0.192276 | 0.101715 | 0.094 | 0.076 | 1 | 2 | Reps1 | 0.018 | Old |
| 0.192496 | 0.317136 | 0.069 | 0.053 | 1 | 2 | Borcs8 | 0.016 | Old |
| 0.192509 | 0.13302 | 0.324 | 0.291 | 1 | 2 | Mkrn1 | 0.033 | Old |
| 0.192513 | 0.719086 | 0.035 | 0.025 | 1 | 2 | Flywch1 | 0.01 | Old |
| 0.193053 | 0.362082 | 0.13 | 0.109 | 1 | 2 | Rras | 0.021 | Old |
| 0.193073 | 0.488723 | 0.03 | 0.02 | 1 | 2 | Cep68 | 0.01 | Old |
| 0.193106 | 0.40371 | 0.112 | 0.094 | 1 | 2 | Acadvl | 0.018 | Old |
| 0.193367 | 0.822138 | 0.04 | 0.029 | 1 | 2 | Sh2b1 | 0.011 | Old |
| 0.193559 | 0.318123 | 0.034 | 0.024 | 1 | 2 | Ftsj1 | 0.01 | Old |
| 0.193595 | 0.741639 | 0.012 | 0.006 | 1 | 2 | Gm13479 | 0.006 | Old |
| 0.193638 | 0.070563 | 0.166 | 0.141 | 1 | 2 | Tmem128 | 0.025 | Old |

| | | | | | | | | |
|---|---|---|---|---|---|---|---|---|
| 0.193983 | 0.540607 | 0.012 | 0.006 | 1 | 2 | Gm5431 | 0.006 | Old |
| 0.194324 | 0.129249 | 0.175 | 0.151 | 1 | 2 | Slc25a37 | 0.024 | Old |
| 0.194473 | 0.300439 | 0.026 | 0.017 | 1 | 2 | Katnal1 | 0.009 | Old |
| 0.1946 | 0.166453 | 0.026 | 0.017 | 1 | 2 | B3galt6 | 0.009 | Old |
| 0.194636 | 0.156137 | 0.293 | 0.265 | 1 | 2 | Crk | 0.028 | Old |
| 0.194727 | 0.215341 | 0.187 | 0.165 | 1 | 2 | Vamp2 | 0.022 | Old |
| 0.194807 | 0.194679 | 0.448 | 0.422 | 1 | 2 | Mcub | 0.026 | Old |
| 0.194975 | 0.154915 | 0.498 | 0.469 | 1 | 2 | Dusp3 | 0.029 | Old |
| 0.195209 | 0.237222 | 0.125 | 0.106 | 1 | 2 | Mrpl3 | 0.019 | Old |
| 0.195267 | 0.136528 | 0.428 | 0.392 | 1 | 2 | Mfsd1 | 0.036 | Old |
| 0.195347 | 0.348547 | 0.012 | 0.006 | 1 | 2 | Sirt5 | 0.006 | Old |
| 0.195347 | 0.228794 | 0.012 | 0.006 | 1 | 2 | AC163354. | 0.006 | Old |
| 0.19538 | 0.128675 | 0.067 | 0.052 | 1 | 2 | Txlng | 0.015 | Old |
| 0.195539 | 0.782193 | 0.04 | 0.029 | 1 | 2 | Hes1 | 0.011 | Old |
| 0.195965 | 0.098141 | 0.511 | 0.464 | 1 | 2 | Abi1 | 0.047 | Old |
| 0.195975 | 0.110518 | 0.101 | 0.082 | 1 | 2 | Gm13212 | 0.019 | Old |
| 0.196012 | 0.043131 | 0.015 | 0.008 | 1 | 2 | Pigb | 0.007 | Old |
| 0.196118 | 0.04635 | 0.03 | 0.02 | 1 | 2 | Scimp | 0.01 | Old |
| 0.196126 | 0.234194 | 0.208 | 0.184 | 1 | 2 | Mcfd2 | 0.024 | Old |
| 0.196355 | 0.408752 | 0.015 | 0.008 | 1 | 2 | Slc9a4 | 0.007 | Old |
| 0.196423 | 0.15957 | 0.142 | 0.121 | 1 | 2 | Cln6 | 0.021 | Old |
| 0.196473 | 0.311544 | 0.048 | 0.036 | 1 | 2 | Wdr7 | 0.012 | Old |
| 0.196574 | 0.280782 | 0.12 | 0.1 | 1 | 2 | Frmd4a | 0.02 | Old |
| 0.196691 | 0.096611 | 0.346 | 0.314 | 1 | 2 | Tnrc18 | 0.032 | Old |
| 0.196698 | 0.548373 | 0.015 | 0.008 | 1 | 2 | Asf1b | 0.007 | Old |
| 0.197355 | 0.154188 | 0.197 | 0.17 | 1 | 2 | Ndufaf8 | 0.027 | Old |
| 0.19742 | 0.414536 | 0.093 | 0.076 | 1 | 2 | Zfp608 | 0.017 | Old |
| 0.197502 | 0.311865 | 0.034 | 0.024 | 1 | 2 | Mthfd1 | 0.01 | Old |
| 0.19751 | 0.301733 | 0.136 | 0.115 | 1 | 2 | Ipmk | 0.021 | Old |
| 0.197512 | 0.040882 | 0.849 | 0.846 | 1 | 2 | Ctsc | 0.003 | Old |
| 0.197768 | 0.53008 | 0.06 | 0.046 | 1 | 2 | Pus1 | 0.014 | Old |
| 0.197847 | 0.144172 | 0.393 | 0.36 | 1 | 2 | Cast | 0.033 | Old |
| 0.198048 | 0.193655 | 0.175 | 0.15 | 1 | 2 | Safb | 0.025 | Old |
| 0.198526 | 0.026417 | 0.072 | 0.056 | 1 | 2 | Ap1m1 | 0.016 | Old |
| 0.198716 | 0.142527 | 0.412 | 0.378 | 1 | 2 | Eif5b | 0.034 | Old |
| 0.198969 | 0.737342 | 0.05 | 0.038 | 1 | 2 | Fbxo45 | 0.012 | Old |
| 0.199091 | 0.627619 | 0.022 | 0.014 | 1 | 2 | Wrap53 | 0.008 | Old |
| 0.199321 | 0.120067 | 0.09 | 0.072 | 1 | 2 | Fam118b | 0.018 | Old |
| 0.199526 | 0.189011 | 0.2 | 0.175 | 1 | 2 | R3hdm2 | 0.025 | Old |
| 0.199622 | 0.171228 | 0.052 | 0.039 | 1 | 2 | Gstp1 | 0.013 | Old |
| 0.19983 | 0.202865 | 0.078 | 0.062 | 1 | 2 | Sh3tc1 | 0.016 | Old |
| 0.199886 | 0.149784 | 0.202 | 0.175 | 1 | 2 | Tmem168 | 0.027 | Old |
| 0.199964 | 0.39678 | 0.133 | 0.114 | 1 | 2 | Mgst1 | 0.019 | Old |
| 0.200333 | 0.69604 | 0.031 | 0.021 | 1 | 2 | Trim23 | 0.01 | Old |
| 0.200341 | 0.496334 | 0.022 | 0.014 | 1 | 2 | B230354K1 | 0.008 | Old |

| | | | | | | | | |
|---|---|---|---|---|---|---|---|---|
| 0.200501 | 0.166984 | 0.163 | 0.14 | 1 | 2 | Cpeb4 | 0.023 | Old |
| 0.20056 | 0.227743 | 0.081 | 0.065 | 1 | 2 | Senp5 | 0.016 | Old |
| 0.200604 | 0.220382 | 0.048 | 0.036 | 1 | 2 | 4930503L1 | 0.012 | Old |
| 0.200845 | 0.144761 | 0.382 | 0.344 | 1 | 2 | Arhgap9 | 0.038 | Old |
| 0.201433 | 0.238871 | 0.04 | 0.029 | 1 | 2 | Pdhx | 0.011 | Old |
| 0.20156 | 0.285548 | 0.09 | 0.074 | 1 | 2 | Phrf1 | 0.016 | Old |
| 0.201772 | 0.186526 | 0.126 | 0.106 | 1 | 2 | Bicd2 | 0.02 | Old |
| 0.201923 | 0.106054 | 0.122 | 0.102 | 1 | 2 | Oxsr1 | 0.02 | Old |
| 0.202053 | 0.02563 | 0.223 | 0.192 | 1 | 2 | Cdk6 | 0.031 | Old |
| 0.202303 | 0.144287 | 0.214 | 0.186 | 1 | 2 | Csnk2a2 | 0.028 | Old |
| 0.202397 | 0.17206 | 0.342 | 0.316 | 1 | 2 | Cers2 | 0.026 | Old |
| 0.202412 | 0.454814 | 0.069 | 0.055 | 1 | 2 | Haghl | 0.014 | Old |
| 0.202946 | 0.361594 | 0.06 | 0.046 | 1 | 2 | Tirap | 0.014 | Old |
| 0.20328 | 0.059858 | 0.022 | 0.014 | 1 | 2 | Mars | 0.008 | Old |
| 0.203421 | 0.309538 | 0.022 | 0.014 | 1 | 2 | Fahd1 | 0.008 | Old |
| 0.203568 | 0.110937 | 0.134 | 0.113 | 1 | 2 | Naip2 | 0.021 | Old |
| 0.20371 | 0.149691 | 0.363 | 0.331 | 1 | 2 | Pigt | 0.032 | Old |
| 0.203912 | 0.422369 | 0.05 | 0.038 | 1 | 2 | Card9 | 0.012 | Old |
| 0.203955 | 0.414827 | 0.103 | 0.086 | 1 | 2 | Wdr43 | 0.017 | Old |
| 0.203987 | 0.571456 | 0.031 | 0.021 | 1 | 2 | Plpp5 | 0.01 | Old |
| 0.204014 | 0.067013 | 0.116 | 0.095 | 1 | 2 | Dpp3 | 0.021 | Old |
| 0.204028 | 0.158275 | 0.378 | 0.347 | 1 | 2 | Ap2m1 | 0.031 | Old |
| 0.204059 | 0.129603 | 0.192 | 0.167 | 1 | 2 | Zdhhc21 | 0.025 | Old |
| 0.204168 | 0.288829 | 0.19 | 0.167 | 1 | 2 | Dpep2 | 0.023 | Old |
| 0.204289 | 0.265343 | 0.211 | 0.186 | 1 | 2 | Nceh1 | 0.025 | Old |
| 0.204467 | 0.243308 | 0.204 | 0.183 | 1 | 2 | C3 | 0.021 | Old |
| 0.20453 | 0.605456 | 0.027 | 0.018 | 1 | 2 | Zfp971 | 0.009 | Old |
| 0.204977 | 0.384175 | 0.049 | 0.037 | 1 | 2 | Itpkc | 0.012 | Old |
| 0.204984 | 0.182202 | 0.338 | 0.303 | 1 | 2 | Mcrip1 | 0.035 | Old |
| 0.205156 | 0.146644 | 0.403 | 0.372 | 1 | 2 | Bsg | 0.031 | Old |
| 0.205607 | 0.254375 | 0.281 | 0.254 | 1 | 2 | Tnfaip2 | 0.027 | Old |
| 0.205658 | 0.215243 | 0.237 | 0.209 | 1 | 2 | Becn1 | 0.028 | Old |
| 0.206018 | 0.19284 | 0.054 | 0.04 | 1 | 2 | Pde12 | 0.014 | Old |
| 0.206057 | 0.113158 | 0.596 | 0.564 | 1 | 2 | Wdr1 | 0.032 | Old |
| 0.206207 | 0.191973 | 0.027 | 0.018 | 1 | 2 | Zscan21 | 0.009 | Old |
| 0.206826 | 0.164074 | 0.32 | 0.29 | 1 | 2 | Cnot1 | 0.03 | Old |
| 0.206868 | 0.393231 | 0.055 | 0.042 | 1 | 2 | Prrg2 | 0.013 | Old |
| 0.207244 | 0.035592 | 0.162 | 0.188 | 1 | 2 | Runx3 | -0.026 | Old |
| 0.20726 | 0.061724 | 0.706 | 0.679 | 1 | 2 | Mob1a | 0.027 | Old |
| 0.207445 | 0.268679 | 0.064 | 0.05 | 1 | 2 | Coasy | 0.014 | Old |
| 0.207633 | 0.110874 | 0.661 | 0.64 | 1 | 2 | Ifngr1 | 0.021 | Old |
| 0.207649 | 0.358437 | 0.096 | 0.08 | 1 | 2 | Ube2l6 | 0.016 | Old |
| 0.207681 | 0.125317 | 0.089 | 0.071 | 1 | 2 | Espl1 | 0.018 | Old |
| 0.207755 | 0.380031 | 0.082 | 0.067 | 1 | 2 | Nol8 | 0.015 | Old |
| 0.208099 | 0.236741 | 0.276 | 0.248 | 1 | 2 | Lair1 | 0.028 | Old |

| | | | | | | | | |
|---|---|---|---|---|---|---|---|---|
| 0.20901 | 0.102447 | 0.722 | 0.71 | 1 | 2 | Hnrnpa3 | 0.012 | Old |
| 0.209057 | 0.092282 | 0.266 | 0.236 | 1 | 2 | Usp15 | 0.03 | Old |
| 0.209282 | 0.084034 | 0.579 | 0.558 | 1 | 2 | Cops9 | 0.021 | Old |
| 0.209899 | 0.177124 | 0.064 | 0.05 | 1 | 2 | E2f1 | 0.014 | Old |
| 0.210032 | 0.274811 | 0.06 | 0.046 | 1 | 2 | Nqo2 | 0.014 | Old |
| 0.21039 | 0.53111 | 0.045 | 0.033 | 1 | 2 | 2300009A( | 0.012 | Old |
| 0.210778 | 0.112729 | 0.188 | 0.164 | 1 | 2 | Pptc7 | 0.024 | Old |
| 0.211622 | 0.053588 | 0.276 | 0.246 | 1 | 2 | Usp4 | 0.03 | Old |
| 0.211625 | 0.517207 | 0.019 | 0.012 | 1 | 2 | Nt5c3b | 0.007 | Old |
| 0.211825 | 0.511416 | 0.046 | 0.034 | 1 | 2 | Tbrg4 | 0.012 | Old |
| 0.211911 | 0.301427 | 0.049 | 0.037 | 1 | 2 | Wdyhv1 | 0.012 | Old |
| 0.211939 | 0.595608 | 0.019 | 0.012 | 1 | 2 | Rsad1 | 0.007 | Old |
| 0.212648 | 0.323094 | 0.162 | 0.14 | 1 | 2 | Plaa | 0.022 | Old |
| 0.213062 | 0.314217 | 0.096 | 0.08 | 1 | 2 | Hras | 0.016 | Old |
| 0.21355 | 0.125708 | 0.34 | 0.304 | 1 | 2 | Arf3 | 0.036 | Old |
| 0.213555 | 0.321129 | 0.036 | 0.026 | 1 | 2 | Ccdc28b | 0.01 | Old |
| 0.213643 | 0.248761 | 0.031 | 0.021 | 1 | 2 | Lyrm9 | 0.01 | Old |
| 0.213703 | 0.141978 | 0.378 | 0.352 | 1 | 2 | Pmaip1 | 0.026 | Old |
| 0.213753 | 0.084848 | 0.116 | 0.096 | 1 | 2 | 3830406C1 | 0.02 | Old |
| 0.213905 | 0.096337 | 0.777 | 0.766 | 1 | 2 | Eif4g2 | 0.011 | Old |
| 0.214081 | 0.106479 | 0.194 | 0.169 | 1 | 2 | St3gal1 | 0.025 | Old |
| 0.214148 | 0.528471 | 0.019 | 0.012 | 1 | 2 | Cd276 | 0.007 | Old |
| 0.214203 | 0.518325 | 0.042 | 0.031 | 1 | 2 | Rfng | 0.011 | Old |
| 0.214292 | 0.38614 | 0.041 | 0.03 | 1 | 2 | Ints8 | 0.011 | Old |
| 0.214292 | 0.385265 | 0.041 | 0.03 | 1 | 2 | Ints10 | 0.011 | Old |
| 0.214417 | 0.57649 | 0.046 | 0.034 | 1 | 2 | Asb3 | 0.012 | Old |
| 0.214674 | 0.135207 | 0.207 | 0.181 | 1 | 2 | Rbck1 | 0.026 | Old |
| 0.214684 | 0.351224 | 0.199 | 0.178 | 1 | 2 | Stx16 | 0.021 | Old |
| 0.214959 | 0.196642 | 0.122 | 0.102 | 1 | 2 | Dld | 0.02 | Old |
| 0.215615 | 0.096842 | 0.317 | 0.283 | 1 | 2 | Tmem14c | 0.034 | Old |
| 0.215717 | 0.688769 | 0.028 | 0.019 | 1 | 2 | Zfp719 | 0.009 | Old |
| 0.215763 | 0.067131 | 0.089 | 0.071 | 1 | 2 | Eif2b5 | 0.018 | Old |
| 0.215966 | 0.081456 | 0.285 | 0.255 | 1 | 2 | Fbxl5 | 0.03 | Old |
| 0.216014 | 0.182634 | 0.324 | 0.295 | 1 | 2 | Strn3 | 0.029 | Old |
| 0.216222 | 0.217924 | 0.178 | 0.154 | 1 | 2 | Rhoq | 0.024 | Old |
| 0.216374 | 0.267773 | 0.019 | 0.012 | 1 | 2 | Gm37305 | 0.007 | Old |
| 0.216627 | 0.21611 | 0.066 | 0.052 | 1 | 2 | Mlx | 0.014 | Old |
| 0.217156 | 0.49395 | 0.028 | 0.019 | 1 | 2 | Ipo4 | 0.009 | Old |
| 0.217264 | 0.120746 | 0.588 | 0.55 | 1 | 2 | Grb2 | 0.038 | Old |
| 0.217425 | 0.149975 | 0.222 | 0.197 | 1 | 2 | Coro1b | 0.025 | Old |
| 0.217436 | 0.309825 | 0.046 | 0.034 | 1 | 2 | Lmf1 | 0.012 | Old |
| 0.218186 | 0.083961 | 0.137 | 0.158 | 1 | 2 | Pmp22 | -0.021 | Old |
| 0.218295 | 0.512056 | 0.024 | 0.015 | 1 | 2 | Xrcc6 | 0.009 | Old |
| 0.218484 | 0.105057 | 0.231 | 0.203 | 1 | 2 | Uhmk1 | 0.028 | Old |
| 0.218508 | 0.031329 | 0.122 | 0.102 | 1 | 2 | Mrps23 | 0.02 | Old |

| | | | | | | | | |
|---|---|---|---|---|---|---|---|---|
| 0.218663 | 0.560923 | 0.032 | 0.023 | 1 | 2 | Btrc | 0.009 | Old |
| 0.218804 | 0.29749 | 0.296 | 0.274 | 1 | 2 | Dusp2 | 0.022 | Old |
| 0.218987 | 0.347681 | 0.089 | 0.072 | 1 | 2 | H2-Q6 | 0.017 | Old |
| 0.219209 | 0.035856 | 0.049 | 0.037 | 1 | 2 | Btg3 | 0.012 | Old |
| 0.219604 | 0.195859 | 0.251 | 0.228 | 1 | 2 | Ggh | 0.023 | Old |
| 0.219622 | 0.321811 | 0.091 | 0.075 | 1 | 2 | Nelfe | 0.016 | Old |
| 0.219899 | 0.222796 | 0.337 | 0.308 | 1 | 2 | Brd4 | 0.029 | Old |
| 0.220603 | 0.223459 | 0.079 | 0.064 | 1 | 2 | Abhd17c | 0.015 | Old |
| 0.220996 | 0.209601 | 0.032 | 0.023 | 1 | 2 | Rusc2 | 0.009 | Old |
| 0.221146 | 0.169452 | 0.467 | 0.445 | 1 | 2 | Ikbkb | 0.022 | Old |
| 0.221201 | 0.518297 | 0.046 | 0.034 | 1 | 2 | Med27 | 0.012 | Old |
| 0.221282 | 0.226306 | 0.162 | 0.139 | 1 | 2 | Acaa2 | 0.023 | Old |
| 0.221798 | 0.036008 | 0.207 | 0.181 | 1 | 2 | Hnrnpll | 0.026 | Old |
| 0.221867 | 0.06376 | 0.416 | 0.378 | 1 | 2 | Eif4h | 0.038 | Old |
| 0.221895 | 0.029308 | 0.206 | 0.177 | 1 | 2 | Zzef1 | 0.029 | Old |
| 0.221915 | 0.534057 | 0.028 | 0.019 | 1 | 2 | Mtg2 | 0.009 | Old |
| 0.222092 | 0.011781 | 0.29 | 0.258 | 1 | 2 | Msr1 | 0.032 | Old |
| 0.222243 | 0.499191 | 0.016 | 0.01 | 1 | 2 | Cep19 | 0.006 | Old |
| 0.22226 | 0.492672 | 0.063 | 0.05 | 1 | 2 | Ctps2 | 0.013 | Old |
| 0.222422 | 0.667338 | 0.016 | 0.01 | 1 | 2 | Snapc4 | 0.006 | Old |
| 0.222567 | 0.357289 | 0.073 | 0.058 | 1 | 2 | Cbll1 | 0.015 | Old |
| 0.222677 | 0.505136 | 0.108 | 0.091 | 1 | 2 | Cog5 | 0.017 | Old |
| 0.222781 | 0.857887 | 0.016 | 0.01 | 1 | 2 | Rab3ip | 0.006 | Old |
| 0.22279 | 0.146121 | 0.232 | 0.209 | 1 | 2 | Smim7 | 0.023 | Old |
| 0.222824 | 0.267431 | 0.084 | 0.068 | 1 | 2 | Cyth2 | 0.016 | Old |
| 0.223126 | 0.063117 | 0.349 | 0.316 | 1 | 2 | Cerk | 0.033 | Old |
| 0.223141 | 0.490837 | 0.016 | 0.01 | 1 | 2 | Tefm | 0.006 | Old |
| 0.223142 | 0.061267 | 0.405 | 0.373 | 1 | 2 | Hmgb2 | 0.032 | Old |
| 0.22325 | 0.332103 | 0.028 | 0.019 | 1 | 2 | Clasrp | 0.009 | Old |
| 0.22342 | 0.435458 | 0.183 | 0.164 | 1 | 2 | Ufsp2 | 0.019 | Old |
| 0.223471 | 0.291065 | 0.057 | 0.044 | 1 | 2 | Nol4l | 0.013 | Old |
| 0.2235 | 0.620234 | 0.016 | 0.01 | 1 | 2 | Itfg2 | 0.006 | Old |
| 0.223631 | 0.161211 | 0.078 | 0.063 | 1 | 2 | Hikeshi | 0.015 | Old |
| 0.223744 | 0.107291 | 0.105 | 0.087 | 1 | 2 | Mms19 | 0.018 | Old |
| 0.223783 | 0.148561 | 0.051 | 0.065 | 1 | 2 | Dclre1b | -0.014 | Old |
| 0.224012 | 0.079122 | 0.125 | 0.105 | 1 | 2 | Mprip | 0.02 | Old |
| 0.224254 | 0.390862 | 0.024 | 0.015 | 1 | 2 | Fpr1 | 0.009 | Old |
| 0.22438 | 0.027993 | 0.102 | 0.083 | 1 | 2 | Ddx39 | 0.019 | Old |
| 0.224384 | 0.12576 | 0.071 | 0.056 | 1 | 2 | Alg5 | 0.015 | Old |
| 0.224554 | 0.48782 | 0.059 | 0.046 | 1 | 2 | Elmod2 | 0.013 | Old |
| 0.224572 | 0.121885 | 0.077 | 0.062 | 1 | 2 | Slc17a5 | 0.015 | Old |
| 0.224582 | 0.21677 | 0.016 | 0.01 | 1 | 2 | Nme6 | 0.006 | Old |
| 0.22462 | 0.499948 | 0.037 | 0.027 | 1 | 2 | Gss | 0.01 | Old |
| 0.22488 | 0.270212 | 0.099 | 0.082 | 1 | 2 | Eef1e1 | 0.017 | Old |
| 0.224977 | 0.17984 | 0.072 | 0.057 | 1 | 2 | Syne3 | 0.015 | Old |

| | | | | | | | | |
|---|---|---|---|---|---|---|---|---|
| 0.225124 | 0.052667 | 0.016 | 0.01 | 1 | 2 | Zswim1 | 0.006 | Old |
| 0.225319 | 0.252894 | 0.13 | 0.109 | 1 | 2 | Ngdn | 0.021 | Old |
| 0.225564 | 0.210747 | 0.073 | 0.058 | 1 | 2 | Fbxo22 | 0.015 | Old |
| 0.225586 | 0.158854 | 0.132 | 0.112 | 1 | 2 | Unc50 | 0.02 | Old |
| 0.225685 | 0.238132 | 0.145 | 0.125 | 1 | 2 | Tdrd7 | 0.02 | Old |
| 0.225839 | 0.152707 | 0.032 | 0.023 | 1 | 2 | Bloc1s1 | 0.009 | Old |
| 0.225855 | 0.022981 | 0.077 | 0.062 | 1 | 2 | Lsm14b | 0.015 | Old |
| 0.226025 | 0.243033 | 0.067 | 0.053 | 1 | 2 | Snx14 | 0.014 | Old |
| 0.226211 | 0.036934 | 0.036 | 0.026 | 1 | 2 | Elovl6 | 0.01 | Old |
| 0.226246 | 0.231954 | 0.116 | 0.097 | 1 | 2 | Stard4 | 0.019 | Old |
| 0.226619 | 1.01998 | 0.01 | 0.005 | 1 | 2 | Rcn1 | 0.005 | Old |
| 0.226619 | 0.942816 | 0.01 | 0.005 | 1 | 2 | Cep152 | 0.005 | Old |
| 0.226619 | 0.727004 | 0.01 | 0.005 | 1 | 2 | Vwf | 0.005 | Old |
| 0.226748 | 0.292871 | 0.123 | 0.105 | 1 | 2 | Slc25a20 | 0.018 | Old |
| 0.226897 | 0.259794 | 0.069 | 0.055 | 1 | 2 | Mtmr2 | 0.014 | Old |
| 0.226906 | 0.203337 | 0.14 | 0.121 | 1 | 2 | Slf2 | 0.019 | Old |
| 0.227039 | 0.18385 | 0.287 | 0.261 | 1 | 2 | Derl1 | 0.026 | Old |
| 0.22705 | 0.239143 | 0.117 | 0.099 | 1 | 2 | Kcnab2 | 0.018 | Old |
| 0.227082 | 0.283955 | 0.042 | 0.031 | 1 | 2 | Lrrc20 | 0.011 | Old |
| 0.227102 | 1.06806 | 0.01 | 0.005 | 1 | 2 | Tedc2 | 0.005 | Old |
| 0.227467 | 0.311354 | 0.073 | 0.058 | 1 | 2 | Itga1 | 0.015 | Old |
| 0.227722 | 0.289028 | 0.032 | 0.023 | 1 | 2 | C330018D2 | 0.009 | Old |
| 0.227827 | 0.951323 | 0.01 | 0.005 | 1 | 2 | Trim68 | 0.005 | Old |
| 0.228311 | 0.681477 | 0.01 | 0.005 | 1 | 2 | Lzts2 | 0.005 | Old |
| 0.228311 | 0.222794 | 0.01 | 0.005 | 1 | 2 | Pcbp4 | 0.005 | Old |
| 0.228411 | 0.243382 | 0.09 | 0.074 | 1 | 2 | Commd2 | 0.016 | Old |
| 0.228554 | 0.6451 | 0.01 | 0.005 | 1 | 2 | Cd5l | 0.005 | Old |
| 0.228648 | 0.116932 | 0.028 | 0.019 | 1 | 2 | Ccdc122 | 0.009 | Old |
| 0.228698 | 0.097813 | 0.349 | 0.318 | 1 | 2 | Hspa9 | 0.031 | Old |
| 0.228785 | 1.241546 | 0.02 | 0.013 | 1 | 2 | Zfp760 | 0.007 | Old |
| 0.228936 | 0.059525 | 0.409 | 0.367 | 1 | 2 | Rab11b | 0.042 | Old |
| 0.229039 | 0.567009 | 0.01 | 0.005 | 1 | 2 | Pkp3 | 0.005 | Old |
| 0.22913 | 0.158592 | 0.016 | 0.01 | 1 | 2 | Them4 | 0.006 | Old |
| 0.229234 | 0.260699 | 0.137 | 0.116 | 1 | 2 | Tmem65 | 0.021 | Old |
| 0.229282 | 0.689465 | 0.01 | 0.005 | 1 | 2 | Apold1 | 0.005 | Old |
| 0.229498 | 0.207713 | 0.251 | 0.226 | 1 | 2 | Ccnl2 | 0.025 | Old |
| 0.229526 | 0.517252 | 0.01 | 0.005 | 1 | 2 | Ptpdc1 | 0.005 | Old |
| 0.229755 | 0.089605 | 0.123 | 0.103 | 1 | 2 | Mettl23 | 0.02 | Old |
| 0.229818 | 0.095391 | 0.071 | 0.056 | 1 | 2 | Pik3r4 | 0.015 | Old |
| 0.229898 | 0.078297 | 0.411 | 0.379 | 1 | 2 | Fam111a | 0.032 | Old |
| 0.229994 | 0.13054 | 0.107 | 0.089 | 1 | 2 | Uxs1 | 0.018 | Old |
| 0.23017 | 0.654728 | 0.013 | 0.007 | 1 | 2 | Eya4 | 0.006 | Old |
| 0.230233 | 1.075404 | 0.02 | 0.013 | 1 | 2 | Rem1 | 0.007 | Old |
| 0.230256 | 0.039432 | 0.01 | 0.005 | 1 | 2 | Il4i1 | 0.005 | Old |
| 0.230377 | 0.450372 | 0.013 | 0.007 | 1 | 2 | Lrrc75a | 0.006 | Old |

| | | | | | | | | |
|---|---|---|---|---|---|---|---|---|
| 0.230518 | 0.099325 | 0.353 | 0.331 | 1 | 2 | Atp6v1a | 0.022 | Old |
| 0.23058 | 0.052809 | 0.136 | 0.114 | 1 | 2 | Naga | 0.022 | Old |
| 0.231067 | 0.027722 | 0.047 | 0.036 | 1 | 2 | B430306N( | 0.011 | Old |
| 0.231383 | 0.227834 | 0.118 | 0.1 | 1 | 2 | Zc3h4 | 0.018 | Old |
| 0.231417 | 0.498814 | 0.013 | 0.007 | 1 | 2 | Tmem26 | 0.006 | Old |
| 0.231577 | 0.263877 | 0.082 | 0.067 | 1 | 2 | Nipsnap2 | 0.015 | Old |
| 0.231625 | 0.603807 | 0.013 | 0.007 | 1 | 2 | Mylk | 0.006 | Old |
| 0.232673 | 0.096538 | 0.109 | 0.091 | 1 | 2 | Ugdh | 0.018 | Old |
| 0.233172 | 0.138926 | 0.111 | 0.093 | 1 | 2 | Uba5 | 0.018 | Old |
| 0.233578 | 0.054255 | 0.106 | 0.088 | 1 | 2 | Cask | 0.018 | Old |
| 0.23362 | 0.100108 | 0.067 | 0.053 | 1 | 2 | Map2k5 | 0.014 | Old |
| 0.233801 | 0.439505 | 0.02 | 0.013 | 1 | 2 | Amer1 | 0.007 | Old |
| 0.234297 | 0.117673 | 0.088 | 0.072 | 1 | 2 | Aldh3a2 | 0.016 | Old |
| 0.234413 | 0.263638 | 0.193 | 0.171 | 1 | 2 | Rgs18 | 0.022 | Old |
| 0.234464 | 0.227713 | 0.128 | 0.11 | 1 | 2 | Rfk | 0.018 | Old |
| 0.234564 | 0.109152 | 0.424 | 0.399 | 1 | 2 | Lcp2 | 0.025 | Old |
| 0.234637 | 0.120015 | 0.293 | 0.267 | 1 | 2 | Arf2 | 0.026 | Old |
| 0.234729 | 0.049398 | 0.066 | 0.052 | 1 | 2 | Vps26b | 0.014 | Old |
| 0.234814 | 0.258521 | 0.086 | 0.07 | 1 | 2 | Cep192 | 0.016 | Old |
| 0.234871 | 1.283242 | 0.025 | 0.017 | 1 | 2 | Lilra5 | 0.008 | Old |
| 0.234961 | 0.187863 | 0.184 | 0.163 | 1 | 2 | Nfatc3 | 0.021 | Old |
| 0.235271 | 0.601079 | 0.02 | 0.013 | 1 | 2 | Tmco4 | 0.007 | Old |
| 0.235599 | 0.444263 | 0.02 | 0.013 | 1 | 2 | Srgap3 | 0.007 | Old |
| 0.235612 | 0.140475 | 0.226 | 0.202 | 1 | 2 | Uqcrc2 | 0.024 | Old |
| 0.235653 | 0.056208 | 0.094 | 0.077 | 1 | 2 | Grpel1 | 0.017 | Old |
| 0.235895 | 0.089162 | 0.061 | 0.048 | 1 | 2 | Nup205 | 0.013 | Old |
| 0.235933 | 0.219133 | 0.138 | 0.12 | 1 | 2 | Plekhm2 | 0.018 | Old |
| 0.236034 | 0.284375 | 0.061 | 0.048 | 1 | 2 | Slco3a1 | 0.013 | Old |
| 0.236255 | 0.690329 | 0.02 | 0.013 | 1 | 2 | Macrod1 | 0.007 | Old |
| 0.237161 | 0.078783 | 0.518 | 0.49 | 1 | 2 | Litaf | 0.028 | Old |
| 0.237184 | 0.261303 | 0.058 | 0.045 | 1 | 2 | Cd82 | 0.013 | Old |
| 0.237369 | 0.149023 | 0.59 | 0.581 | 1 | 2 | Pdia6 | 0.009 | Old |
| 0.237519 | 0.627446 | 0.065 | 0.052 | 1 | 2 | Gpkow | 0.013 | Old |
| 0.237737 | 0.303851 | 0.02 | 0.013 | 1 | 2 | Zfp953 | 0.007 | Old |
| 0.237737 | 0.199401 | 0.02 | 0.013 | 1 | 2 | Kcne3 | 0.007 | Old |
| 0.238009 | 0.370062 | 0.133 | 0.115 | 1 | 2 | Dgat1 | 0.018 | Old |
| 0.238068 | 0.094554 | 0.033 | 0.024 | 1 | 2 | Kctd3 | 0.009 | Old |
| 0.238108 | 0.454072 | 0.102 | 0.086 | 1 | 2 | Washc1 | 0.016 | Old |
| 0.238643 | 0.51523 | 0.029 | 0.02 | 1 | 2 | Diexf | 0.009 | Old |
| 0.238803 | 0.147708 | 0.111 | 0.094 | 1 | 2 | Fem1b | 0.017 | Old |
| 0.238891 | 0.290762 | 0.131 | 0.113 | 1 | 2 | Dars | 0.018 | Old |
| 0.239034 | 0.269866 | 0.194 | 0.172 | 1 | 2 | Ctnnd1 | 0.022 | Old |
| 0.239385 | 0.216563 | 0.118 | 0.1 | 1 | 2 | Akap8 | 0.018 | Old |
| 0.239401 | 0.140218 | 0.322 | 0.295 | 1 | 2 | Rnpep | 0.027 | Old |
| 0.239584 | 0.254932 | 0.07 | 0.056 | 1 | 2 | Cog3 | 0.014 | Old |

| | | | | | | | | |
|---|---|---|---|---|---|---|---|---|
| 0.239621 | 0.034405 | 0.047 | 0.036 | 1 | 2 | Mtmr9 | 0.011 | Old |
| 0.239854 | 0.194648 | 0.201 | 0.178 | 1 | 2 | Pank3 | 0.023 | Old |
| 0.239958 | 0.268741 | 0.152 | 0.133 | 1 | 2 | Ncoa2 | 0.019 | Old |
| 0.240055 | 0.2592 | 0.02 | 0.013 | 1 | 2 | Tnfsfm13 | 0.007 | Old |
| 0.240202 | 0.113149 | 0.095 | 0.078 | 1 | 2 | Kars | 0.017 | Old |
| 0.240799 | 0.060826 | 0.047 | 0.036 | 1 | 2 | Snx25 | 0.011 | Old |
| 0.240821 | 0.435011 | 0.073 | 0.059 | 1 | 2 | Ptpn23 | 0.014 | Old |
| 0.240827 | 0.323782 | 0.025 | 0.017 | 1 | 2 | Ttc27 | 0.008 | Old |
| 0.241181 | 0.41196 | 0.085 | 0.07 | 1 | 2 | 2610002M | 0.015 | Old |
| 0.241373 | 0.422055 | 0.097 | 0.082 | 1 | 2 | Mbd1 | 0.015 | Old |
| 0.241512 | 0.736654 | 0.034 | 0.025 | 1 | 2 | Bnip1 | 0.009 | Old |
| 0.241561 | 0.04681 | 0.463 | 0.496 | 1 | 2 | Gls | -0.033 | Old |
| 0.242001 | 0.119638 | 0.196 | 0.173 | 1 | 2 | Bcl3 | 0.023 | Old |
| 0.242011 | 0.23951 | 0.121 | 0.103 | 1 | 2 | Gas2l1 | 0.018 | Old |
| 0.242054 | 0.025268 | 0.186 | 0.163 | 1 | 2 | Rab3il1 | 0.023 | Old |
| 0.242075 | 0.108208 | 0.232 | 0.204 | 1 | 2 | Hadhb | 0.028 | Old |
| 0.242542 | 0.031102 | 0.167 | 0.144 | 1 | 2 | Glrx | 0.023 | Old |
| 0.242641 | 0.157496 | 0.165 | 0.141 | 1 | 2 | Sarnp | 0.024 | Old |
| 0.242736 | 0.148824 | 0.125 | 0.107 | 1 | 2 | Imp4 | 0.018 | Old |
| 0.242968 | 0.113892 | 0.51 | 0.479 | 1 | 2 | Reep3 | 0.031 | Old |
| 0.243081 | 1.414122 | 0.017 | 0.011 | 1 | 2 | Znrf3 | 0.006 | Old |
| 0.243386 | 0.173155 | 0.107 | 0.09 | 1 | 2 | Taf5l | 0.017 | Old |
| 0.243575 | 0.179816 | 0.08 | 0.065 | 1 | 2 | Prkaa1 | 0.015 | Old |
| 0.244431 | 0.264906 | 0.065 | 0.052 | 1 | 2 | Pam | 0.013 | Old |
| 0.244812 | 0.686719 | 0.034 | 0.025 | 1 | 2 | Mrpl49 | 0.009 | Old |
| 0.245104 | 1.031969 | 0.017 | 0.011 | 1 | 2 | Ppp3cc | 0.006 | Old |
| 0.245288 | 1.196506 | 0.017 | 0.011 | 1 | 2 | Zscan12 | 0.006 | Old |
| 0.245762 | 0.533061 | 0.044 | 0.033 | 1 | 2 | Srrd | 0.011 | Old |
| 0.246582 | 0.863859 | 0.017 | 0.011 | 1 | 2 | Gale | 0.006 | Old |
| 0.246659 | 0.449154 | 0.093 | 0.078 | 1 | 2 | Lcor | 0.015 | Old |
| 0.246731 | 0.603047 | 0.034 | 0.025 | 1 | 2 | Nmral1 | 0.009 | Old |
| 0.246767 | 0.656836 | 0.017 | 0.011 | 1 | 2 | A130010J1 | 0.006 | Old |
| 0.246972 | 0.136146 | 0.343 | 0.314 | 1 | 2 | Sirpb1c | 0.029 | Old |
| 0.246992 | 0.21587 | 0.276 | 0.248 | 1 | 2 | Rab21 | 0.028 | Old |
| 0.247086 | 0.176558 | 0.259 | 0.234 | 1 | 2 | U2surp | 0.025 | Old |
| 0.247342 | 0.063036 | 0.216 | 0.188 | 1 | 2 | Phf14 | 0.028 | Old |
| 0.247528 | 0.19996 | 0.213 | 0.19 | 1 | 2 | Polb | 0.023 | Old |
| 0.247541 | 0.185845 | 0.07 | 0.056 | 1 | 2 | Nt5dc3 | 0.014 | Old |
| 0.247639 | 0.151238 | 0.318 | 0.286 | 1 | 2 | Psmd8 | 0.032 | Old |
| 0.247653 | 0.253837 | 0.223 | 0.201 | 1 | 2 | Cyld | 0.022 | Old |
| 0.248156 | 0.175865 | 0.267 | 0.241 | 1 | 2 | Ube2q1 | 0.026 | Old |
| 0.248179 | 0.213222 | 0.039 | 0.029 | 1 | 2 | Fdx1 | 0.01 | Old |
| 0.248248 | 0.097676 | 0.306 | 0.279 | 1 | 2 | Hnrnpul1 | 0.027 | Old |
| 0.248253 | 1.032732 | 0.017 | 0.011 | 1 | 2 | Inpp1 | 0.006 | Old |
| 0.248265 | 0.385118 | 0.04 | 0.03 | 1 | 2 | Tarm1 | 0.01 | Old |

| | | | | | | | | |
|---|---|---|---|---|---|---|---|---|
| 0.24874 | 0.240728 | 0.077 | 0.063 | 1 | 2 | Oas2 | 0.014 | Old |
| 0.248811 | 0.447945 | 0.017 | 0.011 | 1 | 2 | Polr1e | 0.006 | Old |
| 0.248908 | 0.095659 | 0.097 | 0.081 | 1 | 2 | Slc39a9 | 0.016 | Old |
| 0.249047 | 0.192877 | 0.034 | 0.025 | 1 | 2 | Pradc1 | 0.009 | Old |
| 0.249216 | 0.065608 | 0.716 | 0.69 | 1 | 2 | Cltc | 0.026 | Old |
| 0.249798 | 0.559713 | 0.05 | 0.039 | 1 | 2 | Pkp4 | 0.011 | Old |
| 0.249908 | 0.031985 | 0.145 | 0.125 | 1 | 2 | Gbp7 | 0.02 | Old |
| 0.249977 | 0.177337 | 0.244 | 0.22 | 1 | 2 | Zmynd8 | 0.024 | Old |
| 0.250499 | 0.104545 | 0.324 | 0.293 | 1 | 2 | Myof | 0.031 | Old |
| 0.25068 | 0.583872 | 0.017 | 0.011 | 1 | 2 | Gpsm1 | 0.006 | Old |
| 0.250778 | 0.393686 | 0.172 | 0.154 | 1 | 2 | Cyp4f16 | 0.018 | Old |
| 0.250959 | 0.296626 | 0.095 | 0.08 | 1 | 2 | Ifih1 | 0.015 | Old |
| 0.251026 | 0.237967 | 0.073 | 0.058 | 1 | 2 | Ciz1 | 0.015 | Old |
| 0.251617 | 0.503786 | 0.017 | 0.011 | 1 | 2 | Kbtbd4 | 0.006 | Old |
| 0.251856 | 0.401348 | 0.122 | 0.105 | 1 | 2 | Tpm1 | 0.017 | Old |
| 0.252022 | 0.062785 | 0.076 | 0.062 | 1 | 2 | Cbx5 | 0.014 | Old |
| 0.252369 | 0.318248 | 0.017 | 0.011 | 1 | 2 | Lnx2 | 0.006 | Old |
| 0.252382 | 0.179157 | 0.448 | 0.424 | 1 | 2 | Ube2d2a | 0.024 | Old |
| 0.252458 | 0.06314 | 0.093 | 0.077 | 1 | 2 | Pilrb1 | 0.016 | Old |
| 0.252758 | 0.284545 | 0.165 | 0.145 | 1 | 2 | Med1 | 0.02 | Old |
| 0.252937 | 0.151325 | 0.397 | 0.379 | 1 | 2 | Syncrip | 0.018 | Old |
| 0.253476 | 0.075421 | 0.768 | 0.733 | 1 | 2 | Chchd2 | 0.035 | Old |
| 0.2535 | 0.300031 | 0.017 | 0.011 | 1 | 2 | E2f6 | 0.006 | Old |
| 0.253705 | 0.066267 | 0.397 | 0.362 | 1 | 2 | Rtn3 | 0.035 | Old |
| 0.253747 | 0.175403 | 0.328 | 0.302 | 1 | 2 | Midn | 0.026 | Old |
| 0.25377 | 0.108144 | 0.155 | 0.134 | 1 | 2 | Bcl7c | 0.021 | Old |
| 0.253785 | 0.929258 | 0.021 | 0.014 | 1 | 2 | Nat2 | 0.007 | Old |
| 0.253855 | 0.477102 | 0.034 | 0.025 | 1 | 2 | Rtn4ip1 | 0.009 | Old |
| 0.254247 | 0.098252 | 0.555 | 0.521 | 1 | 2 | Clic1 | 0.034 | Old |
| 0.254511 | 0.386055 | 0.094 | 0.08 | 1 | 2 | Faap20 | 0.014 | Old |
| 0.254961 | 0.373732 | 0.091 | 0.076 | 1 | 2 | Cab39l | 0.015 | Old |
| 0.256 | 0.240184 | 0.131 | 0.113 | 1 | 2 | Napg | 0.018 | Old |
| 0.256041 | 0.658709 | 0.026 | 0.018 | 1 | 2 | Ift172 | 0.008 | Old |
| 0.256124 | 0.479864 | 0.109 | 0.094 | 1 | 2 | Hdac4 | 0.015 | Old |
| 0.256325 | 0.366864 | 0.03 | 0.021 | 1 | 2 | Cpt2 | 0.009 | Old |
| 0.256377 | 0.075592 | 0.168 | 0.146 | 1 | 2 | Atf7 | 0.022 | Old |
| 0.256492 | 0.226392 | 0.163 | 0.143 | 1 | 2 | Pcmt1 | 0.02 | Old |
| 0.256693 | 0.237946 | 0.115 | 0.097 | 1 | 2 | Cdc123 | 0.018 | Old |
| 0.256748 | 0.57038 | 0.03 | 0.021 | 1 | 2 | Letm2 | 0.009 | Old |
| 0.256791 | 0.107294 | 0.482 | 0.448 | 1 | 2 | Slk | 0.034 | Old |
| 0.256876 | 0.113665 | 0.201 | 0.177 | 1 | 2 | Sla | 0.024 | Old |
| 0.257109 | 0.352277 | 0.1 | 0.084 | 1 | 2 | Papss1 | 0.016 | Old |
| 0.257147 | 0.66221 | 0.021 | 0.014 | 1 | 2 | Rfc3 | 0.007 | Old |
| 0.257227 | 0.269639 | 0.1 | 0.083 | 1 | 2 | Epb41l1 | 0.017 | Old |
| 0.257783 | 0.229007 | 0.15 | 0.131 | 1 | 2 | Spopl | 0.019 | Old |

| | | | | | | | | |
|---|---|---|---|---|---|---|---|---|
| 0.257899 | 0.193284 | 0.072 | 0.058 | 1 | 2 | Ddx28 | 0.014 | Old |
| 0.25799 | 0.764132 | 0.041 | 0.031 | 1 | 2 | Zfp943 | 0.01 | Old |
| 0.25804 | 0.274472 | 0.049 | 0.038 | 1 | 2 | Ddx55 | 0.011 | Old |
| 0.258435 | 0.152974 | 0.813 | 0.818 | 1 | 2 | Lgmn | -0.005 | Old |
| 0.258727 | 0.31664 | 0.034 | 0.025 | 1 | 2 | Ttll3 | 0.009 | Old |
| 0.258824 | 0.130627 | 0.224 | 0.2 | 1 | 2 | Fndc3a | 0.024 | Old |
| 0.259157 | 0.269802 | 0.03 | 0.021 | 1 | 2 | Pigf | 0.009 | Old |
| 0.25922 | 0.253602 | 0.186 | 0.166 | 1 | 2 | Tulp4 | 0.02 | Old |
| 0.259418 | 0.349653 | 0.026 | 0.018 | 1 | 2 | Pianp | 0.008 | Old |
| 0.259443 | 0.055473 | 0.055 | 0.043 | 1 | 2 | E130309D( | 0.012 | Old |
| 0.259497 | 0.916297 | 0.014 | 0.008 | 1 | 2 | Bcam | 0.006 | Old |
| 0.259499 | 0.00099 | 0.105 | 0.088 | 1 | 2 | Letm1 | 0.017 | Old |
| 0.259584 | 0.454634 | 0.03 | 0.021 | 1 | 2 | Rps6ka2 | 0.009 | Old |
| 0.259726 | 0.290869 | 0.026 | 0.018 | 1 | 2 | Treml2 | 0.008 | Old |
| 0.259967 | 0.098881 | 0.054 | 0.042 | 1 | 2 | Sh2b2 | 0.012 | Old |
| 0.260615 | 0.22464 | 0.148 | 0.129 | 1 | 2 | Ndufa12 | 0.019 | Old |
| 0.260784 | 0.107468 | 0.239 | 0.216 | 1 | 2 | Tmx3 | 0.023 | Old |
| 0.260957 | 0.325351 | 0.096 | 0.081 | 1 | 2 | Map4k1 | 0.015 | Old |
| 0.260963 | 0.290389 | 0.026 | 0.018 | 1 | 2 | Adck5 | 0.008 | Old |
| 0.261194 | 0.129768 | 0.373 | 0.346 | 1 | 2 | Chmp2a | 0.027 | Old |
| 0.261213 | 0.229836 | 0.085 | 0.07 | 1 | 2 | Nucb2 | 0.015 | Old |
| 0.261922 | 0.29278 | 0.077 | 0.063 | 1 | 2 | Topors | 0.014 | Old |
| 0.262076 | 0.20104 | 0.021 | 0.014 | 1 | 2 | Tango6 | 0.007 | Old |
| 0.262269 | 0.365355 | 0.035 | 0.026 | 1 | 2 | Acd | 0.009 | Old |
| 0.262547 | 0.589156 | 0.041 | 0.031 | 1 | 2 | Panx1 | 0.01 | Old |
| 0.262724 | 0.506666 | 0.014 | 0.008 | 1 | 2 | Usp11 | 0.006 | Old |
| 0.262742 | 0.052768 | 0.218 | 0.194 | 1 | 2 | Rnf111 | 0.024 | Old |
| 0.262825 | 0.164194 | 0.044 | 0.033 | 1 | 2 | Borcs5 | 0.011 | Old |
| 0.263207 | 0.162689 | 0.106 | 0.09 | 1 | 2 | Ilvbl | 0.016 | Old |
| 0.263535 | 0.090816 | 0.128 | 0.109 | 1 | 2 | Abr | 0.019 | Old |
| 0.263619 | 0.062991 | 0.021 | 0.014 | 1 | 2 | Crocc | 0.007 | Old |
| 0.263908 | 0.064848 | 0.049 | 0.038 | 1 | 2 | Ebag9 | 0.011 | Old |
| 0.264055 | 0.063627 | 0.14 | 0.12 | 1 | 2 | Golgb1 | 0.02 | Old |
| 0.264239 | 0.032454 | 0.014 | 0.008 | 1 | 2 | Tgm1 | 0.006 | Old |
| 0.264398 | 0.232433 | 0.073 | 0.059 | 1 | 2 | Hax1 | 0.014 | Old |
| 0.264456 | 0.293912 | 0.014 | 0.008 | 1 | 2 | Trim59 | 0.006 | Old |
| 0.264456 | 0.089427 | 0.014 | 0.008 | 1 | 2 | Sag | 0.006 | Old |
| 0.26456 | 0.034557 | 0.175 | 0.153 | 1 | 2 | Ehmt2 | 0.022 | Old |
| 0.264673 | 0.550005 | 0.014 | 0.008 | 1 | 2 | Slc8a1 | 0.006 | Old |
| 0.264882 | 0.425619 | 0.046 | 0.036 | 1 | 2 | Dus3l | 0.01 | Old |
| 0.265018 | 0.302988 | 0.099 | 0.083 | 1 | 2 | Gle1 | 0.016 | Old |
| 0.265139 | 0.47839 | 0.031 | 0.023 | 1 | 2 | Mast2 | 0.008 | Old |
| 0.265582 | 0.168807 | 0.095 | 0.08 | 1 | 2 | Commd1 | 0.015 | Old |
| 0.265796 | 0.309729 | 0.156 | 0.137 | 1 | 2 | Xpo7 | 0.019 | Old |
| 0.265906 | 0.122386 | 0.148 | 0.128 | 1 | 2 | Aph1c | 0.02 | Old |

| | | | | | | | | |
|---|---|---|---|---|---|---|---|---|
| 0.266181 | 0.118787 | 0.228 | 0.205 | 1 | 2 | Atp6v1c1 | 0.023 | Old |
| 0.266213 | 0.026782 | 0.317 | 0.281 | 1 | 2 | Tbl1xr1 | 0.036 | Old |
| 0.267167 | 0.207067 | 0.103 | 0.087 | 1 | 2 | Dcaf15 | 0.016 | Old |
| 0.267314 | 0.608372 | 0.046 | 0.036 | 1 | 2 | Actr5 | 0.01 | Old |
| 0.267371 | 0.227972 | 0.192 | 0.17 | 1 | 2 | Rap2c | 0.022 | Old |
| 0.267377 | 0.17891 | 0.089 | 0.074 | 1 | 2 | 0610010F0 | 0.015 | Old |
| 0.267429 | 0.373481 | 0.035 | 0.026 | 1 | 2 | Chd1l | 0.009 | Old |
| 0.267497 | 0.268568 | 0.074 | 0.061 | 1 | 2 | Slc35f5 | 0.013 | Old |
| 0.267552 | 0.674244 | 0.031 | 0.023 | 1 | 2 | Acad9 | 0.008 | Old |
| 0.267628 | 0.192937 | 0.102 | 0.086 | 1 | 2 | Ppip5k2 | 0.016 | Old |
| 0.267759 | 0.185816 | 0.157 | 0.138 | 1 | 2 | Casc3 | 0.019 | Old |
| 0.267886 | 0.045077 | 0.388 | 0.354 | 1 | 2 | Psme2 | 0.034 | Old |
| 0.268029 | 0.229712 | 0.035 | 0.026 | 1 | 2 | Ubxn8 | 0.009 | Old |
| 0.269016 | 0.10701 | 0.17 | 0.151 | 1 | 2 | 9030624J0 | 0.019 | Old |
| 0.269142 | 0.046312 | 0.071 | 0.057 | 1 | 2 | Ermp1 | 0.014 | Old |
| 0.269388 | 0.103546 | 0.075 | 0.062 | 1 | 2 | St7l | 0.013 | Old |
| 0.269485 | 0.109044 | 0.33 | 0.302 | 1 | 2 | Slamf9 | 0.028 | Old |
| 0.269551 | 0.064477 | 0.077 | 0.063 | 1 | 2 | Atg9a | 0.014 | Old |
| 0.269828 | 0.97156 | 0.011 | 0.006 | 1 | 2 | Ska2 | 0.005 | Old |
| 0.270225 | 0.193986 | 0.14 | 0.121 | 1 | 2 | Psmd7 | 0.019 | Old |
| 0.27041 | 0.051024 | 0.04 | 0.03 | 1 | 2 | Kremen1 | 0.01 | Old |
| 0.270436 | 0.199692 | 0.166 | 0.146 | 1 | 2 | Trim41 | 0.02 | Old |
| 0.270516 | 0.167731 | 0.168 | 0.147 | 1 | 2 | Paics | 0.021 | Old |
| 0.27059 | 1.055254 | 0.011 | 0.006 | 1 | 2 | Zfp316 | 0.005 | Old |
| 0.270596 | 0.131568 | 0.184 | 0.163 | 1 | 2 | Taf1 | 0.021 | Old |
| 0.270642 | 0.230958 | 0.066 | 0.053 | 1 | 2 | Mycbp | 0.013 | Old |
| 0.270774 | 0.346574 | 0.093 | 0.078 | 1 | 2 | Acsl1 | 0.015 | Old |
| 0.27083 | 1.031721 | 0.018 | 0.012 | 1 | 2 | Tdp1 | 0.006 | Old |
| 0.270844 | 1.089093 | 0.011 | 0.006 | 1 | 2 | C6 | 0.005 | Old |
| 0.270929 | 1.030206 | 0.022 | 0.015 | 1 | 2 | Gm20045 | 0.007 | Old |
| 0.271049 | 0.055959 | 0.077 | 0.063 | 1 | 2 | Gm20696 | 0.014 | Old |
| 0.271243 | 0.73707 | 0.052 | 0.042 | 1 | 2 | Kantr | 0.01 | Old |
| 0.271419 | 0.21275 | 0.095 | 0.08 | 1 | 2 | Rhbdf2 | 0.015 | Old |
| 0.271422 | 0.158597 | 0.24 | 0.217 | 1 | 2 | Thoc2 | 0.023 | Old |
| 0.271565 | 0.235621 | 0.123 | 0.106 | 1 | 2 | Nub1 | 0.017 | Old |
| 0.271592 | 1.037612 | 0.018 | 0.012 | 1 | 2 | Atpaf1 | 0.006 | Old |
| 0.271702 | 0.449433 | 0.031 | 0.023 | 1 | 2 | Trip10 | 0.008 | Old |
| 0.272629 | 1.079586 | 0.011 | 0.006 | 1 | 2 | Numbl | 0.005 | Old |
| 0.272629 | 0.82936 | 0.011 | 0.006 | 1 | 2 | 1110008L1 | 0.005 | Old |
| 0.272737 | 1.054502 | 0.018 | 0.012 | 1 | 2 | Ntmt1 | 0.006 | Old |
| 0.272885 | 0.741736 | 0.011 | 0.006 | 1 | 2 | Tmem62 | 0.005 | Old |
| 0.273064 | 0.115249 | 0.035 | 0.026 | 1 | 2 | Cnot11 | 0.009 | Old |
| 0.273141 | 0.584629 | 0.011 | 0.006 | 1 | 2 | Cep95 | 0.005 | Old |
| 0.273353 | 0.177487 | 0.561 | 0.548 | 1 | 2 | Tnip3 | 0.013 | Old |
| 0.273742 | 0.655914 | 0.027 | 0.019 | 1 | 2 | Gpatch1 | 0.008 | Old |

| | | | | | | | | |
|---|---|---|---|---|---|---|---|---|
| 0.273951 | 0.086349 | 0.712 | 0.732 | 1 | 2 | Pcbp2 | -0.02 | Old |
| 0.274505 | 0.08704 | 0.113 | 0.096 | 1 | 2 | Vps37a | 0.017 | Old |
| 0.274546 | 0.304307 | 0.08 | 0.067 | 1 | 2 | Mrpl19 | 0.013 | Old |
| 0.274648 | 0.23685 | 0.233 | 0.208 | 1 | 2 | Sp3 | 0.025 | Old |
| 0.274679 | 0.554257 | 0.011 | 0.006 | 1 | 2 | Gnal | 0.005 | Old |
| 0.274679 | 0.441428 | 0.011 | 0.006 | 1 | 2 | Arhgap27o | 0.005 | Old |
| 0.274879 | 0.371946 | 0.031 | 0.023 | 1 | 2 | Urgcp | 0.008 | Old |
| 0.274907 | 0.105531 | 0.287 | 0.259 | 1 | 2 | Was | 0.028 | Old |
| 0.275119 | 0.120859 | 0.142 | 0.124 | 1 | 2 | Strn4 | 0.018 | Old |
| 0.275124 | 0.07588 | 0.056 | 0.044 | 1 | 2 | Camta1 | 0.012 | Old |
| 0.275366 | 0.296032 | 0.117 | 0.101 | 1 | 2 | Stard7 | 0.016 | Old |
| 0.275625 | 0.204281 | 0.2 | 0.177 | 1 | 2 | Sike1 | 0.023 | Old |
| 0.275972 | 0.143265 | 0.172 | 0.153 | 1 | 2 | Taz | 0.019 | Old |
| 0.276026 | 0.009395 | 0.193 | 0.17 | 1 | 2 | Mob1b | 0.023 | Old |
| 0.276061 | 0.559249 | 0.042 | 0.032 | 1 | 2 | Fmo5 | 0.01 | Old |
| 0.276203 | 0.024605 | 0.056 | 0.044 | 1 | 2 | Akt1s1 | 0.012 | Old |
| 0.276248 | 0.755069 | 0.027 | 0.019 | 1 | 2 | Tmem241 | 0.008 | Old |
| 0.276579 | 0.547845 | 0.018 | 0.012 | 1 | 2 | 1110004E0 | 0.006 | Old |
| 0.276721 | 0.033755 | 0.119 | 0.101 | 1 | 2 | Seh1l | 0.018 | Old |
| 0.276978 | 0.129617 | 0.216 | 0.192 | 1 | 2 | Map3k8 | 0.024 | Old |
| 0.277176 | 0.070437 | 0.301 | 0.271 | 1 | 2 | Pttg1ip | 0.03 | Old |
| 0.277412 | 0.081061 | 0.035 | 0.026 | 1 | 2 | Hspbap1 | 0.009 | Old |
| 0.27747 | 0.152223 | 0.184 | 0.164 | 1 | 2 | Fam172a | 0.02 | Old |
| 0.277636 | 0.197769 | 0.161 | 0.141 | 1 | 2 | Osbp | 0.02 | Old |
| 0.277768 | 0.156042 | 0.075 | 0.062 | 1 | 2 | Zfp664 | 0.013 | Old |
| 0.278029 | 0.763082 | 0.018 | 0.012 | 1 | 2 | Ankrd33b | 0.006 | Old |
| 0.278246 | 0.071864 | 0.291 | 0.262 | 1 | 2 | Dnajc5 | 0.029 | Old |
| 0.27832 | 0.48941 | 0.018 | 0.012 | 1 | 2 | Gtpbp3 | 0.006 | Old |
| 0.278792 | 0.390424 | 0.064 | 0.052 | 1 | 2 | Eci2 | 0.012 | Old |
| 0.279219 | 0.067762 | 0.234 | 0.21 | 1 | 2 | Sri | 0.024 | Old |
| 0.279293 | 0.262588 | 0.168 | 0.151 | 1 | 2 | Ibtk | 0.017 | Old |
| 0.279343 | 0.383862 | 0.036 | 0.027 | 1 | 2 | Jade2 | 0.009 | Old |
| 0.279414 | 0.07373 | 0.17 | 0.15 | 1 | 2 | Dync1li1 | 0.02 | Old |
| 0.279435 | 0.140571 | 0.315 | 0.291 | 1 | 2 | Pten | 0.024 | Old |
| 0.279771 | 0.131124 | 0.221 | 0.197 | 1 | 2 | Secisbp2l | 0.024 | Old |
| 0.279791 | 0.260457 | 0.148 | 0.131 | 1 | 2 | Tmem43 | 0.017 | Old |
| 0.279933 | 0.472491 | 0.059 | 0.048 | 1 | 2 | Akr7a5 | 0.011 | Old |
| 0.280211 | 0.320583 | 0.022 | 0.015 | 1 | 2 | Slc36a3os | 0.007 | Old |
| 0.280274 | 0.312632 | 0.027 | 0.019 | 1 | 2 | Cox15 | 0.008 | Old |
| 0.280396 | 0.193006 | 0.086 | 0.071 | 1 | 2 | Atp1a3 | 0.015 | Old |
| 0.2804 | 0.067123 | 0.106 | 0.089 | 1 | 2 | Dhx16 | 0.017 | Old |
| 0.280661 | 0.29429 | 0.173 | 0.156 | 1 | 2 | Vat1 | 0.017 | Old |
| 0.280671 | 0.163897 | 0.027 | 0.019 | 1 | 2 | Gm6563 | 0.008 | Old |
| 0.280791 | 0.14603 | 0.315 | 0.289 | 1 | 2 | Ubash3b | 0.026 | Old |
| 0.28111 | 0.13348 | 0.217 | 0.195 | 1 | 2 | Vma21 | 0.022 | Old |

| | | | | | | | | |
|---|---|---|---|---|---|---|---|---|
| 0.28114 | 0.467911 | 0.048 | 0.038 | 1 | 2 | Fkbp5 | 0.01 | Old |
| 0.281472 | 0.10811 | 0.317 | 0.287 | 1 | 2 | Ube2r2 | 0.03 | Old |
| 0.281768 | 0.238041 | 0.147 | 0.129 | 1 | 2 | Rprd2 | 0.018 | Old |
| 0.282653 | 0.4655 | 0.022 | 0.015 | 1 | 2 | Deaf1 | 0.007 | Old |
| 0.28306 | 0.428696 | 0.027 | 0.019 | 1 | 2 | Tmem107 | 0.008 | Old |
| 0.283313 | 0.052497 | 0.09 | 0.075 | 1 | 2 | Wdr5 | 0.015 | Old |
| 0.283318 | 0.370443 | 0.103 | 0.088 | 1 | 2 | Trak1 | 0.015 | Old |
| 0.283815 | 0.080213 | 0.626 | 0.6 | 1 | 2 | Ctsa | 0.026 | Old |
| 0.284113 | 0.038198 | 0.137 | 0.118 | 1 | 2 | Snrnp48 | 0.019 | Old |
| 0.284209 | 0.105397 | 0.216 | 0.195 | 1 | 2 | Ppp6r1 | 0.021 | Old |
| 0.285075 | 0.361179 | 0.032 | 0.024 | 1 | 2 | Zfp143 | 0.008 | Old |
| 0.285136 | 0.151517 | 0.042 | 0.032 | 1 | 2 | Paxx | 0.01 | Old |
| 0.285307 | 0.136192 | 0.031 | 0.023 | 1 | 2 | Pi4kb | 0.008 | Old |
| 0.285446 | 0.21345 | 0.183 | 0.164 | 1 | 2 | Rps27rt | 0.019 | Old |
| 0.285805 | 0.430452 | 0.032 | 0.024 | 1 | 2 | Aar2 | 0.008 | Old |
| 0.285928 | 0.270848 | 0.064 | 0.052 | 1 | 2 | Uchl3 | 0.012 | Old |
| 0.286142 | 0.18507 | 0.036 | 0.027 | 1 | 2 | Atg13 | 0.009 | Old |
| 0.286271 | 0.349187 | 0.048 | 0.038 | 1 | 2 | Tstd2 | 0.01 | Old |
| 0.286312 | 0.067739 | 0.782 | 0.77 | 1 | 2 | Ptp4a2 | 0.012 | Old |
| 0.286564 | 0.11855 | 0.138 | 0.12 | 1 | 2 | Naa20 | 0.018 | Old |
| 0.286814 | 0.196262 | 0.079 | 0.065 | 1 | 2 | Prdx4 | 0.014 | Old |
| 0.287204 | 0.235292 | 0.236 | 0.215 | 1 | 2 | Ypel5 | 0.021 | Old |
| 0.287216 | 0.113354 | 0.063 | 0.051 | 1 | 2 | Dhx58 | 0.012 | Old |
| 0.287223 | 0.065972 | 0.198 | 0.175 | 1 | 2 | Cpsf2 | 0.023 | Old |
| 0.28724 | 0.387569 | 0.036 | 0.027 | 1 | 2 | Tmem242 | 0.009 | Old |
| 0.287543 | 0.260597 | 0.091 | 0.077 | 1 | 2 | Prkag2 | 0.014 | Old |
| 0.287841 | 0.205144 | 0.063 | 0.051 | 1 | 2 | Timm8a1 | 0.012 | Old |
| 0.288035 | 0.149217 | 0.026 | 0.034 | 1 | 2 | Smim8 | -0.008 | Old |
| 0.288782 | 0.27511 | 0.089 | 0.075 | 1 | 2 | Usp32 | 0.014 | Old |
| 0.289011 | 0.528236 | 0.08 | 0.068 | 1 | 2 | Insig2 | 0.012 | Old |
| 0.289376 | 0.444968 | 0.036 | 0.027 | 1 | 2 | Mbip | 0.009 | Old |
| 0.289676 | 0.493321 | 0.082 | 0.069 | 1 | 2 | Timm44 | 0.013 | Old |
| 0.290333 | 0.260431 | 0.313 | 0.293 | 1 | 2 | Gpcpd1 | 0.02 | Old |
| 0.290442 | 0.268301 | 0.275 | 0.255 | 1 | 2 | Pik3r1 | 0.02 | Old |
| 0.290499 | 0.139466 | 0.128 | 0.112 | 1 | 2 | Slc4a2 | 0.016 | Old |
| 0.290927 | 0.02118 | 0.606 | 0.574 | 1 | 2 | Tm6sf1 | 0.032 | Old |
| 0.291038 | 0.314771 | 0.066 | 0.055 | 1 | 2 | Exosc4 | 0.011 | Old |
| 0.291258 | 0.229369 | 0.138 | 0.12 | 1 | 2 | Sh3pxd2b | 0.018 | Old |
| 0.291505 | 0.100984 | 0.606 | 0.571 | 1 | 2 | Wdr26 | 0.035 | Old |
| 0.291762 | 0.160573 | 0.137 | 0.158 | 1 | 2 | Skint3 | -0.021 | Old |
| 0.292146 | 0.196652 | 0.482 | 0.458 | 1 | 2 | Ppp2r5c | 0.024 | Old |
| 0.292267 | 0.18219 | 0.106 | 0.091 | 1 | 2 | Lpcat3 | 0.015 | Old |
| 0.292423 | 0.476726 | 0.079 | 0.067 | 1 | 2 | Galns | 0.012 | Old |
| 0.292637 | 0.012549 | 0.163 | 0.143 | 1 | 2 | Grk6 | 0.02 | Old |
| 0.292953 | 0.358652 | 0.043 | 0.033 | 1 | 2 | Tmem80 | 0.01 | Old |

| | | | | | | | | |
|---|---|---|---|---|---|---|---|---|
| 0.293058 | 0.550602 | 0.056 | 0.045 | 1 | 2 | Ripk2 | 0.011 | Old |
| 0.293836 | 0.338828 | 0.077 | 0.064 | 1 | 2 | Acot9 | 0.013 | Old |
| 0.29386 | 0.049425 | 0.135 | 0.116 | 1 | 2 | Mvb12a | 0.019 | Old |
| 0.293942 | 0.020455 | 0.703 | 0.682 | 1 | 2 | Glud1 | 0.021 | Old |
| 0.294293 | 0.337744 | 0.104 | 0.089 | 1 | 2 | Atp6v0a2 | 0.015 | Old |
| 0.294594 | 0.140527 | 0.086 | 0.071 | 1 | 2 | Kctd2 | 0.015 | Old |
| 0.294696 | 0.190777 | 0.33 | 0.308 | 1 | 2 | Rassf1 | 0.022 | Old |
| 0.295386 | 0.444477 | 0.163 | 0.147 | 1 | 2 | Wdr37 | 0.016 | Old |
| 0.295885 | 0.13564 | 0.059 | 0.048 | 1 | 2 | Orc3 | 0.011 | Old |
| 0.296586 | 0.016498 | 0.07 | 0.057 | 1 | 2 | Fnip2 | 0.013 | Old |
| 0.296595 | 0.186335 | 0.032 | 0.024 | 1 | 2 | Tapt1 | 0.008 | Old |
| 0.29669 | 0.178747 | 0.078 | 0.065 | 1 | 2 | Med17 | 0.013 | Old |
| 0.296754 | 0.111463 | 0.53 | 0.505 | 1 | 2 | Tm9sf3 | 0.025 | Old |
| 0.296918 | 0.612242 | 0.015 | 0.01 | 1 | 2 | Hist1h2ap | 0.005 | Old |
| 0.297128 | 0.361137 | 0.024 | 0.017 | 1 | 2 | Zfp65 | 0.007 | Old |
| 0.29719 | 0.064754 | 0.044 | 0.055 | 1 | 2 | Tmem8 | -0.011 | Old |
| 0.297194 | 0.36124 | 0.032 | 0.024 | 1 | 2 | Ercc1 | 0.008 | Old |
| 0.29748 | 0.842231 | 0.024 | 0.017 | 1 | 2 | Mx1 | 0.007 | Old |
| 0.297895 | 0.241368 | 0.169 | 0.151 | 1 | 2 | Cpsf7 | 0.018 | Old |
| 0.298 | 0.160532 | 0.363 | 0.341 | 1 | 2 | Ubald2 | 0.022 | Old |
| 0.298045 | 0.110889 | 0.11 | 0.094 | 1 | 2 | Relb | 0.016 | Old |
| 0.298069 | 0.403424 | 0.117 | 0.102 | 1 | 2 | Mctp1 | 0.015 | Old |
| 0.298095 | 0.132908 | 0.203 | 0.182 | 1 | 2 | Rraga | 0.021 | Old |
| 0.298222 | 0.084789 | 0.253 | 0.229 | 1 | 2 | Rad23b | 0.024 | Old |
| 0.298368 | 0.113151 | 0.413 | 0.387 | 1 | 2 | Plxnb2 | 0.026 | Old |
| 0.298435 | 0.22246 | 0.091 | 0.077 | 1 | 2 | Ifi35 | 0.014 | Old |
| 0.298454 | 0.512203 | 0.028 | 0.02 | 1 | 2 | Fpgs | 0.008 | Old |
| 0.298488 | 0.753733 | 0.015 | 0.01 | 1 | 2 | Tut1 | 0.005 | Old |
| 0.298488 | 0.627995 | 0.015 | 0.01 | 1 | 2 | Rdh10 | 0.005 | Old |
| 0.298611 | 0.385641 | 0.087 | 0.074 | 1 | 2 | Rab3gap2 | 0.013 | Old |
| 0.298686 | 0.23451 | 0.12 | 0.102 | 1 | 2 | Sar1b | 0.018 | Old |
| 0.29894 | 0.438936 | 0.028 | 0.02 | 1 | 2 | Riox1 | 0.008 | Old |
| 0.299389 | 0.366564 | 0.015 | 0.01 | 1 | 2 | Tmem41a | 0.005 | Old |
| 0.299641 | 0.4807 | 0.055 | 0.044 | 1 | 2 | Tfb2m | 0.011 | Old |
| 0.300065 | 0.289059 | 0.015 | 0.01 | 1 | 2 | BC049352 | 0.005 | Old |
| 0.300122 | 0.229919 | 0.13 | 0.113 | 1 | 2 | Cyb5r1 | 0.017 | Old |
| 0.30029 | 0.872538 | 0.015 | 0.01 | 1 | 2 | Hdac8 | 0.005 | Old |
| 0.300395 | 0.233559 | 0.072 | 0.059 | 1 | 2 | Cog7 | 0.013 | Old |
| 0.300528 | 0.118595 | 0.142 | 0.125 | 1 | 2 | Mysm1 | 0.017 | Old |
| 0.300709 | 0.032871 | 0.11 | 0.094 | 1 | 2 | Tbc1d15 | 0.016 | Old |
| 0.300817 | 0.477427 | 0.094 | 0.081 | 1 | 2 | Timm10 | 0.013 | Old |
| 0.300839 | 0.100917 | 0.392 | 0.361 | 1 | 2 | Ctdsp2 | 0.031 | Old |
| 0.300949 | 0.572758 | 0.09 | 0.077 | 1 | 2 | Rai1 | 0.013 | Old |
| 0.301014 | 0.472546 | 0.024 | 0.017 | 1 | 2 | Fam20a | 0.007 | Old |
| 0.301283 | 0.091072 | 0.297 | 0.273 | 1 | 2 | B4galt6 | 0.024 | Old |

| | | | | | | | | |
|---|---|---|---|---|---|---|---|---|
| 0.301428 | 0.038421 | 0.137 | 0.119 | 1 | 2 | St6galnac6 | 0.018 | Old |
| 0.301457 | 0.180202 | 0.405 | 0.378 | 1 | 2 | Cyp4f18 | 0.027 | Old |
| 0.301487 | 0.003241 | 0.222 | 0.197 | 1 | 2 | Coa5 | 0.025 | Old |
| 0.301637 | 0.306907 | 0.032 | 0.024 | 1 | 2 | Cdkal1 | 0.008 | Old |
| 0.301764 | 0.018419 | 0.117 | 0.1 | 1 | 2 | Mfap3 | 0.017 | Old |
| 0.301952 | 0.275679 | 0.085 | 0.071 | 1 | 2 | Nol11 | 0.014 | Old |
| 0.302017 | 0.315254 | 0.105 | 0.091 | 1 | 2 | Slc35a5 | 0.014 | Old |
| 0.3021 | 0.127141 | 0.015 | 0.01 | 1 | 2 | Lppos | 0.005 | Old |
| 0.302118 | 0.475614 | 0.044 | 0.034 | 1 | 2 | Elk1 | 0.01 | Old |
| 0.302222 | 0.041453 | 0.465 | 0.425 | 1 | 2 | Wipf1 | 0.04 | Old |
| 0.302529 | 0.526393 | 0.019 | 0.013 | 1 | 2 | Cables1 | 0.006 | Old |
| 0.302678 | 0.176271 | 0.36 | 0.333 | 1 | 2 | Mkln1 | 0.027 | Old |
| 0.303039 | 0.199316 | 0.164 | 0.145 | 1 | 2 | Egln2 | 0.019 | Old |
| 0.303212 | 0.239977 | 0.111 | 0.096 | 1 | 2 | Hibadh | 0.015 | Old |
| 0.303504 | 0.25894 | 0.043 | 0.033 | 1 | 2 | Cep85 | 0.01 | Old |
| 0.303718 | 0.486782 | 0.019 | 0.013 | 1 | 2 | Wdr59 | 0.006 | Old |
| 0.303837 | 0.218006 | 0.032 | 0.024 | 1 | 2 | Btbd2 | 0.008 | Old |
| 0.303856 | 0.573053 | 0.033 | 0.025 | 1 | 2 | Rps6kb2 | 0.008 | Old |
| 0.30424 | 0.045404 | 0.213 | 0.189 | 1 | 2 | Plcg2 | 0.024 | Old |
| 0.30455 | 0.101498 | 0.385 | 0.359 | 1 | 2 | Vdac2 | 0.026 | Old |
| 0.304669 | 0.366418 | 0.094 | 0.081 | 1 | 2 | Klhdc4 | 0.013 | Old |
| 0.304717 | 0.147602 | 0.07 | 0.057 | 1 | 2 | Ppp1r7 | 0.013 | Old |
| 0.304915 | 0.236645 | 0.164 | 0.146 | 1 | 2 | Gpatch8 | 0.018 | Old |
| 0.305052 | 0.061455 | 0.571 | 0.524 | 1 | 2 | Foxp1 | 0.047 | Old |
| 0.305242 | 0.02601 | 0.037 | 0.029 | 1 | 2 | Pstk | 0.008 | Old |
| 0.30569 | 0.352449 | 0.111 | 0.097 | 1 | 2 | Tmem251 | 0.014 | Old |
| 0.306378 | 0.048121 | 0.076 | 0.063 | 1 | 2 | S1pr2 | 0.013 | Old |
| 0.306608 | 0.010383 | 0.075 | 0.062 | 1 | 2 | Soga1 | 0.013 | Old |
| 0.307204 | 0.064807 | 0.376 | 0.347 | 1 | 2 | Stx7 | 0.029 | Old |
| 0.307932 | 0.020053 | 0.005 | 0.01 | 1 | 2 | 2610301B2 | -0.005 | Old |
| 0.308023 | 0.134734 | 0.206 | 0.183 | 1 | 2 | Mcmbp | 0.023 | Old |
| 0.308329 | 0.137241 | 0.079 | 0.094 | 1 | 2 | Arhgap11a | -0.015 | Old |
| 0.308501 | 0.185072 | 0.222 | 0.201 | 1 | 2 | Nrd1 | 0.021 | Old |
| 0.309068 | 0.114741 | 0.133 | 0.116 | 1 | 2 | Slc31a1 | 0.017 | Old |
| 0.309096 | 0.035214 | 0.128 | 0.112 | 1 | 2 | Lars | 0.016 | Old |
| 0.309751 | 0.338314 | 0.162 | 0.145 | 1 | 2 | Nhsl2 | 0.017 | Old |
| 0.309773 | 0.40068 | 0.141 | 0.125 | 1 | 2 | Arrdc3 | 0.016 | Old |
| 0.309923 | 0.276482 | 0.249 | 0.233 | 1 | 2 | Pum1 | 0.016 | Old |
| 0.310021 | 0.122348 | 0.005 | 0.01 | 1 | 2 | Gm38394 | -0.005 | Old |
| 0.310179 | 0.20812 | 0.085 | 0.071 | 1 | 2 | Erg28 | 0.014 | Old |
| 0.31066 | 0.057938 | 0.592 | 0.569 | 1 | 2 | Cox5a | 0.023 | Old |
| 0.310804 | 0.186104 | 0.104 | 0.089 | 1 | 2 | Flcn | 0.015 | Old |
| 0.311646 | 0.228963 | 0.112 | 0.097 | 1 | 2 | Esrra | 0.015 | Old |
| 0.31172 | 0.37511 | 0.069 | 0.057 | 1 | 2 | Mtmr14 | 0.012 | Old |
| 0.311851 | 0.093221 | 0.233 | 0.213 | 1 | 2 | Kat2b | 0.02 | Old |

| | | | | | | | | |
|---|---|---|---|---|---|---|---|---|
| 0.312145 | 0.855078 | 0.012 | 0.007 | 1 | 2 | 1700109H( | 0.005 | Old |
| 0.312256 | 0.224469 | 0.039 | 0.03 | 1 | 2 | Mfsd8 | 0.009 | Old |
| 0.312397 | 0.186172 | 0.039 | 0.03 | 1 | 2 | Spsb2 | 0.009 | Old |
| 0.312508 | 0.159226 | 0.055 | 0.044 | 1 | 2 | Nckipsd | 0.011 | Old |
| 0.31255 | 0.141532 | 0.133 | 0.115 | 1 | 2 | Utrn | 0.018 | Old |
| 0.312738 | 0.092574 | 0.181 | 0.162 | 1 | 2 | Capn1 | 0.019 | Old |
| 0.312932 | 0.125395 | 0.048 | 0.038 | 1 | 2 | Ccdc43 | 0.01 | Old |
| 0.313051 | 0.081439 | 0.135 | 0.119 | 1 | 2 | Cdk4 | 0.016 | Old |
| 0.313173 | 0.235777 | 0.17 | 0.152 | 1 | 2 | Idnk | 0.018 | Old |
| 0.313311 | 0.17374 | 0.135 | 0.118 | 1 | 2 | Slmap | 0.017 | Old |
| 0.31358 | 0.127185 | 0.426 | 0.4 | 1 | 2 | Ssb | 0.026 | Old |
| 0.313816 | 0.946991 | 0.025 | 0.018 | 1 | 2 | Rfc4 | 0.007 | Old |
| 0.313824 | 0.089449 | 0.291 | 0.268 | 1 | 2 | Slc12a6 | 0.023 | Old |
| 0.31412 | 0.373587 | 0.134 | 0.119 | 1 | 2 | Dusp6 | 0.015 | Old |
| 0.314385 | 0.197644 | 0.044 | 0.034 | 1 | 2 | Ano10 | 0.01 | Old |
| 0.314424 | 0.390597 | 0.045 | 0.036 | 1 | 2 | Abt1 | 0.009 | Old |
| 0.314491 | 0.422719 | 0.056 | 0.045 | 1 | 2 | Rbm15b | 0.011 | Old |
| 0.31464 | 0.179677 | 0.124 | 0.108 | 1 | 2 | Stx5a | 0.016 | Old |
| 0.314709 | 0.49794 | 0.029 | 0.021 | 1 | 2 | Actr6 | 0.008 | Old |
| 0.31497 | 0.146065 | 0.217 | 0.197 | 1 | 2 | Txnl1 | 0.02 | Old |
| 0.315105 | 0.667583 | 0.034 | 0.026 | 1 | 2 | Vps72 | 0.008 | Old |
| 0.315215 | 0.500957 | 0.052 | 0.043 | 1 | 2 | Tecpr1 | 0.009 | Old |
| 0.315433 | 0.235173 | 0.067 | 0.056 | 1 | 2 | Cyp4f13 | 0.011 | Old |
| 0.315499 | 0.581018 | 0.051 | 0.042 | 1 | 2 | Dis3l2 | 0.009 | Old |
| 0.315543 | 0.047214 | 0.378 | 0.35 | 1 | 2 | Bclaf1 | 0.028 | Old |
| 0.315732 | 0.067274 | 0.244 | 0.217 | 1 | 2 | Ripk1 | 0.027 | Old |
| 0.315747 | 0.135936 | 0.116 | 0.1 | 1 | 2 | Get4 | 0.016 | Old |
| 0.315927 | 0.197842 | 0.039 | 0.03 | 1 | 2 | Guf1 | 0.009 | Old |
| 0.316498 | 0.161534 | 0.191 | 0.171 | 1 | 2 | Chchd1 | 0.02 | Old |
| 0.316672 | 0.238514 | 0.135 | 0.12 | 1 | 2 | BC004004 | 0.015 | Old |
| 0.316935 | 0.83656 | 0.025 | 0.018 | 1 | 2 | Dnajc27 | 0.007 | Old |
| 0.317125 | 1.129488 | 0.016 | 0.011 | 1 | 2 | A4galt | 0.005 | Old |
| 0.31717 | 0.135877 | 0.012 | 0.007 | 1 | 2 | AC139671. | 0.005 | Old |
| 0.317207 | 0.17567 | 0.155 | 0.137 | 1 | 2 | Yipf1 | 0.018 | Old |
| 0.317293 | 0.479203 | 0.088 | 0.075 | 1 | 2 | Chst12 | 0.013 | Old |
| 0.317562 | 0.2694 | 0.244 | 0.223 | 1 | 2 | Tmem164 | 0.021 | Old |
| 0.317731 | 0.152139 | 0.605 | 0.594 | 1 | 2 | Ier2 | 0.011 | Old |
| 0.317968 | 0.652506 | 0.012 | 0.007 | 1 | 2 | Nlrp10 | 0.005 | Old |
| 0.317968 | 0.073947 | 0.012 | 0.007 | 1 | 2 | Aldoc | 0.005 | Old |
| 0.318234 | 0.440316 | 0.012 | 0.007 | 1 | 2 | Tesmin | 0.005 | Old |
| 0.318252 | 0.277101 | 0.05 | 0.04 | 1 | 2 | Cinp | 0.01 | Old |
| 0.318318 | 0.198432 | 0.103 | 0.088 | 1 | 2 | Aoah | 0.015 | Old |
| 0.31915 | 0.187268 | 0.172 | 0.154 | 1 | 2 | Cand1 | 0.018 | Old |
| 0.31948 | 0.02591 | 0.927 | 0.912 | 1 | 2 | Cfl1 | 0.015 | Old |
| 0.319803 | 0.638052 | 0.025 | 0.018 | 1 | 2 | Lyrm4 | 0.007 | Old |

| | | | | | | | | |
|---|---|---|---|---|---|---|---|---|
| 0.319835 | 0.096317 | 0.012 | 0.007 | 1 | 2 | Spry2 | 0.005 | Old |
| 0.319937 | 0.088204 | 0.135 | 0.118 | 1 | 2 | Spg7 | 0.017 | Old |
| 0.319977 | 0.296367 | 0.104 | 0.09 | 1 | 2 | Lipe | 0.014 | Old |
| 0.320001 | 0.215639 | 0.166 | 0.148 | 1 | 2 | Ap3m1 | 0.018 | Old |
| 0.320126 | 0.714546 | 0.046 | 0.037 | 1 | 2 | Fam219a | 0.009 | Old |
| 0.320469 | 0.294333 | 0.112 | 0.099 | 1 | 2 | Cirbp | 0.013 | Old |
| 0.320616 | 0.203801 | 0.201 | 0.183 | 1 | 2 | Psma5 | 0.018 | Old |
| 0.320818 | 0.006336 | 0.12 | 0.103 | 1 | 2 | Tmem170b | 0.017 | Old |
| 0.321428 | 0.166872 | 0.125 | 0.109 | 1 | 2 | Helz | 0.016 | Old |
| 0.321847 | 0.347689 | 0.044 | 0.034 | 1 | 2 | Gramd1b | 0.01 | Old |
| 0.321895 | 0.084055 | 0.382 | 0.363 | 1 | 2 | Susd6 | 0.019 | Old |
| 0.321895 | 0.113319 | 0.332 | 0.31 | 1 | 2 | Ciita | 0.022 | Old |
| 0.321966 | 0.341644 | 0.025 | 0.018 | 1 | 2 | Nudt7 | 0.007 | Old |
| 0.321966 | 0.313645 | 0.025 | 0.018 | 1 | 2 | Bphl | 0.007 | Old |
| 0.322327 | 0.507187 | 0.025 | 0.018 | 1 | 2 | Atp6v0d2 | 0.007 | Old |
| 0.322417 | 0.082683 | 0.454 | 0.43 | 1 | 2 | Calm3 | 0.024 | Old |
| 0.322735 | 0.237789 | 0.062 | 0.051 | 1 | 2 | Tmco3 | 0.011 | Old |
| 0.322827 | 0.111189 | 0.109 | 0.094 | 1 | 2 | Hnrnph3 | 0.015 | Old |
| 0.323032 | 0.6223 | 0.04 | 0.031 | 1 | 2 | Dgcr6 | 0.009 | Old |
| 0.323124 | 0.310853 | 0.05 | 0.04 | 1 | 2 | Slx4ip | 0.01 | Old |
| 0.323234 | 0.195511 | 0.182 | 0.164 | 1 | 2 | Puf60 | 0.018 | Old |
| 0.323342 | 0.175145 | 0.061 | 0.05 | 1 | 2 | Mthfr | 0.011 | Old |
| 0.323594 | 0.127684 | 0.251 | 0.229 | 1 | 2 | Atp8b4 | 0.022 | Old |
| 0.323594 | 0.486761 | 0.025 | 0.018 | 1 | 2 | Ogfod3 | 0.007 | Old |
| 0.323742 | 0.102566 | 0.095 | 0.081 | 1 | 2 | Zeb2os | 0.014 | Old |
| 0.323788 | 0.436753 | 0.064 | 0.053 | 1 | 2 | Ints11 | 0.011 | Old |
| 0.323845 | 0.136653 | 0.26 | 0.241 | 1 | 2 | Sumo1 | 0.019 | Old |
| 0.323911 | 0.252433 | 0.072 | 0.059 | 1 | 2 | Thap2 | 0.013 | Old |
| 0.323996 | 0.102397 | 0.122 | 0.106 | 1 | 2 | Mrps9 | 0.016 | Old |
| 0.324056 | 0.047594 | 0.126 | 0.11 | 1 | 2 | Gng12 | 0.016 | Old |
| 0.324231 | 0.11561 | 0.087 | 0.074 | 1 | 2 | Phka2 | 0.013 | Old |
| 0.324234 | 0.169878 | 0.135 | 0.119 | 1 | 2 | Arhgap39 | 0.016 | Old |
| 0.324517 | 0.354758 | 0.058 | 0.048 | 1 | 2 | Zfp414 | 0.01 | Old |
| 0.32497 | 0.501437 | 0.02 | 0.014 | 1 | 2 | Dgke | 0.006 | Old |
| 0.325025 | 0.177137 | 0.208 | 0.188 | 1 | 2 | Btbd1 | 0.02 | Old |
| 0.325034 | 1.018363 | 0.016 | 0.011 | 1 | 2 | Cd200r4 | 0.005 | Old |
| 0.325322 | 0.162591 | 0.084 | 0.07 | 1 | 2 | 1600012H0 | 0.014 | Old |
| 0.325361 | 0.481832 | 0.059 | 0.049 | 1 | 2 | Ss18l2 | 0.01 | Old |
| 0.325439 | 0.494895 | 0.087 | 0.075 | 1 | 2 | Ccnc | 0.012 | Old |
| 0.325565 | 0.098137 | 0.763 | 0.754 | 1 | 2 | Arhgdib | 0.009 | Old |
| 0.325678 | 0.101589 | 0.342 | 0.312 | 1 | 2 | Ist1 | 0.03 | Old |
| 0.325908 | 0.212009 | 0.115 | 0.1 | 1 | 2 | Eif4enif1 | 0.015 | Old |
| 0.325982 | 0.150151 | 0.11 | 0.095 | 1 | 2 | Cds2 | 0.015 | Old |
| 0.326023 | 0.224937 | 0.192 | 0.175 | 1 | 2 | Odc1 | 0.017 | Old |
| 0.326749 | 0.338341 | 0.046 | 0.037 | 1 | 2 | 2410004B1 | 0.009 | Old |

| | | | | | | | | |
|---|---|---|---|---|---|---|---|---|
| 0.326792 | 0.43768 | 0.076 | 0.064 | 1 | 2 | Trim5 | 0.012 | Old |
| 0.327155 | 0.173268 | 0.04 | 0.031 | 1 | 2 | Ap5z1 | 0.009 | Old |
| 0.327385 | 0.559559 | 0.02 | 0.014 | 1 | 2 | Vcpkmt | 0.006 | Old |
| 0.327483 | 0.225159 | 0.063 | 0.052 | 1 | 2 | Chchd4 | 0.011 | Old |
| 0.327734 | 0.099387 | 0.18 | 0.159 | 1 | 2 | Zdhhc9 | 0.021 | Old |
| 0.327765 | 0.06831 | 0.186 | 0.167 | 1 | 2 | Commd7 | 0.019 | Old |
| 0.327809 | 0.141342 | 0.056 | 0.045 | 1 | 2 | Wdr77 | 0.011 | Old |
| 0.328171 | 0.157919 | 0.36 | 0.336 | 1 | 2 | 1110008F1 | 0.024 | Old |
| 0.328315 | 0.068441 | 0.117 | 0.1 | 1 | 2 | Ddx10 | 0.017 | Old |
| 0.328463 | 0.590255 | 0.016 | 0.011 | 1 | 2 | Plppr3 | 0.005 | Old |
| 0.328463 | 0.573541 | 0.016 | 0.011 | 1 | 2 | Batf2 | 0.005 | Old |
| 0.32876 | 0.219607 | 0.271 | 0.248 | 1 | 2 | Usp8 | 0.023 | Old |
| 0.3288 | 0.647375 | 0.02 | 0.014 | 1 | 2 | Asb1 | 0.006 | Old |
| 0.329043 | 0.021297 | 0.079 | 0.093 | 1 | 2 | Vps9d1 | -0.014 | Old |
| 0.32914 | 0.471311 | 0.081 | 0.07 | 1 | 2 | Calhm2 | 0.011 | Old |
| 0.329151 | 0.41789 | 0.016 | 0.011 | 1 | 2 | Pms2 | 0.005 | Old |
| 0.329233 | 0.519878 | 0.03 | 0.023 | 1 | 2 | Icam2 | 0.007 | Old |
| 0.329339 | 0.070733 | 0.146 | 0.128 | 1 | 2 | Atf6b | 0.018 | Old |
| 0.329409 | 0.144459 | 0.278 | 0.257 | 1 | 2 | Gapvd1 | 0.021 | Old |
| 0.329543 | 0.098066 | 0.361 | 0.336 | 1 | 2 | Atf6 | 0.025 | Old |
| 0.32964 | 0.01236 | 0.752 | 0.733 | 1 | 2 | Fxyd5 | 0.019 | Old |
| 0.329841 | 0.201485 | 0.016 | 0.011 | 1 | 2 | Mettl25 | 0.005 | Old |
| 0.330089 | 0.120647 | 0.149 | 0.131 | 1 | 2 | Sec23b | 0.018 | Old |
| 0.330335 | 0.018313 | 0.045 | 0.036 | 1 | 2 | Mecr | 0.009 | Old |
| 0.330856 | 0.091168 | 0.067 | 0.056 | 1 | 2 | Hsf1 | 0.011 | Old |
| 0.331059 | 0.096193 | 0.294 | 0.271 | 1 | 2 | Tpp1 | 0.023 | Old |
| 0.331097 | 0.005185 | 0.072 | 0.059 | 1 | 2 | Rev3l | 0.013 | Old |
| 0.331222 | 0.392514 | 0.016 | 0.011 | 1 | 2 | Slc9a3r2 | 0.005 | Old |
| 0.331439 | 0.093511 | 0.238 | 0.216 | 1 | 2 | Cmtm3 | 0.022 | Old |
| 0.331472 | 0.588043 | 0.047 | 0.038 | 1 | 2 | Mcm5 | 0.009 | Old |
| 0.332192 | 0.215266 | 0.046 | 0.037 | 1 | 2 | E2f3 | 0.009 | Old |
| 0.332385 | 0.774124 | 0.026 | 0.019 | 1 | 2 | Spice1 | 0.007 | Old |
| 0.332501 | 0.046336 | 0.251 | 0.224 | 1 | 2 | Mib1 | 0.027 | Old |
| 0.332622 | 0.105017 | 0.599 | 0.593 | 1 | 2 | Csnk1a1 | 0.006 | Old |
| 0.332962 | 0.26774 | 0.07 | 0.058 | 1 | 2 | Usp36 | 0.012 | Old |
| 0.333524 | 0.279722 | 0.09 | 0.077 | 1 | 2 | Tuba4a | 0.013 | Old |
| 0.334027 | 0.04911 | 0.315 | 0.287 | 1 | 2 | Baz1a | 0.028 | Old |
| 0.334548 | 0.304115 | 0.03 | 0.023 | 1 | 2 | Rpl36a-ps1 | 0.007 | Old |
| 0.335 | 0.043273 | 0.332 | 0.304 | 1 | 2 | Phip | 0.028 | Old |
| 0.335496 | 0.072475 | 0.318 | 0.29 | 1 | 2 | Bcap31 | 0.028 | Old |
| 0.335633 | 0.940466 | 0.026 | 0.019 | 1 | 2 | Ifi206 | 0.007 | Old |
| 0.335992 | 0.234731 | 0.216 | 0.198 | 1 | 2 | Sorl1 | 0.018 | Old |
| 0.336588 | 0.042546 | 0.224 | 0.202 | 1 | 2 | Tsc22d1 | 0.022 | Old |
| 0.336901 | 0.089846 | 0.019 | 0.026 | 1 | 2 | Stat4 | -0.007 | Old |
| 0.336932 | 0.172984 | 0.035 | 0.027 | 1 | 2 | Osbpl3 | 0.008 | Old |

| | | | | | | | | |
|---|---|---|---|---|---|---|---|---|
| 0.337177 | 0.165662 | 0.233 | 0.21 | 1 | 2 | Pbxip1 | 0.023 | Old |
| 0.337288 | 0.198451 | 0.107 | 0.094 | 1 | 2 | Phf11b | 0.013 | Old |
| 0.337337 | 0.158401 | 0.102 | 0.088 | 1 | 2 | Fnta | 0.014 | Old |
| 0.337369 | 0.061012 | 0.02 | 0.014 | 1 | 2 | Ly6i | 0.006 | Old |
| 0.337514 | 0.114414 | 0.349 | 0.324 | 1 | 2 | Map7d1 | 0.025 | Old |
| 0.33768 | 0.068512 | 0.468 | 0.506 | 1 | 2 | Cx3cr1 | -0.038 | Old |
| 0.338344 | 0.104338 | 0.219 | 0.2 | 1 | 2 | Stk40 | 0.019 | Old |
| 0.338563 | 0.031822 | 0.086 | 0.072 | 1 | 2 | Med11 | 0.014 | Old |
| 0.338806 | 0.078324 | 0.524 | 0.492 | 1 | 2 | Tab2 | 0.032 | Old |
| 0.339026 | 0.368641 | 0.063 | 0.052 | 1 | 2 | Hat1 | 0.011 | Old |
| 0.339145 | 0.194632 | 0.197 | 0.178 | 1 | 2 | Foxn2 | 0.019 | Old |
| 0.339427 | 0.20619 | 0.117 | 0.102 | 1 | 2 | Pofut2 | 0.015 | Old |
| 0.339449 | 0.420773 | 0.077 | 0.065 | 1 | 2 | Brix1 | 0.012 | Old |
| 0.339519 | 0.126728 | 0.272 | 0.251 | 1 | 2 | Nab1 | 0.021 | Old |
| 0.339852 | 0.250681 | 0.198 | 0.183 | 1 | 2 | Rbm6 | 0.015 | Old |
| 0.340082 | 0.007478 | 0.384 | 0.405 | 1 | 2 | Spop | -0.021 | Old |
| 0.340322 | 0.388652 | 0.054 | 0.044 | 1 | 2 | Flt3l | 0.01 | Old |
| 0.341606 | 0.224376 | 0.03 | 0.023 | 1 | 2 | Orc5 | 0.007 | Old |
| 0.341754 | 0.269078 | 0.046 | 0.037 | 1 | 2 | Lsm1 | 0.009 | Old |
| 0.341862 | 0.173972 | 0.214 | 0.195 | 1 | 2 | Mfsd14a | 0.019 | Old |
| 0.342753 | 0.416625 | 0.085 | 0.072 | 1 | 2 | Gpd1l | 0.013 | Old |
| 0.342818 | 0.060066 | 0.107 | 0.093 | 1 | 2 | Ftsj3 | 0.014 | Old |
| 0.342949 | 0.243619 | 0.2 | 0.184 | 1 | 2 | Gorasp2 | 0.016 | Old |
| 0.343098 | 0.272181 | 0.201 | 0.184 | 1 | 2 | Znhit1 | 0.017 | Old |
| 0.343221 | 0.187141 | 0.303 | 0.28 | 1 | 2 | Sgpp1 | 0.023 | Old |
| 0.343833 | 0.736567 | 0.021 | 0.015 | 1 | 2 | Zkscan8 | 0.006 | Old |
| 0.344674 | 0.29942 | 0.065 | 0.055 | 1 | 2 | Ctc1 | 0.01 | Old |
| 0.344948 | 0.180298 | 0.312 | 0.29 | 1 | 2 | Bnip3l | 0.022 | Old |
| 0.345085 | 0.08341 | 0.304 | 0.285 | 1 | 2 | Batf3 | 0.019 | Old |
| 0.345387 | 0.03389 | 0.051 | 0.042 | 1 | 2 | Sugp1 | 0.009 | Old |
| 0.345683 | 0.078858 | 0.276 | 0.252 | 1 | 2 | Vps13c | 0.024 | Old |
| 0.345865 | 0.37088 | 0.026 | 0.019 | 1 | 2 | Ip6k2 | 0.007 | Old |
| 0.346094 | 0.139842 | 0.16 | 0.141 | 1 | 2 | Kidins220 | 0.019 | Old |
| 0.34624 | 0.086728 | 0.618 | 0.612 | 1 | 2 | Ier5 | 0.006 | Old |
| 0.347073 | 0.709785 | 0.021 | 0.015 | 1 | 2 | Zbtb11os1 | 0.006 | Old |
| 0.347371 | 0.229164 | 0.099 | 0.086 | 1 | 2 | Meaf6 | 0.013 | Old |
| 0.347375 | 0.256725 | 0.066 | 0.056 | 1 | 2 | Lmf2 | 0.01 | Old |
| 0.34792 | 0.018224 | 0.091 | 0.077 | 1 | 2 | Faf2 | 0.014 | Old |
| 0.348031 | 0.25881 | 0.054 | 0.044 | 1 | 2 | Kat2a | 0.01 | Old |
| 0.348313 | 0.04962 | 0.082 | 0.095 | 1 | 2 | Rbpms | -0.013 | Old |
| 0.348565 | 0.132212 | 0.494 | 0.477 | 1 | 2 | Ywhah | 0.017 | Old |
| 0.348598 | 0.049368 | 0.195 | 0.175 | 1 | 2 | Chtop | 0.02 | Old |
| 0.348817 | 0.218045 | 0.126 | 0.112 | 1 | 2 | Smarca2 | 0.014 | Old |
| 0.348847 | 0.060328 | 0.028 | 0.036 | 1 | 2 | Scai | -0.008 | Old |
| 0.34901 | 0.084767 | 0.026 | 0.019 | 1 | 2 | Slc25a38 | 0.007 | Old |

| | | | | | | | | |
|---|---|---|---|---|---|---|---|---|
| 0.349374 | 0.483855 | 0.042 | 0.033 | 1 | 2 | Acot13 | 0.009 | Old |
| 0.349379 | 1.070948 | 0.013 | 0.008 | 1 | 2 | Pecam1 | 0.005 | Old |
| 0.349456 | 0.189433 | 0.065 | 0.055 | 1 | 2 | Golim4 | 0.01 | Old |
| 0.349779 | 0.288272 | 0.031 | 0.024 | 1 | 2 | Kbtbd7 | 0.007 | Old |
| 0.349834 | 0.117154 | 0.11 | 0.096 | 1 | 2 | Rab3gap1 | 0.014 | Old |
| 0.349915 | 0.865556 | 0.013 | 0.008 | 1 | 2 | Chst1 | 0.005 | Old |
| 0.350086 | 0.168422 | 0.111 | 0.097 | 1 | 2 | Mboat7 | 0.014 | Old |
| 0.350223 | 0.322342 | 0.078 | 0.067 | 1 | 2 | Psmd9 | 0.011 | Old |
| 0.350581 | 0.447139 | 0.043 | 0.034 | 1 | 2 | Ryr1 | 0.009 | Old |
| 0.350619 | 0.161731 | 0.024 | 0.031 | 1 | 2 | Zfp146 | -0.007 | Old |
| 0.350682 | 0.041288 | 0.27 | 0.246 | 1 | 2 | Vdac1 | 0.024 | Old |
| 0.35084 | 0.149816 | 0.176 | 0.158 | 1 | 2 | Lars2 | 0.018 | Old |
| 0.350941 | 0.220907 | 0.143 | 0.127 | 1 | 2 | Socs4 | 0.016 | Old |
| 0.350965 | 0.110833 | 0.07 | 0.058 | 1 | 2 | Csrp1 | 0.012 | Old |
| 0.351257 | 0.359494 | 0.013 | 0.008 | 1 | 2 | Osgepl1 | 0.005 | Old |
| 0.351561 | 0.565088 | 0.021 | 0.015 | 1 | 2 | Sco1 | 0.006 | Old |
| 0.351976 | 0.397013 | 0.069 | 0.058 | 1 | 2 | Slc4a1ap | 0.011 | Old |
| 0.352054 | 0.022632 | 0.149 | 0.166 | 1 | 2 | Abcc1 | -0.017 | Old |
| 0.352098 | 0.05862 | 0.239 | 0.216 | 1 | 2 | Snrpd1 | 0.023 | Old |
| 0.352125 | 0.322387 | 0.178 | 0.162 | 1 | 2 | Stat5b | 0.016 | Old |
| 0.352306 | 0.073515 | 0.701 | 0.686 | 1 | 2 | Ube2d3 | 0.015 | Old |
| 0.352381 | 0.267547 | 0.021 | 0.015 | 1 | 2 | Vamp1 | 0.006 | Old |
| 0.35242 | 0.045207 | 0.256 | 0.234 | 1 | 2 | Pkn2 | 0.022 | Old |
| 0.352669 | 0.128229 | 0.082 | 0.07 | 1 | 2 | Trim47 | 0.012 | Old |
| 0.352913 | 0.016612 | 0.398 | 0.36 | 1 | 2 | Usp9x | 0.038 | Old |
| 0.353018 | 0.288529 | 0.103 | 0.09 | 1 | 2 | Ikzf5 | 0.013 | Old |
| 0.353141 | 0.465768 | 0.013 | 0.008 | 1 | 2 | Rpusd1 | 0.005 | Old |
| 0.353166 | 0.042856 | 0.101 | 0.087 | 1 | 2 | Gabpa | 0.014 | Old |
| 0.353198 | 0.306555 | 0.169 | 0.153 | 1 | 2 | Sptlc2 | 0.016 | Old |
| 0.353299 | 0.104092 | 0.047 | 0.038 | 1 | 2 | Dbf4 | 0.009 | Old |
| 0.353314 | 0.137118 | 0.47 | 0.439 | 1 | 2 | Creg1 | 0.031 | Old |
| 0.353407 | 0.264437 | 0.021 | 0.015 | 1 | 2 | Zfp52 | 0.006 | Old |
| 0.353411 | 0.88269 | 0.013 | 0.008 | 1 | 2 | Gm27003 | 0.005 | Old |
| 0.353723 | 0.078181 | 0.414 | 0.385 | 1 | 2 | Glul | 0.029 | Old |
| 0.353981 | 0.494789 | 0.054 | 0.044 | 1 | 2 | Cpped1 | 0.01 | Old |
| 0.35402 | 0.049139 | 0.154 | 0.137 | 1 | 2 | Ndufs3 | 0.017 | Old |
| 0.354436 | 0.408948 | 0.021 | 0.015 | 1 | 2 | Prpf3 | 0.006 | Old |
| 0.354742 | 0.370996 | 0.092 | 0.08 | 1 | 2 | Fut8 | 0.012 | Old |
| 0.354761 | 0.516121 | 0.013 | 0.008 | 1 | 2 | Suox | 0.005 | Old |
| 0.354761 | 0.496349 | 0.013 | 0.008 | 1 | 2 | Fbxl8 | 0.005 | Old |
| 0.354875 | 0.088779 | 0.298 | 0.274 | 1 | 2 | Atxn2l | 0.024 | Old |
| 0.35523 | 0.061385 | 0.149 | 0.133 | 1 | 2 | Pqlc3 | 0.016 | Old |
| 0.35526 | 0.090211 | 0.314 | 0.292 | 1 | 2 | Scpep1 | 0.022 | Old |
| 0.355295 | 0.336292 | 0.036 | 0.029 | 1 | 2 | Mettl21a | 0.007 | Old |
| 0.355573 | 0.174597 | 0.013 | 0.008 | 1 | 2 | Mtrf1 | 0.005 | Old |

| | | | | | | | | |
|---|---|---|---|---|---|---|---|---|
| 0.355672 | 0.344117 | 0.021 | 0.015 | 1 | 2 | Zfp81 | 0.006 | Old |
| 0.35568 | 0.118957 | 0.177 | 0.158 | 1 | 2 | Dnajc2 | 0.019 | Old |
| 0.355696 | 0.754807 | 0.017 | 0.012 | 1 | 2 | Car5b | 0.005 | Old |
| 0.355844 | 0.162161 | 0.013 | 0.008 | 1 | 2 | Nck2 | 0.005 | Old |
| 0.356064 | 0.185154 | 0.08 | 0.069 | 1 | 2 | B3galnt2 | 0.011 | Old |
| 0.356178 | 0.375596 | 0.108 | 0.096 | 1 | 2 | Tlk2 | 0.012 | Old |
| 0.35626 | 0.076093 | 0.089 | 0.076 | 1 | 2 | Tax1bp3 | 0.013 | Old |
| 0.356794 | 0.115097 | 0.078 | 0.067 | 1 | 2 | Mrpl37 | 0.011 | Old |
| 0.356928 | 0.300514 | 0.013 | 0.008 | 1 | 2 | Lyrm1 | 0.005 | Old |
| 0.356967 | 0.077238 | 0.275 | 0.249 | 1 | 2 | Eif3b | 0.026 | Old |
| 0.357007 | 0.151445 | 0.111 | 0.097 | 1 | 2 | BC003331 | 0.014 | Old |
| 0.357106 | 0.43994 | 0.081 | 0.07 | 1 | 2 | Slc2a6 | 0.011 | Old |
| 0.3572 | 0.253568 | 0.013 | 0.008 | 1 | 2 | Spryd4 | 0.005 | Old |
| 0.3572 | 0.154117 | 0.013 | 0.008 | 1 | 2 | Zfp553 | 0.005 | Old |
| 0.357323 | 0.630031 | 0.017 | 0.012 | 1 | 2 | Neurl1a | 0.005 | Old |
| 0.35749 | 0.010311 | 0.312 | 0.284 | 1 | 2 | Tnpo1 | 0.028 | Old |
| 0.357686 | 0.417757 | 0.082 | 0.071 | 1 | 2 | Casp6 | 0.011 | Old |
| 0.357788 | 0.523353 | 0.017 | 0.012 | 1 | 2 | Sfxn5 | 0.005 | Old |
| 0.357869 | 0.245959 | 0.092 | 0.08 | 1 | 2 | Phyh | 0.012 | Old |
| 0.357926 | 0.042428 | 0.554 | 0.526 | 1 | 2 | Cbl | 0.028 | Old |
| 0.358361 | 0.055331 | 0.021 | 0.015 | 1 | 2 | Zfp809 | 0.006 | Old |
| 0.35868 | 0.316219 | 0.11 | 0.097 | 1 | 2 | Snrpa1 | 0.013 | Old |
| 0.359025 | 0.044461 | 0.504 | 0.476 | 1 | 2 | Csrnp1 | 0.028 | Old |
| 0.359107 | 0.387903 | 0.055 | 0.045 | 1 | 2 | Ocel1 | 0.01 | Old |
| 0.359352 | 0.161253 | 0.158 | 0.143 | 1 | 2 | Arl1 | 0.015 | Old |
| 0.35946 | 0.3442 | 0.08 | 0.069 | 1 | 2 | Zmat5 | 0.011 | Old |
| 0.359619 | 0.009276 | 0.029 | 0.037 | 1 | 2 | Atp2b4 | -0.008 | Old |
| 0.359872 | 0.000707 | 0.2 | 0.181 | 1 | 2 | Isy1 | 0.019 | Old |
| 0.360022 | 0.201645 | 0.021 | 0.015 | 1 | 2 | Emc9 | 0.006 | Old |
| 0.360089 | 0.183626 | 0.091 | 0.078 | 1 | 2 | Trappc6a | 0.013 | Old |
| 0.360171 | 0.177567 | 0.036 | 0.029 | 1 | 2 | Ift22 | 0.007 | Old |
| 0.360331 | 0.125687 | 0.134 | 0.118 | 1 | 2 | Cetn3 | 0.016 | Old |
| 0.360365 | 0.105356 | 0.891 | 0.9 | 1 | 2 | Klf6 | -0.009 | Old |
| 0.360689 | 0.325172 | 0.074 | 0.063 | 1 | 2 | Rnf167 | 0.011 | Old |
| 0.360737 | 0.094017 | 0.124 | 0.109 | 1 | 2 | Dram1 | 0.015 | Old |
| 0.361016 | 0.238373 | 0.145 | 0.129 | 1 | 2 | Tlnrd1 | 0.016 | Old |
| 0.361037 | 0.145915 | 0.054 | 0.044 | 1 | 2 | Dnajc25 | 0.01 | Old |
| 0.361292 | 0.132061 | 0.017 | 0.012 | 1 | 2 | Polr1b | 0.005 | Old |
| 0.361693 | 0.012709 | 0.045 | 0.055 | 1 | 2 | Mbtps2 | -0.01 | Old |
| 0.361846 | 0.120106 | 0.262 | 0.241 | 1 | 2 | Sltm | 0.021 | Old |
| 0.362107 | 0.27225 | 0.261 | 0.243 | 1 | 2 | Grk3 | 0.018 | Old |
| 0.362229 | 0.08897 | 0.036 | 0.029 | 1 | 2 | Mettl3 | 0.007 | Old |
| 0.36223 | 0.176789 | 0.017 | 0.012 | 1 | 2 | Supv3l1 | 0.005 | Old |
| 0.362291 | 0.178597 | 0.338 | 0.321 | 1 | 2 | Psmb8 | 0.017 | Old |
| 0.362339 | 0.023171 | 0.092 | 0.078 | 1 | 2 | Zfp322a | 0.014 | Old |

| | | | | | | | | |
|---|---|---|---|---|---|---|---|---|
| 0.362391 | 0.023194 | 0.042 | 0.033 | 1 | 2 | Slc35d1 | 0.009 | Old |
| 0.362563 | 0.057085 | 0.089 | 0.076 | 1 | 2 | Tdg | 0.013 | Old |
| 0.363242 | 0.081976 | 0.204 | 0.184 | 1 | 2 | Ocrl | 0.02 | Old |
| 0.363369 | 0.084933 | 0.088 | 0.076 | 1 | 2 | Cd9 | 0.012 | Old |
| 0.363609 | 0.139072 | 0.187 | 0.169 | 1 | 2 | Trappc6b | 0.018 | Old |
| 0.363838 | 0.124228 | 0.164 | 0.146 | 1 | 2 | Ddx42 | 0.018 | Old |
| 0.364026 | 0.44067 | 0.037 | 0.03 | 1 | 2 | Miip | 0.007 | Old |
| 0.364258 | 0.198736 | 0.078 | 0.067 | 1 | 2 | Timm50 | 0.011 | Old |
| 0.364301 | 0.253237 | 0.233 | 0.215 | 1 | 2 | Dcaf12 | 0.018 | Old |
| 0.364537 | 0.215182 | 0.631 | 0.624 | 1 | 2 | Pltp | 0.007 | Old |
| 0.364971 | 0.14605 | 0.086 | 0.074 | 1 | 2 | Phactr4 | 0.012 | Old |
| 0.365197 | 0.256213 | 0.043 | 0.034 | 1 | 2 | Gm12840 | 0.009 | Old |
| 0.365449 | 0.139017 | 0.134 | 0.119 | 1 | 2 | Psen1 | 0.015 | Old |
| 0.365736 | 0.002806 | 0.143 | 0.126 | 1 | 2 | Ppt2 | 0.017 | Old |
| 0.365764 | 0.031147 | 0.032 | 0.04 | 1 | 2 | Sppl2b | -0.008 | Old |
| 0.365932 | 0.199331 | 0.135 | 0.12 | 1 | 2 | Riok1 | 0.015 | Old |
| 0.365996 | 0.074292 | 0.137 | 0.121 | 1 | 2 | Pdzd8 | 0.016 | Old |
| 0.366001 | 0.057512 | 0.091 | 0.078 | 1 | 2 | Nolc1 | 0.013 | Old |
| 0.366329 | 0.253905 | 0.027 | 0.02 | 1 | 2 | Tmem94 | 0.007 | Old |
| 0.366329 | 0.04423 | 0.027 | 0.02 | 1 | 2 | Vamp5 | 0.007 | Old |
| 0.366338 | 0.189008 | 0.16 | 0.143 | 1 | 2 | Tcf20 | 0.017 | Old |
| 0.366346 | 0.0116 | 0.18 | 0.16 | 1 | 2 | Pa2g4 | 0.02 | Old |
| 0.366371 | 0.299631 | 0.043 | 0.034 | 1 | 2 | Zfp64 | 0.009 | Old |
| 0.366385 | 0.252316 | 0.061 | 0.051 | 1 | 2 | Adora2a | 0.01 | Old |
| 0.366738 | 0.191587 | 0.124 | 0.11 | 1 | 2 | Upf1 | 0.014 | Old |
| 0.366875 | 0.072972 | 0.354 | 0.331 | 1 | 2 | Atp5o.1 | 0.023 | Old |
| 0.367045 | 0.0086 | 0.158 | 0.141 | 1 | 2 | Slamf8 | 0.017 | Old |
| 0.367073 | 0.39193 | 0.089 | 0.077 | 1 | 2 | Sympk | 0.012 | Old |
| 0.367252 | 0.133562 | 0.056 | 0.046 | 1 | 2 | Bzw2 | 0.01 | Old |
| 0.367463 | 0.068284 | 0.179 | 0.16 | 1 | 2 | Mtm1 | 0.019 | Old |
| 0.368136 | 0.275904 | 0.043 | 0.034 | 1 | 2 | Spred2 | 0.009 | Old |
| 0.368175 | 0.064266 | 0.14 | 0.125 | 1 | 2 | Nfu1 | 0.015 | Old |
| 0.368229 | 0.228323 | 0.252 | 0.233 | 1 | 2 | Nus1 | 0.019 | Old |
| 0.368248 | 0.089765 | 0.292 | 0.266 | 1 | 2 | AC149090. | 0.026 | Old |
| 0.368467 | 0.091171 | 0.077 | 0.065 | 1 | 2 | Cpq | 0.012 | Old |
| 0.368731 | 0.187111 | 0.123 | 0.109 | 1 | 2 | Map2k4 | 0.014 | Old |
| 0.369519 | 0.005873 | 0.214 | 0.192 | 1 | 2 | Rdx | 0.022 | Old |
| 0.369575 | 0.621199 | 0.051 | 0.043 | 1 | 2 | Dubr | 0.008 | Old |
| 0.369686 | 0.242988 | 0.162 | 0.146 | 1 | 2 | Kmt2d | 0.016 | Old |
| 0.369855 | 0.320098 | 0.124 | 0.112 | 1 | 2 | Copg1 | 0.012 | Old |
| 0.370019 | 0.213154 | 0.118 | 0.132 | 1 | 2 | Il6 | -0.014 | Old |
| 0.370265 | 0.154208 | 0.027 | 0.02 | 1 | 2 | Zfp994 | 0.007 | Old |
| 0.370342 | 0.244193 | 0.088 | 0.076 | 1 | 2 | Tada1 | 0.012 | Old |
| 0.370432 | 0.055091 | 0.118 | 0.103 | 1 | 2 | Mrpl43 | 0.015 | Old |
| 0.370684 | 0.003564 | 0.336 | 0.31 | 1 | 2 | Cnn2 | 0.026 | Old |

| | | | | | | | | |
|---|---|---|---|---|---|---|---|---|
| 0.370749 | 0.469999 | 0.039 | 0.031 | 1 | 2 | Ifi213 | 0.008 | Old |
| 0.371142 | 0.140472 | 0.132 | 0.116 | 1 | 2 | Gtf2b | 0.016 | Old |
| 0.37159 | 0.102055 | 0.175 | 0.158 | 1 | 2 | Srrt | 0.017 | Old |
| 0.37218 | 0.34787 | 0.057 | 0.048 | 1 | 2 | Jrkl | 0.009 | Old |
| 0.372535 | 0.708951 | 0.033 | 0.026 | 1 | 2 | Pithd1 | 0.007 | Old |
| 0.373088 | 0.068791 | 0.398 | 0.376 | 1 | 2 | Taok3 | 0.022 | Old |
| 0.373145 | 0.275675 | 0.056 | 0.046 | 1 | 2 | Wdr81 | 0.01 | Old |
| 0.373719 | 0.071615 | 0.257 | 0.234 | 1 | 2 | Eif1b | 0.023 | Old |
| 0.374355 | 0.197696 | 0.193 | 0.176 | 1 | 2 | Ccnd3 | 0.017 | Old |
| 0.374549 | 0.174355 | 0.246 | 0.228 | 1 | 2 | Ik | 0.018 | Old |
| 0.37455 | 0.168994 | 0.075 | 0.064 | 1 | 2 | Trp53bp2 | 0.011 | Old |
| 0.374821 | 0.964149 | 0.01 | 0.006 | 1 | 2 | Efcab2 | 0.004 | Old |
| 0.37497 | 0.450674 | 0.039 | 0.031 | 1 | 2 | Rnf185 | 0.008 | Old |
| 0.375192 | 0.166962 | 0.078 | 0.067 | 1 | 2 | Thumpd3 | 0.011 | Old |
| 0.37522 | 0.10216 | 0.079 | 0.068 | 1 | 2 | Pno1 | 0.011 | Old |
| 0.375308 | 0.220959 | 0.214 | 0.195 | 1 | 2 | Myd88 | 0.019 | Old |
| 0.375562 | 0.184097 | 0.395 | 0.378 | 1 | 2 | Ube2b | 0.017 | Old |
| 0.375798 | 0.841923 | 0.01 | 0.006 | 1 | 2 | Jcad | 0.004 | Old |
| 0.376018 | 0.245552 | 0.141 | 0.127 | 1 | 2 | 8-Sep | 0.014 | Old |
| 0.376082 | 0.027248 | 0.333 | 0.304 | 1 | 2 | Elavl1 | 0.029 | Old |
| 0.376124 | 0.574915 | 0.01 | 0.006 | 1 | 2 | Gm12089 | 0.004 | Old |
| 0.376451 | 0.77863 | 0.01 | 0.006 | 1 | 2 | Zrsr1 | 0.004 | Old |
| 0.376777 | 0.599437 | 0.01 | 0.006 | 1 | 2 | Trim6 | 0.004 | Old |
| 0.376801 | 0.353783 | 0.056 | 0.046 | 1 | 2 | Mapk9 | 0.01 | Old |
| 0.376934 | 0.076042 | 0.225 | 0.205 | 1 | 2 | Rasa3 | 0.02 | Old |
| 0.376985 | 0.33903 | 0.067 | 0.057 | 1 | 2 | Cln5 | 0.01 | Old |
| 0.377431 | 0.867346 | 0.01 | 0.006 | 1 | 2 | Crybb1 | 0.004 | Old |
| 0.377486 | 0.091626 | 0.302 | 0.28 | 1 | 2 | Mdfic | 0.022 | Old |
| 0.377736 | 0.078178 | 0.128 | 0.113 | 1 | 2 | Cul5 | 0.015 | Old |
| 0.377758 | 0.073793 | 0.074 | 0.063 | 1 | 2 | Ubl7 | 0.011 | Old |
| 0.378005 | 0.123444 | 0.061 | 0.051 | 1 | 2 | Clk2 | 0.01 | Old |
| 0.378162 | 0.006276 | 0.088 | 0.075 | 1 | 2 | Ctbs | 0.013 | Old |
| 0.378387 | 0.193193 | 0.074 | 0.063 | 1 | 2 | Ap3s2 | 0.011 | Old |
| 0.378394 | 0.069813 | 0.07 | 0.081 | 1 | 2 | Snx15 | -0.011 | Old |
| 0.378602 | 0.146949 | 0.46 | 0.444 | 1 | 2 | Pbrm1 | 0.016 | Old |
| 0.378755 | 0.821516 | 0.034 | 0.027 | 1 | 2 | Zfp386 | 0.007 | Old |
| 0.378758 | 0.064686 | 0.113 | 0.1 | 1 | 2 | Adar | 0.013 | Old |
| 0.379068 | 0.889414 | 0.01 | 0.006 | 1 | 2 | Gdpd5 | 0.004 | Old |
| 0.379396 | 0.523529 | 0.01 | 0.006 | 1 | 2 | Zfp213 | 0.004 | Old |
| 0.379456 | 0.007778 | 0.022 | 0.017 | 1 | 2 | Dcun1d2 | 0.005 | Old |
| 0.379724 | 0.60593 | 0.01 | 0.006 | 1 | 2 | Rtel1 | 0.004 | Old |
| 0.379724 | 0.419105 | 0.01 | 0.006 | 1 | 2 | Tmc4 | 0.004 | Old |
| 0.380014 | 0.528068 | 0.051 | 0.043 | 1 | 2 | Blcap | 0.008 | Old |
| 0.380052 | 0.520366 | 0.01 | 0.006 | 1 | 2 | Rad51b | 0.004 | Old |
| 0.380201 | 0.193888 | 0.543 | 0.529 | 1 | 2 | Itga4 | 0.014 | Old |

| | | | | | | | | |
|---|---|---|---|---|---|---|---|---|
| 0.380266 | 0.069603 | 0.08 | 0.069 | 1 | 2 | Nfyc | 0.011 | Old |
| 0.380381 | 0.279653 | 0.01 | 0.006 | 1 | 2 | Tmem218 | 0.004 | Old |
| 0.380411 | 0.060149 | 0.136 | 0.121 | 1 | 2 | Pcif1 | 0.015 | Old |
| 0.380501 | 0.193994 | 0.022 | 0.017 | 1 | 2 | Whamm | 0.005 | Old |
| 0.380548 | 0.067423 | 0.142 | 0.127 | 1 | 2 | Tmem248 | 0.015 | Old |
| 0.380851 | 0.15847 | 0.121 | 0.107 | 1 | 2 | Cnot7 | 0.014 | Old |
| 0.380939 | 0.154321 | 0.043 | 0.034 | 1 | 2 | Zc3h10 | 0.009 | Old |
| 0.381105 | 0.03068 | 0.506 | 0.482 | 1 | 2 | Prrc2c | 0.024 | Old |
| 0.381203 | 0.181806 | 0.056 | 0.046 | 1 | 2 | Tusc2 | 0.01 | Old |
| 0.381263 | 0.083728 | 0.049 | 0.04 | 1 | 2 | Idua | 0.009 | Old |
| 0.381948 | 0.024262 | 0.15 | 0.132 | 1 | 2 | Magoh | 0.018 | Old |
| 0.382003 | 0.072585 | 0.513 | 0.483 | 1 | 2 | Rpl22l1 | 0.03 | Old |
| 0.382026 | 0.375482 | 0.01 | 0.006 | 1 | 2 | Gpc1 | 0.004 | Old |
| 0.382069 | 0.265775 | 0.039 | 0.031 | 1 | 2 | Iffo2 | 0.008 | Old |
| 0.382231 | 1.502343 | 0.014 | 0.01 | 1 | 2 | Gm12166 | 0.004 | Old |
| 0.3825 | 1.438807 | 0.014 | 0.01 | 1 | 2 | Tm7sf2 | 0.004 | Old |
| 0.382651 | 0.125995 | 0.102 | 0.089 | 1 | 2 | Mfsd6 | 0.013 | Old |
| 0.383015 | 0.409414 | 0.01 | 0.006 | 1 | 2 | Pfkfb2 | 0.004 | Old |
| 0.383015 | 0.032008 | 0.01 | 0.006 | 1 | 2 | Meis3 | 0.004 | Old |
| 0.383091 | 0.183584 | 0.245 | 0.229 | 1 | 2 | Rad21 | 0.016 | Old |
| 0.383189 | 0.107502 | 0.461 | 0.441 | 1 | 2 | Etv3 | 0.02 | Old |
| 0.383265 | 0.046342 | 0.204 | 0.186 | 1 | 2 | Plekhg3 | 0.018 | Old |
| 0.384486 | 0.149526 | 0.022 | 0.017 | 1 | 2 | Acsf3 | 0.005 | Old |
| 0.384495 | 0.081491 | 0.223 | 0.201 | 1 | 2 | Plekhj1 | 0.022 | Old |
| 0.384678 | 0.099215 | 0.305 | 0.287 | 1 | 2 | Rab11fip1 | 0.018 | Old |
| 0.385058 | 0.098054 | 0.074 | 0.063 | 1 | 2 | Akr1c13 | 0.011 | Old |
| 0.385556 | 0.036891 | 0.082 | 0.071 | 1 | 2 | Shc1 | 0.011 | Old |
| 0.385587 | 0.116858 | 0.404 | 0.379 | 1 | 2 | Adam17 | 0.025 | Old |
| 0.385806 | 0.083339 | 0.338 | 0.314 | 1 | 2 | Wapl | 0.024 | Old |
| 0.385888 | 0.241955 | 0.028 | 0.021 | 1 | 2 | Ctsf | 0.007 | Old |
| 0.385983 | 0.218313 | 0.158 | 0.141 | 1 | 2 | Psme3 | 0.017 | Old |
| 0.386055 | 0.215177 | 0.1 | 0.088 | 1 | 2 | Cryl1 | 0.012 | Old |
| 0.386491 | 0.495735 | 0.045 | 0.037 | 1 | 2 | Hars2 | 0.008 | Old |
| 0.386598 | 0.069246 | 0.084 | 0.071 | 1 | 2 | Pnpla7 | 0.013 | Old |
| 0.387165 | 0.118162 | 0.179 | 0.163 | 1 | 2 | Ccdc115 | 0.016 | Old |
| 0.387492 | 0.242033 | 0.089 | 0.077 | 1 | 2 | Shkbp1 | 0.012 | Old |
| 0.387504 | 0.097767 | 0.147 | 0.131 | 1 | 2 | Nck1 | 0.016 | Old |
| 0.38761 | 0.136484 | 0.142 | 0.128 | 1 | 2 | Ubr3 | 0.014 | Old |
| 0.387635 | 1.180361 | 0.024 | 0.018 | 1 | 2 | Zfp617 | 0.006 | Old |
| 0.387917 | 0.588462 | 0.014 | 0.01 | 1 | 2 | 2510046G1 | 0.004 | Old |
| 0.388189 | 0.370472 | 0.014 | 0.01 | 1 | 2 | Pigz | 0.004 | Old |
| 0.388229 | 0.467952 | 0.029 | 0.023 | 1 | 2 | Sclt1 | 0.006 | Old |
| 0.38863 | 0.113062 | 0.109 | 0.095 | 1 | 2 | Ddx23 | 0.014 | Old |
| 0.388718 | 0.275496 | 0.018 | 0.013 | 1 | 2 | Thop1 | 0.005 | Old |
| 0.388833 | 0.559131 | 0.091 | 0.081 | 1 | 2 | Gmppa | 0.01 | Old |

| | | | | | | | | |
|---|---|---|---|---|---|---|---|---|
| 0.388863 | 0.294424 | 0.099 | 0.087 | 1 | 2 | Pdcd7 | 0.012 | Old |
| 0.388887 | 0.206718 | 0.49 | 0.488 | 1 | 2 | Sgms1 | 0.002 | Old |
| 0.388955 | 0.424836 | 0.018 | 0.013 | 1 | 2 | Rpgrip1 | 0.005 | Old |
| 0.389006 | 0.113719 | 0.115 | 0.1 | 1 | 2 | Smad1 | 0.015 | Old |
| 0.38902 | 0.04222 | 0.061 | 0.051 | 1 | 2 | Ppard | 0.01 | Old |
| 0.389043 | 0.039125 | 0.142 | 0.126 | 1 | 2 | Mid1ip1 | 0.016 | Old |
| 0.389194 | 0.022909 | 0.281 | 0.254 | 1 | 2 | Samd4b | 0.027 | Old |
| 0.389279 | 0.498164 | 0.014 | 0.01 | 1 | 2 | Spaca9 | 0.004 | Old |
| 0.389503 | 0.207086 | 0.094 | 0.082 | 1 | 2 | Tbcel | 0.012 | Old |
| 0.389552 | 0.668133 | 0.014 | 0.01 | 1 | 2 | Prnp | 0.004 | Old |
| 0.389669 | 0.191708 | 0.157 | 0.143 | 1 | 2 | Esyt2 | 0.014 | Old |
| 0.389824 | 0.537321 | 0.014 | 0.01 | 1 | 2 | Zfp40 | 0.004 | Old |
| 0.389901 | 0.033408 | 0.195 | 0.177 | 1 | 2 | Cebpa | 0.018 | Old |
| 0.390186 | 0.031912 | 0.066 | 0.056 | 1 | 2 | 4930455G( | 0.01 | Old |
| 0.390202 | 0.19527 | 0.084 | 0.072 | 1 | 2 | Tbc1d17 | 0.012 | Old |
| 0.39036 | 0.174821 | 0.191 | 0.176 | 1 | 2 | Cacybp | 0.015 | Old |
| 0.39037 | 0.35202 | 0.014 | 0.01 | 1 | 2 | Dtd1 | 0.004 | Old |
| 0.390375 | 0.350409 | 0.018 | 0.013 | 1 | 2 | Slc25a40 | 0.005 | Old |
| 0.390417 | 0.392869 | 0.065 | 0.056 | 1 | 2 | Nit1 | 0.009 | Old |
| 0.390458 | 0.182821 | 0.252 | 0.236 | 1 | 2 | Ugcg | 0.016 | Old |
| 0.391131 | 0.06426 | 0.316 | 0.291 | 1 | 2 | Lrrc25 | 0.025 | Old |
| 0.39117 | 0.147029 | 0.179 | 0.163 | 1 | 2 | Lpar6 | 0.016 | Old |
| 0.391979 | 0.125809 | 0.148 | 0.133 | 1 | 2 | Eif4e3 | 0.015 | Old |
| 0.391986 | 0.143204 | 0.05 | 0.042 | 1 | 2 | Ring1 | 0.008 | Old |
| 0.392011 | 0.324551 | 0.014 | 0.01 | 1 | 2 | Abcb6 | 0.004 | Old |
| 0.392024 | 0.387802 | 0.051 | 0.043 | 1 | 2 | Snhg9 | 0.008 | Old |
| 0.392036 | 0.116705 | 0.018 | 0.013 | 1 | 2 | Zfp825 | 0.005 | Old |
| 0.392279 | 0.267654 | 0.045 | 0.037 | 1 | 2 | Pcyt2 | 0.008 | Old |
| 0.392578 | 0.051524 | 0.033 | 0.026 | 1 | 2 | Fbxo25 | 0.007 | Old |
| 0.392833 | 0.152642 | 0.014 | 0.01 | 1 | 2 | Gm16124 | 0.004 | Old |
| 0.392896 | 0.116501 | 0.1 | 0.087 | 1 | 2 | Rbmxl1 | 0.013 | Old |
| 0.392964 | 0.020902 | 0.146 | 0.129 | 1 | 2 | Dctn2 | 0.017 | Old |
| 0.393107 | 0.525063 | 0.014 | 0.01 | 1 | 2 | Hist1h1d | 0.004 | Old |
| 0.393144 | 0.249691 | 0.046 | 0.038 | 1 | 2 | Faap100 | 0.008 | Old |
| 0.393381 | 0.111197 | 0.014 | 0.01 | 1 | 2 | Ctu1 | 0.004 | Old |
| 0.393497 | 0.295421 | 0.119 | 0.107 | 1 | 2 | Fbxo3 | 0.012 | Old |
| 0.393506 | 0.21713 | 0.052 | 0.044 | 1 | 2 | Mterf3 | 0.008 | Old |
| 0.393552 | 0.143104 | 0.128 | 0.114 | 1 | 2 | Commd3 | 0.014 | Old |
| 0.393655 | 0.289131 | 0.014 | 0.01 | 1 | 2 | Bora | 0.004 | Old |
| 0.39374 | 0.127518 | 0.182 | 0.164 | 1 | 2 | Ubxn6 | 0.018 | Old |
| 0.393838 | 0.728874 | 0.024 | 0.018 | 1 | 2 | Stradb | 0.006 | Old |
| 0.393944 | 0.212232 | 0.084 | 0.072 | 1 | 2 | Tk2 | 0.012 | Old |
| 0.394271 | 0.01903 | 0.118 | 0.103 | 1 | 2 | Ccdc22 | 0.015 | Old |
| 0.394337 | 0.165797 | 0.092 | 0.081 | 1 | 2 | Dgcr2 | 0.011 | Old |
| 0.394616 | 0.495173 | 0.054 | 0.045 | 1 | 2 | Rnaseh2b | 0.009 | Old |

| | | | | | | | | |
|---|---|---|---|---|---|---|---|---|
| 0.394754 | 0.208792 | 0.014 | 0.01 | 1 | 2 | Ndufaf6 | 0.004 | Old |
| 0.395336 | 0.243481 | 0.072 | 0.062 | 1 | 2 | Mmd | 0.01 | Old |
| 0.395511 | 0.107209 | 0.542 | 0.529 | 1 | 2 | Lnpep | 0.013 | Old |
| 0.395745 | 0.315562 | 0.093 | 0.082 | 1 | 2 | Ccnh | 0.011 | Old |
| 0.395797 | 0.330327 | 0.078 | 0.068 | 1 | 2 | Uckl1 | 0.01 | Old |
| 0.396523 | 0.177592 | 0.192 | 0.175 | 1 | 2 | Uqcrc1 | 0.017 | Old |
| 0.396693 | 0.258155 | 0.182 | 0.169 | 1 | 2 | Dhrs7 | 0.013 | Old |
| 0.396919 | 0.167242 | 0.151 | 0.137 | 1 | 2 | Vrk1 | 0.014 | Old |
| 0.396965 | 0.066553 | 0.545 | 0.524 | 1 | 2 | Atf3 | 0.021 | Old |
| 0.397589 | 0.484937 | 0.024 | 0.018 | 1 | 2 | Slc30a4 | 0.006 | Old |
| 0.397771 | 0.223496 | 0.092 | 0.081 | 1 | 2 | Trak2 | 0.011 | Old |
| 0.397774 | 0.097049 | 0.153 | 0.138 | 1 | 2 | Cdk13 | 0.015 | Old |
| 0.397807 | 0.381167 | 0.08 | 0.07 | 1 | 2 | Nmi | 0.01 | Old |
| 0.398115 | 0.065586 | 0.257 | 0.238 | 1 | 2 | Sdhc | 0.019 | Old |
| 0.398171 | 0.090902 | 0.033 | 0.026 | 1 | 2 | Pcyox1l | 0.007 | Old |
| 0.39821 | 0.165487 | 0.11 | 0.097 | 1 | 2 | F10 | 0.013 | Old |
| 0.398722 | 0.114405 | 0.057 | 0.048 | 1 | 2 | Rabggta | 0.009 | Old |
| 0.399073 | 0.191792 | 0.173 | 0.158 | 1 | 2 | Maz | 0.015 | Old |
| 0.399314 | 0.12753 | 0.136 | 0.121 | 1 | 2 | Ankrd40 | 0.015 | Old |
| 0.39969 | 0.232846 | 0.173 | 0.159 | 1 | 2 | Smu1 | 0.014 | Old |
| 0.399782 | 0.049675 | 0.128 | 0.144 | 1 | 2 | Itsn1 | -0.016 | Old |
| 0.399844 | 0.132804 | 0.358 | 0.338 | 1 | 2 | Lat2 | 0.02 | Old |
| 0.399852 | 0.054189 | 0.3 | 0.278 | 1 | 2 | Gm16286 | 0.022 | Old |
| 0.399924 | 0.117422 | 0.188 | 0.172 | 1 | 2 | Mgrn1 | 0.016 | Old |
| 0.400071 | 0.112976 | 0.05 | 0.042 | 1 | 2 | Plrg1 | 0.008 | Old |
| 0.400198 | 0.026425 | 0.152 | 0.137 | 1 | 2 | Mgat4b | 0.015 | Old |
| 0.400242 | 0.009956 | 0.126 | 0.112 | 1 | 2 | Prelid3b | 0.014 | Old |
| 0.400459 | 0.368345 | 0.058 | 0.049 | 1 | 2 | Memo1 | 0.009 | Old |
| 0.401307 | 0.03078 | 0.32 | 0.295 | 1 | 2 | Nrp1 | 0.025 | Old |
| 0.401352 | 0.069244 | 0.069 | 0.058 | 1 | 2 | Rwdd4a | 0.011 | Old |
| 0.401365 | 0.032671 | 0.767 | 0.76 | 1 | 2 | Gabarap | 0.007 | Old |
| 0.401506 | 0.199293 | 0.191 | 0.175 | 1 | 2 | Inpp4a | 0.016 | Old |
| 0.401783 | 0.216169 | 0.07 | 0.059 | 1 | 2 | Thoc3 | 0.011 | Old |
| 0.401965 | 0.144291 | 0.014 | 0.019 | 1 | 2 | A530064D( | -0.005 | Old |
| 0.40208 | 0.249597 | 0.029 | 0.023 | 1 | 2 | Calml4 | 0.006 | Old |
| 0.402268 | 0.116438 | 0.091 | 0.08 | 1 | 2 | Mpnd | 0.011 | Old |
| 0.402418 | 0.045415 | 0.055 | 0.064 | 1 | 2 | 2310022B( | -0.009 | Old |
| 0.402438 | 0.097385 | 0.193 | 0.176 | 1 | 2 | Ap1s1 | 0.017 | Old |
| 0.402997 | 0.129605 | 0.406 | 0.387 | 1 | 2 | Dnajc3 | 0.019 | Old |
| 0.402999 | 0.014547 | 0.12 | 0.106 | 1 | 2 | Lypla1 | 0.014 | Old |
| 0.403029 | 0.14601 | 0.057 | 0.048 | 1 | 2 | Chpt1 | 0.009 | Old |
| 0.403686 | 0.421184 | 0.095 | 0.084 | 1 | 2 | Crkl | 0.011 | Old |
| 0.403743 | 0.031197 | 0.944 | 0.937 | 1 | 2 | Ptma | 0.007 | Old |
| 0.40377 | 0.083793 | 0.324 | 0.296 | 1 | 2 | Akt1 | 0.028 | Old |
| 0.403945 | 0.195938 | 0.089 | 0.077 | 1 | 2 | Dgkg | 0.012 | Old |

| | | | | | | | | |
|---|---|---|---|---|---|---|---|---|
| 0.404255 | 0.09315 | 0.892 | 0.88 | 1 | 2 | Itm2b | 0.012 | Old |
| 0.404545 | 0.157422 | 0.227 | 0.208 | 1 | 2 | Stk24 | 0.019 | Old |
| 0.404964 | 0.136587 | 0.046 | 0.038 | 1 | 2 | Pccb | 0.008 | Old |
| 0.405212 | 0.405553 | 0.052 | 0.044 | 1 | 2 | Stam | 0.008 | Old |
| 0.405222 | 0.215643 | 0.015 | 0.02 | 1 | 2 | Mgp | -0.005 | Old |
| 0.405396 | 0.326976 | 0.1 | 0.088 | 1 | 2 | Dpp9 | 0.012 | Old |
| 0.405494 | 0.07536 | 0.029 | 0.023 | 1 | 2 | Alg3 | 0.006 | Old |
| 0.405508 | 0.63859 | 0.047 | 0.039 | 1 | 2 | Zfp780b | 0.008 | Old |
| 0.406073 | 0.346687 | 0.04 | 0.032 | 1 | 2 | Hist1h4i | 0.008 | Old |
| 0.406309 | 0.254792 | 0.123 | 0.11 | 1 | 2 | Homer1 | 0.013 | Old |
| 0.40633 | 0.178838 | 0.121 | 0.107 | 1 | 2 | Lrrk1 | 0.014 | Old |
| 0.406595 | 0.236852 | 0.139 | 0.127 | 1 | 2 | Tnks | 0.012 | Old |
| 0.406826 | 0.377967 | 0.041 | 0.033 | 1 | 2 | Cep290 | 0.008 | Old |
| 0.407317 | 0.020874 | 0.064 | 0.055 | 1 | 2 | Zfp422 | 0.009 | Old |
| 0.407684 | 0.166852 | 0.059 | 0.05 | 1 | 2 | Rpp25l | 0.009 | Old |
| 0.407697 | 0.067056 | 0.162 | 0.144 | 1 | 2 | Vbp1 | 0.018 | Old |
| 0.407886 | 0.538071 | 0.054 | 0.045 | 1 | 2 | Ptgir | 0.009 | Old |
| 0.407906 | 0.70781 | 0.019 | 0.014 | 1 | 2 | Polg2 | 0.005 | Old |
| 0.408025 | 0.16273 | 0.058 | 0.049 | 1 | 2 | Anapc10 | 0.009 | Old |
| 0.408071 | 0.107319 | 0.139 | 0.124 | 1 | 2 | Mrpl34 | 0.015 | Old |
| 0.408329 | 0.02091 | 0.124 | 0.11 | 1 | 2 | Mrpl51 | 0.014 | Old |
| 0.408361 | 0.107755 | 0.436 | 0.42 | 1 | 2 | Cflar | 0.016 | Old |
| 0.408377 | 0.76424 | 0.019 | 0.014 | 1 | 2 | Laptm4b | 0.005 | Old |
| 0.408403 | 0.07621 | 0.407 | 0.384 | 1 | 2 | Camk1d | 0.023 | Old |
| 0.408668 | 0.587276 | 0.041 | 0.033 | 1 | 2 | Pyurf | 0.008 | Old |
| 0.408788 | 0.028385 | 0.182 | 0.164 | 1 | 2 | Sec63 | 0.018 | Old |
| 0.408863 | 0.102818 | 0.134 | 0.119 | 1 | 2 | Wdr82 | 0.015 | Old |
| 0.409394 | 0.038436 | 0.443 | 0.413 | 1 | 2 | Pum2 | 0.03 | Old |
| 0.409797 | 0.033108 | 0.141 | 0.126 | 1 | 2 | Ubald1 | 0.015 | Old |
| 0.409798 | 0.211743 | 0.092 | 0.081 | 1 | 2 | Fam120b | 0.011 | Old |
| 0.409973 | 0.154509 | 0.211 | 0.195 | 1 | 2 | Por | 0.016 | Old |
| 0.409986 | 0.266718 | 0.335 | 0.321 | 1 | 2 | Hnrnpdl | 0.014 | Old |
| 0.410156 | 0.014661 | 0.248 | 0.227 | 1 | 2 | Dpm3 | 0.021 | Old |
| 0.410201 | 0.647939 | 0.03 | 0.024 | 1 | 2 | Zc3h3 | 0.006 | Old |
| 0.410204 | 0.094854 | 0.367 | 0.349 | 1 | 2 | Trip12 | 0.018 | Old |
| 0.410272 | 0.0139 | 0.016 | 0.021 | 1 | 2 | Sec22c | -0.005 | Old |
| 0.410377 | 0.193298 | 0.23 | 0.216 | 1 | 2 | Fam126a | 0.014 | Old |
| 0.410515 | 0.383253 | 0.041 | 0.033 | 1 | 2 | Slc35e2 | 0.008 | Old |
| 0.410792 | 0.174562 | 0.04 | 0.032 | 1 | 2 | Scn1b | 0.008 | Old |
| 0.411376 | 0.108357 | 0.185 | 0.198 | 1 | 2 | Ccl7 | -0.013 | Old |
| 0.411581 | 0.103317 | 0.718 | 0.694 | 1 | 2 | Dbi | 0.024 | Old |
| 0.41167 | 0.256315 | 0.121 | 0.108 | 1 | 2 | Rcor1 | 0.013 | Old |
| 0.411702 | 0.053943 | 0.062 | 0.072 | 1 | 2 | Smim19 | -0.01 | Old |
| 0.412184 | 0.370713 | 0.036 | 0.03 | 1 | 2 | Pgm3 | 0.006 | Old |
| 0.412389 | 0.44686 | 0.019 | 0.014 | 1 | 2 | 2610044O1 | 0.005 | Old |

| | | | | | | | | |
|---|---|---|---|---|---|---|---|---|
| 0.412549 | 0.510273 | 0.03 | 0.024 | 1 | 2 | Nprl3 | 0.006 | Old |
| 0.412672 | 0.438254 | 0.042 | 0.034 | 1 | 2 | Slc35b4 | 0.008 | Old |
| 0.412797 | 0.037494 | 0.98 | 0.973 | 1 | 2 | Rpsa | 0.007 | Old |
| 0.413089 | 0.015975 | 0.117 | 0.103 | 1 | 2 | Ptdss1 | 0.014 | Old |
| 0.413113 | 0.127207 | 0.046 | 0.038 | 1 | 2 | Wdfy1 | 0.008 | Old |
| 0.413362 | 0.186015 | 0.082 | 0.072 | 1 | 2 | Slc39a6 | 0.01 | Old |
| 0.413415 | 0.067105 | 0.263 | 0.241 | 1 | 2 | Ubxn4 | 0.022 | Old |
| 0.413692 | 0.159991 | 0.369 | 0.35 | 1 | 2 | Cdc42se1 | 0.019 | Old |
| 0.414297 | 0.132027 | 0.065 | 0.056 | 1 | 2 | Fubp3 | 0.009 | Old |
| 0.414368 | 0.142661 | 0.197 | 0.179 | 1 | 2 | Baz2a | 0.018 | Old |
| 0.414691 | 0.623733 | 0.025 | 0.019 | 1 | 2 | Rfc5 | 0.006 | Old |
| 0.414979 | 0.196961 | 0.079 | 0.069 | 1 | 2 | Vps33a | 0.01 | Old |
| 0.415429 | 0.116566 | 0.331 | 0.309 | 1 | 2 | Rab31 | 0.022 | Old |
| 0.416065 | 0.471309 | 0.048 | 0.04 | 1 | 2 | Trappc2 | 0.008 | Old |
| 0.416218 | 0.027566 | 0.077 | 0.067 | 1 | 2 | Mtmr12 | 0.01 | Old |
| 0.41651 | 0.141019 | 0.194 | 0.177 | 1 | 2 | Stk11 | 0.017 | Old |
| 0.416557 | 0.046466 | 0.412 | 0.393 | 1 | 2 | Ncor1 | 0.019 | Old |
| 0.416692 | 0.075832 | 0.36 | 0.342 | 1 | 2 | Dpysl2 | 0.018 | Old |
| 0.416735 | 0.130861 | 0.092 | 0.081 | 1 | 2 | Heatr1 | 0.011 | Old |
| 0.416749 | 0.043884 | 0.151 | 0.135 | 1 | 2 | Lima1 | 0.016 | Old |
| 0.417158 | 0.074513 | 0.641 | 0.626 | 1 | 2 | Ctsd | 0.015 | Old |
| 0.417232 | 0.458951 | 0.14 | 0.129 | 1 | 2 | Pank2 | 0.011 | Old |
| 0.417352 | 0.126072 | 0.125 | 0.112 | 1 | 2 | Hmg20b | 0.013 | Old |
| 0.418147 | 0.112372 | 0.137 | 0.122 | 1 | 2 | Ola1 | 0.015 | Old |
| 0.418184 | 0.385738 | 0.061 | 0.052 | 1 | 2 | Ttc39c | 0.009 | Old |
| 0.418723 | 0.276886 | 0.117 | 0.105 | 1 | 2 | 2610037D( | 0.012 | Old |
| 0.418735 | 0.051565 | 0.048 | 0.057 | 1 | 2 | Slc38a7 | -0.009 | Old |
| 0.418846 | 0.464057 | 0.056 | 0.048 | 1 | 2 | Txndc12 | 0.008 | Old |
| 0.419041 | 0.1487 | 0.123 | 0.11 | 1 | 2 | Slc35a4 | 0.013 | Old |
| 0.419721 | 0.454814 | 0.025 | 0.019 | 1 | 2 | Wfdc18 | 0.006 | Old |
| 0.419862 | 0.198971 | 0.118 | 0.105 | 1 | 2 | Actl6a | 0.013 | Old |
| 0.419928 | 0.706059 | 0.015 | 0.011 | 1 | 2 | Tmem38a | 0.004 | Old |
| 0.419928 | 0.695235 | 0.015 | 0.011 | 1 | 2 | Pop1 | 0.004 | Old |
| 0.42012 | 0.323805 | 0.082 | 0.072 | 1 | 2 | Apex1 | 0.01 | Old |
| 0.4202 | 0.613017 | 0.015 | 0.011 | 1 | 2 | Hgh1 | 0.004 | Old |
| 0.420337 | 0.657395 | 0.015 | 0.011 | 1 | 2 | Harbi1 | 0.004 | Old |
| 0.420423 | 0.087129 | 0.054 | 0.045 | 1 | 2 | Fam214a | 0.009 | Old |
| 0.420501 | 0.111438 | 0.362 | 0.344 | 1 | 2 | Pgls | 0.018 | Old |
| 0.420614 | 0.146352 | 0.06 | 0.051 | 1 | 2 | Atr | 0.009 | Old |
| 0.420754 | 0.059247 | 0.317 | 0.295 | 1 | 2 | Vapa | 0.022 | Old |
| 0.420868 | 0.133965 | 0.099 | 0.087 | 1 | 2 | Mta1 | 0.012 | Old |
| 0.421085 | 0.015759 | 0.435 | 0.414 | 1 | 2 | Ran | 0.021 | Old |
| 0.421195 | 0.379467 | 0.025 | 0.019 | 1 | 2 | Ctps | 0.006 | Old |
| 0.42139 | 0.06074 | 0.072 | 0.062 | 1 | 2 | Gar1 | 0.01 | Old |
| 0.421531 | 0.141505 | 0.155 | 0.14 | 1 | 2 | Ccng1 | 0.015 | Old |

| | | | | | | | | |
|---|---|---|---|---|---|---|---|---|
| 0.421682 | 0.041274 | 0.019 | 0.014 | 1 | 2 | Ift27 | 0.005 | Old |
| 0.421811 | 0.306017 | 0.075 | 0.065 | 1 | 2 | Tom1 | 0.01 | Old |
| 0.422052 | 0.400906 | 0.075 | 0.065 | 1 | 2 | Zbtb22 | 0.01 | Old |
| 0.422235 | 0.482686 | 0.086 | 0.076 | 1 | 2 | Pmpcb | 0.01 | Old |
| 0.422399 | 0.672447 | 0.035 | 0.043 | 1 | 2 | Gm20186 | -0.008 | Old |
| 0.422756 | 0.169932 | 0.207 | 0.192 | 1 | 2 | Cdc40 | 0.015 | Old |
| 0.422839 | 0.104236 | 0.207 | 0.188 | 1 | 2 | Mff | 0.019 | Old |
| 0.422883 | 0.084786 | 0.019 | 0.014 | 1 | 2 | Qrsl1 | 0.005 | Old |
| 0.423208 | 0.501182 | 0.015 | 0.011 | 1 | 2 | Gpat3 | 0.004 | Old |
| 0.42326 | 0.307018 | 0.03 | 0.024 | 1 | 2 | Tubgcp6 | 0.006 | Old |
| 0.423401 | 0.20422 | 0.101 | 0.089 | 1 | 2 | Dctn5 | 0.012 | Old |
| 0.423482 | 0.181456 | 0.108 | 0.096 | 1 | 2 | Ccdc174 | 0.012 | Old |
| 0.423762 | 0.160015 | 0.08 | 0.07 | 1 | 2 | Celf4 | 0.01 | Old |
| 0.423904 | 0.504203 | 0.071 | 0.062 | 1 | 2 | Nagk | 0.009 | Old |
| 0.423936 | 0.157212 | 0.017 | 0.023 | 1 | 2 | Ankrd28 | -0.006 | Old |
| 0.42403 | 0.374498 | 0.015 | 0.011 | 1 | 2 | Fkbp9 | 0.004 | Old |
| 0.424085 | 0.255807 | 0.112 | 0.101 | 1 | 2 | Ppie | 0.011 | Old |
| 0.424406 | 0.174776 | 0.166 | 0.151 | 1 | 2 | Lats2 | 0.015 | Old |
| 0.425044 | 0.439849 | 0.048 | 0.04 | 1 | 2 | Gtdc1 | 0.008 | Old |
| 0.42512 | 0.078761 | 0.061 | 0.052 | 1 | 2 | Arl10 | 0.009 | Old |
| 0.425164 | 0.036181 | 0.263 | 0.243 | 1 | 2 | Pura | 0.02 | Old |
| 0.4252 | 0.622613 | 0.06 | 0.051 | 1 | 2 | Gdf15 | 0.009 | Old |
| 0.42538 | 0.154354 | 0.035 | 0.029 | 1 | 2 | Zdhhc23 | 0.006 | Old |
| 0.425497 | 0.062244 | 0.419 | 0.388 | 1 | 2 | Evi2a | 0.031 | Old |
| 0.425953 | 0.025772 | 0.015 | 0.011 | 1 | 2 | Zfp518b | 0.004 | Old |
| 0.426228 | 0.241333 | 0.015 | 0.011 | 1 | 2 | Bpgm | 0.004 | Old |
| 0.426322 | 0.256366 | 0.095 | 0.084 | 1 | 2 | Ylpm1 | 0.011 | Old |
| 0.426419 | 0.161373 | 0.06 | 0.051 | 1 | 2 | Fiz1 | 0.009 | Old |
| 0.426419 | 0.110534 | 0.03 | 0.024 | 1 | 2 | Amd2 | 0.006 | Old |
| 0.426426 | 0.447211 | 0.092 | 0.082 | 1 | 2 | Lrrc41 | 0.01 | Old |
| 0.426473 | 0.399208 | 0.011 | 0.007 | 1 | 2 | Zfp512b | 0.004 | Old |
| 0.426473 | 0.344835 | 0.011 | 0.007 | 1 | 2 | Ercc6l | 0.004 | Old |
| 0.426776 | 0.688951 | 0.02 | 0.015 | 1 | 2 | Fastkd1 | 0.005 | Old |
| 0.426778 | 0.265277 | 0.015 | 0.011 | 1 | 2 | Rilp | 0.004 | Old |
| 0.426802 | 0.29742 | 0.011 | 0.007 | 1 | 2 | Nipsnap1 | 0.004 | Old |
| 0.426946 | 0.159301 | 0.041 | 0.033 | 1 | 2 | Plekhn1 | 0.008 | Old |
| 0.427024 | 0.005051 | 0.126 | 0.113 | 1 | 2 | Usp24 | 0.013 | Old |
| 0.427367 | 0.098433 | 0.343 | 0.324 | 1 | 2 | Ehbp1l1 | 0.019 | Old |
| 0.427461 | 0.236906 | 0.011 | 0.007 | 1 | 2 | Polrmt | 0.004 | Old |
| 0.427577 | 0.127931 | 0.101 | 0.089 | 1 | 2 | Rbl2 | 0.012 | Old |
| 0.427665 | 0.390589 | 0.036 | 0.03 | 1 | 2 | Fam114a1 | 0.006 | Old |
| 0.427667 | 0.15054 | 0.03 | 0.024 | 1 | 2 | 5730480H( | 0.006 | Old |
| 0.427791 | 0.237166 | 0.011 | 0.007 | 1 | 2 | Ifit2 | 0.004 | Old |
| 0.427936 | 0.055237 | 0.145 | 0.129 | 1 | 2 | Suds3 | 0.016 | Old |
| 0.428156 | 0.19956 | 0.015 | 0.011 | 1 | 2 | Slc30a6 | 0.004 | Old |

| | | | | | | | | |
|---|---|---|---|---|---|---|---|---|
| 0.428187 | 0.274631 | 0.036 | 0.03 | 1 | 2 | Pdp1 | 0.006 | Old |
| 0.428203 | 0.111047 | 0.14 | 0.126 | 1 | 2 | Rin3 | 0.014 | Old |
| 0.428328 | 0.843748 | 0.026 | 0.02 | 1 | 2 | Dnajc12 | 0.006 | Old |
| 0.428451 | 0.255532 | 0.011 | 0.007 | 1 | 2 | Map7d3 | 0.004 | Old |
| 0.428451 | 0.057054 | 0.011 | 0.007 | 1 | 2 | Pdss1 | 0.004 | Old |
| 0.429112 | 0.100996 | 0.011 | 0.007 | 1 | 2 | Zfyve28 | 0.004 | Old |
| 0.429112 | 0.008038 | 0.011 | 0.007 | 1 | 2 | Slc22a18 | 0.004 | Old |
| 0.429159 | 0.298023 | 0.02 | 0.026 | 1 | 2 | Rint1 | -0.006 | Old |
| 0.429305 | 0.351562 | 0.031 | 0.025 | 1 | 2 | Bcl9 | 0.006 | Old |
| 0.430333 | 0.607787 | 0.044 | 0.037 | 1 | 2 | Zfp26 | 0.007 | Old |
| 0.430487 | 0.058156 | 0.024 | 0.03 | 1 | 2 | Rpap3 | -0.006 | Old |
| 0.430711 | 0.023591 | 0.101 | 0.089 | 1 | 2 | Rfc2 | 0.012 | Old |
| 0.430809 | 0.089196 | 0.126 | 0.113 | 1 | 2 | Kdm5c | 0.013 | Old |
| 0.431383 | 0.4489 | 0.019 | 0.025 | 1 | 2 | Scd1 | -0.006 | Old |
| 0.43235 | 0.296051 | 0.108 | 0.097 | 1 | 2 | Amn1 | 0.011 | Old |
| 0.432368 | 0.204637 | 0.022 | 0.029 | 1 | 2 | Sacs | -0.007 | Old |
| 0.432375 | 0.282429 | 0.036 | 0.03 | 1 | 2 | Fbxl4 | 0.006 | Old |
| 0.43241 | 0.02555 | 0.315 | 0.29 | 1 | 2 | Ttc7 | 0.025 | Old |
| 0.432685 | 0.155085 | 0.131 | 0.118 | 1 | 2 | Dnajc14 | 0.013 | Old |
| 0.43333 | 0.235776 | 0.162 | 0.15 | 1 | 2 | Qdpr | 0.012 | Old |
| 0.433555 | 0.270706 | 0.055 | 0.046 | 1 | 2 | Nup210 | 0.009 | Old |
| 0.43362 | 0.248695 | 0.26 | 0.247 | 1 | 2 | Eif3d | 0.013 | Old |
| 0.433889 | 0.055466 | 0.035 | 0.029 | 1 | 2 | Tmem18 | 0.006 | Old |
| 0.434086 | 0.136426 | 0.128 | 0.116 | 1 | 2 | Eps15l1 | 0.012 | Old |
| 0.434167 | 0.283714 | 0.026 | 0.02 | 1 | 2 | C030034I2 | 0.006 | Old |
| 0.434573 | 0.134063 | 0.284 | 0.267 | 1 | 2 | 9930111J2 | 0.017 | Old |
| 0.434678 | 0.096556 | 0.116 | 0.102 | 1 | 2 | Mrps12 | 0.014 | Old |
| 0.434834 | 0.110389 | 0.191 | 0.173 | 1 | 2 | Gtpbp4 | 0.018 | Old |
| 0.434888 | 0.058263 | 0.08 | 0.091 | 1 | 2 | Matk | -0.011 | Old |
| 0.434937 | 0.519101 | 0.057 | 0.049 | 1 | 2 | Pop7 | 0.008 | Old |
| 0.435062 | 0.261106 | 0.065 | 0.056 | 1 | 2 | Rfwd3 | 0.009 | Old |
| 0.435104 | 0.129732 | 0.688 | 0.665 | 1 | 2 | Fus | 0.023 | Old |
| 0.435912 | 0.081449 | 0.185 | 0.169 | 1 | 2 | Man1c1 | 0.016 | Old |
| 0.435973 | 0.006212 | 0.288 | 0.264 | 1 | 2 | Raly | 0.024 | Old |
| 0.436012 | 0.00272 | 0.094 | 0.083 | 1 | 2 | Cdk5rap3 | 0.011 | Old |
| 0.436115 | 0.224411 | 0.11 | 0.1 | 1 | 2 | Clcc1 | 0.01 | Old |
| 0.436221 | 0.260295 | 0.099 | 0.088 | 1 | 2 | Gpr160 | 0.011 | Old |
| 0.436234 | 0.293643 | 0.036 | 0.03 | 1 | 2 | Lin37 | 0.006 | Old |
| 0.436501 | 0.113745 | 0.048 | 0.04 | 1 | 2 | Ogfod1 | 0.008 | Old |
| 0.436893 | 0.070072 | 0.08 | 0.07 | 1 | 2 | Mrpl38 | 0.01 | Old |
| 0.437202 | 0.04886 | 0.03 | 0.037 | 1 | 2 | Zbtb33 | -0.007 | Old |
| 0.437278 | 0.006185 | 0.031 | 0.025 | 1 | 2 | Rnf25 | 0.006 | Old |
| 0.437878 | 0.063501 | 0.041 | 0.049 | 1 | 2 | Acvr2a | -0.008 | Old |
| 0.438569 | 0.522093 | 0.02 | 0.015 | 1 | 2 | Pex11a | 0.005 | Old |
| 0.439066 | 0.08163 | 0.085 | 0.074 | 1 | 2 | Nfyb | 0.011 | Old |

| | | | | | | | | |
|---|---|---|---|---|---|---|---|---|
| 0.439222 | 0.251582 | 0.021 | 0.027 | 1 | 2 | Trim21 | -0.006 | Old |
| 0.439629 | 0.349017 | 0.026 | 0.02 | 1 | 2 | Gmppb | 0.006 | Old |
| 0.43968 | 0.167953 | 0.049 | 0.042 | 1 | 2 | Lxn | 0.007 | Old |
| 0.439727 | 0.039397 | 0.215 | 0.196 | 1 | 2 | Skp1a | 0.019 | Old |
| 0.440334 | 0.102618 | 0.275 | 0.255 | 1 | 2 | Psmb6 | 0.02 | Old |
| 0.440411 | 0.154017 | 0.028 | 0.034 | 1 | 2 | Tmtc3 | -0.006 | Old |
| 0.440601 | 0.15114 | 0.18 | 0.166 | 1 | 2 | Pcmtd1 | 0.014 | Old |
| 0.440716 | 0.126685 | 0.255 | 0.239 | 1 | 2 | Arl8b | 0.016 | Old |
| 0.440738 | 0.37912 | 0.077 | 0.068 | 1 | 2 | Cabin1 | 0.009 | Old |
| 0.441816 | 0.047358 | 0.139 | 0.126 | 1 | 2 | Tor1aip2 | 0.013 | Old |
| 0.442064 | 0.043189 | 0.261 | 0.241 | 1 | 2 | Ap3s1 | 0.02 | Old |
| 0.442275 | 0.070276 | 0.453 | 0.431 | 1 | 2 | Fubp1 | 0.022 | Old |
| 0.442359 | 0.313087 | 0.107 | 0.096 | 1 | 2 | Zyg11b | 0.011 | Old |
| 0.442428 | 0.066934 | 0.246 | 0.229 | 1 | 2 | Socs6 | 0.017 | Old |
| 0.442555 | 0.141627 | 0.095 | 0.084 | 1 | 2 | Akap10 | 0.011 | Old |
| 0.44301 | 0.378014 | 0.026 | 0.02 | 1 | 2 | Nudt18 | 0.006 | Old |
| 0.443099 | 0.051823 | 0.02 | 0.015 | 1 | 2 | Poc5 | 0.005 | Old |
| 0.443121 | 0.153935 | 0.21 | 0.196 | 1 | 2 | Csnk2a1 | 0.014 | Old |
| 0.44331 | 0.010621 | 0.348 | 0.321 | 1 | 2 | Tle3 | 0.027 | Old |
| 0.443339 | 0.157332 | 0.02 | 0.015 | 1 | 2 | BC017158 | 0.005 | Old |
| 0.443795 | 0.167569 | 0.096 | 0.086 | 1 | 2 | Parg | 0.01 | Old |
| 0.445315 | 0.142731 | 0.049 | 0.042 | 1 | 2 | Bcl2l13 | 0.007 | Old |
| 0.445337 | 0.243507 | 0.111 | 0.1 | 1 | 2 | Plekhf2 | 0.011 | Old |
| 0.44534 | 0.244756 | 0.043 | 0.036 | 1 | 2 | Fam210a | 0.007 | Old |
| 0.445483 | 0.079333 | 0.258 | 0.242 | 1 | 2 | Xrn2 | 0.016 | Old |
| 0.4461 | 0.104011 | 0.282 | 0.261 | 1 | 2 | Ppp1r12c | 0.021 | Old |
| 0.446972 | 0.229771 | 0.153 | 0.14 | 1 | 2 | Dhrs1 | 0.013 | Old |
| 0.447171 | 0.307113 | 0.188 | 0.173 | 1 | 2 | Glrx5 | 0.015 | Old |
| 0.447341 | 0.029484 | 0.31 | 0.286 | 1 | 2 | Pgd | 0.024 | Old |
| 0.447482 | 0.102416 | 0.374 | 0.354 | 1 | 2 | Rilpl2 | 0.02 | Old |
| 0.447559 | 0.181584 | 0.397 | 0.385 | 1 | 2 | Ezr | 0.012 | Old |
| 0.448091 | 0.127538 | 0.388 | 0.372 | 1 | 2 | Nipbl | 0.016 | Old |
| 0.448461 | 0.002381 | 0.043 | 0.036 | 1 | 2 | Polr3f | 0.007 | Old |
| 0.44886 | 0.097439 | 0.02 | 0.015 | 1 | 2 | Nup37 | 0.005 | Old |
| 0.448956 | 0.038831 | 0.09 | 0.08 | 1 | 2 | Rnf41 | 0.01 | Old |
| 0.449007 | 0.090672 | 0.062 | 0.053 | 1 | 2 | Gaa | 0.009 | Old |
| 0.449182 | 0.194105 | 0.036 | 0.03 | 1 | 2 | Incenp | 0.006 | Old |
| 0.449696 | 0.091914 | 0.043 | 0.036 | 1 | 2 | Wdr70 | 0.007 | Old |
| 0.450048 | 0.097087 | 0.423 | 0.405 | 1 | 2 | Xiap | 0.018 | Old |
| 0.450278 | 0.071919 | 0.229 | 0.214 | 1 | 2 | Phb2 | 0.015 | Old |
| 0.450347 | 0.669198 | 0.021 | 0.017 | 1 | 2 | Cdkn2c | 0.004 | Old |
| 0.450638 | 0.008569 | 0.369 | 0.34 | 1 | 2 | Sf1 | 0.029 | Old |
| 0.451392 | 0.259609 | 0.044 | 0.037 | 1 | 2 | Zscan29 | 0.007 | Old |
| 0.451584 | 0.080224 | 0.196 | 0.179 | 1 | 2 | Pik3ca | 0.017 | Old |
| 0.452217 | 0.124059 | 0.228 | 0.211 | 1 | 2 | Adipor2 | 0.017 | Old |

| | | | | | | | | |
|---|---|---|---|---|---|---|---|---|
| 0.452341 | 0.117185 | 0.116 | 0.103 | 1 | 2 | Mertk | 0.013 | Old |
| 0.452451 | 0.064123 | 0.558 | 0.544 | 1 | 2 | Rgs2 | 0.014 | Old |
| 0.452992 | 0.7452 | 0.016 | 0.012 | 1 | 2 | Zfp952 | 0.004 | Old |
| 0.453985 | 0.075773 | 0.413 | 0.386 | 1 | 2 | Degs1 | 0.027 | Old |
| 0.454762 | 0.336178 | 0.056 | 0.048 | 1 | 2 | Nbdy | 0.008 | Old |
| 0.455281 | 0.01698 | 0.146 | 0.159 | 1 | 2 | Abca3 | -0.013 | Old |
| 0.455557 | 0.036001 | 0.13 | 0.116 | 1 | 2 | Kdm6a | 0.014 | Old |
| 0.45566 | 0.610635 | 0.027 | 0.021 | 1 | 2 | Rbm10 | 0.006 | Old |
| 0.456008 | 0.223037 | 0.125 | 0.114 | 1 | 2 | Dyrk1a | 0.011 | Old |
| 0.456134 | 0.001186 | 0.207 | 0.189 | 1 | 2 | Peak1 | 0.018 | Old |
| 0.45614 | 0.223533 | 0.101 | 0.09 | 1 | 2 | Pspc1 | 0.011 | Old |
| 0.456148 | 0.172172 | 0.208 | 0.194 | 1 | 2 | Ctdsp1 | 0.014 | Old |
| 0.456227 | 0.026519 | 0.08 | 0.07 | 1 | 2 | Parp2 | 0.01 | Old |
| 0.456299 | 0.054219 | 0.904 | 0.893 | 1 | 2 | H3f3a | 0.011 | Old |
| 0.456506 | 0.311248 | 0.045 | 0.038 | 1 | 2 | Kif16b | 0.007 | Old |
| 0.456611 | 0.336508 | 0.094 | 0.084 | 1 | 2 | Entpd7 | 0.01 | Old |
| 0.456767 | 0.325951 | 0.089 | 0.08 | 1 | 2 | Med20 | 0.009 | Old |
| 0.457379 | 0.006808 | 0.016 | 0.012 | 1 | 2 | Zfp74 | 0.004 | Old |
| 0.458608 | 0.375974 | 0.027 | 0.021 | 1 | 2 | Dhrs11 | 0.006 | Old |
| 0.458755 | 0.091389 | 0.016 | 0.012 | 1 | 2 | Fadd | 0.004 | Old |
| 0.458991 | 0.339384 | 0.063 | 0.055 | 1 | 2 | Rbbp5 | 0.008 | Old |
| 0.459212 | 0.072025 | 0.125 | 0.113 | 1 | 2 | Renbp | 0.012 | Old |
| 0.459216 | 0.161239 | 0.209 | 0.195 | 1 | 2 | Ssrp1 | 0.014 | Old |
| 0.459617 | 0.144357 | 0.232 | 0.217 | 1 | 2 | Safb2 | 0.015 | Old |
| 0.459962 | 0.025445 | 0.118 | 0.105 | 1 | 2 | Tnfaip1 | 0.013 | Old |
| 0.460133 | 0.033952 | 0.016 | 0.012 | 1 | 2 | Ptcd1 | 0.004 | Old |
| 0.460202 | 0.006276 | 0.139 | 0.125 | 1 | 2 | Dync1li2 | 0.014 | Old |
| 0.460484 | 0.549011 | 0.021 | 0.017 | 1 | 2 | Casz1 | 0.004 | Old |
| 0.460584 | 0.186884 | 0.133 | 0.121 | 1 | 2 | Htra2 | 0.012 | Old |
| 0.460686 | 0.093903 | 0.191 | 0.175 | 1 | 2 | Tia1 | 0.016 | Old |
| 0.460693 | 0.255765 | 0.222 | 0.21 | 1 | 2 | Retreg2 | 0.012 | Old |
| 0.460763 | 0.042026 | 0.117 | 0.105 | 1 | 2 | Gk | 0.012 | Old |
| 0.460869 | 0.002516 | 0.196 | 0.179 | 1 | 2 | Tmem50b | 0.017 | Old |
| 0.460924 | 0.005978 | 0.407 | 0.419 | 1 | 2 | Hnrnpc | -0.012 | Old |
| 0.46096 | 0.125315 | 0.016 | 0.012 | 1 | 2 | Med26 | 0.004 | Old |
| 0.46096 | 0.088169 | 0.016 | 0.012 | 1 | 2 | Gm47754 | 0.004 | Old |
| 0.461266 | 0.078719 | 0.037 | 0.031 | 1 | 2 | Slc41a2 | 0.006 | Old |
| 0.461595 | 0.194336 | 0.261 | 0.245 | 1 | 2 | Arhgef2 | 0.016 | Old |
| 0.461802 | 0.131702 | 0.467 | 0.445 | 1 | 2 | Usf2 | 0.022 | Old |
| 0.461934 | 0.136168 | 0.24 | 0.227 | 1 | 2 | Rexo2 | 0.013 | Old |
| 0.462368 | 0.141905 | 0.056 | 0.048 | 1 | 2 | Lrrc42 | 0.008 | Old |
| 0.462755 | 0.150117 | 0.135 | 0.122 | 1 | 2 | Zfp217 | 0.013 | Old |
| 0.463079 | 0.009683 | 0.139 | 0.127 | 1 | 2 | Gm49336 | 0.012 | Old |
| 0.463104 | 0.845838 | 0.028 | 0.023 | 1 | 2 | Gpn2 | 0.005 | Old |
| 0.463331 | 0.254002 | 0.072 | 0.063 | 1 | 2 | Orc4 | 0.009 | Old |

| | | | | | | | | |
|---|---|---|---|---|---|---|---|---|
| 0.463593 | 0.07836 | 0.16 | 0.146 | 1 | 2 | Parl | 0.014 | Old |
| 0.463996 | 0.188037 | 0.109 | 0.097 | 1 | 2 | Lrrc47 | 0.012 | Old |
| 0.464071 | 0.097322 | 0.11 | 0.099 | 1 | 2 | Zbtb1 | 0.011 | Old |
| 0.464474 | 0.002585 | 0.128 | 0.115 | 1 | 2 | Kansl3 | 0.013 | Old |
| 0.464723 | 0.208608 | 0.057 | 0.049 | 1 | 2 | Pde4dip | 0.008 | Old |
| 0.464835 | 0.110289 | 0.016 | 0.012 | 1 | 2 | Gstm4 | 0.004 | Old |
| 0.464966 | 0.591256 | 0.045 | 0.038 | 1 | 2 | Lrrc8b | 0.007 | Old |
| 0.464969 | 0.453575 | 0.012 | 0.008 | 1 | 2 | Gm6133 | 0.004 | Old |
| 0.465169 | 0.082709 | 0.256 | 0.24 | 1 | 2 | Nin | 0.016 | Old |
| 0.465391 | 0.156026 | 0.1 | 0.088 | 1 | 2 | Senp1 | 0.012 | Old |
| 0.465687 | 0.126862 | 0.101 | 0.09 | 1 | 2 | Glipr1 | 0.011 | Old |
| 0.465853 | 0.117916 | 0.183 | 0.166 | 1 | 2 | Psen2 | 0.017 | Old |
| 0.465949 | 1.143376 | 0.012 | 0.008 | 1 | 2 | Zfp933 | 0.004 | Old |
| 0.466096 | 0.019216 | 0.104 | 0.115 | 1 | 2 | Pi4ka | -0.011 | Old |
| 0.46624 | 0.115737 | 0.027 | 0.021 | 1 | 2 | Gm20604 | 0.006 | Old |
| 0.466436 | 0.097385 | 0.021 | 0.017 | 1 | 2 | Mrpl44 | 0.004 | Old |
| 0.466602 | 0.613802 | 0.012 | 0.008 | 1 | 2 | Tesk2 | 0.004 | Old |
| 0.466738 | 0.164196 | 0.107 | 0.096 | 1 | 2 | Ppfia1 | 0.011 | Old |
| 0.466763 | 0.197751 | 0.073 | 0.064 | 1 | 2 | Trappc13 | 0.009 | Old |
| 0.466929 | 0.940049 | 0.012 | 0.008 | 1 | 2 | Rccd1 | 0.004 | Old |
| 0.467256 | 0.562877 | 0.012 | 0.008 | 1 | 2 | Lamb3 | 0.004 | Old |
| 0.467286 | 0.214885 | 0.071 | 0.062 | 1 | 2 | Ascc1 | 0.009 | Old |
| 0.467583 | 0.159502 | 0.139 | 0.127 | 1 | 2 | Cd80 | 0.012 | Old |
| 0.46766 | 0.01612 | 0.131 | 0.119 | 1 | 2 | Kif3b | 0.012 | Old |
| 0.468442 | 0.033651 | 0.091 | 0.081 | 1 | 2 | Klhl18 | 0.01 | Old |
| 0.468566 | 0.338253 | 0.012 | 0.008 | 1 | 2 | A530040E1 | 0.004 | Old |
| 0.468566 | 0.25579 | 0.012 | 0.008 | 1 | 2 | Nlrp1a | 0.004 | Old |
| 0.468662 | 0.01735 | 0.33 | 0.304 | 1 | 2 | Tmem59 | 0.026 | Old |
| 0.468794 | 0.530525 | 0.006 | 0.01 | 1 | 2 | 4732496C0 | -0.004 | Old |
| 0.468906 | 0.091741 | 0.226 | 0.211 | 1 | 2 | Mydgf | 0.015 | Old |
| 0.468974 | 0.253133 | 0.11 | 0.1 | 1 | 2 | Clasp1 | 0.01 | Old |
| 0.46916 | 0.085423 | 0.449 | 0.437 | 1 | 2 | Aprt | 0.012 | Old |
| 0.469588 | 0.281646 | 0.096 | 0.087 | 1 | 2 | Arpc5l | 0.009 | Old |
| 0.469866 | 0.099106 | 0.087 | 0.077 | 1 | 2 | 1700123O2 | 0.01 | Old |
| 0.469939 | 0.138748 | 0.09 | 0.08 | 1 | 2 | Sae1 | 0.01 | Old |
| 0.470038 | 0.108614 | 0.142 | 0.129 | 1 | 2 | Heatr5a | 0.013 | Old |
| 0.470274 | 0.177004 | 0.234 | 0.222 | 1 | 2 | Osbpl9 | 0.012 | Old |
| 0.470523 | 0.013693 | 0.202 | 0.186 | 1 | 2 | Stag1 | 0.016 | Old |
| 0.471192 | 0.06522 | 0.012 | 0.008 | 1 | 2 | F3 | 0.004 | Old |
| 0.471672 | 0.298066 | 0.052 | 0.045 | 1 | 2 | Usb1 | 0.007 | Old |
| 0.471848 | 0.079406 | 0.48 | 0.464 | 1 | 2 | C5ar1 | 0.016 | Old |
| 0.47185 | 0.481611 | 0.087 | 0.078 | 1 | 2 | Med30 | 0.009 | Old |
| 0.472134 | 0.132844 | 0.544 | 0.531 | 1 | 2 | Adgre1 | 0.013 | Old |
| 0.472508 | 0.028371 | 0.012 | 0.008 | 1 | 2 | Gm42595 | 0.004 | Old |
| 0.472671 | 0.113471 | 0.021 | 0.017 | 1 | 2 | Gm10138 | 0.004 | Old |

| | | | | | | | | |
|---|---|---|---|---|---|---|---|---|
| 0.473703 | 0.214246 | 0.201 | 0.188 | 1 | 2 | Setd1b | 0.013 | Old |
| 0.47402 | 0.934939 | 0.022 | 0.018 | 1 | 2 | Cyp2ab1 | 0.004 | Old |
| 0.474313 | 0.112092 | 0.066 | 0.076 | 1 | 2 | Ints12 | -0.01 | Old |
| 0.474737 | 0.141294 | 0.247 | 0.233 | 1 | 2 | Tnrc6b | 0.014 | Old |
| 0.475016 | 0.378896 | 0.028 | 0.023 | 1 | 2 | Tnfsf14 | 0.005 | Old |
| 0.475222 | 0.089102 | 0.317 | 0.298 | 1 | 2 | Cul3 | 0.019 | Old |
| 0.475648 | 0.362002 | 0.052 | 0.045 | 1 | 2 | Tapbpl | 0.007 | Old |
| 0.475763 | 0.033928 | 0.503 | 0.488 | 1 | 2 | Dok2 | 0.015 | Old |
| 0.475921 | 0.019974 | 0.27 | 0.248 | 1 | 2 | Tex261 | 0.022 | Old |
| 0.476049 | 0.189886 | 0.081 | 0.072 | 1 | 2 | Cluh | 0.009 | Old |
| 0.476482 | 0.125505 | 0.147 | 0.133 | 1 | 2 | Galnt11 | 0.014 | Old |
| 0.476521 | 0.226533 | 0.058 | 0.05 | 1 | 2 | Nbeal2 | 0.008 | Old |
| 0.476534 | 0.035994 | 0.289 | 0.271 | 1 | 2 | Fuca1 | 0.018 | Old |
| 0.477319 | 0.040873 | 0.195 | 0.177 | 1 | 2 | Cycs | 0.018 | Old |
| 0.477505 | 0.349816 | 0.054 | 0.046 | 1 | 2 | Trub2 | 0.008 | Old |
| 0.477548 | 0.142926 | 0.176 | 0.162 | 1 | 2 | Htt | 0.014 | Old |
| 0.478333 | 0.044869 | 0.206 | 0.221 | 1 | 2 | Atg3 | -0.015 | Old |
| 0.478488 | 0.054514 | 0.127 | 0.115 | 1 | 2 | Stk3 | 0.012 | Old |
| 0.478667 | 0.110391 | 0.033 | 0.027 | 1 | 2 | Wdcp | 0.006 | Old |
| 0.478711 | 0.220761 | 0.222 | 0.207 | 1 | 2 | Aldh3b1 | 0.015 | Old |
| 0.478807 | 0.145711 | 0.138 | 0.126 | 1 | 2 | Tbc1d13 | 0.012 | Old |
| 0.478812 | 0.34504 | 0.028 | 0.023 | 1 | 2 | Qtrt2 | 0.005 | Old |
| 0.47948 | 0.047516 | 0.726 | 0.692 | 1 | 2 | Arhgdia | 0.034 | Old |
| 0.479756 | 0.127562 | 0.104 | 0.094 | 1 | 2 | Mcrs1 | 0.01 | Old |
| 0.479913 | 0.014595 | 0.085 | 0.075 | 1 | 2 | Rinl | 0.01 | Old |
| 0.480358 | 0.042338 | 0.46 | 0.493 | 1 | 2 | Egr1 | -0.033 | Old |
| 0.480372 | 0.178895 | 0.094 | 0.106 | 1 | 2 | Usf1 | -0.012 | Old |
| 0.480504 | 0.286885 | 0.028 | 0.023 | 1 | 2 | Rbks | 0.005 | Old |
| 0.481192 | 0.428526 | 0.076 | 0.068 | 1 | 2 | Trim34a | 0.008 | Old |
| 0.481558 | 0.114907 | 0.007 | 0.011 | 1 | 2 | 3300005D( | -0.004 | Old |
| 0.481874 | 0.072156 | 0.691 | 0.659 | 1 | 2 | Sat1 | 0.032 | Old |
| 0.482268 | 0.469595 | 0.034 | 0.029 | 1 | 2 | Slc35d2 | 0.005 | Old |
| 0.482525 | 0.099937 | 0.074 | 0.065 | 1 | 2 | Wdr11 | 0.009 | Old |
| 0.482615 | 0.043636 | 0.281 | 0.259 | 1 | 2 | Ggnbp2 | 0.022 | Old |
| 0.482639 | 0.038012 | 0.193 | 0.176 | 1 | 2 | Stip1 | 0.017 | Old |
| 0.482851 | 0.064769 | 0.597 | 0.58 | 1 | 2 | Itgam | 0.017 | Old |
| 0.482861 | 0.659227 | 0.029 | 0.024 | 1 | 2 | Znhit2 | 0.005 | Old |
| 0.483127 | 0.334878 | 0.033 | 0.027 | 1 | 2 | Aptx | 0.006 | Old |
| 0.483164 | 0.205236 | 0.162 | 0.151 | 1 | 2 | Irf2bpl | 0.011 | Old |
| 0.483677 | 0.259269 | 0.007 | 0.011 | 1 | 2 | Klhl33 | -0.004 | Old |
| 0.483947 | 0.286213 | 0.1 | 0.09 | 1 | 2 | Setdb1 | 0.01 | Old |
| 0.484272 | 0.253695 | 0.11 | 0.1 | 1 | 2 | Smad5 | 0.01 | Old |
| 0.484769 | 0.196009 | 0.104 | 0.094 | 1 | 2 | Gcc2 | 0.01 | Old |
| 0.484859 | 0.040318 | 0.125 | 0.113 | 1 | 2 | Armcx3 | 0.012 | Old |
| 0.484933 | 0.180442 | 0.246 | 0.232 | 1 | 2 | Tspan31 | 0.014 | Old |

| | | | | | | | | |
|---|---|---|---|---|---|---|---|---|
| 0.484943 | 0.341878 | 0.067 | 0.059 | 1 | 2 | Smpd1 | 0.008 | Old |
| 0.485425 | 0.069136 | 0.375 | 0.356 | 1 | 2 | Adam10 | 0.019 | Old |
| 0.485835 | 0.1777 | 0.103 | 0.093 | 1 | 2 | Ckap5 | 0.01 | Old |
| 0.485979 | 0.121639 | 0.046 | 0.039 | 1 | 2 | Fam13b | 0.007 | Old |
| 0.486088 | 0.104325 | 0.105 | 0.095 | 1 | 2 | Tmem106b | 0.01 | Old |
| 0.486252 | 0.326432 | 0.017 | 0.013 | 1 | 2 | 2900076A0 | 0.004 | Old |
| 0.486601 | 0.544141 | 0.058 | 0.051 | 1 | 2 | Neurl4 | 0.007 | Old |
| 0.48677 | 0.388259 | 0.022 | 0.018 | 1 | 2 | Brat1 | 0.004 | Old |
| 0.486816 | 0.352435 | 0.124 | 0.114 | 1 | 2 | Cant1 | 0.01 | Old |
| 0.486995 | 0.217506 | 0.07 | 0.062 | 1 | 2 | Slc9a6 | 0.008 | Old |
| 0.487277 | 0.33209 | 0.152 | 0.143 | 1 | 2 | Irak4 | 0.009 | Old |
| 0.487348 | 0.106827 | 0.017 | 0.013 | 1 | 2 | Aatk | 0.004 | Old |
| 0.487369 | 0.053302 | 0.291 | 0.27 | 1 | 2 | Rbm5 | 0.021 | Old |
| 0.487734 | 0.145545 | 0.363 | 0.347 | 1 | 2 | Gns | 0.016 | Old |
| 0.487777 | 0.313849 | 0.086 | 0.077 | 1 | 2 | Ric1 | 0.009 | Old |
| 0.488249 | 0.040937 | 0.193 | 0.176 | 1 | 2 | Ubr2 | 0.017 | Old |
| 0.488545 | 0.017929 | 0.155 | 0.167 | 1 | 2 | Uqcc3 | -0.012 | Old |
| 0.489171 | 0.080675 | 0.218 | 0.203 | 1 | 2 | Pdia4 | 0.015 | Old |
| 0.48921 | 0.10087 | 0.046 | 0.039 | 1 | 2 | Ino80b | 0.007 | Old |
| 0.489356 | 0.194333 | 0.034 | 0.029 | 1 | 2 | Acot2 | 0.005 | Old |
| 0.489456 | 0.024759 | 0.214 | 0.197 | 1 | 2 | Kmt2a | 0.017 | Old |
| 0.489547 | 0.067529 | 0.198 | 0.183 | 1 | 2 | Chd3 | 0.015 | Old |
| 0.489549 | 0.226781 | 0.034 | 0.029 | 1 | 2 | Fam35a | 0.005 | Old |
| 0.48985 | 0.029889 | 0.092 | 0.082 | 1 | 2 | Smg9 | 0.01 | Old |
| 0.490302 | 0.13392 | 0.009 | 0.012 | 1 | 2 | Fam109b | -0.003 | Old |
| 0.491281 | 0.62014 | 0.217 | 0.235 | 1 | 2 | Ccl4 | -0.018 | Old |
| 0.491974 | 0.115146 | 0.284 | 0.27 | 1 | 2 | Pdlim5 | 0.014 | Old |
| 0.492097 | 0.142924 | 0.138 | 0.126 | 1 | 2 | C330007P0 | 0.012 | Old |
| 0.492161 | 0.294915 | 0.067 | 0.059 | 1 | 2 | Plxnc1 | 0.008 | Old |
| 0.492211 | 0.133052 | 0.266 | 0.254 | 1 | 2 | Fxr1 | 0.012 | Old |
| 0.49232 | 0.052118 | 0.009 | 0.012 | 1 | 2 | Dtwd2 | -0.003 | Old |
| 0.492478 | 0.127844 | 0.198 | 0.184 | 1 | 2 | Rab1b | 0.014 | Old |
| 0.492496 | 0.144436 | 0.022 | 0.018 | 1 | 2 | Acot11 | 0.004 | Old |
| 0.492681 | 0.117589 | 0.273 | 0.259 | 1 | 2 | Ppp4r3b | 0.014 | Old |
| 0.492834 | 0.131048 | 0.051 | 0.059 | 1 | 2 | Afdn | -0.008 | Old |
| 0.492846 | 0.065928 | 0.017 | 0.013 | 1 | 2 | Pard6a | 0.004 | Old |
| 0.493082 | 0.060426 | 0.661 | 0.648 | 1 | 2 | Ywhae | 0.013 | Old |
| 0.493239 | 0.066215 | 0.178 | 0.162 | 1 | 2 | Eml4 | 0.016 | Old |
| 0.493331 | 0.249786 | 0.009 | 0.012 | 1 | 2 | Dbp | -0.003 | Old |
| 0.493487 | 0.205649 | 0.137 | 0.126 | 1 | 2 | 4931406C0 | 0.011 | Old |
| 0.493917 | 0.034284 | 0.394 | 0.374 | 1 | 2 | Atp5d | 0.02 | Old |
| 0.493984 | 0.27139 | 0.034 | 0.029 | 1 | 2 | Psmg1 | 0.005 | Old |
| 0.493998 | 0.005946 | 0.072 | 0.063 | 1 | 2 | Banp | 0.009 | Old |
| 0.494773 | 0.192785 | 0.022 | 0.018 | 1 | 2 | Rabep2 | 0.004 | Old |
| 0.494806 | 0.051738 | 0.228 | 0.213 | 1 | 2 | Pacsin2 | 0.015 | Old |

| | | | | | | | | |
|---|---|---|---|---|---|---|---|---|
| 0.494928 | 0.252547 | 0.113 | 0.103 | 1 | 2 | Setd7 | 0.01 | Old |
| 0.495211 | 0.157212 | 0.029 | 0.024 | 1 | 2 | 1810032O( | 0.005 | Old |
| 0.495468 | 0.083561 | 0.138 | 0.127 | 1 | 2 | Utp3 | 0.011 | Old |
| 0.495469 | 0.104127 | 0.376 | 0.362 | 1 | 2 | Sec62 | 0.014 | Old |
| 0.495495 | 0.380165 | 0.078 | 0.07 | 1 | 2 | Apmap | 0.008 | Old |
| 0.495644 | 0.190997 | 0.097 | 0.088 | 1 | 2 | Smurf2 | 0.009 | Old |
| 0.495648 | 0.004798 | 0.197 | 0.182 | 1 | 2 | BC028528 | 0.015 | Old |
| 0.495701 | 0.440466 | 0.056 | 0.049 | 1 | 2 | Tlr6 | 0.007 | Old |
| 0.495933 | 0.257728 | 0.041 | 0.034 | 1 | 2 | Map3k15 | 0.007 | Old |
| 0.496065 | 0.251787 | 0.208 | 0.196 | 1 | 2 | Ugp2 | 0.012 | Old |
| 0.496073 | 0.230795 | 0.09 | 0.081 | 1 | 2 | Usp33 | 0.009 | Old |
| 0.496189 | 0.0095 | 0.042 | 0.049 | 1 | 2 | 1190002N1 | -0.007 | Old |
| 0.496237 | 0.096384 | 0.041 | 0.048 | 1 | 2 | Zfp746 | -0.007 | Old |
| 0.496245 | 0.073198 | 0.409 | 0.395 | 1 | 2 | Psmd2 | 0.014 | Old |
| 0.496579 | 0.027113 | 0.18 | 0.166 | 1 | 2 | Acbd3 | 0.014 | Old |
| 0.497109 | 0.2921 | 0.029 | 0.024 | 1 | 2 | Mios | 0.005 | Old |
| 0.497837 | 0.084687 | 0.366 | 0.349 | 1 | 2 | Fgfr1op2 | 0.017 | Old |
| 0.498017 | 0.934314 | 0.013 | 0.01 | 1 | 2 | Zfp799 | 0.003 | Old |
| 0.498512 | 0.121125 | 0.227 | 0.213 | 1 | 2 | Akt2 | 0.014 | Old |
| 0.498588 | 0.297622 | 0.029 | 0.024 | 1 | 2 | Retsat | 0.005 | Old |
| 0.498701 | 0.23325 | 0.215 | 0.204 | 1 | 2 | Trpm7 | 0.011 | Old |
| 0.500157 | 0.186771 | 0.125 | 0.114 | 1 | 2 | Ilf2 | 0.011 | Old |
| 0.50055 | 0.062979 | 0.142 | 0.129 | 1 | 2 | Mettl9 | 0.013 | Old |
| 0.500872 | 0.166048 | 0.349 | 0.338 | 1 | 2 | Atp6v0d1 | 0.011 | Old |
| 0.500915 | 0.409789 | 0.013 | 0.01 | 1 | 2 | Platr25 | 0.003 | Old |
| 0.500943 | 0.255552 | 0.042 | 0.036 | 1 | 2 | Gnl3l | 0.006 | Old |
| 0.501262 | 0.073099 | 0.042 | 0.049 | 1 | 2 | Rpa2 | -0.007 | Old |
| 0.501738 | 0.262466 | 0.075 | 0.067 | 1 | 2 | Carnmt1 | 0.008 | Old |
| 0.501824 | 0.106274 | 0.042 | 0.036 | 1 | 2 | Wdr45 | 0.006 | Old |
| 0.502624 | 0.404816 | 0.049 | 0.043 | 1 | 2 | Prr5l | 0.006 | Old |
| 0.502722 | 0.370793 | 0.029 | 0.024 | 1 | 2 | 0610009B2 | 0.005 | Old |
| 0.503176 | 0.579738 | 0.013 | 0.01 | 1 | 2 | Gm45894 | 0.003 | Old |
| 0.503312 | 0.231874 | 0.077 | 0.069 | 1 | 2 | Idh3a | 0.008 | Old |
| 0.503493 | 0.1407 | 0.208 | 0.196 | 1 | 2 | Mnt | 0.012 | Old |
| 0.503583 | 0.239768 | 0.057 | 0.05 | 1 | 2 | Rnaset2a | 0.007 | Old |
| 0.504115 | 0.015313 | 0.106 | 0.096 | 1 | 2 | Nr1h2 | 0.01 | Old |
| 0.504132 | 0.622934 | 0.018 | 0.014 | 1 | 2 | Nxn | 0.004 | Old |
| 0.504263 | 0.009246 | 0.11 | 0.121 | 1 | 2 | Usp1 | -0.011 | Old |
| 0.50447 | 0.311544 | 0.013 | 0.01 | 1 | 2 | Cdc25b | 0.003 | Old |
| 0.504476 | 0.193972 | 0.055 | 0.048 | 1 | 2 | Mtfr1 | 0.007 | Old |
| 0.504536 | 0.046528 | 0.156 | 0.144 | 1 | 2 | Msmo1 | 0.012 | Old |
| 0.504986 | 0.258563 | 0.103 | 0.094 | 1 | 2 | Klf11 | 0.009 | Old |
| 0.505025 | 0.106132 | 0.035 | 0.03 | 1 | 2 | Hebp1 | 0.005 | Old |
| 0.5051 | 0.047625 | 0.464 | 0.439 | 1 | 2 | Larp4b | 0.025 | Old |
| 0.505441 | 0.257114 | 0.013 | 0.01 | 1 | 2 | Mboat1 | 0.003 | Old |

| | | | | | | | | |
|---|---|---|---|---|---|---|---|---|
| 0.505594 | 0.117722 | 0.029 | 0.024 | 1 | 2 | Gm5086 | 0.005 | Old |
| 0.505638 | 0.343427 | 0.024 | 0.019 | 1 | 2 | Fam206a | 0.005 | Old |
| 0.505708 | 0.063197 | 0.293 | 0.278 | 1 | 2 | Hivep2 | 0.015 | Old |
| 0.505766 | 0.163507 | 0.013 | 0.01 | 1 | 2 | Tmem138 | 0.003 | Old |
| 0.50583 | 0.054457 | 0.251 | 0.233 | 1 | 2 | Kpnb1 | 0.018 | Old |
| 0.505875 | 0.262994 | 0.024 | 0.019 | 1 | 2 | Ttf2 | 0.005 | Old |
| 0.506414 | 0.295471 | 0.013 | 0.01 | 1 | 2 | Tubd1 | 0.003 | Old |
| 0.50656 | 0.686124 | 0.018 | 0.014 | 1 | 2 | Akap1 | 0.004 | Old |
| 0.506998 | 0.043736 | 0.41 | 0.391 | 1 | 2 | 11-Sep | 0.019 | Old |
| 0.507055 | 0.014698 | 0.124 | 0.113 | 1 | 2 | Pfdn1 | 0.011 | Old |
| 0.507072 | 0.088916 | 0.088 | 0.078 | 1 | 2 | Parp11 | 0.01 | Old |
| 0.507315 | 0.109103 | 0.219 | 0.205 | 1 | 2 | Arih2 | 0.014 | Old |
| 0.507755 | 0.052254 | 0.106 | 0.095 | 1 | 2 | A630001G2 | 0.011 | Old |
| 0.508101 | 0.232161 | 0.077 | 0.069 | 1 | 2 | Eif1ad | 0.008 | Old |
| 0.508453 | 0.438351 | 0.018 | 0.014 | 1 | 2 | Gm48066 | 0.004 | Old |
| 0.508563 | 0.039939 | 0.569 | 0.557 | 1 | 2 | Vsir | 0.012 | Old |
| 0.508811 | 0.173887 | 0.137 | 0.125 | 1 | 2 | Impad1 | 0.012 | Old |
| 0.508895 | 0.019565 | 0.148 | 0.159 | 1 | 2 | Ap1g1 | -0.011 | Old |
| 0.508912 | 0.081072 | 0.277 | 0.26 | 1 | 2 | Mapk1ip1l | 0.017 | Old |
| 0.509777 | 0.308074 | 0.056 | 0.049 | 1 | 2 | Rab28 | 0.007 | Old |
| 0.510078 | 0.496759 | 0.018 | 0.014 | 1 | 2 | Aqp9 | 0.004 | Old |
| 0.510156 | 0.067107 | 0.214 | 0.201 | 1 | 2 | Hck | 0.013 | Old |
| 0.510169 | 0.142069 | 0.155 | 0.144 | 1 | 2 | Hcfc1 | 0.011 | Old |
| 0.510842 | 0.050852 | 0.101 | 0.11 | 1 | 2 | Bag3 | -0.009 | Old |
| 0.510944 | 0.012366 | 0.991 | 0.998 | 1 | 2 | Rpl37a | -0.007 | Old |
| 0.511099 | 0.257334 | 0.059 | 0.052 | 1 | 2 | Sun1 | 0.007 | Old |
| 0.51113 | 0.179088 | 0.198 | 0.188 | 1 | 2 | Numa1 | 0.01 | Old |
| 0.511237 | 0.410508 | 0.055 | 0.048 | 1 | 2 | Caap1 | 0.007 | Old |
| 0.511392 | 0.061291 | 0.047 | 0.04 | 1 | 2 | Ctdp1 | 0.007 | Old |
| 0.511442 | 0.12784 | 0.233 | 0.22 | 1 | 2 | Il17ra | 0.013 | Old |
| 0.51162 | 0.104299 | 0.199 | 0.184 | 1 | 2 | Elk4 | 0.015 | Old |
| 0.512377 | 0.125127 | 0.192 | 0.179 | 1 | 2 | Rnh1 | 0.013 | Old |
| 0.512832 | 0.005778 | 0.463 | 0.475 | 1 | 2 | Dhrs3 | -0.012 | Old |
| 0.512853 | 0.182006 | 0.108 | 0.099 | 1 | 2 | Klhl28 | 0.009 | Old |
| 0.512871 | 0.026619 | 0.43 | 0.455 | 1 | 2 | Nr4a2 | -0.025 | Old |
| 0.512949 | 0.077498 | 0.218 | 0.202 | 1 | 2 | Emc1 | 0.016 | Old |
| 0.512981 | 0.066629 | 0.466 | 0.444 | 1 | 2 | Capns1 | 0.022 | Old |
| 0.513065 | 0.321481 | 0.018 | 0.014 | 1 | 2 | Gm10521 | 0.004 | Old |
| 0.513204 | 0.010766 | 0.589 | 0.603 | 1 | 2 | Fyb | -0.014 | Old |
| 0.513333 | 0.00028 | 0.119 | 0.108 | 1 | 2 | Hmgcl | 0.011 | Old |
| 0.513507 | 0.03647 | 0.125 | 0.114 | 1 | 2 | Psmc4 | 0.011 | Old |
| 0.513681 | 0.247642 | 0.138 | 0.128 | 1 | 2 | Dock1 | 0.01 | Old |
| 0.513911 | 0.160464 | 0.035 | 0.03 | 1 | 2 | Haus3 | 0.005 | Old |
| 0.51459 | 0.127358 | 0.035 | 0.03 | 1 | 2 | Myg1 | 0.005 | Old |
| 0.514596 | 0.338131 | 0.081 | 0.074 | 1 | 2 | Rnf138 | 0.007 | Old |

| | | | | | | | | |
|---|---|---|---|---|---|---|---|---|
| 0.514663 | 0.092776 | 0.394 | 0.376 | 1 | 2 | Zbtb7a | 0.018 | Old |
| 0.514893 | 0.301216 | 0.052 | 0.061 | 1 | 2 | Tmx2 | -0.009 | Old |
| 0.51497 | 0.53093 | 0.018 | 0.014 | 1 | 2 | Tlr3 | 0.004 | Old |
| 0.515076 | 0.001375 | 0.035 | 0.03 | 1 | 2 | Rmnd5b | 0.005 | Old |
| 0.515367 | 0.05222 | 0.113 | 0.103 | 1 | 2 | Mbd6 | 0.01 | Old |
| 0.516863 | 0.019465 | 0.047 | 0.04 | 1 | 2 | Gfm2 | 0.007 | Old |
| 0.517051 | 0.101617 | 0.087 | 0.077 | 1 | 2 | Frat2 | 0.01 | Old |
| 0.517124 | 0.298013 | 0.03 | 0.025 | 1 | 2 | Txnl4b | 0.005 | Old |
| 0.517679 | 0.165036 | 0.101 | 0.091 | 1 | 2 | Zbtb7b | 0.01 | Old |
| 0.517894 | 0.183894 | 0.036 | 0.031 | 1 | 2 | Dusp12 | 0.005 | Old |
| 0.518007 | 0.015223 | 0.642 | 0.666 | 1 | 2 | Ccnl1 | -0.024 | Old |
| 0.518293 | 0.106563 | 0.238 | 0.224 | 1 | 2 | Nrip1 | 0.014 | Old |
| 0.51872 | 0.135009 | 0.23 | 0.219 | 1 | 2 | Nrbp1 | 0.011 | Old |
| 0.518756 | 0.222554 | 0.049 | 0.043 | 1 | 2 | Zfp398 | 0.006 | Old |
| 0.518795 | 0.142993 | 0.106 | 0.096 | 1 | 2 | Cdk2 | 0.01 | Old |
| 0.519066 | 0.070202 | 0.035 | 0.03 | 1 | 2 | Gm26853 | 0.005 | Old |
| 0.519426 | 0.03825 | 0.662 | 0.638 | 1 | 2 | Eif3a | 0.024 | Old |
| 0.520167 | 0.076664 | 0.055 | 0.048 | 1 | 2 | Ankrd16 | 0.007 | Old |
| 0.520292 | 0.529698 | 0.036 | 0.031 | 1 | 2 | Eif2ak4 | 0.005 | Old |
| 0.520466 | 0.225182 | 0.133 | 0.122 | 1 | 2 | Kctd10 | 0.011 | Old |
| 0.520673 | 0.211718 | 0.018 | 0.023 | 1 | 2 | Zfp429 | -0.005 | Old |
| 0.520905 | 0.11277 | 0.318 | 0.299 | 1 | 2 | 7-Mar | 0.019 | Old |
| 0.520926 | 0.162985 | 0.14 | 0.127 | 1 | 2 | Ahcyl1 | 0.013 | Old |
| 0.521502 | 0.199883 | 0.062 | 0.055 | 1 | 2 | Gtf3c3 | 0.007 | Old |
| 0.521552 | 0.07359 | 0.09 | 0.081 | 1 | 2 | Ruvbl1 | 0.009 | Old |
| 0.521611 | 0.125908 | 0.268 | 0.251 | 1 | 2 | Pkn1 | 0.017 | Old |
| 0.522163 | 0.078843 | 0.048 | 0.042 | 1 | 2 | Dvl3 | 0.006 | Old |
| 0.523113 | 0.030622 | 0.168 | 0.154 | 1 | 2 | Aida | 0.014 | Old |
| 0.5234 | 0.604387 | 0.039 | 0.033 | 1 | 2 | Zfp866 | 0.006 | Old |
| 0.523783 | 0.289177 | 0.089 | 0.081 | 1 | 2 | Ubqln2 | 0.008 | Old |
| 0.523879 | 0.104379 | 0.077 | 0.069 | 1 | 2 | Zbtb4 | 0.008 | Old |
| 0.523901 | 0.266142 | 0.087 | 0.078 | 1 | 2 | Cisd1 | 0.009 | Old |
| 0.524128 | 0.33386 | 0.14 | 0.131 | 1 | 2 | Adrb2 | 0.009 | Old |
| 0.524188 | 0.335764 | 0.014 | 0.018 | 1 | 2 | Afap1 | -0.004 | Old |
| 0.525166 | 0.152672 | 0.145 | 0.157 | 1 | 2 | Serpinb6a | -0.012 | Old |
| 0.526965 | 0.020631 | 0.781 | 0.76 | 1 | 2 | Unc93b1 | 0.021 | Old |
| 0.527107 | 0.161494 | 0.161 | 0.15 | 1 | 2 | Maea | 0.011 | Old |
| 0.527144 | 0.08057 | 0.019 | 0.024 | 1 | 2 | Maoa | -0.005 | Old |
| 0.527317 | 0.111777 | 0.05 | 0.044 | 1 | 2 | Myo1e | 0.006 | Old |
| 0.527427 | 0.096975 | 0.077 | 0.069 | 1 | 2 | Tubb4b | 0.008 | Old |
| 0.527527 | 0.129371 | 0.221 | 0.21 | 1 | 2 | Slc16a3 | 0.011 | Old |
| 0.528121 | 0.212834 | 0.07 | 0.062 | 1 | 2 | Gnptab | 0.008 | Old |
| 0.528283 | 0.225271 | 0.195 | 0.185 | 1 | 2 | Naa15 | 0.01 | Old |
| 0.528873 | 0.025331 | 0.179 | 0.165 | 1 | 2 | Dstyk | 0.014 | Old |
| 0.529179 | 0.142966 | 0.054 | 0.061 | 1 | 2 | Mkl1 | -0.007 | Old |

| | | | | | | | | |
|---|---|---|---|---|---|---|---|---|
| 0.529389 | 0.201195 | 0.024 | 0.029 | 1 | 2 | Dmap1 | -0.005 | Old |
| 0.530666 | 0.358834 | 0.031 | 0.026 | 1 | 2 | Bclaf3 | 0.005 | Old |
| 0.531881 | 0.078161 | 0.21 | 0.198 | 1 | 2 | Psmb9 | 0.012 | Old |
| 0.531985 | 0.214676 | 0.095 | 0.087 | 1 | 2 | Med12 | 0.008 | Old |
| 0.532134 | 0.101533 | 0.395 | 0.378 | 1 | 2 | Eif4ebp2 | 0.017 | Old |
| 0.532364 | 0.410357 | 0.019 | 0.015 | 1 | 2 | Rpl9-ps6 | 0.004 | Old |
| 0.532617 | 0.675347 | 0.026 | 0.021 | 1 | 2 | Plpp6 | 0.005 | Old |
| 0.53266 | 0.354124 | 0.051 | 0.045 | 1 | 2 | Trim11 | 0.006 | Old |
| 0.533029 | 0.028426 | 0.134 | 0.122 | 1 | 2 | Me2 | 0.012 | Old |
| 0.533065 | 0.038878 | 0.073 | 0.081 | 1 | 2 | Akap8l | -0.008 | Old |
| 0.533169 | 0.613299 | 0.019 | 0.015 | 1 | 2 | Crip2 | 0.004 | Old |
| 0.533334 | 0.168442 | 0.236 | 0.223 | 1 | 2 | Ccni | 0.013 | Old |
| 0.533535 | 0.08639 | 0.086 | 0.077 | 1 | 2 | Mrps7 | 0.009 | Old |
| 0.533872 | 0.094146 | 0.237 | 0.222 | 1 | 2 | Cic | 0.015 | Old |
| 0.534308 | 0.081715 | 0.126 | 0.115 | 1 | 2 | Itgb7 | 0.011 | Old |
| 0.534457 | 0.083509 | 0.102 | 0.093 | 1 | 2 | Ash2l | 0.009 | Old |
| 0.534482 | 0.036129 | 0.078 | 0.07 | 1 | 2 | Nup214 | 0.008 | Old |
| 0.534646 | 0.509518 | 0.019 | 0.015 | 1 | 2 | Tctn3 | 0.004 | Old |
| 0.534695 | 0.273448 | 0.025 | 0.02 | 1 | 2 | Gatc | 0.005 | Old |
| 0.534884 | 0.593317 | 0.014 | 0.011 | 1 | 2 | D3Ertd254 | 0.003 | Old |
| 0.534923 | 0.058755 | 0.212 | 0.2 | 1 | 2 | Appbp2 | 0.012 | Old |
| 0.535046 | 0.221294 | 0.097 | 0.089 | 1 | 2 | Rsrc1 | 0.008 | Old |
| 0.535203 | 0.527667 | 0.014 | 0.011 | 1 | 2 | Rnd1 | 0.003 | Old |
| 0.535289 | 0.083805 | 0.151 | 0.139 | 1 | 2 | Cdc26 | 0.012 | Old |
| 0.535318 | 0.239652 | 0.019 | 0.015 | 1 | 2 | Nup43 | 0.004 | Old |
| 0.535492 | 0.047732 | 0.076 | 0.068 | 1 | 2 | Mfsd7a | 0.008 | Old |
| 0.535521 | 0.431317 | 0.014 | 0.011 | 1 | 2 | Katnb1 | 0.003 | Old |
| 0.535644 | 0.112386 | 0.025 | 0.02 | 1 | 2 | 3110001I2 | 0.005 | Old |
| 0.53603 | 0.567779 | 0.07 | 0.063 | 1 | 2 | Topbp1 | 0.007 | Old |
| 0.536428 | 0.113971 | 0.403 | 0.386 | 1 | 2 | Mef2d | 0.017 | Old |
| 0.536561 | 0.311123 | 0.095 | 0.087 | 1 | 2 | Oard1 | 0.008 | Old |
| 0.536825 | 0.224778 | 0.283 | 0.271 | 1 | 2 | Tmem167 | 0.012 | Old |
| 0.536933 | 0.418017 | 0.019 | 0.015 | 1 | 2 | Mterf4 | 0.004 | Old |
| 0.537395 | 0.038057 | 0.543 | 0.524 | 1 | 2 | M6pr | 0.019 | Old |
| 0.537854 | 0.070568 | 0.065 | 0.058 | 1 | 2 | Vps45 | 0.007 | Old |
| 0.538072 | 0.57313 | 0.014 | 0.011 | 1 | 2 | Gm16973 | 0.003 | Old |
| 0.538135 | 0.050594 | 0.337 | 0.315 | 1 | 2 | Sh3kbp1 | 0.022 | Old |
| 0.538142 | 0.250825 | 0.087 | 0.078 | 1 | 2 | Cep250 | 0.009 | Old |
| 0.53882 | 0.272248 | 0.019 | 0.015 | 1 | 2 | Zfp768 | 0.004 | Old |
| 0.539135 | 0.099739 | 0.237 | 0.223 | 1 | 2 | Stat6 | 0.014 | Old |
| 0.53935 | 0.285009 | 0.014 | 0.011 | 1 | 2 | Gm12689 | 0.003 | Old |
| 0.539405 | 0.48584 | 0.037 | 0.032 | 1 | 2 | Tmem175 | 0.005 | Old |
| 0.539547 | 0.109623 | 0.048 | 0.055 | 1 | 2 | Ddx41 | -0.007 | Old |
| 0.53957 | 0.129411 | 0.05 | 0.044 | 1 | 2 | Dvl1 | 0.006 | Old |
| 0.540036 | 0.124501 | 0.044 | 0.038 | 1 | 2 | Net1 | 0.006 | Old |

| | | | | | | | | |
|---|---|---|---|---|---|---|---|---|
| 0.54025 | 0.059091 | 0.451 | 0.468 | 1 | 2 | Myh9 | -0.017 | Old |
| 0.54031 | 0.282451 | 0.014 | 0.011 | 1 | 2 | Efnb1 | 0.003 | Old |
| 0.54031 | 0.273361 | 0.014 | 0.011 | 1 | 2 | Snx33 | 0.003 | Old |
| 0.540596 | 0.070665 | 0.177 | 0.164 | 1 | 2 | Dpy19l1 | 0.013 | Old |
| 0.540981 | 0.054042 | 0.019 | 0.015 | 1 | 2 | Slfn4 | 0.004 | Old |
| 0.541068 | 0.180286 | 0.167 | 0.181 | 1 | 2 | Smarcc2 | -0.014 | Old |
| 0.541329 | 0.05507 | 0.15 | 0.14 | 1 | 2 | Dtx3l | 0.01 | Old |
| 0.54159 | 0.280346 | 0.014 | 0.011 | 1 | 2 | Tfb1m | 0.003 | Old |
| 0.542104 | 0.401193 | 0.06 | 0.053 | 1 | 2 | Tubgcp5 | 0.007 | Old |
| 0.542552 | 0.826001 | 0.014 | 0.011 | 1 | 2 | P2ry10 | 0.003 | Old |
| 0.542552 | 0.206826 | 0.014 | 0.011 | 1 | 2 | Apex2 | 0.003 | Old |
| 0.543021 | 0.045217 | 0.772 | 0.747 | 1 | 2 | Hnrnpk | 0.025 | Old |
| 0.544254 | 0.118789 | 0.139 | 0.127 | 1 | 2 | Synj2bp | 0.012 | Old |
| 0.544254 | 0.013968 | 0.208 | 0.195 | 1 | 2 | Ogfr | 0.013 | Old |
| 0.544477 | 0.18226 | 0.014 | 0.011 | 1 | 2 | Zfp692 | 0.003 | Old |
| 0.5452 | 0.247205 | 0.132 | 0.122 | 1 | 2 | Pygl | 0.01 | Old |
| 0.545316 | 0.133906 | 0.019 | 0.015 | 1 | 2 | 4930430F0 | 0.004 | Old |
| 0.545539 | 0.067136 | 0.124 | 0.114 | 1 | 2 | Ppp2r5e | 0.01 | Old |
| 0.546061 | 0.085803 | 0.177 | 0.165 | 1 | 2 | Rbm17 | 0.012 | Old |
| 0.546063 | 0.022114 | 0.044 | 0.05 | 1 | 2 | Sass6 | -0.006 | Old |
| 0.546129 | 0.023693 | 0.105 | 0.115 | 1 | 2 | Taok2 | -0.01 | Old |
| 0.546237 | 0.037629 | 0.743 | 0.738 | 1 | 2 | Canx | 0.005 | Old |
| 0.546419 | 0.233624 | 0.032 | 0.027 | 1 | 2 | Prdm15 | 0.005 | Old |
| 0.546442 | 0.321442 | 0.039 | 0.033 | 1 | 2 | 2310061I0 | 0.006 | Old |
| 0.546782 | 0.138031 | 0.081 | 0.074 | 1 | 2 | Rhot1 | 0.007 | Old |
| 0.547432 | 0.104175 | 0.154 | 0.144 | 1 | 2 | Mbtps1 | 0.01 | Old |
| 0.547459 | 1.65E-05 | 0.116 | 0.125 | 1 | 2 | Atp9b | -0.009 | Old |
| 0.548303 | 0.278639 | 0.032 | 0.027 | 1 | 2 | Rab2b | 0.005 | Old |
| 0.548392 | 0.098625 | 0.488 | 0.471 | 1 | 2 | Ppp1r18 | 0.017 | Old |
| 0.548656 | 0.081033 | 0.352 | 0.337 | 1 | 2 | Id3 | 0.015 | Old |
| 0.548974 | 0.007803 | 0.511 | 0.537 | 1 | 2 | Rab7b | -0.026 | Old |
| 0.549014 | 0.24206 | 0.087 | 0.078 | 1 | 2 | Nrm | 0.009 | Old |
| 0.549328 | 0.083907 | 0.286 | 0.273 | 1 | 2 | Cdk9 | 0.013 | Old |
| 0.549533 | 0.088012 | 0.522 | 0.505 | 1 | 2 | Emb | 0.017 | Old |
| 0.549642 | 0.102675 | 0.253 | 0.236 | 1 | 2 | Ndufs2 | 0.017 | Old |
| 0.550247 | 0.069335 | 0.368 | 0.354 | 1 | 2 | Ube3a | 0.014 | Old |
| 0.550305 | 0.627475 | 0.047 | 0.042 | 1 | 2 | Ap4m1 | 0.005 | Old |
| 0.55065 | 0.404763 | 0.039 | 0.033 | 1 | 2 | Arl2 | 0.006 | Old |
| 0.550676 | 0.318328 | 0.148 | 0.139 | 1 | 2 | Zzz3 | 0.009 | Old |
| 0.551403 | 0.187813 | 0.075 | 0.068 | 1 | 2 | Herc6 | 0.007 | Old |
| 0.551703 | 0.269585 | 0.127 | 0.119 | 1 | 2 | Ythdf2 | 0.008 | Old |
| 0.551721 | 0.947272 | 0.033 | 0.029 | 1 | 2 | Cd69 | 0.004 | Old |
| 0.551745 | 0.178743 | 0.026 | 0.021 | 1 | 2 | Abca6 | 0.005 | Old |
| 0.551975 | 0.056966 | 0.081 | 0.074 | 1 | 2 | Hps5 | 0.007 | Old |
| 0.552028 | 0.05177 | 0.305 | 0.287 | 1 | 2 | Tiam1 | 0.018 | Old |

| | | | | | | | | |
|---|---|---|---|---|---|---|---|---|
| 0.55229 | 0.148129 | 0.032 | 0.027 | 1 | 2 | 2810402E2 | 0.005 | Old |
| 0.552754 | 0.231902 | 0.165 | 0.154 | 1 | 2 | Tmem131 | 0.011 | Old |
| 0.552871 | 0.172079 | 0.168 | 0.157 | 1 | 2 | Kcnq1ot1 | 0.011 | Old |
| 0.552895 | 0.041957 | 0.306 | 0.287 | 1 | 2 | Med13l | 0.019 | Old |
| 0.552921 | 0.332472 | 0.032 | 0.027 | 1 | 2 | Shmt2 | 0.005 | Old |
| 0.553207 | 0.034629 | 0.36 | 0.34 | 1 | 2 | Plekhb2 | 0.02 | Old |
| 0.553953 | 0.187835 | 0.045 | 0.039 | 1 | 2 | Mrps18b | 0.006 | Old |
| 0.554245 | 0.164185 | 0.169 | 0.158 | 1 | 2 | Mga | 0.011 | Old |
| 0.554833 | 0.059553 | 0.257 | 0.242 | 1 | 2 | Zfand6 | 0.015 | Old |
| 0.555096 | 0.039643 | 0.168 | 0.178 | 1 | 2 | Hcst | -0.01 | Old |
| 0.555172 | 0.03433 | 0.112 | 0.103 | 1 | 2 | Brd3 | 0.009 | Old |
| 0.555672 | 0.235463 | 0.088 | 0.08 | 1 | 2 | Samd1 | 0.008 | Old |
| 0.555737 | 0.01134 | 0.044 | 0.038 | 1 | 2 | Vipas39 | 0.006 | Old |
| 0.555968 | 0.01433 | 0.624 | 0.603 | 1 | 2 | Hexa | 0.021 | Old |
| 0.556291 | 0.148941 | 0.032 | 0.027 | 1 | 2 | Scfd2 | 0.005 | Old |
| 0.556565 | 0.179899 | 0.054 | 0.048 | 1 | 2 | Sephs2 | 0.006 | Old |
| 0.556657 | 0.012264 | 0.089 | 0.081 | 1 | 2 | Cstf2t | 0.008 | Old |
| 0.557478 | 0.06916 | 0.68 | 0.666 | 1 | 2 | Atp5a1 | 0.014 | Old |
| 0.557525 | 0.072655 | 0.369 | 0.349 | 1 | 2 | Kpna4 | 0.02 | Old |
| 0.557819 | 0.029267 | 0.346 | 0.327 | 1 | 2 | Pacs2 | 0.019 | Old |
| 0.558144 | 0.092426 | 0.123 | 0.133 | 1 | 2 | Gab2 | -0.01 | Old |
| 0.558192 | 0.169539 | 0.032 | 0.027 | 1 | 2 | Ercc2 | 0.005 | Old |
| 0.558296 | 0.077522 | 0.218 | 0.204 | 1 | 2 | Snw1 | 0.014 | Old |
| 0.559004 | 0.893114 | 0.01 | 0.007 | 1 | 2 | Cdkn1c | 0.003 | Old |
| 0.559029 | 0.215082 | 0.155 | 0.145 | 1 | 2 | Fam76b | 0.01 | Old |
| 0.559791 | 0.694004 | 0.01 | 0.007 | 1 | 2 | Ankzf1 | 0.003 | Old |
| 0.55998 | 0.109493 | 0.046 | 0.052 | 1 | 2 | Gmeb2 | -0.006 | Old |
| 0.560076 | 0.110889 | 0.245 | 0.232 | 1 | 2 | BC017643 | 0.013 | Old |
| 0.560184 | 0.775854 | 0.01 | 0.007 | 1 | 2 | Slc4a11 | 0.003 | Old |
| 0.560578 | 0.950018 | 0.01 | 0.007 | 1 | 2 | Mill2 | 0.003 | Old |
| 0.560778 | 0.521504 | 0.046 | 0.04 | 1 | 2 | Acot8 | 0.006 | Old |
| 0.561037 | 0.521163 | 0.041 | 0.036 | 1 | 2 | Nup107 | 0.005 | Old |
| 0.56116 | 0.020356 | 0.435 | 0.416 | 1 | 2 | Psmb1 | 0.019 | Old |
| 0.56121 | 0.01885 | 0.084 | 0.075 | 1 | 2 | Nsmaf | 0.009 | Old |
| 0.561359 | 0.127865 | 0.346 | 0.333 | 1 | 2 | Ap1s2 | 0.013 | Old |
| 0.561366 | 0.898998 | 0.01 | 0.007 | 1 | 2 | Oas1c | 0.003 | Old |
| 0.561451 | 0.332675 | 0.062 | 0.056 | 1 | 2 | Akt3 | 0.006 | Old |
| 0.561825 | 0.44788 | 0.072 | 0.065 | 1 | 2 | Crot | 0.007 | Old |
| 0.561902 | 0.044298 | 0.065 | 0.058 | 1 | 2 | Rad17 | 0.007 | Old |
| 0.561911 | 0.173006 | 0.318 | 0.304 | 1 | 2 | Zc3hav1 | 0.014 | Old |
| 0.562154 | 0.198221 | 0.01 | 0.007 | 1 | 2 | Cidec | 0.003 | Old |
| 0.562351 | 0.026817 | 0.88 | 0.893 | 1 | 2 | Pabpc1 | -0.013 | Old |
| 0.562543 | 0.001041 | 0.079 | 0.087 | 1 | 2 | Trip4 | -0.008 | Old |
| 0.562549 | 0.006504 | 0.354 | 0.329 | 1 | 2 | Prpf40a | 0.025 | Old |
| 0.562859 | 0.017533 | 0.026 | 0.021 | 1 | 2 | Ttl | 0.005 | Old |

| | | | | | | | | |
|---|---|---|---|---|---|---|---|---|
| 0.562915 | 0.081256 | 0.138 | 0.127 | 1 | 2 | Rabep1 | 0.011 | Old |
| 0.562943 | 0.604187 | 0.01 | 0.007 | 1 | 2 | Jchain | 0.003 | Old |
| 0.562987 | 0.166749 | 0.16 | 0.15 | 1 | 2 | Il3ra | 0.01 | Old |
| 0.563093 | 0.079878 | 0.708 | 0.704 | 1 | 2 | Tpd52 | 0.004 | Old |
| 0.563134 | 0.18976 | 0.02 | 0.017 | 1 | 2 | Gm15564 | 0.003 | Old |
| 0.563338 | 0.631111 | 0.01 | 0.007 | 1 | 2 | Bbs7 | 0.003 | Old |
| 0.563531 | 0.1212 | 0.067 | 0.061 | 1 | 2 | Nop9 | 0.006 | Old |
| 0.563733 | 0.496754 | 0.01 | 0.007 | 1 | 2 | Zranb3 | 0.003 | Old |
| 0.564176 | 0.15849 | 0.227 | 0.214 | 1 | 2 | Lpxn | 0.013 | Old |
| 0.564358 | 0.261559 | 0.062 | 0.056 | 1 | 2 | Ptgr1 | 0.006 | Old |
| 0.564596 | 0.142184 | 0.074 | 0.067 | 1 | 2 | Wdr91 | 0.007 | Old |
| 0.564628 | 0.083704 | 0.377 | 0.362 | 1 | 2 | Irf5 | 0.015 | Old |
| 0.564666 | 0.025127 | 0.185 | 0.172 | 1 | 2 | Psmd6 | 0.013 | Old |
| 0.564899 | 0.076694 | 0.106 | 0.096 | 1 | 2 | Smim27 | 0.01 | Old |
| 0.566578 | 0.489249 | 0.056 | 0.05 | 1 | 2 | Ccdc93 | 0.006 | Old |
| 0.566595 | 0.055718 | 0.09 | 0.099 | 1 | 2 | Tmem243 | -0.009 | Old |
| 0.566783 | 0.035015 | 0.197 | 0.184 | 1 | 2 | Atxn2 | 0.013 | Old |
| 0.567152 | 0.152951 | 0.02 | 0.017 | 1 | 2 | Vps51 | 0.003 | Old |
| 0.567869 | 0.007893 | 0.238 | 0.223 | 1 | 2 | Kat6a | 0.015 | Old |
| 0.567942 | 0.286359 | 0.015 | 0.012 | 1 | 2 | Herc3 | 0.003 | Old |
| 0.568075 | 0.112946 | 0.058 | 0.051 | 1 | 2 | Borcs7 | 0.007 | Old |
| 0.568321 | 0.498312 | 0.046 | 0.04 | 1 | 2 | Slpi | 0.006 | Old |
| 0.56848 | 0.076272 | 0.01 | 0.007 | 1 | 2 | Mansc1 | 0.003 | Old |
| 0.568509 | 0.180407 | 0.075 | 0.068 | 1 | 2 | Creb3 | 0.007 | Old |
| 0.568522 | 0.019461 | 0.154 | 0.163 | 1 | 2 | Atxn7l1 | -0.009 | Old |
| 0.568877 | 0.049817 | 0.01 | 0.007 | 1 | 2 | Mbnl3 | 0.003 | Old |
| 0.568885 | 0.405114 | 0.015 | 0.012 | 1 | 2 | Slc19a2 | 0.003 | Old |
| 0.569065 | 0.163001 | 0.094 | 0.087 | 1 | 2 | Tfrc | 0.007 | Old |
| 0.569108 | 0.031271 | 0.283 | 0.262 | 1 | 2 | Psmd11 | 0.021 | Old |
| 0.569192 | 0.248575 | 0.052 | 0.046 | 1 | 2 | Mthfsl | 0.006 | Old |
| 0.569504 | 0.29258 | 0.033 | 0.029 | 1 | 2 | Dcakd | 0.004 | Old |
| 0.569983 | 0.055771 | 0.06 | 0.053 | 1 | 2 | Slc44a2 | 0.007 | Old |
| 0.570143 | 0.788953 | 0.015 | 0.012 | 1 | 2 | 1700019L1 | 0.003 | Old |
| 0.570457 | 0.223307 | 0.015 | 0.012 | 1 | 2 | Nme7 | 0.003 | Old |
| 0.571402 | 0.392083 | 0.015 | 0.012 | 1 | 2 | Zfp46 | 0.003 | Old |
| 0.572 | 0.055131 | 0.177 | 0.164 | 1 | 2 | 4930523C0 | 0.013 | Old |
| 0.572032 | 0.317019 | 0.015 | 0.012 | 1 | 2 | Zbtb39 | 0.003 | Old |
| 0.572158 | 0.373963 | 0.047 | 0.042 | 1 | 2 | Prcc | 0.005 | Old |
| 0.572373 | 0.012382 | 0.052 | 0.046 | 1 | 2 | Sqle | 0.006 | Old |
| 0.572998 | 0.184412 | 0.055 | 0.049 | 1 | 2 | Orai2 | 0.006 | Old |
| 0.57304 | 0.185167 | 0.047 | 0.042 | 1 | 2 | Ppargc1b | 0.005 | Old |
| 0.573409 | 0.078285 | 0.027 | 0.023 | 1 | 2 | Supt7l | 0.004 | Old |
| 0.573909 | 0.011028 | 0.153 | 0.14 | 1 | 2 | Hspa14 | 0.013 | Old |
| 0.574389 | 0.099116 | 0.061 | 0.055 | 1 | 2 | Gab1 | 0.006 | Old |
| 0.574556 | 0.144424 | 0.015 | 0.012 | 1 | 2 | Gm7160 | 0.003 | Old |

| | | | | | | | | |
|---|---|---|---|---|---|---|---|---|
| 0.575286 | 0.076152 | 0.027 | 0.023 | 1 | 2 | Ggct | 0.004 | Old |
| 0.575504 | 0.013376 | 0.015 | 0.012 | 1 | 2 | Hrh2 | 0.003 | Old |
| 0.57571 | 0.060392 | 0.39 | 0.371 | 1 | 2 | Ndufa3 | 0.019 | Old |
| 0.575835 | 0.113485 | 0.209 | 0.196 | 1 | 2 | Rap2b | 0.013 | Old |
| 0.576036 | 0.799472 | 0.034 | 0.03 | 1 | 2 | Tada3 | 0.004 | Old |
| 0.576452 | 0.099178 | 0.429 | 0.411 | 1 | 2 | Scp2 | 0.018 | Old |
| 0.576452 | 0.462496 | 0.015 | 0.012 | 1 | 2 | Gatsl3 | 0.003 | Old |
| 0.57656 | 0.126656 | 0.036 | 0.042 | 1 | 2 | Foxo4 | -0.006 | Old |
| 0.576756 | 0.264369 | 0.033 | 0.029 | 1 | 2 | Alkbh8 | 0.004 | Old |
| 0.577628 | 0.043751 | 0.332 | 0.317 | 1 | 2 | Cfh | 0.015 | Old |
| 0.579203 | 0.089614 | 0.109 | 0.101 | 1 | 2 | Aldh16a1 | 0.008 | Old |
| 0.579413 | 0.541921 | 0.021 | 0.018 | 1 | 2 | Mthfd2l | 0.003 | Old |
| 0.579453 | 0.029505 | 0.054 | 0.048 | 1 | 2 | Dcp1a | 0.006 | Old |
| 0.57947 | 0.3053 | 0.042 | 0.037 | 1 | 2 | Cd99l2 | 0.005 | Old |
| 0.579688 | 0.036706 | 0.206 | 0.191 | 1 | 2 | Uba2 | 0.015 | Old |
| 0.579785 | 0.054995 | 0.47 | 0.46 | 1 | 2 | Rab1a | 0.01 | Old |
| 0.580253 | 0.044331 | 0.712 | 0.702 | 1 | 2 | Lgals3 | 0.01 | Old |
| 0.580314 | 0.420819 | 0.096 | 0.089 | 1 | 2 | Hdac3 | 0.007 | Old |
| 0.580398 | 0.168626 | 0.062 | 0.056 | 1 | 2 | Reep4 | 0.006 | Old |
| 0.580411 | 0.394647 | 0.042 | 0.037 | 1 | 2 | Rdh14 | 0.005 | Old |
| 0.580615 | 0.172369 | 0.191 | 0.179 | 1 | 2 | Rab32 | 0.012 | Old |
| 0.581349 | 0.013968 | 0.708 | 0.695 | 1 | 2 | Wnk1 | 0.013 | Old |
| 0.581718 | 0.157771 | 0.115 | 0.107 | 1 | 2 | Slc39a10 | 0.008 | Old |
| 0.581747 | 0.216493 | 0.127 | 0.119 | 1 | 2 | Tiprl | 0.008 | Old |
| 0.58184 | 0.056788 | 0.046 | 0.04 | 1 | 2 | Kat14 | 0.006 | Old |
| 0.582023 | 0.208932 | 0.069 | 0.062 | 1 | 2 | Ttc32 | 0.007 | Old |
| 0.582588 | 0.099817 | 0.084 | 0.076 | 1 | 2 | Smg6 | 0.008 | Old |
| 0.582892 | 0.119501 | 0.192 | 0.181 | 1 | 2 | Stxbp2 | 0.011 | Old |
| 0.583088 | 0.238219 | 0.086 | 0.078 | 1 | 2 | Dera | 0.008 | Old |
| 0.583194 | 0.061382 | 0.074 | 0.081 | 1 | 2 | Mrto4 | -0.007 | Old |
| 0.583284 | 0.083639 | 0.067 | 0.061 | 1 | 2 | Cwc22 | 0.006 | Old |
| 0.584573 | 0.043287 | 0.145 | 0.153 | 1 | 2 | Spata6 | -0.008 | Old |
| 0.584675 | 0.223259 | 0.126 | 0.116 | 1 | 2 | Sos2 | 0.01 | Old |
| 0.584876 | 0.101394 | 0.063 | 0.07 | 1 | 2 | 1700037H( | -0.007 | Old |
| 0.585038 | 0.177705 | 0.041 | 0.036 | 1 | 2 | Unc119b | 0.005 | Old |
| 0.585353 | 0.552727 | 0.021 | 0.018 | 1 | 2 | Phtf1 | 0.003 | Old |
| 0.585363 | 0.030248 | 0.186 | 0.176 | 1 | 2 | Fam32a | 0.01 | Old |
| 0.585517 | 0.295859 | 0.063 | 0.057 | 1 | 2 | Qser1 | 0.006 | Old |
| 0.585856 | 0.140921 | 0.141 | 0.132 | 1 | 2 | Angel2 | 0.009 | Old |
| 0.586179 | 0.131581 | 0.261 | 0.248 | 1 | 2 | Card19 | 0.013 | Old |
| 0.586267 | 0.159918 | 0.047 | 0.042 | 1 | 2 | Tmem37 | 0.005 | Old |
| 0.586391 | 0.008047 | 0.1 | 0.091 | 1 | 2 | Guca1a | 0.009 | Old |
| 0.586427 | 0.087729 | 0.293 | 0.277 | 1 | 2 | Psma1 | 0.016 | Old |
| 0.586697 | 0.193384 | 0.139 | 0.131 | 1 | 2 | Rbm22 | 0.008 | Old |
| 0.586949 | 0.009282 | 0.09 | 0.082 | 1 | 2 | Strn | 0.008 | Old |

| | | | | | | | | |
|---|---|---|---|---|---|---|---|---|
| 0.586962 | 0.277488 | 0.056 | 0.05 | 1 | 2 | Timm9 | 0.006 | Old |
| 0.587141 | 0.258479 | 0.063 | 0.057 | 1 | 2 | Rapgef1 | 0.006 | Old |
| 0.587526 | 0.282444 | 0.066 | 0.061 | 1 | 2 | Ipo11 | 0.005 | Old |
| 0.588209 | 0.063822 | 0.286 | 0.274 | 1 | 2 | Ndufs6 | 0.012 | Old |
| 0.588483 | 0.188511 | 0.072 | 0.065 | 1 | 2 | Ebpl | 0.007 | Old |
| 0.588709 | 0.236968 | 0.122 | 0.114 | 1 | 2 | Snrpa | 0.008 | Old |
| 0.588766 | 0.226698 | 0.063 | 0.057 | 1 | 2 | Skiv2l2 | 0.006 | Old |
| 0.588854 | 0.07907 | 0.309 | 0.299 | 1 | 2 | Vps4b | 0.01 | Old |
| 0.588944 | 0.307114 | 0.034 | 0.03 | 1 | 2 | Sh2d1b1 | 0.004 | Old |
| 0.589215 | 0.000322 | 0.52 | 0.488 | 1 | 2 | Stat3 | 0.032 | Old |
| 0.589304 | 0.074591 | 0.094 | 0.102 | 1 | 2 | Ap4s1 | -0.008 | Old |
| 0.589482 | 0.000903 | 0.343 | 0.327 | 1 | 2 | Tppp3 | 0.016 | Old |
| 0.589524 | 0.087893 | 0.178 | 0.166 | 1 | 2 | Dcaf8 | 0.012 | Old |
| 0.589594 | 0.048461 | 0.021 | 0.018 | 1 | 2 | Gm26518 | 0.003 | Old |
| 0.589774 | 0.037892 | 0.511 | 0.479 | 1 | 2 | Celf1 | 0.032 | Old |
| 0.59018 | 0.094569 | 0.308 | 0.293 | 1 | 2 | Ndufb5 | 0.015 | Old |
| 0.590284 | 0.121013 | 0.092 | 0.084 | 1 | 2 | Fbxo28 | 0.008 | Old |
| 0.590652 | 0.266871 | 0.077 | 0.07 | 1 | 2 | Vac14 | 0.007 | Old |
| 0.591109 | 0.142286 | 0.194 | 0.183 | 1 | 2 | Rps6ka1 | 0.011 | Old |
| 0.591454 | 0.163283 | 0.021 | 0.018 | 1 | 2 | Ppp1r16a | 0.003 | Old |
| 0.591484 | 0.066206 | 0.227 | 0.214 | 1 | 2 | Dock11 | 0.013 | Old |
| 0.591853 | 0.088866 | 0.458 | 0.442 | 1 | 2 | Selenot | 0.016 | Old |
| 0.591947 | 0.067419 | 0.247 | 0.235 | 1 | 2 | Fcgr1 | 0.012 | Old |
| 0.59237 | 0.500473 | 0.016 | 0.013 | 1 | 2 | Shq1 | 0.003 | Old |
| 0.593401 | 0.029536 | 0.34 | 0.321 | 1 | 2 | Rbx1 | 0.019 | Old |
| 0.594456 | 0.037982 | 0.184 | 0.173 | 1 | 2 | Gdap10 | 0.011 | Old |
| 0.594576 | 0.02123 | 0.113 | 0.105 | 1 | 2 | Bbx | 0.008 | Old |
| 0.594845 | 0.156821 | 0.064 | 0.058 | 1 | 2 | Gtpbp6 | 0.006 | Old |
| 0.594997 | 0.20181 | 0.028 | 0.024 | 1 | 2 | Atg10 | 0.004 | Old |
| 0.595635 | 0.429532 | 0.1 | 0.093 | 1 | 2 | Rpl10-ps3 | 0.007 | Old |
| 0.595993 | 0.081838 | 0.133 | 0.122 | 1 | 2 | Ldah | 0.011 | Old |
| 0.596005 | 0.015054 | 0.226 | 0.209 | 1 | 2 | Ube2f | 0.017 | Old |
| 0.596158 | 0.010062 | 0.066 | 0.072 | 1 | 2 | Slc11a2 | -0.006 | Old |
| 0.596308 | 0.12799 | 0.087 | 0.08 | 1 | 2 | Tes | 0.007 | Old |
| 0.596326 | 0.149048 | 0.201 | 0.192 | 1 | 2 | Pik3r5 | 0.009 | Old |
| 0.596688 | 0.227865 | 0.016 | 0.013 | 1 | 2 | Mrps22 | 0.003 | Old |
| 0.596713 | 0.414955 | 0.086 | 0.08 | 1 | 2 | Padi2 | 0.006 | Old |
| 0.596815 | 0.159652 | 0.034 | 0.03 | 1 | 2 | Nol9 | 0.004 | Old |
| 0.596964 | 0.098479 | 0.126 | 0.118 | 1 | 2 | Coro7 | 0.008 | Old |
| 0.597131 | 0.361838 | 0.03 | 0.034 | 1 | 2 | Pomgnt1 | -0.004 | Old |
| 0.597614 | 0.105056 | 0.314 | 0.303 | 1 | 2 | Arl8a | 0.011 | Old |
| 0.597644 | 0.024192 | 0.382 | 0.365 | 1 | 2 | Polr1d | 0.017 | Old |
| 0.598063 | 0.572295 | 0.022 | 0.019 | 1 | 2 | Fam110b | 0.003 | Old |
| 0.598247 | 0.003357 | 0.244 | 0.255 | 1 | 2 | H2afv | -0.011 | Old |
| 0.598639 | 0.005347 | 0.031 | 0.036 | 1 | 2 | Zfp182 | -0.005 | Old |

| | | | | | | | | |
|---|---|---|---|---|---|---|---|---|
| 0.598816 | 0.187978 | 0.034 | 0.03 | 1 | 2 | Fhl3 | 0.004 | Old |
| 0.598853 | 0.446785 | 0.016 | 0.013 | 1 | 2 | Hacd1 | 0.003 | Old |
| 0.599451 | 0.142954 | 0.08 | 0.074 | 1 | 2 | Thumpd1 | 0.006 | Old |
| 0.599903 | 0.138758 | 0.028 | 0.032 | 1 | 2 | Cep104 | -0.004 | Old |
| 0.600091 | 0.191099 | 0.016 | 0.013 | 1 | 2 | Mmp12 | 0.003 | Old |
| 0.600886 | 0.334415 | 0.036 | 0.042 | 1 | 2 | Fbxl12 | -0.006 | Old |
| 0.60098 | 0.179355 | 0.043 | 0.038 | 1 | 2 | Cacna1d | 0.005 | Old |
| 0.601021 | 0.267385 | 0.016 | 0.013 | 1 | 2 | Aldh1b1 | 0.003 | Old |
| 0.601077 | 0.005538 | 0.028 | 0.024 | 1 | 2 | C87436 | 0.004 | Old |
| 0.601106 | 0.548245 | 0.011 | 0.008 | 1 | 2 | Arhgef37 | 0.003 | Old |
| 0.601437 | 0.152278 | 0.049 | 0.044 | 1 | 2 | Slc39a11 | 0.005 | Old |
| 0.601531 | 0.126946 | 0.289 | 0.276 | 1 | 2 | Egr2 | 0.013 | Old |
| 0.601576 | 0.11948 | 0.042 | 0.037 | 1 | 2 | Mad2l1bp | 0.005 | Old |
| 0.601641 | 0.021675 | 0.016 | 0.013 | 1 | 2 | Aurkb | 0.003 | Old |
| 0.602076 | 0.160393 | 0.143 | 0.134 | 1 | 2 | Vps41 | 0.009 | Old |
| 0.602221 | 0.036246 | 0.119 | 0.127 | 1 | 2 | Cops2 | -0.008 | Old |
| 0.602953 | 0.422765 | 0.028 | 0.024 | 1 | 2 | 2610008E1 | 0.004 | Old |
| 0.603192 | 0.202924 | 0.016 | 0.013 | 1 | 2 | Gm29340 | 0.003 | Old |
| 0.603204 | 0.018565 | 0.059 | 0.065 | 1 | 2 | Rae1 | -0.006 | Old |
| 0.603209 | 0.186147 | 0.031 | 0.036 | 1 | 2 | Mynn | -0.005 | Old |
| 0.603367 | 0.113039 | 0.59 | 0.578 | 1 | 2 | Ppp3ca | 0.012 | Old |
| 0.603489 | 0.177531 | 0.122 | 0.114 | 1 | 2 | Ttc39b | 0.008 | Old |
| 0.603786 | 0.190925 | 0.011 | 0.008 | 1 | 2 | Acot1 | 0.003 | Old |
| 0.603946 | 0.032119 | 0.649 | 0.635 | 1 | 2 | Akr1a1 | 0.014 | Old |
| 0.603999 | 0.217745 | 0.188 | 0.181 | 1 | 2 | Man2a2 | 0.007 | Old |
| 0.604124 | 0.114203 | 0.016 | 0.013 | 1 | 2 | 8030462N1 | 0.003 | Old |
| 0.604552 | 0.130069 | 0.011 | 0.008 | 1 | 2 | 4933417E1 | 0.003 | Old |
| 0.604842 | 0.40077 | 0.022 | 0.019 | 1 | 2 | Hdac10 | 0.003 | Old |
| 0.605442 | 0.036816 | 0.173 | 0.162 | 1 | 2 | Chd8 | 0.011 | Old |
| 0.605703 | 0.357224 | 0.011 | 0.008 | 1 | 2 | Gstt3 | 0.003 | Old |
| 0.605832 | 0.080024 | 0.127 | 0.119 | 1 | 2 | Sptlc1 | 0.008 | Old |
| 0.606634 | 0.006196 | 0.276 | 0.262 | 1 | 2 | Huwe1 | 0.014 | Old |
| 0.606935 | 0.109207 | 0.022 | 0.019 | 1 | 2 | 1110002J0 | 0.003 | Old |
| 0.607138 | 0.391109 | 0.044 | 0.039 | 1 | 2 | Utp23 | 0.005 | Old |
| 0.607238 | 0.2394 | 0.011 | 0.008 | 1 | 2 | Rab39 | 0.003 | Old |
| 0.607623 | 0.16276 | 0.011 | 0.008 | 1 | 2 | Ipp | 0.003 | Old |
| 0.608007 | 0.324764 | 0.011 | 0.008 | 1 | 2 | Fitm2 | 0.003 | Old |
| 0.608007 | 0.17878 | 0.011 | 0.008 | 1 | 2 | Zfp7 | 0.003 | Old |
| 0.608007 | 0.066743 | 0.011 | 0.008 | 1 | 2 | Eml6 | 0.003 | Old |
| 0.608719 | 0.086313 | 0.201 | 0.194 | 1 | 2 | Ube2z | 0.007 | Old |
| 0.609545 | 0.130145 | 0.011 | 0.008 | 1 | 2 | 0610040J0 | 0.003 | Old |
| 0.610289 | 0.297751 | 0.029 | 0.025 | 1 | 2 | Abhd10 | 0.004 | Old |
| 0.610424 | 0.046655 | 0.058 | 0.052 | 1 | 2 | Slc27a4 | 0.006 | Old |
| 0.610487 | 0.188166 | 0.075 | 0.069 | 1 | 2 | Zfp407 | 0.006 | Old |
| 0.6107 | 0.027444 | 0.011 | 0.008 | 1 | 2 | Polr3h | 0.003 | Old |

| | | | | | | | | |
|---|---|---|---|---|---|---|---|---|
| 0.610868 | 0.188938 | 0.022 | 0.019 | 1 | 2 | Tsen54 | 0.003 | Old |
| 0.610887 | 0.09986 | 0.05 | 0.045 | 1 | 2 | Ppil3 | 0.005 | Old |
| 0.611765 | 0.070628 | 0.05 | 0.045 | 1 | 2 | R3hcc1l | 0.005 | Old |
| 0.611966 | 0.03265 | 0.207 | 0.194 | 1 | 2 | Mlec | 0.013 | Old |
| 0.612168 | 0.08514 | 0.269 | 0.254 | 1 | 2 | Mpp1 | 0.015 | Old |
| 0.612182 | 0.49526 | 0.022 | 0.019 | 1 | 2 | Nipal3 | 0.003 | Old |
| 0.612248 | 0.024249 | 0.325 | 0.311 | 1 | 2 | Ywhaq | 0.014 | Old |
| 0.613127 | 0.187984 | 0.185 | 0.195 | 1 | 2 | Sppl3 | -0.01 | Old |
| 0.613315 | 0.288447 | 0.075 | 0.069 | 1 | 2 | Golga5 | 0.006 | Old |
| 0.613397 | 0.116412 | 0.051 | 0.057 | 1 | 2 | Zfp639 | -0.006 | Old |
| 0.613599 | 0.160591 | 0.021 | 0.025 | 1 | 2 | Nudcd1 | -0.004 | Old |
| 0.613739 | 0.016395 | 0.025 | 0.029 | 1 | 2 | Xlr4b | -0.004 | Old |
| 0.61375 | 0.280527 | 0.075 | 0.069 | 1 | 2 | Ago1 | 0.006 | Old |
| 0.613775 | 0.04786 | 0.209 | 0.221 | 1 | 2 | Nfix | -0.012 | Old |
| 0.614349 | 0.247456 | 0.226 | 0.219 | 1 | 2 | Zfp148 | 0.007 | Old |
| 0.614376 | 0.005546 | 0.054 | 0.048 | 1 | 2 | E4f1 | 0.006 | Old |
| 0.614672 | 0.072141 | 0.156 | 0.146 | 1 | 2 | Ubp1 | 0.01 | Old |
| 0.614707 | 0.4807 | 0.03 | 0.026 | 1 | 2 | Zfp839 | 0.004 | Old |
| 0.615024 | 0.099306 | 0.148 | 0.138 | 1 | 2 | Spag7 | 0.01 | Old |
| 0.615114 | 0.172218 | 0.061 | 0.068 | 1 | 2 | Rgcc | -0.007 | Old |
| 0.615391 | 0.238409 | 0.13 | 0.122 | 1 | 2 | Gtf2a2 | 0.008 | Old |
| 0.615425 | 0.011719 | 0.887 | 0.906 | 1 | 2 | Oaz1 | -0.019 | Old |
| 0.61585 | 0.192949 | 0.151 | 0.143 | 1 | 2 | Gabpb2 | 0.008 | Old |
| 0.615914 | 0.029936 | 0.128 | 0.119 | 1 | 2 | Tec | 0.009 | Old |
| 0.615947 | 0.174603 | 0.051 | 0.046 | 1 | 2 | Kctd5 | 0.005 | Old |
| 0.616071 | 0.311713 | 0.03 | 0.026 | 1 | 2 | Trmt10c | 0.004 | Old |
| 0.616185 | 0.502505 | 0.03 | 0.026 | 1 | 2 | Qsox2 | 0.004 | Old |
| 0.616258 | 0.080297 | 0.043 | 0.038 | 1 | 2 | Slc7a6 | 0.005 | Old |
| 0.616841 | 0.117553 | 0.148 | 0.14 | 1 | 2 | Arhgap25 | 0.008 | Old |
| 0.617263 | 0.158834 | 0.073 | 0.067 | 1 | 2 | Gtf3a | 0.006 | Old |
| 0.617428 | 0.204769 | 0.131 | 0.124 | 1 | 2 | Tlr7 | 0.007 | Old |
| 0.618211 | 0.448879 | 0.037 | 0.033 | 1 | 2 | Dgka | 0.004 | Old |
| 0.618635 | 0.212711 | 0.029 | 0.025 | 1 | 2 | Zbtb5 | 0.004 | Old |
| 0.618868 | 0.008296 | 0.029 | 0.025 | 1 | 2 | Sirt3 | 0.004 | Old |
| 0.619077 | 0.050562 | 0.099 | 0.09 | 1 | 2 | Adh5 | 0.009 | Old |
| 0.619166 | 0.203628 | 0.044 | 0.039 | 1 | 2 | Rbms2 | 0.005 | Old |
| 0.619182 | 0.034305 | 0.1 | 0.093 | 1 | 2 | Hilpda | 0.007 | Old |
| 0.62002 | 0.597767 | 0.017 | 0.014 | 1 | 2 | Enc1 | 0.003 | Old |
| 0.620455 | 0.133762 | 0.095 | 0.088 | 1 | 2 | Fibp | 0.007 | Old |
| 0.62095 | 0.195497 | 0.085 | 0.078 | 1 | 2 | Plekhg2 | 0.007 | Old |
| 0.621455 | 0.662047 | 0.039 | 0.034 | 1 | 2 | Fpgt | 0.005 | Old |
| 0.621645 | 0.290216 | 0.166 | 0.157 | 1 | 2 | Sun2 | 0.009 | Old |
| 0.621781 | 0.165446 | 0.126 | 0.119 | 1 | 2 | Tspan3 | 0.007 | Old |
| 0.621963 | 0.075932 | 0.158 | 0.148 | 1 | 2 | Rps12-ps3 | 0.01 | Old |
| 0.622446 | 0.00832 | 0.15 | 0.139 | 1 | 2 | Gigyf1 | 0.011 | Old |

| | | | | | | | | |
|---|---|---|---|---|---|---|---|---|
| 0.622474 | 0.095005 | 0.178 | 0.166 | 1 | 2 | Ptpn2 | 0.012 | Old |
| 0.62275 | 0.376132 | 0.017 | 0.014 | 1 | 2 | Crtc1 | 0.003 | Old |
| 0.623332 | 0.080858 | 0.073 | 0.067 | 1 | 2 | Nsmf | 0.006 | Old |
| 0.623535 | 0.329644 | 0.105 | 0.099 | 1 | 2 | Irak3 | 0.006 | Old |
| 0.623609 | 0.178509 | 0.19 | 0.181 | 1 | 2 | Snf8 | 0.009 | Old |
| 0.623683 | 0.326693 | 0.036 | 0.032 | 1 | 2 | Uck1 | 0.004 | Old |
| 0.623685 | 0.111388 | 0.431 | 0.42 | 1 | 2 | Capza1 | 0.011 | Old |
| 0.623703 | 0.133669 | 0.044 | 0.039 | 1 | 2 | Msantd2 | 0.005 | Old |
| 0.623814 | 0.107129 | 0.115 | 0.107 | 1 | 2 | Lrpprc | 0.008 | Old |
| 0.6239 | 0.238936 | 0.051 | 0.046 | 1 | 2 | Gpn1 | 0.005 | Old |
| 0.623908 | 0.06006 | 0.829 | 0.816 | 1 | 2 | Ddx5 | 0.013 | Old |
| 0.624209 | 0.117166 | 0.05 | 0.045 | 1 | 2 | Elac2 | 0.005 | Old |
| 0.625108 | 0.016967 | 0.638 | 0.663 | 1 | 2 | Sh3bgrl | -0.025 | Old |
| 0.625257 | 0.078966 | 0.304 | 0.289 | 1 | 2 | Prex1 | 0.015 | Old |
| 0.625589 | 0.23724 | 0.057 | 0.063 | 1 | 2 | Fas | -0.006 | Old |
| 0.62564 | 0.063566 | 0.231 | 0.217 | 1 | 2 | Rap1gds1 | 0.014 | Old |
| 0.625789 | 0.223968 | 0.017 | 0.014 | 1 | 2 | Acss2 | 0.003 | Old |
| 0.626094 | 0.254193 | 0.017 | 0.014 | 1 | 2 | Poll | 0.003 | Old |
| 0.626094 | 0.246098 | 0.017 | 0.014 | 1 | 2 | Ccdc125 | 0.003 | Old |
| 0.6266 | 0.126872 | 0.073 | 0.067 | 1 | 2 | Stat5a | 0.006 | Old |
| 0.626878 | 0.095438 | 0.075 | 0.069 | 1 | 2 | Mtx1 | 0.006 | Old |
| 0.627305 | 0.097063 | 0.044 | 0.039 | 1 | 2 | Exosc1 | 0.005 | Old |
| 0.627674 | 0.038948 | 0.471 | 0.461 | 1 | 2 | Abracl | 0.01 | Old |
| 0.627689 | 0.415596 | 0.072 | 0.067 | 1 | 2 | Phkb | 0.005 | Old |
| 0.627808 | 0.052037 | 0.149 | 0.14 | 1 | 2 | AI839979 | 0.009 | Old |
| 0.627953 | 0.479732 | 0.03 | 0.026 | 1 | 2 | Serpine1 | 0.004 | Old |
| 0.62807 | 0.118513 | 0.582 | 0.574 | 1 | 2 | Akap13 | 0.008 | Old |
| 0.628225 | 0.350546 | 0.017 | 0.014 | 1 | 2 | Plcxd2 | 0.003 | Old |
| 0.628528 | 0.136496 | 0.589 | 0.583 | 1 | 2 | Gm26917 | 0.006 | Old |
| 0.628835 | 0.145764 | 0.017 | 0.014 | 1 | 2 | Ugt1a6a | 0.003 | Old |
| 0.629271 | 0.107098 | 0.105 | 0.113 | 1 | 2 | Snx29 | -0.008 | Old |
| 0.629311 | 0.188514 | 0.183 | 0.176 | 1 | 2 | Pla2g15 | 0.007 | Old |
| 0.630488 | 0.067827 | 0.064 | 0.058 | 1 | 2 | Gopc | 0.006 | Old |
| 0.630525 | 0.118921 | 0.1 | 0.091 | 1 | 2 | Zcchc17 | 0.009 | Old |
| 0.630555 | 0.024549 | 0.036 | 0.032 | 1 | 2 | Pkdcc | 0.004 | Old |
| 0.630651 | 0.971677 | 0.022 | 0.026 | 1 | 2 | Ets1 | -0.004 | Old |
| 0.630741 | 0.042608 | 0.107 | 0.1 | 1 | 2 | Cir1 | 0.007 | Old |
| 0.631167 | 0.045954 | 0.092 | 0.099 | 1 | 2 | Slc23a2 | -0.007 | Old |
| 0.631428 | 0.258466 | 0.075 | 0.069 | 1 | 2 | 1110059G1 | 0.006 | Old |
| 0.631464 | 0.035433 | 0.047 | 0.052 | 1 | 2 | Ptrhd1 | -0.005 | Old |
| 0.632417 | 0.247595 | 0.079 | 0.074 | 1 | 2 | Tmem19 | 0.005 | Old |
| 0.63249 | 0.63382 | 0.025 | 0.021 | 1 | 2 | Sh3bp5l | 0.004 | Old |
| 0.633722 | 0.131846 | 0.101 | 0.094 | 1 | 2 | Ctr9 | 0.007 | Old |
| 0.634356 | 0.021601 | 0.092 | 0.086 | 1 | 2 | Rhoc | 0.006 | Old |
| 0.634368 | 0.053333 | 0.051 | 0.046 | 1 | 2 | Isyna1 | 0.005 | Old |

| | | | | | | | | |
|---|---|---|---|---|---|---|---|---|
| 0.6344 | 0.098427 | 0.037 | 0.033 | 1 | 2 | Ercc5 | 0.004 | Old |
| 0.634606 | 0.217227 | 0.037 | 0.033 | 1 | 2 | Slc20a2 | 0.004 | Old |
| 0.635123 | 0.360095 | 0.103 | 0.097 | 1 | 2 | Map3k5 | 0.006 | Old |
| 0.635865 | 0.741448 | 0.012 | 0.01 | 1 | 2 | Wdr73 | 0.002 | Old |
| 0.635977 | 0.08604 | 0.195 | 0.183 | 1 | 2 | Rab18 | 0.012 | Old |
| 0.63602 | 0.052662 | 0.232 | 0.222 | 1 | 2 | Ddx21 | 0.01 | Old |
| 0.636063 | 0.762799 | 0.017 | 0.02 | 1 | 2 | Saa3 | -0.003 | Old |
| 0.636236 | 0.421681 | 0.012 | 0.01 | 1 | 2 | Zdhhc1 | 0.002 | Old |
| 0.636931 | 0.049555 | 0.859 | 0.868 | 1 | 2 | Marcks | -0.009 | Old |
| 0.636979 | 0.642737 | 0.012 | 0.01 | 1 | 2 | Tnks1bp1 | 0.002 | Old |
| 0.637058 | 0.068181 | 0.345 | 0.331 | 1 | 2 | Fam129a | 0.014 | Old |
| 0.63735 | 0.941496 | 0.012 | 0.01 | 1 | 2 | Gja1 | 0.002 | Old |
| 0.637489 | 0.293402 | 0.016 | 0.019 | 1 | 2 | 5830432E0 | -0.003 | Old |
| 0.637532 | 0.18064 | 0.061 | 0.056 | 1 | 2 | Kdm5d | 0.005 | Old |
| 0.637938 | 0.128081 | 0.066 | 0.061 | 1 | 2 | Ndufaf4 | 0.005 | Old |
| 0.63842 | 0.176738 | 0.122 | 0.115 | 1 | 2 | Pip4k2b | 0.007 | Old |
| 0.638695 | 0.14165 | 0.186 | 0.177 | 1 | 2 | Cpne3 | 0.009 | Old |
| 0.638903 | 0.038683 | 0.266 | 0.255 | 1 | 2 | Sf3b3 | 0.011 | Old |
| 0.639461 | 0.115821 | 0.085 | 0.078 | 1 | 2 | Acbd6 | 0.007 | Old |
| 0.639776 | 0.071499 | 0.321 | 0.305 | 1 | 2 | Birc6 | 0.016 | Old |
| 0.639835 | 0.100726 | 0.322 | 0.309 | 1 | 2 | Cmtm7 | 0.013 | Old |
| 0.640096 | 0.152239 | 0.127 | 0.12 | 1 | 2 | 1810013L2 | 0.007 | Old |
| 0.64014 | 0.078155 | 0.062 | 0.068 | 1 | 2 | Gtf2f2 | -0.006 | Old |
| 0.640695 | 0.423075 | 0.012 | 0.01 | 1 | 2 | Dsel | 0.002 | Old |
| 0.641004 | 0.131622 | 0.018 | 0.021 | 1 | 2 | 2810001G2 | -0.003 | Old |
| 0.641241 | 0.054942 | 0.024 | 0.02 | 1 | 2 | C1rl | 0.004 | Old |
| 0.64128 | 0.037619 | 0.172 | 0.162 | 1 | 2 | Gpbp1l1 | 0.01 | Old |
| 0.641387 | 0.109158 | 0.019 | 0.023 | 1 | 2 | Dexi | -0.004 | Old |
| 0.64145 | 0.117812 | 0.337 | 0.322 | 1 | 2 | Psma6 | 0.015 | Old |
| 0.641487 | 0.565301 | 0.046 | 0.042 | 1 | 2 | Tmem110 | 0.004 | Old |
| 0.641812 | 0.243121 | 0.012 | 0.01 | 1 | 2 | A230050P2 | 0.002 | Old |
| 0.642147 | 0.027519 | 0.105 | 0.112 | 1 | 2 | Prdx3 | -0.007 | Old |
| 0.642184 | 0.171927 | 0.012 | 0.01 | 1 | 2 | Tubg1 | 0.002 | Old |
| 0.642252 | 0.448069 | 0.014 | 0.017 | 1 | 2 | Tnfaip6 | -0.003 | Old |
| 0.642406 | 0.193019 | 0.054 | 0.049 | 1 | 2 | Brpf3 | 0.005 | Old |
| 0.642622 | 0.063494 | 0.109 | 0.102 | 1 | 2 | Tmem192 | 0.007 | Old |
| 0.642912 | 0.06022 | 0.307 | 0.293 | 1 | 2 | Rnf141 | 0.014 | Old |
| 0.643545 | 0.02599 | 0.115 | 0.107 | 1 | 2 | Mob3b | 0.008 | Old |
| 0.643557 | 0.056315 | 0.139 | 0.147 | 1 | 2 | Hdac1 | -0.008 | Old |
| 0.643711 | 0.095173 | 0.268 | 0.255 | 1 | 2 | Eif4g3 | 0.013 | Old |
| 0.644093 | 0.331602 | 0.031 | 0.027 | 1 | 2 | Taf5 | 0.004 | Old |
| 0.644579 | 0.009415 | 0.164 | 0.153 | 1 | 2 | 0610012G0 | 0.011 | Old |
| 0.644621 | 0.000447 | 0.264 | 0.249 | 1 | 2 | Tmem109 | 0.015 | Old |
| 0.644734 | 0.035894 | 0.139 | 0.132 | 1 | 2 | Tmem87b | 0.007 | Old |
| 0.644737 | 0.113672 | 0.173 | 0.165 | 1 | 2 | Herc2 | 0.008 | Old |

| | | | | | | | | |
|---|---|---|---|---|---|---|---|---|
| 0.644794 | 0.112158 | 0.012 | 0.01 | 1 | 2 | Catip | 0.002 | Old |
| 0.644799 | 0.224423 | 0.016 | 0.019 | 1 | 2 | Zc2hc1a | -0.003 | Old |
| 0.644821 | 0.091286 | 0.007 | 0.01 | 1 | 2 | Akr1b8 | -0.003 | Old |
| 0.644856 | 0.054953 | 0.258 | 0.248 | 1 | 2 | Etfb | 0.01 | Old |
| 0.645167 | 0.271611 | 0.012 | 0.01 | 1 | 2 | Pex11g | 0.002 | Old |
| 0.645169 | 0.051177 | 0.234 | 0.222 | 1 | 2 | Pdcd6 | 0.012 | Old |
| 0.645235 | 0.581921 | 0.018 | 0.015 | 1 | 2 | Mars2 | 0.003 | Old |
| 0.645241 | 0.470545 | 0.099 | 0.093 | 1 | 2 | Mzt1 | 0.006 | Old |
| 0.645686 | 0.181634 | 0.031 | 0.027 | 1 | 2 | Csrp2 | 0.004 | Old |
| 0.646287 | 0.093566 | 0.012 | 0.01 | 1 | 2 | Lgi4 | 0.002 | Old |
| 0.646667 | 0.087712 | 0.242 | 0.233 | 1 | 2 | Pon2 | 0.009 | Old |
| 0.646691 | 0.046359 | 0.18 | 0.169 | 1 | 2 | Frg1 | 0.011 | Old |
| 0.646878 | 0.096792 | 0.151 | 0.143 | 1 | 2 | Tbc1d20 | 0.008 | Old |
| 0.647366 | 0.268172 | 0.009 | 0.011 | 1 | 2 | Tmem14a | -0.002 | Old |
| 0.647408 | 0.213674 | 0.015 | 0.018 | 1 | 2 | Bud13 | -0.003 | Old |
| 0.647627 | 0.208804 | 0.054 | 0.049 | 1 | 2 | Tmem63b | 0.005 | Old |
| 0.647741 | 0.21145 | 0.05 | 0.056 | 1 | 2 | Pmf1 | -0.006 | Old |
| 0.647761 | 0.089275 | 0.009 | 0.011 | 1 | 2 | Tagap1 | -0.002 | Old |
| 0.647785 | 0.164877 | 0.111 | 0.119 | 1 | 2 | Cstf3 | -0.008 | Old |
| 0.647793 | 0.123708 | 0.126 | 0.119 | 1 | 2 | Gsto1 | 0.007 | Old |
| 0.647845 | 0.329565 | 0.039 | 0.034 | 1 | 2 | Dhx35 | 0.005 | Old |
| 0.647852 | 0.019705 | 0.095 | 0.102 | 1 | 2 | Plod1 | -0.007 | Old |
| 0.647966 | 0.093739 | 0.031 | 0.027 | 1 | 2 | Ints4 | 0.004 | Old |
| 0.64798 | 0.141402 | 0.011 | 0.013 | 1 | 2 | Tyms | -0.002 | Old |
| 0.648162 | 0.114591 | 0.082 | 0.076 | 1 | 2 | Cpsf3 | 0.006 | Old |
| 0.648271 | 0.042517 | 0.013 | 0.015 | 1 | 2 | Ddr1 | -0.002 | Old |
| 0.649286 | 0.077868 | 0.18 | 0.17 | 1 | 2 | Mtch2 | 0.01 | Old |
| 0.649786 | 0.37098 | 0.014 | 0.017 | 1 | 2 | Nemp2 | -0.003 | Old |
| 0.650083 | 0.232359 | 0.025 | 0.021 | 1 | 2 | Rpap2 | 0.004 | Old |
| 0.650308 | 0.256078 | 0.018 | 0.015 | 1 | 2 | Izumo4 | 0.003 | Old |
| 0.6504 | 0.074832 | 0.01 | 0.012 | 1 | 2 | Dynlt1c | -0.002 | Old |
| 0.650829 | 0.187507 | 0.011 | 0.013 | 1 | 2 | Mocs3 | -0.002 | Old |
| 0.650923 | 0.368021 | 0.009 | 0.011 | 1 | 2 | Hist2h3c2 | -0.002 | Old |
| 0.650966 | 0.41993 | 0.016 | 0.019 | 1 | 2 | Lmln | -0.003 | Old |
| 0.65123 | 0.050539 | 0.092 | 0.086 | 1 | 2 | Relt | 0.006 | Old |
| 0.651516 | 0.079546 | 0.012 | 0.014 | 1 | 2 | Ankrd37 | -0.002 | Old |
| 0.652273 | 0.239474 | 0.01 | 0.012 | 1 | 2 | Pdia5 | -0.002 | Old |
| 0.652522 | 0.245304 | 0.013 | 0.015 | 1 | 2 | Gm26890 | -0.002 | Old |
| 0.6533 | 0.01182 | 0.018 | 0.015 | 1 | 2 | Top3a | 0.003 | Old |
| 0.653475 | 0.159548 | 0.121 | 0.114 | 1 | 2 | Retreg3 | 0.007 | Old |
| 0.653683 | 0.189955 | 0.011 | 0.013 | 1 | 2 | Ceacam19 | -0.002 | Old |
| 0.653832 | 0.569951 | 0.013 | 0.015 | 1 | 2 | Txnrd3 | -0.002 | Old |
| 0.654122 | 0.005233 | 0.166 | 0.157 | 1 | 2 | Tns3 | 0.009 | Old |
| 0.654558 | 0.006732 | 0.241 | 0.251 | 1 | 2 | Fam208a | -0.01 | Old |
| 0.654639 | 0.000258 | 0.596 | 0.564 | 1 | 2 | Sfpq | 0.032 | Old |

| | | | | | | | | |
|---|---|---|---|---|---|---|---|---|
| 0.654701 | 0.077663 | 0.683 | 0.682 | 1 | 2 | Ly86 | 0.001 | Old |
| 0.655426 | 0.138227 | 0.156 | 0.148 | 1 | 2 | Tmem9b | 0.008 | Old |
| 0.655735 | 0.18227 | 0.025 | 0.021 | 1 | 2 | Eif2d | 0.004 | Old |
| 0.656127 | 0.182266 | 0.013 | 0.015 | 1 | 2 | Naif1 | -0.002 | Old |
| 0.656437 | 0.029192 | 0.227 | 0.215 | 1 | 2 | Fam168a | 0.012 | Old |
| 0.65657 | 0.497583 | 0.034 | 0.031 | 1 | 2 | Nxt2 | 0.003 | Old |
| 0.656597 | 0.080969 | 0.018 | 0.015 | 1 | 2 | Nphp3 | 0.003 | Old |
| 0.656972 | 0.087605 | 0.055 | 0.05 | 1 | 2 | Stx17 | 0.005 | Old |
| 0.657368 | 0.383107 | 0.041 | 0.037 | 1 | 2 | Man2c1 | 0.004 | Old |
| 0.657668 | 0.053088 | 0.123 | 0.131 | 1 | 2 | Tars | -0.008 | Old |
| 0.657772 | 0.159273 | 0.088 | 0.082 | 1 | 2 | Rsbn1 | 0.006 | Old |
| 0.658101 | 0.040731 | 0.191 | 0.179 | 1 | 2 | 2310022A1 | 0.012 | Old |
| 0.65841 | 0.119689 | 0.061 | 0.067 | 1 | 2 | Fech | -0.006 | Old |
| 0.659334 | 0.058395 | 0.281 | 0.291 | 1 | 2 | Cisd2 | -0.01 | Old |
| 0.659709 | 0.1142 | 0.175 | 0.164 | 1 | 2 | Bmt2 | 0.011 | Old |
| 0.659853 | 0.21432 | 0.048 | 0.044 | 1 | 2 | Tinf2 | 0.004 | Old |
| 0.660105 | 0.04518 | 0.091 | 0.084 | 1 | 2 | Kansl2 | 0.007 | Old |
| 0.660124 | 0.168488 | 0.032 | 0.029 | 1 | 2 | Zfp335 | 0.003 | Old |
| 0.660301 | 0.114097 | 0.26 | 0.251 | 1 | 2 | Uggt1 | 0.009 | Old |
| 0.660502 | 0.010643 | 0.018 | 0.015 | 1 | 2 | Socs1 | 0.003 | Old |
| 0.660803 | 0.003494 | 0.018 | 0.015 | 1 | 2 | Lin52 | 0.003 | Old |
| 0.660914 | 0.344081 | 0.032 | 0.029 | 1 | 2 | Rcc1l | 0.003 | Old |
| 0.661347 | 0.015406 | 0.153 | 0.16 | 1 | 2 | Slain2 | -0.007 | Old |
| 0.662043 | 0.044416 | 0.727 | 0.719 | 1 | 2 | Naca | 0.008 | Old |
| 0.662043 | 0.006211 | 0.075 | 0.081 | 1 | 2 | Sap30l | -0.006 | Old |
| 0.662758 | 0.082285 | 0.097 | 0.091 | 1 | 2 | Slc27a1 | 0.006 | Old |
| 0.663303 | 0.035726 | 0.034 | 0.038 | 1 | 2 | Tmem29 | -0.004 | Old |
| 0.663405 | 0.252981 | 0.026 | 0.023 | 1 | 2 | Fam84b | 0.003 | Old |
| 0.664026 | 0.140524 | 0.109 | 0.102 | 1 | 2 | Fbxw7 | 0.007 | Old |
| 0.664595 | 0.141375 | 0.057 | 0.052 | 1 | 2 | Arfgef2 | 0.005 | Old |
| 0.665166 | 0.131507 | 0.268 | 0.28 | 1 | 2 | Tnks2 | -0.012 | Old |
| 0.665215 | 0.324833 | 0.048 | 0.044 | 1 | 2 | Decr1 | 0.004 | Old |
| 0.66544 | 0.247386 | 0.101 | 0.095 | 1 | 2 | Faim | 0.006 | Old |
| 0.665465 | 0.069578 | 0.12 | 0.113 | 1 | 2 | Rnf146 | 0.007 | Old |
| 0.665869 | 0.74311 | 0.013 | 0.011 | 1 | 2 | Tal1 | 0.002 | Old |
| 0.665983 | 0.062382 | 0.476 | 0.455 | 1 | 2 | Bzw1 | 0.021 | Old |
| 0.666148 | 0.512361 | 0.049 | 0.045 | 1 | 2 | Rnf126 | 0.004 | Old |
| 0.666229 | 0.721344 | 0.013 | 0.011 | 1 | 2 | Zfp41 | 0.002 | Old |
| 0.666319 | 0.022773 | 0.145 | 0.135 | 1 | 2 | Zfp326 | 0.01 | Old |
| 0.666366 | 0.200071 | 0.075 | 0.07 | 1 | 2 | Nol12 | 0.005 | Old |
| 0.667001 | 0.000855 | 0.039 | 0.034 | 1 | 2 | Zdhhc12 | 0.005 | Old |
| 0.667786 | 0.10029 | 0.429 | 0.419 | 1 | 2 | Arf4 | 0.01 | Old |
| 0.667794 | 0.241719 | 0.091 | 0.086 | 1 | 2 | Zfp120 | 0.005 | Old |
| 0.668029 | 0.348093 | 0.013 | 0.011 | 1 | 2 | Ciart | 0.002 | Old |
| 0.668082 | 0.02203 | 0.123 | 0.115 | 1 | 2 | Otud4 | 0.008 | Old |

| | | | | | | | | |
|---|---|---|---|---|---|---|---|---|
| 0.668255 | 0.217485 | 0.033 | 0.03 | 1 | 2 | Dclre1a | 0.003 | Old |
| 0.66828 | 0.003115 | 0.337 | 0.315 | 1 | 2 | Mindy2 | 0.022 | Old |
| 0.668597 | 0.128733 | 0.2 | 0.195 | 1 | 2 | Rtf1 | 0.005 | Old |
| 0.668717 | 0.438659 | 0.026 | 0.023 | 1 | 2 | Dgcr14 | 0.003 | Old |
| 0.668766 | 0.031156 | 0.236 | 0.242 | 1 | 2 | Prkcsh | -0.006 | Old |
| 0.669074 | 0.009822 | 0.361 | 0.348 | 1 | 2 | Hnrnpa1 | 0.013 | Old |
| 0.669837 | 0.121872 | 0.112 | 0.106 | 1 | 2 | Atp11c | 0.006 | Old |
| 0.669904 | 0.017232 | 0.039 | 0.034 | 1 | 2 | Pias3 | 0.005 | Old |
| 0.670031 | 0.221654 | 0.062 | 0.057 | 1 | 2 | Cherp | 0.005 | Old |
| 0.670492 | 0.370035 | 0.026 | 0.023 | 1 | 2 | Osbpl7 | 0.003 | Old |
| 0.671 | 0.150069 | 0.026 | 0.023 | 1 | 2 | Zfhx2 | 0.003 | Old |
| 0.671506 | 0.114327 | 0.07 | 0.064 | 1 | 2 | Vkorc1l1 | 0.006 | Old |
| 0.671552 | 0.06613 | 0.223 | 0.211 | 1 | 2 | Slc20a1 | 0.012 | Old |
| 0.671687 | 0.04968 | 0.073 | 0.078 | 1 | 2 | Abi2 | -0.005 | Old |
| 0.671996 | 0.302346 | 0.013 | 0.011 | 1 | 2 | Gm38251 | 0.002 | Old |
| 0.672075 | 0.378486 | 0.019 | 0.017 | 1 | 2 | Pola1 | 0.002 | Old |
| 0.672357 | 0.284688 | 0.013 | 0.011 | 1 | 2 | Lipt2 | 0.002 | Old |
| 0.672357 | 0.250684 | 0.013 | 0.011 | 1 | 2 | Hr | 0.002 | Old |
| 0.672382 | 0.342378 | 0.092 | 0.087 | 1 | 2 | Uri1 | 0.005 | Old |
| 0.672448 | 0.145687 | 0.09 | 0.084 | 1 | 2 | Specc1 | 0.006 | Old |
| 0.672717 | 0.240498 | 0.033 | 0.03 | 1 | 2 | Coq10a | 0.003 | Old |
| 0.672718 | 0.48946 | 0.013 | 0.011 | 1 | 2 | Them6 | 0.002 | Old |
| 0.673079 | 0.077868 | 0.288 | 0.277 | 1 | 2 | Hdlbp | 0.011 | Old |
| 0.673164 | 0.279056 | 0.033 | 0.03 | 1 | 2 | Gm14005 | 0.003 | Old |
| 0.673307 | 0.008388 | 0.196 | 0.184 | 1 | 2 | Abca7 | 0.012 | Old |
| 0.673513 | 0.128456 | 0.263 | 0.255 | 1 | 2 | Ppp2r1a | 0.008 | Old |
| 0.673576 | 0.016769 | 0.082 | 0.077 | 1 | 2 | Uspl1 | 0.005 | Old |
| 0.674164 | 0.328276 | 0.013 | 0.011 | 1 | 2 | Magee1 | 0.002 | Old |
| 0.674654 | 0.003474 | 0.104 | 0.097 | 1 | 2 | Ly96 | 0.007 | Old |
| 0.674709 | 0.228657 | 0.057 | 0.052 | 1 | 2 | Gtpbp1 | 0.005 | Old |
| 0.675972 | 0.482365 | 0.013 | 0.011 | 1 | 2 | Trim7 | 0.002 | Old |
| 0.675972 | 0.325968 | 0.013 | 0.011 | 1 | 2 | Itgb3bp | 0.002 | Old |
| 0.676039 | 0.125442 | 0.171 | 0.162 | 1 | 2 | Cnot4 | 0.009 | Old |
| 0.676119 | 0.023101 | 0.923 | 0.935 | 1 | 2 | H2-Aa | -0.012 | Old |
| 0.676182 | 0.111525 | 0.027 | 0.024 | 1 | 2 | Ankrd39 | 0.003 | Old |
| 0.677058 | 0.01405 | 0.013 | 0.011 | 1 | 2 | Pus7l | 0.002 | Old |
| 0.67718 | 0.083277 | 0.17 | 0.162 | 1 | 2 | Dip2b | 0.008 | Old |
| 0.67742 | 0.159974 | 0.013 | 0.011 | 1 | 2 | Tceanc | 0.002 | Old |
| 0.677527 | 0.10292 | 0.033 | 0.03 | 1 | 2 | Mzt2 | 0.003 | Old |
| 0.67789 | 0.087892 | 0.186 | 0.178 | 1 | 2 | Setx | 0.008 | Old |
| 0.678268 | 0.264561 | 0.115 | 0.109 | 1 | 2 | Sec23ip | 0.006 | Old |
| 0.678976 | 0.249274 | 0.096 | 0.091 | 1 | 2 | Slc35e1 | 0.005 | Old |
| 0.679182 | 0.026388 | 0.208 | 0.201 | 1 | 2 | Osgep | 0.007 | Old |
| 0.679279 | 0.077777 | 0.561 | 0.556 | 1 | 2 | Rpn1 | 0.005 | Old |
| 0.679853 | 0.483007 | 0.042 | 0.038 | 1 | 2 | Prmt5 | 0.004 | Old |

| | | | | | | | | |
|---|---|---|---|---|---|---|---|---|
| 0.6805 | 0.0808 | 0.254 | 0.245 | 1 | 2 | Notch1 | 0.009 | Old |
| 0.680695 | 0.059058 | 0.086 | 0.08 | 1 | 2 | Ahsa2 | 0.006 | Old |
| 0.680793 | 0.246643 | 0.027 | 0.024 | 1 | 2 | Sema4b | 0.003 | Old |
| 0.681368 | 0.164245 | 0.146 | 0.14 | 1 | 2 | Glrx2 | 0.006 | Old |
| 0.681498 | 0.262861 | 0.062 | 0.057 | 1 | 2 | Hdac7 | 0.005 | Old |
| 0.682155 | 0.09917 | 0.532 | 0.517 | 1 | 2 | Itgb1 | 0.015 | Old |
| 0.682347 | 0.182495 | 0.186 | 0.179 | 1 | 2 | Lsm12 | 0.007 | Old |
| 0.682364 | 0.032894 | 0.306 | 0.309 | 1 | 2 | Gmfb | -0.003 | Old |
| 0.682612 | 0.029845 | 0.07 | 0.064 | 1 | 2 | Phb | 0.006 | Old |
| 0.682788 | 0.032055 | 0.072 | 0.067 | 1 | 2 | Dph5 | 0.005 | Old |
| 0.682806 | 0.184838 | 0.056 | 0.051 | 1 | 2 | Pou5f2 | 0.005 | Old |
| 0.682954 | 0.043823 | 0.176 | 0.165 | 1 | 2 | Elf4 | 0.011 | Old |
| 0.683091 | 0.207123 | 0.163 | 0.157 | 1 | 2 | Cdk19 | 0.006 | Old |
| 0.683097 | 0.191798 | 0.082 | 0.077 | 1 | 2 | Atg4d | 0.005 | Old |
| 0.683125 | 0.197938 | 0.05 | 0.046 | 1 | 2 | St3gal2 | 0.004 | Old |
| 0.683257 | 0.168568 | 0.042 | 0.038 | 1 | 2 | Erich1 | 0.004 | Old |
| 0.683373 | 0.359508 | 0.058 | 0.053 | 1 | 2 | Tmem161a | 0.005 | Old |
| 0.683542 | 0.034808 | 0.779 | 0.784 | 1 | 2 | H2afz | -0.005 | Old |
| 0.68384 | 0.05864 | 0.089 | 0.095 | 1 | 2 | Nup54 | -0.006 | Old |
| 0.684147 | 0.137784 | 0.033 | 0.037 | 1 | 2 | Mllt1 | -0.004 | Old |
| 0.684486 | 0.240237 | 0.058 | 0.053 | 1 | 2 | Gm5617 | 0.005 | Old |
| 0.684549 | 0.117369 | 0.165 | 0.157 | 1 | 2 | Acat1 | 0.008 | Old |
| 0.685117 | 0.179608 | 0.049 | 0.045 | 1 | 2 | Smagp | 0.004 | Old |
| 0.68547 | 0.719003 | 0.035 | 0.032 | 1 | 2 | Gpx3 | 0.003 | Old |
| 0.685912 | 0.257718 | 0.116 | 0.11 | 1 | 2 | Slc38a9 | 0.006 | Old |
| 0.686039 | 0.175517 | 0.14 | 0.133 | 1 | 2 | Nup88 | 0.007 | Old |
| 0.686473 | 0.101818 | 0.056 | 0.051 | 1 | 2 | Rbl1 | 0.005 | Old |
| 0.68659 | 0.077703 | 0.418 | 0.405 | 1 | 2 | Rab7 | 0.013 | Old |
| 0.686958 | 0.022739 | 0.115 | 0.121 | 1 | 2 | Micu2 | -0.006 | Old |
| 0.687043 | 0.159204 | 0.027 | 0.024 | 1 | 2 | Trmt61a | 0.003 | Old |
| 0.689124 | 0.148796 | 0.269 | 0.259 | 1 | 2 | Fndc3b | 0.01 | Old |
| 0.68916 | 0.002783 | 0.143 | 0.15 | 1 | 2 | Rnf216 | -0.007 | Old |
| 0.690289 | 0.058803 | 0.158 | 0.151 | 1 | 2 | Cnpy2 | 0.007 | Old |
| 0.690493 | 0.02565 | 0.162 | 0.152 | 1 | 2 | Vcpip1 | 0.01 | Old |
| 0.690533 | 0.023044 | 0.344 | 0.324 | 1 | 2 | Caprin1 | 0.02 | Old |
| 0.690615 | 0.11151 | 0.152 | 0.145 | 1 | 2 | Wbp4 | 0.007 | Old |
| 0.690904 | 0.142108 | 0.095 | 0.09 | 1 | 2 | Parp4 | 0.005 | Old |
| 0.691302 | 0.088119 | 0.04 | 0.036 | 1 | 2 | Parp3 | 0.004 | Old |
| 0.691558 | 0.001602 | 0.335 | 0.338 | 1 | 2 | Samsn1 | -0.003 | Old |
| 0.691803 | 0.010903 | 0.048 | 0.044 | 1 | 2 | Mon1b | 0.004 | Old |
| 0.692059 | 0.098553 | 0.027 | 0.024 | 1 | 2 | Dolk | 0.003 | Old |
| 0.692328 | 0.038799 | 0.145 | 0.138 | 1 | 2 | Wipf2 | 0.007 | Old |
| 0.692484 | 0.106182 | 0.069 | 0.064 | 1 | 2 | Tmem2 | 0.005 | Old |
| 0.692615 | 0.030079 | 0.196 | 0.185 | 1 | 2 | Cfap43 | 0.011 | Old |
| 0.692649 | 0.280767 | 0.048 | 0.044 | 1 | 2 | Nfrkb | 0.004 | Old |

| | | | | | | | | |
|---|---|---|---|---|---|---|---|---|
| 0.692904 | 0.091105 | 0.042 | 0.038 | 1 | 2 | Man2b2 | 0.004 | Old |
| 0.693284 | 0.018988 | 0.12 | 0.126 | 1 | 2 | Ppm1b | -0.006 | Old |
| 0.694222 | 0.356022 | 0.02 | 0.018 | 1 | 2 | Gm4673 | 0.002 | Old |
| 0.694335 | 0.159499 | 0.122 | 0.115 | 1 | 2 | Sh3bp2 | 0.007 | Old |
| 0.694726 | 0.29337 | 0.105 | 0.1 | 1 | 2 | Glod4 | 0.005 | Old |
| 0.695425 | 0.053045 | 0.139 | 0.131 | 1 | 2 | Polr2g | 0.008 | Old |
| 0.695499 | 0.029956 | 0.089 | 0.083 | 1 | 2 | Eef1akmt1 | 0.006 | Old |
| 0.695525 | 0.107404 | 0.042 | 0.038 | 1 | 2 | Naglu | 0.004 | Old |
| 0.695804 | 0.131573 | 0.11 | 0.105 | 1 | 2 | Med10 | 0.005 | Old |
| 0.696366 | 0.001433 | 0.243 | 0.229 | 1 | 2 | Osbpl8 | 0.014 | Old |
| 0.696433 | 0.169539 | 0.05 | 0.046 | 1 | 2 | Cep295 | 0.004 | Old |
| 0.696819 | 0.195381 | 0.02 | 0.018 | 1 | 2 | Ap1s3 | 0.002 | Old |
| 0.696849 | 0.181908 | 0.041 | 0.037 | 1 | 2 | Snx11 | 0.004 | Old |
| 0.696959 | 0.117319 | 0.031 | 0.034 | 1 | 2 | Mthfd1l | -0.003 | Old |
| 0.697048 | 0.015206 | 0.12 | 0.113 | 1 | 2 | Tsnax | 0.007 | Old |
| 0.697117 | 0.064719 | 0.276 | 0.267 | 1 | 2 | Stk4 | 0.009 | Old |
| 0.697362 | 0.24634 | 0.043 | 0.039 | 1 | 2 | Nudt2 | 0.004 | Old |
| 0.697882 | 0.057385 | 0.049 | 0.053 | 1 | 2 | Gfm1 | -0.004 | Old |
| 0.698542 | 0.068378 | 0.122 | 0.115 | 1 | 2 | Casp8ap2 | 0.007 | Old |
| 0.69864 | 0.025045 | 0.558 | 0.54 | 1 | 2 | Mat2a | 0.018 | Old |
| 0.698678 | 0.008296 | 0.413 | 0.397 | 1 | 2 | Fermt3 | 0.016 | Old |
| 0.699232 | 0.225793 | 0.014 | 0.012 | 1 | 2 | Mrps27 | 0.002 | Old |
| 0.700189 | 0.086912 | 0.034 | 0.031 | 1 | 2 | Ptrh2 | 0.003 | Old |
| 0.700634 | 0.398769 | 0.014 | 0.012 | 1 | 2 | Eml2 | 0.002 | Old |
| 0.700917 | 0.17719 | 0.137 | 0.132 | 1 | 2 | Rab24 | 0.005 | Old |
| 0.70138 | 0.02121 | 0.458 | 0.463 | 1 | 2 | Tmem30a | -0.005 | Old |
| 0.701501 | 0.170241 | 0.105 | 0.1 | 1 | 2 | Cspp1 | 0.005 | Old |
| 0.701709 | 0.229435 | 0.124 | 0.119 | 1 | 2 | Ssr2 | 0.005 | Old |
| 0.701751 | 0.312828 | 0.043 | 0.039 | 1 | 2 | Yars | 0.004 | Old |
| 0.70217 | 0.043283 | 0.065 | 0.07 | 1 | 2 | Rpf2 | -0.005 | Old |
| 0.702171 | 0.054854 | 0.102 | 0.096 | 1 | 2 | Zfp62 | 0.006 | Old |
| 0.702249 | 0.250782 | 0.069 | 0.064 | 1 | 2 | Tpra1 | 0.005 | Old |
| 0.702875 | 0.033464 | 0.063 | 0.058 | 1 | 2 | Siah1a | 0.005 | Old |
| 0.703261 | 0.102266 | 0.05 | 0.046 | 1 | 2 | Prr12 | 0.004 | Old |
| 0.703669 | 0.153765 | 0.086 | 0.081 | 1 | 2 | Naa60 | 0.005 | Old |
| 0.704836 | 0.209443 | 0.106 | 0.101 | 1 | 2 | Ube3b | 0.005 | Old |
| 0.704956 | 0.188565 | 0.052 | 0.049 | 1 | 2 | Btbd10 | 0.003 | Old |
| 0.705247 | 0.414584 | 0.037 | 0.034 | 1 | 2 | Coq7 | 0.003 | Old |
| 0.705273 | 0.229939 | 0.029 | 0.032 | 1 | 2 | Umps | -0.003 | Old |
| 0.706174 | 0.023567 | 0.027 | 0.024 | 1 | 2 | Dmkn | 0.003 | Old |
| 0.706239 | 0.010997 | 0.145 | 0.152 | 1 | 2 | Zmym5 | -0.007 | Old |
| 0.70653 | 0.049668 | 0.082 | 0.077 | 1 | 2 | Gm1673 | 0.005 | Old |
| 0.706828 | 0.10917 | 0.122 | 0.116 | 1 | 2 | Sars | 0.006 | Old |
| 0.707039 | 0.101256 | 0.052 | 0.049 | 1 | 2 | Casp9 | 0.003 | Old |
| 0.707091 | 0.252172 | 0.028 | 0.025 | 1 | 2 | C2cd2 | 0.003 | Old |

| | | | | | | | | |
|---|---|---|---|---|---|---|---|---|
| 0.707635 | 0.154633 | 0.103 | 0.097 | 1 | 2 | Ccnk | 0.006 | Old |
| 0.70766 | 0.363117 | 0.014 | 0.012 | 1 | 2 | Dnal4 | 0.002 | Old |
| 0.707917 | 0.066314 | 0.261 | 0.253 | 1 | 2 | Prdx6 | 0.008 | Old |
| 0.707936 | 0.072661 | 0.55 | 0.54 | 1 | 2 | Ppp2ca | 0.01 | Old |
| 0.708722 | 0.227663 | 0.05 | 0.046 | 1 | 2 | Gbe1 | 0.004 | Old |
| 0.709074 | 0.505698 | 0.028 | 0.025 | 1 | 2 | Eif2b1 | 0.003 | Old |
| 0.709522 | 0.003321 | 0.919 | 0.933 | 1 | 2 | S100a6 | -0.014 | Old |
| 0.709729 | 0.027492 | 0.158 | 0.148 | 1 | 2 | Uba52 | 0.01 | Old |
| 0.709968 | 0.116629 | 0.125 | 0.119 | 1 | 2 | Ltb4r1 | 0.006 | Old |
| 0.710125 | 0.007687 | 0.014 | 0.012 | 1 | 2 | Gm15448 | 0.002 | Old |
| 0.710211 | 0.284015 | 0.107 | 0.102 | 1 | 2 | Ubfd1 | 0.005 | Old |
| 0.710806 | 0.001096 | 0.027 | 0.03 | 1 | 2 | Pym1 | -0.003 | Old |
| 0.711375 | 0.187453 | 0.054 | 0.05 | 1 | 2 | Sssca1 | 0.004 | Old |
| 0.711595 | 0.001019 | 0.213 | 0.203 | 1 | 2 | Srpr | 0.01 | Old |
| 0.712367 | 0.073344 | 0.043 | 0.039 | 1 | 2 | Narfl | 0.004 | Old |
| 0.712734 | 0.249828 | 0.094 | 0.09 | 1 | 2 | Zfp330 | 0.004 | Old |
| 0.71276 | 0.242638 | 0.124 | 0.12 | 1 | 2 | Ing1 | 0.004 | Old |
| 0.712785 | 0.052539 | 0.112 | 0.107 | 1 | 2 | Chrac1 | 0.005 | Old |
| 0.713147 | 0.045351 | 0.534 | 0.525 | 1 | 2 | Rab14 | 0.009 | Old |
| 0.713295 | 0.11111 | 0.028 | 0.025 | 1 | 2 | Ncapd3 | 0.003 | Old |
| 0.714155 | 0.004704 | 0.334 | 0.317 | 1 | 2 | Fis1 | 0.017 | Old |
| 0.714287 | 0.199455 | 0.076 | 0.081 | 1 | 2 | Shprh | -0.005 | Old |
| 0.714524 | 0.064013 | 0.63 | 0.61 | 1 | 2 | Cox7a2l | 0.02 | Old |
| 0.714728 | 0.0019 | 0.074 | 0.078 | 1 | 2 | Ttc19 | -0.004 | Old |
| 0.715389 | 0.098153 | 0.08 | 0.075 | 1 | 2 | Ino80 | 0.005 | Old |
| 0.715461 | 0.02566 | 0.202 | 0.21 | 1 | 2 | Dcun1d5 | -0.008 | Old |
| 0.715958 | 0.052795 | 0.238 | 0.229 | 1 | 2 | Ifnar2 | 0.009 | Old |
| 0.715962 | 0.119895 | 0.021 | 0.019 | 1 | 2 | Tti2 | 0.002 | Old |
| 0.716278 | 0.020196 | 0.318 | 0.305 | 1 | 2 | Snrpg | 0.013 | Old |
| 0.716759 | 0.074144 | 0.162 | 0.154 | 1 | 2 | Ncoa1 | 0.008 | Old |
| 0.717123 | 0.115916 | 0.154 | 0.15 | 1 | 2 | Xrn1 | 0.004 | Old |
| 0.717256 | 0.089251 | 0.046 | 0.05 | 1 | 2 | Pgap1 | -0.004 | Old |
| 0.717336 | 0.368922 | 0.029 | 0.026 | 1 | 2 | 2700097O( | 0.003 | Old |
| 0.717693 | 0.050115 | 0.044 | 0.04 | 1 | 2 | Mrpl46 | 0.004 | Old |
| 0.717738 | 0.005179 | 0.058 | 0.053 | 1 | 2 | Atxn1 | 0.005 | Old |
| 0.718119 | 0.191493 | 0.094 | 0.089 | 1 | 2 | Lgals8 | 0.005 | Old |
| 0.718231 | 0.045484 | 0.021 | 0.019 | 1 | 2 | Gm45606 | 0.002 | Old |
| 0.71865 | 0.067548 | 0.539 | 0.551 | 1 | 2 | Psmb4 | -0.012 | Old |
| 0.719483 | 0.025837 | 0.062 | 0.067 | 1 | 2 | Vti1a | -0.005 | Old |
| 0.71965 | 0.259387 | 0.021 | 0.019 | 1 | 2 | Zfp747 | 0.002 | Old |
| 0.719797 | 0.201087 | 0.051 | 0.056 | 1 | 2 | Cnot9 | -0.005 | Old |
| 0.720042 | 0.025581 | 0.738 | 0.729 | 1 | 2 | Btf3 | 0.009 | Old |
| 0.720252 | 0.220474 | 0.046 | 0.043 | 1 | 2 | Fam160a2 | 0.003 | Old |
| 0.720307 | 0.079885 | 0.221 | 0.21 | 1 | 2 | Zfp516 | 0.011 | Old |
| 0.720737 | 0.524378 | 0.046 | 0.043 | 1 | 2 | 3110082I1 | 0.003 | Old |

| | | | | | | | | |
|---|---|---|---|---|---|---|---|---|
| 0.720855 | 0.06389 | 0.249 | 0.239 | 1 | 2 | Amfr | 0.01 | Old |
| 0.720962 | 0.121137 | 0.107 | 0.101 | 1 | 2 | Abraxas2 | 0.006 | Old |
| 0.721045 | 0.234098 | 0.073 | 0.069 | 1 | 2 | Prpf19 | 0.004 | Old |
| 0.721112 | 0.041457 | 0.1 | 0.106 | 1 | 2 | Pag1 | -0.006 | Old |
| 0.721624 | 0.079803 | 0.06 | 0.056 | 1 | 2 | Fuom | 0.004 | Old |
| 0.721624 | 0.037609 | 0.06 | 0.056 | 1 | 2 | Timm17b | 0.004 | Old |
| 0.721678 | 0.012979 | 0.158 | 0.151 | 1 | 2 | Sertad2 | 0.007 | Old |
| 0.721767 | 0.132096 | 0.077 | 0.082 | 1 | 2 | Mfsd12 | -0.005 | Old |
| 0.722417 | 0.181408 | 0.036 | 0.033 | 1 | 2 | Sergef | 0.003 | Old |
| 0.722552 | 0.162848 | 0.054 | 0.05 | 1 | 2 | Ctns | 0.004 | Old |
| 0.722679 | 0.447465 | 0.046 | 0.043 | 1 | 2 | Bpnt1 | 0.003 | Old |
| 0.723346 | 0.119999 | 0.021 | 0.019 | 1 | 2 | Pycrl | 0.002 | Old |
| 0.723462 | 0.063132 | 0.087 | 0.082 | 1 | 2 | Sertad3 | 0.005 | Old |
| 0.723567 | 0.066724 | 0.476 | 0.469 | 1 | 2 | Zcchc6 | 0.007 | Old |
| 0.723715 | 0.026215 | 0.539 | 0.534 | 1 | 2 | Pomp | 0.005 | Old |
| 0.723879 | 0.147901 | 0.037 | 0.034 | 1 | 2 | Gbp5 | 0.003 | Old |
| 0.724273 | 0.036844 | 0.036 | 0.033 | 1 | 2 | Tmem126b | 0.003 | Old |
| 0.724382 | 0.049423 | 0.645 | 0.635 | 1 | 2 | Ncf2 | 0.01 | Old |
| 0.724666 | 0.0262 | 0.365 | 0.353 | 1 | 2 | Hnrnph2 | 0.012 | Old |
| 0.725173 | 0.154904 | 0.024 | 0.026 | 1 | 2 | Ppip5k1 | -0.002 | Old |
| 0.72519 | 0.404089 | 0.03 | 0.027 | 1 | 2 | Cc2d1a | 0.003 | Old |
| 0.725902 | 0.036819 | 0.619 | 0.638 | 1 | 2 | Il6ra | -0.019 | Old |
| 0.726041 | 0.001884 | 0.125 | 0.119 | 1 | 2 | Sec13 | 0.006 | Old |
| 0.726418 | 0.090072 | 0.199 | 0.192 | 1 | 2 | Ppp6c | 0.007 | Old |
| 0.72704 | 0.143027 | 0.206 | 0.198 | 1 | 2 | Usp19 | 0.008 | Old |
| 0.727442 | 0.063595 | 0.089 | 0.094 | 1 | 2 | Dnpep | -0.005 | Old |
| 0.728304 | 0.02604 | 0.103 | 0.097 | 1 | 2 | Tubgcp4 | 0.006 | Old |
| 0.728314 | 0.023557 | 0.796 | 0.799 | 1 | 2 | Efhd2 | -0.003 | Old |
| 0.728316 | 0.653732 | 0.03 | 0.027 | 1 | 2 | Nbas | 0.003 | Old |
| 0.728727 | 0.006781 | 0.2 | 0.191 | 1 | 2 | 2610001J0 | 0.009 | Old |
| 0.72877 | 0.146732 | 0.069 | 0.064 | 1 | 2 | Chpf2 | 0.005 | Old |
| 0.729142 | 0.375641 | 0.015 | 0.013 | 1 | 2 | Acad11 | 0.002 | Old |
| 0.729684 | 0.004795 | 0.05 | 0.046 | 1 | 2 | Rnf157 | 0.004 | Old |
| 0.730125 | 0.231846 | 0.074 | 0.07 | 1 | 2 | Gm19951 | 0.004 | Old |
| 0.730219 | 0.101183 | 0.327 | 0.324 | 1 | 2 | Uvrag | 0.003 | Old |
| 0.730318 | 0.147828 | 0.04 | 0.043 | 1 | 2 | Stmn1 | -0.003 | Old |
| 0.73052 | 0.223614 | 0.054 | 0.05 | 1 | 2 | Slc33a1 | 0.004 | Old |
| 0.730652 | 0.160601 | 0.073 | 0.069 | 1 | 2 | Urm1 | 0.004 | Old |
| 0.73072 | 0.047203 | 0.16 | 0.166 | 1 | 2 | Eif2s3y | -0.006 | Old |
| 0.731081 | 0.192354 | 0.092 | 0.088 | 1 | 2 | Maml3 | 0.004 | Old |
| 0.731144 | 0.055753 | 0.022 | 0.025 | 1 | 2 | Slc2a9 | -0.003 | Old |
| 0.731166 | 0.054102 | 0.036 | 0.033 | 1 | 2 | Fblim1 | 0.003 | Old |
| 0.731885 | 0.037084 | 0.123 | 0.118 | 1 | 2 | Vps13d | 0.005 | Old |
| 0.732182 | 0.170866 | 0.133 | 0.138 | 1 | 2 | Ccl3 | -0.005 | Old |
| 0.732557 | 0.104512 | 0.015 | 0.013 | 1 | 2 | Map3k6 | 0.002 | Old |

| | | | | | | | | |
|---|---|---|---|---|---|---|---|---|
| 0.732653 | 0.025651 | 0.044 | 0.04 | 1 | 2 | Ppp2r1b | 0.004 | Old |
| 0.732692 | 0.044987 | 0.085 | 0.089 | 1 | 2 | Cetn2 | -0.004 | Old |
| 0.73273 | 0.288877 | 0.025 | 0.027 | 1 | 2 | Klhl7 | -0.002 | Old |
| 0.733497 | 0.115963 | 0.042 | 0.045 | 1 | 2 | Patz1 | -0.003 | Old |
| 0.73394 | 0.029743 | 0.228 | 0.219 | 1 | 2 | Cd302 | 0.009 | Old |
| 0.734756 | 0.143647 | 0.131 | 0.125 | 1 | 2 | Actr10 | 0.006 | Old |
| 0.734783 | 0.240945 | 0.045 | 0.042 | 1 | 2 | Havcr2 | 0.003 | Old |
| 0.734861 | 0.338762 | 0.046 | 0.043 | 1 | 2 | Mocs1 | 0.003 | Old |
| 0.735565 | 0.076535 | 0.096 | 0.091 | 1 | 2 | Chm | 0.005 | Old |
| 0.735787 | 0.522548 | 0.03 | 0.027 | 1 | 2 | Wbp1 | 0.003 | Old |
| 0.736213 | 0.055603 | 0.314 | 0.299 | 1 | 2 | Uba1 | 0.015 | Old |
| 0.736549 | 0.543013 | 0.022 | 0.02 | 1 | 2 | Tmem237 | 0.002 | Old |
| 0.736549 | 0.162849 | 0.022 | 0.02 | 1 | 2 | Pdss2 | 0.002 | Old |
| 0.737068 | 0.108071 | 0.135 | 0.129 | 1 | 2 | Ccdc124 | 0.006 | Old |
| 0.737385 | 0.213186 | 0.022 | 0.02 | 1 | 2 | Mon1a | 0.002 | Old |
| 0.737461 | 0.33309 | 0.039 | 0.036 | 1 | 2 | Rbm12b1 | 0.003 | Old |
| 0.73799 | 0.127154 | 0.046 | 0.043 | 1 | 2 | Stambp | 0.003 | Old |
| 0.73878 | 0.034893 | 0.02 | 0.023 | 1 | 2 | Arhgap27 | -0.003 | Old |
| 0.738966 | 0.224584 | 0.093 | 0.089 | 1 | 2 | Zfp280d | 0.004 | Old |
| 0.739388 | 0.172264 | 0.193 | 0.188 | 1 | 2 | Tollip | 0.005 | Old |
| 0.74006 | 0.03623 | 0.147 | 0.138 | 1 | 2 | Pip5k1a | 0.009 | Old |
| 0.740176 | 0.210028 | 0.022 | 0.02 | 1 | 2 | Smyd2 | 0.002 | Old |
| 0.740379 | 0.182662 | 0.03 | 0.027 | 1 | 2 | 1500011B( | 0.003 | Old |
| 0.740814 | 0.062864 | 0.3 | 0.291 | 1 | 2 | Eif1a | 0.009 | Old |
| 0.740937 | 0.013202 | 0.368 | 0.361 | 1 | 2 | Ralb | 0.007 | Old |
| 0.740947 | 0.026076 | 0.366 | 0.354 | 1 | 2 | Zfp706 | 0.012 | Old |
| 0.741076 | 0.141097 | 0.123 | 0.116 | 1 | 2 | Cdc34 | 0.007 | Old |
| 0.741161 | 0.074042 | 0.352 | 0.352 | 1 | 2 | Rassf3 | 0 | Old |
| 0.741572 | 0.058578 | 0.022 | 0.02 | 1 | 2 | Mettl2 | 0.002 | Old |
| 0.741592 | 0.22859 | 0.082 | 0.078 | 1 | 2 | Abl1 | 0.004 | Old |
| 0.742144 | 0.072246 | 0.188 | 0.179 | 1 | 2 | Psmc1 | 0.009 | Old |
| 0.742192 | 0.02737 | 0.077 | 0.072 | 1 | 2 | Lmnb1 | 0.005 | Old |
| 0.742431 | 0.073377 | 0.165 | 0.159 | 1 | 2 | Pbx2 | 0.006 | Old |
| 0.742472 | 0.044845 | 0.096 | 0.091 | 1 | 2 | Tmem57 | 0.005 | Old |
| 0.74262 | 0.089523 | 0.193 | 0.186 | 1 | 2 | Slc25a36 | 0.007 | Old |
| 0.74313 | 0.066235 | 0.212 | 0.22 | 1 | 2 | Acbd5 | -0.008 | Old |
| 0.743757 | 0.248109 | 0.064 | 0.061 | 1 | 2 | Mmgt1 | 0.003 | Old |
| 0.743838 | 0.020295 | 0.024 | 0.026 | 1 | 2 | Mthfsd | -0.002 | Old |
| 0.743954 | 0.416016 | 0.066 | 0.063 | 1 | 2 | Dbt | 0.003 | Old |
| 0.744107 | 0.047819 | 0.514 | 0.501 | 1 | 2 | Dstn | 0.013 | Old |
| 0.744228 | 0.367856 | 0.04 | 0.037 | 1 | 2 | Slc39a13 | 0.003 | Old |
| 0.74438 | 0.006943 | 0.335 | 0.322 | 1 | 2 | Clint1 | 0.013 | Old |
| 0.744863 | 0.03552 | 0.338 | 0.341 | 1 | 2 | Cct4 | -0.003 | Old |
| 0.745321 | 0.074293 | 0.167 | 0.159 | 1 | 2 | Ociad1 | 0.008 | Old |
| 0.745335 | 0.027699 | 0.192 | 0.197 | 1 | 2 | Krcc1 | -0.005 | Old |

| | | | | | | | | |
|---|---|---|---|---|---|---|---|---|
| 0.745436 | 0.12476 | 0.147 | 0.14 | 1 | 2 | Eif4ebp1 | 0.007 | Old |
| 0.745492 | 0.13074 | 0.042 | 0.045 | 1 | 2 | Exoc6 | -0.003 | Old |
| 0.746367 | 0.061287 | 0.037 | 0.034 | 1 | 2 | Gm20707 | 0.003 | Old |
| 0.74669 | 0.047101 | 0.355 | 0.367 | 1 | 2 | Ddx3y | -0.012 | Old |
| 0.747263 | 0.096274 | 0.077 | 0.072 | 1 | 2 | BC005561 | 0.005 | Old |
| 0.747343 | 0.354384 | 0.021 | 0.024 | 1 | 2 | Cks1b | -0.003 | Old |
| 0.747666 | 0.184408 | 0.055 | 0.051 | 1 | 2 | Emilin1 | 0.004 | Old |
| 0.747916 | 0.065069 | 0.258 | 0.248 | 1 | 2 | Ccar1 | 0.01 | Old |
| 0.748073 | 0.036895 | 0.186 | 0.192 | 1 | 2 | Hmox2 | -0.006 | Old |
| 0.748102 | 0.522465 | 0.016 | 0.014 | 1 | 2 | Ick | 0.002 | Old |
| 0.748673 | 0.12907 | 0.472 | 0.49 | 1 | 2 | Mt1 | -0.018 | Old |
| 0.749127 | 0.071094 | 0.02 | 0.023 | 1 | 2 | Trmt10a | -0.003 | Old |
| 0.75024 | 0.195077 | 0.113 | 0.109 | 1 | 2 | Bcap29 | 0.004 | Old |
| 0.750529 | 0.041936 | 0.022 | 0.02 | 1 | 2 | Hnmt | 0.002 | Old |
| 0.750923 | 0.052426 | 0.288 | 0.279 | 1 | 2 | Ghitm | 0.009 | Old |
| 0.751427 | 0.062207 | 0.231 | 0.236 | 1 | 2 | Lilrb4a | -0.005 | Old |
| 0.75146 | 0.006906 | 0.6 | 0.589 | 1 | 2 | Snrnp70 | 0.011 | Old |
| 0.751956 | 0.078727 | 0.292 | 0.287 | 1 | 2 | Ubxn1 | 0.005 | Old |
| 0.752633 | 0.064222 | 0.02 | 0.023 | 1 | 2 | Vwa8 | -0.003 | Old |
| 0.753162 | 0.097963 | 0.041 | 0.044 | 1 | 2 | Cep44 | -0.003 | Old |
| 0.753343 | 0.00686 | 0.188 | 0.181 | 1 | 2 | Emc6 | 0.007 | Old |
| 0.754074 | 0.18812 | 0.016 | 0.014 | 1 | 2 | Als2 | 0.002 | Old |
| 0.754406 | 0.44921 | 0.016 | 0.014 | 1 | 2 | Smim5 | 0.002 | Old |
| 0.754436 | 0.051292 | 0.32 | 0.321 | 1 | 2 | Lipa | -0.001 | Old |
| 0.755009 | 0.115695 | 0.084 | 0.08 | 1 | 2 | Yipf6 | 0.004 | Old |
| 0.755025 | 0.259004 | 0.031 | 0.029 | 1 | 2 | Zer1 | 0.002 | Old |
| 0.755362 | 0.073369 | 0.575 | 0.57 | 1 | 2 | Srsf3 | 0.005 | Old |
| 0.756172 | 0.323527 | 0.025 | 0.023 | 1 | 2 | Gkap1 | 0.002 | Old |
| 0.756348 | 0.050774 | 0.443 | 0.457 | 1 | 2 | Stap1 | -0.014 | Old |
| 0.756719 | 0.001825 | 0.317 | 0.305 | 1 | 2 | Tfe3 | 0.012 | Old |
| 0.757262 | 0.038359 | 0.337 | 0.328 | 1 | 2 | Crebbp | 0.009 | Old |
| 0.757539 | 0.085537 | 0.231 | 0.222 | 1 | 2 | Larp4 | 0.009 | Old |
| 0.757599 | 0.070157 | 0.035 | 0.038 | 1 | 2 | Zfp865 | -0.003 | Old |
| 0.758292 | 0.301965 | 0.062 | 0.058 | 1 | 2 | Psip1 | 0.004 | Old |
| 0.758723 | 0.211354 | 0.109 | 0.106 | 1 | 2 | Pttg1 | 0.003 | Old |
| 0.759061 | 0.118957 | 0.016 | 0.014 | 1 | 2 | Gin1 | 0.002 | Old |
| 0.759394 | 0.097921 | 0.016 | 0.014 | 1 | 2 | Mpp7 | 0.002 | Old |
| 0.759474 | 0.06597 | 0.126 | 0.132 | 1 | 2 | Tspan13 | -0.006 | Old |
| 0.759669 | 0.203872 | 0.094 | 0.09 | 1 | 2 | Fam193a | 0.004 | Old |
| 0.759753 | 0.283211 | 0.041 | 0.038 | 1 | 2 | 1110034G2 | 0.003 | Old |
| 0.759825 | 0.022365 | 0.037 | 0.04 | 1 | 2 | Usp31 | -0.003 | Old |
| 0.759835 | 0.260134 | 0.032 | 0.03 | 1 | 2 | Qpctl | 0.002 | Old |
| 0.760144 | 0.212919 | 0.018 | 0.02 | 1 | 2 | Gm49359 | -0.002 | Old |
| 0.760224 | 0.843035 | 0.01 | 0.008 | 1 | 2 | Brf2 | 0.002 | Old |
| 0.760292 | 0.203251 | 0.086 | 0.082 | 1 | 2 | Mt2 | 0.004 | Old |

| | | | | | | | | |
|---|---|---|---|---|---|---|---|---|
| 0.760393 | 0.142431 | 0.016 | 0.014 | 1 | 2 | Pkd1l2 | 0.002 | Old |
| 0.760683 | 0.046415 | 0.097 | 0.102 | 1 | 2 | Desi2 | -0.005 | Old |
| 0.760736 | 0.058333 | 0.389 | 0.381 | 1 | 2 | Cd68 | 0.008 | Old |
| 0.76082 | 0.021876 | 0.71 | 0.722 | 1 | 2 | Rtn4 | -0.012 | Old |
| 0.761328 | 0.115393 | 0.057 | 0.053 | 1 | 2 | E2f2 | 0.004 | Old |
| 0.761597 | 0.092042 | 0.105 | 0.109 | 1 | 2 | Mapkapk3 | -0.004 | Old |
| 0.761615 | 0.083603 | 0.064 | 0.061 | 1 | 2 | Armc5 | 0.003 | Old |
| 0.761727 | 0.084557 | 0.086 | 0.09 | 1 | 2 | Acadsb | -0.004 | Old |
| 0.76215 | 0.220363 | 0.048 | 0.045 | 1 | 2 | Casp2 | 0.003 | Old |
| 0.762398 | 0.250435 | 0.085 | 0.081 | 1 | 2 | Tcf3 | 0.004 | Old |
| 0.762756 | 0.043426 | 0.057 | 0.053 | 1 | 2 | Sgms2 | 0.004 | Old |
| 0.762855 | 6.87E-06 | 0.35 | 0.338 | 1 | 2 | Ppp1cb | 0.012 | Old |
| 0.762872 | 0.048274 | 0.158 | 0.164 | 1 | 2 | Limd2 | -0.006 | Old |
| 0.762962 | 0.202909 | 0.064 | 0.061 | 1 | 2 | Trappc11 | 0.003 | Old |
| 0.763797 | 0.011935 | 0.223 | 0.216 | 1 | 2 | Gna12 | 0.007 | Old |
| 0.763867 | 0.270062 | 0.067 | 0.064 | 1 | 2 | Hif1an | 0.003 | Old |
| 0.763893 | 0.041162 | 0.327 | 0.316 | 1 | 2 | Adcy7 | 0.011 | Old |
| 0.764249 | 0.354404 | 0.043 | 0.04 | 1 | 2 | Bex3 | 0.003 | Old |
| 0.764623 | 0.09509 | 0.471 | 0.463 | 1 | 2 | Ldha | 0.008 | Old |
| 0.764832 | 0.014988 | 0.052 | 0.056 | 1 | 2 | Akap7 | -0.004 | Old |
| 0.765153 | 0.071951 | 0.1 | 0.095 | 1 | 2 | C1galt1c1 | 0.005 | Old |
| 0.765235 | 0.012546 | 0.338 | 0.333 | 1 | 2 | Lbr | 0.005 | Old |
| 0.765642 | 0.101937 | 0.137 | 0.132 | 1 | 2 | Ube2q2 | 0.005 | Old |
| 0.765836 | 0.318378 | 0.01 | 0.008 | 1 | 2 | Dennd2d | 0.002 | Old |
| 0.76589 | 0.091425 | 0.066 | 0.07 | 1 | 2 | St8sia6 | -0.004 | Old |
| 0.766367 | 0.027872 | 0.037 | 0.04 | 1 | 2 | Dnajc9 | -0.003 | Old |
| 0.766545 | 0.114139 | 0.032 | 0.03 | 1 | 2 | Zkscan6 | 0.002 | Old |
| 0.767133 | 0.48953 | 0.01 | 0.008 | 1 | 2 | Miga1 | 0.002 | Old |
| 0.767492 | 0.055107 | 0.207 | 0.201 | 1 | 2 | Igf1 | 0.006 | Old |
| 0.767565 | 0.22174 | 0.01 | 0.008 | 1 | 2 | Coq8a | 0.002 | Old |
| 0.767744 | 0.147984 | 0.339 | 0.334 | 1 | 2 | Manf | 0.005 | Old |
| 0.768589 | 0.055058 | 0.149 | 0.143 | 1 | 2 | Gba | 0.006 | Old |
| 0.768708 | 0.114646 | 0.04 | 0.037 | 1 | 2 | Pex16 | 0.003 | Old |
| 0.768836 | 0.007188 | 0.366 | 0.372 | 1 | 2 | Lmna | -0.006 | Old |
| 0.768848 | 0.097179 | 0.222 | 0.214 | 1 | 2 | 1110004F1 | 0.008 | Old |
| 0.768863 | 0.123036 | 0.01 | 0.008 | 1 | 2 | Rbfox2 | 0.002 | Old |
| 0.769295 | 0.313432 | 0.01 | 0.008 | 1 | 2 | Plscr4 | 0.002 | Old |
| 0.769295 | 0.112655 | 0.01 | 0.008 | 1 | 2 | Dhrs13 | 0.002 | Old |
| 0.769295 | 0.007987 | 0.01 | 0.008 | 1 | 2 | Nipa1 | 0.002 | Old |
| 0.769387 | 0.114841 | 0.051 | 0.055 | 1 | 2 | Rdh11 | -0.004 | Old |
| 0.769429 | 0.257863 | 0.085 | 0.081 | 1 | 2 | Cd200r1 | 0.004 | Old |
| 0.769716 | 0.057965 | 0.179 | 0.171 | 1 | 2 | Cebpd | 0.008 | Old |
| 0.769974 | 0.172759 | 0.018 | 0.02 | 1 | 2 | 4732440D( | -0.002 | Old |
| 0.770142 | 0.275514 | 0.017 | 0.019 | 1 | 2 | Gramd3 | -0.002 | Old |
| 0.770142 | 0.266522 | 0.017 | 0.019 | 1 | 2 | Samd4 | -0.002 | Old |

| | | | | | | | | |
|---|---|---|---|---|---|---|---|---|
| 0.770161 | 0.249639 | 0.01 | 0.008 | 1 | 2 | Zbtb48 | 0.002 | Old |
| 0.770161 | 0.194592 | 0.01 | 0.008 | 1 | 2 | Tulp3 | 0.002 | Old |
| 0.770588 | 0.007632 | 0.419 | 0.4 | 1 | 2 | G3bp1 | 0.019 | Old |
| 0.77087 | 0.101184 | 0.091 | 0.087 | 1 | 2 | Pex7 | 0.004 | Old |
| 0.770873 | 0.070229 | 0.049 | 0.052 | 1 | 2 | Fyco1 | -0.003 | Old |
| 0.77089 | 0.18467 | 0.366 | 0.384 | 1 | 2 | Ccl2 | -0.018 | Old |
| 0.770898 | 0.125596 | 0.033 | 0.031 | 1 | 2 | Mrgbp | 0.002 | Old |
| 0.77146 | 0.040456 | 0.01 | 0.008 | 1 | 2 | Mical2 | 0.002 | Old |
| 0.771476 | 0.328588 | 0.025 | 0.023 | 1 | 2 | 2700038G2 | 0.002 | Old |
| 0.771506 | 0.14249 | 0.035 | 0.038 | 1 | 2 | Tdp2 | -0.003 | Old |
| 0.771746 | 0.252344 | 0.025 | 0.023 | 1 | 2 | Gorab | 0.002 | Old |
| 0.771973 | 0.42491 | 0.032 | 0.03 | 1 | 2 | Bcar3 | 0.002 | Old |
| 0.771984 | 0.058817 | 0.132 | 0.126 | 1 | 2 | Eed | 0.006 | Old |
| 0.772363 | 0.127943 | 0.018 | 0.02 | 1 | 2 | Ppp1r3d | -0.002 | Old |
| 0.772554 | 0.12539 | 0.025 | 0.023 | 1 | 2 | Sema6b | 0.002 | Old |
| 0.772641 | 0.32795 | 0.034 | 0.032 | 1 | 2 | Mlh1 | 0.002 | Old |
| 0.772733 | 0.09608 | 0.509 | 0.507 | 1 | 2 | Hnrnpm | 0.002 | Old |
| 0.772985 | 0.00895 | 0.966 | 0.97 | 1 | 2 | Rpl38 | -0.004 | Old |
| 0.772988 | 0.329666 | 0.033 | 0.031 | 1 | 2 | Klhl12 | 0.002 | Old |
| 0.773219 | 0.021027 | 0.017 | 0.019 | 1 | 2 | Zfp958 | -0.002 | Old |
| 0.773252 | 0.27419 | 0.051 | 0.049 | 1 | 2 | Megf8 | 0.002 | Old |
| 0.773373 | 0.043496 | 0.017 | 0.019 | 1 | 2 | Phc1 | -0.002 | Old |
| 0.773521 | 0.104106 | 0.105 | 0.11 | 1 | 2 | Taf13 | -0.005 | Old |
| 0.77375 | 0.06208 | 0.049 | 0.046 | 1 | 2 | Pcid2 | 0.003 | Old |
| 0.773914 | 0.597442 | 0.046 | 0.044 | 1 | 2 | Fam60a | 0.002 | Old |
| 0.774101 | 0.478274 | 0.063 | 0.061 | 1 | 2 | Aph1b | 0.002 | Old |
| 0.775213 | 0.228436 | 0.017 | 0.015 | 1 | 2 | Chac2 | 0.002 | Old |
| 0.775426 | 0.011661 | 0.065 | 0.062 | 1 | 2 | Enoph1 | 0.003 | Old |
| 0.77548 | 0.026004 | 0.203 | 0.209 | 1 | 2 | Tnfrsf13b | -0.006 | Old |
| 0.775784 | 0.097633 | 0.051 | 0.055 | 1 | 2 | Tprkb | -0.004 | Old |
| 0.775863 | 0.023018 | 0.482 | 0.464 | 1 | 2 | Skap2 | 0.018 | Old |
| 0.776245 | 0.238992 | 0.049 | 0.046 | 1 | 2 | Rrp36 | 0.003 | Old |
| 0.7766 | 0.085631 | 0.081 | 0.077 | 1 | 2 | Pot1b | 0.004 | Old |
| 0.776668 | 0.113441 | 0.015 | 0.017 | 1 | 2 | Pde4d | -0.002 | Old |
| 0.776916 | 0.402617 | 0.017 | 0.019 | 1 | 2 | Rasal3 | -0.002 | Old |
| 0.777153 | 0.709031 | 0.017 | 0.015 | 1 | 2 | C730034F0 | 0.002 | Old |
| 0.777153 | 0.196224 | 0.017 | 0.015 | 1 | 2 | Ccnyl1 | 0.002 | Old |
| 0.777353 | 0.076698 | 0.088 | 0.084 | 1 | 2 | Scaf4 | 0.004 | Old |
| 0.777649 | 0.054821 | 0.245 | 0.236 | 1 | 2 | Dnajc8 | 0.009 | Old |
| 0.777653 | 0.621068 | 0.026 | 0.024 | 1 | 2 | Fut4 | 0.002 | Old |
| 0.777886 | 0.320374 | 0.064 | 0.062 | 1 | 2 | Dusp7 | 0.002 | Old |
| 0.779542 | 0.087054 | 0.069 | 0.072 | 1 | 2 | Ufd1 | -0.003 | Old |
| 0.780027 | 0.081355 | 0.072 | 0.068 | 1 | 2 | Tex264 | 0.004 | Old |
| 0.780087 | 0.050685 | 0.049 | 0.046 | 1 | 2 | Lztr1 | 0.003 | Old |
| 0.78022 | 0.083015 | 0.127 | 0.122 | 1 | 2 | Gtf3c6 | 0.005 | Old |

| | | | | | | | | |
|---|---|---|---|---|---|---|---|---|
| 0.780647 | 0.043127 | 0.433 | 0.433 | 1 | 2 | Birc3 | 0 | Old |
| 0.780718 | 0.0028 | 0.147 | 0.141 | 1 | 2 | Otud5 | 0.006 | Old |
| 0.780904 | 0.232428 | 0.123 | 0.12 | 1 | 2 | Rlf | 0.003 | Old |
| 0.780955 | 0.012192 | 0.471 | 0.454 | 1 | 2 | Atp6v0e | 0.017 | Old |
| 0.780964 | 0.009464 | 0.291 | 0.297 | 1 | 2 | Irf8 | -0.006 | Old |
| 0.781409 | 0.071099 | 0.531 | 0.527 | 1 | 2 | Tomm7 | 0.004 | Old |
| 0.781857 | 0.067817 | 0.016 | 0.018 | 1 | 2 | Ptgfrn | -0.002 | Old |
| 0.782029 | 0.104368 | 0.052 | 0.056 | 1 | 2 | Yae1d1 | -0.004 | Old |
| 0.78207 | 0.103849 | 0.084 | 0.08 | 1 | 2 | Sec16a | 0.004 | Old |
| 0.782074 | 0.019552 | 0.089 | 0.084 | 1 | 2 | Stx8 | 0.005 | Old |
| 0.783054 | 0.030614 | 0.032 | 0.034 | 1 | 2 | Gpr68 | -0.002 | Old |
| 0.783133 | 0.085001 | 0.016 | 0.018 | 1 | 2 | Chek2 | -0.002 | Old |
| 0.783665 | 0.254251 | 0.071 | 0.068 | 1 | 2 | Kctd12b | 0.003 | Old |
| 0.783922 | 0.138253 | 0.043 | 0.04 | 1 | 2 | Rcbtb1 | 0.003 | Old |
| 0.784132 | 0.019066 | 0.014 | 0.015 | 1 | 2 | Ighmbp2 | -0.001 | Old |
| 0.784759 | 0.137574 | 0.034 | 0.037 | 1 | 2 | Nsun4 | -0.003 | Old |
| 0.784811 | 0.015529 | 0.216 | 0.209 | 1 | 2 | Mcts1 | 0.007 | Old |
| 0.784919 | 0.007596 | 0.05 | 0.048 | 1 | 2 | Gm38399 | 0.002 | Old |
| 0.78501 | 0.233067 | 0.214 | 0.211 | 1 | 2 | Zc3h13 | 0.003 | Old |
| 0.785051 | 0.253531 | 0.026 | 0.024 | 1 | 2 | Kbtbd3 | 0.002 | Old |
| 0.785239 | 0.040424 | 0.015 | 0.017 | 1 | 2 | Zfp628 | -0.002 | Old |
| 0.78525 | 0.055649 | 0.204 | 0.198 | 1 | 2 | Sptbn1 | 0.006 | Old |
| 0.785564 | 0.070187 | 0.13 | 0.125 | 1 | 2 | Srp54a | 0.005 | Old |
| 0.785674 | 0.433236 | 0.062 | 0.059 | 1 | 2 | Cd244 | 0.003 | Old |
| 0.785839 | 0.014167 | 0.236 | 0.233 | 1 | 2 | Grina | 0.003 | Old |
| 0.786529 | 0.266051 | 0.014 | 0.015 | 1 | 2 | Szt2 | -0.001 | Old |
| 0.787227 | 0.277455 | 0.087 | 0.083 | 1 | 2 | Fgfr1op | 0.004 | Old |
| 0.787648 | 0.230167 | 0.125 | 0.122 | 1 | 2 | Bri3bp | 0.003 | Old |
| 0.788186 | 0.013319 | 0.226 | 0.223 | 1 | 2 | Zmat2 | 0.003 | Old |
| 0.788452 | 0.159716 | 0.132 | 0.128 | 1 | 2 | Acp1 | 0.004 | Old |
| 0.788803 | 0.045323 | 0.111 | 0.115 | 1 | 2 | Cacul1 | -0.004 | Old |
| 0.788804 | 0.108669 | 0.048 | 0.051 | 1 | 2 | Top3b | -0.003 | Old |
| 0.789374 | 0.005279 | 0.211 | 0.204 | 1 | 2 | Dusp22 | 0.007 | Old |
| 0.789614 | 0.262143 | 0.014 | 0.015 | 1 | 2 | Rbak | -0.001 | Old |
| 0.790187 | 0.051074 | 0.633 | 0.632 | 1 | 2 | Ddx3x | 0.001 | Old |
| 0.791719 | 0.031499 | 0.047 | 0.05 | 1 | 2 | Smarcal1 | -0.003 | Old |
| 0.79185 | 0.360995 | 0.055 | 0.052 | 1 | 2 | Nxt1 | 0.003 | Old |
| 0.792648 | 0.403743 | 0.018 | 0.017 | 1 | 2 | Prmt6 | 0.001 | Old |
| 0.793037 | 0.120958 | 0.033 | 0.031 | 1 | 2 | Wdr55 | 0.002 | Old |
| 0.793045 | 0.350292 | 0.034 | 0.032 | 1 | 2 | Alkbh3 | 0.002 | Old |
| 0.79312 | 0.115262 | 0.05 | 0.048 | 1 | 2 | Fbxw5 | 0.002 | Old |
| 0.793505 | 0.447883 | 0.015 | 0.017 | 1 | 2 | Aldh7a1 | -0.002 | Old |
| 0.793771 | 0.16922 | 0.222 | 0.217 | 1 | 2 | Pkig | 0.005 | Old |
| 0.793896 | 0.141032 | 0.113 | 0.109 | 1 | 2 | Siah2 | 0.004 | Old |
| 0.793957 | 0.047667 | 0.482 | 0.477 | 1 | 2 | Cnbp | 0.005 | Old |

| | | | | | | | | |
|---|---|---|---|---|---|---|---|---|
| 0.794107 | 0.05289 | 0.105 | 0.109 | 1 | 2 | Trabd | -0.004 | Old |
| 0.794908 | 0.099417 | 0.106 | 0.11 | 1 | 2 | Rabgap1 | -0.004 | Old |
| 0.795206 | 0.122567 | 0.365 | 0.357 | 1 | 2 | Trir | 0.008 | Old |
| 0.795241 | 0.111185 | 0.166 | 0.162 | 1 | 2 | Nudcd3 | 0.004 | Old |
| 0.795242 | 0.069163 | 0.097 | 0.094 | 1 | 2 | Grk5 | 0.003 | Old |
| 0.795296 | 0.373204 | 0.077 | 0.082 | 1 | 2 | Ipo8 | -0.005 | Old |
| 0.79545 | 0.097086 | 0.178 | 0.172 | 1 | 2 | Emsy | 0.006 | Old |
| 0.795915 | 0.014329 | 0.204 | 0.196 | 1 | 2 | Ndufa8 | 0.008 | Old |
| 0.795967 | 0.03283 | 0.319 | 0.31 | 1 | 2 | Cers6 | 0.009 | Old |
| 0.795984 | 0.012179 | 0.444 | 0.441 | 1 | 2 | Selplg | 0.003 | Old |
| 0.796224 | 0.05763 | 0.373 | 0.357 | 1 | 2 | Scamp2 | 0.016 | Old |
| 0.796625 | 1.033448 | 0.011 | 0.01 | 1 | 2 | Ddit4 | 0.001 | Old |
| 0.796625 | 0.136219 | 0.011 | 0.01 | 1 | 2 | 9130221H1 | 0.001 | Old |
| 0.79745 | 0.447788 | 0.011 | 0.01 | 1 | 2 | Tmem17 | 0.001 | Old |
| 0.797513 | 0.086932 | 0.014 | 0.015 | 1 | 2 | Zfp934 | -0.001 | Old |
| 0.79763 | 0.175878 | 0.09 | 0.087 | 1 | 2 | Armc1 | 0.003 | Old |
| 0.79769 | 0.216226 | 0.018 | 0.017 | 1 | 2 | Ulk3 | 0.001 | Old |
| 0.798274 | 0.406005 | 0.011 | 0.01 | 1 | 2 | Nmb | 0.001 | Old |
| 0.798408 | 0.308871 | 0.063 | 0.061 | 1 | 2 | Zbtb37 | 0.002 | Old |
| 0.799117 | 0.008527 | 0.542 | 0.526 | 1 | 2 | Gltp | 0.016 | Old |
| 0.799899 | 0.407202 | 0.018 | 0.017 | 1 | 2 | Nubpl | 0.001 | Old |
| 0.800135 | 0.626082 | 0.015 | 0.017 | 1 | 2 | Sparc | -0.002 | Old |
| 0.80029 | 0.050593 | 0.649 | 0.646 | 1 | 2 | Calr | 0.003 | Old |
| 0.800643 | 0.062002 | 0.093 | 0.089 | 1 | 2 | Fnbp4 | 0.004 | Old |
| 0.800846 | 0.196496 | 0.018 | 0.017 | 1 | 2 | Neurl1b | 0.001 | Old |
| 0.801169 | 0.08612 | 0.306 | 0.299 | 1 | 2 | N4bp1 | 0.007 | Old |
| 0.801346 | 0.093262 | 0.127 | 0.124 | 1 | 2 | Lta4h | 0.003 | Old |
| 0.801793 | 0.214236 | 0.018 | 0.017 | 1 | 2 | Gm7694 | 0.001 | Old |
| 0.801879 | 0.037496 | 0.262 | 0.252 | 1 | 2 | Ids | 0.01 | Old |
| 0.801987 | 0.500367 | 0.014 | 0.015 | 1 | 2 | Cntrob | -0.001 | Old |
| 0.802148 | 0.097815 | 0.034 | 0.032 | 1 | 2 | Kdm1b | 0.002 | Old |
| 0.802245 | 0.615513 | 0.013 | 0.014 | 1 | 2 | Zfp579 | -0.001 | Old |
| 0.802425 | 0.063884 | 0.018 | 0.017 | 1 | 2 | Tmed1 | 0.001 | Old |
| 0.802442 | 0.104531 | 0.141 | 0.138 | 1 | 2 | Hltf | 0.003 | Old |
| 0.802815 | 0.008392 | 0.011 | 0.01 | 1 | 2 | Tmem191c | 0.001 | Old |
| 0.803293 | 0.16731 | 0.074 | 0.071 | 1 | 2 | Dgkh | 0.003 | Old |
| 0.803678 | 0.116677 | 0.013 | 0.014 | 1 | 2 | 1110019D1 | -0.001 | Old |
| 0.803699 | 0.034125 | 0.353 | 0.347 | 1 | 2 | Rab10 | 0.006 | Old |
| 0.804217 | 0.081584 | 0.262 | 0.258 | 1 | 2 | Cpt1a | 0.004 | Old |
| 0.805604 | 0.054525 | 0.067 | 0.064 | 1 | 2 | Gpatch2 | 0.003 | Old |
| 0.805719 | 0.02874 | 0.104 | 0.1 | 1 | 2 | Tmem141 | 0.004 | Old |
| 0.806306 | 0.003412 | 0.122 | 0.118 | 1 | 2 | Mindy3 | 0.004 | Old |
| 0.806488 | 0.131706 | 0.146 | 0.141 | 1 | 2 | Usp14 | 0.005 | Old |
| 0.807115 | 0.008104 | 0.141 | 0.137 | 1 | 2 | Ccdc59 | 0.004 | Old |
| 0.807126 | 0.050041 | 0.202 | 0.196 | 1 | 2 | Mcur1 | 0.006 | Old |

| | | | | | | | | |
|---|---|---|---|---|---|---|---|---|
| 0.807159 | 0.457646 | 0.039 | 0.037 | 1 | 2 | Rhou | 0.002 | Old |
| 0.807519 | 0.486768 | 0.036 | 0.034 | 1 | 2 | Dhx29 | 0.002 | Old |
| 0.808192 | 0.682312 | 0.014 | 0.015 | 1 | 2 | Tfec | -0.001 | Old |
| 0.808295 | 0.088733 | 0.181 | 0.177 | 1 | 2 | Strap | 0.004 | Old |
| 0.808341 | 0.482803 | 0.013 | 0.014 | 1 | 2 | Ehbp1 | -0.001 | Old |
| 0.808478 | 0.034062 | 0.226 | 0.219 | 1 | 2 | Yeats4 | 0.007 | Old |
| 0.808505 | 0.133709 | 0.097 | 0.094 | 1 | 2 | Timm17a | 0.003 | Old |
| 0.809277 | 0.175091 | 0.012 | 0.013 | 1 | 2 | Mtif3 | -0.001 | Old |
| 0.809277 | 0.170798 | 0.012 | 0.013 | 1 | 2 | Coil | -0.001 | Old |
| 0.809824 | 0.187253 | 0.054 | 0.051 | 1 | 2 | Ctnnbl1 | 0.003 | Old |
| 0.81038 | 0.159417 | 0.045 | 0.048 | 1 | 2 | Hectd3 | -0.003 | Old |
| 0.811278 | 0.087276 | 0.052 | 0.05 | 1 | 2 | Tmem198b | 0.002 | Old |
| 0.81135 | 0.699921 | 0.019 | 0.018 | 1 | 2 | Mus81 | 0.001 | Old |
| 0.81135 | 0.581045 | 0.019 | 0.018 | 1 | 2 | Clmp | 0.001 | Old |
| 0.811795 | 0.193009 | 0.084 | 0.081 | 1 | 2 | Hacd3 | 0.003 | Old |
| 0.81182 | 0.116256 | 0.046 | 0.049 | 1 | 2 | Ercc3 | -0.003 | Old |
| 0.812377 | 0.084803 | 0.163 | 0.16 | 1 | 2 | Foxj2 | 0.003 | Old |
| 0.812503 | 0.074161 | 0.029 | 0.031 | 1 | 2 | Mlst8 | -0.002 | Old |
| 0.812907 | 0.193921 | 0.06 | 0.063 | 1 | 2 | Mdc1 | -0.003 | Old |
| 0.812969 | 0.148088 | 0.155 | 0.16 | 1 | 2 | Brox | -0.005 | Old |
| 0.812984 | 0.200334 | 0.108 | 0.105 | 1 | 2 | Jdp2 | 0.003 | Old |
| 0.813233 | 0.001614 | 0.231 | 0.222 | 1 | 2 | Rab5a | 0.009 | Old |
| 0.813778 | 0.564153 | 0.012 | 0.013 | 1 | 2 | 0610040B1 | -0.001 | Old |
| 0.814029 | 0.10256 | 0.132 | 0.127 | 1 | 2 | Ppp2r2d | 0.005 | Old |
| 0.81471 | 0.272255 | 0.103 | 0.1 | 1 | 2 | Nf1 | 0.003 | Old |
| 0.815212 | 0.057597 | 0.088 | 0.091 | 1 | 2 | Tmem120a | -0.003 | Old |
| 0.815262 | 0.026468 | 0.242 | 0.236 | 1 | 2 | Nudt3 | 0.006 | Old |
| 0.815733 | 0.051796 | 0.3 | 0.296 | 1 | 2 | Dlst | 0.004 | Old |
| 0.815785 | 0.256321 | 0.011 | 0.012 | 1 | 2 | Lhfpl2 | -0.001 | Old |
| 0.81604 | 0.036875 | 0.35 | 0.361 | 1 | 2 | Hook3 | -0.011 | Old |
| 0.816115 | 0.184466 | 0.019 | 0.018 | 1 | 2 | Dis3l | 0.001 | Old |
| 0.816203 | 0.073608 | 0.233 | 0.227 | 1 | 2 | Atp6v0a1 | 0.006 | Old |
| 0.816313 | 0.244013 | 0.045 | 0.048 | 1 | 2 | L3mbtl3 | -0.003 | Old |
| 0.816897 | 0.028147 | 0.163 | 0.167 | 1 | 2 | Slc8b1 | -0.004 | Old |
| 0.816934 | 0.01202 | 0.11 | 0.106 | 1 | 2 | Chchd3 | 0.004 | Old |
| 0.816997 | 0.34655 | 0.028 | 0.026 | 1 | 2 | Clpb | 0.002 | Old |
| 0.81753 | 0.080611 | 0.078 | 0.075 | 1 | 2 | Mis12 | 0.003 | Old |
| 0.817754 | 0.050524 | 0.011 | 0.012 | 1 | 2 | Gnb5 | -0.001 | Old |
| 0.818141 | 0.001016 | 0.345 | 0.349 | 1 | 2 | Nras | -0.004 | Old |
| 0.818763 | 0.071574 | 0.167 | 0.164 | 1 | 2 | Ints6l | 0.003 | Old |
| 0.81878 | 0.089459 | 0.297 | 0.29 | 1 | 2 | Slc11a1 | 0.007 | Old |
| 0.818936 | 0.542578 | 0.011 | 0.012 | 1 | 2 | Ikzf2 | -0.001 | Old |
| 0.818966 | 0.195463 | 0.12 | 0.116 | 1 | 2 | Rab3d | 0.004 | Old |
| 0.819253 | 0.096302 | 0.097 | 0.101 | 1 | 2 | Rxrb | -0.004 | Old |
| 0.819263 | 0.162632 | 0.073 | 0.07 | 1 | 2 | Ints1 | 0.003 | Old |

| | | | | | | | | |
|---|---|---|---|---|---|---|---|---|
| 0.819347 | 0.209242 | 0.019 | 0.018 | 1 | 2 | Phka1 | 0.001 | Old |
| 0.82033 | 0.203864 | 0.028 | 0.026 | 1 | 2 | Hsd17b7 | 0.002 | Old |
| 0.820513 | 0.030273 | 0.011 | 0.012 | 1 | 2 | Wdr53 | -0.001 | Old |
| 0.821696 | 1.246679 | 0.011 | 0.012 | 1 | 2 | Lrg1 | -0.001 | Old |
| 0.821927 | 0.082563 | 0.353 | 0.347 | 1 | 2 | Ywhag | 0.006 | Old |
| 0.821955 | 0.151328 | 0.137 | 0.133 | 1 | 2 | Hgsnat | 0.004 | Old |
| 0.822255 | 0.177475 | 0.045 | 0.048 | 1 | 2 | Nat10 | -0.003 | Old |
| 0.822287 | 0.020448 | 0.037 | 0.036 | 1 | 2 | Rbm45 | 0.001 | Old |
| 0.822808 | 0.02063 | 0.503 | 0.512 | 1 | 2 | Ptprj | -0.009 | Old |
| 0.822894 | 0.15316 | 0.046 | 0.044 | 1 | 2 | Polr3d | 0.002 | Old |
| 0.823044 | 0.096863 | 0.019 | 0.018 | 1 | 2 | Med9 | 0.001 | Old |
| 0.823179 | 0.298265 | 0.039 | 0.037 | 1 | 2 | Rmdn3 | 0.002 | Old |
| 0.823661 | 0.100289 | 0.019 | 0.018 | 1 | 2 | Mrm1 | 0.001 | Old |
| 0.823923 | 0.232529 | 0.028 | 0.026 | 1 | 2 | Alg6 | 0.002 | Old |
| 0.82438 | 0.205831 | 0.034 | 0.032 | 1 | 2 | Mrpl1 | 0.002 | Old |
| 0.824503 | 0.07947 | 0.13 | 0.133 | 1 | 2 | Dgkz | -0.003 | Old |
| 0.824826 | 0.079159 | 0.227 | 0.223 | 1 | 2 | Ak2 | 0.004 | Old |
| 0.82493 | 0.039844 | 0.054 | 0.051 | 1 | 2 | Zbtb2 | 0.003 | Old |
| 0.825051 | 0.05952 | 0.243 | 0.239 | 1 | 2 | Abcg1 | 0.004 | Old |
| 0.825203 | 0.127569 | 0.019 | 0.018 | 1 | 2 | Noa1 | 0.001 | Old |
| 0.825616 | 0.078644 | 0.071 | 0.074 | 1 | 2 | Prps1 | -0.003 | Old |
| 0.825792 | 0.214142 | 0.082 | 0.086 | 1 | 2 | Tm7sf3 | -0.004 | Old |
| 0.82655 | 0.060946 | 0.288 | 0.286 | 1 | 2 | Hsd17b11 | 0.002 | Old |
| 0.826911 | 0.051826 | 0.035 | 0.033 | 1 | 2 | Clp1 | 0.002 | Old |
| 0.826957 | 0.037454 | 0.028 | 0.03 | 1 | 2 | Ttc38 | -0.002 | Old |
| 0.827241 | 0.012076 | 0.162 | 0.157 | 1 | 2 | Clock | 0.005 | Old |
| 0.827419 | 0.053888 | 0.012 | 0.011 | 1 | 2 | Surf6 | 0.001 | Old |
| 0.827488 | 0.182176 | 0.01 | 0.011 | 1 | 2 | Gstp3 | -0.001 | Old |
| 0.827617 | 0.631208 | 0.011 | 0.012 | 1 | 2 | Tmod1 | -0.001 | Old |
| 0.828063 | 0.139631 | 0.113 | 0.11 | 1 | 2 | Ehmt1 | 0.003 | Old |
| 0.829 | 0.152405 | 0.012 | 0.011 | 1 | 2 | Uros | 0.001 | Old |
| 0.829533 | 0.04063 | 0.163 | 0.165 | 1 | 2 | Chd9 | -0.002 | Old |
| 0.829569 | 0.414255 | 0.01 | 0.011 | 1 | 2 | Cavin2 | -0.001 | Old |
| 0.829569 | 0.066213 | 0.01 | 0.011 | 1 | 2 | Gpt2 | -0.001 | Old |
| 0.829873 | 0.012247 | 0.078 | 0.081 | 1 | 2 | Sfxn2 | -0.003 | Old |
| 0.830051 | 0.034206 | 0.099 | 0.102 | 1 | 2 | Dpy30 | -0.003 | Old |
| 0.830078 | 0.171007 | 0.17 | 0.167 | 1 | 2 | Prpf39 | 0.003 | Old |
| 0.830091 | 0.150207 | 0.028 | 0.026 | 1 | 2 | Tmem101 | 0.002 | Old |
| 0.830345 | 0.048608 | 0.054 | 0.056 | 1 | 2 | Tsta3 | -0.002 | Old |
| 0.830528 | 0.024248 | 0.157 | 0.152 | 1 | 2 | Fto | 0.005 | Old |
| 0.830533 | 0.02575 | 0.045 | 0.043 | 1 | 2 | Scnm1 | 0.002 | Old |
| 0.831032 | 0.019659 | 0.035 | 0.033 | 1 | 2 | Caml | 0.002 | Old |
| 0.831069 | 0.022407 | 0.019 | 0.018 | 1 | 2 | Mblac2 | 0.001 | Old |
| 0.831077 | 0.009237 | 0.041 | 0.043 | 1 | 2 | Acaca | -0.002 | Old |
| 0.831121 | 0.020845 | 0.036 | 0.034 | 1 | 2 | Gm20342 | 0.002 | Old |

| | | | | | | | | |
|---|---|---|---|---|---|---|---|---|
| 0.831432 | 0.148957 | 0.046 | 0.044 | 1 | 2 | Carm1 | 0.002 | Old |
| 0.832305 | 0.095976 | 0.11 | 0.107 | 1 | 2 | Impa1 | 0.003 | Old |
| 0.832371 | 0.024487 | 0.147 | 0.143 | 1 | 2 | Mien1 | 0.004 | Old |
| 0.833272 | 0.089329 | 0.251 | 0.246 | 1 | 2 | Eif2s3x | 0.005 | Old |
| 0.833319 | 0.397781 | 0.01 | 0.011 | 1 | 2 | Ncapd2 | -0.001 | Old |
| 0.833492 | 0.008715 | 0.308 | 0.298 | 1 | 2 | Aes | 0.01 | Old |
| 0.834281 | 0.072071 | 0.009 | 0.01 | 1 | 2 | Gm10762 | -0.001 | Old |
| 0.834424 | 0.135334 | 0.136 | 0.133 | 1 | 2 | Rabl6 | 0.003 | Old |
| 0.835503 | 0.365596 | 0.02 | 0.019 | 1 | 2 | Rnf135 | 0.001 | Old |
| 0.835888 | 0.039286 | 0.276 | 0.277 | 1 | 2 | Rb1cc1 | -0.001 | Old |
| 0.836063 | 0.008878 | 0.195 | 0.189 | 1 | 2 | Gmip | 0.006 | Old |
| 0.836491 | 0.198371 | 0.009 | 0.01 | 1 | 2 | Csf1 | -0.001 | Old |
| 0.836491 | 0.039028 | 0.009 | 0.01 | 1 | 2 | Mgmt | -0.001 | Old |
| 0.836934 | 0.056367 | 0.009 | 0.01 | 1 | 2 | Dcbld2 | -0.001 | Old |
| 0.837818 | 0.288961 | 0.009 | 0.01 | 1 | 2 | Timm23 | -0.001 | Old |
| 0.838378 | 0.115522 | 0.046 | 0.044 | 1 | 2 | Mvd | 0.002 | Old |
| 0.839258 | 0.006243 | 0.342 | 0.335 | 1 | 2 | Ssr1 | 0.007 | Old |
| 0.839588 | 0.239296 | 0.009 | 0.01 | 1 | 2 | Zfp458 | -0.001 | Old |
| 0.83972 | 0.142909 | 0.258 | 0.257 | 1 | 2 | Zcchc24 | 0.001 | Old |
| 0.839884 | 0.012135 | 0.156 | 0.152 | 1 | 2 | Inpp5k | 0.004 | Old |
| 0.84006 | 0.193196 | 0.037 | 0.036 | 1 | 2 | Zfp84 | 0.001 | Old |
| 0.840292 | 0.05661 | 0.373 | 0.365 | 1 | 2 | Ep300 | 0.008 | Old |
| 0.840395 | 0.037763 | 0.438 | 0.447 | 1 | 2 | Ccdc88a | -0.009 | Old |
| 0.840615 | 0.121463 | 0.143 | 0.147 | 1 | 2 | Eftud2 | -0.004 | Old |
| 0.841085 | 0.043221 | 0.15 | 0.152 | 1 | 2 | Gsdmd | -0.002 | Old |
| 0.841508 | 0.204103 | 0.037 | 0.036 | 1 | 2 | Yaf2 | 0.001 | Old |
| 0.841765 | 0.089806 | 0.093 | 0.09 | 1 | 2 | Fut11 | 0.003 | Old |
| 0.841796 | 0.055277 | 0.111 | 0.108 | 1 | 2 | Sgta | 0.003 | Old |
| 0.841963 | 0.319289 | 0.085 | 0.083 | 1 | 2 | Mesd | 0.002 | Old |
| 0.842408 | 0.034828 | 0.131 | 0.126 | 1 | 2 | Clasp2 | 0.005 | Old |
| 0.842496 | 0.461899 | 0.021 | 0.02 | 1 | 2 | Sgsm2 | 0.001 | Old |
| 0.842531 | 0.081048 | 0.101 | 0.097 | 1 | 2 | Dhdh | 0.004 | Old |
| 0.843339 | 0.045925 | 0.252 | 0.255 | 1 | 2 | Cebpz | -0.003 | Old |
| 0.843375 | 0.194272 | 0.021 | 0.02 | 1 | 2 | Tacc3 | 0.001 | Old |
| 0.84342 | 0.090988 | 0.141 | 0.138 | 1 | 2 | Rnf20 | 0.003 | Old |
| 0.843668 | 0.162173 | 0.021 | 0.02 | 1 | 2 | Zfp959 | 0.001 | Old |
| 0.84372 | 0.001994 | 0.773 | 0.755 | 1 | 2 | Sdcbp | 0.018 | Old |
| 0.843948 | 0.141626 | 0.105 | 0.102 | 1 | 2 | Nbr1 | 0.003 | Old |
| 0.845015 | 0.119923 | 0.162 | 0.158 | 1 | 2 | Plekha1 | 0.004 | Old |
| 0.845855 | 0.042462 | 0.479 | 0.473 | 1 | 2 | Klf3 | 0.006 | Old |
| 0.84602 | 0.58722 | 0.013 | 0.012 | 1 | 2 | Ppp1r9a | 0.001 | Old |
| 0.846264 | 0.002294 | 0.419 | 0.41 | 1 | 2 | H13 | 0.009 | Old |
| 0.846398 | 0.443816 | 0.013 | 0.012 | 1 | 2 | Fbxo46 | 0.001 | Old |
| 0.846399 | 0.014944 | 0.029 | 0.027 | 1 | 2 | Cnn3 | 0.002 | Old |
| 0.8468 | 0.152899 | 0.121 | 0.118 | 1 | 2 | Vrk2 | 0.003 | Old |

| | | | | | | | | |
|---|---|---|---|---|---|---|---|---|
| 0.846889 | 0.128081 | 0.325 | 0.331 | 1 | 2 | Ccr1 | -0.006 | Old |
| 0.847388 | 0.068907 | 0.257 | 0.259 | 1 | 2 | Pias1 | -0.002 | Old |
| 0.847633 | 0.152114 | 0.089 | 0.087 | 1 | 2 | Rnpc3 | 0.002 | Old |
| 0.847639 | 0.096328 | 0.037 | 0.036 | 1 | 2 | Scap | 0.001 | Old |
| 0.848172 | 0.012927 | 0.029 | 0.027 | 1 | 2 | Zfp687 | 0.002 | Old |
| 0.848743 | 0.114627 | 0.051 | 0.053 | 1 | 2 | Phtf2 | -0.002 | Old |
| 0.848747 | 0.226058 | 0.069 | 0.067 | 1 | 2 | Fhod1 | 0.002 | Old |
| 0.849086 | 0.015279 | 0.21 | 0.204 | 1 | 2 | Rap2a | 0.006 | Old |
| 0.849868 | 0.119785 | 0.055 | 0.052 | 1 | 2 | Exoc7 | 0.003 | Old |
| 0.849886 | 0.040067 | 0.122 | 0.125 | 1 | 2 | Ip6k1 | -0.003 | Old |
| 0.850186 | 0.164862 | 0.071 | 0.069 | 1 | 2 | Eya3 | 0.002 | Old |
| 0.850562 | 0.61518 | 0.021 | 0.02 | 1 | 2 | Kctd13 | 0.001 | Old |
| 0.850726 | 0.234831 | 0.125 | 0.124 | 1 | 2 | Pwwp2a | 0.001 | Old |
| 0.851167 | 0.165672 | 0.049 | 0.051 | 1 | 2 | Mrpl27 | -0.002 | Old |
| 0.851279 | 0.012196 | 0.115 | 0.118 | 1 | 2 | Arl6ip6 | -0.003 | Old |
| 0.85188 | 0.044084 | 0.024 | 0.025 | 1 | 2 | Strada | -0.001 | Old |
| 0.852842 | 0.129892 | 0.013 | 0.012 | 1 | 2 | Cmtr2 | 0.001 | Old |
| 0.853958 | 0.003546 | 0.738 | 0.739 | 1 | 2 | Tmbim6 | -0.001 | Old |
| 0.854162 | 0.613599 | 0.031 | 0.03 | 1 | 2 | Pgghg | 0.001 | Old |
| 0.854389 | 0.027353 | 0.305 | 0.296 | 1 | 2 | Dnm2 | 0.009 | Old |
| 0.855409 | 0.213945 | 0.021 | 0.02 | 1 | 2 | Zmym1 | 0.001 | Old |
| 0.856627 | 0.036702 | 0.33 | 0.328 | 1 | 2 | Lifr | 0.002 | Old |
| 0.856819 | 0.110742 | 0.022 | 0.024 | 1 | 2 | Plcg1 | -0.002 | Old |
| 0.857548 | 0.136623 | 0.04 | 0.038 | 1 | 2 | D6Wsu163 | 0.002 | Old |
| 0.857585 | 0.005769 | 0.111 | 0.108 | 1 | 2 | Plgrkt | 0.003 | Old |
| 0.858017 | 0.086168 | 0.106 | 0.103 | 1 | 2 | Ccdc82 | 0.003 | Old |
| 0.858243 | 0.0042 | 0.095 | 0.093 | 1 | 2 | Kdm3b | 0.002 | Old |
| 0.858657 | 0.026417 | 0.133 | 0.129 | 1 | 2 | Fbxl20 | 0.004 | Old |
| 0.858706 | 0.125267 | 0.094 | 0.091 | 1 | 2 | Srek1ip1 | 0.003 | Old |
| 0.859023 | 0.031411 | 0.06 | 0.058 | 1 | 2 | Pigq | 0.002 | Old |
| 0.859437 | 0.070864 | 0.09 | 0.087 | 1 | 2 | Fam98c | 0.003 | Old |
| 0.859573 | 0.239553 | 0.088 | 0.091 | 1 | 2 | Ncbp1 | -0.003 | Old |
| 0.860061 | 0.385044 | 0.039 | 0.037 | 1 | 2 | 6-Sep | 0.002 | Old |
| 0.86061 | 0.234926 | 0.05 | 0.052 | 1 | 2 | Uty | -0.002 | Old |
| 0.860722 | 0.049724 | 0.135 | 0.131 | 1 | 2 | Hlcs | 0.004 | Old |
| 0.861186 | 0.111042 | 0.118 | 0.115 | 1 | 2 | Xpot | 0.003 | Old |
| 0.861763 | 0.054727 | 0.201 | 0.196 | 1 | 2 | Ankrd13a | 0.005 | Old |
| 0.862367 | 0.29896 | 0.042 | 0.04 | 1 | 2 | Slc25a22 | 0.002 | Old |
| 0.86252 | 0.123366 | 0.057 | 0.055 | 1 | 2 | Zbtb16 | 0.002 | Old |
| 0.863328 | 0.131234 | 0.059 | 0.057 | 1 | 2 | Mpv17l2 | 0.002 | Old |
| 0.863429 | 0.098811 | 0.177 | 0.173 | 1 | 2 | Mllt10 | 0.004 | Old |
| 0.86346 | 0.26283 | 0.031 | 0.03 | 1 | 2 | Homer3 | 0.001 | Old |
| 0.863795 | 0.075302 | 0.145 | 0.141 | 1 | 2 | Slfn8 | 0.004 | Old |
| 0.864278 | 0.045406 | 0.192 | 0.185 | 1 | 2 | Trim28 | 0.007 | Old |
| 0.864552 | 0.000257 | 0.134 | 0.129 | 1 | 2 | Dpf2 | 0.005 | Old |

| | | | | | | | | |
|---|---|---|---|---|---|---|---|---|
| 0.864587 | 0.073681 | 0.17 | 0.166 | 1 | 2 | Camta2 | 0.004 | Old |
| 0.86483 | 0.062025 | 0.03 | 0.029 | 1 | 2 | Vps37c | 0.001 | Old |
| 0.864915 | 0.019929 | 0.126 | 0.129 | 1 | 2 | Tpcn1 | -0.003 | Old |
| 0.864968 | 0.062199 | 0.234 | 0.228 | 1 | 2 | Smc1a | 0.006 | Old |
| 0.86497 | 0.848823 | 0.014 | 0.013 | 1 | 2 | C1qtnf12 | 0.001 | Old |
| 0.865121 | 0.032624 | 0.173 | 0.17 | 1 | 2 | Lrp6 | 0.003 | Old |
| 0.866113 | 0.005709 | 0.117 | 0.114 | 1 | 2 | Epb41l4aos | 0.003 | Old |
| 0.866903 | 0.02936 | 0.04 | 0.038 | 1 | 2 | Fam217b | 0.002 | Old |
| 0.867226 | 0.145096 | 0.11 | 0.114 | 1 | 2 | Deptor | -0.004 | Old |
| 0.867265 | 0.034134 | 0.246 | 0.239 | 1 | 2 | Rlim | 0.007 | Old |
| 0.86737 | 0.015308 | 0.522 | 0.525 | 1 | 2 | Ostf1 | -0.003 | Old |
| 0.867499 | 0.181569 | 0.079 | 0.077 | 1 | 2 | Lrif1 | 0.002 | Old |
| 0.867594 | 0.257723 | 0.039 | 0.04 | 1 | 2 | Stxbp1 | -0.001 | Old |
| 0.867886 | 0.43111 | 0.014 | 0.013 | 1 | 2 | Abhd14b | 0.001 | Old |
| 0.867945 | 0.026797 | 0.223 | 0.217 | 1 | 2 | Il13ra1 | 0.006 | Old |
| 0.867992 | 1.28904 | 0.04 | 0.038 | 1 | 2 | Hdc | 0.002 | Old |
| 0.868158 | 0.070695 | 0.652 | 0.673 | 1 | 2 | Plbd1 | -0.021 | Old |
| 0.868279 | 0.033849 | 0.047 | 0.049 | 1 | 2 | Mtap | -0.002 | Old |
| 0.868699 | 0.319635 | 0.07 | 0.072 | 1 | 2 | Vps16 | -0.002 | Old |
| 0.869527 | 0.492151 | 0.014 | 0.013 | 1 | 2 | Amotl2 | 0.001 | Old |
| 0.871795 | 0.014052 | 0.031 | 0.03 | 1 | 2 | Zbtb41 | 0.001 | Old |
| 0.871966 | 0.359111 | 0.022 | 0.021 | 1 | 2 | Srbd1 | 0.001 | Old |
| 0.871982 | 0.075181 | 0.412 | 0.399 | 1 | 2 | Cct8 | 0.013 | Old |
| 0.872201 | 0.006829 | 0.108 | 0.11 | 1 | 2 | Srpk2 | -0.002 | Old |
| 0.872505 | 0.274687 | 0.022 | 0.024 | 1 | 2 | Eogt | -0.002 | Old |
| 0.872563 | 0.070365 | 0.123 | 0.127 | 1 | 2 | 4833439L1 | -0.004 | Old |
| 0.873346 | 0.160119 | 0.124 | 0.122 | 1 | 2 | Tmx1 | 0.002 | Old |
| 0.873696 | 0.026787 | 0.034 | 0.036 | 1 | 2 | Hcar2 | -0.002 | Old |
| 0.873984 | 0.204119 | 0.037 | 0.039 | 1 | 2 | Cep164 | -0.002 | Old |
| 0.874415 | 0.109753 | 0.086 | 0.088 | 1 | 2 | Mtus1 | -0.002 | Old |
| 0.874659 | 0.072701 | 0.139 | 0.137 | 1 | 2 | Cops5 | 0.002 | Old |
| 0.875705 | 0.099223 | 0.301 | 0.306 | 1 | 2 | Ensa | -0.005 | Old |
| 0.875778 | 0.084826 | 0.346 | 0.349 | 1 | 2 | Mtss1 | -0.003 | Old |
| 0.875937 | 0.013118 | 0.293 | 0.286 | 1 | 2 | Dgkd | 0.007 | Old |
| 0.876065 | 0.309318 | 0.032 | 0.031 | 1 | 2 | Cwc27 | 0.001 | Old |
| 0.876393 | 0.065317 | 0.241 | 0.239 | 1 | 2 | Etnk1 | 0.002 | Old |
| 0.876541 | 0.093084 | 0.093 | 0.091 | 1 | 2 | Zpr1 | 0.002 | Old |
| 0.876785 | 0.244016 | 0.084 | 0.082 | 1 | 2 | Nip7 | 0.002 | Old |
| 0.876904 | 0.038431 | 0.196 | 0.191 | 1 | 2 | Fip1l1 | 0.005 | Old |
| 0.877432 | 0.060545 | 0.277 | 0.28 | 1 | 2 | Tnpo3 | -0.003 | Old |
| 0.877795 | 0.187901 | 0.095 | 0.094 | 1 | 2 | Phf2 | 0.001 | Old |
| 0.87788 | 0.441819 | 0.025 | 0.024 | 1 | 2 | Slc2a8 | 0.001 | Old |
| 0.878481 | 0.056678 | 0.08 | 0.082 | 1 | 2 | Gtf2i | -0.002 | Old |
| 0.878686 | 0.066276 | 0.182 | 0.181 | 1 | 2 | Evi5 | 0.001 | Old |
| 0.878699 | 0.564674 | 0.034 | 0.033 | 1 | 2 | Metap1d | 0.001 | Old |

| | | | | | | | | |
|---|---|---|---|---|---|---|---|---|
| 0.879523 | 0.018984 | 0.072 | 0.074 | 1 | 2 | Ngly1 | -0.002 | Old |
| 0.879886 | 0.003523 | 0.041 | 0.039 | 1 | 2 | Snip1 | 0.002 | Old |
| 0.880241 | 0.072666 | 0.139 | 0.135 | 1 | 2 | Filip1l | 0.004 | Old |
| 0.88039 | 0.288263 | 0.033 | 0.032 | 1 | 2 | Zfp513 | 0.001 | Old |
| 0.880964 | 0.094055 | 0.141 | 0.139 | 1 | 2 | Psd3 | 0.002 | Old |
| 0.880987 | 0.02415 | 0.061 | 0.063 | 1 | 2 | Selenoh | -0.002 | Old |
| 0.88103 | 0.023454 | 0.014 | 0.013 | 1 | 2 | Rabl3 | 0.001 | Old |
| 0.881331 | 0.011292 | 0.399 | 0.407 | 1 | 2 | Eif4b | -0.008 | Old |
| 0.881545 | 0.066471 | 0.112 | 0.114 | 1 | 2 | Tbc1d9 | -0.002 | Old |
| 0.882163 | 0.088666 | 0.057 | 0.058 | 1 | 2 | Zfp518a | -0.001 | Old |
| 0.88224 | 0.025744 | 0.751 | 0.742 | 1 | 2 | Picalm | 0.009 | Old |
| 0.882858 | 0.02374 | 0.085 | 0.087 | 1 | 2 | Anapc2 | -0.002 | Old |
| 0.882951 | 0.086018 | 0.221 | 0.215 | 1 | 2 | Dnajc13 | 0.006 | Old |
| 0.88301 | 0.172535 | 0.152 | 0.151 | 1 | 2 | Fam76a | 0.001 | Old |
| 0.883279 | 0.106392 | 0.192 | 0.19 | 1 | 2 | Otub1 | 0.002 | Old |
| 0.883477 | 0.111385 | 0.033 | 0.032 | 1 | 2 | Ifitm5 | 0.001 | Old |
| 0.883547 | 0.010706 | 0.571 | 0.578 | 1 | 2 | Hnrnpa0 | -0.007 | Old |
| 0.883741 | 0.032807 | 0.024 | 0.023 | 1 | 2 | Zbtb24 | 0.001 | Old |
| 0.883835 | 0.081358 | 0.199 | 0.2 | 1 | 2 | Osm | -0.001 | Old |
| 0.883946 | 0.419739 | 0.059 | 0.058 | 1 | 2 | Tigd2 | 0.001 | Old |
| 0.884254 | 0.013886 | 0.97 | 0.974 | 1 | 2 | Rps26 | -0.004 | Old |
| 0.885007 | 0.143135 | 0.092 | 0.094 | 1 | 2 | Scaf8 | -0.002 | Old |
| 0.885172 | 0.076018 | 0.091 | 0.089 | 1 | 2 | Myo18a | 0.002 | Old |
| 0.885781 | 0.371037 | 0.036 | 0.038 | 1 | 2 | Unkl | -0.002 | Old |
| 0.886014 | 0.112573 | 0.087 | 0.089 | 1 | 2 | Mtmr10 | -0.002 | Old |
| 0.886434 | 0.091922 | 0.236 | 0.233 | 1 | 2 | Wdr45b | 0.003 | Old |
| 0.886605 | 0.022402 | 0.323 | 0.324 | 1 | 2 | Dennd5a | -0.001 | Old |
| 0.887089 | 0.11559 | 0.065 | 0.068 | 1 | 2 | Pkd2 | -0.003 | Old |
| 0.88715 | 0.037101 | 0.136 | 0.134 | 1 | 2 | Ythdc2 | 0.002 | Old |
| 0.887215 | 0.048 | 0.059 | 0.057 | 1 | 2 | Tex2 | 0.002 | Old |
| 0.88727 | 0.031688 | 0.058 | 0.056 | 1 | 2 | Fam117a | 0.002 | Old |
| 0.887425 | 0.003652 | 0.057 | 0.058 | 1 | 2 | Pam16 | -0.001 | Old |
| 0.887541 | 0.110841 | 0.044 | 0.043 | 1 | 2 | Nup160 | 0.001 | Old |
| 0.887836 | 0.214818 | 0.034 | 0.036 | 1 | 2 | Ccdc32 | -0.002 | Old |
| 0.888336 | 0.042138 | 0.264 | 0.265 | 1 | 2 | Baz1b | -0.001 | Old |
| 0.888446 | 0.443807 | 0.02 | 0.021 | 1 | 2 | Urb1 | -0.001 | Old |
| 0.888446 | 0.229352 | 0.02 | 0.021 | 1 | 2 | Gga2 | -0.001 | Old |
| 0.888627 | 0.350568 | 0.032 | 0.031 | 1 | 2 | Carf | 0.001 | Old |
| 0.889812 | 0.024359 | 0.321 | 0.323 | 1 | 2 | Git2 | -0.002 | Old |
| 0.890138 | 0.017824 | 0.635 | 0.65 | 1 | 2 | Eif4a1 | -0.015 | Old |
| 0.890254 | 0.043733 | 0.019 | 0.02 | 1 | 2 | Mrrf | -0.001 | Old |
| 0.890283 | 0.243222 | 0.078 | 0.077 | 1 | 2 | Hmg20a | 0.001 | Old |
| 0.890453 | 0.069192 | 0.081 | 0.083 | 1 | 2 | Edem2 | -0.002 | Old |
| 0.891055 | 0.373672 | 0.015 | 0.014 | 1 | 2 | Disc1 | 0.001 | Old |
| 0.891332 | 0.235624 | 0.073 | 0.071 | 1 | 2 | Afg3l2 | 0.002 | Old |

| | | | | | | | | |
|---|---|---|---|---|---|---|---|---|
| 0.891688 | 0.077003 | 0.353 | 0.349 | 1 | 2 | Foxn3 | 0.004 | Old |
| 0.891899 | 0.60831 | 0.044 | 0.043 | 1 | 2 | Cfap36 | 0.001 | Old |
| 0.892266 | 0.069716 | 0.051 | 0.05 | 1 | 2 | Trove2 | 0.001 | Old |
| 0.892681 | 0.256993 | 0.063 | 0.065 | 1 | 2 | Klhdc2 | -0.002 | Old |
| 0.893023 | 0.16488 | 0.139 | 0.137 | 1 | 2 | Pitpnc1 | 0.002 | Old |
| 0.893077 | 0.071719 | 0.033 | 0.034 | 1 | 2 | Taf8 | -0.001 | Old |
| 0.893519 | 0.207841 | 0.051 | 0.05 | 1 | 2 | Fbxw8 | 0.001 | Old |
| 0.893635 | 0.105523 | 0.101 | 0.099 | 1 | 2 | Creb5 | 0.002 | Old |
| 0.893747 | 0.006281 | 0.574 | 0.569 | 1 | 2 | Tmed10 | 0.005 | Old |
| 0.894371 | 0.367203 | 0.025 | 0.024 | 1 | 2 | Ercc8 | 0.001 | Old |
| 0.894491 | 0.03817 | 0.4 | 0.405 | 1 | 2 | S100a13 | -0.005 | Old |
| 0.895123 | 0.092302 | 0.019 | 0.02 | 1 | 2 | Lancl2 | -0.001 | Old |
| 0.895634 | 0.133108 | 0.015 | 0.014 | 1 | 2 | Haao | 0.001 | Old |
| 0.895872 | 0.134837 | 0.116 | 0.118 | 1 | 2 | Sirt7 | -0.002 | Old |
| 0.896214 | 0.089232 | 0.069 | 0.067 | 1 | 2 | Arhgef7 | 0.002 | Old |
| 0.896539 | 0.021619 | 0.198 | 0.195 | 1 | 2 | Gga1 | 0.003 | Old |
| 0.896947 | 0.033106 | 0.067 | 0.069 | 1 | 2 | Mis18a | -0.002 | Old |
| 0.896986 | 0.110414 | 0.111 | 0.109 | 1 | 2 | Ndufs1 | 0.002 | Old |
| 0.897613 | 0.03455 | 0.106 | 0.108 | 1 | 2 | Lsm8 | -0.002 | Old |
| 0.897625 | 0.017994 | 0.511 | 0.508 | 1 | 2 | Mapkapk2 | 0.003 | Old |
| 0.897748 | 0.118621 | 0.015 | 0.014 | 1 | 2 | Rdh13 | 0.001 | Old |
| 0.898132 | 0.165628 | 0.045 | 0.044 | 1 | 2 | Coq5 | 0.001 | Old |
| 0.898344 | 0.410257 | 0.032 | 0.033 | 1 | 2 | Saal1 | -0.001 | Old |
| 0.898399 | 0.238234 | 0.066 | 0.065 | 1 | 2 | Sik3 | 0.001 | Old |
| 0.898453 | 0.157012 | 0.015 | 0.014 | 1 | 2 | Mphosph9 | 0.001 | Old |
| 0.898464 | 0.229927 | 0.033 | 0.034 | 1 | 2 | Trmt2b | -0.001 | Old |
| 0.898575 | 0.002447 | 0.132 | 0.129 | 1 | 2 | Zbtb44 | 0.003 | Old |
| 0.899158 | 0.075244 | 0.015 | 0.014 | 1 | 2 | Ulbp1 | 0.001 | Old |
| 0.899268 | 0.184407 | 0.072 | 0.07 | 1 | 2 | Hdac9 | 0.002 | Old |
| 0.900494 | 0.26988 | 0.062 | 0.064 | 1 | 2 | Klhl21 | -0.002 | Old |
| 0.900959 | 0.033437 | 0.997 | 1 | 1 | 2 | Malat1 | -0.003 | Old |
| 0.901315 | 0.017694 | 0.044 | 0.045 | 1 | 2 | Tsr1 | -0.001 | Old |
| 0.901394 | 0.041474 | 0.034 | 0.033 | 1 | 2 | Rabepk | 0.001 | Old |
| 0.901627 | 0.075811 | 0.015 | 0.014 | 1 | 2 | Ube2e2 | 0.001 | Old |
| 0.902122 | 0.055865 | 0.132 | 0.133 | 1 | 2 | Jmjd6 | -0.001 | Old |
| 0.902296 | 0.241766 | 0.047 | 0.046 | 1 | 2 | Slc25a32 | 0.001 | Old |
| 0.902574 | 0.027831 | 0.141 | 0.143 | 1 | 2 | Pja2 | -0.002 | Old |
| 0.903099 | 0.475051 | 0.047 | 0.046 | 1 | 2 | Prkrip1 | 0.001 | Old |
| 0.903269 | 0.003979 | 0.389 | 0.382 | 1 | 2 | Zfp207 | 0.007 | Old |
| 0.9034 | 0.021316 | 0.281 | 0.276 | 1 | 2 | Casp4 | 0.005 | Old |
| 0.904564 | 0.089006 | 0.025 | 0.024 | 1 | 2 | Fance | 0.001 | Old |
| 0.904739 | 0.208879 | 0.087 | 0.086 | 1 | 2 | Acox1 | 0.001 | Old |
| 0.905036 | 0.068559 | 0.054 | 0.055 | 1 | 2 | Naf1 | -0.001 | Old |
| 0.905136 | 0.031088 | 0.033 | 0.032 | 1 | 2 | Adck1 | 0.001 | Old |
| 0.905865 | 0.023248 | 0.255 | 0.249 | 1 | 2 | Rin2 | 0.006 | Old |

| | | | | | | | | |
|---|---|---|---|---|---|---|---|---|
| 0.906785 | 0.018716 | 0.286 | 0.291 | 1 | 2 | Ctdnep1 | -0.005 | Old |
| 0.907123 | 0.198631 | 0.018 | 0.019 | 1 | 2 | Nsdhl | -0.001 | Old |
| 0.907445 | 0.04546 | 0.065 | 0.067 | 1 | 2 | Abhd16a | -0.002 | Old |
| 0.907583 | 0.067553 | 0.044 | 0.045 | 1 | 2 | 1600014C1 | -0.001 | Old |
| 0.90761 | 0.317267 | 0.057 | 0.058 | 1 | 2 | Mier3 | -0.001 | Old |
| 0.907841 | 0.022609 | 0.307 | 0.309 | 1 | 2 | Lasp1 | -0.002 | Old |
| 0.907918 | 0.048264 | 0.227 | 0.227 | 1 | 2 | Psmc6 | 0 | Old |
| 0.908049 | 0.026241 | 0.09 | 0.088 | 1 | 2 | Gon4l | 0.002 | Old |
| 0.908354 | 0.121328 | 0.272 | 0.272 | 1 | 2 | Id2 | 0 | Old |
| 0.90838 | 0.089746 | 0.27 | 0.265 | 1 | 2 | Slc29a3 | 0.005 | Old |
| 0.908472 | 0.112945 | 0.085 | 0.083 | 1 | 2 | Ralgapa1 | 0.002 | Old |
| 0.908693 | 0.117597 | 0.018 | 0.019 | 1 | 2 | Bcr | -0.001 | Old |
| 0.908821 | 0.079033 | 0.287 | 0.291 | 1 | 2 | Smap2 | -0.004 | Old |
| 0.909363 | 0.086371 | 0.044 | 0.043 | 1 | 2 | Metrn | 0.001 | Old |
| 0.909389 | 0.394089 | 0.016 | 0.015 | 1 | 2 | Gins2 | 0.001 | Old |
| 0.910096 | 0.168907 | 0.054 | 0.055 | 1 | 2 | Traf3ip3 | -0.001 | Old |
| 0.910613 | 0.210375 | 0.156 | 0.156 | 1 | 2 | Efcab14 | 0 | Old |
| 0.911091 | 0.890541 | 0.016 | 0.015 | 1 | 2 | Spic | 0.001 | Old |
| 0.911688 | 0.087528 | 0.017 | 0.018 | 1 | 2 | 4933427D1 | -0.001 | Old |
| 0.913281 | 0.018378 | 0.528 | 0.53 | 1 | 2 | Trib1 | -0.002 | Old |
| 0.914757 | 0.045189 | 0.042 | 0.043 | 1 | 2 | Inafm2 | -0.001 | Old |
| 0.914835 | 0.023637 | 0.611 | 0.626 | 1 | 2 | Cxcl2 | -0.015 | Old |
| 0.915068 | 0.000658 | 0.328 | 0.318 | 1 | 2 | Asap1 | 0.01 | Old |
| 0.91552 | 0.04983 | 0.016 | 0.015 | 1 | 2 | Sap130 | 0.001 | Old |
| 0.915648 | 0.093648 | 0.179 | 0.177 | 1 | 2 | Atg2a | 0.002 | Old |
| 0.915911 | 0.077453 | 0.054 | 0.055 | 1 | 2 | Jade1 | -0.001 | Old |
| 0.915992 | 0.38648 | 0.046 | 0.045 | 1 | 2 | Iffo1 | 0.001 | Old |
| 0.916324 | 0.228782 | 0.09 | 0.089 | 1 | 2 | Zdhhc7 | 0.001 | Old |
| 0.916573 | 0.133525 | 0.089 | 0.087 | 1 | 2 | Pcnx | 0.002 | Old |
| 0.917233 | 0.308745 | 0.057 | 0.056 | 1 | 2 | Epm2aip1 | 0.001 | Old |
| 0.918141 | 0.015893 | 0.35 | 0.343 | 1 | 2 | Sqstm1 | 0.007 | Old |
| 0.918766 | 0.148392 | 0.042 | 0.043 | 1 | 2 | Ptcd3 | -0.001 | Old |
| 0.919494 | 0.048355 | 0.18 | 0.178 | 1 | 2 | Fbxo33 | 0.002 | Old |
| 0.919938 | 0.123394 | 0.034 | 0.033 | 1 | 2 | Ndst1 | 0.001 | Old |
| 0.920146 | 0.156185 | 0.026 | 0.025 | 1 | 2 | Tlr8 | 0.001 | Old |
| 0.920341 | 0.054481 | 0.045 | 0.044 | 1 | 2 | Stx2 | 0.001 | Old |
| 0.920398 | 0.05426 | 0.168 | 0.164 | 1 | 2 | Eif6 | 0.004 | Old |
| 0.920602 | 0.062446 | 0.658 | 0.664 | 1 | 2 | Serp1 | -0.006 | Old |
| 0.921276 | 0.005305 | 0.074 | 0.075 | 1 | 2 | Tgfbrap1 | -0.001 | Old |
| 0.921342 | 0.074243 | 0.117 | 0.115 | 1 | 2 | Bckdk | 0.002 | Old |
| 0.921498 | 0.090547 | 0.026 | 0.025 | 1 | 2 | Etfbkmt | 0.001 | Old |
| 0.922038 | 0.140279 | 0.065 | 0.064 | 1 | 2 | Rbm15 | 0.001 | Old |
| 0.922126 | 0.048404 | 0.147 | 0.144 | 1 | 2 | Shoc2 | 0.003 | Old |
| 0.922154 | 0.011984 | 0.051 | 0.052 | 1 | 2 | Gmcl1 | -0.001 | Old |
| 0.923968 | 0.206492 | 0.081 | 0.081 | 1 | 2 | Nrbf2 | 0 | Old |

| | | | | | | | | |
|---|---|---|---|---|---|---|---|---|
| 0.924453 | 0.172413 | 0.086 | 0.086 | 1 | 2 | Trmt1l | 0 | Old |
| 0.924468 | 0.20659 | 0.042 | 0.043 | 1 | 2 | Bcl11a | -0.001 | Old |
| 0.925392 | 0.029073 | 0.116 | 0.114 | 1 | 2 | Cnpy3 | 0.002 | Old |
| 0.925407 | 0.071199 | 0.198 | 0.197 | 1 | 2 | Etfa | 0.001 | Old |
| 0.925415 | 0.308092 | 0.017 | 0.017 | 1 | 2 | Gm42047 | 0 | Old |
| 0.925486 | 0.025759 | 0.054 | 0.055 | 1 | 2 | Heatr3 | -0.001 | Old |
| 0.926557 | 0.007118 | 0.126 | 0.124 | 1 | 2 | Rcsd1 | 0.002 | Old |
| 0.927184 | 0.29812 | 0.016 | 0.017 | 1 | 2 | Tmem115 | -0.001 | Old |
| 0.928516 | 0.032493 | 0.579 | 0.572 | 1 | 2 | Fcgr2b | 0.007 | Old |
| 0.928826 | 0.379705 | 0.028 | 0.027 | 1 | 2 | Toe1 | 0.001 | Old |
| 0.929116 | 0.312605 | 0.042 | 0.043 | 1 | 2 | Eef2kmt | -0.001 | Old |
| 0.929125 | 0.124293 | 0.03 | 0.031 | 1 | 2 | Slc35a1 | -0.001 | Old |
| 0.930188 | 0.020006 | 0.182 | 0.178 | 1 | 2 | Vps36 | 0.004 | Old |
| 0.930779 | 2.54E-05 | 0.495 | 0.493 | 1 | 2 | Tmcc1 | 0.002 | Old |
| 0.930853 | 0.009267 | 0.423 | 0.42 | 1 | 2 | Tram1 | 0.003 | Old |
| 0.930906 | 0.082802 | 0.108 | 0.109 | 1 | 2 | Srsf4 | -0.001 | Old |
| 0.931902 | 0.090629 | 0.349 | 0.347 | 1 | 2 | Mapre2 | 0.002 | Old |
| 0.932403 | 0.246341 | 0.094 | 0.094 | 1 | 2 | Gemin7 | 0 | Old |
| 0.93242 | 0.048462 | 0.035 | 0.034 | 1 | 2 | Rnf113a2 | 0.001 | Old |
| 0.93324 | 0.343695 | 0.028 | 0.027 | 1 | 2 | Fzd1 | 0.001 | Old |
| 0.933662 | 0.107373 | 0.075 | 0.074 | 1 | 2 | Plekha3 | 0.001 | Old |
| 0.933735 | 0.222279 | 0.077 | 0.078 | 1 | 2 | Ptp4a3 | -0.001 | Old |
| 0.933746 | 0.108268 | 0.037 | 0.037 | 1 | 2 | Gins4 | 0 | Old |
| 0.933837 | 0.148831 | 0.12 | 0.12 | 1 | 2 | Dnttip2 | 0 | Old |
| 0.933974 | 0.181405 | 0.058 | 0.057 | 1 | 2 | Yod1 | 0.001 | Old |
| 0.934496 | 0.336672 | 0.04 | 0.039 | 1 | 2 | 2010315B0 | 0.001 | Old |
| 0.934751 | 0.016681 | 0.18 | 0.178 | 1 | 2 | St3gal5 | 0.002 | Old |
| 0.934912 | 0.022254 | 0.345 | 0.34 | 1 | 2 | Nup98 | 0.005 | Old |
| 0.934993 | 0.196147 | 0.052 | 0.053 | 1 | 2 | Pck2 | -0.001 | Old |
| 0.935278 | 0.107818 | 0.052 | 0.053 | 1 | 2 | Ephx1 | -0.001 | Old |
| 0.935315 | 0.17793 | 0.017 | 0.017 | 1 | 2 | Map3k10 | 0 | Old |
| 0.936193 | 0.057402 | 0.063 | 0.064 | 1 | 2 | B9d2 | -0.001 | Old |
| 0.93696 | 0.103772 | 0.123 | 0.122 | 1 | 2 | Sf3a3 | 0.001 | Old |
| 0.937101 | 0.210434 | 0.049 | 0.049 | 1 | 2 | Pus10 | 0 | Old |
| 0.937135 | 0.042166 | 0.028 | 0.027 | 1 | 2 | B4gat1 | 0.001 | Old |
| 0.93814 | 0.084034 | 0.015 | 0.015 | 1 | 2 | Nup35 | 0 | Old |
| 0.93844 | 0.147713 | 0.081 | 0.081 | 1 | 2 | Ccdc6 | 0 | Old |
| 0.938488 | 0.013017 | 0.015 | 0.015 | 1 | 2 | Endov | 0 | Old |
| 0.938824 | 0.111537 | 0.073 | 0.074 | 1 | 2 | Atrn | -0.001 | Old |
| 0.939039 | 0.074703 | 0.232 | 0.234 | 1 | 2 | Frrs1 | -0.002 | Old |
| 0.939061 | 0.118545 | 0.079 | 0.078 | 1 | 2 | Stk19 | 0.001 | Old |
| 0.940425 | 0.007114 | 0.787 | 0.803 | 1 | 2 | Irf2bp2 | -0.016 | Old |
| 0.941349 | 0.079976 | 0.178 | 0.179 | 1 | 2 | Wdfy4 | -0.001 | Old |
| 0.941383 | 0.021563 | 0.124 | 0.125 | 1 | 2 | Smarcad1 | -0.001 | Old |
| 0.94215 | 0.014208 | 0.137 | 0.135 | 1 | 2 | Pstpip1 | 0.002 | Old |

| | | | | | | | | |
|---|---|---|---|---|---|---|---|---|
| 0.942508 | 0.344687 | 0.028 | 0.029 | 1 | 2 | Fdxacb1 | -0.001 | Old |
| 0.942942 | 0.397564 | 0.103 | 0.105 | 1 | 2 | Osgin2 | -0.002 | Old |
| 0.943458 | 0.251077 | 0.027 | 0.027 | 1 | 2 | Coq9 | 0 | Old |
| 0.943569 | 0.420146 | 0.018 | 0.018 | 1 | 2 | Ice2 | 0 | Old |
| 0.944343 | 0.060395 | 0.076 | 0.075 | 1 | 2 | Snhg12 | 0.001 | Old |
| 0.944377 | 0.028662 | 0.027 | 0.027 | 1 | 2 | Gm43660 | 0 | Old |
| 0.944407 | 0.041588 | 0.407 | 0.4 | 1 | 2 | 7-Sep | 0.007 | Old |
| 0.944461 | 0.076332 | 0.172 | 0.171 | 1 | 2 | Rsbn1l | 0.001 | Old |
| 0.944751 | 0.717852 | 0.015 | 0.015 | 1 | 2 | Faap24 | 0 | Old |
| 0.944797 | 0.082872 | 0.05 | 0.051 | 1 | 2 | Cars2 | -0.001 | Old |
| 0.945055 | 0.027696 | 0.037 | 0.038 | 1 | 2 | Gprasp1 | -0.001 | Old |
| 0.945132 | 0.010638 | 0.329 | 0.328 | 1 | 2 | Irf1 | 0.001 | Old |
| 0.945962 | 0.015795 | 0.084 | 0.082 | 1 | 2 | Ate1 | 0.002 | Old |
| 0.946245 | 0.150548 | 0.119 | 0.118 | 1 | 2 | Zfp385a | 0.001 | Old |
| 0.946259 | 0.199566 | 0.059 | 0.058 | 1 | 2 | Dmxl2 | 0.001 | Old |
| 0.946652 | 0.020028 | 0.522 | 0.51 | 1 | 2 | Pafah1b1 | 0.012 | Old |
| 0.946839 | 0.17818 | 0.015 | 0.015 | 1 | 2 | 1810024B0 | 0 | Old |
| 0.948371 | 0.113759 | 0.018 | 0.018 | 1 | 2 | Tfcp2 | 0 | Old |
| 0.948765 | 0.162485 | 0.014 | 0.014 | 1 | 2 | Kctd9 | 0 | Old |
| 0.948927 | 0.042979 | 0.273 | 0.267 | 1 | 2 | Igsf6 | 0.006 | Old |
| 0.949114 | 0.094826 | 0.051 | 0.052 | 1 | 2 | Cdt1 | -0.001 | Old |
| 0.949236 | 0.30175 | 0.051 | 0.051 | 1 | 2 | Nt5c3 | 0 | Old |
| 0.949271 | 0.26441 | 0.037 | 0.037 | 1 | 2 | Mlycd | 0 | Old |
| 0.949327 | 0.2358 | 0.049 | 0.049 | 1 | 2 | Nuak1 | 0 | Old |
| 0.949624 | 0.448684 | 0.015 | 0.015 | 1 | 2 | Rfx2 | 0 | Old |
| 0.949666 | 0.018375 | 0.118 | 0.119 | 1 | 2 | Aim2 | -0.001 | Old |
| 0.95125 | 0.030553 | 0.521 | 0.517 | 1 | 2 | Gpx4 | 0.004 | Old |
| 0.951254 | 0.175332 | 0.018 | 0.018 | 1 | 2 | Tyw5 | 0 | Old |
| 0.951784 | 0.024785 | 0.149 | 0.146 | 1 | 2 | Yipf5 | 0.003 | Old |
| 0.951816 | 0.15093 | 0.052 | 0.053 | 1 | 2 | Rrn3 | -0.001 | Old |
| 0.952156 | 0.003291 | 0.249 | 0.248 | 1 | 2 | Stom | 0.001 | Old |
| 0.953015 | 0.00035 | 0.155 | 0.157 | 1 | 2 | Tsen34 | -0.002 | Old |
| 0.953386 | 0.567934 | 0.029 | 0.029 | 1 | 2 | Prkdc | 0 | Old |
| 0.95371 | 0.220984 | 0.066 | 0.068 | 1 | 2 | Tmem260 | -0.002 | Old |
| 0.953816 | 0.063269 | 0.018 | 0.018 | 1 | 2 | BC002059 | 0 | Old |
| 0.954017 | 0.202243 | 0.163 | 0.164 | 1 | 2 | Rchy1 | -0.001 | Old |
| 0.954313 | 0.011272 | 0.207 | 0.205 | 1 | 2 | Sgk1 | 0.002 | Old |
| 0.954347 | 0.383033 | 0.041 | 0.042 | 1 | 2 | Habp4 | -0.001 | Old |
| 0.95439 | 0.101099 | 0.077 | 0.078 | 1 | 2 | Ube2g1 | -0.001 | Old |
| 0.954682 | 0.02381 | 0.162 | 0.162 | 1 | 2 | Skiv2l | 0 | Old |
| 0.955024 | 0.285482 | 0.06 | 0.061 | 1 | 2 | Pkib | -0.001 | Old |
| 0.955739 | 0.434546 | 0.018 | 0.018 | 1 | 2 | Tiam2 | 0 | Old |
| 0.957084 | 0.417844 | 0.014 | 0.014 | 1 | 2 | Dynlt1a | 0 | Old |
| 0.957322 | 0.01412 | 0.427 | 0.431 | 1 | 2 | Eps8 | -0.004 | Old |
| 0.957429 | 0.10358 | 0.026 | 0.026 | 1 | 2 | Tdrd3 | 0 | Old |

| | | | | | | | | |
|---|---|---|---|---|---|---|---|---|
| 0.957804 | 0.042557 | 0.183 | 0.183 | 1 | 2 | Med15 | 0 | Old |
| 0.957837 | 0.072189 | 0.187 | 0.186 | 1 | 2 | Mark3 | 0.001 | Old |
| 0.957998 | 0.004838 | 0.334 | 0.329 | 1 | 2 | Psma4 | 0.005 | Old |
| 0.95802 | 0.084345 | 0.133 | 0.132 | 1 | 2 | Tuba1a | 0.001 | Old |
| 0.958315 | 0.358247 | 0.091 | 0.093 | 1 | 2 | Thra | -0.002 | Old |
| 0.958712 | 0.059576 | 0.014 | 0.014 | 1 | 2 | Gorasp1 | 0 | Old |
| 0.958851 | 0.265591 | 0.05 | 0.05 | 1 | 2 | Kif1bp | 0 | Old |
| 0.95901 | 0.158848 | 0.132 | 0.132 | 1 | 2 | Rab12 | 0 | Old |
| 0.959037 | 0.151351 | 0.063 | 0.064 | 1 | 2 | Ubqln4 | -0.001 | Old |
| 0.960089 | 0.033114 | 0.047 | 0.046 | 1 | 2 | Nemp1 | 0.001 | Old |
| 0.961556 | 0.026632 | 0.099 | 0.099 | 1 | 2 | Scamp1 | 0 | Old |
| 0.962407 | 0.079974 | 0.123 | 0.124 | 1 | 2 | 3110043O2 | -0.001 | Old |
| 0.963101 | 0.145268 | 0.019 | 0.019 | 1 | 2 | B930036N1 | 0 | Old |
| 0.963265 | 0.224823 | 0.03 | 0.03 | 1 | 2 | Fam46c | 0 | Old |
| 0.963276 | 0.262325 | 0.115 | 0.114 | 1 | 2 | Trim12a | 0.001 | Old |
| 0.963312 | 0.044476 | 0.123 | 0.124 | 1 | 2 | Sf3a1 | -0.001 | Old |
| 0.963492 | 0.084129 | 0.025 | 0.025 | 1 | 2 | Pex1 | 0 | Old |
| 0.964127 | 0.162514 | 0.383 | 0.382 | 1 | 2 | Cdkn1b | 0.001 | Old |
| 0.964792 | 0.065589 | 0.095 | 0.095 | 1 | 2 | Spr | 0 | Old |
| 0.965059 | 0.247411 | 0.062 | 0.062 | 1 | 2 | Kif13b | 0 | Old |
| 0.965757 | 0.013808 | 0.238 | 0.233 | 1 | 2 | Cep350 | 0.005 | Old |
| 0.96622 | 0.017425 | 0.013 | 0.013 | 1 | 2 | Ptgs2os2 | 0 | Old |
| 0.966472 | 1.362086 | 0.01 | 0.01 | 1 | 2 | Cxcr2 | 0 | Old |
| 0.966554 | 0.019902 | 0.075 | 0.075 | 1 | 2 | Diablo | 0 | Old |
| 0.966689 | 0.003897 | 0.264 | 0.264 | 1 | 2 | Rbm47 | 0 | Old |
| 0.966947 | 0.069828 | 0.179 | 0.178 | 1 | 2 | Bbip1 | 0.001 | Old |
| 0.967109 | 0.672538 | 0.027 | 0.027 | 1 | 2 | Ccdc112 | 0 | Old |
| 0.967187 | 0.085347 | 0.122 | 0.119 | 1 | 2 | Dicer1 | 0.003 | Old |
| 0.96853 | 0.108035 | 0.036 | 0.037 | 1 | 2 | Ccl25 | -0.001 | Old |
| 0.968757 | 0.045466 | 0.059 | 0.058 | 1 | 2 | Bicral | 0.001 | Old |
| 0.96886 | 0.04391 | 0.013 | 0.013 | 1 | 2 | Fer | 0 | Old |
| 0.968962 | 0.56065 | 0.02 | 0.02 | 1 | 2 | Tnfsf13b | 0 | Old |
| 0.969105 | 0.024771 | 0.591 | 0.583 | 1 | 2 | Aplp2 | 0.008 | Old |
| 0.969398 | 0.028413 | 0.123 | 0.121 | 1 | 2 | 5-Mar | 0.002 | Old |
| 0.969526 | 0.188123 | 0.058 | 0.058 | 1 | 2 | Rrm1 | 0 | Old |
| 0.969892 | 0.005811 | 0.048 | 0.049 | 1 | 2 | 2410089E0 | -0.001 | Old |
| 0.969992 | 0.170338 | 0.013 | 0.013 | 1 | 2 | Ddah2 | 0 | Old |
| 0.97012 | 0.27574 | 0.036 | 0.037 | 1 | 2 | Phospho2 | -0.001 | Old |
| 0.970328 | 0.159025 | 0.037 | 0.038 | 1 | 2 | Oxsm | -0.001 | Old |
| 0.970514 | 0.180302 | 0.106 | 0.106 | 1 | 2 | AI413582 | 0 | Old |
| 0.970902 | 0.047575 | 0.09 | 0.09 | 1 | 2 | Asl | 0 | Old |
| 0.971193 | 0.180154 | 0.019 | 0.019 | 1 | 2 | Usp2 | 0 | Old |
| 0.971232 | 0.244368 | 0.02 | 0.02 | 1 | 2 | Il1r2 | 0 | Old |
| 0.971533 | 0.09471 | 0.034 | 0.034 | 1 | 2 | Cd3eap | 0 | Old |
| 0.972605 | 0.444884 | 0.01 | 0.01 | 1 | 2 | Tpmt | 0 | Old |

| | | | | | | | | |
|---|---|---|---|---|---|---|---|---|
| 0.972605 | 0.218249 | 0.01 | 0.01 | 1 | 2 | Agrn | 0 | Old |
| 0.973017 | 0.03177 | 0.078 | 0.078 | 1 | 2 | Ranbp10 | 0 | Old |
| 0.973158 | 0.65227 | 0.027 | 0.027 | 1 | 2 | Rad9a | 0 | Old |
| 0.973372 | 0.018075 | 0.019 | 0.019 | 1 | 2 | Zfp938 | 0 | Old |
| 0.973481 | 0.455069 | 0.01 | 0.01 | 1 | 2 | B3gnt2 | 0 | Old |
| 0.974142 | 0.392262 | 0.013 | 0.013 | 1 | 2 | Asrgl1 | 0 | Old |
| 0.97419 | 0.032079 | 0.05 | 0.05 | 1 | 2 | Oma1 | 0 | Old |
| 0.974258 | 0.209536 | 0.031 | 0.031 | 1 | 2 | Zdhhc14 | 0 | Old |
| 0.974354 | 0.235954 | 0.077 | 0.077 | 1 | 2 | Sigmar1 | 0 | Old |
| 0.974371 | 0.000583 | 0.229 | 0.228 | 1 | 2 | Ndufs7 | 0.001 | Old |
| 0.974457 | 0.501816 | 0.032 | 0.032 | 1 | 2 | Brms1l | 0 | Old |
| 0.974544 | 0.062032 | 0.03 | 0.03 | 1 | 2 | Trpm4 | 0 | Old |
| 0.975224 | 0.183208 | 0.059 | 0.059 | 1 | 2 | Rabgap1l | 0 | Old |
| 0.976095 | 0.179107 | 0.073 | 0.072 | 1 | 2 | AC163018. | 0.001 | Old |
| 0.97611 | 0.185804 | 0.01 | 0.01 | 1 | 2 | Tmem267 | 0 | Old |
| 0.976249 | 0.072845 | 0.084 | 0.083 | 1 | 2 | Fam43a | 0.001 | Old |
| 0.97626 | 0.126382 | 0.207 | 0.203 | 1 | 2 | Psmd1 | 0.004 | Old |
| 0.976299 | 0.247525 | 0.042 | 0.042 | 1 | 2 | Ccs | 0 | Old |
| 0.976549 | 0.088851 | 0.01 | 0.01 | 1 | 2 | Rsl1 | 0 | Old |
| 0.976811 | 0.261053 | 0.105 | 0.107 | 1 | 2 | Ubac2 | -0.002 | Old |
| 0.976863 | 0.244453 | 0.064 | 0.065 | 1 | 2 | AI606181 | -0.001 | Old |
| 0.977242 | 0.059573 | 0.6 | 0.591 | 1 | 2 | Kdm7a | 0.009 | Old |
| 0.977425 | 0.184679 | 0.01 | 0.01 | 1 | 2 | Col3a1 | 0 | Old |
| 0.977863 | 0.090614 | 0.01 | 0.01 | 1 | 2 | Dlg3 | 0 | Old |
| 0.97891 | 0.186497 | 0.057 | 0.057 | 1 | 2 | Gpr155 | 0 | Old |
| 0.979736 | 0.026566 | 0.153 | 0.152 | 1 | 2 | Fuca2 | 0.001 | Old |
| 0.979752 | 0.057243 | 0.103 | 0.103 | 1 | 2 | Snrnp40 | 0 | Old |
| 0.980181 | 0.884591 | 0.013 | 0.013 | 1 | 2 | Arhgap24 | 0 | Old |
| 0.980313 | 0.08814 | 0.089 | 0.089 | 1 | 2 | Ccdc186 | 0 | Old |
| 0.980405 | 0.090447 | 0.279 | 0.278 | 1 | 2 | Hprt | 0.001 | Old |
| 0.981525 | 0.125853 | 0.02 | 0.02 | 1 | 2 | Hist1h1e | 0 | Old |
| 0.981828 | 0.153947 | 0.02 | 0.02 | 1 | 2 | Ghdc | 0 | Old |
| 0.982301 | 0.031611 | 0.775 | 0.771 | 1 | 2 | Grn | 0.004 | Old |
| 0.982478 | 0.001997 | 0.955 | 0.971 | 1 | 2 | Rps28 | -0.016 | Old |
| 0.982823 | 0.956892 | 0.013 | 0.013 | 1 | 2 | Gsdme | 0 | Old |
| 0.982953 | 0.011543 | 0.365 | 0.36 | 1 | 2 | Matr3 | 0.005 | Old |
| 0.983371 | 0.043071 | 0.031 | 0.031 | 1 | 2 | Dtx3 | 0 | Old |
| 0.983599 | 0.111929 | 0.054 | 0.053 | 1 | 2 | Fbxo7 | 0.001 | Old |
| 0.984008 | 0.092977 | 0.012 | 0.012 | 1 | 2 | Cenpj | 0 | Old |
| 0.984886 | 0.136526 | 0.034 | 0.034 | 1 | 2 | Nbn | 0 | Old |
| 0.985587 | 0.164802 | 0.012 | 0.012 | 1 | 2 | Fuz | 0 | Old |
| 0.985866 | 0.069007 | 0.249 | 0.247 | 1 | 2 | Apobec3 | 0.002 | Old |
| 0.985922 | 0.071489 | 0.076 | 0.076 | 1 | 2 | Slc25a12 | 0 | Old |
| 0.985982 | 0.29585 | 0.012 | 0.012 | 1 | 2 | 2010320M | 0 | Old |
| 0.986297 | 0.195827 | 0.112 | 0.112 | 1 | 2 | Vps11 | 0 | Old |

| | | | | | | | | |
|---|---|---|---|---|---|---|---|---|
| 0.986319 | 0.052522 | 0.045 | 0.045 | 1 | 2 | Wdsub1 | 0 | Old |
| 0.986795 | 0.045559 | 0.704 | 0.724 | 1 | 2 | Lsp1 | -0.02 | Old |
| 0.987166 | 0.596291 | 0.012 | 0.012 | 1 | 2 | Inhba | 0 | Old |
| 0.987345 | 0.527326 | 0.011 | 0.011 | 1 | 2 | Aff3 | 0 | Old |
| 0.987345 | 0.452275 | 0.011 | 0.011 | 1 | 2 | E130311K1 | 0 | Old |
| 0.987561 | 0.04262 | 0.012 | 0.012 | 1 | 2 | Pcnx4 | 0 | Old |
| 0.987561 | 0.023669 | 0.012 | 0.012 | 1 | 2 | Exo5 | 0 | Old |
| 0.988084 | 0.014115 | 0.391 | 0.385 | 1 | 2 | Serinc1 | 0.006 | Old |
| 0.988351 | 0.106942 | 0.012 | 0.012 | 1 | 2 | Ms4a14 | 0 | Old |
| 0.988763 | 0.125946 | 0.057 | 0.057 | 1 | 2 | Arhgap10 | 0 | Old |
| 0.989849 | 0.04209 | 0.133 | 0.133 | 1 | 2 | Tm9sf4 | 0 | Old |
| 0.98993 | 0.18762 | 0.012 | 0.012 | 1 | 2 | Gm43672 | 0 | Old |
| 0.990005 | 0.009351 | 0.02 | 0.02 | 1 | 2 | Fbxl19 | 0 | Old |
| 0.990219 | 0.146584 | 0.022 | 0.023 | 1 | 2 | Ms4a4b | -0.001 | Old |
| 0.990524 | 0.088175 | 0.063 | 0.063 | 1 | 2 | D430042O( | 0 | Old |
| 0.990615 | 0.10668 | 0.051 | 0.051 | 1 | 2 | Pcnx3 | 0 | Old |
| 0.990656 | 0.321276 | 0.035 | 0.036 | 1 | 2 | P2ry1 | -0.001 | Old |
| 0.990899 | 0.042455 | 0.281 | 0.28 | 1 | 2 | Necap2 | 0.001 | Old |
| 0.99118 | 0.039326 | 0.123 | 0.122 | 1 | 2 | Stim2 | 0.001 | Old |
| 0.991458 | 0.160615 | 0.042 | 0.042 | 1 | 2 | Slc10a3 | 0 | Old |
| 0.99151 | 0.668864 | 0.012 | 0.012 | 1 | 2 | Fchsd1 | 0 | Old |
| 0.991668 | 0.081277 | 0.052 | 0.052 | 1 | 2 | Atn1 | 0 | Old |
| 0.991707 | 0.277891 | 0.012 | 0.012 | 1 | 2 | Nradd | 0 | Old |
| 0.991716 | 0.031935 | 0.313 | 0.311 | 1 | 2 | Herpud1 | 0.002 | Old |
| 0.992287 | 0.006734 | 0.084 | 0.083 | 1 | 2 | Ergic1 | 0.001 | Old |
| 0.992404 | 0.04277 | 0.335 | 0.327 | 1 | 2 | Fcho2 | 0.008 | Old |
| 0.992746 | 0.037646 | 0.1 | 0.1 | 1 | 2 | AW549877 | 0 | Old |
| 0.993189 | 0.082351 | 0.214 | 0.213 | 1 | 2 | Ndufb4 | 0.001 | Old |
| 0.993713 | 0.029128 | 0.264 | 0.265 | 1 | 2 | Srebf2 | -0.001 | Old |
| 0.993879 | 0.488665 | 0.012 | 0.012 | 1 | 2 | Olfr920 | 0 | Old |
| 0.994024 | 0.152459 | 0.034 | 0.034 | 1 | 2 | Cbx7 | 0 | Old |
| 0.994126 | 0.105936 | 0.224 | 0.224 | 1 | 2 | Rasgrp4 | 0 | Old |
| 0.994247 | 0.132665 | 0.022 | 0.023 | 1 | 2 | Zfp944 | -0.001 | Old |
| 0.994377 | 0.259117 | 0.044 | 0.044 | 1 | 2 | 1600010M | 0 | Old |
| 0.994529 | 0.085877 | 0.625 | 0.622 | 1 | 2 | Pcbp1 | 0.003 | Old |
| 0.995002 | 0.037863 | 0.044 | 0.044 | 1 | 2 | Dvl2 | 0 | Old |
| 0.995138 | 0.054551 | 0.148 | 0.146 | 1 | 2 | Katna1 | 0.002 | Old |
| 0.995564 | 0.032381 | 0.041 | 0.04 | 1 | 2 | Sepsecs | 0.001 | Old |
| 0.995643 | 0.118465 | 0.011 | 0.011 | 1 | 2 | Chst14 | 0 | Old |
| 0.995643 | 0.342822 | 0.011 | 0.011 | 1 | 2 | Alkbh2 | 0 | Old |
| 0.996058 | 0.629484 | 0.011 | 0.011 | 1 | 2 | Tmem140 | 0 | Old |
| 0.99606 | 0.01958 | 0.139 | 0.138 | 1 | 2 | Ubqln1 | 0.001 | Old |
| 0.996099 | 0.078541 | 0.043 | 0.043 | 1 | 2 | Calcoco1 | 0 | Old |
| 0.996105 | 0.087633 | 0.202 | 0.203 | 1 | 2 | Fbxl3 | -0.001 | Old |
| 0.996171 | 0.010712 | 0.154 | 0.153 | 1 | 2 | Atg101 | 0.001 | Old |

| | | | | | | | | |
|---|---|---|---|---|---|---|---|---|
| 0.996247 | 0.001554 | 0.155 | 0.156 | 1 | 2 | Klhdc3 | -0.001 | Old |
| 0.996473 | 0.078266 | 0.011 | 0.011 | 1 | 2 | Gm14326 | 0 | Old |
| 0.997093 | 0.184882 | 0.071 | 0.071 | 1 | 2 | Zadh2 | 0 | Old |
| 0.997183 | 0.015554 | 0.096 | 0.096 | 1 | 2 | D10Jhu81e | 0 | Old |
| 0.997659 | 0.009233 | 0.223 | 0.22 | 1 | 2 | Hnrnpr | 0.003 | Old |
| 0.997718 | 0.044779 | 0.011 | 0.011 | 1 | 2 | Zhx3 | 0 | Old |
| 0.99784 | 0.218859 | 0.045 | 0.045 | 1 | 2 | Daam1 | 0 | Old |
| 0.998282 | 0.025485 | 0.165 | 0.164 | 1 | 2 | Cr1l | 0.001 | Old |
| 0.999378 | 0.195203 | 0.011 | 0.011 | 1 | 2 | Gins1 | 0 | Old |
| 0.999699 | 0.027095 | 0.363 | 0.359 | 1 | 2 | Hnrnpl | 0.004 | Old |
| 5.79E-19 | 0.345838 | 0.954 | 0.928 | 1.80E-14 | 2 | Rpl30 | 0.026 | Young |
| 6.01E-19 | 0.312762 | 0.964 | 0.952 | 1.87E-14 | 2 | Rps12 | 0.012 | Young |
| 7.18E-19 | 0.33513 | 0.96 | 0.95 | 2.23E-14 | 2 | Rps14 | 0.01 | Young |
| 3.38E-17 | 1.972781 | 0.283 | 0.132 | 1.05E-12 | 2 | Hspa1b | 0.151 | Young |
| 1.34E-15 | 0.393307 | 0.907 | 0.86 | 4.16E-11 | 2 | Rpl34 | 0.047 | Young |
| 2.36E-15 | 0.447445 | 0.879 | 0.797 | 7.31E-11 | 2 | Rpl32 | 0.082 | Young |
| 2.99E-15 | 0.251976 | 0.985 | 0.975 | 9.29E-11 | 2 | Rps24 | 0.01 | Young |
| 6.16E-15 | 0.341062 | 0.955 | 0.93 | 1.91E-10 | 2 | Rps16 | 0.025 | Young |
| 8.28E-15 | 0.29445 | 0.964 | 0.936 | 2.57E-10 | 2 | Rps20 | 0.028 | Young |
| 2.35E-14 | 0.243814 | 0.983 | 0.972 | 7.29E-10 | 2 | Rplp1 | 0.011 | Young |
| 1.10E-13 | 0.402013 | 0.888 | 0.808 | 3.42E-09 | 2 | Rpl26 | 0.08 | Young |
| 2.69E-13 | 0.317092 | 0.955 | 0.923 | 8.36E-09 | 2 | Rps10 | 0.032 | Young |
| 1.10E-12 | 0.247854 | 0.983 | 0.974 | 3.41E-08 | 2 | Fau | 0.009 | Young |
| 3.19E-12 | 1.406314 | 0.295 | 0.167 | 9.92E-08 | 2 | Hspa1a | 0.128 | Young |
| 9.67E-12 | 0.312187 | 0.947 | 0.92 | 3.00E-07 | 2 | Rps19 | 0.027 | Young |
| 4.15E-11 | 0.332108 | 0.913 | 0.881 | 1.29E-06 | 2 | Rps13 | 0.032 | Young |
| 3.65E-10 | 0.371082 | 0.815 | 0.771 | 1.13E-05 | 2 | Rpl11 | 0.044 | Young |
| 4.77E-10 | 0.158379 | 0.994 | 0.994 | 1.48E-05 | 2 | Rps29 | 0 | Young |
| 6.74E-10 | 1.067258 | 0.238 | 0.126 | 2.09E-05 | 2 | Cd33 | 0.112 | Young |
| 8.79E-10 | 0.296828 | 0.904 | 0.85 | 2.73E-05 | 2 | Rps15a | 0.054 | Young |
| 1.94E-09 | 0.29158 | 0.92 | 0.897 | 6.04E-05 | 2 | Rpl18 | 0.023 | Young |
| 2.23E-09 | 0.357406 | 0.861 | 0.804 | 6.94E-05 | 2 | Rps6 | 0.057 | Young |
| 2.52E-09 | 1.171533 | 0.234 | 0.132 | 7.81E-05 | 2 | Gas6 | 0.102 | Young |
| 3.14E-09 | 0.296316 | 0.899 | 0.852 | 9.76E-05 | 2 | Rps5 | 0.047 | Young |
| 3.17E-09 | 0.2096 | 0.951 | 0.949 | 9.85E-05 | 2 | Rps23 | 0.002 | Young |
| 7.49E-09 | 0.225739 | 0.964 | 0.938 | 0.000233 | 2 | Rpl13 | 0.026 | Young |
| 1.02E-08 | 0.18543 | 0.983 | 0.974 | 0.000316 | 2 | Rps8 | 0.009 | Young |
| 1.02E-08 | 0.360661 | 0.791 | 0.711 | 0.000318 | 2 | Rpl12 | 0.08 | Young |
| 1.21E-08 | 0.271027 | 0.912 | 0.867 | 0.000377 | 2 | Rps11 | 0.045 | Young |
| 1.25E-08 | 0.274199 | 0.952 | 0.923 | 0.000389 | 2 | Rps4x | 0.029 | Young |
| 1.51E-08 | 0.173469 | 0.985 | 0.984 | 0.000468 | 2 | Rpl35a | 0.001 | Young |
| 1.53E-08 | 0.277342 | 0.916 | 0.885 | 0.000476 | 2 | Rpl18a | 0.031 | Young |
| 3.69E-08 | 1.40208 | 0.103 | 0.037 | 0.001145 | 2 | Slc15a2 | 0.066 | Young |
| 3.86E-08 | 0.392804 | 0.644 | 0.543 | 0.001199 | 2 | Rel | 0.101 | Young |
| 4.03E-08 | 0.397727 | 0.799 | 0.726 | 0.001252 | 2 | Nfkbiz | 0.073 | Young |

| | | | | | | | | |
|---|---|---|---|---|---|---|---|---|
| 1.27E-07 | 0.17869 | 0.974 | 0.961 | 0.00394 | 2 | Rps3a1 | 0.013 | Young |
| 1.68E-07 | 0.225484 | 0.974 | 0.96 | 0.005205 | 2 | Fth1 | 0.014 | Young |
| 1.96E-07 | 0.248528 | 0.891 | 0.854 | 0.006074 | 2 | Rps25 | 0.037 | Young |
| 2.26E-07 | 1.006537 | 0.255 | 0.165 | 0.007009 | 2 | Stab1 | 0.09 | Young |
| 2.32E-07 | 0.216758 | 0.952 | 0.913 | 0.007218 | 2 | Rpl37 | 0.039 | Young |
| 3.55E-07 | 0.853074 | 0.227 | 0.138 | 0.011032 | 2 | Gatm | 0.089 | Young |
| 4.23E-07 | 0.600894 | 0.483 | 0.38 | 0.01313 | 2 | Pf4 | 0.103 | Young |
| 4.89E-07 | 0.298231 | 0.831 | 0.785 | 0.015183 | 2 | Rpl36a | 0.046 | Young |
| 5.08E-07 | 0.226123 | 0.956 | 0.939 | 0.015768 | 2 | Rpl17 | 0.017 | Young |
| 5.17E-07 | 0.266474 | 0.873 | 0.825 | 0.016066 | 2 | Rpl28 | 0.048 | Young |
| 5.26E-07 | 0.226136 | 0.948 | 0.926 | 0.016342 | 2 | Rpl39 | 0.022 | Young |
| 6.71E-07 | 0.231572 | 0.909 | 0.865 | 0.020848 | 2 | Rpl9 | 0.044 | Young |
| 9.45E-07 | 0.230778 | 0.922 | 0.887 | 0.029348 | 2 | Rpl19 | 0.035 | Young |
| 1.14E-06 | 0.19294 | 0.948 | 0.93 | 0.035313 | 2 | Rpl27a | 0.018 | Young |
| 1.21E-06 | 0.005536 | 1 | 1 | 0.037494 | 2 | Gm42418 | 0 | Young |
| 1.48E-06 | 0.264817 | 0.822 | 0.806 | 0.046077 | 2 | Rpl36 | 0.016 | Young |
| 1.53E-06 | 0.397578 | 0.774 | 0.7 | 0.047563 | 2 | Cd14 | 0.074 | Young |
| 1.55E-06 | 1.559762 | 0.106 | 0.047 | 0.048 | 2 | Cd38 | 0.059 | Young |
| 1.63E-06 | 0.267911 | 0.86 | 0.818 | 0.050466 | 2 | Rps7 | 0.042 | Young |
| 1.73E-06 | 0.751934 | 0.327 | 0.236 | 0.053637 | 2 | Bcl2a1b | 0.091 | Young |
| 1.74E-06 | 0.762158 | 0.296 | 0.202 | 0.054055 | 2 | Cd81 | 0.094 | Young |
| 1.78E-06 | 0.342159 | 0.752 | 0.675 | 0.055371 | 2 | Rpl7a | 0.077 | Young |
| 2.06E-06 | 0.274432 | 0.847 | 0.781 | 0.06394 | 2 | Rpl35 | 0.066 | Young |
| 2.16E-06 | 1.488033 | 0.127 | 0.063 | 0.067094 | 2 | Cd209a | 0.064 | Young |
| 2.47E-06 | 0.14662 | 0.988 | 0.98 | 0.076834 | 2 | Rplp2 | 0.008 | Young |
| 2.80E-06 | 0.288995 | 0.8 | 0.756 | 0.086985 | 2 | Rps3 | 0.044 | Young |
| 2.86E-06 | 0.430417 | 0.544 | 0.46 | 0.088802 | 2 | Atpif1 | 0.084 | Young |
| 4.22E-06 | 0.755628 | 0.287 | 0.198 | 0.131034 | 2 | Fcrls | 0.089 | Young |
| 4.73E-06 | 1.610871 | 0.07 | 0.025 | 0.146833 | 2 | Plxdc2 | 0.045 | Young |
| 5.93E-06 | 0.812365 | 0.192 | 0.118 | 0.184244 | 2 | Hpgd | 0.074 | Young |
| 6.94E-06 | 2.563757 | 0.039 | 0.007 | 0.215414 | 2 | AA467197 | 0.032 | Young |
| 7.10E-06 | 0.242145 | 0.827 | 0.768 | 0.22036 | 2 | Rpl21 | 0.059 | Young |
| 7.30E-06 | 0.182803 | 0.937 | 0.914 | 0.226794 | 2 | Hspa8 | 0.023 | Young |
| 8.79E-06 | 0.168612 | 0.973 | 0.948 | 0.273013 | 2 | Cd74 | 0.025 | Young |
| 9.21E-06 | 0.055616 | 0.752 | 0.637 | 0.28593 | 2 | Retnla | 0.115 | Young |
| 1.02E-05 | 0.161003 | 0.979 | 0.961 | 0.315875 | 2 | Rps21 | 0.018 | Young |
| 1.02E-05 | 1.285777 | 0.1 | 0.046 | 0.316664 | 2 | Gpr34 | 0.054 | Young |
| 1.72E-05 | 0.309625 | 0.717 | 0.658 | 0.534036 | 2 | Rpl15 | 0.059 | Young |
| 1.90E-05 | 0.343502 | 0.613 | 0.534 | 0.590035 | 2 | Pnrc1 | 0.079 | Young |
| 1.95E-05 | 0.210237 | 0.743 | 0.688 | 0.605885 | 2 | Selenop | 0.055 | Young |
| 2.00E-05 | 0.162089 | 0.951 | 0.946 | 0.620172 | 2 | Rpl23 | 0.005 | Young |
| 2.60E-05 | 1.722711 | 0.05 | 0.015 | 0.806054 | 2 | Snta1 | 0.035 | Young |
| 2.82E-05 | 0.368759 | 0.575 | 0.484 | 0.875081 | 2 | Tlr2 | 0.091 | Young |
| 3.26E-05 | 1.108459 | 0.08 | 0.034 | 1 | 2 | Itga6 | 0.046 | Young |
| 3.55E-05 | 0.13203 | 0.996 | 0.991 | 1 | 2 | Rps9 | 0.005 | Young |

| | | | | | | | | |
|---|---|---|---|---|---|---|---|---|
| 3.63E-05 | 1.094855 | 0.086 | 0.039 | 1 | 2 | Tanc2 | 0.047 | Young |
| 4.01E-05 | 0.650658 | 0.322 | 0.246 | 1 | 2 | Lamtor2 | 0.076 | Young |
| 4.30E-05 | 0.590931 | 0.279 | 0.2 | 1 | 2 | Igfbp4 | 0.079 | Young |
| 4.85E-05 | 0.387892 | 0.518 | 0.44 | 1 | 2 | Spag9 | 0.078 | Young |
| 4.91E-05 | 0.367007 | 0.884 | 0.855 | 1 | 2 | Apoe | 0.029 | Young |
| 4.93E-05 | 0.26295 | 0.778 | 0.714 | 1 | 2 | Rpl10 | 0.064 | Young |
| 5.31E-05 | 0.126877 | 0.983 | 0.987 | 1 | 2 | Rps27a | -0.004 | Young |
| 5.31E-05 | 0.279248 | 0.689 | 0.604 | 1 | 2 | Gng5 | 0.085 | Young |
| 6.46E-05 | 0.30252 | 0.698 | 0.656 | 1 | 2 | Csf1r | 0.042 | Young |
| 6.76E-05 | 0.181816 | 0.961 | 0.948 | 1 | 2 | Ftl1 | 0.013 | Young |
| 7.99E-05 | 0.234139 | 0.761 | 0.712 | 1 | 2 | Rps18 | 0.049 | Young |
| 8.44E-05 | 0.902859 | 0.061 | 0.024 | 1 | 2 | P2ry12 | 0.037 | Young |
| 8.58E-05 | 1.764847 | 0.053 | 0.019 | 1 | 2 | Serpinf1 | 0.034 | Young |
| 9.87E-05 | 0.184482 | 0.918 | 0.88 | 1 | 2 | Rplp0 | 0.038 | Young |
| 0.000101 | 0.456594 | 0.308 | 0.23 | 1 | 2 | Abca1 | 0.078 | Young |
| 0.000138 | 0.549263 | 0.205 | 0.138 | 1 | 2 | Bin1 | 0.067 | Young |
| 0.000164 | 0.243194 | 0.831 | 0.785 | 1 | 2 | Cd83 | 0.046 | Young |
| 0.000166 | 1.277386 | 0.139 | 0.085 | 1 | 2 | Ltc4s | 0.054 | Young |
| 0.000172 | 0.350416 | 0.57 | 0.499 | 1 | 2 | Pfdn5 | 0.071 | Young |
| 0.000182 | 1.728704 | 0.036 | 0.01 | 1 | 2 | Atf5 | 0.026 | Young |
| 0.000194 | 0.267524 | 0.86 | 0.818 | 1 | 2 | Kctd12 | 0.042 | Young |
| 0.000222 | 0.784273 | 0.164 | 0.107 | 1 | 2 | Cttnbp2nl | 0.057 | Young |
| 0.000235 | 0.266688 | 0.822 | 0.778 | 1 | 2 | Tgfbi | 0.044 | Young |
| 0.000246 | 1.353573 | 0.056 | 0.022 | 1 | 2 | Rtn1 | 0.034 | Young |
| 0.000308 | 0.69063 | 0.065 | 0.029 | 1 | 2 | Cmah | 0.036 | Young |
| 0.000323 | 0.72802 | 0.31 | 0.252 | 1 | 2 | Ndufv3 | 0.058 | Young |
| 0.000343 | 0.233148 | 0.862 | 0.784 | 1 | 2 | Il1b | 0.078 | Young |
| 0.000354 | 0.80984 | 0.27 | 0.206 | 1 | 2 | Lyz1 | 0.064 | Young |
| 0.00038 | 0.342559 | 0.588 | 0.515 | 1 | 2 | Atox1 | 0.073 | Young |
| 0.000399 | 0.831414 | 0.121 | 0.073 | 1 | 2 | Cd163 | 0.048 | Young |
| 0.000403 | 0.437496 | 0.532 | 0.467 | 1 | 2 | Ier3 | 0.065 | Young |
| 0.000408 | 0.575642 | 0.154 | 0.1 | 1 | 2 | Fgfr1 | 0.054 | Young |
| 0.000417 | 0.550847 | 0.338 | 0.271 | 1 | 2 | Cxcl16 | 0.067 | Young |
| 0.000419 | 0.983083 | 0.091 | 0.049 | 1 | 2 | Dcaf5 | 0.042 | Young |
| 0.000433 | 1.009485 | 0.043 | 0.015 | 1 | 2 | Nedd4l | 0.028 | Young |
| 0.000437 | 0.281017 | 0.441 | 0.363 | 1 | 2 | Axl | 0.078 | Young |
| 0.000442 | 0.266482 | 0.583 | 0.507 | 1 | 2 | Srp9 | 0.076 | Young |
| 0.00045 | 0.177299 | 0.9 | 0.855 | 1 | 2 | Tyrobp | 0.045 | Young |
| 0.000455 | 0.321712 | 0.515 | 0.455 | 1 | 2 | Gadd45b | 0.06 | Young |
| 0.000477 | 1.817505 | 0.052 | 0.021 | 1 | 2 | Arg1 | 0.031 | Young |
| 0.000489 | 0.387119 | 0.504 | 0.446 | 1 | 2 | Txn1 | 0.058 | Young |
| 0.000496 | 0.547692 | 0.302 | 0.233 | 1 | 2 | Cited2 | 0.069 | Young |
| 0.000595 | 0.929779 | 0.049 | 0.019 | 1 | 2 | Abcc3 | 0.03 | Young |
| 0.000613 | 1.614393 | 0.025 | 0.005 | 1 | 2 | Chst11 | 0.02 | Young |
| 0.000619 | 0.204746 | 0.837 | 0.814 | 1 | 2 | Rpl10a | 0.023 | Young |

| | | | | | | | | |
|---|---|---|---|---|---|---|---|---|
| 0.00064 | 0.766703 | 0.1 | 0.057 | 1 | 2 | Slco2b1 | 0.043 | Young |
| 0.000652 | 0.689679 | 0.208 | 0.15 | 1 | 2 | Lpgat1 | 0.058 | Young |
| 0.000664 | 0.84746 | 0.076 | 0.039 | 1 | 2 | Chn2 | 0.037 | Young |
| 0.000716 | 1.841772 | 0.029 | 0.007 | 1 | 2 | Inpp5b | 0.022 | Young |
| 0.000727 | 0.183438 | 0.882 | 0.857 | 1 | 2 | Rpl24 | 0.025 | Young |
| 0.000765 | 1.827041 | 0.032 | 0.01 | 1 | 2 | Cfd | 0.022 | Young |
| 0.000808 | 0.287526 | 0.546 | 0.478 | 1 | 2 | Hint1 | 0.068 | Young |
| 0.00086 | 1.206589 | 0.044 | 0.017 | 1 | 2 | Ddx31 | 0.027 | Young |
| 0.000888 | 0.339002 | 0.595 | 0.546 | 1 | 2 | Ptpn18 | 0.049 | Young |
| 0.001034 | 1.327796 | 0.037 | 0.013 | 1 | 2 | Tsen15 | 0.024 | Young |
| 0.001088 | 0.443988 | 0.346 | 0.284 | 1 | 2 | Eif4g1 | 0.062 | Young |
| 0.001159 | 1.618063 | 0.014 | 0.001 | 1 | 2 | Smim10l2a | 0.013 | Young |
| 0.001192 | 0.563498 | 0.232 | 0.176 | 1 | 2 | Nr4a3 | 0.056 | Young |
| 0.001197 | 0.206 | 0.755 | 0.698 | 1 | 2 | Npm1 | 0.057 | Young |
| 0.001238 | 0.982875 | 0.084 | 0.047 | 1 | 2 | Cds1 | 0.037 | Young |
| 0.001295 | 1.28593 | 0.034 | 0.012 | 1 | 2 | Pmepa1 | 0.022 | Young |
| 0.001305 | 1.320298 | 0.031 | 0.01 | 1 | 2 | P3h2 | 0.021 | Young |
| 0.001371 | 0.420327 | 0.45 | 0.384 | 1 | 2 | Sdc4 | 0.066 | Young |
| 0.001459 | 0.284695 | 0.539 | 0.465 | 1 | 2 | Arl4c | 0.074 | Young |
| 0.00147 | 0.56185 | 0.183 | 0.13 | 1 | 2 | Ccl24 | 0.053 | Young |
| 0.001481 | 0.298361 | 0.384 | 0.313 | 1 | 2 | Mgl2 | 0.071 | Young |
| 0.001577 | 0.417874 | 0.385 | 0.327 | 1 | 2 | Ifngr2 | 0.058 | Young |
| 0.001615 | 0.226714 | 0.589 | 0.53 | 1 | 2 | Marcksl1 | 0.059 | Young |
| 0.00162 | 0.290194 | 0.571 | 0.515 | 1 | 2 | Nfe2l2 | 0.056 | Young |
| 0.001648 | 0.451227 | 0.164 | 0.112 | 1 | 2 | Lpin2 | 0.052 | Young |
| 0.001682 | 1.281074 | 0.062 | 0.031 | 1 | 2 | Ap2a1 | 0.031 | Young |
| 0.001693 | 0.186303 | 0.688 | 0.639 | 1 | 2 | Atp5h | 0.049 | Young |
| 0.001814 | 0.193791 | 0.809 | 0.802 | 1 | 2 | Rps17 | 0.007 | Young |
| 0.001856 | 0.180299 | 0.705 | 0.655 | 1 | 2 | Eef2 | 0.05 | Young |
| 0.001971 | 0.596724 | 0.02 | 0.004 | 1 | 2 | Slc40a1 | 0.016 | Young |
| 0.002035 | 0.189757 | 0.871 | 0.836 | 1 | 2 | Rpl3 | 0.035 | Young |
| 0.002114 | 0.154088 | 0.878 | 0.849 | 1 | 2 | Cox4i1 | 0.029 | Young |
| 0.002125 | 0.461828 | 0.39 | 0.337 | 1 | 2 | Nlrp3 | 0.053 | Young |
| 0.002268 | 0.78678 | 0.058 | 0.029 | 1 | 2 | Pak4 | 0.029 | Young |
| 0.002301 | 0.176085 | 0.835 | 0.794 | 1 | 2 | Rps2 | 0.041 | Young |
| 0.002392 | 0.80398 | 0.071 | 0.039 | 1 | 2 | Zfp704 | 0.032 | Young |
| 0.002499 | 0.179211 | 0.847 | 0.8 | 1 | 2 | Uqcrh | 0.047 | Young |
| 0.002538 | 0.12702 | 0.951 | 0.936 | 1 | 2 | Rps15 | 0.015 | Young |
| 0.002715 | 0.677404 | 0.053 | 0.026 | 1 | 2 | Rhobtb1 | 0.027 | Young |
| 0.002795 | 0.331404 | 0.12 | 0.078 | 1 | 2 | Cks2 | 0.042 | Young |
| 0.00281 | 0.834306 | 0.068 | 0.036 | 1 | 2 | Spp1 | 0.032 | Young |
| 0.002823 | 0.414911 | 0.148 | 0.102 | 1 | 2 | 1700017B0 | 0.046 | Young |
| 0.00284 | 1.754187 | 0.036 | 0.014 | 1 | 2 | Arg2 | 0.022 | Young |
| 0.002919 | 1.355412 | 0.021 | 0.005 | 1 | 2 | Colec12 | 0.016 | Young |
| 0.002973 | 0.521462 | 0.304 | 0.249 | 1 | 2 | Tnfsf9 | 0.055 | Young |

| | | | | | | | | |
|---|---|---|---|---|---|---|---|---|
| 0.003011 | 0.514473 | 0.167 | 0.119 | 1 | 2 | St6gal1 | 0.048 | Young |
| 0.003212 | 0.664548 | 0.125 | 0.082 | 1 | 2 | Tagap | 0.043 | Young |
| 0.003222 | 0.919245 | 0.072 | 0.041 | 1 | 2 | Zfp266 | 0.031 | Young |
| 0.003246 | 1.26846 | 0.059 | 0.031 | 1 | 2 | Lyve1 | 0.028 | Young |
| 0.00326 | 0.221367 | 0.751 | 0.717 | 1 | 2 | Plek | 0.034 | Young |
| 0.003266 | 0.074762 | 0.993 | 0.987 | 1 | 2 | Tpt1 | 0.006 | Young |
| 0.003393 | 1.518806 | 0.019 | 0.004 | 1 | 2 | Klhl25 | 0.015 | Young |
| 0.003665 | 1.0767 | 0.089 | 0.055 | 1 | 2 | Nacc1 | 0.034 | Young |
| 0.003844 | 1.368943 | 0.038 | 0.016 | 1 | 2 | Fmnl3 | 0.022 | Young |
| 0.003854 | 1.372011 | 0.056 | 0.029 | 1 | 2 | Car3 | 0.027 | Young |
| 0.003854 | 0.154266 | 0.911 | 0.907 | 1 | 2 | Rpl22 | 0.004 | Young |
| 0.003855 | 1.749666 | 0.023 | 0.006 | 1 | 2 | Tbc1d16 | 0.017 | Young |
| 0.003861 | 0.405681 | 0.017 | 0.003 | 1 | 2 | Sox4 | 0.014 | Young |
| 0.003977 | 0.347604 | 0.513 | 0.445 | 1 | 2 | Prdx5 | 0.068 | Young |
| 0.0041 | 0.121646 | 0.909 | 0.909 | 1 | 2 | Rpl23a | 0 | Young |
| 0.004108 | 0.72085 | 0.133 | 0.091 | 1 | 2 | Rogdi | 0.042 | Young |
| 0.004268 | 0.145605 | 0.78 | 0.715 | 1 | 2 | Atp5e | 0.065 | Young |
| 0.004317 | 0.537927 | 0.309 | 0.264 | 1 | 2 | Cebpg | 0.045 | Young |
| 0.00432 | 0.202422 | 0.733 | 0.684 | 1 | 2 | Ctsh | 0.049 | Young |
| 0.004337 | 0.368662 | 0.5 | 0.452 | 1 | 2 | Mrpl52 | 0.048 | Young |
| 0.004398 | 0.311762 | 0.196 | 0.146 | 1 | 2 | Rmnd5a | 0.05 | Young |
| 0.004515 | 0.324859 | 0.129 | 0.088 | 1 | 2 | Retreg1 | 0.041 | Young |
| 0.004554 | 0.25605 | 0.624 | 0.59 | 1 | 2 | Sh3glb1 | 0.034 | Young |
| 0.004563 | 0.151055 | 0.89 | 0.885 | 1 | 2 | Rpl6 | 0.005 | Young |
| 0.004661 | 1.835312 | 0.02 | 0.005 | 1 | 2 | Grasp | 0.015 | Young |
| 0.004806 | 1.033492 | 0.045 | 0.021 | 1 | 2 | F11r | 0.024 | Young |
| 0.004828 | 0.398202 | 0.159 | 0.113 | 1 | 2 | Arid4b | 0.046 | Young |
| 0.004967 | 0.380862 | 0.292 | 0.237 | 1 | 2 | Med13 | 0.055 | Young |
| 0.005126 | 0.729616 | 0.116 | 0.078 | 1 | 2 | Wdr13 | 0.038 | Young |
| 0.005276 | 0.473722 | 0.279 | 0.229 | 1 | 2 | Dot1l | 0.05 | Young |
| 0.005306 | 0.498693 | 0.243 | 0.196 | 1 | 2 | Tnrc6c | 0.047 | Young |
| 0.005372 | 0.446197 | 0.21 | 0.162 | 1 | 2 | 2410015M | 0.048 | Young |
| 0.005389 | 0.324679 | 0.481 | 0.424 | 1 | 2 | Ncf1 | 0.057 | Young |
| 0.005536 | 0.941272 | 0.046 | 0.022 | 1 | 2 | Xpc | 0.024 | Young |
| 0.005652 | 1.812084 | 0.018 | 0.004 | 1 | 2 | Siglech | 0.014 | Young |
| 0.005714 | 0.55298 | 0.22 | 0.172 | 1 | 2 | Trem2 | 0.048 | Young |
| 0.005809 | 1.01302 | 0.048 | 0.024 | 1 | 2 | Ang | 0.024 | Young |
| 0.005811 | 1.012235 | 0.037 | 0.016 | 1 | 2 | 2900089D1 | 0.021 | Young |
| 0.005935 | 1.030521 | 0.048 | 0.024 | 1 | 2 | Bend4 | 0.024 | Young |
| 0.005956 | 1.279516 | 0.029 | 0.011 | 1 | 2 | Map2k3os | 0.018 | Young |
| 0.005959 | 0.886925 | 0.095 | 0.061 | 1 | 2 | Hpgds | 0.034 | Young |
| 0.006065 | 0.562762 | 0.16 | 0.118 | 1 | 2 | Ptgs1 | 0.042 | Young |
| 0.006151 | 1.414716 | 0.029 | 0.011 | 1 | 2 | Ass1 | 0.018 | Young |
| 0.006166 | 0.228196 | 0.677 | 0.661 | 1 | 2 | Tmem176b | 0.016 | Young |
| 0.006306 | 0.173587 | 0.647 | 0.587 | 1 | 2 | C1qa | 0.06 | Young |

| | | | | | | | | |
|---|---|---|---|---|---|---|---|---|
| 0.006399 | 0.89481 | 0.089 | 0.057 | 1 | 2 | Egr3 | 0.032 | Young |
| 0.006513 | 1.125842 | 0.025 | 0.009 | 1 | 2 | Gpr84 | 0.016 | Young |
| 0.006566 | 0.556009 | 0.163 | 0.12 | 1 | 2 | Sbf2 | 0.043 | Young |
| 0.00668 | 0.36628 | 0.281 | 0.23 | 1 | 2 | Zbtb20 | 0.051 | Young |
| 0.006819 | 1.691328 | 0.011 | 0.001 | 1 | 2 | Uaca | 0.01 | Young |
| 0.006878 | 1.360062 | 0.011 | 0.001 | 1 | 2 | Gm15247 | 0.01 | Young |
| 0.007025 | 1.592591 | 0.03 | 0.012 | 1 | 2 | Ramp3 | 0.018 | Young |
| 0.007077 | 0.292922 | 0.482 | 0.426 | 1 | 2 | Mrfap1 | 0.056 | Young |
| 0.007533 | 0.441265 | 0.151 | 0.109 | 1 | 2 | Esf1 | 0.042 | Young |
| 0.007612 | 0.311819 | 0.468 | 0.412 | 1 | 2 | Smdt1 | 0.056 | Young |
| 0.007731 | 0.12763 | 0.847 | 0.79 | 1 | 2 | Cyba | 0.057 | Young |
| 0.007826 | 1.581799 | 0.019 | 0.005 | 1 | 2 | Dnase1l3 | 0.014 | Young |
| 0.007945 | 0.726901 | 0.134 | 0.096 | 1 | 2 | Sesn1 | 0.038 | Young |
| 0.008213 | 1.436246 | 0.023 | 0.007 | 1 | 2 | Atad5 | 0.016 | Young |
| 0.008266 | 0.468641 | 0.145 | 0.104 | 1 | 2 | Tgfbr1 | 0.041 | Young |
| 0.008485 | 1.074045 | 0.04 | 0.019 | 1 | 2 | Plekhg5 | 0.021 | Young |
| 0.008591 | 1.099706 | 0.064 | 0.037 | 1 | 2 | Mrps34 | 0.027 | Young |
| 0.008726 | 0.27256 | 0.412 | 0.352 | 1 | 2 | Ndufa13 | 0.06 | Young |
| 0.008935 | 0.510327 | 0.196 | 0.151 | 1 | 2 | Rnase4 | 0.045 | Young |
| 0.00912 | 0.271177 | 0.538 | 0.5 | 1 | 2 | Ubb | 0.038 | Young |
| 0.00919 | 0.804118 | 0.062 | 0.035 | 1 | 2 | Fbxo30 | 0.027 | Young |
| 0.009269 | 0.172984 | 0.162 | 0.12 | 1 | 2 | Gclc | 0.042 | Young |
| 0.009274 | 0.737529 | 0.093 | 0.06 | 1 | 2 | Erf | 0.033 | Young |
| 0.009366 | 0.765382 | 0.106 | 0.072 | 1 | 2 | Kin | 0.034 | Young |
| 0.009435 | 0.382006 | 0.29 | 0.239 | 1 | 2 | Icam1 | 0.051 | Young |
| 0.009516 | 1.243164 | 0.017 | 0.004 | 1 | 2 | Pxdc1 | 0.013 | Young |
| 0.00958 | 0.501292 | 0.159 | 0.12 | 1 | 2 | Gabarapl1 | 0.039 | Young |
| 0.009635 | 1.157127 | 0.017 | 0.004 | 1 | 2 | Carmil1 | 0.013 | Young |
| 0.009678 | 0.178287 | 0.669 | 0.62 | 1 | 2 | Rpl31 | 0.049 | Young |
| 0.009697 | 0.229337 | 0.378 | 0.322 | 1 | 2 | Anxa3 | 0.056 | Young |
| 0.009731 | 0.868511 | 0.101 | 0.069 | 1 | 2 | Ssbp2 | 0.032 | Young |
| 0.009785 | 0.4374 | 0.189 | 0.145 | 1 | 2 | Tbl1x | 0.044 | Young |
| 0.010076 | 0.246568 | 0.602 | 0.558 | 1 | 2 | Cox8a | 0.044 | Young |
| 0.010123 | 0.34618 | 0.362 | 0.315 | 1 | 2 | Eef1d | 0.047 | Young |
| 0.010133 | 0.738941 | 0.027 | 0.011 | 1 | 2 | Chst7 | 0.016 | Young |
| 0.010233 | 0.390434 | 0.219 | 0.175 | 1 | 2 | Prkacb | 0.044 | Young |
| 0.010258 | 0.515891 | 0.24 | 0.195 | 1 | 2 | Slc35b2 | 0.045 | Young |
| 0.010314 | 0.31019 | 0.581 | 0.541 | 1 | 2 | Dennd4a | 0.04 | Young |
| 0.01032 | 1.092704 | 0.02 | 0.006 | 1 | 2 | Cxxc5 | 0.014 | Young |
| 0.010426 | 1.570175 | 0.033 | 0.015 | 1 | 2 | Qtrt1 | 0.018 | Young |
| 0.010877 | 0.256885 | 0.486 | 0.435 | 1 | 2 | Dnaja1 | 0.051 | Young |
| 0.011034 | 1.575681 | 0.014 | 0.003 | 1 | 2 | Cd320 | 0.011 | Young |
| 0.011188 | 0.925966 | 0.077 | 0.049 | 1 | 2 | Csad | 0.028 | Young |
| 0.011238 | 0.482031 | 0.094 | 0.062 | 1 | 2 | Ophn1 | 0.032 | Young |
| 0.011239 | 0.774955 | 0.082 | 0.052 | 1 | 2 | Pias4 | 0.03 | Young |

| | | | | | | | | |
|---|---|---|---|---|---|---|---|---|
| 0.011334 | 0.772541 | 0.086 | 0.056 | 1 | 2 | Pik3c3 | 0.03 | Young |
| 0.011388 | 0.672711 | 0.108 | 0.075 | 1 | 2 | Bcl2a1a | 0.033 | Young |
| 0.011615 | 1.290113 | 0.046 | 0.025 | 1 | 2 | St7 | 0.021 | Young |
| 0.011639 | 0.224788 | 0.586 | 0.552 | 1 | 2 | Kdm6b | 0.034 | Young |
| 0.011837 | 0.594718 | 0.102 | 0.07 | 1 | 2 | Klhl6 | 0.032 | Young |
| 0.011897 | 1.681062 | 0.04 | 0.02 | 1 | 2 | Mvb12b | 0.02 | Young |
| 0.012296 | 1.805918 | 0.012 | 0.002 | 1 | 2 | Gm42670 | 0.01 | Young |
| 0.012327 | 0.304062 | 0.373 | 0.323 | 1 | 2 | Itpripl2 | 0.05 | Young |
| 0.012376 | 1.187036 | 0.025 | 0.01 | 1 | 2 | Mrvi1 | 0.015 | Young |
| 0.012392 | 0.493536 | 0.196 | 0.153 | 1 | 2 | Cbr2 | 0.043 | Young |
| 0.012434 | 1.707513 | 0.01 | 0.001 | 1 | 2 | Tnfrsf25 | 0.009 | Young |
| 0.012479 | 1.023991 | 0.012 | 0.002 | 1 | 2 | Phlda3 | 0.01 | Young |
| 0.012513 | 1.618631 | 0.01 | 0.001 | 1 | 2 | Cep112 | 0.009 | Young |
| 0.01272 | 0.103013 | 0.862 | 0.839 | 1 | 2 | Arpc1b | 0.023 | Young |
| 0.012744 | 0.59562 | 0.025 | 0.01 | 1 | 2 | Per3 | 0.015 | Young |
| 0.012753 | 0.233278 | 0.052 | 0.029 | 1 | 2 | Wdr46 | 0.023 | Young |
| 0.013217 | 1.067827 | 0.046 | 0.025 | 1 | 2 | Bmp2 | 0.021 | Young |
| 0.01327 | 0.08013 | 0.986 | 0.982 | 1 | 2 | Eef1a1 | 0.004 | Young |
| 0.013477 | 0.588559 | 0.135 | 0.099 | 1 | 2 | Slc16a6 | 0.036 | Young |
| 0.013893 | 0.877022 | 0.089 | 0.06 | 1 | 2 | Gm46224 | 0.029 | Young |
| 0.013982 | 0.209196 | 0.439 | 0.391 | 1 | 2 | Eif5 | 0.048 | Young |
| 0.014015 | 0.605754 | 0.076 | 0.048 | 1 | 2 | Nelfb | 0.028 | Young |
| 0.014056 | 0.510128 | 0.072 | 0.045 | 1 | 2 | Ralgps2 | 0.027 | Young |
| 0.014092 | 0.330831 | 0.324 | 0.281 | 1 | 2 | Ost4 | 0.043 | Young |
| 0.014276 | 0.706374 | 0.036 | 0.017 | 1 | 2 | Peli2 | 0.019 | Young |
| 0.014303 | 0.122371 | 0.948 | 0.913 | 1 | 2 | Rps27 | 0.035 | Young |
| 0.014329 | 1.090393 | 0.036 | 0.017 | 1 | 2 | Eid3 | 0.019 | Young |
| 0.014368 | 0.453633 | 0.172 | 0.133 | 1 | 2 | Setd5 | 0.039 | Young |
| 0.014427 | 0.735007 | 0.078 | 0.05 | 1 | 2 | Triap1 | 0.028 | Young |
| 0.014505 | 0.90333 | 0.049 | 0.027 | 1 | 2 | Noc3l | 0.022 | Young |
| 0.014629 | 0.938176 | 0.043 | 0.022 | 1 | 2 | Fzd5 | 0.021 | Young |
| 0.014803 | 0.353488 | 0.296 | 0.254 | 1 | 2 | Ccdc12 | 0.042 | Young |
| 0.014962 | 0.315968 | 0.222 | 0.177 | 1 | 2 | Ndufs5 | 0.045 | Young |
| 0.015173 | 1.178267 | 0.044 | 0.024 | 1 | 2 | Flrt3 | 0.02 | Young |
| 0.01553 | 1.083584 | 0.051 | 0.029 | 1 | 2 | Rasgrp3 | 0.022 | Young |
| 0.01554 | 1.638198 | 0.015 | 0.004 | 1 | 2 | Msrb2 | 0.011 | Young |
| 0.016001 | 1.113807 | 0.023 | 0.009 | 1 | 2 | Gstz1 | 0.014 | Young |
| 0.016039 | 1.201733 | 0.023 | 0.009 | 1 | 2 | Cbx6 | 0.014 | Young |
| 0.016112 | 1.012889 | 0.015 | 0.004 | 1 | 2 | 4933433G1 | 0.011 | Young |
| 0.016159 | 0.519501 | 0.195 | 0.155 | 1 | 2 | Pcyt1a | 0.04 | Young |
| 0.016307 | 0.404479 | 0.07 | 0.044 | 1 | 2 | Sh3pxd2a | 0.026 | Young |
| 0.016525 | 0.78699 | 0.064 | 0.04 | 1 | 2 | Med25 | 0.024 | Young |
| 0.01666 | 0.517699 | 0.127 | 0.092 | 1 | 2 | Mylip | 0.035 | Young |
| 0.016725 | 0.31709 | 0.387 | 0.342 | 1 | 2 | Dek | 0.045 | Young |
| 0.016802 | 0.304662 | 0.362 | 0.324 | 1 | 2 | Mier1 | 0.038 | Young |

| | | | | | | | | |
|---|---|---|---|---|---|---|---|---|
| 0.017009 | 0.757033 | 0.102 | 0.072 | 1 | 2 | Gadd45g | 0.03 | Young |
| 0.017193 | 0.442727 | 0.234 | 0.192 | 1 | 2 | Sft2d2 | 0.042 | Young |
| 0.017614 | 0.259421 | 0.504 | 0.448 | 1 | 2 | Gapdh | 0.056 | Young |
| 0.018093 | 0.549893 | 0.144 | 0.108 | 1 | 2 | Adap2 | 0.036 | Young |
| 0.018232 | 1.153192 | 0.058 | 0.035 | 1 | 2 | Cd72 | 0.023 | Young |
| 0.018729 | 1.954256 | 0.013 | 0.003 | 1 | 2 | Gm43936 | 0.01 | Young |
| 0.018789 | 0.237992 | 0.411 | 0.368 | 1 | 2 | Tgoln1 | 0.043 | Young |
| 0.018853 | 0.274948 | 0.375 | 0.339 | 1 | 2 | Rabac1 | 0.036 | Young |
| 0.018882 | 1.637039 | 0.013 | 0.003 | 1 | 2 | Prag1 | 0.01 | Young |
| 0.018943 | 1.326988 | 0.013 | 0.003 | 1 | 2 | Cd28 | 0.01 | Young |
| 0.019036 | 0.907908 | 0.013 | 0.003 | 1 | 2 | Ift88 | 0.01 | Young |
| 0.019221 | 0.358659 | 0.346 | 0.307 | 1 | 2 | Cox14 | 0.039 | Young |
| 0.019535 | 0.791958 | 0.04 | 0.021 | 1 | 2 | Setdb2 | 0.019 | Young |
| 0.019727 | 0.30998 | 0.352 | 0.307 | 1 | 2 | Ndufa1 | 0.045 | Young |
| 0.019866 | 0.328264 | 0.354 | 0.313 | 1 | 2 | Slc15a3 | 0.041 | Young |
| 0.019944 | 0.339875 | 0.232 | 0.188 | 1 | 2 | Tial1 | 0.044 | Young |
| 0.020393 | 0.374253 | 0.203 | 0.163 | 1 | 2 | Bdp1 | 0.04 | Young |
| 0.020555 | 0.209298 | 0.603 | 0.567 | 1 | 2 | Atp5g2 | 0.036 | Young |
| 0.020832 | 0.777034 | 0.071 | 0.046 | 1 | 2 | Lncpint | 0.025 | Young |
| 0.020868 | 0.188118 | 0.548 | 0.504 | 1 | 2 | Snrpe | 0.044 | Young |
| 0.020895 | 0.456103 | 0.101 | 0.071 | 1 | 2 | Mmp14 | 0.03 | Young |
| 0.020935 | 0.418552 | 0.356 | 0.319 | 1 | 2 | Ndufa7 | 0.037 | Young |
| 0.021102 | 0.126192 | 0.831 | 0.79 | 1 | 2 | Serinc3 | 0.041 | Young |
| 0.021187 | 0.44608 | 0.194 | 0.153 | 1 | 2 | Ivns1abp | 0.041 | Young |
| 0.021191 | 0.55537 | 0.178 | 0.141 | 1 | 2 | Maff | 0.037 | Young |
| 0.021244 | 0.850142 | 0.091 | 0.063 | 1 | 2 | Naa30 | 0.028 | Young |
| 0.02134 | 1.16473 | 0.017 | 0.005 | 1 | 2 | Armc9 | 0.012 | Young |
| 0.021549 | 0.210227 | 0.38 | 0.332 | 1 | 2 | Tsc22d3 | 0.048 | Young |
| 0.02162 | 0.350301 | 0.369 | 0.324 | 1 | 2 | Pde4b | 0.045 | Young |
| 0.02167 | 0.150589 | 0.343 | 0.291 | 1 | 2 | Fcgrt | 0.052 | Young |
| 0.021774 | 1.693811 | 0.011 | 0.002 | 1 | 2 | Cdiptos | 0.009 | Young |
| 0.021792 | 1.256753 | 0.025 | 0.011 | 1 | 2 | Hirip3 | 0.014 | Young |
| 0.021866 | 0.265595 | 0.419 | 0.374 | 1 | 2 | Snrpf | 0.045 | Young |
| 0.021878 | 1.050439 | 0.043 | 0.024 | 1 | 2 | Ppcdc | 0.019 | Young |
| 0.021931 | 1.398151 | 0.011 | 0.002 | 1 | 2 | Zfp956 | 0.009 | Young |
| 0.021953 | 0.583497 | 0.09 | 0.062 | 1 | 2 | Ncoa5 | 0.028 | Young |
| 0.022128 | 1.048058 | 0.011 | 0.002 | 1 | 2 | Gm12355 | 0.009 | Young |
| 0.022195 | 0.908569 | 0.025 | 0.011 | 1 | 2 | 2810006K2 | 0.014 | Young |
| 0.022348 | 1.04232 | 0.103 | 0.075 | 1 | 2 | Dnajb4 | 0.028 | Young |
| 0.022444 | 0.269307 | 0.077 | 0.05 | 1 | 2 | Zdhhc17 | 0.027 | Young |
| 0.022622 | 0.439222 | 0.17 | 0.133 | 1 | 2 | Man1a2 | 0.037 | Young |
| 0.022724 | 0.461143 | 0.058 | 0.035 | 1 | 2 | Ctdspl | 0.023 | Young |
| 0.0229 | 0.613117 | 0.036 | 0.018 | 1 | 2 | Commd9 | 0.018 | Young |
| 0.022963 | 0.210649 | 0.558 | 0.52 | 1 | 2 | Mrpl33 | 0.038 | Young |
| 0.022976 | 0.465895 | 0.138 | 0.104 | 1 | 2 | Opa1 | 0.034 | Young |

| | | | | | | | | |
|---|---|---|---|---|---|---|---|---|
| 0.023293 | 0.649786 | 0.12 | 0.089 | 1 | 2 | 9930021J0 | 0.031 | Young |
| 0.023347 | 0.115188 | 0.846 | 0.813 | 1 | 2 | Ppia | 0.033 | Young |
| 0.023492 | 0.157073 | 0.789 | 0.737 | 1 | 2 | Rbpj | 0.052 | Young |
| 0.023504 | 0.901897 | 0.084 | 0.058 | 1 | 2 | Hs2st1 | 0.026 | Young |
| 0.023602 | 0.418303 | 0.279 | 0.238 | 1 | 2 | Fkbp2 | 0.041 | Young |
| 0.023857 | 0.285072 | 0.192 | 0.154 | 1 | 2 | Bhlhe40 | 0.038 | Young |
| 0.023876 | 0.224965 | 0.565 | 0.545 | 1 | 2 | Tob2 | 0.02 | Young |
| 0.023995 | 1.4165 | 0.021 | 0.009 | 1 | 2 | Gm37233 | 0.012 | Young |
| 0.0243 | 1.372854 | 0.026 | 0.012 | 1 | 2 | Slc14a1 | 0.014 | Young |
| 0.024358 | 0.273429 | 0.466 | 0.427 | 1 | 2 | Gpi1 | 0.039 | Young |
| 0.024444 | 1.38225 | 0.021 | 0.009 | 1 | 2 | Slc37a1 | 0.012 | Young |
| 0.024692 | 0.538681 | 0.27 | 0.231 | 1 | 2 | Aebp2 | 0.039 | Young |
| 0.02513 | 0.887124 | 0.021 | 0.009 | 1 | 2 | Adat2 | 0.012 | Young |
| 0.025221 | 1.327461 | 0.032 | 0.016 | 1 | 2 | Eva1b | 0.016 | Young |
| 0.025227 | 0.867667 | 0.076 | 0.051 | 1 | 2 | Rassf8 | 0.025 | Young |
| 0.025567 | 1.361481 | 0.032 | 0.016 | 1 | 2 | Tatdn3 | 0.016 | Young |
| 0.025807 | 1.312795 | 0.039 | 0.021 | 1 | 2 | Alg14 | 0.018 | Young |
| 0.02592 | 0.192894 | 0.555 | 0.514 | 1 | 2 | Rpl29 | 0.041 | Young |
| 0.025952 | 1.564502 | 0.018 | 0.006 | 1 | 2 | Xpo5 | 0.012 | Young |
| 0.025987 | 0.263724 | 0.365 | 0.325 | 1 | 2 | Tgif1 | 0.04 | Young |
| 0.02609 | 0.125299 | 0.635 | 0.572 | 1 | 2 | C1qb | 0.063 | Young |
| 0.026183 | 0.476759 | 0.165 | 0.131 | 1 | 2 | Itga5 | 0.034 | Young |
| 0.026239 | 1.727457 | 0.014 | 0.004 | 1 | 2 | Cfap126 | 0.01 | Young |
| 0.026252 | 1.493882 | 0.018 | 0.006 | 1 | 2 | Lrrc49 | 0.012 | Young |
| 0.026555 | 0.186112 | 0.17 | 0.131 | 1 | 2 | Galc | 0.039 | Young |
| 0.026585 | 1.214141 | 0.014 | 0.004 | 1 | 2 | Osgin1 | 0.01 | Young |
| 0.026662 | 0.777683 | 0.014 | 0.004 | 1 | 2 | Gm15232 | 0.01 | Young |
| 0.0268 | 1.078222 | 0.032 | 0.016 | 1 | 2 | Flt3 | 0.016 | Young |
| 0.026893 | 0.773837 | 0.039 | 0.021 | 1 | 2 | Cep162 | 0.018 | Young |
| 0.026903 | 0.724746 | 0.124 | 0.093 | 1 | 2 | Gla | 0.031 | Young |
| 0.026972 | 0.322374 | 0.296 | 0.256 | 1 | 2 | Pdcd5 | 0.04 | Young |
| 0.027226 | 0.146052 | 0.786 | 0.754 | 1 | 2 | Nfkbia | 0.032 | Young |
| 0.027374 | 0.562558 | 0.018 | 0.006 | 1 | 2 | Zfp810 | 0.012 | Young |
| 0.028323 | 1.118929 | 0.033 | 0.017 | 1 | 2 | Cox6b2 | 0.016 | Young |
| 0.028512 | 0.454065 | 0.198 | 0.161 | 1 | 2 | Errfi1 | 0.037 | Young |
| 0.029389 | 0.100559 | 0.489 | 0.449 | 1 | 2 | Ckb | 0.04 | Young |
| 0.029536 | 0.600372 | 0.033 | 0.017 | 1 | 2 | Ppp1r12b | 0.016 | Young |
| 0.029697 | 1.325096 | 0.029 | 0.014 | 1 | 2 | Dcp1b | 0.015 | Young |
| 0.030085 | 0.694517 | 0.049 | 0.029 | 1 | 2 | Uqcc1 | 0.02 | Young |
| 0.030244 | 0.914611 | 0.05 | 0.03 | 1 | 2 | Sesn3 | 0.02 | Young |
| 0.030355 | 0.241087 | 0.327 | 0.29 | 1 | 2 | Pds5a | 0.037 | Young |
| 0.03039 | 0.123024 | 0.213 | 0.171 | 1 | 2 | Tgfbr2 | 0.042 | Young |
| 0.030407 | 0.209931 | 0.4 | 0.362 | 1 | 2 | Tmem256 | 0.038 | Young |
| 0.030554 | 0.527056 | 0.144 | 0.111 | 1 | 2 | Gatad2a | 0.033 | Young |
| 0.030857 | 1.39686 | 0.019 | 0.007 | 1 | 2 | Det1 | 0.012 | Young |

| | | | | | | | | |
|---|---|---|---|---|---|---|---|---|
| 0.031331 | 1.393876 | 0.055 | 0.034 | 1 | 2 | Hint3 | 0.021 | Young |
| 0.031334 | 0.479822 | 0.049 | 0.029 | 1 | 2 | Akr1e1 | 0.02 | Young |
| 0.031427 | 0.713385 | 0.05 | 0.03 | 1 | 2 | Cd209d | 0.02 | Young |
| 0.031745 | 0.084657 | 0.127 | 0.093 | 1 | 2 | Clec2i | 0.034 | Young |
| 0.031767 | 0.553477 | 0.265 | 0.23 | 1 | 2 | Mif | 0.035 | Young |
| 0.032013 | 1.445173 | 0.019 | 0.007 | 1 | 2 | Evi5l | 0.012 | Young |
| 0.032213 | 1.384776 | 0.012 | 0.003 | 1 | 2 | Gm43569 | 0.009 | Young |
| 0.03272 | 0.583885 | 0.124 | 0.094 | 1 | 2 | Sult1a1 | 0.03 | Young |
| 0.032722 | 0.769325 | 0.012 | 0.003 | 1 | 2 | Iqcc | 0.009 | Young |
| 0.033216 | 0.618619 | 0.115 | 0.087 | 1 | 2 | Tomm40 | 0.028 | Young |
| 0.033469 | 0.961305 | 0.07 | 0.047 | 1 | 2 | Lrwd1 | 0.023 | Young |
| 0.033637 | 0.926797 | 0.055 | 0.034 | 1 | 2 | Xxylt1 | 0.021 | Young |
| 0.033831 | 0.405024 | 0.247 | 0.209 | 1 | 2 | Calu | 0.038 | Young |
| 0.034194 | 0.671533 | 0.108 | 0.08 | 1 | 2 | Fh1 | 0.028 | Young |
| 0.034228 | 1.051024 | 0.057 | 0.036 | 1 | 2 | Fcna | 0.021 | Young |
| 0.034289 | 0.72221 | 0.036 | 0.019 | 1 | 2 | Decr2 | 0.017 | Young |
| 0.03431 | 0.454262 | 0.158 | 0.125 | 1 | 2 | Slc9a9 | 0.033 | Young |
| 0.034582 | 0.391791 | 0.134 | 0.103 | 1 | 2 | Arsb | 0.031 | Young |
| 0.034811 | 0.290709 | 0.381 | 0.345 | 1 | 2 | Ndufc2 | 0.036 | Young |
| 0.035054 | 0.616425 | 0.053 | 0.033 | 1 | 2 | Pigyl | 0.02 | Young |
| 0.035422 | 0.510787 | 0.03 | 0.015 | 1 | 2 | Gtf2h2 | 0.015 | Young |
| 0.03549 | 0.841915 | 0.069 | 0.046 | 1 | 2 | Sik2 | 0.023 | Young |
| 0.035856 | 0.168088 | 0.35 | 0.302 | 1 | 2 | Ptgs2 | 0.048 | Young |
| 0.036231 | 0.285863 | 0.097 | 0.07 | 1 | 2 | Mrpl9 | 0.027 | Young |
| 0.036386 | 0.26478 | 0.398 | 0.362 | 1 | 2 | Wasf2 | 0.036 | Young |
| 0.036423 | 1.236522 | 0.025 | 0.012 | 1 | 2 | Pald1 | 0.013 | Young |
| 0.036438 | 1.348952 | 0.039 | 0.022 | 1 | 2 | Chp2 | 0.017 | Young |
| 0.036565 | 0.839088 | 0.059 | 0.039 | 1 | 2 | Med29 | 0.02 | Young |
| 0.036683 | 0.540445 | 0.09 | 0.064 | 1 | 2 | Aak1 | 0.026 | Young |
| 0.036861 | 0.329597 | 0.182 | 0.143 | 1 | 2 | Sh2b3 | 0.039 | Young |
| 0.036867 | 0.486144 | 0.044 | 0.026 | 1 | 2 | Nat6 | 0.018 | Young |
| 0.036945 | 0.411733 | 0.182 | 0.148 | 1 | 2 | Banf1 | 0.034 | Young |
| 0.037176 | 0.246595 | 0.335 | 0.294 | 1 | 2 | Pea15a | 0.041 | Young |
| 0.037205 | 0.401474 | 0.207 | 0.17 | 1 | 2 | Phf21a | 0.037 | Young |
| 0.037295 | 0.852087 | 0.025 | 0.012 | 1 | 2 | Ankrd13d | 0.013 | Young |
| 0.037457 | 0.368913 | 0.278 | 0.241 | 1 | 2 | Sumo3 | 0.037 | Young |
| 0.037516 | 1.12252 | 0.02 | 0.009 | 1 | 2 | Nmnat3 | 0.011 | Young |
| 0.037658 | 0.606877 | 0.046 | 0.028 | 1 | 2 | Chkb | 0.018 | Young |
| 0.037708 | 0.839672 | 0.031 | 0.016 | 1 | 2 | Snx21 | 0.015 | Young |
| 0.03777 | 0.647117 | 0.087 | 0.062 | 1 | 2 | AU019823 | 0.025 | Young |
| 0.037874 | 0.219121 | 0.482 | 0.446 | 1 | 2 | Mef2c | 0.036 | Young |
| 0.038112 | 0.104163 | 0.861 | 0.81 | 1 | 2 | Serf2 | 0.051 | Young |
| 0.03814 | 0.264045 | 0.373 | 0.335 | 1 | 2 | Sema4a | 0.038 | Young |
| 0.038222 | 0.324552 | 0.025 | 0.012 | 1 | 2 | Dyrk1b | 0.013 | Young |
| 0.038349 | 0.326702 | 0.265 | 0.227 | 1 | 2 | Mrpl23 | 0.038 | Young |

| | | | | | | | | |
|---|---|---|---|---|---|---|---|---|
| 0.038522 | 0.651583 | 0.087 | 0.062 | 1 | 2 | Xpo4 | 0.025 | Young |
| 0.038562 | 1.507234 | 0.01 | 0.002 | 1 | 2 | Tle2 | 0.008 | Young |
| 0.038562 | 1.352859 | 0.01 | 0.002 | 1 | 2 | Zfp341 | 0.008 | Young |
| 0.038629 | 1.345499 | 0.01 | 0.002 | 1 | 2 | H2-M2 | 0.008 | Young |
| 0.038629 | 1.182585 | 0.01 | 0.002 | 1 | 2 | 2210417A( | 0.008 | Young |
| 0.038737 | 0.650004 | 0.031 | 0.016 | 1 | 2 | Ppat | 0.015 | Young |
| 0.038983 | 0.296915 | 0.055 | 0.034 | 1 | 2 | Slc7a11 | 0.021 | Young |
| 0.039051 | 0.353493 | 0.182 | 0.148 | 1 | 2 | Bcl2a1d | 0.034 | Young |
| 0.039275 | 0.174077 | 0.614 | 0.587 | 1 | 2 | Aif1 | 0.027 | Young |
| 0.039331 | 0.302774 | 0.109 | 0.08 | 1 | 2 | Tns1 | 0.029 | Young |
| 0.039434 | 0.707151 | 0.133 | 0.105 | 1 | 2 | Fundc2 | 0.028 | Young |
| 0.039513 | 1.013262 | 0.032 | 0.017 | 1 | 2 | 2-Sep | 0.015 | Young |
| 0.039569 | 0.563014 | 0.076 | 0.052 | 1 | 2 | Kcnk13 | 0.024 | Young |
| 0.039979 | 0.185516 | 0.768 | 0.725 | 1 | 2 | Fn1 | 0.043 | Young |
| 0.04047 | 0.328367 | 0.12 | 0.09 | 1 | 2 | Hspbp1 | 0.03 | Young |
| 0.040729 | 1.103217 | 0.026 | 0.013 | 1 | 2 | Gm38843 | 0.013 | Young |
| 0.040933 | 0.494111 | 0.049 | 0.03 | 1 | 2 | Mtmr4 | 0.019 | Young |
| 0.041047 | 0.155856 | 0.455 | 0.419 | 1 | 2 | Ighm | 0.036 | Young |
| 0.041105 | 1.343567 | 0.017 | 0.006 | 1 | 2 | Lrsam1 | 0.011 | Young |
| 0.041404 | 0.443435 | 0.16 | 0.128 | 1 | 2 | Mtmr3 | 0.032 | Young |
| 0.041447 | 0.507163 | 0.133 | 0.104 | 1 | 2 | Sec24a | 0.029 | Young |
| 0.041711 | 0.997641 | 0.017 | 0.006 | 1 | 2 | Pomt1 | 0.011 | Young |
| 0.041777 | 0.707551 | 0.026 | 0.013 | 1 | 2 | Il18rap | 0.013 | Young |
| 0.042069 | 1.142108 | 0.017 | 0.006 | 1 | 2 | Ptrh1 | 0.011 | Young |
| 0.042206 | 0.116854 | 0.715 | 0.675 | 1 | 2 | mt-Nd4l | 0.04 | Young |
| 0.042217 | 1.477089 | 0.052 | 0.033 | 1 | 2 | Cxcl10 | 0.019 | Young |
| 0.04265 | 0.243239 | 0.512 | 0.482 | 1 | 2 | Fcgr3 | 0.03 | Young |
| 0.042732 | 1.213696 | 0.021 | 0.01 | 1 | 2 | Cd209b | 0.011 | Young |
| 0.042827 | 1.564878 | 0.013 | 0.004 | 1 | 2 | Zfp251 | 0.009 | Young |
| 0.042827 | 1.548375 | 0.013 | 0.004 | 1 | 2 | Gmpr | 0.009 | Young |
| 0.043105 | 0.406057 | 0.075 | 0.051 | 1 | 2 | Rnaseh2a | 0.024 | Young |
| 0.043111 | 0.577875 | 0.084 | 0.06 | 1 | 2 | Anapc4 | 0.024 | Young |
| 0.043117 | 0.829456 | 0.065 | 0.044 | 1 | 2 | Etv1 | 0.021 | Young |
| 0.043128 | 1.922254 | 0.013 | 0.004 | 1 | 2 | Il7r | 0.009 | Young |
| 0.04343 | 1.205336 | 0.027 | 0.014 | 1 | 2 | Praf2 | 0.013 | Young |
| 0.043872 | 0.139927 | 0.576 | 0.544 | 1 | 2 | Purb | 0.032 | Young |
| 0.043948 | 0.622857 | 0.083 | 0.059 | 1 | 2 | Ankrd10 | 0.024 | Young |
| 0.044258 | 0.641598 | 0.069 | 0.047 | 1 | 2 | Fam207a | 0.022 | Young |
| 0.044626 | 1.013258 | 0.027 | 0.014 | 1 | 2 | Mapkapk5 | 0.013 | Young |
| 0.044703 | 0.572574 | 0.072 | 0.05 | 1 | 2 | Trap1 | 0.022 | Young |
| 0.044808 | 0.267102 | 0.293 | 0.254 | 1 | 2 | Cd86 | 0.039 | Young |
| 0.045234 | 0.987623 | 0.027 | 0.014 | 1 | 2 | Tmem86b | 0.013 | Young |
| 0.045309 | 0.940206 | 0.055 | 0.035 | 1 | 2 | Klrk1 | 0.02 | Young |
| 0.045657 | 0.823447 | 0.043 | 0.026 | 1 | 2 | Mical1 | 0.017 | Young |
| 0.045687 | 0.352254 | 0.133 | 0.103 | 1 | 2 | Ulk2 | 0.03 | Young |

| | | | | | | | | |
|---|---|---|---|---|---|---|---|---|
| 0.045973 | 0.738179 | 0.027 | 0.014 | 1 | 2 | Pde1b | 0.013 | Young |
| 0.046077 | 0.401853 | 0.178 | 0.145 | 1 | 2 | Idh3g | 0.033 | Young |
| 0.046183 | 0.347055 | 0.252 | 0.218 | 1 | 2 | Arrb1 | 0.034 | Young |
| 0.046201 | 0.265261 | 0.42 | 0.38 | 1 | 2 | Atp5j | 0.04 | Young |
| 0.046271 | 0.28006 | 0.513 | 0.484 | 1 | 2 | Nfkb1 | 0.029 | Young |
| 0.046624 | 0.670655 | 0.056 | 0.036 | 1 | 2 | Thoc5 | 0.02 | Young |
| 0.046759 | 0.575896 | 0.074 | 0.051 | 1 | 2 | Arntl | 0.023 | Young |
| 0.047463 | 0.991334 | 0.029 | 0.015 | 1 | 2 | Ppp2r3a | 0.014 | Young |
| 0.04761 | 0.898997 | 0.061 | 0.041 | 1 | 2 | Pigc | 0.02 | Young |
| 0.047687 | 0.701288 | 0.044 | 0.027 | 1 | 2 | Mir155hg | 0.017 | Young |
| 0.047975 | 0.4225 | 0.166 | 0.134 | 1 | 2 | 1500011K1 | 0.032 | Young |
| 0.04803 | 0.3111 | 0.412 | 0.381 | 1 | 2 | Hexb | 0.031 | Young |
| 0.048104 | 0.551518 | 0.163 | 0.134 | 1 | 2 | Rnf145 | 0.029 | Young |
| 0.048123 | 0.15601 | 0.609 | 0.585 | 1 | 2 | Tmem176a | 0.024 | Young |
| 0.048127 | 0.261746 | 0.177 | 0.143 | 1 | 2 | Rab11fip5 | 0.034 | Young |
| 0.048498 | 0.747678 | 0.055 | 0.035 | 1 | 2 | Sdhaf1 | 0.02 | Young |
| 0.049168 | 0.984923 | 0.037 | 0.021 | 1 | 2 | 1500015A0 | 0.016 | Young |
| 0.049211 | 0.600121 | 0.056 | 0.036 | 1 | 2 | Nrde2 | 0.02 | Young |
| 0.049386 | 0.0767 | 0.956 | 0.942 | 1 | 2 | Mcl1 | 0.014 | Young |
| 0.049787 | 0.554382 | 0.094 | 0.07 | 1 | 2 | Bambi | 0.024 | Young |
| 0.049838 | 0.298717 | 0.35 | 0.315 | 1 | 2 | Tmem258 | 0.035 | Young |
| 0.050127 | 0.22812 | 0.397 | 0.357 | 1 | 2 | Skil | 0.04 | Young |
| 0.050602 | 0.04801 | 0.386 | 0.343 | 1 | 2 | Ssh2 | 0.043 | Young |
| 0.051757 | 0.802811 | 0.048 | 0.03 | 1 | 2 | Rcn3 | 0.018 | Young |
| 0.051802 | 0.182468 | 0.475 | 0.435 | 1 | 2 | Ets2 | 0.04 | Young |
| 0.051864 | 0.410602 | 0.141 | 0.111 | 1 | 2 | Igf1r | 0.03 | Young |
| 0.052442 | 0.296345 | 0.099 | 0.073 | 1 | 2 | Rfx7 | 0.026 | Young |
| 0.053256 | 1.133536 | 0.039 | 0.024 | 1 | 2 | Zfp715 | 0.015 | Young |
| 0.053519 | 0.844716 | 0.03 | 0.016 | 1 | 2 | Fbxo32 | 0.014 | Young |
| 0.053746 | 1.49662 | 0.014 | 0.005 | 1 | 2 | Usp20 | 0.009 | Young |
| 0.053784 | 0.398972 | 0.057 | 0.037 | 1 | 2 | Rnf40 | 0.02 | Young |
| 0.053874 | 1.687154 | 0.011 | 0.003 | 1 | 2 | Vmac | 0.008 | Young |
| 0.054037 | 1.715664 | 0.011 | 0.003 | 1 | 2 | Nid2 | 0.008 | Young |
| 0.054118 | 1.391421 | 0.011 | 0.003 | 1 | 2 | Slc7a2 | 0.008 | Young |
| 0.054432 | 0.870822 | 0.014 | 0.005 | 1 | 2 | Epn2 | 0.009 | Young |
| 0.054957 | 1.613012 | 0.019 | 0.009 | 1 | 2 | Atp10a | 0.01 | Young |
| 0.055762 | 0.542977 | 0.049 | 0.031 | 1 | 2 | Vps39 | 0.018 | Young |
| 0.055941 | 0.203404 | 0.506 | 0.479 | 1 | 2 | Nsa2 | 0.027 | Young |
| 0.055947 | 0.387403 | 0.138 | 0.109 | 1 | 2 | Pitpnb | 0.029 | Young |
| 0.056083 | 0.076444 | 0.918 | 0.885 | 1 | 2 | Atp2b1 | 0.033 | Young |
| 0.056196 | 0.28303 | 0.346 | 0.316 | 1 | 2 | Ralbp1 | 0.03 | Young |
| 0.056221 | 0.228534 | 0.355 | 0.317 | 1 | 2 | Ppp1r15a | 0.038 | Young |
| 0.056326 | 0.514833 | 0.05 | 0.032 | 1 | 2 | Polr3b | 0.018 | Young |
| 0.056541 | 0.452322 | 0.133 | 0.106 | 1 | 2 | Ern1 | 0.027 | Young |
| 0.05699 | 0.415878 | 0.09 | 0.066 | 1 | 2 | Rpf1 | 0.024 | Young |

| | | | | | | | | |
|---|---|---|---|---|---|---|---|---|
| 0.057374 | 0.101756 | 0.836 | 0.813 | 1 | 2 | Ucp2 | 0.023 | Young |
| 0.057462 | 1.006555 | 0.025 | 0.013 | 1 | 2 | B3galnt1 | 0.012 | Young |
| 0.057511 | 0.78813 | 0.04 | 0.025 | 1 | 2 | Cry2 | 0.015 | Young |
| 0.057775 | 0.868757 | 0.025 | 0.013 | 1 | 2 | Arhgef10 | 0.012 | Young |
| 0.057854 | 0.18257 | 0.501 | 0.464 | 1 | 2 | H2afj | 0.037 | Young |
| 0.057861 | 0.529194 | 0.106 | 0.08 | 1 | 2 | Polr1c | 0.026 | Young |
| 0.057925 | 0.281669 | 0.081 | 0.058 | 1 | 2 | AU020206 | 0.023 | Young |
| 0.057948 | 0.40354 | 0.039 | 0.024 | 1 | 2 | Lclat1 | 0.015 | Young |
| 0.058046 | 0.581159 | 0.089 | 0.066 | 1 | 2 | Usp40 | 0.023 | Young |
| 0.058167 | 0.340041 | 0.154 | 0.125 | 1 | 2 | 5031425E2 | 0.029 | Young |
| 0.058448 | 0.331822 | 0.2 | 0.168 | 1 | 2 | Chd1 | 0.032 | Young |
| 0.05909 | 0.6449 | 0.074 | 0.052 | 1 | 2 | Lsm10 | 0.022 | Young |
| 0.059716 | 0.627895 | 0.094 | 0.071 | 1 | 2 | 2310009B1 | 0.023 | Young |
| 0.059885 | 0.69562 | 0.095 | 0.071 | 1 | 2 | Zc3h14 | 0.024 | Young |
| 0.059925 | 0.614554 | 0.128 | 0.103 | 1 | 2 | Gm47283 | 0.025 | Young |
| 0.059951 | 0.815505 | 0.025 | 0.013 | 1 | 2 | Ssx2ip | 0.012 | Young |
| 0.060154 | 0.179055 | 0.52 | 0.484 | 1 | 2 | Uqcrq | 0.036 | Young |
| 0.060621 | 0.117298 | 0.57 | 0.541 | 1 | 2 | Slfn2 | 0.029 | Young |
| 0.060748 | 0.600031 | 0.075 | 0.054 | 1 | 2 | Mpg | 0.021 | Young |
| 0.060879 | 0.225005 | 0.391 | 0.353 | 1 | 2 | Ap2s1 | 0.038 | Young |
| 0.061179 | 0.415676 | 0.144 | 0.116 | 1 | 2 | Cenpx | 0.028 | Young |
| 0.061182 | 0.61312 | 0.074 | 0.052 | 1 | 2 | Mrps30 | 0.022 | Young |
| 0.061417 | 0.171603 | 0.032 | 0.018 | 1 | 2 | Mapk1ip1 | 0.014 | Young |
| 0.061513 | 0.473316 | 0.032 | 0.018 | 1 | 2 | Plxna4 | 0.014 | Young |
| 0.061636 | 0.263966 | 0.289 | 0.252 | 1 | 2 | Msl2 | 0.037 | Young |
| 0.061761 | 0.214572 | 0.577 | 0.544 | 1 | 2 | Txnip | 0.033 | Young |
| 0.061929 | 0.755674 | 0.042 | 0.026 | 1 | 2 | Grap | 0.016 | Young |
| 0.062121 | 0.276829 | 0.264 | 0.228 | 1 | 2 | Commd8 | 0.036 | Young |
| 0.062522 | 0.158541 | 0.638 | 0.608 | 1 | 2 | Eif3f | 0.03 | Young |
| 0.062609 | 0.550041 | 0.053 | 0.035 | 1 | 2 | Siae | 0.018 | Young |
| 0.062766 | 0.668129 | 0.118 | 0.092 | 1 | 2 | Itpk1 | 0.026 | Young |
| 0.062782 | 0.033951 | 0.739 | 0.702 | 1 | 2 | Mrc1 | 0.037 | Young |
| 0.062843 | 0.686018 | 0.089 | 0.066 | 1 | 2 | Foxj3 | 0.023 | Young |
| 0.062892 | 0.234312 | 0.27 | 0.233 | 1 | 2 | Serpinb8 | 0.037 | Young |
| 0.063248 | 0.306344 | 0.27 | 0.239 | 1 | 2 | Fkbp8 | 0.031 | Young |
| 0.063542 | 0.659446 | 0.057 | 0.039 | 1 | 2 | Bach2 | 0.018 | Young |
| 0.06359 | 0.252123 | 0.362 | 0.333 | 1 | 2 | Txndc17 | 0.029 | Young |
| 0.063864 | 1.834634 | 0.015 | 0.006 | 1 | 2 | Fndc7 | 0.009 | Young |
| 0.06392 | 0.405059 | 0.083 | 0.061 | 1 | 2 | Pld1 | 0.022 | Young |
| 0.063959 | 0.419766 | 0.07 | 0.049 | 1 | 2 | Lrrcc1 | 0.021 | Young |
| 0.064015 | 0.47698 | 0.145 | 0.116 | 1 | 2 | Commd6 | 0.029 | Young |
| 0.064238 | 1.557013 | 0.015 | 0.006 | 1 | 2 | Dnah2 | 0.009 | Young |
| 0.064238 | 1.213637 | 0.015 | 0.006 | 1 | 2 | Ryk | 0.009 | Young |
| 0.064269 | 0.856435 | 0.02 | 0.01 | 1 | 2 | Trim16 | 0.01 | Young |
| 0.064275 | 0.853072 | 0.044 | 0.028 | 1 | 2 | Sp4 | 0.016 | Young |

| | | | | | | | | |
|---|---|---|---|---|---|---|---|---|
| 0.064361 | 0.311341 | 0.121 | 0.094 | 1 | 2 | Gstm1 | 0.027 | Young |
| 0.064829 | 0.587281 | 0.057 | 0.039 | 1 | 2 | Fam110a | 0.018 | Young |
| 0.06489 | 0.293921 | 0.286 | 0.257 | 1 | 2 | Top2b | 0.029 | Young |
| 0.064973 | 0.820099 | 0.069 | 0.049 | 1 | 2 | Serhl | 0.02 | Young |
| 0.06499 | 1.026859 | 0.015 | 0.006 | 1 | 2 | Amigo1 | 0.009 | Young |
| 0.065042 | 0.46141 | 0.02 | 0.01 | 1 | 2 | Adamtsl5 | 0.01 | Young |
| 0.065086 | 0.748695 | 0.065 | 0.046 | 1 | 2 | Pnkd | 0.019 | Young |
| 0.065362 | 0.789923 | 0.081 | 0.06 | 1 | 2 | Thap3 | 0.021 | Young |
| 0.065495 | 0.844746 | 0.07 | 0.05 | 1 | 2 | 2510009E0 | 0.02 | Young |
| 0.065502 | 0.120408 | 0.784 | 0.768 | 1 | 2 | Rpl14 | 0.016 | Young |
| 0.065571 | 0.370308 | 0.169 | 0.139 | 1 | 2 | Pvr | 0.03 | Young |
| 0.065693 | 0.334757 | 0.02 | 0.01 | 1 | 2 | Cald1 | 0.01 | Young |
| 0.065958 | 0.48468 | 0.214 | 0.184 | 1 | 2 | Zcrb1 | 0.03 | Young |
| 0.066043 | 1.419623 | 0.027 | 0.015 | 1 | 2 | Spout1 | 0.012 | Young |
| 0.066081 | 0.383443 | 0.216 | 0.185 | 1 | 2 | Rtf2 | 0.031 | Young |
| 0.066084 | 0.389417 | 0.16 | 0.132 | 1 | 2 | Idh2 | 0.028 | Young |
| 0.066168 | 0.504793 | 0.043 | 0.027 | 1 | 2 | Arhgap19 | 0.016 | Young |
| 0.066209 | 0.516998 | 0.015 | 0.006 | 1 | 2 | Fam109a | 0.009 | Young |
| 0.066415 | 0.337275 | 0.02 | 0.01 | 1 | 2 | Rfesd | 0.01 | Young |
| 0.066439 | 0.827549 | 0.015 | 0.006 | 1 | 2 | D030056L2 | 0.009 | Young |
| 0.067034 | 0.919587 | 0.045 | 0.029 | 1 | 2 | Lig3 | 0.016 | Young |
| 0.067298 | 0.447004 | 0.12 | 0.093 | 1 | 2 | Lcorl | 0.027 | Young |
| 0.067322 | 1.167927 | 0.027 | 0.015 | 1 | 2 | Pank1 | 0.012 | Young |
| 0.067414 | 0.23869 | 0.215 | 0.181 | 1 | 2 | Acsl5 | 0.034 | Young |
| 0.067916 | 0.209902 | 0.356 | 0.32 | 1 | 2 | Use1 | 0.036 | Young |
| 0.06796 | 0.526459 | 0.107 | 0.082 | 1 | 2 | Pum3 | 0.025 | Young |
| 0.06813 | 0.310579 | 0.072 | 0.051 | 1 | 2 | Slc10a7 | 0.021 | Young |
| 0.068185 | 0.417074 | 0.129 | 0.104 | 1 | 2 | Ednrb | 0.025 | Young |
| 0.068387 | 1.336944 | 0.021 | 0.011 | 1 | 2 | Surf2 | 0.01 | Young |
| 0.068838 | 0.462803 | 0.045 | 0.029 | 1 | 2 | Saysd1 | 0.016 | Young |
| 0.068901 | 0.126875 | 0.899 | 0.882 | 1 | 2 | Hsp90ab1 | 0.017 | Young |
| 0.069078 | 0.671493 | 0.027 | 0.015 | 1 | 2 | Oxnad1 | 0.012 | Young |
| 0.069092 | 0.062876 | 0.923 | 0.908 | 1 | 2 | Rpl8 | 0.015 | Young |
| 0.069135 | 0.749689 | 0.027 | 0.015 | 1 | 2 | Arfip2 | 0.012 | Young |
| 0.06928 | 0.210788 | 0.29 | 0.256 | 1 | 2 | Ubtf | 0.034 | Young |
| 0.069581 | 0.1108 | 0.624 | 0.593 | 1 | 2 | Ndufa2 | 0.031 | Young |
| 0.069741 | 0.329086 | 0.24 | 0.21 | 1 | 2 | P2ry6 | 0.03 | Young |
| 0.069787 | 0.276168 | 0.209 | 0.178 | 1 | 2 | Iscu | 0.031 | Young |
| 0.069961 | 1.506942 | 0.012 | 0.004 | 1 | 2 | Pdpn | 0.008 | Young |
| 0.07023 | 1.199062 | 0.021 | 0.011 | 1 | 2 | Dpysl3 | 0.01 | Young |
| 0.070235 | 0.267183 | 0.455 | 0.437 | 1 | 2 | Slc3a2 | 0.018 | Young |
| 0.070358 | 0.292481 | 0.337 | 0.308 | 1 | 2 | Srsf9 | 0.029 | Young |
| 0.070429 | 1.023155 | 0.012 | 0.004 | 1 | 2 | Cdk5r1 | 0.008 | Young |
| 0.070467 | 0.285011 | 0.343 | 0.315 | 1 | 2 | Abca9 | 0.028 | Young |
| 0.070523 | 1.22042 | 0.012 | 0.004 | 1 | 2 | Epor | 0.008 | Young |

| | | | | | | | | |
|---|---|---|---|---|---|---|---|---|
| 0.070523 | 1.194865 | 0.012 | 0.004 | 1 | 2 | Rangrf | 0.008 | Young |
| 0.070831 | 0.994101 | 0.021 | 0.011 | 1 | 2 | 2310057M | 0.01 | Young |
| 0.070888 | 0.481451 | 0.11 | 0.086 | 1 | 2 | Smim3 | 0.024 | Young |
| 0.071184 | 0.86299 | 0.012 | 0.004 | 1 | 2 | Nav2 | 0.008 | Young |
| 0.071393 | 0.205612 | 0.283 | 0.251 | 1 | 2 | Crlf3 | 0.032 | Young |
| 0.071469 | 0.60604 | 0.012 | 0.004 | 1 | 2 | 1810009A1 | 0.008 | Young |
| 0.071695 | 0.149043 | 0.254 | 0.219 | 1 | 2 | Arl5c | 0.035 | Young |
| 0.071733 | 0.286797 | 0.27 | 0.238 | 1 | 2 | Lmo2 | 0.032 | Young |
| 0.071815 | 0.826858 | 0.05 | 0.033 | 1 | 2 | Agpat2 | 0.017 | Young |
| 0.072019 | 0.372799 | 0.162 | 0.133 | 1 | 2 | Rnaseh2c | 0.029 | Young |
| 0.072181 | 0.338885 | 0.083 | 0.061 | 1 | 2 | Txndc16 | 0.022 | Young |
| 0.072233 | 0.446715 | 0.012 | 0.004 | 1 | 2 | Ankrd9 | 0.008 | Young |
| 0.072327 | 0.686433 | 0.05 | 0.033 | 1 | 2 | Rbmx | 0.017 | Young |
| 0.072519 | 0.572835 | 0.05 | 0.033 | 1 | 2 | Ubap2 | 0.017 | Young |
| 0.072519 | 1.830885 | 0.017 | 0.007 | 1 | 2 | Klrb1b | 0.01 | Young |
| 0.072799 | 0.293623 | 0.281 | 0.252 | 1 | 2 | Psmd4 | 0.029 | Young |
| 0.073025 | 0.208879 | 0.413 | 0.383 | 1 | 2 | P2rx4 | 0.03 | Young |
| 0.073685 | 0.263356 | 0.342 | 0.315 | 1 | 2 | Tmem160 | 0.027 | Young |
| 0.07414 | 0.641652 | 0.037 | 0.022 | 1 | 2 | Sema4c | 0.015 | Young |
| 0.074451 | 0.546453 | 0.071 | 0.051 | 1 | 2 | Mcee | 0.02 | Young |
| 0.07468 | 0.404117 | 0.128 | 0.103 | 1 | 2 | Pes1 | 0.025 | Young |
| 0.074983 | 0.957948 | 0.03 | 0.017 | 1 | 2 | Fam151b | 0.013 | Young |
| 0.074994 | 1.004278 | 0.017 | 0.007 | 1 | 2 | Pcolce | 0.01 | Young |
| 0.075191 | 1.694799 | 0.023 | 0.012 | 1 | 2 | Jaml | 0.011 | Young |
| 0.075415 | 0.534318 | 0.05 | 0.033 | 1 | 2 | Cntln | 0.017 | Young |
| 0.076298 | 0.867742 | 0.017 | 0.007 | 1 | 2 | Zfp658 | 0.01 | Young |
| 0.07635 | 0.106492 | 0.803 | 0.783 | 1 | 2 | Atp5l | 0.02 | Young |
| 0.076414 | 0.408345 | 0.152 | 0.125 | 1 | 2 | Baiap2 | 0.027 | Young |
| 0.07654 | 0.415792 | 0.129 | 0.104 | 1 | 2 | Timm8b | 0.025 | Young |
| 0.076709 | 0.599266 | 0.017 | 0.007 | 1 | 2 | C130013H( | 0.01 | Young |
| 0.077046 | 0.692176 | 0.023 | 0.012 | 1 | 2 | Plk4 | 0.011 | Young |
| 0.077184 | 0.12032 | 0.675 | 0.641 | 1 | 2 | Rpl4 | 0.034 | Young |
| 0.077377 | 0.29834 | 0.272 | 0.245 | 1 | 2 | Trmt112 | 0.027 | Young |
| 0.077489 | 0.520818 | 0.145 | 0.119 | 1 | 2 | Naa38 | 0.026 | Young |
| 0.077703 | 0.368581 | 0.186 | 0.155 | 1 | 2 | Zfc3h1 | 0.031 | Young |
| 0.077866 | 0.383419 | 0.162 | 0.135 | 1 | 2 | Slc25a4 | 0.027 | Young |
| 0.078131 | 0.450474 | 0.14 | 0.115 | 1 | 2 | Slc30a7 | 0.025 | Young |
| 0.078132 | 0.286348 | 0.037 | 0.022 | 1 | 2 | Ankrd54 | 0.015 | Young |
| 0.078426 | 0.772556 | 0.055 | 0.037 | 1 | 2 | Cdc42ep4 | 0.018 | Young |
| 0.078444 | 0.832223 | 0.023 | 0.012 | 1 | 2 | 9930104L0 | 0.011 | Young |
| 0.078907 | 0.243143 | 0.336 | 0.3 | 1 | 2 | Eps15 | 0.036 | Young |
| 0.078939 | 0.138811 | 0.6 | 0.555 | 1 | 2 | C1qc | 0.045 | Young |
| 0.079084 | 0.124276 | 0.648 | 0.61 | 1 | 2 | Gnas | 0.038 | Young |
| 0.079335 | 0.350385 | 0.128 | 0.103 | 1 | 2 | Zkscan3 | 0.025 | Young |
| 0.079691 | 0.539488 | 0.114 | 0.09 | 1 | 2 | Nfatc2 | 0.024 | Young |

| | | | | | | | | |
|---|---|---|---|---|---|---|---|---|
| 0.080014 | 0.952496 | 0.042 | 0.027 | 1 | 2 | Snrnp35 | 0.015 | Young |
| 0.080143 | 1.573421 | 0.032 | 0.019 | 1 | 2 | Areg | 0.013 | Young |
| 0.080269 | 0.685102 | 0.055 | 0.037 | 1 | 2 | Cebpzos | 0.018 | Young |
| 0.080484 | 0.523357 | 0.051 | 0.034 | 1 | 2 | Pla2g2d | 0.017 | Young |
| 0.080932 | 0.743707 | 0.062 | 0.044 | 1 | 2 | Jagn1 | 0.018 | Young |
| 0.081402 | 0.882931 | 0.042 | 0.027 | 1 | 2 | Zfp358 | 0.015 | Young |
| 0.081686 | 0.376006 | 0.207 | 0.18 | 1 | 2 | Spg21 | 0.027 | Young |
| 0.082273 | 1.241554 | 0.024 | 0.013 | 1 | 2 | Galnt6 | 0.011 | Young |
| 0.082298 | 0.511413 | 0.075 | 0.055 | 1 | 2 | Ezh2 | 0.02 | Young |
| 0.082333 | 1.342581 | 0.018 | 0.009 | 1 | 2 | Ogg1 | 0.009 | Young |
| 0.082415 | 0.933941 | 0.058 | 0.041 | 1 | 2 | Arl11 | 0.017 | Young |
| 0.0825 | 0.206202 | 0.273 | 0.241 | 1 | 2 | Fam117b | 0.032 | Young |
| 0.082996 | 0.129884 | 0.663 | 0.638 | 1 | 2 | Rps27l | 0.025 | Young |
| 0.083836 | 0.43633 | 0.22 | 0.195 | 1 | 2 | Szrd1 | 0.025 | Young |
| 0.084614 | 0.155665 | 0.234 | 0.202 | 1 | 2 | Iqgap2 | 0.032 | Young |
| 0.084649 | 0.701413 | 0.032 | 0.019 | 1 | 2 | Mcm7 | 0.013 | Young |
| 0.08481 | 0.524339 | 0.099 | 0.077 | 1 | 2 | Virma | 0.022 | Young |
| 0.085312 | 0.124375 | 0.083 | 0.062 | 1 | 2 | Pgm2l1 | 0.021 | Young |
| 0.085424 | 0.402085 | 0.122 | 0.099 | 1 | 2 | Arhgef3 | 0.023 | Young |
| 0.086119 | 0.416115 | 0.089 | 0.067 | 1 | 2 | Nop16 | 0.022 | Young |
| 0.086201 | 0.39223 | 0.23 | 0.202 | 1 | 2 | Fgd2 | 0.028 | Young |
| 0.086445 | 0.341169 | 0.032 | 0.019 | 1 | 2 | Supt3 | 0.013 | Young |
| 0.086787 | 0.195733 | 0.549 | 0.519 | 1 | 2 | Ncl | 0.03 | Young |
| 0.087169 | 0.098462 | 0.583 | 0.551 | 1 | 2 | Mtdh | 0.032 | Young |
| 0.087651 | 0.191754 | 0.429 | 0.398 | 1 | 2 | Sfr1 | 0.031 | Young |
| 0.087668 | 0.643536 | 0.061 | 0.043 | 1 | 2 | Paox | 0.018 | Young |
| 0.087767 | 0.606489 | 0.11 | 0.088 | 1 | 2 | Zfp992 | 0.022 | Young |
| 0.087854 | 1.049788 | 0.025 | 0.014 | 1 | 2 | Gm46430 | 0.011 | Young |
| 0.088295 | 0.340145 | 0.113 | 0.089 | 1 | 2 | Scrib | 0.024 | Young |
| 0.088782 | 0.605792 | 0.032 | 0.019 | 1 | 2 | Cdc23 | 0.013 | Young |
| 0.088948 | 0.268389 | 0.385 | 0.357 | 1 | 2 | Llph | 0.028 | Young |
| 0.089149 | 0.349468 | 0.127 | 0.103 | 1 | 2 | Rnf220 | 0.024 | Young |
| 0.08923 | 0.773187 | 0.121 | 0.1 | 1 | 2 | Adam19 | 0.021 | Young |
| 0.089238 | 0.350356 | 0.084 | 0.063 | 1 | 2 | Gtf3c1 | 0.021 | Young |
| 0.089274 | 0.114212 | 0.711 | 0.677 | 1 | 2 | Ptafr | 0.034 | Young |
| 0.089374 | 0.74695 | 0.045 | 0.03 | 1 | 2 | S1pr1 | 0.015 | Young |
| 0.089923 | 0.170026 | 0.45 | 0.416 | 1 | 2 | Trf | 0.034 | Young |
| 0.090525 | 0.585839 | 0.045 | 0.03 | 1 | 2 | Lin9 | 0.015 | Young |
| 0.090558 | 0.20897 | 0.357 | 0.327 | 1 | 2 | Ifnar1 | 0.03 | Young |
| 0.090564 | 0.322027 | 0.171 | 0.143 | 1 | 2 | Tifa | 0.028 | Young |
| 0.090859 | 1.257024 | 0.037 | 0.024 | 1 | 2 | Tma16 | 0.013 | Young |
| 0.091005 | 1.226295 | 0.01 | 0.003 | 1 | 2 | 4932422M | 0.007 | Young |
| 0.0911 | 0.220323 | 0.121 | 0.096 | 1 | 2 | Fxyd2 | 0.025 | Young |
| 0.091104 | 0.210235 | 0.319 | 0.289 | 1 | 2 | Arhgap18 | 0.03 | Young |
| 0.091136 | 1.102229 | 0.01 | 0.003 | 1 | 2 | Wee1 | 0.007 | Young |

| | | | | | | | | |
|---|---|---|---|---|---|---|---|---|
| 0.091266 | 1.270069 | 0.01 | 0.003 | 1 | 2 | 2900026A( | 0.007 | Young |
| 0.091266 | 1.138046 | 0.01 | 0.003 | 1 | 2 | Socs2 | 0.007 | Young |
| 0.091266 | 0.993415 | 0.01 | 0.003 | 1 | 2 | Ccdc15 | 0.007 | Young |
| 0.091659 | 0.903622 | 0.01 | 0.003 | 1 | 2 | Tex9 | 0.007 | Young |
| 0.091669 | 0.783649 | 0.034 | 0.021 | 1 | 2 | Mfsd13a | 0.013 | Young |
| 0.091791 | 0.912727 | 0.01 | 0.003 | 1 | 2 | Abhd6 | 0.007 | Young |
| 0.091922 | 1.425291 | 0.01 | 0.003 | 1 | 2 | Acy1 | 0.007 | Young |
| 0.091922 | 1.059735 | 0.01 | 0.003 | 1 | 2 | Tmem72 | 0.007 | Young |
| 0.091922 | 0.784511 | 0.01 | 0.003 | 1 | 2 | Oxld1 | 0.007 | Young |
| 0.092223 | 0.440877 | 0.151 | 0.125 | 1 | 2 | Acvrl1 | 0.026 | Young |
| 0.092297 | 1.142501 | 0.026 | 0.015 | 1 | 2 | Twnk | 0.011 | Young |
| 0.092657 | 0.579272 | 0.096 | 0.075 | 1 | 2 | Phf12 | 0.021 | Young |
| 0.092783 | 0.233014 | 0.113 | 0.089 | 1 | 2 | Mb21d1 | 0.024 | Young |
| 0.092806 | 0.426492 | 0.108 | 0.085 | 1 | 2 | Proser1 | 0.023 | Young |
| 0.092979 | 0.671391 | 0.01 | 0.003 | 1 | 2 | Ccnj | 0.007 | Young |
| 0.093068 | 0.171739 | 0.359 | 0.327 | 1 | 2 | Rab2a | 0.032 | Young |
| 0.093196 | 1.016676 | 0.057 | 0.041 | 1 | 2 | Tmem64 | 0.016 | Young |
| 0.093307 | 1.16096 | 0.019 | 0.01 | 1 | 2 | Zfp946 | 0.009 | Young |
| 0.094024 | 0.59622 | 0.076 | 0.057 | 1 | 2 | Apeh | 0.019 | Young |
| 0.094335 | 0.058942 | 0.461 | 0.422 | 1 | 2 | Eid1 | 0.039 | Young |
| 0.094357 | 0.448542 | 0.126 | 0.103 | 1 | 2 | Opa3 | 0.023 | Young |
| 0.094384 | 0.632737 | 0.076 | 0.057 | 1 | 2 | Agl | 0.019 | Young |
| 0.094413 | 0.227034 | 0.232 | 0.202 | 1 | 2 | Ypel3 | 0.03 | Young |
| 0.094465 | 0.596078 | 0.019 | 0.01 | 1 | 2 | Setd4 | 0.009 | Young |
| 0.095033 | 0.160825 | 0.201 | 0.171 | 1 | 2 | Fam91a1 | 0.03 | Young |
| 0.095183 | 0.701155 | 0.019 | 0.01 | 1 | 2 | Csf3 | 0.009 | Young |
| 0.095864 | 0.659834 | 0.049 | 0.033 | 1 | 2 | Ccdc58 | 0.016 | Young |
| 0.096087 | 0.719185 | 0.052 | 0.036 | 1 | 2 | Poldip2 | 0.016 | Young |
| 0.096256 | 0.671968 | 0.026 | 0.015 | 1 | 2 | Cnnm2 | 0.011 | Young |
| 0.096661 | 0.570633 | 0.132 | 0.109 | 1 | 2 | Pdgfb | 0.023 | Young |
| 0.096781 | 0.478079 | 0.068 | 0.049 | 1 | 2 | Uck2 | 0.019 | Young |
| 0.096997 | 0.579637 | 0.048 | 0.032 | 1 | 2 | Prkci | 0.016 | Young |
| 0.097048 | 0.096049 | 0.502 | 0.465 | 1 | 2 | Dab2 | 0.037 | Young |
| 0.097106 | 0.529184 | 0.05 | 0.034 | 1 | 2 | Prr14l | 0.016 | Young |
| 0.097216 | 0.382141 | 0.078 | 0.059 | 1 | 2 | Cd209f | 0.019 | Young |
| 0.097323 | 0.281434 | 0.192 | 0.165 | 1 | 2 | Mark2 | 0.027 | Young |
| 0.097666 | 0.483218 | 0.036 | 0.022 | 1 | 2 | Snx19 | 0.014 | Young |
| 0.097838 | 0.245977 | 0.163 | 0.136 | 1 | 2 | Antxr2 | 0.027 | Young |
| 0.098261 | 0.28707 | 0.305 | 0.278 | 1 | 2 | Eif3i | 0.027 | Young |
| 0.098576 | 1.035449 | 0.014 | 0.006 | 1 | 2 | E130102H2 | 0.008 | Young |
| 0.099122 | 1.266647 | 0.014 | 0.006 | 1 | 2 | Cdyl2 | 0.008 | Young |
| 0.099303 | 0.441226 | 0.158 | 0.134 | 1 | 2 | Blnk | 0.024 | Young |
| 0.099936 | 0.748768 | 0.027 | 0.016 | 1 | 2 | Dimt1 | 0.011 | Young |
| 0.100256 | 0.689395 | 0.055 | 0.039 | 1 | 2 | Mark4 | 0.016 | Young |
| 0.100479 | 0.178815 | 0.248 | 0.222 | 1 | 2 | Spty2d1 | 0.026 | Young |

| | | | | | | | | |
|---|---|---|---|---|---|---|---|---|
| 0.100695 | 0.446935 | 0.108 | 0.086 | 1 | 2 | Exosc5 | 0.022 | Young |
| 0.101002 | 0.378657 | 0.131 | 0.106 | 1 | 2 | Rpp21 | 0.025 | Young |
| 0.101068 | 0.106538 | 0.411 | 0.376 | 1 | 2 | Fkbp1a | 0.035 | Young |
| 0.101263 | 0.151537 | 0.148 | 0.123 | 1 | 2 | Ppfia4 | 0.025 | Young |
| 0.101317 | 0.911247 | 0.02 | 0.011 | 1 | 2 | Rnf26 | 0.009 | Young |
| 0.101664 | 0.646248 | 0.014 | 0.006 | 1 | 2 | Dcaf4 | 0.008 | Young |
| 0.101682 | 1.045915 | 0.02 | 0.011 | 1 | 2 | Scrn3 | 0.009 | Young |
| 0.101956 | 0.766282 | 0.02 | 0.011 | 1 | 2 | Myh10 | 0.009 | Young |
| 0.101961 | 0.333913 | 0.178 | 0.151 | 1 | 2 | Pdpk1 | 0.027 | Young |
| 0.102011 | 0.517141 | 0.027 | 0.016 | 1 | 2 | Gstcd | 0.011 | Young |
| 0.102143 | 0.717886 | 0.038 | 0.025 | 1 | 2 | Gprc5c | 0.013 | Young |
| 0.102188 | 0.305625 | 0.138 | 0.112 | 1 | 2 | Herpud2 | 0.026 | Young |
| 0.102323 | 0.871309 | 0.02 | 0.011 | 1 | 2 | Hepacam2 | 0.009 | Young |
| 0.102529 | 0.315096 | 0.308 | 0.281 | 1 | 2 | Scarb2 | 0.027 | Young |
| 0.102801 | 1.059746 | 0.039 | 0.026 | 1 | 2 | Tmem154 | 0.013 | Young |
| 0.102868 | 0.059336 | 0.753 | 0.709 | 1 | 2 | Cox7c | 0.044 | Young |
| 0.102897 | 0.381549 | 0.156 | 0.131 | 1 | 2 | Tmem173 | 0.025 | Young |
| 0.103095 | 0.390612 | 0.084 | 0.064 | 1 | 2 | Fam193b | 0.02 | Young |
| 0.103158 | 0.178464 | 0.233 | 0.2 | 1 | 2 | Sash1 | 0.033 | Young |
| 0.103219 | 0.058648 | 0.126 | 0.1 | 1 | 2 | Fbxw4 | 0.026 | Young |
| 0.103726 | 0.239227 | 0.324 | 0.299 | 1 | 2 | Bcl10 | 0.025 | Young |
| 0.103883 | 0.338891 | 0.154 | 0.131 | 1 | 2 | Ldb1 | 0.023 | Young |
| 0.104176 | 0.286594 | 0.235 | 0.208 | 1 | 2 | Bod1l | 0.027 | Young |
| 0.104196 | 0.464145 | 0.052 | 0.036 | 1 | 2 | Ints7 | 0.016 | Young |
| 0.104913 | 0.006103 | 0.074 | 0.096 | 1 | 2 | Rras2 | -0.022 | Young |
| 0.10549 | 0.331727 | 0.126 | 0.103 | 1 | 2 | Pcbd2 | 0.023 | Young |
| 0.105606 | 0.161409 | 0.429 | 0.401 | 1 | 2 | Sowahc | 0.028 | Young |
| 0.106298 | 0.35062 | 0.171 | 0.146 | 1 | 2 | Ccdc47 | 0.025 | Young |
| 0.106625 | 1.149806 | 0.021 | 0.012 | 1 | 2 | Slamf6 | 0.009 | Young |
| 0.106716 | 0.61675 | 0.062 | 0.045 | 1 | 2 | Kat6b | 0.017 | Young |
| 0.106754 | 0.378891 | 0.148 | 0.124 | 1 | 2 | Hmgn1 | 0.024 | Young |
| 0.106794 | 0.036593 | 0.049 | 0.033 | 1 | 2 | Kifc3 | 0.016 | Young |
| 0.107137 | 0.434943 | 0.118 | 0.095 | 1 | 2 | Drg1 | 0.023 | Young |
| 0.107284 | 0.174298 | 0.439 | 0.412 | 1 | 2 | Tbca | 0.027 | Young |
| 0.107822 | 0.995785 | 0.021 | 0.012 | 1 | 2 | Gm26778 | 0.009 | Young |
| 0.108047 | 0.620677 | 0.069 | 0.051 | 1 | 2 | Ppa1 | 0.018 | Young |
| 0.108442 | 0.484225 | 0.151 | 0.127 | 1 | 2 | Impdh2 | 0.024 | Young |
| 0.108489 | 0.084157 | 0.89 | 0.876 | 1 | 2 | Rpl5 | 0.014 | Young |
| 0.109337 | 0.211947 | 0.219 | 0.192 | 1 | 2 | Cd164 | 0.027 | Young |
| 0.109473 | 0.677271 | 0.04 | 0.027 | 1 | 2 | Dhx32 | 0.013 | Young |
| 0.109615 | 0.459212 | 0.069 | 0.051 | 1 | 2 | Htatip2 | 0.018 | Young |
| 0.10989 | 0.288245 | 0.391 | 0.37 | 1 | 2 | Smarca5 | 0.021 | Young |
| 0.110002 | 0.233155 | 0.141 | 0.116 | 1 | 2 | Snrk | 0.025 | Young |
| 0.110222 | 0.447837 | 0.203 | 0.181 | 1 | 2 | Igbp1 | 0.022 | Young |
| 0.110246 | 1.184655 | 0.021 | 0.012 | 1 | 2 | 2700049A( | 0.009 | Young |

| | | | | | | | | |
|---|---|---|---|---|---|---|---|---|
| 0.110572 | 0.621127 | 0.091 | 0.073 | 1 | 2 | Fbxo34 | 0.018 | Young |
| 0.110574 | 0.134879 | 0.191 | 0.163 | 1 | 2 | Twistnb | 0.028 | Young |
| 0.110659 | 0.227637 | 0.343 | 0.312 | 1 | 2 | Tiparp | 0.031 | Young |
| 0.110905 | 0.570827 | 0.106 | 0.085 | 1 | 2 | Alox5 | 0.021 | Young |
| 0.110941 | 0.186724 | 0.483 | 0.465 | 1 | 2 | Sec61g | 0.018 | Young |
| 0.111096 | 0.794974 | 0.021 | 0.012 | 1 | 2 | Odf2l | 0.009 | Young |
| 0.111162 | 0.432372 | 0.128 | 0.106 | 1 | 2 | Fam98b | 0.022 | Young |
| 0.111538 | 0.092545 | 0.658 | 0.638 | 1 | 2 | Erp29 | 0.02 | Young |
| 0.111539 | 0.664141 | 0.043 | 0.029 | 1 | 2 | Tex30 | 0.014 | Young |
| 0.111845 | 0.677898 | 0.072 | 0.055 | 1 | 2 | Vps52 | 0.017 | Young |
| 0.112086 | 1.22374 | 0.023 | 0.013 | 1 | 2 | Serf1 | 0.01 | Young |
| 0.112228 | 0.398867 | 0.061 | 0.044 | 1 | 2 | Ggt5 | 0.017 | Young |
| 0.112485 | 0.632069 | 0.043 | 0.029 | 1 | 2 | Adprhl2 | 0.014 | Young |
| 0.112537 | 0.362677 | 0.159 | 0.134 | 1 | 2 | Folr2 | 0.025 | Young |
| 0.112759 | 0.248667 | 0.137 | 0.112 | 1 | 2 | Txndc11 | 0.025 | Young |
| 0.112793 | 0.147179 | 0.352 | 0.322 | 1 | 2 | Zfp106 | 0.03 | Young |
| 0.112906 | 0.604303 | 0.021 | 0.012 | 1 | 2 | Cmtm4 | 0.009 | Young |
| 0.113381 | 1.197153 | 0.011 | 0.004 | 1 | 2 | Zfp951 | 0.007 | Young |
| 0.11345 | 0.791556 | 0.015 | 0.007 | 1 | 2 | Mad2l1 | 0.008 | Young |
| 0.113566 | 0.751739 | 0.015 | 0.007 | 1 | 2 | Col14a1 | 0.008 | Young |
| 0.113566 | 0.716007 | 0.015 | 0.007 | 1 | 2 | Pex12 | 0.008 | Young |
| 0.113644 | 0.149871 | 0.144 | 0.118 | 1 | 2 | Ccnd2 | 0.026 | Young |
| 0.113692 | 0.536366 | 0.124 | 0.102 | 1 | 2 | Ppp1r11 | 0.022 | Young |
| 0.11375 | 0.729962 | 0.107 | 0.087 | 1 | 2 | Ube2w | 0.02 | Young |
| 0.113985 | 0.87916 | 0.031 | 0.019 | 1 | 2 | Haus6 | 0.012 | Young |
| 0.114029 | 0.641107 | 0.015 | 0.007 | 1 | 2 | Zfp58 | 0.008 | Young |
| 0.114146 | 0.467425 | 0.015 | 0.007 | 1 | 2 | 3-Mar | 0.008 | Young |
| 0.114242 | 0.829255 | 0.011 | 0.004 | 1 | 2 | AC174678. | 0.007 | Young |
| 0.114317 | 0.999714 | 0.023 | 0.013 | 1 | 2 | Mamdc2 | 0.01 | Young |
| 0.114378 | 0.432498 | 0.015 | 0.007 | 1 | 2 | Mccc2 | 0.008 | Young |
| 0.114386 | 1.018694 | 0.011 | 0.004 | 1 | 2 | Ppcs | 0.007 | Young |
| 0.114534 | 0.446712 | 0.071 | 0.054 | 1 | 2 | Eif2ak3 | 0.017 | Young |
| 0.114674 | 0.487092 | 0.011 | 0.004 | 1 | 2 | Fntb | 0.007 | Young |
| 0.114702 | 0.654334 | 0.075 | 0.057 | 1 | 2 | Icosl | 0.018 | Young |
| 0.114881 | 0.334944 | 0.058 | 0.042 | 1 | 2 | Ddx49 | 0.016 | Young |
| 0.114963 | 1.12995 | 0.011 | 0.004 | 1 | 2 | S100a9 | 0.007 | Young |
| 0.115041 | 0.379826 | 0.152 | 0.128 | 1 | 2 | Qpct | 0.024 | Young |
| 0.115108 | 0.57467 | 0.011 | 0.004 | 1 | 2 | Prdm9 | 0.007 | Young |
| 0.115429 | 0.306397 | 0.015 | 0.007 | 1 | 2 | Usp35 | 0.008 | Young |
| 0.11604 | 0.740653 | 0.048 | 0.033 | 1 | 2 | Kmo | 0.015 | Young |
| 0.116293 | 0.131134 | 0.499 | 0.473 | 1 | 2 | Rbm25 | 0.026 | Young |
| 0.11632 | 0.778713 | 0.032 | 0.02 | 1 | 2 | Strbp | 0.012 | Young |
| 0.116347 | 0.804409 | 0.051 | 0.036 | 1 | 2 | Tbl3 | 0.015 | Young |
| 0.116665 | 0.252815 | 0.253 | 0.228 | 1 | 2 | Zc3h12a | 0.025 | Young |
| 0.116705 | 0.509555 | 0.12 | 0.099 | 1 | 2 | Mllt6 | 0.021 | Young |

| | | | | | | | | |
|---|---|---|---|---|---|---|---|---|
| 0.117535 | 0.265773 | 0.342 | 0.318 | 1 | 2 | Cggbp1 | 0.024 | Young |
| 0.117559 | 0.455037 | 0.072 | 0.055 | 1 | 2 | Tmem106c | 0.017 | Young |
| 0.117613 | 0.095509 | 0.189 | 0.158 | 1 | 2 | Kmt5a | 0.031 | Young |
| 0.118383 | 0.391389 | 0.061 | 0.044 | 1 | 2 | BC035044 | 0.017 | Young |
| 0.118713 | 0.295698 | 0.139 | 0.115 | 1 | 2 | Mefv | 0.024 | Young |
| 0.118845 | 0.151025 | 0.152 | 0.126 | 1 | 2 | Dapk1 | 0.026 | Young |
| 0.119018 | 0.293599 | 0.112 | 0.09 | 1 | 2 | Mrpl13 | 0.022 | Young |
| 0.119582 | 0.089294 | 0.043 | 0.06 | 1 | 2 | Alg1 | -0.017 | Young |
| 0.120107 | 1.181826 | 0.024 | 0.014 | 1 | 2 | Cd2 | 0.01 | Young |
| 0.120136 | 0.387894 | 0.164 | 0.139 | 1 | 2 | Mcemp1 | 0.025 | Young |
| 0.120353 | 0.375674 | 0.186 | 0.163 | 1 | 2 | Borcs6 | 0.023 | Young |
| 0.120456 | 0.213483 | 0.044 | 0.03 | 1 | 2 | Ofd1 | 0.014 | Young |
| 0.120493 | 0.086615 | 0.007 | 0.015 | 1 | 2 | Cbwd1 | -0.008 | Young |
| 0.120559 | 0.236849 | 0.14 | 0.116 | 1 | 2 | Atg5 | 0.024 | Young |
| 0.120674 | 1.071949 | 0.024 | 0.014 | 1 | 2 | Trp53i13 | 0.01 | Young |
| 0.121091 | 0.780071 | 0.053 | 0.039 | 1 | 2 | Ppfibp1 | 0.014 | Young |
| 0.12138 | 0.375582 | 0.205 | 0.178 | 1 | 2 | Ms4a7 | 0.027 | Young |
| 0.121382 | 1.176435 | 0.017 | 0.009 | 1 | 2 | Taf9b | 0.008 | Young |
| 0.121391 | 0.085164 | 0.715 | 0.672 | 1 | 2 | Rbm39 | 0.043 | Young |
| 0.121468 | 0.141718 | 0.568 | 0.541 | 1 | 2 | Bri3 | 0.027 | Young |
| 0.121497 | 0.452335 | 0.075 | 0.057 | 1 | 2 | Jmy | 0.018 | Young |
| 0.121865 | 0.286686 | 0.11 | 0.089 | 1 | 2 | Ppil4 | 0.021 | Young |
| 0.122575 | 0.425661 | 0.046 | 0.032 | 1 | 2 | Mfng | 0.014 | Young |
| 0.122666 | 1.473616 | 0.017 | 0.009 | 1 | 2 | Grhpr | 0.008 | Young |
| 0.122966 | 0.873179 | 0.024 | 0.014 | 1 | 2 | 0610009O2 | 0.01 | Young |
| 0.123357 | 0.286521 | 0.191 | 0.165 | 1 | 2 | Apc | 0.026 | Young |
| 0.123411 | 0.57282 | 0.09 | 0.072 | 1 | 2 | Dpy19l4 | 0.018 | Young |
| 0.123582 | 0.132486 | 0.253 | 0.224 | 1 | 2 | Myo1g | 0.029 | Young |
| 0.123584 | 0.396976 | 0.046 | 0.032 | 1 | 2 | Mgst3 | 0.014 | Young |
| 0.123738 | 0.801589 | 0.024 | 0.014 | 1 | 2 | Ddx20 | 0.01 | Young |
| 0.124279 | 0.181436 | 0.156 | 0.132 | 1 | 2 | Luzp1 | 0.024 | Young |
| 0.124416 | 0.625699 | 0.024 | 0.014 | 1 | 2 | Ptpn4 | 0.01 | Young |
| 0.124624 | 0.657373 | 0.088 | 0.07 | 1 | 2 | Brap | 0.018 | Young |
| 0.124625 | 0.090988 | 0.418 | 0.389 | 1 | 2 | Dnajb6 | 0.029 | Young |
| 0.124672 | 0.505897 | 0.017 | 0.009 | 1 | 2 | Ccl22 | 0.008 | Young |
| 0.124898 | 0.737941 | 0.059 | 0.044 | 1 | 2 | Wdtc1 | 0.015 | Young |
| 0.125022 | 0.484477 | 0.034 | 0.022 | 1 | 2 | Hectd4 | 0.012 | Young |
| 0.125546 | 0.188045 | 0.178 | 0.151 | 1 | 2 | Hpf1 | 0.027 | Young |
| 0.12563 | 0.190278 | 0.147 | 0.123 | 1 | 2 | Dnajc21 | 0.024 | Young |
| 0.126011 | 0.370113 | 0.132 | 0.11 | 1 | 2 | Smim4 | 0.022 | Young |
| 0.126198 | 0.322486 | 0.162 | 0.137 | 1 | 2 | Prmt1 | 0.025 | Young |
| 0.126342 | 0.452758 | 0.017 | 0.009 | 1 | 2 | Kbtbd8 | 0.008 | Young |
| 0.126939 | 0.958907 | 0.025 | 0.015 | 1 | 2 | Dnajb5 | 0.01 | Young |
| 0.127276 | 0.686879 | 0.058 | 0.043 | 1 | 2 | Eng | 0.015 | Young |
| 0.127655 | 0.553274 | 0.051 | 0.036 | 1 | 2 | Acvr1 | 0.015 | Young |

| | | | | | | | | |
|---|---|---|---|---|---|---|---|---|
| 0.127667 | 0.385527 | 0.017 | 0.009 | 1 | 2 | Zfp770 | 0.008 | Young |
| 0.127711 | 0.168978 | 0.512 | 0.493 | 1 | 2 | Tubb5 | 0.019 | Young |
| 0.128008 | 0.635767 | 0.078 | 0.061 | 1 | 2 | Zfp397 | 0.017 | Young |
| 0.128463 | 0.357702 | 0.09 | 0.072 | 1 | 2 | Mgat5 | 0.018 | Young |
| 0.128496 | 0.171184 | 0.401 | 0.376 | 1 | 2 | Tcf25 | 0.025 | Young |
| 0.128538 | 0.216061 | 0.352 | 0.327 | 1 | 2 | Arhgef6 | 0.025 | Young |
| 0.128554 | 0.592655 | 0.076 | 0.059 | 1 | 2 | Gpr137 | 0.017 | Young |
| 0.128628 | 0.590506 | 0.056 | 0.041 | 1 | 2 | H2-DMb2 | 0.015 | Young |
| 0.128682 | 0.151426 | 0.23 | 0.201 | 1 | 2 | Rnf150 | 0.029 | Young |
| 0.128847 | 0.156778 | 0.075 | 0.057 | 1 | 2 | Cish | 0.018 | Young |
| 0.128886 | 0.209461 | 0.158 | 0.133 | 1 | 2 | Cntrl | 0.025 | Young |
| 0.129053 | 0.364112 | 0.05 | 0.035 | 1 | 2 | Tbc1d4 | 0.015 | Young |
| 0.129145 | 0.312883 | 0.024 | 0.014 | 1 | 2 | Cep70 | 0.01 | Young |
| 0.129733 | 0.318642 | 0.211 | 0.188 | 1 | 2 | Sdhb | 0.023 | Young |
| 0.129742 | 0.20509 | 0.312 | 0.286 | 1 | 2 | Srrm1 | 0.026 | Young |
| 0.129991 | 0.912267 | 0.038 | 0.026 | 1 | 2 | Gtf3c4 | 0.012 | Young |
| 0.130094 | 0.176484 | 0.176 | 0.15 | 1 | 2 | Pip5k1c | 0.026 | Young |
| 0.130227 | 0.130244 | 0.663 | 0.645 | 1 | 2 | Ifrd1 | 0.018 | Young |
| 0.130645 | 0.28272 | 0.083 | 0.064 | 1 | 2 | Fam199x | 0.019 | Young |
| 0.131003 | 0.870144 | 0.059 | 0.044 | 1 | 2 | Slc2a1 | 0.015 | Young |
| 0.131059 | 0.547586 | 0.034 | 0.022 | 1 | 2 | Cables2 | 0.012 | Young |
| 0.131094 | 0.203812 | 0.102 | 0.081 | 1 | 2 | Dock4 | 0.021 | Young |
| 0.131118 | 0.501893 | 0.053 | 0.039 | 1 | 2 | Piga | 0.014 | Young |
| 0.131402 | 1.072342 | 0.012 | 0.005 | 1 | 2 | Synpo | 0.007 | Young |
| 0.131575 | 0.597177 | 0.096 | 0.077 | 1 | 2 | Nmt2 | 0.019 | Young |
| 0.13181 | 0.53169 | 0.051 | 0.036 | 1 | 2 | Etfdh | 0.015 | Young |
| 0.1319 | 1.072604 | 0.026 | 0.016 | 1 | 2 | Gtpbp8 | 0.01 | Young |
| 0.132211 | 1.225271 | 0.018 | 0.01 | 1 | 2 | Pomk | 0.008 | Young |
| 0.132573 | 0.206962 | 0.38 | 0.359 | 1 | 2 | Cox7b | 0.021 | Young |
| 0.132629 | 0.63813 | 0.037 | 0.025 | 1 | 2 | Fra10ac1 | 0.012 | Young |
| 0.132753 | 1.393845 | 0.012 | 0.005 | 1 | 2 | Hspb1 | 0.007 | Young |
| 0.132753 | 1.022965 | 0.012 | 0.005 | 1 | 2 | Cd4 | 0.007 | Young |
| 0.132807 | 1.368157 | 0.018 | 0.01 | 1 | 2 | Sccpdh | 0.008 | Young |
| 0.13303 | 0.213001 | 0.14 | 0.117 | 1 | 2 | Marveld1 | 0.023 | Young |
| 0.133054 | 1.083909 | 0.012 | 0.005 | 1 | 2 | Mmp13 | 0.007 | Young |
| 0.133156 | 0.852758 | 0.061 | 0.045 | 1 | 2 | Smad7 | 0.016 | Young |
| 0.133205 | 0.867512 | 0.012 | 0.005 | 1 | 2 | Akap17b | 0.007 | Young |
| 0.133234 | 0.156399 | 0.082 | 0.063 | 1 | 2 | Itgax | 0.019 | Young |
| 0.133811 | 0.899596 | 0.012 | 0.005 | 1 | 2 | Satb2 | 0.007 | Young |
| 0.134061 | 0.723806 | 0.039 | 0.027 | 1 | 2 | Abhd11 | 0.012 | Young |
| 0.134609 | 0.615913 | 0.018 | 0.01 | 1 | 2 | Cdan1 | 0.008 | Young |
| 0.135027 | 0.267053 | 0.116 | 0.095 | 1 | 2 | Fam214b | 0.021 | Young |
| 0.135129 | 0.334016 | 0.159 | 0.135 | 1 | 2 | Jup | 0.024 | Young |
| 0.135134 | 0.531888 | 0.037 | 0.025 | 1 | 2 | Lix1l | 0.012 | Young |
| 0.135333 | 0.411249 | 0.012 | 0.005 | 1 | 2 | A530013C2 | 0.007 | Young |

| | | | | | | | | |
|---|---|---|---|---|---|---|---|---|
| 0.135727 | 0.843752 | 0.043 | 0.03 | 1 | 2 | Rrp9 | 0.013 | Young |
| 0.135753 | 0.296105 | 0.211 | 0.185 | 1 | 2 | Col4a3bp | 0.026 | Young |
| 0.135884 | 0.845945 | 0.026 | 0.016 | 1 | 2 | Vars2 | 0.01 | Young |
| 0.135941 | 0.170929 | 0.097 | 0.078 | 1 | 2 | Pdcd2l | 0.019 | Young |
| 0.13626 | 0.33292 | 0.115 | 0.095 | 1 | 2 | Bag6 | 0.02 | Young |
| 0.136261 | 0.551899 | 0.006 | 0.013 | 1 | 2 | Tnfrsf23 | -0.007 | Young |
| 0.136956 | 0.53785 | 0.076 | 0.059 | 1 | 2 | Cops7a | 0.017 | Young |
| 0.137248 | 0.045299 | 0.063 | 0.082 | 1 | 2 | Kif13a | -0.019 | Young |
| 0.137357 | 0.101087 | 0.053 | 0.072 | 1 | 2 | Fam222b | -0.019 | Young |
| 0.137939 | 0.583715 | 0.124 | 0.103 | 1 | 2 | Vegfa | 0.021 | Young |
| 0.138095 | 0.062791 | 0.132 | 0.108 | 1 | 2 | Ncbp3 | 0.024 | Young |
| 0.138159 | 0.681335 | 0.026 | 0.016 | 1 | 2 | Maged1 | 0.01 | Young |
| 0.138516 | 1.330627 | 0.029 | 0.018 | 1 | 2 | Mettl26 | 0.011 | Young |
| 0.138883 | 0.40131 | 0.064 | 0.048 | 1 | 2 | Wdr36 | 0.016 | Young |
| 0.139241 | 0.863072 | 0.042 | 0.029 | 1 | 2 | Hdhd2 | 0.013 | Young |
| 0.139275 | 0.604203 | 0.113 | 0.094 | 1 | 2 | Pak1 | 0.019 | Young |
| 0.139358 | 0.613909 | 0.026 | 0.016 | 1 | 2 | Ralgps1 | 0.01 | Young |
| 0.139371 | 0.490343 | 0.076 | 0.059 | 1 | 2 | Upf3b | 0.017 | Young |
| 0.14027 | 0.342098 | 0.101 | 0.081 | 1 | 2 | Trp53inp2 | 0.02 | Young |
| 0.141797 | 0.250253 | 0.057 | 0.042 | 1 | 2 | Ampd3 | 0.015 | Young |
| 0.14185 | 0.219383 | 0.216 | 0.188 | 1 | 2 | Oat | 0.028 | Young |
| 0.14194 | 0.582658 | 0.04 | 0.028 | 1 | 2 | Msh2 | 0.012 | Young |
| 0.141985 | 0.142889 | 0.342 | 0.319 | 1 | 2 | Elob | 0.023 | Young |
| 0.142157 | 0.948263 | 0.029 | 0.018 | 1 | 2 | Mapkbp1 | 0.011 | Young |
| 0.142221 | 0.413859 | 0.095 | 0.076 | 1 | 2 | Med14 | 0.019 | Young |
| 0.142535 | 0.345177 | 0.082 | 0.064 | 1 | 2 | Fam213b | 0.018 | Young |
| 0.142613 | 0.999336 | 0.019 | 0.011 | 1 | 2 | Gem | 0.008 | Young |
| 0.142664 | 0.561894 | 0.04 | 0.028 | 1 | 2 | Armt1 | 0.012 | Young |
| 0.143251 | 0.089073 | 0.6 | 0.577 | 1 | 2 | Prdx1 | 0.023 | Young |
| 0.143255 | 0.717672 | 0.045 | 0.032 | 1 | 2 | Kifap3 | 0.013 | Young |
| 0.143368 | 0.257917 | 0.201 | 0.178 | 1 | 2 | Smndc1 | 0.023 | Young |
| 0.143479 | 0.042622 | 0.988 | 0.986 | 1 | 2 | Rpl41 | 0.002 | Young |
| 0.144128 | 0.266518 | 0.115 | 0.094 | 1 | 2 | Nabp2 | 0.021 | Young |
| 0.144201 | 0.604009 | 0.029 | 0.018 | 1 | 2 | Igfbp6 | 0.011 | Young |
| 0.144301 | 0.332541 | 0.115 | 0.095 | 1 | 2 | Mrps36 | 0.02 | Young |
| 0.144322 | 0.404304 | 0.252 | 0.232 | 1 | 2 | Rp9 | 0.02 | Young |
| 0.144889 | 0.095959 | 0.008 | 0.016 | 1 | 2 | Cfap45 | -0.008 | Young |
| 0.145106 | 0.171134 | 0.058 | 0.043 | 1 | 2 | Nubp2 | 0.015 | Young |
| 0.145128 | 0.256145 | 0.065 | 0.049 | 1 | 2 | Atad2 | 0.016 | Young |
| 0.14517 | 0.565998 | 0.044 | 0.031 | 1 | 2 | Fbxo42 | 0.013 | Young |
| 0.1454 | 0.437571 | 0.113 | 0.093 | 1 | 2 | Usp38 | 0.02 | Young |
| 0.145874 | 0.271797 | 0.217 | 0.192 | 1 | 2 | Chd7 | 0.025 | Young |
| 0.146401 | 0.829336 | 0.048 | 0.034 | 1 | 2 | Zfp369 | 0.014 | Young |
| 0.14674 | 0.225062 | 0.285 | 0.26 | 1 | 2 | Ssr4 | 0.025 | Young |
| 0.147267 | 0.576375 | 0.12 | 0.101 | 1 | 2 | Vgll4 | 0.019 | Young |

| | | | | | | | | |
|---|---|---|---|---|---|---|---|---|
| 0.147858 | 0.420394 | 0.158 | 0.137 | 1 | 2 | Zfp652 | 0.021 | Young |
| 0.147862 | 0.287322 | 0.182 | 0.157 | 1 | 2 | Ccnd1 | 0.025 | Young |
| 0.147885 | 0.282221 | 0.167 | 0.146 | 1 | 2 | Il6st | 0.021 | Young |
| 0.148029 | 0.628441 | 0.019 | 0.011 | 1 | 2 | Dennd2c | 0.008 | Young |
| 0.148179 | 0.308717 | 0.099 | 0.079 | 1 | 2 | Scaf1 | 0.02 | Young |
| 0.148499 | 1.108413 | 0.031 | 0.02 | 1 | 2 | Rbm41 | 0.011 | Young |
| 0.148506 | 0.569717 | 0.029 | 0.018 | 1 | 2 | Rac3 | 0.011 | Young |
| 0.148605 | 0.424669 | 0.132 | 0.111 | 1 | 2 | Luc7l | 0.021 | Young |
| 0.148985 | 1.068689 | 0.03 | 0.019 | 1 | 2 | Fam118a | 0.011 | Young |
| 0.149065 | 0.268154 | 0.167 | 0.145 | 1 | 2 | Ube2v2 | 0.022 | Young |
| 0.149636 | 0.196578 | 0.116 | 0.095 | 1 | 2 | Wdr44 | 0.021 | Young |
| 0.149667 | 0.21451 | 0.143 | 0.121 | 1 | 2 | Anapc16 | 0.022 | Young |
| 0.149686 | 0.373245 | 0.045 | 0.032 | 1 | 2 | Mcm4 | 0.013 | Young |
| 0.149818 | 0.54654 | 0.07 | 0.055 | 1 | 2 | Nde1 | 0.015 | Young |
| 0.150209 | 0.301278 | 0.081 | 0.063 | 1 | 2 | Fem1a | 0.018 | Young |
| 0.150558 | 0.541705 | 0.083 | 0.066 | 1 | 2 | Sin3a | 0.017 | Young |
| 0.150963 | 0.114999 | 0.371 | 0.342 | 1 | 2 | Cnot6l | 0.029 | Young |
| 0.151018 | 0.165384 | 0.487 | 0.47 | 1 | 2 | Chd4 | 0.017 | Young |
| 0.151301 | 0.373989 | 0.097 | 0.078 | 1 | 2 | Nacc2 | 0.019 | Young |
| 0.151311 | 0.165321 | 0.061 | 0.045 | 1 | 2 | Cd177 | 0.016 | Young |
| 0.151312 | 0.778938 | 0.013 | 0.006 | 1 | 2 | Tpbgl | 0.007 | Young |
| 0.151333 | 0.655032 | 0.051 | 0.037 | 1 | 2 | Nup155 | 0.014 | Young |
| 0.151594 | 0.628 | 0.051 | 0.037 | 1 | 2 | Git1 | 0.014 | Young |
| 0.151782 | 0.64689 | 0.013 | 0.006 | 1 | 2 | Prodh | 0.007 | Young |
| 0.152314 | 0.208317 | 0.121 | 0.101 | 1 | 2 | Zbtb18 | 0.02 | Young |
| 0.152569 | 0.454342 | 0.013 | 0.006 | 1 | 2 | Ncs1 | 0.007 | Young |
| 0.152569 | 0.399249 | 0.013 | 0.006 | 1 | 2 | Stbd1 | 0.007 | Young |
| 0.152631 | 0.355288 | 0.08 | 0.062 | 1 | 2 | Olr1 | 0.018 | Young |
| 0.152803 | 0.858761 | 0.02 | 0.012 | 1 | 2 | Adam33 | 0.008 | Young |
| 0.15295 | 0.530879 | 0.065 | 0.05 | 1 | 2 | Styx | 0.015 | Young |
| 0.153023 | 0.129603 | 0.572 | 0.548 | 1 | 2 | Tgfb1 | 0.024 | Young |
| 0.153546 | 0.628748 | 0.03 | 0.019 | 1 | 2 | Cluap1 | 0.011 | Young |
| 0.153764 | 1.277839 | 0.053 | 0.04 | 1 | 2 | Rcan1 | 0.013 | Young |
| 0.153779 | 0.235438 | 0.131 | 0.109 | 1 | 2 | Uap1 | 0.022 | Young |
| 0.153843 | 0.242841 | 0.153 | 0.132 | 1 | 2 | Trim56 | 0.021 | Young |
| 0.154119 | 0.251349 | 0.077 | 0.06 | 1 | 2 | Pgm1 | 0.017 | Young |
| 0.154737 | 0.09263 | 0.178 | 0.151 | 1 | 2 | Irs2 | 0.027 | Young |
| 0.155127 | 0.708717 | 0.099 | 0.08 | 1 | 2 | Il1rl1 | 0.019 | Young |
| 0.156297 | 0.209352 | 0.327 | 0.304 | 1 | 2 | Zfhx3 | 0.023 | Young |
| 0.156319 | 0.173585 | 0.046 | 0.033 | 1 | 2 | Cp | 0.013 | Young |
| 0.156631 | 0.221728 | 0.185 | 0.162 | 1 | 2 | Gch1 | 0.023 | Young |
| 0.157297 | 0.267543 | 0.093 | 0.074 | 1 | 2 | Fam160b2 | 0.019 | Young |
| 0.157433 | 0.243578 | 0.086 | 0.067 | 1 | 2 | Tmem88 | 0.019 | Young |
| 0.157975 | 0.614961 | 0.032 | 0.021 | 1 | 2 | Mib2 | 0.011 | Young |
| 0.157988 | 0.432434 | 0.051 | 0.037 | 1 | 2 | Phf13 | 0.014 | Young |

| | | | | | | | | |
|---|---|---|---|---|---|---|---|---|
| 0.158165 | 0.263098 | 0.253 | 0.227 | 1 | 2 | Ube2i | 0.026 | Young |
| 0.158235 | 0.58537 | 0.071 | 0.056 | 1 | 2 | Hsd17b10 | 0.015 | Young |
| 0.1583 | 0.575616 | 0.031 | 0.02 | 1 | 2 | Tom1l2 | 0.011 | Young |
| 0.158688 | 0.101605 | 0.86 | 0.819 | 1 | 2 | Gnai2 | 0.041 | Young |
| 0.158725 | 0.360629 | 0.251 | 0.231 | 1 | 2 | Bmp2k | 0.02 | Young |
| 0.158884 | 0.220548 | 0.121 | 0.101 | 1 | 2 | Trrap | 0.02 | Young |
| 0.158885 | 0.672099 | 0.052 | 0.039 | 1 | 2 | Edc3 | 0.013 | Young |
| 0.159304 | 0.269754 | 0.291 | 0.266 | 1 | 2 | Glyr1 | 0.025 | Young |
| 0.159629 | 0.415476 | 0.065 | 0.05 | 1 | 2 | Calcrl | 0.015 | Young |
| 0.160238 | 0.573193 | 0.05 | 0.036 | 1 | 2 | Rpp14 | 0.014 | Young |
| 0.160368 | 1.411083 | 0.033 | 0.022 | 1 | 2 | Il1a | 0.011 | Young |
| 0.160438 | 0.387523 | 0.062 | 0.047 | 1 | 2 | Aatf | 0.015 | Young |
| 0.160902 | 0.095177 | 0.209 | 0.183 | 1 | 2 | Polr2m | 0.026 | Young |
| 0.161014 | 0.485629 | 0.138 | 0.118 | 1 | 2 | Acox3 | 0.02 | Young |
| 0.161441 | 0.517795 | 0.033 | 0.022 | 1 | 2 | Tfdp2 | 0.011 | Young |
| 0.162248 | 0.138551 | 0.27 | 0.241 | 1 | 2 | Ppp6r3 | 0.029 | Young |
| 0.162413 | 0.326951 | 0.52 | 0.518 | 1 | 2 | Jun | 0.002 | Young |
| 0.162627 | 0.439564 | 0.158 | 0.136 | 1 | 2 | Prdm1 | 0.022 | Young |
| 0.163021 | 1.472986 | 0.014 | 0.007 | 1 | 2 | Lrmda | 0.007 | Young |
| 0.163081 | 0.478804 | 0.12 | 0.101 | 1 | 2 | Clec4n | 0.019 | Young |
| 0.1638 | 0.715932 | 0.033 | 0.022 | 1 | 2 | AW554918 | 0.011 | Young |
| 0.163892 | 0.417464 | 0.179 | 0.158 | 1 | 2 | Nhp2 | 0.021 | Young |
| 0.163926 | 0.092356 | 0.521 | 0.488 | 1 | 2 | Eif3k | 0.033 | Young |
| 0.164053 | 0.373743 | 0.071 | 0.056 | 1 | 2 | Fmc1 | 0.015 | Young |
| 0.164321 | 0.113274 | 0.053 | 0.071 | 1 | 2 | Golt1b | -0.018 | Young |
| 0.164503 | 0.420883 | 0.069 | 0.054 | 1 | 2 | Rexo4 | 0.015 | Young |
| 0.164553 | 0.493415 | 0.059 | 0.045 | 1 | 2 | Urod | 0.014 | Young |
| 0.164919 | 0.444782 | 0.059 | 0.045 | 1 | 2 | Ttc1 | 0.014 | Young |
| 0.16522 | 0.29789 | 0.147 | 0.126 | 1 | 2 | Tor1b | 0.021 | Young |
| 0.165587 | 0.524894 | 0.033 | 0.022 | 1 | 2 | Fam122b | 0.011 | Young |
| 0.16561 | 0.145961 | 0.012 | 0.02 | 1 | 2 | Gm42031 | -0.008 | Young |
| 0.165863 | 0.909103 | 0.014 | 0.007 | 1 | 2 | Cwf19l1 | 0.007 | Young |
| 0.165974 | 0.261517 | 0.411 | 0.396 | 1 | 2 | Fnbp1 | 0.015 | Young |
| 0.166285 | 0.415179 | 0.033 | 0.022 | 1 | 2 | Ttc28 | 0.011 | Young |
| 0.167408 | 0.078074 | 0.312 | 0.282 | 1 | 2 | Fam102b | 0.03 | Young |
| 0.167457 | 0.575332 | 0.014 | 0.007 | 1 | 2 | Abraxas1 | 0.007 | Young |
| 0.167905 | 0.286095 | 0.254 | 0.231 | 1 | 2 | Ndufb6 | 0.023 | Young |
| 0.167938 | 0.616697 | 0.014 | 0.007 | 1 | 2 | Csrnp2 | 0.007 | Young |
| 0.168022 | 0.307351 | 0.122 | 0.103 | 1 | 2 | Ten1 | 0.019 | Young |
| 0.168124 | 0.432832 | 0.034 | 0.024 | 1 | 2 | Ncdn | 0.01 | Young |
| 0.168208 | 0.099152 | 0.246 | 0.219 | 1 | 2 | Mknk2 | 0.027 | Young |
| 0.168496 | 0.561223 | 0.033 | 0.022 | 1 | 2 | Sema6d | 0.011 | Young |
| 0.16878 | 0.831766 | 0.063 | 0.049 | 1 | 2 | Slc9a1 | 0.014 | Young |
| 0.168868 | 0.024271 | 1 | 0.999 | 1 | 2 | Tmsb4x | 0.001 | Young |
| 0.168902 | 0.413736 | 0.014 | 0.007 | 1 | 2 | Chchd6 | 0.007 | Young |

| | | | | | | | | |
|---|---|---|---|---|---|---|---|---|
| 0.168906 | 0.149338 | 0.264 | 0.239 | 1 | 2 | Gm26740 | 0.025 | Young |
| 0.169215 | 0.111205 | 0.454 | 0.427 | 1 | 2 | Anp32b | 0.027 | Young |
| 0.169327 | 0.236393 | 0.333 | 0.313 | 1 | 2 | Tbxas1 | 0.02 | Young |
| 0.169845 | 0.164579 | 0.118 | 0.097 | 1 | 2 | Lemd3 | 0.021 | Young |
| 0.169868 | 0.366743 | 0.163 | 0.143 | 1 | 2 | Lin7c | 0.02 | Young |
| 0.170032 | 0.411595 | 0.014 | 0.007 | 1 | 2 | Cxcl3 | 0.007 | Young |
| 0.17041 | 1.102478 | 0.024 | 0.015 | 1 | 2 | D2hgdh | 0.009 | Young |
| 0.170828 | 0.19147 | 0.281 | 0.262 | 1 | 2 | Gnai3 | 0.019 | Young |
| 0.171022 | 0.115497 | 0.165 | 0.141 | 1 | 2 | 4933434E2 | 0.024 | Young |
| 0.171041 | 0.297688 | 0.176 | 0.153 | 1 | 2 | Kdm1a | 0.023 | Young |
| 0.171429 | 0.127294 | 0.183 | 0.16 | 1 | 2 | Stard8 | 0.023 | Young |
| 0.171484 | 0.461829 | 0.146 | 0.127 | 1 | 2 | Ndfip2 | 0.019 | Young |
| 0.171676 | 0.472921 | 0.037 | 0.026 | 1 | 2 | Zfp667 | 0.011 | Young |
| 0.171753 | 1.16922 | 0.024 | 0.015 | 1 | 2 | 1810043G( | 0.009 | Young |
| 0.172373 | 0.889273 | 0.023 | 0.014 | 1 | 2 | Cd1d1 | 0.009 | Young |
| 0.172688 | 0.459236 | 0.036 | 0.025 | 1 | 2 | Cep128 | 0.011 | Young |
| 0.174167 | 0.834436 | 0.046 | 0.034 | 1 | 2 | Pwwp2b | 0.012 | Young |
| 0.174416 | 0.350629 | 0.108 | 0.09 | 1 | 2 | Mmadhc | 0.018 | Young |
| 0.174482 | 0.081107 | 0.021 | 0.032 | 1 | 2 | F5 | -0.011 | Young |
| 0.174788 | 0.307012 | 0.023 | 0.014 | 1 | 2 | Mypopos | 0.009 | Young |
| 0.175223 | 0.410868 | 0.226 | 0.207 | 1 | 2 | Krtcap2 | 0.019 | Young |
| 0.17545 | 0.228799 | 0.102 | 0.085 | 1 | 2 | Plk3 | 0.017 | Young |
| 0.175559 | 0.196018 | 0.223 | 0.2 | 1 | 2 | Ank | 0.023 | Young |
| 0.176109 | 0.740729 | 0.039 | 0.028 | 1 | 2 | Pef1 | 0.011 | Young |
| 0.176114 | 0.620857 | 0.038 | 0.027 | 1 | 2 | Lmo1 | 0.011 | Young |
| 0.176374 | 0.23612 | 0.367 | 0.349 | 1 | 2 | Srp14 | 0.018 | Young |
| 0.177876 | 0.426234 | 0.095 | 0.077 | 1 | 2 | Nxpe5 | 0.018 | Young |
| 0.177997 | 1.422513 | 0.01 | 0.004 | 1 | 2 | Shf | 0.006 | Young |
| 0.178418 | 1.51098 | 0.01 | 0.004 | 1 | 2 | Nrg2 | 0.006 | Young |
| 0.178418 | 1.155122 | 0.01 | 0.004 | 1 | 2 | Prkcg | 0.006 | Young |
| 0.178418 | 0.99487 | 0.01 | 0.004 | 1 | 2 | Gm26759 | 0.006 | Young |
| 0.17846 | 0.628998 | 0.024 | 0.015 | 1 | 2 | Cdk20 | 0.009 | Young |
| 0.17857 | 0.095342 | 0.217 | 0.193 | 1 | 2 | Dennd1b | 0.024 | Young |
| 0.178742 | 0.756565 | 0.043 | 0.031 | 1 | 2 | Wipi1 | 0.012 | Young |
| 0.178748 | 0.351796 | 0.108 | 0.09 | 1 | 2 | Jak2 | 0.018 | Young |
| 0.178877 | 0.060781 | 0.733 | 0.717 | 1 | 2 | mt-Nd5 | 0.016 | Young |
| 0.179051 | 0.958376 | 0.01 | 0.004 | 1 | 2 | Cdh5 | 0.006 | Young |
| 0.179051 | 0.947625 | 0.01 | 0.004 | 1 | 2 | Clstn1 | 0.006 | Young |
| 0.179051 | 0.944531 | 0.01 | 0.004 | 1 | 2 | Cep89 | 0.006 | Young |
| 0.179539 | 0.413519 | 0.194 | 0.176 | 1 | 2 | Smim11 | 0.018 | Young |
| 0.179686 | 0.681542 | 0.01 | 0.004 | 1 | 2 | Wdr34 | 0.006 | Young |
| 0.179695 | 0.203894 | 0.316 | 0.292 | 1 | 2 | Eif3j1 | 0.024 | Young |
| 0.179864 | 0.17236 | 0.156 | 0.134 | 1 | 2 | Ttc3 | 0.022 | Young |
| 0.180046 | 0.101267 | 0.57 | 0.545 | 1 | 2 | Cox6b1 | 0.025 | Young |
| 0.180333 | 0.241012 | 0.087 | 0.07 | 1 | 2 | Tango2 | 0.017 | Young |

| | | | | | | | | |
|---|---|---|---|---|---|---|---|---|
| 0.18096 | 0.803595 | 0.044 | 0.032 | 1 | 2 | Selenbp1 | 0.012 | Young |
| 0.181369 | 0.765864 | 0.015 | 0.009 | 1 | 2 | Neil1 | 0.006 | Young |
| 0.181371 | 0.289937 | 0.219 | 0.196 | 1 | 2 | Madd | 0.023 | Young |
| 0.181507 | 0.531449 | 0.099 | 0.082 | 1 | 2 | Ccm2 | 0.017 | Young |
| 0.181511 | 0.544322 | 0.039 | 0.028 | 1 | 2 | Zfp771 | 0.011 | Young |
| 0.181641 | 0.326176 | 0.069 | 0.054 | 1 | 2 | Setd1a | 0.015 | Young |
| 0.181977 | 0.194973 | 0.036 | 0.025 | 1 | 2 | Gab3 | 0.011 | Young |
| 0.182169 | 0.23767 | 0.074 | 0.058 | 1 | 2 | Gm45716 | 0.016 | Young |
| 0.182666 | 0.511006 | 0.015 | 0.009 | 1 | 2 | Tigar | 0.006 | Young |
| 0.182817 | 0.217147 | 0.251 | 0.229 | 1 | 2 | Ube2s | 0.022 | Young |
| 0.182844 | 0.198647 | 0.064 | 0.049 | 1 | 2 | Klhl20 | 0.015 | Young |
| 0.18324 | 0.155725 | 0.653 | 0.636 | 1 | 2 | S100a10 | 0.017 | Young |
| 0.183355 | 0.364788 | 0.279 | 0.262 | 1 | 2 | Smim14 | 0.017 | Young |
| 0.183527 | 0.365712 | 0.095 | 0.078 | 1 | 2 | Suclg2 | 0.017 | Young |
| 0.183539 | 0.55487 | 0.101 | 0.085 | 1 | 2 | Cyp20a1 | 0.016 | Young |
| 0.183778 | 0.61651 | 0.05 | 0.037 | 1 | 2 | Cars | 0.013 | Young |
| 0.183806 | 0.386667 | 0.015 | 0.009 | 1 | 2 | Ifi214 | 0.006 | Young |
| 0.184254 | 0.200721 | 0.12 | 0.101 | 1 | 2 | Rnd3 | 0.019 | Young |
| 0.184427 | 0.061356 | 0.84 | 0.807 | 1 | 2 | Gm10076 | 0.033 | Young |
| 0.184471 | 0.328116 | 0.04 | 0.029 | 1 | 2 | Nos1ap | 0.011 | Young |
| 0.184554 | 0.269399 | 0.039 | 0.028 | 1 | 2 | Accs | 0.011 | Young |
| 0.184943 | 0.329054 | 0.221 | 0.199 | 1 | 2 | Mpc1 | 0.022 | Young |
| 0.184978 | 0.110255 | 0.246 | 0.222 | 1 | 2 | Vapb | 0.024 | Young |
| 0.185116 | 0.465702 | 0.015 | 0.009 | 1 | 2 | Gstt2 | 0.006 | Young |
| 0.185181 | 0.647036 | 0.059 | 0.046 | 1 | 2 | Zfp830 | 0.013 | Young |
| 0.185389 | 0.197326 | 0.521 | 0.501 | 1 | 2 | Nktr | 0.02 | Young |
| 0.185776 | 0.361177 | 0.045 | 0.033 | 1 | 2 | Zfp68 | 0.012 | Young |
| 0.185937 | 0.414177 | 0.088 | 0.072 | 1 | 2 | Ndrg3 | 0.016 | Young |
| 0.186382 | 0.178094 | 0.139 | 0.118 | 1 | 2 | Afg3l1 | 0.021 | Young |
| 0.186404 | 0.0131 | 0.335 | 0.37 | 1 | 2 | Rac2 | -0.035 | Young |
| 0.186734 | 0.310923 | 0.025 | 0.016 | 1 | 2 | Zc3h8 | 0.009 | Young |
| 0.187067 | 0.527437 | 0.072 | 0.058 | 1 | 2 | Plcb3 | 0.014 | Young |
| 0.187215 | 0.134186 | 0.013 | 0.021 | 1 | 2 | Zfp503 | -0.008 | Young |
| 0.187325 | 0.808681 | 0.017 | 0.01 | 1 | 2 | Abhd8 | 0.007 | Young |
| 0.187429 | 0.113699 | 0.141 | 0.12 | 1 | 2 | Hivep1 | 0.021 | Young |
| 0.1878 | 0.66629 | 0.055 | 0.042 | 1 | 2 | Zfp219 | 0.013 | Young |
| 0.18785 | 0.069163 | 0.77 | 0.743 | 1 | 2 | Ybx1 | 0.027 | Young |
| 0.188333 | 0.541336 | 0.046 | 0.034 | 1 | 2 | Gm9493 | 0.012 | Young |
| 0.189393 | 0.472206 | 0.027 | 0.018 | 1 | 2 | Slf1 | 0.009 | Young |
| 0.189572 | 0.529631 | 0.083 | 0.067 | 1 | 2 | Nudt5 | 0.016 | Young |
| 0.189881 | 0.083474 | 0.236 | 0.21 | 1 | 2 | Exoc5 | 0.026 | Young |
| 0.190436 | 0.655784 | 0.026 | 0.017 | 1 | 2 | Kcnj2 | 0.009 | Young |
| 0.190635 | 0.242136 | 0.114 | 0.095 | 1 | 2 | Dirc2 | 0.019 | Young |
| 0.190766 | 0.265547 | 0.196 | 0.173 | 1 | 2 | Dennd6a | 0.023 | Young |
| 0.190831 | 0.741246 | 0.017 | 0.01 | 1 | 2 | Sac3d1 | 0.007 | Young |

| | | | | | | | | |
|---|---|---|---|---|---|---|---|---|
| 0.190911 | 0.162488 | 0.102 | 0.084 | 1 | 2 | Cep120 | 0.018 | Young |
| 0.19149 | 0.015718 | 0.112 | 0.134 | 1 | 2 | Fam129b | -0.022 | Young |
| 0.191606 | 0.546853 | 0.055 | 0.042 | 1 | 2 | Aldh1a2 | 0.013 | Young |
| 0.191832 | 0.188111 | 0.042 | 0.03 | 1 | 2 | Msh6 | 0.012 | Young |
| 0.191885 | 0.168625 | 0.31 | 0.286 | 1 | 2 | Tcea1 | 0.024 | Young |
| 0.191902 | 0.292815 | 0.151 | 0.131 | 1 | 2 | Trp53 | 0.02 | Young |
| 0.19201 | 0.192222 | 0.328 | 0.3 | 1 | 2 | Wac | 0.028 | Young |
| 0.192279 | 0.478346 | 0.017 | 0.01 | 1 | 2 | Wtip | 0.007 | Young |
| 0.192446 | 0.560225 | 0.056 | 0.043 | 1 | 2 | Ppa2 | 0.013 | Young |
| 0.192582 | 0.38942 | 0.099 | 0.081 | 1 | 2 | 2810004N2 | 0.018 | Young |
| 0.193374 | 0.880948 | 0.029 | 0.019 | 1 | 2 | Cdk5rap2 | 0.01 | Young |
| 0.193409 | 0.089108 | 0.025 | 0.016 | 1 | 2 | Mettl17 | 0.009 | Young |
| 0.194094 | 0.561123 | 0.139 | 0.122 | 1 | 2 | Pgrmc1 | 0.017 | Young |
| 0.194105 | 0.165501 | 0.148 | 0.127 | 1 | 2 | Ostm1 | 0.021 | Young |
| 0.195269 | 0.274483 | 0.091 | 0.075 | 1 | 2 | Vkorc1 | 0.016 | Young |
| 0.195509 | 0.028104 | 0.048 | 0.062 | 1 | 2 | Fdft1 | -0.014 | Young |
| 0.19574 | 0.242673 | 0.3 | 0.279 | 1 | 2 | Tmem234 | 0.021 | Young |
| 0.195815 | 0.172831 | 0.342 | 0.324 | 1 | 2 | Rgs10 | 0.018 | Young |
| 0.1961 | 0.059744 | 0.101 | 0.124 | 1 | 2 | Ilkap | -0.023 | Young |
| 0.19627 | 0.456142 | 0.083 | 0.067 | 1 | 2 | Msi2 | 0.016 | Young |
| 0.196368 | 0.617428 | 0.089 | 0.074 | 1 | 2 | Nufip1 | 0.015 | Young |
| 0.196675 | 0.319375 | 0.258 | 0.24 | 1 | 2 | Cdc37 | 0.018 | Young |
| 0.196727 | 0.560194 | 0.067 | 0.052 | 1 | 2 | Cmc2 | 0.015 | Young |
| 0.197071 | 0.654594 | 0.051 | 0.039 | 1 | 2 | Dppa3 | 0.012 | Young |
| 0.197072 | 0.43476 | 0.049 | 0.036 | 1 | 2 | Wdr75 | 0.013 | Young |
| 0.197561 | 0.10205 | 0.196 | 0.171 | 1 | 2 | Rapgef6 | 0.025 | Young |
| 0.197752 | 0.009222 | 0.125 | 0.104 | 1 | 2 | Ncoa4 | 0.021 | Young |
| 0.197865 | 0.118497 | 0.461 | 0.44 | 1 | 2 | Scand1 | 0.021 | Young |
| 0.197872 | 0.626202 | 0.029 | 0.019 | 1 | 2 | Tsr2 | 0.01 | Young |
| 0.198165 | 0.181916 | 0.359 | 0.342 | 1 | 2 | Pycard | 0.017 | Young |
| 0.198616 | 0.046276 | 0.167 | 0.145 | 1 | 2 | Stxbp6 | 0.022 | Young |
| 0.199116 | 0.10018 | 0.485 | 0.456 | 1 | 2 | Snx3 | 0.029 | Young |
| 0.200025 | 0.077045 | 0.594 | 0.565 | 1 | 2 | Qk | 0.029 | Young |
| 0.200268 | 0.032722 | 0.717 | 0.71 | 1 | 2 | F13a1 | 0.007 | Young |
| 0.200768 | 0.397322 | 0.132 | 0.115 | 1 | 2 | Btaf1 | 0.017 | Young |
| 0.200835 | 0.108571 | 0.172 | 0.151 | 1 | 2 | Slc35f6 | 0.021 | Young |
| 0.200931 | 0.198859 | 0.224 | 0.204 | 1 | 2 | Phf20l1 | 0.02 | Young |
| 0.201149 | 1.2811 | 0.011 | 0.005 | 1 | 2 | 2700081O1 | 0.006 | Young |
| 0.20157 | 0.243154 | 0.366 | 0.35 | 1 | 2 | Cct2 | 0.016 | Young |
| 0.201704 | 0.349989 | 0.214 | 0.196 | 1 | 2 | Eif2a | 0.018 | Young |
| 0.201862 | 0.312125 | 0.08 | 0.064 | 1 | 2 | Cblb | 0.016 | Young |
| 0.202217 | 1.292879 | 0.011 | 0.005 | 1 | 2 | Ambp | 0.006 | Young |
| 0.202254 | 0.443751 | 0.051 | 0.039 | 1 | 2 | Klc4 | 0.012 | Young |
| 0.202489 | 0.692372 | 0.018 | 0.011 | 1 | 2 | Serpinb9 | 0.007 | Young |
| 0.202572 | 0.162688 | 0.029 | 0.019 | 1 | 2 | Eef2k | 0.01 | Young |

| | | | | | | | | |
|---|---|---|---|---|---|---|---|---|
| 0.203008 | 0.093845 | 0.479 | 0.46 | 1 | 2 | Dock10 | 0.019 | Young |
| 0.203425 | 0.6578 | 0.031 | 0.021 | 1 | 2 | Nprl2 | 0.01 | Young |
| 0.203617 | 0.57137 | 0.018 | 0.011 | 1 | 2 | Gk5 | 0.007 | Young |
| 0.203935 | 0.822825 | 0.011 | 0.005 | 1 | 2 | Shpk | 0.006 | Young |
| 0.20415 | 0.664073 | 0.011 | 0.005 | 1 | 2 | Gm4285 | 0.006 | Young |
| 0.204482 | 0.153687 | 0.044 | 0.032 | 1 | 2 | Haus2 | 0.012 | Young |
| 0.204651 | 0.161864 | 0.154 | 0.134 | 1 | 2 | Braf | 0.02 | Young |
| 0.204816 | 0.016966 | 0.034 | 0.047 | 1 | 2 | Prr3 | -0.013 | Young |
| 0.204949 | 0.327583 | 0.081 | 0.065 | 1 | 2 | Slc41a1 | 0.016 | Young |
| 0.205014 | 0.630541 | 0.011 | 0.005 | 1 | 2 | A830010M | 0.006 | Young |
| 0.20523 | 0.484136 | 0.011 | 0.005 | 1 | 2 | A730063M | 0.006 | Young |
| 0.205248 | 0.306728 | 0.109 | 0.092 | 1 | 2 | Tbc1d22a | 0.017 | Young |
| 0.205564 | 0.444517 | 0.129 | 0.11 | 1 | 2 | Clec4d | 0.019 | Young |
| 0.205663 | 0.421259 | 0.011 | 0.005 | 1 | 2 | AW011738 | 0.006 | Young |
| 0.206428 | 0.260594 | 0.229 | 0.209 | 1 | 2 | Nfic | 0.02 | Young |
| 0.206528 | 0.517171 | 0.087 | 0.072 | 1 | 2 | Glo1 | 0.015 | Young |
| 0.206531 | 0.607236 | 0.011 | 0.005 | 1 | 2 | Nnt | 0.006 | Young |
| 0.207372 | 0.079428 | 0.505 | 0.49 | 1 | 2 | Tpr | 0.015 | Young |
| 0.207788 | 0.267888 | 0.243 | 0.224 | 1 | 2 | Ptbp1 | 0.019 | Young |
| 0.207839 | 0.030036 | 0.011 | 0.005 | 1 | 2 | 1700056N1 | 0.006 | Young |
| 0.207959 | 0.441496 | 0.135 | 0.118 | 1 | 2 | Bet1 | 0.017 | Young |
| 0.207979 | 0.273636 | 0.1 | 0.084 | 1 | 2 | Lats1 | 0.016 | Young |
| 0.208857 | 0.210602 | 0.368 | 0.351 | 1 | 2 | Rpl13a | 0.017 | Young |
| 0.209157 | 0.126032 | 0.529 | 0.521 | 1 | 2 | Jpt1 | 0.008 | Young |
| 0.209462 | 0.113484 | 0.492 | 0.469 | 1 | 2 | Lgals1 | 0.023 | Young |
| 0.209517 | 0.045755 | 0.391 | 0.364 | 1 | 2 | Laptm4a | 0.027 | Young |
| 0.209968 | 0.311617 | 0.268 | 0.252 | 1 | 2 | Gpbp1 | 0.016 | Young |
| 0.210558 | 0.294079 | 0.241 | 0.222 | 1 | 2 | Atp5g1 | 0.019 | Young |
| 0.210985 | 0.451889 | 0.134 | 0.118 | 1 | 2 | Nudt4 | 0.016 | Young |
| 0.211906 | 0.223399 | 0.311 | 0.292 | 1 | 2 | Prelid1 | 0.019 | Young |
| 0.212114 | 0.48349 | 0.096 | 0.08 | 1 | 2 | Ago4 | 0.016 | Young |
| 0.212122 | 0.2533 | 0.337 | 0.322 | 1 | 2 | Per1 | 0.015 | Young |
| 0.212299 | 0.115475 | 0.229 | 0.207 | 1 | 2 | Npc1 | 0.022 | Young |
| 0.212422 | 0.274425 | 0.16 | 0.142 | 1 | 2 | Tspan32 | 0.018 | Young |
| 0.212683 | 0.961177 | 0.044 | 0.033 | 1 | 2 | Cdk5 | 0.011 | Young |
| 0.213088 | 0.053951 | 0.391 | 0.362 | 1 | 2 | Pnisr | 0.029 | Young |
| 0.213131 | 0.452294 | 0.019 | 0.012 | 1 | 2 | Mfsd3 | 0.007 | Young |
| 0.213312 | 0.473351 | 0.097 | 0.081 | 1 | 2 | Cpd | 0.016 | Young |
| 0.213597 | 0.073398 | 0.648 | 0.633 | 1 | 2 | Sub1 | 0.015 | Young |
| 0.213606 | 0.179334 | 0.055 | 0.042 | 1 | 2 | Abcd4 | 0.013 | Young |
| 0.214116 | 0.484266 | 0.033 | 0.024 | 1 | 2 | 1700021F0 | 0.009 | Young |
| 0.214476 | 0.116791 | 0.018 | 0.027 | 1 | 2 | Tbc1d8b | -0.009 | Young |
| 0.21485 | 0.492325 | 0.032 | 0.022 | 1 | 2 | Megf9 | 0.01 | Young |
| 0.21539 | 0.46829 | 0.019 | 0.012 | 1 | 2 | Arhgap22 | 0.007 | Young |
| 0.21558 | 0.259245 | 0.233 | 0.213 | 1 | 2 | Sec14l1 | 0.02 | Young |

| | | | | | | | | |
|---|---|---|---|---|---|---|---|---|
| 0.215582 | 0.392507 | 0.032 | 0.022 | 1 | 2 | Bloc1s5 | 0.01 | Young |
| 0.216051 | 0.152754 | 0.354 | 0.339 | 1 | 2 | Nop53 | 0.015 | Young |
| 0.216363 | 0.250922 | 0.019 | 0.012 | 1 | 2 | Clybl | 0.007 | Young |
| 0.216538 | 0.105107 | 0.076 | 0.061 | 1 | 2 | Zscan26 | 0.015 | Young |
| 0.216711 | 1.016738 | 0.063 | 0.05 | 1 | 2 | Ifitm1 | 0.013 | Young |
| 0.217014 | 0.299754 | 0.019 | 0.012 | 1 | 2 | Tslp | 0.007 | Young |
| 0.217257 | 0.431536 | 0.188 | 0.171 | 1 | 2 | Uqcc2 | 0.017 | Young |
| 0.217341 | 0.342048 | 0.1 | 0.084 | 1 | 2 | Pcyox1 | 0.016 | Young |
| 0.218045 | 0.766692 | 0.042 | 0.031 | 1 | 2 | Apool | 0.011 | Young |
| 0.218271 | 0.302211 | 0.112 | 0.094 | 1 | 2 | Mmp19 | 0.018 | Young |
| 0.218374 | 0.292442 | 0.162 | 0.141 | 1 | 2 | Dmxl1 | 0.021 | Young |
| 0.218729 | 0.273426 | 0.148 | 0.131 | 1 | 2 | Dnajb1 | 0.017 | Young |
| 0.220002 | 0.551729 | 0.038 | 0.028 | 1 | 2 | Klhl15 | 0.01 | Young |
| 0.220659 | 0.737523 | 0.02 | 0.013 | 1 | 2 | Taf4b | 0.007 | Young |
| 0.221696 | 0.589424 | 0.069 | 0.056 | 1 | 2 | Amz2 | 0.013 | Young |
| 0.221773 | 0.195207 | 0.143 | 0.124 | 1 | 2 | Churc1 | 0.019 | Young |
| 0.221836 | 0.388871 | 0.076 | 0.062 | 1 | 2 | Stau1 | 0.014 | Young |
| 0.222021 | 0.510876 | 0.077 | 0.063 | 1 | 2 | Ak3 | 0.014 | Young |
| 0.222091 | 0.317379 | 0.128 | 0.11 | 1 | 2 | Atad1 | 0.018 | Young |
| 0.222159 | 0.203585 | 0.219 | 0.198 | 1 | 2 | Pabpn1 | 0.021 | Young |
| 0.222404 | 0.052584 | 0.038 | 0.05 | 1 | 2 | Ddt | -0.012 | Young |
| 0.222637 | 0.686164 | 0.042 | 0.031 | 1 | 2 | Mpp5 | 0.011 | Young |
| 0.222773 | 1.170099 | 0.021 | 0.014 | 1 | 2 | 2310040G2 | 0.007 | Young |
| 0.222844 | 0.134403 | 0.406 | 0.385 | 1 | 2 | Romo1 | 0.021 | Young |
| 0.222873 | 0.434908 | 0.082 | 0.067 | 1 | 2 | Ppfibp2 | 0.015 | Young |
| 0.223115 | 0.672371 | 0.039 | 0.029 | 1 | 2 | Cdk7 | 0.01 | Young |
| 0.223627 | 0.320225 | 0.197 | 0.179 | 1 | 2 | Smarcc1 | 0.018 | Young |
| 0.223784 | 0.404635 | 0.099 | 0.082 | 1 | 2 | Pop5 | 0.017 | Young |
| 0.223794 | 0.760131 | 0.012 | 0.006 | 1 | 2 | Poli | 0.006 | Young |
| 0.223839 | 0.185417 | 0.293 | 0.27 | 1 | 2 | B230219D2 | 0.023 | Young |
| 0.223973 | 0.532877 | 0.068 | 0.055 | 1 | 2 | Gm17018 | 0.013 | Young |
| 0.224441 | 0.811351 | 0.012 | 0.006 | 1 | 2 | 9430015G1 | 0.006 | Young |
| 0.224441 | 0.605914 | 0.012 | 0.006 | 1 | 2 | Zbtb12 | 0.006 | Young |
| 0.224657 | 0.702644 | 0.012 | 0.006 | 1 | 2 | Mtmr11 | 0.006 | Young |
| 0.224744 | 0.504872 | 0.045 | 0.034 | 1 | 2 | Trpv4 | 0.011 | Young |
| 0.225021 | 0.309601 | 0.078 | 0.064 | 1 | 2 | Ddx1 | 0.014 | Young |
| 0.225263 | 1.048386 | 0.021 | 0.014 | 1 | 2 | Gm21781 | 0.007 | Young |
| 0.225315 | 0.702734 | 0.042 | 0.031 | 1 | 2 | Rubcn | 0.011 | Young |
| 0.225425 | 0.512895 | 0.042 | 0.031 | 1 | 2 | Ikbkg | 0.011 | Young |
| 0.225717 | 0.275784 | 0.214 | 0.195 | 1 | 2 | Impact | 0.019 | Young |
| 0.22598 | 0.645215 | 0.093 | 0.078 | 1 | 2 | Zfp277 | 0.015 | Young |
| 0.226085 | 0.62765 | 0.042 | 0.031 | 1 | 2 | Parp6 | 0.011 | Young |
| 0.226174 | 0.521962 | 0.012 | 0.006 | 1 | 2 | Tmem202 | 0.006 | Young |
| 0.226571 | 0.491454 | 0.046 | 0.035 | 1 | 2 | Zfp282 | 0.011 | Young |
| 0.226579 | 0.258025 | 0.034 | 0.025 | 1 | 2 | 3110062M | 0.009 | Young |

| | | | | | | | | |
|---|---|---|---|---|---|---|---|---|
| 0.226592 | 0.186234 | 0.219 | 0.196 | 1 | 2 | Dnajc19 | 0.023 | Young |
| 0.227144 | 0.502676 | 0.021 | 0.014 | 1 | 2 | Uevld | 0.007 | Young |
| 0.227377 | 0.518005 | 0.065 | 0.052 | 1 | 2 | Togaram1 | 0.013 | Young |
| 0.228099 | 0.08174 | 0.368 | 0.346 | 1 | 2 | Arhgap15 | 0.022 | Young |
| 0.228205 | 0.382867 | 0.125 | 0.108 | 1 | 2 | Nudcd2 | 0.017 | Young |
| 0.228356 | 0.219091 | 0.188 | 0.167 | 1 | 2 | Zranb1 | 0.021 | Young |
| 0.228404 | 0.646705 | 0.021 | 0.014 | 1 | 2 | Zfp1 | 0.007 | Young |
| 0.228451 | 0.278776 | 0.241 | 0.223 | 1 | 2 | Ssfa2 | 0.018 | Young |
| 0.228648 | 0.048111 | 0.721 | 0.7 | 1 | 2 | Rrbp1 | 0.021 | Young |
| 0.229292 | 0.004361 | 0.043 | 0.056 | 1 | 2 | Mrpl50 | -0.013 | Young |
| 0.229519 | 0.505148 | 0.058 | 0.046 | 1 | 2 | Zbtb43 | 0.012 | Young |
| 0.229565 | 0.22202 | 0.362 | 0.345 | 1 | 2 | Dnajb14 | 0.017 | Young |
| 0.23025 | 0.173885 | 0.238 | 0.215 | 1 | 2 | Diaph2 | 0.023 | Young |
| 0.23085 | 0.30132 | 0.177 | 0.161 | 1 | 2 | Ndufa5 | 0.016 | Young |
| 0.231131 | 0.603442 | 0.053 | 0.042 | 1 | 2 | Psmg2 | 0.011 | Young |
| 0.231335 | 0.237996 | 0.172 | 0.152 | 1 | 2 | Csnk1g3 | 0.02 | Young |
| 0.231601 | 0.298844 | 0.04 | 0.03 | 1 | 2 | Rps6kc1 | 0.01 | Young |
| 0.231606 | 0.118365 | 0.469 | 0.448 | 1 | 2 | Cxcr4 | 0.021 | Young |
| 0.231699 | 0.163592 | 0.476 | 0.468 | 1 | 2 | Rock1 | 0.008 | Young |
| 0.232084 | 0.428648 | 0.042 | 0.031 | 1 | 2 | Crebl2 | 0.011 | Young |
| 0.232671 | 0.346571 | 0.158 | 0.141 | 1 | 2 | B3gat3 | 0.017 | Young |
| 0.232872 | 0.122568 | 0.363 | 0.348 | 1 | 2 | Rsrc2 | 0.015 | Young |
| 0.232889 | 0.140425 | 0.203 | 0.183 | 1 | 2 | Stard9 | 0.02 | Young |
| 0.233368 | 0.22668 | 0.04 | 0.03 | 1 | 2 | Rnf8 | 0.01 | Young |
| 0.233519 | 0.083109 | 0.574 | 0.545 | 1 | 2 | Cyth4 | 0.029 | Young |
| 0.233716 | 0.109665 | 0.007 | 0.013 | 1 | 2 | Atp5s | -0.006 | Young |
| 0.23377 | 0.443515 | 0.042 | 0.031 | 1 | 2 | Gm26782 | 0.011 | Young |
| 0.234373 | 0.419296 | 0.126 | 0.11 | 1 | 2 | Pold4 | 0.016 | Young |
| 0.234481 | 0.119133 | 0.445 | 0.424 | 1 | 2 | Top1 | 0.021 | Young |
| 0.235219 | 0.079064 | 0.065 | 0.081 | 1 | 2 | Plscr3 | -0.016 | Young |
| 0.235506 | 0.561909 | 0.023 | 0.015 | 1 | 2 | Platr17 | 0.008 | Young |
| 0.236001 | 0.044066 | 0.118 | 0.1 | 1 | 2 | Selenon | 0.018 | Young |
| 0.236011 | 0.115323 | 0.375 | 0.36 | 1 | 2 | Nisch | 0.015 | Young |
| 0.236435 | 0.215177 | 0.039 | 0.029 | 1 | 2 | Kank3 | 0.01 | Young |
| 0.236646 | 0.398066 | 0.067 | 0.054 | 1 | 2 | Ccdc86 | 0.013 | Young |
| 0.236759 | 0.518933 | 0.023 | 0.015 | 1 | 2 | Cers4 | 0.008 | Young |
| 0.236913 | 0.180072 | 0.464 | 0.459 | 1 | 2 | Hif1a | 0.005 | Young |
| 0.237073 | 0.317874 | 0.023 | 0.015 | 1 | 2 | Tnni2 | 0.008 | Young |
| 0.237118 | 0.353893 | 0.127 | 0.11 | 1 | 2 | Mrpl4 | 0.017 | Young |
| 0.237498 | 0.195445 | 0.19 | 0.17 | 1 | 2 | Thoc7 | 0.02 | Young |
| 0.237698 | 0.466187 | 0.055 | 0.043 | 1 | 2 | Arhgef11 | 0.012 | Young |
| 0.2377 | 0.497901 | 0.07 | 0.057 | 1 | 2 | Wnt11 | 0.013 | Young |
| 0.237867 | 1.201163 | 0.027 | 0.019 | 1 | 2 | Rmnd1 | 0.008 | Young |
| 0.238084 | 0.066962 | 0.504 | 0.475 | 1 | 2 | Ppp1ca | 0.029 | Young |
| 0.238093 | 0.534133 | 0.063 | 0.05 | 1 | 2 | Mpv17 | 0.013 | Young |

| | | | | | | | | |
|---|---|---|---|---|---|---|---|---|
| 0.238506 | 0.716181 | 0.024 | 0.016 | 1 | 2 | Zmat1 | 0.008 | Young |
| 0.238859 | 0.206775 | 0.153 | 0.135 | 1 | 2 | Tnfsf12 | 0.018 | Young |
| 0.238982 | 0.146991 | 0.088 | 0.073 | 1 | 2 | Rnf168 | 0.015 | Young |
| 0.23914 | 0.360747 | 0.113 | 0.097 | 1 | 2 | Mast3 | 0.016 | Young |
| 0.239471 | 0.008406 | 0.043 | 0.056 | 1 | 2 | Thoc1 | -0.013 | Young |
| 0.239534 | 0.897785 | 0.013 | 0.007 | 1 | 2 | Dhfr | 0.006 | Young |
| 0.23967 | 0.478085 | 0.086 | 0.072 | 1 | 2 | Mphosph8 | 0.014 | Young |
| 0.239682 | 0.41268 | 0.058 | 0.046 | 1 | 2 | Rpa1 | 0.012 | Young |
| 0.24023 | 0.125505 | 0.126 | 0.108 | 1 | 2 | Ccng2 | 0.018 | Young |
| 0.240432 | 0.233048 | 0.075 | 0.061 | 1 | 2 | Taf7 | 0.014 | Young |
| 0.240541 | 0.080844 | 0.58 | 0.567 | 1 | 2 | Tnfaip3 | 0.013 | Young |
| 0.240639 | 0.424867 | 0.061 | 0.048 | 1 | 2 | Rfx1 | 0.013 | Young |
| 0.240817 | 0.507976 | 0.013 | 0.007 | 1 | 2 | Hyal1 | 0.006 | Young |
| 0.240968 | 0.899831 | 0.024 | 0.016 | 1 | 2 | Prtn3 | 0.008 | Young |
| 0.241665 | 0.218259 | 0.125 | 0.107 | 1 | 2 | Snapin | 0.018 | Young |
| 0.242089 | 0.498497 | 0.056 | 0.044 | 1 | 2 | Vill | 0.012 | Young |
| 0.243636 | 0.201895 | 0.044 | 0.033 | 1 | 2 | Zxdc | 0.011 | Young |
| 0.243681 | 0.479661 | 0.024 | 0.016 | 1 | 2 | Isoc2b | 0.008 | Young |
| 0.244755 | 0.737701 | 0.025 | 0.017 | 1 | 2 | Aldh6a1 | 0.008 | Young |
| 0.244786 | 0.036146 | 0.052 | 0.041 | 1 | 2 | Foxred2 | 0.011 | Young |
| 0.244844 | 0.337757 | 0.189 | 0.172 | 1 | 2 | Ankrd13c | 0.017 | Young |
| 0.245694 | 0.501996 | 0.116 | 0.102 | 1 | 2 | Mex3c | 0.014 | Young |
| 0.245796 | 0.892291 | 0.026 | 0.018 | 1 | 2 | Orc6 | 0.008 | Young |
| 0.245825 | 0.06125 | 0.403 | 0.378 | 1 | 2 | Sgpl1 | 0.025 | Young |
| 0.245896 | 0.06062 | 0.186 | 0.166 | 1 | 2 | Crem | 0.02 | Young |
| 0.245996 | 0.101329 | 0.013 | 0.007 | 1 | 2 | Rnf180 | 0.006 | Young |
| 0.24643 | 0.769423 | 0.025 | 0.017 | 1 | 2 | Zc3h6 | 0.008 | Young |
| 0.246663 | 0.381888 | 0.061 | 0.048 | 1 | 2 | Uvssa | 0.013 | Young |
| 0.24669 | 0.273258 | 0.063 | 0.05 | 1 | 2 | 2410002F2 | 0.013 | Young |
| 0.247194 | 0.479673 | 0.025 | 0.017 | 1 | 2 | Gys1 | 0.008 | Young |
| 0.247713 | 0.584386 | 0.077 | 0.064 | 1 | 2 | Tmem203 | 0.013 | Young |
| 0.247732 | 0.629978 | 0.026 | 0.018 | 1 | 2 | Ccdc71 | 0.008 | Young |
| 0.248026 | 0.408811 | 0.097 | 0.082 | 1 | 2 | Tnfaip8l2 | 0.015 | Young |
| 0.248272 | 0.186963 | 0.059 | 0.047 | 1 | 2 | Rcor3 | 0.012 | Young |
| 0.24885 | 0.330752 | 0.056 | 0.044 | 1 | 2 | Spg11 | 0.012 | Young |
| 0.249251 | 1.638078 | 0.014 | 0.009 | 1 | 2 | Pmm1 | 0.005 | Young |
| 0.25022 | 0.229576 | 0.343 | 0.327 | 1 | 2 | Vasp | 0.016 | Young |
| 0.250359 | 0.202053 | 0.024 | 0.016 | 1 | 2 | Gypc | 0.008 | Young |
| 0.250576 | 0.520075 | 0.025 | 0.017 | 1 | 2 | Fads6 | 0.008 | Young |
| 0.250589 | 0.248233 | 0.171 | 0.152 | 1 | 2 | Supt4a | 0.019 | Young |
| 0.250591 | 0.098833 | 0.407 | 0.388 | 1 | 2 | Kmt2e | 0.019 | Young |
| 0.250882 | 0.71509 | 0.026 | 0.018 | 1 | 2 | Nek4 | 0.008 | Young |
| 0.252314 | 0.217608 | 0.277 | 0.263 | 1 | 2 | Sec61a1 | 0.014 | Young |
| 0.252599 | 0.904917 | 0.014 | 0.009 | 1 | 2 | Zfp873 | 0.005 | Young |
| 0.252748 | 0.022251 | 0.441 | 0.414 | 1 | 2 | Neurl3 | 0.027 | Young |

| | | | | | | | | |
|---|---|---|---|---|---|---|---|---|
| 0.252913 | 0.263435 | 0.127 | 0.11 | 1 | 2 | Sptan1 | 0.017 | Young |
| 0.253077 | 0.437299 | 0.186 | 0.171 | 1 | 2 | Cenpb | 0.015 | Young |
| 0.253298 | 0.172225 | 0.059 | 0.047 | 1 | 2 | 9330160F1 | 0.012 | Young |
| 0.253567 | 1.048524 | 0.029 | 0.02 | 1 | 2 | Gfod1 | 0.009 | Young |
| 0.253878 | 0.075482 | 0.011 | 0.017 | 1 | 2 | Gm15859 | -0.006 | Young |
| 0.254074 | 0.9496 | 0.014 | 0.009 | 1 | 2 | Spred3 | 0.005 | Young |
| 0.254211 | 0.43154 | 0.026 | 0.018 | 1 | 2 | Akip1 | 0.008 | Young |
| 0.254708 | 0.814825 | 0.014 | 0.009 | 1 | 2 | Enpp4 | 0.005 | Young |
| 0.255131 | 0.738543 | 0.014 | 0.009 | 1 | 2 | Alg12 | 0.005 | Young |
| 0.255163 | 0.089892 | 0.107 | 0.09 | 1 | 2 | Hsph1 | 0.017 | Young |
| 0.255277 | 0.461609 | 0.026 | 0.018 | 1 | 2 | Tatdn1 | 0.008 | Young |
| 0.25566 | 0.51907 | 0.082 | 0.069 | 1 | 2 | Scyl2 | 0.013 | Young |
| 0.255758 | 0.081911 | 0.424 | 0.406 | 1 | 2 | Tnfaip8 | 0.018 | Young |
| 0.255797 | 0.00549 | 0.166 | 0.188 | 1 | 2 | Ap3b1 | -0.022 | Young |
| 0.256465 | 0.104391 | 0.232 | 0.211 | 1 | 2 | Dhx9 | 0.021 | Young |
| 0.256765 | 0.689418 | 0.029 | 0.02 | 1 | 2 | 2900093K2 | 0.009 | Young |
| 0.257107 | 0.806172 | 0.057 | 0.046 | 1 | 2 | C2cd2l | 0.011 | Young |
| 0.257381 | 0.448967 | 0.027 | 0.019 | 1 | 2 | 2610524H( | 0.008 | Young |
| 0.257514 | 0.050907 | 0.175 | 0.154 | 1 | 2 | Myo9a | 0.021 | Young |
| 0.25798 | 0.483153 | 0.027 | 0.019 | 1 | 2 | Extl2 | 0.008 | Young |
| 0.258024 | 0.219037 | 0.127 | 0.11 | 1 | 2 | Fam114a2 | 0.017 | Young |
| 0.258107 | 0.098575 | 0.014 | 0.009 | 1 | 2 | Arl16 | 0.005 | Young |
| 0.258236 | 0.045877 | 0.705 | 0.675 | 1 | 2 | Morf4l1 | 0.03 | Young |
| 0.25834 | 0.209758 | 0.026 | 0.018 | 1 | 2 | Ahdc1 | 0.008 | Young |
| 0.258865 | 0.09401 | 0.075 | 0.061 | 1 | 2 | Tmem250- | 0.014 | Young |
| 0.258878 | 0.163094 | 0.152 | 0.135 | 1 | 2 | Vti1b | 0.017 | Young |
| 0.25892 | 0.813631 | 0.055 | 0.044 | 1 | 2 | Selenom | 0.011 | Young |
| 0.258962 | 0.365724 | 0.014 | 0.009 | 1 | 2 | Ptch1 | 0.005 | Young |
| 0.259206 | 0.167653 | 0.353 | 0.335 | 1 | 2 | Ndufb10 | 0.018 | Young |
| 0.259784 | 0.126682 | 0.056 | 0.044 | 1 | 2 | Dglucy | 0.012 | Young |
| 0.260596 | 0.237279 | 0.179 | 0.163 | 1 | 2 | Dclre1c | 0.016 | Young |
| 0.260992 | 0.236273 | 0.271 | 0.254 | 1 | 2 | Mxd4 | 0.017 | Young |
| 0.261519 | 0.18738 | 0.126 | 0.109 | 1 | 2 | Ero1lb | 0.017 | Young |
| 0.261903 | 0.319531 | 0.089 | 0.075 | 1 | 2 | Tbck | 0.014 | Young |
| 0.262023 | 0.429923 | 0.115 | 0.101 | 1 | 2 | Lpcat1 | 0.014 | Young |
| 0.262564 | 0.030691 | 0.186 | 0.167 | 1 | 2 | Mospd2 | 0.019 | Young |
| 0.263297 | 0.703558 | 0.044 | 0.034 | 1 | 2 | Vps8 | 0.01 | Young |
| 0.263491 | 0.204475 | 0.261 | 0.243 | 1 | 2 | Mrpl20 | 0.018 | Young |
| 0.264597 | 0.100798 | 0.14 | 0.122 | 1 | 2 | Pcm1 | 0.018 | Young |
| 0.264635 | 0.699099 | 0.027 | 0.019 | 1 | 2 | Esr1 | 0.008 | Young |
| 0.264884 | 0.826634 | 0.049 | 0.039 | 1 | 2 | Anpep | 0.01 | Young |
| 0.265032 | 0.408559 | 0.084 | 0.071 | 1 | 2 | Nsmce2 | 0.013 | Young |
| 0.26515 | 0.397357 | 0.031 | 0.022 | 1 | 2 | 3110040N1 | 0.009 | Young |
| 0.265203 | 0.804243 | 0.039 | 0.03 | 1 | 2 | Med6 | 0.009 | Young |
| 0.265504 | 0.02249 | 0.046 | 0.059 | 1 | 2 | Gnpat | -0.013 | Young |

| | | | | | | | | |
|---|---|---|---|---|---|---|---|---|
| 0.26572 | 0.30457 | 0.113 | 0.097 | 1 | 2 | Siglece | 0.016 | Young |
| 0.265759 | 0.102648 | 0.126 | 0.108 | 1 | 2 | Tmem183a | 0.018 | Young |
| 0.265846 | 0.099614 | 0.154 | 0.136 | 1 | 2 | Ago3 | 0.018 | Young |
| 0.265873 | 0.463236 | 0.03 | 0.021 | 1 | 2 | Nfkbil1 | 0.009 | Young |
| 0.266142 | 0.751108 | 0.015 | 0.01 | 1 | 2 | Nsun5 | 0.005 | Young |
| 0.266288 | 0.057933 | 0.344 | 0.317 | 1 | 2 | Rpl36al | 0.027 | Young |
| 0.266498 | 0.447557 | 0.08 | 0.066 | 1 | 2 | Bcl2l2 | 0.014 | Young |
| 0.266558 | 0.693049 | 0.015 | 0.01 | 1 | 2 | Ift74 | 0.005 | Young |
| 0.266725 | 0.173265 | 0.186 | 0.169 | 1 | 2 | Kdm2b | 0.017 | Young |
| 0.267183 | 0.710835 | 0.015 | 0.01 | 1 | 2 | Pdk2 | 0.005 | Young |
| 0.267199 | 0.140201 | 0.121 | 0.104 | 1 | 2 | Epb41 | 0.017 | Young |
| 0.2676 | 0.683242 | 0.015 | 0.01 | 1 | 2 | Ak8 | 0.005 | Young |
| 0.267882 | 0.228081 | 0.363 | 0.351 | 1 | 2 | Cox5b | 0.012 | Young |
| 0.268018 | 0.759713 | 0.015 | 0.01 | 1 | 2 | Galnt9 | 0.005 | Young |
| 0.268018 | 0.686563 | 0.015 | 0.01 | 1 | 2 | Kdm4b | 0.005 | Young |
| 0.268238 | 0.816835 | 0.039 | 0.03 | 1 | 2 | Dmac2 | 0.009 | Young |
| 0.268368 | 0.243132 | 0.137 | 0.12 | 1 | 2 | Srek1 | 0.017 | Young |
| 0.268617 | 0.061509 | 0.799 | 0.786 | 1 | 2 | Cd52 | 0.013 | Young |
| 0.268919 | 0.280954 | 0.095 | 0.081 | 1 | 2 | Kdm3a | 0.014 | Young |
| 0.269054 | 0.298854 | 0.115 | 0.1 | 1 | 2 | Clcn5 | 0.015 | Young |
| 0.269169 | 0.768181 | 0.05 | 0.04 | 1 | 2 | Nat8l | 0.01 | Young |
| 0.269172 | 0.125851 | 0.278 | 0.258 | 1 | 2 | Sema4d | 0.02 | Young |
| 0.269212 | 0.534044 | 0.033 | 0.025 | 1 | 2 | Hist2h2be | 0.008 | Young |
| 0.269304 | 0.670756 | 0.055 | 0.044 | 1 | 2 | Pdk3 | 0.011 | Young |
| 0.269775 | 0.623604 | 0.052 | 0.042 | 1 | 2 | Rrp15 | 0.01 | Young |
| 0.269921 | 0.269742 | 0.008 | 0.014 | 1 | 2 | Repin1 | -0.006 | Young |
| 0.269927 | 0.414111 | 0.074 | 0.061 | 1 | 2 | Ascc2 | 0.013 | Young |
| 0.270273 | 0.063696 | 0.068 | 0.082 | 1 | 2 | Gart | -0.014 | Young |
| 0.270302 | 0.173106 | 0.025 | 0.034 | 1 | 2 | Polr3a | -0.009 | Young |
| 0.270799 | 0.083571 | 0.095 | 0.08 | 1 | 2 | Polr3k | 0.015 | Young |
| 0.270815 | 0.277629 | 0.094 | 0.079 | 1 | 2 | Plekhm1 | 0.015 | Young |
| 0.271068 | 0.132108 | 0.062 | 0.076 | 1 | 2 | Zdhhc18 | -0.014 | Young |
| 0.271826 | 0.496515 | 0.063 | 0.051 | 1 | 2 | Ppp2r3c | 0.012 | Young |
| 0.272093 | 0.120414 | 0.428 | 0.412 | 1 | 2 | Sf3b2 | 0.016 | Young |
| 0.272391 | 0.386397 | 0.088 | 0.075 | 1 | 2 | Mak16 | 0.013 | Young |
| 0.272571 | 0.387597 | 0.058 | 0.047 | 1 | 2 | Slc25a44 | 0.011 | Young |
| 0.272786 | 0.515081 | 0.037 | 0.028 | 1 | 2 | Fam136a | 0.009 | Young |
| 0.272853 | 0.310888 | 0.032 | 0.024 | 1 | 2 | Mtg1 | 0.008 | Young |
| 0.27286 | 0.362626 | 0.183 | 0.166 | 1 | 2 | Dapp1 | 0.017 | Young |
| 0.272914 | 0.115962 | 0.018 | 0.026 | 1 | 2 | Lsm11 | -0.008 | Young |
| 0.272918 | 0.124541 | 0.7 | 0.693 | 1 | 2 | Cdkn1a | 0.007 | Young |
| 0.272938 | 0.307356 | 0.101 | 0.087 | 1 | 2 | Kbtbd2 | 0.014 | Young |
| 0.273141 | 0.081167 | 0.006 | 0.011 | 1 | 2 | Epop | -0.005 | Young |
| 0.273228 | 0.497289 | 0.044 | 0.034 | 1 | 2 | 6030458C1 | 0.01 | Young |
| 0.273418 | 0.182961 | 0.196 | 0.177 | 1 | 2 | Slc29a1 | 0.019 | Young |

| | | | | | | | | |
|---|---|---|---|---|---|---|---|---|
| 0.273484 | 0.053329 | 0.214 | 0.193 | 1 | 2 | Tnrc6a | 0.021 | Young |
| 0.273594 | 0.833878 | 0.042 | 0.032 | 1 | 2 | Mkrn2 | 0.01 | Young |
| 0.273669 | 0.748879 | 0.053 | 0.043 | 1 | 2 | Uxt | 0.01 | Young |
| 0.274001 | 0.076761 | 0.239 | 0.218 | 1 | 2 | Pnn | 0.021 | Young |
| 0.274225 | 0.012286 | 0.289 | 0.327 | 1 | 2 | Rbm8a | -0.038 | Young |
| 0.275082 | 0.040014 | 0.072 | 0.087 | 1 | 2 | Gapt | -0.015 | Young |
| 0.275202 | 0.363873 | 0.058 | 0.047 | 1 | 2 | Txnrd2 | 0.011 | Young |
| 0.275425 | 0.147182 | 0.163 | 0.145 | 1 | 2 | Ift20 | 0.018 | Young |
| 0.275879 | 0.365125 | 0.151 | 0.135 | 1 | 2 | Slc25a51 | 0.016 | Young |
| 0.276172 | 0.290586 | 0.285 | 0.272 | 1 | 2 | Arl5b | 0.013 | Young |
| 0.27628 | 0.285857 | 0.28 | 0.264 | 1 | 2 | Elmo1 | 0.016 | Young |
| 0.276481 | 0.06795 | 0.006 | 0.011 | 1 | 2 | Trmu | -0.005 | Young |
| 0.27671 | 0.09557 | 0.473 | 0.455 | 1 | 2 | Rock2 | 0.018 | Young |
| 0.276716 | 0.730267 | 0.017 | 0.011 | 1 | 2 | Tlcd2 | 0.006 | Young |
| 0.276759 | 0.287271 | 0.072 | 0.06 | 1 | 2 | Prune2 | 0.012 | Young |
| 0.277039 | 0.203439 | 0.126 | 0.11 | 1 | 2 | Chml | 0.016 | Young |
| 0.277275 | 0.265251 | 0.033 | 0.025 | 1 | 2 | Rprd1a | 0.008 | Young |
| 0.277296 | 0.275003 | 0.201 | 0.183 | 1 | 2 | Lsm5 | 0.018 | Young |
| 0.277324 | 0.325771 | 0.15 | 0.134 | 1 | 2 | Rgl2 | 0.016 | Young |
| 0.27777 | 0.432736 | 0.055 | 0.044 | 1 | 2 | Fam8a1 | 0.011 | Young |
| 0.277781 | 0.35341 | 0.166 | 0.151 | 1 | 2 | Ebp | 0.015 | Young |
| 0.277992 | 0.353252 | 0.075 | 0.062 | 1 | 2 | Mtr | 0.013 | Young |
| 0.278099 | 0.142324 | 0.101 | 0.086 | 1 | 2 | Rasa2 | 0.015 | Young |
| 0.278163 | 0.101948 | 0.456 | 0.436 | 1 | 2 | C3ar1 | 0.02 | Young |
| 0.27848 | 0.32919 | 0.033 | 0.025 | 1 | 2 | Ptges2 | 0.008 | Young |
| 0.278529 | 0.484261 | 0.053 | 0.043 | 1 | 2 | Glt8d1 | 0.01 | Young |
| 0.278646 | 0.175044 | 0.134 | 0.118 | 1 | 2 | Rcbtb2 | 0.016 | Young |
| 0.278803 | 0.556829 | 0.037 | 0.028 | 1 | 2 | Ston2 | 0.009 | Young |
| 0.278825 | 0.161337 | 0.115 | 0.1 | 1 | 2 | Fundc1 | 0.015 | Young |
| 0.278973 | 0.735471 | 0.017 | 0.011 | 1 | 2 | Ttc21b | 0.006 | Young |
| 0.279046 | 0.251676 | 0.366 | 0.354 | 1 | 2 | Eno1 | 0.012 | Young |
| 0.27909 | 0.165618 | 0.051 | 0.064 | 1 | 2 | Bap1 | -0.013 | Young |
| 0.279198 | 0.433592 | 0.061 | 0.049 | 1 | 2 | Atad3a | 0.012 | Young |
| 0.279372 | 0.312753 | 0.175 | 0.161 | 1 | 2 | Anapc13 | 0.014 | Young |
| 0.279683 | 0.312172 | 0.052 | 0.042 | 1 | 2 | Ltbp3 | 0.01 | Young |
| 0.279835 | 0.061521 | 0.252 | 0.23 | 1 | 2 | Chic2 | 0.022 | Young |
| 0.279882 | 0.284056 | 0.037 | 0.028 | 1 | 2 | Jak3 | 0.009 | Young |
| 0.280042 | 0.266561 | 0.127 | 0.112 | 1 | 2 | Ktn1 | 0.015 | Young |
| 0.280162 | 0.165854 | 0.094 | 0.079 | 1 | 2 | Nelfa | 0.015 | Young |
| 0.280191 | 0.498342 | 0.055 | 0.044 | 1 | 2 | Ttc37 | 0.011 | Young |
| 0.280444 | 0.478391 | 0.173 | 0.158 | 1 | 2 | Dna2 | 0.015 | Young |
| 0.280816 | 0.187265 | 0.188 | 0.169 | 1 | 2 | Mfhas1 | 0.019 | Young |
| 0.280987 | 0.188202 | 0.228 | 0.21 | 1 | 2 | Cul1 | 0.018 | Young |
| 0.281288 | 0.395803 | 0.043 | 0.033 | 1 | 2 | Mfge8 | 0.01 | Young |
| 0.28145 | 0.472305 | 0.017 | 0.011 | 1 | 2 | Trub1 | 0.006 | Young |

| | | | | | | | | |
|---|---|---|---|---|---|---|---|---|
| 0.28145 | 0.303279 | 0.017 | 0.011 | 1 | 2 | Kiz | 0.006 | Young |
| 0.281488 | 0.139986 | 0.247 | 0.226 | 1 | 2 | Nek7 | 0.021 | Young |
| 0.282129 | 0.4793 | 0.038 | 0.029 | 1 | 2 | Sgf29 | 0.009 | Young |
| 0.282505 | 0.425383 | 0.04 | 0.031 | 1 | 2 | Dip2a | 0.009 | Young |
| 0.282509 | 0.018568 | 0.09 | 0.107 | 1 | 2 | Ppp4r1 | -0.017 | Young |
| 0.282633 | 0.088267 | 0.108 | 0.093 | 1 | 2 | Hspa13 | 0.015 | Young |
| 0.282764 | 0.286569 | 0.16 | 0.143 | 1 | 2 | Hdac5 | 0.017 | Young |
| 0.283267 | 0.044274 | 0.631 | 0.611 | 1 | 2 | App | 0.02 | Young |
| 0.283787 | 0.153446 | 0.258 | 0.238 | 1 | 2 | Creb1 | 0.02 | Young |
| 0.283803 | 0.178497 | 0.468 | 0.453 | 1 | 2 | Jmjd1c | 0.015 | Young |
| 0.283864 | 0.067529 | 0.403 | 0.382 | 1 | 2 | Prpf38b | 0.021 | Young |
| 0.284114 | 0.559735 | 0.07 | 0.059 | 1 | 2 | Gadd45gip | 0.011 | Young |
| 0.284153 | 0.230418 | 0.065 | 0.054 | 1 | 2 | Pygb | 0.011 | Young |
| 0.284396 | 0.429725 | 0.045 | 0.035 | 1 | 2 | Hbegf | 0.01 | Young |
| 0.284856 | 0.825739 | 0.018 | 0.012 | 1 | 2 | Tmem132a | 0.006 | Young |
| 0.284994 | 0.20628 | 0.036 | 0.027 | 1 | 2 | Tbccd1 | 0.009 | Young |
| 0.285646 | 0.342369 | 0.146 | 0.131 | 1 | 2 | Abcd1 | 0.015 | Young |
| 0.285659 | 0.865474 | 0.018 | 0.012 | 1 | 2 | Mfsd4a | 0.006 | Young |
| 0.285745 | 0.298647 | 0.232 | 0.217 | 1 | 2 | Arpp19 | 0.015 | Young |
| 0.285891 | 0.164811 | 0.096 | 0.082 | 1 | 2 | Tmx4 | 0.014 | Young |
| 0.285944 | 0.356245 | 0.061 | 0.049 | 1 | 2 | Neurl2 | 0.012 | Young |
| 0.286206 | 0.549335 | 0.076 | 0.064 | 1 | 2 | Rab9 | 0.012 | Young |
| 0.286409 | 0.239418 | 0.157 | 0.141 | 1 | 2 | Zhx1 | 0.016 | Young |
| 0.286634 | 0.057594 | 0.07 | 0.085 | 1 | 2 | Slc28a2 | -0.015 | Young |
| 0.28676 | 0.397631 | 0.145 | 0.131 | 1 | 2 | 1700025G( | 0.014 | Young |
| 0.286964 | 0.493327 | 0.038 | 0.029 | 1 | 2 | Dhcr7 | 0.009 | Young |
| 0.28698 | 0.570191 | 0.05 | 0.04 | 1 | 2 | Plpp2 | 0.01 | Young |
| 0.286986 | 0.034914 | 0.076 | 0.091 | 1 | 2 | Nifk | -0.015 | Young |
| 0.287029 | 0.293761 | 0.198 | 0.182 | 1 | 2 | Tmed3 | 0.016 | Young |
| 0.287507 | 0.039434 | 0.232 | 0.211 | 1 | 2 | Ppp3cb | 0.021 | Young |
| 0.287568 | 0.486791 | 0.046 | 0.036 | 1 | 2 | Crls1 | 0.01 | Young |
| 0.287914 | 0.41737 | 0.061 | 0.049 | 1 | 2 | Icmt | 0.012 | Young |
| 0.287979 | 0.166109 | 0.202 | 0.186 | 1 | 2 | Tcf12 | 0.016 | Young |
| 0.288159 | 0.409991 | 0.045 | 0.035 | 1 | 2 | Slc16a1 | 0.01 | Young |
| 0.288466 | 0.212381 | 0.264 | 0.252 | 1 | 2 | Clk4 | 0.012 | Young |
| 0.288531 | 0.376788 | 0.045 | 0.035 | 1 | 2 | Usp21 | 0.01 | Young |
| 0.288851 | 0.10065 | 0.019 | 0.027 | 1 | 2 | Fbf1 | -0.008 | Young |
| 0.289311 | 0.300179 | 0.048 | 0.037 | 1 | 2 | Hps4 | 0.011 | Young |
| 0.29049 | 0.2247 | 0.235 | 0.216 | 1 | 2 | Tubb2a | 0.019 | Young |
| 0.290531 | 0.095189 | 0.327 | 0.306 | 1 | 2 | Zdhhc20 | 0.021 | Young |
| 0.29077 | 0.03037 | 0.099 | 0.084 | 1 | 2 | Rbm4b | 0.015 | Young |
| 0.291245 | 0.231943 | 0.043 | 0.033 | 1 | 2 | Smim24 | 0.01 | Young |
| 0.291296 | 0.202229 | 0.185 | 0.169 | 1 | 2 | Rnf187 | 0.016 | Young |
| 0.291471 | 0.086296 | 0.268 | 0.245 | 1 | 2 | Clptm1l | 0.023 | Young |
| 0.291697 | 0.080246 | 0.159 | 0.184 | 1 | 2 | Eif3g | -0.025 | Young |

| | | | | | | | | |
|---|---|---|---|---|---|---|---|---|
| 0.291938 | 0.04032 | 0.886 | 0.874 | 1 | 2 | H2-Eb1 | 0.012 | Young |
| 0.292548 | 0.387369 | 0.018 | 0.012 | 1 | 2 | Vdr | 0.006 | Young |
| 0.292567 | 0.734261 | 0.071 | 0.06 | 1 | 2 | Sdhaf4 | 0.011 | Young |
| 0.292807 | 1.114034 | 0.019 | 0.013 | 1 | 2 | Ccdc138 | 0.006 | Young |
| 0.292807 | 0.794419 | 0.019 | 0.013 | 1 | 2 | Chd3os | 0.006 | Young |
| 0.292947 | 0.114127 | 0.417 | 0.405 | 1 | 2 | Ehd1 | 0.012 | Young |
| 0.293514 | 0.166677 | 0.309 | 0.299 | 1 | 2 | Ndufb8 | 0.01 | Young |
| 0.29436 | 0.084364 | 0.074 | 0.061 | 1 | 2 | Msra | 0.013 | Young |
| 0.29439 | 0.250809 | 0.165 | 0.15 | 1 | 2 | Psmg4 | 0.015 | Young |
| 0.294793 | 0.272524 | 0.093 | 0.079 | 1 | 2 | Vav2 | 0.014 | Young |
| 0.294846 | 0.160538 | 0.134 | 0.118 | 1 | 2 | Hspa4l | 0.016 | Young |
| 0.294851 | 0.044568 | 0.042 | 0.032 | 1 | 2 | Tmed8 | 0.01 | Young |
| 0.295074 | 0.049574 | 0.184 | 0.165 | 1 | 2 | Pilra | 0.019 | Young |
| 0.295171 | 0.017326 | 0.118 | 0.137 | 1 | 2 | Psmd3 | -0.019 | Young |
| 0.295848 | 0.061932 | 0.027 | 0.036 | 1 | 2 | Dynlt1f | -0.009 | Young |
| 0.295956 | 0.315838 | 0.109 | 0.095 | 1 | 2 | Lsm7 | 0.014 | Young |
| 0.296084 | 0.085371 | 0.078 | 0.065 | 1 | 2 | Cnep1r1 | 0.013 | Young |
| 0.296169 | 0.898336 | 0.019 | 0.013 | 1 | 2 | Ccdc126 | 0.006 | Young |
| 0.29626 | 0.093919 | 0.219 | 0.201 | 1 | 2 | mt-Atp8 | 0.018 | Young |
| 0.296451 | 0.158508 | 0.036 | 0.046 | 1 | 2 | Lpar1 | -0.01 | Young |
| 0.296964 | 0.65596 | 0.019 | 0.013 | 1 | 2 | Wdhd1 | 0.006 | Young |
| 0.297007 | 0.551557 | 0.077 | 0.065 | 1 | 2 | Prmt3 | 0.012 | Young |
| 0.297974 | 0.068828 | 0.419 | 0.401 | 1 | 2 | Bnip2 | 0.018 | Young |
| 0.298114 | 0.179918 | 0.048 | 0.037 | 1 | 2 | Atg2b | 0.011 | Young |
| 0.298182 | 0.28003 | 0.049 | 0.039 | 1 | 2 | Zbtb42 | 0.01 | Young |
| 0.298384 | 0.082911 | 0.067 | 0.055 | 1 | 2 | Ice1 | 0.012 | Young |
| 0.298522 | 0.120835 | 0.486 | 0.481 | 1 | 2 | Nfkbid | 0.005 | Young |
| 0.298529 | 0.168632 | 0.198 | 0.183 | 1 | 2 | Dhx40 | 0.015 | Young |
| 0.29871 | 0.004416 | 0.207 | 0.23 | 1 | 2 | Emp1 | -0.023 | Young |
| 0.29943 | 0.168927 | 0.35 | 0.337 | 1 | 2 | Timm13 | 0.013 | Young |
| 0.299731 | 0.375356 | 0.076 | 0.064 | 1 | 2 | Snrnp25 | 0.012 | Young |
| 0.299847 | 0.372211 | 0.109 | 0.096 | 1 | 2 | Socs7 | 0.013 | Young |
| 0.29998 | 0.183197 | 0.141 | 0.125 | 1 | 2 | Ebi3 | 0.016 | Young |
| 0.300388 | 0.019041 | 0.05 | 0.062 | 1 | 2 | Zcchc8 | -0.012 | Young |
| 0.301502 | 0.068047 | 0.373 | 0.351 | 1 | 2 | Etf1 | 0.022 | Young |
| 0.301843 | 0.271446 | 0.063 | 0.051 | 1 | 2 | Ipo9 | 0.012 | Young |
| 0.302071 | 0.297392 | 0.052 | 0.042 | 1 | 2 | Bmf | 0.01 | Young |
| 0.302276 | 0.21676 | 0.05 | 0.04 | 1 | 2 | Def6 | 0.01 | Young |
| 0.302727 | 0.216147 | 0.186 | 0.171 | 1 | 2 | Pole4 | 0.015 | Young |
| 0.302924 | 0.057393 | 0.451 | 0.425 | 1 | 2 | Eif3c | 0.026 | Young |
| 0.302983 | 0.036663 | 0.126 | 0.145 | 1 | 2 | Amdhd2 | -0.019 | Young |
| 0.303171 | 0.07899 | 0.211 | 0.239 | 1 | 2 | Dctn1 | -0.028 | Young |
| 0.304039 | 0.257801 | 0.145 | 0.13 | 1 | 2 | Crybg3 | 0.015 | Young |
| 0.304142 | 0.074249 | 0.024 | 0.032 | 1 | 2 | Lrrc40 | -0.008 | Young |
| 0.304371 | 0.110395 | 0.134 | 0.154 | 1 | 2 | Vps28 | -0.02 | Young |

| | | | | | | | | |
|---|---|---|---|---|---|---|---|---|
| 0.304668 | 1.046687 | 0.01 | 0.005 | 1 | 2 | Diaph3 | 0.005 | Young |
| 0.305055 | 0.567809 | 0.01 | 0.015 | 1 | 2 | Actn1 | -0.005 | Young |
| 0.30511 | 0.581587 | 0.091 | 0.079 | 1 | 2 | Polr2b | 0.012 | Young |
| 0.305545 | 0.06021 | 0.17 | 0.151 | 1 | 2 | Sh3bp5 | 0.019 | Young |
| 0.305556 | 0.849956 | 0.01 | 0.005 | 1 | 2 | Bgn | 0.005 | Young |
| 0.305763 | 0.35339 | 0.02 | 0.014 | 1 | 2 | Frat1 | 0.006 | Young |
| 0.305852 | 0.489465 | 0.01 | 0.005 | 1 | 2 | Mgll | 0.005 | Young |
| 0.305916 | 0.195185 | 0.163 | 0.148 | 1 | 2 | Cdc73 | 0.015 | Young |
| 0.305994 | 0.118735 | 0.049 | 0.039 | 1 | 2 | Abhd5 | 0.01 | Young |
| 0.30629 | 0.055511 | 0.787 | 0.756 | 1 | 2 | Cotl1 | 0.031 | Young |
| 0.306445 | 0.502621 | 0.01 | 0.005 | 1 | 2 | Smtn | 0.005 | Young |
| 0.307039 | 0.396392 | 0.01 | 0.005 | 1 | 2 | Apoc1 | 0.005 | Young |
| 0.307123 | 0.589401 | 0.061 | 0.05 | 1 | 2 | Mcm6 | 0.011 | Young |
| 0.307358 | 0.279625 | 0.241 | 0.225 | 1 | 2 | Leprotl1 | 0.016 | Young |
| 0.307513 | 0.1093 | 0.081 | 0.096 | 1 | 2 | Rnf14 | -0.015 | Young |
| 0.307634 | 0.331025 | 0.01 | 0.005 | 1 | 2 | Cd209c | 0.005 | Young |
| 0.307658 | 0.361041 | 0.215 | 0.203 | 1 | 2 | B4galt1 | 0.012 | Young |
| 0.307826 | 0.180616 | 0.242 | 0.225 | 1 | 2 | Taf10 | 0.017 | Young |
| 0.307905 | 0.235706 | 0.013 | 0.019 | 1 | 2 | Zfp629 | -0.006 | Young |
| 0.307955 | 0.307288 | 0.115 | 0.101 | 1 | 2 | Mepce | 0.014 | Young |
| 0.308073 | 0.383286 | 0.099 | 0.086 | 1 | 2 | Flot1 | 0.013 | Young |
| 0.308271 | 0.043715 | 0.1 | 0.086 | 1 | 2 | Uchl5 | 0.014 | Young |
| 0.308528 | 0.328606 | 0.01 | 0.005 | 1 | 2 | Fuk | 0.005 | Young |
| 0.309174 | 0.392389 | 0.154 | 0.141 | 1 | 2 | Aff1 | 0.013 | Young |
| 0.309366 | 0.022223 | 0.037 | 0.047 | 1 | 2 | Anapc7 | -0.01 | Young |
| 0.309379 | 0.686229 | 0.021 | 0.015 | 1 | 2 | Nek6 | 0.006 | Young |
| 0.309423 | 0.027052 | 0.01 | 0.005 | 1 | 2 | Map2k6 | 0.005 | Young |
| 0.310515 | 0.954182 | 0.024 | 0.017 | 1 | 2 | Tmem150b | 0.007 | Young |
| 0.31073 | 0.045342 | 0.051 | 0.041 | 1 | 2 | Ypel2 | 0.01 | Young |
| 0.310888 | 0.012859 | 0.02 | 0.014 | 1 | 2 | Cmc4 | 0.006 | Young |
| 0.311243 | 0.343279 | 0.108 | 0.095 | 1 | 2 | Lpcat2 | 0.013 | Young |
| 0.311642 | 0.302316 | 0.094 | 0.081 | 1 | 2 | Mcm3 | 0.013 | Young |
| 0.311681 | 0.382155 | 0.021 | 0.015 | 1 | 2 | Ccdc130 | 0.006 | Young |
| 0.311917 | 0.203664 | 0.208 | 0.191 | 1 | 2 | Rragc | 0.017 | Young |
| 0.312074 | 0.184051 | 0.357 | 0.339 | 1 | 2 | Malt1 | 0.018 | Young |
| 0.312326 | 0.64825 | 0.023 | 0.016 | 1 | 2 | Ankrd26 | 0.007 | Young |
| 0.312625 | 0.233713 | 0.158 | 0.142 | 1 | 2 | Parp8 | 0.016 | Young |
| 0.312875 | 0.217538 | 0.02 | 0.014 | 1 | 2 | Glb1l | 0.006 | Young |
| 0.312879 | 0.509015 | 0.109 | 0.097 | 1 | 2 | Nup62 | 0.012 | Young |
| 0.313513 | 0.841801 | 0.024 | 0.017 | 1 | 2 | 1110012L1 | 0.007 | Young |
| 0.313557 | 0.263665 | 0.096 | 0.082 | 1 | 2 | Wdr3 | 0.014 | Young |
| 0.313704 | 0.265296 | 0.205 | 0.192 | 1 | 2 | Nudt21 | 0.013 | Young |
| 0.313719 | 0.159563 | 0.165 | 0.148 | 1 | 2 | Alkbh1 | 0.017 | Young |
| 0.31426 | 1.167029 | 0.025 | 0.018 | 1 | 2 | Dtd2 | 0.007 | Young |
| 0.314371 | 0.046327 | 0.033 | 0.043 | 1 | 2 | Tbkbp1 | -0.01 | Young |

| | | | | | | | | |
|---|---|---|---|---|---|---|---|---|
| 0.314689 | 0.097157 | 0.153 | 0.176 | 1 | 2 | Atp7a | -0.023 | Young |
| 0.314859 | 0.146441 | 0.296 | 0.279 | 1 | 2 | Ninj1 | 0.017 | Young |
| 0.314927 | 0.065482 | 0.169 | 0.152 | 1 | 2 | Gsk3a | 0.017 | Young |
| 0.315932 | 0.274058 | 0.021 | 0.015 | 1 | 2 | Ippk | 0.006 | Young |
| 0.316531 | 0.326154 | 0.074 | 0.062 | 1 | 2 | Smurf1 | 0.012 | Young |
| 0.316809 | 0.304481 | 0.179 | 0.164 | 1 | 2 | Eri1 | 0.015 | Young |
| 0.316835 | 0.630559 | 0.023 | 0.016 | 1 | 2 | Cdpf1 | 0.007 | Young |
| 0.317076 | 0.35817 | 0.222 | 0.213 | 1 | 2 | Sirt2 | 0.009 | Young |
| 0.317315 | 0.156544 | 0.28 | 0.267 | 1 | 2 | Map2k2 | 0.013 | Young |
| 0.317501 | 0.804554 | 0.026 | 0.019 | 1 | 2 | Kpna2 | 0.007 | Young |
| 0.317614 | 0.085371 | 0.038 | 0.048 | 1 | 2 | Paqr7 | -0.01 | Young |
| 0.318008 | 0.819821 | 0.025 | 0.018 | 1 | 2 | Zfp874a | 0.007 | Young |
| 0.318134 | 0.276199 | 0.228 | 0.214 | 1 | 2 | N4bp2l2 | 0.014 | Young |
| 0.318235 | 0.06122 | 0.034 | 0.044 | 1 | 2 | Adprm | -0.01 | Young |
| 0.319532 | 0.754082 | 0.051 | 0.042 | 1 | 2 | 0610010K1 | 0.009 | Young |
| 0.319542 | 0.142682 | 0.115 | 0.101 | 1 | 2 | Ube4b | 0.014 | Young |
| 0.3196 | 0.016853 | 0.068 | 0.081 | 1 | 2 | Ccdc97 | -0.013 | Young |
| 0.319864 | 0.112385 | 0.023 | 0.016 | 1 | 2 | Lrrc45 | 0.007 | Young |
| 0.320334 | 0.150363 | 0.108 | 0.094 | 1 | 2 | Arhgap5 | 0.014 | Young |
| 0.321137 | 0.403664 | 0.024 | 0.017 | 1 | 2 | Rhobtb2 | 0.007 | Young |
| 0.321481 | 0.296214 | 0.101 | 0.088 | 1 | 2 | Hint2 | 0.013 | Young |
| 0.321497 | 0.000673 | 0.223 | 0.247 | 1 | 2 | Rab8a | -0.024 | Young |
| 0.321584 | 0.091717 | 0.546 | 0.545 | 1 | 2 | Ndufa6 | 0.001 | Young |
| 0.322288 | 0.577033 | 0.059 | 0.049 | 1 | 2 | Zfp329 | 0.01 | Young |
| 0.322435 | 0.511767 | 0.024 | 0.017 | 1 | 2 | Slx1b | 0.007 | Young |
| 0.322439 | 0.849929 | 0.011 | 0.006 | 1 | 2 | Trmt44 | 0.005 | Young |
| 0.32253 | 0.250365 | 0.023 | 0.016 | 1 | 2 | Miga2 | 0.007 | Young |
| 0.322914 | 0.38679 | 0.12 | 0.107 | 1 | 2 | Naa10 | 0.013 | Young |
| 0.323098 | 0.281421 | 0.223 | 0.211 | 1 | 2 | Lsm6 | 0.012 | Young |
| 0.323107 | 0.222134 | 0.181 | 0.166 | 1 | 2 | Rbm33 | 0.015 | Young |
| 0.323251 | 0.144636 | 0.216 | 0.201 | 1 | 2 | Atp13a2 | 0.015 | Young |
| 0.323297 | 1.308187 | 0.011 | 0.006 | 1 | 2 | Gtf2ird2 | 0.005 | Young |
| 0.323473 | 0.160033 | 0.357 | 0.344 | 1 | 2 | Ppig | 0.013 | Young |
| 0.324074 | 0.416842 | 0.09 | 0.078 | 1 | 2 | Arl13b | 0.012 | Young |
| 0.324232 | 0.092633 | 0.457 | 0.441 | 1 | 2 | Tcf4 | 0.016 | Young |
| 0.324255 | 0.732926 | 0.027 | 0.02 | 1 | 2 | Zfyve19 | 0.007 | Young |
| 0.324463 | 0.206154 | 0.286 | 0.273 | 1 | 2 | Hsbp1 | 0.013 | Young |
| 0.324582 | 0.647721 | 0.03 | 0.022 | 1 | 2 | Gna11 | 0.008 | Young |
| 0.324731 | 0.633065 | 0.011 | 0.006 | 1 | 2 | Mr1 | 0.005 | Young |
| 0.325045 | 0.386801 | 0.173 | 0.162 | 1 | 2 | Rheb | 0.011 | Young |
| 0.325306 | 0.779033 | 0.011 | 0.006 | 1 | 2 | 4833445I0 | 0.005 | Young |
| 0.325306 | 0.752064 | 0.011 | 0.006 | 1 | 2 | Map3k12 | 0.005 | Young |
| 0.326266 | 0.099271 | 0.064 | 0.077 | 1 | 2 | Umad1 | -0.013 | Young |
| 0.326457 | 0.455039 | 0.011 | 0.006 | 1 | 2 | Coq4 | 0.005 | Young |
| 0.326507 | 0.604103 | 0.027 | 0.02 | 1 | 2 | Zeb1 | 0.007 | Young |

| | | | | | | | | |
|---|---|---|---|---|---|---|---|---|
| 0.326691 | 0.18804 | 0.178 | 0.162 | 1 | 2 | Tcof1 | 0.016 | Young |
| 0.326746 | 0.345912 | 0.011 | 0.006 | 1 | 2 | Enho | 0.005 | Young |
| 0.326844 | 0.141042 | 0.181 | 0.163 | 1 | 2 | Fam104a | 0.018 | Young |
| 0.326881 | 0.200558 | 0.188 | 0.171 | 1 | 2 | Rabgef1 | 0.017 | Young |
| 0.326903 | 0.962882 | 0.03 | 0.022 | 1 | 2 | Alad | 0.008 | Young |
| 0.327323 | 0.876964 | 0.011 | 0.006 | 1 | 2 | Fn3krp | 0.005 | Young |
| 0.327381 | 0.446452 | 0.107 | 0.095 | 1 | 2 | Rnf10 | 0.012 | Young |
| 0.327612 | 0.487957 | 0.011 | 0.006 | 1 | 2 | Mex3b | 0.005 | Young |
| 0.328094 | 0.211187 | 0.096 | 0.084 | 1 | 2 | Map3k20 | 0.012 | Young |
| 0.328103 | 0.017534 | 0.05 | 0.061 | 1 | 2 | D10Wsu10 | -0.011 | Young |
| 0.328479 | 0.386484 | 0.011 | 0.006 | 1 | 2 | Ttll1 | 0.005 | Young |
| 0.328479 | 0.12897 | 0.011 | 0.006 | 1 | 2 | Col1a2 | 0.005 | Young |
| 0.328538 | 0.261803 | 0.107 | 0.094 | 1 | 2 | Crebzf | 0.013 | Young |
| 0.328622 | 0.040318 | 0.498 | 0.48 | 1 | 2 | Mycbp2 | 0.018 | Young |
| 0.328786 | 0.312636 | 0.071 | 0.06 | 1 | 2 | Med23 | 0.011 | Young |
| 0.328934 | 0.271383 | 0.126 | 0.112 | 1 | 2 | Pom121 | 0.014 | Young |
| 0.329337 | 0.336836 | 0.026 | 0.019 | 1 | 2 | Bloc1s3 | 0.007 | Young |
| 0.329587 | 0.592738 | 0.09 | 0.079 | 1 | 2 | 4931406P1 | 0.011 | Young |
| 0.329645 | 0.15925 | 0.068 | 0.057 | 1 | 2 | Elof1 | 0.011 | Young |
| 0.330111 | 0.533481 | 0.038 | 0.03 | 1 | 2 | Sufu | 0.008 | Young |
| 0.330113 | 0.870831 | 0.04 | 0.032 | 1 | 2 | Numb | 0.008 | Young |
| 0.330348 | 0.012497 | 0.191 | 0.215 | 1 | 2 | Mfsd5 | -0.024 | Young |
| 0.330353 | 0.008982 | 0.159 | 0.181 | 1 | 2 | Myo1c | -0.022 | Young |
| 0.330662 | 0.390862 | 0.101 | 0.089 | 1 | 2 | Ccdc9 | 0.012 | Young |
| 0.330712 | 0.106948 | 0.025 | 0.018 | 1 | 2 | Wdr92 | 0.007 | Young |
| 0.331264 | 0.243124 | 0.025 | 0.018 | 1 | 2 | Klhl36 | 0.007 | Young |
| 0.331354 | 0.609398 | 0.034 | 0.027 | 1 | 2 | Elp3 | 0.007 | Young |
| 0.331591 | 0.488012 | 0.049 | 0.04 | 1 | 2 | Zdhhc4 | 0.009 | Young |
| 0.331668 | 0.074023 | 0.413 | 0.401 | 1 | 2 | Tax1bp1 | 0.012 | Young |
| 0.332201 | 0.424517 | 0.088 | 0.076 | 1 | 2 | Zfyve27 | 0.012 | Young |
| 0.332417 | 0.061108 | 0.267 | 0.246 | 1 | 2 | Cnppd1 | 0.021 | Young |
| 0.332561 | 0.559399 | 0.036 | 0.028 | 1 | 2 | Hddc2 | 0.008 | Young |
| 0.332861 | 0.248989 | 0.124 | 0.11 | 1 | 2 | Oser1 | 0.014 | Young |
| 0.332912 | 0.654299 | 0.038 | 0.03 | 1 | 2 | Rbm19 | 0.008 | Young |
| 0.33338 | 0.16249 | 0.426 | 0.409 | 1 | 2 | Plekho1 | 0.017 | Young |
| 0.333393 | 0.324435 | 0.059 | 0.049 | 1 | 2 | Ppp5c | 0.01 | Young |
| 0.333874 | 0.411251 | 0.05 | 0.041 | 1 | 2 | Arfgap3 | 0.009 | Young |
| 0.333931 | 0.537341 | 0.03 | 0.022 | 1 | 2 | Nr1d1 | 0.008 | Young |
| 0.334384 | 0.23909 | 0.14 | 0.126 | 1 | 2 | Adrm1 | 0.014 | Young |
| 0.334966 | 0.328348 | 0.052 | 0.043 | 1 | 2 | Nat9 | 0.009 | Young |
| 0.335901 | 0.068744 | 0.704 | 0.712 | 1 | 2 | Tln1 | -0.008 | Young |
| 0.336018 | 0.231473 | 0.442 | 0.434 | 1 | 2 | Minos1 | 0.008 | Young |
| 0.336157 | 0.418397 | 0.116 | 0.104 | 1 | 2 | Atic | 0.012 | Young |
| 0.336165 | 0.400774 | 0.048 | 0.039 | 1 | 2 | Frmd6 | 0.009 | Young |
| 0.336485 | 0.544259 | 0.032 | 0.025 | 1 | 2 | Oraov1 | 0.007 | Young |

| | | | | | | | | |
|---|---|---|---|---|---|---|---|---|
| 0.337004 | 0.444795 | 0.043 | 0.034 | 1 | 2 | Zc3hc1 | 0.009 | Young |
| 0.337188 | 0.371886 | 0.059 | 0.049 | 1 | 2 | Rrad | 0.01 | Young |
| 0.337231 | 0.16442 | 0.353 | 0.338 | 1 | 2 | Brd2 | 0.015 | Young |
| 0.337476 | 0.278361 | 0.088 | 0.075 | 1 | 2 | Adora2b | 0.013 | Young |
| 0.337598 | 0.36068 | 0.08 | 0.069 | 1 | 2 | Aen | 0.011 | Young |
| 0.337821 | 0.30332 | 0.03 | 0.022 | 1 | 2 | Dalrd3 | 0.008 | Young |
| 0.337823 | 0.529043 | 0.05 | 0.041 | 1 | 2 | Dnttip1 | 0.009 | Young |
| 0.338147 | 0.292636 | 0.065 | 0.055 | 1 | 2 | Cdc16 | 0.01 | Young |
| 0.338163 | 0.083109 | 0.682 | 0.675 | 1 | 2 | Calm2 | 0.007 | Young |
| 0.338224 | 0.020257 | 0.303 | 0.283 | 1 | 2 | Hipk1 | 0.02 | Young |
| 0.338263 | 0.19658 | 0.026 | 0.019 | 1 | 2 | Npl | 0.007 | Young |
| 0.339128 | 0.324036 | 0.029 | 0.021 | 1 | 2 | Tmem206 | 0.008 | Young |
| 0.339466 | 0.40124 | 0.033 | 0.026 | 1 | 2 | Sprtn | 0.007 | Young |
| 0.339538 | 0.27622 | 0.131 | 0.117 | 1 | 2 | Cnot8 | 0.014 | Young |
| 0.339641 | 0.173513 | 0.416 | 0.41 | 1 | 2 | Cox7a2 | 0.006 | Young |
| 0.339755 | 0.344124 | 0.145 | 0.132 | 1 | 2 | Ahsa1 | 0.013 | Young |
| 0.339997 | 0.405395 | 0.031 | 0.024 | 1 | 2 | Nfatc2ip | 0.007 | Young |
| 0.340265 | 0.391515 | 0.045 | 0.036 | 1 | 2 | 4930453N2 | 0.009 | Young |
| 0.340293 | 0.66676 | 0.012 | 0.007 | 1 | 2 | Gm26601 | 0.005 | Young |
| 0.340293 | 0.486238 | 0.012 | 0.007 | 1 | 2 | Cd101 | 0.005 | Young |
| 0.340571 | 0.946306 | 0.012 | 0.007 | 1 | 2 | Ccdc181 | 0.005 | Young |
| 0.340571 | 0.798765 | 0.012 | 0.007 | 1 | 2 | Pgap3 | 0.005 | Young |
| 0.340639 | 0.007023 | 0.125 | 0.141 | 1 | 2 | Galk2 | -0.016 | Young |
| 0.340668 | 0.497252 | 0.046 | 0.037 | 1 | 2 | Dgcr8 | 0.009 | Young |
| 0.340724 | 0.816824 | 0.044 | 0.035 | 1 | 2 | St3gal6 | 0.009 | Young |
| 0.340754 | 0.106315 | 0.031 | 0.04 | 1 | 2 | Tpgs2 | -0.009 | Young |
| 0.34085 | 0.933159 | 0.012 | 0.007 | 1 | 2 | Gm340 | 0.005 | Young |
| 0.340958 | 0.165939 | 0.132 | 0.118 | 1 | 2 | Atraid | 0.014 | Young |
| 0.340981 | 0.154258 | 0.034 | 0.027 | 1 | 2 | Ydjc | 0.007 | Young |
| 0.341128 | 0.553442 | 0.012 | 0.007 | 1 | 2 | Efemp2 | 0.005 | Young |
| 0.341145 | 0.359244 | 0.037 | 0.029 | 1 | 2 | Sec22a | 0.008 | Young |
| 0.341686 | 0.858776 | 0.012 | 0.007 | 1 | 2 | Cenpv | 0.005 | Young |
| 0.341686 | 0.811563 | 0.012 | 0.007 | 1 | 2 | Pecr | 0.005 | Young |
| 0.341965 | 1.495451 | 0.012 | 0.007 | 1 | 2 | S100a8 | 0.005 | Young |
| 0.342062 | 0.454736 | 0.037 | 0.029 | 1 | 2 | Kdelc2 | 0.008 | Young |
| 0.342117 | 0.016523 | 0.208 | 0.228 | 1 | 2 | Usp16 | -0.02 | Young |
| 0.342148 | 0.490126 | 0.069 | 0.059 | 1 | 2 | Arl4a | 0.01 | Young |
| 0.342152 | 0.279803 | 0.132 | 0.119 | 1 | 2 | Psmd12 | 0.013 | Young |
| 0.342244 | 0.858115 | 0.012 | 0.007 | 1 | 2 | Macrod2 | 0.005 | Young |
| 0.342299 | 0.141188 | 0.058 | 0.048 | 1 | 2 | Crim1 | 0.01 | Young |
| 0.342512 | 0.087293 | 0.452 | 0.437 | 1 | 2 | Rpl27 | 0.015 | Young |
| 0.342569 | 0.517574 | 0.042 | 0.033 | 1 | 2 | Spire1 | 0.009 | Young |
| 0.342577 | 0.309093 | 0.102 | 0.09 | 1 | 2 | Pink1 | 0.012 | Young |
| 0.342987 | 0.408455 | 0.018 | 0.025 | 1 | 2 | Zcchc4 | -0.007 | Young |
| 0.343379 | 0.256371 | 0.051 | 0.042 | 1 | 2 | Sharpin | 0.009 | Young |

| | | | | | | | | |
|---|---|---|---|---|---|---|---|---|
| 0.343584 | 0.329632 | 0.062 | 0.051 | 1 | 2 | Rad54l2 | 0.011 | Young |
| 0.344634 | 0.366028 | 0.039 | 0.031 | 1 | 2 | Upb1 | 0.008 | Young |
| 0.344934 | 0.386411 | 0.048 | 0.039 | 1 | 2 | Stk35 | 0.009 | Young |
| 0.345086 | 0.309889 | 0.05 | 0.041 | 1 | 2 | Zfp90 | 0.009 | Young |
| 0.345188 | 0.24187 | 0.097 | 0.085 | 1 | 2 | Fam53b | 0.012 | Young |
| 0.345292 | 0.131739 | 0.473 | 0.471 | 1 | 2 | 1810037I1 | 0.002 | Young |
| 0.345476 | 0.092121 | 0.238 | 0.222 | 1 | 2 | Ddb1 | 0.016 | Young |
| 0.345607 | 0.09682 | 0.246 | 0.229 | 1 | 2 | Golph3 | 0.017 | Young |
| 0.345838 | 0.119254 | 0.074 | 0.087 | 1 | 2 | Pggt1b | -0.013 | Young |
| 0.345982 | 0.201824 | 0.031 | 0.04 | 1 | 2 | Dedd | -0.009 | Young |
| 0.346351 | 0.290214 | 0.127 | 0.113 | 1 | 2 | Rfc1 | 0.014 | Young |
| 0.346683 | 0.512591 | 0.042 | 0.033 | 1 | 2 | Pdcd2 | 0.009 | Young |
| 0.346844 | 0.138384 | 0.295 | 0.277 | 1 | 2 | Traf1 | 0.018 | Young |
| 0.347771 | 0.100201 | 0.173 | 0.16 | 1 | 2 | Gabarapl2 | 0.013 | Young |
| 0.347916 | 0.311841 | 0.043 | 0.034 | 1 | 2 | Def8 | 0.009 | Young |
| 0.348165 | 0.818247 | 0.07 | 0.06 | 1 | 2 | Cd24a | 0.01 | Young |
| 0.348269 | 0.125225 | 0.019 | 0.026 | 1 | 2 | Lmtk2 | -0.007 | Young |
| 0.348818 | 0.319973 | 0.337 | 0.323 | 1 | 2 | Drap1 | 0.014 | Young |
| 0.348819 | 0.290511 | 0.038 | 0.03 | 1 | 2 | Slc25a33 | 0.008 | Young |
| 0.34897 | 0.128294 | 0.319 | 0.302 | 1 | 2 | Peli1 | 0.017 | Young |
| 0.349076 | 0.205229 | 0.188 | 0.172 | 1 | 2 | Ankhd1 | 0.016 | Young |
| 0.349262 | 0.174163 | 0.189 | 0.173 | 1 | 2 | Atad2b | 0.016 | Young |
| 0.34947 | 0.327297 | 0.124 | 0.111 | 1 | 2 | Coa3 | 0.013 | Young |
| 0.349478 | 0.268001 | 0.036 | 0.028 | 1 | 2 | 2510002D2 | 0.008 | Young |
| 0.349568 | 0.228555 | 0.093 | 0.08 | 1 | 2 | Gse1 | 0.013 | Young |
| 0.35016 | 0.36417 | 0.075 | 0.064 | 1 | 2 | 4930402H2 | 0.011 | Young |
| 0.350259 | 0.012322 | 0.226 | 0.251 | 1 | 2 | Nfkbie | -0.025 | Young |
| 0.350535 | 0.12449 | 0.2 | 0.185 | 1 | 2 | Entpd1 | 0.015 | Young |
| 0.35063 | 0.382022 | 0.076 | 0.065 | 1 | 2 | Gm2000 | 0.011 | Young |
| 0.35072 | 1.037907 | 0.013 | 0.009 | 1 | 2 | Sap18b | 0.004 | Young |
| 0.350896 | 0.180995 | 0.112 | 0.099 | 1 | 2 | Stam2 | 0.013 | Young |
| 0.351459 | 0.293678 | 0.097 | 0.086 | 1 | 2 | Slc39a14 | 0.011 | Young |
| 0.351513 | 0.203566 | 0.342 | 0.328 | 1 | 2 | Arrb2 | 0.014 | Young |
| 0.351837 | 0.042683 | 0.09 | 0.105 | 1 | 2 | Dok3 | -0.015 | Young |
| 0.351905 | 0.102541 | 0.178 | 0.163 | 1 | 2 | Tcp11l2 | 0.015 | Young |
| 0.352477 | 0.092234 | 0.048 | 0.039 | 1 | 2 | Tfam | 0.009 | Young |
| 0.35264 | 0.441857 | 0.036 | 0.028 | 1 | 2 | Pdik1l | 0.008 | Young |
| 0.353107 | 0.135635 | 0.19 | 0.175 | 1 | 2 | Ndufa10 | 0.015 | Young |
| 0.353621 | 0.065131 | 0.283 | 0.266 | 1 | 2 | Tprgl | 0.017 | Young |
| 0.35422 | 0.600484 | 0.013 | 0.009 | 1 | 2 | Ada | 0.004 | Young |
| 0.354491 | 1.38448 | 0.013 | 0.009 | 1 | 2 | P2ry13 | 0.004 | Young |
| 0.354585 | 0.219538 | 0.177 | 0.164 | 1 | 2 | Ube2e3 | 0.013 | Young |
| 0.355302 | 0.499403 | 0.013 | 0.009 | 1 | 2 | Dph2 | 0.004 | Young |
| 0.355844 | 0.676779 | 0.013 | 0.009 | 1 | 2 | Kcng2 | 0.004 | Young |
| 0.356009 | 0.098623 | 0.108 | 0.095 | 1 | 2 | Pfdn4 | 0.013 | Young |

| | | | | | | | | |
|---|---|---|---|---|---|---|---|---|
| 0.356989 | 0.217475 | 0.254 | 0.24 | 1 | 2 | Basp1 | 0.014 | Young |
| 0.35703 | 0.24096 | 0.26 | 0.248 | 1 | 2 | Dynlrb1 | 0.012 | Young |
| 0.357159 | 0.054047 | 0.157 | 0.179 | 1 | 2 | Xpo6 | -0.022 | Young |
| 0.357887 | 0.202542 | 0.229 | 0.215 | 1 | 2 | Ddrgk1 | 0.014 | Young |
| 0.358559 | 0.242444 | 0.013 | 0.009 | 1 | 2 | Ccdc114 | 0.004 | Young |
| 0.358614 | 0.376672 | 0.156 | 0.142 | 1 | 2 | Nfatc1 | 0.014 | Young |
| 0.358653 | 0.114694 | 0.055 | 0.045 | 1 | 2 | Arhgap21 | 0.01 | Young |
| 0.358776 | 0.007936 | 0.015 | 0.021 | 1 | 2 | Aste1 | -0.006 | Young |
| 0.359272 | 0.091529 | 0.439 | 0.43 | 1 | 2 | Dynll1 | 0.009 | Young |
| 0.35932 | 0.348235 | 0.082 | 0.072 | 1 | 2 | Pigp | 0.01 | Young |
| 0.359455 | 0.036789 | 0.076 | 0.064 | 1 | 2 | Fbxo8 | 0.012 | Young |
| 0.360571 | 0.150331 | 0.049 | 0.04 | 1 | 2 | Zfp945 | 0.009 | Young |
| 0.36068 | 0.44125 | 0.204 | 0.194 | 1 | 2 | Ipo7 | 0.01 | Young |
| 0.36098 | 0.602608 | 0.061 | 0.051 | 1 | 2 | Inpp5a | 0.01 | Young |
| 0.36124 | 0.335028 | 0.137 | 0.124 | 1 | 2 | Tmem147 | 0.013 | Young |
| 0.362663 | 0.242444 | 0.094 | 0.082 | 1 | 2 | Cdk16 | 0.012 | Young |
| 0.362935 | 0.405297 | 0.129 | 0.119 | 1 | 2 | Zfp131 | 0.01 | Young |
| 0.363169 | 0.084992 | 0.012 | 0.017 | 1 | 2 | Gm48604 | -0.005 | Young |
| 0.363382 | 0.294905 | 0.144 | 0.132 | 1 | 2 | Sra1 | 0.012 | Young |
| 0.364212 | 0.159592 | 0.172 | 0.158 | 1 | 2 | Lsr | 0.014 | Young |
| 0.36422 | 0.485413 | 0.062 | 0.052 | 1 | 2 | Kbtbd11 | 0.01 | Young |
| 0.364606 | 0.285602 | 0.097 | 0.086 | 1 | 2 | Mdp1 | 0.011 | Young |
| 0.365053 | 0.090159 | 0.088 | 0.076 | 1 | 2 | Pde3b | 0.012 | Young |
| 0.365139 | 0.068299 | 0.038 | 0.047 | 1 | 2 | Fastk | -0.009 | Young |
| 0.365479 | 0.80615 | 0.014 | 0.01 | 1 | 2 | Diras2 | 0.004 | Young |
| 0.365479 | 0.648953 | 0.014 | 0.01 | 1 | 2 | Dgat2 | 0.004 | Young |
| 0.365553 | 0.255686 | 0.176 | 0.163 | 1 | 2 | Acadm | 0.013 | Young |
| 0.365741 | 0.728104 | 0.014 | 0.01 | 1 | 2 | E030030I0( | 0.004 | Young |
| 0.365741 | 0.611393 | 0.014 | 0.01 | 1 | 2 | Zfp472 | 0.004 | Young |
| 0.365913 | 0.053666 | 0.113 | 0.101 | 1 | 2 | Spata2 | 0.012 | Young |
| 0.366713 | 0.21673 | 0.078 | 0.067 | 1 | 2 | Ralgapb | 0.011 | Young |
| 0.36679 | 0.271153 | 0.11 | 0.099 | 1 | 2 | Mrpl15 | 0.011 | Young |
| 0.367207 | 0.074983 | 0.183 | 0.208 | 1 | 2 | Yipf3 | -0.025 | Young |
| 0.36758 | 0.683017 | 0.014 | 0.01 | 1 | 2 | Rita1 | 0.004 | Young |
| 0.367587 | 0.033533 | 0.128 | 0.113 | 1 | 2 | Tlk1 | 0.015 | Young |
| 0.367598 | 0.092825 | 0.499 | 0.487 | 1 | 2 | Selenok | 0.012 | Young |
| 0.368047 | 0.079314 | 0.308 | 0.296 | 1 | 2 | Dusp11 | 0.012 | Young |
| 0.36863 | 0.179081 | 0.158 | 0.145 | 1 | 2 | Eif4a3 | 0.013 | Young |
| 0.368898 | 0.064623 | 0.014 | 0.01 | 1 | 2 | Nxpe3 | 0.004 | Young |
| 0.369157 | 0.211118 | 0.131 | 0.119 | 1 | 2 | Trappc1 | 0.012 | Young |
| 0.369425 | 0.364598 | 0.014 | 0.01 | 1 | 2 | Il1rl2 | 0.004 | Young |
| 0.369524 | 0.054534 | 0.304 | 0.29 | 1 | 2 | Ccrl2 | 0.014 | Young |
| 0.370061 | 0.277024 | 0.09 | 0.079 | 1 | 2 | Ddit3 | 0.011 | Young |
| 0.37011 | 0.196127 | 0.609 | 0.607 | 1 | 2 | Hsp90aa1 | 0.002 | Young |
| 0.370369 | 0.311862 | 0.115 | 0.103 | 1 | 2 | Prkag1 | 0.012 | Young |

| | | | | | | | | |
|---|---|---|---|---|---|---|---|---|
| 0.370708 | 0.079975 | 0.125 | 0.142 | 1 | 2 | Gm8186 | -0.017 | Young |
| 0.37085 | 0.2132 | 0.309 | 0.299 | 1 | 2 | Frmd4b | 0.01 | Young |
| 0.370851 | 0.071463 | 0.11 | 0.126 | 1 | 2 | Mplkip | -0.016 | Young |
| 0.371011 | 0.357668 | 0.014 | 0.01 | 1 | 2 | Cav1 | 0.004 | Young |
| 0.371127 | 0.462271 | 0.087 | 0.076 | 1 | 2 | Tmem11 | 0.011 | Young |
| 0.371276 | 0.967092 | 0.015 | 0.011 | 1 | 2 | Nars2 | 0.004 | Young |
| 0.371276 | 0.172004 | 0.014 | 0.01 | 1 | 2 | Wdr6 | 0.004 | Young |
| 0.371532 | 0.041633 | 0.941 | 0.928 | 1 | 2 | Rack1 | 0.013 | Young |
| 0.372289 | 0.896077 | 0.015 | 0.011 | 1 | 2 | Coq6 | 0.004 | Young |
| 0.372402 | 0.501497 | 0.059 | 0.05 | 1 | 2 | Ormdl3 | 0.009 | Young |
| 0.372561 | 0.09696 | 0.12 | 0.108 | 1 | 2 | Bcor | 0.012 | Young |
| 0.372673 | 0.036474 | 0.038 | 0.047 | 1 | 2 | Hscb | -0.009 | Young |
| 0.372769 | 0.205413 | 0.246 | 0.231 | 1 | 2 | Dyrk2 | 0.015 | Young |
| 0.372776 | 0.188213 | 0.247 | 0.232 | 1 | 2 | Xpr1 | 0.015 | Young |
| 0.372837 | 0.216573 | 0.113 | 0.101 | 1 | 2 | Ube3c | 0.012 | Young |
| 0.373459 | 0.044505 | 0.08 | 0.092 | 1 | 2 | Prkra | -0.012 | Young |
| 0.373664 | 0.096902 | 0.014 | 0.01 | 1 | 2 | Arhgap35 | 0.004 | Young |
| 0.373681 | 0.072258 | 0.449 | 0.427 | 1 | 2 | Timp2 | 0.022 | Young |
| 0.373944 | 0.11912 | 0.228 | 0.213 | 1 | 2 | Heca | 0.015 | Young |
| 0.374279 | 0.162117 | 0.264 | 0.248 | 1 | 2 | Ncoa3 | 0.016 | Young |
| 0.374747 | 0.431011 | 0.091 | 0.08 | 1 | 2 | Polr3gl | 0.011 | Young |
| 0.374897 | 0.137475 | 0.462 | 0.451 | 1 | 2 | Itpkb | 0.011 | Young |
| 0.375041 | 0.296901 | 0.053 | 0.064 | 1 | 2 | D730003I1 | -0.011 | Young |
| 0.375205 | 0.091241 | 0.089 | 0.104 | 1 | 2 | Cfdp1 | -0.015 | Young |
| 0.375372 | 0.198472 | 0.067 | 0.057 | 1 | 2 | Ciapin1 | 0.01 | Young |
| 0.375624 | 0.476261 | 0.078 | 0.069 | 1 | 2 | Utp18 | 0.009 | Young |
| 0.375801 | 0.179049 | 0.19 | 0.176 | 1 | 2 | Slc15a4 | 0.014 | Young |
| 0.375949 | 0.150041 | 0.279 | 0.266 | 1 | 2 | Elf1 | 0.013 | Young |
| 0.376202 | 0.389779 | 0.064 | 0.055 | 1 | 2 | 5430427O1 | 0.009 | Young |
| 0.376344 | 0.04971 | 0.118 | 0.104 | 1 | 2 | Ddhd2 | 0.014 | Young |
| 0.376462 | 0.037727 | 0.321 | 0.303 | 1 | 2 | Il10rb | 0.018 | Young |
| 0.376616 | 0.517509 | 0.015 | 0.011 | 1 | 2 | Adnp | 0.004 | Young |
| 0.376871 | 0.251377 | 0.015 | 0.011 | 1 | 2 | Pfkm | 0.004 | Young |
| 0.377382 | 0.408518 | 0.015 | 0.011 | 1 | 2 | E2f8 | 0.004 | Young |
| 0.37746 | 0.917224 | 0.017 | 0.012 | 1 | 2 | Mms22l | 0.005 | Young |
| 0.377517 | 0.060926 | 0.167 | 0.188 | 1 | 2 | Slc25a39 | -0.021 | Young |
| 0.377668 | 0.231423 | 0.184 | 0.17 | 1 | 2 | Appl1 | 0.014 | Young |
| 0.378045 | 0.158294 | 0.071 | 0.061 | 1 | 2 | Itpa | 0.01 | Young |
| 0.378406 | 0.72444 | 0.015 | 0.011 | 1 | 2 | Kyat1 | 0.004 | Young |
| 0.378443 | 0.767596 | 0.017 | 0.012 | 1 | 2 | Erc1 | 0.005 | Young |
| 0.379405 | 0.198273 | 0.067 | 0.057 | 1 | 2 | Malsu1 | 0.01 | Young |
| 0.380459 | 0.641199 | 0.015 | 0.011 | 1 | 2 | A430046D1 | 0.004 | Young |
| 0.38226 | 0.050764 | 0.015 | 0.011 | 1 | 2 | Fzd4 | 0.004 | Young |
| 0.382832 | 0.382784 | 0.052 | 0.044 | 1 | 2 | Adamts10 | 0.008 | Young |
| 0.383034 | 0.077119 | 0.015 | 0.011 | 1 | 2 | Hyal2 | 0.004 | Young |

| | | | | | | | | |
|---|---|---|---|---|---|---|---|---|
| 0.383348 | 0.218529 | 0.298 | 0.285 | 1 | 2 | Gng2 | 0.013 | Young |
| 0.383607 | 0.047005 | 0.21 | 0.234 | 1 | 2 | Hsd17b12 | -0.024 | Young |
| 0.383762 | 0.476504 | 0.049 | 0.041 | 1 | 2 | Tubgcp2 | 0.008 | Young |
| 0.383827 | 0.08222 | 0.2 | 0.185 | 1 | 2 | Park7 | 0.015 | Young |
| 0.384105 | 0.149188 | 0.189 | 0.175 | 1 | 2 | N4bp2l1 | 0.014 | Young |
| 0.384123 | 0.454324 | 0.017 | 0.012 | 1 | 2 | Mcoln2 | 0.005 | Young |
| 0.38417 | 0.079831 | 0.082 | 0.095 | 1 | 2 | Vta1 | -0.013 | Young |
| 0.384504 | 0.209352 | 0.19 | 0.179 | 1 | 2 | Pik3c2a | 0.011 | Young |
| 0.384708 | 0.054638 | 0.04 | 0.049 | 1 | 2 | Mthfs | -0.009 | Young |
| 0.385009 | 0.201495 | 0.147 | 0.134 | 1 | 2 | Sec24c | 0.013 | Young |
| 0.385051 | 0.184722 | 0.129 | 0.117 | 1 | 2 | Nfkbib | 0.012 | Young |
| 0.385094 | 0.746125 | 0.018 | 0.013 | 1 | 2 | Zfp141 | 0.005 | Young |
| 0.385114 | 0.167843 | 0.28 | 0.272 | 1 | 2 | Zcchc11 | 0.008 | Young |
| 0.385173 | 0.216414 | 0.179 | 0.167 | 1 | 2 | Ptprs | 0.012 | Young |
| 0.385638 | 0.293769 | 0.053 | 0.045 | 1 | 2 | Cib2 | 0.008 | Young |
| 0.385788 | 0.037843 | 0.203 | 0.19 | 1 | 2 | Abcf1 | 0.013 | Young |
| 0.385953 | 0.124985 | 0.17 | 0.156 | 1 | 2 | Nemf | 0.014 | Young |
| 0.386051 | 0.882989 | 0.018 | 0.013 | 1 | 2 | Paqr4 | 0.005 | Young |
| 0.386418 | 0.19309 | 0.26 | 0.243 | 1 | 2 | Phlda1 | 0.017 | Young |
| 0.386442 | 0.383936 | 0.103 | 0.093 | 1 | 2 | Vars | 0.01 | Young |
| 0.38662 | 0.209469 | 0.203 | 0.191 | 1 | 2 | Ccdc71l | 0.012 | Young |
| 0.387055 | 0.114948 | 0.414 | 0.409 | 1 | 2 | Cd300c2 | 0.005 | Young |
| 0.387251 | 0.412596 | 0.018 | 0.013 | 1 | 2 | Zmynd15 | 0.005 | Young |
| 0.38739 | 0.054886 | 0.068 | 0.058 | 1 | 2 | Sos1 | 0.01 | Young |
| 0.387405 | 0.052847 | 0.025 | 0.032 | 1 | 2 | Lzic | -0.007 | Young |
| 0.387924 | 0.075376 | 0.228 | 0.257 | 1 | 2 | Rwdd1 | -0.029 | Young |
| 0.388211 | 0.764863 | 0.018 | 0.013 | 1 | 2 | 5430416N( | 0.005 | Young |
| 0.38835 | 0.163176 | 0.224 | 0.211 | 1 | 2 | Taf15 | 0.013 | Young |
| 0.388478 | 0.100075 | 0.072 | 0.062 | 1 | 2 | Exosc10 | 0.01 | Young |
| 0.388529 | 0.533797 | 0.048 | 0.04 | 1 | 2 | Ubr7 | 0.008 | Young |
| 0.388808 | 0.442499 | 0.07 | 0.061 | 1 | 2 | Trim24 | 0.009 | Young |
| 0.388855 | 0.141521 | 0.017 | 0.012 | 1 | 2 | Zfp113 | 0.005 | Young |
| 0.389287 | 0.666489 | 0.019 | 0.014 | 1 | 2 | D930030I0 | 0.005 | Young |
| 0.389415 | 0.505809 | 0.018 | 0.013 | 1 | 2 | Simc1 | 0.005 | Young |
| 0.389468 | 0.041595 | 0.258 | 0.239 | 1 | 2 | Prkcb | 0.019 | Young |
| 0.389493 | 0.030416 | 0.12 | 0.107 | 1 | 2 | St6galnac4 | 0.013 | Young |
| 0.389752 | 0.794772 | 0.019 | 0.014 | 1 | 2 | Sdhaf3 | 0.005 | Young |
| 0.390099 | 0.240303 | 0.122 | 0.11 | 1 | 2 | Slirp | 0.012 | Young |
| 0.390203 | 0.074378 | 0.395 | 0.381 | 1 | 2 | Tmed7 | 0.014 | Young |
| 0.390411 | 0.146442 | 0.228 | 0.215 | 1 | 2 | Adam8 | 0.013 | Young |
| 0.39062 | 0.563279 | 0.018 | 0.013 | 1 | 2 | Nsl1 | 0.005 | Young |
| 0.391103 | 0.513253 | 0.018 | 0.013 | 1 | 2 | Gm17745 | 0.005 | Young |
| 0.391106 | 0.22681 | 0.204 | 0.192 | 1 | 2 | Hp1bp3 | 0.012 | Young |
| 0.391504 | 0.096672 | 0.213 | 0.195 | 1 | 2 | Higd1a | 0.018 | Young |
| 0.392181 | 0.337936 | 0.057 | 0.048 | 1 | 2 | Prps2 | 0.009 | Young |

| | | | | | | | | |
|---|---|---|---|---|---|---|---|---|
| 0.392236 | 0.15864 | 0.285 | 0.271 | 1 | 2 | Acin1 | 0.014 | Young |
| 0.392312 | 0.517923 | 0.018 | 0.013 | 1 | 2 | Tlr5 | 0.005 | Young |
| 0.392662 | 0.979019 | 0.02 | 0.015 | 1 | 2 | Slc26a6 | 0.005 | Young |
| 0.392697 | 0.342119 | 0.075 | 0.065 | 1 | 2 | Extl3 | 0.01 | Young |
| 0.393063 | 0.244915 | 0.055 | 0.046 | 1 | 2 | Zfp991 | 0.009 | Young |
| 0.393109 | 0.179936 | 0.036 | 0.044 | 1 | 2 | H2-T24 | -0.008 | Young |
| 0.393512 | 0.531052 | 0.046 | 0.039 | 1 | 2 | Ddx19b | 0.007 | Young |
| 0.393529 | 0.225043 | 0.059 | 0.071 | 1 | 2 | Slc25a17 | -0.012 | Young |
| 0.394321 | 0.0609 | 0.026 | 0.033 | 1 | 2 | Pex11b | -0.007 | Young |
| 0.394979 | 0.200876 | 0.018 | 0.013 | 1 | 2 | Taf1c | 0.005 | Young |
| 0.395302 | 0.693089 | 0.042 | 0.034 | 1 | 2 | Zcchc10 | 0.008 | Young |
| 0.395421 | 0.327851 | 0.107 | 0.095 | 1 | 2 | Rbm12 | 0.012 | Young |
| 0.395988 | 0.019323 | 0.102 | 0.116 | 1 | 2 | Paip1 | -0.014 | Young |
| 0.396377 | 0.263947 | 0.186 | 0.176 | 1 | 2 | Stx12 | 0.01 | Young |
| 0.396406 | 0.098935 | 0.01 | 0.014 | 1 | 2 | Rhebl1 | -0.004 | Young |
| 0.396672 | 0.795319 | 0.021 | 0.016 | 1 | 2 | Pik3ap1 | 0.005 | Young |
| 0.396804 | 0.171 | 0.013 | 0.018 | 1 | 2 | Ccdc91 | -0.005 | Young |
| 0.396957 | 0.098131 | 0.01 | 0.014 | 1 | 2 | Acsf2 | -0.004 | Young |
| 0.397704 | 0.65255 | 0.038 | 0.031 | 1 | 2 | Usp42 | 0.007 | Young |
| 0.398183 | 0.375881 | 0.019 | 0.014 | 1 | 2 | Ifrd2 | 0.005 | Young |
| 0.398228 | 0.02377 | 0.648 | 0.625 | 1 | 2 | Clta | 0.023 | Young |
| 0.398332 | 0.076426 | 0.078 | 0.091 | 1 | 2 | Rad50 | -0.013 | Young |
| 0.398379 | 0.647528 | 0.067 | 0.058 | 1 | 2 | Zfp512 | 0.009 | Young |
| 0.398388 | 0.834302 | 0.025 | 0.019 | 1 | 2 | Spink2 | 0.006 | Young |
| 0.398472 | 0.131948 | 0.246 | 0.234 | 1 | 2 | Edf1 | 0.012 | Young |
| 0.398646 | 0.004721 | 0.341 | 0.373 | 1 | 2 | Epb41l2 | -0.032 | Young |
| 0.398682 | 0.068827 | 0.134 | 0.122 | 1 | 2 | Mob3c | 0.012 | Young |
| 0.398825 | 0.042985 | 0.121 | 0.136 | 1 | 2 | Gskip | -0.015 | Young |
| 0.399026 | 0.516143 | 0.02 | 0.015 | 1 | 2 | Zbtb26 | 0.005 | Young |
| 0.399395 | 0.192792 | 0.082 | 0.072 | 1 | 2 | Avl9 | 0.01 | Young |
| 0.399431 | 0.305091 | 0.091 | 0.08 | 1 | 2 | Cyp27a1 | 0.011 | Young |
| 0.399524 | 0.069423 | 0.031 | 0.039 | 1 | 2 | Scly | -0.008 | Young |
| 0.399675 | 0.009205 | 0.951 | 0.924 | 1 | 2 | Ccr2 | 0.027 | Young |
| 0.399843 | 0.05174 | 0.309 | 0.291 | 1 | 2 | Myo5a | 0.018 | Young |
| 0.400071 | 0.301618 | 0.019 | 0.014 | 1 | 2 | Cacna1f | 0.005 | Young |
| 0.400431 | 0.075972 | 0.058 | 0.069 | 1 | 2 | Ankrd27 | -0.011 | Young |
| 0.401231 | 0.153502 | 0.063 | 0.054 | 1 | 2 | Cacfd1 | 0.009 | Young |
| 0.401271 | 0.000462 | 0.087 | 0.076 | 1 | 2 | Bank1 | 0.011 | Young |
| 0.401345 | 0.26365 | 0.112 | 0.101 | 1 | 2 | Atl2 | 0.011 | Young |
| 0.401426 | 0.152311 | 0.166 | 0.153 | 1 | 2 | Pigs | 0.013 | Young |
| 0.401515 | 0.785819 | 0.033 | 0.027 | 1 | 2 | Dpp4 | 0.006 | Young |
| 0.401548 | 0.818458 | 0.037 | 0.03 | 1 | 2 | Fars2 | 0.007 | Young |
| 0.401689 | 0.016441 | 0.093 | 0.106 | 1 | 2 | Trmt1 | -0.013 | Young |
| 0.40192 | 0.599039 | 0.034 | 0.028 | 1 | 2 | Thap1 | 0.006 | Young |
| 0.402056 | 0.198493 | 0.031 | 0.039 | 1 | 2 | Sugp2 | -0.008 | Young |

| | | | | | | | | |
|---|---|---|---|---|---|---|---|---|
| 0.402202 | 0.596562 | 0.024 | 0.018 | 1 | 2 | Ccdc28a | 0.006 | Young |
| 0.40245 | 0.018664 | 0.105 | 0.092 | 1 | 2 | Mpdu1 | 0.013 | Young |
| 0.403017 | 0.097536 | 0.183 | 0.204 | 1 | 2 | Pgap2 | -0.021 | Young |
| 0.403102 | 0.244914 | 0.045 | 0.037 | 1 | 2 | Prmt7 | 0.008 | Young |
| 0.40315 | 0.44029 | 0.02 | 0.015 | 1 | 2 | Pdgfc | 0.005 | Young |
| 0.403227 | 0.181649 | 0.502 | 0.493 | 1 | 2 | Atp6v1g1 | 0.009 | Young |
| 0.403568 | 0.397789 | 0.217 | 0.206 | 1 | 2 | Elmsan1 | 0.011 | Young |
| 0.40361 | 0.296623 | 0.02 | 0.015 | 1 | 2 | Zfp709 | 0.005 | Young |
| 0.403771 | 0.106506 | 0.063 | 0.074 | 1 | 2 | Ech1 | -0.011 | Young |
| 0.403995 | 0.310381 | 0.021 | 0.016 | 1 | 2 | Irgq | 0.005 | Young |
| 0.40407 | 0.373995 | 0.02 | 0.015 | 1 | 2 | Atg4a | 0.005 | Young |
| 0.404218 | 0.471075 | 0.021 | 0.016 | 1 | 2 | Fig4 | 0.005 | Young |
| 0.404782 | 0.329957 | 0.049 | 0.041 | 1 | 2 | Serinc5 | 0.008 | Young |
| 0.404971 | 0.3277 | 0.042 | 0.034 | 1 | 2 | Hmgxb4 | 0.008 | Young |
| 0.405092 | 0.469031 | 0.044 | 0.036 | 1 | 2 | Ercc6l2 | 0.008 | Young |
| 0.405127 | 0.065039 | 0.441 | 0.429 | 1 | 2 | Map4k4 | 0.012 | Young |
| 0.405328 | 0.788631 | 0.032 | 0.026 | 1 | 2 | Coa4 | 0.006 | Young |
| 0.405441 | 0.058856 | 0.153 | 0.171 | 1 | 2 | Pdcl3 | -0.018 | Young |
| 0.40559 | 0.531606 | 0.065 | 0.057 | 1 | 2 | Cd79b | 0.008 | Young |
| 0.4058 | 0.045685 | 0.166 | 0.187 | 1 | 2 | Trip11 | -0.021 | Young |
| 0.405806 | 0.126968 | 0.049 | 0.041 | 1 | 2 | Atg16l2 | 0.008 | Young |
| 0.406059 | 0.483795 | 0.082 | 0.073 | 1 | 2 | Pfkfb3 | 0.009 | Young |
| 0.406175 | 0.174741 | 0.048 | 0.04 | 1 | 2 | Rab40c | 0.008 | Young |
| 0.40623 | 0.334065 | 0.021 | 0.016 | 1 | 2 | Eif5a2 | 0.005 | Young |
| 0.406295 | 0.552517 | 0.033 | 0.027 | 1 | 2 | Mad1l1 | 0.006 | Young |
| 0.406454 | 0.219172 | 0.021 | 0.016 | 1 | 2 | Invs | 0.005 | Young |
| 0.406606 | 0.168809 | 0.02 | 0.015 | 1 | 2 | Cmss1 | 0.005 | Young |
| 0.406785 | 0.18027 | 0.055 | 0.046 | 1 | 2 | Exosc2 | 0.009 | Young |
| 0.406804 | 0.72167 | 0.031 | 0.025 | 1 | 2 | Zbtb40 | 0.006 | Young |
| 0.406944 | 0.130608 | 0.181 | 0.168 | 1 | 2 | Pdcd4 | 0.013 | Young |
| 0.407341 | 0.140776 | 0.179 | 0.166 | 1 | 2 | Hadha | 0.013 | Young |
| 0.407351 | 0.373225 | 0.021 | 0.016 | 1 | 2 | Cbr4 | 0.005 | Young |
| 0.407361 | 0.031621 | 0.126 | 0.142 | 1 | 2 | Tor1a | -0.016 | Young |
| 0.407695 | 0.302411 | 0.024 | 0.018 | 1 | 2 | Tspan33 | 0.006 | Young |
| 0.407695 | 0.028321 | 0.024 | 0.018 | 1 | 2 | Mier2 | 0.006 | Young |
| 0.408024 | 0.345134 | 0.021 | 0.016 | 1 | 2 | Nle1 | 0.005 | Young |
| 0.408075 | 0.626924 | 0.033 | 0.027 | 1 | 2 | Rtl8a | 0.006 | Young |
| 0.408256 | 0.080034 | 0.512 | 0.505 | 1 | 2 | Eif3e | 0.007 | Young |
| 0.408322 | 0.093664 | 0.328 | 0.314 | 1 | 2 | Gatad2b | 0.014 | Young |
| 0.408428 | 0.14954 | 0.106 | 0.094 | 1 | 2 | Pofut1 | 0.012 | Young |
| 0.408844 | 0.031587 | 0.211 | 0.231 | 1 | 2 | Lamtor3 | -0.02 | Young |
| 0.409076 | 0.509487 | 0.025 | 0.019 | 1 | 2 | Wfs1 | 0.006 | Young |
| 0.409307 | 0.07786 | 0.023 | 0.029 | 1 | 2 | Taf6 | -0.006 | Young |
| 0.40955 | 0.430892 | 0.03 | 0.024 | 1 | 2 | Tab1 | 0.006 | Young |
| 0.409928 | 0.395837 | 0.03 | 0.024 | 1 | 2 | Mospd1 | 0.006 | Young |

| | | | | | | | | |
|---|---|---|---|---|---|---|---|---|
| 0.410015 | 0.075577 | 0.818 | 0.82 | 1 | 2 | Eef1b2 | -0.002 | Young |
| 0.41015 | 0.217865 | 0.317 | 0.309 | 1 | 2 | Mapk14 | 0.008 | Young |
| 0.410386 | 0.075754 | 0.046 | 0.039 | 1 | 2 | Gm44751 | 0.007 | Young |
| 0.410497 | 0.579963 | 0.03 | 0.024 | 1 | 2 | Fancg | 0.006 | Young |
| 0.410569 | 0.045299 | 0.266 | 0.291 | 1 | 2 | Daglb | -0.025 | Young |
| 0.411204 | 0.57804 | 0.024 | 0.018 | 1 | 2 | Acad12 | 0.006 | Young |
| 0.411314 | 0.255589 | 0.048 | 0.04 | 1 | 2 | Stx18 | 0.008 | Young |
| 0.411346 | 0.125631 | 0.146 | 0.134 | 1 | 2 | C1galt1 | 0.012 | Young |
| 0.411347 | 0.110947 | 0.302 | 0.287 | 1 | 2 | Mia3 | 0.015 | Young |
| 0.411516 | 0.61207 | 0.029 | 0.022 | 1 | 2 | Zfp367 | 0.007 | Young |
| 0.411586 | 0.22463 | 0.166 | 0.155 | 1 | 2 | Sfswap | 0.011 | Young |
| 0.411894 | 0.224478 | 0.102 | 0.091 | 1 | 2 | Fxr2 | 0.011 | Young |
| 0.412251 | 0.037157 | 0.094 | 0.107 | 1 | 2 | Pfdn6 | -0.013 | Young |
| 0.413207 | 0.372366 | 0.031 | 0.025 | 1 | 2 | Coprs | 0.006 | Young |
| 0.413393 | 0.548671 | 0.031 | 0.025 | 1 | 2 | Kif3a | 0.006 | Young |
| 0.413471 | 0.218928 | 0.045 | 0.055 | 1 | 2 | Pdzd11 | -0.01 | Young |
| 0.413882 | 0.274559 | 0.163 | 0.184 | 1 | 2 | Tmem165 | -0.021 | Young |
| 0.413938 | 0.15883 | 0.399 | 0.395 | 1 | 2 | Il10ra | 0.004 | Young |
| 0.41409 | 0.710238 | 0.059 | 0.051 | 1 | 2 | Ttc17 | 0.008 | Young |
| 0.414303 | 0.179559 | 0.024 | 0.018 | 1 | 2 | Zfp317 | 0.006 | Young |
| 0.414526 | 0.376467 | 0.04 | 0.033 | 1 | 2 | Ppp6r2 | 0.007 | Young |
| 0.414709 | 0.064588 | 0.044 | 0.036 | 1 | 2 | Ndufaf2 | 0.008 | Young |
| 0.415046 | 0.357596 | 0.07 | 0.061 | 1 | 2 | Metap1 | 0.009 | Young |
| 0.415251 | 0.319032 | 0.03 | 0.024 | 1 | 2 | Grpel2 | 0.006 | Young |
| 0.415366 | 0.108531 | 0.27 | 0.259 | 1 | 2 | Tomm5 | 0.011 | Young |
| 0.41573 | 0.148031 | 0.023 | 0.017 | 1 | 2 | Zkscan17 | 0.006 | Young |
| 0.415777 | 0.32521 | 0.072 | 0.063 | 1 | 2 | Tnfrsf12a | 0.009 | Young |
| 0.415915 | 0.320587 | 0.031 | 0.025 | 1 | 2 | Paip2b | 0.006 | Young |
| 0.416233 | 0.166894 | 0.024 | 0.018 | 1 | 2 | Lrrc28 | 0.006 | Young |
| 0.416662 | 0.349727 | 0.024 | 0.018 | 1 | 2 | Iqcb1 | 0.006 | Young |
| 0.416987 | 0.385804 | 0.038 | 0.031 | 1 | 2 | Aamdc | 0.007 | Young |
| 0.417067 | 0.261379 | 0.095 | 0.085 | 1 | 2 | Lcmt1 | 0.01 | Young |
| 0.417209 | 0.415013 | 0.039 | 0.032 | 1 | 2 | Prkar2b | 0.007 | Young |
| 0.41741 | 0.224521 | 0.135 | 0.124 | 1 | 2 | Zfp451 | 0.011 | Young |
| 0.417831 | 0.267559 | 0.025 | 0.019 | 1 | 2 | Npat | 0.006 | Young |
| 0.417953 | 0.085654 | 0.024 | 0.018 | 1 | 2 | Shb | 0.006 | Young |
| 0.418447 | 0.292856 | 0.04 | 0.033 | 1 | 2 | Dph6 | 0.007 | Young |
| 0.418539 | 0.316004 | 0.036 | 0.029 | 1 | 2 | Lrrc61 | 0.007 | Young |
| 0.418608 | 0.093163 | 0.366 | 0.357 | 1 | 2 | Furin | 0.009 | Young |
| 0.418638 | 0.207213 | 0.293 | 0.283 | 1 | 2 | Tubb6 | 0.01 | Young |
| 0.418738 | 0.104403 | 0.347 | 0.338 | 1 | 2 | Txn2 | 0.009 | Young |
| 0.41881 | 0.167903 | 0.014 | 0.019 | 1 | 2 | Zfp346 | -0.005 | Young |
| 0.418939 | 0.088955 | 0.044 | 0.036 | 1 | 2 | Ccdc134 | 0.008 | Young |
| 0.419126 | 0.221367 | 0.023 | 0.029 | 1 | 2 | Blzf1 | -0.006 | Young |
| 0.419159 | 0.228782 | 0.172 | 0.16 | 1 | 2 | Tbc1d5 | 0.012 | Young |

| | | | | | | | | |
|---|---|---|---|---|---|---|---|---|
| 0.420079 | 0.012932 | 0.141 | 0.128 | 1 | 2 | Snx8 | 0.013 | Young |
| 0.420118 | 0.163901 | 0.22 | 0.21 | 1 | 2 | Slbp | 0.01 | Young |
| 0.420204 | 0.18428 | 0.039 | 0.032 | 1 | 2 | Polr2d | 0.007 | Young |
| 0.420209 | 0.067738 | 0.063 | 0.074 | 1 | 2 | Aspscr1 | -0.011 | Young |
| 0.420468 | 0.022408 | 0.057 | 0.067 | 1 | 2 | Spns1 | -0.01 | Young |
| 0.421165 | 0.258571 | 0.196 | 0.185 | 1 | 2 | Tecr | 0.011 | Young |
| 0.421304 | 0.047797 | 0.154 | 0.141 | 1 | 2 | Ing2 | 0.013 | Young |
| 0.421348 | 0.078965 | 0.049 | 0.058 | 1 | 2 | Tpgs1 | -0.009 | Young |
| 0.421694 | 0.251653 | 0.154 | 0.141 | 1 | 2 | Cdc37l1 | 0.013 | Young |
| 0.421977 | 0.295638 | 0.034 | 0.028 | 1 | 2 | Atm | 0.006 | Young |
| 0.422098 | 0.056852 | 0.293 | 0.318 | 1 | 2 | Lamtor1 | -0.025 | Young |
| 0.422117 | 0.185977 | 0.129 | 0.119 | 1 | 2 | Oaz2 | 0.01 | Young |
| 0.422324 | 0.182526 | 0.086 | 0.075 | 1 | 2 | Traf6 | 0.011 | Young |
| 0.422595 | 0.111406 | 0.132 | 0.148 | 1 | 2 | Sms | -0.016 | Young |
| 0.422642 | 0.059987 | 0.014 | 0.019 | 1 | 2 | Fastkd5 | -0.005 | Young |
| 0.42276 | 0.303179 | 0.141 | 0.13 | 1 | 2 | Ssbp3 | 0.011 | Young |
| 0.422825 | 0.836843 | 0.055 | 0.047 | 1 | 2 | Poc1b | 0.008 | Young |
| 0.423017 | 0.381528 | 0.07 | 0.061 | 1 | 2 | Tbc1d8 | 0.009 | Young |
| 0.423943 | 0.257611 | 0.027 | 0.034 | 1 | 2 | Ankrd52 | -0.007 | Young |
| 0.424097 | 0.109445 | 0.069 | 0.06 | 1 | 2 | Trappc9 | 0.009 | Young |
| 0.424324 | 0.19593 | 0.036 | 0.029 | 1 | 2 | Pdgfa | 0.007 | Young |
| 0.424401 | 0.283234 | 0.108 | 0.097 | 1 | 2 | Brwd3 | 0.011 | Young |
| 0.425216 | 0.031253 | 0.175 | 0.192 | 1 | 2 | Camk1 | -0.017 | Young |
| 0.425219 | 0.072044 | 0.026 | 0.02 | 1 | 2 | Hmga1 | 0.006 | Young |
| 0.425602 | 0.105658 | 0.488 | 0.475 | 1 | 2 | Tra2a | 0.013 | Young |
| 0.425739 | 0.165843 | 0.113 | 0.102 | 1 | 2 | Slc25a30 | 0.011 | Young |
| 0.426623 | 0.27306 | 0.037 | 0.03 | 1 | 2 | Mrps2 | 0.007 | Young |
| 0.426797 | 0.199445 | 0.245 | 0.23 | 1 | 2 | Pcnp | 0.015 | Young |
| 0.427166 | 0.115904 | 0.16 | 0.148 | 1 | 2 | Clptm1 | 0.012 | Young |
| 0.427505 | 0.170306 | 0.131 | 0.12 | 1 | 2 | Ric8a | 0.011 | Young |
| 0.427672 | 0.188189 | 0.127 | 0.116 | 1 | 2 | Pdlim4 | 0.011 | Young |
| 0.427889 | 0.006654 | 0.083 | 0.095 | 1 | 2 | Magohb | -0.012 | Young |
| 0.427902 | 0.255593 | 0.182 | 0.171 | 1 | 2 | Gosr2 | 0.011 | Young |
| 0.428084 | 0.184195 | 0.2 | 0.185 | 1 | 2 | Wdfy3 | 0.015 | Young |
| 0.428139 | 0.120254 | 0.129 | 0.117 | 1 | 2 | Xylt1 | 0.012 | Young |
| 0.428358 | 0.297141 | 0.1 | 0.09 | 1 | 2 | Atg12 | 0.01 | Young |
| 0.428426 | 0.011574 | 0.347 | 0.377 | 1 | 2 | Atp2a2 | -0.03 | Young |
| 0.428471 | 0.200903 | 0.031 | 0.025 | 1 | 2 | Hmgxb3 | 0.006 | Young |
| 0.428642 | 0.049925 | 0.62 | 0.615 | 1 | 2 | Trps1 | 0.005 | Young |
| 0.429094 | 0.05069 | 0.577 | 0.564 | 1 | 2 | Ddx6 | 0.013 | Young |
| 0.430397 | 0.231394 | 0.156 | 0.145 | 1 | 2 | Mecp2 | 0.011 | Young |
| 0.430559 | 0.271303 | 0.091 | 0.081 | 1 | 2 | Asnsd1 | 0.01 | Young |
| 0.430575 | 0.079429 | 0.241 | 0.227 | 1 | 2 | Cdk11b | 0.014 | Young |
| 0.430766 | 0.216611 | 0.007 | 0.011 | 1 | 2 | Large1 | -0.004 | Young |
| 0.430766 | 0.07056 | 0.007 | 0.011 | 1 | 2 | Gvin1 | -0.004 | Young |

| | | | | | | | | |
|---|---|---|---|---|---|---|---|---|
| 0.430814 | 0.133345 | 0.259 | 0.249 | 1 | 2 | Adss | 0.01 | Young |
| 0.431098 | 0.144417 | 0.007 | 0.011 | 1 | 2 | Gm9530 | -0.004 | Young |
| 0.43132 | 0.452851 | 0.087 | 0.077 | 1 | 2 | Dhps | 0.01 | Young |
| 0.432432 | 0.239591 | 0.154 | 0.143 | 1 | 2 | Nudc | 0.011 | Young |
| 0.432559 | 0.166057 | 0.078 | 0.069 | 1 | 2 | Polr2j | 0.009 | Young |
| 0.432706 | 0.001182 | 0.116 | 0.105 | 1 | 2 | Nploc4 | 0.011 | Young |
| 0.432768 | 0.572533 | 0.055 | 0.047 | 1 | 2 | Cd226 | 0.008 | Young |
| 0.43378 | 0.047012 | 0.652 | 0.636 | 1 | 2 | Arpc3 | 0.016 | Young |
| 0.434567 | 0.229318 | 0.108 | 0.099 | 1 | 2 | Ctdspl2 | 0.009 | Young |
| 0.434627 | 0.643626 | 0.048 | 0.041 | 1 | 2 | Stk38l | 0.007 | Young |
| 0.434971 | 0.125379 | 0.235 | 0.224 | 1 | 2 | Nfe2l1 | 0.011 | Young |
| 0.435325 | 1.058786 | 0.043 | 0.036 | 1 | 2 | Hvcn1 | 0.007 | Young |
| 0.435374 | 0.015823 | 0.107 | 0.095 | 1 | 2 | Cbx4 | 0.012 | Young |
| 0.435706 | 0.059372 | 0.046 | 0.055 | 1 | 2 | Dnajb12 | -0.009 | Young |
| 0.436055 | 0.263153 | 0.29 | 0.285 | 1 | 2 | Arih1 | 0.005 | Young |
| 0.436154 | 0.108898 | 0.292 | 0.273 | 1 | 2 | Cd63 | 0.019 | Young |
| 0.436654 | 0.008961 | 0.196 | 0.181 | 1 | 2 | Kxd1 | 0.015 | Young |
| 0.436993 | 0.096397 | 0.157 | 0.145 | 1 | 2 | Ufc1 | 0.012 | Young |
| 0.437038 | 0.016321 | 0.093 | 0.105 | 1 | 2 | Ell | -0.012 | Young |
| 0.437293 | 0.448507 | 0.134 | 0.125 | 1 | 2 | Pisd | 0.009 | Young |
| 0.43742 | 0.050684 | 0.47 | 0.461 | 1 | 2 | Fam46a | 0.009 | Young |
| 0.437659 | 0.188745 | 0.198 | 0.19 | 1 | 2 | Ythdf1 | 0.008 | Young |
| 0.437848 | 0.007395 | 0.07 | 0.08 | 1 | 2 | Cyhr1 | -0.01 | Young |
| 0.440448 | 0.133741 | 0.086 | 0.076 | 1 | 2 | Mrps28 | 0.01 | Young |
| 0.440616 | 0.054487 | 0.264 | 0.249 | 1 | 2 | Bax | 0.015 | Young |
| 0.441017 | 0.040143 | 0.43 | 0.452 | 1 | 2 | Ubl5 | -0.022 | Young |
| 0.441326 | 0.037173 | 0.045 | 0.054 | 1 | 2 | Nt5c2 | -0.009 | Young |
| 0.441479 | 0.378559 | 0.056 | 0.048 | 1 | 2 | Tmem159 | 0.008 | Young |
| 0.441878 | 0.053854 | 0.068 | 0.059 | 1 | 2 | Etv5 | 0.009 | Young |
| 0.442199 | 0.311924 | 0.185 | 0.178 | 1 | 2 | Ssbp4 | 0.007 | Young |
| 0.442729 | 0.310137 | 0.113 | 0.103 | 1 | 2 | Rnf5 | 0.01 | Young |
| 0.442734 | 0.311799 | 0.106 | 0.095 | 1 | 2 | Arl6ip4 | 0.011 | Young |
| 0.442873 | 0.088882 | 0.04 | 0.048 | 1 | 2 | Vezt | -0.008 | Young |
| 0.443192 | 0.022861 | 0.878 | 0.854 | 1 | 2 | Npc2 | 0.024 | Young |
| 0.443618 | 0.287703 | 0.407 | 0.407 | 1 | 2 | S100a11 | 0 | Young |
| 0.44369 | 0.030092 | 0.248 | 0.273 | 1 | 2 | Ppp4c | -0.025 | Young |
| 0.443794 | 0.103137 | 0.025 | 0.031 | 1 | 2 | Mul1 | -0.006 | Young |
| 0.443826 | 0.084536 | 0.157 | 0.143 | 1 | 2 | Chtf8 | 0.014 | Young |
| 0.443946 | 0.029179 | 0.08 | 0.07 | 1 | 2 | Thap11 | 0.01 | Young |
| 0.444416 | 0.153844 | 0.209 | 0.198 | 1 | 2 | 4932438A1 | 0.011 | Young |
| 0.444994 | 0.142632 | 0.24 | 0.226 | 1 | 2 | Mpc2 | 0.014 | Young |
| 0.44541 | 0.57039 | 0.052 | 0.045 | 1 | 2 | Ccdc117 | 0.007 | Young |
| 0.445506 | 0.270065 | 0.2 | 0.187 | 1 | 2 | Lrrc8d | 0.013 | Young |
| 0.445696 | 0.068187 | 0.182 | 0.169 | 1 | 2 | Rsl1d1 | 0.013 | Young |
| 0.445734 | 0.191136 | 0.015 | 0.02 | 1 | 2 | Dars2 | -0.005 | Young |

| | | | | | | | | |
|---|---|---|---|---|---|---|---|---|
| 0.44574 | 0.232044 | 0.068 | 0.059 | 1 | 2 | Pign | 0.009 | Young |
| 0.445808 | 0.354873 | 0.115 | 0.106 | 1 | 2 | Fchsd2 | 0.009 | Young |
| 0.445974 | 0.117808 | 0.015 | 0.02 | 1 | 2 | Gstk1 | -0.005 | Young |
| 0.4469 | 0.103686 | 0.334 | 0.325 | 1 | 2 | Tcp1 | 0.009 | Young |
| 0.447106 | 0.062882 | 0.063 | 0.073 | 1 | 2 | Agpat1 | -0.01 | Young |
| 0.447527 | 0.178407 | 0.153 | 0.142 | 1 | 2 | Snx1 | 0.011 | Young |
| 0.447959 | 0.002661 | 0.088 | 0.1 | 1 | 2 | Eif2ak1 | -0.012 | Young |
| 0.448189 | 0.097578 | 0.034 | 0.042 | 1 | 2 | Alg9 | -0.008 | Young |
| 0.448349 | 0.237586 | 0.195 | 0.184 | 1 | 2 | Gm10073 | 0.011 | Young |
| 0.448355 | 0.09105 | 0.106 | 0.095 | 1 | 2 | Rasgrp1 | 0.011 | Young |
| 0.448474 | 0.184205 | 0.245 | 0.234 | 1 | 2 | Parvb | 0.011 | Young |
| 0.449153 | 0.150295 | 0.205 | 0.193 | 1 | 2 | Rab35 | 0.012 | Young |
| 0.449395 | 0.252269 | 0.025 | 0.031 | 1 | 2 | AU022252 | -0.006 | Young |
| 0.44943 | 0.503934 | 0.065 | 0.058 | 1 | 2 | Tesk1 | 0.007 | Young |
| 0.450975 | 0.061915 | 0.299 | 0.285 | 1 | 2 | Srsf1 | 0.014 | Young |
| 0.451037 | 0.036624 | 0.116 | 0.128 | 1 | 2 | Rnf11 | -0.012 | Young |
| 0.451429 | 0.041475 | 0.367 | 0.352 | 1 | 2 | Rps6ka3 | 0.015 | Young |
| 0.452337 | 0.329322 | 0.051 | 0.044 | 1 | 2 | Rtca | 0.007 | Young |
| 0.452966 | 0.271444 | 0.015 | 0.02 | 1 | 2 | Btd | -0.005 | Young |
| 0.453056 | 0.419499 | 0.055 | 0.047 | 1 | 2 | Creb3l2 | 0.008 | Young |
| 0.453097 | 0.026341 | 0.205 | 0.191 | 1 | 2 | Erlin1 | 0.014 | Young |
| 0.453235 | 0.070505 | 0.539 | 0.53 | 1 | 2 | Atf4 | 0.009 | Young |
| 0.453594 | 0.212657 | 0.12 | 0.109 | 1 | 2 | Zbtb11 | 0.011 | Young |
| 0.455663 | 0.522447 | 0.065 | 0.058 | 1 | 2 | Gid4 | 0.007 | Young |
| 0.455908 | 0.012481 | 0.058 | 0.067 | 1 | 2 | Josd2 | -0.009 | Young |
| 0.456113 | 0.320772 | 0.179 | 0.171 | 1 | 2 | Zfr | 0.008 | Young |
| 0.456266 | 0.005924 | 0.081 | 0.091 | 1 | 2 | Wdr33 | -0.01 | Young |
| 0.456301 | 0.20288 | 0.173 | 0.161 | 1 | 2 | Atf1 | 0.012 | Young |
| 0.456317 | 0.122773 | 0.23 | 0.221 | 1 | 2 | Kmt2c | 0.009 | Young |
| 0.456631 | 1.049412 | 0.01 | 0.006 | 1 | 2 | Naprt | 0.004 | Young |
| 0.456652 | 0.051817 | 0.492 | 0.485 | 1 | 2 | Ptms | 0.007 | Young |
| 0.456838 | 0.176344 | 0.209 | 0.198 | 1 | 2 | Snx6 | 0.011 | Young |
| 0.457116 | 0.117087 | 0.243 | 0.231 | 1 | 2 | Klhl9 | 0.012 | Young |
| 0.45731 | 0.36142 | 0.052 | 0.045 | 1 | 2 | Edem3 | 0.007 | Young |
| 0.457448 | 0.330063 | 0.068 | 0.06 | 1 | 2 | Emc4 | 0.008 | Young |
| 0.458283 | 0.143148 | 0.175 | 0.162 | 1 | 2 | Ell2 | 0.013 | Young |
| 0.458384 | 0.359321 | 0.09 | 0.081 | 1 | 2 | Man1b1 | 0.009 | Young |
| 0.459121 | 0.247411 | 0.052 | 0.045 | 1 | 2 | Ubtd1 | 0.007 | Young |
| 0.459348 | 0.115535 | 0.044 | 0.052 | 1 | 2 | Rab11fip2 | -0.008 | Young |
| 0.459563 | 1.106012 | 0.01 | 0.006 | 1 | 2 | Slc27a3 | 0.004 | Young |
| 0.459581 | 0.108698 | 0.012 | 0.016 | 1 | 2 | Blm | -0.004 | Young |
| 0.460298 | 1.144799 | 0.01 | 0.006 | 1 | 2 | Per2 | 0.004 | Young |
| 0.460325 | 0.028177 | 0.606 | 0.597 | 1 | 2 | Sec61b | 0.009 | Young |
| 0.460539 | 0.253327 | 0.242 | 0.237 | 1 | 2 | Arid4a | 0.005 | Young |
| 0.461025 | 0.034194 | 0.222 | 0.208 | 1 | 2 | Pfdn2 | 0.014 | Young |

| | | | | | | | | |
|---|---|---|---|---|---|---|---|---|
| 0.461033 | 0.71889 | 0.01 | 0.006 | 1 | 2 | Gm36839 | 0.004 | Young |
| 0.461401 | 1.011489 | 0.01 | 0.006 | 1 | 2 | Zfp974 | 0.004 | Young |
| 0.461401 | 0.724087 | 0.01 | 0.006 | 1 | 2 | Taf1a | 0.004 | Young |
| 0.461618 | 0.58915 | 0.077 | 0.07 | 1 | 2 | Mtx2 | 0.007 | Young |
| 0.461754 | 0.317711 | 0.046 | 0.04 | 1 | 2 | Tmem214 | 0.006 | Young |
| 0.461861 | 0.153053 | 0.119 | 0.108 | 1 | 2 | Stx6 | 0.011 | Young |
| 0.46223 | 0.003423 | 0.203 | 0.221 | 1 | 2 | Copb2 | -0.018 | Young |
| 0.462243 | 0.097467 | 0.07 | 0.08 | 1 | 2 | Pprc1 | -0.01 | Young |
| 0.46261 | 0.089719 | 0.247 | 0.236 | 1 | 2 | Eprs | 0.011 | Young |
| 0.462667 | 0.020354 | 0.381 | 0.36 | 1 | 2 | Cep170 | 0.021 | Young |
| 0.462875 | 0.484085 | 0.01 | 0.006 | 1 | 2 | Tkfc | 0.004 | Young |
| 0.463123 | 0.157443 | 0.063 | 0.055 | 1 | 2 | Rbsn | 0.008 | Young |
| 0.463612 | 0.877245 | 0.01 | 0.006 | 1 | 2 | Gm30978 | 0.004 | Young |
| 0.463612 | 0.652841 | 0.01 | 0.006 | 1 | 2 | Rhox8 | 0.004 | Young |
| 0.463612 | 0.298034 | 0.01 | 0.006 | 1 | 2 | Gm42722 | 0.004 | Young |
| 0.463878 | 0.089694 | 0.816 | 0.801 | 1 | 2 | Srgn | 0.015 | Young |
| 0.463904 | 0.16343 | 0.163 | 0.151 | 1 | 2 | Cdk12 | 0.012 | Young |
| 0.463981 | 0.360789 | 0.01 | 0.006 | 1 | 2 | Zfp948 | 0.004 | Young |
| 0.464152 | 0.0054 | 0.069 | 0.079 | 1 | 2 | Gtf2e2 | -0.01 | Young |
| 0.464339 | 0.233111 | 0.069 | 0.079 | 1 | 2 | Trim65 | -0.01 | Young |
| 0.464351 | 0.399485 | 0.01 | 0.006 | 1 | 2 | Gm11696 | 0.004 | Young |
| 0.464459 | 0.101621 | 0.433 | 0.413 | 1 | 2 | Ndufb7 | 0.02 | Young |
| 0.464493 | 0.226222 | 0.046 | 0.04 | 1 | 2 | Nlrc4 | 0.006 | Young |
| 0.46472 | 0.281074 | 0.01 | 0.006 | 1 | 2 | Phf19 | 0.004 | Young |
| 0.464925 | 0.243174 | 0.181 | 0.169 | 1 | 2 | Crtc3 | 0.012 | Young |
| 0.46509 | 0.259382 | 0.01 | 0.006 | 1 | 2 | Id1 | 0.004 | Young |
| 0.465459 | 0.493401 | 0.01 | 0.006 | 1 | 2 | Trim13 | 0.004 | Young |
| 0.465459 | 0.301303 | 0.01 | 0.006 | 1 | 2 | Usp49 | 0.004 | Young |
| 0.465827 | 0.195059 | 0.133 | 0.123 | 1 | 2 | Epc2 | 0.01 | Young |
| 0.465852 | 0.13349 | 0.247 | 0.236 | 1 | 2 | Csnk1e | 0.011 | Young |
| 0.465936 | 0.001061 | 0.21 | 0.195 | 1 | 2 | Ankrd12 | 0.015 | Young |
| 0.46657 | 0.265484 | 0.01 | 0.006 | 1 | 2 | Timeless | 0.004 | Young |
| 0.466598 | 0.18897 | 0.2 | 0.19 | 1 | 2 | Flii | 0.01 | Young |
| 0.466772 | 0.0986 | 0.064 | 0.056 | 1 | 2 | Impdh1 | 0.008 | Young |
| 0.466939 | 0.002614 | 0.194 | 0.18 | 1 | 2 | Swap70 | 0.014 | Young |
| 0.46706 | 0.124202 | 0.093 | 0.082 | 1 | 2 | Khnyn | 0.011 | Young |
| 0.467707 | 0.017957 | 0.053 | 0.046 | 1 | 2 | Map3k3 | 0.007 | Young |
| 0.468248 | 0.063783 | 0.249 | 0.237 | 1 | 2 | Usp34 | 0.012 | Young |
| 0.468372 | 0.070457 | 0.186 | 0.204 | 1 | 2 | Mrpl28 | -0.018 | Young |
| 0.468566 | 0.115459 | 0.008 | 0.012 | 1 | 2 | Slc25a53 | -0.004 | Young |
| 0.468606 | 0.060536 | 0.084 | 0.075 | 1 | 2 | 9130401M | 0.009 | Young |
| 0.468693 | 0.095956 | 0.08 | 0.071 | 1 | 2 | Polr2h | 0.009 | Young |
| 0.468728 | 0.459254 | 0.012 | 0.016 | 1 | 2 | Sirt4 | -0.004 | Young |
| 0.468785 | 0.033601 | 0.048 | 0.056 | 1 | 2 | Synj2 | -0.008 | Young |
| 0.469103 | 0.181351 | 0.09 | 0.081 | 1 | 2 | Hbs1l | 0.009 | Young |

| | | | | | | | | |
|---|---|---|---|---|---|---|---|---|
| 0.469222 | 0.106744 | 0.008 | 0.012 | 1 | 2 | Tti1 | -0.004 | Young |
| 0.470491 | 0.513976 | 0.067 | 0.059 | 1 | 2 | Agap3 | 0.008 | Young |
| 0.470814 | 0.183189 | 0.08 | 0.071 | 1 | 2 | Rrp7a | 0.009 | Young |
| 0.471018 | 0.135602 | 0.173 | 0.161 | 1 | 2 | Polr2e | 0.012 | Young |
| 0.4712 | 0.040242 | 0.115 | 0.105 | 1 | 2 | Upf3a | 0.01 | Young |
| 0.47122 | 0.201175 | 0.147 | 0.138 | 1 | 2 | Ide | 0.009 | Young |
| 0.471381 | 0.094339 | 0.114 | 0.127 | 1 | 2 | CAAA0111 | -0.013 | Young |
| 0.471475 | 0.059467 | 0.184 | 0.172 | 1 | 2 | Slc25a11 | 0.012 | Young |
| 0.471503 | 0.005341 | 0.406 | 0.439 | 1 | 2 | Rab5c | -0.033 | Young |
| 0.47159 | 0.276681 | 0.032 | 0.039 | 1 | 2 | Galt | -0.007 | Young |
| 0.472303 | 0.262395 | 0.19 | 0.183 | 1 | 2 | Atxn7 | 0.007 | Young |
| 0.472349 | 0.130031 | 0.049 | 0.042 | 1 | 2 | Fbxl15 | 0.007 | Young |
| 0.473527 | 0.108778 | 0.198 | 0.187 | 1 | 2 | Mrpl57 | 0.011 | Young |
| 0.473942 | 0.089731 | 0.268 | 0.255 | 1 | 2 | Fbl | 0.013 | Young |
| 0.47419 | 0.194831 | 0.153 | 0.142 | 1 | 2 | Gtf3c2 | 0.011 | Young |
| 0.474194 | 0.072406 | 0.439 | 0.429 | 1 | 2 | Apobec1 | 0.01 | Young |
| 0.474206 | 0.298122 | 0.185 | 0.178 | 1 | 2 | Mrpl30 | 0.007 | Young |
| 0.474227 | 0.140009 | 0.107 | 0.097 | 1 | 2 | Dcaf13 | 0.01 | Young |
| 0.474292 | 0.257884 | 0.131 | 0.122 | 1 | 2 | Slc35a3 | 0.009 | Young |
| 0.474579 | 0.203826 | 0.093 | 0.084 | 1 | 2 | Cops3 | 0.009 | Young |
| 0.474834 | 0.142996 | 0.308 | 0.302 | 1 | 2 | Gpsm3 | 0.006 | Young |
| 0.474883 | 0.241799 | 0.011 | 0.007 | 1 | 2 | Slc22a21 | 0.004 | Young |
| 0.474924 | 0.311564 | 0.051 | 0.044 | 1 | 2 | 4921524J1 | 0.007 | Young |
| 0.474946 | 0.048764 | 0.08 | 0.071 | 1 | 2 | Ccl12 | 0.009 | Young |
| 0.475233 | 1.00701 | 0.011 | 0.007 | 1 | 2 | Trim2 | 0.004 | Young |
| 0.475233 | 0.288909 | 0.011 | 0.007 | 1 | 2 | Dnajc17 | 0.004 | Young |
| 0.475566 | 0.368948 | 0.043 | 0.036 | 1 | 2 | Acad8 | 0.007 | Young |
| 0.475583 | 0.431237 | 0.011 | 0.007 | 1 | 2 | Gabbr1 | 0.004 | Young |
| 0.475806 | 0.138351 | 0.017 | 0.021 | 1 | 2 | Mpst | -0.004 | Young |
| 0.475933 | 0.677598 | 0.011 | 0.007 | 1 | 2 | Kif20b | 0.004 | Young |
| 0.476303 | 1.079081 | 0.012 | 0.009 | 1 | 2 | Epas1 | 0.003 | Young |
| 0.476413 | 0.073638 | 0.328 | 0.313 | 1 | 2 | 1700020I1 | 0.015 | Young |
| 0.476546 | 0.074986 | 0.102 | 0.092 | 1 | 2 | Vps4a | 0.01 | Young |
| 0.476635 | 0.735255 | 0.011 | 0.007 | 1 | 2 | Ltb | 0.004 | Young |
| 0.476799 | 0.444383 | 0.008 | 0.012 | 1 | 2 | Tuft1 | -0.004 | Young |
| 0.476846 | 0.070342 | 0.504 | 0.501 | 1 | 2 | Ccdc50 | 0.003 | Young |
| 0.476985 | 0.379988 | 0.011 | 0.007 | 1 | 2 | Pls3 | 0.004 | Young |
| 0.477175 | 0.026406 | 0.044 | 0.051 | 1 | 2 | Zfp942 | -0.007 | Young |
| 0.477333 | 0.022805 | 0.58 | 0.551 | 1 | 2 | Psma7 | 0.029 | Young |
| 0.47751 | 0.106687 | 0.049 | 0.057 | 1 | 2 | Acsl3 | -0.008 | Young |
| 0.478038 | 0.250648 | 0.011 | 0.007 | 1 | 2 | Atp23 | 0.004 | Young |
| 0.478214 | 0.164951 | 0.296 | 0.289 | 1 | 2 | Nop10 | 0.007 | Young |
| 0.478233 | 0.317147 | 0.045 | 0.039 | 1 | 2 | Fgd3 | 0.006 | Young |
| 0.478289 | 1.239297 | 0.012 | 0.009 | 1 | 2 | Hpn | 0.003 | Young |
| 0.478444 | 0.134776 | 0.336 | 0.321 | 1 | 2 | Cndp2 | 0.015 | Young |

| | | | | | | | | |
|---|---|---|---|---|---|---|---|---|
| 0.478456 | 0.127954 | 0.074 | 0.065 | 1 | 2 | Ginm1 | 0.009 | Young |
| 0.478494 | 0.054517 | 0.367 | 0.347 | 1 | 2 | Ndufc1 | 0.02 | Young |
| 0.47856 | 0.127991 | 0.037 | 0.044 | 1 | 2 | Ubac1 | -0.007 | Young |
| 0.478741 | 0.461455 | 0.011 | 0.007 | 1 | 2 | Cyb5rl | 0.004 | Young |
| 0.478741 | 0.133008 | 0.011 | 0.007 | 1 | 2 | Hist2h2aa1 | 0.004 | Young |
| 0.479093 | 0.288709 | 0.011 | 0.007 | 1 | 2 | Cd300ld3 | 0.004 | Young |
| 0.479126 | 0.742524 | 0.034 | 0.029 | 1 | 2 | Prmt2 | 0.005 | Young |
| 0.479156 | 0.33906 | 0.062 | 0.055 | 1 | 2 | Tspan4 | 0.007 | Young |
| 0.47931 | 0.164167 | 0.026 | 0.032 | 1 | 2 | Papss2 | -0.006 | Young |
| 0.479423 | 0.316735 | 0.118 | 0.109 | 1 | 2 | Grsf1 | 0.009 | Young |
| 0.479796 | 0.17966 | 0.011 | 0.007 | 1 | 2 | Zbtb10 | 0.004 | Young |
| 0.479817 | 0.11051 | 0.253 | 0.238 | 1 | 2 | Rbbp6 | 0.015 | Young |
| 0.479948 | 1.079721 | 0.012 | 0.009 | 1 | 2 | 1110002L0 | 0.003 | Young |
| 0.480054 | 0.034195 | 0.134 | 0.123 | 1 | 2 | Cnot2 | 0.011 | Young |
| 0.480945 | 0.859658 | 0.012 | 0.009 | 1 | 2 | Mterf1a | 0.003 | Young |
| 0.48106 | 0.002397 | 0.038 | 0.045 | 1 | 2 | Cd274 | -0.007 | Young |
| 0.481238 | 0.261759 | 0.087 | 0.078 | 1 | 2 | Dolpp1 | 0.009 | Young |
| 0.48151 | 0.210389 | 0.144 | 0.135 | 1 | 2 | Nap1l4 | 0.009 | Young |
| 0.481984 | 0.415785 | 0.039 | 0.033 | 1 | 2 | Spryd3 | 0.006 | Young |
| 0.481992 | 0.162418 | 0.239 | 0.225 | 1 | 2 | Sdf2 | 0.014 | Young |
| 0.482278 | 0.043738 | 0.145 | 0.134 | 1 | 2 | Nfya | 0.011 | Young |
| 0.48244 | 0.420008 | 0.044 | 0.037 | 1 | 2 | Tmem184c | 0.007 | Young |
| 0.482656 | 0.029732 | 0.075 | 0.066 | 1 | 2 | H2-Ke6 | 0.009 | Young |
| 0.482705 | 0.481502 | 0.037 | 0.031 | 1 | 2 | Alkbh6 | 0.006 | Young |
| 0.482835 | 0.280317 | 0.033 | 0.028 | 1 | 2 | Myc | 0.005 | Young |
| 0.483193 | 0.386118 | 0.059 | 0.052 | 1 | 2 | Tpst1 | 0.007 | Young |
| 0.483217 | 0.031422 | 0.051 | 0.044 | 1 | 2 | Rnf170 | 0.007 | Young |
| 0.483275 | 0.426519 | 0.012 | 0.009 | 1 | 2 | Hemk1 | 0.003 | Young |
| 0.483422 | 0.044064 | 0.141 | 0.156 | 1 | 2 | Hipk2 | -0.015 | Young |
| 0.483447 | 0.150678 | 0.192 | 0.182 | 1 | 2 | Chd2 | 0.01 | Young |
| 0.483818 | 1.039096 | 0.013 | 0.01 | 1 | 2 | Bend3 | 0.003 | Young |
| 0.484026 | 0.182831 | 0.069 | 0.061 | 1 | 2 | Strip1 | 0.008 | Young |
| 0.484501 | 0.167444 | 0.083 | 0.074 | 1 | 2 | Bckdha | 0.009 | Young |
| 0.48461 | 0.026292 | 0.144 | 0.157 | 1 | 2 | Prrc2b | -0.013 | Young |
| 0.484837 | 0.016966 | 0.122 | 0.111 | 1 | 2 | Nfxl1 | 0.011 | Young |
| 0.484943 | 0.632294 | 0.012 | 0.009 | 1 | 2 | Hmgn3 | 0.003 | Young |
| 0.484943 | 0.336751 | 0.012 | 0.009 | 1 | 2 | 6330403L0 | 0.003 | Young |
| 0.485231 | 0.396856 | 0.077 | 0.07 | 1 | 2 | Stoml2 | 0.007 | Young |
| 0.485347 | 0.164329 | 0.266 | 0.258 | 1 | 2 | Sod2 | 0.008 | Young |
| 0.485527 | 0.165529 | 0.27 | 0.264 | 1 | 2 | Erbin | 0.006 | Young |
| 0.485642 | 1.056433 | 0.027 | 0.022 | 1 | 2 | Cgnl1 | 0.005 | Young |
| 0.485946 | 0.326202 | 0.012 | 0.009 | 1 | 2 | Nckap5l | 0.003 | Young |
| 0.486038 | 0.076484 | 0.197 | 0.217 | 1 | 2 | Smap1 | -0.02 | Young |
| 0.48628 | 0.159936 | 0.012 | 0.009 | 1 | 2 | Ehf | 0.003 | Young |
| 0.486439 | 0.360174 | 0.061 | 0.054 | 1 | 2 | Tmem126a | 0.007 | Young |

| | | | | | | | | |
|---|---|---|---|---|---|---|---|---|
| 0.486614 | 0.193228 | 0.012 | 0.009 | 1 | 2 | Atp6v0c | 0.003 | Young |
| 0.486651 | 0.498252 | 0.036 | 0.03 | 1 | 2 | Mettl14 | 0.006 | Young |
| 0.486736 | 0.097066 | 0.659 | 0.656 | 1 | 2 | Gdi2 | 0.003 | Young |
| 0.486762 | 0.414294 | 0.037 | 0.044 | 1 | 2 | Phf6 | -0.007 | Young |
| 0.487284 | 0.275889 | 0.012 | 0.009 | 1 | 2 | Nupl2 | 0.003 | Young |
| 0.487309 | 0.175404 | 0.171 | 0.162 | 1 | 2 | Tmcc3 | 0.009 | Young |
| 0.487415 | 0.06871 | 0.354 | 0.349 | 1 | 2 | Eif3h | 0.005 | Young |
| 0.487482 | 0.096016 | 0.044 | 0.051 | 1 | 2 | Acvr1b | -0.007 | Young |
| 0.487619 | 0.306151 | 0.012 | 0.009 | 1 | 2 | Itpr3 | 0.003 | Young |
| 0.487619 | 0.159571 | 0.012 | 0.009 | 1 | 2 | Pou6f1 | 0.003 | Young |
| 0.487625 | 0.708102 | 0.013 | 0.01 | 1 | 2 | Glmn | 0.003 | Young |
| 0.48788 | 1.131459 | 0.025 | 0.02 | 1 | 2 | 1300002E1 | 0.005 | Young |
| 0.487945 | 0.036227 | 0.169 | 0.184 | 1 | 2 | Paf1 | -0.015 | Young |
| 0.487954 | 0.467676 | 0.012 | 0.009 | 1 | 2 | Trip6 | 0.003 | Young |
| 0.488297 | 0.434163 | 0.059 | 0.052 | 1 | 2 | Abcf2 | 0.007 | Young |
| 0.488399 | 0.158051 | 0.056 | 0.064 | 1 | 2 | Cpsf1 | -0.008 | Young |
| 0.488953 | 0.106298 | 0.058 | 0.066 | 1 | 2 | Larp1b | -0.008 | Young |
| 0.489028 | 0.057869 | 0.114 | 0.127 | 1 | 2 | Trim44 | -0.013 | Young |
| 0.489295 | 0.225128 | 0.012 | 0.009 | 1 | 2 | Lmbr1 | 0.003 | Young |
| 0.489721 | 0.135783 | 0.065 | 0.075 | 1 | 2 | Rpl27-ps3 | -0.01 | Young |
| 0.489743 | 0.271582 | 0.069 | 0.061 | 1 | 2 | Mrps5 | 0.008 | Young |
| 0.489798 | 0.12583 | 0.147 | 0.137 | 1 | 2 | Ccdc88b | 0.01 | Young |
| 0.489853 | 0.72511 | 0.013 | 0.01 | 1 | 2 | Ficd | 0.003 | Young |
| 0.490081 | 0.095515 | 0.044 | 0.037 | 1 | 2 | AC165953. | 0.007 | Young |
| 0.490248 | 0.26306 | 0.039 | 0.033 | 1 | 2 | Ankrd50 | 0.006 | Young |
| 0.490335 | 0.118409 | 0.409 | 0.393 | 1 | 2 | Pgk1 | 0.016 | Young |
| 0.490613 | 0.249322 | 0.038 | 0.032 | 1 | 2 | Gm31718 | 0.006 | Young |
| 0.49081 | 0.636013 | 0.013 | 0.01 | 1 | 2 | Ccdc102a | 0.003 | Young |
| 0.49083 | 0.197794 | 0.048 | 0.056 | 1 | 2 | Rfxap | -0.008 | Young |
| 0.491334 | 0.314374 | 0.068 | 0.06 | 1 | 2 | Mrpl45 | 0.008 | Young |
| 0.491647 | 0.057394 | 0.012 | 0.009 | 1 | 2 | Clec2g | 0.003 | Young |
| 0.491829 | 0.291086 | 0.04 | 0.034 | 1 | 2 | Kti12 | 0.006 | Young |
| 0.491859 | 0.166716 | 0.133 | 0.123 | 1 | 2 | Smad2 | 0.01 | Young |
| 0.492149 | 0.383748 | 0.034 | 0.029 | 1 | 2 | Nlrp1b | 0.005 | Young |
| 0.492443 | 0.127065 | 0.023 | 0.028 | 1 | 2 | Endog | -0.005 | Young |
| 0.492455 | 0.525957 | 0.031 | 0.026 | 1 | 2 | Phf7 | 0.005 | Young |
| 0.49248 | 0.02563 | 0.05 | 0.058 | 1 | 2 | Mrs2 | -0.008 | Young |
| 0.492646 | 0.999231 | 0.014 | 0.011 | 1 | 2 | Zfp169 | 0.003 | Young |
| 0.492951 | 0.197186 | 0.014 | 0.011 | 1 | 2 | Rad18 | 0.003 | Young |
| 0.493105 | 0.363835 | 0.037 | 0.031 | 1 | 2 | E130307A1 | 0.006 | Young |
| 0.493501 | 0.13529 | 0.07 | 0.062 | 1 | 2 | Gmeb1 | 0.008 | Young |
| 0.493686 | 0.364989 | 0.013 | 0.01 | 1 | 2 | Gpt | 0.003 | Young |
| 0.493686 | 0.332316 | 0.013 | 0.01 | 1 | 2 | Fam83f | 0.003 | Young |
| 0.493862 | 0.703932 | 0.027 | 0.022 | 1 | 2 | Alkbh7 | 0.005 | Young |
| 0.494006 | 0.348212 | 0.013 | 0.01 | 1 | 2 | Cnrip1 | 0.003 | Young |

| | | | | | | | | |
|---|---|---|---|---|---|---|---|---|
| 0.49404 | 0.02647 | 0.376 | 0.36 | 1 | 2 | Ndufb3 | 0.016 | Young |
| 0.494267 | 0.145921 | 0.323 | 0.316 | 1 | 2 | Tmed9 | 0.007 | Young |
| 0.494756 | 0.380381 | 0.084 | 0.077 | 1 | 2 | Lage3 | 0.007 | Young |
| 0.494804 | 0.331024 | 0.075 | 0.067 | 1 | 2 | Ext2 | 0.008 | Young |
| 0.494846 | 0.171538 | 0.109 | 0.1 | 1 | 2 | Slc12a9 | 0.009 | Young |
| 0.495225 | 0.515657 | 0.093 | 0.086 | 1 | 2 | Elp2 | 0.007 | Young |
| 0.495534 | 0.007221 | 0.12 | 0.132 | 1 | 2 | Tfg | -0.012 | Young |
| 0.495554 | 0.34258 | 0.036 | 0.03 | 1 | 2 | Eif2b3 | 0.006 | Young |
| 0.495618 | 0.717659 | 0.026 | 0.021 | 1 | 2 | Rpap1 | 0.005 | Young |
| 0.495628 | 0.151015 | 0.034 | 0.029 | 1 | 2 | Traf2 | 0.005 | Young |
| 0.496056 | 0.854843 | 0.018 | 0.014 | 1 | 2 | Upp2 | 0.004 | Young |
| 0.49608 | 0.152303 | 0.042 | 0.035 | 1 | 2 | Fasn | 0.007 | Young |
| 0.496083 | 0.285738 | 0.119 | 0.11 | 1 | 2 | Zfp655 | 0.009 | Young |
| 0.496095 | 0.283576 | 0.071 | 0.063 | 1 | 2 | Hjurp | 0.008 | Young |
| 0.496275 | 0.067532 | 0.356 | 0.339 | 1 | 2 | Ndufb11 | 0.017 | Young |
| 0.496286 | 0.231952 | 0.232 | 0.223 | 1 | 2 | Specc1l | 0.009 | Young |
| 0.496287 | 0.903132 | 0.026 | 0.021 | 1 | 2 | Rtl8b | 0.005 | Young |
| 0.496619 | 0.585675 | 0.014 | 0.011 | 1 | 2 | Vegfb | 0.003 | Young |
| 0.497006 | 0.061436 | 0.093 | 0.104 | 1 | 2 | Alas1 | -0.011 | Young |
| 0.497134 | 0.284168 | 0.033 | 0.028 | 1 | 2 | Tjap1 | 0.005 | Young |
| 0.49723 | 0.183136 | 0.048 | 0.056 | 1 | 2 | Gnpda2 | -0.008 | Young |
| 0.497388 | 0.229545 | 0.062 | 0.071 | 1 | 2 | Ttf1 | -0.009 | Young |
| 0.497539 | 0.26249 | 0.014 | 0.011 | 1 | 2 | Zfp764 | 0.003 | Young |
| 0.497552 | 0.088027 | 0.367 | 0.354 | 1 | 2 | Rassf2 | 0.013 | Young |
| 0.497848 | 0.635529 | 0.026 | 0.021 | 1 | 2 | Zfp426 | 0.005 | Young |
| 0.497897 | 0.011682 | 0.07 | 0.079 | 1 | 2 | Ddx27 | -0.009 | Young |
| 0.498152 | 0.130656 | 0.014 | 0.011 | 1 | 2 | Gm11772 | 0.003 | Young |
| 0.498397 | 0.364119 | 0.11 | 0.103 | 1 | 2 | Hadh | 0.007 | Young |
| 0.498738 | 0.305244 | 0.032 | 0.027 | 1 | 2 | Iqce | 0.005 | Young |
| 0.498791 | 0.150353 | 0.214 | 0.206 | 1 | 2 | Eloa | 0.008 | Young |
| 0.499091 | 0.730186 | 0.027 | 0.022 | 1 | 2 | Pqlc2 | 0.005 | Young |
| 0.499581 | 0.462475 | 0.056 | 0.049 | 1 | 2 | Gm48065 | 0.007 | Young |
| 0.499703 | 0.302675 | 0.034 | 0.029 | 1 | 2 | Gm16536 | 0.005 | Young |
| 0.499721 | 0.332806 | 0.031 | 0.026 | 1 | 2 | Bcl9l | 0.005 | Young |
| 0.499976 | 0.146241 | 0.21 | 0.203 | 1 | 2 | Rela | 0.007 | Young |
| 0.500185 | 0.443776 | 0.027 | 0.022 | 1 | 2 | Slc26a11 | 0.005 | Young |
| 0.500287 | 0.382903 | 0.034 | 0.029 | 1 | 2 | Cnpy4 | 0.005 | Young |
| 0.50036 | 0.11708 | 0.049 | 0.042 | 1 | 2 | Itga9 | 0.007 | Young |
| 0.500483 | 0.068871 | 0.43 | 0.419 | 1 | 2 | Clic4 | 0.011 | Young |
| 0.5006 | 0.139474 | 0.07 | 0.062 | 1 | 2 | Fbxo9 | 0.008 | Young |
| 0.500871 | 0.35866 | 0.034 | 0.029 | 1 | 2 | Zdhhc16 | 0.005 | Young |
| 0.500943 | 0.057645 | 0.07 | 0.062 | 1 | 2 | Golph3l | 0.008 | Young |
| 0.500961 | 0.272759 | 0.018 | 0.014 | 1 | 2 | E330009J0 | 0.004 | Young |
| 0.501138 | 0.249057 | 0.109 | 0.101 | 1 | 2 | Acss1 | 0.008 | Young |
| 0.501375 | 0.394742 | 0.015 | 0.012 | 1 | 2 | Ap5b1 | 0.003 | Young |

| | | | | | | | | |
|---|---|---|---|---|---|---|---|---|
| 0.501376 | 0.085607 | 0.281 | 0.271 | 1 | 2 | Orai1 | 0.01 | Young |
| 0.501442 | 0.051342 | 0.521 | 0.509 | 1 | 2 | Ndel1 | 0.012 | Young |
| 0.501635 | 0.792928 | 0.023 | 0.018 | 1 | 2 | Gcdh | 0.005 | Young |
| 0.5018 | 0.348726 | 0.141 | 0.133 | 1 | 2 | Ptov1 | 0.008 | Young |
| 0.501877 | 0.181724 | 0.018 | 0.022 | 1 | 2 | Mxra7 | -0.004 | Young |
| 0.50202 | 0.044593 | 0.222 | 0.213 | 1 | 2 | Psme4 | 0.009 | Young |
| 0.502069 | 0.333075 | 0.057 | 0.05 | 1 | 2 | Ticam1 | 0.007 | Young |
| 0.502156 | 0.688647 | 0.027 | 0.022 | 1 | 2 | Rora | 0.005 | Young |
| 0.502161 | 0.233797 | 0.382 | 0.385 | 1 | 2 | Clec2d | -0.003 | Young |
| 0.502171 | 0.114634 | 0.431 | 0.421 | 1 | 2 | Atp5j2 | 0.01 | Young |
| 0.502196 | 0.330631 | 0.017 | 0.013 | 1 | 2 | Traf3ip1 | 0.004 | Young |
| 0.502335 | 0.745258 | 0.024 | 0.019 | 1 | 2 | Zfp568 | 0.005 | Young |
| 0.502814 | 0.284203 | 0.027 | 0.022 | 1 | 2 | Heatr5b | 0.005 | Young |
| 0.502904 | 0.071771 | 0.159 | 0.173 | 1 | 2 | Dennd4b | -0.014 | Young |
| 0.502966 | 0.637708 | 0.019 | 0.015 | 1 | 2 | Kank2 | 0.004 | Young |
| 0.503208 | 0.535419 | 0.023 | 0.018 | 1 | 2 | Rab26os | 0.005 | Young |
| 0.503233 | 0.135 | 0.439 | 0.435 | 1 | 2 | Capg | 0.004 | Young |
| 0.503442 | 0.14729 | 0.015 | 0.012 | 1 | 2 | Pcgf1 | 0.003 | Young |
| 0.503503 | 0.539187 | 0.025 | 0.02 | 1 | 2 | Alyref2 | 0.005 | Young |
| 0.50367 | 0.493337 | 0.026 | 0.021 | 1 | 2 | Prkd2 | 0.005 | Young |
| 0.503707 | 0.098543 | 0.069 | 0.061 | 1 | 2 | Mre11a | 0.008 | Young |
| 0.504221 | 0.688737 | 0.024 | 0.019 | 1 | 2 | Ccp110 | 0.005 | Young |
| 0.504246 | 0.525078 | 0.018 | 0.014 | 1 | 2 | Mcrip2 | 0.004 | Young |
| 0.504431 | 0.155427 | 0.074 | 0.066 | 1 | 2 | Zfp646 | 0.008 | Young |
| 0.504657 | 0.611977 | 0.018 | 0.014 | 1 | 2 | R3hcc1 | 0.004 | Young |
| 0.504761 | 0.345838 | 0.058 | 0.051 | 1 | 2 | Rab33b | 0.007 | Young |
| 0.504774 | 0.777723 | 0.021 | 0.017 | 1 | 2 | Creld1 | 0.004 | Young |
| 0.504783 | 0.13493 | 0.132 | 0.146 | 1 | 2 | Cyc1 | -0.014 | Young |
| 0.504794 | 0.44469 | 0.018 | 0.014 | 1 | 2 | Ust | 0.004 | Young |
| 0.504884 | 0.442067 | 0.02 | 0.016 | 1 | 2 | Clec16a | 0.004 | Young |
| 0.504899 | 0.558161 | 0.021 | 0.017 | 1 | 2 | Mettl8 | 0.004 | Young |
| 0.504999 | 0.679328 | 0.025 | 0.02 | 1 | 2 | Exd2 | 0.005 | Young |
| 0.50509 | 0.080595 | 0.572 | 0.565 | 1 | 2 | Mef2a | 0.007 | Young |
| 0.505343 | 0.607571 | 0.018 | 0.014 | 1 | 2 | Sumf2 | 0.004 | Young |
| 0.50554 | 0.292257 | 0.062 | 0.055 | 1 | 2 | Sgsh | 0.007 | Young |
| 0.505595 | 0.252017 | 0.031 | 0.026 | 1 | 2 | Timm21 | 0.005 | Young |
| 0.505607 | 0.159568 | 0.017 | 0.013 | 1 | 2 | Fancc | 0.004 | Young |
| 0.505713 | 0.051534 | 0.095 | 0.087 | 1 | 2 | Gpd2 | 0.008 | Young |
| 0.50594 | 0.162413 | 0.226 | 0.219 | 1 | 2 | Ndufa11 | 0.007 | Young |
| 0.506111 | 0.538488 | 0.024 | 0.019 | 1 | 2 | 1600020E0 | 0.005 | Young |
| 0.506177 | 0.78316 | 0.017 | 0.013 | 1 | 2 | Fcor | 0.004 | Young |
| 0.506282 | 0.188551 | 0.315 | 0.307 | 1 | 2 | Ankrd17 | 0.008 | Young |
| 0.506462 | 0.329656 | 0.017 | 0.013 | 1 | 2 | Gm26669 | 0.004 | Young |
| 0.506662 | 0.289959 | 0.03 | 0.025 | 1 | 2 | Ranbp6 | 0.005 | Young |
| 0.506806 | 0.140043 | 0.039 | 0.046 | 1 | 2 | Mcm3ap | -0.007 | Young |

| | | | | | | | | |
|---|---|---|---|---|---|---|---|---|
| 0.507058 | 0.57449 | 0.024 | 0.019 | 1 | 2 | Zfp574 | 0.005 | Young |
| 0.507115 | 0.091891 | 0.077 | 0.07 | 1 | 2 | Cnot10 | 0.007 | Young |
| 0.507164 | 0.320042 | 0.118 | 0.109 | 1 | 2 | Zfp592 | 0.009 | Young |
| 0.507454 | 0.542568 | 0.023 | 0.018 | 1 | 2 | Zfp846 | 0.005 | Young |
| 0.507697 | 0.400901 | 0.023 | 0.018 | 1 | 2 | Etaa1 | 0.005 | Young |
| 0.507709 | 0.046424 | 0.329 | 0.317 | 1 | 2 | Csf2rb | 0.012 | Young |
| 0.507713 | 0.036141 | 0.01 | 0.013 | 1 | 2 | Adgrl2 | -0.003 | Young |
| 0.507817 | 0.395078 | 0.018 | 0.014 | 1 | 2 | Timmdc1 | 0.004 | Young |
| 0.508506 | 0.084118 | 0.045 | 0.052 | 1 | 2 | Tmem268 | -0.007 | Young |
| 0.508512 | 0.153024 | 0.262 | 0.256 | 1 | 2 | Ctnna1 | 0.006 | Young |
| 0.508806 | 0.383891 | 0.019 | 0.015 | 1 | 2 | Zfp777 | 0.004 | Young |
| 0.508906 | 0.04909 | 0.07 | 0.079 | 1 | 2 | Tnip2 | -0.009 | Young |
| 0.509316 | 0.228055 | 0.017 | 0.013 | 1 | 2 | Ddx59 | 0.004 | Young |
| 0.509525 | 0.063281 | 0.477 | 0.465 | 1 | 2 | Lamtor4 | 0.012 | Young |
| 0.509602 | 0.058606 | 0.017 | 0.013 | 1 | 2 | 1700037C1 | 0.004 | Young |
| 0.509973 | 0.019621 | 0.087 | 0.097 | 1 | 2 | Fam208b | -0.01 | Young |
| 0.510189 | 0.068115 | 0.3 | 0.292 | 1 | 2 | Bptf | 0.008 | Young |
| 0.510886 | 0.002572 | 0.106 | 0.096 | 1 | 2 | Maml1 | 0.01 | Young |
| 0.510948 | 0.089672 | 0.305 | 0.293 | 1 | 2 | Vps35 | 0.012 | Young |
| 0.511293 | 0.018515 | 0.01 | 0.013 | 1 | 2 | Setd6 | -0.003 | Young |
| 0.511393 | 0.337803 | 0.055 | 0.048 | 1 | 2 | Mphosph6 | 0.007 | Young |
| 0.511596 | 0.094897 | 0.023 | 0.018 | 1 | 2 | Stk26 | 0.005 | Young |
| 0.511728 | 0.497819 | 0.095 | 0.088 | 1 | 2 | Vrk3 | 0.007 | Young |
| 0.511898 | 0.009876 | 0.201 | 0.19 | 1 | 2 | Rere | 0.011 | Young |
| 0.511903 | 0.003831 | 0.052 | 0.06 | 1 | 2 | Ciao1 | -0.008 | Young |
| 0.512008 | 0.3742 | 0.172 | 0.168 | 1 | 2 | Vps13b | 0.004 | Young |
| 0.51223 | 0.284087 | 0.018 | 0.014 | 1 | 2 | Pfas | 0.004 | Young |
| 0.512257 | 0.008508 | 0.407 | 0.397 | 1 | 2 | Ewsr1 | 0.01 | Young |
| 0.512297 | 0.061824 | 0.055 | 0.062 | 1 | 2 | Timm22 | -0.007 | Young |
| 0.512329 | 0.396181 | 0.023 | 0.018 | 1 | 2 | Cep76 | 0.005 | Young |
| 0.512551 | 0.334655 | 0.029 | 0.024 | 1 | 2 | D930016D( | 0.005 | Young |
| 0.512787 | 0.220418 | 0.021 | 0.017 | 1 | 2 | Fbxo31 | 0.004 | Young |
| 0.513246 | 0.13654 | 0.103 | 0.095 | 1 | 2 | Nhlrc3 | 0.008 | Young |
| 0.513251 | 0.161639 | 0.01 | 0.013 | 1 | 2 | Tmem186 | -0.003 | Young |
| 0.513518 | 0.211295 | 0.1 | 0.091 | 1 | 2 | Rnf139 | 0.009 | Young |
| 0.513576 | 0.02044 | 0.071 | 0.063 | 1 | 2 | Xab2 | 0.008 | Young |
| 0.513939 | 0.196834 | 0.024 | 0.029 | 1 | 2 | Pigv | -0.005 | Young |
| 0.514041 | 0.308251 | 0.023 | 0.018 | 1 | 2 | C1qtnf1 | 0.005 | Young |
| 0.514069 | 0.168598 | 0.496 | 0.495 | 1 | 2 | Cstb | 0.001 | Young |
| 0.514082 | 0.176494 | 0.297 | 0.293 | 1 | 2 | Tmbim1 | 0.004 | Young |
| 0.514674 | 0.020873 | 0.021 | 0.017 | 1 | 2 | Zswim7 | 0.004 | Young |
| 0.514891 | 0.006042 | 0.65 | 0.642 | 1 | 2 | Srrm2 | 0.008 | Young |
| 0.514901 | 0.119957 | 0.024 | 0.019 | 1 | 2 | Rbm48 | 0.005 | Young |
| 0.51505 | 0.057119 | 0.017 | 0.013 | 1 | 2 | Cbarp | 0.004 | Young |
| 0.515052 | 0.214097 | 0.021 | 0.017 | 1 | 2 | Zfp874b | 0.004 | Young |

| | | | | | | | | |
|---|---|---|---|---|---|---|---|---|
| 0.515339 | 0.298209 | 0.173 | 0.167 | 1 | 2 | Rictor | 0.006 | Young |
| 0.515444 | 0.122688 | 0.024 | 0.029 | 1 | 2 | Tysnd1 | -0.005 | Young |
| 0.515445 | 0.119796 | 0.285 | 0.278 | 1 | 2 | Lsm4 | 0.007 | Young |
| 0.515809 | 0.2055 | 0.029 | 0.024 | 1 | 2 | Mcph1 | 0.005 | Young |
| 0.516247 | 0.214573 | 0.023 | 0.018 | 1 | 2 | Fam213a | 0.005 | Young |
| 0.5168 | 0.537482 | 0.114 | 0.107 | 1 | 2 | Fam133b | 0.007 | Young |
| 0.516811 | 0.014189 | 0.019 | 0.024 | 1 | 2 | Tsr3 | -0.005 | Young |
| 0.51707 | 0.192735 | 0.021 | 0.017 | 1 | 2 | Angptl4 | 0.004 | Young |
| 0.517956 | 0.006329 | 0.167 | 0.181 | 1 | 2 | Dpp8 | -0.014 | Young |
| 0.517984 | 0.055539 | 0.341 | 0.33 | 1 | 2 | Aldh2 | 0.011 | Young |
| 0.518126 | 0.276002 | 0.128 | 0.119 | 1 | 2 | Dapk3 | 0.009 | Young |
| 0.518171 | 0.079564 | 0.019 | 0.015 | 1 | 2 | Dync2h1 | 0.004 | Young |
| 0.518175 | 0.332734 | 0.107 | 0.1 | 1 | 2 | Ammecr1l | 0.007 | Young |
| 0.518314 | 0.067758 | 0.118 | 0.131 | 1 | 2 | Hmbox1 | -0.013 | Young |
| 0.518845 | 0.178499 | 0.024 | 0.019 | 1 | 2 | Prelid2 | 0.005 | Young |
| 0.518944 | 0.027066 | 0.317 | 0.346 | 1 | 2 | Metap2 | -0.029 | Young |
| 0.519421 | 0.217129 | 0.209 | 0.201 | 1 | 2 | Psmb7 | 0.008 | Young |
| 0.519498 | 0.593725 | 0.065 | 0.059 | 1 | 2 | Flt1 | 0.006 | Young |
| 0.519613 | 0.025478 | 0.014 | 0.018 | 1 | 2 | Suv39h1 | -0.004 | Young |
| 0.519724 | 0.125024 | 0.133 | 0.125 | 1 | 2 | Zfp740 | 0.008 | Young |
| 0.51983 | 0.067639 | 0.53 | 0.535 | 1 | 2 | Gmfg | -0.005 | Young |
| 0.519973 | 0.151469 | 0.232 | 0.225 | 1 | 2 | Cdc5l | 0.007 | Young |
| 0.520359 | 0.094959 | 0.021 | 0.017 | 1 | 2 | Cdr2 | 0.004 | Young |
| 0.520544 | 0.150822 | 0.077 | 0.07 | 1 | 2 | Dcaf17 | 0.007 | Young |
| 0.520571 | 0.076475 | 0.014 | 0.018 | 1 | 2 | Cmtr1 | -0.004 | Young |
| 0.520948 | 0.025592 | 0.348 | 0.337 | 1 | 2 | Uqcr10 | 0.011 | Young |
| 0.52114 | 0.298606 | 0.151 | 0.142 | 1 | 2 | Exoc4 | 0.009 | Young |
| 0.52178 | 0.140211 | 0.312 | 0.307 | 1 | 2 | Dmac1 | 0.005 | Young |
| 0.522672 | 0.051867 | 0.061 | 0.069 | 1 | 2 | Fkbp1b | -0.008 | Young |
| 0.52273 | 0.216557 | 0.317 | 0.317 | 1 | 2 | Tsn | 0 | Young |
| 0.523904 | 0.059045 | 0.329 | 0.32 | 1 | 2 | Plaur | 0.009 | Young |
| 0.524813 | 0.357153 | 0.08 | 0.073 | 1 | 2 | Rffl | 0.007 | Young |
| 0.524971 | 0.007901 | 0.025 | 0.03 | 1 | 2 | Nup188 | -0.005 | Young |
| 0.525581 | 0.322885 | 0.071 | 0.064 | 1 | 2 | Rpa3 | 0.007 | Young |
| 0.525653 | 0.42749 | 0.014 | 0.018 | 1 | 2 | Zfp748 | -0.004 | Young |
| 0.525653 | 0.127383 | 0.014 | 0.018 | 1 | 2 | Cep78 | -0.004 | Young |
| 0.525797 | 0.146963 | 0.164 | 0.155 | 1 | 2 | Nmt1 | 0.009 | Young |
| 0.525831 | 0.338531 | 0.088 | 0.08 | 1 | 2 | Rnf103 | 0.008 | Young |
| 0.526538 | 0.332328 | 0.069 | 0.062 | 1 | 2 | Klf16 | 0.007 | Young |
| 0.526609 | 0.211031 | 0.141 | 0.133 | 1 | 2 | Epc1 | 0.008 | Young |
| 0.526928 | 0.137427 | 0.103 | 0.095 | 1 | 2 | Slc30a9 | 0.008 | Young |
| 0.527181 | 0.225134 | 0.071 | 0.08 | 1 | 2 | Pold3 | -0.009 | Young |
| 0.527183 | 0.171044 | 0.131 | 0.121 | 1 | 2 | Iws1 | 0.01 | Young |
| 0.527316 | 0.10428 | 0.025 | 0.03 | 1 | 2 | Pigh | -0.005 | Young |
| 0.527488 | 0.389922 | 0.083 | 0.076 | 1 | 2 | Mocs2 | 0.007 | Young |

| | | | | | | | | |
|---|---|---|---|---|---|---|---|---|
| 0.52779 | 0.152215 | 0.072 | 0.065 | 1 | 2 | Utp6 | 0.007 | Young |
| 0.528018 | 0.285002 | 0.05 | 0.044 | 1 | 2 | Hacd2 | 0.006 | Young |
| 0.528502 | 0.263762 | 0.063 | 0.072 | 1 | 2 | Dctn6 | -0.009 | Young |
| 0.529281 | 0.118079 | 0.053 | 0.061 | 1 | 2 | Ikbkap | -0.008 | Young |
| 0.529635 | 0.380512 | 0.048 | 0.042 | 1 | 2 | Sesn2 | 0.006 | Young |
| 0.530638 | 0.008222 | 0.138 | 0.151 | 1 | 2 | Dnajc1 | -0.013 | Young |
| 0.530746 | 0.311446 | 0.096 | 0.089 | 1 | 2 | Ubap1 | 0.007 | Young |
| 0.531201 | 0.047813 | 0.138 | 0.15 | 1 | 2 | Lactb | -0.012 | Young |
| 0.531212 | 0.021746 | 0.223 | 0.24 | 1 | 2 | AI314180 | -0.017 | Young |
| 0.53125 | 0.034873 | 0.134 | 0.123 | 1 | 2 | Spen | 0.011 | Young |
| 0.531852 | 0.01532 | 0.298 | 0.315 | 1 | 2 | Hmox1 | -0.017 | Young |
| 0.531906 | 0.322823 | 0.112 | 0.105 | 1 | 2 | Psmc2 | 0.007 | Young |
| 0.532243 | 0.05238 | 0.09 | 0.101 | 1 | 2 | Exosc3 | -0.011 | Young |
| 0.532384 | 0.290355 | 0.15 | 0.142 | 1 | 2 | Nt5c | 0.008 | Young |
| 0.532565 | 0.629136 | 0.07 | 0.064 | 1 | 2 | Bmyc | 0.006 | Young |
| 0.532711 | 0.467446 | 0.119 | 0.112 | 1 | 2 | Usp45 | 0.007 | Young |
| 0.532834 | 0.387725 | 0.05 | 0.044 | 1 | 2 | Lrp5 | 0.006 | Young |
| 0.532958 | 0.242259 | 0.1 | 0.092 | 1 | 2 | Vhl | 0.008 | Young |
| 0.533568 | 0.044665 | 0.091 | 0.102 | 1 | 2 | Fbxo18 | -0.011 | Young |
| 0.534197 | 0.103167 | 0.109 | 0.101 | 1 | 2 | Ing3 | 0.008 | Young |
| 0.534212 | 0.05009 | 0.321 | 0.31 | 1 | 2 | Dad1 | 0.011 | Young |
| 0.53478 | 0.017091 | 0.121 | 0.132 | 1 | 2 | Dguok | -0.011 | Young |
| 0.534845 | 0.276942 | 0.146 | 0.139 | 1 | 2 | Fam234a | 0.007 | Young |
| 0.534932 | 0.111389 | 0.02 | 0.025 | 1 | 2 | Rttn | -0.005 | Young |
| 0.535051 | 0.199196 | 0.113 | 0.105 | 1 | 2 | Fkbp3 | 0.008 | Young |
| 0.535575 | 0.110193 | 0.189 | 0.179 | 1 | 2 | Pim3 | 0.01 | Young |
| 0.535886 | 0.362749 | 0.025 | 0.03 | 1 | 2 | Ncapg2 | -0.005 | Young |
| 0.535888 | 0.06732 | 0.323 | 0.343 | 1 | 2 | Adipor1 | -0.02 | Young |
| 0.535909 | 0.01824 | 0.058 | 0.051 | 1 | 2 | Khk | 0.007 | Young |
| 0.536094 | 0.056055 | 0.106 | 0.118 | 1 | 2 | Armc8 | -0.012 | Young |
| 0.537148 | 0.40899 | 0.064 | 0.058 | 1 | 2 | Pogk | 0.006 | Young |
| 0.537192 | 0.038366 | 0.495 | 0.481 | 1 | 2 | Tspo | 0.014 | Young |
| 0.537297 | 0.044774 | 0.208 | 0.196 | 1 | 2 | Prkd3 | 0.012 | Young |
| 0.537417 | 0.157614 | 0.259 | 0.253 | 1 | 2 | Mdh1 | 0.006 | Young |
| 0.537513 | 0.101449 | 0.077 | 0.087 | 1 | 2 | Lap3 | -0.01 | Young |
| 0.537687 | 0.24041 | 0.069 | 0.062 | 1 | 2 | Rnf34 | 0.007 | Young |
| 0.537726 | 0.023527 | 0.235 | 0.255 | 1 | 2 | Pan3 | -0.02 | Young |
| 0.53804 | 0.259618 | 0.025 | 0.03 | 1 | 2 | Pwp2 | -0.005 | Young |
| 0.538272 | 0.074702 | 0.436 | 0.438 | 1 | 2 | Atp6v1f | -0.002 | Young |
| 0.538274 | 0.113956 | 0.081 | 0.074 | 1 | 2 | Traf7 | 0.007 | Young |
| 0.538551 | 0.026843 | 0.015 | 0.019 | 1 | 2 | Gtf2h3 | -0.004 | Young |
| 0.539176 | 0.074755 | 0.049 | 0.043 | 1 | 2 | Slc25a16 | 0.006 | Young |
| 0.539291 | 0.134033 | 0.102 | 0.112 | 1 | 2 | Uhrf1bp1l | -0.01 | Young |
| 0.539316 | 0.040424 | 0.507 | 0.489 | 1 | 2 | Slc38a2 | 0.018 | Young |
| 0.539332 | 0.319149 | 0.081 | 0.074 | 1 | 2 | Phf1 | 0.007 | Young |

| | | | | | | | | |
|---|---|---|---|---|---|---|---|---|
| 0.539686 | 0.093557 | 0.031 | 0.036 | 1 | 2 | Plcl1 | -0.005 | Young |
| 0.53989 | 0.079736 | 0.153 | 0.168 | 1 | 2 | Ptpn22 | -0.015 | Young |
| 0.539938 | 0.25073 | 0.195 | 0.186 | 1 | 2 | Ascc3 | 0.009 | Young |
| 0.540666 | 0.023277 | 0.088 | 0.097 | 1 | 2 | Mafk | -0.009 | Young |
| 0.541362 | 0.036603 | 0.166 | 0.155 | 1 | 2 | Tcerg1 | 0.011 | Young |
| 0.541973 | 0.045407 | 0.568 | 0.594 | 1 | 2 | Wfdc17 | -0.026 | Young |
| 0.542075 | 0.086146 | 0.02 | 0.025 | 1 | 2 | Rbm4 | -0.005 | Young |
| 0.542199 | 0.119911 | 0.121 | 0.112 | 1 | 2 | Itpr2 | 0.009 | Young |
| 0.542351 | 0.347457 | 0.046 | 0.041 | 1 | 2 | Gm27010 | 0.005 | Young |
| 0.542577 | 0.237169 | 0.116 | 0.109 | 1 | 2 | Chd6 | 0.007 | Young |
| 0.542624 | 0.010319 | 0.464 | 0.454 | 1 | 2 | Syngr2 | 0.01 | Young |
| 0.543739 | 0.293897 | 0.109 | 0.102 | 1 | 2 | P2rx7 | 0.007 | Young |
| 0.544187 | 0.007695 | 0.077 | 0.07 | 1 | 2 | Zfp260 | 0.007 | Young |
| 0.544393 | 0.191748 | 0.031 | 0.036 | 1 | 2 | Asna1 | -0.005 | Young |
| 0.544798 | 0.587386 | 0.011 | 0.014 | 1 | 2 | Rnf24 | -0.003 | Young |
| 0.544798 | 0.229629 | 0.011 | 0.014 | 1 | 2 | Tmem35b | -0.003 | Young |
| 0.544878 | 0.20109 | 0.046 | 0.041 | 1 | 2 | Cox19 | 0.005 | Young |
| 0.545 | 0.257314 | 0.182 | 0.175 | 1 | 2 | Ddx54 | 0.007 | Young |
| 0.54512 | 0.217048 | 0.011 | 0.014 | 1 | 2 | Mtrr | -0.003 | Young |
| 0.545151 | 0.006814 | 0.316 | 0.299 | 1 | 2 | Lmo4 | 0.017 | Young |
| 0.545286 | 0.027816 | 0.119 | 0.131 | 1 | 2 | Ilf3 | -0.012 | Young |
| 0.545762 | 0.409704 | 0.011 | 0.014 | 1 | 2 | Iigp1 | -0.003 | Young |
| 0.546126 | 0.038567 | 0.106 | 0.117 | 1 | 2 | Ormdl2 | -0.011 | Young |
| 0.546456 | 0.122289 | 0.271 | 0.262 | 1 | 2 | Sptssa | 0.009 | Young |
| 0.546607 | 0.108589 | 0.197 | 0.191 | 1 | 2 | Zfp644 | 0.006 | Young |
| 0.546752 | 0.397396 | 0.114 | 0.108 | 1 | 2 | Sart1 | 0.006 | Young |
| 0.547814 | 0.008638 | 0.146 | 0.16 | 1 | 2 | Ptpn7 | -0.014 | Young |
| 0.548016 | 0.043261 | 0.011 | 0.014 | 1 | 2 | Cptp | -0.003 | Young |
| 0.54818 | 0.11934 | 0.228 | 0.221 | 1 | 2 | Syf2 | 0.007 | Young |
| 0.54866 | 0.109539 | 0.011 | 0.014 | 1 | 2 | 2610021A( | -0.003 | Young |
| 0.548689 | 0.007973 | 0.032 | 0.037 | 1 | 2 | Fam58b | -0.005 | Young |
| 0.548938 | 0.364745 | 0.04 | 0.035 | 1 | 2 | Galnt4 | 0.005 | Young |
| 0.548951 | 0.037814 | 0.778 | 0.76 | 1 | 2 | Ubc | 0.018 | Young |
| 0.549649 | 0.07421 | 0.156 | 0.168 | 1 | 2 | Akna | -0.012 | Young |
| 0.549816 | 0.067094 | 0.1 | 0.091 | 1 | 2 | Casd1 | 0.009 | Young |
| 0.54982 | 0.046639 | 0.181 | 0.169 | 1 | 2 | Ppp1r10 | 0.012 | Young |
| 0.551602 | 0.101994 | 0.172 | 0.163 | 1 | 2 | Maf1 | 0.009 | Young |
| 0.551755 | 0.101834 | 0.096 | 0.089 | 1 | 2 | Atl3 | 0.007 | Young |
| 0.551868 | 0.20622 | 0.207 | 0.198 | 1 | 2 | Gxylt1 | 0.009 | Young |
| 0.551996 | 0.073919 | 0.032 | 0.037 | 1 | 2 | Tbc1d32 | -0.005 | Young |
| 0.552132 | 0.195411 | 0.044 | 0.05 | 1 | 2 | Slc35b3 | -0.006 | Young |
| 0.552351 | 0.311927 | 0.045 | 0.04 | 1 | 2 | Parp10 | 0.005 | Young |
| 0.552454 | 0.028424 | 0.151 | 0.164 | 1 | 2 | Zdhhc3 | -0.013 | Young |
| 0.552827 | 0.223597 | 0.068 | 0.061 | 1 | 2 | Mettl1 | 0.007 | Young |
| 0.553212 | 0.142544 | 0.075 | 0.067 | 1 | 2 | Iars2 | 0.008 | Young |

| | | | | | | | | |
|---|---|---|---|---|---|---|---|---|
| 0.55337 | 0.033045 | 0.026 | 0.031 | 1 | 2 | Sec61a2 | -0.005 | Young |
| 0.554057 | 0.106353 | 0.15 | 0.163 | 1 | 2 | Exoc3 | -0.013 | Young |
| 0.554165 | 0.254845 | 0.061 | 0.055 | 1 | 2 | Desi1 | 0.006 | Young |
| 0.554193 | 0.193323 | 0.037 | 0.043 | 1 | 2 | Cox10 | -0.006 | Young |
| 0.554312 | 0.279471 | 0.063 | 0.057 | 1 | 2 | Ppp2r5d | 0.006 | Young |
| 0.554466 | 0.035946 | 0.568 | 0.555 | 1 | 2 | Uqcr11 | 0.013 | Young |
| 0.554467 | 0.165935 | 0.131 | 0.122 | 1 | 2 | Kcnn4 | 0.009 | Young |
| 0.555159 | 0.038678 | 0.219 | 0.209 | 1 | 2 | Trafd1 | 0.01 | Young |
| 0.555867 | 0.099158 | 0.258 | 0.249 | 1 | 2 | Slc26a2 | 0.009 | Young |
| 0.555899 | 0.333473 | 0.032 | 0.037 | 1 | 2 | Gtpbp10 | -0.005 | Young |
| 0.556384 | 0.103017 | 0.086 | 0.095 | 1 | 2 | Rbm14 | -0.009 | Young |
| 0.556528 | 0.129193 | 0.078 | 0.088 | 1 | 2 | Arsa | -0.01 | Young |
| 0.556695 | 0.281024 | 0.132 | 0.124 | 1 | 2 | Susd3 | 0.008 | Young |
| 0.556797 | 0.31086 | 0.172 | 0.165 | 1 | 2 | Nrp2 | 0.007 | Young |
| 0.557216 | 0.094496 | 0.19 | 0.18 | 1 | 2 | Hbp1 | 0.01 | Young |
| 0.557231 | 0.018273 | 0.23 | 0.217 | 1 | 2 | Kdm5a | 0.013 | Young |
| 0.557492 | 0.00848 | 0.121 | 0.132 | 1 | 2 | Slc7a1 | -0.011 | Young |
| 0.557804 | 0.016217 | 0.45 | 0.437 | 1 | 2 | Tomm20 | 0.013 | Young |
| 0.557848 | 0.310458 | 0.089 | 0.1 | 1 | 2 | Arfrp1 | -0.011 | Young |
| 0.558685 | 0.366675 | 0.088 | 0.081 | 1 | 2 | Snapc5 | 0.007 | Young |
| 0.558722 | 0.068068 | 0.044 | 0.039 | 1 | 2 | Ndufaf7 | 0.005 | Young |
| 0.559302 | 0.074597 | 0.081 | 0.074 | 1 | 2 | G2e3 | 0.007 | Young |
| 0.559487 | 0.012591 | 0.045 | 0.04 | 1 | 2 | Iqsec1 | 0.005 | Young |
| 0.559497 | 0.09197 | 0.125 | 0.116 | 1 | 2 | Mknk1 | 0.009 | Young |
| 0.559624 | 0.156244 | 0.05 | 0.044 | 1 | 2 | Tmem161b | 0.006 | Young |
| 0.559707 | 0.248742 | 0.072 | 0.081 | 1 | 2 | Frmd8 | -0.009 | Young |
| 0.559764 | 0.279574 | 0.084 | 0.078 | 1 | 2 | Batf | 0.006 | Young |
| 0.560048 | 0.153745 | 0.154 | 0.148 | 1 | 2 | Ano6 | 0.006 | Young |
| 0.560143 | 0.0922 | 0.158 | 0.149 | 1 | 2 | Dynll2 | 0.009 | Young |
| 0.56088 | 0.022921 | 0.119 | 0.11 | 1 | 2 | Mrpl18 | 0.009 | Young |
| 0.561288 | 0.166802 | 0.063 | 0.057 | 1 | 2 | Ubr1 | 0.006 | Young |
| 0.561311 | 0.202724 | 0.255 | 0.249 | 1 | 2 | Mpp6 | 0.006 | Young |
| 0.561413 | 0.042339 | 0.217 | 0.209 | 1 | 2 | Tank | 0.008 | Young |
| 0.561434 | 0.062901 | 0.044 | 0.05 | 1 | 2 | Ksr1 | -0.006 | Young |
| 0.562294 | 0.052032 | 0.083 | 0.092 | 1 | 2 | Dpm1 | -0.009 | Young |
| 0.562423 | 0.113078 | 0.211 | 0.206 | 1 | 2 | Ppm1a | 0.005 | Young |
| 0.562789 | 0.006549 | 0.051 | 0.045 | 1 | 2 | Fam219b | 0.006 | Young |
| 0.563084 | 0.064763 | 0.095 | 0.087 | 1 | 2 | Ubxn7 | 0.008 | Young |
| 0.563271 | 0.667325 | 0.052 | 0.047 | 1 | 2 | Rbm38 | 0.005 | Young |
| 0.563321 | 0.048517 | 0.239 | 0.232 | 1 | 2 | Eif4e | 0.007 | Young |
| 0.563325 | 0.321894 | 0.038 | 0.033 | 1 | 2 | Gpr89 | 0.005 | Young |
| 0.563462 | 0.090102 | 0.596 | 0.599 | 1 | 2 | Son | -0.003 | Young |
| 0.563623 | 0.157672 | 0.124 | 0.116 | 1 | 2 | Nasp | 0.008 | Young |
| 0.563777 | 0.242203 | 0.059 | 0.054 | 1 | 2 | Xrcc1 | 0.005 | Young |
| 0.5638 | 0.304828 | 0.049 | 0.056 | 1 | 2 | Rmi1 | -0.007 | Young |

| | | | | | | | | |
|---|---|---|---|---|---|---|---|---|
| 0.563813 | 0.155427 | 0.078 | 0.072 | 1 | 2 | Tssc4 | 0.006 | Young |
| 0.564941 | 0.106752 | 0.043 | 0.037 | 1 | 2 | Appl2 | 0.006 | Young |
| 0.565249 | 0.146266 | 0.062 | 0.07 | 1 | 2 | Eepd1 | -0.008 | Young |
| 0.565624 | 0.410654 | 0.04 | 0.035 | 1 | 2 | Rrnad1 | 0.005 | Young |
| 0.565708 | 0.193663 | 0.007 | 0.01 | 1 | 2 | Tbc1d10c | -0.003 | Young |
| 0.566051 | 0.033174 | 0.062 | 0.07 | 1 | 2 | Dlat | -0.008 | Young |
| 0.566098 | 0.277807 | 0.065 | 0.059 | 1 | 2 | Aga | 0.006 | Young |
| 0.566243 | 0.120014 | 0.296 | 0.291 | 1 | 2 | Snu13 | 0.005 | Young |
| 0.566542 | 0.0809 | 0.172 | 0.185 | 1 | 2 | Rybp | -0.013 | Young |
| 0.566571 | 0.115321 | 0.133 | 0.146 | 1 | 2 | Txndc5 | -0.013 | Young |
| 0.566651 | 0.056985 | 0.202 | 0.219 | 1 | 2 | Aup1 | -0.017 | Young |
| 0.566928 | 0.44951 | 0.037 | 0.032 | 1 | 2 | Tmco6 | 0.005 | Young |
| 0.56747 | 0.082436 | 0.154 | 0.167 | 1 | 2 | Uap1l1 | -0.013 | Young |
| 0.567528 | 0.2203 | 0.039 | 0.034 | 1 | 2 | Dhx34 | 0.005 | Young |
| 0.567697 | 0.466053 | 0.086 | 0.079 | 1 | 2 | Ntpcr | 0.007 | Young |
| 0.568337 | 0.056382 | 0.158 | 0.149 | 1 | 2 | Parp1 | 0.009 | Young |
| 0.568442 | 0.000466 | 0.151 | 0.164 | 1 | 2 | Zfp800 | -0.013 | Young |
| 0.568572 | 0.081393 | 0.091 | 0.084 | 1 | 2 | Plcl2 | 0.007 | Young |
| 0.568985 | 0.055172 | 0.208 | 0.202 | 1 | 2 | Ddx50 | 0.006 | Young |
| 0.569099 | 0.017238 | 0.052 | 0.059 | 1 | 2 | Bola3 | -0.007 | Young |
| 0.569637 | 0.087113 | 0.122 | 0.113 | 1 | 2 | Trim30d | 0.009 | Young |
| 0.569839 | 0.45582 | 0.034 | 0.03 | 1 | 2 | Tbcd | 0.004 | Young |
| 0.569932 | 0.223511 | 0.065 | 0.059 | 1 | 2 | Mrpl16 | 0.006 | Young |
| 0.570275 | 0.088854 | 0.14 | 0.153 | 1 | 2 | P4ha1 | -0.013 | Young |
| 0.570335 | 0.122448 | 0.241 | 0.232 | 1 | 2 | Rc3h1 | 0.009 | Young |
| 0.570499 | 0.000618 | 0.186 | 0.176 | 1 | 2 | Btk | 0.01 | Young |
| 0.570546 | 0.148301 | 0.273 | 0.266 | 1 | 2 | Zfp710 | 0.007 | Young |
| 0.570747 | 0.025469 | 0.048 | 0.042 | 1 | 2 | Pnpo | 0.006 | Young |
| 0.570861 | 0.756736 | 0.007 | 0.01 | 1 | 2 | Fgf11 | -0.003 | Young |
| 0.570936 | 0.035698 | 0.065 | 0.073 | 1 | 2 | Ppp2r3d | -0.008 | Young |
| 0.571216 | 0.053586 | 0.063 | 0.071 | 1 | 2 | Pin1 | -0.008 | Young |
| 0.571256 | 0.14988 | 0.148 | 0.139 | 1 | 2 | Ankle2 | 0.009 | Young |
| 0.571282 | 0.595672 | 0.034 | 0.03 | 1 | 2 | Mtor | 0.004 | Young |
| 0.571323 | 0.187708 | 0.119 | 0.11 | 1 | 2 | Casp3 | 0.009 | Young |
| 0.571415 | 0.130152 | 0.251 | 0.242 | 1 | 2 | Ybx3 | 0.009 | Young |
| 0.571491 | 0.369966 | 0.081 | 0.075 | 1 | 2 | Slc25a25 | 0.006 | Young |
| 0.571827 | 0.187433 | 0.099 | 0.092 | 1 | 2 | Mrpl2 | 0.007 | Young |
| 0.572203 | 0.064958 | 0.124 | 0.135 | 1 | 2 | Ifi209 | -0.011 | Young |
| 0.572311 | 0.279167 | 0.037 | 0.032 | 1 | 2 | Phlpp2 | 0.005 | Young |
| 0.572517 | 0.206268 | 0.057 | 0.051 | 1 | 2 | Crcp | 0.006 | Young |
| 0.573079 | 0.623168 | 0.032 | 0.028 | 1 | 2 | Ube2o | 0.004 | Young |
| 0.573194 | 0.18969 | 0.138 | 0.13 | 1 | 2 | Uimc1 | 0.008 | Young |
| 0.573214 | 0.262813 | 0.063 | 0.057 | 1 | 2 | Zfyve26 | 0.006 | Young |
| 0.573239 | 0.181546 | 0.169 | 0.162 | 1 | 2 | Plekha2 | 0.007 | Young |
| 0.573268 | 0.071124 | 0.08 | 0.073 | 1 | 2 | Mrps18a | 0.007 | Young |

| | | | | | | | | |
|---|---|---|---|---|---|---|---|---|
| 0.573918 | 0.507444 | 0.077 | 0.072 | 1 | 2 | Iars | 0.005 | Young |
| 0.573925 | 0.420421 | 0.012 | 0.015 | 1 | 2 | 1700066M | -0.003 | Young |
| 0.573966 | 0.032221 | 0.144 | 0.155 | 1 | 2 | Blvra | -0.011 | Young |
| 0.57424 | 0.342514 | 0.012 | 0.015 | 1 | 2 | Zfp637 | -0.003 | Young |
| 0.574403 | 0.266696 | 0.036 | 0.031 | 1 | 2 | Rcc1 | 0.005 | Young |
| 0.574662 | 0.148476 | 0.072 | 0.066 | 1 | 2 | Ccdc90b | 0.006 | Young |
| 0.57469 | 0.215422 | 0.017 | 0.02 | 1 | 2 | Aldh4a1 | -0.003 | Young |
| 0.575045 | 0.22162 | 0.058 | 0.052 | 1 | 2 | Nedd1 | 0.006 | Young |
| 0.57515 | 0.192864 | 0.312 | 0.313 | 1 | 2 | Nav1 | -0.001 | Young |
| 0.575587 | 0.097448 | 0.081 | 0.074 | 1 | 2 | Srp68 | 0.007 | Young |
| 0.575611 | 0.057142 | 0.134 | 0.125 | 1 | 2 | Rab5b | 0.009 | Young |
| 0.575705 | 0.190798 | 0.039 | 0.045 | 1 | 2 | Efl1 | -0.006 | Young |
| 0.57582 | 0.221527 | 0.012 | 0.015 | 1 | 2 | Xndc1 | -0.003 | Young |
| 0.575955 | 0.627715 | 0.053 | 0.048 | 1 | 2 | Abhd13 | 0.005 | Young |
| 0.575987 | 0.122368 | 0.42 | 0.422 | 1 | 2 | Eif2s2 | -0.002 | Young |
| 0.576201 | 0.168778 | 0.192 | 0.184 | 1 | 2 | Ep400 | 0.008 | Young |
| 0.576281 | 0.252598 | 0.038 | 0.033 | 1 | 2 | Il21r | 0.005 | Young |
| 0.576294 | 0.349116 | 0.012 | 0.015 | 1 | 2 | H2-Q10 | -0.003 | Young |
| 0.576622 | 0.396048 | 0.09 | 0.085 | 1 | 2 | Plxna1 | 0.005 | Young |
| 0.576862 | 0.110583 | 0.068 | 0.061 | 1 | 2 | Slc38a6 | 0.007 | Young |
| 0.576962 | 0.128067 | 0.216 | 0.21 | 1 | 2 | Rex1bd | 0.006 | Young |
| 0.577138 | 0.125623 | 0.132 | 0.141 | 1 | 2 | Irak2 | -0.009 | Young |
| 0.577402 | 0.018304 | 0.012 | 0.015 | 1 | 2 | Cdc42bpg | -0.003 | Young |
| 0.577451 | 0.211693 | 0.064 | 0.072 | 1 | 2 | Trappc3 | -0.008 | Young |
| 0.577773 | 0.110395 | 0.4 | 0.399 | 1 | 2 | Cd93 | 0.001 | Young |
| 0.577931 | 0.200626 | 0.124 | 0.117 | 1 | 2 | Qars | 0.007 | Young |
| 0.57807 | 0.354182 | 0.067 | 0.075 | 1 | 2 | Itpripl1 | -0.008 | Young |
| 0.578171 | 0.450043 | 0.052 | 0.047 | 1 | 2 | Adcy9 | 0.005 | Young |
| 0.578272 | 0.098421 | 0.19 | 0.182 | 1 | 2 | Ube2h | 0.008 | Young |
| 0.578349 | 0.290648 | 0.106 | 0.099 | 1 | 2 | Mxi1 | 0.007 | Young |
| 0.578471 | 0.002278 | 0.608 | 0.585 | 1 | 2 | Fosb | 0.023 | Young |
| 0.578757 | 0.299723 | 0.118 | 0.111 | 1 | 2 | Slc45a4 | 0.007 | Young |
| 0.579153 | 0.0087 | 0.065 | 0.059 | 1 | 2 | Emc8 | 0.006 | Young |
| 0.579282 | 0.158863 | 0.023 | 0.027 | 1 | 2 | Snupn | -0.004 | Young |
| 0.579429 | 0.010373 | 0.062 | 0.069 | 1 | 2 | Tbrg1 | -0.007 | Young |
| 0.579697 | 0.043742 | 0.67 | 0.652 | 1 | 2 | Slc25a5 | 0.018 | Young |
| 0.579821 | 0.234985 | 0.07 | 0.064 | 1 | 2 | Scoc | 0.006 | Young |
| 0.579947 | 0.277045 | 0.189 | 0.184 | 1 | 2 | Lamtor5 | 0.005 | Young |
| 0.580433 | 0.332383 | 0.038 | 0.033 | 1 | 2 | Kif19a | 0.005 | Young |
| 0.580926 | 0.025119 | 0.17 | 0.185 | 1 | 2 | Ube2e1 | -0.015 | Young |
| 0.581459 | 0.08545 | 0.599 | 0.606 | 1 | 2 | Hnrnpu | -0.007 | Young |
| 0.582121 | 0.251829 | 0.056 | 0.05 | 1 | 2 | Rev1 | 0.006 | Young |
| 0.582385 | 0.10876 | 0.039 | 0.034 | 1 | 2 | Aifm2 | 0.005 | Young |
| 0.582401 | 0.051331 | 0.308 | 0.302 | 1 | 2 | Nol7 | 0.006 | Young |
| 0.582681 | 0.325732 | 0.034 | 0.03 | 1 | 2 | Rabif | 0.004 | Young |

| | | | | | | | | |
|---|---|---|---|---|---|---|---|---|
| 0.583098 | 0.339312 | 0.033 | 0.029 | 1 | 2 | Fen1 | 0.004 | Young |
| 0.583149 | 0.076862 | 0.037 | 0.032 | 1 | 2 | Pik3r2 | 0.005 | Young |
| 0.58324 | 0.165851 | 0.128 | 0.121 | 1 | 2 | Tceal9 | 0.007 | Young |
| 0.584099 | 0.039338 | 0.129 | 0.141 | 1 | 2 | Mrps17 | -0.012 | Young |
| 0.584165 | 0.014955 | 0.046 | 0.052 | 1 | 2 | Knop1 | -0.006 | Young |
| 0.584476 | 0.132416 | 0.065 | 0.059 | 1 | 2 | Tshz1 | 0.006 | Young |
| 0.584681 | 0.080924 | 0.16 | 0.175 | 1 | 2 | Poldip3 | -0.015 | Young |
| 0.584707 | 0.018323 | 0.178 | 0.191 | 1 | 2 | Clk3 | -0.013 | Young |
| 0.584774 | 0.076393 | 0.078 | 0.072 | 1 | 2 | Rif1 | 0.006 | Young |
| 0.585538 | 0.212425 | 0.023 | 0.027 | 1 | 2 | Zxdb | -0.004 | Young |
| 0.586376 | 0.158953 | 0.297 | 0.291 | 1 | 2 | Snap23 | 0.006 | Young |
| 0.586625 | 0.029857 | 0.739 | 0.74 | 1 | 2 | Tpm3 | -0.001 | Young |
| 0.586706 | 0.357337 | 0.033 | 0.029 | 1 | 2 | 10-Sep | 0.004 | Young |
| 0.586904 | 0.03253 | 0.165 | 0.179 | 1 | 2 | Cuedc2 | -0.014 | Young |
| 0.58696 | 0.176776 | 0.099 | 0.092 | 1 | 2 | Rarg | 0.007 | Young |
| 0.587374 | 0.005223 | 0.2 | 0.217 | 1 | 2 | Mbd2 | -0.017 | Young |
| 0.587562 | 0.080605 | 0.606 | 0.606 | 1 | 2 | Rbm3 | 0 | Young |
| 0.587829 | 0.106572 | 0.1 | 0.092 | 1 | 2 | Sumf1 | 0.008 | Young |
| 0.587895 | 0.022935 | 0.046 | 0.052 | 1 | 2 | Mtpap | -0.006 | Young |
| 0.587969 | 0.092742 | 0.211 | 0.201 | 1 | 2 | Chp1 | 0.01 | Young |
| 0.588202 | 0.133637 | 0.074 | 0.067 | 1 | 2 | Wasl | 0.007 | Young |
| 0.589445 | 0.30512 | 0.036 | 0.031 | 1 | 2 | Tspyl2 | 0.005 | Young |
| 0.589449 | 0.068702 | 0.512 | 0.511 | 1 | 2 | Maf | 0.001 | Young |
| 0.589711 | 0.139077 | 0.089 | 0.082 | 1 | 2 | Slc7a7 | 0.007 | Young |
| 0.589859 | 0.056093 | 0.018 | 0.021 | 1 | 2 | Nit2 | -0.003 | Young |
| 0.590303 | 0.09097 | 0.285 | 0.282 | 1 | 2 | Klhl24 | 0.003 | Young |
| 0.590541 | 0.560209 | 0.03 | 0.026 | 1 | 2 | Gpatch4 | 0.004 | Young |
| 0.590658 | 0.049024 | 0.468 | 0.467 | 1 | 2 | Bola2 | 0.001 | Young |
| 0.590689 | 0.275474 | 0.096 | 0.09 | 1 | 2 | Ube4a | 0.006 | Young |
| 0.590844 | 0.043532 | 0.069 | 0.076 | 1 | 2 | Nudt13 | -0.007 | Young |
| 0.59094 | 0.063148 | 0.328 | 0.322 | 1 | 2 | Pebp1 | 0.006 | Young |
| 0.591192 | 0.009059 | 0.11 | 0.12 | 1 | 2 | Pih1d1 | -0.01 | Young |
| 0.591206 | 0.051322 | 0.084 | 0.078 | 1 | 2 | Ccpg1 | 0.006 | Young |
| 0.591492 | 0.213334 | 0.056 | 0.05 | 1 | 2 | Xylt2 | 0.006 | Young |
| 0.592175 | 0.057584 | 0.096 | 0.105 | 1 | 2 | Suclg1 | -0.009 | Young |
| 0.592209 | 0.038911 | 0.053 | 0.06 | 1 | 2 | Gcnt1 | -0.007 | Young |
| 0.592457 | 0.282929 | 0.074 | 0.067 | 1 | 2 | Ttc9c | 0.007 | Young |
| 0.592665 | 0.326441 | 0.067 | 0.061 | 1 | 2 | Isoc1 | 0.006 | Young |
| 0.592718 | 0.064683 | 0.205 | 0.196 | 1 | 2 | Rbm7 | 0.009 | Young |
| 0.592764 | 0.029173 | 0.055 | 0.061 | 1 | 2 | Las1l | -0.006 | Young |
| 0.59292 | 0.01496 | 0.08 | 0.073 | 1 | 2 | Mfap1a | 0.007 | Young |
| 0.59351 | 0.353871 | 0.046 | 0.052 | 1 | 2 | Gzf1 | -0.006 | Young |
| 0.594099 | 0.799081 | 0.027 | 0.024 | 1 | 2 | B4galt7 | 0.003 | Young |
| 0.594715 | 0.136585 | 0.061 | 0.067 | 1 | 2 | Narf | -0.006 | Young |
| 0.595221 | 0.068456 | 0.071 | 0.079 | 1 | 2 | Foxk1 | -0.008 | Young |

| | | | | | | | | |
|---|---|---|---|---|---|---|---|---|
| 0.59526 | 0.174173 | 0.076 | 0.07 | 1 | 2 | Bcl2l1 | 0.006 | Young |
| 0.595401 | 0.081706 | 0.19 | 0.202 | 1 | 2 | Wdr89 | -0.012 | Young |
| 0.595824 | 0.201963 | 0.298 | 0.297 | 1 | 2 | Arhgap45 | 0.001 | Young |
| 0.595958 | 0.054873 | 0.125 | 0.135 | 1 | 2 | Mrpl12 | -0.01 | Young |
| 0.595982 | 0.193171 | 0.018 | 0.021 | 1 | 2 | Psph | -0.003 | Young |
| 0.596545 | 0.065332 | 0.076 | 0.07 | 1 | 2 | Ecd | 0.006 | Young |
| 0.596679 | 0.074796 | 0.169 | 0.181 | 1 | 2 | Cat | -0.012 | Young |
| 0.596913 | 0.022637 | 0.089 | 0.082 | 1 | 2 | Prrc1 | 0.007 | Young |
| 0.597282 | 0.046238 | 0.032 | 0.028 | 1 | 2 | Zw10 | 0.004 | Young |
| 0.597379 | 0.054525 | 0.038 | 0.033 | 1 | 2 | Zbtb17 | 0.005 | Young |
| 0.598092 | 0.057749 | 0.167 | 0.181 | 1 | 2 | Tbc1d14 | -0.014 | Young |
| 0.598121 | 0.478676 | 0.072 | 0.067 | 1 | 2 | Exosc9 | 0.005 | Young |
| 0.598396 | 0.062959 | 0.178 | 0.192 | 1 | 2 | Rad23a | -0.014 | Young |
| 0.598902 | 0.25161 | 0.089 | 0.082 | 1 | 2 | Prpf31 | 0.007 | Young |
| 0.599399 | 0.004006 | 0.099 | 0.091 | 1 | 2 | Irf4 | 0.008 | Young |
| 0.599552 | 0.018134 | 0.926 | 0.93 | 1 | 2 | Cst3 | -0.004 | Young |
| 0.599552 | 0.212791 | 0.099 | 0.092 | 1 | 2 | Mdn1 | 0.007 | Young |
| 0.599713 | 0.240075 | 0.031 | 0.027 | 1 | 2 | Tmem238 | 0.004 | Young |
| 0.599793 | 0.142112 | 0.078 | 0.072 | 1 | 2 | Bid | 0.006 | Young |
| 0.599864 | 0.095219 | 0.175 | 0.167 | 1 | 2 | Klf9 | 0.008 | Young |
| 0.599893 | 0.549484 | 0.045 | 0.041 | 1 | 2 | Ppih | 0.004 | Young |
| 0.600059 | 0.163433 | 0.26 | 0.254 | 1 | 2 | AC160336. | 0.006 | Young |
| 0.600063 | 0.086219 | 0.087 | 0.094 | 1 | 2 | Slc16a7 | -0.007 | Young |
| 0.600075 | 0.087392 | 0.131 | 0.141 | 1 | 2 | Phactr2 | -0.01 | Young |
| 0.600233 | 0.234597 | 0.197 | 0.193 | 1 | 2 | Ramp1 | 0.004 | Young |
| 0.600591 | 0.313655 | 0.051 | 0.046 | 1 | 2 | Usp53 | 0.005 | Young |
| 0.600731 | 0.100808 | 0.304 | 0.3 | 1 | 2 | Gpr65 | 0.004 | Young |
| 0.600961 | 0.275464 | 0.069 | 0.063 | 1 | 2 | Fez2 | 0.006 | Young |
| 0.60124 | 0.344728 | 0.029 | 0.033 | 1 | 2 | Dusp18 | -0.004 | Young |
| 0.601312 | 0.059918 | 0.226 | 0.216 | 1 | 2 | Kansl1 | 0.01 | Young |
| 0.601545 | 0.048553 | 0.024 | 0.028 | 1 | 2 | Drosha | -0.004 | Young |
| 0.601574 | 0.155169 | 0.068 | 0.062 | 1 | 2 | Tceanc2 | 0.006 | Young |
| 0.601636 | 0.070408 | 0.093 | 0.086 | 1 | 2 | Atrnl1 | 0.007 | Young |
| 0.602255 | 0.011325 | 0.205 | 0.222 | 1 | 2 | Ms4a8a | -0.017 | Young |
| 0.602265 | 0.319213 | 0.201 | 0.198 | 1 | 2 | Eea1 | 0.003 | Young |
| 0.602314 | 0.030165 | 0.587 | 0.58 | 1 | 2 | Lrp1 | 0.007 | Young |
| 0.602517 | 0.01972 | 0.267 | 0.258 | 1 | 2 | Arfgef1 | 0.009 | Young |
| 0.60285 | 0.051251 | 0.07 | 0.077 | 1 | 2 | Kat7 | -0.007 | Young |
| 0.603322 | 0.373222 | 0.027 | 0.024 | 1 | 2 | Clip2 | 0.003 | Young |
| 0.603462 | 0.169455 | 0.208 | 0.204 | 1 | 2 | Kif1b | 0.004 | Young |
| 0.603992 | 0.088261 | 0.053 | 0.06 | 1 | 2 | Zfp110 | -0.007 | Young |
| 0.604128 | 0.582886 | 0.046 | 0.042 | 1 | 2 | Agfg2 | 0.004 | Young |
| 0.604853 | 0.641033 | 0.024 | 0.02 | 1 | 2 | Bcorl1 | 0.004 | Young |
| 0.605233 | 0.176109 | 0.17 | 0.163 | 1 | 2 | Pnrc2 | 0.007 | Young |
| 0.60545 | 0.01734 | 0.128 | 0.121 | 1 | 2 | Cdk2ap1 | 0.007 | Young |

| | | | | | | | | |
|---|---|---|---|---|---|---|---|---|
| 0.605697 | 0.172187 | 0.027 | 0.024 | 1 | 2 | Timp1 | 0.003 | Young |
| 0.606154 | 0.23766 | 0.099 | 0.093 | 1 | 2 | Gpr183 | 0.006 | Young |
| 0.606528 | 0.018797 | 0.935 | 0.951 | 1 | 2 | Rpl7 | -0.016 | Young |
| 0.606611 | 0.606041 | 0.026 | 0.022 | 1 | 2 | Btf3l4 | 0.004 | Young |
| 0.606648 | 0.289966 | 0.03 | 0.026 | 1 | 2 | Wrnip1 | 0.004 | Young |
| 0.606653 | 0.063098 | 0.139 | 0.132 | 1 | 2 | Abl2 | 0.007 | Young |
| 0.607674 | 0.248817 | 0.182 | 0.177 | 1 | 2 | Ppp2r2a | 0.005 | Young |
| 0.6077 | 0.30069 | 0.044 | 0.04 | 1 | 2 | Mettl7a1 | 0.004 | Young |
| 0.608169 | 0.083796 | 0.013 | 0.016 | 1 | 2 | 2810454H( | -0.003 | Young |
| 0.608185 | 0.049928 | 0.037 | 0.032 | 1 | 2 | Mrps11 | 0.005 | Young |
| 0.608316 | 0.399992 | 0.075 | 0.07 | 1 | 2 | Ptpn9 | 0.005 | Young |
| 0.608326 | 0.109529 | 0.051 | 0.046 | 1 | 2 | Rhno1 | 0.005 | Young |
| 0.608331 | 0.119233 | 0.17 | 0.163 | 1 | 2 | Cib1 | 0.007 | Young |
| 0.608832 | 0.285748 | 0.029 | 0.025 | 1 | 2 | Zfp729b | 0.004 | Young |
| 0.609075 | 0.039239 | 0.159 | 0.152 | 1 | 2 | Ganab | 0.007 | Young |
| 0.609851 | 0.487784 | 0.025 | 0.021 | 1 | 2 | Zfp619 | 0.004 | Young |
| 0.610096 | 0.063621 | 0.493 | 0.496 | 1 | 2 | Tmed2 | -0.003 | Young |
| 0.610102 | 0.277645 | 0.027 | 0.024 | 1 | 2 | Abcg2 | 0.003 | Young |
| 0.610214 | 0.59785 | 0.024 | 0.02 | 1 | 2 | Mov10 | 0.004 | Young |
| 0.610315 | 0.318583 | 0.008 | 0.011 | 1 | 2 | Spon1 | -0.003 | Young |
| 0.610315 | 0.109457 | 0.008 | 0.011 | 1 | 2 | Tspyl4 | -0.003 | Young |
| 0.610408 | 0.179728 | 0.181 | 0.173 | 1 | 2 | Nbeal1 | 0.008 | Young |
| 0.610564 | 0.25458 | 0.139 | 0.133 | 1 | 2 | Npepl1 | 0.006 | Young |
| 0.610676 | 0.258362 | 0.036 | 0.041 | 1 | 2 | Tmem39b | -0.005 | Young |
| 0.611085 | 0.037337 | 0.008 | 0.011 | 1 | 2 | Gm45871 | -0.003 | Young |
| 0.611186 | 0.143977 | 0.287 | 0.283 | 1 | 2 | Efr3a | 0.004 | Young |
| 0.612518 | 0.477177 | 0.024 | 0.02 | 1 | 2 | Prickle3 | 0.004 | Young |
| 0.612627 | 0.4955 | 0.008 | 0.011 | 1 | 2 | Slc9a7 | -0.003 | Young |
| 0.612651 | 0.108292 | 0.094 | 0.103 | 1 | 2 | Ddx52 | -0.009 | Young |
| 0.612827 | 0.138523 | 0.216 | 0.207 | 1 | 2 | 1-Mar | 0.009 | Young |
| 0.613377 | 0.053747 | 0.35 | 0.337 | 1 | 2 | Pgam1 | 0.013 | Young |
| 0.613599 | 0.372477 | 0.025 | 0.021 | 1 | 2 | Pced1a | 0.004 | Young |
| 0.613823 | 0.168455 | 0.151 | 0.146 | 1 | 2 | Babam2 | 0.005 | Young |
| 0.614022 | 0.125367 | 0.329 | 0.322 | 1 | 2 | Gna13 | 0.007 | Young |
| 0.614251 | 0.713658 | 0.021 | 0.018 | 1 | 2 | Usp28 | 0.003 | Young |
| 0.614302 | 0.007282 | 0.078 | 0.086 | 1 | 2 | Odf2 | -0.008 | Young |
| 0.614811 | 0.353807 | 0.048 | 0.043 | 1 | 2 | Ppm1k | 0.005 | Young |
| 0.615052 | 0.08915 | 0.185 | 0.178 | 1 | 2 | Smad4 | 0.007 | Young |
| 0.61533 | 0.214413 | 0.031 | 0.035 | 1 | 2 | Cd300lf | -0.004 | Young |
| 0.615441 | 0.225072 | 0.083 | 0.077 | 1 | 2 | Pcnt | 0.006 | Young |
| 0.615681 | 0.160762 | 0.252 | 0.247 | 1 | 2 | Ranbp1 | 0.005 | Young |
| 0.615848 | 0.401461 | 0.029 | 0.025 | 1 | 2 | Pacs1 | 0.004 | Young |
| 0.616433 | 0.036058 | 0.122 | 0.116 | 1 | 2 | Akap9 | 0.006 | Young |
| 0.616584 | 0.446298 | 0.075 | 0.07 | 1 | 2 | Lipo3 | 0.005 | Young |
| 0.616719 | 0.309662 | 0.051 | 0.046 | 1 | 2 | Trem1 | 0.005 | Young |

| | | | | | | | | |
|---|---|---|---|---|---|---|---|---|
| 0.616963 | 0.055022 | 0.086 | 0.093 | 1 | 2 | Ift52 | -0.007 | Young |
| 0.61716 | 0.131113 | 0.134 | 0.127 | 1 | 2 | Ipo5 | 0.007 | Young |
| 0.617211 | 0.559088 | 0.019 | 0.016 | 1 | 2 | Mrpl47 | 0.003 | Young |
| 0.617308 | 0.774469 | 0.043 | 0.039 | 1 | 2 | Ttc4 | 0.004 | Young |
| 0.617453 | 0.349383 | 0.05 | 0.045 | 1 | 2 | Riok2 | 0.005 | Young |
| 0.617702 | 0.04617 | 0.084 | 0.078 | 1 | 2 | Gtf2f1 | 0.006 | Young |
| 0.617727 | 0.20715 | 0.181 | 0.176 | 1 | 2 | Blvrb | 0.005 | Young |
| 0.617976 | 0.312054 | 0.023 | 0.019 | 1 | 2 | Ap4b1 | 0.004 | Young |
| 0.61863 | 0.225528 | 0.071 | 0.078 | 1 | 2 | Vcam1 | -0.007 | Young |
| 0.618703 | 0.16115 | 0.026 | 0.022 | 1 | 2 | Prdm10 | 0.004 | Young |
| 0.618714 | 0.389288 | 0.048 | 0.043 | 1 | 2 | Kat5 | 0.005 | Young |
| 0.618764 | 0.349119 | 0.062 | 0.057 | 1 | 2 | Qsox1 | 0.005 | Young |
| 0.618819 | 0.211019 | 0.067 | 0.061 | 1 | 2 | Fam45a | 0.006 | Young |
| 0.619112 | 0.262171 | 0.137 | 0.131 | 1 | 2 | Chmp1b | 0.006 | Young |
| 0.619129 | 0.111391 | 0.082 | 0.09 | 1 | 2 | Hmgn2 | -0.008 | Young |
| 0.619597 | 0.09373 | 0.338 | 0.333 | 1 | 2 | Coro1c | 0.005 | Young |
| 0.619609 | 0.079202 | 0.069 | 0.063 | 1 | 2 | Smn1 | 0.006 | Young |
| 0.619911 | 0.132936 | 0.106 | 0.1 | 1 | 2 | Pou2f1 | 0.006 | Young |
| 0.620001 | 0.171038 | 0.196 | 0.212 | 1 | 2 | Map4 | -0.016 | Young |
| 0.620132 | 0.008335 | 0.721 | 0.708 | 1 | 2 | Rsrp1 | 0.013 | Young |
| 0.620353 | 0.347279 | 0.023 | 0.019 | 1 | 2 | Fmnl2 | 0.004 | Young |
| 0.620544 | 0.025587 | 0.026 | 0.022 | 1 | 2 | Hus1 | 0.004 | Young |
| 0.620562 | 0.126035 | 0.05 | 0.045 | 1 | 2 | Gphn | 0.005 | Young |
| 0.620644 | 0.129403 | 0.131 | 0.124 | 1 | 2 | Pqbp1 | 0.007 | Young |
| 0.621146 | 0.445352 | 0.023 | 0.019 | 1 | 2 | Ano8 | 0.004 | Young |
| 0.621255 | 0.378029 | 0.024 | 0.02 | 1 | 2 | Nt5m | 0.004 | Young |
| 0.621283 | 0.497227 | 0.045 | 0.041 | 1 | 2 | Smc2 | 0.004 | Young |
| 0.621565 | 0.223301 | 0.091 | 0.086 | 1 | 2 | Ppp1r35 | 0.005 | Young |
| 0.621704 | 0.223049 | 0.178 | 0.172 | 1 | 2 | Lrrfip2 | 0.006 | Young |
| 0.621777 | 0.028089 | 0.088 | 0.081 | 1 | 2 | Brwd1 | 0.007 | Young |
| 0.62179 | 0.273416 | 0.061 | 0.056 | 1 | 2 | Nr2f6 | 0.005 | Young |
| 0.62203 | 0.205249 | 0.045 | 0.041 | 1 | 2 | Cd151 | 0.004 | Young |
| 0.62211 | 0.719783 | 0.019 | 0.016 | 1 | 2 | Aldh18a1 | 0.003 | Young |
| 0.622184 | 0.001219 | 0.089 | 0.096 | 1 | 2 | Tmem41b | -0.007 | Young |
| 0.622398 | 0.756935 | 0.019 | 0.016 | 1 | 2 | Snapc2 | 0.003 | Young |
| 0.622469 | 0.305825 | 0.023 | 0.019 | 1 | 2 | Ppox | 0.004 | Young |
| 0.622469 | 0.147811 | 0.019 | 0.022 | 1 | 2 | Msh3 | -0.003 | Young |
| 0.622809 | 0.327668 | 0.048 | 0.043 | 1 | 2 | 0610037L1 | 0.005 | Young |
| 0.623158 | 0.084096 | 0.202 | 0.194 | 1 | 2 | Zfp292 | 0.008 | Young |
| 0.623187 | 0.12979 | 0.049 | 0.044 | 1 | 2 | Mrps10 | 0.005 | Young |
| 0.62321 | 0.154858 | 0.107 | 0.116 | 1 | 2 | Paxbp1 | -0.009 | Young |
| 0.623994 | 0.035414 | 0.025 | 0.029 | 1 | 2 | A530017D2 | -0.004 | Young |
| 0.624009 | 0.092386 | 0.719 | 0.706 | 1 | 2 | H2-DMa | 0.013 | Young |
| 0.624061 | 0.002414 | 0.129 | 0.123 | 1 | 2 | Nr2c2 | 0.006 | Young |
| 0.624104 | 0.050352 | 0.234 | 0.254 | 1 | 2 | Aff4 | -0.02 | Young |

| | | | | | | | | |
|---|---|---|---|---|---|---|---|---|
| 0.624142 | 0.021424 | 0.103 | 0.111 | 1 | 2 | Zfp871 | -0.008 | Young |
| 0.624268 | 0.172145 | 0.049 | 0.044 | 1 | 2 | Hsdl2 | 0.005 | Young |
| 0.624294 | 0.205815 | 0.021 | 0.018 | 1 | 2 | Coa7 | 0.003 | Young |
| 0.624538 | 0.271524 | 0.099 | 0.093 | 1 | 2 | Tfeb | 0.006 | Young |
| 0.624869 | 0.172295 | 0.024 | 0.02 | 1 | 2 | Mpi | 0.004 | Young |
| 0.624877 | 0.285053 | 0.014 | 0.017 | 1 | 2 | Pigw | -0.003 | Young |
| 0.625256 | 0.128729 | 0.038 | 0.043 | 1 | 2 | Bud23 | -0.005 | Young |
| 0.625573 | 0.270162 | 0.062 | 0.057 | 1 | 2 | Dopey2 | 0.005 | Young |
| 0.626279 | 0.158076 | 0.08 | 0.074 | 1 | 2 | Kmt5c | 0.006 | Young |
| 0.626625 | 0.048531 | 0.077 | 0.085 | 1 | 2 | Scfd1 | -0.008 | Young |
| 0.626798 | 0.025685 | 0.341 | 0.365 | 1 | 2 | Sec11c | -0.024 | Young |
| 0.62686 | 0.105614 | 0.17 | 0.182 | 1 | 2 | Nuak2 | -0.012 | Young |
| 0.62691 | 0.036061 | 0.393 | 0.384 | 1 | 2 | Zmiz1 | 0.009 | Young |
| 0.626918 | 0.56716 | 0.018 | 0.015 | 1 | 2 | Tmem9 | 0.003 | Young |
| 0.62698 | 0.034224 | 0.094 | 0.088 | 1 | 2 | Txndc15 | 0.006 | Young |
| 0.627052 | 0.081086 | 0.184 | 0.198 | 1 | 2 | Taf6l | -0.014 | Young |
| 0.627123 | 0.01684 | 0.245 | 0.257 | 1 | 2 | Klf10 | -0.012 | Young |
| 0.627126 | 0.284142 | 0.077 | 0.072 | 1 | 2 | Ppm1m | 0.005 | Young |
| 0.627213 | 0.305566 | 0.059 | 0.055 | 1 | 2 | Ints13 | 0.004 | Young |
| 0.627242 | 0.473766 | 0.023 | 0.019 | 1 | 2 | Yipf2 | 0.004 | Young |
| 0.627242 | 0.256546 | 0.023 | 0.019 | 1 | 2 | Zmym6 | 0.004 | Young |
| 0.627286 | 0.126866 | 0.07 | 0.064 | 1 | 2 | Tmem39a | 0.006 | Young |
| 0.627839 | 0.162195 | 0.021 | 0.018 | 1 | 2 | Pold2 | 0.003 | Young |
| 0.628144 | 0.20793 | 0.062 | 0.057 | 1 | 2 | Pex3 | 0.005 | Young |
| 0.628166 | 0.257187 | 0.058 | 0.054 | 1 | 2 | Terf2 | 0.004 | Young |
| 0.628275 | 0.447685 | 0.094 | 0.089 | 1 | 2 | Carhsp1 | 0.005 | Young |
| 0.628324 | 0.136231 | 0.076 | 0.071 | 1 | 2 | Med7 | 0.005 | Young |
| 0.628498 | 0.292564 | 0.134 | 0.13 | 1 | 2 | Ncor2 | 0.004 | Young |
| 0.628695 | 0.166817 | 0.196 | 0.214 | 1 | 2 | Glmp | -0.018 | Young |
| 0.628932 | 0.187159 | 0.021 | 0.018 | 1 | 2 | Alms1 | 0.003 | Young |
| 0.629138 | 0.315921 | 0.025 | 0.029 | 1 | 2 | Zfp212 | -0.004 | Young |
| 0.629226 | 0.074813 | 0.083 | 0.077 | 1 | 2 | Polg | 0.006 | Young |
| 0.629295 | 0.38761 | 0.02 | 0.017 | 1 | 2 | Urb2 | 0.003 | Young |
| 0.629376 | 0.24489 | 0.087 | 0.081 | 1 | 2 | Lsm2 | 0.006 | Young |
| 0.629602 | 0.035826 | 0.552 | 0.539 | 1 | 2 | Clec12a | 0.013 | Young |
| 0.629776 | 0.64561 | 0.017 | 0.014 | 1 | 2 | Zfp949 | 0.003 | Young |
| 0.629843 | 0.15033 | 0.156 | 0.148 | 1 | 2 | Zfp395 | 0.008 | Young |
| 0.629858 | 0.468166 | 0.02 | 0.017 | 1 | 2 | Fbxo21 | 0.003 | Young |
| 0.629858 | 0.437332 | 0.02 | 0.017 | 1 | 2 | Dis3 | 0.003 | Young |
| 0.629901 | 0.213633 | 0.019 | 0.022 | 1 | 2 | Dnajc4 | -0.003 | Young |
| 0.629919 | 0.02953 | 0.068 | 0.062 | 1 | 2 | Cyp4v3 | 0.006 | Young |
| 0.630051 | 0.036734 | 0.049 | 0.044 | 1 | 2 | Hagh | 0.005 | Young |
| 0.630102 | 0.58472 | 0.057 | 0.052 | 1 | 2 | Ranbp3 | 0.005 | Young |
| 0.63014 | 0.347633 | 0.02 | 0.017 | 1 | 2 | Dnmbp | 0.003 | Young |
| 0.630163 | 0.381946 | 0.059 | 0.055 | 1 | 2 | BC029722 | 0.004 | Young |

| | | | | | | | | |
|---|---|---|---|---|---|---|---|---|
| 0.630213 | 1.137023 | 0.018 | 0.015 | 1 | 2 | Cxcl9 | 0.003 | Young |
| 0.630878 | 0.583228 | 0.042 | 0.037 | 1 | 2 | Aimp2 | 0.005 | Young |
| 0.631114 | 0.535542 | 0.018 | 0.015 | 1 | 2 | Zfp607a | 0.003 | Young |
| 0.63133 | 0.710059 | 0.017 | 0.014 | 1 | 2 | Tgif2 | 0.003 | Young |
| 0.631393 | 0.002035 | 0.021 | 0.018 | 1 | 2 | BC147527 | 0.003 | Young |
| 0.631414 | 0.471723 | 0.018 | 0.015 | 1 | 2 | Card11 | 0.003 | Young |
| 0.631569 | 0.097837 | 0.084 | 0.092 | 1 | 2 | Mfsd10 | -0.008 | Young |
| 0.631604 | 0.146683 | 0.02 | 0.024 | 1 | 2 | Zfp444 | -0.004 | Young |
| 0.631984 | 0.087557 | 0.241 | 0.233 | 1 | 2 | R3hdm4 | 0.008 | Young |
| 0.632215 | 0.101599 | 0.021 | 0.018 | 1 | 2 | Top2a | 0.003 | Young |
| 0.632385 | 0.232834 | 0.131 | 0.124 | 1 | 2 | Fbrs | 0.007 | Young |
| 0.632393 | 0.248245 | 0.02 | 0.017 | 1 | 2 | Dut | 0.003 | Young |
| 0.632626 | 0.112902 | 0.077 | 0.072 | 1 | 2 | Hsdl1 | 0.005 | Young |
| 0.632804 | 0.059664 | 0.014 | 0.017 | 1 | 2 | Nmrk1 | -0.003 | Young |
| 0.633 | 0.651573 | 0.015 | 0.013 | 1 | 2 | Purg | 0.002 | Young |
| 0.633364 | 0.098796 | 0.023 | 0.019 | 1 | 2 | Mrps31 | 0.004 | Young |
| 0.633557 | 0.264108 | 0.087 | 0.081 | 1 | 2 | Cul4a | 0.006 | Young |
| 0.635109 | 0.16658 | 0.096 | 0.106 | 1 | 2 | Qrich1 | -0.01 | Young |
| 0.635181 | 0.173523 | 0.24 | 0.233 | 1 | 2 | Srp72 | 0.007 | Young |
| 0.635251 | 0.182062 | 0.014 | 0.017 | 1 | 2 | Pdlim7 | -0.003 | Young |
| 0.635396 | 0.014169 | 0.382 | 0.381 | 1 | 2 | Selenof | 0.001 | Young |
| 0.635422 | 0.171071 | 0.059 | 0.055 | 1 | 2 | Pop4 | 0.004 | Young |
| 0.635876 | 0.199348 | 0.058 | 0.054 | 1 | 2 | Pitrm1 | 0.004 | Young |
| 0.636236 | 1.133048 | 0.012 | 0.01 | 1 | 2 | Efnb2 | 0.002 | Young |
| 0.636249 | 0.642985 | 0.014 | 0.012 | 1 | 2 | Gcat | 0.002 | Young |
| 0.636313 | 0.450535 | 0.017 | 0.014 | 1 | 2 | Bco2 | 0.003 | Young |
| 0.63656 | 0.384333 | 0.015 | 0.013 | 1 | 2 | Msl3l2 | 0.002 | Young |
| 0.636716 | 0.030922 | 0.093 | 0.1 | 1 | 2 | Znfx1 | -0.007 | Young |
| 0.636826 | 0.160813 | 0.044 | 0.04 | 1 | 2 | Ttll12 | 0.004 | Young |
| 0.637133 | 1.324777 | 0.011 | 0.009 | 1 | 2 | Klrd1 | 0.002 | Young |
| 0.637153 | 0.022612 | 0.021 | 0.018 | 1 | 2 | Osbpl1a | 0.003 | Young |
| 0.637195 | 0.010106 | 0.046 | 0.051 | 1 | 2 | Acot7 | -0.005 | Young |
| 0.637249 | 0.506707 | 0.017 | 0.014 | 1 | 2 | Gm41307 | 0.003 | Young |
| 0.637261 | 0.068051 | 0.169 | 0.18 | 1 | 2 | Ndufb2 | -0.011 | Young |
| 0.637525 | 1.084822 | 0.011 | 0.009 | 1 | 2 | Dusp28 | 0.002 | Young |
| 0.637644 | 0.054173 | 0.173 | 0.185 | 1 | 2 | Gnl3 | -0.012 | Young |
| 0.637857 | 0.459797 | 0.015 | 0.013 | 1 | 2 | Asgr2 | 0.002 | Young |
| 0.637881 | 0.031371 | 0.075 | 0.081 | 1 | 2 | Hk3 | -0.006 | Young |
| 0.638125 | 0.674537 | 0.01 | 0.007 | 1 | 2 | Nhej1 | 0.003 | Young |
| 0.638506 | 0.496569 | 0.015 | 0.013 | 1 | 2 | Trp53rka | 0.002 | Young |
| 0.639133 | 0.271317 | 0.164 | 0.161 | 1 | 2 | Tomm34 | 0.003 | Young |
| 0.639152 | 0.19633 | 0.132 | 0.126 | 1 | 2 | Mitf | 0.006 | Young |
| 0.639436 | 0.285004 | 0.017 | 0.014 | 1 | 2 | Tsen2 | 0.003 | Young |
| 0.639464 | 0.519151 | 0.013 | 0.011 | 1 | 2 | AI480526 | 0.002 | Young |
| 0.639494 | 0.077551 | 0.048 | 0.043 | 1 | 2 | Tfdp1 | 0.005 | Young |

| | | | | | | | | |
|---|---|---|---|---|---|---|---|---|
| 0.639713 | 0.164679 | 0.258 | 0.253 | 1 | 2 | Eif3l | 0.005 | Young |
| 0.639796 | 0.87644 | 0.01 | 0.007 | 1 | 2 | Cebpe | 0.003 | Young |
| 0.639881 | 0.905525 | 0.011 | 0.009 | 1 | 2 | Rsph3b | 0.002 | Young |
| 0.640176 | 0.101282 | 0.099 | 0.107 | 1 | 2 | Mrpl21 | -0.008 | Young |
| 0.640274 | 0.921411 | 0.011 | 0.009 | 1 | 2 | Fam216a | 0.002 | Young |
| 0.640652 | 0.082952 | 0.039 | 0.044 | 1 | 2 | Ankrd46 | -0.005 | Young |
| 0.640667 | 0.51583 | 0.011 | 0.009 | 1 | 2 | 4833418N( | 0.002 | Young |
| 0.640667 | 0.439884 | 0.011 | 0.009 | 1 | 2 | Ccl5 | 0.002 | Young |
| 0.640691 | 0.234077 | 0.078 | 0.073 | 1 | 2 | Nsmce1 | 0.005 | Young |
| 0.640695 | 0.491659 | 0.012 | 0.01 | 1 | 2 | Tbc1d7 | 0.002 | Young |
| 0.641051 | 0.697146 | 0.01 | 0.007 | 1 | 2 | Arvcf | 0.003 | Young |
| 0.64114 | 0.220033 | 0.086 | 0.08 | 1 | 2 | Thap12 | 0.006 | Young |
| 0.641429 | 0.36149 | 0.015 | 0.013 | 1 | 2 | Dus2 | 0.002 | Young |
| 0.641454 | 0.505584 | 0.011 | 0.009 | 1 | 2 | Tmem220 | 0.002 | Young |
| 0.641754 | 0.193344 | 0.015 | 0.013 | 1 | 2 | Ecsit | 0.002 | Young |
| 0.641847 | 0.492327 | 0.011 | 0.009 | 1 | 2 | Ptpn21 | 0.002 | Young |
| 0.641888 | 0.408071 | 0.01 | 0.007 | 1 | 2 | Dennd5b | 0.003 | Young |
| 0.641999 | 0.35379 | 0.014 | 0.012 | 1 | 2 | 1810044D( | 0.002 | Young |
| 0.642187 | 0.213213 | 0.12 | 0.113 | 1 | 2 | Iah1 | 0.007 | Young |
| 0.642544 | 0.182776 | 0.19 | 0.185 | 1 | 2 | Plekhm3 | 0.005 | Young |
| 0.642557 | 0.401704 | 0.012 | 0.01 | 1 | 2 | Lamc1 | 0.002 | Young |
| 0.642795 | 0.10448 | 0.338 | 0.333 | 1 | 2 | Myo9b | 0.005 | Young |
| 0.642929 | 0.002207 | 0.01 | 0.012 | 1 | 2 | Ggcx | -0.002 | Young |
| 0.643145 | 0.292163 | 0.01 | 0.007 | 1 | 2 | 3222401L1 | 0.003 | Young |
| 0.643458 | 0.022167 | 0.262 | 0.256 | 1 | 2 | Ppp1r14b | 0.006 | Young |
| 0.643564 | 0.318099 | 0.01 | 0.007 | 1 | 2 | Tmem221 | 0.003 | Young |
| 0.643675 | 0.254961 | 0.012 | 0.01 | 1 | 2 | Lepr | 0.002 | Young |
| 0.643694 | 0.116164 | 0.014 | 0.012 | 1 | 2 | Hrh1 | 0.002 | Young |
| 0.643806 | 0.261892 | 0.186 | 0.183 | 1 | 2 | Tmem219 | 0.003 | Young |
| 0.644071 | 0.292272 | 0.013 | 0.011 | 1 | 2 | Fam187b | 0.002 | Young |
| 0.644289 | 0.199248 | 0.268 | 0.268 | 1 | 2 | Ap2b1 | 0 | Young |
| 0.644402 | 0.328878 | 0.01 | 0.007 | 1 | 2 | Pex10 | 0.003 | Young |
| 0.644402 | 0.286124 | 0.01 | 0.007 | 1 | 2 | Hells | 0.003 | Young |
| 0.644426 | 0.242069 | 0.013 | 0.011 | 1 | 2 | Car9 | 0.002 | Young |
| 0.644684 | 0.126643 | 0.015 | 0.013 | 1 | 2 | Ccdc171 | 0.002 | Young |
| 0.644712 | 0.511346 | 0.014 | 0.012 | 1 | 2 | Fggy | 0.002 | Young |
| 0.644781 | 0.389619 | 0.013 | 0.011 | 1 | 2 | Hdhd5 | 0.002 | Young |
| 0.644821 | 0.23056 | 0.01 | 0.007 | 1 | 2 | Rnf122 | 0.003 | Young |
| 0.644821 | 0.05286 | 0.01 | 0.007 | 1 | 2 | Pcgf6 | 0.003 | Young |
| 0.644882 | 0.252212 | 0.014 | 0.012 | 1 | 2 | Bckdhb | 0.002 | Young |
| 0.644934 | 0.087481 | 0.249 | 0.242 | 1 | 2 | Ubn1 | 0.007 | Young |
| 0.644998 | 0.3726 | 0.011 | 0.009 | 1 | 2 | Ube2c | 0.002 | Young |
| 0.645097 | 0.136931 | 0.251 | 0.247 | 1 | 2 | Ski | 0.004 | Young |
| 0.645178 | 0.020262 | 0.045 | 0.041 | 1 | 2 | Nadk2 | 0.004 | Young |
| 0.645645 | 0.433907 | 0.055 | 0.05 | 1 | 2 | Cpsf4 | 0.005 | Young |

| | | | | | | | | |
|---|---|---|---|---|---|---|---|---|
| 0.645787 | 0.639856 | 0.011 | 0.009 | 1 | 2 | Zbtb25 | 0.002 | Young |
| 0.645913 | 0.057674 | 0.01 | 0.012 | 1 | 2 | Haus5 | -0.002 | Young |
| 0.646181 | 0.062409 | 0.011 | 0.009 | 1 | 2 | Cdca7l | 0.002 | Young |
| 0.646287 | 0.257629 | 0.012 | 0.01 | 1 | 2 | Mmab | 0.002 | Young |
| 0.646576 | 0.156233 | 0.011 | 0.009 | 1 | 2 | Ubtd2 | 0.002 | Young |
| 0.64668 | 0.19667 | 0.04 | 0.036 | 1 | 2 | Zfp821 | 0.004 | Young |
| 0.646913 | 0.228552 | 0.013 | 0.011 | 1 | 2 | A430078G2 | 0.002 | Young |
| 0.647034 | 0.393463 | 0.012 | 0.01 | 1 | 2 | Gcfc2 | 0.002 | Young |
| 0.647052 | 0.160323 | 0.14 | 0.151 | 1 | 2 | Acaa1a | -0.011 | Young |
| 0.647366 | 0.25695 | 0.011 | 0.009 | 1 | 2 | Zfp707 | 0.002 | Young |
| 0.647676 | 0.239102 | 0.12 | 0.113 | 1 | 2 | Arid5b | 0.007 | Young |
| 0.647761 | 0.008152 | 0.011 | 0.009 | 1 | 2 | Pusl1 | 0.002 | Young |
| 0.647781 | 0.042297 | 0.012 | 0.01 | 1 | 2 | Trp53rkb | 0.002 | Young |
| 0.647785 | 0.184064 | 0.121 | 0.116 | 1 | 2 | Ctnnbip1 | 0.005 | Young |
| 0.648155 | 0.22818 | 0.012 | 0.01 | 1 | 2 | Cby1 | 0.002 | Young |
| 0.648155 | 0.050545 | 0.01 | 0.012 | 1 | 2 | Zc3h11a | -0.002 | Young |
| 0.648212 | 0.095664 | 0.164 | 0.156 | 1 | 2 | Zrsr2 | 0.008 | Young |
| 0.648336 | 0.067754 | 0.013 | 0.011 | 1 | 2 | Mamld1 | 0.002 | Young |
| 0.648529 | 0.48208 | 0.012 | 0.01 | 1 | 2 | Fbxo10 | 0.002 | Young |
| 0.648529 | 0.187495 | 0.012 | 0.01 | 1 | 2 | 2900060B1 | 0.002 | Young |
| 0.648529 | 0.184363 | 0.01 | 0.012 | 1 | 2 | Ubiad1 | -0.002 | Young |
| 0.648551 | 0.241689 | 0.011 | 0.009 | 1 | 2 | Btbd3 | 0.002 | Young |
| 0.648626 | 0.231256 | 0.102 | 0.097 | 1 | 2 | Ado | 0.005 | Young |
| 0.648725 | 0.315951 | 0.039 | 0.035 | 1 | 2 | Utp4 | 0.004 | Young |
| 0.648747 | 0.033249 | 0.853 | 0.85 | 1 | 2 | Lamp1 | 0.003 | Young |
| 0.649251 | 0.115376 | 0.015 | 0.013 | 1 | 2 | Cox7a1 | 0.002 | Young |
| 0.649267 | 0.128501 | 0.09 | 0.085 | 1 | 2 | Axin1 | 0.005 | Young |
| 0.649277 | 0.162615 | 0.012 | 0.01 | 1 | 2 | Mpv17l | 0.002 | Young |
| 0.649277 | 0.124852 | 0.01 | 0.012 | 1 | 2 | Mks1 | -0.002 | Young |
| 0.649341 | 0.089027 | 0.011 | 0.009 | 1 | 2 | Slc5a6 | 0.002 | Young |
| 0.649597 | 0.020887 | 0.121 | 0.115 | 1 | 2 | Pcmtd2 | 0.006 | Young |
| 0.649792 | 0.107277 | 0.027 | 0.031 | 1 | 2 | Map4k3 | -0.004 | Young |
| 0.649966 | 0.057349 | 0.099 | 0.106 | 1 | 2 | Zfp984 | -0.007 | Young |
| 0.649984 | 0.03978 | 0.214 | 0.208 | 1 | 2 | Noc2l | 0.006 | Young |
| 0.650664 | 0.078118 | 0.355 | 0.349 | 1 | 2 | Stxbp3 | 0.006 | Young |
| 0.650757 | 0.083384 | 0.061 | 0.056 | 1 | 2 | Chchd5 | 0.005 | Young |
| 0.651412 | 0.094913 | 0.115 | 0.109 | 1 | 2 | Papd5 | 0.006 | Young |
| 0.651475 | 0.2423 | 0.059 | 0.055 | 1 | 2 | Fopnl | 0.004 | Young |
| 0.651574 | 0.111484 | 0.055 | 0.06 | 1 | 2 | Arsk | -0.005 | Young |
| 0.651609 | 0.153386 | 0.061 | 0.066 | 1 | 2 | As3mt | -0.005 | Young |
| 0.651672 | 0.028467 | 0.279 | 0.293 | 1 | 2 | Rab6a | -0.014 | Young |
| 0.651899 | 0.016231 | 0.013 | 0.011 | 1 | 2 | Gm17586 | 0.002 | Young |
| 0.652157 | 0.337782 | 0.106 | 0.101 | 1 | 2 | Dido1 | 0.005 | Young |
| 0.653197 | 0.60221 | 0.051 | 0.047 | 1 | 2 | Nudt19 | 0.004 | Young |
| 0.653761 | 0.024062 | 0.209 | 0.223 | 1 | 2 | Cmklr1 | -0.014 | Young |

| | | | | | | | | |
|---|---|---|---|---|---|---|---|---|
| 0.653902 | 0.030765 | 0.158 | 0.168 | 1 | 2 | Lrrc59 | -0.01 | Young |
| 0.654245 | 0.099522 | 0.179 | 0.172 | 1 | 2 | Clcn4 | 0.007 | Young |
| 0.654383 | 0.175808 | 0.075 | 0.07 | 1 | 2 | Abcb1b | 0.005 | Young |
| 0.654427 | 0.090264 | 0.114 | 0.108 | 1 | 2 | Ticam2 | 0.006 | Young |
| 0.654444 | 0.06586 | 0.082 | 0.076 | 1 | 2 | Exosc8 | 0.006 | Young |
| 0.654888 | 0.008874 | 0.044 | 0.04 | 1 | 2 | Nt5dc1 | 0.004 | Young |
| 0.654986 | 0.195903 | 0.046 | 0.051 | 1 | 2 | Tor2a | -0.005 | Young |
| 0.655012 | 0.053284 | 0.184 | 0.178 | 1 | 2 | Abce1 | 0.006 | Young |
| 0.655344 | 0.164856 | 0.067 | 0.073 | 1 | 2 | Ccdc34 | -0.006 | Young |
| 0.655501 | 0.004081 | 0.368 | 0.358 | 1 | 2 | Nedd8 | 0.01 | Young |
| 0.655683 | 0.002242 | 0.126 | 0.134 | 1 | 2 | Psme2b | -0.008 | Young |
| 0.655723 | 0.240833 | 0.072 | 0.067 | 1 | 2 | Ssna1 | 0.005 | Young |
| 0.655767 | 0.042978 | 0.16 | 0.171 | 1 | 2 | Uba6 | -0.011 | Young |
| 0.656313 | 0.099602 | 0.103 | 0.097 | 1 | 2 | Ahctf1 | 0.006 | Young |
| 0.656388 | 0.094716 | 0.12 | 0.128 | 1 | 2 | Stx11 | -0.008 | Young |
| 0.656426 | 0.218325 | 0.148 | 0.142 | 1 | 2 | Ncoa7 | 0.006 | Young |
| 0.656613 | 0.032476 | 0.248 | 0.264 | 1 | 2 | Rnf4 | -0.016 | Young |
| 0.656616 | 0.278629 | 0.061 | 0.056 | 1 | 2 | Pdlim2 | 0.005 | Young |
| 0.65678 | 0.014292 | 0.131 | 0.124 | 1 | 2 | Prdm2 | 0.007 | Young |
| 0.657401 | 0.128967 | 0.081 | 0.076 | 1 | 2 | Ap5m1 | 0.005 | Young |
| 0.657931 | 0.216842 | 0.07 | 0.065 | 1 | 2 | Rufy1 | 0.005 | Young |
| 0.658275 | 0.278382 | 0.052 | 0.048 | 1 | 2 | Cog6 | 0.004 | Young |
| 0.658731 | 0.026519 | 0.215 | 0.231 | 1 | 2 | Dnase2a | -0.016 | Young |
| 0.658741 | 0.075489 | 0.327 | 0.338 | 1 | 2 | Ythdc1 | -0.011 | Young |
| 0.658784 | 0.09183 | 0.163 | 0.173 | 1 | 2 | Hipk3 | -0.01 | Young |
| 0.658955 | 0.030414 | 0.021 | 0.025 | 1 | 2 | Selenoo | -0.004 | Young |
| 0.659057 | 0.050323 | 0.075 | 0.081 | 1 | 2 | Nvl | -0.006 | Young |
| 0.659102 | 0.040082 | 0.26 | 0.273 | 1 | 2 | Snx9 | -0.013 | Young |
| 0.659341 | 0.136002 | 0.021 | 0.025 | 1 | 2 | Ift57 | -0.004 | Young |
| 0.659437 | 0.017599 | 0.133 | 0.142 | 1 | 2 | Nsmce4a | -0.009 | Young |
| 0.659604 | 0.196946 | 0.089 | 0.085 | 1 | 2 | Ubxn2a | 0.004 | Young |
| 0.660381 | 0.376721 | 0.034 | 0.031 | 1 | 2 | Cxcl12 | 0.003 | Young |
| 0.660449 | 0.130587 | 0.274 | 0.27 | 1 | 2 | Cysltr1 | 0.004 | Young |
| 0.66106 | 0.287894 | 0.083 | 0.078 | 1 | 2 | Lactb2 | 0.005 | Young |
| 0.661181 | 0.298816 | 0.037 | 0.033 | 1 | 2 | Slc37a4 | 0.004 | Young |
| 0.661291 | 0.216794 | 0.082 | 0.077 | 1 | 2 | Lztfl1 | 0.005 | Young |
| 0.661403 | 0.030566 | 0.312 | 0.312 | 1 | 2 | Bag1 | 0 | Young |
| 0.661405 | 0.197442 | 0.015 | 0.018 | 1 | 2 | Btbd19 | -0.003 | Young |
| 0.662098 | 0.092684 | 0.127 | 0.136 | 1 | 2 | Erap1 | -0.009 | Young |
| 0.662204 | 0.110073 | 0.062 | 0.067 | 1 | 2 | Zfp787 | -0.005 | Young |
| 0.662212 | 0.189511 | 0.188 | 0.183 | 1 | 2 | Slc7a8 | 0.005 | Young |
| 0.662493 | 0.333808 | 0.094 | 0.089 | 1 | 2 | Zdhhc5 | 0.005 | Young |
| 0.663253 | 0.056772 | 0.106 | 0.115 | 1 | 2 | Bcat2 | -0.009 | Young |
| 0.663564 | 0.227481 | 0.064 | 0.071 | 1 | 2 | Mphosph1 | -0.007 | Young |
| 0.664449 | 0.124065 | 0.038 | 0.034 | 1 | 2 | Drg2 | 0.004 | Young |

| | | | | | | | | |
|---|---|---|---|---|---|---|---|---|
| 0.664987 | 0.345598 | 0.037 | 0.033 | 1 | 2 | 5031425F1 | 0.004 | Young |
| 0.665113 | 0.302412 | 0.131 | 0.126 | 1 | 2 | Gan | 0.005 | Young |
| 0.665173 | 0.385878 | 0.059 | 0.055 | 1 | 2 | Il1rap | 0.004 | Young |
| 0.665176 | 0.215413 | 0.082 | 0.089 | 1 | 2 | Tsc1 | -0.007 | Young |
| 0.66535 | 0.058848 | 0.09 | 0.085 | 1 | 2 | Cdk8 | 0.005 | Young |
| 0.665493 | 0.14867 | 0.297 | 0.294 | 1 | 2 | Ttc14 | 0.003 | Young |
| 0.665569 | 0.219323 | 0.086 | 0.093 | 1 | 2 | Tmem86a | -0.007 | Young |
| 0.665736 | 0.080836 | 0.217 | 0.215 | 1 | 2 | Tmf1 | 0.002 | Young |
| 0.665802 | 0.145779 | 0.021 | 0.025 | 1 | 2 | Klhl22 | -0.004 | Young |
| 0.665881 | 0.069216 | 0.433 | 0.424 | 1 | 2 | Cct5 | 0.009 | Young |
| 0.665909 | 0.186071 | 0.038 | 0.034 | 1 | 2 | Tmem70 | 0.004 | Young |
| 0.665916 | 0.009739 | 0.04 | 0.036 | 1 | 2 | Dzip3 | 0.004 | Young |
| 0.665931 | 0.018363 | 0.361 | 0.353 | 1 | 2 | Anp32e | 0.008 | Young |
| 0.66594 | 0.406955 | 0.037 | 0.033 | 1 | 2 | Aktip | 0.004 | Young |
| 0.66595 | 0.443343 | 0.034 | 0.031 | 1 | 2 | Cyb561d2 | 0.003 | Young |
| 0.666109 | 0.072915 | 0.074 | 0.069 | 1 | 2 | Il16 | 0.005 | Young |
| 0.66659 | 0.331581 | 0.099 | 0.094 | 1 | 2 | Fam162a | 0.005 | Young |
| 0.666631 | 0.08071 | 0.048 | 0.052 | 1 | 2 | Usp10 | -0.004 | Young |
| 0.667373 | 0.318595 | 0.034 | 0.031 | 1 | 2 | Zbtb14 | 0.003 | Young |
| 0.667558 | 0.04031 | 0.154 | 0.164 | 1 | 2 | Lrpap1 | -0.01 | Young |
| 0.668105 | 0.011498 | 0.366 | 0.352 | 1 | 2 | Socs3 | 0.014 | Young |
| 0.668229 | 0.0729 | 0.253 | 0.248 | 1 | 2 | Psmc3 | 0.005 | Young |
| 0.668271 | 0.044628 | 0.063 | 0.069 | 1 | 2 | Mnat1 | -0.006 | Young |
| 0.668301 | 0.20793 | 0.056 | 0.051 | 1 | 2 | Sord | 0.005 | Young |
| 0.668395 | 0.027698 | 0.374 | 0.385 | 1 | 2 | Clec4e | -0.011 | Young |
| 0.669011 | 0.162346 | 0.095 | 0.09 | 1 | 2 | Arhgap12 | 0.005 | Young |
| 0.669013 | 0.556618 | 0.057 | 0.054 | 1 | 2 | Smim26 | 0.003 | Young |
| 0.669127 | 0.172497 | 0.057 | 0.062 | 1 | 2 | Sdhaf2 | -0.005 | Young |
| 0.669258 | 0.547968 | 0.033 | 0.03 | 1 | 2 | Depdc5 | 0.003 | Young |
| 0.669419 | 0.013848 | 0.2 | 0.214 | 1 | 2 | Ogfrl1 | -0.014 | Young |
| 0.66953 | 0.148051 | 0.15 | 0.146 | 1 | 2 | Vps13a | 0.004 | Young |
| 0.669574 | 0.110212 | 0.042 | 0.046 | 1 | 2 | Cog4 | -0.004 | Young |
| 0.670426 | 0.108607 | 0.106 | 0.101 | 1 | 2 | Polr3c | 0.005 | Young |
| 0.670564 | 0.113121 | 0.055 | 0.05 | 1 | 2 | Inip | 0.005 | Young |
| 0.670605 | 0.042867 | 0.26 | 0.253 | 1 | 2 | Ptpa | 0.007 | Young |
| 0.670939 | 0.32324 | 0.027 | 0.031 | 1 | 2 | Kptn | -0.004 | Young |
| 0.671041 | 0.114971 | 0.062 | 0.057 | 1 | 2 | Dnajc16 | 0.005 | Young |
| 0.671252 | 0.315143 | 0.039 | 0.035 | 1 | 2 | Tecpr2 | 0.004 | Young |
| 0.671439 | 0.319358 | 0.063 | 0.059 | 1 | 2 | Zfpl1 | 0.004 | Young |
| 0.671673 | 0.100348 | 0.17 | 0.165 | 1 | 2 | Glipr2 | 0.005 | Young |
| 0.671967 | 0.073909 | 0.21 | 0.222 | 1 | 2 | Rc3h2 | -0.012 | Young |
| 0.672158 | 0.130203 | 0.295 | 0.29 | 1 | 2 | Supt16 | 0.005 | Young |
| 0.672577 | 0.89612 | 0.015 | 0.018 | 1 | 2 | Rmdn1 | -0.003 | Young |
| 0.673052 | 0.328703 | 0.033 | 0.03 | 1 | 2 | Gm14548 | 0.003 | Young |
| 0.673072 | 0.438485 | 0.126 | 0.124 | 1 | 2 | 1810022K0 | 0.002 | Young |

| | | | | | | | | |
|---|---|---|---|---|---|---|---|---|
| 0.673218 | 0.164111 | 0.215 | 0.213 | 1 | 2 | Chmp3 | 0.002 | Young |
| 0.673219 | 0.039826 | 0.886 | 0.88 | 1 | 2 | Sem1 | 0.006 | Young |
| 0.673587 | 0.402549 | 0.021 | 0.025 | 1 | 2 | Mlh3 | -0.004 | Young |
| 0.673623 | 0.027104 | 0.072 | 0.067 | 1 | 2 | Cnnm4 | 0.005 | Young |
| 0.67392 | 0.06451 | 0.077 | 0.084 | 1 | 2 | Pdcl | -0.007 | Young |
| 0.674396 | 0.509279 | 0.046 | 0.043 | 1 | 2 | Hes6 | 0.003 | Young |
| 0.674453 | 0.05788 | 0.382 | 0.408 | 1 | 2 | Pepd | -0.026 | Young |
| 0.675236 | 0.472855 | 0.051 | 0.047 | 1 | 2 | Gdpd3 | 0.004 | Young |
| 0.675408 | 0.430981 | 0.021 | 0.025 | 1 | 2 | Tbcc | -0.004 | Young |
| 0.675573 | 0.249814 | 0.023 | 0.026 | 1 | 2 | Mcm2 | -0.003 | Young |
| 0.675741 | 0.196103 | 0.038 | 0.034 | 1 | 2 | Bag5 | 0.004 | Young |
| 0.675798 | 0.045574 | 0.588 | 0.596 | 1 | 2 | Ddx17 | -0.008 | Young |
| 0.676246 | 0.040065 | 0.122 | 0.117 | 1 | 2 | Sf3b4 | 0.005 | Young |
| 0.676338 | 0.025566 | 0.062 | 0.057 | 1 | 2 | Pole3 | 0.005 | Young |
| 0.67682 | 0.046829 | 0.176 | 0.171 | 1 | 2 | Emc3 | 0.005 | Young |
| 0.677138 | 0.383124 | 0.055 | 0.05 | 1 | 2 | Ppp1r3b | 0.005 | Young |
| 0.677181 | 0.091871 | 0.102 | 0.109 | 1 | 2 | Gars | -0.007 | Young |
| 0.677669 | 0.118398 | 0.207 | 0.204 | 1 | 2 | Ecm1 | 0.003 | Young |
| 0.677932 | 0.352734 | 0.029 | 0.032 | 1 | 2 | Tars2 | -0.003 | Young |
| 0.678039 | 0.411404 | 0.048 | 0.044 | 1 | 2 | Slc46a3 | 0.004 | Young |
| 0.67827 | 0.102343 | 0.037 | 0.033 | 1 | 2 | Pgm2 | 0.004 | Young |
| 0.678628 | 0.363326 | 0.067 | 0.062 | 1 | 2 | Eif2b4 | 0.005 | Young |
| 0.679012 | 0.124675 | 0.078 | 0.085 | 1 | 2 | E2f4 | -0.007 | Young |
| 0.679072 | 0.110447 | 0.138 | 0.131 | 1 | 2 | Rbm28 | 0.007 | Young |
| 0.679315 | 0.173316 | 0.051 | 0.047 | 1 | 2 | Zbtb34 | 0.004 | Young |
| 0.679599 | 0.002836 | 0.05 | 0.055 | 1 | 2 | Brcc3 | -0.005 | Young |
| 0.679656 | 0.295226 | 0.033 | 0.03 | 1 | 2 | Clcf1 | 0.003 | Young |
| 0.679677 | 0.115372 | 0.034 | 0.031 | 1 | 2 | Alg13 | 0.003 | Young |
| 0.679957 | 0.376829 | 0.011 | 0.013 | 1 | 2 | Fahd2a | -0.002 | Young |
| 0.679957 | 0.131949 | 0.011 | 0.013 | 1 | 2 | Nmnat1 | -0.002 | Young |
| 0.680683 | 0.406156 | 0.011 | 0.013 | 1 | 2 | Cryz | -0.002 | Young |
| 0.680683 | 0.369323 | 0.011 | 0.013 | 1 | 2 | Gpr157 | -0.002 | Young |
| 0.680716 | 0.062271 | 0.081 | 0.087 | 1 | 2 | Mir142hg | -0.006 | Young |
| 0.680765 | 0.154814 | 0.043 | 0.047 | 1 | 2 | Jpt2 | -0.004 | Young |
| 0.680918 | 0.082813 | 0.425 | 0.413 | 1 | 2 | Man2b1 | 0.012 | Young |
| 0.681045 | 0.11874 | 0.011 | 0.013 | 1 | 2 | Arsi | -0.002 | Young |
| 0.681263 | 0.383671 | 0.031 | 0.028 | 1 | 2 | Zfp160 | 0.003 | Young |
| 0.681424 | 0.189611 | 0.065 | 0.061 | 1 | 2 | Pabpc4 | 0.004 | Young |
| 0.681549 | 0.154278 | 0.122 | 0.117 | 1 | 2 | Utp14a | 0.005 | Young |
| 0.68163 | 0.200344 | 0.156 | 0.152 | 1 | 2 | Slc25a24 | 0.004 | Young |
| 0.682135 | 0.099492 | 0.093 | 0.088 | 1 | 2 | Mta3 | 0.005 | Young |
| 0.682858 | 0.105739 | 0.242 | 0.236 | 1 | 2 | Rcc2 | 0.006 | Young |
| 0.682899 | 0.027138 | 0.062 | 0.057 | 1 | 2 | Scamp5 | 0.005 | Young |
| 0.683047 | 0.503278 | 0.089 | 0.086 | 1 | 2 | Zswim4 | 0.003 | Young |
| 0.68313 | 0.167499 | 0.089 | 0.097 | 1 | 2 | Bahd1 | -0.008 | Young |

| | | | | | | | | |
|---|---|---|---|---|---|---|---|---|
| 0.683693 | 0.021831 | 0.669 | 0.651 | 1 | 2 | Gm2a | 0.018 | Young |
| 0.683742 | 0.26435 | 0.065 | 0.072 | 1 | 2 | Ggps1 | -0.007 | Young |
| 0.683761 | 0.208172 | 0.034 | 0.031 | 1 | 2 | Rab6b | 0.003 | Young |
| 0.683966 | 0.214854 | 0.078 | 0.074 | 1 | 2 | Cdkn2aipnl | 0.004 | Young |
| 0.684147 | 0.17731 | 0.03 | 0.033 | 1 | 2 | Med31 | -0.003 | Young |
| 0.684363 | 0.030636 | 0.057 | 0.052 | 1 | 2 | Rnf19a | 0.005 | Young |
| 0.685046 | 0.157997 | 0.167 | 0.164 | 1 | 2 | Anapc5 | 0.003 | Young |
| 0.685818 | 0.01126 | 0.118 | 0.112 | 1 | 2 | Arid2 | 0.006 | Young |
| 0.685885 | 0.198365 | 0.346 | 0.354 | 1 | 2 | Rnf13 | -0.008 | Young |
| 0.685956 | 0.007638 | 0.139 | 0.133 | 1 | 2 | Slc36a1 | 0.006 | Young |
| 0.68606 | 0.595293 | 0.081 | 0.076 | 1 | 2 | Tgm2 | 0.005 | Young |
| 0.686076 | 0.160014 | 0.189 | 0.187 | 1 | 2 | Emd | 0.002 | Young |
| 0.68673 | 0.136898 | 0.043 | 0.047 | 1 | 2 | Gm45902 | -0.004 | Young |
| 0.687103 | 0.012325 | 0.036 | 0.032 | 1 | 2 | Fam173b | 0.004 | Young |
| 0.687166 | 0.043526 | 0.215 | 0.209 | 1 | 2 | Mrps14 | 0.006 | Young |
| 0.687354 | 0.101961 | 0.05 | 0.055 | 1 | 2 | Timm29 | -0.005 | Young |
| 0.687662 | 0.130105 | 0.088 | 0.084 | 1 | 2 | Znrf1 | 0.004 | Young |
| 0.688086 | 0.03296 | 0.958 | 0.94 | 1 | 2 | Zeb2 | 0.018 | Young |
| 0.688256 | 0.033498 | 0.139 | 0.134 | 1 | 2 | Ncaph2 | 0.005 | Young |
| 0.68838 | 0.162659 | 0.159 | 0.154 | 1 | 2 | Rftn1 | 0.005 | Young |
| 0.688572 | 0.173403 | 0.053 | 0.049 | 1 | 2 | Arhgap6 | 0.004 | Young |
| 0.68861 | 0.174091 | 0.044 | 0.048 | 1 | 2 | Zfp362 | -0.004 | Young |
| 0.688826 | 0.052666 | 0.074 | 0.079 | 1 | 2 | Dnmt1 | -0.005 | Young |
| 0.688918 | 0.006528 | 0.347 | 0.364 | 1 | 2 | Tep1 | -0.017 | Young |
| 0.688927 | 0.00453 | 0.192 | 0.184 | 1 | 2 | Bmpr2 | 0.008 | Young |
| 0.689155 | 0.171676 | 0.215 | 0.211 | 1 | 2 | Saraf | 0.004 | Young |
| 0.689497 | 0.008328 | 0.12 | 0.126 | 1 | 2 | Tmem184b | -0.006 | Young |
| 0.690317 | 0.528777 | 0.03 | 0.027 | 1 | 2 | Gna15 | 0.003 | Young |
| 0.69039 | 0.415664 | 0.023 | 0.026 | 1 | 2 | Zfp868 | -0.003 | Young |
| 0.690538 | 0.183167 | 0.176 | 0.171 | 1 | 2 | Gtpbp2 | 0.005 | Young |
| 0.691018 | 0.161321 | 0.033 | 0.03 | 1 | 2 | Fam122a | 0.003 | Young |
| 0.691197 | 0.045594 | 0.262 | 0.274 | 1 | 2 | Pet100 | -0.012 | Young |
| 0.691637 | 0.088021 | 0.037 | 0.041 | 1 | 2 | Arel1 | -0.004 | Young |
| 0.691786 | 0.156155 | 0.223 | 0.219 | 1 | 2 | Ppp4r2 | 0.004 | Young |
| 0.692328 | 0.178805 | 0.05 | 0.046 | 1 | 2 | Lonrf1 | 0.004 | Young |
| 0.692732 | 0.280097 | 0.058 | 0.055 | 1 | 2 | Cdkn2aip | 0.003 | Young |
| 0.692952 | 0.601309 | 0.052 | 0.049 | 1 | 2 | Tpm3-rs7 | 0.003 | Young |
| 0.692979 | 0.357294 | 0.151 | 0.149 | 1 | 2 | Ccny | 0.002 | Young |
| 0.6935 | 0.066002 | 0.43 | 0.425 | 1 | 2 | Clec4a2 | 0.005 | Young |
| 0.693633 | 0.055357 | 0.245 | 0.24 | 1 | 2 | Ptger4 | 0.005 | Young |
| 0.69398 | 0.130208 | 0.135 | 0.146 | 1 | 2 | Slc1a5 | -0.011 | Young |
| 0.694008 | 0.082229 | 0.077 | 0.073 | 1 | 2 | Stk25 | 0.004 | Young |
| 0.694358 | 0.023571 | 0.058 | 0.054 | 1 | 2 | Abcd3 | 0.004 | Young |
| 0.694605 | 0.0005 | 0.119 | 0.126 | 1 | 2 | Fam103a1 | -0.007 | Young |
| 0.694749 | 0.070369 | 0.355 | 0.35 | 1 | 2 | Epn1 | 0.005 | Young |

| | | | | | | | | |
|---|---|---|---|---|---|---|---|---|
| 0.695439 | 0.006673 | 0.189 | 0.195 | 1 | 2 | Ahr | -0.006 | Young |
| 0.695484 | 0.473017 | 0.045 | 0.042 | 1 | 2 | Cgrrf1 | 0.003 | Young |
| 0.696074 | 0.008756 | 0.281 | 0.299 | 1 | 2 | Kdelr1 | -0.018 | Young |
| 0.696116 | 0.147118 | 0.223 | 0.221 | 1 | 2 | Srsf7 | 0.002 | Young |
| 0.696341 | 0.068146 | 0.107 | 0.111 | 1 | 2 | Utp11 | -0.004 | Young |
| 0.696957 | 0.087306 | 0.154 | 0.164 | 1 | 2 | Hsd17b4 | -0.01 | Young |
| 0.697662 | 0.037962 | 0.127 | 0.134 | 1 | 2 | Arid3a | -0.007 | Young |
| 0.697747 | 0.173507 | 0.109 | 0.105 | 1 | 2 | Slc36a4 | 0.004 | Young |
| 0.698173 | 0.067288 | 0.33 | 0.327 | 1 | 2 | Mrps21 | 0.003 | Young |
| 0.69831 | 0.029103 | 0.045 | 0.049 | 1 | 2 | Hmgn5 | -0.004 | Young |
| 0.699033 | 0.186414 | 0.031 | 0.028 | 1 | 2 | Igip | 0.003 | Young |
| 0.69916 | 0.164969 | 0.273 | 0.276 | 1 | 2 | Hspa4 | -0.003 | Young |
| 0.699973 | 0.258428 | 0.031 | 0.028 | 1 | 2 | Rundc1 | 0.003 | Young |
| 0.700289 | 0.049776 | 0.196 | 0.208 | 1 | 2 | Alkbh5 | -0.012 | Young |
| 0.701074 | 0.361534 | 0.03 | 0.027 | 1 | 2 | Cmpk2 | 0.003 | Young |
| 0.701183 | 0.049996 | 0.038 | 0.042 | 1 | 2 | Ift46 | -0.004 | Young |
| 0.701189 | 0.001181 | 0.232 | 0.224 | 1 | 2 | Uso1 | 0.008 | Young |
| 0.701402 | 0.208125 | 0.196 | 0.195 | 1 | 2 | Ssu72 | 0.001 | Young |
| 0.70176 | 0.012577 | 0.064 | 0.06 | 1 | 2 | Myef2 | 0.004 | Young |
| 0.70198 | 0.053126 | 0.076 | 0.081 | 1 | 2 | Wdr48 | -0.005 | Young |
| 0.702254 | 0.003898 | 0.059 | 0.064 | 1 | 2 | Hinfp | -0.005 | Young |
| 0.703177 | 0.169155 | 0.077 | 0.073 | 1 | 2 | Golga3 | 0.004 | Young |
| 0.703194 | 0.043682 | 0.134 | 0.143 | 1 | 2 | Mrpl48 | -0.009 | Young |
| 0.703393 | 0.052262 | 0.024 | 0.027 | 1 | 2 | Msto1 | -0.003 | Young |
| 0.703728 | 0.343783 | 0.061 | 0.057 | 1 | 2 | Cd209g | 0.004 | Young |
| 0.704183 | 0.068789 | 0.133 | 0.141 | 1 | 2 | Fam50a | -0.008 | Young |
| 0.704542 | 0.070787 | 0.1 | 0.106 | 1 | 2 | Gcn1l1 | -0.006 | Young |
| 0.704579 | 0.0392 | 0.032 | 0.029 | 1 | 2 | Asf1a | 0.003 | Young |
| 0.704639 | 0.25118 | 0.152 | 0.149 | 1 | 2 | Snrnp200 | 0.003 | Young |
| 0.70483 | 0.23527 | 0.107 | 0.103 | 1 | 2 | Igsf8 | 0.004 | Young |
| 0.705223 | 0.117285 | 0.274 | 0.273 | 1 | 2 | Nars | 0.001 | Young |
| 0.705548 | 0.060111 | 0.157 | 0.152 | 1 | 2 | Pdhb | 0.005 | Young |
| 0.706159 | 0.074311 | 0.357 | 0.36 | 1 | 2 | Nucks1 | -0.003 | Young |
| 0.706399 | 0.167127 | 0.14 | 0.136 | 1 | 2 | Clcn3 | 0.004 | Young |
| 0.706749 | 0.218046 | 0.029 | 0.026 | 1 | 2 | Leng1 | 0.003 | Young |
| 0.70693 | 0.003323 | 0.074 | 0.07 | 1 | 2 | Erlin2 | 0.004 | Young |
| 0.707052 | 0.351066 | 0.043 | 0.04 | 1 | 2 | Ubl4a | 0.003 | Young |
| 0.707068 | 0.090976 | 0.143 | 0.151 | 1 | 2 | Nans | -0.008 | Young |
| 0.707893 | 0.012403 | 0.222 | 0.234 | 1 | 2 | Bud31 | -0.012 | Young |
| 0.707947 | 0.281282 | 0.089 | 0.086 | 1 | 2 | Rxra | 0.003 | Young |
| 0.709229 | 0.383664 | 0.026 | 0.024 | 1 | 2 | Dst | 0.002 | Young |
| 0.709294 | 0.115469 | 0.068 | 0.073 | 1 | 2 | Krr1 | -0.005 | Young |
| 0.709322 | 0.003653 | 0.025 | 0.028 | 1 | 2 | Stoml1 | -0.003 | Young |
| 0.709364 | 0.160067 | 0.077 | 0.073 | 1 | 2 | Plcb2 | 0.004 | Young |
| 0.709475 | 0.015123 | 0.201 | 0.192 | 1 | 2 | Sec24b | 0.009 | Young |

| | | | | | | | | |
|---|---|---|---|---|---|---|---|---|
| 0.70963 | 0.090337 | 0.376 | 0.375 | 1 | 2 | Taok1 | 0.001 | Young |
| 0.709692 | 0.15908 | 0.151 | 0.148 | 1 | 2 | Lrrc8c | 0.003 | Young |
| 0.70992 | 0.200372 | 0.1 | 0.096 | 1 | 2 | Smim13 | 0.004 | Young |
| 0.710572 | 0.01909 | 0.105 | 0.11 | 1 | 2 | Tmem230 | -0.005 | Young |
| 0.71083 | 0.288211 | 0.012 | 0.014 | 1 | 2 | Ptk2 | -0.002 | Young |
| 0.710868 | 0.469928 | 0.03 | 0.033 | 1 | 2 | Orc2 | -0.003 | Young |
| 0.711031 | 0.305547 | 0.045 | 0.049 | 1 | 2 | Ccdc85b | -0.004 | Young |
| 0.711073 | 0.042023 | 0.162 | 0.158 | 1 | 2 | Map2k7 | 0.004 | Young |
| 0.711353 | 0.016082 | 0.051 | 0.047 | 1 | 2 | Cracr2b | 0.004 | Young |
| 0.711458 | 0.158303 | 0.141 | 0.138 | 1 | 2 | Snx30 | 0.003 | Young |
| 0.711864 | 0.019968 | 0.671 | 0.69 | 1 | 2 | Atp6v0b | -0.019 | Young |
| 0.712483 | 0.079796 | 0.591 | 0.595 | 1 | 2 | Tma7 | -0.004 | Young |
| 0.71285 | 0.055613 | 0.165 | 0.161 | 1 | 2 | Tcn2 | 0.004 | Young |
| 0.712881 | 0.380116 | 0.062 | 0.058 | 1 | 2 | Taf12 | 0.004 | Young |
| 0.713119 | 0.278872 | 0.087 | 0.082 | 1 | 2 | Dag1 | 0.005 | Young |
| 0.713257 | 0.008871 | 0.278 | 0.272 | 1 | 2 | Dhx15 | 0.006 | Young |
| 0.713374 | 0.007588 | 0.067 | 0.062 | 1 | 2 | Cnnm3 | 0.005 | Young |
| 0.713854 | 0.066595 | 0.057 | 0.061 | 1 | 2 | Psmg3 | -0.004 | Young |
| 0.714005 | 0.266523 | 0.012 | 0.014 | 1 | 2 | A630072M | -0.002 | Young |
| 0.714401 | 0.007984 | 0.051 | 0.047 | 1 | 2 | Cep63 | 0.004 | Young |
| 0.714478 | 0.108683 | 0.308 | 0.302 | 1 | 2 | Got1 | 0.006 | Young |
| 0.71448 | 0.234662 | 0.062 | 0.058 | 1 | 2 | Abtb1 | 0.004 | Young |
| 0.714642 | 0.008581 | 0.799 | 0.798 | 1 | 2 | Sh3bgrl3 | 0.001 | Young |
| 0.714869 | 0.104216 | 0.221 | 0.214 | 1 | 2 | Snx17 | 0.007 | Young |
| 0.714951 | 0.057418 | 0.287 | 0.283 | 1 | 2 | Nsd1 | 0.004 | Young |
| 0.715006 | 0.226903 | 0.084 | 0.08 | 1 | 2 | Brd9 | 0.004 | Young |
| 0.715039 | 0.210702 | 0.086 | 0.091 | 1 | 2 | Tmem63a | -0.005 | Young |
| 0.715085 | 0.269736 | 0.061 | 0.057 | 1 | 2 | Mex3d | 0.004 | Young |
| 0.71539 | 0.39987 | 0.109 | 0.106 | 1 | 2 | Stim1 | 0.003 | Young |
| 0.715392 | 0.361923 | 0.057 | 0.054 | 1 | 2 | Wdr18 | 0.003 | Young |
| 0.715576 | 0.119481 | 0.062 | 0.058 | 1 | 2 | Ambra1 | 0.004 | Young |
| 0.715602 | 0.107734 | 0.071 | 0.067 | 1 | 2 | Usp5 | 0.004 | Young |
| 0.715965 | 0.010405 | 0.922 | 0.915 | 1 | 2 | H2-Ab1 | 0.007 | Young |
| 0.716041 | 0.22584 | 0.055 | 0.051 | 1 | 2 | Pcsk7 | 0.004 | Young |
| 0.716146 | 0.293868 | 0.153 | 0.15 | 1 | 2 | Dazap1 | 0.003 | Young |
| 0.716286 | 0.036319 | 0.276 | 0.292 | 1 | 2 | Tmem189 | -0.016 | Young |
| 0.716349 | 0.213631 | 0.202 | 0.198 | 1 | 2 | Fnip1 | 0.004 | Young |
| 0.716647 | 0.231992 | 0.106 | 0.102 | 1 | 2 | Smg5 | 0.004 | Young |
| 0.716671 | 0.269824 | 0.026 | 0.024 | 1 | 2 | Zfp609 | 0.002 | Young |
| 0.717071 | 0.207774 | 0.027 | 0.025 | 1 | 2 | Grwd1 | 0.002 | Young |
| 0.717778 | 0.020408 | 0.361 | 0.362 | 1 | 2 | Znrf2 | -0.001 | Young |
| 0.717905 | 0.591398 | 0.051 | 0.048 | 1 | 2 | Maip1 | 0.003 | Young |
| 0.718026 | 0.331668 | 0.056 | 0.052 | 1 | 2 | Csf2rb2 | 0.004 | Young |
| 0.718152 | 0.421068 | 0.037 | 0.041 | 1 | 2 | Ruvbl2 | -0.004 | Young |
| 0.718431 | 0.060918 | 0.083 | 0.089 | 1 | 2 | Pmpca | -0.006 | Young |

| | | | | | | | | |
|---|---|---|---|---|---|---|---|---|
| 0.718566 | 0.107685 | 0.1 | 0.096 | 1 | 2 | Clns1a | 0.004 | Young |
| 0.718627 | 0.341976 | 0.067 | 0.063 | 1 | 2 | Gm11808 | 0.004 | Young |
| 0.718703 | 0.009113 | 0.12 | 0.126 | 1 | 2 | Phf8 | -0.006 | Young |
| 0.718729 | 0.338548 | 0.026 | 0.024 | 1 | 2 | Oaf | 0.002 | Young |
| 0.718785 | 0.041637 | 0.099 | 0.093 | 1 | 2 | Trappc5 | 0.006 | Young |
| 0.719144 | 0.140256 | 0.025 | 0.028 | 1 | 2 | Unk | -0.003 | Young |
| 0.719214 | 0.084916 | 0.096 | 0.092 | 1 | 2 | Amz1 | 0.004 | Young |
| 0.719787 | 0.379581 | 0.025 | 0.022 | 1 | 2 | Eefsec | 0.003 | Young |
| 0.719817 | 0.000297 | 0.325 | 0.339 | 1 | 2 | Akr1b3 | -0.014 | Young |
| 0.719933 | 0.093943 | 0.156 | 0.166 | 1 | 2 | Tgs1 | -0.01 | Young |
| 0.720114 | 0.364715 | 0.043 | 0.04 | 1 | 2 | Bcas3 | 0.003 | Young |
| 0.720152 | 0.2329 | 0.087 | 0.082 | 1 | 2 | Mrpl11 | 0.005 | Young |
| 0.720274 | 0.215117 | 0.026 | 0.024 | 1 | 2 | Ttbk2 | 0.002 | Young |
| 0.720516 | 0.244155 | 0.025 | 0.028 | 1 | 2 | Tada2b | -0.003 | Young |
| 0.721197 | 0.074575 | 0.279 | 0.294 | 1 | 2 | Ppib | -0.015 | Young |
| 0.721499 | 0.018317 | 0.032 | 0.035 | 1 | 2 | Abhd18 | -0.003 | Young |
| 0.721692 | 0.139337 | 0.176 | 0.171 | 1 | 2 | Os9 | 0.005 | Young |
| 0.721701 | 0.046416 | 0.21 | 0.206 | 1 | 2 | Blmh | 0.004 | Young |
| 0.72183 | 0.055152 | 0.221 | 0.214 | 1 | 2 | Wdfy2 | 0.007 | Young |
| 0.722047 | 0.009583 | 0.279 | 0.291 | 1 | 2 | Arhgef1 | -0.012 | Young |
| 0.722309 | 0.477014 | 0.065 | 0.062 | 1 | 2 | Dohh | 0.003 | Young |
| 0.722331 | 0.040664 | 0.422 | 0.439 | 1 | 2 | Tardbp | -0.017 | Young |
| 0.722369 | 0.09431 | 0.185 | 0.182 | 1 | 2 | Timm10b | 0.003 | Young |
| 0.723085 | 0.135166 | 0.09 | 0.096 | 1 | 2 | Cltb | -0.006 | Young |
| 0.723516 | 0.198607 | 0.07 | 0.066 | 1 | 2 | Helb | 0.004 | Young |
| 0.724014 | 0.121278 | 0.052 | 0.057 | 1 | 2 | Atxn1l | -0.005 | Young |
| 0.724162 | 0.087561 | 0.323 | 0.342 | 1 | 2 | Myo1f | -0.019 | Young |
| 0.724208 | 0.143199 | 0.119 | 0.125 | 1 | 2 | Pex6 | -0.006 | Young |
| 0.724574 | 0.24028 | 0.058 | 0.055 | 1 | 2 | Slc25a13 | 0.003 | Young |
| 0.724798 | 0.091385 | 0.038 | 0.042 | 1 | 2 | Spsb3 | -0.004 | Young |
| 0.725002 | 0.311032 | 0.059 | 0.064 | 1 | 2 | Coq8b | -0.005 | Young |
| 0.725086 | 0.149571 | 0.166 | 0.162 | 1 | 2 | Zfx | 0.004 | Young |
| 0.725463 | 0.20147 | 0.043 | 0.04 | 1 | 2 | Bola1 | 0.003 | Young |
| 0.726463 | 0.102364 | 0.185 | 0.197 | 1 | 2 | Coro2a | -0.012 | Young |
| 0.726489 | 0.063846 | 0.279 | 0.29 | 1 | 2 | Fam168b | -0.011 | Young |
| 0.726647 | 0.20195 | 0.025 | 0.022 | 1 | 2 | Tgds | 0.003 | Young |
| 0.726648 | 0.149681 | 0.075 | 0.071 | 1 | 2 | Utp20 | 0.004 | Young |
| 0.72726 | 0.388112 | 0.052 | 0.049 | 1 | 2 | Nop14 | 0.003 | Young |
| 0.727455 | 0.210598 | 0.044 | 0.041 | 1 | 2 | Pde4a | 0.003 | Young |
| 0.72774 | 0.057125 | 0.233 | 0.23 | 1 | 2 | Comt | 0.003 | Young |
| 0.727751 | 0.083169 | 0.097 | 0.093 | 1 | 2 | Rbbp8 | 0.004 | Young |
| 0.727841 | 0.171369 | 0.15 | 0.146 | 1 | 2 | Pigx | 0.004 | Young |
| 0.728055 | 0.229919 | 0.112 | 0.108 | 1 | 2 | Bms1 | 0.004 | Young |
| 0.72836 | 0.209392 | 0.226 | 0.223 | 1 | 2 | Txnrd1 | 0.003 | Young |
| 0.728929 | 0.13337 | 0.101 | 0.108 | 1 | 2 | Sucla2 | -0.007 | Young |

| | | | | | | | | |
|---|---|---|---|---|---|---|---|---|
| 0.729013 | 0.051726 | 0.059 | 0.056 | 1 | 2 | Zfp24 | 0.003 | Young |
| 0.729184 | 0.024369 | 0.287 | 0.3 | 1 | 2 | Snrpd3 | -0.013 | Young |
| 0.729387 | 0.200949 | 0.024 | 0.021 | 1 | 2 | Zfp961 | 0.003 | Young |
| 0.72982 | 0.198149 | 0.038 | 0.042 | 1 | 2 | Dhx57 | -0.004 | Young |
| 0.729852 | 0.042944 | 0.044 | 0.041 | 1 | 2 | Zfp654 | 0.003 | Young |
| 0.72988 | 0.057338 | 0.148 | 0.145 | 1 | 2 | Elovl1 | 0.003 | Young |
| 0.7299 | 0.014737 | 0.452 | 0.469 | 1 | 2 | Hmgb1 | -0.017 | Young |
| 0.729932 | 0.194042 | 0.101 | 0.097 | 1 | 2 | Erlec1 | 0.004 | Young |
| 0.730264 | 0.647822 | 0.049 | 0.046 | 1 | 2 | Letmd1 | 0.003 | Young |
| 0.730879 | 0.488787 | 0.025 | 0.022 | 1 | 2 | Dcaf6 | 0.003 | Young |
| 0.731013 | 0.203329 | 0.024 | 0.021 | 1 | 2 | Trmt11 | 0.003 | Young |
| 0.731032 | 0.207254 | 0.11 | 0.106 | 1 | 2 | Clcn7 | 0.004 | Young |
| 0.731264 | 0.028679 | 0.062 | 0.066 | 1 | 2 | Hspb11 | -0.004 | Young |
| 0.731844 | 0.160484 | 0.239 | 0.238 | 1 | 2 | Vamp3 | 0.001 | Young |
| 0.73195 | 0.47576 | 0.038 | 0.035 | 1 | 2 | Cisd3 | 0.003 | Young |
| 0.73203 | 0.058406 | 0.026 | 0.029 | 1 | 2 | Lrrk2 | -0.003 | Young |
| 0.732098 | 0.144739 | 0.024 | 0.021 | 1 | 2 | Tdrkh | 0.003 | Young |
| 0.732176 | 0.116532 | 0.208 | 0.218 | 1 | 2 | Npepps | -0.01 | Young |
| 0.732355 | 0.06908 | 0.112 | 0.108 | 1 | 2 | Zswim8 | 0.004 | Young |
| 0.73237 | 0.048896 | 0.024 | 0.021 | 1 | 2 | Cnksr3 | 0.003 | Young |
| 0.732582 | 0.214591 | 0.04 | 0.037 | 1 | 2 | Rpain | 0.003 | Young |
| 0.733065 | 0.26012 | 0.114 | 0.11 | 1 | 2 | Samd8 | 0.004 | Young |
| 0.733151 | 0.024318 | 0.514 | 0.515 | 1 | 2 | Ehd4 | -0.001 | Young |
| 0.733209 | 0.207539 | 0.042 | 0.039 | 1 | 2 | Riox2 | 0.003 | Young |
| 0.733278 | 0.065978 | 0.112 | 0.118 | 1 | 2 | Trpv2 | -0.006 | Young |
| 0.733286 | 0.093303 | 0.273 | 0.269 | 1 | 2 | Ncf4 | 0.004 | Young |
| 0.733838 | 0.037751 | 0.071 | 0.067 | 1 | 2 | Ndst2 | 0.004 | Young |
| 0.734184 | 0.064918 | 0.019 | 0.021 | 1 | 2 | Dip2c | -0.002 | Young |
| 0.734544 | 0.062173 | 0.179 | 0.177 | 1 | 2 | Agfg1 | 0.002 | Young |
| 0.73469 | 0.042995 | 0.356 | 0.35 | 1 | 2 | Fam120a | 0.006 | Young |
| 0.734756 | 0.191642 | 0.019 | 0.021 | 1 | 2 | Mccc1 | -0.002 | Young |
| 0.734813 | 0.305276 | 0.024 | 0.021 | 1 | 2 | Zbtb45 | 0.003 | Young |
| 0.734932 | 0.151315 | 0.144 | 0.153 | 1 | 2 | Rnmt | -0.009 | Young |
| 0.735329 | 0.07626 | 0.074 | 0.079 | 1 | 2 | Pmvk | -0.005 | Young |
| 0.735368 | 0.17118 | 0.042 | 0.039 | 1 | 2 | Rusc1 | 0.003 | Young |
| 0.735555 | 0.175477 | 0.101 | 0.108 | 1 | 2 | Map1s | -0.007 | Young |
| 0.736024 | 0.188931 | 0.132 | 0.127 | 1 | 2 | Gmps | 0.005 | Young |
| 0.736126 | 0.108568 | 0.076 | 0.072 | 1 | 2 | Tmem55a | 0.004 | Young |
| 0.736186 | 0.80952 | 0.019 | 0.021 | 1 | 2 | Gemin2 | -0.002 | Young |
| 0.736839 | 0.077653 | 0.219 | 0.216 | 1 | 2 | Lrp10 | 0.003 | Young |
| 0.736861 | 0.182477 | 0.042 | 0.045 | 1 | 2 | Snx16 | -0.003 | Young |
| 0.736905 | 0.110145 | 0.258 | 0.259 | 1 | 2 | Prpf4b | -0.001 | Young |
| 0.737004 | 0.317939 | 0.013 | 0.015 | 1 | 2 | Knstrn | -0.002 | Young |
| 0.737004 | 0.18372 | 0.013 | 0.015 | 1 | 2 | Wars2 | -0.002 | Young |
| 0.737013 | 0.053835 | 0.764 | 0.758 | 1 | 2 | Ctsz | 0.006 | Young |

| | | | | | | | | |
|---|---|---|---|---|---|---|---|---|
| 0.737197 | 0.437932 | 0.026 | 0.029 | 1 | 2 | Wdr47 | -0.003 | Young |
| 0.737265 | 0.098233 | 0.265 | 0.262 | 1 | 2 | Ddost | 0.003 | Young |
| 0.738083 | 0.421559 | 0.023 | 0.02 | 1 | 2 | Slc52a2 | 0.003 | Young |
| 0.738083 | 0.290636 | 0.086 | 0.082 | 1 | 2 | Pde7a | 0.004 | Young |
| 0.738222 | 0.244121 | 0.023 | 0.02 | 1 | 2 | 1190007I0 | 0.003 | Young |
| 0.738403 | 0.116442 | 0.375 | 0.381 | 1 | 2 | Bcl2l11 | -0.006 | Young |
| 0.738889 | 0.075941 | 0.013 | 0.015 | 1 | 2 | Ccdc73 | -0.002 | Young |
| 0.739063 | 0.026616 | 0.22 | 0.232 | 1 | 2 | Brd1 | -0.012 | Young |
| 0.739078 | 0.004666 | 0.796 | 0.804 | 1 | 2 | Btg2 | -0.008 | Young |
| 0.739338 | 0.351826 | 0.023 | 0.02 | 1 | 2 | Echdc1 | 0.003 | Young |
| 0.739829 | 0.02287 | 0.488 | 0.479 | 1 | 2 | Hnrnpab | 0.009 | Young |
| 0.739897 | 0.205744 | 0.023 | 0.02 | 1 | 2 | Fam102a | 0.003 | Young |
| 0.740117 | 0.024023 | 0.489 | 0.497 | 1 | 2 | Sod1 | -0.008 | Young |
| 0.740145 | 0.020076 | 0.116 | 0.123 | 1 | 2 | Ykt6 | -0.007 | Young |
| 0.740293 | 0.105409 | 0.094 | 0.1 | 1 | 2 | Lman2l | -0.006 | Young |
| 0.740402 | 0.006239 | 0.359 | 0.372 | 1 | 2 | Cd2ap | -0.013 | Young |
| 0.740581 | 0.004615 | 0.072 | 0.077 | 1 | 2 | Ppid | -0.005 | Young |
| 0.740591 | 0.017039 | 0.406 | 0.407 | 1 | 2 | Ptp4a1 | -0.001 | Young |
| 0.740677 | 0.379471 | 0.037 | 0.034 | 1 | 2 | Rrs1 | 0.003 | Young |
| 0.740734 | 0.125414 | 0.023 | 0.02 | 1 | 2 | F8a | 0.003 | Young |
| 0.740998 | 0.139583 | 0.067 | 0.063 | 1 | 2 | Mfap1b | 0.004 | Young |
| 0.741035 | 0.072505 | 0.464 | 0.475 | 1 | 2 | Snrpb | -0.011 | Young |
| 0.741074 | 0.168257 | 0.171 | 0.168 | 1 | 2 | Xpo1 | 0.003 | Young |
| 0.742112 | 0.096374 | 0.062 | 0.058 | 1 | 2 | Bmi1 | 0.004 | Young |
| 0.742237 | 0.142116 | 0.247 | 0.247 | 1 | 2 | Nxf1 | 0 | Young |
| 0.742381 | 0.045372 | 0.14 | 0.136 | 1 | 2 | Brpf1 | 0.004 | Young |
| 0.742525 | 0.085776 | 0.131 | 0.137 | 1 | 2 | Tatdn2 | -0.006 | Young |
| 0.742527 | 0.042504 | 0.05 | 0.054 | 1 | 2 | Hace1 | -0.004 | Young |
| 0.74255 | 0.061297 | 0.116 | 0.122 | 1 | 2 | Milr1 | -0.006 | Young |
| 0.742576 | 0.039034 | 0.163 | 0.157 | 1 | 2 | Tnf | 0.006 | Young |
| 0.742827 | 0.507042 | 0.048 | 0.045 | 1 | 2 | Aifm1 | 0.003 | Young |
| 0.743085 | 0.002845 | 0.043 | 0.046 | 1 | 2 | Med24 | -0.003 | Young |
| 0.743249 | 0.096345 | 0.024 | 0.021 | 1 | 2 | Mtx3 | 0.003 | Young |
| 0.743406 | 0.173706 | 0.042 | 0.039 | 1 | 2 | Tmem87a | 0.003 | Young |
| 0.743511 | 0.09386 | 0.184 | 0.18 | 1 | 2 | Itgb5 | 0.004 | Young |
| 0.74358 | 0.229659 | 0.063 | 0.06 | 1 | 2 | Commd10 | 0.003 | Young |
| 0.744075 | 0.310927 | 0.055 | 0.059 | 1 | 2 | Stxbp5 | -0.004 | Young |
| 0.744412 | 0.245879 | 0.034 | 0.037 | 1 | 2 | Cep135 | -0.003 | Young |
| 0.744728 | 0.15272 | 0.089 | 0.085 | 1 | 2 | Wipi2 | 0.004 | Young |
| 0.744917 | 0.020908 | 0.987 | 0.99 | 1 | 2 | mt-Nd4 | -0.003 | Young |
| 0.745115 | 0.105234 | 0.125 | 0.132 | 1 | 2 | Gm15987 | -0.007 | Young |
| 0.74565 | 0.28889 | 0.021 | 0.019 | 1 | 2 | Nkiras1 | 0.002 | Young |
| 0.746029 | 0.029721 | 0.336 | 0.344 | 1 | 2 | Cdv3 | -0.008 | Young |
| 0.746291 | 0.019116 | 0.251 | 0.262 | 1 | 2 | Cuta | -0.011 | Young |
| 0.747027 | 0.111816 | 0.171 | 0.168 | 1 | 2 | Cbx1 | 0.003 | Young |

| | | | | | | | | |
|---|---|---|---|---|---|---|---|---|
| 0.747549 | 0.230234 | 0.157 | 0.154 | 1 | 2 | C1qbp | 0.003 | Young |
| 0.747596 | 0.371372 | 0.074 | 0.071 | 1 | 2 | Cd2bp2 | 0.003 | Young |
| 0.747608 | 0.074043 | 0.467 | 0.473 | 1 | 2 | Colgalt1 | -0.006 | Young |
| 0.74773 | 0.059374 | 0.27 | 0.267 | 1 | 2 | Rgl1 | 0.003 | Young |
| 0.747866 | 0.25995 | 0.038 | 0.035 | 1 | 2 | Dctpp1 | 0.003 | Young |
| 0.747991 | 0.105778 | 0.056 | 0.06 | 1 | 2 | Ccser2 | -0.004 | Young |
| 0.74839 | 0.197183 | 0.131 | 0.128 | 1 | 2 | Oxr1 | 0.003 | Young |
| 0.748464 | 0.037818 | 0.072 | 0.069 | 1 | 2 | Plxna4os1 | 0.003 | Young |
| 0.748828 | 0.167259 | 0.138 | 0.135 | 1 | 2 | Rspry1 | 0.003 | Young |
| 0.749118 | 0.532583 | 0.059 | 0.057 | 1 | 2 | Noc4l | 0.002 | Young |
| 0.749275 | 0.085898 | 0.229 | 0.226 | 1 | 2 | Wls | 0.003 | Young |
| 0.749392 | 0.018681 | 0.234 | 0.243 | 1 | 2 | Cnot6 | -0.009 | Young |
| 0.749528 | 0.055017 | 0.064 | 0.061 | 1 | 2 | Stk16 | 0.003 | Young |
| 0.749606 | 0.056202 | 0.182 | 0.191 | 1 | 2 | Samm50 | -0.009 | Young |
| 0.749847 | 0.132526 | 0.034 | 0.037 | 1 | 2 | D17H6S53I | -0.003 | Young |
| 0.749933 | 0.255022 | 0.04 | 0.044 | 1 | 2 | Ripk3 | -0.004 | Young |
| 0.749979 | 0.043147 | 0.221 | 0.234 | 1 | 2 | Cript | -0.013 | Young |
| 0.750076 | 0.027432 | 0.126 | 0.133 | 1 | 2 | Tbc1d1 | -0.007 | Young |
| 0.750234 | 0.650361 | 0.019 | 0.017 | 1 | 2 | 1700096K1 | 0.002 | Young |
| 0.750354 | 0.427678 | 0.048 | 0.045 | 1 | 2 | Fbxo6 | 0.003 | Young |
| 0.750519 | 0.205745 | 0.09 | 0.087 | 1 | 2 | Crnkl1 | 0.003 | Young |
| 0.750562 | 0.161499 | 0.027 | 0.03 | 1 | 2 | Ints5.1 | -0.003 | Young |
| 0.750723 | 0.017086 | 0.036 | 0.039 | 1 | 2 | Rufy2 | -0.003 | Young |
| 0.750868 | 0.299928 | 0.064 | 0.061 | 1 | 2 | Syap1 | 0.003 | Young |
| 0.75106 | 0.103602 | 0.083 | 0.079 | 1 | 2 | Zfyve16 | 0.004 | Young |
| 0.751608 | 0.204482 | 0.039 | 0.036 | 1 | 2 | Anks3 | 0.003 | Young |
| 0.751665 | 0.175388 | 0.05 | 0.054 | 1 | 2 | Rnf121 | -0.004 | Young |
| 0.751908 | 0.286101 | 0.197 | 0.198 | 1 | 2 | Pdlim1 | -0.001 | Young |
| 0.752107 | 0.129967 | 0.126 | 0.123 | 1 | 2 | Ralgds | 0.003 | Young |
| 0.752249 | 0.379972 | 0.071 | 0.069 | 1 | 2 | Mapk8ip3 | 0.002 | Young |
| 0.752434 | 0.228015 | 0.11 | 0.107 | 1 | 2 | Gps1 | 0.003 | Young |
| 0.752599 | 0.161949 | 0.132 | 0.13 | 1 | 2 | Mrps18c | 0.002 | Young |
| 0.752828 | 0.078915 | 0.044 | 0.047 | 1 | 2 | Trmt2a | -0.003 | Young |
| 0.752992 | 0.152502 | 0.027 | 0.03 | 1 | 2 | Nlrx1 | -0.003 | Young |
| 0.753774 | 0.001669 | 0.1 | 0.096 | 1 | 2 | Sipa1l3 | 0.004 | Young |
| 0.754045 | 0.284552 | 0.083 | 0.08 | 1 | 2 | Chchd7 | 0.003 | Young |
| 0.754065 | 0.061319 | 0.065 | 0.07 | 1 | 2 | mt-Nd6 | -0.005 | Young |
| 0.754237 | 0.015062 | 0.297 | 0.291 | 1 | 2 | Zc3h7a | 0.006 | Young |
| 0.754543 | 0.081481 | 0.407 | 0.408 | 1 | 2 | Inpp5d | -0.001 | Young |
| 0.754675 | 0.221219 | 0.119 | 0.116 | 1 | 2 | Ezh1 | 0.003 | Young |
| 0.754769 | 0.042246 | 0.575 | 0.585 | 1 | 2 | Taldo1 | -0.01 | Young |
| 0.754824 | 0.055348 | 0.055 | 0.051 | 1 | 2 | G6pc3 | 0.004 | Young |
| 0.754918 | 0.35609 | 0.037 | 0.034 | 1 | 2 | Hotairm1 | 0.003 | Young |
| 0.755358 | 0.29256 | 0.037 | 0.034 | 1 | 2 | Atg14 | 0.003 | Young |
| 0.755688 | 0.251614 | 0.02 | 0.018 | 1 | 2 | Lmnb2 | 0.002 | Young |

| | | | | | | | | |
|---|---|---|---|---|---|---|---|---|
| 0.755729 | 0.064582 | 0.145 | 0.152 | 1 | 2 | Agtrap | -0.007 | Young |
| 0.755932 | 0.11574 | 0.249 | 0.246 | 1 | 2 | Nfat5 | 0.003 | Young |
| 0.755957 | 0.021443 | 0.362 | 0.358 | 1 | 2 | Ctnnb1 | 0.004 | Young |
| 0.756037 | 0.289024 | 0.019 | 0.017 | 1 | 2 | Rasgrf2 | 0.002 | Young |
| 0.756556 | 0.010259 | 0.601 | 0.592 | 1 | 2 | Cox6c | 0.009 | Young |
| 0.756589 | 0.017051 | 0.126 | 0.133 | 1 | 2 | Trappc10 | -0.007 | Young |
| 0.7567 | 0.058131 | 0.119 | 0.115 | 1 | 2 | Fam3c | 0.004 | Young |
| 0.756732 | 0.02086 | 0.014 | 0.016 | 1 | 2 | Capn5 | -0.002 | Young |
| 0.757248 | 0.134535 | 0.321 | 0.321 | 1 | 2 | Lrrc58 | 0 | Young |
| 0.757371 | 0.162371 | 0.175 | 0.172 | 1 | 2 | Smarca4 | 0.003 | Young |
| 0.757384 | 0.0275 | 0.065 | 0.07 | 1 | 2 | Dpm2 | -0.005 | Young |
| 0.757453 | 0.207293 | 0.083 | 0.088 | 1 | 2 | Igsf9 | -0.005 | Young |
| 0.757867 | 0.194495 | 0.065 | 0.062 | 1 | 2 | Ccdc107 | 0.003 | Young |
| 0.758105 | 0.127052 | 0.027 | 0.03 | 1 | 2 | Cul9 | -0.003 | Young |
| 0.758451 | 0.000599 | 0.204 | 0.201 | 1 | 2 | Slc39a7 | 0.003 | Young |
| 0.758472 | 0.409514 | 0.049 | 0.046 | 1 | 2 | Ppan | 0.003 | Young |
| 0.75855 | 0.190188 | 0.226 | 0.223 | 1 | 2 | Slamf7 | 0.003 | Young |
| 0.759658 | 0.119797 | 0.089 | 0.094 | 1 | 2 | Dcaf11 | -0.005 | Young |
| 0.75969 | 0.212946 | 0.096 | 0.093 | 1 | 2 | Wwc2 | 0.003 | Young |
| 0.759745 | 0.069907 | 0.488 | 0.498 | 1 | 2 | Brk1 | -0.01 | Young |
| 0.759767 | 0.343845 | 0.034 | 0.032 | 1 | 2 | Hs1bp3 | 0.002 | Young |
| 0.760393 | 0.021381 | 0.043 | 0.04 | 1 | 2 | Acyp1 | 0.003 | Young |
| 0.760618 | 0.152377 | 0.093 | 0.089 | 1 | 2 | Oxa1l | 0.004 | Young |
| 0.76072 | 0.064857 | 0.366 | 0.363 | 1 | 2 | Slc38a1 | 0.003 | Young |
| 0.760726 | 0.051039 | 0.014 | 0.016 | 1 | 2 | Fcho1 | -0.002 | Young |
| 0.760934 | 0.006779 | 0.11 | 0.116 | 1 | 2 | Lyar | -0.006 | Young |
| 0.761246 | 0.108076 | 0.099 | 0.105 | 1 | 2 | Trim33 | -0.006 | Young |
| 0.761552 | 0.147052 | 0.182 | 0.18 | 1 | 2 | Chka | 0.002 | Young |
| 0.762015 | 0.120011 | 0.059 | 0.063 | 1 | 2 | Tbc1d31 | -0.004 | Young |
| 0.762196 | 0.038039 | 0.553 | 0.565 | 1 | 2 | Sf3b1 | -0.012 | Young |
| 0.76238 | 0.05888 | 0.343 | 0.343 | 1 | 2 | Nckap1l | 0 | Young |
| 0.762429 | 0.012187 | 0.315 | 0.314 | 1 | 2 | Coq10b | 0.001 | Young |
| 0.762583 | 0.040505 | 0.209 | 0.208 | 1 | 2 | Arl6ip5 | 0.001 | Young |
| 0.763058 | 0.186492 | 0.014 | 0.016 | 1 | 2 | Jmjd8 | -0.002 | Young |
| 0.763058 | 0.158459 | 0.014 | 0.016 | 1 | 2 | Rbbp9 | -0.002 | Young |
| 0.763188 | 0.089166 | 0.245 | 0.242 | 1 | 2 | Neu1 | 0.003 | Young |
| 0.763387 | 0.124383 | 0.019 | 0.017 | 1 | 2 | Chaf1b | 0.002 | Young |
| 0.763416 | 0.158704 | 0.02 | 0.018 | 1 | 2 | Snx7 | 0.002 | Young |
| 0.763481 | 0.602584 | 0.033 | 0.031 | 1 | 2 | Tpk1 | 0.002 | Young |
| 0.763693 | 0.04737 | 0.019 | 0.017 | 1 | 2 | Wrb | 0.002 | Young |
| 0.763901 | 0.095802 | 0.078 | 0.075 | 1 | 2 | Abhd2 | 0.003 | Young |
| 0.764007 | 0.044378 | 0.119 | 0.125 | 1 | 2 | Apopt1 | -0.006 | Young |
| 0.764376 | 0.142562 | 0.335 | 0.339 | 1 | 2 | Aldoa | -0.004 | Young |
| 0.764621 | 0.373496 | 0.129 | 0.127 | 1 | 2 | Cmip | 0.002 | Young |
| 0.764845 | 0.777019 | 0.017 | 0.015 | 1 | 2 | Pstpip2 | 0.002 | Young |

| | | | | | | | | |
|---|---|---|---|---|---|---|---|---|
| 0.76504 | 0.000643 | 0.52 | 0.517 | 1 | 2 | Snx5 | 0.003 | Young |
| 0.765282 | 0.101071 | 0.213 | 0.225 | 1 | 2 | Glrx3 | -0.012 | Young |
| 0.76588 | 0.034845 | 0.057 | 0.061 | 1 | 2 | Nkap | -0.004 | Young |
| 0.766329 | 0.020866 | 0.053 | 0.057 | 1 | 2 | Sp3os | -0.004 | Young |
| 0.766389 | 0.01136 | 0.371 | 0.384 | 1 | 2 | Ptpra | -0.013 | Young |
| 0.766592 | 0.226081 | 0.037 | 0.034 | 1 | 2 | Pwp1 | 0.003 | Young |
| 0.766649 | 0.115116 | 0.251 | 0.248 | 1 | 2 | Mrps33 | 0.003 | Young |
| 0.766882 | 0.341629 | 0.044 | 0.047 | 1 | 2 | Gclm | -0.003 | Young |
| 0.766995 | 0.042303 | 0.755 | 0.739 | 1 | 2 | Tmsb10 | 0.016 | Young |
| 0.767092 | 0.317457 | 0.061 | 0.058 | 1 | 2 | Slc7a6os | 0.003 | Young |
| 0.767631 | 0.672287 | 0.017 | 0.015 | 1 | 2 | Lhpp | 0.002 | Young |
| 0.768316 | 0.165709 | 0.103 | 0.101 | 1 | 2 | Arrdc4 | 0.002 | Young |
| 0.768661 | 0.020803 | 0.09 | 0.095 | 1 | 2 | Tbpl1 | -0.005 | Young |
| 0.769043 | 0.073984 | 0.201 | 0.2 | 1 | 2 | Tnfrsf21 | 0.001 | Young |
| 0.769113 | 0.161622 | 0.051 | 0.055 | 1 | 2 | 1110008P1 | -0.004 | Young |
| 0.769295 | 0.318845 | 0.008 | 0.01 | 1 | 2 | Smyd4 | -0.002 | Young |
| 0.769295 | 0.130675 | 0.008 | 0.01 | 1 | 2 | Rnf128 | -0.002 | Young |
| 0.770013 | 0.152791 | 0.135 | 0.133 | 1 | 2 | Emc2 | 0.002 | Young |
| 0.770105 | 0.016745 | 0.036 | 0.039 | 1 | 2 | Zfp280c | -0.003 | Young |
| 0.770404 | 0.234298 | 0.194 | 0.208 | 1 | 2 | Tox4 | -0.014 | Young |
| 0.770722 | 0.1668 | 0.209 | 0.207 | 1 | 2 | Atp6v1h | 0.002 | Young |
| 0.771027 | 0.085077 | 0.008 | 0.01 | 1 | 2 | 9030025P2 | -0.002 | Young |
| 0.771041 | 0.554771 | 0.018 | 0.016 | 1 | 2 | Ccdc191 | 0.002 | Young |
| 0.771107 | 0.055221 | 0.129 | 0.137 | 1 | 2 | Gbp3 | -0.008 | Young |
| 0.771249 | 0.040988 | 0.257 | 0.255 | 1 | 2 | Ap1b1 | 0.002 | Young |
| 0.771348 | 0.15477 | 0.05 | 0.047 | 1 | 2 | Eif2b2 | 0.003 | Young |
| 0.771676 | 0.302601 | 0.018 | 0.016 | 1 | 2 | Ndufaf5 | 0.002 | Young |
| 0.771713 | 0.042601 | 0.052 | 0.049 | 1 | 2 | Zfp410 | 0.003 | Young |
| 0.772012 | 0.04481 | 0.217 | 0.213 | 1 | 2 | Slc4a7 | 0.004 | Young |
| 0.772169 | 0.197793 | 0.07 | 0.067 | 1 | 2 | Zfp598 | 0.003 | Young |
| 0.772326 | 0.028935 | 0.008 | 0.01 | 1 | 2 | Tmem201 | -0.002 | Young |
| 0.772392 | 0.400138 | 0.017 | 0.015 | 1 | 2 | Hdac6 | 0.002 | Young |
| 0.772488 | 0.097237 | 0.108 | 0.105 | 1 | 2 | Trim35 | 0.003 | Young |
| 0.772564 | 0.119925 | 0.044 | 0.047 | 1 | 2 | Mcu | -0.003 | Young |
| 0.772735 | 0.34298 | 0.07 | 0.067 | 1 | 2 | BC003965 | 0.003 | Young |
| 0.772742 | 0.188234 | 0.065 | 0.062 | 1 | 2 | Zmym4 | 0.003 | Young |
| 0.772782 | 0.139032 | 0.096 | 0.102 | 1 | 2 | Dpagt1 | -0.006 | Young |
| 0.772969 | 0.063245 | 0.226 | 0.225 | 1 | 2 | Pdxk | 0.001 | Young |
| 0.773094 | 0.004369 | 0.133 | 0.139 | 1 | 2 | Smc4 | -0.006 | Young |
| 0.773255 | 0.002231 | 0.061 | 0.064 | 1 | 2 | Lonp2 | -0.003 | Young |
| 0.773378 | 0.177796 | 0.017 | 0.015 | 1 | 2 | Abca2 | 0.002 | Young |
| 0.773809 | 0.171174 | 0.148 | 0.146 | 1 | 2 | Tm2d1 | 0.002 | Young |
| 0.773921 | 0.038163 | 0.34 | 0.343 | 1 | 2 | Ptpro | -0.003 | Young |
| 0.774036 | 0.425923 | 0.017 | 0.015 | 1 | 2 | Grap2 | 0.002 | Young |
| 0.7741 | 0.457154 | 0.032 | 0.03 | 1 | 2 | Cyb561d1 | 0.002 | Young |

| | | | | | | | | |
|---|---|---|---|---|---|---|---|---|
| 0.774279 | 0.010145 | 0.204 | 0.2 | 1 | 2 | Hexim1 | 0.004 | Young |
| 0.774927 | 0.185824 | 0.008 | 0.01 | 1 | 2 | Mtbp | -0.002 | Young |
| 0.774933 | 0.560989 | 0.044 | 0.042 | 1 | 2 | Papolg | 0.002 | Young |
| 0.775273 | 0.092204 | 0.122 | 0.127 | 1 | 2 | Fbxo11 | -0.005 | Young |
| 0.775283 | 0.348724 | 0.071 | 0.069 | 1 | 2 | Nomo1 | 0.002 | Young |
| 0.775351 | 0.39278 | 0.017 | 0.015 | 1 | 2 | Nkrf | 0.002 | Young |
| 0.775351 | 0.346497 | 0.017 | 0.015 | 1 | 2 | Ndufaf1 | 0.002 | Young |
| 0.77546 | 0.070236 | 0.284 | 0.3 | 1 | 2 | Tpi1 | -0.016 | Young |
| 0.77568 | 0.214766 | 0.017 | 0.015 | 1 | 2 | Nop2 | 0.002 | Young |
| 0.776148 | 0.367648 | 0.044 | 0.042 | 1 | 2 | AI837181 | 0.002 | Young |
| 0.776211 | 0.178457 | 0.096 | 0.093 | 1 | 2 | Fam53c | 0.003 | Young |
| 0.776355 | 0.171385 | 0.198 | 0.197 | 1 | 2 | Psmd14 | 0.001 | Young |
| 0.776459 | 0.151175 | 0.132 | 0.128 | 1 | 2 | Brd8 | 0.004 | Young |
| 0.776662 | 0.360489 | 0.008 | 0.01 | 1 | 2 | Loxl3 | -0.002 | Young |
| 0.776719 | 0.00475 | 0.341 | 0.357 | 1 | 2 | Psma3 | -0.016 | Young |
| 0.77708 | 0.092853 | 0.018 | 0.016 | 1 | 2 | Sars2 | 0.002 | Young |
| 0.777434 | 0.227129 | 0.05 | 0.054 | 1 | 2 | Aig1 | -0.004 | Young |
| 0.777597 | 0.236942 | 0.089 | 0.087 | 1 | 2 | Ing5 | 0.002 | Young |
| 0.777635 | 0.607518 | 0.015 | 0.014 | 1 | 2 | Zdhhc24 | 0.001 | Young |
| 0.777662 | 0.160936 | 0.048 | 0.045 | 1 | 2 | Rhot2 | 0.003 | Young |
| 0.777738 | 0.002924 | 0.508 | 0.505 | 1 | 2 | Ogt | 0.003 | Young |
| 0.778123 | 0.059941 | 0.089 | 0.086 | 1 | 2 | AU040320 | 0.003 | Young |
| 0.778205 | 0.11318 | 0.057 | 0.055 | 1 | 2 | Hira | 0.002 | Young |
| 0.778337 | 0.106684 | 0.071 | 0.069 | 1 | 2 | Rab10os | 0.002 | Young |
| 0.778366 | 0.076068 | 0.105 | 0.109 | 1 | 2 | Kdm4a | -0.004 | Young |
| 0.778482 | 0.048688 | 0.219 | 0.212 | 1 | 2 | Crebrf | 0.007 | Young |
| 0.778716 | 0.288669 | 0.057 | 0.061 | 1 | 2 | Med4 | -0.004 | Young |
| 0.778973 | 0.467397 | 0.017 | 0.015 | 1 | 2 | Thada | 0.002 | Young |
| 0.77903 | 0.204173 | 0.158 | 0.167 | 1 | 2 | Tspyl1 | -0.009 | Young |
| 0.779142 | 0.033136 | 0.278 | 0.276 | 1 | 2 | Dnaja2 | 0.002 | Young |
| 0.779267 | 0.392004 | 0.033 | 0.031 | 1 | 2 | Pus7 | 0.002 | Young |
| 0.780198 | 0.459347 | 0.021 | 0.024 | 1 | 2 | Fbxw17 | -0.003 | Young |
| 0.780252 | 0.111576 | 0.146 | 0.142 | 1 | 2 | Atp13a1 | 0.004 | Young |
| 0.780774 | 0.004343 | 0.138 | 0.142 | 1 | 2 | Lyst | -0.004 | Young |
| 0.78095 | 0.172626 | 0.017 | 0.015 | 1 | 2 | Ccdc66 | 0.002 | Young |
| 0.781081 | 0.061936 | 0.181 | 0.178 | 1 | 2 | Pi4k2a | 0.003 | Young |
| 0.78161 | 0.050528 | 0.017 | 0.015 | 1 | 2 | Tnfaip8l1 | 0.002 | Young |
| 0.781737 | 0.390071 | 0.015 | 0.014 | 1 | 2 | Abhd14a | 0.001 | Young |
| 0.781737 | 0.299756 | 0.015 | 0.014 | 1 | 2 | Ier5l | 0.001 | Young |
| 0.781794 | 0.081081 | 0.051 | 0.048 | 1 | 2 | Tbc1d24 | 0.003 | Young |
| 0.781818 | 0.331028 | 0.103 | 0.102 | 1 | 2 | Lrch3 | 0.001 | Young |
| 0.782282 | 0.165426 | 0.286 | 0.292 | 1 | 2 | Irak1 | -0.006 | Young |
| 0.782599 | 0.393793 | 0.017 | 0.015 | 1 | 2 | Dffb | 0.002 | Young |
| 0.782793 | 0.152449 | 0.183 | 0.182 | 1 | 2 | Vps29 | 0.001 | Young |
| 0.783282 | 0.026101 | 0.227 | 0.227 | 1 | 2 | Mgea5 | 0 | Young |

| | | | | | | | | |
|---|---|---|---|---|---|---|---|---|
| 0.783658 | 0.172295 | 0.133 | 0.131 | 1 | 2 | Clip1 | 0.002 | Young |
| 0.783827 | 0.112662 | 0.229 | 0.229 | 1 | 2 | Parp14 | 0 | Young |
| 0.783898 | 0.219163 | 0.067 | 0.064 | 1 | 2 | Mrps6 | 0.003 | Young |
| 0.783927 | 0.194047 | 0.033 | 0.031 | 1 | 2 | Pdpr | 0.002 | Young |
| 0.783957 | 0.100213 | 0.015 | 0.017 | 1 | 2 | Gm15446 | -0.002 | Young |
| 0.784162 | 0.141354 | 0.07 | 0.067 | 1 | 2 | Atg4b | 0.003 | Young |
| 0.784194 | 0.220525 | 0.036 | 0.039 | 1 | 2 | Pnpla6 | -0.003 | Young |
| 0.784212 | 0.145688 | 0.204 | 0.202 | 1 | 2 | Ap3d1 | 0.002 | Young |
| 0.78443 | 0.131565 | 0.034 | 0.032 | 1 | 2 | Rgp1 | 0.002 | Young |
| 0.784549 | 0.24388 | 0.045 | 0.043 | 1 | 2 | Nrg1 | 0.002 | Young |
| 0.784558 | 0.005929 | 0.245 | 0.253 | 1 | 2 | Map2k3 | -0.008 | Young |
| 0.784753 | 0.14155 | 0.361 | 0.365 | 1 | 2 | Fem1c | -0.004 | Young |
| 0.78512 | 0.215861 | 0.063 | 0.06 | 1 | 2 | Mid1 | 0.003 | Young |
| 0.785235 | 0.268273 | 0.03 | 0.032 | 1 | 2 | Pla2g12a | -0.002 | Young |
| 0.785377 | 0.254034 | 0.141 | 0.14 | 1 | 2 | D16Ertd47 | 0.001 | Young |
| 0.785579 | 0.427376 | 0.015 | 0.017 | 1 | 2 | Serpinb1a | -0.002 | Young |
| 0.786009 | 0.364459 | 0.124 | 0.122 | 1 | 2 | Tmem131l | 0.002 | Young |
| 0.786011 | 0.266575 | 0.08 | 0.077 | 1 | 2 | Xpa | 0.003 | Young |
| 0.786195 | 0.152089 | 0.162 | 0.171 | 1 | 2 | Trio | -0.009 | Young |
| 0.786229 | 0.170251 | 0.015 | 0.017 | 1 | 2 | Gm10036 | -0.002 | Young |
| 0.786379 | 0.318636 | 0.119 | 0.118 | 1 | 2 | Mob3a | 0.001 | Young |
| 0.786529 | 0.057485 | 0.015 | 0.014 | 1 | 2 | Mocos | 0.001 | Young |
| 0.786907 | 0.130072 | 0.163 | 0.161 | 1 | 2 | Pqlc1 | 0.002 | Young |
| 0.787059 | 0.08487 | 0.053 | 0.057 | 1 | 2 | Rps19bp1 | -0.004 | Young |
| 0.787126 | 0.045617 | 0.209 | 0.218 | 1 | 2 | Sipa1 | -0.009 | Young |
| 0.78759 | 0.400097 | 0.014 | 0.013 | 1 | 2 | Cenpw | 0.001 | Young |
| 0.787618 | 0.051821 | 0.165 | 0.172 | 1 | 2 | Tpst2 | -0.007 | Young |
| 0.787646 | 0.10735 | 0.172 | 0.169 | 1 | 2 | Ndufs4 | 0.003 | Young |
| 0.788421 | 0.083534 | 0.167 | 0.176 | 1 | 2 | Polr2f | -0.009 | Young |
| 0.788427 | 0.03602 | 0.169 | 0.176 | 1 | 2 | Fam107b | -0.007 | Young |
| 0.788585 | 0.070456 | 0.015 | 0.014 | 1 | 2 | Dcstamp | 0.001 | Young |
| 0.788867 | 0.010606 | 0.219 | 0.228 | 1 | 2 | Ttyh3 | -0.009 | Young |
| 0.788957 | 0.740292 | 0.042 | 0.04 | 1 | 2 | Tjp2 | 0.002 | Young |
| 0.788988 | 0.005656 | 0.147 | 0.142 | 1 | 2 | Rnf166 | 0.005 | Young |
| 0.789006 | 0.012679 | 0.622 | 0.609 | 1 | 2 | Reep5 | 0.013 | Young |
| 0.789192 | 0.001892 | 0.091 | 0.095 | 1 | 2 | Tlr4 | -0.004 | Young |
| 0.789908 | 0.025969 | 0.31 | 0.322 | 1 | 2 | Cwc15 | -0.012 | Young |
| 0.790225 | 0.153767 | 0.032 | 0.03 | 1 | 2 | Aven | 0.002 | Young |
| 0.790939 | 0.253258 | 0.03 | 0.032 | 1 | 2 | Pde6d | -0.002 | Young |
| 0.790939 | 0.140629 | 0.032 | 0.03 | 1 | 2 | Spata13 | 0.002 | Young |
| 0.791085 | 0.184405 | 0.113 | 0.11 | 1 | 2 | Slc38a10 | 0.003 | Young |
| 0.791161 | 0.055538 | 0.46 | 0.458 | 1 | 2 | Gas7 | 0.002 | Young |
| 0.791211 | 0.015279 | 0.767 | 0.794 | 1 | 2 | Gnb2 | -0.027 | Young |
| 0.79125 | 0.134994 | 0.181 | 0.19 | 1 | 2 | Psmc5 | -0.009 | Young |
| 0.791329 | 0.035108 | 0.015 | 0.014 | 1 | 2 | Zgpat | 0.001 | Young |

| | | | | | | | | |
|---|---|---|---|---|---|---|---|---|
| 0.791478 | 0.228095 | 0.071 | 0.069 | 1 | 2 | Wrn | 0.002 | Young |
| 0.791646 | 0.482206 | 0.043 | 0.041 | 1 | 2 | Apip | 0.002 | Young |
| 0.791872 | 0.517171 | 0.014 | 0.013 | 1 | 2 | 5730409E0 | 0.001 | Young |
| 0.791981 | 0.012896 | 0.045 | 0.043 | 1 | 2 | Dlg4 | 0.002 | Young |
| 0.79202 | 0.345684 | 0.023 | 0.025 | 1 | 2 | Phykpl | -0.002 | Young |
| 0.792087 | 0.047921 | 0.062 | 0.065 | 1 | 2 | Syne2 | -0.003 | Young |
| 0.792229 | 0.129664 | 0.014 | 0.013 | 1 | 2 | Sh2d3c | 0.001 | Young |
| 0.792291 | 0.222607 | 0.023 | 0.025 | 1 | 2 | Ddx51 | -0.002 | Young |
| 0.792337 | 0.087171 | 0.046 | 0.044 | 1 | 2 | Fads1 | 0.002 | Young |
| 0.792399 | 0.116991 | 0.052 | 0.056 | 1 | 2 | Psmd5 | -0.004 | Young |
| 0.79247 | 0.135346 | 0.034 | 0.032 | 1 | 2 | Cul4b | 0.002 | Young |
| 0.792668 | 0.016889 | 0.308 | 0.329 | 1 | 2 | Hnrnpul2 | -0.021 | Young |
| 0.792702 | 0.319112 | 0.015 | 0.014 | 1 | 2 | Eid2b | 0.001 | Young |
| 0.792943 | 0.471179 | 0.014 | 0.013 | 1 | 2 | Cul7 | 0.001 | Young |
| 0.793182 | 0.001094 | 0.148 | 0.155 | 1 | 2 | Nfkb2 | -0.007 | Young |
| 0.794152 | 0.227174 | 0.032 | 0.03 | 1 | 2 | Rubcnl | 0.002 | Young |
| 0.794411 | 0.287661 | 0.05 | 0.054 | 1 | 2 | Dcxr | -0.004 | Young |
| 0.794451 | 0.112869 | 0.173 | 0.17 | 1 | 2 | Ergic3 | 0.003 | Young |
| 0.794514 | 0.579038 | 0.029 | 0.027 | 1 | 2 | Kctd11 | 0.002 | Young |
| 0.794682 | 0.62788 | 0.015 | 0.017 | 1 | 2 | Gm4707 | -0.002 | Young |
| 0.794763 | 0.02698 | 0.015 | 0.014 | 1 | 2 | Xlr | 0.001 | Young |
| 0.795184 | 0.236023 | 0.045 | 0.048 | 1 | 2 | Triobp | -0.003 | Young |
| 0.795231 | 0.083669 | 0.328 | 0.329 | 1 | 2 | Sdha | -0.001 | Young |
| 0.795445 | 0.115706 | 0.014 | 0.013 | 1 | 2 | Tmtc4 | 0.001 | Young |
| 0.795888 | 0.057121 | 0.107 | 0.112 | 1 | 2 | Gramd1a | -0.005 | Young |
| 0.796032 | 0.348376 | 0.109 | 0.108 | 1 | 2 | Fmr1 | 0.001 | Young |
| 0.796134 | 0.245136 | 0.096 | 0.102 | 1 | 2 | Gid8 | -0.006 | Young |
| 0.796206 | 0.225437 | 0.07 | 0.067 | 1 | 2 | Ppp1r37 | 0.003 | Young |
| 0.796493 | 0.030271 | 0.308 | 0.305 | 1 | 2 | Ash1l | 0.003 | Young |
| 0.796517 | 0.375363 | 0.014 | 0.013 | 1 | 2 | Rida | 0.001 | Young |
| 0.796538 | 0.058647 | 0.094 | 0.099 | 1 | 2 | Cnp | -0.005 | Young |
| 0.797417 | 0.019449 | 0.045 | 0.043 | 1 | 2 | Tstd3 | 0.002 | Young |
| 0.797483 | 0.027294 | 0.069 | 0.073 | 1 | 2 | Camk2g | -0.004 | Young |
| 0.797678 | 0.72473 | 0.013 | 0.012 | 1 | 2 | Tprn | 0.001 | Young |
| 0.797878 | 0.199028 | 0.059 | 0.057 | 1 | 2 | Poglut1 | 0.002 | Young |
| 0.797879 | 0.065806 | 0.211 | 0.223 | 1 | 2 | Cnih1 | -0.012 | Young |
| 0.798337 | 0.064847 | 0.063 | 0.06 | 1 | 2 | Gfra2 | 0.003 | Young |
| 0.798526 | 0.066342 | 0.084 | 0.088 | 1 | 2 | Reps2 | -0.004 | Young |
| 0.798537 | 0.146038 | 0.132 | 0.131 | 1 | 2 | Synrg | 0.001 | Young |
| 0.798719 | 0.366626 | 0.058 | 0.056 | 1 | 2 | Cav2 | 0.002 | Young |
| 0.798719 | 0.081106 | 0.058 | 0.056 | 1 | 2 | Tbc1d10a | 0.002 | Young |
| 0.799016 | 0.050415 | 0.334 | 0.333 | 1 | 2 | Atxn7l3b | 0.001 | Young |
| 0.799022 | 0.297104 | 0.014 | 0.013 | 1 | 2 | Zfp319 | 0.001 | Young |
| 0.799332 | 0.087476 | 0.132 | 0.127 | 1 | 2 | 1810030O( | 0.005 | Young |
| 0.799512 | 0.199452 | 0.01 | 0.011 | 1 | 2 | Zfp12 | -0.001 | Young |

| | | | | | | | | |
|---|---|---|---|---|---|---|---|---|
| 0.799951 | 0.2692 | 0.051 | 0.055 | 1 | 2 | Ripor2 | -0.004 | Young |
| 0.800137 | 0.018392 | 0.101 | 0.097 | 1 | 2 | Ssbp1 | 0.004 | Young |
| 0.800574 | 0.25917 | 0.059 | 0.063 | 1 | 2 | Zfand2b | -0.004 | Young |
| 0.800602 | 0.049238 | 0.22 | 0.216 | 1 | 2 | Tsc22d2 | 0.004 | Young |
| 0.800681 | 0.061564 | 0.072 | 0.076 | 1 | 2 | Golga2 | -0.004 | Young |
| 0.800886 | 0.115211 | 0.167 | 0.175 | 1 | 2 | Lman1 | -0.008 | Young |
| 0.800941 | 0.227095 | 0.095 | 0.092 | 1 | 2 | Ndrg1 | 0.003 | Young |
| 0.801235 | 0.08469 | 0.024 | 0.026 | 1 | 2 | Zfp276 | -0.002 | Young |
| 0.801423 | 0.102734 | 0.032 | 0.03 | 1 | 2 | Ripor1 | 0.002 | Young |
| 0.801536 | 0.099428 | 0.048 | 0.05 | 1 | 2 | Chmp7 | -0.002 | Young |
| 0.801789 | 0.303243 | 0.013 | 0.012 | 1 | 2 | Thg1l | 0.001 | Young |
| 0.801964 | 0.038341 | 0.116 | 0.113 | 1 | 2 | Micu1 | 0.003 | Young |
| 0.802063 | 0.00507 | 0.059 | 0.062 | 1 | 2 | Fbxl17 | -0.003 | Young |
| 0.802245 | 0.04494 | 0.014 | 0.013 | 1 | 2 | Zfp930 | 0.001 | Young |
| 0.802459 | 0.05121 | 0.081 | 0.086 | 1 | 2 | Nubp1 | -0.005 | Young |
| 0.802483 | 0.154403 | 0.094 | 0.099 | 1 | 2 | Phf20 | -0.005 | Young |
| 0.802563 | 0.02961 | 0.166 | 0.164 | 1 | 2 | Dhx36 | 0.002 | Young |
| 0.802754 | 0.113911 | 0.138 | 0.146 | 1 | 2 | Slc31a2 | -0.008 | Young |
| 0.802887 | 0.059441 | 0.143 | 0.149 | 1 | 2 | Rnf38 | -0.006 | Young |
| 0.803149 | 0.016191 | 0.475 | 0.469 | 1 | 2 | Jak1 | 0.006 | Young |
| 0.803364 | 0.165273 | 0.024 | 0.026 | 1 | 2 | Mad2l2 | -0.002 | Young |
| 0.804589 | 0.051788 | 0.325 | 0.345 | 1 | 2 | Kdm2a | -0.02 | Young |
| 0.804628 | 0.177439 | 0.166 | 0.173 | 1 | 2 | Denr | -0.007 | Young |
| 0.804974 | 0.31778 | 0.045 | 0.048 | 1 | 2 | Exosc7 | -0.003 | Young |
| 0.805295 | 0.097526 | 0.01 | 0.011 | 1 | 2 | Gm16104 | -0.001 | Young |
| 0.805355 | 0.27985 | 0.097 | 0.095 | 1 | 2 | Naxd | 0.002 | Young |
| 0.805508 | 0.05377 | 0.076 | 0.08 | 1 | 2 | Ifi27 | -0.004 | Young |
| 0.805584 | 0.042027 | 0.127 | 0.133 | 1 | 2 | Agtpbp1 | -0.006 | Young |
| 0.805708 | 0.191971 | 0.01 | 0.011 | 1 | 2 | Cep131 | -0.001 | Young |
| 0.806068 | 0.024122 | 0.226 | 0.223 | 1 | 2 | Fes | 0.003 | Young |
| 0.806072 | 0.04416 | 0.044 | 0.042 | 1 | 2 | Zmat3 | 0.002 | Young |
| 0.806114 | 0.027734 | 0.404 | 0.406 | 1 | 2 | Tet3 | -0.002 | Young |
| 0.806125 | 0.272551 | 0.039 | 0.042 | 1 | 2 | Kctd6 | -0.003 | Young |
| 0.806409 | 0.108491 | 0.325 | 0.33 | 1 | 2 | Ptges3 | -0.005 | Young |
| 0.80644 | 0.028221 | 0.032 | 0.03 | 1 | 2 | Spata5 | 0.002 | Young |
| 0.80653 | 0.187687 | 0.064 | 0.062 | 1 | 2 | Ints3 | 0.002 | Young |
| 0.80656 | 0.043427 | 0.024 | 0.026 | 1 | 2 | Fktn | -0.002 | Young |
| 0.806583 | 0.382677 | 0.044 | 0.042 | 1 | 2 | Tsc2 | 0.002 | Young |
| 0.806588 | 0.050039 | 0.265 | 0.263 | 1 | 2 | AI504432 | 0.002 | Young |
| 0.807029 | 0.01921 | 0.013 | 0.012 | 1 | 2 | E130308A1 | 0.001 | Young |
| 0.807334 | 0.330333 | 0.075 | 0.079 | 1 | 2 | Ankrd49 | -0.004 | Young |
| 0.807403 | 0.290832 | 0.013 | 0.012 | 1 | 2 | Zfat | 0.001 | Young |
| 0.807483 | 0.316118 | 0.017 | 0.018 | 1 | 2 | Zfp275 | -0.001 | Young |
| 0.807628 | 0.044581 | 0.226 | 0.234 | 1 | 2 | Alyref | -0.008 | Young |
| 0.807634 | 0.073653 | 0.087 | 0.091 | 1 | 2 | Cul2 | -0.004 | Young |

| | | | | | | | | |
|---|---|---|---|---|---|---|---|---|
| 0.807635 | 0.059012 | 0.083 | 0.087 | 1 | 2 | Usf3 | -0.004 | Young |
| 0.807894 | 0.382299 | 0.029 | 0.027 | 1 | 2 | Mgme1 | 0.002 | Young |
| 0.808116 | 0.296173 | 0.038 | 0.041 | 1 | 2 | Dusp23 | -0.003 | Young |
| 0.808153 | 0.068866 | 0.013 | 0.012 | 1 | 2 | Zfp523 | 0.001 | Young |
| 0.808153 | 0.056533 | 0.013 | 0.012 | 1 | 2 | Eps8l1 | 0.001 | Young |
| 0.808221 | 0.088942 | 0.097 | 0.102 | 1 | 2 | Hivep3 | -0.005 | Young |
| 0.808331 | 0.223751 | 0.042 | 0.04 | 1 | 2 | Sil1 | 0.002 | Young |
| 0.808397 | 0.16217 | 0.043 | 0.041 | 1 | 2 | Scaper | 0.002 | Young |
| 0.808574 | 0.080129 | 0.1 | 0.105 | 1 | 2 | Ethe1 | -0.005 | Young |
| 0.80861 | 0.066832 | 0.056 | 0.059 | 1 | 2 | Ppp1r8 | -0.003 | Young |
| 0.808706 | 0.619382 | 0.012 | 0.011 | 1 | 2 | Ift80 | 0.001 | Young |
| 0.808909 | 0.068427 | 0.394 | 0.398 | 1 | 2 | Atp6v1e1 | -0.004 | Young |
| 0.809203 | 0.060038 | 0.112 | 0.117 | 1 | 2 | Pfkfb4 | -0.005 | Young |
| 0.809382 | 0.585873 | 0.017 | 0.018 | 1 | 2 | Adck2 | -0.001 | Young |
| 0.809421 | 0.290195 | 0.122 | 0.122 | 1 | 2 | Cmc1 | 0 | Young |
| 0.809492 | 0.378762 | 0.012 | 0.011 | 1 | 2 | Prkca | 0.001 | Young |
| 0.809581 | 0.535013 | 0.039 | 0.037 | 1 | 2 | Pdk1 | 0.002 | Young |
| 0.809885 | 0.358072 | 0.012 | 0.011 | 1 | 2 | Sorbs3 | 0.001 | Young |
| 0.810293 | 0.190635 | 0.024 | 0.026 | 1 | 2 | Mrpl39 | -0.002 | Young |
| 0.810617 | 0.008474 | 0.605 | 0.618 | 1 | 2 | Csde1 | -0.013 | Young |
| 0.810649 | 0.155653 | 0.017 | 0.018 | 1 | 2 | Nectin4 | -0.001 | Young |
| 0.810671 | 0.338446 | 0.012 | 0.011 | 1 | 2 | Pigo | 0.001 | Young |
| 0.810698 | 0.031482 | 0.032 | 0.034 | 1 | 2 | Eci1 | -0.002 | Young |
| 0.810796 | 0.282729 | 0.031 | 0.029 | 1 | 2 | Rhbdd2 | 0.002 | Young |
| 0.810966 | 0.164295 | 0.017 | 0.018 | 1 | 2 | Depdc7 | -0.001 | Young |
| 0.81126 | 0.005933 | 0.424 | 0.42 | 1 | 2 | Slc43a2 | 0.004 | Young |
| 0.811927 | 0.012822 | 0.162 | 0.169 | 1 | 2 | Arid1b | -0.007 | Young |
| 0.811944 | 0.236648 | 0.029 | 0.027 | 1 | 2 | Syce2 | 0.002 | Young |
| 0.812244 | 0.269461 | 0.012 | 0.011 | 1 | 2 | Phldb3 | 0.001 | Young |
| 0.81233 | 0.081851 | 0.106 | 0.11 | 1 | 2 | Txlna | -0.004 | Young |
| 0.812434 | 0.064231 | 0.228 | 0.239 | 1 | 2 | 1810058I2 | -0.011 | Young |
| 0.812471 | 0.623257 | 0.038 | 0.036 | 1 | 2 | Ogfod2 | 0.002 | Young |
| 0.812637 | 0.364997 | 0.012 | 0.011 | 1 | 2 | Lrfn1 | 0.001 | Young |
| 0.812928 | 0.014034 | 0.058 | 0.056 | 1 | 2 | Sart3 | 0.002 | Young |
| 0.813259 | 0.064918 | 0.374 | 0.393 | 1 | 2 | Ndufb9 | -0.019 | Young |
| 0.813509 | 0.135406 | 0.03 | 0.028 | 1 | 2 | Fbxl18 | 0.002 | Young |
| 0.813977 | 0.003361 | 0.165 | 0.171 | 1 | 2 | Pcf11 | -0.006 | Young |
| 0.814006 | 0.1985 | 0.03 | 0.028 | 1 | 2 | Tmem185a | 0.002 | Young |
| 0.814404 | 0.119546 | 0.12 | 0.118 | 1 | 2 | Crat | 0.002 | Young |
| 0.814896 | 0.395563 | 0.063 | 0.061 | 1 | 2 | Nfs1 | 0.002 | Young |
| 0.81517 | 0.053082 | 0.2 | 0.196 | 1 | 2 | Sfxn3 | 0.004 | Young |
| 0.815329 | 0.244246 | 0.112 | 0.109 | 1 | 2 | Vps54 | 0.003 | Young |
| 0.815402 | 0.179529 | 0.045 | 0.043 | 1 | 2 | Hps1 | 0.002 | Young |
| 0.815483 | 0.021567 | 0.317 | 0.33 | 1 | 2 | Twf2 | -0.013 | Young |
| 0.815516 | 0.311895 | 0.071 | 0.07 | 1 | 2 | Cenpc1 | 0.001 | Young |

| | | | | | | | | |
|---|---|---|---|---|---|---|---|---|
| 0.815834 | 0.126554 | 0.137 | 0.134 | 1 | 2 | Mtfr1l | 0.003 | Young |
| 0.81597 | 0.004613 | 0.051 | 0.054 | 1 | 2 | Kansl1l | -0.003 | Young |
| 0.816257 | 0.005374 | 0.032 | 0.034 | 1 | 2 | Edrf1 | -0.002 | Young |
| 0.816684 | 0.411827 | 0.011 | 0.01 | 1 | 2 | AW146154 | 0.001 | Young |
| 0.8167 | 0.108285 | 0.056 | 0.054 | 1 | 2 | Endod1 | 0.002 | Young |
| 0.816887 | 0.118196 | 0.057 | 0.055 | 1 | 2 | Plau | 0.002 | Young |
| 0.816891 | 0.523236 | 0.011 | 0.01 | 1 | 2 | Zfp677 | 0.001 | Young |
| 0.816925 | 0.114049 | 0.198 | 0.198 | 1 | 2 | Papd4 | 0 | Young |
| 0.816959 | 0.035722 | 0.593 | 0.597 | 1 | 2 | Luc7l2 | -0.004 | Young |
| 0.817314 | 0.16911 | 0.033 | 0.035 | 1 | 2 | Zc3h12c | -0.002 | Young |
| 0.817532 | 0.104953 | 0.379 | 0.382 | 1 | 2 | Ier3ip1 | -0.003 | Young |
| 0.817603 | 0.058627 | 0.371 | 0.375 | 1 | 2 | Usmg5 | -0.004 | Young |
| 0.817729 | 0.106387 | 0.127 | 0.132 | 1 | 2 | Prpf8 | -0.005 | Young |
| 0.817825 | 0.19271 | 0.063 | 0.061 | 1 | 2 | Rnf215 | 0.002 | Young |
| 0.818312 | 0.100368 | 0.175 | 0.184 | 1 | 2 | Tmem208 | -0.009 | Young |
| 0.819623 | 0.0019 | 0.124 | 0.12 | 1 | 2 | Mospd3 | 0.004 | Young |
| 0.82009 | 0.104748 | 0.14 | 0.139 | 1 | 2 | Prkab2 | 0.001 | Young |
| 0.820536 | 0.18611 | 0.097 | 0.102 | 1 | 2 | Supt20 | -0.005 | Young |
| 0.820765 | 0.109196 | 0.119 | 0.124 | 1 | 2 | BC037034 | -0.005 | Young |
| 0.821107 | 0.067067 | 0.087 | 0.091 | 1 | 2 | Rprd1b | -0.004 | Young |
| 0.821237 | 0.033857 | 0.502 | 0.5 | 1 | 2 | Adam15 | 0.002 | Young |
| 0.82129 | 0.096479 | 0.251 | 0.252 | 1 | 2 | Cbx3 | -0.001 | Young |
| 0.821605 | 0.119399 | 0.072 | 0.07 | 1 | 2 | Lrch1 | 0.002 | Young |
| 0.821619 | 0.173664 | 0.133 | 0.14 | 1 | 2 | Ncoa6 | -0.007 | Young |
| 0.821674 | 0.17198 | 0.076 | 0.074 | 1 | 2 | Akap11 | 0.002 | Young |
| 0.821702 | 0.083172 | 0.088 | 0.086 | 1 | 2 | Bloc1s2 | 0.002 | Young |
| 0.821789 | 0.130855 | 0.184 | 0.195 | 1 | 2 | Cyb5b | -0.011 | Young |
| 0.822082 | 0.380663 | 0.011 | 0.01 | 1 | 2 | Cbx8 | 0.001 | Young |
| 0.822082 | 0.349661 | 0.011 | 0.01 | 1 | 2 | Klf8 | 0.001 | Young |
| 0.822497 | 0.00187 | 0.011 | 0.01 | 1 | 2 | Sh3rf1 | 0.001 | Young |
| 0.822842 | 0.014567 | 0.72 | 0.721 | 1 | 2 | Pitpna | -0.001 | Young |
| 0.823044 | 0.246237 | 0.018 | 0.019 | 1 | 2 | Ssh3 | -0.001 | Young |
| 0.823111 | 0.02831 | 0.045 | 0.043 | 1 | 2 | Scyl3 | 0.002 | Young |
| 0.82343 | 0.536305 | 0.026 | 0.025 | 1 | 2 | Olfr1167 | 0.001 | Young |
| 0.823551 | 0.099946 | 0.233 | 0.234 | 1 | 2 | Mlf2 | -0.001 | Young |
| 0.823902 | 0.080572 | 0.324 | 0.335 | 1 | 2 | Mapk1 | -0.011 | Young |
| 0.82416 | 0.135498 | 0.011 | 0.01 | 1 | 2 | Fam212a | 0.001 | Young |
| 0.824416 | 0.032889 | 0.12 | 0.124 | 1 | 2 | Nab2 | -0.004 | Young |
| 0.824576 | 0.025705 | 0.011 | 0.01 | 1 | 2 | Ddb2 | 0.001 | Young |
| 0.824664 | 0.036688 | 0.722 | 0.722 | 1 | 2 | Tagln2 | 0 | Young |
| 0.824701 | 0.064224 | 0.094 | 0.091 | 1 | 2 | Mtf2 | 0.003 | Young |
| 0.824847 | 0.442971 | 0.039 | 0.037 | 1 | 2 | Pld2 | 0.002 | Young |
| 0.824992 | 0.105785 | 0.011 | 0.01 | 1 | 2 | Tmem263 | 0.001 | Young |
| 0.824992 | 0.020413 | 0.011 | 0.01 | 1 | 2 | Adamtsl4 | 0.001 | Young |
| 0.825113 | 0.26166 | 0.084 | 0.082 | 1 | 2 | Cwc25 | 0.002 | Young |

| | | | | | | | | |
|---|---|---|---|---|---|---|---|---|
| 0.8253 | 0.053978 | 0.089 | 0.087 | 1 | 2 | Arhgef12 | 0.002 | Young |
| 0.825619 | 0.054986 | 0.23 | 0.24 | 1 | 2 | Zfand3 | -0.01 | Young |
| 0.825823 | 0.248324 | 0.011 | 0.01 | 1 | 2 | Inpp4b | 0.001 | Young |
| 0.825855 | 0.008719 | 0.482 | 0.48 | 1 | 2 | Prkar1a | 0.002 | Young |
| 0.825859 | 0.17276 | 0.118 | 0.116 | 1 | 2 | Orai3 | 0.002 | Young |
| 0.826132 | 0.059061 | 0.167 | 0.165 | 1 | 2 | Kctd20 | 0.002 | Young |
| 0.826279 | 0.112792 | 0.185 | 0.182 | 1 | 2 | Fkbp15 | 0.003 | Young |
| 0.826514 | 0.191616 | 0.131 | 0.13 | 1 | 2 | Mau2 | 0.001 | Young |
| 0.826585 | 0.173752 | 0.056 | 0.054 | 1 | 2 | Mfn2 | 0.002 | Young |
| 0.826629 | 0.021209 | 0.011 | 0.012 | 1 | 2 | Cenpo | -0.001 | Young |
| 0.826984 | 0.304694 | 0.025 | 0.027 | 1 | 2 | 1110032A( | -0.002 | Young |
| 0.827216 | 0.449093 | 0.01 | 0.009 | 1 | 2 | Ccdc94 | 0.001 | Young |
| 0.827237 | 0.004712 | 0.224 | 0.222 | 1 | 2 | Copb1 | 0.002 | Young |
| 0.827406 | 0.350908 | 0.026 | 0.025 | 1 | 2 | Mtrf1l | 0.001 | Young |
| 0.827617 | 0.065956 | 0.099 | 0.096 | 1 | 2 | Polr1a | 0.003 | Young |
| 0.827627 | 0.051212 | 0.11 | 0.116 | 1 | 2 | Hdac2 | -0.006 | Young |
| 0.82832 | 0.099031 | 0.011 | 0.01 | 1 | 2 | Bahcc1 | 0.001 | Young |
| 0.82864 | 0.043105 | 0.106 | 0.11 | 1 | 2 | Cops6 | -0.004 | Young |
| 0.828806 | 0.119311 | 0.156 | 0.154 | 1 | 2 | Rapgef2 | 0.002 | Young |
| 0.828906 | 0.119649 | 0.018 | 0.019 | 1 | 2 | Mipep | -0.001 | Young |
| 0.828944 | 0.00603 | 0.158 | 0.164 | 1 | 2 | Map3k2 | -0.006 | Young |
| 0.829002 | 0.128726 | 0.19 | 0.191 | 1 | 2 | Arpc1a | -0.001 | Young |
| 0.829125 | 0.148216 | 0.164 | 0.163 | 1 | 2 | Phc3 | 0.001 | Young |
| 0.829146 | 0.103998 | 0.176 | 0.176 | 1 | 2 | Aimp1 | 0 | Young |
| 0.829377 | 0.117993 | 0.109 | 0.107 | 1 | 2 | Mapkap1 | 0.002 | Young |
| 0.829406 | 0.011612 | 0.165 | 0.162 | 1 | 2 | Usp3 | 0.003 | Young |
| 0.829479 | 0.18763 | 0.131 | 0.13 | 1 | 2 | Usp12 | 0.001 | Young |
| 0.829833 | 0.029736 | 0.018 | 0.019 | 1 | 2 | Nek1 | -0.001 | Young |
| 0.829893 | 0.003463 | 0.087 | 0.084 | 1 | 2 | Pbx3 | 0.003 | Young |
| 0.830006 | 0.073753 | 0.089 | 0.087 | 1 | 2 | Zfp622 | 0.002 | Young |
| 0.830108 | 0.235609 | 0.025 | 0.024 | 1 | 2 | Pold1 | 0.001 | Young |
| 0.830516 | 0.026541 | 0.308 | 0.31 | 1 | 2 | Cmtm6 | -0.002 | Young |
| 0.830747 | 0.464722 | 0.01 | 0.009 | 1 | 2 | Cela1 | 0.001 | Young |
| 0.830862 | 0.079012 | 0.385 | 0.4 | 1 | 2 | Prdx2 | -0.015 | Young |
| 0.831206 | 0.120432 | 0.144 | 0.148 | 1 | 2 | Plpbp | -0.004 | Young |
| 0.831451 | 0.059858 | 0.121 | 0.119 | 1 | 2 | Gtf2h1 | 0.002 | Young |
| 0.831531 | 0.109918 | 0.084 | 0.081 | 1 | 2 | Fzr1 | 0.003 | Young |
| 0.83163 | 0.387898 | 0.01 | 0.009 | 1 | 2 | Hmgb3 | 0.001 | Young |
| 0.83176 | 0.032054 | 0.169 | 0.176 | 1 | 2 | Elovl5 | -0.007 | Young |
| 0.831767 | 0.058014 | 0.011 | 0.012 | 1 | 2 | Gbp4 | -0.001 | Young |
| 0.831909 | 0.226529 | 0.089 | 0.093 | 1 | 2 | Gcsh | -0.004 | Young |
| 0.831973 | 0.519032 | 0.025 | 0.027 | 1 | 2 | Tor4a | -0.002 | Young |
| 0.832164 | 0.029151 | 0.102 | 0.106 | 1 | 2 | Dok1 | -0.004 | Young |
| 0.832435 | 0.118555 | 0.04 | 0.043 | 1 | 2 | H6pd | -0.003 | Young |
| 0.83251 | 0.083039 | 0.076 | 0.074 | 1 | 2 | Zfp236 | 0.002 | Young |

| | | | | | | | | |
|---|---|---|---|---|---|---|---|---|
| 0.832513 | 0.095482 | 0.01 | 0.009 | 1 | 2 | Arsg | 0.001 | Young |
| 0.832522 | 0.488853 | 0.05 | 0.048 | 1 | 2 | Rcan3 | 0.002 | Young |
| 0.832639 | 0.050911 | 0.171 | 0.177 | 1 | 2 | Rab20 | -0.006 | Young |
| 0.832787 | 0.077079 | 0.05 | 0.052 | 1 | 2 | Bcl2l12 | -0.002 | Young |
| 0.832955 | 0.432469 | 0.01 | 0.009 | 1 | 2 | Pkmyt1 | 0.001 | Young |
| 0.832961 | 0.06025 | 0.182 | 0.18 | 1 | 2 | Mta2 | 0.002 | Young |
| 0.832997 | 0.121541 | 0.141 | 0.149 | 1 | 2 | Kdelr2 | -0.008 | Young |
| 0.833231 | 0.124454 | 0.115 | 0.12 | 1 | 2 | Rbbp7 | -0.005 | Young |
| 0.833377 | 0.186516 | 0.034 | 0.036 | 1 | 2 | Ap5s1 | -0.002 | Young |
| 0.833397 | 0.115088 | 0.01 | 0.009 | 1 | 2 | Rwdd2a | 0.001 | Young |
| 0.83394 | 0.041458 | 0.211 | 0.219 | 1 | 2 | Dock7 | -0.008 | Young |
| 0.8341 | 0.150331 | 0.053 | 0.051 | 1 | 2 | Dtx4 | 0.002 | Young |
| 0.834141 | 0.001297 | 0.011 | 0.012 | 1 | 2 | 6330562C2 | -0.001 | Young |
| 0.834223 | 0.081269 | 0.114 | 0.119 | 1 | 2 | Wdr41 | -0.005 | Young |
| 0.83424 | 0.113299 | 0.033 | 0.035 | 1 | 2 | Exoc8 | -0.002 | Young |
| 0.83424 | 0.295459 | 0.033 | 0.035 | 1 | 2 | Ganc | -0.002 | Young |
| 0.834466 | 0.081802 | 0.026 | 0.028 | 1 | 2 | Dbr1 | -0.002 | Young |
| 0.834556 | 0.202324 | 0.086 | 0.084 | 1 | 2 | Upf2 | 0.002 | Young |
| 0.835329 | 0.694104 | 0.011 | 0.012 | 1 | 2 | B130006D( | -0.001 | Young |
| 0.835381 | 0.099513 | 0.115 | 0.121 | 1 | 2 | D5Ertd579 | -0.006 | Young |
| 0.835559 | 0.096262 | 0.304 | 0.305 | 1 | 2 | Scaf11 | -0.001 | Young |
| 0.836104 | 0.061517 | 0.019 | 0.02 | 1 | 2 | Mmp8 | -0.001 | Young |
| 0.83612 | 0.345928 | 0.011 | 0.012 | 1 | 2 | Ms4a4a | -0.001 | Young |
| 0.83612 | 0.088536 | 0.011 | 0.012 | 1 | 2 | Fkbp7 | -0.001 | Young |
| 0.837034 | 0.283092 | 0.025 | 0.024 | 1 | 2 | Cipc | 0.001 | Young |
| 0.837168 | 0.263442 | 0.057 | 0.06 | 1 | 2 | Mavs | -0.003 | Young |
| 0.837547 | 0.257099 | 0.051 | 0.049 | 1 | 2 | Zfp318 | 0.002 | Young |
| 0.837574 | 0.010122 | 0.075 | 0.078 | 1 | 2 | Faf1 | -0.003 | Young |
| 0.837818 | 0.054743 | 0.01 | 0.009 | 1 | 2 | Zfp119b | 0.001 | Young |
| 0.838238 | 0.14291 | 0.375 | 0.38 | 1 | 2 | Prrc2a | -0.005 | Young |
| 0.838564 | 0.033645 | 0.489 | 0.495 | 1 | 2 | Set | -0.006 | Young |
| 0.838622 | 0.025246 | 0.128 | 0.126 | 1 | 2 | Galnt7 | 0.002 | Young |
| 0.838784 | 0.08829 | 0.038 | 0.036 | 1 | 2 | Arap3 | 0.002 | Young |
| 0.8388 | 0.035937 | 0.236 | 0.232 | 1 | 2 | Ppt1 | 0.004 | Young |
| 0.839023 | 0.19555 | 0.034 | 0.036 | 1 | 2 | Nr2c1 | -0.002 | Young |
| 0.839056 | 0.113284 | 0.249 | 0.251 | 1 | 2 | Taf1d | -0.002 | Young |
| 0.839071 | 0.178846 | 0.072 | 0.071 | 1 | 2 | Isca2 | 0.001 | Young |
| 0.839169 | 0.035959 | 0.31 | 0.307 | 1 | 2 | Msl1 | 0.003 | Young |
| 0.839247 | 0.071118 | 0.128 | 0.126 | 1 | 2 | 0610030E2 | 0.002 | Young |
| 0.839253 | 0.150001 | 0.081 | 0.085 | 1 | 2 | Pex19 | -0.004 | Young |
| 0.839397 | 0.01229 | 0.043 | 0.045 | 1 | 2 | Dusp10 | -0.002 | Young |
| 0.839695 | 0.050634 | 0.167 | 0.175 | 1 | 2 | Ftl1-ps1 | -0.008 | Young |
| 0.839928 | 0.125932 | 0.185 | 0.185 | 1 | 2 | Mrpl54 | 0 | Young |
| 0.840077 | 0.188764 | 0.049 | 0.047 | 1 | 2 | Camsap1 | 0.002 | Young |
| 0.840283 | 0.077318 | 0.169 | 0.168 | 1 | 2 | Tomm70a | 0.001 | Young |

| | | | | | | | | |
|---|---|---|---|---|---|---|---|---|
| 0.84037 | 0.041148 | 0.324 | 0.32 | 1 | 2 | Stag2 | 0.004 | Young |
| 0.840783 | 0.016569 | 0.395 | 0.407 | 1 | 2 | Tmem106a | -0.012 | Young |
| 0.841356 | 0.089677 | 0.026 | 0.025 | 1 | 2 | H2-Ob | 0.001 | Young |
| 0.841399 | 0.134739 | 0.091 | 0.09 | 1 | 2 | Gbf1 | 0.001 | Young |
| 0.841584 | 0.289516 | 0.033 | 0.035 | 1 | 2 | Capn15 | -0.002 | Young |
| 0.843 | 0.196811 | 0.105 | 0.108 | 1 | 2 | Daxx | -0.003 | Young |
| 0.843071 | 0.361831 | 0.043 | 0.045 | 1 | 2 | Pramef8 | -0.002 | Young |
| 0.843111 | 0.03433 | 0.486 | 0.496 | 1 | 2 | Hcls1 | -0.01 | Young |
| 0.843298 | 0.216275 | 0.101 | 0.105 | 1 | 2 | Pafah1b2 | -0.004 | Young |
| 0.843429 | 0.132797 | 0.025 | 0.024 | 1 | 2 | Ttll4 | 0.001 | Young |
| 0.843557 | 0.165535 | 0.05 | 0.048 | 1 | 2 | N6amt1 | 0.002 | Young |
| 0.843685 | 0.258179 | 0.067 | 0.065 | 1 | 2 | Dtymk | 0.002 | Young |
| 0.843955 | 0.034459 | 0.076 | 0.079 | 1 | 2 | Zbtb21 | -0.003 | Young |
| 0.844195 | 0.087725 | 0.077 | 0.08 | 1 | 2 | Plcb1 | -0.003 | Young |
| 0.844533 | 0.370596 | 0.024 | 0.022 | 1 | 2 | Gemin5 | 0.002 | Young |
| 0.844875 | 0.020149 | 0.065 | 0.063 | 1 | 2 | Ormdl1 | 0.002 | Young |
| 0.845003 | 0.313846 | 0.057 | 0.06 | 1 | 2 | Prorsd1 | -0.003 | Young |
| 0.845388 | 0.065704 | 0.388 | 0.392 | 1 | 2 | Ubap2l | -0.004 | Young |
| 0.8455 | 0.032894 | 0.037 | 0.035 | 1 | 2 | Alg11 | 0.002 | Young |
| 0.845906 | 0.024278 | 0.029 | 0.03 | 1 | 2 | Acrbp | -0.001 | Young |
| 0.845947 | 0.173534 | 0.138 | 0.145 | 1 | 2 | Cklf | -0.007 | Young |
| 0.846054 | 0.004976 | 0.118 | 0.116 | 1 | 2 | Naa35 | 0.002 | Young |
| 0.846102 | 0.194213 | 0.058 | 0.061 | 1 | 2 | Ttc33 | -0.003 | Young |
| 0.846285 | 0.007811 | 0.026 | 0.025 | 1 | 2 | Agk | 0.001 | Young |
| 0.846412 | 0.094455 | 0.065 | 0.063 | 1 | 2 | Adpgk | 0.002 | Young |
| 0.846456 | 0.119382 | 0.09 | 0.094 | 1 | 2 | Lin54 | -0.004 | Young |
| 0.846659 | 0.049829 | 0.078 | 0.081 | 1 | 2 | Nudt9 | -0.003 | Young |
| 0.847123 | 0.01594 | 0.078 | 0.076 | 1 | 2 | Pbdc1 | 0.002 | Young |
| 0.847133 | 0.078398 | 0.299 | 0.313 | 1 | 2 | U2af2 | -0.014 | Young |
| 0.847412 | 0.088832 | 0.303 | 0.315 | 1 | 2 | Mbtd1 | -0.012 | Young |
| 0.847486 | 0.093772 | 0.039 | 0.037 | 1 | 2 | Ppm1d | 0.002 | Young |
| 0.847855 | 0.177706 | 0.038 | 0.036 | 1 | 2 | Pde8a | 0.002 | Young |
| 0.84788 | 0.152476 | 0.024 | 0.022 | 1 | 2 | Met | 0.002 | Young |
| 0.847937 | 0.101263 | 0.258 | 0.271 | 1 | 2 | Lpp | -0.013 | Young |
| 0.848028 | 0.014041 | 0.131 | 0.135 | 1 | 2 | Tmem135 | -0.004 | Young |
| 0.848421 | 0.058102 | 0.045 | 0.047 | 1 | 2 | Bysl | -0.002 | Young |
| 0.848476 | 0.00051 | 0.932 | 0.926 | 1 | 2 | Eif1 | 0.006 | Young |
| 0.848547 | 0.049875 | 0.411 | 0.409 | 1 | 2 | Sbno2 | 0.002 | Young |
| 0.848975 | 0.03062 | 0.063 | 0.065 | 1 | 2 | Farsb | -0.002 | Young |
| 0.849184 | 0.160245 | 0.038 | 0.036 | 1 | 2 | Zc3h12d | 0.002 | Young |
| 0.849186 | 0.254717 | 0.027 | 0.029 | 1 | 2 | Rrp8 | -0.002 | Young |
| 0.849257 | 0.108185 | 0.075 | 0.073 | 1 | 2 | Tvp23b | 0.002 | Young |
| 0.849464 | 0.009695 | 0.175 | 0.181 | 1 | 2 | Cers5 | -0.006 | Young |
| 0.849533 | 0.12995 | 0.059 | 0.062 | 1 | 2 | Dhdds | -0.003 | Young |
| 0.849807 | 0.032847 | 0.074 | 0.072 | 1 | 2 | Ttc5 | 0.002 | Young |

| | | | | | | | | |
|---|---|---|---|---|---|---|---|---|
| 0.849955 | 0.039681 | 0.192 | 0.19 | 1 | 2 | Rsf1 | 0.002 | Young |
| 0.850515 | 0.024174 | 0.025 | 0.024 | 1 | 2 | Alg2 | 0.001 | Young |
| 0.85096 | 0.165479 | 0.027 | 0.029 | 1 | 2 | Zkscan14 | -0.002 | Young |
| 0.85098 | 0.118466 | 0.234 | 0.233 | 1 | 2 | Vav1 | 0.001 | Young |
| 0.851133 | 0.039012 | 0.207 | 0.206 | 1 | 2 | Wbp11 | 0.001 | Young |
| 0.851311 | 0.148198 | 0.179 | 0.179 | 1 | 2 | Vav3 | 0 | Young |
| 0.851343 | 0.058647 | 0.164 | 0.17 | 1 | 2 | Ndufv2 | -0.006 | Young |
| 0.851346 | 0.208624 | 0.037 | 0.035 | 1 | 2 | Polr3e | 0.002 | Young |
| 0.851455 | 0.204128 | 0.023 | 0.021 | 1 | 2 | Fzd7 | 0.002 | Young |
| 0.851532 | 0.054653 | 0.311 | 0.313 | 1 | 2 | Nr3c1 | -0.002 | Young |
| 0.851636 | 0.199254 | 0.12 | 0.119 | 1 | 2 | Atf2 | 0.001 | Young |
| 0.851848 | 0.015338 | 0.027 | 0.029 | 1 | 2 | Trim37 | -0.002 | Young |
| 0.852365 | 0.426386 | 0.046 | 0.045 | 1 | 2 | Dhrs4 | 0.001 | Young |
| 0.852698 | 0.027012 | 0.025 | 0.024 | 1 | 2 | Gpr85 | 0.001 | Young |
| 0.852927 | 0.575584 | 0.033 | 0.032 | 1 | 2 | Mmgt2 | 0.001 | Young |
| 0.853211 | 0.2179 | 0.059 | 0.058 | 1 | 2 | Kdsr | 0.001 | Young |
| 0.853393 | 0.205497 | 0.067 | 0.065 | 1 | 2 | Dcps | 0.002 | Young |
| 0.85378 | 0.097151 | 0.191 | 0.192 | 1 | 2 | Napa | -0.001 | Young |
| 0.853907 | 0.469623 | 0.08 | 0.078 | 1 | 2 | B3gnt5 | 0.002 | Young |
| 0.853983 | 0.096606 | 0.084 | 0.088 | 1 | 2 | Vamp7 | -0.004 | Young |
| 0.854238 | 0.304377 | 0.067 | 0.065 | 1 | 2 | Dtnbp1 | 0.002 | Young |
| 0.854459 | 0.063262 | 0.053 | 0.056 | 1 | 2 | Ints14 | -0.003 | Young |
| 0.854766 | 0.348888 | 0.051 | 0.054 | 1 | 2 | Yif1a | -0.003 | Young |
| 0.855213 | 0.13963 | 0.061 | 0.059 | 1 | 2 | Cfap20 | 0.002 | Young |
| 0.85525 | 0.261326 | 0.076 | 0.075 | 1 | 2 | Ppme1 | 0.001 | Young |
| 0.855878 | 0.104885 | 0.012 | 0.013 | 1 | 2 | Meis1 | -0.001 | Young |
| 0.856245 | 0.052928 | 0.038 | 0.04 | 1 | 2 | Eml3 | -0.002 | Young |
| 0.857372 | 0.272041 | 0.034 | 0.036 | 1 | 2 | Unc13d | -0.002 | Young |
| 0.857593 | 0.168858 | 0.051 | 0.054 | 1 | 2 | Mcm9 | -0.003 | Young |
| 0.857733 | 0.186287 | 0.195 | 0.197 | 1 | 2 | Srp19 | -0.002 | Young |
| 0.858053 | 0.521274 | 0.034 | 0.036 | 1 | 2 | Psmf1 | -0.002 | Young |
| 0.858191 | 0.193208 | 0.058 | 0.057 | 1 | 2 | Tmem167b | 0.001 | Young |
| 0.858586 | 0.212203 | 0.036 | 0.037 | 1 | 2 | Fancl | -0.001 | Young |
| 0.85876 | 0.278837 | 0.023 | 0.021 | 1 | 2 | Flnb | 0.002 | Young |
| 0.859235 | 0.255151 | 0.043 | 0.045 | 1 | 2 | Rgmb | -0.002 | Young |
| 0.859377 | 0.080794 | 0.061 | 0.059 | 1 | 2 | Brms1 | 0.002 | Young |
| 0.85962 | 0.51674 | 0.019 | 0.02 | 1 | 2 | Dnajc18 | -0.001 | Young |
| 0.859906 | 0.312289 | 0.023 | 0.021 | 1 | 2 | Rad1 | 0.002 | Young |
| 0.859976 | 0.108423 | 0.067 | 0.07 | 1 | 2 | Cass4 | -0.003 | Young |
| 0.860193 | 0.169742 | 0.023 | 0.021 | 1 | 2 | Gm35154 | 0.002 | Young |
| 0.860252 | 0.122602 | 0.213 | 0.211 | 1 | 2 | Mgat1 | 0.002 | Young |
| 0.860436 | 0.039645 | 0.012 | 0.013 | 1 | 2 | Sh3bp4 | -0.001 | Young |
| 0.860682 | 0.031476 | 0.147 | 0.146 | 1 | 2 | Egln1 | 0.001 | Young |
| 0.860816 | 0.274318 | 0.012 | 0.013 | 1 | 2 | Otof | -0.001 | Young |
| 0.860997 | 0.255364 | 0.02 | 0.021 | 1 | 2 | Cadm1 | -0.001 | Young |

| | | | | | | | | |
|---|---|---|---|---|---|---|---|---|
| 0.861298 | 0.079021 | 0.08 | 0.082 | 1 | 2 | 9430038I0: | -0.002 | Young |
| 0.86134 | 0.286705 | 0.023 | 0.021 | 1 | 2 | Padi4 | 0.002 | Young |
| 0.861463 | 0.188324 | 0.107 | 0.106 | 1 | 2 | Supt6 | 0.001 | Young |
| 0.861525 | 0.204869 | 0.07 | 0.069 | 1 | 2 | Fbxl14 | 0.001 | Young |
| 0.861576 | 0.122186 | 0.012 | 0.013 | 1 | 2 | Mvk | -0.001 | Young |
| 0.862431 | 0.270046 | 0.101 | 0.106 | 1 | 2 | Rbm34 | -0.005 | Young |
| 0.862469 | 0.0241 | 0.02 | 0.021 | 1 | 2 | Gng11 | -0.001 | Young |
| 0.862631 | 0.232651 | 0.023 | 0.021 | 1 | 2 | Twsg1 | 0.002 | Young |
| 0.863115 | 0.039198 | 0.024 | 0.022 | 1 | 2 | Wdr4 | 0.002 | Young |
| 0.863184 | 0.003092 | 0.063 | 0.065 | 1 | 2 | Gm15417 | -0.002 | Young |
| 0.863239 | 0.229858 | 0.044 | 0.046 | 1 | 2 | Gnl2 | -0.002 | Young |
| 0.863298 | 0.033327 | 0.109 | 0.107 | 1 | 2 | Ncln | 0.002 | Young |
| 0.863349 | 0.030062 | 0.122 | 0.126 | 1 | 2 | Dlg1 | -0.004 | Young |
| 0.863401 | 0.264679 | 0.027 | 0.029 | 1 | 2 | Zfp101 | -0.002 | Young |
| 0.863772 | 0.022294 | 0.201 | 0.208 | 1 | 2 | Ddx46 | -0.007 | Young |
| 0.863941 | 0.0915 | 0.02 | 0.021 | 1 | 2 | Rfxank | -0.001 | Young |
| 0.864666 | 0.264788 | 0.051 | 0.054 | 1 | 2 | Rrm2b | -0.003 | Young |
| 0.864739 | 0.15108 | 0.131 | 0.131 | 1 | 2 | S100a1 | 0 | Young |
| 0.864784 | 0.174159 | 0.023 | 0.021 | 1 | 2 | Mbd4 | 0.002 | Young |
| 0.865 | 0.009963 | 0.192 | 0.196 | 1 | 2 | Rbm26 | -0.004 | Young |
| 0.865097 | 0.063487 | 0.048 | 0.046 | 1 | 2 | Ipo13 | 0.002 | Young |
| 0.865302 | 0.322666 | 0.1 | 0.1 | 1 | 2 | Asxl1 | 0 | Young |
| 0.865443 | 0.326148 | 0.057 | 0.056 | 1 | 2 | Naa40 | 0.001 | Young |
| 0.865493 | 0.073566 | 0.093 | 0.091 | 1 | 2 | Prkce | 0.002 | Young |
| 0.865511 | 0.066654 | 0.499 | 0.509 | 1 | 2 | Pkm | -0.01 | Young |
| 0.865774 | 0.124148 | 0.217 | 0.221 | 1 | 2 | Cox20 | -0.004 | Young |
| 0.866111 | 0.019371 | 0.096 | 0.1 | 1 | 2 | Fcf1 | -0.004 | Young |
| 0.86652 | 0.488949 | 0.012 | 0.013 | 1 | 2 | Abcc10 | -0.001 | Young |
| 0.866674 | 0.113959 | 0.105 | 0.109 | 1 | 2 | Zfp281 | -0.004 | Young |
| 0.867124 | 0.100005 | 0.037 | 0.035 | 1 | 2 | Smarcd1 | 0.002 | Young |
| 0.867411 | 0.243611 | 0.052 | 0.055 | 1 | 2 | Nr2c2ap | -0.003 | Young |
| 0.868203 | 0.131268 | 0.229 | 0.232 | 1 | 2 | Gspt1 | -0.003 | Young |
| 0.868277 | 0.252493 | 0.2 | 0.204 | 1 | 2 | Ndufab1 | -0.004 | Young |
| 0.868312 | 0.225615 | 0.056 | 0.055 | 1 | 2 | Atg4c | 0.001 | Young |
| 0.869092 | 0.002729 | 0.021 | 0.022 | 1 | 2 | Zfp653 | -0.001 | Young |
| 0.869245 | 0.302193 | 0.021 | 0.02 | 1 | 2 | Abtb2 | 0.001 | Young |
| 0.869464 | 0.290813 | 0.045 | 0.044 | 1 | 2 | Gatsl2 | 0.001 | Young |
| 0.869554 | 0.1654 | 0.259 | 0.271 | 1 | 2 | Fam105a | -0.012 | Young |
| 0.869565 | 0.091719 | 0.036 | 0.034 | 1 | 2 | Skp2 | 0.002 | Young |
| 0.869571 | 0.097038 | 0.242 | 0.245 | 1 | 2 | Kcmf1 | -0.003 | Young |
| 0.869761 | 0.068812 | 0.173 | 0.178 | 1 | 2 | Psmb2 | -0.005 | Young |
| 0.869835 | 0.378166 | 0.021 | 0.02 | 1 | 2 | Pigg | 0.001 | Young |
| 0.869861 | 0.243629 | 0.11 | 0.116 | 1 | 2 | Vps18 | -0.006 | Young |
| 0.870077 | 0.219454 | 0.045 | 0.044 | 1 | 2 | Zfp180 | 0.001 | Young |
| 0.87013 | 0.350509 | 0.02 | 0.021 | 1 | 2 | Tfpi | -0.001 | Young |

| | | | | | | | | |
|---|---|---|---|---|---|---|---|---|
| 0.870246 | 0.008618 | 0.625 | 0.642 | 1 | 2 | Slc6a6 | -0.017 | Young |
| 0.870532 | 0.096347 | 0.088 | 0.087 | 1 | 2 | N4bp2 | 0.001 | Young |
| 0.870768 | 0.093815 | 0.082 | 0.08 | 1 | 2 | Fam126b | 0.002 | Young |
| 0.872165 | 0.244458 | 0.034 | 0.033 | 1 | 2 | Il27ra | 0.001 | Young |
| 0.872266 | 0.263297 | 0.12 | 0.12 | 1 | 2 | Ndufa9 | 0 | Young |
| 0.872488 | 0.168847 | 0.156 | 0.163 | 1 | 2 | Aurkaip1 | -0.007 | Young |
| 0.872823 | 0.036125 | 0.289 | 0.293 | 1 | 2 | Cab39 | -0.004 | Young |
| 0.873028 | 0.073953 | 0.284 | 0.294 | 1 | 2 | Clec4b1 | -0.01 | Young |
| 0.873248 | 0.032884 | 0.405 | 0.406 | 1 | 2 | Tsc22d4 | -0.001 | Young |
| 0.87367 | 0.207762 | 0.021 | 0.02 | 1 | 2 | Acvr2b | 0.001 | Young |
| 0.873767 | 0.293187 | 0.095 | 0.094 | 1 | 2 | Hist3h2a | 0.001 | Young |
| 0.874398 | 0.262192 | 0.056 | 0.055 | 1 | 2 | Fbxl6 | 0.001 | Young |
| 0.87448 | 0.184532 | 0.166 | 0.167 | 1 | 2 | Agpat5 | -0.001 | Young |
| 0.874554 | 0.204003 | 0.021 | 0.022 | 1 | 2 | Ubxn2b | -0.001 | Young |
| 0.874762 | 0.461447 | 0.02 | 0.019 | 1 | 2 | Cbr1 | 0.001 | Young |
| 0.87477 | 0.238317 | 0.11 | 0.116 | 1 | 2 | Mbd3 | -0.006 | Young |
| 0.875065 | 0.392824 | 0.02 | 0.019 | 1 | 2 | Rnf219 | 0.001 | Young |
| 0.875252 | 0.197962 | 0.151 | 0.154 | 1 | 2 | Imp3 | -0.003 | Young |
| 0.875301 | 0.177226 | 0.062 | 0.064 | 1 | 2 | Aars | -0.002 | Young |
| 0.875748 | 0.169725 | 0.075 | 0.078 | 1 | 2 | Trmt6 | -0.003 | Young |
| 0.875843 | 0.291686 | 0.036 | 0.037 | 1 | 2 | Xpnpep3 | -0.001 | Young |
| 0.875896 | 0.176719 | 0.077 | 0.08 | 1 | 2 | Taf2 | -0.003 | Young |
| 0.876173 | 0.029906 | 0.166 | 0.171 | 1 | 2 | Atp6v1d | -0.005 | Young |
| 0.87624 | 0.107719 | 0.324 | 0.332 | 1 | 2 | Mafg | -0.008 | Young |
| 0.876313 | 0.032357 | 0.238 | 0.244 | 1 | 2 | Senp6 | -0.006 | Young |
| 0.876621 | 0.04235 | 0.021 | 0.02 | 1 | 2 | Smg8 | 0.001 | Young |
| 0.876632 | 0.113715 | 0.112 | 0.116 | 1 | 2 | Hlx | -0.004 | Young |
| 0.876999 | 0.087952 | 0.296 | 0.304 | 1 | 2 | Sf3b5 | -0.008 | Young |
| 0.877682 | 0.099806 | 0.086 | 0.084 | 1 | 2 | Edc4 | 0.002 | Young |
| 0.877736 | 0.159901 | 0.132 | 0.131 | 1 | 2 | Map3k7 | 0.001 | Young |
| 0.877784 | 0.138682 | 0.096 | 0.1 | 1 | 2 | Svbp | -0.004 | Young |
| 0.877995 | 0.698067 | 0.019 | 0.018 | 1 | 2 | Arrdc2 | 0.001 | Young |
| 0.878138 | 0.000333 | 0.076 | 0.078 | 1 | 2 | Pgp | -0.002 | Young |
| 0.878248 | 0.216274 | 0.137 | 0.141 | 1 | 2 | Znrd1 | -0.004 | Young |
| 0.878628 | 0.212116 | 0.058 | 0.057 | 1 | 2 | Polk | 0.001 | Young |
| 0.878736 | 0.046923 | 0.236 | 0.237 | 1 | 2 | Add1 | -0.001 | Young |
| 0.878948 | 0.062802 | 0.501 | 0.506 | 1 | 2 | Gpr132 | -0.005 | Young |
| 0.879202 | 0.008907 | 0.013 | 0.014 | 1 | 2 | Ybey | -0.001 | Young |
| 0.879236 | 0.135369 | 0.137 | 0.137 | 1 | 2 | Ergic2 | 0 | Young |
| 0.879637 | 0.120836 | 0.062 | 0.064 | 1 | 2 | Kif21b | -0.002 | Young |
| 0.879958 | 0.095111 | 0.333 | 0.348 | 1 | 2 | Hspd1 | -0.015 | Young |
| 0.880047 | 0.123636 | 0.272 | 0.282 | 1 | 2 | Psenen | -0.01 | Young |
| 0.880132 | 0.221664 | 0.113 | 0.117 | 1 | 2 | Preb | -0.004 | Young |
| 0.880299 | 0.103191 | 0.013 | 0.014 | 1 | 2 | Ppp2r5b | -0.001 | Young |
| 0.881255 | 0.239954 | 0.032 | 0.031 | 1 | 2 | Bod1 | 0.001 | Young |

| | | | | | | | | |
|---|---|---|---|---|---|---|---|---|
| 0.881296 | 0.117079 | 0.038 | 0.04 | 1 | 2 | Nt5dc2 | -0.002 | Young |
| 0.881667 | 0.107363 | 0.048 | 0.049 | 1 | 2 | Pxmp4 | -0.001 | Young |
| 0.88175 | 0.072812 | 0.021 | 0.022 | 1 | 2 | Gtf3c5 | -0.001 | Young |
| 0.881963 | 0.09907 | 0.423 | 0.439 | 1 | 2 | Hspe1 | -0.016 | Young |
| 0.882201 | 0.022159 | 0.353 | 0.357 | 1 | 2 | Ppp1r9b | -0.004 | Young |
| 0.882234 | 0.222222 | 0.112 | 0.111 | 1 | 2 | Got2 | 0.001 | Young |
| 0.882388 | 0.160167 | 0.038 | 0.04 | 1 | 2 | Gon7 | -0.002 | Young |
| 0.882654 | 0.252956 | 0.02 | 0.019 | 1 | 2 | Slc7a5 | 0.001 | Young |
| 0.882709 | 0.19608 | 0.056 | 0.055 | 1 | 2 | Inafm1 | 0.001 | Young |
| 0.882858 | 0.095385 | 0.013 | 0.014 | 1 | 2 | Trmt10b | -0.001 | Young |
| 0.883437 | 0.062494 | 0.048 | 0.049 | 1 | 2 | Fam98a | -0.001 | Young |
| 0.883585 | 0.082481 | 0.164 | 0.163 | 1 | 2 | Snd1 | 0.001 | Young |
| 0.883708 | 0.016511 | 0.24 | 0.238 | 1 | 2 | Sh3bp1 | 0.002 | Young |
| 0.883766 | 0.046491 | 0.159 | 0.165 | 1 | 2 | Dnase1l1 | -0.006 | Young |
| 0.884321 | 0.29728 | 0.013 | 0.014 | 1 | 2 | Trmo | -0.001 | Young |
| 0.884872 | 0.594005 | 0.019 | 0.018 | 1 | 2 | Phldb1 | 0.001 | Young |
| 0.885061 | 0.261349 | 0.063 | 0.062 | 1 | 2 | Gps2 | 0.001 | Young |
| 0.885484 | 0.124543 | 0.139 | 0.145 | 1 | 2 | Tmem5 | -0.006 | Young |
| 0.885552 | 0.194847 | 0.209 | 0.211 | 1 | 2 | Eif1ax | -0.002 | Young |
| 0.88578 | 0.309852 | 0.074 | 0.073 | 1 | 2 | Mrpl58 | 0.001 | Young |
| 0.885786 | 0.083682 | 0.063 | 0.065 | 1 | 2 | Mettl16 | -0.002 | Young |
| 0.886008 | 0.092585 | 0.089 | 0.092 | 1 | 2 | Isg20l2 | -0.003 | Young |
| 0.886039 | 0.009701 | 0.352 | 0.357 | 1 | 2 | Snx18 | -0.005 | Young |
| 0.886151 | 0.397381 | 0.013 | 0.014 | 1 | 2 | Cep57l1 | -0.001 | Young |
| 0.886523 | 0.113594 | 0.133 | 0.133 | 1 | 2 | Pxk | 0 | Young |
| 0.886616 | 0.247795 | 0.053 | 0.052 | 1 | 2 | Rps6ka5 | 0.001 | Young |
| 0.886638 | 0.082462 | 0.105 | 0.107 | 1 | 2 | Mir17hg | -0.002 | Young |
| 0.886661 | 0.349566 | 0.031 | 0.03 | 1 | 2 | Slc13a3 | 0.001 | Young |
| 0.886779 | 0.009258 | 0.266 | 0.269 | 1 | 2 | Phf3 | -0.003 | Young |
| 0.886944 | 0.141274 | 0.125 | 0.123 | 1 | 2 | Trnt1 | 0.002 | Young |
| 0.887066 | 0.275981 | 0.048 | 0.046 | 1 | 2 | Zfp560 | 0.002 | Young |
| 0.887071 | 0.033925 | 0.405 | 0.41 | 1 | 2 | Asah1 | -0.005 | Young |
| 0.887124 | 0.015287 | 0.342 | 0.351 | 1 | 2 | Map1lc3b | -0.009 | Young |
| 0.887217 | 0.003431 | 0.204 | 0.202 | 1 | 2 | Pin4 | 0.002 | Young |
| 0.887352 | 0.190823 | 0.146 | 0.153 | 1 | 2 | Cd40 | -0.007 | Young |
| 0.887423 | 0.073077 | 0.045 | 0.047 | 1 | 2 | Arhgef18 | -0.002 | Young |
| 0.88745 | 0.476262 | 0.106 | 0.107 | 1 | 2 | Snx12 | -0.001 | Young |
| 0.887609 | 0.019925 | 0.171 | 0.17 | 1 | 2 | Dhx38 | 0.001 | Young |
| 0.887662 | 0.129939 | 0.139 | 0.146 | 1 | 2 | Polr2i | -0.007 | Young |
| 0.887797 | 0.177577 | 0.062 | 0.064 | 1 | 2 | Pibf1 | -0.002 | Young |
| 0.88813 | 0.012496 | 0.143 | 0.14 | 1 | 2 | Insr | 0.003 | Young |
| 0.888228 | 0.33399 | 0.045 | 0.047 | 1 | 2 | Nostrin | -0.002 | Young |
| 0.888337 | 0.140364 | 0.082 | 0.081 | 1 | 2 | Eaf1 | 0.001 | Young |
| 0.888578 | 0.477582 | 0.044 | 0.043 | 1 | 2 | Syt11 | 0.001 | Young |
| 0.888592 | 0.669905 | 0.029 | 0.03 | 1 | 2 | Dusp8 | -0.001 | Young |

| | | | | | | | | |
|---|---|---|---|---|---|---|---|---|
| 0.888627 | 0.165744 | 0.019 | 0.018 | 1 | 2 | Gm4070 | 0.001 | Young |
| 0.888644 | 0.105073 | 0.069 | 0.067 | 1 | 2 | Josd1 | 0.002 | Young |
| 0.888792 | 0.210872 | 0.046 | 0.045 | 1 | 2 | Fkrp | 0.001 | Young |
| 0.888825 | 0.167741 | 0.033 | 0.032 | 1 | 2 | Hook2 | 0.001 | Young |
| 0.889079 | 0.131774 | 0.189 | 0.19 | 1 | 2 | Sugt1 | -0.001 | Young |
| 0.889141 | 0.077172 | 0.102 | 0.106 | 1 | 2 | Snapc1 | -0.004 | Young |
| 0.889652 | 0.067288 | 0.172 | 0.173 | 1 | 2 | Rps6kb1 | -0.001 | Young |
| 0.889737 | 0.278657 | 0.03 | 0.031 | 1 | 2 | Zkscan1 | -0.001 | Young |
| 0.890013 | 0.01314 | 0.319 | 0.318 | 1 | 2 | Srsf11 | 0.001 | Young |
| 0.89011 | 0.181285 | 0.021 | 0.022 | 1 | 2 | Zfp758 | -0.001 | Young |
| 0.890491 | 0.031713 | 0.325 | 0.329 | 1 | 2 | Arid1a | -0.004 | Young |
| 0.890517 | 0.277767 | 0.129 | 0.128 | 1 | 2 | Osbpl11 | 0.001 | Young |
| 0.89141 | 0.04307 | 0.606 | 0.628 | 1 | 2 | Eef1g | -0.022 | Young |
| 0.892071 | 0.193212 | 0.019 | 0.018 | 1 | 2 | Arl15 | 0.001 | Young |
| 0.893057 | 0.05298 | 0.095 | 0.097 | 1 | 2 | Pex2 | -0.002 | Young |
| 0.893115 | 0.140572 | 0.076 | 0.075 | 1 | 2 | Bbc3 | 0.001 | Young |
| 0.893212 | 0.050202 | 0.39 | 0.409 | 1 | 2 | St13 | -0.019 | Young |
| 0.893341 | 0.140702 | 0.078 | 0.081 | 1 | 2 | Sacm1l | -0.003 | Young |
| 0.893593 | 0.199903 | 0.042 | 0.041 | 1 | 2 | 5730455P1 | 0.001 | Young |
| 0.893708 | 0.183213 | 0.031 | 0.032 | 1 | 2 | Irf2bp1 | -0.001 | Young |
| 0.894048 | 0.188384 | 0.031 | 0.03 | 1 | 2 | Mki67 | 0.001 | Young |
| 0.894207 | 0.814867 | 0.013 | 0.014 | 1 | 2 | Nif3l1 | -0.001 | Young |
| 0.894246 | 0.038419 | 0.229 | 0.233 | 1 | 2 | Stard3nl | -0.004 | Young |
| 0.894414 | 0.03197 | 0.395 | 0.399 | 1 | 2 | Snx20 | -0.004 | Young |
| 0.894415 | 0.091078 | 0.121 | 0.125 | 1 | 2 | Aco1 | -0.004 | Young |
| 0.894556 | 0.000878 | 0.115 | 0.118 | 1 | 2 | Ranbp9 | -0.003 | Young |
| 0.894719 | 0.122671 | 0.023 | 0.024 | 1 | 2 | Haus4 | -0.001 | Young |
| 0.895151 | 0.054391 | 0.306 | 0.318 | 1 | 2 | Larp1 | -0.012 | Young |
| 0.895458 | 0.06535 | 0.222 | 0.233 | 1 | 2 | Zranb2 | -0.011 | Young |
| 0.895487 | 0.175079 | 0.065 | 0.067 | 1 | 2 | Tmem71 | -0.002 | Young |
| 0.896001 | 0.041933 | 0.208 | 0.214 | 1 | 2 | Lmbrd1 | -0.006 | Young |
| 0.896002 | 0.169504 | 0.278 | 0.293 | 1 | 2 | Gtf2h5 | -0.015 | Young |
| 0.896008 | 0.254775 | 0.032 | 0.031 | 1 | 2 | Tfpt | 0.001 | Young |
| 0.896282 | 0.020652 | 0.049 | 0.05 | 1 | 2 | Srxn1 | -0.001 | Young |
| 0.896545 | 0.009753 | 0.188 | 0.192 | 1 | 2 | Ext1 | -0.004 | Young |
| 0.896894 | 0.104771 | 0.29 | 0.294 | 1 | 2 | Nufip2 | -0.004 | Young |
| 0.896903 | 0.021368 | 0.113 | 0.116 | 1 | 2 | Dda1 | -0.003 | Young |
| 0.89724 | 0.017945 | 0.309 | 0.309 | 1 | 2 | Pxn | 0 | Young |
| 0.897325 | 0.021911 | 0.102 | 0.105 | 1 | 2 | Fam204a | -0.003 | Young |
| 0.897398 | 0.006596 | 0.019 | 0.018 | 1 | 2 | Atrip | 0.001 | Young |
| 0.897599 | 0.017985 | 0.362 | 0.362 | 1 | 2 | Uqcrb | 0 | Young |
| 0.8976 | 0.019903 | 0.215 | 0.224 | 1 | 2 | Rb1 | -0.009 | Young |
| 0.897657 | 0.002982 | 0.071 | 0.07 | 1 | 2 | Usp39 | 0.001 | Young |
| 0.897773 | 0.566633 | 0.018 | 0.017 | 1 | 2 | Pvt1 | 0.001 | Young |
| 0.897881 | 0.18956 | 0.121 | 0.125 | 1 | 2 | Dscr3 | -0.004 | Young |

| | | | | | | | | |
|---|---|---|---|---|---|---|---|---|
| 0.898148 | 0.29442 | 0.052 | 0.051 | 1 | 2 | Ssh1 | 0.001 | Young |
| 0.898382 | 0.046727 | 0.023 | 0.024 | 1 | 2 | Ccdc61 | -0.001 | Young |
| 0.898438 | 0.283239 | 0.039 | 0.041 | 1 | 2 | Nf2 | -0.002 | Young |
| 0.899178 | 0.06792 | 0.032 | 0.033 | 1 | 2 | Nsun3 | -0.001 | Young |
| 0.89927 | 0.06469 | 0.12 | 0.123 | 1 | 2 | Itgb1bp1 | -0.003 | Young |
| 0.899425 | 0.101413 | 0.057 | 0.059 | 1 | 2 | Mob2 | -0.002 | Young |
| 0.899465 | 0.10806 | 0.31 | 0.314 | 1 | 2 | Rab11a | -0.004 | Young |
| 0.900085 | 0.088782 | 0.049 | 0.05 | 1 | 2 | Zbed4 | -0.001 | Young |
| 0.900635 | 0.03853 | 0.055 | 0.054 | 1 | 2 | Slc22a15 | 0.001 | Young |
| 0.90092 | 0.34462 | 0.023 | 0.024 | 1 | 2 | Rnaseh1 | -0.001 | Young |
| 0.901 | 0.069779 | 0.105 | 0.103 | 1 | 2 | Rbm18 | 0.002 | Young |
| 0.901065 | 0.156478 | 0.048 | 0.049 | 1 | 2 | Tmem199 | -0.001 | Young |
| 0.901586 | 0.099622 | 0.334 | 0.339 | 1 | 2 | 9530068E0 | -0.005 | Young |
| 0.901603 | 0.1182 | 0.065 | 0.067 | 1 | 2 | Tmem60 | -0.002 | Young |
| 0.901627 | 0.271948 | 0.014 | 0.015 | 1 | 2 | Zcchc14 | -0.001 | Young |
| 0.901951 | 0.152174 | 0.2 | 0.199 | 1 | 2 | Akirin1 | 0.001 | Young |
| 0.902332 | 0.24507 | 0.014 | 0.015 | 1 | 2 | Trdmt1 | -0.001 | Young |
| 0.902612 | 0.022458 | 0.262 | 0.263 | 1 | 2 | Herc1 | -0.001 | Young |
| 0.902651 | 0.06923 | 0.493 | 0.502 | 1 | 2 | Atp5k | -0.009 | Young |
| 0.902817 | 0.165813 | 0.049 | 0.05 | 1 | 2 | Rpusd4 | -0.001 | Young |
| 0.902939 | 0.22723 | 0.048 | 0.049 | 1 | 2 | Txnl4a | -0.001 | Young |
| 0.903158 | 0.001936 | 0.194 | 0.19 | 1 | 2 | Smc3 | 0.004 | Young |
| 0.90321 | 0.106608 | 0.067 | 0.069 | 1 | 2 | Xpnpep1 | -0.002 | Young |
| 0.903409 | 0.125524 | 0.057 | 0.059 | 1 | 2 | Prdm4 | -0.002 | Young |
| 0.904166 | 0.173912 | 0.039 | 0.041 | 1 | 2 | Foxred1 | -0.002 | Young |
| 0.904368 | 0.107187 | 0.031 | 0.032 | 1 | 2 | Kctd17 | -0.001 | Young |
| 0.904368 | 0.048846 | 0.044 | 0.043 | 1 | 2 | Spryd7 | 0.001 | Young |
| 0.904414 | 0.076985 | 0.31 | 0.321 | 1 | 2 | Rtraf | -0.011 | Young |
| 0.904453 | 0.065182 | 0.154 | 0.153 | 1 | 2 | Mindy1 | 0.001 | Young |
| 0.904534 | 0.208327 | 0.21 | 0.212 | 1 | 2 | Bcl6 | -0.002 | Young |
| 0.90456 | 0.069557 | 0.159 | 0.163 | 1 | 2 | Gnl1 | -0.004 | Young |
| 0.905212 | 0.304881 | 0.018 | 0.017 | 1 | 2 | Slc19a1 | 0.001 | Young |
| 0.905291 | 0.233725 | 0.076 | 0.079 | 1 | 2 | Sphk2 | -0.003 | Young |
| 0.905543 | 0.01336 | 0.122 | 0.12 | 1 | 2 | Rnf2 | 0.002 | Young |
| 0.905613 | 0.099808 | 0.032 | 0.033 | 1 | 2 | Zfyve1 | -0.001 | Young |
| 0.90568 | 0.33322 | 0.04 | 0.04 | 1 | 2 | Fgd6 | 0 | Young |
| 0.905713 | 0.339852 | 0.147 | 0.15 | 1 | 2 | Gpalpp1 | -0.003 | Young |
| 0.905861 | 0.220894 | 0.014 | 0.015 | 1 | 2 | Slc16a13 | -0.001 | Young |
| 0.90603 | 0.017066 | 0.14 | 0.14 | 1 | 2 | Kpna1 | 0 | Young |
| 0.906362 | 0.29276 | 0.093 | 0.096 | 1 | 2 | Irf3 | -0.003 | Young |
| 0.906389 | 0.062307 | 0.042 | 0.041 | 1 | 2 | Med22 | 0.001 | Young |
| 0.906553 | 0.023281 | 0.215 | 0.219 | 1 | 2 | Pnpla8 | -0.004 | Young |
| 0.90661 | 0.073022 | 0.122 | 0.125 | 1 | 2 | Nenf | -0.003 | Young |
| 0.906921 | 0.537406 | 0.038 | 0.037 | 1 | 2 | Bag4 | 0.001 | Young |
| 0.908063 | 0.018385 | 0.605 | 0.586 | 1 | 2 | Pld4 | 0.019 | Young |

| | | | | | | | | |
|---|---|---|---|---|---|---|---|---|
| 0.908207 | 0.202995 | 0.265 | 0.273 | 1 | 2 | Cct3 | -0.008 | Young |
| 0.908226 | 0.24853 | 0.059 | 0.059 | 1 | 2 | Nmd3 | 0 | Young |
| 0.90823 | 0.12908 | 0.107 | 0.109 | 1 | 2 | Mapk8 | -0.002 | Young |
| 0.908265 | 0.107558 | 0.121 | 0.121 | 1 | 2 | Ddx18 | 0 | Young |
| 0.908491 | 0.063918 | 0.032 | 0.031 | 1 | 2 | Runx2 | 0.001 | Young |
| 0.908619 | 0.044564 | 0.173 | 0.178 | 1 | 2 | Yme1l1 | -0.005 | Young |
| 0.908685 | 0.2955 | 0.014 | 0.015 | 1 | 2 | Slc25a26 | -0.001 | Young |
| 0.908701 | 0.010986 | 0.024 | 0.025 | 1 | 2 | Lig1 | -0.001 | Young |
| 0.908863 | 0.1896 | 0.077 | 0.077 | 1 | 2 | Slc25a46 | 0 | Young |
| 0.908905 | 0.084726 | 0.069 | 0.071 | 1 | 2 | Sh3glb2 | -0.002 | Young |
| 0.909038 | 0.431787 | 0.014 | 0.015 | 1 | 2 | Hspa2 | -0.001 | Young |
| 0.909171 | 0.31006 | 0.067 | 0.066 | 1 | 2 | Prep | 0.001 | Young |
| 0.909557 | 0.092284 | 0.146 | 0.146 | 1 | 2 | Lemd2 | 0 | Young |
| 0.909639 | 0.073537 | 0.05 | 0.051 | 1 | 2 | Cdk10 | -0.001 | Young |
| 0.909681 | 0.104026 | 0.065 | 0.067 | 1 | 2 | Smim12 | -0.002 | Young |
| 0.909745 | 0.150987 | 0.018 | 0.017 | 1 | 2 | Slfn9 | 0.001 | Young |
| 0.909749 | 0.137498 | 0.048 | 0.049 | 1 | 2 | Paxip1 | -0.001 | Young |
| 0.909983 | 0.205737 | 0.083 | 0.082 | 1 | 2 | Mrpl35 | 0.001 | Young |
| 0.910114 | 0.109079 | 0.285 | 0.297 | 1 | 2 | Emc7 | -0.012 | Young |
| 0.910197 | 0.514985 | 0.038 | 0.04 | 1 | 2 | Slc12a4 | -0.002 | Young |
| 0.910416 | 0.187123 | 0.039 | 0.039 | 1 | 2 | Rbmx2 | 0 | Young |
| 0.910871 | 0.018546 | 0.056 | 0.055 | 1 | 2 | Alpk1 | 0.001 | Young |
| 0.91094 | 0.067354 | 0.381 | 0.389 | 1 | 2 | Runx1 | -0.008 | Young |
| 0.911384 | 0.16753 | 0.099 | 0.097 | 1 | 2 | B4galt3 | 0.002 | Young |
| 0.911412 | 0.156731 | 0.178 | 0.181 | 1 | 2 | Phf23 | -0.003 | Young |
| 0.911576 | 0.195967 | 0.032 | 0.033 | 1 | 2 | Rbm43 | -0.001 | Young |
| 0.911863 | 0.509764 | 0.014 | 0.015 | 1 | 2 | Lrp4 | -0.001 | Young |
| 0.912113 | 0.251935 | 0.105 | 0.105 | 1 | 2 | Trim26 | 0 | Young |
| 0.912314 | 0.096553 | 0.216 | 0.225 | 1 | 2 | Tmem55b | -0.009 | Young |
| 0.912323 | 0.053489 | 0.068 | 0.07 | 1 | 2 | Srpk1 | -0.002 | Young |
| 0.912635 | 0.176457 | 0.05 | 0.051 | 1 | 2 | Psat1 | -0.001 | Young |
| 0.91266 | 0.174725 | 0.018 | 0.017 | 1 | 2 | Stk11ip | 0.001 | Young |
| 0.912883 | 0.081144 | 0.154 | 0.157 | 1 | 2 | Ube2a | -0.003 | Young |
| 0.912944 | 0.024402 | 0.323 | 0.323 | 1 | 2 | Paip2 | 0 | Young |
| 0.913203 | 0.051473 | 0.112 | 0.115 | 1 | 2 | Cc2d1b | -0.003 | Young |
| 0.913392 | 0.094456 | 0.024 | 0.025 | 1 | 2 | Gm49368 | -0.001 | Young |
| 0.913486 | 0.190795 | 0.032 | 0.033 | 1 | 2 | Entpd6 | -0.001 | Young |
| 0.913542 | 0.05196 | 0.031 | 0.03 | 1 | 2 | Sfmbt1 | 0.001 | Young |
| 0.913702 | 0.133675 | 0.083 | 0.086 | 1 | 2 | Nup50 | -0.003 | Young |
| 0.913946 | 0.043809 | 0.235 | 0.241 | 1 | 2 | Lacc1 | -0.006 | Young |
| 0.91408 | 0.094889 | 0.063 | 0.062 | 1 | 2 | Immp1l | 0.001 | Young |
| 0.914114 | 0.235944 | 0.116 | 0.121 | 1 | 2 | Arhgap1 | -0.005 | Young |
| 0.914136 | 0.007766 | 0.108 | 0.111 | 1 | 2 | Atxn3 | -0.003 | Young |
| 0.914255 | 0.069546 | 0.169 | 0.173 | 1 | 2 | Eny2 | -0.004 | Young |
| 0.914317 | 0.415762 | 0.031 | 0.032 | 1 | 2 | Mto1 | -0.001 | Young |

| | | | | | | | | |
|---|---|---|---|---|---|---|---|---|
| 0.914363 | 0.299572 | 0.063 | 0.065 | 1 | 2 | Mosmo | -0.002 | Young |
| 0.914496 | 0.089363 | 0.024 | 0.025 | 1 | 2 | Pyroxd1 | -0.001 | Young |
| 0.914679 | 0.302421 | 0.032 | 0.033 | 1 | 2 | Trim39 | -0.001 | Young |
| 0.915058 | 0.1986 | 0.076 | 0.078 | 1 | 2 | Tmem205 | -0.002 | Young |
| 0.915309 | 0.118826 | 0.069 | 0.071 | 1 | 2 | Ppwd1 | -0.002 | Young |
| 0.915343 | 0.26896 | 0.051 | 0.05 | 1 | 2 | Lsg1 | 0.001 | Young |
| 0.91552 | 0.081256 | 0.015 | 0.016 | 1 | 2 | Usp30 | -0.001 | Young |
| 0.915708 | 0.093307 | 0.033 | 0.034 | 1 | 2 | Zfp626 | -0.001 | Young |
| 0.915853 | 0.114983 | 0.191 | 0.196 | 1 | 2 | Mxd1 | -0.005 | Young |
| 0.916127 | 0.041915 | 0.385 | 0.388 | 1 | 2 | Glg1 | -0.003 | Young |
| 0.916178 | 0.116768 | 0.033 | 0.034 | 1 | 2 | Gbp9 | -0.001 | Young |
| 0.916458 | 0.170972 | 0.017 | 0.016 | 1 | 2 | Atp6v1g2 | 0.001 | Young |
| 0.916838 | 0.106902 | 0.284 | 0.297 | 1 | 2 | Mapk6 | -0.013 | Young |
| 0.916857 | 0.035573 | 0.135 | 0.138 | 1 | 2 | Ppil2 | -0.003 | Young |
| 0.916982 | 0.286284 | 0.024 | 0.025 | 1 | 2 | Spindoc | -0.001 | Young |
| 0.91699 | 0.097051 | 0.075 | 0.074 | 1 | 2 | Camsap2 | 0.001 | Young |
| 0.917444 | 0.162523 | 0.165 | 0.165 | 1 | 2 | Max | 0 | Young |
| 0.917471 | 0.003928 | 0.051 | 0.05 | 1 | 2 | Rnf31 | 0.001 | Young |
| 0.91863 | 0.00735 | 0.169 | 0.171 | 1 | 2 | Dctn3 | -0.002 | Young |
| 0.91879 | 0.159811 | 0.064 | 0.063 | 1 | 2 | Pik3cb | 0.001 | Young |
| 0.918896 | 0.110412 | 0.125 | 0.128 | 1 | 2 | Dcun1d1 | -0.003 | Young |
| 0.919049 | 0.003352 | 0.647 | 0.643 | 1 | 2 | Prcp | 0.004 | Young |
| 0.919269 | 0.283392 | 0.015 | 0.016 | 1 | 2 | Otud7b | -0.001 | Young |
| 0.919467 | 0.06523 | 0.018 | 0.017 | 1 | 2 | Gemin6 | 0.001 | Young |
| 0.919495 | 0.007552 | 0.454 | 0.456 | 1 | 2 | Lilr4b | -0.002 | Young |
| 0.919509 | 0.165753 | 0.076 | 0.078 | 1 | 2 | Mea1 | -0.002 | Young |
| 0.920007 | 0.146199 | 0.273 | 0.283 | 1 | 2 | Uqcrfs1 | -0.01 | Young |
| 0.92002 | 0.448813 | 0.024 | 0.025 | 1 | 2 | Rad51d | -0.001 | Young |
| 0.92036 | 0.155915 | 0.125 | 0.128 | 1 | 2 | Phf10 | -0.003 | Young |
| 0.920738 | 0.257621 | 0.039 | 0.039 | 1 | 2 | Mrps35 | 0 | Young |
| 0.920854 | 0.02062 | 0.052 | 0.054 | 1 | 2 | Tcp11l1 | -0.002 | Young |
| 0.920869 | 0.233646 | 0.083 | 0.086 | 1 | 2 | AY036118 | -0.003 | Young |
| 0.92099 | 0.190759 | 0.05 | 0.049 | 1 | 2 | Zfp597 | 0.001 | Young |
| 0.921102 | 0.048947 | 0.21 | 0.215 | 1 | 2 | R3hdm1 | -0.005 | Young |
| 0.921141 | 0.012219 | 0.108 | 0.107 | 1 | 2 | Mlxip | 0.001 | Young |
| 0.92138 | 0.644223 | 0.039 | 0.041 | 1 | 2 | Pelp1 | -0.002 | Young |
| 0.921564 | 0.071757 | 0.097 | 0.1 | 1 | 2 | Foxo3 | -0.003 | Young |
| 0.921746 | 0.05497 | 0.064 | 0.063 | 1 | 2 | Pogz | 0.001 | Young |
| 0.921807 | 0.500501 | 0.015 | 0.015 | 1 | 2 | Asap2 | 0 | Young |
| 0.922498 | 0.137488 | 0.109 | 0.109 | 1 | 2 | Nup153 | 0 | Young |
| 0.922607 | 0.133936 | 0.157 | 0.157 | 1 | 2 | Rasa4 | 0 | Young |
| 0.922895 | 0.174096 | 0.198 | 0.207 | 1 | 2 | Jtb | -0.009 | Young |
| 0.922964 | 0.035702 | 0.063 | 0.064 | 1 | 2 | Foxo1 | -0.001 | Young |
| 0.923051 | 0.066732 | 0.064 | 0.063 | 1 | 2 | Tbc1d9b | 0.001 | Young |
| 0.92306 | 0.340882 | 0.024 | 0.025 | 1 | 2 | Ppil1 | -0.001 | Young |

| | | | | | | | | |
|---|---|---|---|---|---|---|---|---|
| 0.923073 | 0.025667 | 0.064 | 0.065 | 1 | 2 | Nfia | -0.001 | Young |
| 0.923245 | 0.019762 | 0.064 | 0.065 | 1 | 2 | Mras | -0.001 | Young |
| 0.923289 | 0.020911 | 0.276 | 0.271 | 1 | 2 | 2810474O1 | 0.005 | Young |
| 0.923335 | 0.062292 | 0.264 | 0.266 | 1 | 2 | Trpc4ap | -0.002 | Young |
| 0.923337 | 0.39031 | 0.024 | 0.025 | 1 | 2 | Pgpep1 | -0.001 | Young |
| 0.923378 | 0.024466 | 0.43 | 0.439 | 1 | 2 | Plxnd1 | -0.009 | Young |
| 0.923486 | 0.099683 | 0.101 | 0.104 | 1 | 2 | Atxn7l3 | -0.003 | Young |
| 0.923487 | 0.134961 | 0.042 | 0.041 | 1 | 2 | Il18 | 0.001 | Young |
| 0.923652 | 0.020972 | 0.059 | 0.061 | 1 | 2 | Atp11a | -0.002 | Young |
| 0.92366 | 0.136667 | 0.11 | 0.11 | 1 | 2 | Flot2 | 0 | Young |
| 0.924124 | 0.243605 | 0.029 | 0.028 | 1 | 2 | Cdc25a | 0.001 | Young |
| 0.924376 | 0.060583 | 0.386 | 0.396 | 1 | 2 | Atp5c1 | -0.01 | Young |
| 0.924442 | 0.12258 | 0.024 | 0.025 | 1 | 2 | Wdr24 | -0.001 | Young |
| 0.92451 | 0.018827 | 0.139 | 0.137 | 1 | 2 | Chmp2b | 0.002 | Young |
| 0.924834 | 0.186385 | 0.083 | 0.082 | 1 | 2 | Hk1 | 0.001 | Young |
| 0.924839 | 0.088282 | 0.146 | 0.15 | 1 | 2 | Cyb5a | -0.004 | Young |
| 0.92493 | 0.237574 | 0.096 | 0.1 | 1 | 2 | Sfxn1 | -0.004 | Young |
| 0.925172 | 0.360614 | 0.017 | 0.016 | 1 | 2 | Zfp995 | 0.001 | Young |
| 0.925322 | 0.066283 | 0.072 | 0.072 | 1 | 2 | Nln | 0 | Young |
| 0.925689 | 0.09559 | 0.306 | 0.316 | 1 | 2 | Eloc | -0.01 | Young |
| 0.925891 | 0.151628 | 0.105 | 0.107 | 1 | 2 | Nfx1 | -0.002 | Young |
| 0.925947 | 0.203977 | 0.042 | 0.043 | 1 | 2 | Morc2a | -0.001 | Young |
| 0.926093 | 0.090788 | 0.119 | 0.119 | 1 | 2 | Ero1l | 0 | Young |
| 0.926272 | 0.287948 | 0.128 | 0.13 | 1 | 2 | Vps37b | -0.002 | Young |
| 0.926848 | 0.010983 | 0.017 | 0.016 | 1 | 2 | Aasdh | 0.001 | Young |
| 0.926971 | 0.525382 | 0.076 | 0.076 | 1 | 2 | Slc5a3 | 0 | Young |
| 0.926978 | 0.007213 | 0.217 | 0.219 | 1 | 2 | Tnfrsf1a | -0.002 | Young |
| 0.927156 | 0.365031 | 0.049 | 0.05 | 1 | 2 | Clec5a | -0.001 | Young |
| 0.927993 | 0.261639 | 0.039 | 0.039 | 1 | 2 | Wiz | 0 | Young |
| 0.928126 | 0.223185 | 0.159 | 0.162 | 1 | 2 | Zmiz2 | -0.003 | Young |
| 0.928359 | 0.122143 | 0.127 | 0.131 | 1 | 2 | Birc2 | -0.004 | Young |
| 0.928379 | 0.02487 | 0.101 | 0.102 | 1 | 2 | C4b | -0.001 | Young |
| 0.928825 | 0.060911 | 0.156 | 0.158 | 1 | 2 | Hk2 | -0.002 | Young |
| 0.928962 | 0.062544 | 0.053 | 0.055 | 1 | 2 | Sdc1 | -0.002 | Young |
| 0.92898 | 0.09436 | 0.153 | 0.156 | 1 | 2 | Tmem33 | -0.003 | Young |
| 0.928998 | 0.078925 | 0.122 | 0.122 | 1 | 2 | Rps6ka4 | 0 | Young |
| 0.929474 | 0.053307 | 0.306 | 0.309 | 1 | 2 | Papola | -0.003 | Young |
| 0.929703 | 0.04246 | 0.108 | 0.11 | 1 | 2 | Cpeb2 | -0.002 | Young |
| 0.929842 | 0.218289 | 0.077 | 0.079 | 1 | 2 | Snap29 | -0.002 | Young |
| 0.930257 | 0.213424 | 0.075 | 0.077 | 1 | 2 | Cpne2 | -0.002 | Young |
| 0.93028 | 0.050653 | 0.252 | 0.256 | 1 | 2 | Baz2b | -0.004 | Young |
| 0.930516 | 0.117518 | 0.049 | 0.048 | 1 | 2 | Pla2g16 | 0.001 | Young |
| 0.930602 | 0.152774 | 0.027 | 0.027 | 1 | 2 | Ercc6 | 0 | Young |
| 0.930817 | 0.019807 | 0.154 | 0.153 | 1 | 2 | Sec22b | 0.001 | Young |
| 0.931148 | 0.023947 | 0.601 | 0.603 | 1 | 2 | Bach1 | -0.002 | Young |

| | | | | | | | | |
|---|---|---|---|---|---|---|---|---|
| 0.931161 | 0.108887 | 0.12 | 0.12 | 1 | 2 | Ctbp2 | 0 | Young |
| 0.931209 | 0.486867 | 0.015 | 0.016 | 1 | 2 | Ccdc84 | -0.001 | Young |
| 0.931227 | 0.072949 | 0.033 | 0.034 | 1 | 2 | Ndc1 | -0.001 | Young |
| 0.931339 | 0.140833 | 0.106 | 0.106 | 1 | 2 | Spast | 0 | Young |
| 0.931638 | 0.388847 | 0.057 | 0.059 | 1 | 2 | Gfer | -0.002 | Young |
| 0.932529 | 0.11153 | 0.245 | 0.245 | 1 | 2 | Csnk1d | 0 | Young |
| 0.932831 | 0.114852 | 0.167 | 0.168 | 1 | 2 | Tsg101 | -0.001 | Young |
| 0.932995 | 0.073049 | 0.038 | 0.037 | 1 | 2 | Pip4k2c | 0.001 | Young |
| 0.932998 | 0.107178 | 0.135 | 0.138 | 1 | 2 | Gpr107 | -0.003 | Young |
| 0.933218 | 0.065144 | 0.138 | 0.14 | 1 | 2 | 2310011J0 | -0.002 | Young |
| 0.933608 | 0.147934 | 0.091 | 0.091 | 1 | 2 | Fam241a | 0 | Young |
| 0.933906 | 0.124024 | 0.334 | 0.346 | 1 | 2 | Zfp91 | -0.012 | Young |
| 0.934072 | 0.269105 | 0.043 | 0.044 | 1 | 2 | Bet1l | -0.001 | Young |
| 0.93433 | 0.261385 | 0.051 | 0.052 | 1 | 2 | Lrp12 | -0.001 | Young |
| 0.934455 | 0.009058 | 0.27 | 0.275 | 1 | 2 | Ube2k | -0.005 | Young |
| 0.934663 | 0.039816 | 0.015 | 0.015 | 1 | 2 | Plxna3 | 0 | Young |
| 0.934698 | 0.420271 | 0.043 | 0.044 | 1 | 2 | Gmpr2 | -0.001 | Young |
| 0.934899 | 0.146494 | 0.064 | 0.065 | 1 | 2 | Rpia | -0.001 | Young |
| 0.934919 | 0.006484 | 0.094 | 0.093 | 1 | 2 | Arl2bp | 0.001 | Young |
| 0.935501 | 0.075456 | 0.129 | 0.13 | 1 | 2 | Plin3 | -0.001 | Young |
| 0.935763 | 0.111653 | 0.081 | 0.082 | 1 | 2 | Patl1 | -0.001 | Young |
| 0.93609 | 0.137083 | 0.192 | 0.193 | 1 | 2 | Dnajb9 | -0.001 | Young |
| 0.936093 | 0.013285 | 0.346 | 0.351 | 1 | 2 | Dync1i2 | -0.005 | Young |
| 0.93612 | 0.302507 | 0.038 | 0.037 | 1 | 2 | Trim3 | 0.001 | Young |
| 0.936683 | 0.019971 | 0.112 | 0.113 | 1 | 2 | Snrnp27 | -0.001 | Young |
| 0.936959 | 0.013955 | 0.309 | 0.306 | 1 | 2 | Sdf4 | 0.003 | Young |
| 0.937018 | 0.172555 | 0.219 | 0.224 | 1 | 2 | Tmem259 | -0.005 | Young |
| 0.937191 | 0.037365 | 0.042 | 0.041 | 1 | 2 | Coa6 | 0.001 | Young |
| 0.937277 | 0.112813 | 0.091 | 0.093 | 1 | 2 | Acads | -0.002 | Young |
| 0.937367 | 0.021365 | 0.096 | 0.096 | 1 | 2 | 8-Mar | 0 | Young |
| 0.9376 | 0.140692 | 0.072 | 0.074 | 1 | 2 | Pex5 | -0.002 | Young |
| 0.937626 | 0.084455 | 0.112 | 0.115 | 1 | 2 | Atg16l1 | -0.003 | Young |
| 0.937934 | 0.02269 | 0.398 | 0.404 | 1 | 2 | D8Ertd738 | -0.006 | Young |
| 0.93814 | 0.011406 | 0.015 | 0.015 | 1 | 2 | Zfp668 | 0 | Young |
| 0.93815 | 0.409771 | 0.086 | 0.087 | 1 | 2 | Zcchc9 | -0.001 | Young |
| 0.938209 | 0.204883 | 0.027 | 0.027 | 1 | 2 | Zfp87 | 0 | Young |
| 0.938617 | 0.198602 | 0.017 | 0.017 | 1 | 2 | Tmem42 | 0 | Young |
| 0.93866 | 0.313376 | 0.026 | 0.026 | 1 | 2 | Zfand1 | 0 | Young |
| 0.938662 | 0.050156 | 0.015 | 0.015 | 1 | 2 | Fxn | 0 | Young |
| 0.938695 | 0.324602 | 0.037 | 0.036 | 1 | 2 | Slc25a10 | 0.001 | Young |
| 0.938753 | 0.06342 | 0.108 | 0.11 | 1 | 2 | Larp7 | -0.002 | Young |
| 0.939058 | 0.100335 | 0.069 | 0.071 | 1 | 2 | Psmd10 | -0.002 | Young |
| 0.939907 | 0.133016 | 0.061 | 0.06 | 1 | 2 | Kpna6 | 0.001 | Young |
| 0.939916 | 0.035143 | 0.527 | 0.549 | 1 | 2 | Vcp | -0.022 | Young |
| 0.940167 | 0.326009 | 0.025 | 0.026 | 1 | 2 | Gpr180 | -0.001 | Young |

| | | | | | | | | |
|---|---|---|---|---|---|---|---|---|
| 0.940268 | 0.139342 | 0.017 | 0.017 | 1 | 2 | A430106G: | 0 | Young |
| 0.940762 | 0.251299 | 0.077 | 0.077 | 1 | 2 | Tle1 | 0 | Young |
| 0.941206 | 0.026665 | 0.179 | 0.182 | 1 | 2 | Dap | -0.003 | Young |
| 0.941347 | 0.055253 | 0.378 | 0.382 | 1 | 2 | Raph1 | -0.004 | Young |
| 0.94221 | 0.185709 | 0.119 | 0.12 | 1 | 2 | Mrps16 | -0.001 | Young |
| 0.94258 | 0.99898 | 0.017 | 0.017 | 1 | 2 | Mfsd4b4 | 0 | Young |
| 0.942978 | 0.060174 | 0.158 | 0.162 | 1 | 2 | Actr1a | -0.004 | Young |
| 0.943038 | 0.032441 | 0.134 | 0.137 | 1 | 2 | Bcl7b | -0.003 | Young |
| 0.943723 | 0.034372 | 0.084 | 0.086 | 1 | 2 | Ints9 | -0.002 | Young |
| 0.943982 | 0.055041 | 0.094 | 0.095 | 1 | 2 | Slc35e4 | -0.001 | Young |
| 0.944071 | 0.058283 | 0.026 | 0.027 | 1 | 2 | Mllt11 | -0.001 | Young |
| 0.944233 | 0.147806 | 0.22 | 0.226 | 1 | 2 | Ppp1cc | -0.006 | Young |
| 0.944239 | 0.097522 | 0.204 | 0.208 | 1 | 2 | Ilk | -0.004 | Young |
| 0.944322 | 0.163953 | 0.125 | 0.125 | 1 | 2 | Prpf18 | 0 | Young |
| 0.944384 | 0.206016 | 0.038 | 0.037 | 1 | 2 | Gde1 | 0.001 | Young |
| 0.944411 | 0.214663 | 0.027 | 0.028 | 1 | 2 | Terf2ip | -0.001 | Young |
| 0.945149 | 0.084688 | 0.014 | 0.014 | 1 | 2 | Zfp931 | 0 | Young |
| 0.945239 | 0.155763 | 0.145 | 0.147 | 1 | 2 | Plbd2 | -0.002 | Young |
| 0.94608 | 0.192713 | 0.205 | 0.21 | 1 | 2 | Hcfc1r1 | -0.005 | Young |
| 0.946704 | 0.341083 | 0.084 | 0.087 | 1 | 2 | Tctex1d2 | -0.003 | Young |
| 0.946957 | 0.221778 | 0.014 | 0.014 | 1 | 2 | Nim1k | 0 | Young |
| 0.947023 | 0.368268 | 0.382 | 0.393 | 1 | 2 | Thbs1 | -0.011 | Young |
| 0.947427 | 0.327718 | 0.069 | 0.069 | 1 | 2 | Pygo2 | 0 | Young |
| 0.947471 | 0.214622 | 0.065 | 0.065 | 1 | 2 | Dkc1 | 0 | Young |
| 0.947627 | 0.278789 | 0.07 | 0.072 | 1 | 2 | Mgat4a | -0.002 | Young |
| 0.947946 | 0.045893 | 0.354 | 0.355 | 1 | 2 | Hacd4 | -0.001 | Young |
| 0.9481 | 0.024712 | 0.069 | 0.069 | 1 | 2 | Prpsap1 | 0 | Young |
| 0.948155 | 0.048178 | 0.115 | 0.115 | 1 | 2 | Lsm3 | 0 | Young |
| 0.948356 | 0.010976 | 0.413 | 0.421 | 1 | 2 | Sumo2 | -0.008 | Young |
| 0.94836 | 0.002374 | 0.361 | 0.367 | 1 | 2 | Gnaq | -0.006 | Young |
| 0.948403 | 0.208529 | 0.014 | 0.014 | 1 | 2 | Bcs1l | 0 | Young |
| 0.948403 | 0.1255 | 0.014 | 0.014 | 1 | 2 | Armc6 | 0 | Young |
| 0.948444 | 0.035256 | 0.58 | 0.576 | 1 | 2 | Cfp | 0.004 | Young |
| 0.949334 | 0.064069 | 0.121 | 0.121 | 1 | 2 | Ankib1 | 0 | Young |
| 0.949425 | 0.012702 | 0.196 | 0.201 | 1 | 2 | Tnfrsf11a | -0.005 | Young |
| 0.949486 | 0.033299 | 0.08 | 0.079 | 1 | 2 | Fam96b | 0.001 | Young |
| 0.949801 | 0.109962 | 0.107 | 0.107 | 1 | 2 | Lpin1 | 0 | Young |
| 0.949918 | 0.203592 | 0.026 | 0.026 | 1 | 2 | Entpd5 | 0 | Young |
| 0.949976 | 0.100629 | 0.096 | 0.096 | 1 | 2 | Cd300lg | 0 | Young |
| 0.950114 | 0.022901 | 0.07 | 0.072 | 1 | 2 | Pfkl | -0.002 | Young |
| 0.950501 | 0.013776 | 0.138 | 0.14 | 1 | 2 | Ube2d1 | -0.002 | Young |
| 0.950842 | 0.051274 | 0.045 | 0.046 | 1 | 2 | Osbpl2 | -0.001 | Young |
| 0.950847 | 0.166288 | 0.037 | 0.037 | 1 | 2 | Rpp30 | 0 | Young |
| 0.951372 | 0.1984 | 0.045 | 0.045 | 1 | 2 | AC127341. | 0 | Young |
| 0.951425 | 0.103297 | 0.163 | 0.163 | 1 | 2 | Marf1 | 0 | Young |

| | | | | | | | | |
|---|---|---|---|---|---|---|---|---|
| 0.951714 | 0.271983 | 0.093 | 0.094 | 1 | 2 | Katnbl1 | -0.001 | Young |
| 0.951835 | 0.518569 | 0.017 | 0.017 | 1 | 2 | Ift140 | 0 | Young |
| 0.951992 | 0.090074 | 0.11 | 0.112 | 1 | 2 | Aaed1 | -0.002 | Young |
| 0.952145 | 0.132996 | 0.063 | 0.064 | 1 | 2 | Comtd1 | -0.001 | Young |
| 0.952347 | 0.108093 | 0.113 | 0.115 | 1 | 2 | Syvn1 | -0.002 | Young |
| 0.952357 | 0.007662 | 0.291 | 0.293 | 1 | 2 | Ubn2 | -0.002 | Young |
| 0.952467 | 0.065041 | 0.131 | 0.13 | 1 | 2 | Fam192a | 0.001 | Young |
| 0.953257 | 0.387258 | 0.024 | 0.024 | 1 | 2 | E2f5 | 0 | Young |
| 0.954033 | 0.016477 | 0.046 | 0.047 | 1 | 2 | Map4k2 | -0.001 | Young |
| 0.95419 | 0.058868 | 0.014 | 0.014 | 1 | 2 | Plekhh3 | 0 | Young |
| 0.954457 | 0.033891 | 0.018 | 0.018 | 1 | 2 | Eef1akmt4 | 0 | Young |
| 0.954551 | 0.009382 | 0.014 | 0.014 | 1 | 2 | Tnnt1 | 0 | Young |
| 0.954684 | 0.403797 | 0.034 | 0.035 | 1 | 2 | Gm20703 | -0.001 | Young |
| 0.954811 | 0.494587 | 0.017 | 0.017 | 1 | 2 | Acp6 | 0 | Young |
| 0.954819 | 0.313198 | 0.026 | 0.027 | 1 | 2 | Smpd4 | -0.001 | Young |
| 0.954829 | 0.007456 | 0.232 | 0.233 | 1 | 2 | Dnajc7 | -0.001 | Young |
| 0.954903 | 0.063001 | 0.422 | 0.433 | 1 | 2 | Spi1 | -0.011 | Young |
| 0.955097 | 0.11938 | 0.048 | 0.047 | 1 | 2 | Rnf214 | 0.001 | Young |
| 0.955136 | 0.021048 | 0.037 | 0.036 | 1 | 2 | Smyd5 | 0.001 | Young |
| 0.955438 | 0.072906 | 0.233 | 0.237 | 1 | 2 | Ctbp1 | -0.004 | Young |
| 0.955497 | 0.005117 | 0.344 | 0.351 | 1 | 2 | Wwp1 | -0.007 | Young |
| 0.955803 | 0.070927 | 0.428 | 0.439 | 1 | 2 | Selenow | -0.011 | Young |
| 0.955835 | 0.013395 | 0.69 | 0.676 | 1 | 2 | Emp3 | 0.014 | Young |
| 0.955849 | 0.05299 | 0.046 | 0.047 | 1 | 2 | Nosip | -0.001 | Young |
| 0.955946 | 0.106508 | 0.125 | 0.125 | 1 | 2 | Anapc1 | 0 | Young |
| 0.956162 | 0.237051 | 0.08 | 0.08 | 1 | 2 | Snhg6 | 0 | Young |
| 0.956314 | 0.111982 | 0.186 | 0.19 | 1 | 2 | Zfp445 | -0.004 | Young |
| 0.9567 | 0.186465 | 0.018 | 0.018 | 1 | 2 | Tacc2 | 0 | Young |
| 0.956968 | 0.055167 | 0.126 | 0.127 | 1 | 2 | Mrpl42 | -0.001 | Young |
| 0.957169 | 0.531543 | 0.013 | 0.013 | 1 | 2 | P3h1 | 0 | Young |
| 0.957171 | 0.033951 | 0.188 | 0.192 | 1 | 2 | Il4ra | -0.004 | Young |
| 0.957176 | 0.251295 | 0.108 | 0.111 | 1 | 2 | Elp5 | -0.003 | Young |
| 0.957462 | 0.203807 | 0.062 | 0.063 | 1 | 2 | Map3k4 | -0.001 | Young |
| 0.957511 | 0.074189 | 0.195 | 0.195 | 1 | 2 | Rnasel | 0 | Young |
| 0.958023 | 0.356513 | 0.075 | 0.077 | 1 | 2 | Mink1 | -0.002 | Young |
| 0.95809 | 0.036011 | 0.118 | 0.119 | 1 | 2 | Arfgap2 | -0.001 | Young |
| 0.958487 | 0.015458 | 0.152 | 0.154 | 1 | 2 | Ube2g2 | -0.002 | Young |
| 0.958929 | 0.067852 | 0.089 | 0.091 | 1 | 2 | Ulk1 | -0.002 | Young |
| 0.959226 | 0.225911 | 0.1 | 0.101 | 1 | 2 | Zwint | -0.001 | Young |
| 0.959511 | 0.196145 | 0.029 | 0.029 | 1 | 2 | Armc7 | 0 | Young |
| 0.959919 | 0.130696 | 0.128 | 0.132 | 1 | 2 | Slc44a1 | -0.004 | Young |
| 0.960053 | 0.042706 | 0.2 | 0.2 | 1 | 2 | Rala | 0 | Young |
| 0.960056 | 0.124287 | 0.099 | 0.099 | 1 | 2 | Ap1ar | 0 | Young |
| 0.960474 | 0.024261 | 0.025 | 0.025 | 1 | 2 | Micu3 | 0 | Young |
| 0.96059 | 0.236911 | 0.055 | 0.056 | 1 | 2 | Arhgap23 | -0.001 | Young |

| | | | | | | | | |
|---|---|---|---|---|---|---|---|---|
| 0.960997 | 0.320025 | 0.034 | 0.034 | 1 | 2 | Ndufaf3 | 0 | Young |
| 0.961013 | 0.000496 | 0.099 | 0.1 | 1 | 2 | Ino80e | -0.001 | Young |
| 0.961067 | 0.080814 | 0.083 | 0.084 | 1 | 2 | Gt(ROSA)2( | -0.001 | Young |
| 0.961157 | 0.025625 | 0.257 | 0.257 | 1 | 2 | Il1rn | 0 | Young |
| 0.961571 | 0.151917 | 0.025 | 0.025 | 1 | 2 | Pus3 | 0 | Young |
| 0.961875 | 0.000913 | 0.088 | 0.089 | 1 | 2 | Mfsd11 | -0.001 | Young |
| 0.962123 | 0.001213 | 0.048 | 0.047 | 1 | 2 | Arfgap1 | 0.001 | Young |
| 0.962469 | 0.269627 | 0.069 | 0.07 | 1 | 2 | Mtmr1 | -0.001 | Young |
| 0.962599 | 0.103954 | 0.154 | 0.16 | 1 | 2 | 2510039O1 | -0.006 | Young |
| 0.962825 | 0.065435 | 0.322 | 0.328 | 1 | 2 | Wdr83os | -0.006 | Young |
| 0.963093 | 0.063897 | 0.148 | 0.15 | 1 | 2 | Tug1 | -0.002 | Young |
| 0.963808 | 0.16011 | 0.201 | 0.208 | 1 | 2 | Bfar | -0.007 | Young |
| 0.963818 | 0.073482 | 0.167 | 0.17 | 1 | 2 | Tm2d2 | -0.003 | Young |
| 0.964105 | 0.279245 | 0.029 | 0.029 | 1 | 2 | Tab3 | 0 | Young |
| 0.964151 | 0.183747 | 0.151 | 0.153 | 1 | 2 | Dynlt3 | -0.002 | Young |
| 0.964216 | 0.060125 | 0.037 | 0.036 | 1 | 2 | Cog2 | 0.001 | Young |
| 0.964233 | 0.053081 | 0.029 | 0.029 | 1 | 2 | Zmym3 | 0 | Young |
| 0.964239 | 0.016418 | 0.486 | 0.488 | 1 | 2 | Itm2c | -0.002 | Young |
| 0.964457 | 0.032435 | 0.094 | 0.094 | 1 | 2 | Pdxdc1 | 0 | Young |
| 0.964814 | 0.195158 | 0.125 | 0.125 | 1 | 2 | Msl3 | 0 | Young |
| 0.964846 | 0.104929 | 0.056 | 0.057 | 1 | 2 | Zfp672 | -0.001 | Young |
| 0.964863 | 0.030911 | 0.025 | 0.025 | 1 | 2 | Syne1 | 0 | Young |
| 0.96563 | 0.033805 | 0.048 | 0.048 | 1 | 2 | Mbd5 | 0 | Young |
| 0.965796 | 0.064349 | 0.039 | 0.04 | 1 | 2 | Nudt14 | -0.001 | Young |
| 0.966253 | 0.036094 | 0.227 | 0.229 | 1 | 2 | Rnf7 | -0.002 | Young |
| 0.966992 | 0.034969 | 0.075 | 0.076 | 1 | 2 | Phax | -0.001 | Young |
| 0.967234 | 0.061359 | 0.146 | 0.148 | 1 | 2 | Eif4e2 | -0.002 | Young |
| 0.967399 | 0.166051 | 0.045 | 0.045 | 1 | 2 | Eri3 | 0 | Young |
| 0.967846 | 0.012147 | 0.048 | 0.047 | 1 | 2 | Cxxc1 | 0.001 | Young |
| 0.968077 | 0.609749 | 0.023 | 0.022 | 1 | 2 | Dffa | 0.001 | Young |
| 0.96842 | 0.184269 | 0.064 | 0.064 | 1 | 2 | Tarbp2 | 0 | Young |
| 0.968584 | 0.137446 | 0.135 | 0.136 | 1 | 2 | Nop58 | -0.001 | Young |
| 0.968838 | 0.062638 | 0.045 | 0.045 | 1 | 2 | Frs2 | 0 | Young |
| 0.969057 | 0.011145 | 0.071 | 0.072 | 1 | 2 | Pycr2 | -0.001 | Young |
| 0.969201 | 0.470207 | 0.018 | 0.018 | 1 | 2 | Rhbdf1 | 0 | Young |
| 0.96922 | 0.126506 | 0.162 | 0.165 | 1 | 2 | Vezf1 | -0.003 | Young |
| 0.969382 | 0.01574 | 0.147 | 0.148 | 1 | 2 | Rbfa | -0.001 | Young |
| 0.969438 | 0.028344 | 0.31 | 0.314 | 1 | 2 | Snx10 | -0.004 | Young |
| 0.970201 | 0.056079 | 0.077 | 0.077 | 1 | 2 | Lrrc8a | 0 | Young |
| 0.970398 | 0.456729 | 0.039 | 0.04 | 1 | 2 | Dnajb2 | -0.001 | Young |
| 0.970558 | 0.060426 | 0.156 | 0.156 | 1 | 2 | Samd9l | 0 | Young |
| 0.971102 | 0.013138 | 0.284 | 0.289 | 1 | 2 | Cd48 | -0.005 | Young |
| 0.971104 | 0.078924 | 0.059 | 0.06 | 1 | 2 | Nr1d2 | -0.001 | Young |
| 0.97112 | 0.218868 | 0.109 | 0.112 | 1 | 2 | Prr14 | -0.003 | Young |
| 0.971208 | 0.085906 | 0.103 | 0.105 | 1 | 2 | Tmem68 | -0.002 | Young |

| | | | | | | | | |
|---|---|---|---|---|---|---|---|---|
| 0.971376 | 0.755555 | 0.012 | 0.012 | 1 | 2 | Acat2 | 0 | Young |
| 0.97152 | 0.194462 | 0.063 | 0.064 | 1 | 2 | Nagpa | -0.001 | Young |
| 0.971651 | 0.011316 | 0.985 | 0.976 | 1 | 2 | Lyz2 | 0.009 | Young |
| 0.971654 | 0.024089 | 0.077 | 0.078 | 1 | 2 | Rnf181 | -0.001 | Young |
| 0.971785 | 0.296681 | 0.033 | 0.033 | 1 | 2 | Prim2 | 0 | Young |
| 0.971858 | 0.270625 | 0.043 | 0.043 | 1 | 2 | Cops7b | 0 | Young |
| 0.972395 | 0.070368 | 0.245 | 0.251 | 1 | 2 | Snrpc | -0.006 | Young |
| 0.972458 | 0.436793 | 0.053 | 0.055 | 1 | 2 | Ralgapa2 | -0.002 | Young |
| 0.972905 | 0.165622 | 0.027 | 0.028 | 1 | 2 | Otud1 | -0.001 | Young |
| 0.972927 | 0.211973 | 0.082 | 0.084 | 1 | 2 | Uba3 | -0.002 | Young |
| 0.973078 | 0.071297 | 0.122 | 0.124 | 1 | 2 | Chordc1 | -0.002 | Young |
| 0.973318 | 0.144004 | 0.101 | 0.103 | 1 | 2 | Nop56 | -0.002 | Young |
| 0.973551 | 0.013335 | 0.235 | 0.24 | 1 | 2 | Svil | -0.005 | Young |
| 0.9741 | 0.230726 | 0.122 | 0.125 | 1 | 2 | Sap18 | -0.003 | Young |
| 0.97414 | 0.102997 | 0.133 | 0.136 | 1 | 2 | 2210016F1 | -0.003 | Young |
| 0.974188 | 0.13205 | 0.14 | 0.142 | 1 | 2 | Wwp2 | -0.002 | Young |
| 0.974294 | 0.104491 | 0.03 | 0.03 | 1 | 2 | Fam210b | 0 | Young |
| 0.974302 | 0.028109 | 0.024 | 0.024 | 1 | 2 | Ercc4 | 0 | Young |
| 0.974425 | 0.126487 | 0.057 | 0.058 | 1 | 2 | Rtl5 | -0.001 | Young |
| 0.974576 | 0.190351 | 0.048 | 0.047 | 1 | 2 | Asb7 | 0.001 | Young |
| 0.974859 | 0.043891 | 0.057 | 0.057 | 1 | 2 | Ampd2 | 0 | Young |
| 0.974864 | 0.101342 | 0.024 | 0.024 | 1 | 2 | Mtfmt | 0 | Young |
| 0.974928 | 0.486891 | 0.012 | 0.012 | 1 | 2 | Tmem81 | 0 | Young |
| 0.975011 | 0.208021 | 0.159 | 0.162 | 1 | 2 | Ube2m | -0.003 | Young |
| 0.9752 | 0.039959 | 0.175 | 0.177 | 1 | 2 | Arl5a | -0.002 | Young |
| 0.975247 | 0.258033 | 0.125 | 0.128 | 1 | 2 | Nrf1 | -0.003 | Young |
| 0.975296 | 0.059011 | 0.03 | 0.03 | 1 | 2 | Kif23 | 0 | Young |
| 0.975518 | 0.001151 | 0.048 | 0.048 | 1 | 2 | Cep57 | 0 | Young |
| 0.975845 | 0.032847 | 0.121 | 0.122 | 1 | 2 | Avpi1 | -0.001 | Young |
| 0.976043 | 0.015298 | 0.372 | 0.376 | 1 | 2 | Fli1 | -0.004 | Young |
| 0.976204 | 0.347503 | 0.101 | 0.103 | 1 | 2 | Wsb2 | -0.002 | Young |
| 0.976209 | 0.086954 | 0.097 | 0.099 | 1 | 2 | Ltv1 | -0.002 | Young |
| 0.976314 | 0.053657 | 0.175 | 0.177 | 1 | 2 | Setd3 | -0.002 | Young |
| 0.976449 | 0.005191 | 0.924 | 0.916 | 1 | 2 | Fcer1g | 0.008 | Young |
| 0.976619 | 0.527566 | 0.029 | 0.029 | 1 | 2 | Asb13 | 0 | Young |
| 0.976892 | 0.001335 | 0.374 | 0.378 | 1 | 2 | 2900097C1 | -0.004 | Young |
| 0.977108 | 0.042697 | 0.019 | 0.019 | 1 | 2 | Cad | 0 | Young |
| 0.977425 | 0.020426 | 0.01 | 0.01 | 1 | 2 | Gm39469 | 0 | Young |
| 0.977852 | 0.307492 | 0.023 | 0.022 | 1 | 2 | Casp7 | 0.001 | Young |
| 0.977898 | 0.067879 | 0.214 | 0.219 | 1 | 2 | Anxa6 | -0.005 | Young |
| 0.978015 | 0.08883 | 0.154 | 0.156 | 1 | 2 | Cnot3 | -0.002 | Young |
| 0.978086 | 0.496735 | 0.012 | 0.012 | 1 | 2 | Wdr62 | 0 | Young |
| 0.978172 | 0.129523 | 0.132 | 0.135 | 1 | 2 | Ei24 | -0.003 | Young |
| 0.978481 | 0.579144 | 0.012 | 0.012 | 1 | 2 | Ptchd1 | 0 | Young |
| 0.978567 | 0.089198 | 0.143 | 0.143 | 1 | 2 | Abhd17a | 0 | Young |

| | | | | | | | | |
|---|---|---|---|---|---|---|---|---|
| 0.978887 | 0.010328 | 0.802 | 0.804 | 1 | 2 | Pid1 | -0.002 | Young |
| 0.978977 | 0.282142 | 0.019 | 0.019 | 1 | 2 | Spaca6 | 0 | Young |
| 0.979054 | 0.01985 | 0.112 | 0.113 | 1 | 2 | Fbrsl1 | -0.001 | Young |
| 0.979122 | 0.164191 | 0.036 | 0.035 | 1 | 2 | Etfrf1 | 0.001 | Young |
| 0.979234 | 0.312744 | 0.043 | 0.043 | 1 | 2 | Ints2 | 0 | Young |
| 0.979589 | 0.076664 | 0.124 | 0.124 | 1 | 2 | Zfp950 | 0 | Young |
| 0.980055 | 0.195227 | 0.01 | 0.01 | 1 | 2 | Prdm11 | 0 | Young |
| 0.980149 | 0.061311 | 0.065 | 0.065 | 1 | 2 | Tsga10 | 0 | Young |
| 0.980331 | 0.223913 | 0.118 | 0.12 | 1 | 2 | Smox | -0.002 | Young |
| 0.980433 | 0.203467 | 0.139 | 0.14 | 1 | 2 | Mybbp1a | -0.001 | Young |
| 0.980455 | 0.165561 | 0.012 | 0.012 | 1 | 2 | Gm4951 | 0 | Young |
| 0.980545 | 0.121681 | 0.122 | 0.123 | 1 | 2 | Ndufs8 | -0.001 | Young |
| 0.980721 | 0.029225 | 0.166 | 0.168 | 1 | 2 | Nupr1 | -0.002 | Young |
| 0.981244 | 0.213299 | 0.012 | 0.012 | 1 | 2 | Ccdc77 | 0 | Young |
| 0.981244 | 0.18274 | 0.012 | 0.012 | 1 | 2 | Gm10642 | 0 | Young |
| 0.981369 | 0.451276 | 0.01 | 0.01 | 1 | 2 | Spata1 | 0 | Young |
| 0.981369 | 0.274585 | 0.01 | 0.01 | 1 | 2 | Cdca8 | 0 | Young |
| 0.981537 | 0.019996 | 0.201 | 0.202 | 1 | 2 | Zc3h15 | -0.001 | Young |
| 0.981601 | 0.109869 | 0.055 | 0.055 | 1 | 2 | Tmem104 | 0 | Young |
| 0.981761 | 0.624522 | 0.027 | 0.028 | 1 | 2 | Prmt9 | -0.001 | Young |
| 0.982149 | 0.019698 | 0.056 | 0.056 | 1 | 2 | Asb8 | 0 | Young |
| 0.982684 | 0.497089 | 0.01 | 0.01 | 1 | 2 | Hexdc | 0 | Young |
| 0.982684 | 0.390863 | 0.01 | 0.01 | 1 | 2 | Flrt2 | 0 | Young |
| 0.982704 | 0.185035 | 0.049 | 0.049 | 1 | 2 | Nup85 | 0 | Young |
| 0.982741 | 0.324546 | 0.023 | 0.022 | 1 | 2 | Pick1 | 0.001 | Young |
| 0.982745 | 0.675587 | 0.021 | 0.021 | 1 | 2 | 1110065P2 | 0 | Young |
| 0.983007 | 0.067557 | 0.024 | 0.024 | 1 | 2 | Serpinb2 | 0 | Young |
| 0.983074 | 0.09941 | 0.19 | 0.192 | 1 | 2 | Akirin2 | -0.002 | Young |
| 0.983218 | 0.282019 | 0.012 | 0.012 | 1 | 2 | Trim32 | 0 | Young |
| 0.983307 | 0.176227 | 0.146 | 0.15 | 1 | 2 | Nsun2 | -0.004 | Young |
| 0.98345 | 0.096082 | 0.043 | 0.043 | 1 | 2 | Agpat3 | 0 | Young |
| 0.983561 | 0.457598 | 0.01 | 0.01 | 1 | 2 | Wdr90 | 0 | Young |
| 0.983673 | 0.193223 | 0.099 | 0.1 | 1 | 2 | Rnf169 | -0.001 | Young |
| 0.983891 | 0.027336 | 0.023 | 0.022 | 1 | 2 | Cenpl | 0.001 | Young |
| 0.983999 | 0.469678 | 0.01 | 0.01 | 1 | 2 | Arap2 | 0 | Young |
| 0.984036 | 0.244449 | 0.051 | 0.051 | 1 | 2 | Trim36 | 0 | Young |
| 0.984879 | 0.038934 | 0.033 | 0.033 | 1 | 2 | Tram2 | 0 | Young |
| 0.985072 | 0.038598 | 0.27 | 0.272 | 1 | 2 | Emg1 | -0.002 | Young |
| 0.985412 | 0.228542 | 0.07 | 0.071 | 1 | 2 | Ak6 | -0.001 | Young |
| 0.985773 | 0.075779 | 0.16 | 0.164 | 1 | 2 | Mrpl36 | -0.004 | Young |
| 0.985849 | 0.261839 | 0.071 | 0.072 | 1 | 2 | Pias2 | -0.001 | Young |
| 0.986157 | 0.218076 | 0.057 | 0.058 | 1 | 2 | Tbce | -0.001 | Young |
| 0.986191 | 0.50033 | 0.01 | 0.01 | 1 | 2 | B3galt4 | 0 | Young |
| 0.986273 | 0.141541 | 0.241 | 0.247 | 1 | 2 | Sar1a | -0.006 | Young |
| 0.986584 | 0.676886 | 0.029 | 0.029 | 1 | 2 | Zfand2a | 0 | Young |

| | | | | | | | | |
|---|---|---|---|---|---|---|---|---|
| 0.987464 | 0.418278 | 0.021 | 0.021 | 1 | 2 | Trim14 | 0 | Young |
| 0.987712 | 0.263646 | 0.051 | 0.051 | 1 | 2 | Akr1b10 | 0 | Young |
| 0.988027 | 0.083614 | 0.094 | 0.094 | 1 | 2 | Prpf38a | 0 | Young |
| 0.988206 | 0.067366 | 0.023 | 0.022 | 1 | 2 | Coq3 | 0.001 | Young |
| 0.988437 | 0.069471 | 0.309 | 0.316 | 1 | 2 | Rp2 | -0.007 | Young |
| 0.988664 | 0.359867 | 0.101 | 0.103 | 1 | 2 | Enox2 | -0.002 | Young |
| 0.98868 | 0.016599 | 0.051 | 0.051 | 1 | 2 | Champ1 | 0 | Young |
| 0.98878 | 0.492618 | 0.046 | 0.047 | 1 | 2 | Arl3 | -0.001 | Young |
| 0.988821 | 0.521349 | 0.01 | 0.01 | 1 | 2 | Cfb | 0 | Young |
| 0.988831 | 0.465438 | 0.049 | 0.049 | 1 | 2 | Mri1 | 0 | Young |
| 0.988875 | 0.024927 | 0.219 | 0.221 | 1 | 2 | Washc2 | -0.002 | Young |
| 0.988939 | 0.058902 | 0.242 | 0.248 | 1 | 2 | Mvp | -0.006 | Young |
| 0.989152 | 0.061889 | 0.189 | 0.192 | 1 | 2 | Rasgef1b | -0.003 | Young |
| 0.989469 | 0.289998 | 0.061 | 0.061 | 1 | 2 | Bnip3 | 0 | Young |
| 0.989471 | 0.081597 | 0.063 | 0.063 | 1 | 2 | Thyn1 | 0 | Young |
| 0.989698 | 0.786094 | 0.01 | 0.01 | 1 | 2 | Mtcp1 | 0 | Young |
| 0.989932 | 0.178569 | 0.023 | 0.022 | 1 | 2 | Haus7 | 0.001 | Young |
| 0.989982 | 0.071074 | 0.125 | 0.126 | 1 | 2 | Idh3b | -0.001 | Young |
| 0.990251 | 0.065385 | 0.077 | 0.077 | 1 | 2 | Tusc3 | 0 | Young |
| 0.990677 | 0.264424 | 0.032 | 0.032 | 1 | 2 | Rdm1 | 0 | Young |
| 0.99077 | 0.314132 | 0.089 | 0.091 | 1 | 2 | Tufm | -0.002 | Young |
| 0.990894 | 0.125673 | 0.055 | 0.055 | 1 | 2 | Mettl6 | 0 | Young |
| 0.99117 | 0.074415 | 0.204 | 0.209 | 1 | 2 | Rufy3 | -0.005 | Young |
| 0.991381 | 0.110465 | 0.127 | 0.127 | 1 | 2 | Ppm1g | 0 | Young |
| 0.99184 | 0.131066 | 0.063 | 0.063 | 1 | 2 | Surf1 | 0 | Young |
| 0.991982 | 0.094708 | 0.05 | 0.05 | 1 | 2 | Unc45a | 0 | Young |
| 0.992034 | 0.20219 | 0.126 | 0.13 | 1 | 2 | Cldnd1 | -0.004 | Young |
| 0.992281 | 0.066952 | 0.09 | 0.091 | 1 | 2 | Zbtb8os | -0.001 | Young |
| 0.992953 | 0.039279 | 0.042 | 0.042 | 1 | 2 | Dpp7 | 0 | Young |
| 0.993337 | 0.135509 | 0.176 | 0.179 | 1 | 2 | Ubr5 | -0.003 | Young |
| 0.993569 | 0.633932 | 0.011 | 0.011 | 1 | 2 | Bub1b | 0 | Young |
| 0.993658 | 0.050969 | 0.021 | 0.021 | 1 | 2 | Cox11 | 0 | Young |
| 0.993718 | 0.516519 | 0.031 | 0.031 | 1 | 2 | Lancl1 | 0 | Young |
| 0.993833 | 0.085141 | 0.04 | 0.041 | 1 | 2 | Snx24 | -0.001 | Young |
| 0.994099 | 0.042237 | 0.159 | 0.162 | 1 | 2 | Btbd7 | -0.003 | Young |
| 0.994254 | 0.070656 | 0.102 | 0.104 | 1 | 2 | Smarcb1 | -0.002 | Young |
| 0.994431 | 0.152777 | 0.032 | 0.032 | 1 | 2 | Sephs1 | 0 | Young |
| 0.994457 | 0.38488 | 0.031 | 0.031 | 1 | 2 | Clpp | 0 | Young |
| 0.994764 | 0.212143 | 0.26 | 0.27 | 1 | 2 | Man1a | -0.01 | Young |
| 0.994813 | 0.533066 | 0.011 | 0.011 | 1 | 2 | Rab38 | 0 | Young |
| 0.995196 | 0.197413 | 0.031 | 0.031 | 1 | 2 | Ap3m2 | 0 | Young |
| 0.996018 | 0.026372 | 0.021 | 0.021 | 1 | 2 | Gm15445 | 0 | Young |
| 0.996162 | 0.176237 | 0.067 | 0.067 | 1 | 2 | Rce1 | 0 | Young |
| 0.996303 | 0.054996 | 0.072 | 0.073 | 1 | 2 | Smim20 | -0.001 | Young |
| 0.996399 | 0.083454 | 0.184 | 0.186 | 1 | 2 | Mrpl24 | -0.002 | Young |

| | | | | | | | | |
|---|---|---|---|---|---|---|---|---|
| 0.996473 | 0.193955 | 0.011 | 0.011 | 1 | 2 | Amot | 0 | Young |
| 0.996474 | 0.01699 | 0.084 | 0.085 | 1 | 2 | Abcg3 | -0.001 | Young |
| 0.997295 | 0.205883 | 0.04 | 0.041 | 1 | 2 | 2310039H( | -0.001 | Young |
| 0.997771 | 0.004163 | 0.251 | 0.254 | 1 | 2 | Acp2 | -0.003 | Young |
| 0.997772 | 0.181346 | 0.112 | 0.112 | 1 | 2 | Zufsp | 0 | Young |
| 0.997906 | 0.25782 | 0.031 | 0.031 | 1 | 2 | Znhit6 | 0 | Young |
| 0.998133 | 0.573609 | 0.011 | 0.011 | 1 | 2 | Acpp | 0 | Young |
| 0.998353 | 0.079476 | 0.23 | 0.236 | 1 | 2 | Ahcyl2 | -0.006 | Young |
| 0.998548 | 0.208661 | 0.011 | 0.011 | 1 | 2 | Gm47675 | 0 | Young |
| 0.998548 | 0.038344 | 0.011 | 0.011 | 1 | 2 | Copz2 | 0 | Young |
| 0.998963 | 0.121672 | 0.011 | 0.011 | 1 | 2 | Ckap2l | 0 | Young |
| 0.998968 | 0.105736 | 0.021 | 0.021 | 1 | 2 | Thap4 | 0 | Young |
| 0.999036 | 0.136878 | 0.166 | 0.17 | 1 | 2 | 2810403A( | -0.004 | Young |
| 0.999178 | 0.048191 | 0.709 | 0.741 | 1 | 2 | Zfp36l1 | -0.032 | Young |
| 0.999193 | 0.008873 | 0.093 | 0.093 | 1 | 2 | Swt1 | 0 | Young |
| 0.999229 | 0.107245 | 0.102 | 0.104 | 1 | 2 | Tpd52l2 | -0.002 | Young |
| 0.999231 | 0.038085 | 0.103 | 0.104 | 1 | 2 | Casp1 | -0.001 | Young |
| 0.999292 | 0.153483 | 0.1 | 0.102 | 1 | 2 | Sp140 | -0.002 | Young |
| 0.999413 | 0.628636 | 0.05 | 0.05 | 1 | 2 | Mettl13 | 0 | Young |
| 0.999793 | 0.137024 | 0.011 | 0.011 | 1 | 2 | Rab23 | 0 | Young |
| 3.52E-15 | 0.213883 | 1 | 1 | 1.09E-10 | 3 | mt-Co1 | 0 | Old |
| 2.14E-14 | 0.38603 | 0.976 | 0.932 | 6.66E-10 | 3 | B2m | 0.044 | Old |
| 3.56E-14 | 3.804452 | 0.086 | 0.006 | 1.11E-09 | 3 | Cxcl13 | 0.08 | Old |
| 7.23E-14 | 0.19462 | 0.999 | 0.997 | 2.24E-09 | 3 | Actb | 0.002 | Old |
| 2.83E-13 | 0.300421 | 0.958 | 0.934 | 8.78E-09 | 3 | Gda | 0.024 | Old |
| 6.99E-13 | 0.214012 | 0.999 | 0.995 | 2.17E-08 | 3 | Actg1 | 0.004 | Old |
| 9.16E-12 | 0.74903 | 0.471 | 0.312 | 2.84E-07 | 3 | Coro2a | 0.159 | Old |
| 1.75E-11 | 0.35375 | 0.869 | 0.755 | 5.44E-07 | 3 | Tnip3 | 0.114 | Old |
| 4.31E-11 | 0.697733 | 0.548 | 0.389 | 1.34E-06 | 3 | Rgs1 | 0.159 | Old |
| 1.12E-10 | 0.196755 | 1 | 1 | 3.48E-06 | 3 | mt-Co3 | 0 | Old |
| 1.77E-09 | 0.347324 | 0.828 | 0.717 | 5.48E-05 | 3 | Lamp2 | 0.111 | Old |
| 1.90E-09 | 0.204836 | 0.986 | 0.961 | 5.90E-05 | 3 | Cfl1 | 0.025 | Old |
| 2.57E-09 | 0.2547 | 0.957 | 0.923 | 7.98E-05 | 3 | Actr3 | 0.034 | Old |
| 2.64E-09 | 0.437005 | 0.662 | 0.499 | 8.19E-05 | 3 | Cstb | 0.163 | Old |
| 2.84E-09 | 0.147078 | 0.999 | 0.997 | 8.83E-05 | 3 | Psap | 0.002 | Old |
| 3.28E-09 | 0.276257 | 0.94 | 0.862 | 0.000102 | 3 | Hnrnpa2b1 | 0.078 | Old |
| 3.57E-09 | 0.216856 | 0.946 | 0.908 | 0.000111 | 3 | Capza2 | 0.038 | Old |
| 5.26E-09 | 0.169969 | 0.992 | 0.984 | 0.000163 | 3 | Gpx1 | 0.008 | Old |
| 1.83E-08 | 2.061249 | 0.075 | 0.016 | 0.000568 | 3 | Igkc | 0.059 | Old |
| 1.88E-08 | 0.250829 | 0.982 | 0.967 | 0.000583 | 3 | Hsp90b1 | 0.015 | Old |
| 1.95E-08 | 0.232218 | 0.941 | 0.9 | 0.000606 | 3 | Arpc5 | 0.041 | Old |
| 2.04E-08 | 0.37787 | 0.778 | 0.672 | 0.000634 | 3 | Atp5g3 | 0.106 | Old |
| 2.05E-08 | 0.261491 | 0.967 | 0.934 | 0.000636 | 3 | Ywhaz | 0.033 | Old |
| 2.43E-08 | 0.184618 | 0.995 | 0.989 | 0.000754 | 3 | Ccl9 | 0.006 | Old |
| 2.45E-08 | 0.457193 | 0.717 | 0.612 | 0.000761 | 3 | Ifi27l2a | 0.105 | Old |

| | | | | | | | | |
|---|---|---|---|---|---|---|---|---|
| 2.84E-08 | 0.263995 | 0.904 | 0.846 | 0.000883 | 3 | Sdcbp | 0.058 | Old |
| 2.95E-08 | 0.286818 | 0.858 | 0.773 | 0.000915 | 3 | Wdr1 | 0.085 | Old |
| 3.75E-08 | 0.925734 | 0.164 | 0.073 | 0.001165 | 3 | Dusp10 | 0.091 | Old |
| 7.38E-08 | 0.39993 | 0.72 | 0.609 | 0.002291 | 3 | Cd36 | 0.111 | Old |
| 7.73E-08 | 0.347363 | 0.748 | 0.638 | 0.0024 | 3 | Ccr5 | 0.11 | Old |
| 8.74E-08 | 0.309053 | 0.814 | 0.688 | 0.002713 | 3 | Ssr3 | 0.126 | Old |
| 9.42E-08 | 0.308593 | 0.848 | 0.775 | 0.002925 | 3 | Slc25a5 | 0.073 | Old |
| 9.72E-08 | 0.303423 | 0.951 | 0.931 | 0.003018 | 3 | H3f3b | 0.02 | Old |
| 1.19E-07 | 0.725249 | 0.251 | 0.145 | 0.003682 | 3 | F10 | 0.106 | Old |
| 1.24E-07 | 0.205009 | 0.987 | 0.989 | 0.003836 | 3 | mt-Cytb | -0.002 | Old |
| 1.64E-07 | 0.617232 | 0.311 | 0.196 | 0.005078 | 3 | Ctsl | 0.115 | Old |
| 2.07E-07 | 0.255628 | 0.957 | 0.957 | 0.006431 | 3 | Ctss | 0 | Old |
| 2.20E-07 | 0.548175 | 0.351 | 0.232 | 0.00682 | 3 | Man2a1 | 0.119 | Old |
| 2.20E-07 | 0.371211 | 0.761 | 0.676 | 0.006836 | 3 | Rpn1 | 0.085 | Old |
| 2.42E-07 | 0.216893 | 0.938 | 0.893 | 0.007528 | 3 | Actr2 | 0.045 | Old |
| 2.60E-07 | 0.248828 | 0.931 | 0.847 | 0.008074 | 3 | Pim1 | 0.084 | Old |
| 3.01E-07 | 0.269976 | 0.91 | 0.865 | 0.009349 | 3 | Sdc3 | 0.045 | Old |
| 3.58E-07 | 0.518872 | 0.376 | 0.258 | 0.011112 | 3 | Psmd6 | 0.118 | Old |
| 5.50E-07 | 0.390901 | 0.57 | 0.431 | 0.017092 | 3 | Tcp1 | 0.139 | Old |
| 7.56E-07 | 0.256749 | 0.9 | 0.824 | 0.023469 | 3 | Lcp1 | 0.076 | Old |
| 7.85E-07 | 0.326065 | 0.781 | 0.679 | 0.02437 | 3 | Ldha | 0.102 | Old |
| 7.97E-07 | 0.327246 | 0.736 | 0.595 | 0.024747 | 3 | Prkar1a | 0.141 | Old |
| 8.29E-07 | 0.53978 | 0.418 | 0.304 | 0.025758 | 3 | Klf7 | 0.114 | Old |
| 9.67E-07 | 0.237968 | 0.896 | 0.854 | 0.030016 | 3 | Cd53 | 0.042 | Old |
| 1.05E-06 | 0.162369 | 1 | 0.997 | 0.032535 | 3 | mt-Atp6 | 0.003 | Old |
| 1.19E-06 | 0.4903 | 0.442 | 0.323 | 0.036906 | 3 | Ccnd2 | 0.119 | Old |
| 1.27E-06 | 0.297733 | 0.932 | 0.88 | 0.039471 | 3 | Samhd1 | 0.052 | Old |
| 1.32E-06 | 1.238723 | 0.134 | 0.062 | 0.041103 | 3 | Ccl8 | 0.072 | Old |
| 1.43E-06 | 0.366747 | 0.692 | 0.588 | 0.044415 | 3 | Cmpk1 | 0.104 | Old |
| 1.74E-06 | 1.776416 | 0.061 | 0.015 | 0.053896 | 3 | Gm26870 | 0.046 | Old |
| 2.05E-06 | 0.409566 | 0.767 | 0.687 | 0.063602 | 3 | Plin2 | 0.08 | Old |
| 2.44E-06 | 0.519574 | 0.326 | 0.216 | 0.075649 | 3 | Snx4 | 0.11 | Old |
| 2.49E-06 | 1.707781 | 0.06 | 0.015 | 0.077454 | 3 | Cd300lf | 0.045 | Old |
| 2.60E-06 | 0.158479 | 0.992 | 0.966 | 0.080889 | 3 | Junb | 0.026 | Old |
| 2.88E-06 | 0.200672 | 0.919 | 0.884 | 0.089418 | 3 | Gm10076 | 0.035 | Old |
| 3.27E-06 | 0.826924 | 0.155 | 0.08 | 0.101522 | 3 | Igf2r | 0.075 | Old |
| 3.74E-06 | 0.187062 | 0.984 | 0.968 | 0.116009 | 3 | Ptma | 0.016 | Old |
| 3.82E-06 | 0.388127 | 0.449 | 0.335 | 0.118671 | 3 | Dusp5 | 0.114 | Old |
| 4.05E-06 | 0.245428 | 0.918 | 0.899 | 0.125795 | 3 | Ctsd | 0.019 | Old |
| 4.16E-06 | 0.621874 | 0.214 | 0.125 | 0.129088 | 3 | Eef1e1 | 0.089 | Old |
| 5.13E-06 | 0.134452 | 0.999 | 1 | 0.159178 | 3 | mt-Co2 | -0.001 | Old |
| 5.38E-06 | 0.567826 | 0.37 | 0.268 | 0.1672 | 3 | Palld | 0.102 | Old |
| 5.44E-06 | 0.269954 | 0.902 | 0.87 | 0.168908 | 3 | Ctsb | 0.032 | Old |
| 5.61E-06 | 0.263293 | 0.846 | 0.799 | 0.174176 | 3 | Hnrnpa3 | 0.047 | Old |
| 5.68E-06 | 1.929207 | 0.074 | 0.025 | 0.176385 | 3 | Irgm1 | 0.049 | Old |

| | | | | | | | | |
|---|---|---|---|---|---|---|---|---|
| 6.05E-06 | 0.333655 | 0.667 | 0.549 | 0.187858 | 3 | Pgam1 | 0.118 | Old |
| 6.21E-06 | 0.580162 | 0.346 | 0.239 | 0.192892 | 3 | Uqcrc1 | 0.107 | Old |
| 6.32E-06 | 0.367304 | 0.297 | 0.192 | 0.196113 | 3 | Slc7a1 | 0.105 | Old |
| 6.67E-06 | 0.395435 | 0.612 | 0.491 | 0.20701 | 3 | Atp6ap2 | 0.121 | Old |
| 7.09E-06 | 0.506609 | 0.644 | 0.544 | 0.220137 | 3 | Dab2 | 0.1 | Old |
| 7.22E-06 | 0.352513 | 0.429 | 0.314 | 0.224243 | 3 | Trim8 | 0.115 | Old |
| 7.80E-06 | 0.329554 | 0.789 | 0.699 | 0.242172 | 3 | Ly86 | 0.09 | Old |
| 8.04E-06 | 0.423385 | 0.676 | 0.609 | 0.249563 | 3 | Egr1 | 0.067 | Old |
| 9.15E-06 | 0.470043 | 0.437 | 0.328 | 0.28414 | 3 | Nsf | 0.109 | Old |
| 9.28E-06 | 0.147682 | 0.947 | 0.932 | 0.288146 | 3 | mt-Nd3 | 0.015 | Old |
| 9.61E-06 | 0.334663 | 0.448 | 0.331 | 0.298568 | 3 | Mfsd1 | 0.117 | Old |
| 1.00E-05 | 1.10593 | 0.081 | 0.03 | 0.311955 | 3 | Gmppb | 0.051 | Old |
| 1.08E-05 | 0.384491 | 0.351 | 0.246 | 0.336068 | 3 | Noct | 0.105 | Old |
| 1.20E-05 | 0.333642 | 0.551 | 0.436 | 0.372816 | 3 | 9530068E0 | 0.115 | Old |
| 1.23E-05 | 0.321502 | 0.565 | 0.443 | 0.383188 | 3 | Larp1 | 0.122 | Old |
| 1.26E-05 | 0.186498 | 0.995 | 0.994 | 0.389883 | 3 | Jund | 0.001 | Old |
| 1.28E-05 | 0.136984 | 0.992 | 0.979 | 0.396315 | 3 | Rps26 | 0.013 | Old |
| 1.28E-05 | 0.285437 | 0.712 | 0.604 | 0.396889 | 3 | Pgk1 | 0.108 | Old |
| 1.33E-05 | 0.680579 | 0.152 | 0.081 | 0.412498 | 3 | Ergic1 | 0.071 | Old |
| 1.38E-05 | 0.248258 | 0.624 | 0.499 | 0.428525 | 3 | Plaur | 0.125 | Old |
| 1.47E-05 | 0.720278 | 0.268 | 0.182 | 0.455085 | 3 | Ccnd1 | 0.086 | Old |
| 1.48E-05 | 0.368008 | 0.513 | 0.4 | 0.458311 | 3 | Pld3 | 0.113 | Old |
| 1.48E-05 | 0.65758 | 0.211 | 0.129 | 0.460822 | 3 | Slc39a14 | 0.082 | Old |
| 1.57E-05 | 0.241596 | 0.873 | 0.812 | 0.486385 | 3 | Atp5a1 | 0.061 | Old |
| 1.64E-05 | 0.410256 | 0.449 | 0.342 | 0.508901 | 3 | Ptprs | 0.107 | Old |
| 1.76E-05 | 0.136504 | 0.974 | 0.97 | 0.546367 | 3 | Arpc2 | 0.004 | Old |
| 2.01E-05 | 0.259424 | 0.839 | 0.74 | 0.623792 | 3 | Atp5f1 | 0.099 | Old |
| 2.22E-05 | 0.169748 | 0.901 | 0.855 | 0.689857 | 3 | Cd44 | 0.046 | Old |
| 2.29E-05 | 0.246084 | 0.805 | 0.713 | 0.712401 | 3 | Xbp1 | 0.092 | Old |
| 2.53E-05 | 0.865125 | 0.149 | 0.082 | 0.785002 | 3 | Pde12 | 0.067 | Old |
| 2.66E-05 | 0.218588 | 0.914 | 0.854 | 0.826498 | 3 | Hnrnpf | 0.06 | Old |
| 2.73E-05 | 0.619977 | 0.239 | 0.154 | 0.847215 | 3 | Hilpda | 0.085 | Old |
| 2.79E-05 | 0.644359 | 0.155 | 0.086 | 0.865657 | 3 | Nup93 | 0.069 | Old |
| 2.82E-05 | 0.362607 | 0.547 | 0.441 | 0.876251 | 3 | Eif1a | 0.106 | Old |
| 3.14E-05 | 0.138876 | 0.974 | 0.965 | 0.97597 | 3 | Myl6 | 0.009 | Old |
| 3.48E-05 | 0.166548 | 0.946 | 0.894 | 1 | 3 | Eif4g2 | 0.052 | Old |
| 3.93E-05 | 0.274438 | 0.658 | 0.548 | 1 | 3 | Clec4a2 | 0.11 | Old |
| 4.20E-05 | 0.374568 | 0.593 | 0.484 | 1 | 3 | Clec4a3 | 0.109 | Old |
| 4.20E-05 | 0.291712 | 0.535 | 0.427 | 1 | 3 | Nfil3 | 0.108 | Old |
| 4.28E-05 | 0.279543 | 0.704 | 0.606 | 1 | 3 | Hnrnpm | 0.098 | Old |
| 4.39E-05 | 0.432965 | 0.332 | 0.232 | 1 | 3 | Ccdc71l | 0.1 | Old |
| 4.46E-05 | 0.745111 | 0.232 | 0.153 | 1 | 3 | Parp14 | 0.079 | Old |
| 4.60E-05 | 0.983762 | 0.125 | 0.066 | 1 | 3 | Jagn1 | 0.059 | Old |
| 4.72E-05 | 0.998338 | 0.129 | 0.068 | 1 | 3 | Cdc42ep2 | 0.061 | Old |
| 4.92E-05 | 0.419993 | 0.404 | 0.301 | 1 | 3 | Sdhc | 0.103 | Old |

| | | | | | | | | |
|---|---|---|---|---|---|---|---|---|
| 5.03E-05 | 0.850716 | 0.226 | 0.146 | 1 | 3 | Snrpa | 0.08 | Old |
| 5.38E-05 | 0.151251 | 0.992 | 0.992 | 1 | 3 | Ccl6 | 0 | Old |
| 5.54E-05 | 1.307138 | 0.075 | 0.03 | 1 | 3 | Grwd1 | 0.045 | Old |
| 5.85E-05 | 0.409493 | 0.593 | 0.482 | 1 | 3 | Lgmn | 0.111 | Old |
| 6.04E-05 | 0.342593 | 0.269 | 0.178 | 1 | 3 | Odc1 | 0.091 | Old |
| 6.05E-05 | 0.539659 | 0.235 | 0.154 | 1 | 3 | Lpxn | 0.081 | Old |
| 6.39E-05 | 0.35171 | 0.521 | 0.414 | 1 | 3 | Ppp1r15b | 0.107 | Old |
| 6.48E-05 | 0.264457 | 0.789 | 0.713 | 1 | 3 | Metrnl | 0.076 | Old |
| 6.58E-05 | 0.369443 | 0.418 | 0.321 | 1 | 3 | Mcur1 | 0.097 | Old |
| 6.59E-05 | 0.443773 | 0.251 | 0.165 | 1 | 3 | Chchd3 | 0.086 | Old |
| 7.03E-05 | 0.343503 | 0.419 | 0.316 | 1 | 3 | Etfa | 0.103 | Old |
| 7.17E-05 | 0.42562 | 0.544 | 0.453 | 1 | 3 | Atp2a2 | 0.091 | Old |
| 7.47E-05 | 0.254758 | 0.812 | 0.73 | 1 | 3 | Ier2 | 0.082 | Old |
| 7.63E-05 | 0.701718 | 0.173 | 0.106 | 1 | 3 | Tmem185b | 0.067 | Old |
| 7.67E-05 | 0.251625 | 0.848 | 0.766 | 1 | 3 | 2010107E0 | 0.082 | Old |
| 7.75E-05 | 0.372729 | 0.254 | 0.169 | 1 | 3 | Bet1 | 0.085 | Old |
| 7.77E-05 | 0.237391 | 0.889 | 0.835 | 1 | 3 | Hspa5 | 0.054 | Old |
| 8.02E-05 | 0.263112 | 0.638 | 0.52 | 1 | 3 | Jarid2 | 0.118 | Old |
| 8.16E-05 | 0.910365 | 0.16 | 0.095 | 1 | 3 | Slfn5 | 0.065 | Old |
| 8.25E-05 | 0.453822 | 0.263 | 0.178 | 1 | 3 | Ap3m1 | 0.085 | Old |
| 8.37E-05 | 0.2178 | 0.925 | 0.869 | 1 | 3 | H2-DMb1 | 0.056 | Old |
| 8.64E-05 | 0.260805 | 0.949 | 0.937 | 1 | 3 | Lpl | 0.012 | Old |
| 8.71E-05 | 0.353781 | 0.441 | 0.342 | 1 | 3 | Mif | 0.099 | Old |
| 8.96E-05 | 0.918199 | 0.178 | 0.111 | 1 | 3 | Mpdu1 | 0.067 | Old |
| 9.01E-05 | 0.251926 | 0.761 | 0.657 | 1 | 3 | Psmb4 | 0.104 | Old |
| 9.04E-05 | 0.278599 | 0.795 | 0.722 | 1 | 3 | Pdia6 | 0.073 | Old |
| 9.17E-05 | 0.393597 | 0.578 | 0.48 | 1 | 3 | Sppl2a | 0.098 | Old |
| 9.40E-05 | 0.2632 | 0.459 | 0.354 | 1 | 3 | Api5 | 0.105 | Old |
| 9.63E-05 | 0.441212 | 0.185 | 0.112 | 1 | 3 | Ado | 0.073 | Old |
| 0.000102 | 0.293282 | 0.522 | 0.409 | 1 | 3 | Vapa | 0.113 | Old |
| 0.00011 | 0.356116 | 0.553 | 0.432 | 1 | 3 | Il1rn | 0.121 | Old |
| 0.000112 | 2.001376 | 0.025 | 0.003 | 1 | 3 | Csprs | 0.022 | Old |
| 0.000117 | 0.29349 | 0.343 | 0.246 | 1 | 3 | Frrs1 | 0.097 | Old |
| 0.000128 | 0.544956 | 0.249 | 0.169 | 1 | 3 | Cdk17 | 0.08 | Old |
| 0.00013 | 0.25642 | 0.648 | 0.54 | 1 | 3 | Trim25 | 0.108 | Old |
| 0.000141 | 0.14057 | 0.967 | 0.949 | 1 | 3 | Anxa2 | 0.018 | Old |
| 0.000143 | 0.353749 | 0.331 | 0.237 | 1 | 3 | B4galt5 | 0.094 | Old |
| 0.000144 | 0.724376 | 0.16 | 0.098 | 1 | 3 | Vdr | 0.062 | Old |
| 0.000145 | 0.321135 | 0.449 | 0.348 | 1 | 3 | Dnaja2 | 0.101 | Old |
| 0.000149 | 0.325844 | 0.258 | 0.174 | 1 | 3 | Utp18 | 0.084 | Old |
| 0.000151 | 0.430316 | 0.251 | 0.17 | 1 | 3 | Etnk1 | 0.081 | Old |
| 0.000152 | 0.382007 | 0.272 | 0.188 | 1 | 3 | Suz12 | 0.084 | Old |
| 0.000155 | 0.787579 | 0.142 | 0.083 | 1 | 3 | Fam220a | 0.059 | Old |
| 0.000158 | 0.272394 | 0.688 | 0.583 | 1 | 3 | Larp4b | 0.105 | Old |
| 0.00016 | 0.341456 | 0.225 | 0.146 | 1 | 3 | Ndufs1 | 0.079 | Old |

| | | | | | | | | |
|---|---|---|---|---|---|---|---|---|
| 0.000163 | 0.278227 | 0.56 | 0.466 | 1 | 3 | Tmem258 | 0.094 | Old |
| 0.000165 | 0.360904 | 0.435 | 0.342 | 1 | 3 | B4galt6 | 0.093 | Old |
| 0.000172 | 0.467156 | 0.275 | 0.192 | 1 | 3 | Cyc1 | 0.083 | Old |
| 0.000174 | 1.548501 | 0.056 | 0.02 | 1 | 3 | Gngt2 | 0.036 | Old |
| 0.000175 | 0.211853 | 0.857 | 0.77 | 1 | 3 | Tkt | 0.087 | Old |
| 0.000175 | 0.429288 | 0.326 | 0.236 | 1 | 3 | Mmp19 | 0.09 | Old |
| 0.000175 | 0.331662 | 0.7 | 0.629 | 1 | 3 | Bhlhe40 | 0.071 | Old |
| 0.000181 | 0.191318 | 0.866 | 0.798 | 1 | 3 | Ywhab | 0.068 | Old |
| 0.000181 | 0.554695 | 0.196 | 0.125 | 1 | 3 | Rhoc | 0.071 | Old |
| 0.000181 | 0.288714 | 0.568 | 0.462 | 1 | 3 | Lat2 | 0.106 | Old |
| 0.000187 | 0.265419 | 0.356 | 0.255 | 1 | 3 | Pnp | 0.101 | Old |
| 0.000189 | 0.180825 | 0.836 | 0.775 | 1 | 3 | Srsf5 | 0.061 | Old |
| 0.000194 | 0.338598 | 0.504 | 0.409 | 1 | 3 | Sdhd | 0.095 | Old |
| 0.000196 | 1.423004 | 0.063 | 0.025 | 1 | 3 | F7 | 0.038 | Old |
| 0.000197 | 0.29894 | 0.466 | 0.361 | 1 | 3 | Riok3 | 0.105 | Old |
| 0.0002 | 0.39082 | 0.605 | 0.53 | 1 | 3 | Anxa1 | 0.075 | Old |
| 0.000202 | 0.451755 | 0.509 | 0.417 | 1 | 3 | Naa50 | 0.092 | Old |
| 0.000204 | 0.137678 | 0.956 | 0.908 | 1 | 3 | BC005537 | 0.048 | Old |
| 0.000206 | 0.175477 | 0.892 | 0.843 | 1 | 3 | Btf3 | 0.049 | Old |
| 0.000218 | 0.325218 | 0.249 | 0.169 | 1 | 3 | Ogfr | 0.08 | Old |
| 0.000219 | 0.322436 | 0.274 | 0.192 | 1 | 3 | Tomm34 | 0.082 | Old |
| 0.000222 | 0.295243 | 0.372 | 0.271 | 1 | 3 | Ppp1r21 | 0.101 | Old |
| 0.000225 | 0.287117 | 0.737 | 0.66 | 1 | 3 | Srsf3 | 0.077 | Old |
| 0.000225 | 0.283557 | 0.489 | 0.395 | 1 | 3 | Bcap31 | 0.094 | Old |
| 0.000228 | 0.261289 | 0.681 | 0.59 | 1 | 3 | Taldo1 | 0.091 | Old |
| 0.00023 | 0.826821 | 0.137 | 0.081 | 1 | 3 | Abhd2 | 0.056 | Old |
| 0.000238 | 1.590694 | 0.047 | 0.015 | 1 | 3 | Oasl2 | 0.032 | Old |
| 0.000243 | 0.447537 | 0.258 | 0.179 | 1 | 3 | Creld2 | 0.079 | Old |
| 0.000244 | 0.882518 | 0.11 | 0.059 | 1 | 3 | Mtmr2 | 0.051 | Old |
| 0.00026 | 1.583973 | 0.059 | 0.023 | 1 | 3 | Cald1 | 0.036 | Old |
| 0.000268 | 0.306119 | 0.411 | 0.316 | 1 | 3 | Tmem164 | 0.095 | Old |
| 0.000282 | 0.311865 | 0.563 | 0.473 | 1 | 3 | Kcnn4 | 0.09 | Old |
| 0.00029 | 0.443553 | 0.329 | 0.246 | 1 | 3 | Cyp51 | 0.083 | Old |
| 0.000291 | 0.25511 | 0.603 | 0.495 | 1 | 3 | Rab7 | 0.108 | Old |
| 0.000292 | 0.137182 | 0.956 | 0.932 | 1 | 3 | Capzb | 0.024 | Old |
| 0.000295 | 0.237084 | 0.345 | 0.255 | 1 | 3 | Strap | 0.09 | Old |
| 0.000298 | 0.238608 | 0.673 | 0.568 | 1 | 3 | Khdrbs1 | 0.105 | Old |
| 0.000301 | 1.345732 | 0.114 | 0.064 | 1 | 3 | Ifi47 | 0.05 | Old |
| 0.000304 | 0.239096 | 0.658 | 0.557 | 1 | 3 | Vdac2 | 0.101 | Old |
| 0.000307 | 0.914636 | 0.128 | 0.074 | 1 | 3 | Crtap | 0.054 | Old |
| 0.000311 | 0.258522 | 0.352 | 0.263 | 1 | 3 | Dnajb11 | 0.089 | Old |
| 0.000313 | 0.532533 | 0.179 | 0.115 | 1 | 3 | 1700123O2 | 0.064 | Old |
| 0.000315 | 0.349047 | 0.315 | 0.229 | 1 | 3 | Znhit1 | 0.086 | Old |
| 0.000321 | 0.289489 | 0.33 | 0.24 | 1 | 3 | Smarcd2 | 0.09 | Old |
| 0.000326 | 0.500606 | 0.166 | 0.104 | 1 | 3 | Rpf2 | 0.062 | Old |

| | | | | | | | | |
|---|---|---|---|---|---|---|---|---|
| 0.000327 | 0.337772 | 0.211 | 0.14 | 1 | 3 | Fbxw11 | 0.071 | Old |
| 0.000333 | 0.522863 | 0.204 | 0.136 | 1 | 3 | Nploc4 | 0.068 | Old |
| 0.000336 | 0.409531 | 0.442 | 0.348 | 1 | 3 | Atp6v1a | 0.094 | Old |
| 0.000342 | 0.307956 | 0.202 | 0.13 | 1 | 3 | Tnfrsf11a | 0.072 | Old |
| 0.000347 | 0.288637 | 0.553 | 0.446 | 1 | 3 | Rab11a | 0.107 | Old |
| 0.000348 | 0.304375 | 0.516 | 0.431 | 1 | 3 | Got1 | 0.085 | Old |
| 0.000348 | 0.683407 | 0.166 | 0.105 | 1 | 3 | Polr3k | 0.061 | Old |
| 0.000348 | 0.327814 | 0.708 | 0.634 | 1 | 3 | Klf4 | 0.074 | Old |
| 0.000359 | 0.749291 | 0.194 | 0.129 | 1 | 3 | Ly6e | 0.065 | Old |
| 0.000363 | 0.766818 | 0.151 | 0.092 | 1 | 3 | Slc27a1 | 0.059 | Old |
| 0.000363 | 0.474375 | 0.406 | 0.321 | 1 | 3 | Ap3s1 | 0.085 | Old |
| 0.000363 | 0.205163 | 0.687 | 0.59 | 1 | 3 | Ran | 0.097 | Old |
| 0.00037 | 0.182831 | 0.853 | 0.758 | 1 | 3 | Olfm1 | 0.095 | Old |
| 0.000372 | 1.575462 | 0.037 | 0.01 | 1 | 3 | Nck2 | 0.027 | Old |
| 0.000373 | 0.31355 | 0.381 | 0.292 | 1 | 3 | Cyb5r4 | 0.089 | Old |
| 0.000377 | 0.23662 | 0.627 | 0.547 | 1 | 3 | Tpd52 | 0.08 | Old |
| 0.000388 | 0.591929 | 0.127 | 0.073 | 1 | 3 | Pgm1 | 0.054 | Old |
| 0.000389 | 0.248924 | 0.325 | 0.236 | 1 | 3 | Kpna1 | 0.089 | Old |
| 0.000394 | 1.93632 | 0.035 | 0.009 | 1 | 3 | Chac2 | 0.026 | Old |
| 0.000395 | 0.283687 | 0.584 | 0.497 | 1 | 3 | Cdv3 | 0.087 | Old |
| 0.000398 | 1.6247 | 0.019 | 0.001 | 1 | 3 | A530040E1 | 0.018 | Old |
| 0.000401 | 2.008314 | 0.027 | 0.005 | 1 | 3 | Cd55 | 0.022 | Old |
| 0.000415 | 0.259722 | 0.644 | 0.543 | 1 | 3 | Gm26917 | 0.101 | Old |
| 0.000415 | 0.556121 | 0.191 | 0.124 | 1 | 3 | Commd2 | 0.067 | Old |
| 0.000432 | 1.873736 | 0.016 | 0 | 1 | 3 | Dmpk | 0.016 | Old |
| 0.000464 | 0.102204 | 0.994 | 0.99 | 1 | 3 | Laptm5 | 0.004 | Old |
| 0.000477 | 0.323134 | 0.409 | 0.318 | 1 | 3 | Psmd14 | 0.091 | Old |
| 0.00048 | 0.158938 | 0.956 | 0.927 | 1 | 3 | Btg1 | 0.029 | Old |
| 0.000487 | 0.548214 | 0.534 | 0.457 | 1 | 3 | Ccl2 | 0.077 | Old |
| 0.000518 | 0.531853 | 0.179 | 0.117 | 1 | 3 | Ppp1r7 | 0.062 | Old |
| 0.000519 | 0.202553 | 0.841 | 0.774 | 1 | 3 | Fam49b | 0.067 | Old |
| 0.000524 | 0.122836 | 0.391 | 0.289 | 1 | 3 | Ube2q1 | 0.102 | Old |
| 0.00054 | 0.492429 | 0.226 | 0.157 | 1 | 3 | Ccdc22 | 0.069 | Old |
| 0.00054 | 0.246442 | 0.711 | 0.601 | 1 | 3 | Kif5b | 0.11 | Old |
| 0.000559 | 0.401891 | 0.43 | 0.346 | 1 | 3 | Mfhas1 | 0.084 | Old |
| 0.000567 | 0.244486 | 0.626 | 0.533 | 1 | 3 | Ywhag | 0.093 | Old |
| 0.000569 | 0.28422 | 0.427 | 0.34 | 1 | 3 | Socs3 | 0.087 | Old |
| 0.000584 | 0.18914 | 0.762 | 0.67 | 1 | 3 | Hnrnpab | 0.092 | Old |
| 0.000589 | 0.21946 | 0.734 | 0.643 | 1 | 3 | Cap1 | 0.091 | Old |
| 0.000607 | 0.338993 | 0.344 | 0.265 | 1 | 3 | Rbm47 | 0.079 | Old |
| 0.000615 | 0.413907 | 0.33 | 0.25 | 1 | 3 | Hist1h1c | 0.08 | Old |
| 0.000616 | 0.581222 | 0.159 | 0.1 | 1 | 3 | Pspc1 | 0.059 | Old |
| 0.000618 | 0.245267 | 0.53 | 0.418 | 1 | 3 | Rbx1 | 0.112 | Old |
| 0.000632 | 0.267571 | 0.452 | 0.356 | 1 | 3 | Hnrnpdl | 0.096 | Old |
| 0.00064 | 1.687284 | 0.026 | 0.005 | 1 | 3 | Gng11 | 0.021 | Old |

| | | | | | | | | |
|---|---|---|---|---|---|---|---|---|
| 0.000643 | 0.142183 | 0.324 | 0.24 | 1 | 3 | Mt2 | 0.084 | Old |
| 0.000666 | 0.987889 | 0.033 | 0.009 | 1 | 3 | Hps6 | 0.024 | Old |
| 0.000667 | 0.2385 | 0.607 | 0.519 | 1 | 3 | St8sia4 | 0.088 | Old |
| 0.000672 | 0.331878 | 0.643 | 0.559 | 1 | 3 | Rnf149 | 0.084 | Old |
| 0.000676 | 0.250887 | 0.528 | 0.438 | 1 | 3 | Tmed7 | 0.09 | Old |
| 0.000679 | 0.226013 | 0.786 | 0.71 | 1 | 3 | Nme1 | 0.076 | Old |
| 0.000691 | 0.17791 | 0.749 | 0.659 | 1 | 3 | Gls | 0.09 | Old |
| 0.000691 | 0.704634 | 0.078 | 0.038 | 1 | 3 | Ikbke | 0.04 | Old |
| 0.000704 | 0.238968 | 0.431 | 0.348 | 1 | 3 | Cers6 | 0.083 | Old |
| 0.000706 | 0.345535 | 0.499 | 0.418 | 1 | 3 | Fgr | 0.081 | Old |
| 0.000707 | 1.32329 | 0.057 | 0.024 | 1 | 3 | Cep104 | 0.033 | Old |
| 0.000717 | 0.221777 | 0.737 | 0.669 | 1 | 3 | Cops9 | 0.068 | Old |
| 0.000728 | 0.237901 | 0.498 | 0.403 | 1 | 3 | Cd200r1 | 0.095 | Old |
| 0.000731 | 0.544271 | 0.194 | 0.131 | 1 | 3 | Cox16 | 0.063 | Old |
| 0.00075 | 0.226366 | 0.522 | 0.423 | 1 | 3 | Gnaq | 0.099 | Old |
| 0.000751 | 0.502422 | 0.202 | 0.138 | 1 | 3 | Cxcl1 | 0.064 | Old |
| 0.000759 | 0.543592 | 0.164 | 0.106 | 1 | 3 | Nufip1 | 0.058 | Old |
| 0.000763 | 0.238243 | 0.658 | 0.54 | 1 | 3 | Arf4 | 0.118 | Old |
| 0.00077 | 0.680454 | 0.111 | 0.063 | 1 | 3 | Nt5dc3 | 0.048 | Old |
| 0.00077 | 0.432736 | 0.297 | 0.221 | 1 | 3 | Bst2 | 0.076 | Old |
| 0.00077 | 0.645937 | 0.134 | 0.082 | 1 | 3 | Nup214 | 0.052 | Old |
| 0.000782 | 0.246201 | 0.569 | 0.475 | 1 | 3 | Fam168a | 0.094 | Old |
| 0.000784 | 0.274127 | 0.323 | 0.239 | 1 | 3 | Agps | 0.084 | Old |
| 0.00079 | 0.311759 | 0.444 | 0.356 | 1 | 3 | Ddb1 | 0.088 | Old |
| 0.000805 | 0.379635 | 0.241 | 0.169 | 1 | 3 | Ireb2 | 0.072 | Old |
| 0.00082 | 0.249553 | 0.626 | 0.532 | 1 | 3 | Tmed5 | 0.094 | Old |
| 0.000826 | 0.41246 | 0.186 | 0.124 | 1 | 3 | Lrp12 | 0.062 | Old |
| 0.000838 | 0.485692 | 0.112 | 0.064 | 1 | 3 | Dkc1 | 0.048 | Old |
| 0.000842 | 0.355787 | 0.211 | 0.146 | 1 | 3 | Utp3 | 0.065 | Old |
| 0.000856 | 0.244471 | 0.33 | 0.25 | 1 | 3 | Arf6 | 0.08 | Old |
| 0.00086 | 0.690281 | 0.09 | 0.047 | 1 | 3 | Fam198b | 0.043 | Old |
| 0.000863 | 0.251176 | 0.581 | 0.47 | 1 | 3 | Arf3 | 0.111 | Old |
| 0.000864 | 0.346669 | 0.381 | 0.301 | 1 | 3 | Vcl | 0.08 | Old |
| 0.000878 | 0.405672 | 0.24 | 0.169 | 1 | 3 | Gtf2h1 | 0.071 | Old |
| 0.00088 | 0.35441 | 0.536 | 0.442 | 1 | 3 | Psmd8 | 0.094 | Old |
| 0.000884 | 0.185226 | 0.24 | 0.169 | 1 | 3 | Agpat5 | 0.071 | Old |
| 0.000898 | 0.417165 | 0.36 | 0.284 | 1 | 3 | Cnih1 | 0.076 | Old |
| 0.00091 | 0.293786 | 0.521 | 0.442 | 1 | 3 | Bak1 | 0.079 | Old |
| 0.000911 | 0.138554 | 0.924 | 0.893 | 1 | 3 | Slc25a3 | 0.031 | Old |
| 0.000923 | 0.185695 | 0.799 | 0.708 | 1 | 3 | Pdia3 | 0.091 | Old |
| 0.000924 | 0.214788 | 0.686 | 0.602 | 1 | 3 | Rab8b | 0.084 | Old |
| 0.000936 | 0.279023 | 0.72 | 0.659 | 1 | 3 | Id2 | 0.061 | Old |
| 0.000947 | 0.248925 | 0.601 | 0.511 | 1 | 3 | Atp6v1e1 | 0.09 | Old |
| 0.000971 | 0.746194 | 0.085 | 0.044 | 1 | 3 | Bcar3 | 0.041 | Old |
| 0.000973 | 0.118181 | 0.98 | 0.955 | 1 | 3 | Pfn1 | 0.025 | Old |

| | | | | | | | | |
|---|---|---|---|---|---|---|---|---|
| 0.000974 | 0.182072 | 0.35 | 0.264 | 1 | 3 | Arih2 | 0.086 | Old |
| 0.000981 | 0.329239 | 0.306 | 0.232 | 1 | 3 | Prelid3b | 0.074 | Old |
| 0.000984 | 0.894173 | 0.035 | 0.01 | 1 | 3 | Prmt6 | 0.025 | Old |
| 0.001012 | 1.191758 | 0.041 | 0.014 | 1 | 3 | Cmtr2 | 0.027 | Old |
| 0.001015 | 0.618041 | 0.088 | 0.047 | 1 | 3 | Aacs | 0.041 | Old |
| 0.001027 | 0.296955 | 0.468 | 0.374 | 1 | 3 | Vdac3 | 0.094 | Old |
| 0.00103 | 0.342077 | 0.314 | 0.236 | 1 | 3 | Cpsf2 | 0.078 | Old |
| 0.001046 | 0.385235 | 0.217 | 0.153 | 1 | 3 | Mbtps1 | 0.064 | Old |
| 0.001066 | 0.676 | 0.081 | 0.042 | 1 | 3 | Tpcn2 | 0.039 | Old |
| 0.001073 | 0.514058 | 0.088 | 0.047 | 1 | 3 | Leo1 | 0.041 | Old |
| 0.001083 | 1.301376 | 0.065 | 0.03 | 1 | 3 | C3 | 0.035 | Old |
| 0.001106 | 0.265887 | 0.524 | 0.432 | 1 | 3 | Tmem106a | 0.092 | Old |
| 0.001116 | 0.203399 | 0.716 | 0.633 | 1 | 3 | Arf1 | 0.083 | Old |
| 0.001122 | 0.186311 | 0.726 | 0.631 | 1 | 3 | Tpm4 | 0.095 | Old |
| 0.001123 | 0.290701 | 0.529 | 0.434 | 1 | 3 | Arcn1 | 0.095 | Old |
| 0.001125 | 0.402904 | 0.231 | 0.163 | 1 | 3 | Mecp2 | 0.068 | Old |
| 0.00113 | 0.202601 | 0.625 | 0.537 | 1 | 3 | Capza1 | 0.088 | Old |
| 0.001136 | 0.245378 | 0.424 | 0.327 | 1 | 3 | Eif2s3x | 0.097 | Old |
| 0.001139 | 0.338038 | 0.336 | 0.261 | 1 | 3 | Rsl1d1 | 0.075 | Old |
| 0.001144 | 1.305883 | 0.057 | 0.025 | 1 | 3 | Fcgr4 | 0.032 | Old |
| 0.001177 | 0.3393 | 0.124 | 0.073 | 1 | 3 | Mthfsl | 0.051 | Old |
| 0.00118 | 0.382046 | 0.197 | 0.135 | 1 | 3 | Ykt6 | 0.062 | Old |
| 0.001188 | 0.281057 | 0.368 | 0.29 | 1 | 3 | Becn1 | 0.078 | Old |
| 0.00119 | 0.88628 | 0.088 | 0.048 | 1 | 3 | Wdr7 | 0.04 | Old |
| 0.001203 | 0.714344 | 0.044 | 0.016 | 1 | 3 | Sgcb | 0.028 | Old |
| 0.001213 | 0.444441 | 0.268 | 0.198 | 1 | 3 | Rab43 | 0.07 | Old |
| 0.001216 | 0.199064 | 0.679 | 0.605 | 1 | 3 | Atp6ap1 | 0.074 | Old |
| 0.001223 | 0.274606 | 0.695 | 0.621 | 1 | 3 | Atp5k | 0.074 | Old |
| 0.001225 | 0.32399 | 0.429 | 0.342 | 1 | 3 | Prkcsh | 0.087 | Old |
| 0.001234 | 0.101468 | 0.958 | 0.942 | 1 | 3 | Cotl1 | 0.016 | Old |
| 0.001237 | 0.137995 | 0.957 | 0.934 | 1 | 3 | Arf5 | 0.023 | Old |
| 0.001266 | 0.19296 | 0.588 | 0.501 | 1 | 3 | Psmd13 | 0.087 | Old |
| 0.001282 | 0.874168 | 0.103 | 0.059 | 1 | 3 | Eepd1 | 0.044 | Old |
| 0.001286 | 0.685565 | 0.128 | 0.078 | 1 | 3 | Mapk9 | 0.05 | Old |
| 0.001322 | 1.079982 | 0.106 | 0.063 | 1 | 3 | Lgals3bp | 0.043 | Old |
| 0.001326 | 0.331714 | 0.269 | 0.198 | 1 | 3 | Hnrnpll | 0.071 | Old |
| 0.001348 | 0.668671 | 0.124 | 0.076 | 1 | 3 | Usp39 | 0.048 | Old |
| 0.001366 | 0.152651 | 0.93 | 0.91 | 1 | 3 | Nfkbia | 0.02 | Old |
| 0.00138 | 0.508194 | 0.178 | 0.12 | 1 | 3 | Sulf2 | 0.058 | Old |
| 0.001383 | 0.383653 | 0.233 | 0.167 | 1 | 3 | Snx1 | 0.066 | Old |
| 0.001398 | 0.438493 | 0.213 | 0.149 | 1 | 3 | Nrp1 | 0.064 | Old |
| 0.001423 | 0.501811 | 0.178 | 0.122 | 1 | 3 | Basp1 | 0.056 | Old |
| 0.001441 | 0.384453 | 0.253 | 0.181 | 1 | 3 | Zmiz2 | 0.072 | Old |
| 0.001453 | 0.679859 | 0.085 | 0.045 | 1 | 3 | Foxred1 | 0.04 | Old |
| 0.001462 | 0.173411 | 0.756 | 0.683 | 1 | 3 | Ostf1 | 0.073 | Old |

| | | | | | | | | |
|---|---|---|---|---|---|---|---|---|
| 0.001487 | 0.222486 | 0.699 | 0.622 | 1 | 3 | Ppp1r18 | 0.077 | Old |
| 0.001511 | 0.410251 | 0.278 | 0.206 | 1 | 3 | 4931406CC | 0.072 | Old |
| 0.001534 | 0.090711 | 0.99 | 0.989 | 1 | 3 | Rpl41 | 0.001 | Old |
| 0.001548 | 0.310005 | 0.398 | 0.323 | 1 | 3 | Usp4 | 0.075 | Old |
| 0.001549 | 0.228357 | 0.583 | 0.491 | 1 | 3 | Surf4 | 0.092 | Old |
| 0.001551 | 1.507333 | 0.024 | 0.005 | 1 | 3 | Efemp2 | 0.019 | Old |
| 0.001552 | 0.246116 | 0.616 | 0.545 | 1 | 3 | Degs1 | 0.071 | Old |
| 0.001568 | 0.605838 | 0.096 | 0.053 | 1 | 3 | Cherp | 0.043 | Old |
| 0.001576 | 0.43597 | 0.167 | 0.111 | 1 | 3 | Pitpnm1 | 0.056 | Old |
| 0.001585 | 0.30201 | 0.403 | 0.319 | 1 | 3 | Uso1 | 0.084 | Old |
| 0.001587 | 0.638404 | 0.098 | 0.056 | 1 | 3 | Dram1 | 0.042 | Old |
| 0.001598 | 0.33336 | 0.459 | 0.378 | 1 | 3 | Sec61a1 | 0.081 | Old |
| 0.0016 | 0.487239 | 0.159 | 0.105 | 1 | 3 | Gas2l1 | 0.054 | Old |
| 0.0016 | 0.280962 | 0.548 | 0.456 | 1 | 3 | Plekhb2 | 0.092 | Old |
| 0.00161 | 0.332606 | 0.318 | 0.247 | 1 | 3 | Mgat2 | 0.071 | Old |
| 0.001613 | 0.368356 | 0.41 | 0.33 | 1 | 3 | Erp44 | 0.08 | Old |
| 0.001619 | 0.179608 | 0.57 | 0.476 | 1 | 3 | Slc39a1 | 0.094 | Old |
| 0.001621 | 0.215468 | 0.611 | 0.519 | 1 | 3 | Rab7b | 0.092 | Old |
| 0.001642 | 1.375144 | 0.024 | 0.005 | 1 | 3 | Spef1 | 0.019 | Old |
| 0.001645 | 0.257598 | 0.389 | 0.308 | 1 | 3 | Tcirg1 | 0.081 | Old |
| 0.001682 | 0.192569 | 0.839 | 0.784 | 1 | 3 | Cox5a | 0.055 | Old |
| 0.001725 | 0.238262 | 0.588 | 0.497 | 1 | 3 | Ssr1 | 0.091 | Old |
| 0.001751 | 0.207056 | 0.599 | 0.514 | 1 | 3 | Cct2 | 0.085 | Old |
| 0.001765 | 0.17929 | 0.859 | 0.813 | 1 | 3 | Ptbp3 | 0.046 | Old |
| 0.001779 | 0.21754 | 0.621 | 0.548 | 1 | 3 | Selenos | 0.073 | Old |
| 0.001814 | 0.248106 | 0.389 | 0.308 | 1 | 3 | Derl1 | 0.081 | Old |
| 0.001822 | 0.420792 | 0.161 | 0.106 | 1 | 3 | Traf3 | 0.055 | Old |
| 0.001853 | 1.007408 | 0.016 | 0.001 | 1 | 3 | Npl | 0.015 | Old |
| 0.00187 | 0.724921 | 0.127 | 0.08 | 1 | 3 | Fubp3 | 0.047 | Old |
| 0.001883 | 0.28095 | 0.394 | 0.309 | 1 | 3 | 1110004F1 | 0.085 | Old |
| 0.001915 | 0.171167 | 0.863 | 0.795 | 1 | 3 | Eif4a1 | 0.068 | Old |
| 0.001927 | 0.120737 | 0.944 | 0.922 | 1 | 3 | Efhd2 | 0.022 | Old |
| 0.001937 | 0.145962 | 0.92 | 0.896 | 1 | 3 | Flna | 0.024 | Old |
| 0.001942 | 0.298 | 0.832 | 0.746 | 1 | 3 | Fos | 0.086 | Old |
| 0.001943 | 0.167013 | 0.392 | 0.307 | 1 | 3 | Gorasp2 | 0.085 | Old |
| 0.001946 | 0.060918 | 0.999 | 0.994 | 1 | 3 | Tmsb4x | 0.005 | Old |
| 0.001951 | 0.292526 | 0.351 | 0.266 | 1 | 3 | Atp6v0a1 | 0.085 | Old |
| 0.001952 | 0.621091 | 0.102 | 0.059 | 1 | 3 | P2ry1 | 0.043 | Old |
| 0.001972 | 0.355628 | 0.228 | 0.165 | 1 | 3 | Ccndbp1 | 0.063 | Old |
| 0.002031 | 0.941283 | 0.054 | 0.024 | 1 | 3 | Slc36a3os | 0.03 | Old |
| 0.002034 | 0.915776 | 0.057 | 0.027 | 1 | 3 | Akr1b10 | 0.03 | Old |
| 0.002063 | 0.92975 | 0.081 | 0.044 | 1 | 3 | Prkci | 0.037 | Old |
| 0.002071 | 0.269746 | 0.565 | 0.491 | 1 | 3 | Hspa9 | 0.074 | Old |
| 0.002099 | 0.277891 | 0.263 | 0.193 | 1 | 3 | Gpalpp1 | 0.07 | Old |
| 0.002113 | 0.459374 | 0.143 | 0.092 | 1 | 3 | Insig2 | 0.051 | Old |

| | | | | | | | | |
|---|---|---|---|---|---|---|---|---|
| 0.002132 | 0.302733 | 0.35 | 0.275 | 1 | 3 | Ascc3 | 0.075 | Old |
| 0.002142 | 0.31113 | 0.19 | 0.131 | 1 | 3 | Plekhm2 | 0.059 | Old |
| 0.002205 | 0.443803 | 0.207 | 0.149 | 1 | 3 | Fabp5 | 0.058 | Old |
| 0.002209 | 0.256712 | 0.761 | 0.674 | 1 | 3 | Arpc4 | 0.087 | Old |
| 0.002232 | 0.140415 | 0.443 | 0.359 | 1 | 3 | Lgals1 | 0.084 | Old |
| 0.002241 | 0.50539 | 0.122 | 0.076 | 1 | 3 | Rrp15 | 0.046 | Old |
| 0.002251 | 0.288548 | 0.321 | 0.249 | 1 | 3 | Ncstn | 0.072 | Old |
| 0.002283 | 1.616071 | 0.018 | 0.003 | 1 | 3 | Ubxn11 | 0.015 | Old |
| 0.00233 | 0.288281 | 0.454 | 0.379 | 1 | 3 | Sbno1 | 0.075 | Old |
| 0.002346 | 0.579555 | 0.076 | 0.04 | 1 | 3 | Ddx28 | 0.036 | Old |
| 0.00235 | 0.330022 | 0.262 | 0.196 | 1 | 3 | Nfu1 | 0.066 | Old |
| 0.002371 | 0.192399 | 0.728 | 0.648 | 1 | 3 | Cdk2ap2 | 0.08 | Old |
| 0.002432 | 0.307593 | 0.333 | 0.261 | 1 | 3 | Ptpn2 | 0.072 | Old |
| 0.002436 | 0.527842 | 0.158 | 0.106 | 1 | 3 | Psat1 | 0.052 | Old |
| 0.002437 | 0.187503 | 0.565 | 0.489 | 1 | 3 | Metap2 | 0.076 | Old |
| 0.00246 | 0.527423 | 0.136 | 0.087 | 1 | 3 | Stk16 | 0.049 | Old |
| 0.002475 | 0.757285 | 0.073 | 0.038 | 1 | 3 | Tex2 | 0.035 | Old |
| 0.002491 | 0.437744 | 0.311 | 0.241 | 1 | 3 | Wdr89 | 0.07 | Old |
| 0.002497 | 0.342367 | 0.17 | 0.114 | 1 | 3 | Dennd1a | 0.056 | Old |
| 0.0025 | 0.499784 | 0.18 | 0.126 | 1 | 3 | Zfp280d | 0.054 | Old |
| 0.002509 | 0.47932 | 0.17 | 0.116 | 1 | 3 | Pofut2 | 0.054 | Old |
| 0.002525 | 0.867158 | 0.055 | 0.025 | 1 | 3 | Fzd1 | 0.03 | Old |
| 0.002525 | 0.550895 | 0.027 | 0.008 | 1 | 3 | Adnp | 0.019 | Old |
| 0.002549 | 0.288749 | 0.151 | 0.1 | 1 | 3 | A430005L1 | 0.051 | Old |
| 0.002549 | 0.570506 | 0.099 | 0.058 | 1 | 3 | Snx25 | 0.041 | Old |
| 0.002557 | 0.407792 | 0.164 | 0.111 | 1 | 3 | Larp7 | 0.053 | Old |
| 0.002609 | 0.272106 | 0.597 | 0.508 | 1 | 3 | Morf4l2 | 0.089 | Old |
| 0.002629 | 0.206931 | 0.375 | 0.295 | 1 | 3 | Ilk | 0.08 | Old |
| 0.002629 | 0.300481 | 0.321 | 0.247 | 1 | 3 | Rad21 | 0.074 | Old |
| 0.002635 | 0.182727 | 0.465 | 0.374 | 1 | 3 | Supt16 | 0.091 | Old |
| 0.002643 | 0.366424 | 0.407 | 0.333 | 1 | 3 | Aldh9a1 | 0.074 | Old |
| 0.002645 | 0.395759 | 0.202 | 0.145 | 1 | 3 | Hivep1 | 0.057 | Old |
| 0.002649 | 0.355886 | 0.166 | 0.112 | 1 | 3 | Rbm14 | 0.054 | Old |
| 0.002656 | 0.430324 | 0.209 | 0.149 | 1 | 3 | Hdac2 | 0.06 | Old |
| 0.002677 | 0.465381 | 0.084 | 0.047 | 1 | 3 | Pus7 | 0.037 | Old |
| 0.002683 | 0.707464 | 0.125 | 0.08 | 1 | 3 | Zfp407 | 0.045 | Old |
| 0.002698 | 0.439131 | 0.222 | 0.164 | 1 | 3 | Cd63 | 0.058 | Old |
| 0.00271 | 0.378987 | 0.422 | 0.34 | 1 | 3 | Sod2 | 0.082 | Old |
| 0.002742 | 0.165922 | 0.569 | 0.467 | 1 | 3 | Ogdh | 0.102 | Old |
| 0.002775 | 1.323866 | 0.039 | 0.015 | 1 | 3 | Slc25a33 | 0.024 | Old |
| 0.002776 | 0.197089 | 0.384 | 0.302 | 1 | 3 | Rab21 | 0.082 | Old |
| 0.002796 | 0.307416 | 0.495 | 0.41 | 1 | 3 | Ddost | 0.085 | Old |
| 0.002837 | 0.344091 | 0.45 | 0.371 | 1 | 3 | Cndp2 | 0.079 | Old |
| 0.00288 | 0.25962 | 0.532 | 0.447 | 1 | 3 | Smpdl3a | 0.085 | Old |
| 0.002905 | 0.434952 | 0.288 | 0.221 | 1 | 3 | Dpep2 | 0.067 | Old |

| | | | | | | | | |
|---|---|---|---|---|---|---|---|---|
| 0.002943 | 0.188484 | 0.386 | 0.309 | 1 | 3 | Noc2l | 0.077 | Old |
| 0.002948 | 0.377199 | 0.154 | 0.102 | 1 | 3 | Lsm2 | 0.052 | Old |
| 0.002962 | 0.75406 | 0.1 | 0.059 | 1 | 3 | Bysl | 0.041 | Old |
| 0.002964 | 0.137072 | 0.882 | 0.83 | 1 | 3 | Sirpa | 0.052 | Old |
| 0.003003 | 0.246403 | 0.516 | 0.431 | 1 | 3 | Spcs1 | 0.085 | Old |
| 0.003004 | 0.527775 | 0.159 | 0.107 | 1 | 3 | Parp9 | 0.052 | Old |
| 0.003009 | 0.168181 | 0.802 | 0.739 | 1 | 3 | Atp1a1 | 0.063 | Old |
| 0.003029 | 0.239646 | 0.707 | 0.626 | 1 | 3 | Ptpre | 0.081 | Old |
| 0.003033 | 0.278871 | 0.271 | 0.203 | 1 | 3 | Ppdpf | 0.068 | Old |
| 0.003082 | 0.402166 | 0.16 | 0.106 | 1 | 3 | Prkaca | 0.054 | Old |
| 0.00309 | 0.192268 | 0.528 | 0.444 | 1 | 3 | Pigt | 0.084 | Old |
| 0.00312 | 0.948763 | 0.076 | 0.042 | 1 | 3 | Bcl2l12 | 0.034 | Old |
| 0.003125 | 0.378079 | 0.361 | 0.292 | 1 | 3 | Lta4h | 0.069 | Old |
| 0.003159 | 0.303274 | 0.419 | 0.34 | 1 | 3 | Lsm6 | 0.079 | Old |
| 0.003166 | 0.127394 | 0.4 | 0.312 | 1 | 3 | Ggh | 0.088 | Old |
| 0.003203 | 0.255236 | 0.432 | 0.352 | 1 | 3 | Esd | 0.08 | Old |
| 0.003251 | 0.123766 | 0.926 | 0.899 | 1 | 3 | Coro1a | 0.027 | Old |
| 0.00326 | 0.816348 | 0.047 | 0.02 | 1 | 3 | Inpp5b | 0.027 | Old |
| 0.003283 | 0.311888 | 0.329 | 0.258 | 1 | 3 | Tspan31 | 0.071 | Old |
| 0.003299 | 0.241536 | 0.313 | 0.242 | 1 | 3 | Eif6 | 0.071 | Old |
| 0.003299 | 0.23328 | 0.631 | 0.544 | 1 | 3 | Adam10 | 0.087 | Old |
| 0.003305 | 0.551293 | 0.153 | 0.104 | 1 | 3 | Ap1g2 | 0.049 | Old |
| 0.003312 | 0.888154 | 0.13 | 0.086 | 1 | 3 | Dgat1 | 0.044 | Old |
| 0.003331 | 0.530856 | 0.191 | 0.138 | 1 | 3 | Klra2 | 0.053 | Old |
| 0.003334 | 0.906657 | 0.037 | 0.014 | 1 | 3 | B4galt4 | 0.023 | Old |
| 0.003362 | 0.152544 | 0.91 | 0.859 | 1 | 3 | Hnrnpk | 0.051 | Old |
| 0.003398 | 0.167124 | 0.168 | 0.112 | 1 | 3 | Cacna1d | 0.056 | Old |
| 0.00341 | 0.207444 | 0.59 | 0.511 | 1 | 3 | Sec11c | 0.079 | Old |
| 0.00342 | 0.086399 | 0.962 | 0.958 | 1 | 3 | Sem1 | 0.004 | Old |
| 0.003428 | 0.664041 | 0.05 | 0.023 | 1 | 3 | Ctps | 0.027 | Old |
| 0.003438 | 0.23543 | 0.357 | 0.282 | 1 | 3 | Pum1 | 0.075 | Old |
| 0.003444 | 0.166594 | 0.683 | 0.583 | 1 | 3 | Atf3 | 0.1 | Old |
| 0.003444 | 1.343216 | 0.011 | 0 | 1 | 3 | Fuk | 0.011 | Old |
| 0.003453 | 0.201649 | 0.726 | 0.626 | 1 | 3 | Stat3 | 0.1 | Old |
| 0.003456 | 0.332077 | 0.284 | 0.216 | 1 | 3 | Nbeal1 | 0.068 | Old |
| 0.003478 | 0.156908 | 0.62 | 0.535 | 1 | 3 | G3bp1 | 0.085 | Old |
| 0.003493 | 0.30441 | 0.244 | 0.183 | 1 | 3 | Gtf3c6 | 0.061 | Old |
| 0.003505 | 0.353611 | 0.438 | 0.355 | 1 | 3 | Cs | 0.083 | Old |
| 0.003506 | 0.176381 | 0.614 | 0.535 | 1 | 3 | Ssb | 0.079 | Old |
| 0.003511 | 0.208976 | 0.358 | 0.279 | 1 | 3 | Sec63 | 0.079 | Old |
| 0.003528 | 0.193408 | 0.3 | 0.23 | 1 | 3 | Gpat4 | 0.07 | Old |
| 0.003538 | 0.264391 | 0.405 | 0.327 | 1 | 3 | Alyref | 0.078 | Old |
| 0.003564 | 0.749106 | 0.069 | 0.037 | 1 | 3 | Prmt7 | 0.032 | Old |
| 0.003606 | 0.808067 | 0.066 | 0.034 | 1 | 3 | Ms4a6d | 0.032 | Old |
| 0.003643 | 0.751912 | 0.114 | 0.072 | 1 | 3 | Fech | 0.042 | Old |

| | | | | | | | | |
|---|---|---|---|---|---|---|---|---|
| 0.00368 | 0.34849 | 0.387 | 0.317 | 1 | 3 | Psmc5 | 0.07 | Old |
| 0.003708 | 0.033191 | 0.548 | 0.444 | 1 | 3 | Tuba1c | 0.104 | Old |
| 0.003728 | 0.398148 | 0.102 | 0.061 | 1 | 3 | Milr1 | 0.041 | Old |
| 0.003756 | 0.202853 | 0.552 | 0.467 | 1 | 3 | Anxa7 | 0.085 | Old |
| 0.003799 | 0.151836 | 0.362 | 0.285 | 1 | 3 | Lpar1 | 0.077 | Old |
| 0.003838 | 0.375787 | 0.276 | 0.217 | 1 | 3 | Sarnp | 0.059 | Old |
| 0.003852 | 0.151221 | 0.613 | 0.529 | 1 | 3 | Rab1a | 0.084 | Old |
| 0.003912 | 0.171347 | 0.668 | 0.567 | 1 | 3 | Rpn2 | 0.101 | Old |
| 0.003914 | 0.537367 | 0.17 | 0.119 | 1 | 3 | Abcc5 | 0.051 | Old |
| 0.00393 | 0.324923 | 0.37 | 0.294 | 1 | 3 | Rnf7 | 0.076 | Old |
| 0.003934 | 0.196244 | 0.814 | 0.758 | 1 | 3 | Rbm3 | 0.056 | Old |
| 0.003956 | 0.305233 | 0.507 | 0.426 | 1 | 3 | Copz1 | 0.081 | Old |
| 0.003971 | 0.337943 | 0.311 | 0.239 | 1 | 3 | Ugcg | 0.072 | Old |
| 0.003987 | 0.15329 | 0.341 | 0.264 | 1 | 3 | Glrx | 0.077 | Old |
| 0.004002 | 0.297757 | 0.231 | 0.17 | 1 | 3 | Sptlc2 | 0.061 | Old |
| 0.004005 | 0.370975 | 0.202 | 0.148 | 1 | 3 | Naa20 | 0.054 | Old |
| 0.004015 | 1.269449 | 0.019 | 0.004 | 1 | 3 | Phpt1 | 0.015 | Old |
| 0.00404 | 1.221919 | 0.056 | 0.028 | 1 | 3 | Fen1 | 0.028 | Old |
| 0.004045 | 0.240673 | 0.315 | 0.242 | 1 | 3 | Ppp3cb | 0.073 | Old |
| 0.004058 | 1.04847 | 0.038 | 0.015 | 1 | 3 | Slfn1 | 0.023 | Old |
| 0.004066 | 1.588285 | 0.022 | 0.005 | 1 | 3 | Tubd1 | 0.017 | Old |
| 0.004086 | 0.350283 | 0.585 | 0.509 | 1 | 3 | Lbh | 0.076 | Old |
| 0.004087 | 0.297915 | 0.166 | 0.116 | 1 | 3 | Ero1lb | 0.05 | Old |
| 0.004098 | 0.129965 | 0.907 | 0.865 | 1 | 3 | Cybb | 0.042 | Old |
| 0.004129 | 1.51962 | 0.032 | 0.011 | 1 | 3 | Gbp4 | 0.021 | Old |
| 0.004151 | 0.180089 | 0.841 | 0.818 | 1 | 3 | Itgam | 0.023 | Old |
| 0.00417 | 0.121359 | 0.133 | 0.086 | 1 | 3 | Wdr77 | 0.047 | Old |
| 0.004177 | 0.401568 | 0.174 | 0.121 | 1 | 3 | Amdhd2 | 0.053 | Old |
| 0.004193 | 0.454818 | 0.114 | 0.071 | 1 | 3 | Pi16 | 0.043 | Old |
| 0.00422 | 0.148212 | 0.851 | 0.774 | 1 | 3 | Ywhae | 0.077 | Old |
| 0.004226 | 1.265062 | 0.042 | 0.018 | 1 | 3 | Smpd3 | 0.024 | Old |
| 0.00425 | 0.055093 | 0.994 | 0.992 | 1 | 3 | mt-Nd1 | 0.002 | Old |
| 0.00426 | 0.059752 | 0.239 | 0.177 | 1 | 3 | Ccl7 | 0.062 | Old |
| 0.004263 | 0.152069 | 0.453 | 0.367 | 1 | 3 | Gm16286 | 0.086 | Old |
| 0.004279 | 0.19835 | 0.385 | 0.308 | 1 | 3 | Pim3 | 0.077 | Old |
| 0.004289 | 0.963198 | 0.06 | 0.03 | 1 | 3 | Orc6 | 0.03 | Old |
| 0.004302 | 0.17165 | 0.611 | 0.534 | 1 | 3 | Hspd1 | 0.077 | Old |
| 0.004349 | 0.704762 | 0.129 | 0.085 | 1 | 3 | Thumpd3 | 0.044 | Old |
| 0.004523 | 0.218879 | 0.519 | 0.434 | 1 | 3 | Lman2 | 0.085 | Old |
| 0.004542 | 0.44969 | 0.25 | 0.189 | 1 | 3 | Rab20 | 0.061 | Old |
| 0.004563 | 0.324715 | 0.375 | 0.302 | 1 | 3 | Abce1 | 0.073 | Old |
| 0.004607 | 0.433851 | 0.13 | 0.086 | 1 | 3 | Timm22 | 0.044 | Old |
| 0.004653 | 0.535144 | 0.076 | 0.043 | 1 | 3 | Tnfaip1 | 0.033 | Old |
| 0.004678 | 0.249807 | 0.411 | 0.335 | 1 | 3 | Ppp4c | 0.076 | Old |
| 0.00468 | 0.085484 | 0.933 | 0.913 | 1 | 3 | Neat1 | 0.02 | Old |

| | | | | | | | | |
|---|---|---|---|---|---|---|---|---|
| 0.004684 | 0.113385 | 0.354 | 0.275 | 1 | 3 | Tomm70a | 0.079 | Old |
| 0.00473 | 0.287006 | 0.251 | 0.186 | 1 | 3 | Clcn7 | 0.065 | Old |
| 0.004731 | 0.489479 | 0.147 | 0.1 | 1 | 3 | Secisbp2 | 0.047 | Old |
| 0.004762 | 0.626271 | 0.183 | 0.131 | 1 | 3 | Idh3a | 0.052 | Old |
| 0.004787 | 1.036705 | 0.066 | 0.035 | 1 | 3 | Kat2a | 0.031 | Old |
| 0.004802 | 0.652898 | 0.098 | 0.061 | 1 | 3 | Sort1 | 0.037 | Old |
| 0.004827 | 0.183014 | 0.779 | 0.705 | 1 | 3 | Wdr26 | 0.074 | Old |
| 0.004832 | 0.337158 | 0.235 | 0.177 | 1 | 3 | Fyttd1 | 0.058 | Old |
| 0.004841 | 0.279662 | 0.43 | 0.357 | 1 | 3 | Smg1 | 0.073 | Old |
| 0.004925 | 0.834371 | 0.049 | 0.023 | 1 | 3 | Trim37 | 0.026 | Old |
| 0.004926 | 0.302808 | 0.533 | 0.466 | 1 | 3 | Actn4 | 0.067 | Old |
| 0.004967 | 0.212097 | 0.179 | 0.124 | 1 | 3 | Brwd3 | 0.055 | Old |
| 0.005028 | 0.210889 | 0.583 | 0.514 | 1 | 3 | Gusb | 0.069 | Old |
| 0.005032 | 0.243272 | 0.41 | 0.332 | 1 | 3 | Psma1 | 0.078 | Old |
| 0.005044 | 0.186354 | 0.247 | 0.183 | 1 | 3 | Cpeb4 | 0.064 | Old |
| 0.005086 | 0.436055 | 0.173 | 0.122 | 1 | 3 | Nifk | 0.051 | Old |
| 0.005093 | 0.375318 | 0.186 | 0.134 | 1 | 3 | Slc23a2 | 0.052 | Old |
| 0.005175 | 0.129472 | 0.481 | 0.405 | 1 | 3 | Psme2 | 0.076 | Old |
| 0.005218 | 0.03616 | 0.725 | 0.643 | 1 | 3 | Prdx1 | 0.082 | Old |
| 0.00522 | 0.299432 | 0.179 | 0.128 | 1 | 3 | Tmem120a | 0.051 | Old |
| 0.005236 | 0.17376 | 0.748 | 0.698 | 1 | 3 | Ezr | 0.05 | Old |
| 0.005245 | 0.241652 | 0.232 | 0.172 | 1 | 3 | Pgap2 | 0.06 | Old |
| 0.005245 | 0.604836 | 0.097 | 0.059 | 1 | 3 | Ly9 | 0.038 | Old |
| 0.005248 | 0.431288 | 0.253 | 0.192 | 1 | 3 | Man1a | 0.061 | Old |
| 0.00525 | 0.305151 | 0.515 | 0.443 | 1 | 3 | St3gal5 | 0.072 | Old |
| 0.005269 | 0.234676 | 0.857 | 0.84 | 1 | 3 | Mrc1 | 0.017 | Old |
| 0.005276 | 0.246209 | 0.652 | 0.586 | 1 | 3 | Rgs2 | 0.066 | Old |
| 0.005312 | 0.350426 | 0.284 | 0.222 | 1 | 3 | Flvcr1 | 0.062 | Old |
| 0.005322 | 0.302548 | 0.28 | 0.217 | 1 | 3 | Atp1a3 | 0.063 | Old |
| 0.005337 | 0.284295 | 0.339 | 0.271 | 1 | 3 | Hexb | 0.068 | Old |
| 0.005338 | 0.277178 | 0.164 | 0.114 | 1 | 3 | Ufl1 | 0.05 | Old |
| 0.005342 | 0.383567 | 0.265 | 0.203 | 1 | 3 | Rnft1 | 0.062 | Old |
| 0.005406 | 2.024859 | 0.025 | 0.008 | 1 | 3 | Oas3 | 0.017 | Old |
| 0.00541 | 0.355631 | 0.286 | 0.222 | 1 | 3 | Taf1 | 0.064 | Old |
| 0.005435 | 0.329087 | 0.308 | 0.242 | 1 | 3 | Acadl | 0.066 | Old |
| 0.005451 | 0.068491 | 0.998 | 0.995 | 1 | 3 | Crip1 | 0.003 | Old |
| 0.005452 | 0.541358 | 0.079 | 0.045 | 1 | 3 | Trim34a | 0.034 | Old |
| 0.005492 | 0.217387 | 0.32 | 0.25 | 1 | 3 | Ifngr2 | 0.07 | Old |
| 0.005526 | 0.384618 | 0.354 | 0.285 | 1 | 3 | Stt3a | 0.069 | Old |
| 0.005566 | 0.559446 | 0.087 | 0.052 | 1 | 3 | Slc27a4 | 0.035 | Old |
| 0.005571 | 0.467757 | 0.198 | 0.145 | 1 | 3 | Fam129b | 0.053 | Old |
| 0.005574 | 0.405557 | 0.096 | 0.058 | 1 | 3 | Crbn | 0.038 | Old |
| 0.005585 | 0.307973 | 0.228 | 0.17 | 1 | 3 | Tsnax | 0.058 | Old |
| 0.005639 | 0.176063 | 0.474 | 0.403 | 1 | 3 | Stk10 | 0.071 | Old |
| 0.005726 | 0.240717 | 0.472 | 0.388 | 1 | 3 | Vps4b | 0.084 | Old |

| | | | | | | | | |
|---|---|---|---|---|---|---|---|---|
| 0.005768 | 0.516475 | 0.147 | 0.101 | 1 | 3 | Rpa3 | 0.046 | Old |
| 0.005768 | 0.51815 | 0.165 | 0.116 | 1 | 3 | Hbs1l | 0.049 | Old |
| 0.005779 | 0.260746 | 0.559 | 0.476 | 1 | 3 | Edem1 | 0.083 | Old |
| 0.005836 | 1.650026 | 0.01 | 0 | 1 | 3 | Akap12 | 0.01 | Old |
| 0.005836 | 1.560206 | 0.01 | 0 | 1 | 3 | Iglc2 | 0.01 | Old |
| 0.005836 | 1.354469 | 0.01 | 0 | 1 | 3 | Tpx2 | 0.01 | Old |
| 0.005856 | 0.22158 | 0.282 | 0.215 | 1 | 3 | Nsfl1c | 0.067 | Old |
| 0.005866 | 0.722969 | 0.08 | 0.047 | 1 | 3 | Kif1bp | 0.033 | Old |
| 0.005893 | 0.259723 | 0.41 | 0.335 | 1 | 3 | Tnfrsf21 | 0.075 | Old |
| 0.005897 | 0.212693 | 0.509 | 0.431 | 1 | 3 | Azin1 | 0.078 | Old |
| 0.005986 | 0.216308 | 0.662 | 0.592 | 1 | 3 | M6pr | 0.07 | Old |
| 0.005989 | 0.120925 | 0.692 | 0.597 | 1 | 3 | Atp5b | 0.095 | Old |
| 0.005993 | 0.563839 | 0.057 | 0.029 | 1 | 3 | 3110040N1 | 0.028 | Old |
| 0.006002 | 0.288217 | 0.399 | 0.33 | 1 | 3 | Dhx9 | 0.069 | Old |
| 0.006014 | 0.508168 | 0.165 | 0.116 | 1 | 3 | Slc28a2 | 0.049 | Old |
| 0.00604 | 0.095959 | 0.964 | 0.957 | 1 | 3 | Calm1 | 0.007 | Old |
| 0.00605 | 0.183383 | 0.453 | 0.366 | 1 | 3 | Usp25 | 0.087 | Old |
| 0.00606 | 0.307488 | 0.394 | 0.324 | 1 | 3 | Mlf2 | 0.07 | Old |
| 0.006078 | 0.195766 | 0.571 | 0.484 | 1 | 3 | Eif4h | 0.087 | Old |
| 0.006106 | 0.41765 | 0.363 | 0.302 | 1 | 3 | Rhob | 0.061 | Old |
| 0.006109 | 1.12353 | 0.016 | 0.003 | 1 | 3 | Ehd2 | 0.013 | Old |
| 0.006138 | 0.399115 | 0.105 | 0.066 | 1 | 3 | Phb | 0.039 | Old |
| 0.006149 | 0.348894 | 0.222 | 0.167 | 1 | 3 | Camk2g | 0.055 | Old |
| 0.006198 | 0.331275 | 0.155 | 0.107 | 1 | 3 | Gpd2 | 0.048 | Old |
| 0.006258 | 0.082713 | 0.965 | 0.97 | 1 | 3 | Iqgap1 | -0.005 | Old |
| 0.006345 | 1.449609 | 0.018 | 0.004 | 1 | 3 | Gm42595 | 0.014 | Old |
| 0.006347 | 0.285946 | 0.208 | 0.153 | 1 | 3 | Ric8a | 0.055 | Old |
| 0.006388 | 0.117421 | 0.895 | 0.864 | 1 | 3 | Prkcd | 0.031 | Old |
| 0.006389 | 0.7116 | 0.073 | 0.042 | 1 | 3 | Ctdp1 | 0.031 | Old |
| 0.006402 | 0.330193 | 0.299 | 0.234 | 1 | 3 | Rbm17 | 0.065 | Old |
| 0.006462 | 0.283359 | 0.282 | 0.218 | 1 | 3 | Fam91a1 | 0.064 | Old |
| 0.006548 | 0.632117 | 0.067 | 0.037 | 1 | 3 | Slc12a7 | 0.03 | Old |
| 0.006587 | 0.703555 | 0.076 | 0.044 | 1 | 3 | Ubap2 | 0.032 | Old |
| 0.006633 | 0.222182 | 0.753 | 0.684 | 1 | 3 | Ywhah | 0.069 | Old |
| 0.006637 | 0.248293 | 0.314 | 0.245 | 1 | 3 | Lsm5 | 0.069 | Old |
| 0.006681 | 0.635498 | 0.06 | 0.032 | 1 | 3 | Dhrs9 | 0.028 | Old |
| 0.006709 | 0.191003 | 0.554 | 0.472 | 1 | 3 | Arhgef6 | 0.082 | Old |
| 0.006777 | 0.097949 | 0.367 | 0.289 | 1 | 3 | Psmb7 | 0.078 | Old |
| 0.006849 | 0.416882 | 0.085 | 0.051 | 1 | 3 | Rai14 | 0.034 | Old |
| 0.006866 | 0.230528 | 0.459 | 0.369 | 1 | 3 | Smchd1 | 0.09 | Old |
| 0.006871 | 0.26001 | 0.412 | 0.347 | 1 | 3 | Epb41l2 | 0.065 | Old |
| 0.006895 | 0.129866 | 0.803 | 0.75 | 1 | 3 | Nrros | 0.053 | Old |
| 0.006896 | 0.402614 | 0.215 | 0.162 | 1 | 3 | Xpot | 0.053 | Old |
| 0.006912 | 0.155177 | 0.614 | 0.524 | 1 | 3 | Malt1 | 0.09 | Old |
| 0.00695 | 0.143544 | 0.26 | 0.197 | 1 | 3 | Dpp8 | 0.063 | Old |

| | | | | | | | | |
|---|---|---|---|---|---|---|---|---|
| 0.00697 | 0.388958 | 0.179 | 0.129 | 1 | 3 | Pmpca | 0.05 | Old |
| 0.006985 | 0.56155 | 0.145 | 0.101 | 1 | 3 | Aars | 0.044 | Old |
| 0.00701 | 0.398909 | 0.257 | 0.199 | 1 | 3 | Xpo1 | 0.058 | Old |
| 0.007026 | 0.215398 | 0.228 | 0.17 | 1 | 3 | Hras | 0.058 | Old |
| 0.007087 | 0.363027 | 0.329 | 0.27 | 1 | 3 | Atp6v1c1 | 0.059 | Old |
| 0.007105 | 0.641075 | 0.082 | 0.049 | 1 | 3 | Arfgap1 | 0.033 | Old |
| 0.007132 | 0.199922 | 0.527 | 0.453 | 1 | 3 | Tpi1 | 0.074 | Old |
| 0.007136 | 0.46464 | 0.164 | 0.117 | 1 | 3 | 9430038I0 | 0.047 | Old |
| 0.007143 | 0.169097 | 0.622 | 0.54 | 1 | 3 | Tomm20 | 0.082 | Old |
| 0.007145 | 0.279389 | 0.44 | 0.366 | 1 | 3 | Net1 | 0.074 | Old |
| 0.007241 | 0.579216 | 0.145 | 0.101 | 1 | 3 | Atg4b | 0.044 | Old |
| 0.007251 | 0.486184 | 0.108 | 0.069 | 1 | 3 | Fbxo9 | 0.039 | Old |
| 0.007333 | 0.527304 | 0.134 | 0.092 | 1 | 3 | Ndst2 | 0.042 | Old |
| 0.00734 | 0.319473 | 0.418 | 0.343 | 1 | 3 | Ube2k | 0.075 | Old |
| 0.007354 | 0.457117 | 0.115 | 0.076 | 1 | 3 | Mtmr1 | 0.039 | Old |
| 0.007361 | 0.320346 | 0.295 | 0.234 | 1 | 3 | Tcf4 | 0.061 | Old |
| 0.007371 | 0.637548 | 0.128 | 0.086 | 1 | 3 | Rrp12 | 0.042 | Old |
| 0.007374 | 0.481323 | 0.133 | 0.09 | 1 | 3 | Pparg | 0.043 | Old |
| 0.00738 | 0.174427 | 0.802 | 0.726 | 1 | 3 | Pcbp1 | 0.076 | Old |
| 0.007412 | 0.221625 | 0.484 | 0.413 | 1 | 3 | Cd300ld | 0.071 | Old |
| 0.007468 | 0.262458 | 0.281 | 0.217 | 1 | 3 | Plxna1 | 0.064 | Old |
| 0.007491 | 0.095104 | 0.969 | 0.937 | 1 | 3 | Mbnl1 | 0.032 | Old |
| 0.00751 | 0.284346 | 0.438 | 0.365 | 1 | 3 | Nudt3 | 0.073 | Old |
| 0.007516 | 0.382222 | 0.246 | 0.189 | 1 | 3 | Ola1 | 0.057 | Old |
| 0.007565 | 0.238156 | 0.251 | 0.191 | 1 | 3 | Dennd4c | 0.06 | Old |
| 0.007632 | 0.220956 | 0.461 | 0.385 | 1 | 3 | Psmd11 | 0.076 | Old |
| 0.007636 | 0.690655 | 0.049 | 0.024 | 1 | 3 | Rasal3 | 0.025 | Old |
| 0.007699 | 0.263317 | 0.079 | 0.045 | 1 | 3 | Tbl3 | 0.034 | Old |
| 0.007769 | 0.161025 | 0.811 | 0.745 | 1 | 3 | Msn | 0.066 | Old |
| 0.007786 | 0.651834 | 0.092 | 0.057 | 1 | 3 | Elac2 | 0.035 | Old |
| 0.007797 | 0.389533 | 0.139 | 0.095 | 1 | 3 | Scaf4 | 0.044 | Old |
| 0.007836 | 0.214119 | 0.718 | 0.652 | 1 | 3 | Minos1 | 0.066 | Old |
| 0.007878 | 0.257268 | 0.211 | 0.157 | 1 | 3 | Fryl | 0.054 | Old |
| 0.007884 | 0.425894 | 0.173 | 0.125 | 1 | 3 | Parg | 0.048 | Old |
| 0.007892 | 0.129762 | 0.93 | 0.885 | 1 | 3 | Btg2 | 0.045 | Old |
| 0.007898 | 0.187153 | 0.162 | 0.114 | 1 | 3 | Sipa1l3 | 0.048 | Old |
| 0.007914 | 0.189518 | 0.318 | 0.247 | 1 | 3 | Gatad2a | 0.071 | Old |
| 0.007918 | 0.179478 | 0.578 | 0.506 | 1 | 3 | Ppp1r14b | 0.072 | Old |
| 0.007961 | 0.337387 | 0.188 | 0.135 | 1 | 3 | Cab39l | 0.053 | Old |
| 0.007989 | 0.331007 | 0.249 | 0.189 | 1 | 3 | Bcat2 | 0.06 | Old |
| 0.008026 | 0.503136 | 0.112 | 0.073 | 1 | 3 | Fam53b | 0.039 | Old |
| 0.008074 | 0.229048 | 0.232 | 0.174 | 1 | 3 | Mrps12 | 0.058 | Old |
| 0.008083 | 0.223586 | 0.091 | 0.056 | 1 | 3 | Mthfs | 0.035 | Old |
| 0.008125 | 0.310594 | 0.189 | 0.141 | 1 | 3 | Slc16a10 | 0.048 | Old |
| 0.008162 | 0.582234 | 0.263 | 0.207 | 1 | 3 | Fabp4 | 0.056 | Old |

| | | | | | | | | |
|---|---|---|---|---|---|---|---|---|
| 0.008263 | 0.051245 | 0.335 | 0.261 | 1 | 3 | Elmo1 | 0.074 | Old |
| 0.008307 | 0.249203 | 0.331 | 0.269 | 1 | 3 | Sumo3 | 0.062 | Old |
| 0.008366 | 0.38383 | 0.122 | 0.081 | 1 | 3 | Sacm1l | 0.041 | Old |
| 0.008439 | 0.278799 | 0.314 | 0.25 | 1 | 3 | Klhl9 | 0.064 | Old |
| 0.008469 | 0.232953 | 0.544 | 0.467 | 1 | 3 | Manf | 0.077 | Old |
| 0.008503 | 1.571179 | 0.012 | 0.001 | 1 | 3 | Hpn | 0.011 | Old |
| 0.008533 | 0.161561 | 0.651 | 0.572 | 1 | 3 | Rab11fip1 | 0.079 | Old |
| 0.008542 | 0.859548 | 0.059 | 0.032 | 1 | 3 | Prune1 | 0.027 | Old |
| 0.008577 | 1.864214 | 0.012 | 0.001 | 1 | 3 | Apold1 | 0.011 | Old |
| 0.008615 | 1.02001 | 0.012 | 0.001 | 1 | 3 | Itpr3 | 0.011 | Old |
| 0.008618 | 0.102153 | 0.839 | 0.785 | 1 | 3 | mt-Nd4l | 0.054 | Old |
| 0.008647 | 0.146998 | 0.753 | 0.702 | 1 | 3 | Uqcr11 | 0.051 | Old |
| 0.008653 | 0.26689 | 0.146 | 0.101 | 1 | 3 | Sumf1 | 0.045 | Old |
| 0.008655 | 0.482592 | 0.18 | 0.134 | 1 | 3 | Lypla1 | 0.046 | Old |
| 0.008675 | 0.167576 | 0.746 | 0.693 | 1 | 3 | Litaf | 0.053 | Old |
| 0.008689 | 0.100734 | 0.122 | 0.08 | 1 | 3 | Rexo4 | 0.042 | Old |
| 0.008694 | 0.224153 | 0.243 | 0.184 | 1 | 3 | Cope | 0.059 | Old |
| 0.008825 | 0.282555 | 0.106 | 0.068 | 1 | 3 | Prkaa1 | 0.038 | Old |
| 0.008846 | 0.150149 | 0.49 | 0.422 | 1 | 3 | Tmco1 | 0.068 | Old |
| 0.008872 | 0.334914 | 0.339 | 0.278 | 1 | 3 | Ifi203 | 0.061 | Old |
| 0.008901 | 0.137415 | 0.452 | 0.361 | 1 | 3 | B4galnt1 | 0.091 | Old |
| 0.008929 | 0.409719 | 0.264 | 0.208 | 1 | 3 | Pgrmc1 | 0.056 | Old |
| 0.008973 | 0.289039 | 0.228 | 0.173 | 1 | 3 | 1500011K1 | 0.055 | Old |
| 0.009012 | 0.188762 | 0.306 | 0.241 | 1 | 3 | Dr1 | 0.065 | Old |
| 0.009061 | 0.941278 | 0.047 | 0.023 | 1 | 3 | Xaf1 | 0.024 | Old |
| 0.009079 | 0.568268 | 0.124 | 0.085 | 1 | 3 | Slc39a11 | 0.039 | Old |
| 0.009105 | 0.252648 | 0.614 | 0.539 | 1 | 3 | Chmp4b | 0.075 | Old |
| 0.009126 | 1.036623 | 0.03 | 0.011 | 1 | 3 | Car6 | 0.019 | Old |
| 0.009172 | 0.456811 | 0.205 | 0.158 | 1 | 3 | Ubr3 | 0.047 | Old |
| 0.009209 | 0.24623 | 0.186 | 0.135 | 1 | 3 | Fgd6 | 0.051 | Old |
| 0.009237 | 1.100115 | 0.022 | 0.006 | 1 | 3 | Fitm2 | 0.016 | Old |
| 0.009249 | 0.532405 | 0.105 | 0.068 | 1 | 3 | Smim3 | 0.037 | Old |
| 0.009453 | 0.331255 | 0.134 | 0.091 | 1 | 3 | Slc25a25 | 0.043 | Old |
| 0.009505 | 0.18767 | 0.391 | 0.324 | 1 | 3 | Gtpbp4 | 0.067 | Old |
| 0.009564 | 0.233157 | 0.146 | 0.101 | 1 | 3 | Ube2l6 | 0.045 | Old |
| 0.009634 | 0.181228 | 0.817 | 0.766 | 1 | 3 | Sh3bgrl | 0.051 | Old |
| 0.009653 | 1.421703 | 0.037 | 0.016 | 1 | 3 | Rtp4 | 0.021 | Old |
| 0.009659 | 0.309846 | 0.168 | 0.121 | 1 | 3 | 5-Mar | 0.047 | Old |
| 0.009663 | 0.146527 | 0.524 | 0.441 | 1 | 3 | Arhgap18 | 0.083 | Old |
| 0.009671 | 0.15335 | 0.713 | 0.646 | 1 | 3 | Ptpn1 | 0.067 | Old |
| 0.009691 | 0.91744 | 0.065 | 0.037 | 1 | 3 | Phf11d | 0.028 | Old |
| 0.009701 | 0.247173 | 0.208 | 0.154 | 1 | 3 | Smarcad1 | 0.054 | Old |
| 0.009741 | 0.075254 | 0.973 | 0.958 | 1 | 3 | Cdc42 | 0.015 | Old |
| 0.009762 | 0.208669 | 0.317 | 0.251 | 1 | 3 | Med28 | 0.066 | Old |
| 0.009774 | 0.128451 | 0.895 | 0.876 | 1 | 3 | Ear2 | 0.019 | Old |

| | | | | | | | | |
|---|---|---|---|---|---|---|---|---|
| 0.00979 | 0.291875 | 0.104 | 0.067 | 1 | 3 | Sec24d | 0.037 | Old |
| 0.009793 | 0.438323 | 0.116 | 0.077 | 1 | 3 | Gsr | 0.039 | Old |
| 0.009796 | 0.216095 | 0.337 | 0.27 | 1 | 3 | Eif4e | 0.067 | Old |
| 0.009806 | 0.276668 | 0.243 | 0.188 | 1 | 3 | Ube2v2 | 0.055 | Old |
| 0.009837 | 1.320383 | 0.014 | 0.003 | 1 | 3 | Intu | 0.011 | Old |
| 0.009841 | 0.34359 | 0.208 | 0.155 | 1 | 3 | Commd6 | 0.053 | Old |
| 0.009869 | 0.237116 | 0.247 | 0.191 | 1 | 3 | Ocrl | 0.056 | Old |
| 0.009871 | 0.464989 | 0.055 | 0.029 | 1 | 3 | Rabepk | 0.026 | Old |
| 0.009921 | 0.62234 | 0.043 | 0.02 | 1 | 3 | Psph | 0.023 | Old |
| 0.009964 | 0.582589 | 0.09 | 0.056 | 1 | 3 | Oma1 | 0.034 | Old |
| 0.010014 | 0.399312 | 0.17 | 0.124 | 1 | 3 | Babam1 | 0.046 | Old |
| 0.010044 | 0.355699 | 0.198 | 0.149 | 1 | 3 | Uba5 | 0.049 | Old |
| 0.010049 | 0.1085 | 0.183 | 0.133 | 1 | 3 | Stard7 | 0.05 | Old |
| 0.010063 | 0.186356 | 0.337 | 0.266 | 1 | 3 | Ufm1 | 0.071 | Old |
| 0.01007 | 0.37968 | 0.184 | 0.134 | 1 | 3 | Lap3 | 0.05 | Old |
| 0.010239 | 0.770817 | 0.079 | 0.048 | 1 | 3 | Lpar6 | 0.031 | Old |
| 0.010265 | 0.282052 | 0.272 | 0.211 | 1 | 3 | Tap1 | 0.061 | Old |
| 0.010285 | 0.182263 | 0.234 | 0.176 | 1 | 3 | Elp5 | 0.058 | Old |
| 0.010321 | 0.18016 | 0.244 | 0.187 | 1 | 3 | Tmem248 | 0.057 | Old |
| 0.010347 | 0.235024 | 0.344 | 0.282 | 1 | 3 | Bcas2 | 0.062 | Old |
| 0.010394 | 0.119248 | 0.311 | 0.244 | 1 | 3 | Ythdf1 | 0.067 | Old |
| 0.010405 | 0.241853 | 0.338 | 0.271 | 1 | 3 | Abcf1 | 0.067 | Old |
| 0.010437 | 0.247222 | 0.366 | 0.299 | 1 | 3 | Rlim | 0.067 | Old |
| 0.010438 | 0.129903 | 0.272 | 0.211 | 1 | 3 | Hs6st1 | 0.061 | Old |
| 0.010442 | 0.847043 | 0.044 | 0.021 | 1 | 3 | Coq3 | 0.023 | Old |
| 0.010465 | 0.727049 | 0.104 | 0.068 | 1 | 3 | Zdhhc6 | 0.036 | Old |
| 0.010482 | 0.216979 | 0.413 | 0.341 | 1 | 3 | Sar1a | 0.072 | Old |
| 0.010509 | 0.186438 | 0.501 | 0.42 | 1 | 3 | Snx9 | 0.081 | Old |
| 0.010537 | 0.205874 | 0.539 | 0.465 | 1 | 3 | Tle3 | 0.074 | Old |
| 0.010578 | 0.141221 | 0.796 | 0.747 | 1 | 3 | Eif5a | 0.049 | Old |
| 0.010603 | 0.453757 | 0.168 | 0.124 | 1 | 3 | Tmx4 | 0.044 | Old |
| 0.010619 | 0.177645 | 0.664 | 0.602 | 1 | 3 | Mtpn | 0.062 | Old |
| 0.010641 | 0.316901 | 0.147 | 0.104 | 1 | 3 | Slu7 | 0.043 | Old |
| 0.010667 | 0.434165 | 0.244 | 0.194 | 1 | 3 | Atp13a1 | 0.05 | Old |
| 0.010679 | 1.05485 | 0.044 | 0.021 | 1 | 3 | Helz2 | 0.023 | Old |
| 0.010687 | 0.076887 | 0.594 | 0.497 | 1 | 3 | Sbno2 | 0.097 | Old |
| 0.010692 | 0.281372 | 0.422 | 0.357 | 1 | 3 | Siva1 | 0.065 | Old |
| 0.010711 | 0.904337 | 0.037 | 0.016 | 1 | 3 | Pus7l | 0.021 | Old |
| 0.010787 | 0.391154 | 0.124 | 0.085 | 1 | 3 | Wrn | 0.039 | Old |
| 0.010815 | 0.735532 | 0.105 | 0.068 | 1 | 3 | Rars2 | 0.037 | Old |
| 0.010822 | 0.777408 | 0.078 | 0.047 | 1 | 3 | Dcn | 0.031 | Old |
| 0.010842 | 0.469716 | 0.16 | 0.116 | 1 | 3 | Rnf213 | 0.044 | Old |
| 0.010854 | 0.041257 | 0.385 | 0.301 | 1 | 3 | Clec4e | 0.084 | Old |
| 0.010858 | 0.358473 | 0.27 | 0.216 | 1 | 3 | Cept1 | 0.054 | Old |
| 0.01086 | 0.056037 | 0.485 | 0.405 | 1 | 3 | Traf1 | 0.08 | Old |

| | | | | | | | | |
|---|---|---|---|---|---|---|---|---|
| 0.01086 | 0.223735 | 0.188 | 0.139 | 1 | 3 | Ifi207 | 0.049 | Old |
| 0.01087 | 0.269084 | 0.429 | 0.361 | 1 | 3 | Snrpb2 | 0.068 | Old |
| 0.010893 | 0.183974 | 0.441 | 0.372 | 1 | 3 | Acsl4 | 0.069 | Old |
| 0.010932 | 0.397266 | 0.072 | 0.042 | 1 | 3 | Haus2 | 0.03 | Old |
| 0.010967 | 0.255677 | 0.338 | 0.277 | 1 | 3 | Ddx24 | 0.061 | Old |
| 0.010975 | 0.119155 | 0.571 | 0.484 | 1 | 3 | Uqcrb | 0.087 | Old |
| 0.010994 | 0.278045 | 0.232 | 0.177 | 1 | 3 | Tars | 0.055 | Old |
| 0.010997 | 0.117781 | 0.369 | 0.293 | 1 | 3 | Cox20 | 0.076 | Old |
| 0.011017 | 0.269304 | 0.276 | 0.217 | 1 | 3 | Cops4 | 0.059 | Old |
| 0.011028 | 0.140543 | 0.884 | 0.835 | 1 | 3 | Naaa | 0.049 | Old |
| 0.011118 | 0.809197 | 0.084 | 0.052 | 1 | 3 | Il1rap | 0.032 | Old |
| 0.01116 | 0.387569 | 0.327 | 0.268 | 1 | 3 | Tspan3 | 0.059 | Old |
| 0.011179 | 0.362116 | 0.188 | 0.143 | 1 | 3 | Tdg | 0.045 | Old |
| 0.011182 | 0.088098 | 0.556 | 0.468 | 1 | 3 | Ptpa | 0.088 | Old |
| 0.011239 | 0.524926 | 0.125 | 0.086 | 1 | 3 | Nop9 | 0.039 | Old |
| 0.011297 | 0.262815 | 0.32 | 0.256 | 1 | 3 | Prkx | 0.064 | Old |
| 0.011363 | 0.419293 | 0.079 | 0.048 | 1 | 3 | Tradd | 0.031 | Old |
| 0.011363 | 0.979681 | 0.042 | 0.02 | 1 | 3 | Zmym3 | 0.022 | Old |
| 0.011434 | 0.276862 | 0.215 | 0.165 | 1 | 3 | Eml4 | 0.05 | Old |
| 0.011451 | 0.781659 | 0.053 | 0.028 | 1 | 3 | F8a | 0.025 | Old |
| 0.011524 | 0.332125 | 0.098 | 0.063 | 1 | 3 | Wdr18 | 0.035 | Old |
| 0.011548 | 0.478753 | 0.155 | 0.114 | 1 | 3 | Rab3il1 | 0.041 | Old |
| 0.011613 | 0.372773 | 0.069 | 0.04 | 1 | 3 | Ccdc167 | 0.029 | Old |
| 0.011624 | 0.10006 | 0.393 | 0.314 | 1 | 3 | Eif3d | 0.079 | Old |
| 0.011639 | 0.217931 | 0.456 | 0.39 | 1 | 3 | Pdia4 | 0.066 | Old |
| 0.011682 | 0.551068 | 0.093 | 0.061 | 1 | 3 | Lias | 0.032 | Old |
| 0.011792 | 0.146194 | 0.305 | 0.236 | 1 | 3 | Usp19 | 0.069 | Old |
| 0.011796 | 0.096339 | 0.413 | 0.336 | 1 | 3 | Pdcd6 | 0.077 | Old |
| 0.011819 | 0.544105 | 0.068 | 0.039 | 1 | 3 | Engase | 0.029 | Old |
| 0.011867 | 0.242806 | 0.222 | 0.169 | 1 | 3 | Trio | 0.053 | Old |
| 0.011906 | 0.126468 | 0.671 | 0.588 | 1 | 3 | Brk1 | 0.083 | Old |
| 0.011941 | 0.120119 | 0.646 | 0.571 | 1 | 3 | Arhgap30 | 0.075 | Old |
| 0.012083 | 0.217239 | 0.399 | 0.33 | 1 | 3 | Cnot6 | 0.069 | Old |
| 0.012115 | 0.416929 | 0.257 | 0.205 | 1 | 3 | Hyou1 | 0.052 | Old |
| 0.012159 | 0.328077 | 0.152 | 0.109 | 1 | 3 | Sdad1 | 0.043 | Old |
| 0.012184 | 0.398705 | 0.074 | 0.044 | 1 | 3 | Fuom | 0.03 | Old |
| 0.012255 | 0.244623 | 0.137 | 0.096 | 1 | 3 | Sipa1l1 | 0.041 | Old |
| 0.012293 | 0.129001 | 0.763 | 0.726 | 1 | 3 | Tm6sf1 | 0.037 | Old |
| 0.012372 | 0.400501 | 0.143 | 0.102 | 1 | 3 | Dctn5 | 0.041 | Old |
| 0.012409 | 0.108522 | 0.892 | 0.842 | 1 | 3 | Tmem50a | 0.05 | Old |
| 0.012415 | 0.471341 | 0.155 | 0.114 | 1 | 3 | Nudt5 | 0.041 | Old |
| 0.01243 | 0.600046 | 0.092 | 0.059 | 1 | 3 | Kti12 | 0.033 | Old |
| 0.012482 | 0.365365 | 0.104 | 0.069 | 1 | 3 | Mcoln1 | 0.035 | Old |
| 0.01249 | 0.099208 | 0.367 | 0.297 | 1 | 3 | Piezo1 | 0.07 | Old |
| 0.012516 | 0.481818 | 0.266 | 0.215 | 1 | 3 | Lrrc59 | 0.051 | Old |

| | | | | | | | | |
|---|---|---|---|---|---|---|---|---|
| 0.012521 | 0.211931 | 0.27 | 0.211 | 1 | 3 | Ttyh3 | 0.059 | Old |
| 0.012556 | 0.127466 | 0.357 | 0.284 | 1 | 3 | Sec24b | 0.073 | Old |
| 0.012589 | 0.349379 | 0.284 | 0.225 | 1 | 3 | Uba6 | 0.059 | Old |
| 0.012626 | 0.417051 | 0.134 | 0.093 | 1 | 3 | Prpf19 | 0.041 | Old |
| 0.012791 | 0.442138 | 0.249 | 0.199 | 1 | 3 | Otulin | 0.05 | Old |
| 0.012842 | 0.72496 | 0.081 | 0.051 | 1 | 3 | Pcid2 | 0.03 | Old |
| 0.012879 | 0.762783 | 0.072 | 0.043 | 1 | 3 | Zdhhc16 | 0.029 | Old |
| 0.012882 | 0.290645 | 0.257 | 0.202 | 1 | 3 | Ndufs8 | 0.055 | Old |
| 0.01294 | 0.367699 | 0.114 | 0.077 | 1 | 3 | Ascc2 | 0.037 | Old |
| 0.012954 | 0.488902 | 0.118 | 0.081 | 1 | 3 | Phkg2 | 0.037 | Old |
| 0.01311 | 0.146592 | 0.432 | 0.359 | 1 | 3 | Stx4a | 0.073 | Old |
| 0.013122 | 1.007083 | 0.063 | 0.037 | 1 | 3 | Golga1 | 0.026 | Old |
| 0.013167 | 0.12233 | 0.321 | 0.255 | 1 | 3 | Mff | 0.066 | Old |
| 0.013168 | 0.245016 | 0.355 | 0.292 | 1 | 3 | Fam174a | 0.063 | Old |
| 0.013181 | 0.9876 | 0.023 | 0.008 | 1 | 3 | Tlr3 | 0.015 | Old |
| 0.013297 | 0.102961 | 0.185 | 0.134 | 1 | 3 | Rasgrp1 | 0.051 | Old |
| 0.013311 | 0.064409 | 0.341 | 0.273 | 1 | 3 | Bcl2l11 | 0.068 | Old |
| 0.013357 | 0.160295 | 0.652 | 0.596 | 1 | 3 | Sec61g | 0.056 | Old |
| 0.013396 | 0.562154 | 0.225 | 0.177 | 1 | 3 | Wnt11 | 0.048 | Old |
| 0.013503 | 0.245776 | 0.321 | 0.26 | 1 | 3 | Smc1a | 0.061 | Old |
| 0.013577 | 0.137799 | 0.216 | 0.162 | 1 | 3 | Srek1 | 0.054 | Old |
| 0.013632 | 0.270949 | 0.165 | 0.12 | 1 | 3 | Zfp330 | 0.045 | Old |
| 0.013754 | 0.033274 | 0.137 | 0.095 | 1 | 3 | Atp7a | 0.042 | Old |
| 0.013794 | 0.134291 | 0.627 | 0.552 | 1 | 3 | Acer3 | 0.075 | Old |
| 0.013804 | 0.242456 | 0.249 | 0.196 | 1 | 3 | Sec23b | 0.053 | Old |
| 0.014048 | 0.129769 | 0.662 | 0.6 | 1 | 3 | Vmp1 | 0.062 | Old |
| 0.014065 | 0.452066 | 0.05 | 0.027 | 1 | 3 | Timm21 | 0.023 | Old |
| 0.014156 | 1.126892 | 0.043 | 0.021 | 1 | 3 | Oas2 | 0.022 | Old |
| 0.014202 | 1.035806 | 0.039 | 0.019 | 1 | 3 | Zbtb11os1 | 0.02 | Old |
| 0.014216 | 0.467513 | 0.184 | 0.139 | 1 | 3 | Pom121 | 0.045 | Old |
| 0.014234 | 0.374256 | 0.188 | 0.141 | 1 | 3 | Plk2 | 0.047 | Old |
| 0.014248 | 1.415813 | 0.011 | 0.001 | 1 | 3 | 4933406I1 | 0.01 | Old |
| 0.014258 | 0.42687 | 0.059 | 0.033 | 1 | 3 | Edrf1 | 0.026 | Old |
| 0.014273 | 0.348051 | 0.094 | 0.061 | 1 | 3 | Ttc33 | 0.033 | Old |
| 0.014279 | 1.238444 | 0.011 | 0.001 | 1 | 3 | Lpcat4 | 0.01 | Old |
| 0.014324 | 0.176567 | 0.657 | 0.571 | 1 | 3 | Creg1 | 0.086 | Old |
| 0.014329 | 0.226274 | 0.195 | 0.146 | 1 | 3 | D5Ertd579 | 0.049 | Old |
| 0.014345 | 0.217935 | 0.362 | 0.297 | 1 | 3 | Vma21 | 0.065 | Old |
| 0.01445 | 0.333623 | 0.167 | 0.124 | 1 | 3 | Lrrc41 | 0.043 | Old |
| 0.014465 | 1.033156 | 0.02 | 0.006 | 1 | 3 | Eml6 | 0.014 | Old |
| 0.01448 | 0.179592 | 0.133 | 0.091 | 1 | 3 | Rnf103 | 0.042 | Old |
| 0.014507 | 1.077708 | 0.02 | 0.006 | 1 | 3 | Ccdc137 | 0.014 | Old |
| 0.014559 | 0.305095 | 0.128 | 0.09 | 1 | 3 | Nfyb | 0.038 | Old |
| 0.014563 | 0.461483 | 0.076 | 0.047 | 1 | 3 | Smad3 | 0.029 | Old |
| 0.014638 | 0.209597 | 0.278 | 0.22 | 1 | 3 | Atf1 | 0.058 | Old |

| | | | | | | | | |
|---|---|---|---|---|---|---|---|---|
| 0.014695 | 0.191685 | 0.521 | 0.444 | 1 | 3 | Sidt2 | 0.077 | Old |
| 0.014702 | 0.252302 | 0.48 | 0.412 | 1 | 3 | Acly | 0.068 | Old |
| 0.014773 | 0.212934 | 0.458 | 0.393 | 1 | 3 | Aco2 | 0.065 | Old |
| 0.014801 | 0.295438 | 0.143 | 0.101 | 1 | 3 | Atmin | 0.042 | Old |
| 0.014881 | 0.275053 | 0.094 | 0.061 | 1 | 3 | Gpatch11 | 0.033 | Old |
| 0.014914 | 0.122833 | 0.509 | 0.427 | 1 | 3 | Ranbp1 | 0.082 | Old |
| 0.014946 | 0.393709 | 0.225 | 0.174 | 1 | 3 | Osgin2 | 0.051 | Old |
| 0.015056 | 0.286099 | 0.382 | 0.314 | 1 | 3 | Mob4 | 0.068 | Old |
| 0.01511 | 0.236056 | 0.341 | 0.277 | 1 | 3 | Srp19 | 0.064 | Old |
| 0.015124 | 0.219827 | 0.398 | 0.335 | 1 | 3 | Gmfb | 0.063 | Old |
| 0.01518 | 0.504317 | 0.111 | 0.076 | 1 | 3 | Chid1 | 0.035 | Old |
| 0.015267 | 0.187276 | 0.085 | 0.053 | 1 | 3 | Wdr75 | 0.032 | Old |
| 0.015287 | 0.217737 | 0.219 | 0.168 | 1 | 3 | Spg7 | 0.051 | Old |
| 0.015318 | 0.170237 | 0.51 | 0.437 | 1 | 3 | Anp32e | 0.073 | Old |
| 0.015328 | 0.286105 | 0.251 | 0.199 | 1 | 3 | Atic | 0.052 | Old |
| 0.015345 | 0.21021 | 0.195 | 0.145 | 1 | 3 | Srd5a3 | 0.05 | Old |
| 0.015506 | 0.212509 | 0.108 | 0.072 | 1 | 3 | Slc9a8 | 0.036 | Old |
| 0.015558 | 0.357025 | 0.148 | 0.107 | 1 | 3 | Pmm2 | 0.041 | Old |
| 0.015572 | 0.290004 | 0.269 | 0.213 | 1 | 3 | Zdhhc21 | 0.056 | Old |
| 0.015601 | 0.473297 | 0.177 | 0.131 | 1 | 3 | Ddx39 | 0.046 | Old |
| 0.015628 | 0.132243 | 0.542 | 0.484 | 1 | 3 | Ccr1 | 0.058 | Old |
| 0.015667 | 0.994005 | 0.018 | 0.005 | 1 | 3 | Tmem218 | 0.013 | Old |
| 0.015686 | 0.804697 | 0.026 | 0.01 | 1 | 3 | Prr15 | 0.016 | Old |
| 0.015716 | 0.316894 | 0.204 | 0.154 | 1 | 3 | Galk2 | 0.05 | Old |
| 0.015736 | 0.215729 | 0.509 | 0.441 | 1 | 3 | Snx2 | 0.068 | Old |
| 0.015839 | 0.25924 | 0.137 | 0.097 | 1 | 3 | Senp5 | 0.04 | Old |
| 0.015854 | 0.175929 | 0.545 | 0.472 | 1 | 3 | Csk | 0.073 | Old |
| 0.015879 | 0.200961 | 0.149 | 0.107 | 1 | 3 | Cln5 | 0.042 | Old |
| 0.015909 | 0.269248 | 0.352 | 0.289 | 1 | 3 | Ndufab1 | 0.063 | Old |
| 0.015919 | 1.476633 | 0.013 | 0.003 | 1 | 3 | Vasn | 0.01 | Old |
| 0.015991 | 0.256296 | 0.245 | 0.191 | 1 | 3 | Rnps1 | 0.054 | Old |
| 0.016025 | 0.235203 | 0.102 | 0.067 | 1 | 3 | Slc30a1 | 0.035 | Old |
| 0.016069 | 1.280156 | 0.013 | 0.003 | 1 | 3 | Gm14296 | 0.01 | Old |
| 0.016069 | 1.070169 | 0.013 | 0.003 | 1 | 3 | Usp2 | 0.01 | Old |
| 0.016081 | 0.149379 | 0.755 | 0.686 | 1 | 3 | Serp1 | 0.069 | Old |
| 0.016097 | 0.278078 | 0.205 | 0.158 | 1 | 3 | Evi2 | 0.047 | Old |
| 0.016111 | 1.283839 | 0.016 | 0.004 | 1 | 3 | Chst7 | 0.012 | Old |
| 0.016162 | 0.310189 | 0.276 | 0.222 | 1 | 3 | Pkig | 0.054 | Old |
| 0.016192 | 1.458905 | 0.016 | 0.004 | 1 | 3 | Prdm9 | 0.012 | Old |
| 0.016202 | 0.061648 | 0.211 | 0.158 | 1 | 3 | Esf1 | 0.053 | Old |
| 0.01625 | 0.963692 | 0.013 | 0.003 | 1 | 3 | Olfr920 | 0.01 | Old |
| 0.016398 | 0.126896 | 0.507 | 0.429 | 1 | 3 | Cers2 | 0.078 | Old |
| 0.016432 | 0.252877 | 0.519 | 0.448 | 1 | 3 | Fbl | 0.071 | Old |
| 0.016441 | 0.992899 | 0.03 | 0.013 | 1 | 3 | Gm14286 | 0.017 | Old |
| 0.016501 | 0.484331 | 0.066 | 0.039 | 1 | 3 | Acad9 | 0.027 | Old |

| | | | | | | | | |
|---|---|---|---|---|---|---|---|---|
| 0.016527 | 0.073392 | 0.183 | 0.134 | 1 | 3 | Erh | 0.049 | Old |
| 0.016553 | 0.181426 | 0.388 | 0.318 | 1 | 3 | Snrpc | 0.07 | Old |
| 0.016633 | 0.072636 | 0.337 | 0.27 | 1 | 3 | Dcaf7 | 0.067 | Old |
| 0.016656 | 0.41389 | 0.211 | 0.162 | 1 | 3 | Cops2 | 0.049 | Old |
| 0.016746 | 0.116028 | 0.237 | 0.186 | 1 | 3 | Tnip2 | 0.051 | Old |
| 0.016748 | 0.084674 | 0.507 | 0.427 | 1 | 3 | Elmo2 | 0.08 | Old |
| 0.016759 | 0.125821 | 0.667 | 0.591 | 1 | 3 | Dync1h1 | 0.076 | Old |
| 0.016765 | 0.268168 | 0.198 | 0.15 | 1 | 3 | Oxsr1 | 0.048 | Old |
| 0.016771 | 0.137612 | 0.63 | 0.559 | 1 | 3 | Cct8 | 0.071 | Old |
| 0.016817 | 0.139079 | 0.312 | 0.25 | 1 | 3 | Smim10l1 | 0.062 | Old |
| 0.016824 | 0.460455 | 0.075 | 0.047 | 1 | 3 | Cd40 | 0.028 | Old |
| 0.016824 | 0.241265 | 0.407 | 0.343 | 1 | 3 | Plcg2 | 0.064 | Old |
| 0.016832 | 0.315105 | 0.259 | 0.211 | 1 | 3 | Hibadh | 0.048 | Old |
| 0.016841 | 0.475887 | 0.172 | 0.13 | 1 | 3 | Slc25a30 | 0.042 | Old |
| 0.016846 | 0.270589 | 0.252 | 0.199 | 1 | 3 | Adam9 | 0.053 | Old |
| 0.016961 | 0.507049 | 0.14 | 0.1 | 1 | 3 | Xdh | 0.04 | Old |
| 0.017046 | 1.03251 | 0.078 | 0.049 | 1 | 3 | Mex3d | 0.029 | Old |
| 0.017069 | 0.232109 | 0.283 | 0.227 | 1 | 3 | Ccdc115 | 0.056 | Old |
| 0.017111 | 0.639471 | 0.066 | 0.039 | 1 | 3 | Fmo5 | 0.027 | Old |
| 0.017115 | 0.244847 | 0.264 | 0.212 | 1 | 3 | Dgkz | 0.052 | Old |
| 0.017179 | 0.309651 | 0.24 | 0.191 | 1 | 3 | Dhcr24 | 0.049 | Old |
| 0.01723 | 0.698255 | 0.062 | 0.037 | 1 | 3 | Gmnn | 0.025 | Old |
| 0.017239 | 0.272524 | 0.189 | 0.141 | 1 | 3 | Ranbp9 | 0.048 | Old |
| 0.017244 | 0.083937 | 0.818 | 0.756 | 1 | 3 | Mcub | 0.062 | Old |
| 0.017298 | 0.193628 | 0.474 | 0.402 | 1 | 3 | Eloc | 0.072 | Old |
| 0.017329 | 0.465996 | 0.131 | 0.095 | 1 | 3 | Commd10 | 0.036 | Old |
| 0.017452 | 0.296993 | 0.323 | 0.269 | 1 | 3 | Arl5a | 0.054 | Old |
| 0.017499 | 0.65806 | 0.033 | 0.015 | 1 | 3 | Smug1 | 0.018 | Old |
| 0.017503 | 0.345931 | 0.134 | 0.096 | 1 | 3 | Tfrc | 0.038 | Old |
| 0.017518 | 0.729916 | 0.033 | 0.015 | 1 | 3 | Deaf1 | 0.018 | Old |
| 0.01753 | 0.89811 | 0.037 | 0.018 | 1 | 3 | Tgm2 | 0.019 | Old |
| 0.017595 | 0.237962 | 0.364 | 0.306 | 1 | 3 | Eif1b | 0.058 | Old |
| 0.017657 | 0.336882 | 0.251 | 0.202 | 1 | 3 | Zfp263 | 0.049 | Old |
| 0.017728 | 1.011064 | 0.024 | 0.009 | 1 | 3 | Jag1 | 0.015 | Old |
| 0.017778 | 0.162219 | 0.246 | 0.191 | 1 | 3 | Polr2c | 0.055 | Old |
| 0.017827 | 0.291083 | 0.417 | 0.359 | 1 | 3 | Cpsf6 | 0.058 | Old |
| 0.01783 | 0.159353 | 0.438 | 0.367 | 1 | 3 | Dennd5a | 0.071 | Old |
| 0.017891 | 0.414164 | 0.191 | 0.146 | 1 | 3 | Ss18l2 | 0.045 | Old |
| 0.018015 | 0.474931 | 0.13 | 0.092 | 1 | 3 | Slc41a2 | 0.038 | Old |
| 0.018102 | 0.521289 | 0.037 | 0.018 | 1 | 3 | Gm38843 | 0.019 | Old |
| 0.018103 | 0.620579 | 0.041 | 0.02 | 1 | 3 | Gtf2h2 | 0.021 | Old |
| 0.018119 | 0.132508 | 0.593 | 0.514 | 1 | 3 | Eif5b | 0.079 | Old |
| 0.018153 | 0.301427 | 0.344 | 0.287 | 1 | 3 | Cyb5b | 0.057 | Old |
| 0.018242 | 0.164059 | 0.466 | 0.396 | 1 | 3 | Glipr2 | 0.07 | Old |
| 0.018302 | 0.11123 | 0.603 | 0.515 | 1 | 3 | 2900097C1 | 0.088 | Old |

| | | | | | | | | |
|---|---|---|---|---|---|---|---|---|
| 0.018384 | 0.341702 | 0.136 | 0.097 | 1 | 3 | Mphosph1 | 0.039 | Old |
| 0.018472 | 0.706089 | 0.062 | 0.037 | 1 | 3 | Ppat | 0.025 | Old |
| 0.01856 | 0.282925 | 0.21 | 0.163 | 1 | 3 | Lyar | 0.047 | Old |
| 0.018584 | 0.386113 | 0.217 | 0.17 | 1 | 3 | Txndc5 | 0.047 | Old |
| 0.018589 | 0.464356 | 0.07 | 0.043 | 1 | 3 | Pwp1 | 0.027 | Old |
| 0.01859 | 0.37973 | 0.189 | 0.145 | 1 | 3 | Lyst | 0.044 | Old |
| 0.018591 | 0.305961 | 0.239 | 0.188 | 1 | 3 | Gskip | 0.051 | Old |
| 0.018635 | 0.414231 | 0.2 | 0.155 | 1 | 3 | Dnm1l | 0.045 | Old |
| 0.018669 | 0.109682 | 0.127 | 0.087 | 1 | 3 | Nop14 | 0.04 | Old |
| 0.018767 | 0.173541 | 0.47 | 0.396 | 1 | 3 | Sdf4 | 0.074 | Old |
| 0.018812 | 0.193558 | 0.593 | 0.528 | 1 | 3 | Cd24a | 0.065 | Old |
| 0.018831 | 0.522406 | 0.131 | 0.095 | 1 | 3 | Slc31a1 | 0.036 | Old |
| 0.018832 | 0.331326 | 0.116 | 0.081 | 1 | 3 | Golga3 | 0.035 | Old |
| 0.018862 | 0.747659 | 0.072 | 0.044 | 1 | 3 | Etfrf1 | 0.028 | Old |
| 0.018938 | 0.512329 | 0.147 | 0.107 | 1 | 3 | Dera | 0.04 | Old |
| 0.018956 | 0.208683 | 0.287 | 0.231 | 1 | 3 | Mogs | 0.056 | Old |
| 0.018958 | 0.43801 | 0.092 | 0.061 | 1 | 3 | Tmem161a | 0.031 | Old |
| 0.018966 | 0.159892 | 0.529 | 0.453 | 1 | 3 | Bag1 | 0.076 | Old |
| 0.019008 | 1.883806 | 0.022 | 0.008 | 1 | 3 | Ccnb2 | 0.014 | Old |
| 0.019112 | 0.478733 | 0.176 | 0.133 | 1 | 3 | Wdr61 | 0.043 | Old |
| 0.019131 | 0.432542 | 0.149 | 0.11 | 1 | 3 | Wdr5 | 0.039 | Old |
| 0.019137 | 0.256659 | 0.44 | 0.383 | 1 | 3 | Rp2 | 0.057 | Old |
| 0.01916 | 0.178473 | 0.492 | 0.414 | 1 | 3 | Ralb | 0.078 | Old |
| 0.01923 | 0.184553 | 0.491 | 0.424 | 1 | 3 | Smap2 | 0.067 | Old |
| 0.019256 | 0.197518 | 0.231 | 0.181 | 1 | 3 | Mindy1 | 0.05 | Old |
| 0.019359 | 0.204498 | 0.564 | 0.491 | 1 | 3 | Gsn | 0.073 | Old |
| 0.019386 | 0.150603 | 0.229 | 0.173 | 1 | 3 | 1810013L2 | 0.056 | Old |
| 0.019466 | 0.199717 | 0.546 | 0.47 | 1 | 3 | Sf3b6 | 0.076 | Old |
| 0.019484 | 0.302101 | 0.225 | 0.177 | 1 | 3 | Inppl1 | 0.048 | Old |
| 0.019491 | 0.596114 | 0.096 | 0.064 | 1 | 3 | Las1l | 0.032 | Old |
| 0.019491 | 0.285468 | 0.145 | 0.105 | 1 | 3 | Chchd4 | 0.04 | Old |
| 0.019495 | 0.114626 | 0.327 | 0.264 | 1 | 3 | Tmem51 | 0.063 | Old |
| 0.019592 | 0.161077 | 0.196 | 0.149 | 1 | 3 | Gatm | 0.047 | Old |
| 0.019913 | 0.356194 | 0.295 | 0.247 | 1 | 3 | Nop58 | 0.048 | Old |
| 0.019972 | 0.792737 | 0.074 | 0.047 | 1 | 3 | Cyb561d2 | 0.027 | Old |
| 0.020073 | 0.522089 | 0.139 | 0.101 | 1 | 3 | Ift52 | 0.038 | Old |
| 0.020078 | 0.574674 | 0.111 | 0.077 | 1 | 3 | Memo1 | 0.034 | Old |
| 0.020144 | 0.767775 | 0.022 | 0.008 | 1 | 3 | Tnfsfm13 | 0.014 | Old |
| 0.020197 | 0.323158 | 0.296 | 0.246 | 1 | 3 | Senp2 | 0.05 | Old |
| 0.020224 | 0.509239 | 0.042 | 0.021 | 1 | 3 | Ifrd2 | 0.021 | Old |
| 0.020253 | 0.250367 | 0.123 | 0.087 | 1 | 3 | Hck | 0.036 | Old |
| 0.020272 | 0.299539 | 0.152 | 0.111 | 1 | 3 | Iars | 0.041 | Old |
| 0.020282 | 0.199803 | 0.744 | 0.662 | 1 | 3 | Furin | 0.082 | Old |
| 0.020325 | 0.2037 | 0.239 | 0.186 | 1 | 3 | Lin7c | 0.053 | Old |
| 0.020329 | 0.071908 | 0.919 | 0.898 | 1 | 3 | Srgn | 0.021 | Old |

| | | | | | | | | |
|---|---|---|---|---|---|---|---|---|
| 0.020381 | 0.222287 | 0.164 | 0.122 | 1 | 3 | Gan | 0.042 | Old |
| 0.020437 | 0.13639 | 0.342 | 0.28 | 1 | 3 | Tulp4 | 0.062 | Old |
| 0.020502 | 0.315118 | 0.417 | 0.356 | 1 | 3 | Gm26532 | 0.061 | Old |
| 0.020519 | 0.498142 | 0.062 | 0.037 | 1 | 3 | Cd274 | 0.025 | Old |
| 0.020554 | 0.216794 | 0.152 | 0.111 | 1 | 3 | Prrc1 | 0.041 | Old |
| 0.020571 | 0.194703 | 0.318 | 0.261 | 1 | 3 | C1qbp | 0.057 | Old |
| 0.020699 | 0.061931 | 0.897 | 0.855 | 1 | 3 | Gnb2 | 0.042 | Old |
| 0.020715 | 0.138202 | 0.244 | 0.186 | 1 | 3 | Dctn2 | 0.058 | Old |
| 0.020726 | 0.415407 | 0.141 | 0.102 | 1 | 3 | Cnot10 | 0.039 | Old |
| 0.020727 | 0.497199 | 0.084 | 0.054 | 1 | 3 | Ppih | 0.03 | Old |
| 0.02079 | 0.796523 | 0.035 | 0.016 | 1 | 3 | B3galt6 | 0.019 | Old |
| 0.020851 | 0.714799 | 0.038 | 0.019 | 1 | 3 | Rpgrip1 | 0.019 | Old |
| 0.02087 | 0.053408 | 0.154 | 0.112 | 1 | 3 | Pex2 | 0.042 | Old |
| 0.021029 | 0.348737 | 0.127 | 0.09 | 1 | 3 | Srm | 0.037 | Old |
| 0.021048 | 0.805901 | 0.031 | 0.014 | 1 | 3 | Gm4673 | 0.017 | Old |
| 0.021083 | 0.437558 | 0.068 | 0.042 | 1 | 3 | Cstf2 | 0.026 | Old |
| 0.021088 | 0.662276 | 0.035 | 0.016 | 1 | 3 | Pdss1 | 0.019 | Old |
| 0.02109 | 0.160339 | 0.79 | 0.723 | 1 | 3 | Sfpq | 0.067 | Old |
| 0.02116 | 0.090886 | 0.594 | 0.533 | 1 | 3 | Ubash3b | 0.061 | Old |
| 0.02124 | 0.074293 | 0.294 | 0.234 | 1 | 3 | Yme1l1 | 0.06 | Old |
| 0.021285 | 0.329319 | 0.201 | 0.154 | 1 | 3 | Preb | 0.047 | Old |
| 0.021304 | 0.162591 | 0.339 | 0.277 | 1 | 3 | Rap2a | 0.062 | Old |
| 0.02132 | 0.283003 | 0.461 | 0.383 | 1 | 3 | Ids | 0.078 | Old |
| 0.021396 | 0.098691 | 0.845 | 0.788 | 1 | 3 | Calr | 0.057 | Old |
| 0.021465 | 0.184959 | 0.361 | 0.289 | 1 | 3 | Commd7 | 0.072 | Old |
| 0.021469 | 0.273065 | 0.198 | 0.153 | 1 | 3 | Cluh | 0.045 | Old |
| 0.021554 | 0.176278 | 0.271 | 0.218 | 1 | 3 | Tmbim4 | 0.053 | Old |
| 0.021573 | 0.161216 | 0.33 | 0.273 | 1 | 3 | Gnl1 | 0.057 | Old |
| 0.021606 | 0.746151 | 0.038 | 0.019 | 1 | 3 | Rabl3 | 0.019 | Old |
| 0.02168 | 0.260688 | 0.14 | 0.102 | 1 | 3 | Mrpl19 | 0.038 | Old |
| 0.021909 | 0.19383 | 0.201 | 0.155 | 1 | 3 | Slc35a3 | 0.046 | Old |
| 0.022072 | 0.256829 | 0.294 | 0.237 | 1 | 3 | Dnajc13 | 0.057 | Old |
| 0.022076 | 0.236438 | 0.059 | 0.034 | 1 | 3 | Pml | 0.025 | Old |
| 0.022142 | 0.222684 | 0.188 | 0.14 | 1 | 3 | Plekhf2 | 0.048 | Old |
| 0.022208 | 0.16646 | 0.545 | 0.47 | 1 | 3 | Spcs3 | 0.075 | Old |
| 0.022242 | 0.292605 | 0.215 | 0.168 | 1 | 3 | Zc3h18 | 0.047 | Old |
| 0.022246 | 0.325597 | 0.127 | 0.091 | 1 | 3 | Trip4 | 0.036 | Old |
| 0.022409 | 0.069207 | 0.348 | 0.284 | 1 | 3 | Irf4 | 0.064 | Old |
| 0.022513 | 0.086666 | 0.165 | 0.122 | 1 | 3 | Bop1 | 0.043 | Old |
| 0.022577 | 0.282579 | 0.207 | 0.16 | 1 | 3 | Sar1b | 0.047 | Old |
| 0.022593 | 0.117092 | 0.843 | 0.761 | 1 | 3 | Tnfaip3 | 0.082 | Old |
| 0.022609 | 0.15178 | 0.686 | 0.615 | 1 | 3 | Hdgf | 0.071 | Old |
| 0.022654 | 0.279494 | 0.131 | 0.095 | 1 | 3 | Slc35f5 | 0.036 | Old |
| 0.022665 | 0.248854 | 0.26 | 0.21 | 1 | 3 | Frg1 | 0.05 | Old |
| 0.022758 | 0.115124 | 0.571 | 0.491 | 1 | 3 | Rras2 | 0.08 | Old |

| | | | | | | | | |
|---|---|---|---|---|---|---|---|---|
| 0.022795 | 0.319504 | 0.213 | 0.169 | 1 | 3 | Nudcd2 | 0.044 | Old |
| 0.022863 | 0.122688 | 0.97 | 0.952 | 1 | 3 | H2-K1 | 0.018 | Old |
| 0.022875 | 0.218823 | 0.13 | 0.092 | 1 | 3 | Cd200r4 | 0.038 | Old |
| 0.022947 | 0.845291 | 0.07 | 0.044 | 1 | 3 | AC127341. | 0.026 | Old |
| 0.023094 | 0.135814 | 0.557 | 0.484 | 1 | 3 | Sf1 | 0.073 | Old |
| 0.023103 | 0.666166 | 0.025 | 0.01 | 1 | 3 | Slc4a11 | 0.015 | Old |
| 0.02311 | 0.090925 | 0.492 | 0.419 | 1 | 3 | Wdr83os | 0.073 | Old |
| 0.023113 | 0.157447 | 0.296 | 0.244 | 1 | 3 | Secisbp2l | 0.052 | Old |
| 0.023141 | 0.348509 | 0.217 | 0.173 | 1 | 3 | Cdk4 | 0.044 | Old |
| 0.023159 | 0.277389 | 0.139 | 0.101 | 1 | 3 | Pik3cg | 0.038 | Old |
| 0.023166 | 0.303056 | 0.315 | 0.261 | 1 | 3 | Ppp6c | 0.054 | Old |
| 0.023174 | 0.485171 | 0.109 | 0.076 | 1 | 3 | Heatr6 | 0.033 | Old |
| 0.023222 | 0.215312 | 0.173 | 0.13 | 1 | 3 | Zmym2 | 0.043 | Old |
| 0.023289 | 0.774754 | 0.036 | 0.018 | 1 | 3 | Fancg | 0.018 | Old |
| 0.023303 | 0.3543 | 0.168 | 0.128 | 1 | 3 | Zbtb7b | 0.04 | Old |
| 0.023337 | 0.102492 | 0.375 | 0.306 | 1 | 3 | Psmc1 | 0.069 | Old |
| 0.023398 | 0.166149 | 0.468 | 0.404 | 1 | 3 | Ghitm | 0.064 | Old |
| 0.023424 | 0.171621 | 0.173 | 0.131 | 1 | 3 | Otud6b | 0.042 | Old |
| 0.023458 | 0.550015 | 0.105 | 0.073 | 1 | 3 | Ankrd27 | 0.032 | Old |
| 0.023523 | 0.276993 | 0.133 | 0.096 | 1 | 3 | Irak3 | 0.037 | Old |
| 0.023622 | 0.468893 | 0.06 | 0.035 | 1 | 3 | Piwil2 | 0.025 | Old |
| 0.023736 | 0.157274 | 0.25 | 0.198 | 1 | 3 | Emc2 | 0.052 | Old |
| 0.023838 | 1.241021 | 0.01 | 0.001 | 1 | 3 | Pdzd4 | 0.009 | Old |
| 0.02389 | 0.164625 | 0.481 | 0.417 | 1 | 3 | Mrpl20 | 0.064 | Old |
| 0.023999 | 0.740194 | 0.051 | 0.029 | 1 | 3 | Entpd6 | 0.022 | Old |
| 0.024003 | 0.181442 | 0.272 | 0.218 | 1 | 3 | Adprh | 0.054 | Old |
| 0.024018 | 0.19958 | 0.226 | 0.179 | 1 | 3 | Vps13a | 0.047 | Old |
| 0.024021 | 1.013068 | 0.017 | 0.005 | 1 | 3 | Chchd10 | 0.012 | Old |
| 0.024026 | 0.983487 | 0.053 | 0.03 | 1 | 3 | Phlpp1 | 0.023 | Old |
| 0.024086 | 0.377718 | 0.129 | 0.093 | 1 | 3 | Ppid | 0.036 | Old |
| 0.024101 | 0.254598 | 0.176 | 0.133 | 1 | 3 | Bag6 | 0.043 | Old |
| 0.024217 | 0.211684 | 0.176 | 0.133 | 1 | 3 | Wdr48 | 0.043 | Old |
| 0.024249 | 0.180892 | 0.01 | 0.001 | 1 | 3 | C6 | 0.009 | Old |
| 0.024314 | 1.063651 | 0.017 | 0.005 | 1 | 3 | Gm38251 | 0.012 | Old |
| 0.024354 | 0.253447 | 0.288 | 0.239 | 1 | 3 | Rnf6 | 0.049 | Old |
| 0.024461 | 0.843545 | 0.017 | 0.005 | 1 | 3 | Slc52a3 | 0.012 | Old |
| 0.024498 | 1.282516 | 0.017 | 0.005 | 1 | 3 | Rgs5 | 0.012 | Old |
| 0.024529 | 0.939131 | 0.062 | 0.038 | 1 | 3 | Ccdc112 | 0.024 | Old |
| 0.024596 | 0.795134 | 0.029 | 0.013 | 1 | 3 | Zfp933 | 0.016 | Old |
| 0.024602 | 0.572914 | 0.056 | 0.033 | 1 | 3 | Lyrm4 | 0.023 | Old |
| 0.024714 | 0.372738 | 0.149 | 0.111 | 1 | 3 | Cirbp | 0.038 | Old |
| 0.02473 | 0.142517 | 0.471 | 0.394 | 1 | 3 | Notch2 | 0.077 | Old |
| 0.024807 | 0.197782 | 0.587 | 0.521 | 1 | 3 | Hpcal1 | 0.066 | Old |
| 0.02489 | 0.168415 | 0.614 | 0.551 | 1 | 3 | Ube2d2a | 0.063 | Old |
| 0.025012 | 0.331715 | 0.118 | 0.083 | 1 | 3 | Mrps30 | 0.035 | Old |

| | | | | | | | | |
|---|---|---|---|---|---|---|---|---|
| 0.02506 | 0.698635 | 0.044 | 0.024 | 1 | 3 | Cdc23 | 0.02 | Old |
| 0.025189 | 0.369055 | 0.097 | 0.066 | 1 | 3 | Aifm2 | 0.031 | Old |
| 0.025209 | 0.445114 | 0.133 | 0.096 | 1 | 3 | Prps1 | 0.037 | Old |
| 0.025259 | 0.1499 | 0.29 | 0.234 | 1 | 3 | Sppl3 | 0.056 | Old |
| 0.025272 | 0.25701 | 0.243 | 0.192 | 1 | 3 | Slc25a51 | 0.051 | Old |
| 0.025338 | 0.366605 | 0.103 | 0.071 | 1 | 3 | Inpp5a | 0.032 | Old |
| 0.025363 | 0.203496 | 0.017 | 0.005 | 1 | 3 | Nemp2 | 0.012 | Old |
| 0.025386 | 0.353597 | 0.179 | 0.138 | 1 | 3 | Dusp2 | 0.041 | Old |
| 0.025469 | 0.208065 | 0.252 | 0.201 | 1 | 3 | Sh3pxd2b | 0.051 | Old |
| 0.025519 | 0.217946 | 0.217 | 0.17 | 1 | 3 | Lars | 0.047 | Old |
| 0.025547 | 0.287638 | 0.262 | 0.212 | 1 | 3 | Ccdc88b | 0.05 | Old |
| 0.025614 | 0.24883 | 0.066 | 0.04 | 1 | 3 | Pstk | 0.026 | Old |
| 0.025658 | 1.387854 | 0.014 | 0.004 | 1 | 3 | Arap2 | 0.01 | Old |
| 0.025783 | 0.21619 | 0.229 | 0.177 | 1 | 3 | Stim2 | 0.052 | Old |
| 0.025797 | 0.327022 | 0.036 | 0.018 | 1 | 3 | Nle1 | 0.018 | Old |
| 0.025877 | 0.355502 | 0.205 | 0.163 | 1 | 3 | Ccz1 | 0.042 | Old |
| 0.025888 | 0.23976 | 0.136 | 0.1 | 1 | 3 | Brd9 | 0.036 | Old |
| 0.025988 | 0.087909 | 0.706 | 0.621 | 1 | 3 | Rab14 | 0.085 | Old |
| 0.02604 | 0.928103 | 0.014 | 0.004 | 1 | 3 | Nutf2 | 0.01 | Old |
| 0.026063 | 0.221065 | 0.159 | 0.119 | 1 | 3 | Sympk | 0.04 | Old |
| 0.026076 | 0.144578 | 0.319 | 0.263 | 1 | 3 | Aprt | 0.056 | Old |
| 0.026076 | 1.303427 | 0.026 | 0.011 | 1 | 3 | Il18rap | 0.015 | Old |
| 0.026127 | 0.106188 | 0.231 | 0.179 | 1 | 3 | Zdhhc3 | 0.052 | Old |
| 0.026154 | 0.39353 | 0.1 | 0.069 | 1 | 3 | Mre11a | 0.031 | Old |
| 0.026168 | 0.899455 | 0.014 | 0.004 | 1 | 3 | Gm15853 | 0.01 | Old |
| 0.026174 | 0.53955 | 0.131 | 0.097 | 1 | 3 | Ifi204 | 0.034 | Old |
| 0.026211 | 1.089128 | 0.014 | 0.004 | 1 | 3 | Ttyh2 | 0.01 | Old |
| 0.026226 | 0.106102 | 0.821 | 0.75 | 1 | 3 | Eif3a | 0.071 | Old |
| 0.026296 | 0.213658 | 0.26 | 0.21 | 1 | 3 | 9030624J0 | 0.05 | Old |
| 0.026309 | 0.212136 | 0.437 | 0.367 | 1 | 3 | Cd2ap | 0.07 | Old |
| 0.026313 | 0.481749 | 0.149 | 0.112 | 1 | 3 | Nol11 | 0.037 | Old |
| 0.026321 | 0.40116 | 0.056 | 0.033 | 1 | 3 | Tapt1 | 0.023 | Old |
| 0.026331 | 0.303689 | 0.14 | 0.102 | 1 | 3 | Pla2g4a | 0.038 | Old |
| 0.026349 | 0.174379 | 0.483 | 0.42 | 1 | 3 | Dync1i2 | 0.063 | Old |
| 0.026376 | 0.025703 | 0.179 | 0.133 | 1 | 3 | Get4 | 0.046 | Old |
| 0.026432 | 0.2415 | 0.136 | 0.098 | 1 | 3 | Usp50 | 0.038 | Old |
| 0.026474 | 0.23398 | 0.204 | 0.159 | 1 | 3 | Manba | 0.045 | Old |
| 0.026567 | 0.127633 | 0.272 | 0.215 | 1 | 3 | Fam172a | 0.057 | Old |
| 0.026588 | 0.22266 | 0.099 | 0.067 | 1 | 3 | Fam98a | 0.032 | Old |
| 0.026594 | 0.815215 | 0.012 | 0.003 | 1 | 3 | 3222401L1 | 0.009 | Old |
| 0.026644 | 0.151851 | 0.228 | 0.181 | 1 | 3 | Actl6a | 0.047 | Old |
| 0.026668 | 0.024508 | 0.244 | 0.189 | 1 | 3 | Stat5b | 0.055 | Old |
| 0.026739 | 0.101243 | 0.387 | 0.319 | 1 | 3 | Psmd1 | 0.068 | Old |
| 0.026858 | 0.200342 | 0.527 | 0.457 | 1 | 3 | Gm21188 | 0.07 | Old |
| 0.02686 | 0.215606 | 0.119 | 0.086 | 1 | 3 | Poglut1 | 0.033 | Old |

| | | | | | | | | |
|---|---|---|---|---|---|---|---|---|
| 0.026862 | 0.263172 | 0.562 | 0.496 | 1 | 3 | Dock10 | 0.066 | Old |
| 0.026971 | 0.143912 | 0.553 | 0.485 | 1 | 3 | Fam120a | 0.068 | Old |
| 0.02699 | 0.160577 | 0.588 | 0.51 | 1 | 3 | Selenot | 0.078 | Old |
| 0.027134 | 0.136264 | 0.87 | 0.803 | 1 | 3 | Ctsc | 0.067 | Old |
| 0.027277 | 0.138813 | 0.751 | 0.682 | 1 | 3 | Zfand5 | 0.069 | Old |
| 0.027279 | 0.34705 | 0.295 | 0.245 | 1 | 3 | Zfp148 | 0.05 | Old |
| 0.027285 | 0.363238 | 0.179 | 0.138 | 1 | 3 | Tmem131 | 0.041 | Old |
| 0.027333 | 0.094671 | 0.257 | 0.206 | 1 | 3 | Hpf1 | 0.051 | Old |
| 0.027362 | 0.242943 | 0.376 | 0.319 | 1 | 3 | Lfng | 0.057 | Old |
| 0.027371 | 0.142023 | 0.397 | 0.337 | 1 | 3 | Mia3 | 0.06 | Old |
| 0.027408 | 0.26104 | 0.117 | 0.083 | 1 | 3 | Vhl | 0.034 | Old |
| 0.027424 | 0.374734 | 0.084 | 0.056 | 1 | 3 | Ttll12 | 0.028 | Old |
| 0.027444 | 0.344678 | 0.105 | 0.074 | 1 | 3 | Fbxo6 | 0.031 | Old |
| 0.027457 | 0.155087 | 0.447 | 0.378 | 1 | 3 | Tbl1xr1 | 0.069 | Old |
| 0.027633 | 0.22159 | 0.417 | 0.35 | 1 | 3 | Irak1 | 0.067 | Old |
| 0.027651 | 0.025876 | 0.333 | 0.27 | 1 | 3 | Slc26a2 | 0.063 | Old |
| 0.027726 | 0.184434 | 0.314 | 0.258 | 1 | 3 | Ptpn22 | 0.056 | Old |
| 0.027907 | 0.494638 | 0.096 | 0.066 | 1 | 3 | Ddx58 | 0.03 | Old |
| 0.027916 | 0.141477 | 0.484 | 0.405 | 1 | 3 | Zfp91 | 0.079 | Old |
| 0.02794 | 0.791499 | 0.037 | 0.019 | 1 | 3 | Gm12589 | 0.018 | Old |
| 0.02804 | 0.252462 | 0.234 | 0.184 | 1 | 3 | Mospd2 | 0.05 | Old |
| 0.028095 | 0.206797 | 0.294 | 0.242 | 1 | 3 | U2af1 | 0.052 | Old |
| 0.028119 | 0.400178 | 0.118 | 0.085 | 1 | 3 | Rtca | 0.033 | Old |
| 0.028355 | 0.174697 | 0.335 | 0.278 | 1 | 3 | Mrpl57 | 0.057 | Old |
| 0.028377 | 0.365906 | 0.098 | 0.068 | 1 | 3 | Wdr11 | 0.03 | Old |
| 0.028383 | 0.280852 | 0.14 | 0.102 | 1 | 3 | Zbtb18 | 0.038 | Old |
| 0.028464 | 0.324187 | 0.174 | 0.135 | 1 | 3 | Ccdc186 | 0.039 | Old |
| 0.028545 | 0.373885 | 0.153 | 0.115 | 1 | 3 | Rnf14 | 0.038 | Old |
| 0.028617 | 0.512822 | 0.044 | 0.024 | 1 | 3 | Nkrf | 0.02 | Old |
| 0.028695 | 0.180233 | 0.563 | 0.499 | 1 | 3 | Rilpl2 | 0.064 | Old |
| 0.028774 | 0.372163 | 0.171 | 0.131 | 1 | 3 | Rbm18 | 0.04 | Old |
| 0.028835 | 0.250487 | 0.086 | 0.057 | 1 | 3 | Mrps34 | 0.029 | Old |
| 0.028891 | 0.202798 | 0.177 | 0.136 | 1 | 3 | Dhrs1 | 0.041 | Old |
| 0.028919 | 0.664638 | 0.059 | 0.035 | 1 | 3 | Numb | 0.024 | Old |
| 0.029003 | 0.174976 | 0.522 | 0.458 | 1 | 3 | Ptges3 | 0.064 | Old |
| 0.029081 | 0.078133 | 0.458 | 0.388 | 1 | 3 | Aebp2 | 0.07 | Old |
| 0.029104 | 0.145816 | 0.458 | 0.39 | 1 | 3 | Arap1 | 0.068 | Old |
| 0.029171 | 0.133384 | 0.556 | 0.486 | 1 | 3 | Ppp3r1 | 0.07 | Old |
| 0.029176 | 0.116812 | 0.349 | 0.287 | 1 | 3 | Taf1d | 0.062 | Old |
| 0.029229 | 0.331579 | 0.141 | 0.105 | 1 | 3 | Slc39a6 | 0.036 | Old |
| 0.029264 | 0.242879 | 0.3 | 0.246 | 1 | 3 | Taf15 | 0.054 | Old |
| 0.029299 | 0.488552 | 0.154 | 0.117 | 1 | 3 | Snrnp25 | 0.037 | Old |
| 0.029325 | 0.674222 | 0.03 | 0.014 | 1 | 3 | Mov10 | 0.016 | Old |
| 0.029342 | 0.031507 | 0.033 | 0.016 | 1 | 3 | Twsg1 | 0.017 | Old |
| 0.029423 | 0.041568 | 0.215 | 0.168 | 1 | 3 | Ago3 | 0.047 | Old |

| | | | | | | | | |
|---|---|---|---|---|---|---|---|---|
| 0.029425 | 0.292492 | 0.19 | 0.146 | 1 | 3 | Tbc1d1 | 0.044 | Old |
| 0.029436 | 0.146733 | 0.227 | 0.178 | 1 | 3 | Mtfr1l | 0.049 | Old |
| 0.029446 | 0.903223 | 0.02 | 0.008 | 1 | 3 | Cmtr1 | 0.012 | Old |
| 0.029457 | 0.558347 | 0.033 | 0.016 | 1 | 3 | Ppp2r3a | 0.017 | Old |
| 0.029475 | 0.584923 | 0.084 | 0.056 | 1 | 3 | Bcl2 | 0.028 | Old |
| 0.029486 | 0.799921 | 0.055 | 0.033 | 1 | 3 | Nlrc5 | 0.022 | Old |
| 0.0295 | 0.223521 | 0.223 | 0.178 | 1 | 3 | Mettl9 | 0.045 | Old |
| 0.029551 | 0.112932 | 0.816 | 0.76 | 1 | 3 | Cox6a1 | 0.056 | Old |
| 0.02958 | 0.309216 | 0.278 | 0.23 | 1 | 3 | Trim28 | 0.048 | Old |
| 0.029604 | 0.250202 | 0.454 | 0.407 | 1 | 3 | Il1rl1 | 0.047 | Old |
| 0.029649 | 0.188041 | 0.274 | 0.221 | 1 | 3 | Mrpl17 | 0.053 | Old |
| 0.029702 | 0.43467 | 0.122 | 0.088 | 1 | 3 | Cdca4 | 0.034 | Old |
| 0.029776 | 0.143102 | 0.473 | 0.404 | 1 | 3 | Tcea1 | 0.069 | Old |
| 0.029782 | 0.727522 | 0.081 | 0.054 | 1 | 3 | Nrm | 0.027 | Old |
| 0.029815 | 0.448199 | 0.096 | 0.066 | 1 | 3 | Tmem260 | 0.03 | Old |
| 0.02982 | 0.394991 | 0.09 | 0.061 | 1 | 3 | Rrp36 | 0.029 | Old |
| 0.029865 | 0.345927 | 0.16 | 0.12 | 1 | 3 | Tusc1 | 0.04 | Old |
| 0.029906 | 0.311058 | 0.131 | 0.096 | 1 | 3 | Dubr | 0.035 | Old |
| 0.030137 | 0.180447 | 0.313 | 0.258 | 1 | 3 | Gm6377 | 0.055 | Old |
| 0.030172 | 0.222355 | 0.211 | 0.169 | 1 | 3 | Uap1l1 | 0.042 | Old |
| 0.030203 | 0.053449 | 0.243 | 0.196 | 1 | 3 | Ubp1 | 0.047 | Old |
| 0.030205 | 0.365422 | 0.094 | 0.064 | 1 | 3 | Efl1 | 0.03 | Old |
| 0.030248 | 0.069343 | 0.295 | 0.235 | 1 | 3 | Avpi1 | 0.06 | Old |
| 0.03038 | 0.369592 | 0.244 | 0.201 | 1 | 3 | Nceh1 | 0.043 | Old |
| 0.030583 | 0.407271 | 0.2 | 0.16 | 1 | 3 | Exosc3 | 0.04 | Old |
| 0.030628 | 0.167788 | 0.137 | 0.101 | 1 | 3 | Dnajc11 | 0.036 | Old |
| 0.030647 | 0.264765 | 0.165 | 0.125 | 1 | 3 | Gtf2f1 | 0.04 | Old |
| 0.030729 | 0.504294 | 0.131 | 0.098 | 1 | 3 | Eipr1 | 0.033 | Old |
| 0.030756 | 0.288497 | 0.226 | 0.183 | 1 | 3 | Sec22b | 0.043 | Old |
| 0.030963 | 0.204712 | 0.173 | 0.131 | 1 | 3 | Gadd45g | 0.042 | Old |
| 0.031002 | 0.558292 | 0.05 | 0.029 | 1 | 3 | Mettl26 | 0.021 | Old |
| 0.031032 | 0.117766 | 0.166 | 0.125 | 1 | 3 | Kcnk13 | 0.041 | Old |
| 0.031035 | 0.137145 | 0.202 | 0.157 | 1 | 3 | Anapc1 | 0.045 | Old |
| 0.031059 | 0.172066 | 0.22 | 0.173 | 1 | 3 | Sptlc1 | 0.047 | Old |
| 0.0311 | 0.455761 | 0.085 | 0.057 | 1 | 3 | Pccb | 0.028 | Old |
| 0.0311 | 0.350703 | 0.226 | 0.178 | 1 | 3 | Rnmt | 0.048 | Old |
| 0.031133 | 0.119412 | 0.904 | 0.864 | 1 | 3 | Atp5l | 0.04 | Old |
| 0.03115 | 0.355509 | 0.133 | 0.097 | 1 | 3 | Mrpl38 | 0.036 | Old |
| 0.031154 | 0.18617 | 0.266 | 0.212 | 1 | 3 | Tmem165 | 0.054 | Old |
| 0.031227 | 0.109776 | 0.157 | 0.116 | 1 | 3 | Eif2b5 | 0.041 | Old |
| 0.031386 | 0.075956 | 0.245 | 0.193 | 1 | 3 | Psmd12 | 0.052 | Old |
| 0.0315 | 0.864474 | 0.056 | 0.034 | 1 | 3 | Pde4dip | 0.022 | Old |
| 0.031547 | 0.22831 | 0.382 | 0.328 | 1 | 3 | Srp72 | 0.054 | Old |
| 0.03157 | 0.198364 | 0.275 | 0.223 | 1 | 3 | Parl | 0.052 | Old |
| 0.031619 | 0.121623 | 0.307 | 0.251 | 1 | 3 | Gga1 | 0.056 | Old |

| | | | | | | | | |
|---|---|---|---|---|---|---|---|---|
| 0.031732 | 0.287965 | 0.121 | 0.087 | 1 | 3 | Copg2 | 0.034 | Old |
| 0.03178 | 0.958964 | 0.047 | 0.027 | 1 | 3 | Tbc1d24 | 0.02 | Old |
| 0.031796 | 0.291273 | 0.136 | 0.1 | 1 | 3 | Smim4 | 0.036 | Old |
| 0.031817 | 0.271453 | 0.384 | 0.33 | 1 | 3 | Eif1ax | 0.054 | Old |
| 0.031834 | 0.18111 | 0.221 | 0.174 | 1 | 3 | Nipsnap3b | 0.047 | Old |
| 0.031891 | 0.117602 | 0.646 | 0.596 | 1 | 3 | Adam8 | 0.05 | Old |
| 0.031944 | 0.355093 | 0.142 | 0.106 | 1 | 3 | Aph1b | 0.036 | Old |
| 0.031954 | 0.626875 | 0.092 | 0.063 | 1 | 3 | Smcr8 | 0.029 | Old |
| 0.031994 | 0.149977 | 0.232 | 0.182 | 1 | 3 | Yrdc | 0.05 | Old |
| 0.031999 | 0.238551 | 0.235 | 0.188 | 1 | 3 | Btbd7 | 0.047 | Old |
| 0.032054 | 0.108334 | 0.297 | 0.245 | 1 | 3 | Alkbh5 | 0.052 | Old |
| 0.032077 | 0.069395 | 0.934 | 0.912 | 1 | 3 | Ctsz | 0.022 | Old |
| 0.03209 | 0.235599 | 0.3 | 0.247 | 1 | 3 | Otub1 | 0.053 | Old |
| 0.032169 | 0.454205 | 0.086 | 0.058 | 1 | 3 | Prps2 | 0.028 | Old |
| 0.032221 | 0.061697 | 0.349 | 0.287 | 1 | 3 | Polr2k | 0.062 | Old |
| 0.032302 | 0.77329 | 0.043 | 0.024 | 1 | 3 | Tgds | 0.019 | Old |
| 0.032322 | 0.497697 | 0.081 | 0.054 | 1 | 3 | Alg8 | 0.027 | Old |
| 0.032421 | 0.297344 | 0.122 | 0.09 | 1 | 3 | Mfn2 | 0.032 | Old |
| 0.032561 | 0.274928 | 0.284 | 0.235 | 1 | 3 | BC031181 | 0.049 | Old |
| 0.032666 | 0.795017 | 0.024 | 0.01 | 1 | 3 | BC024978 | 0.014 | Old |
| 0.032757 | 0.1865 | 0.388 | 0.335 | 1 | 3 | Ak2 | 0.053 | Old |
| 0.032777 | 0.517154 | 0.067 | 0.043 | 1 | 3 | Eif2b3 | 0.024 | Old |
| 0.03286 | 0.084436 | 0.405 | 0.345 | 1 | 3 | Usp34 | 0.06 | Old |
| 0.033011 | 0.753606 | 0.024 | 0.01 | 1 | 3 | B3gntl1 | 0.014 | Old |
| 0.033095 | 0.335531 | 0.295 | 0.245 | 1 | 3 | Azi2 | 0.05 | Old |
| 0.033152 | 0.189193 | 0.448 | 0.386 | 1 | 3 | Snrpd3 | 0.062 | Old |
| 0.033228 | 0.238059 | 0.343 | 0.285 | 1 | 3 | Vamp3 | 0.058 | Old |
| 0.033374 | 0.912914 | 0.048 | 0.028 | 1 | 3 | Mx1 | 0.02 | Old |
| 0.033383 | 0.108002 | 0.861 | 0.822 | 1 | 3 | Mob1a | 0.039 | Old |
| 0.03347 | 0.433496 | 0.104 | 0.073 | 1 | 3 | Zfp277 | 0.031 | Old |
| 0.033486 | 0.941587 | 0.073 | 0.048 | 1 | 3 | Polr3d | 0.025 | Old |
| 0.03357 | 0.187532 | 0.235 | 0.188 | 1 | 3 | Ebna1bp2 | 0.047 | Old |
| 0.033615 | 0.161351 | 0.341 | 0.287 | 1 | 3 | Nhp2 | 0.054 | Old |
| 0.033665 | 1.020307 | 0.018 | 0.006 | 1 | 3 | Zmat1 | 0.012 | Old |
| 0.033812 | 0.919723 | 0.027 | 0.013 | 1 | 3 | Bpgm | 0.014 | Old |
| 0.034036 | 0.717393 | 0.018 | 0.006 | 1 | 3 | Mboat1 | 0.012 | Old |
| 0.034178 | 0.413476 | 0.081 | 0.054 | 1 | 3 | Tbcd | 0.027 | Old |
| 0.034232 | 0.396292 | 0.082 | 0.056 | 1 | 3 | Rnf40 | 0.026 | Old |
| 0.03427 | 0.128553 | 0.303 | 0.253 | 1 | 3 | Cbx1 | 0.05 | Old |
| 0.034343 | 0.211565 | 0.315 | 0.26 | 1 | 3 | 2610001J0 | 0.055 | Old |
| 0.034376 | 0.456136 | 0.07 | 0.045 | 1 | 3 | Xrcc4 | 0.025 | Old |
| 0.0344 | 0.237502 | 0.065 | 0.04 | 1 | 3 | Zdhhc12 | 0.025 | Old |
| 0.034421 | 0.070492 | 0.982 | 0.976 | 1 | 3 | Rpl38 | 0.006 | Old |
| 0.03444 | 0.372002 | 0.11 | 0.078 | 1 | 3 | Bloc1s4 | 0.032 | Old |
| 0.034446 | 0.483385 | 0.066 | 0.042 | 1 | 3 | Vcam1 | 0.024 | Old |

| | | | | | | | | |
|---|---|---|---|---|---|---|---|---|
| 0.034537 | 0.102795 | 0.283 | 0.23 | 1 | 3 | Mta2 | 0.053 | Old |
| 0.034537 | 0.082246 | 0.886 | 0.856 | 1 | 3 | Canx | 0.03 | Old |
| 0.034607 | 0.281949 | 0.146 | 0.109 | 1 | 3 | Sec16a | 0.037 | Old |
| 0.034689 | 0.091921 | 0.698 | 0.638 | 1 | 3 | Itgb2 | 0.06 | Old |
| 0.034723 | 0.216194 | 0.143 | 0.106 | 1 | 3 | Mcrs1 | 0.037 | Old |
| 0.03476 | 0.142735 | 0.423 | 0.36 | 1 | 3 | Cuta | 0.063 | Old |
| 0.034785 | 0.359977 | 0.143 | 0.109 | 1 | 3 | Ttc13 | 0.034 | Old |
| 0.03491 | 0.11061 | 0.595 | 0.515 | 1 | 3 | Syncrip | 0.08 | Old |
| 0.034966 | 0.185255 | 0.517 | 0.437 | 1 | 3 | Prpf40a | 0.08 | Old |
| 0.034969 | 0.126826 | 0.281 | 0.229 | 1 | 3 | Nsun2 | 0.052 | Old |
| 0.034986 | 0.157831 | 0.48 | 0.412 | 1 | 3 | Synj1 | 0.068 | Old |
| 0.034991 | 0.382356 | 0.123 | 0.091 | 1 | 3 | Ripk2 | 0.032 | Old |
| 0.035001 | 0.231685 | 0.214 | 0.169 | 1 | 3 | Galnt7 | 0.045 | Old |
| 0.035024 | 0.464059 | 0.081 | 0.054 | 1 | 3 | Tmem70 | 0.027 | Old |
| 0.035073 | 0.406082 | 0.165 | 0.126 | 1 | 3 | Rnf5 | 0.039 | Old |
| 0.035085 | 0.525483 | 0.057 | 0.035 | 1 | 3 | Isca1 | 0.022 | Old |
| 0.035146 | 0.529252 | 0.121 | 0.09 | 1 | 3 | Rab10os | 0.031 | Old |
| 0.035261 | 0.74492 | 0.039 | 0.021 | 1 | 3 | Dcakd | 0.018 | Old |
| 0.035422 | 0.454552 | 0.108 | 0.077 | 1 | 3 | 1600012H( | 0.031 | Old |
| 0.035587 | 0.169894 | 0.375 | 0.316 | 1 | 3 | Stt3b | 0.059 | Old |
| 0.035589 | 0.592066 | 0.043 | 0.024 | 1 | 3 | Mdga1 | 0.019 | Old |
| 0.03562 | 0.073282 | 0.888 | 0.855 | 1 | 3 | Gabarap | 0.033 | Old |
| 0.035638 | 0.445023 | 0.165 | 0.128 | 1 | 3 | Ms4a6c | 0.037 | Old |
| 0.035682 | 0.293268 | 0.235 | 0.189 | 1 | 3 | Ergic2 | 0.046 | Old |
| 0.035929 | 0.47991 | 0.076 | 0.051 | 1 | 3 | Champ1 | 0.025 | Old |
| 0.036022 | 0.38822 | 0.073 | 0.048 | 1 | 3 | Pdhx | 0.025 | Old |
| 0.036025 | 0.639558 | 0.065 | 0.042 | 1 | 3 | Eef1akmt2 | 0.023 | Old |
| 0.036039 | 0.126899 | 0.757 | 0.711 | 1 | 3 | Alcam | 0.046 | Old |
| 0.036143 | 0.286857 | 0.093 | 0.064 | 1 | 3 | Xab2 | 0.029 | Old |
| 0.036151 | 0.145718 | 0.446 | 0.383 | 1 | 3 | Emc7 | 0.063 | Old |
| 0.036177 | 0.14377 | 0.354 | 0.294 | 1 | 3 | Acsl5 | 0.06 | Old |
| 0.036344 | 0.331221 | 0.166 | 0.128 | 1 | 3 | Mif4gd | 0.038 | Old |
| 0.036354 | 0.15983 | 0.366 | 0.309 | 1 | 3 | Ythdf3 | 0.057 | Old |
| 0.036372 | 0.321339 | 0.217 | 0.177 | 1 | 3 | Rabl6 | 0.04 | Old |
| 0.036447 | 0.333592 | 0.096 | 0.067 | 1 | 3 | Nt5c3 | 0.029 | Old |
| 0.036556 | 0.07933 | 0.906 | 0.898 | 1 | 3 | Lamp1 | 0.008 | Old |
| 0.036615 | 0.232484 | 0.329 | 0.278 | 1 | 3 | Gfpt1 | 0.051 | Old |
| 0.036818 | 0.148432 | 0.321 | 0.271 | 1 | 3 | Rtf2 | 0.05 | Old |
| 0.036906 | 1.155647 | 0.016 | 0.005 | 1 | 3 | Atp8b5 | 0.011 | Old |
| 0.036914 | 0.005126 | 0.265 | 0.213 | 1 | 3 | Prpf39 | 0.052 | Old |
| 0.036962 | 0.305096 | 0.057 | 0.035 | 1 | 3 | Kctd18 | 0.022 | Old |
| 0.037075 | 0.231469 | 0.145 | 0.107 | 1 | 3 | Fbxo38 | 0.038 | Old |
| 0.037114 | 0.386238 | 0.127 | 0.093 | 1 | 3 | Nol8 | 0.034 | Old |
| 0.037123 | 1.20513 | 0.016 | 0.005 | 1 | 3 | Gnb1l | 0.011 | Old |
| 0.037174 | 0.384868 | 0.048 | 0.028 | 1 | 3 | Zfp729a | 0.02 | Old |

| | | | | | | | | |
|---|---|---|---|---|---|---|---|---|
| 0.037442 | 0.145705 | 0.809 | 0.76 | 1 | 3 | Atp6v1b2 | 0.049 | Old |
| 0.037471 | 0.172737 | 0.312 | 0.26 | 1 | 3 | Mrpl28 | 0.052 | Old |
| 0.037471 | 0.267161 | 0.25 | 0.206 | 1 | 3 | Ap1g1 | 0.044 | Old |
| 0.037563 | 0.10474 | 0.409 | 0.346 | 1 | 3 | Xrn2 | 0.063 | Old |
| 0.03764 | 0.457073 | 0.119 | 0.088 | 1 | 3 | Polr3gl | 0.031 | Old |
| 0.037718 | 0.212557 | 0.493 | 0.428 | 1 | 3 | Nono | 0.065 | Old |
| 0.037824 | 0.108048 | 0.153 | 0.115 | 1 | 3 | Exoc6b | 0.038 | Old |
| 0.037878 | 0.133532 | 0.544 | 0.475 | 1 | 3 | Atp6v0d1 | 0.069 | Old |
| 0.037881 | 0.071734 | 0.698 | 0.606 | 1 | 3 | Cnbp | 0.092 | Old |
| 0.037886 | 0.296989 | 0.186 | 0.148 | 1 | 3 | Lman2l | 0.038 | Old |
| 0.037896 | 0.08447 | 0.545 | 0.462 | 1 | 3 | Hivep2 | 0.083 | Old |
| 0.037903 | 0.433332 | 0.08 | 0.054 | 1 | 3 | Mcm6 | 0.026 | Old |
| 0.037923 | 0.130943 | 0.283 | 0.229 | 1 | 3 | Pdcl3 | 0.054 | Old |
| 0.037935 | 0.668191 | 0.065 | 0.042 | 1 | 3 | 2010111I0 | 0.023 | Old |
| 0.037988 | 0.186925 | 0.407 | 0.35 | 1 | 3 | Cct7 | 0.057 | Old |
| 0.03803 | 0.749461 | 0.016 | 0.005 | 1 | 3 | Car9 | 0.011 | Old |
| 0.038175 | 0.839306 | 0.032 | 0.016 | 1 | 3 | Mospd1 | 0.016 | Old |
| 0.038333 | 0.64668 | 0.022 | 0.009 | 1 | 3 | Uhrf1 | 0.013 | Old |
| 0.038359 | 0.299926 | 0.16 | 0.124 | 1 | 3 | AW112010 | 0.036 | Old |
| 0.038376 | 0.145422 | 0.315 | 0.265 | 1 | 3 | Dnajc15 | 0.05 | Old |
| 0.038383 | 0.21804 | 0.343 | 0.287 | 1 | 3 | Smim7 | 0.056 | Old |
| 0.038396 | 0.443735 | 0.103 | 0.073 | 1 | 3 | C4b | 0.03 | Old |
| 0.038544 | 0.10129 | 0.676 | 0.611 | 1 | 3 | Sh3glb1 | 0.065 | Old |
| 0.038559 | 0.157313 | 0.223 | 0.178 | 1 | 3 | Mrpl18 | 0.045 | Old |
| 0.038622 | 0.163774 | 0.18 | 0.14 | 1 | 3 | Rgl1 | 0.04 | Old |
| 0.038739 | 0.242978 | 0.232 | 0.188 | 1 | 3 | Adrm1 | 0.044 | Old |
| 0.038853 | 0.821499 | 0.041 | 0.023 | 1 | 3 | Polr1e | 0.018 | Old |
| 0.038953 | 0.196087 | 0.327 | 0.274 | 1 | 3 | Sirt2 | 0.053 | Old |
| 0.039013 | 0.253258 | 0.117 | 0.086 | 1 | 3 | Timm9 | 0.031 | Old |
| 0.039021 | 0.099839 | 0.124 | 0.09 | 1 | 3 | Zfyve16 | 0.034 | Old |
| 0.03909 | 0.172034 | 0.133 | 0.098 | 1 | 3 | Immp1l | 0.035 | Old |
| 0.0391 | 0.018061 | 0.178 | 0.139 | 1 | 3 | Ten1 | 0.039 | Old |
| 0.039113 | 0.093499 | 0.379 | 0.317 | 1 | 3 | Lmbrd1 | 0.062 | Old |
| 0.039139 | 0.219582 | 0.51 | 0.449 | 1 | 3 | Lasp1 | 0.061 | Old |
| 0.039164 | 0.269414 | 0.24 | 0.193 | 1 | 3 | Dcp2 | 0.047 | Old |
| 0.03925 | 0.148267 | 0.368 | 0.311 | 1 | 3 | Ppp1r2 | 0.057 | Old |
| 0.03931 | 1.221878 | 0.025 | 0.011 | 1 | 3 | Ifi211 | 0.014 | Old |
| 0.039311 | 0.356884 | 0.049 | 0.029 | 1 | 3 | Prim2 | 0.02 | Old |
| 0.039339 | 0.244353 | 0.123 | 0.091 | 1 | 3 | Sos1 | 0.032 | Old |
| 0.039357 | 0.042659 | 0.305 | 0.246 | 1 | 3 | Tsg101 | 0.059 | Old |
| 0.03941 | 0.542949 | 0.044 | 0.025 | 1 | 3 | Gtf3c5 | 0.019 | Old |
| 0.039416 | 0.133229 | 0.171 | 0.131 | 1 | 3 | Fam98c | 0.04 | Old |
| 0.039442 | 0.176912 | 0.233 | 0.187 | 1 | 3 | Gigyf2 | 0.046 | Old |
| 0.039569 | 0.49096 | 0.036 | 0.019 | 1 | 3 | Hs1bp3 | 0.017 | Old |
| 0.039683 | 0.235345 | 0.146 | 0.11 | 1 | 3 | Necap1 | 0.036 | Old |

| | | | | | | | | |
|---|---|---|---|---|---|---|---|---|
| 0.03969 | 0.240063 | 0.281 | 0.232 | 1 | 3 | Chd3 | 0.049 | Old |
| 0.039691 | 0.4157 | 0.125 | 0.093 | 1 | 3 | Itpa | 0.032 | Old |
| 0.039759 | 0.384795 | 0.192 | 0.153 | 1 | 3 | Sfxn1 | 0.039 | Old |
| 0.039785 | 0.195615 | 0.477 | 0.417 | 1 | 3 | Spcs2 | 0.06 | Old |
| 0.039786 | 0.823918 | 0.029 | 0.014 | 1 | 3 | C1qtnf6 | 0.015 | Old |
| 0.039939 | 0.170146 | 0.166 | 0.126 | 1 | 3 | Gm36161 | 0.04 | Old |
| 0.039999 | 0.871869 | 0.045 | 0.027 | 1 | 3 | Zfp346 | 0.018 | Old |
| 0.040016 | 0.328634 | 0.197 | 0.155 | 1 | 3 | Atrnl1 | 0.042 | Old |
| 0.040157 | 0.056672 | 0.342 | 0.285 | 1 | 3 | Spred1 | 0.057 | Old |
| 0.04022 | 0.355319 | 0.305 | 0.256 | 1 | 3 | Ube2m | 0.049 | Old |
| 0.04025 | 0.437542 | 0.14 | 0.106 | 1 | 3 | Gm17018 | 0.034 | Old |
| 0.040309 | 0.309794 | 0.188 | 0.146 | 1 | 3 | Oxr1 | 0.042 | Old |
| 0.040312 | 0.124816 | 0.748 | 0.701 | 1 | 3 | H2afy | 0.047 | Old |
| 0.040331 | 0.103164 | 0.916 | 0.895 | 1 | 3 | Arhgdib | 0.021 | Old |
| 0.04038 | 0.366656 | 0.063 | 0.04 | 1 | 3 | Zfp422 | 0.023 | Old |
| 0.04039 | 0.64095 | 0.079 | 0.053 | 1 | 3 | Fkbp5 | 0.026 | Old |
| 0.040433 | 0.484497 | 0.059 | 0.037 | 1 | 3 | Sgf29 | 0.022 | Old |
| 0.040473 | 0.29708 | 0.209 | 0.168 | 1 | 3 | Impad1 | 0.041 | Old |
| 0.040548 | 0.250414 | 0.196 | 0.154 | 1 | 3 | Polr3c | 0.042 | Old |
| 0.040582 | 0.182225 | 0.55 | 0.487 | 1 | 3 | Psme1 | 0.063 | Old |
| 0.040584 | 0.11319 | 0.136 | 0.102 | 1 | 3 | Apmap | 0.034 | Old |
| 0.040691 | 0.176172 | 0.637 | 0.583 | 1 | 3 | Glg1 | 0.054 | Old |
| 0.040748 | 0.203058 | 0.476 | 0.419 | 1 | 3 | Psma2 | 0.057 | Old |
| 0.04087 | 0.134268 | 0.395 | 0.332 | 1 | 3 | Cbfb | 0.063 | Old |
| 0.040995 | 0.157787 | 0.319 | 0.264 | 1 | 3 | Atp6v0a2 | 0.055 | Old |
| 0.041018 | 0.803852 | 0.066 | 0.043 | 1 | 3 | Zfp746 | 0.023 | Old |
| 0.04105 | 0.47311 | 0.167 | 0.131 | 1 | 3 | Ralgapa1 | 0.036 | Old |
| 0.041155 | 0.364752 | 0.073 | 0.048 | 1 | 3 | Fnip2 | 0.025 | Old |
| 0.04121 | 0.32904 | 0.097 | 0.068 | 1 | 3 | Banp | 0.029 | Old |
| 0.041337 | 0.113146 | 0.229 | 0.182 | 1 | 3 | Tlnrd1 | 0.047 | Old |
| 0.041377 | 0.313623 | 0.166 | 0.129 | 1 | 3 | Ipo8 | 0.037 | Old |
| 0.041378 | 0.363233 | 0.079 | 0.053 | 1 | 3 | Recql | 0.026 | Old |
| 0.041405 | 0.174021 | 0.351 | 0.297 | 1 | 3 | Lrrfip2 | 0.054 | Old |
| 0.041406 | 0.061573 | 0.499 | 0.413 | 1 | 3 | Map2k3 | 0.086 | Old |
| 0.041419 | 0.246432 | 0.201 | 0.159 | 1 | 3 | Fam78b | 0.042 | Old |
| 0.041442 | 0.220272 | 0.153 | 0.116 | 1 | 3 | Nagk | 0.037 | Old |
| 0.041506 | 0.181039 | 0.331 | 0.275 | 1 | 3 | Brd7 | 0.056 | Old |
| 0.041559 | 0.199649 | 0.271 | 0.221 | 1 | 3 | Dock1 | 0.05 | Old |
| 0.041615 | 0.203055 | 0.305 | 0.251 | 1 | 3 | Ccny | 0.054 | Old |
| 0.041627 | 0.059088 | 0.208 | 0.164 | 1 | 3 | Dync1li2 | 0.044 | Old |
| 0.041731 | 0.265572 | 0.084 | 0.057 | 1 | 3 | Mcm4 | 0.027 | Old |
| 0.041934 | 0.384647 | 0.217 | 0.178 | 1 | 3 | Spag7 | 0.039 | Old |
| 0.041988 | 0.2638 | 0.158 | 0.121 | 1 | 3 | Fam160b2 | 0.037 | Old |
| 0.042016 | 0.09561 | 0.496 | 0.423 | 1 | 3 | Cysltr1 | 0.073 | Old |
| 0.042056 | 0.099257 | 0.489 | 0.423 | 1 | 3 | Rad23b | 0.066 | Old |

| | | | | | | | | |
|---|---|---|---|---|---|---|---|---|
| 0.042095 | 0.185332 | 0.434 | 0.378 | 1 | 3 | Ddx21 | 0.056 | Old |
| 0.042115 | 0.235934 | 0.154 | 0.119 | 1 | 3 | Scd2 | 0.035 | Old |
| 0.042193 | 0.167647 | 0.385 | 0.322 | 1 | 3 | Rab35 | 0.063 | Old |
| 0.042238 | 0.398598 | 0.049 | 0.029 | 1 | 3 | Nprl2 | 0.02 | Old |
| 0.042331 | 0.111357 | 0.228 | 0.181 | 1 | 3 | Arnt | 0.047 | Old |
| 0.042652 | 0.01605 | 0.738 | 0.658 | 1 | 3 | Ppp2ca | 0.08 | Old |
| 0.042664 | 0.367266 | 0.041 | 0.023 | 1 | 3 | Fastkd1 | 0.018 | Old |
| 0.042723 | 0.45005 | 0.256 | 0.211 | 1 | 3 | Nampt | 0.045 | Old |
| 0.042732 | 1.105679 | 0.011 | 0.003 | 1 | 3 | Tfec | 0.008 | Old |
| 0.042806 | 0.404403 | 0.054 | 0.033 | 1 | 3 | Mrrf | 0.021 | Old |
| 0.042938 | 0.119323 | 0.693 | 0.625 | 1 | 3 | Cytip | 0.068 | Old |
| 0.042962 | 0.225704 | 0.186 | 0.148 | 1 | 3 | Nup54 | 0.038 | Old |
| 0.043111 | 0.152821 | 0.385 | 0.328 | 1 | 3 | Lamtor3 | 0.057 | Old |
| 0.043115 | 0.166769 | 0.143 | 0.107 | 1 | 3 | Tmem251 | 0.036 | Old |
| 0.04314 | 0.198821 | 0.176 | 0.138 | 1 | 3 | Trak2 | 0.038 | Old |
| 0.043157 | 0.337249 | 0.086 | 0.059 | 1 | 3 | Acap3 | 0.027 | Old |
| 0.043191 | 0.943307 | 0.011 | 0.003 | 1 | 3 | Myo10 | 0.008 | Old |
| 0.043199 | 0.257197 | 0.245 | 0.201 | 1 | 3 | Rps12-ps3 | 0.044 | Old |
| 0.043225 | 0.633552 | 0.045 | 0.027 | 1 | 3 | Mios | 0.018 | Old |
| 0.043241 | 0.417343 | 0.115 | 0.085 | 1 | 3 | Exosc9 | 0.03 | Old |
| 0.043259 | 0.809059 | 0.045 | 0.027 | 1 | 3 | D17H6S53I | 0.018 | Old |
| 0.043268 | 0.680233 | 0.011 | 0.003 | 1 | 3 | 1700030K0 | 0.008 | Old |
| 0.043268 | 0.941452 | 0.011 | 0.003 | 1 | 3 | Gm17435 | 0.008 | Old |
| 0.043428 | 0.385916 | 0.118 | 0.086 | 1 | 3 | Washc5 | 0.032 | Old |
| 0.043488 | 0.156478 | 0.222 | 0.178 | 1 | 3 | Scamp3 | 0.044 | Old |
| 0.043506 | 0.157611 | 0.289 | 0.24 | 1 | 3 | Emd | 0.049 | Old |
| 0.043584 | 0.091544 | 0.118 | 0.086 | 1 | 3 | Smim12 | 0.032 | Old |
| 0.043787 | 0.133321 | 0.306 | 0.254 | 1 | 3 | Ndufb2 | 0.052 | Old |
| 0.043807 | 0.116577 | 0.409 | 0.342 | 1 | 3 | Nxf1 | 0.067 | Old |
| 0.043809 | 0.647704 | 0.072 | 0.048 | 1 | 3 | Bpnt1 | 0.024 | Old |
| 0.043894 | 0.143665 | 0.41 | 0.359 | 1 | 3 | Ndufb6 | 0.051 | Old |
| 0.043924 | 0.238912 | 0.231 | 0.188 | 1 | 3 | Cdc26 | 0.043 | Old |
| 0.044044 | 0.448844 | 0.104 | 0.076 | 1 | 3 | Etv5 | 0.028 | Old |
| 0.044056 | 0.17329 | 0.556 | 0.484 | 1 | 3 | Atp13a3 | 0.072 | Old |
| 0.044102 | 0.622849 | 0.05 | 0.03 | 1 | 3 | Nbdy | 0.02 | Old |
| 0.044147 | 0.166873 | 0.361 | 0.307 | 1 | 3 | Smarcc1 | 0.054 | Old |
| 0.044195 | 1.037092 | 0.019 | 0.008 | 1 | 3 | Gm26601 | 0.011 | Old |
| 0.044203 | 0.115149 | 0.329 | 0.274 | 1 | 3 | Myd88 | 0.055 | Old |
| 0.04433 | 0.190243 | 0.559 | 0.513 | 1 | 3 | Snrpf | 0.046 | Old |
| 0.04453 | 0.810561 | 0.019 | 0.008 | 1 | 3 | Jchain | 0.011 | Old |
| 0.044537 | 0.295085 | 0.153 | 0.117 | 1 | 3 | Scamp1 | 0.036 | Old |
| 0.044564 | 0.388221 | 0.065 | 0.042 | 1 | 3 | Polr3e | 0.023 | Old |
| 0.04475 | 0.919463 | 0.033 | 0.018 | 1 | 3 | Gpx3 | 0.015 | Old |
| 0.044802 | 0.133179 | 0.781 | 0.735 | 1 | 3 | Fosb | 0.046 | Old |
| 0.044892 | 0.364013 | 0.119 | 0.088 | 1 | 3 | Pygl | 0.031 | Old |

| | | | | | | | | |
|---|---|---|---|---|---|---|---|---|
| 0.044919 | 0.161394 | 0.225 | 0.181 | 1 | 3 | 4933434E2 | 0.044 | Old |
| 0.045036 | 0.391564 | 0.165 | 0.13 | 1 | 3 | Paip1 | 0.035 | Old |
| 0.045317 | 0.261522 | 0.096 | 0.068 | 1 | 3 | Jpt2 | 0.028 | Old |
| 0.04547 | 0.320951 | 0.13 | 0.098 | 1 | 3 | Txnl4a | 0.032 | Old |
| 0.045662 | 0.110398 | 0.262 | 0.215 | 1 | 3 | Max | 0.047 | Old |
| 0.045667 | 0.320756 | 0.055 | 0.034 | 1 | 3 | Nxn | 0.021 | Old |
| 0.0457 | 0.272017 | 0.143 | 0.109 | 1 | 3 | Ube4a | 0.034 | Old |
| 0.045729 | 0.155485 | 0.799 | 0.735 | 1 | 3 | Sgms1 | 0.064 | Old |
| 0.045743 | 0.085328 | 0.686 | 0.598 | 1 | 3 | Oxct1 | 0.088 | Old |
| 0.046053 | 0.255604 | 0.284 | 0.237 | 1 | 3 | Psme3 | 0.047 | Old |
| 0.046077 | 0.161821 | 0.176 | 0.135 | 1 | 3 | Pes1 | 0.041 | Old |
| 0.046077 | 0.235898 | 0.183 | 0.145 | 1 | 3 | Rars | 0.038 | Old |
| 0.046148 | 0.238465 | 0.153 | 0.117 | 1 | 3 | Med21 | 0.036 | Old |
| 0.046247 | 0.320293 | 0.168 | 0.133 | 1 | 3 | Fem1b | 0.035 | Old |
| 0.046374 | 0.212572 | 0.479 | 0.419 | 1 | 3 | Tmem123 | 0.06 | Old |
| 0.046402 | 0.736636 | 0.115 | 0.086 | 1 | 3 | Clpx | 0.029 | Old |
| 0.046488 | 0.173214 | 0.257 | 0.205 | 1 | 3 | Anxa11 | 0.052 | Old |
| 0.046555 | 0.255624 | 0.176 | 0.139 | 1 | 3 | Helz | 0.037 | Old |
| 0.046636 | 0.274802 | 0.076 | 0.052 | 1 | 3 | Diexf | 0.024 | Old |
| 0.046786 | 0.376926 | 0.066 | 0.043 | 1 | 3 | Rnf157 | 0.023 | Old |
| 0.046831 | 0.093257 | 0.861 | 0.84 | 1 | 3 | Arhgdia | 0.021 | Old |
| 0.04685 | 0.296716 | 0.134 | 0.101 | 1 | 3 | Pggt1b | 0.033 | Old |
| 0.04687 | 0.064821 | 0.557 | 0.48 | 1 | 3 | Ptpra | 0.077 | Old |
| 0.046893 | 0.925864 | 0.063 | 0.042 | 1 | 3 | Nop2 | 0.021 | Old |
| 0.046925 | 0.261916 | 0.21 | 0.169 | 1 | 3 | Lsm7 | 0.041 | Old |
| 0.046944 | 0.14707 | 0.219 | 0.176 | 1 | 3 | Slc30a5 | 0.043 | Old |
| 0.046964 | 0.977512 | 0.038 | 0.021 | 1 | 3 | Ippk | 0.017 | Old |
| 0.047277 | 0.115313 | 0.432 | 0.367 | 1 | 3 | Gspt1 | 0.065 | Old |
| 0.047306 | 0.150701 | 0.454 | 0.396 | 1 | 3 | Psmb8 | 0.058 | Old |
| 0.047336 | 0.05711 | 0.228 | 0.184 | 1 | 3 | Slc25a4 | 0.044 | Old |
| 0.047364 | 0.095727 | 0.183 | 0.143 | 1 | 3 | Clns1a | 0.04 | Old |
| 0.047393 | 0.290017 | 0.092 | 0.064 | 1 | 3 | Rbm15b | 0.028 | Old |
| 0.047394 | 0.209753 | 0.357 | 0.304 | 1 | 3 | Nus1 | 0.053 | Old |
| 0.047475 | 0.232912 | 0.245 | 0.201 | 1 | 3 | Pdhb | 0.044 | Old |
| 0.047485 | 0.420266 | 0.159 | 0.124 | 1 | 3 | Sf3a2 | 0.035 | Old |
| 0.047536 | 0.181511 | 0.492 | 0.442 | 1 | 3 | Mapk1 | 0.05 | Old |
| 0.047595 | 0.199768 | 0.459 | 0.404 | 1 | 3 | Arid1a | 0.055 | Old |
| 0.047684 | 0.128777 | 0.763 | 0.71 | 1 | 3 | Tuba1b | 0.053 | Old |
| 0.047847 | 0.789773 | 0.094 | 0.068 | 1 | 3 | Aatf | 0.026 | Old |
| 0.04793 | 0.093929 | 0.318 | 0.261 | 1 | 3 | Lsm14a | 0.057 | Old |
| 0.048026 | 0.044735 | 0.949 | 0.96 | 1 | 3 | Rbpj | -0.011 | Old |
| 0.04811 | 0.15826 | 0.042 | 0.024 | 1 | 3 | Acp5 | 0.018 | Old |
| 0.048127 | 0.236297 | 0.293 | 0.245 | 1 | 3 | Ddx54 | 0.048 | Old |
| 0.048178 | 0.1595 | 0.302 | 0.256 | 1 | 3 | Rbck1 | 0.046 | Old |
| 0.048192 | 0.133031 | 0.149 | 0.115 | 1 | 3 | Pdgfb | 0.034 | Old |

| | | | | | | | | |
|---|---|---|---|---|---|---|---|---|
| 0.048566 | 0.662781 | 0.038 | 0.021 | 1 | 3 | Sema6b | 0.017 | Old |
| 0.048577 | 0.199291 | 0.264 | 0.22 | 1 | 3 | Dram2 | 0.044 | Old |
| 0.048733 | 0.185773 | 0.154 | 0.119 | 1 | 3 | Zfp384 | 0.035 | Old |
| 0.04903 | 0.104636 | 0.361 | 0.307 | 1 | 3 | Serpinb6a | 0.054 | Old |
| 0.049054 | 0.216171 | 0.282 | 0.234 | 1 | 3 | Slc7a8 | 0.048 | Old |
| 0.04945 | 0.475903 | 0.053 | 0.033 | 1 | 3 | Noa1 | 0.02 | Old |
| 0.04945 | 0.420442 | 0.161 | 0.126 | 1 | 3 | Zfp655 | 0.035 | Old |
| 0.049483 | 0.312745 | 0.149 | 0.115 | 1 | 3 | Emp1 | 0.034 | Old |
| 0.04949 | 0.15746 | 0.52 | 0.46 | 1 | 3 | Adssl1 | 0.06 | Old |
| 0.049588 | 0.29082 | 0.103 | 0.074 | 1 | 3 | Cep192 | 0.029 | Old |
| 0.049592 | 0.031884 | 0.814 | 0.761 | 1 | 3 | Rtn4 | 0.053 | Old |
| 0.049686 | 0.443415 | 0.118 | 0.088 | 1 | 3 | Impdh1 | 0.03 | Old |
| 0.049717 | 0.124566 | 0.417 | 0.364 | 1 | 3 | Soat1 | 0.053 | Old |
| 0.049729 | 0.070169 | 0.196 | 0.155 | 1 | 3 | Sav1 | 0.041 | Old |
| 0.04974 | 0.16553 | 0.343 | 0.287 | 1 | 3 | Mydgf | 0.056 | Old |
| 0.049771 | 0.057986 | 0.394 | 0.333 | 1 | 3 | Sdc1 | 0.061 | Old |
| 0.049923 | 0.628617 | 0.096 | 0.069 | 1 | 3 | Coq7 | 0.027 | Old |
| 0.050088 | 0.253977 | 0.268 | 0.223 | 1 | 3 | Mybbp1a | 0.045 | Old |
| 0.050098 | 0.148869 | 0.189 | 0.15 | 1 | 3 | Ncbp3 | 0.039 | Old |
| 0.050234 | 0.264988 | 0.082 | 0.057 | 1 | 3 | Ebag9 | 0.025 | Old |
| 0.05033 | 0.215695 | 0.302 | 0.254 | 1 | 3 | Chp1 | 0.048 | Old |
| 0.050455 | 0.368083 | 0.139 | 0.106 | 1 | 3 | Prpf38a | 0.033 | Old |
| 0.050465 | 0.257247 | 0.358 | 0.304 | 1 | 3 | Ube2i | 0.054 | Old |
| 0.050496 | 0.19458 | 0.108 | 0.078 | 1 | 3 | Llgl1 | 0.03 | Old |
| 0.050628 | 0.188759 | 0.491 | 0.443 | 1 | 3 | Pros1 | 0.048 | Old |
| 0.050639 | 0.286714 | 0.154 | 0.121 | 1 | 3 | Slc39a9 | 0.033 | Old |
| 0.050688 | 0.073192 | 0.533 | 0.465 | 1 | 3 | Pgls | 0.068 | Old |
| 0.051031 | 0.074404 | 0.708 | 0.641 | 1 | 3 | Tomm7 | 0.067 | Old |
| 0.051057 | 0.621542 | 0.043 | 0.025 | 1 | 3 | Riox1 | 0.018 | Old |
| 0.051063 | 0.228747 | 0.062 | 0.04 | 1 | 3 | Clcn6 | 0.022 | Old |
| 0.051206 | 0.091858 | 0.983 | 0.994 | 1 | 3 | Fn1 | -0.011 | Old |
| 0.051243 | 0.519387 | 0.069 | 0.047 | 1 | 3 | Tipin | 0.022 | Old |
| 0.051262 | 0.113757 | 0.278 | 0.229 | 1 | 3 | Eif3g | 0.049 | Old |
| 0.051321 | 0.245794 | 0.148 | 0.114 | 1 | 3 | Fibp | 0.034 | Old |
| 0.051511 | 0.323995 | 0.099 | 0.072 | 1 | 3 | Kctd5 | 0.027 | Old |
| 0.051526 | 0.15155 | 0.319 | 0.269 | 1 | 3 | Guk1 | 0.05 | Old |
| 0.051566 | 0.198465 | 0.375 | 0.317 | 1 | 3 | Rnf4 | 0.058 | Old |
| 0.051567 | 0.280463 | 0.203 | 0.165 | 1 | 3 | Ctnnd1 | 0.038 | Old |
| 0.051607 | 0.093992 | 0.64 | 0.567 | 1 | 3 | Hnrnpc | 0.073 | Old |
| 0.051721 | 0.003453 | 0.131 | 0.097 | 1 | 3 | Eif2ak2 | 0.034 | Old |
| 0.051758 | 0.113833 | 0.357 | 0.303 | 1 | 3 | Ppp4r2 | 0.054 | Old |
| 0.051784 | 0.128481 | 0.281 | 0.234 | 1 | 3 | Prmt1 | 0.047 | Old |
| 0.051794 | 0.162861 | 0.256 | 0.215 | 1 | 3 | Vav1 | 0.041 | Old |
| 0.051949 | 0.330032 | 0.133 | 0.101 | 1 | 3 | Mettl1 | 0.032 | Old |
| 0.052231 | 0.272476 | 0.114 | 0.083 | 1 | 3 | Ctnnbip1 | 0.031 | Old |

| | | | | | | | | |
|---|---|---|---|---|---|---|---|---|
| 0.052406 | 0.089808 | 0.217 | 0.176 | 1 | 3 | Zfp451 | 0.041 | Old |
| 0.052547 | 0.147829 | 0.611 | 0.563 | 1 | 3 | Tram1 | 0.048 | Old |
| 0.052605 | 0.032792 | 0.38 | 0.323 | 1 | 3 | Aff4 | 0.057 | Old |
| 0.052614 | 0.314181 | 0.211 | 0.17 | 1 | 3 | Ube3b | 0.041 | Old |
| 0.05262 | 0.246732 | 0.192 | 0.154 | 1 | 3 | Hps5 | 0.038 | Old |
| 0.052725 | 0.141825 | 0.13 | 0.098 | 1 | 3 | Chm | 0.032 | Old |
| 0.052779 | 0.445312 | 0.115 | 0.086 | 1 | 3 | Med27 | 0.029 | Old |
| 0.053254 | 0.186201 | 0.128 | 0.096 | 1 | 3 | Cfl2 | 0.032 | Old |
| 0.053389 | 0.272648 | 0.121 | 0.091 | 1 | 3 | Dusp23 | 0.03 | Old |
| 0.05351 | 1.147882 | 0.02 | 0.009 | 1 | 3 | Pfkfb2 | 0.011 | Old |
| 0.053517 | 0.155236 | 0.364 | 0.312 | 1 | 3 | Copb2 | 0.052 | Old |
| 0.053599 | 0.000247 | 0.477 | 0.409 | 1 | 3 | Lypla2 | 0.068 | Old |
| 0.053655 | 0.19913 | 0.085 | 0.059 | 1 | 3 | Rrs1 | 0.026 | Old |
| 0.053909 | 0.318343 | 0.162 | 0.128 | 1 | 3 | Gm2000 | 0.034 | Old |
| 0.053971 | 0.138334 | 0.35 | 0.294 | 1 | 3 | Map3k1 | 0.056 | Old |
| 0.054179 | 0.173938 | 0.054 | 0.034 | 1 | 3 | Kctd6 | 0.02 | Old |
| 0.054199 | 0.707571 | 0.078 | 0.054 | 1 | 3 | Edem3 | 0.024 | Old |
| 0.054225 | 0.192987 | 0.229 | 0.189 | 1 | 3 | Kitl | 0.04 | Old |
| 0.05429 | 0.2251 | 0.116 | 0.086 | 1 | 3 | Mief1 | 0.03 | Old |
| 0.054361 | 0.031306 | 0.17 | 0.131 | 1 | 3 | Trak1 | 0.039 | Old |
| 0.054424 | 0.248225 | 0.162 | 0.128 | 1 | 3 | Riok1 | 0.034 | Old |
| 0.054442 | 0.181843 | 0.301 | 0.253 | 1 | 3 | Pdap1 | 0.048 | Old |
| 0.054452 | 0.10937 | 0.148 | 0.112 | 1 | 3 | Trappc9 | 0.036 | Old |
| 0.054455 | 0.465269 | 0.06 | 0.039 | 1 | 3 | C130050O1 | 0.021 | Old |
| 0.054489 | 0.167243 | 0.335 | 0.287 | 1 | 3 | Nfat5 | 0.048 | Old |
| 0.054609 | 0.449886 | 0.133 | 0.102 | 1 | 3 | Yipf6 | 0.031 | Old |
| 0.054679 | 0.045327 | 0.621 | 0.553 | 1 | 3 | Txn2 | 0.068 | Old |
| 0.054819 | 0.149144 | 0.277 | 0.229 | 1 | 3 | Tspan5 | 0.048 | Old |
| 0.054878 | 0.244963 | 0.252 | 0.21 | 1 | 3 | Cops5 | 0.042 | Old |
| 0.054886 | 0.747031 | 0.054 | 0.034 | 1 | 3 | Ggta1 | 0.02 | Old |
| 0.054932 | 0.149384 | 0.866 | 0.833 | 1 | 3 | Grn | 0.033 | Old |
| 0.055124 | 0.167985 | 0.252 | 0.207 | 1 | 3 | Phf5a | 0.045 | Old |
| 0.055204 | 0.256817 | 0.146 | 0.112 | 1 | 3 | Nlk | 0.034 | Old |
| 0.055451 | 0.070644 | 0.566 | 0.501 | 1 | 3 | Srsf6 | 0.065 | Old |
| 0.055452 | 0.671491 | 0.027 | 0.014 | 1 | 3 | Rnf24 | 0.013 | Old |
| 0.055479 | 0.060238 | 0.221 | 0.178 | 1 | 3 | Yipf5 | 0.043 | Old |
| 0.055488 | 0.233799 | 0.257 | 0.21 | 1 | 3 | Srp54a | 0.047 | Old |
| 0.055538 | 0.197708 | 0.309 | 0.259 | 1 | 3 | Mfsd5 | 0.05 | Old |
| 0.055907 | 0.064456 | 0.358 | 0.297 | 1 | 3 | Rad23a | 0.061 | Old |
| 0.055944 | 0.100533 | 0.536 | 0.475 | 1 | 3 | Ewsr1 | 0.061 | Old |
| 0.056015 | 0.271077 | 0.148 | 0.115 | 1 | 3 | Enoph1 | 0.033 | Old |
| 0.056092 | 0.93586 | 0.024 | 0.011 | 1 | 3 | Dagla | 0.013 | Old |
| 0.056092 | 0.215651 | 0.278 | 0.234 | 1 | 3 | Polr2g | 0.044 | Old |
| 0.056253 | 0.638023 | 0.02 | 0.009 | 1 | 3 | Nupl2 | 0.011 | Old |
| 0.05638 | 0.752241 | 0.044 | 0.027 | 1 | 3 | Mbip | 0.017 | Old |

| | | | | | | | | |
|---|---|---|---|---|---|---|---|---|
| 0.056398 | 0.428745 | 0.155 | 0.121 | 1 | 3 | Vps54 | 0.034 | Old |
| 0.056455 | 0.293732 | 0.031 | 0.016 | 1 | 3 | Bst1 | 0.015 | Old |
| 0.056505 | 0.12196 | 0.707 | 0.644 | 1 | 3 | Tmed2 | 0.063 | Old |
| 0.05661 | 0.247355 | 0.031 | 0.016 | 1 | 3 | C130026I2 | 0.015 | Old |
| 0.056678 | 0.280261 | 0.049 | 0.03 | 1 | 3 | Sft2d3 | 0.019 | Old |
| 0.056796 | 0.069054 | 0.235 | 0.192 | 1 | 3 | Mark3 | 0.043 | Old |
| 0.056814 | 0.052283 | 0.142 | 0.107 | 1 | 3 | Il17ra | 0.035 | Old |
| 0.056957 | 0.148675 | 0.166 | 0.13 | 1 | 3 | Lpcat3 | 0.036 | Old |
| 0.057028 | 0.131648 | 0.542 | 0.465 | 1 | 3 | Dnmt3a | 0.077 | Old |
| 0.057035 | 0.157859 | 0.332 | 0.275 | 1 | 3 | Rnpep | 0.057 | Old |
| 0.057088 | 0.198098 | 0.122 | 0.091 | 1 | 3 | Phrf1 | 0.031 | Old |
| 0.057246 | 0.284428 | 0.054 | 0.034 | 1 | 3 | Mul1 | 0.02 | Old |
| 0.057265 | 0.20894 | 0.292 | 0.245 | 1 | 3 | Psmd7 | 0.047 | Old |
| 0.057268 | 0.142276 | 0.364 | 0.309 | 1 | 3 | Bud31 | 0.055 | Old |
| 0.05727 | 0.730474 | 0.041 | 0.024 | 1 | 3 | Pold2 | 0.017 | Old |
| 0.057317 | 0.15643 | 0.173 | 0.138 | 1 | 3 | Nip7 | 0.035 | Old |
| 0.057319 | 0.325919 | 0.171 | 0.134 | 1 | 3 | Usp22 | 0.037 | Old |
| 0.057352 | 0.125144 | 0.313 | 0.263 | 1 | 3 | Rbm27 | 0.05 | Old |
| 0.057485 | 0.50491 | 0.059 | 0.038 | 1 | 3 | Tmem80 | 0.021 | Old |
| 0.057582 | 0.270038 | 0.241 | 0.202 | 1 | 3 | Tspyl1 | 0.039 | Old |
| 0.05761 | 0.41056 | 0.014 | 0.005 | 1 | 3 | Rbpms2 | 0.009 | Old |
| 0.057615 | 0.339085 | 0.054 | 0.034 | 1 | 3 | Mrpl47 | 0.02 | Old |
| 0.057782 | 0.246714 | 0.241 | 0.198 | 1 | 3 | Syvn1 | 0.043 | Old |
| 0.057883 | 0.139467 | 0.435 | 0.378 | 1 | 3 | Map7d1 | 0.057 | Old |
| 0.057923 | 0.115861 | 0.742 | 0.682 | 1 | 3 | Slfn2 | 0.06 | Old |
| 0.057986 | 0.393317 | 0.143 | 0.112 | 1 | 3 | Uspl1 | 0.031 | Old |
| 0.058099 | 0.811971 | 0.014 | 0.005 | 1 | 3 | D130040H: | 0.009 | Old |
| 0.058174 | 0.30149 | 0.06 | 0.039 | 1 | 3 | Wrap73 | 0.021 | Old |
| 0.058289 | 0.119316 | 0.608 | 0.54 | 1 | 3 | Pycard | 0.068 | Old |
| 0.058316 | 0.289201 | 0.044 | 0.027 | 1 | 3 | Tubgcp6 | 0.017 | Old |
| 0.058335 | 0.20784 | 0.305 | 0.259 | 1 | 3 | Eea1 | 0.046 | Old |
| 0.058344 | 0.891826 | 0.014 | 0.005 | 1 | 3 | Met | 0.009 | Old |
| 0.058381 | 0.167062 | 0.183 | 0.145 | 1 | 3 | Ldlrap1 | 0.038 | Old |
| 0.058509 | 0.530914 | 0.014 | 0.005 | 1 | 3 | Spc25 | 0.009 | Old |
| 0.058533 | 0.065828 | 0.99 | 0.985 | 1 | 3 | Rpsa | 0.005 | Old |
| 0.05855 | 0.290843 | 0.179 | 0.141 | 1 | 3 | Hccs | 0.038 | Old |
| 0.058615 | 0.092311 | 0.054 | 0.034 | 1 | 3 | Rfc4 | 0.02 | Old |
| 0.058621 | 0.054404 | 0.456 | 0.388 | 1 | 3 | Evl | 0.068 | Old |
| 0.058713 | 0.212201 | 0.332 | 0.285 | 1 | 3 | U2surp | 0.047 | Old |
| 0.058751 | 0.27621 | 0.065 | 0.043 | 1 | 3 | Shb | 0.022 | Old |
| 0.058798 | 0.361695 | 0.109 | 0.081 | 1 | 3 | Mrpl37 | 0.028 | Old |
| 0.05884 | 0.713392 | 0.136 | 0.105 | 1 | 3 | Dck | 0.031 | Old |
| 0.058982 | 0.053052 | 0.381 | 0.322 | 1 | 3 | Glrx5 | 0.059 | Old |
| 0.058983 | 0.15189 | 0.225 | 0.179 | 1 | 3 | Acvrl1 | 0.046 | Old |
| 0.059002 | 0.618244 | 0.036 | 0.02 | 1 | 3 | Pola2 | 0.016 | Old |

| | | | | | | | | |
|---|---|---|---|---|---|---|---|---|
| 0.059016 | 0.04523 | 0.157 | 0.121 | 1 | 3 | Cog5 | 0.036 | Old |
| 0.059064 | 0.158139 | 0.252 | 0.206 | 1 | 3 | Tollip | 0.046 | Old |
| 0.059076 | 0.072788 | 0.591 | 0.529 | 1 | 3 | Fermt3 | 0.062 | Old |
| 0.059083 | 0.087642 | 0.219 | 0.177 | 1 | 3 | Smad2 | 0.042 | Old |
| 0.059108 | 0.543285 | 0.061 | 0.04 | 1 | 3 | Gpr180 | 0.021 | Old |
| 0.059269 | 0.272321 | 0.116 | 0.087 | 1 | 3 | Rpf1 | 0.029 | Old |
| 0.0593 | 0.484958 | 0.079 | 0.056 | 1 | 3 | Card9 | 0.023 | Old |
| 0.05932 | 0.267619 | 0.301 | 0.255 | 1 | 3 | Ivns1abp | 0.046 | Old |
| 0.059386 | 0.110299 | 0.317 | 0.269 | 1 | 3 | Akirin1 | 0.048 | Old |
| 0.059426 | 0.199123 | 0.177 | 0.141 | 1 | 3 | Mrpl13 | 0.036 | Old |
| 0.05945 | 0.361922 | 0.115 | 0.086 | 1 | 3 | Tle1 | 0.029 | Old |
| 0.059593 | 0.283 | 0.179 | 0.143 | 1 | 3 | Hdac4 | 0.036 | Old |
| 0.059607 | 0.306023 | 0.142 | 0.111 | 1 | 3 | Erlin2 | 0.031 | Old |
| 0.059623 | 0.115117 | 0.732 | 0.674 | 1 | 3 | 5031439G( | 0.058 | Old |
| 0.059793 | 0.015912 | 0.384 | 0.323 | 1 | 3 | Larp4 | 0.061 | Old |
| 0.059839 | 0.151665 | 0.232 | 0.188 | 1 | 3 | Dmxl1 | 0.044 | Old |
| 0.059862 | 0.222755 | 0.464 | 0.407 | 1 | 3 | Tnfrsf1b | 0.057 | Old |
| 0.05993 | 0.139695 | 0.432 | 0.379 | 1 | 3 | Ryr1 | 0.053 | Old |
| 0.060144 | 0.11052 | 0.366 | 0.313 | 1 | 3 | Ap1b1 | 0.053 | Old |
| 0.06032 | 0.368343 | 0.072 | 0.049 | 1 | 3 | Mum1 | 0.023 | Old |
| 0.060392 | 0.106327 | 0.785 | 0.723 | 1 | 3 | Mpeg1 | 0.062 | Old |
| 0.060411 | 0.158814 | 0.544 | 0.482 | 1 | 3 | Stx7 | 0.062 | Old |
| 0.060502 | 0.121813 | 0.282 | 0.234 | 1 | 3 | Anapc5 | 0.048 | Old |
| 0.060533 | 0.1893 | 0.669 | 0.609 | 1 | 3 | Myadm | 0.06 | Old |
| 0.060571 | 0.342943 | 0.091 | 0.066 | 1 | 3 | Cgrrf1 | 0.025 | Old |
| 0.060627 | 0.134876 | 0.204 | 0.164 | 1 | 3 | Tbc1d2b | 0.04 | Old |
| 0.060647 | 0.343388 | 0.116 | 0.087 | 1 | 3 | Bad | 0.029 | Old |
| 0.060819 | 0.17701 | 0.351 | 0.302 | 1 | 3 | Dcun1d5 | 0.049 | Old |
| 0.060891 | 0.188488 | 0.148 | 0.116 | 1 | 3 | Smox | 0.032 | Old |
| 0.060907 | 0.19958 | 0.392 | 0.336 | 1 | 3 | Cdipt | 0.056 | Old |
| 0.061031 | 0.072966 | 0.713 | 0.634 | 1 | 3 | Lmna | 0.079 | Old |
| 0.061152 | 0.197458 | 0.143 | 0.111 | 1 | 3 | Myo1e | 0.032 | Old |
| 0.06132 | 0.260178 | 0.055 | 0.035 | 1 | 3 | Rps6kb2 | 0.02 | Old |
| 0.061325 | 0.202726 | 0.164 | 0.129 | 1 | 3 | 2810004N2 | 0.035 | Old |
| 0.061447 | 0.39158 | 0.078 | 0.054 | 1 | 3 | Syn1 | 0.024 | Old |
| 0.061612 | 0.085936 | 0.723 | 0.672 | 1 | 3 | Ifitm2 | 0.051 | Old |
| 0.061684 | 0.131855 | 0.441 | 0.394 | 1 | 3 | Gpsm3 | 0.047 | Old |
| 0.061736 | 1.252249 | 0.025 | 0.013 | 1 | 3 | Gm15564 | 0.012 | Old |
| 0.061812 | 0.565924 | 0.041 | 0.024 | 1 | 3 | Pgpep1 | 0.017 | Old |
| 0.061948 | 0.245255 | 0.069 | 0.047 | 1 | 3 | Bcl2l13 | 0.022 | Old |
| 0.062173 | 0.223881 | 0.185 | 0.146 | 1 | 3 | Atf6b | 0.039 | Old |
| 0.062205 | 0.243796 | 0.186 | 0.15 | 1 | 3 | Rab5b | 0.036 | Old |
| 0.062255 | 0.111418 | 0.305 | 0.255 | 1 | 3 | Pi4k2a | 0.05 | Old |
| 0.06227 | 0.100694 | 0.122 | 0.092 | 1 | 3 | Mrps5 | 0.03 | Old |
| 0.062285 | 0.064544 | 0.967 | 0.967 | 1 | 3 | Ptprc | 0 | Old |

| | | | | | | | | |
|---|---|---|---|---|---|---|---|---|
| 0.062317 | 0.268697 | 0.127 | 0.096 | 1 | 3 | Mtmr12 | 0.031 | Old |
| 0.062338 | 0.017306 | 0.303 | 0.25 | 1 | 3 | Wdr45b | 0.053 | Old |
| 0.062484 | 0.21482 | 0.041 | 0.024 | 1 | 3 | Trim30b | 0.017 | Old |
| 0.06251 | 0.168962 | 0.29 | 0.246 | 1 | 3 | Pigx | 0.044 | Old |
| 0.062571 | 0.146566 | 0.105 | 0.078 | 1 | 3 | Cwc22 | 0.027 | Old |
| 0.062572 | 0.103282 | 0.848 | 0.778 | 1 | 3 | Colgalt1 | 0.07 | Old |
| 0.062683 | 0.22553 | 0.094 | 0.068 | 1 | 3 | Rnf121 | 0.026 | Old |
| 0.062746 | 0.027159 | 0.276 | 0.227 | 1 | 3 | C5ar1 | 0.049 | Old |
| 0.062781 | 0.100383 | 0.258 | 0.213 | 1 | 3 | Cd300lb | 0.045 | Old |
| 0.062839 | 0.138952 | 0.514 | 0.441 | 1 | 3 | Uqcrfs1 | 0.073 | Old |
| 0.062852 | 0.18343 | 0.251 | 0.208 | 1 | 3 | Smu1 | 0.043 | Old |
| 0.062873 | 0.51469 | 0.056 | 0.037 | 1 | 3 | Flrt2 | 0.019 | Old |
| 0.062988 | 1.811085 | 0.012 | 0.004 | 1 | 3 | Mettl15 | 0.008 | Old |
| 0.063017 | 0.23724 | 0.332 | 0.287 | 1 | 3 | Snx17 | 0.045 | Old |
| 0.063037 | 0.191848 | 0.2 | 0.159 | 1 | 3 | Elk3 | 0.041 | Old |
| 0.063228 | 0.384521 | 0.088 | 0.064 | 1 | 3 | E130309D( | 0.024 | Old |
| 0.063402 | 0.136135 | 0.397 | 0.337 | 1 | 3 | Atg3 | 0.06 | Old |
| 0.063438 | 0.26468 | 0.088 | 0.063 | 1 | 3 | Fam222b | 0.025 | Old |
| 0.063469 | 0.222331 | 0.136 | 0.105 | 1 | 3 | 0610010F0 | 0.031 | Old |
| 0.0635 | 0.140895 | 0.732 | 0.667 | 1 | 3 | Tm9sf3 | 0.065 | Old |
| 0.063554 | 0.162539 | 0.348 | 0.295 | 1 | 3 | Sp100 | 0.053 | Old |
| 0.063569 | 0.4399 | 0.105 | 0.078 | 1 | 3 | Krr1 | 0.027 | Old |
| 0.063587 | 0.150303 | 0.066 | 0.044 | 1 | 3 | Vps8 | 0.022 | Old |
| 0.063648 | 0.228978 | 0.131 | 0.101 | 1 | 3 | Ipcef1 | 0.03 | Old |
| 0.063669 | 1.159508 | 0.012 | 0.004 | 1 | 3 | Mgll | 0.008 | Old |
| 0.063674 | 0.161804 | 0.501 | 0.443 | 1 | 3 | Ndufs6 | 0.058 | Old |
| 0.063794 | 0.137913 | 0.722 | 0.66 | 1 | 3 | Wfdc17 | 0.062 | Old |
| 0.063837 | 0.464479 | 0.092 | 0.067 | 1 | 3 | Tmem104 | 0.025 | Old |
| 0.063862 | 0.107842 | 0.274 | 0.226 | 1 | 3 | Mrpl51 | 0.048 | Old |
| 0.064088 | 0.197044 | 0.319 | 0.275 | 1 | 3 | Arl1 | 0.044 | Old |
| 0.064154 | 0.324063 | 0.234 | 0.197 | 1 | 3 | H2-M3 | 0.037 | Old |
| 0.064161 | 0.150612 | 0.356 | 0.306 | 1 | 3 | Skp1a | 0.05 | Old |
| 0.064231 | 0.07139 | 0.131 | 0.101 | 1 | 3 | Klhl2 | 0.03 | Old |
| 0.064235 | 0.208155 | 0.281 | 0.237 | 1 | 3 | Plod3 | 0.044 | Old |
| 0.06424 | 0.337818 | 0.152 | 0.117 | 1 | 3 | Dph5 | 0.035 | Old |
| 0.064243 | 0.060479 | 0.276 | 0.227 | 1 | 3 | Fnip1 | 0.049 | Old |
| 0.064258 | 0.896781 | 0.012 | 0.004 | 1 | 3 | Gpr21 | 0.008 | Old |
| 0.064281 | 0.823263 | 0.018 | 0.008 | 1 | 3 | Rab27a | 0.01 | Old |
| 0.06436 | 0.635147 | 0.029 | 0.015 | 1 | 3 | Vwa8 | 0.014 | Old |
| 0.06443 | 0.105739 | 0.671 | 0.593 | 1 | 3 | Psma4 | 0.078 | Old |
| 0.064455 | 0.793049 | 0.012 | 0.004 | 1 | 3 | Gm10800 | 0.008 | Old |
| 0.064477 | 0.128664 | 0.307 | 0.26 | 1 | 3 | Etv6 | 0.047 | Old |
| 0.064491 | 0.306501 | 0.124 | 0.096 | 1 | 3 | Taf4 | 0.028 | Old |
| 0.064576 | 0.12473 | 0.104 | 0.076 | 1 | 3 | Ccdc34 | 0.028 | Old |
| 0.064645 | 0.205177 | 0.454 | 0.399 | 1 | 3 | Emc10 | 0.055 | Old |

| | | | | | | | | |
|---|---|---|---|---|---|---|---|---|
| 0.064777 | 0.067201 | 0.962 | 0.938 | 1 | 3 | Cebpb | 0.024 | Old |
| 0.064924 | 0.219832 | 0.11 | 0.082 | 1 | 3 | Gpr146 | 0.028 | Old |
| 0.06493 | 0.330575 | 0.099 | 0.072 | 1 | 3 | Pop4 | 0.027 | Old |
| 0.064953 | 0.089639 | 0.18 | 0.144 | 1 | 3 | Snx12 | 0.036 | Old |
| 0.064984 | 0.768178 | 0.018 | 0.008 | 1 | 3 | Adgrl1 | 0.01 | Old |
| 0.065388 | 0.219552 | 0.079 | 0.056 | 1 | 3 | Nostrin | 0.023 | Old |
| 0.065449 | 0.55802 | 0.012 | 0.004 | 1 | 3 | Mcts2 | 0.008 | Old |
| 0.065477 | 0.371432 | 0.117 | 0.09 | 1 | 3 | Tmem39a | 0.027 | Old |
| 0.065549 | 0.296972 | 0.012 | 0.004 | 1 | 3 | Oasl1 | 0.008 | Old |
| 0.065565 | 0.112695 | 0.487 | 0.427 | 1 | 3 | Lrmp | 0.06 | Old |
| 0.065617 | 0.256731 | 0.194 | 0.155 | 1 | 3 | Tufm | 0.039 | Old |
| 0.065642 | 0.173799 | 0.274 | 0.229 | 1 | 3 | Chka | 0.045 | Old |
| 0.065643 | 0.01334 | 0.338 | 0.285 | 1 | 3 | Prkacb | 0.053 | Old |
| 0.065717 | 0.184937 | 0.356 | 0.308 | 1 | 3 | Slc25a24 | 0.048 | Old |
| 0.065744 | 1.059268 | 0.022 | 0.01 | 1 | 3 | Zfp202 | 0.012 | Old |
| 0.065774 | 0.16426 | 0.173 | 0.138 | 1 | 3 | Rab29 | 0.035 | Old |
| 0.065999 | 0.176282 | 0.252 | 0.213 | 1 | 3 | Zfp800 | 0.039 | Old |
| 0.066048 | 0.184859 | 0.337 | 0.284 | 1 | 3 | Csgalnact2 | 0.053 | Old |
| 0.066384 | 0.172699 | 0.245 | 0.199 | 1 | 3 | Polr2l | 0.046 | Old |
| 0.066669 | 0.022368 | 0.455 | 0.393 | 1 | 3 | Rbm8a | 0.062 | Old |
| 0.066905 | 0.143628 | 0.122 | 0.092 | 1 | 3 | Brms1 | 0.03 | Old |
| 0.067001 | 0.212512 | 0.131 | 0.101 | 1 | 3 | Thyn1 | 0.03 | Old |
| 0.067011 | 0.180994 | 0.203 | 0.165 | 1 | 3 | Synj2bp | 0.038 | Old |
| 0.067085 | 0.202799 | 0.147 | 0.115 | 1 | 3 | Mtf2 | 0.032 | Old |
| 0.067138 | 0.314801 | 0.185 | 0.152 | 1 | 3 | Ppt1 | 0.033 | Old |
| 0.067171 | 0.456461 | 0.056 | 0.037 | 1 | 3 | Tlr6 | 0.019 | Old |
| 0.067255 | 0.729746 | 0.022 | 0.01 | 1 | 3 | Fam187b | 0.012 | Old |
| 0.067461 | 0.191617 | 0.213 | 0.173 | 1 | 3 | Actr10 | 0.04 | Old |
| 0.067534 | 0.195606 | 0.253 | 0.21 | 1 | 3 | Aco1 | 0.043 | Old |
| 0.067546 | 0.799537 | 0.022 | 0.01 | 1 | 3 | Trmt5 | 0.012 | Old |
| 0.067573 | 0.107342 | 0.157 | 0.122 | 1 | 3 | Atp8a1 | 0.035 | Old |
| 0.067594 | 0.394489 | 0.166 | 0.133 | 1 | 3 | Gid8 | 0.033 | Old |
| 0.067764 | 0.109136 | 0.172 | 0.135 | 1 | 3 | Nmt2 | 0.037 | Old |
| 0.067779 | 0.092699 | 0.897 | 0.866 | 1 | 3 | Chchd2 | 0.031 | Old |
| 0.067868 | 0.318378 | 0.025 | 0.013 | 1 | 3 | 4632404H1 | 0.012 | Old |
| 0.067949 | 0.12169 | 0.738 | 0.67 | 1 | 3 | Pkm | 0.068 | Old |
| 0.06801 | 0.323407 | 0.078 | 0.054 | 1 | 3 | Gmeb2 | 0.024 | Old |
| 0.068192 | 0.221962 | 0.319 | 0.273 | 1 | 3 | Arf2 | 0.046 | Old |
| 0.068204 | 0.145253 | 0.424 | 0.378 | 1 | 3 | Med13l | 0.046 | Old |
| 0.068291 | 0.101086 | 0.56 | 0.504 | 1 | 3 | Dnajc3 | 0.056 | Old |
| 0.068325 | 0.177472 | 0.091 | 0.066 | 1 | 3 | Ggps1 | 0.025 | Old |
| 0.068627 | 0.068329 | 0.51 | 0.452 | 1 | 3 | Scamp2 | 0.058 | Old |
| 0.068717 | 0.133465 | 0.257 | 0.215 | 1 | 3 | Atxn2 | 0.042 | Old |
| 0.068877 | 0.32204 | 0.112 | 0.085 | 1 | 3 | Trp53inp2 | 0.027 | Old |
| 0.068963 | 0.069832 | 0.108 | 0.08 | 1 | 3 | Pphln1 | 0.028 | Old |

| | | | | | | | | |
|---|---|---|---|---|---|---|---|---|
| 0.069044 | 0.087317 | 0.079 | 0.056 | 1 | 3 | Zfp410 | 0.023 | Old |
| 0.069055 | 0.219924 | 0.473 | 0.427 | 1 | 3 | Baz1a | 0.046 | Old |
| 0.069084 | 0.386432 | 0.025 | 0.013 | 1 | 3 | E130308A1 | 0.012 | Old |
| 0.069213 | 0.145862 | 0.314 | 0.266 | 1 | 3 | Fbxl3 | 0.048 | Old |
| 0.069235 | 0.950119 | 0.01 | 0.003 | 1 | 3 | 4632428C0 | 0.007 | Old |
| 0.069253 | 0.183904 | 0.076 | 0.053 | 1 | 3 | Tspyl2 | 0.023 | Old |
| 0.069376 | 0.954202 | 0.03 | 0.016 | 1 | 3 | Slc30a6 | 0.014 | Old |
| 0.069408 | 0.105971 | 0.147 | 0.115 | 1 | 3 | Gtf2f2 | 0.032 | Old |
| 0.069474 | 0.870893 | 0.01 | 0.003 | 1 | 3 | Rdh5 | 0.007 | Old |
| 0.069713 | 0.818409 | 0.01 | 0.003 | 1 | 3 | Penk | 0.007 | Old |
| 0.06982 | 0.299391 | 0.195 | 0.158 | 1 | 3 | Cul5 | 0.037 | Old |
| 0.069835 | 0.307468 | 0.1 | 0.074 | 1 | 3 | Lxn | 0.026 | Old |
| 0.069916 | 0.039418 | 0.464 | 0.41 | 1 | 3 | Slc20a1 | 0.054 | Old |
| 0.069921 | 0.407205 | 0.033 | 0.019 | 1 | 3 | Lppos | 0.014 | Old |
| 0.069954 | 0.401778 | 0.01 | 0.003 | 1 | 3 | Setmar | 0.007 | Old |
| 0.069981 | 0.082698 | 0.208 | 0.167 | 1 | 3 | Tor1aip2 | 0.041 | Old |
| 0.070126 | 0.409742 | 0.153 | 0.121 | 1 | 3 | Naxd | 0.032 | Old |
| 0.070158 | 0.520675 | 0.059 | 0.039 | 1 | 3 | Tmem175 | 0.02 | Old |
| 0.07016 | 0.224907 | 0.299 | 0.256 | 1 | 3 | Chd1 | 0.043 | Old |
| 0.070315 | 0.569629 | 0.01 | 0.003 | 1 | 3 | Dst | 0.007 | Old |
| 0.070315 | 0.376572 | 0.01 | 0.003 | 1 | 3 | Slfn9 | 0.007 | Old |
| 0.070363 | 0.319449 | 0.121 | 0.092 | 1 | 3 | Rce1 | 0.029 | Old |
| 0.070437 | 0.068371 | 0.472 | 0.405 | 1 | 3 | D17Wsu92 | 0.067 | Old |
| 0.070531 | 0.141084 | 0.082 | 0.058 | 1 | 3 | Trub2 | 0.024 | Old |
| 0.070551 | 0.3481 | 0.073 | 0.051 | 1 | 3 | Lig3 | 0.022 | Old |
| 0.070747 | 0.23447 | 0.264 | 0.222 | 1 | 3 | Grk6 | 0.042 | Old |
| 0.070787 | 0.474071 | 0.048 | 0.03 | 1 | 3 | Gemin6 | 0.018 | Old |
| 0.070795 | 0.123039 | 0.46 | 0.407 | 1 | 3 | Ostc | 0.053 | Old |
| 0.070936 | 0.922984 | 0.03 | 0.016 | 1 | 3 | Trmo | 0.014 | Old |
| 0.070938 | 0.253954 | 0.197 | 0.16 | 1 | 3 | Mmadhc | 0.037 | Old |
| 0.070945 | 0.263491 | 0.142 | 0.111 | 1 | 3 | Ate1 | 0.031 | Old |
| 0.071205 | 0.094608 | 0.579 | 0.515 | 1 | 3 | Lcp2 | 0.064 | Old |
| 0.071205 | 0.353289 | 0.216 | 0.181 | 1 | 3 | Sp110 | 0.035 | Old |
| 0.071246 | 0.182475 | 0.134 | 0.102 | 1 | 3 | Mertk | 0.032 | Old |
| 0.071475 | 0.599204 | 0.059 | 0.039 | 1 | 3 | AI597479 | 0.02 | Old |
| 0.071844 | 0.410205 | 0.152 | 0.121 | 1 | 3 | 1110038F1 | 0.031 | Old |
| 0.071917 | 0.036914 | 0.177 | 0.139 | 1 | 3 | Rab3gap1 | 0.038 | Old |
| 0.071926 | 0.004043 | 0.245 | 0.198 | 1 | 3 | Cdk13 | 0.047 | Old |
| 0.072138 | 0.203299 | 0.158 | 0.124 | 1 | 3 | Nomo1 | 0.034 | Old |
| 0.07242 | 0.031367 | 0.603 | 0.529 | 1 | 3 | Emb | 0.074 | Old |
| 0.072483 | 0.174217 | 0.229 | 0.191 | 1 | 3 | Rcn2 | 0.038 | Old |
| 0.072583 | 0.220049 | 0.159 | 0.125 | 1 | 3 | Ddx1 | 0.034 | Old |
| 0.072641 | 0.112407 | 0.288 | 0.241 | 1 | 3 | Rnf145 | 0.047 | Old |
| 0.072713 | 0.38176 | 0.069 | 0.048 | 1 | 3 | Rasa4 | 0.021 | Old |
| 0.072756 | 0.194639 | 0.162 | 0.129 | 1 | 3 | Commd1 | 0.033 | Old |

| | | | | | | | | |
|---|---|---|---|---|---|---|---|---|
| 0.07296 | 0.174097 | 0.053 | 0.034 | 1 | 3 | Ipo4 | 0.019 | Old |
| 0.073016 | 0.439752 | 0.127 | 0.098 | 1 | 3 | Thoc3 | 0.029 | Old |
| 0.073377 | 0.111251 | 0.18 | 0.144 | 1 | 3 | Tmem131l | 0.036 | Old |
| 0.073629 | 0.280784 | 0.053 | 0.034 | 1 | 3 | Asb7 | 0.019 | Old |
| 0.073678 | 0.08036 | 0.189 | 0.152 | 1 | 3 | Nup153 | 0.037 | Old |
| 0.073731 | 0.280305 | 0.092 | 0.067 | 1 | 3 | Coa6 | 0.025 | Old |
| 0.073732 | 0.596839 | 0.096 | 0.071 | 1 | 3 | Ppp2r1b | 0.025 | Old |
| 0.07378 | 0.391689 | 0.084 | 0.061 | 1 | 3 | Triobp | 0.023 | Old |
| 0.073873 | 0.022176 | 0.069 | 0.048 | 1 | 3 | Lclat1 | 0.021 | Old |
| 0.073881 | 0.376222 | 0.03 | 0.016 | 1 | 3 | Zcchc14 | 0.014 | Old |
| 0.073954 | 0.364684 | 0.174 | 0.143 | 1 | 3 | Cyp20a1 | 0.031 | Old |
| 0.073981 | 0.320005 | 0.143 | 0.114 | 1 | 3 | Srp68 | 0.029 | Old |
| 0.074052 | 0.672219 | 0.063 | 0.043 | 1 | 3 | Akr1e1 | 0.02 | Old |
| 0.074086 | 0.138632 | 0.111 | 0.083 | 1 | 3 | Sec23a | 0.028 | Old |
| 0.074192 | 0.142882 | 0.376 | 0.326 | 1 | 3 | Ptpn12 | 0.05 | Old |
| 0.074342 | 0.627932 | 0.06 | 0.04 | 1 | 3 | Elovl6 | 0.02 | Old |
| 0.07435 | 0.130762 | 0.08 | 0.057 | 1 | 3 | Pyurf | 0.023 | Old |
| 0.074421 | 0.506178 | 0.059 | 0.039 | 1 | 3 | Wdr4 | 0.02 | Old |
| 0.074502 | 0.061235 | 0.244 | 0.201 | 1 | 3 | Ptgir | 0.043 | Old |
| 0.07452 | 0.108681 | 0.812 | 0.754 | 1 | 3 | Atp1b3 | 0.058 | Old |
| 0.07486 | 0.08353 | 0.297 | 0.247 | 1 | 3 | Mrps15 | 0.05 | Old |
| 0.074867 | 0.825033 | 0.016 | 0.006 | 1 | 3 | Gm31718 | 0.01 | Old |
| 0.075058 | 0.791953 | 0.016 | 0.006 | 1 | 3 | Epop | 0.01 | Old |
| 0.075065 | 0.685937 | 0.026 | 0.014 | 1 | 3 | Ankdd1a | 0.012 | Old |
| 0.075119 | 0.093139 | 0.504 | 0.448 | 1 | 3 | Ms4a8a | 0.056 | Old |
| 0.075158 | 0.263592 | 0.135 | 0.106 | 1 | 3 | Tsfm | 0.029 | Old |
| 0.07521 | 0.280869 | 0.221 | 0.186 | 1 | 3 | Prdx3 | 0.035 | Old |
| 0.075212 | 0.150222 | 0.53 | 0.482 | 1 | 3 | Pik3cd | 0.048 | Old |
| 0.075422 | 0.179048 | 0.294 | 0.247 | 1 | 3 | R3hdm4 | 0.047 | Old |
| 0.075469 | 0.387027 | 0.086 | 0.063 | 1 | 3 | Scoc | 0.023 | Old |
| 0.07555 | 0.096587 | 0.783 | 0.74 | 1 | 3 | Slc6a6 | 0.043 | Old |
| 0.075604 | 0.095288 | 0.287 | 0.24 | 1 | 3 | Tcf12 | 0.047 | Old |
| 0.075613 | 0.087521 | 0.762 | 0.718 | 1 | 3 | Ndufa4 | 0.044 | Old |
| 0.075631 | 0.734438 | 0.016 | 0.006 | 1 | 3 | 9130230L2 | 0.01 | Old |
| 0.075857 | 0.162124 | 0.283 | 0.241 | 1 | 3 | Acat1 | 0.042 | Old |
| 0.07586 | 0.07908 | 0.789 | 0.712 | 1 | 3 | Arpc3 | 0.077 | Old |
| 0.076011 | 0.408478 | 0.105 | 0.08 | 1 | 3 | Vegfa | 0.025 | Old |
| 0.076031 | 0.06941 | 0.264 | 0.221 | 1 | 3 | Cavin1 | 0.043 | Old |
| 0.076111 | 0.135342 | 0.33 | 0.279 | 1 | 3 | Pira2 | 0.051 | Old |
| 0.076193 | 0.054278 | 0.435 | 0.37 | 1 | 3 | Mapk1ip1l | 0.065 | Old |
| 0.076341 | 0.526347 | 0.055 | 0.037 | 1 | 3 | Phtf1 | 0.018 | Old |
| 0.076461 | 0.499667 | 0.039 | 0.024 | 1 | 3 | Kpna2 | 0.015 | Old |
| 0.076544 | 0.244356 | 0.18 | 0.146 | 1 | 3 | Tm9sf4 | 0.034 | Old |
| 0.076555 | 0.095101 | 0.509 | 0.442 | 1 | 3 | Nras | 0.067 | Old |
| 0.07656 | 0.076166 | 0.539 | 0.484 | 1 | 3 | Gns | 0.055 | Old |

| | | | | | | | | |
|---|---|---|---|---|---|---|---|---|
| 0.07668 | 0.187993 | 0.182 | 0.145 | 1 | 3 | Dlg1 | 0.037 | Old |
| 0.076761 | 0.165898 | 0.225 | 0.188 | 1 | 3 | BC004004 | 0.037 | Old |
| 0.076799 | 0.102317 | 0.904 | 0.842 | 1 | 3 | Dazap2 | 0.062 | Old |
| 0.076856 | 0.302396 | 0.115 | 0.087 | 1 | 3 | Rae1 | 0.028 | Old |
| 0.076895 | 0.243438 | 0.179 | 0.145 | 1 | 3 | Cpq | 0.034 | Old |
| 0.076927 | 0.004579 | 0.235 | 0.194 | 1 | 3 | Cse1l | 0.041 | Old |
| 0.076982 | 0.355817 | 0.016 | 0.006 | 1 | 3 | Kcng2 | 0.01 | Old |
| 0.077038 | 0.731197 | 0.039 | 0.024 | 1 | 3 | Leng9 | 0.015 | Old |
| 0.077372 | 0.317342 | 0.219 | 0.183 | 1 | 3 | Cmc1 | 0.036 | Old |
| 0.077514 | 0.149393 | 0.299 | 0.251 | 1 | 3 | Rtf1 | 0.048 | Old |
| 0.077819 | 0.171189 | 0.102 | 0.076 | 1 | 3 | Gpatch2l | 0.026 | Old |
| 0.077829 | 0.015205 | 0.552 | 0.49 | 1 | 3 | Vasp | 0.062 | Old |
| 0.078008 | 0.554565 | 0.026 | 0.014 | 1 | 3 | Mkks | 0.012 | Old |
| 0.078009 | 0.585765 | 0.079 | 0.057 | 1 | 3 | Tspan13 | 0.022 | Old |
| 0.078088 | 0.130247 | 0.301 | 0.251 | 1 | 3 | Cib1 | 0.05 | Old |
| 0.078196 | 0.824466 | 0.031 | 0.018 | 1 | 3 | Zfp609 | 0.013 | Old |
| 0.07825 | 0.331032 | 0.103 | 0.077 | 1 | 3 | Riox2 | 0.026 | Old |
| 0.078279 | 0.13884 | 0.616 | 0.537 | 1 | 3 | Ube2l3 | 0.079 | Old |
| 0.078315 | 0.111044 | 0.588 | 0.525 | 1 | 3 | Zfp706 | 0.063 | Old |
| 0.078374 | 0.339817 | 0.026 | 0.014 | 1 | 3 | Megf9 | 0.012 | Old |
| 0.078395 | 0.038652 | 0.143 | 0.111 | 1 | 3 | Ppa1 | 0.032 | Old |
| 0.078405 | 0.856604 | 0.019 | 0.009 | 1 | 3 | Ptgs2os2 | 0.01 | Old |
| 0.078406 | 0.41941 | 0.039 | 0.024 | 1 | 3 | Mtx3 | 0.015 | Old |
| 0.078498 | 0.014691 | 0.51 | 0.443 | 1 | 3 | Usp9x | 0.067 | Old |
| 0.078594 | 0.360483 | 0.026 | 0.014 | 1 | 3 | Shank1 | 0.012 | Old |
| 0.078612 | 0.157932 | 0.072 | 0.051 | 1 | 3 | Frs2 | 0.021 | Old |
| 0.07864 | 0.240359 | 0.06 | 0.04 | 1 | 3 | Nek1 | 0.02 | Old |
| 0.078674 | 0.112021 | 0.229 | 0.188 | 1 | 3 | Ece1 | 0.041 | Old |
| 0.078788 | 0.283548 | 0.104 | 0.078 | 1 | 3 | Fez2 | 0.026 | Old |
| 0.078912 | 0.127285 | 0.397 | 0.341 | 1 | 3 | Immt | 0.056 | Old |
| 0.078937 | 0.07673 | 0.875 | 0.819 | 1 | 3 | Rbms1 | 0.056 | Old |
| 0.079037 | 0.307638 | 0.092 | 0.068 | 1 | 3 | 1700017B( | 0.024 | Old |
| 0.079118 | 0.161814 | 0.308 | 0.268 | 1 | 3 | Psme4 | 0.04 | Old |
| 0.079143 | 0.0345 | 0.478 | 0.417 | 1 | 3 | Uba1 | 0.061 | Old |
| 0.079424 | 0.111112 | 0.579 | 0.524 | 1 | 3 | Prkar2a | 0.055 | Old |
| 0.079536 | 0.195881 | 0.308 | 0.265 | 1 | 3 | Naa15 | 0.043 | Old |
| 0.079548 | 0.667599 | 0.019 | 0.009 | 1 | 3 | Zfp810 | 0.01 | Old |
| 0.07984 | 0.084509 | 0.466 | 0.408 | 1 | 3 | AI467606 | 0.058 | Old |
| 0.07994 | 0.7893 | 0.031 | 0.018 | 1 | 3 | Gpsm2 | 0.013 | Old |
| 0.080016 | 0.53465 | 0.073 | 0.052 | 1 | 3 | Ncdn | 0.021 | Old |
| 0.08011 | 0.104754 | 0.55 | 0.491 | 1 | 3 | Mapk3 | 0.059 | Old |
| 0.08015 | 0.108072 | 0.17 | 0.135 | 1 | 3 | Drg1 | 0.035 | Old |
| 0.080158 | 0.214335 | 0.268 | 0.23 | 1 | 3 | Psmb2 | 0.038 | Old |
| 0.080193 | 0.370539 | 0.066 | 0.045 | 1 | 3 | Hmces | 0.021 | Old |
| 0.080214 | 0.639987 | 0.05 | 0.033 | 1 | 3 | Setdb2 | 0.017 | Old |

| | | | | | | | | |
|---|---|---|---|---|---|---|---|---|
| 0.08029 | 0.122585 | 0.313 | 0.265 | 1 | 3 | Emc6 | 0.048 | Old |
| 0.080323 | 0.184032 | 0.437 | 0.383 | 1 | 3 | Pon2 | 0.054 | Old |
| 0.080438 | 0.169627 | 0.127 | 0.098 | 1 | 3 | Sash1 | 0.029 | Old |
| 0.080526 | 0.63699 | 0.019 | 0.009 | 1 | 3 | Cd72 | 0.01 | Old |
| 0.080526 | 0.574151 | 0.019 | 0.009 | 1 | 3 | Gcnt2 | 0.01 | Old |
| 0.080543 | 0.235422 | 0.281 | 0.241 | 1 | 3 | Lgals9 | 0.04 | Old |
| 0.080549 | 0.096013 | 0.24 | 0.197 | 1 | 3 | Arhgap10 | 0.043 | Old |
| 0.08091 | 0.313105 | 0.152 | 0.121 | 1 | 3 | Dtx3l | 0.031 | Old |
| 0.080961 | 0.661664 | 0.031 | 0.018 | 1 | 3 | Pdk1 | 0.013 | Old |
| 0.080994 | 0.109371 | 0.729 | 0.67 | 1 | 3 | Abracl | 0.059 | Old |
| 0.081053 | 0.428636 | 0.039 | 0.024 | 1 | 3 | Dffa | 0.015 | Old |
| 0.081116 | 0.130305 | 0.324 | 0.278 | 1 | 3 | Fndc3b | 0.046 | Old |
| 0.081177 | 0.181852 | 0.118 | 0.091 | 1 | 3 | Ciapin1 | 0.027 | Old |
| 0.081303 | 0.156772 | 0.204 | 0.169 | 1 | 3 | 1810030O( | 0.035 | Old |
| 0.081536 | 0.401671 | 0.05 | 0.033 | 1 | 3 | Cipc | 0.017 | Old |
| 0.081556 | 0.134089 | 0.241 | 0.202 | 1 | 3 | Ube2j1 | 0.039 | Old |
| 0.081631 | 0.184439 | 0.351 | 0.302 | 1 | 3 | Pa2g4 | 0.049 | Old |
| 0.081662 | 0.340722 | 0.139 | 0.11 | 1 | 3 | Psmd9 | 0.029 | Old |
| 0.081723 | 0.348625 | 0.111 | 0.086 | 1 | 3 | Sqle | 0.025 | Old |
| 0.081773 | 0.235378 | 0.054 | 0.035 | 1 | 3 | Hmgxb4 | 0.019 | Old |
| 0.08178 | 0.12616 | 0.338 | 0.294 | 1 | 3 | Cd300a | 0.044 | Old |
| 0.081896 | 0.775312 | 0.062 | 0.043 | 1 | 3 | Alkbh6 | 0.019 | Old |
| 0.081911 | 0.266538 | 0.131 | 0.104 | 1 | 3 | Mrps7 | 0.027 | Old |
| 0.081954 | 0.050172 | 0.053 | 0.034 | 1 | 3 | Mgat5 | 0.019 | Old |
| 0.082121 | 0.078604 | 0.183 | 0.146 | 1 | 3 | Atl3 | 0.037 | Old |
| 0.082162 | 0.189705 | 0.112 | 0.086 | 1 | 3 | Slc10a7 | 0.026 | Old |
| 0.082175 | 0.199385 | 0.271 | 0.231 | 1 | 3 | Pole4 | 0.04 | Old |
| 0.082201 | 0.042544 | 0.384 | 0.331 | 1 | 3 | Cycs | 0.053 | Old |
| 0.082296 | 0.118294 | 0.146 | 0.114 | 1 | 3 | Cul4a | 0.032 | Old |
| 0.082362 | 0.295456 | 0.08 | 0.058 | 1 | 3 | Zfp266 | 0.022 | Old |
| 0.082536 | 0.267227 | 0.044 | 0.028 | 1 | 3 | Cep162 | 0.016 | Old |
| 0.082619 | 0.310545 | 0.097 | 0.072 | 1 | 3 | Hs2st1 | 0.025 | Old |
| 0.08271 | 0.121438 | 0.146 | 0.114 | 1 | 3 | Heatr1 | 0.032 | Old |
| 0.083047 | 0.084951 | 0.287 | 0.242 | 1 | 3 | Stxbp2 | 0.045 | Old |
| 0.083075 | 0.092114 | 0.327 | 0.279 | 1 | 3 | Hk2 | 0.048 | Old |
| 0.083105 | 0.531596 | 0.031 | 0.018 | 1 | 3 | Itgb5 | 0.013 | Old |
| 0.083142 | 0.171647 | 0.283 | 0.245 | 1 | 3 | Zfr | 0.038 | Old |
| 0.083188 | 0.071403 | 0.405 | 0.354 | 1 | 3 | Mtch1 | 0.051 | Old |
| 0.08336 | 0.380211 | 0.111 | 0.086 | 1 | 3 | Apeh | 0.025 | Old |
| 0.083374 | 0.227646 | 0.131 | 0.102 | 1 | 3 | Kpna6 | 0.029 | Old |
| 0.083401 | 0.107008 | 0.503 | 0.457 | 1 | 3 | Ddx39b | 0.046 | Old |
| 0.083482 | 0.439524 | 0.093 | 0.069 | 1 | 3 | Nup160 | 0.024 | Old |
| 0.083538 | 0.434741 | 0.073 | 0.052 | 1 | 3 | Nt5dc1 | 0.021 | Old |
| 0.083552 | 0.043626 | 0.459 | 0.402 | 1 | 3 | Psmc3 | 0.057 | Old |
| 0.083735 | 0.223925 | 0.171 | 0.138 | 1 | 3 | Tbc1d22a | 0.033 | Old |

| | | | | | | | | |
|---|---|---|---|---|---|---|---|---|
| 0.083873 | 0.208327 | 0.084 | 0.061 | 1 | 3 | Tpst1 | 0.023 | Old |
| 0.083902 | 0.243122 | 0.164 | 0.131 | 1 | 3 | Hdac9 | 0.033 | Old |
| 0.083909 | 0.054298 | 0.994 | 0.989 | 1 | 3 | mt-Nd4 | 0.005 | Old |
| 0.083913 | 0.23468 | 0.208 | 0.172 | 1 | 3 | Slc36a4 | 0.036 | Old |
| 0.083994 | 0.127726 | 0.591 | 0.533 | 1 | 3 | Mdh2 | 0.058 | Old |
| 0.084012 | 0.231478 | 0.441 | 0.393 | 1 | 3 | Clec10a | 0.048 | Old |
| 0.084137 | 0.432308 | 0.157 | 0.126 | 1 | 3 | Cnnm4 | 0.031 | Old |
| 0.084162 | 0.14261 | 0.53 | 0.486 | 1 | 3 | Irf1 | 0.044 | Old |
| 0.084169 | 0.215512 | 0.237 | 0.196 | 1 | 3 | Cnot4 | 0.041 | Old |
| 0.084418 | 0.09962 | 0.339 | 0.288 | 1 | 3 | Pfdn2 | 0.051 | Old |
| 0.084418 | 0.067654 | 0.358 | 0.301 | 1 | 3 | Aftph | 0.057 | Old |
| 0.084435 | 0.086465 | 0.296 | 0.25 | 1 | 3 | Mob1b | 0.046 | Old |
| 0.084454 | 0.129751 | 0.519 | 0.468 | 1 | 3 | Snx10 | 0.051 | Old |
| 0.084512 | 0.127349 | 0.496 | 0.449 | 1 | 3 | Magt1 | 0.047 | Old |
| 0.084591 | 0.117917 | 0.339 | 0.288 | 1 | 3 | Usp8 | 0.051 | Old |
| 0.084744 | 0.120308 | 0.257 | 0.216 | 1 | 3 | Dctn4 | 0.041 | Old |
| 0.085088 | 0.166649 | 0.151 | 0.119 | 1 | 3 | Vps11 | 0.032 | Old |
| 0.085685 | 0.366411 | 0.045 | 0.029 | 1 | 3 | Alg11 | 0.016 | Old |
| 0.085791 | 0.070996 | 0.155 | 0.121 | 1 | 3 | Nudt9 | 0.034 | Old |
| 0.085806 | 0.654221 | 0.078 | 0.057 | 1 | 3 | Rhot2 | 0.021 | Old |
| 0.085979 | 0.55475 | 0.081 | 0.059 | 1 | 3 | Itgal | 0.022 | Old |
| 0.086051 | 0.23619 | 0.105 | 0.08 | 1 | 3 | Flot1 | 0.025 | Old |
| 0.086056 | 0.202496 | 0.374 | 0.324 | 1 | 3 | Psma5 | 0.05 | Old |
| 0.086139 | 0.687514 | 0.037 | 0.023 | 1 | 3 | Tigar | 0.014 | Old |
| 0.086193 | 0.120512 | 0.369 | 0.323 | 1 | 3 | Golph3 | 0.046 | Old |
| 0.086208 | 0.078678 | 0.336 | 0.283 | 1 | 3 | Pik3r5 | 0.053 | Old |
| 0.086345 | 0.138448 | 0.535 | 0.49 | 1 | 3 | 1110008F1 | 0.045 | Old |
| 0.086391 | 0.09942 | 0.239 | 0.198 | 1 | 3 | Suds3 | 0.041 | Old |
| 0.086393 | 0.067033 | 0.287 | 0.237 | 1 | 3 | Rbm26 | 0.05 | Old |
| 0.086423 | 0.113414 | 0.084 | 0.061 | 1 | 3 | Lztr1 | 0.023 | Old |
| 0.086677 | 0.088932 | 0.241 | 0.201 | 1 | 3 | Tle4 | 0.04 | Old |
| 0.086682 | 0.997814 | 0.013 | 0.005 | 1 | 3 | Zrsr1 | 0.008 | Old |
| 0.086682 | 0.675478 | 0.013 | 0.005 | 1 | 3 | Vegfb | 0.008 | Old |
| 0.086741 | 0.187503 | 0.098 | 0.073 | 1 | 3 | Slc2a1 | 0.025 | Old |
| 0.086799 | 0.975513 | 0.013 | 0.005 | 1 | 3 | Bbs12 | 0.008 | Old |
| 0.086805 | 0.249424 | 0.16 | 0.13 | 1 | 3 | Rbmxl1 | 0.03 | Old |
| 0.086917 | 0.005819 | 0.124 | 0.154 | 1 | 3 | Cd209a | -0.03 | Old |
| 0.087199 | 0.122565 | 0.705 | 0.643 | 1 | 3 | Plekho2 | 0.062 | Old |
| 0.087386 | 0.289547 | 0.201 | 0.17 | 1 | 3 | Plbd2 | 0.031 | Old |
| 0.08739 | 0.073479 | 0.118 | 0.091 | 1 | 3 | Dhrs7b | 0.027 | Old |
| 0.087437 | 0.226795 | 0.183 | 0.149 | 1 | 3 | Letm1 | 0.034 | Old |
| 0.087438 | 0.12255 | 0.164 | 0.129 | 1 | 3 | Magohb | 0.035 | Old |
| 0.087445 | 0.102744 | 0.282 | 0.239 | 1 | 3 | Eif2s1 | 0.043 | Old |
| 0.087614 | 0.276195 | 0.21 | 0.176 | 1 | 3 | Erap1 | 0.034 | Old |
| 0.087639 | 0.152151 | 0.162 | 0.129 | 1 | 3 | Nolc1 | 0.033 | Old |

| | | | | | | | | |
|---|---|---|---|---|---|---|---|---|
| 0.087728 | 0.172832 | 0.545 | 0.475 | 1 | 3 | Sowahc | 0.07 | Old |
| 0.087856 | 0.569532 | 0.032 | 0.019 | 1 | 3 | Sars2 | 0.013 | Old |
| 0.087883 | 0.212626 | 0.162 | 0.13 | 1 | 3 | Uri1 | 0.032 | Old |
| 0.087906 | 0.064721 | 0.249 | 0.21 | 1 | 3 | Dyrk2 | 0.039 | Old |
| 0.087953 | 0.156331 | 0.091 | 0.067 | 1 | 3 | Nelfcd | 0.024 | Old |
| 0.087979 | 0.866452 | 0.037 | 0.023 | 1 | 3 | Tcta | 0.014 | Old |
| 0.088035 | 0.104253 | 0.151 | 0.121 | 1 | 3 | Nqo2 | 0.03 | Old |
| 0.088238 | 0.034983 | 0.33 | 0.28 | 1 | 3 | Abca7 | 0.05 | Old |
| 0.08824 | 0.063201 | 0.508 | 0.448 | 1 | 3 | Lrrc58 | 0.06 | Old |
| 0.088295 | 0.100458 | 0.424 | 0.369 | 1 | 3 | Pdcd10 | 0.055 | Old |
| 0.088398 | 0.489132 | 0.108 | 0.082 | 1 | 3 | Ptgr1 | 0.026 | Old |
| 0.088453 | 0.61002 | 0.013 | 0.005 | 1 | 3 | Hist1h4h | 0.008 | Old |
| 0.088479 | 0.087081 | 0.507 | 0.438 | 1 | 3 | Gnai3 | 0.069 | Old |
| 0.088672 | 0.187586 | 0.238 | 0.201 | 1 | 3 | Klhdc3 | 0.037 | Old |
| 0.088682 | 0.181416 | 0.198 | 0.164 | 1 | 3 | Ddx10 | 0.034 | Old |
| 0.088692 | 0.41078 | 0.013 | 0.005 | 1 | 3 | Adat1 | 0.008 | Old |
| 0.088982 | 0.064346 | 0.822 | 0.799 | 1 | 3 | App | 0.023 | Old |
| 0.088986 | 0.0414 | 0.235 | 0.196 | 1 | 3 | Cat | 0.039 | Old |
| 0.089072 | 0.082079 | 0.612 | 0.551 | 1 | 3 | Pum2 | 0.061 | Old |
| 0.089209 | 0.339961 | 0.112 | 0.087 | 1 | 3 | Pik3cb | 0.025 | Old |
| 0.089313 | 0.696742 | 0.037 | 0.023 | 1 | 3 | Zc3h8 | 0.014 | Old |
| 0.089373 | 0.175513 | 0.355 | 0.306 | 1 | 3 | Tank | 0.049 | Old |
| 0.089461 | 0.089913 | 0.216 | 0.178 | 1 | 3 | Slc25a11 | 0.038 | Old |
| 0.089473 | 0.213626 | 0.178 | 0.145 | 1 | 3 | Dpp9 | 0.033 | Old |
| 0.089798 | 0.245835 | 0.146 | 0.116 | 1 | 3 | Smad5 | 0.03 | Old |
| 0.090042 | 0.051929 | 0.287 | 0.246 | 1 | 3 | Dennd4b | 0.041 | Old |
| 0.090114 | 0.245924 | 0.145 | 0.115 | 1 | 3 | Dpy30 | 0.03 | Old |
| 0.090152 | 0.12137 | 0.582 | 0.514 | 1 | 3 | Mrpl52 | 0.068 | Old |
| 0.090375 | 0.020588 | 0.151 | 0.119 | 1 | 3 | Rinl | 0.032 | Old |
| 0.090447 | 0.838756 | 0.024 | 0.013 | 1 | 3 | Dhrs11 | 0.011 | Old |
| 0.090481 | 0.099351 | 0.485 | 0.428 | 1 | 3 | Copa | 0.057 | Old |
| 0.090518 | 0.248888 | 0.041 | 0.025 | 1 | 3 | Lonrf1 | 0.016 | Old |
| 0.090718 | 0.062728 | 0.235 | 0.192 | 1 | 3 | Cln8 | 0.043 | Old |
| 0.090783 | 0.110908 | 0.376 | 0.328 | 1 | 3 | Mdh1 | 0.048 | Old |
| 0.090879 | 0.668781 | 0.024 | 0.013 | 1 | 3 | Polrmt | 0.011 | Old |
| 0.090963 | 0.049341 | 0.577 | 0.509 | 1 | 3 | Txndc17 | 0.068 | Old |
| 0.091049 | 0.152969 | 0.462 | 0.418 | 1 | 3 | Sf3b5 | 0.044 | Old |
| 0.091062 | 0.040499 | 1 | 0.999 | 1 | 3 | Rpl37a | 0.001 | Old |
| 0.091193 | 0.068842 | 0.564 | 0.5 | 1 | 3 | Selenof | 0.064 | Old |
| 0.091209 | 0.240395 | 0.051 | 0.034 | 1 | 3 | Hist1h4i | 0.017 | Old |
| 0.091211 | 0.049652 | 0.284 | 0.237 | 1 | 3 | Setd1b | 0.047 | Old |
| 0.091258 | 0.035136 | 0.24 | 0.203 | 1 | 3 | Ell2 | 0.037 | Old |
| 0.091277 | 1.108997 | 0.017 | 0.008 | 1 | 3 | C920006O[illegible] | 0.009 | Old |
| 0.091361 | 0.039963 | 0.712 | 0.646 | 1 | 3 | Csrnp1 | 0.066 | Old |
| 0.091387 | 0.36298 | 0.099 | 0.076 | 1 | 3 | Strip1 | 0.023 | Old |

| | | | | | | | | |
|---|---|---|---|---|---|---|---|---|
| 0.091416 | 0.086958 | 0.642 | 0.58 | 1 | 3 | Romo1 | 0.062 | Old |
| 0.091702 | 1.061777 | 0.017 | 0.008 | 1 | 3 | Otud3 | 0.009 | Old |
| 0.091792 | 0.019202 | 0.211 | 0.173 | 1 | 3 | Efcab14 | 0.038 | Old |
| 0.091889 | 0.671117 | 0.043 | 0.028 | 1 | 3 | Alad | 0.015 | Old |
| 0.091913 | 0.121093 | 0.097 | 0.072 | 1 | 3 | Zswim4 | 0.025 | Old |
| 0.091947 | 0.813839 | 0.029 | 0.016 | 1 | 3 | Rubcnl | 0.013 | Old |
| 0.092086 | 0.081358 | 0.112 | 0.086 | 1 | 3 | Mrpl40 | 0.026 | Old |
| 0.092183 | 0.790452 | 0.02 | 0.01 | 1 | 3 | Zfp951 | 0.01 | Old |
| 0.09228 | 0.043006 | 0.53 | 0.471 | 1 | 3 | Prrc2a | 0.059 | Old |
| 0.092304 | 0.34425 | 0.1 | 0.077 | 1 | 3 | Ikbip | 0.023 | Old |
| 0.092388 | 0.480566 | 0.09 | 0.067 | 1 | 3 | Rufy1 | 0.023 | Old |
| 0.0924 | 0.005582 | 0.437 | 0.374 | 1 | 3 | Cab39 | 0.063 | Old |
| 0.092658 | 0.150389 | 0.116 | 0.088 | 1 | 3 | Nom1 | 0.028 | Old |
| 0.092707 | 0.315392 | 0.129 | 0.102 | 1 | 3 | F2rl2 | 0.027 | Old |
| 0.092951 | 0.598716 | 0.02 | 0.01 | 1 | 3 | Gm5150 | 0.01 | Old |
| 0.093026 | 0.0526 | 0.406 | 0.352 | 1 | 3 | Gnpda1 | 0.054 | Old |
| 0.0932 | 0.604104 | 0.017 | 0.008 | 1 | 3 | Gm20513 | 0.009 | Old |
| 0.093292 | 0.093803 | 0.594 | 0.533 | 1 | 3 | Bsg | 0.061 | Old |
| 0.093325 | 0.110068 | 0.324 | 0.274 | 1 | 3 | Nudt21 | 0.05 | Old |
| 0.093328 | 0.166751 | 0.388 | 0.345 | 1 | 3 | Uqcrc2 | 0.043 | Old |
| 0.093381 | 0.074299 | 0.091 | 0.067 | 1 | 3 | E4f1 | 0.024 | Old |
| 0.093496 | 0.074907 | 0.284 | 0.245 | 1 | 3 | Bcl3 | 0.039 | Old |
| 0.093638 | 0.21336 | 0.178 | 0.144 | 1 | 3 | Slc25a20 | 0.034 | Old |
| 0.093657 | 0.234 | 0.171 | 0.138 | 1 | 3 | St6galnac4 | 0.033 | Old |
| 0.093702 | 0.112966 | 0.823 | 0.756 | 1 | 3 | Serbp1 | 0.067 | Old |
| 0.093722 | 0.278425 | 0.133 | 0.105 | 1 | 3 | Hikeshi | 0.028 | Old |
| 0.093912 | 0.102888 | 0.785 | 0.735 | 1 | 3 | Zyx | 0.05 | Old |
| 0.094165 | 0.307271 | 0.103 | 0.078 | 1 | 3 | Rnf41 | 0.025 | Old |
| 0.09423 | 0.261824 | 0.209 | 0.177 | 1 | 3 | Ftsj3 | 0.032 | Old |
| 0.094472 | 0.243143 | 0.176 | 0.144 | 1 | 3 | Ppfibp1 | 0.032 | Old |
| 0.09451 | 0.280738 | 0.075 | 0.054 | 1 | 3 | Ngrn | 0.021 | Old |
| 0.094523 | 0.037644 | 0.319 | 0.274 | 1 | 3 | Ddrgk1 | 0.045 | Old |
| 0.094541 | 0.138419 | 0.205 | 0.169 | 1 | 3 | Prkce | 0.036 | Old |
| 0.094548 | 0.117551 | 0.477 | 0.414 | 1 | 3 | Cisd2 | 0.063 | Old |
| 0.094613 | 0.718374 | 0.049 | 0.033 | 1 | 3 | Gtpbp10 | 0.016 | Old |
| 0.09475 | 0.147951 | 0.341 | 0.294 | 1 | 3 | Ssrp1 | 0.047 | Old |
| 0.094784 | 0.221551 | 0.368 | 0.326 | 1 | 3 | Uhmk1 | 0.042 | Old |
| 0.094815 | 0.124886 | 0.205 | 0.169 | 1 | 3 | Ldah | 0.036 | Old |
| 0.094863 | 0.106203 | 0.209 | 0.177 | 1 | 3 | Usp6nl | 0.032 | Old |
| 0.095034 | 0.058198 | 0.123 | 0.096 | 1 | 3 | Cstf2t | 0.027 | Old |
| 0.095046 | 0.1438 | 0.074 | 0.053 | 1 | 3 | 4930453N2 | 0.021 | Old |
| 0.095099 | 0.151834 | 0.128 | 0.101 | 1 | 3 | Hmg20a | 0.027 | Old |
| 0.095105 | 0.0639 | 0.33 | 0.288 | 1 | 3 | Dcstamp | 0.042 | Old |
| 0.095192 | 0.11759 | 0.349 | 0.302 | 1 | 3 | Napa | 0.047 | Old |
| 0.095291 | 0.362719 | 0.127 | 0.1 | 1 | 3 | Igsf8 | 0.027 | Old |

| | | | | | | | | |
|---|---|---|---|---|---|---|---|---|
| 0.095401 | 0.099531 | 0.086 | 0.063 | 1 | 3 | Cyth3 | 0.023 | Old |
| 0.095634 | 0.106618 | 0.638 | 0.58 | 1 | 3 | Tapbp | 0.058 | Old |
| 0.095714 | 0.261384 | 0.141 | 0.111 | 1 | 3 | Pdxdc1 | 0.03 | Old |
| 0.095769 | 0.147823 | 0.106 | 0.081 | 1 | 3 | Mrps18b | 0.025 | Old |
| 0.095937 | 0.116861 | 0.797 | 0.73 | 1 | 3 | Hnrnpa0 | 0.067 | Old |
| 0.096075 | 0.252187 | 0.075 | 0.054 | 1 | 3 | Rbl1 | 0.021 | Old |
| 0.096089 | 0.29433 | 0.067 | 0.047 | 1 | 3 | Cdk7 | 0.02 | Old |
| 0.09609 | 0.222414 | 0.195 | 0.163 | 1 | 3 | Ptbp2 | 0.032 | Old |
| 0.096174 | 0.416522 | 0.065 | 0.045 | 1 | 3 | Serhl | 0.02 | Old |
| 0.09618 | 0.236137 | 0.153 | 0.122 | 1 | 3 | B9d2 | 0.031 | Old |
| 0.096203 | 0.3768 | 0.082 | 0.061 | 1 | 3 | Btg3 | 0.021 | Old |
| 0.096418 | 0.200006 | 0.063 | 0.044 | 1 | 3 | Dpp7 | 0.019 | Old |
| 0.09649 | 0.753462 | 0.055 | 0.038 | 1 | 3 | Actr5 | 0.017 | Old |
| 0.096546 | 0.12309 | 0.73 | 0.669 | 1 | 3 | Far1 | 0.061 | Old |
| 0.096598 | 0.095728 | 0.535 | 0.489 | 1 | 3 | Stxbp3 | 0.046 | Old |
| 0.096646 | 0.193239 | 0.215 | 0.178 | 1 | 3 | Exosc5 | 0.037 | Old |
| 0.096742 | 0.359556 | 0.076 | 0.056 | 1 | 3 | Rcc1 | 0.02 | Old |
| 0.096789 | 0.169515 | 0.241 | 0.203 | 1 | 3 | Sars | 0.038 | Old |
| 0.096797 | 0.392157 | 0.139 | 0.11 | 1 | 3 | Apex1 | 0.029 | Old |
| 0.096913 | 0.575841 | 0.1 | 0.077 | 1 | 3 | Gpn1 | 0.023 | Old |
| 0.097049 | 0.348716 | 0.09 | 0.067 | 1 | 3 | Cdk6 | 0.023 | Old |
| 0.097056 | 0.323375 | 0.109 | 0.085 | 1 | 3 | Tbrg4 | 0.024 | Old |
| 0.097127 | 0.392726 | 0.07 | 0.051 | 1 | 3 | Pus10 | 0.019 | Old |
| 0.097429 | 0.607473 | 0.029 | 0.016 | 1 | 3 | Cox11 | 0.013 | Old |
| 0.097582 | 0.188585 | 0.366 | 0.323 | 1 | 3 | Atp8b4 | 0.043 | Old |
| 0.097691 | 0.561534 | 0.067 | 0.048 | 1 | 3 | Ap5z1 | 0.019 | Old |
| 0.097786 | 0.19766 | 0.075 | 0.054 | 1 | 3 | Gss | 0.021 | Old |
| 0.097969 | 0.152905 | 0.177 | 0.144 | 1 | 3 | Dda1 | 0.033 | Old |
| 0.098078 | 0.113747 | 0.139 | 0.11 | 1 | 3 | Nvl | 0.029 | Old |
| 0.09809 | 0.535634 | 0.029 | 0.016 | 1 | 3 | Nmnat1 | 0.013 | Old |
| 0.098195 | 0.487591 | 0.038 | 0.024 | 1 | 3 | Irgm2 | 0.014 | Old |
| 0.098231 | 0.043707 | 0.262 | 0.221 | 1 | 3 | Ociad1 | 0.041 | Old |
| 0.098241 | 0.080693 | 0.881 | 0.861 | 1 | 3 | Rrbp1 | 0.02 | Old |
| 0.098275 | 0.081799 | 0.466 | 0.396 | 1 | 3 | Arhgap17 | 0.07 | Old |
| 0.098301 | 0.26593 | 0.179 | 0.148 | 1 | 3 | Cish | 0.031 | Old |
| 0.098339 | 0.536606 | 0.029 | 0.016 | 1 | 3 | B930036N1 | 0.013 | Old |
| 0.098521 | 0.299449 | 0.136 | 0.109 | 1 | 3 | Nipsnap2 | 0.027 | Old |
| 0.098592 | 0.038625 | 0.133 | 0.104 | 1 | 3 | 2610002M | 0.029 | Old |
| 0.098922 | 0.040345 | 0.748 | 0.683 | 1 | 3 | Jpt1 | 0.065 | Old |
| 0.098936 | 0.083469 | 0.677 | 0.604 | 1 | 3 | Itpkb | 0.073 | Old |
| 0.098975 | 0.329246 | 0.084 | 0.062 | 1 | 3 | Cd244 | 0.022 | Old |
| 0.099001 | 0.361889 | 0.125 | 0.098 | 1 | 3 | Plekha3 | 0.027 | Old |
| 0.099078 | 0.18079 | 0.256 | 0.218 | 1 | 3 | Tap2 | 0.038 | Old |
| 0.09909 | 0.604001 | 0.029 | 0.016 | 1 | 3 | 1110008L1 | 0.013 | Old |
| 0.099149 | 0.260555 | 0.128 | 0.1 | 1 | 3 | Slc35a5 | 0.028 | Old |

| | | | | | | | | |
|---|---|---|---|---|---|---|---|---|
| 0.099162 | 0.95313 | 0.011 | 0.004 | 1 | 3 | Gm26825 | 0.007 | Old |
| 0.099386 | 0.557238 | 0.055 | 0.038 | 1 | 3 | Exd2 | 0.017 | Old |
| 0.099501 | 0.30833 | 0.215 | 0.183 | 1 | 3 | Stat1 | 0.032 | Old |
| 0.0996 | 1.05456 | 0.011 | 0.004 | 1 | 3 | 2700046G( | 0.007 | Old |
| 0.099806 | 0.188111 | 0.393 | 0.354 | 1 | 3 | Atf6 | 0.039 | Old |
| 0.099844 | 0.097243 | 0.238 | 0.198 | 1 | 3 | Sae1 | 0.04 | Old |
| 0.10004 | 1.225139 | 0.011 | 0.004 | 1 | 3 | Gm4631 | 0.007 | Old |
| 0.100053 | 0.290723 | 0.085 | 0.063 | 1 | 3 | Nudt13 | 0.022 | Old |
| 0.100157 | 0.076905 | 0.394 | 0.341 | 1 | 3 | Mdfic | 0.053 | Old |
| 0.100286 | 0.076737 | 0.283 | 0.241 | 1 | 3 | Neu1 | 0.042 | Old |
| 0.10029 | 0.130505 | 0.511 | 0.467 | 1 | 3 | Ikzf1 | 0.044 | Old |
| 0.100305 | 0.142791 | 0.252 | 0.216 | 1 | 3 | Rictor | 0.036 | Old |
| 0.100334 | 0.892496 | 0.011 | 0.004 | 1 | 3 | Gprc5b | 0.007 | Old |
| 0.10036 | 0.061134 | 0.877 | 0.826 | 1 | 3 | Irf2bp2 | 0.051 | Old |
| 0.100481 | 0.917952 | 0.011 | 0.004 | 1 | 3 | Zfp975 | 0.007 | Old |
| 0.10053 | 0.154937 | 0.157 | 0.125 | 1 | 3 | Renbp | 0.032 | Old |
| 0.100629 | 0.656458 | 0.011 | 0.004 | 1 | 3 | Dpf3 | 0.007 | Old |
| 0.100685 | 0.10962 | 0.444 | 0.398 | 1 | 3 | Cwc15 | 0.046 | Old |
| 0.100704 | 0.030684 | 0.254 | 0.216 | 1 | 3 | Maff | 0.038 | Old |
| 0.100808 | 0.045679 | 0.346 | 0.297 | 1 | 3 | Tpp2 | 0.049 | Old |
| 0.100834 | 0.060449 | 0.231 | 0.189 | 1 | 3 | Mon2 | 0.042 | Old |
| 0.100945 | 0.107926 | 0.245 | 0.206 | 1 | 3 | Magoh | 0.039 | Old |
| 0.10095 | 0.229257 | 0.205 | 0.172 | 1 | 3 | Tatdn2 | 0.033 | Old |
| 0.100959 | 0.425673 | 0.068 | 0.049 | 1 | 3 | Rnaset2b | 0.019 | Old |
| 0.100975 | 0.393267 | 0.054 | 0.037 | 1 | 3 | Hars2 | 0.017 | Old |
| 0.101039 | 0.204548 | 0.29 | 0.25 | 1 | 3 | Slc35b2 | 0.04 | Old |
| 0.101097 | 0.359775 | 0.134 | 0.107 | 1 | 3 | Stoml2 | 0.027 | Old |
| 0.101101 | 0.053678 | 0.806 | 0.759 | 1 | 3 | Vcp | 0.047 | Old |
| 0.101187 | 0.077216 | 0.173 | 0.14 | 1 | 3 | Sucla2 | 0.033 | Old |
| 0.101267 | 0.23796 | 0.097 | 0.073 | 1 | 3 | Mri1 | 0.024 | Old |
| 0.101285 | 0.050869 | 0.331 | 0.284 | 1 | 3 | Chtop | 0.047 | Old |
| 0.101404 | 0.211392 | 0.253 | 0.216 | 1 | 3 | H1f0 | 0.037 | Old |
| 0.101464 | 0.033502 | 0.189 | 0.154 | 1 | 3 | Mapkapk3 | 0.035 | Old |
| 0.10147 | 0.316256 | 0.068 | 0.049 | 1 | 3 | Nxpe4 | 0.019 | Old |
| 0.101517 | 0.517573 | 0.011 | 0.004 | 1 | 3 | Gpr18 | 0.007 | Old |
| 0.101638 | 0.366373 | 0.06 | 0.042 | 1 | 3 | Pot1a | 0.018 | Old |
| 0.101665 | 0.772302 | 0.011 | 0.004 | 1 | 3 | Erbb3 | 0.007 | Old |
| 0.101784 | 0.013638 | 0.226 | 0.191 | 1 | 3 | Esyt2 | 0.035 | Old |
| 0.101794 | 0.302321 | 0.029 | 0.016 | 1 | 3 | Casp7 | 0.013 | Old |
| 0.101926 | 0.052274 | 0.033 | 0.02 | 1 | 3 | BC002059 | 0.013 | Old |
| 0.101953 | 0.161391 | 0.554 | 0.509 | 1 | 3 | Sgpl1 | 0.045 | Old |
| 0.102112 | 0.167215 | 0.011 | 0.004 | 1 | 3 | Fosl1 | 0.007 | Old |
| 0.102365 | 0.060504 | 0.184 | 0.152 | 1 | 3 | 8-Mar | 0.032 | Old |
| 0.102461 | 0.319056 | 0.141 | 0.114 | 1 | 3 | Elp2 | 0.027 | Old |
| 0.102477 | 0.483699 | 0.033 | 0.02 | 1 | 3 | Zfp653 | 0.013 | Old |

| | | | | | | | | |
|---|---|---|---|---|---|---|---|---|
| 0.1026 | 0.143583 | 0.541 | 0.482 | 1 | 3 | Trir | 0.059 | Old |
| 0.102622 | 0.089765 | 0.104 | 0.08 | 1 | 3 | Srprb | 0.024 | Old |
| 0.102638 | 0.093838 | 0.106 | 0.081 | 1 | 3 | Pop7 | 0.025 | Old |
| 0.102674 | 0.159031 | 0.589 | 0.53 | 1 | 3 | Psmd2 | 0.059 | Old |
| 0.102747 | 0.526755 | 0.093 | 0.071 | 1 | 3 | Sssca1 | 0.022 | Old |
| 0.102977 | 0.309002 | 0.086 | 0.064 | 1 | 3 | Chmp7 | 0.022 | Old |
| 0.102994 | 0.10839 | 0.054 | 0.037 | 1 | 3 | Mgme1 | 0.017 | Old |
| 0.103068 | 0.247234 | 0.184 | 0.153 | 1 | 3 | Pitpnb | 0.031 | Old |
| 0.103309 | 0.415912 | 0.033 | 0.02 | 1 | 3 | Zscan21 | 0.013 | Old |
| 0.103547 | 0.271947 | 0.059 | 0.04 | 1 | 3 | Zfp64 | 0.019 | Old |
| 0.103616 | 0.15374 | 0.19 | 0.157 | 1 | 3 | Sec24a | 0.033 | Old |
| 0.103621 | 0.155988 | 0.344 | 0.303 | 1 | 3 | Ap1s1 | 0.041 | Old |
| 0.103711 | 0.086614 | 0.619 | 0.578 | 1 | 3 | Cox7a2 | 0.041 | Old |
| 0.103802 | 0.128373 | 0.441 | 0.386 | 1 | 3 | Svil | 0.055 | Old |
| 0.103829 | 0.034764 | 0.498 | 0.438 | 1 | 3 | Tgif1 | 0.06 | Old |
| 0.103837 | 0.438351 | 0.049 | 0.033 | 1 | 3 | Zfp971 | 0.016 | Old |
| 0.103893 | 0.340763 | 0.025 | 0.014 | 1 | 3 | Tgfbr3 | 0.011 | Old |
| 0.103939 | 0.050899 | 0.196 | 0.162 | 1 | 3 | Cnn3 | 0.034 | Old |
| 0.103983 | 0.18525 | 0.177 | 0.145 | 1 | 3 | Rpl7l1 | 0.032 | Old |
| 0.103987 | 0.793752 | 0.025 | 0.014 | 1 | 3 | Mcoln2 | 0.011 | Old |
| 0.10406 | 0.206935 | 0.502 | 0.451 | 1 | 3 | Hnrnph2 | 0.051 | Old |
| 0.104161 | 0.241884 | 0.245 | 0.208 | 1 | 3 | Cenpb | 0.037 | Old |
| 0.104181 | 0.144995 | 0.41 | 0.361 | 1 | 3 | Psmb5 | 0.049 | Old |
| 0.104661 | 0.042471 | 0.32 | 0.275 | 1 | 3 | Elk4 | 0.045 | Old |
| 0.104978 | 0.100609 | 0.342 | 0.301 | 1 | 3 | Ipo7 | 0.041 | Old |
| 0.105051 | 0.382366 | 0.043 | 0.028 | 1 | 3 | Trim30c | 0.015 | Old |
| 0.105144 | 0.205328 | 0.146 | 0.117 | 1 | 3 | Synj2 | 0.029 | Old |
| 0.105354 | 0.308022 | 0.038 | 0.024 | 1 | 3 | Ttll4 | 0.014 | Old |
| 0.105396 | 0.20809 | 0.065 | 0.045 | 1 | 3 | Med31 | 0.02 | Old |
| 0.105529 | 0.073518 | 0.419 | 0.366 | 1 | 3 | Cmtm3 | 0.053 | Old |
| 0.105554 | 0.146085 | 0.452 | 0.407 | 1 | 3 | Tpp1 | 0.045 | Old |
| 0.105628 | 0.129523 | 0.18 | 0.146 | 1 | 3 | Ddx23 | 0.034 | Old |
| 0.105787 | 0.167775 | 0.195 | 0.162 | 1 | 3 | Arhgef3 | 0.033 | Old |
| 0.106006 | 0.293292 | 0.135 | 0.109 | 1 | 3 | Tango2 | 0.026 | Old |
| 0.106528 | 0.902936 | 0.03 | 0.018 | 1 | 3 | Blm | 0.012 | Old |
| 0.106583 | 0.421808 | 0.039 | 0.025 | 1 | 3 | Trmt61a | 0.014 | Old |
| 0.106615 | 0.084315 | 0.252 | 0.215 | 1 | 3 | Eif4a3 | 0.037 | Old |
| 0.10685 | 0.357578 | 0.062 | 0.044 | 1 | 3 | Svbp | 0.018 | Old |
| 0.10689 | 0.152876 | 0.114 | 0.088 | 1 | 3 | Slfn8 | 0.026 | Old |
| 0.10689 | 0.151946 | 0.072 | 0.052 | 1 | 3 | Dhx33 | 0.02 | Old |
| 0.106917 | 0.415754 | 0.056 | 0.039 | 1 | 3 | Fam46c | 0.017 | Old |
| 0.107065 | 0.123636 | 0.718 | 0.665 | 1 | 3 | Ap2a2 | 0.053 | Old |
| 0.107331 | 0.108012 | 0.681 | 0.602 | 1 | 3 | Nptn | 0.079 | Old |
| 0.10736 | 0.573216 | 0.035 | 0.021 | 1 | 3 | BC051226 | 0.014 | Old |
| 0.107394 | 0.506358 | 0.03 | 0.018 | 1 | 3 | Nr1h3 | 0.012 | Old |

| | | | | | | | | |
|---|---|---|---|---|---|---|---|---|
| 0.107598 | 0.265246 | 0.129 | 0.102 | 1 | 3 | E2f1 | 0.027 | Old |
| 0.107763 | 0.200246 | 0.129 | 0.102 | 1 | 3 | Stau1 | 0.027 | Old |
| 0.108274 | 0.084414 | 0.735 | 0.681 | 1 | 3 | Pdcd6ip | 0.054 | Old |
| 0.108288 | 0.973424 | 0.022 | 0.011 | 1 | 3 | N4bp3 | 0.011 | Old |
| 0.108394 | 0.189294 | 0.13 | 0.102 | 1 | 3 | Crebzf | 0.028 | Old |
| 0.108414 | 0.445797 | 0.093 | 0.071 | 1 | 3 | Cd151 | 0.022 | Old |
| 0.10845 | 0.157803 | 0.406 | 0.367 | 1 | 3 | Ubald2 | 0.039 | Old |
| 0.10851 | 0.291341 | 0.061 | 0.043 | 1 | 3 | Zfp414 | 0.018 | Old |
| 0.108513 | 0.174658 | 0.578 | 0.516 | 1 | 3 | Bnip2 | 0.062 | Old |
| 0.108531 | 0.340826 | 0.039 | 0.025 | 1 | 3 | Zfp276 | 0.014 | Old |
| 0.108563 | 0.078828 | 0.287 | 0.244 | 1 | 3 | H2-Q7 | 0.043 | Old |
| 0.108652 | 0.285373 | 0.044 | 0.029 | 1 | 3 | Sgsm3 | 0.015 | Old |
| 0.109092 | 0.120077 | 0.209 | 0.176 | 1 | 3 | Keap1 | 0.033 | Old |
| 0.109109 | 0.215861 | 0.185 | 0.154 | 1 | 3 | Ppie | 0.031 | Old |
| 0.109143 | 0.185426 | 0.17 | 0.139 | 1 | 3 | Rasa3 | 0.031 | Old |
| 0.109387 | 0.342417 | 0.049 | 0.033 | 1 | 3 | Ap5s1 | 0.016 | Old |
| 0.109405 | 0.507667 | 0.091 | 0.069 | 1 | 3 | Cpsf1 | 0.022 | Old |
| 0.109446 | 0.060262 | 0.777 | 0.73 | 1 | 3 | Tma7 | 0.047 | Old |
| 0.109482 | 0.34773 | 0.102 | 0.078 | 1 | 3 | Il7r | 0.024 | Old |
| 0.109502 | 0.144564 | 0.066 | 0.047 | 1 | 3 | Slc37a4 | 0.019 | Old |
| 0.10971 | 0.342156 | 0.141 | 0.116 | 1 | 3 | Ddx56 | 0.025 | Old |
| 0.109742 | 0.355361 | 0.275 | 0.236 | 1 | 3 | Erlin1 | 0.039 | Old |
| 0.109866 | 0.492981 | 0.05 | 0.034 | 1 | 3 | Ubxn2b | 0.016 | Old |
| 0.110052 | 1.11056 | 0.014 | 0.006 | 1 | 3 | Ifit1 | 0.008 | Old |
| 0.110093 | 0.054887 | 0.276 | 0.235 | 1 | 3 | Acaa1a | 0.041 | Old |
| 0.11027 | 0.102504 | 0.357 | 0.314 | 1 | 3 | Slamf8 | 0.043 | Old |
| 0.11028 | 0.295248 | 0.055 | 0.038 | 1 | 3 | Zbtb17 | 0.017 | Old |
| 0.110294 | 0.207611 | 0.253 | 0.221 | 1 | 3 | Hars | 0.032 | Old |
| 0.110319 | 0.704462 | 0.014 | 0.006 | 1 | 3 | Brf2 | 0.008 | Old |
| 0.110373 | 0.414761 | 0.119 | 0.096 | 1 | 3 | Reps1 | 0.023 | Old |
| 0.110416 | 0.157594 | 0.215 | 0.179 | 1 | 3 | Atp2c1 | 0.036 | Old |
| 0.110474 | 0.161915 | 0.478 | 0.429 | 1 | 3 | Nop10 | 0.049 | Old |
| 0.110743 | 0.315591 | 0.129 | 0.104 | 1 | 3 | Psmd10 | 0.025 | Old |
| 0.110854 | 0.327984 | 0.014 | 0.006 | 1 | 3 | Enkur | 0.008 | Old |
| 0.111009 | 0.145584 | 0.184 | 0.15 | 1 | 3 | Haus8 | 0.034 | Old |
| 0.111315 | 0.214554 | 0.093 | 0.071 | 1 | 3 | Zfp991 | 0.022 | Old |
| 0.111331 | 0.102286 | 0.324 | 0.278 | 1 | 3 | Dnajc2 | 0.046 | Old |
| 0.111404 | 0.170652 | 0.084 | 0.062 | 1 | 3 | P2rx7 | 0.022 | Old |
| 0.111536 | 0.126084 | 0.637 | 0.587 | 1 | 3 | Myl12a | 0.05 | Old |
| 0.111557 | 0.098118 | 0.195 | 0.16 | 1 | 3 | Ero1l | 0.035 | Old |
| 0.111672 | 0.224239 | 0.158 | 0.129 | 1 | 3 | Cul2 | 0.029 | Old |
| 0.111707 | 0.097198 | 0.235 | 0.197 | 1 | 3 | Twistnb | 0.038 | Old |
| 0.111779 | 0.05276 | 0.325 | 0.273 | 1 | 3 | Pcmt1 | 0.052 | Old |
| 0.111804 | 0.213415 | 0.129 | 0.104 | 1 | 3 | Smim19 | 0.025 | Old |
| 0.112029 | 0.23631 | 0.14 | 0.111 | 1 | 3 | Rhot1 | 0.029 | Old |

| | | | | | | | | |
|---|---|---|---|---|---|---|---|---|
| 0.112063 | 0.188266 | 0.25 | 0.211 | 1 | 3 | Phc2 | 0.039 | Old |
| 0.112185 | 0.557734 | 0.065 | 0.047 | 1 | 3 | Mlh1 | 0.018 | Old |
| 0.11232 | 0.579211 | 0.069 | 0.051 | 1 | 3 | Dnaja3 | 0.018 | Old |
| 0.112336 | 0.532993 | 0.014 | 0.006 | 1 | 3 | Tnk2 | 0.008 | Old |
| 0.112437 | 0.327455 | 0.03 | 0.018 | 1 | 3 | 2700062C0 | 0.012 | Old |
| 0.112499 | 0.333549 | 0.092 | 0.069 | 1 | 3 | Wrnip1 | 0.023 | Old |
| 0.112545 | 0.706255 | 0.018 | 0.009 | 1 | 3 | Neil1 | 0.009 | Old |
| 0.112556 | 0.178829 | 0.182 | 0.15 | 1 | 3 | Ncbp2 | 0.032 | Old |
| 0.112578 | 0.071734 | 0.639 | 0.577 | 1 | 3 | Usf2 | 0.062 | Old |
| 0.112596 | 0.071733 | 0.476 | 0.42 | 1 | 3 | Dhx15 | 0.056 | Old |
| 0.112649 | 0.691971 | 0.022 | 0.011 | 1 | 3 | Nlrp10 | 0.011 | Old |
| 0.112671 | 0.247991 | 0.085 | 0.063 | 1 | 3 | Atr | 0.022 | Old |
| 0.1127 | 0.05985 | 0.216 | 0.178 | 1 | 3 | Tmem183a | 0.038 | Old |
| 0.112743 | 0.599805 | 0.014 | 0.006 | 1 | 3 | Ocstamp | 0.008 | Old |
| 0.112784 | 0.925718 | 0.018 | 0.009 | 1 | 3 | Jade3 | 0.009 | Old |
| 0.112879 | 0.494082 | 0.014 | 0.006 | 1 | 3 | Aldh1l1 | 0.008 | Old |
| 0.113015 | 0.579304 | 0.014 | 0.006 | 1 | 3 | Ccdc126 | 0.008 | Old |
| 0.113016 | 0.039624 | 0.424 | 0.366 | 1 | 3 | Adam17 | 0.058 | Old |
| 0.113117 | 0.249805 | 0.091 | 0.069 | 1 | 3 | Zfp672 | 0.022 | Old |
| 0.113234 | 0.104582 | 0.325 | 0.283 | 1 | 3 | Cul1 | 0.042 | Old |
| 0.113245 | 0.148428 | 0.102 | 0.078 | 1 | 3 | 2510009E0 | 0.024 | Old |
| 0.113453 | 0.177305 | 0.143 | 0.116 | 1 | 3 | Ell | 0.027 | Old |
| 0.113503 | 0.708014 | 0.018 | 0.009 | 1 | 3 | Homez | 0.009 | Old |
| 0.113661 | 0.126236 | 0.179 | 0.148 | 1 | 3 | Ap4s1 | 0.031 | Old |
| 0.113797 | 0.158799 | 0.358 | 0.314 | 1 | 3 | Cop1 | 0.044 | Old |
| 0.113801 | 0.151834 | 0.102 | 0.078 | 1 | 3 | Tfdp1 | 0.024 | Old |
| 0.113865 | 0.737909 | 0.018 | 0.009 | 1 | 3 | Epas1 | 0.009 | Old |
| 0.114136 | 0.216891 | 0.184 | 0.152 | 1 | 3 | Ankrd40 | 0.032 | Old |
| 0.114169 | 0.357977 | 0.022 | 0.011 | 1 | 3 | Vcan | 0.011 | Old |
| 0.114285 | 0.622576 | 0.041 | 0.027 | 1 | 3 | Entpd5 | 0.014 | Old |
| 0.114347 | 0.442018 | 0.106 | 0.085 | 1 | 3 | Wasl | 0.021 | Old |
| 0.114464 | 0.256759 | 0.067 | 0.048 | 1 | 3 | Carm1 | 0.019 | Old |
| 0.114832 | 0.344686 | 0.018 | 0.009 | 1 | 3 | Rom1 | 0.009 | Old |
| 0.114845 | 0.136711 | 0.22 | 0.183 | 1 | 3 | Rap2b | 0.037 | Old |
| 0.115231 | 0.052785 | 0.584 | 0.524 | 1 | 3 | Shtn1 | 0.06 | Old |
| 0.11529 | 0.884268 | 0.026 | 0.015 | 1 | 3 | Tmem186 | 0.011 | Old |
| 0.115498 | 0.627961 | 0.053 | 0.037 | 1 | 3 | Alg14 | 0.016 | Old |
| 0.115719 | 0.229193 | 0.275 | 0.242 | 1 | 3 | Abhd17a | 0.033 | Old |
| 0.115739 | 0.083284 | 0.341 | 0.294 | 1 | 3 | Cdk14 | 0.047 | Old |
| 0.115907 | 0.692483 | 0.053 | 0.037 | 1 | 3 | Parp10 | 0.016 | Old |
| 0.116065 | 0.069985 | 0.446 | 0.395 | 1 | 3 | Acin1 | 0.051 | Old |
| 0.116068 | 0.873931 | 0.036 | 0.023 | 1 | 3 | Cdc7 | 0.013 | Old |
| 0.116195 | 0.198467 | 0.249 | 0.216 | 1 | 3 | Cebpe | 0.033 | Old |
| 0.116197 | 0.057603 | 0.583 | 0.53 | 1 | 3 | Rnf130 | 0.053 | Old |
| 0.116257 | 0.128231 | 0.274 | 0.234 | 1 | 3 | Blmh | 0.04 | Old |

| | | | | | | | | |
|---|---|---|---|---|---|---|---|---|
| 0.116428 | 0.109952 | 0.123 | 0.097 | 1 | 3 | Mrpl16 | 0.026 | Old |
| 0.116534 | 0.244136 | 0.207 | 0.173 | 1 | 3 | Acot7 | 0.034 | Old |
| 0.116777 | 0.236179 | 0.115 | 0.091 | 1 | 3 | Cmc2 | 0.024 | Old |
| 0.117107 | 0.236181 | 0.214 | 0.183 | 1 | 3 | Dlgap4 | 0.031 | Old |
| 0.11713 | 0.093617 | 0.281 | 0.241 | 1 | 3 | Tnfrsf1a | 0.04 | Old |
| 0.117218 | 0.069986 | 0.219 | 0.182 | 1 | 3 | Bmpr2 | 0.037 | Old |
| 0.117399 | 0.057465 | 0.926 | 0.918 | 1 | 3 | Tagln2 | 0.008 | Old |
| 0.117481 | 0.128562 | 0.114 | 0.088 | 1 | 3 | Gcsh | 0.026 | Old |
| 0.117703 | 0.09246 | 0.379 | 0.328 | 1 | 3 | Sel1l | 0.051 | Old |
| 0.117874 | 0.026858 | 0.256 | 0.215 | 1 | 3 | Slc9a3r1 | 0.041 | Old |
| 0.118009 | 0.119165 | 0.19 | 0.155 | 1 | 3 | Ncln | 0.035 | Old |
| 0.118285 | 0.192894 | 0.074 | 0.054 | 1 | 3 | Ndufaf7 | 0.02 | Old |
| 0.118349 | 0.066667 | 0.216 | 0.181 | 1 | 3 | Ahcyl1 | 0.035 | Old |
| 0.1185 | 0.135913 | 0.067 | 0.048 | 1 | 3 | Endog | 0.019 | Old |
| 0.118541 | 0.278908 | 0.197 | 0.167 | 1 | 3 | Plgrkt | 0.03 | Old |
| 0.118722 | 0.547759 | 0.031 | 0.019 | 1 | 3 | Hmga1 | 0.012 | Old |
| 0.118756 | 0.175537 | 0.325 | 0.283 | 1 | 3 | Arl6ip5 | 0.042 | Old |
| 0.11877 | 0.273714 | 0.026 | 0.015 | 1 | 3 | Nsun5 | 0.011 | Old |
| 0.118904 | 0.139303 | 0.098 | 0.074 | 1 | 3 | Cbll1 | 0.024 | Old |
| 0.118904 | 0.067582 | 0.474 | 0.408 | 1 | 3 | Camk1d | 0.066 | Old |
| 0.118927 | 0.116099 | 0.252 | 0.216 | 1 | 3 | Ndufa12 | 0.036 | Old |
| 0.118993 | 0.098635 | 0.777 | 0.726 | 1 | 3 | Cyth4 | 0.051 | Old |
| 0.119048 | 0.1777 | 0.36 | 0.318 | 1 | 3 | Dennd1b | 0.042 | Old |
| 0.11926 | 0.161722 | 0.182 | 0.15 | 1 | 3 | Cltb | 0.032 | Old |
| 0.119274 | 0.241928 | 0.166 | 0.136 | 1 | 3 | Fam98b | 0.03 | Old |
| 0.119374 | 0.449799 | 0.026 | 0.015 | 1 | 3 | Tlr13 | 0.011 | Old |
| 0.119668 | 0.08236 | 0.79 | 0.739 | 1 | 3 | Tacc1 | 0.051 | Old |
| 0.119756 | 0.290087 | 0.051 | 0.035 | 1 | 3 | Tdrkh | 0.016 | Old |
| 0.120004 | 0.151796 | 0.373 | 0.327 | 1 | 3 | Commd8 | 0.046 | Old |
| 0.120281 | 0.148594 | 0.16 | 0.131 | 1 | 3 | Fmr1 | 0.029 | Old |
| 0.12056 | 0.398842 | 0.036 | 0.023 | 1 | 3 | B4gat1 | 0.013 | Old |
| 0.120778 | 0.263863 | 0.047 | 0.032 | 1 | 3 | Tarm1 | 0.015 | Old |
| 0.12086 | 0.086099 | 0.424 | 0.367 | 1 | 3 | Ndufb4 | 0.057 | Old |
| 0.120872 | 0.139256 | 0.104 | 0.081 | 1 | 3 | Gpatch2 | 0.023 | Old |
| 0.120932 | 0.051138 | 0.784 | 0.721 | 1 | 3 | Luc7l2 | 0.063 | Old |
| 0.12103 | 0.045486 | 0.428 | 0.38 | 1 | 3 | Srsf1 | 0.048 | Old |
| 0.121079 | 0.158297 | 0.123 | 0.098 | 1 | 3 | Tmem198b | 0.025 | Old |
| 0.12139 | 0.256993 | 0.053 | 0.037 | 1 | 3 | Ralgapa2 | 0.016 | Old |
| 0.121467 | 0.016517 | 0.241 | 0.201 | 1 | 3 | Elovl1 | 0.04 | Old |
| 0.12151 | 0.438839 | 0.026 | 0.015 | 1 | 3 | Efnb1 | 0.011 | Old |
| 0.121517 | 0.143943 | 0.075 | 0.056 | 1 | 3 | Snip1 | 0.019 | Old |
| 0.121544 | 0.157768 | 0.259 | 0.222 | 1 | 3 | Appbp2 | 0.037 | Old |
| 0.121862 | 0.11363 | 0.173 | 0.143 | 1 | 3 | Fam76b | 0.03 | Old |
| 0.121889 | 0.125672 | 0.516 | 0.468 | 1 | 3 | Ago2 | 0.048 | Old |
| 0.121978 | 0.040168 | 0.062 | 0.044 | 1 | 3 | Snhg9 | 0.018 | Old |

| | | | | | | | | |
|---|---|---|---|---|---|---|---|---|
| 0.122118 | 0.38062 | 0.088 | 0.068 | 1 | 3 | Gtf2e1 | 0.02 | Old |
| 0.12236 | 0.473324 | 0.054 | 0.038 | 1 | 3 | Nup107 | 0.016 | Old |
| 0.122387 | 0.202168 | 0.208 | 0.174 | 1 | 3 | Itgav | 0.034 | Old |
| 0.12239 | 0.709875 | 0.048 | 0.033 | 1 | 3 | Ddx31 | 0.015 | Old |
| 0.122536 | 0.308897 | 0.087 | 0.067 | 1 | 3 | Tmem209 | 0.02 | Old |
| 0.122555 | 0.170627 | 0.141 | 0.115 | 1 | 3 | Kri1 | 0.026 | Old |
| 0.12258 | 0.055056 | 0.455 | 0.4 | 1 | 3 | Tmbim1 | 0.055 | Old |
| 0.122593 | 0.027382 | 0.202 | 0.167 | 1 | 3 | Epc1 | 0.035 | Old |
| 0.122623 | 1.294073 | 0.023 | 0.013 | 1 | 3 | Irf7 | 0.01 | Old |
| 0.122675 | 0.028371 | 0.097 | 0.074 | 1 | 3 | Gm34084 | 0.023 | Old |
| 0.122712 | 0.146912 | 0.105 | 0.082 | 1 | 3 | Plekhm1 | 0.023 | Old |
| 0.12288 | 0.559119 | 0.042 | 0.028 | 1 | 3 | Fzd7 | 0.014 | Old |
| 0.123095 | 0.196137 | 0.157 | 0.128 | 1 | 3 | Cdk2 | 0.029 | Old |
| 0.123197 | 0.007022 | 0.116 | 0.09 | 1 | 3 | Zbp1 | 0.026 | Old |
| 0.123253 | 0.193169 | 0.272 | 0.237 | 1 | 3 | Kansl1 | 0.035 | Old |
| 0.123478 | 0.007677 | 0.339 | 0.289 | 1 | 3 | Dpm3 | 0.05 | Old |
| 0.123664 | 0.596982 | 0.054 | 0.038 | 1 | 3 | Mkrn2 | 0.016 | Old |
| 0.123742 | 0.689007 | 0.023 | 0.013 | 1 | 3 | Oas1c | 0.01 | Old |
| 0.123879 | 0.826709 | 0.049 | 0.034 | 1 | 3 | Map3k15 | 0.015 | Old |
| 0.123992 | 0.81677 | 0.037 | 0.024 | 1 | 3 | Tmem38b | 0.013 | Old |
| 0.123993 | 0.23456 | 0.082 | 0.062 | 1 | 3 | Habp4 | 0.02 | Old |
| 0.123995 | 0.245754 | 0.201 | 0.168 | 1 | 3 | Tm2d1 | 0.033 | Old |
| 0.124352 | 0.211402 | 0.313 | 0.273 | 1 | 3 | Zmat2 | 0.04 | Old |
| 0.124364 | 0.16117 | 0.191 | 0.16 | 1 | 3 | Opa1 | 0.031 | Old |
| 0.124554 | 0.377568 | 0.125 | 0.101 | 1 | 3 | Ifi35 | 0.024 | Old |
| 0.124697 | 0.081324 | 0.349 | 0.299 | 1 | 3 | Csnk1g2 | 0.05 | Old |
| 0.124719 | 0.189693 | 0.102 | 0.08 | 1 | 3 | Malsu1 | 0.022 | Old |
| 0.124746 | 0.010518 | 0.246 | 0.205 | 1 | 3 | Lrpap1 | 0.041 | Old |
| 0.124814 | 0.227128 | 0.22 | 0.187 | 1 | 3 | Acp2 | 0.033 | Old |
| 0.125136 | 0.160646 | 0.385 | 0.341 | 1 | 3 | Glrx3 | 0.044 | Old |
| 0.125483 | 0.474237 | 0.111 | 0.09 | 1 | 3 | Vps33a | 0.021 | Old |
| 0.12565 | 0.332201 | 0.091 | 0.069 | 1 | 3 | Mtpap | 0.022 | Old |
| 0.126003 | 0.190321 | 0.135 | 0.11 | 1 | 3 | Pign | 0.025 | Old |
| 0.126196 | 0.281666 | 0.164 | 0.138 | 1 | 3 | Gatd1 | 0.026 | Old |
| 0.126414 | 0.014989 | 0.271 | 0.227 | 1 | 3 | Cand1 | 0.044 | Old |
| 0.126465 | 0.141139 | 0.047 | 0.032 | 1 | 3 | Ppif | 0.015 | Old |
| 0.126469 | 0.077062 | 0.094 | 0.072 | 1 | 3 | Casp6 | 0.022 | Old |
| 0.126599 | 0.156566 | 0.413 | 0.365 | 1 | 3 | Scaf11 | 0.048 | Old |
| 0.126657 | 0.25677 | 0.289 | 0.255 | 1 | 3 | Chfr | 0.034 | Old |
| 0.127179 | 0.12111 | 0.103 | 0.08 | 1 | 3 | Ppil3 | 0.023 | Old |
| 0.127275 | 0.101816 | 0.307 | 0.263 | 1 | 3 | Ypel5 | 0.044 | Old |
| 0.127338 | 0.226906 | 0.084 | 0.063 | 1 | 3 | Phkb | 0.021 | Old |
| 0.127466 | 0.190734 | 0.258 | 0.223 | 1 | 3 | Spty2d1 | 0.035 | Old |
| 0.127606 | 0.347061 | 0.023 | 0.013 | 1 | 3 | Arl16 | 0.01 | Old |
| 0.127634 | 0.124753 | 0.258 | 0.222 | 1 | 3 | Ube2a | 0.036 | Old |

| | | | | | | | | |
|---|---|---|---|---|---|---|---|---|
| 0.12796 | 0.006097 | 0.225 | 0.187 | 1 | 3 | Angel2 | 0.038 | Old |
| 0.128203 | 0.057403 | 0.566 | 0.496 | 1 | 3 | Egr2 | 0.07 | Old |
| 0.128482 | 0.026024 | 0.747 | 0.682 | 1 | 3 | Pak2 | 0.065 | Old |
| 0.128691 | 0.19764 | 0.213 | 0.179 | 1 | 3 | Bcap29 | 0.034 | Old |
| 0.128873 | 0.052999 | 0.726 | 0.688 | 1 | 3 | Tspo | 0.038 | Old |
| 0.129042 | 0.22836 | 0.117 | 0.093 | 1 | 3 | Daxx | 0.024 | Old |
| 0.129078 | 0.516994 | 0.032 | 0.02 | 1 | 3 | Simc1 | 0.012 | Old |
| 0.129151 | 0.025693 | 0.226 | 0.189 | 1 | 3 | Fnbp1l | 0.037 | Old |
| 0.129214 | 0.839725 | 0.019 | 0.01 | 1 | 3 | Trdmt1 | 0.009 | Old |
| 0.129269 | 0.567877 | 0.032 | 0.02 | 1 | 3 | Fbxw17 | 0.012 | Old |
| 0.129342 | 0.781794 | 0.019 | 0.01 | 1 | 3 | Mfsd4b4 | 0.009 | Old |
| 0.129461 | 0.100692 | 0.64 | 0.586 | 1 | 3 | Myof | 0.054 | Old |
| 0.129511 | 0.124833 | 0.263 | 0.223 | 1 | 3 | BC028528 | 0.04 | Old |
| 0.12978 | 0.078874 | 0.059 | 0.042 | 1 | 3 | Bclaf3 | 0.017 | Old |
| 0.129903 | 0.273829 | 0.142 | 0.116 | 1 | 3 | Itpr2 | 0.026 | Old |
| 0.129941 | 0.049145 | 0.503 | 0.434 | 1 | 3 | Dpysl2 | 0.069 | Old |
| 0.130149 | 0.468801 | 0.055 | 0.039 | 1 | 3 | Ubxn8 | 0.016 | Old |
| 0.130345 | 0.179131 | 0.067 | 0.049 | 1 | 3 | Siah1a | 0.018 | Old |
| 0.130387 | 0.033206 | 0.276 | 0.239 | 1 | 3 | Iscu | 0.037 | Old |
| 0.130434 | 0.34832 | 0.027 | 0.016 | 1 | 3 | Mmgt2 | 0.011 | Old |
| 0.130482 | 0.271156 | 0.158 | 0.133 | 1 | 3 | Pdcd11 | 0.025 | Old |
| 0.130809 | 0.536626 | 0.032 | 0.02 | 1 | 3 | Rsad2 | 0.012 | Old |
| 0.130938 | 0.880954 | 0.012 | 0.005 | 1 | 3 | Dnph1 | 0.007 | Old |
| 0.131024 | 0.638335 | 0.055 | 0.039 | 1 | 3 | Rbm43 | 0.016 | Old |
| 0.131036 | 0.219482 | 0.086 | 0.066 | 1 | 3 | Atg4c | 0.02 | Old |
| 0.131039 | 0.117017 | 0.037 | 0.024 | 1 | 3 | Zkscan6 | 0.013 | Old |
| 0.131129 | 0.358019 | 0.037 | 0.024 | 1 | 3 | Fut7 | 0.013 | Old |
| 0.131462 | 0.419464 | 0.061 | 0.044 | 1 | 3 | Anapc7 | 0.017 | Old |
| 0.131888 | 0.111528 | 0.562 | 0.494 | 1 | 3 | Etf1 | 0.068 | Old |
| 0.131962 | 0.075426 | 0.449 | 0.396 | 1 | 3 | Ppp1r9b | 0.053 | Old |
| 0.132119 | 0.09127 | 0.247 | 0.21 | 1 | 3 | Mbd2 | 0.037 | Old |
| 0.132335 | 0.215581 | 0.08 | 0.061 | 1 | 3 | Cyp4v3 | 0.019 | Old |
| 0.132352 | 0.214844 | 0.091 | 0.069 | 1 | 3 | Pex3 | 0.022 | Old |
| 0.132485 | 0.119566 | 0.037 | 0.024 | 1 | 3 | Actr6 | 0.013 | Old |
| 0.132635 | 0.114769 | 0.235 | 0.201 | 1 | 3 | Sec24c | 0.034 | Old |
| 0.132821 | 0.39002 | 0.019 | 0.01 | 1 | 3 | Zbtb39 | 0.009 | Old |
| 0.132834 | 0.240569 | 0.092 | 0.071 | 1 | 3 | Ttc32 | 0.021 | Old |
| 0.132978 | 0.428124 | 0.027 | 0.016 | 1 | 3 | Aig1 | 0.011 | Old |
| 0.13302 | 0.012322 | 0.474 | 0.412 | 1 | 3 | Cflar | 0.062 | Old |
| 0.133082 | 0.535608 | 0.019 | 0.01 | 1 | 3 | Sirt5 | 0.009 | Old |
| 0.133133 | 0.88433 | 0.044 | 0.03 | 1 | 3 | Rbm12b2 | 0.014 | Old |
| 0.133252 | 0.926331 | 0.016 | 0.008 | 1 | 3 | 2610301B2 | 0.008 | Old |
| 0.133437 | 0.318867 | 0.127 | 0.102 | 1 | 3 | Mycbp | 0.025 | Old |
| 0.133554 | 0.305212 | 0.196 | 0.164 | 1 | 3 | Sgk1 | 0.032 | Old |
| 0.133617 | 0.101193 | 0.18 | 0.149 | 1 | 3 | Abhd17c | 0.031 | Old |

| | | | | | | | | |
|---|---|---|---|---|---|---|---|---|
| 0.133669 | 0.256857 | 0.037 | 0.024 | 1 | 3 | Lrrc61 | 0.013 | Old |
| 0.133819 | 0.311707 | 0.012 | 0.005 | 1 | 3 | E430018J2 | 0.007 | Old |
| 0.133894 | 0.116331 | 0.326 | 0.283 | 1 | 3 | Stat6 | 0.043 | Old |
| 0.133966 | 0.364065 | 0.043 | 0.029 | 1 | 3 | Sufu | 0.014 | Old |
| 0.133974 | 0.11362 | 0.384 | 0.337 | 1 | 3 | Sec11a | 0.047 | Old |
| 0.13401 | 0.253538 | 0.152 | 0.125 | 1 | 3 | Brix1 | 0.027 | Old |
| 0.134028 | 0.053148 | 0.07 | 0.052 | 1 | 3 | Pdzd11 | 0.018 | Old |
| 0.13407 | 0.517084 | 0.049 | 0.034 | 1 | 3 | Lrrc57 | 0.015 | Old |
| 0.134134 | 0.799323 | 0.016 | 0.008 | 1 | 3 | Cttn | 0.008 | Old |
| 0.13414 | 0.007884 | 0.357 | 0.311 | 1 | 3 | Stk40 | 0.046 | Old |
| 0.134144 | 0.106408 | 0.584 | 0.535 | 1 | 3 | Eno1 | 0.049 | Old |
| 0.134221 | 0.354539 | 0.114 | 0.091 | 1 | 3 | Farsa | 0.023 | Old |
| 0.134428 | 0.92151 | 0.016 | 0.008 | 1 | 3 | Oxld1 | 0.008 | Old |
| 0.134428 | 0.847971 | 0.016 | 0.008 | 1 | 3 | Trmt44 | 0.008 | Old |
| 0.134463 | 0.126616 | 0.161 | 0.131 | 1 | 3 | Mepce | 0.03 | Old |
| 0.134482 | 0.155222 | 0.147 | 0.117 | 1 | 3 | Aspscr1 | 0.03 | Old |
| 0.134633 | 0.290521 | 0.076 | 0.057 | 1 | 3 | Lrig2 | 0.019 | Old |
| 0.134653 | 0.020988 | 0.338 | 0.289 | 1 | 3 | Usp15 | 0.049 | Old |
| 0.134755 | 0.055429 | 0.165 | 0.136 | 1 | 3 | Ino80e | 0.029 | Old |
| 0.134829 | 0.011124 | 0.299 | 0.255 | 1 | 3 | Vps36 | 0.044 | Old |
| 0.13494 | 0.021792 | 0.238 | 0.201 | 1 | 3 | Tbc1d14 | 0.037 | Old |
| 0.135019 | 0.580085 | 0.016 | 0.008 | 1 | 3 | Senp8 | 0.008 | Old |
| 0.135038 | 0.22448 | 0.073 | 0.054 | 1 | 3 | Hotairm1 | 0.019 | Old |
| 0.135217 | 0.379223 | 0.032 | 0.02 | 1 | 3 | Tprn | 0.012 | Old |
| 0.13523 | 0.164304 | 0.037 | 0.024 | 1 | 3 | Sfmbt1 | 0.013 | Old |
| 0.135243 | 0.486495 | 0.063 | 0.047 | 1 | 3 | Sec22c | 0.016 | Old |
| 0.135298 | 0.099779 | 0.073 | 0.054 | 1 | 3 | Il1a | 0.019 | Old |
| 0.135362 | 0.095301 | 0.172 | 0.143 | 1 | 3 | Cops8 | 0.029 | Old |
| 0.135636 | 0.121311 | 0.262 | 0.223 | 1 | 3 | Bcl7b | 0.039 | Old |
| 0.135734 | 0.032831 | 0.565 | 0.504 | 1 | 3 | Pde4b | 0.061 | Old |
| 0.135758 | 0.09808 | 0.659 | 0.609 | 1 | 3 | Tra2b | 0.05 | Old |
| 0.135761 | 0.738874 | 0.016 | 0.008 | 1 | 3 | Ahnak2 | 0.008 | Old |
| 0.135761 | 0.531146 | 0.016 | 0.008 | 1 | 3 | BC052040 | 0.008 | Old |
| 0.135848 | 0.105552 | 0.497 | 0.433 | 1 | 3 | Zmiz1 | 0.064 | Old |
| 0.135922 | 0.471976 | 0.074 | 0.056 | 1 | 3 | Pon3 | 0.018 | Old |
| 0.136091 | 0.1795 | 0.663 | 0.629 | 1 | 3 | Rbm25 | 0.034 | Old |
| 0.136314 | 0.237314 | 0.188 | 0.157 | 1 | 3 | Atg5 | 0.031 | Old |
| 0.136423 | 0.032236 | 0.309 | 0.266 | 1 | 3 | Usp14 | 0.043 | Old |
| 0.136657 | 0.22396 | 0.11 | 0.088 | 1 | 3 | Dcaf1 | 0.022 | Old |
| 0.136721 | 0.095399 | 0.332 | 0.292 | 1 | 3 | Mnt | 0.04 | Old |
| 0.136768 | 0.052882 | 0.46 | 0.409 | 1 | 3 | Eif3l | 0.051 | Old |
| 0.136861 | 0.058378 | 0.072 | 0.053 | 1 | 3 | Mecr | 0.019 | Old |
| 0.136898 | 0.72156 | 0.044 | 0.03 | 1 | 3 | Xrcc6 | 0.014 | Old |
| 0.137008 | 0.430896 | 0.139 | 0.115 | 1 | 3 | Ttc1 | 0.024 | Old |
| 0.137403 | 0.348571 | 0.016 | 0.008 | 1 | 3 | Trim6 | 0.008 | Old |

| | | | | | | | | |
|---|---|---|---|---|---|---|---|---|
| 0.137726 | 0.355949 | 0.08 | 0.061 | 1 | 3 | Tmem184c | 0.019 | Old |
| 0.137742 | 0.049994 | 0.128 | 0.102 | 1 | 3 | Nrbf2 | 0.026 | Old |
| 0.137773 | 0.240126 | 0.13 | 0.106 | 1 | 3 | Trap1 | 0.024 | Old |
| 0.137856 | 0.015632 | 0.41 | 0.359 | 1 | 3 | Elovl5 | 0.051 | Old |
| 0.137915 | 0.087108 | 0.317 | 0.278 | 1 | 3 | Trim30a | 0.039 | Old |
| 0.138019 | 0.386899 | 0.061 | 0.044 | 1 | 3 | Bex6 | 0.017 | Old |
| 0.13827 | 0.40919 | 0.093 | 0.073 | 1 | 3 | Gm20186 | 0.02 | Old |
| 0.13849 | 0.351295 | 0.056 | 0.04 | 1 | 3 | Tmem110 | 0.016 | Old |
| 0.138523 | 0.134907 | 0.091 | 0.069 | 1 | 3 | Dnajc16 | 0.022 | Old |
| 0.13858 | 0.041781 | 0.297 | 0.263 | 1 | 3 | Ppm1a | 0.034 | Old |
| 0.138854 | 0.063572 | 0.154 | 0.128 | 1 | 3 | Tlk2 | 0.026 | Old |
| 0.139032 | 0.567226 | 0.05 | 0.035 | 1 | 3 | Dusp18 | 0.015 | Old |
| 0.139102 | 0.173731 | 0.209 | 0.176 | 1 | 3 | Msmo1 | 0.033 | Old |
| 0.13928 | 0.161983 | 0.344 | 0.304 | 1 | 3 | Tsc22d1 | 0.04 | Old |
| 0.139417 | 0.055749 | 0.306 | 0.263 | 1 | 3 | Aup1 | 0.043 | Old |
| 0.139679 | 0.380265 | 0.087 | 0.067 | 1 | 3 | Ptcd2 | 0.02 | Old |
| 0.139833 | 0.370631 | 0.067 | 0.049 | 1 | 3 | Gaa | 0.018 | Old |
| 0.14004 | 0.069488 | 0.26 | 0.22 | 1 | 3 | Meaf6 | 0.04 | Old |
| 0.14009 | 0.112535 | 0.756 | 0.691 | 1 | 3 | Reep5 | 0.065 | Old |
| 0.140391 | 0.157413 | 0.332 | 0.293 | 1 | 3 | Ik | 0.039 | Old |
| 0.140432 | 0.290893 | 0.155 | 0.13 | 1 | 3 | Ppme1 | 0.025 | Old |
| 0.140519 | 0.214204 | 0.076 | 0.058 | 1 | 3 | Gnptab | 0.018 | Old |
| 0.14058 | 0.151809 | 0.161 | 0.134 | 1 | 3 | Pigm | 0.027 | Old |
| 0.140714 | 0.063768 | 0.689 | 0.622 | 1 | 3 | Mapre1 | 0.067 | Old |
| 0.140993 | 0.716812 | 0.024 | 0.014 | 1 | 3 | Wee1 | 0.01 | Old |
| 0.141366 | 0.307411 | 0.096 | 0.074 | 1 | 3 | Mgat4a | 0.022 | Old |
| 0.141474 | 0.071774 | 0.271 | 0.231 | 1 | 3 | Pdpk1 | 0.04 | Old |
| 0.141491 | 0.081519 | 0.097 | 0.074 | 1 | 3 | Tfb2m | 0.023 | Old |
| 0.141652 | 0.216511 | 0.067 | 0.049 | 1 | 3 | Slc35e3 | 0.018 | Old |
| 0.142013 | 0.201887 | 0.079 | 0.059 | 1 | 3 | Dnttip1 | 0.02 | Old |
| 0.142074 | 0.084869 | 0.067 | 0.049 | 1 | 3 | Zc3hc1 | 0.018 | Old |
| 0.142505 | 0.081617 | 0.068 | 0.051 | 1 | 3 | Sertad3 | 0.017 | Old |
| 0.142642 | 0.275385 | 0.182 | 0.153 | 1 | 3 | Utp11 | 0.029 | Old |
| 0.143026 | 0.437258 | 0.186 | 0.163 | 1 | 3 | Tnfsf9 | 0.023 | Old |
| 0.143074 | 0.067946 | 0.219 | 0.186 | 1 | 3 | Atp13a2 | 0.033 | Old |
| 0.143519 | 0.196449 | 0.505 | 0.465 | 1 | 3 | Pla2g7 | 0.04 | Old |
| 0.143564 | 0.097235 | 0.09 | 0.069 | 1 | 3 | Wdr46 | 0.021 | Old |
| 0.14362 | 0.271968 | 0.186 | 0.16 | 1 | 3 | Polr2h | 0.026 | Old |
| 0.143943 | 0.291731 | 0.076 | 0.058 | 1 | 3 | C2cd2l | 0.018 | Old |
| 0.143962 | 0.194034 | 0.038 | 0.025 | 1 | 3 | Mtrr | 0.013 | Old |
| 0.144153 | 0.01003 | 0.303 | 0.265 | 1 | 3 | Abcg1 | 0.038 | Old |
| 0.144209 | 0.571755 | 0.065 | 0.048 | 1 | 3 | Gatsl2 | 0.017 | Old |
| 0.144246 | 0.191846 | 0.231 | 0.196 | 1 | 3 | Anks1 | 0.035 | Old |
| 0.14428 | 0.107997 | 0.114 | 0.09 | 1 | 3 | Fasn | 0.024 | Old |
| 0.14445 | 0.240003 | 0.159 | 0.133 | 1 | 3 | Washc3 | 0.026 | Old |

| | | | | | | | | |
|---|---|---|---|---|---|---|---|---|
| 0.144536 | 0.181417 | 0.074 | 0.056 | 1 | 3 | Hectd3 | 0.018 | Old |
| 0.144564 | 0.107484 | 0.266 | 0.23 | 1 | 3 | Hsd17b4 | 0.036 | Old |
| 0.144681 | 0.106932 | 0.223 | 0.189 | 1 | 3 | Vps26a | 0.034 | Old |
| 0.144715 | 0.484508 | 0.029 | 0.018 | 1 | 3 | Acat2 | 0.011 | Old |
| 0.144909 | 0.089192 | 0.547 | 0.504 | 1 | 3 | Mbtd1 | 0.043 | Old |
| 0.144919 | 0.171123 | 0.222 | 0.188 | 1 | 3 | 6430548M | 0.034 | Old |
| 0.145022 | 0.292834 | 0.055 | 0.039 | 1 | 3 | Rrp1b | 0.016 | Old |
| 0.145587 | 0.197663 | 0.14 | 0.115 | 1 | 3 | Med16 | 0.025 | Old |
| 0.145603 | 0.463908 | 0.088 | 0.069 | 1 | 3 | Tmem106c | 0.019 | Old |
| 0.145617 | 0.00782 | 0.528 | 0.453 | 1 | 3 | Emilin2 | 0.075 | Old |
| 0.14565 | 0.2065 | 0.431 | 0.398 | 1 | 3 | Nabp1 | 0.033 | Old |
| 0.145681 | 0.049497 | 0.209 | 0.176 | 1 | 3 | Trim12c | 0.033 | Old |
| 0.145805 | 0.083458 | 0.652 | 0.6 | 1 | 3 | Eif2s2 | 0.052 | Old |
| 0.146214 | 0.205558 | 0.075 | 0.057 | 1 | 3 | Spryd7 | 0.018 | Old |
| 0.146237 | 0.25201 | 0.1 | 0.08 | 1 | 3 | Snx14 | 0.02 | Old |
| 0.146285 | 0.43498 | 0.039 | 0.027 | 1 | 3 | Panx1 | 0.012 | Old |
| 0.146327 | 0.134885 | 0.049 | 0.034 | 1 | 3 | Fzd5 | 0.015 | Old |
| 0.14653 | 0.161405 | 0.207 | 0.177 | 1 | 3 | Tmem128 | 0.03 | Old |
| 0.146692 | 0.110087 | 0.544 | 0.492 | 1 | 3 | Matr3 | 0.052 | Old |
| 0.146734 | 0.153104 | 0.104 | 0.082 | 1 | 3 | Fbxl6 | 0.022 | Old |
| 0.147344 | 0.17348 | 0.049 | 0.034 | 1 | 3 | Ppp1r16b | 0.015 | Old |
| 0.147407 | 0.178492 | 0.467 | 0.428 | 1 | 3 | Cep170 | 0.039 | Old |
| 0.147411 | 0.21084 | 0.033 | 0.021 | 1 | 3 | Aste1 | 0.012 | Old |
| 0.147641 | 0.096363 | 0.208 | 0.177 | 1 | 3 | Map3k7 | 0.031 | Old |
| 0.147741 | 0.085313 | 0.317 | 0.275 | 1 | 3 | Snf8 | 0.042 | Old |
| 0.147754 | 0.696044 | 0.068 | 0.051 | 1 | 3 | Klhl12 | 0.017 | Old |
| 0.147788 | 0.193693 | 0.112 | 0.091 | 1 | 3 | Gm11808 | 0.021 | Old |
| 0.147797 | 0.066009 | 0.191 | 0.16 | 1 | 3 | Dnttip2 | 0.031 | Old |
| 0.147917 | 0.019217 | 0.099 | 0.077 | 1 | 3 | Mtap | 0.022 | Old |
| 0.148221 | 0.010301 | 0.226 | 0.191 | 1 | 3 | Wwp2 | 0.035 | Old |
| 0.148236 | 0.100031 | 0.281 | 0.242 | 1 | 3 | Gosr2 | 0.039 | Old |
| 0.148343 | 0.179192 | 0.25 | 0.215 | 1 | 3 | Sik1 | 0.035 | Old |
| 0.148498 | 0.018649 | 0.123 | 0.097 | 1 | 3 | Nos1ap | 0.026 | Old |
| 0.148515 | 0.079847 | 0.968 | 0.928 | 1 | 3 | Zfp36 | 0.04 | Old |
| 0.148801 | 0.111803 | 0.056 | 0.04 | 1 | 3 | Mocs1 | 0.016 | Old |
| 0.148878 | 0.111156 | 0.501 | 0.458 | 1 | 3 | Csf2rb | 0.043 | Old |
| 0.148949 | 0.052996 | 0.164 | 0.138 | 1 | 3 | Med14 | 0.026 | Old |
| 0.149016 | 0.028608 | 0.075 | 0.057 | 1 | 3 | Mtor | 0.018 | Old |
| 0.149041 | 0.071647 | 0.232 | 0.197 | 1 | 3 | Hdac1 | 0.035 | Old |
| 0.149064 | 0.115705 | 0.143 | 0.117 | 1 | 3 | Fam20c | 0.026 | Old |
| 0.14919 | 0.37011 | 0.07 | 0.053 | 1 | 3 | Prcc | 0.017 | Old |
| 0.14928 | 0.946938 | 0.02 | 0.011 | 1 | 3 | Ifi214 | 0.009 | Old |
| 0.149301 | 0.228594 | 0.081 | 0.062 | 1 | 3 | Gbe1 | 0.019 | Old |
| 0.149417 | 0.594934 | 0.02 | 0.011 | 1 | 3 | Pex12 | 0.009 | Old |
| 0.149467 | 0.257606 | 0.122 | 0.1 | 1 | 3 | Nelfe | 0.022 | Old |

| | | | | | | | | |
|---|---|---|---|---|---|---|---|---|
| 0.149516 | 0.210917 | 0.207 | 0.176 | 1 | 3 | Dpagt1 | 0.031 | Old |
| 0.149639 | 0.141878 | 0.073 | 0.054 | 1 | 3 | 2310001H1 | 0.019 | Old |
| 0.149641 | 0.437174 | 0.065 | 0.048 | 1 | 3 | Pgm3 | 0.017 | Old |
| 0.150327 | 0.048659 | 0.128 | 0.102 | 1 | 3 | Abcf2 | 0.026 | Old |
| 0.150366 | 0.090843 | 0.133 | 0.107 | 1 | 3 | Stk19 | 0.026 | Old |
| 0.150498 | 0.102658 | 0.364 | 0.319 | 1 | 3 | Ube2e1 | 0.045 | Old |
| 0.150806 | 0.400897 | 0.029 | 0.018 | 1 | 3 | Suv39h1 | 0.011 | Old |
| 0.15084 | 0.358484 | 0.093 | 0.073 | 1 | 3 | Bap1 | 0.02 | Old |
| 0.150947 | 0.111169 | 0.256 | 0.222 | 1 | 3 | Npm3 | 0.034 | Old |
| 0.150989 | 0.012082 | 0.719 | 0.669 | 1 | 3 | Atp6v0e | 0.05 | Old |
| 0.151303 | 0.314062 | 0.081 | 0.062 | 1 | 3 | Dclre1b | 0.019 | Old |
| 0.151319 | 0.108286 | 0.192 | 0.162 | 1 | 3 | Commd4 | 0.03 | Old |
| 0.151382 | 0.175416 | 0.104 | 0.082 | 1 | 3 | Mlx | 0.022 | Old |
| 0.151624 | 0.65096 | 0.116 | 0.095 | 1 | 3 | Plp2 | 0.021 | Old |
| 0.151714 | 0.184252 | 0.152 | 0.125 | 1 | 3 | Kif13a | 0.027 | Old |
| 0.151741 | 0.124032 | 0.039 | 0.027 | 1 | 3 | Irgq | 0.012 | Old |
| 0.152253 | 0.150904 | 0.458 | 0.422 | 1 | 3 | Eif3i | 0.036 | Old |
| 0.152282 | 0.190521 | 0.098 | 0.078 | 1 | 3 | Borcs5 | 0.02 | Old |
| 0.152337 | 0.338421 | 0.063 | 0.047 | 1 | 3 | Scyl3 | 0.016 | Old |
| 0.152433 | 0.323451 | 0.062 | 0.045 | 1 | 3 | Lmtk2 | 0.017 | Old |
| 0.15257 | 0.326505 | 0.057 | 0.042 | 1 | 3 | Rbsn | 0.015 | Old |
| 0.152915 | 0.258979 | 0.136 | 0.114 | 1 | 3 | Slc44a1 | 0.022 | Old |
| 0.153012 | 0.748125 | 0.02 | 0.011 | 1 | 3 | Mmaa | 0.009 | Old |
| 0.153025 | 0.151808 | 0.14 | 0.114 | 1 | 3 | Polr1c | 0.026 | Old |
| 0.15324 | 0.06906 | 0.416 | 0.366 | 1 | 3 | Mndal | 0.05 | Old |
| 0.153309 | 0.103215 | 0.272 | 0.236 | 1 | 3 | Rps6ka4 | 0.036 | Old |
| 0.153365 | 0.329893 | 0.103 | 0.082 | 1 | 3 | Ctnnbl1 | 0.021 | Old |
| 0.153664 | 0.331454 | 0.116 | 0.095 | 1 | 3 | Rab28 | 0.021 | Old |
| 0.153708 | 0.130564 | 0.194 | 0.162 | 1 | 3 | Kdm6a | 0.032 | Old |
| 0.153764 | 0.020739 | 0.228 | 0.192 | 1 | 3 | Zzz3 | 0.036 | Old |
| 0.153858 | 0.268383 | 0.057 | 0.042 | 1 | 3 | Armt1 | 0.015 | Old |
| 0.153924 | 0.29576 | 0.105 | 0.083 | 1 | 3 | Fbxo8 | 0.022 | Old |
| 0.153985 | 0.055515 | 0.196 | 0.167 | 1 | 3 | Tifa | 0.029 | Old |
| 0.154018 | 0.085737 | 0.294 | 0.254 | 1 | 3 | Ugp2 | 0.04 | Old |
| 0.15421 | 0.502245 | 0.035 | 0.023 | 1 | 3 | Fam151b | 0.012 | Old |
| 0.154467 | 0.139664 | 0.106 | 0.083 | 1 | 3 | Sdhaf4 | 0.023 | Old |
| 0.155329 | 0.293355 | 0.115 | 0.093 | 1 | 3 | Lactb2 | 0.022 | Old |
| 0.155344 | 0.528719 | 0.082 | 0.064 | 1 | 3 | Dvl1 | 0.018 | Old |
| 0.155347 | 0.05748 | 0.22 | 0.188 | 1 | 3 | Zfp385a | 0.032 | Old |
| 0.155472 | 0.629347 | 0.01 | 0.004 | 1 | 3 | Ctsk | 0.006 | Old |
| 0.155472 | 0.574347 | 0.01 | 0.004 | 1 | 3 | Rtl6 | 0.006 | Old |
| 0.155592 | 0.112138 | 0.127 | 0.102 | 1 | 3 | Pfkl | 0.025 | Old |
| 0.155756 | 0.071662 | 0.935 | 0.924 | 1 | 3 | Alox5ap | 0.011 | Old |
| 0.15585 | 0.043316 | 0.106 | 0.083 | 1 | 3 | Anapc10 | 0.023 | Old |
| 0.155905 | 0.773205 | 0.01 | 0.004 | 1 | 3 | Htr7 | 0.006 | Old |

| | | | | | | | | |
|---|---|---|---|---|---|---|---|---|
| 0.156074 | 0.043353 | 0.214 | 0.181 | 1 | 3 | Rnf20 | 0.033 | Old |
| 0.156201 | 0.208765 | 0.025 | 0.015 | 1 | 3 | Ogg1 | 0.01 | Old |
| 0.156263 | 0.077669 | 0.157 | 0.129 | 1 | 3 | Plagl2 | 0.028 | Old |
| 0.156338 | 0.458113 | 0.01 | 0.004 | 1 | 3 | C920021L1 | 0.006 | Old |
| 0.156417 | 0.236229 | 0.108 | 0.086 | 1 | 3 | Eaf1 | 0.022 | Old |
| 0.156556 | 0.723528 | 0.01 | 0.004 | 1 | 3 | Osbpl3 | 0.006 | Old |
| 0.156556 | 0.21694 | 0.01 | 0.004 | 1 | 3 | Trmt12 | 0.006 | Old |
| 0.156643 | 0.341353 | 0.116 | 0.093 | 1 | 3 | Prpsap1 | 0.023 | Old |
| 0.156773 | 0.420121 | 0.01 | 0.004 | 1 | 3 | Inmt | 0.006 | Old |
| 0.156773 | 0.381254 | 0.01 | 0.004 | 1 | 3 | Col6a2 | 0.006 | Old |
| 0.156921 | 0.166837 | 0.143 | 0.117 | 1 | 3 | Plk3 | 0.026 | Old |
| 0.156959 | 0.185196 | 0.025 | 0.015 | 1 | 3 | Fahd1 | 0.01 | Old |
| 0.156991 | 0.231913 | 0.01 | 0.004 | 1 | 3 | Gm14698 | 0.006 | Old |
| 0.157137 | 0.054509 | 0.583 | 0.524 | 1 | 3 | Ank | 0.059 | Old |
| 0.157295 | 0.182378 | 0.085 | 0.066 | 1 | 3 | Isoc1 | 0.019 | Old |
| 0.157353 | 0.06178 | 0.959 | 0.958 | 1 | 3 | Klf6 | 0.001 | Old |
| 0.157357 | 0.143638 | 0.375 | 0.333 | 1 | 3 | Fam168b | 0.042 | Old |
| 0.15751 | 0.352798 | 0.053 | 0.038 | 1 | 3 | Pop1 | 0.015 | Old |
| 0.157528 | 0.109471 | 0.186 | 0.157 | 1 | 3 | Cgnl1 | 0.029 | Old |
| 0.157631 | 0.976659 | 0.017 | 0.009 | 1 | 3 | Gm45716 | 0.008 | Old |
| 0.157863 | 0.10635 | 0.22 | 0.188 | 1 | 3 | Rsbn1l | 0.032 | Old |
| 0.158107 | 0.469956 | 0.017 | 0.009 | 1 | 3 | Gm29340 | 0.008 | Old |
| 0.15814 | 0.136537 | 0.142 | 0.117 | 1 | 3 | Akap10 | 0.025 | Old |
| 0.158457 | 0.101775 | 0.228 | 0.193 | 1 | 3 | Irs2 | 0.035 | Old |
| 0.158738 | 0.151006 | 0.162 | 0.135 | 1 | 3 | Zcchc17 | 0.027 | Old |
| 0.158868 | 0.047554 | 0.36 | 0.311 | 1 | 3 | Fam126a | 0.049 | Old |
| 0.158903 | 0.614393 | 0.017 | 0.009 | 1 | 3 | Cul7 | 0.008 | Old |
| 0.158985 | 0.244119 | 0.131 | 0.109 | 1 | 3 | Fbrs | 0.022 | Old |
| 0.159029 | 0.041979 | 0.121 | 0.097 | 1 | 3 | Sphk2 | 0.024 | Old |
| 0.159065 | 0.009653 | 0.374 | 0.323 | 1 | 3 | Sf3b3 | 0.051 | Old |
| 0.159405 | 0.581955 | 0.03 | 0.019 | 1 | 3 | Mgst1 | 0.011 | Old |
| 0.159405 | 0.118045 | 0.03 | 0.019 | 1 | 3 | 9830107B1 | 0.011 | Old |
| 0.159696 | 0.663308 | 0.036 | 0.024 | 1 | 3 | I830077J02 | 0.012 | Old |
| 0.159702 | 0.093538 | 0.017 | 0.009 | 1 | 3 | Ksr2 | 0.008 | Old |
| 0.160098 | 0.17984 | 0.152 | 0.125 | 1 | 3 | Pycr2 | 0.027 | Old |
| 0.160264 | 0.210459 | 0.065 | 0.048 | 1 | 3 | Plekhg1 | 0.017 | Old |
| 0.160533 | 0.249895 | 0.025 | 0.015 | 1 | 3 | Rwdd2b | 0.01 | Old |
| 0.160648 | 0.010968 | 0.241 | 0.208 | 1 | 3 | Fyn | 0.033 | Old |
| 0.160826 | 0.45167 | 0.017 | 0.009 | 1 | 3 | Bub1b | 0.008 | Old |
| 0.161325 | 0.076859 | 0.196 | 0.165 | 1 | 3 | Pum3 | 0.031 | Old |
| 0.161384 | 0.210227 | 0.123 | 0.1 | 1 | 3 | Tmem181a | 0.023 | Old |
| 0.161565 | 0.11759 | 0.025 | 0.015 | 1 | 3 | Msl3l2 | 0.01 | Old |
| 0.16165 | 0.881276 | 0.013 | 0.006 | 1 | 3 | 1110002L0 | 0.007 | Old |
| 0.161666 | 0.215154 | 0.165 | 0.139 | 1 | 3 | Thoc6 | 0.026 | Old |
| 0.161705 | 0.044259 | 0.283 | 0.246 | 1 | 3 | Snx18 | 0.037 | Old |

| | | | | | | | | |
|---|---|---|---|---|---|---|---|---|
| 0.161724 | 0.033163 | 0.973 | 0.968 | 1 | 3 | Mcl1 | 0.005 | Old |
| 0.161833 | 0.126183 | 0.157 | 0.13 | 1 | 3 | Slc25a1 | 0.027 | Old |
| 0.162089 | 0.067049 | 0.553 | 0.494 | 1 | 3 | Atp5o.1 | 0.059 | Old |
| 0.16214 | 0.037657 | 0.988 | 0.989 | 1 | 3 | mt-Nd2 | -0.001 | Old |
| 0.162236 | 0.433225 | 0.069 | 0.053 | 1 | 3 | Tmem2 | 0.016 | Old |
| 0.162243 | 0.060595 | 0.195 | 0.167 | 1 | 3 | Usp24 | 0.028 | Old |
| 0.162391 | 0.597315 | 0.013 | 0.006 | 1 | 3 | 1700001L0 | 0.007 | Old |
| 0.162593 | 0.070685 | 0.652 | 0.614 | 1 | 3 | Srsf2 | 0.038 | Old |
| 0.162762 | 0.864025 | 0.013 | 0.006 | 1 | 3 | Spry1 | 0.007 | Old |
| 0.162894 | 0.056745 | 0.665 | 0.61 | 1 | 3 | Prrc2c | 0.055 | Old |
| 0.163262 | 0.024094 | 0.194 | 0.163 | 1 | 3 | Lsm8 | 0.031 | Old |
| 0.163423 | 0.107382 | 0.164 | 0.135 | 1 | 3 | St6gal1 | 0.029 | Old |
| 0.163432 | 0.208924 | 0.195 | 0.168 | 1 | 3 | Msl3 | 0.027 | Old |
| 0.163481 | 0.102248 | 0.442 | 0.395 | 1 | 3 | Srrm1 | 0.047 | Old |
| 0.163645 | 0.307976 | 0.152 | 0.128 | 1 | 3 | Ubqln2 | 0.024 | Old |
| 0.16367 | 0.127238 | 0.162 | 0.135 | 1 | 3 | Cnot2 | 0.027 | Old |
| 0.163757 | 0.041764 | 0.217 | 0.186 | 1 | 3 | Sec13 | 0.031 | Old |
| 0.163885 | 0.156716 | 0.072 | 0.054 | 1 | 3 | Ikbkg | 0.018 | Old |
| 0.164147 | 0.344223 | 0.069 | 0.053 | 1 | 3 | Pigc | 0.016 | Old |
| 0.164151 | 0.031356 | 0.407 | 0.361 | 1 | 3 | Tmcc3 | 0.046 | Old |
| 0.164187 | 0.130553 | 0.229 | 0.197 | 1 | 3 | Gabarapl2 | 0.032 | Old |
| 0.164254 | 0.773404 | 0.013 | 0.006 | 1 | 3 | Tarbp1 | 0.007 | Old |
| 0.164254 | 0.424084 | 0.013 | 0.006 | 1 | 3 | C030006K1 | 0.007 | Old |
| 0.164443 | 0.368197 | 0.067 | 0.051 | 1 | 3 | Uqcc1 | 0.016 | Old |
| 0.164473 | 0.325195 | 0.068 | 0.052 | 1 | 3 | Mcu | 0.016 | Old |
| 0.164593 | 0.093465 | 0.262 | 0.232 | 1 | 3 | Tmem208 | 0.03 | Old |
| 0.164609 | 0.095371 | 0.428 | 0.389 | 1 | 3 | Crebbp | 0.039 | Old |
| 0.164628 | 0.417272 | 0.013 | 0.006 | 1 | 3 | Dpt | 0.007 | Old |
| 0.164816 | 0.402235 | 0.013 | 0.006 | 1 | 3 | Enc1 | 0.007 | Old |
| 0.164816 | 0.300447 | 0.013 | 0.006 | 1 | 3 | 9130011E1 | 0.007 | Old |
| 0.164816 | 0.279133 | 0.013 | 0.006 | 1 | 3 | Rhof | 0.007 | Old |
| 0.164847 | 0.083341 | 0.139 | 0.114 | 1 | 3 | Exoc1 | 0.025 | Old |
| 0.165312 | 0.244476 | 0.106 | 0.086 | 1 | 3 | Pik3c3 | 0.02 | Old |
| 0.165412 | 0.016799 | 0.411 | 0.367 | 1 | 3 | Aph1a | 0.044 | Old |
| 0.165455 | 0.118421 | 0.152 | 0.126 | 1 | 3 | Parp4 | 0.026 | Old |
| 0.165665 | 0.573772 | 0.022 | 0.013 | 1 | 3 | Sell | 0.009 | Old |
| 0.165858 | 0.358432 | 0.048 | 0.034 | 1 | 3 | Cnr2 | 0.014 | Old |
| 0.166046 | 0.155402 | 0.22 | 0.189 | 1 | 3 | Acox3 | 0.031 | Old |
| 0.166348 | 0.017806 | 0.119 | 0.096 | 1 | 3 | Creb3 | 0.023 | Old |
| 0.166404 | 0.017117 | 0.176 | 0.146 | 1 | 3 | Nup210 | 0.03 | Old |
| 0.166673 | 0.160931 | 0.308 | 0.269 | 1 | 3 | Osgep | 0.039 | Old |
| 0.166704 | 0.045604 | 0.155 | 0.13 | 1 | 3 | Mbd6 | 0.025 | Old |
| 0.166834 | 0.221712 | 0.202 | 0.174 | 1 | 3 | Bccip | 0.028 | Old |
| 0.166874 | 0.217855 | 0.178 | 0.152 | 1 | 3 | Cnp | 0.026 | Old |
| 0.166886 | 0.16037 | 0.082 | 0.064 | 1 | 3 | Ccs | 0.018 | Old |

| | | | | | | | | |
|---|---|---|---|---|---|---|---|---|
| 0.166955 | 0.204499 | 0.179 | 0.157 | 1 | 3 | Ube4b | 0.022 | Old |
| 0.167092 | 0.084926 | 0.575 | 0.528 | 1 | 3 | Ppp2r5a | 0.047 | Old |
| 0.167458 | 0.375473 | 0.082 | 0.064 | 1 | 3 | Nr1d2 | 0.018 | Old |
| 0.167581 | 0.039037 | 0.501 | 0.457 | 1 | 3 | Dbnl | 0.044 | Old |
| 0.167819 | 0.637271 | 0.022 | 0.013 | 1 | 3 | Btbd3 | 0.009 | Old |
| 0.167893 | 0.446742 | 0.031 | 0.02 | 1 | 3 | Mettl8 | 0.011 | Old |
| 0.167925 | 0.038587 | 0.933 | 0.902 | 1 | 3 | Npc2 | 0.031 | Old |
| 0.168148 | 0.262199 | 0.252 | 0.222 | 1 | 3 | Chuk | 0.03 | Old |
| 0.168211 | 0.092106 | 0.214 | 0.183 | 1 | 3 | AI504432 | 0.031 | Old |
| 0.168473 | 0.649009 | 0.049 | 0.035 | 1 | 3 | D2hgdh | 0.014 | Old |
| 0.168489 | 0.788963 | 0.026 | 0.016 | 1 | 3 | Serpine1 | 0.01 | Old |
| 0.1685 | 0.133441 | 0.125 | 0.102 | 1 | 3 | Adh5 | 0.023 | Old |
| 0.168541 | 0.591229 | 0.022 | 0.013 | 1 | 3 | Slc5a6 | 0.009 | Old |
| 0.168554 | 0.069197 | 0.67 | 0.615 | 1 | 3 | Bola2 | 0.055 | Old |
| 0.168752 | 0.10741 | 0.116 | 0.095 | 1 | 3 | Mtx1 | 0.021 | Old |
| 0.168842 | 0.30824 | 0.031 | 0.02 | 1 | 3 | Tssk6 | 0.011 | Old |
| 0.168939 | 0.071016 | 0.22 | 0.187 | 1 | 3 | Ano6 | 0.033 | Old |
| 0.169251 | 0.100677 | 0.397 | 0.345 | 1 | 3 | Me2 | 0.052 | Old |
| 0.169379 | 0.024907 | 0.195 | 0.163 | 1 | 3 | Suco | 0.032 | Old |
| 0.16965 | 0.051465 | 0.424 | 0.372 | 1 | 3 | U2af2 | 0.052 | Old |
| 0.169954 | 0.202735 | 0.216 | 0.188 | 1 | 3 | Psmc4 | 0.028 | Old |
| 0.17007 | 0.26066 | 0.114 | 0.092 | 1 | 3 | Pprc1 | 0.022 | Old |
| 0.17019 | 0.115846 | 0.615 | 0.572 | 1 | 3 | Psma3 | 0.043 | Old |
| 0.170315 | 0.008442 | 0.171 | 0.143 | 1 | 3 | Il27ra | 0.028 | Old |
| 0.170399 | 0.014992 | 0.362 | 0.316 | 1 | 3 | Rsu1 | 0.046 | Old |
| 0.170407 | 0.552186 | 0.037 | 0.025 | 1 | 3 | Ift27 | 0.012 | Old |
| 0.170467 | 0.412831 | 0.1 | 0.081 | 1 | 3 | Slc35b3 | 0.019 | Old |
| 0.170678 | 0.135349 | 0.166 | 0.139 | 1 | 3 | Ncoa4 | 0.027 | Old |
| 0.170773 | 0.035457 | 0.249 | 0.211 | 1 | 3 | Prelid2 | 0.038 | Old |
| 0.170945 | 0.237186 | 0.128 | 0.106 | 1 | 3 | Pdcd2l | 0.022 | Old |
| 0.170966 | 0.028371 | 0.406 | 0.356 | 1 | 3 | Fes | 0.05 | Old |
| 0.171136 | 0.075091 | 0.13 | 0.106 | 1 | 3 | Lnpk | 0.024 | Old |
| 0.171148 | 0.311525 | 0.115 | 0.093 | 1 | 3 | Pus1 | 0.022 | Old |
| 0.171326 | 0.003734 | 0.055 | 0.04 | 1 | 3 | Asf1a | 0.015 | Old |
| 0.171502 | 0.252283 | 0.096 | 0.076 | 1 | 3 | Dis3l2 | 0.02 | Old |
| 0.171503 | 0.035328 | 0.931 | 0.874 | 1 | 3 | Nr4a1 | 0.057 | Old |
| 0.171988 | 0.037038 | 0.17 | 0.14 | 1 | 3 | Usp1 | 0.03 | Old |
| 0.172215 | 0.073417 | 0.179 | 0.15 | 1 | 3 | Pla2g15 | 0.029 | Old |
| 0.17223 | 0.088849 | 0.663 | 0.606 | 1 | 3 | Prr13 | 0.057 | Old |
| 0.172483 | 0.640039 | 0.022 | 0.013 | 1 | 3 | Wfdc18 | 0.009 | Old |
| 0.172546 | 0.354668 | 0.026 | 0.016 | 1 | 3 | Zfp518b | 0.01 | Old |
| 0.172697 | 0.132824 | 0.068 | 0.052 | 1 | 3 | Zfp830 | 0.016 | Old |
| 0.172703 | 0.466731 | 0.062 | 0.047 | 1 | 3 | Pdcd2 | 0.015 | Old |
| 0.172773 | 0.018088 | 0.713 | 0.653 | 1 | 3 | Lnpep | 0.06 | Old |
| 0.172798 | 0.328893 | 0.031 | 0.02 | 1 | 3 | Pag1 | 0.011 | Old |

| | | | | | | | | |
|---|---|---|---|---|---|---|---|---|
| 0.17308 | 0.247078 | 0.185 | 0.158 | 1 | 3 | Hcfc1 | 0.027 | Old |
| 0.17325 | 0.159057 | 0.253 | 0.22 | 1 | 3 | Pcna | 0.033 | Old |
| 0.173271 | 0.182406 | 0.264 | 0.231 | 1 | 3 | Por | 0.033 | Old |
| 0.173352 | 0.13439 | 0.042 | 0.029 | 1 | 3 | Wdr83 | 0.013 | Old |
| 0.173452 | 0.163389 | 0.09 | 0.071 | 1 | 3 | Stam | 0.019 | Old |
| 0.173532 | 0.464635 | 0.093 | 0.074 | 1 | 3 | Zfpl1 | 0.019 | Old |
| 0.173699 | 0.136118 | 0.654 | 0.624 | 1 | 3 | Ctsa | 0.03 | Old |
| 0.173799 | 0.088075 | 0.37 | 0.323 | 1 | 3 | Gpr35 | 0.047 | Old |
| 0.173807 | 0.157344 | 0.142 | 0.117 | 1 | 3 | Tax1bp3 | 0.025 | Old |
| 0.173893 | 0.08587 | 0.244 | 0.21 | 1 | 3 | Herc2 | 0.034 | Old |
| 0.173919 | 0.452285 | 0.037 | 0.025 | 1 | 3 | Zfp874a | 0.012 | Old |
| 0.174161 | 0.16244 | 0.148 | 0.122 | 1 | 3 | Anpep | 0.026 | Old |
| 0.174309 | 0.280327 | 0.195 | 0.167 | 1 | 3 | Rpe | 0.028 | Old |
| 0.174355 | 0.544893 | 0.086 | 0.068 | 1 | 3 | Slc25a22 | 0.018 | Old |
| 0.174361 | 0.0629 | 0.244 | 0.212 | 1 | 3 | Ssr2 | 0.032 | Old |
| 0.174367 | 0.17937 | 0.43 | 0.388 | 1 | 3 | Sin3b | 0.042 | Old |
| 0.174379 | 0.129604 | 0.397 | 0.36 | 1 | 3 | Usp7 | 0.037 | Old |
| 0.17445 | 0.160896 | 0.201 | 0.173 | 1 | 3 | Lcmt1 | 0.028 | Old |
| 0.174564 | 0.237543 | 0.203 | 0.176 | 1 | 3 | Mrpl32 | 0.027 | Old |
| 0.175018 | 0.068994 | 0.258 | 0.221 | 1 | 3 | Med15 | 0.037 | Old |
| 0.175069 | 0.081079 | 0.228 | 0.197 | 1 | 3 | Derl2 | 0.031 | Old |
| 0.175289 | 0.032024 | 0.667 | 0.609 | 1 | 3 | Gpr132 | 0.058 | Old |
| 0.175465 | 0.250998 | 0.055 | 0.04 | 1 | 3 | Mrgbp | 0.015 | Old |
| 0.175967 | 0.488884 | 0.056 | 0.042 | 1 | 3 | Nr2c1 | 0.014 | Old |
| 0.17601 | 0.17621 | 0.357 | 0.326 | 1 | 3 | Stub1 | 0.031 | Old |
| 0.176253 | 0.061505 | 0.294 | 0.256 | 1 | 3 | Ppp4r3a | 0.038 | Old |
| 0.176358 | 0.209097 | 0.078 | 0.061 | 1 | 3 | BC035044 | 0.017 | Old |
| 0.176513 | 0.042384 | 0.407 | 0.359 | 1 | 3 | Dnajb14 | 0.048 | Old |
| 0.176952 | 0.086322 | 0.209 | 0.179 | 1 | 3 | Vps37a | 0.03 | Old |
| 0.176962 | 0.113267 | 0.472 | 0.437 | 1 | 3 | Snu13 | 0.035 | Old |
| 0.177116 | 0.272875 | 0.088 | 0.069 | 1 | 3 | Cdc16 | 0.019 | Old |
| 0.177323 | 0.052754 | 0.269 | 0.229 | 1 | 3 | Myo1c | 0.04 | Old |
| 0.177341 | 0.155444 | 0.227 | 0.197 | 1 | 3 | Pip5k1a | 0.03 | Old |
| 0.177364 | 0.14168 | 0.227 | 0.197 | 1 | 3 | 4833439L1 | 0.03 | Old |
| 0.177437 | 0.385511 | 0.167 | 0.145 | 1 | 3 | BC003331 | 0.022 | Old |
| 0.177519 | 0.221095 | 0.228 | 0.202 | 1 | 3 | Gba | 0.026 | Old |
| 0.177554 | 0.107812 | 0.133 | 0.109 | 1 | 3 | BC037034 | 0.024 | Old |
| 0.177559 | 0.032839 | 0.116 | 0.093 | 1 | 3 | Ralgds | 0.023 | Old |
| 0.177586 | 0.099696 | 0.358 | 0.323 | 1 | 3 | Vav3 | 0.035 | Old |
| 0.177888 | 0.115057 | 0.026 | 0.016 | 1 | 3 | Rnf26 | 0.01 | Old |
| 0.17791 | 0.079138 | 0.331 | 0.29 | 1 | 3 | Btbd1 | 0.041 | Old |
| 0.177911 | 0.189628 | 0.286 | 0.254 | 1 | 3 | Ubr5 | 0.032 | Old |
| 0.177955 | 0.020387 | 0.238 | 0.202 | 1 | 3 | Skiv2l | 0.036 | Old |
| 0.178268 | 0.083741 | 0.569 | 0.513 | 1 | 3 | Slc3a2 | 0.056 | Old |
| 0.178429 | 0.093839 | 0.026 | 0.016 | 1 | 3 | Gm26982 | 0.01 | Old |

| | | | | | | | | |
|---|---|---|---|---|---|---|---|---|
| 0.178503 | 0.642267 | 0.102 | 0.082 | 1 | 3 | Nagpa | 0.02 | Old |
| 0.178507 | 0.140159 | 0.323 | 0.292 | 1 | 3 | Golga4 | 0.031 | Old |
| 0.178569 | 0.089121 | 0.654 | 0.604 | 1 | 3 | Cnn2 | 0.05 | Old |
| 0.178928 | 0.13071 | 0.17 | 0.141 | 1 | 3 | Wdr43 | 0.029 | Old |
| 0.179215 | 0.072576 | 0.297 | 0.261 | 1 | 3 | Smarca4 | 0.036 | Old |
| 0.179218 | 0.120572 | 0.453 | 0.412 | 1 | 3 | Vat1 | 0.041 | Old |
| 0.179412 | 0.001346 | 0.319 | 0.275 | 1 | 3 | Nucb1 | 0.044 | Old |
| 0.179576 | 0.206288 | 0.244 | 0.215 | 1 | 3 | Arhgap25 | 0.029 | Old |
| 0.179674 | 0.13994 | 0.208 | 0.177 | 1 | 3 | Samd9l | 0.031 | Old |
| 0.179687 | 0.059002 | 0.043 | 0.03 | 1 | 3 | Dut | 0.013 | Old |
| 0.179741 | 0.041421 | 0.229 | 0.199 | 1 | 3 | Peak1 | 0.03 | Old |
| 0.179744 | 0.772178 | 0.018 | 0.01 | 1 | 3 | Atp6v0d2 | 0.008 | Old |
| 0.179757 | 0.103061 | 0.496 | 0.438 | 1 | 3 | Eid1 | 0.058 | Old |
| 0.179799 | 0.171104 | 0.061 | 0.045 | 1 | 3 | Ankmy2 | 0.016 | Old |
| 0.180701 | 0.051777 | 0.406 | 0.365 | 1 | 3 | Gna13 | 0.041 | Old |
| 0.180706 | 0.258981 | 0.139 | 0.114 | 1 | 3 | Foxj3 | 0.025 | Old |
| 0.180713 | 0.019611 | 0.222 | 0.188 | 1 | 3 | Eif2s3y | 0.034 | Old |
| 0.180918 | 0.70799 | 0.018 | 0.01 | 1 | 3 | Gm32856 | 0.008 | Old |
| 0.181138 | 0.095189 | 0.096 | 0.076 | 1 | 3 | Nkap | 0.02 | Old |
| 0.181152 | 0.27659 | 0.038 | 0.027 | 1 | 3 | Snupn | 0.011 | Old |
| 0.181255 | 0.263486 | 0.018 | 0.01 | 1 | 3 | Tmem41a | 0.008 | Old |
| 0.181375 | 0.374013 | 0.056 | 0.042 | 1 | 3 | Foxp4 | 0.014 | Old |
| 0.181739 | 0.044014 | 0.502 | 0.452 | 1 | 3 | Ctnnb1 | 0.05 | Old |
| 0.18192 | 0.036074 | 0.222 | 0.191 | 1 | 3 | Exoc3 | 0.031 | Old |
| 0.182002 | 0.243979 | 0.076 | 0.059 | 1 | 3 | Pym1 | 0.017 | Old |
| 0.182045 | 0.07647 | 0.164 | 0.139 | 1 | 3 | Pcyt1a | 0.025 | Old |
| 0.182051 | 0.243109 | 0.168 | 0.143 | 1 | 3 | Ccdc9 | 0.025 | Old |
| 0.182054 | 0.093477 | 0.418 | 0.37 | 1 | 3 | Kdelr1 | 0.048 | Old |
| 0.182101 | 0.221715 | 0.074 | 0.057 | 1 | 3 | Specc1 | 0.017 | Old |
| 0.182122 | 0.671492 | 0.027 | 0.018 | 1 | 3 | Wars2 | 0.009 | Old |
| 0.182124 | 0.220208 | 0.102 | 0.082 | 1 | 3 | Alg10b | 0.02 | Old |
| 0.18227 | 0.015532 | 0.391 | 0.337 | 1 | 3 | Trpc4ap | 0.054 | Old |
| 0.182314 | 0.167259 | 0.3 | 0.266 | 1 | 3 | Fgl2 | 0.034 | Old |
| 0.182415 | 0.281027 | 0.08 | 0.063 | 1 | 3 | Mettl6 | 0.017 | Old |
| 0.182436 | 0.519708 | 0.018 | 0.01 | 1 | 3 | Rbak | 0.008 | Old |
| 0.182734 | 0.096706 | 0.161 | 0.136 | 1 | 3 | Cdkn2d | 0.025 | Old |
| 0.182925 | 0.182809 | 0.292 | 0.26 | 1 | 3 | Osbpl9 | 0.032 | Old |
| 0.182952 | 0.065706 | 0.405 | 0.352 | 1 | 3 | Cnih4 | 0.053 | Old |
| 0.183081 | 0.227056 | 0.062 | 0.047 | 1 | 3 | Wfs1 | 0.015 | Old |
| 0.183287 | 0.001954 | 0.205 | 0.174 | 1 | 3 | Upf1 | 0.031 | Old |
| 0.183463 | 0.244028 | 0.049 | 0.035 | 1 | 3 | Usp42 | 0.014 | Old |
| 0.183566 | 0.264634 | 0.139 | 0.117 | 1 | 3 | Naip5 | 0.022 | Old |
| 0.183966 | 0.34128 | 0.038 | 0.027 | 1 | 3 | BC147527 | 0.011 | Old |
| 0.184219 | 0.114761 | 0.161 | 0.136 | 1 | 3 | Aldh16a1 | 0.025 | Old |
| 0.184378 | 0.19643 | 0.306 | 0.27 | 1 | 3 | Thoc7 | 0.036 | Old |

| | | | | | | | | |
|---|---|---|---|---|---|---|---|---|
| 0.184437 | 0.099361 | 0.136 | 0.112 | 1 | 3 | Rhoq | 0.024 | Old |
| 0.185116 | 0.237901 | 0.106 | 0.086 | 1 | 3 | Zmym4 | 0.02 | Old |
| 0.185181 | 0.29457 | 0.146 | 0.124 | 1 | 3 | 4833420G1 | 0.022 | Old |
| 0.185329 | 0.086051 | 0.018 | 0.01 | 1 | 3 | Nectin2 | 0.008 | Old |
| 0.185419 | 0.146151 | 0.105 | 0.085 | 1 | 3 | Ddx49 | 0.02 | Old |
| 0.185517 | 0.178979 | 0.349 | 0.313 | 1 | 3 | St3gal1 | 0.036 | Old |
| 0.185774 | 0.054068 | 0.195 | 0.165 | 1 | 3 | Hk1 | 0.03 | Old |
| 0.185967 | 0.382762 | 0.108 | 0.088 | 1 | 3 | Clcn5 | 0.02 | Old |
| 0.186284 | 0.036766 | 0.228 | 0.197 | 1 | 3 | Vps41 | 0.031 | Old |
| 0.186325 | 0.148806 | 0.09 | 0.071 | 1 | 3 | Fam3a | 0.019 | Old |
| 0.186354 | 0.670663 | 0.038 | 0.027 | 1 | 3 | Dtx4 | 0.011 | Old |
| 0.18648 | 0.132975 | 0.143 | 0.117 | 1 | 3 | Mtx2 | 0.026 | Old |
| 0.186755 | 0.302686 | 0.098 | 0.08 | 1 | 3 | Stxbp5 | 0.018 | Old |
| 0.186817 | 0.172671 | 0.24 | 0.21 | 1 | 3 | Chordc1 | 0.03 | Old |
| 0.186898 | 0.458201 | 0.124 | 0.105 | 1 | 3 | Gon4l | 0.019 | Old |
| 0.187239 | 0.026232 | 0.669 | 0.622 | 1 | 3 | Ehd1 | 0.047 | Old |
| 0.187597 | 0.071018 | 0.811 | 0.798 | 1 | 3 | Csde1 | 0.013 | Old |
| 0.187655 | 0.041641 | 0.204 | 0.176 | 1 | 3 | 5031425E2 | 0.028 | Old |
| 0.188231 | 0.046452 | 0.055 | 0.04 | 1 | 3 | Acd | 0.015 | Old |
| 0.18829 | 0.250779 | 0.023 | 0.014 | 1 | 3 | Rbfox2 | 0.009 | Old |
| 0.188837 | 0.011528 | 0.719 | 0.66 | 1 | 3 | Rap1a | 0.059 | Old |
| 0.188917 | 0.130154 | 0.416 | 0.372 | 1 | 3 | Psmb10 | 0.044 | Old |
| 0.189125 | 0.117203 | 0.082 | 0.064 | 1 | 3 | Dusp7 | 0.018 | Old |
| 0.18961 | 1.175664 | 0.029 | 0.019 | 1 | 3 | Gpatch3 | 0.01 | Old |
| 0.189708 | 0.037093 | 0.891 | 0.862 | 1 | 3 | Rap1b | 0.029 | Old |
| 0.189814 | 0.536551 | 0.023 | 0.014 | 1 | 3 | Atp23 | 0.009 | Old |
| 0.190225 | 0.091561 | 0.082 | 0.064 | 1 | 3 | Inip | 0.018 | Old |
| 0.190392 | 0.065406 | 0.326 | 0.287 | 1 | 3 | Rnf166 | 0.039 | Old |
| 0.190527 | 0.097817 | 0.545 | 0.481 | 1 | 3 | Wsb1 | 0.064 | Old |
| 0.190569 | 0.056877 | 0.301 | 0.266 | 1 | 3 | Bub3 | 0.035 | Old |
| 0.190615 | 0.199999 | 0.18 | 0.153 | 1 | 3 | Pfdn4 | 0.027 | Old |
| 0.190771 | 0.454994 | 0.033 | 0.023 | 1 | 3 | Mars2 | 0.01 | Old |
| 0.190842 | 0.36549 | 0.05 | 0.037 | 1 | 3 | Slc35d2 | 0.013 | Old |
| 0.190976 | 0.162774 | 0.294 | 0.264 | 1 | 3 | Dnajb9 | 0.03 | Old |
| 0.191039 | 0.693975 | 0.014 | 0.008 | 1 | 3 | P3h1 | 0.006 | Old |
| 0.191279 | 0.075046 | 0.352 | 0.317 | 1 | 3 | Mef2c | 0.035 | Old |
| 0.19133 | 0.19917 | 0.1 | 0.081 | 1 | 3 | Spns1 | 0.019 | Old |
| 0.191354 | 0.136721 | 0.063 | 0.048 | 1 | 3 | Eif2b1 | 0.015 | Old |
| 0.191536 | 0.062373 | 0.742 | 0.705 | 1 | 3 | Ddx3x | 0.037 | Old |
| 0.191571 | 0.117811 | 0.275 | 0.246 | 1 | 3 | Ufsp2 | 0.029 | Old |
| 0.191817 | 0.265139 | 0.174 | 0.15 | 1 | 3 | Tomm40 | 0.024 | Old |
| 0.192011 | 0.098907 | 0.348 | 0.308 | 1 | 3 | Ckap4 | 0.04 | Old |
| 0.192389 | 0.322723 | 0.075 | 0.059 | 1 | 3 | Cinp | 0.016 | Old |
| 0.192572 | 0.05674 | 0.341 | 0.304 | 1 | 3 | Plekhj1 | 0.037 | Old |
| 0.192581 | 0.003049 | 0.622 | 0.573 | 1 | 3 | Tnfaip8 | 0.049 | Old |

| | | | | | | | | |
|---|---|---|---|---|---|---|---|---|
| 0.192823 | 0.804253 | 0.014 | 0.008 | 1 | 3 | Cep131 | 0.006 | Old |
| 0.193022 | 0.47015 | 0.014 | 0.008 | 1 | 3 | 0610040B1 | 0.006 | Old |
| 0.193112 | 0.100224 | 0.614 | 0.568 | 1 | 3 | Lims1 | 0.046 | Old |
| 0.193221 | 0.553494 | 0.014 | 0.008 | 1 | 3 | Fam189b | 0.006 | Old |
| 0.193293 | 0.141353 | 0.201 | 0.173 | 1 | 3 | Nudt4 | 0.028 | Old |
| 0.19333 | 0.13843 | 0.209 | 0.179 | 1 | 3 | Nub1 | 0.03 | Old |
| 0.19366 | 0.629371 | 0.074 | 0.058 | 1 | 3 | Tmem199 | 0.016 | Old |
| 0.193805 | 0.058825 | 0.478 | 0.422 | 1 | 3 | Ranbp2 | 0.056 | Old |
| 0.193819 | 0.363894 | 0.014 | 0.008 | 1 | 3 | Kcnip4 | 0.006 | Old |
| 0.19417 | 0.131327 | 0.269 | 0.237 | 1 | 3 | Tcof1 | 0.032 | Old |
| 0.194252 | 0.000639 | 0.114 | 0.092 | 1 | 3 | Adar | 0.022 | Old |
| 0.194297 | 0.711503 | 0.053 | 0.039 | 1 | 3 | P2ry10 | 0.014 | Old |
| 0.19445 | 1.190504 | 0.011 | 0.005 | 1 | 3 | Cidec | 0.006 | Old |
| 0.194618 | 0.110416 | 0.087 | 0.069 | 1 | 3 | Itpkc | 0.018 | Old |
| 0.195392 | 0.663137 | 0.011 | 0.005 | 1 | 3 | Treml2 | 0.006 | Old |
| 0.19546 | 0.098055 | 0.146 | 0.122 | 1 | 3 | Kdm3b | 0.024 | Old |
| 0.195717 | 0.096588 | 0.091 | 0.072 | 1 | 3 | Ptrhd1 | 0.019 | Old |
| 0.195864 | 0.455582 | 0.011 | 0.005 | 1 | 3 | Slc22a5 | 0.006 | Old |
| 0.196006 | 0.01572 | 0.221 | 0.187 | 1 | 3 | Casc3 | 0.034 | Old |
| 0.196432 | 0.086594 | 0.568 | 0.514 | 1 | 3 | Ier3ip1 | 0.054 | Old |
| 0.196681 | 0.161932 | 0.104 | 0.085 | 1 | 3 | Hax1 | 0.019 | Old |
| 0.196787 | 0.445911 | 0.082 | 0.066 | 1 | 3 | Arhgap5 | 0.016 | Old |
| 0.196802 | 0.007778 | 0.21 | 0.179 | 1 | 3 | Mpp1 | 0.031 | Old |
| 0.196948 | 0.04841 | 0.731 | 0.663 | 1 | 3 | Tmed10 | 0.068 | Old |
| 0.196997 | 0.042514 | 0.568 | 0.524 | 1 | 3 | Cox17 | 0.044 | Old |
| 0.197022 | 0.089868 | 0.455 | 0.415 | 1 | 3 | Eif3b | 0.04 | Old |
| 0.197049 | 0.51602 | 0.011 | 0.005 | 1 | 3 | Gm48420 | 0.006 | Old |
| 0.197056 | 0.309959 | 0.121 | 0.101 | 1 | 3 | AU019823 | 0.02 | Old |
| 0.197104 | 0.185562 | 0.118 | 0.098 | 1 | 3 | Nob1 | 0.02 | Old |
| 0.197285 | 0.301174 | 0.141 | 0.119 | 1 | 3 | Timm50 | 0.022 | Old |
| 0.197286 | 0.668442 | 0.011 | 0.005 | 1 | 3 | Ubox5 | 0.006 | Old |
| 0.197312 | 0.082387 | 0.797 | 0.769 | 1 | 3 | Sec61b | 0.028 | Old |
| 0.197334 | 0.17096 | 0.062 | 0.047 | 1 | 3 | Creb5 | 0.015 | Old |
| 0.19746 | 0.052743 | 0.472 | 0.429 | 1 | 3 | Cct3 | 0.043 | Old |
| 0.197762 | 0.351689 | 0.011 | 0.005 | 1 | 3 | Gm28707 | 0.006 | Old |
| 0.197808 | 0.199827 | 0.22 | 0.193 | 1 | 3 | Ncaph2 | 0.027 | Old |
| 0.197839 | 0.650875 | 0.041 | 0.029 | 1 | 3 | Igtp | 0.012 | Old |
| 0.198159 | 0.801436 | 0.019 | 0.011 | 1 | 3 | Arsi | 0.008 | Old |
| 0.198238 | 0.313871 | 0.011 | 0.005 | 1 | 3 | Rflnb | 0.006 | Old |
| 0.198449 | 0.101267 | 0.346 | 0.308 | 1 | 3 | Atp6v1h | 0.038 | Old |
| 0.198477 | 0.206876 | 0.011 | 0.005 | 1 | 3 | Rsph3a | 0.006 | Old |
| 0.199026 | 0.300114 | 0.125 | 0.105 | 1 | 3 | Tada1 | 0.02 | Old |
| 0.199409 | 0.473018 | 0.054 | 0.04 | 1 | 3 | Slc16a1 | 0.014 | Old |
| 0.199441 | 0.109726 | 0.074 | 0.057 | 1 | 3 | Pex14 | 0.017 | Old |
| 0.199718 | 0.242049 | 0.074 | 0.058 | 1 | 3 | Txndc12 | 0.016 | Old |

| | | | | | | | | |
|---|---|---|---|---|---|---|---|---|
| 0.199728 | 0.253091 | 0.066 | 0.051 | 1 | 3 | Ptcd3 | 0.015 | Old |
| 0.199759 | 0.079014 | 0.575 | 0.523 | 1 | 3 | Cct4 | 0.052 | Old |
| 0.19989 | 0.775213 | 0.019 | 0.011 | 1 | 3 | Gm15446 | 0.008 | Old |
| 0.199913 | 0.06038 | 0.011 | 0.005 | 1 | 3 | Ablim1 | 0.006 | Old |
| 0.200015 | 0.125564 | 0.124 | 0.102 | 1 | 3 | Gm46224 | 0.022 | Old |
| 0.200308 | 0.11124 | 0.281 | 0.244 | 1 | 3 | Gatad1 | 0.037 | Old |
| 0.200528 | 0.029167 | 0.722 | 0.669 | 1 | 3 | Itgb1 | 0.053 | Old |
| 0.200836 | 0.006153 | 0.215 | 0.184 | 1 | 3 | Trnt1 | 0.031 | Old |
| 0.200959 | 0.206075 | 0.066 | 0.051 | 1 | 3 | Zxdc | 0.015 | Old |
| 0.200967 | 0.010614 | 0.22 | 0.189 | 1 | 3 | Coa3 | 0.031 | Old |
| 0.201189 | 0.656461 | 0.024 | 0.015 | 1 | 3 | Map7 | 0.009 | Old |
| 0.201192 | 0.016882 | 0.296 | 0.254 | 1 | 3 | Usp47 | 0.042 | Old |
| 0.201397 | 0.03632 | 0.305 | 0.266 | 1 | 3 | Mgea5 | 0.039 | Old |
| 0.201546 | 0.022365 | 0.142 | 0.117 | 1 | 3 | Mrps26 | 0.025 | Old |
| 0.201553 | 0.238233 | 0.143 | 0.12 | 1 | 3 | AI839979 | 0.023 | Old |
| 0.201758 | 0.142121 | 0.096 | 0.076 | 1 | 3 | E2f3 | 0.02 | Old |
| 0.201974 | 0.058464 | 0.639 | 0.593 | 1 | 3 | Fubp1 | 0.046 | Old |
| 0.202007 | 0.06077 | 0.515 | 0.468 | 1 | 3 | Prelid1 | 0.047 | Old |
| 0.202037 | 0.057584 | 0.07 | 0.054 | 1 | 3 | Nol9 | 0.016 | Old |
| 0.202133 | 0.4529 | 0.054 | 0.04 | 1 | 3 | 3110062M | 0.014 | Old |
| 0.202298 | 0.062745 | 0.243 | 0.207 | 1 | 3 | Pja2 | 0.036 | Old |
| 0.202527 | 0.074434 | 0.142 | 0.119 | 1 | 3 | Abcb7 | 0.023 | Old |
| 0.202718 | 0.085279 | 0.386 | 0.35 | 1 | 3 | Emg1 | 0.036 | Old |
| 0.202951 | 0.144875 | 0.174 | 0.149 | 1 | 3 | Ccdc174 | 0.025 | Old |
| 0.20305 | 0.589514 | 0.024 | 0.015 | 1 | 3 | Fastkd5 | 0.009 | Old |
| 0.203212 | 0.273696 | 0.139 | 0.117 | 1 | 3 | Slc39a10 | 0.022 | Old |
| 0.203386 | 0.085817 | 0.651 | 0.602 | 1 | 3 | Atp5j2 | 0.049 | Old |
| 0.203665 | 0.048151 | 0.289 | 0.251 | 1 | 3 | Cd302 | 0.038 | Old |
| 0.203679 | 0.371451 | 0.029 | 0.019 | 1 | 3 | Dntt | 0.01 | Old |
| 0.203721 | 0.059332 | 0.514 | 0.466 | 1 | 3 | Ndufb3 | 0.048 | Old |
| 0.203799 | 0.336604 | 0.13 | 0.109 | 1 | 3 | Gpaa1 | 0.021 | Old |
| 0.203815 | 0.121784 | 0.341 | 0.306 | 1 | 3 | Yeats4 | 0.035 | Old |
| 0.20382 | 0.231302 | 0.18 | 0.154 | 1 | 3 | Slc25a46 | 0.026 | Old |
| 0.203916 | 0.037455 | 0.057 | 0.043 | 1 | 3 | Dph6 | 0.014 | Old |
| 0.204018 | 0.18969 | 0.178 | 0.152 | 1 | 3 | Ugdh | 0.026 | Old |
| 0.204066 | 0.238298 | 0.054 | 0.04 | 1 | 3 | Zfyve26 | 0.014 | Old |
| 0.204104 | 0.339261 | 0.029 | 0.019 | 1 | 3 | Nme7 | 0.01 | Old |
| 0.204199 | 0.033795 | 0.372 | 0.33 | 1 | 3 | Txnl1 | 0.042 | Old |
| 0.20445 | 0.22807 | 0.202 | 0.176 | 1 | 3 | Kdm5c | 0.026 | Old |
| 0.204654 | 0.3353 | 0.047 | 0.034 | 1 | 3 | Igip | 0.013 | Old |
| 0.205278 | 0.172769 | 0.172 | 0.146 | 1 | 3 | Slc4a2 | 0.026 | Old |
| 0.205617 | 0.17153 | 0.382 | 0.343 | 1 | 3 | Epn1 | 0.039 | Old |
| 0.205799 | 0.104526 | 0.066 | 0.051 | 1 | 3 | Rarb | 0.015 | Old |
| 0.20624 | 0.072788 | 0.232 | 0.201 | 1 | 3 | Klhl5 | 0.031 | Old |
| 0.206258 | 0.335192 | 0.035 | 0.024 | 1 | 3 | Tfcp2 | 0.011 | Old |

| | | | | | | | | |
|---|---|---|---|---|---|---|---|---|
| 0.20655 | 0.053077 | 0.479 | 0.434 | 1 | 3 | 1700020I14 | 0.045 | Old |
| 0.206621 | 0.17415 | 0.116 | 0.096 | 1 | 3 | Ints1 | 0.02 | Old |
| 0.206681 | 0.459557 | 0.054 | 0.04 | 1 | 3 | Tctex1d2 | 0.014 | Old |
| 0.206793 | 0.028735 | 0.112 | 0.092 | 1 | 3 | Cdk10 | 0.02 | Old |
| 0.206809 | 0.315615 | 0.024 | 0.015 | 1 | 3 | Gm10642 | 0.009 | Old |
| 0.207082 | 0.077881 | 0.243 | 0.211 | 1 | 3 | Ss18 | 0.032 | Old |
| 0.207189 | 0.081797 | 0.366 | 0.326 | 1 | 3 | Leng8 | 0.04 | Old |
| 0.207219 | 0.223442 | 0.041 | 0.029 | 1 | 3 | Tbc1d22b | 0.012 | Old |
| 0.207547 | 0.105336 | 0.146 | 0.125 | 1 | 3 | Hspa13 | 0.021 | Old |
| 0.208081 | 0.100898 | 0.442 | 0.399 | 1 | 3 | Clec4a1 | 0.043 | Old |
| 0.208232 | 0.003614 | 0.386 | 0.348 | 1 | 3 | Sgk3 | 0.038 | Old |
| 0.208401 | 0.319932 | 0.087 | 0.071 | 1 | 3 | Smyd5 | 0.016 | Old |
| 0.208413 | 0.280211 | 0.041 | 0.029 | 1 | 3 | Tor4a | 0.012 | Old |
| 0.208507 | 0.068174 | 0.536 | 0.479 | 1 | 3 | P2rx4 | 0.057 | Old |
| 0.208653 | 0.209635 | 0.041 | 0.029 | 1 | 3 | Tecpr2 | 0.012 | Old |
| 0.208787 | 0.245469 | 0.053 | 0.039 | 1 | 3 | Fastkd2 | 0.014 | Old |
| 0.208952 | 0.059666 | 0.534 | 0.485 | 1 | 3 | Ap2s1 | 0.049 | Old |
| 0.209069 | 0.087416 | 0.019 | 0.011 | 1 | 3 | Fndc7 | 0.008 | Old |
| 0.209086 | 0.047829 | 0.521 | 0.46 | 1 | 3 | Ube2n | 0.061 | Old |
| 0.209387 | 0.325382 | 0.048 | 0.035 | 1 | 3 | Pcca | 0.013 | Old |
| 0.209392 | 0.067117 | 0.582 | 0.524 | 1 | 3 | Eif4b | 0.058 | Old |
| 0.209516 | 0.061801 | 0.165 | 0.141 | 1 | 3 | Ddhd1 | 0.024 | Old |
| 0.209547 | 0.342729 | 0.146 | 0.124 | 1 | 3 | Mrpl41 | 0.022 | Old |
| 0.20975 | 0.290844 | 0.061 | 0.047 | 1 | 3 | Sco2 | 0.014 | Old |
| 0.209862 | 0.058039 | 0.712 | 0.662 | 1 | 3 | Hspe1 | 0.05 | Old |
| 0.21143 | 0.554759 | 0.03 | 0.02 | 1 | 3 | Ttc38 | 0.01 | Old |
| 0.212024 | 0.302919 | 0.085 | 0.068 | 1 | 3 | Pkd2 | 0.017 | Old |
| 0.212246 | 0.140018 | 0.152 | 0.129 | 1 | 3 | Phf2 | 0.023 | Old |
| 0.212425 | 0.610968 | 0.03 | 0.02 | 1 | 3 | Ccdc191 | 0.01 | Old |
| 0.212576 | 0.130722 | 0.1 | 0.082 | 1 | 3 | Ccdc97 | 0.018 | Old |
| 0.212785 | 0.09984 | 0.306 | 0.273 | 1 | 3 | Ap3d1 | 0.033 | Old |
| 0.212796 | 0.343452 | 0.061 | 0.047 | 1 | 3 | Zw10 | 0.014 | Old |
| 0.213077 | 0.601791 | 0.036 | 0.025 | 1 | 3 | Lrrc14 | 0.011 | Old |
| 0.213083 | 0.056717 | 0.356 | 0.318 | 1 | 3 | Sdf2l1 | 0.038 | Old |
| 0.213163 | 0.227762 | 0.075 | 0.059 | 1 | 3 | Acot11 | 0.016 | Old |
| 0.213174 | 0.078781 | 0.342 | 0.303 | 1 | 3 | Elf1 | 0.039 | Old |
| 0.213298 | 0.090114 | 0.167 | 0.143 | 1 | 3 | Vopp1 | 0.024 | Old |
| 0.213595 | 0.011111 | 0.311 | 0.273 | 1 | 3 | Ccnt2 | 0.038 | Old |
| 0.213664 | 0.376317 | 0.042 | 0.03 | 1 | 3 | Rundc1 | 0.012 | Old |
| 0.214028 | 0.103783 | 0.221 | 0.192 | 1 | 3 | Sra1 | 0.029 | Old |
| 0.214035 | 0.054341 | 0.21 | 0.182 | 1 | 3 | Chtf8 | 0.028 | Old |
| 0.214037 | 0.013984 | 0.324 | 0.284 | 1 | 3 | Srsf7 | 0.04 | Old |
| 0.214139 | 0.995295 | 0.03 | 0.02 | 1 | 3 | Ccl22 | 0.01 | Old |
| 0.215026 | 0.229056 | 0.194 | 0.169 | 1 | 3 | Ngdn | 0.025 | Old |
| 0.21581 | 0.005587 | 0.324 | 0.283 | 1 | 3 | Yipf3 | 0.041 | Old |

| | | | | | | | | |
|---|---|---|---|---|---|---|---|---|
| 0.215826 | 0.183252 | 0.142 | 0.12 | 1 | 3 | Cetn2 | 0.022 | Old |
| 0.215887 | 0.00324 | 0.191 | 0.164 | 1 | 3 | Mrpl21 | 0.027 | Old |
| 0.215958 | 0.1348 | 0.259 | 0.225 | 1 | 3 | Clk3 | 0.034 | Old |
| 0.216088 | 0.030363 | 0.593 | 0.533 | 1 | 3 | Cerk | 0.06 | Old |
| 0.216093 | 0.150222 | 0.136 | 0.115 | 1 | 3 | Med30 | 0.021 | Old |
| 0.216246 | 0.041818 | 0.515 | 0.466 | 1 | 3 | Stap1 | 0.049 | Old |
| 0.216325 | 0.166771 | 0.305 | 0.271 | 1 | 3 | Crybg1 | 0.034 | Old |
| 0.216679 | 0.038948 | 0.196 | 0.169 | 1 | 3 | Dph3 | 0.027 | Old |
| 0.216814 | 0.333852 | 0.073 | 0.058 | 1 | 3 | Arhgef11 | 0.015 | Old |
| 0.216909 | 0.060878 | 0.281 | 0.245 | 1 | 3 | Glrx2 | 0.036 | Old |
| 0.216931 | 0.150951 | 0.139 | 0.117 | 1 | 3 | Tmem11 | 0.022 | Old |
| 0.216999 | 0.130138 | 0.386 | 0.348 | 1 | 3 | Lactb | 0.038 | Old |
| 0.217838 | 0.293016 | 0.207 | 0.184 | 1 | 3 | Cdc123 | 0.023 | Old |
| 0.217962 | 0.063904 | 0.13 | 0.11 | 1 | 3 | Tbcel | 0.02 | Old |
| 0.218316 | 0.472216 | 0.042 | 0.03 | 1 | 3 | Arfip2 | 0.012 | Old |
| 0.218317 | 0.076017 | 0.055 | 0.042 | 1 | 3 | Cds1 | 0.013 | Old |
| 0.218322 | 0.249004 | 0.03 | 0.02 | 1 | 3 | Gm10138 | 0.01 | Old |
| 0.218631 | 0.137713 | 0.079 | 0.063 | 1 | 3 | Fkbp1b | 0.016 | Old |
| 0.218871 | 0.162535 | 0.063 | 0.049 | 1 | 3 | Rttn | 0.014 | Old |
| 0.218889 | 0.219527 | 0.148 | 0.126 | 1 | 3 | Mesd | 0.022 | Old |
| 0.219231 | 0.389893 | 0.042 | 0.03 | 1 | 3 | Ap4e1 | 0.012 | Old |
| 0.219242 | 0.194441 | 0.254 | 0.229 | 1 | 3 | Dgkd | 0.025 | Old |
| 0.219361 | 0.107769 | 0.093 | 0.076 | 1 | 3 | Msra | 0.017 | Old |
| 0.219617 | 0.120352 | 0.284 | 0.249 | 1 | 3 | Tmem55b | 0.035 | Old |
| 0.220173 | 0.136826 | 0.115 | 0.096 | 1 | 3 | Crls1 | 0.019 | Old |
| 0.220244 | 0.403369 | 0.025 | 0.016 | 1 | 3 | Cetn4 | 0.009 | Old |
| 0.220258 | 0.018881 | 0.22 | 0.189 | 1 | 3 | Cacul1 | 0.031 | Old |
| 0.220558 | 0.015639 | 0.37 | 0.327 | 1 | 3 | Clec2d | 0.043 | Old |
| 0.220577 | 0.20758 | 0.042 | 0.03 | 1 | 3 | Gstcd | 0.012 | Old |
| 0.220955 | 0.416805 | 0.07 | 0.056 | 1 | 3 | Sap30 | 0.014 | Old |
| 0.220956 | 1.144751 | 0.02 | 0.013 | 1 | 3 | Cenpj | 0.007 | Old |
| 0.220983 | 0.021204 | 0.088 | 0.071 | 1 | 3 | Hsdl1 | 0.017 | Old |
| 0.221146 | 0.492213 | 0.109 | 0.091 | 1 | 3 | Ppp1r8 | 0.018 | Old |
| 0.22121 | 0.108151 | 0.222 | 0.196 | 1 | 3 | Apobec3 | 0.026 | Old |
| 0.22122 | 0.236158 | 0.128 | 0.107 | 1 | 3 | AU040320 | 0.021 | Old |
| 0.2213 | 0.055869 | 0.466 | 0.415 | 1 | 3 | Hspa4 | 0.051 | Old |
| 0.221311 | 0.105117 | 0.069 | 0.054 | 1 | 3 | Nckipsd | 0.015 | Old |
| 0.22135 | 0.666032 | 0.016 | 0.009 | 1 | 3 | Spata13 | 0.007 | Old |
| 0.22144 | 0.015545 | 0.708 | 0.649 | 1 | 3 | Grk2 | 0.059 | Old |
| 0.221456 | 0.182403 | 0.049 | 0.037 | 1 | 3 | B4galt7 | 0.012 | Old |
| 0.221528 | 0.377033 | 0.025 | 0.016 | 1 | 3 | Kbtbd3 | 0.009 | Old |
| 0.221556 | 0.217562 | 0.036 | 0.025 | 1 | 3 | Spice1 | 0.011 | Old |
| 0.221563 | 0.089014 | 0.409 | 0.372 | 1 | 3 | Ndufs2 | 0.037 | Old |
| 0.221659 | 0.134415 | 0.141 | 0.12 | 1 | 3 | Snhg6 | 0.021 | Old |
| 0.221756 | 0.168878 | 0.036 | 0.025 | 1 | 3 | Sec22a | 0.011 | Old |

| | | | | | | | | |
|---|---|---|---|---|---|---|---|---|
| 0.222186 | 0.170866 | 0.016 | 0.009 | 1 | 3 | Gm30025 | 0.007 | Old |
| 0.222335 | 0.559574 | 0.037 | 0.027 | 1 | 3 | Zbtb24 | 0.01 | Old |
| 0.22235 | 0.092818 | 0.185 | 0.16 | 1 | 3 | Atad1 | 0.025 | Old |
| 0.222395 | 0.492016 | 0.016 | 0.009 | 1 | 3 | Ace | 0.007 | Old |
| 0.222395 | 0.328572 | 0.016 | 0.009 | 1 | 3 | Amacr | 0.007 | Old |
| 0.222806 | 0.095663 | 0.086 | 0.069 | 1 | 3 | Camta1 | 0.017 | Old |
| 0.222989 | 0.125486 | 0.203 | 0.177 | 1 | 3 | Nedd9 | 0.026 | Old |
| 0.223024 | 0.244082 | 0.016 | 0.009 | 1 | 3 | Sap18b | 0.007 | Old |
| 0.223225 | 0.314808 | 0.07 | 0.056 | 1 | 3 | Unkl | 0.014 | Old |
| 0.223408 | 0.006419 | 0.602 | 0.54 | 1 | 3 | Parvb | 0.062 | Old |
| 0.223829 | 0.045813 | 0.416 | 0.376 | 1 | 3 | Nipa2 | 0.04 | Old |
| 0.223878 | 0.207063 | 0.085 | 0.068 | 1 | 3 | Prkrip1 | 0.017 | Old |
| 0.224872 | 0.099399 | 0.45 | 0.414 | 1 | 3 | Hnrnpd | 0.036 | Old |
| 0.225104 | 0.032587 | 0.352 | 0.311 | 1 | 3 | Sptssa | 0.041 | Old |
| 0.225484 | 0.090793 | 0.177 | 0.152 | 1 | 3 | Plekhg3 | 0.025 | Old |
| 0.225533 | 0.183961 | 0.306 | 0.277 | 1 | 3 | Tspan14 | 0.029 | Old |
| 0.225572 | 0.086477 | 0.092 | 0.074 | 1 | 3 | Setd1a | 0.018 | Old |
| 0.225627 | 0.023808 | 0.239 | 0.208 | 1 | 3 | Araf | 0.031 | Old |
| 0.225973 | 0.381459 | 0.134 | 0.114 | 1 | 3 | Ctdspl2 | 0.02 | Old |
| 0.226157 | 0.101865 | 0.102 | 0.083 | 1 | 3 | Plekhg2 | 0.019 | Old |
| 0.226174 | 0.221334 | 0.125 | 0.106 | 1 | 3 | Ammecr1 | 0.019 | Old |
| 0.226579 | 0.683107 | 0.02 | 0.013 | 1 | 3 | AU041133 | 0.007 | Old |
| 0.226652 | 0.049849 | 0.569 | 0.504 | 1 | 3 | Abi1 | 0.065 | Old |
| 0.227247 | 0.212097 | 0.171 | 0.148 | 1 | 3 | Atp2a3 | 0.023 | Old |
| 0.227385 | 0.189186 | 0.082 | 0.067 | 1 | 3 | Papd7 | 0.015 | Old |
| 0.227471 | 0.050008 | 0.048 | 0.035 | 1 | 3 | 2010315B0 | 0.013 | Old |
| 0.228146 | 0.188795 | 0.056 | 0.043 | 1 | 3 | Ogfod2 | 0.013 | Old |
| 0.228194 | 0.004551 | 0.185 | 0.158 | 1 | 3 | Pop5 | 0.027 | Old |
| 0.228287 | 0.136565 | 0.196 | 0.17 | 1 | 3 | Ing1 | 0.026 | Old |
| 0.228369 | 0.272966 | 0.098 | 0.081 | 1 | 3 | Ppan | 0.017 | Old |
| 0.228452 | 0.314372 | 0.223 | 0.201 | 1 | 3 | Hspa14 | 0.022 | Old |
| 0.228599 | 0.193963 | 0.02 | 0.013 | 1 | 3 | Cep95 | 0.007 | Old |
| 0.229051 | 0.589876 | 0.059 | 0.045 | 1 | 3 | Gfm2 | 0.014 | Old |
| 0.229254 | 0.085788 | 0.694 | 0.654 | 1 | 3 | Qk | 0.04 | Old |
| 0.229299 | 0.04216 | 0.171 | 0.145 | 1 | 3 | Klhdc4 | 0.026 | Old |
| 0.229335 | 0.049699 | 0.197 | 0.17 | 1 | 3 | Adgrg6 | 0.027 | Old |
| 0.229644 | 0.284254 | 0.117 | 0.098 | 1 | 3 | Eef1akmt1 | 0.019 | Old |
| 0.229809 | 0.055913 | 0.36 | 0.317 | 1 | 3 | 2610507B1 | 0.043 | Old |
| 0.230076 | 0.26842 | 0.02 | 0.013 | 1 | 3 | Arl6 | 0.007 | Old |
| 0.230106 | 0.19548 | 0.143 | 0.124 | 1 | 3 | Wdr33 | 0.019 | Old |
| 0.230196 | 0.071667 | 0.108 | 0.088 | 1 | 3 | Dcps | 0.02 | Old |
| 0.23028 | 0.123795 | 0.122 | 0.102 | 1 | 3 | Pkd1 | 0.02 | Old |
| 0.230421 | 0.180604 | 0.099 | 0.082 | 1 | 3 | Chmp6 | 0.017 | Old |
| 0.230461 | 0.040887 | 0.067 | 0.052 | 1 | 3 | 0610010K1 | 0.015 | Old |
| 0.230559 | 0.072813 | 0.272 | 0.241 | 1 | 3 | Pmp22 | 0.031 | Old |

| | | | | | | | | |
|---|---|---|---|---|---|---|---|---|
| 0.230677 | 0.080217 | 0.412 | 0.375 | 1 | 3 | Slc15a3 | 0.037 | Old |
| 0.230716 | 0.14535 | 0.16 | 0.136 | 1 | 3 | Tenm4 | 0.024 | Old |
| 0.231314 | 0.055882 | 0.18 | 0.153 | 1 | 3 | G6pdx | 0.027 | Old |
| 0.23154 | 0.122818 | 0.436 | 0.404 | 1 | 3 | Zc3hav1 | 0.032 | Old |
| 0.23168 | 0.15076 | 0.043 | 0.032 | 1 | 3 | Sh2b2 | 0.011 | Old |
| 0.232077 | 0.438328 | 0.051 | 0.039 | 1 | 3 | Fbxl18 | 0.012 | Old |
| 0.232279 | 0.01488 | 0.413 | 0.381 | 1 | 3 | Tnpo3 | 0.032 | Old |
| 0.232343 | 0.816374 | 0.012 | 0.006 | 1 | 3 | Tmem72 | 0.006 | Old |
| 0.232439 | 0.201824 | 0.072 | 0.057 | 1 | 3 | Slc11a2 | 0.015 | Old |
| 0.232637 | 0.074718 | 0.192 | 0.167 | 1 | 3 | Raf1 | 0.025 | Old |
| 0.232939 | 0.135967 | 0.183 | 0.159 | 1 | 3 | Cdc34 | 0.024 | Old |
| 0.232992 | 0.02346 | 0.692 | 0.636 | 1 | 3 | Jun | 0.056 | Old |
| 0.23301 | 0.156163 | 0.118 | 0.1 | 1 | 3 | Rbm15 | 0.018 | Old |
| 0.233138 | 0.177402 | 0.037 | 0.027 | 1 | 3 | Tmem86b | 0.01 | Old |
| 0.233255 | 0.047402 | 0.799 | 0.729 | 1 | 3 | Cox6b1 | 0.07 | Old |
| 0.233341 | 1.243276 | 0.012 | 0.006 | 1 | 3 | Iigp1 | 0.006 | Old |
| 0.233341 | 0.655062 | 0.012 | 0.006 | 1 | 3 | Cacna1b | 0.006 | Old |
| 0.233506 | 0.11573 | 0.072 | 0.057 | 1 | 3 | Dus1l | 0.015 | Old |
| 0.234593 | 0.446817 | 0.012 | 0.006 | 1 | 3 | Sparcl1 | 0.006 | Old |
| 0.234722 | 0.141272 | 0.489 | 0.448 | 1 | 3 | Hdlbp | 0.041 | Old |
| 0.234843 | 0.814908 | 0.012 | 0.006 | 1 | 3 | A4galt | 0.006 | Old |
| 0.23491 | 0.026414 | 0.099 | 0.081 | 1 | 3 | Bcl2l2 | 0.018 | Old |
| 0.234981 | 0.075673 | 0.243 | 0.215 | 1 | 3 | Safb | 0.028 | Old |
| 0.235282 | 0.231033 | 0.146 | 0.125 | 1 | 3 | Serpinb1a | 0.021 | Old |
| 0.235597 | 0.475056 | 0.012 | 0.006 | 1 | 3 | Zfp692 | 0.006 | Old |
| 0.235623 | 0.027814 | 0.055 | 0.042 | 1 | 3 | Ttl | 0.013 | Old |
| 0.235705 | 0.349998 | 0.043 | 0.032 | 1 | 3 | Sv2b | 0.011 | Old |
| 0.23578 | 0.256563 | 0.05 | 0.038 | 1 | 3 | Zfp386 | 0.012 | Old |
| 0.236543 | 0.084319 | 0.179 | 0.155 | 1 | 3 | Cd300c2 | 0.024 | Old |
| 0.236545 | 0.03324 | 0.143 | 0.121 | 1 | 3 | Clasp1 | 0.022 | Old |
| 0.236611 | 0.250304 | 0.085 | 0.069 | 1 | 3 | Zfp24 | 0.016 | Old |
| 0.236815 | 0.592532 | 0.038 | 0.028 | 1 | 3 | Polr1b | 0.01 | Old |
| 0.236857 | 0.504333 | 0.012 | 0.006 | 1 | 3 | Myo1b | 0.006 | Old |
| 0.236943 | 0.399122 | 0.069 | 0.056 | 1 | 3 | Alkbh7 | 0.013 | Old |
| 0.237168 | 0.235979 | 0.031 | 0.021 | 1 | 3 | Fxn | 0.01 | Old |
| 0.237319 | 0.248717 | 0.031 | 0.021 | 1 | 3 | Zxdb | 0.01 | Old |
| 0.237616 | 0.190744 | 0.012 | 0.006 | 1 | 3 | Il17rc | 0.006 | Old |
| 0.237717 | 0.104425 | 0.051 | 0.039 | 1 | 3 | Ermard | 0.012 | Old |
| 0.2379 | 0.114203 | 0.491 | 0.453 | 1 | 3 | Gramd4 | 0.038 | Old |
| 0.238267 | 0.053774 | 0.094 | 0.077 | 1 | 3 | Prdm4 | 0.017 | Old |
| 0.238376 | 0.037057 | 0.012 | 0.006 | 1 | 3 | Sirpb1a | 0.006 | Old |
| 0.23875 | 0.105176 | 0.226 | 0.197 | 1 | 3 | Slc13a3 | 0.029 | Old |
| 0.23907 | 0.043093 | 0.274 | 0.24 | 1 | 3 | Cyb5a | 0.034 | Old |
| 0.239105 | 0.399206 | 0.09 | 0.073 | 1 | 3 | Unc45a | 0.017 | Old |
| 0.239203 | 0.285968 | 0.151 | 0.129 | 1 | 3 | Pgp | 0.022 | Old |

| | | | | | | | | |
|---|---|---|---|---|---|---|---|---|
| 0.239226 | 0.050132 | 0.092 | 0.076 | 1 | 3 | Tmem55a | 0.016 | Old |
| 0.239434 | 0.037527 | 0.07 | 0.056 | 1 | 3 | Tars2 | 0.014 | Old |
| 0.239534 | 0.288409 | 0.067 | 0.053 | 1 | 3 | Dzip3 | 0.014 | Old |
| 0.23964 | 0.301097 | 0.081 | 0.066 | 1 | 3 | Coq2 | 0.015 | Old |
| 0.239714 | 0.013218 | 0.037 | 0.027 | 1 | 3 | Ears2 | 0.01 | Old |
| 0.239993 | 0.124421 | 0.13 | 0.109 | 1 | 3 | mt-Nd6 | 0.021 | Old |
| 0.240005 | 0.344827 | 0.143 | 0.124 | 1 | 3 | Mfsd7a | 0.019 | Old |
| 0.2403 | 0.115762 | 0.182 | 0.159 | 1 | 3 | Tmem141 | 0.023 | Old |
| 0.240439 | 0.031455 | 0.266 | 0.237 | 1 | 3 | Ints6 | 0.029 | Old |
| 0.240614 | 0.168024 | 0.026 | 0.018 | 1 | 3 | Amer1 | 0.008 | Old |
| 0.240892 | 0.110293 | 0.032 | 0.023 | 1 | 3 | Zfp429 | 0.009 | Old |
| 0.241057 | 0.118654 | 0.297 | 0.263 | 1 | 3 | Itpr1 | 0.034 | Old |
| 0.241278 | 0.766297 | 0.087 | 0.073 | 1 | 3 | Ankrd16 | 0.014 | Old |
| 0.241334 | 0.349442 | 0.051 | 0.039 | 1 | 3 | Cox18 | 0.012 | Old |
| 0.241377 | 0.250547 | 0.119 | 0.101 | 1 | 3 | Rpia | 0.018 | Old |
| 0.241443 | 0.113994 | 0.103 | 0.085 | 1 | 3 | Tex10 | 0.018 | Old |
| 0.241491 | 0.014068 | 0.278 | 0.247 | 1 | 3 | Zfp445 | 0.031 | Old |
| 0.24161 | 0.021972 | 0.476 | 0.436 | 1 | 3 | Fli1 | 0.04 | Old |
| 0.241716 | 0.049643 | 0.114 | 0.095 | 1 | 3 | BC003965 | 0.019 | Old |
| 0.242065 | 0.128451 | 0.134 | 0.114 | 1 | 3 | Zfp593 | 0.02 | Old |
| 0.242109 | 0.485964 | 0.038 | 0.028 | 1 | 3 | Alyref2 | 0.01 | Old |
| 0.242366 | 0.092044 | 0.21 | 0.184 | 1 | 3 | Slc35b1 | 0.026 | Old |
| 0.242579 | 0.090507 | 0.327 | 0.294 | 1 | 3 | Htt | 0.033 | Old |
| 0.243008 | 0.245494 | 0.164 | 0.144 | 1 | 3 | Mrpl35 | 0.02 | Old |
| 0.243051 | 0.040359 | 0.227 | 0.198 | 1 | 3 | Ndufa9 | 0.029 | Old |
| 0.243211 | 0.10814 | 0.535 | 0.482 | 1 | 3 | Tor1aip1 | 0.053 | Old |
| 0.243234 | 0.403482 | 0.066 | 0.052 | 1 | 3 | Zbtb42 | 0.014 | Old |
| 0.243423 | 0.340257 | 0.131 | 0.112 | 1 | 3 | Tex264 | 0.019 | Old |
| 0.243755 | 0.086493 | 0.228 | 0.201 | 1 | 3 | Dscr3 | 0.027 | Old |
| 0.243847 | 0.625444 | 0.022 | 0.014 | 1 | 3 | Atg9b | 0.008 | Old |
| 0.244029 | 0.242554 | 0.038 | 0.028 | 1 | 3 | Zfp943 | 0.01 | Old |
| 0.244396 | 0.196507 | 0.124 | 0.105 | 1 | 3 | Ppp1r35 | 0.019 | Old |
| 0.244478 | 0.249093 | 0.119 | 0.1 | 1 | 3 | Nelfa | 0.019 | Old |
| 0.244618 | 0.077499 | 0.026 | 0.018 | 1 | 3 | Cep19 | 0.008 | Old |
| 0.244796 | 0.007193 | 0.195 | 0.167 | 1 | 3 | Wipf2 | 0.028 | Old |
| 0.244925 | 0.035313 | 0.544 | 0.489 | 1 | 3 | Xiap | 0.055 | Old |
| 0.244935 | 0.540257 | 0.032 | 0.023 | 1 | 3 | Diaph3 | 0.009 | Old |
| 0.245018 | 0.296697 | 0.017 | 0.01 | 1 | 3 | Cav1 | 0.007 | Old |
| 0.245241 | 0.078255 | 0.087 | 0.071 | 1 | 3 | Slc25a19 | 0.016 | Old |
| 0.245269 | 0.212535 | 0.038 | 0.028 | 1 | 3 | Dcaf6 | 0.01 | Old |
| 0.245322 | 0.16757 | 0.059 | 0.045 | 1 | 3 | Mettl5 | 0.014 | Old |
| 0.245496 | 0.092175 | 0.094 | 0.078 | 1 | 3 | Nt5c2 | 0.016 | Old |
| 0.245525 | 0.242299 | 0.102 | 0.085 | 1 | 3 | Rapgef1 | 0.017 | Old |
| 0.245575 | 0.067147 | 0.281 | 0.25 | 1 | 3 | Isy1 | 0.031 | Old |
| 0.245626 | 0.094659 | 0.05 | 0.038 | 1 | 3 | Zdhhc14 | 0.012 | Old |

| | | | | | | | | |
|---|---|---|---|---|---|---|---|---|
| 0.24566 | 0.53797 | 0.017 | 0.01 | 1 | 3 | Trem1 | 0.007 | Old |
| 0.245689 | 0.233517 | 0.032 | 0.023 | 1 | 3 | Spout1 | 0.009 | Old |
| 0.245999 | 0.019646 | 0.166 | 0.141 | 1 | 3 | Grpel1 | 0.025 | Old |
| 0.246028 | 0.499277 | 0.039 | 0.029 | 1 | 3 | Cdc25a | 0.01 | Old |
| 0.246152 | 0.422029 | 0.103 | 0.086 | 1 | 3 | Slc25a28 | 0.017 | Old |
| 0.246403 | 0.109947 | 0.109 | 0.091 | 1 | 3 | Ern1 | 0.018 | Old |
| 0.246435 | 0.207946 | 0.059 | 0.045 | 1 | 3 | Pan2 | 0.014 | Old |
| 0.246519 | 0.565463 | 0.017 | 0.01 | 1 | 3 | 1700066M | 0.007 | Old |
| 0.246596 | 0.167624 | 0.032 | 0.023 | 1 | 3 | Zfp729b | 0.009 | Old |
| 0.246763 | 0.098884 | 0.235 | 0.21 | 1 | 3 | Tug1 | 0.025 | Old |
| 0.246826 | 0.207824 | 0.068 | 0.054 | 1 | 3 | Nes | 0.014 | Old |
| 0.24689 | 0.010366 | 0.087 | 0.071 | 1 | 3 | Tom1 | 0.016 | Old |
| 0.24692 | 0.352999 | 0.06 | 0.047 | 1 | 3 | Msh2 | 0.013 | Old |
| 0.247441 | 0.099855 | 0.262 | 0.227 | 1 | 3 | Twf1 | 0.035 | Old |
| 0.247603 | 0.050477 | 0.229 | 0.199 | 1 | 3 | Med1 | 0.03 | Old |
| 0.247642 | 0.010735 | 0.066 | 0.052 | 1 | 3 | Stk35 | 0.014 | Old |
| 0.247925 | 0.15322 | 0.277 | 0.255 | 1 | 3 | Rbm6 | 0.022 | Old |
| 0.248026 | 0.389815 | 0.017 | 0.01 | 1 | 3 | Sh3d19 | 0.007 | Old |
| 0.248111 | 0.146565 | 0.059 | 0.045 | 1 | 3 | Sp3os | 0.014 | Old |
| 0.248175 | 0.011578 | 0.127 | 0.107 | 1 | 3 | Uhrf1bp1l | 0.02 | Old |
| 0.248447 | 0.229946 | 0.059 | 0.045 | 1 | 3 | Aamdc | 0.014 | Old |
| 0.248668 | 0.227872 | 0.076 | 0.062 | 1 | 3 | Exoc7 | 0.014 | Old |
| 0.248718 | 0.017094 | 0.841 | 0.806 | 1 | 3 | Dok2 | 0.035 | Old |
| 0.248804 | 0.122575 | 0.611 | 0.559 | 1 | 3 | Myl12b | 0.052 | Old |
| 0.248834 | 0.023138 | 0.189 | 0.163 | 1 | 3 | Slc29a1 | 0.026 | Old |
| 0.249119 | 0.15656 | 0.345 | 0.317 | 1 | 3 | Thoc2 | 0.028 | Old |
| 0.250169 | 0.131319 | 0.222 | 0.194 | 1 | 3 | Atp11b | 0.028 | Old |
| 0.250303 | 0.024314 | 0.854 | 0.857 | 1 | 3 | Rac1 | -0.003 | Old |
| 0.250423 | 0.399359 | 0.022 | 0.014 | 1 | 3 | Nadsyn1 | 0.008 | Old |
| 0.250688 | 0.173792 | 0.092 | 0.076 | 1 | 3 | Snapc1 | 0.016 | Old |
| 0.251061 | 0.273867 | 0.017 | 0.01 | 1 | 3 | Zfp799 | 0.007 | Old |
| 0.251112 | 0.353014 | 0.047 | 0.035 | 1 | 3 | Rdm1 | 0.012 | Old |
| 0.251716 | 0.0414 | 0.233 | 0.203 | 1 | 3 | Arid1b | 0.03 | Old |
| 0.251716 | 0.218767 | 0.053 | 0.04 | 1 | 3 | Sirpb1c | 0.013 | Old |
| 0.2518 | 0.279563 | 0.087 | 0.072 | 1 | 3 | Faap100 | 0.015 | Old |
| 0.25187 | 0.042233 | 0.35 | 0.307 | 1 | 3 | Stard3nl | 0.043 | Old |
| 0.251955 | 0.034249 | 0.053 | 0.04 | 1 | 3 | Slc37a2 | 0.013 | Old |
| 0.252186 | 0.041257 | 0.228 | 0.199 | 1 | 3 | Spink2 | 0.029 | Old |
| 0.252386 | 0.228964 | 0.114 | 0.096 | 1 | 3 | Tmem19 | 0.018 | Old |
| 0.252427 | 0.009816 | 0.284 | 0.251 | 1 | 3 | Zc3h13 | 0.033 | Old |
| 0.252906 | 0.013866 | 0.337 | 0.297 | 1 | 3 | Arhgef2 | 0.04 | Old |
| 0.252959 | 0.100268 | 0.059 | 0.045 | 1 | 3 | Polr3b | 0.014 | Old |
| 0.253113 | 0.072728 | 0.061 | 0.048 | 1 | 3 | Gfod1 | 0.013 | Old |
| 0.253216 | 0.038131 | 0.136 | 0.116 | 1 | 3 | Kcnq1ot1 | 0.02 | Old |
| 0.253236 | 0.130841 | 0.153 | 0.134 | 1 | 3 | Acot9 | 0.019 | Old |

| | | | | | | | | |
|---|---|---|---|---|---|---|---|---|
| 0.253469 | 0.081717 | 0.164 | 0.14 | 1 | 3 | Ythdc2 | 0.024 | Old |
| 0.253546 | 0.024287 | 0.139 | 0.117 | 1 | 3 | Mrpl10 | 0.022 | Old |
| 0.253718 | 0.356742 | 0.06 | 0.047 | 1 | 3 | Prmt9 | 0.013 | Old |
| 0.253801 | 0.311433 | 0.103 | 0.086 | 1 | 3 | Trappc11 | 0.017 | Old |
| 0.254121 | 0.206655 | 0.017 | 0.01 | 1 | 3 | Cenps | 0.007 | Old |
| 0.254266 | 0.131019 | 0.145 | 0.124 | 1 | 3 | Mut | 0.021 | Old |
| 0.254402 | 0.06116 | 0.244 | 0.216 | 1 | 3 | Eapp | 0.028 | Old |
| 0.254588 | 0.207935 | 0.045 | 0.034 | 1 | 3 | Kctd13 | 0.011 | Old |
| 0.254784 | 0.187924 | 0.061 | 0.048 | 1 | 3 | Cog2 | 0.013 | Old |
| 0.25499 | 0.25908 | 0.062 | 0.049 | 1 | 3 | Pnpt1 | 0.013 | Old |
| 0.255203 | 0.259842 | 0.054 | 0.042 | 1 | 3 | Rhbdd2 | 0.012 | Old |
| 0.255268 | 0.224927 | 0.179 | 0.158 | 1 | 3 | Rprd2 | 0.021 | Old |
| 0.255405 | 0.035373 | 0.164 | 0.139 | 1 | 3 | Qars | 0.025 | Old |
| 0.255677 | 0.2666 | 0.075 | 0.061 | 1 | 3 | Mtmr9 | 0.014 | Old |
| 0.255952 | 0.335523 | 0.027 | 0.019 | 1 | 3 | Zmym6 | 0.008 | Old |
| 0.256065 | 0.102261 | 0.344 | 0.309 | 1 | 3 | Slc1a5 | 0.035 | Old |
| 0.25612 | 0.471511 | 0.027 | 0.019 | 1 | 3 | Zdhhc1 | 0.008 | Old |
| 0.25623 | 0.064423 | 0.033 | 0.024 | 1 | 3 | Otud1 | 0.009 | Old |
| 0.256433 | 0.138393 | 0.162 | 0.139 | 1 | 3 | Uchl5 | 0.023 | Old |
| 0.256439 | 0.11409 | 0.111 | 0.092 | 1 | 3 | Gucd1 | 0.019 | Old |
| 0.256583 | 0.125224 | 0.171 | 0.148 | 1 | 3 | Zmpste24 | 0.023 | Old |
| 0.257292 | 0.059553 | 0.642 | 0.587 | 1 | 3 | Rpl27 | 0.055 | Old |
| 0.257428 | 0.496056 | 0.047 | 0.035 | 1 | 3 | Haus6 | 0.012 | Old |
| 0.257499 | 0.061378 | 0.084 | 0.068 | 1 | 3 | Gm49368 | 0.016 | Old |
| 0.257632 | 0.051422 | 0.039 | 0.029 | 1 | 3 | Lmnb2 | 0.01 | Old |
| 0.258092 | 0.0204 | 0.012 | 0.019 | 1 | 3 | Cep57l1 | -0.007 | Old |
| 0.258103 | 0.00959 | 0.045 | 0.034 | 1 | 3 | Lrrc40 | 0.011 | Old |
| 0.258192 | 0.177663 | 0.094 | 0.078 | 1 | 3 | Akr7a5 | 0.016 | Old |
| 0.258277 | 0.097798 | 0.246 | 0.217 | 1 | 3 | Idi1 | 0.029 | Old |
| 0.258321 | 0.350919 | 0.027 | 0.019 | 1 | 3 | Cenpw | 0.008 | Old |
| 0.258877 | 0.191212 | 0.124 | 0.106 | 1 | 3 | Btbd17 | 0.018 | Old |
| 0.258889 | 0.207332 | 0.045 | 0.034 | 1 | 3 | Serf1 | 0.011 | Old |
| 0.259193 | 0.249252 | 0.134 | 0.115 | 1 | 3 | Nhlrc2 | 0.019 | Old |
| 0.259231 | 0.049353 | 0.066 | 0.052 | 1 | 3 | Atg2b | 0.014 | Old |
| 0.259489 | 0.047726 | 0.133 | 0.112 | 1 | 3 | Arfip1 | 0.021 | Old |
| 0.259597 | 0.09104 | 0.576 | 0.529 | 1 | 3 | Atp5c1 | 0.047 | Old |
| 0.259613 | 0.126403 | 0.268 | 0.241 | 1 | 3 | Srrt | 0.027 | Old |
| 0.259813 | 0.362303 | 0.07 | 0.057 | 1 | 3 | Lmnb1 | 0.013 | Old |
| 0.259852 | 0.230588 | 0.027 | 0.019 | 1 | 3 | Ice2 | 0.008 | Old |
| 0.260142 | 0.000437 | 0.268 | 0.232 | 1 | 3 | Cr1l | 0.036 | Old |
| 0.260255 | 0.131375 | 0.158 | 0.138 | 1 | 3 | Fam103a1 | 0.02 | Old |
| 0.260295 | 0.348354 | 0.069 | 0.056 | 1 | 3 | Ddx41 | 0.013 | Old |
| 0.260338 | 0.057297 | 0.906 | 0.87 | 1 | 3 | Cd83 | 0.036 | Old |
| 0.260451 | 0.05321 | 0.444 | 0.402 | 1 | 3 | Tmem14c | 0.042 | Old |
| 0.260961 | 0.153759 | 0.143 | 0.122 | 1 | 3 | Rrp7a | 0.021 | Old |

| | | | | | | | | |
|---|---|---|---|---|---|---|---|---|
| 0.260992 | 0.448495 | 0.048 | 0.037 | 1 | 3 | Cenpl | 0.011 | Old |
| 0.261286 | 0.285987 | 0.022 | 0.03 | 1 | 3 | Ier5l | -0.008 | Old |
| 0.261476 | 0.247269 | 0.063 | 0.051 | 1 | 3 | Zbtb2 | 0.012 | Old |
| 0.261984 | 0.00059 | 0.051 | 0.039 | 1 | 3 | Zbtb6 | 0.012 | Old |
| 0.262099 | 0.442745 | 0.023 | 0.015 | 1 | 3 | Pigo | 0.008 | Old |
| 0.262212 | 0.322707 | 0.069 | 0.056 | 1 | 3 | Zdhhc4 | 0.013 | Old |
| 0.262427 | 0.399233 | 0.072 | 0.058 | 1 | 3 | Cox10 | 0.014 | Old |
| 0.262506 | 1.32331 | 0.013 | 0.008 | 1 | 3 | Zfp169 | 0.005 | Old |
| 0.262815 | 0.018029 | 0.511 | 0.466 | 1 | 3 | Itpripl2 | 0.045 | Old |
| 0.262848 | 0.193174 | 0.085 | 0.069 | 1 | 3 | Ttf1 | 0.016 | Old |
| 0.262893 | 0.082118 | 0.097 | 0.081 | 1 | 3 | Zdhhc18 | 0.016 | Old |
| 0.262918 | 0.281046 | 0.048 | 0.037 | 1 | 3 | Cnpy4 | 0.011 | Old |
| 0.263198 | 0.060228 | 0.108 | 0.09 | 1 | 3 | Ginm1 | 0.018 | Old |
| 0.263201 | 0.372012 | 0.035 | 0.025 | 1 | 3 | Vsig8 | 0.01 | Old |
| 0.263499 | 0.044576 | 0.184 | 0.162 | 1 | 3 | Mrpl15 | 0.022 | Old |
| 0.263537 | 0.150792 | 0.166 | 0.144 | 1 | 3 | Armc1 | 0.022 | Old |
| 0.263771 | 0.017985 | 0.381 | 0.342 | 1 | 3 | Gapvd1 | 0.039 | Old |
| 0.263874 | 0.127482 | 0.449 | 0.414 | 1 | 3 | Plxnb2 | 0.035 | Old |
| 0.263952 | 0.268636 | 0.072 | 0.058 | 1 | 3 | Usp40 | 0.014 | Old |
| 0.264081 | 0.205016 | 0.146 | 0.124 | 1 | 3 | Usp5 | 0.022 | Old |
| 0.264138 | 0.126345 | 0.027 | 0.019 | 1 | 3 | Zfp932 | 0.008 | Old |
| 0.26433 | 0.081417 | 0.552 | 0.515 | 1 | 3 | Mfsd14b | 0.037 | Old |
| 0.264379 | 0.607618 | 0.023 | 0.015 | 1 | 3 | 3300005D( | 0.008 | Old |
| 0.264418 | 0.383885 | 0.035 | 0.025 | 1 | 3 | Pkd1l2 | 0.01 | Old |
| 0.264687 | 0.19297 | 0.208 | 0.184 | 1 | 3 | Gm49336 | 0.024 | Old |
| 0.26487 | 0.028087 | 0.798 | 0.769 | 1 | 3 | Calm2 | 0.029 | Old |
| 0.264951 | 0.503223 | 0.023 | 0.015 | 1 | 3 | Fuz | 0.008 | Old |
| 0.265085 | 0.096948 | 0.085 | 0.069 | 1 | 3 | Nol6 | 0.016 | Old |
| 0.265123 | 0.03955 | 0.148 | 0.126 | 1 | 3 | Lipe | 0.022 | Old |
| 0.265605 | 0.130805 | 0.434 | 0.4 | 1 | 3 | Stag2 | 0.034 | Old |
| 0.265619 | 0.125128 | 0.146 | 0.125 | 1 | 3 | Acbd6 | 0.021 | Old |
| 0.26564 | 0.203793 | 0.035 | 0.025 | 1 | 3 | Zfand1 | 0.01 | Old |
| 0.26568 | 0.130806 | 0.229 | 0.206 | 1 | 3 | C1d | 0.023 | Old |
| 0.265888 | 0.075563 | 0.047 | 0.035 | 1 | 3 | Mus81 | 0.012 | Old |
| 0.266127 | 0.117556 | 0.146 | 0.126 | 1 | 3 | Hnrnph3 | 0.02 | Old |
| 0.266366 | 0.918382 | 0.029 | 0.02 | 1 | 3 | Ptcd1 | 0.009 | Old |
| 0.266438 | 0.103508 | 0.118 | 0.1 | 1 | 3 | Med17 | 0.018 | Old |
| 0.266592 | 0.152456 | 0.067 | 0.053 | 1 | 3 | Ncs1 | 0.014 | Old |
| 0.266678 | 0.013399 | 0.446 | 0.399 | 1 | 3 | Cd164 | 0.047 | Old |
| 0.266726 | 0.108899 | 0.174 | 0.152 | 1 | 3 | Psen2 | 0.022 | Old |
| 0.266866 | 0.760089 | 0.013 | 0.008 | 1 | 3 | Chst14 | 0.005 | Old |
| 0.267315 | 0.102998 | 0.142 | 0.121 | 1 | 3 | Vkorc1l1 | 0.021 | Old |
| 0.267891 | 0.607418 | 0.029 | 0.02 | 1 | 3 | Dppa3 | 0.009 | Old |
| 0.267899 | 0.534656 | 0.013 | 0.008 | 1 | 3 | Gpr150 | 0.005 | Old |
| 0.268194 | 0.015641 | 0.288 | 0.258 | 1 | 3 | Polr2f | 0.03 | Old |

| | | | | | | | | |
|---|---|---|---|---|---|---|---|---|
| 0.268231 | 0.553319 | 0.029 | 0.02 | 1 | 3 | Pdlim7 | 0.009 | Old |
| 0.268659 | 1.126388 | 0.018 | 0.011 | 1 | 3 | Zfp579 | 0.007 | Old |
| 0.26898 | 0.478485 | 0.023 | 0.015 | 1 | 3 | Serpinb9 | 0.008 | Old |
| 0.269047 | 0.251341 | 0.007 | 0.013 | 1 | 3 | Rab11fip3 | -0.006 | Old |
| 0.269081 | 0.571286 | 0.029 | 0.02 | 1 | 3 | Coa7 | 0.009 | Old |
| 0.26917 | 0.041711 | 0.146 | 0.125 | 1 | 3 | Fam160b1 | 0.021 | Old |
| 0.269343 | 0.059414 | 0.366 | 0.332 | 1 | 3 | Uhrf2 | 0.034 | Old |
| 0.269454 | 0.512789 | 0.013 | 0.008 | 1 | 3 | Gm15050 | 0.005 | Old |
| 0.269473 | 0.176774 | 0.188 | 0.167 | 1 | 3 | Asxl2 | 0.021 | Old |
| 0.26954 | 0.031661 | 0.26 | 0.227 | 1 | 3 | Coro1b | 0.033 | Old |
| 0.269986 | 0.256788 | 0.062 | 0.049 | 1 | 3 | Cd1d1 | 0.013 | Old |
| 0.270144 | 0.079481 | 0.069 | 0.056 | 1 | 3 | Ublcp1 | 0.013 | Old |
| 0.270148 | 0.044023 | 0.179 | 0.154 | 1 | 3 | Slc50a1 | 0.025 | Old |
| 0.270333 | 0.050625 | 0.307 | 0.27 | 1 | 3 | Rb1 | 0.037 | Old |
| 0.270412 | 0.397605 | 0.018 | 0.011 | 1 | 3 | Fancm | 0.007 | Old |
| 0.270433 | 0.033495 | 0.358 | 0.322 | 1 | 3 | Trmt112 | 0.036 | Old |
| 0.270754 | 0.487086 | 0.013 | 0.008 | 1 | 3 | Gm29994 | 0.005 | Old |
| 0.271057 | 0.271423 | 0.063 | 0.051 | 1 | 3 | Tmem29 | 0.012 | Old |
| 0.271085 | 0.210295 | 0.078 | 0.063 | 1 | 3 | Nit1 | 0.015 | Old |
| 0.27129 | 0.055185 | 0.321 | 0.29 | 1 | 3 | Wdfy4 | 0.031 | Old |
| 0.271536 | 0.329374 | 0.013 | 0.008 | 1 | 3 | Cysltr2 | 0.005 | Old |
| 0.271746 | 0.146533 | 0.102 | 0.085 | 1 | 3 | Pla2g16 | 0.017 | Old |
| 0.271843 | 0.126237 | 0.115 | 0.096 | 1 | 3 | Tnip1 | 0.019 | Old |
| 0.271945 | 0.221973 | 0.062 | 0.049 | 1 | 3 | Rpa2 | 0.013 | Old |
| 0.272261 | 0.082315 | 0.357 | 0.319 | 1 | 3 | Cacybp | 0.038 | Old |
| 0.272658 | 0.07122 | 0.277 | 0.245 | 1 | 3 | Hnrnpr | 0.032 | Old |
| 0.272901 | 0.162842 | 0.076 | 0.062 | 1 | 3 | Stx18 | 0.014 | Old |
| 0.273105 | 0.019184 | 0.013 | 0.008 | 1 | 3 | Gstp2 | 0.005 | Old |
| 0.273275 | 0.51969 | 0.018 | 0.011 | 1 | 3 | Rab38 | 0.007 | Old |
| 0.273492 | 0.177876 | 0.254 | 0.229 | 1 | 3 | Rnf44 | 0.025 | Old |
| 0.273555 | 0.07833 | 0.444 | 0.402 | 1 | 3 | Tmed9 | 0.042 | Old |
| 0.273871 | 0.377356 | 0.06 | 0.048 | 1 | 3 | Pskh1 | 0.012 | Old |
| 0.273885 | 0.060435 | 0.129 | 0.11 | 1 | 3 | Polg | 0.019 | Old |
| 0.274155 | 0.053485 | 0.013 | 0.008 | 1 | 3 | Fastkd3 | 0.005 | Old |
| 0.274172 | 0.139998 | 0.264 | 0.236 | 1 | 3 | Tmem33 | 0.028 | Old |
| 0.274357 | 0.08655 | 0.125 | 0.107 | 1 | 3 | Kctd17 | 0.018 | Old |
| 0.274923 | 0.341731 | 0.067 | 0.054 | 1 | 3 | Rcc1l | 0.013 | Old |
| 0.275495 | 0.055699 | 0.246 | 0.218 | 1 | 3 | Spata6 | 0.028 | Old |
| 0.275536 | 0.279614 | 0.122 | 0.105 | 1 | 3 | Gtf2e2 | 0.017 | Old |
| 0.275592 | 0.194294 | 0.079 | 0.064 | 1 | 3 | Gpr85 | 0.015 | Old |
| 0.275615 | 0.238246 | 0.186 | 0.167 | 1 | 3 | Ifnar2 | 0.019 | Old |
| 0.275766 | 0.230295 | 0.049 | 0.038 | 1 | 3 | Abcb10 | 0.011 | Old |
| 0.27583 | 0.219091 | 0.152 | 0.133 | 1 | 3 | Mapkap1 | 0.019 | Old |
| 0.2759 | 0.026975 | 0.364 | 0.327 | 1 | 3 | Supt5 | 0.037 | Old |
| 0.276077 | 0.125881 | 0.108 | 0.091 | 1 | 3 | Gmeb1 | 0.017 | Old |

| | | | | | | | | |
|---|---|---|---|---|---|---|---|---|
| 0.276116 | 0.055776 | 0.324 | 0.292 | 1 | 3 | Ndufa10 | 0.032 | Old |
| 0.276383 | 0.264792 | 0.018 | 0.011 | 1 | 3 | Capn10 | 0.007 | Old |
| 0.276692 | 0.113443 | 0.277 | 0.251 | 1 | 3 | Acbd3 | 0.026 | Old |
| 0.276774 | 0.008363 | 0.092 | 0.076 | 1 | 3 | Gpkow | 0.016 | Old |
| 0.27682 | 0.056873 | 0.029 | 0.02 | 1 | 3 | Fam206a | 0.009 | Old |
| 0.27689 | 0.095477 | 0.21 | 0.189 | 1 | 3 | Ncoa6 | 0.021 | Old |
| 0.277282 | 0.079745 | 0.477 | 0.441 | 1 | 3 | Lima1 | 0.036 | Old |
| 0.278346 | 0.06483 | 0.662 | 0.609 | 1 | 3 | Taok3 | 0.053 | Old |
| 0.278579 | 0.084124 | 0.13 | 0.111 | 1 | 3 | Arid3a | 0.019 | Old |
| 0.278746 | 0.010162 | 0.194 | 0.17 | 1 | 3 | Smarcb1 | 0.024 | Old |
| 0.278784 | 0.09759 | 0.151 | 0.131 | 1 | 3 | Trim27 | 0.02 | Old |
| 0.279742 | 0.012113 | 0.128 | 0.109 | 1 | 3 | Rbm34 | 0.019 | Old |
| 0.28017 | 0.247456 | 0.056 | 0.044 | 1 | 3 | Pmf1 | 0.012 | Old |
| 0.28068 | 0.252044 | 0.066 | 0.053 | 1 | 3 | Fam60a | 0.013 | Old |
| 0.281212 | 0.136227 | 0.364 | 0.33 | 1 | 3 | Zfand6 | 0.034 | Old |
| 0.28147 | 0.557774 | 0.024 | 0.016 | 1 | 3 | Mad2l1 | 0.008 | Old |
| 0.281497 | 0.222139 | 0.112 | 0.096 | 1 | 3 | Yif1a | 0.016 | Old |
| 0.282762 | 0.205411 | 0.098 | 0.083 | 1 | 3 | Smc2 | 0.015 | Old |
| 0.282764 | 0.405401 | 0.057 | 0.045 | 1 | 3 | Tdrd3 | 0.012 | Old |
| 0.282876 | 0.106092 | 0.227 | 0.203 | 1 | 3 | Snx8 | 0.024 | Old |
| 0.282912 | 0.122709 | 0.036 | 0.027 | 1 | 3 | Dmap1 | 0.009 | Old |
| 0.282943 | 0.035176 | 0.333 | 0.302 | 1 | 3 | Rap1gds1 | 0.031 | Old |
| 0.283049 | 0.027737 | 0.314 | 0.285 | 1 | 3 | Papd4 | 0.029 | Old |
| 0.283402 | 0.401567 | 0.067 | 0.054 | 1 | 3 | Mterf3 | 0.013 | Old |
| 0.28349 | 0.185292 | 0.106 | 0.09 | 1 | 3 | St7l | 0.016 | Old |
| 0.283619 | 0.051753 | 0.446 | 0.396 | 1 | 3 | Wapl | 0.05 | Old |
| 0.283695 | 0.485937 | 0.036 | 0.027 | 1 | 3 | Mcph1 | 0.009 | Old |
| 0.283787 | 0.029821 | 0.078 | 0.063 | 1 | 3 | Nat10 | 0.015 | Old |
| 0.284184 | 0.011393 | 0.124 | 0.105 | 1 | 3 | Fam241a | 0.019 | Old |
| 0.284203 | 0.092007 | 0.076 | 0.062 | 1 | 3 | Dnajc24 | 0.014 | Old |
| 0.284374 | 0.493641 | 0.024 | 0.016 | 1 | 3 | Ift172 | 0.008 | Old |
| 0.284456 | 0.028759 | 0.486 | 0.442 | 1 | 3 | Mapre2 | 0.044 | Old |
| 0.284958 | 0.734781 | 0.024 | 0.016 | 1 | 3 | Trub1 | 0.008 | Old |
| 0.285048 | 0.956843 | 0.01 | 0.005 | 1 | 3 | Pkmyt1 | 0.005 | Old |
| 0.28514 | 0.050652 | 0.111 | 0.093 | 1 | 3 | Synpo | 0.018 | Old |
| 0.285209 | 0.065156 | 0.535 | 0.491 | 1 | 3 | Fem1c | 0.044 | Old |
| 0.28531 | 0.256978 | 0.03 | 0.021 | 1 | 3 | Gale | 0.009 | Old |
| 0.28534 | 0.213909 | 0.231 | 0.208 | 1 | 3 | Atxn7l1 | 0.023 | Old |
| 0.285736 | 0.152314 | 0.024 | 0.016 | 1 | 3 | Acad12 | 0.008 | Old |
| 0.285901 | 0.051192 | 0.497 | 0.451 | 1 | 3 | Prkd3 | 0.046 | Old |
| 0.285929 | 0.008171 | 0.097 | 0.081 | 1 | 3 | Dhdds | 0.016 | Old |
| 0.285931 | 0.711378 | 0.024 | 0.016 | 1 | 3 | Zfp953 | 0.008 | Old |
| 0.285955 | 0.225661 | 0.043 | 0.033 | 1 | 3 | Prpf3 | 0.01 | Old |
| 0.286033 | 0.034546 | 0.612 | 0.563 | 1 | 3 | St13 | 0.049 | Old |
| 0.286069 | 0.026828 | 0.434 | 0.389 | 1 | 3 | Vps35 | 0.045 | Old |

| | | | | | | | | |
|---|---|---|---|---|---|---|---|---|
| 0.286321 | 0.469784 | 0.024 | 0.016 | 1 | 3 | Plscr1 | 0.008 | Old |
| 0.286333 | 0.678447 | 0.01 | 0.005 | 1 | 3 | Zbtb10 | 0.005 | Old |
| 0.286356 | 0.055502 | 0.399 | 0.362 | 1 | 3 | Calu | 0.037 | Old |
| 0.286771 | 0.461067 | 0.079 | 0.066 | 1 | 3 | Mfap1b | 0.013 | Old |
| 0.286903 | 0.086288 | 0.226 | 0.201 | 1 | 3 | Rps27rt | 0.025 | Old |
| 0.286968 | 0.006267 | 0.716 | 0.668 | 1 | 3 | Slk | 0.048 | Old |
| 0.286976 | 0.666903 | 0.01 | 0.005 | 1 | 3 | 1700102PC | 0.005 | Old |
| 0.287016 | 0.00111 | 0.352 | 0.321 | 1 | 3 | Mcts1 | 0.031 | Old |
| 0.287069 | 0.234549 | 0.161 | 0.14 | 1 | 3 | Stim1 | 0.021 | Old |
| 0.287218 | 0.400029 | 0.03 | 0.021 | 1 | 3 | Clec16a | 0.009 | Old |
| 0.287218 | 0.34346 | 0.03 | 0.021 | 1 | 3 | Kif3a | 0.009 | Old |
| 0.287289 | 0.207092 | 0.068 | 0.056 | 1 | 3 | Sorl1 | 0.012 | Old |
| 0.287393 | 0.332602 | 0.043 | 0.033 | 1 | 3 | Cacna1f | 0.01 | Old |
| 0.287621 | 0.625955 | 0.01 | 0.005 | 1 | 3 | Zfp28 | 0.005 | Old |
| 0.287621 | 0.541346 | 0.01 | 0.005 | 1 | 3 | 1200007C1 | 0.005 | Old |
| 0.287625 | 0.247032 | 0.061 | 0.049 | 1 | 3 | Rmdn3 | 0.012 | Old |
| 0.287929 | 0.016355 | 0.232 | 0.205 | 1 | 3 | Gm8186 | 0.027 | Old |
| 0.287944 | 0.60342 | 0.01 | 0.005 | 1 | 3 | Bmpr1a | 0.005 | Old |
| 0.287944 | 0.409876 | 0.01 | 0.005 | 1 | 3 | Phf11c | 0.005 | Old |
| 0.28821 | 0.210441 | 0.037 | 0.028 | 1 | 3 | Sirpb1b | 0.009 | Old |
| 0.28859 | 0.417429 | 0.01 | 0.005 | 1 | 3 | Zfp60 | 0.005 | Old |
| 0.288592 | 0.189497 | 0.243 | 0.22 | 1 | 3 | Mcfd2 | 0.023 | Old |
| 0.288678 | 0.205202 | 0.102 | 0.086 | 1 | 3 | Car5b | 0.016 | Old |
| 0.288751 | 0.482629 | 0.01 | 0.005 | 1 | 3 | Gm48529 | 0.005 | Old |
| 0.288913 | 0.344001 | 0.01 | 0.005 | 1 | 3 | Gm4951 | 0.005 | Old |
| 0.289187 | 0.180148 | 0.066 | 0.053 | 1 | 3 | Uvssa | 0.013 | Old |
| 0.289237 | 0.182301 | 0.01 | 0.005 | 1 | 3 | Gm17767 | 0.005 | Old |
| 0.289783 | 0.025947 | 0.616 | 0.573 | 1 | 3 | Ap2m1 | 0.043 | Old |
| 0.290209 | 0.227326 | 0.01 | 0.005 | 1 | 3 | Gpr137c | 0.005 | Old |
| 0.290217 | 0.001519 | 0.201 | 0.177 | 1 | 3 | Ahctf1 | 0.024 | Old |
| 0.290534 | 0.251657 | 0.01 | 0.005 | 1 | 3 | Nudt8.1 | 0.005 | Old |
| 0.290544 | 0.642173 | 0.059 | 0.047 | 1 | 3 | Oxsm | 0.012 | Old |
| 0.290798 | 0.043842 | 0.079 | 0.064 | 1 | 3 | Cebpzos | 0.015 | Old |
| 0.290859 | 0.280131 | 0.01 | 0.005 | 1 | 3 | Dus4l | 0.005 | Old |
| 0.290923 | 0.035259 | 0.479 | 0.428 | 1 | 3 | Hook3 | 0.051 | Old |
| 0.291128 | 0.026739 | 0.277 | 0.249 | 1 | 3 | Ebp | 0.028 | Old |
| 0.291238 | 0.127807 | 0.168 | 0.148 | 1 | 3 | Acads | 0.02 | Old |
| 0.291423 | 0.16465 | 0.115 | 0.097 | 1 | 3 | Abhd16a | 0.018 | Old |
| 0.291795 | 1.027909 | 0.019 | 0.013 | 1 | 3 | Csf3 | 0.006 | Old |
| 0.291805 | 0.521188 | 0.037 | 0.028 | 1 | 3 | Scrn3 | 0.009 | Old |
| 0.292161 | 0.004504 | 0.01 | 0.005 | 1 | 3 | Aplp1 | 0.005 | Old |
| 0.292177 | 0.109698 | 0.164 | 0.145 | 1 | 3 | Fbxw7 | 0.019 | Old |
| 0.292288 | 0.363694 | 0.069 | 0.057 | 1 | 3 | Nudt14 | 0.012 | Old |
| 0.292858 | 0.183091 | 0.051 | 0.04 | 1 | 3 | Rbm10 | 0.011 | Old |
| 0.292881 | 0.262423 | 0.074 | 0.061 | 1 | 3 | Myc | 0.013 | Old |

| | | | | | | | | |
|---|---|---|---|---|---|---|---|---|
| 0.292934 | 0.262916 | 0.086 | 0.072 | 1 | 3 | Prdm1 | 0.014 | Old |
| 0.292957 | 0.479489 | 0.054 | 0.043 | 1 | 3 | Cnot11 | 0.011 | Old |
| 0.293041 | 0.019842 | 0.301 | 0.268 | 1 | 3 | Smndc1 | 0.033 | Old |
| 0.293077 | 0.051567 | 0.265 | 0.237 | 1 | 3 | Mtch2 | 0.028 | Old |
| 0.29308 | 0.172447 | 0.219 | 0.196 | 1 | 3 | Tmem127 | 0.023 | Old |
| 0.293876 | 0.469896 | 0.03 | 0.021 | 1 | 3 | AA467197 | 0.009 | Old |
| 0.293959 | 0.035978 | 0.142 | 0.122 | 1 | 3 | Vps52 | 0.02 | Old |
| 0.294094 | 0.044412 | 0.184 | 0.162 | 1 | 3 | Plpp2 | 0.022 | Old |
| 0.294199 | 0.026446 | 0.56 | 0.516 | 1 | 3 | Daglb | 0.044 | Old |
| 0.294431 | 0.00383 | 0.075 | 0.062 | 1 | 3 | Mavs | 0.013 | Old |
| 0.294804 | 0.361845 | 0.062 | 0.051 | 1 | 3 | Stmn1 | 0.011 | Old |
| 0.29493 | 0.364206 | 0.053 | 0.042 | 1 | 3 | Tbc1d10a | 0.011 | Old |
| 0.295096 | 0.011224 | 0.043 | 0.033 | 1 | 3 | Rusc2 | 0.01 | Old |
| 0.295568 | 0.066028 | 0.336 | 0.306 | 1 | 3 | Nrd1 | 0.03 | Old |
| 0.295733 | 0.164255 | 0.096 | 0.081 | 1 | 3 | Sp2 | 0.015 | Old |
| 0.295945 | 0.080541 | 0.174 | 0.154 | 1 | 3 | Trim35 | 0.02 | Old |
| 0.296192 | 0.125581 | 0.024 | 0.016 | 1 | 3 | Atad5 | 0.008 | Old |
| 0.296195 | 0.46679 | 0.073 | 0.061 | 1 | 3 | Ercc3 | 0.012 | Old |
| 0.296712 | 0.630729 | 0.019 | 0.013 | 1 | 3 | D3Ertd254 | 0.006 | Old |
| 0.296777 | 0.084492 | 0.044 | 0.034 | 1 | 3 | Mad2l2 | 0.01 | Old |
| 0.296888 | 0.124583 | 0.03 | 0.021 | 1 | 3 | Sntg1 | 0.009 | Old |
| 0.296901 | 0.225911 | 0.105 | 0.09 | 1 | 3 | Ubtd1 | 0.015 | Old |
| 0.297063 | 0.159538 | 0.194 | 0.174 | 1 | 3 | Ltbp3 | 0.02 | Old |
| 0.297299 | 0.248225 | 0.059 | 0.047 | 1 | 3 | Qtrt1 | 0.012 | Old |
| 0.297412 | 0.10232 | 0.037 | 0.028 | 1 | 3 | Wdr92 | 0.009 | Old |
| 0.297588 | 0.036709 | 0.571 | 0.514 | 1 | 3 | Aamp | 0.057 | Old |
| 0.29781 | 0.110211 | 0.037 | 0.028 | 1 | 3 | Hes1 | 0.009 | Old |
| 0.297909 | 0.270762 | 0.06 | 0.048 | 1 | 3 | Nol10 | 0.012 | Old |
| 0.297943 | 0.363854 | 0.061 | 0.049 | 1 | 3 | Ripk3 | 0.012 | Old |
| 0.298177 | 0.008445 | 0.211 | 0.188 | 1 | 3 | Hypk | 0.023 | Old |
| 0.298483 | 0.102765 | 0.238 | 0.212 | 1 | 3 | Rchy1 | 0.026 | Old |
| 0.298574 | 0.152986 | 0.127 | 0.11 | 1 | 3 | Mfsd12 | 0.017 | Old |
| 0.298758 | 0.508442 | 0.025 | 0.018 | 1 | 3 | Smim24 | 0.007 | Old |
| 0.298861 | 0.085135 | 0.345 | 0.308 | 1 | 3 | Dnase2a | 0.037 | Old |
| 0.299067 | 0.046495 | 0.293 | 0.263 | 1 | 3 | Emc3 | 0.03 | Old |
| 0.299147 | 0.110088 | 0.234 | 0.207 | 1 | 3 | Gpr108 | 0.027 | Old |
| 0.29919 | 0.353249 | 0.019 | 0.013 | 1 | 3 | Akap17b | 0.006 | Old |
| 0.299269 | 0.018067 | 0.456 | 0.419 | 1 | 3 | Adss | 0.037 | Old |
| 0.299462 | 0.185836 | 0.067 | 0.054 | 1 | 3 | Nrde2 | 0.013 | Old |
| 0.300057 | 0.178288 | 0.323 | 0.295 | 1 | 3 | Emc1 | 0.028 | Old |
| 0.300161 | 0.050002 | 0.11 | 0.093 | 1 | 3 | Rogdi | 0.017 | Old |
| 0.300548 | 0.190384 | 0.019 | 0.013 | 1 | 3 | Tut1 | 0.006 | Old |
| 0.300548 | 0.053003 | 0.019 | 0.013 | 1 | 3 | Gm17586 | 0.006 | Old |
| 0.300634 | 0.173248 | 0.09 | 0.074 | 1 | 3 | Hira | 0.016 | Old |
| 0.300684 | 0.037606 | 0.177 | 0.154 | 1 | 3 | Rlf | 0.023 | Old |

| | | | | | | | | |
|---|---|---|---|---|---|---|---|---|
| 0.301075 | 0.032315 | 0.533 | 0.499 | 1 | 3 | Trip12 | 0.034 | Old |
| 0.301318 | 0.387991 | 0.014 | 0.009 | 1 | 3 | Prc1 | 0.005 | Old |
| 0.301469 | 0.173451 | 0.031 | 0.023 | 1 | 3 | Zfp574 | 0.008 | Old |
| 0.301862 | 0.01358 | 0.293 | 0.265 | 1 | 3 | Fip1l1 | 0.028 | Old |
| 0.301865 | 0.159594 | 0.182 | 0.162 | 1 | 3 | Dnajc14 | 0.02 | Old |
| 0.302118 | 0.183168 | 0.133 | 0.114 | 1 | 3 | Lrrk1 | 0.019 | Old |
| 0.302182 | 0.038581 | 0.288 | 0.259 | 1 | 3 | Epsti1 | 0.029 | Old |
| 0.302635 | 0.029419 | 0.385 | 0.345 | 1 | 3 | Msrb1 | 0.04 | Old |
| 0.302817 | 0.200758 | 0.133 | 0.115 | 1 | 3 | Eif1ad | 0.018 | Old |
| 0.302898 | 0.072529 | 0.221 | 0.198 | 1 | 3 | Suclg1 | 0.023 | Old |
| 0.302931 | 0.183259 | 0.06 | 0.048 | 1 | 3 | Mettl14 | 0.012 | Old |
| 0.302946 | 0.302103 | 0.159 | 0.141 | 1 | 3 | Cbx4 | 0.018 | Old |
| 0.302952 | 0.226649 | 0.081 | 0.068 | 1 | 3 | Lrch1 | 0.013 | Old |
| 0.30353 | 0.03833 | 0.41 | 0.371 | 1 | 3 | Ccar1 | 0.039 | Old |
| 0.303662 | 0.462761 | 0.025 | 0.018 | 1 | 3 | Rnf219 | 0.007 | Old |
| 0.303923 | 0.051246 | 0.363 | 0.331 | 1 | 3 | Lilrb4a | 0.032 | Old |
| 0.304418 | 0.058977 | 0.019 | 0.013 | 1 | 3 | Mtif3 | 0.006 | Old |
| 0.30451 | 0.319904 | 0.014 | 0.009 | 1 | 3 | Tada2a | 0.005 | Old |
| 0.305319 | 0.053665 | 0.909 | 0.891 | 1 | 3 | Marcks | 0.018 | Old |
| 0.305578 | 0.192843 | 0.014 | 0.009 | 1 | 3 | Trip13 | 0.005 | Old |
| 0.305578 | 0.09717 | 0.014 | 0.009 | 1 | 3 | Tmem138 | 0.005 | Old |
| 0.305965 | 0.292897 | 0.079 | 0.066 | 1 | 3 | Mcm3ap | 0.013 | Old |
| 0.306114 | 0.236433 | 0.014 | 0.009 | 1 | 3 | Sfxn5 | 0.005 | Old |
| 0.306114 | 0.074011 | 0.014 | 0.009 | 1 | 3 | Acbd4 | 0.005 | Old |
| 0.30617 | 0.235017 | 0.066 | 0.053 | 1 | 3 | Zfp654 | 0.013 | Old |
| 0.306234 | 0.386854 | 0.025 | 0.018 | 1 | 3 | Ston2 | 0.007 | Old |
| 0.306307 | 0.063532 | 0.806 | 0.758 | 1 | 3 | Sub1 | 0.048 | Old |
| 0.306382 | 0.415239 | 0.014 | 0.009 | 1 | 3 | Slc25a43 | 0.005 | Old |
| 0.306784 | 0.308943 | 0.047 | 0.037 | 1 | 3 | Tmem126b | 0.01 | Old |
| 0.306812 | 0.077948 | 0.147 | 0.129 | 1 | 3 | Glb1 | 0.018 | Old |
| 0.306829 | 0.296771 | 0.025 | 0.018 | 1 | 3 | 1600014C1 | 0.007 | Old |
| 0.306904 | 0.062072 | 0.168 | 0.146 | 1 | 3 | Smg9 | 0.022 | Old |
| 0.306909 | 0.044912 | 0.524 | 0.473 | 1 | 3 | Eif4g1 | 0.051 | Old |
| 0.306969 | 0.072201 | 0.338 | 0.308 | 1 | 3 | Csnk2a1 | 0.03 | Old |
| 0.307129 | 0.333811 | 0.069 | 0.057 | 1 | 3 | Tpgs2 | 0.012 | Old |
| 0.307326 | 0.783432 | 0.026 | 0.019 | 1 | 3 | Gstk1 | 0.007 | Old |
| 0.308375 | 0.02199 | 0.216 | 0.189 | 1 | 3 | Zbtb1 | 0.027 | Old |
| 0.308692 | 0.034736 | 0.406 | 0.361 | 1 | 3 | Ctbp1 | 0.045 | Old |
| 0.308993 | 0.043197 | 0.051 | 0.04 | 1 | 3 | Mapk7 | 0.011 | Old |
| 0.309148 | 0.198737 | 0.106 | 0.091 | 1 | 3 | Irf3 | 0.015 | Old |
| 0.309329 | 0.051669 | 0.434 | 0.393 | 1 | 3 | Mrps33 | 0.041 | Old |
| 0.309441 | 0.39891 | 0.084 | 0.071 | 1 | 3 | Tmem97 | 0.013 | Old |
| 0.309449 | 0.031751 | 0.073 | 0.059 | 1 | 3 | Mfn1 | 0.014 | Old |
| 0.309679 | 0.037595 | 0.539 | 0.499 | 1 | 3 | Cdc42se2 | 0.04 | Old |
| 0.30973 | 0.010113 | 0.148 | 0.129 | 1 | 3 | Mea1 | 0.019 | Old |

| | | | | | | | | |
|---|---|---|---|---|---|---|---|---|
| 0.309791 | 0.18434 | 0.088 | 0.074 | 1 | 3 | Wdr12 | 0.014 | Old |
| 0.309901 | 0.401008 | 0.038 | 0.029 | 1 | 3 | Usp30 | 0.009 | Old |
| 0.30998 | 0.05387 | 0.495 | 0.444 | 1 | 3 | Tbcb | 0.051 | Old |
| 0.31023 | 0.090597 | 0.381 | 0.341 | 1 | 3 | Hadhb | 0.04 | Old |
| 0.311305 | 0.008718 | 0.793 | 0.739 | 1 | 3 | Vamp8 | 0.054 | Old |
| 0.311374 | 0.101247 | 0.076 | 0.063 | 1 | 3 | Dcp1a | 0.013 | Old |
| 0.311675 | 0.13342 | 0.115 | 0.098 | 1 | 3 | Ticam1 | 0.017 | Old |
| 0.312027 | 0.27448 | 0.128 | 0.111 | 1 | 3 | Isg20l2 | 0.017 | Old |
| 0.312045 | 0.033445 | 0.523 | 0.457 | 1 | 3 | Gfra2 | 0.066 | Old |
| 0.312196 | 0.02852 | 0.443 | 0.412 | 1 | 3 | Pdlim5 | 0.031 | Old |
| 0.312196 | 0.019291 | 0.127 | 0.109 | 1 | 3 | Nubp1 | 0.018 | Old |
| 0.312441 | 0.23951 | 0.088 | 0.074 | 1 | 3 | Il13ra1 | 0.014 | Old |
| 0.312725 | 0.063062 | 0.25 | 0.22 | 1 | 3 | Kmt2d | 0.03 | Old |
| 0.312746 | 0.06328 | 0.683 | 0.628 | 1 | 3 | Dynll1 | 0.055 | Old |
| 0.313941 | 0.115308 | 0.168 | 0.148 | 1 | 3 | Esco1 | 0.02 | Old |
| 0.313959 | 0.182653 | 0.1 | 0.086 | 1 | 3 | Tlr4 | 0.014 | Old |
| 0.314031 | 0.023013 | 0.099 | 0.083 | 1 | 3 | Alpk1 | 0.016 | Old |
| 0.314291 | 0.052113 | 0.4 | 0.369 | 1 | 3 | Fam96a | 0.031 | Old |
| 0.314344 | 0.052499 | 0.076 | 0.063 | 1 | 3 | Abr | 0.013 | Old |
| 0.314438 | 0.138846 | 0.053 | 0.042 | 1 | 3 | Mamld1 | 0.011 | Old |
| 0.315008 | 0.021191 | 0.031 | 0.023 | 1 | 3 | Taf9b | 0.008 | Old |
| 0.315581 | 0.154129 | 0.11 | 0.095 | 1 | 3 | Nr2c2ap | 0.015 | Old |
| 0.315668 | 0.077751 | 0.045 | 0.035 | 1 | 3 | Gbp9 | 0.01 | Old |
| 0.315685 | 0.39026 | 0.02 | 0.014 | 1 | 3 | Sh3pxd2a | 0.006 | Old |
| 0.315848 | 0.054054 | 0.078 | 0.064 | 1 | 3 | Fbxo25 | 0.014 | Old |
| 0.316238 | 0.178565 | 0.136 | 0.12 | 1 | 3 | Washc1 | 0.016 | Old |
| 0.316314 | 0.05524 | 0.685 | 0.643 | 1 | 3 | Macf1 | 0.042 | Old |
| 0.316653 | 0.155437 | 0.153 | 0.135 | 1 | 3 | Mrpl11 | 0.018 | Old |
| 0.316698 | 0.103023 | 0.534 | 0.499 | 1 | 3 | Rab10 | 0.035 | Old |
| 0.316842 | 0.017875 | 0.283 | 0.253 | 1 | 3 | Slc2a6 | 0.03 | Old |
| 0.316919 | 0.069587 | 0.271 | 0.241 | 1 | 3 | Jak2 | 0.03 | Old |
| 0.317221 | 0.280823 | 0.063 | 0.052 | 1 | 3 | Prkch | 0.011 | Old |
| 0.317379 | 0.235134 | 0.073 | 0.061 | 1 | 3 | Fam217b | 0.012 | Old |
| 0.317932 | 0.176462 | 0.186 | 0.168 | 1 | 3 | 3830406C1 | 0.018 | Old |
| 0.318204 | 0.549992 | 0.069 | 0.057 | 1 | 3 | Oaf | 0.012 | Old |
| 0.318231 | 0.03448 | 0.275 | 0.247 | 1 | 3 | Vamp4 | 0.028 | Old |
| 0.318321 | 0.109358 | 0.131 | 0.114 | 1 | 3 | Ctps2 | 0.017 | Old |
| 0.31869 | 0.11545 | 0.239 | 0.213 | 1 | 3 | Pank3 | 0.026 | Old |
| 0.318856 | 0.422066 | 0.02 | 0.014 | 1 | 3 | Gm10036 | 0.006 | Old |
| 0.319179 | 0.091072 | 0.032 | 0.024 | 1 | 3 | Urb2 | 0.008 | Old |
| 0.319523 | 0.244386 | 0.096 | 0.081 | 1 | 3 | Dnajb12 | 0.015 | Old |
| 0.319537 | 0.156983 | 0.189 | 0.169 | 1 | 3 | Slc35a4 | 0.02 | Old |
| 0.319538 | 0.45888 | 0.02 | 0.014 | 1 | 3 | Cntrob | 0.006 | Old |
| 0.319656 | 0.028233 | 0.392 | 0.357 | 1 | 3 | Samd4b | 0.035 | Old |
| 0.319821 | 0.02778 | 0.682 | 0.634 | 1 | 3 | Nktr | 0.048 | Old |

| | | | | | | | | |
|---|---|---|---|---|---|---|---|---|
| 0.319891 | 0.034964 | 0.257 | 0.231 | 1 | 3 | Tbc1d20 | 0.026 | Old |
| 0.319908 | 0.509144 | 0.041 | 0.032 | 1 | 3 | Tacc3 | 0.009 | Old |
| 0.319992 | 0.021354 | 0.165 | 0.143 | 1 | 3 | Mak16 | 0.022 | Old |
| 0.320116 | 0.120749 | 0.097 | 0.082 | 1 | 3 | BC005561 | 0.015 | Old |
| 0.320214 | 0.036198 | 0.4 | 0.367 | 1 | 3 | Rer1 | 0.033 | Old |
| 0.320677 | 0.393631 | 0.02 | 0.014 | 1 | 3 | Tnfsf13os | 0.006 | Old |
| 0.321022 | 0.076971 | 0.26 | 0.235 | 1 | 3 | Atad2b | 0.025 | Old |
| 0.321233 | 0.040282 | 0.743 | 0.691 | 1 | 3 | Rpl22l1 | 0.052 | Old |
| 0.321238 | 0.108278 | 0.054 | 0.043 | 1 | 3 | Vps33b | 0.011 | Old |
| 0.321863 | 0.441858 | 0.032 | 0.024 | 1 | 3 | Usp18 | 0.008 | Old |
| 0.322464 | 0.448449 | 0.026 | 0.019 | 1 | 3 | Zfp512b | 0.007 | Old |
| 0.322897 | 0.219461 | 0.192 | 0.172 | 1 | 3 | Gramd1a | 0.02 | Old |
| 0.323422 | 0.204964 | 0.02 | 0.014 | 1 | 3 | Paqr4 | 0.006 | Old |
| 0.324008 | 0.123849 | 0.054 | 0.043 | 1 | 3 | Gm10521 | 0.011 | Old |
| 0.3242 | 0.034524 | 0.209 | 0.186 | 1 | 3 | Dusp16 | 0.023 | Old |
| 0.324418 | 0.032165 | 0.395 | 0.359 | 1 | 3 | Arfgef1 | 0.036 | Old |
| 0.325062 | 0.028267 | 0.89 | 0.871 | 1 | 3 | Ucp2 | 0.019 | Old |
| 0.325067 | 0.193306 | 0.026 | 0.019 | 1 | 3 | C2cd2 | 0.007 | Old |
| 0.325163 | 0.251929 | 0.041 | 0.032 | 1 | 3 | Cep68 | 0.009 | Old |
| 0.325198 | 0.110557 | 0.118 | 0.101 | 1 | 3 | Lztfl1 | 0.017 | Old |
| 0.325286 | 0.004212 | 0.186 | 0.164 | 1 | 3 | Atf2 | 0.022 | Old |
| 0.325695 | 0.236007 | 0.06 | 0.048 | 1 | 3 | 4930512J1 | 0.012 | Old |
| 0.325825 | 0.061889 | 0.384 | 0.342 | 1 | 3 | Ankrd13a | 0.042 | Old |
| 0.325912 | 0.392088 | 0.076 | 0.064 | 1 | 3 | Ptgfrn | 0.012 | Old |
| 0.325952 | 0.326139 | 0.02 | 0.014 | 1 | 3 | Tmem132a | 0.006 | Old |
| 0.326767 | 0.302836 | 0.041 | 0.032 | 1 | 3 | Oas1a | 0.009 | Old |
| 0.326779 | 0.20255 | 0.048 | 0.038 | 1 | 3 | Slc25a38 | 0.01 | Old |
| 0.327017 | 0.136118 | 0.146 | 0.129 | 1 | 3 | Pmpcb | 0.017 | Old |
| 0.327053 | 0.227114 | 0.055 | 0.044 | 1 | 3 | Kat14 | 0.011 | Old |
| 0.327516 | 0.023438 | 0.149 | 0.13 | 1 | 3 | A530064D( | 0.019 | Old |
| 0.327831 | 0.296209 | 0.087 | 0.073 | 1 | 3 | Naa40 | 0.014 | Old |
| 0.328007 | 0.196082 | 0.123 | 0.107 | 1 | 3 | Abraxas2 | 0.016 | Old |
| 0.328332 | 0.131505 | 0.228 | 0.206 | 1 | 3 | Rbbp7 | 0.022 | Old |
| 0.328447 | 0.276424 | 0.091 | 0.077 | 1 | 3 | Ubr1 | 0.014 | Old |
| 0.329341 | 0.678044 | 0.027 | 0.02 | 1 | 3 | Pdk4 | 0.007 | Old |
| 0.329341 | 0.07533 | 0.315 | 0.283 | 1 | 3 | Dip2b | 0.032 | Old |
| 0.329399 | 0.26989 | 0.74 | 0.726 | 1 | 3 | Klf2 | 0.014 | Old |
| 0.330118 | 0.352031 | 0.011 | 0.006 | 1 | 3 | Zfp455 | 0.005 | Old |
| 0.330206 | 0.077357 | 0.176 | 0.155 | 1 | 3 | Dnase1l1 | 0.021 | Old |
| 0.330366 | 0.117659 | 0.146 | 0.128 | 1 | 3 | Ppt2 | 0.018 | Old |
| 0.330614 | 0.658334 | 0.035 | 0.027 | 1 | 3 | Mms22l | 0.008 | Old |
| 0.330671 | 0.107298 | 0.369 | 0.34 | 1 | 3 | Hsd17b11 | 0.029 | Old |
| 0.330698 | 0.229039 | 0.016 | 0.01 | 1 | 3 | Slc17a9 | 0.006 | Old |
| 0.330769 | 0.491581 | 0.042 | 0.033 | 1 | 3 | Brms1l | 0.009 | Old |
| 0.330922 | 0.04756 | 0.554 | 0.51 | 1 | 3 | Selenow | 0.044 | Old |

| | | | | | | | | |
|---|---|---|---|---|---|---|---|---|
| 0.330977 | 0.065998 | 0.086 | 0.072 | 1 | 3 | Slc39a3 | 0.014 | Old |
| 0.330993 | 0.163102 | 0.417 | 0.393 | 1 | 3 | Inpp5d | 0.024 | Old |
| 0.33103 | 0.361695 | 0.088 | 0.076 | 1 | 3 | Tm2d3 | 0.012 | Old |
| 0.331059 | 0.120511 | 0.033 | 0.043 | 1 | 3 | Rora | -0.01 | Old |
| 0.331234 | 0.552615 | 0.016 | 0.01 | 1 | 3 | Tubb2b | 0.006 | Old |
| 0.331329 | 0.125507 | 0.026 | 0.019 | 1 | 3 | Slc25a53 | 0.007 | Old |
| 0.331757 | 0.580963 | 0.011 | 0.006 | 1 | 3 | Cyp2s1 | 0.005 | Old |
| 0.331859 | 0.099095 | 0.079 | 0.066 | 1 | 3 | Timm29 | 0.013 | Old |
| 0.33194 | 0.135064 | 0.142 | 0.124 | 1 | 3 | Taf5l | 0.018 | Old |
| 0.332043 | 0.076489 | 0.114 | 0.097 | 1 | 3 | Fbxl17 | 0.017 | Old |
| 0.332071 | 0.261957 | 0.099 | 0.086 | 1 | 3 | Dohh | 0.013 | Old |
| 0.332188 | 0.037035 | 0.369 | 0.337 | 1 | 3 | Aldh3b1 | 0.032 | Old |
| 0.332377 | 0.07707 | 0.245 | 0.221 | 1 | 3 | Mien1 | 0.024 | Old |
| 0.332414 | 0.447466 | 0.011 | 0.006 | 1 | 3 | Cd46 | 0.005 | Old |
| 0.332414 | 0.331291 | 0.011 | 0.006 | 1 | 3 | Cpeb3 | 0.005 | Old |
| 0.33247 | 0.143989 | 0.458 | 0.424 | 1 | 3 | Mgl2 | 0.034 | Old |
| 0.332622 | 0.022315 | 0.532 | 0.499 | 1 | 3 | 2810474O1 | 0.033 | Old |
| 0.332806 | 0.601707 | 0.027 | 0.02 | 1 | 3 | Ropn1l | 0.007 | Old |
| 0.333402 | 0.514459 | 0.011 | 0.006 | 1 | 3 | 5430403G1 | 0.005 | Old |
| 0.333402 | 0.212667 | 0.011 | 0.006 | 1 | 3 | Cdiptos | 0.005 | Old |
| 0.333766 | 0.445545 | 0.078 | 0.066 | 1 | 3 | Iffo2 | 0.012 | Old |
| 0.333802 | 0.084698 | 0.28 | 0.251 | 1 | 3 | Dctn3 | 0.029 | Old |
| 0.333995 | 0.024036 | 0.379 | 0.342 | 1 | 3 | Lsm12 | 0.037 | Old |
| 0.334061 | 0.317067 | 0.011 | 0.006 | 1 | 3 | Il15ra | 0.005 | Old |
| 0.334236 | 0.009119 | 0.088 | 0.074 | 1 | 3 | Ppp5c | 0.014 | Old |
| 0.334409 | 0.161467 | 0.145 | 0.128 | 1 | 3 | Rai1 | 0.017 | Old |
| 0.33462 | 0.240947 | 0.063 | 0.052 | 1 | 3 | Spred2 | 0.011 | Old |
| 0.334698 | 0.206518 | 0.041 | 0.032 | 1 | 3 | Abhd18 | 0.009 | Old |
| 0.334721 | 0.303814 | 0.011 | 0.006 | 1 | 3 | Rtkn | 0.005 | Old |
| 0.334721 | 0.300856 | 0.011 | 0.006 | 1 | 3 | Zbtb32 | 0.005 | Old |
| 0.334847 | 0.03412 | 0.568 | 0.518 | 1 | 3 | Coro1c | 0.05 | Old |
| 0.334916 | 0.394735 | 0.079 | 0.067 | 1 | 3 | Trim5 | 0.012 | Old |
| 0.33519 | 0.037956 | 0.318 | 0.283 | 1 | 3 | Slc4a7 | 0.035 | Old |
| 0.335382 | 0.021428 | 0.011 | 0.006 | 1 | 3 | Dock4 | 0.005 | Old |
| 0.335723 | 0.12857 | 0.093 | 0.078 | 1 | 3 | Creb3l2 | 0.015 | Old |
| 0.335803 | 0.027467 | 0.082 | 0.069 | 1 | 3 | Pgm2l1 | 0.013 | Old |
| 0.335876 | 0.024977 | 0.729 | 0.684 | 1 | 3 | Hif1a | 0.045 | Old |
| 0.335913 | 0.043626 | 0.442 | 0.407 | 1 | 3 | Ensa | 0.035 | Old |
| 0.336023 | 0.011971 | 0.424 | 0.386 | 1 | 3 | Lamtor1 | 0.038 | Old |
| 0.336037 | 0.04656 | 0.13 | 0.114 | 1 | 3 | Tmem65 | 0.016 | Old |
| 0.336088 | 0.192143 | 0.016 | 0.01 | 1 | 3 | Birc5 | 0.006 | Old |
| 0.336194 | 0.194831 | 0.027 | 0.02 | 1 | 3 | Arhgap27 | 0.007 | Old |
| 0.336449 | 0.009841 | 0.265 | 0.236 | 1 | 3 | Tcerg1 | 0.029 | Old |
| 0.33678 | 0.22051 | 0.063 | 0.052 | 1 | 3 | Zmat3 | 0.011 | Old |
| 0.33688 | 0.261492 | 0.142 | 0.126 | 1 | 3 | Zc3h4 | 0.016 | Old |

| | | | | | | | | |
|---|---|---|---|---|---|---|---|---|
| 0.3369 | 0.124789 | 0.033 | 0.025 | 1 | 3 | Zfp667 | 0.008 | Old |
| 0.337314 | 0.002224 | 0.211 | 0.189 | 1 | 3 | Zbtb11 | 0.022 | Old |
| 0.337654 | 0.069763 | 0.613 | 0.567 | 1 | 3 | Uqcr10 | 0.046 | Old |
| 0.337656 | 0.106845 | 0.125 | 0.109 | 1 | 3 | Itsn1 | 0.016 | Old |
| 0.338669 | 0.341674 | 0.122 | 0.106 | 1 | 3 | Trim30d | 0.016 | Old |
| 0.338781 | 0.225738 | 0.081 | 0.068 | 1 | 3 | Mrpl22 | 0.013 | Old |
| 0.338817 | 0.062677 | 0.117 | 0.101 | 1 | 3 | Thap11 | 0.016 | Old |
| 0.33919 | 0.037615 | 0.182 | 0.16 | 1 | 3 | Fam114a2 | 0.022 | Old |
| 0.339413 | 0.240294 | 0.207 | 0.188 | 1 | 3 | Strn4 | 0.019 | Old |
| 0.339431 | 0.182775 | 0.116 | 0.101 | 1 | 3 | Cpox | 0.015 | Old |
| 0.339445 | 0.020979 | 0.363 | 0.335 | 1 | 3 | Gnl3 | 0.028 | Old |
| 0.33964 | 0.024757 | 0.039 | 0.03 | 1 | 3 | Gm26853 | 0.009 | Old |
| 0.339858 | 0.035469 | 0.374 | 0.34 | 1 | 3 | AB124611 | 0.034 | Old |
| 0.340209 | 0.217431 | 0.027 | 0.02 | 1 | 3 | Hist2h3c2 | 0.007 | Old |
| 0.340221 | 0.181872 | 0.108 | 0.093 | 1 | 3 | Ctu2 | 0.015 | Old |
| 0.340928 | 0.271211 | 0.155 | 0.14 | 1 | 3 | Slc25a12 | 0.015 | Old |
| 0.340997 | 0.194334 | 0.288 | 0.26 | 1 | 3 | Kmt2a | 0.028 | Old |
| 0.341104 | 0.202087 | 0.115 | 0.1 | 1 | 3 | Arhgef7 | 0.015 | Old |
| 0.341233 | 0.21195 | 0.123 | 0.107 | 1 | 3 | Mnat1 | 0.016 | Old |
| 0.341255 | 0.019887 | 0.252 | 0.226 | 1 | 3 | Lman1 | 0.026 | Old |
| 0.341419 | 0.195003 | 0.027 | 0.02 | 1 | 3 | Nlrp1a | 0.007 | Old |
| 0.341795 | 0.02411 | 0.756 | 0.727 | 1 | 3 | Ppp3ca | 0.029 | Old |
| 0.341993 | 0.172504 | 0.033 | 0.025 | 1 | 3 | Ulbp1 | 0.008 | Old |
| 0.342025 | 0.223266 | 0.027 | 0.02 | 1 | 3 | Card11 | 0.007 | Old |
| 0.342499 | 0.160133 | 0.142 | 0.125 | 1 | 3 | Ppp2r3d | 0.017 | Old |
| 0.342511 | 0.268645 | 0.096 | 0.082 | 1 | 3 | Cfap20 | 0.014 | Old |
| 0.342624 | 0.091646 | 0.498 | 0.475 | 1 | 3 | Pilra | 0.023 | Old |
| 0.342683 | 0.047754 | 0.17 | 0.149 | 1 | 3 | Ppm1m | 0.021 | Old |
| 0.342828 | 0.059359 | 0.297 | 0.268 | 1 | 3 | Hmgb2 | 0.029 | Old |
| 0.343247 | 0.576318 | 0.022 | 0.015 | 1 | 3 | Lysmd1 | 0.007 | Old |
| 0.343297 | 0.019749 | 0.135 | 0.119 | 1 | 3 | Map3k5 | 0.016 | Old |
| 0.343681 | 0.125991 | 0.388 | 0.356 | 1 | 3 | Snrpd1 | 0.032 | Old |
| 0.343689 | 0.046209 | 0.211 | 0.191 | 1 | 3 | Pigs | 0.02 | Old |
| 0.343693 | 0.044479 | 0.022 | 0.029 | 1 | 3 | Smad1 | -0.007 | Old |
| 0.344041 | 0.443374 | 0.061 | 0.051 | 1 | 3 | Metrn | 0.01 | Old |
| 0.344472 | 0.098225 | 0.339 | 0.311 | 1 | 3 | Ahr | 0.028 | Old |
| 0.344628 | 0.219898 | 0.123 | 0.109 | 1 | 3 | Pin1 | 0.014 | Old |
| 0.344636 | 0.132961 | 0.124 | 0.109 | 1 | 3 | Xist | 0.015 | Old |
| 0.345191 | 0.223289 | 0.035 | 0.027 | 1 | 3 | Tmem62 | 0.008 | Old |
| 0.345351 | 0.122225 | 0.088 | 0.074 | 1 | 3 | Micall1 | 0.014 | Old |
| 0.34558 | 0.386008 | 0.066 | 0.054 | 1 | 3 | Smim8 | 0.012 | Old |
| 0.345627 | 0.043112 | 0.26 | 0.234 | 1 | 3 | Dazap1 | 0.026 | Old |
| 0.345788 | 0.600535 | 0.022 | 0.015 | 1 | 3 | Klrb1b | 0.007 | Old |
| 0.345874 | 0.34766 | 0.029 | 0.021 | 1 | 3 | Odf2l | 0.008 | Old |
| 0.346162 | 0.105781 | 0.049 | 0.039 | 1 | 3 | Acot13 | 0.01 | Old |

| | | | | | | | | |
|---|---|---|---|---|---|---|---|---|
| 0.346189 | 0.054899 | 0.136 | 0.12 | 1 | 3 | Ccdc6 | 0.016 | Old |
| 0.346432 | 0.571188 | 0.012 | 0.018 | 1 | 3 | Pigl | -0.006 | Old |
| 0.34649 | 0.027013 | 0.216 | 0.194 | 1 | 3 | Lpin1 | 0.022 | Old |
| 0.346668 | 0.018781 | 0.547 | 0.5 | 1 | 3 | Uvrag | 0.047 | Old |
| 0.346892 | 0.229531 | 0.067 | 0.056 | 1 | 3 | Pfkfb3 | 0.011 | Old |
| 0.348132 | 0.023827 | 0.471 | 0.428 | 1 | 3 | Ost4 | 0.043 | Old |
| 0.348137 | 0.027578 | 0.159 | 0.14 | 1 | 3 | Dirc2 | 0.019 | Old |
| 0.348901 | 0.088049 | 0.131 | 0.114 | 1 | 3 | Ap1m1 | 0.017 | Old |
| 0.34901 | 0.000246 | 0.153 | 0.135 | 1 | 3 | Il11ra1 | 0.018 | Old |
| 0.349969 | 0.061353 | 0.027 | 0.02 | 1 | 3 | Zfp747 | 0.007 | Old |
| 0.350049 | 0.222817 | 0.096 | 0.082 | 1 | 3 | Cfh | 0.014 | Old |
| 0.350204 | 0.030023 | 0.022 | 0.015 | 1 | 3 | Pex26 | 0.007 | Old |
| 0.350988 | 0.17572 | 0.115 | 0.1 | 1 | 3 | Ssbp1 | 0.015 | Old |
| 0.351016 | 0.264941 | 0.081 | 0.069 | 1 | 3 | Txlng | 0.012 | Old |
| 0.351511 | 0.03853 | 0.851 | 0.803 | 1 | 3 | Ccnl1 | 0.048 | Old |
| 0.352545 | 0.071827 | 0.539 | 0.495 | 1 | 3 | Srsf9 | 0.044 | Old |
| 0.352812 | 0.307742 | 0.07 | 0.059 | 1 | 3 | Gm38399 | 0.011 | Old |
| 0.352938 | 0.185881 | 0.11 | 0.095 | 1 | 3 | Rnf2 | 0.015 | Old |
| 0.3533 | 0.022052 | 0.203 | 0.183 | 1 | 3 | Wdr82 | 0.02 | Old |
| 0.353559 | 0.141344 | 0.185 | 0.165 | 1 | 3 | Med8 | 0.02 | Old |
| 0.353622 | 0.473718 | 0.006 | 0.01 | 1 | 3 | Gm17745 | -0.004 | Old |
| 0.35383 | 0.092492 | 0.616 | 0.586 | 1 | 3 | Atp5j | 0.03 | Old |
| 0.354024 | 0.091011 | 0.247 | 0.223 | 1 | 3 | Shoc2 | 0.024 | Old |
| 0.354051 | 0.10174 | 0.644 | 0.601 | 1 | 3 | Bzw1 | 0.043 | Old |
| 0.354419 | 0.406962 | 0.043 | 0.034 | 1 | 3 | Ankrd52 | 0.009 | Old |
| 0.354453 | 0.035292 | 0.179 | 0.159 | 1 | 3 | Abi2 | 0.02 | Old |
| 0.355116 | 0.202263 | 0.07 | 0.059 | 1 | 3 | Trmt2a | 0.011 | Old |
| 0.355497 | 0.039611 | 0.029 | 0.021 | 1 | 3 | Coro2b | 0.008 | Old |
| 0.355623 | 0.075976 | 0.282 | 0.255 | 1 | 3 | Chchd1 | 0.027 | Old |
| 0.355703 | 0.525859 | 0.017 | 0.011 | 1 | 3 | Dhodh | 0.006 | Old |
| 0.355837 | 0.050591 | 0.579 | 0.533 | 1 | 3 | Birc3 | 0.046 | Old |
| 0.355858 | 0.065834 | 0.099 | 0.085 | 1 | 3 | Nsun6 | 0.014 | Old |
| 0.355977 | 0.028911 | 0.969 | 0.929 | 1 | 3 | Dusp1 | 0.04 | Old |
| 0.356086 | 0.075198 | 0.094 | 0.08 | 1 | 3 | Tirap | 0.014 | Old |
| 0.356308 | 0.017708 | 0.059 | 0.048 | 1 | 3 | Hook2 | 0.011 | Old |
| 0.356665 | 0.078505 | 0.221 | 0.198 | 1 | 3 | Cldnd1 | 0.023 | Old |
| 0.356779 | 0.87866 | 0.017 | 0.011 | 1 | 3 | Gm26759 | 0.006 | Old |
| 0.356848 | 0.518036 | 0.055 | 0.045 | 1 | 3 | Isg15 | 0.01 | Old |
| 0.357226 | 0.569942 | 0.036 | 0.028 | 1 | 3 | Mtrf1l | 0.008 | Old |
| 0.357271 | 0.241945 | 0.079 | 0.067 | 1 | 3 | Ipo9 | 0.012 | Old |
| 0.357768 | 0.789736 | 0.036 | 0.028 | 1 | 3 | Qrsl1 | 0.008 | Old |
| 0.358011 | 0.464406 | 0.045 | 0.037 | 1 | 3 | Cdt1 | 0.008 | Old |
| 0.358074 | 0.070659 | 0.168 | 0.15 | 1 | 3 | Esrra | 0.018 | Old |
| 0.358126 | 0.47052 | 0.017 | 0.011 | 1 | 3 | Cacnb1 | 0.006 | Old |
| 0.358196 | 0.132255 | 0.106 | 0.092 | 1 | 3 | Med11 | 0.014 | Old |

| | | | | | | | | |
|---|---|---|---|---|---|---|---|---|
| 0.358493 | 0.432819 | 0.023 | 0.016 | 1 | 3 | Nudt15 | 0.007 | Old |
| 0.359336 | 0.121741 | 0.057 | 0.047 | 1 | 3 | Ints2 | 0.01 | Old |
| 0.359354 | 0.245011 | 0.165 | 0.148 | 1 | 3 | Arpc5l | 0.017 | Old |
| 0.359748 | 0.081435 | 0.104 | 0.09 | 1 | 3 | Flnb | 0.014 | Old |
| 0.359909 | 0.001176 | 0.386 | 0.35 | 1 | 3 | Dnajc8 | 0.036 | Old |
| 0.360349 | 0.136389 | 0.061 | 0.051 | 1 | 3 | Alg1 | 0.01 | Old |
| 0.360488 | 0.128543 | 0.061 | 0.051 | 1 | 3 | 1700021F0 | 0.01 | Old |
| 0.360559 | 0.38424 | 0.017 | 0.011 | 1 | 3 | Dnm1 | 0.006 | Old |
| 0.360559 | 0.35146 | 0.017 | 0.011 | 1 | 3 | 2610021A0 | 0.006 | Old |
| 0.36083 | 0.051293 | 0.017 | 0.011 | 1 | 3 | Tmem268 | 0.006 | Old |
| 0.360845 | 0.043881 | 0.05 | 0.04 | 1 | 3 | Scn2b | 0.01 | Old |
| 0.361146 | 0.102967 | 0.099 | 0.086 | 1 | 3 | Lmf1 | 0.013 | Old |
| 0.361321 | 0.028856 | 0.147 | 0.13 | 1 | 3 | Dopey1 | 0.017 | Old |
| 0.3615 | 0.031705 | 0.056 | 0.045 | 1 | 3 | Irf2bp1 | 0.011 | Old |
| 0.361643 | 0.080216 | 0.017 | 0.011 | 1 | 3 | Pou5f1 | 0.006 | Old |
| 0.361715 | 0.297141 | 0.023 | 0.016 | 1 | 3 | Gm20300 | 0.007 | Old |
| 0.361928 | 0.107886 | 0.043 | 0.034 | 1 | 3 | Fcho1 | 0.009 | Old |
| 0.362204 | 0.007913 | 0.084 | 0.071 | 1 | 3 | Mrps10 | 0.013 | Old |
| 0.362639 | 0.261304 | 0.023 | 0.016 | 1 | 3 | Txnrd3 | 0.007 | Old |
| 0.363198 | 0.054915 | 0.297 | 0.271 | 1 | 3 | Pin4 | 0.026 | Old |
| 0.363274 | 0.206798 | 0.017 | 0.011 | 1 | 3 | Tbc1d19 | 0.006 | Old |
| 0.36363 | 0.26815 | 0.09 | 0.077 | 1 | 3 | Prorsd1 | 0.013 | Old |
| 0.36365 | 0.386977 | 0.037 | 0.029 | 1 | 3 | Cdkal1 | 0.008 | Old |
| 0.36365 | 0.205101 | 0.037 | 0.029 | 1 | 3 | Hdac10 | 0.008 | Old |
| 0.363964 | 0.055734 | 0.535 | 0.492 | 1 | 3 | Tax1bp1 | 0.043 | Old |
| 0.364043 | 0.016828 | 0.241 | 0.216 | 1 | 3 | Ndufs3 | 0.025 | Old |
| 0.364091 | 0.262497 | 0.017 | 0.011 | 1 | 3 | Fam84b | 0.006 | Old |
| 0.36429 | 1.188452 | 0.012 | 0.008 | 1 | 3 | Gm26510 | 0.004 | Old |
| 0.364569 | 0.0259 | 0.135 | 0.119 | 1 | 3 | C1galt1c1 | 0.016 | Old |
| 0.364616 | 0.412472 | 0.045 | 0.037 | 1 | 3 | Kdm1b | 0.008 | Old |
| 0.364636 | 0.050445 | 0.017 | 0.011 | 1 | 3 | Erc1 | 0.006 | Old |
| 0.364816 | 0.066022 | 0.762 | 0.711 | 1 | 3 | Akr1a1 | 0.051 | Old |
| 0.364954 | 0.193947 | 0.023 | 0.016 | 1 | 3 | Txnl4b | 0.007 | Old |
| 0.36514 | 0.100324 | 0.09 | 0.077 | 1 | 3 | Cars2 | 0.013 | Old |
| 0.365805 | 0.000791 | 0.27 | 0.242 | 1 | 3 | Wbp1l | 0.028 | Old |
| 0.366031 | 0.416276 | 0.03 | 0.023 | 1 | 3 | Ppcs | 0.007 | Old |
| 0.366165 | 0.049145 | 0.203 | 0.184 | 1 | 3 | Ppm1b | 0.019 | Old |
| 0.366227 | 0.13722 | 0.026 | 0.034 | 1 | 3 | Ift57 | -0.008 | Old |
| 0.366251 | 0.736397 | 0.012 | 0.008 | 1 | 3 | Kif22 | 0.004 | Old |
| 0.366581 | 0.034202 | 0.023 | 0.016 | 1 | 3 | Fadd | 0.007 | Old |
| 0.366724 | 0.147156 | 0.044 | 0.035 | 1 | 3 | Umps | 0.009 | Old |
| 0.366892 | 0.034651 | 0.178 | 0.158 | 1 | 3 | Cass4 | 0.02 | Old |
| 0.367004 | 0.145143 | 0.09 | 0.077 | 1 | 3 | Smarcd1 | 0.013 | Old |
| 0.367047 | 0.359497 | 0.023 | 0.016 | 1 | 3 | Snapc4 | 0.007 | Old |
| 0.367171 | 0.012691 | 0.205 | 0.183 | 1 | 3 | Ei24 | 0.022 | Old |

| | | | | | | | | |
|---|---|---|---|---|---|---|---|---|
| 0.367281 | 0.056319 | 0.078 | 0.066 | 1 | 3 | Exosc2 | 0.012 | Old |
| 0.367306 | 0.111558 | 0.299 | 0.274 | 1 | 3 | Rgs18 | 0.025 | Old |
| 0.367385 | 0.134458 | 0.154 | 0.138 | 1 | 3 | Zfp592 | 0.016 | Old |
| 0.367519 | 0.00678 | 0.173 | 0.153 | 1 | 3 | Jmjd6 | 0.02 | Old |
| 0.367561 | 0.725252 | 0.012 | 0.008 | 1 | 3 | Sirt6 | 0.004 | Old |
| 0.367779 | 0.03917 | 0.945 | 0.938 | 1 | 3 | Vim | 0.007 | Old |
| 0.368218 | 0.572929 | 0.012 | 0.008 | 1 | 3 | Pole2 | 0.004 | Old |
| 0.368322 | 0.203538 | 0.051 | 0.042 | 1 | 3 | AI987944 | 0.009 | Old |
| 0.36835 | 0.047927 | 0.519 | 0.482 | 1 | 3 | Jmjd1c | 0.037 | Old |
| 0.368533 | 0.070429 | 0.271 | 0.245 | 1 | 3 | Ift20 | 0.026 | Old |
| 0.368679 | 0.111953 | 0.023 | 0.016 | 1 | 3 | AC130815. | 0.007 | Old |
| 0.368913 | 0.044306 | 0.023 | 0.016 | 1 | 3 | Fkbp7 | 0.007 | Old |
| 0.369204 | 0.496953 | 0.012 | 0.008 | 1 | 3 | Nphp3 | 0.004 | Old |
| 0.369388 | 0.0505 | 0.136 | 0.121 | 1 | 3 | Ssh1 | 0.015 | Old |
| 0.369474 | 0.378111 | 0.038 | 0.03 | 1 | 3 | Sprtn | 0.008 | Old |
| 0.369603 | 0.016447 | 0.108 | 0.093 | 1 | 3 | Cog3 | 0.015 | Old |
| 0.370032 | 0.422239 | 0.031 | 0.024 | 1 | 3 | Rpap2 | 0.007 | Old |
| 0.370191 | 0.702608 | 0.012 | 0.008 | 1 | 3 | Gdpgp1 | 0.004 | Old |
| 0.370191 | 0.529955 | 0.012 | 0.008 | 1 | 3 | Hist1h2be | 0.004 | Old |
| 0.370194 | 0.030495 | 0.191 | 0.17 | 1 | 3 | Trim56 | 0.021 | Old |
| 0.370419 | 0.173382 | 0.162 | 0.146 | 1 | 3 | Tmem222 | 0.016 | Old |
| 0.37078 | 0.021505 | 0.044 | 0.035 | 1 | 3 | Ddx51 | 0.009 | Old |
| 0.37083 | 0.025885 | 0.452 | 0.419 | 1 | 3 | Dmac1 | 0.033 | Old |
| 0.370851 | 0.832433 | 0.012 | 0.008 | 1 | 3 | Papss2 | 0.004 | Old |
| 0.370851 | 0.398845 | 0.012 | 0.008 | 1 | 3 | Adamts1 | 0.004 | Old |
| 0.371349 | 0.023017 | 0.557 | 0.506 | 1 | 3 | Timm13 | 0.051 | Old |
| 0.371511 | 0.576612 | 0.012 | 0.008 | 1 | 3 | Bcl2a1c | 0.004 | Old |
| 0.371841 | 0.303381 | 0.012 | 0.008 | 1 | 3 | Trip6 | 0.004 | Old |
| 0.372053 | 0.008213 | 0.036 | 0.028 | 1 | 3 | Tnfrsf18 | 0.008 | Old |
| 0.372145 | 0.306876 | 0.037 | 0.029 | 1 | 3 | Zfp868 | 0.008 | Old |
| 0.372273 | 0.107717 | 0.084 | 0.072 | 1 | 3 | Hmgn1 | 0.012 | Old |
| 0.372758 | 0.043335 | 0.959 | 0.951 | 1 | 3 | Oaz1 | 0.008 | Old |
| 0.372826 | 0.090915 | 0.705 | 0.673 | 1 | 3 | Hnrnph1 | 0.032 | Old |
| 0.37341 | 0.208313 | 0.075 | 0.064 | 1 | 3 | Gnpnat1 | 0.011 | Old |
| 0.37341 | 0.087962 | 0.051 | 0.042 | 1 | 3 | Trmt11 | 0.009 | Old |
| 0.373431 | 0.023022 | 0.045 | 0.037 | 1 | 3 | Cdpf1 | 0.008 | Old |
| 0.373642 | 0.519076 | 0.024 | 0.018 | 1 | 3 | Lrrk2 | 0.006 | Old |
| 0.373817 | 0.033998 | 0.108 | 0.093 | 1 | 3 | Mrs2 | 0.015 | Old |
| 0.374158 | 0.219761 | 0.012 | 0.008 | 1 | 3 | Mfsd13a | 0.004 | Old |
| 0.374193 | 0.566655 | 0.056 | 0.047 | 1 | 3 | Dpp4 | 0.009 | Old |
| 0.374194 | 0.093237 | 0.086 | 0.073 | 1 | 3 | Med4 | 0.013 | Old |
| 0.375306 | 0.149315 | 0.205 | 0.187 | 1 | 3 | Cetn3 | 0.018 | Old |
| 0.376036 | 0.053677 | 0.822 | 0.808 | 1 | 3 | Eef1g | 0.014 | Old |
| 0.376115 | 0.035115 | 0.892 | 0.88 | 1 | 3 | Pid1 | 0.012 | Old |
| 0.376262 | 0.244876 | 0.062 | 0.052 | 1 | 3 | Med7 | 0.01 | Old |

| | | | | | | | | |
|---|---|---|---|---|---|---|---|---|
| 0.376486 | 0.33653 | 0.03 | 0.023 | 1 | 3 | 9930104L0 | 0.007 | Old |
| 0.376539 | 0.152908 | 0.044 | 0.035 | 1 | 3 | Prkar2b | 0.009 | Old |
| 0.376905 | 0.087964 | 0.252 | 0.229 | 1 | 3 | Kcnab2 | 0.023 | Old |
| 0.377101 | 0.023977 | 0.03 | 0.023 | 1 | 3 | Chrna5 | 0.007 | Old |
| 0.377432 | 0.045237 | 0.068 | 0.057 | 1 | 3 | Stard5 | 0.011 | Old |
| 0.377885 | 0.408665 | 0.024 | 0.018 | 1 | 3 | Zfp275 | 0.006 | Old |
| 0.378126 | 0.019045 | 0.14 | 0.122 | 1 | 3 | Trim12a | 0.018 | Old |
| 0.378127 | 0.086794 | 0.03 | 0.023 | 1 | 3 | Nmral1 | 0.007 | Old |
| 0.378565 | 0.027085 | 0.473 | 0.434 | 1 | 3 | Ppp1r15a | 0.039 | Old |
| 0.378819 | 0.305483 | 0.012 | 0.008 | 1 | 3 | Tcea2 | 0.004 | Old |
| 0.378891 | 0.144879 | 0.044 | 0.035 | 1 | 3 | D930030I0 | 0.009 | Old |
| 0.379077 | 0.068165 | 0.364 | 0.336 | 1 | 3 | Kif1b | 0.028 | Old |
| 0.379096 | 0.00175 | 0.054 | 0.064 | 1 | 3 | Vps39 | -0.01 | Old |
| 0.379614 | 0.069599 | 0.024 | 0.018 | 1 | 3 | Tmem129 | 0.006 | Old |
| 0.379845 | 0.188549 | 0.024 | 0.018 | 1 | 3 | Tspan4 | 0.006 | Old |
| 0.379855 | 0.000828 | 0.532 | 0.485 | 1 | 3 | Dnajb6 | 0.047 | Old |
| 0.380596 | 0.248579 | 0.03 | 0.023 | 1 | 3 | Tmem202 | 0.007 | Old |
| 0.381057 | 0.02991 | 0.223 | 0.202 | 1 | 3 | Wbp4 | 0.021 | Old |
| 0.381071 | 0.163117 | 0.207 | 0.187 | 1 | 3 | Ptgs2 | 0.02 | Old |
| 0.381152 | 0.484161 | 0.018 | 0.013 | 1 | 3 | Zfp658 | 0.005 | Old |
| 0.381468 | 0.062586 | 0.098 | 0.085 | 1 | 3 | Noc4l | 0.013 | Old |
| 0.381537 | 0.229093 | 0.092 | 0.08 | 1 | 3 | Caap1 | 0.012 | Old |
| 0.381727 | 0.01488 | 0.381 | 0.355 | 1 | 3 | Psmc6 | 0.026 | Old |
| 0.381748 | 0.084236 | 0.066 | 0.077 | 1 | 3 | Ampd2 | -0.011 | Old |
| 0.381839 | 0.20434 | 0.031 | 0.024 | 1 | 3 | Izumo4 | 0.007 | Old |
| 0.382156 | 0.075546 | 0.205 | 0.184 | 1 | 3 | Mrpl3 | 0.021 | Old |
| 0.382185 | 0.188325 | 0.065 | 0.054 | 1 | 3 | Klc4 | 0.011 | Old |
| 0.382314 | 0.1816 | 0.145 | 0.129 | 1 | 3 | Nup50 | 0.016 | Old |
| 0.382499 | 0.808116 | 0.018 | 0.013 | 1 | 3 | Pld2 | 0.005 | Old |
| 0.38285 | 0.537175 | 0.031 | 0.024 | 1 | 3 | Etaa1 | 0.007 | Old |
| 0.382854 | 0.314721 | 0.024 | 0.018 | 1 | 3 | Gbp8 | 0.006 | Old |
| 0.383309 | 0.492714 | 0.018 | 0.013 | 1 | 3 | Ahcy | 0.005 | Old |
| 0.383759 | 0.21788 | 0.104 | 0.091 | 1 | 3 | Ascc1 | 0.013 | Old |
| 0.384119 | 0.80256 | 0.018 | 0.013 | 1 | 3 | Ift80 | 0.005 | Old |
| 0.384127 | 0.121071 | 0.127 | 0.112 | 1 | 3 | Tcn2 | 0.015 | Old |
| 0.384632 | 0.128322 | 0.155 | 0.139 | 1 | 3 | Arid2 | 0.016 | Old |
| 0.384814 | 0.093632 | 0.069 | 0.058 | 1 | 3 | Exosc1 | 0.011 | Old |
| 0.38491 | 0.025875 | 0.999 | 0.995 | 1 | 3 | H2-Aa | 0.004 | Old |
| 0.386358 | 0.332134 | 0.038 | 0.03 | 1 | 3 | Qtrt2 | 0.008 | Old |
| 0.386416 | 0.17311 | 0.174 | 0.157 | 1 | 3 | Batf | 0.017 | Old |
| 0.387101 | 0.532743 | 0.018 | 0.013 | 1 | 3 | Mast4 | 0.005 | Old |
| 0.388298 | 0.137603 | 0.196 | 0.181 | 1 | 3 | Snrpa1 | 0.015 | Old |
| 0.388797 | 0.061417 | 0.599 | 0.552 | 1 | 3 | Sumo2 | 0.047 | Old |
| 0.389224 | 0.035399 | 0.519 | 0.471 | 1 | 3 | Ncor1 | 0.048 | Old |
| 0.389371 | 0.019328 | 0.092 | 0.08 | 1 | 3 | Ly96 | 0.012 | Old |

| | | | | | | | | |
|---|---|---|---|---|---|---|---|---|
| 0.38955 | 0.177382 | 0.018 | 0.013 | 1 | 3 | Rtn4rl1 | 0.005 | Old |
| 0.389905 | 0.10173 | 0.137 | 0.122 | 1 | 3 | Chd6 | 0.015 | Old |
| 0.390369 | 0.10732 | 0.018 | 0.013 | 1 | 3 | Rccd1 | 0.005 | Old |
| 0.390373 | 0.066421 | 0.158 | 0.139 | 1 | 3 | Lrrc47 | 0.019 | Old |
| 0.390383 | 0.248353 | 0.031 | 0.024 | 1 | 3 | 2310057M | 0.007 | Old |
| 0.391123 | 0.078313 | 0.115 | 0.1 | 1 | 3 | Sbf1 | 0.015 | Old |
| 0.391407 | 0.310762 | 0.031 | 0.024 | 1 | 3 | Armc6 | 0.007 | Old |
| 0.391635 | 0.02618 | 0.431 | 0.395 | 1 | 3 | Laptm4a | 0.036 | Old |
| 0.391736 | 0.147281 | 0.018 | 0.013 | 1 | 3 | Abtb2 | 0.005 | Old |
| 0.391859 | 0.066178 | 0.11 | 0.096 | 1 | 3 | Agl | 0.014 | Old |
| 0.39201 | 0.070451 | 0.018 | 0.013 | 1 | 3 | Mgmt | 0.005 | Old |
| 0.392963 | 0.056737 | 0.092 | 0.08 | 1 | 3 | Rabggta | 0.012 | Old |
| 0.393203 | 0.002752 | 0.08 | 0.068 | 1 | 3 | Fbxw5 | 0.012 | Old |
| 0.393319 | 0.0276 | 0.131 | 0.116 | 1 | 3 | Lage3 | 0.015 | Old |
| 0.394212 | 0.077006 | 0.048 | 0.039 | 1 | 3 | Aptx | 0.009 | Old |
| 0.394878 | 0.018738 | 0.155 | 0.14 | 1 | 3 | Cnot8 | 0.015 | Old |
| 0.3956 | 0.142564 | 0.076 | 0.066 | 1 | 3 | Morc2a | 0.01 | Old |
| 0.395634 | 0.130294 | 0.115 | 0.1 | 1 | 3 | Trpv2 | 0.015 | Old |
| 0.396176 | 0.049159 | 0.527 | 0.481 | 1 | 3 | Ndufa3 | 0.046 | Old |
| 0.396494 | 0.092459 | 0.489 | 0.461 | 1 | 3 | Tomm5 | 0.028 | Old |
| 0.396752 | 0.272987 | 0.039 | 0.032 | 1 | 3 | Srrd | 0.007 | Old |
| 0.396899 | 0.063717 | 0.535 | 0.495 | 1 | 3 | Prdx2 | 0.04 | Old |
| 0.396921 | 1.320829 | 0.013 | 0.009 | 1 | 3 | Limk2 | 0.004 | Old |
| 0.396935 | 0.207731 | 0.039 | 0.032 | 1 | 3 | Meis3 | 0.007 | Old |
| 0.397383 | 0.039164 | 0.143 | 0.128 | 1 | 3 | Ing5 | 0.015 | Old |
| 0.397876 | 0.241572 | 0.114 | 0.1 | 1 | 3 | Vamp7 | 0.014 | Old |
| 0.398767 | 0.06413 | 0.039 | 0.032 | 1 | 3 | Dgcr14 | 0.007 | Old |
| 0.398864 | 0.186798 | 0.088 | 0.077 | 1 | 3 | Ermp1 | 0.011 | Old |
| 0.399289 | 0.081374 | 0.135 | 0.121 | 1 | 3 | Lmbrd2 | 0.014 | Old |
| 0.399502 | 0.126733 | 0.039 | 0.032 | 1 | 3 | Btf3l4 | 0.007 | Old |
| 0.400471 | 0.050002 | 0.087 | 0.076 | 1 | 3 | Tpra1 | 0.011 | Old |
| 0.40067 | 0.104355 | 0.032 | 0.025 | 1 | 3 | Miga2 | 0.007 | Old |
| 0.400944 | 0.101322 | 0.155 | 0.138 | 1 | 3 | Tssc4 | 0.017 | Old |
| 0.401071 | 0.015876 | 0.416 | 0.384 | 1 | 3 | Clptm1l | 0.032 | Old |
| 0.401086 | 0.032963 | 0.297 | 0.271 | 1 | 3 | Slain2 | 0.026 | Old |
| 0.401642 | 0.135193 | 0.192 | 0.176 | 1 | 3 | Mrps9 | 0.016 | Old |
| 0.402085 | 0.04591 | 0.286 | 0.261 | 1 | 3 | Ddx46 | 0.025 | Old |
| 0.402426 | 0.022891 | 0.581 | 0.532 | 1 | 3 | Itm2c | 0.049 | Old |
| 0.402954 | 0.088969 | 0.295 | 0.269 | 1 | 3 | Btk | 0.026 | Old |
| 0.403007 | 0.003169 | 0.237 | 0.212 | 1 | 3 | Nmt1 | 0.025 | Old |
| 0.40315 | 0.067905 | 0.447 | 0.422 | 1 | 3 | Icam1 | 0.025 | Old |
| 0.40325 | 0.280977 | 0.093 | 0.081 | 1 | 3 | Mob2 | 0.012 | Old |
| 0.403254 | 0.14715 | 0.205 | 0.189 | 1 | 3 | Nfkbib | 0.016 | Old |
| 0.403766 | 0.007986 | 0.096 | 0.083 | 1 | 3 | Fem1a | 0.013 | Old |
| 0.403932 | 0.013615 | 0.458 | 0.423 | 1 | 3 | Itsn2 | 0.035 | Old |

| | | | | | | | | |
|---|---|---|---|---|---|---|---|---|
| 0.403937 | 0.281001 | 0.032 | 0.025 | 1 | 3 | Zfp53 | 0.007 | Old |
| 0.404208 | 0.029994 | 0.048 | 0.039 | 1 | 3 | Cbfa2t2 | 0.009 | Old |
| 0.404518 | 0.106464 | 0.075 | 0.063 | 1 | 3 | Phf13 | 0.012 | Old |
| 0.404548 | 0.092 | 0.094 | 0.082 | 1 | 3 | Hat1 | 0.012 | Old |
| 0.40458 | 0.095931 | 0.536 | 0.497 | 1 | 3 | Fis1 | 0.039 | Old |
| 0.40471 | 0.059591 | 0.295 | 0.269 | 1 | 3 | Samm50 | 0.026 | Old |
| 0.404752 | 0.806241 | 0.013 | 0.009 | 1 | 3 | A530013C2 | 0.004 | Old |
| 0.404897 | 0.367831 | 0.05 | 0.042 | 1 | 3 | Bnip1 | 0.008 | Old |
| 0.405082 | 0.091058 | 0.183 | 0.167 | 1 | 3 | Spr | 0.016 | Old |
| 0.405105 | 0.007185 | 0.354 | 0.326 | 1 | 3 | Myo9a | 0.028 | Old |
| 0.405211 | 0.051963 | 0.123 | 0.107 | 1 | 3 | Fbxw4 | 0.016 | Old |
| 0.405409 | 0.526682 | 0.013 | 0.009 | 1 | 3 | Dph2 | 0.004 | Old |
| 0.405576 | 0.128022 | 0.032 | 0.025 | 1 | 3 | Cox15 | 0.007 | Old |
| 0.405663 | 0.166177 | 0.078 | 0.067 | 1 | 3 | Rbmx | 0.011 | Old |
| 0.405664 | 0.063476 | 0.208 | 0.188 | 1 | 3 | Dhx38 | 0.02 | Old |
| 0.405925 | 0.306634 | 0.103 | 0.091 | 1 | 3 | Aak1 | 0.012 | Old |
| 0.406067 | 0.247176 | 0.013 | 0.009 | 1 | 3 | Eno3 | 0.004 | Old |
| 0.406395 | 0.378395 | 0.013 | 0.009 | 1 | 3 | Pla2g6 | 0.004 | Old |
| 0.406947 | 0.041027 | 0.105 | 0.092 | 1 | 3 | Utp6 | 0.013 | Old |
| 0.406963 | 0.411315 | 0.06 | 0.051 | 1 | 3 | Naf1 | 0.009 | Old |
| 0.40773 | 0.008618 | 0.571 | 0.524 | 1 | 3 | Ikbkb | 0.047 | Old |
| 0.408042 | 0.383027 | 0.013 | 0.009 | 1 | 3 | Svip | 0.004 | Old |
| 0.408079 | 0.240501 | 0.117 | 0.104 | 1 | 3 | Golga5 | 0.013 | Old |
| 0.408372 | 0.446197 | 0.013 | 0.009 | 1 | 3 | Fkbp9 | 0.004 | Old |
| 0.408735 | 0.188104 | 0.081 | 0.071 | 1 | 3 | Dhrs4 | 0.01 | Old |
| 0.408919 | 0.140792 | 0.05 | 0.042 | 1 | 3 | Sephs1 | 0.008 | Old |
| 0.40909 | 0.0101 | 0.228 | 0.206 | 1 | 3 | Fkbp15 | 0.022 | Old |
| 0.409164 | 0.228489 | 0.019 | 0.014 | 1 | 3 | Lrrc32 | 0.005 | Old |
| 0.409363 | 0.191674 | 0.013 | 0.009 | 1 | 3 | Porcn | 0.004 | Old |
| 0.409401 | 0.309231 | 0.067 | 0.057 | 1 | 3 | Mrpl46 | 0.01 | Old |
| 0.409422 | 0.26007 | 0.092 | 0.081 | 1 | 3 | Abcf3 | 0.011 | Old |
| 0.409443 | 0.038494 | 0.154 | 0.138 | 1 | 3 | Gsap | 0.016 | Old |
| 0.410344 | 0.243291 | 0.125 | 0.112 | 1 | 3 | Polr2b | 0.013 | Old |
| 0.410478 | 0.493067 | 0.117 | 0.106 | 1 | 3 | Cep120 | 0.011 | Old |
| 0.411331 | 0.487351 | 0.019 | 0.014 | 1 | 3 | Cd99l2 | 0.005 | Old |
| 0.412757 | 0.025489 | 0.265 | 0.24 | 1 | 3 | Xpr1 | 0.025 | Old |
| 0.413233 | 0.156126 | 0.019 | 0.014 | 1 | 3 | Ighmbp2 | 0.005 | Old |
| 0.413285 | 0.061067 | 0.097 | 0.083 | 1 | 3 | Rdh11 | 0.014 | Old |
| 0.413689 | 0.119256 | 0.09 | 0.078 | 1 | 3 | Mroh1 | 0.012 | Old |
| 0.414047 | 0.122893 | 0.026 | 0.02 | 1 | 3 | Cdkn2c | 0.006 | Old |
| 0.414164 | 0.018787 | 0.682 | 0.643 | 1 | 3 | Ppp2r5c | 0.039 | Old |
| 0.414322 | 0.080965 | 0.019 | 0.014 | 1 | 3 | Tmed1 | 0.005 | Old |
| 0.414374 | 0.118658 | 0.082 | 0.071 | 1 | 3 | Gpn3 | 0.011 | Old |
| 0.414547 | 0.107888 | 0.272 | 0.251 | 1 | 3 | Rtcb | 0.021 | Old |
| 0.414739 | 0.329078 | 0.026 | 0.02 | 1 | 3 | Zfp961 | 0.006 | Old |

| | | | | | | | | |
|---|---|---|---|---|---|---|---|---|
| 0.414901 | 0.044015 | 0.75 | 0.713 | 1 | 3 | Ddx6 | 0.037 | Old |
| 0.415353 | 0.016096 | 0.13 | 0.116 | 1 | 3 | Nf1 | 0.014 | Old |
| 0.415477 | 0.241083 | 0.105 | 0.093 | 1 | 3 | Tubgcp4 | 0.012 | Old |
| 0.41597 | 0.076114 | 0.614 | 0.573 | 1 | 3 | Snrpb | 0.041 | Old |
| 0.416041 | 0.009945 | 0.176 | 0.157 | 1 | 3 | Dcun1d1 | 0.019 | Old |
| 0.416185 | 0.235281 | 0.102 | 0.09 | 1 | 3 | Rhbdf2 | 0.012 | Old |
| 0.416374 | 0.14188 | 0.202 | 0.187 | 1 | 3 | Mkl2 | 0.015 | Old |
| 0.417115 | 0.025858 | 0.127 | 0.112 | 1 | 3 | Gpr160 | 0.015 | Old |
| 0.417286 | 0.072733 | 0.258 | 0.236 | 1 | 3 | Zmynd11 | 0.022 | Old |
| 0.417592 | 0.179867 | 0.033 | 0.027 | 1 | 3 | Dhx32 | 0.006 | Old |
| 0.417628 | 0.20915 | 0.098 | 0.087 | 1 | 3 | Cnot9 | 0.011 | Old |
| 0.417713 | 0.103151 | 0.157 | 0.14 | 1 | 3 | Taf13 | 0.017 | Old |
| 0.417727 | 0.136552 | 0.174 | 0.159 | 1 | 3 | Rnf220 | 0.015 | Old |
| 0.417841 | 0.081189 | 0.292 | 0.27 | 1 | 3 | Slc25a37 | 0.022 | Old |
| 0.418311 | 0.008862 | 0.389 | 0.36 | 1 | 3 | Sec31a | 0.029 | Old |
| 0.418695 | 0.027674 | 0.019 | 0.014 | 1 | 3 | Cdk5r1 | 0.005 | Old |
| 0.41905 | 0.114556 | 0.311 | 0.287 | 1 | 3 | Kdm2b | 0.024 | Old |
| 0.419366 | 0.226551 | 0.026 | 0.02 | 1 | 3 | Lnx2 | 0.006 | Old |
| 0.419373 | 0.304708 | 0.093 | 0.082 | 1 | 3 | Fam193b | 0.011 | Old |
| 0.419789 | 0.083387 | 0.252 | 0.234 | 1 | 3 | Gak | 0.018 | Old |
| 0.41993 | 0.052235 | 0.067 | 0.057 | 1 | 3 | Frat2 | 0.01 | Old |
| 0.42022 | 0.197764 | 0.042 | 0.034 | 1 | 3 | Otud7b | 0.008 | Old |
| 0.420384 | 0.156549 | 0.155 | 0.14 | 1 | 3 | Alas1 | 0.015 | Old |
| 0.420526 | 0.175831 | 0.162 | 0.146 | 1 | 3 | Tpd52l2 | 0.016 | Old |
| 0.42096 | 0.152217 | 0.05 | 0.042 | 1 | 3 | Nudt18 | 0.008 | Old |
| 0.420993 | 0.276232 | 0.026 | 0.02 | 1 | 3 | Mfsd6l | 0.006 | Old |
| 0.421039 | 0.324125 | 0.051 | 0.043 | 1 | 3 | Mad2l1bp | 0.008 | Old |
| 0.422164 | 0.549229 | 0.036 | 0.029 | 1 | 3 | Ints4 | 0.007 | Old |
| 0.423428 | 0.23738 | 0.136 | 0.124 | 1 | 3 | Rwdd4a | 0.012 | Old |
| 0.423662 | 0.367773 | 0.074 | 0.064 | 1 | 3 | Amz1 | 0.01 | Old |
| 0.423939 | 0.077975 | 0.134 | 0.12 | 1 | 3 | Cenpc1 | 0.014 | Old |
| 0.424048 | 0.685527 | 0.02 | 0.015 | 1 | 3 | Nuak1 | 0.005 | Old |
| 0.4243 | 0.005621 | 0.467 | 0.422 | 1 | 3 | Chn2 | 0.045 | Old |
| 0.424462 | 0.171712 | 0.128 | 0.115 | 1 | 3 | Crnkl1 | 0.013 | Old |
| 0.424848 | 0.525797 | 0.02 | 0.015 | 1 | 3 | Smco4 | 0.005 | Old |
| 0.425295 | 0.055176 | 0.099 | 0.087 | 1 | 3 | Hdac7 | 0.012 | Old |
| 0.425351 | 0.031549 | 0.355 | 0.322 | 1 | 3 | Dctn1 | 0.033 | Old |
| 0.425565 | 0.096755 | 0.097 | 0.085 | 1 | 3 | Ppargc1b | 0.012 | Old |
| 0.425681 | 0.046864 | 0.434 | 0.403 | 1 | 3 | Mapk6 | 0.031 | Old |
| 0.426891 | 0.012893 | 0.398 | 0.366 | 1 | 3 | Lbr | 0.032 | Old |
| 0.427391 | 0.036743 | 0.493 | 0.456 | 1 | 3 | Rtraf | 0.037 | Old |
| 0.428116 | 0.06102 | 0.191 | 0.174 | 1 | 3 | Coro7 | 0.017 | Old |
| 0.428203 | 0.174598 | 0.078 | 0.067 | 1 | 3 | Hhex | 0.011 | Old |
| 0.428378 | 0.017842 | 0.423 | 0.394 | 1 | 3 | Huwe1 | 0.029 | Old |
| 0.428428 | 0.100205 | 0.158 | 0.141 | 1 | 3 | Gemin7 | 0.017 | Old |

| | | | | | | | | |
|---|---|---|---|---|---|---|---|---|
| 0.428571 | 0.006597 | 0.321 | 0.292 | 1 | 3 | Lrrc25 | 0.029 | Old |
| 0.428639 | 0.046144 | 0.108 | 0.095 | 1 | 3 | Alg5 | 0.013 | Old |
| 0.429323 | 0.151095 | 0.106 | 0.095 | 1 | 3 | Zfp524 | 0.011 | Old |
| 0.429336 | 0.474629 | 0.036 | 0.029 | 1 | 3 | Haus7 | 0.007 | Old |
| 0.429434 | 0.050243 | 0.411 | 0.383 | 1 | 3 | Coa5 | 0.028 | Old |
| 0.429435 | 0.091791 | 0.142 | 0.126 | 1 | 3 | Cpsf3 | 0.016 | Old |
| 0.430422 | 0.293929 | 0.114 | 0.101 | 1 | 3 | Nol12 | 0.013 | Old |
| 0.430691 | 0.040777 | 0.45 | 0.422 | 1 | 3 | Vdac1 | 0.028 | Old |
| 0.430864 | 0.021718 | 0.011 | 0.015 | 1 | 3 | Klk13 | -0.004 | Old |
| 0.431015 | 0.402588 | 0.02 | 0.015 | 1 | 3 | Tsen2 | 0.005 | Old |
| 0.431553 | 0.400699 | 0.02 | 0.015 | 1 | 3 | Prepl | 0.005 | Old |
| 0.432712 | 0.654424 | 0.037 | 0.03 | 1 | 3 | Dus2 | 0.007 | Old |
| 0.432884 | 1.098551 | 0.014 | 0.01 | 1 | 3 | Chst1 | 0.004 | Old |
| 0.43299 | 0.270691 | 0.178 | 0.163 | 1 | 3 | Nsmce4a | 0.015 | Old |
| 0.433015 | 0.132971 | 0.035 | 0.028 | 1 | 3 | E2f5 | 0.007 | Old |
| 0.433153 | 0.036784 | 0.535 | 0.499 | 1 | 3 | Hnrnpa1 | 0.036 | Old |
| 0.433173 | 0.029545 | 0.085 | 0.073 | 1 | 3 | Hsf1 | 0.012 | Old |
| 0.433284 | 0.067483 | 0.134 | 0.119 | 1 | 3 | Ube2w | 0.015 | Old |
| 0.433493 | 0.136105 | 0.117 | 0.104 | 1 | 3 | Htatip2 | 0.013 | Old |
| 0.433718 | 0.057921 | 0.027 | 0.021 | 1 | 3 | Mfsd3 | 0.006 | Old |
| 0.433955 | 0.264738 | 0.036 | 0.029 | 1 | 3 | Zc3h7b | 0.007 | Old |
| 0.434863 | 0.084452 | 0.172 | 0.157 | 1 | 3 | Snapin | 0.015 | Old |
| 0.435482 | 0.848601 | 0.014 | 0.01 | 1 | 3 | Ficd | 0.004 | Old |
| 0.435606 | 0.204119 | 0.02 | 0.015 | 1 | 3 | Zfp52 | 0.005 | Old |
| 0.435952 | 0.282324 | 0.097 | 0.085 | 1 | 3 | Pvr | 0.012 | Old |
| 0.436158 | 0.052627 | 0.231 | 0.211 | 1 | 3 | Mrps17 | 0.02 | Old |
| 0.436456 | 0.018306 | 0.233 | 0.212 | 1 | 3 | Psen1 | 0.021 | Old |
| 0.436784 | 0.380187 | 0.014 | 0.01 | 1 | 3 | Zcwpw1 | 0.004 | Old |
| 0.437037 | 0.147462 | 0.084 | 0.073 | 1 | 3 | Entpd7 | 0.011 | Old |
| 0.438305 | 0.099549 | 0.162 | 0.149 | 1 | 3 | Mob3b | 0.013 | Old |
| 0.43832 | 0.26753 | 0.02 | 0.015 | 1 | 3 | Hnmt | 0.005 | Old |
| 0.438637 | 0.271759 | 0.165 | 0.152 | 1 | 3 | Brd3 | 0.013 | Old |
| 0.438858 | 0.054725 | 0.189 | 0.172 | 1 | 3 | Agtrap | 0.017 | Old |
| 0.43889 | 0.061492 | 0.09 | 0.078 | 1 | 3 | Neurl4 | 0.012 | Old |
| 0.439118 | 0.081909 | 0.121 | 0.107 | 1 | 3 | Lemd3 | 0.014 | Old |
| 0.439136 | 0.07385 | 0.02 | 0.015 | 1 | 3 | Fbxo46 | 0.005 | Old |
| 0.439694 | 0.288697 | 0.088 | 0.078 | 1 | 3 | Sntb2 | 0.01 | Old |
| 0.439986 | 0.065967 | 0.069 | 0.059 | 1 | 3 | Utp4 | 0.01 | Old |
| 0.440049 | 0.258148 | 0.014 | 0.01 | 1 | 3 | Gstt3 | 0.004 | Old |
| 0.440376 | 0.029035 | 0.014 | 0.01 | 1 | 3 | Asgr2 | 0.004 | Old |
| 0.440703 | 0.250047 | 0.014 | 0.01 | 1 | 3 | D11Wsu47 | 0.004 | Old |
| 0.440703 | 0.151636 | 0.014 | 0.01 | 1 | 3 | Lysmd4 | 0.004 | Old |
| 0.440831 | 0.157516 | 0.044 | 0.037 | 1 | 3 | Gna11 | 0.007 | Old |
| 0.44113 | 0.283062 | 0.057 | 0.049 | 1 | 3 | Fyco1 | 0.008 | Old |
| 0.441163 | 0.201382 | 0.07 | 0.061 | 1 | 3 | Sugp1 | 0.009 | Old |

| | | | | | | | | |
|---|---|---|---|---|---|---|---|---|
| 0.441201 | 0.065543 | 0.221 | 0.203 | 1 | 3 | Psme2b | 0.018 | Old |
| 0.441359 | 0.240095 | 0.014 | 0.01 | 1 | 3 | Mief2 | 0.004 | Old |
| 0.44175 | 0.257678 | 0.036 | 0.029 | 1 | 3 | Tdp1 | 0.007 | Old |
| 0.442241 | 0.189956 | 0.054 | 0.045 | 1 | 3 | Snhg20 | 0.009 | Old |
| 0.442503 | 0.059188 | 0.339 | 0.317 | 1 | 3 | Cript | 0.022 | Old |
| 0.442673 | 0.141062 | 0.102 | 0.114 | 1 | 3 | Folr2 | -0.012 | Old |
| 0.44298 | 0.047319 | 0.277 | 0.251 | 1 | 3 | Hexim1 | 0.026 | Old |
| 0.443228 | 0.062585 | 0.158 | 0.144 | 1 | 3 | Cnot7 | 0.014 | Old |
| 0.443694 | 0.015283 | 0.676 | 0.63 | 1 | 3 | Tbxas1 | 0.046 | Old |
| 0.443723 | 0.058537 | 0.072 | 0.062 | 1 | 3 | Rgp1 | 0.01 | Old |
| 0.443856 | 0.39493 | 0.037 | 0.03 | 1 | 3 | Alg6 | 0.007 | Old |
| 0.44452 | 0.06765 | 0.149 | 0.135 | 1 | 3 | Foxk2 | 0.014 | Old |
| 0.444588 | 0.244263 | 0.178 | 0.163 | 1 | 3 | Mindy3 | 0.015 | Old |
| 0.444598 | 0.384866 | 0.029 | 0.023 | 1 | 3 | Ppp1r12b | 0.006 | Old |
| 0.445056 | 0.155434 | 0.029 | 0.023 | 1 | 3 | Arl15 | 0.006 | Old |
| 0.446307 | 0.186023 | 0.112 | 0.101 | 1 | 3 | Lsm14b | 0.011 | Old |
| 0.448716 | 0.067316 | 0.131 | 0.117 | 1 | 3 | Usf1 | 0.014 | Old |
| 0.448807 | 0.020796 | 0.108 | 0.095 | 1 | 3 | Rps19bp1 | 0.013 | Old |
| 0.448955 | 0.232723 | 0.029 | 0.023 | 1 | 3 | Socs1 | 0.006 | Old |
| 0.449668 | 0.683476 | 0.03 | 0.024 | 1 | 3 | Wrap53 | 0.006 | Old |
| 0.449679 | 0.04642 | 0.729 | 0.689 | 1 | 3 | 9-Sep | 0.04 | Old |
| 0.44985 | 0.010796 | 0.524 | 0.489 | 1 | 3 | Necap2 | 0.035 | Old |
| 0.449862 | 0.081089 | 0.022 | 0.016 | 1 | 3 | Map4k3 | 0.006 | Old |
| 0.450396 | 0.438769 | 0.022 | 0.016 | 1 | 3 | Il7 | 0.006 | Old |
| 0.450603 | 0.018911 | 0.648 | 0.616 | 1 | 3 | 7-Sep | 0.032 | Old |
| 0.451477 | 0.008659 | 0.123 | 0.11 | 1 | 3 | Dnpep | 0.013 | Old |
| 0.451653 | 0.005312 | 0.072 | 0.062 | 1 | 3 | Hdhd2 | 0.01 | Old |
| 0.452736 | 0.12453 | 0.055 | 0.047 | 1 | 3 | Zfp398 | 0.008 | Old |
| 0.452967 | 0.02348 | 0.091 | 0.08 | 1 | 3 | Rnaseh2b | 0.011 | Old |
| 0.453341 | 0.229917 | 0.022 | 0.016 | 1 | 3 | Dusp19 | 0.006 | Old |
| 0.453352 | 0.033696 | 0.074 | 0.064 | 1 | 3 | Hspbap1 | 0.01 | Old |
| 0.45361 | 0.490122 | 0.022 | 0.016 | 1 | 3 | Rhebl1 | 0.006 | Old |
| 0.453729 | 0.097764 | 0.087 | 0.077 | 1 | 3 | Slc25a32 | 0.01 | Old |
| 0.453739 | 0.289677 | 0.037 | 0.03 | 1 | 3 | Thap1 | 0.007 | Old |
| 0.453791 | 0.051522 | 0.096 | 0.085 | 1 | 3 | Fbxo45 | 0.011 | Old |
| 0.453891 | 0.154098 | 0.069 | 0.059 | 1 | 3 | Chst11 | 0.01 | Old |
| 0.454032 | 0.024675 | 0.308 | 0.284 | 1 | 3 | Sugt1 | 0.024 | Old |
| 0.454068 | 0.111229 | 0.287 | 0.263 | 1 | 3 | Ube2h | 0.024 | Old |
| 0.454574 | 0.142643 | 0.036 | 0.029 | 1 | 3 | Rnf25 | 0.007 | Old |
| 0.454769 | 0.002603 | 0.094 | 0.083 | 1 | 3 | Slamf7 | 0.011 | Old |
| 0.454813 | 0.108773 | 0.227 | 0.21 | 1 | 3 | Wdfy3 | 0.017 | Old |
| 0.455052 | 0.10923 | 0.172 | 0.158 | 1 | 3 | Arhgef10l | 0.014 | Old |
| 0.455138 | 0.021762 | 0.053 | 0.044 | 1 | 3 | Leng1 | 0.009 | Old |
| 0.455489 | 0.113104 | 0.091 | 0.081 | 1 | 3 | Cog8 | 0.01 | Old |
| 0.455553 | 0.070164 | 0.012 | 0.016 | 1 | 3 | Opn3 | -0.004 | Old |

| | | | | | | | | |
|---|---|---|---|---|---|---|---|---|
| 0.455621 | 0.062512 | 0.076 | 0.067 | 1 | 3 | Mcm5 | 0.009 | Old |
| 0.455749 | 0.460907 | 0.048 | 0.04 | 1 | 3 | Gtdc1 | 0.008 | Old |
| 0.455759 | 0.187669 | 0.022 | 0.016 | 1 | 3 | G430095P1 | 0.006 | Old |
| 0.456028 | 0.257485 | 0.022 | 0.016 | 1 | 3 | Pusl1 | 0.006 | Old |
| 0.456071 | 0.075112 | 0.03 | 0.037 | 1 | 3 | H6pd | -0.007 | Old |
| 0.456094 | 0.069723 | 0.051 | 0.043 | 1 | 3 | Tmem237 | 0.008 | Old |
| 0.456382 | 0.222005 | 0.037 | 0.03 | 1 | 3 | Mthfd2l | 0.007 | Old |
| 0.456448 | 0.109288 | 0.117 | 0.105 | 1 | 3 | Ap5m1 | 0.012 | Old |
| 0.456486 | 0.044337 | 0.227 | 0.208 | 1 | 3 | N4bp1 | 0.019 | Old |
| 0.456597 | 0.033898 | 0.07 | 0.061 | 1 | 3 | Ccdc86 | 0.009 | Old |
| 0.45705 | 0.036093 | 0.644 | 0.614 | 1 | 3 | Sh3kbp1 | 0.03 | Old |
| 0.4573 | 1.233692 | 0.01 | 0.006 | 1 | 3 | Zfp623 | 0.004 | Old |
| 0.457374 | 0.420005 | 0.038 | 0.032 | 1 | 3 | Yae1d1 | 0.006 | Old |
| 0.457488 | 0.60776 | 0.023 | 0.018 | 1 | 3 | Mapkbp1 | 0.005 | Old |
| 0.457852 | 0.017507 | 0.968 | 0.962 | 1 | 3 | Pabpc1 | 0.006 | Old |
| 0.457991 | 0.035898 | 0.932 | 0.918 | 1 | 3 | Ptp4a2 | 0.014 | Old |
| 0.458256 | 0.187658 | 0.048 | 0.04 | 1 | 3 | Cpt2 | 0.008 | Old |
| 0.458453 | 0.106114 | 0.022 | 0.016 | 1 | 3 | Nipa1 | 0.006 | Old |
| 0.458706 | 0.02367 | 0.062 | 0.053 | 1 | 3 | Parp12 | 0.009 | Old |
| 0.458904 | 0.422473 | 0.029 | 0.023 | 1 | 3 | Ndufaf6 | 0.006 | Old |
| 0.459357 | 0.099679 | 0.065 | 0.056 | 1 | 3 | Naa16 | 0.009 | Old |
| 0.459572 | 0.101655 | 0.074 | 0.064 | 1 | 3 | Pex16 | 0.01 | Old |
| 0.459619 | 0.069983 | 0.299 | 0.273 | 1 | 3 | Cd180 | 0.026 | Old |
| 0.459953 | 0.036994 | 0.685 | 0.644 | 1 | 3 | Pomp | 0.041 | Old |
| 0.461033 | 0.374117 | 0.01 | 0.006 | 1 | 3 | Cbx6 | 0.004 | Old |
| 0.461205 | 0.228485 | 0.094 | 0.085 | 1 | 3 | Gnpat | 0.009 | Old |
| 0.461346 | 0.554478 | 0.016 | 0.011 | 1 | 3 | Focad | 0.005 | Old |
| 0.461376 | 0.026805 | 0.219 | 0.199 | 1 | 3 | Sipa1 | 0.02 | Old |
| 0.461449 | 0.338081 | 0.01 | 0.006 | 1 | 3 | Nnt | 0.004 | Old |
| 0.461651 | 0.065041 | 0.179 | 0.16 | 1 | 3 | Atxn10 | 0.019 | Old |
| 0.461987 | 0.870717 | 0.016 | 0.011 | 1 | 3 | L2hgdh | 0.005 | Old |
| 0.462273 | 0.136034 | 0.369 | 0.348 | 1 | 3 | Dap | 0.021 | Old |
| 0.462282 | 0.416447 | 0.01 | 0.006 | 1 | 3 | Polh | 0.004 | Old |
| 0.462722 | 0.254411 | 0.056 | 0.048 | 1 | 3 | Smyd2 | 0.008 | Old |
| 0.462828 | 0.031589 | 0.099 | 0.087 | 1 | 3 | Tmem205 | 0.012 | Old |
| 0.46313 | 0.170242 | 0.037 | 0.03 | 1 | 3 | Telo2 | 0.007 | Old |
| 0.463671 | 0.106086 | 0.129 | 0.116 | 1 | 3 | Farsb | 0.013 | Old |
| 0.463812 | 0.193196 | 0.038 | 0.032 | 1 | 3 | Ppp1r16a | 0.006 | Old |
| 0.463912 | 0.120972 | 0.111 | 0.1 | 1 | 3 | Ccnk | 0.011 | Old |
| 0.463949 | 0.190149 | 0.01 | 0.006 | 1 | 3 | Fam124a | 0.004 | Old |
| 0.464083 | 0.066773 | 0.192 | 0.176 | 1 | 3 | Gas2l3 | 0.016 | Old |
| 0.464141 | 0.035722 | 0.022 | 0.016 | 1 | 3 | Tmem267 | 0.006 | Old |
| 0.464515 | 0.118473 | 0.067 | 0.058 | 1 | 3 | Lrrc8b | 0.009 | Old |
| 0.465202 | 0.490247 | 0.016 | 0.011 | 1 | 3 | Cep89 | 0.005 | Old |
| 0.465597 | 0.008748 | 0.037 | 0.03 | 1 | 3 | Surf6 | 0.007 | Old |

| | | | | | | | | |
|---|---|---|---|---|---|---|---|---|
| 0.465847 | 0.295248 | 0.016 | 0.011 | 1 | 3 | Dnmbp | 0.005 | Old |
| 0.465942 | 0.077548 | 0.317 | 0.293 | 1 | 3 | Pnn | 0.024 | Old |
| 0.466038 | 0.53059 | 0.01 | 0.006 | 1 | 3 | Mybpc3 | 0.004 | Old |
| 0.466038 | 0.009878 | 0.01 | 0.006 | 1 | 3 | Cdk18 | 0.004 | Old |
| 0.466169 | 0.159126 | 0.016 | 0.011 | 1 | 3 | Cdh24 | 0.005 | Old |
| 0.466968 | 0.243057 | 0.061 | 0.053 | 1 | 3 | Psmg1 | 0.008 | Old |
| 0.467036 | 0.043007 | 0.122 | 0.109 | 1 | 3 | Psmg3 | 0.013 | Old |
| 0.46709 | 0.206137 | 0.048 | 0.04 | 1 | 3 | Trpm2 | 0.008 | Old |
| 0.46746 | 0.323148 | 0.016 | 0.011 | 1 | 3 | 3-Sep | 0.005 | Old |
| 0.46785 | 0.117561 | 0.047 | 0.039 | 1 | 3 | Gprasp1 | 0.008 | Old |
| 0.468106 | 0.484911 | 0.016 | 0.011 | 1 | 3 | Klri2 | 0.005 | Old |
| 0.468106 | 0.191907 | 0.016 | 0.011 | 1 | 3 | Crocc | 0.005 | Old |
| 0.468354 | 0.044005 | 0.223 | 0.205 | 1 | 3 | Gapt | 0.018 | Old |
| 0.469073 | 0.574209 | 0.023 | 0.018 | 1 | 3 | Tti1 | 0.005 | Old |
| 0.46951 | 0.12489 | 0.039 | 0.033 | 1 | 3 | Zkscan14 | 0.006 | Old |
| 0.469735 | 0.082261 | 0.014 | 0.019 | 1 | 3 | Zfp873 | -0.005 | Old |
| 0.471175 | 0.233862 | 0.037 | 0.03 | 1 | 3 | Gemin2 | 0.007 | Old |
| 0.471546 | 0.0892 | 0.105 | 0.095 | 1 | 3 | Coq5 | 0.01 | Old |
| 0.471597 | 0.118692 | 0.066 | 0.057 | 1 | 3 | Zfp738 | 0.009 | Old |
| 0.472081 | 0.32266 | 0.014 | 0.019 | 1 | 3 | Exo5 | -0.005 | Old |
| 0.472536 | 0.201363 | 0.049 | 0.042 | 1 | 3 | Rpap1 | 0.007 | Old |
| 0.472668 | 0.052558 | 0.014 | 0.019 | 1 | 3 | Hvcn1 | -0.005 | Old |
| 0.472806 | 0.517951 | 0.066 | 0.058 | 1 | 3 | Apool | 0.008 | Old |
| 0.473132 | 0.02889 | 0.26 | 0.24 | 1 | 3 | Hmox2 | 0.02 | Old |
| 0.473282 | 0.213923 | 0.092 | 0.082 | 1 | 3 | Lsg1 | 0.01 | Old |
| 0.474029 | 0.457961 | 0.031 | 0.025 | 1 | 3 | Dimt1 | 0.006 | Old |
| 0.474259 | 0.064447 | 0.228 | 0.213 | 1 | 3 | Nfya | 0.015 | Old |
| 0.474292 | 0.146232 | 0.085 | 0.074 | 1 | 3 | Nosip | 0.011 | Old |
| 0.474343 | 0.010927 | 0.246 | 0.226 | 1 | 3 | Dnajc21 | 0.02 | Old |
| 0.475521 | 0.28917 | 0.041 | 0.034 | 1 | 3 | Cbr4 | 0.007 | Old |
| 0.475623 | 0.122817 | 0.102 | 0.09 | 1 | 3 | Irf8 | 0.012 | Old |
| 0.475706 | 0.142757 | 0.059 | 0.051 | 1 | 3 | Ptges2 | 0.008 | Old |
| 0.475872 | 0.168414 | 0.059 | 0.051 | 1 | 3 | 3110002H1 | 0.008 | Old |
| 0.476231 | 0.041136 | 0.105 | 0.095 | 1 | 3 | Shprh | 0.01 | Old |
| 0.476328 | 0.053871 | 0.061 | 0.053 | 1 | 3 | Nadk2 | 0.008 | Old |
| 0.476539 | 0.166641 | 0.05 | 0.043 | 1 | 3 | Wwox | 0.007 | Old |
| 0.477321 | 0.060383 | 0.117 | 0.104 | 1 | 3 | Dnajc25 | 0.013 | Old |
| 0.477668 | 0.207846 | 0.048 | 0.04 | 1 | 3 | Traf2 | 0.008 | Old |
| 0.478254 | 0.226147 | 0.067 | 0.077 | 1 | 3 | Mta3 | -0.01 | Old |
| 0.479327 | 0.03834 | 0.047 | 0.039 | 1 | 3 | Rtn1 | 0.008 | Old |
| 0.479406 | 0.010466 | 0.263 | 0.244 | 1 | 3 | Mrpl14 | 0.019 | Old |
| 0.480025 | 0.001801 | 0.178 | 0.163 | 1 | 3 | Galnt11 | 0.015 | Old |
| 0.480134 | 0.022539 | 0.167 | 0.153 | 1 | 3 | Btaf1 | 0.014 | Old |
| 0.480142 | 0.036538 | 0.55 | 0.513 | 1 | 3 | Rab11b | 0.037 | Old |
| 0.480284 | 0.083304 | 0.331 | 0.301 | 1 | 3 | Syf2 | 0.03 | Old |

| | | | | | | | | |
|---|---|---|---|---|---|---|---|---|
| 0.480464 | 0.073312 | 0.286 | 0.266 | 1 | 3 | Eloa | 0.02 | Old |
| 0.480577 | 0.388726 | 0.057 | 0.049 | 1 | 3 | Tecpr1 | 0.008 | Old |
| 0.480871 | 0.09432 | 0.166 | 0.153 | 1 | 3 | Ddx47 | 0.013 | Old |
| 0.481398 | 0.037633 | 0.484 | 0.453 | 1 | 3 | Birc6 | 0.031 | Old |
| 0.481695 | 0.030432 | 0.296 | 0.271 | 1 | 3 | Cep350 | 0.025 | Old |
| 0.481994 | 0.348281 | 0.032 | 0.027 | 1 | 3 | Twnk | 0.005 | Old |
| 0.482234 | 0.014718 | 0.485 | 0.455 | 1 | 3 | Cbfa2t3 | 0.03 | Old |
| 0.482579 | 0.026641 | 0.229 | 0.211 | 1 | 3 | Echs1 | 0.018 | Old |
| 0.483006 | 0.066774 | 0.081 | 0.072 | 1 | 3 | Nup85 | 0.009 | Old |
| 0.483353 | 0.116975 | 0.161 | 0.146 | 1 | 3 | Abca1 | 0.015 | Old |
| 0.483406 | 0.034048 | 0.221 | 0.203 | 1 | 3 | Runx3 | 0.018 | Old |
| 0.483419 | 0.283669 | 0.035 | 0.042 | 1 | 3 | Nudt7 | -0.007 | Old |
| 0.484322 | 0.102274 | 0.116 | 0.104 | 1 | 3 | Asb8 | 0.012 | Old |
| 0.484486 | 0.140843 | 0.041 | 0.034 | 1 | 3 | Pla2g12a | 0.007 | Old |
| 0.484603 | 0.339519 | 0.13 | 0.121 | 1 | 3 | Zfp236 | 0.009 | Old |
| 0.485743 | 0.206533 | 0.117 | 0.105 | 1 | 3 | Dym | 0.012 | Old |
| 0.485864 | 0.142759 | 0.18 | 0.167 | 1 | 3 | Hmgcl | 0.013 | Old |
| 0.485908 | 0.082632 | 0.038 | 0.032 | 1 | 3 | Thap7 | 0.006 | Old |
| 0.485999 | 0.040752 | 0.26 | 0.239 | 1 | 3 | Dync1li1 | 0.021 | Old |
| 0.486527 | 0.020805 | 0.341 | 0.309 | 1 | 3 | Stip1 | 0.032 | Old |
| 0.487523 | 0.026584 | 0.648 | 0.612 | 1 | 3 | Wipf1 | 0.036 | Old |
| 0.487774 | 0.136799 | 0.038 | 0.032 | 1 | 3 | 1810024B0 | 0.006 | Old |
| 0.488004 | 0.059607 | 0.048 | 0.04 | 1 | 3 | Rdh14 | 0.008 | Old |
| 0.488224 | 0.256633 | 0.016 | 0.02 | 1 | 3 | Gstm4 | -0.004 | Old |
| 0.488683 | 0.012892 | 0.315 | 0.292 | 1 | 3 | Banf1 | 0.023 | Old |
| 0.48882 | 0.058119 | 0.49 | 0.462 | 1 | 3 | Tomm22 | 0.028 | Old |
| 0.489355 | 0.154619 | 0.031 | 0.025 | 1 | 3 | Nkiras1 | 0.006 | Old |
| 0.48938 | 0.136358 | 0.016 | 0.02 | 1 | 3 | Adck2 | -0.004 | Old |
| 0.489696 | 0.079639 | 0.238 | 0.217 | 1 | 3 | Vbp1 | 0.021 | Old |
| 0.490161 | 0.439887 | 0.017 | 0.013 | 1 | 3 | Ppm1f | 0.004 | Old |
| 0.490237 | 0.003793 | 0.237 | 0.218 | 1 | 3 | Hlcs | 0.019 | Old |
| 0.490553 | 0.011743 | 0.032 | 0.027 | 1 | 3 | Olfr1167 | 0.005 | Old |
| 0.490562 | 0.278299 | 0.116 | 0.105 | 1 | 3 | Tbck | 0.011 | Old |
| 0.490796 | 0.543609 | 0.017 | 0.013 | 1 | 3 | Tgm1 | 0.004 | Old |
| 0.490796 | 0.469007 | 0.017 | 0.013 | 1 | 3 | Chst15 | 0.004 | Old |
| 0.491006 | 0.058385 | 0.032 | 0.027 | 1 | 3 | Slc26a6 | 0.005 | Old |
| 0.491188 | 0.106195 | 0.049 | 0.042 | 1 | 3 | Gfi1b | 0.007 | Old |
| 0.49192 | 0.154779 | 0.088 | 0.08 | 1 | 3 | Fam3c | 0.008 | Old |
| 0.492068 | 0.298632 | 0.017 | 0.013 | 1 | 3 | Gm16845 | 0.004 | Old |
| 0.492167 | 0.015185 | 0.45 | 0.432 | 1 | 3 | Il6 | 0.018 | Old |
| 0.493239 | 0.118117 | 0.147 | 0.135 | 1 | 3 | Prep | 0.012 | Old |
| 0.494905 | 0.825066 | 0.011 | 0.008 | 1 | 3 | Zfp553 | 0.003 | Old |
| 0.495314 | 0.18014 | 0.042 | 0.035 | 1 | 3 | Tjap1 | 0.007 | Old |
| 0.495685 | 0.173789 | 0.061 | 0.053 | 1 | 3 | Tcp11l1 | 0.008 | Old |
| 0.495828 | 0.180448 | 0.082 | 0.073 | 1 | 3 | Armc5 | 0.009 | Old |

| | | | | | | | | |
|---|---|---|---|---|---|---|---|---|
| 0.496984 | 0.300509 | 0.053 | 0.045 | 1 | 3 | Sgsm2 | 0.008 | Old |
| 0.497209 | 0.056153 | 0.024 | 0.019 | 1 | 3 | Adora3 | 0.005 | Old |
| 0.497788 | 0.184927 | 0.216 | 0.202 | 1 | 3 | Eif4e3 | 0.014 | Old |
| 0.497944 | 0.149312 | 0.069 | 0.061 | 1 | 3 | Manbal | 0.008 | Old |
| 0.498014 | 0.01136 | 0.811 | 0.782 | 1 | 3 | Cltc | 0.029 | Old |
| 0.498026 | 0.08409 | 0.906 | 0.881 | 1 | 3 | Anxa5 | 0.025 | Old |
| 0.498259 | 0.399229 | 0.025 | 0.02 | 1 | 3 | Nars2 | 0.005 | Old |
| 0.498358 | 0.161942 | 0.115 | 0.104 | 1 | 3 | F3 | 0.011 | Old |
| 0.498512 | 0.24197 | 0.042 | 0.035 | 1 | 3 | Klhl36 | 0.007 | Old |
| 0.498576 | 0.010066 | 0.186 | 0.168 | 1 | 3 | Sgta | 0.018 | Old |
| 0.498585 | 0.039341 | 0.039 | 0.033 | 1 | 3 | Ctsf | 0.006 | Old |
| 0.498944 | 0.170079 | 0.079 | 0.069 | 1 | 3 | Fam219a | 0.01 | Old |
| 0.499345 | 0.672165 | 0.011 | 0.008 | 1 | 3 | Iqgap3 | 0.003 | Old |
| 0.499989 | 0.549169 | 0.033 | 0.028 | 1 | 3 | Tmem241 | 0.005 | Old |
| 0.500108 | 0.110817 | 0.032 | 0.027 | 1 | 3 | Mipep | 0.005 | Old |
| 0.500154 | 0.46238 | 0.011 | 0.008 | 1 | 3 | Cdkl2 | 0.003 | Old |
| 0.500182 | 0.02213 | 0.3 | 0.277 | 1 | 3 | Nenf | 0.023 | Old |
| 0.500237 | 0.029879 | 0.055 | 0.047 | 1 | 3 | Coprs | 0.008 | Old |
| 0.500704 | 0.210804 | 0.177 | 0.164 | 1 | 3 | Tsen34 | 0.013 | Old |
| 0.501092 | 0.008766 | 0.048 | 0.04 | 1 | 3 | Med6 | 0.008 | Old |
| 0.501113 | 0.012737 | 0.025 | 0.02 | 1 | 3 | Tnfsf14 | 0.005 | Old |
| 0.501785 | 0.020824 | 0.135 | 0.121 | 1 | 3 | Mir17hg | 0.014 | Old |
| 0.501871 | 0.005297 | 0.515 | 0.475 | 1 | 3 | Tet3 | 0.04 | Old |
| 0.501932 | 0.090149 | 0.078 | 0.068 | 1 | 3 | Mrpl27 | 0.01 | Old |
| 0.502248 | 0.055739 | 0.336 | 0.318 | 1 | 3 | Apaf1 | 0.018 | Old |
| 0.502341 | 0.1203 | 0.061 | 0.053 | 1 | 3 | Pgghg | 0.008 | Old |
| 0.502367 | 0.235178 | 0.054 | 0.047 | 1 | 3 | Zfp142 | 0.007 | Old |
| 0.502527 | 0.338525 | 0.053 | 0.045 | 1 | 3 | Qsox2 | 0.008 | Old |
| 0.502626 | 0.210219 | 0.032 | 0.027 | 1 | 3 | Setd6 | 0.005 | Old |
| 0.50263 | 0.032471 | 0.866 | 0.824 | 1 | 3 | Celf2 | 0.042 | Old |
| 0.503805 | 0.332803 | 0.011 | 0.008 | 1 | 3 | Kif20b | 0.003 | Old |
| 0.503805 | 0.150458 | 0.011 | 0.008 | 1 | 3 | Tpmt | 0.003 | Old |
| 0.504143 | 0.043349 | 0.487 | 0.462 | 1 | 3 | Jup | 0.025 | Old |
| 0.504619 | 0.198511 | 0.011 | 0.008 | 1 | 3 | Rpp38 | 0.003 | Old |
| 0.504683 | 0.050706 | 0.019 | 0.024 | 1 | 3 | Large1 | -0.005 | Old |
| 0.505017 | 0.119718 | 0.112 | 0.102 | 1 | 3 | Epb41 | 0.01 | Old |
| 0.505527 | 1.06054 | 0.018 | 0.014 | 1 | 3 | Zfp930 | 0.004 | Old |
| 0.505839 | 0.040307 | 0.498 | 0.468 | 1 | 3 | Ybx3 | 0.03 | Old |
| 0.50584 | 0.171994 | 0.011 | 0.008 | 1 | 3 | Itgb3 | 0.003 | Old |
| 0.50584 | 0.007565 | 0.011 | 0.008 | 1 | 3 | Gm4117 | 0.003 | Old |
| 0.50662 | 0.158561 | 0.051 | 0.044 | 1 | 3 | Coq9 | 0.007 | Old |
| 0.506942 | 0.150599 | 0.11 | 0.1 | 1 | 3 | Cryl1 | 0.01 | Old |
| 0.507088 | 0.032961 | 0.319 | 0.299 | 1 | 3 | Nans | 0.02 | Old |
| 0.507552 | 0.060273 | 0.092 | 0.082 | 1 | 3 | D6Wsu163 | 0.01 | Old |
| 0.508156 | 0.077041 | 0.137 | 0.125 | 1 | 3 | Ube2g1 | 0.012 | Old |

| | | | | | | | | |
|---|---|---|---|---|---|---|---|---|
| 0.508544 | 0.129765 | 0.033 | 0.028 | 1 | 3 | Kat8 | 0.005 | Old |
| 0.509185 | 0.035837 | 0.557 | 0.518 | 1 | 3 | Sf3b2 | 0.039 | Old |
| 0.51023 | 0.015292 | 0.339 | 0.317 | 1 | 3 | Senp6 | 0.022 | Old |
| 0.510541 | 0.334204 | 0.057 | 0.051 | 1 | 3 | Mpp5 | 0.006 | Old |
| 0.510831 | 0.066096 | 0.167 | 0.152 | 1 | 3 | Xpnpep1 | 0.015 | Old |
| 0.511882 | 0.051973 | 0.044 | 0.038 | 1 | 3 | Trip10 | 0.006 | Old |
| 0.512089 | 0.079663 | 0.025 | 0.02 | 1 | 3 | Lyrm1 | 0.005 | Old |
| 0.51239 | 0.09364 | 0.085 | 0.076 | 1 | 3 | Bola1 | 0.009 | Old |
| 0.513146 | 0.029931 | 0.276 | 0.259 | 1 | 3 | Setx | 0.017 | Old |
| 0.513952 | 0.018179 | 0.06 | 0.052 | 1 | 3 | Ints8 | 0.008 | Old |
| 0.513988 | 0.117835 | 0.033 | 0.028 | 1 | 3 | Spidr | 0.005 | Old |
| 0.514272 | 0.118598 | 0.116 | 0.104 | 1 | 3 | Zcchc9 | 0.012 | Old |
| 0.514694 | 0.220041 | 0.085 | 0.076 | 1 | 3 | Ipo13 | 0.009 | Old |
| 0.515385 | 0.185995 | 0.086 | 0.077 | 1 | 3 | Cfap36 | 0.009 | Old |
| 0.516137 | 0.225486 | 0.018 | 0.014 | 1 | 3 | Gm45871 | 0.004 | Old |
| 0.516137 | 0.179528 | 0.018 | 0.014 | 1 | 3 | Hist1h1e | 0.004 | Old |
| 0.516155 | 0.044114 | 0.444 | 0.404 | 1 | 3 | Pds5a | 0.04 | Old |
| 0.516493 | 0.014901 | 0.184 | 0.169 | 1 | 3 | Txlna | 0.015 | Old |
| 0.516787 | 0.622355 | 0.026 | 0.021 | 1 | 3 | Paox | 0.005 | Old |
| 0.517655 | 0.013933 | 0.097 | 0.086 | 1 | 3 | Pibf1 | 0.011 | Old |
| 0.517755 | 0.040322 | 0.613 | 0.578 | 1 | 3 | Atp6v1g1 | 0.035 | Old |
| 0.518021 | 0.114707 | 0.018 | 0.014 | 1 | 3 | Tfb1m | 0.004 | Old |
| 0.518637 | 0.13686 | 0.104 | 0.095 | 1 | 3 | Agpat1 | 0.009 | Old |
| 0.518964 | 0.240012 | 0.018 | 0.014 | 1 | 3 | Nxpe5 | 0.004 | Old |
| 0.519259 | 0.06021 | 0.173 | 0.159 | 1 | 3 | Casp8ap2 | 0.014 | Old |
| 0.519682 | 0.024989 | 0.191 | 0.176 | 1 | 3 | Rbm22 | 0.015 | Old |
| 0.520069 | 0.12416 | 0.062 | 0.054 | 1 | 3 | Hsdl2 | 0.008 | Old |
| 0.520103 | 0.034591 | 0.296 | 0.279 | 1 | 3 | Ssfa2 | 0.017 | Old |
| 0.520538 | 0.02656 | 0.018 | 0.014 | 1 | 3 | Dnaaf3 | 0.004 | Old |
| 0.522115 | 0.24911 | 0.018 | 0.014 | 1 | 3 | Hells | 0.004 | Old |
| 0.522465 | 0.051419 | 0.123 | 0.111 | 1 | 3 | Tmem63a | 0.012 | Old |
| 0.522549 | 0.094415 | 0.137 | 0.126 | 1 | 3 | Gadd45gip | 0.011 | Old |
| 0.522657 | 0.107193 | 0.327 | 0.302 | 1 | 3 | Ripk1 | 0.025 | Old |
| 0.522922 | 0.036114 | 0.025 | 0.02 | 1 | 3 | Dffb | 0.005 | Old |
| 0.523062 | 0.23919 | 0.018 | 0.014 | 1 | 3 | Pcyt1b | 0.004 | Old |
| 0.523226 | 0.036297 | 0.49 | 0.455 | 1 | 3 | Myo9b | 0.035 | Old |
| 0.52376 | 0.178904 | 0.026 | 0.021 | 1 | 3 | Gm37305 | 0.005 | Old |
| 0.524947 | 0.035698 | 0.007 | 0.01 | 1 | 3 | Rpusd1 | -0.003 | Old |
| 0.525276 | 0.017502 | 0.018 | 0.014 | 1 | 3 | Ccdc181 | 0.004 | Old |
| 0.52535 | 0.37199 | 0.078 | 0.069 | 1 | 3 | Zbtb43 | 0.009 | Old |
| 0.525713 | 0.088918 | 0.092 | 0.083 | 1 | 3 | Plau | 0.009 | Old |
| 0.525937 | 0.34216 | 0.036 | 0.03 | 1 | 3 | Ptrh1 | 0.006 | Old |
| 0.526145 | 0.458322 | 0.045 | 0.039 | 1 | 3 | 1190007I0 | 0.006 | Old |
| 0.526158 | 0.282284 | 0.036 | 0.03 | 1 | 3 | Pex11g | 0.006 | Old |
| 0.526277 | 0.266584 | 0.065 | 0.057 | 1 | 3 | Ppp6r2 | 0.008 | Old |

| | | | | | | | | |
|---|---|---|---|---|---|---|---|---|
| 0.526565 | 0.254092 | 0.117 | 0.106 | 1 | 3 | Fam43a | 0.011 | Old |
| 0.526782 | 0.124807 | 0.265 | 0.246 | 1 | 3 | Smc3 | 0.019 | Old |
| 0.527934 | 0.344727 | 0.036 | 0.03 | 1 | 3 | Pigv | 0.006 | Old |
| 0.528042 | 0.059079 | 0.072 | 0.063 | 1 | 3 | Med23 | 0.009 | Old |
| 0.528824 | 0.108516 | 0.362 | 0.34 | 1 | 3 | Cdkn1b | 0.022 | Old |
| 0.528921 | 0.002897 | 0.474 | 0.443 | 1 | 3 | Anxa4 | 0.031 | Old |
| 0.529798 | 0.344364 | 0.045 | 0.039 | 1 | 3 | Uba7 | 0.006 | Old |
| 0.529998 | 0.172997 | 0.026 | 0.021 | 1 | 3 | Fmn1 | 0.005 | Old |
| 0.530356 | 0.238711 | 0.057 | 0.051 | 1 | 3 | Ndufaf3 | 0.006 | Old |
| 0.530365 | 1.214831 | 0.012 | 0.009 | 1 | 3 | Acod1 | 0.003 | Old |
| 0.531613 | 0.03518 | 0.127 | 0.115 | 1 | 3 | Mvb12a | 0.012 | Old |
| 0.531667 | 0.141168 | 0.116 | 0.128 | 1 | 3 | Eif2b4 | -0.012 | Old |
| 0.531786 | 0.13956 | 0.044 | 0.038 | 1 | 3 | Katnb1 | 0.006 | Old |
| 0.531857 | 0.143383 | 0.216 | 0.203 | 1 | 3 | Pilrb1 | 0.013 | Old |
| 0.532188 | 0.148663 | 0.044 | 0.038 | 1 | 3 | Btrc | 0.006 | Old |
| 0.532625 | 0.053336 | 0.569 | 0.538 | 1 | 3 | Usmg5 | 0.031 | Old |
| 0.532744 | 0.035036 | 0.122 | 0.111 | 1 | 3 | Trim26 | 0.011 | Old |
| 0.532877 | 0.159739 | 0.131 | 0.121 | 1 | 3 | Stx8 | 0.01 | Old |
| 0.532913 | 0.024939 | 0.356 | 0.33 | 1 | 3 | Thrap3 | 0.026 | Old |
| 0.53303 | 0.028749 | 0.085 | 0.076 | 1 | 3 | Gstp1 | 0.009 | Old |
| 0.533149 | 0.074396 | 0.26 | 0.242 | 1 | 3 | Wbp11 | 0.018 | Old |
| 0.533305 | 0.330747 | 0.059 | 0.052 | 1 | 3 | Mtg1 | 0.007 | Old |
| 0.533367 | 0.112728 | 0.119 | 0.11 | 1 | 3 | Gab2 | 0.009 | Old |
| 0.533393 | 0.093164 | 0.026 | 0.021 | 1 | 3 | Gstz1 | 0.005 | Old |
| 0.533414 | 0.466836 | 0.019 | 0.015 | 1 | 3 | Snap47 | 0.004 | Old |
| 0.533509 | 0.25406 | 0.012 | 0.009 | 1 | 3 | Tsix | 0.003 | Old |
| 0.53407 | 0.020265 | 0.03 | 0.035 | 1 | 3 | Ccdc61 | -0.005 | Old |
| 0.534296 | 0.28618 | 0.012 | 0.009 | 1 | 3 | Mterf1a | 0.003 | Old |
| 0.534384 | 0.006483 | 0.638 | 0.59 | 1 | 3 | Dusp3 | 0.048 | Old |
| 0.534572 | 0.100289 | 0.142 | 0.13 | 1 | 3 | Idh2 | 0.012 | Old |
| 0.534869 | 0.050661 | 0.222 | 0.206 | 1 | 3 | Babam2 | 0.016 | Old |
| 0.535175 | 0.017201 | 0.152 | 0.139 | 1 | 3 | Cdip1 | 0.013 | Old |
| 0.53548 | 0.019249 | 0.343 | 0.319 | 1 | 3 | Smim15 | 0.024 | Old |
| 0.536239 | 0.3288 | 0.027 | 0.023 | 1 | 3 | Cmss1 | 0.004 | Old |
| 0.536661 | 0.845791 | 0.012 | 0.009 | 1 | 3 | Tspyl4 | 0.003 | Old |
| 0.536877 | 0.081802 | 0.287 | 0.266 | 1 | 3 | 2-Mar | 0.021 | Old |
| 0.537056 | 0.572741 | 0.012 | 0.009 | 1 | 3 | A530032D1 | 0.003 | Old |
| 0.537171 | 0.322459 | 0.025 | 0.03 | 1 | 3 | Cluap1 | -0.005 | Old |
| 0.537451 | 0.108193 | 0.012 | 0.009 | 1 | 3 | Isoc2a | 0.003 | Old |
| 0.537885 | 0.009102 | 0.628 | 0.58 | 1 | 3 | P4hb | 0.048 | Old |
| 0.538773 | 0.077564 | 0.286 | 0.268 | 1 | 3 | Rap2c | 0.018 | Old |
| 0.539249 | 0.043678 | 0.044 | 0.038 | 1 | 3 | Chd1l | 0.006 | Old |
| 0.539895 | 0.17189 | 0.019 | 0.015 | 1 | 3 | Rnase2a | 0.004 | Old |
| 0.540772 | 0.073047 | 0.213 | 0.198 | 1 | 3 | Acp1 | 0.015 | Old |
| 0.541466 | 0.220078 | 0.032 | 0.038 | 1 | 3 | Cracr2b | -0.006 | Old |

| | | | | | | | | |
|---|---|---|---|---|---|---|---|---|
| 0.541627 | 0.127522 | 0.027 | 0.023 | 1 | 3 | Tchp | 0.004 | Old |
| 0.541753 | 0.040198 | 0.019 | 0.015 | 1 | 3 | Mrvi1 | 0.004 | Old |
| 0.541993 | 0.261801 | 0.055 | 0.048 | 1 | 3 | Csf2rb2 | 0.007 | Old |
| 0.542353 | 0.201486 | 0.243 | 0.229 | 1 | 3 | Ddx42 | 0.014 | Old |
| 0.543187 | 0.019733 | 0.008 | 0.011 | 1 | 3 | Farp2 | -0.003 | Old |
| 0.545115 | 0.076672 | 0.068 | 0.061 | 1 | 3 | Cep57 | 0.007 | Old |
| 0.545517 | 0.042936 | 0.209 | 0.194 | 1 | 3 | Dap3 | 0.015 | Old |
| 0.545794 | 0.36697 | 0.061 | 0.054 | 1 | 3 | L1cam | 0.007 | Old |
| 0.546247 | 0.027395 | 0.195 | 0.181 | 1 | 3 | Prpf8 | 0.014 | Old |
| 0.546629 | 0.093726 | 0.742 | 0.718 | 1 | 3 | Nap1l1 | 0.024 | Old |
| 0.546635 | 0.048508 | 0.631 | 0.6 | 1 | 3 | Snx3 | 0.031 | Old |
| 0.54666 | 0.195396 | 0.072 | 0.064 | 1 | 3 | Gpatch4 | 0.008 | Old |
| 0.546676 | 0.00926 | 0.441 | 0.398 | 1 | 3 | Psmb3 | 0.043 | Old |
| 0.546866 | 0.117556 | 0.097 | 0.087 | 1 | 3 | Arl4a | 0.01 | Old |
| 0.547033 | 0.019485 | 0.047 | 0.04 | 1 | 3 | Gm20342 | 0.007 | Old |
| 0.547423 | 0.061494 | 0.161 | 0.149 | 1 | 3 | Snx30 | 0.012 | Old |
| 0.548494 | 0.33488 | 0.026 | 0.032 | 1 | 3 | Smg8 | -0.006 | Old |
| 0.549607 | 0.012333 | 0.345 | 0.326 | 1 | 3 | Lrrfip1 | 0.019 | Old |
| 0.549813 | 0.145168 | 0.047 | 0.04 | 1 | 3 | Prdm10 | 0.007 | Old |
| 0.550656 | 0.058933 | 0.201 | 0.187 | 1 | 3 | Gabpb2 | 0.014 | Old |
| 0.550757 | 0.689909 | 0.02 | 0.016 | 1 | 3 | Gga2 | 0.004 | Old |
| 0.551568 | 0.276766 | 0.029 | 0.024 | 1 | 3 | Mrps27 | 0.005 | Old |
| 0.551599 | 0.067957 | 0.209 | 0.196 | 1 | 3 | Mrps25 | 0.013 | Old |
| 0.551875 | 0.003196 | 0.055 | 0.048 | 1 | 3 | Nit2 | 0.007 | Old |
| 0.552731 | 0.046173 | 0.388 | 0.365 | 1 | 3 | Lpp | 0.023 | Old |
| 0.552895 | 0.03582 | 0.129 | 0.117 | 1 | 3 | Proser1 | 0.012 | Old |
| 0.553091 | 0.359364 | 0.029 | 0.024 | 1 | 3 | Eefsec | 0.005 | Old |
| 0.553586 | 0.438996 | 0.05 | 0.044 | 1 | 3 | Myg1 | 0.006 | Old |
| 0.553885 | 0.219544 | 0.061 | 0.054 | 1 | 3 | Tmco6 | 0.007 | Old |
| 0.554389 | 0.146177 | 0.02 | 0.016 | 1 | 3 | Gm16701.: | 0.004 | Old |
| 0.554863 | 0.321543 | 0.103 | 0.093 | 1 | 3 | Rnf138 | 0.01 | Old |
| 0.555288 | 0.209377 | 0.049 | 0.043 | 1 | 3 | Sema4c | 0.006 | Old |
| 0.555602 | 0.244915 | 0.02 | 0.016 | 1 | 3 | Phka1 | 0.004 | Old |
| 0.555644 | 0.030987 | 0.214 | 0.198 | 1 | 3 | Rnf114 | 0.016 | Old |
| 0.556043 | 0.056306 | 0.429 | 0.403 | 1 | 3 | Trappc4 | 0.026 | Old |
| 0.55655 | 0.041355 | 0.108 | 0.097 | 1 | 3 | Smn1 | 0.011 | Old |
| 0.556571 | 0.067295 | 0.421 | 0.402 | 1 | 3 | Kmt2e | 0.019 | Old |
| 0.556674 | 0.185145 | 0.037 | 0.032 | 1 | 3 | Zbtb5 | 0.005 | Old |
| 0.556816 | 0.143643 | 0.067 | 0.059 | 1 | 3 | Lrwd1 | 0.008 | Old |
| 0.557278 | 0.108125 | 0.074 | 0.066 | 1 | 3 | N4bp2 | 0.008 | Old |
| 0.557433 | 0.601998 | 0.03 | 0.025 | 1 | 3 | Srbd1 | 0.005 | Old |
| 0.557679 | 0.023298 | 0.069 | 0.062 | 1 | 3 | Hps1 | 0.007 | Old |
| 0.558229 | 0.2018 | 0.038 | 0.033 | 1 | 3 | Lancl2 | 0.005 | Old |
| 0.558643 | 0.150023 | 0.092 | 0.083 | 1 | 3 | Coasy | 0.009 | Old |
| 0.559559 | 0.075169 | 0.164 | 0.152 | 1 | 3 | Tnpo2 | 0.012 | Old |

| | | | | | | | | |
|---|---|---|---|---|---|---|---|---|
| 0.559707 | 0.127586 | 0.151 | 0.139 | 1 | 3 | Atxn7l3 | 0.012 | Old |
| 0.560426 | 0.15838 | 0.133 | 0.122 | 1 | 3 | Stk3 | 0.011 | Old |
| 0.560437 | 0.221981 | 0.122 | 0.114 | 1 | 3 | Timm44 | 0.008 | Old |
| 0.561098 | 0.068239 | 0.121 | 0.111 | 1 | 3 | Ddx19a | 0.01 | Old |
| 0.561522 | 0.259725 | 0.096 | 0.087 | 1 | 3 | Atad3a | 0.009 | Old |
| 0.563162 | 0.006683 | 0.122 | 0.112 | 1 | 3 | Orai3 | 0.01 | Old |
| 0.564106 | 0.236379 | 0.07 | 0.063 | 1 | 3 | Nup133 | 0.007 | Old |
| 0.5646 | 0.357418 | 0.013 | 0.01 | 1 | 3 | Pecr | 0.003 | Old |
| 0.564676 | 0.242956 | 0.01 | 0.013 | 1 | 3 | Gm31597 | -0.003 | Old |
| 0.565366 | 0.422279 | 0.013 | 0.01 | 1 | 3 | Naip1 | 0.003 | Old |
| 0.565404 | 0.032368 | 0.202 | 0.187 | 1 | 3 | Utp14a | 0.015 | Old |
| 0.565416 | 0.070762 | 0.16 | 0.146 | 1 | 3 | Tdrd7 | 0.014 | Old |
| 0.565661 | 0.041643 | 0.02 | 0.016 | 1 | 3 | Trmt10b | 0.004 | Old |
| 0.565753 | 0.154669 | 0.011 | 0.014 | 1 | 3 | Tnfaip6 | -0.003 | Old |
| 0.566117 | 0.202339 | 0.207 | 0.194 | 1 | 3 | Chsy1 | 0.013 | Old |
| 0.566137 | 0.04671 | 0.276 | 0.259 | 1 | 3 | Fam32a | 0.017 | Old |
| 0.566837 | 0.129474 | 0.27 | 0.254 | 1 | 3 | Naga | 0.016 | Old |
| 0.567541 | 0.052675 | 0.041 | 0.047 | 1 | 3 | Gtf3c4 | -0.006 | Old |
| 0.568436 | 0.588583 | 0.013 | 0.01 | 1 | 3 | Cadm1 | 0.003 | Old |
| 0.568592 | 0.068262 | 0.271 | 0.254 | 1 | 3 | Eri1 | 0.017 | Old |
| 0.568821 | 0.210359 | 0.013 | 0.01 | 1 | 3 | Zbtb49 | 0.003 | Old |
| 0.569 | 0.03748 | 0.06 | 0.053 | 1 | 3 | Mfsd8 | 0.007 | Old |
| 0.569173 | 0.355517 | 0.022 | 0.018 | 1 | 3 | Tagap1 | 0.004 | Old |
| 0.56919 | 0.065483 | 0.137 | 0.129 | 1 | 3 | Ints9 | 0.008 | Old |
| 0.569746 | 0.06791 | 0.271 | 0.254 | 1 | 3 | Bscl2 | 0.017 | Old |
| 0.569872 | 0.400951 | 0.011 | 0.014 | 1 | 3 | Ciart | -0.003 | Old |
| 0.569903 | 0.107064 | 0.157 | 0.146 | 1 | 3 | Rnf146 | 0.011 | Old |
| 0.570053 | 0.329378 | 0.075 | 0.068 | 1 | 3 | Slc27a3 | 0.007 | Old |
| 0.570331 | 0.006596 | 0.489 | 0.467 | 1 | 3 | Esyt1 | 0.022 | Old |
| 0.570471 | 0.10815 | 0.03 | 0.025 | 1 | 3 | Dars2 | 0.005 | Old |
| 0.570818 | 0.223259 | 0.174 | 0.164 | 1 | 3 | Ptger2 | 0.01 | Old |
| 0.570998 | 0.022859 | 0.011 | 0.014 | 1 | 3 | Akr1c12 | -0.003 | Old |
| 0.571228 | 0.144176 | 0.03 | 0.025 | 1 | 3 | Cd200r3 | 0.005 | Old |
| 0.571614 | 0.025188 | 0.483 | 0.446 | 1 | 3 | Ets2 | 0.037 | Old |
| 0.571838 | 0.003225 | 0.045 | 0.039 | 1 | 3 | Clybl | 0.006 | Old |
| 0.572851 | 0.63691 | 0.031 | 0.027 | 1 | 3 | Lss | 0.004 | Old |
| 0.573047 | 0.080057 | 0.022 | 0.018 | 1 | 3 | Gm16701 | 0.004 | Old |
| 0.573733 | 0.00791 | 0.508 | 0.463 | 1 | 3 | Camkk2 | 0.045 | Old |
| 0.574064 | 0.049532 | 0.68 | 0.654 | 1 | 3 | Trps1 | 0.026 | Old |
| 0.574248 | 0.170378 | 0.104 | 0.096 | 1 | 3 | Fam126b | 0.008 | Old |
| 0.574348 | 0.037164 | 0.219 | 0.203 | 1 | 3 | Tor1b | 0.016 | Old |
| 0.574407 | 0.279988 | 0.041 | 0.035 | 1 | 3 | Plekhn1 | 0.006 | Old |
| 0.575085 | 0.363839 | 0.012 | 0.015 | 1 | 3 | Tcea3 | -0.003 | Old |
| 0.575185 | 0.039797 | 0.141 | 0.131 | 1 | 3 | Mcm3 | 0.01 | Old |
| 0.57578 | 0.230124 | 0.053 | 0.047 | 1 | 3 | Atm | 0.006 | Old |

| | | | | | | | | |
|---|---|---|---|---|---|---|---|---|
| 0.575833 | 0.043148 | 0.091 | 0.083 | 1 | 3 | Fam173b | 0.008 | Old |
| 0.576114 | 0.079568 | 0.223 | 0.211 | 1 | 3 | Chd9 | 0.012 | Old |
| 0.576582 | 0.003637 | 0.219 | 0.199 | 1 | 3 | Tbc1d13 | 0.02 | Old |
| 0.577655 | 0.028784 | 0.072 | 0.064 | 1 | 3 | Zbtb4 | 0.008 | Old |
| 0.577983 | 0.015817 | 0.149 | 0.14 | 1 | 3 | Ccl4 | 0.009 | Old |
| 0.578164 | 0.010903 | 0.194 | 0.182 | 1 | 3 | Nup88 | 0.012 | Old |
| 0.578477 | 0.099718 | 0.048 | 0.054 | 1 | 3 | Mcm7 | -0.006 | Old |
| 0.5785 | 0.024934 | 0.042 | 0.037 | 1 | 3 | Impa2 | 0.005 | Old |
| 0.578527 | 0.0146 | 0.14 | 0.129 | 1 | 3 | Rnf139 | 0.011 | Old |
| 0.579064 | 0.055778 | 0.099 | 0.09 | 1 | 3 | Med12 | 0.009 | Old |
| 0.579116 | 0.155447 | 0.039 | 0.034 | 1 | 3 | Zfp839 | 0.005 | Old |
| 0.57963 | 0.366804 | 0.022 | 0.018 | 1 | 3 | Hhat | 0.004 | Old |
| 0.579862 | 0.048841 | 0.128 | 0.117 | 1 | 3 | Rsbn1 | 0.011 | Old |
| 0.58046 | 0.061152 | 0.072 | 0.064 | 1 | 3 | Xrcc1 | 0.008 | Old |
| 0.580737 | 0.014328 | 0.594 | 0.568 | 1 | 3 | Top1 | 0.026 | Old |
| 0.580843 | 0.010087 | 0.228 | 0.212 | 1 | 3 | Impdh2 | 0.016 | Old |
| 0.580924 | 0.083676 | 0.041 | 0.035 | 1 | 3 | Nedd4l | 0.006 | Old |
| 0.58179 | 0.114619 | 0.1 | 0.092 | 1 | 3 | Pole3 | 0.008 | Old |
| 0.582058 | 0.142598 | 0.086 | 0.078 | 1 | 3 | Mtif2 | 0.008 | Old |
| 0.5826 | 0.00879 | 0.265 | 0.247 | 1 | 3 | Sys1 | 0.018 | Old |
| 0.583434 | 0.059018 | 0.71 | 0.67 | 1 | 3 | Jak1 | 0.04 | Old |
| 0.583524 | 0.184441 | 0.11 | 0.102 | 1 | 3 | Nectin1 | 0.008 | Old |
| 0.584376 | 0.165888 | 0.065 | 0.058 | 1 | 3 | Exosc7 | 0.007 | Old |
| 0.585939 | 0.165565 | 0.053 | 0.047 | 1 | 3 | Dedd | 0.006 | Old |
| 0.586877 | 0.231418 | 0.042 | 0.037 | 1 | 3 | 1110004E0 | 0.005 | Old |
| 0.587058 | 0.06949 | 0.013 | 0.016 | 1 | 3 | Zscan12 | -0.003 | Old |
| 0.587154 | 0.09556 | 0.022 | 0.018 | 1 | 3 | Alg12 | 0.004 | Old |
| 0.587939 | 0.366282 | 0.078 | 0.071 | 1 | 3 | Ocel1 | 0.007 | Old |
| 0.588409 | 0.084284 | 0.031 | 0.027 | 1 | 3 | Slc30a4 | 0.004 | Old |
| 0.588874 | 0.024651 | 0.143 | 0.134 | 1 | 3 | Ltbr | 0.009 | Old |
| 0.589016 | 0.044034 | 0.149 | 0.139 | 1 | 3 | Traf5 | 0.01 | Old |
| 0.589048 | 0.105948 | 0.194 | 0.182 | 1 | 3 | Rhoh | 0.012 | Old |
| 0.589276 | 0.33082 | 0.014 | 0.011 | 1 | 3 | Hyal2 | 0.003 | Old |
| 0.58958 | 0.122542 | 0.066 | 0.059 | 1 | 3 | Bud23 | 0.007 | Old |
| 0.590092 | 0.086157 | 0.104 | 0.095 | 1 | 3 | Oard1 | 0.009 | Old |
| 0.590393 | 0.363318 | 0.014 | 0.011 | 1 | 3 | Rnf122 | 0.003 | Old |
| 0.590548 | 0.012887 | 0.14 | 0.13 | 1 | 3 | Pgrmc2 | 0.01 | Old |
| 0.590826 | 0.008823 | 0.706 | 0.66 | 1 | 3 | Anp32b | 0.046 | Old |
| 0.591918 | 0.031388 | 0.128 | 0.119 | 1 | 3 | Ddx52 | 0.009 | Old |
| 0.592599 | 0.211333 | 0.022 | 0.018 | 1 | 3 | Krt10 | 0.004 | Old |
| 0.592827 | 0.038937 | 0.428 | 0.388 | 1 | 3 | Comt | 0.04 | Old |
| 0.59313 | 0.166666 | 0.128 | 0.119 | 1 | 3 | Ruvbl1 | 0.009 | Old |
| 0.593854 | 0.057189 | 0.053 | 0.047 | 1 | 3 | Mettl3 | 0.006 | Old |
| 0.594226 | 0.122122 | 0.042 | 0.037 | 1 | 3 | Dhx37 | 0.005 | Old |
| 0.594263 | 0.069029 | 0.066 | 0.059 | 1 | 3 | Scai | 0.007 | Old |

| | | | | | | | | |
|---|---|---|---|---|---|---|---|---|
| 0.594302 | 0.245563 | 0.023 | 0.019 | 1 | 3 | B230118H( | 0.004 | Old |
| 0.594434 | 0.153456 | 0.031 | 0.027 | 1 | 3 | Mxra7 | 0.004 | Old |
| 0.594622 | 0.038121 | 0.055 | 0.049 | 1 | 3 | Adck1 | 0.006 | Old |
| 0.594975 | 0.024616 | 0.876 | 0.845 | 1 | 3 | Naca | 0.031 | Old |
| 0.595687 | 0.124754 | 0.033 | 0.029 | 1 | 3 | Dhx58 | 0.004 | Old |
| 0.595882 | 0.239041 | 0.08 | 0.073 | 1 | 3 | Etfdh | 0.007 | Old |
| 0.596512 | 0.271894 | 0.032 | 0.028 | 1 | 3 | Iba57 | 0.004 | Old |
| 0.596707 | 0.316376 | 0.091 | 0.085 | 1 | 3 | Gm14548 | 0.006 | Old |
| 0.596738 | 0.213552 | 0.014 | 0.011 | 1 | 3 | Gatsl3 | 0.003 | Old |
| 0.596738 | 0.19135 | 0.014 | 0.011 | 1 | 3 | Gm39469 | 0.003 | Old |
| 0.596794 | 0.239481 | 0.044 | 0.039 | 1 | 3 | Extl2 | 0.005 | Old |
| 0.596957 | 0.314817 | 0.023 | 0.019 | 1 | 3 | Hes2 | 0.004 | Old |
| 0.597458 | 0.037851 | 0.031 | 0.027 | 1 | 3 | Mon1a | 0.004 | Old |
| 0.597525 | 0.07796 | 0.079 | 0.072 | 1 | 3 | Plscr3 | 0.007 | Old |
| 0.597689 | 0.258431 | 0.045 | 0.04 | 1 | 3 | Clpb | 0.005 | Old |
| 0.59786 | 0.062675 | 0.014 | 0.011 | 1 | 3 | Hist2h3c1 | 0.003 | Old |
| 0.597928 | 0.168294 | 0.043 | 0.038 | 1 | 3 | Ankrd37 | 0.005 | Old |
| 0.598211 | 0.225385 | 0.014 | 0.018 | 1 | 3 | Zfp472 | -0.004 | Old |
| 0.59874 | 0.051865 | 0.106 | 0.098 | 1 | 3 | Fam214a | 0.008 | Old |
| 0.599035 | 0.432045 | 0.024 | 0.02 | 1 | 3 | Icam2 | 0.004 | Old |
| 0.599426 | 0.003732 | 0.278 | 0.259 | 1 | 3 | Nudcd3 | 0.019 | Old |
| 0.599913 | 0.21941 | 0.076 | 0.069 | 1 | 3 | Cops7b | 0.007 | Old |
| 0.599985 | 0.002118 | 0.467 | 0.441 | 1 | 3 | Dlst | 0.026 | Old |
| 0.601655 | 0.117831 | 0.044 | 0.039 | 1 | 3 | Rfx5 | 0.005 | Old |
| 0.601697 | 0.211182 | 0.131 | 0.122 | 1 | 3 | Zfp560 | 0.009 | Old |
| 0.601732 | 0.149287 | 0.105 | 0.097 | 1 | 3 | Tubgcp3 | 0.008 | Old |
| 0.601986 | 0.395059 | 0.014 | 0.011 | 1 | 3 | Mapre3 | 0.003 | Old |
| 0.60234 | 0.048997 | 0.102 | 0.093 | 1 | 3 | Nup205 | 0.009 | Old |
| 0.603469 | 0.129319 | 0.023 | 0.019 | 1 | 3 | Mpp7 | 0.004 | Old |
| 0.603799 | 0.066392 | 0.042 | 0.037 | 1 | 3 | Mcm2 | 0.005 | Old |
| 0.603865 | 0.187458 | 0.014 | 0.011 | 1 | 3 | Cp | 0.003 | Old |
| 0.604247 | 0.014219 | 0.889 | 0.852 | 1 | 3 | Wnk1 | 0.037 | Old |
| 0.604821 | 0.09769 | 0.016 | 0.019 | 1 | 3 | Zfp770 | -0.003 | Old |
| 0.604873 | 0.022184 | 0.051 | 0.045 | 1 | 3 | 1700037H( | 0.006 | Old |
| 0.605844 | 0.025255 | 0.023 | 0.019 | 1 | 3 | 3110009E1 | 0.004 | Old |
| 0.606293 | 0.137028 | 0.056 | 0.051 | 1 | 3 | Mynn | 0.005 | Old |
| 0.608023 | 0.113788 | 0.044 | 0.039 | 1 | 3 | Hmgxb3 | 0.005 | Old |
| 0.609449 | 0.10787 | 0.076 | 0.069 | 1 | 3 | Gm48065 | 0.007 | Old |
| 0.609778 | 0.040306 | 0.167 | 0.155 | 1 | 3 | Sec23ip | 0.012 | Old |
| 0.610008 | 0.250786 | 0.024 | 0.02 | 1 | 3 | Ccdc66 | 0.004 | Old |
| 0.610723 | 0.053902 | 0.099 | 0.091 | 1 | 3 | Enox2 | 0.008 | Old |
| 0.611364 | 0.007382 | 0.148 | 0.138 | 1 | 3 | Mrps28 | 0.01 | Old |
| 0.611579 | 0.059944 | 0.116 | 0.107 | 1 | 3 | Phf12 | 0.009 | Old |
| 0.612534 | 0.084809 | 0.133 | 0.124 | 1 | 3 | Rps6kc1 | 0.009 | Old |
| 0.613845 | 0.36687 | 0.016 | 0.013 | 1 | 3 | Kbtbd8 | 0.003 | Old |

| | | | | | | | | |
|---|---|---|---|---|---|---|---|---|
| 0.614388 | 0.018182 | 0.211 | 0.197 | 1 | 3 | Ing2 | 0.014 | Old |
| 0.614399 | 0.170721 | 0.072 | 0.066 | 1 | 3 | Tmem87a | 0.006 | Old |
| 0.614732 | 0.102931 | 0.093 | 0.086 | 1 | 3 | Ttc39c | 0.007 | Old |
| 0.614932 | 0.340697 | 0.016 | 0.013 | 1 | 3 | Pgap3 | 0.003 | Old |
| 0.615097 | 0.466218 | 0.025 | 0.021 | 1 | 3 | Palm | 0.004 | Old |
| 0.615223 | 0.148254 | 0.018 | 0.021 | 1 | 3 | Traf3ip1 | -0.003 | Old |
| 0.615504 | 0.290608 | 0.045 | 0.04 | 1 | 3 | Ercc8 | 0.005 | Old |
| 0.615636 | 0.053043 | 0.029 | 0.033 | 1 | 3 | Rtn4ip1 | -0.004 | Old |
| 0.615774 | 0.046777 | 0.056 | 0.051 | 1 | 3 | Tom1l2 | 0.005 | Old |
| 0.61605 | 0.311785 | 0.035 | 0.03 | 1 | 3 | Ercc1 | 0.005 | Old |
| 0.616367 | 0.038329 | 0.608 | 0.567 | 1 | 3 | Lyn | 0.041 | Old |
| 0.616471 | 0.004998 | 0.191 | 0.199 | 1 | 3 | Cdc37l1 | -0.008 | Old |
| 0.616559 | 0.141737 | 0.045 | 0.04 | 1 | 3 | Amd1 | 0.005 | Old |
| 0.616689 | 0.253523 | 0.047 | 0.042 | 1 | 3 | Papolg | 0.005 | Old |
| 0.617552 | 0.003079 | 0.233 | 0.221 | 1 | 3 | Mtmr6 | 0.012 | Old |
| 0.618662 | 0.334653 | 0.07 | 0.064 | 1 | 3 | Zc3h12c | 0.006 | Old |
| 0.6187 | 0.097357 | 0.055 | 0.049 | 1 | 3 | Alg9 | 0.006 | Old |
| 0.618727 | 0.013751 | 0.037 | 0.042 | 1 | 3 | Thap4 | -0.005 | Old |
| 0.618889 | 0.030707 | 0.811 | 0.783 | 1 | 3 | Cd84 | 0.028 | Old |
| 0.619196 | 0.04096 | 0.401 | 0.38 | 1 | 3 | Snap23 | 0.021 | Old |
| 0.619354 | 0.236547 | 0.025 | 0.021 | 1 | 3 | 1110019D1 | 0.004 | Old |
| 0.620762 | 0.250676 | 0.078 | 0.071 | 1 | 3 | Sap30bp | 0.007 | Old |
| 0.621345 | 0.406834 | 0.025 | 0.021 | 1 | 3 | Vcpkmt | 0.004 | Old |
| 0.621361 | 0.039359 | 0.024 | 0.02 | 1 | 3 | Rsph3b | 0.004 | Old |
| 0.621594 | 0.009919 | 0.055 | 0.049 | 1 | 3 | Eci1 | 0.006 | Old |
| 0.622478 | 0.06268 | 0.057 | 0.052 | 1 | 3 | Zbtb34 | 0.005 | Old |
| 0.62253 | 0.05417 | 0.024 | 0.02 | 1 | 3 | R3hcc1 | 0.004 | Old |
| 0.623036 | 0.028461 | 0.192 | 0.181 | 1 | 3 | Insr | 0.011 | Old |
| 0.623218 | 0.031212 | 0.067 | 0.061 | 1 | 3 | Spire1 | 0.006 | Old |
| 0.624507 | 0.056181 | 0.154 | 0.143 | 1 | 3 | Copg1 | 0.011 | Old |
| 0.625485 | 0.07412 | 0.016 | 0.013 | 1 | 3 | Sirt4 | 0.003 | Old |
| 0.626041 | 0.128278 | 0.114 | 0.106 | 1 | 3 | Diablo | 0.008 | Old |
| 0.626193 | 0.026767 | 0.025 | 0.021 | 1 | 3 | Jmjd8 | 0.004 | Old |
| 0.626216 | 0.110658 | 0.016 | 0.013 | 1 | 3 | Gm36445 | 0.003 | Old |
| 0.626627 | 0.030981 | 0.64 | 0.616 | 1 | 3 | Clic1 | 0.024 | Old |
| 0.626642 | 0.332702 | 0.037 | 0.033 | 1 | 3 | Pde6d | 0.004 | Old |
| 0.626846 | 0.013619 | 0.06 | 0.054 | 1 | 3 | Gtpbp1 | 0.006 | Old |
| 0.626929 | 0.055791 | 0.092 | 0.085 | 1 | 3 | Acsl1 | 0.007 | Old |
| 0.626973 | 0.053158 | 0.036 | 0.032 | 1 | 3 | Creld1 | 0.004 | Old |
| 0.627312 | 0.045488 | 0.016 | 0.013 | 1 | 3 | Ctif | 0.003 | Old |
| 0.627444 | 0.030253 | 0.145 | 0.135 | 1 | 3 | Atp11a | 0.01 | Old |
| 0.627519 | 0.05085 | 0.154 | 0.144 | 1 | 3 | Ncbp1 | 0.01 | Old |
| 0.627563 | 0.226552 | 0.047 | 0.042 | 1 | 3 | Mllt1 | 0.005 | Old |
| 0.627702 | 0.405127 | 0.053 | 0.048 | 1 | 3 | Acy1 | 0.005 | Old |
| 0.627732 | 0.504434 | 0.039 | 0.035 | 1 | 3 | Dcxr | 0.004 | Old |

| | | | | | | | | |
|---|---|---|---|---|---|---|---|---|
| 0.628471 | 0.13294 | 0.037 | 0.042 | 1 | 3 | Toe1 | -0.005 | Old |
| 0.629133 | 0.073256 | 0.109 | 0.101 | 1 | 3 | Slc9a4 | 0.008 | Old |
| 0.630466 | 0.68021 | 0.017 | 0.014 | 1 | 3 | Cmbl | 0.003 | Old |
| 0.630535 | 0.151355 | 0.139 | 0.13 | 1 | 3 | Gla | 0.009 | Old |
| 0.630665 | 0.040394 | 0.177 | 0.167 | 1 | 3 | Tmem230 | 0.01 | Old |
| 0.630686 | 0.045178 | 0.069 | 0.063 | 1 | 3 | Zbed4 | 0.006 | Old |
| 0.630997 | 0.097807 | 0.08 | 0.073 | 1 | 3 | Dvl3 | 0.007 | Old |
| 0.631028 | 0.141123 | 0.189 | 0.181 | 1 | 3 | Klhl6 | 0.008 | Old |
| 0.631585 | 0.374955 | 0.057 | 0.052 | 1 | 3 | Ddx19b | 0.005 | Old |
| 0.631874 | 0.421903 | 0.017 | 0.014 | 1 | 3 | Gpsm1 | 0.003 | Old |
| 0.632216 | 0.102347 | 0.151 | 0.141 | 1 | 3 | Cops3 | 0.01 | Old |
| 0.6326 | 0.017743 | 0.999 | 0.999 | 1 | 3 | H2-Ab1 | 0 | Old |
| 0.632903 | 0.002661 | 0.134 | 0.125 | 1 | 3 | Mfap3 | 0.009 | Old |
| 0.633829 | 0.010952 | 0.068 | 0.062 | 1 | 3 | Miip | 0.006 | Old |
| 0.635649 | 0.071732 | 0.025 | 0.021 | 1 | 3 | Fam110a | 0.004 | Old |
| 0.635752 | 0.730829 | 0.017 | 0.014 | 1 | 3 | Cd320 | 0.003 | Old |
| 0.635848 | 0.001401 | 0.538 | 0.539 | 1 | 3 | Eif4ebp2 | -0.001 | Old |
| 0.635945 | 0.174596 | 0.166 | 0.157 | 1 | 3 | Gm28875 | 0.009 | Old |
| 0.636118 | 0.068205 | 0.663 | 0.649 | 1 | 3 | Bri3 | 0.014 | Old |
| 0.636436 | 0.166935 | 0.265 | 0.256 | 1 | 3 | Ppp1cc | 0.009 | Old |
| 0.637382 | 0.096696 | 0.029 | 0.033 | 1 | 3 | 2810001G2 | -0.004 | Old |
| 0.637768 | 0.107626 | 0.153 | 0.144 | 1 | 3 | Mrto4 | 0.009 | Old |
| 0.638005 | 0.035635 | 0.121 | 0.112 | 1 | 3 | Gse1 | 0.009 | Old |
| 0.638579 | 0.142256 | 0.017 | 0.014 | 1 | 3 | Ctu1 | 0.003 | Old |
| 0.638587 | 0.06443 | 0.109 | 0.101 | 1 | 3 | Vps45 | 0.008 | Old |
| 0.63943 | 0.009886 | 0.079 | 0.072 | 1 | 3 | Naa25 | 0.007 | Old |
| 0.64003 | 0.207964 | 0.081 | 0.076 | 1 | 3 | Trim36 | 0.005 | Old |
| 0.641057 | 0.130188 | 0.139 | 0.13 | 1 | 3 | Minpp1 | 0.009 | Old |
| 0.64109 | 0.024739 | 0.628 | 0.605 | 1 | 3 | 1810037I1 | 0.023 | Old |
| 0.64145 | 0.198279 | 0.079 | 0.086 | 1 | 3 | Tomm6 | -0.007 | Old |
| 0.641763 | 0.037827 | 0.076 | 0.083 | 1 | 3 | Rbbp5 | -0.007 | Old |
| 0.642052 | 0.128933 | 0.084 | 0.077 | 1 | 3 | Cdc27 | 0.007 | Old |
| 0.643125 | 0.00052 | 0.38 | 0.36 | 1 | 3 | Tm9sf2 | 0.02 | Old |
| 0.643178 | 0.288468 | 0.026 | 0.023 | 1 | 3 | Fbxo32 | 0.003 | Old |
| 0.643538 | 0.221285 | 0.017 | 0.014 | 1 | 3 | Pigb | 0.003 | Old |
| 0.644272 | 0.271369 | 0.158 | 0.149 | 1 | 3 | Ltn1 | 0.009 | Old |
| 0.644486 | 0.133975 | 0.057 | 0.052 | 1 | 3 | Adsl | 0.005 | Old |
| 0.644835 | 0.036438 | 0.129 | 0.121 | 1 | 3 | Rexo1 | 0.008 | Old |
| 0.646237 | 0.3044 | 0.048 | 0.043 | 1 | 3 | Rbmx2 | 0.005 | Old |
| 0.646579 | 0.178233 | 0.037 | 0.042 | 1 | 3 | Aldh18a1 | -0.005 | Old |
| 0.647026 | 0.08188 | 0.151 | 0.143 | 1 | 3 | Lrpprc | 0.008 | Old |
| 0.647028 | 0.058343 | 0.097 | 0.09 | 1 | 3 | Bckdha | 0.007 | Old |
| 0.647198 | 0.066707 | 0.037 | 0.033 | 1 | 3 | 1110032A0 | 0.004 | Old |
| 0.647706 | 0.093093 | 0.08 | 0.074 | 1 | 3 | Clpp | 0.006 | Old |
| 0.648157 | 0.076107 | 0.017 | 0.014 | 1 | 3 | Wdr6 | 0.003 | Old |

| | | | | | | | | |
|---|---|---|---|---|---|---|---|---|
| 0.648506 | 0.330921 | 0.065 | 0.059 | 1 | 3 | Map4k2 | 0.006 | Old |
| 0.648937 | 0.053921 | 0.073 | 0.067 | 1 | 3 | Mtmr4 | 0.006 | Old |
| 0.64916 | 0.017657 | 0.049 | 0.044 | 1 | 3 | Donson | 0.005 | Old |
| 0.650293 | 0.202666 | 0.037 | 0.033 | 1 | 3 | Rmnd1 | 0.004 | Old |
| 0.65037 | 0.108899 | 0.158 | 0.149 | 1 | 3 | Arhgap4 | 0.009 | Old |
| 0.650519 | 0.476973 | 0.05 | 0.045 | 1 | 3 | Plekhg5 | 0.005 | Old |
| 0.651854 | 0.118167 | 0.11 | 0.102 | 1 | 3 | Pak1 | 0.008 | Old |
| 0.652021 | 0.002026 | 0.068 | 0.062 | 1 | 3 | Msi2 | 0.006 | Old |
| 0.653227 | 0.222457 | 0.065 | 0.059 | 1 | 3 | Asb4 | 0.006 | Old |
| 0.653473 | 0.043626 | 0.643 | 0.595 | 1 | 3 | Tmem30a | 0.048 | Old |
| 0.653622 | 0.034777 | 0.188 | 0.176 | 1 | 3 | Xpo7 | 0.012 | Old |
| 0.654099 | 0.301028 | 0.039 | 0.035 | 1 | 3 | Ptdss2 | 0.004 | Old |
| 0.655545 | 0.278997 | 0.037 | 0.033 | 1 | 3 | Mex3b | 0.004 | Old |
| 0.656141 | 0.9224 | 0.01 | 0.008 | 1 | 3 | Arhgef4 | 0.002 | Old |
| 0.65615 | 0.01726 | 0.139 | 0.129 | 1 | 3 | Samd1 | 0.01 | Old |
| 0.656695 | 0.315282 | 0.063 | 0.058 | 1 | 3 | Cd209g | 0.005 | Old |
| 0.656815 | 0.016803 | 0.203 | 0.189 | 1 | 3 | Mau2 | 0.014 | Old |
| 0.657247 | 0.189485 | 0.063 | 0.058 | 1 | 3 | 6030458C1 | 0.005 | Old |
| 0.657462 | 0.53865 | 0.018 | 0.015 | 1 | 3 | AC163032. | 0.003 | Old |
| 0.657763 | 0.177388 | 0.098 | 0.092 | 1 | 3 | Emc4 | 0.006 | Old |
| 0.657827 | 0.096047 | 0.103 | 0.096 | 1 | 3 | Senp7 | 0.007 | Old |
| 0.658841 | 0.340477 | 0.018 | 0.015 | 1 | 3 | Ppp1r3d | 0.003 | Old |
| 0.659876 | 0.294036 | 0.018 | 0.015 | 1 | 3 | Aldoc | 0.003 | Old |
| 0.660188 | 0.073861 | 0.05 | 0.045 | 1 | 3 | Abt1 | 0.005 | Old |
| 0.660433 | 0.454013 | 0.01 | 0.008 | 1 | 3 | Trim45 | 0.002 | Old |
| 0.660567 | 0.240765 | 0.018 | 0.015 | 1 | 3 | Cdk20 | 0.003 | Old |
| 0.660911 | 0.675898 | 0.01 | 0.008 | 1 | 3 | Ddx59 | 0.002 | Old |
| 0.660911 | 0.392595 | 0.01 | 0.008 | 1 | 3 | Rgmb | 0.002 | Old |
| 0.661261 | 0.108388 | 0.027 | 0.024 | 1 | 3 | Ube2e2 | 0.003 | Old |
| 0.661385 | 0.241397 | 0.096 | 0.09 | 1 | 3 | Exosc10 | 0.006 | Old |
| 0.661388 | 1.100553 | 0.01 | 0.008 | 1 | 3 | Zfp438 | 0.002 | Old |
| 0.661603 | 0.127684 | 0.018 | 0.015 | 1 | 3 | Znhit3 | 0.003 | Old |
| 0.661866 | 0.459373 | 0.01 | 0.008 | 1 | 3 | Znrf3 | 0.002 | Old |
| 0.662344 | 0.440752 | 0.01 | 0.008 | 1 | 3 | Gpr84 | 0.002 | Old |
| 0.662622 | 0.045933 | 0.073 | 0.067 | 1 | 3 | Pgm2 | 0.006 | Old |
| 0.662822 | 0.479411 | 0.01 | 0.008 | 1 | 3 | Dner | 0.002 | Old |
| 0.662975 | 0.021646 | 0.037 | 0.033 | 1 | 3 | Atg4a | 0.004 | Old |
| 0.663301 | 0.28857 | 0.01 | 0.008 | 1 | 3 | Fndc10 | 0.002 | Old |
| 0.663437 | 0.145765 | 0.053 | 0.048 | 1 | 3 | 1500011B0 | 0.005 | Old |
| 0.663684 | 0.011451 | 0.311 | 0.29 | 1 | 3 | Acaa2 | 0.021 | Old |
| 0.663779 | 0.26984 | 0.01 | 0.008 | 1 | 3 | Gm9856 | 0.002 | Old |
| 0.663779 | 0.039787 | 0.01 | 0.008 | 1 | 3 | Mss51 | 0.002 | Old |
| 0.664258 | 0.605427 | 0.01 | 0.008 | 1 | 3 | Zbtb48 | 0.002 | Old |
| 0.664369 | 0.059303 | 0.018 | 0.015 | 1 | 3 | Repin1 | 0.003 | Old |
| 0.664526 | 0.046527 | 0.038 | 0.034 | 1 | 3 | Ccdc28a | 0.004 | Old |

| | | | | | | | | |
|---|---|---|---|---|---|---|---|---|
| 0.664893 | 0.017529 | 0.168 | 0.158 | 1 | 3 | Slc25a45 | 0.01 | Old |
| 0.665215 | 0.26189 | 0.01 | 0.008 | 1 | 3 | Serpinh1 | 0.002 | Old |
| 0.665305 | 0.042272 | 0.08 | 0.073 | 1 | 3 | Abhd13 | 0.007 | Old |
| 0.665694 | 0.155426 | 0.01 | 0.008 | 1 | 3 | Gm10320 | 0.002 | Old |
| 0.665694 | 0.11114 | 0.01 | 0.008 | 1 | 3 | Gm40304 | 0.002 | Old |
| 0.666428 | 0.055391 | 0.054 | 0.049 | 1 | 3 | Zfp568 | 0.005 | Old |
| 0.667132 | 0.074452 | 0.01 | 0.008 | 1 | 3 | Tmlhe | 0.002 | Old |
| 0.667132 | 0.001942 | 0.01 | 0.008 | 1 | 3 | Gm12355 | 0.002 | Old |
| 0.66741 | 0.028111 | 0.121 | 0.128 | 1 | 3 | Cxcl10 | -0.007 | Old |
| 0.667491 | 0.00765 | 0.767 | 0.76 | 1 | 3 | Aplp2 | 0.007 | Old |
| 0.668224 | 0.018542 | 0.159 | 0.149 | 1 | 3 | Dmtf1 | 0.01 | Old |
| 0.668571 | 0.006993 | 0.01 | 0.008 | 1 | 3 | Aplf | 0.002 | Old |
| 0.669293 | 0.045427 | 0.07 | 0.076 | 1 | 3 | Noc3l | -0.006 | Old |
| 0.669548 | 0.302401 | 0.039 | 0.035 | 1 | 3 | Trim23 | 0.004 | Old |
| 0.669938 | 0.267346 | 0.051 | 0.047 | 1 | 3 | Dhx57 | 0.004 | Old |
| 0.67112 | 0.223488 | 0.029 | 0.025 | 1 | 3 | Tepsin | 0.004 | Old |
| 0.672048 | 0.10621 | 0.352 | 0.335 | 1 | 3 | Ube2z | 0.017 | Old |
| 0.672151 | 0.140547 | 0.041 | 0.037 | 1 | 3 | Thop1 | 0.004 | Old |
| 0.672194 | 0.157227 | 0.108 | 0.101 | 1 | 3 | Desi1 | 0.007 | Old |
| 0.672756 | 0.688412 | 0.019 | 0.016 | 1 | 3 | Lrsam1 | 0.003 | Old |
| 0.673091 | 0.379064 | 0.019 | 0.016 | 1 | 3 | E2f8 | 0.003 | Old |
| 0.673871 | 0.253327 | 0.039 | 0.035 | 1 | 3 | Pigw | 0.004 | Old |
| 0.674338 | 0.114402 | 0.041 | 0.037 | 1 | 3 | Tfip11 | 0.004 | Old |
| 0.674407 | 0.096329 | 0.029 | 0.025 | 1 | 3 | Nek4 | 0.004 | Old |
| 0.675176 | 0.137324 | 0.081 | 0.087 | 1 | 3 | Thoc1 | -0.006 | Old |
| 0.67555 | 0.010743 | 0.188 | 0.177 | 1 | 3 | Cds2 | 0.011 | Old |
| 0.676316 | 0.00864 | 0.061 | 0.056 | 1 | 3 | Ddr1 | 0.005 | Old |
| 0.677248 | 0.022243 | 0.959 | 0.968 | 1 | 3 | H2-D1 | -0.009 | Old |
| 0.67826 | 0.007635 | 0.041 | 0.037 | 1 | 3 | Rmdn1 | 0.004 | Old |
| 0.6792 | 0.082455 | 0.191 | 0.182 | 1 | 3 | Gsdmd | 0.009 | Old |
| 0.679475 | 0.480548 | 0.043 | 0.039 | 1 | 3 | Slc7a7 | 0.004 | Old |
| 0.679484 | 0.051495 | 0.019 | 0.016 | 1 | 3 | Fntb | 0.003 | Old |
| 0.679833 | 0.180559 | 0.147 | 0.14 | 1 | 3 | Thbs1 | 0.007 | Old |
| 0.679877 | 0.406289 | 0.054 | 0.049 | 1 | 3 | Eva1b | 0.005 | Old |
| 0.680083 | 0.00569 | 0.476 | 0.446 | 1 | 3 | Ndufc2 | 0.03 | Old |
| 0.680381 | 0.130538 | 0.054 | 0.049 | 1 | 3 | Rev1 | 0.005 | Old |
| 0.680689 | 0.068949 | 0.063 | 0.058 | 1 | 3 | Med24 | 0.005 | Old |
| 0.680916 | 0.076338 | 0.065 | 0.059 | 1 | 3 | Tbc1d8 | 0.006 | Old |
| 0.681248 | 0.302336 | 0.03 | 0.027 | 1 | 3 | Fads6 | 0.003 | Old |
| 0.68134 | 0.365225 | 0.031 | 0.028 | 1 | 3 | Ptpn4 | 0.003 | Old |
| 0.681553 | 0.167474 | 0.062 | 0.057 | 1 | 3 | Osbpl2 | 0.005 | Old |
| 0.681845 | 0.204716 | 0.019 | 0.016 | 1 | 3 | Grk4 | 0.003 | Old |
| 0.682448 | 0.042555 | 0.114 | 0.105 | 1 | 3 | Ccdc127 | 0.009 | Old |
| 0.68262 | 0.278981 | 0.043 | 0.039 | 1 | 3 | Aurkb | 0.004 | Old |
| 0.683196 | 0.092417 | 0.019 | 0.016 | 1 | 3 | Lhpp | 0.003 | Old |

| | | | | | | | | |
|---|---|---|---|---|---|---|---|---|
| 0.683791 | 0.000381 | 0.573 | 0.543 | 1 | 3 | Nucks1 | 0.03 | Old |
| 0.684223 | 0.040465 | 0.22 | 0.205 | 1 | 3 | Emsy | 0.015 | Old |
| 0.684494 | 0.050802 | 0.171 | 0.16 | 1 | 3 | Pold4 | 0.011 | Old |
| 0.685161 | 0.535919 | 0.02 | 0.018 | 1 | 3 | Nubpl | 0.002 | Old |
| 0.685642 | 0.186678 | 0.042 | 0.038 | 1 | 3 | Dgkh | 0.004 | Old |
| 0.685642 | 0.111003 | 0.042 | 0.038 | 1 | 3 | Wdr55 | 0.004 | Old |
| 0.68578 | 0.049981 | 0.112 | 0.105 | 1 | 3 | Pttg1 | 0.007 | Old |
| 0.686229 | 0.019729 | 0.209 | 0.193 | 1 | 3 | Fam20b | 0.016 | Old |
| 0.686673 | 0.103177 | 0.105 | 0.098 | 1 | 3 | Cep250 | 0.007 | Old |
| 0.687571 | 0.017625 | 0.64 | 0.602 | 1 | 3 | Ubl5 | 0.038 | Old |
| 0.688269 | 0.084455 | 0.019 | 0.016 | 1 | 3 | Gm16287 | 0.003 | Old |
| 0.688669 | 0.006261 | 0.582 | 0.558 | 1 | 3 | Dock8 | 0.024 | Old |
| 0.690818 | 0.006043 | 0.153 | 0.144 | 1 | 3 | Psd4 | 0.009 | Old |
| 0.690861 | 0.020493 | 0.458 | 0.434 | 1 | 3 | Mcrip1 | 0.024 | Old |
| 0.690966 | 0.091547 | 0.102 | 0.096 | 1 | 3 | Carnmt1 | 0.006 | Old |
| 0.691062 | 0.414097 | 0.02 | 0.018 | 1 | 3 | Pcgf6 | 0.002 | Old |
| 0.691351 | 0.214334 | 0.06 | 0.056 | 1 | 3 | Caml | 0.004 | Old |
| 0.692302 | 0.077377 | 0.227 | 0.217 | 1 | 3 | Wdr37 | 0.01 | Old |
| 0.692729 | 0.350853 | 0.011 | 0.009 | 1 | 3 | Gm43672 | 0.002 | Old |
| 0.693064 | 0.202759 | 0.124 | 0.117 | 1 | 3 | Hectd4 | 0.007 | Old |
| 0.695365 | 0.07922 | 0.088 | 0.093 | 1 | 3 | Crot | -0.005 | Old |
| 0.695463 | 0.270995 | 0.011 | 0.009 | 1 | 3 | Hist2h2aa1 | 0.002 | Old |
| 0.695663 | 0.54421 | 0.02 | 0.018 | 1 | 3 | Mmab | 0.002 | Old |
| 0.695663 | 0.15364 | 0.02 | 0.018 | 1 | 3 | Kmo | 0.002 | Old |
| 0.695983 | 0.020371 | 0.276 | 0.265 | 1 | 3 | Gtf2a1 | 0.011 | Old |
| 0.696225 | 0.038957 | 0.153 | 0.143 | 1 | 3 | 3110043O2 | 0.01 | Old |
| 0.696322 | 0.022056 | 0.02 | 0.018 | 1 | 3 | Tnnt1 | 0.002 | Old |
| 0.696884 | 0.019307 | 0.227 | 0.215 | 1 | 3 | Trim41 | 0.012 | Old |
| 0.697134 | 0.156727 | 0.119 | 0.114 | 1 | 3 | Arhgap11a | 0.005 | Old |
| 0.697244 | 0.371393 | 0.075 | 0.071 | 1 | 3 | Pcyox1l | 0.004 | Old |
| 0.698326 | 0.166444 | 0.143 | 0.136 | 1 | 3 | Timm8a1 | 0.007 | Old |
| 0.698844 | 0.005311 | 0.343 | 0.327 | 1 | 3 | Pcnp | 0.016 | Old |
| 0.700368 | 0.303748 | 0.084 | 0.078 | 1 | 3 | Mmp12 | 0.006 | Old |
| 0.700402 | 0.238583 | 0.032 | 0.029 | 1 | 3 | Mpi | 0.003 | Old |
| 0.701084 | 0.088744 | 0.048 | 0.052 | 1 | 3 | Cul4b | -0.004 | Old |
| 0.702053 | 0.045704 | 0.043 | 0.039 | 1 | 3 | Slc16a6 | 0.004 | Old |
| 0.702221 | 0.023797 | 0.053 | 0.048 | 1 | 3 | Rpp30 | 0.005 | Old |
| 0.702487 | 0.241771 | 0.059 | 0.054 | 1 | 3 | Eif2d | 0.005 | Old |
| 0.703653 | 0.011576 | 0.719 | 0.669 | 1 | 3 | Syk | 0.05 | Old |
| 0.70367 | 0.146907 | 0.044 | 0.04 | 1 | 3 | Gm44751 | 0.004 | Old |
| 0.704118 | 0.009369 | 0.044 | 0.04 | 1 | 3 | Pwwp2b | 0.004 | Old |
| 0.704145 | 0.194178 | 0.011 | 0.009 | 1 | 3 | Ly6a | 0.002 | Old |
| 0.705559 | 0.094718 | 0.057 | 0.053 | 1 | 3 | Bnip3 | 0.004 | Old |
| 0.706716 | 0.103701 | 0.069 | 0.064 | 1 | 3 | Poc1b | 0.005 | Old |
| 0.707067 | 0.019414 | 0.344 | 0.331 | 1 | 3 | Rab1b | 0.013 | Old |

| | | | | | | | | |
|---|---|---|---|---|---|---|---|---|
| 0.707216 | 0.041987 | 0.104 | 0.097 | 1 | 3 | Slc35c1 | 0.007 | Old |
| 0.707872 | 0.181372 | 0.13 | 0.124 | 1 | 3 | Wdr41 | 0.006 | Old |
| 0.707933 | 0.021647 | 0.044 | 0.04 | 1 | 3 | Bloc1s3 | 0.004 | Old |
| 0.708183 | 0.025551 | 0.161 | 0.154 | 1 | 3 | Spen | 0.007 | Old |
| 0.708482 | 0.201689 | 0.13 | 0.124 | 1 | 3 | Gcc1 | 0.006 | Old |
| 0.709063 | 0.107914 | 0.022 | 0.019 | 1 | 3 | Wdr73 | 0.003 | Old |
| 0.710065 | 0.012246 | 0.716 | 0.691 | 1 | 3 | Set | 0.025 | Old |
| 0.710471 | 0.074494 | 0.096 | 0.091 | 1 | 3 | Bid | 0.005 | Old |
| 0.710596 | 0.138712 | 0.237 | 0.23 | 1 | 3 | Osbpl8 | 0.007 | Old |
| 0.710857 | 0.171582 | 0.031 | 0.028 | 1 | 3 | Strbp | 0.003 | Old |
| 0.71119 | 0.160651 | 0.055 | 0.051 | 1 | 3 | Trp53bp1 | 0.004 | Old |
| 0.711825 | 0.028539 | 0.397 | 0.372 | 1 | 3 | Pten | 0.025 | Old |
| 0.712764 | 0.055242 | 0.521 | 0.496 | 1 | 3 | Stk17b | 0.025 | Old |
| 0.713193 | 0.018647 | 0.836 | 0.812 | 1 | 3 | Unc93b1 | 0.024 | Old |
| 0.713269 | 0.046823 | 0.06 | 0.064 | 1 | 3 | Ppcdc | -0.004 | Old |
| 0.714206 | 0.018545 | 0.751 | 0.729 | 1 | 3 | H13 | 0.022 | Old |
| 0.71428 | 0.017547 | 0.886 | 0.87 | 1 | 3 | Tmsb10 | 0.016 | Old |
| 0.71488 | 0.529436 | 0.008 | 0.01 | 1 | 3 | Tbc1d16 | -0.002 | Old |
| 0.715009 | 0.110983 | 0.07 | 0.066 | 1 | 3 | Scn1b | 0.004 | Old |
| 0.715239 | 0.279322 | 0.023 | 0.02 | 1 | 3 | Ssx2ip | 0.003 | Old |
| 0.716307 | 0.081497 | 0.008 | 0.01 | 1 | 3 | Ccr7 | -0.002 | Old |
| 0.716783 | 0.058773 | 0.008 | 0.01 | 1 | 3 | Zscan22 | -0.002 | Old |
| 0.717171 | 0.057819 | 0.131 | 0.139 | 1 | 3 | Psip1 | -0.008 | Old |
| 0.717686 | 0.05761 | 0.105 | 0.1 | 1 | 3 | Erf | 0.005 | Old |
| 0.717921 | 0.015522 | 0.05 | 0.054 | 1 | 3 | Zbtb14 | -0.004 | Old |
| 0.717992 | 0.061009 | 0.012 | 0.01 | 1 | 3 | Zfp507 | 0.002 | Old |
| 0.718083 | 0.196166 | 0.022 | 0.019 | 1 | 3 | BC017158 | 0.003 | Old |
| 0.719733 | 0.23833 | 0.012 | 0.01 | 1 | 3 | Ftx | 0.002 | Old |
| 0.720519 | 0.055595 | 0.087 | 0.082 | 1 | 3 | Gosr1 | 0.005 | Old |
| 0.720604 | 0.105548 | 0.012 | 0.01 | 1 | 3 | Gm42670 | 0.002 | Old |
| 0.721459 | 0.070369 | 0.029 | 0.032 | 1 | 3 | Cnnm2 | -0.003 | Old |
| 0.721475 | 0.086296 | 0.012 | 0.01 | 1 | 3 | Mtus1 | 0.002 | Old |
| 0.721556 | 0.272945 | 0.048 | 0.044 | 1 | 3 | Zfp467 | 0.004 | Old |
| 0.721911 | 0.334118 | 0.012 | 0.01 | 1 | 3 | Tyms | 0.002 | Old |
| 0.723256 | 0.00183 | 0.341 | 0.323 | 1 | 3 | Slc11a1 | 0.018 | Old |
| 0.723326 | 0.015741 | 0.029 | 0.032 | 1 | 3 | Slc25a10 | -0.003 | Old |
| 0.724006 | 0.001832 | 0.025 | 0.028 | 1 | 3 | Gm35154 | -0.003 | Old |
| 0.724091 | 0.198281 | 0.012 | 0.01 | 1 | 3 | Csf1 | 0.002 | Old |
| 0.724358 | 0.367615 | 0.01 | 0.011 | 1 | 3 | Ldlrad3 | -0.001 | Old |
| 0.725838 | 0.044371 | 0.012 | 0.01 | 1 | 3 | 1810055G( | 0.002 | Old |
| 0.725974 | 0.09317 | 0.059 | 0.054 | 1 | 3 | Zfp26 | 0.005 | Old |
| 0.726276 | 0.114108 | 0.06 | 0.056 | 1 | 3 | Hacd2 | 0.004 | Old |
| 0.726755 | 0.074076 | 0.078 | 0.073 | 1 | 3 | Lsm1 | 0.005 | Old |
| 0.727352 | 0.004901 | 0.06 | 0.056 | 1 | 3 | Snapc3 | 0.004 | Old |
| 0.72779 | 0.070644 | 0.022 | 0.019 | 1 | 3 | Rbks | 0.003 | Old |

| | | | | | | | | |
|---|---|---|---|---|---|---|---|---|
| 0.728121 | 0.267785 | 0.023 | 0.02 | 1 | 3 | AW209491 | 0.003 | Old |
| 0.728464 | 0.103682 | 0.105 | 0.098 | 1 | 3 | Gk | 0.007 | Old |
| 0.729175 | 0.039323 | 0.033 | 0.03 | 1 | 3 | Gm9844 | 0.003 | Old |
| 0.729315 | 0.08401 | 0.109 | 0.114 | 1 | 3 | Gm15417 | -0.005 | Old |
| 0.729378 | 0.374321 | 0.011 | 0.013 | 1 | 3 | Chd3os | -0.002 | Old |
| 0.729445 | 0.019606 | 0.707 | 0.673 | 1 | 3 | Myh9 | 0.034 | Old |
| 0.730207 | 0.323074 | 0.037 | 0.034 | 1 | 3 | Ddx60 | 0.003 | Old |
| 0.730771 | 0.026712 | 0.141 | 0.135 | 1 | 3 | Trim33 | 0.006 | Old |
| 0.731328 | 0.10448 | 0.035 | 0.032 | 1 | 3 | Zfp143 | 0.003 | Old |
| 0.731824 | 0.011977 | 0.102 | 0.096 | 1 | 3 | Khnyn | 0.006 | Old |
| 0.731986 | 0.01796 | 0.012 | 0.014 | 1 | 3 | Fer | -0.002 | Old |
| 0.732222 | 0.147489 | 0.023 | 0.02 | 1 | 3 | Acvr2a | 0.003 | Old |
| 0.73367 | 0.024158 | 0.254 | 0.261 | 1 | 3 | Wars | -0.007 | Old |
| 0.734176 | 0.126112 | 0.013 | 0.015 | 1 | 3 | AI480526 | -0.002 | Old |
| 0.735642 | 0.169636 | 0.012 | 0.014 | 1 | 3 | Ttc28 | -0.002 | Old |
| 0.736119 | 0.039713 | 0.013 | 0.015 | 1 | 3 | Atxn7l2 | -0.002 | Old |
| 0.736362 | 0.043276 | 0.016 | 0.018 | 1 | 3 | Ogfod3 | -0.002 | Old |
| 0.736878 | 0.120458 | 0.179 | 0.172 | 1 | 3 | Agtpbp1 | 0.007 | Old |
| 0.73704 | 0.183473 | 0.036 | 0.039 | 1 | 3 | Mrpl49 | -0.003 | Old |
| 0.73728 | 0.000263 | 0.023 | 0.02 | 1 | 3 | Map3k10 | 0.003 | Old |
| 0.738151 | 0.189173 | 0.054 | 0.051 | 1 | 3 | Ccl25 | 0.003 | Old |
| 0.738165 | 0.007618 | 0.07 | 0.074 | 1 | 3 | Tsta3 | -0.004 | Old |
| 0.738393 | 0.649061 | 0.013 | 0.011 | 1 | 3 | Zswim1 | 0.002 | Old |
| 0.738432 | 0.100845 | 0.032 | 0.035 | 1 | 3 | Map3k6 | -0.003 | Old |
| 0.738455 | 0.014763 | 0.33 | 0.328 | 1 | 3 | Krt80 | 0.002 | Old |
| 0.738543 | 0.12065 | 0.168 | 0.162 | 1 | 3 | Limd2 | 0.006 | Old |
| 0.73855 | 0.286391 | 0.038 | 0.035 | 1 | 3 | Mccc1 | 0.003 | Old |
| 0.739598 | 0.015255 | 0.016 | 0.018 | 1 | 3 | Csrp2 | -0.002 | Old |
| 0.739644 | 0.285807 | 0.013 | 0.011 | 1 | 3 | Cdk5rap1 | 0.002 | Old |
| 0.740236 | 0.058445 | 0.637 | 0.619 | 1 | 3 | Aldoa | 0.018 | Old |
| 0.740478 | 0.083473 | 0.013 | 0.011 | 1 | 3 | Nfib | 0.002 | Old |
| 0.741482 | 0.112664 | 0.065 | 0.061 | 1 | 3 | Rnf34 | 0.004 | Old |
| 0.741603 | 0.07948 | 0.018 | 0.02 | 1 | 3 | Dtx3 | -0.002 | Old |
| 0.74173 | 0.317956 | 0.013 | 0.011 | 1 | 3 | Uhrf1bp1 | 0.002 | Old |
| 0.741881 | 0.039296 | 0.563 | 0.535 | 1 | 3 | G3bp2 | 0.028 | Old |
| 0.741917 | 0.108227 | 0.256 | 0.244 | 1 | 3 | 4930455G( | 0.012 | Old |
| 0.743288 | 0.021431 | 0.207 | 0.199 | 1 | 3 | Synrg | 0.008 | Old |
| 0.744225 | 0.147544 | 0.022 | 0.024 | 1 | 3 | Mypopos | -0.002 | Old |
| 0.744266 | 0.086423 | 0.017 | 0.019 | 1 | 3 | Ybey | -0.002 | Old |
| 0.744614 | 0.000448 | 0.017 | 0.019 | 1 | 3 | Cptp | -0.002 | Old |
| 0.74497 | 0.483509 | 0.054 | 0.051 | 1 | 3 | Galnt4 | 0.003 | Old |
| 0.745188 | 0.075722 | 0.202 | 0.194 | 1 | 3 | Psmc2 | 0.008 | Old |
| 0.745491 | 0.196102 | 0.013 | 0.011 | 1 | 3 | Dtl | 0.002 | Old |
| 0.745795 | 0.128078 | 0.031 | 0.034 | 1 | 3 | Pick1 | -0.003 | Old |
| 0.746195 | 0.074248 | 0.094 | 0.09 | 1 | 3 | Fbxw8 | 0.004 | Old |

| | | | | | | | | |
|---|---|---|---|---|---|---|---|---|
| 0.746698 | 0.076669 | 0.024 | 0.021 | 1 | 3 | Dnajb5 | 0.003 | Old |
| 0.74755 | 0.097315 | 0.037 | 0.034 | 1 | 3 | Sec16b | 0.003 | Old |
| 0.74859 | 0.017115 | 0.073 | 0.068 | 1 | 3 | Blcap | 0.005 | Old |
| 0.748686 | 0.218649 | 0.038 | 0.035 | 1 | 3 | Paip2b | 0.003 | Old |
| 0.748983 | 0.033015 | 0.053 | 0.049 | 1 | 3 | Zfp715 | 0.004 | Old |
| 0.750044 | 0.305888 | 0.025 | 0.023 | 1 | 3 | C130013H( | 0.002 | Old |
| 0.750794 | 0.003856 | 0.209 | 0.212 | 1 | 3 | Agfg1 | -0.003 | Old |
| 0.751023 | 0.075628 | 0.068 | 0.064 | 1 | 3 | Mbd5 | 0.004 | Old |
| 0.751087 | 0.006233 | 0.023 | 0.02 | 1 | 3 | Hpgds | 0.003 | Old |
| 0.752436 | 0.032248 | 0.08 | 0.076 | 1 | 3 | Senp3 | 0.004 | Old |
| 0.752779 | 0.018744 | 0.037 | 0.034 | 1 | 3 | Tlr7 | 0.003 | Old |
| 0.753028 | 0.036928 | 0.037 | 0.034 | 1 | 3 | Msh6 | 0.003 | Old |
| 0.753537 | 0.123901 | 0.162 | 0.157 | 1 | 3 | Akna | 0.005 | Old |
| 0.755059 | 0.502205 | 0.038 | 0.035 | 1 | 3 | Zswim7 | 0.003 | Old |
| 0.755353 | 0.007814 | 0.216 | 0.22 | 1 | 3 | Fcgrt | -0.004 | Old |
| 0.756176 | 0.24795 | 0.042 | 0.039 | 1 | 3 | Zmynd19 | 0.003 | Old |
| 0.758044 | 0.06522 | 0.1 | 0.096 | 1 | 3 | Txndc16 | 0.004 | Old |
| 0.759443 | 0.049884 | 0.098 | 0.093 | 1 | 3 | Neurl2 | 0.005 | Old |
| 0.759615 | 0.043199 | 0.331 | 0.319 | 1 | 3 | Srpr | 0.012 | Old |
| 0.762282 | 0.137957 | 0.108 | 0.102 | 1 | 3 | Trappc3 | 0.006 | Old |
| 0.762451 | 0.002451 | 0.025 | 0.023 | 1 | 3 | Uros | 0.002 | Old |
| 0.762754 | 0.09299 | 0.025 | 0.023 | 1 | 3 | Gatc | 0.002 | Old |
| 0.763632 | 0.246167 | 0.014 | 0.013 | 1 | 3 | Adamts6 | 0.001 | Old |
| 0.764894 | 0.131426 | 0.026 | 0.024 | 1 | 3 | Dpy19l3 | 0.002 | Old |
| 0.766525 | 0.067591 | 0.177 | 0.169 | 1 | 3 | Clasp2 | 0.008 | Old |
| 0.767956 | 0.161153 | 0.027 | 0.025 | 1 | 3 | Mlh3 | 0.002 | Old |
| 0.768426 | 0.095621 | 0.067 | 0.063 | 1 | 3 | Acad8 | 0.004 | Old |
| 0.76846 | 0.294109 | 0.014 | 0.013 | 1 | 3 | 4930481A1 | 0.001 | Old |
| 0.768863 | 0.03078 | 0.014 | 0.013 | 1 | 3 | Rida | 0.001 | Old |
| 0.770276 | 0.490587 | 0.027 | 0.025 | 1 | 3 | Depdc7 | 0.002 | Old |
| 0.770911 | 0.014561 | 0.091 | 0.086 | 1 | 3 | Prdx4 | 0.005 | Old |
| 0.771182 | 0.01831 | 0.146 | 0.138 | 1 | 3 | Glo1 | 0.008 | Old |
| 0.771281 | 0.234241 | 0.014 | 0.013 | 1 | 3 | Mmachc | 0.001 | Old |
| 0.771559 | 0.107444 | 0.082 | 0.087 | 1 | 3 | Smpd1 | -0.005 | Old |
| 0.771744 | 0.07653 | 0.097 | 0.101 | 1 | 3 | Kin | -0.004 | Old |
| 0.773415 | 0.035936 | 0.172 | 0.164 | 1 | 3 | Ppfia1 | 0.008 | Old |
| 0.773418 | 0.304012 | 0.078 | 0.073 | 1 | 3 | 2410004B1 | 0.005 | Old |
| 0.773906 | 0.17084 | 0.038 | 0.035 | 1 | 3 | H2-Ob | 0.003 | Old |
| 0.774243 | 0.045005 | 0.344 | 0.327 | 1 | 3 | Rell1 | 0.017 | Old |
| 0.775258 | 0.285781 | 0.056 | 0.053 | 1 | 3 | Pdpr | 0.003 | Old |
| 0.776375 | 0.044353 | 0.088 | 0.085 | 1 | 3 | Usp10 | 0.003 | Old |
| 0.77659 | 0.102783 | 0.056 | 0.053 | 1 | 3 | Saal1 | 0.003 | Old |
| 0.776694 | 0.175662 | 0.099 | 0.095 | 1 | 3 | Tbc1d17 | 0.004 | Old |
| 0.777229 | 0.182323 | 0.073 | 0.069 | 1 | 3 | Stx17 | 0.004 | Old |
| 0.777559 | 0.036823 | 0.055 | 0.052 | 1 | 3 | Rab40c | 0.003 | Old |

| | | | | | | | | |
|---|---|---|---|---|---|---|---|---|
| 0.779288 | 0.001374 | 0.938 | 0.939 | 1 | 3 | Lsp1 | -0.001 | Old |
| 0.779538 | 0.162691 | 0.091 | 0.087 | 1 | 3 | Trappc2 | 0.004 | Old |
| 0.780112 | 0.004472 | 0.204 | 0.198 | 1 | 3 | Slc36a1 | 0.006 | Old |
| 0.780244 | 0.422578 | 0.016 | 0.014 | 1 | 3 | Trim59 | 0.002 | Old |
| 0.781051 | 0.184557 | 0.1 | 0.096 | 1 | 3 | Pik3r4 | 0.004 | Old |
| 0.781791 | 0.145696 | 0.044 | 0.042 | 1 | 3 | Tab1 | 0.002 | Old |
| 0.78244 | 0.082414 | 0.238 | 0.229 | 1 | 3 | Ctcf | 0.009 | Old |
| 0.7835 | 0.230967 | 0.045 | 0.043 | 1 | 3 | Rab2b | 0.002 | Old |
| 0.784276 | 0.141744 | 0.142 | 0.136 | 1 | 3 | Oaz2 | 0.006 | Old |
| 0.785048 | 0.26646 | 0.136 | 0.134 | 1 | 3 | Zfp62 | 0.002 | Old |
| 0.785276 | 0.062818 | 0.029 | 0.027 | 1 | 3 | Det1 | 0.002 | Old |
| 0.786812 | 0.112796 | 0.051 | 0.054 | 1 | 3 | Fdxr | -0.003 | Old |
| 0.787274 | 0.298124 | 0.029 | 0.027 | 1 | 3 | Mki67 | 0.002 | Old |
| 0.787275 | 0.224084 | 0.056 | 0.053 | 1 | 3 | Flad1 | 0.003 | Old |
| 0.787742 | 0.331079 | 0.027 | 0.025 | 1 | 3 | Aldh4a1 | 0.002 | Old |
| 0.787821 | 0.213576 | 0.063 | 0.061 | 1 | 3 | Trem2 | 0.002 | Old |
| 0.78857 | 0.01983 | 0.09 | 0.086 | 1 | 3 | Ramp3 | 0.004 | Old |
| 0.789494 | 0.145206 | 0.027 | 0.025 | 1 | 3 | Rfxank | 0.002 | Old |
| 0.789683 | 0.036867 | 0.094 | 0.09 | 1 | 3 | Hacd3 | 0.004 | Old |
| 0.790023 | 0.109371 | 0.072 | 0.068 | 1 | 3 | Pigyl | 0.004 | Old |
| 0.790101 | 0.166294 | 0.045 | 0.043 | 1 | 3 | Dexi | 0.002 | Old |
| 0.792174 | 0.263578 | 0.044 | 0.042 | 1 | 3 | 2510002D2 | 0.002 | Old |
| 0.792426 | 0.019 | 0.056 | 0.053 | 1 | 3 | Nek6 | 0.003 | Old |
| 0.793564 | 0.095516 | 0.029 | 0.027 | 1 | 3 | Ranbp6 | 0.002 | Old |
| 0.793587 | 0.084449 | 0.084 | 0.08 | 1 | 3 | Flt3l | 0.004 | Old |
| 0.794281 | 0.002648 | 0.085 | 0.081 | 1 | 3 | C2cd3 | 0.004 | Old |
| 0.794904 | 0.178881 | 0.074 | 0.071 | 1 | 3 | Siae | 0.003 | Old |
| 0.79587 | 0.12525 | 0.057 | 0.054 | 1 | 3 | Utp23 | 0.003 | Old |
| 0.796459 | 0.071619 | 0.289 | 0.293 | 1 | 3 | Pdxk | -0.004 | Old |
| 0.796762 | 0.236955 | 0.017 | 0.015 | 1 | 3 | Samd10 | 0.002 | Old |
| 0.797611 | 0.043036 | 0.08 | 0.083 | 1 | 3 | Aga | -0.003 | Old |
| 0.798497 | 0.001279 | 0.375 | 0.357 | 1 | 3 | Hip1 | 0.018 | Old |
| 0.79898 | 0.025234 | 0.042 | 0.039 | 1 | 3 | Tmcc2 | 0.003 | Old |
| 0.800511 | 0.123246 | 0.017 | 0.015 | 1 | 3 | Nup43 | 0.002 | Old |
| 0.800676 | 0.417095 | 0.147 | 0.143 | 1 | 3 | Gbp2 | 0.004 | Old |
| 0.801013 | 0.127798 | 0.03 | 0.028 | 1 | 3 | Chchd6 | 0.002 | Old |
| 0.801014 | 0.004707 | 0.029 | 0.027 | 1 | 3 | Mcf2l | 0.002 | Old |
| 0.801504 | 0.021677 | 0.06 | 0.063 | 1 | 3 | As3mt | -0.003 | Old |
| 0.801637 | 0.251826 | 0.017 | 0.015 | 1 | 3 | 9130008F2 | 0.002 | Old |
| 0.803731 | 0.021173 | 0.373 | 0.359 | 1 | 3 | Il10ra | 0.014 | Old |
| 0.804504 | 0.027435 | 0.048 | 0.051 | 1 | 3 | Snx19 | -0.003 | Old |
| 0.805768 | 0.455301 | 0.017 | 0.015 | 1 | 3 | Thumpd2 | 0.002 | Old |
| 0.806896 | 0.021039 | 0.017 | 0.015 | 1 | 3 | Gm30074 | 0.002 | Old |
| 0.807272 | 0.026463 | 0.017 | 0.015 | 1 | 3 | Gpt2 | 0.002 | Old |
| 0.808879 | 0.0383 | 0.579 | 0.557 | 1 | 3 | Prpf38b | 0.022 | Old |

| | | | | | | | | |
|---|---|---|---|---|---|---|---|---|
| 0.809305 | 0.044901 | 0.045 | 0.043 | 1 | 3 | Cd3eap | 0.002 | Old |
| 0.809649 | 0.074633 | 0.087 | 0.083 | 1 | 3 | Fbxo30 | 0.004 | Old |
| 0.810228 | 0.025133 | 0.625 | 0.62 | 1 | 3 | Pirb | 0.005 | Old |
| 0.810408 | 0.351567 | 0.018 | 0.016 | 1 | 3 | Szt2 | 0.002 | Old |
| 0.811154 | 0.088792 | 0.121 | 0.116 | 1 | 3 | Pkib | 0.005 | Old |
| 0.811366 | 0.015907 | 0.891 | 0.857 | 1 | 3 | Pcbp2 | 0.034 | Old |
| 0.811433 | 0.146582 | 0.057 | 0.061 | 1 | 3 | Thap3 | -0.004 | Old |
| 0.812646 | 0.080003 | 0.116 | 0.119 | 1 | 3 | Vac14 | -0.003 | Old |
| 0.812779 | 0.180536 | 0.031 | 0.029 | 1 | 3 | Dcp1b | 0.002 | Old |
| 0.814003 | 0.270541 | 0.032 | 0.03 | 1 | 3 | Dnajc12 | 0.002 | Old |
| 0.814766 | 0.007681 | 0.018 | 0.016 | 1 | 3 | Atp6v1g2 | 0.002 | Old |
| 0.814915 | 0.049992 | 0.102 | 0.097 | 1 | 3 | Kat7 | 0.005 | Old |
| 0.815144 | 0.048579 | 0.044 | 0.042 | 1 | 3 | Map3k20 | 0.002 | Old |
| 0.815803 | 0.066589 | 0.097 | 0.092 | 1 | 3 | Spn | 0.005 | Old |
| 0.817534 | 0.021602 | 0.145 | 0.139 | 1 | 3 | Lcorl | 0.006 | Old |
| 0.819151 | 0.0423 | 0.094 | 0.091 | 1 | 3 | Dnajc30 | 0.003 | Old |
| 0.819857 | 0.22709 | 0.018 | 0.016 | 1 | 3 | Dnaaf5 | 0.002 | Old |
| 0.822328 | 0.038861 | 0.079 | 0.076 | 1 | 3 | Ikbkap | 0.003 | Old |
| 0.822509 | 0.047265 | 0.232 | 0.234 | 1 | 3 | Areg | -0.002 | Old |
| 0.822652 | 0.170304 | 0.035 | 0.033 | 1 | 3 | Rabep2 | 0.002 | Old |
| 0.822958 | 0.069116 | 0.151 | 0.146 | 1 | 3 | Tnks | 0.005 | Old |
| 0.823069 | 0.292697 | 0.05 | 0.048 | 1 | 3 | Kctd3 | 0.002 | Old |
| 0.823316 | 0.051832 | 0.171 | 0.167 | 1 | 3 | Arhgef12 | 0.004 | Old |
| 0.823846 | 0.082834 | 0.121 | 0.116 | 1 | 3 | Edc4 | 0.005 | Old |
| 0.824731 | 0.125597 | 0.066 | 0.063 | 1 | 3 | Cog4 | 0.003 | Old |
| 0.824792 | 0.190306 | 0.036 | 0.038 | 1 | 3 | Pola1 | -0.002 | Old |
| 0.825774 | 0.057145 | 0.399 | 0.385 | 1 | 3 | Ppp1r12a | 0.014 | Old |
| 0.826371 | 0.077224 | 0.19 | 0.184 | 1 | 3 | Rab32 | 0.006 | Old |
| 0.828415 | 0.099582 | 0.349 | 0.337 | 1 | 3 | Chmp5 | 0.012 | Old |
| 0.829738 | 0.453335 | 0.02 | 0.019 | 1 | 3 | Gm47230 | 0.001 | Old |
| 0.830763 | 0.110543 | 0.02 | 0.019 | 1 | 3 | Polg2 | 0.001 | Old |
| 0.831465 | 0.018142 | 0.875 | 0.874 | 1 | 3 | Pitpna | 0.001 | Old |
| 0.832855 | 0.060791 | 0.094 | 0.091 | 1 | 3 | Vti1a | 0.003 | Old |
| 0.833505 | 0.179369 | 0.036 | 0.034 | 1 | 3 | Oxnad1 | 0.002 | Old |
| 0.835934 | 0.087282 | 0.033 | 0.032 | 1 | 3 | Zhx3 | 0.001 | Old |
| 0.836923 | 0.289868 | 0.02 | 0.019 | 1 | 3 | Mppe1 | 0.001 | Old |
| 0.837433 | 0.025442 | 0.303 | 0.29 | 1 | 3 | Rdx | 0.013 | Old |
| 0.838995 | 0.103079 | 0.174 | 0.179 | 1 | 3 | Ubald1 | -0.005 | Old |
| 0.839193 | 0.106354 | 0.036 | 0.034 | 1 | 3 | Rdh10 | 0.002 | Old |
| 0.839663 | 0.70009 | 0.02 | 0.019 | 1 | 3 | Scarf1 | 0.001 | Old |
| 0.840184 | 0.056962 | 0.066 | 0.063 | 1 | 3 | Endod1 | 0.003 | Old |
| 0.840791 | 0.252641 | 0.031 | 0.033 | 1 | 3 | Pex11b | -0.002 | Old |
| 0.841572 | 0.089904 | 0.306 | 0.302 | 1 | 3 | Scarb2 | 0.004 | Old |
| 0.841605 | 0.174823 | 0.099 | 0.096 | 1 | 3 | Zeb2os | 0.003 | Old |
| 0.842686 | 0.050059 | 0.031 | 0.033 | 1 | 3 | Ankrd39 | -0.002 | Old |

| | | | | | | | | |
|---|---|---|---|---|---|---|---|---|
| 0.842817 | 0.264429 | 0.036 | 0.034 | 1 | 3 | Tbp | 0.002 | Old |
| 0.843478 | 0.13128 | 0.053 | 0.051 | 1 | 3 | Gm26782 | 0.002 | Old |
| 0.844962 | 0.05081 | 0.167 | 0.162 | 1 | 3 | Ifi209 | 0.005 | Old |
| 0.845049 | 0.098888 | 0.033 | 0.032 | 1 | 3 | Eef1akmt4 | 0.001 | Old |
| 0.845144 | 0.014161 | 0.528 | 0.492 | 1 | 3 | Nedd8 | 0.036 | Old |
| 0.845905 | 0.026603 | 0.1 | 0.097 | 1 | 3 | Stard9 | 0.003 | Old |
| 0.846478 | 0.002345 | 0.031 | 0.033 | 1 | 3 | Slc2a9 | -0.002 | Old |
| 0.846582 | 0.019739 | 0.033 | 0.035 | 1 | 3 | Trit1 | -0.002 | Old |
| 0.849413 | 0.127244 | 0.061 | 0.059 | 1 | 3 | Elp3 | 0.002 | Old |
| 0.8496 | 0.006192 | 0.145 | 0.14 | 1 | 3 | Pknox1 | 0.005 | Old |
| 0.850036 | 0.112903 | 0.115 | 0.111 | 1 | 3 | Tsr1 | 0.004 | Old |
| 0.852293 | 0.066241 | 0.061 | 0.063 | 1 | 3 | Elmod2 | -0.002 | Old |
| 0.852613 | 0.161146 | 0.07 | 0.068 | 1 | 3 | Gnptg | 0.002 | Old |
| 0.854141 | 0.056643 | 0.088 | 0.086 | 1 | 3 | Stx2 | 0.002 | Old |
| 0.854605 | 0.029583 | 0.168 | 0.165 | 1 | 3 | Zhx1 | 0.003 | Old |
| 0.854748 | 0.182033 | 0.036 | 0.034 | 1 | 3 | Fam210b | 0.002 | Old |
| 0.855024 | 0.240947 | 0.023 | 0.021 | 1 | 3 | A130010J1 | 0.002 | Old |
| 0.855349 | 0.269189 | 0.023 | 0.021 | 1 | 3 | Tsen54 | 0.002 | Old |
| 0.856096 | 0.087089 | 0.123 | 0.119 | 1 | 3 | Dhx30 | 0.004 | Old |
| 0.856648 | 0.282687 | 0.023 | 0.021 | 1 | 3 | Fahd2a | 0.002 | Old |
| 0.857679 | 0.186402 | 0.042 | 0.04 | 1 | 3 | Cstf1 | 0.002 | Old |
| 0.858254 | 0.064404 | 0.085 | 0.082 | 1 | 3 | P2ry6 | 0.003 | Old |
| 0.858746 | 0.012291 | 0.207 | 0.201 | 1 | 3 | Pfdn1 | 0.006 | Old |
| 0.859843 | 0.010516 | 0.087 | 0.085 | 1 | 3 | 2210016L2 | 0.002 | Old |
| 0.860442 | 0.178042 | 0.026 | 0.028 | 1 | 3 | 6330418K0 | -0.002 | Old |
| 0.861827 | 0.232614 | 0.03 | 0.032 | 1 | 3 | Cep76 | -0.002 | Old |
| 0.862258 | 0.078752 | 0.103 | 0.1 | 1 | 3 | Prr14l | 0.003 | Old |
| 0.863514 | 0.25888 | 0.024 | 0.023 | 1 | 3 | Gm4070 | 0.001 | Old |
| 0.864217 | 0.336272 | 0.025 | 0.027 | 1 | 3 | 2010320M | -0.002 | Old |
| 0.866455 | 0.660416 | 0.01 | 0.009 | 1 | 3 | Gm9774 | 0.001 | Old |
| 0.867938 | 0.01115 | 0.024 | 0.025 | 1 | 3 | Dpys | -0.001 | Old |
| 0.868916 | 0.049671 | 0.057 | 0.056 | 1 | 3 | Fhl3 | 0.001 | Old |
| 0.869021 | 0.398103 | 0.025 | 0.024 | 1 | 3 | 1810011H1 | 0.001 | Old |
| 0.871024 | 0.049622 | 0.02 | 0.021 | 1 | 3 | Pbx1 | -0.001 | Old |
| 0.87129 | 0.078613 | 0.022 | 0.023 | 1 | 3 | Pdia5 | -0.001 | Old |
| 0.871694 | 0.285312 | 0.02 | 0.021 | 1 | 3 | Smyd3 | -0.001 | Old |
| 0.871762 | 0.013888 | 0.119 | 0.121 | 1 | 3 | Bzw2 | -0.002 | Old |
| 0.871968 | 0.024295 | 0.038 | 0.037 | 1 | 3 | Dnajc4 | 0.001 | Old |
| 0.872029 | 0.176657 | 0.066 | 0.064 | 1 | 3 | Socs5 | 0.002 | Old |
| 0.872472 | 0.255678 | 0.01 | 0.009 | 1 | 3 | Vmac | 0.001 | Old |
| 0.873626 | 0.027122 | 0.018 | 0.019 | 1 | 3 | Pstpip2 | -0.001 | Old |
| 0.873918 | 0.049019 | 0.137 | 0.135 | 1 | 3 | Mrpl2 | 0.002 | Old |
| 0.873982 | 0.106254 | 0.024 | 0.023 | 1 | 3 | Plpp5 | 0.001 | Old |
| 0.874026 | 0.004784 | 0.897 | 0.884 | 1 | 3 | Glud1 | 0.013 | Old |
| 0.875398 | 0.779565 | 0.026 | 0.025 | 1 | 3 | Cutc | 0.001 | Old |

| | | | | | | | | |
|---|---|---|---|---|---|---|---|---|
| 0.876181 | 0.104182 | 0.022 | 0.023 | 1 | 3 | Rbm41 | -0.001 | Old |
| 0.876523 | 0.066948 | 0.024 | 0.023 | 1 | 3 | Spred3 | 0.001 | Old |
| 0.87707 | 0.091583 | 0.053 | 0.051 | 1 | 3 | Ahdc1 | 0.002 | Old |
| 0.877463 | 0.029736 | 0.412 | 0.396 | 1 | 3 | Fkbp8 | 0.016 | Old |
| 0.877692 | 0.175593 | 0.025 | 0.024 | 1 | 3 | Tm9sf1 | 0.001 | Old |
| 0.878496 | 0.141681 | 0.01 | 0.009 | 1 | 3 | Rhou | 0.001 | Old |
| 0.878998 | 0.258906 | 0.01 | 0.009 | 1 | 3 | Fbxo10 | 0.001 | Old |
| 0.879501 | 0.087449 | 0.01 | 0.009 | 1 | 3 | Gm26699 | 0.001 | Old |
| 0.880172 | 0.135808 | 0.025 | 0.024 | 1 | 3 | Coil | 0.001 | Old |
| 0.88031 | 0.001474 | 0.501 | 0.48 | 1 | 3 | Tmem256 | 0.021 | Old |
| 0.880506 | 0.05483 | 0.01 | 0.009 | 1 | 3 | Ptpdc1 | 0.001 | Old |
| 0.88075 | 0.307768 | 0.022 | 0.023 | 1 | 3 | Rangrf | -0.001 | Old |
| 0.881292 | 0.117351 | 0.024 | 0.023 | 1 | 3 | Slc25a14 | 0.001 | Old |
| 0.881511 | 0.075674 | 0.01 | 0.009 | 1 | 3 | Zfp532 | 0.001 | Old |
| 0.882243 | 0.028195 | 0.881 | 0.856 | 1 | 3 | Lgals3 | 0.025 | Old |
| 0.882864 | 0.070556 | 0.016 | 0.016 | 1 | 3 | Zfp607a | 0 | Old |
| 0.883495 | 0.005731 | 0.43 | 0.405 | 1 | 3 | Tnpo1 | 0.025 | Old |
| 0.884029 | 0.185119 | 0.094 | 0.093 | 1 | 3 | Zfp120 | 0.001 | Old |
| 0.884169 | 0.015407 | 0.045 | 0.047 | 1 | 3 | Map4k5 | -0.002 | Old |
| 0.884931 | 0.058185 | 0.06 | 0.058 | 1 | 3 | Gm9493 | 0.002 | Old |
| 0.885304 | 0.271334 | 0.051 | 0.053 | 1 | 3 | Hsd17b7 | -0.002 | Old |
| 0.886067 | 0.031895 | 0.025 | 0.024 | 1 | 3 | Zfp65 | 0.001 | Old |
| 0.886697 | 0.052824 | 0.129 | 0.126 | 1 | 3 | Tbpl1 | 0.003 | Old |
| 0.886758 | 0.025119 | 0.049 | 0.051 | 1 | 3 | Mapkapk5 | -0.002 | Old |
| 0.886833 | 0.437293 | 0.018 | 0.019 | 1 | 3 | Gm3448 | -0.001 | Old |
| 0.888506 | 0.125847 | 0.017 | 0.018 | 1 | 3 | Lins1 | -0.001 | Old |
| 0.890051 | 0.046125 | 0.018 | 0.019 | 1 | 3 | Tmem201 | -0.001 | Old |
| 0.891749 | 0.012214 | 0.014 | 0.015 | 1 | 3 | Ptchd1 | -0.001 | Old |
| 0.892064 | 0.278889 | 0.026 | 0.025 | 1 | 3 | Ncapg2 | 0.001 | Old |
| 0.892131 | 0.314629 | 0.011 | 0.01 | 1 | 3 | Itgb3bp | 0.001 | Old |
| 0.892547 | 0.113909 | 0.014 | 0.015 | 1 | 3 | Prune2 | -0.001 | Old |
| 0.893109 | 0.215351 | 0.019 | 0.02 | 1 | 3 | Rwdd2a | -0.001 | Old |
| 0.893551 | 0.221237 | 0.011 | 0.01 | 1 | 3 | Dnah2 | 0.001 | Old |
| 0.893987 | 0.227631 | 0.016 | 0.016 | 1 | 3 | Shpk | 0 | Old |
| 0.894169 | 0.458372 | 0.027 | 0.027 | 1 | 3 | Gys1 | 0 | Old |
| 0.894424 | 0.059186 | 0.185 | 0.181 | 1 | 3 | Anapc16 | 0.004 | Old |
| 0.894498 | 0.186123 | 0.011 | 0.01 | 1 | 3 | Zfp341 | 0.001 | Old |
| 0.894702 | 1.396249 | 0.018 | 0.019 | 1 | 3 | Inhba | -0.001 | Old |
| 0.89601 | 0.050596 | 0.026 | 0.025 | 1 | 3 | Ift140 | 0.001 | Old |
| 0.896683 | 0.066708 | 0.129 | 0.124 | 1 | 3 | Nudt19 | 0.005 | Old |
| 0.898288 | 0.080905 | 0.011 | 0.01 | 1 | 3 | Nedd4 | 0.001 | Old |
| 0.898596 | 0.275526 | 0.016 | 0.016 | 1 | 3 | Bend3 | 0 | Old |
| 0.899235 | 0.000198 | 0.011 | 0.01 | 1 | 3 | Mrm3 | 0.001 | Old |
| 0.899709 | 0.101894 | 0.011 | 0.01 | 1 | 3 | Tmc4 | 0.001 | Old |
| 0.900375 | 0.145507 | 0.013 | 0.014 | 1 | 3 | Tceanc | -0.001 | Old |

| | | | | | | | | |
|---|---|---|---|---|---|---|---|---|
| 0.901246 | 0.005205 | 0.918 | 0.894 | 1 | 3 | Eef1b2 | 0.024 | Old |
| 0.903913 | 0.380759 | 0.012 | 0.013 | 1 | 3 | Itgb8 | -0.001 | Old |
| 0.904403 | 0.050011 | 0.087 | 0.085 | 1 | 3 | Ccdc43 | 0.002 | Old |
| 0.904796 | 0.090089 | 0.011 | 0.011 | 1 | 3 | B630019K0 | 0 | Old |
| 0.904831 | 0.015405 | 0.14 | 0.136 | 1 | 3 | Mms19 | 0.004 | Old |
| 0.905226 | 0.049548 | 0.012 | 0.013 | 1 | 3 | 9030025P2 | -0.001 | Old |
| 0.905802 | 0.194473 | 0.013 | 0.014 | 1 | 3 | Tkfc | -0.001 | Old |
| 0.907415 | 0.489858 | 0.012 | 0.013 | 1 | 3 | Tedc2 | -0.001 | Old |
| 0.907468 | 0.076769 | 0.029 | 0.028 | 1 | 3 | Pfas | 0.001 | Old |
| 0.908428 | 0.002118 | 0.264 | 0.259 | 1 | 3 | Bri3bp | 0.005 | Old |
| 0.908728 | 0.744068 | 0.012 | 0.013 | 1 | 3 | Ppef2 | -0.001 | Old |
| 0.908827 | 0.059066 | 0.139 | 0.141 | 1 | 3 | Akap8 | -0.002 | Old |
| 0.908947 | 0.001637 | 0.011 | 0.011 | 1 | 3 | 4930477G0 | 0 | Old |
| 0.910819 | 0.078052 | 0.045 | 0.044 | 1 | 3 | Zbtb33 | 0.001 | Old |
| 0.911034 | 0.046635 | 0.173 | 0.174 | 1 | 3 | Pde3b | -0.001 | Old |
| 0.911563 | 0.48787 | 0.012 | 0.011 | 1 | 3 | Wdr74 | 0.001 | Old |
| 0.912012 | 0.099391 | 0.012 | 0.011 | 1 | 3 | 2510046G1 | 0.001 | Old |
| 0.912127 | 0.225427 | 0.029 | 0.028 | 1 | 3 | Bcorl1 | 0.001 | Old |
| 0.913649 | 0.05244 | 0.044 | 0.043 | 1 | 3 | Calcoco1 | 0.001 | Old |
| 0.91381 | 0.59152 | 0.012 | 0.011 | 1 | 3 | Xlr3b | 0.001 | Old |
| 0.913886 | 0.725096 | 0.013 | 0.013 | 1 | 3 | Il1r2 | 0 | Old |
| 0.914514 | 0.1315 | 0.066 | 0.064 | 1 | 3 | Xlr4b | 0.002 | Old |
| 0.914563 | 0.224424 | 0.072 | 0.073 | 1 | 3 | Il18 | -0.001 | Old |
| 0.914757 | 0.093114 | 0.01 | 0.01 | 1 | 3 | Ankrd9 | 0 | Old |
| 0.914824 | 0.072305 | 0.397 | 0.385 | 1 | 3 | Apobec1 | 0.012 | Old |
| 0.915124 | 0.099047 | 0.124 | 0.126 | 1 | 3 | Tgfbr2 | -0.002 | Old |
| 0.916025 | 0.583543 | 0.013 | 0.013 | 1 | 3 | Hdac6 | 0 | Old |
| 0.917256 | 0.363858 | 0.011 | 0.011 | 1 | 3 | Klhl42 | 0 | Old |
| 0.917627 | 0.095925 | 0.03 | 0.029 | 1 | 3 | Zfp821 | 0.001 | Old |
| 0.917657 | 0.038714 | 0.027 | 0.027 | 1 | 3 | Slc25a40 | 0 | Old |
| 0.919187 | 0.037618 | 0.081 | 0.08 | 1 | 3 | Med29 | 0.001 | Old |
| 0.920738 | 0.21607 | 0.103 | 0.106 | 1 | 3 | Rpl27-ps3 | -0.003 | Old |
| 0.922055 | 0.018178 | 0.078 | 0.076 | 1 | 3 | Ctso | 0.002 | Old |
| 0.922366 | 0.105331 | 0.044 | 0.045 | 1 | 3 | D230025D1 | -0.001 | Old |
| 0.923155 | 0.044089 | 0.068 | 0.069 | 1 | 3 | Eef2kmt | -0.001 | Old |
| 0.924369 | 0.093428 | 0.029 | 0.028 | 1 | 3 | Pus3 | 0.001 | Old |
| 0.924623 | 0.041905 | 0.109 | 0.109 | 1 | 3 | H2-Q6 | 0 | Old |
| 0.92539 | 0.140016 | 0.029 | 0.028 | 1 | 3 | Zbtb46 | 0.001 | Old |
| 0.925697 | 0.157989 | 0.032 | 0.032 | 1 | 3 | Hacd1 | 0 | Old |
| 0.926001 | 0.34329 | 0.027 | 0.027 | 1 | 3 | Gm42715 | 0 | Old |
| 0.926303 | 0.116796 | 0.013 | 0.013 | 1 | 3 | Afap1 | 0 | Old |
| 0.927547 | 0.053623 | 0.401 | 0.396 | 1 | 3 | Tnrc18 | 0.005 | Old |
| 0.928852 | 0.039164 | 0.062 | 0.061 | 1 | 3 | Slc38a7 | 0.001 | Old |
| 0.929724 | 0.019123 | 0.049 | 0.048 | 1 | 3 | Dip2a | 0.001 | Old |
| 0.930063 | 0.511318 | 0.055 | 0.054 | 1 | 3 | Uck2 | 0.001 | Old |

| | | | | | | | | |
|---|---|---|---|---|---|---|---|---|
| 0.930067 | 0.024202 | 0.263 | 0.258 | 1 | 3 | Wls | 0.005 | Old |
| 0.930299 | 0.020744 | 0.332 | 0.324 | 1 | 3 | Srsf10 | 0.008 | Old |
| 0.930409 | 0.184942 | 0.069 | 0.068 | 1 | 3 | Zfp329 | 0.001 | Old |
| 0.932561 | 0.050624 | 0.375 | 0.369 | 1 | 3 | Phip | 0.006 | Old |
| 0.932696 | 0.400874 | 0.037 | 0.037 | 1 | 3 | Bcl11a | 0 | Old |
| 0.932715 | 0.021201 | 0.217 | 0.212 | 1 | 3 | Pih1d1 | 0.005 | Old |
| 0.934087 | 0.093286 | 0.031 | 0.03 | 1 | 3 | Creb3l1 | 0.001 | Old |
| 0.935735 | 0.583502 | 0.035 | 0.034 | 1 | 3 | Zcchc4 | 0.001 | Old |
| 0.936343 | 0.099881 | 0.114 | 0.114 | 1 | 3 | Topbp1 | 0 | Old |
| 0.936741 | 0.020684 | 0.084 | 0.085 | 1 | 3 | Ndc1 | -0.001 | Old |
| 0.938375 | 0.06946 | 0.042 | 0.043 | 1 | 3 | C87436 | -0.001 | Old |
| 0.939407 | 0.039455 | 0.085 | 0.086 | 1 | 3 | Nelfb | -0.001 | Old |
| 0.940003 | 0.007346 | 0.032 | 0.032 | 1 | 3 | Bex3 | 0 | Old |
| 0.94041 | 0.038658 | 0.033 | 0.033 | 1 | 3 | Lrrc1 | 0 | Old |
| 0.941518 | 0.021374 | 0.129 | 0.128 | 1 | 3 | Vkorc1 | 0.001 | Old |
| 0.941548 | 0.064106 | 0.035 | 0.035 | 1 | 3 | Ggct | 0 | Old |
| 0.941626 | 0.038728 | 0.014 | 0.014 | 1 | 3 | Fam102a | 0 | Old |
| 0.942071 | 0.025411 | 0.13 | 0.128 | 1 | 3 | Gmppa | 0.002 | Old |
| 0.942078 | 0.115781 | 0.031 | 0.032 | 1 | 3 | Pdgfa | -0.001 | Old |
| 0.942811 | 0.094262 | 0.076 | 0.076 | 1 | 3 | Cox19 | 0 | Old |
| 0.942822 | 0.07165 | 0.125 | 0.124 | 1 | 3 | Sp140 | 0.001 | Old |
| 0.943393 | 0.092176 | 0.352 | 0.346 | 1 | 3 | B230219D2 | 0.006 | Old |
| 0.944562 | 0.284169 | 0.016 | 0.015 | 1 | 3 | Gm45894 | 0.001 | Old |
| 0.945214 | 0.030914 | 0.313 | 0.306 | 1 | 3 | Klf9 | 0.007 | Old |
| 0.945331 | 0.033157 | 0.161 | 0.159 | 1 | 3 | Utp20 | 0.002 | Old |
| 0.9455 | 0.064915 | 0.035 | 0.035 | 1 | 3 | Fam118a | 0 | Old |
| 0.946267 | 0.161074 | 0.036 | 0.037 | 1 | 3 | Dusp12 | -0.001 | Old |
| 0.946945 | 0.022975 | 0.103 | 0.101 | 1 | 3 | Mmgt1 | 0.002 | Old |
| 0.948783 | 0.141563 | 0.054 | 0.053 | 1 | 3 | Prr12 | 0.001 | Old |
| 0.94885 | 0.009819 | 0.416 | 0.4 | 1 | 3 | Cd48 | 0.016 | Old |
| 0.949594 | 0.018431 | 0.281 | 0.278 | 1 | 3 | Ap3b1 | 0.003 | Old |
| 0.950635 | 0.009512 | 0.087 | 0.086 | 1 | 3 | Ap4m1 | 0.001 | Old |
| 0.952162 | 0.017275 | 0.508 | 0.496 | 1 | 3 | Tmem59 | 0.012 | Old |
| 0.952837 | 0.047292 | 0.016 | 0.015 | 1 | 3 | Zkscan17 | 0.001 | Old |
| 0.953232 | 0.199558 | 0.016 | 0.015 | 1 | 3 | 4632415L0 | 0.001 | Old |
| 0.954305 | 0.016644 | 0.871 | 0.848 | 1 | 3 | H2afz | 0.023 | Old |
| 0.955537 | 0.36074 | 0.017 | 0.016 | 1 | 3 | Fmnl3 | 0.001 | Old |
| 0.955601 | 0.194688 | 0.036 | 0.037 | 1 | 3 | Ccdc71 | -0.001 | Old |
| 0.95564 | 0.055048 | 0.078 | 0.077 | 1 | 3 | Trim11 | 0.001 | Old |
| 0.955929 | 0.083102 | 0.074 | 0.073 | 1 | 3 | Dgkg | 0.001 | Old |
| 0.960009 | 0.004554 | 0.17 | 0.167 | 1 | 3 | Kansl3 | 0.003 | Old |
| 0.960357 | 0.034371 | 0.093 | 0.093 | 1 | 3 | Rrp9 | 0 | Old |
| 0.960391 | 0.039357 | 0.186 | 0.183 | 1 | 3 | Swap70 | 0.003 | Old |
| 0.961006 | 0.162357 | 0.068 | 0.069 | 1 | 3 | Pou2f2 | -0.001 | Old |
| 0.961652 | 0.480495 | 0.018 | 0.018 | 1 | 3 | Elp6 | 0 | Old |

| | | | | | | | | |
|---|---|---|---|---|---|---|---|---|
| 0.962361 | 0.113192 | 0.03 | 0.03 | 1 | 3 | Mtg2 | 0 | Old |
| 0.96293 | 0.454647 | 0.033 | 0.034 | 1 | 3 | Lrba | -0.001 | Old |
| 0.964586 | 0.081053 | 0.018 | 0.018 | 1 | 3 | Zfp874b | 0 | Old |
| 0.964651 | 0.007135 | 0.017 | 0.016 | 1 | 3 | Gm11423 | 0.001 | Old |
| 0.965686 | 0.094413 | 0.018 | 0.018 | 1 | 3 | Chp2 | 0 | Old |
| 0.96666 | 0.0385 | 0.039 | 0.039 | 1 | 3 | Tmtc2 | 0 | Old |
| 0.967659 | 0.036622 | 0.319 | 0.321 | 1 | 3 | Pikfyve | -0.002 | Old |
| 0.968254 | 0.265558 | 0.018 | 0.018 | 1 | 3 | Agpat3 | 0 | Old |
| 0.968534 | 0.001065 | 0.829 | 0.787 | 1 | 3 | Gnb1 | 0.042 | Old |
| 0.969659 | 0.005433 | 0.176 | 0.178 | 1 | 3 | Pabpc4 | -0.002 | Old |
| 0.969962 | 0.105709 | 0.08 | 0.081 | 1 | 3 | Lrrcc1 | -0.001 | Old |
| 0.970839 | 0.043909 | 0.115 | 0.115 | 1 | 3 | Tmem192 | 0 | Old |
| 0.971555 | 0.036738 | 0.018 | 0.018 | 1 | 3 | Nup35 | 0 | Old |
| 0.972632 | 0.0306 | 0.165 | 0.165 | 1 | 3 | Pbx3 | 0 | Old |
| 0.972852 | 0.016183 | 0.117 | 0.116 | 1 | 3 | Cd2bp2 | 0.001 | Old |
| 0.97396 | 0.021808 | 0.547 | 0.537 | 1 | 3 | Eif4a2 | 0.01 | Old |
| 0.974647 | 0.125971 | 0.042 | 0.042 | 1 | 3 | Ppard | 0 | Old |
| 0.975987 | 0.235224 | 0.025 | 0.025 | 1 | 3 | Rad9a | 0 | Old |
| 0.976632 | 0.081023 | 0.096 | 0.095 | 1 | 3 | Arsa | 0.001 | Old |
| 0.976823 | 0.252776 | 0.08 | 0.081 | 1 | 3 | Bicral | -0.001 | Old |
| 0.97777 | 0.111904 | 0.039 | 0.039 | 1 | 3 | Sesn2 | 0 | Old |
| 0.977975 | 0.205877 | 0.019 | 0.019 | 1 | 3 | Carmil1 | 0 | Old |
| 0.97833 | 0.033729 | 0.019 | 0.019 | 1 | 3 | Zfp113 | 0 | Old |
| 0.979099 | 0.183851 | 0.031 | 0.032 | 1 | 3 | Invs | -0.001 | Old |
| 0.979203 | 0.063844 | 0.041 | 0.04 | 1 | 3 | Ercc4 | 0.001 | Old |
| 0.979287 | 0.005239 | 0.229 | 0.23 | 1 | 3 | Mtmr3 | -0.001 | Old |
| 0.980106 | 0.228706 | 0.019 | 0.019 | 1 | 3 | Gm15232 | 0 | Old |
| 0.980142 | 0.010458 | 0.039 | 0.039 | 1 | 3 | Hspa2 | 0 | Old |
| 0.982098 | 0.057148 | 0.09 | 0.09 | 1 | 3 | Plcb3 | 0 | Old |
| 0.9825 | 0.020613 | 0.444 | 0.436 | 1 | 3 | Tiparp | 0.008 | Old |
| 0.983403 | 0.025913 | 0.125 | 0.125 | 1 | 3 | Arrdc4 | 0 | Old |
| 0.983452 | 0.039226 | 0.061 | 0.061 | 1 | 3 | Tomm40l | 0 | Old |
| 0.983651 | 0.044352 | 0.026 | 0.027 | 1 | 3 | Osbpl7 | -0.001 | Old |
| 0.985044 | 0.042951 | 0.047 | 0.047 | 1 | 3 | Tmem8 | 0 | Old |
| 0.985083 | 0.063126 | 0.068 | 0.068 | 1 | 3 | Pcnx3 | 0 | Old |
| 0.985778 | 0.02204 | 0.065 | 0.064 | 1 | 3 | 4930503L1 | 0.001 | Old |
| 0.987019 | 0.118302 | 0.042 | 0.042 | 1 | 3 | Piga | 0 | Old |
| 0.987999 | 0.00019 | 0.045 | 0.045 | 1 | 3 | Rnaseh1 | 0 | Old |
| 0.988953 | 0.10687 | 0.022 | 0.021 | 1 | 3 | Wdr53 | 0.001 | Old |
| 0.991631 | 0.024763 | 0.022 | 0.021 | 1 | 3 | 9430015G1 | 0.001 | Old |
| 0.991744 | 0.017781 | 0.024 | 0.024 | 1 | 3 | 5430427O1 | 0 | Old |
| 0.992 | 0.066055 | 0.041 | 0.04 | 1 | 3 | Top2a | 0.001 | Old |
| 0.992062 | 0.025705 | 0.024 | 0.024 | 1 | 3 | Gorasp1 | 0 | Old |
| 0.992296 | 0.08977 | 0.109 | 0.109 | 1 | 3 | Acox1 | 0 | Old |
| 0.992408 | 0.337019 | 0.025 | 0.025 | 1 | 3 | Gadd45a | 0 | Old |

| | | | | | | | | |
|---|---|---|---|---|---|---|---|---|
| 0.993182 | 0.056925 | 0.048 | 0.048 | 1 | 3 | Fhod1 | 0 | Old |
| 0.993559 | 0.192603 | 0.082 | 0.082 | 1 | 3 | Bcl2l1 | 0 | Old |
| 0.993692 | 0.040218 | 0.081 | 0.081 | 1 | 3 | Rrn3 | 0 | Old |
| 0.993974 | 0.424841 | 0.022 | 0.021 | 1 | 3 | Coq4 | 0.001 | Old |
| 0.994309 | 0.123527 | 0.022 | 0.021 | 1 | 3 | Ss18l1 | 0.001 | Old |
| 0.994465 | 0.016304 | 0.06 | 0.059 | 1 | 3 | 2300009A( | 0.001 | Old |
| 0.99477 | 0.076901 | 0.062 | 0.062 | 1 | 3 | Stk39 | 0 | Old |
| 0.994865 | 0.101897 | 0.081 | 0.082 | 1 | 3 | Tusc2 | -0.001 | Old |
| 0.997416 | 0.477324 | 0.02 | 0.02 | 1 | 3 | Ggt5 | 0 | Old |
| 0.997487 | 0.046149 | 0.043 | 0.043 | 1 | 3 | Usp46 | 0 | Old |
| 0.998025 | 0.14487 | 0.065 | 0.064 | 1 | 3 | Micu3 | 0.001 | Old |
| 0.998105 | 0.003655 | 0.315 | 0.308 | 1 | 3 | Smc4 | 0.007 | Old |
| 0.998977 | 0.269803 | 0.048 | 0.048 | 1 | 3 | Zcchc10 | 0 | Old |
| 0.999162 | 0.245666 | 0.043 | 0.043 | 1 | 3 | Zfp595 | 0 | Old |
| 0.999235 | 0.011999 | 0.173 | 0.173 | 1 | 3 | Paxbp1 | 0 | Old |
| 7.67E-19 | 0.68796 | 0.784 | 0.625 | 2.38E-14 | 3 | Lyz1 | 0.159 | Young |
| 6.32E-18 | 1.014672 | 0.42 | 0.223 | 1.96E-13 | 3 | Hspa1a | 0.197 | Young |
| 6.42E-15 | 1.018229 | 0.431 | 0.262 | 1.99E-10 | 3 | Hspa1b | 0.169 | Young |
| 8.34E-14 | 0.127887 | 1 | 1 | 2.59E-09 | 3 | Gm42418 | 0 | Young |
| 4.75E-13 | 0.258868 | 0.977 | 0.984 | 1.48E-08 | 3 | Rps14 | -0.007 | Young |
| 1.53E-12 | 0.380686 | 0.876 | 0.873 | 4.76E-08 | 3 | Rps3 | 0.003 | Young |
| 4.89E-11 | 0.352932 | 0.995 | 0.996 | 1.52E-06 | 3 | Malat1 | -0.001 | Young |
| 6.28E-11 | 0.178223 | 0.999 | 0.998 | 1.95E-06 | 3 | Cd74 | 0.001 | Young |
| 4.14E-10 | 0.597372 | 0.655 | 0.551 | 1.29E-05 | 3 | Ecm1 | 0.104 | Young |
| 1.19E-09 | 0.277745 | 0.976 | 0.971 | 3.68E-05 | 3 | Ftl1 | 0.005 | Young |
| 1.42E-09 | 0.19339 | 0.99 | 0.987 | 4.40E-05 | 3 | Fau | 0.003 | Young |
| 1.91E-09 | 0.263283 | 0.957 | 0.947 | 5.92E-05 | 3 | Rps13 | 0.01 | Young |
| 3.29E-09 | 0.259141 | 0.942 | 0.939 | 0.000102 | 3 | Rps5 | 0.003 | Young |
| 5.98E-09 | 0.802579 | 0.532 | 0.405 | 0.000186 | 3 | Apoe | 0.127 | Young |
| 7.88E-09 | 0.269667 | 0.914 | 0.916 | 0.000245 | 3 | Rpl26 | -0.002 | Young |
| 2.90E-08 | 0.213756 | 0.951 | 0.956 | 0.000902 | 3 | Rps19 | -0.005 | Young |
| 2.99E-08 | 0.306152 | 0.857 | 0.829 | 0.000928 | 3 | Rpl10 | 0.028 | Young |
| 3.57E-08 | 0.256437 | 0.933 | 0.928 | 0.001109 | 3 | Rpl18a | 0.005 | Young |
| 7.76E-08 | 0.325474 | 0.835 | 0.787 | 0.00241 | 3 | Rsrp1 | 0.048 | Young |
| 6.51E-07 | 0.178872 | 0.975 | 0.976 | 0.020215 | 3 | Rpl13 | -0.001 | Young |
| 6.66E-07 | 0.237582 | 0.937 | 0.922 | 0.020683 | 3 | Rpl28 | 0.015 | Young |
| 7.98E-07 | 0.137862 | 0.991 | 1 | 0.024783 | 3 | Rps9 | -0.009 | Young |
| 2.64E-06 | 0.221155 | 0.917 | 0.908 | 0.08186 | 3 | Fcrls | 0.009 | Young |
| 3.20E-06 | 0.749391 | 0.298 | 0.208 | 0.099299 | 3 | Plxdc2 | 0.09 | Young |
| 3.86E-06 | 0.187156 | 0.976 | 0.961 | 0.12 | 3 | Hspa8 | 0.015 | Young |
| 3.95E-06 | 0.368804 | 0.708 | 0.669 | 0.122786 | 3 | Ubb | 0.039 | Young |
| 6.15E-06 | 0.206222 | 0.889 | 0.902 | 0.190952 | 3 | Rpl21 | -0.013 | Young |
| 8.76E-06 | 0.554699 | 0.508 | 0.424 | 0.272051 | 3 | Nr4a3 | 0.084 | Young |
| 1.28E-05 | 0.263839 | 0.883 | 0.891 | 0.398561 | 3 | Rpl11 | -0.008 | Young |
| 1.29E-05 | 0.222624 | 0.902 | 0.91 | 0.399944 | 3 | Rps6 | -0.008 | Young |

| | | | | | | | | |
|---|---|---|---|---|---|---|---|---|
| 2.01E-05 | 0.257117 | 0.876 | 0.897 | 0.624185 | 3 | Rpl12 | -0.021 | Young |
| 2.42E-05 | 0.151769 | 0.966 | 0.971 | 0.750941 | 3 | Rps20 | -0.005 | Young |
| 3.09E-05 | 0.237975 | 0.995 | 0.99 | 0.959018 | 3 | Retnla | 0.005 | Young |
| 3.90E-05 | 0.2337 | 0.912 | 0.908 | 1 | 3 | S100a10 | 0.004 | Young |
| 4.24E-05 | 0.182702 | 0.956 | 0.965 | 1 | 3 | Rps10 | -0.009 | Young |
| 4.85E-05 | 0.223389 | 0.838 | 0.815 | 1 | 3 | Eef2 | 0.023 | Young |
| 5.35E-05 | 0.161291 | 0.946 | 0.927 | 1 | 3 | H2-DMa | 0.019 | Young |
| 6.21E-05 | 0.581909 | 0.386 | 0.302 | 1 | 3 | Pf4 | 0.084 | Young |
| 6.59E-05 | 0.54895 | 0.438 | 0.368 | 1 | 3 | Arhgap45 | 0.07 | Young |
| 7.25E-05 | 0.16119 | 0.946 | 0.947 | 1 | 3 | Rplp0 | -0.001 | Young |
| 7.31E-05 | 0.161164 | 0.967 | 0.967 | 1 | 3 | Rpl23 | 0 | Young |
| 7.60E-05 | 1.939829 | 0.048 | 0.014 | 1 | 3 | Hpgd | 0.034 | Young |
| 8.22E-05 | 0.125665 | 0.984 | 0.988 | 1 | 3 | Rps24 | -0.004 | Young |
| 8.32E-05 | 0.165836 | 0.957 | 0.965 | 1 | 3 | Rpl30 | -0.008 | Young |
| 8.98E-05 | 0.181347 | 0.951 | 0.959 | 1 | 3 | Rpl19 | -0.008 | Young |
| 9.14E-05 | 0.286301 | 0.813 | 0.76 | 1 | 3 | F13a1 | 0.053 | Young |
| 9.73E-05 | 0.376618 | 0.756 | 0.722 | 1 | 3 | Fam46a | 0.034 | Young |
| 0.000105 | 0.139703 | 0.972 | 0.969 | 1 | 3 | Rps12 | 0.003 | Young |
| 0.00011 | 0.234369 | 0.857 | 0.839 | 1 | 3 | Tln1 | 0.018 | Young |
| 0.000116 | 0.635922 | 0.312 | 0.238 | 1 | 3 | Anxa6 | 0.074 | Young |
| 0.00012 | 0.159948 | 0.943 | 0.94 | 1 | 3 | Rps27 | 0.003 | Young |
| 0.000123 | 0.587119 | 0.424 | 0.351 | 1 | 3 | Ccl24 | 0.073 | Young |
| 0.000124 | 0.180698 | 0.926 | 0.927 | 1 | 3 | Tyrobp | -0.001 | Young |
| 0.000146 | 0.139969 | 0.956 | 0.962 | 1 | 3 | Rpl27a | -0.006 | Young |
| 0.000152 | 0.222812 | 0.942 | 0.916 | 1 | 3 | Atp2b1 | 0.026 | Young |
| 0.000159 | 0.724965 | 0.17 | 0.108 | 1 | 3 | Hepacam2 | 0.062 | Young |
| 0.000162 | 0.155881 | 0.967 | 0.965 | 1 | 3 | Rps16 | 0.002 | Young |
| 0.000192 | 0.324202 | 0.699 | 0.671 | 1 | 3 | Tmem176b | 0.028 | Young |
| 0.000194 | 0.198176 | 0.913 | 0.907 | 1 | 3 | Rpl36 | 0.006 | Young |
| 0.000229 | 0.151521 | 0.96 | 0.973 | 1 | 3 | Rpl17 | -0.013 | Young |
| 0.000287 | 0.531197 | 0.41 | 0.348 | 1 | 3 | Tsc22d3 | 0.062 | Young |
| 0.000373 | 0.203031 | 0.917 | 0.938 | 1 | 3 | Rpl34 | -0.021 | Young |
| 0.000376 | 0.243984 | 0.814 | 0.827 | 1 | 3 | Rpl15 | -0.013 | Young |
| 0.00047 | 1.609874 | 0.043 | 0.014 | 1 | 3 | Gm46430 | 0.029 | Young |
| 0.000473 | 0.143253 | 0.97 | 0.98 | 1 | 3 | Rps4x | -0.01 | Young |
| 0.000475 | 1.344666 | 0.054 | 0.022 | 1 | 3 | Car3 | 0.032 | Young |
| 0.000488 | 0.32542 | 0.663 | 0.607 | 1 | 3 | Itga4 | 0.056 | Young |
| 0.000494 | 2.006811 | 0.018 | 0.001 | 1 | 3 | 1700056N1 | 0.017 | Young |
| 0.000636 | 0.212516 | 0.896 | 0.915 | 1 | 3 | Nfkbiz | -0.019 | Young |
| 0.000651 | 1.064857 | 0.109 | 0.063 | 1 | 3 | Slc15a2 | 0.046 | Young |
| 0.000675 | 0.58191 | 0.487 | 0.44 | 1 | 3 | Tppp3 | 0.047 | Young |
| 0.000679 | 1.537239 | 0.038 | 0.012 | 1 | 3 | Sparc | 0.026 | Young |
| 0.000751 | 0.800266 | 0.176 | 0.119 | 1 | 3 | Otof | 0.057 | Young |
| 0.000758 | 0.133125 | 0.955 | 0.953 | 1 | 3 | Rpl23a | 0.002 | Young |
| 0.00077 | 0.770799 | 0.112 | 0.066 | 1 | 3 | Cd209d | 0.046 | Young |

| | | | | | | | | |
|---|---|---|---|---|---|---|---|---|
| 0.000797 | 0.424975 | 0.51 | 0.458 | 1 | 3 | Ptpn18 | 0.052 | Young |
| 0.000838 | 0.663245 | 0.196 | 0.136 | 1 | 3 | Tmem88 | 0.06 | Young |
| 0.000845 | 0.922152 | 0.235 | 0.179 | 1 | 3 | Dnajb1 | 0.056 | Young |
| 0.000851 | 0.153952 | 0.929 | 0.915 | 1 | 3 | Sh3bgrl3 | 0.014 | Young |
| 0.000851 | 0.50069 | 0.408 | 0.356 | 1 | 3 | Stxbp6 | 0.052 | Young |
| 0.000858 | 0.314653 | 0.682 | 0.638 | 1 | 3 | Cd81 | 0.044 | Young |
| 0.000888 | 0.234081 | 0.701 | 0.68 | 1 | 3 | Gmfg | 0.021 | Young |
| 0.001125 | 0.696741 | 0.287 | 0.228 | 1 | 3 | Heca | 0.059 | Young |
| 0.001189 | 0.236761 | 0.818 | 0.822 | 1 | 3 | Tgfbi | -0.004 | Young |
| 0.00137 | 0.111738 | 0.947 | 0.943 | 1 | 3 | Arpc1b | 0.004 | Young |
| 0.001407 | 0.15341 | 0.953 | 0.939 | 1 | 3 | Rpl24 | 0.014 | Young |
| 0.00145 | 0.126102 | 0.99 | 0.989 | 1 | 3 | S100a6 | 0.001 | Young |
| 0.001486 | 0.612955 | 0.135 | 0.087 | 1 | 3 | 6-Sep | 0.048 | Young |
| 0.001507 | 0.523531 | 0.426 | 0.373 | 1 | 3 | Baiap2 | 0.053 | Young |
| 0.001525 | 0.183685 | 0.937 | 0.949 | 1 | 3 | Rpl18 | -0.012 | Young |
| 0.001583 | 0.201794 | 0.923 | 0.922 | 1 | 3 | Rpl32 | 0.001 | Young |
| 0.001778 | 0.391397 | 0.504 | 0.456 | 1 | 3 | Abca9 | 0.048 | Young |
| 0.001793 | 0.097503 | 0.984 | 0.99 | 1 | 3 | Rps8 | -0.006 | Young |
| 0.0018 | 0.122903 | 0.996 | 0.996 | 1 | 3 | Ahnak | 0 | Young |
| 0.001858 | 0.645233 | 0.169 | 0.118 | 1 | 3 | Igfbp4 | 0.051 | Young |
| 0.001929 | 0.300652 | 0.639 | 0.633 | 1 | 3 | Ndufa13 | 0.006 | Young |
| 0.001993 | 2.018216 | 0.011 | 0 | 1 | 3 | 3830403N1 | 0.011 | Young |
| 0.002118 | 0.176384 | 0.941 | 0.924 | 1 | 3 | Cfp | 0.017 | Young |
| 0.002234 | 0.187136 | 0.831 | 0.817 | 1 | 3 | Atp5h | 0.014 | Young |
| 0.002305 | 0.456327 | 0.396 | 0.343 | 1 | 3 | Rb1cc1 | 0.053 | Young |
| 0.002668 | 0.135933 | 0.929 | 0.938 | 1 | 3 | Rpl9 | -0.009 | Young |
| 0.00275 | 1.81686 | 0.014 | 0.001 | 1 | 3 | Serpinf1 | 0.013 | Young |
| 0.002944 | 1.033583 | 0.057 | 0.027 | 1 | 3 | Rasgrf2 | 0.03 | Young |
| 0.002991 | 0.458079 | 0.426 | 0.381 | 1 | 3 | Mier1 | 0.045 | Young |
| 0.003042 | 0.082435 | 0.995 | 0.988 | 1 | 3 | Lyz2 | 0.007 | Young |
| 0.003212 | 0.082443 | 0.986 | 0.976 | 1 | 3 | Rps21 | 0.01 | Young |
| 0.003277 | 1.449237 | 0.035 | 0.013 | 1 | 3 | Zc2hc1a | 0.022 | Young |
| 0.003408 | 1.428724 | 0.044 | 0.019 | 1 | 3 | Mblac2 | 0.025 | Young |
| 0.003466 | 0.593024 | 0.143 | 0.097 | 1 | 3 | 4930402H2 | 0.046 | Young |
| 0.00377 | 1.272017 | 0.04 | 0.017 | 1 | 3 | Tlcd2 | 0.023 | Young |
| 0.003822 | 0.221805 | 0.585 | 0.538 | 1 | 3 | Adgre5 | 0.047 | Young |
| 0.003828 | 0.424834 | 0.446 | 0.409 | 1 | 3 | Nop53 | 0.037 | Young |
| 0.003987 | 0.104248 | 0.965 | 0.973 | 1 | 3 | Cst3 | -0.008 | Young |
| 0.004211 | 0.06638 | 0.987 | 0.996 | 1 | 3 | Tpt1 | -0.009 | Young |
| 0.004607 | 1.125938 | 0.052 | 0.025 | 1 | 3 | Tanc2 | 0.027 | Young |
| 0.004692 | 0.640071 | 0.178 | 0.13 | 1 | 3 | Relb | 0.048 | Young |
| 0.004791 | 0.087711 | 0.966 | 0.977 | 1 | 3 | Rps23 | -0.011 | Young |
| 0.004833 | 0.967057 | 0.144 | 0.103 | 1 | 3 | Ubac2 | 0.041 | Young |
| 0.004867 | 0.261932 | 0.587 | 0.554 | 1 | 3 | D8Ertd738 | 0.033 | Young |
| 0.004894 | 0.125846 | 0.966 | 0.969 | 1 | 3 | Hsp90ab1 | -0.003 | Young |

| | | | | | | | | |
|---|---|---|---|---|---|---|---|---|
| 0.004911 | 1.474494 | 0.03 | 0.011 | 1 | 3 | Csf3r | 0.019 | Young |
| 0.004944 | 1.494171 | 0.03 | 0.011 | 1 | 3 | Bcl9l | 0.019 | Young |
| 0.00496 | 0.54896 | 0.241 | 0.192 | 1 | 3 | Supt4a | 0.049 | Young |
| 0.005084 | 1.455935 | 0.018 | 0.004 | 1 | 3 | E130102H2 | 0.014 | Young |
| 0.005228 | 0.564355 | 0.29 | 0.235 | 1 | 3 | Atf7ip | 0.055 | Young |
| 0.005356 | 0.272032 | 0.643 | 0.624 | 1 | 3 | Csf1r | 0.019 | Young |
| 0.005907 | 0.294521 | 0.687 | 0.691 | 1 | 3 | Gadd45b | -0.004 | Young |
| 0.006008 | 0.348773 | 0.169 | 0.121 | 1 | 3 | Ddhd2 | 0.048 | Young |
| 0.006056 | 0.305668 | 0.497 | 0.466 | 1 | 3 | Ralbp1 | 0.031 | Young |
| 0.006103 | 0.382487 | 0.432 | 0.394 | 1 | 3 | Tmem234 | 0.038 | Young |
| 0.006104 | 1.164548 | 0.062 | 0.033 | 1 | 3 | 2-Sep | 0.029 | Young |
| 0.006499 | 0.773287 | 0.14 | 0.099 | 1 | 3 | Ethe1 | 0.041 | Young |
| 0.006566 | 1.010253 | 0.051 | 0.025 | 1 | 3 | Dgkq | 0.026 | Young |
| 0.006661 | 0.773981 | 0.093 | 0.059 | 1 | 3 | Pygb | 0.034 | Young |
| 0.006758 | 1.744178 | 0.021 | 0.006 | 1 | 3 | Tubb3 | 0.015 | Young |
| 0.00684 | 0.933908 | 0.111 | 0.074 | 1 | 3 | Tmem63b | 0.037 | Young |
| 0.007207 | 0.161057 | 0.903 | 0.914 | 1 | 3 | Rps7 | -0.011 | Young |
| 0.007258 | 0.639495 | 0.091 | 0.056 | 1 | 3 | Sesn1 | 0.035 | Young |
| 0.007593 | 0.263558 | 0.626 | 0.597 | 1 | 3 | Tmem176a | 0.029 | Young |
| 0.007633 | 0.163653 | 0.83 | 0.821 | 1 | 3 | Rpl4 | 0.009 | Young |
| 0.007821 | 1.361772 | 0.038 | 0.017 | 1 | 3 | Cfd | 0.021 | Young |
| 0.007952 | 0.224168 | 0.699 | 0.679 | 1 | 3 | Tpr | 0.02 | Young |
| 0.008545 | 1.844211 | 0.016 | 0.004 | 1 | 3 | Hopxos | 0.012 | Young |
| 0.008576 | 0.286421 | 0.667 | 0.636 | 1 | 3 | Tmcc1 | 0.031 | Young |
| 0.008654 | 1.566433 | 0.011 | 0.001 | 1 | 3 | Gm16638 | 0.01 | Young |
| 0.008654 | 1.359424 | 0.011 | 0.001 | 1 | 3 | Angptl6 | 0.01 | Young |
| 0.008781 | 0.994605 | 0.016 | 0.004 | 1 | 3 | A530010L1 | 0.012 | Young |
| 0.009104 | 0.234145 | 0.702 | 0.7 | 1 | 3 | Mrpl33 | 0.002 | Young |
| 0.009123 | 0.438282 | 0.408 | 0.369 | 1 | 3 | Zcchc11 | 0.039 | Young |
| 0.00921 | 0.161035 | 0.848 | 0.84 | 1 | 3 | Rpl7a | 0.008 | Young |
| 0.009385 | 1.339632 | 0.027 | 0.01 | 1 | 3 | Magi2 | 0.017 | Young |
| 0.009436 | 0.977044 | 0.039 | 0.018 | 1 | 3 | Lrrc28 | 0.021 | Young |
| 0.009669 | 1.30026 | 0.058 | 0.032 | 1 | 3 | Rbm12b1 | 0.026 | Young |
| 0.010013 | 0.026374 | 0.259 | 0.337 | 1 | 3 | Ube2j2 | -0.078 | Young |
| 0.01006 | 0.942753 | 0.083 | 0.053 | 1 | 3 | Grap2 | 0.03 | Young |
| 0.010114 | 0.731569 | 0.149 | 0.108 | 1 | 3 | Kdm3a | 0.041 | Young |
| 0.010395 | 1.027105 | 0.035 | 0.016 | 1 | 3 | Kdelc1 | 0.019 | Young |
| 0.01078 | 0.393223 | 0.303 | 0.259 | 1 | 3 | Espn | 0.044 | Young |
| 0.010849 | 0.665248 | 0.122 | 0.086 | 1 | 3 | Atxn1 | 0.036 | Young |
| 0.010999 | 1.175577 | 0.02 | 0.006 | 1 | 3 | 1-Mar | 0.014 | Young |
| 0.011147 | 0.227029 | 0.696 | 0.673 | 1 | 3 | Hint1 | 0.023 | Young |
| 0.011274 | 0.2035 | 0.434 | 0.38 | 1 | 3 | Neurl3 | 0.054 | Young |
| 0.011317 | 0.205296 | 0.783 | 0.76 | 1 | 3 | Rassf4 | 0.023 | Young |
| 0.011441 | 0.077841 | 0.977 | 0.983 | 1 | 3 | Rpl7 | -0.006 | Young |
| 0.011471 | 0.659692 | 0.128 | 0.09 | 1 | 3 | Siglece | 0.038 | Young |

| | | | | | | | | |
|---|---|---|---|---|---|---|---|---|
| 0.011486 | 0.216338 | 0.842 | 0.852 | 1 | 3 | Rps18 | -0.01 | Young |
| 0.012133 | 0.397024 | 0.423 | 0.388 | 1 | 3 | Raly | 0.035 | Young |
| 0.012252 | 0.333502 | 0.446 | 0.403 | 1 | 3 | Ash1l | 0.043 | Young |
| 0.012417 | 0.239061 | 0.654 | 0.626 | 1 | 3 | Fcgr2b | 0.028 | Young |
| 0.012444 | 0.905983 | 0.042 | 0.02 | 1 | 3 | Ccdc102a | 0.022 | Young |
| 0.012534 | 0.559037 | 0.302 | 0.264 | 1 | 3 | Cd80 | 0.038 | Young |
| 0.012609 | 0.078133 | 0.991 | 0.995 | 1 | 3 | H2-Eb1 | -0.004 | Young |
| 0.013025 | 0.434393 | 0.331 | 0.288 | 1 | 3 | Skint3 | 0.043 | Young |
| 0.013178 | 0.124499 | 0.909 | 0.924 | 1 | 3 | Rpl10a | -0.015 | Young |
| 0.013286 | 0.487978 | 0.249 | 0.203 | 1 | 3 | Psmg4 | 0.046 | Young |
| 0.013526 | 0.614562 | 0.172 | 0.133 | 1 | 3 | Ccm2 | 0.039 | Young |
| 0.01354 | 1.191287 | 0.043 | 0.022 | 1 | 3 | Eif5a2 | 0.021 | Young |
| 0.014017 | 0.611697 | 0.126 | 0.091 | 1 | 3 | Stab1 | 0.035 | Young |
| 0.014161 | 0.439203 | 0.141 | 0.102 | 1 | 3 | Wipi2 | 0.039 | Young |
| 0.01417 | 0.563427 | 0.192 | 0.148 | 1 | 3 | Kcnq1 | 0.044 | Young |
| 0.014187 | 0.002498 | 0.254 | 0.325 | 1 | 3 | Rnf19b | -0.071 | Young |
| 0.014286 | 1.403915 | 0.021 | 0.007 | 1 | 3 | Harbi1 | 0.014 | Young |
| 0.014347 | 1.332685 | 0.021 | 0.007 | 1 | 3 | Hacl1 | 0.014 | Young |
| 0.014508 | 1.134364 | 0.015 | 0.004 | 1 | 3 | 2900060B1 | 0.011 | Young |
| 0.014727 | 1.243614 | 0.025 | 0.01 | 1 | 3 | Fam110b | 0.015 | Young |
| 0.014926 | 0.255618 | 0.486 | 0.444 | 1 | 3 | Slc43a2 | 0.042 | Young |
| 0.015005 | 0.152437 | 0.751 | 0.744 | 1 | 3 | Cox7a2l | 0.007 | Young |
| 0.015105 | 0.664059 | 0.064 | 0.038 | 1 | 3 | 9530036M | 0.026 | Young |
| 0.015208 | 0.561316 | 0.102 | 0.069 | 1 | 3 | Pex5 | 0.033 | Young |
| 0.015303 | 1.671662 | 0.01 | 0.001 | 1 | 3 | Dact3 | 0.009 | Young |
| 0.016074 | 0.18203 | 0.74 | 0.734 | 1 | 3 | Pnrc1 | 0.006 | Young |
| 0.016127 | 0.937364 | 0.066 | 0.039 | 1 | 3 | Vipas39 | 0.027 | Young |
| 0.016417 | 1.331635 | 0.047 | 0.025 | 1 | 3 | Reps2 | 0.022 | Young |
| 0.01759 | 0.93576 | 0.059 | 0.035 | 1 | 3 | Scd1 | 0.024 | Young |
| 0.017772 | 0.395679 | 0.332 | 0.288 | 1 | 3 | Hcfc1r1 | 0.044 | Young |
| 0.018514 | 0.940356 | 0.053 | 0.03 | 1 | 3 | Dedd2 | 0.023 | Young |
| 0.018829 | 0.820228 | 0.097 | 0.067 | 1 | 3 | Akr1c18 | 0.03 | Young |
| 0.019269 | 0.253518 | 0.562 | 0.523 | 1 | 3 | Selenop | 0.039 | Young |
| 0.019293 | 0.483335 | 0.068 | 0.042 | 1 | 3 | Rnase4 | 0.026 | Young |
| 0.019374 | 0.076609 | 0.979 | 0.977 | 1 | 3 | Eif1 | 0.002 | Young |
| 0.019522 | 0.806098 | 0.21 | 0.17 | 1 | 3 | Bank1 | 0.04 | Young |
| 0.019682 | 0.43113 | 0.129 | 0.093 | 1 | 3 | Cd163 | 0.036 | Young |
| 0.019697 | 0.772892 | 0.107 | 0.075 | 1 | 3 | Gstm1 | 0.032 | Young |
| 0.019829 | 0.829525 | 0.032 | 0.014 | 1 | 3 | Ip6k2 | 0.018 | Young |
| 0.020231 | 0.680627 | 0.073 | 0.047 | 1 | 3 | Ctc1 | 0.026 | Young |
| 0.020637 | 0.099486 | 0.008 | 0.022 | 1 | 3 | Top1mt | -0.014 | Young |
| 0.020698 | 1.04183 | 0.086 | 0.057 | 1 | 3 | Actr8 | 0.029 | Young |
| 0.020935 | 1.087616 | 0.068 | 0.043 | 1 | 3 | 1700006J1 | 0.025 | Young |
| 0.021149 | 0.643351 | 0.21 | 0.172 | 1 | 3 | Uimc1 | 0.038 | Young |
| 0.021165 | 0.999535 | 0.016 | 0.005 | 1 | 3 | Negr1 | 0.011 | Young |

| | | | | | | | | |
|---|---|---|---|---|---|---|---|---|
| 0.021183 | 0.093452 | 0.951 | 0.951 | 1 | 3 | Rpl6 | 0 | Young |
| 0.02136 | 0.553398 | 0.054 | 0.031 | 1 | 3 | Pck2 | 0.023 | Young |
| 0.021996 | 0.859111 | 0.064 | 0.039 | 1 | 3 | Rps6ka5 | 0.025 | Young |
| 0.022184 | 0.399272 | 0.225 | 0.182 | 1 | 3 | Map3k2 | 0.043 | Young |
| 0.022229 | 1.256772 | 0.024 | 0.01 | 1 | 3 | Inpp4b | 0.014 | Young |
| 0.022339 | 0.633529 | 0.139 | 0.104 | 1 | 3 | Hdac5 | 0.035 | Young |
| 0.022489 | 0.616774 | 0.152 | 0.117 | 1 | 3 | Xpa | 0.035 | Young |
| 0.022563 | 0.839554 | 0.038 | 0.019 | 1 | 3 | Tfpi | 0.019 | Young |
| 0.022889 | 0.154302 | 0.797 | 0.783 | 1 | 3 | Capg | 0.014 | Young |
| 0.022911 | 0.302562 | 0.439 | 0.403 | 1 | 3 | Stk38 | 0.036 | Young |
| 0.022954 | 0.879418 | 0.02 | 0.007 | 1 | 3 | P2ry12 | 0.013 | Young |
| 0.023144 | 0.55998 | 0.02 | 0.007 | 1 | 3 | Inpp1 | 0.013 | Young |
| 0.023149 | 0.211439 | 0.564 | 0.517 | 1 | 3 | Aes | 0.047 | Young |
| 0.023909 | 0.895725 | 0.039 | 0.02 | 1 | 3 | Gk5 | 0.019 | Young |
| 0.023953 | 1.351638 | 0.014 | 0.004 | 1 | 3 | Zbtb12 | 0.01 | Young |
| 0.023994 | 1.432098 | 0.014 | 0.004 | 1 | 3 | Nipsnap1 | 0.01 | Young |
| 0.02419 | 0.205475 | 0.554 | 0.526 | 1 | 3 | Aldh1a2 | 0.028 | Young |
| 0.024201 | 1.388302 | 0.014 | 0.004 | 1 | 3 | Bend6 | 0.01 | Young |
| 0.024243 | 1.050473 | 0.014 | 0.004 | 1 | 3 | D130017N( | 0.01 | Young |
| 0.024257 | 0.104032 | 0.914 | 0.921 | 1 | 3 | Rpl3 | -0.007 | Young |
| 0.024284 | 1.270427 | 0.014 | 0.004 | 1 | 3 | A930033H1 | 0.01 | Young |
| 0.024299 | 1.302888 | 0.047 | 0.026 | 1 | 3 | Dpysl3 | 0.021 | Young |
| 0.024367 | 1.287293 | 0.014 | 0.004 | 1 | 3 | Tle2 | 0.01 | Young |
| 0.024378 | 0.380172 | 0.289 | 0.246 | 1 | 3 | Trafd1 | 0.043 | Young |
| 0.024406 | 1.17979 | 0.029 | 0.013 | 1 | 3 | Gm16867 | 0.016 | Young |
| 0.024577 | 1.064454 | 0.014 | 0.004 | 1 | 3 | 4933433G1 | 0.01 | Young |
| 0.024621 | 0.681103 | 0.102 | 0.072 | 1 | 3 | Faap20 | 0.03 | Young |
| 0.02503 | 0.629385 | 0.146 | 0.111 | 1 | 3 | Tbrg1 | 0.035 | Young |
| 0.025106 | 0.265455 | 0.553 | 0.517 | 1 | 3 | Zcchc6 | 0.036 | Young |
| 0.025219 | 0.366964 | 0.412 | 0.375 | 1 | 3 | Pdcd4 | 0.037 | Young |
| 0.026409 | 0.492335 | 0.324 | 0.289 | 1 | 3 | Arpp19 | 0.035 | Young |
| 0.026842 | 0.277517 | 0.534 | 0.515 | 1 | 3 | Wasf2 | 0.019 | Young |
| 0.027392 | 0.287851 | 0.612 | 0.603 | 1 | 3 | Raph1 | 0.009 | Young |
| 0.027824 | 0.898901 | 0.146 | 0.115 | 1 | 3 | Atg16l1 | 0.031 | Young |
| 0.027882 | 0.257268 | 0.664 | 0.676 | 1 | 3 | H2afj | -0.012 | Young |
| 0.028085 | 0.366603 | 0.096 | 0.066 | 1 | 3 | Zbtb16 | 0.03 | Young |
| 0.028295 | 0.790714 | 0.03 | 0.014 | 1 | 3 | Gpr34 | 0.016 | Young |
| 0.028778 | 0.739185 | 0.042 | 0.023 | 1 | 3 | P3h2 | 0.019 | Young |
| 0.02884 | 0.271522 | 0.456 | 0.434 | 1 | 3 | Nfe2l1 | 0.022 | Young |
| 0.02923 | 0.53932 | 0.202 | 0.167 | 1 | 3 | Tns1 | 0.035 | Young |
| 0.029233 | 0.457788 | 0.188 | 0.151 | 1 | 3 | Mid1ip1 | 0.037 | Young |
| 0.030435 | 1.098951 | 0.037 | 0.019 | 1 | 3 | Acvr2b | 0.018 | Young |
| 0.030975 | 1.134257 | 0.027 | 0.012 | 1 | 3 | Atp2b4 | 0.015 | Young |
| 0.031299 | 0.848744 | 0.037 | 0.019 | 1 | 3 | Cdan1 | 0.018 | Young |
| 0.031587 | 0.177395 | 0.655 | 0.643 | 1 | 3 | Chd4 | 0.012 | Young |

| | | | | | | | | |
|---|---|---|---|---|---|---|---|---|
| 0.031849 | 0.362403 | 0.072 | 0.047 | 1 | 3 | Mrpl1 | 0.025 | Young |
| 0.032125 | 0.727601 | 0.058 | 0.036 | 1 | 3 | Rpain | 0.022 | Young |
| 0.032157 | 0.252362 | 0.525 | 0.495 | 1 | 3 | Dot1l | 0.03 | Young |
| 0.032706 | 0.122146 | 0.864 | 0.857 | 1 | 3 | Npm1 | 0.007 | Young |
| 0.0329 | 0.22647 | 0.735 | 0.76 | 1 | 3 | Akap13 | -0.025 | Young |
| 0.033089 | 0.573221 | 0.043 | 0.024 | 1 | 3 | 1500015A( | 0.019 | Young |
| 0.033529 | 1.49265 | 0.015 | 0.005 | 1 | 3 | Maged2 | 0.01 | Young |
| 0.033657 | 4.790365 | 0.023 | 0.01 | 1 | 3 | Spp1 | 0.013 | Young |
| 0.033787 | 1.066928 | 0.015 | 0.005 | 1 | 3 | Ispd | 0.01 | Young |
| 0.034571 | 0.713933 | 0.015 | 0.005 | 1 | 3 | Gm42031 | 0.01 | Young |
| 0.034903 | 0.898265 | 0.045 | 0.026 | 1 | 3 | Wipi1 | 0.019 | Young |
| 0.035248 | 0.751431 | 0.115 | 0.085 | 1 | 3 | Smg6 | 0.03 | Young |
| 0.035292 | 0.268787 | 0.63 | 0.63 | 1 | 3 | Tob2 | 0 | Young |
| 0.035643 | 0.304399 | 0.471 | 0.447 | 1 | 3 | Ankrd44 | 0.024 | Young |
| 0.035675 | 0.734002 | 0.133 | 0.102 | 1 | 3 | Anapc2 | 0.031 | Young |
| 0.035761 | 1.046771 | 0.028 | 0.013 | 1 | 3 | Rnf135 | 0.015 | Young |
| 0.035775 | 0.177333 | 0.76 | 0.759 | 1 | 3 | Purb | 0.001 | Young |
| 0.03584 | 0.318527 | 0.359 | 0.327 | 1 | 3 | Ski | 0.032 | Young |
| 0.036654 | 0.128171 | 0.811 | 0.816 | 1 | 3 | Ctsh | -0.005 | Young |
| 0.037613 | 1.700457 | 0.024 | 0.011 | 1 | 3 | Pmepa1 | 0.013 | Young |
| 0.037764 | 0.929565 | 0.047 | 0.027 | 1 | 3 | Iqce | 0.02 | Young |
| 0.038157 | 1.203452 | 0.029 | 0.014 | 1 | 3 | Ldlrad1 | 0.015 | Young |
| 0.038462 | 0.651314 | 0.097 | 0.07 | 1 | 3 | Hebp2 | 0.027 | Young |
| 0.038634 | 0.283002 | 0.414 | 0.382 | 1 | 3 | Armc8 | 0.032 | Young |
| 0.038804 | 0.400868 | 0.201 | 0.164 | 1 | 3 | Foxj2 | 0.037 | Young |
| 0.038933 | 0.180704 | 0.124 | 0.164 | 1 | 3 | Cd9 | -0.04 | Young |
| 0.040059 | 1.501943 | 0.013 | 0.004 | 1 | 3 | Dhrs13 | 0.009 | Young |
| 0.040072 | 0.26294 | 0.622 | 0.619 | 1 | 3 | Nfkb1 | 0.003 | Young |
| 0.040097 | 0.103673 | 0.894 | 0.892 | 1 | 3 | Rpl35 | 0.002 | Young |
| 0.040325 | 1.153579 | 0.013 | 0.004 | 1 | 3 | Phtf1os | 0.009 | Young |
| 0.040409 | 0.072379 | 0.985 | 0.992 | 1 | 3 | Rplp1 | -0.007 | Young |
| 0.040552 | 0.861701 | 0.048 | 0.029 | 1 | 3 | Tmem37 | 0.019 | Young |
| 0.040782 | 0.486737 | 0.319 | 0.29 | 1 | 3 | Ssh2 | 0.029 | Young |
| 0.040913 | 0.736186 | 0.024 | 0.011 | 1 | 3 | Rhobtb2 | 0.013 | Young |
| 0.041064 | 0.445366 | 0.013 | 0.004 | 1 | 3 | Gm48604 | 0.009 | Young |
| 0.041758 | 0.251936 | 0.494 | 0.471 | 1 | 3 | Fam111a | 0.023 | Young |
| 0.042664 | 0.948666 | 0.035 | 0.019 | 1 | 3 | Sirt3 | 0.016 | Young |
| 0.043069 | 0.335505 | 0.35 | 0.32 | 1 | 3 | Upb1 | 0.03 | Young |
| 0.04319 | 0.55006 | 0.183 | 0.152 | 1 | 3 | Hmg20b | 0.031 | Young |
| 0.043213 | 0.027929 | 0.096 | 0.13 | 1 | 3 | Smim13 | -0.034 | Young |
| 0.043338 | 0.644945 | 0.09 | 0.063 | 1 | 3 | Rnf152 | 0.027 | Young |
| 0.043392 | 0.457678 | 0.076 | 0.051 | 1 | 3 | Edc3 | 0.025 | Young |
| 0.043741 | 0.707582 | 0.02 | 0.008 | 1 | 3 | Hoxb4 | 0.012 | Young |
| 0.044184 | 1.132996 | 0.016 | 0.006 | 1 | 3 | Rsad1 | 0.01 | Young |
| 0.044937 | 0.031632 | 0.178 | 0.226 | 1 | 3 | Rasa1 | -0.048 | Young |

| | | | | | | | | |
|---|---|---|---|---|---|---|---|---|
| 0.045188 | 0.391989 | 0.389 | 0.367 | 1 | 3 | 9930111J2 | 0.022 | Young |
| 0.045243 | 0.909193 | 0.037 | 0.02 | 1 | 3 | Abcb6 | 0.017 | Young |
| 0.046629 | 0.075574 | 0.131 | 0.171 | 1 | 3 | Bcor | -0.04 | Young |
| 0.046784 | 0.757482 | 0.01 | 0.002 | 1 | 3 | Slc1a4 | 0.008 | Young |
| 0.04687 | 0.785516 | 0.01 | 0.002 | 1 | 3 | Nat14 | 0.008 | Young |
| 0.046956 | 0.653158 | 0.01 | 0.002 | 1 | 3 | Tmem79 | 0.008 | Young |
| 0.047191 | 0.474696 | 0.203 | 0.171 | 1 | 3 | Blvrb | 0.032 | Young |
| 0.047352 | 0.025992 | 0.056 | 0.081 | 1 | 3 | Slc33a1 | -0.025 | Young |
| 0.047419 | 0.326486 | 0.333 | 0.3 | 1 | 3 | Bcl2a1a | 0.033 | Young |
| 0.047665 | 0.096297 | 0.936 | 0.939 | 1 | 3 | Cyba | -0.003 | Young |
| 0.04769 | 0.595171 | 0.143 | 0.112 | 1 | 3 | Usp37 | 0.031 | Young |
| 0.047972 | 0.64939 | 0.088 | 0.063 | 1 | 3 | Prpsap2 | 0.025 | Young |
| 0.048411 | 0.491333 | 0.037 | 0.02 | 1 | 3 | Tamm41 | 0.017 | Young |
| 0.048762 | 0.500894 | 0.164 | 0.131 | 1 | 3 | Gm12840 | 0.033 | Young |
| 0.04879 | 0.253345 | 0.307 | 0.264 | 1 | 3 | Tnrc6b | 0.043 | Young |
| 0.048932 | 0.283765 | 0.465 | 0.443 | 1 | 3 | Lamtor2 | 0.022 | Young |
| 0.049486 | 0.43166 | 0.126 | 0.097 | 1 | 3 | Pak4 | 0.029 | Young |
| 0.049859 | 0.632964 | 0.159 | 0.129 | 1 | 3 | Crim1 | 0.03 | Young |
| 0.049946 | 0.232022 | 0.458 | 0.444 | 1 | 3 | Cox14 | 0.014 | Young |
| 0.050286 | 0.4306 | 0.316 | 0.29 | 1 | 3 | Iqgap2 | 0.026 | Young |
| 0.050473 | 0.002189 | 0.254 | 0.312 | 1 | 3 | R3hdm1 | -0.058 | Young |
| 0.050642 | 0.002357 | 0.133 | 0.171 | 1 | 3 | Unc50 | -0.038 | Young |
| 0.050906 | 0.514473 | 0.128 | 0.099 | 1 | 3 | Acsl3 | 0.029 | Young |
| 0.051343 | 0.175643 | 0.553 | 0.53 | 1 | 3 | Ndufb10 | 0.023 | Young |
| 0.051509 | 0.394225 | 0.313 | 0.28 | 1 | 3 | Rab8a | 0.033 | Young |
| 0.05163 | 0.154502 | 0.768 | 0.771 | 1 | 3 | Cox8a | -0.003 | Young |
| 0.05167 | 0.456253 | 0.306 | 0.272 | 1 | 3 | Smim11 | 0.034 | Young |
| 0.051762 | 0.107597 | 0.902 | 0.908 | 1 | 3 | Uqcrh | -0.006 | Young |
| 0.052138 | 0.428781 | 0.222 | 0.188 | 1 | 3 | Rapgef6 | 0.034 | Young |
| 0.052567 | 0.570602 | 0.199 | 0.166 | 1 | 3 | Fbxl20 | 0.033 | Young |
| 0.052571 | 1.358968 | 0.018 | 0.007 | 1 | 3 | Cwf19l1 | 0.011 | Young |
| 0.053014 | 0.469445 | 0.095 | 0.068 | 1 | 3 | Mir142hg | 0.027 | Young |
| 0.053042 | 0.087111 | 0.917 | 0.925 | 1 | 3 | Pltp | -0.008 | Young |
| 0.053427 | 0.036861 | 0.354 | 0.416 | 1 | 3 | Eprs | -0.062 | Young |
| 0.053857 | 0.813964 | 0.056 | 0.036 | 1 | 3 | Zfp362 | 0.02 | Young |
| 0.054633 | 1.157072 | 0.014 | 0.005 | 1 | 3 | Ccdc80 | 0.009 | Young |
| 0.054714 | 0.844582 | 0.014 | 0.005 | 1 | 3 | Cryz | 0.009 | Young |
| 0.054874 | 0.394265 | 0.014 | 0.005 | 1 | 3 | Samd11 | 0.009 | Young |
| 0.055821 | 0.923435 | 0.057 | 0.037 | 1 | 3 | Clec4n | 0.02 | Young |
| 0.056383 | 0.166446 | 0.677 | 0.675 | 1 | 3 | S100a13 | 0.002 | Young |
| 0.056463 | 1.010057 | 0.058 | 0.038 | 1 | 3 | Casp2 | 0.02 | Young |
| 0.056828 | 0.682102 | 0.098 | 0.073 | 1 | 3 | Ezh2 | 0.025 | Young |
| 0.057155 | 0.653471 | 0.028 | 0.014 | 1 | 3 | Cyb5d2 | 0.014 | Young |
| 0.057317 | 0.00596 | 0.433 | 0.509 | 1 | 3 | Rnf141 | -0.076 | Young |
| 0.05741 | 0.182164 | 0.682 | 0.673 | 1 | 3 | Txnip | 0.009 | Young |

| | | | | | | | | |
|---|---|---|---|---|---|---|---|---|
| 0.057761 | 0.249586 | 0.051 | 0.075 | 1 | 3 | Scmh1 | -0.024 | Young |
| 0.057784 | 0.061124 | 0.977 | 0.974 | 1 | 3 | Zeb2 | 0.003 | Young |
| 0.057848 | 0.617474 | 0.117 | 0.091 | 1 | 3 | Smim26 | 0.026 | Young |
| 0.058414 | 1.116329 | 0.023 | 0.011 | 1 | 3 | Mbd4 | 0.012 | Young |
| 0.058491 | 0.04051 | 0.148 | 0.185 | 1 | 3 | Atp8b2 | -0.037 | Young |
| 0.058532 | 0.426907 | 0.155 | 0.125 | 1 | 3 | Dok1 | 0.03 | Young |
| 0.058971 | 0.288599 | 0.056 | 0.036 | 1 | 3 | Dus3l | 0.02 | Young |
| 0.059004 | 0.12918 | 0.005 | 0.014 | 1 | 3 | Acsbg1 | -0.009 | Young |
| 0.059982 | 0.205432 | 0.619 | 0.618 | 1 | 3 | Cast | 0.001 | Young |
| 0.060032 | 0.304833 | 0.529 | 0.514 | 1 | 3 | Cyp4f18 | 0.015 | Young |
| 0.060284 | 0.640158 | 0.104 | 0.078 | 1 | 3 | Plxna4os1 | 0.026 | Young |
| 0.060419 | 0.066591 | 0.938 | 0.95 | 1 | 3 | Rpl5 | -0.012 | Young |
| 0.061011 | 0.435715 | 0.247 | 0.217 | 1 | 3 | Cdk12 | 0.03 | Young |
| 0.061279 | 0.088458 | 0.91 | 0.895 | 1 | 3 | Rps17 | 0.015 | Young |
| 0.062368 | 0.283609 | 0.359 | 0.32 | 1 | 3 | Maf | 0.039 | Young |
| 0.062377 | 0.54901 | 0.172 | 0.142 | 1 | 3 | Clip1 | 0.03 | Young |
| 0.063034 | 0.990224 | 0.056 | 0.037 | 1 | 3 | Ccdc170 | 0.019 | Young |
| 0.063235 | 1.043992 | 0.029 | 0.016 | 1 | 3 | Mpv17l | 0.013 | Young |
| 0.063499 | 1.007035 | 0.019 | 0.008 | 1 | 3 | Gstt2 | 0.011 | Young |
| 0.063649 | 1.082438 | 0.019 | 0.008 | 1 | 3 | 1810009A1 | 0.011 | Young |
| 0.06374 | 0.288661 | 0.196 | 0.162 | 1 | 3 | Apopt1 | 0.034 | Young |
| 0.063754 | 0.098559 | 0.058 | 0.038 | 1 | 3 | Spata5 | 0.02 | Young |
| 0.063787 | 0.212265 | 0.578 | 0.57 | 1 | 3 | Scand1 | 0.008 | Young |
| 0.064103 | 0.956515 | 0.019 | 0.008 | 1 | 3 | Btbd6 | 0.011 | Young |
| 0.064103 | 0.846111 | 0.019 | 0.008 | 1 | 3 | Setbp1 | 0.011 | Young |
| 0.064573 | 0.515802 | 0.133 | 0.104 | 1 | 3 | Oxa1l | 0.029 | Young |
| 0.065181 | 0.016371 | 0.105 | 0.139 | 1 | 3 | Zfp984 | -0.034 | Young |
| 0.065212 | 0.275603 | 0.21 | 0.176 | 1 | 3 | Rnf216 | 0.034 | Young |
| 0.065408 | 1.486572 | 0.011 | 0.004 | 1 | 3 | Thbs3 | 0.007 | Young |
| 0.065565 | 0.962761 | 0.057 | 0.038 | 1 | 3 | Dnajb2 | 0.019 | Young |
| 0.06572 | 1.232383 | 0.011 | 0.004 | 1 | 3 | Gemin8 | 0.007 | Young |
| 0.065737 | 0.348798 | 0.024 | 0.041 | 1 | 3 | Gdf15 | -0.017 | Young |
| 0.065777 | 0.308813 | 0.168 | 0.136 | 1 | 3 | Tmem135 | 0.032 | Young |
| 0.065788 | 0.450288 | 0.09 | 0.066 | 1 | 3 | Zfp322a | 0.024 | Young |
| 0.065829 | 0.115504 | 0.811 | 0.82 | 1 | 3 | Ndufa6 | -0.009 | Young |
| 0.065929 | 1.159682 | 0.011 | 0.004 | 1 | 3 | AC162302. | 0.007 | Young |
| 0.065939 | 0.72214 | 0.08 | 0.057 | 1 | 3 | Ogfod1 | 0.023 | Young |
| 0.066052 | 0.316867 | 0.413 | 0.397 | 1 | 3 | Arrb1 | 0.016 | Young |
| 0.066253 | 0.810925 | 0.024 | 0.012 | 1 | 3 | A430078G2 | 0.012 | Young |
| 0.066771 | 0.871247 | 0.011 | 0.004 | 1 | 3 | Gm19412 | 0.007 | Young |
| 0.066838 | 0.854286 | 0.037 | 0.022 | 1 | 3 | Tmem115 | 0.015 | Young |
| 0.066855 | 0.370204 | 0.124 | 0.096 | 1 | 3 | Fam219b | 0.028 | Young |
| 0.066877 | 0.555607 | 0.011 | 0.004 | 1 | 3 | Fam234b | 0.007 | Young |
| 0.067089 | 0.536247 | 0.011 | 0.004 | 1 | 3 | Zswim3 | 0.007 | Young |
| 0.06766 | 0.061263 | 0.955 | 0.957 | 1 | 3 | Ccr2 | -0.002 | Young |

| | | | | | | | | |
|---|---|---|---|---|---|---|---|---|
| 0.068751 | 1.099958 | 0.058 | 0.039 | 1 | 3 | Rbms2 | 0.019 | Young |
| 0.06887 | 0.156966 | 0.355 | 0.323 | 1 | 3 | Lmo4 | 0.032 | Young |
| 0.06906 | 0.571188 | 0.077 | 0.055 | 1 | 3 | Agfg2 | 0.022 | Young |
| 0.069161 | 0.155823 | 0.726 | 0.73 | 1 | 3 | Add3 | -0.004 | Young |
| 0.069398 | 0.163618 | 0.028 | 0.045 | 1 | 3 | Btbd19 | -0.017 | Young |
| 0.069708 | 0.281324 | 0.37 | 0.338 | 1 | 3 | Ppp1r12c | 0.032 | Young |
| 0.069734 | 0.03705 | 0.157 | 0.196 | 1 | 3 | Ceacam1 | -0.039 | Young |
| 0.070103 | 0.90932 | 0.047 | 0.03 | 1 | 3 | Xpnpep3 | 0.017 | Young |
| 0.070347 | 0.310852 | 0.376 | 0.348 | 1 | 3 | Maz | 0.028 | Young |
| 0.071803 | 0.002729 | 0.129 | 0.162 | 1 | 3 | Pofut1 | -0.033 | Young |
| 0.071956 | 0.524373 | 0.168 | 0.141 | 1 | 3 | Igf1r | 0.027 | Young |
| 0.072322 | 0.517516 | 0.045 | 0.029 | 1 | 3 | Dalrd3 | 0.016 | Young |
| 0.072421 | 1.035388 | 0.02 | 0.01 | 1 | 3 | Hdac8 | 0.01 | Young |
| 0.072683 | 0.457804 | 0.056 | 0.037 | 1 | 3 | Mfng | 0.019 | Young |
| 0.072705 | 0.291542 | 0.157 | 0.127 | 1 | 3 | Kbtbd11 | 0.03 | Young |
| 0.072819 | 0.56641 | 0.08 | 0.057 | 1 | 3 | Trove2 | 0.023 | Young |
| 0.073394 | 0.230598 | 0.48 | 0.458 | 1 | 3 | Arl5b | 0.022 | Young |
| 0.073584 | 0.746495 | 0.059 | 0.041 | 1 | 3 | Ttll3 | 0.018 | Young |
| 0.074117 | 0.20976 | 0.61 | 0.619 | 1 | 3 | Vsir | -0.009 | Young |
| 0.074286 | 1.036063 | 0.02 | 0.01 | 1 | 3 | Zfp12 | 0.01 | Young |
| 0.074672 | 0.089779 | 0.913 | 0.922 | 1 | 3 | Serf2 | -0.009 | Young |
| 0.074795 | 0.194128 | 0.664 | 0.661 | 1 | 3 | Pfdn5 | 0.003 | Young |
| 0.075123 | 0.236408 | 0.342 | 0.312 | 1 | 3 | Retreg2 | 0.03 | Young |
| 0.075856 | 0.330938 | 0.02 | 0.01 | 1 | 3 | Faap24 | 0.01 | Young |
| 0.076532 | 0.192079 | 0.664 | 0.673 | 1 | 3 | Uqcrq | -0.009 | Young |
| 0.076883 | 1.140386 | 0.04 | 0.025 | 1 | 3 | Zfp944 | 0.015 | Young |
| 0.076903 | 0.663588 | 0.039 | 0.024 | 1 | 3 | Tmem101 | 0.015 | Young |
| 0.077388 | 0.510404 | 0.157 | 0.129 | 1 | 3 | Rps6ka2 | 0.028 | Young |
| 0.077756 | 0.109792 | 0.942 | 0.933 | 1 | 3 | Rps25 | 0.009 | Young |
| 0.077784 | 0.396823 | 0.199 | 0.174 | 1 | 3 | Stx6 | 0.025 | Young |
| 0.077865 | 0.170284 | 0.521 | 0.503 | 1 | 3 | Pdlim1 | 0.018 | Young |
| 0.0785 | 0.325597 | 0.109 | 0.084 | 1 | 3 | Rtl5 | 0.025 | Young |
| 0.078592 | 0.681897 | 0.114 | 0.088 | 1 | 3 | Rnf215 | 0.026 | Young |
| 0.078784 | 0.325882 | 0.167 | 0.135 | 1 | 3 | Trappc5 | 0.032 | Young |
| 0.078839 | 0.000307 | 0.229 | 0.272 | 1 | 3 | Tyk2 | -0.043 | Young |
| 0.079163 | 0.607692 | 0.136 | 0.111 | 1 | 3 | Capn5 | 0.025 | Young |
| 0.080059 | 0.627048 | 0.073 | 0.053 | 1 | 3 | Capn15 | 0.02 | Young |
| 0.080334 | 1.389133 | 0.016 | 0.007 | 1 | 3 | Zfp41 | 0.009 | Young |
| 0.080829 | 0.43617 | 0.278 | 0.253 | 1 | 3 | Fam208a | 0.025 | Young |
| 0.081066 | 0.564796 | 0.187 | 0.16 | 1 | 3 | Erlec1 | 0.027 | Young |
| 0.081121 | 1.190777 | 0.016 | 0.007 | 1 | 3 | Bbs7 | 0.009 | Young |
| 0.081508 | 0.304286 | 0.639 | 0.663 | 1 | 3 | Dennd4a | -0.024 | Young |
| 0.081517 | 0.748171 | 0.016 | 0.007 | 1 | 3 | Ass1 | 0.009 | Young |
| 0.081586 | 0.724254 | 0.098 | 0.076 | 1 | 3 | Topors | 0.022 | Young |
| 0.081809 | 0.299424 | 0.143 | 0.115 | 1 | 3 | Pcmtd2 | 0.028 | Young |

| | | | | | | | | |
|---|---|---|---|---|---|---|---|---|
| 0.081947 | 0.29526 | 0.489 | 0.468 | 1 | 3 | Rabac1 | 0.021 | Young |
| 0.082116 | 0.582064 | 0.092 | 0.069 | 1 | 3 | Cnnm3 | 0.023 | Young |
| 0.082205 | 0.83841 | 0.034 | 0.02 | 1 | 3 | Ccdc163 | 0.014 | Young |
| 0.082276 | 0.390066 | 0.14 | 0.114 | 1 | 3 | Hmgn2 | 0.026 | Young |
| 0.082662 | 0.712323 | 0.063 | 0.044 | 1 | 3 | Adap2 | 0.019 | Young |
| 0.082889 | 0.334993 | 0.215 | 0.183 | 1 | 3 | Akap9 | 0.032 | Young |
| 0.083825 | 0.0731 | 0.956 | 0.968 | 1 | 3 | Rpl37 | -0.012 | Young |
| 0.084179 | 0.675269 | 0.064 | 0.045 | 1 | 3 | Abhd10 | 0.019 | Young |
| 0.084423 | 0.561096 | 0.162 | 0.135 | 1 | 3 | Ppp4r1 | 0.027 | Young |
| 0.084551 | 0.881952 | 0.043 | 0.027 | 1 | 3 | Cdc42bpg | 0.016 | Young |
| 0.084876 | 0.320571 | 0.146 | 0.118 | 1 | 3 | Fnta | 0.028 | Young |
| 0.085086 | 0.786391 | 0.067 | 0.048 | 1 | 3 | Atpaf2 | 0.019 | Young |
| 0.085493 | 1.216542 | 0.028 | 0.016 | 1 | 3 | Maged1 | 0.012 | Young |
| 0.085563 | 0.45254 | 0.021 | 0.011 | 1 | 3 | Tex14 | 0.01 | Young |
| 0.085843 | 0.460865 | 0.12 | 0.093 | 1 | 3 | Nbr1 | 0.027 | Young |
| 0.086173 | 0.629157 | 0.066 | 0.047 | 1 | 3 | Fbxo31 | 0.019 | Young |
| 0.086513 | 0.7085 | 0.013 | 0.005 | 1 | 3 | Tnfrsf14 | 0.008 | Young |
| 0.086781 | 0.384364 | 0.107 | 0.082 | 1 | 3 | Flcn | 0.025 | Young |
| 0.087119 | 0.642002 | 0.013 | 0.005 | 1 | 3 | Lcmt2 | 0.008 | Young |
| 0.087467 | 0.859001 | 0.057 | 0.039 | 1 | 3 | Nfatc2ip | 0.018 | Young |
| 0.087678 | 0.486587 | 0.059 | 0.041 | 1 | 3 | Pigf | 0.018 | Young |
| 0.087683 | 0.325794 | 0.338 | 0.317 | 1 | 3 | Higd2a | 0.021 | Young |
| 0.087772 | 0.043672 | 0.172 | 0.21 | 1 | 3 | Brd8 | -0.038 | Young |
| 0.087971 | 0.250346 | 0.379 | 0.354 | 1 | 3 | Klhl24 | 0.025 | Young |
| 0.088367 | 0.378957 | 0.192 | 0.165 | 1 | 3 | Atp11c | 0.027 | Young |
| 0.088376 | 0.001262 | 0.1 | 0.129 | 1 | 3 | Stk25 | -0.029 | Young |
| 0.088547 | 0.000692 | 0.261 | 0.311 | 1 | 3 | Tes | -0.05 | Young |
| 0.088722 | 0.43441 | 0.336 | 0.321 | 1 | 3 | Smap1 | 0.015 | Young |
| 0.088758 | 0.55412 | 0.035 | 0.022 | 1 | 3 | Gabbr1 | 0.013 | Young |
| 0.089126 | 0.100291 | 0.029 | 0.045 | 1 | 3 | Tsr3 | -0.016 | Young |
| 0.089148 | 0.36783 | 0.053 | 0.036 | 1 | 3 | Ift46 | 0.017 | Young |
| 0.090658 | 0.349999 | 0.369 | 0.345 | 1 | 3 | Ier3 | 0.024 | Young |
| 0.090734 | 0.819208 | 0.057 | 0.039 | 1 | 3 | Bckdhb | 0.018 | Young |
| 0.090889 | 0.163705 | 0.677 | 0.663 | 1 | 3 | Batf3 | 0.014 | Young |
| 0.091485 | 0.088891 | 0.928 | 0.944 | 1 | 3 | Rps11 | -0.016 | Young |
| 0.09239 | 0.684003 | 0.044 | 0.029 | 1 | 3 | Vps72 | 0.015 | Young |
| 0.09251 | 0.549936 | 0.348 | 0.321 | 1 | 3 | Egr3 | 0.027 | Young |
| 0.093142 | 0.66344 | 0.071 | 0.051 | 1 | 3 | Cd33 | 0.02 | Young |
| 0.09346 | 0.832238 | 0.023 | 0.012 | 1 | 3 | Ccdc138 | 0.011 | Young |
| 0.093962 | 0.357326 | 0.374 | 0.357 | 1 | 3 | Pkn1 | 0.017 | Young |
| 0.094264 | 1.066774 | 0.018 | 0.008 | 1 | 3 | Nxpe3 | 0.01 | Young |
| 0.094477 | 0.958562 | 0.018 | 0.008 | 1 | 3 | Trp53rkb | 0.01 | Young |
| 0.094604 | 0.640618 | 0.057 | 0.039 | 1 | 3 | Cry1 | 0.018 | Young |
| 0.094737 | 0.700255 | 0.029 | 0.017 | 1 | 3 | Fam109b | 0.012 | Young |
| 0.095766 | 0.061439 | 0.173 | 0.211 | 1 | 3 | Smg5 | -0.038 | Young |

| | | | | | | | | |
|---|---|---|---|---|---|---|---|---|
| 0.095813 | 0.363949 | 0.343 | 0.32 | 1 | 3 | Cxcr4 | 0.023 | Young |
| 0.095967 | 0.311619 | 0.299 | 0.274 | 1 | 3 | Bod1l | 0.025 | Young |
| 0.095975 | 0.801069 | 0.018 | 0.008 | 1 | 3 | Wdr62 | 0.01 | Young |
| 0.096097 | 0.064948 | 0.158 | 0.194 | 1 | 3 | P2ry10b | -0.036 | Young |
| 0.096194 | 0.028397 | 0.064 | 0.087 | 1 | 3 | Cttnbp2nl | -0.023 | Young |
| 0.096292 | 0.578373 | 0.188 | 0.164 | 1 | 3 | Rxra | 0.024 | Young |
| 0.096312 | 0.598069 | 0.11 | 0.087 | 1 | 3 | Cdyl | 0.023 | Young |
| 0.097039 | 0.56198 | 0.12 | 0.096 | 1 | 3 | Arrdc3 | 0.024 | Young |
| 0.097118 | 0.054725 | 0.241 | 0.283 | 1 | 3 | Tox4 | -0.042 | Young |
| 0.097364 | 0.315643 | 0.098 | 0.076 | 1 | 3 | Rabgap1l | 0.022 | Young |
| 0.097398 | 0.487055 | 0.126 | 0.102 | 1 | 3 | Bbc3 | 0.024 | Young |
| 0.099124 | 0.188947 | 0.692 | 0.701 | 1 | 3 | Sf3b1 | -0.009 | Young |
| 0.09921 | 0.866782 | 0.024 | 0.013 | 1 | 3 | Osbpl1a | 0.011 | Young |
| 0.099365 | 0.414636 | 0.182 | 0.154 | 1 | 3 | Snrnp48 | 0.028 | Young |
| 0.099537 | 0.227389 | 0.235 | 0.205 | 1 | 3 | Rem1 | 0.03 | Young |
| 0.099834 | 0.25601 | 0.473 | 0.456 | 1 | 3 | Klf13 | 0.017 | Young |
| 0.10037 | 0.446131 | 0.059 | 0.042 | 1 | 3 | Efnb2 | 0.017 | Young |
| 0.100576 | 0.018956 | 0.115 | 0.146 | 1 | 3 | Ino80c | -0.031 | Young |
| 0.101287 | 0.044792 | 0.024 | 0.038 | 1 | 3 | Nudcd1 | -0.014 | Young |
| 0.101718 | 0.104331 | 0.851 | 0.851 | 1 | 3 | Ube2d3 | 0 | Young |
| 0.102561 | 0.548047 | 0.004 | 0.011 | 1 | 3 | Zswim9 | -0.007 | Young |
| 0.102957 | 0.400314 | 0.093 | 0.072 | 1 | 3 | Zfp180 | 0.021 | Young |
| 0.103097 | 0.644233 | 0.093 | 0.072 | 1 | 3 | Prkca | 0.021 | Young |
| 0.10331 | 0.785028 | 0.087 | 0.067 | 1 | 3 | Jade1 | 0.02 | Young |
| 0.103564 | 1.353241 | 0.014 | 0.006 | 1 | 3 | Trim16 | 0.008 | Young |
| 0.103564 | 1.281397 | 0.014 | 0.006 | 1 | 3 | Amigo3 | 0.008 | Young |
| 0.10361 | 0.267036 | 0.468 | 0.456 | 1 | 3 | Adcy7 | 0.012 | Young |
| 0.10364 | 0.449995 | 0.038 | 0.024 | 1 | 3 | 5730480H( | 0.014 | Young |
| 0.103894 | 0.713191 | 0.039 | 0.025 | 1 | 3 | Slc25a15 | 0.014 | Young |
| 0.1039 | 0.650681 | 0.059 | 0.042 | 1 | 3 | Arel1 | 0.017 | Young |
| 0.10432 | 0.687252 | 0.024 | 0.013 | 1 | 3 | Ift43 | 0.011 | Young |
| 0.104494 | 0.342535 | 0.375 | 0.361 | 1 | 3 | Cnppd1 | 0.014 | Young |
| 0.105068 | 0.431441 | 0.03 | 0.018 | 1 | 3 | Clmp | 0.012 | Young |
| 0.105489 | 0.792038 | 0.032 | 0.019 | 1 | 3 | Pik3ap1 | 0.013 | Young |
| 0.10575 | 0.354769 | 0.11 | 0.087 | 1 | 3 | Mfap1a | 0.023 | Young |
| 0.105912 | 0.054298 | 0.982 | 0.99 | 1 | 3 | Rps3a1 | -0.008 | Young |
| 0.105931 | 0.290332 | 0.29 | 0.268 | 1 | 3 | Fam129a | 0.022 | Young |
| 0.106027 | 0.644548 | 0.064 | 0.047 | 1 | 3 | Fas | 0.017 | Young |
| 0.106103 | 0.831579 | 0.049 | 0.033 | 1 | 3 | A530017D: | 0.016 | Young |
| 0.106276 | 0.353268 | 0.409 | 0.392 | 1 | 3 | Ubn2 | 0.017 | Young |
| 0.106319 | 0.967154 | 0.053 | 0.037 | 1 | 3 | Zbtb40 | 0.016 | Young |
| 0.106852 | 0.436977 | 0.231 | 0.204 | 1 | 3 | Slc41a3 | 0.027 | Young |
| 0.107012 | 1.233883 | 0.01 | 0.004 | 1 | 3 | Stox2 | 0.006 | Young |
| 0.107222 | 0.210207 | 0.259 | 0.227 | 1 | 3 | Cnpy2 | 0.032 | Young |
| 0.107375 | 0.486867 | 0.082 | 0.062 | 1 | 3 | Zfp704 | 0.02 | Young |

| | | | | | | | | |
|---|---|---|---|---|---|---|---|---|
| 0.107639 | 0.188699 | 0.548 | 0.547 | 1 | 3 | Apbb1ip | 0.001 | Young |
| 0.107661 | 1.421528 | 0.01 | 0.004 | 1 | 3 | Pir | 0.006 | Young |
| 0.107676 | 0.077196 | 0.014 | 0.025 | 1 | 3 | Ggcx | -0.011 | Young |
| 0.107823 | 0.870451 | 0.01 | 0.004 | 1 | 3 | Smkr-ps | 0.006 | Young |
| 0.10789 | 0.357721 | 0.192 | 0.165 | 1 | 3 | Gtf3c2 | 0.027 | Young |
| 0.107904 | 0.317353 | 0.182 | 0.154 | 1 | 3 | Lmo1 | 0.028 | Young |
| 0.107955 | 0.079017 | 0.014 | 0.006 | 1 | 3 | Bicd1 | 0.008 | Young |
| 0.108312 | 0.87074 | 0.01 | 0.004 | 1 | 3 | Fam212b | 0.006 | Young |
| 0.108803 | 0.660625 | 0.01 | 0.004 | 1 | 3 | Gm49207 | 0.006 | Young |
| 0.109296 | 0.764744 | 0.01 | 0.004 | 1 | 3 | Zscan20 | 0.006 | Young |
| 0.10946 | 0.651552 | 0.01 | 0.004 | 1 | 3 | 9330020H( | 0.006 | Young |
| 0.109625 | 0.516915 | 0.01 | 0.004 | 1 | 3 | Xpo5 | 0.006 | Young |
| 0.109885 | 0.522585 | 0.071 | 0.053 | 1 | 3 | Abl1 | 0.018 | Young |
| 0.110028 | 0.528749 | 0.078 | 0.059 | 1 | 3 | F630028O1 | 0.019 | Young |
| 0.11012 | 0.505687 | 0.01 | 0.004 | 1 | 3 | 4732471J0 | 0.006 | Young |
| 0.110284 | 0.230722 | 0.393 | 0.369 | 1 | 3 | Phb2 | 0.024 | Young |
| 0.11036 | 0.080915 | 0.121 | 0.152 | 1 | 3 | Xylt1 | -0.031 | Young |
| 0.110397 | 0.171225 | 0.069 | 0.05 | 1 | 3 | Mtfr1 | 0.019 | Young |
| 0.110432 | 0.402041 | 0.145 | 0.121 | 1 | 3 | Susd3 | 0.024 | Young |
| 0.110452 | 0.130919 | 0.01 | 0.004 | 1 | 3 | Dbndd2 | 0.006 | Young |
| 0.110716 | 0.393468 | 0.126 | 0.103 | 1 | 3 | Tshz1 | 0.023 | Young |
| 0.111016 | 0.283924 | 0.438 | 0.418 | 1 | 3 | Git2 | 0.02 | Young |
| 0.111148 | 0.396747 | 0.091 | 0.07 | 1 | 3 | Tpgs1 | 0.021 | Young |
| 0.111271 | 0.906442 | 0.033 | 0.02 | 1 | 3 | Gm12089 | 0.013 | Young |
| 0.111616 | 1.039337 | 0.043 | 0.029 | 1 | 3 | Gstp3 | 0.014 | Young |
| 0.112072 | 0.443111 | 0.264 | 0.24 | 1 | 3 | Tbc1d23 | 0.024 | Young |
| 0.112077 | 0.535217 | 0.213 | 0.189 | 1 | 3 | Gm47283 | 0.024 | Young |
| 0.112259 | 0.510238 | 0.186 | 0.161 | 1 | 3 | 8-Sep | 0.025 | Young |
| 0.112329 | 0.049651 | 0.996 | 0.994 | 1 | 3 | Rps29 | 0.002 | Young |
| 0.112388 | 0.659134 | 0.068 | 0.05 | 1 | 3 | Scimp | 0.018 | Young |
| 0.113237 | 0.535548 | 0.157 | 0.131 | 1 | 3 | Ranbp10 | 0.026 | Young |
| 0.113328 | 0.322808 | 0.247 | 0.221 | 1 | 3 | Aff1 | 0.026 | Young |
| 0.113539 | 0.287651 | 0.141 | 0.117 | 1 | 3 | Ufd1 | 0.024 | Young |
| 0.113569 | 0.349274 | 0.188 | 0.161 | 1 | 3 | Antxr2 | 0.027 | Young |
| 0.113641 | 0.540088 | 0.133 | 0.109 | 1 | 3 | Parp1 | 0.024 | Young |
| 0.114112 | 0.502763 | 0.225 | 0.202 | 1 | 3 | Mob3a | 0.023 | Young |
| 0.114603 | 0.465688 | 0.101 | 0.08 | 1 | 3 | Gtpbp6 | 0.021 | Young |
| 0.114832 | 0.016367 | 0.093 | 0.121 | 1 | 3 | Gm5547 | -0.028 | Young |
| 0.11485 | 0.084933 | 0.087 | 0.114 | 1 | 3 | Selenon | -0.027 | Young |
| 0.114862 | 0.289237 | 0.253 | 0.222 | 1 | 3 | Prkab2 | 0.031 | Young |
| 0.114898 | 1.150969 | 0.027 | 0.016 | 1 | 3 | Tchh | 0.011 | Young |
| 0.115957 | 0.729593 | 0.034 | 0.022 | 1 | 3 | Prkd2 | 0.012 | Young |
| 0.11702 | 0.213608 | 0.726 | 0.762 | 1 | 3 | Rel | -0.036 | Young |
| 0.11714 | 0.39563 | 0.159 | 0.134 | 1 | 3 | Smim27 | 0.025 | Young |
| 0.117308 | 0.70676 | 0.044 | 0.03 | 1 | 3 | 1700096K1 | 0.014 | Young |

| | | | | | | | | |
|---|---|---|---|---|---|---|---|---|
| 0.119049 | 0.98141 | 0.02 | 0.011 | 1 | 3 | Nme6 | 0.009 | Young |
| 0.119172 | 0.073957 | 0.015 | 0.026 | 1 | 3 | Wdr59 | -0.011 | Young |
| 0.119675 | 0.164139 | 0.678 | 0.7 | 1 | 3 | Nfam1 | -0.022 | Young |
| 0.119944 | 0.088708 | 0.725 | 0.725 | 1 | 3 | Plbd1 | 0 | Young |
| 0.119951 | 0.254951 | 0.509 | 0.502 | 1 | 3 | Bcl10 | 0.007 | Young |
| 0.119957 | 0.371757 | 0.16 | 0.134 | 1 | 3 | Cir1 | 0.026 | Young |
| 0.120091 | 0.307422 | 0.288 | 0.26 | 1 | 3 | Fam49a | 0.028 | Young |
| 0.120325 | 0.43376 | 0.034 | 0.022 | 1 | 3 | Lyve1 | 0.012 | Young |
| 0.12068 | 0.635848 | 0.034 | 0.022 | 1 | 3 | Prickle3 | 0.012 | Young |
| 0.120784 | 0.445482 | 0.056 | 0.039 | 1 | 3 | Gde1 | 0.017 | Young |
| 0.120928 | 0.280352 | 0.479 | 0.467 | 1 | 3 | Zbtb20 | 0.012 | Young |
| 0.121327 | 0.076004 | 0.936 | 0.949 | 1 | 3 | Fcer1g | -0.013 | Young |
| 0.12142 | 0.072665 | 0.841 | 0.849 | 1 | 3 | Emp3 | -0.008 | Young |
| 0.121498 | 0.538223 | 0.092 | 0.072 | 1 | 3 | Extl3 | 0.02 | Young |
| 0.121649 | 0.580972 | 0.02 | 0.011 | 1 | 3 | Exog | 0.009 | Young |
| 0.12165 | 1.039351 | 0.015 | 0.007 | 1 | 3 | C1qtnf1 | 0.008 | Young |
| 0.12192 | 0.463875 | 0.027 | 0.016 | 1 | 3 | Zfp948 | 0.011 | Young |
| 0.121946 | 0.534515 | 0.043 | 0.029 | 1 | 3 | Selenoo | 0.014 | Young |
| 0.122028 | 0.499643 | 0.097 | 0.076 | 1 | 3 | Igf1 | 0.021 | Young |
| 0.122259 | 0.781586 | 0.058 | 0.042 | 1 | 3 | Pkdcc | 0.016 | Young |
| 0.122433 | 0.999933 | 0.037 | 0.024 | 1 | 3 | Nsun4 | 0.013 | Young |
| 0.122466 | 0.544059 | 0.034 | 0.022 | 1 | 3 | Zkscan8 | 0.012 | Young |
| 0.122634 | 0.93775 | 0.015 | 0.007 | 1 | 3 | Fgfbp3 | 0.008 | Young |
| 0.123137 | 0.110284 | 0.338 | 0.311 | 1 | 3 | Mknk2 | 0.027 | Young |
| 0.123632 | 0.978942 | 0.028 | 0.017 | 1 | 3 | Ophn1 | 0.011 | Young |
| 0.124673 | 0.240983 | 0.293 | 0.266 | 1 | 3 | Pik3r1 | 0.027 | Young |
| 0.124678 | 0.171955 | 0.527 | 0.516 | 1 | 3 | Elob | 0.011 | Young |
| 0.12469 | 0.477976 | 0.076 | 0.057 | 1 | 3 | Hjurp | 0.019 | Young |
| 0.124978 | 0.030597 | 0.015 | 0.007 | 1 | 3 | Nat8f1 | 0.008 | Young |
| 0.125653 | 0.798667 | 0.076 | 0.057 | 1 | 3 | Ciao1 | 0.019 | Young |
| 0.125729 | 0.911989 | 0.047 | 0.032 | 1 | 3 | Mthfr | 0.015 | Young |
| 0.125771 | 1.038941 | 0.051 | 0.036 | 1 | 3 | Nsdhl | 0.015 | Young |
| 0.1261 | 0.01616 | 0.092 | 0.117 | 1 | 3 | Wdr20 | -0.025 | Young |
| 0.126278 | 0.510534 | 0.028 | 0.017 | 1 | 3 | Pcgf1 | 0.011 | Young |
| 0.126294 | 0.809407 | 0.066 | 0.049 | 1 | 3 | Kif16b | 0.017 | Young |
| 0.126709 | 0.134384 | 0.042 | 0.059 | 1 | 3 | Frmd4a | -0.017 | Young |
| 0.126823 | 0.026849 | 0.277 | 0.326 | 1 | 3 | Hsd17b12 | -0.049 | Young |
| 0.127165 | 0.016426 | 0.163 | 0.197 | 1 | 3 | Aagab | -0.034 | Young |
| 0.127415 | 0.475988 | 0.077 | 0.059 | 1 | 3 | Spryd3 | 0.018 | Young |
| 0.127547 | 0.043321 | 0.064 | 0.086 | 1 | 3 | Nxt1 | -0.022 | Young |
| 0.127721 | 0.74605 | 0.028 | 0.017 | 1 | 3 | Naif1 | 0.011 | Young |
| 0.127969 | 0.147378 | 0.054 | 0.038 | 1 | 3 | S1pr2 | 0.016 | Young |
| 0.128084 | 0.216043 | 0.244 | 0.217 | 1 | 3 | Cebpa | 0.027 | Young |
| 0.128244 | 0.696481 | 0.048 | 0.033 | 1 | 3 | Fam122a | 0.015 | Young |
| 0.128542 | 0.332873 | 0.26 | 0.235 | 1 | 3 | Pip5k1c | 0.025 | Young |

| | | | | | | | | |
|---|---|---|---|---|---|---|---|---|
| 0.128761 | 0.380846 | 0.144 | 0.121 | 1 | 3 | Tagap | 0.023 | Young |
| 0.129281 | 0.798723 | 0.028 | 0.017 | 1 | 3 | Syce2 | 0.011 | Young |
| 0.130296 | 0.261157 | 0.412 | 0.401 | 1 | 3 | Gatad2b | 0.011 | Young |
| 0.130623 | 0.077191 | 0.848 | 0.853 | 1 | 3 | Cox7c | -0.005 | Young |
| 0.130738 | 0.846732 | 0.051 | 0.036 | 1 | 3 | Lsm11 | 0.015 | Young |
| 0.13079 | 0.541539 | 0.061 | 0.044 | 1 | 3 | Ankrd54 | 0.017 | Young |
| 0.13104 | 0.041255 | 0.128 | 0.158 | 1 | 3 | Cdkn2aipnl | -0.03 | Young |
| 0.131248 | 0.510969 | 0.021 | 0.012 | 1 | 3 | Rsl1 | 0.009 | Young |
| 0.131368 | 0.246614 | 0.46 | 0.438 | 1 | 3 | Tuba1a | 0.022 | Young |
| 0.132347 | 0.698432 | 0.021 | 0.012 | 1 | 3 | Nr6a1 | 0.009 | Young |
| 0.132961 | 0.369637 | 0.021 | 0.012 | 1 | 3 | Capsl | 0.009 | Young |
| 0.133012 | 1.08315 | 0.011 | 0.005 | 1 | 3 | Hs3st3b1 | 0.006 | Young |
| 0.133049 | 0.007673 | 0.342 | 0.395 | 1 | 3 | Clec12a | -0.053 | Young |
| 0.133335 | 0.085638 | 0.894 | 0.912 | 1 | 3 | Rpl36a | -0.018 | Young |
| 0.133371 | 0.007249 | 0.205 | 0.247 | 1 | 3 | Aif1 | -0.042 | Young |
| 0.133849 | 0.125683 | 0.13 | 0.106 | 1 | 3 | Usp36 | 0.024 | Young |
| 0.133945 | 0.541439 | 0.111 | 0.09 | 1 | 3 | Pias4 | 0.021 | Young |
| 0.134252 | 1.371059 | 0.011 | 0.005 | 1 | 3 | Ankrd28 | 0.006 | Young |
| 0.13443 | 0.882741 | 0.011 | 0.005 | 1 | 3 | Itga6 | 0.006 | Young |
| 0.134496 | 0.499924 | 0.141 | 0.118 | 1 | 3 | Vps53 | 0.023 | Young |
| 0.134786 | 1.346798 | 0.011 | 0.005 | 1 | 3 | Zfp111 | 0.006 | Young |
| 0.134965 | 0.6258 | 0.011 | 0.005 | 1 | 3 | Steap4 | 0.006 | Young |
| 0.135501 | 1.262048 | 0.011 | 0.005 | 1 | 3 | Zfp784 | 0.006 | Young |
| 0.135543 | 0.66523 | 0.064 | 0.048 | 1 | 3 | Fam35a | 0.016 | Young |
| 0.135647 | 0.435258 | 0.17 | 0.147 | 1 | 3 | Asxl1 | 0.023 | Young |
| 0.135739 | 0.059948 | 0.101 | 0.129 | 1 | 3 | Tmub2 | -0.028 | Young |
| 0.135843 | 0.262005 | 0.114 | 0.091 | 1 | 3 | Mrpl45 | 0.023 | Young |
| 0.136358 | 0.864539 | 0.057 | 0.042 | 1 | 3 | Mylip | 0.015 | Young |
| 0.136374 | 0.100876 | 0.811 | 0.826 | 1 | 3 | Psma7 | -0.015 | Young |
| 0.136683 | 0.003204 | 0.048 | 0.066 | 1 | 3 | Csad | -0.018 | Young |
| 0.136717 | 0.509888 | 0.106 | 0.086 | 1 | 3 | Map3k4 | 0.02 | Young |
| 0.136759 | 0.63482 | 0.011 | 0.005 | 1 | 3 | Kyat3 | 0.006 | Young |
| 0.136773 | 0.643137 | 0.038 | 0.025 | 1 | 3 | Fbxl4 | 0.013 | Young |
| 0.137019 | 0.135339 | 0.657 | 0.654 | 1 | 3 | Ogt | 0.003 | Young |
| 0.137902 | 0.365999 | 0.292 | 0.274 | 1 | 3 | Arrb2 | 0.018 | Young |
| 0.137975 | 0.646735 | 0.038 | 0.025 | 1 | 3 | Akap1 | 0.013 | Young |
| 0.138029 | 0.802681 | 0.016 | 0.008 | 1 | 3 | Tesmin | 0.008 | Young |
| 0.138624 | 0.653395 | 0.072 | 0.055 | 1 | 3 | Apip | 0.017 | Young |
| 0.138954 | 0.857001 | 0.04 | 0.027 | 1 | 3 | C030034I2 | 0.013 | Young |
| 0.139137 | 0.467746 | 0.029 | 0.018 | 1 | 3 | Zfp825 | 0.011 | Young |
| 0.139203 | 0.397977 | 0.115 | 0.093 | 1 | 3 | Pam16 | 0.022 | Young |
| 0.139353 | 0.580569 | 0.029 | 0.018 | 1 | 3 | Atrip | 0.011 | Young |
| 0.139371 | 0.585709 | 0.038 | 0.025 | 1 | 3 | Cbx7 | 0.013 | Young |
| 0.139561 | 0.834892 | 0.08 | 0.062 | 1 | 3 | Nfrkb | 0.018 | Young |
| 0.139746 | 0.618435 | 0.038 | 0.025 | 1 | 3 | Ttf2 | 0.013 | Young |

| | | | | | | | | |
|---|---|---|---|---|---|---|---|---|
| 0.14052 | 0.049392 | 0.341 | 0.392 | 1 | 3 | Susd6 | -0.051 | Young |
| 0.140929 | 0.561324 | 0.063 | 0.047 | 1 | 3 | D730003I1 | 0.016 | Young |
| 0.141094 | 0.037315 | 0.157 | 0.188 | 1 | 3 | Rabggtb | -0.031 | Young |
| 0.141638 | 0.010172 | 0.058 | 0.078 | 1 | 3 | Wdsub1 | -0.02 | Young |
| 0.141663 | 0.411166 | 0.162 | 0.14 | 1 | 3 | Gcn1l1 | 0.022 | Young |
| 0.141808 | 0.648503 | 0.023 | 0.013 | 1 | 3 | 1110051M | 0.01 | Young |
| 0.14205 | 0.151815 | 0.114 | 0.141 | 1 | 3 | Gpd1l | -0.027 | Young |
| 0.142179 | 0.352834 | 0.186 | 0.16 | 1 | 3 | Sh2d1b1 | 0.026 | Young |
| 0.142269 | 0.324058 | 0.27 | 0.247 | 1 | 3 | Polb | 0.023 | Young |
| 0.142614 | 0.451986 | 0.163 | 0.141 | 1 | 3 | Mrpl9 | 0.022 | Young |
| 0.142668 | 0.322767 | 0.105 | 0.084 | 1 | 3 | Gtf3c1 | 0.021 | Young |
| 0.142787 | 0.041939 | 0.074 | 0.097 | 1 | 3 | Cars | -0.023 | Young |
| 0.142798 | 0.713195 | 0.039 | 0.026 | 1 | 3 | Serinc5 | 0.013 | Young |
| 0.14339 | 0.549931 | 0.016 | 0.008 | 1 | 3 | Gm26809 | 0.008 | Young |
| 0.143501 | 0.594713 | 0.023 | 0.013 | 1 | 3 | Apex2 | 0.01 | Young |
| 0.14352 | 0.205759 | 0.313 | 0.288 | 1 | 3 | Skil | 0.025 | Young |
| 0.143722 | 0.333817 | 0.356 | 0.336 | 1 | 3 | St3gal4 | 0.02 | Young |
| 0.143969 | 0.087524 | 0.856 | 0.858 | 1 | 3 | Dock2 | -0.002 | Young |
| 0.144681 | 0.235437 | 0.313 | 0.292 | 1 | 3 | Ccdc88c | 0.021 | Young |
| 0.145399 | 0.500145 | 0.023 | 0.013 | 1 | 3 | Tbc1d8b | 0.01 | Young |
| 0.146016 | 0.13567 | 0.131 | 0.162 | 1 | 3 | Ppm1h | -0.031 | Young |
| 0.146334 | 0.582747 | 0.074 | 0.057 | 1 | 3 | Psmf1 | 0.017 | Young |
| 0.14634 | 0.179308 | 0.184 | 0.221 | 1 | 3 | Katna1 | -0.037 | Young |
| 0.146412 | 0.772048 | 0.032 | 0.02 | 1 | 3 | Acp6 | 0.012 | Young |
| 0.147364 | 0.293666 | 0.275 | 0.253 | 1 | 3 | Uqcc3 | 0.022 | Young |
| 0.147537 | 0.75183 | 0.042 | 0.029 | 1 | 3 | Zfp866 | 0.013 | Young |
| 0.147794 | 0.60865 | 0.04 | 0.027 | 1 | 3 | Zfp809 | 0.013 | Young |
| 0.14854 | 0.485111 | 0.072 | 0.055 | 1 | 3 | Rab33b | 0.017 | Young |
| 0.148682 | 0.057407 | 0.255 | 0.3 | 1 | 3 | Bfar | -0.045 | Young |
| 0.148741 | 0.01925 | 0.15 | 0.18 | 1 | 3 | Rap1gap2 | -0.03 | Young |
| 0.148793 | 0.190327 | 0.034 | 0.049 | 1 | 3 | Epg5 | -0.015 | Young |
| 0.148957 | 0.011954 | 0.436 | 0.492 | 1 | 3 | Ppig | -0.056 | Young |
| 0.149689 | 0.510316 | 0.04 | 0.027 | 1 | 3 | Mfsd6 | 0.013 | Young |
| 0.150105 | 0.277585 | 0.274 | 0.253 | 1 | 3 | Dstyk | 0.021 | Young |
| 0.150612 | 0.26257 | 0.313 | 0.296 | 1 | 3 | Brd1 | 0.017 | Young |
| 0.150649 | 0.064719 | 0.157 | 0.186 | 1 | 3 | Nae1 | -0.029 | Young |
| 0.150688 | 0.215481 | 0.139 | 0.17 | 1 | 3 | Slc30a9 | -0.031 | Young |
| 0.150948 | 0.728073 | 0.066 | 0.05 | 1 | 3 | Phf6 | 0.016 | Young |
| 0.151109 | 0.369694 | 0.091 | 0.072 | 1 | 3 | 1600010M | 0.019 | Young |
| 0.152302 | 0.100802 | 0.196 | 0.232 | 1 | 3 | Egln2 | -0.036 | Young |
| 0.152381 | 0.332502 | 0.091 | 0.072 | 1 | 3 | Suclg2 | 0.019 | Young |
| 0.152599 | 0.397825 | 0.058 | 0.043 | 1 | 3 | Zfp719 | 0.015 | Young |
| 0.153034 | 0.460884 | 0.091 | 0.073 | 1 | 3 | Gnpda2 | 0.018 | Young |
| 0.153348 | 0.38198 | 0.331 | 0.314 | 1 | 3 | Inpp4a | 0.017 | Young |
| 0.153966 | 0.025784 | 0.04 | 0.056 | 1 | 3 | Pogk | -0.016 | Young |

| | | | | | | | | |
|---|---|---|---|---|---|---|---|---|
| 0.154147 | 0.68099 | 0.018 | 0.01 | 1 | 3 | Haus5 | 0.008 | Young |
| 0.15426 | 0.846493 | 0.044 | 0.031 | 1 | 3 | Acsf3 | 0.013 | Young |
| 0.154829 | 0.87434 | 0.024 | 0.014 | 1 | 3 | Kif23 | 0.01 | Young |
| 0.154923 | 0.277796 | 0.186 | 0.161 | 1 | 3 | Cramp1l | 0.025 | Young |
| 0.154949 | 0.306589 | 0.21 | 0.186 | 1 | 3 | Stard8 | 0.024 | Young |
| 0.155523 | 0.351426 | 0.018 | 0.01 | 1 | 3 | Pfkm | 0.008 | Young |
| 0.155798 | 0.659628 | 0.063 | 0.048 | 1 | 3 | Dnajc9 | 0.015 | Young |
| 0.155984 | 0.577975 | 0.018 | 0.01 | 1 | 3 | Chek2 | 0.008 | Young |
| 0.156908 | 0.669614 | 0.018 | 0.01 | 1 | 3 | Mtcp1 | 0.008 | Young |
| 0.157008 | 0.010699 | 0.236 | 0.272 | 1 | 3 | Zfp292 | -0.036 | Young |
| 0.157238 | 1.015739 | 0.013 | 0.006 | 1 | 3 | Bbs2 | 0.007 | Young |
| 0.157322 | 0.342016 | 0.234 | 0.211 | 1 | 3 | Mrpl48 | 0.023 | Young |
| 0.157433 | 0.040511 | 0.596 | 0.656 | 1 | 3 | Bach1 | -0.06 | Young |
| 0.157645 | 0.317434 | 0.004 | 0.01 | 1 | 3 | 4932422M | -0.006 | Young |
| 0.157799 | 0.955759 | 0.013 | 0.006 | 1 | 3 | Zfp764 | 0.007 | Young |
| 0.157987 | 1.387377 | 0.013 | 0.006 | 1 | 3 | AC163040. | 0.007 | Young |
| 0.158458 | 0.703101 | 0.018 | 0.01 | 1 | 3 | Ccl17 | 0.008 | Young |
| 0.158857 | 0.883401 | 0.034 | 0.023 | 1 | 3 | 2900093K2 | 0.011 | Young |
| 0.159088 | 0.58343 | 0.044 | 0.031 | 1 | 3 | Fdxacb1 | 0.013 | Young |
| 0.159171 | 0.16078 | 0.638 | 0.624 | 1 | 3 | Timp2 | 0.014 | Young |
| 0.159603 | 0.186436 | 0.125 | 0.152 | 1 | 3 | Spata2 | -0.027 | Young |
| 0.15965 | 0.004828 | 0.335 | 0.389 | 1 | 3 | Prex1 | -0.054 | Young |
| 0.159728 | 0.686008 | 0.034 | 0.023 | 1 | 3 | Klhl7 | 0.011 | Young |
| 0.159732 | 0.433257 | 0.112 | 0.093 | 1 | 3 | Lyrm2 | 0.019 | Young |
| 0.160147 | 1.003995 | 0.025 | 0.016 | 1 | 3 | Ralgps1 | 0.009 | Young |
| 0.160489 | 0.067671 | 0.908 | 0.912 | 1 | 3 | Rpl14 | -0.004 | Young |
| 0.160496 | 0.365726 | 0.082 | 0.065 | 1 | 3 | Terf2 | 0.017 | Young |
| 0.160542 | 0.504098 | 0.043 | 0.03 | 1 | 3 | 0610009B2 | 0.013 | Young |
| 0.160554 | 0.346596 | 0.184 | 0.161 | 1 | 3 | Pcbd2 | 0.023 | Young |
| 0.161004 | 0.047302 | 0.197 | 0.232 | 1 | 3 | Crtc3 | -0.035 | Young |
| 0.16116 | 0.557129 | 0.136 | 0.117 | 1 | 3 | Slc9a9 | 0.019 | Young |
| 0.1612 | 0.494084 | 0.013 | 0.006 | 1 | 3 | E030030I0 | 0.007 | Young |
| 0.161329 | 0.467916 | 0.184 | 0.162 | 1 | 3 | Mthfd1 | 0.022 | Young |
| 0.161554 | 0.258455 | 0.255 | 0.231 | 1 | 3 | Phc3 | 0.024 | Young |
| 0.161615 | 0.047611 | 0.188 | 0.221 | 1 | 3 | Setd5 | -0.033 | Young |
| 0.16168 | 0.211335 | 0.359 | 0.339 | 1 | 3 | Ptpn7 | 0.02 | Young |
| 0.162346 | 0.288615 | 0.013 | 0.006 | 1 | 3 | Naprt | 0.007 | Young |
| 0.162394 | 0.122801 | 0.144 | 0.118 | 1 | 3 | Mafk | 0.026 | Young |
| 0.162577 | 0.512862 | 0.061 | 0.045 | 1 | 3 | Gm42047 | 0.016 | Young |
| 0.162716 | 0.005874 | 0.054 | 0.072 | 1 | 3 | Cpne2 | -0.018 | Young |
| 0.162993 | 0.672929 | 0.025 | 0.016 | 1 | 3 | Dclk2 | 0.009 | Young |
| 0.16316 | 0.363486 | 0.181 | 0.158 | 1 | 3 | Vps37b | 0.023 | Young |
| 0.163451 | 0.063249 | 0.193 | 0.232 | 1 | 3 | Faim | -0.039 | Young |
| 0.163491 | 0.781437 | 0.035 | 0.024 | 1 | 3 | Gm12689 | 0.011 | Young |
| 0.163625 | 1.189057 | 0.019 | 0.011 | 1 | 3 | B130006D0 | 0.008 | Young |

| | | | | | | | | |
|---|---|---|---|---|---|---|---|---|
| 0.16387 | 0.554675 | 0.083 | 0.066 | 1 | 3 | Dnlz | 0.017 | Young |
| 0.164506 | 0.655562 | 0.048 | 0.035 | 1 | 3 | Accs | 0.013 | Young |
| 0.165347 | 0.810571 | 0.035 | 0.024 | 1 | 3 | Ppip5k1 | 0.011 | Young |
| 0.16539 | 0.262988 | 0.203 | 0.178 | 1 | 3 | Nabp2 | 0.025 | Young |
| 0.165975 | 0.059955 | 0.133 | 0.16 | 1 | 3 | Plxnc1 | -0.027 | Young |
| 0.16647 | 0.0277 | 0.111 | 0.136 | 1 | 3 | Brwd1 | -0.025 | Young |
| 0.166685 | 0.515662 | 0.104 | 0.086 | 1 | 3 | Pnkd | 0.018 | Young |
| 0.167011 | 0.136242 | 0.571 | 0.565 | 1 | 3 | Rac2 | 0.006 | Young |
| 0.167198 | 0.609758 | 0.115 | 0.097 | 1 | 3 | Arhgef18 | 0.018 | Young |
| 0.167207 | 0.012427 | 0.178 | 0.21 | 1 | 3 | Abhd17b | -0.032 | Young |
| 0.167805 | 0.114841 | 0.199 | 0.174 | 1 | 3 | Lst1 | 0.025 | Young |
| 0.168264 | 0.370528 | 0.025 | 0.016 | 1 | 3 | Lamb3 | 0.009 | Young |
| 0.168632 | 0.025861 | 0.256 | 0.293 | 1 | 3 | Plekha1 | -0.037 | Young |
| 0.169132 | 0.063719 | 0.176 | 0.21 | 1 | 3 | Ptdss1 | -0.034 | Young |
| 0.16987 | 0.136878 | 0.015 | 0.025 | 1 | 3 | Ndufaf1 | -0.01 | Young |
| 0.169954 | 0.316576 | 0.332 | 0.311 | 1 | 3 | Kmt2c | 0.021 | Young |
| 0.170745 | 0.053058 | 0.072 | 0.092 | 1 | 3 | Rad17 | -0.02 | Young |
| 0.170929 | 0.225187 | 0.355 | 0.336 | 1 | 3 | Mrpl23 | 0.019 | Young |
| 0.171295 | 0.404399 | 0.109 | 0.09 | 1 | 3 | Cdc14b | 0.019 | Young |
| 0.171391 | 0.027257 | 0.33 | 0.375 | 1 | 3 | Cebpz | -0.045 | Young |
| 0.171887 | 0.084223 | 0.082 | 0.104 | 1 | 3 | Abcd3 | -0.022 | Young |
| 0.172003 | 0.055429 | 0.912 | 0.921 | 1 | 3 | Gnai2 | -0.009 | Young |
| 0.172033 | 0.048485 | 0.471 | 0.532 | 1 | 3 | Drap1 | -0.061 | Young |
| 0.172205 | 0.177118 | 0.708 | 0.705 | 1 | 3 | Il6ra | 0.003 | Young |
| 0.172901 | 0.081372 | 0.197 | 0.233 | 1 | 3 | Ubr2 | -0.036 | Young |
| 0.1732 | 0.494441 | 0.15 | 0.13 | 1 | 3 | Tuba4a | 0.02 | Young |
| 0.173382 | 0.737423 | 0.092 | 0.074 | 1 | 3 | 9530059O1 | 0.018 | Young |
| 0.173389 | 0.042876 | 0.986 | 0.986 | 1 | 3 | Fth1 | 0 | Young |
| 0.173879 | 0.021128 | 0.183 | 0.215 | 1 | 3 | Man1c1 | -0.032 | Young |
| 0.174231 | 0.107112 | 0.793 | 0.821 | 1 | 3 | Erp29 | -0.028 | Young |
| 0.174407 | 0.153491 | 0.403 | 0.388 | 1 | 3 | H2-DMb2 | 0.015 | Young |
| 0.17443 | 0.069983 | 0.146 | 0.177 | 1 | 3 | Rab12 | -0.031 | Young |
| 0.175621 | 0.119928 | 0.092 | 0.074 | 1 | 3 | Rhobtb1 | 0.018 | Young |
| 0.17578 | 0.544966 | 0.092 | 0.075 | 1 | 3 | Pelp1 | 0.017 | Young |
| 0.176195 | 0.111737 | 0.013 | 0.022 | 1 | 3 | Apoo | -0.009 | Young |
| 0.176484 | 0.552799 | 0.085 | 0.068 | 1 | 3 | Nsl1 | 0.017 | Young |
| 0.177223 | 0.931207 | 0.02 | 0.012 | 1 | 3 | Cd101 | 0.008 | Young |
| 0.177229 | 0.027686 | 0.269 | 0.308 | 1 | 3 | Sp3 | -0.039 | Young |
| 0.177265 | 0.379543 | 0.225 | 0.203 | 1 | 3 | Cnot3 | 0.022 | Young |
| 0.177574 | 0.056603 | 0.168 | 0.197 | 1 | 3 | Timm17a | -0.029 | Young |
| 0.178099 | 0.413338 | 0.284 | 0.274 | 1 | 3 | Fgd2 | 0.01 | Young |
| 0.178527 | 1.112745 | 0.014 | 0.007 | 1 | 3 | Rassf7 | 0.007 | Young |
| 0.17858 | 0.350317 | 0.183 | 0.161 | 1 | 3 | Snrk | 0.022 | Young |
| 0.178806 | 0.037241 | 0.029 | 0.042 | 1 | 3 | Tlr1 | -0.013 | Young |
| 0.178923 | 0.016558 | 0.045 | 0.061 | 1 | 3 | Tbc1d31 | -0.016 | Young |

| | | | | | | | | |
|---|---|---|---|---|---|---|---|---|
| 0.179304 | 0.519682 | 0.083 | 0.067 | 1 | 3 | Togaram1 | 0.016 | Young |
| 0.179369 | 0.304251 | 0.049 | 0.036 | 1 | 3 | Cep290 | 0.013 | Young |
| 0.179912 | 0.254381 | 0.01 | 0.018 | 1 | 3 | Tmem206 | -0.008 | Young |
| 0.179976 | 0.551799 | 0.052 | 0.038 | 1 | 3 | Zfyve1 | 0.014 | Young |
| 0.180468 | 0.886714 | 0.014 | 0.007 | 1 | 3 | Gm10031 | 0.007 | Young |
| 0.180673 | 0.005032 | 0.313 | 0.358 | 1 | 3 | Dnajc7 | -0.045 | Young |
| 0.180676 | 0.742965 | 0.02 | 0.012 | 1 | 3 | 8030462N1 | 0.008 | Young |
| 0.181053 | 0.78703 | 0.014 | 0.007 | 1 | 3 | AW011738 | 0.007 | Young |
| 0.181444 | 0.620082 | 0.014 | 0.007 | 1 | 3 | Mr1 | 0.007 | Young |
| 0.181671 | 0.613529 | 0.056 | 0.042 | 1 | 3 | Cd93 | 0.014 | Young |
| 0.18171 | 0.510768 | 0.082 | 0.066 | 1 | 3 | Fam120b | 0.016 | Young |
| 0.18212 | 0.164517 | 0.428 | 0.423 | 1 | 3 | Ramp1 | 0.005 | Young |
| 0.182253 | 0.891283 | 0.039 | 0.027 | 1 | 3 | Rrnad1 | 0.012 | Young |
| 0.182347 | 0.542327 | 0.051 | 0.037 | 1 | 3 | Sipa1l2 | 0.014 | Young |
| 0.18239 | 0.437142 | 0.028 | 0.018 | 1 | 3 | Ptar1 | 0.01 | Young |
| 0.182581 | 0.655499 | 0.02 | 0.012 | 1 | 3 | Pdk2 | 0.008 | Young |
| 0.182639 | 0.56207 | 0.054 | 0.041 | 1 | 3 | Plpp6 | 0.013 | Young |
| 0.183381 | 0.02175 | 0.245 | 0.287 | 1 | 3 | Poldip3 | -0.042 | Young |
| 0.183699 | 0.681556 | 0.02 | 0.012 | 1 | 3 | Pglyrp1 | 0.008 | Young |
| 0.184198 | 0.66044 | 0.014 | 0.007 | 1 | 3 | Mcc | 0.007 | Young |
| 0.185129 | 0.48594 | 0.169 | 0.148 | 1 | 3 | Nfatc2 | 0.021 | Young |
| 0.185242 | 0.050459 | 0.025 | 0.037 | 1 | 3 | Zmym1 | -0.012 | Young |
| 0.185282 | 0.251601 | 0.306 | 0.283 | 1 | 3 | Tmem154 | 0.023 | Young |
| 0.185532 | 0.050198 | 0.16 | 0.189 | 1 | 3 | Tmem41b | -0.029 | Young |
| 0.185666 | 0.037452 | 0.99 | 0.992 | 1 | 3 | Rpl35a | -0.002 | Young |
| 0.185946 | 0.026036 | 0.186 | 0.217 | 1 | 3 | Ube2d1 | -0.031 | Young |
| 0.18609 | 0.693164 | 0.058 | 0.044 | 1 | 3 | Gdpd3 | 0.014 | Young |
| 0.186273 | 0.508042 | 0.02 | 0.012 | 1 | 3 | Itga2 | 0.008 | Young |
| 0.186785 | 0.764169 | 0.042 | 0.03 | 1 | 3 | Abca2 | 0.012 | Young |
| 0.186802 | 0.274798 | 0.125 | 0.105 | 1 | 3 | Pmvk | 0.02 | Young |
| 0.186855 | 0.170748 | 0.428 | 0.422 | 1 | 3 | Herc1 | 0.006 | Young |
| 0.187344 | 0.61461 | 0.168 | 0.152 | 1 | 3 | Rcsd1 | 0.016 | Young |
| 0.187428 | 0.613711 | 0.058 | 0.044 | 1 | 3 | Wiz | 0.014 | Young |
| 0.187557 | 0.26464 | 0.263 | 0.244 | 1 | 3 | Apc | 0.019 | Young |
| 0.187808 | 0.230174 | 0.124 | 0.104 | 1 | 3 | Chchd7 | 0.02 | Young |
| 0.188408 | 0.824815 | 0.04 | 0.029 | 1 | 3 | Slc7a6 | 0.011 | Young |
| 0.188561 | 0.032541 | 0.376 | 0.437 | 1 | 3 | Strn3 | -0.061 | Young |
| 0.188634 | 0.091872 | 0.139 | 0.167 | 1 | 3 | Erg28 | -0.028 | Young |
| 0.188654 | 0.793732 | 0.069 | 0.055 | 1 | 3 | Syne1 | 0.014 | Young |
| 0.188686 | 0.560545 | 0.054 | 0.041 | 1 | 3 | Oraov1 | 0.013 | Young |
| 0.188849 | 0.036937 | 0.966 | 0.971 | 1 | 3 | Rps15 | -0.005 | Young |
| 0.188977 | 0.370298 | 0.229 | 0.211 | 1 | 3 | Idnk | 0.018 | Young |
| 0.1891 | 1.174338 | 0.021 | 0.013 | 1 | 3 | Spaca6 | 0.008 | Young |
| 0.189768 | 0.353713 | 0.371 | 0.36 | 1 | 3 | Gpr65 | 0.011 | Young |
| 0.190428 | 0.554752 | 0.086 | 0.069 | 1 | 3 | Phf1 | 0.017 | Young |

| | | | | | | | | |
|---|---|---|---|---|---|---|---|---|
| 0.190938 | 0.169972 | 0.03 | 0.043 | 1 | 3 | Nprl3 | -0.013 | Young |
| 0.191009 | 0.134712 | 0.158 | 0.135 | 1 | 3 | Akt3 | 0.023 | Young |
| 0.191143 | 0.1407 | 0.595 | 0.585 | 1 | 3 | Nfe2l2 | 0.01 | Young |
| 0.191636 | 0.980413 | 0.021 | 0.013 | 1 | 3 | Mfge8 | 0.008 | Young |
| 0.192096 | 0.579091 | 0.064 | 0.05 | 1 | 3 | Alkbh3 | 0.014 | Young |
| 0.192234 | 0.061215 | 0.905 | 0.916 | 1 | 3 | Ddx5 | -0.011 | Young |
| 0.192393 | 0.178176 | 0.391 | 0.378 | 1 | 3 | Mapk14 | 0.013 | Young |
| 0.193403 | 0.023894 | 0.126 | 0.152 | 1 | 3 | Pex7 | -0.026 | Young |
| 0.193555 | 0.47054 | 0.057 | 0.043 | 1 | 3 | 3110082I1 | 0.014 | Young |
| 0.193876 | 0.31693 | 0.029 | 0.019 | 1 | 3 | Tldc1 | 0.01 | Young |
| 0.193994 | 0.135083 | 0.039 | 0.027 | 1 | 3 | Vps51 | 0.012 | Young |
| 0.19411 | 0.247069 | 0.235 | 0.216 | 1 | 3 | Ankhd1 | 0.019 | Young |
| 0.194357 | 0.721844 | 0.021 | 0.013 | 1 | 3 | Dusp28 | 0.008 | Young |
| 0.194419 | 0.089968 | 0.008 | 0.014 | 1 | 3 | Timp3 | -0.006 | Young |
| 0.194552 | 0.347449 | 0.192 | 0.172 | 1 | 3 | Sh2b3 | 0.02 | Young |
| 0.194743 | 0.189092 | 0.225 | 0.201 | 1 | 3 | Mbd3 | 0.024 | Young |
| 0.194826 | 0.095863 | 0.117 | 0.097 | 1 | 3 | Ptger4 | 0.02 | Young |
| 0.196038 | 0.015283 | 0.237 | 0.274 | 1 | 3 | Ube2f | -0.037 | Young |
| 0.196491 | 0.012665 | 0.027 | 0.038 | 1 | 3 | 2700097O0 | -0.011 | Young |
| 0.196634 | 0.69595 | 0.03 | 0.02 | 1 | 3 | Supv3l1 | 0.01 | Young |
| 0.197259 | 0.538438 | 0.043 | 0.031 | 1 | 3 | Nod2 | 0.012 | Young |
| 0.197521 | 0.536896 | 0.23 | 0.215 | 1 | 3 | Gpbp1l1 | 0.015 | Young |
| 0.197545 | 0.152194 | 0.6 | 0.614 | 1 | 3 | Syngr2 | -0.014 | Young |
| 0.197594 | 0.465939 | 0.021 | 0.013 | 1 | 3 | Gm9938 | 0.008 | Young |
| 0.197796 | 0.004343 | 0.043 | 0.057 | 1 | 3 | Slc35a1 | -0.014 | Young |
| 0.197811 | 0.070967 | 0.804 | 0.816 | 1 | 3 | Rbm39 | -0.012 | Young |
| 0.198285 | 0.476387 | 0.085 | 0.068 | 1 | 3 | Sharpin | 0.017 | Young |
| 0.198376 | 0.060787 | 0.13 | 0.157 | 1 | 3 | Atraid | -0.027 | Young |
| 0.198421 | 0.780769 | 0.015 | 0.008 | 1 | 3 | Bcr | 0.007 | Young |
| 0.199277 | 0.023793 | 0.157 | 0.185 | 1 | 3 | Akr1c13 | -0.028 | Young |
| 0.199433 | 0.255526 | 0.187 | 0.164 | 1 | 3 | Pcm1 | 0.023 | Young |
| 0.19953 | 0.673758 | 0.044 | 0.032 | 1 | 3 | Dcun1d4 | 0.012 | Young |
| 0.199676 | 0.086133 | 0.038 | 0.051 | 1 | 3 | Zfp780b | -0.013 | Young |
| 0.199809 | 1.036832 | 0.015 | 0.008 | 1 | 3 | Per2 | 0.007 | Young |
| 0.199809 | 0.991545 | 0.015 | 0.008 | 1 | 3 | 1810034E1 | 0.007 | Young |
| 0.200174 | 0.051628 | 0.971 | 0.969 | 1 | 3 | Kctd12 | 0.002 | Young |
| 0.200186 | 0.276288 | 0.211 | 0.189 | 1 | 3 | Ankle2 | 0.022 | Young |
| 0.200592 | 0.360522 | 0.167 | 0.147 | 1 | 3 | Ndufv1 | 0.02 | Young |
| 0.201004 | 0.79537 | 0.015 | 0.008 | 1 | 3 | Poli | 0.007 | Young |
| 0.201466 | 0.261856 | 0.131 | 0.112 | 1 | 3 | Mtmr14 | 0.019 | Young |
| 0.201662 | 0.628711 | 0.071 | 0.056 | 1 | 3 | Pik3r2 | 0.015 | Young |
| 0.202405 | 1.087466 | 0.015 | 0.008 | 1 | 3 | Klhl11 | 0.007 | Young |
| 0.202527 | 0.37225 | 0.106 | 0.088 | 1 | 3 | Klhl21 | 0.018 | Young |
| 0.202806 | 0.627812 | 0.015 | 0.008 | 1 | 3 | Gm29367 | 0.007 | Young |
| 0.202815 | 0.922254 | 0.023 | 0.014 | 1 | 3 | Per3 | 0.009 | Young |

| | | | | | | | | |
|---|---|---|---|---|---|---|---|---|
| 0.203007 | 0.594217 | 0.015 | 0.008 | 1 | 3 | Ndor1 | 0.007 | Young |
| 0.203317 | 0.549563 | 0.116 | 0.099 | 1 | 3 | Trmt6 | 0.017 | Young |
| 0.203385 | 0.511328 | 0.011 | 0.019 | 1 | 3 | Cxcl9 | -0.008 | Young |
| 0.203425 | 0.509276 | 0.059 | 0.045 | 1 | 3 | Fbxo42 | 0.014 | Young |
| 0.203528 | 0.107394 | 0.059 | 0.075 | 1 | 3 | B3glct | -0.016 | Young |
| 0.20369 | 0.31 | 0.093 | 0.076 | 1 | 3 | Pld1 | 0.017 | Young |
| 0.203756 | 0.514286 | 0.095 | 0.078 | 1 | 3 | Mcee | 0.017 | Young |
| 0.203995 | 0.300202 | 0.042 | 0.03 | 1 | 3 | Wdcp | 0.012 | Young |
| 0.204419 | 0.634646 | 0.015 | 0.008 | 1 | 3 | Map2k6 | 0.007 | Young |
| 0.204482 | 0.029365 | 0.172 | 0.201 | 1 | 3 | Cdk19 | -0.029 | Young |
| 0.204589 | 0.511539 | 0.044 | 0.032 | 1 | 3 | Gm16536 | 0.012 | Young |
| 0.20482 | 0.12886 | 0.369 | 0.342 | 1 | 3 | Bmt2 | 0.027 | Young |
| 0.205233 | 0.733985 | 0.023 | 0.014 | 1 | 3 | Ipp | 0.009 | Young |
| 0.20527 | 0.439374 | 0.229 | 0.21 | 1 | 3 | Smg7 | 0.019 | Young |
| 0.205292 | 0.223603 | 0.263 | 0.246 | 1 | 3 | Paics | 0.017 | Young |
| 0.205591 | 0.290208 | 0.396 | 0.384 | 1 | 3 | Nsd3 | 0.012 | Young |
| 0.205876 | 0.201609 | 0.455 | 0.449 | 1 | 3 | Ptpn6 | 0.006 | Young |
| 0.205979 | 0.881121 | 0.054 | 0.042 | 1 | 3 | Bmyc | 0.012 | Young |
| 0.206485 | 1.162491 | 0.01 | 0.005 | 1 | 3 | Slc12a5 | 0.005 | Young |
| 0.206693 | 0.466387 | 0.023 | 0.014 | 1 | 3 | Slc2a3 | 0.009 | Young |
| 0.20672 | 0.331948 | 0.058 | 0.044 | 1 | 3 | Fdx1 | 0.014 | Young |
| 0.206882 | 0.029575 | 0.197 | 0.232 | 1 | 3 | Rala | -0.035 | Young |
| 0.206962 | 0.349256 | 0.154 | 0.135 | 1 | 3 | Ryk | 0.019 | Young |
| 0.207504 | 0.27707 | 0.438 | 0.432 | 1 | 3 | Ddx3y | 0.006 | Young |
| 0.207759 | 0.233863 | 0.346 | 0.324 | 1 | 3 | Creb1 | 0.022 | Young |
| 0.207776 | 0.302542 | 0.381 | 0.37 | 1 | 3 | Coq10b | 0.011 | Young |
| 0.207902 | 0.482383 | 0.069 | 0.055 | 1 | 3 | Cep295 | 0.014 | Young |
| 0.208184 | 0.899538 | 0.054 | 0.042 | 1 | 3 | Ampd3 | 0.012 | Young |
| 0.208537 | 0.315684 | 0.01 | 0.005 | 1 | 3 | Tmem14a | 0.005 | Young |
| 0.20855 | 0.154164 | 0.302 | 0.272 | 1 | 3 | Itpk1 | 0.03 | Young |
| 0.20869 | 0.009973 | 0.259 | 0.3 | 1 | 3 | Arpc1a | -0.041 | Young |
| 0.209053 | 0.789273 | 0.01 | 0.005 | 1 | 3 | 2510017J1 | 0.005 | Young |
| 0.209053 | 0.644261 | 0.01 | 0.005 | 1 | 3 | Wdr27 | 0.005 | Young |
| 0.209059 | 0.247319 | 0.165 | 0.145 | 1 | 3 | Dcaf15 | 0.02 | Young |
| 0.209175 | 0.253804 | 0.318 | 0.3 | 1 | 3 | Arl4c | 0.018 | Young |
| 0.209311 | 0.914778 | 0.01 | 0.005 | 1 | 3 | Ccdc114 | 0.005 | Young |
| 0.209374 | 0.221572 | 0.191 | 0.171 | 1 | 3 | Cep83 | 0.02 | Young |
| 0.209552 | 0.199325 | 0.654 | 0.687 | 1 | 3 | Txn1 | -0.033 | Young |
| 0.209632 | 0.324261 | 0.237 | 0.215 | 1 | 3 | Dhx40 | 0.022 | Young |
| 0.20979 | 0.343844 | 0.164 | 0.143 | 1 | 3 | Ak6 | 0.021 | Young |
| 0.210086 | 0.616559 | 0.01 | 0.005 | 1 | 3 | Stk26 | 0.005 | Young |
| 0.210086 | 0.523778 | 0.01 | 0.005 | 1 | 3 | Gm26826 | 0.005 | Young |
| 0.210345 | 0.44808 | 0.01 | 0.005 | 1 | 3 | Gm39459 | 0.005 | Young |
| 0.210345 | 0.406011 | 0.01 | 0.005 | 1 | 3 | Nthl1 | 0.005 | Young |
| 0.210345 | 0.392125 | 0.01 | 0.005 | 1 | 3 | Ift88 | 0.005 | Young |

| | | | | | | | | |
|---|---|---|---|---|---|---|---|---|
| 0.210637 | 0.285707 | 0.116 | 0.098 | 1 | 3 | Slc31a2 | 0.018 | Young |
| 0.210767 | 0.654163 | 0.049 | 0.037 | 1 | 3 | Vps37c | 0.012 | Young |
| 0.211051 | 0.00618 | 0.408 | 0.456 | 1 | 3 | Galnt1 | -0.048 | Young |
| 0.211124 | 0.565171 | 0.045 | 0.033 | 1 | 3 | Rcbtb1 | 0.012 | Young |
| 0.211451 | 0.116112 | 0.707 | 0.712 | 1 | 3 | Gapdh | -0.005 | Young |
| 0.211989 | 0.165267 | 0.485 | 0.478 | 1 | 3 | Ube2b | 0.007 | Young |
| 0.212144 | 0.023064 | 0.158 | 0.186 | 1 | 3 | Dpp3 | -0.028 | Young |
| 0.212262 | 0.352546 | 0.109 | 0.092 | 1 | 3 | Gid4 | 0.017 | Young |
| 0.212288 | 0.519871 | 0.149 | 0.131 | 1 | 3 | Mknk1 | 0.018 | Young |
| 0.212426 | 0.053926 | 0.225 | 0.258 | 1 | 3 | Fbxl5 | -0.033 | Young |
| 0.212649 | 0.041452 | 0.134 | 0.16 | 1 | 3 | Dag1 | -0.026 | Young |
| 0.212747 | 0.029494 | 0.391 | 0.44 | 1 | 3 | Efr3a | -0.049 | Young |
| 0.213602 | 0.109392 | 0.138 | 0.164 | 1 | 3 | Strn | -0.026 | Young |
| 0.213843 | 0.355874 | 0.097 | 0.08 | 1 | 3 | Rad54l2 | 0.017 | Young |
| 0.213992 | 0.025051 | 0.095 | 0.116 | 1 | 3 | Ranbp3 | -0.021 | Young |
| 0.214217 | 0.144433 | 0.032 | 0.022 | 1 | 3 | Xlr | 0.01 | Young |
| 0.214787 | 0.469534 | 0.092 | 0.075 | 1 | 3 | Txnrd2 | 0.017 | Young |
| 0.21495 | 0.419851 | 0.218 | 0.2 | 1 | 3 | Celf4 | 0.018 | Young |
| 0.215365 | 0.029458 | 0.063 | 0.08 | 1 | 3 | Ppp2r5d | -0.017 | Young |
| 0.215545 | 0.48626 | 0.033 | 0.023 | 1 | 3 | D030056L2 | 0.01 | Young |
| 0.215696 | 0.051858 | 0.222 | 0.259 | 1 | 3 | Kif19a | -0.037 | Young |
| 0.215785 | 0.651137 | 0.035 | 0.025 | 1 | 3 | Cnst | 0.01 | Young |
| 0.216274 | 0.222245 | 0.463 | 0.461 | 1 | 3 | Pdcd5 | 0.002 | Young |
| 0.216392 | 0.960383 | 0.016 | 0.01 | 1 | 3 | Zc3h11a | 0.006 | Young |
| 0.21648 | 0.235641 | 0.439 | 0.45 | 1 | 3 | Mbp | -0.011 | Young |
| 0.216688 | 0.029054 | 0.444 | 0.509 | 1 | 3 | Slc38a1 | -0.065 | Young |
| 0.21709 | 0.068587 | 0.141 | 0.167 | 1 | 3 | Snrnp40 | -0.026 | Young |
| 0.217104 | 0.428425 | 0.183 | 0.165 | 1 | 3 | Nr2c2 | 0.018 | Young |
| 0.217321 | 0.224907 | 0.427 | 0.422 | 1 | 3 | Ssr4 | 0.005 | Young |
| 0.217501 | 0.028491 | 0.182 | 0.211 | 1 | 3 | Dld | -0.029 | Young |
| 0.217534 | 0.102711 | 0.308 | 0.358 | 1 | 3 | Cdk11b | -0.05 | Young |
| 0.217822 | 0.054199 | 0.263 | 0.299 | 1 | 3 | Vps29 | -0.036 | Young |
| 0.218332 | 0.286902 | 0.277 | 0.262 | 1 | 3 | Srgap2 | 0.015 | Young |
| 0.218403 | 0.795568 | 0.016 | 0.01 | 1 | 3 | Hist1h4d | 0.006 | Young |
| 0.218511 | 0.160251 | 0.039 | 0.053 | 1 | 3 | Mettl13 | -0.014 | Young |
| 0.218569 | 0.324344 | 0.316 | 0.307 | 1 | 3 | Dnajc19 | 0.009 | Young |
| 0.218652 | 0.21596 | 0.335 | 0.319 | 1 | 3 | Cd2 | 0.016 | Young |
| 0.218953 | 0.08382 | 0.838 | 0.834 | 1 | 3 | Morf4l1 | 0.004 | Young |
| 0.219985 | 0.496822 | 0.034 | 0.024 | 1 | 3 | Spryd4 | 0.01 | Young |
| 0.219998 | 0.151078 | 0.1 | 0.122 | 1 | 3 | Rasa2 | -0.022 | Young |
| 0.22 | 0.048868 | 0.427 | 0.479 | 1 | 3 | Nup98 | -0.052 | Young |
| 0.220224 | 0.528051 | 0.016 | 0.01 | 1 | 3 | Gm26522 | 0.006 | Young |
| 0.220365 | 0.349964 | 0.063 | 0.049 | 1 | 3 | Znhit6 | 0.014 | Young |
| 0.221037 | 0.885157 | 0.016 | 0.01 | 1 | 3 | Mis18bp1 | 0.006 | Young |
| 0.221064 | 0.263107 | 0.251 | 0.233 | 1 | 3 | Phf23 | 0.018 | Young |

| | | | | | | | | |
|---|---|---|---|---|---|---|---|---|
| 0.221207 | 0.856924 | 0.025 | 0.017 | 1 | 3 | Gm33973 | 0.008 | Young |
| 0.221921 | 0.028814 | 0.082 | 0.1 | 1 | 3 | Isca2 | -0.018 | Young |
| 0.222166 | 0.046281 | 0.097 | 0.119 | 1 | 3 | Adap1 | -0.022 | Young |
| 0.222184 | 0.372698 | 0.024 | 0.016 | 1 | 3 | Pomt2 | 0.008 | Young |
| 0.222261 | 0.48712 | 0.016 | 0.01 | 1 | 3 | Gins1 | 0.006 | Young |
| 0.222287 | 0.120677 | 0.119 | 0.141 | 1 | 3 | Sc5d | -0.022 | Young |
| 0.222333 | 0.764703 | 0.025 | 0.017 | 1 | 3 | Rcn3 | 0.008 | Young |
| 0.223606 | 0.026277 | 0.348 | 0.395 | 1 | 3 | Baz1b | -0.047 | Young |
| 0.223682 | 0.13074 | 0.024 | 0.016 | 1 | 3 | Irgc1 | 0.008 | Young |
| 0.224425 | 0.007129 | 0.287 | 0.324 | 1 | 3 | Rheb | -0.037 | Young |
| 0.224667 | 0.169704 | 0.034 | 0.024 | 1 | 3 | Spindoc | 0.01 | Young |
| 0.224732 | 0.165915 | 0.581 | 0.589 | 1 | 3 | Qpct | -0.008 | Young |
| 0.224979 | 0.307751 | 0.083 | 0.068 | 1 | 3 | N6amt1 | 0.015 | Young |
| 0.225212 | 0.276681 | 0.091 | 0.074 | 1 | 3 | Rnf169 | 0.017 | Young |
| 0.22526 | 0.014845 | 0.078 | 0.097 | 1 | 3 | Zbtb38 | -0.019 | Young |
| 0.225328 | 0.557159 | 0.059 | 0.047 | 1 | 3 | Cxcl12 | 0.012 | Young |
| 0.225864 | 0.011476 | 0.295 | 0.337 | 1 | 3 | Csnk1e | -0.042 | Young |
| 0.22621 | 0.031857 | 0.091 | 0.111 | 1 | 3 | Chpf2 | -0.02 | Young |
| 0.226741 | 0.050947 | 0.121 | 0.145 | 1 | 3 | Taf2 | -0.024 | Young |
| 0.227064 | 0.173142 | 0.085 | 0.068 | 1 | 3 | Trappc13 | 0.017 | Young |
| 0.227079 | 0.640069 | 0.058 | 0.045 | 1 | 3 | Fra10ac1 | 0.013 | Young |
| 0.227132 | 0.627705 | 0.038 | 0.027 | 1 | 3 | Mrps31 | 0.011 | Young |
| 0.227209 | 0.549401 | 0.025 | 0.017 | 1 | 3 | Arhgap19 | 0.008 | Young |
| 0.227373 | 0.584467 | 0.025 | 0.017 | 1 | 3 | 4732440D( | 0.008 | Young |
| 0.2275 | 0.368399 | 0.077 | 0.062 | 1 | 3 | Scfd2 | 0.015 | Young |
| 0.227604 | 0.067585 | 0.187 | 0.213 | 1 | 3 | Pigk | -0.026 | Young |
| 0.228148 | 0.372403 | 0.17 | 0.151 | 1 | 3 | Gtf2i | 0.019 | Young |
| 0.22835 | 0.642352 | 0.061 | 0.048 | 1 | 3 | Slc10a3 | 0.013 | Young |
| 0.228397 | 0.367559 | 0.126 | 0.109 | 1 | 3 | Ckap5 | 0.017 | Young |
| 0.228547 | 0.368621 | 0.199 | 0.179 | 1 | 3 | Zfp652 | 0.02 | Young |
| 0.228561 | 0.663327 | 0.112 | 0.097 | 1 | 3 | Klhdc2 | 0.015 | Young |
| 0.228612 | 0.098751 | 0.198 | 0.232 | 1 | 3 | Gna12 | -0.034 | Young |
| 0.228743 | 0.059797 | 0.081 | 0.099 | 1 | 3 | Wdfy1 | -0.018 | Young |
| 0.228976 | 0.842159 | 0.072 | 0.059 | 1 | 3 | Recql5 | 0.013 | Young |
| 0.229199 | 1.158169 | 0.027 | 0.018 | 1 | 3 | Kiz | 0.009 | Young |
| 0.229237 | 0.387717 | 0.121 | 0.104 | 1 | 3 | Dgcr8 | 0.017 | Young |
| 0.229413 | 0.48724 | 0.02 | 0.03 | 1 | 3 | Aasdh | -0.01 | Young |
| 0.229448 | 0.382366 | 0.078 | 0.063 | 1 | 3 | Epm2aip1 | 0.015 | Young |
| 0.22953 | 0.03602 | 0.042 | 0.055 | 1 | 3 | Usp28 | -0.013 | Young |
| 0.229842 | 0.458963 | 0.058 | 0.045 | 1 | 3 | Cactin | 0.013 | Young |
| 0.23016 | 0.002369 | 0.458 | 0.516 | 1 | 3 | Hnrnpul2 | -0.058 | Young |
| 0.230275 | 0.04095 | 0.074 | 0.092 | 1 | 3 | Arhgap23 | -0.018 | Young |
| 0.230338 | 0.733775 | 0.039 | 0.029 | 1 | 3 | Slc44a2 | 0.01 | Young |
| 0.230797 | 0.037024 | 0.782 | 0.804 | 1 | 3 | Mt1 | -0.022 | Young |
| 0.231234 | 0.144578 | 0.62 | 0.615 | 1 | 3 | Fcgr3 | 0.005 | Young |

| | | | | | | | | |
|---|---|---|---|---|---|---|---|---|
| 0.23165 | 1.336416 | 0.018 | 0.011 | 1 | 3 | Usp35 | 0.007 | Young |
| 0.232316 | 0.252305 | 0.17 | 0.152 | 1 | 3 | Cyp4f16 | 0.018 | Young |
| 0.232891 | 0.241069 | 0.187 | 0.166 | 1 | 3 | Prdm2 | 0.021 | Young |
| 0.233069 | 0.668703 | 0.027 | 0.018 | 1 | 3 | Abhd14b | 0.009 | Young |
| 0.233412 | 0.041934 | 0.395 | 0.441 | 1 | 3 | Tmod3 | -0.046 | Young |
| 0.233482 | 0.369496 | 0.082 | 0.067 | 1 | 3 | Zfp110 | 0.015 | Young |
| 0.233486 | 0.281722 | 0.241 | 0.222 | 1 | 3 | Zfc3h1 | 0.019 | Young |
| 0.233639 | 0.293366 | 0.265 | 0.251 | 1 | 3 | Ppp2r2a | 0.014 | Young |
| 0.23381 | 0.348463 | 0.167 | 0.148 | 1 | 3 | Sult1a1 | 0.019 | Young |
| 0.234316 | 0.135367 | 0.53 | 0.526 | 1 | 3 | Eef1d | 0.004 | Young |
| 0.235699 | 0.466842 | 0.018 | 0.011 | 1 | 3 | Gm6133 | 0.007 | Young |
| 0.235832 | 0.113983 | 0.032 | 0.043 | 1 | 3 | Thrb | -0.011 | Young |
| 0.235862 | 0.686661 | 0.122 | 0.108 | 1 | 3 | Mapk8ip3 | 0.014 | Young |
| 0.236618 | 0.866194 | 0.011 | 0.006 | 1 | 3 | Lrp5 | 0.005 | Young |
| 0.236879 | 0.852639 | 0.011 | 0.006 | 1 | 3 | Fam57a | 0.005 | Young |
| 0.237229 | 0.436891 | 0.064 | 0.051 | 1 | 3 | Asb3 | 0.013 | Young |
| 0.237402 | 0.624313 | 0.011 | 0.006 | 1 | 3 | Bahcc1 | 0.005 | Young |
| 0.237664 | 0.656612 | 0.011 | 0.006 | 1 | 3 | 4931414P1 | 0.005 | Young |
| 0.237732 | 0.422003 | 0.117 | 0.102 | 1 | 3 | Nfatc1 | 0.015 | Young |
| 0.2379 | 0.095077 | 0.111 | 0.094 | 1 | 3 | Gas6 | 0.017 | Young |
| 0.237901 | 0.505633 | 0.098 | 0.084 | 1 | 3 | Vezt | 0.014 | Young |
| 0.238363 | 0.009821 | 0.03 | 0.042 | 1 | 3 | Acot2 | -0.012 | Young |
| 0.238458 | 0.531565 | 0.083 | 0.069 | 1 | 3 | Ormdl3 | 0.014 | Young |
| 0.238469 | 0.488772 | 0.145 | 0.128 | 1 | 3 | Acpp | 0.017 | Young |
| 0.238713 | 0.514085 | 0.011 | 0.006 | 1 | 3 | Ang | 0.005 | Young |
| 0.238813 | 0.131459 | 0.761 | 0.755 | 1 | 3 | Gng5 | 0.006 | Young |
| 0.238948 | 0.934449 | 0.043 | 0.032 | 1 | 3 | Rft1 | 0.011 | Young |
| 0.238954 | 0.304451 | 0.038 | 0.027 | 1 | 3 | Heatr5b | 0.011 | Young |
| 0.239317 | 0.53784 | 0.071 | 0.057 | 1 | 3 | Rprd1a | 0.014 | Young |
| 0.239413 | 0.503363 | 0.107 | 0.091 | 1 | 3 | Thra | 0.016 | Young |
| 0.239766 | 0.462538 | 0.011 | 0.006 | 1 | 3 | Ckap2 | 0.005 | Young |
| 0.240307 | 0.046317 | 0.061 | 0.076 | 1 | 3 | Arfgef2 | -0.015 | Young |
| 0.240386 | 0.40846 | 0.04 | 0.03 | 1 | 3 | Tgif2 | 0.01 | Young |
| 0.240437 | 0.137565 | 0.49 | 0.479 | 1 | 3 | Kpna4 | 0.011 | Young |
| 0.240557 | 0.44435 | 0.011 | 0.006 | 1 | 3 | Pald1 | 0.005 | Young |
| 0.240738 | 0.091892 | 0.086 | 0.105 | 1 | 3 | Tor3a | -0.019 | Young |
| 0.242013 | 0.349229 | 0.213 | 0.197 | 1 | 3 | Aqp9 | 0.016 | Young |
| 0.242085 | 0.212985 | 0.126 | 0.109 | 1 | 3 | Ago1 | 0.017 | Young |
| 0.242239 | 0.052255 | 0.085 | 0.103 | 1 | 3 | Sik3 | -0.018 | Young |
| 0.242246 | 0.053937 | 0.681 | 0.7 | 1 | 3 | Ahcyl2 | -0.019 | Young |
| 0.2423 | 0.073219 | 0.078 | 0.096 | 1 | 3 | Lncpint | -0.018 | Young |
| 0.242599 | 0.234572 | 0.443 | 0.443 | 1 | 3 | Mxd4 | 0 | Young |
| 0.242692 | 0.080765 | 0.056 | 0.07 | 1 | 3 | Acyp1 | -0.014 | Young |
| 0.242758 | 0.000956 | 0.136 | 0.161 | 1 | 3 | Sh3tc1 | -0.025 | Young |
| 0.243197 | 0.125505 | 0.078 | 0.097 | 1 | 3 | Ppp1r37 | -0.019 | Young |

| | | | | | | | | |
|---|---|---|---|---|---|---|---|---|
| 0.243411 | 0.258979 | 0.264 | 0.249 | 1 | 3 | Ighm | 0.015 | Young |
| 0.243499 | 0.306183 | 0.054 | 0.042 | 1 | 3 | Ms4a7 | 0.012 | Young |
| 0.243861 | 0.567315 | 0.066 | 0.053 | 1 | 3 | 1110059E2 | 0.013 | Young |
| 0.244047 | 0.828772 | 0.029 | 0.02 | 1 | 3 | Plcxd2 | 0.009 | Young |
| 0.245399 | 0.827602 | 0.019 | 0.012 | 1 | 3 | 2900076A( | 0.007 | Young |
| 0.24541 | 0.408212 | 0.112 | 0.097 | 1 | 3 | Vps26b | 0.015 | Young |
| 0.245672 | 0.24107 | 0.297 | 0.28 | 1 | 3 | Ttc3 | 0.017 | Young |
| 0.246 | 0.031874 | 0.16 | 0.186 | 1 | 3 | Hmbox1 | -0.026 | Young |
| 0.246248 | 0.19561 | 0.436 | 0.423 | 1 | 3 | Ankrd17 | 0.013 | Young |
| 0.246254 | 0.339074 | 0.216 | 0.2 | 1 | 3 | Mctp1 | 0.016 | Young |
| 0.246311 | 0.47417 | 0.043 | 0.032 | 1 | 3 | Ankrd50 | 0.011 | Young |
| 0.24695 | 0.124997 | 0.208 | 0.243 | 1 | 3 | Egln1 | -0.035 | Young |
| 0.246978 | 0.057593 | 0.471 | 0.515 | 1 | 3 | Cnot6l | -0.044 | Young |
| 0.247369 | 0.04637 | 0.194 | 0.222 | 1 | 3 | Gripap1 | -0.028 | Young |
| 0.247557 | 0.479689 | 0.071 | 0.057 | 1 | 3 | Gon7 | 0.014 | Young |
| 0.248293 | 0.545045 | 0.042 | 0.031 | 1 | 3 | Sh3bp5 | 0.011 | Young |
| 0.248521 | 0.342717 | 0.073 | 0.06 | 1 | 3 | Gnl2 | 0.013 | Young |
| 0.248581 | 0.318857 | 0.181 | 0.165 | 1 | 3 | Sart1 | 0.016 | Young |
| 0.24862 | 0.517317 | 0.167 | 0.152 | 1 | 3 | Lats2 | 0.015 | Young |
| 0.248715 | 0.199191 | 0.715 | 0.73 | 1 | 3 | Dnaja1 | -0.015 | Young |
| 0.249117 | 0.047392 | 0.143 | 0.166 | 1 | 3 | Snapc5 | -0.023 | Young |
| 0.250325 | 0.475479 | 0.044 | 0.033 | 1 | 3 | Padi4 | 0.011 | Young |
| 0.250346 | 0.065478 | 0.869 | 0.9 | 1 | 3 | Fxyd5 | -0.031 | Young |
| 0.250499 | 0.225474 | 0.039 | 0.029 | 1 | 3 | Ano8 | 0.01 | Young |
| 0.250683 | 0.73249 | 0.03 | 0.022 | 1 | 3 | Bmf | 0.008 | Young |
| 0.250796 | 0.157358 | 0.644 | 0.663 | 1 | 3 | Spag9 | -0.019 | Young |
| 0.251012 | 0.169686 | 0.146 | 0.128 | 1 | 3 | Arl3 | 0.018 | Young |
| 0.251802 | 0.057544 | 0.154 | 0.182 | 1 | 3 | Irf2bpl | -0.028 | Young |
| 0.251804 | 0.127535 | 0.098 | 0.118 | 1 | 3 | Nfyc | -0.02 | Young |
| 0.2521 | 0.59495 | 0.019 | 0.012 | 1 | 3 | Abcg2 | 0.007 | Young |
| 0.252205 | 0.379676 | 0.24 | 0.221 | 1 | 3 | Smad4 | 0.019 | Young |
| 0.25263 | 0.292286 | 0.261 | 0.244 | 1 | 3 | Dnajc1 | 0.017 | Young |
| 0.253003 | 0.435118 | 0.064 | 0.051 | 1 | 3 | Adamts10 | 0.013 | Young |
| 0.253014 | 0.092992 | 0.028 | 0.038 | 1 | 3 | Lgi3 | -0.01 | Young |
| 0.253156 | 0.989038 | 0.059 | 0.048 | 1 | 3 | Lmbr1l | 0.011 | Young |
| 0.253166 | 0.173504 | 0.375 | 0.356 | 1 | 3 | Sgpp1 | 0.019 | Young |
| 0.253253 | 0.138035 | 0.056 | 0.043 | 1 | 3 | Tab3 | 0.013 | Young |
| 0.253328 | 0.316567 | 0.064 | 0.051 | 1 | 3 | Clk2 | 0.013 | Young |
| 0.253596 | 0.661349 | 0.053 | 0.042 | 1 | 3 | Btbd2 | 0.011 | Young |
| 0.254329 | 0.131091 | 0.034 | 0.045 | 1 | 3 | 1810043G( | -0.011 | Young |
| 0.254543 | 0.46843 | 0.076 | 0.062 | 1 | 3 | Erich1 | 0.014 | Young |
| 0.254548 | 0.764304 | 0.052 | 0.041 | 1 | 3 | Katnal1 | 0.011 | Young |
| 0.25546 | 0.17125 | 0.475 | 0.468 | 1 | 3 | Ube2r2 | 0.007 | Young |
| 0.255529 | 0.337018 | 0.029 | 0.02 | 1 | 3 | Dyrk1b | 0.009 | Young |
| 0.255559 | 0.23522 | 0.341 | 0.323 | 1 | 3 | Rgs10 | 0.018 | Young |

| | | | | | | | | |
|---|---|---|---|---|---|---|---|---|
| 0.25557 | 0.266427 | 0.042 | 0.031 | 1 | 3 | Zfp87 | 0.011 | Young |
| 0.255629 | 1.08786 | 0.02 | 0.013 | 1 | 3 | 4930430F0 | 0.007 | Young |
| 0.255656 | 0.036867 | 0.179 | 0.205 | 1 | 3 | Pnrc2 | -0.026 | Young |
| 0.256111 | 0.233961 | 0.553 | 0.564 | 1 | 3 | Znrf2 | -0.011 | Young |
| 0.256187 | 0.255104 | 0.419 | 0.415 | 1 | 3 | Cggbp1 | 0.004 | Young |
| 0.256308 | 0.282839 | 0.249 | 0.232 | 1 | 3 | 4932438A1 | 0.017 | Young |
| 0.256355 | 0.106788 | 0.044 | 0.033 | 1 | 3 | Smad7 | 0.011 | Young |
| 0.256929 | 0.009375 | 0.196 | 0.225 | 1 | 3 | Kif3b | -0.029 | Young |
| 0.25711 | 0.111326 | 0.173 | 0.202 | 1 | 3 | Impa1 | -0.029 | Young |
| 0.257703 | 0.271755 | 0.284 | 0.274 | 1 | 3 | Hcst | 0.01 | Young |
| 0.257877 | 0.248241 | 0.504 | 0.516 | 1 | 3 | Nav1 | -0.012 | Young |
| 0.258273 | 0.747525 | 0.032 | 0.023 | 1 | 3 | Ick | 0.009 | Young |
| 0.258376 | 0.384404 | 0.302 | 0.293 | 1 | 3 | Chd2 | 0.009 | Young |
| 0.258687 | 0.49482 | 0.03 | 0.022 | 1 | 3 | Trim21 | 0.008 | Young |
| 0.258798 | 0.509781 | 0.121 | 0.106 | 1 | 3 | Axin1 | 0.015 | Young |
| 0.258801 | 0.046132 | 0.169 | 0.197 | 1 | 3 | Mrps23 | -0.028 | Young |
| 0.259126 | 0.330821 | 0.042 | 0.031 | 1 | 3 | Usp53 | 0.011 | Young |
| 0.259292 | 0.198131 | 0.109 | 0.092 | 1 | 3 | Aff3 | 0.017 | Young |
| 0.259431 | 0.022233 | 0.073 | 0.09 | 1 | 3 | Slc7a6os | -0.017 | Young |
| 0.259602 | 0.187756 | 0.029 | 0.039 | 1 | 3 | Ints5.1 | -0.01 | Young |
| 0.260073 | 0.056828 | 0.144 | 0.17 | 1 | 3 | Nup62 | -0.026 | Young |
| 0.26024 | 0.506532 | 0.02 | 0.013 | 1 | 3 | Abhd8 | 0.007 | Young |
| 0.260258 | 0.915625 | 0.033 | 0.024 | 1 | 3 | Foxo4 | 0.009 | Young |
| 0.260356 | 0.298746 | 0.402 | 0.394 | 1 | 3 | Ccdc12 | 0.008 | Young |
| 0.260484 | 0.005818 | 0.266 | 0.299 | 1 | 3 | Leprotl1 | -0.033 | Young |
| 0.260588 | 0.971331 | 0.013 | 0.007 | 1 | 3 | Mrm2 | 0.006 | Young |
| 0.260704 | 0.407558 | 0.076 | 0.062 | 1 | 3 | Smim5 | 0.014 | Young |
| 0.261048 | 0.675305 | 0.02 | 0.013 | 1 | 3 | Dtwd1 | 0.007 | Young |
| 0.261142 | 0.137164 | 0.722 | 0.763 | 1 | 3 | Eif3f | -0.041 | Young |
| 0.261416 | 0.618621 | 0.074 | 0.061 | 1 | 3 | Ypel2 | 0.013 | Young |
| 0.261453 | 0.761239 | 0.02 | 0.013 | 1 | 3 | D830025C0 | 0.007 | Young |
| 0.261587 | 0.443899 | 0.098 | 0.084 | 1 | 3 | Gm20696 | 0.014 | Young |
| 0.26216 | 0.629579 | 0.013 | 0.007 | 1 | 3 | Rhox8 | 0.006 | Young |
| 0.262628 | 0.301517 | 0.12 | 0.103 | 1 | 3 | Traf7 | 0.017 | Young |
| 0.262728 | 0.39023 | 0.138 | 0.121 | 1 | 3 | Ip6k1 | 0.017 | Young |
| 0.263157 | 0.210127 | 0.068 | 0.055 | 1 | 3 | Tyw1 | 0.013 | Young |
| 0.263211 | 0.721201 | 0.013 | 0.007 | 1 | 3 | Gba2 | 0.006 | Young |
| 0.263211 | 0.467623 | 0.013 | 0.007 | 1 | 3 | Rcn1 | 0.006 | Young |
| 0.263475 | 0.574949 | 0.013 | 0.007 | 1 | 3 | Cracr2a | 0.006 | Young |
| 0.263482 | 0.020652 | 0.256 | 0.287 | 1 | 3 | Nfatc3 | -0.031 | Young |
| 0.263859 | 0.073652 | 0.283 | 0.318 | 1 | 3 | Spg21 | -0.035 | Young |
| 0.264002 | 0.672711 | 0.013 | 0.007 | 1 | 3 | Dbp | 0.006 | Young |
| 0.264104 | 0.200719 | 0.043 | 0.032 | 1 | 3 | Nudt1 | 0.011 | Young |
| 0.264235 | 0.466619 | 0.138 | 0.121 | 1 | 3 | Ccpg1 | 0.017 | Young |
| 0.264339 | 0.06746 | 0.335 | 0.372 | 1 | 3 | Homer1 | -0.037 | Young |

| | | | | | | | | |
|---|---|---|---|---|---|---|---|---|
| 0.264389 | 0.284247 | 0.119 | 0.103 | 1 | 3 | Padi2 | 0.016 | Young |
| 0.265035 | 0.124478 | 0.619 | 0.637 | 1 | 3 | S100a11 | -0.018 | Young |
| 0.265273 | 0.266554 | 0.067 | 0.054 | 1 | 3 | Gprc5a | 0.013 | Young |
| 0.265279 | 0.007974 | 0.091 | 0.109 | 1 | 3 | Mrpl58 | -0.018 | Young |
| 0.266915 | 0.265614 | 0.013 | 0.007 | 1 | 3 | Ada | 0.006 | Young |
| 0.267498 | 0.004029 | 0.494 | 0.556 | 1 | 3 | Tmem160 | -0.062 | Young |
| 0.267915 | 0.270636 | 0.067 | 0.054 | 1 | 3 | Cep44 | 0.013 | Young |
| 0.26794 | 0.110159 | 0.217 | 0.246 | 1 | 3 | Bag3 | -0.029 | Young |
| 0.267989 | 0.516909 | 0.051 | 0.039 | 1 | 3 | Mcm9 | 0.012 | Young |
| 0.268595 | 0.50712 | 0.047 | 0.036 | 1 | 3 | Ttc27 | 0.011 | Young |
| 0.268628 | 0.341556 | 0.077 | 0.063 | 1 | 3 | Bet1l | 0.014 | Young |
| 0.26868 | 0.10082 | 0.237 | 0.271 | 1 | 3 | Hmgcr | -0.034 | Young |
| 0.269634 | 0.064934 | 0.211 | 0.241 | 1 | 3 | Naa38 | -0.03 | Young |
| 0.269701 | 0.731863 | 0.054 | 0.043 | 1 | 3 | Sp4 | 0.011 | Young |
| 0.269942 | 0.111804 | 0.178 | 0.207 | 1 | 3 | Aph1c | -0.029 | Young |
| 0.269996 | 0.316073 | 0.045 | 0.035 | 1 | 3 | Rufy2 | 0.01 | Young |
| 0.270181 | 0.5747 | 0.033 | 0.024 | 1 | 3 | Aldh7a1 | 0.009 | Young |
| 0.270441 | 0.388743 | 0.202 | 0.184 | 1 | 3 | Cstf3 | 0.018 | Young |
| 0.270453 | 0.238662 | 0.495 | 0.497 | 1 | 3 | Plec | -0.002 | Young |
| 0.270542 | 0.282248 | 0.215 | 0.198 | 1 | 3 | Carhsp1 | 0.017 | Young |
| 0.270905 | 0.273732 | 0.12 | 0.103 | 1 | 3 | 2610037D( | 0.017 | Young |
| 0.271128 | 0.419067 | 0.125 | 0.11 | 1 | 3 | Golga2 | 0.015 | Young |
| 0.271207 | 0.230945 | 0.177 | 0.158 | 1 | 3 | Nsd2 | 0.019 | Young |
| 0.271558 | 0.387715 | 0.234 | 0.217 | 1 | 3 | Cited2 | 0.017 | Young |
| 0.271953 | 0.046789 | 0.112 | 0.133 | 1 | 3 | Kbtbd2 | -0.021 | Young |
| 0.272576 | 0.685207 | 0.037 | 0.027 | 1 | 3 | Zeb1 | 0.01 | Young |
| 0.272582 | 0.173782 | 0.51 | 0.505 | 1 | 3 | Ubap2l | 0.005 | Young |
| 0.272804 | 0.300718 | 0.304 | 0.299 | 1 | 3 | Nt5c | 0.005 | Young |
| 0.272877 | 0.595324 | 0.069 | 0.057 | 1 | 3 | Trpv4 | 0.012 | Young |
| 0.27363 | 0.393965 | 0.008 | 0.013 | 1 | 3 | Fabp7 | -0.005 | Young |
| 0.274137 | 0.006228 | 0.138 | 0.159 | 1 | 3 | 9930021J0 | -0.021 | Young |
| 0.274796 | 0.103457 | 0.311 | 0.293 | 1 | 3 | Hp1bp3 | 0.018 | Young |
| 0.274846 | 0.604735 | 0.034 | 0.025 | 1 | 3 | Nipal3 | 0.009 | Young |
| 0.275188 | 0.150565 | 0.063 | 0.079 | 1 | 3 | Slc17a5 | -0.016 | Young |
| 0.275609 | 0.127171 | 0.542 | 0.522 | 1 | 3 | Cox5b | 0.02 | Young |
| 0.275726 | 0.508436 | 0.021 | 0.014 | 1 | 3 | Thada | 0.007 | Young |
| 0.275897 | 0.184766 | 0.178 | 0.204 | 1 | 3 | Nop56 | -0.026 | Young |
| 0.277066 | 0.46947 | 0.034 | 0.025 | 1 | 3 | Tmco4 | 0.009 | Young |
| 0.27738 | 0.029877 | 0.404 | 0.46 | 1 | 3 | Ist1 | -0.056 | Young |
| 0.277558 | 1.027721 | 0.023 | 0.016 | 1 | 3 | Cdyl2 | 0.007 | Young |
| 0.277701 | 0.152086 | 0.544 | 0.552 | 1 | 3 | Mycbp2 | -0.008 | Young |
| 0.277739 | 0.091476 | 0.342 | 0.388 | 1 | 3 | Mkln1 | -0.046 | Young |
| 0.278028 | 0.237467 | 0.239 | 0.223 | 1 | 3 | Blvra | 0.016 | Young |
| 0.278186 | 0.339557 | 0.035 | 0.047 | 1 | 3 | Gin1 | -0.012 | Young |
| 0.278285 | 0.012152 | 0.087 | 0.104 | 1 | 3 | Filip1l | -0.017 | Young |

| | | | | | | | | |
|---|---|---|---|---|---|---|---|---|
| 0.278396 | 0.541961 | 0.098 | 0.085 | 1 | 3 | Trappc12 | 0.013 | Young |
| 0.278574 | 0.501643 | 0.021 | 0.014 | 1 | 3 | E2f2 | 0.007 | Young |
| 0.27968 | 0.454125 | 0.052 | 0.041 | 1 | 3 | Terf1 | 0.011 | Young |
| 0.279707 | 0.48627 | 0.033 | 0.024 | 1 | 3 | Ckap2l | 0.009 | Young |
| 0.279886 | 0.00227 | 0.155 | 0.178 | 1 | 3 | Dhps | -0.023 | Young |
| 0.280109 | 0.026909 | 0.984 | 0.992 | 1 | 3 | Rplp2 | -0.008 | Young |
| 0.280242 | 0.157639 | 0.413 | 0.407 | 1 | 3 | Nrp2 | 0.006 | Young |
| 0.28039 | 0.34699 | 0.096 | 0.081 | 1 | 3 | Ints13 | 0.015 | Young |
| 0.280581 | 0.140587 | 0.186 | 0.166 | 1 | 3 | Arl2bp | 0.02 | Young |
| 0.280642 | 0.146266 | 0.051 | 0.063 | 1 | 3 | Utp15 | -0.012 | Young |
| 0.280979 | 0.182284 | 0.405 | 0.395 | 1 | 3 | Ltb4r1 | 0.01 | Young |
| 0.281138 | 0.287372 | 0.225 | 0.207 | 1 | 3 | Ptov1 | 0.018 | Young |
| 0.281145 | 0.066407 | 0.073 | 0.088 | 1 | 3 | Usb1 | -0.015 | Young |
| 0.281226 | 0.737106 | 0.063 | 0.051 | 1 | 3 | Gbp2b | 0.012 | Young |
| 0.281355 | 0.475636 | 0.054 | 0.043 | 1 | 3 | Strada | 0.011 | Young |
| 0.281667 | 0.425013 | 0.179 | 0.164 | 1 | 3 | Zbtb44 | 0.015 | Young |
| 0.28194 | 0.026205 | 0.867 | 0.866 | 1 | 3 | S100a4 | 0.001 | Young |
| 0.282251 | 0.37225 | 0.048 | 0.037 | 1 | 3 | Rfc5 | 0.011 | Young |
| 0.282769 | 0.65791 | 0.014 | 0.008 | 1 | 3 | Ssc4d | 0.006 | Young |
| 0.282873 | 0.177921 | 0.258 | 0.238 | 1 | 3 | Rps6ka1 | 0.02 | Young |
| 0.282913 | 0.795525 | 0.023 | 0.016 | 1 | 3 | Cenpv | 0.007 | Young |
| 0.283093 | 0.123092 | 0.053 | 0.066 | 1 | 3 | Ercc6 | -0.013 | Young |
| 0.283144 | 0.01566 | 0.254 | 0.289 | 1 | 3 | Themis2 | -0.035 | Young |
| 0.283901 | 0.160504 | 0.087 | 0.105 | 1 | 3 | Glt8d1 | -0.018 | Young |
| 0.284075 | 0.797628 | 0.014 | 0.008 | 1 | 3 | Zfp46 | 0.006 | Young |
| 0.284904 | 0.063502 | 0.206 | 0.234 | 1 | 3 | Wsb2 | -0.028 | Young |
| 0.284913 | 0.549408 | 0.023 | 0.016 | 1 | 3 | Plppr4 | 0.007 | Young |
| 0.284913 | 0.236599 | 0.023 | 0.016 | 1 | 3 | Nmrk1 | 0.007 | Young |
| 0.284921 | 0.455387 | 0.102 | 0.087 | 1 | 3 | Ttc5 | 0.015 | Young |
| 0.285123 | 0.516404 | 0.014 | 0.008 | 1 | 3 | Gm13431 | 0.006 | Young |
| 0.285157 | 0.43675 | 0.124 | 0.109 | 1 | 3 | Dcaf10 | 0.015 | Young |
| 0.285206 | 0.193502 | 0.46 | 0.456 | 1 | 3 | Pea15a | 0.004 | Young |
| 0.285212 | 0.137868 | 0.466 | 0.461 | 1 | 3 | Dek | 0.005 | Young |
| 0.285648 | 0.618819 | 0.014 | 0.008 | 1 | 3 | Zfyve9 | 0.006 | Young |
| 0.286262 | 0.029218 | 0.819 | 0.832 | 1 | 3 | Csf2ra | -0.013 | Young |
| 0.286537 | 0.238323 | 0.254 | 0.237 | 1 | 3 | Hmgcs1 | 0.017 | Young |
| 0.286595 | 0.146507 | 0.109 | 0.129 | 1 | 3 | Srek1ip1 | -0.02 | Young |
| 0.286776 | 0.303467 | 0.256 | 0.243 | 1 | 3 | Rasgrp4 | 0.013 | Young |
| 0.287353 | 0.258945 | 0.37 | 0.36 | 1 | 3 | Gng2 | 0.01 | Young |
| 0.287354 | 0.252064 | 0.086 | 0.072 | 1 | 3 | Pnpla7 | 0.014 | Young |
| 0.287377 | 0.327275 | 0.119 | 0.103 | 1 | 3 | Ticam2 | 0.016 | Young |
| 0.28749 | 0.252887 | 0.014 | 0.008 | 1 | 3 | Hdhd5 | 0.006 | Young |
| 0.288018 | 0.378342 | 0.014 | 0.008 | 1 | 3 | Dph7 | 0.006 | Young |
| 0.288035 | 0.170403 | 0.037 | 0.048 | 1 | 3 | Ercc2 | -0.011 | Young |
| 0.288132 | 0.087878 | 0.754 | 0.785 | 1 | 3 | Clta | -0.031 | Young |

| | | | | | | | | |
|---|---|---|---|---|---|---|---|---|
| 0.288539 | 0.253148 | 0.023 | 0.016 | 1 | 3 | Elmod3 | 0.007 | Young |
| 0.288813 | 0.108773 | 0.049 | 0.062 | 1 | 3 | Rbm19 | -0.013 | Young |
| 0.288863 | 0.154729 | 0.213 | 0.245 | 1 | 3 | Gpatch8 | -0.032 | Young |
| 0.289566 | 0.049205 | 0.208 | 0.237 | 1 | 3 | Nemf | -0.029 | Young |
| 0.290137 | 0.100437 | 0.107 | 0.092 | 1 | 3 | Tnfaip8l2 | 0.015 | Young |
| 0.290192 | 0.440618 | 0.037 | 0.027 | 1 | 3 | Nim1k | 0.01 | Young |
| 0.290767 | 0.163478 | 0.28 | 0.325 | 1 | 3 | Ubn1 | -0.045 | Young |
| 0.290917 | 0.192039 | 0.25 | 0.239 | 1 | 3 | Tnfsf12 | 0.011 | Young |
| 0.29106 | 0.126414 | 0.037 | 0.048 | 1 | 3 | Orc5 | -0.011 | Young |
| 0.291283 | 0.076066 | 0.258 | 0.289 | 1 | 3 | Slc39a7 | -0.031 | Young |
| 0.291309 | 0.253606 | 0.405 | 0.405 | 1 | 3 | Yy1 | 0 | Young |
| 0.291399 | 0.243205 | 0.379 | 0.37 | 1 | 3 | Sbds | 0.009 | Young |
| 0.291821 | 0.761321 | 0.024 | 0.017 | 1 | 3 | Slc25a26 | 0.007 | Young |
| 0.292768 | 0.018707 | 0.322 | 0.36 | 1 | 3 | Cyld | -0.038 | Young |
| 0.292782 | 0.33229 | 0.143 | 0.129 | 1 | 3 | Dcaf11 | 0.014 | Young |
| 0.292975 | 0.410005 | 0.102 | 0.088 | 1 | 3 | Tceanc2 | 0.014 | Young |
| 0.293163 | 0.658977 | 0.039 | 0.03 | 1 | 3 | Ccdc180 | 0.009 | Young |
| 0.293488 | 0.39389 | 0.071 | 0.059 | 1 | 3 | Zfp626 | 0.012 | Young |
| 0.293764 | 0.201875 | 0.563 | 0.576 | 1 | 3 | Unc119 | -0.013 | Young |
| 0.293882 | 0.048828 | 0.284 | 0.315 | 1 | 3 | Rnf111 | -0.031 | Young |
| 0.293989 | 0.255121 | 0.035 | 0.026 | 1 | 3 | Iqsec2 | 0.009 | Young |
| 0.294148 | 0.12074 | 0.194 | 0.176 | 1 | 3 | Dguok | 0.018 | Young |
| 0.294176 | 0.432833 | 0.124 | 0.109 | 1 | 3 | Hgsnat | 0.015 | Young |
| 0.294305 | 0.472442 | 0.131 | 0.117 | 1 | 3 | E2f4 | 0.014 | Young |
| 0.294373 | 0.57529 | 0.087 | 0.075 | 1 | 3 | H2afx | 0.012 | Young |
| 0.294899 | 0.413056 | 0.087 | 0.074 | 1 | 3 | Adcy9 | 0.013 | Young |
| 0.29498 | 0.392231 | 0.067 | 0.055 | 1 | 3 | Rmi1 | 0.012 | Young |
| 0.295106 | 0.305794 | 0.263 | 0.247 | 1 | 3 | Nhsl2 | 0.016 | Young |
| 0.295279 | 0.368565 | 0.124 | 0.108 | 1 | 3 | Faf2 | 0.016 | Young |
| 0.295968 | 0.187227 | 0.106 | 0.125 | 1 | 3 | Pdlim4 | -0.019 | Young |
| 0.296561 | 0.076311 | 0.087 | 0.073 | 1 | 3 | Rnaset2a | 0.014 | Young |
| 0.296839 | 0.376295 | 0.063 | 0.051 | 1 | 3 | Agmo | 0.012 | Young |
| 0.296864 | 0.108172 | 0.549 | 0.545 | 1 | 3 | Ehd4 | 0.004 | Young |
| 0.29703 | 0.001389 | 0.253 | 0.284 | 1 | 3 | Acbd5 | -0.031 | Young |
| 0.297291 | 0.178672 | 0.427 | 0.427 | 1 | 3 | Nufip2 | 0 | Young |
| 0.297425 | 0.42022 | 0.038 | 0.029 | 1 | 3 | Polr3a | 0.009 | Young |
| 0.297759 | 0.100855 | 0.712 | 0.738 | 1 | 3 | Atf4 | -0.026 | Young |
| 0.298003 | 0.760643 | 0.042 | 0.032 | 1 | 3 | Stk11ip | 0.01 | Young |
| 0.298355 | 0.01293 | 0.234 | 0.264 | 1 | 3 | B3gat3 | -0.03 | Young |
| 0.298383 | 0.554502 | 0.177 | 0.164 | 1 | 3 | Ipmk | 0.013 | Young |
| 0.298464 | 0.05811 | 0.244 | 0.275 | 1 | 3 | Cmas | -0.031 | Young |
| 0.298629 | 0.607444 | 0.043 | 0.033 | 1 | 3 | Rpl9-ps6 | 0.01 | Young |
| 0.298788 | 0.38104 | 0.14 | 0.124 | 1 | 3 | Rbpms | 0.016 | Young |
| 0.299966 | 0.354746 | 0.141 | 0.127 | 1 | 3 | Sap30l | 0.014 | Young |
| 0.300013 | 0.062999 | 0.197 | 0.225 | 1 | 3 | Fto | -0.028 | Young |

| | | | | | | | | |
|---|---|---|---|---|---|---|---|---|
| 0.300516 | 0.29467 | 0.367 | 0.366 | 1 | 3 | Mfsd14a | 0.001 | Young |
| 0.300623 | 1.118954 | 0.015 | 0.01 | 1 | 3 | Rasal2 | 0.005 | Young |
| 0.30101 | 1.008468 | 0.015 | 0.01 | 1 | 3 | Clec5a | 0.005 | Young |
| 0.301017 | 0.019186 | 0.376 | 0.421 | 1 | 3 | Hipk1 | -0.045 | Young |
| 0.301132 | 0.025007 | 0.148 | 0.17 | 1 | 3 | Tmem87b | -0.022 | Young |
| 0.302326 | 0.34718 | 0.208 | 0.195 | 1 | 3 | Zcchc24 | 0.013 | Young |
| 0.302692 | 0.684204 | 0.015 | 0.01 | 1 | 3 | Gcat | 0.005 | Young |
| 0.302877 | 0.463724 | 0.04 | 0.031 | 1 | 3 | Nbas | 0.009 | Young |
| 0.303282 | 0.101624 | 0.032 | 0.042 | 1 | 3 | Gcdh | -0.01 | Young |
| 0.303587 | 0.090284 | 0.139 | 0.16 | 1 | 3 | Arfrp1 | -0.021 | Young |
| 0.30373 | 0.766556 | 0.015 | 0.01 | 1 | 3 | Mical3 | 0.005 | Young |
| 0.303751 | 0.724742 | 0.053 | 0.043 | 1 | 3 | Trim3 | 0.01 | Young |
| 0.303859 | 0.149337 | 0.018 | 0.025 | 1 | 3 | Eps8l1 | -0.007 | Young |
| 0.303989 | 0.661198 | 0.015 | 0.01 | 1 | 3 | Zfp605 | 0.005 | Young |
| 0.30451 | 0.579676 | 0.015 | 0.01 | 1 | 3 | Gm19325 | 0.005 | Young |
| 0.3046 | 0.233804 | 0.193 | 0.176 | 1 | 3 | Gtpbp2 | 0.017 | Young |
| 0.305082 | 0.221748 | 0.234 | 0.219 | 1 | 3 | Tpm1 | 0.015 | Young |
| 0.305194 | 0.444107 | 0.04 | 0.031 | 1 | 3 | Asb13 | 0.009 | Young |
| 0.305266 | 0.099222 | 0.133 | 0.154 | 1 | 3 | Fam204a | -0.021 | Young |
| 0.305291 | 0.37964 | 0.015 | 0.01 | 1 | 3 | Gprc5c | 0.005 | Young |
| 0.305305 | 0.306148 | 0.112 | 0.098 | 1 | 3 | Mosmo | 0.014 | Young |
| 0.305441 | 0.508337 | 0.025 | 0.018 | 1 | 3 | Gm16973 | 0.007 | Young |
| 0.305812 | 0.461211 | 0.015 | 0.01 | 1 | 3 | Adamtsl5 | 0.005 | Young |
| 0.307288 | 0.037475 | 0.153 | 0.176 | 1 | 3 | Znrd1 | -0.023 | Young |
| 0.307402 | 0.0645 | 0.345 | 0.326 | 1 | 3 | Polr2a | 0.019 | Young |
| 0.307425 | 0.474391 | 0.025 | 0.018 | 1 | 3 | Dennd2d | 0.007 | Young |
| 0.307592 | 0.033138 | 0.072 | 0.087 | 1 | 3 | Kdm5d | -0.015 | Young |
| 0.307784 | 0.389977 | 0.23 | 0.221 | 1 | 3 | Cdc73 | 0.009 | Young |
| 0.307907 | 0.826362 | 0.027 | 0.019 | 1 | 3 | Runx2 | 0.008 | Young |
| 0.308005 | 0.270242 | 0.164 | 0.148 | 1 | 3 | Hltf | 0.016 | Young |
| 0.308316 | 0.018207 | 0.146 | 0.17 | 1 | 3 | Arl6ip6 | -0.024 | Young |
| 0.308676 | 0.643798 | 0.062 | 0.051 | 1 | 3 | Cyb561d1 | 0.011 | Young |
| 0.308683 | 0.155388 | 0.027 | 0.019 | 1 | 3 | 5430416N( | 0.008 | Young |
| 0.309037 | 0.060246 | 0.14 | 0.162 | 1 | 3 | Gar1 | -0.022 | Young |
| 0.309168 | 0.011566 | 0.112 | 0.13 | 1 | 3 | Atg4d | -0.018 | Young |
| 0.309249 | 0.154225 | 0.612 | 0.634 | 1 | 3 | Prdx5 | -0.022 | Young |
| 0.30939 | 0.306089 | 0.042 | 0.032 | 1 | 3 | Pdik1l | 0.01 | Young |
| 0.309544 | 0.7937 | 0.059 | 0.049 | 1 | 3 | Zfp90 | 0.01 | Young |
| 0.309654 | 0.498679 | 0.027 | 0.019 | 1 | 3 | Thg1l | 0.008 | Young |
| 0.309893 | 0.462251 | 0.067 | 0.055 | 1 | 3 | Adam19 | 0.012 | Young |
| 0.309899 | 0.027401 | 0.197 | 0.225 | 1 | 3 | Klrk1 | -0.028 | Young |
| 0.310206 | 0.291498 | 0.189 | 0.174 | 1 | 3 | Gabarapl1 | 0.015 | Young |
| 0.310608 | 0.087363 | 0.088 | 0.105 | 1 | 3 | Scfd1 | -0.017 | Young |
| 0.310835 | 0.204264 | 0.451 | 0.465 | 1 | 3 | Ppib | -0.014 | Young |
| 0.311492 | 0.222534 | 0.189 | 0.173 | 1 | 3 | Fundc1 | 0.016 | Young |

| | | | | | | | | |
|---|---|---|---|---|---|---|---|---|
| 0.31168 | 0.052416 | 0.676 | 0.723 | 1 | 3 | Lilr4b | -0.047 | Young |
| 0.311717 | 0.007771 | 0.625 | 0.663 | 1 | 3 | H2-T23 | -0.038 | Young |
| 0.312227 | 0.02285 | 0.088 | 0.105 | 1 | 3 | Heg1 | -0.017 | Young |
| 0.31288 | 0.190389 | 0.152 | 0.174 | 1 | 3 | Kcnk6 | -0.022 | Young |
| 0.312939 | 0.539741 | 0.042 | 0.032 | 1 | 3 | Zfp994 | 0.01 | Young |
| 0.312968 | 0.013119 | 0.399 | 0.442 | 1 | 3 | Leprot | -0.043 | Young |
| 0.313343 | 0.003925 | 0.139 | 0.16 | 1 | 3 | Acadvl | -0.021 | Young |
| 0.313697 | 0.37795 | 0.154 | 0.14 | 1 | 3 | Gcc2 | 0.014 | Young |
| 0.31391 | 0.498135 | 0.044 | 0.035 | 1 | 3 | Dnal4 | 0.009 | Young |
| 0.314076 | 0.898162 | 0.029 | 0.022 | 1 | 3 | Zfp958 | 0.007 | Young |
| 0.314722 | 0.278807 | 0.043 | 0.033 | 1 | 3 | Tmed8 | 0.01 | Young |
| 0.314871 | 0.00974 | 0.189 | 0.213 | 1 | 3 | Mrpl42 | -0.024 | Young |
| 0.315003 | 0.251109 | 0.11 | 0.096 | 1 | 3 | Pdcd7 | 0.014 | Young |
| 0.315721 | 0.954525 | 0.027 | 0.019 | 1 | 3 | Taf5 | 0.008 | Young |
| 0.316201 | 0.790218 | 0.016 | 0.011 | 1 | 3 | Btd | 0.005 | Young |
| 0.316241 | 0.019578 | 0.615 | 0.659 | 1 | 3 | Dhrs3 | -0.044 | Young |
| 0.31633 | 0.132182 | 0.105 | 0.123 | 1 | 3 | Mtmr10 | -0.018 | Young |
| 0.316384 | 0.052979 | 0.061 | 0.074 | 1 | 3 | Rpp14 | -0.013 | Young |
| 0.316388 | 0.6119 | 0.028 | 0.02 | 1 | 3 | Zfp995 | 0.008 | Young |
| 0.31658 | 0.178163 | 0.443 | 0.443 | 1 | 3 | Asah1 | 0 | Young |
| 0.316904 | 0.295792 | 0.115 | 0.103 | 1 | 3 | Agap3 | 0.012 | Young |
| 0.317158 | 0.160487 | 0.115 | 0.1 | 1 | 3 | Ednrb | 0.015 | Young |
| 0.317545 | 0.297121 | 0.023 | 0.031 | 1 | 3 | Tcf7l2 | -0.008 | Young |
| 0.31757 | 0.028944 | 0.14 | 0.16 | 1 | 3 | Txndc15 | -0.02 | Young |
| 0.317731 | 0.890798 | 0.016 | 0.011 | 1 | 3 | Gm15859 | 0.005 | Young |
| 0.317878 | 0.251192 | 0.128 | 0.112 | 1 | 3 | Mis18a | 0.016 | Young |
| 0.318199 | 0.243227 | 0.385 | 0.386 | 1 | 3 | Arid4a | -0.001 | Young |
| 0.318859 | 0.651437 | 0.052 | 0.042 | 1 | 3 | Dhx34 | 0.01 | Young |
| 0.318971 | 0.006959 | 0.199 | 0.226 | 1 | 3 | Itgb1bp1 | -0.027 | Young |
| 0.319 | 0.390859 | 0.091 | 0.078 | 1 | 3 | Cdc42bpb | 0.013 | Young |
| 0.319005 | 0.330293 | 0.154 | 0.139 | 1 | 3 | Adrb2 | 0.015 | Young |
| 0.319478 | 0.099005 | 0.019 | 0.026 | 1 | 3 | Timp1 | -0.007 | Young |
| 0.320032 | 0.075641 | 0.538 | 0.533 | 1 | 3 | Mia2 | 0.005 | Young |
| 0.320183 | 0.02424 | 0.319 | 0.354 | 1 | 3 | Myo5a | -0.035 | Young |
| 0.320185 | 0.526378 | 0.051 | 0.041 | 1 | 3 | Stradb | 0.01 | Young |
| 0.321168 | 0.129726 | 0.027 | 0.019 | 1 | 3 | Fbxo21 | 0.008 | Young |
| 0.32118 | 0.027707 | 0.057 | 0.069 | 1 | 3 | Kat5 | -0.012 | Young |
| 0.321412 | 0.004258 | 0.104 | 0.121 | 1 | 3 | Lrch3 | -0.017 | Young |
| 0.321423 | 0.012473 | 0.418 | 0.465 | 1 | 3 | Ap2b1 | -0.047 | Young |
| 0.321698 | 0.501169 | 0.047 | 0.037 | 1 | 3 | Nlrc4 | 0.01 | Young |
| 0.321863 | 0.052135 | 0.187 | 0.209 | 1 | 3 | Rraga | -0.022 | Young |
| 0.322224 | 0.072663 | 0.064 | 0.053 | 1 | 3 | Cdca8 | 0.011 | Young |
| 0.322626 | 0.562979 | 0.125 | 0.111 | 1 | 3 | Cdc14a | 0.014 | Young |
| 0.322864 | 0.276977 | 0.027 | 0.019 | 1 | 3 | Slc46a3 | 0.008 | Young |
| 0.323084 | 0.310614 | 0.206 | 0.191 | 1 | 3 | Hist1h2bc | 0.015 | Young |

| | | | | | | | | |
|---|---|---|---|---|---|---|---|---|
| 0.323362 | 0.166289 | 0.179 | 0.161 | 1 | 3 | Fam192a | 0.018 | Young |
| 0.323541 | 0.191165 | 0.107 | 0.125 | 1 | 3 | Srxn1 | -0.018 | Young |
| 0.323562 | 0.041248 | 0.318 | 0.363 | 1 | 3 | Eif4g3 | -0.045 | Young |
| 0.323863 | 0.137022 | 0.184 | 0.211 | 1 | 3 | Tmpo | -0.027 | Young |
| 0.323989 | 0.328108 | 0.186 | 0.17 | 1 | 3 | Gigyf1 | 0.016 | Young |
| 0.324003 | 0.234822 | 0.288 | 0.269 | 1 | 3 | Pabpn1 | 0.019 | Young |
| 0.324074 | 0.550832 | 0.029 | 0.022 | 1 | 3 | Zfp319 | 0.007 | Young |
| 0.324184 | 0.381876 | 0.093 | 0.08 | 1 | 3 | Pip4k2c | 0.013 | Young |
| 0.324339 | 0.029216 | 0.256 | 0.286 | 1 | 3 | Kat2b | -0.03 | Young |
| 0.324465 | 0.111452 | 0.019 | 0.026 | 1 | 3 | Nradd | -0.007 | Young |
| 0.32455 | 0.564907 | 0.029 | 0.022 | 1 | 3 | Gm21781 | 0.007 | Young |
| 0.324781 | 0.375369 | 0.211 | 0.198 | 1 | 3 | Mlxip | 0.013 | Young |
| 0.324958 | 0.312392 | 0.183 | 0.17 | 1 | 3 | Ksr1 | 0.013 | Young |
| 0.325043 | 0.145517 | 0.082 | 0.098 | 1 | 3 | Coq8b | -0.016 | Young |
| 0.325198 | 0.062679 | 0.234 | 0.268 | 1 | 3 | Igsf6 | -0.034 | Young |
| 0.32571 | 0.400962 | 0.016 | 0.011 | 1 | 3 | Gtf2ird2 | 0.005 | Young |
| 0.326169 | 0.010493 | 0.088 | 0.104 | 1 | 3 | D10Wsu10 | -0.016 | Young |
| 0.326252 | 0.328618 | 0.044 | 0.035 | 1 | 3 | Haus3 | 0.009 | Young |
| 0.327874 | 0.204545 | 0.107 | 0.093 | 1 | 3 | Fdps | 0.014 | Young |
| 0.327968 | 0.021813 | 0.427 | 0.468 | 1 | 3 | Cd226 | -0.041 | Young |
| 0.328024 | 0.163157 | 0.496 | 0.502 | 1 | 3 | Rpl13a | -0.006 | Young |
| 0.328572 | 0.386309 | 0.016 | 0.011 | 1 | 3 | Trp53i13 | 0.005 | Young |
| 0.328933 | 0.178595 | 0.083 | 0.098 | 1 | 3 | Mrpl50 | -0.015 | Young |
| 0.329266 | 0.01813 | 0.051 | 0.062 | 1 | 3 | Bambi | -0.011 | Young |
| 0.329559 | 0.079729 | 0.154 | 0.177 | 1 | 3 | C330007P0 | -0.023 | Young |
| 0.329639 | 0.182884 | 0.23 | 0.213 | 1 | 3 | Nudc | 0.017 | Young |
| 0.330001 | 0.00947 | 0.239 | 0.266 | 1 | 3 | Ube2e3 | -0.027 | Young |
| 0.330084 | 0.150721 | 0.048 | 0.06 | 1 | 3 | Fancl | -0.012 | Young |
| 0.330311 | 0.101486 | 0.169 | 0.192 | 1 | 3 | Eftud2 | -0.023 | Young |
| 0.330366 | 0.008582 | 0.15 | 0.171 | 1 | 3 | Kif2a | -0.021 | Young |
| 0.330648 | 0.002973 | 0.249 | 0.276 | 1 | 3 | Sdhb | -0.027 | Young |
| 0.330786 | 0.311972 | 0.173 | 0.157 | 1 | 3 | Apba1 | 0.016 | Young |
| 0.331417 | 0.326192 | 0.047 | 0.059 | 1 | 3 | St7 | -0.012 | Young |
| 0.331607 | 0.339543 | 0.085 | 0.072 | 1 | 3 | Nfxl1 | 0.013 | Young |
| 0.331724 | 0.043221 | 0.114 | 0.131 | 1 | 3 | Rnpc3 | -0.017 | Young |
| 0.331949 | 0.024762 | 0.174 | 0.197 | 1 | 3 | Arhgap1 | -0.023 | Young |
| 0.332185 | 0.499112 | 0.032 | 0.024 | 1 | 3 | Pcnx4 | 0.008 | Young |
| 0.332517 | 0.148336 | 0.126 | 0.147 | 1 | 3 | Bahd1 | -0.021 | Young |
| 0.332792 | 0.354405 | 0.181 | 0.165 | 1 | 3 | Slc38a10 | 0.016 | Young |
| 0.333168 | 0.302998 | 0.053 | 0.043 | 1 | 3 | Vill | 0.01 | Young |
| 0.333493 | 0.322738 | 0.052 | 0.042 | 1 | 3 | 2410089E0 | 0.01 | Young |
| 0.334119 | 0.12825 | 0.415 | 0.404 | 1 | 3 | Mkrn1 | 0.011 | Young |
| 0.334159 | 0.607407 | 0.018 | 0.012 | 1 | 3 | Evi5l | 0.006 | Young |
| 0.334297 | 0.004603 | 0.266 | 0.296 | 1 | 3 | Ctdsp1 | -0.03 | Young |
| 0.334599 | 0.050829 | 0.542 | 0.6 | 1 | 3 | Nr4a2 | -0.058 | Young |

| | | | | | | | | |
|---|---|---|---|---|---|---|---|---|
| 0.334721 | 0.043539 | 0.006 | 0.011 | 1 | 3 | Zfp316 | -0.005 | Young |
| 0.334867 | 0.215463 | 0.199 | 0.185 | 1 | 3 | Seh1l | 0.014 | Young |
| 0.334917 | 0.830648 | 0.018 | 0.012 | 1 | 3 | Sccpdh | 0.006 | Young |
| 0.335001 | 0.125555 | 0.473 | 0.468 | 1 | 3 | C1qa | 0.005 | Young |
| 0.335054 | 0.215859 | 0.029 | 0.022 | 1 | 3 | Tm7sf2 | 0.007 | Young |
| 0.335382 | 0.001445 | 0.006 | 0.011 | 1 | 3 | Plac8 | -0.005 | Young |
| 0.335541 | 0.362953 | 0.205 | 0.192 | 1 | 3 | Marf1 | 0.013 | Young |
| 0.335548 | 0.481994 | 0.061 | 0.05 | 1 | 3 | AC165953. | 0.011 | Young |
| 0.335574 | 0.379266 | 0.106 | 0.093 | 1 | 3 | Kif21b | 0.013 | Young |
| 0.335713 | 0.047391 | 0.006 | 0.011 | 1 | 3 | Ercc6l | -0.005 | Young |
| 0.335832 | 0.037208 | 0.21 | 0.235 | 1 | 3 | Man1a2 | -0.025 | Young |
| 0.335862 | 0.060862 | 0.052 | 0.063 | 1 | 3 | AC163018. | -0.011 | Young |
| 0.335868 | 0.043869 | 0.239 | 0.268 | 1 | 3 | Rufy3 | -0.029 | Young |
| 0.33592 | 0.072827 | 0.109 | 0.127 | 1 | 3 | Golt1b | -0.018 | Young |
| 0.336285 | 0.043065 | 0.059 | 0.072 | 1 | 3 | Fam136a | -0.013 | Young |
| 0.336375 | 0.189607 | 0.006 | 0.011 | 1 | 3 | Gm4610 | -0.005 | Young |
| 0.336719 | 0.019988 | 0.236 | 0.259 | 1 | 3 | Stom | -0.023 | Young |
| 0.336854 | 0.359057 | 0.117 | 0.104 | 1 | 3 | Papss1 | 0.013 | Young |
| 0.337236 | 0.317168 | 0.311 | 0.305 | 1 | 3 | Aimp1 | 0.006 | Young |
| 0.337455 | 0.55451 | 0.018 | 0.012 | 1 | 3 | Tppp | 0.006 | Young |
| 0.337989 | 0.251496 | 0.01 | 0.016 | 1 | 3 | Calml4 | -0.006 | Young |
| 0.338575 | 0.101128 | 0.225 | 0.256 | 1 | 3 | Rps6kb1 | -0.031 | Young |
| 0.338796 | 0.181817 | 0.176 | 0.159 | 1 | 3 | Htatsf1 | 0.017 | Young |
| 0.338807 | 0.357253 | 0.124 | 0.109 | 1 | 3 | Uchl3 | 0.015 | Young |
| 0.339208 | 0.28537 | 0.03 | 0.023 | 1 | 3 | Nudt22 | 0.007 | Young |
| 0.339466 | 0.222555 | 0.187 | 0.172 | 1 | 3 | Fam234a | 0.015 | Young |
| 0.339637 | 0.019636 | 0.095 | 0.111 | 1 | 3 | Zfp646 | -0.016 | Young |
| 0.339749 | 0.334639 | 0.018 | 0.012 | 1 | 3 | Platr17 | 0.006 | Young |
| 0.339836 | 0.350524 | 0.109 | 0.096 | 1 | 3 | Taf3 | 0.013 | Young |
| 0.339864 | 0.415822 | 0.107 | 0.094 | 1 | 3 | Fam117a | 0.013 | Young |
| 0.33999 | 0.430757 | 0.216 | 0.205 | 1 | 3 | Phf10 | 0.011 | Young |
| 0.340568 | 0.019433 | 0.338 | 0.372 | 1 | 3 | Ciita | -0.034 | Young |
| 0.340585 | 0.251186 | 0.203 | 0.189 | 1 | 3 | Cks2 | 0.014 | Young |
| 0.340917 | 0.116064 | 0.1 | 0.086 | 1 | 3 | Ccdc85b | 0.014 | Young |
| 0.341201 | 0.388533 | 0.203 | 0.191 | 1 | 3 | Tnfrsf13b | 0.012 | Young |
| 0.341218 | 0.08352 | 0.242 | 0.276 | 1 | 3 | Il2rg | -0.034 | Young |
| 0.341422 | 0.149766 | 0.567 | 0.579 | 1 | 3 | Man2b1 | -0.012 | Young |
| 0.341613 | 0.475511 | 0.106 | 0.094 | 1 | 3 | Ciz1 | 0.012 | Young |
| 0.341875 | 0.356543 | 0.23 | 0.222 | 1 | 3 | Sap18 | 0.008 | Young |
| 0.342051 | 0.07183 | 0.184 | 0.207 | 1 | 3 | 2210016F1 | -0.023 | Young |
| 0.342086 | 0.451003 | 0.064 | 0.054 | 1 | 3 | Fkrp | 0.01 | Young |
| 0.342716 | 0.173644 | 0.12 | 0.106 | 1 | 3 | Camsap2 | 0.014 | Young |
| 0.343085 | 0.05556 | 0.309 | 0.346 | 1 | 3 | Mrps6 | -0.037 | Young |
| 0.343378 | 0.013108 | 0.066 | 0.079 | 1 | 3 | Aifm1 | -0.013 | Young |
| 0.343834 | 0.432677 | 0.062 | 0.051 | 1 | 3 | Pacs1 | 0.011 | Young |

| | | | | | | | | |
|---|---|---|---|---|---|---|---|---|
| 0.34426 | 0.223545 | 0.032 | 0.024 | 1 | 3 | Smpd2 | 0.008 | Young |
| 0.344829 | 0.345959 | 0.032 | 0.024 | 1 | 3 | Fig4 | 0.008 | Young |
| 0.344859 | 0.178061 | 0.335 | 0.33 | 1 | 3 | Hacd4 | 0.005 | Young |
| 0.345011 | 0.03977 | 0.159 | 0.182 | 1 | 3 | Timm10 | -0.023 | Young |
| 0.345048 | 0.120446 | 0.12 | 0.14 | 1 | 3 | Usp32 | -0.02 | Young |
| 0.345726 | 0.051013 | 0.899 | 0.926 | 1 | 3 | Rps2 | -0.027 | Young |
| 0.345831 | 0.100016 | 0.144 | 0.162 | 1 | 3 | Vps9d1 | -0.018 | Young |
| 0.346187 | 0.035993 | 0.189 | 0.213 | 1 | 3 | Dicer1 | -0.024 | Young |
| 0.346409 | 0.084868 | 0.032 | 0.041 | 1 | 3 | Etfbkmt | -0.009 | Young |
| 0.346442 | 0.20272 | 0.235 | 0.219 | 1 | 3 | Trp53 | 0.016 | Young |
| 0.347465 | 0.034549 | 0.381 | 0.423 | 1 | 3 | Ifnar1 | -0.042 | Young |
| 0.347822 | 0.590073 | 0.034 | 0.026 | 1 | 3 | Dis3l | 0.008 | Young |
| 0.347822 | 0.45551 | 0.034 | 0.026 | 1 | 3 | Adat2 | 0.008 | Young |
| 0.348324 | 0.668726 | 0.01 | 0.006 | 1 | 3 | Fcor | 0.004 | Young |
| 0.348676 | 0.942509 | 0.01 | 0.006 | 1 | 3 | Gm47926 | 0.004 | Young |
| 0.348699 | 0.09722 | 0.081 | 0.096 | 1 | 3 | Bend4 | -0.015 | Young |
| 0.349028 | 0.60677 | 0.01 | 0.006 | 1 | 3 | Rad9b | 0.004 | Young |
| 0.349028 | 0.468775 | 0.01 | 0.006 | 1 | 3 | Ascl2 | 0.004 | Young |
| 0.349287 | 0.345742 | 0.034 | 0.026 | 1 | 3 | Mphosph9 | 0.008 | Young |
| 0.349353 | 0.257321 | 0.039 | 0.049 | 1 | 3 | Zkscan1 | -0.01 | Young |
| 0.34938 | 0.603936 | 0.01 | 0.006 | 1 | 3 | Gm38832 | 0.004 | Young |
| 0.349479 | 0.185611 | 0.17 | 0.195 | 1 | 3 | Papd5 | -0.025 | Young |
| 0.349821 | 0.229571 | 0.117 | 0.104 | 1 | 3 | Cwf19l2 | 0.013 | Young |
| 0.349838 | 0.274003 | 0.034 | 0.026 | 1 | 3 | Mtfmt | 0.008 | Young |
| 0.349979 | 0.297734 | 0.066 | 0.055 | 1 | 3 | Rnf214 | 0.011 | Young |
| 0.350159 | 0.430049 | 0.019 | 0.013 | 1 | 3 | Zfp74 | 0.006 | Young |
| 0.350247 | 0.498267 | 0.088 | 0.076 | 1 | 3 | Ctdspl | 0.012 | Young |
| 0.350259 | 0.100654 | 0.455 | 0.514 | 1 | 3 | Ptp4a1 | -0.059 | Young |
| 0.350689 | 0.770307 | 0.038 | 0.03 | 1 | 3 | Zfp426 | 0.008 | Young |
| 0.351045 | 0.070853 | 0.111 | 0.128 | 1 | 3 | Cryzl1 | -0.017 | Young |
| 0.351335 | 0.367254 | 0.126 | 0.114 | 1 | 3 | AY036118 | 0.012 | Young |
| 0.351404 | 0.179218 | 0.237 | 0.223 | 1 | 3 | Gbp3 | 0.014 | Young |
| 0.351491 | 0.150091 | 0.105 | 0.092 | 1 | 3 | Il1rl2 | 0.013 | Young |
| 0.35185 | 0.586013 | 0.01 | 0.006 | 1 | 3 | Gm44777 | 0.004 | Young |
| 0.351851 | 0.163902 | 0.229 | 0.219 | 1 | 3 | Snrnp200 | 0.01 | Young |
| 0.351861 | 0.574946 | 0.034 | 0.026 | 1 | 3 | Mad1l1 | 0.008 | Young |
| 0.351882 | 0.341002 | 0.128 | 0.114 | 1 | 3 | Klhl20 | 0.014 | Young |
| 0.351895 | 0.345051 | 0.033 | 0.025 | 1 | 3 | Ankrd26 | 0.008 | Young |
| 0.352091 | 0.1527 | 0.131 | 0.153 | 1 | 3 | Flot2 | -0.022 | Young |
| 0.352256 | 0.546055 | 0.037 | 0.029 | 1 | 3 | Zfp444 | 0.008 | Young |
| 0.352283 | 0.390615 | 0.083 | 0.072 | 1 | 3 | Sema4d | 0.011 | Young |
| 0.352349 | 0.334302 | 0.247 | 0.238 | 1 | 3 | Baz2a | 0.009 | Young |
| 0.352466 | 0.013108 | 0.115 | 0.133 | 1 | 3 | Ahsa2 | -0.018 | Young |
| 0.352544 | 0.082146 | 0.29 | 0.329 | 1 | 3 | Prpf4b | -0.039 | Young |
| 0.352558 | 0.723505 | 0.01 | 0.006 | 1 | 3 | Arv1 | 0.004 | Young |

| | | | | | | | | |
|---|---|---|---|---|---|---|---|---|
| 0.352739 | 0.19868 | 0.167 | 0.153 | 1 | 3 | Brpf1 | 0.014 | Young |
| 0.352946 | 0.064167 | 0.33 | 0.368 | 1 | 3 | Mtm1 | -0.038 | Young |
| 0.353232 | 0.028823 | 0.311 | 0.339 | 1 | 3 | Was | -0.028 | Young |
| 0.353261 | 0.208592 | 0.083 | 0.098 | 1 | 3 | 4931406P1 | -0.015 | Young |
| 0.353267 | 0.092443 | 0.01 | 0.006 | 1 | 3 | Gm17259 | 0.004 | Young |
| 0.353565 | 0.581649 | 0.038 | 0.03 | 1 | 3 | Agk | 0.008 | Young |
| 0.353681 | 0.030022 | 0.24 | 0.271 | 1 | 3 | Uba2 | -0.031 | Young |
| 0.353879 | 0.038803 | 0.332 | 0.372 | 1 | 3 | Nisch | -0.04 | Young |
| 0.354013 | 0.121634 | 0.206 | 0.19 | 1 | 3 | Nuak2 | 0.016 | Young |
| 0.354073 | 0.185252 | 0.405 | 0.404 | 1 | 3 | Sfxn3 | 0.001 | Young |
| 0.354251 | 0.362536 | 0.144 | 0.13 | 1 | 3 | Taok2 | 0.014 | Young |
| 0.354255 | 0.655408 | 0.043 | 0.035 | 1 | 3 | Cep164 | 0.008 | Young |
| 0.354379 | 0.677102 | 0.02 | 0.014 | 1 | 3 | Cbr3 | 0.006 | Young |
| 0.354756 | 0.442425 | 0.225 | 0.214 | 1 | 3 | Rmnd5a | 0.011 | Young |
| 0.354825 | 0.342408 | 0.107 | 0.094 | 1 | 3 | Gclm | 0.013 | Young |
| 0.354842 | 0.653315 | 0.077 | 0.067 | 1 | 3 | Urod | 0.01 | Young |
| 0.355101 | 0.349697 | 0.033 | 0.025 | 1 | 3 | Siglecf | 0.008 | Young |
| 0.355177 | 0.108983 | 0.101 | 0.117 | 1 | 3 | Slc35e4 | -0.016 | Young |
| 0.355251 | 0.002659 | 0.216 | 0.239 | 1 | 3 | Yipf4 | -0.023 | Young |
| 0.355306 | 0.841507 | 0.042 | 0.033 | 1 | 3 | Pced1b | 0.009 | Young |
| 0.355466 | 0.682231 | 0.095 | 0.084 | 1 | 3 | Tnfrsf12a | 0.011 | Young |
| 0.355517 | 0.72961 | 0.059 | 0.05 | 1 | 3 | Mark4 | 0.009 | Young |
| 0.35595 | 0.216776 | 0.02 | 0.027 | 1 | 3 | Drosha | -0.007 | Young |
| 0.355969 | 0.163164 | 0.193 | 0.179 | 1 | 3 | Mrpl43 | 0.014 | Young |
| 0.356061 | 0.089492 | 0.131 | 0.149 | 1 | 3 | Fam105a | -0.018 | Young |
| 0.356285 | 0.032327 | 0.045 | 0.056 | 1 | 3 | Cdr2 | -0.011 | Young |
| 0.356307 | 0.289125 | 0.021 | 0.029 | 1 | 3 | Atat1 | -0.008 | Young |
| 0.356381 | 0.348835 | 0.177 | 0.165 | 1 | 3 | Slc45a4 | 0.012 | Young |
| 0.356512 | 0.132317 | 0.378 | 0.372 | 1 | 3 | Ndufa11 | 0.006 | Young |
| 0.356658 | 0.287474 | 0.122 | 0.11 | 1 | 3 | Smurf2 | 0.012 | Young |
| 0.35671 | 0.437191 | 0.033 | 0.025 | 1 | 3 | Ccdc84 | 0.008 | Young |
| 0.356756 | 0.57331 | 0.098 | 0.087 | 1 | 3 | Hlx | 0.011 | Young |
| 0.356802 | 0.215697 | 0.259 | 0.249 | 1 | 3 | Nfic | 0.01 | Young |
| 0.356834 | 0.081094 | 0.598 | 0.605 | 1 | 3 | Phlda1 | -0.007 | Young |
| 0.357191 | 0.32348 | 0.088 | 0.076 | 1 | 3 | Gmcl1 | 0.012 | Young |
| 0.357561 | 0.079568 | 0.371 | 0.415 | 1 | 3 | Zc3h12a | -0.044 | Young |
| 0.357868 | 0.105875 | 0.607 | 0.624 | 1 | 3 | Gas7 | -0.017 | Young |
| 0.358284 | 0.537219 | 0.02 | 0.014 | 1 | 3 | Herc3 | 0.006 | Young |
| 0.359468 | 0.141479 | 0.42 | 0.409 | 1 | 3 | Dusp11 | 0.011 | Young |
| 0.35951 | 0.921957 | 0.02 | 0.014 | 1 | 3 | Gm8113 | 0.006 | Young |
| 0.359616 | 0.006486 | 0.027 | 0.035 | 1 | 3 | Stambpl1 | -0.008 | Young |
| 0.359779 | 0.146259 | 0.058 | 0.048 | 1 | 3 | Stoml1 | 0.01 | Young |
| 0.359966 | 0.094501 | 0.587 | 0.596 | 1 | 3 | Spop | -0.009 | Young |
| 0.359983 | 0.045542 | 0.188 | 0.211 | 1 | 3 | Sf3b4 | -0.023 | Young |
| 0.360069 | 0.33264 | 0.124 | 0.111 | 1 | 3 | Ppm1k | 0.013 | Young |

| | | | | | | | | |
|---|---|---|---|---|---|---|---|---|
| 0.36038 | 0.162796 | 0.345 | 0.339 | 1 | 3 | AC149090. | 0.006 | Young |
| 0.36066 | 0.809363 | 0.045 | 0.037 | 1 | 3 | Men1 | 0.008 | Young |
| 0.360778 | 0.108627 | 0.058 | 0.048 | 1 | 3 | Tef | 0.01 | Young |
| 0.361044 | 0.381028 | 0.184 | 0.174 | 1 | 3 | Snap29 | 0.01 | Young |
| 0.361566 | 0.206177 | 0.033 | 0.025 | 1 | 3 | Batf2 | 0.008 | Young |
| 0.361621 | 0.196708 | 0.254 | 0.238 | 1 | 3 | Otud5 | 0.016 | Young |
| 0.361724 | 0.418911 | 0.069 | 0.059 | 1 | 3 | Spg11 | 0.01 | Young |
| 0.361807 | 0.149048 | 0.319 | 0.306 | 1 | 3 | Chmp2b | 0.013 | Young |
| 0.362052 | 0.188689 | 0.417 | 0.407 | 1 | 3 | Zfhx3 | 0.01 | Young |
| 0.362113 | 0.077113 | 0.265 | 0.288 | 1 | 3 | Clcn4 | -0.023 | Young |
| 0.362737 | 0.359297 | 0.203 | 0.192 | 1 | 3 | Trmt1 | 0.011 | Young |
| 0.362772 | 0.335251 | 0.266 | 0.256 | 1 | 3 | Zdhhc9 | 0.01 | Young |
| 0.36295 | 0.809184 | 0.021 | 0.016 | 1 | 3 | Tcaim | 0.005 | Young |
| 0.362957 | 0.485367 | 0.02 | 0.014 | 1 | 3 | Gm16104 | 0.006 | Young |
| 0.363117 | 0.053797 | 0.154 | 0.174 | 1 | 3 | Exoc2 | -0.02 | Young |
| 0.363295 | 0.44621 | 0.095 | 0.084 | 1 | 3 | Ppp2r3c | 0.011 | Young |
| 0.363551 | 0.272191 | 0.324 | 0.319 | 1 | 3 | Safb2 | 0.005 | Young |
| 0.363941 | 0.101472 | 0.155 | 0.174 | 1 | 3 | Nudt16l1 | -0.019 | Young |
| 0.364027 | 0.43625 | 0.016 | 0.023 | 1 | 3 | Ubiad1 | -0.007 | Young |
| 0.364224 | 0.582328 | 0.077 | 0.067 | 1 | 3 | Fiz1 | 0.01 | Young |
| 0.364563 | 0.13494 | 0.061 | 0.05 | 1 | 3 | Cacfd1 | 0.011 | Young |
| 0.364701 | 0.358579 | 0.26 | 0.25 | 1 | 3 | Ehmt2 | 0.01 | Young |
| 0.364763 | 0.53389 | 0.08 | 0.069 | 1 | 3 | Snx29 | 0.011 | Young |
| 0.364947 | 0.252634 | 0.071 | 0.06 | 1 | 3 | Il16 | 0.011 | Young |
| 0.364994 | 0.202402 | 0.378 | 0.374 | 1 | 3 | Cebpg | 0.004 | Young |
| 0.365431 | 0.062492 | 0.02 | 0.014 | 1 | 3 | Plxna3 | 0.006 | Young |
| 0.365513 | 0.180126 | 0.1 | 0.086 | 1 | 3 | Pja1 | 0.014 | Young |
| 0.365533 | 0.134167 | 0.162 | 0.183 | 1 | 3 | Tmem57 | -0.021 | Young |
| 0.365549 | 0.100322 | 0.093 | 0.109 | 1 | 3 | Xpo4 | -0.016 | Young |
| 0.365909 | 0.314723 | 0.037 | 0.029 | 1 | 3 | Slc14a1 | 0.008 | Young |
| 0.366606 | 0.096228 | 0.434 | 0.48 | 1 | 3 | Sec62 | -0.046 | Young |
| 0.366645 | 0.153961 | 0.524 | 0.527 | 1 | 3 | Bcl2a1b | -0.003 | Young |
| 0.366738 | 0.365987 | 0.136 | 0.124 | 1 | 3 | Polr2j | 0.012 | Young |
| 0.36675 | 0.052223 | 0.176 | 0.196 | 1 | 3 | Kctd10 | -0.02 | Young |
| 0.36728 | 0.171848 | 0.016 | 0.023 | 1 | 3 | H2-Q10 | -0.007 | Young |
| 0.367554 | 0.006592 | 0.021 | 0.029 | 1 | 3 | 2610044O1 | -0.008 | Young |
| 0.367994 | 0.550343 | 0.122 | 0.111 | 1 | 3 | Mbd1 | 0.011 | Young |
| 0.368225 | 0.991001 | 0.021 | 0.016 | 1 | 3 | Wdr76 | 0.005 | Young |
| 0.368225 | 0.475586 | 0.021 | 0.016 | 1 | 3 | Top3a | 0.005 | Young |
| 0.368586 | 0.117999 | 0.021 | 0.016 | 1 | 3 | Gtpbp3 | 0.005 | Young |
| 0.368969 | 0.196989 | 0.045 | 0.056 | 1 | 3 | Ccdc32 | -0.011 | Young |
| 0.369533 | 0.39628 | 0.231 | 0.22 | 1 | 3 | Rnf38 | 0.011 | Young |
| 0.370883 | 0.012294 | 0.074 | 0.088 | 1 | 3 | Gm43848 | -0.014 | Young |
| 0.370916 | 0.106007 | 0.593 | 0.595 | 1 | 3 | Ccdc50 | -0.002 | Young |
| 0.371297 | 0.32864 | 0.25 | 0.239 | 1 | 3 | Hsph1 | 0.011 | Young |

| | | | | | | | | |
|---|---|---|---|---|---|---|---|---|
| 0.371489 | 0.784018 | 0.011 | 0.007 | 1 | 3 | B230217C1 | 0.004 | Young |
| 0.371662 | 0.500799 | 0.078 | 0.068 | 1 | 3 | Cntln | 0.01 | Young |
| 0.371781 | 0.036883 | 0.029 | 0.037 | 1 | 3 | 2610008E1 | -0.008 | Young |
| 0.37183 | 1.036922 | 0.011 | 0.007 | 1 | 3 | Rnd1 | 0.004 | Young |
| 0.37183 | 0.750976 | 0.011 | 0.007 | 1 | 3 | Gm12166 | 0.004 | Young |
| 0.372343 | 0.029367 | 0.109 | 0.123 | 1 | 3 | Dennd1c | -0.014 | Young |
| 0.372777 | 0.263179 | 0.044 | 0.036 | 1 | 3 | Zfp709 | 0.008 | Young |
| 0.372967 | 0.022852 | 0.404 | 0.454 | 1 | 3 | Csnk2b | -0.05 | Young |
| 0.373041 | 0.057397 | 0.063 | 0.053 | 1 | 3 | Retreg1 | 0.01 | Young |
| 0.374222 | 0.45149 | 0.011 | 0.007 | 1 | 3 | Sardh | 0.004 | Young |
| 0.37431 | 0.276954 | 0.016 | 0.023 | 1 | 3 | 5730409E0 | -0.007 | Young |
| 0.374429 | 0.24355 | 0.136 | 0.122 | 1 | 3 | Gm41307 | 0.014 | Young |
| 0.374454 | 0.16298 | 0.22 | 0.204 | 1 | 3 | 1810022K0 | 0.016 | Young |
| 0.37449 | 0.104773 | 0.008 | 0.012 | 1 | 3 | Tmem140 | -0.004 | Young |
| 0.374831 | 0.24784 | 0.158 | 0.146 | 1 | 3 | Mboat7 | 0.012 | Young |
| 0.374908 | 0.53138 | 0.011 | 0.007 | 1 | 3 | Uprt | 0.004 | Young |
| 0.375104 | 0.065478 | 0.664 | 0.67 | 1 | 3 | Pafah1b1 | -0.006 | Young |
| 0.375231 | 0.272637 | 0.04 | 0.032 | 1 | 3 | Slco2b1 | 0.008 | Young |
| 0.375241 | 0.157348 | 0.47 | 0.47 | 1 | 3 | Tubb2a | 0 | Young |
| 0.375251 | 0.7638 | 0.011 | 0.007 | 1 | 3 | Them6 | 0.004 | Young |
| 0.375251 | 0.300482 | 0.011 | 0.007 | 1 | 3 | Spata1 | 0.004 | Young |
| 0.375423 | 0.225466 | 0.105 | 0.093 | 1 | 3 | Arl10 | 0.012 | Young |
| 0.375736 | 0.723373 | 0.066 | 0.056 | 1 | 3 | Parp6 | 0.01 | Young |
| 0.375918 | 0.174916 | 0.059 | 0.049 | 1 | 3 | Crebl2 | 0.01 | Young |
| 0.376281 | 0.417926 | 0.011 | 0.007 | 1 | 3 | Ccl27a | 0.004 | Young |
| 0.376331 | 0.055374 | 0.895 | 0.895 | 1 | 3 | Rhoa | 0 | Young |
| 0.376486 | 0.115536 | 0.023 | 0.03 | 1 | 3 | Hectd2 | -0.007 | Young |
| 0.376968 | 0.126349 | 0.011 | 0.007 | 1 | 3 | Pter | 0.004 | Young |
| 0.377093 | 0.055592 | 0.307 | 0.341 | 1 | 3 | Mdm2 | -0.034 | Young |
| 0.377346 | 0.503029 | 0.096 | 0.085 | 1 | 3 | Cpped1 | 0.011 | Young |
| 0.377476 | 0.223513 | 0.235 | 0.221 | 1 | 3 | Cdc40 | 0.014 | Young |
| 0.377482 | 0.049539 | 0.304 | 0.337 | 1 | 3 | Map4 | -0.033 | Young |
| 0.377494 | 1.279231 | 0.025 | 0.019 | 1 | 3 | Them4 | 0.006 | Young |
| 0.377619 | 0.427492 | 0.023 | 0.017 | 1 | 3 | Clec2g | 0.006 | Young |
| 0.378164 | 0.638672 | 0.052 | 0.043 | 1 | 3 | Gm26518 | 0.009 | Young |
| 0.378238 | 0.089455 | 0.559 | 0.562 | 1 | 3 | Tcf25 | -0.003 | Young |
| 0.378522 | 0.162206 | 0.241 | 0.269 | 1 | 3 | Alkbh1 | -0.028 | Young |
| 0.378568 | 0.444953 | 0.023 | 0.017 | 1 | 3 | Bphl | 0.006 | Young |
| 0.378691 | 0.191152 | 0.011 | 0.007 | 1 | 3 | Cdkl5 | 0.004 | Young |
| 0.379043 | 0.497037 | 0.023 | 0.017 | 1 | 3 | Dyrk3 | 0.006 | Young |
| 0.379145 | 0.104202 | 0.218 | 0.243 | 1 | 3 | Kdelr2 | -0.025 | Young |
| 0.379488 | 0.33382 | 0.008 | 0.012 | 1 | 3 | Gm34006 | -0.004 | Young |
| 0.379633 | 0.093336 | 0.039 | 0.031 | 1 | 3 | Tango6 | 0.008 | Young |
| 0.379872 | 0.590027 | 0.052 | 0.043 | 1 | 3 | Ppp1r13b | 0.009 | Young |
| 0.379994 | 0.418228 | 0.023 | 0.017 | 1 | 3 | Phlda3 | 0.006 | Young |

| | | | | | | | | |
|---|---|---|---|---|---|---|---|---|
| 0.380201 | 0.103137 | 0.439 | 0.479 | 1 | 3 | Lsm4 | -0.04 | Young |
| 0.380232 | 0.161615 | 0.023 | 0.017 | 1 | 3 | Copz2 | 0.006 | Young |
| 0.380273 | 0.166921 | 0.117 | 0.135 | 1 | 3 | Kmt2b | -0.018 | Young |
| 0.380317 | 0.381147 | 0.188 | 0.177 | 1 | 3 | Rgs14 | 0.011 | Young |
| 0.38047 | 0.454193 | 0.023 | 0.017 | 1 | 3 | Glb1l | 0.006 | Young |
| 0.380484 | 0.048825 | 0.128 | 0.145 | 1 | 3 | Gart | -0.017 | Young |
| 0.381211 | 0.070561 | 0.129 | 0.147 | 1 | 3 | Vta1 | -0.018 | Young |
| 0.381423 | 0.656472 | 0.023 | 0.017 | 1 | 3 | Zfhx2 | 0.006 | Young |
| 0.381457 | 0.13672 | 0.011 | 0.007 | 1 | 3 | Slc26a11 | 0.004 | Young |
| 0.381577 | 0.557169 | 0.068 | 0.059 | 1 | 3 | Qser1 | 0.009 | Young |
| 0.381658 | 0.472159 | 0.062 | 0.053 | 1 | 3 | Depdc5 | 0.009 | Young |
| 0.381837 | 0.104513 | 0.199 | 0.223 | 1 | 3 | Kidins220 | -0.024 | Young |
| 0.381959 | 0.648562 | 0.045 | 0.037 | 1 | 3 | Pold1 | 0.008 | Young |
| 0.382074 | 0.064442 | 0.313 | 0.346 | 1 | 3 | Rragc | -0.033 | Young |
| 0.382173 | 0.439926 | 0.048 | 0.039 | 1 | 3 | Haao | 0.009 | Young |
| 0.382189 | 0.19281 | 0.095 | 0.082 | 1 | 3 | Rnf168 | 0.013 | Young |
| 0.382363 | 0.458868 | 0.071 | 0.061 | 1 | 3 | Zdhhc17 | 0.01 | Young |
| 0.38243 | 0.034381 | 0.963 | 0.976 | 1 | 3 | Rack1 | -0.013 | Young |
| 0.382634 | 0.462514 | 0.043 | 0.035 | 1 | 3 | Coa4 | 0.008 | Young |
| 0.382673 | 0.131665 | 0.097 | 0.086 | 1 | 3 | Rev3l | 0.011 | Young |
| 0.382832 | 0.096637 | 0.067 | 0.079 | 1 | 3 | Zfp945 | -0.012 | Young |
| 0.38289 | 0.065042 | 0.029 | 0.037 | 1 | 3 | Slc7a5 | -0.008 | Young |
| 0.383081 | 0.122433 | 0.128 | 0.147 | 1 | 3 | Vps16 | -0.019 | Young |
| 0.383156 | 0.271512 | 0.212 | 0.202 | 1 | 3 | Pip4k2b | 0.01 | Young |
| 0.383284 | 0.340125 | 0.035 | 0.044 | 1 | 3 | Slc12a2 | -0.009 | Young |
| 0.38329 | 0.083003 | 0.095 | 0.11 | 1 | 3 | Ints11 | -0.015 | Young |
| 0.383298 | 0.013408 | 0.135 | 0.153 | 1 | 3 | Slc35e1 | -0.018 | Young |
| 0.38335 | 0.284122 | 0.04 | 0.032 | 1 | 3 | Pced1a | 0.008 | Young |
| 0.383488 | 0.232998 | 0.023 | 0.03 | 1 | 3 | Polr3h | -0.007 | Young |
| 0.383551 | 0.516081 | 0.024 | 0.018 | 1 | 3 | Galnt6 | 0.006 | Young |
| 0.383712 | 0.020159 | 0.076 | 0.088 | 1 | 3 | Pnpo | -0.012 | Young |
| 0.384034 | 0.681611 | 0.104 | 0.094 | 1 | 3 | Odf2 | 0.01 | Young |
| 0.384835 | 0.066639 | 0.179 | 0.2 | 1 | 3 | Ndfip2 | -0.021 | Young |
| 0.38516 | 0.02533 | 0.134 | 0.152 | 1 | 3 | Mzt1 | -0.018 | Young |
| 0.38555 | 0.077473 | 0.729 | 0.751 | 1 | 3 | Serinc3 | -0.022 | Young |
| 0.386783 | 0.230992 | 0.124 | 0.142 | 1 | 3 | Ppp2cb | -0.018 | Young |
| 0.387024 | 0.267178 | 0.316 | 0.308 | 1 | 3 | Ddx50 | 0.008 | Young |
| 0.387318 | 0.064248 | 0.024 | 0.031 | 1 | 3 | Rdh13 | -0.007 | Young |
| 0.387928 | 0.412993 | 0.053 | 0.044 | 1 | 3 | Rnf8 | 0.009 | Young |
| 0.388682 | 0.50308 | 0.024 | 0.018 | 1 | 3 | Atp6v0c | 0.006 | Young |
| 0.389346 | 0.056467 | 0.198 | 0.223 | 1 | 3 | Gsto1 | -0.025 | Young |
| 0.389366 | 0.057902 | 0.044 | 0.054 | 1 | 3 | Rbm45 | -0.01 | Young |
| 0.390035 | 0.438868 | 0.091 | 0.08 | 1 | 3 | Appl2 | 0.011 | Young |
| 0.390427 | 0.399594 | 0.096 | 0.086 | 1 | 3 | Dnmt1 | 0.01 | Young |
| 0.390544 | 0.030627 | 0.438 | 0.489 | 1 | 3 | Ubxn1 | -0.051 | Young |

| | | | | | | | | |
|---|---|---|---|---|---|---|---|---|
| 0.390757 | 0.357525 | 0.168 | 0.155 | 1 | 3 | Rit1 | 0.013 | Young |
| 0.391121 | 0.075125 | 0.342 | 0.386 | 1 | 3 | Nars | -0.044 | Young |
| 0.391164 | 0.318145 | 0.235 | 0.222 | 1 | 3 | Elf2 | 0.013 | Young |
| 0.391183 | 0.110019 | 0.116 | 0.133 | 1 | 3 | Naa30 | -0.017 | Young |
| 0.391343 | 0.066223 | 0.347 | 0.391 | 1 | 3 | Kcmf1 | -0.044 | Young |
| 0.391391 | 0.033537 | 0.271 | 0.303 | 1 | 3 | Sun2 | -0.032 | Young |
| 0.391642 | 0.672795 | 0.069 | 0.06 | 1 | 3 | Gramd1b | 0.009 | Young |
| 0.391645 | 0.511607 | 0.052 | 0.043 | 1 | 3 | Parn | 0.009 | Young |
| 0.391733 | 0.201442 | 0.381 | 0.372 | 1 | 3 | Fam102b | 0.009 | Young |
| 0.39182 | 0.008101 | 0.035 | 0.044 | 1 | 3 | Zfp938 | -0.009 | Young |
| 0.391833 | 0.210628 | 0.309 | 0.311 | 1 | 3 | Crlf3 | -0.002 | Young |
| 0.39206 | 0.629714 | 0.027 | 0.02 | 1 | 3 | Nmnat3 | 0.007 | Young |
| 0.392114 | 0.006494 | 0.253 | 0.278 | 1 | 3 | Ndufaf8 | -0.025 | Young |
| 0.392189 | 0.555118 | 0.061 | 0.051 | 1 | 3 | Slc39a13 | 0.01 | Young |
| 0.39219 | 0.241606 | 0.064 | 0.076 | 1 | 3 | Gpr137 | -0.012 | Young |
| 0.39244 | 0.265074 | 0.018 | 0.024 | 1 | 3 | Taf4b | -0.006 | Young |
| 0.392513 | 0.046002 | 0.14 | 0.158 | 1 | 3 | Rnd3 | -0.018 | Young |
| 0.3926 | 0.338894 | 0.239 | 0.232 | 1 | 3 | Mdm4 | 0.007 | Young |
| 0.392672 | 0.131377 | 0.149 | 0.134 | 1 | 3 | Clcc1 | 0.015 | Young |
| 0.392993 | 0.039279 | 0.443 | 0.483 | 1 | 3 | Gsk3b | -0.04 | Young |
| 0.393106 | 0.072751 | 0.013 | 0.018 | 1 | 3 | Ccne1 | -0.005 | Young |
| 0.393269 | 0.356007 | 0.234 | 0.228 | 1 | 3 | Lars2 | 0.006 | Young |
| 0.393403 | 0.632714 | 0.013 | 0.008 | 1 | 3 | Swsap1 | 0.005 | Young |
| 0.393489 | 0.000206 | 0.501 | 0.539 | 1 | 3 | Snx20 | -0.038 | Young |
| 0.393735 | 0.612836 | 0.013 | 0.008 | 1 | 3 | Btc | 0.005 | Young |
| 0.394067 | 0.92456 | 0.013 | 0.008 | 1 | 3 | Lsmem1 | 0.005 | Young |
| 0.394067 | 0.744054 | 0.013 | 0.008 | 1 | 3 | Zfp287 | 0.005 | Young |
| 0.394067 | 0.48961 | 0.013 | 0.008 | 1 | 3 | Zfp566 | 0.005 | Young |
| 0.394142 | 0.147595 | 0.217 | 0.243 | 1 | 3 | Mga | -0.026 | Young |
| 0.394208 | 0.203736 | 0.068 | 0.059 | 1 | 3 | Cd209f | 0.009 | Young |
| 0.394217 | 0.157308 | 0.1 | 0.115 | 1 | 3 | Slc35a2 | -0.015 | Young |
| 0.3944 | 0.541166 | 0.013 | 0.008 | 1 | 3 | Gm42722 | 0.005 | Young |
| 0.394564 | 0.065468 | 0.024 | 0.018 | 1 | 3 | Gdf3 | 0.006 | Young |
| 0.394698 | 0.053756 | 0.024 | 0.031 | 1 | 3 | Gm20703 | -0.007 | Young |
| 0.394747 | 0.28075 | 0.047 | 0.038 | 1 | 3 | Asna1 | 0.009 | Young |
| 0.395112 | 0.020829 | 0.274 | 0.307 | 1 | 3 | Ncoa3 | -0.033 | Young |
| 0.395186 | 0.500287 | 0.054 | 0.045 | 1 | 3 | Sesn3 | 0.009 | Young |
| 0.395305 | 0.288739 | 0.105 | 0.093 | 1 | 3 | Rnf126 | 0.012 | Young |
| 0.395398 | 0.486755 | 0.013 | 0.008 | 1 | 3 | Rab39 | 0.005 | Young |
| 0.395577 | 0.443008 | 0.087 | 0.076 | 1 | 3 | Tmem250- | 0.011 | Young |
| 0.39581 | 0.009757 | 0.186 | 0.207 | 1 | 3 | Tmem50b | -0.021 | Young |
| 0.395829 | 0.255047 | 0.179 | 0.205 | 1 | 3 | Socs4 | -0.026 | Young |
| 0.396092 | 0.097905 | 0.231 | 0.257 | 1 | 3 | Pqlc3 | -0.026 | Young |
| 0.396269 | 0.009881 | 0.501 | 0.548 | 1 | 3 | Spi1 | -0.047 | Young |
| 0.396454 | 0.089454 | 0.106 | 0.094 | 1 | 3 | Scarb1 | 0.012 | Young |

| | | | | | | | | |
|---|---|---|---|---|---|---|---|---|
| 0.396568 | 0.187964 | 0.081 | 0.094 | 1 | 3 | Gpr68 | -0.013 | Young |
| 0.396616 | 0.377129 | 0.212 | 0.202 | 1 | 3 | Plpbp | 0.01 | Young |
| 0.39681 | 0.195863 | 0.067 | 0.08 | 1 | 3 | Def6 | -0.013 | Young |
| 0.39865 | 0.606094 | 0.025 | 0.019 | 1 | 3 | Tslp | 0.006 | Young |
| 0.398772 | 0.130454 | 0.4 | 0.4 | 1 | 3 | Use1 | 0 | Young |
| 0.39914 | 0.317604 | 0.101 | 0.09 | 1 | 3 | Trp53bp2 | 0.011 | Young |
| 0.399339 | 0.076091 | 0.125 | 0.141 | 1 | 3 | Dhdh | -0.016 | Young |
| 0.399405 | 0.144352 | 0.013 | 0.008 | 1 | 3 | Dynlt1a | 0.005 | Young |
| 0.399487 | 0.250512 | 0.215 | 0.203 | 1 | 3 | Arid4b | 0.012 | Young |
| 0.399641 | 0.257449 | 0.091 | 0.08 | 1 | 3 | Eci2 | 0.011 | Young |
| 0.399787 | 0.369563 | 0.08 | 0.069 | 1 | 3 | Acot8 | 0.011 | Young |
| 0.400039 | 0.284389 | 0.025 | 0.019 | 1 | 3 | 0610009O2 | 0.006 | Young |
| 0.400098 | 0.00068 | 0.823 | 0.854 | 1 | 3 | 11-Sep | -0.031 | Young |
| 0.400271 | 0.415568 | 0.025 | 0.019 | 1 | 3 | Ankzf1 | 0.006 | Young |
| 0.400515 | 0.481884 | 0.076 | 0.066 | 1 | 3 | Stambp | 0.01 | Young |
| 0.400814 | 0.61334 | 0.027 | 0.02 | 1 | 3 | Cenpq | 0.007 | Young |
| 0.400816 | 0.266228 | 0.053 | 0.044 | 1 | 3 | Ift22 | 0.009 | Young |
| 0.401062 | 0.28868 | 0.062 | 0.053 | 1 | 3 | Ints7 | 0.009 | Young |
| 0.401573 | 0.125018 | 0.067 | 0.057 | 1 | 3 | Man2c1 | 0.01 | Young |
| 0.401846 | 0.27224 | 0.076 | 0.066 | 1 | 3 | Elk1 | 0.01 | Young |
| 0.401895 | 0.236538 | 0.025 | 0.019 | 1 | 3 | Ift74 | 0.006 | Young |
| 0.402034 | 0.151932 | 0.129 | 0.146 | 1 | 3 | Sft2d1 | -0.017 | Young |
| 0.40229 | 0.322792 | 0.153 | 0.141 | 1 | 3 | Zswim8 | 0.012 | Young |
| 0.402426 | 0.110647 | 0.013 | 0.008 | 1 | 3 | Ccdc149 | 0.005 | Young |
| 0.402448 | 0.177896 | 0.048 | 0.039 | 1 | 3 | Zfp182 | 0.009 | Young |
| 0.402567 | 0.082826 | 0.295 | 0.327 | 1 | 3 | Csnk2a2 | -0.032 | Young |
| 0.402746 | 0.16037 | 0.117 | 0.134 | 1 | 3 | Mospd3 | -0.017 | Young |
| 0.402762 | 0.116204 | 0.013 | 0.008 | 1 | 3 | Spata2l | 0.005 | Young |
| 0.403098 | 0.006676 | 0.691 | 0.749 | 1 | 3 | Dstn | -0.058 | Young |
| 0.4032 | 0.104299 | 0.112 | 0.128 | 1 | 3 | Mpv17l2 | -0.016 | Young |
| 0.403281 | 0.135824 | 0.053 | 0.044 | 1 | 3 | Ifitm1 | 0.009 | Young |
| 0.403516 | 0.016167 | 0.218 | 0.243 | 1 | 3 | BC005624 | -0.025 | Young |
| 0.403545 | 0.040103 | 0.126 | 0.142 | 1 | 3 | Eed | -0.016 | Young |
| 0.40361 | 1.052691 | 0.035 | 0.029 | 1 | 3 | Sumf2 | 0.006 | Young |
| 0.40399 | 0.585839 | 0.025 | 0.019 | 1 | 3 | Zfp846 | 0.006 | Young |
| 0.404163 | 0.196746 | 0.351 | 0.345 | 1 | 3 | Trf | 0.006 | Young |
| 0.40472 | 0.119131 | 0.122 | 0.139 | 1 | 3 | Ogfrl1 | -0.017 | Young |
| 0.404831 | 0.082068 | 0.282 | 0.305 | 1 | 3 | Col4a3bp | -0.023 | Young |
| 0.404853 | 0.001854 | 0.903 | 0.918 | 1 | 3 | Tpm3 | -0.015 | Young |
| 0.404942 | 0.155996 | 0.129 | 0.117 | 1 | 3 | Hadh | 0.012 | Young |
| 0.40513 | 0.057248 | 0.116 | 0.133 | 1 | 3 | Asap2 | -0.017 | Young |
| 0.40547 | 0.009608 | 0.097 | 0.085 | 1 | 3 | Tjp2 | 0.012 | Young |
| 0.405479 | 0.951181 | 0.014 | 0.01 | 1 | 3 | Hgf | 0.004 | Young |
| 0.405681 | 0.329001 | 0.294 | 0.292 | 1 | 3 | Herc4 | 0.002 | Young |
| 0.405816 | 0.054184 | 0.179 | 0.2 | 1 | 3 | Ythdf2 | -0.021 | Young |

| | | | | | | | | |
|---|---|---|---|---|---|---|---|---|
| 0.405818 | 0.248667 | 0.106 | 0.094 | 1 | 3 | G6pc3 | 0.012 | Young |
| 0.406117 | 0.000471 | 0.76 | 0.8 | 1 | 3 | Hnrnpu | -0.04 | Young |
| 0.406295 | 0.285222 | 0.181 | 0.17 | 1 | 3 | Ube2q2 | 0.011 | Young |
| 0.406375 | 0.241054 | 0.052 | 0.062 | 1 | 3 | Fcna | -0.01 | Young |
| 0.406859 | 0.434432 | 0.13 | 0.121 | 1 | 3 | Mrps18a | 0.009 | Young |
| 0.406917 | 0.043949 | 0.098 | 0.112 | 1 | 3 | Gopc | -0.014 | Young |
| 0.40738 | 0.028025 | 0.154 | 0.172 | 1 | 3 | Cmklr1 | -0.018 | Young |
| 0.407395 | 0.250582 | 0.024 | 0.031 | 1 | 3 | Kbtbd7 | -0.007 | Young |
| 0.407605 | 0.160533 | 0.024 | 0.031 | 1 | 3 | E2f6 | -0.007 | Young |
| 0.408208 | 0.344929 | 0.083 | 0.073 | 1 | 3 | Abhd5 | 0.01 | Young |
| 0.408349 | 0.184694 | 0.381 | 0.381 | 1 | 3 | Arhgef1 | 0 | Young |
| 0.409268 | 0.000595 | 0.174 | 0.196 | 1 | 3 | Ktn1 | -0.022 | Young |
| 0.409326 | 0.002847 | 0.068 | 0.08 | 1 | 3 | Srgap3 | -0.012 | Young |
| 0.409418 | 0.051828 | 0.152 | 0.17 | 1 | 3 | Arl13b | -0.018 | Young |
| 0.409455 | 0.213588 | 0.027 | 0.02 | 1 | 3 | Tfap4 | 0.007 | Young |
| 0.409843 | 0.126917 | 0.025 | 0.019 | 1 | 3 | Dcaf4 | 0.006 | Young |
| 0.409899 | 0.259695 | 0.083 | 0.098 | 1 | 3 | Fam207a | -0.015 | Young |
| 0.410214 | 0.775658 | 0.03 | 0.024 | 1 | 3 | Neurl1b | 0.006 | Young |
| 0.410234 | 0.284906 | 0.061 | 0.051 | 1 | 3 | Mettl7a1 | 0.01 | Young |
| 0.4106 | 0.423773 | 0.027 | 0.02 | 1 | 3 | Txk | 0.007 | Young |
| 0.410627 | 0.039158 | 0.056 | 0.066 | 1 | 3 | Znrf1 | -0.01 | Young |
| 0.410991 | 0.146567 | 0.125 | 0.112 | 1 | 3 | Zfp664 | 0.013 | Young |
| 0.411151 | 0.510605 | 0.086 | 0.076 | 1 | 3 | Jmy | 0.01 | Young |
| 0.411233 | 0.199936 | 0.096 | 0.111 | 1 | 3 | Tmem238 | -0.015 | Young |
| 0.411676 | 0.049529 | 0.078 | 0.092 | 1 | 3 | Tmed4 | -0.014 | Young |
| 0.411724 | 0.253796 | 0.244 | 0.235 | 1 | 3 | Chd8 | 0.009 | Young |
| 0.411769 | 0.246566 | 0.227 | 0.256 | 1 | 3 | Lacc1 | -0.029 | Young |
| 0.411833 | 0.227264 | 0.082 | 0.072 | 1 | 3 | Pfkfb4 | 0.01 | Young |
| 0.411907 | 0.584737 | 0.014 | 0.01 | 1 | 3 | Afg1l | 0.004 | Young |
| 0.412497 | 0.24208 | 0.65 | 0.673 | 1 | 3 | Cdkn1a | -0.023 | Young |
| 0.412553 | 0.603613 | 0.014 | 0.01 | 1 | 3 | Cavin2 | 0.004 | Young |
| 0.412675 | 0.234266 | 0.009 | 0.013 | 1 | 3 | Gnb5 | -0.004 | Young |
| 0.413269 | 0.014774 | 0.24 | 0.265 | 1 | 3 | Imp3 | -0.025 | Young |
| 0.41333 | 0.089569 | 0.259 | 0.288 | 1 | 3 | Atg2a | -0.029 | Young |
| 0.413516 | 0.156486 | 0.024 | 0.031 | 1 | 3 | Fbxw9 | -0.007 | Young |
| 0.413531 | 0.07624 | 0.71 | 0.753 | 1 | 3 | Pip4k2a | -0.043 | Young |
| 0.413606 | 0.698658 | 0.08 | 0.07 | 1 | 3 | Smarcal1 | 0.01 | Young |
| 0.414212 | 0.25096 | 0.189 | 0.176 | 1 | 3 | Opa3 | 0.013 | Young |
| 0.41438 | 0.082537 | 0.28 | 0.313 | 1 | 3 | Copb1 | -0.033 | Young |
| 0.414495 | 0.525988 | 0.014 | 0.01 | 1 | 3 | Pard3b | 0.004 | Young |
| 0.414819 | 0.505248 | 0.014 | 0.01 | 1 | 3 | Cryab | 0.004 | Young |
| 0.415455 | 0.095328 | 0.158 | 0.177 | 1 | 3 | Pafah1b2 | -0.019 | Young |
| 0.41553 | 0.177822 | 0.074 | 0.065 | 1 | 3 | Fxyd2 | 0.009 | Young |
| 0.416191 | 0.045066 | 0.361 | 0.392 | 1 | 3 | Anxa3 | -0.031 | Young |
| 0.416676 | 0.067092 | 0.237 | 0.26 | 1 | 3 | Srf | -0.023 | Young |

| | | | | | | | | |
|---|---|---|---|---|---|---|---|---|
| 0.417051 | 0.034817 | 0.08 | 0.092 | 1 | 3 | Poldip2 | -0.012 | Young |
| 0.417105 | 0.05123 | 0.162 | 0.148 | 1 | 3 | Tmem173 | 0.014 | Young |
| 0.417448 | 0.212011 | 0.211 | 0.237 | 1 | 3 | Pik3ca | -0.026 | Young |
| 0.417745 | 0.095914 | 0.059 | 0.07 | 1 | 3 | Mettl21a | -0.011 | Young |
| 0.418408 | 0.026004 | 0.101 | 0.088 | 1 | 3 | Espl1 | 0.013 | Young |
| 0.418438 | 0.118696 | 0.02 | 0.026 | 1 | 3 | Kremen1 | -0.006 | Young |
| 0.418446 | 0.013167 | 0.538 | 0.596 | 1 | 3 | Mapkapk2 | -0.058 | Young |
| 0.418721 | 0.213503 | 0.014 | 0.01 | 1 | 3 | Zfp946 | 0.004 | Young |
| 0.418721 | 0.141446 | 0.014 | 0.01 | 1 | 3 | Klk8 | 0.004 | Young |
| 0.418745 | 0.243899 | 0.085 | 0.098 | 1 | 3 | Ap2a1 | -0.013 | Young |
| 0.418969 | 0.497345 | 0.029 | 0.023 | 1 | 3 | Ccdc130 | 0.006 | Young |
| 0.419499 | 0.125859 | 0.777 | 0.803 | 1 | 3 | Srrm2 | -0.026 | Young |
| 0.419654 | 0.024083 | 0.22 | 0.241 | 1 | 3 | 2310011J0 | -0.021 | Young |
| 0.419663 | 0.082601 | 0.658 | 0.687 | 1 | 3 | Clk1 | -0.029 | Young |
| 0.4197 | 0.316931 | 0.014 | 0.01 | 1 | 3 | Hyal1 | 0.004 | Young |
| 0.420741 | 0.033745 | 0.047 | 0.038 | 1 | 3 | Gm26778 | 0.009 | Young |
| 0.421109 | 0.213901 | 0.311 | 0.352 | 1 | 3 | Ppp6r3 | -0.041 | Young |
| 0.421279 | 0.039917 | 0.115 | 0.103 | 1 | 3 | Fam199x | 0.012 | Young |
| 0.421339 | 0.209063 | 0.112 | 0.129 | 1 | 3 | Slc25a13 | -0.017 | Young |
| 0.42137 | 0.465959 | 0.035 | 0.029 | 1 | 3 | Pcolce | 0.006 | Young |
| 0.421458 | 0.861292 | 0.015 | 0.011 | 1 | 3 | Upp2 | 0.004 | Young |
| 0.421576 | 0.010995 | 0.135 | 0.151 | 1 | 3 | Gabpa | -0.016 | Young |
| 0.421598 | 0.754592 | 0.063 | 0.055 | 1 | 3 | Stk32c | 0.008 | Young |
| 0.421704 | 0.031875 | 0.426 | 0.461 | 1 | 3 | Ctdnep1 | -0.035 | Young |
| 0.421767 | 0.016056 | 0.177 | 0.195 | 1 | 3 | Mex3c | -0.018 | Young |
| 0.421804 | 0.109624 | 0.119 | 0.134 | 1 | 3 | A630001G2 | -0.015 | Young |
| 0.421813 | 0.036376 | 0.265 | 0.289 | 1 | 3 | Kmt5a | -0.024 | Young |
| 0.421829 | 0.184587 | 0.167 | 0.155 | 1 | 3 | Larp1b | 0.012 | Young |
| 0.422409 | 0.007342 | 0.144 | 0.16 | 1 | 3 | Tcf3 | -0.016 | Young |
| 0.422624 | 0.328199 | 0.096 | 0.086 | 1 | 3 | Dpm2 | 0.01 | Young |
| 0.42323 | 0.07524 | 0.062 | 0.073 | 1 | 3 | Sil1 | -0.011 | Young |
| 0.42329 | 0.210877 | 0.372 | 0.372 | 1 | 3 | Cdk9 | 0 | Young |
| 0.423721 | 0.10568 | 0.197 | 0.219 | 1 | 3 | Tmem109 | -0.022 | Young |
| 0.423754 | 0.011456 | 0.082 | 0.094 | 1 | 3 | Arhgap21 | -0.012 | Young |
| 0.42463 | 0.328126 | 0.234 | 0.226 | 1 | 3 | Ilf3 | 0.008 | Young |
| 0.424876 | 0.184698 | 0.035 | 0.029 | 1 | 3 | Suox | 0.006 | Young |
| 0.424893 | 1.270296 | 0.015 | 0.011 | 1 | 3 | Prdm11 | 0.004 | Young |
| 0.425031 | 0.015374 | 0.482 | 0.527 | 1 | 3 | Fgfr1op2 | -0.045 | Young |
| 0.425206 | 0.625145 | 0.015 | 0.011 | 1 | 3 | Zfat | 0.004 | Young |
| 0.425486 | 0.023183 | 0.128 | 0.116 | 1 | 3 | Slf2 | 0.012 | Young |
| 0.426036 | 0.139838 | 0.121 | 0.136 | 1 | 3 | Nde1 | -0.015 | Young |
| 0.426338 | 0.21278 | 0.361 | 0.358 | 1 | 3 | Cd86 | 0.003 | Young |
| 0.426459 | 0.564563 | 0.015 | 0.011 | 1 | 3 | Cd209b | 0.004 | Young |
| 0.426479 | 0.04172 | 0.034 | 0.042 | 1 | 3 | Asb6 | -0.008 | Young |
| 0.426675 | 0.42896 | 0.093 | 0.084 | 1 | 3 | Lsm10 | 0.009 | Young |

| | | | | | | | | |
|---|---|---|---|---|---|---|---|---|
| 0.426738 | 0.556947 | 0.058 | 0.05 | 1 | 3 | Selenom | 0.008 | Young |
| 0.426773 | 0.503453 | 0.015 | 0.011 | 1 | 3 | C330027C( | 0.004 | Young |
| 0.427065 | 0.099136 | 0.783 | 0.8 | 1 | 3 | Adam15 | -0.017 | Young |
| 0.42716 | 0.075654 | 0.114 | 0.129 | 1 | 3 | Cln6 | -0.015 | Young |
| 0.427605 | 0.049135 | 0.061 | 0.051 | 1 | 3 | Ttc17 | 0.01 | Young |
| 0.427715 | 1.250374 | 0.015 | 0.011 | 1 | 3 | Wdr25 | 0.004 | Young |
| 0.428081 | 0.603447 | 0.034 | 0.027 | 1 | 3 | Jaml | 0.007 | Young |
| 0.428141 | 0.067262 | 0.083 | 0.096 | 1 | 3 | Syne3 | -0.013 | Young |
| 0.428382 | 0.334439 | 0.143 | 0.133 | 1 | 3 | Nrf1 | 0.01 | Young |
| 0.429201 | 0.047362 | 0.321 | 0.355 | 1 | 3 | Casp4 | -0.034 | Young |
| 0.429258 | 0.639998 | 0.042 | 0.035 | 1 | 3 | Dsel | 0.007 | Young |
| 0.429476 | 0.229344 | 0.056 | 0.066 | 1 | 3 | Kcne3 | -0.01 | Young |
| 0.42976 | 0.04763 | 0.698 | 0.71 | 1 | 3 | Son | -0.012 | Young |
| 0.429784 | 0.471647 | 0.167 | 0.157 | 1 | 3 | Slc7a11 | 0.01 | Young |
| 0.430153 | 0.173879 | 0.468 | 0.483 | 1 | 3 | Arhgap15 | -0.015 | Young |
| 0.430181 | 0.266462 | 0.207 | 0.197 | 1 | 3 | Commd3 | 0.01 | Young |
| 0.430347 | 0.38628 | 0.211 | 0.202 | 1 | 3 | P4ha1 | 0.009 | Young |
| 0.430596 | 0.308788 | 0.201 | 0.188 | 1 | 3 | Vrk1 | 0.013 | Young |
| 0.430649 | 0.124828 | 0.102 | 0.091 | 1 | 3 | Phf20 | 0.011 | Young |
| 0.430864 | 0.256089 | 0.015 | 0.011 | 1 | 3 | Rpgr | 0.004 | Young |
| 0.431053 | 0.015838 | 0.141 | 0.157 | 1 | 3 | Mrpl4 | -0.016 | Young |
| 0.431109 | 0.004946 | 0.595 | 0.649 | 1 | 3 | Hmgb1 | -0.054 | Young |
| 0.431284 | 0.23267 | 0.015 | 0.02 | 1 | 3 | 6430531B1 | -0.005 | Young |
| 0.431754 | 0.048619 | 0.282 | 0.307 | 1 | 3 | Ppp6r1 | -0.025 | Young |
| 0.431999 | 0.014857 | 0.135 | 0.151 | 1 | 3 | Casd1 | -0.016 | Young |
| 0.432116 | 0.175423 | 0.518 | 0.519 | 1 | 3 | Brd2 | -0.001 | Young |
| 0.432246 | 0.124533 | 0.02 | 0.026 | 1 | 3 | Armcx5 | -0.006 | Young |
| 0.432755 | 0.018177 | 0.676 | 0.72 | 1 | 3 | Ncl | -0.044 | Young |
| 0.43276 | 0.364637 | 0.015 | 0.011 | 1 | 3 | Tmem67 | 0.004 | Young |
| 0.43291 | 0.301763 | 0.049 | 0.042 | 1 | 3 | Tnfsf13b | 0.007 | Young |
| 0.433188 | 0.246673 | 0.275 | 0.269 | 1 | 3 | Trappc2l | 0.006 | Young |
| 0.433393 | 0.107784 | 0.015 | 0.011 | 1 | 3 | Nap1l2 | 0.004 | Young |
| 0.4334 | 0.248349 | 0.106 | 0.096 | 1 | 3 | Snhg12 | 0.01 | Young |
| 0.433507 | 0.086322 | 0.367 | 0.41 | 1 | 3 | Arl8a | -0.043 | Young |
| 0.433599 | 0.303084 | 0.283 | 0.274 | 1 | 3 | Evi5 | 0.009 | Young |
| 0.433913 | 0.079601 | 0.069 | 0.081 | 1 | 3 | Zdhhc13 | -0.012 | Young |
| 0.433957 | 0.552046 | 0.039 | 0.032 | 1 | 3 | Prr7 | 0.007 | Young |
| 0.433963 | 0.155068 | 0.034 | 0.027 | 1 | 3 | Wrb | 0.007 | Young |
| 0.434065 | 0.093356 | 0.035 | 0.043 | 1 | 3 | Zbtb41 | -0.008 | Young |
| 0.434071 | 0.418321 | 0.035 | 0.029 | 1 | 3 | Isoc2b | 0.006 | Young |
| 0.434485 | 0.061434 | 0.143 | 0.158 | 1 | 3 | Wdr13 | -0.015 | Young |
| 0.434496 | 0.561971 | 0.033 | 0.026 | 1 | 3 | Rubcn | 0.007 | Young |
| 0.434982 | 0.401139 | 0.136 | 0.125 | 1 | 3 | Anapc15 | 0.011 | Young |
| 0.435064 | 0.026299 | 0.318 | 0.357 | 1 | 3 | Rc3h1 | -0.039 | Young |
| 0.435125 | 0.069297 | 0.121 | 0.137 | 1 | 3 | Taf11 | -0.016 | Young |

| | | | | | | | | |
|---|---|---|---|---|---|---|---|---|
| 0.435295 | 0.328367 | 0.015 | 0.011 | 1 | 3 | Bdh1 | 0.004 | Young |
| 0.435341 | 0.607856 | 0.057 | 0.049 | 1 | 3 | Tmem107 | 0.008 | Young |
| 0.435341 | 0.198179 | 0.269 | 0.256 | 1 | 3 | Acss1 | 0.013 | Young |
| 0.435623 | 0.042494 | 0.088 | 0.102 | 1 | 3 | Gnl3l | -0.014 | Young |
| 0.436069 | 0.271497 | 0.104 | 0.093 | 1 | 3 | Rgcc | 0.011 | Young |
| 0.436815 | 0.527265 | 0.054 | 0.047 | 1 | 3 | Qpctl | 0.007 | Young |
| 0.437082 | 0.645145 | 0.016 | 0.012 | 1 | 3 | Mocs3 | 0.004 | Young |
| 0.437552 | 0.064511 | 0.318 | 0.351 | 1 | 3 | Card19 | -0.033 | Young |
| 0.437764 | 0.016469 | 0.154 | 0.173 | 1 | 3 | Cdk5rap3 | -0.019 | Young |
| 0.438186 | 0.040666 | 0.125 | 0.141 | 1 | 3 | Ndufaf4 | -0.016 | Young |
| 0.438783 | 0.514589 | 0.042 | 0.035 | 1 | 3 | Tbcc | 0.007 | Young |
| 0.438909 | 0.737345 | 0.016 | 0.012 | 1 | 3 | Gprin3 | 0.004 | Young |
| 0.438931 | 0.012859 | 0.316 | 0.348 | 1 | 3 | Jtb | -0.032 | Young |
| 0.438966 | 0.18364 | 0.035 | 0.029 | 1 | 3 | A630072M | 0.006 | Young |
| 0.439005 | 0.457313 | 0.037 | 0.03 | 1 | 3 | Gm43660 | 0.007 | Young |
| 0.439113 | 0.132483 | 0.015 | 0.011 | 1 | 3 | Ccsap | 0.004 | Young |
| 0.43922 | 0.197466 | 0.311 | 0.303 | 1 | 3 | Cmtm7 | 0.008 | Young |
| 0.439408 | 0.038116 | 0.015 | 0.02 | 1 | 3 | Nptxr | -0.005 | Young |
| 0.439608 | 0.325898 | 0.037 | 0.03 | 1 | 3 | Aldh1b1 | 0.007 | Young |
| 0.439912 | 0.265227 | 0.034 | 0.042 | 1 | 3 | Zfp101 | -0.008 | Young |
| 0.440204 | 0.28626 | 0.034 | 0.027 | 1 | 3 | Cenpt | 0.007 | Young |
| 0.440372 | 0.148555 | 0.176 | 0.16 | 1 | 3 | Polr2i | 0.016 | Young |
| 0.440426 | 0.327463 | 0.187 | 0.179 | 1 | 3 | Ppp2r2d | 0.008 | Young |
| 0.440436 | 0.502179 | 0.016 | 0.012 | 1 | 3 | Chaf1a | 0.004 | Young |
| 0.440518 | 0.029017 | 0.169 | 0.186 | 1 | 3 | Crtc2 | -0.017 | Young |
| 0.440713 | 0.0539 | 0.338 | 0.37 | 1 | 3 | Ptbp1 | -0.032 | Young |
| 0.440761 | 0.060716 | 0.207 | 0.228 | 1 | 3 | Fbxw2 | -0.021 | Young |
| 0.440773 | 0.058624 | 0.36 | 0.345 | 1 | 3 | Csnk1d | 0.015 | Young |
| 0.441047 | 0.505902 | 0.016 | 0.012 | 1 | 3 | Zfp456 | 0.004 | Young |
| 0.44108 | 0.30446 | 0.047 | 0.039 | 1 | 3 | Armc7 | 0.008 | Young |
| 0.441228 | 0.395747 | 0.073 | 0.065 | 1 | 3 | Rcor3 | 0.008 | Young |
| 0.441268 | 0.159205 | 0.133 | 0.148 | 1 | 3 | Ssna1 | -0.015 | Young |
| 0.441547 | 0.58069 | 0.068 | 0.06 | 1 | 3 | Spg20 | 0.008 | Young |
| 0.441635 | 0.111097 | 0.115 | 0.129 | 1 | 3 | Ctr9 | -0.014 | Young |
| 0.441744 | 0.111664 | 0.336 | 0.333 | 1 | 3 | Glul | 0.003 | Young |
| 0.44197 | 0.289496 | 0.182 | 0.171 | 1 | 3 | Ppp2r5e | 0.011 | Young |
| 0.442088 | 0.010433 | 0.028 | 0.035 | 1 | 3 | Phf7 | -0.007 | Young |
| 0.442343 | 0.310142 | 0.01 | 0.014 | 1 | 3 | Trim32 | -0.004 | Young |
| 0.442369 | 0.383907 | 0.086 | 0.076 | 1 | 3 | Zdhhc8 | 0.01 | Young |
| 0.442577 | 0.394335 | 0.016 | 0.012 | 1 | 3 | Zfp677 | 0.004 | Young |
| 0.442955 | 0.044885 | 0.015 | 0.02 | 1 | 3 | Abcb1b | -0.005 | Young |
| 0.442975 | 0.166368 | 0.093 | 0.084 | 1 | 3 | Tmem243 | 0.009 | Young |
| 0.443011 | 0.044209 | 0.36 | 0.395 | 1 | 3 | Cnot1 | -0.035 | Young |
| 0.443024 | 0.333789 | 0.042 | 0.035 | 1 | 3 | Nfe2 | 0.007 | Young |
| 0.443271 | 0.326594 | 0.039 | 0.032 | 1 | 3 | Rrad | 0.007 | Young |

| | | | | | | | | |
|---|---|---|---|---|---|---|---|---|
| 0.443328 | 0.156166 | 0.01 | 0.014 | 1 | 3 | Zfp35 | -0.004 | Young |
| 0.443726 | 0.063878 | 0.223 | 0.245 | 1 | 3 | Yipf1 | -0.022 | Young |
| 0.443804 | 0.565742 | 0.016 | 0.012 | 1 | 3 | Fchsd1 | 0.004 | Young |
| 0.444177 | 0.023303 | 0.444 | 0.489 | 1 | 3 | Nckap1l | -0.045 | Young |
| 0.444479 | 0.055137 | 0.181 | 0.198 | 1 | 3 | Fbxo4 | -0.017 | Young |
| 0.444671 | 0.010614 | 0.21 | 0.232 | 1 | 3 | Rnf115 | -0.022 | Young |
| 0.444826 | 0.099871 | 0.249 | 0.278 | 1 | 3 | Elf4 | -0.029 | Young |
| 0.445194 | 0.150298 | 0.157 | 0.174 | 1 | 3 | Dtymk | -0.017 | Young |
| 0.445284 | 0.268919 | 0.197 | 0.185 | 1 | 3 | Pdzd8 | 0.012 | Young |
| 0.445417 | 0.098782 | 0.037 | 0.044 | 1 | 3 | Pam | -0.007 | Young |
| 0.445619 | 0.170496 | 0.038 | 0.031 | 1 | 3 | Rpap3 | 0.007 | Young |
| 0.445751 | 0.368439 | 0.15 | 0.142 | 1 | 3 | Naip2 | 0.008 | Young |
| 0.445767 | 0.070717 | 0.114 | 0.102 | 1 | 3 | Zscan26 | 0.012 | Young |
| 0.445773 | 0.063043 | 0.742 | 0.778 | 1 | 3 | Gdi2 | -0.036 | Young |
| 0.445896 | 0.157741 | 0.12 | 0.136 | 1 | 3 | Nacc2 | -0.016 | Young |
| 0.445944 | 0.200321 | 0.332 | 0.323 | 1 | 3 | Utrn | 0.009 | Young |
| 0.446102 | 0.060282 | 0.705 | 0.706 | 1 | 3 | Atpif1 | -0.001 | Young |
| 0.446263 | 0.144973 | 0.016 | 0.012 | 1 | 3 | Gcfc2 | 0.004 | Young |
| 0.446291 | 0.314098 | 0.01 | 0.014 | 1 | 3 | Hspb1 | -0.004 | Young |
| 0.446399 | 0.22585 | 0.064 | 0.075 | 1 | 3 | Styx | -0.011 | Young |
| 0.446571 | 0.218095 | 0.016 | 0.012 | 1 | 3 | Itga9 | 0.004 | Young |
| 0.446623 | 0.0295 | 0.073 | 0.084 | 1 | 3 | Lmf2 | -0.011 | Young |
| 0.446631 | 0.081088 | 0.152 | 0.168 | 1 | 3 | Fam133b | -0.016 | Young |
| 0.44687 | 0.135634 | 0.617 | 0.649 | 1 | 3 | Ptms | -0.032 | Young |
| 0.446979 | 0.344492 | 0.129 | 0.118 | 1 | 3 | Ilvbl | 0.011 | Young |
| 0.447196 | 0.305687 | 0.035 | 0.029 | 1 | 3 | Ppox | 0.006 | Young |
| 0.447354 | 0.414308 | 0.049 | 0.042 | 1 | 3 | Ap1s3 | 0.007 | Young |
| 0.447419 | 0.164297 | 0.251 | 0.243 | 1 | 3 | Os9 | 0.008 | Young |
| 0.447458 | 0.17754 | 0.441 | 0.495 | 1 | 3 | Mindy2 | -0.054 | Young |
| 0.447479 | 0.047128 | 0.731 | 0.756 | 1 | 3 | Pld4 | -0.025 | Young |
| 0.44756 | 0.026411 | 0.943 | 0.958 | 1 | 3 | Ppia | -0.015 | Young |
| 0.448051 | 0.557621 | 0.054 | 0.047 | 1 | 3 | Pramef8 | 0.007 | Young |
| 0.448417 | 0.199935 | 0.048 | 0.057 | 1 | 3 | Zfp282 | -0.009 | Young |
| 0.44861 | 0.506956 | 0.051 | 0.043 | 1 | 3 | Vamp5 | 0.008 | Young |
| 0.448713 | 0.036445 | 0.266 | 0.289 | 1 | 3 | Puf60 | -0.023 | Young |
| 0.448806 | 0.322131 | 0.102 | 0.092 | 1 | 3 | Fbxl14 | 0.01 | Young |
| 0.448832 | 0.025393 | 0.131 | 0.147 | 1 | 3 | Tmco3 | -0.016 | Young |
| 0.448948 | 0.443119 | 0.04 | 0.033 | 1 | 3 | Nup188 | 0.007 | Young |
| 0.449113 | 0.904152 | 0.043 | 0.036 | 1 | 3 | Adnp2 | 0.007 | Young |
| 0.449448 | 0.2571 | 0.078 | 0.09 | 1 | 3 | Afg3l2 | -0.012 | Young |
| 0.44956 | 0.10303 | 0.213 | 0.235 | 1 | 3 | Mitf | -0.022 | Young |
| 0.449595 | 0.011678 | 0.292 | 0.314 | 1 | 3 | Rab18 | -0.022 | Young |
| 0.449806 | 0.191013 | 0.279 | 0.27 | 1 | 3 | Ep400 | 0.009 | Young |
| 0.44986 | 0.032086 | 0.328 | 0.357 | 1 | 3 | Ndufs7 | -0.029 | Young |
| 0.449916 | 0.081539 | 0.04 | 0.033 | 1 | 3 | Mlst8 | 0.007 | Young |

| | | | | | | | | |
|---|---|---|---|---|---|---|---|---|
| 0.45024 | 0.428377 | 0.043 | 0.036 | 1 | 3 | 2700038G2 | 0.007 | Young |
| 0.450338 | 0.478818 | 0.063 | 0.055 | 1 | 3 | Rab11fip2 | 0.008 | Young |
| 0.450341 | 0.352954 | 0.037 | 0.03 | 1 | 3 | Msr1 | 0.007 | Young |
| 0.451727 | 0.540849 | 0.018 | 0.013 | 1 | 3 | Xlr4a | 0.005 | Young |
| 0.452007 | 0.512458 | 0.057 | 0.049 | 1 | 3 | Trp53inp1 | 0.008 | Young |
| 0.452082 | 0.056378 | 0.732 | 0.79 | 1 | 3 | Picalm | -0.058 | Young |
| 0.45234 | 0.049612 | 0.194 | 0.215 | 1 | 3 | Cuedc2 | -0.021 | Young |
| 0.452444 | 0.048885 | 0.016 | 0.012 | 1 | 3 | Neurl1a | 0.004 | Young |
| 0.452648 | 0.263013 | 0.121 | 0.111 | 1 | 3 | Swt1 | 0.01 | Young |
| 0.452728 | 0.221215 | 0.047 | 0.039 | 1 | 3 | Tmem185a | 0.008 | Young |
| 0.452874 | 0.237234 | 0.035 | 0.043 | 1 | 3 | Tada2b | -0.008 | Young |
| 0.452925 | 0.009271 | 0.596 | 0.627 | 1 | 3 | Ndel1 | -0.031 | Young |
| 0.452949 | 0.051202 | 0.182 | 0.201 | 1 | 3 | Clcn3 | -0.019 | Young |
| 0.452975 | 0.451928 | 0.059 | 0.051 | 1 | 3 | Arhgef10 | 0.008 | Young |
| 0.45356 | 0.132029 | 0.15 | 0.167 | 1 | 3 | Cntrl | -0.017 | Young |
| 0.453812 | 0.58294 | 0.018 | 0.013 | 1 | 3 | Eid2b | 0.005 | Young |
| 0.453997 | 0.019715 | 0.016 | 0.012 | 1 | 3 | Zfp790 | 0.004 | Young |
| 0.454125 | 0.096863 | 0.324 | 0.357 | 1 | 3 | Sdf2 | -0.033 | Young |
| 0.454193 | 0.348134 | 0.049 | 0.042 | 1 | 3 | Fpgt | 0.007 | Young |
| 0.454477 | 0.044269 | 0.177 | 0.197 | 1 | 3 | Slirp | -0.02 | Young |
| 0.45472 | 0.46161 | 0.052 | 0.044 | 1 | 3 | Mrpl44 | 0.008 | Young |
| 0.454935 | 0.716121 | 0.019 | 0.014 | 1 | 3 | Gm45238 | 0.005 | Young |
| 0.454986 | 0.214622 | 0.277 | 0.27 | 1 | 3 | Mcmbp | 0.007 | Young |
| 0.455127 | 0.002653 | 0.356 | 0.381 | 1 | 3 | Ubxn4 | -0.025 | Young |
| 0.455135 | 0.156082 | 0.616 | 0.657 | 1 | 3 | Slc38a2 | -0.041 | Young |
| 0.455448 | 0.359886 | 0.081 | 0.072 | 1 | 3 | Tpcn1 | 0.009 | Young |
| 0.455469 | 0.316031 | 0.039 | 0.032 | 1 | 3 | Supt7l | 0.007 | Young |
| 0.455472 | 0.133892 | 0.207 | 0.228 | 1 | 3 | Gsk3a | -0.021 | Young |
| 0.456112 | 0.009482 | 0.202 | 0.219 | 1 | 3 | Pdha1 | -0.017 | Young |
| 0.456133 | 0.208331 | 0.059 | 0.069 | 1 | 3 | Trim47 | -0.01 | Young |
| 0.456424 | 0.07546 | 0.678 | 0.706 | 1 | 3 | Ccdc88a | -0.028 | Young |
| 0.456521 | 0.185237 | 0.124 | 0.14 | 1 | 3 | Rxrb | -0.016 | Young |
| 0.4566 | 0.31081 | 0.097 | 0.088 | 1 | 3 | Naxe | 0.009 | Young |
| 0.456626 | 0.254101 | 0.073 | 0.065 | 1 | 3 | Xpc | 0.008 | Young |
| 0.457139 | 0.384867 | 0.066 | 0.057 | 1 | 3 | Nbeal2 | 0.009 | Young |
| 0.457595 | 0.302869 | 0.045 | 0.038 | 1 | 3 | Dcbld2 | 0.007 | Young |
| 0.458105 | 0.758228 | 0.019 | 0.014 | 1 | 3 | Snx24 | 0.005 | Young |
| 0.458195 | 0.394951 | 0.08 | 0.07 | 1 | 3 | Orc3 | 0.01 | Young |
| 0.458319 | 0.052538 | 0.165 | 0.184 | 1 | 3 | Snx15 | -0.019 | Young |
| 0.458615 | 0.371124 | 0.097 | 0.087 | 1 | 3 | Rrm2b | 0.01 | Young |
| 0.458765 | 0.344595 | 0.061 | 0.053 | 1 | 3 | Mzt2 | 0.008 | Young |
| 0.458948 | 0.250673 | 0.218 | 0.21 | 1 | 3 | Nkiras2 | 0.008 | Young |
| 0.459352 | 0.069087 | 0.319 | 0.351 | 1 | 3 | Add1 | -0.032 | Young |
| 0.459718 | 0.212667 | 0.187 | 0.174 | 1 | 3 | Trrap | 0.013 | Young |
| 0.459916 | 0.196532 | 0.116 | 0.105 | 1 | 3 | Smurf1 | 0.011 | Young |

| | | | | | | | | |
|---|---|---|---|---|---|---|---|---|
| 0.460074 | 0.141461 | 0.016 | 0.022 | 1 | 3 | Disc1 | -0.006 | Young |
| 0.460358 | 0.032226 | 0.174 | 0.191 | 1 | 3 | Sfswap | -0.017 | Young |
| 0.460509 | 0.224084 | 0.042 | 0.035 | 1 | 3 | Zfp146 | 0.007 | Young |
| 0.460534 | 0.199004 | 0.058 | 0.068 | 1 | 3 | Nbn | -0.01 | Young |
| 0.460683 | 0.0094 | 0.812 | 0.84 | 1 | 3 | Diaph1 | -0.028 | Young |
| 0.460702 | 0.191375 | 0.018 | 0.013 | 1 | 3 | Sfn | 0.005 | Young |
| 0.460979 | 0.183157 | 0.131 | 0.147 | 1 | 3 | Kpna3 | -0.016 | Young |
| 0.46099 | 0.207624 | 0.058 | 0.05 | 1 | 3 | Scaper | 0.008 | Young |
| 0.461003 | 0.501096 | 0.018 | 0.013 | 1 | 3 | Tmem81 | 0.005 | Young |
| 0.461042 | 0.324364 | 0.16 | 0.151 | 1 | 3 | Supt20 | 0.009 | Young |
| 0.461093 | 0.017613 | 0.218 | 0.24 | 1 | 3 | Ide | -0.022 | Young |
| 0.461178 | 0.003097 | 0.183 | 0.201 | 1 | 3 | Vps13d | -0.018 | Young |
| 0.461287 | 0.597191 | 0.019 | 0.014 | 1 | 3 | Chaf1b | 0.005 | Young |
| 0.46131 | 0.089041 | 0.705 | 0.735 | 1 | 3 | Atox1 | -0.03 | Young |
| 0.461388 | 0.242048 | 0.093 | 0.108 | 1 | 3 | Tsga10 | -0.015 | Young |
| 0.461419 | 0.129778 | 0.178 | 0.196 | 1 | 3 | Kif1c | -0.018 | Young |
| 0.461421 | 0.0135 | 0.071 | 0.081 | 1 | 3 | Clcf1 | -0.01 | Young |
| 0.461965 | 0.163938 | 0.063 | 0.055 | 1 | 3 | Wdr45 | 0.008 | Young |
| 0.462241 | 0.92414 | 0.021 | 0.017 | 1 | 3 | Mccc2 | 0.004 | Young |
| 0.462516 | 0.170865 | 0.157 | 0.147 | 1 | 3 | Myo7a | 0.01 | Young |
| 0.462713 | 0.12495 | 0.037 | 0.044 | 1 | 3 | Spaca9 | -0.007 | Young |
| 0.462735 | 0.23966 | 0.122 | 0.114 | 1 | 3 | Cyhr1 | 0.008 | Young |
| 0.463008 | 0.136258 | 0.178 | 0.167 | 1 | 3 | Zbtb8os | 0.011 | Young |
| 0.463056 | 0.261999 | 0.056 | 0.048 | 1 | 3 | Mcemp1 | 0.008 | Young |
| 0.463065 | 0.487383 | 0.083 | 0.075 | 1 | 3 | Borcs7 | 0.008 | Young |
| 0.463147 | 0.071662 | 0.245 | 0.27 | 1 | 3 | Eif2a | -0.025 | Young |
| 0.463853 | 0.263536 | 0.29 | 0.284 | 1 | 3 | Tecr | 0.006 | Young |
| 0.464803 | 0.155803 | 0.251 | 0.245 | 1 | 3 | Kdm1a | 0.006 | Young |
| 0.464862 | 0.563475 | 0.02 | 0.016 | 1 | 3 | S1pr1 | 0.004 | Young |
| 0.46503 | 0.49095 | 0.057 | 0.049 | 1 | 3 | Hmgn5 | 0.008 | Young |
| 0.465463 | 0.340799 | 0.187 | 0.174 | 1 | 3 | Pi4k2b | 0.013 | Young |
| 0.465605 | 0.045777 | 0.289 | 0.276 | 1 | 3 | Dennd6a | 0.013 | Young |
| 0.465756 | 0.136466 | 0.289 | 0.28 | 1 | 3 | 4930523C0 | 0.009 | Young |
| 0.465938 | 0.359737 | 0.019 | 0.014 | 1 | 3 | 2700049A0 | 0.005 | Young |
| 0.466049 | 0.102615 | 0.427 | 0.423 | 1 | 3 | Srebf2 | 0.004 | Young |
| 0.467305 | 0.083001 | 0.924 | 0.927 | 1 | 3 | Cd52 | -0.003 | Young |
| 0.467397 | 0.170883 | 0.077 | 0.088 | 1 | 3 | Tmem167b | -0.011 | Young |
| 0.467594 | 0.146085 | 0.042 | 0.05 | 1 | 3 | Ndrg1 | -0.008 | Young |
| 0.467634 | 0.186169 | 0.048 | 0.041 | 1 | 3 | Ripor1 | 0.007 | Young |
| 0.46769 | 0.236752 | 0.059 | 0.051 | 1 | 3 | Gpr89 | 0.008 | Young |
| 0.467811 | 0.042159 | 0.534 | 0.583 | 1 | 3 | Ccrl2 | -0.049 | Young |
| 0.467863 | 0.193998 | 0.256 | 0.25 | 1 | 3 | Ube2g2 | 0.006 | Young |
| 0.468552 | 0.164924 | 0.006 | 0.01 | 1 | 3 | Kif3c | -0.004 | Young |
| 0.468863 | 0.133182 | 0.316 | 0.306 | 1 | 3 | Arhgap31 | 0.01 | Young |
| 0.469129 | 0.425978 | 0.044 | 0.037 | 1 | 3 | Xylt2 | 0.007 | Young |

| | | | | | | | | |
|---|---|---|---|---|---|---|---|---|
| 0.469391 | 0.253925 | 0.006 | 0.01 | 1 | 3 | Cldn10 | -0.004 | Young |
| 0.469735 | 0.329644 | 0.019 | 0.014 | 1 | 3 | Zbtb7c | 0.005 | Young |
| 0.469834 | 0.130533 | 0.294 | 0.329 | 1 | 3 | Ccni | -0.035 | Young |
| 0.469932 | 0.015001 | 0.03 | 0.037 | 1 | 3 | Gemin5 | -0.007 | Young |
| 0.470219 | 0.644444 | 0.02 | 0.016 | 1 | 3 | Zfp788 | 0.004 | Young |
| 0.470502 | 0.46499 | 0.02 | 0.016 | 1 | 3 | Atpaf1 | 0.004 | Young |
| 0.470524 | 0.020354 | 0.302 | 0.33 | 1 | 3 | Ndufv2 | -0.028 | Young |
| 0.470592 | 0.29782 | 0.236 | 0.225 | 1 | 3 | Smarcc2 | 0.011 | Young |
| 0.470611 | 0.05445 | 0.318 | 0.349 | 1 | 3 | Borcs6 | -0.031 | Young |
| 0.470777 | 0.345958 | 0.078 | 0.069 | 1 | 3 | Pianp | 0.009 | Young |
| 0.470807 | 0.046095 | 0.605 | 0.648 | 1 | 3 | Tra2a | -0.043 | Young |
| 0.471147 | 0.009678 | 0.102 | 0.115 | 1 | 3 | Ubl4a | -0.013 | Young |
| 0.471231 | 0.024443 | 0.13 | 0.147 | 1 | 3 | Rgl2 | -0.017 | Young |
| 0.471713 | 0.27607 | 0.163 | 0.153 | 1 | 3 | St3gal3 | 0.01 | Young |
| 0.472201 | 0.306115 | 0.02 | 0.016 | 1 | 3 | 2810454H( | 0.004 | Young |
| 0.472216 | 0.052283 | 0.074 | 0.085 | 1 | 3 | Ivd | -0.011 | Young |
| 0.472529 | 0.337281 | 0.114 | 0.104 | 1 | 3 | Ubl7 | 0.01 | Young |
| 0.472548 | 0.265322 | 0.158 | 0.148 | 1 | 3 | Tubb4b | 0.01 | Young |
| 0.472649 | 0.201554 | 0.109 | 0.099 | 1 | 3 | Paqr7 | 0.01 | Young |
| 0.473065 | 0.210622 | 0.054 | 0.047 | 1 | 3 | Lzic | 0.007 | Young |
| 0.473071 | 0.009301 | 0.093 | 0.105 | 1 | 3 | Cxcl16 | -0.012 | Young |
| 0.473146 | 0.018699 | 0.131 | 0.147 | 1 | 3 | Samd8 | -0.016 | Young |
| 0.473148 | 0.3063 | 0.146 | 0.136 | 1 | 3 | Eif2ak1 | 0.01 | Young |
| 0.474449 | 0.048304 | 0.037 | 0.044 | 1 | 3 | Zfp942 | -0.007 | Young |
| 0.474451 | 0.113616 | 0.216 | 0.207 | 1 | 3 | Irak4 | 0.009 | Young |
| 0.474856 | 0.040509 | 0.338 | 0.363 | 1 | 3 | Rcc2 | -0.025 | Young |
| 0.474922 | 0.080318 | 0.011 | 0.016 | 1 | 3 | Ppm1l | -0.005 | Young |
| 0.475023 | 0.115804 | 0.08 | 0.09 | 1 | 3 | Dcaf17 | -0.01 | Young |
| 0.475083 | 0.715332 | 0.021 | 0.017 | 1 | 3 | Klhl26 | 0.004 | Young |
| 0.475123 | 0.061508 | 0.03 | 0.037 | 1 | 3 | Zgpat | -0.007 | Young |
| 0.475139 | 0.138772 | 0.073 | 0.065 | 1 | 3 | Ddx55 | 0.008 | Young |
| 0.475326 | 0.311679 | 0.02 | 0.016 | 1 | 3 | Ttbk2 | 0.004 | Young |
| 0.476168 | 0.11447 | 0.134 | 0.149 | 1 | 3 | Epb41l4ao: | -0.015 | Young |
| 0.47646 | 0.553871 | 0.021 | 0.017 | 1 | 3 | Aaas | 0.004 | Young |
| 0.476618 | 0.548563 | 0.081 | 0.073 | 1 | 3 | Qsox1 | 0.008 | Young |
| 0.476644 | 0.210725 | 0.223 | 0.211 | 1 | 3 | Ccdc47 | 0.012 | Young |
| 0.476945 | 0.114705 | 0.506 | 0.526 | 1 | 3 | Bclaf1 | -0.02 | Young |
| 0.476986 | 0.241103 | 0.236 | 0.226 | 1 | 3 | Tmem170b | 0.01 | Young |
| 0.47734 | 0.117192 | 0.114 | 0.127 | 1 | 3 | Pgs1 | -0.013 | Young |
| 0.477394 | 0.305631 | 0.167 | 0.159 | 1 | 3 | Pink1 | 0.008 | Young |
| 0.477499 | 0.173824 | 0.092 | 0.105 | 1 | 3 | Prpf6 | -0.013 | Young |
| 0.477583 | 0.09277 | 0.342 | 0.378 | 1 | 3 | Exoc5 | -0.036 | Young |
| 0.477608 | 0.219592 | 0.133 | 0.123 | 1 | 3 | Ltc4s | 0.01 | Young |
| 0.477649 | 0.236048 | 0.299 | 0.292 | 1 | 3 | Gm26740 | 0.007 | Young |
| 0.477821 | 0.029504 | 0.024 | 0.03 | 1 | 3 | Phykpl | -0.006 | Young |

| | | | | | | | | |
|---|---|---|---|---|---|---|---|---|
| 0.478156 | 0.30745 | 0.312 | 0.312 | 1 | 3 | Cic | 0 | Young |
| 0.478763 | 0.472727 | 0.102 | 0.093 | 1 | 3 | Brip1os | 0.009 | Young |
| 0.479222 | 0.648494 | 0.021 | 0.017 | 1 | 3 | Fggy | 0.004 | Young |
| 0.479575 | 0.056667 | 0.234 | 0.257 | 1 | 3 | Mgat4b | -0.023 | Young |
| 0.479837 | 0.158936 | 0.58 | 0.601 | 1 | 3 | Asap1 | -0.021 | Young |
| 0.479927 | 0.152136 | 0.22 | 0.246 | 1 | 3 | Rcor1 | -0.026 | Young |
| 0.480281 | 0.544403 | 0.023 | 0.018 | 1 | 3 | Ap4b1 | 0.005 | Young |
| 0.480407 | 0.03315 | 0.164 | 0.18 | 1 | 3 | Apobr | -0.016 | Young |
| 0.480616 | 0.380739 | 0.247 | 0.244 | 1 | 3 | Crebrf | 0.003 | Young |
| 0.480747 | 0.041656 | 0.527 | 0.568 | 1 | 3 | Cox7b | -0.041 | Young |
| 0.480791 | 0.003562 | 0.284 | 0.305 | 1 | 3 | Actr1a | -0.021 | Young |
| 0.480818 | 0.823785 | 0.023 | 0.018 | 1 | 3 | Trim7 | 0.005 | Young |
| 0.480842 | 0.183792 | 0.275 | 0.265 | 1 | 3 | 2810403A( | 0.01 | Young |
| 0.480952 | 0.024687 | 0.236 | 0.258 | 1 | 3 | 1-Mar | -0.022 | Young |
| 0.481249 | 0.085992 | 0.043 | 0.036 | 1 | 3 | Retsat | 0.007 | Young |
| 0.481281 | 0.184153 | 0.066 | 0.075 | 1 | 3 | Mier2 | -0.009 | Young |
| 0.481329 | 0.245841 | 0.27 | 0.269 | 1 | 3 | Gm10073 | 0.001 | Young |
| 0.481438 | 0.331061 | 0.021 | 0.017 | 1 | 3 | Ppp2r5b | 0.004 | Young |
| 0.481603 | 0.236336 | 0.054 | 0.063 | 1 | 3 | F5 | -0.009 | Young |
| 0.481625 | 1.08194 | 0.023 | 0.018 | 1 | 3 | Ube2c | 0.005 | Young |
| 0.482225 | 0.169156 | 0.379 | 0.378 | 1 | 3 | Ptpn11 | 0.001 | Young |
| 0.48227 | 0.541499 | 0.021 | 0.017 | 1 | 3 | Fam177a | 0.004 | Young |
| 0.482301 | 0.190719 | 0.144 | 0.133 | 1 | 3 | Narf | 0.011 | Young |
| 0.482408 | 0.228688 | 0.169 | 0.158 | 1 | 3 | Rras | 0.011 | Young |
| 0.482599 | 0.045495 | 0.032 | 0.038 | 1 | 3 | Polm | -0.006 | Young |
| 0.483199 | 0.02128 | 0.076 | 0.086 | 1 | 3 | Siglecg | -0.01 | Young |
| 0.483545 | 0.092554 | 0.41 | 0.45 | 1 | 3 | Zdhhc20 | -0.04 | Young |
| 0.483955 | 0.192847 | 0.074 | 0.066 | 1 | 3 | Abtb1 | 0.008 | Young |
| 0.484502 | 0.038141 | 0.426 | 0.459 | 1 | 3 | Midn | -0.033 | Young |
| 0.485372 | 0.082531 | 0.216 | 0.24 | 1 | 3 | Psmd3 | -0.024 | Young |
| 0.485409 | 0.204089 | 0.04 | 0.048 | 1 | 3 | Med9 | -0.008 | Young |
| 0.48648 | 0.105418 | 0.018 | 0.023 | 1 | 3 | Zfp1 | -0.005 | Young |
| 0.487578 | 0.27685 | 0.342 | 0.333 | 1 | 3 | Sdc4 | 0.009 | Young |
| 0.487759 | 0.121426 | 0.057 | 0.049 | 1 | 3 | Tatdn3 | 0.008 | Young |
| 0.487867 | 0.039144 | 0.12 | 0.131 | 1 | 3 | Fh1 | -0.011 | Young |
| 0.489468 | 0.008846 | 0.809 | 0.814 | 1 | 3 | Hexa | -0.005 | Young |
| 0.48948 | 0.189314 | 0.095 | 0.085 | 1 | 3 | Akap8l | 0.01 | Young |
| 0.48951 | 0.002595 | 0.145 | 0.161 | 1 | 3 | Chrac1 | -0.016 | Young |
| 0.489516 | 0.044076 | 0.021 | 0.017 | 1 | 3 | A430046D: | 0.004 | Young |
| 0.489778 | 0.840971 | 0.029 | 0.024 | 1 | 3 | Zfp523 | 0.005 | Young |
| 0.490546 | 0.494646 | 0.023 | 0.018 | 1 | 3 | Ssh3 | 0.005 | Young |
| 0.490767 | 0.150567 | 0.077 | 0.068 | 1 | 3 | Dctpp1 | 0.009 | Young |
| 0.490912 | 0.217136 | 0.182 | 0.172 | 1 | 3 | Imp4 | 0.01 | Young |
| 0.491361 | 0.395068 | 0.023 | 0.018 | 1 | 3 | Mtrf1 | 0.005 | Young |
| 0.491402 | 0.194967 | 0.069 | 0.061 | 1 | 3 | Golph3l | 0.008 | Young |

| | | | | | | | | |
|---|---|---|---|---|---|---|---|---|
| 0.49142 | 0.037506 | 0.063 | 0.072 | 1 | 3 | Naip6 | -0.009 | Young |
| 0.491422 | 0.026309 | 0.402 | 0.447 | 1 | 3 | Rassf2 | -0.045 | Young |
| 0.491506 | 0.176958 | 0.186 | 0.178 | 1 | 3 | Vcpip1 | 0.008 | Young |
| 0.491549 | 0.600916 | 0.025 | 0.02 | 1 | 3 | Taf1a | 0.005 | Young |
| 0.491633 | 0.350266 | 0.018 | 0.023 | 1 | 3 | Srfbp1 | -0.005 | Young |
| 0.491642 | 0.66409 | 0.024 | 0.019 | 1 | 3 | Agrn | 0.005 | Young |
| 0.491907 | 0.590056 | 0.024 | 0.019 | 1 | 3 | Gm45606 | 0.005 | Young |
| 0.492349 | 0.263684 | 0.189 | 0.182 | 1 | 3 | Ppp1r11 | 0.007 | Young |
| 0.492582 | 0.098246 | 0.066 | 0.075 | 1 | 3 | Sdhaf1 | -0.009 | Young |
| 0.492597 | 0.000778 | 0.21 | 0.229 | 1 | 3 | Tor1a | -0.019 | Young |
| 0.492829 | 0.499101 | 0.107 | 0.1 | 1 | 3 | Tsc1 | 0.007 | Young |
| 0.49297 | 0.138633 | 0.412 | 0.405 | 1 | 3 | Rassf3 | 0.007 | Young |
| 0.493556 | 0.047607 | 0.283 | 0.311 | 1 | 3 | Mras | -0.028 | Young |
| 0.493633 | 0.053139 | 0.186 | 0.2 | 1 | 3 | Snx27 | -0.014 | Young |
| 0.494076 | 0.018512 | 0.148 | 0.164 | 1 | 3 | Bckdk | -0.016 | Young |
| 0.494115 | 0.141089 | 0.177 | 0.194 | 1 | 3 | Galnt10 | -0.017 | Young |
| 0.494149 | 0.097642 | 0.578 | 0.594 | 1 | 3 | Snx5 | -0.016 | Young |
| 0.494361 | 0.075535 | 0.131 | 0.146 | 1 | 3 | Map3k14 | -0.015 | Young |
| 0.4944 | 0.610295 | 0.027 | 0.022 | 1 | 3 | Kbtbd4 | 0.005 | Young |
| 0.494848 | 0.014037 | 0.021 | 0.017 | 1 | 3 | Sap130 | 0.004 | Young |
| 0.495576 | 0.204506 | 0.203 | 0.195 | 1 | 3 | Ap1ar | 0.008 | Young |
| 0.495577 | 0.695966 | 0.035 | 0.03 | 1 | 3 | Eral1 | 0.005 | Young |
| 0.495593 | 0.172217 | 0.032 | 0.038 | 1 | 3 | Dbf4 | -0.006 | Young |
| 0.495673 | 0.345331 | 0.025 | 0.02 | 1 | 3 | Fcgr1 | 0.005 | Young |
| 0.49599 | 0.214003 | 0.241 | 0.233 | 1 | 3 | Relt | 0.008 | Young |
| 0.496001 | 0.917351 | 0.029 | 0.024 | 1 | 3 | Rbm48 | 0.005 | Young |
| 0.496414 | 0.614381 | 0.027 | 0.022 | 1 | 3 | Peli2 | 0.005 | Young |
| 0.496531 | 0.271702 | 0.053 | 0.062 | 1 | 3 | Mefv | -0.009 | Young |
| 0.49661 | 0.022347 | 0.427 | 0.461 | 1 | 3 | Hectd1 | -0.034 | Young |
| 0.496699 | 0.256348 | 0.14 | 0.13 | 1 | 3 | Cdk8 | 0.01 | Young |
| 0.496746 | 0.186866 | 0.13 | 0.147 | 1 | 3 | Zfp608 | -0.017 | Young |
| 0.496763 | 0.249111 | 0.285 | 0.278 | 1 | 3 | Sms | 0.007 | Young |
| 0.49681 | 0.045972 | 0.019 | 0.024 | 1 | 3 | Hgh1 | -0.005 | Young |
| 0.496858 | 0.123128 | 0.141 | 0.159 | 1 | 3 | Nr1h2 | -0.018 | Young |
| 0.496912 | 0.050708 | 0.027 | 0.032 | 1 | 3 | Mfsd4a | -0.005 | Young |
| 0.496967 | 0.042614 | 0.241 | 0.262 | 1 | 3 | Plaa | -0.021 | Young |
| 0.497001 | 0.003407 | 0.025 | 0.031 | 1 | 3 | Ftsj1 | -0.006 | Young |
| 0.497111 | 0.148219 | 0.467 | 0.468 | 1 | 3 | Marcksl1 | -0.001 | Young |
| 0.497124 | 0.009928 | 0.428 | 0.464 | 1 | 3 | Swi5 | -0.036 | Young |
| 0.497763 | 0.003475 | 0.117 | 0.13 | 1 | 3 | Rabgap1 | -0.013 | Young |
| 0.497977 | 0.213165 | 0.12 | 0.134 | 1 | 3 | Nacc1 | -0.014 | Young |
| 0.498179 | 0.67847 | 0.027 | 0.022 | 1 | 3 | Cbx8 | 0.005 | Young |
| 0.49904 | 0.803377 | 0.048 | 0.042 | 1 | 3 | Zfp160 | 0.006 | Young |
| 0.499128 | 0.760288 | 0.029 | 0.024 | 1 | 3 | Lin9 | 0.005 | Young |
| 0.499303 | 0.461435 | 0.102 | 0.094 | 1 | 3 | Tacc2 | 0.008 | Young |

| | | | | | | | | |
|---|---|---|---|---|---|---|---|---|
| 0.499563 | 0.056382 | 0.347 | 0.384 | 1 | 3 | Smc6 | -0.037 | Young |
| 0.499737 | 0.185582 | 0.013 | 0.017 | 1 | 3 | Mst1r | -0.004 | Young |
| 0.499772 | 0.035971 | 0.866 | 0.89 | 1 | 3 | Gm2a | -0.024 | Young |
| 0.500775 | 0.022123 | 0.187 | 0.205 | 1 | 3 | Tmem147 | -0.018 | Young |
| 0.500819 | 0.02809 | 0.553 | 0.597 | 1 | 3 | Tiam1 | -0.044 | Young |
| 0.501112 | 0.063508 | 0.25 | 0.272 | 1 | 3 | Tmem9b | -0.022 | Young |
| 0.501363 | 0.240712 | 0.225 | 0.22 | 1 | 3 | Tnrc6c | 0.005 | Young |
| 0.50174 | 0.299032 | 0.024 | 0.019 | 1 | 3 | Eogt | 0.005 | Young |
| 0.501818 | 0.010208 | 0.318 | 0.308 | 1 | 3 | Zcrb1 | 0.01 | Young |
| 0.501865 | 0.052412 | 0.085 | 0.075 | 1 | 3 | Rhbdd1 | 0.01 | Young |
| 0.502175 | 0.087012 | 0.177 | 0.195 | 1 | 3 | Slc8b1 | -0.018 | Young |
| 0.502224 | 0.193505 | 0.456 | 0.483 | 1 | 3 | Per1 | -0.027 | Young |
| 0.502307 | 0.024982 | 0.013 | 0.017 | 1 | 3 | Tbc1d25 | -0.004 | Young |
| 0.50251 | 0.423707 | 0.154 | 0.147 | 1 | 3 | Ezh1 | 0.007 | Young |
| 0.502726 | 0.070454 | 0.033 | 0.039 | 1 | 3 | 1300002E1 | -0.006 | Young |
| 0.502813 | 0.146421 | 0.066 | 0.075 | 1 | 3 | B4galt3 | -0.009 | Young |
| 0.502994 | 0.145518 | 0.304 | 0.332 | 1 | 3 | Phf20l1 | -0.028 | Young |
| 0.503 | 0.086969 | 0.198 | 0.215 | 1 | 3 | Ldb1 | -0.017 | Young |
| 0.503087 | 0.624667 | 0.034 | 0.029 | 1 | 3 | Dgka | 0.005 | Young |
| 0.503109 | 0.033556 | 0.337 | 0.364 | 1 | 3 | Zfand3 | -0.027 | Young |
| 0.503192 | 0.240423 | 0.212 | 0.205 | 1 | 3 | Ftl1-ps1 | 0.007 | Young |
| 0.503505 | 0.081794 | 0.246 | 0.233 | 1 | 3 | Mrpl30 | 0.013 | Young |
| 0.503654 | 0.446312 | 0.069 | 0.062 | 1 | 3 | Cib2 | 0.007 | Young |
| 0.503681 | 0.026734 | 0.222 | 0.241 | 1 | 3 | Dolpp1 | -0.019 | Young |
| 0.503697 | 0.03744 | 0.135 | 0.149 | 1 | 3 | Qrich1 | -0.014 | Young |
| 0.503733 | 0.075555 | 0.087 | 0.098 | 1 | 3 | Ormdl1 | -0.011 | Young |
| 0.504017 | 0.012934 | 0.068 | 0.078 | 1 | 3 | Hic2 | -0.01 | Young |
| 0.50432 | 0.106292 | 0.034 | 0.041 | 1 | 3 | Alkbh2 | -0.007 | Young |
| 0.504487 | 0.035653 | 0.196 | 0.213 | 1 | 3 | Gdap10 | -0.017 | Young |
| 0.504688 | 0.040092 | 0.114 | 0.127 | 1 | 3 | Thap2 | -0.013 | Young |
| 0.504929 | 0.388975 | 0.096 | 0.088 | 1 | 3 | Cblb | 0.008 | Young |
| 0.505025 | 0.049396 | 0.008 | 0.011 | 1 | 3 | Rap1gap | -0.003 | Young |
| 0.505274 | 0.271757 | 0.027 | 0.022 | 1 | 3 | L3mbtl2 | 0.005 | Young |
| 0.505414 | 0.488776 | 0.029 | 0.024 | 1 | 3 | Cnbd2 | 0.005 | Young |
| 0.505429 | 0.035443 | 0.206 | 0.222 | 1 | 3 | Polr1a | -0.016 | Young |
| 0.5056 | 0.256055 | 0.172 | 0.164 | 1 | 3 | Fkbp3 | 0.008 | Young |
| 0.50584 | 0.188931 | 0.008 | 0.011 | 1 | 3 | Dnmt3aos | -0.003 | Young |
| 0.506016 | 0.140883 | 0.074 | 0.085 | 1 | 3 | Limk1 | -0.011 | Young |
| 0.506655 | 0.085594 | 0.008 | 0.011 | 1 | 3 | Timm23 | -0.003 | Young |
| 0.50686 | 0.237503 | 0.356 | 0.357 | 1 | 3 | Ppp4r3b | -0.001 | Young |
| 0.506915 | 0.756461 | 0.045 | 0.039 | 1 | 3 | Pmm1 | 0.006 | Young |
| 0.507069 | 0.086631 | 0.093 | 0.105 | 1 | 3 | Adgre4 | -0.012 | Young |
| 0.507104 | 0.107106 | 0.244 | 0.231 | 1 | 3 | Usp3 | 0.013 | Young |
| 0.507506 | 0.068554 | 0.162 | 0.151 | 1 | 3 | Socs7 | 0.011 | Young |
| 0.507851 | 0.095219 | 0.705 | 0.729 | 1 | 3 | Mtdh | -0.024 | Young |

| | | | | | | | | |
|---|---|---|---|---|---|---|---|---|
| 0.508004 | 0.432786 | 0.064 | 0.057 | 1 | 3 | Rfxap | 0.007 | Young |
| 0.508081 | 0.082452 | 0.049 | 0.057 | 1 | 3 | Gm45902 | -0.008 | Young |
| 0.508412 | 0.281109 | 0.187 | 0.176 | 1 | 3 | Tiprl | 0.011 | Young |
| 0.509333 | 0.141074 | 0.139 | 0.154 | 1 | 3 | Ccdc82 | -0.015 | Young |
| 0.509427 | 0.126337 | 0.124 | 0.112 | 1 | 3 | Yeats2 | 0.012 | Young |
| 0.50966 | 0.196234 | 0.119 | 0.11 | 1 | 3 | Pex19 | 0.009 | Young |
| 0.510494 | 0.1312 | 0.082 | 0.073 | 1 | 3 | Ganc | 0.009 | Young |
| 0.510618 | 0.346703 | 0.091 | 0.082 | 1 | 3 | Tns4 | 0.009 | Young |
| 0.51068 | 0.172871 | 0.074 | 0.066 | 1 | 3 | Selenoh | 0.008 | Young |
| 0.510721 | 0.267256 | 0.09 | 0.081 | 1 | 3 | Prr3 | 0.009 | Young |
| 0.510846 | 0.030808 | 0.785 | 0.834 | 1 | 3 | Fosl2 | -0.049 | Young |
| 0.510877 | 0.293264 | 0.106 | 0.097 | 1 | 3 | Nfs1 | 0.009 | Young |
| 0.511198 | 0.591628 | 0.04 | 0.035 | 1 | 3 | Ripor2 | 0.005 | Young |
| 0.5112 | 0.059123 | 0.207 | 0.196 | 1 | 3 | Kctd20 | 0.011 | Young |
| 0.51125 | 0.310097 | 0.205 | 0.196 | 1 | 3 | Rab11fip5 | 0.009 | Young |
| 0.511253 | 0.348504 | 0.029 | 0.024 | 1 | 3 | Gm15445 | 0.005 | Young |
| 0.51139 | 0.026639 | 0.409 | 0.444 | 1 | 3 | Dnm2 | -0.035 | Young |
| 0.511687 | 0.375504 | 0.13 | 0.122 | 1 | 3 | Fbxo28 | 0.008 | Young |
| 0.511692 | 0.011655 | 0.062 | 0.07 | 1 | 3 | Fam210a | -0.008 | Young |
| 0.511919 | 0.71569 | 0.052 | 0.045 | 1 | 3 | Snx16 | 0.007 | Young |
| 0.51209 | 0.370879 | 0.064 | 0.057 | 1 | 3 | Chkb | 0.007 | Young |
| 0.512195 | 0.021753 | 0.508 | 0.545 | 1 | 3 | Tmem189 | -0.037 | Young |
| 0.512462 | 0.203395 | 0.241 | 0.232 | 1 | 3 | Rere | 0.009 | Young |
| 0.512614 | 0.031215 | 0.088 | 0.099 | 1 | 3 | Gfm1 | -0.011 | Young |
| 0.51268 | 0.244626 | 0.129 | 0.118 | 1 | 3 | Frmd4b | 0.011 | Young |
| 0.512759 | 0.073063 | 0.256 | 0.284 | 1 | 3 | Sertad2 | -0.028 | Young |
| 0.512999 | 0.015566 | 0.114 | 0.103 | 1 | 3 | Plod1 | 0.011 | Young |
| 0.513029 | 0.098638 | 0.232 | 0.256 | 1 | 3 | Wbp2 | -0.024 | Young |
| 0.513237 | 0.463728 | 0.032 | 0.026 | 1 | 3 | Gm19951 | 0.006 | Young |
| 0.513427 | 0.235383 | 0.153 | 0.145 | 1 | 3 | Senp1 | 0.008 | Young |
| 0.513704 | 0.251306 | 0.027 | 0.022 | 1 | 3 | 2310040G2 | 0.005 | Young |
| 0.51399 | 0.00344 | 0.458 | 0.492 | 1 | 3 | Mrps21 | -0.034 | Young |
| 0.514019 | 0.16717 | 0.293 | 0.293 | 1 | 3 | Dpy19l1 | 0 | Young |
| 0.514118 | 0.236092 | 0.085 | 0.076 | 1 | 3 | Rfwd3 | 0.009 | Young |
| 0.514155 | 0.289302 | 0.196 | 0.189 | 1 | 3 | Rabgef1 | 0.007 | Young |
| 0.514375 | 0.18608 | 0.043 | 0.05 | 1 | 3 | Ofd1 | -0.007 | Young |
| 0.51467 | 0.177193 | 0.028 | 0.033 | 1 | 3 | Npat | -0.005 | Young |
| 0.514824 | 0.082178 | 0.077 | 0.087 | 1 | 3 | Prpf4 | -0.01 | Young |
| 0.515184 | 0.24129 | 0.115 | 0.106 | 1 | 3 | Zfp740 | 0.009 | Young |
| 0.515208 | 0.240223 | 0.279 | 0.277 | 1 | 3 | Cers5 | 0.002 | Young |
| 0.515338 | 0.043994 | 0.189 | 0.205 | 1 | 3 | Fuca2 | -0.016 | Young |
| 0.515414 | 0.370497 | 0.051 | 0.044 | 1 | 3 | Prkdc | 0.007 | Young |
| 0.515963 | 0.699788 | 0.01 | 0.007 | 1 | 3 | Ska2 | 0.003 | Young |
| 0.51608 | 0.088516 | 0.504 | 0.508 | 1 | 3 | Taok1 | -0.004 | Young |
| 0.516439 | 0.596059 | 0.076 | 0.068 | 1 | 3 | Lrrc20 | 0.008 | Young |

| | | | | | | | | |
|---|---|---|---|---|---|---|---|---|
| 0.516692 | 0.098657 | 0.116 | 0.128 | 1 | 3 | Pdrg1 | -0.012 | Young |
| 0.516774 | 0.429545 | 0.037 | 0.031 | 1 | 3 | Rab3ip | 0.006 | Young |
| 0.516794 | 0.038931 | 0.342 | 0.37 | 1 | 3 | Ddi2 | -0.028 | Young |
| 0.516833 | 0.082774 | 0.255 | 0.245 | 1 | 3 | Slc25a36 | 0.01 | Young |
| 0.517146 | 0.168393 | 0.343 | 0.341 | 1 | 3 | Mpc1 | 0.002 | Young |
| 0.517514 | 0.056086 | 0.703 | 0.72 | 1 | 3 | Ncf1 | -0.017 | Young |
| 0.5176 | 0.3804 | 0.066 | 0.075 | 1 | 3 | Ankrd49 | -0.009 | Young |
| 0.517698 | 0.158869 | 0.139 | 0.13 | 1 | 3 | Casp1 | 0.009 | Young |
| 0.517892 | 0.52805 | 0.033 | 0.027 | 1 | 3 | Thbd | 0.006 | Young |
| 0.5179 | 0.370957 | 0.034 | 0.029 | 1 | 3 | Gm49359 | 0.005 | Young |
| 0.517986 | 0.021525 | 0.211 | 0.232 | 1 | 3 | Ndufs4 | -0.021 | Young |
| 0.518095 | 0.475106 | 0.01 | 0.007 | 1 | 3 | 4833418N( | 0.003 | Young |
| 0.518578 | 0.278621 | 0.034 | 0.029 | 1 | 3 | Dip2c | 0.005 | Young |
| 0.518681 | 0.061371 | 0.02 | 0.025 | 1 | 3 | Sac3d1 | -0.005 | Young |
| 0.519065 | 0.44467 | 0.109 | 0.1 | 1 | 3 | Edem2 | 0.009 | Young |
| 0.519069 | 0.00704 | 0.14 | 0.154 | 1 | 3 | Gbf1 | -0.014 | Young |
| 0.519093 | 0.047633 | 0.029 | 0.024 | 1 | 3 | Cd38 | 0.005 | Young |
| 0.519206 | 0.085473 | 0.16 | 0.177 | 1 | 3 | Lpcat1 | -0.017 | Young |
| 0.519308 | 0.380131 | 0.051 | 0.044 | 1 | 3 | Mto1 | 0.007 | Young |
| 0.5194 | 0.068559 | 0.278 | 0.305 | 1 | 3 | Snx6 | -0.027 | Young |
| 0.51958 | 0.313541 | 0.187 | 0.18 | 1 | 3 | Sepsecs | 0.007 | Young |
| 0.51962 | 0.114872 | 0.069 | 0.079 | 1 | 3 | Srcap | -0.01 | Young |
| 0.519622 | 0.156997 | 0.168 | 0.185 | 1 | 3 | Pank2 | -0.017 | Young |
| 0.519804 | 0.399712 | 0.01 | 0.007 | 1 | 3 | Usp49 | 0.003 | Young |
| 0.519984 | 0.319001 | 0.064 | 0.057 | 1 | 3 | Sord | 0.007 | Young |
| 0.520139 | 0.136138 | 0.638 | 0.648 | 1 | 3 | Pepd | -0.01 | Young |
| 0.520367 | 0.114821 | 0.235 | 0.229 | 1 | 3 | Ephx1 | 0.006 | Young |
| 0.520492 | 0.371791 | 0.044 | 0.038 | 1 | 3 | Acrbp | 0.006 | Young |
| 0.520659 | 0.392604 | 0.01 | 0.007 | 1 | 3 | Zbtb8a | 0.003 | Young |
| 0.521052 | 0.283319 | 0.188 | 0.18 | 1 | 3 | Smarca2 | 0.008 | Young |
| 0.521231 | 0.280859 | 0.032 | 0.026 | 1 | 3 | Mrps22 | 0.006 | Young |
| 0.521501 | 0.387746 | 0.062 | 0.055 | 1 | 3 | Agpat2 | 0.007 | Young |
| 0.521515 | 0.244122 | 0.072 | 0.065 | 1 | 3 | Rmnd5b | 0.007 | Young |
| 0.52177 | 0.219788 | 0.068 | 0.078 | 1 | 3 | AI837181 | -0.01 | Young |
| 0.521822 | 0.248816 | 0.341 | 0.344 | 1 | 3 | Ndufs5 | -0.003 | Young |
| 0.521909 | 0.023702 | 0.494 | 0.527 | 1 | 3 | Eif3j1 | -0.033 | Young |
| 0.522187 | 0.080436 | 0.249 | 0.237 | 1 | 3 | Xpo6 | 0.012 | Young |
| 0.522591 | 0.272251 | 0.294 | 0.293 | 1 | 3 | Lrp10 | 0.001 | Young |
| 0.522633 | 0.252169 | 0.035 | 0.03 | 1 | 3 | Zer1 | 0.005 | Young |
| 0.522657 | 0.180258 | 0.02 | 0.025 | 1 | 3 | Echdc1 | -0.005 | Young |
| 0.522798 | 0.058808 | 0.667 | 0.687 | 1 | 3 | Arl6ip1 | -0.02 | Young |
| 0.522801 | 0.358063 | 0.01 | 0.007 | 1 | 3 | Lekr1 | 0.003 | Young |
| 0.522817 | 0.084512 | 0.236 | 0.259 | 1 | 3 | Hipk3 | -0.023 | Young |
| 0.523213 | 0.056717 | 0.168 | 0.182 | 1 | 3 | Wdr91 | -0.014 | Young |
| 0.523612 | 0.155489 | 0.053 | 0.061 | 1 | 3 | Dcun1d3 | -0.008 | Young |

| | | | | | | | | |
|---|---|---|---|---|---|---|---|---|
| 0.523972 | 0.288246 | 0.035 | 0.03 | 1 | 3 | Lig1 | 0.005 | Young |
| 0.52419 | 0.384441 | 0.043 | 0.05 | 1 | 3 | Tmem18 | -0.007 | Young |
| 0.524254 | 0.111735 | 0.16 | 0.176 | 1 | 3 | Zpr1 | -0.016 | Young |
| 0.524438 | 0.082727 | 0.53 | 0.542 | 1 | 3 | Scp2 | -0.012 | Young |
| 0.524517 | 0.230545 | 0.01 | 0.007 | 1 | 3 | Klri1 | 0.003 | Young |
| 0.524643 | 0.305658 | 0.014 | 0.018 | 1 | 3 | Letm2 | -0.004 | Young |
| 0.52469 | 0.202263 | 0.148 | 0.139 | 1 | 3 | Zfp992 | 0.009 | Young |
| 0.524987 | 0.542184 | 0.043 | 0.037 | 1 | 3 | Ydjc | 0.006 | Young |
| 0.525377 | 0.266729 | 0.01 | 0.007 | 1 | 3 | Zfp324 | 0.003 | Young |
| 0.525618 | 0.321939 | 0.034 | 0.029 | 1 | 3 | Gypc | 0.005 | Young |
| 0.525957 | 0.111634 | 0.1 | 0.11 | 1 | 3 | Lonp2 | -0.01 | Young |
| 0.525988 | 0.227326 | 0.197 | 0.188 | 1 | 3 | Il4ra | 0.009 | Young |
| 0.526198 | 0.000884 | 0.187 | 0.203 | 1 | 3 | Mprip | -0.016 | Young |
| 0.526331 | 0.009114 | 0.442 | 0.476 | 1 | 3 | Sqstm1 | -0.034 | Young |
| 0.526443 | 0.158294 | 0.266 | 0.26 | 1 | 3 | Tcp11l2 | 0.006 | Young |
| 0.526656 | 0.127903 | 0.255 | 0.24 | 1 | 3 | Arid5a | 0.015 | Young |
| 0.526875 | 0.559394 | 0.04 | 0.035 | 1 | 3 | Inafm2 | 0.005 | Young |
| 0.526994 | 0.100283 | 0.054 | 0.062 | 1 | 3 | Ccdc107 | -0.008 | Young |
| 0.527016 | 0.358309 | 0.12 | 0.112 | 1 | 3 | Syap1 | 0.008 | Young |
| 0.527381 | 0.068867 | 0.085 | 0.094 | 1 | 3 | Rngtt | -0.009 | Young |
| 0.527445 | 0.07502 | 0.159 | 0.174 | 1 | 3 | Upf3a | -0.015 | Young |
| 0.527669 | 0.100704 | 0.078 | 0.088 | 1 | 3 | Gm1673 | -0.01 | Young |
| 0.527673 | 0.055572 | 0.836 | 0.851 | 1 | 3 | Plxnd1 | -0.015 | Young |
| 0.527727 | 0.279894 | 0.165 | 0.157 | 1 | 3 | Ulk2 | 0.008 | Young |
| 0.527775 | 0.482979 | 0.035 | 0.03 | 1 | 3 | Gm37233 | 0.005 | Young |
| 0.527789 | 0.207333 | 0.319 | 0.315 | 1 | 3 | Rc3h2 | 0.004 | Young |
| 0.527827 | 0.623873 | 0.043 | 0.037 | 1 | 3 | Smo | 0.006 | Young |
| 0.528113 | 0.035562 | 0.184 | 0.201 | 1 | 3 | Dse | -0.017 | Young |
| 0.528382 | 0.021587 | 0.537 | 0.588 | 1 | 3 | Pbrm1 | -0.051 | Young |
| 0.528394 | 0.541969 | 0.038 | 0.032 | 1 | 3 | Tiam2 | 0.006 | Young |
| 0.528629 | 0.316792 | 0.105 | 0.097 | 1 | 3 | Itpripl1 | 0.008 | Young |
| 0.528922 | 0.2711 | 0.152 | 0.142 | 1 | 3 | Rffl | 0.01 | Young |
| 0.529219 | 0.063085 | 0.149 | 0.165 | 1 | 3 | Irak2 | -0.016 | Young |
| 0.529259 | 0.054588 | 0.173 | 0.188 | 1 | 3 | Prkcb | -0.015 | Young |
| 0.529364 | 1.102089 | 0.011 | 0.008 | 1 | 3 | Tmem150k | 0.003 | Young |
| 0.529437 | 0.130839 | 0.28 | 0.276 | 1 | 3 | Map3k11 | 0.004 | Young |
| 0.529485 | 0.638279 | 0.037 | 0.031 | 1 | 3 | Clasrp | 0.006 | Young |
| 0.529609 | 0.158451 | 0.03 | 0.025 | 1 | 3 | Nat6 | 0.005 | Young |
| 0.529913 | 0.237416 | 0.04 | 0.035 | 1 | 3 | Tsen15 | 0.005 | Young |
| 0.530112 | 0.317164 | 0.081 | 0.073 | 1 | 3 | Akt1s1 | 0.008 | Young |
| 0.530282 | 0.284659 | 0.229 | 0.223 | 1 | 3 | Rarg | 0.006 | Young |
| 0.530409 | 0.196895 | 0.22 | 0.237 | 1 | 3 | Gclc | -0.017 | Young |
| 0.530744 | 0.184992 | 0.087 | 0.079 | 1 | 3 | Tgfbrap1 | 0.008 | Young |
| 0.530889 | 0.150016 | 0.087 | 0.079 | 1 | 3 | Sigirr | 0.008 | Young |
| 0.530963 | 0.391755 | 0.072 | 0.065 | 1 | 3 | Zfp639 | 0.007 | Young |

| | | | | | | | | |
|---|---|---|---|---|---|---|---|---|
| 0.531549 | 0.028753 | 0.818 | 0.818 | 1 | 3 | Gnas | 0 | Young |
| 0.532069 | 0.444821 | 0.052 | 0.045 | 1 | 3 | Mpg | 0.007 | Young |
| 0.532135 | 0.198936 | 0.037 | 0.031 | 1 | 3 | Snx21 | 0.006 | Young |
| 0.532287 | 0.035217 | 0.107 | 0.117 | 1 | 3 | Ccnh | -0.01 | Young |
| 0.532546 | 0.103806 | 0.482 | 0.487 | 1 | 3 | Eif3h | -0.005 | Young |
| 0.532916 | 0.388169 | 0.091 | 0.084 | 1 | 3 | Pcyox1 | 0.007 | Young |
| 0.533575 | 0.2408 | 0.14 | 0.154 | 1 | 3 | Rfc1 | -0.014 | Young |
| 0.533742 | 0.332462 | 0.109 | 0.102 | 1 | 3 | Josd2 | 0.007 | Young |
| 0.533798 | 0.022909 | 0.038 | 0.044 | 1 | 3 | Nxt2 | -0.006 | Young |
| 0.533816 | 0.504005 | 0.011 | 0.008 | 1 | 3 | Tmem156 | 0.003 | Young |
| 0.533984 | 0.189716 | 0.09 | 0.081 | 1 | 3 | Thoc5 | 0.009 | Young |
| 0.534272 | 0.01976 | 0.561 | 0.601 | 1 | 3 | Ptprj | -0.04 | Young |
| 0.534354 | 0.471586 | 0.216 | 0.214 | 1 | 3 | Wwp1 | 0.002 | Young |
| 0.534387 | 0.121851 | 0.513 | 0.521 | 1 | 3 | Rab5c | -0.008 | Young |
| 0.53501 | 0.138141 | 0.058 | 0.051 | 1 | 3 | D430042O( | 0.007 | Young |
| 0.535053 | 0.235264 | 0.045 | 0.039 | 1 | 3 | Lrch4 | 0.006 | Young |
| 0.535062 | 0.018828 | 0.402 | 0.443 | 1 | 3 | Brd4 | -0.041 | Young |
| 0.53514 | 0.004968 | 0.049 | 0.056 | 1 | 3 | Gtf2h4 | -0.007 | Young |
| 0.535414 | 0.053867 | 0.299 | 0.323 | 1 | 3 | Lpgat1 | -0.024 | Young |
| 0.53544 | 0.371803 | 0.011 | 0.008 | 1 | 3 | Fam92a | 0.003 | Young |
| 0.535621 | 0.200268 | 0.064 | 0.057 | 1 | 3 | Scnm1 | 0.007 | Young |
| 0.535797 | 0.304835 | 0.212 | 0.205 | 1 | 3 | Pcgf5 | 0.007 | Young |
| 0.535846 | 0.509841 | 0.011 | 0.008 | 1 | 3 | Zfp119a | 0.003 | Young |
| 0.535846 | 0.50503 | 0.011 | 0.008 | 1 | 3 | Plscr4 | 0.003 | Young |
| 0.536432 | 0.069564 | 0.029 | 0.035 | 1 | 3 | 2810006K2 | -0.006 | Young |
| 0.536656 | 0.047454 | 0.16 | 0.173 | 1 | 3 | Irf9 | -0.013 | Young |
| 0.536712 | 0.436238 | 0.048 | 0.042 | 1 | 3 | Kansl1l | 0.006 | Young |
| 0.53683 | 0.400095 | 0.087 | 0.079 | 1 | 3 | Herc6 | 0.008 | Young |
| 0.536847 | 0.187363 | 0.164 | 0.177 | 1 | 3 | Stx11 | -0.013 | Young |
| 0.537247 | 0.267466 | 0.202 | 0.195 | 1 | 3 | Aqr | 0.007 | Young |
| 0.537286 | 0.397011 | 0.052 | 0.045 | 1 | 3 | Ptp4a3 | 0.007 | Young |
| 0.537408 | 0.202544 | 0.035 | 0.042 | 1 | 3 | Atg10 | -0.007 | Young |
| 0.537656 | 0.034268 | 0.384 | 0.412 | 1 | 3 | 7-Mar | -0.028 | Young |
| 0.537925 | 0.110898 | 0.242 | 0.262 | 1 | 3 | Nlrp3 | -0.02 | Young |
| 0.538238 | 0.003109 | 0.039 | 0.045 | 1 | 3 | Trim39 | -0.006 | Young |
| 0.538374 | 0.077089 | 0.462 | 0.508 | 1 | 3 | Cmtm6 | -0.046 | Young |
| 0.538376 | 0.044045 | 0.201 | 0.221 | 1 | 3 | Mrps18c | -0.02 | Young |
| 0.538695 | 0.242704 | 0.011 | 0.008 | 1 | 3 | Zfp606 | 0.003 | Young |
| 0.538713 | 0.121888 | 0.725 | 0.757 | 1 | 3 | Kdm6b | -0.032 | Young |
| 0.538824 | 0.313297 | 0.1 | 0.092 | 1 | 3 | Cpd | 0.008 | Young |
| 0.539284 | 0.07185 | 0.136 | 0.151 | 1 | 3 | Rnf181 | -0.015 | Young |
| 0.539396 | 0.307798 | 0.15 | 0.143 | 1 | 3 | Ints6l | 0.007 | Young |
| 0.53951 | 0.175113 | 0.011 | 0.008 | 1 | 3 | Ubtd2 | 0.003 | Young |
| 0.539702 | 0.011068 | 0.431 | 0.416 | 1 | 3 | C1qc | 0.015 | Young |
| 0.53985 | 0.038785 | 0.468 | 0.526 | 1 | 3 | Uggt1 | -0.058 | Young |

| | | | | | | | | |
|---|---|---|---|---|---|---|---|---|
| 0.540036 | 0.126484 | 0.135 | 0.125 | 1 | 3 | Aen | 0.01 | Young |
| 0.540108 | 0.187568 | 0.072 | 0.081 | 1 | 3 | Abhd11 | -0.009 | Young |
| 0.540173 | 0.016491 | 0.534 | 0.581 | 1 | 3 | Ndufb11 | -0.047 | Young |
| 0.540371 | 0.250652 | 0.038 | 0.032 | 1 | 3 | Fktn | 0.006 | Young |
| 0.54099 | 0.156549 | 0.061 | 0.054 | 1 | 3 | Rgs3 | 0.007 | Young |
| 0.541126 | 0.091158 | 0.626 | 0.667 | 1 | 3 | Cbl | -0.041 | Young |
| 0.541143 | 0.517815 | 0.011 | 0.008 | 1 | 3 | A630089N( | 0.003 | Young |
| 0.54137 | 0.107001 | 0.042 | 0.036 | 1 | 3 | Fbxl12 | 0.006 | Young |
| 0.541408 | 0.133828 | 0.009 | 0.012 | 1 | 3 | Dnmt3b | -0.003 | Young |
| 0.541508 | 0.096663 | 0.433 | 0.429 | 1 | 3 | 2-Mar | 0.004 | Young |
| 0.541551 | 0.189512 | 0.011 | 0.008 | 1 | 3 | Gm15247 | 0.003 | Young |
| 0.541813 | 0.003791 | 0.17 | 0.188 | 1 | 3 | Otud4 | -0.018 | Young |
| 0.541891 | 0.022047 | 0.948 | 0.943 | 1 | 3 | H3f3a | 0.005 | Young |
| 0.54196 | 0.077817 | 0.011 | 0.008 | 1 | 3 | Zc4h2 | 0.003 | Young |
| 0.542343 | 0.295561 | 0.038 | 0.032 | 1 | 3 | Ms4a6b | 0.006 | Young |
| 0.542598 | 0.23832 | 0.009 | 0.012 | 1 | 3 | Zfp652os | -0.003 | Young |
| 0.542964 | 0.198397 | 0.136 | 0.128 | 1 | 3 | Znfx1 | 0.008 | Young |
| 0.543086 | 0.170661 | 0.149 | 0.165 | 1 | 3 | Atg12 | -0.016 | Young |
| 0.543227 | 0.131911 | 0.121 | 0.112 | 1 | 3 | Guca1a | 0.009 | Young |
| 0.543251 | 0.27846 | 0.035 | 0.042 | 1 | 3 | Gmds | -0.007 | Young |
| 0.543388 | 0.084763 | 0.679 | 0.7 | 1 | 3 | Eif3e | -0.021 | Young |
| 0.543596 | 0.258274 | 0.011 | 0.008 | 1 | 3 | Gvin1 | 0.003 | Young |
| 0.544075 | 0.01057 | 0.133 | 0.146 | 1 | 3 | Fcf1 | -0.013 | Young |
| 0.544187 | 0.015957 | 0.009 | 0.012 | 1 | 3 | Prps1l3 | -0.003 | Young |
| 0.544236 | 0.165516 | 0.015 | 0.019 | 1 | 3 | Zfp688 | -0.004 | Young |
| 0.544247 | 0.237767 | 0.073 | 0.066 | 1 | 3 | Cx3cr1 | 0.007 | Young |
| 0.544571 | 0.115564 | 0.138 | 0.129 | 1 | 3 | Smc5 | 0.009 | Young |
| 0.544841 | 0.072511 | 0.676 | 0.725 | 1 | 3 | Tgfb1 | -0.049 | Young |
| 0.544959 | 0.320556 | 0.092 | 0.102 | 1 | 3 | Brf1 | -0.01 | Young |
| 0.545088 | 0.020308 | 0.068 | 0.076 | 1 | 3 | Exosc4 | -0.008 | Young |
| 0.545185 | 0.225594 | 0.364 | 0.372 | 1 | 3 | Lemd2 | -0.008 | Young |
| 0.545379 | 0.278219 | 0.069 | 0.062 | 1 | 3 | Maml3 | 0.007 | Young |
| 0.54544 | 0.325091 | 0.303 | 0.302 | 1 | 3 | St6galnac6 | 0.001 | Young |
| 0.545493 | 0.405528 | 0.021 | 0.026 | 1 | 3 | Gm20604 | -0.005 | Young |
| 0.545778 | 0.098553 | 0.009 | 0.012 | 1 | 3 | Gm27003 | -0.003 | Young |
| 0.54602 | 0.208171 | 0.068 | 0.061 | 1 | 3 | Tsc2 | 0.007 | Young |
| 0.546058 | 0.400837 | 0.176 | 0.171 | 1 | 3 | C1galt1 | 0.005 | Young |
| 0.546114 | 0.023746 | 0.17 | 0.183 | 1 | 3 | Ubfd1 | -0.013 | Young |
| 0.546332 | 0.909713 | 0.013 | 0.01 | 1 | 3 | Ccdc94 | 0.003 | Young |
| 0.546719 | 0.496088 | 0.013 | 0.01 | 1 | 3 | Lrp4 | 0.003 | Young |
| 0.546741 | 0.335827 | 0.052 | 0.06 | 1 | 3 | Bag5 | -0.008 | Young |
| 0.546761 | 0.145423 | 0.178 | 0.197 | 1 | 3 | Ak3 | -0.019 | Young |
| 0.546918 | 0.220795 | 0.08 | 0.09 | 1 | 3 | Pitrm1 | -0.01 | Young |
| 0.546995 | 0.145088 | 0.08 | 0.072 | 1 | 3 | Ccdc90b | 0.008 | Young |
| 0.547003 | 0.256907 | 0.138 | 0.129 | 1 | 3 | Mast3 | 0.009 | Young |

| | | | | | | | | |
|---|---|---|---|---|---|---|---|---|
| 0.547027 | 0.392866 | 0.105 | 0.098 | 1 | 3 | Ndrg3 | 0.007 | Young |
| 0.547053 | 0.254298 | 0.073 | 0.082 | 1 | 3 | Triap1 | -0.009 | Young |
| 0.547498 | 0.17787 | 0.249 | 0.243 | 1 | 3 | Anapc13 | 0.006 | Young |
| 0.547584 | 0.179022 | 0.068 | 0.076 | 1 | 3 | Fam53a | -0.008 | Young |
| 0.547769 | 0.250928 | 0.153 | 0.146 | 1 | 3 | Zfp395 | 0.007 | Young |
| 0.54777 | 0.267407 | 0.009 | 0.012 | 1 | 3 | Cenpk | -0.003 | Young |
| 0.548101 | 0.278321 | 0.129 | 0.121 | 1 | 3 | Scrib | 0.008 | Young |
| 0.54826 | 0.0965 | 0.039 | 0.033 | 1 | 3 | Incenp | 0.006 | Young |
| 0.548415 | 0.073796 | 0.054 | 0.048 | 1 | 3 | Tysnd1 | 0.006 | Young |
| 0.548466 | 0.16391 | 0.129 | 0.119 | 1 | 3 | AC163354. | 0.01 | Young |
| 0.54852 | 0.144778 | 0.087 | 0.079 | 1 | 3 | Hint3 | 0.008 | Young |
| 0.548658 | 0.490716 | 0.013 | 0.01 | 1 | 3 | Fbxl22 | 0.003 | Young |
| 0.548779 | 0.044757 | 0.122 | 0.134 | 1 | 3 | Aldh3a2 | -0.012 | Young |
| 0.549046 | 0.781913 | 0.013 | 0.01 | 1 | 3 | Tha1 | 0.003 | Young |
| 0.54928 | 0.182483 | 0.029 | 0.035 | 1 | 3 | Zfp617 | -0.006 | Young |
| 0.549296 | 0.320746 | 0.076 | 0.068 | 1 | 3 | Prkab1 | 0.008 | Young |
| 0.549434 | 0.61452 | 0.013 | 0.01 | 1 | 3 | Tmod1 | 0.003 | Young |
| 0.549434 | 0.401747 | 0.013 | 0.01 | 1 | 3 | Cdkn1c | 0.003 | Young |
| 0.549655 | 0.485159 | 0.091 | 0.085 | 1 | 3 | Dpy19l4 | 0.006 | Young |
| 0.549736 | 0.051001 | 0.361 | 0.395 | 1 | 3 | Ube2s | -0.034 | Young |
| 0.549828 | 0.009514 | 0.038 | 0.032 | 1 | 3 | Zfp952 | 0.006 | Young |
| 0.549844 | 0.139374 | 0.015 | 0.019 | 1 | 3 | Gm47675 | -0.004 | Young |
| 0.550226 | 0.275707 | 0.208 | 0.202 | 1 | 3 | Lyl1 | 0.006 | Young |
| 0.550339 | 0.525426 | 0.09 | 0.082 | 1 | 3 | Atrn | 0.008 | Young |
| 0.550408 | 0.31757 | 0.023 | 0.027 | 1 | 3 | Natd1 | -0.004 | Young |
| 0.550627 | 0.034412 | 0.067 | 0.06 | 1 | 3 | Eif2ak4 | 0.007 | Young |
| 0.550851 | 0.178288 | 0.178 | 0.168 | 1 | 3 | Jkamp | 0.01 | Young |
| 0.550908 | 0.184589 | 0.16 | 0.153 | 1 | 3 | AI413582 | 0.007 | Young |
| 0.551192 | 0.095385 | 0.049 | 0.056 | 1 | 3 | Tigd2 | -0.007 | Young |
| 0.551478 | 0.000979 | 0.039 | 0.045 | 1 | 3 | Wdtc1 | -0.006 | Young |
| 0.551613 | 0.119597 | 0.26 | 0.282 | 1 | 3 | Tm2d2 | -0.022 | Young |
| 0.551656 | 0.267697 | 0.191 | 0.184 | 1 | 3 | Sema4b | 0.007 | Young |
| 0.551694 | 0.094544 | 0.049 | 0.043 | 1 | 3 | Dmac2 | 0.006 | Young |
| 0.551765 | 0.701903 | 0.013 | 0.01 | 1 | 3 | Lmln | 0.003 | Young |
| 0.551765 | 0.351697 | 0.013 | 0.01 | 1 | 3 | Zfp7 | 0.003 | Young |
| 0.551871 | 0.113361 | 0.038 | 0.044 | 1 | 3 | Homer3 | -0.006 | Young |
| 0.55192 | 0.312514 | 0.124 | 0.116 | 1 | 3 | Mfsd10 | 0.008 | Young |
| 0.551934 | 0.132418 | 0.08 | 0.088 | 1 | 3 | Mtf1 | -0.008 | Young |
| 0.552018 | 0.125226 | 0.217 | 0.207 | 1 | 3 | Atp9b | 0.01 | Young |
| 0.552063 | 0.35584 | 0.045 | 0.053 | 1 | 3 | Fam213b | -0.008 | Young |
| 0.552083 | 0.188777 | 0.206 | 0.196 | 1 | 3 | Afg3l1 | 0.01 | Young |
| 0.552154 | 0.433821 | 0.013 | 0.01 | 1 | 3 | Trim2 | 0.003 | Young |
| 0.552154 | 0.337793 | 0.013 | 0.01 | 1 | 3 | Zcchc3 | 0.003 | Young |
| 0.552248 | 0.183333 | 0.282 | 0.277 | 1 | 3 | Dusp22 | 0.005 | Young |
| 0.55228 | 0.049356 | 0.067 | 0.075 | 1 | 3 | Maip1 | -0.008 | Young |

| | | | | | | | | |
|---|---|---|---|---|---|---|---|---|
| 0.552488 | 0.454668 | 0.078 | 0.072 | 1 | 3 | Wdr81 | 0.006 | Young |
| 0.552544 | 0.242078 | 0.013 | 0.01 | 1 | 3 | Arhgap22 | 0.003 | Young |
| 0.552611 | 0.105711 | 0.374 | 0.373 | 1 | 3 | Mlec | 0.001 | Young |
| 0.552731 | 0.026963 | 0.293 | 0.317 | 1 | 3 | Arrdc1 | -0.024 | Young |
| 0.553004 | 0.188944 | 0.023 | 0.027 | 1 | 3 | Msh3 | -0.004 | Young |
| 0.553084 | 0.044037 | 0.515 | 0.536 | 1 | 3 | Mtss1 | -0.021 | Young |
| 0.553297 | 0.062744 | 0.047 | 0.041 | 1 | 3 | Dtd2 | 0.006 | Young |
| 0.553455 | 0.12932 | 0.057 | 0.05 | 1 | 3 | Diras2 | 0.007 | Young |
| 0.553596 | 0.013145 | 0.04 | 0.047 | 1 | 3 | Fam149b | -0.007 | Young |
| 0.554035 | 0.318695 | 0.052 | 0.045 | 1 | 3 | Phospho2 | 0.007 | Young |
| 0.554637 | 0.303568 | 0.054 | 0.062 | 1 | 3 | Gab1 | -0.008 | Young |
| 0.554899 | 0.063404 | 0.365 | 0.357 | 1 | 3 | Cd37 | 0.008 | Young |
| 0.555097 | 0.400465 | 0.23 | 0.226 | 1 | 3 | Bsdc1 | 0.004 | Young |
| 0.555582 | 0.195694 | 0.073 | 0.066 | 1 | 3 | Map3k3 | 0.007 | Young |
| 0.555986 | 0.1901 | 0.181 | 0.172 | 1 | 3 | Vrk2 | 0.009 | Young |
| 0.556366 | 0.11889 | 0.206 | 0.226 | 1 | 3 | Trnau1ap | -0.02 | Young |
| 0.556369 | 0.019721 | 0.093 | 0.103 | 1 | 3 | Grap | -0.01 | Young |
| 0.556443 | 0.017334 | 0.013 | 0.01 | 1 | 3 | Pim2 | 0.003 | Young |
| 0.557187 | 0.690189 | 0.014 | 0.011 | 1 | 3 | Rtel1 | 0.003 | Young |
| 0.557225 | 0.534358 | 0.013 | 0.01 | 1 | 3 | Adarb1 | 0.003 | Young |
| 0.557334 | 0.21731 | 0.051 | 0.044 | 1 | 3 | Cwc27 | 0.007 | Young |
| 0.557654 | 0.088044 | 0.107 | 0.119 | 1 | 3 | Tprkb | -0.012 | Young |
| 0.557721 | 0.070663 | 0.044 | 0.038 | 1 | 3 | Tmem242 | 0.006 | Young |
| 0.558259 | 0.218619 | 0.093 | 0.086 | 1 | 3 | Ssbp2 | 0.007 | Young |
| 0.558449 | 0.394316 | 0.081 | 0.088 | 1 | 3 | Ccl3 | -0.007 | Young |
| 0.558937 | 0.028992 | 0.289 | 0.311 | 1 | 3 | Hadha | -0.022 | Young |
| 0.559044 | 1.239872 | 0.014 | 0.011 | 1 | 3 | Gm26756 | 0.003 | Young |
| 0.559349 | 0.172345 | 0.323 | 0.321 | 1 | 3 | Tmf1 | 0.002 | Young |
| 0.559816 | 0.163552 | 0.032 | 0.037 | 1 | 3 | Urb1 | -0.005 | Young |
| 0.560036 | 0.208575 | 0.077 | 0.086 | 1 | 3 | Usp31 | -0.009 | Young |
| 0.560187 | 0.119408 | 0.177 | 0.191 | 1 | 3 | Atxn3 | -0.014 | Young |
| 0.560397 | 0.085045 | 0.04 | 0.047 | 1 | 3 | 1110034G2 | -0.007 | Young |
| 0.560928 | 0.050348 | 0.091 | 0.1 | 1 | 3 | Dtnbp1 | -0.009 | Young |
| 0.560951 | 0.16643 | 0.566 | 0.596 | 1 | 3 | Samsn1 | -0.03 | Young |
| 0.56108 | 0.293879 | 0.139 | 0.131 | 1 | 3 | Lysmd3 | 0.008 | Young |
| 0.562701 | 0.06901 | 0.173 | 0.189 | 1 | 3 | Rnaseh2c | -0.016 | Young |
| 0.562802 | 0.020566 | 0.181 | 0.197 | 1 | 3 | Lsm3 | -0.016 | Young |
| 0.562813 | 0.005809 | 0.073 | 0.081 | 1 | 3 | Zcchc8 | -0.008 | Young |
| 0.563127 | 0.141845 | 0.083 | 0.092 | 1 | 3 | Tmem71 | -0.009 | Young |
| 0.563147 | 0.209839 | 0.104 | 0.096 | 1 | 3 | Camsap1 | 0.008 | Young |
| 0.563402 | 0.070147 | 0.149 | 0.159 | 1 | 3 | Htra2 | -0.01 | Young |
| 0.563568 | 0.069567 | 0.592 | 0.644 | 1 | 3 | Klf3 | -0.052 | Young |
| 0.563784 | 0.122617 | 0.163 | 0.178 | 1 | 3 | Pxk | -0.015 | Young |
| 0.564003 | 0.062623 | 0.133 | 0.146 | 1 | 3 | Pde7a | -0.013 | Young |
| 0.564355 | 0.046736 | 0.153 | 0.166 | 1 | 3 | Cisd1 | -0.013 | Young |

| | | | | | | | | |
|---|---|---|---|---|---|---|---|---|
| 0.564709 | 0.223588 | 0.188 | 0.18 | 1 | 3 | Map2k4 | 0.008 | Young |
| 0.564743 | 0.089982 | 0.016 | 0.02 | 1 | 3 | Zfp619 | -0.004 | Young |
| 0.565038 | 0.046062 | 0.581 | 0.609 | 1 | 3 | Tbca | -0.028 | Young |
| 0.565195 | 0.009922 | 0.598 | 0.657 | 1 | 3 | Atp6v1f | -0.059 | Young |
| 0.565456 | 0.022132 | 0.061 | 0.054 | 1 | 3 | Atg14 | 0.007 | Young |
| 0.565595 | 0.15665 | 0.318 | 0.356 | 1 | 3 | mt-Atp8 | -0.038 | Young |
| 0.565713 | 0.018483 | 0.034 | 0.039 | 1 | 3 | Med19 | -0.005 | Young |
| 0.56585 | 0.11013 | 0.634 | 0.657 | 1 | 3 | Shisa5 | -0.023 | Young |
| 0.565854 | 0.067034 | 0.598 | 0.613 | 1 | 3 | Tep1 | -0.015 | Young |
| 0.566095 | 0.048577 | 0.979 | 0.99 | 1 | 3 | Rps28 | -0.011 | Young |
| 0.566489 | 0.639572 | 0.015 | 0.012 | 1 | 3 | Cfap45 | 0.003 | Young |
| 0.566756 | 0.231806 | 0.091 | 0.084 | 1 | 3 | Decr1 | 0.007 | Young |
| 0.566875 | 0.156289 | 0.014 | 0.011 | 1 | 3 | Lamc1 | 0.003 | Young |
| 0.567022 | 0.054642 | 0.42 | 0.452 | 1 | 3 | Notch1 | -0.032 | Young |
| 0.567443 | 0.136775 | 0.182 | 0.196 | 1 | 3 | Ilf2 | -0.014 | Young |
| 0.56756 | 0.699463 | 0.015 | 0.012 | 1 | 3 | AW146154 | 0.003 | Young |
| 0.567623 | 0.206641 | 0.014 | 0.011 | 1 | 3 | Ppp1r18os | 0.003 | Young |
| 0.567892 | 0.046155 | 0.087 | 0.097 | 1 | 3 | Rrm1 | -0.01 | Young |
| 0.567991 | 0.001044 | 0.492 | 0.527 | 1 | 3 | Llph | -0.035 | Young |
| 0.567998 | 0.523334 | 0.014 | 0.011 | 1 | 3 | Gm48066 | 0.003 | Young |
| 0.568058 | 0.509979 | 0.085 | 0.079 | 1 | 3 | Aimp2 | 0.006 | Young |
| 0.568199 | 0.449083 | 0.145 | 0.14 | 1 | 3 | Map1s | 0.005 | Young |
| 0.568427 | 0.24504 | 0.139 | 0.133 | 1 | 3 | AW549877 | 0.006 | Young |
| 0.568632 | 0.529106 | 0.015 | 0.012 | 1 | 3 | Cep78 | 0.003 | Young |
| 0.569187 | 0.068367 | 0.08 | 0.088 | 1 | 3 | Pkp4 | -0.008 | Young |
| 0.569603 | 0.061092 | 0.357 | 0.387 | 1 | 3 | Man2a2 | -0.03 | Young |
| 0.569653 | 0.324076 | 0.088 | 0.081 | 1 | 3 | Slc9a1 | 0.007 | Young |
| 0.569672 | 0.461084 | 0.064 | 0.059 | 1 | 3 | Mthfd1l | 0.005 | Young |
| 0.569975 | 0.206286 | 0.024 | 0.029 | 1 | 3 | Sphk1 | -0.005 | Young |
| 0.570263 | 0.162569 | 0.176 | 0.166 | 1 | 3 | Bloc1s2 | 0.01 | Young |
| 0.570331 | 0.460037 | 0.157 | 0.152 | 1 | 3 | Micu1 | 0.005 | Young |
| 0.570674 | 0.240545 | 0.04 | 0.047 | 1 | 3 | Calcrl | -0.007 | Young |
| 0.570737 | 0.004736 | 0.059 | 0.067 | 1 | 3 | Hagh | -0.008 | Young |
| 0.570876 | 0.415615 | 0.053 | 0.061 | 1 | 3 | Lyrm9 | -0.008 | Young |
| 0.570883 | 0.198518 | 0.149 | 0.162 | 1 | 3 | Pex6 | -0.013 | Young |
| 0.570954 | 0.176489 | 0.066 | 0.059 | 1 | 3 | Slc35b4 | 0.007 | Young |
| 0.571011 | 0.136546 | 0.174 | 0.164 | 1 | 3 | Ppfibp2 | 0.01 | Young |
| 0.571146 | 0.485962 | 0.082 | 0.076 | 1 | 3 | Rnf19a | 0.006 | Young |
| 0.571247 | 0.0155 | 0.186 | 0.2 | 1 | 3 | Vps28 | -0.014 | Young |
| 0.571702 | 0.325818 | 0.077 | 0.07 | 1 | 3 | Wwc2 | 0.007 | Young |
| 0.571749 | 0.272229 | 0.014 | 0.011 | 1 | 3 | Gm527 | 0.003 | Young |
| 0.571842 | 0.51314 | 0.071 | 0.065 | 1 | 3 | AL590144.: | 0.006 | Young |
| 0.57203 | 0.186808 | 0.062 | 0.069 | 1 | 3 | Sfi1 | -0.007 | Young |
| 0.57213 | 0.147851 | 0.129 | 0.141 | 1 | 3 | Stam2 | -0.012 | Young |
| 0.572177 | 0.095247 | 0.253 | 0.247 | 1 | 3 | Hps3 | 0.006 | Young |

| | | | | | | | | |
|---|---|---|---|---|---|---|---|---|
| 0.57272 | 0.379882 | 0.119 | 0.112 | 1 | 3 | Cdc42ep3 | 0.007 | Young |
| 0.57312 | 0.30907 | 0.083 | 0.076 | 1 | 3 | Zfp518a | 0.007 | Young |
| 0.57377 | 0.006591 | 0.374 | 0.4 | 1 | 3 | Stk4 | -0.026 | Young |
| 0.57382 | 0.140085 | 0.21 | 0.228 | 1 | 3 | Sla | -0.018 | Young |
| 0.574011 | 0.215996 | 0.449 | 0.472 | 1 | 3 | Ythdc1 | -0.023 | Young |
| 0.574335 | 0.190707 | 0.121 | 0.115 | 1 | 3 | Gm13212 | 0.006 | Young |
| 0.574733 | 0.006071 | 0.138 | 0.148 | 1 | 3 | Shc1 | -0.01 | Young |
| 0.574742 | 0.189251 | 0.364 | 0.368 | 1 | 3 | Aldh2 | -0.004 | Young |
| 0.574757 | 0.04475 | 0.379 | 0.404 | 1 | 3 | Hnrnpul1 | -0.025 | Young |
| 0.57487 | 0.114193 | 0.069 | 0.078 | 1 | 3 | Hinfp | -0.009 | Young |
| 0.574966 | 0.151783 | 0.121 | 0.133 | 1 | 3 | Psd3 | -0.012 | Young |
| 0.575085 | 0.378095 | 0.015 | 0.012 | 1 | 3 | 1700008J0 | 0.003 | Young |
| 0.575373 | 0.14022 | 0.01 | 0.013 | 1 | 3 | Yes1 | -0.003 | Young |
| 0.575373 | 0.051848 | 0.01 | 0.013 | 1 | 3 | Ugt1a7c | -0.003 | Young |
| 0.57556 | 0.331083 | 0.14 | 0.134 | 1 | 3 | Gtf3a | 0.006 | Young |
| 0.57576 | 0.331674 | 0.01 | 0.013 | 1 | 3 | BC048403 | -0.003 | Young |
| 0.575929 | 0.123983 | 0.138 | 0.129 | 1 | 3 | Stx5a | 0.009 | Young |
| 0.576163 | 0.454114 | 0.015 | 0.012 | 1 | 3 | Cdc6 | 0.003 | Young |
| 0.576422 | 0.005343 | 0.116 | 0.127 | 1 | 3 | C2cd5 | -0.011 | Young |
| 0.57724 | 0.781561 | 0.039 | 0.035 | 1 | 3 | Ulk3 | 0.004 | Young |
| 0.577243 | 0.066403 | 0.015 | 0.012 | 1 | 3 | Pomk | 0.003 | Young |
| 0.577663 | 0.191057 | 0.196 | 0.19 | 1 | 3 | Epb41l1 | 0.006 | Young |
| 0.577699 | 0.102202 | 0.249 | 0.271 | 1 | 3 | Bmp2k | -0.022 | Young |
| 0.578027 | 0.104415 | 0.417 | 0.41 | 1 | 3 | Fuca1 | 0.007 | Young |
| 0.578082 | 0.260035 | 0.01 | 0.013 | 1 | 3 | Trmu | -0.003 | Young |
| 0.578324 | 0.287402 | 0.015 | 0.012 | 1 | 3 | Ddah2 | 0.003 | Young |
| 0.578324 | 0.049354 | 0.015 | 0.012 | 1 | 3 | Dnajc17 | 0.003 | Young |
| 0.578396 | 0.321364 | 0.016 | 0.013 | 1 | 3 | Dtd1 | 0.003 | Young |
| 0.578814 | 0.211177 | 0.196 | 0.216 | 1 | 3 | Bcam | -0.02 | Young |
| 0.579161 | 0.423541 | 0.077 | 0.07 | 1 | 3 | Ube2o | 0.007 | Young |
| 0.579433 | 0.066749 | 0.196 | 0.189 | 1 | 3 | Thap12 | 0.007 | Young |
| 0.579463 | 0.251533 | 0.215 | 0.209 | 1 | 3 | Fam162a | 0.006 | Young |
| 0.579597 | 0.008447 | 0.217 | 0.237 | 1 | 3 | Map2k1 | -0.02 | Young |
| 0.579985 | 0.080352 | 0.096 | 0.105 | 1 | 3 | Ttc19 | -0.009 | Young |
| 0.580124 | 0.999334 | 0.016 | 0.013 | 1 | 3 | Cep152 | 0.003 | Young |
| 0.580127 | 0.036544 | 0.015 | 0.012 | 1 | 3 | Stat4 | 0.003 | Young |
| 0.580388 | 0.250979 | 0.102 | 0.096 | 1 | 3 | Kdm5b | 0.006 | Young |
| 0.580957 | 0.253222 | 0.205 | 0.201 | 1 | 3 | Tmem64 | 0.004 | Young |
| 0.581182 | 0.201722 | 0.295 | 0.289 | 1 | 3 | N4bp2l2 | 0.006 | Young |
| 0.581198 | 0.103222 | 0.359 | 0.354 | 1 | 3 | Crat | 0.005 | Young |
| 0.581241 | 0.160287 | 0.467 | 0.479 | 1 | 3 | Tsc22d4 | -0.012 | Young |
| 0.581351 | 0.063009 | 0.044 | 0.05 | 1 | 3 | Arfgap3 | -0.006 | Young |
| 0.581451 | 0.205169 | 0.295 | 0.292 | 1 | 3 | Zfp644 | 0.003 | Young |
| 0.58166 | 0.186739 | 0.236 | 0.258 | 1 | 3 | Rin2 | -0.022 | Young |
| 0.581797 | 0.557944 | 0.018 | 0.014 | 1 | 3 | Bcs1l | 0.004 | Young |

| | | | | | | | | |
|---|---|---|---|---|---|---|---|---|
| 0.581921 | 0.024369 | 0.739 | 0.79 | 1 | 3 | Fus | -0.051 | Young |
| 0.582287 | 0.141375 | 0.074 | 0.084 | 1 | 3 | Alg13 | -0.01 | Young |
| 0.582428 | 0.04393 | 0.237 | 0.252 | 1 | 3 | Rsl24d1 | -0.015 | Young |
| 0.582546 | 0.657356 | 0.016 | 0.013 | 1 | 3 | Mgp | 0.003 | Young |
| 0.582684 | 0.066063 | 0.071 | 0.079 | 1 | 3 | Nat9 | -0.008 | Young |
| 0.58275 | 0.073401 | 0.052 | 0.059 | 1 | 3 | Ttc4 | -0.007 | Young |
| 0.583314 | 0.297781 | 0.245 | 0.243 | 1 | 3 | Nck1 | 0.002 | Young |
| 0.583413 | 0.014213 | 0.122 | 0.133 | 1 | 3 | Prpf31 | -0.011 | Young |
| 0.583586 | 0.056575 | 0.016 | 0.013 | 1 | 3 | Asf1b | 0.003 | Young |
| 0.583643 | 0.087399 | 0.109 | 0.118 | 1 | 3 | Dcaf5 | -0.009 | Young |
| 0.584333 | 0.03966 | 0.396 | 0.429 | 1 | 3 | Snrpd2 | -0.033 | Young |
| 0.584353 | 0.340796 | 0.146 | 0.143 | 1 | 3 | Scyl2 | 0.003 | Young |
| 0.584376 | 0.322547 | 0.064 | 0.059 | 1 | 3 | Adprhl2 | 0.005 | Young |
| 0.58445 | 0.043585 | 0.461 | 0.513 | 1 | 3 | Arih1 | -0.052 | Young |
| 0.584478 | 0.078825 | 0.629 | 0.656 | 1 | 3 | Lamtor4 | -0.027 | Young |
| 0.584817 | 0.125892 | 0.081 | 0.074 | 1 | 3 | Cyb5r1 | 0.007 | Young |
| 0.58511 | 0.267987 | 0.08 | 0.073 | 1 | 3 | Med22 | 0.007 | Young |
| 0.585463 | 0.469823 | 0.018 | 0.014 | 1 | 3 | Cers4 | 0.004 | Young |
| 0.586766 | 0.259778 | 0.293 | 0.314 | 1 | 3 | Slc5a3 | -0.021 | Young |
| 0.586811 | 0.121372 | 0.174 | 0.166 | 1 | 3 | Arsb | 0.008 | Young |
| 0.586966 | 0.211219 | 0.066 | 0.073 | 1 | 3 | Elac1 | -0.007 | Young |
| 0.587526 | 0.044834 | 0.283 | 0.303 | 1 | 3 | Fndc3a | -0.02 | Young |
| 0.58759 | 0.177154 | 0.081 | 0.09 | 1 | 3 | Ahi1 | -0.009 | Young |
| 0.587849 | 0.329671 | 0.098 | 0.092 | 1 | 3 | Scaf1 | 0.006 | Young |
| 0.587908 | 0.222785 | 0.143 | 0.135 | 1 | 3 | Fmc1 | 0.008 | Young |
| 0.588136 | 0.274915 | 0.018 | 0.014 | 1 | 3 | Cxxc5 | 0.004 | Young |
| 0.588501 | 0.103225 | 0.273 | 0.302 | 1 | 3 | Nfkbie | -0.029 | Young |
| 0.588569 | 0.721836 | 0.023 | 0.019 | 1 | 3 | Map3k12 | 0.004 | Young |
| 0.588798 | 0.12383 | 0.016 | 0.013 | 1 | 3 | Kctd7 | 0.003 | Young |
| 0.588799 | 0.047362 | 0.653 | 0.693 | 1 | 3 | Rpl29 | -0.04 | Young |
| 0.588858 | 0.02018 | 0.804 | 0.848 | 1 | 3 | mt-Nd5 | -0.044 | Young |
| 0.58914 | 0.278181 | 0.018 | 0.014 | 1 | 3 | Gm26624 | 0.004 | Young |
| 0.589158 | 0.097426 | 0.102 | 0.112 | 1 | 3 | Chchd5 | -0.01 | Young |
| 0.589301 | 0.267331 | 0.277 | 0.276 | 1 | 3 | Zcchc7 | 0.001 | Young |
| 0.589321 | 0.102003 | 0.034 | 0.039 | 1 | 3 | Adra2a | -0.005 | Young |
| 0.589439 | 0.532231 | 0.131 | 0.125 | 1 | 3 | Zufsp | 0.006 | Young |
| 0.589979 | 0.381794 | 0.056 | 0.05 | 1 | 3 | Zfp367 | 0.006 | Young |
| 0.590432 | 0.106835 | 0.302 | 0.299 | 1 | 3 | Tifab | 0.003 | Young |
| 0.59048 | 0.64686 | 0.018 | 0.014 | 1 | 3 | Sag | 0.004 | Young |
| 0.590625 | 0.201017 | 0.398 | 0.403 | 1 | 3 | Bcl2a1d | -0.005 | Young |
| 0.590662 | 0.555856 | 0.025 | 0.022 | 1 | 3 | Eif3j2 | 0.003 | Young |
| 0.590885 | 0.007172 | 0.114 | 0.105 | 1 | 3 | Ttc9c | 0.009 | Young |
| 0.591339 | 0.295485 | 0.058 | 0.066 | 1 | 3 | Jade2 | -0.008 | Young |
| 0.591486 | 0.147556 | 0.018 | 0.014 | 1 | 3 | Pard6a | 0.004 | Young |
| 0.591638 | 0.37343 | 0.059 | 0.054 | 1 | 3 | Atn1 | 0.005 | Young |

| | | | | | | | | |
|---|---|---|---|---|---|---|---|---|
| 0.591817 | 0.431621 | 0.019 | 0.016 | 1 | 3 | Mocos | 0.003 | Young |
| 0.591989 | 0.24518 | 0.018 | 0.014 | 1 | 3 | Zfp40 | 0.004 | Young |
| 0.592024 | 0.615085 | 0.038 | 0.033 | 1 | 3 | Atg16l2 | 0.005 | Young |
| 0.593217 | 0.458744 | 0.02 | 0.017 | 1 | 3 | Gm9530 | 0.003 | Young |
| 0.59352 | 0.080402 | 0.263 | 0.286 | 1 | 3 | Bdp1 | -0.023 | Young |
| 0.593739 | 0.006875 | 0.745 | 0.767 | 1 | 3 | Sat1 | -0.022 | Young |
| 0.593913 | 0.15886 | 0.085 | 0.093 | 1 | 3 | Fopnl | -0.008 | Young |
| 0.594302 | 0.002909 | 0.019 | 0.023 | 1 | 3 | Scamp5 | -0.004 | Young |
| 0.594746 | 0.63017 | 0.061 | 0.055 | 1 | 3 | 6720427I0 | 0.006 | Young |
| 0.595104 | 0.218316 | 0.13 | 0.123 | 1 | 3 | Bola3 | 0.007 | Young |
| 0.595248 | 0.227163 | 0.102 | 0.096 | 1 | 3 | Avl9 | 0.006 | Young |
| 0.59538 | 0.015651 | 0.019 | 0.016 | 1 | 3 | Gm10762 | 0.003 | Young |
| 0.595673 | 0.065353 | 0.121 | 0.131 | 1 | 3 | Mxd1 | -0.01 | Young |
| 0.595693 | 0.385941 | 0.027 | 0.031 | 1 | 3 | Cxcl3 | -0.004 | Young |
| 0.595707 | 0.561595 | 0.035 | 0.031 | 1 | 3 | Purg | 0.004 | Young |
| 0.595788 | 0.594143 | 0.043 | 0.038 | 1 | 3 | Ankrd13b | 0.005 | Young |
| 0.59622 | 0.33097 | 0.047 | 0.042 | 1 | 3 | Hps4 | 0.005 | Young |
| 0.596267 | 0.169056 | 0.078 | 0.087 | 1 | 3 | Slc25a44 | -0.009 | Young |
| 0.596396 | 0.089196 | 0.037 | 0.042 | 1 | 3 | Slc52a2 | -0.005 | Young |
| 0.596426 | 0.782678 | 0.047 | 0.042 | 1 | 3 | Jrkl | 0.005 | Young |
| 0.596666 | 0.019356 | 0.043 | 0.049 | 1 | 3 | Fblim1 | -0.006 | Young |
| 0.59685 | 0.32404 | 0.018 | 0.022 | 1 | 3 | Adck5 | -0.004 | Young |
| 0.596877 | 0.258516 | 0.114 | 0.108 | 1 | 3 | Foxo3 | 0.006 | Young |
| 0.596931 | 0.043867 | 0.306 | 0.331 | 1 | 3 | Hprt | -0.025 | Young |
| 0.597051 | 0.650494 | 0.03 | 0.026 | 1 | 3 | Haus4 | 0.004 | Young |
| 0.597123 | 0.162922 | 0.111 | 0.123 | 1 | 3 | Mettl16 | -0.012 | Young |
| 0.59719 | 0.218269 | 0.23 | 0.228 | 1 | 3 | Il3ra | 0.002 | Young |
| 0.597653 | 0.280529 | 0.019 | 0.016 | 1 | 3 | Alms1 | 0.003 | Young |
| 0.597653 | 0.090303 | 0.019 | 0.016 | 1 | 3 | Rad18 | 0.003 | Young |
| 0.597816 | 0.082478 | 0.107 | 0.117 | 1 | 3 | Pygo2 | -0.01 | Young |
| 0.598005 | 0.084732 | 0.273 | 0.299 | 1 | 3 | Nab1 | -0.026 | Young |
| 0.598028 | 0.254021 | 0.194 | 0.19 | 1 | 3 | Trabd | 0.004 | Young |
| 0.599035 | 0.576778 | 0.024 | 0.02 | 1 | 3 | Trim14 | 0.004 | Young |
| 0.599493 | 0.294809 | 0.02 | 0.017 | 1 | 3 | 4833445I0 | 0.003 | Young |
| 0.599493 | 0.092564 | 0.02 | 0.017 | 1 | 3 | Pdgfc | 0.003 | Young |
| 0.599505 | 0.011533 | 0.152 | 0.164 | 1 | 3 | Eif4enif1 | -0.012 | Young |
| 0.599737 | 0.038449 | 0.21 | 0.227 | 1 | 3 | Fkbp4 | -0.017 | Young |
| 0.599809 | 0.346343 | 0.1 | 0.094 | 1 | 3 | Ankra2 | 0.006 | Young |
| 0.599834 | 0.238658 | 0.092 | 0.086 | 1 | 3 | Golim4 | 0.006 | Young |
| 0.599895 | 0.550851 | 0.021 | 0.018 | 1 | 3 | D3Ertd751 | 0.003 | Young |
| 0.599971 | 0.501459 | 0.09 | 0.084 | 1 | 3 | Zfp954 | 0.006 | Young |
| 0.600014 | 0.017315 | 0.66 | 0.688 | 1 | 3 | Nfkbid | -0.028 | Young |
| 0.60017 | 0.535426 | 0.028 | 0.024 | 1 | 3 | 1600020E0 | 0.004 | Young |
| 0.600208 | 0.019873 | 0.447 | 0.489 | 1 | 3 | Wac | -0.042 | Young |
| 0.600806 | 0.217658 | 0.133 | 0.125 | 1 | 3 | Ikzf5 | 0.008 | Young |

| | | | | | | | | |
|---|---|---|---|---|---|---|---|---|
| 0.600859 | 0.320336 | 0.011 | 0.014 | 1 | 3 | Il21r | -0.003 | Young |
| 0.600891 | 0.120183 | 0.105 | 0.115 | 1 | 3 | Ralgapb | -0.01 | Young |
| 0.601558 | 0.202695 | 0.019 | 0.016 | 1 | 3 | Fancc | 0.003 | Young |
| 0.601626 | 0.486522 | 0.024 | 0.02 | 1 | 3 | Cbr1 | 0.004 | Young |
| 0.601686 | 0.299444 | 0.199 | 0.21 | 1 | 3 | Tnf | -0.011 | Young |
| 0.601697 | 0.127208 | 0.356 | 0.354 | 1 | 3 | Edf1 | 0.002 | Young |
| 0.601756 | 0.293302 | 0.057 | 0.051 | 1 | 3 | Bloc1s5 | 0.006 | Young |
| 0.602173 | 0.213233 | 0.168 | 0.161 | 1 | 3 | Ifi27 | 0.007 | Young |
| 0.602335 | 0.348904 | 0.021 | 0.018 | 1 | 3 | Arvcf | 0.003 | Young |
| 0.602511 | 0.116872 | 0.063 | 0.07 | 1 | 3 | Ercc6l2 | -0.007 | Young |
| 0.602628 | 0.035329 | 0.105 | 0.114 | 1 | 3 | Zfp281 | -0.009 | Young |
| 0.602869 | 0.1927 | 0.071 | 0.065 | 1 | 3 | Frmd6 | 0.006 | Young |
| 0.603161 | 0.197198 | 0.163 | 0.155 | 1 | 3 | H2-Q4 | 0.008 | Young |
| 0.603215 | 0.251703 | 0.047 | 0.042 | 1 | 3 | Zscan29 | 0.005 | Young |
| 0.603241 | 0.160551 | 0.298 | 0.321 | 1 | 3 | Tnks2 | -0.023 | Young |
| 0.603525 | 0.423434 | 0.047 | 0.042 | 1 | 3 | Kptn | 0.005 | Young |
| 0.603525 | 0.020918 | 0.027 | 0.031 | 1 | 3 | St3gal6 | -0.004 | Young |
| 0.603526 | 0.466485 | 0.025 | 0.022 | 1 | 3 | Slc22a18 | 0.003 | Young |
| 0.603588 | 0.20278 | 0.336 | 0.339 | 1 | 3 | 2410015M | -0.003 | Young |
| 0.603647 | 0.300622 | 0.058 | 0.053 | 1 | 3 | Cisd3 | 0.005 | Young |
| 0.603933 | 0.518731 | 0.024 | 0.02 | 1 | 3 | Mapk1ip1 | 0.004 | Young |
| 0.604029 | 0.345166 | 0.066 | 0.06 | 1 | 3 | 2610524H( | 0.006 | Young |
| 0.604169 | 0.399108 | 0.021 | 0.018 | 1 | 3 | Usp20 | 0.003 | Young |
| 0.604169 | 0.177602 | 0.021 | 0.018 | 1 | 3 | Platr25 | 0.003 | Young |
| 0.604241 | 0.07026 | 0.011 | 0.014 | 1 | 3 | E230032D2 | -0.003 | Young |
| 0.604331 | 0.524016 | 0.039 | 0.035 | 1 | 3 | Gpatch1 | 0.004 | Young |
| 0.604491 | 0.350362 | 0.054 | 0.049 | 1 | 3 | Uck1 | 0.005 | Young |
| 0.604508 | 0.290627 | 0.116 | 0.128 | 1 | 3 | Zbtb21 | -0.012 | Young |
| 0.60455 | 0.020234 | 0.378 | 0.401 | 1 | 3 | Ptpro | -0.023 | Young |
| 0.604606 | 0.416517 | 0.148 | 0.143 | 1 | 3 | Ddx27 | 0.005 | Young |
| 0.60465 | 0.397458 | 0.025 | 0.022 | 1 | 3 | Scml4 | 0.003 | Young |
| 0.6047 | 0.246379 | 0.056 | 0.05 | 1 | 3 | Man2b2 | 0.006 | Young |
| 0.605087 | 0.217509 | 0.021 | 0.018 | 1 | 3 | Hus1 | 0.003 | Young |
| 0.605087 | 0.122096 | 0.021 | 0.018 | 1 | 3 | Rbbp9 | 0.003 | Young |
| 0.605547 | 0.41241 | 0.023 | 0.019 | 1 | 3 | Ppp1r3b | 0.004 | Young |
| 0.605618 | 0.139294 | 0.605 | 0.622 | 1 | 3 | Gpx4 | -0.017 | Young |
| 0.606061 | 0.514024 | 0.027 | 0.031 | 1 | 3 | Zfp157 | -0.004 | Young |
| 0.606065 | 0.14819 | 0.266 | 0.262 | 1 | 3 | Ankrd13c | 0.004 | Young |
| 0.606924 | 0.072536 | 0.021 | 0.018 | 1 | 3 | Tlr8 | 0.003 | Young |
| 0.607151 | 0.193159 | 0.096 | 0.09 | 1 | 3 | Ing3 | 0.006 | Young |
| 0.607517 | 0.277358 | 0.096 | 0.09 | 1 | 3 | Zfp511 | 0.006 | Young |
| 0.607652 | 0.374277 | 0.051 | 0.045 | 1 | 3 | Nfkbil1 | 0.006 | Young |
| 0.60769 | 0.35894 | 0.024 | 0.02 | 1 | 3 | Gm340 | 0.004 | Young |
| 0.60812 | 0.300405 | 0.056 | 0.05 | 1 | 3 | Spsb2 | 0.006 | Young |
| 0.609506 | 0.00083 | 0.269 | 0.287 | 1 | 3 | Sertad1 | -0.018 | Young |

| | | | | | | | | |
|---|---|---|---|---|---|---|---|---|
| 0.609583 | 0.047147 | 0.287 | 0.309 | 1 | 3 | Morc3 | -0.022 | Young |
| 0.609669 | 0.146601 | 0.049 | 0.055 | 1 | 3 | Gpank1 | -0.006 | Young |
| 0.609717 | 0.197855 | 0.167 | 0.161 | 1 | 3 | Rabep1 | 0.006 | Young |
| 0.609723 | 0.222728 | 0.025 | 0.022 | 1 | 3 | Cbarp | 0.003 | Young |
| 0.609913 | 0.008528 | 0.02 | 0.017 | 1 | 3 | Nme2 | 0.003 | Young |
| 0.609925 | 0.094669 | 0.102 | 0.111 | 1 | 3 | Gm30978 | -0.009 | Young |
| 0.610008 | 0.358319 | 0.024 | 0.02 | 1 | 3 | Slf1 | 0.004 | Young |
| 0.610126 | 0.128961 | 0.074 | 0.082 | 1 | 3 | Thsd4 | -0.008 | Young |
| 0.610517 | 0.083261 | 0.213 | 0.228 | 1 | 3 | Grina | -0.015 | Young |
| 0.610639 | 0.165734 | 0.419 | 0.422 | 1 | 3 | Ttc7 | -0.003 | Young |
| 0.610782 | 0.175025 | 0.295 | 0.292 | 1 | 3 | Abcc1 | 0.003 | Young |
| 0.61091 | 0.033885 | 0.11 | 0.121 | 1 | 3 | Tmem68 | -0.011 | Young |
| 0.611113 | 0.191838 | 0.116 | 0.127 | 1 | 3 | Ino80 | -0.011 | Young |
| 0.611751 | 0.441118 | 0.044 | 0.039 | 1 | 3 | Rtl8b | 0.005 | Young |
| 0.611877 | 0.140928 | 0.069 | 0.063 | 1 | 3 | Yaf2 | 0.006 | Young |
| 0.611888 | 0.254609 | 0.256 | 0.251 | 1 | 3 | Bin1 | 0.005 | Young |
| 0.612462 | 0.335983 | 0.229 | 0.227 | 1 | 3 | Slmap | 0.002 | Young |
| 0.61248 | 0.152695 | 0.121 | 0.115 | 1 | 3 | Bbx | 0.006 | Young |
| 0.612505 | 0.419673 | 0.117 | 0.111 | 1 | 3 | Gt(ROSA)2( | 0.006 | Young |
| 0.612629 | 0.425524 | 0.032 | 0.027 | 1 | 3 | Sh3bp5l | 0.005 | Young |
| 0.612638 | 0.393421 | 0.048 | 0.043 | 1 | 3 | Timmdc1 | 0.005 | Young |
| 0.613268 | 0.234445 | 0.139 | 0.152 | 1 | 3 | Med10 | -0.013 | Young |
| 0.613274 | 0.561464 | 0.034 | 0.03 | 1 | 3 | Kdm4b | 0.004 | Young |
| 0.613498 | 0.2213 | 0.206 | 0.202 | 1 | 3 | Atg101 | 0.004 | Young |
| 0.613616 | 0.19888 | 0.122 | 0.134 | 1 | 3 | Scaf8 | -0.012 | Young |
| 0.613849 | 0.254728 | 0.22 | 0.215 | 1 | 3 | Pilrb2 | 0.005 | Young |
| 0.613854 | 0.18514 | 0.028 | 0.024 | 1 | 3 | Wdhd1 | 0.004 | Young |
| 0.613892 | 0.157999 | 0.032 | 0.027 | 1 | 3 | Unk | 0.005 | Young |
| 0.613989 | 0.173268 | 0.172 | 0.184 | 1 | 3 | Arl5c | -0.012 | Young |
| 0.614002 | 0.361926 | 0.101 | 0.096 | 1 | 3 | Upf2 | 0.005 | Young |
| 0.614113 | 0.148245 | 0.117 | 0.111 | 1 | 3 | Pctp | 0.006 | Young |
| 0.614642 | 0.148572 | 0.051 | 0.045 | 1 | 3 | Metap1d | 0.006 | Young |
| 0.614712 | 0.066253 | 0.25 | 0.269 | 1 | 3 | Trappc6b | -0.019 | Young |
| 0.614734 | 0.123133 | 0.336 | 0.357 | 1 | 3 | Chic2 | -0.021 | Young |
| 0.61474 | 0.173224 | 0.038 | 0.043 | 1 | 3 | Zfyve21 | -0.005 | Young |
| 0.614894 | 0.065993 | 0.107 | 0.116 | 1 | 3 | Tspan17 | -0.009 | Young |
| 0.614942 | 0.027287 | 0.41 | 0.448 | 1 | 3 | Slc29a3 | -0.038 | Young |
| 0.615021 | 0.130257 | 0.268 | 0.289 | 1 | 3 | Specc1l | -0.021 | Young |
| 0.615315 | 0.28331 | 0.052 | 0.047 | 1 | 3 | Ino80b | 0.005 | Young |
| 0.615389 | 0.039198 | 0.208 | 0.226 | 1 | 3 | Mrps16 | -0.018 | Young |
| 0.61539 | 0.219855 | 0.158 | 0.151 | 1 | 3 | Icosl | 0.007 | Young |
| 0.615656 | 0.146869 | 0.122 | 0.134 | 1 | 3 | Naa60 | -0.012 | Young |
| 0.615847 | 0.250858 | 0.071 | 0.065 | 1 | 3 | Cxxc1 | 0.006 | Young |
| 0.616255 | 0.501958 | 0.029 | 0.025 | 1 | 3 | Vwf | 0.004 | Young |
| 0.616461 | 0.227117 | 0.259 | 0.253 | 1 | 3 | 2310035C2 | 0.006 | Young |

| | | | | | | | | |
|---|---|---|---|---|---|---|---|---|
| 0.616652 | 0.01139 | 0.201 | 0.214 | 1 | 3 | Rest | -0.013 | Young |
| 0.616707 | 0.207811 | 0.056 | 0.062 | 1 | 3 | Tada3 | -0.006 | Young |
| 0.616892 | 0.159267 | 0.158 | 0.17 | 1 | 3 | Vps50 | -0.012 | Young |
| 0.617003 | 0.063101 | 0.245 | 0.237 | 1 | 3 | Ergic3 | 0.008 | Young |
| 0.617234 | 0.197726 | 0.039 | 0.035 | 1 | 3 | Tbc1d4 | 0.004 | Young |
| 0.617277 | 0.326425 | 0.122 | 0.135 | 1 | 3 | Adk | -0.013 | Young |
| 0.617446 | 0.085093 | 0.28 | 0.274 | 1 | 3 | Sp1 | 0.006 | Young |
| 0.617543 | 0.194523 | 0.143 | 0.155 | 1 | 3 | Aggf1 | -0.012 | Young |
| 0.617556 | 0.1161 | 0.271 | 0.271 | 1 | 3 | Kdm5a | 0 | Young |
| 0.617672 | 0.324789 | 0.188 | 0.183 | 1 | 3 | Oser1 | 0.005 | Young |
| 0.617678 | 0.130898 | 0.051 | 0.056 | 1 | 3 | Tubgcp2 | -0.005 | Young |
| 0.618098 | 0.102937 | 0.119 | 0.129 | 1 | 3 | Rsrc1 | -0.01 | Young |
| 0.618289 | 0.17836 | 0.134 | 0.128 | 1 | 3 | Crybg3 | 0.006 | Young |
| 0.618311 | 0.18001 | 0.027 | 0.023 | 1 | 3 | Cmc4 | 0.004 | Young |
| 0.618392 | 0.351549 | 0.076 | 0.07 | 1 | 3 | Fam8a1 | 0.006 | Young |
| 0.618618 | 0.003371 | 0.021 | 0.018 | 1 | 3 | Uevld | 0.003 | Young |
| 0.618965 | 0.164615 | 0.297 | 0.319 | 1 | 3 | Tnrc6a | -0.022 | Young |
| 0.618985 | 0.482948 | 0.019 | 0.023 | 1 | 3 | Ptch1 | -0.004 | Young |
| 0.619294 | 0.131135 | 0.515 | 0.529 | 1 | 3 | Ndufb9 | -0.014 | Young |
| 0.619486 | 0.071299 | 0.177 | 0.168 | 1 | 3 | Supt6 | 0.009 | Young |
| 0.619672 | 0.163639 | 0.072 | 0.08 | 1 | 3 | Naglu | -0.008 | Young |
| 0.619947 | 0.329737 | 0.035 | 0.031 | 1 | 3 | Ankrd66 | 0.004 | Young |
| 0.62018 | 0.623162 | 0.082 | 0.091 | 1 | 3 | Msantd4 | -0.009 | Young |
| 0.620319 | 0.042179 | 0.698 | 0.734 | 1 | 3 | Plek | -0.036 | Young |
| 0.620341 | 0.290289 | 0.028 | 0.024 | 1 | 3 | Eml2 | 0.004 | Young |
| 0.62042 | 0.188795 | 0.124 | 0.116 | 1 | 3 | Ulk1 | 0.008 | Young |
| 0.620494 | 0.085504 | 0.037 | 0.042 | 1 | 3 | Ccdc91 | -0.005 | Young |
| 0.620607 | 0.27621 | 0.043 | 0.038 | 1 | 3 | 1110012L1 | 0.005 | Young |
| 0.6208 | 0.094691 | 0.543 | 0.557 | 1 | 3 | Mpp6 | -0.014 | Young |
| 0.621069 | 0.076066 | 0.067 | 0.074 | 1 | 3 | Gfer | -0.007 | Young |
| 0.621134 | 0.168603 | 0.126 | 0.119 | 1 | 3 | Sdhaf2 | 0.007 | Young |
| 0.62151 | 0.149815 | 0.34 | 0.344 | 1 | 3 | Tec | -0.004 | Young |
| 0.621561 | 0.237939 | 0.034 | 0.03 | 1 | 3 | Taf6 | 0.004 | Young |
| 0.621655 | 0.028923 | 0.247 | 0.265 | 1 | 3 | Krcc1 | -0.018 | Young |
| 0.621737 | 0.201636 | 0.03 | 0.026 | 1 | 3 | Vamp1 | 0.004 | Young |
| 0.621748 | 0.187278 | 0.032 | 0.027 | 1 | 3 | Ddx20 | 0.005 | Young |
| 0.621961 | 0.048233 | 0.266 | 0.283 | 1 | 3 | Rnf187 | -0.017 | Young |
| 0.622033 | 0.039601 | 0.769 | 0.791 | 1 | 3 | Ifrd1 | -0.022 | Young |
| 0.622132 | 0.016144 | 0.048 | 0.054 | 1 | 3 | Tmem263 | -0.006 | Young |
| 0.622178 | 0.04485 | 0.183 | 0.195 | 1 | 3 | Slc15a4 | -0.012 | Young |
| 0.622238 | 0.051013 | 0.02 | 0.024 | 1 | 3 | Rnf217 | -0.004 | Young |
| 0.622465 | 0.116792 | 0.157 | 0.17 | 1 | 3 | H2-T22 | -0.013 | Young |
| 0.622899 | 0.033904 | 0.193 | 0.207 | 1 | 3 | Pqbp1 | -0.014 | Young |
| 0.623133 | 0.018249 | 0.08 | 0.087 | 1 | 3 | Hes6 | -0.007 | Young |
| 0.623331 | 0.150749 | 0.091 | 0.099 | 1 | 3 | Flrt3 | -0.008 | Young |

| | | | | | | | | |
|---|---|---|---|---|---|---|---|---|
| 0.623454 | 0.074893 | 0.187 | 0.202 | 1 | 3 | Hspa4l | -0.015 | Young |
| 0.623734 | 0.028396 | 0.359 | 0.388 | 1 | 3 | Fkbp2 | -0.029 | Young |
| 0.624034 | 0.024919 | 0.202 | 0.216 | 1 | 3 | Napg | -0.014 | Young |
| 0.624046 | 0.042655 | 0.138 | 0.148 | 1 | 3 | Nop16 | -0.01 | Young |
| 0.624396 | 0.159509 | 0.226 | 0.223 | 1 | 3 | Nab2 | 0.003 | Young |
| 0.624504 | 0.26443 | 0.143 | 0.136 | 1 | 3 | Nudt16 | 0.007 | Young |
| 0.624531 | 0.283283 | 0.039 | 0.035 | 1 | 3 | Zfp358 | 0.004 | Young |
| 0.624532 | 0.08907 | 0.038 | 0.043 | 1 | 3 | Abcb8 | -0.005 | Young |
| 0.624622 | 0.244664 | 0.355 | 0.364 | 1 | 3 | Lmo2 | -0.009 | Young |
| 0.624698 | 0.251465 | 0.037 | 0.042 | 1 | 3 | Zfp84 | -0.005 | Young |
| 0.624755 | 0.361614 | 0.013 | 0.016 | 1 | 3 | Casz1 | -0.003 | Young |
| 0.625101 | 0.179612 | 0.038 | 0.033 | 1 | 3 | Serpinb2 | 0.005 | Young |
| 0.625113 | 0.108834 | 0.03 | 0.026 | 1 | 3 | Tmc8 | 0.004 | Young |
| 0.625499 | 0.367728 | 0.148 | 0.142 | 1 | 3 | Man1b1 | 0.006 | Young |
| 0.625677 | 0.098403 | 0.287 | 0.308 | 1 | 3 | Lrrc8c | -0.021 | Young |
| 0.625998 | 0.335647 | 0.16 | 0.157 | 1 | 3 | Rpp21 | 0.003 | Young |
| 0.626308 | 0.087249 | 0.208 | 0.225 | 1 | 3 | Cklf | -0.017 | Young |
| 0.62711 | 0.243533 | 0.159 | 0.153 | 1 | 3 | Zfp871 | 0.006 | Young |
| 0.627119 | 0.222319 | 0.049 | 0.044 | 1 | 3 | Tmem39b | 0.005 | Young |
| 0.627358 | 0.140847 | 0.112 | 0.105 | 1 | 3 | Pdk3 | 0.007 | Young |
| 0.627365 | 0.306081 | 0.09 | 0.084 | 1 | 3 | Phax | 0.006 | Young |
| 0.62739 | 0.199315 | 0.112 | 0.106 | 1 | 3 | Thumpd1 | 0.006 | Young |
| 0.627812 | 0.473508 | 0.038 | 0.043 | 1 | 3 | Ccnyl1 | -0.005 | Young |
| 0.628062 | 0.056426 | 0.241 | 0.253 | 1 | 3 | Dynlt3 | -0.012 | Young |
| 0.628251 | 0.31318 | 0.148 | 0.143 | 1 | 3 | Dnajb4 | 0.005 | Young |
| 0.628318 | 0.21385 | 0.019 | 0.023 | 1 | 3 | Fsd1l | -0.004 | Young |
| 0.628327 | 0.140724 | 0.061 | 0.067 | 1 | 3 | Tubgcp5 | -0.006 | Young |
| 0.628362 | 0.128456 | 0.095 | 0.103 | 1 | 3 | Fam13b | -0.008 | Young |
| 0.628602 | 0.413067 | 0.085 | 0.079 | 1 | 3 | Eif2ak3 | 0.006 | Young |
| 0.628651 | 0.223075 | 0.102 | 0.096 | 1 | 3 | Aasdhppt | 0.006 | Young |
| 0.629609 | 0.132466 | 0.119 | 0.112 | 1 | 3 | Eya3 | 0.007 | Young |
| 0.629899 | 0.064421 | 0.092 | 0.099 | 1 | 3 | Orc4 | -0.007 | Young |
| 0.629918 | 0.102058 | 0.162 | 0.172 | 1 | 3 | Zfp217 | -0.01 | Young |
| 0.630132 | 0.070774 | 0.093 | 0.102 | 1 | 3 | G2e3 | -0.009 | Young |
| 0.630144 | 0.10489 | 0.131 | 0.124 | 1 | 3 | Akap11 | 0.007 | Young |
| 0.630233 | 0.159588 | 0.064 | 0.059 | 1 | 3 | Mir155hg | 0.005 | Young |
| 0.630236 | 0.405491 | 0.086 | 0.08 | 1 | 3 | Slc12a9 | 0.006 | Young |
| 0.63032 | 0.053472 | 0.029 | 0.025 | 1 | 3 | Ncapd2 | 0.004 | Young |
| 0.630599 | 0.071612 | 0.033 | 0.029 | 1 | 3 | E130307A1 | 0.004 | Young |
| 0.631092 | 0.267121 | 0.04 | 0.036 | 1 | 3 | Ccar2 | 0.004 | Young |
| 0.631093 | 0.199975 | 0.091 | 0.085 | 1 | 3 | Elof1 | 0.006 | Young |
| 0.631093 | 0.357159 | 0.234 | 0.233 | 1 | 3 | Cfap43 | 0.001 | Young |
| 0.631121 | 0.106238 | 0.029 | 0.025 | 1 | 3 | Inpp5e | 0.004 | Young |
| 0.631503 | 0.05515 | 0.037 | 0.032 | 1 | 3 | Apba3 | 0.005 | Young |
| 0.631555 | 0.135259 | 0.104 | 0.097 | 1 | 3 | Pcnt | 0.007 | Young |

| | | | | | | | | |
|---|---|---|---|---|---|---|---|---|
| 0.631648 | 0.166727 | 0.125 | 0.137 | 1 | 3 | Hspbp1 | -0.012 | Young |
| 0.632003 | 0.256598 | 0.239 | 0.241 | 1 | 3 | Sft2d2 | -0.002 | Young |
| 0.632379 | 0.16926 | 0.412 | 0.423 | 1 | 3 | Kdm2a | -0.011 | Young |
| 0.632438 | 0.165906 | 0.504 | 0.508 | 1 | 3 | Rsrc2 | -0.004 | Young |
| 0.633384 | 0.088928 | 0.148 | 0.139 | 1 | 3 | Tbc1d15 | 0.009 | Young |
| 0.634275 | 0.209828 | 0.111 | 0.105 | 1 | 3 | Chst12 | 0.006 | Young |
| 0.634323 | 0.04919 | 0.187 | 0.203 | 1 | 3 | Il6st | -0.016 | Young |
| 0.634444 | 0.007541 | 0.511 | 0.544 | 1 | 3 | Caprin1 | -0.033 | Young |
| 0.634581 | 0.140627 | 0.28 | 0.3 | 1 | 3 | Pnpla8 | -0.02 | Young |
| 0.634816 | 0.019753 | 0.264 | 0.284 | 1 | 3 | Ssbp4 | -0.02 | Young |
| 0.635211 | 0.115846 | 0.051 | 0.045 | 1 | 3 | Gpn2 | 0.006 | Young |
| 0.635391 | 0.068819 | 0.379 | 0.409 | 1 | 3 | Klf10 | -0.03 | Young |
| 0.63566 | 0.548086 | 0.136 | 0.133 | 1 | 3 | Sestd1 | 0.003 | Young |
| 0.6362 | 0.114602 | 0.335 | 0.333 | 1 | 3 | Med13 | 0.002 | Young |
| 0.636242 | 0.053986 | 0.275 | 0.293 | 1 | 3 | Setd3 | -0.018 | Young |
| 0.63629 | 0.047302 | 0.783 | 0.814 | 1 | 3 | Zfp36l2 | -0.031 | Young |
| 0.636513 | 0.019024 | 0.571 | 0.616 | 1 | 3 | Clic4 | -0.045 | Young |
| 0.636823 | 0.225791 | 0.039 | 0.044 | 1 | 3 | Gm20707 | -0.005 | Young |
| 0.637949 | 0.24338 | 0.255 | 0.256 | 1 | 3 | Tia1 | -0.001 | Young |
| 0.63812 | 0.122696 | 0.193 | 0.186 | 1 | 3 | Ipo5 | 0.007 | Young |
| 0.638238 | 0.008136 | 0.021 | 0.025 | 1 | 3 | B230354K1 | -0.004 | Young |
| 0.63824 | 0.107488 | 0.282 | 0.277 | 1 | 3 | Adipor2 | 0.005 | Young |
| 0.638545 | 0.190429 | 0.087 | 0.096 | 1 | 3 | Map2k5 | -0.009 | Young |
| 0.638548 | 0.14693 | 0.323 | 0.354 | 1 | 3 | Rp9 | -0.031 | Young |
| 0.638797 | 0.02737 | 0.672 | 0.72 | 1 | 3 | Cct5 | -0.048 | Young |
| 0.638797 | 0.004279 | 0.086 | 0.08 | 1 | 3 | Scap | 0.006 | Young |
| 0.639342 | 0.04999 | 0.876 | 0.907 | 1 | 3 | Atp5e | -0.031 | Young |
| 0.639458 | 0.071832 | 0.253 | 0.269 | 1 | 3 | Ccdc124 | -0.016 | Young |
| 0.639465 | 0.046724 | 0.49 | 0.539 | 1 | 3 | Rtn3 | -0.049 | Young |
| 0.639562 | 0.033022 | 0.051 | 0.056 | 1 | 3 | Cep128 | -0.005 | Young |
| 0.639852 | 0.115356 | 0.049 | 0.044 | 1 | 3 | Acvr1b | 0.005 | Young |
| 0.63997 | 0.122143 | 0.04 | 0.045 | 1 | 3 | Tnni2 | -0.005 | Young |
| 0.640147 | 0.074621 | 0.269 | 0.262 | 1 | 3 | Snw1 | 0.007 | Young |
| 0.640515 | 0.047087 | 0.332 | 0.36 | 1 | 3 | Dock7 | -0.028 | Young |
| 0.640646 | 0.07105 | 0.119 | 0.112 | 1 | 3 | Fbrsl1 | 0.007 | Young |
| 0.641351 | 0.118962 | 0.282 | 0.301 | 1 | 3 | Ttc39b | -0.019 | Young |
| 0.64147 | 0.13328 | 0.13 | 0.141 | 1 | 3 | Hsd17b10 | -0.011 | Young |
| 0.642107 | 0.042152 | 0.558 | 0.593 | 1 | 3 | Twf2 | -0.035 | Young |
| 0.642952 | 0.197456 | 0.188 | 0.203 | 1 | 3 | Aim2 | -0.015 | Young |
| 0.643118 | 0.078021 | 0.616 | 0.631 | 1 | 3 | Rps6ka3 | -0.015 | Young |
| 0.643122 | 0.19085 | 0.259 | 0.263 | 1 | 3 | Timm10b | -0.004 | Young |
| 0.643312 | 0.1559 | 0.126 | 0.136 | 1 | 3 | Hspb11 | -0.01 | Young |
| 0.643451 | 0.032163 | 0.261 | 0.282 | 1 | 3 | Smarce1 | -0.021 | Young |
| 0.643527 | 0.162449 | 0.074 | 0.081 | 1 | 3 | Rcbtb2 | -0.007 | Young |
| 0.643656 | 0.395335 | 0.096 | 0.091 | 1 | 3 | Megf8 | 0.005 | Young |

| | | | | | | | | |
|---|---|---|---|---|---|---|---|---|
| 0.643741 | 0.233408 | 0.511 | 0.536 | 1 | 3 | Hmox1 | -0.025 | Young |
| 0.644024 | 0.163415 | 0.032 | 0.036 | 1 | 3 | Pycrl | -0.004 | Young |
| 0.644555 | 0.154414 | 0.03 | 0.035 | 1 | 3 | Slx1b | -0.005 | Young |
| 0.644987 | 0.439803 | 0.246 | 0.247 | 1 | 3 | Tmem219 | -0.001 | Young |
| 0.64547 | 0.056806 | 0.133 | 0.145 | 1 | 3 | Pcif1 | -0.012 | Young |
| 0.645499 | 0.179761 | 0.076 | 0.082 | 1 | 3 | Mbtps2 | -0.006 | Young |
| 0.645561 | 0.136105 | 0.316 | 0.317 | 1 | 3 | Nin | -0.001 | Young |
| 0.645824 | 0.226557 | 0.337 | 0.345 | 1 | 3 | Stk24 | -0.008 | Young |
| 0.645835 | 0.094382 | 0.223 | 0.24 | 1 | 3 | Nap1l4 | -0.017 | Young |
| 0.646023 | 0.174089 | 0.014 | 0.017 | 1 | 3 | Gm10053 | -0.003 | Young |
| 0.646299 | 0.526518 | 0.12 | 0.117 | 1 | 3 | Cabin1 | 0.003 | Young |
| 0.646419 | 0.205647 | 0.265 | 0.265 | 1 | 3 | Mrps24 | 0 | Young |
| 0.646668 | 0.295058 | 0.11 | 0.105 | 1 | 3 | Mmp9 | 0.005 | Young |
| 0.647008 | 0.029704 | 0.104 | 0.112 | 1 | 3 | Usp33 | -0.008 | Young |
| 0.647462 | 0.000637 | 0.093 | 0.102 | 1 | 3 | Adpgk | -0.009 | Young |
| 0.647581 | 0.036106 | 0.663 | 0.674 | 1 | 3 | Capns1 | -0.011 | Young |
| 0.647801 | 0.11576 | 0.014 | 0.017 | 1 | 3 | Setd4 | -0.003 | Young |
| 0.648157 | 0.081026 | 0.014 | 0.017 | 1 | 3 | Zfp141 | -0.003 | Young |
| 0.648254 | 0.119954 | 0.116 | 0.11 | 1 | 3 | Zadh2 | 0.006 | Young |
| 0.648513 | 0.125242 | 0.014 | 0.017 | 1 | 3 | Grhpr | -0.003 | Young |
| 0.648825 | 0.10467 | 0.097 | 0.091 | 1 | 3 | Borcs8 | 0.006 | Young |
| 0.648869 | 0.047226 | 0.014 | 0.017 | 1 | 3 | Cbwd1 | -0.003 | Young |
| 0.649637 | 0.093275 | 0.494 | 0.495 | 1 | 3 | Idh1 | -0.001 | Young |
| 0.649689 | 0.082961 | 0.25 | 0.271 | 1 | 3 | Map2k7 | -0.021 | Young |
| 0.6497 | 0.1878 | 0.073 | 0.08 | 1 | 3 | Ifih1 | -0.007 | Young |
| 0.649778 | 0.248857 | 0.067 | 0.062 | 1 | 3 | Cfap97 | 0.005 | Young |
| 0.649938 | 0.061215 | 0.529 | 0.556 | 1 | 3 | Pfkp | -0.027 | Young |
| 0.65016 | 0.230204 | 0.207 | 0.207 | 1 | 3 | Chmp1b | 0 | Young |
| 0.650472 | 0.114858 | 0.174 | 0.185 | 1 | 3 | Csnk1g1 | -0.011 | Young |
| 0.650677 | 0.012811 | 0.396 | 0.423 | 1 | 3 | Tet2 | -0.027 | Young |
| 0.650841 | 0.309183 | 0.04 | 0.045 | 1 | 3 | Rnf113a2 | -0.005 | Young |
| 0.651082 | 0.007043 | 0.703 | 0.747 | 1 | 3 | Smdt1 | -0.044 | Young |
| 0.65109 | 0.062371 | 0.24 | 0.254 | 1 | 3 | Hbp1 | -0.014 | Young |
| 0.651439 | 0.20097 | 0.242 | 0.26 | 1 | 3 | Uap1 | -0.018 | Young |
| 0.651981 | 0.287101 | 0.078 | 0.073 | 1 | 3 | Dusp6 | 0.005 | Young |
| 0.652076 | 0.301898 | 0.014 | 0.017 | 1 | 3 | Pecam1 | -0.003 | Young |
| 0.65211 | 0.27169 | 0.042 | 0.047 | 1 | 3 | Unc13d | -0.005 | Young |
| 0.652127 | 0.053986 | 0.155 | 0.166 | 1 | 3 | Tlk1 | -0.011 | Young |
| 0.652498 | 0.090889 | 0.096 | 0.104 | 1 | 3 | Chpf | -0.008 | Young |
| 0.654156 | 0.411279 | 0.021 | 0.025 | 1 | 3 | 1700056E2 | -0.004 | Young |
| 0.654204 | 0.072329 | 0.023 | 0.026 | 1 | 3 | Ghdc | -0.003 | Young |
| 0.654571 | 0.198558 | 0.186 | 0.179 | 1 | 3 | Gtf2a2 | 0.007 | Young |
| 0.654886 | 0.016487 | 0.066 | 0.072 | 1 | 3 | Pou5f2 | -0.006 | Young |
| 0.655411 | 0.190051 | 0.217 | 0.211 | 1 | 3 | Cyb5r3 | 0.006 | Young |
| 0.655415 | 0.06459 | 0.226 | 0.243 | 1 | 3 | Tfg | -0.017 | Young |

| | | | | | | | | |
|---|---|---|---|---|---|---|---|---|
| 0.655447 | 0.069825 | 0.26 | 0.253 | 1 | 3 | Trip11 | 0.007 | Young |
| 0.6555 | 0.110426 | 0.255 | 0.251 | 1 | 3 | Snd1 | 0.004 | Young |
| 0.655709 | 0.008013 | 0.14 | 0.152 | 1 | 3 | Phf11b | -0.012 | Young |
| 0.656191 | 0.191431 | 0.182 | 0.196 | 1 | 3 | Gps1 | -0.014 | Young |
| 0.656255 | 0.15303 | 0.549 | 0.585 | 1 | 3 | Nsa2 | -0.036 | Young |
| 0.656733 | 0.173661 | 0.104 | 0.112 | 1 | 3 | Lrrc42 | -0.008 | Young |
| 0.657078 | 0.229572 | 0.014 | 0.017 | 1 | 3 | Gas8 | -0.003 | Young |
| 0.657197 | 0.238956 | 0.069 | 0.065 | 1 | 3 | 4921524J1 | 0.004 | Young |
| 0.657283 | 0.196005 | 0.068 | 0.075 | 1 | 3 | Pcyt2 | -0.007 | Young |
| 0.657455 | 0.037187 | 0.769 | 0.798 | 1 | 3 | Cox6c | -0.029 | Young |
| 0.657547 | 0.121048 | 0.144 | 0.155 | 1 | 3 | Mdp1 | -0.011 | Young |
| 0.657612 | 0.154157 | 0.023 | 0.026 | 1 | 3 | Iqcb1 | -0.003 | Young |
| 0.657749 | 0.093964 | 0.332 | 0.333 | 1 | 3 | Capn1 | -0.001 | Young |
| 0.657772 | 0.146833 | 0.154 | 0.165 | 1 | 3 | 0610030E2 | -0.011 | Young |
| 0.658499 | 0.071705 | 0.301 | 0.325 | 1 | 3 | Nrbp1 | -0.024 | Young |
| 0.659083 | 0.096554 | 0.043 | 0.048 | 1 | 3 | Rbm38 | -0.005 | Young |
| 0.659085 | 0.074542 | 0.117 | 0.125 | 1 | 3 | Fam53c | -0.008 | Young |
| 0.659136 | 0.045617 | 0.033 | 0.037 | 1 | 3 | Bcl9 | -0.004 | Young |
| 0.659727 | 0.087658 | 0.287 | 0.307 | 1 | 3 | Ccng1 | -0.02 | Young |
| 0.660113 | 0.02178 | 0.691 | 0.713 | 1 | 3 | Grb2 | -0.022 | Young |
| 0.660163 | 0.051799 | 0.918 | 0.938 | 1 | 3 | Rps15a | -0.02 | Young |
| 0.660489 | 0.01531 | 0.295 | 0.317 | 1 | 3 | Akirin2 | -0.022 | Young |
| 0.661074 | 0.20567 | 0.068 | 0.063 | 1 | 3 | Mpv17 | 0.005 | Young |
| 0.66113 | 0.115618 | 0.295 | 0.292 | 1 | 3 | Ccnd3 | 0.003 | Young |
| 0.661267 | 0.094523 | 0.23 | 0.246 | 1 | 3 | Ubqln1 | -0.016 | Young |
| 0.661415 | 0.087611 | 0.265 | 0.263 | 1 | 3 | Paf1 | 0.002 | Young |
| 0.661539 | 0.140777 | 0.024 | 0.027 | 1 | 3 | Gm15448 | -0.003 | Young |
| 0.661893 | 0.016209 | 0.593 | 0.634 | 1 | 3 | Plekho1 | -0.041 | Young |
| 0.662277 | 0.038513 | 0.097 | 0.105 | 1 | 3 | Osm | -0.008 | Young |
| 0.662294 | 0.001067 | 0.015 | 0.018 | 1 | 3 | Zfp768 | -0.003 | Young |
| 0.662294 | 0.059141 | 0.015 | 0.018 | 1 | 3 | Gramd3 | -0.003 | Young |
| 0.662374 | 0.089305 | 0.024 | 0.027 | 1 | 3 | Rnf123 | -0.003 | Young |
| 0.662715 | 0.137387 | 0.242 | 0.262 | 1 | 3 | Maea | -0.02 | Young |
| 0.662962 | 0.302047 | 0.165 | 0.164 | 1 | 3 | Kank3 | 0.001 | Young |
| 0.663491 | 0.102327 | 0.705 | 0.736 | 1 | 3 | Ddx17 | -0.031 | Young |
| 0.663602 | 0.213827 | 0.063 | 0.069 | 1 | 3 | Trmt13 | -0.006 | Young |
| 0.663626 | 0.134188 | 0.168 | 0.182 | 1 | 3 | Inf2 | -0.014 | Young |
| 0.663711 | 0.184353 | 0.213 | 0.209 | 1 | 3 | Luc7l | 0.004 | Young |
| 0.66373 | 0.186316 | 0.109 | 0.103 | 1 | 3 | Gpr155 | 0.006 | Young |
| 0.663891 | 0.007494 | 0.139 | 0.148 | 1 | 3 | Glod4 | -0.009 | Young |
| 0.663931 | 0.314694 | 0.133 | 0.128 | 1 | 3 | Ppil4 | 0.005 | Young |
| 0.664092 | 0.533786 | 0.1 | 0.094 | 1 | 3 | Il15 | 0.006 | Young |
| 0.664638 | 0.099482 | 0.208 | 0.223 | 1 | 3 | Dynll2 | -0.015 | Young |
| 0.664794 | 0.001657 | 0.064 | 0.07 | 1 | 3 | Tfam | -0.006 | Young |
| 0.664865 | 0.481267 | 0.057 | 0.053 | 1 | 3 | Guf1 | 0.004 | Young |

| | | | | | | | | |
|---|---|---|---|---|---|---|---|---|
| 0.665103 | 0.188795 | 0.35 | 0.356 | 1 | 3 | Rbm5 | -0.006 | Young |
| 0.665173 | 0.100402 | 0.042 | 0.047 | 1 | 3 | Pyroxd1 | -0.005 | Young |
| 0.665266 | 0.265242 | 0.091 | 0.086 | 1 | 3 | Ppwd1 | 0.005 | Young |
| 0.665408 | 0.083795 | 0.015 | 0.018 | 1 | 3 | Pms2 | -0.003 | Young |
| 0.665442 | 0.214384 | 0.164 | 0.159 | 1 | 3 | Dpf2 | 0.005 | Young |
| 0.665615 | 0.153134 | 0.277 | 0.299 | 1 | 3 | Cdc5l | -0.022 | Young |
| 0.665719 | 0.042541 | 0.024 | 0.027 | 1 | 3 | Tmem42 | -0.003 | Young |
| 0.665952 | 0.056592 | 0.226 | 0.244 | 1 | 3 | Inpp5f | -0.018 | Young |
| 0.666168 | 0.12321 | 0.332 | 0.325 | 1 | 3 | Smim14 | 0.007 | Young |
| 0.66622 | 0.108619 | 0.372 | 0.404 | 1 | 3 | Fcho2 | -0.032 | Young |
| 0.666272 | 0.115224 | 0.106 | 0.1 | 1 | 3 | Klhl18 | 0.006 | Young |
| 0.666653 | 0.07823 | 0.008 | 0.01 | 1 | 3 | Taco1 | -0.002 | Young |
| 0.666676 | 0.171417 | 0.172 | 0.189 | 1 | 3 | Cmip | -0.017 | Young |
| 0.666681 | 0.028595 | 0.396 | 0.419 | 1 | 3 | Herpud1 | -0.023 | Young |
| 0.666755 | 0.21518 | 0.15 | 0.146 | 1 | 3 | Dhx16 | 0.004 | Young |
| 0.667343 | 0.030968 | 0.293 | 0.309 | 1 | 3 | Slbp | -0.016 | Young |
| 0.667359 | 0.206317 | 0.056 | 0.062 | 1 | 3 | B230307C2 | -0.006 | Young |
| 0.667493 | 0.170022 | 0.229 | 0.226 | 1 | 3 | Taf6l | 0.003 | Young |
| 0.667714 | 0.14493 | 0.077 | 0.072 | 1 | 3 | Arl2 | 0.005 | Young |
| 0.667795 | 0.013282 | 0.499 | 0.501 | 1 | 3 | Lipa | -0.002 | Young |
| 0.668445 | 0.252302 | 0.176 | 0.173 | 1 | 3 | Tmem43 | 0.003 | Young |
| 0.668571 | 0.151886 | 0.008 | 0.01 | 1 | 3 | Wwc1 | -0.002 | Young |
| 0.668709 | 0.162107 | 0.106 | 0.115 | 1 | 3 | Pdcl | -0.009 | Young |
| 0.668929 | 0.049107 | 0.307 | 0.33 | 1 | 3 | Rassf5 | -0.023 | Young |
| 0.669012 | 0.103047 | 0.242 | 0.239 | 1 | 3 | Numa1 | 0.003 | Young |
| 0.669075 | 0.333363 | 0.11 | 0.105 | 1 | 3 | Uckl1 | 0.005 | Young |
| 0.669393 | 0.1394 | 0.169 | 0.182 | 1 | 3 | Rybp | -0.013 | Young |
| 0.66946 | 0.30578 | 0.148 | 0.145 | 1 | 3 | Ypel3 | 0.003 | Young |
| 0.669633 | 0.010852 | 0.076 | 0.07 | 1 | 3 | Ust | 0.006 | Young |
| 0.669735 | 0.148415 | 0.247 | 0.241 | 1 | 3 | Idh3g | 0.006 | Young |
| 0.669909 | 0.008922 | 0.048 | 0.053 | 1 | 3 | Fars2 | -0.005 | Young |
| 0.669955 | 0.018737 | 0.069 | 0.075 | 1 | 3 | Ccdc69 | -0.006 | Young |
| 0.670011 | 0.129754 | 0.008 | 0.01 | 1 | 3 | Gm9796 | -0.002 | Young |
| 0.670011 | 0.020535 | 0.008 | 0.01 | 1 | 3 | Hist1h2ae | -0.002 | Young |
| 0.670041 | 0.228778 | 0.073 | 0.08 | 1 | 3 | Casp9 | -0.007 | Young |
| 0.670321 | 0.097603 | 0.28 | 0.299 | 1 | 3 | Ibtk | -0.019 | Young |
| 0.670577 | 0.044388 | 0.58 | 0.616 | 1 | 3 | Eif5 | -0.036 | Young |
| 0.670871 | 0.041575 | 0.616 | 0.658 | 1 | 3 | Gpi1 | -0.042 | Young |
| 0.670883 | 0.179438 | 0.124 | 0.133 | 1 | 3 | Hcfc2 | -0.009 | Young |
| 0.670972 | 0.34962 | 0.008 | 0.01 | 1 | 3 | Gm6710 | -0.002 | Young |
| 0.671305 | 0.104065 | 0.015 | 0.018 | 1 | 3 | Mmp14 | -0.003 | Young |
| 0.672496 | 0.150181 | 0.068 | 0.063 | 1 | 3 | Iffo1 | 0.005 | Young |
| 0.672697 | 0.124386 | 0.09 | 0.084 | 1 | 3 | Klrb1a | 0.006 | Young |
| 0.672765 | 0.019419 | 0.511 | 0.523 | 1 | 3 | Ankrd11 | -0.012 | Young |
| 0.67282 | 0.07443 | 0.284 | 0.301 | 1 | 3 | Marveld1 | -0.017 | Young |

| | | | | | | | | |
|---|---|---|---|---|---|---|---|---|
| 0.672965 | 0.160945 | 0.135 | 0.146 | 1 | 3 | Asnsd1 | -0.011 | Young |
| 0.673203 | 0.092357 | 0.265 | 0.287 | 1 | 3 | Golga7 | -0.022 | Young |
| 0.673278 | 0.086118 | 0.125 | 0.134 | 1 | 3 | Unc119b | -0.009 | Young |
| 0.673297 | 0.286215 | 0.052 | 0.048 | 1 | 3 | Cryzl2 | 0.004 | Young |
| 0.673341 | 0.309051 | 0.21 | 0.205 | 1 | 3 | Arhgap6 | 0.005 | Young |
| 0.673391 | 0.181695 | 0.015 | 0.018 | 1 | 3 | Zfp949 | -0.003 | Young |
| 0.673457 | 0.115499 | 0.155 | 0.152 | 1 | 3 | Zfp131 | 0.003 | Young |
| 0.673639 | 0.093636 | 0.48 | 0.481 | 1 | 3 | Ctdsp2 | -0.001 | Young |
| 0.673679 | 0.017866 | 0.506 | 0.538 | 1 | 3 | Sdha | -0.032 | Young |
| 0.673813 | 0.127007 | 0.261 | 0.278 | 1 | 3 | Prr5l | -0.017 | Young |
| 0.673897 | 0.223203 | 0.023 | 0.026 | 1 | 3 | Bud13 | -0.003 | Young |
| 0.674317 | 0.335488 | 0.138 | 0.134 | 1 | 3 | Hcar2 | 0.004 | Young |
| 0.674383 | 0.050428 | 0.072 | 0.067 | 1 | 3 | Hbegf | 0.005 | Young |
| 0.674606 | 0.431146 | 0.034 | 0.038 | 1 | 3 | Mlkl | -0.004 | Young |
| 0.674713 | 0.201351 | 0.077 | 0.084 | 1 | 3 | 2410002F2 | -0.007 | Young |
| 0.674944 | 0.259457 | 0.117 | 0.128 | 1 | 3 | Iars2 | -0.011 | Young |
| 0.67501 | 0.178227 | 0.165 | 0.177 | 1 | 3 | Vps18 | -0.012 | Young |
| 0.675141 | 0.001617 | 0.487 | 0.516 | 1 | 3 | Blnk | -0.029 | Young |
| 0.675471 | 0.031028 | 0.064 | 0.06 | 1 | 3 | Fads1 | 0.004 | Young |
| 0.675758 | 0.066058 | 0.386 | 0.425 | 1 | 3 | 1810058I2 | -0.039 | Young |
| 0.675965 | 0.19582 | 0.1 | 0.109 | 1 | 3 | Uba3 | -0.009 | Young |
| 0.676 | 0.17136 | 0.138 | 0.147 | 1 | 3 | Vars | -0.009 | Young |
| 0.676185 | 0.048866 | 0.092 | 0.099 | 1 | 3 | Lipo3 | -0.007 | Young |
| 0.676446 | 0.289821 | 0.035 | 0.039 | 1 | 3 | Tfpt | -0.004 | Young |
| 0.676533 | 0.034612 | 0.355 | 0.37 | 1 | 3 | Sumo1 | -0.015 | Young |
| 0.676739 | 0.025441 | 0.364 | 0.388 | 1 | 3 | Arl8b | -0.024 | Young |
| 0.676814 | 0.014642 | 0.222 | 0.233 | 1 | 3 | Atp6v1d | -0.011 | Young |
| 0.67694 | 0.166514 | 0.074 | 0.081 | 1 | 3 | Cep63 | -0.007 | Young |
| 0.676972 | 0.506276 | 0.048 | 0.044 | 1 | 3 | Lix1l | 0.004 | Young |
| 0.677018 | 0.071793 | 0.679 | 0.689 | 1 | 3 | Fgfr1 | -0.01 | Young |
| 0.67717 | 0.038649 | 0.482 | 0.515 | 1 | 3 | Zfp207 | -0.033 | Young |
| 0.677375 | 0.463485 | 0.051 | 0.047 | 1 | 3 | Galt | 0.004 | Young |
| 0.677398 | 0.11648 | 0.117 | 0.125 | 1 | 3 | Adora2a | -0.008 | Young |
| 0.677471 | 0.311864 | 0.198 | 0.216 | 1 | 3 | Slc38a9 | -0.018 | Young |
| 0.678088 | 0.037085 | 0.17 | 0.183 | 1 | 3 | Pisd | -0.013 | Young |
| 0.678362 | 0.156307 | 0.434 | 0.444 | 1 | 3 | Pnisr | -0.01 | Young |
| 0.678683 | 0.15741 | 0.145 | 0.158 | 1 | 3 | 2310009B1 | -0.013 | Young |
| 0.678937 | 0.117086 | 0.254 | 0.271 | 1 | 3 | Mrpl12 | -0.017 | Young |
| 0.678953 | 0.389682 | 0.111 | 0.108 | 1 | 3 | Fzr1 | 0.003 | Young |
| 0.678964 | 0.028254 | 0.824 | 0.845 | 1 | 3 | Ubl3 | -0.021 | Young |
| 0.678981 | 0.124239 | 0.326 | 0.327 | 1 | 3 | Luzp1 | -0.001 | Young |
| 0.679254 | 0.021187 | 0.35 | 0.372 | 1 | 3 | Asph | -0.022 | Young |
| 0.679379 | 0.18363 | 0.054 | 0.06 | 1 | 3 | Tstd2 | -0.006 | Young |
| 0.679768 | 0.180084 | 0.292 | 0.289 | 1 | 3 | Qdpr | 0.003 | Young |
| 0.68005 | 0.200961 | 0.052 | 0.048 | 1 | 3 | Tinf2 | 0.004 | Young |

| | | | | | | | | |
|---|---|---|---|---|---|---|---|---|
| 0.680088 | 0.063377 | 0.114 | 0.122 | 1 | 3 | Maml1 | -0.008 | Young |
| 0.680094 | 0.387449 | 0.057 | 0.053 | 1 | 3 | Ints10 | 0.004 | Young |
| 0.680388 | 0.056725 | 0.432 | 0.432 | 1 | 3 | Nol7 | 0 | Young |
| 0.68042 | 0.070292 | 0.395 | 0.389 | 1 | 3 | Taf10 | 0.006 | Young |
| 0.680587 | 0.129558 | 0.096 | 0.103 | 1 | 3 | Traf6 | -0.007 | Young |
| 0.680719 | 0.259436 | 0.098 | 0.093 | 1 | 3 | Tbce | 0.005 | Young |
| 0.680833 | 0.265566 | 0.016 | 0.019 | 1 | 3 | Gpt | -0.003 | Young |
| 0.680977 | 0.079393 | 0.087 | 0.093 | 1 | 3 | Sbk1 | -0.006 | Young |
| 0.681016 | 0.289551 | 0.081 | 0.076 | 1 | 3 | Anks3 | 0.005 | Young |
| 0.681552 | 0.118585 | 0.312 | 0.331 | 1 | 3 | Pkn2 | -0.019 | Young |
| 0.681846 | 0.123829 | 0.371 | 0.399 | 1 | 3 | Nsd1 | -0.028 | Young |
| 0.68265 | 0.279921 | 0.025 | 0.029 | 1 | 3 | Stbd1 | -0.004 | Young |
| 0.682842 | 0.227683 | 0.121 | 0.116 | 1 | 3 | Fam96b | 0.005 | Young |
| 0.683898 | 0.213842 | 0.179 | 0.174 | 1 | 3 | Bin3 | 0.005 | Young |
| 0.684243 | 0.391368 | 0.049 | 0.045 | 1 | 3 | Tmem94 | 0.004 | Young |
| 0.684303 | 0.002442 | 0.025 | 0.029 | 1 | 3 | Akip1 | -0.004 | Young |
| 0.684591 | 0.156992 | 0.278 | 0.302 | 1 | 3 | Plekhm3 | -0.024 | Young |
| 0.684619 | 0.185705 | 0.064 | 0.07 | 1 | 3 | Ints14 | -0.006 | Young |
| 0.685109 | 0.126268 | 0.241 | 0.263 | 1 | 3 | 0610012G( | -0.022 | Young |
| 0.685343 | 0.184453 | 0.302 | 0.302 | 1 | 3 | Slc12a6 | 0 | Young |
| 0.686977 | 0.115888 | 0.302 | 0.321 | 1 | 3 | Slc16a3 | -0.019 | Young |
| 0.686995 | 0.254563 | 0.067 | 0.073 | 1 | 3 | Surf1 | -0.006 | Young |
| 0.687612 | 0.443981 | 0.039 | 0.036 | 1 | 3 | Zfp959 | 0.003 | Young |
| 0.687755 | 0.026962 | 0.907 | 0.92 | 1 | 3 | Ybx1 | -0.013 | Young |
| 0.688349 | 0.113313 | 0.066 | 0.072 | 1 | 3 | Coq10a | -0.006 | Young |
| 0.68835 | 0.095872 | 0.062 | 0.057 | 1 | 3 | Zfp597 | 0.005 | Young |
| 0.688367 | 0.241776 | 0.331 | 0.344 | 1 | 3 | Phf3 | -0.013 | Young |
| 0.688872 | 0.067052 | 0.386 | 0.391 | 1 | 3 | Psmb6 | -0.005 | Young |
| 0.689124 | 0.128702 | 0.114 | 0.123 | 1 | 3 | Shkbp1 | -0.009 | Young |
| 0.689145 | 0.083173 | 0.035 | 0.039 | 1 | 3 | Tbc1d32 | -0.004 | Young |
| 0.689185 | 0.006867 | 0.061 | 0.066 | 1 | 3 | Tnfaip8l1 | -0.005 | Young |
| 0.689285 | 0.309325 | 0.078 | 0.074 | 1 | 3 | Slx4ip | 0.004 | Young |
| 0.689365 | 0.123391 | 0.355 | 0.358 | 1 | 3 | Mvp | -0.003 | Young |
| 0.689365 | 0.192721 | 0.116 | 0.11 | 1 | 3 | Pold3 | 0.006 | Young |
| 0.689562 | 0.081093 | 0.278 | 0.299 | 1 | 3 | Polr2m | -0.021 | Young |
| 0.690146 | 0.204294 | 0.116 | 0.124 | 1 | 3 | Fut8 | -0.008 | Young |
| 0.690581 | 0.049278 | 0.045 | 0.05 | 1 | 3 | Cep85 | -0.005 | Young |
| 0.690937 | 0.44346 | 0.035 | 0.032 | 1 | 3 | Grpel2 | 0.003 | Young |
| 0.691151 | 0.136103 | 0.072 | 0.078 | 1 | 3 | Polr2d | -0.006 | Young |
| 0.691285 | 0.88492 | 0.032 | 0.029 | 1 | 3 | Surf2 | 0.003 | Young |
| 0.691443 | 0.411062 | 0.074 | 0.07 | 1 | 3 | Tmub1 | 0.004 | Young |
| 0.691746 | 0.211769 | 0.097 | 0.104 | 1 | 3 | Stat2 | -0.007 | Young |
| 0.691778 | 0.200195 | 0.076 | 0.082 | 1 | 3 | Uty | -0.006 | Young |
| 0.692119 | 0.075587 | 0.174 | 0.186 | 1 | 3 | Got2 | -0.012 | Young |
| 0.692533 | 0.369823 | 0.054 | 0.05 | 1 | 3 | 1190002N1 | 0.004 | Young |

| | | | | | | | | |
|---|---|---|---|---|---|---|---|---|
| 0.692585 | 0.332259 | 0.096 | 0.104 | 1 | 3 | Patl1 | -0.008 | Young |
| 0.692604 | 0.088614 | 0.293 | 0.311 | 1 | 3 | Ndufa5 | -0.018 | Young |
| 0.692757 | 0.029915 | 0.076 | 0.07 | 1 | 3 | Traf3ip3 | 0.006 | Young |
| 0.693102 | 0.350153 | 0.043 | 0.039 | 1 | 3 | Slc22a15 | 0.004 | Young |
| 0.693458 | 0.03994 | 0.109 | 0.103 | 1 | 3 | H2-Ke6 | 0.006 | Young |
| 0.693663 | 0.284264 | 0.082 | 0.088 | 1 | 3 | Mis12 | -0.006 | Young |
| 0.693777 | 0.104566 | 0.117 | 0.125 | 1 | 3 | Zdhhc7 | -0.008 | Young |
| 0.693801 | 0.141485 | 0.037 | 0.041 | 1 | 3 | Dcun1d2 | -0.004 | Young |
| 0.694115 | 0.011482 | 0.038 | 0.042 | 1 | 3 | Eng | -0.004 | Young |
| 0.6944 | 0.098742 | 0.086 | 0.081 | 1 | 3 | Phtf2 | 0.005 | Young |
| 0.694525 | 0.012813 | 0.867 | 0.895 | 1 | 3 | Ubc | -0.028 | Young |
| 0.694645 | 0.129536 | 0.135 | 0.145 | 1 | 3 | Zcchc2 | -0.01 | Young |
| 0.694764 | 0.105951 | 0.027 | 0.03 | 1 | 3 | Pigh | -0.003 | Young |
| 0.695396 | 0.330457 | 0.016 | 0.019 | 1 | 3 | Fam26f | -0.003 | Young |
| 0.695528 | 0.342769 | 0.098 | 0.094 | 1 | 3 | Sirt1 | 0.004 | Young |
| 0.695578 | 0.233535 | 0.027 | 0.03 | 1 | 3 | Mllt11 | -0.003 | Young |
| 0.696022 | 0.471722 | 0.034 | 0.038 | 1 | 3 | A430106G1 | -0.004 | Young |
| 0.696715 | 0.429551 | 0.04 | 0.037 | 1 | 3 | Dbr1 | 0.003 | Young |
| 0.69683 | 0.180099 | 0.098 | 0.093 | 1 | 3 | Kmt5c | 0.005 | Young |
| 0.696906 | 0.652915 | 0.035 | 0.032 | 1 | 3 | Plekha8 | 0.003 | Young |
| 0.69757 | 0.035868 | 0.158 | 0.152 | 1 | 3 | Myo18a | 0.006 | Young |
| 0.697964 | 0.320078 | 0.203 | 0.203 | 1 | 3 | Sec14l1 | 0 | Young |
| 0.698115 | 0.052197 | 0.436 | 0.46 | 1 | 3 | Tmem167 | -0.024 | Young |
| 0.698196 | 0.390645 | 0.13 | 0.127 | 1 | 3 | Cbx5 | 0.003 | Young |
| 0.6982 | 0.026356 | 0.009 | 0.011 | 1 | 3 | Gm47815 | -0.002 | Young |
| 0.698294 | 0.323074 | 0.095 | 0.091 | 1 | 3 | Kansl2 | 0.004 | Young |
| 0.698566 | 0.067945 | 0.027 | 0.03 | 1 | 3 | Gatb | -0.003 | Young |
| 0.698726 | 0.081133 | 0.607 | 0.627 | 1 | 3 | Rock2 | -0.02 | Young |
| 0.699036 | 0.027686 | 0.109 | 0.115 | 1 | 3 | Ankrd10 | -0.006 | Young |
| 0.699304 | 0.158747 | 0.044 | 0.041 | 1 | 3 | Tbccd1 | 0.003 | Young |
| 0.699572 | 0.042765 | 0.389 | 0.405 | 1 | 3 | Sltm | -0.016 | Young |
| 0.699781 | 0.166191 | 0.331 | 0.333 | 1 | 3 | Fmnl1 | -0.002 | Young |
| 0.699922 | 0.028486 | 0.311 | 0.331 | 1 | 3 | Zranb2 | -0.02 | Young |
| 0.700054 | 0.467314 | 0.043 | 0.048 | 1 | 3 | Msto1 | -0.005 | Young |
| 0.700078 | 0.240242 | 0.258 | 0.259 | 1 | 3 | Ino80d | -0.001 | Young |
| 0.700662 | 0.102815 | 0.193 | 0.208 | 1 | 3 | Zdhhc5 | -0.015 | Young |
| 0.700728 | 0.165024 | 0.466 | 0.495 | 1 | 3 | Nipbl | -0.029 | Young |
| 0.7009 | 0.180539 | 0.057 | 0.053 | 1 | 3 | Pot1b | 0.004 | Young |
| 0.701177 | 0.36308 | 0.059 | 0.065 | 1 | 3 | Urgcp | -0.006 | Young |
| 0.701422 | 0.169043 | 0.101 | 0.097 | 1 | 3 | Nsmce3 | 0.004 | Young |
| 0.70153 | 0.196732 | 0.045 | 0.05 | 1 | 3 | Ndufaf5 | -0.005 | Young |
| 0.701534 | 0.040616 | 0.102 | 0.097 | 1 | 3 | Klf11 | 0.005 | Young |
| 0.701857 | 0.107496 | 0.009 | 0.011 | 1 | 3 | Pomt1 | -0.002 | Young |
| 0.701925 | 0.103369 | 0.059 | 0.065 | 1 | 3 | Fbxo7 | -0.006 | Young |
| 0.702103 | 0.309136 | 0.027 | 0.03 | 1 | 3 | Nif3l1 | -0.003 | Young |

| | | | | | | | | |
|---|---|---|---|---|---|---|---|---|
| 0.702208 | 0.107314 | 0.548 | 0.578 | 1 | 3 | Pttg1ip | -0.03 | Young |
| 0.702224 | 0.224719 | 0.237 | 0.237 | 1 | 3 | Pcf11 | 0 | Young |
| 0.702362 | 0.146686 | 0.312 | 0.321 | 1 | 3 | Impact | -0.009 | Young |
| 0.702437 | 0.158251 | 0.056 | 0.051 | 1 | 3 | Alg3 | 0.005 | Young |
| 0.702565 | 0.085049 | 0.059 | 0.055 | 1 | 3 | Hscb | 0.004 | Young |
| 0.703173 | 0.012298 | 0.345 | 0.372 | 1 | 3 | AC160336. | -0.027 | Young |
| 0.70323 | 0.119082 | 0.009 | 0.011 | 1 | 3 | Pnpla1 | -0.002 | Young |
| 0.703472 | 0.024934 | 0.288 | 0.301 | 1 | 3 | Map3k8 | -0.013 | Young |
| 0.704135 | 0.022933 | 0.284 | 0.301 | 1 | 3 | Rrp1 | -0.017 | Young |
| 0.704349 | 0.07001 | 0.476 | 0.486 | 1 | 3 | Ndufa7 | -0.01 | Young |
| 0.704607 | 0.390115 | 0.11 | 0.106 | 1 | 3 | Lgals8 | 0.004 | Young |
| 0.704651 | 0.125856 | 0.413 | 0.438 | 1 | 3 | Zfp106 | -0.025 | Young |
| 0.70475 | 0.00232 | 0.074 | 0.08 | 1 | 3 | Ptgr2 | -0.006 | Young |
| 0.704768 | 0.443207 | 0.035 | 0.032 | 1 | 3 | Zbtb45 | 0.003 | Young |
| 0.705097 | 0.017383 | 0.58 | 0.606 | 1 | 3 | Trib1 | -0.026 | Young |
| 0.705232 | 0.447675 | 0.018 | 0.02 | 1 | 3 | Ccp110 | -0.002 | Young |
| 0.705385 | 0.046758 | 0.062 | 0.067 | 1 | 3 | Eid3 | -0.005 | Young |
| 0.705683 | 0.20771 | 0.082 | 0.078 | 1 | 3 | Rif1 | 0.004 | Young |
| 0.705742 | 0.428346 | 0.069 | 0.066 | 1 | 3 | Bach2 | 0.003 | Young |
| 0.705971 | 0.190196 | 0.042 | 0.038 | 1 | 3 | Ccdc122 | 0.004 | Young |
| 0.705975 | 0.210542 | 0.193 | 0.191 | 1 | 3 | Herpud2 | 0.002 | Young |
| 0.706304 | 0.358491 | 0.04 | 0.037 | 1 | 3 | Mvk | 0.003 | Young |
| 0.706346 | 0.075073 | 0.381 | 0.407 | 1 | 3 | Zfp703 | -0.026 | Young |
| 0.70639 | 0.038055 | 0.682 | 0.698 | 1 | 3 | Socs6 | -0.016 | Young |
| 0.706559 | 0.035232 | 0.503 | 0.54 | 1 | 3 | Hnrnpl | -0.037 | Young |
| 0.706575 | 0.150003 | 0.265 | 0.288 | 1 | 3 | Rangap1 | -0.023 | Young |
| 0.706653 | 0.255554 | 0.092 | 0.087 | 1 | 3 | Kdm4c | 0.005 | Young |
| 0.706814 | 0.435624 | 0.01 | 0.008 | 1 | 3 | Ehf | 0.002 | Young |
| 0.707493 | 0.192813 | 0.322 | 0.331 | 1 | 3 | Baz2b | -0.009 | Young |
| 0.707579 | 0.009093 | 0.335 | 0.341 | 1 | 3 | Krit1 | -0.006 | Young |
| 0.707761 | 0.521132 | 0.01 | 0.008 | 1 | 3 | Mblac1 | 0.002 | Young |
| 0.708305 | 0.269449 | 0.109 | 0.104 | 1 | 3 | Atad2 | 0.005 | Young |
| 0.708337 | 0.215893 | 0.21 | 0.208 | 1 | 3 | Gars | 0.002 | Young |
| 0.708709 | 0.439185 | 0.01 | 0.008 | 1 | 3 | 4930579G2 | 0.002 | Young |
| 0.708855 | 0.254595 | 0.077 | 0.073 | 1 | 3 | Mical1 | 0.004 | Young |
| 0.708949 | 0.007855 | 0.989 | 0.996 | 1 | 3 | Eef1a1 | -0.007 | Young |
| 0.709183 | 0.326929 | 0.01 | 0.008 | 1 | 3 | Prim1 | 0.002 | Young |
| 0.709269 | 0.029723 | 0.196 | 0.208 | 1 | 3 | Camta2 | -0.012 | Young |
| 0.709349 | 0.107402 | 0.288 | 0.309 | 1 | 3 | Npepps | -0.021 | Young |
| 0.709439 | 0.805666 | 0.024 | 0.022 | 1 | 3 | Tspan33 | 0.002 | Young |
| 0.709597 | 0.76759 | 0.011 | 0.01 | 1 | 3 | Cdca7l | 0.001 | Young |
| 0.709835 | 0.148366 | 0.15 | 0.146 | 1 | 3 | Zfp950 | 0.004 | Young |
| 0.709895 | 0.532971 | 0.024 | 0.022 | 1 | 3 | Brat1 | 0.002 | Young |
| 0.709917 | 0.518683 | 0.03 | 0.027 | 1 | 3 | Pdss2 | 0.003 | Young |
| 0.710017 | 0.214137 | 0.027 | 0.03 | 1 | 3 | Gtf2h3 | -0.003 | Young |

| | | | | | | | | |
|---|---|---|---|---|---|---|---|---|
| 0.710131 | 0.407353 | 0.01 | 0.008 | 1 | 3 | Gpr183 | 0.002 | Young |
| 0.710296 | 0.148097 | 0.117 | 0.112 | 1 | 3 | Setdb1 | 0.005 | Young |
| 0.710391 | 0.073723 | 0.326 | 0.344 | 1 | 3 | Rin3 | -0.018 | Young |
| 0.710513 | 0.029704 | 0.402 | 0.431 | 1 | 3 | Atp5g1 | -0.029 | Young |
| 0.710526 | 0.45906 | 0.033 | 0.03 | 1 | 3 | Yars2 | 0.003 | Young |
| 0.710863 | 0.074783 | 0.018 | 0.02 | 1 | 3 | Glmn | -0.002 | Young |
| 0.711568 | 0.692918 | 0.024 | 0.022 | 1 | 3 | Coq6 | 0.002 | Young |
| 0.711629 | 0.036305 | 0.258 | 0.275 | 1 | 3 | Kxd1 | -0.017 | Young |
| 0.711724 | 0.124379 | 0.235 | 0.251 | 1 | 3 | Phf14 | -0.016 | Young |
| 0.711781 | 0.225066 | 0.069 | 0.066 | 1 | 3 | Eml3 | 0.003 | Young |
| 0.711792 | 0.306451 | 0.101 | 0.098 | 1 | 3 | Mllt6 | 0.003 | Young |
| 0.712029 | 0.361118 | 0.01 | 0.008 | 1 | 3 | Glis2 | 0.002 | Young |
| 0.712114 | 0.120099 | 0.059 | 0.055 | 1 | 3 | Mink1 | 0.004 | Young |
| 0.712238 | 0.288487 | 0.148 | 0.146 | 1 | 3 | Kctd2 | 0.002 | Young |
| 0.712273 | 0.517255 | 0.011 | 0.01 | 1 | 3 | Ttll1 | 0.001 | Young |
| 0.712504 | 0.297292 | 0.01 | 0.008 | 1 | 3 | Zfp235 | 0.002 | Young |
| 0.712504 | 0.29012 | 0.01 | 0.008 | 1 | 3 | Sgms2 | 0.002 | Young |
| 0.712542 | 0.327947 | 0.037 | 0.033 | 1 | 3 | Mllt3 | 0.004 | Young |
| 0.712629 | 0.565206 | 0.025 | 0.023 | 1 | 3 | Gab3 | 0.002 | Young |
| 0.712729 | 0.334631 | 0.027 | 0.024 | 1 | 3 | C1qtnf12 | 0.003 | Young |
| 0.713007 | 0.000109 | 0.062 | 0.057 | 1 | 3 | Lair1 | 0.005 | Young |
| 0.713019 | 0.668085 | 0.027 | 0.024 | 1 | 3 | Cry2 | 0.003 | Young |
| 0.713321 | 0.093104 | 0.432 | 0.452 | 1 | 3 | Map2k2 | -0.02 | Young |
| 0.713454 | 0.140679 | 0.01 | 0.008 | 1 | 3 | Ildr1 | 0.002 | Young |
| 0.713454 | 0.131077 | 0.01 | 0.008 | 1 | 3 | Gm4285 | 0.002 | Young |
| 0.713557 | 0.089928 | 0.144 | 0.153 | 1 | 3 | Dclre1c | -0.009 | Young |
| 0.713612 | 0.455872 | 0.011 | 0.01 | 1 | 3 | Cacnb4 | 0.001 | Young |
| 0.713678 | 0.050465 | 0.062 | 0.067 | 1 | 3 | Zc3h10 | -0.005 | Young |
| 0.713679 | 0.508731 | 0.087 | 0.084 | 1 | 3 | Deptor | 0.003 | Young |
| 0.71368 | 0.105374 | 0.176 | 0.173 | 1 | 3 | Tmem5 | 0.003 | Young |
| 0.713929 | 0.236201 | 0.01 | 0.008 | 1 | 3 | Rab4b | 0.002 | Young |
| 0.714058 | 0.806017 | 0.011 | 0.01 | 1 | 3 | Pdpn | 0.001 | Young |
| 0.714399 | 0.29198 | 0.11 | 0.117 | 1 | 3 | Hk3 | -0.007 | Young |
| 0.71462 | 0.157098 | 0.072 | 0.078 | 1 | 3 | Daam1 | -0.006 | Young |
| 0.714679 | 0.15544 | 0.059 | 0.065 | 1 | 3 | Nmi | -0.006 | Young |
| 0.714731 | 0.1075 | 0.11 | 0.104 | 1 | 3 | Pxmp4 | 0.006 | Young |
| 0.71475 | 0.181846 | 0.261 | 0.263 | 1 | 3 | Sike1 | -0.002 | Young |
| 0.714793 | 0.410706 | 0.03 | 0.027 | 1 | 3 | Zdhhc23 | 0.003 | Young |
| 0.715164 | 0.018531 | 0.187 | 0.197 | 1 | 3 | Zkscan3 | -0.01 | Young |
| 0.715175 | 0.30918 | 0.035 | 0.032 | 1 | 3 | Carf | 0.003 | Young |
| 0.715201 | 0.281418 | 0.032 | 0.029 | 1 | 3 | Mpst | 0.003 | Young |
| 0.715287 | 0.349531 | 0.082 | 0.078 | 1 | 3 | Zfyve27 | 0.004 | Young |
| 0.71537 | 0.900663 | 0.013 | 0.011 | 1 | 3 | Ica1l | 0.002 | Young |
| 0.715399 | 0.59897 | 0.011 | 0.01 | 1 | 3 | 1700037C1 | 0.001 | Young |
| 0.715399 | 0.323702 | 0.011 | 0.01 | 1 | 3 | Catip | 0.001 | Young |

| | | | | | | | | |
|---|---|---|---|---|---|---|---|---|
| 0.715493 | 0.10568 | 0.489 | 0.501 | 1 | 3 | Sema4a | -0.012 | Young |
| 0.715758 | 0.43567 | 0.034 | 0.031 | 1 | 3 | Zfp212 | 0.003 | Young |
| 0.715823 | 0.136903 | 0.211 | 0.205 | 1 | 3 | Zranb1 | 0.006 | Young |
| 0.715831 | 0.006214 | 0.01 | 0.008 | 1 | 3 | Hibch | 0.002 | Young |
| 0.715852 | 0.014671 | 0.869 | 0.864 | 1 | 3 | Ifitm3 | 0.005 | Young |
| 0.716091 | 0.349393 | 0.048 | 0.053 | 1 | 3 | Fam20a | -0.005 | Young |
| 0.716146 | 0.448916 | 0.021 | 0.019 | 1 | 3 | AU022793 | 0.002 | Young |
| 0.716213 | 1.002614 | 0.016 | 0.014 | 1 | 3 | Zfp707 | 0.002 | Young |
| 0.716224 | 0.179936 | 0.028 | 0.031 | 1 | 3 | Ano10 | -0.003 | Young |
| 0.716273 | 0.14068 | 0.083 | 0.079 | 1 | 3 | Galnt9 | 0.004 | Young |
| 0.716293 | 0.354798 | 0.011 | 0.01 | 1 | 3 | Ccdc152 | 0.001 | Young |
| 0.716872 | 0.034797 | 0.039 | 0.043 | 1 | 3 | Sclt1 | -0.004 | Young |
| 0.716954 | 0.062169 | 0.038 | 0.035 | 1 | 3 | Fam58b | 0.003 | Young |
| 0.716961 | 0.271362 | 0.106 | 0.102 | 1 | 3 | B3galnt2 | 0.004 | Young |
| 0.717063 | 0.831666 | 0.013 | 0.011 | 1 | 3 | Mks1 | 0.002 | Young |
| 0.71717 | 0.207999 | 0.213 | 0.213 | 1 | 3 | CAAA0111ξ | 0 | Young |
| 0.717188 | 0.231038 | 0.011 | 0.01 | 1 | 3 | Tmem38a | 0.001 | Young |
| 0.717244 | 0.022025 | 0.61 | 0.622 | 1 | 3 | Psmb1 | -0.012 | Young |
| 0.717352 | 0.136301 | 0.343 | 0.373 | 1 | 3 | Glyr1 | -0.03 | Young |
| 0.717366 | 0.065299 | 0.736 | 0.757 | 1 | 3 | Tubb5 | -0.021 | Young |
| 0.717557 | 0.08349 | 0.01 | 0.012 | 1 | 3 | 1110002J0 | -0.002 | Young |
| 0.71762 | 0.134465 | 0.597 | 0.619 | 1 | 3 | Mrfap1 | -0.022 | Young |
| 0.717635 | 0.241633 | 0.011 | 0.01 | 1 | 3 | Ugt1a6a | 0.001 | Young |
| 0.717769 | 0.088576 | 0.097 | 0.093 | 1 | 3 | Kat6b | 0.004 | Young |
| 0.717807 | 0.298644 | 0.034 | 0.031 | 1 | 3 | Kctd11 | 0.003 | Young |
| 0.717959 | 0.147724 | 0.28 | 0.302 | 1 | 3 | Sqor | -0.022 | Young |
| 0.718083 | 0.353033 | 0.011 | 0.01 | 1 | 3 | Tmem158 | 0.001 | Young |
| 0.718131 | 0.246922 | 0.038 | 0.042 | 1 | 3 | E330009J0 | -0.004 | Young |
| 0.718421 | 0.075439 | 0.086 | 0.092 | 1 | 3 | Helb | -0.006 | Young |
| 0.718581 | 0.030286 | 0.145 | 0.152 | 1 | 3 | Braf | -0.007 | Young |
| 0.718887 | 0.203518 | 0.177 | 0.173 | 1 | 3 | Slc35f6 | 0.004 | Young |
| 0.718978 | 0.292924 | 0.011 | 0.01 | 1 | 3 | Kif5c | 0.001 | Young |
| 0.718978 | 0.031795 | 0.169 | 0.179 | 1 | 3 | Gtf2b | -0.01 | Young |
| 0.719181 | 0.310999 | 0.013 | 0.011 | 1 | 3 | Mbnl3 | 0.002 | Young |
| 0.719213 | 0.684225 | 0.021 | 0.019 | 1 | 3 | Mterf4 | 0.002 | Young |
| 0.719229 | 0.146995 | 0.101 | 0.097 | 1 | 3 | 2310033P0 | 0.004 | Young |
| 0.719295 | 0.256522 | 0.309 | 0.312 | 1 | 3 | Entpd1 | -0.003 | Young |
| 0.719605 | 0.156492 | 0.013 | 0.011 | 1 | 3 | Caprin2 | 0.002 | Young |
| 0.719679 | 0.060611 | 0.207 | 0.202 | 1 | 3 | Clock | 0.005 | Young |
| 0.719779 | 0.000388 | 0.513 | 0.541 | 1 | 3 | Pebp1 | -0.028 | Young |
| 0.719846 | 0.398554 | 0.014 | 0.012 | 1 | 3 | Mical2 | 0.002 | Young |
| 0.720453 | 0.519625 | 0.013 | 0.011 | 1 | 3 | Rmdn2 | 0.002 | Young |
| 0.720453 | 0.247458 | 0.013 | 0.011 | 1 | 3 | Slc22a21 | 0.002 | Young |
| 0.720539 | 0.0643 | 0.294 | 0.312 | 1 | 3 | Appl1 | -0.018 | Young |
| 0.720849 | 0.281014 | 0.038 | 0.035 | 1 | 3 | Cdk5rap2 | 0.003 | Young |

| | | | | | | | | |
|---|---|---|---|---|---|---|---|---|
| 0.720901 | 0.209636 | 0.109 | 0.105 | 1 | 3 | Stx3 | 0.004 | Young |
| 0.721004 | 0.020331 | 0.889 | 0.891 | 1 | 3 | Dbi | -0.002 | Young |
| 0.721032 | 0.177566 | 0.298 | 0.315 | 1 | 3 | Mat2b | -0.017 | Young |
| 0.721461 | 0.327162 | 0.014 | 0.012 | 1 | 3 | Efcab2 | 0.002 | Young |
| 0.72157 | 0.057473 | 0.775 | 0.777 | 1 | 3 | Eps8 | -0.002 | Young |
| 0.721579 | 0.127545 | 0.347 | 0.367 | 1 | 3 | Ctnna1 | -0.02 | Young |
| 0.721764 | 0.215901 | 0.102 | 0.11 | 1 | 3 | Mag | -0.008 | Young |
| 0.721836 | 0.046291 | 0.047 | 0.043 | 1 | 3 | Plcl1 | 0.004 | Young |
| 0.721974 | 0.068882 | 0.035 | 0.032 | 1 | 3 | Prdm15 | 0.003 | Young |
| 0.722143 | 0.186147 | 0.029 | 0.026 | 1 | 3 | Acyp2 | 0.003 | Young |
| 0.722346 | 0.28384 | 0.155 | 0.166 | 1 | 3 | Rftn1 | -0.011 | Young |
| 0.722444 | 0.361546 | 0.019 | 0.017 | 1 | 3 | Xndc1 | 0.002 | Young |
| 0.722695 | 0.148208 | 0.207 | 0.223 | 1 | 3 | Csnk1g3 | -0.016 | Young |
| 0.722731 | 0.376023 | 0.035 | 0.032 | 1 | 3 | Bcdin3d | 0.003 | Young |
| 0.722824 | 0.31631 | 0.069 | 0.075 | 1 | 3 | Tmem203 | -0.006 | Young |
| 0.722828 | 0.15462 | 0.361 | 0.37 | 1 | 3 | Eps15 | -0.009 | Young |
| 0.722962 | 0.113109 | 0.144 | 0.14 | 1 | 3 | Mta1 | 0.004 | Young |
| 0.722999 | 0.203458 | 0.013 | 0.011 | 1 | 3 | 4930524J0 | 0.002 | Young |
| 0.723063 | 0.345446 | 0.016 | 0.014 | 1 | 3 | Macrod2 | 0.002 | Young |
| 0.723077 | 0.504634 | 0.014 | 0.012 | 1 | 3 | Fam149a | 0.002 | Young |
| 0.723172 | 0.307093 | 0.02 | 0.018 | 1 | 3 | Lpcat2 | 0.002 | Young |
| 0.723213 | 0.029802 | 0.308 | 0.325 | 1 | 3 | Fam104a | -0.017 | Young |
| 0.723257 | 0.176845 | 0.247 | 0.245 | 1 | 3 | Clptm1 | 0.002 | Young |
| 0.72346 | 0.137136 | 0.011 | 0.01 | 1 | 3 | Gm10384 | 0.001 | Young |
| 0.723478 | 0.254221 | 0.019 | 0.017 | 1 | 3 | Mcat | 0.002 | Young |
| 0.723481 | 0.47128 | 0.014 | 0.012 | 1 | 3 | Lrrc56 | 0.002 | Young |
| 0.723481 | 0.243617 | 0.014 | 0.012 | 1 | 3 | Ankrd33b | 0.002 | Young |
| 0.723587 | 0.221483 | 0.129 | 0.127 | 1 | 3 | Pds5b | 0.002 | Young |
| 0.723636 | 0.053253 | 0.059 | 0.065 | 1 | 3 | Tceal8 | -0.006 | Young |
| 0.723749 | 0.011457 | 0.624 | 0.649 | 1 | 3 | Calm3 | -0.025 | Young |
| 0.723763 | 0.356749 | 0.027 | 0.024 | 1 | 3 | Gins2 | 0.003 | Young |
| 0.72409 | 0.676778 | 0.029 | 0.026 | 1 | 3 | Tyw5 | 0.003 | Young |
| 0.724136 | 0.045722 | 0.179 | 0.189 | 1 | 3 | Cops6 | -0.01 | Young |
| 0.724225 | 0.054303 | 0.109 | 0.116 | 1 | 3 | Sdccag8 | -0.007 | Young |
| 0.724273 | 0.172013 | 0.013 | 0.011 | 1 | 3 | Gm47754 | 0.002 | Young |
| 0.724289 | 0.663057 | 0.014 | 0.012 | 1 | 3 | Rab26os | 0.002 | Young |
| 0.724333 | 0.251256 | 0.057 | 0.054 | 1 | 3 | Ric8b | 0.003 | Young |
| 0.724384 | 0.38247 | 0.067 | 0.063 | 1 | 3 | Dhx35 | 0.004 | Young |
| 0.724657 | 0.070569 | 0.484 | 0.493 | 1 | 3 | Runx1 | -0.009 | Young |
| 0.724895 | 0.327462 | 0.018 | 0.016 | 1 | 3 | Vars2 | 0.002 | Young |
| 0.724945 | 0.097692 | 0.511 | 0.529 | 1 | 3 | Pgd | -0.018 | Young |
| 0.725098 | 0.366787 | 0.014 | 0.012 | 1 | 3 | Arrdc2 | 0.002 | Young |
| 0.725098 | 0.187645 | 0.014 | 0.012 | 1 | 3 | Gpr141 | 0.002 | Young |
| 0.725105 | 0.227254 | 0.172 | 0.17 | 1 | 3 | Iws1 | 0.002 | Young |
| 0.725159 | 0.106887 | 0.077 | 0.073 | 1 | 3 | Sugp2 | 0.004 | Young |

| | | | | | | | | |
|---|---|---|---|---|---|---|---|---|
| 0.725167 | 0.04615 | 0.311 | 0.327 | 1 | 3 | Usp16 | -0.016 | Young |
| 0.725175 | 0.712632 | 0.018 | 0.02 | 1 | 3 | Polr3g | -0.002 | Young |
| 0.725253 | 0.824864 | 0.018 | 0.016 | 1 | 3 | Jpx | 0.002 | Young |
| 0.725287 | 0.379676 | 0.02 | 0.023 | 1 | 3 | Tubg1 | -0.003 | Young |
| 0.725306 | 0.063427 | 0.23 | 0.245 | 1 | 3 | Zrsr2 | -0.015 | Young |
| 0.725548 | 0.128202 | 0.013 | 0.011 | 1 | 3 | Syp | 0.002 | Young |
| 0.725761 | 0.127939 | 0.246 | 0.262 | 1 | 3 | Rnpepl1 | -0.016 | Young |
| 0.725895 | 0.227165 | 0.074 | 0.08 | 1 | 3 | 9330160F1 | -0.006 | Young |
| 0.726264 | 0.384455 | 0.032 | 0.029 | 1 | 3 | Crtc1 | 0.003 | Young |
| 0.726425 | 0.05545 | 0.139 | 0.148 | 1 | 3 | Usp38 | -0.009 | Young |
| 0.726558 | 0.533991 | 0.105 | 0.102 | 1 | 3 | Sun1 | 0.003 | Young |
| 0.726561 | 0.017679 | 0.236 | 0.249 | 1 | 3 | Glmp | -0.013 | Young |
| 0.726711 | 0.406974 | 0.09 | 0.097 | 1 | 3 | Mier3 | -0.007 | Young |
| 0.726971 | 0.088378 | 0.053 | 0.049 | 1 | 3 | Msantd2 | 0.004 | Young |
| 0.727019 | 0.043132 | 0.11 | 0.116 | 1 | 3 | Ecd | -0.006 | Young |
| 0.727146 | 0.27892 | 0.016 | 0.014 | 1 | 3 | Celsr3 | 0.002 | Young |
| 0.727242 | 0.17434 | 0.24 | 0.257 | 1 | 3 | Mllt10 | -0.017 | Young |
| 0.727494 | 0.353399 | 0.106 | 0.103 | 1 | 3 | Tm7sf3 | 0.003 | Young |
| 0.727543 | 0.216305 | 0.109 | 0.116 | 1 | 3 | Nupr1 | -0.007 | Young |
| 0.727569 | 0.333708 | 0.034 | 0.031 | 1 | 3 | Snapc2 | 0.003 | Young |
| 0.727583 | 0.087379 | 0.015 | 0.013 | 1 | 3 | Zfp628 | 0.002 | Young |
| 0.727598 | 0.154498 | 0.068 | 0.065 | 1 | 3 | Ubac1 | 0.003 | Young |
| 0.727692 | 0.101563 | 0.134 | 0.13 | 1 | 3 | Slc9a6 | 0.004 | Young |
| 0.727729 | 0.241715 | 0.216 | 0.234 | 1 | 3 | Pitpnc1 | -0.018 | Young |
| 0.727753 | 0.128562 | 0.054 | 0.059 | 1 | 3 | Ttc7b | -0.005 | Young |
| 0.72779 | 0.313232 | 0.021 | 0.019 | 1 | 3 | C730034F0 | 0.002 | Young |
| 0.728145 | 0.026256 | 0.215 | 0.226 | 1 | 3 | Dhx36 | -0.011 | Young |
| 0.728225 | 0.182319 | 0.192 | 0.202 | 1 | 3 | Siah2 | -0.01 | Young |
| 0.728228 | 0.333016 | 0.053 | 0.057 | 1 | 3 | Mgst3 | -0.004 | Young |
| 0.728248 | 0.015293 | 0.668 | 0.708 | 1 | 3 | Ppp1ca | -0.04 | Young |
| 0.728351 | 0.199698 | 0.093 | 0.099 | 1 | 3 | Lrrc8a | -0.006 | Young |
| 0.728381 | 0.145359 | 0.524 | 0.539 | 1 | 3 | Itch | -0.015 | Young |
| 0.72839 | 0.080704 | 0.356 | 0.379 | 1 | 3 | Crk | -0.023 | Young |
| 0.728718 | 0.27038 | 0.027 | 0.024 | 1 | 3 | Ankrd13d | 0.003 | Young |
| 0.72925 | 0.125572 | 0.062 | 0.067 | 1 | 3 | Uxt | -0.005 | Young |
| 0.729672 | 0.084709 | 0.624 | 0.633 | 1 | 3 | Ndufb7 | -0.009 | Young |
| 0.729736 | 0.461431 | 0.021 | 0.019 | 1 | 3 | Slc37a1 | 0.002 | Young |
| 0.729834 | 0.185301 | 0.1 | 0.096 | 1 | 3 | Parp3 | 0.004 | Young |
| 0.729845 | 0.190233 | 0.259 | 0.276 | 1 | 3 | Eif4e2 | -0.017 | Young |
| 0.729864 | 0.064577 | 0.388 | 0.413 | 1 | 3 | Rexo2 | -0.025 | Young |
| 0.730027 | 0.131028 | 0.111 | 0.118 | 1 | 3 | Lrif1 | -0.007 | Young |
| 0.730164 | 0.176408 | 0.288 | 0.289 | 1 | 3 | Zfp516 | -0.001 | Young |
| 0.730241 | 0.06133 | 0.232 | 0.233 | 1 | 3 | Dcaf8 | -0.001 | Young |
| 0.730273 | 0.240392 | 0.028 | 0.025 | 1 | 3 | Macrod1 | 0.003 | Young |
| 0.730387 | 0.042622 | 0.841 | 0.858 | 1 | 3 | Tmbim6 | -0.017 | Young |

| | | | | | | | | |
|---|---|---|---|---|---|---|---|---|
| 0.730585 | 0.210657 | 0.03 | 0.027 | 1 | 3 | Gsdme | 0.003 | Young |
| 0.730705 | 0.201155 | 0.11 | 0.118 | 1 | 3 | Jdp2 | -0.008 | Young |
| 0.730992 | 0.037845 | 0.117 | 0.111 | 1 | 3 | Ints3 | 0.006 | Young |
| 0.731071 | 0.106527 | 0.015 | 0.013 | 1 | 3 | Klhl25 | 0.002 | Young |
| 0.731082 | 0.038931 | 0.013 | 0.011 | 1 | 3 | Thnsl1 | 0.002 | Young |
| 0.731122 | 0.09173 | 0.386 | 0.412 | 1 | 3 | Myo1f | -0.026 | Young |
| 0.73123 | 0.12836 | 0.072 | 0.078 | 1 | 3 | Tmem159 | -0.006 | Young |
| 0.731583 | 0.260783 | 0.073 | 0.07 | 1 | 3 | Cops7a | 0.003 | Young |
| 0.731853 | 0.099823 | 0.116 | 0.112 | 1 | 3 | Srpk1 | 0.004 | Young |
| 0.732429 | 0.445232 | 0.015 | 0.013 | 1 | 3 | Gm28187 | 0.002 | Young |
| 0.732495 | 0.202586 | 0.208 | 0.223 | 1 | 3 | Mrpl34 | -0.015 | Young |
| 0.732516 | 0.229062 | 0.024 | 0.022 | 1 | 3 | Slc19a2 | 0.002 | Young |
| 0.732536 | 0.297473 | 0.02 | 0.018 | 1 | 3 | Poll | 0.002 | Young |
| 0.732576 | 0.110857 | 0.162 | 0.171 | 1 | 3 | Trappc10 | -0.009 | Young |
| 0.732628 | 0.161441 | 0.253 | 0.254 | 1 | 3 | Stx16 | -0.001 | Young |
| 0.732715 | 0.187094 | 0.102 | 0.098 | 1 | 3 | Tmem60 | 0.004 | Young |
| 0.732759 | 0.241249 | 0.033 | 0.03 | 1 | 3 | Primpol | 0.003 | Young |
| 0.733458 | 0.065126 | 0.066 | 0.07 | 1 | 3 | Arntl | -0.004 | Young |
| 0.733601 | 0.052112 | 0.03 | 0.033 | 1 | 3 | Chpt1 | -0.003 | Young |
| 0.7337 | 0.016172 | 0.028 | 0.025 | 1 | 3 | Zfp637 | 0.003 | Young |
| 0.733761 | 0.19248 | 0.032 | 0.029 | 1 | 3 | Slc2a8 | 0.003 | Young |
| 0.733766 | 0.166524 | 0.088 | 0.085 | 1 | 3 | Tmem126a | 0.003 | Young |
| 0.733788 | 0.077755 | 0.015 | 0.013 | 1 | 3 | Pgap1 | 0.002 | Young |
| 0.733914 | 0.030602 | 0.168 | 0.178 | 1 | 3 | Usp12 | -0.01 | Young |
| 0.734017 | 0.110571 | 0.125 | 0.121 | 1 | 3 | Soga1 | 0.004 | Young |
| 0.73434 | 0.200873 | 0.159 | 0.17 | 1 | 3 | Birc2 | -0.011 | Young |
| 0.734541 | 0.030018 | 0.019 | 0.017 | 1 | 3 | Hirip3 | 0.002 | Young |
| 0.734565 | 0.022926 | 0.015 | 0.013 | 1 | 3 | Magee1 | 0.002 | Young |
| 0.734589 | 0.304621 | 0.01 | 0.012 | 1 | 3 | Strip2 | -0.002 | Young |
| 0.734793 | 0.001547 | 0.319 | 0.338 | 1 | 3 | Sypl | -0.019 | Young |
| 0.734834 | 0.295096 | 0.032 | 0.029 | 1 | 3 | Pank4 | 0.003 | Young |
| 0.734844 | 0.047924 | 0.683 | 0.7 | 1 | 3 | Slamf9 | -0.017 | Young |
| 0.734975 | 0.014654 | 0.047 | 0.043 | 1 | 3 | Kctd9 | 0.004 | Young |
| 0.735101 | 0.306929 | 0.066 | 0.062 | 1 | 3 | Ring1 | 0.004 | Young |
| 0.735284 | 0.050352 | 0.018 | 0.016 | 1 | 3 | Cby1 | 0.002 | Young |
| 0.735342 | 0.196871 | 0.015 | 0.013 | 1 | 3 | Slc16a13 | 0.002 | Young |
| 0.735382 | 0.056762 | 0.102 | 0.098 | 1 | 3 | 2310022B0 | 0.004 | Young |
| 0.735432 | 0.037433 | 0.122 | 0.13 | 1 | 3 | Itgb7 | -0.008 | Young |
| 0.735631 | 0.108102 | 0.222 | 0.235 | 1 | 3 | Ankfy1 | -0.013 | Young |
| 0.735969 | 0.17141 | 0.217 | 0.216 | 1 | 3 | Rab22a | 0.001 | Young |
| 0.73599 | 0.035121 | 0.061 | 0.065 | 1 | 3 | Wdr70 | -0.004 | Young |
| 0.73599 | 0.116597 | 0.08 | 0.075 | 1 | 3 | Bicra | 0.005 | Young |
| 0.736082 | 0.155089 | 0.034 | 0.031 | 1 | 3 | Rfc3 | 0.003 | Young |
| 0.736082 | 0.03747 | 0.016 | 0.014 | 1 | 3 | Rpgrip1l | 0.002 | Young |
| 0.736083 | 0.059989 | 0.154 | 0.162 | 1 | 3 | Ipo11 | -0.008 | Young |

| | | | | | | | | |
|---|---|---|---|---|---|---|---|---|
| 0.736238 | 0.233987 | 0.021 | 0.019 | 1 | 3 | Cenpo | 0.002 | Young |
| 0.736556 | 0.050114 | 0.039 | 0.036 | 1 | 3 | Phlpp2 | 0.003 | Young |
| 0.736828 | 0.132364 | 0.016 | 0.014 | 1 | 3 | Fam109a | 0.002 | Young |
| 0.736845 | 0.209654 | 0.042 | 0.045 | 1 | 3 | Dlg4 | -0.003 | Young |
| 0.736897 | 0.159848 | 0.02 | 0.018 | 1 | 3 | Ndst1 | 0.002 | Young |
| 0.736931 | 0.024589 | 0.048 | 0.044 | 1 | 3 | Galns | 0.004 | Young |
| 0.737268 | 0.066914 | 0.085 | 0.09 | 1 | 3 | Fam45a | -0.005 | Young |
| 0.73727 | 0.328727 | 0.058 | 0.055 | 1 | 3 | Slc9a3r2 | 0.003 | Young |
| 0.73728 | 0.083126 | 0.023 | 0.02 | 1 | 3 | Fbxl8 | 0.003 | Young |
| 0.737328 | 0.070037 | 0.335 | 0.361 | 1 | 3 | Pik3r6 | -0.026 | Young |
| 0.737684 | 0.142589 | 0.172 | 0.17 | 1 | 3 | Vps4a | 0.002 | Young |
| 0.738145 | 0.27605 | 0.096 | 0.092 | 1 | 3 | Acaca | 0.004 | Young |
| 0.738237 | 0.016651 | 0.633 | 0.667 | 1 | 3 | Atrx | -0.034 | Young |
| 0.738763 | 0.113807 | 0.673 | 0.704 | 1 | 3 | Snrpe | -0.031 | Young |
| 0.73882 | 0.050598 | 0.284 | 0.287 | 1 | 3 | Rbm7 | -0.003 | Young |
| 0.738981 | 0.118146 | 0.121 | 0.129 | 1 | 3 | Vgll4 | -0.008 | Young |
| 0.739089 | 0.146821 | 0.304 | 0.302 | 1 | 3 | Pcmtd1 | 0.002 | Young |
| 0.73949 | 0.034077 | 0.08 | 0.085 | 1 | 3 | Dok3 | -0.005 | Young |
| 0.739584 | 0.129132 | 0.02 | 0.018 | 1 | 3 | Zbtb25 | 0.002 | Young |
| 0.73988 | 0.092167 | 0.119 | 0.114 | 1 | 3 | Ccnc | 0.005 | Young |
| 0.739886 | 0.011633 | 0.139 | 0.145 | 1 | 3 | Nsmaf | -0.006 | Young |
| 0.739978 | 0.058556 | 0.093 | 0.099 | 1 | 3 | Hist3h2a | -0.006 | Young |
| 0.740148 | 0.052354 | 0.021 | 0.019 | 1 | 3 | Tasp1 | 0.002 | Young |
| 0.740171 | 0.008497 | 0.03 | 0.027 | 1 | 3 | Smpdl3b | 0.003 | Young |
| 0.740247 | 0.138559 | 0.384 | 0.398 | 1 | 3 | Cpt1a | -0.014 | Young |
| 0.740585 | 0.043068 | 0.761 | 0.786 | 1 | 3 | Atp6v0b | -0.025 | Young |
| 0.741209 | 0.072774 | 0.058 | 0.062 | 1 | 3 | Tdp2 | -0.004 | Young |
| 0.74143 | 0.003573 | 0.192 | 0.202 | 1 | 3 | Camk1 | -0.01 | Young |
| 0.741454 | 0.224342 | 0.078 | 0.084 | 1 | 3 | Yod1 | -0.006 | Young |
| 0.741534 | 0.011037 | 0.317 | 0.337 | 1 | 3 | Map4k1 | -0.02 | Young |
| 0.741558 | 0.125166 | 0.184 | 0.196 | 1 | 3 | Sash3 | -0.012 | Young |
| 0.741633 | 0.055472 | 0.095 | 0.1 | 1 | 3 | 1110059G1 | -0.005 | Young |
| 0.741774 | 0.19443 | 0.098 | 0.105 | 1 | 3 | Nsmce2 | -0.007 | Young |
| 0.742075 | 0.274806 | 0.043 | 0.047 | 1 | 3 | 1810032O( | -0.004 | Young |
| 0.742349 | 0.058541 | 0.357 | 0.357 | 1 | 3 | Tmed3 | 0 | Young |
| 0.742817 | 0.220455 | 0.22 | 0.221 | 1 | 3 | Tob1 | -0.001 | Young |
| 0.742832 | 0.181293 | 0.08 | 0.085 | 1 | 3 | Cdc42ep4 | -0.005 | Young |
| 0.742988 | 0.076162 | 0.098 | 0.105 | 1 | 3 | Camk2d | -0.007 | Young |
| 0.743005 | 0.003272 | 0.345 | 0.363 | 1 | 3 | Ttc14 | -0.018 | Young |
| 0.743298 | 0.170914 | 0.311 | 0.313 | 1 | 3 | Mrpl54 | -0.002 | Young |
| 0.743534 | 0.180713 | 0.198 | 0.195 | 1 | 3 | Dhrs7 | 0.003 | Young |
| 0.743618 | 0.206128 | 0.02 | 0.023 | 1 | 3 | Wdr24 | -0.003 | Young |
| 0.743676 | 0.459739 | 0.086 | 0.082 | 1 | 3 | Ndufaf2 | 0.004 | Young |
| 0.743732 | 0.079145 | 0.351 | 0.354 | 1 | 3 | Lilra6 | -0.003 | Young |
| 0.743873 | 0.069478 | 0.246 | 0.262 | 1 | 3 | Vti1b | -0.016 | Young |

| | | | | | | | | |
|---|---|---|---|---|---|---|---|---|
| 0.743965 | 0.084223 | 0.619 | 0.648 | 1 | 3 | Etv3 | -0.029 | Young |
| 0.744105 | 0.116228 | 0.433 | 0.453 | 1 | 3 | Tsn | -0.02 | Young |
| 0.744728 | 0.181045 | 0.179 | 0.192 | 1 | 3 | Ostm1 | -0.013 | Young |
| 0.745259 | 0.132709 | 0.259 | 0.257 | 1 | 3 | Park7 | 0.002 | Young |
| 0.745267 | 0.064121 | 0.107 | 0.115 | 1 | 3 | Brap | -0.008 | Young |
| 0.745288 | 0.307541 | 0.063 | 0.06 | 1 | 3 | Gm5617 | 0.003 | Young |
| 0.745354 | 0.064626 | 0.068 | 0.073 | 1 | 3 | Pgam5 | -0.005 | Young |
| 0.745523 | 0.070935 | 0.02 | 0.023 | 1 | 3 | Ntmt1 | -0.003 | Young |
| 0.745613 | 0.423261 | 0.061 | 0.057 | 1 | 3 | Nemp1 | 0.004 | Young |
| 0.745664 | 0.067207 | 0.241 | 0.239 | 1 | 3 | Pik3c2a | 0.002 | Young |
| 0.746519 | 0.075266 | 0.47 | 0.486 | 1 | 3 | Pxn | -0.016 | Young |
| 0.746795 | 0.126789 | 0.117 | 0.124 | 1 | 3 | Socs2 | -0.007 | Young |
| 0.746884 | 0.092969 | 0.324 | 0.329 | 1 | 3 | Chmp3 | -0.005 | Young |
| 0.746996 | 0.090209 | 0.062 | 0.067 | 1 | 3 | R3hcc1l | -0.005 | Young |
| 0.747141 | 0.425286 | 0.194 | 0.197 | 1 | 3 | Vav2 | -0.003 | Young |
| 0.747178 | 0.189413 | 0.064 | 0.061 | 1 | 3 | Cav2 | 0.003 | Young |
| 0.747341 | 0.125694 | 0.111 | 0.118 | 1 | 3 | Tmem229b | -0.007 | Young |
| 0.747781 | 0.370427 | 0.059 | 0.063 | 1 | 3 | Ubr7 | -0.004 | Young |
| 0.748002 | 0.215899 | 0.011 | 0.013 | 1 | 3 | Il18bp | -0.002 | Young |
| 0.748004 | 0.006102 | 0.032 | 0.035 | 1 | 3 | Sergef | -0.003 | Young |
| 0.748806 | 0.121317 | 0.321 | 0.342 | 1 | 3 | Pan3 | -0.021 | Young |
| 0.748885 | 0.077653 | 0.216 | 0.227 | 1 | 3 | Ddx18 | -0.011 | Young |
| 0.749531 | 0.115896 | 0.372 | 0.38 | 1 | 3 | Cul3 | -0.008 | Young |
| 0.749606 | 0.186519 | 0.155 | 0.166 | 1 | 3 | Zfp326 | -0.011 | Young |
| 0.749685 | 0.101574 | 0.042 | 0.045 | 1 | 3 | 2310061I04 | -0.003 | Young |
| 0.750519 | 0.182369 | 0.221 | 0.22 | 1 | 3 | Sbf2 | 0.001 | Young |
| 0.751031 | 0.105316 | 0.021 | 0.024 | 1 | 3 | Tarsl2 | -0.003 | Young |
| 0.751096 | 0.250372 | 0.302 | 0.313 | 1 | 3 | Pias1 | -0.011 | Young |
| 0.751216 | 0.350256 | 0.051 | 0.048 | 1 | 3 | Idua | 0.003 | Young |
| 0.751217 | 0.326028 | 0.067 | 0.073 | 1 | 3 | Eif2b2 | -0.006 | Young |
| 0.751507 | 0.054547 | 0.069 | 0.074 | 1 | 3 | Ctns | -0.005 | Young |
| 0.751865 | 0.756126 | 0.029 | 0.032 | 1 | 3 | Stxbp1 | -0.003 | Young |
| 0.75214 | 0.311048 | 0.154 | 0.162 | 1 | 3 | Sh3bp2 | -0.008 | Young |
| 0.752439 | 0.02243 | 0.122 | 0.128 | 1 | 3 | Klhl28 | -0.006 | Young |
| 0.752507 | 0.038835 | 0.468 | 0.497 | 1 | 3 | Rab2a | -0.029 | Young |
| 0.752611 | 0.31593 | 0.011 | 0.013 | 1 | 3 | Cmpk2 | -0.002 | Young |
| 0.752611 | 0.209529 | 0.011 | 0.013 | 1 | 3 | Clmn | -0.002 | Young |
| 0.752685 | 0.0009 | 0.179 | 0.19 | 1 | 3 | Ammecr1l | -0.011 | Young |
| 0.752753 | 0.001271 | 0.085 | 0.09 | 1 | 3 | Bmi1 | -0.005 | Young |
| 0.752962 | 0.142034 | 0.145 | 0.157 | 1 | 3 | Naa10 | -0.012 | Young |
| 0.753029 | 0.032416 | 0.163 | 0.17 | 1 | 3 | Steap3 | -0.007 | Young |
| 0.753821 | 0.37066 | 0.021 | 0.024 | 1 | 3 | H2-T24 | -0.003 | Young |
| 0.753864 | 0.038941 | 0.336 | 0.357 | 1 | 3 | Rnh1 | -0.021 | Young |
| 0.754363 | 0.195341 | 0.068 | 0.065 | 1 | 3 | Haghl | 0.003 | Young |
| 0.754426 | 0.071169 | 0.653 | 0.679 | 1 | 3 | Celf1 | -0.026 | Young |

| | | | | | | | | |
|---|---|---|---|---|---|---|---|---|
| 0.754428 | 0.085542 | 0.275 | 0.28 | 1 | 3 | R3hdm2 | -0.005 | Young |
| 0.754627 | 0.089438 | 0.297 | 0.308 | 1 | 3 | Acadm | -0.011 | Young |
| 0.7548 | 0.062691 | 0.111 | 0.108 | 1 | 3 | Hdgfl2 | 0.003 | Young |
| 0.754842 | 0.422689 | 0.12 | 0.117 | 1 | 3 | Parp8 | 0.003 | Young |
| 0.755123 | 0.003056 | 0.491 | 0.527 | 1 | 3 | Srp14 | -0.036 | Young |
| 0.755129 | 0.312189 | 0.011 | 0.013 | 1 | 3 | Lgi4 | -0.002 | Young |
| 0.755176 | 0.061385 | 0.36 | 0.384 | 1 | 3 | Ndufa8 | -0.024 | Young |
| 0.755549 | 0.29012 | 0.011 | 0.013 | 1 | 3 | A730063M | -0.002 | Young |
| 0.755717 | 0.041688 | 0.083 | 0.079 | 1 | 3 | Psmg2 | 0.004 | Young |
| 0.756476 | 0.350888 | 0.091 | 0.087 | 1 | 3 | Klhdc10 | 0.004 | Young |
| 0.756591 | 0.103305 | 0.172 | 0.18 | 1 | 3 | Ppm1g | -0.008 | Young |
| 0.756824 | 0.071055 | 0.093 | 0.099 | 1 | 3 | Ak8 | -0.006 | Young |
| 0.757459 | 0.127515 | 0.186 | 0.183 | 1 | 3 | Eif4ebp1 | 0.003 | Young |
| 0.757549 | 0.051667 | 0.032 | 0.035 | 1 | 3 | Cradd | -0.003 | Young |
| 0.757708 | 0.010722 | 0.41 | 0.423 | 1 | 3 | Psmd4 | -0.013 | Young |
| 0.757725 | 0.323003 | 0.191 | 0.191 | 1 | 3 | B4galt1 | 0 | Young |
| 0.75773 | 0.265264 | 0.138 | 0.146 | 1 | 3 | Pno1 | -0.008 | Young |
| 0.757801 | 0.052477 | 0.641 | 0.681 | 1 | 3 | Eif3k | -0.04 | Young |
| 0.757966 | 0.014227 | 0.194 | 0.189 | 1 | 3 | Prkag1 | 0.005 | Young |
| 0.758009 | 0.062621 | 0.22 | 0.231 | 1 | 3 | Fam118b | -0.011 | Young |
| 0.758087 | 0.177695 | 0.231 | 0.246 | 1 | 3 | Rreb1 | -0.015 | Young |
| 0.758583 | 0.296867 | 0.169 | 0.183 | 1 | 3 | Mysm1 | -0.014 | Young |
| 0.758798 | 0.264604 | 0.043 | 0.047 | 1 | 3 | Galk1 | -0.004 | Young |
| 0.758897 | 0.202931 | 0.162 | 0.171 | 1 | 3 | Ehmt1 | -0.009 | Young |
| 0.758937 | 0.176436 | 0.109 | 0.116 | 1 | 3 | Upf3b | -0.007 | Young |
| 0.759732 | 0.275362 | 0.058 | 0.062 | 1 | 3 | Zfand2a | -0.004 | Young |
| 0.759849 | 0.027418 | 0.081 | 0.086 | 1 | 3 | Zfp512 | -0.005 | Young |
| 0.760173 | 0.031401 | 0.343 | 0.364 | 1 | 3 | Gpbp1 | -0.021 | Young |
| 0.760631 | 0.149266 | 0.351 | 0.376 | 1 | 3 | Dcaf12 | -0.025 | Young |
| 0.760661 | 0.069178 | 0.074 | 0.079 | 1 | 3 | Tmx2 | -0.005 | Young |
| 0.760918 | 0.085029 | 0.763 | 0.767 | 1 | 3 | Il1b | -0.004 | Young |
| 0.762307 | 0.013213 | 0.462 | 0.503 | 1 | 3 | Elavl1 | -0.041 | Young |
| 0.762497 | 0.322533 | 0.098 | 0.096 | 1 | 3 | Fdft1 | 0.002 | Young |
| 0.762635 | 0.005145 | 0.048 | 0.051 | 1 | 3 | Nt5m | -0.003 | Young |
| 0.762763 | 0.04898 | 0.65 | 0.675 | 1 | 3 | Reep3 | -0.025 | Young |
| 0.762975 | 0.038456 | 0.091 | 0.087 | 1 | 3 | Rpa1 | 0.004 | Young |
| 0.764267 | 0.155298 | 0.23 | 0.231 | 1 | 3 | Cpne3 | -0.001 | Young |
| 0.764879 | 0.104888 | 0.023 | 0.025 | 1 | 3 | Epha4 | -0.002 | Young |
| 0.764968 | 0.042842 | 0.107 | 0.114 | 1 | 3 | Nmd3 | -0.007 | Young |
| 0.765033 | 0.026536 | 0.069 | 0.074 | 1 | 3 | Slc12a4 | -0.005 | Young |
| 0.765267 | 0.032654 | 0.739 | 0.765 | 1 | 3 | Atp5g2 | -0.026 | Young |
| 0.765643 | 0.291305 | 0.013 | 0.014 | 1 | 3 | Tmem191c | -0.001 | Young |
| 0.765871 | 0.16204 | 0.085 | 0.09 | 1 | 3 | Zfp598 | -0.005 | Young |
| 0.766164 | 0.166672 | 0.037 | 0.039 | 1 | 3 | Hmgn3 | -0.002 | Young |
| 0.766637 | 0.226846 | 0.082 | 0.079 | 1 | 3 | Fastk | 0.003 | Young |

| | | | | | | | | |
|---|---|---|---|---|---|---|---|---|
| 0.766699 | 0.161449 | 0.12 | 0.116 | 1 | 3 | Ice1 | 0.004 | Young |
| 0.766797 | 0.010473 | 0.062 | 0.066 | 1 | 3 | Zfp771 | -0.004 | Young |
| 0.767195 | 0.395138 | 0.021 | 0.024 | 1 | 3 | Praf2 | -0.003 | Young |
| 0.767808 | 0.255312 | 0.077 | 0.082 | 1 | 3 | Tesk1 | -0.005 | Young |
| 0.767832 | 0.245678 | 0.082 | 0.079 | 1 | 3 | Ccdc117 | 0.003 | Young |
| 0.768601 | 0.156824 | 0.27 | 0.283 | 1 | 3 | Rela | -0.013 | Young |
| 0.768876 | 0.119999 | 0.443 | 0.459 | 1 | 3 | Slc48a1 | -0.016 | Young |
| 0.76918 | 0.060146 | 0.063 | 0.067 | 1 | 3 | Gm7694 | -0.004 | Young |
| 0.769266 | 0.046369 | 0.013 | 0.014 | 1 | 3 | Abraxas1 | -0.001 | Young |
| 0.769383 | 0.341875 | 0.064 | 0.069 | 1 | 3 | 1600002K0 | -0.005 | Young |
| 0.769394 | 0.138402 | 0.153 | 0.149 | 1 | 3 | Rfk | 0.004 | Young |
| 0.769462 | 0.041726 | 0.88 | 0.91 | 1 | 3 | Itm2b | -0.03 | Young |
| 0.769477 | 0.093354 | 0.167 | 0.177 | 1 | 3 | Ptk2b | -0.01 | Young |
| 0.769766 | 0.149546 | 0.086 | 0.091 | 1 | 3 | Zfp397 | -0.005 | Young |
| 0.770467 | 0.190701 | 0.193 | 0.203 | 1 | 3 | Auh | -0.01 | Young |
| 0.770478 | 0.01024 | 0.673 | 0.683 | 1 | 3 | Sod1 | -0.01 | Young |
| 0.770601 | 0.250154 | 0.057 | 0.054 | 1 | 3 | Omp | 0.003 | Young |
| 0.770622 | 0.045529 | 0.052 | 0.055 | 1 | 3 | Amd2 | -0.003 | Young |
| 0.770805 | 0.144601 | 0.057 | 0.054 | 1 | 3 | Aven | 0.003 | Young |
| 0.770906 | 0.297321 | 0.072 | 0.069 | 1 | 3 | Orc2 | 0.003 | Young |
| 0.771011 | 0.126371 | 0.303 | 0.324 | 1 | 3 | Crlf2 | -0.021 | Young |
| 0.771021 | 0.128114 | 0.366 | 0.391 | 1 | 3 | Kpnb1 | -0.025 | Young |
| 0.771087 | 0.163405 | 0.24 | 0.257 | 1 | 3 | 1700025G0 | -0.017 | Young |
| 0.771187 | 0.137413 | 0.275 | 0.277 | 1 | 3 | Itgax | -0.002 | Young |
| 0.771284 | 0.094666 | 0.091 | 0.087 | 1 | 3 | Slc41a1 | 0.004 | Young |
| 0.771429 | 0.142679 | 0.08 | 0.085 | 1 | 3 | Kdsr | -0.005 | Young |
| 0.771435 | 0.130754 | 0.035 | 0.038 | 1 | 3 | Decr2 | -0.003 | Young |
| 0.771452 | 0.175098 | 0.061 | 0.057 | 1 | 3 | Pithd1 | 0.004 | Young |
| 0.771499 | 0.020393 | 0.168 | 0.176 | 1 | 3 | Cc2d1b | -0.008 | Young |
| 0.771568 | 0.029667 | 0.023 | 0.025 | 1 | 3 | Taf1c | -0.002 | Young |
| 0.771785 | 0.035205 | 0.076 | 0.08 | 1 | 3 | Pdp2 | -0.004 | Young |
| 0.773069 | 0.076125 | 0.304 | 0.323 | 1 | 3 | Mrps14 | -0.019 | Young |
| 0.773506 | 0.13174 | 0.058 | 0.055 | 1 | 3 | Rcl1 | 0.003 | Young |
| 0.773829 | 0.079886 | 0.059 | 0.063 | 1 | 3 | Nol4l | -0.004 | Young |
| 0.77469 | 0.156515 | 0.404 | 0.429 | 1 | 3 | Vapb | -0.025 | Young |
| 0.774837 | 0.155937 | 0.067 | 0.063 | 1 | 3 | Ddt | 0.004 | Young |
| 0.774989 | 0.351018 | 0.024 | 0.026 | 1 | 3 | Rab13 | -0.002 | Young |
| 0.774989 | 0.07705 | 0.024 | 0.026 | 1 | 3 | Elp4 | -0.002 | Young |
| 0.775061 | 0.35973 | 0.068 | 0.066 | 1 | 3 | Pogz | 0.002 | Young |
| 0.775258 | 0.149911 | 0.247 | 0.265 | 1 | 3 | Rgs19 | -0.018 | Young |
| 0.775332 | 0.09427 | 0.111 | 0.117 | 1 | 3 | Bcl6 | -0.006 | Young |
| 0.775639 | 0.098172 | 0.096 | 0.102 | 1 | 3 | Mitd1 | -0.006 | Young |
| 0.77572 | 0.574686 | 0.042 | 0.039 | 1 | 3 | Dolk | 0.003 | Young |
| 0.775784 | 0.171829 | 0.076 | 0.081 | 1 | 3 | Foxred2 | -0.005 | Young |
| 0.776041 | 0.073717 | 0.053 | 0.05 | 1 | 3 | Patz1 | 0.003 | Young |

| | | | | | | | | |
|---|---|---|---|---|---|---|---|---|
| 0.776429 | 0.567816 | 0.042 | 0.039 | 1 | 3 | Rhbdd3 | 0.003 | Young |
| 0.77646 | 0.178257 | 0.045 | 0.043 | 1 | 3 | Ppm1d | 0.002 | Young |
| 0.7765 | 0.117917 | 0.194 | 0.204 | 1 | 3 | Brox | -0.01 | Young |
| 0.776524 | 0.072861 | 0.215 | 0.222 | 1 | 3 | Cux1 | -0.007 | Young |
| 0.776527 | 0.361323 | 0.013 | 0.014 | 1 | 3 | Trim46 | -0.001 | Young |
| 0.776718 | 0.291817 | 0.192 | 0.191 | 1 | 3 | D16Ertd47: | 0.001 | Young |
| 0.77748 | 0.159066 | 0.095 | 0.099 | 1 | 3 | Pigu | -0.004 | Young |
| 0.777681 | 0.027961 | 0.585 | 0.619 | 1 | 3 | Selenok | -0.034 | Young |
| 0.777849 | 0.399596 | 0.066 | 0.063 | 1 | 3 | Tex30 | 0.003 | Young |
| 0.778157 | 0.058138 | 0.14 | 0.147 | 1 | 3 | Klf16 | -0.007 | Young |
| 0.77869 | 0.112402 | 0.278 | 0.296 | 1 | 3 | Khsrp | -0.018 | Young |
| 0.778835 | 0.317603 | 0.034 | 0.037 | 1 | 3 | Mettl2 | -0.003 | Young |
| 0.778856 | 0.335734 | 0.035 | 0.038 | 1 | 3 | Snx7 | -0.003 | Young |
| 0.779037 | 0.170301 | 0.114 | 0.119 | 1 | 3 | Metap1 | -0.005 | Young |
| 0.779232 | 0.315351 | 0.13 | 0.129 | 1 | 3 | Mphosph8 | 0.001 | Young |
| 0.779406 | 0.159011 | 0.155 | 0.164 | 1 | 3 | Zwint | -0.009 | Young |
| 0.779712 | 0.025568 | 0.426 | 0.458 | 1 | 3 | Cd68 | -0.032 | Young |
| 0.780535 | 0.094915 | 0.092 | 0.097 | 1 | 3 | Hif1an | -0.005 | Young |
| 0.780647 | 0.089909 | 0.51 | 0.542 | 1 | 3 | Serinc1 | -0.032 | Young |
| 0.780819 | 0.072701 | 0.063 | 0.06 | 1 | 3 | Scly | 0.003 | Young |
| 0.780925 | 0.481365 | 0.042 | 0.039 | 1 | 3 | Ptk2 | 0.003 | Young |
| 0.781508 | 0.390929 | 0.051 | 0.048 | 1 | 3 | Fam160a2 | 0.003 | Young |
| 0.781746 | 0.078226 | 0.399 | 0.409 | 1 | 3 | Rex1bd | -0.01 | Young |
| 0.781862 | 0.337585 | 0.047 | 0.05 | 1 | 3 | 5031425F1 | -0.003 | Young |
| 0.781875 | 0.045412 | 0.051 | 0.054 | 1 | 3 | Cep85l | -0.003 | Young |
| 0.781952 | 0.166266 | 0.082 | 0.079 | 1 | 3 | Dock5 | 0.003 | Young |
| 0.78197 | 0.072263 | 0.194 | 0.205 | 1 | 3 | Ilkap | -0.011 | Young |
| 0.782364 | 0.059477 | 0.419 | 0.437 | 1 | 3 | Dynlrb1 | -0.018 | Young |
| 0.782557 | 0.568928 | 0.04 | 0.038 | 1 | 3 | Nat8l | 0.002 | Young |
| 0.782578 | 0.122505 | 0.183 | 0.195 | 1 | 3 | Trappc1 | -0.012 | Young |
| 0.782583 | 0.405321 | 0.042 | 0.039 | 1 | 3 | Tbc1d12 | 0.003 | Young |
| 0.782583 | 0.014379 | 0.039 | 0.042 | 1 | 3 | Mars | -0.003 | Young |
| 0.783407 | 0.115657 | 0.086 | 0.082 | 1 | 3 | Rcan3 | 0.004 | Young |
| 0.783662 | 0.222667 | 0.071 | 0.068 | 1 | 3 | Zfp260 | 0.003 | Young |
| 0.78373 | 0.195801 | 0.227 | 0.225 | 1 | 3 | Grk5 | 0.002 | Young |
| 0.783767 | 0.270462 | 0.213 | 0.216 | 1 | 3 | Tnfaip2 | -0.003 | Young |
| 0.783873 | 0.164221 | 0.069 | 0.074 | 1 | 3 | Zfp787 | -0.005 | Young |
| 0.783886 | 0.111627 | 0.051 | 0.048 | 1 | 3 | Dvl2 | 0.003 | Young |
| 0.784604 | 0.111552 | 0.221 | 0.233 | 1 | 3 | Churc1 | -0.012 | Young |
| 0.784624 | 0.191828 | 0.134 | 0.131 | 1 | 3 | Nhlrc3 | 0.003 | Young |
| 0.785243 | 0.607982 | 0.061 | 0.059 | 1 | 3 | Gmpr2 | 0.002 | Young |
| 0.785282 | 0.187971 | 0.037 | 0.039 | 1 | 3 | Itga1 | -0.002 | Young |
| 0.78557 | 0.239536 | 0.047 | 0.044 | 1 | 3 | Pinx1 | 0.003 | Young |
| 0.785778 | 0.243561 | 0.12 | 0.117 | 1 | 3 | Tbc1d9 | 0.003 | Young |
| 0.786042 | 0.006959 | 0.394 | 0.411 | 1 | 3 | Casp8 | -0.017 | Young |

| | | | | | | | | |
|---|---|---|---|---|---|---|---|---|
| 0.786396 | 0.119473 | 0.251 | 0.254 | 1 | 3 | Kmt5b | -0.003 | Young |
| 0.787063 | 0.089313 | 0.048 | 0.051 | 1 | 3 | Pwp2 | -0.003 | Young |
| 0.787222 | 0.07618 | 0.37 | 0.395 | 1 | 3 | Ndfip1 | -0.025 | Young |
| 0.787488 | 0.268619 | 0.092 | 0.099 | 1 | 3 | Cyp4f13 | -0.007 | Young |
| 0.787576 | 0.127217 | 0.134 | 0.129 | 1 | 3 | Srsf4 | 0.005 | Young |
| 0.788128 | 0.093875 | 0.201 | 0.21 | 1 | 3 | Ankib1 | -0.009 | Young |
| 0.788136 | 0.331232 | 0.076 | 0.08 | 1 | 3 | Foxo1 | -0.004 | Young |
| 0.788366 | 0.109024 | 0.174 | 0.182 | 1 | 3 | Alox5 | -0.008 | Young |
| 0.788642 | 0.193523 | 0.066 | 0.069 | 1 | 3 | Gamt | -0.003 | Young |
| 0.788657 | 0.050527 | 0.735 | 0.767 | 1 | 3 | Cd14 | -0.032 | Young |
| 0.789162 | 0.009452 | 0.202 | 0.209 | 1 | 3 | Atf7 | -0.007 | Young |
| 0.789417 | 0.250132 | 0.045 | 0.043 | 1 | 3 | Blzf1 | 0.002 | Young |
| 0.789468 | 0.190929 | 0.313 | 0.335 | 1 | 3 | Foxn2 | -0.022 | Young |
| 0.789893 | 0.284727 | 0.1 | 0.106 | 1 | 3 | Scyl1 | -0.006 | Young |
| 0.78995 | 0.361495 | 0.014 | 0.016 | 1 | 3 | Gm26890 | -0.002 | Young |
| 0.790122 | 0.060889 | 0.744 | 0.771 | 1 | 3 | Cxcl2 | -0.027 | Young |
| 0.790162 | 0.058058 | 0.434 | 0.47 | 1 | 3 | Papola | -0.036 | Young |
| 0.790295 | 0.39114 | 0.101 | 0.099 | 1 | 3 | Parp11 | 0.002 | Young |
| 0.790979 | 0.235873 | 0.169 | 0.17 | 1 | 3 | Mob3c | -0.001 | Young |
| 0.791248 | 0.094309 | 0.025 | 0.027 | 1 | 3 | Dmxl2 | -0.002 | Young |
| 0.791436 | 0.136589 | 0.059 | 0.063 | 1 | 3 | Pafah1b3 | -0.004 | Young |
| 0.791496 | 0.647635 | 0.086 | 0.084 | 1 | 3 | Phyh | 0.002 | Young |
| 0.79151 | 0.105787 | 0.223 | 0.235 | 1 | 3 | Dars | -0.012 | Young |
| 0.791699 | 0.323435 | 0.024 | 0.026 | 1 | 3 | Rilp | -0.002 | Young |
| 0.791716 | 0.0885 | 0.058 | 0.055 | 1 | 3 | Ccdc134 | 0.003 | Young |
| 0.792616 | 0.062563 | 0.453 | 0.49 | 1 | 3 | Mef2a | -0.037 | Young |
| 0.792752 | 0.040802 | 0.092 | 0.097 | 1 | 3 | Arhgap12 | -0.005 | Young |
| 0.792841 | 0.014215 | 0.066 | 0.069 | 1 | 3 | Snx11 | -0.003 | Young |
| 0.793017 | 0.041509 | 0.856 | 0.894 | 1 | 3 | Prcp | -0.038 | Young |
| 0.793059 | 0.280864 | 0.112 | 0.111 | 1 | 3 | Btbd9 | 0.001 | Young |
| 0.793131 | 0.063063 | 0.051 | 0.054 | 1 | 3 | 2810402E2 | -0.003 | Young |
| 0.793189 | 0.285565 | 0.198 | 0.198 | 1 | 3 | Tusc3 | 0 | Young |
| 0.793274 | 0.416631 | 0.105 | 0.111 | 1 | 3 | Gm37494 | -0.006 | Young |
| 0.793504 | 0.106777 | 0.039 | 0.042 | 1 | 3 | Flywch1 | -0.003 | Young |
| 0.793626 | 0.124461 | 0.052 | 0.049 | 1 | 3 | Acvr1 | 0.003 | Young |
| 0.7938 | 0.045665 | 0.499 | 0.515 | 1 | 3 | Mbnl2 | -0.016 | Young |
| 0.794094 | 0.206772 | 0.311 | 0.319 | 1 | 3 | Zmynd8 | -0.008 | Young |
| 0.794606 | 0.28553 | 0.04 | 0.038 | 1 | 3 | Cmah | 0.002 | Young |
| 0.794657 | 0.108479 | 0.048 | 0.045 | 1 | 3 | 5830432E0 | 0.003 | Young |
| 0.795031 | 0.123066 | 0.207 | 0.207 | 1 | 3 | Uxs1 | 0 | Young |
| 0.795055 | 0.35978 | 0.13 | 0.131 | 1 | 3 | Setd7 | -0.001 | Young |
| 0.795729 | 0.103004 | 0.242 | 0.243 | 1 | 3 | Ganab | -0.001 | Young |
| 0.795763 | 0.0041 | 0.484 | 0.519 | 1 | 3 | Ehbp1l1 | -0.035 | Young |
| 0.79581 | 0.277254 | 0.047 | 0.05 | 1 | 3 | Usp21 | -0.003 | Young |
| 0.795954 | 0.089997 | 0.092 | 0.097 | 1 | 3 | Rab9 | -0.005 | Young |

| | | | | | | | | |
|---|---|---|---|---|---|---|---|---|
| 0.79619 | 0.190583 | 0.051 | 0.054 | 1 | 3 | Nsun3 | -0.003 | Young |
| 0.796315 | 0.217838 | 0.097 | 0.094 | 1 | 3 | Tarbp2 | 0.003 | Young |
| 0.796745 | 0.203978 | 0.351 | 0.379 | 1 | 3 | Luc7l3 | -0.028 | Young |
| 0.797139 | 0.017731 | 0.292 | 0.294 | 1 | 3 | Kat6a | -0.002 | Young |
| 0.797254 | 0.074462 | 0.432 | 0.464 | 1 | 3 | Bnip3l | -0.032 | Young |
| 0.797432 | 0.152535 | 0.052 | 0.055 | 1 | 3 | Trmt10c | -0.003 | Young |
| 0.79755 | 0.129814 | 0.048 | 0.045 | 1 | 3 | Paxx | 0.003 | Young |
| 0.797615 | 0.088589 | 0.333 | 0.329 | 1 | 3 | Vps13c | 0.004 | Young |
| 0.797691 | 0.175641 | 0.104 | 0.1 | 1 | 3 | Mamdc2 | 0.004 | Young |
| 0.797877 | 0.185502 | 0.505 | 0.54 | 1 | 3 | Tgoln1 | -0.035 | Young |
| 0.797948 | 0.216993 | 0.047 | 0.044 | 1 | 3 | Exoc8 | 0.003 | Young |
| 0.797986 | 0.046376 | 0.128 | 0.133 | 1 | 3 | Cspp1 | -0.005 | Young |
| 0.798433 | 0.129218 | 0.158 | 0.154 | 1 | 3 | Tbc1d5 | 0.004 | Young |
| 0.799032 | 0.062215 | 0.058 | 0.055 | 1 | 3 | Tapbpl | 0.003 | Young |
| 0.799451 | 0.39581 | 0.146 | 0.146 | 1 | 3 | Srebf1 | 0 | Young |
| 0.799517 | 0.127986 | 0.312 | 0.313 | 1 | 3 | Anapc11 | -0.001 | Young |
| 0.799651 | 0.162453 | 0.269 | 0.289 | 1 | 3 | Hr | -0.02 | Young |
| 0.799966 | 0.310128 | 0.253 | 0.259 | 1 | 3 | Txndc9 | -0.006 | Young |
| 0.800202 | 0.175613 | 0.051 | 0.054 | 1 | 3 | Rnf144b | -0.003 | Young |
| 0.80023 | 0.187982 | 0.295 | 0.313 | 1 | 3 | Prdx6 | -0.018 | Young |
| 0.80027 | 0.140788 | 0.107 | 0.104 | 1 | 3 | Tns3 | 0.003 | Young |
| 0.80069 | 0.147496 | 0.157 | 0.153 | 1 | 3 | Cpeb2 | 0.004 | Young |
| 0.801434 | 0.194281 | 0.044 | 0.042 | 1 | 3 | 5730455P1 | 0.002 | Young |
| 0.801478 | 0.260006 | 0.351 | 0.368 | 1 | 3 | Tprgl | -0.017 | Young |
| 0.801522 | 0.011704 | 0.369 | 0.378 | 1 | 3 | Hsbp1 | -0.009 | Young |
| 0.801592 | 0.054442 | 0.148 | 0.157 | 1 | 3 | Epc2 | -0.009 | Young |
| 0.801616 | 0.065641 | 0.407 | 0.444 | 1 | 3 | Gtf2h5 | -0.037 | Young |
| 0.801637 | 0.149748 | 0.015 | 0.017 | 1 | 3 | Crybb1 | -0.002 | Young |
| 0.80229 | 0.241403 | 0.071 | 0.068 | 1 | 3 | Sacs | 0.003 | Young |
| 0.802648 | 0.112379 | 0.052 | 0.049 | 1 | 3 | Cc2d1a | 0.003 | Young |
| 0.803485 | 0.219922 | 0.095 | 0.092 | 1 | 3 | Hmbs | 0.003 | Young |
| 0.803514 | 0.083734 | 0.015 | 0.017 | 1 | 3 | Zdhhc24 | -0.002 | Young |
| 0.803597 | 0.126517 | 0.027 | 0.029 | 1 | 3 | Dnajc18 | -0.002 | Young |
| 0.803758 | 0.064062 | 0.206 | 0.216 | 1 | 3 | Arhgap39 | -0.01 | Young |
| 0.804127 | 0.506469 | 0.037 | 0.035 | 1 | 3 | Selenbp1 | 0.002 | Young |
| 0.804265 | 0.451105 | 0.015 | 0.017 | 1 | 3 | Lrp8 | -0.002 | Young |
| 0.804665 | 0.075083 | 0.347 | 0.375 | 1 | 3 | Mpc2 | -0.028 | Young |
| 0.804696 | 0.199635 | 0.071 | 0.068 | 1 | 3 | Rab6b | 0.003 | Young |
| 0.805802 | 0.238482 | 0.062 | 0.06 | 1 | 3 | Gzf1 | 0.002 | Young |
| 0.805898 | 0.020601 | 0.039 | 0.042 | 1 | 3 | Pradc1 | -0.003 | Young |
| 0.806553 | 0.046288 | 0.153 | 0.16 | 1 | 3 | Fundc2 | -0.007 | Young |
| 0.806619 | 0.271882 | 0.134 | 0.134 | 1 | 3 | Knop1 | 0 | Young |
| 0.807177 | 0.348693 | 0.107 | 0.105 | 1 | 3 | Rasgef1b | 0.002 | Young |
| 0.80718 | 0.184951 | 0.236 | 0.238 | 1 | 3 | Gdi1 | -0.002 | Young |
| 0.807206 | 0.050184 | 0.179 | 0.19 | 1 | 3 | Rab24 | -0.011 | Young |

| | | | | | | | | |
|---|---|---|---|---|---|---|---|---|
| 0.80748 | 0.125475 | 0.028 | 0.03 | 1 | 3 | Tbkbp1 | -0.002 | Young |
| 0.807592 | 0.033567 | 0.129 | 0.127 | 1 | 3 | Osbpl11 | 0.002 | Young |
| 0.808024 | 0.020395 | 0.015 | 0.017 | 1 | 3 | Tnfrsf13c | -0.002 | Young |
| 0.808094 | 0.124795 | 0.134 | 0.139 | 1 | 3 | Rab3gap2 | -0.005 | Young |
| 0.808105 | 0.067577 | 0.21 | 0.22 | 1 | 3 | Phf21a | -0.01 | Young |
| 0.808346 | 0.126761 | 0.366 | 0.375 | 1 | 3 | Nfix | -0.009 | Young |
| 0.8084 | 0.207463 | 0.015 | 0.017 | 1 | 3 | Cyb5rl | -0.002 | Young |
| 0.808476 | 0.133268 | 0.096 | 0.093 | 1 | 3 | Zbtb22 | 0.003 | Young |
| 0.808617 | 0.032058 | 0.106 | 0.11 | 1 | 3 | Rprd1b | -0.004 | Young |
| 0.808703 | 0.220446 | 0.167 | 0.166 | 1 | 3 | Map1lc3a | 0.001 | Young |
| 0.808792 | 0.047394 | 0.047 | 0.044 | 1 | 3 | Etv1 | 0.003 | Young |
| 0.809063 | 0.194877 | 0.119 | 0.116 | 1 | 3 | Rfx7 | 0.003 | Young |
| 0.809528 | 0.127648 | 0.015 | 0.017 | 1 | 3 | Srr | -0.002 | Young |
| 0.809631 | 0.005206 | 0.771 | 0.791 | 1 | 3 | Cd47 | -0.02 | Young |
| 0.809639 | 0.42217 | 0.035 | 0.033 | 1 | 3 | Zfp687 | 0.002 | Young |
| 0.809649 | 0.245826 | 0.072 | 0.069 | 1 | 3 | Abcc4 | 0.003 | Young |
| 0.809905 | 0.493552 | 0.015 | 0.017 | 1 | 3 | Arhgap35 | -0.002 | Young |
| 0.810281 | 0.541918 | 0.015 | 0.017 | 1 | 3 | 1700109H( | -0.002 | Young |
| 0.810405 | 0.48392 | 0.054 | 0.053 | 1 | 3 | Atg13 | 0.001 | Young |
| 0.81086 | 0.24837 | 0.028 | 0.03 | 1 | 3 | Lin52 | -0.002 | Young |
| 0.810958 | 0.146408 | 0.04 | 0.043 | 1 | 3 | Tatdn1 | -0.003 | Young |
| 0.811034 | 0.243159 | 0.015 | 0.017 | 1 | 3 | Nek8 | -0.002 | Young |
| 0.811253 | 0.016504 | 0.08 | 0.076 | 1 | 3 | Armc10 | 0.004 | Young |
| 0.811666 | 0.00468 | 0.437 | 0.46 | 1 | 3 | Etfb | -0.023 | Young |
| 0.812002 | 0.403717 | 0.058 | 0.056 | 1 | 3 | Rasgrp2 | 0.002 | Young |
| 0.812191 | 0.198524 | 0.152 | 0.16 | 1 | 3 | Tpst2 | -0.008 | Young |
| 0.812238 | 0.102868 | 0.097 | 0.102 | 1 | 3 | Umad1 | -0.005 | Young |
| 0.812407 | 0.072509 | 0.306 | 0.318 | 1 | 3 | Ankrd12 | -0.012 | Young |
| 0.812474 | 0.085664 | 0.333 | 0.349 | 1 | 3 | Pet100 | -0.016 | Young |
| 0.81254 | 0.252244 | 0.015 | 0.017 | 1 | 3 | Adgrg5 | -0.002 | Young |
| 0.81258 | 0.032126 | 0.11 | 0.114 | 1 | 3 | Lpin2 | -0.004 | Young |
| 0.812685 | 0.022809 | 0.241 | 0.25 | 1 | 3 | Rbm42 | -0.009 | Young |
| 0.812695 | 0.208578 | 0.187 | 0.197 | 1 | 3 | Rspry1 | -0.01 | Young |
| 0.813451 | 0.121489 | 0.08 | 0.076 | 1 | 3 | Ptpn23 | 0.004 | Young |
| 0.813464 | 0.082012 | 0.139 | 0.137 | 1 | 3 | Zyg11b | 0.002 | Young |
| 0.81354 | 0.095427 | 0.38 | 0.404 | 1 | 3 | Ep300 | -0.024 | Young |
| 0.813691 | 0.007269 | 0.527 | 0.562 | 1 | 3 | Rnf13 | -0.035 | Young |
| 0.813841 | 0.099319 | 0.067 | 0.065 | 1 | 3 | Paxip1 | 0.002 | Young |
| 0.814067 | 0.156906 | 0.369 | 0.379 | 1 | 3 | Kras | -0.01 | Young |
| 0.814749 | 0.138466 | 0.203 | 0.202 | 1 | 3 | Idh3b | 0.001 | Young |
| 0.815027 | 0.233395 | 0.034 | 0.032 | 1 | 3 | Il10 | 0.002 | Young |
| 0.815441 | 0.13191 | 0.367 | 0.407 | 1 | 3 | 6-Mar | -0.04 | Young |
| 0.815676 | 0.060833 | 0.258 | 0.265 | 1 | 3 | Sde2 | -0.007 | Young |
| 0.815694 | 0.020691 | 0.713 | 0.75 | 1 | 3 | Ncf2 | -0.037 | Young |
| 0.815834 | 0.138016 | 0.302 | 0.314 | 1 | 3 | Foxn3 | -0.012 | Young |

| | | | | | | | | |
|---|---|---|---|---|---|---|---|---|
| 0.815865 | 0.086661 | 0.437 | 0.462 | 1 | 3 | Ptgs1 | -0.025 | Young |
| 0.815951 | 0.053259 | 0.077 | 0.08 | 1 | 3 | Cdadc1 | -0.003 | Young |
| 0.816087 | 0.307112 | 0.037 | 0.035 | 1 | 3 | Poc5 | 0.002 | Young |
| 0.816292 | 0.054896 | 0.097 | 0.094 | 1 | 3 | Isyna1 | 0.003 | Young |
| 0.816347 | 0.178637 | 0.184 | 0.196 | 1 | 3 | Xrn1 | -0.012 | Young |
| 0.816349 | 0.062687 | 0.111 | 0.108 | 1 | 3 | Nsrp1 | 0.003 | Young |
| 0.816782 | 0.169507 | 0.292 | 0.299 | 1 | 3 | Dock11 | -0.007 | Young |
| 0.81697 | 0.104851 | 0.179 | 0.178 | 1 | 3 | Usp48 | 0.001 | Young |
| 0.817218 | 0.274928 | 0.061 | 0.059 | 1 | 3 | Mdc1 | 0.002 | Young |
| 0.817348 | 0.107542 | 0.116 | 0.121 | 1 | 3 | Crkl | -0.005 | Young |
| 0.818038 | 0.150329 | 0.016 | 0.018 | 1 | 3 | Tmem143 | -0.002 | Young |
| 0.818153 | 0.173559 | 0.347 | 0.357 | 1 | 3 | Ccnl2 | -0.01 | Young |
| 0.818358 | 0.303344 | 0.102 | 0.1 | 1 | 3 | Mtr | 0.002 | Young |
| 0.818379 | 0.008039 | 0.413 | 0.423 | 1 | 3 | Paip2 | -0.01 | Young |
| 0.818461 | 0.064439 | 0.326 | 0.339 | 1 | 3 | Washc2 | -0.013 | Young |
| 0.818511 | 0.000554 | 0.038 | 0.036 | 1 | 3 | Arid3b | 0.002 | Young |
| 0.818829 | 0.138168 | 0.085 | 0.088 | 1 | 3 | Gdap2 | -0.003 | Young |
| 0.818887 | 0.060223 | 0.309 | 0.307 | 1 | 3 | Sptbn1 | 0.002 | Young |
| 0.819274 | 0.541902 | 0.049 | 0.048 | 1 | 3 | Ccl12 | 0.001 | Young |
| 0.819332 | 0.061607 | 0.495 | 0.54 | 1 | 3 | Pacs2 | -0.045 | Young |
| 0.819415 | 0.293617 | 0.061 | 0.059 | 1 | 3 | Ptrh2 | 0.002 | Young |
| 0.819649 | 0.197098 | 0.116 | 0.114 | 1 | 3 | Fchsd2 | 0.002 | Young |
| 0.820294 | 0.063584 | 0.39 | 0.41 | 1 | 3 | Rassf1 | -0.02 | Young |
| 0.820379 | 0.361112 | 0.062 | 0.066 | 1 | 3 | Ttc37 | -0.004 | Young |
| 0.82145 | 0.470893 | 0.027 | 0.029 | 1 | 3 | Gkap1 | -0.002 | Young |
| 0.821563 | 0.025939 | 0.949 | 0.947 | 1 | 3 | Rpl22 | 0.002 | Young |
| 0.822359 | 0.152299 | 0.141 | 0.14 | 1 | 3 | Pdlim2 | 0.001 | Young |
| 0.822671 | 0.144524 | 0.26 | 0.263 | 1 | 3 | Ufc1 | -0.003 | Young |
| 0.823214 | 0.019717 | 0.13 | 0.135 | 1 | 3 | Ddit3 | -0.005 | Young |
| 0.823313 | 0.061143 | 0.528 | 0.546 | 1 | 3 | C1qb | -0.018 | Young |
| 0.823573 | 0.025521 | 0.029 | 0.031 | 1 | 3 | Ccdc171 | -0.002 | Young |
| 0.82385 | 0.14807 | 0.029 | 0.031 | 1 | 3 | Mcrip2 | -0.002 | Young |
| 0.823872 | 0.136107 | 0.203 | 0.198 | 1 | 3 | Abca3 | 0.005 | Young |
| 0.823952 | 0.139516 | 0.194 | 0.194 | 1 | 3 | Tmem184b | 0 | Young |
| 0.824032 | 0.136293 | 0.27 | 0.274 | 1 | 3 | Mthfd2 | -0.004 | Young |
| 0.824046 | 0.41495 | 0.028 | 0.026 | 1 | 3 | Stac2 | 0.002 | Young |
| 0.824148 | 0.217109 | 0.101 | 0.099 | 1 | 3 | Pigq | 0.002 | Young |
| 0.824711 | 0.111627 | 0.063 | 0.061 | 1 | 3 | Fam78a | 0.002 | Young |
| 0.824945 | 0.078675 | 0.157 | 0.154 | 1 | 3 | Arfgap2 | 0.003 | Young |
| 0.825316 | 0.040213 | 0.169 | 0.166 | 1 | 3 | Cebpd | 0.003 | Young |
| 0.825462 | 0.072604 | 0.035 | 0.033 | 1 | 3 | Esr1 | 0.002 | Young |
| 0.825482 | 0.12567 | 0.167 | 0.165 | 1 | 3 | Cfdp1 | 0.002 | Young |
| 0.825553 | 0.219235 | 0.162 | 0.171 | 1 | 3 | Snrnp27 | -0.009 | Young |
| 0.82582 | 0.180445 | 0.253 | 0.253 | 1 | 3 | Ppp1r10 | 0 | Young |
| 0.825974 | 0.116881 | 0.149 | 0.149 | 1 | 3 | Cnpy3 | 0 | Young |

| | | | | | | | | |
|---|---|---|---|---|---|---|---|---|
| 0.826049 | 0.230108 | 0.016 | 0.018 | 1 | 3 | Rita1 | -0.002 | Young |
| 0.826156 | 0.070917 | 0.086 | 0.084 | 1 | 3 | Rnf170 | 0.002 | Young |
| 0.826616 | 0.052029 | 0.049 | 0.047 | 1 | 3 | Adprm | 0.002 | Young |
| 0.82663 | 0.176058 | 0.073 | 0.076 | 1 | 3 | Abcd4 | -0.003 | Young |
| 0.826693 | 0.025355 | 0.76 | 0.78 | 1 | 3 | Rpl31 | -0.02 | Young |
| 0.826736 | 0.084918 | 0.441 | 0.455 | 1 | 3 | Uqcc2 | -0.014 | Young |
| 0.826847 | 0.180617 | 0.247 | 0.251 | 1 | 3 | Fam107b | -0.004 | Young |
| 0.827141 | 0.056821 | 0.463 | 0.486 | 1 | 3 | Psma6 | -0.023 | Young |
| 0.82748 | 0.086021 | 0.179 | 0.189 | 1 | 3 | Sos2 | -0.01 | Young |
| 0.827667 | 0.159968 | 0.12 | 0.118 | 1 | 3 | Pcgf3 | 0.002 | Young |
| 0.827709 | 0.081635 | 0.13 | 0.128 | 1 | 3 | Ptpn9 | 0.002 | Young |
| 0.827733 | 0.177766 | 0.145 | 0.152 | 1 | 3 | Olr1 | -0.007 | Young |
| 0.827943 | 0.437063 | 0.072 | 0.07 | 1 | 3 | Tpm3-rs7 | 0.002 | Young |
| 0.82809 | 0.135782 | 0.037 | 0.035 | 1 | 3 | Abca6 | 0.002 | Young |
| 0.828188 | 0.161425 | 0.208 | 0.21 | 1 | 3 | Tbl1x | -0.002 | Young |
| 0.828237 | 0.244194 | 0.016 | 0.018 | 1 | 3 | Msrb2 | -0.002 | Young |
| 0.828797 | 0.057075 | 0.444 | 0.471 | 1 | 3 | Nadk | -0.027 | Young |
| 0.829261 | 0.058277 | 0.396 | 0.419 | 1 | 3 | Msl1 | -0.023 | Young |
| 0.829455 | 0.184936 | 0.117 | 0.114 | 1 | 3 | Prmt3 | 0.003 | Young |
| 0.829631 | 0.067713 | 0.391 | 0.412 | 1 | 3 | Sri | -0.021 | Young |
| 0.829714 | 0.086069 | 0.201 | 0.2 | 1 | 3 | Fbxo11 | 0.001 | Young |
| 0.830096 | 0.61972 | 0.023 | 0.022 | 1 | 3 | Gm20045 | 0.001 | Young |
| 0.830128 | 0.279295 | 0.183 | 0.183 | 1 | 3 | Ppil2 | 0 | Young |
| 0.830629 | 0.106778 | 0.379 | 0.393 | 1 | 3 | Dnajc5 | -0.014 | Young |
| 0.830648 | 0.128154 | 0.282 | 0.293 | 1 | 3 | Ubxn6 | -0.011 | Young |
| 0.830806 | 0.459875 | 0.052 | 0.055 | 1 | 3 | Clec4d | -0.003 | Young |
| 0.831208 | 0.000927 | 0.039 | 0.037 | 1 | 3 | AU022252 | 0.002 | Young |
| 0.831216 | 0.469661 | 0.03 | 0.029 | 1 | 3 | 3110001I2 | 0.001 | Young |
| 0.831547 | 0.22058 | 0.198 | 0.2 | 1 | 3 | Sptan1 | -0.002 | Young |
| 0.83209 | 0.086061 | 0.086 | 0.09 | 1 | 3 | Prmt5 | -0.004 | Young |
| 0.832216 | 0.087108 | 0.03 | 0.032 | 1 | 3 | Mrm1 | -0.002 | Young |
| 0.832252 | 0.605811 | 0.016 | 0.018 | 1 | 3 | Zfp934 | -0.002 | Young |
| 0.832324 | 0.154142 | 0.131 | 0.139 | 1 | 3 | S100pbp | -0.008 | Young |
| 0.832396 | 0.153444 | 0.072 | 0.075 | 1 | 3 | Kif13b | -0.003 | Young |
| 0.832447 | 0.079722 | 0.379 | 0.385 | 1 | 3 | Tex261 | -0.006 | Young |
| 0.832477 | 0.154867 | 0.215 | 0.217 | 1 | 3 | N4bp2l1 | -0.002 | Young |
| 0.832529 | 0.036593 | 0.598 | 0.608 | 1 | 3 | Hopx | -0.01 | Young |
| 0.832755 | 0.197412 | 0.169 | 0.168 | 1 | 3 | Prr14 | 0.001 | Young |
| 0.832764 | 0.152948 | 0.128 | 0.125 | 1 | 3 | Ebi3 | 0.003 | Young |
| 0.83301 | 0.206359 | 0.043 | 0.045 | 1 | 3 | Arl14ep | -0.002 | Young |
| 0.833168 | 0.178085 | 0.018 | 0.019 | 1 | 3 | Tctn3 | -0.001 | Young |
| 0.833316 | 0.184681 | 0.36 | 0.364 | 1 | 3 | Ext1 | -0.004 | Young |
| 0.834013 | 0.045671 | 0.433 | 0.46 | 1 | 3 | Map4k4 | -0.027 | Young |
| 0.834109 | 0.096002 | 0.432 | 0.473 | 1 | 3 | Arhgap9 | -0.041 | Young |
| 0.834791 | 0.100668 | 0.059 | 0.062 | 1 | 3 | Gtf3c3 | -0.003 | Young |

| | | | | | | | | |
|---|---|---|---|---|---|---|---|---|
| 0.835392 | 0.195317 | 0.12 | 0.125 | 1 | 3 | Smim20 | -0.005 | Young |
| 0.835562 | 0.092881 | 0.212 | 0.22 | 1 | 3 | Ccdc59 | -0.008 | Young |
| 0.835577 | 0.215008 | 0.032 | 0.03 | 1 | 3 | Pde1b | 0.002 | Young |
| 0.835733 | 0.140205 | 0.177 | 0.176 | 1 | 3 | Ssbp3 | 0.001 | Young |
| 0.83595 | 0.224757 | 0.121 | 0.127 | 1 | 3 | Emc8 | -0.006 | Young |
| 0.836038 | 0.785697 | 0.025 | 0.024 | 1 | 3 | Clip2 | 0.001 | Young |
| 0.836096 | 0.136203 | 0.096 | 0.093 | 1 | 3 | Rnaseh2a | 0.003 | Young |
| 0.836144 | 0.124458 | 0.168 | 0.177 | 1 | 3 | Plcl2 | -0.009 | Young |
| 0.836303 | 0.124173 | 0.096 | 0.1 | 1 | 3 | Ccser2 | -0.004 | Young |
| 0.836345 | 0.607705 | 0.025 | 0.024 | 1 | 3 | Cmtm4 | 0.001 | Young |
| 0.837615 | 0.024741 | 0.717 | 0.771 | 1 | 3 | Snrnp70 | -0.054 | Young |
| 0.837698 | 0.257773 | 0.071 | 0.074 | 1 | 3 | Myef2 | -0.003 | Young |
| 0.837763 | 0.486957 | 0.045 | 0.044 | 1 | 3 | Ncapd3 | 0.001 | Young |
| 0.837823 | 0.239871 | 0.054 | 0.057 | 1 | 3 | Bcas3 | -0.003 | Young |
| 0.837963 | 0.303279 | 0.104 | 0.102 | 1 | 3 | Taf12 | 0.002 | Young |
| 0.838155 | 0.378269 | 0.029 | 0.031 | 1 | 3 | Lipt2 | -0.002 | Young |
| 0.838175 | 0.083635 | 0.062 | 0.065 | 1 | 3 | Narfl | -0.003 | Young |
| 0.838197 | 0.719055 | 0.048 | 0.047 | 1 | 3 | Supt3 | 0.001 | Young |
| 0.838276 | 0.332547 | 0.101 | 0.104 | 1 | 3 | Hivep3 | -0.003 | Young |
| 0.838703 | 0.177371 | 0.278 | 0.286 | 1 | 3 | Il10rb | -0.008 | Young |
| 0.838771 | 0.135254 | 0.152 | 0.161 | 1 | 3 | Bicd2 | -0.009 | Young |
| 0.839895 | 0.300378 | 0.235 | 0.24 | 1 | 3 | Exoc4 | -0.005 | Young |
| 0.840013 | 0.250953 | 0.082 | 0.08 | 1 | 3 | Mxi1 | 0.002 | Young |
| 0.84005 | 0.142885 | 0.342 | 0.352 | 1 | 3 | Zfp710 | -0.01 | Young |
| 0.840349 | 0.341222 | 0.063 | 0.061 | 1 | 3 | Trim65 | 0.002 | Young |
| 0.840438 | 0.287035 | 0.027 | 0.025 | 1 | 3 | Knstrn | 0.002 | Young |
| 0.84084 | 0.164825 | 0.217 | 0.231 | 1 | 3 | Crem | -0.014 | Young |
| 0.84085 | 0.010671 | 0.307 | 0.317 | 1 | 3 | Ncf4 | -0.01 | Young |
| 0.841025 | 0.146343 | 0.032 | 0.033 | 1 | 3 | L3mbtl3 | -0.001 | Young |
| 0.841048 | 0.232053 | 0.146 | 0.154 | 1 | 3 | Wdr44 | -0.008 | Young |
| 0.841169 | 0.09684 | 0.543 | 0.582 | 1 | 3 | Dapk1 | -0.039 | Young |
| 0.841351 | 0.096085 | 0.128 | 0.135 | 1 | 3 | Rbbp8 | -0.007 | Young |
| 0.841548 | 0.212482 | 0.073 | 0.072 | 1 | 3 | Riok2 | 0.001 | Young |
| 0.841995 | 0.198477 | 0.051 | 0.049 | 1 | 3 | Tpk1 | 0.002 | Young |
| 0.84229 | 0.237018 | 0.226 | 0.243 | 1 | 3 | Gmps | -0.017 | Young |
| 0.842537 | 0.303859 | 0.095 | 0.093 | 1 | 3 | Fgfr1op | 0.002 | Young |
| 0.842634 | 0.157595 | 0.032 | 0.033 | 1 | 3 | Snta1 | -0.001 | Young |
| 0.842658 | 0.252691 | 0.029 | 0.027 | 1 | 3 | 2410131K1 | 0.002 | Young |
| 0.842858 | 0.192654 | 0.057 | 0.06 | 1 | 3 | Tma16 | -0.003 | Young |
| 0.84315 | 0.048542 | 0.033 | 0.035 | 1 | 3 | Plk4 | -0.002 | Young |
| 0.844213 | 0.343308 | 0.196 | 0.205 | 1 | 3 | Cpsf7 | -0.009 | Young |
| 0.844915 | 0.137424 | 0.176 | 0.174 | 1 | 3 | Iah1 | 0.002 | Young |
| 0.844965 | 0.11352 | 0.054 | 0.053 | 1 | 3 | Snrnp35 | 0.001 | Young |
| 0.844975 | 0.079618 | 0.725 | 0.769 | 1 | 3 | Ndufa2 | -0.044 | Young |
| 0.845013 | 0.233396 | 0.203 | 0.205 | 1 | 3 | Gmip | -0.002 | Young |

| | | | | | | | | |
|---|---|---|---|---|---|---|---|---|
| 0.845032 | 0.12805 | 0.235 | 0.237 | 1 | 3 | Fbxo33 | -0.002 | Young |
| 0.845061 | 0.080188 | 0.194 | 0.195 | 1 | 3 | Pstpip1 | -0.001 | Young |
| 0.845104 | 0.141192 | 0.106 | 0.111 | 1 | 3 | Mrpl55 | -0.005 | Young |
| 0.84519 | 0.124014 | 0.047 | 0.049 | 1 | 3 | Dclre1a | -0.002 | Young |
| 0.845373 | 0.12292 | 0.173 | 0.171 | 1 | 3 | Zfp318 | 0.002 | Young |
| 0.845412 | 0.09935 | 0.177 | 0.176 | 1 | 3 | Txndc11 | 0.001 | Young |
| 0.845425 | 0.02219 | 0.223 | 0.229 | 1 | 3 | Bbip1 | -0.006 | Young |
| 0.845495 | 0.072666 | 0.679 | 0.716 | 1 | 3 | Mat2a | -0.037 | Young |
| 0.845528 | 0.161674 | 0.139 | 0.139 | 1 | 3 | Chml | 0 | Young |
| 0.845984 | 0.160025 | 0.223 | 0.239 | 1 | 3 | Cd300lg | -0.016 | Young |
| 0.846017 | 0.010095 | 0.336 | 0.35 | 1 | 3 | Mib1 | -0.014 | Young |
| 0.84602 | 0.145359 | 0.028 | 0.026 | 1 | 3 | Ralgps2 | 0.002 | Young |
| 0.846107 | 0.120874 | 0.196 | 0.197 | 1 | 3 | Ncor2 | -0.001 | Young |
| 0.846524 | 0.028201 | 0.019 | 0.02 | 1 | 3 | Phc1 | -0.001 | Young |
| 0.846909 | 0.162396 | 0.17 | 0.172 | 1 | 3 | Adora2b | -0.002 | Young |
| 0.846913 | 0.616813 | 0.023 | 0.022 | 1 | 3 | Fzd4 | 0.001 | Young |
| 0.846961 | 0.110452 | 0.03 | 0.029 | 1 | 3 | Rad1 | 0.001 | Young |
| 0.847246 | 0.348832 | 0.066 | 0.065 | 1 | 3 | Nedd1 | 0.001 | Young |
| 0.847643 | 0.516253 | 0.047 | 0.049 | 1 | 3 | D930016D( | -0.002 | Young |
| 0.847839 | 0.005943 | 0.237 | 0.235 | 1 | 3 | Hipk2 | 0.002 | Young |
| 0.848062 | 0.237613 | 0.311 | 0.33 | 1 | 3 | Nek7 | -0.019 | Young |
| 0.848109 | 0.1241 | 0.153 | 0.159 | 1 | 3 | Lats1 | -0.006 | Young |
| 0.848115 | 0.136702 | 0.191 | 0.2 | 1 | 3 | 2310022A1 | -0.009 | Young |
| 0.848156 | 0.21872 | 0.073 | 0.072 | 1 | 3 | Hsf2 | 0.001 | Young |
| 0.848272 | 0.117458 | 0.058 | 0.056 | 1 | 3 | Gabpb1 | 0.002 | Young |
| 0.848299 | 0.292694 | 0.086 | 0.085 | 1 | 3 | Icmt | 0.001 | Young |
| 0.848445 | 0.590304 | 0.052 | 0.05 | 1 | 3 | Sh2b1 | 0.002 | Young |
| 0.848602 | 0.040028 | 0.03 | 0.032 | 1 | 3 | Rfesd | -0.002 | Young |
| 0.848646 | 0.043918 | 0.033 | 0.031 | 1 | 3 | Sco1 | 0.002 | Young |
| 0.848776 | 0.117021 | 0.239 | 0.253 | 1 | 3 | Pi4ka | -0.014 | Young |
| 0.848945 | 0.173201 | 0.237 | 0.241 | 1 | 3 | Casp3 | -0.004 | Young |
| 0.849257 | 0.184624 | 0.13 | 0.135 | 1 | 3 | Heatr3 | -0.005 | Young |
| 0.849359 | 0.240226 | 0.059 | 0.057 | 1 | 3 | Polr3f | 0.002 | Young |
| 0.849745 | 0.055329 | 0.052 | 0.05 | 1 | 3 | Zfp865 | 0.002 | Young |
| 0.850054 | 0.205672 | 0.027 | 0.025 | 1 | 3 | Tspyl3 | 0.002 | Young |
| 0.850541 | 0.406897 | 0.024 | 0.023 | 1 | 3 | Sdhaf3 | 0.001 | Young |
| 0.850575 | 0.261283 | 0.148 | 0.148 | 1 | 3 | Retreg3 | 0 | Young |
| 0.850989 | 0.376551 | 0.019 | 0.02 | 1 | 3 | Zc3h6 | -0.001 | Young |
| 0.851039 | 0.048233 | 0.299 | 0.313 | 1 | 3 | Gbp7 | -0.014 | Young |
| 0.851063 | 0.25035 | 0.033 | 0.035 | 1 | 3 | Sgsh | -0.002 | Young |
| 0.851104 | 0.092458 | 0.153 | 0.151 | 1 | 3 | Nln | 0.002 | Young |
| 0.851202 | 0.153083 | 0.11 | 0.109 | 1 | 3 | Mfsd11 | 0.001 | Young |
| 0.851288 | 0.222035 | 0.37 | 0.4 | 1 | 3 | Mef2d | -0.03 | Young |
| 0.851855 | 0.322132 | 0.051 | 0.049 | 1 | 3 | Gna15 | 0.002 | Young |
| 0.852147 | 0.234736 | 0.187 | 0.196 | 1 | 3 | Phf8 | -0.009 | Young |

| | | | | | | | | |
|---|---|---|---|---|---|---|---|---|
| 0.85215 | 0.392426 | 0.069 | 0.073 | 1 | 3 | Prrg2 | -0.004 | Young |
| 0.852181 | 0.062882 | 0.484 | 0.52 | 1 | 3 | Akr1b3 | -0.036 | Young |
| 0.85223 | 0.149623 | 0.229 | 0.238 | 1 | 3 | Tgs1 | -0.009 | Young |
| 0.852648 | 0.139941 | 0.033 | 0.035 | 1 | 3 | Rint1 | -0.002 | Young |
| 0.852798 | 0.036018 | 0.12 | 0.124 | 1 | 3 | Dnajc10 | -0.004 | Young |
| 0.852925 | 0.243682 | 0.268 | 0.282 | 1 | 3 | Fry | -0.014 | Young |
| 0.852994 | 0.687619 | 0.019 | 0.018 | 1 | 3 | Mturn | 0.001 | Young |
| 0.853635 | 0.027106 | 0.414 | 0.438 | 1 | 3 | Bax | -0.024 | Young |
| 0.853768 | 0.364459 | 0.049 | 0.048 | 1 | 3 | Trp53rka | 0.001 | Young |
| 0.85379 | 0.234165 | 0.047 | 0.045 | 1 | 3 | Lancl1 | 0.002 | Young |
| 0.854018 | 0.198168 | 0.024 | 0.023 | 1 | 3 | Tmem53 | 0.001 | Young |
| 0.854083 | 0.244102 | 0.019 | 0.02 | 1 | 3 | B3gnt5 | -0.001 | Young |
| 0.854152 | 0.133337 | 0.122 | 0.121 | 1 | 3 | Atg9a | 0.001 | Young |
| 0.854276 | 0.251548 | 0.221 | 0.228 | 1 | 3 | Cenpx | -0.007 | Young |
| 0.854374 | 0.129463 | 0.112 | 0.117 | 1 | 3 | Rptor | -0.005 | Young |
| 0.854497 | 0.018132 | 0.033 | 0.035 | 1 | 3 | Pqlc2 | -0.002 | Young |
| 0.854779 | 0.293671 | 0.194 | 0.207 | 1 | 3 | Ccng2 | -0.013 | Young |
| 0.8549 | 0.056618 | 0.069 | 0.067 | 1 | 3 | Gps2 | 0.002 | Young |
| 0.85507 | 0.097068 | 0.384 | 0.394 | 1 | 3 | Atxn2l | -0.01 | Young |
| 0.855122 | 0.180068 | 0.198 | 0.205 | 1 | 3 | Mrps36 | -0.007 | Young |
| 0.855213 | 0.008033 | 0.989 | 0.99 | 1 | 3 | Rps27a | -0.001 | Young |
| 0.855225 | 0.025401 | 0.139 | 0.136 | 1 | 3 | Ube3c | 0.003 | Young |
| 0.855319 | 0.213741 | 0.029 | 0.027 | 1 | 3 | AW554918 | 0.002 | Young |
| 0.855349 | 0.395156 | 0.023 | 0.022 | 1 | 3 | Klrd1 | 0.001 | Young |
| 0.85601 | 0.030123 | 0.126 | 0.13 | 1 | 3 | Trmt1l | -0.004 | Young |
| 0.856128 | 0.232195 | 0.223 | 0.238 | 1 | 3 | Ncoa7 | -0.015 | Young |
| 0.856347 | 0.201228 | 0.033 | 0.035 | 1 | 3 | Fbf1 | -0.002 | Young |
| 0.856352 | 0.257968 | 0.025 | 0.024 | 1 | 3 | Bivm | 0.001 | Young |
| 0.856787 | 0.146435 | 0.145 | 0.147 | 1 | 3 | Abl2 | -0.002 | Young |
| 0.856823 | 0.133955 | 0.25 | 0.266 | 1 | 3 | Sh3bp1 | -0.016 | Young |
| 0.856973 | 0.248452 | 0.023 | 0.022 | 1 | 3 | Phf11a | 0.001 | Young |
| 0.85715 | 0.195663 | 0.236 | 0.241 | 1 | 3 | Vamp2 | -0.005 | Young |
| 0.85725 | 0.215576 | 0.242 | 0.26 | 1 | 3 | Mrpl36 | -0.018 | Young |
| 0.85732 | 0.096605 | 0.48 | 0.504 | 1 | 3 | Ube3a | -0.024 | Young |
| 0.857469 | 0.263603 | 0.095 | 0.093 | 1 | 3 | Skiv2l2 | 0.002 | Young |
| 0.857582 | 0.139799 | 0.027 | 0.025 | 1 | 3 | Fam213a | 0.002 | Young |
| 0.857685 | 0.185472 | 0.098 | 0.103 | 1 | 3 | Tvp23b | -0.005 | Young |
| 0.857847 | 0.184287 | 0.375 | 0.399 | 1 | 3 | Plin3 | -0.024 | Young |
| 0.857866 | 0.303006 | 0.034 | 0.036 | 1 | 3 | Lrrc45 | -0.002 | Young |
| 0.857883 | 0.19012 | 0.027 | 0.025 | 1 | 3 | Smagp | 0.002 | Young |
| 0.858248 | 0.142457 | 0.082 | 0.08 | 1 | 3 | Akap7 | 0.002 | Young |
| 0.858288 | 0.104423 | 0.143 | 0.148 | 1 | 3 | Ylpm1 | -0.005 | Young |
| 0.858379 | 0.006461 | 0.028 | 0.026 | 1 | 3 | Clstn1 | 0.002 | Young |
| 0.858547 | 0.131005 | 0.105 | 0.103 | 1 | 3 | Zmat5 | 0.002 | Young |
| 0.858599 | 0.182109 | 0.053 | 0.051 | 1 | 3 | Cyp2ab1 | 0.002 | Young |

| | | | | | | | | |
|---|---|---|---|---|---|---|---|---|
| 0.858687 | 0.015401 | 0.111 | 0.115 | 1 | 3 | Srpk2 | -0.004 | Young |
| 0.858923 | 0.086691 | 0.247 | 0.259 | 1 | 3 | Serpinb8 | -0.012 | Young |
| 0.85902 | 0.31507 | 0.197 | 0.198 | 1 | 3 | Taz | -0.001 | Young |
| 0.85903 | 0.568252 | 0.032 | 0.033 | 1 | 3 | Sec61a2 | -0.001 | Young |
| 0.859126 | 0.114907 | 0.405 | 0.416 | 1 | 3 | Fam117b | -0.011 | Young |
| 0.859245 | 0.376144 | 0.019 | 0.02 | 1 | 3 | Nat2 | -0.001 | Young |
| 0.859644 | 0.163794 | 0.02 | 0.022 | 1 | 3 | Aoah | -0.002 | Young |
| 0.860152 | 0.046941 | 0.097 | 0.096 | 1 | 3 | Gga3 | 0.001 | Young |
| 0.860569 | 0.119305 | 0.035 | 0.037 | 1 | 3 | Dnajc27 | -0.002 | Young |
| 0.860647 | 0.35734 | 0.021 | 0.02 | 1 | 3 | Tmem69 | 0.001 | Young |
| 0.861081 | 0.264591 | 0.047 | 0.045 | 1 | 3 | Glce | 0.002 | Young |
| 0.861083 | 0.228795 | 0.277 | 0.287 | 1 | 3 | 2510039O1 | -0.01 | Young |
| 0.861112 | 0.096359 | 0.149 | 0.154 | 1 | 3 | Fam208b | -0.005 | Young |
| 0.861159 | 0.143064 | 0.314 | 0.32 | 1 | 3 | Klc1 | -0.006 | Young |
| 0.861274 | 0.109162 | 0.085 | 0.082 | 1 | 3 | Dhx29 | 0.003 | Young |
| 0.861326 | 0.310707 | 0.049 | 0.048 | 1 | 3 | Kcnd1 | 0.001 | Young |
| 0.861793 | 0.054893 | 0.141 | 0.139 | 1 | 3 | Hdac3 | 0.002 | Young |
| 0.861959 | 0.292154 | 0.076 | 0.074 | 1 | 3 | Nudt2 | 0.002 | Young |
| 0.862122 | 0.258704 | 0.165 | 0.168 | 1 | 3 | Prpf18 | -0.003 | Young |
| 0.862319 | 0.098351 | 0.02 | 0.022 | 1 | 3 | Emilin1 | -0.002 | Young |
| 0.862405 | 0.127082 | 0.062 | 0.06 | 1 | 3 | Ankrd46 | 0.002 | Young |
| 0.862439 | 0.256514 | 0.157 | 0.158 | 1 | 3 | Stard4 | -0.001 | Young |
| 0.862532 | 0.370081 | 0.049 | 0.051 | 1 | 3 | Aktip | -0.002 | Young |
| 0.862567 | 0.317401 | 0.101 | 0.105 | 1 | 3 | Tk2 | -0.004 | Young |
| 0.862988 | 0.224166 | 0.02 | 0.022 | 1 | 3 | Mettl17 | -0.002 | Young |
| 0.863071 | 0.320076 | 0.254 | 0.274 | 1 | 3 | Igbp1 | -0.02 | Young |
| 0.863145 | 0.217101 | 0.192 | 0.195 | 1 | 3 | Ahsa1 | -0.003 | Young |
| 0.863164 | 0.241946 | 0.153 | 0.153 | 1 | 3 | Pigp | 0 | Young |
| 0.863177 | 0.164194 | 0.178 | 0.185 | 1 | 3 | Nucb2 | -0.007 | Young |
| 0.863266 | 0.229197 | 0.152 | 0.153 | 1 | 3 | Bms1 | -0.001 | Young |
| 0.863354 | 0.127398 | 0.227 | 0.228 | 1 | 3 | Abcd1 | -0.001 | Young |
| 0.863515 | 0.029621 | 0.823 | 0.86 | 1 | 3 | Gltp | -0.037 | Young |
| 0.863593 | 0.344909 | 0.198 | 0.203 | 1 | 3 | Phactr2 | -0.005 | Young |
| 0.863637 | 0.119638 | 0.054 | 0.053 | 1 | 3 | Fgd3 | 0.001 | Young |
| 0.863724 | 0.663194 | 0.019 | 0.02 | 1 | 3 | Klf8 | -0.001 | Young |
| 0.863991 | 0.310045 | 0.021 | 0.02 | 1 | 3 | Mdm1 | 0.001 | Young |
| 0.86408 | 0.25284 | 0.165 | 0.173 | 1 | 3 | Slc30a7 | -0.008 | Young |
| 0.864132 | 0.241188 | 0.111 | 0.116 | 1 | 3 | Cdk16 | -0.005 | Young |
| 0.864413 | 0.519007 | 0.02 | 0.019 | 1 | 3 | Pigg | 0.001 | Young |
| 0.864874 | 0.145233 | 0.12 | 0.124 | 1 | 3 | Slc25a17 | -0.004 | Young |
| 0.864995 | 0.010017 | 0.02 | 0.022 | 1 | 3 | Tmtc4 | -0.002 | Young |
| 0.865053 | 0.05705 | 0.395 | 0.416 | 1 | 3 | Ppp2r1a | -0.021 | Young |
| 0.865157 | 0.338697 | 0.049 | 0.051 | 1 | 3 | Pdp1 | -0.002 | Young |
| 0.865554 | 0.111299 | 0.088 | 0.092 | 1 | 3 | Zfand2b | -0.004 | Young |
| 0.865706 | 0.282528 | 0.187 | 0.196 | 1 | 3 | Rnf150 | -0.009 | Young |

| | | | | | | | | |
|---|---|---|---|---|---|---|---|---|
| 0.865918 | 0.133535 | 0.097 | 0.102 | 1 | 3 | Cask | -0.005 | Young |
| 0.866 | 0.200538 | 0.02 | 0.022 | 1 | 3 | Sox12 | -0.002 | Young |
| 0.866078 | 0.407217 | 0.061 | 0.06 | 1 | 3 | Spsb3 | 0.001 | Young |
| 0.866469 | 0.29169 | 0.049 | 0.051 | 1 | 3 | Bag4 | -0.002 | Young |
| 0.866705 | 0.068977 | 0.518 | 0.546 | 1 | 3 | Evi2a | -0.028 | Young |
| 0.866984 | 0.355753 | 0.066 | 0.065 | 1 | 3 | Pkp3 | 0.001 | Young |
| 0.867004 | 0.148183 | 0.021 | 0.02 | 1 | 3 | Pcdh7 | 0.001 | Young |
| 0.867139 | 0.204675 | 0.088 | 0.092 | 1 | 3 | Amz2 | -0.004 | Young |
| 0.867593 | 0.147037 | 0.069 | 0.072 | 1 | 3 | Cep135 | -0.003 | Young |
| 0.867648 | 0.181387 | 0.223 | 0.226 | 1 | 3 | Timm8b | -0.003 | Young |
| 0.86772 | 0.049133 | 0.163 | 0.168 | 1 | 3 | Sigmar1 | -0.005 | Young |
| 0.867724 | 0.092122 | 0.128 | 0.127 | 1 | 3 | Tstd3 | 0.001 | Young |
| 0.868088 | 0.018978 | 0.657 | 0.691 | 1 | 3 | Kdm7a | -0.034 | Young |
| 0.868327 | 0.21099 | 0.197 | 0.208 | 1 | 3 | Psmb9 | -0.011 | Young |
| 0.868489 | 0.1098 | 0.341 | 0.35 | 1 | 3 | Cbx3 | -0.009 | Young |
| 0.868536 | 0.143865 | 0.034 | 0.036 | 1 | 3 | Zfp68 | -0.002 | Young |
| 0.868633 | 0.167655 | 0.159 | 0.159 | 1 | 3 | Capn7 | 0 | Young |
| 0.868684 | 0.210099 | 0.023 | 0.022 | 1 | 3 | Sema6d | 0.001 | Young |
| 0.868732 | 0.221806 | 0.184 | 0.186 | 1 | 3 | Inpp5k | -0.002 | Young |
| 0.868938 | 0.016872 | 0.141 | 0.139 | 1 | 3 | Katnbl1 | 0.002 | Young |
| 0.869071 | 0.168599 | 0.247 | 0.262 | 1 | 3 | Slc35c2 | -0.015 | Young |
| 0.869116 | 0.457619 | 0.042 | 0.041 | 1 | 3 | Ppil1 | 0.001 | Young |
| 0.869263 | 0.018176 | 0.734 | 0.757 | 1 | 3 | Csnk1a1 | -0.023 | Young |
| 0.869309 | 0.48773 | 0.018 | 0.017 | 1 | 3 | Zfp758 | 0.001 | Young |
| 0.869661 | 0.170999 | 0.023 | 0.022 | 1 | 3 | Tti2 | 0.001 | Young |
| 0.869683 | 0.330675 | 0.021 | 0.02 | 1 | 3 | Bfsp1 | 0.001 | Young |
| 0.869706 | 0.281302 | 0.019 | 0.018 | 1 | 3 | Cox6b2 | 0.001 | Young |
| 0.869841 | 0.052669 | 0.375 | 0.394 | 1 | 3 | Higd1a | -0.019 | Young |
| 0.869879 | 0.024059 | 0.427 | 0.446 | 1 | 3 | Srsf11 | -0.019 | Young |
| 0.870001 | 0.209044 | 0.153 | 0.154 | 1 | 3 | Ncoa1 | -0.001 | Young |
| 0.870187 | 0.048697 | 0.462 | 0.474 | 1 | 3 | Ywhaq | -0.012 | Young |
| 0.870239 | 0.254451 | 0.056 | 0.054 | 1 | 3 | Stn1 | 0.002 | Young |
| 0.870497 | 0.413916 | 0.076 | 0.074 | 1 | 3 | Zbtb37 | 0.002 | Young |
| 0.870503 | 0.207497 | 0.26 | 0.265 | 1 | 3 | Gxylt1 | -0.005 | Young |
| 0.870623 | 0.179854 | 0.02 | 0.019 | 1 | 3 | Zfp760 | 0.001 | Young |
| 0.871071 | 0.136751 | 0.136 | 0.141 | 1 | 3 | Fut11 | -0.005 | Young |
| 0.871131 | 0.22963 | 0.019 | 0.018 | 1 | 3 | Bora | 0.001 | Young |
| 0.871707 | 0.030823 | 0.202 | 0.21 | 1 | 3 | Rbm28 | -0.008 | Young |
| 0.871801 | 0.099242 | 0.086 | 0.084 | 1 | 3 | Ap3s2 | 0.002 | Young |
| 0.872124 | 0.007792 | 0.587 | 0.619 | 1 | 3 | Tab2 | -0.032 | Young |
| 0.872362 | 0.16821 | 0.045 | 0.044 | 1 | 3 | Acss2 | 0.001 | Young |
| 0.872694 | 0.714887 | 0.02 | 0.019 | 1 | 3 | 10-Sep | 0.001 | Young |
| 0.87279 | 0.267051 | 0.054 | 0.053 | 1 | 3 | Wdyhv1 | 0.001 | Young |
| 0.872911 | 0.434089 | 0.016 | 0.016 | 1 | 3 | Flt1 | 0 | Young |
| 0.872934 | 0.065953 | 0.357 | 0.363 | 1 | 3 | Rbbp6 | -0.006 | Young |

| | | | | | | | | |
|---|---|---|---|---|---|---|---|---|
| 0.87303 | 0.013127 | 0.023 | 0.024 | 1 | 3 | Slx4 | -0.001 | Young |
| 0.873307 | 0.01928 | 0.109 | 0.111 | 1 | 3 | Vrk3 | -0.002 | Young |
| 0.873416 | 0.547824 | 0.015 | 0.014 | 1 | 3 | Zfp446 | 0.001 | Young |
| 0.873656 | 0.042655 | 0.14 | 0.139 | 1 | 3 | Ubxn2a | 0.001 | Young |
| 0.87384 | 0.409301 | 0.044 | 0.043 | 1 | 3 | Gorab | 0.001 | Young |
| 0.873894 | 0.03525 | 0.398 | 0.407 | 1 | 3 | Akt2 | -0.009 | Young |
| 0.873904 | 0.460893 | 0.043 | 0.042 | 1 | 3 | Ifitm5 | 0.001 | Young |
| 0.874441 | 0.56862 | 0.016 | 0.016 | 1 | 3 | Als2 | 0 | Young |
| 0.874791 | 0.032807 | 0.038 | 0.039 | 1 | 3 | C5ar2 | -0.001 | Young |
| 0.875019 | 0.359127 | 0.096 | 0.096 | 1 | 3 | Taf7 | 0 | Young |
| 0.875583 | 0.192523 | 0.128 | 0.128 | 1 | 3 | Frmd8 | 0 | Young |
| 0.876053 | 0.218484 | 0.021 | 0.02 | 1 | 3 | Nup37 | 0.001 | Young |
| 0.876161 | 0.047768 | 0.193 | 0.201 | 1 | 3 | Sf3a1 | -0.008 | Young |
| 0.876815 | 0.029452 | 0.134 | 0.137 | 1 | 3 | Grsf1 | -0.003 | Young |
| 0.876843 | 0.099195 | 0.331 | 0.346 | 1 | 3 | Pura | -0.015 | Young |
| 0.877131 | 0.626563 | 0.04 | 0.039 | 1 | 3 | Whamm | 0.001 | Young |
| 0.877477 | 0.031837 | 0.024 | 0.023 | 1 | 3 | Plcg1 | 0.001 | Young |
| 0.877877 | 0.011315 | 0.02 | 0.019 | 1 | 3 | Dtwd2 | 0.001 | Young |
| 0.878139 | 0.174009 | 0.021 | 0.023 | 1 | 3 | Gm14326 | -0.002 | Young |
| 0.878595 | 0.07502 | 0.083 | 0.082 | 1 | 3 | Tmem161b | 0.001 | Young |
| 0.878656 | 0.233738 | 0.066 | 0.068 | 1 | 3 | Ccdc93 | -0.002 | Young |
| 0.878747 | 0.165712 | 0.342 | 0.348 | 1 | 3 | Npepl1 | -0.006 | Young |
| 0.879335 | 0.150917 | 0.119 | 0.118 | 1 | 3 | Slco3a1 | 0.001 | Young |
| 0.879446 | 0.271903 | 0.101 | 0.1 | 1 | 3 | Sin3a | 0.001 | Young |
| 0.879568 | 0.152711 | 0.383 | 0.398 | 1 | 3 | Bptf | -0.015 | Young |
| 0.879969 | 0.031346 | 0.616 | 0.645 | 1 | 3 | Skap2 | -0.029 | Young |
| 0.880171 | 0.23629 | 0.264 | 0.28 | 1 | 3 | Lsr | -0.016 | Young |
| 0.88034 | 0.085816 | 0.053 | 0.051 | 1 | 3 | Ap3m2 | 0.002 | Young |
| 0.880342 | 0.095268 | 0.568 | 0.599 | 1 | 3 | Irf2 | -0.031 | Young |
| 0.880506 | 0.019979 | 0.009 | 0.01 | 1 | 3 | Selp | -0.001 | Young |
| 0.880535 | 0.328005 | 0.034 | 0.036 | 1 | 3 | Zfyve28 | -0.002 | Young |
| 0.880761 | 0.138773 | 0.019 | 0.018 | 1 | 3 | Lrrc49 | 0.001 | Young |
| 0.880809 | 0.340823 | 0.043 | 0.042 | 1 | 3 | Slc38a6 | 0.001 | Young |
| 0.881007 | 0.300672 | 0.138 | 0.139 | 1 | 3 | Afdn | -0.001 | Young |
| 0.88109 | 0.493053 | 0.043 | 0.045 | 1 | 3 | Emc9 | -0.002 | Young |
| 0.881247 | 0.143657 | 0.13 | 0.135 | 1 | 3 | Ext2 | -0.005 | Young |
| 0.881511 | 0.052581 | 0.009 | 0.01 | 1 | 3 | Prkcg | -0.001 | Young |
| 0.881796 | 0.013069 | 0.9 | 0.924 | 1 | 3 | Ptafr | -0.024 | Young |
| 0.881929 | 0.154894 | 0.163 | 0.165 | 1 | 3 | Dcaf13 | -0.002 | Young |
| 0.882014 | 0.310196 | 0.009 | 0.01 | 1 | 3 | Pclaf | -0.001 | Young |
| 0.882014 | 0.07716 | 0.009 | 0.01 | 1 | 3 | Sdr42e1 | -0.001 | Young |
| 0.882331 | 0.048343 | 0.129 | 0.134 | 1 | 3 | Tmem245 | -0.005 | Young |
| 0.882345 | 0.116783 | 0.174 | 0.178 | 1 | 3 | Dyrk1a | -0.004 | Young |
| 0.882355 | 0.127586 | 0.064 | 0.067 | 1 | 3 | Plrg1 | -0.003 | Young |
| 0.8824 | 0.192193 | 0.095 | 0.094 | 1 | 3 | Clec2i | 0.001 | Young |

| | | | | | | | | |
|---|---|---|---|---|---|---|---|---|
| 0.882753 | 0.576996 | 0.039 | 0.038 | 1 | 3 | Tbl2 | 0.001 | Young |
| 0.882852 | 0.24225 | 0.086 | 0.085 | 1 | 3 | Kifc3 | 0.001 | Young |
| 0.882864 | 0.201507 | 0.016 | 0.016 | 1 | 3 | Gm5431 | 0 | Young |
| 0.882948 | 0.190949 | 0.177 | 0.177 | 1 | 3 | Micu2 | 0 | Young |
| 0.883247 | 0.102674 | 0.016 | 0.016 | 1 | 3 | C1rl | 0 | Young |
| 0.883503 | 0.012185 | 0.371 | 0.38 | 1 | 3 | Cyth1 | -0.009 | Young |
| 0.883522 | 0.263527 | 0.009 | 0.01 | 1 | 3 | Loxl3 | -0.001 | Young |
| 0.88377 | 0.202042 | 0.015 | 0.014 | 1 | 3 | Zbtb26 | 0.001 | Young |
| 0.883846 | 0.037501 | 0.317 | 0.327 | 1 | 3 | Tmx3 | -0.01 | Young |
| 0.884008 | 0.129305 | 0.231 | 0.237 | 1 | 3 | Id3 | -0.006 | Young |
| 0.884025 | 0.187777 | 0.009 | 0.01 | 1 | 3 | Bbs9 | -0.001 | Young |
| 0.88403 | 0.40411 | 0.188 | 0.201 | 1 | 3 | Ing4 | -0.013 | Young |
| 0.884104 | 0.111491 | 0.237 | 0.247 | 1 | 3 | Fam173a | -0.01 | Young |
| 0.884169 | 0.171315 | 0.015 | 0.014 | 1 | 3 | Sorcs2 | 0.001 | Young |
| 0.884449 | 0.087394 | 0.083 | 0.086 | 1 | 3 | Pvt1 | -0.003 | Young |
| 0.884527 | 0.294602 | 0.009 | 0.01 | 1 | 3 | Pcbp3 | -0.001 | Young |
| 0.88478 | 0.193871 | 0.016 | 0.016 | 1 | 3 | 1810044D( | 0 | Young |
| 0.884955 | 0.444988 | 0.014 | 0.013 | 1 | 3 | Mvb12b | 0.001 | Young |
| 0.885703 | 0.260497 | 0.035 | 0.037 | 1 | 3 | Wdr47 | -0.002 | Young |
| 0.885787 | 0.405749 | 0.014 | 0.013 | 1 | 3 | Atp5s | 0.001 | Young |
| 0.885996 | 0.763065 | 0.013 | 0.012 | 1 | 3 | Mir9-3hg | 0.001 | Young |
| 0.886208 | 0.075527 | 0.119 | 0.123 | 1 | 3 | Dbt | -0.004 | Young |
| 0.886295 | 0.204011 | 0.044 | 0.043 | 1 | 3 | Lin37 | 0.001 | Young |
| 0.886304 | 0.221474 | 0.021 | 0.023 | 1 | 3 | Zfp931 | -0.002 | Young |
| 0.886314 | 0.129858 | 0.016 | 0.016 | 1 | 3 | Cd79b | 0 | Young |
| 0.886358 | 0.036412 | 0.399 | 0.41 | 1 | 3 | Gpcpd1 | -0.011 | Young |
| 0.886361 | 0.163799 | 0.167 | 0.173 | 1 | 3 | Ltv1 | -0.006 | Young |
| 0.886507 | 0.107163 | 0.045 | 0.044 | 1 | 3 | Ccdc28b | 0.001 | Young |
| 0.886738 | 0.541999 | 0.038 | 0.039 | 1 | 3 | Exoc6 | -0.001 | Young |
| 0.886941 | 0.191693 | 0.04 | 0.039 | 1 | 3 | Wbp1 | 0.001 | Young |
| 0.887008 | 0.104356 | 0.15 | 0.151 | 1 | 3 | Rbl2 | -0.001 | Young |
| 0.887174 | 0.154455 | 0.058 | 0.06 | 1 | 3 | Terf2ip | -0.002 | Young |
| 0.887486 | 0.191594 | 0.24 | 0.244 | 1 | 3 | Nrip1 | -0.004 | Young |
| 0.887764 | 0.155127 | 0.067 | 0.069 | 1 | 3 | Cks1b | -0.002 | Young |
| 0.88802 | 0.181754 | 0.139 | 0.142 | 1 | 3 | Lcor | -0.003 | Young |
| 0.888165 | 0.031339 | 0.278 | 0.293 | 1 | 3 | Gch1 | -0.015 | Young |
| 0.888552 | 0.485175 | 0.009 | 0.01 | 1 | 3 | Mpzl3 | -0.001 | Young |
| 0.888631 | 0.182801 | 0.11 | 0.112 | 1 | 3 | Nsmf | -0.002 | Young |
| 0.888902 | 0.114891 | 0.037 | 0.038 | 1 | 3 | Pde4a | -0.001 | Young |
| 0.888942 | 0.07405 | 0.386 | 0.409 | 1 | 3 | Rab31 | -0.023 | Young |
| 0.889093 | 0.280589 | 0.086 | 0.09 | 1 | 3 | Inafm1 | -0.004 | Young |
| 0.889535 | 0.256278 | 0.014 | 0.013 | 1 | 3 | Iqcc | 0.001 | Young |
| 0.88995 | 0.085463 | 0.255 | 0.265 | 1 | 3 | Npc1 | -0.01 | Young |
| 0.889951 | 0.256794 | 0.014 | 0.013 | 1 | 3 | Nhej1 | 0.001 | Young |
| 0.890062 | 0.796566 | 0.009 | 0.01 | 1 | 3 | Zfp251 | -0.001 | Young |

| | | | | | | | | |
|---|---|---|---|---|---|---|---|---|
| 0.890111 | 0.179048 | 0.181 | 0.183 | 1 | 3 | Mettl23 | -0.002 | Young |
| 0.890131 | 0.291386 | 0.091 | 0.091 | 1 | 3 | Rfx1 | 0 | Young |
| 0.890174 | 0.019648 | 0.251 | 0.254 | 1 | 3 | Slc25a39 | -0.003 | Young |
| 0.890204 | 0.501777 | 0.037 | 0.036 | 1 | 3 | Dgke | 0.001 | Young |
| 0.890368 | 0.281282 | 0.014 | 0.013 | 1 | 3 | Nkapl | 0.001 | Young |
| 0.890376 | 0.526683 | 0.042 | 0.041 | 1 | 3 | Mthfsd | 0.001 | Young |
| 0.890584 | 0.221512 | 0.176 | 0.178 | 1 | 3 | Trappc6a | -0.002 | Young |
| 0.890785 | 0.41996 | 0.014 | 0.013 | 1 | 3 | Snx33 | 0.001 | Young |
| 0.890798 | 0.591288 | 0.013 | 0.012 | 1 | 3 | Siah1b | 0.001 | Young |
| 0.890798 | 0.533751 | 0.013 | 0.012 | 1 | 3 | Spsb4 | 0.001 | Young |
| 0.890869 | 0.052174 | 0.04 | 0.039 | 1 | 3 | Klhl15 | 0.001 | Young |
| 0.89093 | 0.032513 | 0.135 | 0.14 | 1 | 3 | Tbk1 | -0.005 | Young |
| 0.890995 | 0.163869 | 0.266 | 0.271 | 1 | 3 | Plcb1 | -0.005 | Young |
| 0.8911 | 0.374016 | 0.039 | 0.038 | 1 | 3 | Rnf185 | 0.001 | Young |
| 0.891205 | 0.010077 | 0.136 | 0.14 | 1 | 3 | Rpl10-ps3 | -0.004 | Young |
| 0.891671 | 0.225602 | 0.013 | 0.012 | 1 | 3 | 1700052K1 | 0.001 | Young |
| 0.891673 | 0.105002 | 0.372 | 0.398 | 1 | 3 | Abhd12 | -0.026 | Young |
| 0.891826 | 0.096418 | 0.343 | 0.363 | 1 | 3 | Tfe3 | -0.02 | Young |
| 0.891829 | 0.220442 | 0.042 | 0.041 | 1 | 3 | Fance | 0.001 | Young |
| 0.891897 | 0.132184 | 0.078 | 0.081 | 1 | 3 | Cpsf4 | -0.003 | Young |
| 0.891951 | 0.193223 | 0.48 | 0.532 | 1 | 3 | Akt1 | -0.052 | Young |
| 0.892285 | 0.090296 | 0.519 | 0.536 | 1 | 3 | Zbtb7a | -0.017 | Young |
| 0.892448 | 0.332786 | 0.037 | 0.038 | 1 | 3 | Shq1 | -0.001 | Young |
| 0.892594 | 0.033148 | 0.052 | 0.054 | 1 | 3 | Tgfbr1 | -0.002 | Young |
| 0.892609 | 0.277223 | 0.091 | 0.094 | 1 | 3 | 0610037L1 | -0.003 | Young |
| 0.892704 | 0.047275 | 0.061 | 0.062 | 1 | 3 | Def8 | -0.001 | Young |
| 0.892767 | 0.083781 | 0.441 | 0.468 | 1 | 3 | Snrpg | -0.027 | Young |
| 0.892947 | 0.083706 | 0.015 | 0.014 | 1 | 3 | Tctn1 | 0.001 | Young |
| 0.893122 | 0.197693 | 0.044 | 0.043 | 1 | 3 | Nr1d1 | 0.001 | Young |
| 0.893129 | 0.227201 | 0.279 | 0.296 | 1 | 3 | Plekha2 | -0.017 | Young |
| 0.893335 | 0.080827 | 0.069 | 0.072 | 1 | 3 | Nf2 | -0.003 | Young |
| 0.893346 | 0.064088 | 0.015 | 0.014 | 1 | 3 | Trim68 | 0.001 | Young |
| 0.893698 | 0.087734 | 0.136 | 0.141 | 1 | 3 | Mocs2 | -0.005 | Young |
| 0.893708 | 0.030861 | 0.072 | 0.074 | 1 | 3 | Mrps11 | -0.002 | Young |
| 0.89372 | 0.098934 | 0.116 | 0.121 | 1 | 3 | Mb21d1 | -0.005 | Young |
| 0.893877 | 0.216431 | 0.112 | 0.112 | 1 | 3 | Stard3 | 0 | Young |
| 0.893891 | 0.101126 | 0.179 | 0.184 | 1 | 3 | Bin2 | -0.005 | Young |
| 0.893968 | 0.164791 | 0.14 | 0.146 | 1 | 3 | Mid1 | -0.006 | Young |
| 0.894119 | 0.466142 | 0.014 | 0.013 | 1 | 3 | Fut4 | 0.001 | Young |
| 0.894343 | 0.225995 | 0.039 | 0.038 | 1 | 3 | Skp2 | 0.001 | Young |
| 0.894498 | 0.367281 | 0.01 | 0.011 | 1 | 3 | Gcnt1 | -0.001 | Young |
| 0.89453 | 0.084826 | 0.452 | 0.464 | 1 | 3 | Ndufb5 | -0.012 | Young |
| 0.894766 | 0.160228 | 0.024 | 0.025 | 1 | 3 | Nt5c3b | -0.001 | Young |
| 0.894888 | 0.106324 | 0.407 | 0.436 | 1 | 3 | Mafg | -0.029 | Young |
| 0.894905 | 0.016934 | 0.138 | 0.136 | 1 | 3 | Fbxo22 | 0.002 | Young |

| | | | | | | | | |
|---|---|---|---|---|---|---|---|---|
| 0.894931 | 0.074874 | 0.107 | 0.11 | 1 | 3 | Grcc10 | -0.003 | Young |
| 0.895272 | 0.098718 | 0.179 | 0.185 | 1 | 3 | Ctbp2 | -0.006 | Young |
| 0.895603 | 0.223246 | 0.013 | 0.012 | 1 | 3 | Zfp408 | 0.001 | Young |
| 0.895665 | 0.208755 | 0.168 | 0.176 | 1 | 3 | Sf3a3 | -0.008 | Young |
| 0.895699 | 0.191839 | 0.063 | 0.066 | 1 | 3 | Kifap3 | -0.003 | Young |
| 0.895765 | 0.056636 | 0.145 | 0.145 | 1 | 3 | Ccdc25 | 0 | Young |
| 0.89604 | 0.54163 | 0.013 | 0.012 | 1 | 3 | Zhx2 | 0.001 | Young |
| 0.896074 | 0.237094 | 0.081 | 0.08 | 1 | 3 | Trim24 | 0.001 | Young |
| 0.896153 | 0.24328 | 0.04 | 0.039 | 1 | 3 | Pex1 | 0.001 | Young |
| 0.896477 | 0.60848 | 0.013 | 0.012 | 1 | 3 | B3galt2 | 0.001 | Young |
| 0.896594 | 0.229327 | 0.206 | 0.209 | 1 | 3 | Wdfy2 | -0.003 | Young |
| 0.8969 | 0.296409 | 0.064 | 0.067 | 1 | 3 | Gphn | -0.003 | Young |
| 0.897037 | 0.315394 | 0.014 | 0.013 | 1 | 3 | Zfp748 | 0.001 | Young |
| 0.897096 | 0.038665 | 0.91 | 0.943 | 1 | 3 | Cox4i1 | -0.033 | Young |
| 0.897132 | 0.315146 | 0.066 | 0.065 | 1 | 3 | Rnf31 | 0.001 | Young |
| 0.897165 | 0.058628 | 0.076 | 0.078 | 1 | 3 | Pnkp | -0.002 | Young |
| 0.89735 | 0.023829 | 0.116 | 0.119 | 1 | 3 | Csrp1 | -0.003 | Young |
| 0.897352 | 0.136163 | 0.013 | 0.012 | 1 | 3 | Gm10269 | 0.001 | Young |
| 0.89739 | 0.261916 | 0.054 | 0.056 | 1 | 3 | Rfng | -0.002 | Young |
| 0.897423 | 0.459542 | 0.011 | 0.011 | 1 | 3 | Gm20632 | 0 | Young |
| 0.897521 | 0.065292 | 0.04 | 0.042 | 1 | 3 | Zfp317 | -0.002 | Young |
| 0.897572 | 0.240644 | 0.197 | 0.202 | 1 | 3 | Diaph2 | -0.005 | Young |
| 0.897597 | 0.107705 | 0.268 | 0.276 | 1 | 3 | Rbbp4 | -0.008 | Young |
| 0.897717 | 0.218371 | 0.255 | 0.263 | 1 | 3 | Trpm7 | -0.008 | Young |
| 0.897772 | 0.046912 | 0.092 | 0.096 | 1 | 3 | Sik2 | -0.004 | Young |
| 0.897841 | 0.207036 | 0.081 | 0.084 | 1 | 3 | Lonp1 | -0.003 | Young |
| 0.897871 | 0.085846 | 0.014 | 0.013 | 1 | 3 | Ffar4 | 0.001 | Young |
| 0.897989 | 0.081506 | 0.087 | 0.09 | 1 | 3 | Brcc3 | -0.003 | Young |
| 0.898156 | 0.097397 | 0.09 | 0.092 | 1 | 3 | Sh3gl1 | -0.002 | Young |
| 0.898288 | 0.133323 | 0.01 | 0.011 | 1 | 3 | Coq8a | -0.001 | Young |
| 0.898663 | 0.145913 | 0.013 | 0.012 | 1 | 3 | 4930522L1 | 0.001 | Young |
| 0.89877 | 0.063794 | 0.496 | 0.523 | 1 | 3 | Cdc42se1 | -0.027 | Young |
| 0.898805 | 0.290657 | 0.011 | 0.011 | 1 | 3 | Adal | 0 | Young |
| 0.899101 | 0.04598 | 0.013 | 0.012 | 1 | 3 | B3galnt1 | 0.001 | Young |
| 0.899235 | 0.065801 | 0.01 | 0.011 | 1 | 3 | Eef2k | -0.001 | Young |
| 0.899265 | 0.180071 | 0.011 | 0.011 | 1 | 3 | Cep41 | 0 | Young |
| 0.899495 | 0.062477 | 0.182 | 0.184 | 1 | 3 | Tmem106b | -0.002 | Young |
| 0.89954 | 0.096863 | 0.014 | 0.013 | 1 | 3 | Smyd4 | 0.001 | Young |
| 0.899604 | 0.233301 | 0.043 | 0.044 | 1 | 3 | Gm27010 | -0.001 | Young |
| 0.899668 | 0.089899 | 0.066 | 0.065 | 1 | 3 | Dtx2 | 0.001 | Young |
| 0.899709 | 0.630965 | 0.01 | 0.011 | 1 | 3 | Slc25a42 | -0.001 | Young |
| 0.900085 | 0.034659 | 0.843 | 0.861 | 1 | 3 | Rps27l | -0.018 | Young |
| 0.900183 | 0.072013 | 0.348 | 0.363 | 1 | 3 | Acap2 | -0.015 | Young |
| 0.900183 | 0.082353 | 0.01 | 0.011 | 1 | 3 | Frat1 | -0.001 | Young |
| 0.900187 | 0.250734 | 0.011 | 0.011 | 1 | 3 | Tnks1bp1 | 0 | Young |

| | | | | | | | | |
|---|---|---|---|---|---|---|---|---|
| 0.900226 | 0.1863 | 0.274 | 0.287 | 1 | 3 | Zfp638 | -0.013 | Young |
| 0.900577 | 0.034328 | 0.59 | 0.624 | 1 | 3 | Eif3c | -0.034 | Young |
| 0.900699 | 0.354873 | 0.058 | 0.057 | 1 | 3 | Nup155 | 0.001 | Young |
| 0.90085 | 0.118732 | 0.013 | 0.012 | 1 | 3 | 6330562C2 | 0.001 | Young |
| 0.90085 | 0.096683 | 0.013 | 0.012 | 1 | 3 | Lmbr1 | 0.001 | Young |
| 0.90085 | 0.041644 | 0.013 | 0.012 | 1 | 3 | Gm38394 | 0.001 | Young |
| 0.900859 | 0.097859 | 0.043 | 0.042 | 1 | 3 | Cad | 0.001 | Young |
| 0.900917 | 0.200556 | 0.082 | 0.081 | 1 | 3 | Med25 | 0.001 | Young |
| 0.900922 | 0.074008 | 0.067 | 0.066 | 1 | 3 | Hddc2 | 0.001 | Young |
| 0.901136 | 0.264748 | 0.23 | 0.238 | 1 | 3 | Lrp6 | -0.008 | Young |
| 0.901271 | 0.104226 | 0.194 | 0.195 | 1 | 3 | Plxna4 | -0.001 | Young |
| 0.901288 | 0.130501 | 0.013 | 0.012 | 1 | 3 | Suv39h2 | 0.001 | Young |
| 0.901868 | 0.083548 | 0.081 | 0.084 | 1 | 3 | Mlycd | -0.003 | Young |
| 0.901983 | 0.075394 | 0.146 | 0.151 | 1 | 3 | Fbxo3 | -0.005 | Young |
| 0.902079 | 0.340392 | 0.01 | 0.011 | 1 | 3 | Dpep3 | -0.001 | Young |
| 0.902447 | 0.026648 | 0.054 | 0.056 | 1 | 3 | Gbp5 | -0.002 | Young |
| 0.902491 | 0.161201 | 0.011 | 0.011 | 1 | 3 | Ap5b1 | 0 | Young |
| 0.902552 | 0.122473 | 0.423 | 0.447 | 1 | 3 | Ap1s2 | -0.024 | Young |
| 0.9026 | 0.072596 | 0.013 | 0.012 | 1 | 3 | Bco2 | 0.001 | Young |
| 0.90267 | 0.03674 | 0.043 | 0.044 | 1 | 3 | Gm6563 | -0.001 | Young |
| 0.902754 | 0.05514 | 0.273 | 0.274 | 1 | 3 | Saraf | -0.001 | Young |
| 0.903031 | 0.120906 | 0.056 | 0.057 | 1 | 3 | Nlrp1b | -0.001 | Young |
| 0.903174 | 0.219704 | 0.093 | 0.093 | 1 | 3 | Ppfia4 | 0 | Young |
| 0.903461 | 0.15828 | 0.569 | 0.606 | 1 | 3 | Mafb | -0.037 | Young |
| 0.903509 | 0.592592 | 0.01 | 0.01 | 1 | 3 | Slc22a17 | 0 | Young |
| 0.90361 | 0.024269 | 0.042 | 0.043 | 1 | 3 | Bmp2 | -0.001 | Young |
| 0.903717 | 0.166201 | 0.225 | 0.235 | 1 | 3 | Rbm33 | -0.01 | Young |
| 0.903841 | 0.096431 | 0.47 | 0.507 | 1 | 3 | Amfr | -0.037 | Young |
| 0.903957 | 0.102454 | 0.303 | 0.319 | 1 | 3 | Axl | -0.016 | Young |
| 0.903998 | 0.481665 | 0.01 | 0.01 | 1 | 3 | Il18r1 | 0 | Young |
| 0.904004 | 0.033244 | 0.428 | 0.458 | 1 | 3 | Rhog | -0.03 | Young |
| 0.904601 | 0.068756 | 0.566 | 0.591 | 1 | 3 | Ldlr | -0.025 | Young |
| 0.904796 | 0.14717 | 0.011 | 0.011 | 1 | 3 | Angptl4 | 0 | Young |
| 0.905075 | 0.058495 | 0.289 | 0.29 | 1 | 3 | Mark2 | -0.001 | Young |
| 0.905226 | 0.135516 | 0.013 | 0.012 | 1 | 3 | Gm14005 | 0.001 | Young |
| 0.905236 | 0.06288 | 0.287 | 0.301 | 1 | 3 | Setd2 | -0.014 | Young |
| 0.905287 | 0.048266 | 0.24 | 0.247 | 1 | 3 | Madd | -0.007 | Young |
| 0.905438 | 0.041628 | 0.048 | 0.047 | 1 | 3 | Taf1b | 0.001 | Young |
| 0.905802 | 0.176481 | 0.407 | 0.425 | 1 | 3 | Cdc37 | -0.018 | Young |
| 0.905953 | 1.030616 | 0.01 | 0.01 | 1 | 3 | Cdkl3 | 0 | Young |
| 0.90598 | 0.632372 | 0.052 | 0.051 | 1 | 3 | Mrpl39 | 0.001 | Young |
| 0.906165 | 0.562812 | 0.034 | 0.033 | 1 | 3 | Cnksr3 | 0.001 | Young |
| 0.906179 | 0.361926 | 0.011 | 0.011 | 1 | 3 | A230050P2 | 0 | Young |
| 0.906195 | 0.230508 | 0.246 | 0.251 | 1 | 3 | S100a1 | -0.005 | Young |
| 0.90678 | 0.064291 | 0.139 | 0.137 | 1 | 3 | 9130401M | 0.002 | Young |

| | | | | | | | | |
|---|---|---|---|---|---|---|---|---|
| 0.906822 | 0.116697 | 0.01 | 0.011 | 1 | 3 | Magi3 | -0.001 | Young |
| 0.906956 | 0.031546 | 0.206 | 0.209 | 1 | 3 | Matk | -0.003 | Young |
| 0.907102 | 0.004574 | 0.011 | 0.011 | 1 | 3 | Gm4707 | 0 | Young |
| 0.907112 | 0.155707 | 0.158 | 0.158 | 1 | 3 | Comtd1 | 0 | Young |
| 0.907419 | 0.500327 | 0.01 | 0.01 | 1 | 3 | Ankrd24 | 0 | Young |
| 0.907419 | 0.304669 | 0.01 | 0.01 | 1 | 3 | Proser3 | 0 | Young |
| 0.907442 | 0.245439 | 0.062 | 0.061 | 1 | 3 | Ercc5 | 0.001 | Young |
| 0.907509 | 0.044347 | 0.104 | 0.106 | 1 | 3 | Anapc4 | -0.002 | Young |
| 0.907613 | 0.087532 | 0.293 | 0.309 | 1 | 3 | Rnasel | -0.016 | Young |
| 0.907653 | 0.363638 | 0.077 | 0.076 | 1 | 3 | Crcp | 0.001 | Young |
| 0.908137 | 0.009039 | 0.082 | 0.084 | 1 | 3 | Rpp25l | -0.002 | Young |
| 0.908329 | 0.099904 | 0.313 | 0.317 | 1 | 3 | Vwa5a | -0.004 | Young |
| 0.908397 | 0.292473 | 0.01 | 0.01 | 1 | 3 | Gm16124 | 0 | Young |
| 0.908886 | 0.147606 | 0.01 | 0.01 | 1 | 3 | Napepld | 0 | Young |
| 0.909257 | 0.231791 | 0.098 | 0.098 | 1 | 3 | Psmd5 | 0 | Young |
| 0.90933 | 0.275853 | 0.039 | 0.038 | 1 | 3 | Pomgnt1 | 0.001 | Young |
| 0.909332 | 0.212261 | 0.08 | 0.079 | 1 | 3 | Ntpcr | 0.001 | Young |
| 0.909494 | 0.157325 | 0.133 | 0.131 | 1 | 3 | Gyg | 0.002 | Young |
| 0.909825 | 0.032548 | 0.698 | 0.728 | 1 | 3 | Srp9 | -0.03 | Young |
| 0.909864 | 0.249642 | 0.01 | 0.01 | 1 | 3 | Ly6c1 | 0 | Young |
| 0.909864 | 0.142575 | 0.01 | 0.01 | 1 | 3 | Tmem220 | 0 | Young |
| 0.910332 | 0.12328 | 0.038 | 0.037 | 1 | 3 | 4933417E1 | 0.001 | Young |
| 0.910353 | 0.663078 | 0.01 | 0.01 | 1 | 3 | 6330403L0 | 0 | Young |
| 0.910469 | 0.025617 | 0.058 | 0.06 | 1 | 3 | Mib2 | -0.002 | Young |
| 0.910507 | 0.333724 | 0.064 | 0.067 | 1 | 3 | Dusp8 | -0.003 | Young |
| 0.911332 | 0.239082 | 0.081 | 0.084 | 1 | 3 | AI606181 | -0.003 | Young |
| 0.911332 | 0.208033 | 0.01 | 0.01 | 1 | 3 | Zfp968 | 0 | Young |
| 0.911512 | 0.114429 | 0.321 | 0.337 | 1 | 3 | Pacsin2 | -0.016 | Young |
| 0.911544 | 0.102816 | 0.028 | 0.029 | 1 | 3 | 4933427D1 | -0.001 | Young |
| 0.911715 | 0.122721 | 0.011 | 0.011 | 1 | 3 | Kyat1 | 0 | Young |
| 0.911821 | 0.064269 | 0.01 | 0.01 | 1 | 3 | Zkscan5 | 0 | Young |
| 0.911821 | 0.029525 | 0.01 | 0.01 | 1 | 3 | Tnfrsf22 | 0 | Young |
| 0.912141 | 0.280877 | 0.066 | 0.068 | 1 | 3 | Nlrx1 | -0.002 | Young |
| 0.912272 | 0.224169 | 0.09 | 0.09 | 1 | 3 | Bloc1s6 | 0 | Young |
| 0.91247 | 0.115637 | 0.133 | 0.134 | 1 | 3 | Ppip5k2 | -0.001 | Young |
| 0.912632 | 0.045328 | 0.159 | 0.165 | 1 | 3 | Lin54 | -0.006 | Young |
| 0.912899 | 0.145442 | 0.201 | 0.202 | 1 | 3 | Fam69a | -0.001 | Young |
| 0.912911 | 0.046658 | 0.011 | 0.012 | 1 | 3 | Ddit4 | -0.001 | Young |
| 0.913268 | 0.018126 | 0.088 | 0.09 | 1 | 3 | Slc25a16 | -0.002 | Young |
| 0.913289 | 0.089236 | 0.01 | 0.01 | 1 | 3 | Isg20 | 0 | Young |
| 0.913386 | 0.121509 | 0.04 | 0.039 | 1 | 3 | Aar2 | 0.001 | Young |
| 0.913433 | 0.068296 | 0.523 | 0.545 | 1 | 3 | Clint1 | -0.022 | Young |
| 0.913572 | 0.041866 | 0.157 | 0.16 | 1 | 3 | Pbx2 | -0.003 | Young |
| 0.913778 | 0.177904 | 0.01 | 0.01 | 1 | 3 | Mettl4 | 0 | Young |
| 0.913778 | 0.031203 | 0.01 | 0.01 | 1 | 3 | Elovl7 | 0 | Young |

| | | | | | | | | |
|---|---|---|---|---|---|---|---|---|
| 0.914028 | 0.282728 | 0.098 | 0.099 | 1 | 3 | Arid5b | -0.001 | Young |
| 0.914339 | 0.41531 | 0.058 | 0.057 | 1 | 3 | Cables2 | 0.001 | Young |
| 0.914382 | 0.285765 | 0.027 | 0.027 | 1 | 3 | Ppp1r3e | 0 | Young |
| 0.91467 | 0.049032 | 0.596 | 0.614 | 1 | 3 | Polr1d | -0.018 | Young |
| 0.914672 | 0.415146 | 0.12 | 0.121 | 1 | 3 | Errfi1 | -0.001 | Young |
| 0.914687 | 0.046331 | 0.432 | 0.455 | 1 | 3 | Atxn7l3b | -0.023 | Young |
| 0.914797 | 0.244016 | 0.205 | 0.207 | 1 | 3 | Rnf10 | -0.002 | Young |
| 0.914936 | 0.118282 | 0.064 | 0.063 | 1 | 3 | Orai2 | 0.001 | Young |
| 0.915088 | 0.086222 | 0.295 | 0.309 | 1 | 3 | Washc4 | -0.014 | Young |
| 0.915105 | 0.126677 | 0.154 | 0.154 | 1 | 3 | Ech1 | 0 | Young |
| 0.915221 | 0.225243 | 0.043 | 0.042 | 1 | 3 | Pnpla6 | 0.001 | Young |
| 0.915686 | 0.144562 | 0.189 | 0.197 | 1 | 3 | Rnf11 | -0.008 | Young |
| 0.915736 | 0.202154 | 0.01 | 0.01 | 1 | 3 | Ets1 | 0 | Young |
| 0.915906 | 0.203775 | 0.077 | 0.08 | 1 | 3 | Dhcr7 | -0.003 | Young |
| 0.915928 | 0.097164 | 0.258 | 0.264 | 1 | 3 | Stag1 | -0.006 | Young |
| 0.916507 | 0.304666 | 0.011 | 0.012 | 1 | 3 | Laptm4b | -0.001 | Young |
| 0.916696 | 0.150869 | 0.04 | 0.042 | 1 | 3 | Tmem86a | -0.002 | Young |
| 0.916795 | 0.156226 | 0.054 | 0.056 | 1 | 3 | Cdk5 | -0.002 | Young |
| 0.916858 | 0.125874 | 0.395 | 0.407 | 1 | 3 | Arhgap26 | -0.012 | Young |
| 0.916883 | 0.350061 | 0.052 | 0.051 | 1 | 3 | Zfp335 | 0.001 | Young |
| 0.917056 | 0.245047 | 0.184 | 0.19 | 1 | 3 | Spopl | -0.006 | Young |
| 0.917407 | 0.161483 | 0.011 | 0.012 | 1 | 3 | Clspn | -0.001 | Young |
| 0.918063 | 0.222395 | 0.058 | 0.057 | 1 | 3 | Shmt2 | 0.001 | Young |
| 0.918441 | 0.025266 | 0.04 | 0.039 | 1 | 3 | Fam114a1 | 0.001 | Young |
| 0.918598 | 0.176858 | 0.101 | 0.104 | 1 | 3 | Ebpl | -0.003 | Young |
| 0.918603 | 0.362831 | 0.121 | 0.125 | 1 | 3 | Amn1 | -0.004 | Young |
| 0.918996 | 0.062283 | 0.13 | 0.133 | 1 | 3 | Zfp622 | -0.003 | Young |
| 0.919108 | 0.273882 | 0.086 | 0.088 | 1 | 3 | Gipc1 | -0.002 | Young |
| 0.919206 | 0.555714 | 0.011 | 0.012 | 1 | 3 | Ppp3cc | -0.001 | Young |
| 0.919291 | 0.139164 | 0.13 | 0.129 | 1 | 3 | Fbxo18 | 0.001 | Young |
| 0.91933 | 0.210094 | 0.08 | 0.079 | 1 | 3 | Hace1 | 0.001 | Young |
| 0.919438 | 0.044602 | 0.39 | 0.405 | 1 | 3 | Eif3m | -0.015 | Young |
| 0.919512 | 0.300252 | 0.058 | 0.057 | 1 | 3 | Gem | 0.001 | Young |
| 0.919682 | 0.333482 | 0.044 | 0.045 | 1 | 3 | Trmt2b | -0.001 | Young |
| 0.920382 | 0.122573 | 0.037 | 0.036 | 1 | 3 | Arl11 | 0.001 | Young |
| 0.92076 | 0.134274 | 0.035 | 0.035 | 1 | 3 | Adi1 | 0 | Young |
| 0.921066 | 0.398565 | 0.033 | 0.032 | 1 | 3 | Rfx2 | 0.001 | Young |
| 0.921352 | 0.067806 | 0.277 | 0.281 | 1 | 3 | Aida | -0.004 | Young |
| 0.921903 | 0.53466 | 0.053 | 0.053 | 1 | 3 | Kazald1 | 0 | Young |
| 0.921905 | 0.607537 | 0.011 | 0.012 | 1 | 3 | Wdr90 | -0.001 | Young |
| 0.921958 | 0.068527 | 0.513 | 0.526 | 1 | 3 | Ndufc1 | -0.013 | Young |
| 0.922103 | 0.338593 | 0.054 | 0.054 | 1 | 3 | 1110065P2 | 0 | Young |
| 0.922355 | 0.330931 | 0.011 | 0.012 | 1 | 3 | Maoa | -0.001 | Young |
| 0.922914 | 0.047131 | 0.439 | 0.456 | 1 | 3 | Dad1 | -0.017 | Young |
| 0.922959 | 0.379023 | 0.091 | 0.091 | 1 | 3 | Mphosph6 | 0 | Young |

| | | | | | | | | |
|---|---|---|---|---|---|---|---|---|
| 0.923009 | 0.237337 | 0.071 | 0.073 | 1 | 3 | Crip2 | -0.002 | Young |
| 0.923135 | 0.13187 | 0.234 | 0.244 | 1 | 3 | Tbc1d10b | -0.01 | Young |
| 0.923436 | 0.130975 | 0.208 | 0.213 | 1 | 3 | Pts | -0.005 | Young |
| 0.923889 | 0.04191 | 0.282 | 0.289 | 1 | 3 | Adipor1 | -0.007 | Young |
| 0.924318 | 0.163169 | 0.098 | 0.098 | 1 | 3 | Urm1 | 0 | Young |
| 0.924465 | 0.205952 | 0.159 | 0.161 | 1 | 3 | Nasp | -0.002 | Young |
| 0.92508 | 0.123697 | 0.038 | 0.037 | 1 | 3 | Rabif | 0.001 | Young |
| 0.925121 | 0.167622 | 0.184 | 0.186 | 1 | 3 | Sirt7 | -0.002 | Young |
| 0.925447 | 0.020908 | 0.612 | 0.625 | 1 | 3 | Rock1 | -0.013 | Young |
| 0.925568 | 0.207414 | 0.096 | 0.099 | 1 | 3 | Pcnx | -0.003 | Young |
| 0.925603 | 0.197946 | 0.039 | 0.038 | 1 | 3 | Slc19a1 | 0.001 | Young |
| 0.925694 | 0.121548 | 0.288 | 0.3 | 1 | 3 | Krtcap2 | -0.012 | Young |
| 0.925717 | 0.062779 | 0.146 | 0.146 | 1 | 3 | Zc3h14 | 0 | Young |
| 0.926124 | 0.017223 | 0.114 | 0.116 | 1 | 3 | Atl2 | -0.002 | Young |
| 0.92684 | 0.015698 | 0.686 | 0.691 | 1 | 3 | Ifitm6 | -0.005 | Young |
| 0.927028 | 0.18853 | 0.225 | 0.228 | 1 | 3 | Ninj1 | -0.003 | Young |
| 0.927056 | 0.096994 | 0.16 | 0.162 | 1 | 3 | Atg7 | -0.002 | Young |
| 0.92757 | 0.311173 | 0.042 | 0.043 | 1 | 3 | Zc3h3 | -0.001 | Young |
| 0.927898 | 0.478681 | 0.128 | 0.131 | 1 | 3 | Ash2l | -0.003 | Young |
| 0.92821 | 0.117311 | 0.141 | 0.145 | 1 | 3 | Nfx1 | -0.004 | Young |
| 0.928388 | 0.316153 | 0.064 | 0.065 | 1 | 3 | Mast2 | -0.001 | Young |
| 0.928949 | 0.168502 | 0.056 | 0.055 | 1 | 3 | Sephs2 | 0.001 | Young |
| 0.929543 | 0.063771 | 0.134 | 0.133 | 1 | 3 | Mpnd | 0.001 | Young |
| 0.929701 | 0.126217 | 0.035 | 0.035 | 1 | 3 | Rbm4 | 0 | Young |
| 0.929856 | 0.111278 | 0.37 | 0.393 | 1 | 3 | Ggnbp2 | -0.023 | Young |
| 0.930187 | 0.01007 | 0.348 | 0.356 | 1 | 3 | Zfp36l1 | -0.008 | Young |
| 0.930563 | 0.242154 | 0.043 | 0.044 | 1 | 3 | Smpd4 | -0.001 | Young |
| 0.930694 | 0.133376 | 0.17 | 0.173 | 1 | 3 | Abcg3 | -0.003 | Young |
| 0.93108 | 0.039074 | 0.028 | 0.029 | 1 | 3 | Mmd | -0.001 | Young |
| 0.93111 | 0.093894 | 0.226 | 0.229 | 1 | 3 | Golgb1 | -0.003 | Young |
| 0.931152 | 0.090826 | 0.105 | 0.104 | 1 | 3 | Racgap1 | 0.001 | Young |
| 0.931349 | 0.158779 | 0.109 | 0.108 | 1 | 3 | Kctd12b | 0.001 | Young |
| 0.931759 | 0.082063 | 0.561 | 0.608 | 1 | 3 | Adgre1 | -0.047 | Young |
| 0.931815 | 0.005167 | 0.115 | 0.117 | 1 | 3 | Ric1 | -0.002 | Young |
| 0.931826 | 0.306413 | 0.033 | 0.032 | 1 | 3 | Letmd1 | 0.001 | Young |
| 0.932069 | 0.03082 | 0.035 | 0.035 | 1 | 3 | Cd82 | 0 | Young |
| 0.93259 | 0.363956 | 0.03 | 0.03 | 1 | 3 | Cul9 | 0 | Young |
| 0.933161 | 0.287056 | 0.013 | 0.013 | 1 | 3 | Amotl2 | 0 | Young |
| 0.933169 | 0.011121 | 0.102 | 0.104 | 1 | 3 | Ubqln4 | -0.002 | Young |
| 0.933253 | 0.131779 | 0.035 | 0.035 | 1 | 3 | Hpse | 0 | Young |
| 0.933468 | 0.072392 | 0.54 | 0.577 | 1 | 3 | Tardbp | -0.037 | Young |
| 0.933504 | 0.301951 | 0.063 | 0.065 | 1 | 3 | Pef1 | -0.002 | Young |
| 0.933571 | 0.413041 | 0.053 | 0.053 | 1 | 3 | Clp1 | 0 | Young |
| 0.93359 | 0.261386 | 0.013 | 0.013 | 1 | 3 | Usp11 | 0 | Young |
| 0.933601 | 0.06201 | 0.138 | 0.141 | 1 | 3 | Fnbp4 | -0.003 | Young |

| | | | | | | | | |
|---|---|---|---|---|---|---|---|---|
| 0.933606 | 0.216983 | 0.261 | 0.27 | 1 | 3 | Vps13b | -0.009 | Young |
| 0.933868 | 0.280876 | 0.193 | 0.198 | 1 | 3 | Tcf20 | -0.005 | Young |
| 0.933913 | 0.244785 | 0.17 | 0.174 | 1 | 3 | Hint2 | -0.004 | Young |
| 0.934007 | 0.432621 | 0.033 | 0.032 | 1 | 3 | Kdelc2 | 0.001 | Young |
| 0.934019 | 0.202256 | 0.013 | 0.013 | 1 | 3 | Zfp458 | 0 | Young |
| 0.934037 | 0.049224 | 0.083 | 0.082 | 1 | 3 | Trpm4 | 0.001 | Young |
| 0.934087 | 0.228526 | 0.03 | 0.031 | 1 | 3 | Tram2 | -0.001 | Young |
| 0.934448 | 0.267176 | 0.013 | 0.013 | 1 | 3 | Mettl22 | 0 | Young |
| 0.934458 | 0.099016 | 0.059 | 0.059 | 1 | 3 | Znhit2 | 0 | Young |
| 0.93446 | 0.033891 | 0.159 | 0.16 | 1 | 3 | Spast | -0.001 | Young |
| 0.934565 | 0.001031 | 0.085 | 0.084 | 1 | 3 | Phactr4 | 0.001 | Young |
| 0.934584 | 0.359829 | 0.028 | 0.029 | 1 | 3 | Zfp81 | -0.001 | Young |
| 0.934728 | 0.037518 | 0.076 | 0.075 | 1 | 3 | Actr1b | 0.001 | Young |
| 0.934773 | 0.267201 | 0.032 | 0.032 | 1 | 3 | Pi4kb | 0 | Young |
| 0.934939 | 0.009023 | 0.663 | 0.699 | 1 | 3 | Sfr1 | -0.036 | Young |
| 0.935062 | 0.100569 | 0.014 | 0.014 | 1 | 3 | Gm36266 | 0 | Young |
| 0.93527 | 0.046668 | 0.275 | 0.287 | 1 | 3 | Szrd1 | -0.012 | Young |
| 0.935495 | 0.172141 | 0.061 | 0.06 | 1 | 3 | Top3b | 0.001 | Young |
| 0.935679 | 0.029094 | 0.654 | 0.671 | 1 | 3 | Cyfip1 | -0.017 | Young |
| 0.935716 | 0.044681 | 0.494 | 0.516 | 1 | 3 | Rpl36al | -0.022 | Young |
| 0.935733 | 0.18688 | 0.071 | 0.073 | 1 | 3 | Slc35d1 | -0.002 | Young |
| 0.935901 | 0.135116 | 0.062 | 0.061 | 1 | 3 | Utp14b | 0.001 | Young |
| 0.936269 | 0.064097 | 0.047 | 0.048 | 1 | 3 | Zfyve19 | -0.001 | Young |
| 0.936298 | 0.177581 | 0.181 | 0.186 | 1 | 3 | Ncoa2 | -0.005 | Young |
| 0.936592 | 0.392302 | 0.013 | 0.013 | 1 | 3 | Hebp1 | 0 | Young |
| 0.936985 | 0.151114 | 0.424 | 0.452 | 1 | 3 | Ubtf | -0.028 | Young |
| 0.937126 | 0.589979 | 0.059 | 0.06 | 1 | 3 | Iqsec1 | -0.001 | Young |
| 0.937149 | 0.072133 | 0.792 | 0.818 | 1 | 3 | Hsp90aa1 | -0.026 | Young |
| 0.937153 | 0.050187 | 0.178 | 0.182 | 1 | 3 | Ormdl2 | -0.004 | Young |
| 0.93745 | 0.310855 | 0.013 | 0.013 | 1 | 3 | Ccno | 0 | Young |
| 0.937512 | 0.259367 | 0.061 | 0.062 | 1 | 3 | Golm1 | -0.001 | Young |
| 0.937636 | 0.145801 | 0.029 | 0.03 | 1 | 3 | Taf8 | -0.001 | Young |
| 0.937984 | 0.091014 | 0.068 | 0.068 | 1 | 3 | Tmem214 | 0 | Young |
| 0.938324 | 0.027467 | 0.179 | 0.183 | 1 | 3 | Acadsb | -0.004 | Young |
| 0.938423 | 0.071657 | 0.124 | 0.127 | 1 | 3 | Cnep1r1 | -0.003 | Young |
| 0.938747 | 0.047501 | 0.176 | 0.176 | 1 | 3 | Fam193a | 0 | Young |
| 0.938818 | 0.105488 | 0.473 | 0.491 | 1 | 3 | Lifr | -0.018 | Young |
| 0.938904 | 0.163556 | 0.192 | 0.201 | 1 | 3 | Fam76a | -0.009 | Young |
| 0.939054 | 0.112156 | 0.145 | 0.148 | 1 | 3 | Syne2 | -0.003 | Young |
| 0.939236 | 0.176211 | 0.172 | 0.172 | 1 | 3 | Prrc2b | 0 | Young |
| 0.939375 | 0.122408 | 0.327 | 0.335 | 1 | 3 | Zc3h15 | -0.008 | Young |
| 0.939638 | 0.428383 | 0.054 | 0.056 | 1 | 3 | Rtl8a | -0.002 | Young |
| 0.939808 | 0.1463 | 0.134 | 0.134 | 1 | 3 | Sfxn2 | 0 | Young |
| 0.93985 | 0.316246 | 0.083 | 0.084 | 1 | 3 | Rnf167 | -0.001 | Young |
| 0.940284 | 0.208026 | 0.048 | 0.048 | 1 | 3 | Rad51d | 0 | Young |

| | | | | | | | | |
|---|---|---|---|---|---|---|---|---|
| 0.94038 | 0.207872 | 0.117 | 0.117 | 1 | 3 | Stat5a | 0 | Young |
| 0.94058 | 0.070904 | 0.617 | 0.659 | 1 | 3 | Pmaip1 | -0.042 | Young |
| 0.940816 | 0.005487 | 0.951 | 0.973 | 1 | 3 | Rpl8 | -0.022 | Young |
| 0.941072 | 0.179311 | 0.112 | 0.114 | 1 | 3 | Usf3 | -0.002 | Young |
| 0.941129 | 0.057436 | 0.091 | 0.093 | 1 | 3 | Mrps35 | -0.002 | Young |
| 0.941595 | 0.954995 | 0.028 | 0.029 | 1 | 3 | Dynlt1f | -0.001 | Young |
| 0.941779 | 0.246462 | 0.106 | 0.109 | 1 | 3 | Parp2 | -0.003 | Young |
| 0.941816 | 0.147934 | 0.153 | 0.16 | 1 | 3 | Ttpal | -0.007 | Young |
| 0.942556 | 0.138682 | 0.309 | 0.326 | 1 | 3 | Scpep1 | -0.017 | Young |
| 0.942937 | 0.06185 | 0.102 | 0.104 | 1 | 3 | Dctn6 | -0.002 | Young |
| 0.942992 | 0.180081 | 0.087 | 0.087 | 1 | 3 | Med20 | 0 | Young |
| 0.943048 | 0.149826 | 0.316 | 0.321 | 1 | 3 | Dna2 | -0.005 | Young |
| 0.943132 | 0.146194 | 0.146 | 0.152 | 1 | 3 | Fam214b | -0.006 | Young |
| 0.943395 | 0.291203 | 0.148 | 0.153 | 1 | 3 | Mplkip | -0.005 | Young |
| 0.944533 | 0.251222 | 0.096 | 0.098 | 1 | 3 | Snx13 | -0.002 | Young |
| 0.944893 | 0.331475 | 0.091 | 0.092 | 1 | 3 | Slc4a1ap | -0.001 | Young |
| 0.944957 | 0.101922 | 0.294 | 0.307 | 1 | 3 | Ndnf | -0.013 | Young |
| 0.944975 | 0.257513 | 0.091 | 0.093 | 1 | 3 | BC029722 | -0.002 | Young |
| 0.944976 | 0.118695 | 0.216 | 0.219 | 1 | 3 | Selenoi | -0.003 | Young |
| 0.944985 | 0.133846 | 0.136 | 0.141 | 1 | 3 | Faf1 | -0.005 | Young |
| 0.945112 | 0.17655 | 0.312 | 0.33 | 1 | 3 | H2afv | -0.018 | Young |
| 0.94522 | 0.054187 | 0.708 | 0.736 | 1 | 3 | Ier5 | -0.028 | Young |
| 0.945754 | 0.067409 | 0.119 | 0.121 | 1 | 3 | Mapk8 | -0.002 | Young |
| 0.945998 | 0.047336 | 0.285 | 0.288 | 1 | 3 | Mgrn1 | -0.003 | Young |
| 0.946215 | 0.036073 | 0.124 | 0.124 | 1 | 3 | Glipr1 | 0 | Young |
| 0.946232 | 0.137288 | 0.17 | 0.177 | 1 | 3 | Fxr2 | -0.007 | Young |
| 0.946319 | 0.029584 | 0.198 | 0.204 | 1 | 3 | Dapk3 | -0.006 | Young |
| 0.946416 | 0.352882 | 0.056 | 0.055 | 1 | 3 | Slc20a2 | 0.001 | Young |
| 0.946486 | 0.196267 | 0.066 | 0.066 | 1 | 3 | Ruvbl2 | 0 | Young |
| 0.946564 | 0.106417 | 0.034 | 0.033 | 1 | 3 | Dis3 | 0.001 | Young |
| 0.946681 | 0.217089 | 0.066 | 0.066 | 1 | 3 | Tfdp2 | 0 | Young |
| 0.946861 | 0.139117 | 0.27 | 0.286 | 1 | 3 | Lamtor5 | -0.016 | Young |
| 0.946876 | 0.302595 | 0.155 | 0.159 | 1 | 3 | Dopey2 | -0.004 | Young |
| 0.947372 | 0.446425 | 0.014 | 0.014 | 1 | 3 | Fam216a | 0 | Young |
| 0.947656 | 0.276314 | 0.027 | 0.026 | 1 | 3 | Alg2 | 0.001 | Young |
| 0.947801 | 0.191119 | 0.049 | 0.05 | 1 | 3 | Bloc1s1 | -0.001 | Young |
| 0.947877 | 0.378854 | 0.078 | 0.081 | 1 | 3 | Cog6 | -0.003 | Young |
| 0.948052 | 0.011246 | 0.49 | 0.515 | 1 | 3 | Limd1 | -0.025 | Young |
| 0.948377 | 0.136889 | 0.254 | 0.266 | 1 | 3 | Osbp | -0.012 | Young |
| 0.948465 | 0.131648 | 0.196 | 0.201 | 1 | 3 | Naa35 | -0.005 | Young |
| 0.94868 | 0.283482 | 0.03 | 0.031 | 1 | 3 | 2900089D1 | -0.001 | Young |
| 0.948752 | 0.227728 | 0.032 | 0.031 | 1 | 3 | Ecsit | 0.001 | Young |
| 0.949036 | 0.157382 | 0.115 | 0.116 | 1 | 3 | Dpm1 | -0.001 | Young |
| 0.949051 | 0.284596 | 0.133 | 0.136 | 1 | 3 | Armcx3 | -0.003 | Young |
| 0.949153 | 0.36529 | 0.072 | 0.074 | 1 | 3 | St8sia6 | -0.002 | Young |

| | | | | | | | | |
|---|---|---|---|---|---|---|---|---|
| 0.949269 | 0.279473 | 0.047 | 0.048 | 1 | 3 | Gm13373 | -0.001 | Young |
| 0.949444 | 0.114245 | 0.25 | 0.254 | 1 | 3 | Lrrc8d | -0.004 | Young |
| 0.949756 | 0.106462 | 0.188 | 0.19 | 1 | 3 | Ntan1 | -0.002 | Young |
| 0.949773 | 0.104249 | 0.556 | 0.578 | 1 | 3 | Ndufa1 | -0.022 | Young |
| 0.949839 | 0.136358 | 0.08 | 0.081 | 1 | 3 | Rrp8 | -0.001 | Young |
| 0.949871 | 0.215777 | 0.086 | 0.086 | 1 | 3 | Syt11 | 0 | Young |
| 0.949981 | 0.117688 | 0.476 | 0.504 | 1 | 3 | Fgd4 | -0.028 | Young |
| 0.950029 | 0.171154 | 0.049 | 0.05 | 1 | 3 | Zfp513 | -0.001 | Young |
| 0.950153 | 0.196698 | 0.197 | 0.203 | 1 | 3 | Nek9 | -0.006 | Young |
| 0.950408 | 0.289194 | 0.102 | 0.104 | 1 | 3 | Prkra | -0.002 | Young |
| 0.950453 | 0.09605 | 0.4 | 0.416 | 1 | 3 | Tlr2 | -0.016 | Young |
| 0.950681 | 0.138808 | 0.083 | 0.085 | 1 | 3 | Ubxn7 | -0.002 | Young |
| 0.95079 | 0.224861 | 0.176 | 0.18 | 1 | 3 | Virma | -0.004 | Young |
| 0.950872 | 0.238771 | 0.051 | 0.05 | 1 | 3 | Zfp369 | 0.001 | Young |
| 0.951264 | 0.275227 | 0.051 | 0.051 | 1 | 3 | Gtpbp8 | 0 | Young |
| 0.951455 | 0.035674 | 0.173 | 0.177 | 1 | 3 | Tmx1 | -0.004 | Young |
| 0.951478 | 0.556519 | 0.014 | 0.014 | 1 | 3 | Usp54 | 0 | Young |
| 0.951478 | 0.531773 | 0.014 | 0.014 | 1 | 3 | 1700019L1 | 0 | Young |
| 0.951655 | 0.073263 | 0.015 | 0.016 | 1 | 3 | Tesk2 | -0.001 | Young |
| 0.951889 | 0.780176 | 0.014 | 0.014 | 1 | 3 | Phf19 | 0 | Young |
| 0.951889 | 0.574922 | 0.014 | 0.014 | 1 | 3 | Tefm | 0 | Young |
| 0.951889 | 0.424965 | 0.014 | 0.014 | 1 | 3 | Numbl | 0 | Young |
| 0.951927 | 0.26092 | 0.056 | 0.055 | 1 | 3 | Saysd1 | 0.001 | Young |
| 0.952139 | 0.385177 | 0.045 | 0.045 | 1 | 3 | Commd9 | 0 | Young |
| 0.9523 | 0.754118 | 0.014 | 0.014 | 1 | 3 | Gm31243 | 0 | Young |
| 0.952494 | 0.195575 | 0.069 | 0.069 | 1 | 3 | St3gal2 | 0 | Young |
| 0.952821 | 0.138478 | 0.078 | 0.078 | 1 | 3 | Ppa2 | 0 | Young |
| 0.953006 | 0.380206 | 0.104 | 0.106 | 1 | 3 | Phka2 | -0.002 | Young |
| 0.953082 | 0.164179 | 0.361 | 0.384 | 1 | 3 | Denr | -0.023 | Young |
| 0.953199 | 0.232013 | 0.056 | 0.055 | 1 | 3 | Kantr | 0.001 | Young |
| 0.953214 | 0.008295 | 0.556 | 0.576 | 1 | 3 | Ifngr1 | -0.02 | Young |
| 0.9535 | 0.045321 | 0.564 | 0.615 | 1 | 3 | Fkbp1a | -0.051 | Young |
| 0.953512 | 0.199454 | 0.268 | 0.276 | 1 | 3 | Flii | -0.008 | Young |
| 0.953623 | 0.146021 | 0.319 | 0.336 | 1 | 3 | Nr3c1 | -0.017 | Young |
| 0.953863 | 0.08359 | 0.307 | 0.324 | 1 | 3 | Zzef1 | -0.017 | Young |
| 0.954002 | 0.476858 | 0.042 | 0.043 | 1 | 3 | Rpl36a-ps1 | -0.001 | Young |
| 0.954202 | 0.196349 | 0.112 | 0.114 | 1 | 3 | Cyth2 | -0.002 | Young |
| 0.954354 | 0.319299 | 0.014 | 0.014 | 1 | 3 | Wtip | 0 | Young |
| 0.954472 | 0.161094 | 0.198 | 0.203 | 1 | 3 | Atxn7 | -0.005 | Young |
| 0.954506 | 0.300586 | 0.048 | 0.049 | 1 | 3 | Zfp280c | -0.001 | Young |
| 0.955026 | 0.000187 | 0.282 | 0.289 | 1 | 3 | Gm15987 | -0.007 | Young |
| 0.955092 | 0.035452 | 0.128 | 0.129 | 1 | 3 | Kars | -0.001 | Young |
| 0.955138 | 0.200957 | 0.048 | 0.048 | 1 | 3 | Nepro | 0 | Young |
| 0.955217 | 0.231277 | 0.101 | 0.102 | 1 | 3 | Cbr2 | -0.001 | Young |
| 0.955269 | 0.347153 | 0.047 | 0.047 | 1 | 3 | Sppl2b | 0 | Young |

| | | | | | | | | |
|---|---|---|---|---|---|---|---|---|
| 0.955332 | 0.193204 | 0.295 | 0.307 | 1 | 3 | Insig1 | -0.012 | Young |
| 0.955477 | 0.154146 | 0.064 | 0.066 | 1 | 3 | Git1 | -0.002 | Young |
| 0.955863 | 0.554053 | 0.025 | 0.025 | 1 | 3 | Endov | 0 | Young |
| 0.955997 | 1.002095 | 0.014 | 0.014 | 1 | 3 | Ereg | 0 | Young |
| 0.956236 | 0.018037 | 0.029 | 0.029 | 1 | 3 | Dync2h1 | 0 | Young |
| 0.956337 | 0.27039 | 0.167 | 0.171 | 1 | 3 | Pak1ip1 | -0.004 | Young |
| 0.956343 | 0.029592 | 0.078 | 0.078 | 1 | 3 | B430306N( | 0 | Young |
| 0.956386 | 0.131256 | 0.015 | 0.016 | 1 | 3 | Ccdc77 | -0.001 | Young |
| 0.956467 | 0.234353 | 0.073 | 0.075 | 1 | 3 | Mon1b | -0.002 | Young |
| 0.956746 | 0.056416 | 0.456 | 0.477 | 1 | 3 | Ndufb8 | -0.021 | Young |
| 0.956929 | 0.161191 | 0.09 | 0.092 | 1 | 3 | Cdkn2aip | -0.002 | Young |
| 0.956931 | 0.679392 | 0.044 | 0.044 | 1 | 3 | Dglucy | 0 | Young |
| 0.957179 | 0.013753 | 0.958 | 0.962 | 1 | 3 | Rpl39 | -0.004 | Young |
| 0.95719 | 0.139202 | 0.122 | 0.124 | 1 | 3 | Rbm12 | -0.002 | Young |
| 0.958022 | 0.089191 | 0.327 | 0.337 | 1 | 3 | BC017643 | -0.01 | Young |
| 0.958775 | 0.220673 | 0.073 | 0.075 | 1 | 3 | Yars | -0.002 | Young |
| 0.959529 | 0.352771 | 0.104 | 0.105 | 1 | 3 | Mkl1 | -0.001 | Young |
| 0.959585 | 0.149053 | 0.095 | 0.097 | 1 | 3 | Ints12 | -0.002 | Young |
| 0.959602 | 0.122225 | 0.547 | 0.583 | 1 | 3 | Selplg | -0.036 | Young |
| 0.959616 | 0.227408 | 0.179 | 0.185 | 1 | 3 | Rbfa | -0.006 | Young |
| 0.960027 | 0.143021 | 0.033 | 0.033 | 1 | 3 | Armc3 | 0 | Young |
| 0.960069 | 0.261435 | 0.177 | 0.185 | 1 | 3 | Aaed1 | -0.008 | Young |
| 0.960375 | 0.101457 | 0.03 | 0.03 | 1 | 3 | Sh3rf1 | 0 | Young |
| 0.960446 | 0.230538 | 0.082 | 0.084 | 1 | 3 | Pcsk7 | -0.002 | Young |
| 0.960571 | 0.191471 | 0.076 | 0.078 | 1 | 3 | Prmt2 | -0.002 | Young |
| 0.960651 | 0.208195 | 0.158 | 0.16 | 1 | 3 | Gle1 | -0.002 | Young |
| 0.960724 | 0.249361 | 0.015 | 0.016 | 1 | 3 | Ube2v1 | -0.001 | Young |
| 0.961005 | 0.132856 | 0.365 | 0.382 | 1 | 3 | Orai1 | -0.017 | Young |
| 0.961268 | 0.458611 | 0.059 | 0.06 | 1 | 3 | Cog1 | -0.001 | Young |
| 0.961409 | 0.048257 | 0.242 | 0.247 | 1 | 3 | Bcl7c | -0.005 | Young |
| 0.961498 | 0.374862 | 0.032 | 0.032 | 1 | 3 | Bod1 | 0 | Young |
| 0.961615 | 0.200372 | 0.129 | 0.131 | 1 | 3 | Prkag2 | -0.002 | Young |
| 0.961738 | 0.009127 | 0.029 | 0.029 | 1 | 3 | Tmtc3 | 0 | Young |
| 0.961761 | 0.098876 | 0.256 | 0.264 | 1 | 3 | Rnase6 | -0.008 | Young |
| 0.961867 | 0.125096 | 0.162 | 0.165 | 1 | 3 | Pwwp2a | -0.003 | Young |
| 0.962373 | 0.008173 | 0.569 | 0.594 | 1 | 3 | Atp5d | -0.025 | Young |
| 0.962438 | 0.144961 | 0.058 | 0.059 | 1 | 3 | Khk | -0.001 | Young |
| 0.962469 | 0.275023 | 0.062 | 0.063 | 1 | 3 | Brpf3 | -0.001 | Young |
| 0.962718 | 0.040009 | 0.621 | 0.645 | 1 | 3 | Foxp1 | -0.024 | Young |
| 0.962719 | 0.265919 | 0.114 | 0.116 | 1 | 3 | Ubap1 | -0.002 | Young |
| 0.963491 | 0.099967 | 0.383 | 0.398 | 1 | 3 | Top2b | -0.015 | Young |
| 0.963566 | 0.15429 | 0.135 | 0.136 | 1 | 3 | Abcd2 | -0.001 | Young |
| 0.963604 | 0.346323 | 0.107 | 0.111 | 1 | 3 | Dlat | -0.004 | Young |
| 0.963688 | 0.317606 | 0.08 | 0.08 | 1 | 3 | Manea | 0 | Young |
| 0.964062 | 0.046717 | 0.119 | 0.119 | 1 | 3 | Foxk1 | 0 | Young |

| | | | | | | | | |
|---|---|---|---|---|---|---|---|---|
| 0.964142 | 0.136602 | 0.146 | 0.148 | 1 | 3 | Wdr3 | -0.002 | Young |
| 0.964416 | 0.078494 | 0.293 | 0.303 | 1 | 3 | Mrpl24 | -0.01 | Young |
| 0.96448 | 0.113978 | 0.273 | 0.284 | 1 | 3 | Eny2 | -0.011 | Young |
| 0.964508 | 0.030994 | 0.352 | 0.36 | 1 | 3 | Tmem168 | -0.008 | Young |
| 0.964516 | 0.083506 | 0.309 | 0.319 | 1 | 3 | Kdm4a | -0.01 | Young |
| 0.964745 | 0.017922 | 0.058 | 0.057 | 1 | 3 | Cog7 | 0.001 | Young |
| 0.964821 | 0.063551 | 0.074 | 0.075 | 1 | 3 | Polk | -0.001 | Young |
| 0.96492 | 0.106135 | 0.264 | 0.275 | 1 | 3 | Aurkaip1 | -0.011 | Young |
| 0.964962 | 0.142015 | 0.256 | 0.259 | 1 | 3 | Tial1 | -0.003 | Young |
| 0.964989 | 0.278702 | 0.13 | 0.134 | 1 | 3 | Rbm4b | -0.004 | Young |
| 0.965257 | 0.135121 | 0.051 | 0.051 | 1 | 3 | Tor2a | 0 | Young |
| 0.965444 | 0.016034 | 0.321 | 0.329 | 1 | 3 | Tspan32 | -0.008 | Young |
| 0.965852 | 0.671938 | 0.015 | 0.016 | 1 | 3 | Zfp974 | -0.001 | Young |
| 0.965956 | 0.119694 | 0.235 | 0.241 | 1 | 3 | Rassf8 | -0.006 | Young |
| 0.966146 | 0.048299 | 0.37 | 0.379 | 1 | 3 | Rwdd1 | -0.009 | Young |
| 0.966197 | 0.213985 | 0.202 | 0.209 | 1 | 3 | Fam50a | -0.007 | Young |
| 0.966353 | 0.149674 | 0.37 | 0.391 | 1 | 3 | Txnrd1 | -0.021 | Young |
| 0.966393 | 0.025442 | 0.082 | 0.082 | 1 | 3 | Pex13 | 0 | Young |
| 0.966479 | 0.132644 | 0.11 | 0.11 | 1 | 3 | Nsmce1 | 0 | Young |
| 0.966549 | 0.148553 | 0.052 | 0.051 | 1 | 3 | Tsr2 | 0.001 | Young |
| 0.966723 | 0.201173 | 0.104 | 0.104 | 1 | 3 | Pias2 | 0 | Young |
| 0.967138 | 0.214205 | 0.035 | 0.036 | 1 | 3 | Fam122b | -0.001 | Young |
| 0.967372 | 0.369886 | 0.042 | 0.042 | 1 | 3 | Mrps2 | 0 | Young |
| 0.96769 | 0.196158 | 0.016 | 0.017 | 1 | 3 | Lig4 | -0.001 | Young |
| 0.967905 | 0.097349 | 0.125 | 0.127 | 1 | 3 | Dido1 | -0.002 | Young |
| 0.967913 | 0.373912 | 0.027 | 0.026 | 1 | 3 | Yipf2 | 0.001 | Young |
| 0.968254 | 0.524893 | 0.024 | 0.024 | 1 | 3 | Gm26881 | 0 | Young |
| 0.968355 | 0.254969 | 0.139 | 0.142 | 1 | 3 | Grk3 | -0.003 | Young |
| 0.968472 | 0.196224 | 0.159 | 0.161 | 1 | 3 | Nr2f6 | -0.002 | Young |
| 0.968804 | 0.126386 | 0.24 | 0.243 | 1 | 3 | Rsf1 | -0.003 | Young |
| 0.969166 | 0.079636 | 0.1 | 0.1 | 1 | 3 | Sart3 | 0 | Young |
| 0.969343 | 0.226195 | 0.102 | 0.105 | 1 | 3 | Timm17b | -0.003 | Young |
| 0.969433 | 0.049632 | 0.068 | 0.069 | 1 | 3 | Eri3 | -0.001 | Young |
| 0.969493 | 0.094502 | 0.035 | 0.036 | 1 | 3 | Fbxl19 | -0.001 | Young |
| 0.969564 | 0.169302 | 0.134 | 0.137 | 1 | 3 | Galc | -0.003 | Young |
| 0.96959 | 0.366912 | 0.016 | 0.017 | 1 | 3 | Nudt6 | -0.001 | Young |
| 0.969725 | 0.043881 | 0.153 | 0.153 | 1 | 3 | Pnpla2 | 0 | Young |
| 0.970267 | 0.183542 | 0.189 | 0.196 | 1 | 3 | Heatr5a | -0.007 | Young |
| 0.970519 | 0.038358 | 0.58 | 0.613 | 1 | 3 | Smarca5 | -0.033 | Young |
| 0.970729 | 0.307639 | 0.016 | 0.017 | 1 | 3 | Abhd14a | -0.001 | Young |
| 0.970863 | 0.201813 | 0.17 | 0.176 | 1 | 3 | D10Jhu81e | -0.006 | Young |
| 0.97148 | 0.043245 | 0.346 | 0.362 | 1 | 3 | Pqlc1 | -0.016 | Young |
| 0.971497 | 0.093211 | 0.39 | 0.403 | 1 | 3 | Arglu1 | -0.013 | Young |
| 0.971503 | 0.207065 | 0.196 | 0.201 | 1 | 3 | Itga5 | -0.005 | Young |
| 0.971804 | 0.06171 | 0.254 | 0.257 | 1 | 3 | Chmp1a | -0.003 | Young |

| | | | | | | | | |
|---|---|---|---|---|---|---|---|---|
| 0.97206 | 0.222136 | 0.053 | 0.054 | 1 | 3 | Drg2 | -0.001 | Young |
| 0.972232 | 0.088541 | 0.245 | 0.246 | 1 | 3 | Hfe | -0.001 | Young |
| 0.972238 | 0.223195 | 0.106 | 0.109 | 1 | 3 | Btbd10 | -0.003 | Young |
| 0.972375 | 0.157065 | 0.158 | 0.161 | 1 | 3 | Itfg1 | -0.003 | Young |
| 0.972383 | 0.010489 | 0.143 | 0.143 | 1 | 3 | Cant1 | 0 | Young |
| 0.972823 | 0.163333 | 0.148 | 0.149 | 1 | 3 | Usp45 | -0.001 | Young |
| 0.972894 | 0.310702 | 0.097 | 0.099 | 1 | 3 | Yif1b | -0.002 | Young |
| 0.973033 | 0.063414 | 0.558 | 0.576 | 1 | 3 | Irf5 | -0.018 | Young |
| 0.973108 | 0.208685 | 0.135 | 0.14 | 1 | 3 | Pou2f1 | -0.005 | Young |
| 0.9735 | 0.105294 | 0.106 | 0.108 | 1 | 3 | Tceal9 | -0.002 | Young |
| 0.973927 | 0.052881 | 0.051 | 0.05 | 1 | 3 | Fbxl15 | 0.001 | Young |
| 0.97452 | 0.071632 | 0.665 | 0.691 | 1 | 3 | Lrp1 | -0.026 | Young |
| 0.974564 | 0.080054 | 0.073 | 0.073 | 1 | 3 | 2310039H( | 0 | Young |
| 0.9746 | 0.116388 | 0.283 | 0.293 | 1 | 3 | Chd7 | -0.01 | Young |
| 0.974858 | 0.145861 | 0.097 | 0.098 | 1 | 3 | Slc16a7 | -0.001 | Young |
| 0.975077 | 0.047589 | 0.385 | 0.393 | 1 | 3 | Psenen | -0.008 | Young |
| 0.975118 | 0.265572 | 0.048 | 0.048 | 1 | 3 | Slc35e2 | 0 | Young |
| 0.975262 | 0.125268 | 0.167 | 0.171 | 1 | 3 | Nfkb2 | -0.004 | Young |
| 0.975368 | 0.090759 | 0.025 | 0.025 | 1 | 3 | E130311K1 | 0 | Young |
| 0.975523 | 0.385009 | 0.096 | 0.098 | 1 | 3 | Cwc25 | -0.002 | Young |
| 0.975537 | 0.06827 | 0.153 | 0.154 | 1 | 3 | Asl | -0.001 | Young |
| 0.975684 | 0.008509 | 0.294 | 0.302 | 1 | 3 | Oat | -0.008 | Young |
| 0.975774 | 0.167612 | 0.076 | 0.076 | 1 | 3 | Tbc1d9b | 0 | Young |
| 0.975784 | 0.120831 | 0.386 | 0.401 | 1 | 3 | Msl2 | -0.015 | Young |
| 0.975871 | 0.048309 | 0.23 | 0.235 | 1 | 3 | Stk11 | -0.005 | Young |
| 0.975958 | 0.14029 | 0.047 | 0.047 | 1 | 3 | Asb1 | 0 | Young |
| 0.976212 | 0.172156 | 0.112 | 0.114 | 1 | 3 | Plcb2 | -0.002 | Young |
| 0.976606 | 0.222457 | 0.098 | 0.1 | 1 | 3 | Wdr36 | -0.002 | Young |
| 0.976823 | 0.171529 | 0.024 | 0.024 | 1 | 3 | Tmem35b | 0 | Young |
| 0.976916 | 0.198816 | 0.025 | 0.025 | 1 | 3 | C330018D2 | 0 | Young |
| 0.976983 | 0.316949 | 0.044 | 0.044 | 1 | 3 | Pla2g2d | 0 | Young |
| 0.977226 | 0.158705 | 0.025 | 0.025 | 1 | 3 | Fpgs | 0 | Young |
| 0.97725 | 0.158744 | 0.28 | 0.289 | 1 | 3 | Tsc22d2 | -0.009 | Young |
| 0.977294 | 0.13049 | 0.027 | 0.026 | 1 | 3 | Itfg2 | 0.001 | Young |
| 0.977463 | 0.060879 | 0.481 | 0.496 | 1 | 3 | Anp32a | -0.015 | Young |
| 0.977495 | 0.289262 | 0.051 | 0.051 | 1 | 3 | Trmt10a | 0 | Young |
| 0.97753 | 0.168005 | 0.052 | 0.053 | 1 | 3 | Atf5 | -0.001 | Young |
| 0.97754 | 0.378058 | 0.058 | 0.059 | 1 | 3 | Gpr137b | -0.001 | Young |
| 0.978375 | 0.071811 | 0.232 | 0.237 | 1 | 3 | Wtap | -0.005 | Young |
| 0.978485 | 0.102351 | 0.404 | 0.421 | 1 | 3 | Rab6a | -0.017 | Young |
| 0.978662 | 0.302293 | 0.023 | 0.023 | 1 | 3 | Adamtsl4 | 0 | Young |
| 0.978686 | 0.09118 | 0.317 | 0.324 | 1 | 3 | AI314180 | -0.007 | Young |
| 0.978727 | 0.219361 | 0.024 | 0.024 | 1 | 3 | Tbc1d7 | 0 | Young |
| 0.978757 | 0.566629 | 0.03 | 0.031 | 1 | 3 | AU020206 | -0.001 | Young |
| 0.979041 | 0.123473 | 0.019 | 0.019 | 1 | 3 | Map2k3os | 0 | Young |

| | | | | | | | | |
|---|---|---|---|---|---|---|---|---|
| 0.979063 | 0.174516 | 0.053 | 0.053 | 1 | 3 | Phldb1 | 0 | Young |
| 0.979081 | 0.038042 | 0.321 | 0.333 | 1 | 3 | Ccnt1 | -0.012 | Young |
| 0.979133 | 0.041513 | 0.249 | 0.25 | 1 | 3 | Cdk2ap1 | -0.001 | Young |
| 0.979501 | 0.261891 | 0.02 | 0.02 | 1 | 3 | Med26 | 0 | Young |
| 0.97951 | 0.074412 | 0.169 | 0.171 | 1 | 3 | Pbdc1 | -0.002 | Young |
| 0.979571 | 0.386377 | 0.063 | 0.063 | 1 | 3 | Dgcr6 | 0 | Young |
| 0.979701 | 0.063657 | 0.09 | 0.091 | 1 | 3 | 1110008P1 | -0.001 | Young |
| 0.979768 | 0.230363 | 0.107 | 0.11 | 1 | 3 | Josd1 | -0.003 | Young |
| 0.979837 | 0.133591 | 0.038 | 0.038 | 1 | 3 | Rcan1 | 0 | Young |
| 0.979997 | 0.243827 | 0.024 | 0.024 | 1 | 3 | Ccdc73 | 0 | Young |
| 0.980099 | 0.092645 | 0.356 | 0.372 | 1 | 3 | Fxr1 | -0.016 | Young |
| 0.980191 | 0.073292 | 0.173 | 0.177 | 1 | 3 | Aip | -0.004 | Young |
| 0.980349 | 0.110622 | 0.083 | 0.085 | 1 | 3 | Zfp219 | -0.002 | Young |
| 0.980529 | 0.108789 | 0.293 | 0.305 | 1 | 3 | Polr2e | -0.012 | Young |
| 0.980534 | 0.044099 | 0.02 | 0.02 | 1 | 3 | Gm49339 | 0 | Young |
| 0.980546 | 0.102655 | 0.083 | 0.084 | 1 | 3 | Ctbs | -0.001 | Young |
| 0.981022 | 0.114626 | 0.333 | 0.357 | 1 | 3 | Rab5a | -0.024 | Young |
| 0.981296 | 0.169313 | 0.194 | 0.2 | 1 | 3 | Eps15l1 | -0.006 | Young |
| 0.981476 | 0.108839 | 0.24 | 0.244 | 1 | 3 | Rara | -0.004 | Young |
| 0.981527 | 0.027611 | 0.019 | 0.019 | 1 | 3 | Grasp | 0 | Young |
| 0.981968 | 0.20201 | 0.095 | 0.097 | 1 | 3 | Dhx8 | -0.002 | Young |
| 0.98197 | 0.143198 | 0.207 | 0.211 | 1 | 3 | Stx12 | -0.004 | Young |
| 0.982055 | 0.402155 | 0.125 | 0.129 | 1 | 3 | Exosc8 | -0.004 | Young |
| 0.982061 | 0.065183 | 0.134 | 0.135 | 1 | 3 | Trappc8 | -0.001 | Young |
| 0.982365 | 0.204356 | 0.212 | 0.22 | 1 | 3 | Vezf1 | -0.008 | Young |
| 0.982445 | 0.16679 | 0.178 | 0.183 | 1 | 3 | Pfdn6 | -0.005 | Young |
| 0.982493 | 0.120007 | 0.025 | 0.025 | 1 | 3 | Zfp777 | 0 | Young |
| 0.982531 | 0.06372 | 0.586 | 0.613 | 1 | 3 | Capn2 | -0.027 | Young |
| 0.982883 | 0.151994 | 0.044 | 0.044 | 1 | 3 | Ccdc58 | 0 | Young |
| 0.982889 | 0.141207 | 0.313 | 0.332 | 1 | 3 | Ssu72 | -0.019 | Young |
| 0.983023 | 0.029564 | 0.15 | 0.153 | 1 | 3 | Cenpa | -0.003 | Young |
| 0.983064 | 0.040015 | 0.426 | 0.437 | 1 | 3 | Ndufv3 | -0.011 | Young |
| 0.983241 | 0.378178 | 0.078 | 0.079 | 1 | 3 | Calhm2 | -0.001 | Young |
| 0.983556 | 0.149563 | 0.173 | 0.177 | 1 | 3 | Tmc6 | -0.004 | Young |
| 0.983658 | 0.393986 | 0.019 | 0.019 | 1 | 3 | Rad52 | 0 | Young |
| 0.984199 | 0.269022 | 0.023 | 0.023 | 1 | 3 | Acad11 | 0 | Young |
| 0.984451 | 0.539886 | 0.097 | 0.099 | 1 | 3 | Arsk | -0.002 | Young |
| 0.984565 | 0.255945 | 0.119 | 0.121 | 1 | 3 | Ago4 | -0.002 | Young |
| 0.984589 | 0.33616 | 0.11 | 0.112 | 1 | 3 | Dgcr2 | -0.002 | Young |
| 0.984989 | 0.001276 | 0.202 | 0.204 | 1 | 3 | Desi2 | -0.002 | Young |
| 0.985088 | 0.009002 | 0.625 | 0.645 | 1 | 3 | Hcls1 | -0.02 | Young |
| 0.985563 | 0.14533 | 0.314 | 0.33 | 1 | 3 | Pptc7 | -0.016 | Young |
| 0.985631 | 0.018659 | 0.098 | 0.099 | 1 | 3 | Ncoa5 | -0.001 | Young |
| 0.98566 | 0.177785 | 0.117 | 0.118 | 1 | 3 | Cln3 | -0.001 | Young |
| 0.985867 | 0.380964 | 0.018 | 0.018 | 1 | 3 | Ovca2 | 0 | Young |

| | | | | | | | | |
|---|---|---|---|---|---|---|---|---|
| 0.985927 | 0.08885 | 0.277 | 0.289 | 1 | 3 | Mgat1 | -0.012 | Young |
| 0.986172 | 0.305573 | 0.193 | 0.203 | 1 | 3 | Trim44 | -0.01 | Young |
| 0.986549 | 0.18003 | 0.388 | 0.409 | 1 | 3 | Zc3h7a | -0.021 | Young |
| 0.986568 | 0.118544 | 0.434 | 0.468 | 1 | 3 | Map1lc3b | -0.034 | Young |
| 0.986613 | 0.052511 | 0.591 | 0.632 | 1 | 3 | Ckb | -0.041 | Young |
| 0.986623 | 0.108255 | 0.412 | 0.434 | 1 | 3 | Fyb | -0.022 | Young |
| 0.986713 | 0.120356 | 0.231 | 0.234 | 1 | 3 | Uba52 | -0.003 | Young |
| 0.987079 | 0.575318 | 0.02 | 0.02 | 1 | 3 | Haus1 | 0 | Young |
| 0.98712 | 0.110267 | 0.217 | 0.222 | 1 | 3 | Zmym5 | -0.005 | Young |
| 0.987311 | 0.256234 | 0.287 | 0.301 | 1 | 3 | Elmsan1 | -0.014 | Young |
| 0.987404 | 0.322704 | 0.054 | 0.055 | 1 | 3 | Dxo | -0.001 | Young |
| 0.987415 | 0.04612 | 0.487 | 0.496 | 1 | 3 | Ppp1cb | -0.009 | Young |
| 0.987462 | 0.247264 | 0.21 | 0.219 | 1 | 3 | Rapgef2 | -0.009 | Young |
| 0.98756 | 0.036981 | 0.202 | 0.204 | 1 | 3 | Zfx | -0.002 | Young |
| 0.987599 | 0.039331 | 0.482 | 0.502 | 1 | 3 | Tubb6 | -0.02 | Young |
| 0.987635 | 0.170238 | 0.039 | 0.039 | 1 | 3 | Pias3 | 0 | Young |
| 0.987829 | 0.040925 | 0.311 | 0.315 | 1 | 3 | Rab3d | -0.004 | Young |
| 0.987999 | 0.18504 | 0.114 | 0.116 | 1 | 3 | Rad50 | -0.002 | Young |
| 0.988108 | 0.271559 | 0.023 | 0.023 | 1 | 3 | Hemk1 | 0 | Young |
| 0.988113 | 0.240431 | 0.02 | 0.02 | 1 | 3 | Gm34590 | 0 | Young |
| 0.988292 | 0.23056 | 0.231 | 0.239 | 1 | 3 | C3ar1 | -0.008 | Young |
| 0.988473 | 0.02249 | 0.235 | 0.237 | 1 | 3 | Ubr4 | -0.002 | Young |
| 0.988543 | 0.294233 | 0.064 | 0.065 | 1 | 3 | Atxn1l | -0.001 | Young |
| 0.988769 | 0.269462 | 0.102 | 0.104 | 1 | 3 | Reep4 | -0.002 | Young |
| 0.988987 | 0.25045 | 0.019 | 0.019 | 1 | 3 | Aldh6a1 | 0 | Young |
| 0.98908 | 0.102274 | 0.096 | 0.097 | 1 | 3 | Nubp2 | -0.001 | Young |
| 0.989265 | 0.300375 | 0.287 | 0.296 | 1 | 3 | Peli1 | -0.009 | Young |
| 0.989619 | 0.085928 | 0.025 | 0.025 | 1 | 3 | Cox7a1 | 0 | Young |
| 0.98986 | 0.110952 | 0.301 | 0.313 | 1 | 3 | Erbin | -0.012 | Young |
| 0.990292 | 0.393518 | 0.021 | 0.022 | 1 | 3 | Klhl22 | -0.001 | Young |
| 0.990373 | 0.212994 | 0.058 | 0.059 | 1 | 3 | Xxylt1 | -0.001 | Young |
| 0.990571 | 0.15891 | 0.149 | 0.149 | 1 | 3 | Arl6ip4 | 0 | Young |
| 0.990639 | 0.450784 | 0.018 | 0.018 | 1 | 3 | Susd1 | 0 | Young |
| 0.990679 | 0.312488 | 0.082 | 0.084 | 1 | 3 | Ambra1 | -0.002 | Young |
| 0.990948 | 0.12068 | 0.071 | 0.072 | 1 | 3 | Gins4 | -0.001 | Young |
| 0.990973 | 0.241247 | 0.083 | 0.084 | 1 | 3 | Rusc1 | -0.001 | Young |
| 0.991085 | 0.119362 | 0.331 | 0.349 | 1 | 3 | Agpat4 | -0.018 | Young |
| 0.99126 | 0.185152 | 0.078 | 0.079 | 1 | 3 | Pde8a | -0.001 | Young |
| 0.991567 | 0.02587 | 0.206 | 0.208 | 1 | 3 | Sdccag3 | -0.002 | Young |
| 0.991629 | 0.142644 | 0.042 | 0.042 | 1 | 3 | Jak3 | 0 | Young |
| 0.991692 | 0.120445 | 0.023 | 0.023 | 1 | 3 | Zfp668 | 0 | Young |
| 0.991829 | 0.346795 | 0.019 | 0.019 | 1 | 3 | Mettl25 | 0 | Young |
| 0.991903 | 0.071376 | 0.02 | 0.02 | 1 | 3 | Cd5l | 0 | Young |
| 0.992151 | 0.167868 | 0.218 | 0.227 | 1 | 3 | Maf1 | -0.009 | Young |
| 0.992336 | 0.055541 | 0.12 | 0.121 | 1 | 3 | Gng12 | -0.001 | Young |

| | | | | | | | | |
|---|---|---|---|---|---|---|---|---|
| 0.992464 | 0.134432 | 0.17 | 0.174 | 1 | 3 | Gpr107 | -0.004 | Young |
| 0.992517 | 0.113832 | 0.169 | 0.174 | 1 | 3 | Igsf9 | -0.005 | Young |
| 0.992845 | 0.268832 | 0.333 | 0.351 | 1 | 3 | Clk4 | -0.018 | Young |
| 0.992941 | 0.32548 | 0.034 | 0.035 | 1 | 3 | Rfx3 | -0.001 | Young |
| 0.992978 | 0.290419 | 0.121 | 0.123 | 1 | 3 | Rpusd4 | -0.002 | Young |
| 0.993185 | 0.138137 | 0.126 | 0.129 | 1 | 3 | Tfeb | -0.003 | Young |
| 0.993216 | 0.044413 | 0.487 | 0.514 | 1 | 3 | Chmp2a | -0.027 | Young |
| 0.993337 | 0.070886 | 0.025 | 0.025 | 1 | 3 | Cep70 | 0 | Young |
| 0.993502 | 0.229594 | 0.073 | 0.074 | 1 | 3 | Agap1 | -0.001 | Young |
| 0.993672 | 0.061619 | 0.047 | 0.047 | 1 | 3 | Stk38l | 0 | Young |
| 0.993681 | 0.192017 | 0.143 | 0.147 | 1 | 3 | Sh3glb2 | -0.004 | Young |
| 0.994142 | 0.011973 | 0.593 | 0.618 | 1 | 3 | Clec4b1 | -0.025 | Young |
| 0.994256 | 0.024424 | 0.298 | 0.306 | 1 | 3 | Tmem259 | -0.008 | Young |
| 0.994539 | 0.218696 | 0.173 | 0.177 | 1 | 3 | Mdn1 | -0.004 | Young |
| 0.995268 | 0.174637 | 0.042 | 0.042 | 1 | 3 | Alkbh8 | 0 | Young |
| 0.995328 | 0.10885 | 0.327 | 0.335 | 1 | 3 | Napsa | -0.008 | Young |
| 0.995487 | 0.1209 | 0.112 | 0.114 | 1 | 3 | Rfc2 | -0.002 | Young |
| 0.995722 | 0.250427 | 0.069 | 0.07 | 1 | 3 | Rhno1 | -0.001 | Young |
| 0.996382 | 0.147646 | 0.02 | 0.02 | 1 | 3 | Hist2h2be | 0 | Young |
| 0.996786 | 0.237982 | 0.09 | 0.091 | 1 | 3 | Fbxo34 | -0.001 | Young |
| 0.996897 | 0.120296 | 0.192 | 0.197 | 1 | 3 | D1Ertd622 | -0.005 | Young |
| 0.997014 | 0.180801 | 0.234 | 0.241 | 1 | 3 | Dapp1 | -0.007 | Young |
| 0.99711 | 0.084746 | 0.063 | 0.063 | 1 | 3 | Sass6 | 0 | Young |
| 0.997209 | 0.126414 | 0.116 | 0.117 | 1 | 3 | Ngly1 | -0.001 | Young |
| 0.997236 | 0.044797 | 0.409 | 0.431 | 1 | 3 | Myo1g | -0.022 | Young |
| 0.998618 | 0.043962 | 0.346 | 0.354 | 1 | 3 | Pbxip1 | -0.008 | Young |
| 0.998685 | 0.22436 | 0.083 | 0.084 | 1 | 3 | Abhd4 | -0.001 | Young |
| 0.998823 | 0.198173 | 0.122 | 0.124 | 1 | 3 | Nfia | -0.002 | Young |
| 0.998945 | 0.219467 | 0.135 | 0.139 | 1 | 3 | Spin1 | -0.004 | Young |
| 0.9993 | 0.177113 | 0.138 | 0.14 | 1 | 3 | Mvd | -0.002 | Young |
| 0.999393 | 0.229921 | 0.042 | 0.042 | 1 | 3 | Tmem229a | 0 | Young |
| 0.999645 | 0.418953 | 0.019 | 0.019 | 1 | 3 | Cyp27a1 | 0 | Young |
| 0.999826 | 0.010615 | 0.587 | 0.622 | 1 | 3 | Fnbp1 | -0.035 | Young |
| 7.04E-09 | 1.426931 | 0.809 | 0.537 | 0.000218 | 4 | Cxcl13 | 0.272 | Old |
| 8.14E-09 | 1.257778 | 0.722 | 0.45 | 0.000253 | 4 | Gda | 0.272 | Old |
| 1.13E-08 | 1.482285 | 0.67 | 0.375 | 0.00035 | 4 | C3 | 0.295 | Old |
| 1.76E-08 | 1.470404 | 0.657 | 0.362 | 0.000547 | 4 | Cd55 | 0.295 | Old |
| 5.08E-08 | 0.700901 | 0.925 | 0.825 | 0.001577 | 4 | Ccl6 | 0.1 | Old |
| 8.42E-07 | 0.638099 | 0.961 | 0.95 | 0.026148 | 4 | Crip1 | 0.011 | Old |
| 1.02E-06 | 0.700703 | 0.887 | 0.8 | 0.031805 | 4 | Ccl9 | 0.087 | Old |
| 2.99E-06 | 0.57499 | 0.924 | 0.825 | 0.092721 | 4 | Tmem176a | 0.099 | Old |
| 4.25E-06 | 0.504979 | 0.953 | 0.9 | 0.131922 | 4 | Ctss | 0.053 | Old |
| 4.40E-06 | 1.242273 | 0.539 | 0.312 | 0.136481 | 4 | S100a10 | 0.227 | Old |
| 7.67E-06 | 0.902072 | 0.716 | 0.512 | 0.238253 | 4 | Cstb | 0.204 | Old |
| 1.01E-05 | 0.492929 | 0.883 | 0.762 | 0.314731 | 4 | Cfp | 0.121 | Old |

| | | | | | | | | |
|---|---|---|---|---|---|---|---|---|
| 1.10E-05 | 0.63981 | 0.856 | 0.688 | 0.340506 | 4 | Sdc3 | 0.168 | Old |
| 1.26E-05 | 0.450636 | 0.964 | 0.963 | 0.392756 | 4 | Ahnak | 0.001 | Old |
| 1.67E-05 | 0.914313 | 0.727 | 0.588 | 0.517971 | 4 | Dstn | 0.139 | Old |
| 2.29E-05 | 0.618709 | 0.814 | 0.65 | 0.710592 | 4 | Apobec3 | 0.164 | Old |
| 3.04E-05 | 1.816789 | 0.387 | 0.175 | 0.944335 | 4 | Armc8 | 0.212 | Old |
| 3.92E-05 | 1.246151 | 0.611 | 0.425 | 1 | 4 | Fcna | 0.186 | Old |
| 5.43E-05 | 2.502844 | 0.265 | 0.062 | 1 | 4 | Colq | 0.203 | Old |
| 0.000118 | 1.21699 | 0.852 | 0.825 | 1 | 4 | Retnla | 0.027 | Old |
| 0.000118 | 0.701001 | 0.771 | 0.675 | 1 | 4 | Cd47 | 0.096 | Old |
| 0.000124 | 1.106536 | 0.497 | 0.3 | 1 | 4 | Anxa2 | 0.197 | Old |
| 0.000165 | 0.769686 | 0.557 | 0.35 | 1 | 4 | Notch2 | 0.207 | Old |
| 0.000184 | 0.289873 | 0.997 | 1 | 1 | 4 | Psap | -0.003 | Old |
| 0.000189 | 0.434493 | 0.955 | 0.875 | 1 | 4 | Timp2 | 0.08 | Old |
| 0.000198 | 0.459524 | 0.937 | 0.9 | 1 | 4 | Gpx1 | 0.037 | Old |
| 0.000201 | 0.90862 | 0.649 | 0.475 | 1 | 4 | Fxyd2 | 0.174 | Old |
| 0.000224 | 0.436127 | 0.988 | 0.988 | 1 | 4 | Mafb | 0 | Old |
| 0.000323 | 1.167848 | 0.35 | 0.15 | 1 | 4 | Emp1 | 0.2 | Old |
| 0.000327 | 1.532371 | 0.323 | 0.138 | 1 | 4 | Ldlr | 0.185 | Old |
| 0.000339 | 0.520327 | 0.896 | 0.787 | 1 | 4 | Anxa5 | 0.109 | Old |
| 0.000422 | 1.24569 | 0.433 | 0.238 | 1 | 4 | Anxa1 | 0.195 | Old |
| 0.000435 | 0.601756 | 0.775 | 0.7 | 1 | 4 | Sulf2 | 0.075 | Old |
| 0.000529 | 0.880368 | 0.586 | 0.45 | 1 | 4 | Tspan3 | 0.136 | Old |
| 0.000535 | 0.431455 | 0.936 | 0.925 | 1 | 4 | Hsp90b1 | 0.011 | Old |
| 0.000685 | 0.657051 | 0.663 | 0.5 | 1 | 4 | Serpinb6a | 0.163 | Old |
| 0.001021 | 1.457039 | 0.293 | 0.125 | 1 | 4 | Smpdl3a | 0.168 | Old |
| 0.001027 | 2.364014 | 0.209 | 0.062 | 1 | 4 | Mrpl14 | 0.147 | Old |
| 0.001376 | 0.182368 | 1 | 1 | 1 | 4 | Actb | 0 | Old |
| 0.001433 | 0.382453 | 0.921 | 0.95 | 1 | 4 | Tmem176b | -0.029 | Old |
| 0.001451 | 0.372007 | 0.913 | 0.863 | 1 | 4 | Ppia | 0.05 | Old |
| 0.001625 | 0.455891 | 0.829 | 0.787 | 1 | 4 | Sirpa | 0.042 | Old |
| 0.001748 | 1.150658 | 0.504 | 0.375 | 1 | 4 | S100a6 | 0.129 | Old |
| 0.001769 | 1.009252 | 0.384 | 0.225 | 1 | 4 | Apobec1 | 0.159 | Old |
| 0.00187 | 0.339183 | 0.977 | 0.975 | 1 | 4 | Actg1 | 0.002 | Old |
| 0.001992 | 0.584009 | 0.768 | 0.6 | 1 | 4 | Ier2 | 0.168 | Old |
| 0.002042 | 0.688119 | 0.642 | 0.55 | 1 | 4 | Ssr3 | 0.092 | Old |
| 0.002082 | 4.799676 | 0.126 | 0.013 | 1 | 4 | Ttn | 0.113 | Old |
| 0.00236 | 3.839415 | 0.125 | 0.013 | 1 | 4 | Gprc5b | 0.112 | Old |
| 0.002424 | 0.430839 | 0.826 | 0.75 | 1 | 4 | Ninj1 | 0.076 | Old |
| 0.002654 | 0.409039 | 0.866 | 0.787 | 1 | 4 | Vim | 0.079 | Old |
| 0.002845 | 1.04701 | 0.386 | 0.238 | 1 | 4 | Wls | 0.148 | Old |
| 0.002994 | 0.317026 | 0.955 | 0.975 | 1 | 4 | Ly6e | -0.02 | Old |
| 0.003315 | 5.324354 | 0.098 | 0 | 1 | 4 | Sh2d4a | 0.098 | Old |
| 0.003354 | 0.44141 | 0.861 | 0.762 | 1 | 4 | Cfh | 0.099 | Old |
| 0.003903 | 0.454172 | 0.817 | 0.738 | 1 | 4 | Iqgap1 | 0.079 | Old |
| 0.004164 | 2.548425 | 0.117 | 0.013 | 1 | 4 | Camk2a | 0.104 | Old |

| | | | | | | | | |
|---|---|---|---|---|---|---|---|---|
| 0.004203 | 0.512145 | 0.679 | 0.562 | 1 | 4 | H2-T23 | 0.117 | Old |
| 0.004659 | 0.497286 | 0.728 | 0.675 | 1 | 4 | Sh3bgrl | 0.053 | Old |
| 0.004665 | 3.257869 | 0.113 | 0.013 | 1 | 4 | Sh3bgrl2 | 0.1 | Old |
| 0.004736 | 1.903076 | 0.185 | 0.062 | 1 | 4 | Hmgcs1 | 0.123 | Old |
| 0.004807 | 0.278913 | 0.963 | 0.95 | 1 | 4 | Lamp1 | 0.013 | Old |
| 0.004907 | 1.145724 | 0.327 | 0.188 | 1 | 4 | Ms4a8a | 0.139 | Old |
| 0.005078 | 0.281468 | 0.974 | 0.938 | 1 | 4 | Ptma | 0.036 | Old |
| 0.005268 | 0.603339 | 0.457 | 0.312 | 1 | 4 | Hdgf | 0.145 | Old |
| 0.005458 | 0.402743 | 0.871 | 0.85 | 1 | 4 | Rap1b | 0.021 | Old |
| 0.005559 | 0.481485 | 0.685 | 0.55 | 1 | 4 | Plbd1 | 0.135 | Old |
| 0.005765 | 0.662699 | 0.607 | 0.5 | 1 | 4 | Emp3 | 0.107 | Old |
| 0.005924 | 0.555594 | 0.587 | 0.462 | 1 | 4 | Tubb5 | 0.125 | Old |
| 0.006146 | 1.171267 | 0.313 | 0.175 | 1 | 4 | Hpcal1 | 0.138 | Old |
| 0.006807 | 0.924638 | 0.388 | 0.262 | 1 | 4 | Zbtb7a | 0.126 | Old |
| 0.006965 | 0.83363 | 0.429 | 0.3 | 1 | 4 | Tkt | 0.129 | Old |
| 0.007602 | 4.840336 | 0.082 | 0 | 1 | 4 | Apba1 | 0.082 | Old |
| 0.00778 | 0.543656 | 0.659 | 0.562 | 1 | 4 | Pea15a | 0.097 | Old |
| 0.008144 | 0.762947 | 0.497 | 0.375 | 1 | 4 | Fam43a | 0.122 | Old |
| 0.008146 | 0.461125 | 0.802 | 0.688 | 1 | 4 | BC005537 | 0.114 | Old |
| 0.008982 | 1.048416 | 0.335 | 0.212 | 1 | 4 | Plec | 0.123 | Old |
| 0.008995 | 0.40269 | 0.741 | 0.675 | 1 | 4 | Morf4l1 | 0.066 | Old |
| 0.009077 | 2.692949 | 0.101 | 0.013 | 1 | 4 | Cbx5 | 0.088 | Old |
| 0.009519 | 0.540304 | 0.644 | 0.525 | 1 | 4 | Rgl1 | 0.119 | Old |
| 0.009768 | 1.963223 | 0.154 | 0.05 | 1 | 4 | Fundc2 | 0.104 | Old |
| 0.010001 | 4.42741 | 0.077 | 0 | 1 | 4 | Gm28875 | 0.077 | Old |
| 0.010433 | 1.250844 | 0.273 | 0.15 | 1 | 4 | Slamf9 | 0.123 | Old |
| 0.010679 | 0.358195 | 0.916 | 0.938 | 1 | 4 | Pltp | -0.022 | Old |
| 0.010994 | 0.657147 | 0.526 | 0.425 | 1 | 4 | Mfsd1 | 0.101 | Old |
| 0.011541 | 0.676612 | 0.501 | 0.388 | 1 | 4 | 2900097C1 | 0.113 | Old |
| 0.011792 | 0.809624 | 0.362 | 0.238 | 1 | 4 | Lfng | 0.124 | Old |
| 0.011838 | 0.857651 | 0.241 | 0.112 | 1 | 4 | Birc2 | 0.129 | Old |
| 0.011866 | 4.671867 | 0.074 | 0 | 1 | 4 | Spib | 0.074 | Old |
| 0.011887 | 1.841456 | 0.135 | 0.038 | 1 | 4 | Map1s | 0.097 | Old |
| 0.011952 | 0.433145 | 0.764 | 0.713 | 1 | 4 | Rhoa | 0.051 | Old |
| 0.011988 | 2.455249 | 0.131 | 0.038 | 1 | 4 | Kif2c | 0.093 | Old |
| 0.012052 | 1.854871 | 0.117 | 0.025 | 1 | 4 | Nudcd2 | 0.092 | Old |
| 0.012273 | 0.881298 | 0.334 | 0.212 | 1 | 4 | Anp32b | 0.122 | Old |
| 0.012278 | 4.734762 | 0.073 | 0 | 1 | 4 | Ppp2cb | 0.073 | Old |
| 0.012316 | 0.584257 | 0.493 | 0.388 | 1 | 4 | Lmna | 0.105 | Old |
| 0.012431 | 0.383402 | 0.753 | 0.725 | 1 | 4 | Samhd1 | 0.028 | Old |
| 0.012583 | 0.771596 | 0.35 | 0.225 | 1 | 4 | Txn2 | 0.125 | Old |
| 0.01287 | 0.900874 | 0.364 | 0.238 | 1 | 4 | Psmd2 | 0.126 | Old |
| 0.012949 | 2.805255 | 0.094 | 0.013 | 1 | 4 | Abcd2 | 0.081 | Old |
| 0.012971 | 1.128758 | 0.313 | 0.2 | 1 | 4 | Swi5 | 0.113 | Old |
| 0.013398 | 1.555545 | 0.134 | 0.038 | 1 | 4 | Atf2 | 0.096 | Old |

| | | | | | | | | |
|---|---|---|---|---|---|---|---|---|
| 0.013727 | 0.473831 | 0.852 | 0.762 | 1 | 4 | Klf2 | 0.09 | Old |
| 0.013977 | 0.303976 | 0.851 | 0.775 | 1 | 4 | Cd63 | 0.076 | Old |
| 0.014432 | 0.383854 | 0.874 | 0.838 | 1 | 4 | Egr1 | 0.036 | Old |
| 0.014506 | 0.784683 | 0.421 | 0.3 | 1 | 4 | Tbca | 0.121 | Old |
| 0.014935 | 0.219623 | 0.989 | 0.963 | 1 | 4 | B2m | 0.026 | Old |
| 0.014973 | 2.204815 | 0.111 | 0.025 | 1 | 4 | Smim20 | 0.086 | Old |
| 0.015068 | 4.847613 | 0.069 | 0 | 1 | 4 | Gm26870 | 0.069 | Old |
| 0.015216 | 0.551289 | 0.505 | 0.438 | 1 | 4 | Pak2 | 0.067 | Old |
| 0.015597 | 0.643134 | 0.582 | 0.475 | 1 | 4 | Clec4a3 | 0.107 | Old |
| 0.015694 | 0.966156 | 0.165 | 0.062 | 1 | 4 | Bet1 | 0.103 | Old |
| 0.016143 | 0.467235 | 0.681 | 0.575 | 1 | 4 | Cyfip1 | 0.106 | Old |
| 0.016553 | 1.449166 | 0.206 | 0.1 | 1 | 4 | Trappc10 | 0.106 | Old |
| 0.016991 | 1.265056 | 0.221 | 0.112 | 1 | 4 | Snrpd2 | 0.109 | Old |
| 0.017231 | 0.932426 | 0.416 | 0.325 | 1 | 4 | Necap2 | 0.091 | Old |
| 0.018573 | 0.6112 | 0.605 | 0.537 | 1 | 4 | Gas7 | 0.068 | Old |
| 0.018753 | 1.399952 | 0.203 | 0.1 | 1 | 4 | Ube2z | 0.103 | Old |
| 0.018936 | 1.008153 | 0.311 | 0.2 | 1 | 4 | Gfpt1 | 0.111 | Old |
| 0.019789 | 4.083203 | 0.064 | 0 | 1 | 4 | Bola3 | 0.064 | Old |
| 0.020554 | 1.497025 | 0.142 | 0.05 | 1 | 4 | Stat2 | 0.092 | Old |
| 0.021099 | 0.581724 | 0.516 | 0.425 | 1 | 4 | Prkar1a | 0.091 | Old |
| 0.021321 | 1.142959 | 0.255 | 0.15 | 1 | 4 | Polr2k | 0.105 | Old |
| 0.021625 | 2.473044 | 0.119 | 0.038 | 1 | 4 | Gng11 | 0.081 | Old |
| 0.021635 | 2.179807 | 0.121 | 0.038 | 1 | 4 | Vldlr | 0.083 | Old |
| 0.021807 | 2.347908 | 0.103 | 0.025 | 1 | 4 | Il1rn | 0.078 | Old |
| 0.021865 | 0.546357 | 0.625 | 0.525 | 1 | 4 | Pcbp1 | 0.1 | Old |
| 0.021877 | 1.306894 | 0.285 | 0.188 | 1 | 4 | Tspan32 | 0.097 | Old |
| 0.021964 | 2.87262 | 0.084 | 0.013 | 1 | 4 | Rabgap1l | 0.071 | Old |
| 0.022036 | 1.155804 | 0.175 | 0.075 | 1 | 4 | Aida | 0.1 | Old |
| 0.022138 | 1.568455 | 0.209 | 0.112 | 1 | 4 | Ckap4 | 0.097 | Old |
| 0.022279 | 0.921867 | 0.205 | 0.1 | 1 | 4 | Dscr3 | 0.105 | Old |
| 0.022528 | 0.387391 | 0.729 | 0.65 | 1 | 4 | Ifitm3 | 0.079 | Old |
| 0.022547 | 2.169993 | 0.103 | 0.025 | 1 | 4 | Tmem38b | 0.078 | Old |
| 0.022603 | 0.104612 | 1 | 1 | 1 | 4 | mt-Co1 | 0 | Old |
| 0.02302 | 3.875427 | 0.082 | 0.013 | 1 | 4 | Dnm1 | 0.069 | Old |
| 0.023056 | 3.285502 | 0.082 | 0.013 | 1 | 4 | Cdh6 | 0.069 | Old |
| 0.02316 | 1.260993 | 0.24 | 0.138 | 1 | 4 | Gpr146 | 0.102 | Old |
| 0.023776 | 0.662981 | 0.52 | 0.4 | 1 | 4 | Cd209d | 0.12 | Old |
| 0.025212 | 0.737791 | 0.391 | 0.262 | 1 | 4 | Il2rg | 0.129 | Old |
| 0.02539 | 0.332644 | 0.728 | 0.662 | 1 | 4 | Gltp | 0.066 | Old |
| 0.025984 | 4.267336 | 0.059 | 0 | 1 | 4 | Rxrg | 0.059 | Old |
| 0.026156 | 2.255386 | 0.146 | 0.062 | 1 | 4 | Ifitm6 | 0.084 | Old |
| 0.026262 | 0.977778 | 0.265 | 0.162 | 1 | 4 | Gga1 | 0.103 | Old |
| 0.02698 | 1.299806 | 0.121 | 0.038 | 1 | 4 | Flvcr1 | 0.083 | Old |
| 0.027815 | 3.95773 | 0.057 | 0 | 1 | 4 | Batf | 0.057 | Old |
| 0.027916 | 0.520138 | 0.542 | 0.45 | 1 | 4 | Grk2 | 0.092 | Old |

| | | | | | | | | |
|---|---|---|---|---|---|---|---|---|
| 0.027985 | 0.127738 | 0.958 | 0.95 | 1 | 4 | H3f3b | 0.008 | Old |
| 0.028551 | 1.090351 | 0.305 | 0.212 | 1 | 4 | 11-Sep | 0.093 | Old |
| 0.029514 | 0.465289 | 0.752 | 0.662 | 1 | 4 | Mtss1 | 0.09 | Old |
| 0.029642 | 1.099858 | 0.212 | 0.112 | 1 | 4 | Arpc1a | 0.1 | Old |
| 0.029668 | 0.264512 | 0.843 | 0.787 | 1 | 4 | Wwp1 | 0.056 | Old |
| 0.029782 | 1.365714 | 0.1 | 0.025 | 1 | 4 | 1810022K0 | 0.075 | Old |
| 0.030247 | 0.68966 | 0.322 | 0.212 | 1 | 4 | Mapre2 | 0.11 | Old |
| 0.030704 | 0.759867 | 0.499 | 0.425 | 1 | 4 | Spred1 | 0.074 | Old |
| 0.0309 | 1.448339 | 0.19 | 0.1 | 1 | 4 | Etfa | 0.09 | Old |
| 0.031864 | 1.148312 | 0.136 | 0.05 | 1 | 4 | Bst1 | 0.086 | Old |
| 0.031992 | 1.531591 | 0.176 | 0.088 | 1 | 4 | Espl1 | 0.088 | Old |
| 0.032012 | 0.547085 | 0.614 | 0.575 | 1 | 4 | Cox5a | 0.039 | Old |
| 0.032175 | 1.411973 | 0.161 | 0.075 | 1 | 4 | Ahctf1 | 0.086 | Old |
| 0.032462 | 1.454261 | 0.162 | 0.075 | 1 | 4 | Nfil3 | 0.087 | Old |
| 0.033534 | 0.384813 | 0.62 | 0.588 | 1 | 4 | Clk1 | 0.032 | Old |
| 0.034102 | 2.820955 | 0.109 | 0.038 | 1 | 4 | Ryk | 0.071 | Old |
| 0.03418 | 0.397815 | 0.727 | 0.7 | 1 | 4 | Ywhaz | 0.027 | Old |
| 0.034995 | 0.723833 | 0.286 | 0.175 | 1 | 4 | Inpp4a | 0.111 | Old |
| 0.035165 | 1.473461 | 0.13 | 0.05 | 1 | 4 | Papss1 | 0.08 | Old |
| 0.036021 | 1.565684 | 0.076 | 0.013 | 1 | 4 | Ceacam1 | 0.063 | Old |
| 0.036151 | 1.349373 | 0.175 | 0.088 | 1 | 4 | Etv6 | 0.087 | Old |
| 0.036533 | 4.285933 | 0.052 | 0 | 1 | 4 | Osbpl5 | 0.052 | Old |
| 0.037112 | 1.177341 | 0.131 | 0.05 | 1 | 4 | Slc35a4 | 0.081 | Old |
| 0.037169 | 1.02449 | 0.326 | 0.238 | 1 | 4 | Arcn1 | 0.088 | Old |
| 0.0372 | 1.733039 | 0.142 | 0.062 | 1 | 4 | Idi1 | 0.08 | Old |
| 0.037224 | 1.080928 | 0.179 | 0.088 | 1 | 4 | Cbfa2t3 | 0.091 | Old |
| 0.037233 | 0.35656 | 0.78 | 0.787 | 1 | 4 | Capzb | -0.007 | Old |
| 0.037302 | 0.220019 | 0.996 | 1 | 1 | 4 | Jund | -0.004 | Old |
| 0.037547 | 0.666166 | 0.419 | 0.325 | 1 | 4 | Fam111a | 0.094 | Old |
| 0.037802 | 4.251831 | 0.051 | 0 | 1 | 4 | Spint2 | 0.051 | Old |
| 0.038125 | 0.833422 | 0.13 | 0.05 | 1 | 4 | Kdelc2 | 0.08 | Old |
| 0.038762 | 0.391489 | 0.682 | 0.625 | 1 | 4 | Ywhae | 0.057 | Old |
| 0.039017 | 0.936957 | 0.318 | 0.225 | 1 | 4 | Strn3 | 0.093 | Old |
| 0.039034 | 0.434499 | 0.698 | 0.65 | 1 | 4 | Arpc5 | 0.048 | Old |
| 0.039205 | 0.347903 | 0.653 | 0.512 | 1 | 4 | Btf3 | 0.141 | Old |
| 0.03925 | 0.853671 | 0.267 | 0.175 | 1 | 4 | Epb41l1 | 0.092 | Old |
| 0.039416 | 0.585686 | 0.447 | 0.35 | 1 | 4 | Dync1h1 | 0.097 | Old |
| 0.039868 | 0.648308 | 0.509 | 0.412 | 1 | 4 | Ifi204 | 0.097 | Old |
| 0.040391 | 1.842339 | 0.092 | 0.025 | 1 | 4 | Ppp1r7 | 0.067 | Old |
| 0.040473 | 3.977027 | 0.05 | 0 | 1 | 4 | Tns4 | 0.05 | Old |
| 0.040518 | 0.271439 | 0.881 | 0.863 | 1 | 4 | Cdc42 | 0.018 | Old |
| 0.040522 | 0.512514 | 0.379 | 0.275 | 1 | 4 | Anxa4 | 0.104 | Old |
| 0.040963 | 0.491562 | 0.594 | 0.537 | 1 | 4 | 1810037I1 | 0.057 | Old |
| 0.041386 | 2.039783 | 0.091 | 0.025 | 1 | 4 | Kifc3 | 0.066 | Old |
| 0.041659 | 1.313859 | 0.111 | 0.038 | 1 | 4 | Polr2l | 0.073 | Old |

| | | | | | | | | |
|---|---|---|---|---|---|---|---|---|
| 0.042279 | 0.358707 | 0.765 | 0.713 | 1 | 4 | Rbpj | 0.052 | Old |
| 0.042778 | 0.891987 | 0.358 | 0.275 | 1 | 4 | Jarid2 | 0.083 | Old |
| 0.042828 | 1.050424 | 0.231 | 0.138 | 1 | 4 | Dhx9 | 0.093 | Old |
| 0.042921 | 3.459208 | 0.07 | 0.013 | 1 | 4 | Brwd1 | 0.057 | Old |
| 0.043419 | 0.260457 | 0.762 | 0.688 | 1 | 4 | Tagln2 | 0.074 | Old |
| 0.044123 | 2.87907 | 0.088 | 0.025 | 1 | 4 | Nr1h3 | 0.063 | Old |
| 0.04417 | 1.562586 | 0.072 | 0.013 | 1 | 4 | Gpr68 | 0.059 | Old |
| 0.044565 | 0.408517 | 0.684 | 0.588 | 1 | 4 | Pepd | 0.096 | Old |
| 0.044955 | 0.394694 | 0.685 | 0.512 | 1 | 4 | Cd209g | 0.173 | Old |
| 0.045289 | 0.974982 | 0.191 | 0.1 | 1 | 4 | Sema6a | 0.091 | Old |
| 0.04531 | 1.513539 | 0.125 | 0.05 | 1 | 4 | Fbxw7 | 0.075 | Old |
| 0.045657 | 0.969145 | 0.243 | 0.15 | 1 | 4 | 1110004F1 | 0.093 | Old |
| 0.046405 | 3.909386 | 0.047 | 0 | 1 | 4 | Tmem198b | 0.047 | Old |
| 0.046993 | 2.536816 | 0.103 | 0.038 | 1 | 4 | Yrdc | 0.065 | Old |
| 0.047093 | 0.909937 | 0.219 | 0.125 | 1 | 4 | Lman1 | 0.094 | Old |
| 0.047252 | 0.823362 | 0.26 | 0.162 | 1 | 4 | Pdlim4 | 0.098 | Old |
| 0.047256 | 1.102946 | 0.225 | 0.138 | 1 | 4 | Smc6 | 0.087 | Old |
| 0.047386 | 0.689783 | 0.395 | 0.312 | 1 | 4 | Metrnl | 0.083 | Old |
| 0.047578 | 2.455385 | 0.103 | 0.038 | 1 | 4 | Ak3 | 0.065 | Old |
| 0.047625 | 0.674496 | 0.495 | 0.412 | 1 | 4 | Scpep1 | 0.083 | Old |
| 0.047756 | 0.272565 | 0.595 | 0.462 | 1 | 4 | Lyve1 | 0.133 | Old |
| 0.048022 | 3.952486 | 0.047 | 0 | 1 | 4 | Mast2 | 0.047 | Old |
| 0.049729 | 0.393986 | 0.646 | 0.575 | 1 | 4 | Stard8 | 0.071 | Old |
| 0.050538 | 1.322632 | 0.167 | 0.088 | 1 | 4 | Dmpk | 0.079 | Old |
| 0.050737 | 0.688623 | 0.331 | 0.25 | 1 | 4 | Psmb1 | 0.081 | Old |
| 0.050993 | 0.876021 | 0.343 | 0.262 | 1 | 4 | Ap3s1 | 0.081 | Old |
| 0.051429 | 3.770777 | 0.045 | 0 | 1 | 4 | Oas1a | 0.045 | Old |
| 0.052493 | 2.8552 | 0.067 | 0.013 | 1 | 4 | Naip5 | 0.054 | Old |
| 0.052852 | 0.410561 | 0.608 | 0.562 | 1 | 4 | Gnb1 | 0.046 | Old |
| 0.053224 | 3.830825 | 0.045 | 0 | 1 | 4 | Cacna1a | 0.045 | Old |
| 0.053224 | 3.721923 | 0.045 | 0 | 1 | 4 | Fmc1 | 0.045 | Old |
| 0.05386 | 0.260505 | 0.89 | 0.838 | 1 | 4 | Folr2 | 0.052 | Old |
| 0.053939 | 1.35921 | 0.179 | 0.1 | 1 | 4 | Ubqln2 | 0.079 | Old |
| 0.054039 | 1.759104 | 0.119 | 0.05 | 1 | 4 | Eftud2 | 0.069 | Old |
| 0.054158 | 2.152165 | 0.085 | 0.025 | 1 | 4 | Sms | 0.06 | Old |
| 0.054196 | 0.693203 | 0.447 | 0.375 | 1 | 4 | Rp2 | 0.072 | Old |
| 0.054254 | 0.741964 | 0.273 | 0.188 | 1 | 4 | Arrdc4 | 0.085 | Old |
| 0.054288 | 0.526981 | 0.51 | 0.425 | 1 | 4 | Socs3 | 0.085 | Old |
| 0.054311 | 1.918786 | 0.086 | 0.025 | 1 | 4 | Arl4a | 0.061 | Old |
| 0.054453 | 2.187481 | 0.085 | 0.025 | 1 | 4 | Ear2 | 0.06 | Old |
| 0.054496 | 0.787731 | 0.355 | 0.275 | 1 | 4 | Ndufb7 | 0.08 | Old |
| 0.054595 | 2.739115 | 0.084 | 0.025 | 1 | 4 | Tmem71 | 0.059 | Old |
| 0.054695 | 0.900784 | 0.272 | 0.188 | 1 | 4 | Psma6 | 0.084 | Old |
| 0.054876 | 1.830013 | 0.102 | 0.038 | 1 | 4 | Calhm2 | 0.064 | Old |
| 0.055342 | 1.258412 | 0.138 | 0.062 | 1 | 4 | Lamtor5 | 0.076 | Old |

| | | | | | | | | |
|---|---|---|---|---|---|---|---|---|
| 0.055502 | 2.55792 | 0.099 | 0.038 | 1 | 4 | Mum1 | 0.061 | Old |
| 0.05563 | 1.309361 | 0.165 | 0.088 | 1 | 4 | B4galt5 | 0.077 | Old |
| 0.055648 | 0.591578 | 0.364 | 0.275 | 1 | 4 | Add3 | 0.089 | Old |
| 0.055969 | 0.759629 | 0.212 | 0.125 | 1 | 4 | Tmem167 | 0.087 | Old |
| 0.055978 | 1.205628 | 0.137 | 0.062 | 1 | 4 | Rpl7l1 | 0.075 | Old |
| 0.056046 | 1.826079 | 0.085 | 0.025 | 1 | 4 | Msantd4 | 0.06 | Old |
| 0.056369 | 0.626656 | 0.439 | 0.362 | 1 | 4 | Cdk2ap2 | 0.077 | Old |
| 0.057009 | 3.789928 | 0.043 | 0 | 1 | 4 | Iqsec2 | 0.043 | Old |
| 0.057213 | 1.770974 | 0.117 | 0.05 | 1 | 4 | Kif13b | 0.067 | Old |
| 0.057746 | 2.841813 | 0.065 | 0.013 | 1 | 4 | Gfm1 | 0.052 | Old |
| 0.057894 | 0.133327 | 1 | 1 | 1 | 4 | Rpl37a | 0 | Old |
| 0.057905 | 1.827596 | 0.084 | 0.025 | 1 | 4 | Pmvk | 0.059 | Old |
| 0.058385 | 0.746958 | 0.282 | 0.188 | 1 | 4 | Nus1 | 0.094 | Old |
| 0.058759 | 2.015438 | 0.084 | 0.025 | 1 | 4 | Mta3 | 0.059 | Old |
| 0.059304 | 1.326458 | 0.084 | 0.025 | 1 | 4 | Gucd1 | 0.059 | Old |
| 0.059484 | 2.243132 | 0.083 | 0.025 | 1 | 4 | Rae1 | 0.058 | Old |
| 0.059592 | 1.399794 | 0.119 | 0.05 | 1 | 4 | Pi4ka | 0.069 | Old |
| 0.05963 | 0.4329 | 0.673 | 0.65 | 1 | 4 | Rhob | 0.023 | Old |
| 0.059896 | 2.135238 | 0.065 | 0.013 | 1 | 4 | Prep | 0.052 | Old |
| 0.059977 | 2.133405 | 0.065 | 0.013 | 1 | 4 | Mkl2 | 0.052 | Old |
| 0.060978 | 1.340098 | 0.175 | 0.1 | 1 | 4 | Ube2d1 | 0.075 | Old |
| 0.063212 | 3.690754 | 0.042 | 0 | 1 | 4 | St3gal3 | 0.042 | Old |
| 0.063261 | 0.878302 | 0.309 | 0.225 | 1 | 4 | Tppp3 | 0.084 | Old |
| 0.063499 | 1.213221 | 0.149 | 0.075 | 1 | 4 | Otulin | 0.074 | Old |
| 0.063527 | 3.241138 | 0.063 | 0.013 | 1 | 4 | Ttc37 | 0.05 | Old |
| 0.063999 | 1.383725 | 0.101 | 0.038 | 1 | 4 | Dtx2 | 0.063 | Old |
| 0.064828 | 0.730883 | 0.324 | 0.25 | 1 | 4 | Rps6ka3 | 0.074 | Old |
| 0.064866 | 8.74E-05 | 0.007 | 0.025 | 1 | 4 | Gm10602 | -0.018 | Old |
| 0.065431 | 3.663354 | 0.041 | 0 | 1 | 4 | Ctu2 | 0.041 | Old |
| 0.065431 | 3.638667 | 0.041 | 0 | 1 | 4 | Zfp715 | 0.041 | Old |
| 0.065437 | 0.175951 | 0.956 | 0.975 | 1 | 4 | Rps28 | -0.019 | Old |
| 0.065622 | 2.379812 | 0.063 | 0.013 | 1 | 4 | Nop56 | 0.05 | Old |
| 0.065721 | 0.00988 | 0.026 | 0.062 | 1 | 4 | Natd1 | -0.036 | Old |
| 0.065793 | 2.996476 | 0.08 | 0.025 | 1 | 4 | Serpinb1a | 0.055 | Old |
| 0.065848 | 3.119915 | 0.062 | 0.013 | 1 | 4 | Itpa | 0.049 | Old |
| 0.067303 | 0.107987 | 0.002 | 0.013 | 1 | 4 | Podn | -0.011 | Old |
| 0.067303 | 0.065945 | 0.002 | 0.013 | 1 | 4 | 2410080I0: | -0.011 | Old |
| 0.06773 | 3.61393 | 0.04 | 0 | 1 | 4 | Spsb2 | 0.04 | Old |
| 0.067893 | 1.145301 | 0.204 | 0.125 | 1 | 4 | Rbm6 | 0.079 | Old |
| 0.068457 | 1.122528 | 0.227 | 0.15 | 1 | 4 | St8sia6 | 0.077 | Old |
| 0.068751 | 0.246214 | 0.95 | 0.938 | 1 | 4 | Rpsa | 0.012 | Old |
| 0.069758 | 1.773391 | 0.141 | 0.075 | 1 | 4 | Adgrg1 | 0.066 | Old |
| 0.069953 | 1.685429 | 0.097 | 0.038 | 1 | 4 | Pam | 0.059 | Old |
| 0.070112 | 4.534224 | 0.04 | 0 | 1 | 4 | Fzd8 | 0.04 | Old |
| 0.070112 | 3.927998 | 0.04 | 0 | 1 | 4 | Poc5 | 0.04 | Old |

| | | | | | | | | |
|---|---|---|---|---|---|---|---|---|
| 0.070112 | 3.674801 | 0.04 | 0 | 1 | 4 | Alad | 0.04 | Old |
| 0.070177 | 0.223162 | 0.969 | 0.963 | 1 | 4 | Clec10a | 0.006 | Old |
| 0.070563 | 0.294362 | 0.804 | 0.825 | 1 | 4 | Wnk1 | -0.021 | Old |
| 0.070765 | 0.962972 | 0.243 | 0.162 | 1 | 4 | Appl1 | 0.081 | Old |
| 0.070876 | 1.036494 | 0.224 | 0.15 | 1 | 4 | Tial1 | 0.074 | Old |
| 0.071038 | 0.596362 | 0.117 | 0.05 | 1 | 4 | Cyp51 | 0.067 | Old |
| 0.071059 | 0.274639 | 0.549 | 0.475 | 1 | 4 | Arap1 | 0.074 | Old |
| 0.071516 | 2.709344 | 0.061 | 0.013 | 1 | 4 | Adam8 | 0.048 | Old |
| 0.071559 | 1.034779 | 0.208 | 0.125 | 1 | 4 | Tnfsf9 | 0.083 | Old |
| 0.071851 | 0.840167 | 0.275 | 0.188 | 1 | 4 | Kat2b | 0.087 | Old |
| 0.072016 | 0.954085 | 0.206 | 0.125 | 1 | 4 | Dram1 | 0.081 | Old |
| 0.072582 | 3.578873 | 0.039 | 0 | 1 | 4 | Recql | 0.039 | Old |
| 0.072738 | 1.343008 | 0.129 | 0.062 | 1 | 4 | Lat2 | 0.067 | Old |
| 0.07339 | 0.285868 | 0.629 | 0.512 | 1 | 4 | Eif3a | 0.117 | Old |
| 0.073704 | 1.253138 | 0.115 | 0.05 | 1 | 4 | Hdac3 | 0.065 | Old |
| 0.07398 | 2.266552 | 0.095 | 0.038 | 1 | 4 | Cfap36 | 0.057 | Old |
| 0.074174 | 1.304835 | 0.159 | 0.088 | 1 | 4 | Gcnt1 | 0.071 | Old |
| 0.074176 | 0.54023 | 0.372 | 0.288 | 1 | 4 | Trim30a | 0.084 | Old |
| 0.074514 | 1.387706 | 0.128 | 0.062 | 1 | 4 | Stard4 | 0.066 | Old |
| 0.074744 | 0.528404 | 0.439 | 0.362 | 1 | 4 | Prr13 | 0.077 | Old |
| 0.074836 | 2.405715 | 0.078 | 0.025 | 1 | 4 | Mrpl19 | 0.053 | Old |
| 0.075141 | 3.776653 | 0.038 | 0 | 1 | 4 | Cpeb1 | 0.038 | Old |
| 0.075141 | 3.759935 | 0.038 | 0 | 1 | 4 | Rev3l | 0.038 | Old |
| 0.075141 | 3.656025 | 0.038 | 0 | 1 | 4 | Dip2a | 0.038 | Old |
| 0.075141 | 3.523934 | 0.038 | 0 | 1 | 4 | Tdrkh | 0.038 | Old |
| 0.075707 | 1.158393 | 0.115 | 0.05 | 1 | 4 | Mrpl2 | 0.065 | Old |
| 0.076005 | 0.369552 | 0.457 | 0.362 | 1 | 4 | Ddx3y | 0.095 | Old |
| 0.076118 | 1.543537 | 0.126 | 0.062 | 1 | 4 | Arsk | 0.064 | Old |
| 0.076398 | 0.678624 | 0.287 | 0.2 | 1 | 4 | Raly | 0.087 | Old |
| 0.076858 | 1.263444 | 0.08 | 0.025 | 1 | 4 | Hilpda | 0.055 | Old |
| 0.077183 | 1.464554 | 0.061 | 0.013 | 1 | 4 | Ivd | 0.048 | Old |
| 0.07732 | 3.201582 | 0.059 | 0.013 | 1 | 4 | Slc39a6 | 0.046 | Old |
| 0.077445 | 1.236632 | 0.196 | 0.125 | 1 | 4 | Insr | 0.071 | Old |
| 0.077795 | 3.555311 | 0.038 | 0 | 1 | 4 | Ptgfrn | 0.038 | Old |
| 0.077795 | 3.495682 | 0.038 | 0 | 1 | 4 | Rtn4rl1 | 0.038 | Old |
| 0.077795 | 3.464231 | 0.038 | 0 | 1 | 4 | Znhit6 | 0.038 | Old |
| 0.077804 | 1.171428 | 0.257 | 0.188 | 1 | 4 | S100a4 | 0.069 | Old |
| 0.078111 | 0.960564 | 0.189 | 0.112 | 1 | 4 | Tug1 | 0.077 | Old |
| 0.078359 | 0.801745 | 0.294 | 0.212 | 1 | 4 | Arhgap9 | 0.082 | Old |
| 0.078478 | 1.442775 | 0.096 | 0.038 | 1 | 4 | Zbtb46 | 0.058 | Old |
| 0.078725 | 0.598298 | 0.397 | 0.325 | 1 | 4 | Cnot6l | 0.072 | Old |
| 0.078813 | 1.382158 | 0.096 | 0.038 | 1 | 4 | 4930455G( | 0.058 | Old |
| 0.079151 | 0.426633 | 0.615 | 0.55 | 1 | 4 | Frmd4b | 0.065 | Old |
| 0.079542 | 1.175272 | 0.113 | 0.05 | 1 | 4 | Parp9 | 0.063 | Old |
| 0.079824 | 0.427788 | 0.482 | 0.4 | 1 | 4 | Srp9 | 0.082 | Old |

| | | | | | | | | |
|---|---|---|---|---|---|---|---|---|
| 0.080256 | 2.308081 | 0.093 | 0.038 | 1 | 4 | S1pr2 | 0.055 | Old |
| 0.080294 | 0.509163 | 0.493 | 0.45 | 1 | 4 | Ostf1 | 0.043 | Old |
| 0.080546 | 3.760906 | 0.037 | 0 | 1 | 4 | Dpf3 | 0.037 | Old |
| 0.080546 | 3.606117 | 0.037 | 0 | 1 | 4 | Abcb10 | 0.037 | Old |
| 0.080546 | 3.502125 | 0.037 | 0 | 1 | 4 | Nat10 | 0.037 | Old |
| 0.080568 | 2.077592 | 0.077 | 0.025 | 1 | 4 | Ccdc122 | 0.052 | Old |
| 0.081016 | 2.852101 | 0.058 | 0.013 | 1 | 4 | Amz2 | 0.045 | Old |
| 0.081383 | 0.366642 | 0.217 | 0.138 | 1 | 4 | Atp6v1h | 0.079 | Old |
| 0.081795 | 1.098986 | 0.096 | 0.038 | 1 | 4 | Tiam1 | 0.058 | Old |
| 0.081882 | 0.070128 | 0.096 | 0.162 | 1 | 4 | Ergic1 | -0.066 | Old |
| 0.082322 | 0.781724 | 0.24 | 0.162 | 1 | 4 | Tmem165 | 0.078 | Old |
| 0.082358 | 1.265469 | 0.079 | 0.025 | 1 | 4 | Dusp5 | 0.054 | Old |
| 0.082365 | 0.747439 | 0.245 | 0.162 | 1 | 4 | Eno1 | 0.083 | Old |
| 0.082712 | 2.599343 | 0.092 | 0.038 | 1 | 4 | Pot1a | 0.054 | Old |
| 0.082831 | 1.820198 | 0.094 | 0.038 | 1 | 4 | Alcam | 0.056 | Old |
| 0.083398 | 3.523997 | 0.036 | 0 | 1 | 4 | Dhx34 | 0.036 | Old |
| 0.083398 | 3.435797 | 0.036 | 0 | 1 | 4 | Rint1 | 0.036 | Old |
| 0.083902 | 0.42734 | 0.494 | 0.412 | 1 | 4 | Fkbp1a | 0.082 | Old |
| 0.084037 | 0.797004 | 0.216 | 0.138 | 1 | 4 | Ube2f | 0.078 | Old |
| 0.084041 | 0.181095 | 0.838 | 0.825 | 1 | 4 | Fcgrt | 0.013 | Old |
| 0.084122 | 0.872505 | 0.242 | 0.162 | 1 | 4 | Rap2b | 0.08 | Old |
| 0.084638 | 0.898611 | 0.262 | 0.188 | 1 | 4 | Dapk1 | 0.074 | Old |
| 0.085319 | 2.392201 | 0.075 | 0.025 | 1 | 4 | Xaf1 | 0.05 | Old |
| 0.08535 | 0.420458 | 0.623 | 0.575 | 1 | 4 | Ehd1 | 0.048 | Old |
| 0.085485 | 0.713326 | 0.282 | 0.2 | 1 | 4 | Abcc1 | 0.082 | Old |
| 0.085902 | 0.265029 | 0.811 | 0.825 | 1 | 4 | Aplp2 | -0.014 | Old |
| 0.086028 | 2.225108 | 0.057 | 0.013 | 1 | 4 | Carnmt1 | 0.044 | Old |
| 0.086356 | 3.714218 | 0.036 | 0 | 1 | 4 | Dpyd | 0.036 | Old |
| 0.086356 | 3.476657 | 0.036 | 0 | 1 | 4 | 1700021F0 | 0.036 | Old |
| 0.086356 | 3.403061 | 0.036 | 0 | 1 | 4 | Mrps18b | 0.036 | Old |
| 0.086356 | 3.317892 | 0.036 | 0 | 1 | 4 | Polr1a | 0.036 | Old |
| 0.086356 | 3.317627 | 0.036 | 0 | 1 | 4 | Rcc1l | 0.036 | Old |
| 0.086356 | 3.292687 | 0.036 | 0 | 1 | 4 | Lrwd1 | 0.036 | Old |
| 0.08638 | 0.847782 | 0.059 | 0.013 | 1 | 4 | Tmem209 | 0.046 | Old |
| 0.087891 | 0.247675 | 0.677 | 0.6 | 1 | 4 | Zcchc6 | 0.077 | Old |
| 0.088332 | 1.049395 | 0.11 | 0.05 | 1 | 4 | Ell2 | 0.06 | Old |
| 0.088783 | 0.367553 | 0.549 | 0.488 | 1 | 4 | Naaa | 0.061 | Old |
| 0.088974 | 0.655014 | 0.283 | 0.2 | 1 | 4 | Card19 | 0.083 | Old |
| 0.089308 | 0.21595 | 0.829 | 0.812 | 1 | 4 | Dazap2 | 0.017 | Old |
| 0.089881 | 1.148823 | 0.094 | 0.038 | 1 | 4 | Stat5a | 0.056 | Old |
| 0.090005 | 2.011735 | 0.092 | 0.038 | 1 | 4 | Cytip | 0.054 | Old |
| 0.090554 | 1.060298 | 0.126 | 0.062 | 1 | 4 | Zfp62 | 0.064 | Old |
| 0.09088 | 0.562227 | 0.329 | 0.25 | 1 | 4 | Supt16 | 0.079 | Old |
| 0.091369 | 0.964227 | 0.171 | 0.1 | 1 | 4 | Ptbp1 | 0.071 | Old |
| 0.091655 | 1.656593 | 0.122 | 0.062 | 1 | 4 | Crybg1 | 0.06 | Old |

| | | | | | | | | |
|---|---|---|---|---|---|---|---|---|
| 0.091859 | 0.342445 | 0.713 | 0.675 | 1 | 4 | Ifi207 | 0.038 | Old |
| 0.092695 | 1.229729 | 0.167 | 0.1 | 1 | 4 | Abcf1 | 0.067 | Old |
| 0.093084 | 1.753599 | 0.091 | 0.038 | 1 | 4 | Stk25 | 0.053 | Old |
| 0.093092 | 1.496943 | 0.137 | 0.075 | 1 | 4 | Stk11 | 0.062 | Old |
| 0.093223 | 1.19346 | 0.093 | 0.038 | 1 | 4 | Pgpep1 | 0.055 | Old |
| 0.093289 | 2.675894 | 0.055 | 0.013 | 1 | 4 | Fiz1 | 0.042 | Old |
| 0.0937 | 2.343691 | 0.073 | 0.025 | 1 | 4 | Slc25a25 | 0.048 | Old |
| 0.093747 | 1.242044 | 0.138 | 0.075 | 1 | 4 | Rnps1 | 0.063 | Old |
| 0.094186 | 0.445805 | 0.161 | 0.088 | 1 | 4 | Vamp4 | 0.073 | Old |
| 0.094371 | 0.508769 | 0.131 | 0.062 | 1 | 4 | Chfr | 0.069 | Old |
| 0.094716 | 0.903564 | 0.172 | 0.1 | 1 | 4 | Ireb2 | 0.072 | Old |
| 0.094986 | 1.508105 | 0.107 | 0.05 | 1 | 4 | Tmem55a | 0.057 | Old |
| 0.095909 | 3.511563 | 0.034 | 0 | 1 | 4 | Hectd4 | 0.034 | Old |
| 0.095909 | 3.418492 | 0.034 | 0 | 1 | 4 | Etaa1 | 0.034 | Old |
| 0.095909 | 3.417215 | 0.034 | 0 | 1 | 4 | Ccdc61 | 0.034 | Old |
| 0.095909 | 3.208633 | 0.034 | 0 | 1 | 4 | Mrpl1 | 0.034 | Old |
| 0.096223 | 0.129936 | 1 | 1 | 1 | 4 | C1qa | 0 | Old |
| 0.09688 | 1.010829 | 0.14 | 0.075 | 1 | 4 | Dctn3 | 0.065 | Old |
| 0.097221 | 1.866305 | 0.055 | 0.013 | 1 | 4 | Fbxw8 | 0.042 | Old |
| 0.097223 | 2.532726 | 0.055 | 0.013 | 1 | 4 | Slc2a8 | 0.042 | Old |
| 0.09725 | 0.400717 | 0.466 | 0.388 | 1 | 4 | Myadm | 0.078 | Old |
| 0.09725 | 0.582564 | 0.221 | 0.15 | 1 | 4 | Fn1 | 0.071 | Old |
| 0.09731 | 0.56197 | 0.414 | 0.338 | 1 | 4 | Dip2b | 0.076 | Old |
| 0.09735 | 1.014999 | 0.125 | 0.062 | 1 | 4 | Avpi1 | 0.063 | Old |
| 0.097436 | 0.305862 | 0.709 | 0.675 | 1 | 4 | Chchd2 | 0.034 | Old |
| 0.097871 | 0.402563 | 0.484 | 0.388 | 1 | 4 | Nipa2 | 0.096 | Old |
| 0.097885 | 0.502229 | 0.371 | 0.275 | 1 | 4 | Coro1c | 0.096 | Old |
| 0.097937 | 1.972602 | 0.073 | 0.025 | 1 | 4 | Wipi2 | 0.048 | Old |
| 0.098657 | 2.227852 | 0.055 | 0.013 | 1 | 4 | Slc9a3r2 | 0.042 | Old |
| 0.098786 | 3.042466 | 0.054 | 0.013 | 1 | 4 | 2310022B0 | 0.041 | Old |
| 0.099058 | 0.228356 | 0.77 | 0.762 | 1 | 4 | Srsf5 | 0.008 | Old |
| 0.099335 | 3.594841 | 0.033 | 0 | 1 | 4 | Tchp | 0.033 | Old |
| 0.099335 | 3.254355 | 0.033 | 0 | 1 | 4 | Adsl | 0.033 | Old |
| 0.099335 | 3.140621 | 0.033 | 0 | 1 | 4 | Phf6 | 0.033 | Old |
| 0.099845 | 1.95495 | 0.089 | 0.038 | 1 | 4 | Dcaf5 | 0.051 | Old |
| 0.100053 | 0.852829 | 0.169 | 0.1 | 1 | 4 | Ssbp3 | 0.069 | Old |
| 0.100158 | 0.55381 | 0.27 | 0.188 | 1 | 4 | Aph1a | 0.082 | Old |
| 0.100533 | 0.586702 | 0.375 | 0.3 | 1 | 4 | Macf1 | 0.075 | Old |
| 0.100575 | 1.654149 | 0.073 | 0.025 | 1 | 4 | Tmem167b | 0.048 | Old |
| 0.100652 | 0.25556 | 0.533 | 0.45 | 1 | 4 | Bach1 | 0.083 | Old |
| 0.100756 | 0.130857 | 0.999 | 1 | 1 | 4 | Malat1 | -0.001 | Old |
| 0.100908 | 0.867266 | 0.294 | 0.225 | 1 | 4 | Gas2l1 | 0.069 | Old |
| 0.10185 | 0.622633 | 0.364 | 0.3 | 1 | 4 | Aes | 0.064 | Old |
| 0.102603 | 0.191072 | 0.873 | 0.787 | 1 | 4 | Mmp9 | 0.086 | Old |
| 0.102607 | 1.346088 | 0.136 | 0.075 | 1 | 4 | Vps36 | 0.061 | Old |

| | | | | | | | | |
|---|---|---|---|---|---|---|---|---|
| 0.102702 | 0.629191 | 0.261 | 0.188 | 1 | 4 | Psma4 | 0.073 | Old |
| 0.102891 | 3.58693 | 0.032 | 0 | 1 | 4 | Dis3l | 0.032 | Old |
| 0.102891 | 3.353221 | 0.032 | 0 | 1 | 4 | Grpel2 | 0.032 | Old |
| 0.102894 | 0.941849 | 0.092 | 0.038 | 1 | 4 | Xxylt1 | 0.054 | Old |
| 0.102996 | 0.001484 | 0.09 | 0.15 | 1 | 4 | Gtf2b | -0.06 | Old |
| 0.103742 | 0.821276 | 0.187 | 0.112 | 1 | 4 | Pfkfb3 | 0.075 | Old |
| 0.103747 | 1.683219 | 0.072 | 0.025 | 1 | 4 | Amn1 | 0.047 | Old |
| 0.103825 | 0.48314 | 0.393 | 0.3 | 1 | 4 | A4galt | 0.093 | Old |
| 0.104198 | 1.110927 | 0.123 | 0.062 | 1 | 4 | Ash2l | 0.061 | Old |
| 0.104476 | 0.437108 | 0.289 | 0.2 | 1 | 4 | Cmtm3 | 0.089 | Old |
| 0.104554 | 1.395438 | 0.161 | 0.1 | 1 | 4 | Cript | 0.061 | Old |
| 0.104676 | 1.254479 | 0.121 | 0.062 | 1 | 4 | Phf2 | 0.059 | Old |
| 0.104896 | 3.411977 | 0.053 | 0.013 | 1 | 4 | Zc3h7b | 0.04 | Old |
| 0.105317 | 1.420277 | 0.09 | 0.038 | 1 | 4 | Pcnt | 0.052 | Old |
| 0.105451 | 1.374862 | 0.135 | 0.075 | 1 | 4 | Tmem65 | 0.06 | Old |
| 0.105616 | 0.85345 | 0.165 | 0.1 | 1 | 4 | Traf3 | 0.065 | Old |
| 0.106331 | 0.487457 | 0.202 | 0.125 | 1 | 4 | Aldh9a1 | 0.077 | Old |
| 0.106582 | 3.399136 | 0.032 | 0 | 1 | 4 | Itga2 | 0.032 | Old |
| 0.106582 | 3.348368 | 0.032 | 0 | 1 | 4 | Actr6 | 0.032 | Old |
| 0.10668 | 0.796969 | 0.197 | 0.125 | 1 | 4 | Btk | 0.072 | Old |
| 0.107151 | 0.750245 | 0.091 | 0.038 | 1 | 4 | Fam208b | 0.053 | Old |
| 0.107233 | 1.276036 | 0.105 | 0.05 | 1 | 4 | Rhoh | 0.055 | Old |
| 0.107618 | 0.338696 | 0.222 | 0.138 | 1 | 4 | Ctdsp1 | 0.084 | Old |
| 0.10862 | 0.393826 | 0.557 | 0.488 | 1 | 4 | Tm9sf3 | 0.069 | Old |
| 0.109587 | 1.227318 | 0.162 | 0.1 | 1 | 4 | Dag1 | 0.062 | Old |
| 0.110217 | 1.357887 | 0.134 | 0.075 | 1 | 4 | Gm15675 | 0.059 | Old |
| 0.110926 | 0.808705 | 0.256 | 0.188 | 1 | 4 | Hexim1 | 0.068 | Old |
| 0.111138 | 0.279437 | 0.356 | 0.288 | 1 | 4 | Agtrap | 0.068 | Old |
| 0.111267 | 0.761238 | 0.167 | 0.1 | 1 | 4 | Mark2 | 0.067 | Old |
| 0.111307 | 1.907443 | 0.087 | 0.038 | 1 | 4 | Snrnp40 | 0.049 | Old |
| 0.111451 | 1.265442 | 0.134 | 0.075 | 1 | 4 | Mrpl17 | 0.059 | Old |
| 0.111531 | 2.529262 | 0.052 | 0.013 | 1 | 4 | Srxn1 | 0.039 | Old |
| 0.112192 | 0.686082 | 0.254 | 0.188 | 1 | 4 | Baz1b | 0.066 | Old |
| 0.112443 | 0.147342 | 0.014 | 0.038 | 1 | 4 | Olfr1033 | -0.024 | Old |
| 0.112841 | 0.004424 | 0.008 | 0.025 | 1 | 4 | Nfib | -0.017 | Old |
| 0.113296 | 0.606186 | 0.141 | 0.075 | 1 | 4 | Clk3 | 0.066 | Old |
| 0.113324 | 0.940626 | 0.136 | 0.075 | 1 | 4 | Rnf2 | 0.061 | Old |
| 0.114392 | 3.490929 | 0.03 | 0 | 1 | 4 | Pcnx4 | 0.03 | Old |
| 0.114392 | 3.397242 | 0.03 | 0 | 1 | 4 | Mamld1 | 0.03 | Old |
| 0.114392 | 3.312346 | 0.03 | 0 | 1 | 4 | Ap4b1 | 0.03 | Old |
| 0.114666 | 1.929916 | 0.07 | 0.025 | 1 | 4 | Arl2bp | 0.045 | Old |
| 0.114954 | 0.684572 | 0.297 | 0.238 | 1 | 4 | Ago2 | 0.059 | Old |
| 0.115482 | 0.598941 | 0.236 | 0.162 | 1 | 4 | Cd209b | 0.074 | Old |
| 0.115538 | 0.837319 | 0.181 | 0.112 | 1 | 4 | Mrps14 | 0.069 | Old |
| 0.115935 | 0.306239 | 0.663 | 0.637 | 1 | 4 | Rac1 | 0.026 | Old |

| | | | | | | | | |
|---|---|---|---|---|---|---|---|---|
| 0.11595 | 0.581082 | 0.4 | 0.35 | 1 | 4 | Hnrnpc | 0.05 | Old |
| 0.117371 | 0.12333 | 0.106 | 0.05 | 1 | 4 | Hcfc2 | 0.056 | Old |
| 0.117461 | 0.364204 | 0.598 | 0.55 | 1 | 4 | Adam15 | 0.048 | Old |
| 0.117954 | 1.316745 | 0.07 | 0.025 | 1 | 4 | Ube2l6 | 0.045 | Old |
| 0.118206 | 0.602461 | 0.404 | 0.35 | 1 | 4 | Slc39a1 | 0.054 | Old |
| 0.118524 | 3.285529 | 0.03 | 0 | 1 | 4 | Ulbp1 | 0.03 | Old |
| 0.118524 | 3.162884 | 0.03 | 0 | 1 | 4 | Ttc27 | 0.03 | Old |
| 0.118524 | 3.124885 | 0.03 | 0 | 1 | 4 | Wdr83 | 0.03 | Old |
| 0.118761 | 2.714367 | 0.051 | 0.013 | 1 | 4 | Tenm4 | 0.038 | Old |
| 0.11884 | 0.733997 | 0.244 | 0.175 | 1 | 4 | Srebf2 | 0.069 | Old |
| 0.119025 | 1.86935 | 0.069 | 0.025 | 1 | 4 | Hagh | 0.044 | Old |
| 0.119211 | 1.855672 | 0.086 | 0.038 | 1 | 4 | Cdk8 | 0.048 | Old |
| 0.119906 | 0.696475 | 0.052 | 0.013 | 1 | 4 | Esr1 | 0.039 | Old |
| 0.120131 | 1.536331 | 0.051 | 0.013 | 1 | 4 | Megf8 | 0.038 | Old |
| 0.120156 | 1.126733 | 0.136 | 0.075 | 1 | 4 | Rgs1 | 0.061 | Old |
| 0.120364 | 0.814361 | 0.089 | 0.038 | 1 | 4 | Ino80e | 0.051 | Old |
| 0.120667 | 0.726932 | 0.071 | 0.025 | 1 | 4 | Cluh | 0.046 | Old |
| 0.120876 | 0.497747 | 0.211 | 0.138 | 1 | 4 | Fyttd1 | 0.073 | Old |
| 0.121059 | 0.530997 | 0.505 | 0.462 | 1 | 4 | Fgfr1 | 0.043 | Old |
| 0.121862 | 1.042253 | 0.104 | 0.05 | 1 | 4 | Tor3a | 0.054 | Old |
| 0.122494 | 0.238433 | 0.848 | 0.838 | 1 | 4 | Sem1 | 0.01 | Old |
| 0.122534 | 1.333618 | 0.102 | 0.05 | 1 | 4 | Commd2 | 0.052 | Old |
| 0.122816 | 3.123174 | 0.029 | 0 | 1 | 4 | Ccdc28a | 0.029 | Old |
| 0.122816 | 3.263591 | 0.029 | 0 | 1 | 4 | Alkbh7 | 0.029 | Old |
| 0.122816 | 3.042109 | 0.029 | 0 | 1 | 4 | Rem1 | 0.029 | Old |
| 0.122816 | 2.884938 | 0.029 | 0 | 1 | 4 | Lilra6 | 0.029 | Old |
| 0.122826 | 1.744711 | 0.051 | 0.013 | 1 | 4 | Zbp1 | 0.038 | Old |
| 0.124206 | 0.417214 | 0.183 | 0.112 | 1 | 4 | Rnd3 | 0.071 | Old |
| 0.124305 | 0.236915 | 0.708 | 0.675 | 1 | 4 | Ptbp3 | 0.033 | Old |
| 0.124405 | 0.351063 | 0.57 | 0.525 | 1 | 4 | Rab8b | 0.045 | Old |
| 0.124704 | 1.237585 | 0.131 | 0.075 | 1 | 4 | Kif2a | 0.056 | Old |
| 0.124894 | 1.051143 | 0.119 | 0.062 | 1 | 4 | Sestd1 | 0.057 | Old |
| 0.125141 | 3.107428 | 0.049 | 0.013 | 1 | 4 | Tom1l2 | 0.036 | Old |
| 0.12516 | 0.742445 | 0.316 | 0.25 | 1 | 4 | Stag2 | 0.066 | Old |
| 0.125166 | 1.192968 | 0.146 | 0.088 | 1 | 4 | Vps28 | 0.058 | Old |
| 0.125296 | 0.117088 | 0.003 | 0.013 | 1 | 4 | Cyp1b1 | -0.01 | Old |
| 0.125296 | 0.021391 | 0.003 | 0.013 | 1 | 4 | Dnah12 | -0.01 | Old |
| 0.125334 | 1.367934 | 0.101 | 0.05 | 1 | 4 | Gtf3c6 | 0.051 | Old |
| 0.125703 | 0.872929 | 0.134 | 0.075 | 1 | 4 | Zscan26 | 0.059 | Old |
| 0.125915 | 0.581999 | 0.408 | 0.362 | 1 | 4 | Reps2 | 0.046 | Old |
| 0.12621 | 1.79006 | 0.084 | 0.038 | 1 | 4 | AI837181 | 0.046 | Old |
| 0.126248 | 1.773511 | 0.05 | 0.013 | 1 | 4 | Fam241a | 0.037 | Old |
| 0.126398 | 0.914327 | 0.119 | 0.062 | 1 | 4 | Tcof1 | 0.057 | Old |
| 0.126664 | 0.382754 | 0.617 | 0.575 | 1 | 4 | Tns1 | 0.042 | Old |
| 0.127276 | 3.389697 | 0.028 | 0 | 1 | 4 | Bmpr1a | 0.028 | Old |

| | | | | | | | | |
|---|---|---|---|---|---|---|---|---|
| 0.127276 | 3.329776 | 0.028 | 0 | 1 | 4 | A130010J1 | 0.028 | Old |
| 0.127276 | 3.301107 | 0.028 | 0 | 1 | 4 | Hoxb4 | 0.028 | Old |
| 0.127276 | 3.288333 | 0.028 | 0 | 1 | 4 | Zfp821 | 0.028 | Old |
| 0.127276 | 3.178379 | 0.028 | 0 | 1 | 4 | Atrip | 0.028 | Old |
| 0.127276 | 3.177079 | 0.028 | 0 | 1 | 4 | E130311K1 | 0.028 | Old |
| 0.127508 | 1.021169 | 0.133 | 0.075 | 1 | 4 | Aurkaip1 | 0.058 | Old |
| 0.128008 | 0.377703 | 0.603 | 0.562 | 1 | 4 | Rap1a | 0.041 | Old |
| 0.128572 | 0.66217 | 0.378 | 0.325 | 1 | 4 | Kdelr1 | 0.053 | Old |
| 0.128703 | 0.228129 | 0.881 | 0.838 | 1 | 4 | Myl6 | 0.043 | Old |
| 0.128892 | 0.627921 | 0.396 | 0.35 | 1 | 4 | Hif1a | 0.046 | Old |
| 0.129026 | 1.623145 | 0.068 | 0.025 | 1 | 4 | Vps39 | 0.043 | Old |
| 0.129145 | 0.457706 | 0.298 | 0.225 | 1 | 4 | Smim10l1 | 0.073 | Old |
| 0.129234 | 3.153012 | 0.049 | 0.013 | 1 | 4 | 1500011B0 | 0.036 | Old |
| 0.129465 | 1.695153 | 0.068 | 0.025 | 1 | 4 | Ikzf2 | 0.043 | Old |
| 0.129698 | 1.337688 | 0.069 | 0.025 | 1 | 4 | Trim11 | 0.044 | Old |
| 0.130019 | 0.724991 | 0.198 | 0.138 | 1 | 4 | Csnk2b | 0.06 | Old |
| 0.130562 | 0.365558 | 0.612 | 0.562 | 1 | 4 | Dbi | 0.05 | Old |
| 0.131911 | 3.350567 | 0.028 | 0 | 1 | 4 | Rcl1 | 0.028 | Old |
| 0.131911 | 3.094959 | 0.028 | 0 | 1 | 4 | Spn | 0.028 | Old |
| 0.131963 | 1.169569 | 0.157 | 0.1 | 1 | 4 | Mif4gd | 0.057 | Old |
| 0.132724 | 1.400864 | 0.115 | 0.062 | 1 | 4 | Tbc1d15 | 0.053 | Old |
| 0.133122 | 0.470429 | 0.299 | 0.225 | 1 | 4 | Luc7l3 | 0.074 | Old |
| 0.133218 | 1.085687 | 0.086 | 0.038 | 1 | 4 | Pygo2 | 0.048 | Old |
| 0.134033 | 0.421803 | 0.572 | 0.512 | 1 | 4 | Serbp1 | 0.06 | Old |
| 0.134627 | 1.837559 | 0.049 | 0.013 | 1 | 4 | Mcm9 | 0.036 | Old |
| 0.134841 | 0.249044 | 0.827 | 0.8 | 1 | 4 | Ap2a2 | 0.027 | Old |
| 0.134862 | 0.25723 | 0.831 | 0.85 | 1 | 4 | Ptafr | -0.019 | Old |
| 0.13501 | 2.318577 | 0.066 | 0.025 | 1 | 4 | Ddah2 | 0.041 | Old |
| 0.135063 | 2.760209 | 0.048 | 0.013 | 1 | 4 | Fam210a | 0.035 | Old |
| 0.135566 | 0.500484 | 0.588 | 0.575 | 1 | 4 | Eps8 | 0.013 | Old |
| 0.135653 | 0.385665 | 0.345 | 0.275 | 1 | 4 | Fam120a | 0.07 | Old |
| 0.136728 | 3.215198 | 0.027 | 0 | 1 | 4 | Tfpt | 0.027 | Old |
| 0.136728 | 3.361518 | 0.027 | 0 | 1 | 4 | Trim68 | 0.027 | Old |
| 0.136728 | 3.175331 | 0.027 | 0 | 1 | 4 | B3gnt8 | 0.027 | Old |
| 0.136728 | 3.145995 | 0.027 | 0 | 1 | 4 | Dnajb5 | 0.027 | Old |
| 0.136728 | 2.868755 | 0.027 | 0 | 1 | 4 | Slc16a3 | 0.027 | Old |
| 0.136899 | 0.549318 | 0.276 | 0.2 | 1 | 4 | Rhoq | 0.076 | Old |
| 0.137225 | 0.513762 | 0.353 | 0.288 | 1 | 4 | Ctnna1 | 0.065 | Old |
| 0.137621 | 1.243371 | 0.143 | 0.088 | 1 | 4 | Unc50 | 0.055 | Old |
| 0.137644 | 0.407134 | 0.576 | 0.55 | 1 | 4 | Tmem106a | 0.026 | Old |
| 0.137787 | 0.24137 | 0.057 | 0.1 | 1 | 4 | Rrn3 | -0.043 | Old |
| 0.13812 | 2.172084 | 0.048 | 0.013 | 1 | 4 | N6amt1 | 0.035 | Old |
| 0.138602 | 0.527597 | 0.219 | 0.15 | 1 | 4 | C5ar2 | 0.069 | Old |
| 0.138685 | 0.779377 | 0.204 | 0.138 | 1 | 4 | Usp16 | 0.066 | Old |
| 0.138995 | 0.428503 | 0.386 | 0.312 | 1 | 4 | Romo1 | 0.074 | Old |

| | | | | | | | | |
|---|---|---|---|---|---|---|---|---|
| 0.139266 | 1.474943 | 0.126 | 0.075 | 1 | 4 | Atad1 | 0.051 | Old |
| 0.139473 | 1.028731 | 0.157 | 0.1 | 1 | 4 | Phf10 | 0.057 | Old |
| 0.139901 | 0.234566 | 0.485 | 0.412 | 1 | 4 | Ap2m1 | 0.073 | Old |
| 0.140154 | 1.177715 | 0.155 | 0.1 | 1 | 4 | Snrnp200 | 0.055 | Old |
| 0.141738 | 3.236834 | 0.026 | 0 | 1 | 4 | Tmem39b | 0.026 | Old |
| 0.141738 | 3.14405 | 0.026 | 0 | 1 | 4 | 2810006K2 | 0.026 | Old |
| 0.141738 | 3.075831 | 0.026 | 0 | 1 | 4 | Glis2 | 0.026 | Old |
| 0.141738 | 3.066244 | 0.026 | 0 | 1 | 4 | Wdr55 | 0.026 | Old |
| 0.141738 | 2.915207 | 0.026 | 0 | 1 | 4 | Aacs | 0.026 | Old |
| 0.141882 | 1.604357 | 0.066 | 0.025 | 1 | 4 | Lsm14b | 0.041 | Old |
| 0.142845 | 0.468316 | 0.023 | 0.05 | 1 | 4 | Lamc1 | -0.027 | Old |
| 0.143205 | 1.984421 | 0.047 | 0.013 | 1 | 4 | Gimap6 | 0.034 | Old |
| 0.143434 | 1.168725 | 0.142 | 0.088 | 1 | 4 | Mesd | 0.054 | Old |
| 0.143438 | 0.764148 | 0.176 | 0.112 | 1 | 4 | Uqcc3 | 0.064 | Old |
| 0.144021 | 1.190995 | 0.048 | 0.013 | 1 | 4 | Parn | 0.035 | Old |
| 0.144253 | 2.971185 | 0.047 | 0.013 | 1 | 4 | Gm16104 | 0.034 | Old |
| 0.144538 | 1.819192 | 0.065 | 0.025 | 1 | 4 | Slc13a3 | 0.04 | Old |
| 0.144597 | 1.117359 | 0.099 | 0.05 | 1 | 4 | Mfap1a | 0.049 | Old |
| 0.14464 | 1.052715 | 0.115 | 0.062 | 1 | 4 | Ndrg3 | 0.053 | Old |
| 0.144966 | 0.915368 | 0.13 | 0.075 | 1 | 4 | Cetn3 | 0.055 | Old |
| 0.145093 | 1.752132 | 0.047 | 0.013 | 1 | 4 | Rad17 | 0.034 | Old |
| 0.145289 | 1.109851 | 0.128 | 0.075 | 1 | 4 | Anapc1 | 0.053 | Old |
| 0.145341 | 0.534399 | 0.363 | 0.3 | 1 | 4 | Eif4ebp2 | 0.063 | Old |
| 0.145972 | 0.361935 | 0.513 | 0.438 | 1 | 4 | Atp6v1b2 | 0.075 | Old |
| 0.146724 | 0.954527 | 0.13 | 0.075 | 1 | 4 | Tars | 0.055 | Old |
| 0.14674 | 2.315802 | 0.047 | 0.013 | 1 | 4 | Stx3 | 0.034 | Old |
| 0.146947 | 3.125513 | 0.026 | 0 | 1 | 4 | Trmt10c | 0.026 | Old |
| 0.146947 | 3.092551 | 0.026 | 0 | 1 | 4 | Mctp2 | 0.026 | Old |
| 0.146947 | 3.039945 | 0.026 | 0 | 1 | 4 | Rrp1b | 0.026 | Old |
| 0.146947 | 2.985568 | 0.026 | 0 | 1 | 4 | Ercc2 | 0.026 | Old |
| 0.147144 | 0.872182 | 0.2 | 0.138 | 1 | 4 | Map3k8 | 0.062 | Old |
| 0.14722 | 0.741698 | 0.115 | 0.062 | 1 | 4 | Pofut1 | 0.053 | Old |
| 0.14812 | 0.651534 | 0.143 | 0.088 | 1 | 4 | Chtf8 | 0.055 | Old |
| 0.148407 | 1.893704 | 0.065 | 0.025 | 1 | 4 | Rasa2 | 0.04 | Old |
| 0.148501 | 0.743203 | 0.285 | 0.225 | 1 | 4 | Sptbn1 | 0.06 | Old |
| 0.149174 | 0.723663 | 0.128 | 0.075 | 1 | 4 | Zmpste24 | 0.053 | Old |
| 0.14948 | 0.493975 | 0.321 | 0.25 | 1 | 4 | Cd37 | 0.071 | Old |
| 0.149565 | 1.112724 | 0.154 | 0.1 | 1 | 4 | Rps12-ps3 | 0.054 | Old |
| 0.14958 | 2.902364 | 0.046 | 0.013 | 1 | 4 | Fkrp | 0.033 | Old |
| 0.149645 | 0.424549 | 0.375 | 0.288 | 1 | 4 | Ncor1 | 0.087 | Old |
| 0.149916 | 1.639949 | 0.096 | 0.05 | 1 | 4 | Poc1b | 0.046 | Old |
| 0.150008 | 0.857496 | 0.199 | 0.138 | 1 | 4 | Cept1 | 0.061 | Old |
| 0.150764 | 2.510346 | 0.046 | 0.013 | 1 | 4 | Gnptab | 0.033 | Old |
| 0.150875 | 0.546139 | 0.256 | 0.188 | 1 | 4 | Sgms1 | 0.068 | Old |
| 0.151113 | 0.240899 | 0.708 | 0.688 | 1 | 4 | Pcbp2 | 0.02 | Old |

| | | | | | | | | |
|---|---|---|---|---|---|---|---|---|
| 0.151185 | 0.207395 | 0.896 | 0.85 | 1 | 4 | Glul | 0.046 | Old |
| 0.151221 | 1.730146 | 0.047 | 0.013 | 1 | 4 | Coro2a | 0.034 | Old |
| 0.151559 | 1.315383 | 0.065 | 0.025 | 1 | 4 | Ssna1 | 0.04 | Old |
| 0.151877 | 0.48913 | 0.404 | 0.35 | 1 | 4 | Ppp1r18 | 0.054 | Old |
| 0.151879 | 1.30797 | 0.082 | 0.038 | 1 | 4 | Sp3os | 0.044 | Old |
| 0.152182 | 0.391173 | 0.27 | 0.2 | 1 | 4 | Camkk2 | 0.07 | Old |
| 0.152229 | 0.727876 | 0.279 | 0.225 | 1 | 4 | Psmb10 | 0.054 | Old |
| 0.152367 | 3.288766 | 0.025 | 0 | 1 | 4 | Hspa12a | 0.025 | Old |
| 0.152367 | 3.258144 | 0.025 | 0 | 1 | 4 | Zbtb42 | 0.025 | Old |
| 0.152367 | 3.245292 | 0.025 | 0 | 1 | 4 | D930016D( | 0.025 | Old |
| 0.152367 | 3.175801 | 0.025 | 0 | 1 | 4 | Eps8l1 | 0.025 | Old |
| 0.152367 | 3.14806 | 0.025 | 0 | 1 | 4 | Nmnat3 | 0.025 | Old |
| 0.152367 | 3.03803 | 0.025 | 0 | 1 | 4 | Acpp | 0.025 | Old |
| 0.152367 | 3.037827 | 0.025 | 0 | 1 | 4 | Sumf2 | 0.025 | Old |
| 0.152367 | 3.024279 | 0.025 | 0 | 1 | 4 | Mtrr | 0.025 | Old |
| 0.152394 | 0.851511 | 0.144 | 0.088 | 1 | 4 | Sarnp | 0.056 | Old |
| 0.153569 | 0.15021 | 0.847 | 0.787 | 1 | 4 | Pfn1 | 0.06 | Old |
| 0.153628 | 0.650057 | 0.298 | 0.238 | 1 | 4 | Epb41l2 | 0.06 | Old |
| 0.15371 | 0.958218 | 0.22 | 0.162 | 1 | 4 | Chic2 | 0.058 | Old |
| 0.153799 | 1.270838 | 0.047 | 0.013 | 1 | 4 | Plpp6 | 0.034 | Old |
| 0.153801 | 1.431174 | 0.065 | 0.025 | 1 | 4 | Slc7a1 | 0.04 | Old |
| 0.153987 | 0.455964 | 0.352 | 0.288 | 1 | 4 | Adipor1 | 0.064 | Old |
| 0.154047 | 0.003488 | 0.105 | 0.162 | 1 | 4 | Slirp | -0.057 | Old |
| 0.154377 | 0.169517 | 0.81 | 0.738 | 1 | 4 | Cd209f | 0.072 | Old |
| 0.155071 | 0.578421 | 0.171 | 0.112 | 1 | 4 | Casc3 | 0.059 | Old |
| 0.155924 | 2.648306 | 0.045 | 0.013 | 1 | 4 | Rars2 | 0.032 | Old |
| 0.156173 | 0.555688 | 0.163 | 0.1 | 1 | 4 | Chd3 | 0.063 | Old |
| 0.156567 | 0.443896 | 0.433 | 0.362 | 1 | 4 | Clic4 | 0.071 | Old |
| 0.15698 | 0.641208 | 0.084 | 0.038 | 1 | 4 | Kctd12b | 0.046 | Old |
| 0.157069 | 1.321087 | 0.081 | 0.038 | 1 | 4 | Hdac4 | 0.043 | Old |
| 0.157758 | 1.510549 | 0.11 | 0.062 | 1 | 4 | Malt1 | 0.048 | Old |
| 0.158008 | 2.943071 | 0.024 | 0 | 1 | 4 | St5 | 0.024 | Old |
| 0.158008 | 3.172278 | 0.024 | 0 | 1 | 4 | Zfp574 | 0.024 | Old |
| 0.158008 | 3.008892 | 0.024 | 0 | 1 | 4 | Lzic | 0.024 | Old |
| 0.158008 | 2.918419 | 0.024 | 0 | 1 | 4 | Flad1 | 0.024 | Old |
| 0.158008 | 2.841106 | 0.024 | 0 | 1 | 4 | 3110040N1 | 0.024 | Old |
| 0.158008 | 2.746455 | 0.024 | 0 | 1 | 4 | Bcorl1 | 0.024 | Old |
| 0.158008 | 2.552453 | 0.024 | 0 | 1 | 4 | Plpp2 | 0.024 | Old |
| 0.158204 | 1.622492 | 0.123 | 0.075 | 1 | 4 | Gaa | 0.048 | Old |
| 0.158521 | 1.846796 | 0.079 | 0.038 | 1 | 4 | Traf2 | 0.041 | Old |
| 0.158656 | 1.024566 | 0.082 | 0.038 | 1 | 4 | Wtip | 0.044 | Old |
| 0.158806 | 2.010941 | 0.045 | 0.013 | 1 | 4 | Aen | 0.032 | Old |
| 0.159078 | 1.158071 | 0.111 | 0.062 | 1 | 4 | Hacd3 | 0.049 | Old |
| 0.159259 | 1.335159 | 0.11 | 0.062 | 1 | 4 | Fech | 0.048 | Old |
| 0.159282 | 1.180491 | 0.111 | 0.062 | 1 | 4 | Fam204a | 0.049 | Old |

| | | | | | | | | |
|---|---|---|---|---|---|---|---|---|
| 0.159308 | 0.656189 | 0.341 | 0.288 | 1 | 4 | Gfra2 | 0.053 | Old |
| 0.159316 | 1.350395 | 0.064 | 0.025 | 1 | 4 | Nkap | 0.039 | Old |
| 0.159401 | 2.289042 | 0.063 | 0.025 | 1 | 4 | Smim26 | 0.038 | Old |
| 0.160278 | 0.477409 | 0.098 | 0.05 | 1 | 4 | Ngdn | 0.048 | Old |
| 0.161106 | 0.665309 | 0.065 | 0.025 | 1 | 4 | Dtx4 | 0.04 | Old |
| 0.161164 | 1.089947 | 0.081 | 0.038 | 1 | 4 | Gatsl2 | 0.043 | Old |
| 0.161325 | 0.695276 | 0.2 | 0.138 | 1 | 4 | Slc26a2 | 0.062 | Old |
| 0.161742 | 0.823691 | 0.046 | 0.013 | 1 | 4 | Gm15417 | 0.033 | Old |
| 0.161799 | 1.345165 | 0.064 | 0.025 | 1 | 4 | Elavl4 | 0.039 | Old |
| 0.161987 | 0.705909 | 0.233 | 0.175 | 1 | 4 | AI467606 | 0.058 | Old |
| 0.162059 | 0.296792 | 0.725 | 0.713 | 1 | 4 | App | 0.012 | Old |
| 0.162164 | 2.02309 | 0.078 | 0.038 | 1 | 4 | Fam160b2 | 0.04 | Old |
| 0.162174 | 0.959639 | 0.097 | 0.05 | 1 | 4 | 2010315B0 | 0.047 | Old |
| 0.162264 | 1.021988 | 0.112 | 0.062 | 1 | 4 | Ccdc88b | 0.05 | Old |
| 0.162337 | 0.022885 | 0.167 | 0.238 | 1 | 4 | Il10 | -0.071 | Old |
| 0.162359 | 1.474776 | 0.045 | 0.013 | 1 | 4 | 1110008P1 | 0.032 | Old |
| 0.162468 | 0.231795 | 0.641 | 0.625 | 1 | 4 | Ifi27l2a | 0.016 | Old |
| 0.162834 | 0.339035 | 0.133 | 0.075 | 1 | 4 | Cib1 | 0.058 | Old |
| 0.16334 | 1.905388 | 0.079 | 0.038 | 1 | 4 | Ero1l | 0.041 | Old |
| 0.163389 | 2.296479 | 0.045 | 0.013 | 1 | 4 | Pwp1 | 0.032 | Old |
| 0.163424 | 0.856965 | 0.221 | 0.162 | 1 | 4 | Acadm | 0.059 | Old |
| 0.163751 | 1.346445 | 0.063 | 0.025 | 1 | 4 | Lias | 0.038 | Old |
| 0.163821 | 0.481799 | 0.48 | 0.425 | 1 | 4 | St13 | 0.055 | Old |
| 0.163879 | 2.771156 | 0.024 | 0 | 1 | 4 | Amd2 | 0.024 | Old |
| 0.163879 | 3.349021 | 0.024 | 0 | 1 | 4 | Zfp932 | 0.024 | Old |
| 0.163879 | 3.135766 | 0.024 | 0 | 1 | 4 | Zfp934 | 0.024 | Old |
| 0.163879 | 3.111651 | 0.024 | 0 | 1 | 4 | Gm49359 | 0.024 | Old |
| 0.163879 | 3.071135 | 0.024 | 0 | 1 | 4 | Urb1 | 0.024 | Old |
| 0.163879 | 3.050883 | 0.024 | 0 | 1 | 4 | Mvk | 0.024 | Old |
| 0.163879 | 3.024229 | 0.024 | 0 | 1 | 4 | Svip | 0.024 | Old |
| 0.163879 | 3.02117 | 0.024 | 0 | 1 | 4 | Zswim1 | 0.024 | Old |
| 0.163879 | 2.978622 | 0.024 | 0 | 1 | 4 | Zc3h3 | 0.024 | Old |
| 0.163879 | 2.936281 | 0.024 | 0 | 1 | 4 | Oasl1 | 0.024 | Old |
| 0.163879 | 2.818457 | 0.024 | 0 | 1 | 4 | Nme7 | 0.024 | Old |
| 0.163879 | 2.790368 | 0.024 | 0 | 1 | 4 | Btf3l4 | 0.024 | Old |
| 0.163879 | 2.742239 | 0.024 | 0 | 1 | 4 | Ppp3cc | 0.024 | Old |
| 0.165276 | 0.96424 | 0.191 | 0.138 | 1 | 4 | Slmap | 0.053 | Old |
| 0.165316 | 1.813325 | 0.045 | 0.013 | 1 | 4 | Zfand2b | 0.032 | Old |
| 0.165488 | 0.501281 | 0.255 | 0.188 | 1 | 4 | Nsf | 0.067 | Old |
| 0.165747 | 1.909611 | 0.045 | 0.013 | 1 | 4 | Stk16 | 0.032 | Old |
| 0.16629 | 0.637279 | 0.252 | 0.188 | 1 | 4 | Sec31a | 0.064 | Old |
| 0.166364 | 2.728602 | 0.044 | 0.013 | 1 | 4 | Rai14 | 0.031 | Old |
| 0.166502 | 0.434224 | 0.314 | 0.25 | 1 | 4 | Ppp1r12c | 0.064 | Old |
| 0.166615 | 0.937119 | 0.045 | 0.013 | 1 | 4 | Zfp942 | 0.032 | Old |
| 0.166663 | 0.817195 | 0.196 | 0.138 | 1 | 4 | Usp48 | 0.058 | Old |

| | | | | | | | | |
|---|---|---|---|---|---|---|---|---|
| 0.167108 | 0.220482 | 0.393 | 0.338 | 1 | 4 | Slfn2 | 0.055 | Old |
| 0.16711 | 0.121148 | 0.983 | 0.975 | 1 | 4 | Rps8 | 0.008 | Old |
| 0.167684 | 1.831187 | 0.121 | 0.075 | 1 | 4 | Fam20c | 0.046 | Old |
| 0.167876 | 2.570682 | 0.061 | 0.025 | 1 | 4 | C130026I2 | 0.036 | Old |
| 0.167914 | 2.01657 | 0.062 | 0.025 | 1 | 4 | Tmem87a | 0.037 | Old |
| 0.16804 | 0.993952 | 0.225 | 0.175 | 1 | 4 | Usp4 | 0.05 | Old |
| 0.168124 | 0.137959 | 0.059 | 0.1 | 1 | 4 | Gtf3c4 | -0.041 | Old |
| 0.169142 | 0.517133 | 0.239 | 0.175 | 1 | 4 | Rybp | 0.064 | Old |
| 0.169158 | 0.988336 | 0.187 | 0.138 | 1 | 4 | Themis2 | 0.049 | Old |
| 0.169993 | 3.082536 | 0.023 | 0 | 1 | 4 | Galm | 0.023 | Old |
| 0.169993 | 3.078944 | 0.023 | 0 | 1 | 4 | A930033H1 | 0.023 | Old |
| 0.169993 | 2.961894 | 0.023 | 0 | 1 | 4 | Rnf122 | 0.023 | Old |
| 0.169993 | 2.94735 | 0.023 | 0 | 1 | 4 | Tmem129 | 0.023 | Old |
| 0.169993 | 2.926394 | 0.023 | 0 | 1 | 4 | Gemin2 | 0.023 | Old |
| 0.169993 | 2.697069 | 0.023 | 0 | 1 | 4 | Syne3 | 0.023 | Old |
| 0.169993 | 2.537968 | 0.023 | 0 | 1 | 4 | Prr5l | 0.023 | Old |
| 0.170069 | 0.377783 | 0.377 | 0.312 | 1 | 4 | Ndufb3 | 0.065 | Old |
| 0.170158 | 0.768205 | 0.222 | 0.162 | 1 | 4 | Pgam1 | 0.06 | Old |
| 0.170211 | 0.307541 | 0.37 | 0.288 | 1 | 4 | Ssb | 0.082 | Old |
| 0.17037 | 0.496971 | 0.157 | 0.1 | 1 | 4 | Rbbp8 | 0.057 | Old |
| 0.170538 | 2.00461 | 0.044 | 0.013 | 1 | 4 | Stmn1 | 0.031 | Old |
| 0.170879 | 0.794422 | 0.097 | 0.05 | 1 | 4 | Vac14 | 0.047 | Old |
| 0.171078 | 2.117572 | 0.061 | 0.025 | 1 | 4 | Agpat4 | 0.036 | Old |
| 0.171494 | 0.192561 | 0.579 | 0.488 | 1 | 4 | Uqcr11 | 0.091 | Old |
| 0.17165 | 1.712848 | 0.044 | 0.013 | 1 | 4 | Zfp954 | 0.031 | Old |
| 0.171773 | 1.163095 | 0.152 | 0.1 | 1 | 4 | Lars2 | 0.052 | Old |
| 0.171865 | 0.98021 | 0.045 | 0.013 | 1 | 4 | Rrp8 | 0.032 | Old |
| 0.171878 | 0.601057 | 0.016 | 0.038 | 1 | 4 | Cavin2 | -0.022 | Old |
| 0.172079 | 2.482033 | 0.043 | 0.013 | 1 | 4 | Chn2 | 0.03 | Old |
| 0.172079 | 2.415079 | 0.043 | 0.013 | 1 | 4 | Qpct | 0.03 | Old |
| 0.17291 | 0.21174 | 0.016 | 0.038 | 1 | 4 | Fgf11 | -0.022 | Old |
| 0.173066 | 0.643129 | 0.112 | 0.062 | 1 | 4 | Hmgcr | 0.05 | Old |
| 0.173215 | 1.612438 | 0.044 | 0.013 | 1 | 4 | Dhx35 | 0.031 | Old |
| 0.174687 | 0.369633 | 0.633 | 0.6 | 1 | 4 | Gm26917 | 0.033 | Old |
| 0.175243 | 1.397397 | 0.044 | 0.013 | 1 | 4 | Nat6 | 0.031 | Old |
| 0.175646 | 0.645774 | 0.347 | 0.3 | 1 | 4 | Mndal | 0.047 | Old |
| 0.175856 | 2.10046 | 0.061 | 0.025 | 1 | 4 | Gm15987 | 0.036 | Old |
| 0.176363 | 3.006573 | 0.022 | 0 | 1 | 4 | Shq1 | 0.022 | Old |
| 0.176363 | 3.062816 | 0.022 | 0 | 1 | 4 | Fam222a | 0.022 | Old |
| 0.176363 | 2.873595 | 0.022 | 0 | 1 | 4 | Fig4 | 0.022 | Old |
| 0.176363 | 2.826092 | 0.022 | 0 | 1 | 4 | Tspan13 | 0.022 | Old |
| 0.176363 | 2.714504 | 0.022 | 0 | 1 | 4 | Gstm4 | 0.022 | Old |
| 0.176363 | 2.625545 | 0.022 | 0 | 1 | 4 | Inf2 | 0.022 | Old |
| 0.176963 | 0.215824 | 0.711 | 0.662 | 1 | 4 | Mgl2 | 0.049 | Old |
| 0.177079 | 1.776032 | 0.043 | 0.013 | 1 | 4 | Mapk7 | 0.03 | Old |

| | | | | | | | | |
|---|---|---|---|---|---|---|---|---|
| 0.177079 | 1.660327 | 0.043 | 0.013 | 1 | 4 | Orc2 | 0.03 | Old |
| 0.177108 | 1.404596 | 0.159 | 0.112 | 1 | 4 | Egln1 | 0.047 | Old |
| 0.178039 | 0.402396 | 0.353 | 0.275 | 1 | 4 | Rab20 | 0.078 | Old |
| 0.178062 | 1.070362 | 0.236 | 0.188 | 1 | 4 | Marveld1 | 0.048 | Old |
| 0.178137 | 0.433219 | 0.553 | 0.5 | 1 | 4 | Cops9 | 0.053 | Old |
| 0.178188 | 0.658149 | 0.127 | 0.075 | 1 | 4 | Ilkap | 0.052 | Old |
| 0.178482 | 0.792394 | 0.094 | 0.05 | 1 | 4 | Mpnd | 0.044 | Old |
| 0.178692 | 1.52566 | 0.043 | 0.013 | 1 | 4 | Tubgcp2 | 0.03 | Old |
| 0.178697 | 2.572071 | 0.043 | 0.013 | 1 | 4 | Wrap73 | 0.03 | Old |
| 0.17926 | 0.600379 | 0.096 | 0.05 | 1 | 4 | Ndst2 | 0.046 | Old |
| 0.179262 | 0.806446 | 0.167 | 0.112 | 1 | 4 | Plxna1 | 0.055 | Old |
| 0.179755 | 0.494048 | 0.297 | 0.238 | 1 | 4 | Fam208a | 0.059 | Old |
| 0.179921 | 0.295744 | 0.422 | 0.362 | 1 | 4 | Taok1 | 0.06 | Old |
| 0.180369 | 2.399617 | 0.06 | 0.025 | 1 | 4 | Prrg2 | 0.035 | Old |
| 0.180961 | 1.368288 | 0.077 | 0.038 | 1 | 4 | Atg4c | 0.039 | Old |
| 0.181155 | 1.194548 | 0.061 | 0.025 | 1 | 4 | Lsm1 | 0.036 | Old |
| 0.181368 | 2.085845 | 0.06 | 0.025 | 1 | 4 | Ebag9 | 0.035 | Old |
| 0.181489 | 0.201268 | 0.671 | 0.613 | 1 | 4 | Atp5a1 | 0.058 | Old |
| 0.181863 | 0.462502 | 0.129 | 0.075 | 1 | 4 | Snx1 | 0.054 | Old |
| 0.18269 | 1.835593 | 0.043 | 0.013 | 1 | 4 | Fbxo25 | 0.03 | Old |
| 0.182847 | 0.089873 | 0.127 | 0.188 | 1 | 4 | Pgm2l1 | -0.061 | Old |
| 0.18299 | 1.268127 | 0.078 | 0.038 | 1 | 4 | Sfxn1 | 0.04 | Old |
| 0.183 | 2.918104 | 0.022 | 0 | 1 | 4 | Lrsam1 | 0.022 | Old |
| 0.183 | 2.853294 | 0.022 | 0 | 1 | 4 | Lmbr1 | 0.022 | Old |
| 0.183 | 2.739107 | 0.022 | 0 | 1 | 4 | Ppp2r5b | 0.022 | Old |
| 0.183005 | 1.779779 | 0.105 | 0.062 | 1 | 4 | Sacm1l | 0.043 | Old |
| 0.183832 | 0.913183 | 0.138 | 0.088 | 1 | 4 | Fam53b | 0.05 | Old |
| 0.183859 | 0.549604 | 0.095 | 0.05 | 1 | 4 | Trip4 | 0.045 | Old |
| 0.183969 | 1.457588 | 0.146 | 0.1 | 1 | 4 | Lsp1 | 0.046 | Old |
| 0.184996 | 1.939525 | 0.06 | 0.025 | 1 | 4 | Krr1 | 0.035 | Old |
| 0.185493 | 0.540632 | 0.043 | 0.013 | 1 | 4 | Slc10a3 | 0.03 | Old |
| 0.185528 | 0.504972 | 0.308 | 0.25 | 1 | 4 | Slc7a8 | 0.058 | Old |
| 0.185956 | 0.439534 | 0.412 | 0.35 | 1 | 4 | Nadk | 0.062 | Old |
| 0.186095 | 1.062576 | 0.108 | 0.062 | 1 | 4 | Asb8 | 0.046 | Old |
| 0.186228 | 0.122871 | 0.042 | 0.075 | 1 | 4 | Rbms2 | -0.033 | Old |
| 0.186422 | 2.011092 | 0.06 | 0.025 | 1 | 4 | Stoml2 | 0.035 | Old |
| 0.187951 | 0.111971 | 0.988 | 0.975 | 1 | 4 | H2-Aa | 0.013 | Old |
| 0.188203 | 0.423222 | 0.387 | 0.338 | 1 | 4 | 7-Sep | 0.049 | Old |
| 0.188244 | 2.416084 | 0.042 | 0.013 | 1 | 4 | Evi2 | 0.029 | Old |
| 0.188288 | 2.030497 | 0.059 | 0.025 | 1 | 4 | Irf7 | 0.034 | Old |
| 0.188469 | 0.266254 | 0.798 | 0.762 | 1 | 4 | Lamp2 | 0.036 | Old |
| 0.189173 | 0.45446 | 0.359 | 0.312 | 1 | 4 | Cap1 | 0.047 | Old |
| 0.189214 | 0.88932 | 0.078 | 0.038 | 1 | 4 | Rufy1 | 0.04 | Old |
| 0.189562 | 0.605552 | 0.341 | 0.288 | 1 | 4 | 1700020I14 | 0.053 | Old |
| 0.189572 | 0.197949 | 0.428 | 0.35 | 1 | 4 | Bzw1 | 0.078 | Old |

| | | | | | | | | |
|---|---|---|---|---|---|---|---|---|
| 0.18992 | 3.086685 | 0.021 | 0 | 1 | 4 | Hemk1 | 0.021 | Old |
| 0.18992 | 3.023702 | 0.021 | 0 | 1 | 4 | Scrn3 | 0.021 | Old |
| 0.18992 | 2.958817 | 0.021 | 0 | 1 | 4 | Cyb5d2 | 0.021 | Old |
| 0.18992 | 2.937309 | 0.021 | 0 | 1 | 4 | Tatdn3 | 0.021 | Old |
| 0.18992 | 2.930795 | 0.021 | 0 | 1 | 4 | Nmrk1 | 0.021 | Old |
| 0.18992 | 2.91236 | 0.021 | 0 | 1 | 4 | Noa1 | 0.021 | Old |
| 0.18992 | 2.84444 | 0.021 | 0 | 1 | 4 | Cep57l1 | 0.021 | Old |
| 0.18992 | 2.8439 | 0.021 | 0 | 1 | 4 | Cish | 0.021 | Old |
| 0.18992 | 2.779226 | 0.021 | 0 | 1 | 4 | Dcstamp | 0.021 | Old |
| 0.18992 | 2.760811 | 0.021 | 0 | 1 | 4 | Psat1 | 0.021 | Old |
| 0.18992 | 2.744631 | 0.021 | 0 | 1 | 4 | Chchd6 | 0.021 | Old |
| 0.18992 | 2.714804 | 0.021 | 0 | 1 | 4 | Gm3448 | 0.021 | Old |
| 0.18992 | 2.714153 | 0.021 | 0 | 1 | 4 | Spred3 | 0.021 | Old |
| 0.18992 | 2.618734 | 0.021 | 0 | 1 | 4 | Unc119b | 0.021 | Old |
| 0.18992 | 2.595368 | 0.021 | 0 | 1 | 4 | Aff3 | 0.021 | Old |
| 0.18992 | 2.553987 | 0.021 | 0 | 1 | 4 | Ift57 | 0.021 | Old |
| 0.18992 | 2.489089 | 0.021 | 0 | 1 | 4 | Klf8 | 0.021 | Old |
| 0.18992 | 2.462695 | 0.021 | 0 | 1 | 4 | Ptgir | 0.021 | Old |
| 0.190286 | 1.038556 | 0.061 | 0.025 | 1 | 4 | Qser1 | 0.036 | Old |
| 0.190485 | 2.836863 | 0.042 | 0.013 | 1 | 4 | Ezh2 | 0.029 | Old |
| 0.190628 | 0.742873 | 0.043 | 0.013 | 1 | 4 | Cd226 | 0.03 | Old |
| 0.190655 | 0.678415 | 0.256 | 0.2 | 1 | 4 | Il3ra | 0.056 | Old |
| 0.190762 | 0.792944 | 0.094 | 0.05 | 1 | 4 | Mzt1 | 0.044 | Old |
| 0.190915 | 1.008669 | 0.077 | 0.038 | 1 | 4 | Me2 | 0.039 | Old |
| 0.191179 | 0.100597 | 0.003 | 0.013 | 1 | 4 | Mrm3 | -0.01 | Old |
| 0.191284 | 0.952173 | 0.077 | 0.038 | 1 | 4 | Rpf1 | 0.039 | Old |
| 0.191409 | 1.435224 | 0.119 | 0.075 | 1 | 4 | Washc3 | 0.044 | Old |
| 0.191727 | 2.596592 | 0.042 | 0.013 | 1 | 4 | Zdhhc8 | 0.029 | Old |
| 0.191859 | 0.541666 | 0.2 | 0.138 | 1 | 4 | Spag7 | 0.062 | Old |
| 0.191973 | 1.02417 | 0.221 | 0.175 | 1 | 4 | Fam32a | 0.046 | Old |
| 0.191977 | 0.347152 | 0.003 | 0.013 | 1 | 4 | Akap12 | -0.01 | Old |
| 0.191977 | 0.256903 | 0.003 | 0.013 | 1 | 4 | Gm10382 | -0.01 | Old |
| 0.191977 | 0.187733 | 0.003 | 0.013 | 1 | 4 | Gm31410 | -0.01 | Old |
| 0.191977 | 0.156379 | 0.003 | 0.013 | 1 | 4 | AC174678. | -0.01 | Old |
| 0.191977 | 0.145727 | 0.003 | 0.013 | 1 | 4 | Bicc1 | -0.01 | Old |
| 0.191977 | 0.037516 | 0.003 | 0.013 | 1 | 4 | Fancf | -0.01 | Old |
| 0.192425 | 0.93745 | 0.2 | 0.15 | 1 | 4 | Slc36a1 | 0.05 | Old |
| 0.192877 | 0.777698 | 0.109 | 0.062 | 1 | 4 | Mrps25 | 0.047 | Old |
| 0.193042 | 1.419112 | 0.06 | 0.025 | 1 | 4 | Taf12 | 0.035 | Old |
| 0.193121 | 2.421276 | 0.059 | 0.025 | 1 | 4 | Stx11 | 0.034 | Old |
| 0.194042 | 0.895867 | 0.137 | 0.088 | 1 | 4 | Ube2v2 | 0.049 | Old |
| 0.194354 | 0.322518 | 0.217 | 0.15 | 1 | 4 | Gtf2a1 | 0.067 | Old |
| 0.194388 | 1.156301 | 0.16 | 0.112 | 1 | 4 | Ufsp2 | 0.048 | Old |
| 0.194811 | 0.894302 | 0.248 | 0.2 | 1 | 4 | Vps13a | 0.048 | Old |
| 0.19498 | 1.001668 | 0.15 | 0.1 | 1 | 4 | Cnp | 0.05 | Old |

| | | | | | | | | |
|---|---|---|---|---|---|---|---|---|
| 0.195085 | 1.579982 | 0.104 | 0.062 | 1 | 4 | Pqlc3 | 0.042 | Old |
| 0.195191 | 1.896351 | 0.074 | 0.038 | 1 | 4 | Elmod2 | 0.036 | Old |
| 0.19539 | 0.709028 | 0.285 | 0.238 | 1 | 4 | Fam213b | 0.047 | Old |
| 0.195456 | 1.118609 | 0.148 | 0.1 | 1 | 4 | Ahcyl2 | 0.048 | Old |
| 0.195503 | 0.687971 | 0.109 | 0.062 | 1 | 4 | Glo1 | 0.047 | Old |
| 0.195687 | 0.478827 | 0.017 | 0.038 | 1 | 4 | Nkiras1 | -0.021 | Old |
| 0.195739 | 0.135736 | 0.94 | 1 | 1 | 4 | Zeb2 | -0.06 | Old |
| 0.195879 | 0.210635 | 0.079 | 0.125 | 1 | 4 | Yipf6 | -0.046 | Old |
| 0.196755 | 1.649564 | 0.042 | 0.013 | 1 | 4 | Gpatch11 | 0.029 | Old |
| 0.196811 | 2.909523 | 0.041 | 0.013 | 1 | 4 | Slc41a1 | 0.028 | Old |
| 0.197138 | 2.533744 | 0.02 | 0 | 1 | 4 | Ppcs | 0.02 | Old |
| 0.197138 | 2.981777 | 0.02 | 0 | 1 | 4 | Ercc8 | 0.02 | Old |
| 0.197138 | 2.965277 | 0.02 | 0 | 1 | 4 | Xlr4a | 0.02 | Old |
| 0.197138 | 2.945798 | 0.02 | 0 | 1 | 4 | Tgfbr3 | 0.02 | Old |
| 0.197138 | 2.939909 | 0.02 | 0 | 1 | 4 | Rab23 | 0.02 | Old |
| 0.197138 | 2.847347 | 0.02 | 0 | 1 | 4 | Nbeal2 | 0.02 | Old |
| 0.197138 | 2.826161 | 0.02 | 0 | 1 | 4 | Ncapd2 | 0.02 | Old |
| 0.197138 | 2.791288 | 0.02 | 0 | 1 | 4 | Tmem238 | 0.02 | Old |
| 0.197138 | 2.761512 | 0.02 | 0 | 1 | 4 | Zfp316 | 0.02 | Old |
| 0.197138 | 2.595022 | 0.02 | 0 | 1 | 4 | Klhl36 | 0.02 | Old |
| 0.197138 | 2.507804 | 0.02 | 0 | 1 | 4 | Car5b | 0.02 | Old |
| 0.197171 | 0.332708 | 0.241 | 0.175 | 1 | 4 | Phc3 | 0.066 | Old |
| 0.197344 | 1.247734 | 0.091 | 0.05 | 1 | 4 | Senp5 | 0.041 | Old |
| 0.197646 | 0.477169 | 0.327 | 0.275 | 1 | 4 | Stard3nl | 0.052 | Old |
| 0.197747 | 0.693314 | 0.205 | 0.15 | 1 | 4 | Tuba1a | 0.055 | Old |
| 0.197776 | 1.210636 | 0.076 | 0.038 | 1 | 4 | Topbp1 | 0.038 | Old |
| 0.197904 | 1.01179 | 0.076 | 0.038 | 1 | 4 | Sec23a | 0.038 | Old |
| 0.19807 | 1.077133 | 0.185 | 0.138 | 1 | 4 | Map4 | 0.047 | Old |
| 0.198093 | 2.788509 | 0.041 | 0.013 | 1 | 4 | Flrt2 | 0.028 | Old |
| 0.198217 | 0.722335 | 0.166 | 0.112 | 1 | 4 | Lrpap1 | 0.054 | Old |
| 0.198236 | 0.833664 | 0.108 | 0.062 | 1 | 4 | Utp14a | 0.046 | Old |
| 0.198538 | 0.658121 | 0.206 | 0.15 | 1 | 4 | Coa5 | 0.056 | Old |
| 0.198607 | 2.630731 | 0.041 | 0.013 | 1 | 4 | Ccdc112 | 0.028 | Old |
| 0.198976 | 0.503154 | 0.214 | 0.15 | 1 | 4 | Rad23a | 0.064 | Old |
| 0.198979 | 1.197447 | 0.106 | 0.062 | 1 | 4 | Mrpl16 | 0.044 | Old |
| 0.199117 | 1.522698 | 0.059 | 0.025 | 1 | 4 | Nufip1 | 0.034 | Old |
| 0.199151 | 1.01327 | 0.198 | 0.15 | 1 | 4 | Apc | 0.048 | Old |
| 0.199607 | 0.374029 | 0.111 | 0.062 | 1 | 4 | 6430548M | 0.049 | Old |
| 0.199692 | 1.461952 | 0.104 | 0.062 | 1 | 4 | Dda1 | 0.042 | Old |
| 0.200157 | 0.324739 | 0.44 | 0.375 | 1 | 4 | Eif4b | 0.065 | Old |
| 0.20016 | 1.163883 | 0.091 | 0.05 | 1 | 4 | Pkd2 | 0.041 | Old |
| 0.200476 | 2.17494 | 0.058 | 0.025 | 1 | 4 | Fbxo9 | 0.033 | Old |
| 0.202449 | 0.807679 | 0.177 | 0.125 | 1 | 4 | Lrmp | 0.052 | Old |
| 0.202466 | 0.531367 | 0.139 | 0.088 | 1 | 4 | Map2k1 | 0.051 | Old |
| 0.202716 | 0.219174 | 0.109 | 0.062 | 1 | 4 | Rbm12 | 0.047 | Old |

| | | | | | | | | |
|---|---|---|---|---|---|---|---|---|
| 0.203643 | 0.422424 | 0.14 | 0.088 | 1 | 4 | Gm34084 | 0.052 | Old |
| 0.203725 | 0.30009 | 0.715 | 0.713 | 1 | 4 | Tuba1b | 0.002 | Old |
| 0.20436 | 1.701335 | 0.074 | 0.038 | 1 | 4 | Rrp36 | 0.036 | Old |
| 0.204429 | 0.211218 | 0.75 | 0.812 | 1 | 4 | Mob1a | -0.062 | Old |
| 0.20467 | 2.932972 | 0.02 | 0 | 1 | 4 | BC017158 | 0.02 | Old |
| 0.20467 | 2.889169 | 0.02 | 0 | 1 | 4 | Zbtb8a | 0.02 | Old |
| 0.20467 | 2.849321 | 0.02 | 0 | 1 | 4 | Mrm2 | 0.02 | Old |
| 0.20467 | 2.801952 | 0.02 | 0 | 1 | 4 | Magi1 | 0.02 | Old |
| 0.20467 | 2.794075 | 0.02 | 0 | 1 | 4 | Gem | 0.02 | Old |
| 0.20467 | 2.773948 | 0.02 | 0 | 1 | 4 | Pcbp4 | 0.02 | Old |
| 0.20467 | 2.77174 | 0.02 | 0 | 1 | 4 | Wdr60 | 0.02 | Old |
| 0.20467 | 2.755093 | 0.02 | 0 | 1 | 4 | Map2k3os | 0.02 | Old |
| 0.20467 | 2.684709 | 0.02 | 0 | 1 | 4 | Tacc3 | 0.02 | Old |
| 0.204738 | 0.644277 | 0.042 | 0.013 | 1 | 4 | Mapkapk3 | 0.029 | Old |
| 0.205317 | 0.647052 | 0.094 | 0.05 | 1 | 4 | Irgm2 | 0.044 | Old |
| 0.205695 | 0.738491 | 0.121 | 0.075 | 1 | 4 | Kcnab2 | 0.046 | Old |
| 0.205745 | 0.550714 | 0.01 | 0.025 | 1 | 4 | Padi4 | -0.015 | Old |
| 0.20584 | 0.137556 | 0.026 | 0.05 | 1 | 4 | Nlrx1 | -0.024 | Old |
| 0.205932 | 1.189607 | 0.158 | 0.112 | 1 | 4 | Crtap | 0.046 | Old |
| 0.206011 | 2.325855 | 0.04 | 0.013 | 1 | 4 | Nr2c2ap | 0.027 | Old |
| 0.206146 | 1.922493 | 0.074 | 0.038 | 1 | 4 | Ttc32 | 0.036 | Old |
| 0.206181 | 0.547567 | 0.244 | 0.188 | 1 | 4 | Cnih1 | 0.056 | Old |
| 0.206278 | 2.401313 | 0.04 | 0.013 | 1 | 4 | Lmf1 | 0.027 | Old |
| 0.206452 | 0.620007 | 0.32 | 0.275 | 1 | 4 | Fam126a | 0.045 | Old |
| 0.20658 | 1.063111 | 0.059 | 0.025 | 1 | 4 | Cfap97 | 0.034 | Old |
| 0.206595 | 0.429189 | 0.449 | 0.4 | 1 | 4 | Pbrm1 | 0.049 | Old |
| 0.207609 | 1.907908 | 0.073 | 0.038 | 1 | 4 | Tbc1d10a | 0.035 | Old |
| 0.207615 | 2.289875 | 0.04 | 0.013 | 1 | 4 | Igf2r | 0.027 | Old |
| 0.207908 | 1.131444 | 0.105 | 0.062 | 1 | 4 | Hsdl1 | 0.043 | Old |
| 0.208036 | 1.128971 | 0.041 | 0.013 | 1 | 4 | E130309D( | 0.028 | Old |
| 0.208218 | 0.529881 | 0.168 | 0.112 | 1 | 4 | Rab32 | 0.056 | Old |
| 0.208432 | 0.589275 | 0.139 | 0.088 | 1 | 4 | Slamf8 | 0.051 | Old |
| 0.208689 | 2.017466 | 0.04 | 0.013 | 1 | 4 | H2-T24 | 0.027 | Old |
| 0.208761 | 1.497905 | 0.089 | 0.05 | 1 | 4 | BC029722 | 0.039 | Old |
| 0.209398 | 0.550351 | 0.254 | 0.2 | 1 | 4 | Cd300a | 0.054 | Old |
| 0.209454 | 0.162841 | 0.06 | 0.025 | 1 | 4 | Pctp | 0.035 | Old |
| 0.209559 | 0.347595 | 0.522 | 0.462 | 1 | 4 | Rell1 | 0.06 | Old |
| 0.209824 | 0.62106 | 0.313 | 0.262 | 1 | 4 | Apbb1ip | 0.051 | Old |
| 0.21014 | 1.899932 | 0.057 | 0.025 | 1 | 4 | Hopx | 0.032 | Old |
| 0.211297 | 0.355427 | 0.368 | 0.312 | 1 | 4 | Ppp2r5c | 0.056 | Old |
| 0.211801 | 0.481955 | 0.37 | 0.325 | 1 | 4 | Pias1 | 0.045 | Old |
| 0.21206 | 0.553364 | 0.059 | 0.025 | 1 | 4 | Nostrin | 0.034 | Old |
| 0.212326 | 2.531617 | 0.04 | 0.013 | 1 | 4 | Ddt | 0.027 | Old |
| 0.212472 | 1.623053 | 0.073 | 0.038 | 1 | 4 | Rgs14 | 0.035 | Old |
| 0.212533 | 2.735395 | 0.019 | 0 | 1 | 4 | Gm27010 | 0.019 | Old |

| | | | | | | | | |
|---|---|---|---|---|---|---|---|---|
| 0.212533 | 2.710265 | 0.019 | 0 | 1 | 4 | Abhd14a | 0.019 | Old |
| 0.212533 | 2.668573 | 0.019 | 0 | 1 | 4 | Spindoc | 0.019 | Old |
| 0.212533 | 2.648728 | 0.019 | 0 | 1 | 4 | Cep41 | 0.019 | Old |
| 0.212533 | 2.647935 | 0.019 | 0 | 1 | 4 | Cep131 | 0.019 | Old |
| 0.212533 | 2.623777 | 0.019 | 0 | 1 | 4 | AI427809 | 0.019 | Old |
| 0.212533 | 2.496475 | 0.019 | 0 | 1 | 4 | Gpsm1 | 0.019 | Old |
| 0.212533 | 2.420859 | 0.019 | 0 | 1 | 4 | Tbc1d8b | 0.019 | Old |
| 0.21265 | 0.059985 | 0.987 | 0.963 | 1 | 4 | Eef1a1 | 0.024 | Old |
| 0.213426 | 2.284431 | 0.04 | 0.013 | 1 | 4 | Bod1 | 0.027 | Old |
| 0.214297 | 0.128024 | 0.999 | 1 | 1 | 4 | Selenop | -0.001 | Old |
| 0.214865 | 0.765013 | 0.091 | 0.05 | 1 | 4 | Pigm | 0.041 | Old |
| 0.215084 | 2.070819 | 0.04 | 0.013 | 1 | 4 | Htr1b | 0.027 | Old |
| 0.215361 | 2.209755 | 0.04 | 0.013 | 1 | 4 | Trim37 | 0.027 | Old |
| 0.216029 | 0.976558 | 0.105 | 0.062 | 1 | 4 | Ndufa9 | 0.043 | Old |
| 0.216043 | 1.180998 | 0.04 | 0.013 | 1 | 4 | Smyd2 | 0.027 | Old |
| 0.216268 | 0.064434 | 0.967 | 0.975 | 1 | 4 | H2-Ab1 | -0.008 | Old |
| 0.216536 | 0.39805 | 0.059 | 0.025 | 1 | 4 | AC127341. | 0.034 | Old |
| 0.216762 | 0.155727 | 0.825 | 0.825 | 1 | 4 | Mbnl1 | 0 | Old |
| 0.217167 | 1.722662 | 0.057 | 0.025 | 1 | 4 | Ciao1 | 0.032 | Old |
| 0.217345 | 0.759842 | 0.209 | 0.162 | 1 | 4 | Prkx | 0.047 | Old |
| 0.217426 | 0.915253 | 0.04 | 0.013 | 1 | 4 | Ip6k3 | 0.027 | Old |
| 0.217475 | 0.801921 | 0.075 | 0.038 | 1 | 4 | Asgr2 | 0.037 | Old |
| 0.218261 | 0.564628 | 0.283 | 0.225 | 1 | 4 | Tsen34 | 0.058 | Old |
| 0.218263 | 0.879392 | 0.119 | 0.075 | 1 | 4 | Aim2 | 0.044 | Old |
| 0.218519 | 0.678711 | 0.106 | 0.062 | 1 | 4 | Trim65 | 0.044 | Old |
| 0.218552 | 1.051682 | 0.074 | 0.038 | 1 | 4 | Kdm5b | 0.036 | Old |
| 0.218683 | 1.329704 | 0.057 | 0.025 | 1 | 4 | Slc35e4 | 0.032 | Old |
| 0.218771 | 0.462564 | 0.181 | 0.125 | 1 | 4 | Irf2bpl | 0.056 | Old |
| 0.219519 | 1.581627 | 0.057 | 0.025 | 1 | 4 | Mief1 | 0.032 | Old |
| 0.219628 | 0.861201 | 0.209 | 0.162 | 1 | 4 | Taf15 | 0.047 | Old |
| 0.21988 | 0.820793 | 0.074 | 0.038 | 1 | 4 | B3galnt2 | 0.036 | Old |
| 0.220113 | 1.419847 | 0.04 | 0.013 | 1 | 4 | Plekha8 | 0.027 | Old |
| 0.220233 | 0.723385 | 0.121 | 0.075 | 1 | 4 | Enpep | 0.046 | Old |
| 0.220415 | 0.548462 | 0.162 | 0.112 | 1 | 4 | Lsm5 | 0.05 | Old |
| 0.220747 | 2.790471 | 0.018 | 0 | 1 | 4 | Tmem206 | 0.018 | Old |
| 0.220747 | 2.774933 | 0.018 | 0 | 1 | 4 | Tldc1 | 0.018 | Old |
| 0.220747 | 2.703548 | 0.018 | 0 | 1 | 4 | Thop1 | 0.018 | Old |
| 0.220747 | 2.700415 | 0.018 | 0 | 1 | 4 | Zfp53 | 0.018 | Old |
| 0.220747 | 2.697899 | 0.018 | 0 | 1 | 4 | Tbc1d19 | 0.018 | Old |
| 0.220747 | 2.631699 | 0.018 | 0 | 1 | 4 | Nt5c3b | 0.018 | Old |
| 0.220747 | 2.604195 | 0.018 | 0 | 1 | 4 | Tdp1 | 0.018 | Old |
| 0.220747 | 2.596912 | 0.018 | 0 | 1 | 4 | Eefsec | 0.018 | Old |
| 0.220747 | 2.590416 | 0.018 | 0 | 1 | 4 | Naprt | 0.018 | Old |
| 0.220747 | 2.523228 | 0.018 | 0 | 1 | 4 | Fbxl8 | 0.018 | Old |
| 0.220759 | 1.237779 | 0.073 | 0.038 | 1 | 4 | Cbll1 | 0.035 | Old |

| | | | | | | | | |
|---|---|---|---|---|---|---|---|---|
| 0.220834 | 2.646453 | 0.039 | 0.013 | 1 | 4 | Vwa8 | 0.026 | Old |
| 0.221074 | 1.285136 | 0.073 | 0.038 | 1 | 4 | Relt | 0.035 | Old |
| 0.221128 | 0.378355 | 0.525 | 0.5 | 1 | 4 | Vsir | 0.025 | Old |
| 0.221248 | 0.340015 | 0.076 | 0.038 | 1 | 4 | Nup93 | 0.038 | Old |
| 0.222091 | 1.123884 | 0.04 | 0.013 | 1 | 4 | Mvd | 0.027 | Old |
| 0.222763 | 0.095037 | 0.997 | 1 | 1 | 4 | Tpt1 | -0.003 | Old |
| 0.222779 | 2.170217 | 0.071 | 0.038 | 1 | 4 | Incenp | 0.033 | Old |
| 0.223156 | 0.634108 | 0.3 | 0.25 | 1 | 4 | Psmd13 | 0.05 | Old |
| 0.223797 | 1.003147 | 0.04 | 0.013 | 1 | 4 | Cep85 | 0.027 | Old |
| 0.223801 | 1.15165 | 0.103 | 0.062 | 1 | 4 | Ilvbl | 0.041 | Old |
| 0.224226 | 0.157561 | 0.178 | 0.25 | 1 | 4 | P3h2 | -0.072 | Old |
| 0.224274 | 0.692486 | 0.074 | 0.038 | 1 | 4 | Mindy3 | 0.036 | Old |
| 0.224368 | 1.050373 | 0.04 | 0.013 | 1 | 4 | Dym | 0.027 | Old |
| 0.224441 | 0.460447 | 0.443 | 0.412 | 1 | 4 | Klhl9 | 0.031 | Old |
| 0.224845 | 1.985334 | 0.039 | 0.013 | 1 | 4 | Pdp2 | 0.026 | Old |
| 0.225027 | 0.01042 | 0.072 | 0.112 | 1 | 4 | Fbxo8 | -0.04 | Old |
| 0.225633 | 1.232783 | 0.153 | 0.112 | 1 | 4 | Dmwd | 0.041 | Old |
| 0.225816 | 0.973401 | 0.089 | 0.05 | 1 | 4 | Rarg | 0.039 | Old |
| 0.226151 | 0.905172 | 0.057 | 0.025 | 1 | 4 | Cdc27 | 0.032 | Old |
| 0.226388 | 0.655687 | 0.074 | 0.038 | 1 | 4 | Bicra | 0.036 | Old |
| 0.226581 | 0.834208 | 0.133 | 0.088 | 1 | 4 | H2-T22 | 0.045 | Old |
| 0.226754 | 2.829279 | 0.038 | 0.013 | 1 | 4 | Pou6f1 | 0.025 | Old |
| 0.22716 | 1.417723 | 0.039 | 0.013 | 1 | 4 | Card9 | 0.026 | Old |
| 0.227232 | 2.171272 | 0.055 | 0.025 | 1 | 4 | Med4 | 0.03 | Old |
| 0.227276 | 1.121633 | 0.117 | 0.075 | 1 | 4 | Htatsf1 | 0.042 | Old |
| 0.227558 | 0.342297 | 0.181 | 0.125 | 1 | 4 | Zkscan3 | 0.056 | Old |
| 0.227632 | 2.662404 | 0.038 | 0.013 | 1 | 4 | Uckl1 | 0.025 | Old |
| 0.227713 | 0.382423 | 0.329 | 0.262 | 1 | 4 | Hspd1 | 0.067 | Old |
| 0.227795 | 1.048241 | 0.117 | 0.075 | 1 | 4 | Slc4a2 | 0.042 | Old |
| 0.228741 | 0.186238 | 0.775 | 0.75 | 1 | 4 | Cd81 | 0.025 | Old |
| 0.229333 | 2.606493 | 0.018 | 0 | 1 | 4 | Hirip3 | 0.018 | Old |
| 0.229333 | 2.622582 | 0.018 | 0 | 1 | 4 | Pex12 | 0.018 | Old |
| 0.229333 | 2.621496 | 0.018 | 0 | 1 | 4 | Nckap5l | 0.018 | Old |
| 0.229333 | 2.60723 | 0.018 | 0 | 1 | 4 | Snap47 | 0.018 | Old |
| 0.229333 | 2.512355 | 0.018 | 0 | 1 | 4 | Rad1 | 0.018 | Old |
| 0.229333 | 2.364202 | 0.018 | 0 | 1 | 4 | Slc2a6 | 0.018 | Old |
| 0.229333 | 2.349519 | 0.018 | 0 | 1 | 4 | Riox1 | 0.018 | Old |
| 0.229333 | 2.278513 | 0.018 | 0 | 1 | 4 | Gorab | 0.018 | Old |
| 0.229333 | 2.231029 | 0.018 | 0 | 1 | 4 | Ercc1 | 0.018 | Old |
| 0.229333 | 2.071484 | 0.018 | 0 | 1 | 4 | Kctd17 | 0.018 | Old |
| 0.229692 | 2.324345 | 0.038 | 0.013 | 1 | 4 | Ap5s1 | 0.025 | Old |
| 0.229702 | 0.275192 | 0.153 | 0.1 | 1 | 4 | Arhgap1 | 0.053 | Old |
| 0.229775 | 0.44485 | 0.167 | 0.112 | 1 | 4 | Tcp11l2 | 0.055 | Old |
| 0.229891 | 0.062385 | 0.054 | 0.088 | 1 | 4 | Casp9 | -0.034 | Old |
| 0.230442 | 1.039923 | 0.13 | 0.088 | 1 | 4 | Gypc | 0.042 | Old |

| | | | | | | | | |
|---|---|---|---|---|---|---|---|---|
| 0.230578 | 2.212383 | 0.038 | 0.013 | 1 | 4 | Rad51d | 0.025 | Old |
| 0.230892 | 0.818949 | 0.118 | 0.075 | 1 | 4 | Rnf20 | 0.043 | Old |
| 0.230968 | 0.239184 | 0.253 | 0.2 | 1 | 4 | Pnp | 0.053 | Old |
| 0.231011 | 0.860116 | 0.171 | 0.125 | 1 | 4 | Ganab | 0.046 | Old |
| 0.231269 | 0.566721 | 0.074 | 0.038 | 1 | 4 | Pdlim2 | 0.036 | Old |
| 0.231785 | 0.614098 | 0.317 | 0.275 | 1 | 4 | Nop10 | 0.042 | Old |
| 0.231988 | 1.229014 | 0.056 | 0.025 | 1 | 4 | Gtpbp6 | 0.031 | Old |
| 0.232431 | 1.014822 | 0.039 | 0.013 | 1 | 4 | Tmem237 | 0.026 | Old |
| 0.232476 | 1.045819 | 0.072 | 0.038 | 1 | 4 | Ankrd46 | 0.034 | Old |
| 0.232483 | 1.150573 | 0.056 | 0.025 | 1 | 4 | Slc35b3 | 0.031 | Old |
| 0.233172 | 1.336019 | 0.101 | 0.062 | 1 | 4 | Cog5 | 0.039 | Old |
| 0.233253 | 1.670985 | 0.038 | 0.013 | 1 | 4 | Zkscan6 | 0.025 | Old |
| 0.233351 | 0.304786 | 0.454 | 0.388 | 1 | 4 | Fam174a | 0.066 | Old |
| 0.233512 | 1.444526 | 0.086 | 0.05 | 1 | 4 | Srebf1 | 0.036 | Old |
| 0.233656 | 0.660911 | 0.074 | 0.038 | 1 | 4 | Atg7 | 0.036 | Old |
| 0.235112 | 0.055769 | 0.093 | 0.138 | 1 | 4 | Frmd6 | -0.045 | Old |
| 0.235262 | 2.692362 | 0.038 | 0.013 | 1 | 4 | Pex14 | 0.025 | Old |
| 0.235396 | 0.188092 | 0.039 | 0.013 | 1 | 4 | Ttl | 0.026 | Old |
| 0.235813 | 0.735329 | 0.073 | 0.038 | 1 | 4 | Ccser2 | 0.035 | Old |
| 0.235949 | 1.3692 | 0.038 | 0.013 | 1 | 4 | Prkdc | 0.025 | Old |
| 0.23625 | 1.479363 | 0.038 | 0.013 | 1 | 4 | Wdr12 | 0.025 | Old |
| 0.236391 | 1.107985 | 0.072 | 0.038 | 1 | 4 | Ncbp1 | 0.034 | Old |
| 0.236392 | 0.450674 | 0.415 | 0.362 | 1 | 4 | Lcp2 | 0.053 | Old |
| 0.236591 | 0.829788 | 0.157 | 0.112 | 1 | 4 | Thra | 0.045 | Old |
| 0.236649 | 0.832652 | 0.103 | 0.062 | 1 | 4 | Ube3b | 0.041 | Old |
| 0.236877 | 0.295922 | 0.532 | 0.512 | 1 | 4 | Atp5g3 | 0.02 | Old |
| 0.237266 | 0.285576 | 0.341 | 0.275 | 1 | 4 | Capza1 | 0.066 | Old |
| 0.237422 | 1.464048 | 0.055 | 0.025 | 1 | 4 | Nfs1 | 0.03 | Old |
| 0.238092 | 0.404761 | 0.285 | 0.238 | 1 | 4 | Sh3bp1 | 0.047 | Old |
| 0.238092 | 0.383743 | 0.011 | 0.025 | 1 | 4 | Wdpcp | -0.014 | Old |
| 0.238312 | 2.887575 | 0.017 | 0 | 1 | 4 | Trmt61a | 0.017 | Old |
| 0.238312 | 2.801158 | 0.017 | 0 | 1 | 4 | Tnfsf15 | 0.017 | Old |
| 0.238312 | 2.744891 | 0.017 | 0 | 1 | 4 | Gca | 0.017 | Old |
| 0.238312 | 2.704133 | 0.017 | 0 | 1 | 4 | Cryz | 0.017 | Old |
| 0.238312 | 2.602534 | 0.017 | 0 | 1 | 4 | Fntb | 0.017 | Old |
| 0.238312 | 2.596035 | 0.017 | 0 | 1 | 4 | Snn | 0.017 | Old |
| 0.238312 | 2.545583 | 0.017 | 0 | 1 | 4 | Zscan12 | 0.017 | Old |
| 0.238312 | 2.47018 | 0.017 | 0 | 1 | 4 | Eno3 | 0.017 | Old |
| 0.238312 | 2.441716 | 0.017 | 0 | 1 | 4 | Arhgap35 | 0.017 | Old |
| 0.238312 | 2.31162 | 0.017 | 0 | 1 | 4 | Cdca8 | 0.017 | Old |
| 0.238312 | 2.187449 | 0.017 | 0 | 1 | 4 | Oas3 | 0.017 | Old |
| 0.239145 | 0.807366 | 0.21 | 0.162 | 1 | 4 | Zfc3h1 | 0.048 | Old |
| 0.23922 | 1.963258 | 0.038 | 0.013 | 1 | 4 | Ttbk2 | 0.025 | Old |
| 0.239724 | 2.360743 | 0.069 | 0.038 | 1 | 4 | Dennd2a | 0.031 | Old |
| 0.239754 | 0.529022 | 0.189 | 0.138 | 1 | 4 | Cyb5b | 0.051 | Old |

| | | | | | | | | |
|---|---|---|---|---|---|---|---|---|
| 0.239801 | 0.468212 | 0.19 | 0.138 | 1 | 4 | Ptpn11 | 0.052 | Old |
| 0.240263 | 0.414861 | 0.011 | 0.025 | 1 | 4 | Tmem177 | -0.014 | Old |
| 0.240295 | 0.42782 | 0.018 | 0.038 | 1 | 4 | Kif5c | -0.02 | Old |
| 0.240733 | 0.294067 | 0.289 | 0.238 | 1 | 4 | Trpc4ap | 0.051 | Old |
| 0.241052 | 0.351971 | 0.484 | 0.45 | 1 | 4 | Hnrnpm | 0.034 | Old |
| 0.241063 | 0.750767 | 0.118 | 0.075 | 1 | 4 | Cmc1 | 0.043 | Old |
| 0.241102 | 0.941575 | 0.038 | 0.013 | 1 | 4 | Ttc19 | 0.025 | Old |
| 0.241354 | 0.392442 | 0.011 | 0.025 | 1 | 4 | Tyw3 | -0.014 | Old |
| 0.241581 | 0.405613 | 0.286 | 0.225 | 1 | 4 | Drap1 | 0.061 | Old |
| 0.241679 | 1.574224 | 0.038 | 0.013 | 1 | 4 | Fam122b | 0.025 | Old |
| 0.242085 | 1.792211 | 0.07 | 0.038 | 1 | 4 | Golt1b | 0.032 | Old |
| 0.242215 | 1.762847 | 0.055 | 0.025 | 1 | 4 | Fam58b | 0.03 | Old |
| 0.242308 | 0.331727 | 0.502 | 0.45 | 1 | 4 | Jpt1 | 0.052 | Old |
| 0.242632 | 0.000757 | 0.166 | 0.225 | 1 | 4 | Rbfa | -0.059 | Old |
| 0.243826 | 0.658625 | 0.072 | 0.038 | 1 | 4 | Cdk5rap3 | 0.034 | Old |
| 0.244539 | 1.458589 | 0.055 | 0.025 | 1 | 4 | Gm16536 | 0.03 | Old |
| 0.244731 | 2.357318 | 0.037 | 0.013 | 1 | 4 | Il4i1 | 0.024 | Old |
| 0.244778 | 1.298402 | 0.038 | 0.013 | 1 | 4 | Supt7l | 0.025 | Old |
| 0.244897 | 0.153614 | 0.027 | 0.05 | 1 | 4 | Mcph1 | -0.023 | Old |
| 0.245017 | 0.840307 | 0.182 | 0.138 | 1 | 4 | Mllt10 | 0.044 | Old |
| 0.245058 | 1.288593 | 0.055 | 0.025 | 1 | 4 | Kat14 | 0.03 | Old |
| 0.245162 | 0.682782 | 0.281 | 0.238 | 1 | 4 | Ppp2r1a | 0.043 | Old |
| 0.245996 | 0.9947 | 0.087 | 0.05 | 1 | 4 | Tmem245 | 0.037 | Old |
| 0.245997 | 0.440543 | 0.233 | 0.175 | 1 | 4 | Sfxn3 | 0.058 | Old |
| 0.24633 | 0.696784 | 0.186 | 0.138 | 1 | 4 | Tmem219 | 0.048 | Old |
| 0.246337 | 1.126583 | 0.038 | 0.013 | 1 | 4 | Prdm15 | 0.025 | Old |
| 0.246934 | 2.113082 | 0.037 | 0.013 | 1 | 4 | Gm20707 | 0.024 | Old |
| 0.247067 | 0.501455 | 0.304 | 0.25 | 1 | 4 | Naa50 | 0.054 | Old |
| 0.24759 | 1.086532 | 0.038 | 0.013 | 1 | 4 | Wdyhv1 | 0.025 | Old |
| 0.247709 | 2.755589 | 0.016 | 0 | 1 | 4 | Zfp763 | 0.016 | Old |
| 0.247709 | 2.654937 | 0.016 | 0 | 1 | 4 | Rbak | 0.016 | Old |
| 0.247709 | 2.564694 | 0.016 | 0 | 1 | 4 | Zbtb12 | 0.016 | Old |
| 0.247709 | 2.564382 | 0.016 | 0 | 1 | 4 | Zdhhc24 | 0.016 | Old |
| 0.247709 | 2.531405 | 0.016 | 0 | 1 | 4 | Efcab2 | 0.016 | Old |
| 0.247709 | 2.465165 | 0.016 | 0 | 1 | 4 | Sars2 | 0.016 | Old |
| 0.247709 | 2.453176 | 0.016 | 0 | 1 | 4 | Zfp747 | 0.016 | Old |
| 0.247709 | 2.433899 | 0.016 | 0 | 1 | 4 | A530032D1 | 0.016 | Old |
| 0.247709 | 2.366474 | 0.016 | 0 | 1 | 4 | Nup210 | 0.016 | Old |
| 0.247709 | 2.360432 | 0.016 | 0 | 1 | 4 | Cttn | 0.016 | Old |
| 0.247709 | 2.345987 | 0.016 | 0 | 1 | 4 | Cenpt | 0.016 | Old |
| 0.247709 | 2.345179 | 0.016 | 0 | 1 | 4 | Ankzf1 | 0.016 | Old |
| 0.247709 | 2.023523 | 0.016 | 0 | 1 | 4 | Upb1 | 0.016 | Old |
| 0.247857 | 1.447858 | 0.07 | 0.038 | 1 | 4 | Calml4 | 0.032 | Old |
| 0.248532 | 0.967198 | 0.038 | 0.013 | 1 | 4 | Fyn | 0.025 | Old |
| 0.248989 | 1.334584 | 0.085 | 0.05 | 1 | 4 | Tada1 | 0.035 | Old |

| | | | | | | | | |
|---|---|---|---|---|---|---|---|---|
| 0.249122 | 0.619636 | 0.117 | 0.075 | 1 | 4 | Pex16 | 0.042 | Old |
| 0.249481 | 0.161338 | 0.547 | 0.512 | 1 | 4 | Pdcd6ip | 0.035 | Old |
| 0.249967 | 0.257309 | 0.686 | 0.675 | 1 | 4 | Alox5ap | 0.011 | Old |
| 0.250541 | 1.595832 | 0.054 | 0.025 | 1 | 4 | Nlrc4 | 0.029 | Old |
| 0.250617 | 0.676323 | 0.234 | 0.188 | 1 | 4 | Sp3 | 0.046 | Old |
| 0.250657 | 0.905157 | 0.169 | 0.125 | 1 | 4 | Puf60 | 0.044 | Old |
| 0.250744 | 1.622216 | 0.037 | 0.013 | 1 | 4 | Parp3 | 0.024 | Old |
| 0.251448 | 0.789803 | 0.248 | 0.212 | 1 | 4 | Xpr1 | 0.036 | Old |
| 0.251549 | 0.715604 | 0.208 | 0.162 | 1 | 4 | Rrp1 | 0.046 | Old |
| 0.252023 | 1.375652 | 0.037 | 0.013 | 1 | 4 | Mad2l1bp | 0.024 | Old |
| 0.252106 | 1.034296 | 0.086 | 0.05 | 1 | 4 | Hps5 | 0.036 | Old |
| 0.252811 | 0.441639 | 0.036 | 0.062 | 1 | 4 | Tomm6 | -0.026 | Old |
| 0.25295 | 0.189892 | 0.762 | 0.713 | 1 | 4 | Actr3 | 0.049 | Old |
| 0.252966 | 0.535792 | 0.279 | 0.238 | 1 | 4 | Snx27 | 0.041 | Old |
| 0.253266 | 0.043533 | 0.074 | 0.112 | 1 | 4 | Yif1b | -0.038 | Old |
| 0.253359 | 0.472521 | 0.088 | 0.05 | 1 | 4 | Commd1 | 0.038 | Old |
| 0.253384 | 0.745943 | 0.117 | 0.075 | 1 | 4 | Irf4 | 0.042 | Old |
| 0.253665 | 0.680226 | 0.264 | 0.225 | 1 | 4 | Chmp3 | 0.039 | Old |
| 0.253735 | 1.323266 | 0.054 | 0.025 | 1 | 4 | Ndufaf2 | 0.029 | Old |
| 0.254102 | 0.407196 | 0.34 | 0.3 | 1 | 4 | Tep1 | 0.04 | Old |
| 0.254259 | 0.841344 | 0.055 | 0.025 | 1 | 4 | Ankrd16 | 0.03 | Old |
| 0.254272 | 1.139244 | 0.037 | 0.013 | 1 | 4 | Pcyox1l | 0.024 | Old |
| 0.25428 | 0.012127 | 0.144 | 0.2 | 1 | 4 | Agpat3 | -0.056 | Old |
| 0.254538 | 1.370805 | 0.054 | 0.025 | 1 | 4 | Ipo13 | 0.029 | Old |
| 0.256971 | 0.733423 | 0.101 | 0.062 | 1 | 4 | Itpr2 | 0.039 | Old |
| 0.257552 | 2.993621 | 0.016 | 0 | 1 | 4 | Ttc8 | 0.016 | Old |
| 0.257552 | 2.737081 | 0.016 | 0 | 1 | 4 | Osgepl1 | 0.016 | Old |
| 0.257552 | 2.596938 | 0.016 | 0 | 1 | 4 | Adap2os | 0.016 | Old |
| 0.257552 | 2.595965 | 0.016 | 0 | 1 | 4 | Lss | 0.016 | Old |
| 0.257552 | 2.558623 | 0.016 | 0 | 1 | 4 | Meis1 | 0.016 | Old |
| 0.257552 | 2.539389 | 0.016 | 0 | 1 | 4 | Zfp41 | 0.016 | Old |
| 0.257552 | 2.537024 | 0.016 | 0 | 1 | 4 | Zfp976 | 0.016 | Old |
| 0.257552 | 2.524521 | 0.016 | 0 | 1 | 4 | Ndufaf6 | 0.016 | Old |
| 0.257552 | 2.500926 | 0.016 | 0 | 1 | 4 | Homez | 0.016 | Old |
| 0.257552 | 2.492312 | 0.016 | 0 | 1 | 4 | Tmem62 | 0.016 | Old |
| 0.257552 | 2.466916 | 0.016 | 0 | 1 | 4 | Afg1l | 0.016 | Old |
| 0.257552 | 2.463601 | 0.016 | 0 | 1 | 4 | Zbtb39 | 0.016 | Old |
| 0.257552 | 2.43713 | 0.016 | 0 | 1 | 4 | Zfp775 | 0.016 | Old |
| 0.257552 | 2.430701 | 0.016 | 0 | 1 | 4 | Tmlhe | 0.016 | Old |
| 0.257552 | 2.420277 | 0.016 | 0 | 1 | 4 | Gm35154 | 0.016 | Old |
| 0.257552 | 2.37865 | 0.016 | 0 | 1 | 4 | Ahi1 | 0.016 | Old |
| 0.257552 | 2.361563 | 0.016 | 0 | 1 | 4 | Ubiad1 | 0.016 | Old |
| 0.257552 | 2.336055 | 0.016 | 0 | 1 | 4 | Gm4070 | 0.016 | Old |
| 0.257552 | 2.253186 | 0.016 | 0 | 1 | 4 | Zgpat | 0.016 | Old |
| 0.257552 | 2.241967 | 0.016 | 0 | 1 | 4 | Scn2b | 0.016 | Old |

| | | | | | | | | |
|---|---|---|---|---|---|---|---|---|
| 0.257552 | 1.993376 | 0.016 | 0 | 1 | 4 | Spink2 | 0.016 | Old |
| 0.257552 | 1.934794 | 0.016 | 0 | 1 | 4 | Itgb7 | 0.016 | Old |
| 0.257552 | 1.908438 | 0.016 | 0 | 1 | 4 | Sirpb1c | 0.016 | Old |
| 0.257597 | 0.596317 | 0.086 | 0.05 | 1 | 4 | Mrpl21 | 0.036 | Old |
| 0.257677 | 0.551332 | 0.298 | 0.25 | 1 | 4 | Capg | 0.048 | Old |
| 0.25817 | 1.502991 | 0.111 | 0.075 | 1 | 4 | Aldh16a1 | 0.036 | Old |
| 0.258208 | 1.151403 | 0.113 | 0.075 | 1 | 4 | Edem2 | 0.038 | Old |
| 0.258725 | 0.156715 | 0.587 | 0.55 | 1 | 4 | Sfpq | 0.037 | Old |
| 0.258918 | 0.69957 | 0.117 | 0.075 | 1 | 4 | Smc5 | 0.042 | Old |
| 0.259122 | 0.057118 | 0.135 | 0.188 | 1 | 4 | Flcn | -0.053 | Old |
| 0.259193 | 0.557514 | 0.07 | 0.038 | 1 | 4 | Pgp | 0.032 | Old |
| 0.259265 | 0.042544 | 0.037 | 0.062 | 1 | 4 | Wdr75 | -0.025 | Old |
| 0.259301 | 0.345733 | 0.203 | 0.15 | 1 | 4 | Mfhas1 | 0.053 | Old |
| 0.259394 | 0.936558 | 0.054 | 0.025 | 1 | 4 | Snhg20 | 0.029 | Old |
| 0.259609 | 0.102237 | 0.028 | 0.05 | 1 | 4 | Ttf2 | -0.022 | Old |
| 0.259938 | 0.913359 | 0.054 | 0.025 | 1 | 4 | Zfp746 | 0.029 | Old |
| 0.26006 | 0.447154 | 0.321 | 0.275 | 1 | 4 | Lima1 | 0.046 | Old |
| 0.26063 | 0.743459 | 0.07 | 0.038 | 1 | 4 | Mkl1 | 0.032 | Old |
| 0.260727 | 0.825697 | 0.004 | 0.013 | 1 | 4 | Crisp1 | -0.009 | Old |
| 0.260784 | 1.506679 | 0.053 | 0.025 | 1 | 4 | Pno1 | 0.028 | Old |
| 0.26087 | 0.79569 | 0.07 | 0.038 | 1 | 4 | Tubgcp3 | 0.032 | Old |
| 0.261112 | 0.3692 | 0.35 | 0.3 | 1 | 4 | Srsf6 | 0.05 | Old |
| 0.261424 | 0.61669 | 0.235 | 0.188 | 1 | 4 | Ncstn | 0.047 | Old |
| 0.26165 | 0.2222 | 0.004 | 0.013 | 1 | 4 | Traf3ip3 | -0.009 | Old |
| 0.26165 | 0.087394 | 0.004 | 0.013 | 1 | 4 | Npr1 | -0.009 | Old |
| 0.26165 | 0.08641 | 0.004 | 0.013 | 1 | 4 | 1700010I1 | -0.009 | Old |
| 0.26165 | 0.026872 | 0.004 | 0.013 | 1 | 4 | Meox2 | -0.009 | Old |
| 0.26167 | 1.377574 | 0.111 | 0.075 | 1 | 4 | Slc41a2 | 0.036 | Old |
| 0.262574 | 0.301327 | 0.004 | 0.013 | 1 | 4 | Rad9b | -0.009 | Old |
| 0.262574 | 0.195398 | 0.004 | 0.013 | 1 | 4 | Cdk5rap1 | -0.009 | Old |
| 0.262574 | 0.188912 | 0.004 | 0.013 | 1 | 4 | Fbxo5 | -0.009 | Old |
| 0.262574 | 0.111156 | 0.004 | 0.013 | 1 | 4 | Tctex1d4 | -0.009 | Old |
| 0.263209 | 0.311558 | 0.163 | 0.112 | 1 | 4 | 4933434E2 | 0.051 | Old |
| 0.264111 | 1.023335 | 0.176 | 0.138 | 1 | 4 | Ankhd1 | 0.038 | Old |
| 0.264118 | 1.63194 | 0.053 | 0.025 | 1 | 4 | Memo1 | 0.028 | Old |
| 0.264807 | 0.302758 | 0.31 | 0.25 | 1 | 4 | Ids | 0.06 | Old |
| 0.264876 | 0.415903 | 0.204 | 0.15 | 1 | 4 | Ccr2 | 0.054 | Old |
| 0.265191 | 0.246411 | 0.215 | 0.162 | 1 | 4 | Rex1bd | 0.053 | Old |
| 0.265449 | 0.12036 | 0.019 | 0.038 | 1 | 4 | BC049352 | -0.019 | Old |
| 0.265449 | 0.055947 | 0.019 | 0.038 | 1 | 4 | Slc25a53 | -0.019 | Old |
| 0.265486 | 1.020957 | 0.036 | 0.013 | 1 | 4 | Brms1l | 0.023 | Old |
| 0.265533 | 1.161978 | 0.069 | 0.038 | 1 | 4 | Rtp4 | 0.031 | Old |
| 0.265821 | 1.019153 | 0.036 | 0.013 | 1 | 4 | Zbtb41 | 0.023 | Old |
| 0.265847 | 0.725178 | 0.182 | 0.138 | 1 | 4 | Il11ra1 | 0.044 | Old |
| 0.266213 | 1.838791 | 0.036 | 0.013 | 1 | 4 | Tbcc | 0.023 | Old |

| | | | | | | | | |
|---|---|---|---|---|---|---|---|---|
| 0.266263 | 1.179447 | 0.084 | 0.05 | 1 | 4 | Golph3l | 0.034 | Old |
| 0.266292 | 0.158379 | 0.065 | 0.1 | 1 | 4 | Mrps7 | -0.035 | Old |
| 0.266334 | 0.791585 | 0.181 | 0.138 | 1 | 4 | Kxd1 | 0.043 | Old |
| 0.26699 | 0.402506 | 0.055 | 0.025 | 1 | 4 | Grcc10 | 0.03 | Old |
| 0.267008 | 1.068486 | 0.069 | 0.038 | 1 | 4 | Exosc9 | 0.031 | Old |
| 0.267585 | 0.360707 | 0.275 | 0.225 | 1 | 4 | Olfm1 | 0.05 | Old |
| 0.267868 | 2.673089 | 0.015 | 0 | 1 | 4 | Cldn34c1 | 0.015 | Old |
| 0.267868 | 2.654905 | 0.015 | 0 | 1 | 4 | Magee1 | 0.015 | Old |
| 0.267868 | 2.646547 | 0.015 | 0 | 1 | 4 | Shmt1 | 0.015 | Old |
| 0.267868 | 2.561322 | 0.015 | 0 | 1 | 4 | Wrap53 | 0.015 | Old |
| 0.267868 | 2.514172 | 0.015 | 0 | 1 | 4 | Cystm1 | 0.015 | Old |
| 0.267868 | 2.507182 | 0.015 | 0 | 1 | 4 | Gm13373 | 0.015 | Old |
| 0.267868 | 2.480553 | 0.015 | 0 | 1 | 4 | Traf3ip1 | 0.015 | Old |
| 0.267868 | 2.469575 | 0.015 | 0 | 1 | 4 | Phactr1 | 0.015 | Old |
| 0.267868 | 2.466681 | 0.015 | 0 | 1 | 4 | Tmem69 | 0.015 | Old |
| 0.267868 | 2.431042 | 0.015 | 0 | 1 | 4 | Ahnak2 | 0.015 | Old |
| 0.267868 | 2.39952 | 0.015 | 0 | 1 | 4 | Rad18 | 0.015 | Old |
| 0.267868 | 2.361293 | 0.015 | 0 | 1 | 4 | Insl6 | 0.015 | Old |
| 0.267868 | 2.332322 | 0.015 | 0 | 1 | 4 | Med26 | 0.015 | Old |
| 0.267868 | 2.33127 | 0.015 | 0 | 1 | 4 | Gtf2ird2 | 0.015 | Old |
| 0.267868 | 2.305462 | 0.015 | 0 | 1 | 4 | Cdiptos | 0.015 | Old |
| 0.267868 | 2.223363 | 0.015 | 0 | 1 | 4 | Abcb8 | 0.015 | Old |
| 0.267868 | 2.160168 | 0.015 | 0 | 1 | 4 | Slc25a38 | 0.015 | Old |
| 0.267905 | 1.003735 | 0.099 | 0.062 | 1 | 4 | Gkap1 | 0.037 | Old |
| 0.26805 | 0.036038 | 0.328 | 0.425 | 1 | 4 | Sdhc | -0.097 | Old |
| 0.268528 | 0.260045 | 0.64 | 0.637 | 1 | 4 | Pitpna | 0.003 | Old |
| 0.268593 | 1.401054 | 0.053 | 0.025 | 1 | 4 | Phkg2 | 0.028 | Old |
| 0.268817 | 0.987065 | 0.053 | 0.025 | 1 | 4 | Rps19bp1 | 0.028 | Old |
| 0.269462 | 0.391438 | 0.15 | 0.1 | 1 | 4 | Fabp5 | 0.05 | Old |
| 0.269987 | 0.700945 | 0.274 | 0.238 | 1 | 4 | Emc7 | 0.036 | Old |
| 0.270139 | 0.824934 | 0.191 | 0.15 | 1 | 4 | Mdm4 | 0.041 | Old |
| 0.270142 | 1.526895 | 0.082 | 0.05 | 1 | 4 | Ube2g1 | 0.032 | Old |
| 0.270288 | 0.984313 | 0.153 | 0.112 | 1 | 4 | Stxbp2 | 0.041 | Old |
| 0.270469 | 0.981008 | 0.069 | 0.038 | 1 | 4 | Daxx | 0.031 | Old |
| 0.270561 | 2.111124 | 0.067 | 0.038 | 1 | 4 | 3110082I1 | 0.029 | Old |
| 0.270775 | 0.458912 | 0.336 | 0.288 | 1 | 4 | Nme1 | 0.048 | Old |
| 0.270881 | 0.296104 | 0.101 | 0.062 | 1 | 4 | Ubap1 | 0.039 | Old |
| 0.270981 | 1.26314 | 0.036 | 0.013 | 1 | 4 | Dexi | 0.023 | Old |
| 0.271038 | 2.484712 | 0.035 | 0.013 | 1 | 4 | Zbtb45 | 0.022 | Old |
| 0.271145 | 0.450858 | 0.318 | 0.275 | 1 | 4 | Ssr1 | 0.043 | Old |
| 0.271261 | 0.211979 | 0.437 | 0.388 | 1 | 4 | Raph1 | 0.049 | Old |
| 0.272077 | 2.275155 | 0.035 | 0.013 | 1 | 4 | Park2 | 0.022 | Old |
| 0.272127 | 0.534632 | 0.037 | 0.062 | 1 | 4 | Msantd2 | -0.025 | Old |
| 0.272424 | 2.252551 | 0.035 | 0.013 | 1 | 4 | Atp8b4 | 0.022 | Old |
| 0.272515 | 0.363824 | 0.054 | 0.025 | 1 | 4 | Cdc16 | 0.029 | Old |

| | | | | | | | | |
|---|---|---|---|---|---|---|---|---|
| 0.273042 | 1.073102 | 0.036 | 0.013 | 1 | 4 | Ints8 | 0.023 | Old |
| 0.273085 | 0.793379 | 0.181 | 0.138 | 1 | 4 | Cdk6 | 0.043 | Old |
| 0.273118 | 2.545462 | 0.035 | 0.013 | 1 | 4 | Fkbp7 | 0.022 | Old |
| 0.273356 | 0.601789 | 0.198 | 0.15 | 1 | 4 | Mcts1 | 0.048 | Old |
| 0.273732 | 1.030005 | 0.036 | 0.013 | 1 | 4 | AI597479 | 0.023 | Old |
| 0.273829 | 1.050091 | 0.139 | 0.1 | 1 | 4 | Mtfr1l | 0.039 | Old |
| 0.274 | 0.97808 | 0.098 | 0.062 | 1 | 4 | Minpp1 | 0.036 | Old |
| 0.274163 | 2.251969 | 0.035 | 0.013 | 1 | 4 | Sergef | 0.022 | Old |
| 0.274163 | 2.027579 | 0.035 | 0.013 | 1 | 4 | Atp8b2 | 0.022 | Old |
| 0.274307 | 1.549804 | 0.096 | 0.062 | 1 | 4 | Tmem19 | 0.034 | Old |
| 0.275209 | 1.889475 | 0.035 | 0.013 | 1 | 4 | Mrps2 | 0.022 | Old |
| 0.275562 | 0.495698 | 0.362 | 0.325 | 1 | 4 | Tnrc18 | 0.037 | Old |
| 0.276028 | 0.129774 | 0.773 | 0.713 | 1 | 4 | Rps17 | 0.06 | Old |
| 0.276142 | 0.436534 | 0.522 | 0.512 | 1 | 4 | Arf1 | 0.01 | Old |
| 0.276288 | 0.857461 | 0.214 | 0.175 | 1 | 4 | Mif | 0.039 | Old |
| 0.27631 | 0.11463 | 0.076 | 0.112 | 1 | 4 | Tmem64 | -0.036 | Old |
| 0.276364 | 0.564268 | 0.306 | 0.262 | 1 | 4 | Kif1b | 0.044 | Old |
| 0.277353 | 0.428626 | 0.146 | 0.1 | 1 | 4 | Mrpl36 | 0.046 | Old |
| 0.277426 | 0.033174 | 0.228 | 0.3 | 1 | 4 | Csnk1e | -0.072 | Old |
| 0.278273 | 1.071804 | 0.097 | 0.062 | 1 | 4 | Zcchc17 | 0.035 | Old |
| 0.27828 | 0.281907 | 0.656 | 0.662 | 1 | 4 | Ptms | -0.006 | Old |
| 0.278517 | 0.198227 | 0.619 | 0.537 | 1 | 4 | H2afz | 0.082 | Old |
| 0.278692 | 2.615954 | 0.014 | 0 | 1 | 4 | Gm29340 | 0.014 | Old |
| 0.278692 | 2.529223 | 0.014 | 0 | 1 | 4 | Dclk1 | 0.014 | Old |
| 0.278692 | 2.492276 | 0.014 | 0 | 1 | 4 | Dzip1 | 0.014 | Old |
| 0.278692 | 2.490846 | 0.014 | 0 | 1 | 4 | Phlpp1 | 0.014 | Old |
| 0.278692 | 2.442252 | 0.014 | 0 | 1 | 4 | Nup37 | 0.014 | Old |
| 0.278692 | 2.414738 | 0.014 | 0 | 1 | 4 | Tigar | 0.014 | Old |
| 0.278692 | 2.386332 | 0.014 | 0 | 1 | 4 | Phldb1 | 0.014 | Old |
| 0.278692 | 2.386308 | 0.014 | 0 | 1 | 4 | Rab3ip | 0.014 | Old |
| 0.278692 | 2.376954 | 0.014 | 0 | 1 | 4 | Fam20a | 0.014 | Old |
| 0.278692 | 2.345091 | 0.014 | 0 | 1 | 4 | Siah1b | 0.014 | Old |
| 0.278692 | 2.303234 | 0.014 | 0 | 1 | 4 | Rabl3 | 0.014 | Old |
| 0.278692 | 2.298318 | 0.014 | 0 | 1 | 4 | Tsen54 | 0.014 | Old |
| 0.278692 | 2.290536 | 0.014 | 0 | 1 | 4 | Nemp2 | 0.014 | Old |
| 0.278692 | 2.272171 | 0.014 | 0 | 1 | 4 | Morn2 | 0.014 | Old |
| 0.278692 | 2.268931 | 0.014 | 0 | 1 | 4 | Zfp764 | 0.014 | Old |
| 0.278692 | 2.256242 | 0.014 | 0 | 1 | 4 | Abca6 | 0.014 | Old |
| 0.278692 | 2.185873 | 0.014 | 0 | 1 | 4 | Chek2 | 0.014 | Old |
| 0.278692 | 2.078783 | 0.014 | 0 | 1 | 4 | Asb4 | 0.014 | Old |
| 0.278692 | 1.929695 | 0.014 | 0 | 1 | 4 | Ptger2 | 0.014 | Old |
| 0.278904 | 0.567704 | 0.248 | 0.2 | 1 | 4 | Gpcpd1 | 0.048 | Old |
| 0.279041 | 0.174774 | 0.465 | 0.412 | 1 | 4 | Colgalt1 | 0.053 | Old |
| 0.279065 | 0.446077 | 0.269 | 0.225 | 1 | 4 | Papd4 | 0.044 | Old |
| 0.279085 | 0.120728 | 0.258 | 0.2 | 1 | 4 | Tbl1xr1 | 0.058 | Old |

| | | | | | | | | |
|---|---|---|---|---|---|---|---|---|
| 0.279359 | 0.31262 | 0.472 | 0.45 | 1 | 4 | Tma7 | 0.022 | Old |
| 0.279532 | 1.281353 | 0.052 | 0.025 | 1 | 4 | G6pc3 | 0.027 | Old |
| 0.279777 | 1.487539 | 0.035 | 0.013 | 1 | 4 | Qsox2 | 0.022 | Old |
| 0.279845 | 0.522115 | 0.224 | 0.175 | 1 | 4 | Pfdn2 | 0.049 | Old |
| 0.280482 | 1.168825 | 0.111 | 0.075 | 1 | 4 | Lsm7 | 0.036 | Old |
| 0.280697 | 1.201117 | 0.052 | 0.025 | 1 | 4 | Ints14 | 0.027 | Old |
| 0.280869 | 0.866044 | 0.099 | 0.062 | 1 | 4 | Fibp | 0.037 | Old |
| 0.280983 | 0.063496 | 0.038 | 0.062 | 1 | 4 | Lin9 | -0.024 | Old |
| 0.281047 | 0.400335 | 0.291 | 0.25 | 1 | 4 | Fem1c | 0.041 | Old |
| 0.281193 | 1.190279 | 0.035 | 0.013 | 1 | 4 | Zdhhc12 | 0.022 | Old |
| 0.281469 | 0.602168 | 0.113 | 0.075 | 1 | 4 | Hint2 | 0.038 | Old |
| 0.281659 | 2.65228 | 0.034 | 0.013 | 1 | 4 | Ttll4 | 0.021 | Old |
| 0.281858 | 0.540394 | 0.255 | 0.212 | 1 | 4 | Cnn2 | 0.043 | Old |
| 0.282614 | 0.464502 | 0.241 | 0.188 | 1 | 4 | Eif3m | 0.053 | Old |
| 0.282736 | 0.98526 | 0.083 | 0.05 | 1 | 4 | Umad1 | 0.033 | Old |
| 0.283094 | 2.116092 | 0.034 | 0.013 | 1 | 4 | Nr1d1 | 0.021 | Old |
| 0.283149 | 0.1147 | 0.706 | 0.725 | 1 | 4 | Efhd2 | -0.019 | Old |
| 0.283419 | 0.332874 | 0.391 | 0.338 | 1 | 4 | Tram1 | 0.053 | Old |
| 0.283578 | 0.646709 | 0.221 | 0.175 | 1 | 4 | Galnt7 | 0.046 | Old |
| 0.284064 | 0.669167 | 0.205 | 0.162 | 1 | 4 | Mgat4b | 0.043 | Old |
| 0.284173 | 2.237003 | 0.034 | 0.013 | 1 | 4 | Tab1 | 0.021 | Old |
| 0.284206 | 0.77521 | 0.179 | 0.138 | 1 | 4 | Commd7 | 0.041 | Old |
| 0.284231 | 0.573859 | 0.115 | 0.075 | 1 | 4 | Naa60 | 0.04 | Old |
| 0.284362 | 0.8619 | 0.097 | 0.062 | 1 | 4 | Atl3 | 0.035 | Old |
| 0.284763 | 0.087594 | 0.158 | 0.112 | 1 | 4 | Pcmtd2 | 0.046 | Old |
| 0.28482 | 0.43905 | 0.167 | 0.125 | 1 | 4 | Eif2s3y | 0.042 | Old |
| 0.285176 | 0.763466 | 0.181 | 0.138 | 1 | 4 | AB124611 | 0.043 | Old |
| 0.285308 | 0.41072 | 0.028 | 0.05 | 1 | 4 | Trmt11 | -0.022 | Old |
| 0.286501 | 0.576067 | 0.258 | 0.212 | 1 | 4 | Tspan14 | 0.046 | Old |
| 0.28716 | 0.92354 | 0.188 | 0.15 | 1 | 4 | Madd | 0.038 | Old |
| 0.287264 | 0.676128 | 0.035 | 0.013 | 1 | 4 | Cdc42ep2 | 0.022 | Old |
| 0.287293 | 1.218777 | 0.051 | 0.025 | 1 | 4 | Zfp943 | 0.026 | Old |
| 0.287623 | 0.689101 | 0.035 | 0.013 | 1 | 4 | Zc2hc1a | 0.022 | Old |
| 0.2877 | 0.876675 | 0.083 | 0.05 | 1 | 4 | Secisbp2 | 0.033 | Old |
| 0.287704 | 1.938613 | 0.051 | 0.025 | 1 | 4 | Sh2b1 | 0.026 | Old |
| 0.28805 | 0.546366 | 0.247 | 0.2 | 1 | 4 | Nbeal1 | 0.047 | Old |
| 0.288285 | 0.308263 | 0.353 | 0.3 | 1 | 4 | Eif1a | 0.053 | Old |
| 0.288363 | 0.116997 | 0.106 | 0.15 | 1 | 4 | Kdm5c | -0.044 | Old |
| 0.289409 | 0.582922 | 0.084 | 0.05 | 1 | 4 | Asl | 0.034 | Old |
| 0.289476 | 0.187912 | 0.726 | 0.725 | 1 | 4 | Itm2c | 0.001 | Old |
| 0.289883 | 0.634192 | 0.084 | 0.05 | 1 | 4 | Mrpl32 | 0.034 | Old |
| 0.289934 | 0.337261 | 0.298 | 0.262 | 1 | 4 | Zmynd8 | 0.036 | Old |
| 0.290015 | 0.913908 | 0.111 | 0.075 | 1 | 4 | Agap1 | 0.036 | Old |
| 0.290015 | 0.034626 | 0.107 | 0.15 | 1 | 4 | Zfp592 | -0.043 | Old |
| 0.290057 | 2.365326 | 0.014 | 0 | 1 | 4 | Tmem81 | 0.014 | Old |

| | | | | | | | | |
|---|---|---|---|---|---|---|---|---|
| 0.290057 | 2.787548 | 0.014 | 0 | 1 | 4 | Pcdhb22 | 0.014 | Old |
| 0.290057 | 2.638738 | 0.014 | 0 | 1 | 4 | Gm38251 | 0.014 | Old |
| 0.290057 | 2.628846 | 0.014 | 0 | 1 | 4 | Timp3 | 0.014 | Old |
| 0.290057 | 2.553086 | 0.014 | 0 | 1 | 4 | Cavin3 | 0.014 | Old |
| 0.290057 | 2.496377 | 0.014 | 0 | 1 | 4 | Tmem51os | 0.014 | Old |
| 0.290057 | 2.443422 | 0.014 | 0 | 1 | 4 | Klhl25 | 0.014 | Old |
| 0.290057 | 2.329456 | 0.014 | 0 | 1 | 4 | Usp20 | 0.014 | Old |
| 0.290057 | 2.325364 | 0.014 | 0 | 1 | 4 | Cdyl2 | 0.014 | Old |
| 0.290057 | 2.305244 | 0.014 | 0 | 1 | 4 | Zfp677 | 0.014 | Old |
| 0.290057 | 2.251715 | 0.014 | 0 | 1 | 4 | Acot7 | 0.014 | Old |
| 0.290057 | 2.229399 | 0.014 | 0 | 1 | 4 | Cacna1f | 0.014 | Old |
| 0.290057 | 2.226941 | 0.014 | 0 | 1 | 4 | Nmnat1 | 0.014 | Old |
| 0.290057 | 2.189615 | 0.014 | 0 | 1 | 4 | Gm17025 | 0.014 | Old |
| 0.290057 | 2.175829 | 0.014 | 0 | 1 | 4 | Lamb2 | 0.014 | Old |
| 0.290057 | 2.1684 | 0.014 | 0 | 1 | 4 | Zfp59 | 0.014 | Old |
| 0.290057 | 2.118443 | 0.014 | 0 | 1 | 4 | Tmem191c | 0.014 | Old |
| 0.290057 | 2.045647 | 0.014 | 0 | 1 | 4 | Ccl27a | 0.014 | Old |
| 0.290057 | 2.03305 | 0.014 | 0 | 1 | 4 | Mocos | 0.014 | Old |
| 0.290057 | 2.008915 | 0.014 | 0 | 1 | 4 | Tmem42 | 0.014 | Old |
| 0.290057 | 1.986568 | 0.014 | 0 | 1 | 4 | Ccdc69 | 0.014 | Old |
| 0.290057 | 1.951988 | 0.014 | 0 | 1 | 4 | Prdm11 | 0.014 | Old |
| 0.290702 | 0.648965 | 0.099 | 0.062 | 1 | 4 | Pld2 | 0.037 | Old |
| 0.290817 | 0.029307 | 0.2 | 0.262 | 1 | 4 | Tet2 | -0.062 | Old |
| 0.291072 | 0.541685 | 0.084 | 0.05 | 1 | 4 | Celf4 | 0.034 | Old |
| 0.29193 | 0.221214 | 0.238 | 0.175 | 1 | 4 | Ecm1 | 0.063 | Old |
| 0.291934 | 0.743244 | 0.153 | 0.112 | 1 | 4 | Lrch3 | 0.041 | Old |
| 0.291972 | 2.214229 | 0.034 | 0.013 | 1 | 4 | Ramp1 | 0.021 | Old |
| 0.292252 | 0.592245 | 0.099 | 0.062 | 1 | 4 | Pdcl | 0.037 | Old |
| 0.292343 | 2.366876 | 0.034 | 0.013 | 1 | 4 | Aldh1a2 | 0.021 | Old |
| 0.292532 | 1.45921 | 0.051 | 0.025 | 1 | 4 | Cops7b | 0.026 | Old |
| 0.292628 | 0.345669 | 0.227 | 0.188 | 1 | 4 | Ergic3 | 0.039 | Old |
| 0.292713 | 2.127997 | 0.034 | 0.013 | 1 | 4 | Ppp2r3a | 0.021 | Old |
| 0.292835 | 1.467984 | 0.051 | 0.025 | 1 | 4 | Prr3 | 0.026 | Old |
| 0.292842 | 1.354371 | 0.095 | 0.062 | 1 | 4 | Virma | 0.033 | Old |
| 0.292917 | 0.724743 | 0.177 | 0.138 | 1 | 4 | Ankrd40 | 0.039 | Old |
| 0.293398 | 1.07351 | 0.082 | 0.05 | 1 | 4 | Lrig2 | 0.032 | Old |
| 0.293464 | 0.270326 | 0.602 | 0.588 | 1 | 4 | B4galt6 | 0.014 | Old |
| 0.293828 | 2.266699 | 0.034 | 0.013 | 1 | 4 | Orai2 | 0.021 | Old |
| 0.293854 | 0.093149 | 0.824 | 0.812 | 1 | 4 | Arpc2 | 0.012 | Old |
| 0.293899 | 0.911917 | 0.051 | 0.025 | 1 | 4 | Nop9 | 0.026 | Old |
| 0.294151 | 0.899506 | 0.082 | 0.05 | 1 | 4 | Tmem203 | 0.032 | Old |
| 0.294572 | 2.185638 | 0.034 | 0.013 | 1 | 4 | Pnpla6 | 0.021 | Old |
| 0.294572 | 2.059831 | 0.034 | 0.013 | 1 | 4 | Gpr180 | 0.021 | Old |
| 0.294808 | 0.921382 | 0.051 | 0.025 | 1 | 4 | Dvl2 | 0.026 | Old |
| 0.295132 | 0.333647 | 0.435 | 0.388 | 1 | 4 | Khdrbs1 | 0.047 | Old |

| | | | | | | | | |
|---|---|---|---|---|---|---|---|---|
| 0.295984 | 0.469519 | 0.099 | 0.062 | 1 | 4 | Mbp | 0.037 | Old |
| 0.296056 | 0.879816 | 0.285 | 0.262 | 1 | 4 | Rassf3 | 0.023 | Old |
| 0.296105 | 0.156198 | 0.416 | 0.375 | 1 | 4 | Snrpb | 0.041 | Old |
| 0.296216 | 0.380367 | 0.403 | 0.362 | 1 | 4 | Dync1i2 | 0.041 | Old |
| 0.296418 | 0.330189 | 0.225 | 0.175 | 1 | 4 | 6-Mar | 0.05 | Old |
| 0.29724 | 0.765314 | 0.051 | 0.025 | 1 | 4 | Rnf31 | 0.026 | Old |
| 0.297426 | 0.421733 | 0.209 | 0.162 | 1 | 4 | 4932438A1 | 0.047 | Old |
| 0.297568 | 2.14011 | 0.05 | 0.025 | 1 | 4 | Ppil3 | 0.025 | Old |
| 0.29811 | 0.331846 | 0.247 | 0.2 | 1 | 4 | Ubash3b | 0.047 | Old |
| 0.298335 | 0.24558 | 0.729 | 0.725 | 1 | 4 | Hnrnpk | 0.004 | Old |
| 0.298461 | 0.654196 | 0.034 | 0.013 | 1 | 4 | Zfp141 | 0.021 | Old |
| 0.29869 | 1.46509 | 0.034 | 0.013 | 1 | 4 | Gdf3 | 0.021 | Old |
| 0.299268 | 0.41743 | 0.393 | 0.35 | 1 | 4 | Rab11a | 0.043 | Old |
| 0.299442 | 1.452958 | 0.034 | 0.013 | 1 | 4 | Tysnd1 | 0.021 | Old |
| 0.299693 | 0.174049 | 0.291 | 0.225 | 1 | 4 | Gm26740 | 0.066 | Old |
| 0.299819 | 1.566584 | 0.034 | 0.013 | 1 | 4 | Ano8 | 0.021 | Old |
| 0.299872 | 0.009923 | 0.138 | 0.188 | 1 | 4 | Ift20 | -0.05 | Old |
| 0.300169 | 0.945028 | 0.067 | 0.038 | 1 | 4 | Zscan29 | 0.029 | Old |
| 0.300759 | 0.431415 | 0.358 | 0.312 | 1 | 4 | Tmem256 | 0.046 | Old |
| 0.300966 | 1.744846 | 0.065 | 0.038 | 1 | 4 | St7l | 0.027 | Old |
| 0.301614 | 0.945032 | 0.136 | 0.1 | 1 | 4 | Pigp | 0.036 | Old |
| 0.302005 | 2.625961 | 0.013 | 0 | 1 | 4 | Ggnbp1 | 0.013 | Old |
| 0.302005 | 2.598548 | 0.013 | 0 | 1 | 4 | Cadm3 | 0.013 | Old |
| 0.302005 | 2.540338 | 0.013 | 0 | 1 | 4 | Tmem132c | 0.013 | Old |
| 0.302005 | 2.444067 | 0.013 | 0 | 1 | 4 | Ccdc138 | 0.013 | Old |
| 0.302005 | 2.397941 | 0.013 | 0 | 1 | 4 | Tnfaip8l1 | 0.013 | Old |
| 0.302005 | 2.366383 | 0.013 | 0 | 1 | 4 | Sgms2 | 0.013 | Old |
| 0.302005 | 2.3544 | 0.013 | 0 | 1 | 4 | Gm15564 | 0.013 | Old |
| 0.302005 | 2.333253 | 0.013 | 0 | 1 | 4 | Hgh1 | 0.013 | Old |
| 0.302005 | 2.319542 | 0.013 | 0 | 1 | 4 | Tmem202 | 0.013 | Old |
| 0.302005 | 2.306035 | 0.013 | 0 | 1 | 4 | 4931414P1 | 0.013 | Old |
| 0.302005 | 2.276204 | 0.013 | 0 | 1 | 4 | Trim16 | 0.013 | Old |
| 0.302005 | 2.25875 | 0.013 | 0 | 1 | 4 | Slc25a42 | 0.013 | Old |
| 0.302005 | 2.257704 | 0.013 | 0 | 1 | 4 | Trp53rkb | 0.013 | Old |
| 0.302005 | 2.220932 | 0.013 | 0 | 1 | 4 | Tcaim | 0.013 | Old |
| 0.302005 | 2.213912 | 0.013 | 0 | 1 | 4 | Fam185a | 0.013 | Old |
| 0.302005 | 2.203963 | 0.013 | 0 | 1 | 4 | Vwf | 0.013 | Old |
| 0.302005 | 2.19874 | 0.013 | 0 | 1 | 4 | Srpx | 0.013 | Old |
| 0.302005 | 2.197999 | 0.013 | 0 | 1 | 4 | Gm14326 | 0.013 | Old |
| 0.302005 | 2.181773 | 0.013 | 0 | 1 | 4 | Plac8 | 0.013 | Old |
| 0.302005 | 2.178657 | 0.013 | 0 | 1 | 4 | A430106G1 | 0.013 | Old |
| 0.302005 | 2.116728 | 0.013 | 0 | 1 | 4 | Apex2 | 0.013 | Old |
| 0.302005 | 2.096726 | 0.013 | 0 | 1 | 4 | BC051226 | 0.013 | Old |
| 0.302005 | 2.094291 | 0.013 | 0 | 1 | 4 | Plcxd2 | 0.013 | Old |
| 0.302005 | 2.086028 | 0.013 | 0 | 1 | 4 | Acsf3 | 0.013 | Old |

| | | | | | | | | |
|---|---|---|---|---|---|---|---|---|
| 0.302005 | 2.066431 | 0.013 | 0 | 1 | 4 | Dph2 | 0.013 | Old |
| 0.302005 | 2.056574 | 0.013 | 0 | 1 | 4 | Dhrs11 | 0.013 | Old |
| 0.302005 | 1.985251 | 0.013 | 0 | 1 | 4 | Taf4b | 0.013 | Old |
| 0.302005 | 1.966904 | 0.013 | 0 | 1 | 4 | Pianp | 0.013 | Old |
| 0.302005 | 1.72588 | 0.013 | 0 | 1 | 4 | Atp2a3 | 0.013 | Old |
| 0.302366 | 0.682333 | 0.097 | 0.062 | 1 | 4 | Ppp4r1 | 0.035 | Old |
| 0.302519 | 1.023246 | 0.096 | 0.062 | 1 | 4 | Cyhr1 | 0.034 | Old |
| 0.302681 | 0.560737 | 0.261 | 0.225 | 1 | 4 | Ap2b1 | 0.036 | Old |
| 0.30273 | 0.399755 | 0.125 | 0.088 | 1 | 4 | Noct | 0.037 | Old |
| 0.302889 | 0.494953 | 0.312 | 0.275 | 1 | 4 | Mcub | 0.037 | Old |
| 0.30311 | 0.115386 | 0.73 | 0.688 | 1 | 4 | Rrbp1 | 0.042 | Old |
| 0.303449 | 1.506212 | 0.08 | 0.05 | 1 | 4 | Tcim | 0.03 | Old |
| 0.303454 | 2.327241 | 0.033 | 0.013 | 1 | 4 | Bloc1s3 | 0.02 | Old |
| 0.303838 | 2.469819 | 0.033 | 0.013 | 1 | 4 | Tns2 | 0.02 | Old |
| 0.304075 | 0.515368 | 0.209 | 0.162 | 1 | 4 | Uba2 | 0.047 | Old |
| 0.304153 | 0.50087 | 0.196 | 0.15 | 1 | 4 | Dnaaf3 | 0.046 | Old |
| 0.304222 | 2.206825 | 0.033 | 0.013 | 1 | 4 | Strada | 0.02 | Old |
| 0.304251 | 1.047794 | 0.096 | 0.062 | 1 | 4 | B4galnt1 | 0.034 | Old |
| 0.304263 | 0.233579 | 0.129 | 0.088 | 1 | 4 | Smim27 | 0.041 | Old |
| 0.305232 | 1.088703 | 0.066 | 0.038 | 1 | 4 | Mapk8 | 0.028 | Old |
| 0.306073 | 0.264614 | 0.52 | 0.5 | 1 | 4 | M6pr | 0.02 | Old |
| 0.306206 | 1.411139 | 0.065 | 0.038 | 1 | 4 | Kif16b | 0.027 | Old |
| 0.306238 | 0.588885 | 0.111 | 0.075 | 1 | 4 | Eed | 0.036 | Old |
| 0.306912 | 0.191155 | 0.684 | 0.65 | 1 | 4 | Vamp8 | 0.034 | Old |
| 0.307201 | 0.383278 | 0.248 | 0.2 | 1 | 4 | Saraf | 0.048 | Old |
| 0.307339 | 1.03759 | 0.08 | 0.05 | 1 | 4 | Chst15 | 0.03 | Old |
| 0.307422 | 0.680517 | 0.034 | 0.013 | 1 | 4 | Mlh3 | 0.021 | Old |
| 0.307677 | 0.639946 | 0.179 | 0.138 | 1 | 4 | Srsf10 | 0.041 | Old |
| 0.307677 | 0.598475 | 0.179 | 0.138 | 1 | 4 | Pigk | 0.041 | Old |
| 0.308085 | 0.283486 | 0.394 | 0.362 | 1 | 4 | Pikfyve | 0.032 | Old |
| 0.308736 | 0.330948 | 0.366 | 0.35 | 1 | 4 | Lgals3 | 0.016 | Old |
| 0.309016 | 1.435344 | 0.107 | 0.075 | 1 | 4 | Mex3c | 0.032 | Old |
| 0.309105 | 0.94885 | 0.066 | 0.038 | 1 | 4 | Lmf2 | 0.028 | Old |
| 0.309443 | 0.034244 | 0.02 | 0.038 | 1 | 4 | Plxdc2 | -0.018 | Old |
| 0.309484 | 0.519835 | 0.14 | 0.1 | 1 | 4 | Arih2 | 0.04 | Old |
| 0.309753 | 0.592801 | 0.167 | 0.125 | 1 | 4 | Ipo7 | 0.042 | Old |
| 0.309999 | 0.345915 | 0.263 | 0.212 | 1 | 4 | Uhmk1 | 0.051 | Old |
| 0.310002 | 0.401869 | 0.154 | 0.112 | 1 | 4 | Rbm17 | 0.042 | Old |
| 0.310313 | 0.653015 | 0.165 | 0.125 | 1 | 4 | Tdrd7 | 0.04 | Old |
| 0.311191 | 1.283231 | 0.033 | 0.013 | 1 | 4 | Dis3 | 0.02 | Old |
| 0.311191 | 1.175592 | 0.033 | 0.013 | 1 | 4 | Rnf126 | 0.02 | Old |
| 0.311804 | 0.828225 | 0.123 | 0.088 | 1 | 4 | Cul2 | 0.035 | Old |
| 0.311908 | 1.418427 | 0.049 | 0.025 | 1 | 4 | Xrcc4 | 0.024 | Old |
| 0.312363 | 1.132622 | 0.033 | 0.013 | 1 | 4 | Hibch | 0.02 | Old |
| 0.312433 | 0.701392 | 0.123 | 0.088 | 1 | 4 | Tlk2 | 0.035 | Old |

| | | | | | | | | |
|---|---|---|---|---|---|---|---|---|
| 0.31245 | 0.543157 | 0.066 | 0.038 | 1 | 4 | Pin1 | 0.028 | Old |
| 0.312458 | 0.737451 | 0.096 | 0.062 | 1 | 4 | Arhgap11a | 0.034 | Old |
| 0.313016 | 0.097128 | 0.088 | 0.125 | 1 | 4 | Mpv17 | -0.037 | Old |
| 0.313145 | 1.182466 | 0.033 | 0.013 | 1 | 4 | Ncdn | 0.02 | Old |
| 0.313929 | 1.130903 | 0.033 | 0.013 | 1 | 4 | Rnf25 | 0.02 | Old |
| 0.314 | 0.590835 | 0.082 | 0.05 | 1 | 4 | Tusc3 | 0.032 | Old |
| 0.31422 | 2.546039 | 0.032 | 0.013 | 1 | 4 | Alyref2 | 0.019 | Old |
| 0.31422 | 2.299953 | 0.032 | 0.013 | 1 | 4 | Anln | 0.019 | Old |
| 0.314322 | 1.034623 | 0.033 | 0.013 | 1 | 4 | Ttc38 | 0.02 | Old |
| 0.314457 | 0.696461 | 0.096 | 0.062 | 1 | 4 | Flot1 | 0.034 | Old |
| 0.314541 | 0.412758 | 0.126 | 0.088 | 1 | 4 | Slu7 | 0.038 | Old |
| 0.314579 | 2.821772 | 0.013 | 0 | 1 | 4 | Ccl22 | 0.013 | Old |
| 0.314579 | 2.698933 | 0.013 | 0 | 1 | 4 | Barx2 | 0.013 | Old |
| 0.314579 | 2.391648 | 0.013 | 0 | 1 | 4 | Gpr27 | 0.013 | Old |
| 0.314579 | 2.387291 | 0.013 | 0 | 1 | 4 | AI480526 | 0.013 | Old |
| 0.314579 | 2.381431 | 0.013 | 0 | 1 | 4 | Ltbp4 | 0.013 | Old |
| 0.314579 | 2.346718 | 0.013 | 0 | 1 | 4 | Fastkd5 | 0.013 | Old |
| 0.314579 | 2.324998 | 0.013 | 0 | 1 | 4 | Thumpd2 | 0.013 | Old |
| 0.314579 | 2.321769 | 0.013 | 0 | 1 | 4 | Ahrr | 0.013 | Old |
| 0.314579 | 2.319072 | 0.013 | 0 | 1 | 4 | Mfsd9 | 0.013 | Old |
| 0.314579 | 2.304387 | 0.013 | 0 | 1 | 4 | Rab26os | 0.013 | Old |
| 0.314579 | 2.291297 | 0.013 | 0 | 1 | 4 | Tmem17 | 0.013 | Old |
| 0.314579 | 2.282629 | 0.013 | 0 | 1 | 4 | Gm42047 | 0.013 | Old |
| 0.314579 | 2.269182 | 0.013 | 0 | 1 | 4 | Tm7sf2 | 0.013 | Old |
| 0.314579 | 2.261284 | 0.013 | 0 | 1 | 4 | 9130011E1 | 0.013 | Old |
| 0.314579 | 2.199261 | 0.013 | 0 | 1 | 4 | Ttll5 | 0.013 | Old |
| 0.314579 | 2.182469 | 0.013 | 0 | 1 | 4 | Uros | 0.013 | Old |
| 0.314579 | 2.179652 | 0.013 | 0 | 1 | 4 | Tmem41a | 0.013 | Old |
| 0.314579 | 2.138396 | 0.013 | 0 | 1 | 4 | Mphosph9 | 0.013 | Old |
| 0.314579 | 2.121228 | 0.013 | 0 | 1 | 4 | Wdr4 | 0.013 | Old |
| 0.314579 | 2.111802 | 0.013 | 0 | 1 | 4 | Rps6ka2 | 0.013 | Old |
| 0.314579 | 2.093842 | 0.013 | 0 | 1 | 4 | Ptrh1 | 0.013 | Old |
| 0.314579 | 2.077684 | 0.013 | 0 | 1 | 4 | Ncs1 | 0.013 | Old |
| 0.314579 | 2.035534 | 0.013 | 0 | 1 | 4 | Bco2 | 0.013 | Old |
| 0.314579 | 2.013271 | 0.013 | 0 | 1 | 4 | 4732440D( | 0.013 | Old |
| 0.314579 | 1.971373 | 0.013 | 0 | 1 | 4 | Synpo | 0.013 | Old |
| 0.314579 | 1.939275 | 0.013 | 0 | 1 | 4 | Snapc4 | 0.013 | Old |
| 0.314579 | 1.931733 | 0.013 | 0 | 1 | 4 | Pradc1 | 0.013 | Old |
| 0.314579 | 1.92893 | 0.013 | 0 | 1 | 4 | Zc3h11a | 0.013 | Old |
| 0.314579 | 1.905683 | 0.013 | 0 | 1 | 4 | Adora2a | 0.013 | Old |
| 0.314616 | 2.202822 | 0.032 | 0.013 | 1 | 4 | Dhcr24 | 0.019 | Old |
| 0.314616 | 1.980116 | 0.032 | 0.013 | 1 | 4 | C2cd2 | 0.019 | Old |
| 0.314715 | 1.014818 | 0.033 | 0.013 | 1 | 4 | Dnajc27 | 0.02 | Old |
| 0.31483 | 0.499915 | 0.168 | 0.125 | 1 | 4 | Arid1b | 0.043 | Old |
| 0.314996 | 0.703644 | 0.202 | 0.162 | 1 | 4 | Ogfr | 0.04 | Old |

| | | | | | | | | |
|---|---|---|---|---|---|---|---|---|
| 0.315013 | 2.224742 | 0.032 | 0.013 | 1 | 4 | Ccdc88c | 0.019 | Old |
| 0.315029 | 1.643241 | 0.049 | 0.025 | 1 | 4 | Zmym1 | 0.024 | Old |
| 0.315118 | 0.283386 | 0.267 | 0.212 | 1 | 4 | Supt5 | 0.055 | Old |
| 0.315525 | 0.096027 | 0.937 | 0.912 | 1 | 4 | Atp2b1 | 0.025 | Old |
| 0.315551 | 0.430644 | 0.176 | 0.138 | 1 | 4 | Ntan1 | 0.038 | Old |
| 0.315661 | 0.474501 | 0.138 | 0.1 | 1 | 4 | Tpd52l2 | 0.038 | Old |
| 0.315696 | 0.849586 | 0.136 | 0.1 | 1 | 4 | Dpf2 | 0.036 | Old |
| 0.315824 | 0.274129 | 0.453 | 0.412 | 1 | 4 | Atp5k | 0.041 | Old |
| 0.316206 | 1.860052 | 0.032 | 0.013 | 1 | 4 | Lzts2 | 0.019 | Old |
| 0.316289 | 0.941733 | 0.033 | 0.013 | 1 | 4 | Zhx3 | 0.02 | Old |
| 0.316598 | 0.294794 | 0.02 | 0.038 | 1 | 4 | Klhl13 | -0.018 | Old |
| 0.317002 | 2.121064 | 0.032 | 0.013 | 1 | 4 | Zfp317 | 0.019 | Old |
| 0.317473 | 0.75405 | 0.033 | 0.013 | 1 | 4 | Pnpo | 0.02 | Old |
| 0.317601 | 0.700122 | 0.15 | 0.112 | 1 | 4 | Il16 | 0.038 | Old |
| 0.3178 | 2.048139 | 0.032 | 0.013 | 1 | 4 | Rbm19 | 0.019 | Old |
| 0.317891 | 0.090943 | 0.65 | 0.787 | 1 | 4 | Mat2a | -0.137 | Old |
| 0.318081 | 0.06669 | 0.31 | 0.4 | 1 | 4 | Specc1l | -0.09 | Old |
| 0.318229 | 0.687795 | 0.138 | 0.1 | 1 | 4 | Rnf38 | 0.038 | Old |
| 0.318242 | 0.581927 | 0.096 | 0.062 | 1 | 4 | Metap1 | 0.034 | Old |
| 0.318276 | 0.980529 | 0.108 | 0.075 | 1 | 4 | Slc7a7 | 0.033 | Old |
| 0.318362 | 0.398699 | 0.066 | 0.038 | 1 | 4 | Mrpl35 | 0.028 | Old |
| 0.318499 | 0.864225 | 0.122 | 0.088 | 1 | 4 | Arhgap10 | 0.034 | Old |
| 0.318573 | 0.491487 | 0.139 | 0.1 | 1 | 4 | Nr2f6 | 0.039 | Old |
| 0.318941 | 1.761624 | 0.049 | 0.025 | 1 | 4 | Gtf2e1 | 0.024 | Old |
| 0.319339 | 1.301737 | 0.065 | 0.038 | 1 | 4 | Hspb11 | 0.027 | Old |
| 0.319414 | 0.036919 | 0.049 | 0.075 | 1 | 4 | Ddx55 | -0.026 | Old |
| 0.31993 | 0.44865 | 0.155 | 0.112 | 1 | 4 | Ddx58 | 0.043 | Old |
| 0.320202 | 1.447415 | 0.032 | 0.013 | 1 | 4 | AW554918 | 0.019 | Old |
| 0.320344 | 0.336283 | 0.039 | 0.062 | 1 | 4 | Prkce | -0.023 | Old |
| 0.320528 | 0.381029 | 0.323 | 0.275 | 1 | 4 | Plxnd1 | 0.048 | Old |
| 0.320603 | 1.35328 | 0.032 | 0.013 | 1 | 4 | Haus7 | 0.019 | Old |
| 0.320853 | 0.922216 | 0.065 | 0.038 | 1 | 4 | Exosc4 | 0.027 | Old |
| 0.321005 | 1.366504 | 0.032 | 0.013 | 1 | 4 | Slc12a5 | 0.019 | Old |
| 0.321006 | 0.522729 | 0.05 | 0.025 | 1 | 4 | Noc3l | 0.025 | Old |
| 0.321407 | 1.30179 | 0.032 | 0.013 | 1 | 4 | Rabif | 0.019 | Old |
| 0.321481 | 1.047555 | 0.094 | 0.062 | 1 | 4 | Pold3 | 0.032 | Old |
| 0.321968 | 0.411826 | 0.233 | 0.188 | 1 | 4 | Clptm1l | 0.045 | Old |
| 0.322277 | 0.893104 | 0.049 | 0.025 | 1 | 4 | Polr3f | 0.024 | Old |
| 0.322885 | 0.62338 | 0.258 | 0.225 | 1 | 4 | Atp5o.1 | 0.033 | Old |
| 0.322886 | 0.346002 | 0.02 | 0.038 | 1 | 4 | Jaml | -0.018 | Old |
| 0.323077 | 1.228365 | 0.065 | 0.038 | 1 | 4 | Gart | 0.027 | Old |
| 0.323422 | 1.053148 | 0.032 | 0.013 | 1 | 4 | Tbc1d16 | 0.019 | Old |
| 0.323529 | 0.568401 | 0.179 | 0.138 | 1 | 4 | Cdc40 | 0.041 | Old |
| 0.323712 | 0.926707 | 0.108 | 0.075 | 1 | 4 | Rbpms | 0.033 | Old |
| 0.323935 | 0.374452 | 0.142 | 0.1 | 1 | 4 | Trim30d | 0.042 | Old |

| | | | | | | | | |
|---|---|---|---|---|---|---|---|---|
| 0.324082 | 0.667242 | 0.214 | 0.175 | 1 | 4 | Hbp1 | 0.039 | Old |
| 0.324513 | 0.172363 | 0.705 | 0.725 | 1 | 4 | Rbms1 | -0.02 | Old |
| 0.324778 | 0.43675 | 0.341 | 0.3 | 1 | 4 | Tbc1d14 | 0.041 | Old |
| 0.325409 | 0.487346 | 0.05 | 0.025 | 1 | 4 | Smc2 | 0.025 | Old |
| 0.32545 | 0.841585 | 0.065 | 0.038 | 1 | 4 | Cavin1 | 0.027 | Old |
| 0.326385 | 0.346646 | 0.42 | 0.388 | 1 | 4 | Set | 0.032 | Old |
| 0.326563 | 0.656508 | 0.11 | 0.075 | 1 | 4 | Lsm8 | 0.035 | Old |
| 0.327039 | 2.151806 | 0.032 | 0.013 | 1 | 4 | Rcc1 | 0.019 | Old |
| 0.327327 | 0.09543 | 0.149 | 0.2 | 1 | 4 | Yipf4 | -0.051 | Old |
| 0.327459 | 0.546115 | 0.298 | 0.262 | 1 | 4 | Bag1 | 0.036 | Old |
| 0.327631 | 0.406787 | 0.362 | 0.312 | 1 | 4 | Ccdc88a | 0.05 | Old |
| 0.32783 | 2.088159 | 0.012 | 0 | 1 | 4 | Cep112 | 0.012 | Old |
| 0.32783 | 2.624119 | 0.012 | 0 | 1 | 4 | Clec2g | 0.012 | Old |
| 0.32783 | 2.405991 | 0.012 | 0 | 1 | 4 | Zfp707 | 0.012 | Old |
| 0.32783 | 2.372063 | 0.012 | 0 | 1 | 4 | Frg2f1 | 0.012 | Old |
| 0.32783 | 2.350165 | 0.012 | 0 | 1 | 4 | 1600014C1 | 0.012 | Old |
| 0.32783 | 2.344447 | 0.012 | 0 | 1 | 4 | AC168977. | 0.012 | Old |
| 0.32783 | 2.310211 | 0.012 | 0 | 1 | 4 | Zfp975 | 0.012 | Old |
| 0.32783 | 2.270717 | 0.012 | 0 | 1 | 4 | Ss18l1 | 0.012 | Old |
| 0.32783 | 2.219083 | 0.012 | 0 | 1 | 4 | Hoxb3 | 0.012 | Old |
| 0.32783 | 2.188139 | 0.012 | 0 | 1 | 4 | Kif20b | 0.012 | Old |
| 0.32783 | 2.18588 | 0.012 | 0 | 1 | 4 | Ankrd66 | 0.012 | Old |
| 0.32783 | 2.183648 | 0.012 | 0 | 1 | 4 | Cox6a2 | 0.012 | Old |
| 0.32783 | 2.154986 | 0.012 | 0 | 1 | 4 | Sacs | 0.012 | Old |
| 0.32783 | 2.109088 | 0.012 | 0 | 1 | 4 | Nr6a1 | 0.012 | Old |
| 0.32783 | 2.102588 | 0.012 | 0 | 1 | 4 | Fkbpl | 0.012 | Old |
| 0.32783 | 2.09342 | 0.012 | 0 | 1 | 4 | Dock6 | 0.012 | Old |
| 0.32783 | 2.067368 | 0.012 | 0 | 1 | 4 | Simc1 | 0.012 | Old |
| 0.32783 | 1.995903 | 0.012 | 0 | 1 | 4 | Timp1 | 0.012 | Old |
| 0.32783 | 1.96381 | 0.012 | 0 | 1 | 4 | Oas1g | 0.012 | Old |
| 0.32783 | 1.958738 | 0.012 | 0 | 1 | 4 | Grhpr | 0.012 | Old |
| 0.32783 | 1.942068 | 0.012 | 0 | 1 | 4 | Zbtb25 | 0.012 | Old |
| 0.32783 | 1.931315 | 0.012 | 0 | 1 | 4 | Yars2 | 0.012 | Old |
| 0.32783 | 1.867313 | 0.012 | 0 | 1 | 4 | Nim1k | 0.012 | Old |
| 0.32783 | 1.792848 | 0.012 | 0 | 1 | 4 | Tyms | 0.012 | Old |
| 0.32783 | 1.677923 | 0.012 | 0 | 1 | 4 | Clcf1 | 0.012 | Old |
| 0.32783 | 1.634274 | 0.012 | 0 | 1 | 4 | Sh2d1b1 | 0.012 | Old |
| 0.32783 | 1.508384 | 0.012 | 0 | 1 | 4 | 6-Sep | 0.012 | Old |
| 0.32783 | 1.420759 | 0.012 | 0 | 1 | 4 | Slc2a1 | 0.012 | Old |
| 0.328114 | 0.70434 | 0.255 | 0.225 | 1 | 4 | Pum1 | 0.03 | Old |
| 0.328558 | 1.023295 | 0.133 | 0.1 | 1 | 4 | Slc38a9 | 0.033 | Old |
| 0.32879 | 0.170776 | 0.439 | 0.4 | 1 | 4 | Selenot | 0.039 | Old |
| 0.32919 | 0.178051 | 0.05 | 0.025 | 1 | 4 | Lsg1 | 0.025 | Old |
| 0.329518 | 0.431594 | 0.032 | 0.013 | 1 | 4 | Bloc1s5 | 0.019 | Old |
| 0.329527 | 1.164787 | 0.049 | 0.025 | 1 | 4 | Tmem199 | 0.024 | Old |

| | | | | | | | | |
|---|---|---|---|---|---|---|---|---|
| 0.329651 | 0.500463 | 0.238 | 0.2 | 1 | 4 | Srpr | 0.038 | Old |
| 0.329846 | 0.12484 | 0.159 | 0.212 | 1 | 4 | Cstf2t | -0.053 | Old |
| 0.329898 | 0.738249 | 0.198 | 0.162 | 1 | 4 | Setx | 0.036 | Old |
| 0.330157 | 0.725828 | 0.095 | 0.062 | 1 | 4 | Nabp2 | 0.033 | Old |
| 0.330381 | 0.521119 | 0.065 | 0.038 | 1 | 4 | Dhx8 | 0.027 | Old |
| 0.330531 | 1.005592 | 0.049 | 0.025 | 1 | 4 | 9130401M | 0.024 | Old |
| 0.330671 | 1.140532 | 0.078 | 0.05 | 1 | 4 | Dctn6 | 0.028 | Old |
| 0.331161 | 0.499833 | 0.109 | 0.075 | 1 | 4 | Usp3 | 0.034 | Old |
| 0.331396 | 1.060253 | 0.106 | 0.075 | 1 | 4 | Pes1 | 0.031 | Old |
| 0.331468 | 0.450824 | 0.151 | 0.112 | 1 | 4 | Zcchc2 | 0.039 | Old |
| 0.331971 | 0.203963 | 0.687 | 0.675 | 1 | 4 | Pmp22 | 0.012 | Old |
| 0.332043 | 0.091119 | 0.005 | 0.013 | 1 | 4 | Tnni2 | -0.008 | Old |
| 0.332534 | 0.544789 | 0.049 | 0.025 | 1 | 4 | Rab13 | 0.024 | Old |
| 0.332823 | 1.407342 | 0.032 | 0.013 | 1 | 4 | Hsd17b14 | 0.019 | Old |
| 0.333058 | 0.362539 | 0.005 | 0.013 | 1 | 4 | Lzts3 | -0.008 | Old |
| 0.333058 | 0.310548 | 0.005 | 0.013 | 1 | 4 | Pcdhgb5 | -0.008 | Old |
| 0.333058 | 0.133712 | 0.005 | 0.013 | 1 | 4 | Piwil2 | -0.008 | Old |
| 0.333115 | 0.971193 | 0.121 | 0.088 | 1 | 4 | Ankle2 | 0.033 | Old |
| 0.33353 | 0.198327 | 0.588 | 0.575 | 1 | 4 | Plin2 | 0.013 | Old |
| 0.333532 | 0.746883 | 0.269 | 0.238 | 1 | 4 | H1f0 | 0.031 | Old |
| 0.33354 | 1.051927 | 0.064 | 0.038 | 1 | 4 | Desi1 | 0.026 | Old |
| 0.334075 | 0.41822 | 0.005 | 0.013 | 1 | 4 | Adgrl4 | -0.008 | Old |
| 0.334075 | 0.408405 | 0.005 | 0.013 | 1 | 4 | Ace | -0.008 | Old |
| 0.334075 | 0.256196 | 0.005 | 0.013 | 1 | 4 | Gm28874 | -0.008 | Old |
| 0.334075 | 0.040379 | 0.005 | 0.013 | 1 | 4 | AC163354. | -0.008 | Old |
| 0.334488 | 1.089567 | 0.032 | 0.013 | 1 | 4 | Tgds | 0.019 | Old |
| 0.334604 | 0.603963 | 0.258 | 0.225 | 1 | 4 | Irak1 | 0.033 | Old |
| 0.334727 | 0.353728 | 0.272 | 0.225 | 1 | 4 | Zmiz2 | 0.047 | Old |
| 0.334769 | 0.484986 | 0.165 | 0.125 | 1 | 4 | Kdelr2 | 0.04 | Old |
| 0.334788 | 0.811822 | 0.184 | 0.15 | 1 | 4 | Fip1l1 | 0.034 | Old |
| 0.335166 | 0.821462 | 0.134 | 0.1 | 1 | 4 | Fam160b1 | 0.034 | Old |
| 0.335426 | 0.57842 | 0.096 | 0.062 | 1 | 4 | Braf | 0.034 | Old |
| 0.33547 | 1.179044 | 0.143 | 0.112 | 1 | 4 | Pip4k2b | 0.031 | Old |
| 0.335803 | 0.44268 | 0.248 | 0.212 | 1 | 4 | Acsl4 | 0.036 | Old |
| 0.3365 | 0.236895 | 0.717 | 0.725 | 1 | 4 | Tpm3 | -0.008 | Old |
| 0.336592 | 0.255977 | 0.097 | 0.062 | 1 | 4 | Hk1 | 0.035 | Old |
| 0.336827 | 0.692889 | 0.094 | 0.062 | 1 | 4 | Sart1 | 0.032 | Old |
| 0.337251 | 0.924695 | 0.169 | 0.138 | 1 | 4 | Hivep1 | 0.031 | Old |
| 0.337431 | 2.360452 | 0.031 | 0.013 | 1 | 4 | Klhl8 | 0.018 | Old |
| 0.337436 | 0.13509 | 0.377 | 0.338 | 1 | 4 | Rhog | 0.039 | Old |
| 0.33756 | 1.541722 | 0.09 | 0.062 | 1 | 4 | Aga | 0.028 | Old |
| 0.337963 | 0.027356 | 0.09 | 0.125 | 1 | 4 | Zfp664 | -0.035 | Old |
| 0.33804 | 0.344858 | 0.294 | 0.25 | 1 | 4 | Gapdh | 0.044 | Old |
| 0.338051 | 0.517936 | 0.135 | 0.1 | 1 | 4 | Acss1 | 0.035 | Old |
| 0.338253 | 0.84554 | 0.032 | 0.013 | 1 | 4 | Gstt3 | 0.019 | Old |

| | | | | | | | | |
|---|---|---|---|---|---|---|---|---|
| 0.338585 | 0.197418 | 0.597 | 0.55 | 1 | 4 | Ap1b1 | 0.047 | Old |
| 0.338986 | 0.06121 | 0.13 | 0.175 | 1 | 4 | Exoc6b | -0.045 | Old |
| 0.339748 | 1.071118 | 0.105 | 0.075 | 1 | 4 | Pex7 | 0.03 | Old |
| 0.339935 | 0.872736 | 0.032 | 0.013 | 1 | 4 | Coq9 | 0.019 | Old |
| 0.33995 | 0.687696 | 0.065 | 0.038 | 1 | 4 | Gm5617 | 0.027 | Old |
| 0.340221 | 0.332078 | 0.111 | 0.075 | 1 | 4 | Ipo5 | 0.036 | Old |
| 0.340263 | 0.522195 | 0.227 | 0.188 | 1 | 4 | Pts | 0.039 | Old |
| 0.340272 | 0.654788 | 0.221 | 0.188 | 1 | 4 | Rexo2 | 0.033 | Old |
| 0.340447 | 0.158485 | 0.079 | 0.112 | 1 | 4 | Cabin1 | -0.033 | Old |
| 0.340458 | 0.552219 | 0.224 | 0.188 | 1 | 4 | Apaf1 | 0.036 | Old |
| 0.341064 | 0.925704 | 0.048 | 0.025 | 1 | 4 | Mettl21a | 0.023 | Old |
| 0.341083 | 1.902931 | 0.047 | 0.025 | 1 | 4 | Nop16 | 0.022 | Old |
| 0.341581 | 0.919399 | 0.048 | 0.025 | 1 | 4 | Nup155 | 0.023 | Old |
| 0.341682 | 1.929378 | 0.031 | 0.013 | 1 | 4 | Ppox | 0.018 | Old |
| 0.341813 | 2.375639 | 0.011 | 0 | 1 | 4 | Gadd45a | 0.011 | Old |
| 0.341813 | 2.36242 | 0.011 | 0 | 1 | 4 | Gm17435 | 0.011 | Old |
| 0.341813 | 2.345592 | 0.011 | 0 | 1 | 4 | Atp1a2 | 0.011 | Old |
| 0.341813 | 2.228958 | 0.011 | 0 | 1 | 4 | Ajuba | 0.011 | Old |
| 0.341813 | 2.216853 | 0.011 | 0 | 1 | 4 | Adgra3 | 0.011 | Old |
| 0.341813 | 2.210173 | 0.011 | 0 | 1 | 4 | Mettl2 | 0.011 | Old |
| 0.341813 | 2.210066 | 0.011 | 0 | 1 | 4 | Ptges3l | 0.011 | Old |
| 0.341813 | 2.200981 | 0.011 | 0 | 1 | 4 | 1700030K0 | 0.011 | Old |
| 0.341813 | 2.200842 | 0.011 | 0 | 1 | 4 | Vcpkmt | 0.011 | Old |
| 0.341813 | 2.198686 | 0.011 | 0 | 1 | 4 | Parp16 | 0.011 | Old |
| 0.341813 | 2.198032 | 0.011 | 0 | 1 | 4 | Ccdc51 | 0.011 | Old |
| 0.341813 | 2.195312 | 0.011 | 0 | 1 | 4 | Gdf9 | 0.011 | Old |
| 0.341813 | 2.190343 | 0.011 | 0 | 1 | 4 | D130020L0 | 0.011 | Old |
| 0.341813 | 2.160854 | 0.011 | 0 | 1 | 4 | Abcc10 | 0.011 | Old |
| 0.341813 | 2.140222 | 0.011 | 0 | 1 | 4 | Carmil1 | 0.011 | Old |
| 0.341813 | 2.131005 | 0.011 | 0 | 1 | 4 | Dach1 | 0.011 | Old |
| 0.341813 | 2.127738 | 0.011 | 0 | 1 | 4 | Ift88 | 0.011 | Old |
| 0.341813 | 2.101474 | 0.011 | 0 | 1 | 4 | Depdc7 | 0.011 | Old |
| 0.341813 | 2.069563 | 0.011 | 0 | 1 | 4 | Senp8 | 0.011 | Old |
| 0.341813 | 2.06569 | 0.011 | 0 | 1 | 4 | Wdr25 | 0.011 | Old |
| 0.341813 | 2.053509 | 0.011 | 0 | 1 | 4 | Adamts14 | 0.011 | Old |
| 0.341813 | 2.048191 | 0.011 | 0 | 1 | 4 | Ppp1r3d | 0.011 | Old |
| 0.341813 | 2.040777 | 0.011 | 0 | 1 | 4 | Rwdd2b | 0.011 | Old |
| 0.341813 | 2.03192 | 0.011 | 0 | 1 | 4 | Cbwd1 | 0.011 | Old |
| 0.341813 | 2.014825 | 0.011 | 0 | 1 | 4 | Mfsd3 | 0.011 | Old |
| 0.341813 | 2.000999 | 0.011 | 0 | 1 | 4 | Pnpla1 | 0.011 | Old |
| 0.341813 | 1.993642 | 0.011 | 0 | 1 | 4 | Hddc3 | 0.011 | Old |
| 0.341813 | 1.992591 | 0.011 | 0 | 1 | 4 | Rbks | 0.011 | Old |
| 0.341813 | 1.970195 | 0.011 | 0 | 1 | 4 | Ftx | 0.011 | Old |
| 0.341813 | 1.968254 | 0.011 | 0 | 1 | 4 | Nlrp1a | 0.011 | Old |
| 0.341813 | 1.961374 | 0.011 | 0 | 1 | 4 | Ssx2ip | 0.011 | Old |

| | | | | | | | | |
|---|---|---|---|---|---|---|---|---|
| 0.341813 | 1.95622 | 0.011 | 0 | 1 | 4 | Nupl2 | 0.011 | Old |
| 0.341813 | 1.901087 | 0.011 | 0 | 1 | 4 | Neurl1b | 0.011 | Old |
| 0.341813 | 1.900719 | 0.011 | 0 | 1 | 4 | Fsd1l | 0.011 | Old |
| 0.341813 | 1.889498 | 0.011 | 0 | 1 | 4 | Fam198b | 0.011 | Old |
| 0.341813 | 1.82891 | 0.011 | 0 | 1 | 4 | Cfb | 0.011 | Old |
| 0.341813 | 1.635368 | 0.011 | 0 | 1 | 4 | Cnn3 | 0.011 | Old |
| 0.341813 | 1.290694 | 0.011 | 0 | 1 | 4 | Cd2 | 0.011 | Old |
| 0.342391 | 0.475431 | 0.049 | 0.025 | 1 | 4 | Utp18 | 0.024 | Old |
| 0.342559 | 0.22845 | 0.03 | 0.05 | 1 | 4 | Clpb | -0.02 | Old |
| 0.342978 | 0.008724 | 0.03 | 0.05 | 1 | 4 | Phka1 | -0.02 | Old |
| 0.343011 | 0.236456 | 0.09 | 0.125 | 1 | 4 | Kdm3a | -0.035 | Old |
| 0.343275 | 0.292601 | 0.082 | 0.05 | 1 | 4 | Socs5 | 0.032 | Old |
| 0.343419 | 0.515908 | 0.095 | 0.062 | 1 | 4 | Rnpc3 | 0.033 | Old |
| 0.343818 | 0.085431 | 0.049 | 0.075 | 1 | 4 | Guf1 | -0.026 | Old |
| 0.343821 | 1.510591 | 0.031 | 0.013 | 1 | 4 | Thap7 | 0.018 | Old |
| 0.343824 | 0.482073 | 0.138 | 0.1 | 1 | 4 | Spg20 | 0.038 | Old |
| 0.343989 | 0.640235 | 0.233 | 0.2 | 1 | 4 | Med28 | 0.033 | Old |
| 0.344324 | 0.570647 | 0.289 | 0.262 | 1 | 4 | Nol7 | 0.027 | Old |
| 0.344394 | 0.933868 | 0.132 | 0.1 | 1 | 4 | Slc39a9 | 0.032 | Old |
| 0.344452 | 0.193651 | 0.408 | 0.362 | 1 | 4 | Eif3e | 0.046 | Old |
| 0.344519 | 1.482248 | 0.076 | 0.05 | 1 | 4 | Rwdd4a | 0.026 | Old |
| 0.344637 | 1.090293 | 0.13 | 0.1 | 1 | 4 | Gm36161 | 0.03 | Old |
| 0.3447 | 1.089244 | 0.092 | 0.062 | 1 | 4 | Polr3k | 0.03 | Old |
| 0.344731 | 0.155198 | 0.887 | 0.863 | 1 | 4 | Oaz1 | 0.024 | Old |
| 0.344869 | 0.671371 | 0.108 | 0.075 | 1 | 4 | Hmg20b | 0.033 | Old |
| 0.344912 | 0.899481 | 0.064 | 0.038 | 1 | 4 | Thrb | 0.026 | Old |
| 0.345003 | 0.40379 | 0.096 | 0.062 | 1 | 4 | Jade2 | 0.034 | Old |
| 0.345313 | 0.204089 | 0.493 | 0.45 | 1 | 4 | Plxnb2 | 0.043 | Old |
| 0.345349 | 0.914566 | 0.157 | 0.125 | 1 | 4 | Pik3r5 | 0.032 | Old |
| 0.345538 | 1.103688 | 0.031 | 0.013 | 1 | 4 | Slc35e2 | 0.018 | Old |
| 0.346348 | 0.227978 | 0.638 | 0.6 | 1 | 4 | Naca | 0.038 | Old |
| 0.346398 | 1.048882 | 0.031 | 0.013 | 1 | 4 | Cutc | 0.018 | Old |
| 0.346437 | 0.461827 | 0.304 | 0.262 | 1 | 4 | Tex261 | 0.042 | Old |
| 0.346704 | 0.353394 | 0.065 | 0.038 | 1 | 4 | Jagn1 | 0.027 | Old |
| 0.346812 | 0.145943 | 0.013 | 0.025 | 1 | 4 | 2610035D1 | -0.012 | Old |
| 0.347056 | 0.045182 | 0.08 | 0.112 | 1 | 4 | Usp38 | -0.032 | Old |
| 0.347364 | 0.873689 | 0.092 | 0.062 | 1 | 4 | Nvl | 0.03 | Old |
| 0.347458 | 0.380721 | 0.013 | 0.025 | 1 | 4 | Slc37a1 | -0.012 | Old |
| 0.347636 | 0.432429 | 0.272 | 0.225 | 1 | 4 | Plekhg5 | 0.047 | Old |
| 0.347963 | 1.094037 | 0.189 | 0.162 | 1 | 4 | Ak2 | 0.027 | Old |
| 0.348082 | 0.252679 | 0.342 | 0.3 | 1 | 4 | Itch | 0.042 | Old |
| 0.348223 | 0.000829 | 0.251 | 0.312 | 1 | 4 | Map2k2 | -0.061 | Old |
| 0.348274 | 0.14684 | 0.06 | 0.088 | 1 | 4 | Eif2b4 | -0.028 | Old |
| 0.348755 | 1.148212 | 0.047 | 0.025 | 1 | 4 | Trub2 | 0.022 | Old |
| 0.348898 | 0.120807 | 0.286 | 0.362 | 1 | 4 | Myof | -0.076 | Old |

| | | | | | | | | |
|---|---|---|---|---|---|---|---|---|
| 0.349183 | 0.474829 | 0.344 | 0.312 | 1 | 4 | Eef1d | 0.032 | Old |
| 0.349259 | 0.525941 | 0.175 | 0.138 | 1 | 4 | Tbc1d20 | 0.037 | Old |
| 0.349508 | 1.636431 | 0.047 | 0.025 | 1 | 4 | Cfap45 | 0.022 | Old |
| 0.349583 | 0.246119 | 0.65 | 0.637 | 1 | 4 | Npm1 | 0.013 | Old |
| 0.351219 | 1.063017 | 0.047 | 0.025 | 1 | 4 | Pak4 | 0.022 | Old |
| 0.351355 | 1.038195 | 0.063 | 0.038 | 1 | 4 | Helb | 0.025 | Old |
| 0.35159 | 0.857469 | 0.031 | 0.013 | 1 | 4 | Iqcb1 | 0.018 | Old |
| 0.351666 | 0.635738 | 0.161 | 0.125 | 1 | 4 | Rfc1 | 0.036 | Old |
| 0.351842 | 0.378112 | 0.166 | 0.125 | 1 | 4 | Rsf1 | 0.041 | Old |
| 0.352425 | 0.331769 | 0.327 | 0.288 | 1 | 4 | Leng8 | 0.039 | Old |
| 0.352576 | 1.784451 | 0.03 | 0.013 | 1 | 4 | Tubgcp6 | 0.017 | Old |
| 0.352631 | 0.509512 | 0.294 | 0.262 | 1 | 4 | Dmac1 | 0.032 | Old |
| 0.352872 | 0.739394 | 0.107 | 0.075 | 1 | 4 | Tyk2 | 0.032 | Old |
| 0.353025 | 0.352762 | 0.424 | 0.388 | 1 | 4 | Cdc42se2 | 0.036 | Old |
| 0.353159 | 0.083423 | 0.05 | 0.075 | 1 | 4 | Atg2b | -0.025 | Old |
| 0.354193 | 1.055652 | 0.077 | 0.05 | 1 | 4 | Foxk2 | 0.027 | Old |
| 0.354609 | 0.060165 | 0.121 | 0.162 | 1 | 4 | Prdm2 | -0.041 | Old |
| 0.354756 | 1.053927 | 0.077 | 0.05 | 1 | 4 | Kat7 | 0.027 | Old |
| 0.354945 | 0.891234 | 0.169 | 0.138 | 1 | 4 | Cdc73 | 0.031 | Old |
| 0.355037 | 0.711245 | 0.063 | 0.038 | 1 | 4 | Nt5dc3 | 0.025 | Old |
| 0.355578 | 0.46062 | 0.364 | 0.338 | 1 | 4 | Morf4l2 | 0.026 | Old |
| 0.355791 | 0.265723 | 0.47 | 0.488 | 1 | 4 | Plekhb2 | -0.018 | Old |
| 0.355885 | 1.112492 | 0.077 | 0.05 | 1 | 4 | Ccdc9 | 0.027 | Old |
| 0.356079 | 0.255421 | 0.729 | 0.713 | 1 | 4 | Cd53 | 0.016 | Old |
| 0.356122 | 0.801246 | 0.206 | 0.175 | 1 | 4 | Uso1 | 0.031 | Old |
| 0.356187 | 0.48101 | 0.24 | 0.2 | 1 | 4 | Riok3 | 0.04 | Old |
| 0.356509 | 0.350619 | 0.416 | 0.388 | 1 | 4 | Mycbp2 | 0.028 | Old |
| 0.356549 | 1.637788 | 0.03 | 0.013 | 1 | 4 | AI839979 | 0.017 | Old |
| 0.356594 | 2.771431 | 0.011 | 0 | 1 | 4 | Gm26802 | 0.011 | Old |
| 0.356594 | 2.283147 | 0.011 | 0 | 1 | 4 | Frk | 0.011 | Old |
| 0.356594 | 2.251439 | 0.011 | 0 | 1 | 4 | Mex3b | 0.011 | Old |
| 0.356594 | 2.195749 | 0.011 | 0 | 1 | 4 | Gpnmb | 0.011 | Old |
| 0.356594 | 2.188942 | 0.011 | 0 | 1 | 4 | 1810034E1 | 0.011 | Old |
| 0.356594 | 2.183707 | 0.011 | 0 | 1 | 4 | Ispd | 0.011 | Old |
| 0.356594 | 2.165087 | 0.011 | 0 | 1 | 4 | Stxbp4 | 0.011 | Old |
| 0.356594 | 2.159259 | 0.011 | 0 | 1 | 4 | Flrt3 | 0.011 | Old |
| 0.356594 | 2.146839 | 0.011 | 0 | 1 | 4 | Zranb3 | 0.011 | Old |
| 0.356594 | 2.128788 | 0.011 | 0 | 1 | 4 | Tmem132a | 0.011 | Old |
| 0.356594 | 2.128493 | 0.011 | 0 | 1 | 4 | Bahcc1 | 0.011 | Old |
| 0.356594 | 2.122598 | 0.011 | 0 | 1 | 4 | Hoxb7 | 0.011 | Old |
| 0.356594 | 2.104486 | 0.011 | 0 | 1 | 4 | Pla2g6 | 0.011 | Old |
| 0.356594 | 2.096561 | 0.011 | 0 | 1 | 4 | Ugt1a7c | 0.011 | Old |
| 0.356594 | 2.07987 | 0.011 | 0 | 1 | 4 | Zfp623 | 0.011 | Old |
| 0.356594 | 2.048688 | 0.011 | 0 | 1 | 4 | Mterf1a | 0.011 | Old |
| 0.356594 | 2.047768 | 0.011 | 0 | 1 | 4 | Cebpe | 0.011 | Old |

| | | | | | | | | |
|---|---|---|---|---|---|---|---|---|
| 0.356594 | 2.040836 | 0.011 | 0 | 1 | 4 | Bbs4 | 0.011 | Old |
| 0.356594 | 2.035665 | 0.011 | 0 | 1 | 4 | Slc6a8 | 0.011 | Old |
| 0.356594 | 2.033447 | 0.011 | 0 | 1 | 4 | Srfbp1 | 0.011 | Old |
| 0.356594 | 2.033401 | 0.011 | 0 | 1 | 4 | Zfp324 | 0.011 | Old |
| 0.356594 | 2.015339 | 0.011 | 0 | 1 | 4 | Ska2 | 0.011 | Old |
| 0.356594 | 2.011217 | 0.011 | 0 | 1 | 4 | Zfp712 | 0.011 | Old |
| 0.356594 | 2.008746 | 0.011 | 0 | 1 | 4 | AC129328. | 0.011 | Old |
| 0.356594 | 2.001728 | 0.011 | 0 | 1 | 4 | Cd320 | 0.011 | Old |
| 0.356594 | 1.957255 | 0.011 | 0 | 1 | 4 | Ak8 | 0.011 | Old |
| 0.356594 | 1.942322 | 0.011 | 0 | 1 | 4 | Fastkd1 | 0.011 | Old |
| 0.356594 | 1.926747 | 0.011 | 0 | 1 | 4 | Stab2 | 0.011 | Old |
| 0.356594 | 1.906914 | 0.011 | 0 | 1 | 4 | Mlkl | 0.011 | Old |
| 0.356594 | 1.877747 | 0.011 | 0 | 1 | 4 | Tmprss4 | 0.011 | Old |
| 0.356594 | 1.855663 | 0.011 | 0 | 1 | 4 | Syt3 | 0.011 | Old |
| 0.356594 | 1.855102 | 0.011 | 0 | 1 | 4 | Pigo | 0.011 | Old |
| 0.356594 | 1.846939 | 0.011 | 0 | 1 | 4 | A430046D1 | 0.011 | Old |
| 0.356594 | 1.790539 | 0.011 | 0 | 1 | 4 | C130050O1 | 0.011 | Old |
| 0.356594 | 1.789713 | 0.011 | 0 | 1 | 4 | Plcb1 | 0.011 | Old |
| 0.356594 | 1.783362 | 0.011 | 0 | 1 | 4 | Nsl1 | 0.011 | Old |
| 0.356594 | 1.772019 | 0.011 | 0 | 1 | 4 | Nicn1 | 0.011 | Old |
| 0.356594 | 1.720282 | 0.011 | 0 | 1 | 4 | Crip2 | 0.011 | Old |
| 0.356594 | 1.70424 | 0.011 | 0 | 1 | 4 | Ifrd2 | 0.011 | Old |
| 0.356594 | 1.662909 | 0.011 | 0 | 1 | 4 | 1700056E2 | 0.011 | Old |
| 0.356594 | 1.635651 | 0.011 | 0 | 1 | 4 | Mefv | 0.011 | Old |
| 0.356833 | 0.596582 | 0.269 | 0.238 | 1 | 4 | Sf3b6 | 0.031 | Old |
| 0.356873 | 0.546113 | 0.148 | 0.112 | 1 | 4 | Mrps17 | 0.036 | Old |
| 0.357151 | 0.748423 | 0.157 | 0.125 | 1 | 4 | Nsmce4a | 0.032 | Old |
| 0.357296 | 0.589328 | 0.385 | 0.375 | 1 | 4 | Cd44 | 0.01 | Old |
| 0.357298 | 0.247198 | 0.294 | 0.25 | 1 | 4 | R3hdm4 | 0.044 | Old |
| 0.357554 | 0.069832 | 1 | 1 | 1 | 4 | mt-Co3 | 0 | Old |
| 0.357605 | 1.197865 | 0.062 | 0.038 | 1 | 4 | Zdhhc6 | 0.024 | Old |
| 0.357805 | 1.208039 | 0.141 | 0.112 | 1 | 4 | Slfn8 | 0.029 | Old |
| 0.358014 | 0.249321 | 0.125 | 0.088 | 1 | 4 | Tbcel | 0.037 | Old |
| 0.358251 | 1.160084 | 0.09 | 0.062 | 1 | 4 | Zdhhc21 | 0.028 | Old |
| 0.358312 | 0.522088 | 0.254 | 0.212 | 1 | 4 | Pitpnc1 | 0.042 | Old |
| 0.358318 | 0.314477 | 0.047 | 0.025 | 1 | 4 | Mycbp | 0.022 | Old |
| 0.358452 | 0.152416 | 0.063 | 0.038 | 1 | 4 | Urgcp | 0.025 | Old |
| 0.359214 | 1.027696 | 0.03 | 0.013 | 1 | 4 | Sema6b | 0.017 | Old |
| 0.359527 | 0.56423 | 0.16 | 0.125 | 1 | 4 | Hipk3 | 0.035 | Old |
| 0.359659 | 1.100337 | 0.03 | 0.013 | 1 | 4 | Alg13 | 0.017 | Old |
| 0.359867 | 1.086476 | 0.047 | 0.025 | 1 | 4 | Ublcp1 | 0.022 | Old |
| 0.359898 | 0.135137 | 0.198 | 0.15 | 1 | 4 | Ccni | 0.048 | Old |
| 0.359899 | 0.47361 | 0.251 | 0.212 | 1 | 4 | Micu1 | 0.039 | Old |
| 0.36043 | 0.743777 | 0.132 | 0.1 | 1 | 4 | Utrn | 0.032 | Old |
| 0.360589 | 0.982412 | 0.047 | 0.025 | 1 | 4 | Zbtb24 | 0.022 | Old |

| | | | | | | | | |
|---|---|---|---|---|---|---|---|---|
| 0.360631 | 0.503075 | 0.108 | 0.075 | 1 | 4 | Lta4h | 0.033 | Old |
| 0.36099 | 0.618196 | 0.077 | 0.05 | 1 | 4 | Map3k20 | 0.027 | Old |
| 0.362598 | 0.96038 | 0.118 | 0.088 | 1 | 4 | Ppfibp2 | 0.03 | Old |
| 0.362726 | 0.115682 | 0.093 | 0.062 | 1 | 4 | Creld2 | 0.031 | Old |
| 0.362869 | 1.064734 | 0.076 | 0.05 | 1 | 4 | Snrnp25 | 0.026 | Old |
| 0.362874 | 1.450667 | 0.046 | 0.025 | 1 | 4 | Fam207a | 0.021 | Old |
| 0.362956 | 2.498495 | 0.03 | 0.013 | 1 | 4 | Trim30c | 0.017 | Old |
| 0.363032 | 0.41634 | 0.215 | 0.175 | 1 | 4 | Cyth1 | 0.04 | Old |
| 0.363234 | 0.829498 | 0.03 | 0.013 | 1 | 4 | Zfp386 | 0.017 | Old |
| 0.363381 | 0.57727 | 0.134 | 0.1 | 1 | 4 | Ndufb2 | 0.034 | Old |
| 0.363567 | 0.611326 | 0.198 | 0.162 | 1 | 4 | Gorasp2 | 0.036 | Old |
| 0.364315 | 2.239202 | 0.03 | 0.013 | 1 | 4 | Cd9 | 0.017 | Old |
| 0.364508 | 0.216557 | 0.514 | 0.475 | 1 | 4 | Dnmt3a | 0.039 | Old |
| 0.36477 | 0.693837 | 0.133 | 0.1 | 1 | 4 | Cdk13 | 0.033 | Old |
| 0.36525 | 0.427157 | 0.21 | 0.175 | 1 | 4 | Cic | 0.035 | Old |
| 0.365263 | 0.726777 | 0.092 | 0.062 | 1 | 4 | Slc36a4 | 0.03 | Old |
| 0.365356 | 0.015819 | 0.022 | 0.038 | 1 | 4 | Intu | -0.016 | Old |
| 0.36548 | 0.40784 | 0.03 | 0.013 | 1 | 4 | Ubxn2b | 0.017 | Old |
| 0.365667 | 0.66496 | 0.133 | 0.1 | 1 | 4 | 2310022A1 | 0.033 | Old |
| 0.366239 | 0.644432 | 0.078 | 0.05 | 1 | 4 | Mdn1 | 0.028 | Old |
| 0.36638 | 0.305418 | 0.03 | 0.013 | 1 | 4 | Fam217b | 0.017 | Old |
| 0.366799 | 0.661969 | 0.195 | 0.162 | 1 | 4 | Nuak1 | 0.033 | Old |
| 0.366878 | 0.518813 | 0.136 | 0.1 | 1 | 4 | Hck | 0.036 | Old |
| 0.367295 | 0.023427 | 0.957 | 0.95 | 1 | 4 | H2-Eb1 | 0.007 | Old |
| 0.367824 | 0.487023 | 0.237 | 0.2 | 1 | 4 | Psme4 | 0.037 | Old |
| 0.367829 | 0.525331 | 0.316 | 0.288 | 1 | 4 | Bola2 | 0.028 | Old |
| 0.368255 | 0.41805 | 0.165 | 0.125 | 1 | 4 | Tomm70a | 0.04 | Old |
| 0.368452 | 0.846603 | 0.119 | 0.088 | 1 | 4 | Hdac2 | 0.031 | Old |
| 0.368585 | 0.668118 | 0.047 | 0.025 | 1 | 4 | Ccs | 0.022 | Old |
| 0.368619 | 0.207659 | 0.281 | 0.238 | 1 | 4 | Ist1 | 0.043 | Old |
| 0.369196 | 0.782777 | 0.155 | 0.125 | 1 | 4 | Lrrc8c | 0.03 | Old |
| 0.369571 | 0.504216 | 0.248 | 0.212 | 1 | 4 | Batf3 | 0.036 | Old |
| 0.370703 | 1.286562 | 0.03 | 0.013 | 1 | 4 | 1700096K1 | 0.017 | Old |
| 0.370703 | 1.27481 | 0.03 | 0.013 | 1 | 4 | Stn1 | 0.017 | Old |
| 0.370785 | 0.577388 | 0.047 | 0.025 | 1 | 4 | Stambp | 0.022 | Old |
| 0.371162 | 1.352285 | 0.03 | 0.013 | 1 | 4 | Rrp12 | 0.017 | Old |
| 0.371621 | 1.227298 | 0.03 | 0.013 | 1 | 4 | Snx21 | 0.017 | Old |
| 0.371945 | 0.51747 | 0.308 | 0.275 | 1 | 4 | Lbr | 0.033 | Old |
| 0.372052 | 0.184916 | 0.347 | 0.3 | 1 | 4 | Txndc17 | 0.047 | Old |
| 0.372169 | 0.275421 | 0.302 | 0.262 | 1 | 4 | Etnk1 | 0.04 | Old |
| 0.372246 | 2.377866 | 0.01 | 0 | 1 | 4 | Btc | 0.01 | Old |
| 0.372246 | 2.323792 | 0.01 | 0 | 1 | 4 | Adm | 0.01 | Old |
| 0.372246 | 2.304293 | 0.01 | 0 | 1 | 4 | Kcnt2 | 0.01 | Old |
| 0.372246 | 2.22407 | 0.01 | 0 | 1 | 4 | Tcf7l1 | 0.01 | Old |
| 0.372246 | 2.138533 | 0.01 | 0 | 1 | 4 | Proser3 | 0.01 | Old |

| | | | | | | | | |
|---|---|---|---|---|---|---|---|---|
| 0.372246 | 2.124264 | 0.01 | 0 | 1 | 4 | Gm10800 | 0.01 | Old |
| 0.372246 | 2.121554 | 0.01 | 0 | 1 | 4 | Gm2814 | 0.01 | Old |
| 0.372246 | 2.119592 | 0.01 | 0 | 1 | 4 | Zfp867 | 0.01 | Old |
| 0.372246 | 2.118101 | 0.01 | 0 | 1 | 4 | Dhfr | 0.01 | Old |
| 0.372246 | 2.061593 | 0.01 | 0 | 1 | 4 | Hs3st3a1 | 0.01 | Old |
| 0.372246 | 2.02404 | 0.01 | 0 | 1 | 4 | Gins1 | 0.01 | Old |
| 0.372246 | 2.012729 | 0.01 | 0 | 1 | 4 | Wdr27 | 0.01 | Old |
| 0.372246 | 1.978282 | 0.01 | 0 | 1 | 4 | Acyp2 | 0.01 | Old |
| 0.372246 | 1.946845 | 0.01 | 0 | 1 | 4 | Mpp7 | 0.01 | Old |
| 0.372246 | 1.933098 | 0.01 | 0 | 1 | 4 | Pcdh15 | 0.01 | Old |
| 0.372246 | 1.917935 | 0.01 | 0 | 1 | 4 | Gm26890 | 0.01 | Old |
| 0.372246 | 1.91067 | 0.01 | 0 | 1 | 4 | Ciart | 0.01 | Old |
| 0.372246 | 1.882055 | 0.01 | 0 | 1 | 4 | Eef1akmt4 | 0.01 | Old |
| 0.372246 | 1.866042 | 0.01 | 0 | 1 | 4 | Gm7967 | 0.01 | Old |
| 0.372246 | 1.822969 | 0.01 | 0 | 1 | 4 | Arhgap29 | 0.01 | Old |
| 0.372246 | 1.819916 | 0.01 | 0 | 1 | 4 | Tyw5 | 0.01 | Old |
| 0.372246 | 1.797261 | 0.01 | 0 | 1 | 4 | Mrnip | 0.01 | Old |
| 0.372246 | 1.796808 | 0.01 | 0 | 1 | 4 | Phlda3 | 0.01 | Old |
| 0.372246 | 1.751926 | 0.01 | 0 | 1 | 4 | Hp | 0.01 | Old |
| 0.372246 | 1.751913 | 0.01 | 0 | 1 | 4 | Ubox5 | 0.01 | Old |
| 0.372246 | 1.720989 | 0.01 | 0 | 1 | 4 | Zfp446 | 0.01 | Old |
| 0.372246 | 1.719921 | 0.01 | 0 | 1 | 4 | C130013H( | 0.01 | Old |
| 0.372246 | 1.716753 | 0.01 | 0 | 1 | 4 | Dnajb13 | 0.01 | Old |
| 0.372246 | 1.710851 | 0.01 | 0 | 1 | 4 | Cox6b2 | 0.01 | Old |
| 0.372246 | 1.6917 | 0.01 | 0 | 1 | 4 | Impa2 | 0.01 | Old |
| 0.372246 | 1.584154 | 0.01 | 0 | 1 | 4 | Cd244 | 0.01 | Old |
| 0.37251 | 0.530348 | 0.264 | 0.225 | 1 | 4 | Slc20a1 | 0.039 | Old |
| 0.372883 | 0.853595 | 0.131 | 0.1 | 1 | 4 | Cldnd1 | 0.031 | Old |
| 0.372913 | 0.337739 | 0.179 | 0.138 | 1 | 4 | Tmbim4 | 0.041 | Old |
| 0.37305 | 0.889125 | 0.13 | 0.1 | 1 | 4 | Snupn | 0.03 | Old |
| 0.373081 | 0.65414 | 0.077 | 0.05 | 1 | 4 | Nln | 0.027 | Old |
| 0.373137 | 0.158675 | 0.147 | 0.112 | 1 | 4 | Hpf1 | 0.035 | Old |
| 0.373478 | 0.47169 | 0.238 | 0.2 | 1 | 4 | Agfg1 | 0.038 | Old |
| 0.373573 | 0.776482 | 0.119 | 0.088 | 1 | 4 | Snrk | 0.031 | Old |
| 0.374043 | 0.289336 | 0.269 | 0.225 | 1 | 4 | Ggt5 | 0.044 | Old |
| 0.374232 | 0.378863 | 0.385 | 0.35 | 1 | 4 | Aamp | 0.035 | Old |
| 0.374384 | 0.974369 | 0.03 | 0.013 | 1 | 4 | Tsr2 | 0.017 | Old |
| 0.374649 | 0.462272 | 0.022 | 0.038 | 1 | 4 | Cradd | -0.016 | Old |
| 0.374753 | 0.680061 | 0.076 | 0.05 | 1 | 4 | Rpa1 | 0.026 | Old |
| 0.374846 | 0.841861 | 0.03 | 0.013 | 1 | 4 | Chst14 | 0.017 | Old |
| 0.375321 | 0.359634 | 0.383 | 0.35 | 1 | 4 | Cmpk1 | 0.033 | Old |
| 0.37577 | 0.864825 | 0.03 | 0.013 | 1 | 4 | Optn | 0.017 | Old |
| 0.37577 | 0.625881 | 0.03 | 0.013 | 1 | 4 | Psd4 | 0.017 | Old |
| 0.376103 | 0.631555 | 0.062 | 0.038 | 1 | 4 | Tmem63b | 0.024 | Old |
| 0.376159 | 1.111852 | 0.102 | 0.075 | 1 | 4 | Cox16 | 0.027 | Old |

| | | | | | | | | |
|---|---|---|---|---|---|---|---|---|
| 0.376524 | 0.488177 | 0.284 | 0.25 | 1 | 4 | Cdipt | 0.034 | Old |
| 0.37669 | 0.138785 | 0.256 | 0.325 | 1 | 4 | Ppp1r14b | -0.069 | Old |
| 0.376902 | 0.819156 | 0.046 | 0.025 | 1 | 4 | Exosc7 | 0.021 | Old |
| 0.37692 | 1.213582 | 0.089 | 0.062 | 1 | 4 | Mill2 | 0.027 | Old |
| 0.377342 | 0.496409 | 0.248 | 0.212 | 1 | 4 | Acer3 | 0.036 | Old |
| 0.377459 | 0.372014 | 0.077 | 0.05 | 1 | 4 | Aspa | 0.027 | Old |
| 0.377742 | 1.132639 | 0.089 | 0.062 | 1 | 4 | Bmi1 | 0.027 | Old |
| 0.377938 | 0.67703 | 0.219 | 0.188 | 1 | 4 | 2310035C2 | 0.031 | Old |
| 0.378837 | 1.778338 | 0.029 | 0.013 | 1 | 4 | Dsel | 0.016 | Old |
| 0.379017 | 0.549325 | 0.03 | 0.013 | 1 | 4 | Cox15 | 0.017 | Old |
| 0.379044 | 1.498886 | 0.06 | 0.038 | 1 | 4 | Gpr157 | 0.022 | Old |
| 0.379308 | 1.947447 | 0.029 | 0.013 | 1 | 4 | Yipf2 | 0.016 | Old |
| 0.379589 | 0.169493 | 0.308 | 0.262 | 1 | 4 | Casp8 | 0.046 | Old |
| 0.37968 | 0.303949 | 0.131 | 0.175 | 1 | 4 | Nans | -0.044 | Old |
| 0.379795 | 0.76804 | 0.103 | 0.075 | 1 | 4 | Vamp7 | 0.028 | Old |
| 0.37991 | 0.00348 | 0.093 | 0.125 | 1 | 4 | Slc35e3 | -0.032 | Old |
| 0.379948 | 0.386512 | 0.03 | 0.013 | 1 | 4 | Mus81 | 0.017 | Old |
| 0.38046 | 0.656356 | 0.046 | 0.025 | 1 | 4 | Ring1 | 0.021 | Old |
| 0.380785 | 0.25781 | 0.047 | 0.025 | 1 | 4 | Batf2 | 0.022 | Old |
| 0.380806 | 0.572012 | 0.147 | 0.112 | 1 | 4 | Ube2e3 | 0.035 | Old |
| 0.380894 | 0.445936 | 0.356 | 0.325 | 1 | 4 | Dusp6 | 0.031 | Old |
| 0.381233 | 0.336043 | 0.374 | 0.338 | 1 | 4 | Smim15 | 0.036 | Old |
| 0.38164 | 0.022551 | 0.093 | 0.125 | 1 | 4 | Cog3 | -0.032 | Old |
| 0.381663 | 1.647098 | 0.029 | 0.013 | 1 | 4 | Ift27 | 0.016 | Old |
| 0.381828 | 0.640285 | 0.229 | 0.2 | 1 | 4 | Atf6 | 0.029 | Old |
| 0.381928 | 0.419974 | 0.348 | 0.325 | 1 | 4 | Elk3 | 0.023 | Old |
| 0.382378 | 1.147562 | 0.045 | 0.025 | 1 | 4 | Micu3 | 0.02 | Old |
| 0.382607 | 0.053963 | 0.082 | 0.112 | 1 | 4 | Ankrd27 | -0.03 | Old |
| 0.382609 | 0.515324 | 0.134 | 0.1 | 1 | 4 | Cab39l | 0.034 | Old |
| 0.382793 | 0.548852 | 0.246 | 0.212 | 1 | 4 | Gpr160 | 0.034 | Old |
| 0.3829 | 0.617464 | 0.061 | 0.038 | 1 | 4 | Thoc3 | 0.023 | Old |
| 0.383169 | 0.828819 | 0.103 | 0.075 | 1 | 4 | Ptpn9 | 0.028 | Old |
| 0.383397 | 0.414613 | 0.368 | 0.338 | 1 | 4 | Ptpro | 0.03 | Old |
| 0.383403 | 0.057313 | 0.114 | 0.15 | 1 | 4 | Frrs1 | -0.036 | Old |
| 0.383404 | 0.229581 | 0.047 | 0.025 | 1 | 4 | Ubac1 | 0.022 | Old |
| 0.383423 | 0.722371 | 0.061 | 0.038 | 1 | 4 | Mettl14 | 0.023 | Old |
| 0.383554 | 1.27486 | 0.029 | 0.013 | 1 | 4 | Dbr1 | 0.016 | Old |
| 0.383601 | 0.614345 | 0.119 | 0.088 | 1 | 4 | Med10 | 0.031 | Old |
| 0.383753 | 0.666959 | 0.061 | 0.038 | 1 | 4 | Zfp991 | 0.023 | Old |
| 0.383771 | 0.415493 | 0.148 | 0.112 | 1 | 4 | Gab1 | 0.036 | Old |
| 0.383823 | 0.562884 | 0.119 | 0.088 | 1 | 4 | Naga | 0.031 | Old |
| 0.384191 | 0.406076 | 0.213 | 0.175 | 1 | 4 | Lgals9 | 0.038 | Old |
| 0.384201 | 0.050195 | 0.013 | 0.025 | 1 | 4 | Prepl | -0.012 | Old |
| 0.384313 | 1.401972 | 0.074 | 0.05 | 1 | 4 | Hif1an | 0.024 | Old |
| 0.385385 | 0.939648 | 0.061 | 0.038 | 1 | 4 | Mapk8ip3 | 0.023 | Old |

| | | | | | | | | |
|---|---|---|---|---|---|---|---|---|
| 0.385618 | 0.057187 | 0.285 | 0.35 | 1 | 4 | Eif4g1 | -0.065 | Old |
| 0.385681 | 0.667888 | 0.076 | 0.05 | 1 | 4 | Siae | 0.026 | Old |
| 0.385706 | 0.006906 | 0.104 | 0.138 | 1 | 4 | Bcl7c | -0.034 | Old |
| 0.38573 | 0.015269 | 0.032 | 0.05 | 1 | 4 | Zc3h12c | -0.018 | Old |
| 0.386243 | 0.166984 | 0.112 | 0.15 | 1 | 4 | Maml1 | -0.038 | Old |
| 0.386977 | 0.028327 | 0.047 | 0.025 | 1 | 4 | Zfp369 | 0.022 | Old |
| 0.387182 | 1.708526 | 0.045 | 0.025 | 1 | 4 | Eepd1 | 0.02 | Old |
| 0.387575 | 0.504003 | 0.013 | 0.025 | 1 | 4 | Cped1 | -0.012 | Old |
| 0.387742 | 0.347433 | 0.356 | 0.325 | 1 | 4 | Dock7 | 0.031 | Old |
| 0.387951 | 0.232646 | 0.406 | 0.375 | 1 | 4 | Ifi203 | 0.031 | Old |
| 0.38817 | 2.374059 | 0.044 | 0.025 | 1 | 4 | C2cd2l | 0.019 | Old |
| 0.388592 | 0.0096 | 0.072 | 0.1 | 1 | 4 | Eif1ad | -0.028 | Old |
| 0.388862 | 0.690194 | 0.105 | 0.075 | 1 | 4 | Abhd16a | 0.03 | Old |
| 0.388993 | 0.166789 | 0.072 | 0.1 | 1 | 4 | Slc35b1 | -0.028 | Old |
| 0.389034 | 0.326934 | 0.357 | 0.325 | 1 | 4 | Spcs1 | 0.032 | Old |
| 0.389244 | 0.745686 | 0.045 | 0.025 | 1 | 4 | Elac2 | 0.02 | Old |
| 0.38985 | 0.969959 | 0.165 | 0.138 | 1 | 4 | Rit1 | 0.027 | Old |
| 0.389851 | 0.119061 | 0.383 | 0.325 | 1 | 4 | Tnfrsf1b | 0.058 | Old |
| 0.390007 | 0.601971 | 0.184 | 0.15 | 1 | 4 | Sod2 | 0.034 | Old |
| 0.390396 | 0.750253 | 0.045 | 0.025 | 1 | 4 | Gar1 | 0.02 | Old |
| 0.390553 | 0.839375 | 0.061 | 0.038 | 1 | 4 | Bcl9 | 0.023 | Old |
| 0.39063 | 0.800652 | 0.117 | 0.088 | 1 | 4 | Bak1 | 0.029 | Old |
| 0.391152 | 0.170029 | 0.106 | 0.075 | 1 | 4 | Timm8b | 0.031 | Old |
| 0.391165 | 0.773002 | 0.045 | 0.025 | 1 | 4 | Slc20a2 | 0.02 | Old |
| 0.391767 | 0.751664 | 0.075 | 0.05 | 1 | 4 | Adgrl2 | 0.025 | Old |
| 0.391861 | 0.527158 | 0.09 | 0.062 | 1 | 4 | Pmpca | 0.028 | Old |
| 0.392044 | 0.075584 | 0.124 | 0.162 | 1 | 4 | Tmem131 | -0.038 | Old |
| 0.392185 | 0.334389 | 0.336 | 0.288 | 1 | 4 | Taok3 | 0.048 | Old |
| 0.392615 | 0.397965 | 0.029 | 0.013 | 1 | 4 | Evi5l | 0.016 | Old |
| 0.392699 | 2.050677 | 0.059 | 0.038 | 1 | 4 | Mrps26 | 0.021 | Old |
| 0.392862 | 0.660664 | 0.117 | 0.088 | 1 | 4 | Arfgap2 | 0.029 | Old |
| 0.39288 | 0.106665 | 0.077 | 0.05 | 1 | 4 | Hrh1 | 0.027 | Old |
| 0.392946 | 0.062698 | 0.98 | 0.963 | 1 | 4 | H2-K1 | 0.017 | Old |
| 0.392993 | 0.65968 | 0.09 | 0.062 | 1 | 4 | Trak1 | 0.028 | Old |
| 0.393002 | 1.788363 | 0.028 | 0.013 | 1 | 4 | Cc2d1a | 0.015 | Old |
| 0.393074 | 1.027197 | 0.06 | 0.038 | 1 | 4 | Ap1g2 | 0.022 | Old |
| 0.393918 | 1.357386 | 0.073 | 0.05 | 1 | 4 | Gtf2e2 | 0.023 | Old |
| 0.394079 | 0.449679 | 0.076 | 0.05 | 1 | 4 | 0610010F0 | 0.026 | Old |
| 0.394281 | 0.320256 | 0.203 | 0.162 | 1 | 4 | Igsf6 | 0.041 | Old |
| 0.394344 | 0.568885 | 0.091 | 0.062 | 1 | 4 | Samsn1 | 0.029 | Old |
| 0.394381 | 0.72377 | 0.181 | 0.15 | 1 | 4 | Btbd7 | 0.031 | Old |
| 0.394539 | 0.355589 | 0.029 | 0.013 | 1 | 4 | 1500015A0 | 0.016 | Old |
| 0.394931 | 0.108093 | 0.022 | 0.038 | 1 | 4 | Zfp40 | -0.016 | Old |
| 0.394948 | 1.276987 | 0.028 | 0.013 | 1 | 4 | Tmem218 | 0.015 | Old |
| 0.395021 | 0.33867 | 0.029 | 0.013 | 1 | 4 | Ttll3 | 0.016 | Old |

| | | | | | | | | |
|---|---|---|---|---|---|---|---|---|
| 0.395155 | 0.061879 | 0.221 | 0.275 | 1 | 4 | Asap1 | -0.054 | Old |
| 0.395316 | 0.959445 | 0.045 | 0.025 | 1 | 4 | Slc9a1 | 0.02 | Old |
| 0.395503 | 0.118757 | 0.029 | 0.013 | 1 | 4 | Msh6 | 0.016 | Old |
| 0.395715 | 0.881709 | 0.116 | 0.088 | 1 | 4 | Tec | 0.028 | Old |
| 0.396085 | 0.256923 | 0.454 | 0.425 | 1 | 4 | Mtpn | 0.029 | Old |
| 0.396184 | 0.086302 | 0.968 | 0.95 | 1 | 4 | Rps27a | 0.018 | Old |
| 0.396326 | 0.18845 | 0.697 | 0.688 | 1 | 4 | Arhgdia | 0.009 | Old |
| 0.396386 | 1.565459 | 0.073 | 0.05 | 1 | 4 | Tmem60 | 0.023 | Old |
| 0.396561 | 0.197028 | 0.582 | 0.6 | 1 | 4 | AC149090. | -0.018 | Old |
| 0.39675 | 0.39543 | 0.316 | 0.288 | 1 | 4 | Ap1s2 | 0.028 | Old |
| 0.3969 | 1.739607 | 0.028 | 0.013 | 1 | 4 | Fzd7 | 0.015 | Old |
| 0.39697 | 1.538001 | 0.086 | 0.062 | 1 | 4 | Tbc1d1 | 0.024 | Old |
| 0.397108 | 0.173773 | 0.061 | 0.088 | 1 | 4 | Gbe1 | -0.027 | Old |
| 0.397381 | 0.984304 | 0.074 | 0.05 | 1 | 4 | Cln5 | 0.024 | Old |
| 0.397389 | 1.377871 | 0.028 | 0.013 | 1 | 4 | Csprs | 0.015 | Old |
| 0.397531 | 0.487514 | 0.276 | 0.25 | 1 | 4 | Ocrl | 0.026 | Old |
| 0.397553 | 1.580069 | 0.044 | 0.025 | 1 | 4 | Zdhhc16 | 0.019 | Old |
| 0.397879 | 1.013392 | 0.028 | 0.013 | 1 | 4 | Casp7 | 0.015 | Old |
| 0.398075 | 0.030327 | 0.36 | 0.312 | 1 | 4 | Kitl | 0.048 | Old |
| 0.398368 | 1.094125 | 0.028 | 0.013 | 1 | 4 | Eid3 | 0.015 | Old |
| 0.398384 | 0.850265 | 0.128 | 0.1 | 1 | 4 | B4galt3 | 0.028 | Old |
| 0.398858 | 1.237138 | 0.028 | 0.013 | 1 | 4 | Kyat1 | 0.015 | Old |
| 0.399081 | 0.229814 | 0.379 | 0.35 | 1 | 4 | Prkcb | 0.029 | Old |
| 0.399191 | 0.664015 | 0.154 | 0.125 | 1 | 4 | Brd3 | 0.029 | Old |
| 0.399483 | 1.289216 | 0.073 | 0.05 | 1 | 4 | Tsga10 | 0.023 | Old |
| 0.399823 | 1.088182 | 0.074 | 0.05 | 1 | 4 | Icmt | 0.024 | Old |
| 0.40001 | 0.829041 | 0.045 | 0.025 | 1 | 4 | Ndc1 | 0.02 | Old |
| 0.400315 | 1.444018 | 0.044 | 0.025 | 1 | 4 | Atg14 | 0.019 | Old |
| 0.400465 | 0.363347 | 0.186 | 0.15 | 1 | 4 | Pgrmc1 | 0.036 | Old |
| 0.40047 | 0.119232 | 0.207 | 0.162 | 1 | 4 | Tmem106b | 0.045 | Old |
| 0.400711 | 1.320369 | 0.044 | 0.025 | 1 | 4 | Xpnpep3 | 0.019 | Old |
| 0.400711 | 1.311627 | 0.044 | 0.025 | 1 | 4 | Mrto4 | 0.019 | Old |
| 0.400939 | 0.413055 | 0.156 | 0.125 | 1 | 4 | Ing4 | 0.031 | Old |
| 0.401117 | 0.033137 | 0.052 | 0.075 | 1 | 4 | Mfsd7a | -0.023 | Old |
| 0.401126 | 1.130049 | 0.126 | 0.1 | 1 | 4 | Map3k7 | 0.026 | Old |
| 0.401434 | 0.175558 | 0.005 | 0.013 | 1 | 4 | Angel1 | -0.008 | Old |
| 0.402294 | 0.29497 | 0.105 | 0.075 | 1 | 4 | Dusp7 | 0.03 | Old |
| 0.402306 | 0.529088 | 0.359 | 0.35 | 1 | 4 | H13 | 0.009 | Old |
| 0.402511 | 0.306121 | 0.005 | 0.013 | 1 | 4 | Atxn7l2 | -0.008 | Old |
| 0.402511 | 0.229766 | 0.005 | 0.013 | 1 | 4 | S100a16 | -0.008 | Old |
| 0.402511 | 0.111777 | 0.005 | 0.013 | 1 | 4 | Gm48420 | -0.008 | Old |
| 0.40258 | 0.347748 | 0.257 | 0.212 | 1 | 4 | Ago3 | 0.045 | Old |
| 0.4026 | 0.828176 | 0.06 | 0.038 | 1 | 4 | Fbxo34 | 0.022 | Old |
| 0.402692 | 1.395606 | 0.044 | 0.025 | 1 | 4 | Psmg2 | 0.019 | Old |
| 0.402884 | 0.879646 | 0.115 | 0.088 | 1 | 4 | Clcc1 | 0.027 | Old |

| | | | | | | | | |
|---|---|---|---|---|---|---|---|---|
| 0.403192 | 0.505182 | 0.042 | 0.062 | 1 | 4 | Zer1 | -0.02 | Old |
| 0.403489 | 0.575835 | 0.231 | 0.2 | 1 | 4 | Usp34 | 0.031 | Old |
| 0.40359 | 0.150331 | 0.005 | 0.013 | 1 | 4 | Bgn | -0.008 | Old |
| 0.40359 | 0.341005 | 0.005 | 0.013 | 1 | 4 | Gm43614 | -0.008 | Old |
| 0.40359 | 0.318696 | 0.005 | 0.013 | 1 | 4 | Amigo3 | -0.008 | Old |
| 0.403792 | 2.077545 | 0.043 | 0.025 | 1 | 4 | Polr3b | 0.018 | Old |
| 0.404118 | 0.381609 | 0.076 | 0.05 | 1 | 4 | Ttc9c | 0.026 | Old |
| 0.404281 | 1.284341 | 0.044 | 0.025 | 1 | 4 | Mamdc2 | 0.019 | Old |
| 0.404362 | 0.089566 | 0.822 | 0.838 | 1 | 4 | Ctsh | -0.016 | Old |
| 0.404587 | 0.485143 | 0.155 | 0.125 | 1 | 4 | Hspa14 | 0.03 | Old |
| 0.404671 | 0.552798 | 0.005 | 0.013 | 1 | 4 | Gm11772 | -0.008 | Old |
| 0.404671 | 0.383346 | 0.005 | 0.013 | 1 | 4 | Fignl1 | -0.008 | Old |
| 0.404671 | 0.298045 | 0.005 | 0.013 | 1 | 4 | Lca5 | -0.008 | Old |
| 0.404922 | 0.059788 | 0.18 | 0.138 | 1 | 4 | Appbp2 | 0.042 | Old |
| 0.405107 | 0.886388 | 0.101 | 0.075 | 1 | 4 | Terf2 | 0.026 | Old |
| 0.405183 | 0.450151 | 0.104 | 0.075 | 1 | 4 | 4931406P1 | 0.029 | Old |
| 0.405237 | 2.338212 | 0.028 | 0.013 | 1 | 4 | Fbxo21 | 0.015 | Old |
| 0.405414 | 0.973222 | 0.074 | 0.05 | 1 | 4 | Frs2 | 0.024 | Old |
| 0.405528 | 0.715521 | 0.117 | 0.088 | 1 | 4 | Ezr | 0.029 | Old |
| 0.406155 | 0.118969 | 0.052 | 0.075 | 1 | 4 | Epor | -0.023 | Old |
| 0.406265 | 1.246652 | 0.059 | 0.038 | 1 | 4 | Trmt6 | 0.021 | Old |
| 0.406734 | 0.16223 | 0.045 | 0.025 | 1 | 4 | Med9 | 0.02 | Old |
| 0.406742 | 0.737751 | 0.244 | 0.225 | 1 | 4 | Smarcd2 | 0.019 | Old |
| 0.406898 | 0.36476 | 0.37 | 0.338 | 1 | 4 | Gsn | 0.032 | Old |
| 0.406918 | 0.66466 | 0.155 | 0.125 | 1 | 4 | Stom | 0.03 | Old |
| 0.406968 | 0.127458 | 0.104 | 0.138 | 1 | 4 | Rbm18 | -0.034 | Old |
| 0.406973 | 0.264289 | 0.123 | 0.162 | 1 | 4 | Taf11 | -0.039 | Old |
| 0.407258 | 0.143902 | 0.072 | 0.1 | 1 | 4 | Ahsa2 | -0.028 | Old |
| 0.407988 | 0.103256 | 0.294 | 0.362 | 1 | 4 | Crlf3 | -0.068 | Old |
| 0.407999 | 0.894714 | 0.059 | 0.038 | 1 | 4 | Zfyve1 | 0.021 | Old |
| 0.408238 | 0.361372 | 0.028 | 0.013 | 1 | 4 | Paip2b | 0.015 | Old |
| 0.408497 | 1.74046 | 0.028 | 0.013 | 1 | 4 | Gmeb2 | 0.015 | Old |
| 0.408613 | 1.908294 | 0.043 | 0.025 | 1 | 4 | Magohb | 0.018 | Old |
| 0.408722 | 0.696946 | 0.116 | 0.088 | 1 | 4 | Rnmt | 0.028 | Old |
| 0.409252 | 1.637182 | 0.028 | 0.013 | 1 | 4 | Mettl8 | 0.015 | Old |
| 0.409252 | 1.617734 | 0.028 | 0.013 | 1 | 4 | 5730480H( | 0.015 | Old |
| 0.409252 | 1.59796 | 0.028 | 0.013 | 1 | 4 | Tle1 | 0.015 | Old |
| 0.409519 | 0.386136 | 0.31 | 0.275 | 1 | 4 | Ddi2 | 0.035 | Old |
| 0.409756 | 1.566934 | 0.028 | 0.013 | 1 | 4 | Gstt2 | 0.015 | Old |
| 0.410408 | 0.379145 | 0.175 | 0.138 | 1 | 4 | Acadl | 0.037 | Old |
| 0.410873 | 0.293883 | 0.2 | 0.162 | 1 | 4 | Sowahc | 0.038 | Old |
| 0.411068 | 0.057528 | 0.711 | 0.7 | 1 | 4 | Sh3glb1 | 0.011 | Old |
| 0.411655 | 0.525599 | 0.299 | 0.275 | 1 | 4 | Actn4 | 0.024 | Old |
| 0.411685 | 0.0231 | 0.32 | 0.388 | 1 | 4 | Pkig | -0.068 | Old |
| 0.411774 | 1.36694 | 0.028 | 0.013 | 1 | 4 | Skint3 | 0.015 | Old |

| | | | | | | | | |
|---|---|---|---|---|---|---|---|---|
| 0.411845 | 1.444671 | 0.043 | 0.025 | 1 | 4 | Dcps | 0.018 | Old |
| 0.412118 | 0.347227 | 0.082 | 0.112 | 1 | 4 | Fdx1 | -0.03 | Old |
| 0.41225 | 1.558248 | 0.043 | 0.025 | 1 | 4 | Tbc1d12 | 0.018 | Old |
| 0.412406 | 0.08186 | 0.25 | 0.212 | 1 | 4 | Nck1 | 0.038 | Old |
| 0.412772 | 0.466954 | 0.302 | 0.275 | 1 | 4 | Ndufa1 | 0.027 | Old |
| 0.412785 | 1.126209 | 0.028 | 0.013 | 1 | 4 | Acrbp | 0.015 | Old |
| 0.412975 | 0.255966 | 0.106 | 0.075 | 1 | 4 | Armc5 | 0.031 | Old |
| 0.413089 | 0.878336 | 0.044 | 0.025 | 1 | 4 | Ccnd2 | 0.019 | Old |
| 0.413181 | 0.156797 | 0.094 | 0.125 | 1 | 4 | Rrad | -0.031 | Old |
| 0.413294 | 0.007223 | 0.596 | 0.65 | 1 | 4 | Ighm | -0.054 | Old |
| 0.413582 | 0.580772 | 0.215 | 0.188 | 1 | 4 | Ppp3cb | 0.027 | Old |
| 0.413798 | 1.070631 | 0.028 | 0.013 | 1 | 4 | Acad12 | 0.015 | Old |
| 0.413859 | 0.567099 | 0.263 | 0.238 | 1 | 4 | Rad21 | 0.025 | Old |
| 0.414045 | 0.250184 | 0.106 | 0.075 | 1 | 4 | Bccip | 0.031 | Old |
| 0.414065 | 0.307908 | 0.061 | 0.038 | 1 | 4 | Dnajb12 | 0.023 | Old |
| 0.414172 | 0.026585 | 0.084 | 0.112 | 1 | 4 | Ncoa5 | -0.028 | Old |
| 0.414611 | 0.873286 | 0.087 | 0.062 | 1 | 4 | Ccdc25 | 0.025 | Old |
| 0.414857 | 0.23562 | 0.32 | 0.275 | 1 | 4 | H2-M3 | 0.045 | Old |
| 0.415012 | 0.157849 | 0.125 | 0.162 | 1 | 4 | Armc1 | -0.037 | Old |
| 0.415106 | 0.712872 | 0.044 | 0.025 | 1 | 4 | Ints4 | 0.019 | Old |
| 0.41524 | 0.386388 | 0.235 | 0.2 | 1 | 4 | Yeats4 | 0.035 | Old |
| 0.415296 | 0.823981 | 0.042 | 0.062 | 1 | 4 | Klhl7 | -0.02 | Old |
| 0.415463 | 0.471677 | 0.185 | 0.15 | 1 | 4 | Piezo1 | 0.035 | Old |
| 0.41564 | 0.648111 | 0.309 | 0.3 | 1 | 4 | Rad23b | 0.009 | Old |
| 0.416187 | 0.40258 | 0.146 | 0.112 | 1 | 4 | Hlx | 0.034 | Old |
| 0.416878 | 0.081927 | 0.073 | 0.1 | 1 | 4 | Ciz1 | -0.027 | Old |
| 0.417014 | 0.067825 | 0.156 | 0.2 | 1 | 4 | Smad5 | -0.044 | Old |
| 0.417436 | 0.728463 | 0.059 | 0.038 | 1 | 4 | Atr | 0.021 | Old |
| 0.417552 | 0.26397 | 0.173 | 0.138 | 1 | 4 | Stk3 | 0.035 | Old |
| 0.418121 | 0.234366 | 0.52 | 0.512 | 1 | 4 | H2-DMb1 | 0.008 | Old |
| 0.418126 | 0.240185 | 0.171 | 0.138 | 1 | 4 | Dock1 | 0.033 | Old |
| 0.418346 | 0.361024 | 0.042 | 0.062 | 1 | 4 | Smpd4 | -0.02 | Old |
| 0.4185 | 0.16677 | 0.771 | 0.775 | 1 | 4 | Sat1 | -0.004 | Old |
| 0.418938 | 0.384879 | 0.277 | 0.238 | 1 | 4 | Rcc2 | 0.039 | Old |
| 0.418947 | 2.290242 | 0.027 | 0.013 | 1 | 4 | Stradb | 0.014 | Old |
| 0.419161 | 1.49408 | 0.043 | 0.025 | 1 | 4 | Arel1 | 0.018 | Old |
| 0.419175 | 0.466542 | 0.219 | 0.188 | 1 | 4 | Dock11 | 0.031 | Old |
| 0.419651 | 0.097404 | 0.053 | 0.075 | 1 | 4 | Cdca4 | -0.022 | Old |
| 0.419966 | 0.864913 | 0.073 | 0.05 | 1 | 4 | Zfp524 | 0.023 | Old |
| 0.419992 | 1.108767 | 0.043 | 0.025 | 1 | 4 | Rdh11 | 0.018 | Old |
| 0.420488 | 0.49438 | 0.074 | 0.05 | 1 | 4 | Psip1 | 0.024 | Old |
| 0.420497 | 2.191166 | 0.027 | 0.013 | 1 | 4 | Kdm4b | 0.014 | Old |
| 0.420497 | 2.06529 | 0.027 | 0.013 | 1 | 4 | Pold1 | 0.014 | Old |
| 0.420632 | 0.021459 | 0.023 | 0.038 | 1 | 4 | Chpf | -0.015 | Old |
| 0.421188 | 1.208808 | 0.058 | 0.038 | 1 | 4 | Elp3 | 0.02 | Old |

| | | | | | | | | |
|---|---|---|---|---|---|---|---|---|
| 0.421593 | 0.084779 | 0.246 | 0.2 | 1 | 4 | Mindy1 | 0.046 | Old |
| 0.421719 | 0.097592 | 0.023 | 0.038 | 1 | 4 | Pmepa1 | -0.015 | Old |
| 0.422009 | 0.56136 | 0.042 | 0.062 | 1 | 4 | Ptcd2 | -0.02 | Old |
| 0.422184 | 0.45788 | 0.372 | 0.362 | 1 | 4 | Dpysl2 | 0.01 | Old |
| 0.422188 | 0.400673 | 0.082 | 0.112 | 1 | 4 | Decr1 | -0.03 | Old |
| 0.422264 | 0.143124 | 0.023 | 0.038 | 1 | 4 | Wdr59 | -0.015 | Old |
| 0.422335 | 0.502831 | 0.244 | 0.212 | 1 | 4 | Kat6a | 0.032 | Old |
| 0.42257 | 1.792067 | 0.027 | 0.013 | 1 | 4 | Ccdc130 | 0.014 | Old |
| 0.422709 | 0.31171 | 0.133 | 0.1 | 1 | 4 | Ankib1 | 0.033 | Old |
| 0.422885 | 0.89958 | 0.101 | 0.075 | 1 | 4 | Ankra2 | 0.026 | Old |
| 0.422978 | 0.22387 | 0.028 | 0.013 | 1 | 4 | Ifit1 | 0.015 | Old |
| 0.423017 | 1.153404 | 0.086 | 0.062 | 1 | 4 | Cnot7 | 0.024 | Old |
| 0.42331 | 0.150784 | 0.405 | 0.375 | 1 | 4 | Ubl5 | 0.03 | Old |
| 0.423499 | 0.422361 | 0.103 | 0.075 | 1 | 4 | Phf8 | 0.028 | Old |
| 0.423537 | 0.578315 | 0.206 | 0.175 | 1 | 4 | Atxn2 | 0.031 | Old |
| 0.424236 | 0.211394 | 0.511 | 0.5 | 1 | 4 | Vcp | 0.011 | Old |
| 0.424501 | 0.004795 | 0.116 | 0.15 | 1 | 4 | Akap8 | -0.034 | Old |
| 0.424513 | 0.648338 | 0.043 | 0.025 | 1 | 4 | Atg10 | 0.018 | Old |
| 0.424649 | 1.605229 | 0.027 | 0.013 | 1 | 4 | Lekr1 | 0.014 | Old |
| 0.425169 | 1.230574 | 0.027 | 0.013 | 1 | 4 | Gm26601 | 0.014 | Old |
| 0.425339 | 0.823039 | 0.043 | 0.025 | 1 | 4 | Dnajc25 | 0.018 | Old |
| 0.425629 | 0.039029 | 0.35 | 0.412 | 1 | 4 | Scand1 | -0.062 | Old |
| 0.425674 | 0.016076 | 0.17 | 0.212 | 1 | 4 | Cdc26 | -0.042 | Old |
| 0.42628 | 0.201208 | 0.202 | 0.162 | 1 | 4 | Ddx50 | 0.04 | Old |
| 0.426516 | 0.153668 | 0.063 | 0.088 | 1 | 4 | Qsox1 | -0.025 | Old |
| 0.426658 | 0.413298 | 0.21 | 0.175 | 1 | 4 | Sema6d | 0.035 | Old |
| 0.426701 | 0.615524 | 0.217 | 0.188 | 1 | 4 | Clock | 0.029 | Old |
| 0.426714 | 0.717906 | 0.073 | 0.05 | 1 | 4 | Cfl2 | 0.023 | Old |
| 0.427066 | 0.668586 | 0.128 | 0.1 | 1 | 4 | Dicer1 | 0.028 | Old |
| 0.427137 | 0.269235 | 0.449 | 0.438 | 1 | 4 | Litaf | 0.011 | Old |
| 0.427148 | 0.01952 | 0.126 | 0.162 | 1 | 4 | Cul5 | -0.036 | Old |
| 0.427329 | 0.809118 | 0.126 | 0.1 | 1 | 4 | Sfswap | 0.026 | Old |
| 0.427529 | 0.162974 | 0.268 | 0.225 | 1 | 4 | U2surp | 0.043 | Old |
| 0.427551 | 0.185296 | 0.105 | 0.138 | 1 | 4 | Tnip2 | -0.033 | Old |
| 0.427809 | 0.089125 | 0.033 | 0.05 | 1 | 4 | Tsen15 | -0.017 | Old |
| 0.427891 | 2.348417 | 0.042 | 0.025 | 1 | 4 | Ghdc | 0.017 | Old |
| 0.428062 | 0.107989 | 0.084 | 0.112 | 1 | 4 | Dmxl2 | -0.028 | Old |
| 0.428704 | 1.396143 | 0.043 | 0.025 | 1 | 4 | Ikbkg | 0.018 | Old |
| 0.428824 | 1.112752 | 0.027 | 0.013 | 1 | 4 | Tmem242 | 0.014 | Old |
| 0.429058 | 0.022447 | 0.096 | 0.125 | 1 | 4 | Mrpl43 | -0.029 | Old |
| 0.429457 | 0.358742 | 0.118 | 0.088 | 1 | 4 | Phyh | 0.03 | Old |
| 0.429478 | 0.63243 | 0.043 | 0.025 | 1 | 4 | BC005561 | 0.018 | Old |
| 0.430151 | 0.038785 | 0.592 | 0.713 | 1 | 4 | Fam49b | -0.121 | Old |
| 0.430206 | 1.218243 | 0.113 | 0.088 | 1 | 4 | Tnip3 | 0.025 | Old |
| 0.430214 | 0.55058 | 0.276 | 0.25 | 1 | 4 | Anxa6 | 0.026 | Old |

| | | | | | | | | |
|---|---|---|---|---|---|---|---|---|
| 0.431218 | 0.225257 | 0.55 | 0.537 | 1 | 4 | Rock2 | 0.013 | Old |
| 0.431823 | 0.415847 | 0.128 | 0.1 | 1 | 4 | Psmd3 | 0.028 | Old |
| 0.432279 | 0.526744 | 0.088 | 0.062 | 1 | 4 | Crnkl1 | 0.026 | Old |
| 0.432502 | 0.458797 | 0.117 | 0.088 | 1 | 4 | Cops8 | 0.029 | Old |
| 0.432728 | 0.633637 | 0.059 | 0.038 | 1 | 4 | Nup133 | 0.021 | Old |
| 0.433189 | 1.621163 | 0.057 | 0.038 | 1 | 4 | Mterf3 | 0.019 | Old |
| 0.433485 | 0.67892 | 0.128 | 0.1 | 1 | 4 | Rab3gap2 | 0.028 | Old |
| 0.433528 | 1.717823 | 0.07 | 0.05 | 1 | 4 | Med11 | 0.02 | Old |
| 0.43385 | 0.626432 | 0.073 | 0.05 | 1 | 4 | Tom1 | 0.023 | Old |
| 0.434116 | 0.534734 | 0.191 | 0.162 | 1 | 4 | Azi2 | 0.029 | Old |
| 0.434381 | 0.411536 | 0.157 | 0.125 | 1 | 4 | Strap | 0.032 | Old |
| 0.434544 | 0.236794 | 0.09 | 0.062 | 1 | 4 | Rbm14 | 0.028 | Old |
| 0.434978 | 0.122886 | 0.538 | 0.512 | 1 | 4 | Reep3 | 0.026 | Old |
| 0.434994 | 0.939091 | 0.043 | 0.025 | 1 | 4 | Zfp180 | 0.018 | Old |
| 0.435236 | 0.199862 | 0.043 | 0.062 | 1 | 4 | Cactin | -0.019 | Old |
| 0.435363 | 0.227735 | 0.132 | 0.1 | 1 | 4 | Sdccag3 | 0.032 | Old |
| 0.435587 | 0.596904 | 0.153 | 0.125 | 1 | 4 | Ndufs3 | 0.028 | Old |
| 0.436114 | 0.709848 | 0.354 | 0.325 | 1 | 4 | Lpl | 0.029 | Old |
| 0.436342 | 1.529326 | 0.042 | 0.025 | 1 | 4 | E2f3 | 0.017 | Old |
| 0.436717 | 0.454899 | 0.027 | 0.013 | 1 | 4 | Sdhaf3 | 0.014 | Old |
| 0.436746 | 0.554456 | 0.101 | 0.075 | 1 | 4 | Mosmo | 0.026 | Old |
| 0.437166 | 0.124401 | 0.163 | 0.125 | 1 | 4 | Scn1b | 0.038 | Old |
| 0.437609 | 0.661228 | 0.222 | 0.2 | 1 | 4 | Usp19 | 0.022 | Old |
| 0.438077 | 0.063132 | 0.053 | 0.075 | 1 | 4 | Cpsf4 | -0.022 | Old |
| 0.438305 | 0.398553 | 0.027 | 0.013 | 1 | 4 | Gm20300 | 0.014 | Old |
| 0.438489 | 0.943271 | 0.072 | 0.05 | 1 | 4 | Coa6 | 0.022 | Old |
| 0.438531 | 1.65517 | 0.026 | 0.013 | 1 | 4 | Mageh1 | 0.013 | Old |
| 0.439218 | 0.797793 | 0.043 | 0.025 | 1 | 4 | Snx19 | 0.018 | Old |
| 0.439547 | 0.058591 | 0.3 | 0.25 | 1 | 4 | Prpf40a | 0.05 | Old |
| 0.439605 | 1.680202 | 0.026 | 0.013 | 1 | 4 | Irgq | 0.013 | Old |
| 0.440065 | 0.819534 | 0.043 | 0.025 | 1 | 4 | Prkca | 0.018 | Old |
| 0.440142 | 1.863009 | 0.026 | 0.013 | 1 | 4 | Fdxr | 0.013 | Old |
| 0.440377 | 0.729385 | 0.058 | 0.038 | 1 | 4 | Prkaa1 | 0.02 | Old |
| 0.440651 | 0.2749 | 0.264 | 0.238 | 1 | 4 | Ripk1 | 0.026 | Old |
| 0.440832 | 0.855783 | 0.086 | 0.062 | 1 | 4 | Uty | 0.024 | Old |
| 0.441969 | 0.860745 | 0.072 | 0.05 | 1 | 4 | Gadd45gip | 0.022 | Old |
| 0.442068 | 0.180923 | 0.084 | 0.112 | 1 | 4 | Kdsr | -0.028 | Old |
| 0.442078 | 0.741472 | 0.126 | 0.1 | 1 | 4 | Rpe | 0.026 | Old |
| 0.443235 | 0.687399 | 0.126 | 0.1 | 1 | 4 | Acox1 | 0.026 | Old |
| 0.443311 | 0.443663 | 0.167 | 0.138 | 1 | 4 | Fez2 | 0.029 | Old |
| 0.443713 | 0.296832 | 0.377 | 0.35 | 1 | 4 | Gusb | 0.027 | Old |
| 0.443845 | 0.062622 | 0.064 | 0.088 | 1 | 4 | Rprd1a | -0.024 | Old |
| 0.44395 | 0.185312 | 0.059 | 0.038 | 1 | 4 | Ercc3 | 0.021 | Old |
| 0.444083 | 0.976261 | 0.057 | 0.038 | 1 | 4 | Nomo1 | 0.019 | Old |
| 0.444623 | 0.362693 | 0.246 | 0.212 | 1 | 4 | Dst | 0.034 | Old |

| | | | | | | | | |
|---|---|---|---|---|---|---|---|---|
| 0.444857 | 0.504633 | 0.331 | 0.312 | 1 | 4 | U2af2 | 0.019 | Old |
| 0.444996 | 0.022511 | 0.043 | 0.025 | 1 | 4 | 0610009B2 | 0.018 | Old |
| 0.445425 | 1.034927 | 0.071 | 0.05 | 1 | 4 | Kdm5d | 0.021 | Old |
| 0.4455 | 0.044582 | 0.204 | 0.25 | 1 | 4 | Aebp2 | -0.046 | Old |
| 0.445593 | 1.865649 | 0.056 | 0.038 | 1 | 4 | Glce | 0.018 | Old |
| 0.445674 | 0.508342 | 0.073 | 0.05 | 1 | 4 | Slc16a1 | 0.023 | Old |
| 0.446344 | 1.698816 | 0.056 | 0.038 | 1 | 4 | Nectin2 | 0.018 | Old |
| 0.44661 | 0.957486 | 0.042 | 0.025 | 1 | 4 | Usp31 | 0.017 | Old |
| 0.44672 | 0.441601 | 0.18 | 0.15 | 1 | 4 | Spata6 | 0.03 | Old |
| 0.447166 | 0.063079 | 0.074 | 0.05 | 1 | 4 | Aspscr1 | 0.024 | Old |
| 0.447892 | 0.821137 | 0.086 | 0.062 | 1 | 4 | Tpra1 | 0.024 | Old |
| 0.448097 | 0.051728 | 0.096 | 0.125 | 1 | 4 | Pkp4 | -0.029 | Old |
| 0.448203 | 1.05078 | 0.057 | 0.038 | 1 | 4 | Ryr1 | 0.019 | Old |
| 0.44872 | 0.907725 | 0.16 | 0.138 | 1 | 4 | Nmt1 | 0.022 | Old |
| 0.448782 | 0.909266 | 0.071 | 0.05 | 1 | 4 | Strip1 | 0.021 | Old |
| 0.448837 | 0.54481 | 0.18 | 0.15 | 1 | 4 | Trio | 0.03 | Old |
| 0.449401 | 0.138467 | 0.064 | 0.088 | 1 | 4 | Src | -0.024 | Old |
| 0.4497 | 0.561774 | 0.248 | 0.225 | 1 | 4 | Arhgef3 | 0.023 | Old |
| 0.449921 | 0.373021 | 0.482 | 0.488 | 1 | 4 | Ppp2ca | -0.006 | Old |
| 0.449943 | 0.123972 | 0.088 | 0.062 | 1 | 4 | Tatdn2 | 0.026 | Old |
| 0.450424 | 0.556801 | 0.026 | 0.013 | 1 | 4 | Map3k15 | 0.013 | Old |
| 0.450538 | 0.064206 | 0.246 | 0.3 | 1 | 4 | Pgls | -0.054 | Old |
| 0.450652 | 0.728409 | 0.072 | 0.05 | 1 | 4 | Sos1 | 0.022 | Old |
| 0.450657 | 1.577773 | 0.042 | 0.025 | 1 | 4 | Ccdc86 | 0.017 | Old |
| 0.451149 | 0.839875 | 0.085 | 0.062 | 1 | 4 | Gsr | 0.023 | Old |
| 0.451285 | 1.186119 | 0.07 | 0.05 | 1 | 4 | Usp45 | 0.02 | Old |
| 0.451312 | 1.370425 | 0.042 | 0.025 | 1 | 4 | Atm | 0.017 | Old |
| 0.451618 | 0.53789 | 0.153 | 0.125 | 1 | 4 | Mrpl38 | 0.028 | Old |
| 0.452374 | 0.181997 | 0.729 | 0.775 | 1 | 4 | Actr2 | -0.046 | Old |
| 0.452411 | 0.908731 | 0.148 | 0.125 | 1 | 4 | Smurf1 | 0.023 | Old |
| 0.452567 | 0.572747 | 0.165 | 0.138 | 1 | 4 | Srrt | 0.027 | Old |
| 0.45284 | 1.453655 | 0.042 | 0.025 | 1 | 4 | Gramd4 | 0.017 | Old |
| 0.453129 | 0.171517 | 0.47 | 0.438 | 1 | 4 | Sgpp1 | 0.032 | Old |
| 0.453137 | 0.58667 | 0.127 | 0.1 | 1 | 4 | Zyg11b | 0.027 | Old |
| 0.453324 | 0.293882 | 0.073 | 0.05 | 1 | 4 | Flnb | 0.023 | Old |
| 0.453414 | 0.078642 | 0.152 | 0.188 | 1 | 4 | Man2a2 | -0.036 | Old |
| 0.453717 | 0.18463 | 0.486 | 0.462 | 1 | 4 | Ncf2 | 0.024 | Old |
| 0.453752 | 0.354207 | 0.043 | 0.025 | 1 | 4 | Steap4 | 0.018 | Old |
| 0.453818 | 1.543274 | 0.083 | 0.062 | 1 | 4 | Mrps6 | 0.021 | Old |
| 0.454192 | 0.92695 | 0.098 | 0.075 | 1 | 4 | Smarcb1 | 0.023 | Old |
| 0.454249 | 0.236475 | 0.327 | 0.288 | 1 | 4 | Hnrnpl | 0.039 | Old |
| 0.4544 | 0.661097 | 0.042 | 0.025 | 1 | 4 | Fbxl19 | 0.017 | Old |
| 0.45489 | 0.236744 | 0.024 | 0.038 | 1 | 4 | Glb1l | -0.014 | Old |
| 0.455461 | 0.330495 | 0.115 | 0.088 | 1 | 4 | Npepl1 | 0.027 | Old |
| 0.456439 | 0.012807 | 0.026 | 0.013 | 1 | 4 | Dcun1d2 | 0.013 | Old |

| | | | | | | | | |
|---|---|---|---|---|---|---|---|---|
| 0.456607 | 0.596346 | 0.223 | 0.2 | 1 | 4 | Sptlc2 | 0.023 | Old |
| 0.456734 | 0.189477 | 0.034 | 0.05 | 1 | 4 | Nrm | -0.016 | Old |
| 0.456745 | 0.103959 | 0.343 | 0.288 | 1 | 4 | Surf4 | 0.055 | Old |
| 0.45697 | 0.807846 | 0.098 | 0.075 | 1 | 4 | Med16 | 0.023 | Old |
| 0.457377 | 1.700965 | 0.026 | 0.013 | 1 | 4 | Pik3ap1 | 0.013 | Old |
| 0.457577 | 0.751659 | 0.071 | 0.05 | 1 | 4 | Agl | 0.021 | Old |
| 0.457714 | 0.885 | 0.097 | 0.075 | 1 | 4 | Sec24d | 0.022 | Old |
| 0.457748 | 0.401167 | 0.142 | 0.112 | 1 | 4 | Cdc42ep3 | 0.03 | Old |
| 0.458937 | 0.208216 | 0.065 | 0.088 | 1 | 4 | Rpl10-ps3 | -0.023 | Old |
| 0.459032 | 0.081083 | 0.014 | 0.025 | 1 | 4 | Pde1b | -0.011 | Old |
| 0.459177 | 0.365416 | 0.163 | 0.138 | 1 | 4 | Slc25a24 | 0.025 | Old |
| 0.459827 | 0.464844 | 0.255 | 0.225 | 1 | 4 | Rnf7 | 0.03 | Old |
| 0.460162 | 1.084348 | 0.026 | 0.013 | 1 | 4 | Rfx2 | 0.013 | Old |
| 0.460269 | 0.516165 | 0.086 | 0.062 | 1 | 4 | Sc5d | 0.024 | Old |
| 0.460443 | 0.001237 | 0.119 | 0.15 | 1 | 4 | Katna1 | -0.031 | Old |
| 0.46073 | 0.127 | 0.274 | 0.338 | 1 | 4 | Cpne3 | -0.064 | Old |
| 0.460749 | 1.04153 | 0.042 | 0.025 | 1 | 4 | Ston2 | 0.017 | Old |
| 0.460846 | 0.323943 | 0.126 | 0.162 | 1 | 4 | Itfg1 | -0.036 | Old |
| 0.461182 | 0.246511 | 0.014 | 0.025 | 1 | 4 | Cep95 | -0.011 | Old |
| 0.461182 | 0.180271 | 0.014 | 0.025 | 1 | 4 | Zc3h12d | -0.011 | Old |
| 0.461307 | 0.299568 | 0.382 | 0.362 | 1 | 4 | Hspe1 | 0.02 | Old |
| 0.462102 | 0.072895 | 0.065 | 0.088 | 1 | 4 | Sipa1l1 | -0.023 | Old |
| 0.462326 | 0.0058 | 0.406 | 0.475 | 1 | 4 | Acp2 | -0.069 | Old |
| 0.462746 | 0.050913 | 0.055 | 0.075 | 1 | 4 | Pfkl | -0.02 | Old |
| 0.462811 | 0.381882 | 0.169 | 0.138 | 1 | 4 | Cdc37l1 | 0.031 | Old |
| 0.463024 | 0.148583 | 0.118 | 0.088 | 1 | 4 | Khnyn | 0.03 | Old |
| 0.463233 | 0.81042 | 0.07 | 0.05 | 1 | 4 | Zfp24 | 0.02 | Old |
| 0.463403 | 1.050498 | 0.042 | 0.025 | 1 | 4 | Ticam1 | 0.017 | Old |
| 0.464399 | 0.480518 | 0.203 | 0.175 | 1 | 4 | Trappc4 | 0.028 | Old |
| 0.46473 | 0.24255 | 0.142 | 0.112 | 1 | 4 | Nfu1 | 0.03 | Old |
| 0.465213 | 0.696079 | 0.098 | 0.075 | 1 | 4 | Clasp1 | 0.023 | Old |
| 0.465351 | 0.439351 | 0.128 | 0.1 | 1 | 4 | Ranbp9 | 0.028 | Old |
| 0.465499 | 0.6089 | 0.014 | 0.025 | 1 | 4 | Fndc4 | -0.011 | Old |
| 0.466505 | 0.364312 | 0.469 | 0.475 | 1 | 4 | Zmiz1 | -0.006 | Old |
| 0.466794 | 0.983789 | 0.034 | 0.05 | 1 | 4 | Spsb3 | -0.016 | Old |
| 0.466954 | 0.79139 | 0.042 | 0.025 | 1 | 4 | Map4k5 | 0.017 | Old |
| 0.466985 | 1.467159 | 0.055 | 0.038 | 1 | 4 | Nectin1 | 0.017 | Old |
| 0.467075 | 0.520469 | 0.198 | 0.175 | 1 | 4 | Ap3b1 | 0.023 | Old |
| 0.467272 | 0.000931 | 0.231 | 0.275 | 1 | 4 | Dynlrb1 | -0.044 | Old |
| 0.467833 | 0.040199 | 0.241 | 0.288 | 1 | 4 | Rasgrp4 | -0.047 | Old |
| 0.46822 | 0.397297 | 0.359 | 0.35 | 1 | 4 | Klf13 | 0.009 | Old |
| 0.468365 | 0.330621 | 0.184 | 0.15 | 1 | 4 | Ddx42 | 0.034 | Old |
| 0.46847 | 0.156826 | 0.065 | 0.088 | 1 | 4 | Crot | -0.023 | Old |
| 0.468631 | 0.171342 | 0.159 | 0.2 | 1 | 4 | Emc2 | -0.041 | Old |
| 0.468911 | 0.324499 | 0.057 | 0.038 | 1 | 4 | Mthfd2 | 0.019 | Old |

| | | | | | | | | |
|---|---|---|---|---|---|---|---|---|
| 0.468951 | 0.088152 | 0.044 | 0.062 | 1 | 4 | Lrrc40 | -0.018 | Old |
| 0.469121 | 0.190401 | 0.485 | 0.45 | 1 | 4 | Psmb4 | 0.035 | Old |
| 0.470073 | 0.625868 | 0.042 | 0.025 | 1 | 4 | Riox2 | 0.017 | Old |
| 0.470121 | 0.249276 | 0.206 | 0.175 | 1 | 4 | Rfk | 0.031 | Old |
| 0.470267 | 0.495885 | 0.026 | 0.013 | 1 | 4 | Cstf2 | 0.013 | Old |
| 0.47052 | 0.618252 | 0.042 | 0.025 | 1 | 4 | Polr2d | 0.017 | Old |
| 0.470558 | 0.079653 | 0.076 | 0.1 | 1 | 4 | Dhx16 | -0.024 | Old |
| 0.470671 | 0.550147 | 0.113 | 0.088 | 1 | 4 | Rab29 | 0.025 | Old |
| 0.47108 | 0.026476 | 0.006 | 0.013 | 1 | 4 | Palb2 | -0.007 | Old |
| 0.471308 | 0.04674 | 0.292 | 0.35 | 1 | 4 | Brd2 | -0.058 | Old |
| 0.47133 | 1.914761 | 0.025 | 0.013 | 1 | 4 | Fam117a | 0.012 | Old |
| 0.47146 | 0.090904 | 0.721 | 0.662 | 1 | 4 | Eif4g2 | 0.059 | Old |
| 0.471527 | 0.707636 | 0.07 | 0.05 | 1 | 4 | Ankrd10 | 0.02 | Old |
| 0.47216 | 1.02188 | 0.041 | 0.025 | 1 | 4 | Diras2 | 0.016 | Old |
| 0.4722 | 0.399445 | 0.006 | 0.013 | 1 | 4 | Napepld | -0.007 | Old |
| 0.4722 | 0.228382 | 0.006 | 0.013 | 1 | 4 | 3300005D( | -0.007 | Old |
| 0.4722 | 0.018862 | 0.006 | 0.013 | 1 | 4 | Mettl15 | -0.007 | Old |
| 0.472476 | 1.726258 | 0.025 | 0.013 | 1 | 4 | Gmds | 0.012 | Old |
| 0.472544 | 0.32752 | 0.154 | 0.125 | 1 | 4 | 1810030O( | 0.029 | Old |
| 0.473081 | 0.01758 | 0.119 | 0.15 | 1 | 4 | Ptprs | -0.031 | Old |
| 0.473323 | 1.230696 | 0.006 | 0.013 | 1 | 4 | Dlc1 | -0.007 | Old |
| 0.473323 | 0.729173 | 0.006 | 0.013 | 1 | 4 | Slc16a9 | -0.007 | Old |
| 0.473323 | 0.427324 | 0.006 | 0.013 | 1 | 4 | Pced1b | -0.007 | Old |
| 0.473323 | 0.360362 | 0.006 | 0.013 | 1 | 4 | Zfp273 | -0.007 | Old |
| 0.473328 | 1.779819 | 0.055 | 0.038 | 1 | 4 | Nol12 | 0.017 | Old |
| 0.473401 | 0.301928 | 0.152 | 0.125 | 1 | 4 | Nfya | 0.027 | Old |
| 0.473466 | 0.399046 | 0.072 | 0.05 | 1 | 4 | Coq2 | 0.022 | Old |
| 0.473513 | 1.110212 | 0.041 | 0.025 | 1 | 4 | Zfp865 | 0.016 | Old |
| 0.473994 | 0.045345 | 0.375 | 0.438 | 1 | 4 | Mrpl52 | -0.063 | Old |
| 0.474197 | 1.537858 | 0.025 | 0.013 | 1 | 4 | Prim2 | 0.012 | Old |
| 0.474227 | 0.4234 | 0.056 | 0.038 | 1 | 4 | Thoc1 | 0.018 | Old |
| 0.474231 | 0.23972 | 0.128 | 0.1 | 1 | 4 | Eri1 | 0.028 | Old |
| 0.474427 | 0.129143 | 0.46 | 0.412 | 1 | 4 | Atp1b3 | 0.048 | Old |
| 0.474508 | 0.1909 | 0.034 | 0.05 | 1 | 4 | Phf11d | -0.016 | Old |
| 0.474696 | 0.000473 | 0.372 | 0.45 | 1 | 4 | Atp5j2 | -0.078 | Old |
| 0.474772 | 1.306947 | 0.025 | 0.013 | 1 | 4 | Zfp111 | 0.012 | Old |
| 0.475004 | 0.669743 | 0.056 | 0.038 | 1 | 4 | Mrpl40 | 0.018 | Old |
| 0.475083 | 0.209809 | 0.425 | 0.4 | 1 | 4 | Tfrc | 0.025 | Old |
| 0.475266 | 0.306035 | 0.142 | 0.112 | 1 | 4 | Skiv2l | 0.03 | Old |
| 0.475837 | 0.164617 | 0.721 | 0.75 | 1 | 4 | Arf5 | -0.029 | Old |
| 0.475894 | 0.589471 | 0.112 | 0.088 | 1 | 4 | Mdp1 | 0.024 | Old |
| 0.475922 | 1.203823 | 0.025 | 0.013 | 1 | 4 | Pkib | 0.012 | Old |
| 0.47609 | 0.449663 | 0.26 | 0.238 | 1 | 4 | Arfgef1 | 0.022 | Old |
| 0.476175 | 0.420698 | 0.307 | 0.288 | 1 | 4 | Sin3b | 0.019 | Old |
| 0.4767 | 0.021139 | 0.908 | 0.912 | 1 | 4 | mt-Nd3 | -0.004 | Old |

| | | | | | | | | |
|---|---|---|---|---|---|---|---|---|
| 0.476842 | 0.489452 | 0.189 | 0.162 | 1 | 4 | Polr2e | 0.027 | Old |
| 0.47689 | 0.768047 | 0.084 | 0.062 | 1 | 4 | Copg2 | 0.022 | Old |
| 0.476922 | 0.177179 | 0.034 | 0.05 | 1 | 4 | Ccdc28b | -0.016 | Old |
| 0.47699 | 1.712641 | 0.04 | 0.025 | 1 | 4 | Zfp617 | 0.015 | Old |
| 0.477052 | 0.022662 | 0.109 | 0.138 | 1 | 4 | Lrrc41 | -0.029 | Old |
| 0.477104 | 0.104901 | 0.638 | 0.637 | 1 | 4 | Pdia3 | 0.001 | Old |
| 0.477284 | 0.4438 | 0.084 | 0.062 | 1 | 4 | Tmem120a | 0.022 | Old |
| 0.477302 | 0.005197 | 0.233 | 0.275 | 1 | 4 | Sri | -0.042 | Old |
| 0.477493 | 0.556958 | 0.254 | 0.238 | 1 | 4 | Ppp1r12a | 0.016 | Old |
| 0.477552 | 0.326933 | 0.168 | 0.138 | 1 | 4 | Rest | 0.03 | Old |
| 0.478119 | 0.60619 | 0.056 | 0.038 | 1 | 4 | Mtpap | 0.018 | Old |
| 0.478517 | 0.896332 | 0.122 | 0.1 | 1 | 4 | Srf | 0.022 | Old |
| 0.47874 | 0.214553 | 0.456 | 0.438 | 1 | 4 | Ube2d2a | 0.018 | Old |
| 0.478805 | 0.921022 | 0.025 | 0.013 | 1 | 4 | Rfxank | 0.012 | Old |
| 0.4789 | 0.692696 | 0.056 | 0.038 | 1 | 4 | Cd24a | 0.018 | Old |
| 0.479291 | 0.845524 | 0.096 | 0.075 | 1 | 4 | Nifk | 0.021 | Old |
| 0.47931 | 0.040041 | 0.045 | 0.062 | 1 | 4 | Gga2 | -0.017 | Old |
| 0.479436 | 0.084277 | 0.874 | 0.863 | 1 | 4 | Rpl22 | 0.011 | Old |
| 0.4795 | 0.240061 | 0.042 | 0.025 | 1 | 4 | Farsa | 0.017 | Old |
| 0.479677 | 0.46205 | 0.113 | 0.088 | 1 | 4 | Adar | 0.025 | Old |
| 0.479736 | 1.573651 | 0.04 | 0.025 | 1 | 4 | Xlr4b | 0.015 | Old |
| 0.479914 | 0.363981 | 0.126 | 0.1 | 1 | 4 | Dnajc14 | 0.026 | Old |
| 0.479952 | 0.249215 | 0.042 | 0.025 | 1 | 4 | Ubxn8 | 0.017 | Old |
| 0.480019 | 0.3898 | 0.23 | 0.2 | 1 | 4 | Srsf9 | 0.03 | Old |
| 0.480409 | 0.142902 | 0.087 | 0.112 | 1 | 4 | Snrnp48 | -0.025 | Old |
| 0.480629 | 1.04691 | 0.082 | 0.062 | 1 | 4 | Tusc2 | 0.02 | Old |
| 0.481118 | 0.754396 | 0.025 | 0.013 | 1 | 4 | Ttc21b | 0.012 | Old |
| 0.481118 | 0.577691 | 0.025 | 0.013 | 1 | 4 | Rbm41 | 0.012 | Old |
| 0.481432 | 0.789643 | 0.055 | 0.038 | 1 | 4 | Mpv17l2 | 0.017 | Old |
| 0.481442 | 0.608933 | 0.056 | 0.038 | 1 | 4 | Mir142hg | 0.018 | Old |
| 0.481585 | 0.343107 | 0.261 | 0.225 | 1 | 4 | Pdcd5 | 0.036 | Old |
| 0.481637 | 0.139637 | 0.024 | 0.038 | 1 | 4 | Sgsm2 | -0.014 | Old |
| 0.481995 | 0.424619 | 0.153 | 0.125 | 1 | 4 | Atp9b | 0.028 | Old |
| 0.48203 | 1.455692 | 0.04 | 0.025 | 1 | 4 | Pop4 | 0.015 | Old |
| 0.482298 | 0.299462 | 0.142 | 0.112 | 1 | 4 | Pank2 | 0.03 | Old |
| 0.482419 | 0.026021 | 0.252 | 0.3 | 1 | 4 | Tcp1 | -0.048 | Old |
| 0.4825 | 0.29989 | 0.403 | 0.388 | 1 | 4 | Arhgap30 | 0.015 | Old |
| 0.482571 | 0.54599 | 0.258 | 0.238 | 1 | 4 | Adam9 | 0.02 | Old |
| 0.482605 | 0.226788 | 0.057 | 0.038 | 1 | 4 | Trap1 | 0.019 | Old |
| 0.482728 | 0.211525 | 0.239 | 0.3 | 1 | 4 | Pet100 | -0.061 | Old |
| 0.483049 | 0.753886 | 0.041 | 0.025 | 1 | 4 | Scd2 | 0.016 | Old |
| 0.483064 | 0.362922 | 0.07 | 0.05 | 1 | 4 | Ap3s2 | 0.02 | Old |
| 0.48316 | 0.050095 | 0.045 | 0.062 | 1 | 4 | Ints11 | -0.017 | Old |
| 0.483634 | 0.874849 | 0.158 | 0.138 | 1 | 4 | Foxn2 | 0.02 | Old |
| 0.483766 | 0.718027 | 0.123 | 0.1 | 1 | 4 | Otud4 | 0.023 | Old |

| | | | | | | | | |
|---|---|---|---|---|---|---|---|---|
| 0.483806 | 0.238391 | 0.178 | 0.15 | 1 | 4 | Bcl7b | 0.028 | Old |
| 0.483929 | 0.409262 | 0.024 | 0.038 | 1 | 4 | Spef1 | -0.014 | Old |
| 0.484131 | 0.078677 | 0.055 | 0.075 | 1 | 4 | Arl11 | -0.02 | Old |
| 0.484315 | 0.101049 | 0.228 | 0.275 | 1 | 4 | Ostc | -0.047 | Old |
| 0.484598 | 0.549169 | 0.025 | 0.013 | 1 | 4 | Gemin5 | 0.012 | Old |
| 0.484738 | 0.438281 | 0.111 | 0.088 | 1 | 4 | Atp11b | 0.023 | Old |
| 0.484764 | 0.104747 | 0.173 | 0.212 | 1 | 4 | Tmem51 | -0.039 | Old |
| 0.484884 | 0.390751 | 0.071 | 0.05 | 1 | 4 | Trim2 | 0.021 | Old |
| 0.485022 | 1.22525 | 0.04 | 0.025 | 1 | 4 | Nemp1 | 0.015 | Old |
| 0.48518 | 0.471249 | 0.025 | 0.013 | 1 | 4 | Dtx3 | 0.012 | Old |
| 0.485652 | 0.266214 | 0.024 | 0.038 | 1 | 4 | Amer1 | -0.014 | Old |
| 0.485714 | 1.278452 | 0.04 | 0.025 | 1 | 4 | Cep44 | 0.015 | Old |
| 0.485719 | 0.136236 | 0.542 | 0.525 | 1 | 4 | Gnas | 0.017 | Old |
| 0.485825 | 0.821959 | 0.11 | 0.088 | 1 | 4 | Dhx30 | 0.022 | Old |
| 0.486406 | 1.197711 | 0.04 | 0.025 | 1 | 4 | Atpaf1 | 0.015 | Old |
| 0.486469 | 0.257502 | 0.57 | 0.588 | 1 | 4 | Rpn1 | -0.018 | Old |
| 0.486608 | 0.239951 | 0.356 | 0.325 | 1 | 4 | Map7d1 | 0.031 | Old |
| 0.486635 | 0.627165 | 0.211 | 0.188 | 1 | 4 | Irf9 | 0.023 | Old |
| 0.486926 | 0.167155 | 0.025 | 0.013 | 1 | 4 | Gk5 | 0.012 | Old |
| 0.487099 | 1.14233 | 0.04 | 0.025 | 1 | 4 | Ints7 | 0.015 | Old |
| 0.487342 | 0.397941 | 0.106 | 0.138 | 1 | 4 | Cog7 | -0.032 | Old |
| 0.487745 | 0.220748 | 0.111 | 0.088 | 1 | 4 | Fgr | 0.023 | Old |
| 0.487846 | 0.527662 | 0.098 | 0.075 | 1 | 4 | Acads | 0.023 | Old |
| 0.488004 | 0.497949 | 0.07 | 0.05 | 1 | 4 | Arhgap4 | 0.02 | Old |
| 0.488093 | 0.348871 | 0.025 | 0.013 | 1 | 4 | Zfp868 | 0.012 | Old |
| 0.488145 | 0.058703 | 0.121 | 0.15 | 1 | 4 | Uqcc2 | -0.029 | Old |
| 0.488556 | 0.647098 | 0.097 | 0.075 | 1 | 4 | Edc4 | 0.022 | Old |
| 0.488676 | 0.107078 | 0.025 | 0.013 | 1 | 4 | Osbpl1a | 0.012 | Old |
| 0.488683 | 0.057073 | 0.291 | 0.35 | 1 | 4 | Dad1 | -0.059 | Old |
| 0.488775 | 0.218108 | 0.115 | 0.088 | 1 | 4 | Rsl24d1 | 0.027 | Old |
| 0.48915 | 1.272921 | 0.094 | 0.075 | 1 | 4 | Dpp3 | 0.019 | Old |
| 0.490067 | 0.632886 | 0.123 | 0.1 | 1 | 4 | Zwint | 0.023 | Old |
| 0.490348 | 0.400899 | 0.113 | 0.088 | 1 | 4 | Pou2f1 | 0.025 | Old |
| 0.491119 | 1.436029 | 0.024 | 0.013 | 1 | 4 | Aste1 | 0.011 | Old |
| 0.491158 | 1.043559 | 0.055 | 0.038 | 1 | 4 | Klhl2 | 0.017 | Old |
| 0.491959 | 1.050362 | 0.055 | 0.038 | 1 | 4 | Eipr1 | 0.017 | Old |
| 0.492028 | 0.013806 | 0.243 | 0.288 | 1 | 4 | Ddb1 | -0.045 | Old |
| 0.492127 | 0.684051 | 0.055 | 0.038 | 1 | 4 | Col16a1 | 0.017 | Old |
| 0.492486 | 0.56079 | 0.083 | 0.062 | 1 | 4 | Slc38a1 | 0.021 | Old |
| 0.492761 | 0.90562 | 0.055 | 0.038 | 1 | 4 | Naf1 | 0.017 | Old |
| 0.493026 | 0.11816 | 0.13 | 0.162 | 1 | 4 | Cnot8 | -0.032 | Old |
| 0.493522 | 0.597506 | 0.055 | 0.038 | 1 | 4 | 4921524J1 | 0.017 | Old |
| 0.493904 | 0.022891 | 0.177 | 0.212 | 1 | 4 | Srp19 | -0.035 | Old |
| 0.494025 | 0.037772 | 0.863 | 0.825 | 1 | 4 | Calm1 | 0.038 | Old |
| 0.494029 | 0.004456 | 0.099 | 0.125 | 1 | 4 | Ttc33 | -0.026 | Old |

| | | | | | | | | |
|---|---|---|---|---|---|---|---|---|
| 0.494162 | 0.485402 | 0.305 | 0.288 | 1 | 4 | S1pr1 | 0.017 | Old |
| 0.494326 | 0.793184 | 0.069 | 0.05 | 1 | 4 | Unc5a | 0.019 | Old |
| 0.494546 | 0.316973 | 0.337 | 0.325 | 1 | 4 | Pik3cd | 0.012 | Old |
| 0.494683 | 1.200694 | 0.024 | 0.013 | 1 | 4 | Ptges2 | 0.011 | Old |
| 0.494938 | 1.296393 | 0.068 | 0.05 | 1 | 4 | Mob2 | 0.018 | Old |
| 0.495083 | 0.019495 | 0.099 | 0.125 | 1 | 4 | Suclg2 | -0.026 | Old |
| 0.495279 | 1.051532 | 0.024 | 0.013 | 1 | 4 | Srr | 0.011 | Old |
| 0.495596 | 1.359304 | 0.067 | 0.05 | 1 | 4 | Zfp236 | 0.017 | Old |
| 0.496095 | 0.012183 | 0.345 | 0.412 | 1 | 4 | Fermt3 | -0.067 | Old |
| 0.496268 | 0.034151 | 0.186 | 0.225 | 1 | 4 | Tox4 | -0.039 | Old |
| 0.496376 | 0.873373 | 0.055 | 0.038 | 1 | 4 | Sigmar1 | 0.017 | Old |
| 0.496384 | 0.512037 | 0.187 | 0.162 | 1 | 4 | Nsfl1c | 0.025 | Old |
| 0.496471 | 0.932915 | 0.024 | 0.013 | 1 | 4 | Elmod3 | 0.011 | Old |
| 0.496496 | 0.026421 | 0.158 | 0.125 | 1 | 4 | Ethe1 | 0.033 | Old |
| 0.49676 | 0.886659 | 0.082 | 0.062 | 1 | 4 | Taf7 | 0.02 | Old |
| 0.497199 | 0.092376 | 0.098 | 0.125 | 1 | 4 | Ikzf5 | -0.027 | Old |
| 0.497233 | 0.528854 | 0.07 | 0.05 | 1 | 4 | Sik3 | 0.02 | Old |
| 0.497441 | 0.67129 | 0.083 | 0.062 | 1 | 4 | Utp6 | 0.021 | Old |
| 0.497586 | 0.280014 | 0.015 | 0.025 | 1 | 4 | Ccdc106 | -0.01 | Old |
| 0.497586 | 0.169096 | 0.015 | 0.025 | 1 | 4 | Katnb1 | -0.01 | Old |
| 0.497613 | 0.005197 | 0.484 | 0.438 | 1 | 4 | Srsf2 | 0.046 | Old |
| 0.497744 | 0.868528 | 0.108 | 0.088 | 1 | 4 | Ric1 | 0.02 | Old |
| 0.498123 | 0.476115 | 0.055 | 0.038 | 1 | 4 | Dubr | 0.017 | Old |
| 0.498714 | 1.430213 | 0.04 | 0.025 | 1 | 4 | Ccdc85b | 0.015 | Old |
| 0.498959 | 0.620912 | 0.04 | 0.025 | 1 | 4 | Dclre1a | 0.015 | Old |
| 0.499035 | 0.224334 | 0.596 | 0.6 | 1 | 4 | Dse | -0.004 | Old |
| 0.499053 | 0.243224 | 0.015 | 0.025 | 1 | 4 | Pfkfb2 | -0.01 | Old |
| 0.499053 | 0.100435 | 0.015 | 0.025 | 1 | 4 | Gm10138 | -0.01 | Old |
| 0.499197 | 0.796825 | 0.055 | 0.038 | 1 | 4 | 0610010K1 | 0.017 | Old |
| 0.499326 | 0.361579 | 0.055 | 0.038 | 1 | 4 | Anapc10 | 0.017 | Old |
| 0.499569 | 0.541081 | 0.07 | 0.05 | 1 | 4 | C2cd3 | 0.02 | Old |
| 0.499788 | 0.300213 | 0.015 | 0.025 | 1 | 4 | Acsf2 | -0.01 | Old |
| 0.49981 | 0.25093 | 0.108 | 0.138 | 1 | 4 | Mvb12a | -0.03 | Old |
| 0.500326 | 0.484212 | 0.11 | 0.088 | 1 | 4 | Prkag2 | 0.022 | Old |
| 0.500523 | 0.31118 | 0.015 | 0.025 | 1 | 4 | Bend6 | -0.01 | Old |
| 0.500619 | 0.00017 | 0.387 | 0.462 | 1 | 4 | Hspa9 | -0.075 | Old |
| 0.500638 | 0.106519 | 0.056 | 0.038 | 1 | 4 | Prcc | 0.018 | Old |
| 0.500655 | 0.696095 | 0.024 | 0.013 | 1 | 4 | Zfp653 | 0.011 | Old |
| 0.500813 | 0.667869 | 0.055 | 0.038 | 1 | 4 | Zfp771 | 0.017 | Old |
| 0.501143 | 0.046926 | 0.165 | 0.2 | 1 | 4 | Mrpl51 | -0.035 | Old |
| 0.501156 | 0.667044 | 0.069 | 0.05 | 1 | 4 | Zfp398 | 0.019 | Old |
| 0.501254 | 0.487374 | 0.024 | 0.013 | 1 | 4 | Ncapd3 | 0.011 | Old |
| 0.501259 | 0.279353 | 0.015 | 0.025 | 1 | 4 | Tesk2 | -0.01 | Old |
| 0.501336 | 0.384117 | 0.055 | 0.038 | 1 | 4 | Timm17b | 0.017 | Old |
| 0.502323 | 0.380247 | 0.14 | 0.112 | 1 | 4 | Fkbp4 | 0.028 | Old |

| | | | | | | | | |
|---|---|---|---|---|---|---|---|---|
| 0.502594 | 0.844223 | 0.155 | 0.138 | 1 | 4 | Vps11 | 0.017 | Old |
| 0.502652 | 0.456503 | 0.188 | 0.162 | 1 | 4 | Eif1ax | 0.026 | Old |
| 0.502902 | 0.109737 | 0.878 | 0.887 | 1 | 4 | Rack1 | -0.009 | Old |
| 0.502968 | 1.009992 | 0.04 | 0.025 | 1 | 4 | Pdcd2 | 0.015 | Old |
| 0.50303 | 0.123895 | 0.045 | 0.062 | 1 | 4 | Cxxc1 | -0.017 | Old |
| 0.503054 | 0.513603 | 0.024 | 0.013 | 1 | 4 | Amotl2 | 0.011 | Old |
| 0.503118 | 0.955961 | 0.054 | 0.038 | 1 | 4 | Acap3 | 0.016 | Old |
| 0.503245 | 0.176591 | 0.416 | 0.388 | 1 | 4 | Vwa5a | 0.028 | Old |
| 0.503258 | 0.007423 | 0.133 | 0.162 | 1 | 4 | Cramp1l | -0.029 | Old |
| 0.503352 | 0.574165 | 0.069 | 0.05 | 1 | 4 | Nfyc | 0.019 | Old |
| 0.503564 | 0.281684 | 0.245 | 0.212 | 1 | 4 | Zc3hav1 | 0.033 | Old |
| 0.503704 | 0.722294 | 0.069 | 0.05 | 1 | 4 | Pigq | 0.019 | Old |
| 0.504207 | 1.080873 | 0.015 | 0.025 | 1 | 4 | Peg10 | -0.01 | Old |
| 0.504246 | 0.441833 | 0.227 | 0.2 | 1 | 4 | Hps3 | 0.027 | Old |
| 0.504334 | 0.291783 | 0.14 | 0.112 | 1 | 4 | Dpy19l4 | 0.028 | Old |
| 0.504828 | 0.366064 | 0.04 | 0.025 | 1 | 4 | Cox10 | 0.015 | Old |
| 0.506059 | 0.319686 | 0.263 | 0.238 | 1 | 4 | Snx18 | 0.025 | Old |
| 0.506259 | 0.048125 | 0.067 | 0.088 | 1 | 4 | Fbxl4 | -0.021 | Old |
| 0.506279 | 0.755067 | 0.132 | 0.112 | 1 | 4 | Nup62 | 0.02 | Old |
| 0.506286 | 1.940313 | 0.024 | 0.013 | 1 | 4 | Gm5431 | 0.011 | Old |
| 0.506308 | 1.962861 | 0.039 | 0.025 | 1 | 4 | Men1 | 0.014 | Old |
| 0.506552 | 0.233907 | 0.594 | 0.65 | 1 | 4 | Cflar | -0.056 | Old |
| 0.506897 | 2.022391 | 0.024 | 0.013 | 1 | 4 | 2610044O1 | 0.011 | Old |
| 0.506932 | 0.495335 | 0.248 | 0.225 | 1 | 4 | Il21r | 0.023 | Old |
| 0.507076 | 0.295064 | 0.294 | 0.262 | 1 | 4 | Rtraf | 0.032 | Old |
| 0.507095 | 0.625181 | 0.083 | 0.062 | 1 | 4 | Il15 | 0.021 | Old |
| 0.507692 | 0.403082 | 0.138 | 0.112 | 1 | 4 | Fut8 | 0.026 | Old |
| 0.508293 | 0.517221 | 0.137 | 0.112 | 1 | 4 | Oaz2 | 0.025 | Old |
| 0.508841 | 1.022243 | 0.054 | 0.038 | 1 | 4 | Papd7 | 0.016 | Old |
| 0.508858 | 0.043383 | 0.586 | 0.512 | 1 | 4 | Lrp6 | 0.074 | Old |
| 0.509142 | 1.252366 | 0.053 | 0.038 | 1 | 4 | Polr2h | 0.015 | Old |
| 0.509181 | 1.21152 | 0.067 | 0.05 | 1 | 4 | Scyl1 | 0.017 | Old |
| 0.509319 | 0.523977 | 0.163 | 0.138 | 1 | 4 | Dstyk | 0.025 | Old |
| 0.509344 | 1.515132 | 0.024 | 0.013 | 1 | 4 | Rhoj | 0.011 | Old |
| 0.509623 | 0.712489 | 0.04 | 0.025 | 1 | 4 | Slc25a19 | 0.015 | Old |
| 0.510011 | 0.407237 | 0.284 | 0.262 | 1 | 4 | Hectd1 | 0.022 | Old |
| 0.510021 | 0.51254 | 0.11 | 0.088 | 1 | 4 | Nckipsd | 0.022 | Old |
| 0.510569 | 1.383859 | 0.024 | 0.013 | 1 | 4 | Ifi206 | 0.011 | Old |
| 0.510578 | 0.758948 | 0.04 | 0.025 | 1 | 4 | Tomm40l | 0.015 | Old |
| 0.511055 | 0.744437 | 0.04 | 0.025 | 1 | 4 | Rpl27-ps3 | 0.015 | Old |
| 0.511183 | 1.459957 | 0.024 | 0.013 | 1 | 4 | Wdr24 | 0.011 | Old |
| 0.511873 | 0.106317 | 0.175 | 0.15 | 1 | 4 | Dcp2 | 0.025 | Old |
| 0.512074 | 1.410969 | 0.039 | 0.025 | 1 | 4 | Pwp2 | 0.014 | Old |
| 0.512117 | 0.284385 | 0.219 | 0.188 | 1 | 4 | Egln2 | 0.031 | Old |
| 0.512242 | 0.270363 | 0.385 | 0.362 | 1 | 4 | Tmco1 | 0.023 | Old |

| | | | | | | | | |
|---|---|---|---|---|---|---|---|---|
| 0.512312 | 0.046305 | 0.288 | 0.338 | 1 | 4 | Adap2 | -0.05 | Old |
| 0.512411 | 1.053724 | 0.024 | 0.013 | 1 | 4 | Orc6 | 0.011 | Old |
| 0.51333 | 0.464371 | 0.15 | 0.125 | 1 | 4 | Ythdf2 | 0.025 | Old |
| 0.51343 | 0.104389 | 0.078 | 0.1 | 1 | 4 | Fbxl17 | -0.022 | Old |
| 0.513695 | 0.315035 | 0.403 | 0.388 | 1 | 4 | Ube3a | 0.015 | Old |
| 0.514004 | 1.301021 | 0.039 | 0.025 | 1 | 4 | Pomk | 0.014 | Old |
| 0.514035 | 0.571314 | 0.252 | 0.238 | 1 | 4 | Tpcn1 | 0.014 | Old |
| 0.514147 | 0.435954 | 0.083 | 0.062 | 1 | 4 | Faim | 0.021 | Old |
| 0.514186 | 0.748442 | 0.054 | 0.038 | 1 | 4 | Ppard | 0.016 | Old |
| 0.514298 | 0.105903 | 0.067 | 0.088 | 1 | 4 | Rai1 | -0.021 | Old |
| 0.514904 | 0.567219 | 0.161 | 0.138 | 1 | 4 | Pole4 | 0.023 | Old |
| 0.515936 | 0.153377 | 0.796 | 0.787 | 1 | 4 | Rtn4 | 0.009 | Old |
| 0.516267 | 0.192529 | 0.155 | 0.125 | 1 | 4 | Bdp1 | 0.03 | Old |
| 0.516534 | 0.080732 | 0.036 | 0.05 | 1 | 4 | Dxo | -0.014 | Old |
| 0.516603 | 0.62129 | 0.082 | 0.062 | 1 | 4 | Ankrd50 | 0.02 | Old |
| 0.516691 | 0.217988 | 0.43 | 0.425 | 1 | 4 | Tmed2 | 0.005 | Old |
| 0.51683 | 0.339673 | 0.239 | 0.212 | 1 | 4 | Nfkb2 | 0.027 | Old |
| 0.51684 | 0.434946 | 0.368 | 0.375 | 1 | 4 | Gm2a | -0.007 | Old |
| 0.516958 | 0.464188 | 0.15 | 0.125 | 1 | 4 | Acaa2 | 0.025 | Old |
| 0.516999 | 0.937938 | 0.118 | 0.1 | 1 | 4 | Ephx1 | 0.018 | Old |
| 0.517202 | 0.005095 | 0.321 | 0.362 | 1 | 4 | Idh2 | -0.041 | Old |
| 0.517285 | 0.555876 | 0.04 | 0.025 | 1 | 4 | Gnpnat1 | 0.015 | Old |
| 0.517578 | 0.591457 | 0.208 | 0.188 | 1 | 4 | Tulp4 | 0.02 | Old |
| 0.517955 | 0.690097 | 0.024 | 0.013 | 1 | 4 | Pick1 | 0.011 | Old |
| 0.518196 | 0.058528 | 0.046 | 0.062 | 1 | 4 | Rpain | -0.016 | Old |
| 0.518765 | 0.590489 | 0.069 | 0.05 | 1 | 4 | Cd2bp2 | 0.019 | Old |
| 0.519186 | 0.017025 | 0.098 | 0.075 | 1 | 4 | Mettl23 | 0.023 | Old |
| 0.519429 | 1.270813 | 0.067 | 0.05 | 1 | 4 | Bcam | 0.017 | Old |
| 0.51959 | 1.000686 | 0.067 | 0.05 | 1 | 4 | Pias4 | 0.017 | Old |
| 0.51976 | 0.569313 | 0.229 | 0.212 | 1 | 4 | Rnf213 | 0.017 | Old |
| 0.519802 | 0.361424 | 0.055 | 0.038 | 1 | 4 | Sap30bp | 0.017 | Old |
| 0.520179 | 0.137173 | 0.131 | 0.162 | 1 | 4 | Grk6 | -0.031 | Old |
| 0.520401 | 2.088941 | 0.038 | 0.025 | 1 | 4 | Slc37a4 | 0.013 | Old |
| 0.520401 | 1.928506 | 0.038 | 0.025 | 1 | 4 | Plp2 | 0.013 | Old |
| 0.520406 | 0.333073 | 0.308 | 0.288 | 1 | 4 | Mdh1 | 0.02 | Old |
| 0.520782 | 0.162522 | 0.233 | 0.288 | 1 | 4 | Snrpg | -0.055 | Old |
| 0.520891 | 0.095238 | 0.057 | 0.075 | 1 | 4 | Mphosph6 | -0.018 | Old |
| 0.521382 | 2.02321 | 0.038 | 0.025 | 1 | 4 | Recql5 | 0.013 | Old |
| 0.521936 | 0.463897 | 0.162 | 0.138 | 1 | 4 | Pogk | 0.024 | Old |
| 0.522504 | 0.130581 | 0.036 | 0.05 | 1 | 4 | 2300009A( | -0.014 | Old |
| 0.522513 | 0.016928 | 0.078 | 0.1 | 1 | 4 | Nde1 | -0.022 | Old |
| 0.522906 | 0.724662 | 0.068 | 0.05 | 1 | 4 | Etfrf1 | 0.018 | Old |
| 0.52291 | 0.319838 | 0.024 | 0.013 | 1 | 4 | Fn3krp | 0.011 | Old |
| 0.52291 | 0.257383 | 0.024 | 0.013 | 1 | 4 | Clec4d | 0.011 | Old |
| 0.52326 | 0.633622 | 0.082 | 0.062 | 1 | 4 | Klhdc2 | 0.02 | Old |

| | | | | | | | | |
|---|---|---|---|---|---|---|---|---|
| 0.523521 | 0.815565 | 0.067 | 0.05 | 1 | 4 | Gphn | 0.017 | Old |
| 0.523639 | 0.151475 | 0.285 | 0.338 | 1 | 4 | Snu13 | -0.053 | Old |
| 0.524104 | 0.917457 | 0.053 | 0.038 | 1 | 4 | Syap1 | 0.015 | Old |
| 0.524152 | 0.076019 | 0.024 | 0.013 | 1 | 4 | Abhd11 | 0.011 | Old |
| 0.524459 | 1.005085 | 0.067 | 0.05 | 1 | 4 | Ap4m1 | 0.017 | Old |
| 0.524649 | 0.372053 | 0.083 | 0.062 | 1 | 4 | Lnpk | 0.021 | Old |
| 0.524774 | 0.132159 | 0.024 | 0.013 | 1 | 4 | Pigh | 0.011 | Old |
| 0.524774 | 0.056378 | 0.024 | 0.013 | 1 | 4 | Ftsj1 | 0.011 | Old |
| 0.525328 | 1.102989 | 0.08 | 0.062 | 1 | 4 | Ddx47 | 0.018 | Old |
| 0.52615 | 0.754489 | 0.039 | 0.025 | 1 | 4 | Tfap4 | 0.014 | Old |
| 0.526227 | 0.175504 | 0.404 | 0.4 | 1 | 4 | Rab31 | 0.004 | Old |
| 0.526292 | 0.370174 | 0.272 | 0.25 | 1 | 4 | Mlf2 | 0.022 | Old |
| 0.526489 | 0.706272 | 0.169 | 0.15 | 1 | 4 | Xrn1 | 0.019 | Old |
| 0.5267 | 0.14435 | 0.185 | 0.225 | 1 | 4 | Clk4 | -0.04 | Old |
| 0.52679 | 1.812909 | 0.038 | 0.025 | 1 | 4 | Dtd2 | 0.013 | Old |
| 0.52687 | 0.200229 | 0.566 | 0.575 | 1 | 4 | Sub1 | -0.009 | Old |
| 0.527183 | 0.364377 | 0.097 | 0.075 | 1 | 4 | Pex13 | 0.022 | Old |
| 0.527459 | 0.705551 | 0.053 | 0.038 | 1 | 4 | N4bp2 | 0.015 | Old |
| 0.52784 | 0.611401 | 0.132 | 0.112 | 1 | 4 | Rheb | 0.02 | Old |
| 0.528104 | 0.282594 | 0.4 | 0.388 | 1 | 4 | Matr3 | 0.012 | Old |
| 0.528106 | 0.588249 | 0.039 | 0.025 | 1 | 4 | Ints2 | 0.014 | Old |
| 0.528334 | 1.891722 | 0.023 | 0.013 | 1 | 4 | Ipo4 | 0.01 | Old |
| 0.528402 | 0.323361 | 0.083 | 0.062 | 1 | 4 | Pop5 | 0.021 | Old |
| 0.528484 | 0.366805 | 0.285 | 0.262 | 1 | 4 | Vma21 | 0.023 | Old |
| 0.528619 | 0.33056 | 0.308 | 0.288 | 1 | 4 | Comt | 0.02 | Old |
| 0.529126 | 1.077939 | 0.053 | 0.038 | 1 | 4 | Rnf121 | 0.015 | Old |
| 0.52914 | 0.654648 | 0.053 | 0.038 | 1 | 4 | Cav2 | 0.015 | Old |
| 0.529353 | 0.156873 | 0.226 | 0.262 | 1 | 4 | Egr2 | -0.036 | Old |
| 0.529849 | 0.552014 | 0.096 | 0.075 | 1 | 4 | Eif2s1 | 0.021 | Old |
| 0.529873 | 0.504464 | 0.149 | 0.125 | 1 | 4 | Ipo11 | 0.024 | Old |
| 0.529925 | 0.121749 | 0.111 | 0.138 | 1 | 4 | Mrps18c | -0.027 | Old |
| 0.530082 | 0.093064 | 0.134 | 0.162 | 1 | 4 | Lpin2 | -0.028 | Old |
| 0.530234 | 1.507109 | 0.023 | 0.013 | 1 | 4 | Zbtb40 | 0.01 | Old |
| 0.530246 | 1.502549 | 0.038 | 0.025 | 1 | 4 | Btrc | 0.013 | Old |
| 0.530317 | 0.076474 | 0.155 | 0.188 | 1 | 4 | Pdpk1 | -0.033 | Old |
| 0.530546 | 0.154174 | 0.098 | 0.075 | 1 | 4 | Chchd3 | 0.023 | Old |
| 0.530797 | 0.298561 | 0.083 | 0.062 | 1 | 4 | Bag5 | 0.021 | Old |
| 0.530812 | 0.563728 | 0.096 | 0.075 | 1 | 4 | Sar1b | 0.021 | Old |
| 0.531237 | 0.047104 | 0.309 | 0.362 | 1 | 4 | Prdx2 | -0.053 | Old |
| 0.531288 | 0.244805 | 0.246 | 0.212 | 1 | 4 | Sirt2 | 0.034 | Old |
| 0.531319 | 0.301301 | 0.067 | 0.088 | 1 | 4 | Exosc8 | -0.021 | Old |
| 0.531322 | 0.173282 | 0.04 | 0.025 | 1 | 4 | Rnaset2b | 0.015 | Old |
| 0.53134 | 0.596209 | 0.082 | 0.062 | 1 | 4 | Ccng2 | 0.02 | Old |
| 0.531652 | 0.319756 | 0.054 | 0.038 | 1 | 4 | Ero1lb | 0.016 | Old |
| 0.531779 | 0.02508 | 0.146 | 0.175 | 1 | 4 | Tm2d1 | -0.029 | Old |

| | | | | | | | | |
|---|---|---|---|---|---|---|---|---|
| 0.532152 | 1.312764 | 0.066 | 0.05 | 1 | 4 | Trim3 | 0.016 | Old |
| 0.532367 | 0.010168 | 0.09 | 0.112 | 1 | 4 | Msi2 | -0.022 | Old |
| 0.532538 | 0.131801 | 0.79 | 0.787 | 1 | 4 | Canx | 0.003 | Old |
| 0.532772 | 0.926165 | 0.023 | 0.013 | 1 | 4 | Def8 | 0.01 | Old |
| 0.533037 | 0.560742 | 0.107 | 0.088 | 1 | 4 | Tnfrsf21 | 0.019 | Old |
| 0.533338 | 0.032987 | 0.079 | 0.1 | 1 | 4 | Rab28 | -0.021 | Old |
| 0.533725 | 0.0932 | 0.179 | 0.212 | 1 | 4 | Fam3c | -0.033 | Old |
| 0.533765 | 0.169307 | 0.084 | 0.062 | 1 | 4 | Tax1bp3 | 0.022 | Old |
| 0.534057 | 0.175599 | 0.219 | 0.262 | 1 | 4 | Tmem168 | -0.043 | Old |
| 0.534515 | 0.616786 | 0.159 | 0.138 | 1 | 4 | Fnbp1l | 0.021 | Old |
| 0.534542 | 0.297953 | 0.016 | 0.025 | 1 | 4 | Bhlhe41 | -0.009 | Old |
| 0.534783 | 0.375023 | 0.309 | 0.288 | 1 | 4 | Nfix | 0.021 | Old |
| 0.534826 | 0.090689 | 0.09 | 0.112 | 1 | 4 | Hhex | -0.022 | Old |
| 0.534884 | 0.062669 | 0.007 | 0.013 | 1 | 4 | H2-Q10 | -0.006 | Old |
| 0.535288 | 0.327958 | 0.016 | 0.025 | 1 | 4 | Sag | -0.009 | Old |
| 0.535476 | 0.010043 | 0.098 | 0.075 | 1 | 4 | Pbx1 | 0.023 | Old |
| 0.535559 | 0.891578 | 0.036 | 0.05 | 1 | 4 | Alkbh3 | -0.014 | Old |
| 0.535596 | 0.685882 | 0.081 | 0.062 | 1 | 4 | Ndufaf4 | 0.019 | Old |
| 0.535753 | 0.378952 | 0.099 | 0.125 | 1 | 4 | Rad50 | -0.026 | Old |
| 0.535787 | 0.008452 | 0.211 | 0.25 | 1 | 4 | Ube2k | -0.039 | Old |
| 0.535805 | 0.061115 | 0.156 | 0.188 | 1 | 4 | Deptor | -0.032 | Old |
| 0.535858 | 0.031733 | 0.101 | 0.125 | 1 | 4 | Galnt10 | -0.024 | Old |
| 0.535952 | 0.807107 | 0.023 | 0.013 | 1 | 4 | Klhl12 | 0.01 | Old |
| 0.535952 | 0.750434 | 0.023 | 0.013 | 1 | 4 | Gtf3c5 | 0.01 | Old |
| 0.536041 | 0.056211 | 0.047 | 0.062 | 1 | 4 | Ppa2 | -0.015 | Old |
| 0.53659 | 0.840996 | 0.023 | 0.013 | 1 | 4 | Tepsin | 0.01 | Old |
| 0.536735 | 0.624256 | 0.132 | 0.112 | 1 | 4 | Adgrg6 | 0.02 | Old |
| 0.536755 | 0.403538 | 0.175 | 0.15 | 1 | 4 | Ascc3 | 0.025 | Old |
| 0.53682 | 0.216815 | 0.241 | 0.3 | 1 | 4 | Rbm8a | -0.059 | Old |
| 0.537171 | 0.770285 | 0.007 | 0.013 | 1 | 4 | Aatk | -0.006 | Old |
| 0.537171 | 0.746573 | 0.007 | 0.013 | 1 | 4 | 1700056N1 | -0.006 | Old |
| 0.537171 | 0.248696 | 0.007 | 0.013 | 1 | 4 | Rsph3b | -0.006 | Old |
| 0.537171 | 0.149753 | 0.007 | 0.013 | 1 | 4 | 9130230L2 | -0.006 | Old |
| 0.537171 | 0.079259 | 0.007 | 0.013 | 1 | 4 | Slc2a3 | -0.006 | Old |
| 0.537227 | 0.828437 | 0.023 | 0.013 | 1 | 4 | Klra2 | 0.01 | Old |
| 0.537323 | 0.700927 | 0.107 | 0.088 | 1 | 4 | Prkra | 0.019 | Old |
| 0.537417 | 0.555931 | 0.249 | 0.238 | 1 | 4 | Rmnd5a | 0.011 | Old |
| 0.537818 | 0.130119 | 0.069 | 0.05 | 1 | 4 | Ino80c | 0.019 | Old |
| 0.538069 | 1.321133 | 0.052 | 0.038 | 1 | 4 | Aamdc | 0.014 | Old |
| 0.538317 | 0.553132 | 0.007 | 0.013 | 1 | 4 | Ercc6 | -0.006 | Old |
| 0.538317 | 0.352205 | 0.007 | 0.013 | 1 | 4 | Itga6 | -0.006 | Old |
| 0.538504 | 0.822754 | 0.023 | 0.013 | 1 | 4 | Itgax | 0.01 | Old |
| 0.538895 | 0.434419 | 0.026 | 0.038 | 1 | 4 | Aasdh | -0.012 | Old |
| 0.538909 | 1.882465 | 0.038 | 0.025 | 1 | 4 | Pstk | 0.013 | Old |
| 0.538909 | 0.081835 | 0.036 | 0.05 | 1 | 4 | Dalrd3 | -0.014 | Old |

| | | | | | | | | |
|---|---|---|---|---|---|---|---|---|
| 0.539186 | 1.006356 | 0.038 | 0.025 | 1 | 4 | Mcm4 | 0.013 | Old |
| 0.53943 | 0.361241 | 0.039 | 0.025 | 1 | 4 | Snhg9 | 0.014 | Old |
| 0.539463 | 0.721303 | 0.007 | 0.013 | 1 | 4 | Znrf3 | -0.006 | Old |
| 0.539675 | 0.64225 | 0.094 | 0.075 | 1 | 4 | Suclg1 | 0.019 | Old |
| 0.539779 | 0.575479 | 0.016 | 0.025 | 1 | 4 | Suv39h1 | -0.009 | Old |
| 0.539915 | 1.757693 | 0.038 | 0.025 | 1 | 4 | Mtx3 | 0.013 | Old |
| 0.539915 | 0.094411 | 0.036 | 0.05 | 1 | 4 | Asf1a | -0.014 | Old |
| 0.540014 | 0.47018 | 0.234 | 0.212 | 1 | 4 | Evi5 | 0.022 | Old |
| 0.540184 | 0.942971 | 0.038 | 0.025 | 1 | 4 | Gtf2h4 | 0.013 | Old |
| 0.540476 | 0.572704 | 0.081 | 0.062 | 1 | 4 | Got2 | 0.019 | Old |
| 0.540529 | 0.646138 | 0.016 | 0.025 | 1 | 4 | Gm26982 | -0.009 | Old |
| 0.540674 | 0.384731 | 0.026 | 0.038 | 1 | 4 | Rfx3 | -0.012 | Old |
| 0.541295 | 0.478447 | 0.08 | 0.062 | 1 | 4 | Uba3 | 0.018 | Old |
| 0.542433 | 1.662512 | 0.038 | 0.025 | 1 | 4 | Plpp5 | 0.013 | Old |
| 0.543116 | 0.411015 | 0.236 | 0.212 | 1 | 4 | Emg1 | 0.024 | Old |
| 0.543625 | 0.17507 | 0.023 | 0.013 | 1 | 4 | Uevld | 0.01 | Old |
| 0.544307 | 0.737524 | 0.067 | 0.05 | 1 | 4 | Plaur | 0.017 | Old |
| 0.544556 | 0.330853 | 0.057 | 0.075 | 1 | 4 | Arl3 | -0.018 | Old |
| 0.544742 | 0.668456 | 0.12 | 0.1 | 1 | 4 | Ibtk | 0.02 | Old |
| 0.544824 | 0.640742 | 0.157 | 0.138 | 1 | 4 | Elmo2 | 0.019 | Old |
| 0.545214 | 0.403196 | 0.067 | 0.088 | 1 | 4 | Gpn3 | -0.021 | Old |
| 0.545253 | 0.028188 | 0.101 | 0.125 | 1 | 4 | Sec24a | -0.024 | Old |
| 0.54545 | 0.480958 | 0.148 | 0.125 | 1 | 4 | Prpf18 | 0.023 | Old |
| 0.545764 | 0.19072 | 0.069 | 0.05 | 1 | 4 | Cep250 | 0.019 | Old |
| 0.546349 | 0.051546 | 0.069 | 0.088 | 1 | 4 | Bid | -0.019 | Old |
| 0.546451 | 0.347096 | 0.089 | 0.112 | 1 | 4 | Fam219b | -0.023 | Old |
| 0.546533 | 0.291033 | 0.053 | 0.038 | 1 | 4 | Ltbp3 | 0.015 | Old |
| 0.546694 | 0.539268 | 0.038 | 0.025 | 1 | 4 | Gm20696 | 0.013 | Old |
| 0.546783 | 0.147275 | 0.231 | 0.2 | 1 | 4 | Ndufa5 | 0.031 | Old |
| 0.546825 | 2.168312 | 0.022 | 0.013 | 1 | 4 | Zfp46 | 0.009 | Old |
| 0.546891 | 0.049008 | 0.201 | 0.238 | 1 | 4 | Rab8a | -0.037 | Old |
| 0.547048 | 0.477408 | 0.053 | 0.038 | 1 | 4 | Ogfod1 | 0.015 | Old |
| 0.547155 | 0.052143 | 0.524 | 0.6 | 1 | 4 | Celf1 | -0.076 | Old |
| 0.547196 | 0.37301 | 0.038 | 0.025 | 1 | 4 | Cdc25b | 0.013 | Old |
| 0.54763 | 0.68959 | 0.08 | 0.062 | 1 | 4 | Kars | 0.018 | Old |
| 0.548131 | 1.716039 | 0.022 | 0.013 | 1 | 4 | Slx4 | 0.009 | Old |
| 0.548131 | 1.914196 | 0.022 | 0.013 | 1 | 4 | Pdia5 | 0.009 | Old |
| 0.548131 | 1.811757 | 0.022 | 0.013 | 1 | 4 | Areg | 0.009 | Old |
| 0.548131 | 1.211093 | 0.022 | 0.013 | 1 | 4 | Prelid3a | 0.009 | Old |
| 0.54814 | 0.848483 | 0.118 | 0.1 | 1 | 4 | Tmem43 | 0.018 | Old |
| 0.548701 | 1.356632 | 0.051 | 0.038 | 1 | 4 | Dhx57 | 0.013 | Old |
| 0.548749 | 0.569593 | 0.134 | 0.112 | 1 | 4 | Nr2c2 | 0.022 | Old |
| 0.548784 | 1.463943 | 0.022 | 0.013 | 1 | 4 | Taf5 | 0.009 | Old |
| 0.548895 | 0.074768 | 0.113 | 0.138 | 1 | 4 | Pik3r6 | -0.025 | Old |
| 0.548928 | 0.400853 | 0.067 | 0.05 | 1 | 4 | Ambra1 | 0.017 | Old |

| | | | | | | | | |
|---|---|---|---|---|---|---|---|---|
| 0.549438 | 1.642868 | 0.022 | 0.013 | 1 | 4 | 2010111I0: | 0.009 | Old |
| 0.549653 | 0.562576 | 0.132 | 0.112 | 1 | 4 | Sh3bp2 | 0.02 | Old |
| 0.549691 | 0.026448 | 0.091 | 0.112 | 1 | 4 | Uchl5 | -0.021 | Old |
| 0.550106 | 0.307726 | 0.161 | 0.138 | 1 | 4 | Chchd7 | 0.023 | Old |
| 0.550157 | 0.351914 | 0.15 | 0.125 | 1 | 4 | Dnajc10 | 0.025 | Old |
| 0.55047 | 0.842389 | 0.117 | 0.1 | 1 | 4 | Srsf4 | 0.017 | Old |
| 0.550473 | 0.420093 | 0.067 | 0.05 | 1 | 4 | 4930402H2 | 0.017 | Old |
| 0.550719 | 0.569137 | 0.038 | 0.025 | 1 | 4 | Otud1 | 0.013 | Old |
| 0.550747 | 1.330531 | 0.022 | 0.013 | 1 | 4 | Isoc2b | 0.009 | Old |
| 0.550986 | 0.555751 | 0.121 | 0.1 | 1 | 4 | Ccz1 | 0.021 | Old |
| 0.551015 | 0.8474 | 0.153 | 0.138 | 1 | 4 | Cln3 | 0.015 | Old |
| 0.551699 | 0.210387 | 0.057 | 0.075 | 1 | 4 | Tfam | -0.018 | Old |
| 0.55216 | 0.081509 | 0.163 | 0.138 | 1 | 4 | Bbip1 | 0.025 | Old |
| 0.552253 | 0.087073 | 0.08 | 0.1 | 1 | 4 | Ctnnbip1 | -0.02 | Old |
| 0.552713 | 1.390949 | 0.022 | 0.013 | 1 | 4 | Ccdc91 | 0.009 | Old |
| 0.552824 | 0.484248 | 0.182 | 0.162 | 1 | 4 | Eloa | 0.02 | Old |
| 0.553076 | 1.029296 | 0.038 | 0.025 | 1 | 4 | Tbl3 | 0.013 | Old |
| 0.5532 | 0.106122 | 0.068 | 0.05 | 1 | 4 | Mbtps2 | 0.018 | Old |
| 0.553672 | 0.002577 | 0.331 | 0.388 | 1 | 4 | Znrf2 | -0.057 | Old |
| 0.553774 | 0.45584 | 0.094 | 0.075 | 1 | 4 | Abraxas2 | 0.019 | Old |
| 0.553822 | 0.179222 | 0.46 | 0.45 | 1 | 4 | Prpf38b | 0.01 | Old |
| 0.553952 | 0.515668 | 0.253 | 0.238 | 1 | 4 | Ndufb5 | 0.015 | Old |
| 0.554194 | 0.345601 | 0.287 | 0.262 | 1 | 4 | Ensa | 0.025 | Old |
| 0.554587 | 0.165022 | 0.611 | 0.588 | 1 | 4 | C3ar1 | 0.023 | Old |
| 0.554688 | 0.132256 | 0.053 | 0.038 | 1 | 4 | Cks2 | 0.015 | Old |
| 0.554765 | 0.636718 | 0.094 | 0.075 | 1 | 4 | Tubb4b | 0.019 | Old |
| 0.555076 | 0.998996 | 0.065 | 0.05 | 1 | 4 | Dcaf11 | 0.015 | Old |
| 0.555781 | 0.360218 | 0.081 | 0.062 | 1 | 4 | Ubl7 | 0.019 | Old |
| 0.555822 | 0.017998 | 0.567 | 0.625 | 1 | 4 | Mrfap1 | -0.058 | Old |
| 0.555946 | 0.654396 | 0.08 | 0.062 | 1 | 4 | A430005L1 | 0.018 | Old |
| 0.556294 | 0.042431 | 0.124 | 0.15 | 1 | 4 | Smarca4 | -0.026 | Old |
| 0.556458 | 0.282649 | 0.175 | 0.15 | 1 | 4 | Tiprl | 0.025 | Old |
| 0.556475 | 1.77775 | 0.037 | 0.025 | 1 | 4 | Rarb | 0.012 | Old |
| 0.556661 | 0.001412 | 0.963 | 0.963 | 1 | 4 | Rpl38 | 0 | Old |
| 0.557157 | 0.451108 | 0.036 | 0.05 | 1 | 4 | Nudt18 | -0.014 | Old |
| 0.557336 | 0.066718 | 0.976 | 0.988 | 1 | 4 | Junb | -0.012 | Old |
| 0.557374 | 0.053956 | 0.08 | 0.062 | 1 | 4 | Nom1 | 0.018 | Old |
| 0.557432 | 0.120383 | 0.068 | 0.05 | 1 | 4 | Gtf2f1 | 0.018 | Old |
| 0.557452 | 0.261512 | 0.287 | 0.262 | 1 | 4 | Sf3b5 | 0.025 | Old |
| 0.557879 | 0.382498 | 0.148 | 0.125 | 1 | 4 | Prpf39 | 0.023 | Old |
| 0.557973 | 0.579483 | 0.022 | 0.013 | 1 | 4 | Zfp628 | 0.009 | Old |
| 0.558423 | 0.2443 | 0.188 | 0.162 | 1 | 4 | Cnppd1 | 0.026 | Old |
| 0.558682 | 0.51496 | 0.057 | 0.075 | 1 | 4 | Samd1 | -0.018 | Old |
| 0.558866 | 0.633302 | 0.052 | 0.038 | 1 | 4 | Morc2a | 0.014 | Old |
| 0.559337 | 0.824772 | 0.079 | 0.062 | 1 | 4 | Necap1 | 0.017 | Old |

| | | | | | | | | |
|---|---|---|---|---|---|---|---|---|
| 0.559696 | 0.921757 | 0.079 | 0.062 | 1 | 4 | Lsm2 | 0.017 | Old |
| 0.559813 | 0.121254 | 0.221 | 0.262 | 1 | 4 | Cr1l | -0.041 | Old |
| 0.559887 | 0.586164 | 0.106 | 0.088 | 1 | 4 | Atxn3 | 0.018 | Old |
| 0.559951 | 0.498613 | 0.022 | 0.013 | 1 | 4 | Gm1673 | 0.009 | Old |
| 0.559969 | 0.222252 | 0.419 | 0.4 | 1 | 4 | Aftph | 0.019 | Old |
| 0.560175 | 0.470196 | 0.067 | 0.05 | 1 | 4 | Iffo2 | 0.017 | Old |
| 0.560227 | 0.736998 | 0.038 | 0.025 | 1 | 4 | Akt3 | 0.013 | Old |
| 0.560303 | 0.174289 | 0.858 | 0.887 | 1 | 4 | Txnip | -0.029 | Old |
| 0.560603 | 0.247686 | 0.188 | 0.162 | 1 | 4 | Srd5a3 | 0.026 | Old |
| 0.560629 | 0.034559 | 0.57 | 0.637 | 1 | 4 | Atp6ap1 | -0.067 | Old |
| 0.56074 | 0.519165 | 0.038 | 0.025 | 1 | 4 | Zfp65 | 0.013 | Old |
| 0.561272 | 0.346618 | 0.022 | 0.013 | 1 | 4 | Coa4 | 0.009 | Old |
| 0.561354 | 0.140432 | 0.038 | 0.025 | 1 | 4 | Rnaseh2b | 0.013 | Old |
| 0.561409 | 0.281701 | 0.176 | 0.15 | 1 | 4 | Kmt5b | 0.026 | Old |
| 0.561578 | 0.040314 | 0.408 | 0.388 | 1 | 4 | Shisa5 | 0.02 | Old |
| 0.561765 | 0.689841 | 0.038 | 0.025 | 1 | 4 | Ruvbl2 | 0.013 | Old |
| 0.561842 | 1.193831 | 0.078 | 0.062 | 1 | 4 | Wdr48 | 0.016 | Old |
| 0.561912 | 0.168164 | 0.619 | 0.625 | 1 | 4 | Dhrs3 | -0.006 | Old |
| 0.561983 | 0.41136 | 0.057 | 0.075 | 1 | 4 | Pmf1 | -0.018 | Old |
| 0.562042 | 0.192807 | 0.124 | 0.1 | 1 | 4 | Napg | 0.024 | Old |
| 0.562202 | 0.987773 | 0.127 | 0.112 | 1 | 4 | Lats1 | 0.015 | Old |
| 0.562205 | 1.616955 | 0.051 | 0.038 | 1 | 4 | Mkrn2 | 0.013 | Old |
| 0.562256 | 0.326515 | 0.212 | 0.188 | 1 | 4 | Map2k7 | 0.024 | Old |
| 0.562292 | 0.041609 | 0.082 | 0.062 | 1 | 4 | Rab6b | 0.02 | Old |
| 0.563081 | 0.282364 | 0.274 | 0.25 | 1 | 4 | Tmem14c | 0.024 | Old |
| 0.563256 | 0.358289 | 0.134 | 0.112 | 1 | 4 | Nagk | 0.022 | Old |
| 0.563274 | 0.01554 | 0.069 | 0.088 | 1 | 4 | Slc33a1 | -0.019 | Old |
| 0.563984 | 0.436502 | 0.08 | 0.062 | 1 | 4 | Zdhhc17 | 0.018 | Old |
| 0.564283 | 0.83566 | 0.051 | 0.038 | 1 | 4 | Alpk1 | 0.013 | Old |
| 0.564298 | 0.408922 | 0.121 | 0.1 | 1 | 4 | Dhx38 | 0.021 | Old |
| 0.564335 | 0.134881 | 0.15 | 0.125 | 1 | 4 | Aldh3b1 | 0.025 | Old |
| 0.564611 | 0.181593 | 0.026 | 0.038 | 1 | 4 | Lig3 | -0.012 | Old |
| 0.565166 | 0.09879 | 0.068 | 0.05 | 1 | 4 | Ppargc1b | 0.018 | Old |
| 0.565332 | 0.191629 | 0.51 | 0.512 | 1 | 4 | Ncl | -0.002 | Old |
| 0.565531 | 0.264897 | 0.123 | 0.15 | 1 | 4 | Cdk14 | -0.027 | Old |
| 0.565645 | 0.136357 | 0.814 | 0.812 | 1 | 4 | Gabarap | 0.002 | Old |
| 0.565696 | 0.819507 | 0.065 | 0.05 | 1 | 4 | Arfgap3 | 0.015 | Old |
| 0.566325 | 0.58234 | 0.052 | 0.038 | 1 | 4 | Pdhx | 0.014 | Old |
| 0.566453 | 0.128122 | 0.037 | 0.05 | 1 | 4 | C1rl | -0.013 | Old |
| 0.566473 | 0.248284 | 0.168 | 0.2 | 1 | 4 | Gas2l3 | -0.032 | Old |
| 0.566589 | 0.241061 | 0.24 | 0.212 | 1 | 4 | Herc4 | 0.028 | Old |
| 0.566725 | 0.286219 | 0.161 | 0.138 | 1 | 4 | Cyb5r4 | 0.023 | Old |
| 0.566754 | 2.241924 | 0.022 | 0.013 | 1 | 4 | L2hgdh | 0.009 | Old |
| 0.567068 | 0.2387 | 0.11 | 0.088 | 1 | 4 | Rabggtb | 0.022 | Old |
| 0.567427 | 1.941042 | 0.022 | 0.013 | 1 | 4 | Hdac6 | 0.009 | Old |

| | | | | | | | | |
|---|---|---|---|---|---|---|---|---|
| 0.568471 | 1.390024 | 0.091 | 0.075 | 1 | 4 | Il6 | 0.016 | Old |
| 0.568715 | 0.680886 | 0.079 | 0.062 | 1 | 4 | Mrpl9 | 0.017 | Old |
| 0.568776 | 1.547722 | 0.022 | 0.013 | 1 | 4 | Ccdc181 | 0.009 | Old |
| 0.568776 | 1.357469 | 0.022 | 0.013 | 1 | 4 | 2610021A0 | 0.009 | Old |
| 0.569084 | 0.405935 | 0.159 | 0.138 | 1 | 4 | Capn1 | 0.021 | Old |
| 0.569889 | 0.513525 | 0.094 | 0.075 | 1 | 4 | Get4 | 0.019 | Old |
| 0.570035 | 0.252815 | 0.037 | 0.05 | 1 | 4 | Arhgap6 | -0.013 | Old |
| 0.570042 | 0.639794 | 0.051 | 0.038 | 1 | 4 | Smad3 | 0.013 | Old |
| 0.570125 | 1.179829 | 0.022 | 0.013 | 1 | 4 | Ablim1 | 0.009 | Old |
| 0.570219 | 1.314922 | 0.051 | 0.038 | 1 | 4 | Nol9 | 0.013 | Old |
| 0.570374 | 0.826296 | 0.078 | 0.062 | 1 | 4 | Fkbp5 | 0.016 | Old |
| 0.570447 | 0.946741 | 0.065 | 0.05 | 1 | 4 | Ikbke | 0.015 | Old |
| 0.570463 | 0.917517 | 0.037 | 0.025 | 1 | 4 | Actr5 | 0.012 | Old |
| 0.570931 | 0.753192 | 0.051 | 0.038 | 1 | 4 | Mark4 | 0.013 | Old |
| 0.571477 | 1.234006 | 0.022 | 0.013 | 1 | 4 | Syp | 0.009 | Old |
| 0.571568 | 0.203358 | 0.094 | 0.075 | 1 | 4 | Tmem88 | 0.019 | Old |
| 0.571993 | 0.041153 | 0.227 | 0.262 | 1 | 4 | Dynlt3 | -0.035 | Old |
| 0.572153 | 1.421555 | 0.022 | 0.013 | 1 | 4 | Mthfd1l | 0.009 | Old |
| 0.572153 | 1.375429 | 0.022 | 0.013 | 1 | 4 | Bckdhb | 0.009 | Old |
| 0.572153 | 1.346326 | 0.022 | 0.013 | 1 | 4 | Paqr7 | 0.009 | Old |
| 0.572153 | 1.125312 | 0.022 | 0.013 | 1 | 4 | Mdm1 | 0.009 | Old |
| 0.572321 | 0.149908 | 0.134 | 0.112 | 1 | 4 | Wdr36 | 0.022 | Old |
| 0.572535 | 0.51278 | 0.26 | 0.25 | 1 | 4 | Grk3 | 0.01 | Old |
| 0.572765 | 0.016284 | 0.375 | 0.425 | 1 | 4 | Gpr132 | -0.05 | Old |
| 0.57283 | 1.409661 | 0.022 | 0.013 | 1 | 4 | Top2a | 0.009 | Old |
| 0.573315 | 0.109616 | 0.167 | 0.2 | 1 | 4 | Glb1 | -0.033 | Old |
| 0.573552 | 0.548343 | 0.144 | 0.125 | 1 | 4 | Ndufv2 | 0.019 | Old |
| 0.573586 | 0.012665 | 0.081 | 0.1 | 1 | 4 | Iqsec1 | -0.019 | Old |
| 0.573602 | 0.067098 | 0.053 | 0.038 | 1 | 4 | Adamts10 | 0.015 | Old |
| 0.573999 | 0.929497 | 0.103 | 0.088 | 1 | 4 | Adamtsl5 | 0.015 | Old |
| 0.574074 | 0.055864 | 0.408 | 0.388 | 1 | 4 | Ubap2l | 0.02 | Old |
| 0.574512 | 0.127345 | 0.262 | 0.238 | 1 | 4 | Psma3 | 0.024 | Old |
| 0.574584 | 2.104473 | 0.036 | 0.025 | 1 | 4 | Lrrc14 | 0.011 | Old |
| 0.574639 | 0.772795 | 0.037 | 0.025 | 1 | 4 | Clpp | 0.012 | Old |
| 0.574745 | 0.011083 | 0.565 | 0.662 | 1 | 4 | Sgpl1 | -0.097 | Old |
| 0.574946 | 0.646082 | 0.092 | 0.075 | 1 | 4 | Prmt2 | 0.017 | Old |
| 0.57502 | 0.312983 | 0.016 | 0.025 | 1 | 4 | Hyal1 | -0.009 | Old |
| 0.57502 | 0.102999 | 0.016 | 0.025 | 1 | 4 | Tmco6 | -0.009 | Old |
| 0.575086 | 0.109554 | 0.687 | 0.675 | 1 | 4 | Rbm39 | 0.012 | Old |
| 0.57518 | 0.012612 | 0.115 | 0.138 | 1 | 4 | Psen2 | -0.023 | Old |
| 0.575337 | 1.585465 | 0.05 | 0.038 | 1 | 4 | Sqle | 0.012 | Old |
| 0.575383 | 0.743943 | 0.051 | 0.038 | 1 | 4 | Mrps35 | 0.013 | Old |
| 0.575923 | 0.433804 | 0.08 | 0.062 | 1 | 4 | Fbxl14 | 0.018 | Old |
| 0.575927 | 0.363649 | 0.145 | 0.125 | 1 | 4 | Shoc2 | 0.02 | Old |
| 0.57595 | 0.963551 | 0.116 | 0.1 | 1 | 4 | Ppm1h | 0.016 | Old |

| | | | | | | | | |
|---|---|---|---|---|---|---|---|---|
| 0.57599 | 0.112757 | 0.403 | 0.375 | 1 | 4 | Dnase2a | 0.028 | Old |
| 0.576129 | 0.28841 | 0.25 | 0.225 | 1 | 4 | Eif3h | 0.025 | Old |
| 0.576219 | 0.900484 | 0.022 | 0.013 | 1 | 4 | Prmt6 | 0.009 | Old |
| 0.576652 | 0.456421 | 0.196 | 0.175 | 1 | 4 | Vars | 0.021 | Old |
| 0.576685 | 0.273261 | 0.134 | 0.112 | 1 | 4 | Htra2 | 0.022 | Old |
| 0.576871 | 0.141635 | 0.312 | 0.275 | 1 | 4 | Ppp1r15b | 0.037 | Old |
| 0.576898 | 0.692739 | 0.022 | 0.013 | 1 | 4 | Mpv17l | 0.009 | Old |
| 0.577307 | 0.258201 | 0.08 | 0.062 | 1 | 4 | Trappc11 | 0.018 | Old |
| 0.577438 | 0.642426 | 0.079 | 0.062 | 1 | 4 | Nagpa | 0.017 | Old |
| 0.577614 | 0.320065 | 0.058 | 0.075 | 1 | 4 | Nfrkb | -0.017 | Old |
| 0.577733 | 0.484975 | 0.066 | 0.05 | 1 | 4 | Dcxr | 0.016 | Old |
| 0.578257 | 0.692635 | 0.022 | 0.013 | 1 | 4 | 4933427D1 | 0.009 | Old |
| 0.578281 | 1.66918 | 0.036 | 0.025 | 1 | 4 | Polr3d | 0.011 | Old |
| 0.578361 | 0.419966 | 0.197 | 0.175 | 1 | 4 | Slc15a4 | 0.022 | Old |
| 0.579167 | 0.681152 | 0.13 | 0.112 | 1 | 4 | Epc1 | 0.018 | Old |
| 0.579355 | 0.449841 | 0.037 | 0.025 | 1 | 4 | Bcl2a1d | 0.012 | Old |
| 0.579562 | 0.215775 | 0.134 | 0.112 | 1 | 4 | St6galnac6 | 0.022 | Old |
| 0.579617 | 0.576474 | 0.022 | 0.013 | 1 | 4 | Gm26778 | 0.009 | Old |
| 0.579721 | 0.379936 | 0.066 | 0.05 | 1 | 4 | Psmd5 | 0.016 | Old |
| 0.579854 | 0.468258 | 0.093 | 0.075 | 1 | 4 | Rrm1 | 0.018 | Old |
| 0.580047 | 0.339372 | 0.184 | 0.162 | 1 | 4 | Eif2a | 0.022 | Old |
| 0.580243 | 0.36883 | 0.12 | 0.1 | 1 | 4 | Sgsh | 0.02 | Old |
| 0.580405 | 0.575231 | 0.037 | 0.025 | 1 | 4 | Fmn1 | 0.012 | Old |
| 0.581459 | 1.505804 | 0.036 | 0.025 | 1 | 4 | Myc | 0.011 | Old |
| 0.581489 | 0.756133 | 0.065 | 0.05 | 1 | 4 | Rab40c | 0.015 | Old |
| 0.581545 | 0.129892 | 0.252 | 0.225 | 1 | 4 | Adcy7 | 0.027 | Old |
| 0.581636 | 0.798836 | 0.104 | 0.088 | 1 | 4 | Pigx | 0.016 | Old |
| 0.582057 | 0.511331 | 0.105 | 0.088 | 1 | 4 | Socs7 | 0.017 | Old |
| 0.582183 | 0.302378 | 0.273 | 0.25 | 1 | 4 | Tmod3 | 0.023 | Old |
| 0.582424 | 0.05105 | 0.11 | 0.088 | 1 | 4 | Palld | 0.022 | Old |
| 0.583025 | 0.366999 | 0.022 | 0.013 | 1 | 4 | Sh3bp5l | 0.009 | Old |
| 0.583231 | 0.144773 | 0.136 | 0.162 | 1 | 4 | Rnf146 | -0.026 | Old |
| 0.583252 | 0.682241 | 0.19 | 0.175 | 1 | 4 | Dctn4 | 0.015 | Old |
| 0.58327 | 0.536805 | 0.182 | 0.162 | 1 | 4 | Nupr1 | 0.02 | Old |
| 0.583382 | 1.0059 | 0.064 | 0.05 | 1 | 4 | Trip10 | 0.014 | Old |
| 0.583697 | 0.907595 | 0.051 | 0.038 | 1 | 4 | Dnajc9 | 0.013 | Old |
| 0.583784 | 0.014821 | 0.366 | 0.4 | 1 | 4 | Zdhhc20 | -0.034 | Old |
| 0.583934 | 0.019704 | 0.091 | 0.112 | 1 | 4 | Mrps16 | -0.021 | Old |
| 0.584042 | 0.105659 | 0.85 | 0.812 | 1 | 4 | Cbr2 | 0.038 | Old |
| 0.584113 | 1.077466 | 0.036 | 0.025 | 1 | 4 | Dcaf6 | 0.011 | Old |
| 0.584133 | 0.079261 | 0.352 | 0.325 | 1 | 4 | Pld3 | 0.027 | Old |
| 0.584816 | 0.096442 | 0.027 | 0.038 | 1 | 4 | Setdb2 | -0.011 | Old |
| 0.584819 | 0.596223 | 0.092 | 0.075 | 1 | 4 | Usf3 | 0.017 | Old |
| 0.5849 | 0.405276 | 0.316 | 0.312 | 1 | 4 | Cct8 | 0.004 | Old |
| 0.585384 | 0.141921 | 0.067 | 0.05 | 1 | 4 | Alg5 | 0.017 | Old |

| | | | | | | | | |
|---|---|---|---|---|---|---|---|---|
| 0.585388 | 0.029302 | 0.038 | 0.05 | 1 | 4 | Tmem94 | -0.012 | Old |
| 0.585551 | 0.324843 | 0.145 | 0.125 | 1 | 4 | Rabgap1 | 0.02 | Old |
| 0.58641 | 0.666343 | 0.051 | 0.038 | 1 | 4 | Hps1 | 0.013 | Old |
| 0.586441 | 0.129989 | 0.07 | 0.088 | 1 | 4 | Zfp945 | -0.018 | Old |
| 0.586661 | 0.236186 | 0.091 | 0.112 | 1 | 4 | Fam219a | -0.021 | Old |
| 0.586677 | 0.019582 | 0.138 | 0.162 | 1 | 4 | Prkd3 | -0.024 | Old |
| 0.587103 | 0.166207 | 0.135 | 0.112 | 1 | 4 | Tbc1d13 | 0.023 | Old |
| 0.587127 | 0.042812 | 0.022 | 0.013 | 1 | 4 | Vamp1 | 0.009 | Old |
| 0.587536 | 0.657263 | 0.065 | 0.05 | 1 | 4 | Milr1 | 0.015 | Old |
| 0.587541 | 0.503554 | 0.237 | 0.225 | 1 | 4 | Bcas2 | 0.012 | Old |
| 0.587586 | 0.007488 | 0.082 | 0.1 | 1 | 4 | Pcgf3 | -0.018 | Old |
| 0.588027 | 0.255945 | 0.147 | 0.125 | 1 | 4 | Rabgef1 | 0.022 | Old |
| 0.588165 | 0.013263 | 0.369 | 0.412 | 1 | 4 | Zcchc24 | -0.043 | Old |
| 0.588226 | 1.742573 | 0.021 | 0.013 | 1 | 4 | Nol10 | 0.008 | Old |
| 0.588226 | 2.028124 | 0.021 | 0.013 | 1 | 4 | Slc44a2 | 0.008 | Old |
| 0.588438 | 0.104815 | 0.027 | 0.038 | 1 | 4 | Fignl2 | -0.011 | Old |
| 0.588616 | 0.781657 | 0.078 | 0.062 | 1 | 4 | Mb21d1 | 0.016 | Old |
| 0.588742 | 0.248667 | 0.359 | 0.338 | 1 | 4 | Mdfic | 0.021 | Old |
| 0.588797 | 0.300961 | 0.079 | 0.062 | 1 | 4 | Brwd3 | 0.017 | Old |
| 0.588846 | 0.476358 | 0.217 | 0.2 | 1 | 4 | Ppp4c | 0.017 | Old |
| 0.588921 | 2.148145 | 0.021 | 0.013 | 1 | 4 | Ddx28 | 0.008 | Old |
| 0.588921 | 1.962533 | 0.021 | 0.013 | 1 | 4 | AW146154 | 0.008 | Old |
| 0.588957 | 0.490433 | 0.1 | 0.125 | 1 | 4 | Golga2 | -0.025 | Old |
| 0.589042 | 0.118436 | 0.027 | 0.038 | 1 | 4 | Clasrp | -0.011 | Old |
| 0.589166 | 0.123047 | 0.125 | 0.15 | 1 | 4 | Pde4dip | -0.025 | Old |
| 0.58937 | 0.309302 | 0.037 | 0.025 | 1 | 4 | Ptpn4 | 0.012 | Old |
| 0.589423 | 0.142856 | 0.236 | 0.275 | 1 | 4 | Tbl1x | -0.039 | Old |
| 0.589439 | 1.031428 | 0.036 | 0.025 | 1 | 4 | Sbk1 | 0.011 | Old |
| 0.589466 | 0.901375 | 0.064 | 0.05 | 1 | 4 | Wdtc1 | 0.014 | Old |
| 0.589617 | 1.831416 | 0.021 | 0.013 | 1 | 4 | Elp4 | 0.008 | Old |
| 0.58991 | 0.833817 | 0.151 | 0.138 | 1 | 4 | Samd9l | 0.013 | Old |
| 0.589934 | 0.297065 | 0.376 | 0.362 | 1 | 4 | Trir | 0.014 | Old |
| 0.589973 | 0.904503 | 0.036 | 0.025 | 1 | 4 | Aifm1 | 0.011 | Old |
| 0.590111 | 0.195432 | 0.066 | 0.05 | 1 | 4 | Zbtb21 | 0.016 | Old |
| 0.590376 | 0.908387 | 0.077 | 0.062 | 1 | 4 | Bloc1s4 | 0.015 | Old |
| 0.590761 | 0.306374 | 0.059 | 0.075 | 1 | 4 | Pitrm1 | -0.016 | Old |
| 0.59101 | 1.993302 | 0.021 | 0.013 | 1 | 4 | 6330403L0 | 0.008 | Old |
| 0.59101 | 1.41873 | 0.021 | 0.013 | 1 | 4 | Pgap3 | 0.008 | Old |
| 0.591039 | 0.28787 | 0.381 | 0.375 | 1 | 4 | Cdc42se1 | 0.006 | Old |
| 0.591178 | 0.607365 | 0.065 | 0.05 | 1 | 4 | Rap1gap2 | 0.015 | Old |
| 0.591206 | 0.156224 | 0.406 | 0.4 | 1 | 4 | Sdha | 0.006 | Old |
| 0.59131 | 0.225814 | 0.209 | 0.188 | 1 | 4 | Rp9 | 0.021 | Old |
| 0.591545 | 0.297322 | 0.051 | 0.038 | 1 | 4 | Kctd5 | 0.013 | Old |
| 0.591707 | 1.427265 | 0.021 | 0.013 | 1 | 4 | Lancl2 | 0.008 | Old |
| 0.59207 | 0.210728 | 0.027 | 0.038 | 1 | 4 | Hist1h4i | -0.011 | Old |

| | | | | | | | | |
|---|---|---|---|---|---|---|---|---|
| 0.59207 | 0.152164 | 0.027 | 0.038 | 1 | 4 | Atg4a | -0.011 | Old |
| 0.592104 | 0.096915 | 0.095 | 0.075 | 1 | 4 | Nsmce3 | 0.02 | Old |
| 0.592619 | 0.156568 | 0.464 | 0.462 | 1 | 4 | Gna12 | 0.002 | Old |
| 0.592645 | 0.980957 | 0.036 | 0.025 | 1 | 4 | Txlng | 0.011 | Old |
| 0.592697 | 0.164684 | 0.103 | 0.125 | 1 | 4 | Ccdc124 | -0.022 | Old |
| 0.59289 | 0.238004 | 0.121 | 0.1 | 1 | 4 | Ylpm1 | 0.021 | Old |
| 0.593644 | 1.140379 | 0.076 | 0.062 | 1 | 4 | G6pdx | 0.014 | Old |
| 0.593715 | 0.612244 | 0.036 | 0.025 | 1 | 4 | Tnfsfm13 | 0.011 | Old |
| 0.593799 | 0.210995 | 0.368 | 0.35 | 1 | 4 | Arl8a | 0.018 | Old |
| 0.593894 | 0.172141 | 0.235 | 0.212 | 1 | 4 | St6galnac4 | 0.023 | Old |
| 0.593989 | 0.205847 | 0.161 | 0.138 | 1 | 4 | Xpot | 0.023 | Old |
| 0.594372 | 0.085814 | 0.052 | 0.038 | 1 | 4 | Smim24 | 0.014 | Old |
| 0.5945 | 1.463561 | 0.021 | 0.013 | 1 | 4 | Trub1 | 0.008 | Old |
| 0.5945 | 1.02317 | 0.021 | 0.013 | 1 | 4 | 2900093K2 | 0.008 | Old |
| 0.594655 | 0.296917 | 0.107 | 0.088 | 1 | 4 | Bag6 | 0.019 | Old |
| 0.594898 | 0.848878 | 0.05 | 0.038 | 1 | 4 | Map2k5 | 0.012 | Old |
| 0.59509 | 0.170182 | 0.059 | 0.075 | 1 | 4 | Gamt | -0.016 | Old |
| 0.595171 | 0.870827 | 0.064 | 0.05 | 1 | 4 | Prpf4 | 0.014 | Old |
| 0.595712 | 0.334358 | 0.027 | 0.038 | 1 | 4 | Nck2 | -0.011 | Old |
| 0.595952 | 0.404817 | 0.051 | 0.038 | 1 | 4 | Sepsecs | 0.013 | Old |
| 0.596395 | 0.675072 | 0.036 | 0.025 | 1 | 4 | A530017D2 | 0.011 | Old |
| 0.596395 | 0.674262 | 0.036 | 0.025 | 1 | 4 | Zfp637 | 0.011 | Old |
| 0.596503 | 0.699641 | 0.09 | 0.075 | 1 | 4 | Steap3 | 0.015 | Old |
| 0.596708 | 0.10137 | 0.65 | 0.688 | 1 | 4 | Birc3 | -0.038 | Old |
| 0.596784 | 0.54253 | 0.129 | 0.112 | 1 | 4 | Hnrnph3 | 0.017 | Old |
| 0.597036 | 1.635128 | 0.036 | 0.025 | 1 | 4 | Smg8 | 0.011 | Old |
| 0.59718 | 0.930509 | 0.126 | 0.112 | 1 | 4 | 3830406C1 | 0.014 | Old |
| 0.597182 | 0.077941 | 0.059 | 0.075 | 1 | 4 | Abhd13 | -0.016 | Old |
| 0.597578 | 1.323718 | 0.036 | 0.025 | 1 | 4 | Pdpr | 0.011 | Old |
| 0.597679 | 0.23672 | 0.287 | 0.262 | 1 | 4 | Ndufs6 | 0.025 | Old |
| 0.597868 | 0.116995 | 0.253 | 0.225 | 1 | 4 | Rasa1 | 0.028 | Old |
| 0.59819 | 0.068136 | 0.007 | 0.013 | 1 | 4 | Gm6133 | -0.006 | Old |
| 0.598275 | 0.871681 | 0.047 | 0.062 | 1 | 4 | Tdp2 | -0.015 | Old |
| 0.598699 | 0.730552 | 0.021 | 0.013 | 1 | 4 | Nudt1 | 0.008 | Old |
| 0.598765 | 1.036184 | 0.169 | 0.162 | 1 | 4 | Rnft1 | 0.007 | Old |
| 0.598813 | 0.399836 | 0.144 | 0.125 | 1 | 4 | BC004004 | 0.019 | Old |
| 0.598899 | 0.621792 | 0.117 | 0.1 | 1 | 4 | Lrrfip1 | 0.017 | Old |
| 0.599229 | 0.194141 | 0.156 | 0.188 | 1 | 4 | Med1 | -0.032 | Old |
| 0.599334 | 0.466465 | 0.215 | 0.2 | 1 | 4 | Slc27a1 | 0.015 | Old |
| 0.5994 | 0.753654 | 0.021 | 0.013 | 1 | 4 | Fam136a | 0.008 | Old |
| 0.5994 | 0.712783 | 0.021 | 0.013 | 1 | 4 | 3110001I2 | 0.008 | Old |
| 0.599414 | 0.578897 | 0.078 | 0.062 | 1 | 4 | Naxe | 0.016 | Old |
| 0.600102 | 0.748084 | 0.021 | 0.013 | 1 | 4 | Cbx8 | 0.008 | Old |
| 0.600102 | 0.528019 | 0.021 | 0.013 | 1 | 4 | Mrpl49 | 0.008 | Old |
| 0.600103 | 0.153265 | 0.486 | 0.462 | 1 | 4 | Mapk3 | 0.024 | Old |

| | | | | | | | | |
|---|---|---|---|---|---|---|---|---|
| 0.600286 | 0.202786 | 0.455 | 0.438 | 1 | 4 | Sumo2 | 0.017 | Old |
| 0.600289 | 1.240124 | 0.036 | 0.025 | 1 | 4 | Ddx51 | 0.011 | Old |
| 0.600501 | 0.477439 | 0.007 | 0.013 | 1 | 4 | Cd27 | -0.006 | Old |
| 0.600804 | 0.583304 | 0.021 | 0.013 | 1 | 4 | Dars2 | 0.008 | Old |
| 0.60097 | 0.278078 | 0.065 | 0.05 | 1 | 4 | Lrrcc1 | 0.015 | Old |
| 0.601044 | 0.296501 | 0.133 | 0.112 | 1 | 4 | Cln6 | 0.021 | Old |
| 0.601081 | 0.079525 | 0.713 | 0.738 | 1 | 4 | Eef1b2 | -0.025 | Old |
| 0.601376 | 0.217015 | 0.065 | 0.05 | 1 | 4 | Gla | 0.015 | Old |
| 0.601506 | 0.581197 | 0.021 | 0.013 | 1 | 4 | Rbm48 | 0.008 | Old |
| 0.601634 | 0.029806 | 0.353 | 0.4 | 1 | 4 | Sfr1 | -0.047 | Old |
| 0.601658 | 0.896036 | 0.007 | 0.013 | 1 | 4 | Ptpdc1 | -0.006 | Old |
| 0.601658 | 0.879897 | 0.007 | 0.013 | 1 | 4 | Serping1 | -0.006 | Old |
| 0.601658 | 0.831422 | 0.007 | 0.013 | 1 | 4 | Zfp119b | -0.006 | Old |
| 0.601658 | 0.634934 | 0.007 | 0.013 | 1 | 4 | Kif9 | -0.006 | Old |
| 0.601658 | 0.609394 | 0.007 | 0.013 | 1 | 4 | Platr17 | -0.006 | Old |
| 0.601658 | 0.493668 | 0.007 | 0.013 | 1 | 4 | Dock5 | -0.006 | Old |
| 0.601658 | 0.4095 | 0.007 | 0.013 | 1 | 4 | Retn | -0.006 | Old |
| 0.60184 | 0.141953 | 0.121 | 0.1 | 1 | 4 | Elp2 | 0.021 | Old |
| 0.602253 | 0.161665 | 0.049 | 0.062 | 1 | 4 | Tbcd | -0.013 | Old |
| 0.602272 | 0.271557 | 0.106 | 0.088 | 1 | 4 | Nt5c2 | 0.018 | Old |
| 0.602499 | 1.05242 | 0.063 | 0.05 | 1 | 4 | Med23 | 0.013 | Old |
| 0.602816 | 0.480931 | 0.007 | 0.013 | 1 | 4 | Rgsl1 | -0.006 | Old |
| 0.602912 | 0.353601 | 0.021 | 0.013 | 1 | 4 | Plekhg1 | 0.008 | Old |
| 0.603004 | 0.288059 | 0.065 | 0.05 | 1 | 4 | Champ1 | 0.015 | Old |
| 0.603135 | 0.627207 | 0.128 | 0.112 | 1 | 4 | Fam20b | 0.016 | Old |
| 0.603411 | 0.28083 | 0.065 | 0.05 | 1 | 4 | Tctex1d2 | 0.015 | Old |
| 0.603448 | 0.227478 | 0.185 | 0.162 | 1 | 4 | Rasa3 | 0.023 | Old |
| 0.603572 | 0.418064 | 0.078 | 0.062 | 1 | 4 | Kif1bp | 0.016 | Old |
| 0.603758 | 0.360848 | 0.344 | 0.338 | 1 | 4 | Ehbp1l1 | 0.006 | Old |
| 0.604135 | 0.174457 | 0.133 | 0.112 | 1 | 4 | Kank2 | 0.021 | Old |
| 0.604182 | 0.281268 | 0.181 | 0.162 | 1 | 4 | Mcmbp | 0.019 | Old |
| 0.604664 | 0.117804 | 0.254 | 0.225 | 1 | 4 | Llph | 0.029 | Old |
| 0.604759 | 0.138956 | 0.409 | 0.388 | 1 | 4 | Rab7 | 0.021 | Old |
| 0.604776 | 0.014912 | 0.632 | 0.713 | 1 | 4 | Clec2d | -0.081 | Old |
| 0.605024 | 0.374513 | 0.021 | 0.013 | 1 | 4 | Slc35d1 | 0.008 | Old |
| 0.605024 | 0.225979 | 0.021 | 0.013 | 1 | 4 | 5830416I1! | 0.008 | Old |
| 0.605067 | 0.075832 | 0.078 | 0.062 | 1 | 4 | Uri1 | 0.016 | Old |
| 0.605094 | 0.392609 | 0.248 | 0.238 | 1 | 4 | Parvb | 0.01 | Old |
| 0.605416 | 0.220627 | 0.308 | 0.288 | 1 | 4 | Ywhag | 0.02 | Old |
| 0.605496 | 0.089022 | 0.171 | 0.2 | 1 | 4 | Trappc2l | -0.029 | Old |
| 0.605502 | 0.174005 | 0.119 | 0.1 | 1 | 4 | Phax | 0.019 | Old |
| 0.605946 | 0.500288 | 0.05 | 0.038 | 1 | 4 | Hinfp | 0.012 | Old |
| 0.606243 | 0.62633 | 0.064 | 0.05 | 1 | 4 | Ngrn | 0.014 | Old |
| 0.606258 | 0.447413 | 0.065 | 0.05 | 1 | 4 | Cpne8 | 0.015 | Old |
| 0.60664 | 0.654573 | 0.09 | 0.075 | 1 | 4 | Prr14 | 0.015 | Old |

| | | | | | | | | |
|---|---|---|---|---|---|---|---|---|
| 0.606808 | 0.390477 | 0.092 | 0.075 | 1 | 4 | Nsmf | 0.017 | Old |
| 0.607057 | 0.105803 | 0.169 | 0.2 | 1 | 4 | Mdm2 | -0.031 | Old |
| 0.60723 | 0.386468 | 0.13 | 0.112 | 1 | 4 | Ilf3 | 0.018 | Old |
| 0.607658 | 0.094291 | 0.249 | 0.288 | 1 | 4 | Copb1 | -0.039 | Old |
| 0.607844 | 0.0231 | 0.021 | 0.013 | 1 | 4 | Il15ra | 0.008 | Old |
| 0.607909 | 1.032393 | 0.036 | 0.025 | 1 | 4 | 1190007I0 | 0.011 | Old |
| 0.608027 | 0.400488 | 0.294 | 0.288 | 1 | 4 | Sgk3 | 0.006 | Old |
| 0.608029 | 0.150067 | 0.538 | 0.537 | 1 | 4 | Stxbp3 | 0.001 | Old |
| 0.60869 | 1.078042 | 0.076 | 0.062 | 1 | 4 | Hax1 | 0.014 | Old |
| 0.608895 | 0.681405 | 0.059 | 0.075 | 1 | 4 | Tfb2m | -0.016 | Old |
| 0.609094 | 1.033558 | 0.063 | 0.05 | 1 | 4 | Mfn1 | 0.013 | Old |
| 0.609095 | 0.072383 | 0.017 | 0.025 | 1 | 4 | Tmem115 | -0.008 | Old |
| 0.609861 | 0.512605 | 0.017 | 0.025 | 1 | 4 | Hspa2 | -0.008 | Old |
| 0.610521 | 0.290638 | 0.16 | 0.138 | 1 | 4 | Ing2 | 0.022 | Old |
| 0.610614 | 0.90572 | 0.138 | 0.125 | 1 | 4 | Pisd | 0.013 | Old |
| 0.611056 | 0.172973 | 0.526 | 0.525 | 1 | 4 | Tomm7 | 0.001 | Old |
| 0.611187 | 0.911693 | 0.036 | 0.025 | 1 | 4 | Syce2 | 0.011 | Old |
| 0.611272 | 0.273726 | 0.049 | 0.062 | 1 | 4 | Msra | -0.013 | Old |
| 0.611339 | 0.356383 | 0.291 | 0.275 | 1 | 4 | Atp5d | 0.016 | Old |
| 0.611394 | 0.232485 | 0.017 | 0.025 | 1 | 4 | Limk2 | -0.008 | Old |
| 0.612009 | 0.076644 | 0.065 | 0.05 | 1 | 4 | Senp3 | 0.015 | Old |
| 0.61283 | 0.838054 | 0.036 | 0.025 | 1 | 4 | Rfc5 | 0.011 | Old |
| 0.612929 | 0.292264 | 0.017 | 0.025 | 1 | 4 | Spout1 | -0.008 | Old |
| 0.613513 | 1.52408 | 0.02 | 0.013 | 1 | 4 | Syn1 | 0.007 | Old |
| 0.613697 | 0.392611 | 0.017 | 0.025 | 1 | 4 | Tprn | -0.008 | Old |
| 0.613806 | 0.123009 | 0.534 | 0.512 | 1 | 4 | Rpn2 | 0.022 | Old |
| 0.614466 | 0.384573 | 0.017 | 0.025 | 1 | 4 | Nnt | -0.008 | Old |
| 0.614537 | 0.27529 | 0.093 | 0.112 | 1 | 4 | Nhp2 | -0.019 | Old |
| 0.614814 | 0.626593 | 0.076 | 0.062 | 1 | 4 | Dok1 | 0.014 | Old |
| 0.614884 | 0.205813 | 0.174 | 0.15 | 1 | 4 | Vrk2 | 0.024 | Old |
| 0.614952 | 1.391066 | 0.02 | 0.013 | 1 | 4 | Gstk1 | 0.007 | Old |
| 0.615256 | 1.562323 | 0.035 | 0.025 | 1 | 4 | Zfp319 | 0.01 | Old |
| 0.615472 | 0.214572 | 1 | 1 | 1 | 4 | Gm42418 | 0 | Old |
| 0.615673 | 1.676762 | 0.02 | 0.013 | 1 | 4 | Fuz | 0.007 | Old |
| 0.615891 | 0.317852 | 0.118 | 0.1 | 1 | 4 | Ddx10 | 0.018 | Old |
| 0.616016 | 0.026135 | 0.242 | 0.275 | 1 | 4 | Myo1f | -0.033 | Old |
| 0.616223 | 0.018296 | 0.107 | 0.088 | 1 | 4 | Dnm1l | 0.019 | Old |
| 0.616394 | 1.55549 | 0.02 | 0.013 | 1 | 4 | Det1 | 0.007 | Old |
| 0.616394 | 1.283371 | 0.02 | 0.013 | 1 | 4 | A930007I1 | 0.007 | Old |
| 0.616912 | 0.114383 | 0.132 | 0.112 | 1 | 4 | Mrpl13 | 0.02 | Old |
| 0.617076 | 0.04355 | 0.094 | 0.075 | 1 | 4 | Mcm6 | 0.019 | Old |
| 0.617087 | 0.177118 | 0.7 | 0.7 | 1 | 4 | Klf4 | 0 | Old |
| 0.617116 | 0.226332 | 0.221 | 0.2 | 1 | 4 | Cyb5a | 0.021 | Old |
| 0.617331 | 0.297181 | 0.178 | 0.212 | 1 | 4 | Ppp6r1 | -0.034 | Old |
| 0.617484 | 0.888322 | 0.076 | 0.062 | 1 | 4 | Atg4d | 0.014 | Old |

| | | | | | | | | |
|---|---|---|---|---|---|---|---|---|
| 0.617551 | 0.183803 | 0.049 | 0.062 | 1 | 4 | Inip | -0.013 | Old |
| 0.618118 | 0.010998 | 0.353 | 0.4 | 1 | 4 | Magt1 | -0.047 | Old |
| 0.618189 | 0.250099 | 0.19 | 0.225 | 1 | 4 | Lmo4 | -0.035 | Old |
| 0.618483 | 0.09645 | 0.216 | 0.25 | 1 | 4 | Sf3b3 | -0.034 | Old |
| 0.618558 | 0.891759 | 0.02 | 0.013 | 1 | 4 | Pstpip2 | 0.007 | Old |
| 0.618581 | 1.54996 | 0.035 | 0.025 | 1 | 4 | Psmg1 | 0.01 | Old |
| 0.618951 | 0.092243 | 0.051 | 0.038 | 1 | 4 | Clec4e | 0.013 | Old |
| 0.619452 | 0.059554 | 0.387 | 0.438 | 1 | 4 | Gpi1 | -0.051 | Old |
| 0.619874 | 0.764421 | 0.049 | 0.038 | 1 | 4 | Sbf1 | 0.011 | Old |
| 0.619957 | 0.207318 | 0.271 | 0.25 | 1 | 4 | Calu | 0.021 | Old |
| 0.620399 | 0.374372 | 0.13 | 0.112 | 1 | 4 | Hibadh | 0.018 | Old |
| 0.620492 | 0.156054 | 0.352 | 0.338 | 1 | 4 | Caprin1 | 0.014 | Old |
| 0.620652 | 0.117887 | 0.184 | 0.162 | 1 | 4 | Gba | 0.022 | Old |
| 0.621071 | 0.555314 | 0.128 | 0.112 | 1 | 4 | Hnrnpll | 0.016 | Old |
| 0.621106 | 0.073504 | 0.082 | 0.1 | 1 | 4 | Asnsd1 | -0.018 | Old |
| 0.621696 | 0.491566 | 0.059 | 0.075 | 1 | 4 | Mbd5 | -0.016 | Old |
| 0.621791 | 0.019188 | 0.409 | 0.462 | 1 | 4 | Synj1 | -0.053 | Old |
| 0.621894 | 0.243411 | 0.145 | 0.125 | 1 | 4 | Gabarapl2 | 0.02 | Old |
| 0.62202 | 0.275452 | 0.038 | 0.05 | 1 | 4 | Ermp1 | -0.012 | Old |
| 0.62247 | 0.122512 | 0.212 | 0.188 | 1 | 4 | Sidt2 | 0.024 | Old |
| 0.622587 | 0.283618 | 0.357 | 0.35 | 1 | 4 | Uqcrb | 0.007 | Old |
| 0.62277 | 0.363963 | 0.26 | 0.25 | 1 | 4 | Lpin1 | 0.01 | Old |
| 0.622911 | 0.105447 | 0.625 | 0.613 | 1 | 4 | Tln1 | 0.012 | Old |
| 0.623519 | 0.035068 | 0.078 | 0.062 | 1 | 4 | Mis18a | 0.016 | Old |
| 0.623546 | 0.641148 | 0.09 | 0.075 | 1 | 4 | Pvr | 0.015 | Old |
| 0.623583 | 0.387706 | 0.077 | 0.062 | 1 | 4 | Taf5l | 0.015 | Old |
| 0.623624 | 0.144193 | 0.05 | 0.038 | 1 | 4 | Kcnn4 | 0.012 | Old |
| 0.623648 | 0.315511 | 0.092 | 0.075 | 1 | 4 | Gm49336 | 0.017 | Old |
| 0.623859 | 0.800406 | 0.089 | 0.075 | 1 | 4 | Nup50 | 0.014 | Old |
| 0.623949 | 0.012259 | 0.13 | 0.15 | 1 | 4 | Dlgap4 | -0.02 | Old |
| 0.624216 | 0.724748 | 0.089 | 0.075 | 1 | 4 | Mrpl15 | 0.014 | Old |
| 0.624253 | 0.025137 | 0.179 | 0.15 | 1 | 4 | Lpxn | 0.029 | Old |
| 0.624327 | 0.332509 | 0.091 | 0.075 | 1 | 4 | Ccdc174 | 0.016 | Old |
| 0.624332 | 0.118001 | 0.105 | 0.125 | 1 | 4 | 5-Mar | -0.02 | Old |
| 0.6244 | 0.077591 | 0.117 | 0.138 | 1 | 4 | Lipo3 | -0.021 | Old |
| 0.624612 | 0.057631 | 0.162 | 0.188 | 1 | 4 | Csrp1 | -0.026 | Old |
| 0.624698 | 1.09756 | 0.035 | 0.025 | 1 | 4 | 1110012L1 | 0.01 | Old |
| 0.624844 | 0.000879 | 0.312 | 0.275 | 1 | 4 | Spag9 | 0.037 | Old |
| 0.62485 | 0.248012 | 0.132 | 0.112 | 1 | 4 | Snapin | 0.02 | Old |
| 0.62496 | 0.50162 | 0.063 | 0.05 | 1 | 4 | Josd2 | 0.013 | Old |
| 0.624973 | 0.574344 | 0.028 | 0.038 | 1 | 4 | Smyd5 | -0.01 | Old |
| 0.625072 | 0.542109 | 0.02 | 0.013 | 1 | 4 | Stau2 | 0.007 | Old |
| 0.625138 | 0.233336 | 0.558 | 0.588 | 1 | 4 | Atp5f1 | -0.03 | Old |
| 0.625587 | 0.561265 | 0.028 | 0.038 | 1 | 4 | Heatr5b | -0.01 | Old |
| 0.625587 | 0.284023 | 0.028 | 0.038 | 1 | 4 | Gm47675 | -0.01 | Old |

| | | | | | | | | |
|---|---|---|---|---|---|---|---|---|
| 0.626756 | 0.334542 | 0.24 | 0.225 | 1 | 4 | Tcirg1 | 0.015 | Old |
| 0.62725 | 0.297835 | 0.02 | 0.013 | 1 | 4 | I830077J02 | 0.007 | Old |
| 0.627374 | 0.384486 | 0.049 | 0.062 | 1 | 4 | Nod1 | -0.013 | Old |
| 0.627486 | 0.844587 | 0.035 | 0.025 | 1 | 4 | Coq10a | 0.01 | Old |
| 0.627685 | 0.474482 | 0.153 | 0.138 | 1 | 4 | Parp1 | 0.015 | Old |
| 0.627976 | 0.293548 | 0.144 | 0.125 | 1 | 4 | Setdb1 | 0.019 | Old |
| 0.628006 | 0.16999 | 0.127 | 0.15 | 1 | 4 | Sgta | -0.023 | Old |
| 0.628209 | 0.516312 | 0.09 | 0.075 | 1 | 4 | Patl1 | 0.015 | Old |
| 0.628253 | 0.020119 | 0.092 | 0.075 | 1 | 4 | Cpne2 | 0.017 | Old |
| 0.628297 | 0.023749 | 0.246 | 0.275 | 1 | 4 | Pcyt1a | -0.029 | Old |
| 0.628407 | 0.162332 | 0.192 | 0.225 | 1 | 4 | Zc3h13 | -0.033 | Old |
| 0.628528 | 0.523414 | 0.063 | 0.05 | 1 | 4 | Mrps30 | 0.013 | Old |
| 0.628638 | 0.184032 | 0.082 | 0.1 | 1 | 4 | Stk19 | -0.018 | Old |
| 0.628646 | 0.485677 | 0.128 | 0.112 | 1 | 4 | Akr1c13 | 0.016 | Old |
| 0.628704 | 0.19438 | 0.02 | 0.013 | 1 | 4 | Suox | 0.007 | Old |
| 0.628902 | 0.32915 | 0.216 | 0.2 | 1 | 4 | Uqcrc2 | 0.016 | Old |
| 0.629008 | 0.248193 | 0.284 | 0.262 | 1 | 4 | Tmem258 | 0.022 | Old |
| 0.629266 | 0.085697 | 0.457 | 0.525 | 1 | 4 | Gls | -0.068 | Old |
| 0.62939 | 0.460067 | 0.19 | 0.175 | 1 | 4 | Fam96a | 0.015 | Old |
| 0.629404 | 0.684048 | 0.076 | 0.062 | 1 | 4 | Emc8 | 0.014 | Old |
| 0.629729 | 0.216528 | 0.115 | 0.138 | 1 | 4 | Slc25a46 | -0.023 | Old |
| 0.629895 | 0.593926 | 0.028 | 0.038 | 1 | 4 | Mrps31 | -0.01 | Old |
| 0.629917 | 0.053753 | 0.039 | 0.05 | 1 | 4 | Abtb1 | -0.011 | Old |
| 0.630421 | 0.514872 | 0.063 | 0.05 | 1 | 4 | 9430038I0 | 0.013 | Old |
| 0.63045 | 0.203719 | 0.362 | 0.338 | 1 | 4 | Rnf141 | 0.024 | Old |
| 0.630535 | 0.362328 | 0.218 | 0.2 | 1 | 4 | Stag1 | 0.018 | Old |
| 0.630665 | 0.147319 | 0.082 | 0.1 | 1 | 4 | Phactr4 | -0.018 | Old |
| 0.630665 | 0.020933 | 0.077 | 0.062 | 1 | 4 | Fbxo30 | 0.015 | Old |
| 0.630866 | 0.116849 | 0.26 | 0.238 | 1 | 4 | Mib1 | 0.022 | Old |
| 0.630887 | 0.076655 | 0.02 | 0.013 | 1 | 4 | Olfr920 | 0.007 | Old |
| 0.630938 | 0.391107 | 0.135 | 0.162 | 1 | 4 | Erlec1 | -0.027 | Old |
| 0.631139 | 0.084216 | 0.14 | 0.162 | 1 | 4 | Fam192a | -0.022 | Old |
| 0.631403 | 0.387284 | 0.329 | 0.325 | 1 | 4 | Sdf4 | 0.004 | Old |
| 0.631501 | 0.676594 | 0.038 | 0.05 | 1 | 4 | Sp4 | -0.012 | Old |
| 0.631921 | 0.675112 | 0.101 | 0.088 | 1 | 4 | Tbc1d22a | 0.013 | Old |
| 0.631963 | 0.27042 | 0.091 | 0.075 | 1 | 4 | Ptk2b | 0.016 | Old |
| 0.632058 | 0.181306 | 0.094 | 0.112 | 1 | 4 | Arfip1 | -0.018 | Old |
| 0.632496 | 0.252573 | 0.218 | 0.2 | 1 | 4 | Api5 | 0.018 | Old |
| 0.632519 | 0.569456 | 0.035 | 0.025 | 1 | 4 | 1110034G2 | 0.01 | Old |
| 0.632533 | 0.527989 | 0.175 | 0.162 | 1 | 4 | Clptm1 | 0.013 | Old |
| 0.632863 | 0.184321 | 0.3 | 0.35 | 1 | 4 | Amfr | -0.05 | Old |
| 0.63322 | 0.448975 | 0.076 | 0.062 | 1 | 4 | Gtf3c1 | 0.014 | Old |
| 0.633589 | 0.010473 | 0.039 | 0.05 | 1 | 4 | Fbxo42 | -0.011 | Old |
| 0.633607 | 0.358406 | 0.135 | 0.162 | 1 | 4 | Fblim1 | -0.027 | Old |
| 0.633902 | 0.27353 | 0.204 | 0.188 | 1 | 4 | Mob4 | 0.016 | Old |

| | | | | | | | | |
|---|---|---|---|---|---|---|---|---|
| 0.634063 | 0.601562 | 0.127 | 0.112 | 1 | 4 | Dlg1 | 0.015 | Old |
| 0.634094 | 0.452298 | 0.09 | 0.075 | 1 | 4 | Commd4 | 0.015 | Old |
| 0.634201 | 0.682932 | 0.035 | 0.025 | 1 | 4 | Taf8 | 0.01 | Old |
| 0.634528 | 0.188652 | 0.064 | 0.05 | 1 | 4 | Gdap2 | 0.014 | Old |
| 0.634738 | 0.142837 | 0.064 | 0.05 | 1 | 4 | Fktn | 0.014 | Old |
| 0.634745 | 0.458088 | 0.128 | 0.112 | 1 | 4 | Coa3 | 0.016 | Old |
| 0.634796 | 0.218759 | 0.116 | 0.1 | 1 | 4 | Nae1 | 0.016 | Old |
| 0.634814 | 0.181926 | 0.116 | 0.138 | 1 | 4 | Chst7 | -0.022 | Old |
| 0.635884 | 0.524246 | 0.035 | 0.025 | 1 | 4 | Txnl4a | 0.01 | Old |
| 0.636063 | 0.291731 | 0.091 | 0.075 | 1 | 4 | Pnpla2 | 0.016 | Old |
| 0.636209 | 0.246643 | 0.157 | 0.138 | 1 | 4 | Sf3a1 | 0.019 | Old |
| 0.636267 | 1.824621 | 0.02 | 0.013 | 1 | 4 | Gm13431 | 0.007 | Old |
| 0.636382 | 0.957652 | 0.099 | 0.088 | 1 | 4 | Cetn2 | 0.011 | Old |
| 0.636625 | 0.1074 | 0.077 | 0.062 | 1 | 4 | Selenom | 0.015 | Old |
| 0.636728 | 0.298086 | 0.13 | 0.112 | 1 | 4 | Tnip1 | 0.018 | Old |
| 0.637589 | 0.385335 | 0.103 | 0.088 | 1 | 4 | Bnip3 | 0.015 | Old |
| 0.637716 | 0.270896 | 0.14 | 0.125 | 1 | 4 | Zfp451 | 0.015 | Old |
| 0.637752 | 1.569052 | 0.02 | 0.013 | 1 | 4 | Rfc4 | 0.007 | Old |
| 0.638004 | 0.791799 | 0.171 | 0.162 | 1 | 4 | Hltf | 0.009 | Old |
| 0.638132 | 0.227152 | 0.035 | 0.025 | 1 | 4 | Mrps10 | 0.01 | Old |
| 0.63831 | 0.385923 | 0.049 | 0.038 | 1 | 4 | Mcm3ap | 0.011 | Old |
| 0.638409 | 0.095062 | 0.764 | 0.775 | 1 | 4 | Hnrnpf | -0.011 | Old |
| 0.638634 | 0.535368 | 0.115 | 0.1 | 1 | 4 | Dgkz | 0.015 | Old |
| 0.638779 | 0.28847 | 0.071 | 0.088 | 1 | 4 | Cpped1 | -0.017 | Old |
| 0.638939 | 0.308244 | 0.077 | 0.062 | 1 | 4 | Parp2 | 0.015 | Old |
| 0.639355 | 0.420091 | 0.049 | 0.038 | 1 | 4 | Slc18b1 | 0.011 | Old |
| 0.639405 | 0.639871 | 0.063 | 0.05 | 1 | 4 | Pus1 | 0.013 | Old |
| 0.639407 | 0.384552 | 0.076 | 0.062 | 1 | 4 | Sap30 | 0.014 | Old |
| 0.639882 | 0.04917 | 0.265 | 0.3 | 1 | 4 | Thrap3 | -0.035 | Old |
| 0.639983 | 1.478693 | 0.02 | 0.013 | 1 | 4 | Snapc2 | 0.007 | Old |
| 0.639983 | 1.434058 | 0.02 | 0.013 | 1 | 4 | Brat1 | 0.007 | Old |
| 0.639983 | 1.125946 | 0.02 | 0.013 | 1 | 4 | Sirt5 | 0.007 | Old |
| 0.639998 | 0.139831 | 0.064 | 0.05 | 1 | 4 | Ccdc107 | 0.014 | Old |
| 0.640366 | 0.208034 | 0.259 | 0.238 | 1 | 4 | Arf2 | 0.021 | Old |
| 0.640431 | 0.246931 | 0.039 | 0.05 | 1 | 4 | Stk35 | -0.011 | Old |
| 0.640673 | 0.626217 | 0.174 | 0.162 | 1 | 4 | Zfr | 0.012 | Old |
| 0.64101 | 0.643103 | 0.138 | 0.125 | 1 | 4 | Ckap5 | 0.013 | Old |
| 0.641108 | 0.165162 | 0.063 | 0.05 | 1 | 4 | 2310033P0 | 0.013 | Old |
| 0.641243 | 0.072112 | 0.134 | 0.112 | 1 | 4 | Atxn1 | 0.022 | Old |
| 0.641449 | 0.165605 | 0.16 | 0.188 | 1 | 4 | Glrx3 | -0.028 | Old |
| 0.641472 | 0.955146 | 0.02 | 0.013 | 1 | 4 | Sall2 | 0.007 | Old |
| 0.64187 | 0.904865 | 0.062 | 0.05 | 1 | 4 | Angptl4 | 0.012 | Old |
| 0.64196 | 0.083075 | 0.063 | 0.05 | 1 | 4 | Mcc | 0.013 | Old |
| 0.642016 | 0.017309 | 0.715 | 0.725 | 1 | 4 | Prcp | -0.01 | Old |
| 0.642045 | 0.063292 | 0.208 | 0.238 | 1 | 4 | Atg3 | -0.03 | Old |

| | | | | | | | | |
|---|---|---|---|---|---|---|---|---|
| 0.642084 | 0.784065 | 0.062 | 0.05 | 1 | 4 | Fdft1 | 0.012 | Old |
| 0.642828 | 0.211925 | 0.072 | 0.088 | 1 | 4 | Zfp318 | -0.016 | Old |
| 0.643037 | 0.013678 | 0.028 | 0.038 | 1 | 4 | 4930503L1 | -0.01 | Old |
| 0.643472 | 0.076952 | 0.018 | 0.025 | 1 | 4 | Pold2 | -0.007 | Old |
| 0.643472 | 0.008458 | 0.018 | 0.025 | 1 | 4 | Mtfmt | -0.007 | Old |
| 0.643652 | 0.160068 | 0.028 | 0.038 | 1 | 4 | Fam149b | -0.01 | Old |
| 0.643709 | 1.081382 | 0.02 | 0.013 | 1 | 4 | Dut | 0.007 | Old |
| 0.643709 | 1.006846 | 0.02 | 0.013 | 1 | 4 | Spon1 | 0.007 | Old |
| 0.643715 | 0.606499 | 0.126 | 0.112 | 1 | 4 | Kdm4a | 0.014 | Old |
| 0.643746 | 0.65932 | 0.101 | 0.088 | 1 | 4 | Chsy1 | 0.013 | Old |
| 0.643973 | 0.0303 | 0.745 | 0.787 | 1 | 4 | Atp6v0b | -0.042 | Old |
| 0.64405 | 0.048407 | 0.05 | 0.062 | 1 | 4 | Prdm4 | -0.012 | Old |
| 0.644091 | 0.246165 | 0.063 | 0.05 | 1 | 4 | Pold4 | 0.013 | Old |
| 0.644126 | 0.357217 | 0.167 | 0.15 | 1 | 4 | Cers6 | 0.017 | Old |
| 0.644243 | 0.200733 | 0.018 | 0.025 | 1 | 4 | Mapk1ip1 | -0.007 | Old |
| 0.644455 | 0.537235 | 0.02 | 0.013 | 1 | 4 | Zfp35 | 0.007 | Old |
| 0.644573 | 0.210298 | 0.33 | 0.312 | 1 | 4 | Fgfr1op2 | 0.018 | Old |
| 0.645014 | 0.473967 | 0.018 | 0.025 | 1 | 4 | Coa7 | -0.007 | Old |
| 0.64525 | 0.334434 | 0.155 | 0.138 | 1 | 4 | Frmd4a | 0.017 | Old |
| 0.645459 | 0.143224 | 0.035 | 0.025 | 1 | 4 | Pomgnt1 | 0.01 | Old |
| 0.64551 | 0.999557 | 0.074 | 0.062 | 1 | 4 | Cdk10 | 0.012 | Old |
| 0.645986 | 0.060393 | 0.072 | 0.088 | 1 | 4 | Hivep3 | -0.016 | Old |
| 0.646131 | 0.054336 | 0.196 | 0.225 | 1 | 4 | Ccdc47 | -0.029 | Old |
| 0.646511 | 0.936425 | 0.034 | 0.025 | 1 | 4 | Jak3 | 0.009 | Old |
| 0.646558 | 0.475859 | 0.018 | 0.025 | 1 | 4 | Rgcc | -0.007 | Old |
| 0.646696 | 0.642929 | 0.02 | 0.013 | 1 | 4 | 1110065P2 | 0.007 | Old |
| 0.646696 | 0.622818 | 0.02 | 0.013 | 1 | 4 | Rnf180 | 0.007 | Old |
| 0.647071 | 1.407694 | 0.061 | 0.05 | 1 | 4 | Cenpc1 | 0.011 | Old |
| 0.647076 | 0.233894 | 0.446 | 0.45 | 1 | 4 | Pros1 | -0.004 | Old |
| 0.647295 | 0.297729 | 0.094 | 0.112 | 1 | 4 | Wdr3 | -0.018 | Old |
| 0.647333 | 0.280925 | 0.178 | 0.162 | 1 | 4 | Senp6 | 0.016 | Old |
| 0.647721 | 0.129474 | 0.035 | 0.025 | 1 | 4 | Pdss2 | 0.01 | Old |
| 0.647765 | 1.470286 | 0.047 | 0.038 | 1 | 4 | 2410002F2 | 0.009 | Old |
| 0.64778 | 0.026035 | 0.147 | 0.125 | 1 | 4 | Rictor | 0.022 | Old |
| 0.64809 | 0.349921 | 0.061 | 0.075 | 1 | 4 | B4gat1 | -0.014 | Old |
| 0.648159 | 0.28009 | 0.168 | 0.2 | 1 | 4 | Rgl2 | -0.032 | Old |
| 0.648191 | 0.41837 | 0.02 | 0.013 | 1 | 4 | Pus3 | 0.007 | Old |
| 0.648583 | 0.131639 | 0.028 | 0.038 | 1 | 4 | Zcchc4 | -0.01 | Old |
| 0.648876 | 0.828582 | 0.018 | 0.025 | 1 | 4 | Vsig10 | -0.007 | Old |
| 0.649688 | 0.441211 | 0.02 | 0.013 | 1 | 4 | Nipa1 | 0.007 | Old |
| 0.649816 | 0.158612 | 0.107 | 0.125 | 1 | 4 | Ube2a | -0.018 | Old |
| 0.649998 | 0.025961 | 0.246 | 0.225 | 1 | 4 | Pacsin2 | 0.021 | Old |
| 0.650033 | 0.019982 | 0.142 | 0.162 | 1 | 4 | Ndufa12 | -0.02 | Old |
| 0.650504 | 0.453813 | 0.063 | 0.05 | 1 | 4 | Utp20 | 0.013 | Old |
| 0.650552 | 0.040623 | 0.035 | 0.025 | 1 | 4 | Cnksr3 | 0.01 | Old |

| | | | | | | | | |
|---|---|---|---|---|---|---|---|---|
| 0.650571 | 0.24526 | 0.3 | 0.288 | 1 | 4 | Eloc | 0.012 | Old |
| 0.650834 | 0.713118 | 0.049 | 0.062 | 1 | 4 | 2510002D2 | -0.013 | Old |
| 0.651054 | 0.357405 | 0.028 | 0.038 | 1 | 4 | Flywch1 | -0.01 | Old |
| 0.651084 | 0.388605 | 0.246 | 0.238 | 1 | 4 | Glyr1 | 0.008 | Old |
| 0.651182 | 0.003582 | 0.478 | 0.537 | 1 | 4 | Tspo | -0.059 | Old |
| 0.651863 | 0.064816 | 0.375 | 0.412 | 1 | 4 | Map3k1 | -0.037 | Old |
| 0.651973 | 0.665023 | 0.018 | 0.025 | 1 | 4 | Haus1 | -0.007 | Old |
| 0.65256 | 0.827082 | 0.061 | 0.05 | 1 | 4 | Dpysl3 | 0.011 | Old |
| 0.652588 | 0.066844 | 0.387 | 0.438 | 1 | 4 | Ctnnd1 | -0.051 | Old |
| 0.65261 | 0.020432 | 0.073 | 0.088 | 1 | 4 | Ermard | -0.015 | Old |
| 0.652798 | 0.589603 | 0.034 | 0.025 | 1 | 4 | Sgf29 | 0.009 | Old |
| 0.652871 | 0.196393 | 0.17 | 0.15 | 1 | 4 | Pex6 | 0.02 | Old |
| 0.652891 | 0.15113 | 0.063 | 0.05 | 1 | 4 | Rps6ka5 | 0.013 | Old |
| 0.652909 | 0.287483 | 0.265 | 0.25 | 1 | 4 | Acly | 0.015 | Old |
| 0.653018 | 0.17973 | 0.13 | 0.112 | 1 | 4 | Esyt1 | 0.018 | Old |
| 0.653264 | 0.484808 | 0.174 | 0.162 | 1 | 4 | Esyt2 | 0.012 | Old |
| 0.653348 | 0.064176 | 0.153 | 0.175 | 1 | 4 | Luzp1 | -0.022 | Old |
| 0.653502 | 0.301025 | 0.286 | 0.275 | 1 | 4 | Cers2 | 0.011 | Old |
| 0.653528 | 0.737147 | 0.028 | 0.038 | 1 | 4 | Bud23 | -0.01 | Old |
| 0.653774 | 0.25967 | 0.117 | 0.138 | 1 | 4 | Retreg3 | -0.021 | Old |
| 0.653853 | 0.271959 | 0.103 | 0.088 | 1 | 4 | Ulk1 | 0.015 | Old |
| 0.653944 | 0.705793 | 0.034 | 0.025 | 1 | 4 | Dusp10 | 0.009 | Old |
| 0.654118 | 0.588327 | 0.062 | 0.05 | 1 | 4 | Gnptg | 0.012 | Old |
| 0.654192 | 0.44422 | 0.114 | 0.1 | 1 | 4 | Nedd9 | 0.014 | Old |
| 0.65471 | 0.412497 | 0.203 | 0.188 | 1 | 4 | Cyld | 0.015 | Old |
| 0.654895 | 0.142329 | 0.161 | 0.188 | 1 | 4 | Gpbp1l1 | -0.027 | Old |
| 0.654909 | 0.223144 | 0.169 | 0.15 | 1 | 4 | Eapp | 0.019 | Old |
| 0.654979 | 0.179164 | 0.106 | 0.125 | 1 | 4 | Acadsb | -0.019 | Old |
| 0.655271 | 0.780444 | 0.039 | 0.05 | 1 | 4 | Aifm2 | -0.011 | Old |
| 0.655586 | 1.118731 | 0.047 | 0.038 | 1 | 4 | Lrrc42 | 0.009 | Old |
| 0.655775 | 0.702422 | 0.074 | 0.062 | 1 | 4 | Ksr1 | 0.012 | Old |
| 0.655778 | 0.516059 | 0.163 | 0.15 | 1 | 4 | Txnrd1 | 0.013 | Old |
| 0.655907 | 0.552889 | 0.048 | 0.038 | 1 | 4 | Gtpbp10 | 0.01 | Old |
| 0.656392 | 0.2498 | 0.154 | 0.138 | 1 | 4 | Xpo7 | 0.016 | Old |
| 0.656709 | 0.464466 | 0.062 | 0.05 | 1 | 4 | AL590144.: | 0.012 | Old |
| 0.656764 | 0.245462 | 0.05 | 0.062 | 1 | 4 | Maml2 | -0.012 | Old |
| 0.656803 | 0.522558 | 0.114 | 0.1 | 1 | 4 | Cdk16 | 0.014 | Old |
| 0.656868 | 0.027439 | 0.966 | 0.963 | 1 | 4 | Rps26 | 0.003 | Old |
| 0.657241 | 0.21351 | 0.572 | 0.613 | 1 | 4 | Grb2 | -0.041 | Old |
| 0.657307 | 0.06695 | 0.263 | 0.288 | 1 | 4 | Dnajb9 | -0.025 | Old |
| 0.657735 | 0.112271 | 0.084 | 0.1 | 1 | 4 | Timd4 | -0.016 | Old |
| 0.657803 | 0.027958 | 0.078 | 0.062 | 1 | 4 | Zfp639 | 0.016 | Old |
| 0.658154 | 0.675519 | 0.074 | 0.062 | 1 | 4 | Ints1 | 0.012 | Old |
| 0.658304 | 0.370858 | 0.153 | 0.138 | 1 | 4 | Tmem5 | 0.015 | Old |
| 0.658438 | 0.522604 | 0.062 | 0.05 | 1 | 4 | Nqo2 | 0.012 | Old |

| | | | | | | | | |
|---|---|---|---|---|---|---|---|---|
| 0.658582 | 0.013428 | 0.078 | 0.062 | 1 | 4 | Atad2 | 0.016 | Old |
| 0.658649 | 0.075066 | 0.008 | 0.013 | 1 | 4 | Gins3 | -0.005 | Old |
| 0.658649 | 0.027237 | 0.008 | 0.013 | 1 | 4 | Adarb1 | -0.005 | Old |
| 0.658922 | 0.120583 | 0.244 | 0.275 | 1 | 4 | Ikzf1 | -0.031 | Old |
| 0.65893 | 0.122618 | 0.163 | 0.188 | 1 | 4 | Arhgap12 | -0.025 | Old |
| 0.659176 | 0.405621 | 0.104 | 0.125 | 1 | 4 | Cstf3 | -0.021 | Old |
| 0.659232 | 0.048631 | 0.153 | 0.175 | 1 | 4 | Ap3m1 | -0.022 | Old |
| 0.659805 | 0.408848 | 0.008 | 0.013 | 1 | 4 | Gm11423 | -0.005 | Old |
| 0.659805 | 0.14567 | 0.008 | 0.013 | 1 | 4 | Gm45871 | -0.005 | Old |
| 0.660274 | 1.21142 | 0.034 | 0.025 | 1 | 4 | Rgp1 | 0.009 | Old |
| 0.660291 | 0.216879 | 0.192 | 0.175 | 1 | 4 | Hpse | 0.017 | Old |
| 0.660716 | 0.219173 | 0.218 | 0.2 | 1 | 4 | Atf7ip | 0.018 | Old |
| 0.660739 | 0.910603 | 0.047 | 0.038 | 1 | 4 | Mrs2 | 0.009 | Old |
| 0.660854 | 1.080086 | 0.034 | 0.025 | 1 | 4 | Cdc42bpa | 0.009 | Old |
| 0.660879 | 0.132082 | 0.04 | 0.05 | 1 | 4 | Slc35d2 | -0.01 | Old |
| 0.660963 | 0.521897 | 0.008 | 0.013 | 1 | 4 | Spdl1 | -0.005 | Old |
| 0.660963 | 0.319561 | 0.008 | 0.013 | 1 | 4 | Xlr4c | -0.005 | Old |
| 0.661145 | 0.43819 | 0.101 | 0.088 | 1 | 4 | Trmt1l | 0.013 | Old |
| 0.661447 | 1.160548 | 0.061 | 0.05 | 1 | 4 | Eef1e1 | 0.011 | Old |
| 0.661526 | 0.783333 | 0.087 | 0.075 | 1 | 4 | Gmppa | 0.012 | Old |
| 0.661966 | 0.468109 | 0.14 | 0.125 | 1 | 4 | Plin3 | 0.015 | Old |
| 0.662121 | 0.849282 | 0.008 | 0.013 | 1 | 4 | Slc22a5 | -0.005 | Old |
| 0.662121 | 0.743562 | 0.008 | 0.013 | 1 | 4 | B3galt4 | -0.005 | Old |
| 0.662121 | 0.637213 | 0.008 | 0.013 | 1 | 4 | 9130023H2 | -0.005 | Old |
| 0.662121 | 0.476786 | 0.008 | 0.013 | 1 | 4 | Gcat | -0.005 | Old |
| 0.662279 | 0.594551 | 0.173 | 0.162 | 1 | 4 | Ei24 | 0.011 | Old |
| 0.662541 | 0.126087 | 0.646 | 0.675 | 1 | 4 | Cotl1 | -0.029 | Old |
| 0.662742 | 0.079965 | 0.062 | 0.075 | 1 | 4 | Akr7a5 | -0.013 | Old |
| 0.663 | 1.256113 | 0.049 | 0.038 | 1 | 4 | Igkc | 0.011 | Old |
| 0.663024 | 0.24639 | 0.116 | 0.1 | 1 | 4 | Bag3 | 0.016 | Old |
| 0.663128 | 0.168318 | 0.466 | 0.45 | 1 | 4 | Anp32a | 0.016 | Old |
| 0.663237 | 0.440767 | 0.164 | 0.15 | 1 | 4 | Nup88 | 0.014 | Old |
| 0.663263 | 0.181414 | 0.205 | 0.238 | 1 | 4 | Vapb | -0.033 | Old |
| 0.66328 | 0.997601 | 0.008 | 0.013 | 1 | 4 | Ttc25 | -0.005 | Old |
| 0.66328 | 0.700825 | 0.008 | 0.013 | 1 | 4 | Atl1 | -0.005 | Old |
| 0.66328 | 0.648179 | 0.008 | 0.013 | 1 | 4 | 1810011H1 | -0.005 | Old |
| 0.66328 | 0.369507 | 0.008 | 0.013 | 1 | 4 | Slc39a8 | -0.005 | Old |
| 0.663558 | 0.009543 | 0.202 | 0.225 | 1 | 4 | Ndufs5 | -0.023 | Old |
| 0.663671 | 0.342241 | 0.226 | 0.212 | 1 | 4 | Sertad1 | 0.014 | Old |
| 0.66386 | 0.621291 | 0.061 | 0.05 | 1 | 4 | Rpia | 0.011 | Old |
| 0.663895 | 0.356769 | 0.274 | 0.262 | 1 | 4 | Stk38 | 0.012 | Old |
| 0.663946 | 0.08872 | 0.221 | 0.2 | 1 | 4 | Eif3i | 0.021 | Old |
| 0.664053 | 0.4158 | 0.257 | 0.25 | 1 | 4 | Zmat2 | 0.007 | Old |
| 0.664492 | 0.168884 | 0.128 | 0.15 | 1 | 4 | Ppp2r5e | -0.022 | Old |
| 0.664872 | 0.23443 | 0.117 | 0.138 | 1 | 4 | Dennd1c | -0.021 | Old |

| | | | | | | | | |
|---|---|---|---|---|---|---|---|---|
| 0.664971 | 0.345021 | 0.14 | 0.125 | 1 | 4 | Gpr108 | 0.015 | Old |
| 0.66504 | 0.441251 | 0.188 | 0.175 | 1 | 4 | Klf9 | 0.013 | Old |
| 0.665393 | 1.514372 | 0.019 | 0.013 | 1 | 4 | Hus1 | 0.006 | Old |
| 0.665403 | 0.137603 | 0.161 | 0.188 | 1 | 4 | Cpq | -0.027 | Old |
| 0.665509 | 0.180698 | 0.073 | 0.088 | 1 | 4 | Cpox | -0.015 | Old |
| 0.665614 | 0.435299 | 0.21 | 0.2 | 1 | 4 | Rtf2 | 0.01 | Old |
| 0.666798 | 0.048897 | 0.294 | 0.325 | 1 | 4 | Washc2 | -0.031 | Old |
| 0.666931 | 0.948104 | 0.019 | 0.013 | 1 | 4 | Btbd6 | 0.006 | Old |
| 0.667056 | 0.01134 | 0.029 | 0.038 | 1 | 4 | Crcp | -0.009 | Old |
| 0.667166 | 0.420774 | 0.164 | 0.15 | 1 | 4 | Imp3 | 0.014 | Old |
| 0.66721 | 0.123937 | 0.119 | 0.138 | 1 | 4 | Fnta | -0.019 | Old |
| 0.667354 | 0.620711 | 0.137 | 0.125 | 1 | 4 | Ncor2 | 0.012 | Old |
| 0.667534 | 0.209513 | 0.103 | 0.088 | 1 | 4 | Abcb7 | 0.015 | Old |
| 0.667544 | 0.127242 | 0.243 | 0.225 | 1 | 4 | Slc8b1 | 0.018 | Old |
| 0.667837 | 0.830762 | 0.034 | 0.025 | 1 | 4 | Abhd10 | 0.009 | Old |
| 0.667922 | 0.306675 | 0.347 | 0.35 | 1 | 4 | Metap2 | -0.003 | Old |
| 0.6681 | 0.515148 | 0.074 | 0.062 | 1 | 4 | Pitpnm1 | 0.012 | Old |
| 0.668128 | 0.431325 | 0.048 | 0.038 | 1 | 4 | Yae1d1 | 0.01 | Old |
| 0.668376 | 0.830935 | 0.047 | 0.038 | 1 | 4 | Anapc7 | 0.009 | Old |
| 0.668565 | 0.073198 | 0.119 | 0.138 | 1 | 4 | Med17 | -0.019 | Old |
| 0.669004 | 0.863884 | 0.034 | 0.025 | 1 | 4 | Drosha | 0.009 | Old |
| 0.66954 | 0.182405 | 0.096 | 0.112 | 1 | 4 | Mrps23 | -0.016 | Old |
| 0.669912 | 0.04488 | 0.074 | 0.088 | 1 | 4 | Zfpl1 | -0.014 | Old |
| 0.670411 | 0.247973 | 0.202 | 0.188 | 1 | 4 | Rhobtb1 | 0.014 | Old |
| 0.670531 | 0.054585 | 0.2 | 0.225 | 1 | 4 | Mknk2 | -0.025 | Old |
| 0.67079 | 0.361993 | 0.088 | 0.075 | 1 | 4 | Isoc1 | 0.013 | Old |
| 0.670945 | 0.427548 | 0.231 | 0.225 | 1 | 4 | H2-Q7 | 0.006 | Old |
| 0.671055 | 0.325332 | 0.089 | 0.075 | 1 | 4 | BC037034 | 0.014 | Old |
| 0.671331 | 0.240624 | 0.183 | 0.212 | 1 | 4 | Rbm7 | -0.029 | Old |
| 0.67134 | 0.575049 | 0.034 | 0.025 | 1 | 4 | Accs | 0.009 | Old |
| 0.671384 | 0.121536 | 0.298 | 0.288 | 1 | 4 | Nars | 0.01 | Old |
| 0.671551 | 0.667311 | 0.019 | 0.013 | 1 | 4 | Fut10 | 0.006 | Old |
| 0.671728 | 0.093901 | 0.64 | 0.738 | 1 | 4 | Foxp1 | -0.098 | Old |
| 0.671937 | 0.016864 | 0.109 | 0.125 | 1 | 4 | Lypla1 | -0.016 | Old |
| 0.672509 | 0.715355 | 0.034 | 0.025 | 1 | 4 | Acvr1b | 0.009 | Old |
| 0.672655 | 0.228952 | 0.203 | 0.238 | 1 | 4 | Asxl2 | -0.035 | Old |
| 0.673094 | 0.494817 | 0.019 | 0.013 | 1 | 4 | Gm43936 | 0.006 | Old |
| 0.673098 | 0.305792 | 0.075 | 0.062 | 1 | 4 | Fyco1 | 0.013 | Old |
| 0.673245 | 0.127042 | 0.029 | 0.038 | 1 | 4 | Zfp81 | -0.009 | Old |
| 0.673345 | 0.077998 | 0.498 | 0.5 | 1 | 4 | Purb | -0.002 | Old |
| 0.673385 | 0.218184 | 0.384 | 0.388 | 1 | 4 | Gm21188 | -0.004 | Old |
| 0.673573 | 0.384631 | 0.126 | 0.15 | 1 | 4 | Cracr2b | -0.024 | Old |
| 0.673729 | 0.337954 | 0.128 | 0.112 | 1 | 4 | Fbxo22 | 0.016 | Old |
| 0.673787 | 1.385202 | 0.033 | 0.025 | 1 | 4 | Numb | 0.008 | Old |
| 0.673802 | 0.340633 | 0.212 | 0.2 | 1 | 4 | Tmem164 | 0.012 | Old |

| | | | | | | | | |
|---|---|---|---|---|---|---|---|---|
| 0.673968 | 0.451714 | 0.083 | 0.1 | 1 | 4 | Trappc13 | -0.017 | Old |
| 0.674252 | 0.396148 | 0.019 | 0.013 | 1 | 4 | Mast4 | 0.006 | Old |
| 0.674287 | 0.196786 | 0.204 | 0.238 | 1 | 4 | Ubr5 | -0.034 | Old |
| 0.674486 | 0.324456 | 0.029 | 0.038 | 1 | 4 | Gemin6 | -0.009 | Old |
| 0.674605 | 0.20595 | 0.14 | 0.125 | 1 | 4 | Commd3 | 0.015 | Old |
| 0.674639 | 0.36055 | 0.019 | 0.013 | 1 | 4 | Macrod2 | 0.006 | Old |
| 0.674684 | 0.144146 | 0.076 | 0.062 | 1 | 4 | Wrn | 0.014 | Old |
| 0.674733 | 0.132357 | 0.104 | 0.088 | 1 | 4 | Tmem135 | 0.016 | Old |
| 0.674738 | 0.205138 | 0.261 | 0.25 | 1 | 4 | Sdhd | 0.011 | Old |
| 0.674887 | 0.925784 | 0.06 | 0.05 | 1 | 4 | Sgce | 0.01 | Old |
| 0.675107 | 0.092091 | 0.029 | 0.038 | 1 | 4 | Acyp1 | -0.009 | Old |
| 0.675236 | 0.433892 | 0.061 | 0.05 | 1 | 4 | Trappc2 | 0.011 | Old |
| 0.675325 | 0.104394 | 0.259 | 0.238 | 1 | 4 | Fndc3a | 0.021 | Old |
| 0.675411 | 0.420431 | 0.019 | 0.013 | 1 | 4 | Polr1b | 0.006 | Old |
| 0.676424 | 0.202874 | 0.226 | 0.212 | 1 | 4 | Stard9 | 0.014 | Old |
| 0.676501 | 0.254635 | 0.073 | 0.088 | 1 | 4 | Alg10b | -0.015 | Old |
| 0.676598 | 0.607592 | 0.094 | 0.112 | 1 | 4 | Tmem41b | -0.018 | Old |
| 0.677504 | 0.140055 | 0.073 | 0.088 | 1 | 4 | Mtx2 | -0.015 | Old |
| 0.678118 | 0.054261 | 0.018 | 0.025 | 1 | 4 | Ip6k2 | -0.007 | Old |
| 0.678118 | 0.024613 | 0.018 | 0.025 | 1 | 4 | Endov | -0.007 | Old |
| 0.678121 | 0.812731 | 0.073 | 0.062 | 1 | 4 | Ddx56 | 0.011 | Old |
| 0.678384 | 0.020404 | 0.488 | 0.525 | 1 | 4 | Ywhah | -0.037 | Old |
| 0.678464 | 0.199029 | 0.142 | 0.125 | 1 | 4 | Txndc5 | 0.017 | Old |
| 0.678605 | 0.299961 | 0.154 | 0.138 | 1 | 4 | Polb | 0.016 | Old |
| 0.678777 | 0.459873 | 0.047 | 0.038 | 1 | 4 | Tyw1 | 0.009 | Old |
| 0.679081 | 0.964366 | 0.047 | 0.038 | 1 | 4 | Miip | 0.009 | Old |
| 0.679189 | 0.603473 | 0.087 | 0.075 | 1 | 4 | Tuft1 | 0.012 | Old |
| 0.679667 | 0.133705 | 0.018 | 0.025 | 1 | 4 | Spaca9 | -0.007 | Old |
| 0.679703 | 1.358866 | 0.033 | 0.025 | 1 | 4 | Isyna1 | 0.008 | Old |
| 0.679703 | 1.271453 | 0.033 | 0.025 | 1 | 4 | Mthfs | 0.008 | Old |
| 0.67974 | 0.745112 | 0.073 | 0.062 | 1 | 4 | Ccdc93 | 0.011 | Old |
| 0.679806 | 0.056305 | 0.287 | 0.325 | 1 | 4 | Lman2 | -0.038 | Old |
| 0.680055 | 0.112715 | 0.019 | 0.013 | 1 | 4 | Slc16a12 | 0.006 | Old |
| 0.680368 | 0.629617 | 0.061 | 0.05 | 1 | 4 | Ddx39 | 0.011 | Old |
| 0.680702 | 0.485783 | 0.029 | 0.038 | 1 | 4 | Pus7 | -0.009 | Old |
| 0.680889 | 1.15504 | 0.033 | 0.025 | 1 | 4 | Ctnnbl1 | 0.008 | Old |
| 0.681217 | 0.0058 | 0.018 | 0.025 | 1 | 4 | Cntrob | -0.007 | Old |
| 0.681226 | 0.184946 | 0.393 | 0.388 | 1 | 4 | S100a13 | 0.005 | Old |
| 0.681786 | 0.113668 | 0.04 | 0.05 | 1 | 4 | Tpst1 | -0.01 | Old |
| 0.682075 | 1.009024 | 0.033 | 0.025 | 1 | 4 | Gorasp1 | 0.008 | Old |
| 0.682075 | 0.94626 | 0.033 | 0.025 | 1 | 4 | Zfp280c | 0.008 | Old |
| 0.682173 | 0.336252 | 0.255 | 0.25 | 1 | 4 | Eif3j1 | 0.005 | Old |
| 0.682213 | 0.635419 | 0.232 | 0.238 | 1 | 4 | Capn2 | -0.006 | Old |
| 0.682769 | 0.173435 | 0.018 | 0.025 | 1 | 4 | Izumo4 | -0.007 | Old |
| 0.682914 | 0.529781 | 0.1 | 0.088 | 1 | 4 | Fam69a | 0.012 | Old |

| | | | | | | | | |
|---|---|---|---|---|---|---|---|---|
| 0.683055 | 0.496212 | 0.062 | 0.075 | 1 | 4 | Rtl5 | -0.013 | Old |
| 0.683102 | 0.110569 | 0.197 | 0.225 | 1 | 4 | Khsrp | -0.028 | Old |
| 0.683139 | 0.143941 | 0.279 | 0.325 | 1 | 4 | Srrm1 | -0.046 | Old |
| 0.683319 | 0.232524 | 0.38 | 0.375 | 1 | 4 | Eif4h | 0.005 | Old |
| 0.683385 | 0.184479 | 0.04 | 0.05 | 1 | 4 | Entpd5 | -0.01 | Old |
| 0.683545 | 0.2398 | 0.018 | 0.025 | 1 | 4 | Glmn | -0.007 | Old |
| 0.683856 | 0.839791 | 0.033 | 0.025 | 1 | 4 | Pola2 | 0.008 | Old |
| 0.683924 | 0.284447 | 0.062 | 0.075 | 1 | 4 | Fam78a | -0.013 | Old |
| 0.684015 | 0.070473 | 0.177 | 0.2 | 1 | 4 | Cop1 | -0.023 | Old |
| 0.684061 | 0.016267 | 0.259 | 0.288 | 1 | 4 | Ankrd12 | -0.029 | Old |
| 0.684239 | 0.136553 | 0.177 | 0.162 | 1 | 4 | Dck | 0.015 | Old |
| 0.684279 | 0.088905 | 0.051 | 0.062 | 1 | 4 | Crebl2 | -0.011 | Old |
| 0.684322 | 0.309292 | 0.018 | 0.025 | 1 | 4 | Zfp456 | -0.007 | Old |
| 0.684805 | 0.482106 | 0.087 | 0.075 | 1 | 4 | Crebzf | 0.012 | Old |
| 0.684937 | 0.409617 | 0.199 | 0.188 | 1 | 4 | Rtcb | 0.011 | Old |
| 0.685418 | 0.923026 | 0.073 | 0.062 | 1 | 4 | Polr2j | 0.011 | Old |
| 0.685826 | 0.242239 | 0.181 | 0.162 | 1 | 4 | Kbtbd11 | 0.019 | Old |
| 0.685854 | 0.919917 | 0.072 | 0.062 | 1 | 4 | Fam45a | 0.01 | Old |
| 0.685876 | 0.275722 | 0.018 | 0.025 | 1 | 4 | Zfp825 | -0.007 | Old |
| 0.685902 | 0.056687 | 0.074 | 0.088 | 1 | 4 | Rps6kc1 | -0.014 | Old |
| 0.685978 | 0.243376 | 0.167 | 0.15 | 1 | 4 | Usp47 | 0.017 | Old |
| 0.686053 | 0.280069 | 0.04 | 0.05 | 1 | 4 | Gprasp1 | -0.01 | Old |
| 0.686448 | 0.078119 | 0.074 | 0.088 | 1 | 4 | Csf2rb2 | -0.014 | Old |
| 0.686862 | 0.571003 | 0.099 | 0.088 | 1 | 4 | Plekha3 | 0.011 | Old |
| 0.686948 | 0.030563 | 0.775 | 0.8 | 1 | 4 | Uqcrh | -0.025 | Old |
| 0.687129 | 0.382301 | 0.061 | 0.05 | 1 | 4 | Dgat1 | 0.011 | Old |
| 0.687212 | 1.852792 | 0.018 | 0.013 | 1 | 4 | 1810032O( | 0.005 | Old |
| 0.687386 | 0.286831 | 0.128 | 0.15 | 1 | 4 | Zdhhc14 | -0.022 | Old |
| 0.687564 | 0.68905 | 0.029 | 0.038 | 1 | 4 | Cyb561d1 | -0.009 | Old |
| 0.688003 | 1.943055 | 0.018 | 0.013 | 1 | 4 | 4930481A1 | 0.005 | Old |
| 0.688003 | 1.896988 | 0.018 | 0.013 | 1 | 4 | Zfp606 | 0.005 | Old |
| 0.68805 | 0.308206 | 0.345 | 0.338 | 1 | 4 | Ndufc1 | 0.007 | Old |
| 0.688057 | 0.034745 | 0.167 | 0.188 | 1 | 4 | Mtmr3 | -0.021 | Old |
| 0.688177 | 0.56367 | 0.135 | 0.125 | 1 | 4 | Ptbp2 | 0.01 | Old |
| 0.688182 | 0.470262 | 0.087 | 0.075 | 1 | 4 | Bcl2l13 | 0.012 | Old |
| 0.688189 | 0.701576 | 0.029 | 0.038 | 1 | 4 | Fen1 | -0.009 | Old |
| 0.688662 | 0.240962 | 0.509 | 0.537 | 1 | 4 | Bsg | -0.028 | Old |
| 0.68868 | 0.72478 | 0.06 | 0.05 | 1 | 4 | Hectd3 | 0.01 | Old |
| 0.688795 | 1.776153 | 0.018 | 0.013 | 1 | 4 | Slc7a2 | 0.005 | Old |
| 0.688988 | 0.713819 | 0.018 | 0.025 | 1 | 4 | Alg3 | -0.007 | Old |
| 0.689237 | 0.136637 | 0.366 | 0.412 | 1 | 4 | Lipa | -0.046 | Old |
| 0.689273 | 0.020652 | 0.361 | 0.4 | 1 | 4 | Elf1 | -0.039 | Old |
| 0.689587 | 1.456103 | 0.018 | 0.013 | 1 | 4 | Sdc1 | 0.005 | Old |
| 0.689668 | 0.075482 | 0.202 | 0.225 | 1 | 4 | Syf2 | -0.023 | Old |
| 0.689778 | 0.378728 | 0.126 | 0.112 | 1 | 4 | Akap9 | 0.014 | Old |

| | | | | | | | | |
|---|---|---|---|---|---|---|---|---|
| 0.690021 | 0.819957 | 0.06 | 0.05 | 1 | 4 | D16Ertd47: | 0.01 | Old |
| 0.690332 | 0.463275 | 0.161 | 0.15 | 1 | 4 | Ddhd2 | 0.011 | Old |
| 0.690464 | 0.00889 | 0.088 | 0.075 | 1 | 4 | Inpp5f | 0.013 | Old |
| 0.691201 | 0.852867 | 0.146 | 0.138 | 1 | 4 | Prdx6 | 0.008 | Old |
| 0.691288 | 0.170459 | 0.182 | 0.162 | 1 | 4 | Poldip3 | 0.02 | Old |
| 0.691326 | 0.772668 | 0.018 | 0.025 | 1 | 4 | Slc17a9 | -0.007 | Old |
| 0.691398 | 0.066618 | 0.611 | 0.637 | 1 | 4 | Hmox1 | -0.026 | Old |
| 0.691526 | 0.597586 | 0.124 | 0.112 | 1 | 4 | Mbd6 | 0.012 | Old |
| 0.691657 | 0.101807 | 0.076 | 0.062 | 1 | 4 | Pspc1 | 0.014 | Old |
| 0.69219 | 0.710142 | 0.033 | 0.025 | 1 | 4 | Wrnip1 | 0.008 | Old |
| 0.69219 | 0.668479 | 0.033 | 0.025 | 1 | 4 | Prpf3 | 0.008 | Old |
| 0.692445 | 1.534442 | 0.032 | 0.025 | 1 | 4 | Usp42 | 0.007 | Old |
| 0.692804 | 0.011429 | 0.403 | 0.412 | 1 | 4 | Dnajb6 | -0.009 | Old |
| 0.693244 | 0.333423 | 0.061 | 0.05 | 1 | 4 | Pxmp4 | 0.011 | Old |
| 0.693391 | 1.421504 | 0.059 | 0.05 | 1 | 4 | Tbce | 0.009 | Old |
| 0.693563 | 0.28495 | 0.063 | 0.075 | 1 | 4 | Tppp | -0.012 | Old |
| 0.693704 | 0.245051 | 0.101 | 0.088 | 1 | 4 | Afg3l1 | 0.013 | Old |
| 0.693984 | 0.083151 | 0.109 | 0.125 | 1 | 4 | Plekhm3 | -0.016 | Old |
| 0.694059 | 0.277857 | 0.257 | 0.25 | 1 | 4 | Fkbp2 | 0.007 | Old |
| 0.694345 | 0.826828 | 0.018 | 0.013 | 1 | 4 | Homer2 | 0.005 | Old |
| 0.694663 | 0.196219 | 0.154 | 0.175 | 1 | 4 | Rgs3 | -0.021 | Old |
| 0.695042 | 0.503776 | 0.073 | 0.088 | 1 | 4 | Pum3 | -0.015 | Old |
| 0.695154 | 0.437982 | 0.047 | 0.038 | 1 | 4 | Fbxo6 | 0.009 | Old |
| 0.695689 | 0.206663 | 0.195 | 0.175 | 1 | 4 | Atg5 | 0.02 | Old |
| 0.695742 | 0.175164 | 0.393 | 0.388 | 1 | 4 | Eif3k | 0.005 | Old |
| 0.695934 | 0.77598 | 0.018 | 0.013 | 1 | 4 | Zswim9 | 0.005 | Old |
| 0.696091 | 0.069815 | 0.097 | 0.112 | 1 | 4 | Appl2 | -0.015 | Old |
| 0.696143 | 0.101086 | 0.132 | 0.15 | 1 | 4 | Iah1 | -0.018 | Old |
| 0.696371 | 0.568712 | 0.033 | 0.025 | 1 | 4 | Pik3cb | 0.008 | Old |
| 0.696729 | 0.541909 | 0.018 | 0.013 | 1 | 4 | Inpp1 | 0.005 | Old |
| 0.697446 | 0.571706 | 0.099 | 0.088 | 1 | 4 | Tmem192 | 0.011 | Old |
| 0.697583 | 0.31183 | 0.074 | 0.088 | 1 | 4 | Glt8d1 | -0.014 | Old |
| 0.697662 | 0.218123 | 0.336 | 0.325 | 1 | 4 | Selplg | 0.011 | Old |
| 0.697761 | 0.519906 | 0.161 | 0.15 | 1 | 4 | Dync1li1 | 0.011 | Old |
| 0.697846 | 0.518275 | 0.149 | 0.138 | 1 | 4 | Msmo1 | 0.011 | Old |
| 0.698093 | 0.325781 | 0.321 | 0.312 | 1 | 4 | Dnm2 | 0.009 | Old |
| 0.698143 | 0.56284 | 0.149 | 0.138 | 1 | 4 | Manba | 0.011 | Old |
| 0.698283 | 0.054336 | 0.341 | 0.325 | 1 | 4 | Snx9 | 0.016 | Old |
| 0.69832 | 0.745659 | 0.018 | 0.013 | 1 | 4 | Trmt10a | 0.005 | Old |
| 0.69832 | 0.606764 | 0.018 | 0.013 | 1 | 4 | Abcg3 | 0.005 | Old |
| 0.698741 | 0.792453 | 0.061 | 0.075 | 1 | 4 | Rhbdd1 | -0.014 | Old |
| 0.698765 | 0.388913 | 0.033 | 0.025 | 1 | 4 | Hddc2 | 0.008 | Old |
| 0.6988 | 0.222222 | 0.153 | 0.175 | 1 | 4 | Smarce1 | -0.022 | Old |
| 0.698896 | 0.456903 | 0.084 | 0.1 | 1 | 4 | Zeb2os | -0.016 | Old |
| 0.699076 | 0.323076 | 0.2 | 0.188 | 1 | 4 | Psmc1 | 0.012 | Old |

| | | | | | | | | |
|---|---|---|---|---|---|---|---|---|
| 0.699081 | 0.147331 | 0.185 | 0.212 | 1 | 4 | Mfsd6 | -0.027 | Old |
| 0.699652 | 0.024366 | 0.191 | 0.212 | 1 | 4 | Plod1 | -0.021 | Old |
| 0.699739 | 0.032775 | 0.524 | 0.588 | 1 | 4 | Rps27l | -0.064 | Old |
| 0.699912 | 0.4098 | 0.018 | 0.013 | 1 | 4 | Vegfa | 0.005 | Old |
| 0.700131 | 1.100862 | 0.046 | 0.038 | 1 | 4 | Tjp2 | 0.008 | Old |
| 0.70033 | 0.507838 | 0.06 | 0.05 | 1 | 4 | Nrde2 | 0.01 | Old |
| 0.700366 | 0.051809 | 0.097 | 0.112 | 1 | 4 | Tpm1 | -0.015 | Old |
| 0.70053 | 0.803452 | 0.04 | 0.05 | 1 | 4 | Phlpp2 | -0.01 | Old |
| 0.700562 | 0.242428 | 0.033 | 0.025 | 1 | 4 | Nudt7 | 0.008 | Old |
| 0.701161 | 0.277396 | 0.033 | 0.025 | 1 | 4 | Nup107 | 0.008 | Old |
| 0.701167 | 0.948896 | 0.059 | 0.05 | 1 | 4 | Arl5c | 0.009 | Old |
| 0.701364 | 0.264128 | 0.101 | 0.088 | 1 | 4 | Svil | 0.013 | Old |
| 0.701503 | 1.361468 | 0.032 | 0.025 | 1 | 4 | Mcm5 | 0.007 | Old |
| 0.701539 | 0.073631 | 0.041 | 0.05 | 1 | 4 | Clp1 | -0.009 | Old |
| 0.701947 | 0.674201 | 0.097 | 0.088 | 1 | 4 | Luc7l | 0.009 | Old |
| 0.701981 | 0.301281 | 0.129 | 0.15 | 1 | 4 | Parl | -0.021 | Old |
| 0.702087 | 0.484431 | 0.159 | 0.188 | 1 | 4 | Sh2b3 | -0.029 | Old |
| 0.702199 | 0.048324 | 0.111 | 0.1 | 1 | 4 | Vps4a | 0.011 | Old |
| 0.702361 | 0.327694 | 0.033 | 0.025 | 1 | 4 | Mcm2 | 0.008 | Old |
| 0.702451 | 0.240252 | 0.113 | 0.1 | 1 | 4 | Cops5 | 0.013 | Old |
| 0.702483 | 0.05896 | 0.169 | 0.188 | 1 | 4 | Emsy | -0.019 | Old |
| 0.703053 | 0.045513 | 0.538 | 0.512 | 1 | 4 | Eif3f | 0.026 | Old |
| 0.7031 | 0.102597 | 0.018 | 0.013 | 1 | 4 | Slc25a23 | 0.005 | Old |
| 0.703155 | 0.003265 | 0.121 | 0.138 | 1 | 4 | Gcc2 | -0.017 | Old |
| 0.703944 | 0.12011 | 0.03 | 0.038 | 1 | 4 | Tnk2 | -0.008 | Old |
| 0.705109 | 0.161934 | 0.326 | 0.3 | 1 | 4 | Snap23 | 0.026 | Old |
| 0.705154 | 0.296074 | 0.119 | 0.138 | 1 | 4 | Mrpl34 | -0.019 | Old |
| 0.705258 | 0.364584 | 0.107 | 0.125 | 1 | 4 | Prorsd1 | -0.018 | Old |
| 0.705274 | 0.364677 | 0.099 | 0.088 | 1 | 4 | Atp13a1 | 0.011 | Old |
| 0.705638 | 0.749546 | 0.109 | 0.1 | 1 | 4 | Ube2w | 0.009 | Old |
| 0.706356 | 0.04186 | 0.041 | 0.05 | 1 | 4 | Foxo4 | -0.009 | Old |
| 0.706442 | 0.441613 | 0.03 | 0.038 | 1 | 4 | 1810043G( | -0.008 | Old |
| 0.706442 | 0.328527 | 0.03 | 0.038 | 1 | 4 | Ccdc126 | -0.008 | Old |
| 0.706622 | 0.037377 | 0.213 | 0.238 | 1 | 4 | Ythdf1 | -0.025 | Old |
| 0.707066 | 0.736962 | 0.097 | 0.088 | 1 | 4 | B3galnt1 | 0.009 | Old |
| 0.707332 | 0.257919 | 0.315 | 0.3 | 1 | 4 | Cmah | 0.015 | Old |
| 0.70754 | 0.828669 | 0.04 | 0.05 | 1 | 4 | Fbxl12 | -0.01 | Old |
| 0.707562 | 0.061865 | 0.167 | 0.188 | 1 | 4 | Rbbp7 | -0.021 | Old |
| 0.707687 | 0.5968 | 0.147 | 0.138 | 1 | 4 | Srp54a | 0.009 | Old |
| 0.708293 | 0.57423 | 0.046 | 0.038 | 1 | 4 | D10Wsu10 | 0.008 | Old |
| 0.708317 | 0.30293 | 0.03 | 0.038 | 1 | 4 | Gabbr1 | -0.008 | Old |
| 0.708587 | 0.406489 | 0.149 | 0.138 | 1 | 4 | Ppp2r2d | 0.011 | Old |
| 0.708793 | 0.239883 | 0.052 | 0.062 | 1 | 4 | Narf | -0.01 | Old |
| 0.708872 | 0.540804 | 0.096 | 0.112 | 1 | 4 | Wdr61 | -0.016 | Old |
| 0.709316 | 0.527255 | 0.046 | 0.038 | 1 | 4 | Hars2 | 0.008 | Old |

| | | | | | | | | |
|---|---|---|---|---|---|---|---|---|
| 0.709994 | 1.006341 | 0.032 | 0.025 | 1 | 4 | Casp2 | 0.007 | Old |
| 0.710001 | 0.432641 | 0.221 | 0.212 | 1 | 4 | Ube2q1 | 0.009 | Old |
| 0.710339 | 0.442574 | 0.148 | 0.138 | 1 | 4 | Hebp1 | 0.01 | Old |
| 0.711111 | 0.494508 | 0.072 | 0.062 | 1 | 4 | Tmem185b | 0.01 | Old |
| 0.7114 | 1.265152 | 0.083 | 0.075 | 1 | 4 | Abca7 | 0.008 | Old |
| 0.711427 | 0.004491 | 0.19 | 0.212 | 1 | 4 | Smagp | -0.022 | Old |
| 0.711857 | 0.125288 | 0.109 | 0.125 | 1 | 4 | Arl6ip6 | -0.016 | Old |
| 0.71213 | 0.005957 | 0.296 | 0.275 | 1 | 4 | Mgea5 | 0.021 | Old |
| 0.712198 | 0.513769 | 0.111 | 0.1 | 1 | 4 | Seh1l | 0.011 | Old |
| 0.712239 | 2.082514 | 0.032 | 0.025 | 1 | 4 | Cmpk2 | 0.007 | Old |
| 0.712351 | 0.571787 | 0.072 | 0.062 | 1 | 4 | Vkorc1l1 | 0.01 | Old |
| 0.7127 | 0.66818 | 0.03 | 0.038 | 1 | 4 | Ada | -0.008 | Old |
| 0.71276 | 0.185387 | 0.702 | 0.725 | 1 | 4 | Snx2 | -0.023 | Old |
| 0.712853 | 1.904619 | 0.032 | 0.025 | 1 | 4 | Prkd2 | 0.007 | Old |
| 0.712956 | 0.592249 | 0.147 | 0.138 | 1 | 4 | Cyb5r1 | 0.009 | Old |
| 0.713198 | 0.121787 | 0.783 | 0.775 | 1 | 4 | Ptp4a2 | 0.008 | Old |
| 0.713212 | 0.785142 | 0.097 | 0.088 | 1 | 4 | Acbd6 | 0.009 | Old |
| 0.713235 | 0.126467 | 0.221 | 0.2 | 1 | 4 | Nudt3 | 0.021 | Old |
| 0.713597 | 0.012725 | 0.171 | 0.188 | 1 | 4 | Inpp5k | -0.017 | Old |
| 0.713925 | 1.095001 | 0.045 | 0.038 | 1 | 4 | Nup205 | 0.007 | Old |
| 0.714268 | 0.240552 | 0.153 | 0.175 | 1 | 4 | Hadh | -0.022 | Old |
| 0.714448 | 0.332949 | 0.386 | 0.4 | 1 | 4 | Tcn2 | -0.014 | Old |
| 0.714542 | 0.218239 | 0.052 | 0.062 | 1 | 4 | Abcf2 | -0.01 | Old |
| 0.714612 | 0.016326 | 0.251 | 0.275 | 1 | 4 | Sar1a | -0.024 | Old |
| 0.714807 | 0.597119 | 0.191 | 0.188 | 1 | 4 | Acbd3 | 0.003 | Old |
| 0.714858 | 1.776405 | 0.018 | 0.013 | 1 | 4 | Fbxo46 | 0.005 | Old |
| 0.714928 | 0.073828 | 0.047 | 0.038 | 1 | 4 | Nos1ap | 0.009 | Old |
| 0.715105 | 0.402595 | 0.009 | 0.013 | 1 | 4 | Mttp | -0.004 | Old |
| 0.715105 | 0.149968 | 0.009 | 0.013 | 1 | 4 | Gm10521 | -0.004 | Old |
| 0.715473 | 0.605768 | 0.032 | 0.025 | 1 | 4 | Rrnad1 | 0.007 | Old |
| 0.715608 | 0.325116 | 0.112 | 0.1 | 1 | 4 | Imp4 | 0.012 | Old |
| 0.715675 | 1.505747 | 0.018 | 0.013 | 1 | 4 | Dpt | 0.005 | Old |
| 0.715755 | 0.945686 | 0.108 | 0.1 | 1 | 4 | Abhd17a | 0.008 | Old |
| 0.715857 | 0.105931 | 0.587 | 0.55 | 1 | 4 | Hnrnpa0 | 0.037 | Old |
| 0.71599 | 1.106127 | 0.045 | 0.038 | 1 | 4 | Galns | 0.007 | Old |
| 0.716012 | 0.06134 | 0.22 | 0.2 | 1 | 4 | Cct7 | 0.02 | Old |
| 0.716056 | 0.03147 | 0.191 | 0.212 | 1 | 4 | Plekhg3 | -0.021 | Old |
| 0.716255 | 0.210511 | 0.009 | 0.013 | 1 | 4 | Bcl11a | -0.004 | Old |
| 0.716255 | 0.141587 | 0.009 | 0.013 | 1 | 4 | Aaas | -0.004 | Old |
| 0.716489 | 0.349268 | 0.046 | 0.038 | 1 | 4 | Pira2 | 0.008 | Old |
| 0.716491 | 1.674949 | 0.018 | 0.013 | 1 | 4 | Cby1 | 0.005 | Old |
| 0.716565 | 0.367336 | 0.041 | 0.05 | 1 | 4 | Atg9a | -0.009 | Old |
| 0.716883 | 0.099228 | 0.019 | 0.025 | 1 | 4 | Lyrm4 | -0.006 | Old |
| 0.717309 | 1.711633 | 0.018 | 0.013 | 1 | 4 | Znhit3 | 0.005 | Old |
| 0.717332 | 0.28582 | 0.152 | 0.175 | 1 | 4 | Anpep | -0.023 | Old |

| | | | | | | | | |
|---|---|---|---|---|---|---|---|---|
| 0.717406 | 0.501854 | 0.009 | 0.013 | 1 | 4 | Cpt1c | -0.004 | Old |
| 0.717406 | 0.39673 | 0.009 | 0.013 | 1 | 4 | Havcr2 | -0.004 | Old |
| 0.717406 | 0.172827 | 0.009 | 0.013 | 1 | 4 | Cysltr2 | -0.004 | Old |
| 0.717406 | 0.134374 | 0.009 | 0.013 | 1 | 4 | Gm45051 | -0.004 | Old |
| 0.717571 | 0.310076 | 0.211 | 0.2 | 1 | 4 | Arrdc1 | 0.011 | Old |
| 0.71766 | 0.434944 | 0.019 | 0.025 | 1 | 4 | Mettl25 | -0.006 | Old |
| 0.717731 | 0.485925 | 0.171 | 0.162 | 1 | 4 | BC031181 | 0.009 | Old |
| 0.718099 | 0.292093 | 0.175 | 0.2 | 1 | 4 | Txnl1 | -0.025 | Old |
| 0.718383 | 0.249355 | 0.052 | 0.062 | 1 | 4 | Decr2 | -0.01 | Old |
| 0.718522 | 0.485424 | 0.032 | 0.025 | 1 | 4 | Erc1 | 0.007 | Old |
| 0.718557 | 0.526231 | 0.009 | 0.013 | 1 | 4 | 9130019O2 | -0.004 | Old |
| 0.718557 | 0.426208 | 0.009 | 0.013 | 1 | 4 | Gm12689 | -0.004 | Old |
| 0.71861 | 0.294564 | 0.073 | 0.062 | 1 | 4 | Stx8 | 0.011 | Old |
| 0.718707 | 0.126525 | 0.245 | 0.225 | 1 | 4 | Stk4 | 0.02 | Old |
| 0.718944 | 1.240959 | 0.018 | 0.013 | 1 | 4 | Tsr3 | 0.005 | Old |
| 0.719113 | 0.126051 | 0.214 | 0.238 | 1 | 4 | Dnajc19 | -0.024 | Old |
| 0.719196 | 0.012901 | 0.211 | 0.238 | 1 | 4 | Bmpr2 | -0.027 | Old |
| 0.719216 | 0.55558 | 0.019 | 0.025 | 1 | 4 | Adora2b | -0.006 | Old |
| 0.719216 | 0.386655 | 0.019 | 0.025 | 1 | 4 | Rab34 | -0.006 | Old |
| 0.719232 | 1.072347 | 0.07 | 0.062 | 1 | 4 | Tnfsf13os | 0.008 | Old |
| 0.71945 | 0.169134 | 0.1 | 0.088 | 1 | 4 | Nsmaf | 0.012 | Old |
| 0.719484 | 0.06758 | 0.25 | 0.275 | 1 | 4 | Cpt1a | -0.025 | Old |
| 0.719553 | 0.068198 | 0.122 | 0.138 | 1 | 4 | Trrap | -0.016 | Old |
| 0.719709 | 0.891061 | 0.009 | 0.013 | 1 | 4 | Gm36266 | -0.004 | Old |
| 0.719709 | 0.77872 | 0.009 | 0.013 | 1 | 4 | Col1a2 | -0.004 | Old |
| 0.719744 | 0.088395 | 0.032 | 0.025 | 1 | 4 | Gm20703 | 0.007 | Old |
| 0.719941 | 0.034592 | 0.202 | 0.188 | 1 | 4 | Pdgfc | 0.014 | Old |
| 0.719995 | 0.274645 | 0.019 | 0.025 | 1 | 4 | Nop2 | -0.006 | Old |
| 0.720055 | 0.44915 | 0.111 | 0.1 | 1 | 4 | Zfp704 | 0.011 | Old |
| 0.720206 | 0.322603 | 0.289 | 0.288 | 1 | 4 | Snx30 | 0.001 | Old |
| 0.72061 | 0.204744 | 0.219 | 0.25 | 1 | 4 | Snx4 | -0.031 | Old |
| 0.720663 | 0.28585 | 0.099 | 0.088 | 1 | 4 | Csnk1g1 | 0.011 | Old |
| 0.720736 | 0.060759 | 0.447 | 0.488 | 1 | 4 | Serinc1 | -0.041 | Old |
| 0.720747 | 1.531173 | 0.057 | 0.05 | 1 | 4 | 2900089D1 | 0.007 | Old |
| 0.720861 | 1.02907 | 0.009 | 0.013 | 1 | 4 | Zscan22 | -0.004 | Old |
| 0.720861 | 0.671048 | 0.009 | 0.013 | 1 | 4 | Pde8b | -0.004 | Old |
| 0.720966 | 0.365018 | 0.032 | 0.025 | 1 | 4 | Eif2b1 | 0.007 | Old |
| 0.720966 | 0.260405 | 0.032 | 0.025 | 1 | 4 | Prmt9 | 0.007 | Old |
| 0.721162 | 0.804197 | 0.045 | 0.038 | 1 | 4 | Commd9 | 0.007 | Old |
| 0.721371 | 0.293273 | 0.046 | 0.038 | 1 | 4 | Sik2 | 0.008 | Old |
| 0.7214 | 1.166744 | 0.018 | 0.013 | 1 | 4 | Chd1l | 0.005 | Old |
| 0.721513 | 0.989777 | 0.083 | 0.075 | 1 | 4 | Eya3 | 0.008 | Old |
| 0.721577 | 0.229074 | 0.032 | 0.025 | 1 | 4 | Exd2 | 0.007 | Old |
| 0.721577 | 0.220859 | 0.032 | 0.025 | 1 | 4 | Atpaf2 | 0.007 | Old |
| 0.722275 | 0.418516 | 0.231 | 0.225 | 1 | 4 | Insig1 | 0.006 | Old |

| | | | | | | | | |
|---|---|---|---|---|---|---|---|---|
| 0.722472 | 0.632718 | 0.052 | 0.062 | 1 | 4 | Mtmr9 | -0.01 | Old |
| 0.722976 | 0.429537 | 0.059 | 0.05 | 1 | 4 | Tmco3 | 0.009 | Old |
| 0.72302 | 0.163365 | 0.356 | 0.338 | 1 | 4 | Pttg1ip | 0.018 | Old |
| 0.723172 | 0.011942 | 0.046 | 0.038 | 1 | 4 | Rdh14 | 0.008 | Old |
| 0.72323 | 0.281236 | 0.211 | 0.2 | 1 | 4 | Arf6 | 0.011 | Old |
| 0.723269 | 0.092078 | 0.087 | 0.1 | 1 | 4 | Rif1 | -0.013 | Old |
| 0.723474 | 0.477118 | 0.123 | 0.112 | 1 | 4 | Pdgfa | 0.011 | Old |
| 0.723612 | 0.024494 | 0.357 | 0.388 | 1 | 4 | Idh1 | -0.031 | Old |
| 0.723661 | 0.259193 | 0.172 | 0.162 | 1 | 4 | Nfatc2 | 0.01 | Old |
| 0.723858 | 0.495141 | 0.018 | 0.013 | 1 | 4 | Cdan1 | 0.005 | Old |
| 0.724293 | 0.16031 | 0.06 | 0.05 | 1 | 4 | Sass6 | 0.01 | Old |
| 0.724308 | 0.197947 | 0.086 | 0.1 | 1 | 4 | Dph3 | -0.014 | Old |
| 0.72444 | 0.281195 | 0.341 | 0.338 | 1 | 4 | Spop | 0.003 | Old |
| 0.72446 | 0.314041 | 0.046 | 0.038 | 1 | 4 | Pigu | 0.008 | Old |
| 0.724564 | 1.090935 | 0.032 | 0.025 | 1 | 4 | Ddx19b | 0.007 | Old |
| 0.724881 | 0.500682 | 0.052 | 0.062 | 1 | 4 | Faap100 | -0.01 | Old |
| 0.724912 | 0.443088 | 0.098 | 0.088 | 1 | 4 | Gcn1l1 | 0.01 | Old |
| 0.725092 | 0.002323 | 0.424 | 0.45 | 1 | 4 | Susd6 | -0.026 | Old |
| 0.725452 | 0.914065 | 0.019 | 0.025 | 1 | 4 | Mpst | -0.006 | Old |
| 0.725499 | 0.619005 | 0.018 | 0.013 | 1 | 4 | Zfp518b | 0.005 | Old |
| 0.725794 | 0.059508 | 0.337 | 0.375 | 1 | 4 | Ranbp2 | -0.038 | Old |
| 0.7258 | 1.011959 | 0.032 | 0.025 | 1 | 4 | Tmem241 | 0.007 | Old |
| 0.725816 | 0.530329 | 0.204 | 0.2 | 1 | 4 | Glrx5 | 0.004 | Old |
| 0.726075 | 0.005927 | 0.306 | 0.325 | 1 | 4 | Cerk | -0.019 | Old |
| 0.726278 | 0.24083 | 0.312 | 0.3 | 1 | 4 | Efr3a | 0.012 | Old |
| 0.726335 | 0.045352 | 0.03 | 0.038 | 1 | 4 | Osbpl7 | -0.008 | Old |
| 0.726451 | 0.408351 | 0.148 | 0.138 | 1 | 4 | Abcd1 | 0.01 | Old |
| 0.726556 | 0.237241 | 0.042 | 0.05 | 1 | 4 | Ripor1 | -0.008 | Old |
| 0.727118 | 0.122569 | 0.208 | 0.188 | 1 | 4 | Mbd2 | 0.02 | Old |
| 0.727164 | 0.080651 | 0.123 | 0.138 | 1 | 4 | Mms19 | -0.015 | Old |
| 0.727234 | 0.282777 | 0.15 | 0.138 | 1 | 4 | Rbm22 | 0.012 | Old |
| 0.727281 | 0.257804 | 0.12 | 0.138 | 1 | 4 | Hspbp1 | -0.018 | Old |
| 0.727336 | 0.549543 | 0.085 | 0.075 | 1 | 4 | Akr1b10 | 0.01 | Old |
| 0.727499 | 0.196597 | 0.364 | 0.35 | 1 | 4 | Tns3 | 0.014 | Old |
| 0.727549 | 0.508914 | 0.085 | 0.1 | 1 | 4 | Zfp397 | -0.015 | Old |
| 0.727638 | 0.175456 | 0.086 | 0.075 | 1 | 4 | Pfdn4 | 0.011 | Old |
| 0.727792 | 0.434396 | 0.275 | 0.275 | 1 | 4 | Klc1 | 0 | Old |
| 0.72786 | 0.242375 | 0.098 | 0.112 | 1 | 4 | Anapc2 | -0.014 | Old |
| 0.727933 | 0.055838 | 0.086 | 0.075 | 1 | 4 | Thap2 | 0.011 | Old |
| 0.727962 | 0.412909 | 0.018 | 0.013 | 1 | 4 | Trmt10b | 0.005 | Old |
| 0.727962 | 0.324941 | 0.018 | 0.013 | 1 | 4 | Tram2 | 0.005 | Old |
| 0.727962 | 0.305943 | 0.018 | 0.013 | 1 | 4 | Smyd4 | 0.005 | Old |
| 0.728098 | 0.684574 | 0.071 | 0.062 | 1 | 4 | Ormdl1 | 0.009 | Old |
| 0.728103 | 0.074788 | 0.19 | 0.175 | 1 | 4 | Rbbp4 | 0.015 | Old |
| 0.72821 | 0.336959 | 0.03 | 0.038 | 1 | 4 | Krt10 | -0.008 | Old |

| | | | | | | | | |
|---|---|---|---|---|---|---|---|---|
| 0.728333 | 0.336592 | 0.248 | 0.238 | 1 | 4 | Ptpn12 | 0.01 | Old |
| 0.728638 | 1.572396 | 0.057 | 0.05 | 1 | 4 | Vps9d1 | 0.007 | Old |
| 0.728898 | 0.402857 | 0.098 | 0.088 | 1 | 4 | Pfkfb4 | 0.01 | Old |
| 0.728904 | 0.259474 | 0.112 | 0.1 | 1 | 4 | Ybx3 | 0.012 | Old |
| 0.728916 | 0.485314 | 0.204 | 0.2 | 1 | 4 | Anapc11 | 0.004 | Old |
| 0.728938 | 0.589702 | 0.071 | 0.062 | 1 | 4 | Gosr1 | 0.009 | Old |
| 0.729122 | 0.029707 | 0.099 | 0.088 | 1 | 4 | Hbs1l | 0.011 | Old |
| 0.729181 | 0.042814 | 0.087 | 0.075 | 1 | 4 | Prpf38a | 0.012 | Old |
| 0.729461 | 0.084614 | 0.03 | 0.038 | 1 | 4 | Abhd6 | -0.008 | Old |
| 0.729507 | 0.383496 | 0.174 | 0.162 | 1 | 4 | Otub1 | 0.012 | Old |
| 0.729514 | 0.901807 | 0.032 | 0.025 | 1 | 4 | Acaca | 0.007 | Old |
| 0.729782 | 0.195496 | 0.042 | 0.05 | 1 | 4 | Lin52 | -0.008 | Old |
| 0.730578 | 0.181556 | 0.1 | 0.088 | 1 | 4 | Camsap2 | 0.012 | Old |
| 0.730652 | 0.360404 | 0.211 | 0.2 | 1 | 4 | Sec61a1 | 0.011 | Old |
| 0.730933 | 0.324994 | 0.193 | 0.225 | 1 | 4 | Zfx | -0.032 | Old |
| 0.731189 | 0.133685 | 0.377 | 0.362 | 1 | 4 | Ddx39b | 0.015 | Old |
| 0.731416 | 0.213646 | 0.296 | 0.338 | 1 | 4 | Mfsd5 | -0.042 | Old |
| 0.73158 | 0.135285 | 0.154 | 0.175 | 1 | 4 | Clcn3 | -0.021 | Old |
| 0.732051 | 0.054577 | 0.154 | 0.138 | 1 | 4 | Pxk | 0.016 | Old |
| 0.732239 | 0.152128 | 0.121 | 0.138 | 1 | 4 | Aggf1 | -0.017 | Old |
| 0.733234 | 0.79785 | 0.032 | 0.025 | 1 | 4 | Rpap1 | 0.007 | Old |
| 0.733234 | 0.652015 | 0.032 | 0.025 | 1 | 4 | Lrrc8b | 0.007 | Old |
| 0.733518 | 0.311905 | 0.248 | 0.238 | 1 | 4 | Tbcb | 0.01 | Old |
| 0.73375 | 0.089534 | 0.366 | 0.4 | 1 | 4 | Camk1 | -0.034 | Old |
| 0.734171 | 0.201584 | 0.243 | 0.275 | 1 | 4 | Mrpl23 | -0.032 | Old |
| 0.7345 | 0.085968 | 0.216 | 0.238 | 1 | 4 | Chtop | -0.022 | Old |
| 0.73458 | 0.323202 | 0.136 | 0.125 | 1 | 4 | Sec24c | 0.011 | Old |
| 0.735128 | 0.663488 | 0.109 | 0.1 | 1 | 4 | Ext2 | 0.009 | Old |
| 0.735277 | 0.095194 | 0.122 | 0.138 | 1 | 4 | Cbx4 | -0.016 | Old |
| 0.735329 | 0.215003 | 0.352 | 0.362 | 1 | 4 | Cast | -0.01 | Old |
| 0.735386 | 0.188646 | 0.148 | 0.138 | 1 | 4 | Aqr | 0.01 | Old |
| 0.735664 | 0.109699 | 0.271 | 0.262 | 1 | 4 | Arglu1 | 0.009 | Old |
| 0.735718 | 0.865867 | 0.032 | 0.025 | 1 | 4 | Espn | 0.007 | Old |
| 0.735987 | 0.076836 | 0.223 | 0.238 | 1 | 4 | Id3 | -0.015 | Old |
| 0.736084 | 0.267993 | 0.262 | 0.25 | 1 | 4 | Psma2 | 0.012 | Old |
| 0.736097 | 0.078293 | 0.111 | 0.125 | 1 | 4 | Ltn1 | -0.014 | Old |
| 0.736101 | 0.168175 | 0.177 | 0.162 | 1 | 4 | C330007P0 | 0.015 | Old |
| 0.736415 | 0.31832 | 0.27 | 0.262 | 1 | 4 | Fndc3b | 0.008 | Old |
| 0.736505 | 0.1211 | 0.327 | 0.312 | 1 | 4 | Ythdc1 | 0.015 | Old |
| 0.736689 | 0.012998 | 0.072 | 0.062 | 1 | 4 | Smn1 | 0.01 | Old |
| 0.737025 | 1.181244 | 0.057 | 0.05 | 1 | 4 | Pilrb1 | 0.007 | Old |
| 0.737441 | 0.077073 | 0.087 | 0.075 | 1 | 4 | Ddx1 | 0.012 | Old |
| 0.7378 | 0.103681 | 0.099 | 0.088 | 1 | 4 | Stx5a | 0.011 | Old |
| 0.737801 | 0.318221 | 0.097 | 0.088 | 1 | 4 | Pak1ip1 | 0.009 | Old |
| 0.738316 | 0.053873 | 0.099 | 0.112 | 1 | 4 | Zfp110 | -0.013 | Old |

| | | | | | | | | |
|---|---|---|---|---|---|---|---|---|
| 0.73844 | 0.225927 | 0.2 | 0.188 | 1 | 4 | Cbx7 | 0.012 | Old |
| 0.738499 | 0.115792 | 0.123 | 0.112 | 1 | 4 | Sde2 | 0.011 | Old |
| 0.738713 | 0.510055 | 0.097 | 0.088 | 1 | 4 | Nelfe | 0.009 | Old |
| 0.738732 | 0.099926 | 0.122 | 0.138 | 1 | 4 | Lrrc59 | -0.016 | Old |
| 0.738948 | 0.129815 | 0.163 | 0.15 | 1 | 4 | Yipf3 | 0.013 | Old |
| 0.739107 | 0.371703 | 0.075 | 0.088 | 1 | 4 | Scfd1 | -0.013 | Old |
| 0.739113 | 0.136818 | 0.252 | 0.238 | 1 | 4 | Twf2 | 0.014 | Old |
| 0.739493 | 0.614312 | 0.03 | 0.038 | 1 | 4 | Fdps | -0.008 | Old |
| 0.739545 | 0.352395 | 0.374 | 0.388 | 1 | 4 | Csf2ra | -0.014 | Old |
| 0.740071 | 0.343755 | 0.032 | 0.025 | 1 | 4 | Ppip5k1 | 0.007 | Old |
| 0.74026 | 0.075071 | 0.134 | 0.15 | 1 | 4 | Ehmt1 | -0.016 | Old |
| 0.740383 | 0.388943 | 0.086 | 0.1 | 1 | 4 | Hmg20a | -0.014 | Old |
| 0.740505 | 0.089403 | 0.654 | 0.662 | 1 | 4 | Ppp3ca | -0.008 | Old |
| 0.740571 | 0.546603 | 0.042 | 0.05 | 1 | 4 | Fbxw17 | -0.008 | Old |
| 0.740707 | 0.216211 | 0.099 | 0.088 | 1 | 4 | Mfap1b | 0.011 | Old |
| 0.74081 | 0.385769 | 0.123 | 0.112 | 1 | 4 | Babam1 | 0.011 | Old |
| 0.740972 | 0.042982 | 0.053 | 0.062 | 1 | 4 | Rfng | -0.009 | Old |
| 0.74118 | 0.102846 | 0.113 | 0.1 | 1 | 4 | Ola1 | 0.013 | Old |
| 0.741752 | 0.172507 | 0.138 | 0.125 | 1 | 4 | Eif6 | 0.013 | Old |
| 0.741939 | 0.427407 | 0.032 | 0.025 | 1 | 4 | Nubpl | 0.007 | Old |
| 0.742199 | 0.444605 | 0.109 | 0.1 | 1 | 4 | Casp1 | 0.009 | Old |
| 0.742389 | 0.330208 | 0.072 | 0.062 | 1 | 4 | Fry | 0.01 | Old |
| 0.742403 | 0.099764 | 0.376 | 0.388 | 1 | 4 | Tomm20 | -0.012 | Old |
| 0.742421 | 0.204271 | 0.319 | 0.312 | 1 | 4 | Arl8b | 0.007 | Old |
| 0.742658 | 0.261201 | 0.15 | 0.138 | 1 | 4 | Ndufa10 | 0.012 | Old |
| 0.743155 | 0.029934 | 0.146 | 0.162 | 1 | 4 | Abca3 | -0.016 | Old |
| 0.743266 | 0.872293 | 0.03 | 0.038 | 1 | 4 | Mcu | -0.008 | Old |
| 0.743424 | 0.00418 | 0.065 | 0.075 | 1 | 4 | Pbdc1 | -0.01 | Old |
| 0.743523 | 0.442317 | 0.11 | 0.1 | 1 | 4 | Pnpla7 | 0.01 | Old |
| 0.74373 | 0.721162 | 0.045 | 0.038 | 1 | 4 | AC165953. | 0.007 | Old |
| 0.743771 | 0.386605 | 0.059 | 0.05 | 1 | 4 | Oxsm | 0.009 | Old |
| 0.744257 | 0.460895 | 0.045 | 0.038 | 1 | 4 | Prr14l | 0.007 | Old |
| 0.744641 | 1.813036 | 0.017 | 0.013 | 1 | 4 | Lrrc45 | 0.004 | Old |
| 0.744641 | 1.634744 | 0.017 | 0.013 | 1 | 4 | Surf6 | 0.004 | Old |
| 0.744911 | 0.403894 | 0.11 | 0.1 | 1 | 4 | Gsdmd | 0.01 | Old |
| 0.744976 | 0.135503 | 0.02 | 0.025 | 1 | 4 | Emc9 | -0.005 | Old |
| 0.74498 | 0.420605 | 0.108 | 0.125 | 1 | 4 | Ssr2 | -0.017 | Old |
| 0.745339 | 0.35686 | 0.072 | 0.062 | 1 | 4 | Ell | 0.01 | Old |
| 0.745371 | 0.005556 | 0.16 | 0.175 | 1 | 4 | Nfatc3 | -0.015 | Old |
| 0.745785 | 0.923332 | 0.03 | 0.038 | 1 | 4 | Ube2o | -0.008 | Old |
| 0.746019 | 0.177649 | 0.302 | 0.3 | 1 | 4 | Anxa7 | 0.002 | Old |
| 0.746103 | 0.537421 | 0.192 | 0.188 | 1 | 4 | Gigyf1 | 0.004 | Old |
| 0.746326 | 1.272579 | 0.017 | 0.013 | 1 | 4 | Btnl2 | 0.004 | Old |
| 0.746326 | 1.254499 | 0.017 | 0.013 | 1 | 4 | Hdac7 | 0.004 | Old |
| 0.746326 | 1.217153 | 0.017 | 0.013 | 1 | 4 | Pdlim7 | 0.004 | Old |

| | | | | | | | | |
|---|---|---|---|---|---|---|---|---|
| 0.74633 | 0.425202 | 0.084 | 0.075 | 1 | 4 | Pign | 0.009 | Old |
| 0.746416 | 1.010316 | 0.031 | 0.025 | 1 | 4 | Zfp212 | 0.006 | Old |
| 0.746494 | 0.441698 | 0.064 | 0.075 | 1 | 4 | Sephs2 | -0.011 | Old |
| 0.74717 | 1.273663 | 0.017 | 0.013 | 1 | 4 | Ndnf | 0.004 | Old |
| 0.747195 | 0.159319 | 0.304 | 0.338 | 1 | 4 | Ppp2r5a | -0.034 | Old |
| 0.74729 | 0.513785 | 0.097 | 0.112 | 1 | 4 | Nsd2 | -0.015 | Old |
| 0.747737 | 0.391104 | 0.337 | 0.35 | 1 | 4 | Vat1 | -0.013 | Old |
| 0.747748 | 0.440894 | 0.206 | 0.2 | 1 | 4 | Polr2m | 0.006 | Old |
| 0.747821 | 0.722844 | 0.119 | 0.112 | 1 | 4 | Pml | 0.007 | Old |
| 0.748013 | 1.263216 | 0.017 | 0.013 | 1 | 4 | Caprin2 | 0.004 | Old |
| 0.748013 | 1.532133 | 0.017 | 0.013 | 1 | 4 | Zfp930 | 0.004 | Old |
| 0.748187 | 0.140303 | 0.053 | 0.062 | 1 | 4 | Tmtc3 | -0.009 | Old |
| 0.748781 | 0.147193 | 0.045 | 0.038 | 1 | 4 | Spryd7 | 0.007 | Old |
| 0.748857 | 1.017146 | 0.017 | 0.013 | 1 | 4 | AC162302. | 0.004 | Old |
| 0.748857 | 0.97866 | 0.017 | 0.013 | 1 | 4 | AC163032. | 0.004 | Old |
| 0.748857 | 0.958256 | 0.017 | 0.013 | 1 | 4 | Ildr2 | 0.004 | Old |
| 0.749299 | 0.304927 | 0.283 | 0.275 | 1 | 4 | Nenf | 0.008 | Old |
| 0.749426 | 0.16843 | 0.032 | 0.025 | 1 | 4 | Amigo1 | 0.007 | Old |
| 0.749597 | 0.60471 | 0.086 | 0.1 | 1 | 4 | Slc25a17 | -0.014 | Old |
| 0.749702 | 1.352758 | 0.017 | 0.013 | 1 | 4 | Vars2 | 0.004 | Old |
| 0.750377 | 0.144496 | 0.189 | 0.175 | 1 | 4 | Kcmf1 | 0.014 | Old |
| 0.75041 | 0.088597 | 0.02 | 0.025 | 1 | 4 | Tcaf1 | -0.005 | Old |
| 0.750546 | 1.087687 | 0.017 | 0.013 | 1 | 4 | Zfhx2 | 0.004 | Old |
| 0.750682 | 0.385971 | 0.065 | 0.075 | 1 | 4 | Nudt5 | -0.01 | Old |
| 0.750746 | 0.20014 | 0.352 | 0.35 | 1 | 4 | Copa | 0.002 | Old |
| 0.750832 | 0.982112 | 0.031 | 0.025 | 1 | 4 | Tada2b | 0.006 | Old |
| 0.751391 | 0.562849 | 0.017 | 0.013 | 1 | 4 | Miga2 | 0.004 | Old |
| 0.751513 | 0.005751 | 0.211 | 0.225 | 1 | 4 | Uqcrfs1 | -0.014 | Old |
| 0.751614 | 0.074334 | 0.031 | 0.038 | 1 | 4 | Lmbr1l | -0.007 | Old |
| 0.751964 | 0.162009 | 0.02 | 0.025 | 1 | 4 | Kif23 | -0.005 | Old |
| 0.751985 | 0.402435 | 0.12 | 0.138 | 1 | 4 | Psmg4 | -0.018 | Old |
| 0.752177 | 0.470477 | 0.084 | 0.075 | 1 | 4 | Med12 | 0.009 | Old |
| 0.752188 | 0.080143 | 0.228 | 0.212 | 1 | 4 | Larp4 | 0.016 | Old |
| 0.752207 | 0.033282 | 0.372 | 0.4 | 1 | 4 | Cd302 | -0.028 | Old |
| 0.752236 | 0.035359 | 0.125 | 0.138 | 1 | 4 | Prpsap2 | -0.013 | Old |
| 0.752635 | 0.005083 | 0.2 | 0.188 | 1 | 4 | Igf1r | 0.012 | Old |
| 0.752693 | 0.447831 | 0.122 | 0.112 | 1 | 4 | Arl6ip5 | 0.01 | Old |
| 0.752727 | 1.033242 | 0.03 | 0.038 | 1 | 4 | Mocs1 | -0.008 | Old |
| 0.752958 | 0.042279 | 0.274 | 0.3 | 1 | 4 | Il6st | -0.026 | Old |
| 0.753024 | 0.197773 | 0.085 | 0.075 | 1 | 4 | Zpr1 | 0.01 | Old |
| 0.753107 | 0.119844 | 0.088 | 0.1 | 1 | 4 | Ago1 | -0.012 | Old |
| 0.75313 | 0.004724 | 0.24 | 0.262 | 1 | 4 | Rala | -0.022 | Old |
| 0.753221 | 0.488475 | 0.168 | 0.162 | 1 | 4 | Stim2 | 0.006 | Old |
| 0.753222 | 0.16898 | 0.076 | 0.088 | 1 | 4 | Phf11b | -0.012 | Old |
| 0.753316 | 0.201402 | 0.111 | 0.125 | 1 | 4 | Desi2 | -0.014 | Old |

| | | | | | | | | |
|---|---|---|---|---|---|---|---|---|
| 0.753359 | 0.817678 | 0.031 | 0.025 | 1 | 4 | Zfp444 | 0.006 | Old |
| 0.753371 | 0.459133 | 0.169 | 0.162 | 1 | 4 | Ep400 | 0.007 | Old |
| 0.753685 | 0.053628 | 0.207 | 0.225 | 1 | 4 | Gpbp1 | -0.018 | Old |
| 0.753787 | 0.29211 | 0.072 | 0.062 | 1 | 4 | Odf2 | 0.01 | Old |
| 0.754281 | 0.98485 | 0.105 | 0.1 | 1 | 4 | Pggt1b | 0.005 | Old |
| 0.754298 | 0.222469 | 0.02 | 0.025 | 1 | 4 | Tbc1d30 | -0.005 | Old |
| 0.754298 | 0.484779 | 0.02 | 0.025 | 1 | 4 | Whamm | -0.005 | Old |
| 0.754988 | 0.396264 | 0.121 | 0.112 | 1 | 4 | Utp3 | 0.009 | Old |
| 0.755077 | 0.219268 | 0.02 | 0.025 | 1 | 4 | Ccnyl1 | -0.005 | Old |
| 0.755256 | 0.584223 | 0.031 | 0.025 | 1 | 4 | Mier2 | 0.006 | Old |
| 0.755575 | 0.353815 | 0.071 | 0.062 | 1 | 4 | Tigd2 | 0.009 | Old |
| 0.75561 | 0.484736 | 0.045 | 0.038 | 1 | 4 | Fbxl6 | 0.007 | Old |
| 0.755831 | 0.400042 | 0.169 | 0.162 | 1 | 4 | Baz1a | 0.007 | Old |
| 0.756082 | 0.215161 | 0.286 | 0.312 | 1 | 4 | Mt2 | -0.026 | Old |
| 0.756579 | 0.247342 | 0.158 | 0.15 | 1 | 4 | Atxn7l1 | 0.008 | Old |
| 0.756659 | 0.099368 | 0.636 | 0.662 | 1 | 4 | Zfand5 | -0.026 | Old |
| 0.756666 | 1.096 | 0.044 | 0.038 | 1 | 4 | Kank3 | 0.006 | Old |
| 0.756915 | 0.072224 | 0.304 | 0.288 | 1 | 4 | Vdac2 | 0.016 | Old |
| 0.757185 | 0.017207 | 0.267 | 0.3 | 1 | 4 | Nucb1 | -0.033 | Old |
| 0.757373 | 0.570877 | 0.07 | 0.062 | 1 | 4 | Rnf215 | 0.008 | Old |
| 0.757446 | 0.022407 | 0.397 | 0.412 | 1 | 4 | Mafg | -0.015 | Old |
| 0.757599 | 0.256856 | 0.072 | 0.062 | 1 | 4 | Parp11 | 0.01 | Old |
| 0.757861 | 0.029281 | 0.487 | 0.475 | 1 | 4 | Dnajc3 | 0.012 | Old |
| 0.757899 | 0.303346 | 0.378 | 0.388 | 1 | 4 | Ogdh | -0.01 | Old |
| 0.758009 | 0.167906 | 0.202 | 0.225 | 1 | 4 | Csnk2a1 | -0.023 | Old |
| 0.758152 | 0.024904 | 0.125 | 0.138 | 1 | 4 | Rprd2 | -0.013 | Old |
| 0.758491 | 0.033795 | 0.506 | 0.512 | 1 | 4 | Itgam | -0.006 | Old |
| 0.758964 | 0.31407 | 0.11 | 0.125 | 1 | 4 | Mob3b | -0.015 | Old |
| 0.759008 | 0.198107 | 0.017 | 0.013 | 1 | 4 | Arpin | 0.004 | Old |
| 0.759143 | 0.607113 | 0.108 | 0.1 | 1 | 4 | Gramd1a | 0.008 | Old |
| 0.759224 | 0.853456 | 0.082 | 0.075 | 1 | 4 | Gm43848 | 0.007 | Old |
| 0.759338 | 0.324503 | 0.181 | 0.175 | 1 | 4 | Armcx3 | 0.006 | Old |
| 0.759673 | 0.362921 | 0.084 | 0.075 | 1 | 4 | Sh3tc1 | 0.009 | Old |
| 0.759856 | 0.174155 | 0.017 | 0.013 | 1 | 4 | Dph7 | 0.004 | Old |
| 0.759927 | 0.230283 | 0.286 | 0.325 | 1 | 4 | Oxct1 | -0.039 | Old |
| 0.760353 | 0.204806 | 0.316 | 0.3 | 1 | 4 | Pdlim5 | 0.016 | Old |
| 0.760401 | 0.845682 | 0.044 | 0.038 | 1 | 4 | Tusc1 | 0.006 | Old |
| 0.760401 | 0.811866 | 0.044 | 0.038 | 1 | 4 | Mgmt | 0.006 | Old |
| 0.760407 | 0.076114 | 0.539 | 0.588 | 1 | 4 | Srsf3 | -0.049 | Old |
| 0.760956 | 0.411806 | 0.031 | 0.025 | 1 | 4 | Nprl3 | 0.006 | Old |
| 0.760956 | 0.379087 | 0.031 | 0.025 | 1 | 4 | Ndufaf3 | 0.006 | Old |
| 0.761086 | 0.346364 | 0.144 | 0.162 | 1 | 4 | Eml4 | -0.018 | Old |
| 0.761613 | 0.636327 | 0.131 | 0.125 | 1 | 4 | Cops2 | 0.006 | Old |
| 0.76192 | 0.889539 | 0.106 | 0.1 | 1 | 4 | Nsrp1 | 0.006 | Old |
| 0.762272 | 0.313375 | 0.031 | 0.038 | 1 | 4 | Cnr2 | -0.007 | Old |

| | | | | | | | | |
|---|---|---|---|---|---|---|---|---|
| 0.762732 | 0.021623 | 0.054 | 0.062 | 1 | 4 | Pskh1 | -0.008 | Old |
| 0.762859 | 0.276843 | 0.031 | 0.025 | 1 | 4 | Haus4 | 0.006 | Old |
| 0.762859 | 0.240582 | 0.031 | 0.025 | 1 | 4 | Hotairm1 | 0.006 | Old |
| 0.763016 | 0.179971 | 0.516 | 0.525 | 1 | 4 | Irf8 | -0.009 | Old |
| 0.763185 | 0.059082 | 0.962 | 0.963 | 1 | 4 | Grn | -0.001 | Old |
| 0.763607 | 0.526794 | 0.044 | 0.038 | 1 | 4 | Coq7 | 0.006 | Old |
| 0.763755 | 0.152127 | 0.291 | 0.275 | 1 | 4 | Dgkd | 0.016 | Old |
| 0.764456 | 0.938363 | 0.069 | 0.062 | 1 | 4 | Ubr1 | 0.007 | Old |
| 0.764514 | 0.10408 | 0.077 | 0.088 | 1 | 4 | Srp68 | -0.011 | Old |
| 0.764764 | 0.32318 | 0.031 | 0.025 | 1 | 4 | Tnfrsf12a | 0.006 | Old |
| 0.76505 | 0.03247 | 0.113 | 0.125 | 1 | 4 | Glod4 | -0.012 | Old |
| 0.76528 | 0.075779 | 0.21 | 0.238 | 1 | 4 | Stt3a | -0.028 | Old |
| 0.765335 | 0.104361 | 0.329 | 0.312 | 1 | 4 | Wac | 0.017 | Old |
| 0.765388 | 0.241345 | 0.123 | 0.112 | 1 | 4 | Prdx3 | 0.011 | Old |
| 0.765653 | 0.003923 | 0.265 | 0.288 | 1 | 4 | Sdf2 | -0.023 | Old |
| 0.765741 | 0.154332 | 0.077 | 0.088 | 1 | 4 | Sdhaf1 | -0.011 | Old |
| 0.765984 | 0.18324 | 0.157 | 0.175 | 1 | 4 | Cyb5r3 | -0.018 | Old |
| 0.766038 | 0.779558 | 0.053 | 0.062 | 1 | 4 | Msh2 | -0.009 | Old |
| 0.766197 | 0.380353 | 0.237 | 0.238 | 1 | 4 | Sp1 | -0.001 | Old |
| 0.766211 | 0.591707 | 0.083 | 0.075 | 1 | 4 | Pex5 | 0.008 | Old |
| 0.766296 | 0.18195 | 0.065 | 0.075 | 1 | 4 | Xrcc1 | -0.01 | Old |
| 0.76637 | 0.53359 | 0.042 | 0.05 | 1 | 4 | Pcnx | -0.008 | Old |
| 0.766372 | 0.375201 | 0.058 | 0.05 | 1 | 4 | Zfyve27 | 0.008 | Old |
| 0.766619 | 0.584601 | 0.057 | 0.05 | 1 | 4 | 2310009B1 | 0.007 | Old |
| 0.76667 | 0.153618 | 0.031 | 0.025 | 1 | 4 | Ttc7b | 0.006 | Old |
| 0.76667 | 0.074 | 0.031 | 0.025 | 1 | 4 | Parp6 | 0.006 | Old |
| 0.766791 | 0.419904 | 0.07 | 0.062 | 1 | 4 | Tshz1 | 0.008 | Old |
| 0.766842 | 0.370687 | 0.058 | 0.05 | 1 | 4 | Zfp101 | 0.008 | Old |
| 0.766993 | 0.399801 | 0.084 | 0.075 | 1 | 4 | Wdr5 | 0.009 | Old |
| 0.767191 | 0.328197 | 0.084 | 0.075 | 1 | 4 | Paxbp1 | 0.009 | Old |
| 0.767222 | 0.247837 | 0.123 | 0.138 | 1 | 4 | Ube2j1 | -0.015 | Old |
| 0.76726 | 0.28489 | 0.184 | 0.175 | 1 | 4 | Yme1l1 | 0.009 | Old |
| 0.767464 | 0.015123 | 0.113 | 0.125 | 1 | 4 | Tmem131l | -0.012 | Old |
| 0.767483 | 0.87982 | 0.069 | 0.062 | 1 | 4 | Gan | 0.007 | Old |
| 0.767727 | 0.470006 | 0.042 | 0.05 | 1 | 4 | Chchd4 | -0.008 | Old |
| 0.768253 | 0.226589 | 0.058 | 0.05 | 1 | 4 | Med29 | 0.008 | Old |
| 0.76849 | 0.019638 | 0.3 | 0.325 | 1 | 4 | Rreb1 | -0.025 | Old |
| 0.768746 | 0.631872 | 0.075 | 0.088 | 1 | 4 | Polk | -0.013 | Old |
| 0.768793 | 0.00414 | 0.905 | 0.938 | 1 | 4 | Gas6 | -0.033 | Old |
| 0.768795 | 0.30812 | 0.009 | 0.013 | 1 | 4 | Gm26510 | -0.004 | Old |
| 0.768795 | 0.224566 | 0.009 | 0.013 | 1 | 4 | Poli | -0.004 | Old |
| 0.768839 | 0.393586 | 0.213 | 0.212 | 1 | 4 | Lpp | 0.001 | Old |
| 0.768987 | 0.118708 | 0.112 | 0.125 | 1 | 4 | Wdr33 | -0.013 | Old |
| 0.769056 | 0.256828 | 0.11 | 0.125 | 1 | 4 | Gga3 | -0.015 | Old |
| 0.769192 | 0.948579 | 0.031 | 0.038 | 1 | 4 | Zw10 | -0.007 | Old |

| | | | | | | | | |
|---|---|---|---|---|---|---|---|---|
| 0.769204 | 0.321011 | 0.144 | 0.138 | 1 | 4 | Selenbp1 | 0.006 | Old |
| 0.769464 | 0.037405 | 0.248 | 0.275 | 1 | 4 | Tnfrsf1a | -0.027 | Old |
| 0.76954 | 0.09622 | 0.302 | 0.325 | 1 | 4 | Derl1 | -0.023 | Old |
| 0.769822 | 0.588145 | 0.031 | 0.038 | 1 | 4 | Nynrin | -0.007 | Old |
| 0.770036 | 0.068269 | 0.124 | 0.138 | 1 | 4 | Mcm3 | -0.014 | Old |
| 0.770208 | 0.101859 | 0.045 | 0.038 | 1 | 4 | Scyl3 | 0.007 | Old |
| 0.770377 | 0.125443 | 0.043 | 0.05 | 1 | 4 | Slf1 | -0.007 | Old |
| 0.770751 | 0.129626 | 0.192 | 0.212 | 1 | 4 | Fam76a | -0.02 | Old |
| 0.771059 | 0.00879 | 0.351 | 0.388 | 1 | 4 | Slc29a1 | -0.037 | Old |
| 0.77107 | 0.627799 | 0.009 | 0.013 | 1 | 4 | Tagap1 | -0.004 | Old |
| 0.77107 | 0.403131 | 0.009 | 0.013 | 1 | 4 | Swsap1 | -0.004 | Old |
| 0.77107 | 0.189438 | 0.009 | 0.013 | 1 | 4 | Lins1 | -0.004 | Old |
| 0.77107 | 0.031903 | 0.009 | 0.013 | 1 | 4 | Rasal3 | -0.004 | Old |
| 0.771179 | 0.36054 | 0.186 | 0.212 | 1 | 4 | Hmbox1 | -0.026 | Old |
| 0.771249 | 0.048403 | 0.746 | 0.75 | 1 | 4 | Arpc1b | -0.004 | Old |
| 0.771363 | 0.139382 | 0.241 | 0.262 | 1 | 4 | Tmem141 | -0.021 | Old |
| 0.771416 | 0.011901 | 0.055 | 0.062 | 1 | 4 | Cep164 | -0.007 | Old |
| 0.771456 | 0.972621 | 0.043 | 0.038 | 1 | 4 | Depdc5 | 0.005 | Old |
| 0.771456 | 0.117274 | 0.043 | 0.05 | 1 | 4 | Tipin | -0.007 | Old |
| 0.771812 | 0.735111 | 0.069 | 0.062 | 1 | 4 | Tmem159 | 0.007 | Old |
| 0.771824 | 0.05846 | 0.101 | 0.112 | 1 | 4 | Polr3c | -0.011 | Old |
| 0.771862 | 0.16795 | 0.135 | 0.15 | 1 | 4 | Xdh | -0.015 | Old |
| 0.772103 | 0.926649 | 0.03 | 0.025 | 1 | 4 | Rnf217 | 0.005 | Old |
| 0.772209 | 0.52008 | 0.009 | 0.013 | 1 | 4 | Zbtb49 | -0.004 | Old |
| 0.772209 | 0.35923 | 0.009 | 0.013 | 1 | 4 | Ccdc94 | -0.004 | Old |
| 0.772279 | 0.096012 | 0.188 | 0.175 | 1 | 4 | Arl5b | 0.013 | Old |
| 0.772396 | 0.054559 | 0.031 | 0.025 | 1 | 4 | Zfp142 | 0.006 | Old |
| 0.772474 | 0.126283 | 0.262 | 0.288 | 1 | 4 | Baz2b | -0.026 | Old |
| 0.77295 | 0.141581 | 0.206 | 0.225 | 1 | 4 | Pkn2 | -0.019 | Old |
| 0.772962 | 0.129827 | 0.058 | 0.05 | 1 | 4 | Ppp2r5d | 0.008 | Old |
| 0.773092 | 0.330357 | 0.243 | 0.238 | 1 | 4 | Pnpla8 | 0.005 | Old |
| 0.773248 | 0.070411 | 0.4 | 0.438 | 1 | 4 | Psma7 | -0.038 | Old |
| 0.773285 | 0.033371 | 0.144 | 0.162 | 1 | 4 | Chmp2b | -0.018 | Old |
| 0.773348 | 0.55788 | 0.009 | 0.013 | 1 | 4 | B3gnt2 | -0.004 | Old |
| 0.773348 | 0.499179 | 0.009 | 0.013 | 1 | 4 | Slc25a14 | -0.004 | Old |
| 0.773348 | 0.291198 | 0.009 | 0.013 | 1 | 4 | Pinx1 | -0.004 | Old |
| 0.773376 | 0.467066 | 0.083 | 0.075 | 1 | 4 | Otud6b | 0.008 | Old |
| 0.773485 | 0.137763 | 0.363 | 0.4 | 1 | 4 | Klf7 | -0.037 | Old |
| 0.773693 | 0.422679 | 0.096 | 0.088 | 1 | 4 | Nlk | 0.008 | Old |
| 0.773707 | 0.633772 | 0.042 | 0.05 | 1 | 4 | Fam35a | -0.008 | Old |
| 0.774015 | 0.157053 | 0.226 | 0.25 | 1 | 4 | Hfe | -0.024 | Old |
| 0.774133 | 0.193272 | 0.4 | 0.412 | 1 | 4 | Flna | -0.012 | Old |
| 0.77442 | 0.321228 | 0.208 | 0.2 | 1 | 4 | Tap2 | 0.008 | Old |
| 0.774488 | 1.024555 | 0.009 | 0.013 | 1 | 4 | Smim1 | -0.004 | Old |
| 0.774488 | 1.003139 | 0.009 | 0.013 | 1 | 4 | Cdk1 | -0.004 | Old |

| | | | | | | | | |
|---|---|---|---|---|---|---|---|---|
| 0.774488 | 0.441628 | 0.009 | 0.013 | 1 | 4 | Vill | -0.004 | Old |
| 0.774537 | 0.034325 | 0.566 | 0.588 | 1 | 4 | Rab1a | -0.022 | Old |
| 0.774771 | 0.467113 | 0.083 | 0.075 | 1 | 4 | Selenon | 0.008 | Old |
| 0.774951 | 0.534112 | 0.057 | 0.05 | 1 | 4 | Pdk3 | 0.007 | Old |
| 0.774977 | 1.304923 | 0.016 | 0.013 | 1 | 4 | Zfp931 | 0.003 | Old |
| 0.775061 | 0.383293 | 0.122 | 0.138 | 1 | 4 | Hars | -0.016 | Old |
| 0.775069 | 0.02751 | 0.248 | 0.262 | 1 | 4 | Zyx | -0.014 | Old |
| 0.775085 | 0.176515 | 0.084 | 0.075 | 1 | 4 | C1galt1c1 | 0.009 | Old |
| 0.775239 | 1.03813 | 0.043 | 0.038 | 1 | 4 | Ubap2 | 0.005 | Old |
| 0.775628 | 0.965596 | 0.009 | 0.013 | 1 | 4 | Cald1 | -0.004 | Old |
| 0.775846 | 1.369067 | 0.016 | 0.013 | 1 | 4 | Tgif2 | 0.003 | Old |
| 0.776046 | 0.697189 | 0.082 | 0.075 | 1 | 4 | Smg5 | 0.007 | Old |
| 0.776091 | 0.119095 | 0.182 | 0.2 | 1 | 4 | Slc25a51 | -0.018 | Old |
| 0.776229 | 0.490834 | 0.107 | 0.1 | 1 | 4 | Scoc | 0.007 | Old |
| 0.776416 | 0.611023 | 0.076 | 0.088 | 1 | 4 | Adk | -0.012 | Old |
| 0.776609 | 0.925078 | 0.03 | 0.025 | 1 | 4 | Rmnd1 | 0.005 | Old |
| 0.776849 | 0.781962 | 0.082 | 0.075 | 1 | 4 | Lemd3 | 0.007 | Old |
| 0.777344 | 0.383566 | 0.096 | 0.088 | 1 | 4 | Usp32 | 0.008 | Old |
| 0.777659 | 0.194172 | 0.066 | 0.075 | 1 | 4 | Nup85 | -0.009 | Old |
| 0.778279 | 0.116123 | 0.173 | 0.162 | 1 | 4 | Smarcc1 | 0.011 | Old |
| 0.778485 | 0.77537 | 0.043 | 0.038 | 1 | 4 | Ap5z1 | 0.005 | Old |
| 0.778602 | 0.043904 | 0.09 | 0.1 | 1 | 4 | Usp12 | -0.01 | Old |
| 0.778739 | 0.068355 | 0.206 | 0.225 | 1 | 4 | Mga | -0.019 | Old |
| 0.778862 | 0.262123 | 0.077 | 0.088 | 1 | 4 | Vps37b | -0.011 | Old |
| 0.77889 | 0.021585 | 0.7 | 0.762 | 1 | 4 | mt-Nd4l | -0.062 | Old |
| 0.779059 | 0.198216 | 0.097 | 0.088 | 1 | 4 | Gpd1l | 0.009 | Old |
| 0.779096 | 0.190455 | 0.123 | 0.112 | 1 | 4 | Plagl2 | 0.011 | Old |
| 0.779183 | 0.374607 | 0.291 | 0.3 | 1 | 4 | Ttc7 | -0.009 | Old |
| 0.779306 | 0.041329 | 0.191 | 0.175 | 1 | 4 | Mtmr12 | 0.016 | Old |
| 0.779325 | 1.207315 | 0.016 | 0.013 | 1 | 4 | Polg2 | 0.003 | Old |
| 0.779325 | 1.056523 | 0.016 | 0.013 | 1 | 4 | Klhl33 | 0.003 | Old |
| 0.779695 | 0.160386 | 0.044 | 0.038 | 1 | 4 | Utp23 | 0.006 | Old |
| 0.779833 | 0.680269 | 0.03 | 0.025 | 1 | 4 | Gde1 | 0.005 | Old |
| 0.780109 | 0.866993 | 0.043 | 0.038 | 1 | 4 | Zscan21 | 0.005 | Old |
| 0.780198 | 0.622991 | 0.057 | 0.05 | 1 | 4 | Ss18l2 | 0.007 | Old |
| 0.780402 | 0.070746 | 0.032 | 0.038 | 1 | 4 | Asah2 | -0.006 | Old |
| 0.780597 | 0.028366 | 0.541 | 0.575 | 1 | 4 | Creg1 | -0.034 | Old |
| 0.781028 | 0.232451 | 0.032 | 0.038 | 1 | 4 | Nphp3 | -0.006 | Old |
| 0.781066 | 1.260573 | 0.016 | 0.013 | 1 | 4 | Exog | 0.003 | Old |
| 0.781066 | 0.710851 | 0.016 | 0.013 | 1 | 4 | Clec1a | 0.003 | Old |
| 0.781222 | 0.094061 | 0.058 | 0.05 | 1 | 4 | AI987944 | 0.008 | Old |
| 0.781361 | 0.009389 | 0.102 | 0.112 | 1 | 4 | Ddx19a | -0.01 | Old |
| 0.781585 | 0.293173 | 0.158 | 0.15 | 1 | 4 | Vezf1 | 0.008 | Old |
| 0.782147 | 0.559848 | 0.095 | 0.088 | 1 | 4 | Alkbh1 | 0.007 | Old |
| 0.782256 | 0.204485 | 0.119 | 0.112 | 1 | 4 | Ptgr1 | 0.007 | Old |

| | | | | | | | | |
|---|---|---|---|---|---|---|---|---|
| 0.782261 | 0.340587 | 0.057 | 0.05 | 1 | 4 | Ak6 | 0.007 | Old |
| 0.782281 | 0.278493 | 0.096 | 0.088 | 1 | 4 | Rxrb | 0.008 | Old |
| 0.782414 | 0.47837 | 0.03 | 0.025 | 1 | 4 | Atf5 | 0.005 | Old |
| 0.782473 | 0.352292 | 0.07 | 0.062 | 1 | 4 | Mrps28 | 0.008 | Old |
| 0.78253 | 0.213101 | 0.097 | 0.088 | 1 | 4 | Cd99l2 | 0.009 | Old |
| 0.782808 | 0.750965 | 0.016 | 0.013 | 1 | 4 | Gm14296 | 0.003 | Old |
| 0.78285 | 0.090431 | 0.234 | 0.25 | 1 | 4 | Snx17 | -0.016 | Old |
| 0.783652 | 0.550407 | 0.095 | 0.088 | 1 | 4 | Swt1 | 0.007 | Old |
| 0.784002 | 0.339819 | 0.07 | 0.062 | 1 | 4 | Gramd3 | 0.008 | Old |
| 0.784176 | 0.263448 | 0.089 | 0.1 | 1 | 4 | Nol11 | -0.011 | Old |
| 0.784829 | 0.354118 | 0.17 | 0.162 | 1 | 4 | Maea | 0.008 | Old |
| 0.784872 | 0.133176 | 0.02 | 0.025 | 1 | 4 | Ift140 | -0.005 | Old |
| 0.784872 | 0.015106 | 0.02 | 0.025 | 1 | 4 | Odf2l | -0.005 | Old |
| 0.784964 | 0.091933 | 0.173 | 0.188 | 1 | 4 | Plekha1 | -0.015 | Old |
| 0.785046 | 0.313267 | 0.262 | 0.262 | 1 | 4 | Msl2 | 0 | Old |
| 0.785419 | 0.236916 | 0.032 | 0.038 | 1 | 4 | Ercc4 | -0.006 | Old |
| 0.785532 | 0.335271 | 0.043 | 0.05 | 1 | 4 | Edc3 | -0.007 | Old |
| 0.785618 | 0.029907 | 0.044 | 0.038 | 1 | 4 | Ranbp6 | 0.006 | Old |
| 0.785635 | 0.126837 | 0.082 | 0.075 | 1 | 4 | Pi4k2b | 0.007 | Old |
| 0.785648 | 0.098925 | 0.02 | 0.025 | 1 | 4 | Gnb4 | -0.005 | Old |
| 0.785738 | 0.135431 | 0.57 | 0.613 | 1 | 4 | Glud1 | -0.043 | Old |
| 0.785934 | 0.507685 | 0.057 | 0.05 | 1 | 4 | Trp53inp1 | 0.007 | Old |
| 0.785941 | 0.065093 | 0.238 | 0.225 | 1 | 4 | Pcnp | 0.013 | Old |
| 0.786296 | 0.172504 | 0.016 | 0.013 | 1 | 4 | Pcyt1b | 0.003 | Old |
| 0.786366 | 0.417973 | 0.107 | 0.1 | 1 | 4 | Chml | 0.007 | Old |
| 0.786424 | 0.063735 | 0.02 | 0.025 | 1 | 4 | Smug1 | -0.005 | Old |
| 0.786424 | 0.267733 | 0.02 | 0.025 | 1 | 4 | 1110046J0 | -0.005 | Old |
| 0.786665 | 0.457468 | 0.095 | 0.088 | 1 | 4 | Cd151 | 0.007 | Old |
| 0.787169 | 0.392002 | 0.016 | 0.013 | 1 | 4 | Pyroxd2 | 0.003 | Old |
| 0.787417 | 1.690293 | 0.03 | 0.025 | 1 | 4 | Ampd2 | 0.005 | Old |
| 0.787611 | 0.299107 | 0.099 | 0.112 | 1 | 4 | Ppt2 | -0.013 | Old |
| 0.787748 | 0.195206 | 0.223 | 0.212 | 1 | 4 | Smim11 | 0.011 | Old |
| 0.787871 | 0.0826 | 0.122 | 0.112 | 1 | 4 | Gch1 | 0.01 | Old |
| 0.78793 | 0.478314 | 0.032 | 0.038 | 1 | 4 | Zbtb5 | -0.006 | Old |
| 0.78793 | 0.330761 | 0.032 | 0.038 | 1 | 4 | 5730409E0 | -0.006 | Old |
| 0.788042 | 0.077493 | 0.016 | 0.013 | 1 | 4 | Efna2 | 0.003 | Old |
| 0.788248 | 0.361006 | 0.121 | 0.112 | 1 | 4 | Pttg1 | 0.009 | Old |
| 0.788448 | 0.182404 | 0.057 | 0.05 | 1 | 4 | Phf13 | 0.007 | Old |
| 0.788539 | 0.058088 | 0.187 | 0.2 | 1 | 4 | Nabp1 | -0.013 | Old |
| 0.788559 | 0.292551 | 0.032 | 0.038 | 1 | 4 | Pycrl | -0.006 | Old |
| 0.788754 | 0.391305 | 0.02 | 0.025 | 1 | 4 | Zfp12 | -0.005 | Old |
| 0.78879 | 0.356482 | 0.043 | 0.05 | 1 | 4 | Rdh10 | -0.007 | Old |
| 0.788982 | 0.192404 | 0.184 | 0.175 | 1 | 4 | Pin4 | 0.009 | Old |
| 0.789161 | 0.029525 | 0.458 | 0.488 | 1 | 4 | Rtn3 | -0.03 | Old |
| 0.789271 | 0.040331 | 0.708 | 0.688 | 1 | 4 | Gnb2 | 0.02 | Old |

| | | | | | | | | |
|---|---|---|---|---|---|---|---|---|
| 0.789272 | 0.195044 | 0.055 | 0.062 | 1 | 4 | Agfg2 | -0.007 | Old |
| 0.789333 | 0.505114 | 0.043 | 0.038 | 1 | 4 | Rnf113a2 | 0.005 | Old |
| 0.789664 | 0.058654 | 0.043 | 0.05 | 1 | 4 | Ubl4a | -0.007 | Old |
| 0.789766 | 0.01981 | 1 | 1 | 1 | 4 | mt-Co2 | 0 | Old |
| 0.789893 | 0.078039 | 0.913 | 0.938 | 1 | 4 | Cfl1 | -0.025 | Old |
| 0.790173 | 0.153796 | 0.03 | 0.025 | 1 | 4 | Sh3d19 | 0.005 | Old |
| 0.790532 | 0.127781 | 0.136 | 0.15 | 1 | 4 | Slc9a3r1 | -0.014 | Old |
| 0.790762 | 0.016752 | 0.111 | 0.1 | 1 | 4 | Mcoln1 | 0.011 | Old |
| 0.790976 | 0.057933 | 0.084 | 0.075 | 1 | 4 | Pla2g16 | 0.009 | Old |
| 0.791073 | 0.611809 | 0.032 | 0.038 | 1 | 4 | Cog2 | -0.006 | Old |
| 0.791862 | 0.597528 | 0.02 | 0.025 | 1 | 4 | Taf1a | -0.005 | Old |
| 0.791868 | 0.252701 | 0.101 | 0.112 | 1 | 4 | Cnep1r1 | -0.011 | Old |
| 0.792087 | 0.333123 | 0.219 | 0.212 | 1 | 4 | Psma5 | 0.007 | Old |
| 0.792174 | 0.087903 | 0.722 | 0.725 | 1 | 4 | Bri3 | -0.003 | Old |
| 0.792177 | 0.44064 | 0.143 | 0.162 | 1 | 4 | Trim12a | -0.019 | Old |
| 0.792319 | 0.087591 | 0.161 | 0.175 | 1 | 4 | Hvcn1 | -0.014 | Old |
| 0.79243 | 0.073895 | 0.107 | 0.1 | 1 | 4 | Fryl | 0.007 | Old |
| 0.792904 | 0.068648 | 0.043 | 0.05 | 1 | 4 | Tmem184c | -0.007 | Old |
| 0.793132 | 0.207807 | 0.225 | 0.212 | 1 | 4 | Impact | 0.013 | Old |
| 0.793168 | 0.050214 | 0.084 | 0.075 | 1 | 4 | Nelfa | 0.009 | Old |
| 0.79339 | 0.320823 | 0.055 | 0.062 | 1 | 4 | Med22 | -0.007 | Old |
| 0.793421 | 0.294719 | 0.12 | 0.112 | 1 | 4 | Pwwp2a | 0.008 | Old |
| 0.793445 | 0.097803 | 0.043 | 0.05 | 1 | 4 | Zfp157 | -0.007 | Old |
| 0.79359 | 0.877789 | 0.032 | 0.038 | 1 | 4 | Klc4 | -0.006 | Old |
| 0.79359 | 0.629808 | 0.032 | 0.038 | 1 | 4 | Pgm3 | -0.006 | Old |
| 0.79394 | 0.076819 | 0.684 | 0.688 | 1 | 4 | Rnf130 | -0.004 | Old |
| 0.794315 | 0.069017 | 0.186 | 0.2 | 1 | 4 | Tmpo | -0.014 | Old |
| 0.794333 | 0.042685 | 0.265 | 0.288 | 1 | 4 | Rere | -0.023 | Old |
| 0.794509 | 0.220375 | 0.184 | 0.175 | 1 | 4 | Ndufa8 | 0.009 | Old |
| 0.794637 | 0.054061 | 0.324 | 0.3 | 1 | 4 | Pdia4 | 0.024 | Old |
| 0.794653 | 0.199048 | 0.383 | 0.425 | 1 | 4 | Arhgap17 | -0.042 | Old |
| 0.794675 | 0.182231 | 0.084 | 0.075 | 1 | 4 | Fam162a | 0.009 | Old |
| 0.794709 | 0.047835 | 0.03 | 0.025 | 1 | 4 | Foxp4 | 0.005 | Old |
| 0.794722 | 0.377986 | 0.066 | 0.075 | 1 | 4 | 5031425F1 | -0.009 | Old |
| 0.795075 | 0.212086 | 0.109 | 0.1 | 1 | 4 | Naa35 | 0.009 | Old |
| 0.795542 | 0.211258 | 0.322 | 0.312 | 1 | 4 | Ugcg | 0.01 | Old |
| 0.795669 | 0.071325 | 0.124 | 0.138 | 1 | 4 | Snrnp27 | -0.014 | Old |
| 0.795694 | 0.253656 | 0.121 | 0.112 | 1 | 4 | Nub1 | 0.009 | Old |
| 0.795862 | 0.314818 | 0.043 | 0.038 | 1 | 4 | Mgst3 | 0.005 | Old |
| 0.796152 | 0.077044 | 0.055 | 0.062 | 1 | 4 | Zfp654 | -0.007 | Old |
| 0.796198 | 0.567388 | 0.054 | 0.062 | 1 | 4 | Usp39 | -0.008 | Old |
| 0.796383 | 0.941698 | 0.043 | 0.038 | 1 | 4 | Fam214a | 0.005 | Old |
| 0.796417 | 0.763728 | 0.08 | 0.075 | 1 | 4 | Zswim4 | 0.005 | Old |
| 0.796527 | 0.187777 | 0.181 | 0.2 | 1 | 4 | Keap1 | -0.019 | Old |
| 0.796585 | 0.282071 | 0.333 | 0.338 | 1 | 4 | Cct5 | -0.005 | Old |

| | | | | | | | | |
|---|---|---|---|---|---|---|---|---|
| 0.796648 | 0.09745 | 0.381 | 0.375 | 1 | 4 | Midn | 0.006 | Old |
| 0.796727 | 0.172789 | 0.194 | 0.212 | 1 | 4 | Tmf1 | -0.018 | Old |
| 0.796903 | 0.055263 | 0.067 | 0.075 | 1 | 4 | Lman2l | -0.008 | Old |
| 0.796993 | 0.258532 | 0.134 | 0.125 | 1 | 4 | Lpgat1 | 0.009 | Old |
| 0.797271 | 0.841168 | 0.055 | 0.05 | 1 | 4 | Parp10 | 0.005 | Old |
| 0.798325 | 0.501766 | 0.056 | 0.05 | 1 | 4 | Zfp626 | 0.006 | Old |
| 0.798485 | 0.385462 | 0.055 | 0.062 | 1 | 4 | Lrp12 | -0.007 | Old |
| 0.798523 | 0.053991 | 0.269 | 0.262 | 1 | 4 | Tmem160 | 0.007 | Old |
| 0.798769 | 0.288551 | 0.078 | 0.088 | 1 | 4 | Cfap20 | -0.01 | Old |
| 0.798983 | 0.262305 | 0.07 | 0.062 | 1 | 4 | Sap30l | 0.008 | Old |
| 0.799084 | 0.139611 | 0.208 | 0.2 | 1 | 4 | Bcl3 | 0.008 | Old |
| 0.799292 | 0.557011 | 0.056 | 0.05 | 1 | 4 | Fuom | 0.006 | Old |
| 0.799684 | 0.071154 | 0.763 | 0.775 | 1 | 4 | Tcf4 | -0.012 | Old |
| 0.799716 | 0.290726 | 0.399 | 0.438 | 1 | 4 | Dok2 | -0.039 | Old |
| 0.799851 | 0.095763 | 0.861 | 0.838 | 1 | 4 | Trf | 0.023 | Old |
| 0.800072 | 0.356701 | 0.095 | 0.088 | 1 | 4 | Mnat1 | 0.007 | Old |
| 0.800226 | 0.677123 | 0.043 | 0.038 | 1 | 4 | Extl2 | 0.005 | Old |
| 0.800479 | 0.271559 | 0.043 | 0.05 | 1 | 4 | Exoc8 | -0.007 | Old |
| 0.80066 | 0.06425 | 0.111 | 0.1 | 1 | 4 | Cdk2 | 0.011 | Old |
| 0.800994 | 0.074719 | 0.451 | 0.488 | 1 | 4 | Sod1 | -0.037 | Old |
| 0.80119 | 0.268957 | 0.157 | 0.15 | 1 | 4 | Xpo6 | 0.007 | Old |
| 0.801269 | 0.654072 | 0.069 | 0.062 | 1 | 4 | Lxn | 0.007 | Old |
| 0.8015 | 0.331529 | 0.215 | 0.212 | 1 | 4 | Tcf12 | 0.003 | Old |
| 0.801533 | 0.224688 | 0.173 | 0.162 | 1 | 4 | Anxa11 | 0.011 | Old |
| 0.801654 | 0.100993 | 0.148 | 0.138 | 1 | 4 | Gabpb2 | 0.01 | Old |
| 0.80175 | 0.164547 | 0.078 | 0.088 | 1 | 4 | Naa40 | -0.01 | Old |
| 0.801829 | 0.786102 | 0.03 | 0.025 | 1 | 4 | Trip6 | 0.005 | Old |
| 0.80186 | 0.188089 | 0.043 | 0.038 | 1 | 4 | Ccdc71 | 0.005 | Old |
| 0.802129 | 0.760991 | 0.055 | 0.05 | 1 | 4 | Irak2 | 0.005 | Old |
| 0.802158 | 0.173732 | 0.26 | 0.288 | 1 | 4 | Plbd2 | -0.028 | Old |
| 0.802235 | 0.58303 | 0.162 | 0.188 | 1 | 4 | Sptlc1 | -0.026 | Old |
| 0.802787 | 0.014328 | 0.126 | 0.138 | 1 | 4 | Coro7 | -0.012 | Old |
| 0.802825 | 0.338553 | 0.144 | 0.138 | 1 | 4 | Tsnax | 0.006 | Old |
| 0.803552 | 0.049795 | 0.287 | 0.262 | 1 | 4 | Fgd2 | 0.025 | Old |
| 0.804001 | 0.163044 | 0.564 | 0.562 | 1 | 4 | Tpp1 | 0.002 | Old |
| 0.804153 | 0.084733 | 0.197 | 0.188 | 1 | 4 | Napa | 0.009 | Old |
| 0.804364 | 0.061206 | 0.126 | 0.138 | 1 | 4 | Snrpa | -0.012 | Old |
| 0.804599 | 0.166644 | 0.278 | 0.3 | 1 | 4 | C6 | -0.022 | Old |
| 0.804748 | 0.396814 | 0.111 | 0.125 | 1 | 4 | Prmt1 | -0.014 | Old |
| 0.805095 | 0.411147 | 0.056 | 0.05 | 1 | 4 | Naip6 | 0.006 | Old |
| 0.805184 | 0.055705 | 0.296 | 0.288 | 1 | 4 | Fam102b | 0.008 | Old |
| 0.805233 | 0.076403 | 0.462 | 0.5 | 1 | 4 | Sec61b | -0.038 | Old |
| 0.805465 | 0.505327 | 0.081 | 0.075 | 1 | 4 | Cdadc1 | 0.006 | Old |
| 0.805704 | 0.100026 | 0.599 | 0.588 | 1 | 4 | Arl6ip1 | 0.011 | Old |
| 0.805857 | 0.253711 | 0.364 | 0.412 | 1 | 4 | Fubp1 | -0.048 | Old |

| | | | | | | | | |
|---|---|---|---|---|---|---|---|---|
| 0.805869 | 1.759589 | 0.016 | 0.013 | 1 | 4 | Rab11fip3 | 0.003 | Old |
| 0.806188 | 0.26823 | 0.157 | 0.15 | 1 | 4 | Tbc1d10b | 0.007 | Old |
| 0.806429 | 0.661111 | 0.03 | 0.025 | 1 | 4 | Xrcc6 | 0.005 | Old |
| 0.806429 | 0.41236 | 0.03 | 0.025 | 1 | 4 | Slc39a11 | 0.005 | Old |
| 0.806765 | 1.851076 | 0.016 | 0.013 | 1 | 4 | Psph | 0.003 | Old |
| 0.806765 | 1.766682 | 0.016 | 0.013 | 1 | 4 | Pard6a | 0.003 | Old |
| 0.806858 | 0.026738 | 0.083 | 0.075 | 1 | 4 | Eaf1 | 0.008 | Old |
| 0.807322 | 0.178318 | 0.043 | 0.038 | 1 | 4 | Nsun6 | 0.005 | Old |
| 0.807417 | 1.313418 | 0.042 | 0.038 | 1 | 4 | Fbxo45 | 0.004 | Old |
| 0.807527 | 0.010815 | 0.188 | 0.2 | 1 | 4 | Lsm6 | -0.012 | Old |
| 0.807531 | 0.377428 | 0.043 | 0.05 | 1 | 4 | Mbip | -0.007 | Old |
| 0.80766 | 1.280768 | 0.016 | 0.013 | 1 | 4 | Wdr90 | 0.003 | Old |
| 0.80777 | 0.068829 | 0.148 | 0.138 | 1 | 4 | Cope | 0.01 | Old |
| 0.807931 | 0.395462 | 0.208 | 0.238 | 1 | 4 | Usp15 | -0.03 | Old |
| 0.80799 | 0.335187 | 0.193 | 0.188 | 1 | 4 | Vps29 | 0.005 | Old |
| 0.808213 | 0.442586 | 0.055 | 0.062 | 1 | 4 | Erh | -0.007 | Old |
| 0.808419 | 0.246265 | 0.204 | 0.225 | 1 | 4 | Smad4 | -0.021 | Old |
| 0.808556 | 1.402446 | 0.016 | 0.013 | 1 | 4 | Zfp820 | 0.003 | Old |
| 0.808885 | 0.276325 | 0.169 | 0.188 | 1 | 4 | Nudt21 | -0.019 | Old |
| 0.809067 | 0.387802 | 0.131 | 0.125 | 1 | 4 | Vps13d | 0.006 | Old |
| 0.809411 | 0.208357 | 0.159 | 0.175 | 1 | 4 | Clcn4 | -0.016 | Old |
| 0.809431 | 0.785727 | 0.055 | 0.05 | 1 | 4 | Apip | 0.005 | Old |
| 0.809453 | 1.442513 | 0.016 | 0.013 | 1 | 4 | Thnsl1 | 0.003 | Old |
| 0.809453 | 1.347172 | 0.016 | 0.013 | 1 | 4 | Zfp1 | 0.003 | Old |
| 0.809578 | 0.599961 | 0.043 | 0.038 | 1 | 4 | Hpn | 0.005 | Old |
| 0.809634 | 0.52637 | 0.069 | 0.062 | 1 | 4 | Acot9 | 0.007 | Old |
| 0.809664 | 0.237387 | 0.148 | 0.162 | 1 | 4 | Rbm33 | -0.014 | Old |
| 0.809921 | 0.008186 | 0.41 | 0.45 | 1 | 4 | Uqcrq | -0.04 | Old |
| 0.80994 | 0.508668 | 0.056 | 0.05 | 1 | 4 | Yif1a | 0.006 | Old |
| 0.810349 | 0.938078 | 0.016 | 0.013 | 1 | 4 | Phpt1 | 0.003 | Old |
| 0.810349 | 0.872273 | 0.016 | 0.013 | 1 | 4 | Fam216a | 0.003 | Old |
| 0.810378 | 0.194727 | 0.03 | 0.025 | 1 | 4 | Ifi205 | 0.005 | Old |
| 0.810378 | 0.139578 | 0.03 | 0.025 | 1 | 4 | Fancl | 0.005 | Old |
| 0.810422 | 1.468182 | 0.029 | 0.025 | 1 | 4 | Hmgn5 | 0.004 | Old |
| 0.810802 | 0.145362 | 0.032 | 0.038 | 1 | 4 | Wdr92 | -0.006 | Old |
| 0.81091 | 0.373526 | 0.056 | 0.05 | 1 | 4 | Slc39a13 | 0.006 | Old |
| 0.810962 | 0.187008 | 0.374 | 0.375 | 1 | 4 | Trip12 | -0.001 | Old |
| 0.81134 | 0.042879 | 0.556 | 0.575 | 1 | 4 | Itgb1 | -0.019 | Old |
| 0.811432 | 0.220996 | 0.146 | 0.138 | 1 | 4 | Ppil2 | 0.008 | Old |
| 0.811753 | 0.001522 | 0.294 | 0.312 | 1 | 4 | Rbx1 | -0.018 | Old |
| 0.81188 | 0.376437 | 0.055 | 0.062 | 1 | 4 | Patz1 | -0.007 | Old |
| 0.812005 | 0.189441 | 0.121 | 0.112 | 1 | 4 | Trove2 | 0.009 | Old |
| 0.812092 | 0.347011 | 0.1 | 0.112 | 1 | 4 | Gle1 | -0.012 | Old |
| 0.812143 | 0.932551 | 0.016 | 0.013 | 1 | 4 | Il17rd | 0.003 | Old |
| 0.812143 | 0.887126 | 0.016 | 0.013 | 1 | 4 | Dnajc4 | 0.003 | Old |

| | | | | | | | | |
|---|---|---|---|---|---|---|---|---|
| 0.812466 | 0.939933 | 0.067 | 0.062 | 1 | 4 | Tspan5 | 0.005 | Old |
| 0.812597 | 0.735395 | 0.068 | 0.062 | 1 | 4 | Pabpc4 | 0.006 | Old |
| 0.81304 | 0.898062 | 0.016 | 0.013 | 1 | 4 | Zfp667 | 0.003 | Old |
| 0.81304 | 0.845228 | 0.016 | 0.013 | 1 | 4 | P2ry1 | 0.003 | Old |
| 0.81304 | 0.769081 | 0.016 | 0.013 | 1 | 4 | Pecr | 0.003 | Old |
| 0.813153 | 0.479305 | 0.106 | 0.1 | 1 | 4 | Dnajb4 | 0.006 | Old |
| 0.813245 | 0.207313 | 0.308 | 0.312 | 1 | 4 | Smarca5 | -0.004 | Old |
| 0.813252 | 0.189153 | 0.082 | 0.075 | 1 | 4 | Pex3 | 0.007 | Old |
| 0.813463 | 0.574839 | 0.163 | 0.162 | 1 | 4 | Dguok | 0.001 | Old |
| 0.813471 | 0.158887 | 0.285 | 0.312 | 1 | 4 | Fnip1 | -0.027 | Old |
| 0.813683 | 0.065189 | 0.173 | 0.162 | 1 | 4 | Cnot4 | 0.011 | Old |
| 0.813746 | 0.762918 | 0.08 | 0.075 | 1 | 4 | Sp140 | 0.005 | Old |
| 0.813762 | 0.07075 | 0.363 | 0.362 | 1 | 4 | St8sia4 | 0.001 | Old |
| 0.814057 | 0.063499 | 0.056 | 0.062 | 1 | 4 | Gm17018 | -0.006 | Old |
| 0.814331 | 0.198855 | 0.03 | 0.025 | 1 | 4 | Pigg | 0.005 | Old |
| 0.81454 | 0.396434 | 0.043 | 0.038 | 1 | 4 | Taf1b | 0.005 | Old |
| 0.814772 | 0.106424 | 0.208 | 0.225 | 1 | 4 | Ube2i | -0.017 | Old |
| 0.814835 | 0.655387 | 0.016 | 0.013 | 1 | 4 | Efnb1 | 0.003 | Old |
| 0.815277 | 1.46218 | 0.042 | 0.038 | 1 | 4 | Scai | 0.004 | Old |
| 0.815499 | 0.106666 | 0.23 | 0.25 | 1 | 4 | Ankrd13a | -0.02 | Old |
| 0.815649 | 0.312205 | 0.03 | 0.025 | 1 | 4 | Fam60a | 0.005 | Old |
| 0.8157 | 0.086794 | 0.269 | 0.262 | 1 | 4 | Pgk1 | 0.007 | Old |
| 0.815733 | 0.372636 | 0.016 | 0.013 | 1 | 4 | Gmpr | 0.003 | Old |
| 0.815733 | 0.267334 | 0.016 | 0.013 | 1 | 4 | Vps51 | 0.003 | Old |
| 0.815789 | 0.03333 | 0.152 | 0.162 | 1 | 4 | Ubxn6 | -0.01 | Old |
| 0.815812 | 0.060616 | 0.068 | 0.075 | 1 | 4 | Banp | -0.007 | Old |
| 0.816022 | 0.939664 | 0.055 | 0.05 | 1 | 4 | Gtf3c3 | 0.005 | Old |
| 0.816029 | 0.453072 | 0.067 | 0.075 | 1 | 4 | Sp2 | -0.008 | Old |
| 0.816108 | 0.132346 | 0.195 | 0.188 | 1 | 4 | Snrpd1 | 0.007 | Old |
| 0.816196 | 0.353297 | 0.043 | 0.038 | 1 | 4 | Enoph1 | 0.005 | Old |
| 0.816278 | 0.02321 | 0.08 | 0.088 | 1 | 4 | Egfr | -0.008 | Old |
| 0.816287 | 0.146304 | 0.221 | 0.212 | 1 | 4 | Ndufs2 | 0.009 | Old |
| 0.816519 | 0.355413 | 0.107 | 0.1 | 1 | 4 | Actl6a | 0.007 | Old |
| 0.81675 | 0.544664 | 0.055 | 0.05 | 1 | 4 | Tlr3 | 0.005 | Old |
| 0.816842 | 0.21942 | 0.3 | 0.3 | 1 | 4 | Snrpf | 0 | Old |
| 0.817041 | 0.429594 | 0.188 | 0.188 | 1 | 4 | Pdlim1 | 0 | Old |
| 0.817082 | 1.238384 | 0.029 | 0.025 | 1 | 4 | Ppm1f | 0.004 | Old |
| 0.817143 | 0.063002 | 0.561 | 0.562 | 1 | 4 | Atp6v1g1 | -0.001 | Old |
| 0.817318 | 0.337756 | 0.157 | 0.15 | 1 | 4 | Gstm1 | 0.007 | Old |
| 0.817532 | 0.222466 | 0.095 | 0.088 | 1 | 4 | Mpdu1 | 0.007 | Old |
| 0.817968 | 0.09267 | 0.393 | 0.425 | 1 | 4 | Casp4 | -0.032 | Old |
| 0.817983 | 0.042474 | 0.211 | 0.225 | 1 | 4 | Tollip | -0.014 | Old |
| 0.81828 | 0.179809 | 0.29 | 0.325 | 1 | 4 | Wdfy2 | -0.035 | Old |
| 0.818336 | 0.40816 | 0.032 | 0.038 | 1 | 4 | Colca2 | -0.006 | Old |
| 0.818569 | 0.159919 | 0.15 | 0.162 | 1 | 4 | Bsdc1 | -0.012 | Old |

| | | | | | | | | |
|---|---|---|---|---|---|---|---|---|
| 0.818708 | 0.056365 | 0.01 | 0.013 | 1 | 4 | Aqp9 | -0.003 | Old |
| 0.818862 | 0.444213 | 0.211 | 0.212 | 1 | 4 | Dcaf7 | -0.001 | Old |
| 0.818931 | 0.04124 | 0.125 | 0.138 | 1 | 4 | Oser1 | -0.013 | Old |
| 0.818969 | 0.28469 | 0.206 | 0.2 | 1 | 4 | Atraid | 0.006 | Old |
| 0.818989 | 0.246954 | 0.196 | 0.188 | 1 | 4 | Safb2 | 0.008 | Old |
| 0.819017 | 0.09958 | 0.269 | 0.262 | 1 | 4 | Tmed3 | 0.007 | Old |
| 0.819084 | 0.332637 | 0.09 | 0.1 | 1 | 4 | AY036118 | -0.01 | Old |
| 0.819327 | 0.369957 | 0.016 | 0.013 | 1 | 4 | Epas1 | 0.003 | Old |
| 0.819327 | 0.141051 | 0.016 | 0.013 | 1 | 4 | Ifit3 | 0.003 | Old |
| 0.81935 | 0.00522 | 0.64 | 0.688 | 1 | 4 | Celf2 | -0.048 | Old |
| 0.819446 | 0.299769 | 0.144 | 0.138 | 1 | 4 | Adss | 0.006 | Old |
| 0.819459 | 0.077313 | 0.021 | 0.025 | 1 | 4 | Acss2 | -0.004 | Old |
| 0.81951 | 0.057373 | 0.043 | 0.038 | 1 | 4 | Cdk5 | 0.005 | Old |
| 0.819999 | 0.124488 | 0.091 | 0.1 | 1 | 4 | Snx14 | -0.009 | Old |
| 0.820222 | 0.51704 | 0.032 | 0.038 | 1 | 4 | Slc5a6 | -0.006 | Old |
| 0.820223 | 0.173886 | 0.233 | 0.225 | 1 | 4 | Cox20 | 0.008 | Old |
| 0.82043 | 0.314629 | 0.094 | 0.088 | 1 | 4 | Cdc14b | 0.006 | Old |
| 0.820802 | 0.500634 | 0.123 | 0.138 | 1 | 4 | Mocs2 | -0.015 | Old |
| 0.821003 | 0.302098 | 0.094 | 0.088 | 1 | 4 | Nrf1 | 0.006 | Old |
| 0.821006 | 0.240537 | 0.021 | 0.025 | 1 | 4 | Gimap1 | -0.004 | Old |
| 0.821125 | 0.083389 | 0.016 | 0.013 | 1 | 4 | Phldb3 | 0.003 | Old |
| 0.821358 | 0.429179 | 0.298 | 0.312 | 1 | 4 | Jup | -0.014 | Old |
| 0.821532 | 0.973747 | 0.078 | 0.075 | 1 | 4 | Ocel1 | 0.003 | Old |
| 0.82161 | 0.068011 | 0.128 | 0.138 | 1 | 4 | Atf1 | -0.01 | Old |
| 0.821866 | 0.612674 | 0.042 | 0.038 | 1 | 4 | Pan2 | 0.004 | Old |
| 0.821947 | 0.541376 | 0.078 | 0.088 | 1 | 4 | Gcc1 | -0.01 | Old |
| 0.822071 | 0.037462 | 0.01 | 0.013 | 1 | 4 | Sac3d1 | -0.003 | Old |
| 0.822186 | 0.006544 | 0.586 | 0.55 | 1 | 4 | Ywhab | 0.036 | Old |
| 0.82231 | 0.006754 | 0.426 | 0.462 | 1 | 4 | Rabac1 | -0.036 | Old |
| 0.822733 | 0.118919 | 0.208 | 0.2 | 1 | 4 | Nrbp1 | 0.008 | Old |
| 0.822738 | 0.742625 | 0.032 | 0.038 | 1 | 4 | Kctd18 | -0.006 | Old |
| 0.82298 | 0.717749 | 0.042 | 0.038 | 1 | 4 | Mospd1 | 0.004 | Old |
| 0.823172 | 0.070334 | 0.103 | 0.112 | 1 | 4 | Sphk2 | -0.009 | Old |
| 0.823192 | 0.3929 | 0.01 | 0.013 | 1 | 4 | Slc19a2 | -0.003 | Old |
| 0.823228 | 0.096545 | 0.08 | 0.088 | 1 | 4 | Ift52 | -0.008 | Old |
| 0.823694 | 0.577046 | 0.068 | 0.062 | 1 | 4 | Uba7 | 0.006 | Old |
| 0.82401 | 0.085623 | 0.288 | 0.275 | 1 | 4 | Timm13 | 0.013 | Old |
| 0.824138 | 0.264846 | 0.194 | 0.188 | 1 | 4 | Plekhj1 | 0.006 | Old |
| 0.824314 | 0.808082 | 0.01 | 0.013 | 1 | 4 | Arl16 | -0.003 | Old |
| 0.824314 | 0.531042 | 0.01 | 0.013 | 1 | 4 | Sh3rf1 | -0.003 | Old |
| 0.824728 | 0.068965 | 0.123 | 0.112 | 1 | 4 | Mrpl30 | 0.011 | Old |
| 0.824875 | 0.424669 | 0.021 | 0.025 | 1 | 4 | Frat2 | -0.004 | Old |
| 0.825058 | 0.118802 | 0.186 | 0.175 | 1 | 4 | Rchy1 | 0.011 | Old |
| 0.825398 | 0.80155 | 0.067 | 0.062 | 1 | 4 | Polr1c | 0.005 | Old |
| 0.825437 | 1.022138 | 0.01 | 0.013 | 1 | 4 | Zfp7 | -0.003 | Old |

| | | | | | | | | |
|---|---|---|---|---|---|---|---|---|
| 0.825437 | 0.894417 | 0.01 | 0.013 | 1 | 4 | Cldn10 | -0.003 | Old |
| 0.825437 | 0.446389 | 0.01 | 0.013 | 1 | 4 | Ipcef1 | -0.003 | Old |
| 0.825638 | 0.053692 | 0.045 | 0.05 | 1 | 4 | Tma16 | -0.005 | Old |
| 0.825886 | 0.971158 | 0.032 | 0.038 | 1 | 4 | Map4k3 | -0.006 | Old |
| 0.826045 | 0.568152 | 0.042 | 0.038 | 1 | 4 | Zfp414 | 0.004 | Old |
| 0.82652 | 0.017567 | 0.033 | 0.038 | 1 | 4 | Wdfy1 | -0.005 | Old |
| 0.826559 | 0.973277 | 0.01 | 0.013 | 1 | 4 | Cacnb4 | -0.003 | Old |
| 0.826559 | 0.961984 | 0.01 | 0.013 | 1 | 4 | AC151836. | -0.003 | Old |
| 0.826559 | 0.889648 | 0.01 | 0.013 | 1 | 4 | Snx32 | -0.003 | Old |
| 0.826559 | 0.834884 | 0.01 | 0.013 | 1 | 4 | Plpp1 | -0.003 | Old |
| 0.826881 | 0.547879 | 0.042 | 0.038 | 1 | 4 | Gm44751 | 0.004 | Old |
| 0.827006 | 0.190146 | 0.102 | 0.112 | 1 | 4 | Ddhd1 | -0.01 | Old |
| 0.82705 | 0.487843 | 0.093 | 0.088 | 1 | 4 | Dcaf13 | 0.005 | Old |
| 0.827098 | 0.815962 | 0.029 | 0.025 | 1 | 4 | Dhx37 | 0.004 | Old |
| 0.827199 | 0.585764 | 0.021 | 0.025 | 1 | 4 | Fam167b | -0.004 | Old |
| 0.827241 | 0.044848 | 0.177 | 0.188 | 1 | 4 | Kif1c | -0.011 | Old |
| 0.827468 | 0.180179 | 0.391 | 0.438 | 1 | 4 | Calm3 | -0.047 | Old |
| 0.827683 | 1.226641 | 0.01 | 0.013 | 1 | 4 | Cdkn2a | -0.003 | Old |
| 0.827683 | 1.166079 | 0.01 | 0.013 | 1 | 4 | Fzd4 | -0.003 | Old |
| 0.827683 | 0.956516 | 0.01 | 0.013 | 1 | 4 | Gm340 | -0.003 | Old |
| 0.827766 | 0.622746 | 0.029 | 0.025 | 1 | 4 | Wdr47 | 0.004 | Old |
| 0.828209 | 0.378254 | 0.269 | 0.275 | 1 | 4 | Sel1l | -0.006 | Old |
| 0.828404 | 0.413826 | 0.165 | 0.162 | 1 | 4 | Phb2 | 0.003 | Old |
| 0.828824 | 0.063077 | 0.309 | 0.338 | 1 | 4 | Ubald2 | -0.029 | Old |
| 0.828919 | 0.002247 | 0.069 | 0.075 | 1 | 4 | Smg6 | -0.006 | Old |
| 0.828927 | 0.221784 | 0.219 | 0.212 | 1 | 4 | Ino80d | 0.007 | Old |
| 0.829035 | 0.296717 | 0.132 | 0.125 | 1 | 4 | Rbck1 | 0.007 | Old |
| 0.829061 | 0.116729 | 0.056 | 0.062 | 1 | 4 | Trmt2b | -0.006 | Old |
| 0.829381 | 0.21274 | 0.081 | 0.075 | 1 | 4 | Usp40 | 0.006 | Old |
| 0.829524 | 0.856868 | 0.021 | 0.025 | 1 | 4 | Rpl36a-ps1 | -0.004 | Old |
| 0.829689 | 0.125913 | 0.196 | 0.188 | 1 | 4 | Maz | 0.008 | Old |
| 0.829751 | 0.308021 | 0.167 | 0.162 | 1 | 4 | Phf14 | 0.005 | Old |
| 0.830072 | 0.204455 | 0.121 | 0.112 | 1 | 4 | Gatd1 | 0.009 | Old |
| 0.830201 | 0.323688 | 0.264 | 0.262 | 1 | 4 | Creb1 | 0.002 | Old |
| 0.830443 | 0.484827 | 0.029 | 0.025 | 1 | 4 | Bloc1s1 | 0.004 | Old |
| 0.83079 | 0.118165 | 0.286 | 0.275 | 1 | 4 | 1-Mar | 0.011 | Old |
| 0.831029 | 0.695488 | 0.055 | 0.05 | 1 | 4 | Ddx27 | 0.005 | Old |
| 0.831038 | 0.100056 | 0.184 | 0.175 | 1 | 4 | Psmd6 | 0.009 | Old |
| 0.831043 | 0.032463 | 0.621 | 0.713 | 1 | 4 | Xbp1 | -0.092 | Old |
| 0.831251 | 0.41876 | 0.08 | 0.075 | 1 | 4 | Dap3 | 0.005 | Old |
| 0.831551 | 0.840413 | 0.042 | 0.038 | 1 | 4 | Ankmy2 | 0.004 | Old |
| 0.831781 | 0.389197 | 0.029 | 0.025 | 1 | 4 | Fbxw9 | 0.004 | Old |
| 0.832304 | 0.44568 | 0.187 | 0.188 | 1 | 4 | Ncf4 | -0.001 | Old |
| 0.832314 | 0.23885 | 0.25 | 0.275 | 1 | 4 | Mrps33 | -0.025 | Old |
| 0.832723 | 0.913169 | 0.113 | 0.112 | 1 | 4 | Hspa13 | 0.001 | Old |

| | | | | | | | | |
|---|---|---|---|---|---|---|---|---|
| 0.832978 | 0.597959 | 0.092 | 0.088 | 1 | 4 | Psmc2 | 0.004 | Old |
| 0.833116 | 0.101071 | 0.069 | 0.062 | 1 | 4 | Sdad1 | 0.007 | Old |
| 0.833173 | 0.495728 | 0.044 | 0.05 | 1 | 4 | Fpgt | -0.006 | Old |
| 0.833935 | 0.095078 | 0.416 | 0.412 | 1 | 4 | Selenos | 0.004 | Old |
| 0.83446 | 0.435048 | 0.029 | 0.025 | 1 | 4 | B3gnt3 | 0.004 | Old |
| 0.83446 | 0.404816 | 0.029 | 0.025 | 1 | 4 | Zfp346 | 0.004 | Old |
| 0.834606 | 0.052094 | 0.806 | 0.875 | 1 | 4 | Rpl5 | -0.069 | Old |
| 0.83487 | 0.279276 | 0.055 | 0.05 | 1 | 4 | Zfp26 | 0.005 | Old |
| 0.834976 | 0.573746 | 0.055 | 0.05 | 1 | 4 | Rnaseh2a | 0.005 | Old |
| 0.83502 | 0.33378 | 0.105 | 0.1 | 1 | 4 | Ube3c | 0.005 | Old |
| 0.83565 | 0.3329 | 0.235 | 0.238 | 1 | 4 | Usp8 | -0.003 | Old |
| 0.835895 | 0.084881 | 0.033 | 0.038 | 1 | 4 | Zmym6 | -0.005 | Old |
| 0.835917 | 1.088621 | 0.078 | 0.075 | 1 | 4 | Brix1 | 0.003 | Old |
| 0.836036 | 0.411464 | 0.32 | 0.35 | 1 | 4 | Irf2 | -0.03 | Old |
| 0.836164 | 0.448562 | 0.09 | 0.1 | 1 | 4 | Stx6 | -0.01 | Old |
| 0.836233 | 0.592776 | 0.139 | 0.138 | 1 | 4 | Ccndbp1 | 0.001 | Old |
| 0.83632 | 0.305613 | 0.142 | 0.138 | 1 | 4 | Fkbp3 | 0.004 | Old |
| 0.836375 | 0.281771 | 0.042 | 0.038 | 1 | 4 | Borcs5 | 0.004 | Old |
| 0.836375 | 0.231067 | 0.042 | 0.038 | 1 | 4 | Pnpt1 | 0.004 | Old |
| 0.836471 | 0.249116 | 0.029 | 0.025 | 1 | 4 | Kremen1 | 0.004 | Old |
| 0.836487 | 0.174913 | 0.365 | 0.362 | 1 | 4 | Ikbkb | 0.003 | Old |
| 0.836516 | 0.176194 | 0.219 | 0.238 | 1 | 4 | Alkbh5 | -0.019 | Old |
| 0.836939 | 0.281871 | 0.16 | 0.175 | 1 | 4 | Plod3 | -0.015 | Old |
| 0.837243 | 0.199042 | 0.16 | 0.175 | 1 | 4 | Arntl | -0.015 | Old |
| 0.837445 | 0.474516 | 0.055 | 0.05 | 1 | 4 | 4930453N2 | 0.005 | Old |
| 0.838007 | 0.133288 | 0.08 | 0.088 | 1 | 4 | Mphosph8 | -0.008 | Old |
| 0.838715 | 0.412031 | 0.136 | 0.15 | 1 | 4 | B3gat3 | -0.014 | Old |
| 0.838949 | 0.292586 | 0.367 | 0.375 | 1 | 4 | Ptpn6 | -0.008 | Old |
| 0.839153 | 0.126111 | 0.029 | 0.025 | 1 | 4 | Gm46430 | 0.004 | Old |
| 0.839323 | 0.214145 | 0.107 | 0.1 | 1 | 4 | Tmem128 | 0.007 | Old |
| 0.839403 | 0.324523 | 0.221 | 0.225 | 1 | 4 | Rnf4 | -0.004 | Old |
| 0.839483 | 0.161808 | 0.095 | 0.088 | 1 | 4 | Dnajc21 | 0.007 | Old |
| 0.839485 | 1.514437 | 0.028 | 0.025 | 1 | 4 | Lrrc28 | 0.003 | Old |
| 0.839824 | 0.032321 | 0.029 | 0.025 | 1 | 4 | Bclaf3 | 0.004 | Old |
| 0.839997 | 0.695825 | 0.042 | 0.038 | 1 | 4 | Fbxl15 | 0.004 | Old |
| 0.840068 | 0.034686 | 0.093 | 0.1 | 1 | 4 | Sfi1 | -0.007 | Old |
| 0.840091 | 1.72453 | 0.015 | 0.013 | 1 | 4 | Tmem144 | 0.002 | Old |
| 0.840091 | 1.672164 | 0.015 | 0.013 | 1 | 4 | Fadd | 0.002 | Old |
| 0.84015 | 0.052331 | 0.082 | 0.088 | 1 | 4 | Mis12 | -0.006 | Old |
| 0.840164 | 0.311761 | 0.154 | 0.15 | 1 | 4 | Map3k2 | 0.004 | Old |
| 0.84041 | 0.125977 | 0.368 | 0.362 | 1 | 4 | Hmgb2 | 0.006 | Old |
| 0.840664 | 0.825292 | 0.113 | 0.112 | 1 | 4 | Lcorl | 0.001 | Old |
| 0.840761 | 0.447758 | 0.101 | 0.112 | 1 | 4 | Polg | -0.011 | Old |
| 0.841013 | 1.64968 | 0.015 | 0.013 | 1 | 4 | Lyrm1 | 0.002 | Old |
| 0.841013 | 1.412382 | 0.015 | 0.013 | 1 | 4 | Fam206a | 0.002 | Old |

| | | | | | | | | |
|---|---|---|---|---|---|---|---|---|
| 0.841101 | 0.176367 | 0.132 | 0.125 | 1 | 4 | S100a11 | 0.007 | Old |
| 0.841322 | 0.060383 | 0.233 | 0.25 | 1 | 4 | Csnk1d | -0.017 | Old |
| 0.841749 | 0.168581 | 0.055 | 0.05 | 1 | 4 | Rhbdf1 | 0.005 | Old |
| 0.841837 | 0.114306 | 0.029 | 0.025 | 1 | 4 | Dppa3 | 0.004 | Old |
| 0.841936 | 1.271076 | 0.015 | 0.013 | 1 | 4 | P3h3 | 0.002 | Old |
| 0.841936 | 1.178983 | 0.015 | 0.013 | 1 | 4 | Brca2 | 0.002 | Old |
| 0.842404 | 0.558652 | 0.104 | 0.1 | 1 | 4 | Dopey1 | 0.004 | Old |
| 0.842413 | 0.125531 | 0.132 | 0.125 | 1 | 4 | Ccdc71l | 0.007 | Old |
| 0.842735 | 0.096 | 0.526 | 0.562 | 1 | 4 | Mtdh | -0.036 | Old |
| 0.842774 | 0.224203 | 0.236 | 0.262 | 1 | 4 | Rc3h1 | -0.026 | Old |
| 0.842852 | 0.328561 | 0.102 | 0.112 | 1 | 4 | Zfp384 | -0.01 | Old |
| 0.842859 | 1.364989 | 0.015 | 0.013 | 1 | 4 | 4632404H1 | 0.002 | Old |
| 0.842859 | 1.242397 | 0.015 | 0.013 | 1 | 4 | Nkrf | 0.002 | Old |
| 0.842859 | 0.970132 | 0.015 | 0.013 | 1 | 4 | Gm42715 | 0.002 | Old |
| 0.843402 | 0.120946 | 0.176 | 0.188 | 1 | 4 | Rnf187 | -0.012 | Old |
| 0.843617 | 0.139126 | 0.388 | 0.4 | 1 | 4 | Dennd5a | -0.012 | Old |
| 0.84365 | 0.203146 | 0.042 | 0.038 | 1 | 4 | Zbtb33 | 0.004 | Old |
| 0.843678 | 0.017574 | 0.915 | 0.938 | 1 | 4 | Rpl37 | -0.023 | Old |
| 0.843728 | 0.117574 | 0.247 | 0.262 | 1 | 4 | Ncoa3 | -0.015 | Old |
| 0.844092 | 0.287384 | 0.155 | 0.15 | 1 | 4 | Sec24b | 0.005 | Old |
| 0.844706 | 0.738255 | 0.015 | 0.013 | 1 | 4 | Tmem221 | 0.002 | Old |
| 0.844706 | 0.544654 | 0.015 | 0.013 | 1 | 4 | Nr3c2 | 0.002 | Old |
| 0.844796 | 0.058334 | 0.652 | 0.688 | 1 | 4 | Il6ra | -0.036 | Old |
| 0.845018 | 0.743407 | 0.054 | 0.05 | 1 | 4 | Trp53bp1 | 0.004 | Old |
| 0.845085 | 0.316933 | 0.245 | 0.25 | 1 | 4 | Oat | -0.005 | Old |
| 0.84563 | 0.643282 | 0.015 | 0.013 | 1 | 4 | Nt5m | 0.002 | Old |
| 0.845639 | 0.558714 | 0.042 | 0.038 | 1 | 4 | Endod1 | 0.004 | Old |
| 0.845843 | 0.234511 | 0.126 | 0.138 | 1 | 4 | Opa1 | -0.012 | Old |
| 0.846239 | 0.078272 | 0.3 | 0.325 | 1 | 4 | Mia2 | -0.025 | Old |
| 0.846375 | 0.132624 | 0.17 | 0.162 | 1 | 4 | Naa15 | 0.008 | Old |
| 0.846416 | 0.162565 | 0.145 | 0.138 | 1 | 4 | Ptdss1 | 0.007 | Old |
| 0.84651 | 1.156947 | 0.041 | 0.038 | 1 | 4 | Rbm12b2 | 0.003 | Old |
| 0.846553 | 0.526319 | 0.015 | 0.013 | 1 | 4 | Mrps27 | 0.002 | Old |
| 0.846594 | 0.150423 | 0.289 | 0.288 | 1 | 4 | Slk | 0.001 | Old |
| 0.847176 | 0.605039 | 0.033 | 0.038 | 1 | 4 | Zfp426 | -0.005 | Old |
| 0.847186 | 0.062252 | 0.081 | 0.088 | 1 | 4 | Ppdpf | -0.007 | Old |
| 0.847217 | 0.02365 | 0.022 | 0.025 | 1 | 4 | Ctu1 | -0.003 | Old |
| 0.847478 | 0.907894 | 0.015 | 0.013 | 1 | 4 | Inpp5b | 0.002 | Old |
| 0.848402 | 0.430588 | 0.015 | 0.013 | 1 | 4 | Gm13091 | 0.002 | Old |
| 0.848402 | 0.290947 | 0.015 | 0.013 | 1 | 4 | Mccc2 | 0.002 | Old |
| 0.84847 | 0.464468 | 0.115 | 0.112 | 1 | 4 | Ctbp2 | 0.003 | Old |
| 0.848587 | 0.054215 | 0.159 | 0.15 | 1 | 4 | Htt | 0.009 | Old |
| 0.848678 | 0.327621 | 0.093 | 0.088 | 1 | 4 | Foxo1 | 0.005 | Old |
| 0.848746 | 0.187886 | 0.247 | 0.275 | 1 | 4 | Vdac1 | -0.028 | Old |
| 0.848766 | 0.088308 | 0.838 | 0.863 | 1 | 4 | Pabpc1 | -0.025 | Old |

| | | | | | | | | |
|---|---|---|---|---|---|---|---|---|
| 0.849327 | 0.48162 | 0.015 | 0.013 | 1 | 4 | Acp6 | 0.002 | Old |
| 0.849327 | 0.342096 | 0.015 | 0.013 | 1 | 4 | Eral1 | 0.002 | Old |
| 0.849388 | 0.423868 | 0.104 | 0.1 | 1 | 4 | Nrbf2 | 0.004 | Old |
| 0.849566 | 0.405231 | 0.055 | 0.05 | 1 | 4 | Caap1 | 0.005 | Old |
| 0.849592 | 0.611142 | 0.042 | 0.038 | 1 | 4 | Cblb | 0.004 | Old |
| 0.849636 | 0.059753 | 0.385 | 0.388 | 1 | 4 | Capns1 | -0.003 | Old |
| 0.849725 | 0.271509 | 0.202 | 0.2 | 1 | 4 | Fam49a | 0.002 | Old |
| 0.850259 | 0.025324 | 0.142 | 0.15 | 1 | 4 | Smad2 | -0.008 | Old |
| 0.850503 | 0.934897 | 0.067 | 0.075 | 1 | 4 | Zfp410 | -0.008 | Old |
| 0.851143 | 0.184463 | 0.15 | 0.162 | 1 | 4 | Msl3 | -0.012 | Old |
| 0.851176 | 0.171441 | 0.015 | 0.013 | 1 | 4 | A630072M | 0.002 | Old |
| 0.851176 | 0.153839 | 0.015 | 0.013 | 1 | 4 | Abcc4 | 0.002 | Old |
| 0.851285 | 0.053719 | 0.4 | 0.412 | 1 | 4 | Atxn7l3b | -0.012 | Old |
| 0.85136 | 0.058865 | 0.108 | 0.1 | 1 | 4 | Txndc16 | 0.008 | Old |
| 0.851583 | 0.359722 | 0.045 | 0.05 | 1 | 4 | Med6 | -0.005 | Old |
| 0.851584 | 0.019801 | 0.092 | 0.1 | 1 | 4 | Mavs | -0.008 | Old |
| 0.851943 | 0.200852 | 0.273 | 0.275 | 1 | 4 | Snrpd3 | -0.002 | Old |
| 0.851957 | 0.004593 | 0.034 | 0.038 | 1 | 4 | Rbm45 | -0.004 | Old |
| 0.852101 | 0.194618 | 0.015 | 0.013 | 1 | 4 | Agrn | 0.002 | Old |
| 0.852125 | 0.403708 | 0.045 | 0.05 | 1 | 4 | Rpa2 | -0.005 | Old |
| 0.853029 | 0.328101 | 0.217 | 0.225 | 1 | 4 | Phf20 | -0.008 | Old |
| 0.853038 | 0.389352 | 0.08 | 0.075 | 1 | 4 | Oma1 | 0.005 | Old |
| 0.85307 | 0.519736 | 0.028 | 0.025 | 1 | 4 | Ggcx | 0.003 | Old |
| 0.853157 | 0.227552 | 0.068 | 0.062 | 1 | 4 | Cyp2j6 | 0.006 | Old |
| 0.853899 | 0.221628 | 0.057 | 0.062 | 1 | 4 | Tmem39a | -0.005 | Old |
| 0.853915 | 0.138144 | 0.312 | 0.312 | 1 | 4 | Wdr83os | 0 | Old |
| 0.854267 | 0.064759 | 0.644 | 0.65 | 1 | 4 | Eps15 | -0.006 | Old |
| 0.854484 | 0.71369 | 0.054 | 0.05 | 1 | 4 | Prkch | 0.004 | Old |
| 0.854484 | 0.544438 | 0.054 | 0.05 | 1 | 4 | Zbtb6 | 0.004 | Old |
| 0.854837 | 0.039692 | 0.449 | 0.425 | 1 | 4 | Cndp2 | 0.024 | Old |
| 0.85492 | 0.305628 | 0.225 | 0.225 | 1 | 4 | Zc3h7a | 0 | Old |
| 0.855673 | 0.243134 | 0.022 | 0.025 | 1 | 4 | Zfp280b | -0.003 | Old |
| 0.855727 | 0.258569 | 0.08 | 0.088 | 1 | 4 | Cacna1d | -0.008 | Old |
| 0.855779 | 0.139462 | 0.045 | 0.05 | 1 | 4 | Dnlz | -0.005 | Old |
| 0.85582 | 0.118202 | 0.081 | 0.088 | 1 | 4 | Grk5 | -0.007 | Old |
| 0.856255 | 0.035622 | 0.141 | 0.15 | 1 | 4 | Pih1d1 | -0.009 | Old |
| 0.856461 | 0.038688 | 0.08 | 0.075 | 1 | 4 | Trappc6a | 0.005 | Old |
| 0.856533 | 0.569806 | 0.173 | 0.175 | 1 | 4 | Cant1 | -0.002 | Old |
| 0.85691 | 0.349872 | 0.154 | 0.15 | 1 | 4 | Yipf1 | 0.004 | Old |
| 0.857041 | 0.367166 | 0.117 | 0.112 | 1 | 4 | Stard5 | 0.005 | Old |
| 0.857084 | 0.009263 | 0.226 | 0.238 | 1 | 4 | Mdh2 | -0.012 | Old |
| 0.857154 | 0.375598 | 0.028 | 0.025 | 1 | 4 | Gm17745 | 0.003 | Old |
| 0.857441 | 0.067814 | 0.13 | 0.138 | 1 | 4 | Wbp4 | -0.008 | Old |
| 0.857485 | 0.313003 | 0.201 | 0.2 | 1 | 4 | Tecr | 0.001 | Old |
| 0.857497 | 0.443597 | 0.08 | 0.088 | 1 | 4 | Gsap | -0.008 | Old |

| | | | | | | | | |
|---|---|---|---|---|---|---|---|---|
| 0.857688 | 0.225405 | 0.577 | 0.625 | 1 | 4 | Trps1 | -0.048 | Old |
| 0.85769 | 0.129923 | 0.105 | 0.112 | 1 | 4 | Tlr1 | -0.007 | Old |
| 0.857699 | 0.838856 | 0.053 | 0.05 | 1 | 4 | Hps4 | 0.003 | Old |
| 0.857793 | 0.066211 | 0.081 | 0.075 | 1 | 4 | Cpsf3 | 0.006 | Old |
| 0.85783 | 0.092812 | 0.093 | 0.1 | 1 | 4 | Pon3 | -0.007 | Old |
| 0.857835 | 0.279157 | 0.028 | 0.025 | 1 | 4 | D3Ertd751 | 0.003 | Old |
| 0.857873 | 0.067834 | 0.069 | 0.075 | 1 | 4 | Rfwd3 | -0.006 | Old |
| 0.857976 | 0.293934 | 0.054 | 0.05 | 1 | 4 | Fra10ac1 | 0.004 | Old |
| 0.858077 | 0.303433 | 0.042 | 0.038 | 1 | 4 | Unk | 0.004 | Old |
| 0.858516 | 0.384422 | 0.028 | 0.025 | 1 | 4 | Slc22a15 | 0.003 | Old |
| 0.858671 | 0.486412 | 0.067 | 0.062 | 1 | 4 | Asb7 | 0.005 | Old |
| 0.859042 | 0.863896 | 0.041 | 0.038 | 1 | 4 | Uck1 | 0.003 | Old |
| 0.859123 | 0.462773 | 0.067 | 0.062 | 1 | 4 | Fcf1 | 0.005 | Old |
| 0.859342 | 0.18294 | 0.106 | 0.1 | 1 | 4 | Copg1 | 0.006 | Old |
| 0.859624 | 0.184611 | 0.236 | 0.238 | 1 | 4 | Brd1 | -0.002 | Old |
| 0.859815 | 0.093223 | 0.429 | 0.462 | 1 | 4 | Zfp706 | -0.033 | Old |
| 0.860059 | 0.481637 | 0.034 | 0.038 | 1 | 4 | Pcca | -0.004 | Old |
| 0.860071 | 0.122653 | 0.667 | 0.688 | 1 | 4 | Ddx17 | -0.021 | Old |
| 0.860544 | 0.187993 | 0.401 | 0.425 | 1 | 4 | Itpripl2 | -0.024 | Old |
| 0.860571 | 0.0263 | 0.18 | 0.188 | 1 | 4 | Cpsf2 | -0.008 | Old |
| 0.860827 | 0.048641 | 0.19 | 0.188 | 1 | 4 | Psmb5 | 0.002 | Old |
| 0.860896 | 0.162618 | 0.144 | 0.138 | 1 | 4 | Spty2d1 | 0.006 | Old |
| 0.861682 | 0.184149 | 0.132 | 0.125 | 1 | 4 | Twistnb | 0.007 | Old |
| 0.861694 | 0.740954 | 0.056 | 0.062 | 1 | 4 | Aktip | -0.006 | Old |
| 0.861722 | 0.215525 | 0.055 | 0.05 | 1 | 4 | Fam160a2 | 0.005 | Old |
| 0.861899 | 0.470265 | 0.045 | 0.05 | 1 | 4 | Dhx58 | -0.005 | Old |
| 0.862556 | 0.236154 | 0.034 | 0.038 | 1 | 4 | Gm7694 | -0.004 | Old |
| 0.862604 | 0.989092 | 0.022 | 0.025 | 1 | 4 | Gpank1 | -0.003 | Old |
| 0.86318 | 0.116507 | 0.509 | 0.562 | 1 | 4 | Selenok | -0.053 | Old |
| 0.8632 | 0.335069 | 0.092 | 0.1 | 1 | 4 | Upf3b | -0.008 | Old |
| 0.863211 | 0.026791 | 0.332 | 0.362 | 1 | 4 | Rnpep | -0.03 | Old |
| 0.863285 | 0.113758 | 0.153 | 0.162 | 1 | 4 | Sdhb | -0.009 | Old |
| 0.863287 | 0.060404 | 0.028 | 0.025 | 1 | 4 | Map4k2 | 0.003 | Old |
| 0.863287 | 0.054934 | 0.028 | 0.025 | 1 | 4 | Clybl | 0.003 | Old |
| 0.863736 | 0.747856 | 0.066 | 0.062 | 1 | 4 | Tmem97 | 0.004 | Old |
| 0.863859 | 0.115359 | 0.08 | 0.075 | 1 | 4 | Ifih1 | 0.005 | Old |
| 0.863969 | 0.031531 | 0.028 | 0.025 | 1 | 4 | Mblac2 | 0.003 | Old |
| 0.86418 | 0.082541 | 0.158 | 0.15 | 1 | 4 | Tor1b | 0.008 | Old |
| 0.864309 | 0.242424 | 0.042 | 0.038 | 1 | 4 | Hsd17b7 | 0.004 | Old |
| 0.864351 | 0.165599 | 0.157 | 0.15 | 1 | 4 | Chmp1b | 0.007 | Old |
| 0.864586 | 0.016672 | 0.279 | 0.275 | 1 | 4 | Lypla2 | 0.004 | Old |
| 0.864876 | 0.166239 | 0.042 | 0.038 | 1 | 4 | Abcg2 | 0.004 | Old |
| 0.864876 | 0.127991 | 0.042 | 0.038 | 1 | 4 | Acp5 | 0.004 | Old |
| 0.864957 | 0.149691 | 0.045 | 0.05 | 1 | 4 | Kantr | -0.005 | Old |
| 0.86499 | 1.385391 | 0.053 | 0.05 | 1 | 4 | Vhl | 0.003 | Old |

| | | | | | | | | |
|---|---|---|---|---|---|---|---|---|
| 0.86511 | 0.328883 | 0.08 | 0.075 | 1 | 4 | Btbd9 | 0.005 | Old |
| 0.865201 | 0.108661 | 0.055 | 0.05 | 1 | 4 | Ccdc43 | 0.005 | Old |
| 0.865248 | 1.276018 | 0.028 | 0.025 | 1 | 4 | Tspyl4 | 0.003 | Old |
| 0.865336 | 0.183893 | 0.309 | 0.312 | 1 | 4 | Clint1 | -0.003 | Old |
| 0.865497 | 0.240081 | 0.045 | 0.05 | 1 | 4 | Dctpp1 | -0.005 | Old |
| 0.865737 | 0.658657 | 0.053 | 0.05 | 1 | 4 | Rfx5 | 0.003 | Old |
| 0.865887 | 0.384727 | 0.194 | 0.212 | 1 | 4 | Scamp3 | -0.018 | Old |
| 0.865893 | 0.605818 | 0.041 | 0.038 | 1 | 4 | Cables2 | 0.003 | Old |
| 0.866044 | 0.059076 | 0.584 | 0.625 | 1 | 4 | Sema4a | -0.041 | Old |
| 0.866097 | 0.833798 | 0.056 | 0.062 | 1 | 4 | Zfp729a | -0.006 | Old |
| 0.866101 | 0.243347 | 0.118 | 0.112 | 1 | 4 | Mcoln2 | 0.006 | Old |
| 0.866302 | 0.299197 | 0.034 | 0.038 | 1 | 4 | Zfp758 | -0.004 | Old |
| 0.866494 | 0.192759 | 0.057 | 0.062 | 1 | 4 | Fam110a | -0.005 | Old |
| 0.866524 | 0.328673 | 0.184 | 0.2 | 1 | 4 | Frg1 | -0.016 | Old |
| 0.866562 | 0.759676 | 0.065 | 0.062 | 1 | 4 | Zdhhc18 | 0.003 | Old |
| 0.866628 | 1.168503 | 0.028 | 0.025 | 1 | 4 | Galt | 0.003 | Old |
| 0.866651 | 0.559218 | 0.078 | 0.075 | 1 | 4 | Vta1 | 0.003 | Old |
| 0.866944 | 0.384766 | 0.164 | 0.162 | 1 | 4 | Ip6k1 | 0.002 | Old |
| 0.86706 | 0.217525 | 0.186 | 0.2 | 1 | 4 | Pi4k2a | -0.014 | Old |
| 0.867122 | 0.578165 | 0.102 | 0.1 | 1 | 4 | Fam199x | 0.002 | Old |
| 0.867204 | 0.18097 | 0.186 | 0.2 | 1 | 4 | Vamp2 | -0.014 | Old |
| 0.867233 | 0.16417 | 0.011 | 0.013 | 1 | 4 | Trmt44 | -0.002 | Old |
| 0.867349 | 0.412704 | 0.114 | 0.125 | 1 | 4 | Ccnh | -0.011 | Old |
| 0.867418 | 0.483382 | 0.09 | 0.088 | 1 | 4 | Qars | 0.002 | Old |
| 0.867748 | 0.767663 | 0.053 | 0.05 | 1 | 4 | Tmem70 | 0.003 | Old |
| 0.867765 | 0.389975 | 0.14 | 0.138 | 1 | 4 | Cwf19l2 | 0.002 | Old |
| 0.867895 | 0.499919 | 0.163 | 0.162 | 1 | 4 | Psmb7 | 0.001 | Old |
| 0.868151 | 0.180459 | 0.105 | 0.1 | 1 | 4 | Tmem184b | 0.005 | Old |
| 0.868227 | 0.653069 | 0.09 | 0.088 | 1 | 4 | Ppm1k | 0.002 | Old |
| 0.86828 | 0.12881 | 0.042 | 0.038 | 1 | 4 | Wwox | 0.004 | Old |
| 0.868333 | 0.096594 | 0.011 | 0.013 | 1 | 4 | Ifi214 | -0.002 | Old |
| 0.868333 | 0.068092 | 0.011 | 0.013 | 1 | 4 | Cwf19l1 | -0.002 | Old |
| 0.868667 | 0.136834 | 0.301 | 0.325 | 1 | 4 | Aldoa | -0.024 | Old |
| 0.86869 | 0.30895 | 0.248 | 0.25 | 1 | 4 | Emc1 | -0.002 | Old |
| 0.868739 | 0.054373 | 0.676 | 0.7 | 1 | 4 | Cox7c | -0.024 | Old |
| 0.869258 | 0.526877 | 0.053 | 0.05 | 1 | 4 | Serinc5 | 0.003 | Old |
| 0.869377 | 0.343946 | 0.08 | 0.088 | 1 | 4 | Jak2 | -0.008 | Old |
| 0.869434 | 0.132357 | 0.011 | 0.013 | 1 | 4 | Isg20 | -0.002 | Old |
| 0.869474 | 0.140005 | 0.167 | 0.162 | 1 | 4 | Sec23b | 0.005 | Old |
| 0.869918 | 0.207958 | 0.431 | 0.462 | 1 | 4 | Sumo3 | -0.031 | Old |
| 0.870008 | 0.140887 | 0.283 | 0.288 | 1 | 4 | Phf3 | -0.005 | Old |
| 0.870536 | 0.35712 | 0.011 | 0.013 | 1 | 4 | Btbd19 | -0.002 | Old |
| 0.870536 | 0.182821 | 0.011 | 0.013 | 1 | 4 | Lyplal1 | -0.002 | Old |
| 0.870536 | 0.058584 | 0.011 | 0.013 | 1 | 4 | Bbs5 | -0.002 | Old |
| 0.870795 | 0.598235 | 0.066 | 0.062 | 1 | 4 | Klhl21 | 0.004 | Old |

| | | | | | | | | |
|---|---|---|---|---|---|---|---|---|
| 0.871002 | 0.158483 | 0.092 | 0.088 | 1 | 4 | Tmem63a | 0.004 | Old |
| 0.871002 | 0.263969 | 0.092 | 0.088 | 1 | 4 | Vps54 | 0.004 | Old |
| 0.871038 | 0.814354 | 0.041 | 0.038 | 1 | 4 | Map4k1 | 0.003 | Old |
| 0.871394 | 0.295517 | 0.092 | 0.1 | 1 | 4 | Vps33a | -0.008 | Old |
| 0.871637 | 0.762688 | 0.011 | 0.013 | 1 | 4 | Acbd4 | -0.002 | Old |
| 0.871637 | 0.4364 | 0.011 | 0.013 | 1 | 4 | Zfp51 | -0.002 | Old |
| 0.871637 | 0.093374 | 0.011 | 0.013 | 1 | 4 | 4833418N( | -0.002 | Old |
| 0.871941 | 0.02353 | 0.838 | 0.8 | 1 | 4 | Hexa | 0.038 | Old |
| 0.872101 | 1.204927 | 0.04 | 0.038 | 1 | 4 | Dtnb | 0.002 | Old |
| 0.872164 | 0.338924 | 0.126 | 0.138 | 1 | 4 | Galnt11 | -0.012 | Old |
| 0.872182 | 0.383416 | 0.041 | 0.038 | 1 | 4 | Orc5 | 0.003 | Old |
| 0.872302 | 0.064787 | 0.248 | 0.262 | 1 | 4 | Setd2 | -0.014 | Old |
| 0.872401 | 0.475041 | 0.126 | 0.125 | 1 | 4 | Ndufs4 | 0.001 | Old |
| 0.872528 | 0.453661 | 0.045 | 0.05 | 1 | 4 | Cep83 | -0.005 | Old |
| 0.872541 | 0.046599 | 0.23 | 0.225 | 1 | 4 | Ndufab1 | 0.005 | Old |
| 0.872625 | 0.008382 | 0.117 | 0.112 | 1 | 4 | Mcrs1 | 0.005 | Old |
| 0.872739 | 0.810937 | 0.011 | 0.013 | 1 | 4 | Pde2a | -0.002 | Old |
| 0.872739 | 0.278656 | 0.011 | 0.013 | 1 | 4 | Gm26522 | -0.002 | Old |
| 0.873039 | 0.260159 | 0.188 | 0.188 | 1 | 4 | Vav1 | 0 | Old |
| 0.873604 | 0.029797 | 0.082 | 0.088 | 1 | 4 | Josd1 | -0.006 | Old |
| 0.873841 | 0.380115 | 0.011 | 0.013 | 1 | 4 | Dnaaf5 | -0.002 | Old |
| 0.873871 | 0.570195 | 0.079 | 0.088 | 1 | 4 | Mrpl45 | -0.009 | Old |
| 0.873914 | 0.177947 | 0.301 | 0.3 | 1 | 4 | Nek7 | 0.001 | Old |
| 0.874152 | 0.494263 | 0.045 | 0.05 | 1 | 4 | Ppp1r3b | -0.005 | Old |
| 0.874218 | 0.601637 | 0.078 | 0.075 | 1 | 4 | Dnajc16 | 0.003 | Old |
| 0.874255 | 0.189096 | 0.079 | 0.075 | 1 | 4 | Rras2 | 0.004 | Old |
| 0.874778 | 0.416364 | 0.175 | 0.175 | 1 | 4 | Fam172a | 0 | Old |
| 0.874944 | 0.916529 | 0.011 | 0.013 | 1 | 4 | Zfp958 | -0.002 | Old |
| 0.874944 | 0.748671 | 0.011 | 0.013 | 1 | 4 | Dact3 | -0.002 | Old |
| 0.874944 | 0.695187 | 0.011 | 0.013 | 1 | 4 | Ficd | -0.002 | Old |
| 0.875047 | 1.626594 | 0.014 | 0.013 | 1 | 4 | Gpatch3 | 0.001 | Old |
| 0.875047 | 1.529247 | 0.014 | 0.013 | 1 | 4 | Scn3b | 0.001 | Old |
| 0.875047 | 1.212795 | 0.014 | 0.013 | 1 | 4 | Capn10 | 0.001 | Old |
| 0.875333 | 0.187115 | 0.417 | 0.438 | 1 | 4 | Kif5b | -0.021 | Old |
| 0.875372 | 0.088284 | 0.467 | 0.488 | 1 | 4 | Nap1l1 | -0.021 | Old |
| 0.875416 | 0.171216 | 0.257 | 0.262 | 1 | 4 | Csk | -0.005 | Old |
| 0.875588 | 0.37788 | 0.182 | 0.2 | 1 | 4 | St3gal1 | -0.018 | Old |
| 0.875668 | 0.057005 | 0.358 | 0.375 | 1 | 4 | Srp14 | -0.017 | Old |
| 0.87569 | 0.249905 | 0.092 | 0.088 | 1 | 4 | Ftsj3 | 0.004 | Old |
| 0.875997 | 1.720516 | 0.014 | 0.013 | 1 | 4 | Jchain | 0.001 | Old |
| 0.875997 | 1.092737 | 0.014 | 0.013 | 1 | 4 | Plppr4 | 0.001 | Old |
| 0.876029 | 0.1504 | 0.354 | 0.388 | 1 | 4 | Gnpda1 | -0.034 | Old |
| 0.876035 | 0.181683 | 0.046 | 0.05 | 1 | 4 | St3gal5 | -0.004 | Old |
| 0.876046 | 1.326192 | 0.011 | 0.013 | 1 | 4 | Rcn1 | -0.002 | Old |
| 0.876046 | 1.065699 | 0.011 | 0.013 | 1 | 4 | Rgs5 | -0.002 | Old |

| | | | | | | | | |
|---|---|---|---|---|---|---|---|---|
| 0.876046 | 0.689404 | 0.011 | 0.013 | 1 | 4 | Prr7 | -0.002 | Old |
| 0.876046 | 0.551881 | 0.011 | 0.013 | 1 | 4 | Mtcp1 | -0.002 | Old |
| 0.876137 | 0.926418 | 0.04 | 0.038 | 1 | 4 | Gpr89 | 0.002 | Old |
| 0.876137 | 0.914331 | 0.04 | 0.038 | 1 | 4 | Fam151b | 0.002 | Old |
| 0.876276 | 0.197589 | 0.092 | 0.088 | 1 | 4 | Hlcs | 0.004 | Old |
| 0.876297 | 0.636837 | 0.028 | 0.025 | 1 | 4 | Lsm11 | 0.003 | Old |
| 0.87662 | 0.194745 | 0.379 | 0.388 | 1 | 4 | Man2a1 | -0.009 | Old |
| 0.876725 | 0.254838 | 0.209 | 0.225 | 1 | 4 | Hs6st1 | -0.016 | Old |
| 0.876947 | 1.180083 | 0.014 | 0.013 | 1 | 4 | Atp6v0c | 0.001 | Old |
| 0.876947 | 0.923894 | 0.014 | 0.013 | 1 | 4 | Atp6v1g2 | 0.001 | Old |
| 0.877149 | 1.415195 | 0.011 | 0.013 | 1 | 4 | Rhebl1 | -0.002 | Old |
| 0.877179 | 0.04119 | 0.262 | 0.262 | 1 | 4 | Ndufs7 | 0 | Old |
| 0.877315 | 0.425111 | 0.053 | 0.05 | 1 | 4 | Rusc2 | 0.003 | Old |
| 0.877513 | 0.096774 | 0.13 | 0.125 | 1 | 4 | Map3k5 | 0.005 | Old |
| 0.87768 | 0.642519 | 0.028 | 0.025 | 1 | 4 | Racgap1 | 0.003 | Old |
| 0.878297 | 0.004708 | 0.513 | 0.525 | 1 | 4 | Tgfbr2 | -0.012 | Old |
| 0.878303 | 0.026194 | 0.854 | 0.85 | 1 | 4 | Lcp1 | 0.004 | Old |
| 0.878479 | 0.283121 | 0.041 | 0.038 | 1 | 4 | Ralgapa2 | 0.003 | Old |
| 0.878756 | 0.088942 | 0.383 | 0.4 | 1 | 4 | Lbh | -0.017 | Old |
| 0.878847 | 0.974136 | 0.014 | 0.013 | 1 | 4 | Polrmt | 0.001 | Old |
| 0.878847 | 0.882628 | 0.014 | 0.013 | 1 | 4 | Sgsm3 | 0.001 | Old |
| 0.879063 | 0.745977 | 0.028 | 0.025 | 1 | 4 | Tjap1 | 0.003 | Old |
| 0.879063 | 0.506945 | 0.028 | 0.025 | 1 | 4 | Fam210b | 0.003 | Old |
| 0.879122 | 0.601343 | 0.069 | 0.075 | 1 | 4 | Mmp19 | -0.006 | Old |
| 0.879265 | 0.181149 | 0.046 | 0.05 | 1 | 4 | Eif2b2 | -0.004 | Old |
| 0.879331 | 0.390527 | 0.053 | 0.05 | 1 | 4 | Fmo5 | 0.003 | Old |
| 0.879569 | 0.410079 | 0.045 | 0.05 | 1 | 4 | Pigv | -0.005 | Old |
| 0.879625 | 0.166915 | 0.041 | 0.038 | 1 | 4 | Mfge8 | 0.003 | Old |
| 0.879798 | 0.797923 | 0.014 | 0.013 | 1 | 4 | Nxn | 0.001 | Old |
| 0.879798 | 0.681472 | 0.014 | 0.013 | 1 | 4 | A130077B1 | 0.001 | Old |
| 0.879798 | 0.391083 | 0.014 | 0.013 | 1 | 4 | Trmt5 | 0.001 | Old |
| 0.879959 | 0.289594 | 0.081 | 0.088 | 1 | 4 | Sorbs3 | -0.007 | Old |
| 0.880173 | 0.257935 | 0.117 | 0.125 | 1 | 4 | Ddx18 | -0.008 | Old |
| 0.880541 | 0.017571 | 0.948 | 0.963 | 1 | 4 | H2-D1 | -0.015 | Old |
| 0.880598 | 0.078233 | 0.167 | 0.162 | 1 | 4 | Dtx3l | 0.005 | Old |
| 0.880609 | 0.141901 | 0.348 | 0.388 | 1 | 4 | Copz1 | -0.04 | Old |
| 0.880748 | 0.360789 | 0.014 | 0.013 | 1 | 4 | Hdac8 | 0.001 | Old |
| 0.880868 | 0.127485 | 0.338 | 0.362 | 1 | 4 | Limd1 | -0.024 | Old |
| 0.881143 | 0.015478 | 0.437 | 0.462 | 1 | 4 | Rbm25 | -0.025 | Old |
| 0.881206 | 0.243573 | 0.405 | 0.425 | 1 | 4 | Prrc2a | -0.02 | Old |
| 0.881245 | 0.108683 | 0.619 | 0.688 | 1 | 4 | Ms4a6c | -0.069 | Old |
| 0.88153 | 0.145081 | 0.238 | 0.238 | 1 | 4 | Dbnl | 0 | Old |
| 0.881546 | 0.20778 | 0.336 | 0.375 | 1 | 4 | Vps35 | -0.039 | Old |
| 0.881699 | 0.54269 | 0.014 | 0.013 | 1 | 4 | Il1a | 0.001 | Old |
| 0.881699 | 0.865679 | 0.014 | 0.013 | 1 | 4 | Lmo1 | 0.001 | Old |

| | | | | | | | | |
|---|---|---|---|---|---|---|---|---|
| 0.881809 | 0.199974 | 0.14 | 0.138 | 1 | 4 | Pnrc2 | 0.002 | Old |
| 0.88183 | 0.521637 | 0.028 | 0.025 | 1 | 4 | Ehf | 0.003 | Old |
| 0.882173 | 0.308635 | 0.092 | 0.088 | 1 | 4 | Hdac9 | 0.004 | Old |
| 0.882261 | 0.072024 | 0.406 | 0.4 | 1 | 4 | Pla2g15 | 0.006 | Old |
| 0.882522 | 0.400417 | 0.028 | 0.025 | 1 | 4 | Gss | 0.003 | Old |
| 0.88265 | 0.42395 | 0.014 | 0.013 | 1 | 4 | Itfg2 | 0.001 | Old |
| 0.882696 | 0.418409 | 0.092 | 0.1 | 1 | 4 | Map3k4 | -0.008 | Old |
| 0.882933 | 0.112267 | 0.105 | 0.1 | 1 | 4 | Cdc42bpb | 0.005 | Old |
| 0.883301 | 0.087893 | 0.484 | 0.5 | 1 | 4 | Mapre1 | -0.016 | Old |
| 0.883329 | 0.272243 | 0.141 | 0.138 | 1 | 4 | Pfkp | 0.003 | Old |
| 0.883489 | 0.159631 | 0.318 | 0.35 | 1 | 4 | Pten | -0.032 | Old |
| 0.883575 | 0.176539 | 0.046 | 0.05 | 1 | 4 | Slc25a32 | -0.004 | Old |
| 0.883601 | 0.421008 | 0.014 | 0.013 | 1 | 4 | Rabl2 | 0.001 | Old |
| 0.883601 | 0.197169 | 0.014 | 0.013 | 1 | 4 | Pms2 | 0.001 | Old |
| 0.883738 | 0.143589 | 0.243 | 0.238 | 1 | 4 | Pja2 | 0.005 | Old |
| 0.884552 | 0.213272 | 0.014 | 0.013 | 1 | 4 | Urb2 | 0.001 | Old |
| 0.884567 | 0.156421 | 0.117 | 0.112 | 1 | 4 | Magoh | 0.005 | Old |
| 0.884862 | 0.145127 | 0.403 | 0.45 | 1 | 4 | Nsa2 | -0.047 | Old |
| 0.8849 | 0.290521 | 0.105 | 0.112 | 1 | 4 | Smim19 | -0.007 | Old |
| 0.885504 | 0.433338 | 0.014 | 0.013 | 1 | 4 | Praf2 | 0.001 | Old |
| 0.885504 | 0.224651 | 0.014 | 0.013 | 1 | 4 | Smco4 | 0.001 | Old |
| 0.885632 | 0.017078 | 0.095 | 0.1 | 1 | 4 | Fam120b | -0.005 | Old |
| 0.885733 | 0.123668 | 0.13 | 0.138 | 1 | 4 | Tceal8 | -0.008 | Old |
| 0.88589 | 0.237695 | 0.053 | 0.05 | 1 | 4 | Letmd1 | 0.003 | Old |
| 0.886087 | 0.312635 | 0.066 | 0.062 | 1 | 4 | Dcaf1 | 0.004 | Old |
| 0.886111 | 0.01295 | 0.645 | 0.625 | 1 | 4 | Pid1 | 0.02 | Old |
| 0.886168 | 0.149076 | 0.322 | 0.35 | 1 | 4 | Ptges3 | -0.028 | Old |
| 0.886432 | 0.41692 | 0.078 | 0.075 | 1 | 4 | Rpp25l | 0.003 | Old |
| 0.886455 | 0.196607 | 0.014 | 0.013 | 1 | 4 | Ggct | 0.001 | Old |
| 0.886547 | 0.237244 | 0.24 | 0.25 | 1 | 4 | Nampt | -0.01 | Old |
| 0.886821 | 0.006207 | 0.107 | 0.112 | 1 | 4 | Prdm1 | -0.005 | Old |
| 0.886926 | 0.037042 | 0.976 | 0.975 | 1 | 4 | Lgmn | 0.001 | Old |
| 0.886969 | 0.113188 | 0.106 | 0.112 | 1 | 4 | Hyou1 | -0.006 | Old |
| 0.887036 | 1.109337 | 0.057 | 0.062 | 1 | 4 | Pygl | -0.005 | Old |
| 0.887279 | 0.010861 | 0.329 | 0.35 | 1 | 4 | Zfp207 | -0.021 | Old |
| 0.887469 | 0.094295 | 0.35 | 0.35 | 1 | 4 | Ndufb9 | 0 | Old |
| 0.887706 | 0.598611 | 0.045 | 0.05 | 1 | 4 | Susd1 | -0.005 | Old |
| 0.887868 | 0.047705 | 0.497 | 0.512 | 1 | 4 | Nisch | -0.015 | Old |
| 0.887892 | 0.146009 | 0.246 | 0.262 | 1 | 4 | Rabep1 | -0.016 | Old |
| 0.888062 | 0.292909 | 0.028 | 0.025 | 1 | 4 | Il27ra | 0.003 | Old |
| 0.888089 | 0.585045 | 0.101 | 0.1 | 1 | 4 | Mob3a | 0.001 | Old |
| 0.888602 | 0.196636 | 0.092 | 0.088 | 1 | 4 | Arhgap23 | 0.004 | Old |
| 0.888612 | 0.245461 | 0.117 | 0.125 | 1 | 4 | Dcun1d1 | -0.008 | Old |
| 0.88886 | 0.689697 | 0.022 | 0.025 | 1 | 4 | Zfp946 | -0.003 | Old |
| 0.888884 | 0.354538 | 0.034 | 0.038 | 1 | 4 | Trmt13 | -0.004 | Old |

| | | | | | | | | |
|---|---|---|---|---|---|---|---|---|
| 0.888951 | 0.012459 | 0.229 | 0.225 | 1 | 4 | Slc35b2 | 0.004 | Old |
| 0.889059 | 0.009432 | 0.24 | 0.238 | 1 | 4 | Arih1 | 0.002 | Old |
| 0.889091 | 0.166352 | 0.13 | 0.138 | 1 | 4 | Pdxdc1 | -0.008 | Old |
| 0.889146 | 0.071569 | 0.143 | 0.138 | 1 | 4 | Thoc7 | 0.005 | Old |
| 0.889471 | 0.287614 | 0.233 | 0.238 | 1 | 4 | Dlst | -0.005 | Old |
| 0.88959 | 0.519863 | 0.09 | 0.088 | 1 | 4 | Nup214 | 0.002 | Old |
| 0.889625 | 0.477879 | 0.022 | 0.025 | 1 | 4 | Gpatch4 | -0.003 | Old |
| 0.88966 | 0.307505 | 0.082 | 0.088 | 1 | 4 | Zfp518a | -0.006 | Old |
| 0.889982 | 0.031128 | 0.428 | 0.425 | 1 | 4 | Eif2s2 | 0.003 | Old |
| 0.890389 | 0.260536 | 0.022 | 0.025 | 1 | 4 | Acy1 | -0.003 | Old |
| 0.890421 | 0.729228 | 0.045 | 0.05 | 1 | 4 | Cacfd1 | -0.005 | Old |
| 0.890522 | 0.003284 | 0.041 | 0.038 | 1 | 4 | Kctd11 | 0.003 | Old |
| 0.890747 | 0.07222 | 0.393 | 0.425 | 1 | 4 | Mkln1 | -0.032 | Old |
| 0.89096 | 0.015238 | 0.344 | 0.338 | 1 | 4 | Ndufa3 | 0.006 | Old |
| 0.891287 | 0.015665 | 0.054 | 0.05 | 1 | 4 | Zfp738 | 0.004 | Old |
| 0.891348 | 0.468871 | 0.053 | 0.05 | 1 | 4 | Reep4 | 0.003 | Old |
| 0.891814 | 0.087651 | 0.047 | 0.05 | 1 | 4 | Ldlrap1 | -0.003 | Old |
| 0.892133 | 0.275391 | 0.094 | 0.1 | 1 | 4 | Ing3 | -0.006 | Old |
| 0.892419 | 1.392516 | 0.027 | 0.025 | 1 | 4 | Rttn | 0.002 | Old |
| 0.893 | 0.723313 | 0.113 | 0.112 | 1 | 4 | Elovl5 | 0.001 | Old |
| 0.893105 | 0.085829 | 0.105 | 0.1 | 1 | 4 | Senp7 | 0.005 | Old |
| 0.893239 | 0.326012 | 0.034 | 0.038 | 1 | 4 | Rpp30 | -0.004 | Old |
| 0.893567 | 0.394489 | 0.15 | 0.15 | 1 | 4 | Stk24 | 0 | Old |
| 0.894277 | 0.116997 | 0.059 | 0.062 | 1 | 4 | Slc10a7 | -0.003 | Old |
| 0.894351 | 0.024454 | 0.181 | 0.175 | 1 | 4 | Rabl6 | 0.006 | Old |
| 0.894502 | 0.382204 | 0.344 | 0.375 | 1 | 4 | Sbf2 | -0.031 | Old |
| 0.894523 | 1.126073 | 0.027 | 0.025 | 1 | 4 | Zfp74 | 0.002 | Old |
| 0.894627 | 0.363824 | 0.04 | 0.038 | 1 | 4 | Cry2 | 0.002 | Old |
| 0.894977 | 0.730649 | 0.022 | 0.025 | 1 | 4 | Cenpl | -0.003 | Old |
| 0.895069 | 0.135382 | 0.131 | 0.138 | 1 | 4 | Ptpn2 | -0.007 | Old |
| 0.895514 | 0.40102 | 0.126 | 0.125 | 1 | 4 | Dctn2 | 0.001 | Old |
| 0.895644 | 0.222044 | 0.366 | 0.375 | 1 | 4 | Myl12b | -0.009 | Old |
| 0.895983 | 0.403599 | 0.046 | 0.05 | 1 | 4 | Ddx49 | -0.004 | Old |
| 0.896055 | 0.003495 | 0.446 | 0.462 | 1 | 4 | Atp6v1a | -0.016 | Old |
| 0.896206 | 0.029879 | 0.059 | 0.062 | 1 | 4 | Tmem110 | -0.003 | Old |
| 0.896525 | 0.312538 | 0.184 | 0.2 | 1 | 4 | Atp5g1 | -0.016 | Old |
| 0.896579 | 0.220156 | 0.142 | 0.15 | 1 | 4 | Por | -0.008 | Old |
| 0.896612 | 0.072196 | 0.119 | 0.125 | 1 | 4 | Rsbn1 | -0.006 | Old |
| 0.896898 | 0.317392 | 0.058 | 0.062 | 1 | 4 | Panx1 | -0.004 | Old |
| 0.896901 | 0.149789 | 0.314 | 0.325 | 1 | 4 | Hnrnpul1 | -0.011 | Old |
| 0.897013 | 1.022655 | 0.087 | 0.088 | 1 | 4 | Agtpbp1 | -0.001 | Old |
| 0.897062 | 0.304657 | 0.117 | 0.125 | 1 | 4 | Raf1 | -0.008 | Old |
| 0.897329 | 0.88163 | 0.027 | 0.025 | 1 | 4 | Tbkbp1 | 0.002 | Old |
| 0.897619 | 0.53642 | 0.16 | 0.162 | 1 | 4 | Tmem33 | -0.002 | Old |
| 0.897747 | 0.140589 | 0.067 | 0.062 | 1 | 4 | Lrrc57 | 0.005 | Old |

| | | | | | | | | |
|---|---|---|---|---|---|---|---|---|
| 0.897803 | 0.528385 | 0.089 | 0.088 | 1 | 4 | Mrpl18 | 0.001 | Old |
| 0.898373 | 0.185429 | 0.224 | 0.225 | 1 | 4 | Elmsan1 | -0.001 | Old |
| 0.898922 | 0.172633 | 0.201 | 0.212 | 1 | 4 | Erp44 | -0.011 | Old |
| 0.898987 | 0.056273 | 0.08 | 0.075 | 1 | 4 | 1700123O2 | 0.005 | Old |
| 0.899048 | 0.31289 | 0.103 | 0.1 | 1 | 4 | Zfp281 | 0.003 | Old |
| 0.899283 | 0.011984 | 0.352 | 0.35 | 1 | 4 | Pmaip1 | 0.002 | Old |
| 0.899413 | 0.267361 | 0.14 | 0.15 | 1 | 4 | Fbxo18 | -0.01 | Old |
| 0.899682 | 0.497811 | 0.069 | 0.075 | 1 | 4 | Pigc | -0.006 | Old |
| 0.900334 | 1.215708 | 0.022 | 0.025 | 1 | 4 | Nat2 | -0.003 | Old |
| 0.900616 | 0.411575 | 0.183 | 0.188 | 1 | 4 | Stat5b | -0.005 | Old |
| 0.900749 | 0.0673 | 0.269 | 0.288 | 1 | 4 | Nup98 | -0.019 | Old |
| 0.900754 | 0.22419 | 0.355 | 0.375 | 1 | 4 | Fchsd2 | -0.02 | Old |
| 0.900918 | 0.137418 | 0.284 | 0.288 | 1 | 4 | Eea1 | -0.004 | Old |
| 0.901213 | 0.45192 | 0.137 | 0.138 | 1 | 4 | Fbxw11 | -0.001 | Old |
| 0.901262 | 0.394017 | 0.058 | 0.062 | 1 | 4 | Fhl3 | -0.004 | Old |
| 0.901361 | 0.405538 | 0.093 | 0.1 | 1 | 4 | Cnot10 | -0.007 | Old |
| 0.901461 | 0.110852 | 0.748 | 0.725 | 1 | 4 | C5ar1 | 0.023 | Old |
| 0.901541 | 0.718169 | 0.027 | 0.025 | 1 | 4 | Dus3l | 0.002 | Old |
| 0.902012 | 0.178987 | 0.047 | 0.05 | 1 | 4 | Eml3 | -0.003 | Old |
| 0.902156 | 0.066667 | 0.04 | 0.038 | 1 | 4 | Pi4kb | 0.002 | Old |
| 0.903266 | 0.145105 | 0.389 | 0.425 | 1 | 4 | Akr1b3 | -0.036 | Old |
| 0.903639 | 0.150327 | 0.365 | 0.362 | 1 | 4 | Atp5b | 0.003 | Old |
| 0.903869 | 0.12514 | 0.262 | 0.262 | 1 | 4 | Slc23a2 | 0 | Old |
| 0.9039 | 0.154262 | 0.226 | 0.238 | 1 | 4 | Vps26a | -0.012 | Old |
| 0.904163 | 0.170412 | 0.308 | 0.312 | 1 | 4 | Ankfy1 | -0.004 | Old |
| 0.90435 | 0.563831 | 0.027 | 0.025 | 1 | 4 | Dffa | 0.002 | Old |
| 0.90435 | 0.491518 | 0.027 | 0.025 | 1 | 4 | Nup35 | 0.002 | Old |
| 0.904685 | 0.012751 | 0.198 | 0.2 | 1 | 4 | Plk2 | -0.002 | Old |
| 0.90488 | 0.232299 | 0.071 | 0.075 | 1 | 4 | Cdk2ap1 | -0.004 | Old |
| 0.905053 | 0.573388 | 0.027 | 0.025 | 1 | 4 | Zfp335 | 0.002 | Old |
| 0.905053 | 0.498288 | 0.027 | 0.025 | 1 | 4 | Lrmda | 0.002 | Old |
| 0.905085 | 0.124836 | 0.118 | 0.125 | 1 | 4 | Ranbp10 | -0.007 | Old |
| 0.905413 | 0.025431 | 0.993 | 0.988 | 1 | 4 | Rps9 | 0.005 | Old |
| 0.905756 | 0.880229 | 0.027 | 0.025 | 1 | 4 | Gpn2 | 0.002 | Old |
| 0.905756 | 0.682852 | 0.027 | 0.025 | 1 | 4 | Msto1 | 0.002 | Old |
| 0.906091 | 0.48432 | 0.204 | 0.212 | 1 | 4 | Alyref | -0.008 | Old |
| 0.906746 | 0.396657 | 0.102 | 0.1 | 1 | 4 | Epn2 | 0.002 | Old |
| 0.906928 | 0.001071 | 0.097 | 0.1 | 1 | 4 | Tmx2 | -0.003 | Old |
| 0.906966 | 0.745306 | 0.04 | 0.038 | 1 | 4 | Neurl2 | 0.002 | Old |
| 0.907021 | 0.001322 | 0.072 | 0.075 | 1 | 4 | Rsad2 | -0.003 | Old |
| 0.907166 | 0.025547 | 0.258 | 0.262 | 1 | 4 | Kmt2c | -0.004 | Old |
| 0.90744 | 0.181103 | 0.035 | 0.038 | 1 | 4 | Naa16 | -0.003 | Old |
| 0.90744 | 0.086906 | 0.035 | 0.038 | 1 | 4 | Cnnm4 | -0.003 | Old |
| 0.907463 | 0.163719 | 0.213 | 0.225 | 1 | 4 | Golga4 | -0.012 | Old |
| 0.907864 | 0.660409 | 0.027 | 0.025 | 1 | 4 | Mettl3 | 0.002 | Old |

| | | | | | | | | |
|---|---|---|---|---|---|---|---|---|
| 0.907864 | 0.434462 | 0.027 | 0.025 | 1 | 4 | Rtn4ip1 | 0.002 | Old |
| 0.908069 | 0.089395 | 0.066 | 0.062 | 1 | 4 | Il18 | 0.004 | Old |
| 0.908252 | 0.221017 | 0.234 | 0.238 | 1 | 4 | Uhrf2 | -0.004 | Old |
| 0.908412 | 0.101497 | 0.071 | 0.075 | 1 | 4 | Dhps | -0.004 | Old |
| 0.908605 | 0.276955 | 0.32 | 0.35 | 1 | 4 | Rasa4 | -0.03 | Old |
| 0.908649 | 0.187949 | 0.053 | 0.05 | 1 | 4 | Gclm | 0.003 | Old |
| 0.908779 | 0.464985 | 0.093 | 0.1 | 1 | 4 | Cdc34 | -0.007 | Old |
| 0.909086 | 0.613579 | 0.076 | 0.075 | 1 | 4 | Ccdc97 | 0.001 | Old |
| 0.909296 | 0.065523 | 0.035 | 0.038 | 1 | 4 | Uxt | -0.003 | Old |
| 0.909732 | 0.137937 | 0.104 | 0.1 | 1 | 4 | Agmo | 0.004 | Old |
| 0.909733 | 0.37909 | 0.065 | 0.062 | 1 | 4 | Timm50 | 0.003 | Old |
| 0.909878 | 0.464702 | 0.148 | 0.15 | 1 | 4 | Vps13b | -0.002 | Old |
| 0.910182 | 0.499535 | 0.052 | 0.05 | 1 | 4 | Tada3 | 0.002 | Old |
| 0.910456 | 0.059725 | 0.108 | 0.112 | 1 | 4 | Naxd | -0.004 | Old |
| 0.910473 | 0.571395 | 0.04 | 0.038 | 1 | 4 | Gm37233 | 0.002 | Old |
| 0.910664 | 0.245098 | 0.152 | 0.15 | 1 | 4 | Abcc5 | 0.002 | Old |
| 0.910758 | 1.545903 | 0.014 | 0.013 | 1 | 4 | Dennd2d | 0.001 | Old |
| 0.910758 | 1.420018 | 0.014 | 0.013 | 1 | 4 | Alg12 | 0.001 | Old |
| 0.91089 | 0.146269 | 0.508 | 0.562 | 1 | 4 | Wdr26 | -0.054 | Old |
| 0.910996 | 0.052455 | 0.121 | 0.125 | 1 | 4 | Gtf2i | -0.004 | Old |
| 0.911379 | 0.494428 | 0.027 | 0.025 | 1 | 4 | Nectin4 | 0.002 | Old |
| 0.911379 | 0.279766 | 0.027 | 0.025 | 1 | 4 | Ick | 0.002 | Old |
| 0.911501 | 0.142014 | 0.192 | 0.188 | 1 | 4 | Emc3 | 0.004 | Old |
| 0.911617 | 0.215054 | 0.14 | 0.138 | 1 | 4 | Ncoa7 | 0.002 | Old |
| 0.911734 | 1.673957 | 0.014 | 0.013 | 1 | 4 | Fam135a | 0.001 | Old |
| 0.911771 | 0.27166 | 0.035 | 0.038 | 1 | 4 | Mul1 | -0.003 | Old |
| 0.912082 | 0.224269 | 0.027 | 0.025 | 1 | 4 | Plk4 | 0.002 | Old |
| 0.912239 | 0.066368 | 0.023 | 0.025 | 1 | 4 | Clec16a | -0.002 | Old |
| 0.912365 | 0.060983 | 0.12 | 0.125 | 1 | 4 | Slc35a3 | -0.005 | Old |
| 0.912372 | 0.092059 | 0.133 | 0.138 | 1 | 4 | Mtr | -0.005 | Old |
| 0.912692 | 0.151464 | 0.403 | 0.425 | 1 | 4 | Sec61g | -0.022 | Old |
| 0.913149 | 0.370242 | 0.114 | 0.112 | 1 | 4 | Meaf6 | 0.002 | Old |
| 0.913263 | 0.400861 | 0.101 | 0.1 | 1 | 4 | Capn7 | 0.001 | Old |
| 0.913274 | 0.275541 | 0.163 | 0.175 | 1 | 4 | Bicd2 | -0.012 | Old |
| 0.913688 | 1.174437 | 0.014 | 0.013 | 1 | 4 | Zfp276 | 0.001 | Old |
| 0.913688 | 0.993173 | 0.014 | 0.013 | 1 | 4 | Kcnj2 | 0.001 | Old |
| 0.913688 | 0.729839 | 0.014 | 0.013 | 1 | 4 | Atp1a3 | 0.001 | Old |
| 0.913805 | 0.466272 | 0.07 | 0.075 | 1 | 4 | Fopnl | -0.005 | Old |
| 0.913846 | 0.10053 | 0.626 | 0.688 | 1 | 4 | Nrp2 | -0.062 | Old |
| 0.913846 | 0.264066 | 0.117 | 0.125 | 1 | 4 | Taok2 | -0.008 | Old |
| 0.914119 | 0.022872 | 0.779 | 0.8 | 1 | 4 | Hnrnpa2b1 | -0.021 | Old |
| 0.914193 | 0.107276 | 0.027 | 0.025 | 1 | 4 | Fance | 0.002 | Old |
| 0.914442 | 0.148236 | 0.011 | 0.013 | 1 | 4 | Gas8 | -0.002 | Old |
| 0.914442 | 0.103004 | 0.011 | 0.013 | 1 | 4 | Heg1 | -0.002 | Old |
| 0.914509 | 0.227212 | 0.023 | 0.025 | 1 | 4 | Ms4a4c | -0.002 | Old |

| | | | | | | | | |
|---|---|---|---|---|---|---|---|---|
| 0.914573 | 0.158227 | 0.385 | 0.412 | 1 | 4 | Wsb1 | -0.027 | Old |
| 0.914579 | 0.319567 | 0.114 | 0.112 | 1 | 4 | Fkbp9 | 0.002 | Old |
| 0.914665 | 0.785344 | 0.014 | 0.013 | 1 | 4 | AU041133 | 0.001 | Old |
| 0.914828 | 0.001096 | 0.121 | 0.125 | 1 | 4 | Trnt1 | -0.004 | Old |
| 0.914853 | 0.193881 | 0.13 | 0.138 | 1 | 4 | Dazap1 | -0.008 | Old |
| 0.914977 | 0.320076 | 0.163 | 0.162 | 1 | 4 | Zfp263 | 0.001 | Old |
| 0.915265 | 0.08159 | 0.096 | 0.1 | 1 | 4 | Cd72 | -0.004 | Old |
| 0.915266 | 0.076618 | 0.023 | 0.025 | 1 | 4 | Amd1 | -0.002 | Old |
| 0.915486 | 0.223741 | 0.083 | 0.088 | 1 | 4 | Exosc5 | -0.005 | Old |
| 0.915642 | 0.937732 | 0.014 | 0.013 | 1 | 4 | Coq4 | 0.001 | Old |
| 0.915642 | 0.526181 | 0.014 | 0.013 | 1 | 4 | Tmem138 | 0.001 | Old |
| 0.915696 | 0.008431 | 0.319 | 0.325 | 1 | 4 | Klf10 | -0.006 | Old |
| 0.915724 | 0.078301 | 0.107 | 0.112 | 1 | 4 | AU040320 | -0.005 | Old |
| 0.915811 | 0.289909 | 0.196 | 0.2 | 1 | 4 | Srsf7 | -0.004 | Old |
| 0.915875 | 0.417295 | 0.138 | 0.15 | 1 | 4 | Tbc1d17 | -0.012 | Old |
| 0.91609 | 0.125625 | 0.107 | 0.112 | 1 | 4 | Rnf139 | -0.005 | Old |
| 0.9166 | 0.214511 | 0.011 | 0.013 | 1 | 4 | 4833445I0 | -0.002 | Old |
| 0.9166 | 0.15461 | 0.011 | 0.013 | 1 | 4 | 9430015G1 | -0.002 | Old |
| 0.9166 | 0.095231 | 0.011 | 0.013 | 1 | 4 | Tubd1 | -0.002 | Old |
| 0.91662 | 0.79441 | 0.014 | 0.013 | 1 | 4 | Trmt12 | 0.001 | Old |
| 0.916661 | 0.128583 | 0.167 | 0.175 | 1 | 4 | Nrd1 | -0.008 | Old |
| 0.916989 | 0.057297 | 0.536 | 0.575 | 1 | 4 | Wdr1 | -0.039 | Old |
| 0.917069 | 0.293362 | 0.047 | 0.05 | 1 | 4 | Arsg | -0.003 | Old |
| 0.917112 | 0.163597 | 0.065 | 0.062 | 1 | 4 | Zfyve26 | 0.003 | Old |
| 0.917194 | 0.329966 | 0.197 | 0.2 | 1 | 4 | Gpsm3 | -0.003 | Old |
| 0.917361 | 0.169408 | 0.052 | 0.05 | 1 | 4 | 2410089E0 | 0.002 | Old |
| 0.917461 | 0.574418 | 0.059 | 0.062 | 1 | 4 | Hbegf | -0.003 | Old |
| 0.917532 | 0.327946 | 0.077 | 0.075 | 1 | 4 | Bcl2l2 | 0.002 | Old |
| 0.917538 | 0.143749 | 0.023 | 0.025 | 1 | 4 | Bzw2 | -0.002 | Old |
| 0.917604 | 0.018251 | 0.092 | 0.088 | 1 | 4 | Fam118b | 0.004 | Old |
| 0.91764 | 0.059195 | 0.615 | 0.637 | 1 | 4 | Ogt | -0.022 | Old |
| 0.917764 | 0.074263 | 0.269 | 0.262 | 1 | 4 | BC017643 | 0.007 | Old |
| 0.917804 | 0.148321 | 0.231 | 0.225 | 1 | 4 | Rer1 | 0.006 | Old |
| 0.917961 | 0.305418 | 0.035 | 0.038 | 1 | 4 | Pot1b | -0.003 | Old |
| 0.918078 | 0.359715 | 0.152 | 0.162 | 1 | 4 | Ncbp2 | -0.01 | Old |
| 0.918487 | 0.053238 | 0.263 | 0.275 | 1 | 4 | Hipk1 | -0.012 | Old |
| 0.918574 | 0.436456 | 0.014 | 0.013 | 1 | 4 | Creb3l1 | 0.001 | Old |
| 0.91858 | 0.542167 | 0.035 | 0.038 | 1 | 4 | Bambi | -0.003 | Old |
| 0.918684 | 0.253781 | 0.047 | 0.05 | 1 | 4 | Prkab1 | -0.003 | Old |
| 0.91875 | 1.038918 | 0.075 | 0.075 | 1 | 4 | Trappc12 | 0 | Old |
| 0.918759 | 0.775279 | 0.011 | 0.013 | 1 | 4 | Zbtb26 | -0.002 | Old |
| 0.918759 | 0.368301 | 0.011 | 0.013 | 1 | 4 | Polr1e | -0.002 | Old |
| 0.918937 | 0.490105 | 0.076 | 0.075 | 1 | 4 | Rbm34 | 0.001 | Old |
| 0.918963 | 0.048842 | 0.536 | 0.575 | 1 | 4 | Atp1a1 | -0.039 | Old |
| 0.91913 | 0.137347 | 0.091 | 0.088 | 1 | 4 | Sf3a2 | 0.003 | Old |

| | | | | | | | | |
|---|---|---|---|---|---|---|---|---|
| 0.919279 | 0.2764 | 0.095 | 0.1 | 1 | 4 | Polr3gl | -0.005 | Old |
| 0.919288 | 0.362513 | 0.065 | 0.062 | 1 | 4 | Rab33b | 0.003 | Old |
| 0.919413 | 0.341614 | 0.052 | 0.05 | 1 | 4 | Lrch1 | 0.002 | Old |
| 0.919552 | 0.331271 | 0.014 | 0.013 | 1 | 4 | Tnfrsf26 | 0.001 | Old |
| 0.919552 | 0.260361 | 0.014 | 0.013 | 1 | 4 | Dyrk1b | 0.001 | Old |
| 0.919832 | 0.127498 | 0.04 | 0.038 | 1 | 4 | Zfp579 | 0.002 | Old |
| 0.919838 | 0.58987 | 0.011 | 0.013 | 1 | 4 | Dynlt1a | -0.002 | Old |
| 0.919838 | 0.464009 | 0.011 | 0.013 | 1 | 4 | Gm16867 | -0.002 | Old |
| 0.91988 | 0.345141 | 0.059 | 0.062 | 1 | 4 | Akr1c12 | -0.003 | Old |
| 0.920206 | 0.887879 | 0.11 | 0.112 | 1 | 4 | Crtc2 | -0.002 | Old |
| 0.920263 | 0.010054 | 0.047 | 0.05 | 1 | 4 | Acot13 | -0.003 | Old |
| 0.920378 | 0.100387 | 0.053 | 0.05 | 1 | 4 | Tpk1 | 0.003 | Old |
| 0.920473 | 0.025426 | 0.638 | 0.65 | 1 | 4 | Ccl7 | -0.012 | Old |
| 0.920521 | 0.300243 | 0.244 | 0.25 | 1 | 4 | Huwe1 | -0.006 | Old |
| 0.92053 | 0.742345 | 0.014 | 0.013 | 1 | 4 | Mmab | 0.001 | Old |
| 0.92053 | 0.212085 | 0.014 | 0.013 | 1 | 4 | Nphp1 | 0.001 | Old |
| 0.920748 | 0.443646 | 0.035 | 0.038 | 1 | 4 | Carf | -0.003 | Old |
| 0.920763 | 0.092682 | 0.186 | 0.188 | 1 | 4 | Myo9a | -0.002 | Old |
| 0.920766 | 0.210471 | 0.201 | 0.2 | 1 | 4 | Sec63 | 0.001 | Old |
| 0.920918 | 1.07814 | 0.011 | 0.013 | 1 | 4 | Ift43 | -0.002 | Old |
| 0.920918 | 0.860429 | 0.011 | 0.013 | 1 | 4 | Nsun5 | -0.002 | Old |
| 0.920918 | 0.732231 | 0.011 | 0.013 | 1 | 4 | Slc7a6 | -0.002 | Old |
| 0.920918 | 0.324231 | 0.011 | 0.013 | 1 | 4 | Gm9796 | -0.002 | Old |
| 0.92104 | 0.24328 | 0.103 | 0.1 | 1 | 4 | Kansl3 | 0.003 | Old |
| 0.921325 | 0.503698 | 0.023 | 0.025 | 1 | 4 | Rbfox2 | -0.002 | Old |
| 0.92134 | 0.268854 | 0.246 | 0.25 | 1 | 4 | D17Wsu92 | -0.004 | Old |
| 0.921388 | 0.288379 | 0.102 | 0.1 | 1 | 4 | Senp1 | 0.002 | Old |
| 0.921396 | 0.683731 | 0.115 | 0.125 | 1 | 4 | Supt4a | -0.01 | Old |
| 0.921506 | 0.169682 | 0.594 | 0.625 | 1 | 4 | G3bp2 | -0.031 | Old |
| 0.921508 | 0.218961 | 0.014 | 0.013 | 1 | 4 | Armc9 | 0.001 | Old |
| 0.921508 | 0.178431 | 0.014 | 0.013 | 1 | 4 | Dgke | 0.001 | Old |
| 0.921998 | 0.961852 | 0.011 | 0.013 | 1 | 4 | Amacr | -0.002 | Old |
| 0.921998 | 0.925861 | 0.011 | 0.013 | 1 | 4 | Cyp2j9 | -0.002 | Old |
| 0.921998 | 0.863529 | 0.011 | 0.013 | 1 | 4 | 2510046G1 | -0.002 | Old |
| 0.922174 | 0.220491 | 0.04 | 0.038 | 1 | 4 | Zfp687 | 0.002 | Old |
| 0.922304 | 0.262036 | 0.279 | 0.288 | 1 | 4 | Tnfaip2 | -0.009 | Old |
| 0.922841 | 0.747135 | 0.023 | 0.025 | 1 | 4 | Ripk3 | -0.002 | Old |
| 0.92287 | 1.078874 | 0.063 | 0.062 | 1 | 4 | Skiv2l2 | 0.001 | Old |
| 0.92294 | 0.026772 | 0.047 | 0.05 | 1 | 4 | Prmt7 | -0.003 | Old |
| 0.923078 | 1.049595 | 0.011 | 0.013 | 1 | 4 | Helq | -0.002 | Old |
| 0.923078 | 0.955156 | 0.011 | 0.013 | 1 | 4 | Focad | -0.002 | Old |
| 0.923078 | 0.910831 | 0.011 | 0.013 | 1 | 4 | Zfp455 | -0.002 | Old |
| 0.923464 | 0.002557 | 0.014 | 0.013 | 1 | 4 | Mical1 | 0.001 | Old |
| 0.923552 | 0.652427 | 0.099 | 0.1 | 1 | 4 | Manea | -0.001 | Old |
| 0.924113 | 0.777048 | 0.051 | 0.05 | 1 | 4 | Alg1 | 0.001 | Old |

| | | | | | | | | |
|---|---|---|---|---|---|---|---|---|
| 0.924158 | 0.925581 | 0.011 | 0.013 | 1 | 4 | Palm | -0.002 | Old |
| 0.92428 | 0.275358 | 0.267 | 0.275 | 1 | 4 | Srsf1 | -0.008 | Old |
| 0.924552 | 0.26306 | 0.211 | 0.225 | 1 | 4 | Slbp | -0.014 | Old |
| 0.924609 | 0.525396 | 0.047 | 0.05 | 1 | 4 | Kdm1b | -0.003 | Old |
| 0.92463 | 0.307859 | 0.196 | 0.2 | 1 | 4 | Dcaf12 | -0.004 | Old |
| 0.924982 | 0.017075 | 0.097 | 0.1 | 1 | 4 | Timm17a | -0.003 | Old |
| 0.925036 | 0.057839 | 0.09 | 0.088 | 1 | 4 | Vcl | 0.002 | Old |
| 0.925115 | 0.705719 | 0.023 | 0.025 | 1 | 4 | Ankrd33b | -0.002 | Old |
| 0.925148 | 0.507894 | 0.047 | 0.05 | 1 | 4 | Bpnt1 | -0.003 | Old |
| 0.925237 | 0.260299 | 0.188 | 0.2 | 1 | 4 | Nudcd3 | -0.012 | Old |
| 0.92546 | 0.0146 | 0.06 | 0.062 | 1 | 4 | Nol8 | -0.002 | Old |
| 0.925764 | 0.052619 | 0.54 | 0.6 | 1 | 4 | Atox1 | -0.06 | Old |
| 0.925789 | 0.082625 | 0.153 | 0.15 | 1 | 4 | Spin1 | 0.003 | Old |
| 0.926086 | 0.459358 | 0.287 | 0.312 | 1 | 4 | Edem1 | -0.025 | Old |
| 0.926152 | 0.121003 | 0.047 | 0.05 | 1 | 4 | Clk2 | -0.003 | Old |
| 0.926643 | 0.040833 | 0.036 | 0.038 | 1 | 4 | Gmppb | -0.002 | Old |
| 0.926971 | 0.071363 | 0.355 | 0.362 | 1 | 4 | Cln8 | -0.007 | Old |
| 0.927256 | 0.66678 | 0.035 | 0.038 | 1 | 4 | Mlh1 | -0.003 | Old |
| 0.927389 | 0.872193 | 0.023 | 0.025 | 1 | 4 | Zfp952 | -0.002 | Old |
| 0.927487 | 0.044575 | 0.756 | 0.812 | 1 | 4 | Calm2 | -0.056 | Old |
| 0.927729 | 0.518174 | 0.051 | 0.05 | 1 | 4 | Lyrm2 | 0.001 | Old |
| 0.927874 | 0.054423 | 0.152 | 0.15 | 1 | 4 | Mrpl48 | 0.002 | Old |
| 0.927916 | 0.41431 | 0.194 | 0.212 | 1 | 4 | Uggt1 | -0.018 | Old |
| 0.928101 | 0.559303 | 0.088 | 0.088 | 1 | 4 | Rgmb | 0 | Old |
| 0.928147 | 0.766646 | 0.023 | 0.025 | 1 | 4 | Armt1 | -0.002 | Old |
| 0.928815 | 0.261572 | 0.14 | 0.138 | 1 | 4 | Dapp1 | 0.002 | Old |
| 0.929116 | 0.583404 | 0.035 | 0.038 | 1 | 4 | Cbr3 | -0.003 | Old |
| 0.929155 | 0.107435 | 0.084 | 0.088 | 1 | 4 | Pogz | -0.004 | Old |
| 0.92948 | 0.174102 | 0.072 | 0.075 | 1 | 4 | Cwc22 | -0.003 | Old |
| 0.929737 | 0.88469 | 0.035 | 0.038 | 1 | 4 | Pex11a | -0.003 | Old |
| 0.930136 | 0.024501 | 0.203 | 0.212 | 1 | 4 | Rnf44 | -0.009 | Old |
| 0.930303 | 0.69736 | 0.026 | 0.025 | 1 | 4 | Ms4a14 | 0.001 | Old |
| 0.930577 | 0.379486 | 0.137 | 0.138 | 1 | 4 | Gnl1 | -0.001 | Old |
| 0.930999 | 0.09804 | 0.084 | 0.088 | 1 | 4 | Poglut1 | -0.004 | Old |
| 0.931051 | 0.803794 | 0.081 | 0.088 | 1 | 4 | Rab9 | -0.007 | Old |
| 0.931695 | 0.099507 | 0.06 | 0.062 | 1 | 4 | Gm47754 | -0.002 | Old |
| 0.931746 | 0.022442 | 0.217 | 0.212 | 1 | 4 | Tnfrsf11a | 0.005 | Old |
| 0.932218 | 0.998511 | 0.035 | 0.038 | 1 | 4 | Alkbh6 | -0.003 | Old |
| 0.932485 | 0.153954 | 0.036 | 0.038 | 1 | 4 | Bend4 | -0.002 | Old |
| 0.932695 | 0.711367 | 0.047 | 0.05 | 1 | 4 | Pramef8 | -0.003 | Old |
| 0.932766 | 0.037354 | 0.052 | 0.05 | 1 | 4 | Sdccag8 | 0.002 | Old |
| 0.932897 | 0.436936 | 0.051 | 0.05 | 1 | 4 | Nxt1 | 0.001 | Old |
| 0.933025 | 0.116101 | 0.146 | 0.15 | 1 | 4 | Rnf115 | -0.004 | Old |
| 0.933611 | 0.118948 | 0.102 | 0.1 | 1 | 4 | Zfyve16 | 0.002 | Old |
| 0.933671 | 0.674225 | 0.115 | 0.125 | 1 | 4 | Dok3 | -0.01 | Old |

| | | | | | | | | |
|---|---|---|---|---|---|---|---|---|
| 0.934007 | 0.388386 | 0.1 | 0.1 | 1 | 4 | Prkci | 0 | Old |
| 0.934269 | 0.485249 | 0.039 | 0.038 | 1 | 4 | Ift22 | 0.001 | Old |
| 0.934773 | 0.086905 | 0.072 | 0.075 | 1 | 4 | Gpatch2l | -0.003 | Old |
| 0.934828 | 0.025962 | 0.29 | 0.288 | 1 | 4 | 1110008F1 | 0.002 | Old |
| 0.935261 | 0.344695 | 0.047 | 0.05 | 1 | 4 | Rbl1 | -0.003 | Old |
| 0.935324 | 0.329727 | 0.153 | 0.162 | 1 | 4 | Slc50a1 | -0.009 | Old |
| 0.935425 | 0.048412 | 0.117 | 0.112 | 1 | 4 | Tex264 | 0.005 | Old |
| 0.935514 | 0.231179 | 0.126 | 0.125 | 1 | 4 | Cfdp1 | 0.001 | Old |
| 0.935561 | 0.186075 | 0.036 | 0.038 | 1 | 4 | Zfp846 | -0.002 | Old |
| 0.935569 | 0.112997 | 0.096 | 0.1 | 1 | 4 | Fnbp4 | -0.004 | Old |
| 0.935777 | 0.203931 | 0.143 | 0.15 | 1 | 4 | Kif3b | -0.007 | Old |
| 0.935797 | 0.53108 | 0.047 | 0.05 | 1 | 4 | Rab2b | -0.003 | Old |
| 0.936365 | 0.172901 | 0.052 | 0.05 | 1 | 4 | Zfp593 | 0.002 | Old |
| 0.936475 | 0.113095 | 0.217 | 0.225 | 1 | 4 | Fam91a1 | -0.008 | Old |
| 0.936723 | 0.365719 | 0.026 | 0.025 | 1 | 4 | Usp49 | 0.001 | Old |
| 0.936723 | 0.065452 | 0.191 | 0.188 | 1 | 4 | Mgat2 | 0.003 | Old |
| 0.937026 | 0.316714 | 0.123 | 0.125 | 1 | 4 | Psme3 | -0.002 | Old |
| 0.937045 | 0.089596 | 0.175 | 0.175 | 1 | 4 | Lyl1 | 0 | Old |
| 0.937182 | 0.897175 | 0.035 | 0.038 | 1 | 4 | Adprhl2 | -0.003 | Old |
| 0.937344 | 0.049762 | 0.452 | 0.45 | 1 | 4 | Atrx | 0.002 | Old |
| 0.937436 | 0.35907 | 0.026 | 0.025 | 1 | 4 | Thap4 | 0.001 | Old |
| 0.937436 | 0.498042 | 0.026 | 0.025 | 1 | 4 | Smpd2 | 0.001 | Old |
| 0.937475 | 0.149344 | 0.229 | 0.238 | 1 | 4 | Smc1a | -0.009 | Old |
| 0.93756 | 0.127193 | 0.409 | 0.438 | 1 | 4 | Ralbp1 | -0.029 | Old |
| 0.93772 | 0.323773 | 0.233 | 0.238 | 1 | 4 | Stat1 | -0.005 | Old |
| 0.937784 | 0.055713 | 0.252 | 0.262 | 1 | 4 | Dennd1a | -0.01 | Old |
| 0.937818 | 0.112161 | 0.341 | 0.338 | 1 | 4 | Ndufb11 | 0.003 | Old |
| 0.93787 | 0.338201 | 0.23 | 0.25 | 1 | 4 | Etfb | -0.02 | Old |
| 0.937928 | 0.296652 | 0.138 | 0.138 | 1 | 4 | Ccdc59 | 0 | Old |
| 0.93815 | 0.360736 | 0.026 | 0.025 | 1 | 4 | Polr3a | 0.001 | Old |
| 0.938182 | 0.281349 | 0.1 | 0.1 | 1 | 4 | Fam214b | 0 | Old |
| 0.938475 | 0.164497 | 0.077 | 0.075 | 1 | 4 | Cib2 | 0.002 | Old |
| 0.938863 | 0.073464 | 0.026 | 0.025 | 1 | 4 | Napsa | 0.001 | Old |
| 0.939401 | 0.167832 | 0.084 | 0.088 | 1 | 4 | Specc1 | -0.004 | Old |
| 0.939588 | 0.624024 | 0.039 | 0.038 | 1 | 4 | Sharpin | 0.001 | Old |
| 0.939602 | 0.01745 | 0.086 | 0.088 | 1 | 4 | Slc38a7 | -0.002 | Old |
| 0.939744 | 0.068291 | 0.452 | 0.462 | 1 | 4 | Syngr2 | -0.01 | Old |
| 0.939943 | 0.113267 | 0.065 | 0.062 | 1 | 4 | Ssbp2 | 0.003 | Old |
| 0.939985 | 0.137704 | 0.322 | 0.338 | 1 | 4 | 8-Sep | -0.016 | Old |
| 0.940021 | 0.149064 | 0.139 | 0.138 | 1 | 4 | Wwp2 | 0.001 | Old |
| 0.940153 | 0.241251 | 0.221 | 0.225 | 1 | 4 | Ap3d1 | -0.004 | Old |
| 0.940291 | 0.257845 | 0.026 | 0.025 | 1 | 4 | Tpbgl | 0.001 | Old |
| 0.940309 | 0.079054 | 0.217 | 0.225 | 1 | 4 | Zfp395 | -0.008 | Old |
| 0.940484 | 0.393612 | 0.036 | 0.038 | 1 | 4 | Cbr1 | -0.002 | Old |
| 0.940576 | 0.167335 | 0.102 | 0.1 | 1 | 4 | Dpp9 | 0.002 | Old |

| | | | | | | | | |
|---|---|---|---|---|---|---|---|---|
| 0.940783 | 0.036375 | 0.159 | 0.162 | 1 | 4 | Dirc2 | -0.003 | Old |
| 0.9411 | 0.450763 | 0.036 | 0.038 | 1 | 4 | Rgl3 | -0.002 | Old |
| 0.941201 | 1.422847 | 0.038 | 0.038 | 1 | 4 | Psmf1 | 0 | Old |
| 0.941447 | 0.229718 | 0.155 | 0.162 | 1 | 4 | Dhdh | -0.007 | Old |
| 0.941474 | 0.802766 | 0.082 | 0.088 | 1 | 4 | Tmem11 | -0.006 | Old |
| 0.941718 | 0.037315 | 0.026 | 0.025 | 1 | 4 | Stk38l | 0.001 | Old |
| 0.941787 | 0.278525 | 0.119 | 0.125 | 1 | 4 | Ergic2 | -0.006 | Old |
| 0.941875 | 0.017124 | 0.26 | 0.275 | 1 | 4 | Tle3 | -0.015 | Old |
| 0.941877 | 0.756587 | 0.074 | 0.075 | 1 | 4 | Neurl4 | -0.001 | Old |
| 0.942252 | 0.728942 | 0.07 | 0.075 | 1 | 4 | Prpf6 | -0.005 | Old |
| 0.942499 | 0.105267 | 0.065 | 0.062 | 1 | 4 | Uspl1 | 0.003 | Old |
| 0.942618 | 0.158499 | 0.211 | 0.212 | 1 | 4 | Rcn3 | -0.001 | Old |
| 0.943136 | 0.399047 | 0.039 | 0.038 | 1 | 4 | Zc3h8 | 0.001 | Old |
| 0.943146 | 0.276712 | 0.137 | 0.138 | 1 | 4 | Ccdc22 | -0.001 | Old |
| 0.943307 | 0.227757 | 0.024 | 0.025 | 1 | 4 | Tspan7 | -0.001 | Old |
| 0.94338 | 0.037791 | 0.128 | 0.125 | 1 | 4 | Wdr82 | 0.003 | Old |
| 0.943387 | 0.187683 | 0.072 | 0.075 | 1 | 4 | Rpa3 | -0.003 | Old |
| 0.943665 | 0.248399 | 0.173 | 0.175 | 1 | 4 | Prkar2a | -0.002 | Old |
| 0.94386 | 0.093093 | 0.026 | 0.025 | 1 | 4 | Letm2 | 0.001 | Old |
| 0.944318 | 0.254717 | 0.039 | 0.038 | 1 | 4 | Hscb | 0.001 | Old |
| 0.944408 | 0.331703 | 0.088 | 0.088 | 1 | 4 | AU019823 | 0 | Old |
| 0.944427 | 0.150717 | 0.534 | 0.575 | 1 | 4 | Pdia6 | -0.041 | Old |
| 0.944512 | 0.179009 | 0.164 | 0.162 | 1 | 4 | Csgalnact2 | 0.002 | Old |
| 0.944523 | 0.521142 | 0.123 | 0.125 | 1 | 4 | Ykt6 | -0.002 | Old |
| 0.944568 | 0.089144 | 0.389 | 0.412 | 1 | 4 | Rab10 | -0.023 | Old |
| 0.94504 | 0.227085 | 0.237 | 0.25 | 1 | 4 | Psmd4 | -0.013 | Old |
| 0.945046 | 0.158563 | 0.463 | 0.475 | 1 | 4 | Cdkn1a | -0.012 | Old |
| 0.945167 | 0.069017 | 0.061 | 0.062 | 1 | 4 | Mak16 | -0.001 | Old |
| 0.94538 | 0.160252 | 0.389 | 0.425 | 1 | 4 | Dek | -0.036 | Old |
| 0.945514 | 0.581789 | 0.051 | 0.05 | 1 | 4 | Tlr6 | 0.001 | Old |
| 0.945657 | 0.355338 | 0.205 | 0.212 | 1 | 4 | Pdcd10 | -0.007 | Old |
| 0.945712 | 0.107076 | 0.177 | 0.175 | 1 | 4 | S100a1 | 0.002 | Old |
| 0.945724 | 0.40705 | 0.063 | 0.062 | 1 | 4 | Zfyve21 | 0.001 | Old |
| 0.946093 | 0.170419 | 0.039 | 0.038 | 1 | 4 | Dkc1 | 0.001 | Old |
| 0.946104 | 0.416903 | 0.169 | 0.175 | 1 | 4 | Atp11c | -0.006 | Old |
| 0.946365 | 0.03216 | 0.134 | 0.138 | 1 | 4 | Plaa | -0.004 | Old |
| 0.946516 | 0.252246 | 0.136 | 0.138 | 1 | 4 | mt-Nd6 | -0.002 | Old |
| 0.946537 | 0.153596 | 0.535 | 0.575 | 1 | 4 | Pirb | -0.04 | Old |
| 0.946686 | 0.310879 | 0.101 | 0.1 | 1 | 4 | E2f1 | 0.001 | Old |
| 0.94672 | 0.239799 | 0.339 | 0.375 | 1 | 4 | Mcrip1 | -0.036 | Old |
| 0.946927 | 0.067236 | 0.097 | 0.1 | 1 | 4 | Sec23ip | -0.003 | Old |
| 0.947016 | 0.312086 | 0.084 | 0.088 | 1 | 4 | Larp7 | -0.004 | Old |
| 0.947484 | 0.319913 | 0.187 | 0.2 | 1 | 4 | Mpc2 | -0.013 | Old |
| 0.947799 | 0.121782 | 0.22 | 0.225 | 1 | 4 | R3hdm2 | -0.005 | Old |
| 0.94786 | 0.546801 | 0.088 | 0.088 | 1 | 4 | Gyg | 0 | Old |

| | | | | | | | | |
|---|---|---|---|---|---|---|---|---|
| 0.947942 | 0.062269 | 0.155 | 0.15 | 1 | 4 | Rcn2 | 0.005 | Old |
| 0.947944 | 0.050926 | 0.246 | 0.25 | 1 | 4 | Psmd8 | -0.004 | Old |
| 0.948167 | 0.316427 | 0.096 | 0.1 | 1 | 4 | Hccs | -0.004 | Old |
| 0.948211 | 0.132345 | 0.11 | 0.112 | 1 | 4 | Lactb | -0.002 | Old |
| 0.948255 | 0.154101 | 0.371 | 0.4 | 1 | 4 | Bcl10 | -0.029 | Old |
| 0.948384 | 0.152731 | 0.246 | 0.25 | 1 | 4 | Hadhb | -0.004 | Old |
| 0.94853 | 0.164573 | 0.169 | 0.175 | 1 | 4 | Derl2 | -0.006 | Old |
| 0.948891 | 0.299787 | 0.183 | 0.188 | 1 | 4 | Ugp2 | -0.005 | Old |
| 0.948989 | 0.026644 | 0.048 | 0.05 | 1 | 4 | Scaper | -0.002 | Old |
| 0.949078 | 0.225935 | 0.126 | 0.125 | 1 | 4 | Znrd1 | 0.001 | Old |
| 0.949083 | 0.241274 | 0.221 | 0.225 | 1 | 4 | Sik1 | -0.004 | Old |
| 0.949165 | 0.183697 | 0.36 | 0.375 | 1 | 4 | Atp6v0d1 | -0.015 | Old |
| 0.949398 | 0.014384 | 0.089 | 0.088 | 1 | 4 | Aph1c | 0.001 | Old |
| 0.949555 | 0.246699 | 0.21 | 0.225 | 1 | 4 | Rbm26 | -0.015 | Old |
| 0.949917 | 0.147429 | 0.286 | 0.288 | 1 | 4 | Use1 | -0.002 | Old |
| 0.949944 | 0.203864 | 0.064 | 0.062 | 1 | 4 | Lonp1 | 0.002 | Old |
| 0.950056 | 0.30935 | 0.024 | 0.025 | 1 | 4 | Myo10 | -0.001 | Old |
| 0.950085 | 0.135921 | 0.144 | 0.15 | 1 | 4 | Mospd3 | -0.006 | Old |
| 0.950258 | 1.442405 | 0.013 | 0.013 | 1 | 4 | Mfsd4a | 0 | Old |
| 0.950258 | 1.128044 | 0.013 | 0.013 | 1 | 4 | Ttc12 | 0 | Old |
| 0.950258 | 0.910329 | 0.013 | 0.013 | 1 | 4 | Gm15445 | 0 | Old |
| 0.950424 | 1.024981 | 0.026 | 0.025 | 1 | 4 | Mon1a | 0.001 | Old |
| 0.950478 | 0.213118 | 0.246 | 0.25 | 1 | 4 | Zfp652 | -0.004 | Old |
| 0.950806 | 0.493023 | 0.024 | 0.025 | 1 | 4 | Zfp87 | -0.001 | Old |
| 0.950806 | 0.339645 | 0.024 | 0.025 | 1 | 4 | Ncapg2 | -0.001 | Old |
| 0.950869 | 0.081554 | 0.065 | 0.062 | 1 | 4 | Crls1 | 0.003 | Old |
| 0.950881 | 0.126492 | 0.096 | 0.1 | 1 | 4 | Rnf167 | -0.004 | Old |
| 0.950954 | 0.720152 | 0.036 | 0.038 | 1 | 4 | Pyroxd1 | -0.002 | Old |
| 0.951211 | 0.383028 | 0.185 | 0.2 | 1 | 4 | Herpud2 | -0.015 | Old |
| 0.951234 | 0.226359 | 0.101 | 0.1 | 1 | 4 | Knop1 | 0.001 | Old |
| 0.951253 | 0.783732 | 0.074 | 0.075 | 1 | 4 | Caml | -0.001 | Old |
| 0.951365 | 0.139195 | 0.329 | 0.338 | 1 | 4 | Ncoa4 | -0.009 | Old |
| 0.951382 | 0.300235 | 0.076 | 0.075 | 1 | 4 | Map3k14 | 0.001 | Old |
| 0.951391 | 0.349305 | 0.153 | 0.162 | 1 | 4 | Baz2a | -0.009 | Old |
| 0.951518 | 0.287684 | 0.265 | 0.275 | 1 | 4 | Rapgef6 | -0.01 | Old |
| 0.951818 | 0.477329 | 0.123 | 0.125 | 1 | 4 | Atg2a | -0.002 | Old |
| 0.951825 | 0.398 | 0.063 | 0.062 | 1 | 4 | Nudt13 | 0.001 | Old |
| 0.952265 | 0.884328 | 0.013 | 0.013 | 1 | 4 | Hr | 0 | Old |
| 0.952283 | 0.546124 | 0.051 | 0.05 | 1 | 4 | Arfgap1 | 0.001 | Old |
| 0.952306 | 0.373307 | 0.024 | 0.025 | 1 | 4 | Echdc1 | -0.001 | Old |
| 0.952528 | 0.68351 | 0.038 | 0.038 | 1 | 4 | Zfp760 | 0 | Old |
| 0.952593 | 1.040084 | 0.026 | 0.025 | 1 | 4 | Tgm1 | 0.001 | Old |
| 0.952659 | 0.049712 | 0.199 | 0.2 | 1 | 4 | Ap1s1 | -0.001 | Old |
| 0.952755 | 0.434248 | 0.122 | 0.125 | 1 | 4 | Crkl | -0.003 | Old |
| 0.952771 | 0.367162 | 0.108 | 0.112 | 1 | 4 | Fam133b | -0.004 | Old |

| | | | | | | | | |
|---|---|---|---|---|---|---|---|---|
| 0.952794 | 0.929079 | 0.062 | 0.062 | 1 | 4 | 1110059E2 | 0 | Old |
| 0.952807 | 0.120963 | 0.089 | 0.088 | 1 | 4 | B3glct | 0.001 | Old |
| 0.952895 | 0.096221 | 0.085 | 0.088 | 1 | 4 | Mboat7 | -0.003 | Old |
| 0.953269 | 0.775336 | 0.013 | 0.013 | 1 | 4 | Nmb | 0 | Old |
| 0.953473 | 0.277489 | 0.167 | 0.175 | 1 | 4 | Mrpl20 | -0.008 | Old |
| 0.953503 | 0.755939 | 0.047 | 0.05 | 1 | 4 | Xpc | -0.003 | Old |
| 0.954273 | 0.74089 | 0.013 | 0.013 | 1 | 4 | Gm30411 | 0 | Old |
| 0.954273 | 0.311064 | 0.013 | 0.013 | 1 | 4 | Inmt | 0 | Old |
| 0.954404 | 0.307955 | 0.076 | 0.075 | 1 | 4 | Zfp512 | 0.001 | Old |
| 0.954557 | 0.702316 | 0.024 | 0.025 | 1 | 4 | Slc52a2 | -0.001 | Old |
| 0.954633 | 0.126172 | 0.165 | 0.162 | 1 | 4 | Cpsf7 | 0.003 | Old |
| 0.954635 | 0.018045 | 0.514 | 0.55 | 1 | 4 | Sf3b1 | -0.036 | Old |
| 0.955021 | 0.119844 | 0.277 | 0.275 | 1 | 4 | Dock10 | 0.002 | Old |
| 0.955044 | 0.069116 | 0.65 | 0.725 | 1 | 4 | Cox7a2l | -0.075 | Old |
| 0.955229 | 0.340336 | 0.158 | 0.162 | 1 | 4 | Ly9 | -0.004 | Old |
| 0.955282 | 0.630745 | 0.098 | 0.1 | 1 | 4 | Plcg2 | -0.002 | Old |
| 0.955352 | 0.244764 | 0.132 | 0.138 | 1 | 4 | Slc31a1 | -0.006 | Old |
| 0.955423 | 0.052615 | 0.413 | 0.438 | 1 | 4 | Selenow | -0.025 | Old |
| 0.955499 | 0.368007 | 0.244 | 0.262 | 1 | 4 | Lacc1 | -0.018 | Old |
| 0.955567 | 0.263857 | 0.441 | 0.5 | 1 | 4 | Fcho2 | -0.059 | Old |
| 0.955825 | 0.02299 | 0.289 | 0.3 | 1 | 4 | Larp1 | -0.011 | Old |
| 0.95614 | 0.129115 | 0.121 | 0.125 | 1 | 4 | Cycs | -0.004 | Old |
| 0.956204 | 0.187865 | 0.437 | 0.475 | 1 | 4 | Fli1 | -0.038 | Old |
| 0.956246 | 0.131846 | 0.096 | 0.1 | 1 | 4 | Kctd10 | -0.004 | Old |
| 0.95628 | 0.482225 | 0.013 | 0.013 | 1 | 4 | Trim6 | 0 | Old |
| 0.95628 | 0.479659 | 0.013 | 0.013 | 1 | 4 | Wdhd1 | 0 | Old |
| 0.95628 | 0.362617 | 0.013 | 0.013 | 1 | 4 | Pdk2 | 0 | Old |
| 0.95628 | 0.24183 | 0.013 | 0.013 | 1 | 4 | 5033421B0 | 0 | Old |
| 0.956647 | 0.243461 | 0.061 | 0.062 | 1 | 4 | Gmcl1 | -0.001 | Old |
| 0.9568 | 0.212685 | 0.1 | 0.1 | 1 | 4 | Esrra | 0 | Old |
| 0.956808 | 0.800361 | 0.024 | 0.025 | 1 | 4 | Gm9493 | -0.001 | Old |
| 0.956968 | 0.560276 | 0.144 | 0.15 | 1 | 4 | Usp22 | -0.006 | Old |
| 0.957285 | 0.410409 | 0.013 | 0.013 | 1 | 4 | Gpr85 | 0 | Old |
| 0.957285 | 0.265765 | 0.013 | 0.013 | 1 | 4 | Tha1 | 0 | Old |
| 0.957285 | 0.245032 | 0.013 | 0.013 | 1 | 4 | Zc4h2 | 0 | Old |
| 0.957285 | 0.01264 | 0.013 | 0.013 | 1 | 4 | Card10 | 0 | Old |
| 0.95757 | 0.540958 | 0.071 | 0.075 | 1 | 4 | Arhgap21 | -0.004 | Old |
| 0.957654 | 0.686779 | 0.026 | 0.025 | 1 | 4 | Ahdc1 | 0.001 | Old |
| 0.958102 | 0.313481 | 0.113 | 0.112 | 1 | 4 | Dusp23 | 0.001 | Old |
| 0.958153 | 0.065515 | 0.26 | 0.275 | 1 | 4 | Smim14 | -0.015 | Old |
| 0.958288 | 0.049635 | 0.012 | 0.013 | 1 | 4 | Dtd1 | -0.001 | Old |
| 0.958673 | 0.735538 | 0.109 | 0.112 | 1 | 4 | Impdh1 | -0.003 | Old |
| 0.958747 | 0.263175 | 0.12 | 0.125 | 1 | 4 | Snx24 | -0.005 | Old |
| 0.95912 | 0.093004 | 0.262 | 0.275 | 1 | 4 | Elf2 | -0.013 | Old |
| 0.95926 | 0.044555 | 0.281 | 0.288 | 1 | 4 | Uba1 | -0.007 | Old |

| | | | | | | | | |
|---|---|---|---|---|---|---|---|---|
| 0.959293 | 0.18603 | 0.013 | 0.013 | 1 | 4 | Cep89 | 0 | Old |
| 0.959343 | 0.247199 | 0.012 | 0.013 | 1 | 4 | Amz1 | -0.001 | Old |
| 0.959614 | 0.215712 | 0.036 | 0.038 | 1 | 4 | Rtl8b | -0.002 | Old |
| 0.959625 | 0.671493 | 0.07 | 0.075 | 1 | 4 | AI606181 | -0.005 | Old |
| 0.959739 | 0.169567 | 0.076 | 0.075 | 1 | 4 | Cd209e | 0.001 | Old |
| 0.959948 | 1.127854 | 0.047 | 0.05 | 1 | 4 | Ece1 | -0.003 | Old |
| 0.959991 | 0.714385 | 0.084 | 0.088 | 1 | 4 | Adssl1 | -0.004 | Old |
| 0.960144 | 0.503572 | 0.074 | 0.075 | 1 | 4 | Dpm1 | -0.001 | Old |
| 0.960196 | 0.401194 | 0.048 | 0.05 | 1 | 4 | Spred2 | -0.002 | Old |
| 0.960229 | 0.171856 | 0.302 | 0.312 | 1 | 4 | Acin1 | -0.01 | Old |
| 0.960297 | 0.030381 | 0.013 | 0.013 | 1 | 4 | Agbl5 | 0 | Old |
| 0.960302 | 0.309093 | 0.099 | 0.1 | 1 | 4 | Zfp280d | -0.001 | Old |
| 0.960361 | 0.496733 | 0.169 | 0.175 | 1 | 4 | Mmd | -0.006 | Old |
| 0.960398 | 0.30863 | 0.012 | 0.013 | 1 | 4 | Rad52 | -0.001 | Old |
| 0.960398 | 0.175544 | 0.012 | 0.013 | 1 | 4 | Elp6 | -0.001 | Old |
| 0.960547 | 0.568922 | 0.026 | 0.025 | 1 | 4 | Myg1 | 0.001 | Old |
| 0.960561 | 0.972356 | 0.024 | 0.025 | 1 | 4 | Umps | -0.001 | Old |
| 0.96088 | 0.434772 | 0.038 | 0.038 | 1 | 4 | Rtca | 0 | Old |
| 0.960911 | 0.727222 | 0.05 | 0.05 | 1 | 4 | Ercc6l2 | 0 | Old |
| 0.961048 | 0.129972 | 0.109 | 0.112 | 1 | 4 | Acadvl | -0.003 | Old |
| 0.96118 | 0.117471 | 0.401 | 0.412 | 1 | 4 | Kdm2a | -0.011 | Old |
| 0.961271 | 0.591323 | 0.026 | 0.025 | 1 | 4 | 2010007H( | 0.001 | Old |
| 0.961302 | 0.003287 | 0.013 | 0.013 | 1 | 4 | Olr1 | 0 | Old |
| 0.961311 | 1.41402 | 0.024 | 0.025 | 1 | 4 | Knstrn | -0.001 | Old |
| 0.961453 | 0.202844 | 0.012 | 0.013 | 1 | 4 | Ccdc73 | -0.001 | Old |
| 0.961453 | 0.063935 | 0.012 | 0.013 | 1 | 4 | Zfp287 | -0.001 | Old |
| 0.961471 | 1.084525 | 0.038 | 0.038 | 1 | 4 | Mgat5 | 0 | Old |
| 0.961658 | 0.199535 | 0.414 | 0.438 | 1 | 4 | Etv3 | -0.024 | Old |
| 0.961785 | 0.018051 | 0.19 | 0.188 | 1 | 4 | Mrpl24 | 0.002 | Old |
| 0.961797 | 0.366934 | 0.048 | 0.05 | 1 | 4 | Abt1 | -0.002 | Old |
| 0.961951 | 0.304499 | 0.108 | 0.112 | 1 | 4 | Kmt2b | -0.004 | Old |
| 0.962031 | 0.002193 | 0.975 | 0.988 | 1 | 4 | Dusp1 | -0.013 | Old |
| 0.962074 | 0.447836 | 0.038 | 0.038 | 1 | 4 | Sco2 | 0 | Old |
| 0.962258 | 0.067305 | 0.304 | 0.312 | 1 | 4 | Adam10 | -0.008 | Old |
| 0.962509 | 0.536194 | 0.012 | 0.013 | 1 | 4 | Disc1 | -0.001 | Old |
| 0.962509 | 0.355705 | 0.012 | 0.013 | 1 | 4 | Mllt3 | -0.001 | Old |
| 0.962672 | 0.326795 | 0.036 | 0.038 | 1 | 4 | Tspyl2 | -0.002 | Old |
| 0.962676 | 0.409276 | 0.123 | 0.125 | 1 | 4 | Mrpl12 | -0.002 | Old |
| 0.962739 | 0.205917 | 0.097 | 0.1 | 1 | 4 | Mxi1 | -0.003 | Old |
| 0.962794 | 0.114813 | 0.113 | 0.112 | 1 | 4 | Esf1 | 0.001 | Old |
| 0.963092 | 0.148267 | 0.049 | 0.05 | 1 | 4 | Igsf9 | -0.001 | Old |
| 0.963096 | 0.168251 | 0.401 | 0.425 | 1 | 4 | Atp6v0a1 | -0.024 | Old |
| 0.963262 | 0.124754 | 0.2 | 0.2 | 1 | 4 | Sppl3 | 0 | Old |
| 0.963441 | 0.400076 | 0.026 | 0.025 | 1 | 4 | Zhx2 | 0.001 | Old |
| 0.963441 | 0.265764 | 0.026 | 0.025 | 1 | 4 | Qpctl | 0.001 | Old |

| | | | | | | | | |
|---|---|---|---|---|---|---|---|---|
| 0.963564 | 0.871589 | 0.012 | 0.013 | 1 | 4 | D930030I0 | -0.001 | Old |
| 0.963564 | 0.740215 | 0.012 | 0.013 | 1 | 4 | Col27a1 | -0.001 | Old |
| 0.963714 | 0.013595 | 0.185 | 0.188 | 1 | 4 | Ube2h | -0.003 | Old |
| 0.963734 | 0.154521 | 0.26 | 0.262 | 1 | 4 | Tsc22d2 | -0.002 | Old |
| 0.964206 | 0.321257 | 0.099 | 0.1 | 1 | 4 | Fzr1 | -0.001 | Old |
| 0.964506 | 0.323578 | 0.036 | 0.038 | 1 | 4 | Ccar2 | -0.002 | Old |
| 0.964619 | 0.695862 | 0.012 | 0.013 | 1 | 4 | Usp54 | -0.001 | Old |
| 0.964757 | 0.293628 | 0.155 | 0.162 | 1 | 4 | Man1a2 | -0.007 | Old |
| 0.964789 | 0.251649 | 0.051 | 0.05 | 1 | 4 | Pcyt2 | 0.001 | Old |
| 0.965065 | 1.062176 | 0.024 | 0.025 | 1 | 4 | Ift74 | -0.001 | Old |
| 0.96508 | 1.167932 | 0.038 | 0.038 | 1 | 4 | 1700025G( | 0 | Old |
| 0.965255 | 0.054319 | 0.705 | 0.738 | 1 | 4 | Sh3bgrl3 | -0.033 | Old |
| 0.965485 | 0.186284 | 0.358 | 0.388 | 1 | 4 | Smdt1 | -0.03 | Old |
| 0.965675 | 1.090955 | 0.012 | 0.013 | 1 | 4 | Abcb9 | -0.001 | Old |
| 0.965675 | 1.06824 | 0.012 | 0.013 | 1 | 4 | Fzd1 | -0.001 | Old |
| 0.965675 | 0.692045 | 0.012 | 0.013 | 1 | 4 | Pex26 | -0.001 | Old |
| 0.965721 | 0.422765 | 0.087 | 0.088 | 1 | 4 | Cmc2 | -0.001 | Old |
| 0.965943 | 0.155718 | 0.088 | 0.088 | 1 | 4 | Lancl1 | 0 | Old |
| 0.966309 | 0.284086 | 0.182 | 0.188 | 1 | 4 | Plekhm2 | -0.006 | Old |
| 0.966335 | 0.187733 | 0.026 | 0.025 | 1 | 4 | Lrrc20 | 0.001 | Old |
| 0.966341 | 0.264321 | 0.036 | 0.038 | 1 | 4 | Mbd4 | -0.002 | Old |
| 0.966346 | 0.143342 | 0.088 | 0.088 | 1 | 4 | Ttc5 | 0 | Old |
| 0.966368 | 0.190767 | 0.163 | 0.162 | 1 | 4 | Ythdc2 | 0.001 | Old |
| 0.966464 | 0.069816 | 0.234 | 0.238 | 1 | 4 | Vps13c | -0.004 | Old |
| 0.9667 | 0.298538 | 0.061 | 0.062 | 1 | 4 | Ptrhd1 | -0.001 | Old |
| 0.966725 | 0.586215 | 0.161 | 0.175 | 1 | 4 | Lsm12 | -0.014 | Old |
| 0.96673 | 1.179383 | 0.012 | 0.013 | 1 | 4 | Hist2h2aa1 | -0.001 | Old |
| 0.966881 | 0.114294 | 0.304 | 0.312 | 1 | 4 | Bnip3l | -0.008 | Old |
| 0.967058 | 0.390804 | 0.026 | 0.025 | 1 | 4 | Cdc6 | 0.001 | Old |
| 0.967058 | 0.133124 | 0.026 | 0.025 | 1 | 4 | Rida | 0.001 | Old |
| 0.967279 | 0.334164 | 0.108 | 0.112 | 1 | 4 | Rnf10 | -0.004 | Old |
| 0.967327 | 0.225461 | 0.137 | 0.138 | 1 | 4 | Dido1 | -0.001 | Old |
| 0.967473 | 0.340614 | 0.145 | 0.15 | 1 | 4 | Tmem127 | -0.005 | Old |
| 0.967554 | 0.181007 | 0.088 | 0.088 | 1 | 4 | Tsc1 | 0 | Old |
| 0.96773 | 0.029778 | 0.298 | 0.288 | 1 | 4 | Gnai3 | 0.01 | Old |
| 0.967756 | 0.110054 | 0.163 | 0.162 | 1 | 4 | Hcst | 0.001 | Old |
| 0.967782 | 0.217445 | 0.026 | 0.025 | 1 | 4 | Bcr | 0.001 | Old |
| 0.967786 | 1.121527 | 0.012 | 0.013 | 1 | 4 | Gng7 | -0.001 | Old |
| 0.968091 | 0.072547 | 0.711 | 0.738 | 1 | 4 | Tmem50a | -0.027 | Old |
| 0.968378 | 0.273353 | 0.191 | 0.2 | 1 | 4 | 8-Mar | -0.009 | Old |
| 0.968481 | 0.358574 | 0.072 | 0.075 | 1 | 4 | Ppid | -0.003 | Old |
| 0.968788 | 0.478865 | 0.036 | 0.038 | 1 | 4 | Ppp1r16a | -0.002 | Old |
| 0.968796 | 0.354246 | 0.217 | 0.225 | 1 | 4 | Smarcc2 | -0.008 | Old |
| 0.968842 | 1.166631 | 0.012 | 0.013 | 1 | 4 | Alkbh4 | -0.001 | Old |
| 0.969144 | 0.119651 | 0.163 | 0.162 | 1 | 4 | Ubr4 | 0.001 | Old |

| | | | | | | | | |
|---|---|---|---|---|---|---|---|---|
| 0.969417 | 0.1133 | 0.185 | 0.188 | 1 | 4 | Ube2e1 | -0.003 | Old |
| 0.969439 | 0.084127 | 0.101 | 0.1 | 1 | 4 | Ormdl2 | 0.001 | Old |
| 0.970285 | 0.115816 | 0.206 | 0.212 | 1 | 4 | Ctcf | -0.006 | Old |
| 0.970724 | 0.106323 | 0.329 | 0.35 | 1 | 4 | Hsd17b11 | -0.021 | Old |
| 0.970728 | 0.192749 | 0.169 | 0.175 | 1 | 4 | Mecp2 | -0.006 | Old |
| 0.970808 | 0.299293 | 0.166 | 0.175 | 1 | 4 | Pbx3 | -0.009 | Old |
| 0.970911 | 0.224797 | 0.061 | 0.062 | 1 | 4 | Zfp780b | -0.001 | Old |
| 0.971041 | 0.039233 | 0.758 | 0.762 | 1 | 4 | Cox4i1 | -0.004 | Old |
| 0.971198 | 0.00069 | 0.14 | 0.138 | 1 | 4 | Banf1 | 0.002 | Old |
| 0.971444 | 0.172385 | 0.15 | 0.15 | 1 | 4 | Aagab | 0 | Old |
| 0.971527 | 0.003289 | 0.347 | 0.35 | 1 | 4 | Mier1 | -0.003 | Old |
| 0.971834 | 0.26763 | 0.318 | 0.338 | 1 | 4 | AC160336. | -0.02 | Old |
| 0.971841 | 0.027369 | 0.989 | 0.988 | 1 | 4 | Rpl41 | 0.001 | Old |
| 0.971864 | 0.054556 | 0.061 | 0.062 | 1 | 4 | Uvssa | -0.001 | Old |
| 0.971958 | 0.191756 | 0.354 | 0.375 | 1 | 4 | Sptssa | -0.021 | Old |
| 0.972048 | 0.14817 | 0.379 | 0.4 | 1 | 4 | Epn1 | -0.021 | Old |
| 0.972209 | 0.362946 | 0.087 | 0.088 | 1 | 4 | E2f5 | -0.001 | Old |
| 0.973101 | 0.05123 | 0.074 | 0.075 | 1 | 4 | Ap1ar | -0.001 | Old |
| 0.973272 | 0.019495 | 0.377 | 0.412 | 1 | 4 | Slfn5 | -0.035 | Old |
| 0.973682 | 0.739305 | 0.036 | 0.038 | 1 | 4 | Saysd1 | -0.002 | Old |
| 0.973682 | 0.178369 | 0.354 | 0.375 | 1 | 4 | Sp100 | -0.021 | Old |
| 0.973829 | 0.196241 | 0.219 | 0.225 | 1 | 4 | Mcur1 | -0.006 | Old |
| 0.973984 | 1.337097 | 0.025 | 0.025 | 1 | 4 | F830016B0 | 0 | Old |
| 0.974014 | 0.142583 | 0.038 | 0.038 | 1 | 4 | Rbm38 | 0 | Old |
| 0.974175 | 0.147625 | 0.051 | 0.05 | 1 | 4 | Gtdc1 | 0.001 | Old |
| 0.974239 | 0.259046 | 0.049 | 0.05 | 1 | 4 | Ppa1 | -0.001 | Old |
| 0.974363 | 0.438823 | 0.061 | 0.062 | 1 | 4 | Brap | -0.001 | Old |
| 0.974395 | 0.778159 | 0.049 | 0.05 | 1 | 4 | Cul9 | -0.001 | Old |
| 0.974403 | 0.058252 | 0.024 | 0.025 | 1 | 4 | Zfp788 | -0.001 | Old |
| 0.974413 | 0.090731 | 0.074 | 0.075 | 1 | 4 | 0610037L1 | -0.001 | Old |
| 0.97455 | 0.092113 | 0.321 | 0.338 | 1 | 4 | Pigt | -0.017 | Old |
| 0.975014 | 0.158111 | 0.29 | 0.3 | 1 | 4 | Elavl1 | -0.01 | Old |
| 0.975145 | 0.126574 | 0.024 | 0.025 | 1 | 4 | Tstd3 | -0.001 | Old |
| 0.975147 | 0.334142 | 0.144 | 0.15 | 1 | 4 | Cers5 | -0.006 | Old |
| 0.975159 | 0.147496 | 0.173 | 0.175 | 1 | 4 | Dpp8 | -0.002 | Old |
| 0.975218 | 0.174142 | 0.051 | 0.05 | 1 | 4 | Smim8 | 0.001 | Old |
| 0.975243 | 0.161282 | 0.289 | 0.3 | 1 | 4 | Ankrd17 | -0.011 | Old |
| 0.975324 | 0.264422 | 0.133 | 0.138 | 1 | 4 | 1810013L2 | -0.005 | Old |
| 0.97533 | 0.014177 | 0.813 | 0.838 | 1 | 4 | Cybb | -0.025 | Old |
| 0.975517 | 0.663123 | 0.036 | 0.038 | 1 | 4 | Isg15 | -0.002 | Old |
| 0.975556 | 0.627612 | 0.062 | 0.062 | 1 | 4 | Nosip | 0 | Old |
| 0.975689 | 0.096256 | 0.171 | 0.175 | 1 | 4 | Kidins220 | -0.004 | Old |
| 0.975725 | 0.077895 | 0.462 | 0.475 | 1 | 4 | Dusp3 | -0.013 | Old |
| 0.976379 | 0.392735 | 0.063 | 0.062 | 1 | 4 | Lin54 | 0.001 | Old |
| 0.976405 | 0.008117 | 0.296 | 0.3 | 1 | 4 | Papola | -0.004 | Old |

| | | | | | | | | |
|---|---|---|---|---|---|---|---|---|
| 0.976915 | 0.400888 | 0.265 | 0.288 | 1 | 4 | Hp1bp3 | -0.023 | Old |
| 0.977191 | 0.057333 | 0.026 | 0.025 | 1 | 4 | Ifi213 | 0.001 | Old |
| 0.977809 | 0.204481 | 0.364 | 0.388 | 1 | 4 | Hmgn1 | -0.024 | Old |
| 0.978112 | 0.249423 | 0.024 | 0.025 | 1 | 4 | Snx33 | -0.001 | Old |
| 0.978514 | 0.489916 | 0.188 | 0.2 | 1 | 4 | Trpm7 | -0.012 | Old |
| 0.978638 | 0.308133 | 0.132 | 0.138 | 1 | 4 | Entpd7 | -0.006 | Old |
| 0.978916 | 0.093912 | 0.325 | 0.338 | 1 | 4 | Fam168b | -0.013 | Old |
| 0.979018 | 0.478915 | 0.049 | 0.05 | 1 | 4 | Camsap1 | -0.001 | Old |
| 0.979042 | 0.01252 | 0.279 | 0.275 | 1 | 4 | Mfsd14b | 0.004 | Old |
| 0.979058 | 0.507068 | 0.072 | 0.075 | 1 | 4 | Dhdds | -0.003 | Old |
| 0.979108 | 0.03292 | 0.281 | 0.275 | 1 | 4 | Osbpl11 | 0.006 | Old |
| 0.979238 | 0.173698 | 0.213 | 0.225 | 1 | 4 | Spg21 | -0.012 | Old |
| 0.979374 | 0.16447 | 0.086 | 0.088 | 1 | 4 | Vezt | -0.002 | Old |
| 0.979411 | 0.172417 | 0.35 | 0.375 | 1 | 4 | Grina | -0.025 | Old |
| 0.979454 | 0.037859 | 0.151 | 0.15 | 1 | 4 | Zbtb38 | 0.001 | Old |
| 0.979492 | 0.21359 | 0.061 | 0.062 | 1 | 4 | Rhot2 | -0.001 | Old |
| 0.979683 | 0.005286 | 0.136 | 0.138 | 1 | 4 | Smg7 | -0.002 | Old |
| 0.980056 | 0.212907 | 0.144 | 0.15 | 1 | 4 | Echs1 | -0.006 | Old |
| 0.980607 | 0.013164 | 0.098 | 0.1 | 1 | 4 | Exoc2 | -0.002 | Old |
| 0.980697 | 0.108109 | 0.337 | 0.362 | 1 | 4 | Ywhaq | -0.025 | Old |
| 0.980805 | 0.206197 | 0.312 | 0.325 | 1 | 4 | Cdk9 | -0.013 | Old |
| 0.981019 | 0.539262 | 0.175 | 0.188 | 1 | 4 | Yipf5 | -0.013 | Old |
| 0.981396 | 0.154804 | 0.088 | 0.088 | 1 | 4 | Uba52 | 0 | Old |
| 0.981538 | 0.237562 | 0.087 | 0.088 | 1 | 4 | Cdkn2d | -0.001 | Old |
| 0.981651 | 0.284297 | 0.18 | 0.188 | 1 | 4 | Maf1 | -0.008 | Old |
| 0.981658 | 0.092503 | 0.088 | 0.088 | 1 | 4 | Sympk | 0 | Old |
| 0.981791 | 0.202083 | 0.3 | 0.312 | 1 | 4 | Anxa3 | -0.012 | Old |
| 0.981893 | 0.477486 | 0.05 | 0.05 | 1 | 4 | Rfx1 | 0 | Old |
| 0.982074 | 0.312722 | 0.123 | 0.125 | 1 | 4 | Tmem248 | -0.002 | Old |
| 0.982118 | 0.237609 | 0.11 | 0.112 | 1 | 4 | Spen | -0.002 | Old |
| 0.982541 | 0.035186 | 0.518 | 0.537 | 1 | 4 | Hint1 | -0.019 | Old |
| 0.982762 | 0.054274 | 0.42 | 0.425 | 1 | 4 | Nav1 | -0.005 | Old |
| 0.98284 | 0.715453 | 0.049 | 0.05 | 1 | 4 | Foxred1 | -0.001 | Old |
| 0.982942 | 0.410535 | 0.05 | 0.05 | 1 | 4 | Cars | 0 | Old |
| 0.982976 | 0.062065 | 0.112 | 0.112 | 1 | 4 | Kcnk13 | 0 | Old |
| 0.983305 | 0.662885 | 0.024 | 0.025 | 1 | 4 | 2700038G2 | -0.001 | Old |
| 0.983307 | 0.311502 | 0.061 | 0.062 | 1 | 4 | Mrpl37 | -0.001 | Old |
| 0.983471 | 0.205688 | 0.312 | 0.338 | 1 | 4 | Vapa | -0.026 | Old |
| 0.983515 | 0.405573 | 0.132 | 0.138 | 1 | 4 | Gigyf2 | -0.006 | Old |
| 0.983624 | 0.474004 | 0.062 | 0.062 | 1 | 4 | Ttf1 | 0 | Old |
| 0.983719 | 0.274577 | 0.168 | 0.175 | 1 | 4 | Angel2 | -0.007 | Old |
| 0.983833 | 0.180164 | 0.261 | 0.275 | 1 | 4 | Trmt112 | -0.014 | Old |
| 0.983912 | 0.214843 | 0.037 | 0.038 | 1 | 4 | Wdcp | -0.001 | Old |
| 0.984047 | 0.858918 | 0.024 | 0.025 | 1 | 4 | Zfp709 | -0.001 | Old |
| 0.984328 | 0.406825 | 0.146 | 0.15 | 1 | 4 | Mylk | -0.004 | Old |

| | | | | | | | | |
|---|---|---|---|---|---|---|---|---|
| 0.984404 | 0.028825 | 0.35 | 0.35 | 1 | 4 | Twf1 | 0 | Old |
| 0.984435 | 0.242058 | 0.259 | 0.275 | 1 | 4 | Dpm3 | -0.016 | Old |
| 0.98453 | 0.412745 | 0.086 | 0.088 | 1 | 4 | Exosc3 | -0.002 | Old |
| 0.98465 | 0.13275 | 0.076 | 0.075 | 1 | 4 | Trappc9 | 0.001 | Old |
| 0.984694 | 0.120782 | 0.265 | 0.275 | 1 | 4 | Atxn2l | -0.01 | Old |
| 0.984784 | 0.113467 | 0.087 | 0.088 | 1 | 4 | Fam3a | -0.001 | Old |
| 0.984975 | 0.728884 | 0.025 | 0.025 | 1 | 4 | Mcrip2 | 0 | Old |
| 0.985037 | 0.391642 | 0.11 | 0.112 | 1 | 4 | Ncln | -0.002 | Old |
| 0.985222 | 0.005331 | 0.159 | 0.162 | 1 | 4 | Kctd20 | -0.003 | Old |
| 0.985289 | 0.03425 | 0.54 | 0.562 | 1 | 4 | Cox6b1 | -0.022 | Old |
| 0.985403 | 0.02665 | 0.074 | 0.075 | 1 | 4 | Aph1b | -0.001 | Old |
| 0.985531 | 0.390234 | 0.024 | 0.025 | 1 | 4 | Bpgm | -0.001 | Old |
| 0.985547 | 0.359988 | 0.038 | 0.038 | 1 | 4 | Dpep2 | 0 | Old |
| 0.985595 | 0.149712 | 0.087 | 0.088 | 1 | 4 | BC003965 | -0.001 | Old |
| 0.985672 | 0.451746 | 0.176 | 0.188 | 1 | 4 | Ppp4r3a | -0.012 | Old |
| 0.985923 | 0.615738 | 0.049 | 0.05 | 1 | 4 | Yars | -0.001 | Old |
| 0.985942 | 0.048775 | 0.113 | 0.112 | 1 | 4 | Dld | 0.001 | Old |
| 0.986055 | 0.73836 | 0.072 | 0.075 | 1 | 4 | Siah1a | -0.003 | Old |
| 0.986119 | 0.012647 | 0.138 | 0.138 | 1 | 4 | Tlnrd1 | 0 | Old |
| 0.986221 | 0.273971 | 0.074 | 0.075 | 1 | 4 | Atn1 | -0.001 | Old |
| 0.986328 | 0.119069 | 0.198 | 0.2 | 1 | 4 | Cbfb | -0.002 | Old |
| 0.986751 | 0.229125 | 0.038 | 0.038 | 1 | 4 | Tbp | 0 | Old |
| 0.986776 | 0.192426 | 0.424 | 0.45 | 1 | 4 | Tm9sf2 | -0.026 | Old |
| 0.986875 | 0.064851 | 0.114 | 0.112 | 1 | 4 | Emb | 0.002 | Old |
| 0.987174 | 0.742818 | 0.025 | 0.025 | 1 | 4 | Mturn | 0 | Old |
| 0.987327 | 0.150625 | 0.098 | 0.1 | 1 | 4 | Pmpcb | -0.002 | Old |
| 0.98744 | 0.202701 | 0.161 | 0.162 | 1 | 4 | Ebp | -0.001 | Old |
| 0.98747 | 0.056798 | 0.429 | 0.438 | 1 | 4 | Arpc4 | -0.009 | Old |
| 0.98754 | 0.582101 | 0.025 | 0.025 | 1 | 4 | Nprl2 | 0 | Old |
| 0.987599 | 0.28836 | 0.061 | 0.062 | 1 | 4 | Wiz | -0.001 | Old |
| 0.987813 | 0.213254 | 0.239 | 0.25 | 1 | 4 | Tmx3 | -0.011 | Old |
| 0.987907 | 0.496457 | 0.025 | 0.025 | 1 | 4 | Lmln | 0 | Old |
| 0.988076 | 0.355997 | 0.061 | 0.062 | 1 | 4 | Tex2 | -0.001 | Old |
| 0.988498 | 0.993585 | 0.024 | 0.025 | 1 | 4 | Gm20186 | -0.001 | Old |
| 0.988541 | 0.066978 | 0.076 | 0.075 | 1 | 4 | Htatip2 | 0.001 | Old |
| 0.988639 | 0.556692 | 0.025 | 0.025 | 1 | 4 | Bcl7a | 0 | Old |
| 0.989127 | 0.229533 | 0.238 | 0.25 | 1 | 4 | Cep350 | -0.012 | Old |
| 0.989285 | 0.405423 | 0.177 | 0.188 | 1 | 4 | Wbp1l | -0.011 | Old |
| 0.989381 | 0.350569 | 0.169 | 0.175 | 1 | 4 | Ss18 | -0.006 | Old |
| 0.989514 | 0.26568 | 0.264 | 0.275 | 1 | 4 | Dusp11 | -0.011 | Old |
| 0.989679 | 0.581283 | 0.073 | 0.075 | 1 | 4 | Wdr7 | -0.002 | Old |
| 0.9897 | 1.419406 | 0.013 | 0.013 | 1 | 4 | Gm48653 | 0 | Old |
| 0.990105 | 0.525213 | 0.025 | 0.025 | 1 | 4 | Mad2l2 | 0 | Old |
| 0.990198 | 0.129718 | 0.375 | 0.388 | 1 | 4 | 9930111J2 | -0.013 | Old |
| 0.990282 | 0.239823 | 0.134 | 0.138 | 1 | 4 | Gpr107 | -0.004 | Old |

| | | | | | | | | |
|---|---|---|---|---|---|---|---|---|
| 0.990364 | 0.060473 | 0.038 | 0.038 | 1 | 4 | Zfp719 | 0 | Old |
| 0.990467 | 0.140807 | 0.3 | 0.312 | 1 | 4 | Chmp5 | -0.012 | Old |
| 0.990518 | 0.104737 | 0.318 | 0.325 | 1 | 4 | Mcfd2 | -0.007 | Old |
| 0.990838 | 0.455833 | 0.025 | 0.025 | 1 | 4 | Lrrc49 | 0 | Old |
| 0.99124 | 0.294661 | 0.238 | 0.25 | 1 | 4 | Dcaf8 | -0.012 | Old |
| 0.99176 | 0.765574 | 0.013 | 0.013 | 1 | 4 | Gm4631 | 0 | Old |
| 0.991939 | 0.214293 | 0.074 | 0.075 | 1 | 4 | Stam | -0.001 | Old |
| 0.991966 | 8.05E-05 | 0.405 | 0.412 | 1 | 4 | Rab3il1 | -0.007 | Old |
| 0.992054 | 0.413626 | 0.144 | 0.15 | 1 | 4 | Rbm28 | -0.006 | Old |
| 0.992404 | 0.100361 | 0.075 | 0.075 | 1 | 4 | Agpat1 | 0 | Old |
| 0.992411 | 0.252885 | 0.037 | 0.038 | 1 | 4 | Pphln1 | -0.001 | Old |
| 0.992522 | 0.024324 | 0.601 | 0.637 | 1 | 4 | Csde1 | -0.036 | Old |
| 0.992664 | 0.450369 | 0.109 | 0.112 | 1 | 4 | Drg1 | -0.003 | Old |
| 0.99279 | 1.267293 | 0.013 | 0.013 | 1 | 4 | Fam177a | 0 | Old |
| 0.99279 | 0.786002 | 0.013 | 0.013 | 1 | 4 | Hacl1 | 0 | Old |
| 0.99279 | 0.541684 | 0.013 | 0.013 | 1 | 4 | Trmo | 0 | Old |
| 0.992872 | 0.268359 | 0.049 | 0.05 | 1 | 4 | Gstp1 | -0.001 | Old |
| 0.99295 | 1.264032 | 0.024 | 0.025 | 1 | 4 | Fastkd2 | -0.001 | Old |
| 0.993215 | 0.612518 | 0.13 | 0.138 | 1 | 4 | Trim5 | -0.008 | Old |
| 0.993458 | 0.131097 | 0.442 | 0.462 | 1 | 4 | Mpp1 | -0.02 | Old |
| 0.994077 | 0.40506 | 0.086 | 0.088 | 1 | 4 | Rbm15 | -0.002 | Old |
| 0.994636 | 0.791277 | 0.084 | 0.088 | 1 | 4 | Stxbp1 | -0.004 | Old |
| 0.994716 | 0.136512 | 0.425 | 0.462 | 1 | 4 | Tor1aip1 | -0.037 | Old |
| 0.994798 | 0.339286 | 0.235 | 0.25 | 1 | 4 | Krcc1 | -0.015 | Old |
| 0.994839 | 0.646718 | 0.037 | 0.038 | 1 | 4 | Camta1 | -0.001 | Old |
| 0.99485 | 0.422569 | 0.013 | 0.013 | 1 | 4 | Rmdn2 | 0 | Old |
| 0.994932 | 0.211761 | 0.335 | 0.362 | 1 | 4 | Itsn2 | -0.027 | Old |
| 0.994984 | 0.225536 | 0.049 | 0.05 | 1 | 4 | Rpusd4 | -0.001 | Old |
| 0.995138 | 0.219975 | 0.169 | 0.175 | 1 | 4 | C1qtnf1 | -0.006 | Old |
| 0.995182 | 0.021248 | 0.038 | 0.038 | 1 | 4 | Bud13 | 0 | Old |
| 0.995291 | 0.113785 | 0.112 | 0.112 | 1 | 4 | Med8 | 0 | Old |
| 0.99533 | 0.135358 | 0.497 | 0.512 | 1 | 4 | Lgals1 | -0.015 | Old |
| 0.995752 | 0.338869 | 0.169 | 0.175 | 1 | 4 | Mtmr10 | -0.006 | Old |
| 0.995786 | 0.311875 | 0.158 | 0.162 | 1 | 4 | Socs6 | -0.004 | Old |
| 0.99588 | 0.696485 | 0.013 | 0.013 | 1 | 4 | Rnf24 | 0 | Old |
| 0.99588 | 0.464398 | 0.013 | 0.013 | 1 | 4 | Tceanc | 0 | Old |
| 0.99588 | 0.443048 | 0.013 | 0.013 | 1 | 4 | Ppp1r18os | 0 | Old |
| 0.99588 | 0.360117 | 0.013 | 0.013 | 1 | 4 | Adal | 0 | Old |
| 0.99588 | 0.253949 | 0.013 | 0.013 | 1 | 4 | BC147527 | 0 | Old |
| 0.995919 | 1.304311 | 0.024 | 0.025 | 1 | 4 | Txnrd3 | -0.001 | Old |
| 0.995929 | 0.28392 | 0.086 | 0.088 | 1 | 4 | Ifi211 | -0.002 | Old |
| 0.99644 | 0.144457 | 0.062 | 0.062 | 1 | 4 | Clmp | 0 | Old |
| 0.99691 | 0.424058 | 0.013 | 0.013 | 1 | 4 | Tubb2b | 0 | Old |
| 0.99691 | 0.125708 | 0.013 | 0.013 | 1 | 4 | Nlgn2 | 0 | Old |
| 0.996989 | 0.11677 | 0.038 | 0.038 | 1 | 4 | Fam114a1 | 0 | Old |

| | | | | | | | | |
|---|---|---|---|---|---|---|---|---|
| 0.99703 | 0.19807 | 0.227 | 0.238 | 1 | 4 | Nfia | -0.011 | Old |
| 0.997435 | 0.056654 | 0.025 | 0.025 | 1 | 4 | Rab3a | 0 | Old |
| 0.997435 | 0.021182 | 0.025 | 0.025 | 1 | 4 | Klhl15 | 0 | Old |
| 0.99746 | 0.02242 | 0.31 | 0.312 | 1 | 4 | Mapk1 | -0.002 | Old |
| 0.997599 | 0.36249 | 0.202 | 0.212 | 1 | 4 | Osgep | -0.01 | Old |
| 0.997668 | 0.23167 | 0.348 | 0.362 | 1 | 4 | Degs1 | -0.014 | Old |
| 0.99794 | 0.504069 | 0.013 | 0.013 | 1 | 4 | Zfp235 | 0 | Old |
| 0.99794 | 0.273225 | 0.013 | 0.013 | 1 | 4 | Mrm1 | 0 | Old |
| 0.998039 | 0.267396 | 0.074 | 0.075 | 1 | 4 | Cd80 | -0.001 | Old |
| 0.998039 | 0.236744 | 0.074 | 0.075 | 1 | 4 | Fbxo4 | -0.001 | Old |
| 0.998331 | 0.155246 | 0.169 | 0.175 | 1 | 4 | Aaed1 | -0.006 | Old |
| 0.9985 | 0.02202 | 0.75 | 0.725 | 1 | 4 | Slc6a6 | 0.025 | Old |
| 0.998848 | 0.319471 | 0.098 | 0.1 | 1 | 4 | Acox3 | -0.002 | Old |
| 0.998906 | 0.404977 | 0.074 | 0.075 | 1 | 4 | Med21 | -0.001 | Old |
| 0.99897 | 0.009219 | 0.013 | 0.013 | 1 | 4 | Gm15446 | 0 | Old |
| 0.99913 | 0.198387 | 0.123 | 0.125 | 1 | 4 | Sars | -0.002 | Old |
| 0.999266 | 0.389524 | 0.109 | 0.112 | 1 | 4 | Slc35f5 | -0.003 | Old |
| 0.999893 | 0.144675 | 0.481 | 0.55 | 1 | 4 | Ndufa6 | -0.069 | Old |
| 1.61E-29 | 4.567186 | 0.175 | 0.009 | 5.00E-25 | 4 | Tln2 | 0.166 | Young |
| 1.15E-24 | 1.248213 | 1 | 0.967 | 3.57E-20 | 4 | Apoe | 0.033 | Young |
| 1.58E-18 | 3.498149 | 0.112 | 0.006 | 4.90E-14 | 4 | Xist | 0.106 | Young |
| 4.53E-15 | 2.661048 | 0.288 | 0.061 | 1.41E-10 | 4 | Cx3cr1 | 0.227 | Young |
| 1.30E-11 | 0.586176 | 1 | 0.978 | 4.05E-07 | 4 | Cst3 | 0.022 | Young |
| 1.01E-10 | 2.946664 | 0.125 | 0.016 | 3.13E-06 | 4 | Inpp4b | 0.109 | Young |
| 1.34E-10 | 3.45728 | 0.038 | 0.001 | 4.16E-06 | 4 | Lrrc75a | 0.037 | Young |
| 1.34E-10 | 2.851703 | 0.038 | 0.001 | 4.16E-06 | 4 | Ager | 0.037 | Young |
| 1.34E-10 | 2.650142 | 0.038 | 0.001 | 4.16E-06 | 4 | Marco | 0.037 | Young |
| 2.49E-10 | 1.622352 | 0.275 | 0.073 | 7.75E-06 | 4 | Clec4n | 0.202 | Young |
| 4.02E-10 | 3.366587 | 0.075 | 0.006 | 1.25E-05 | 4 | Igfbp6 | 0.069 | Young |
| 6.25E-10 | 3.445532 | 0.062 | 0.004 | 1.94E-05 | 4 | 1700052K1 | 0.058 | Young |
| 7.26E-10 | 3.615977 | 0.025 | 0 | 2.26E-05 | 4 | Syngr3 | 0.025 | Young |
| 7.26E-10 | 3.115075 | 0.025 | 0 | 2.26E-05 | 4 | A430073D: | 0.025 | Young |
| 7.26E-10 | 1.574556 | 0.025 | 0 | 2.26E-05 | 4 | Gm15283 | 0.025 | Young |
| 9.60E-10 | 2.489055 | 0.188 | 0.04 | 2.98E-05 | 4 | Mtus1 | 0.148 | Young |
| 3.37E-09 | 0.705663 | 0.988 | 0.918 | 0.000105 | 4 | Hspa8 | 0.07 | Young |
| 7.40E-09 | 1.61192 | 0.325 | 0.111 | 0.00023 | 4 | Sh3bp5 | 0.214 | Young |
| 1.63E-08 | 3.046333 | 0.038 | 0.001 | 0.000506 | 4 | Tagln | 0.037 | Young |
| 1.65E-08 | 3.191928 | 0.038 | 0.001 | 0.000514 | 4 | Jag2 | 0.037 | Young |
| 3.86E-08 | 1.659606 | 0.162 | 0.035 | 0.001197 | 4 | Tmem9 | 0.127 | Young |
| 4.37E-08 | 1.498181 | 0.412 | 0.178 | 0.001357 | 4 | Gatm | 0.234 | Young |
| 5.06E-08 | 1.410132 | 0.438 | 0.194 | 0.001571 | 4 | Hpgd | 0.244 | Young |
| 5.64E-08 | 2.701948 | 0.125 | 0.023 | 0.001751 | 4 | Mettl7a1 | 0.102 | Young |
| 1.92E-07 | 0.639506 | 0.925 | 0.875 | 0.005976 | 4 | Csf1r | 0.05 | Young |
| 2.52E-07 | 2.529369 | 0.125 | 0.025 | 0.007813 | 4 | Ggta1 | 0.1 | Young |
| 3.89E-07 | 2.021254 | 0.125 | 0.025 | 0.012088 | 4 | Slc40a1 | 0.1 | Young |

| | | | | | | | | |
|---|---|---|---|---|---|---|---|---|
| 4.20E-07 | 2.571112 | 0.038 | 0.002 | 0.013058 | 4 | Mlxipl | 0.036 | Young |
| 4.78E-07 | 1.896091 | 0.35 | 0.146 | 0.014831 | 4 | Dnajb1 | 0.204 | Young |
| 6.74E-07 | 2.050725 | 0.075 | 0.01 | 0.02094 | 4 | Patj | 0.065 | Young |
| 7.17E-07 | 0.354087 | 0.988 | 0.988 | 0.022255 | 4 | Rplp1 | 0 | Young |
| 7.88E-07 | 1.748662 | 0.5 | 0.277 | 0.024462 | 4 | Hspa1a | 0.223 | Young |
| 9.07E-07 | 2.095633 | 0.238 | 0.08 | 0.02818 | 4 | Osm | 0.158 | Young |
| 9.48E-07 | 3.507284 | 0.025 | 0.001 | 0.029444 | 4 | Des | 0.024 | Young |
| 9.64E-07 | 2.368058 | 0.025 | 0.001 | 0.029946 | 4 | Gpat3 | 0.024 | Young |
| 9.64E-07 | 1.594228 | 0.025 | 0.001 | 0.029946 | 4 | Rapsn | 0.024 | Young |
| 9.81E-07 | 2.164597 | 0.025 | 0.001 | 0.030455 | 4 | Gm12708 | 0.024 | Young |
| 1.26E-06 | 4.080544 | 0.075 | 0.011 | 0.03911 | 4 | Cyr61 | 0.064 | Young |
| 1.27E-06 | 0.742763 | 0.925 | 0.815 | 0.039356 | 4 | Cd83 | 0.11 | Young |
| 1.43E-06 | 2.5743 | 0.075 | 0.011 | 0.044264 | 4 | Mgp | 0.064 | Young |
| 1.67E-06 | 2.21294 | 0.1 | 0.018 | 0.051901 | 4 | Gm42031 | 0.082 | Young |
| 1.74E-06 | 1.858565 | 0.188 | 0.055 | 0.054059 | 4 | Retreg1 | 0.133 | Young |
| 1.87E-06 | 1.394721 | 0.65 | 0.43 | 0.058002 | 4 | Cxcl2 | 0.22 | Young |
| 2.46E-06 | 0.383996 | 1 | 0.989 | 0.076481 | 4 | Mrc1 | 0.011 | Young |
| 2.93E-06 | 1.871309 | 0.188 | 0.058 | 0.091052 | 4 | Fscn1 | 0.13 | Young |
| 4.15E-06 | 1.166222 | 0.325 | 0.139 | 0.128777 | 4 | Limd2 | 0.186 | Young |
| 5.89E-06 | 1.338512 | 0.262 | 0.099 | 0.182755 | 4 | Tbc1d9b | 0.163 | Young |
| 6.10E-06 | 2.909173 | 0.05 | 0.005 | 0.189538 | 4 | Fhit | 0.045 | Young |
| 6.58E-06 | 1.834223 | 0.188 | 0.059 | 0.204478 | 4 | Las1l | 0.129 | Young |
| 8.00E-06 | 0.569363 | 0.9 | 0.786 | 0.248572 | 4 | Tyrobp | 0.114 | Young |
| 8.03E-06 | 1.063198 | 0.575 | 0.362 | 0.249356 | 4 | Gadd45b | 0.213 | Young |
| 9.34E-06 | 0.402745 | 0.938 | 0.912 | 0.290124 | 4 | Fcer1g | 0.026 | Young |
| 1.05E-05 | 1.399933 | 0.225 | 0.081 | 0.325956 | 4 | Naa30 | 0.144 | Young |
| 1.08E-05 | 1.267577 | 0.212 | 0.073 | 0.336494 | 4 | Mgat4a | 0.139 | Young |
| 1.10E-05 | 0.62888 | 0.975 | 0.906 | 0.340911 | 4 | Dab2 | 0.069 | Young |
| 1.27E-05 | 0.819949 | 0.55 | 0.314 | 0.393856 | 4 | Clec12a | 0.236 | Young |
| 1.34E-05 | 3.285014 | 0.013 | 0 | 0.416696 | 4 | H2afy2 | 0.013 | Young |
| 1.34E-05 | 3.138752 | 0.013 | 0 | 0.416696 | 4 | 1700040D1 | 0.013 | Young |
| 1.34E-05 | 3.065276 | 0.013 | 0 | 0.416696 | 4 | Gm5426 | 0.013 | Young |
| 1.34E-05 | 3.065276 | 0.013 | 0 | 0.416696 | 4 | Mlip | 0.013 | Young |
| 1.34E-05 | 2.84746 | 0.013 | 0 | 0.416696 | 4 | Gm17501 | 0.013 | Young |
| 1.34E-05 | 2.84746 | 0.013 | 0 | 0.416696 | 4 | 3632451O( | 0.013 | Young |
| 1.34E-05 | 2.84746 | 0.013 | 0 | 0.416696 | 4 | Hist1h2br | 0.013 | Young |
| 1.34E-05 | 2.833386 | 0.013 | 0 | 0.416696 | 4 | Tbx21 | 0.013 | Young |
| 1.34E-05 | 2.830351 | 0.013 | 0 | 0.416696 | 4 | Gm12925 | 0.013 | Young |
| 1.34E-05 | 2.830351 | 0.013 | 0 | 0.416696 | 4 | Papln | 0.013 | Young |
| 1.34E-05 | 2.829744 | 0.013 | 0 | 0.416696 | 4 | Plekha6 | 0.013 | Young |
| 1.34E-05 | 2.821896 | 0.013 | 0 | 0.416696 | 4 | Wasf3 | 0.013 | Young |
| 1.34E-05 | 2.821896 | 0.013 | 0 | 0.416696 | 4 | Gjc2 | 0.013 | Young |
| 1.34E-05 | 2.792812 | 0.013 | 0 | 0.416696 | 4 | Hmmr | 0.013 | Young |
| 1.34E-05 | 2.778848 | 0.013 | 0 | 0.416696 | 4 | 4930553M | 0.013 | Young |
| 1.34E-05 | 2.778848 | 0.013 | 0 | 0.416696 | 4 | Spock2 | 0.013 | Young |

| | | | | | | | | |
|---|---|---|---|---|---|---|---|---|
| 1.34E-05 | 2.778848 | 0.013 | 0 | 0.416696 | 4 | Cnih2 | 0.013 | Young |
| 1.34E-05 | 2.764492 | 0.013 | 0 | 0.416696 | 4 | Afmid | 0.013 | Young |
| 1.34E-05 | 2.662835 | 0.013 | 0 | 0.416696 | 4 | Foxd1 | 0.013 | Young |
| 1.34E-05 | 2.651495 | 0.013 | 0 | 0.416696 | 4 | Ddx43 | 0.013 | Young |
| 1.34E-05 | 2.64638 | 0.013 | 0 | 0.416696 | 4 | Teddm2 | 0.013 | Young |
| 1.34E-05 | 2.64485 | 0.013 | 0 | 0.416696 | 4 | Foxc2 | 0.013 | Young |
| 1.34E-05 | 2.64485 | 0.013 | 0 | 0.416696 | 4 | Clic5 | 0.013 | Young |
| 1.34E-05 | 2.63471 | 0.013 | 0 | 0.416696 | 4 | Gpam | 0.013 | Young |
| 1.34E-05 | 2.633198 | 0.013 | 0 | 0.416696 | 4 | Krt222 | 0.013 | Young |
| 1.34E-05 | 2.62717 | 0.013 | 0 | 0.416696 | 4 | Flnc | 0.013 | Young |
| 1.34E-05 | 2.561664 | 0.013 | 0 | 0.416696 | 4 | Gm10069 | 0.013 | Young |
| 1.34E-05 | 2.561664 | 0.013 | 0 | 0.416696 | 4 | Vtn | 0.013 | Young |
| 1.34E-05 | 2.510772 | 0.013 | 0 | 0.416696 | 4 | Cyb5d1 | 0.013 | Young |
| 1.34E-05 | 2.501424 | 0.013 | 0 | 0.416696 | 4 | Ltbp1 | 0.013 | Young |
| 1.34E-05 | 2.38882 | 0.013 | 0 | 0.416696 | 4 | B3gnt4 | 0.013 | Young |
| 1.34E-05 | 2.380938 | 0.013 | 0 | 0.416696 | 4 | Olfr1385 | 0.013 | Young |
| 1.34E-05 | 2.375459 | 0.013 | 0 | 0.416696 | 4 | Scn7a | 0.013 | Young |
| 1.34E-05 | 2.375459 | 0.013 | 0 | 0.416696 | 4 | Slc39a4 | 0.013 | Young |
| 1.34E-05 | 2.293545 | 0.013 | 0 | 0.416696 | 4 | 1700034P1 | 0.013 | Young |
| 1.34E-05 | 2.183991 | 0.013 | 0 | 0.416696 | 4 | Pnck | 0.013 | Young |
| 1.34E-05 | 2.113113 | 0.013 | 0 | 0.416696 | 4 | Col9a3 | 0.013 | Young |
| 1.34E-05 | 2.113113 | 0.013 | 0 | 0.416696 | 4 | Gm45184 | 0.013 | Young |
| 1.34E-05 | 2.113113 | 0.013 | 0 | 0.416696 | 4 | Eci3 | 0.013 | Young |
| 1.34E-05 | 2.111933 | 0.013 | 0 | 0.416696 | 4 | Kcne4 | 0.013 | Young |
| 1.34E-05 | 2.104009 | 0.013 | 0 | 0.416696 | 4 | Mylk4 | 0.013 | Young |
| 1.34E-05 | 2.095575 | 0.013 | 0 | 0.416696 | 4 | Olfr624 | 0.013 | Young |
| 1.34E-05 | 2.069616 | 0.013 | 0 | 0.416696 | 4 | Olfr1160 | 0.013 | Young |
| 1.34E-05 | 2.069616 | 0.013 | 0 | 0.416696 | 4 | Ppil6 | 0.013 | Young |
| 1.34E-05 | 2.029493 | 0.013 | 0 | 0.416696 | 4 | Gm29264 | 0.013 | Young |
| 1.34E-05 | 2.028151 | 0.013 | 0 | 0.416696 | 4 | Loxl1 | 0.013 | Young |
| 1.34E-05 | 1.923927 | 0.013 | 0 | 0.416696 | 4 | Zg16 | 0.013 | Young |
| 1.34E-05 | 1.860157 | 0.013 | 0 | 0.416696 | 4 | Gm15612 | 0.013 | Young |
| 1.34E-05 | 1.860157 | 0.013 | 0 | 0.416696 | 4 | Syt12 | 0.013 | Young |
| 1.34E-05 | 1.830458 | 0.013 | 0 | 0.416696 | 4 | Ifitm10 | 0.013 | Young |
| 1.34E-05 | 1.798541 | 0.013 | 0 | 0.416696 | 4 | Gm44148 | 0.013 | Young |
| 1.34E-05 | 1.745141 | 0.013 | 0 | 0.416696 | 4 | 4933411E0 | 0.013 | Young |
| 1.34E-05 | 1.689472 | 0.013 | 0 | 0.416696 | 4 | Gm26541 | 0.013 | Young |
| 1.34E-05 | 1.689472 | 0.013 | 0 | 0.416696 | 4 | Cnn1 | 0.013 | Young |
| 1.34E-05 | 1.689472 | 0.013 | 0 | 0.416696 | 4 | Prss30 | 0.013 | Young |
| 1.34E-05 | 1.624653 | 0.013 | 0 | 0.416696 | 4 | C130073E2 | 0.013 | Young |
| 1.34E-05 | 1.577614 | 0.013 | 0 | 0.416696 | 4 | Gm19434 | 0.013 | Young |
| 1.34E-05 | 1.577614 | 0.013 | 0 | 0.416696 | 4 | Nrxn3 | 0.013 | Young |
| 1.34E-05 | 1.455313 | 0.013 | 0 | 0.416696 | 4 | Zfp819 | 0.013 | Young |
| 1.34E-05 | 1.309825 | 0.013 | 0 | 0.416696 | 4 | Cox8b | 0.013 | Young |
| 1.34E-05 | 1.309825 | 0.013 | 0 | 0.416696 | 4 | Prr36 | 0.013 | Young |

| | | | | | | | | |
|---|---|---|---|---|---|---|---|---|
| 1.34E-05 | 1.228226 | 0.013 | 0 | 0.416696 | 4 | Il23a | 0.013 | Young |
| 1.34E-05 | 1.187926 | 0.013 | 0 | 0.416696 | 4 | Gm19265 | 0.013 | Young |
| 1.34E-05 | 1.187926 | 0.013 | 0 | 0.416696 | 4 | Pcdhb19 | 0.013 | Young |
| 1.34E-05 | 1.187926 | 0.013 | 0 | 0.416696 | 4 | Pcdhga1 | 0.013 | Young |
| 1.34E-05 | 1.147312 | 0.013 | 0 | 0.416696 | 4 | Ak4 | 0.013 | Young |
| 1.34E-05 | 1.074103 | 0.013 | 0 | 0.416696 | 4 | Pbld1 | 0.013 | Young |
| 1.34E-05 | 1.074103 | 0.013 | 0 | 0.416696 | 4 | Mns1 | 0.013 | Young |
| 1.34E-05 | 1.074103 | 0.013 | 0 | 0.416696 | 4 | Pygo1 | 0.013 | Young |
| 1.34E-05 | 1.074103 | 0.013 | 0 | 0.416696 | 4 | Apol9a | 0.013 | Young |
| 1.34E-05 | 0.905372 | 0.013 | 0 | 0.416696 | 4 | Angpt4 | 0.013 | Young |
| 1.34E-05 | 0.905372 | 0.013 | 0 | 0.416696 | 4 | Lama5 | 0.013 | Young |
| 1.34E-05 | 0.905372 | 0.013 | 0 | 0.416696 | 4 | Ssc5d | 0.013 | Young |
| 1.37E-05 | 2.25005 | 0.4 | 0.219 | 0.424938 | 4 | Hspa1b | 0.181 | Young |
| 1.81E-05 | 2.948022 | 0.05 | 0.006 | 0.560915 | 4 | Gas1 | 0.044 | Young |
| 1.99E-05 | 1.965126 | 0.062 | 0.009 | 0.619052 | 4 | C2 | 0.053 | Young |
| 2.28E-05 | 1.214384 | 0.275 | 0.115 | 0.707945 | 4 | Siglech | 0.16 | Young |
| 2.30E-05 | 1.768161 | 0.15 | 0.045 | 0.714085 | 4 | Sesn3 | 0.105 | Young |
| 2.33E-05 | 1.822711 | 0.162 | 0.051 | 0.724388 | 4 | Rcan1 | 0.111 | Young |
| 2.36E-05 | 1.206427 | 0.362 | 0.175 | 0.73223 | 4 | Tmem8 | 0.187 | Young |
| 2.47E-05 | 0.455537 | 0.938 | 0.808 | 0.766297 | 4 | Cebpb | 0.13 | Young |
| 2.51E-05 | 1.993434 | 0.038 | 0.003 | 0.780708 | 4 | Snhg18 | 0.035 | Young |
| 2.56E-05 | 1.831348 | 0.038 | 0.003 | 0.794894 | 4 | Armc2 | 0.035 | Young |
| 2.69E-05 | 0.923994 | 0.625 | 0.385 | 0.833928 | 4 | Fcrls | 0.24 | Young |
| 2.82E-05 | 0.887238 | 0.4 | 0.196 | 0.874291 | 4 | Cebpa | 0.204 | Young |
| 2.82E-05 | 1.593145 | 0.088 | 0.018 | 0.874318 | 4 | Stk39 | 0.07 | Young |
| 2.95E-05 | 0.846885 | 0.575 | 0.358 | 0.914607 | 4 | Rassf4 | 0.217 | Young |
| 3.10E-05 | 0.968951 | 0.525 | 0.308 | 0.961809 | 4 | Pou2f2 | 0.217 | Young |
| 3.21E-05 | 1.03065 | 0.1 | 0.022 | 0.99557 | 4 | Slc16a13 | 0.078 | Young |
| 3.28E-05 | 1.271994 | 0.262 | 0.108 | 1 | 4 | Chrac1 | 0.154 | Young |
| 3.64E-05 | 2.276655 | 0.025 | 0.001 | 1 | 4 | Col6a5 | 0.024 | Young |
| 3.69E-05 | 1.627976 | 0.025 | 0.001 | 1 | 4 | Gm12185 | 0.024 | Young |
| 3.74E-05 | 2.532345 | 0.025 | 0.001 | 1 | 4 | Gm10516 | 0.024 | Young |
| 3.74E-05 | 1.597929 | 0.025 | 0.001 | 1 | 4 | Gm10384 | 0.024 | Young |
| 3.74E-05 | 1.318764 | 0.025 | 0.001 | 1 | 4 | Kcnab1 | 0.024 | Young |
| 3.78E-05 | 0.912077 | 0.537 | 0.333 | 1 | 4 | Cd33 | 0.204 | Young |
| 3.84E-05 | 0.655935 | 0.912 | 0.781 | 1 | 4 | Btg2 | 0.131 | Young |
| 4.18E-05 | 1.921335 | 0.125 | 0.034 | 1 | 4 | Dglucy | 0.091 | Young |
| 4.19E-05 | 0.898834 | 0.725 | 0.543 | 1 | 4 | Ier3 | 0.182 | Young |
| 4.82E-05 | 1.904436 | 0.1 | 0.024 | 1 | 4 | Zmynd15 | 0.076 | Young |
| 5.34E-05 | 1.026538 | 0.2 | 0.072 | 1 | 4 | Mroh1 | 0.128 | Young |
| 5.47E-05 | 2.483692 | 0.075 | 0.014 | 1 | 4 | Lpcat2 | 0.061 | Young |
| 5.69E-05 | 1.344638 | 0.188 | 0.067 | 1 | 4 | Gm45716 | 0.121 | Young |
| 7.22E-05 | 1.167626 | 0.262 | 0.115 | 1 | 4 | Cyp27a1 | 0.147 | Young |
| 7.30E-05 | 1.112854 | 0.325 | 0.159 | 1 | 4 | Cd4 | 0.166 | Young |
| 7.68E-05 | 1.749773 | 0.062 | 0.011 | 1 | 4 | Pafah2 | 0.051 | Young |

| | | | | | | | | |
|---|---|---|---|---|---|---|---|---|
| 7.81E-05 | 1.733072 | 0.112 | 0.03 | 1 | 4 | Vsig4 | 0.082 | Young |
| 8.10E-05 | 2.139287 | 0.088 | 0.02 | 1 | 4 | Mrvi1 | 0.068 | Young |
| 8.27E-05 | 1.571062 | 0.2 | 0.078 | 1 | 4 | Itgb5 | 0.122 | Young |
| 8.63E-05 | 1.231322 | 0.225 | 0.092 | 1 | 4 | Fuca2 | 0.133 | Young |
| 8.76E-05 | 1.031851 | 0.325 | 0.159 | 1 | 4 | Dhx36 | 0.166 | Young |
| 8.97E-05 | 0.782094 | 0.575 | 0.357 | 1 | 4 | Cd86 | 0.218 | Young |
| 9.39E-05 | 2.373234 | 0.038 | 0.004 | 1 | 4 | 2810021J2 | 0.034 | Young |
| 9.62E-05 | 2.250583 | 0.038 | 0.004 | 1 | 4 | Egfl7 | 0.034 | Young |
| 0.000101 | 0.814876 | 0.488 | 0.283 | 1 | 4 | Rel | 0.205 | Young |
| 0.000105 | 0.664268 | 0.8 | 0.601 | 1 | 4 | Mef2c | 0.199 | Young |
| 0.000107 | 1.331222 | 0.188 | 0.069 | 1 | 4 | Txndc12 | 0.119 | Young |
| 0.000108 | 1.006965 | 0.55 | 0.369 | 1 | 4 | Cd38 | 0.181 | Young |
| 0.000108 | 2.809714 | 0.05 | 0.007 | 1 | 4 | Spry2 | 0.043 | Young |
| 0.000112 | 1.428464 | 0.188 | 0.07 | 1 | 4 | Adpgk | 0.118 | Young |
| 0.00012 | 2.464702 | 0.062 | 0.011 | 1 | 4 | Osbpl3 | 0.051 | Young |
| 0.000134 | 1.197215 | 0.225 | 0.094 | 1 | 4 | Rhbdf2 | 0.131 | Young |
| 0.000138 | 0.774335 | 0.662 | 0.465 | 1 | 4 | Slc3a2 | 0.197 | Young |
| 0.000138 | 1.49366 | 0.162 | 0.057 | 1 | 4 | Mrps5 | 0.105 | Young |
| 0.000151 | 1.000776 | 0.3 | 0.143 | 1 | 4 | Bcl2l11 | 0.157 | Young |
| 0.000154 | 1.036096 | 0.3 | 0.144 | 1 | 4 | Prpf8 | 0.156 | Young |
| 0.000154 | 0.794191 | 0.475 | 0.277 | 1 | 4 | Nfxl1 | 0.198 | Young |
| 0.000174 | 0.684644 | 0.812 | 0.628 | 1 | 4 | Nr4a1 | 0.184 | Young |
| 0.000193 | 1.37675 | 0.262 | 0.119 | 1 | 4 | Nlrp3 | 0.143 | Young |
| 0.000204 | 0.66648 | 0.85 | 0.649 | 1 | 4 | Atf3 | 0.201 | Young |
| 0.000208 | 2.050406 | 0.062 | 0.012 | 1 | 4 | C030034I2 | 0.05 | Young |
| 0.000211 | 0.759899 | 0.75 | 0.585 | 1 | 4 | Cd14 | 0.165 | Young |
| 0.000213 | 2.055884 | 0.075 | 0.016 | 1 | 4 | 1600020E0 | 0.059 | Young |
| 0.00022 | 2.650369 | 0.05 | 0.008 | 1 | 4 | Dnal1 | 0.042 | Young |
| 0.000225 | 1.746201 | 0.075 | 0.016 | 1 | 4 | Cfd | 0.059 | Young |
| 0.000226 | 2.342548 | 0.05 | 0.008 | 1 | 4 | Mief2 | 0.042 | Young |
| 0.000239 | 0.900427 | 0.375 | 0.209 | 1 | 4 | Hnrnpr | 0.166 | Young |
| 0.000245 | 1.559121 | 0.1 | 0.027 | 1 | 4 | Cxxc5 | 0.073 | Young |
| 0.000246 | 0.620359 | 0.738 | 0.532 | 1 | 4 | Tab2 | 0.206 | Young |
| 0.000267 | 1.223768 | 0.3 | 0.151 | 1 | 4 | Lrp10 | 0.149 | Young |
| 0.000273 | 2.378416 | 0.088 | 0.022 | 1 | 4 | Taf6 | 0.066 | Young |
| 0.000275 | 1.838933 | 0.112 | 0.033 | 1 | 4 | Ly6a | 0.079 | Young |
| 0.000286 | 1.492811 | 0.038 | 0.005 | 1 | 4 | 2610301B2 | 0.033 | Young |
| 0.000308 | 1.087053 | 0.425 | 0.267 | 1 | 4 | Coq10b | 0.158 | Young |
| 0.000318 | 0.810737 | 0.462 | 0.287 | 1 | 4 | Runx1 | 0.175 | Young |
| 0.000319 | 3.40705 | 0.025 | 0.002 | 1 | 4 | Gm26513 | 0.023 | Young |
| 0.000323 | 2.611181 | 0.025 | 0.002 | 1 | 4 | Gm13710 | 0.023 | Young |
| 0.000326 | 1.411928 | 0.025 | 0.002 | 1 | 4 | 1810021B2 | 0.023 | Young |
| 0.000329 | 2.466027 | 0.025 | 0.002 | 1 | 4 | Bik | 0.023 | Young |
| 0.000329 | 2.451114 | 0.025 | 0.002 | 1 | 4 | Sox13 | 0.023 | Young |
| 0.000329 | 2.153999 | 0.025 | 0.002 | 1 | 4 | Tlcd1 | 0.023 | Young |

| | | | | | | | | |
|---|---|---|---|---|---|---|---|---|
| 0.000332 | 2.473588 | 0.025 | 0.002 | 1 | 4 | Gm26826 | 0.023 | Young |
| 0.000332 | 1.87045 | 0.025 | 0.002 | 1 | 4 | Nat14 | 0.023 | Young |
| 0.000335 | 1.535998 | 0.025 | 0.002 | 1 | 4 | Plppr3 | 0.023 | Young |
| 0.000346 | 1.834108 | 0.1 | 0.028 | 1 | 4 | Zfp595 | 0.072 | Young |
| 0.000368 | 0.622907 | 0.8 | 0.627 | 1 | 4 | Ptpn18 | 0.173 | Young |
| 0.000378 | 0.86829 | 0.412 | 0.241 | 1 | 4 | Gpr34 | 0.171 | Young |
| 0.000397 | 0.507972 | 0.787 | 0.607 | 1 | 4 | Rpl12 | 0.18 | Young |
| 0.000412 | 2.110366 | 0.075 | 0.018 | 1 | 4 | Osgin1 | 0.057 | Young |
| 0.000416 | 0.609574 | 0.675 | 0.525 | 1 | 4 | Ubb | 0.15 | Young |
| 0.000438 | 2.617734 | 0.05 | 0.009 | 1 | 4 | Lrg1 | 0.041 | Young |
| 0.000448 | 1.121093 | 0.212 | 0.093 | 1 | 4 | Irak3 | 0.119 | Young |
| 0.000468 | 0.5943 | 0.488 | 0.289 | 1 | 4 | Nufip2 | 0.199 | Young |
| 0.000489 | 1.438046 | 0.1 | 0.029 | 1 | 4 | Fars2 | 0.071 | Young |
| 0.000514 | 2.215117 | 0.062 | 0.013 | 1 | 4 | Pde4d | 0.049 | Young |
| 0.000523 | 0.928871 | 0.388 | 0.224 | 1 | 4 | Cd93 | 0.164 | Young |
| 0.000531 | 1.105917 | 0.262 | 0.13 | 1 | 4 | Ago4 | 0.132 | Young |
| 0.000544 | 1.094845 | 0.088 | 0.023 | 1 | 4 | Pald1 | 0.065 | Young |
| 0.000547 | 1.010951 | 0.25 | 0.119 | 1 | 4 | Rnf11 | 0.131 | Young |
| 0.000551 | 1.373402 | 0.2 | 0.088 | 1 | 4 | Wdr44 | 0.112 | Young |
| 0.000564 | 0.685219 | 0.388 | 0.213 | 1 | 4 | Ifnar2 | 0.175 | Young |
| 0.000598 | 1.44491 | 0.112 | 0.036 | 1 | 4 | Colec12 | 0.076 | Young |
| 0.000607 | 1.972923 | 0.075 | 0.018 | 1 | 4 | Per2 | 0.057 | Young |
| 0.000654 | 2.754543 | 0.038 | 0.005 | 1 | 4 | Tcea3 | 0.033 | Young |
| 0.000673 | 1.555934 | 0.125 | 0.043 | 1 | 4 | Ap2a1 | 0.082 | Young |
| 0.000675 | 2.359923 | 0.038 | 0.005 | 1 | 4 | B130006D( | 0.033 | Young |
| 0.000675 | 2.3276 | 0.038 | 0.005 | 1 | 4 | Sox4 | 0.033 | Young |
| 0.000675 | 0.875604 | 0.125 | 0.042 | 1 | 4 | Pqlc2 | 0.083 | Young |
| 0.000683 | 1.913633 | 0.038 | 0.005 | 1 | 4 | Fmo2 | 0.033 | Young |
| 0.000685 | 0.600433 | 0.613 | 0.446 | 1 | 4 | Lst1 | 0.167 | Young |
| 0.000717 | 2.441068 | 0.05 | 0.009 | 1 | 4 | Fam109a | 0.041 | Young |
| 0.000723 | 1.209279 | 0.038 | 0.005 | 1 | 4 | A530013C2 | 0.033 | Young |
| 0.000725 | 1.228439 | 0.325 | 0.188 | 1 | 4 | Dennd4a | 0.137 | Young |
| 0.000733 | 0.661742 | 0.038 | 0.005 | 1 | 4 | Ccl11 | 0.033 | Young |
| 0.000737 | 0.538162 | 0.038 | 0.005 | 1 | 4 | Cd22 | 0.033 | Young |
| 0.000739 | 2.412779 | 0.05 | 0.009 | 1 | 4 | Atp23 | 0.041 | Young |
| 0.000753 | 1.708824 | 0.088 | 0.024 | 1 | 4 | Laptm4b | 0.064 | Young |
| 0.000761 | 2.070668 | 0.05 | 0.009 | 1 | 4 | Ms4a4a | 0.041 | Young |
| 0.000773 | 0.820177 | 0.212 | 0.093 | 1 | 4 | Sla | 0.119 | Young |
| 0.000778 | 1.702805 | 0.088 | 0.024 | 1 | 4 | Tapt1 | 0.064 | Young |
| 0.000784 | 1.837998 | 0.05 | 0.009 | 1 | 4 | Serpina3g | 0.041 | Young |
| 0.0008 | 1.314212 | 0.25 | 0.125 | 1 | 4 | Atp7a | 0.125 | Young |
| 0.000805 | 1.947139 | 0.062 | 0.014 | 1 | 4 | Nup188 | 0.048 | Young |
| 0.000805 | 1.866624 | 0.062 | 0.014 | 1 | 4 | Rpgrip1 | 0.048 | Young |
| 0.000848 | 2.291141 | 0.112 | 0.038 | 1 | 4 | Ccl3 | 0.074 | Young |
| 0.000876 | 0.829751 | 0.238 | 0.11 | 1 | 4 | Spg7 | 0.128 | Young |

| | | | | | | | | |
|---|---|---|---|---|---|---|---|---|
| 0.000925 | 1.125803 | 0.138 | 0.05 | 1 | 4 | Hook2 | 0.088 | Young |
| 0.000937 | 1.214336 | 0.262 | 0.133 | 1 | 4 | Hist3h2a | 0.129 | Young |
| 0.000941 | 0.594164 | 0.525 | 0.356 | 1 | 4 | Neurl3 | 0.169 | Young |
| 0.000942 | 2.090264 | 0.162 | 0.07 | 1 | 4 | Nr4a3 | 0.092 | Young |
| 0.001045 | 0.846964 | 0.4 | 0.248 | 1 | 4 | Prdx5 | 0.152 | Young |
| 0.001088 | 0.524011 | 0.675 | 0.481 | 1 | 4 | Tsc22d3 | 0.194 | Young |
| 0.001089 | 1.552593 | 0.188 | 0.083 | 1 | 4 | Arsb | 0.105 | Young |
| 0.001106 | 1.533581 | 0.125 | 0.045 | 1 | 4 | Epg5 | 0.08 | Young |
| 0.001114 | 2.004831 | 0.062 | 0.014 | 1 | 4 | Rsad1 | 0.048 | Young |
| 0.001126 | 0.350996 | 0.887 | 0.783 | 1 | 4 | Rps15a | 0.104 | Young |
| 0.001137 | 1.106406 | 0.175 | 0.074 | 1 | 4 | Selenoh | 0.101 | Young |
| 0.001137 | 0.861081 | 0.388 | 0.229 | 1 | 4 | Arhgap31 | 0.159 | Young |
| 0.001139 | 1.64479 | 0.125 | 0.045 | 1 | 4 | Mettl6 | 0.08 | Young |
| 0.001169 | 0.372875 | 0.875 | 0.81 | 1 | 4 | Rpl26 | 0.065 | Young |
| 0.001177 | 1.456037 | 0.212 | 0.103 | 1 | 4 | Ccrl2 | 0.109 | Young |
| 0.00119 | 0.617616 | 0.35 | 0.192 | 1 | 4 | Bscl2 | 0.158 | Young |
| 0.0012 | 0.943272 | 0.15 | 0.058 | 1 | 4 | Rnaset2a | 0.092 | Young |
| 0.001204 | 1.352636 | 0.138 | 0.052 | 1 | 4 | Tsc2 | 0.086 | Young |
| 0.001243 | 2.123364 | 0.05 | 0.01 | 1 | 4 | Tmem220 | 0.04 | Young |
| 0.00126 | 1.874615 | 0.05 | 0.01 | 1 | 4 | Kcng2 | 0.04 | Young |
| 0.001278 | 1.468258 | 0.075 | 0.02 | 1 | 4 | Slc25a33 | 0.055 | Young |
| 0.001278 | 1.841879 | 0.05 | 0.01 | 1 | 4 | Ccdc114 | 0.04 | Young |
| 0.001284 | 2.023989 | 0.05 | 0.01 | 1 | 4 | Mb21d2 | 0.04 | Young |
| 0.001295 | 1.579802 | 0.175 | 0.074 | 1 | 4 | Gt(ROSA)2( | 0.101 | Young |
| 0.001302 | 1.066001 | 0.15 | 0.059 | 1 | 4 | Serpinf1 | 0.091 | Young |
| 0.001332 | 1.597029 | 0.05 | 0.01 | 1 | 4 | BC030499 | 0.04 | Young |
| 0.001351 | 0.462281 | 0.762 | 0.557 | 1 | 4 | Rps18 | 0.205 | Young |
| 0.001355 | 0.758806 | 0.4 | 0.244 | 1 | 4 | Irf1 | 0.156 | Young |
| 0.001366 | 0.524971 | 0.725 | 0.567 | 1 | 4 | Fcgr3 | 0.158 | Young |
| 0.001371 | 1.659507 | 0.1 | 0.032 | 1 | 4 | Vps72 | 0.068 | Young |
| 0.001375 | 0.727869 | 0.388 | 0.231 | 1 | 4 | Psmd11 | 0.157 | Young |
| 0.001379 | 0.755805 | 0.562 | 0.389 | 1 | 4 | Inpp5d | 0.173 | Young |
| 0.001397 | 2.57404 | 0.038 | 0.006 | 1 | 4 | Mok | 0.032 | Young |
| 0.001407 | 1.067591 | 0.238 | 0.117 | 1 | 4 | Ang | 0.121 | Young |
| 0.001433 | 2.360104 | 0.025 | 0.003 | 1 | 4 | Pomgnt2 | 0.022 | Young |
| 0.001444 | 2.649186 | 0.025 | 0.003 | 1 | 4 | Eda | 0.022 | Young |
| 0.001444 | 2.618826 | 0.025 | 0.003 | 1 | 4 | Gm2415 | 0.022 | Young |
| 0.001444 | 2.178031 | 0.025 | 0.003 | 1 | 4 | Abcb4 | 0.022 | Young |
| 0.001444 | 1.818625 | 0.025 | 0.003 | 1 | 4 | Hist1h2ae | 0.022 | Young |
| 0.001456 | 2.091766 | 0.025 | 0.003 | 1 | 4 | Col1a1 | 0.022 | Young |
| 0.001456 | 1.740137 | 0.025 | 0.003 | 1 | 4 | Tpx2 | 0.022 | Young |
| 0.001456 | 1.38552 | 0.025 | 0.003 | 1 | 4 | Arg1 | 0.022 | Young |
| 0.001468 | 1.869572 | 0.025 | 0.003 | 1 | 4 | Gpr137c | 0.022 | Young |
| 0.001468 | 1.656469 | 0.025 | 0.003 | 1 | 4 | Txk | 0.022 | Young |
| 0.001471 | 1.598119 | 0.038 | 0.006 | 1 | 4 | Tmem120b | 0.032 | Young |

| | | | | | | | | |
|---|---|---|---|---|---|---|---|---|
| 0.001474 | 2.512262 | 0.062 | 0.015 | 1 | 4 | Pxdc1 | 0.047 | Young |
| 0.001511 | 0.443015 | 0.863 | 0.717 | 1 | 4 | Adgre1 | 0.146 | Young |
| 0.001514 | 0.973005 | 0.038 | 0.006 | 1 | 4 | Gne | 0.032 | Young |
| 0.001525 | 1.389275 | 0.15 | 0.061 | 1 | 4 | Pstpip1 | 0.089 | Young |
| 0.001527 | 1.383355 | 0.138 | 0.053 | 1 | 4 | Tgfbr1 | 0.085 | Young |
| 0.00153 | 0.689603 | 0.512 | 0.343 | 1 | 4 | Slc9a9 | 0.169 | Young |
| 0.001536 | 2.23322 | 0.062 | 0.015 | 1 | 4 | Fmnl3 | 0.047 | Young |
| 0.001546 | 0.799131 | 0.175 | 0.074 | 1 | 4 | Trp53inp2 | 0.101 | Young |
| 0.001558 | 0.262341 | 0.038 | 0.006 | 1 | 4 | Ldlrad4 | 0.032 | Young |
| 0.001563 | 1.380725 | 0.225 | 0.114 | 1 | 4 | Nxpe5 | 0.111 | Young |
| 0.001566 | 0.239095 | 0.988 | 0.991 | 1 | 4 | Ftl1 | -0.003 | Young |
| 0.001599 | 0.781763 | 0.475 | 0.339 | 1 | 4 | Per1 | 0.136 | Young |
| 0.001624 | 0.200814 | 1 | 0.995 | 1 | 4 | Fth1 | 0.005 | Young |
| 0.001717 | 1.129741 | 0.262 | 0.139 | 1 | 4 | Acp1 | 0.123 | Young |
| 0.001723 | 0.886589 | 0.238 | 0.117 | 1 | 4 | Cnot2 | 0.121 | Young |
| 0.001778 | 0.311772 | 1 | 0.985 | 1 | 4 | Pf4 | 0.015 | Young |
| 0.001802 | 0.774389 | 0.425 | 0.266 | 1 | 4 | Ypel3 | 0.159 | Young |
| 0.001811 | 2.632606 | 0.05 | 0.011 | 1 | 4 | Gm5086 | 0.039 | Young |
| 0.001884 | 1.004532 | 0.288 | 0.155 | 1 | 4 | Rab3gap1 | 0.133 | Young |
| 0.001899 | 0.417218 | 0.975 | 0.932 | 1 | 4 | Kctd12 | 0.043 | Young |
| 0.001959 | 1.80728 | 0.05 | 0.011 | 1 | 4 | Wars2 | 0.039 | Young |
| 0.001964 | 1.661653 | 0.075 | 0.021 | 1 | 4 | 4930430F0 | 0.054 | Young |
| 0.001985 | 0.606683 | 0.162 | 0.067 | 1 | 4 | Sash3 | 0.095 | Young |
| 0.00199 | 1.337212 | 0.15 | 0.063 | 1 | 4 | Tnf | 0.087 | Young |
| 0.002007 | 0.537997 | 0.425 | 0.286 | 1 | 4 | Rgs2 | 0.139 | Young |
| 0.002007 | 0.281015 | 1 | 0.966 | 1 | 4 | Rps12 | 0.034 | Young |
| 0.002019 | 1.73867 | 0.05 | 0.011 | 1 | 4 | Oplah | 0.039 | Young |
| 0.00202 | 1.040778 | 0.275 | 0.148 | 1 | 4 | Bckdk | 0.127 | Young |
| 0.002062 | 1.451827 | 0.088 | 0.027 | 1 | 4 | Lhpp | 0.061 | Young |
| 0.002167 | 0.685185 | 0.462 | 0.301 | 1 | 4 | Tmed9 | 0.161 | Young |
| 0.002167 | 1.064446 | 0.288 | 0.165 | 1 | 4 | Axl | 0.123 | Young |
| 0.002192 | 0.445776 | 0.875 | 0.754 | 1 | 4 | Marcksl1 | 0.121 | Young |
| 0.002204 | 1.243622 | 0.238 | 0.122 | 1 | 4 | F630028O1 | 0.116 | Young |
| 0.002205 | 1.051756 | 0.15 | 0.061 | 1 | 4 | Soga1 | 0.089 | Young |
| 0.002206 | 0.982373 | 0.112 | 0.04 | 1 | 4 | Edem3 | 0.072 | Young |
| 0.002218 | 1.65691 | 0.062 | 0.016 | 1 | 4 | Klhl26 | 0.046 | Young |
| 0.00226 | 0.633993 | 0.775 | 0.676 | 1 | 4 | Abca1 | 0.099 | Young |
| 0.00237 | 0.449832 | 0.838 | 0.717 | 1 | 4 | Rpl32 | 0.121 | Young |
| 0.00248 | 0.994211 | 0.238 | 0.121 | 1 | 4 | Eif2ak2 | 0.117 | Young |
| 0.002481 | 1.076119 | 0.125 | 0.047 | 1 | 4 | Retsat | 0.078 | Young |
| 0.002489 | 0.71003 | 0.3 | 0.167 | 1 | 4 | Rac2 | 0.133 | Young |
| 0.002568 | 1.64618 | 0.075 | 0.022 | 1 | 4 | Cd3eap | 0.053 | Young |
| 0.002594 | 0.469697 | 0.212 | 0.099 | 1 | 4 | Lap3 | 0.113 | Young |
| 0.002629 | 0.451393 | 0.662 | 0.495 | 1 | 4 | Nfkbid | 0.167 | Young |
| 0.002702 | 1.721182 | 0.038 | 0.007 | 1 | 4 | P3h1 | 0.031 | Young |

| | | | | | | | | |
|---|---|---|---|---|---|---|---|---|
| 0.002716 | 1.912592 | 0.038 | 0.007 | 1 | 4 | Gm13822 | 0.031 | Young |
| 0.002726 | 1.389854 | 0.075 | 0.022 | 1 | 4 | Rad9a | 0.053 | Young |
| 0.002759 | 1.714802 | 0.038 | 0.007 | 1 | 4 | Gm15859 | 0.031 | Young |
| 0.002884 | 1.343209 | 0.15 | 0.064 | 1 | 4 | Prps1 | 0.086 | Young |
| 0.002921 | 0.822065 | 0.288 | 0.157 | 1 | 4 | Stt3b | 0.131 | Young |
| 0.002928 | 0.868982 | 0.088 | 0.028 | 1 | 4 | Gatc | 0.06 | Young |
| 0.002965 | 1.160606 | 0.1 | 0.034 | 1 | 4 | Fbf1 | 0.066 | Young |
| 0.002992 | 1.116074 | 0.138 | 0.056 | 1 | 4 | Mier3 | 0.082 | Young |
| 0.002999 | 1.020342 | 0.475 | 0.329 | 1 | 4 | Il1b | 0.146 | Young |
| 0.003037 | 1.787606 | 0.088 | 0.028 | 1 | 4 | Trpm2 | 0.06 | Young |
| 0.003099 | 0.972143 | 0.238 | 0.126 | 1 | 4 | Slc35a5 | 0.112 | Young |
| 0.00314 | 0.417243 | 0.838 | 0.772 | 1 | 4 | Rps5 | 0.066 | Young |
| 0.003244 | 0.649667 | 0.45 | 0.291 | 1 | 4 | Hpgds | 0.159 | Young |
| 0.003265 | 0.86741 | 0.262 | 0.144 | 1 | 4 | Zdhhc9 | 0.118 | Young |
| 0.003376 | 0.853994 | 0.212 | 0.106 | 1 | 4 | Cklf | 0.106 | Young |
| 0.003442 | 0.747147 | 0.162 | 0.071 | 1 | 4 | Car3 | 0.091 | Young |
| 0.003514 | 3.151417 | 0.013 | 0.001 | 1 | 4 | Alox15 | 0.012 | Young |
| 0.003514 | 3.041213 | 0.013 | 0.001 | 1 | 4 | Plin1 | 0.012 | Young |
| 0.003514 | 2.7134 | 0.013 | 0.001 | 1 | 4 | Itga2b | 0.012 | Young |
| 0.003514 | 2.709564 | 0.013 | 0.001 | 1 | 4 | Hoxb8 | 0.012 | Young |
| 0.003514 | 2.657289 | 0.013 | 0.001 | 1 | 4 | Abi3bp | 0.012 | Young |
| 0.003514 | 2.64825 | 0.013 | 0.001 | 1 | 4 | Gm26664 | 0.012 | Young |
| 0.003514 | 2.604215 | 0.013 | 0.001 | 1 | 4 | Tspan9 | 0.012 | Young |
| 0.003514 | 2.565812 | 0.013 | 0.001 | 1 | 4 | 1110020A2 | 0.012 | Young |
| 0.003514 | 2.553127 | 0.013 | 0.001 | 1 | 4 | Bicd1 | 0.012 | Young |
| 0.003514 | 2.533589 | 0.013 | 0.001 | 1 | 4 | Defb42 | 0.012 | Young |
| 0.003514 | 2.511918 | 0.013 | 0.001 | 1 | 4 | Psd2 | 0.012 | Young |
| 0.003514 | 2.487567 | 0.013 | 0.001 | 1 | 4 | Sfrp2 | 0.012 | Young |
| 0.003514 | 2.458178 | 0.013 | 0.001 | 1 | 4 | Sod3 | 0.012 | Young |
| 0.003514 | 2.453994 | 0.013 | 0.001 | 1 | 4 | Enpp2 | 0.012 | Young |
| 0.003514 | 2.376771 | 0.013 | 0.001 | 1 | 4 | Tcf24 | 0.012 | Young |
| 0.003514 | 2.360608 | 0.013 | 0.001 | 1 | 4 | Rsph3a | 0.012 | Young |
| 0.003514 | 2.338388 | 0.013 | 0.001 | 1 | 4 | Rtkn2 | 0.012 | Young |
| 0.003514 | 2.33692 | 0.013 | 0.001 | 1 | 4 | Adamts2 | 0.012 | Young |
| 0.003514 | 2.313968 | 0.013 | 0.001 | 1 | 4 | Serpinf2 | 0.012 | Young |
| 0.003514 | 2.288105 | 0.013 | 0.001 | 1 | 4 | Gm15651 | 0.012 | Young |
| 0.003514 | 2.275623 | 0.013 | 0.001 | 1 | 4 | Gm11127 | 0.012 | Young |
| 0.003514 | 2.250717 | 0.013 | 0.001 | 1 | 4 | Klrg2 | 0.012 | Young |
| 0.003514 | 1.985976 | 0.013 | 0.001 | 1 | 4 | Fgd5 | 0.012 | Young |
| 0.003514 | 1.850414 | 0.013 | 0.001 | 1 | 4 | Flt3 | 0.012 | Young |
| 0.003514 | 1.845231 | 0.013 | 0.001 | 1 | 4 | Gm31812 | 0.012 | Young |
| 0.003514 | 1.844834 | 0.013 | 0.001 | 1 | 4 | Il12rb1 | 0.012 | Young |
| 0.003514 | 1.776585 | 0.013 | 0.001 | 1 | 4 | Ar | 0.012 | Young |
| 0.003514 | 1.714917 | 0.013 | 0.001 | 1 | 4 | Mkrn3 | 0.012 | Young |
| 0.003514 | 1.38649 | 0.013 | 0.001 | 1 | 4 | P4ha2 | 0.012 | Young |

| | | | | | | | | |
|---|---|---|---|---|---|---|---|---|
| 0.003514 | 1.133817 | 0.013 | 0.001 | 1 | 4 | AK157302 | 0.012 | Young |
| 0.003514 | 0.841492 | 0.013 | 0.001 | 1 | 4 | Epha4 | 0.012 | Young |
| 0.003514 | 0.828459 | 0.013 | 0.001 | 1 | 4 | Gm7467 | 0.012 | Young |
| 0.003524 | 1.098966 | 0.1 | 0.035 | 1 | 4 | Afap1l1 | 0.065 | Young |
| 0.003529 | 0.528129 | 0.65 | 0.499 | 1 | 4 | Ifrd1 | 0.151 | Young |
| 0.003552 | 0.730085 | 0.275 | 0.152 | 1 | 4 | Itga4 | 0.123 | Young |
| 0.003559 | 0.718657 | 0.25 | 0.131 | 1 | 4 | Jdp2 | 0.119 | Young |
| 0.00356 | 2.219201 | 0.013 | 0.001 | 1 | 4 | Ankrd35 | 0.012 | Young |
| 0.00356 | 2.179136 | 0.013 | 0.001 | 1 | 4 | Pcdhb14 | 0.012 | Young |
| 0.00356 | 2.178903 | 0.013 | 0.001 | 1 | 4 | Gm26734 | 0.012 | Young |
| 0.00356 | 2.12202 | 0.013 | 0.001 | 1 | 4 | Arx | 0.012 | Young |
| 0.00356 | 2.070301 | 0.013 | 0.001 | 1 | 4 | Paqr6 | 0.012 | Young |
| 0.00356 | 2.056903 | 0.013 | 0.001 | 1 | 4 | Msh4 | 0.012 | Young |
| 0.00356 | 2.04886 | 0.013 | 0.001 | 1 | 4 | Hook1 | 0.012 | Young |
| 0.00356 | 1.950904 | 0.013 | 0.001 | 1 | 4 | Gpr18 | 0.012 | Young |
| 0.00356 | 1.925525 | 0.013 | 0.001 | 1 | 4 | Tcf15 | 0.012 | Young |
| 0.00356 | 1.901633 | 0.013 | 0.001 | 1 | 4 | Hif3a | 0.012 | Young |
| 0.00356 | 1.84698 | 0.013 | 0.001 | 1 | 4 | Rab40b | 0.012 | Young |
| 0.00356 | 1.717156 | 0.013 | 0.001 | 1 | 4 | Itgb8 | 0.012 | Young |
| 0.00356 | 1.660384 | 0.013 | 0.001 | 1 | 4 | Ptcra | 0.012 | Young |
| 0.00356 | 1.647246 | 0.013 | 0.001 | 1 | 4 | Pard3 | 0.012 | Young |
| 0.00356 | 1.633642 | 0.013 | 0.001 | 1 | 4 | Ccdc159 | 0.012 | Young |
| 0.00356 | 1.559306 | 0.013 | 0.001 | 1 | 4 | Gsdmc4 | 0.012 | Young |
| 0.00356 | 1.55052 | 0.013 | 0.001 | 1 | 4 | Gm45669 | 0.012 | Young |
| 0.00356 | 1.523191 | 0.013 | 0.001 | 1 | 4 | Nr2f1 | 0.012 | Young |
| 0.00356 | 1.436353 | 0.013 | 0.001 | 1 | 4 | Hoxa5 | 0.012 | Young |
| 0.00356 | 1.411753 | 0.013 | 0.001 | 1 | 4 | Adgrg5 | 0.012 | Young |
| 0.00356 | 1.41038 | 0.013 | 0.001 | 1 | 4 | Mapk13 | 0.012 | Young |
| 0.00356 | 1.362562 | 0.013 | 0.001 | 1 | 4 | H2-Eb2 | 0.012 | Young |
| 0.00356 | 1.326566 | 0.013 | 0.001 | 1 | 4 | Gm568 | 0.012 | Young |
| 0.00356 | 1.230247 | 0.013 | 0.001 | 1 | 4 | Adamts15 | 0.012 | Young |
| 0.00356 | 1.207292 | 0.013 | 0.001 | 1 | 4 | Med9os | 0.012 | Young |
| 0.00356 | 1.206331 | 0.013 | 0.001 | 1 | 4 | Gm13594 | 0.012 | Young |
| 0.00356 | 1.128495 | 0.013 | 0.001 | 1 | 4 | Ckap2 | 0.012 | Young |
| 0.00356 | 1.084443 | 0.013 | 0.001 | 1 | 4 | Prx | 0.012 | Young |
| 0.00356 | 1.060571 | 0.013 | 0.001 | 1 | 4 | Elf3 | 0.012 | Young |
| 0.00356 | 1.043719 | 0.013 | 0.001 | 1 | 4 | Tpm2 | 0.012 | Young |
| 0.00356 | 1.012762 | 0.013 | 0.001 | 1 | 4 | Spsb1 | 0.012 | Young |
| 0.00356 | 0.997777 | 0.013 | 0.001 | 1 | 4 | Ampd1 | 0.012 | Young |
| 0.00356 | 0.982029 | 0.013 | 0.001 | 1 | 4 | Capn11 | 0.012 | Young |
| 0.00356 | 0.951499 | 0.013 | 0.001 | 1 | 4 | 1700088E0 | 0.012 | Young |
| 0.00356 | 0.901353 | 0.013 | 0.001 | 1 | 4 | Filip1 | 0.012 | Young |
| 0.00356 | 0.875863 | 0.013 | 0.001 | 1 | 4 | Pacsin3 | 0.012 | Young |
| 0.00356 | 0.840309 | 0.013 | 0.001 | 1 | 4 | Gm17196 | 0.012 | Young |
| 0.00356 | 0.833277 | 0.013 | 0.001 | 1 | 4 | Gm49027 | 0.012 | Young |

| | | | | | | | | |
|---|---|---|---|---|---|---|---|---|
| 0.00356 | 0.824975 | 0.013 | 0.001 | 1 | 4 | 5033403F0 | 0.012 | Young |
| 0.00356 | 0.789053 | 0.013 | 0.001 | 1 | 4 | Hist1h4k | 0.012 | Young |
| 0.00356 | 0.786048 | 0.013 | 0.001 | 1 | 4 | Mfap4 | 0.012 | Young |
| 0.00356 | 0.776574 | 0.013 | 0.001 | 1 | 4 | Gm28086 | 0.012 | Young |
| 0.00356 | 0.754304 | 0.013 | 0.001 | 1 | 4 | 6030443J0 | 0.012 | Young |
| 0.00356 | 0.752607 | 0.013 | 0.001 | 1 | 4 | Zfp811 | 0.012 | Young |
| 0.00356 | 0.747035 | 0.013 | 0.001 | 1 | 4 | Nhlrc1 | 0.012 | Young |
| 0.00356 | 0.705501 | 0.013 | 0.001 | 1 | 4 | Phkg1 | 0.012 | Young |
| 0.00356 | 0.679614 | 0.013 | 0.001 | 1 | 4 | Ptpn14 | 0.012 | Young |
| 0.00356 | 0.618972 | 0.013 | 0.001 | 1 | 4 | Tmtc1 | 0.012 | Young |
| 0.00356 | 0.60209 | 0.013 | 0.001 | 1 | 4 | D030007L0 | 0.012 | Young |
| 0.00356 | 0.591446 | 0.013 | 0.001 | 1 | 4 | Slc35g2 | 0.012 | Young |
| 0.00356 | 0.569736 | 0.013 | 0.001 | 1 | 4 | Tmem44 | 0.012 | Young |
| 0.00356 | 0.568541 | 0.013 | 0.001 | 1 | 4 | P2rx6 | 0.012 | Young |
| 0.00356 | 0.494411 | 0.013 | 0.001 | 1 | 4 | Fhl1 | 0.012 | Young |
| 0.00356 | 0.456721 | 0.013 | 0.001 | 1 | 4 | Gm12279 | 0.012 | Young |
| 0.00356 | 0.414027 | 0.013 | 0.001 | 1 | 4 | Gm13889 | 0.012 | Young |
| 0.003577 | 0.692695 | 0.45 | 0.298 | 1 | 4 | Daglb | 0.152 | Young |
| 0.003646 | 0.312875 | 0.85 | 0.685 | 1 | 4 | Qk | 0.165 | Young |
| 0.003665 | 0.688362 | 0.625 | 0.516 | 1 | 4 | Tob2 | 0.109 | Young |
| 0.003666 | 0.661215 | 0.462 | 0.32 | 1 | 4 | Mpeg1 | 0.142 | Young |
| 0.003725 | 0.671884 | 0.412 | 0.262 | 1 | 4 | Pde4b | 0.15 | Young |
| 0.003758 | 1.034569 | 0.075 | 0.022 | 1 | 4 | Asna1 | 0.053 | Young |
| 0.003808 | 0.805488 | 0.262 | 0.144 | 1 | 4 | Emd | 0.118 | Young |
| 0.003853 | 0.619606 | 0.312 | 0.18 | 1 | 4 | Cttnbp2nl | 0.132 | Young |
| 0.003941 | 2.60394 | 0.062 | 0.017 | 1 | 4 | Fdxacb1 | 0.045 | Young |
| 0.004006 | 0.762792 | 0.262 | 0.141 | 1 | 4 | Tlr7 | 0.121 | Young |
| 0.004037 | 0.468714 | 0.2 | 0.094 | 1 | 4 | Wdr13 | 0.106 | Young |
| 0.004079 | 2.929637 | 0.025 | 0.003 | 1 | 4 | Olfr373 | 0.022 | Young |
| 0.004115 | 1.037789 | 0.188 | 0.092 | 1 | 4 | Pde3b | 0.096 | Young |
| 0.004135 | 2.485731 | 0.025 | 0.003 | 1 | 4 | Gnb1l | 0.022 | Young |
| 0.004158 | 0.526857 | 0.65 | 0.489 | 1 | 4 | Pfdn5 | 0.161 | Young |
| 0.004164 | 2.364966 | 0.025 | 0.003 | 1 | 4 | Tlr12 | 0.022 | Young |
| 0.004167 | 0.434494 | 0.35 | 0.203 | 1 | 4 | Ly96 | 0.147 | Young |
| 0.004193 | 2.038056 | 0.025 | 0.003 | 1 | 4 | Cyp2u1 | 0.022 | Young |
| 0.00425 | 1.6268 | 0.025 | 0.003 | 1 | 4 | Kifc2 | 0.022 | Young |
| 0.004293 | 1.404741 | 0.062 | 0.017 | 1 | 4 | Ptdss2 | 0.045 | Young |
| 0.004308 | 0.461414 | 0.613 | 0.431 | 1 | 4 | Evi2a | 0.182 | Young |
| 0.004309 | 1.003853 | 0.025 | 0.003 | 1 | 4 | 1700030J2 | 0.022 | Young |
| 0.004345 | 2.381464 | 0.05 | 0.012 | 1 | 4 | Hspb1 | 0.038 | Young |
| 0.004364 | 0.349545 | 0.838 | 0.701 | 1 | 4 | Rpl36 | 0.137 | Young |
| 0.004366 | 1.41502 | 0.075 | 0.023 | 1 | 4 | Zfand1 | 0.052 | Young |
| 0.004444 | 1.122547 | 0.125 | 0.051 | 1 | 4 | Ttll12 | 0.074 | Young |
| 0.004507 | 0.229467 | 0.988 | 0.986 | 1 | 4 | mt-Nd2 | 0.002 | Young |
| 0.004529 | 0.450962 | 0.762 | 0.618 | 1 | 4 | Nrp1 | 0.144 | Young |

| | | | | | | | | |
|---|---|---|---|---|---|---|---|---|
| 0.004563 | 1.095383 | 0.162 | 0.074 | 1 | 4 | Ttpal | 0.088 | Young |
| 0.0046 | 2.099476 | 0.038 | 0.007 | 1 | 4 | Ppif | 0.031 | Young |
| 0.004619 | 0.454534 | 0.812 | 0.727 | 1 | 4 | Rpl21 | 0.085 | Young |
| 0.004622 | 1.99094 | 0.038 | 0.007 | 1 | 4 | Zfp652os | 0.031 | Young |
| 0.004628 | 0.915118 | 0.112 | 0.043 | 1 | 4 | Dhx32 | 0.069 | Young |
| 0.004654 | 1.183402 | 0.112 | 0.043 | 1 | 4 | Zmym3 | 0.069 | Young |
| 0.004676 | 1.332424 | 0.05 | 0.012 | 1 | 4 | Zfp873 | 0.038 | Young |
| 0.004689 | 1.867955 | 0.038 | 0.007 | 1 | 4 | Sdr42e1 | 0.031 | Young |
| 0.004689 | 1.862898 | 0.038 | 0.007 | 1 | 4 | Farp2 | 0.031 | Young |
| 0.004709 | 0.757177 | 0.275 | 0.152 | 1 | 4 | Xpo1 | 0.123 | Young |
| 0.004804 | 0.604444 | 0.288 | 0.157 | 1 | 4 | Spopl | 0.131 | Young |
| 0.004867 | 1.125921 | 0.2 | 0.102 | 1 | 4 | Hmgcl | 0.098 | Young |
| 0.004876 | 0.353571 | 0.887 | 0.806 | 1 | 4 | Rpl34 | 0.081 | Young |
| 0.00489 | 0.906272 | 0.312 | 0.184 | 1 | 4 | Phc2 | 0.128 | Young |
| 0.004924 | 1.068486 | 0.15 | 0.067 | 1 | 4 | Mettl16 | 0.083 | Young |
| 0.004925 | 0.88149 | 0.312 | 0.196 | 1 | 4 | Id2 | 0.116 | Young |
| 0.005 | 2.017209 | 0.062 | 0.018 | 1 | 4 | Ttll1 | 0.044 | Young |
| 0.005008 | 0.519498 | 0.4 | 0.24 | 1 | 4 | Icam1 | 0.16 | Young |
| 0.005035 | 1.004084 | 0.088 | 0.03 | 1 | 4 | Vps33b | 0.058 | Young |
| 0.00515 | 0.839363 | 0.325 | 0.209 | 1 | 4 | Fcgr1 | 0.116 | Young |
| 0.005167 | 0.674306 | 0.2 | 0.097 | 1 | 4 | Cbx6 | 0.103 | Young |
| 0.005189 | 0.560834 | 0.537 | 0.382 | 1 | 4 | Map1lc3b | 0.155 | Young |
| 0.005235 | 0.616461 | 0.425 | 0.282 | 1 | 4 | Chmp2a | 0.143 | Young |
| 0.005262 | 0.862943 | 0.275 | 0.157 | 1 | 4 | Rasgef1b | 0.118 | Young |
| 0.005303 | 0.6102 | 0.425 | 0.274 | 1 | 4 | Arl4c | 0.151 | Young |
| 0.005378 | 1.81179 | 0.062 | 0.018 | 1 | 4 | Mettl17 | 0.044 | Young |
| 0.005461 | 0.993363 | 0.125 | 0.051 | 1 | 4 | Pithd1 | 0.074 | Young |
| 0.005484 | 0.754028 | 0.412 | 0.275 | 1 | 4 | Rab6a | 0.137 | Young |
| 0.005493 | 0.76684 | 0.188 | 0.091 | 1 | 4 | Creb3l2 | 0.097 | Young |
| 0.005548 | 0.906185 | 0.212 | 0.109 | 1 | 4 | AW549877 | 0.103 | Young |
| 0.005581 | 1.347747 | 0.088 | 0.03 | 1 | 4 | Sppl2b | 0.058 | Young |
| 0.005582 | 1.134853 | 0.162 | 0.077 | 1 | 4 | Rnf181 | 0.085 | Young |
| 0.00559 | 0.56231 | 0.388 | 0.24 | 1 | 4 | Ilk | 0.148 | Young |
| 0.005623 | 0.4121 | 0.85 | 0.785 | 1 | 4 | Nfkbia | 0.065 | Young |
| 0.005626 | 0.620277 | 0.512 | 0.37 | 1 | 4 | Nfe2l2 | 0.142 | Young |
| 0.005664 | 1.156222 | 0.088 | 0.03 | 1 | 4 | Icosl | 0.058 | Young |
| 0.005725 | 0.359817 | 0.875 | 0.821 | 1 | 4 | Rps25 | 0.054 | Young |
| 0.005738 | 1.108198 | 0.3 | 0.19 | 1 | 4 | Slc11a1 | 0.11 | Young |
| 0.005804 | 0.938044 | 0.25 | 0.138 | 1 | 4 | Mark3 | 0.112 | Young |
| 0.005893 | 1.189344 | 0.112 | 0.045 | 1 | 4 | Nt5dc1 | 0.067 | Young |
| 0.005919 | 0.546699 | 0.462 | 0.31 | 1 | 4 | Lair1 | 0.152 | Young |
| 0.005927 | 1.01985 | 0.15 | 0.068 | 1 | 4 | Vegfb | 0.082 | Young |
| 0.005966 | 1.93551 | 0.075 | 0.024 | 1 | 4 | Sco1 | 0.051 | Young |
| 0.005975 | 1.731199 | 0.05 | 0.013 | 1 | 4 | Xlr | 0.037 | Young |
| 0.00601 | 0.77996 | 0.275 | 0.159 | 1 | 4 | Mink1 | 0.116 | Young |

| | | | | | | | | |
|---|---|---|---|---|---|---|---|---|
| 0.006034 | 0.275518 | 0.975 | 0.926 | 1 | 4 | Rpl30 | 0.049 | Young |
| 0.006198 | 1.389802 | 0.05 | 0.013 | 1 | 4 | Ncaph | 0.037 | Young |
| 0.006299 | 1.134885 | 0.125 | 0.053 | 1 | 4 | Ccdc134 | 0.072 | Young |
| 0.006346 | 0.750773 | 0.175 | 0.083 | 1 | 4 | Nup54 | 0.092 | Young |
| 0.006448 | 0.991776 | 0.162 | 0.076 | 1 | 4 | Gpt2 | 0.086 | Young |
| 0.006453 | 0.328002 | 0.938 | 0.883 | 1 | 4 | Rps16 | 0.055 | Young |
| 0.00649 | 0.707487 | 0.188 | 0.092 | 1 | 4 | Arpc5l | 0.096 | Young |
| 0.006547 | 1.238325 | 0.05 | 0.013 | 1 | 4 | Gpx7 | 0.037 | Young |
| 0.006617 | 1.185337 | 0.088 | 0.031 | 1 | 4 | Sft2d3 | 0.057 | Young |
| 0.006706 | 1.75407 | 0.062 | 0.018 | 1 | 4 | Gm8369 | 0.044 | Young |
| 0.006709 | 1.469382 | 0.162 | 0.08 | 1 | 4 | Ubxn2a | 0.082 | Young |
| 0.006919 | 1.169621 | 0.1 | 0.038 | 1 | 4 | Slc35c1 | 0.062 | Young |
| 0.006952 | 0.979569 | 0.175 | 0.086 | 1 | 4 | Tmem229b | 0.089 | Young |
| 0.007044 | 1.624271 | 0.088 | 0.032 | 1 | 4 | Bmf | 0.056 | Young |
| 0.007104 | 1.342576 | 0.062 | 0.018 | 1 | 4 | Armcx5 | 0.044 | Young |
| 0.007193 | 1.172841 | 0.1 | 0.038 | 1 | 4 | Ino80b | 0.062 | Young |
| 0.007205 | 2.295935 | 0.038 | 0.008 | 1 | 4 | Tesmin | 0.03 | Young |
| 0.007297 | 0.636557 | 0.3 | 0.177 | 1 | 4 | Phf23 | 0.123 | Young |
| 0.007307 | 1.166133 | 0.138 | 0.062 | 1 | 4 | Csf3r | 0.076 | Young |
| 0.007334 | 2.069039 | 0.038 | 0.008 | 1 | 4 | Lrp4 | 0.03 | Young |
| 0.00735 | 1.127058 | 0.1 | 0.038 | 1 | 4 | 5430427O1 | 0.062 | Young |
| 0.007354 | 0.568261 | 0.4 | 0.267 | 1 | 4 | Prune2 | 0.133 | Young |
| 0.007399 | 1.667897 | 0.038 | 0.008 | 1 | 4 | Kif3c | 0.03 | Young |
| 0.007415 | 1.212286 | 0.188 | 0.096 | 1 | 4 | 0610030E2 | 0.092 | Young |
| 0.007455 | 0.512122 | 0.4 | 0.255 | 1 | 4 | Rbbp6 | 0.145 | Young |
| 0.00747 | 1.12061 | 0.188 | 0.098 | 1 | 4 | Pcif1 | 0.09 | Young |
| 0.007543 | 0.665191 | 0.412 | 0.279 | 1 | 4 | Il4ra | 0.133 | Young |
| 0.007665 | 0.77734 | 0.125 | 0.053 | 1 | 4 | Bcl9l | 0.072 | Young |
| 0.007701 | 1.57941 | 0.038 | 0.008 | 1 | 4 | Cd276 | 0.03 | Young |
| 0.007804 | 1.146948 | 0.038 | 0.008 | 1 | 4 | Gm16973 | 0.03 | Young |
| 0.007873 | 0.881781 | 0.038 | 0.008 | 1 | 4 | Trim13 | 0.03 | Young |
| 0.007881 | 1.068058 | 0.262 | 0.154 | 1 | 4 | Crem | 0.108 | Young |
| 0.007973 | 2.139559 | 0.05 | 0.013 | 1 | 4 | Gm7160 | 0.037 | Young |
| 0.008004 | 1.061653 | 0.188 | 0.097 | 1 | 4 | Fmr1 | 0.091 | Young |
| 0.008077 | 0.688155 | 0.1 | 0.038 | 1 | 4 | Exosc1 | 0.062 | Young |
| 0.008221 | 0.520296 | 0.725 | 0.571 | 1 | 4 | Vmp1 | 0.154 | Young |
| 0.008244 | 0.681185 | 0.325 | 0.202 | 1 | 4 | Snx13 | 0.123 | Young |
| 0.008333 | 0.696723 | 0.438 | 0.319 | 1 | 4 | Tlr2 | 0.119 | Young |
| 0.008468 | 0.646614 | 0.162 | 0.078 | 1 | 4 | Dcn | 0.084 | Young |
| 0.008472 | 1.077145 | 0.1 | 0.039 | 1 | 4 | Rnf8 | 0.061 | Young |
| 0.008506 | 0.461987 | 0.775 | 0.671 | 1 | 4 | Picalm | 0.104 | Young |
| 0.008578 | 1.529886 | 0.05 | 0.013 | 1 | 4 | Enho | 0.037 | Young |
| 0.008616 | 1.048687 | 0.15 | 0.071 | 1 | 4 | Adcy3 | 0.079 | Young |
| 0.008627 | 1.680334 | 0.062 | 0.019 | 1 | 4 | Pfas | 0.043 | Young |
| 0.008704 | 0.555485 | 0.3 | 0.175 | 1 | 4 | Leprotl1 | 0.125 | Young |

| | | | | | | | | |
|---|---|---|---|---|---|---|---|---|
| 0.008721 | 0.486424 | 0.238 | 0.128 | 1 | 4 | Ndufs1 | 0.11 | Young |
| 0.008756 | 0.504091 | 0.537 | 0.4 | 1 | 4 | Rpl29 | 0.137 | Young |
| 0.008819 | 1.234318 | 0.05 | 0.013 | 1 | 4 | Zrsr1 | 0.037 | Young |
| 0.008905 | 0.729774 | 0.2 | 0.103 | 1 | 4 | Tbk1 | 0.097 | Young |
| 0.008914 | 0.430255 | 0.462 | 0.312 | 1 | 4 | Foxn3 | 0.15 | Young |
| 0.008927 | 1.268985 | 0.088 | 0.032 | 1 | 4 | Mthfd1 | 0.056 | Young |
| 0.008927 | 0.523115 | 0.588 | 0.447 | 1 | 4 | Stat3 | 0.141 | Young |
| 0.008967 | 1.284012 | 0.088 | 0.032 | 1 | 4 | Acd | 0.056 | Young |
| 0.008971 | 0.657511 | 0.425 | 0.303 | 1 | 4 | Ccl12 | 0.122 | Young |
| 0.009005 | 1.149461 | 0.188 | 0.099 | 1 | 4 | Cenpa | 0.089 | Young |
| 0.009082 | 0.444788 | 0.6 | 0.471 | 1 | 4 | Jmjd1c | 0.129 | Young |
| 0.009112 | 0.346763 | 0.85 | 0.742 | 1 | 4 | Rpl28 | 0.108 | Young |
| 0.009144 | 0.224798 | 1 | 0.972 | 1 | 4 | Ctsc | 0.028 | Young |
| 0.009161 | 3.088674 | 0.025 | 0.004 | 1 | 4 | E130208F1 | 0.021 | Young |
| 0.009181 | 0.731204 | 0.212 | 0.115 | 1 | 4 | Fbxo11 | 0.097 | Young |
| 0.00921 | 0.502044 | 0.388 | 0.244 | 1 | 4 | Nrip1 | 0.144 | Young |
| 0.009215 | 2.350534 | 0.025 | 0.004 | 1 | 4 | Gm15013 | 0.021 | Young |
| 0.009266 | 1.130402 | 0.125 | 0.055 | 1 | 4 | Usp21 | 0.07 | Young |
| 0.00927 | 1.923901 | 0.025 | 0.004 | 1 | 4 | A630089N( | 0.021 | Young |
| 0.009271 | 0.689904 | 0.262 | 0.153 | 1 | 4 | Myo7a | 0.109 | Young |
| 0.009305 | 1.169245 | 0.062 | 0.019 | 1 | 4 | Art2b | 0.043 | Young |
| 0.009436 | 2.072407 | 0.025 | 0.004 | 1 | 4 | Bicdl1 | 0.021 | Young |
| 0.009451 | 1.065944 | 0.3 | 0.188 | 1 | 4 | Eprs | 0.112 | Young |
| 0.009456 | 0.629864 | 0.175 | 0.086 | 1 | 4 | Xylt2 | 0.089 | Young |
| 0.009478 | 0.329641 | 0.7 | 0.52 | 1 | 4 | Rbm3 | 0.18 | Young |
| 0.009496 | 0.845362 | 0.15 | 0.07 | 1 | 4 | Phb | 0.08 | Young |
| 0.00952 | 1.47622 | 0.025 | 0.004 | 1 | 4 | 2510017J1 | 0.021 | Young |
| 0.009548 | 1.670167 | 0.025 | 0.004 | 1 | 4 | Gm44777 | 0.021 | Young |
| 0.009601 | 0.952212 | 0.088 | 0.032 | 1 | 4 | Sephs1 | 0.056 | Young |
| 0.009605 | 1.214525 | 0.025 | 0.004 | 1 | 4 | Ccr7 | 0.021 | Young |
| 0.009658 | 0.507691 | 0.138 | 0.061 | 1 | 4 | R3hcc1l | 0.077 | Young |
| 0.00966 | 1.072482 | 0.1 | 0.04 | 1 | 4 | Ddx31 | 0.06 | Young |
| 0.009725 | 1.241756 | 0.075 | 0.026 | 1 | 4 | Eif2b3 | 0.049 | Young |
| 0.009778 | 1.11565 | 0.112 | 0.047 | 1 | 4 | Zbtb2 | 0.065 | Young |
| 0.009794 | 1.397505 | 0.175 | 0.093 | 1 | 4 | Dnase1l3 | 0.082 | Young |
| 0.009825 | 1.345973 | 0.112 | 0.048 | 1 | 4 | Hdhd2 | 0.064 | Young |
| 0.009881 | 0.537201 | 0.675 | 0.574 | 1 | 4 | Scamp2 | 0.101 | Young |
| 0.009897 | 0.356151 | 0.15 | 0.069 | 1 | 4 | Mtf1 | 0.081 | Young |
| 0.009933 | 0.889117 | 0.125 | 0.055 | 1 | 4 | Fpr2 | 0.07 | Young |
| 0.009959 | 0.962132 | 0.162 | 0.08 | 1 | 4 | Afg3l2 | 0.082 | Young |
| 0.01006 | 1.155116 | 0.112 | 0.047 | 1 | 4 | Dnajc24 | 0.065 | Young |
| 0.010068 | 0.439326 | 0.262 | 0.147 | 1 | 4 | Stip1 | 0.115 | Young |
| 0.01017 | 0.97138 | 0.1 | 0.04 | 1 | 4 | Fbxl18 | 0.06 | Young |
| 0.010468 | 0.887033 | 0.2 | 0.108 | 1 | 4 | Cyth3 | 0.092 | Young |
| 0.010527 | 1.586406 | 0.075 | 0.026 | 1 | 4 | Cry1 | 0.049 | Young |

| | | | | | | | | |
|---|---|---|---|---|---|---|---|---|
| 0.010578 | 0.95313 | 0.1 | 0.04 | 1 | 4 | Itgal | 0.06 | Young |
| 0.01072 | 0.779408 | 0.762 | 0.704 | 1 | 4 | Wfdc17 | 0.058 | Young |
| 0.01073 | 1.142019 | 0.112 | 0.048 | 1 | 4 | Scmh1 | 0.064 | Young |
| 0.010915 | 0.671363 | 0.312 | 0.196 | 1 | 4 | Park7 | 0.116 | Young |
| 0.010943 | 0.426809 | 0.162 | 0.078 | 1 | 4 | Pias2 | 0.084 | Young |
| 0.01095 | 0.523758 | 0.425 | 0.289 | 1 | 4 | Ms4a6d | 0.136 | Young |
| 0.010977 | 1.577644 | 0.075 | 0.026 | 1 | 4 | Gm16638 | 0.049 | Young |
| 0.011031 | 1.546773 | 0.075 | 0.026 | 1 | 4 | Kctd13 | 0.049 | Young |
| 0.011203 | 0.64797 | 0.425 | 0.304 | 1 | 4 | Bptf | 0.121 | Young |
| 0.01122 | 0.785292 | 0.175 | 0.089 | 1 | 4 | Thap12 | 0.086 | Young |
| 0.011222 | 1.375088 | 0.075 | 0.026 | 1 | 4 | Invs | 0.049 | Young |
| 0.01124 | 0.618092 | 0.438 | 0.292 | 1 | 4 | Top2b | 0.146 | Young |
| 0.011265 | 0.648863 | 0.212 | 0.115 | 1 | 4 | Gbp3 | 0.097 | Young |
| 0.011364 | 0.972527 | 0.138 | 0.064 | 1 | 4 | Tfeb | 0.074 | Young |
| 0.011411 | 1.432537 | 0.05 | 0.014 | 1 | 4 | Numbl | 0.036 | Young |
| 0.011445 | 1.545947 | 0.088 | 0.034 | 1 | 4 | Tab3 | 0.054 | Young |
| 0.011528 | 1.145738 | 0.062 | 0.02 | 1 | 4 | Ppil1 | 0.042 | Young |
| 0.011573 | 0.237047 | 0.938 | 0.962 | 1 | 4 | Marcks | -0.024 | Young |
| 0.011781 | 0.777371 | 0.25 | 0.145 | 1 | 4 | Slc14a1 | 0.105 | Young |
| 0.011789 | 1.072907 | 0.038 | 0.009 | 1 | 4 | Zfp169 | 0.029 | Young |
| 0.011831 | 1.11959 | 0.05 | 0.014 | 1 | 4 | Cenpj | 0.036 | Young |
| 0.011837 | 0.711778 | 0.038 | 0.009 | 1 | 4 | Nudt6 | 0.029 | Young |
| 0.011856 | 0.504559 | 0.6 | 0.448 | 1 | 4 | Kdm6b | 0.152 | Young |
| 0.011886 | 0.670916 | 0.038 | 0.009 | 1 | 4 | 1700109H( | 0.029 | Young |
| 0.011962 | 1.032728 | 0.15 | 0.073 | 1 | 4 | Ptp4a3 | 0.077 | Young |
| 0.012187 | 0.445676 | 0.575 | 0.431 | 1 | 4 | Pla2g7 | 0.144 | Young |
| 0.012329 | 0.457693 | 0.312 | 0.186 | 1 | 4 | Setd3 | 0.126 | Young |
| 0.012403 | 0.419011 | 0.275 | 0.161 | 1 | 4 | Chka | 0.114 | Young |
| 0.012515 | 0.597531 | 0.525 | 0.397 | 1 | 4 | Aldh2 | 0.128 | Young |
| 0.012548 | 0.71495 | 0.2 | 0.108 | 1 | 4 | Tmem57 | 0.092 | Young |
| 0.012564 | 0.231404 | 0.062 | 0.02 | 1 | 4 | Mov10 | 0.042 | Young |
| 0.012564 | 0.670159 | 0.138 | 0.064 | 1 | 4 | Foxj3 | 0.074 | Young |
| 0.012689 | 1.651776 | 0.075 | 0.027 | 1 | 4 | Gys1 | 0.048 | Young |
| 0.012763 | 0.76325 | 0.15 | 0.072 | 1 | 4 | Smcr8 | 0.078 | Young |
| 0.012871 | 0.965105 | 0.1 | 0.041 | 1 | 4 | Cep70 | 0.059 | Young |
| 0.012916 | 0.700771 | 0.125 | 0.056 | 1 | 4 | Haghl | 0.069 | Young |
| 0.012919 | 0.637587 | 0.175 | 0.089 | 1 | 4 | Brip1os | 0.086 | Young |
| 0.013013 | 0.204218 | 1 | 0.993 | 1 | 4 | Lyz2 | 0.007 | Young |
| 0.013319 | 0.491914 | 0.55 | 0.416 | 1 | 4 | Rassf2 | 0.134 | Young |
| 0.013329 | 0.817547 | 0.212 | 0.117 | 1 | 4 | Trim44 | 0.095 | Young |
| 0.013331 | 1.555488 | 0.062 | 0.02 | 1 | 4 | Gm6563 | 0.042 | Young |
| 0.013376 | 0.454493 | 0.588 | 0.48 | 1 | 4 | Srgn | 0.108 | Young |
| 0.013518 | 0.691825 | 0.212 | 0.117 | 1 | 4 | Idnk | 0.095 | Young |
| 0.01372 | 1.774142 | 0.05 | 0.014 | 1 | 4 | Zfp748 | 0.036 | Young |
| 0.01373 | 0.631367 | 0.35 | 0.231 | 1 | 4 | Mob3c | 0.119 | Young |

| | | | | | | | | |
|---|---|---|---|---|---|---|---|---|
| 0.013775 | 1.120956 | 0.188 | 0.102 | 1 | 4 | Lrrc8d | 0.086 | Young |
| 0.013785 | 0.786379 | 0.088 | 0.034 | 1 | 4 | Unc45a | 0.054 | Young |
| 0.013919 | 1.3609 | 0.062 | 0.02 | 1 | 4 | Csrp2 | 0.042 | Young |
| 0.013936 | 1.416833 | 0.112 | 0.051 | 1 | 4 | Rcbtb1 | 0.061 | Young |
| 0.014032 | 1.402362 | 0.062 | 0.02 | 1 | 4 | Wdr6 | 0.042 | Young |
| 0.014101 | 0.841021 | 0.262 | 0.16 | 1 | 4 | Tbc1d9 | 0.102 | Young |
| 0.014133 | 0.441748 | 0.488 | 0.345 | 1 | 4 | Ckb | 0.143 | Young |
| 0.014146 | 0.531637 | 0.475 | 0.337 | 1 | 4 | Ube2l3 | 0.138 | Young |
| 0.014404 | 0.263107 | 0.988 | 0.949 | 1 | 4 | Rps23 | 0.039 | Young |
| 0.014443 | 1.527762 | 0.075 | 0.028 | 1 | 4 | Nit2 | 0.047 | Young |
| 0.014507 | 1.138318 | 0.125 | 0.057 | 1 | 4 | Pde4a | 0.068 | Young |
| 0.014569 | 0.872655 | 0.225 | 0.13 | 1 | 4 | Higd1a | 0.095 | Young |
| 0.014578 | 0.630559 | 0.462 | 0.327 | 1 | 4 | Peli1 | 0.135 | Young |
| 0.014587 | 0.868655 | 0.325 | 0.215 | 1 | 4 | Gpr65 | 0.11 | Young |
| 0.014614 | 1.414986 | 0.05 | 0.014 | 1 | 4 | Bivm | 0.036 | Young |
| 0.014713 | 0.234937 | 0.95 | 0.935 | 1 | 4 | Maf | 0.015 | Young |
| 0.014733 | 0.705485 | 0.075 | 0.027 | 1 | 4 | Atp6v0d2 | 0.048 | Young |
| 0.014734 | 0.908344 | 0.112 | 0.049 | 1 | 4 | Thyn1 | 0.063 | Young |
| 0.01474 | 0.294632 | 0.775 | 0.663 | 1 | 4 | Stab1 | 0.112 | Young |
| 0.014777 | 0.379 | 0.2 | 0.105 | 1 | 4 | Zbtb8os | 0.095 | Young |
| 0.014922 | 0.613428 | 0.613 | 0.549 | 1 | 4 | C4b | 0.064 | Young |
| 0.015064 | 1.013618 | 0.088 | 0.034 | 1 | 4 | Arl4d | 0.054 | Young |
| 0.01512 | 0.200612 | 0.988 | 0.994 | 1 | 4 | C1qc | -0.006 | Young |
| 0.015123 | 0.416193 | 0.375 | 0.242 | 1 | 4 | mt-Atp8 | 0.133 | Young |
| 0.015174 | 0.472637 | 0.362 | 0.24 | 1 | 4 | Higd2a | 0.122 | Young |
| 0.015304 | 0.55071 | 0.462 | 0.337 | 1 | 4 | Abhd12 | 0.125 | Young |
| 0.015501 | 1.315599 | 0.1 | 0.043 | 1 | 4 | Dus1l | 0.057 | Young |
| 0.015547 | 0.398717 | 0.412 | 0.276 | 1 | 4 | Sft2d2 | 0.136 | Young |
| 0.015567 | 0.46015 | 0.512 | 0.37 | 1 | 4 | Lrp1 | 0.142 | Young |
| 0.015649 | 0.310624 | 0.775 | 0.74 | 1 | 4 | Rsrp1 | 0.035 | Young |
| 0.015651 | 0.473792 | 0.325 | 0.208 | 1 | 4 | Cdc37 | 0.117 | Young |
| 0.015662 | 0.759573 | 0.25 | 0.15 | 1 | 4 | Tk2 | 0.1 | Young |
| 0.015713 | 1.019316 | 0.175 | 0.096 | 1 | 4 | Ppp1r9a | 0.079 | Young |
| 0.015774 | 1.21583 | 0.075 | 0.028 | 1 | 4 | Gm38843 | 0.047 | Young |
| 0.015815 | 0.680825 | 0.288 | 0.18 | 1 | 4 | Gmip | 0.108 | Young |
| 0.01588 | 1.357142 | 0.125 | 0.059 | 1 | 4 | Pgs1 | 0.066 | Young |
| 0.016052 | 0.897692 | 0.162 | 0.084 | 1 | 4 | Fam98c | 0.078 | Young |
| 0.016064 | 1.486426 | 0.125 | 0.06 | 1 | 4 | Slc1a5 | 0.065 | Young |
| 0.016099 | 0.597569 | 0.275 | 0.166 | 1 | 4 | Elmo1 | 0.109 | Young |
| 0.016373 | 0.827533 | 0.2 | 0.112 | 1 | 4 | Aldh7a1 | 0.088 | Young |
| 0.016375 | 0.485632 | 0.45 | 0.314 | 1 | 4 | Cysltr1 | 0.136 | Young |
| 0.016495 | 1.32185 | 0.1 | 0.043 | 1 | 4 | Mecr | 0.057 | Young |
| 0.016528 | 0.405385 | 0.762 | 0.663 | 1 | 4 | Ly86 | 0.099 | Young |
| 0.016572 | 0.293794 | 0.95 | 0.899 | 1 | 4 | Rpl27a | 0.051 | Young |
| 0.016652 | 1.338856 | 0.038 | 0.009 | 1 | 4 | Zfp251 | 0.029 | Young |

| | | | | | | | | |
|---|---|---|---|---|---|---|---|---|
| 0.016672 | 0.786002 | 0.15 | 0.076 | 1 | 4 | Bcl2l1 | 0.074 | Young |
| 0.0167 | 1.199472 | 0.125 | 0.059 | 1 | 4 | Zbtb34 | 0.066 | Young |
| 0.016759 | 1.262832 | 0.1 | 0.043 | 1 | 4 | Extl3 | 0.057 | Young |
| 0.016822 | 0.910446 | 0.075 | 0.028 | 1 | 4 | Tspan33 | 0.047 | Young |
| 0.016959 | 1.544388 | 0.075 | 0.028 | 1 | 4 | Entpd6 | 0.047 | Young |
| 0.017086 | 2.705213 | 0.025 | 0.005 | 1 | 4 | Tgfa | 0.02 | Young |
| 0.017265 | 2.837952 | 0.025 | 0.005 | 1 | 4 | Zfp27 | 0.02 | Young |
| 0.017265 | 2.427484 | 0.025 | 0.005 | 1 | 4 | 1700029I1! | 0.02 | Young |
| 0.017281 | 1.317919 | 0.175 | 0.097 | 1 | 4 | Oard1 | 0.078 | Young |
| 0.017334 | 1.340416 | 0.125 | 0.059 | 1 | 4 | Ttc1 | 0.066 | Young |
| 0.017355 | 2.972434 | 0.025 | 0.005 | 1 | 4 | H2-M2 | 0.02 | Young |
| 0.01743 | 0.795903 | 0.062 | 0.021 | 1 | 4 | Neil1 | 0.041 | Young |
| 0.017445 | 2.292654 | 0.025 | 0.005 | 1 | 4 | Rpp40 | 0.02 | Young |
| 0.017445 | 2.076381 | 0.025 | 0.005 | 1 | 4 | Otub2 | 0.02 | Young |
| 0.017536 | 2.968602 | 0.025 | 0.005 | 1 | 4 | Id1 | 0.02 | Young |
| 0.017536 | 2.016284 | 0.025 | 0.005 | 1 | 4 | Slc27a3 | 0.02 | Young |
| 0.017536 | 1.976371 | 0.025 | 0.005 | 1 | 4 | Kif15 | 0.02 | Young |
| 0.017628 | 1.97469 | 0.025 | 0.005 | 1 | 4 | Mpzl1 | 0.02 | Young |
| 0.017628 | 1.363302 | 0.025 | 0.005 | 1 | 4 | 1700113A1 | 0.02 | Young |
| 0.017777 | 0.68853 | 0.238 | 0.142 | 1 | 4 | Arl1 | 0.096 | Young |
| 0.017812 | 1.76514 | 0.025 | 0.005 | 1 | 4 | Ndor1 | 0.02 | Young |
| 0.017834 | 0.732011 | 0.375 | 0.263 | 1 | 4 | Nop53 | 0.112 | Young |
| 0.017904 | 1.012965 | 0.025 | 0.005 | 1 | 4 | Ankrd9 | 0.02 | Young |
| 0.017913 | 0.275599 | 0.975 | 0.943 | 1 | 4 | Rpl23 | 0.032 | Young |
| 0.017938 | 0.523137 | 0.412 | 0.292 | 1 | 4 | Blnk | 0.12 | Young |
| 0.017967 | 0.911814 | 0.138 | 0.068 | 1 | 4 | Ormdl3 | 0.07 | Young |
| 0.017999 | 1.64977 | 0.125 | 0.061 | 1 | 4 | Trem2 | 0.064 | Young |
| 0.018026 | 0.92683 | 0.188 | 0.105 | 1 | 4 | Cd300c2 | 0.083 | Young |
| 0.018091 | 1.144367 | 0.025 | 0.005 | 1 | 4 | 4930524J0 | 0.02 | Young |
| 0.018241 | 0.475148 | 0.425 | 0.294 | 1 | 4 | Ash1l | 0.131 | Young |
| 0.018312 | 0.215947 | 0.975 | 0.953 | 1 | 4 | C1qb | 0.022 | Young |
| 0.018319 | 0.811973 | 0.262 | 0.162 | 1 | 4 | Irak4 | 0.1 | Young |
| 0.018474 | 1.220623 | 0.075 | 0.028 | 1 | 4 | Ice2 | 0.047 | Young |
| 0.018598 | 1.39901 | 0.05 | 0.015 | 1 | 4 | Plscr1 | 0.035 | Young |
| 0.01871 | 1.476975 | 0.05 | 0.015 | 1 | 4 | Jmjd8 | 0.035 | Young |
| 0.018902 | 1.332927 | 0.112 | 0.052 | 1 | 4 | Orc4 | 0.06 | Young |
| 0.018972 | 0.288075 | 0.925 | 0.884 | 1 | 4 | Rps27 | 0.041 | Young |
| 0.019048 | 1.283633 | 0.05 | 0.015 | 1 | 4 | Mmp14 | 0.035 | Young |
| 0.019089 | 0.684038 | 0.212 | 0.121 | 1 | 4 | Cryl1 | 0.091 | Young |
| 0.019125 | 0.54286 | 0.275 | 0.167 | 1 | 4 | Rbm27 | 0.108 | Young |
| 0.019185 | 0.564608 | 0.175 | 0.093 | 1 | 4 | Nmd3 | 0.082 | Young |
| 0.01922 | 1.140908 | 0.05 | 0.015 | 1 | 4 | Platr25 | 0.035 | Young |
| 0.019288 | 0.796139 | 0.162 | 0.086 | 1 | 4 | Mbd1 | 0.076 | Young |
| 0.019381 | 0.351779 | 0.6 | 0.457 | 1 | 4 | Pafah1b1 | 0.143 | Young |
| 0.019453 | 0.684732 | 0.2 | 0.113 | 1 | 4 | Siglece | 0.087 | Young |

| | | | | | | | | |
|---|---|---|---|---|---|---|---|---|
| 0.019555 | 0.79455 | 0.188 | 0.105 | 1 | 4 | Tbrg1 | 0.083 | Young |
| 0.01958 | 0.772824 | 0.112 | 0.051 | 1 | 4 | Tpgs1 | 0.061 | Young |
| 0.019683 | 0.620899 | 0.05 | 0.015 | 1 | 4 | R3hcc1 | 0.035 | Young |
| 0.019766 | 0.913981 | 0.112 | 0.051 | 1 | 4 | Coprs | 0.061 | Young |
| 0.019956 | 1.134446 | 0.125 | 0.061 | 1 | 4 | Pip4k2c | 0.064 | Young |
| 0.020065 | 0.374354 | 0.125 | 0.058 | 1 | 4 | Frmd8 | 0.067 | Young |
| 0.020199 | 0.4095 | 0.5 | 0.361 | 1 | 4 | Cd84 | 0.139 | Young |
| 0.020204 | 0.807708 | 0.112 | 0.051 | 1 | 4 | Skp2 | 0.061 | Young |
| 0.020216 | 0.638428 | 0.05 | 0.015 | 1 | 4 | Zfp146 | 0.035 | Young |
| 0.020515 | 0.750635 | 0.25 | 0.151 | 1 | 4 | 2810403A( | 0.099 | Young |
| 0.020613 | 0.365732 | 0.675 | 0.555 | 1 | 4 | Ms4a7 | 0.12 | Young |
| 0.020661 | 0.637322 | 0.262 | 0.165 | 1 | 4 | Scamp1 | 0.097 | Young |
| 0.020909 | 0.463232 | 0.175 | 0.092 | 1 | 4 | Cep120 | 0.083 | Young |
| 0.020913 | 0.917008 | 0.062 | 0.022 | 1 | 4 | Il1rap | 0.04 | Young |
| 0.021185 | 0.674794 | 0.2 | 0.113 | 1 | 4 | Gtf2a2 | 0.087 | Young |
| 0.021288 | 0.695625 | 0.2 | 0.113 | 1 | 4 | Akap10 | 0.087 | Young |
| 0.021317 | 1.869934 | 0.062 | 0.022 | 1 | 4 | Slc25a13 | 0.04 | Young |
| 0.021377 | 0.840453 | 0.138 | 0.069 | 1 | 4 | Mlx | 0.069 | Young |
| 0.021469 | 1.066849 | 0.088 | 0.036 | 1 | 4 | Hacd2 | 0.052 | Young |
| 0.021567 | 1.476542 | 0.1 | 0.044 | 1 | 4 | Acvr2a | 0.056 | Young |
| 0.021601 | 0.646617 | 0.125 | 0.059 | 1 | 4 | Gpkow | 0.066 | Young |
| 0.021632 | 1.168636 | 0.075 | 0.029 | 1 | 4 | Snx7 | 0.046 | Young |
| 0.021666 | 0.783983 | 0.2 | 0.115 | 1 | 4 | Suds3 | 0.085 | Young |
| 0.021698 | 1.839083 | 0.038 | 0.01 | 1 | 4 | Megf9 | 0.028 | Young |
| 0.021909 | 0.713335 | 0.2 | 0.114 | 1 | 4 | Prdx4 | 0.086 | Young |
| 0.021923 | 0.808665 | 0.25 | 0.156 | 1 | 4 | Ube2j2 | 0.094 | Young |
| 0.021926 | 0.589576 | 0.062 | 0.022 | 1 | 4 | Slc2a9 | 0.04 | Young |
| 0.021926 | 0.5023 | 0.2 | 0.111 | 1 | 4 | Dhrs7b | 0.089 | Young |
| 0.02196 | 0.771159 | 0.2 | 0.115 | 1 | 4 | Dars | 0.085 | Young |
| 0.021964 | 1.083938 | 0.075 | 0.029 | 1 | 4 | Utp4 | 0.046 | Young |
| 0.022089 | 1.974739 | 0.05 | 0.016 | 1 | 4 | Runx3 | 0.034 | Young |
| 0.02209 | 1.664977 | 0.038 | 0.01 | 1 | 4 | Tm4sf5 | 0.028 | Young |
| 0.022098 | 0.475498 | 0.388 | 0.26 | 1 | 4 | Smg1 | 0.128 | Young |
| 0.022143 | 0.406754 | 0.25 | 0.148 | 1 | 4 | Lilrb4a | 0.102 | Young |
| 0.022169 | 1.86129 | 0.038 | 0.01 | 1 | 4 | C030006K1 | 0.028 | Young |
| 0.022228 | 0.308362 | 0.863 | 0.736 | 1 | 4 | Clta | 0.127 | Young |
| 0.022265 | 0.477209 | 0.325 | 0.209 | 1 | 4 | Rc3h2 | 0.116 | Young |
| 0.022296 | 0.877957 | 0.275 | 0.181 | 1 | 4 | Ebi3 | 0.094 | Young |
| 0.022333 | 1.574882 | 0.075 | 0.03 | 1 | 4 | Arid3b | 0.045 | Young |
| 0.022448 | 0.51965 | 0.525 | 0.409 | 1 | 4 | Tra2b | 0.116 | Young |
| 0.022608 | 1.45586 | 0.062 | 0.022 | 1 | 4 | 2510009E( | 0.04 | Young |
| 0.02262 | 0.269095 | 0.925 | 0.878 | 1 | 4 | Rpl39 | 0.047 | Young |
| 0.022649 | 1.581685 | 0.038 | 0.01 | 1 | 4 | Lhfpl2 | 0.028 | Young |
| 0.022683 | 0.436275 | 0.5 | 0.372 | 1 | 4 | Usp9x | 0.128 | Young |
| 0.022811 | 1.397519 | 0.038 | 0.01 | 1 | 4 | Zfp11 | 0.028 | Young |

| | | | | | | | | |
|---|---|---|---|---|---|---|---|---|
| 0.022823 | 0.274518 | 0.662 | 0.501 | 1 | 4 | Cbl | 0.161 | Young |
| 0.022909 | 0.812743 | 0.125 | 0.061 | 1 | 4 | Bckdha | 0.064 | Young |
| 0.023087 | 0.648557 | 0.212 | 0.125 | 1 | 4 | Ing1 | 0.087 | Young |
| 0.023418 | 1.013991 | 0.088 | 0.037 | 1 | 4 | B4galt7 | 0.051 | Young |
| 0.023439 | 0.62513 | 0.162 | 0.086 | 1 | 4 | Hk2 | 0.076 | Young |
| 0.02347 | 0.858111 | 0.038 | 0.01 | 1 | 4 | Ndufaf5 | 0.028 | Young |
| 0.023471 | 0.349065 | 0.462 | 0.335 | 1 | 4 | Kmt2e | 0.127 | Young |
| 0.023506 | 1.294875 | 0.062 | 0.022 | 1 | 4 | Prr5 | 0.04 | Young |
| 0.02352 | 0.423841 | 0.65 | 0.511 | 1 | 4 | Myo5a | 0.139 | Young |
| 0.023561 | 0.560109 | 0.537 | 0.42 | 1 | 4 | Pum2 | 0.117 | Young |
| 0.023654 | 0.781264 | 0.162 | 0.086 | 1 | 4 | Tcf7l2 | 0.076 | Young |
| 0.023668 | 0.620957 | 0.088 | 0.036 | 1 | 4 | Zfp513 | 0.052 | Young |
| 0.023704 | 0.533201 | 0.6 | 0.499 | 1 | 4 | Ccl24 | 0.101 | Young |
| 0.02385 | 1.159411 | 0.062 | 0.022 | 1 | 4 | Acvr2b | 0.04 | Young |
| 0.023973 | 0.765114 | 0.2 | 0.115 | 1 | 4 | Gpr137b | 0.085 | Young |
| 0.024117 | 1.253903 | 0.075 | 0.03 | 1 | 4 | Borcs8 | 0.045 | Young |
| 0.024165 | 0.257294 | 0.85 | 0.69 | 1 | 4 | Fus | 0.16 | Young |
| 0.024263 | 2.760252 | 0.013 | 0.001 | 1 | 4 | Mex3a | 0.012 | Young |
| 0.024263 | 2.544161 | 0.013 | 0.001 | 1 | 4 | Capn13 | 0.012 | Young |
| 0.024263 | 2.425681 | 0.013 | 0.001 | 1 | 4 | Slc2a12 | 0.012 | Young |
| 0.024263 | 2.41868 | 0.013 | 0.001 | 1 | 4 | Kcnj10 | 0.012 | Young |
| 0.024263 | 2.41012 | 0.013 | 0.001 | 1 | 4 | Gramd1c | 0.012 | Young |
| 0.024263 | 2.399864 | 0.013 | 0.001 | 1 | 4 | Kcnip3 | 0.012 | Young |
| 0.024263 | 2.383048 | 0.013 | 0.001 | 1 | 4 | Zfp354b | 0.012 | Young |
| 0.024263 | 2.365697 | 0.013 | 0.001 | 1 | 4 | 9430037G( | 0.012 | Young |
| 0.024263 | 2.351714 | 0.013 | 0.001 | 1 | 4 | Vit | 0.012 | Young |
| 0.024263 | 2.340262 | 0.013 | 0.001 | 1 | 4 | Lrrc4 | 0.012 | Young |
| 0.024263 | 2.2926 | 0.013 | 0.001 | 1 | 4 | Zfp334 | 0.012 | Young |
| 0.024263 | 2.229179 | 0.013 | 0.001 | 1 | 4 | Hoxc8 | 0.012 | Young |
| 0.024263 | 2.162294 | 0.013 | 0.001 | 1 | 4 | Rbms3 | 0.012 | Young |
| 0.024263 | 2.062364 | 0.013 | 0.001 | 1 | 4 | Fam178b | 0.012 | Young |
| 0.024263 | 2.036078 | 0.013 | 0.001 | 1 | 4 | Gen1 | 0.012 | Young |
| 0.024263 | 2.012684 | 0.013 | 0.001 | 1 | 4 | Meig1 | 0.012 | Young |
| 0.024263 | 1.889185 | 0.013 | 0.001 | 1 | 4 | 9430078K2 | 0.012 | Young |
| 0.024263 | 1.869588 | 0.013 | 0.001 | 1 | 4 | Myo19 | 0.012 | Young |
| 0.024263 | 1.801146 | 0.013 | 0.001 | 1 | 4 | Col7a1 | 0.012 | Young |
| 0.024263 | 1.773333 | 0.013 | 0.001 | 1 | 4 | Gm13075 | 0.012 | Young |
| 0.024263 | 1.567396 | 0.013 | 0.001 | 1 | 4 | Hist1h1b | 0.012 | Young |
| 0.024263 | 0.773119 | 0.013 | 0.001 | 1 | 4 | Zfyve28 | 0.012 | Young |
| 0.024336 | 1.145868 | 0.1 | 0.045 | 1 | 4 | Snx11 | 0.055 | Young |
| 0.024469 | 0.215489 | 0.95 | 0.874 | 1 | 4 | Rps19 | 0.076 | Young |
| 0.024473 | 1.913918 | 0.013 | 0.001 | 1 | 4 | Col14a1 | 0.012 | Young |
| 0.024473 | 1.884913 | 0.013 | 0.001 | 1 | 4 | A430105I1 | 0.012 | Young |
| 0.024473 | 1.841565 | 0.013 | 0.001 | 1 | 4 | B230208H1 | 0.012 | Young |
| 0.024473 | 1.752291 | 0.013 | 0.001 | 1 | 4 | Xkr6 | 0.012 | Young |

| | | | | | | | | |
|---|---|---|---|---|---|---|---|---|
| 0.024473 | 1.722389 | 0.013 | 0.001 | 1 | 4 | Mxra8 | 0.012 | Young |
| 0.024473 | 1.663543 | 0.013 | 0.001 | 1 | 4 | Slc9a7 | 0.012 | Young |
| 0.024473 | 1.622432 | 0.013 | 0.001 | 1 | 4 | Melk | 0.012 | Young |
| 0.024473 | 1.403935 | 0.013 | 0.001 | 1 | 4 | Lbp | 0.012 | Young |
| 0.024473 | 1.401296 | 0.013 | 0.001 | 1 | 4 | Gm14966 | 0.012 | Young |
| 0.024473 | 1.378126 | 0.013 | 0.001 | 1 | 4 | Hoxd8 | 0.012 | Young |
| 0.024473 | 1.342658 | 0.013 | 0.001 | 1 | 4 | Mettl4 | 0.012 | Young |
| 0.024473 | 1.274801 | 0.013 | 0.001 | 1 | 4 | Pcgf2 | 0.012 | Young |
| 0.024473 | 1.188477 | 0.013 | 0.001 | 1 | 4 | Prima1 | 0.012 | Young |
| 0.024473 | 1.166347 | 0.013 | 0.001 | 1 | 4 | C530050E1 | 0.012 | Young |
| 0.024473 | 0.949418 | 0.013 | 0.001 | 1 | 4 | Fabp7 | 0.012 | Young |
| 0.024473 | 0.933962 | 0.013 | 0.001 | 1 | 4 | AA986860 | 0.012 | Young |
| 0.024473 | 0.887421 | 0.013 | 0.001 | 1 | 4 | 4930426I2 | 0.012 | Young |
| 0.024473 | 0.865719 | 0.013 | 0.001 | 1 | 4 | Lynx1 | 0.012 | Young |
| 0.024473 | 0.86344 | 0.013 | 0.001 | 1 | 4 | Gm17484 | 0.012 | Young |
| 0.024473 | 0.790094 | 0.013 | 0.001 | 1 | 4 | Casc4 | 0.012 | Young |
| 0.024473 | 0.642149 | 0.013 | 0.001 | 1 | 4 | Dpys | 0.012 | Young |
| 0.024513 | 0.500015 | 0.2 | 0.113 | 1 | 4 | Dgcr2 | 0.087 | Young |
| 0.024654 | 0.829274 | 0.112 | 0.053 | 1 | 4 | Txnrd2 | 0.059 | Young |
| 0.024666 | 0.955139 | 0.175 | 0.099 | 1 | 4 | Rogdi | 0.076 | Young |
| 0.024684 | 2.077551 | 0.013 | 0.001 | 1 | 4 | 5330426L2 | 0.012 | Young |
| 0.024684 | 1.757852 | 0.013 | 0.001 | 1 | 4 | 8430408G2 | 0.012 | Young |
| 0.024684 | 1.577884 | 0.013 | 0.001 | 1 | 4 | Zfp575 | 0.012 | Young |
| 0.024684 | 1.535066 | 0.013 | 0.001 | 1 | 4 | Cdkn3 | 0.012 | Young |
| 0.024684 | 1.412608 | 0.013 | 0.001 | 1 | 4 | Gm26620 | 0.012 | Young |
| 0.024684 | 1.357276 | 0.013 | 0.001 | 1 | 4 | 4933429H1 | 0.012 | Young |
| 0.024684 | 1.159525 | 0.013 | 0.001 | 1 | 4 | Pck1 | 0.012 | Young |
| 0.024684 | 1.06416 | 0.013 | 0.001 | 1 | 4 | Gm26981 | 0.012 | Young |
| 0.024684 | 1.037693 | 0.013 | 0.001 | 1 | 4 | Gm16253 | 0.012 | Young |
| 0.024684 | 0.95791 | 0.013 | 0.001 | 1 | 4 | Eml1 | 0.012 | Young |
| 0.024684 | 0.948972 | 0.013 | 0.001 | 1 | 4 | Dab2ip | 0.012 | Young |
| 0.024684 | 0.905337 | 0.013 | 0.001 | 1 | 4 | Chst2 | 0.012 | Young |
| 0.024684 | 0.902397 | 0.013 | 0.001 | 1 | 4 | Tbx3 | 0.012 | Young |
| 0.024684 | 0.900145 | 0.013 | 0.001 | 1 | 4 | 9930012K1 | 0.012 | Young |
| 0.024684 | 0.897098 | 0.013 | 0.001 | 1 | 4 | Zfp558 | 0.012 | Young |
| 0.024684 | 0.839789 | 0.013 | 0.001 | 1 | 4 | Gm16028 | 0.012 | Young |
| 0.024684 | 0.827047 | 0.013 | 0.001 | 1 | 4 | Gm45620 | 0.012 | Young |
| 0.024684 | 0.816606 | 0.013 | 0.001 | 1 | 4 | Gm15510 | 0.012 | Young |
| 0.024684 | 0.740552 | 0.013 | 0.001 | 1 | 4 | Gm11361 | 0.012 | Young |
| 0.024684 | 0.730799 | 0.013 | 0.001 | 1 | 4 | Pard3b | 0.012 | Young |
| 0.024684 | 0.693967 | 0.013 | 0.001 | 1 | 4 | Igfbp3 | 0.012 | Young |
| 0.024684 | 0.685731 | 0.013 | 0.001 | 1 | 4 | Gm11032 | 0.012 | Young |
| 0.024684 | 0.661543 | 0.013 | 0.001 | 1 | 4 | Npr3 | 0.012 | Young |
| 0.024684 | 0.644836 | 0.013 | 0.001 | 1 | 4 | Mmp23 | 0.012 | Young |
| 0.024684 | 0.63174 | 0.013 | 0.001 | 1 | 4 | 4921516A0 | 0.012 | Young |

| | | | | | | | | |
|---|---|---|---|---|---|---|---|---|
| 0.024684 | 0.597525 | 0.013 | 0.001 | 1 | 4 | Clec9a | 0.012 | Young |
| 0.024684 | 0.504534 | 0.013 | 0.001 | 1 | 4 | Vash1 | 0.012 | Young |
| 0.024684 | 0.414115 | 0.013 | 0.001 | 1 | 4 | Gm45151 | 0.012 | Young |
| 0.024684 | 0.405786 | 0.013 | 0.001 | 1 | 4 | Scarf2 | 0.012 | Young |
| 0.024684 | 0.401764 | 0.013 | 0.001 | 1 | 4 | Spsb4 | 0.012 | Young |
| 0.024684 | 0.399926 | 0.013 | 0.001 | 1 | 4 | Gm12500 | 0.012 | Young |
| 0.024731 | 0.952975 | 0.062 | 0.022 | 1 | 4 | 1700066M | 0.04 | Young |
| 0.024823 | 0.827875 | 0.162 | 0.088 | 1 | 4 | Axin1 | 0.074 | Young |
| 0.02485 | 1.048071 | 0.062 | 0.022 | 1 | 4 | Slc37a3 | 0.04 | Young |
| 0.024943 | 0.699122 | 0.2 | 0.116 | 1 | 4 | Ripor2 | 0.084 | Young |
| 0.024948 | 1.229533 | 0.075 | 0.03 | 1 | 4 | Svbp | 0.045 | Young |
| 0.02555 | 0.72789 | 0.238 | 0.145 | 1 | 4 | Tnfaip8l2 | 0.093 | Young |
| 0.025865 | 0.662231 | 0.225 | 0.135 | 1 | 4 | Rassf5 | 0.09 | Young |
| 0.026045 | 1.116739 | 0.112 | 0.054 | 1 | 4 | Nacc1 | 0.058 | Young |
| 0.026172 | 0.534195 | 0.312 | 0.209 | 1 | 4 | Eif4e | 0.103 | Young |
| 0.026285 | 0.623528 | 0.338 | 0.239 | 1 | 4 | Itpkb | 0.099 | Young |
| 0.02643 | 0.416907 | 0.262 | 0.16 | 1 | 4 | Mrpl3 | 0.102 | Young |
| 0.02663 | 0.83245 | 0.075 | 0.03 | 1 | 4 | Scfd2 | 0.045 | Young |
| 0.026667 | 0.615881 | 0.325 | 0.223 | 1 | 4 | Med13l | 0.102 | Young |
| 0.026704 | 0.180479 | 1 | 0.995 | 1 | 4 | Ctsb | 0.005 | Young |
| 0.026776 | 0.727795 | 0.225 | 0.14 | 1 | 4 | Zzz3 | 0.085 | Young |
| 0.026777 | 0.461363 | 0.362 | 0.244 | 1 | 4 | Rpl13a | 0.118 | Young |
| 0.026866 | 0.902051 | 0.162 | 0.09 | 1 | 4 | Rcbtb2 | 0.072 | Young |
| 0.026987 | 1.030745 | 0.1 | 0.045 | 1 | 4 | Klhdc10 | 0.055 | Young |
| 0.027016 | 0.501214 | 0.238 | 0.143 | 1 | 4 | Fam76b | 0.095 | Young |
| 0.027025 | 0.7221 | 0.238 | 0.145 | 1 | 4 | Ext1 | 0.093 | Young |
| 0.027046 | 0.213754 | 1 | 0.978 | 1 | 4 | Rps24 | 0.022 | Young |
| 0.027063 | 0.512976 | 0.25 | 0.149 | 1 | 4 | Idh3g | 0.101 | Young |
| 0.027326 | 0.764268 | 0.138 | 0.07 | 1 | 4 | Herc6 | 0.068 | Young |
| 0.027327 | 0.653911 | 0.338 | 0.231 | 1 | 4 | Commd8 | 0.107 | Young |
| 0.027392 | 0.690649 | 0.25 | 0.157 | 1 | 4 | Ccdc82 | 0.093 | Young |
| 0.027539 | 0.842359 | 0.225 | 0.138 | 1 | 4 | Zrsr2 | 0.087 | Young |
| 0.027618 | 0.554615 | 0.238 | 0.146 | 1 | 4 | Slc31a2 | 0.092 | Young |
| 0.02765 | 0.364051 | 0.288 | 0.178 | 1 | 4 | Pik3c2a | 0.11 | Young |
| 0.027719 | 0.502953 | 0.488 | 0.354 | 1 | 4 | Rnf150 | 0.134 | Young |
| 0.027733 | 1.358904 | 0.062 | 0.023 | 1 | 4 | Kbtbd3 | 0.039 | Young |
| 0.028016 | 0.302329 | 0.338 | 0.227 | 1 | 4 | Fabp4 | 0.111 | Young |
| 0.02803 | 1.234395 | 0.088 | 0.038 | 1 | 4 | Cdc23 | 0.05 | Young |
| 0.02819 | 0.444998 | 0.488 | 0.363 | 1 | 4 | Lamtor4 | 0.125 | Young |
| 0.028305 | 0.571079 | 0.338 | 0.236 | 1 | 4 | Trafd1 | 0.102 | Young |
| 0.028322 | 0.361107 | 0.762 | 0.663 | 1 | 4 | Rpl11 | 0.099 | Young |
| 0.02842 | 0.21197 | 0.938 | 0.868 | 1 | 4 | Rpl23a | 0.07 | Young |
| 0.028492 | 0.487575 | 0.575 | 0.458 | 1 | 4 | Rab5c | 0.117 | Young |
| 0.028497 | 2.272232 | 0.025 | 0.005 | 1 | 4 | Pxmp2 | 0.02 | Young |
| 0.028519 | 0.616521 | 0.2 | 0.117 | 1 | 4 | Lars | 0.083 | Young |

| | | | | | | | | |
|---|---|---|---|---|---|---|---|---|
| 0.028523 | 1.227032 | 0.062 | 0.023 | 1 | 4 | Ighd | 0.039 | Young |
| 0.028616 | 0.871516 | 0.088 | 0.038 | 1 | 4 | Gipc1 | 0.05 | Young |
| 0.028636 | 1.597602 | 0.05 | 0.016 | 1 | 4 | Nradd | 0.034 | Young |
| 0.028748 | 0.752808 | 0.3 | 0.199 | 1 | 4 | Maoa | 0.101 | Young |
| 0.028761 | 1.646289 | 0.025 | 0.005 | 1 | 4 | B430306N( | 0.02 | Young |
| 0.028782 | 0.552998 | 0.262 | 0.165 | 1 | 4 | Flii | 0.097 | Young |
| 0.028829 | 0.761446 | 0.225 | 0.137 | 1 | 4 | Tcf3 | 0.088 | Young |
| 0.02887 | 1.004899 | 0.075 | 0.03 | 1 | 4 | Tmem38a | 0.045 | Young |
| 0.028895 | 1.803079 | 0.025 | 0.005 | 1 | 4 | 1110002L0 | 0.02 | Young |
| 0.028896 | 0.689378 | 0.1 | 0.045 | 1 | 4 | Zbtb43 | 0.055 | Young |
| 0.028926 | 0.852331 | 0.175 | 0.099 | 1 | 4 | Adcy9 | 0.076 | Young |
| 0.029025 | 0.556332 | 0.15 | 0.079 | 1 | 4 | Sdhaf4 | 0.071 | Young |
| 0.029036 | 0.368223 | 0.3 | 0.192 | 1 | 4 | Clasp2 | 0.108 | Young |
| 0.02906 | 0.657095 | 0.062 | 0.023 | 1 | 4 | Ccdc167 | 0.039 | Young |
| 0.029239 | 0.504937 | 0.175 | 0.097 | 1 | 4 | Fgd6 | 0.078 | Young |
| 0.029345 | 2.789582 | 0.038 | 0.011 | 1 | 4 | Kctd21 | 0.027 | Young |
| 0.029345 | 1.874315 | 0.038 | 0.011 | 1 | 4 | H2-Ob | 0.027 | Young |
| 0.029405 | 0.547858 | 0.262 | 0.167 | 1 | 4 | Heatr5a | 0.095 | Young |
| 0.029469 | 0.76113 | 0.062 | 0.023 | 1 | 4 | Atp1b1 | 0.039 | Young |
| 0.029579 | 1.383438 | 0.125 | 0.065 | 1 | 4 | Erf | 0.06 | Young |
| 0.029602 | 1.145422 | 0.212 | 0.132 | 1 | 4 | Slc15a3 | 0.08 | Young |
| 0.029632 | 0.459028 | 0.238 | 0.144 | 1 | 4 | Nbr1 | 0.094 | Young |
| 0.029642 | 1.794716 | 0.038 | 0.011 | 1 | 4 | Fxn | 0.027 | Young |
| 0.029644 | 0.936302 | 0.138 | 0.072 | 1 | 4 | Vgll4 | 0.066 | Young |
| 0.029655 | 0.675645 | 0.312 | 0.213 | 1 | 4 | Kcnk6 | 0.099 | Young |
| 0.029675 | 0.763907 | 0.062 | 0.023 | 1 | 4 | Hexdc | 0.039 | Young |
| 0.029682 | 0.127679 | 0.988 | 0.993 | 1 | 4 | mt-Nd4 | -0.005 | Young |
| 0.029753 | 0.712186 | 0.212 | 0.128 | 1 | 4 | Foxred2 | 0.084 | Young |
| 0.029833 | 0.708964 | 0.138 | 0.071 | 1 | 4 | Cyp4v3 | 0.067 | Young |
| 0.030032 | 1.490375 | 0.088 | 0.039 | 1 | 4 | Tlr13 | 0.049 | Young |
| 0.030191 | 0.365929 | 0.5 | 0.379 | 1 | 4 | Tbxas1 | 0.121 | Young |
| 0.030231 | 0.774411 | 0.075 | 0.03 | 1 | 4 | Peli2 | 0.045 | Young |
| 0.03033 | 0.182236 | 0.188 | 0.107 | 1 | 4 | Lyz1 | 0.081 | Young |
| 0.030501 | 0.349088 | 0.462 | 0.339 | 1 | 4 | Zfp106 | 0.123 | Young |
| 0.030772 | 1.307941 | 0.075 | 0.031 | 1 | 4 | Btg3 | 0.044 | Young |
| 0.030787 | 0.874308 | 0.088 | 0.038 | 1 | 4 | Alg2 | 0.05 | Young |
| 0.030853 | 1.143928 | 0.038 | 0.011 | 1 | 4 | Slc25a26 | 0.027 | Young |
| 0.030958 | 0.255543 | 0.95 | 0.901 | 1 | 4 | Rps15 | 0.049 | Young |
| 0.030965 | 1.079174 | 0.175 | 0.102 | 1 | 4 | Mapkap1 | 0.073 | Young |
| 0.031059 | 1.054896 | 0.038 | 0.011 | 1 | 4 | Vsig10l | 0.027 | Young |
| 0.031175 | 0.705066 | 0.388 | 0.28 | 1 | 4 | Snx8 | 0.108 | Young |
| 0.031267 | 0.595442 | 0.038 | 0.011 | 1 | 4 | Smad7 | 0.027 | Young |
| 0.03129 | 0.291741 | 0.963 | 0.932 | 1 | 4 | Zfp36l1 | 0.031 | Young |
| 0.031379 | 0.433249 | 0.55 | 0.416 | 1 | 4 | Itsn1 | 0.134 | Young |
| 0.031397 | 1.243642 | 0.075 | 0.031 | 1 | 4 | Fzd5 | 0.044 | Young |

| | | | | | | | | |
|---|---|---|---|---|---|---|---|---|
| 0.031453 | 1.425673 | 0.112 | 0.056 | 1 | 4 | Slc41a3 | 0.056 | Young |
| 0.031659 | 0.570582 | 0.188 | 0.107 | 1 | 4 | Igsf8 | 0.081 | Young |
| 0.031892 | 0.661529 | 0.188 | 0.109 | 1 | 4 | Gatad2a | 0.079 | Young |
| 0.032226 | 1.344427 | 0.062 | 0.024 | 1 | 4 | Selenoo | 0.038 | Young |
| 0.03241 | 0.454018 | 0.2 | 0.115 | 1 | 4 | Vps37a | 0.085 | Young |
| 0.032488 | 1.124451 | 0.1 | 0.047 | 1 | 4 | Rbmx | 0.053 | Young |
| 0.032503 | 0.424162 | 0.388 | 0.274 | 1 | 4 | Cul3 | 0.114 | Young |
| 0.032629 | 0.697779 | 0.15 | 0.081 | 1 | 4 | Ap4s1 | 0.069 | Young |
| 0.032685 | 0.301886 | 0.45 | 0.32 | 1 | 4 | Syncrip | 0.13 | Young |
| 0.032787 | 0.374405 | 0.5 | 0.397 | 1 | 4 | Hcls1 | 0.103 | Young |
| 0.03281 | 0.79502 | 0.2 | 0.12 | 1 | 4 | Crim1 | 0.08 | Young |
| 0.032812 | 0.528617 | 0.4 | 0.294 | 1 | 4 | P2ry6 | 0.106 | Young |
| 0.032977 | 0.591455 | 0.325 | 0.222 | 1 | 4 | Psmc3 | 0.103 | Young |
| 0.033024 | 0.767066 | 0.188 | 0.111 | 1 | 4 | Mrps12 | 0.077 | Young |
| 0.033045 | 1.329728 | 0.138 | 0.076 | 1 | 4 | Manbal | 0.062 | Young |
| 0.033071 | 0.35942 | 0.225 | 0.134 | 1 | 4 | Pdhb | 0.091 | Young |
| 0.033103 | 0.43062 | 0.462 | 0.362 | 1 | 4 | Clec4a2 | 0.1 | Young |
| 0.033156 | 0.698775 | 0.238 | 0.15 | 1 | 4 | Abce1 | 0.088 | Young |
| 0.033229 | 0.773603 | 0.225 | 0.143 | 1 | 4 | Rps6ka1 | 0.082 | Young |
| 0.033393 | 0.478724 | 0.325 | 0.221 | 1 | 4 | Kpnb1 | 0.104 | Young |
| 0.033504 | 0.992475 | 0.175 | 0.1 | 1 | 4 | Aldh3a2 | 0.075 | Young |
| 0.03368 | 1.197687 | 0.1 | 0.047 | 1 | 4 | Paxip1 | 0.053 | Young |
| 0.033712 | 0.625355 | 0.088 | 0.038 | 1 | 4 | Tsr1 | 0.05 | Young |
| 0.033834 | 0.833072 | 0.1 | 0.047 | 1 | 4 | Lrp5 | 0.053 | Young |
| 0.034056 | 0.667076 | 0.325 | 0.231 | 1 | 4 | Eif4g3 | 0.094 | Young |
| 0.034311 | 1.491615 | 0.05 | 0.017 | 1 | 4 | Rnf26 | 0.033 | Young |
| 0.034451 | 0.377484 | 0.312 | 0.205 | 1 | 4 | Srp72 | 0.107 | Young |
| 0.03464 | 0.827839 | 0.062 | 0.024 | 1 | 4 | Kiz | 0.038 | Young |
| 0.034803 | 0.401248 | 0.262 | 0.164 | 1 | 4 | Wtap | 0.098 | Young |
| 0.034843 | 0.562288 | 0.375 | 0.268 | 1 | 4 | Ndufb8 | 0.107 | Young |
| 0.034916 | 0.349289 | 0.5 | 0.36 | 1 | 4 | Hsbp1 | 0.14 | Young |
| 0.035581 | 1.174645 | 0.075 | 0.032 | 1 | 4 | Pdp1 | 0.043 | Young |
| 0.035604 | 0.340059 | 0.688 | 0.59 | 1 | 4 | Cd163 | 0.098 | Young |
| 0.035626 | 0.836363 | 0.088 | 0.039 | 1 | 4 | Cyb561d2 | 0.049 | Young |
| 0.035663 | 0.46173 | 0.062 | 0.024 | 1 | 4 | Kptn | 0.038 | Young |
| 0.035724 | 1.044998 | 0.088 | 0.04 | 1 | 4 | Rfxap | 0.048 | Young |
| 0.035909 | 0.782489 | 0.15 | 0.083 | 1 | 4 | Chd7 | 0.067 | Young |
| 0.036067 | 0.465059 | 0.35 | 0.236 | 1 | 4 | Fam234a | 0.114 | Young |
| 0.036337 | 0.783332 | 0.05 | 0.017 | 1 | 4 | Zbtb10 | 0.033 | Young |
| 0.036835 | 0.693013 | 0.125 | 0.064 | 1 | 4 | Dgcr6 | 0.061 | Young |
| 0.036906 | 0.805117 | 0.225 | 0.142 | 1 | 4 | Polr2f | 0.083 | Young |
| 0.036926 | 0.374456 | 0.738 | 0.646 | 1 | 4 | Rps3 | 0.092 | Young |
| 0.037198 | 0.384625 | 0.375 | 0.255 | 1 | 4 | Ppp6r3 | 0.12 | Young |
| 0.037375 | 0.880264 | 0.3 | 0.208 | 1 | 4 | Adrb2 | 0.092 | Young |
| 0.037739 | 0.639642 | 0.188 | 0.11 | 1 | 4 | Mau2 | 0.078 | Young |

| | | | | | | | | |
|---|---|---|---|---|---|---|---|---|
| 0.037921 | 1.165501 | 0.088 | 0.04 | 1 | 4 | Wdr45 | 0.048 | Young |
| 0.038011 | 0.697952 | 0.2 | 0.122 | 1 | 4 | Klhdc3 | 0.078 | Young |
| 0.038348 | 0.516381 | 0.138 | 0.072 | 1 | 4 | Med7 | 0.066 | Young |
| 0.038423 | 0.752101 | 0.238 | 0.152 | 1 | 4 | Rxra | 0.086 | Young |
| 0.038493 | 0.629737 | 0.338 | 0.244 | 1 | 4 | Snx29 | 0.094 | Young |
| 0.038622 | 1.517616 | 0.075 | 0.032 | 1 | 4 | Tacc2 | 0.043 | Young |
| 0.038746 | 1.523638 | 0.038 | 0.011 | 1 | 4 | Msrb2 | 0.027 | Young |
| 0.038788 | 0.349717 | 0.3 | 0.196 | 1 | 4 | Rap2c | 0.104 | Young |
| 0.038808 | 1.191169 | 0.138 | 0.075 | 1 | 4 | Wdr81 | 0.063 | Young |
| 0.038854 | 0.64575 | 0.262 | 0.177 | 1 | 4 | Psmb3 | 0.085 | Young |
| 0.038926 | 0.919596 | 0.088 | 0.04 | 1 | 4 | Cog4 | 0.048 | Young |
| 0.038944 | 0.645923 | 0.188 | 0.111 | 1 | 4 | Mfap3 | 0.077 | Young |
| 0.03904 | 0.463746 | 0.55 | 0.425 | 1 | 4 | Rnf13 | 0.125 | Young |
| 0.03914 | 0.538004 | 0.2 | 0.119 | 1 | 4 | Agpat5 | 0.081 | Young |
| 0.039323 | 0.767774 | 0.212 | 0.132 | 1 | 4 | Slc16a6 | 0.08 | Young |
| 0.039359 | 1.298273 | 0.038 | 0.011 | 1 | 4 | Mansc1 | 0.027 | Young |
| 0.039377 | 0.423649 | 0.475 | 0.358 | 1 | 4 | Ppp1r15a | 0.117 | Young |
| 0.039416 | 0.608583 | 0.162 | 0.092 | 1 | 4 | Bcl2a1b | 0.07 | Young |
| 0.039463 | 0.518407 | 0.175 | 0.1 | 1 | 4 | Slc12a9 | 0.075 | Young |
| 0.039482 | 1.380681 | 0.038 | 0.011 | 1 | 4 | Pomt1 | 0.027 | Young |
| 0.039485 | 0.787643 | 0.162 | 0.091 | 1 | 4 | Abcg1 | 0.071 | Young |
| 0.039562 | 0.89024 | 0.212 | 0.134 | 1 | 4 | Tfg | 0.078 | Young |
| 0.039622 | 1.470127 | 0.05 | 0.018 | 1 | 4 | Slc19a1 | 0.032 | Young |
| 0.039855 | 1.446434 | 0.038 | 0.011 | 1 | 4 | Ap5b1 | 0.027 | Young |
| 0.039895 | 0.683397 | 0.075 | 0.032 | 1 | 4 | Cep135 | 0.043 | Young |
| 0.03998 | 1.21653 | 0.038 | 0.011 | 1 | 4 | Tsku | 0.027 | Young |
| 0.040063 | 0.586078 | 0.412 | 0.303 | 1 | 4 | Mon2 | 0.109 | Young |
| 0.040083 | 0.481095 | 0.188 | 0.11 | 1 | 4 | Plxnc1 | 0.078 | Young |
| 0.040139 | 0.928219 | 0.15 | 0.084 | 1 | 4 | Esco1 | 0.066 | Young |
| 0.040247 | 0.433424 | 0.775 | 0.689 | 1 | 4 | Stk17b | 0.086 | Young |
| 0.040327 | 0.604594 | 0.312 | 0.212 | 1 | 4 | Ulk2 | 0.1 | Young |
| 0.040327 | 2.102983 | 0.05 | 0.018 | 1 | 4 | C030034L1 | 0.032 | Young |
| 0.040356 | 0.699004 | 0.038 | 0.011 | 1 | 4 | Arhgef18 | 0.027 | Young |
| 0.040413 | 0.505013 | 0.138 | 0.073 | 1 | 4 | Itpk1 | 0.065 | Young |
| 0.040541 | 0.421028 | 0.475 | 0.351 | 1 | 4 | Git2 | 0.124 | Young |
| 0.04057 | 1.012558 | 0.088 | 0.04 | 1 | 4 | Clcn6 | 0.048 | Young |
| 0.040802 | 0.647212 | 0.162 | 0.093 | 1 | 4 | Zcchc14 | 0.069 | Young |
| 0.040954 | 0.381032 | 0.212 | 0.126 | 1 | 4 | Cnpy3 | 0.086 | Young |
| 0.04099 | 1.132584 | 0.125 | 0.066 | 1 | 4 | Exosc10 | 0.059 | Young |
| 0.041003 | 0.666773 | 0.088 | 0.04 | 1 | 4 | 2310039H( | 0.048 | Young |
| 0.041049 | 0.382106 | 0.225 | 0.137 | 1 | 4 | 2210016F1 | 0.088 | Young |
| 0.041159 | 1.095243 | 0.138 | 0.075 | 1 | 4 | Mrps18a | 0.063 | Young |
| 0.041208 | 1.008345 | 0.075 | 0.032 | 1 | 4 | Elk1 | 0.043 | Young |
| 0.041298 | 0.31626 | 0.112 | 0.055 | 1 | 4 | Exoc7 | 0.057 | Young |
| 0.041464 | 0.788596 | 0.238 | 0.156 | 1 | 4 | Pink1 | 0.082 | Young |

| | | | | | | | | |
|---|---|---|---|---|---|---|---|---|
| 0.041505 | 0.216702 | 0.688 | 0.542 | 1 | 4 | Lifr | 0.146 | Young |
| 0.0416 | 0.810595 | 0.1 | 0.048 | 1 | 4 | Naa25 | 0.052 | Young |
| 0.041742 | 0.993048 | 0.15 | 0.085 | 1 | 4 | Pdcd7 | 0.065 | Young |
| 0.041936 | 0.357966 | 0.175 | 0.099 | 1 | 4 | Ccdc6 | 0.076 | Young |
| 0.041966 | 0.435751 | 0.312 | 0.211 | 1 | 4 | Tanc2 | 0.101 | Young |
| 0.042075 | 0.968925 | 0.075 | 0.032 | 1 | 4 | Ttyh2 | 0.043 | Young |
| 0.042367 | 0.522886 | 0.088 | 0.04 | 1 | 4 | Rfc3 | 0.048 | Young |
| 0.042373 | 0.649009 | 0.15 | 0.083 | 1 | 4 | Tmem183a | 0.067 | Young |
| 0.042793 | 1.072174 | 0.1 | 0.049 | 1 | 4 | Nf2 | 0.051 | Young |
| 0.042864 | 0.805731 | 0.1 | 0.049 | 1 | 4 | Dusp8 | 0.051 | Young |
| 0.04288 | 2.278338 | 0.025 | 0.006 | 1 | 4 | Gpr141 | 0.019 | Young |
| 0.043466 | 1.139016 | 0.175 | 0.108 | 1 | 4 | Akna | 0.067 | Young |
| 0.043598 | 2.020333 | 0.025 | 0.006 | 1 | 4 | Hes7 | 0.019 | Young |
| 0.043738 | 0.522028 | 0.275 | 0.182 | 1 | 4 | Bfar | 0.093 | Young |
| 0.04378 | 2.063358 | 0.025 | 0.006 | 1 | 4 | Tedc2 | 0.019 | Young |
| 0.043789 | 0.23957 | 0.625 | 0.473 | 1 | 4 | Brk1 | 0.152 | Young |
| 0.043838 | 0.56775 | 0.288 | 0.194 | 1 | 4 | Rraga | 0.094 | Young |
| 0.04389 | 0.444007 | 0.45 | 0.339 | 1 | 4 | Bin1 | 0.111 | Young |
| 0.044013 | 0.91848 | 0.05 | 0.018 | 1 | 4 | Neurl1a | 0.032 | Young |
| 0.044013 | 0.764173 | 0.05 | 0.018 | 1 | 4 | Ptcd1 | 0.032 | Young |
| 0.044133 | 0.223207 | 0.188 | 0.107 | 1 | 4 | Rhot1 | 0.081 | Young |
| 0.044282 | 1.097319 | 0.088 | 0.041 | 1 | 4 | Kat2a | 0.047 | Young |
| 0.044327 | 1.872951 | 0.025 | 0.006 | 1 | 4 | Tle2 | 0.019 | Young |
| 0.044391 | 0.423446 | 0.362 | 0.253 | 1 | 4 | Srsf11 | 0.109 | Young |
| 0.04451 | 1.723342 | 0.025 | 0.006 | 1 | 4 | Oxld1 | 0.019 | Young |
| 0.04451 | 1.647341 | 0.025 | 0.006 | 1 | 4 | Zfp119a | 0.019 | Young |
| 0.04451 | 1.51973 | 0.025 | 0.006 | 1 | 4 | Ropn1l | 0.019 | Young |
| 0.044606 | 0.361117 | 0.4 | 0.289 | 1 | 4 | Hprt | 0.111 | Young |
| 0.044627 | 1.561895 | 0.075 | 0.033 | 1 | 4 | Dolk | 0.042 | Young |
| 0.044676 | 0.290836 | 0.175 | 0.099 | 1 | 4 | Sae1 | 0.076 | Young |
| 0.044695 | 1.392884 | 0.025 | 0.006 | 1 | 4 | Bbs10 | 0.019 | Young |
| 0.044695 | 1.287933 | 0.025 | 0.006 | 1 | 4 | Brdt | 0.019 | Young |
| 0.044749 | 0.25056 | 0.887 | 0.871 | 1 | 4 | Rps10 | 0.016 | Young |
| 0.044988 | 0.822881 | 0.112 | 0.058 | 1 | 4 | Shkbp1 | 0.054 | Young |
| 0.045065 | 1.393491 | 0.025 | 0.006 | 1 | 4 | 2700046G( | 0.019 | Young |
| 0.045438 | 0.952919 | 0.025 | 0.006 | 1 | 4 | 0610040J0 | 0.019 | Young |
| 0.045438 | 0.569601 | 0.025 | 0.006 | 1 | 4 | Tmem53 | 0.019 | Young |
| 0.045567 | 1.405227 | 0.088 | 0.042 | 1 | 4 | Exosc2 | 0.046 | Young |
| 0.045604 | 0.40251 | 0.3 | 0.204 | 1 | 4 | Osbpl8 | 0.096 | Young |
| 0.045742 | 0.54039 | 0.175 | 0.102 | 1 | 4 | Nsmce1 | 0.073 | Young |
| 0.045871 | 0.71291 | 0.4 | 0.29 | 1 | 4 | Ppp1cc | 0.11 | Young |
| 0.046054 | 1.134865 | 0.112 | 0.059 | 1 | 4 | Tbc1d8 | 0.053 | Young |
| 0.046156 | 0.731007 | 0.238 | 0.155 | 1 | 4 | Gosr2 | 0.083 | Young |
| 0.046193 | 0.688002 | 0.025 | 0.006 | 1 | 4 | Gvin1 | 0.019 | Young |
| 0.046592 | 0.625369 | 0.438 | 0.343 | 1 | 4 | Ndel1 | 0.095 | Young |

| | | | | | | | | |
|---|---|---|---|---|---|---|---|---|
| 0.046617 | 0.299141 | 0.438 | 0.321 | 1 | 4 | Dnajc13 | 0.117 | Young |
| 0.046671 | 0.816882 | 0.1 | 0.049 | 1 | 4 | Copz2 | 0.051 | Young |
| 0.046767 | 0.354222 | 0.35 | 0.244 | 1 | 4 | Cct3 | 0.106 | Young |
| 0.046771 | 1.15621 | 0.238 | 0.155 | 1 | 4 | Fbxw4 | 0.083 | Young |
| 0.046894 | 1.491789 | 0.062 | 0.026 | 1 | 4 | Gdpd3 | 0.036 | Young |
| 0.046913 | 1.045376 | 0.075 | 0.033 | 1 | 4 | Vipas39 | 0.042 | Young |
| 0.047321 | 0.793165 | 0.188 | 0.115 | 1 | 4 | Hs2st1 | 0.073 | Young |
| 0.047444 | 0.660282 | 0.438 | 0.336 | 1 | 4 | Rpl27 | 0.102 | Young |
| 0.047505 | 0.892333 | 0.162 | 0.096 | 1 | 4 | Gpr183 | 0.066 | Young |
| 0.047538 | 0.536884 | 0.225 | 0.144 | 1 | 4 | Dync1li2 | 0.081 | Young |
| 0.047567 | 0.063958 | 0.112 | 0.056 | 1 | 4 | Tecpr1 | 0.056 | Young |
| 0.047794 | 0.589606 | 0.1 | 0.049 | 1 | 4 | Noc4l | 0.051 | Young |
| 0.047872 | 0.988377 | 0.075 | 0.033 | 1 | 4 | Mtfr1 | 0.042 | Young |
| 0.047874 | 1.315319 | 0.062 | 0.026 | 1 | 4 | Acad11 | 0.036 | Young |
| 0.048014 | 0.802804 | 0.125 | 0.067 | 1 | 4 | Slc5a3 | 0.058 | Young |
| 0.048054 | 0.862196 | 0.125 | 0.067 | 1 | 4 | Tmem2 | 0.058 | Young |
| 0.048062 | 0.903904 | 0.088 | 0.041 | 1 | 4 | Sufu | 0.047 | Young |
| 0.048181 | 0.593735 | 0.15 | 0.084 | 1 | 4 | Gbf1 | 0.066 | Young |
| 0.04825 | 0.734726 | 0.25 | 0.166 | 1 | 4 | Ints6l | 0.084 | Young |
| 0.048715 | 0.35674 | 0.35 | 0.246 | 1 | 4 | Stx12 | 0.104 | Young |
| 0.048777 | 0.425508 | 0.238 | 0.148 | 1 | 4 | Map1lc3a | 0.09 | Young |
| 0.049078 | 0.89204 | 0.15 | 0.087 | 1 | 4 | Clns1a | 0.063 | Young |
| 0.04938 | 0.418288 | 0.362 | 0.254 | 1 | 4 | Tmem234 | 0.108 | Young |
| 0.049491 | 0.546785 | 0.338 | 0.252 | 1 | 4 | Cox5b | 0.086 | Young |
| 0.049515 | 0.807174 | 0.2 | 0.123 | 1 | 4 | Eif4enif1 | 0.077 | Young |
| 0.04954 | 0.650839 | 0.1 | 0.049 | 1 | 4 | Slc35a1 | 0.051 | Young |
| 0.04954 | 0.727554 | 0.112 | 0.058 | 1 | 4 | Cep295 | 0.054 | Young |
| 0.049661 | 1.389041 | 0.038 | 0.012 | 1 | 4 | Ets1 | 0.026 | Young |
| 0.049681 | 0.322756 | 0.462 | 0.35 | 1 | 4 | Lims1 | 0.112 | Young |
| 0.049723 | 0.60983 | 0.175 | 0.103 | 1 | 4 | Stk10 | 0.072 | Young |
| 0.049869 | 0.592177 | 0.475 | 0.379 | 1 | 4 | Sash1 | 0.096 | Young |
| 0.050102 | 1.372018 | 0.038 | 0.012 | 1 | 4 | Trp53i13 | 0.026 | Young |
| 0.050102 | 1.12864 | 0.038 | 0.012 | 1 | 4 | Szt2 | 0.026 | Young |
| 0.05031 | 1.028527 | 0.088 | 0.042 | 1 | 4 | Cisd3 | 0.046 | Young |
| 0.050398 | 1.163575 | 0.038 | 0.012 | 1 | 4 | Tnks1bp1 | 0.026 | Young |
| 0.050402 | 0.463272 | 0.288 | 0.198 | 1 | 4 | Map3k11 | 0.09 | Young |
| 0.050426 | 0.486686 | 0.5 | 0.395 | 1 | 4 | Rab7b | 0.105 | Young |
| 0.050475 | 1.097601 | 0.088 | 0.042 | 1 | 4 | Itpkc | 0.046 | Young |
| 0.050547 | 1.132399 | 0.038 | 0.012 | 1 | 4 | Fam46c | 0.026 | Young |
| 0.050621 | 1.173016 | 0.038 | 0.012 | 1 | 4 | Car9 | 0.026 | Young |
| 0.050668 | 0.465635 | 0.162 | 0.094 | 1 | 4 | Psmd7 | 0.068 | Young |
| 0.050825 | 0.561732 | 0.25 | 0.166 | 1 | 4 | Slc38a6 | 0.084 | Young |
| 0.050937 | 0.521643 | 0.362 | 0.266 | 1 | 4 | Tnrc6b | 0.096 | Young |
| 0.051071 | 0.817653 | 0.138 | 0.077 | 1 | 4 | H2-Q6 | 0.061 | Young |
| 0.051339 | 0.486683 | 0.412 | 0.304 | 1 | 4 | Mef2d | 0.108 | Young |

| | | | | | | | | |
|---|---|---|---|---|---|---|---|---|
| 0.051349 | 1.062546 | 0.05 | 0.018 | 1 | 4 | Tbl2 | 0.032 | Young |
| 0.051453 | 0.376317 | 0.225 | 0.142 | 1 | 4 | Kctd2 | 0.083 | Young |
| 0.051503 | 0.460588 | 0.65 | 0.535 | 1 | 4 | Plek | 0.115 | Young |
| 0.051529 | 0.645069 | 0.138 | 0.076 | 1 | 4 | Pja1 | 0.062 | Young |
| 0.051559 | 0.498464 | 0.2 | 0.122 | 1 | 4 | Cdip1 | 0.078 | Young |
| 0.051644 | 1.574731 | 0.075 | 0.034 | 1 | 4 | Fancg | 0.041 | Young |
| 0.051721 | 0.243001 | 0.537 | 0.421 | 1 | 4 | Nptn | 0.116 | Young |
| 0.051779 | 0.662083 | 0.05 | 0.018 | 1 | 4 | Rfesd | 0.032 | Young |
| 0.051803 | 0.608732 | 0.25 | 0.168 | 1 | 4 | Tbc1d4 | 0.082 | Young |
| 0.051853 | 0.486777 | 0.088 | 0.041 | 1 | 4 | Zdhhc4 | 0.047 | Young |
| 0.051958 | 0.336539 | 0.338 | 0.235 | 1 | 4 | Ggh | 0.103 | Young |
| 0.052052 | 0.649871 | 0.038 | 0.012 | 1 | 4 | Hcar2 | 0.026 | Young |
| 0.052179 | 0.897931 | 0.062 | 0.026 | 1 | 4 | Zfand2a | 0.036 | Young |
| 0.052338 | 1.284009 | 0.1 | 0.05 | 1 | 4 | Acad8 | 0.05 | Young |
| 0.052512 | 1.087722 | 0.125 | 0.07 | 1 | 4 | Fam8a1 | 0.055 | Young |
| 0.052632 | 0.350644 | 0.075 | 0.033 | 1 | 4 | Zcchc10 | 0.042 | Young |
| 0.052729 | 0.908673 | 0.1 | 0.05 | 1 | 4 | Tlr5 | 0.05 | Young |
| 0.052747 | 0.815739 | 0.088 | 0.042 | 1 | 4 | Ptrh2 | 0.046 | Young |
| 0.052818 | 0.151886 | 0.038 | 0.012 | 1 | 4 | Fmnl2 | 0.026 | Young |
| 0.052899 | 0.515637 | 0.05 | 0.018 | 1 | 4 | Top3a | 0.032 | Young |
| 0.052942 | 1.149343 | 0.088 | 0.042 | 1 | 4 | Mzt2 | 0.046 | Young |
| 0.053806 | 0.332066 | 0.25 | 0.163 | 1 | 4 | Atp8a1 | 0.087 | Young |
| 0.053896 | 0.405281 | 0.138 | 0.075 | 1 | 4 | Thumpd3 | 0.063 | Young |
| 0.053973 | 0.864449 | 0.15 | 0.088 | 1 | 4 | Papd5 | 0.062 | Young |
| 0.054091 | 0.27791 | 0.45 | 0.35 | 1 | 4 | Cited2 | 0.1 | Young |
| 0.054135 | 0.571998 | 0.188 | 0.116 | 1 | 4 | Arid2 | 0.072 | Young |
| 0.054217 | 0.747028 | 0.188 | 0.117 | 1 | 4 | Pom121 | 0.071 | Young |
| 0.054296 | 1.312461 | 0.062 | 0.026 | 1 | 4 | Srrd | 0.036 | Young |
| 0.054307 | 0.601673 | 0.238 | 0.153 | 1 | 4 | Gsk3a | 0.085 | Young |
| 0.05438 | 0.563951 | 0.112 | 0.059 | 1 | 4 | Fam107b | 0.053 | Young |
| 0.054398 | 0.491908 | 0.225 | 0.145 | 1 | 4 | Zfp950 | 0.08 | Young |
| 0.054613 | 0.542369 | 0.162 | 0.094 | 1 | 4 | Ntpcr | 0.068 | Young |
| 0.054624 | 0.986377 | 0.138 | 0.078 | 1 | 4 | Tmem37 | 0.06 | Young |
| 0.05466 | 0.253838 | 0.8 | 0.639 | 1 | 4 | Rpl35 | 0.161 | Young |
| 0.054743 | 0.602866 | 0.275 | 0.186 | 1 | 4 | Mff | 0.089 | Young |
| 0.054853 | 0.528693 | 0.375 | 0.27 | 1 | 4 | Rbm5 | 0.105 | Young |
| 0.054881 | 0.230416 | 0.887 | 0.826 | 1 | 4 | Rps11 | 0.061 | Young |
| 0.055 | 0.43622 | 0.2 | 0.121 | 1 | 4 | Rasgrp3 | 0.079 | Young |
| 0.055023 | 0.788977 | 0.2 | 0.13 | 1 | 4 | Otud5 | 0.07 | Young |
| 0.055242 | 1.384719 | 0.1 | 0.051 | 1 | 4 | Ganc | 0.049 | Young |
| 0.055289 | 0.864784 | 0.088 | 0.042 | 1 | 4 | Kat5 | 0.046 | Young |
| 0.055296 | 1.007487 | 0.112 | 0.06 | 1 | 4 | Cherp | 0.052 | Young |
| 0.055357 | 0.852521 | 0.112 | 0.059 | 1 | 4 | Tarbp2 | 0.053 | Young |
| 0.055514 | 0.591796 | 0.362 | 0.274 | 1 | 4 | Rgs18 | 0.088 | Young |
| 0.05562 | 0.242137 | 0.8 | 0.686 | 1 | 4 | Ier5 | 0.114 | Young |

| | | | | | | | | |
|---|---|---|---|---|---|---|---|---|
| 0.055639 | 0.623079 | 0.275 | 0.193 | 1 | 4 | Atf7 | 0.082 | Young |
| 0.055768 | 1.808675 | 0.05 | 0.019 | 1 | 4 | Als2 | 0.031 | Young |
| 0.055889 | 0.747013 | 0.112 | 0.059 | 1 | 4 | Bap1 | 0.053 | Young |
| 0.055957 | 1.035071 | 0.175 | 0.111 | 1 | 4 | Ppil4 | 0.064 | Young |
| 0.0561 | 0.3787 | 0.688 | 0.589 | 1 | 4 | Tnfaip3 | 0.099 | Young |
| 0.056109 | 0.263719 | 0.912 | 0.889 | 1 | 4 | Hsp90ab1 | 0.023 | Young |
| 0.05614 | 0.54392 | 0.138 | 0.076 | 1 | 4 | Pank1 | 0.062 | Young |
| 0.056215 | 0.2565 | 0.188 | 0.111 | 1 | 4 | Wsb2 | 0.077 | Young |
| 0.05631 | 0.515127 | 0.15 | 0.086 | 1 | 4 | Ate1 | 0.064 | Young |
| 0.056378 | 0.596423 | 0.138 | 0.077 | 1 | 4 | Prpf19 | 0.061 | Young |
| 0.056481 | 0.453349 | 0.2 | 0.124 | 1 | 4 | Crtc3 | 0.076 | Young |
| 0.05681 | 1.487878 | 0.05 | 0.019 | 1 | 4 | C1qtnf12 | 0.031 | Young |
| 0.056843 | 1.085797 | 0.062 | 0.026 | 1 | 4 | Micall1 | 0.036 | Young |
| 0.057516 | 0.329748 | 0.262 | 0.171 | 1 | 4 | Gpatch8 | 0.091 | Young |
| 0.057524 | 1.080517 | 0.062 | 0.026 | 1 | 4 | 2-Sep | 0.036 | Young |
| 0.05754 | 0.972127 | 0.088 | 0.042 | 1 | 4 | Mccc1 | 0.046 | Young |
| 0.05778 | 0.216873 | 0.4 | 0.283 | 1 | 4 | Ddost | 0.117 | Young |
| 0.058241 | 1.343022 | 0.1 | 0.052 | 1 | 4 | Brpf3 | 0.048 | Young |
| 0.058302 | 0.362093 | 0.688 | 0.59 | 1 | 4 | Gng5 | 0.098 | Young |
| 0.058691 | 1.256251 | 0.138 | 0.08 | 1 | 4 | Tmem260 | 0.058 | Young |
| 0.058942 | 1.15853 | 0.05 | 0.019 | 1 | 4 | Cmss1 | 0.031 | Young |
| 0.059031 | 0.663106 | 0.1 | 0.051 | 1 | 4 | Pprc1 | 0.049 | Young |
| 0.059113 | 0.666455 | 0.088 | 0.042 | 1 | 4 | Homer3 | 0.046 | Young |
| 0.059207 | 0.72108 | 0.088 | 0.042 | 1 | 4 | Borcs7 | 0.046 | Young |
| 0.059349 | 1.130684 | 0.05 | 0.019 | 1 | 4 | Zfp951 | 0.031 | Young |
| 0.059501 | 1.957834 | 0.038 | 0.013 | 1 | 4 | Dnajc28 | 0.025 | Young |
| 0.059623 | 0.686877 | 0.138 | 0.078 | 1 | 4 | Chpf2 | 0.06 | Young |
| 0.059642 | 0.758133 | 0.2 | 0.13 | 1 | 4 | Rps27rt | 0.07 | Young |
| 0.059668 | 1.805924 | 0.038 | 0.013 | 1 | 4 | Gm21781 | 0.025 | Young |
| 0.059727 | 0.716551 | 0.162 | 0.097 | 1 | 4 | Suco | 0.065 | Young |
| 0.059728 | 0.360259 | 0.188 | 0.115 | 1 | 4 | Epb41 | 0.073 | Young |
| 0.059881 | 0.552318 | 0.288 | 0.2 | 1 | 4 | Chd9 | 0.088 | Young |
| 0.060033 | 1.341928 | 0.05 | 0.019 | 1 | 4 | Slfn1 | 0.031 | Young |
| 0.060033 | 0.785805 | 0.05 | 0.019 | 1 | 4 | Pced1a | 0.031 | Young |
| 0.060128 | 0.145829 | 0.088 | 0.042 | 1 | 4 | Dvl1 | 0.046 | Young |
| 0.060283 | 1.666263 | 0.062 | 0.027 | 1 | 4 | Gm12840 | 0.035 | Young |
| 0.060358 | 0.879471 | 0.112 | 0.061 | 1 | 4 | Phtf2 | 0.051 | Young |
| 0.060623 | 0.780999 | 0.125 | 0.069 | 1 | 4 | Ino80 | 0.056 | Young |
| 0.060725 | 0.502845 | 0.15 | 0.086 | 1 | 4 | Ccnk | 0.064 | Young |
| 0.060826 | 0.389935 | 0.175 | 0.103 | 1 | 4 | Nmt2 | 0.072 | Young |
| 0.061298 | 1.121967 | 0.138 | 0.081 | 1 | 4 | Mxra7 | 0.057 | Young |
| 0.061329 | 2.296186 | 0.025 | 0.007 | 1 | 4 | Tshz3 | 0.018 | Young |
| 0.061346 | 1.363151 | 0.062 | 0.027 | 1 | 4 | Gatb | 0.035 | Young |
| 0.061527 | 1.383869 | 0.038 | 0.013 | 1 | 4 | Qrsl1 | 0.025 | Young |
| 0.06156 | 2.118018 | 0.025 | 0.007 | 1 | 4 | Dnajc17 | 0.018 | Young |

| | | | | | | | | |
|---|---|---|---|---|---|---|---|---|
| 0.061784 | 0.424399 | 0.288 | 0.201 | 1 | 4 | Ralgps2 | 0.087 | Young |
| 0.061791 | 2.011128 | 0.025 | 0.007 | 1 | 4 | Arap3 | 0.018 | Young |
| 0.062214 | 1.311363 | 0.038 | 0.013 | 1 | 4 | Zfp202 | 0.025 | Young |
| 0.062214 | 1.105065 | 0.038 | 0.013 | 1 | 4 | 2700062C0 | 0.025 | Young |
| 0.062256 | 1.835568 | 0.025 | 0.007 | 1 | 4 | Asrgl1 | 0.018 | Young |
| 0.062331 | 1.33817 | 0.075 | 0.035 | 1 | 4 | Exoc6 | 0.04 | Young |
| 0.062358 | 0.280439 | 0.062 | 0.026 | 1 | 4 | Clip2 | 0.036 | Young |
| 0.06253 | 0.471468 | 0.112 | 0.059 | 1 | 4 | Vps16 | 0.053 | Young |
| 0.062676 | 0.466254 | 0.175 | 0.105 | 1 | 4 | Slc35c2 | 0.07 | Young |
| 0.062724 | 1.74214 | 0.025 | 0.007 | 1 | 4 | Gm30329 | 0.018 | Young |
| 0.062724 | 1.513242 | 0.025 | 0.007 | 1 | 4 | Hist2h3c1 | 0.018 | Young |
| 0.062784 | 0.83458 | 0.188 | 0.119 | 1 | 4 | 3110043O2 | 0.069 | Young |
| 0.062908 | 1.024399 | 0.038 | 0.013 | 1 | 4 | Chac2 | 0.025 | Young |
| 0.062994 | 0.902743 | 0.088 | 0.043 | 1 | 4 | Ctdp1 | 0.045 | Young |
| 0.063056 | 0.839167 | 0.1 | 0.051 | 1 | 4 | Ccdc152 | 0.049 | Young |
| 0.06312 | 0.610658 | 0.225 | 0.148 | 1 | 4 | Slc35f6 | 0.077 | Young |
| 0.063151 | 0.363689 | 0.512 | 0.402 | 1 | 4 | Man1c1 | 0.11 | Young |
| 0.063397 | 1.145594 | 0.062 | 0.027 | 1 | 4 | Nfatc2ip | 0.035 | Young |
| 0.063535 | 0.252813 | 0.35 | 0.239 | 1 | 4 | Kmt2a | 0.111 | Young |
| 0.063542 | 0.525599 | 0.362 | 0.268 | 1 | 4 | Npc1 | 0.094 | Young |
| 0.06357 | 0.56185 | 0.162 | 0.097 | 1 | 4 | Arhgap39 | 0.065 | Young |
| 0.063733 | 1.083189 | 0.075 | 0.035 | 1 | 4 | 2610008E1 | 0.04 | Young |
| 0.063785 | 0.781364 | 0.038 | 0.013 | 1 | 4 | Fbxo10 | 0.025 | Young |
| 0.064115 | 0.908987 | 0.075 | 0.035 | 1 | 4 | Pafah1b3 | 0.04 | Young |
| 0.064139 | 0.351392 | 0.112 | 0.059 | 1 | 4 | Elof1 | 0.053 | Young |
| 0.064312 | 0.655297 | 0.188 | 0.119 | 1 | 4 | Rcsd1 | 0.069 | Young |
| 0.064418 | 1.239895 | 0.1 | 0.053 | 1 | 4 | Tex10 | 0.047 | Young |
| 0.064435 | 0.436965 | 0.288 | 0.198 | 1 | 4 | 0610012G0 | 0.09 | Young |
| 0.06445 | 0.388827 | 0.212 | 0.135 | 1 | 4 | Psen1 | 0.077 | Young |
| 0.064615 | 0.717865 | 0.138 | 0.078 | 1 | 4 | Mcee | 0.06 | Young |
| 0.064721 | 0.409759 | 0.162 | 0.096 | 1 | 4 | Impdh2 | 0.066 | Young |
| 0.064734 | 0.798405 | 0.1 | 0.052 | 1 | 4 | Terf1 | 0.048 | Young |
| 0.064796 | 1.383888 | 0.075 | 0.036 | 1 | 4 | Tti2 | 0.039 | Young |
| 0.064866 | 0.771582 | 0.025 | 0.007 | 1 | 4 | Noxo1 | 0.018 | Young |
| 0.064866 | 0.606449 | 0.025 | 0.007 | 1 | 4 | Rwdd3 | 0.018 | Young |
| 0.065073 | 0.448311 | 0.15 | 0.088 | 1 | 4 | Hmgn2 | 0.062 | Young |
| 0.065108 | 0.175337 | 0.025 | 0.007 | 1 | 4 | Gdpgp1 | 0.018 | Young |
| 0.065173 | 0.643406 | 0.162 | 0.096 | 1 | 4 | Eif2ak1 | 0.066 | Young |
| 0.065215 | 0.449728 | 0.175 | 0.105 | 1 | 4 | Bms1 | 0.07 | Young |
| 0.065271 | 0.803358 | 0.112 | 0.061 | 1 | 4 | Per3 | 0.051 | Young |
| 0.065484 | 0.346875 | 0.1 | 0.051 | 1 | 4 | Dnaja3 | 0.049 | Young |
| 0.065512 | 0.392902 | 0.35 | 0.252 | 1 | 4 | Orai1 | 0.098 | Young |
| 0.065771 | 0.915171 | 0.225 | 0.153 | 1 | 4 | Guk1 | 0.072 | Young |
| 0.065967 | 0.361294 | 0.537 | 0.449 | 1 | 4 | Selenof | 0.088 | Young |
| 0.066024 | 4.56985 | 0.013 | 0.002 | 1 | 4 | Slpi | 0.011 | Young |

| | | | | | | | | |
|---|---|---|---|---|---|---|---|---|
| 0.066024 | 3.273761 | 0.013 | 0.002 | 1 | 4 | Lrrd1 | 0.011 | Young |
| 0.066024 | 2.676874 | 0.013 | 0.002 | 1 | 4 | Zfp651 | 0.011 | Young |
| 0.066024 | 2.59993 | 0.013 | 0.002 | 1 | 4 | 5430416N( | 0.011 | Young |
| 0.066024 | 2.515853 | 0.013 | 0.002 | 1 | 4 | Shc4 | 0.011 | Young |
| 0.066024 | 2.469001 | 0.013 | 0.002 | 1 | 4 | Gm10282 | 0.011 | Young |
| 0.066024 | 2.316369 | 0.013 | 0.002 | 1 | 4 | Sfmbt2 | 0.011 | Young |
| 0.066024 | 2.298575 | 0.013 | 0.002 | 1 | 4 | Tspoap1 | 0.011 | Young |
| 0.066024 | 2.220563 | 0.013 | 0.002 | 1 | 4 | Fbln2 | 0.011 | Young |
| 0.066024 | 2.202955 | 0.013 | 0.002 | 1 | 4 | Hap1 | 0.011 | Young |
| 0.066024 | 2.152568 | 0.013 | 0.002 | 1 | 4 | Spats2l | 0.011 | Young |
| 0.066024 | 2.129326 | 0.013 | 0.002 | 1 | 4 | Gstm2 | 0.011 | Young |
| 0.066024 | 2.01491 | 0.013 | 0.002 | 1 | 4 | Akap5 | 0.011 | Young |
| 0.066024 | 1.991841 | 0.013 | 0.002 | 1 | 4 | AC125351. | 0.011 | Young |
| 0.066024 | 1.985994 | 0.013 | 0.002 | 1 | 4 | Ccdc170 | 0.011 | Young |
| 0.066024 | 1.936365 | 0.013 | 0.002 | 1 | 4 | Gdpd5 | 0.011 | Young |
| 0.066024 | 1.770141 | 0.013 | 0.002 | 1 | 4 | Gsg1 | 0.011 | Young |
| 0.066024 | 1.757315 | 0.013 | 0.002 | 1 | 4 | Rab4b | 0.011 | Young |
| 0.06604 | 0.483825 | 0.138 | 0.078 | 1 | 4 | Pofut2 | 0.06 | Young |
| 0.066151 | 0.533295 | 0.125 | 0.069 | 1 | 4 | Kansl2 | 0.056 | Young |
| 0.066448 | 1.991086 | 0.013 | 0.002 | 1 | 4 | Hspa1l | 0.011 | Young |
| 0.066448 | 1.971872 | 0.013 | 0.002 | 1 | 4 | A130051J0 | 0.011 | Young |
| 0.066448 | 1.817085 | 0.013 | 0.002 | 1 | 4 | Gm9465 | 0.011 | Young |
| 0.066448 | 1.70871 | 0.013 | 0.002 | 1 | 4 | Cyp26b1 | 0.011 | Young |
| 0.066448 | 1.61247 | 0.013 | 0.002 | 1 | 4 | Rragd | 0.011 | Young |
| 0.066448 | 1.492703 | 0.013 | 0.002 | 1 | 4 | Grin2d | 0.011 | Young |
| 0.066448 | 1.478659 | 0.013 | 0.002 | 1 | 4 | Vangl2 | 0.011 | Young |
| 0.066448 | 1.2271 | 0.013 | 0.002 | 1 | 4 | Zfp428 | 0.011 | Young |
| 0.066448 | 1.175666 | 0.013 | 0.002 | 1 | 4 | Gbp8 | 0.011 | Young |
| 0.066448 | 1.106297 | 0.013 | 0.002 | 1 | 4 | Gm47204 | 0.011 | Young |
| 0.06645 | 1.249752 | 0.05 | 0.02 | 1 | 4 | Nek6 | 0.03 | Young |
| 0.06666 | 0.13686 | 0.112 | 0.059 | 1 | 4 | Rexo4 | 0.053 | Young |
| 0.066661 | 1.00173 | 0.013 | 0.002 | 1 | 4 | Spef2 | 0.011 | Young |
| 0.066769 | 0.614062 | 0.062 | 0.027 | 1 | 4 | Upp2 | 0.035 | Young |
| 0.066875 | 1.572618 | 0.013 | 0.002 | 1 | 4 | Fabp1 | 0.011 | Young |
| 0.066875 | 1.51515 | 0.013 | 0.002 | 1 | 4 | 5430402O: | 0.011 | Young |
| 0.066875 | 1.500452 | 0.013 | 0.002 | 1 | 4 | Gas2 | 0.011 | Young |
| 0.066875 | 1.429644 | 0.013 | 0.002 | 1 | 4 | Ccdc63 | 0.011 | Young |
| 0.066875 | 1.385022 | 0.013 | 0.002 | 1 | 4 | Ltb | 0.011 | Young |
| 0.066875 | 1.319263 | 0.013 | 0.002 | 1 | 4 | Cox4i2 | 0.011 | Young |
| 0.066875 | 1.178319 | 0.013 | 0.002 | 1 | 4 | Tmem116 | 0.011 | Young |
| 0.066875 | 1.020109 | 0.013 | 0.002 | 1 | 4 | Prrx1 | 0.011 | Young |
| 0.066875 | 1.016344 | 0.013 | 0.002 | 1 | 4 | A930029G: | 0.011 | Young |
| 0.066875 | 0.936875 | 0.013 | 0.002 | 1 | 4 | Dmd | 0.011 | Young |
| 0.066875 | 0.931167 | 0.013 | 0.002 | 1 | 4 | Mfap5 | 0.011 | Young |
| 0.066875 | 0.785519 | 0.013 | 0.002 | 1 | 4 | Tnfrsf13c | 0.011 | Young |

| | | | | | | | | |
|---|---|---|---|---|---|---|---|---|
| 0.066875 | 0.591092 | 0.013 | 0.002 | 1 | 4 | D130017N( | 0.011 | Young |
| 0.066913 | 0.711944 | 0.125 | 0.07 | 1 | 4 | Naa10 | 0.055 | Young |
| 0.066965 | 0.53337 | 0.325 | 0.236 | 1 | 4 | Clcn5 | 0.089 | Young |
| 0.067007 | 0.309406 | 0.6 | 0.495 | 1 | 4 | Man1a | 0.105 | Young |
| 0.067111 | 0.212595 | 0.45 | 0.329 | 1 | 4 | Smap2 | 0.121 | Young |
| 0.06716 | 0.220225 | 0.863 | 0.806 | 1 | 4 | Gnai2 | 0.057 | Young |
| 0.067178 | 0.879489 | 0.138 | 0.081 | 1 | 4 | Zfp598 | 0.057 | Young |
| 0.067242 | 1.231708 | 0.075 | 0.036 | 1 | 4 | Dnmbp | 0.039 | Young |
| 0.067303 | 1.124766 | 0.013 | 0.002 | 1 | 4 | Klf15 | 0.011 | Young |
| 0.067303 | 1.071037 | 0.013 | 0.002 | 1 | 4 | Defb28 | 0.011 | Young |
| 0.067303 | 1.001321 | 0.013 | 0.002 | 1 | 4 | C130074G1 | 0.011 | Young |
| 0.067303 | 0.947452 | 0.013 | 0.002 | 1 | 4 | Hal | 0.011 | Young |
| 0.067303 | 0.658267 | 0.013 | 0.002 | 1 | 4 | Hspb8 | 0.011 | Young |
| 0.067303 | 0.565143 | 0.013 | 0.002 | 1 | 4 | Omp | 0.011 | Young |
| 0.067303 | 0.555245 | 0.013 | 0.002 | 1 | 4 | Gm43672 | 0.011 | Young |
| 0.067303 | 0.547412 | 0.013 | 0.002 | 1 | 4 | Gm43625 | 0.011 | Young |
| 0.067303 | 0.544217 | 0.013 | 0.002 | 1 | 4 | Car11 | 0.011 | Young |
| 0.067303 | 0.471354 | 0.013 | 0.002 | 1 | 4 | H1fx | 0.011 | Young |
| 0.067303 | 0.451118 | 0.013 | 0.002 | 1 | 4 | Nudt8.1 | 0.011 | Young |
| 0.067303 | 0.337358 | 0.013 | 0.002 | 1 | 4 | Epha2 | 0.011 | Young |
| 0.067303 | 0.30371 | 0.013 | 0.002 | 1 | 4 | 2210011C2 | 0.011 | Young |
| 0.067303 | 0.25876 | 0.013 | 0.002 | 1 | 4 | Tgfbr3l | 0.011 | Young |
| 0.067303 | 0.19255 | 0.013 | 0.002 | 1 | 4 | Zbtb3 | 0.011 | Young |
| 0.067303 | 0.187049 | 0.013 | 0.002 | 1 | 4 | Zscan25 | 0.011 | Young |
| 0.067404 | 0.746942 | 0.238 | 0.163 | 1 | 4 | Zranb1 | 0.075 | Young |
| 0.067468 | 1.167577 | 0.075 | 0.036 | 1 | 4 | Rundc1 | 0.039 | Young |
| 0.067762 | 0.79451 | 0.15 | 0.088 | 1 | 4 | 1600012H( | 0.062 | Young |
| 0.067888 | 0.75939 | 0.125 | 0.069 | 1 | 4 | Casp6 | 0.056 | Young |
| 0.067895 | 0.375378 | 0.212 | 0.136 | 1 | 4 | Fam118a | 0.076 | Young |
| 0.068025 | 0.966427 | 0.162 | 0.099 | 1 | 4 | 2610037D( | 0.063 | Young |
| 0.068034 | 0.380581 | 0.2 | 0.126 | 1 | 4 | Ldah | 0.074 | Young |
| 0.068152 | 0.629069 | 0.2 | 0.129 | 1 | 4 | Mlxip | 0.071 | Young |
| 0.068566 | 0.75749 | 0.088 | 0.043 | 1 | 4 | Ubtd1 | 0.045 | Young |
| 0.068617 | 0.357266 | 0.325 | 0.229 | 1 | 4 | Rpl36al | 0.096 | Young |
| 0.068634 | 0.779135 | 0.162 | 0.099 | 1 | 4 | Abhd4 | 0.063 | Young |
| 0.069174 | 0.847423 | 0.125 | 0.071 | 1 | 4 | Eng | 0.054 | Young |
| 0.069251 | 0.347227 | 0.488 | 0.379 | 1 | 4 | Pura | 0.109 | Young |
| 0.06945 | 0.664973 | 0.1 | 0.052 | 1 | 4 | 1190002N1 | 0.048 | Young |
| 0.06953 | 1.266761 | 0.075 | 0.036 | 1 | 4 | Spats2 | 0.039 | Young |
| 0.069578 | 0.472892 | 0.15 | 0.088 | 1 | 4 | Ddit3 | 0.062 | Young |
| 0.069652 | 0.375434 | 0.275 | 0.189 | 1 | 4 | Trpv4 | 0.086 | Young |
| 0.069849 | 0.820016 | 0.138 | 0.08 | 1 | 4 | Lpar6 | 0.058 | Young |
| 0.069983 | 0.409114 | 0.4 | 0.306 | 1 | 4 | Stub1 | 0.094 | Young |
| 0.070062 | 0.374979 | 0.3 | 0.206 | 1 | 4 | Crlf2 | 0.094 | Young |
| 0.070087 | 0.604699 | 0.2 | 0.129 | 1 | 4 | Pfdn1 | 0.071 | Young |

| | | | | | | | | |
|---|---|---|---|---|---|---|---|---|
| 0.07016 | 0.348604 | 0.362 | 0.255 | 1 | 4 | Dnaja2 | 0.107 | Young |
| 0.070196 | 0.651486 | 0.162 | 0.099 | 1 | 4 | Aak1 | 0.063 | Young |
| 0.070348 | 1.183292 | 0.125 | 0.072 | 1 | 4 | Mrpl10 | 0.053 | Young |
| 0.070511 | 0.289013 | 0.338 | 0.242 | 1 | 4 | Mkrn1 | 0.096 | Young |
| 0.070849 | 0.220544 | 0.575 | 0.482 | 1 | 4 | Rgs10 | 0.093 | Young |
| 0.070917 | 0.681075 | 0.125 | 0.07 | 1 | 4 | Mdc1 | 0.055 | Young |
| 0.071174 | 0.948195 | 0.138 | 0.081 | 1 | 4 | Pdrg1 | 0.057 | Young |
| 0.071458 | 0.59609 | 0.05 | 0.02 | 1 | 4 | Plekhh3 | 0.03 | Young |
| 0.071493 | 1.048231 | 0.088 | 0.044 | 1 | 4 | Gps2 | 0.044 | Young |
| 0.071615 | 0.790754 | 0.05 | 0.02 | 1 | 4 | Mars | 0.03 | Young |
| 0.071726 | 1.749227 | 0.038 | 0.013 | 1 | 4 | Cenpv | 0.025 | Young |
| 0.071769 | 0.182169 | 0.6 | 0.468 | 1 | 4 | Klf3 | 0.132 | Young |
| 0.071827 | 0.58712 | 0.238 | 0.161 | 1 | 4 | Smad1 | 0.077 | Young |
| 0.071999 | 0.664299 | 0.125 | 0.07 | 1 | 4 | Utp14b | 0.055 | Young |
| 0.072121 | 0.902942 | 0.075 | 0.036 | 1 | 4 | Cep104 | 0.039 | Young |
| 0.072219 | 0.231552 | 0.075 | 0.035 | 1 | 4 | Habp4 | 0.04 | Young |
| 0.072255 | 0.181285 | 0.875 | 0.756 | 1 | 4 | Tmbim6 | 0.119 | Young |
| 0.072578 | 0.362983 | 0.2 | 0.128 | 1 | 4 | Arid5b | 0.072 | Young |
| 0.072629 | 0.55027 | 0.288 | 0.208 | 1 | 4 | Arid4a | 0.08 | Young |
| 0.072782 | 0.410718 | 0.212 | 0.138 | 1 | 4 | Ano6 | 0.074 | Young |
| 0.072841 | 0.773632 | 0.075 | 0.036 | 1 | 4 | Tmem35b | 0.039 | Young |
| 0.07291 | 1.049958 | 0.138 | 0.083 | 1 | 4 | Mtf2 | 0.055 | Young |
| 0.073036 | 1.092373 | 0.125 | 0.071 | 1 | 4 | Snhg12 | 0.054 | Young |
| 0.073446 | 0.715277 | 0.075 | 0.036 | 1 | 4 | Oraov1 | 0.039 | Young |
| 0.073537 | 0.716647 | 0.25 | 0.177 | 1 | 4 | Golga7 | 0.073 | Young |
| 0.073701 | 0.391882 | 0.25 | 0.168 | 1 | 4 | Crybg3 | 0.082 | Young |
| 0.073737 | 0.965394 | 0.138 | 0.08 | 1 | 4 | Phrf1 | 0.058 | Young |
| 0.074289 | 0.455556 | 0.275 | 0.19 | 1 | 4 | Ppp4r2 | 0.085 | Young |
| 0.07456 | 0.665194 | 0.138 | 0.081 | 1 | 4 | Zdhhc7 | 0.057 | Young |
| 0.074632 | 0.375991 | 0.613 | 0.505 | 1 | 4 | Cox8a | 0.108 | Young |
| 0.074713 | 0.491868 | 0.238 | 0.163 | 1 | 4 | Cux1 | 0.075 | Young |
| 0.074973 | 1.499584 | 0.062 | 0.028 | 1 | 4 | Abcc3 | 0.034 | Young |
| 0.075297 | 0.637211 | 0.25 | 0.169 | 1 | 4 | Pcmt1 | 0.081 | Young |
| 0.075555 | 0.6337 | 0.15 | 0.09 | 1 | 4 | Tmed4 | 0.06 | Young |
| 0.075876 | 1.142367 | 0.075 | 0.036 | 1 | 4 | Dedd | 0.039 | Young |
| 0.075912 | 0.619615 | 0.2 | 0.132 | 1 | 4 | Smc4 | 0.068 | Young |
| 0.075978 | 0.604589 | 0.125 | 0.071 | 1 | 4 | Zcchc9 | 0.054 | Young |
| 0.076241 | 0.734916 | 0.225 | 0.159 | 1 | 4 | Pbx2 | 0.066 | Young |
| 0.076275 | 0.371011 | 0.575 | 0.507 | 1 | 4 | Man2b1 | 0.068 | Young |
| 0.076379 | 0.49904 | 0.25 | 0.173 | 1 | 4 | P2ry12 | 0.077 | Young |
| 0.076505 | 0.639797 | 0.188 | 0.122 | 1 | 4 | Wipf2 | 0.066 | Young |
| 0.076662 | 0.516734 | 0.25 | 0.17 | 1 | 4 | D1Ertd622 | 0.08 | Young |
| 0.076686 | 0.207901 | 0.7 | 0.567 | 1 | 4 | Slc25a5 | 0.133 | Young |
| 0.076814 | 0.193111 | 0.412 | 0.29 | 1 | 4 | Ubtf | 0.122 | Young |
| 0.076947 | 0.280144 | 0.212 | 0.136 | 1 | 4 | Rubcn | 0.076 | Young |

| | | | | | | | | |
|---|---|---|---|---|---|---|---|---|
| 0.077002 | 1.251363 | 0.05 | 0.02 | 1 | 4 | Rbbp9 | 0.03 | Young |
| 0.077008 | 0.968649 | 0.038 | 0.013 | 1 | 4 | 2810001G2 | 0.025 | Young |
| 0.077169 | 0.534876 | 0.1 | 0.053 | 1 | 4 | Tmem126b | 0.047 | Young |
| 0.077209 | 0.443044 | 0.038 | 0.013 | 1 | 4 | Mcemp1 | 0.025 | Young |
| 0.077221 | 0.138822 | 0.388 | 0.291 | 1 | 4 | Fgl2 | 0.097 | Young |
| 0.078172 | 0.81931 | 0.138 | 0.083 | 1 | 4 | Bmyc | 0.055 | Young |
| 0.078561 | 0.628026 | 0.162 | 0.101 | 1 | 4 | Gclc | 0.061 | Young |
| 0.078632 | 0.539287 | 0.038 | 0.013 | 1 | 4 | Asf1b | 0.025 | Young |
| 0.078652 | 0.311912 | 0.525 | 0.426 | 1 | 4 | Ctnnb1 | 0.099 | Young |
| 0.078837 | 0.192036 | 0.038 | 0.013 | 1 | 4 | Aldh18a1 | 0.025 | Young |
| 0.079075 | 0.42376 | 0.25 | 0.166 | 1 | 4 | Zcchc7 | 0.084 | Young |
| 0.079088 | 0.349003 | 0.388 | 0.289 | 1 | 4 | Acat1 | 0.099 | Young |
| 0.079104 | 0.845371 | 0.088 | 0.045 | 1 | 4 | Rnf185 | 0.043 | Young |
| 0.079163 | 1.056871 | 0.1 | 0.054 | 1 | 4 | Slc35b4 | 0.046 | Young |
| 0.079553 | 0.285621 | 0.138 | 0.08 | 1 | 4 | Psmc4 | 0.058 | Young |
| 0.079662 | 0.144234 | 0.038 | 0.013 | 1 | 4 | Cenpw | 0.025 | Young |
| 0.0798 | 0.505855 | 0.162 | 0.1 | 1 | 4 | Chm | 0.062 | Young |
| 0.079919 | 0.329688 | 0.488 | 0.387 | 1 | 4 | Gsk3b | 0.101 | Young |
| 0.08014 | 0.847072 | 0.175 | 0.113 | 1 | 4 | Pex2 | 0.062 | Young |
| 0.080166 | 1.025051 | 0.05 | 0.02 | 1 | 4 | Rilp | 0.03 | Young |
| 0.080264 | 0.118091 | 0.6 | 0.472 | 1 | 4 | Fosl2 | 0.128 | Young |
| 0.080338 | 0.326343 | 0.262 | 0.179 | 1 | 4 | Zfp516 | 0.083 | Young |
| 0.080657 | 0.423812 | 0.362 | 0.275 | 1 | 4 | Elob | 0.087 | Young |
| 0.080679 | 0.632168 | 0.225 | 0.153 | 1 | 4 | Mgst1 | 0.072 | Young |
| 0.080712 | 0.409283 | 0.588 | 0.505 | 1 | 4 | Zfhx3 | 0.083 | Young |
| 0.080811 | 0.363743 | 0.138 | 0.08 | 1 | 4 | Tm2d3 | 0.058 | Young |
| 0.080917 | 0.675647 | 0.112 | 0.063 | 1 | 4 | Dis3l2 | 0.049 | Young |
| 0.081346 | 0.614517 | 0.075 | 0.036 | 1 | 4 | Fam53a | 0.039 | Young |
| 0.081358 | 0.867809 | 0.05 | 0.02 | 1 | 4 | Thada | 0.03 | Young |
| 0.081389 | 0.421548 | 0.162 | 0.099 | 1 | 4 | Nr1h2 | 0.063 | Young |
| 0.081449 | 0.589154 | 0.15 | 0.09 | 1 | 4 | Exoc1 | 0.06 | Young |
| 0.081503 | 0.544025 | 0.3 | 0.223 | 1 | 4 | Med13 | 0.077 | Young |
| 0.081545 | 0.394487 | 0.312 | 0.223 | 1 | 4 | Zfp638 | 0.089 | Young |
| 0.081598 | 0.27372 | 0.65 | 0.574 | 1 | 4 | Atp5h | 0.076 | Young |
| 0.081637 | 0.371236 | 0.488 | 0.383 | 1 | 4 | Snx6 | 0.105 | Young |
| 0.081663 | 0.933863 | 0.062 | 0.028 | 1 | 4 | D030056L2 | 0.034 | Young |
| 0.081717 | 0.408591 | 0.275 | 0.193 | 1 | 4 | Ccdc12 | 0.082 | Young |
| 0.081777 | 0.177295 | 0.175 | 0.107 | 1 | 4 | Sirt7 | 0.068 | Young |
| 0.081852 | 0.523543 | 0.25 | 0.175 | 1 | 4 | Exoc4 | 0.075 | Young |
| 0.082037 | 0.265373 | 0.875 | 0.819 | 1 | 4 | Igfbp4 | 0.056 | Young |
| 0.082126 | 0.65341 | 0.112 | 0.063 | 1 | 4 | Gid4 | 0.049 | Young |
| 0.082203 | 0.453133 | 0.375 | 0.292 | 1 | 4 | Slc12a6 | 0.083 | Young |
| 0.082211 | 0.227028 | 0.662 | 0.516 | 1 | 4 | Tm6sf1 | 0.146 | Young |
| 0.082245 | 0.806168 | 0.125 | 0.073 | 1 | 4 | Shc1 | 0.052 | Young |
| 0.082252 | 0.418976 | 0.15 | 0.09 | 1 | 4 | Vav2 | 0.06 | Young |

| | | | | | | | | |
|---|---|---|---|---|---|---|---|---|
| 0.082495 | 0.29213 | 0.762 | 0.708 | 1 | 4 | Ctsa | 0.054 | Young |
| 0.082539 | 0.526203 | 0.162 | 0.098 | 1 | 4 | Wdr11 | 0.064 | Young |
| 0.082565 | 0.571099 | 0.05 | 0.02 | 1 | 4 | Mtif3 | 0.03 | Young |
| 0.082906 | 1.06465 | 0.1 | 0.055 | 1 | 4 | Sipa1l2 | 0.045 | Young |
| 0.083045 | 0.84663 | 0.175 | 0.114 | 1 | 4 | Abl1 | 0.061 | Young |
| 0.083328 | 0.353543 | 0.425 | 0.341 | 1 | 4 | Vps4b | 0.084 | Young |
| 0.083515 | 0.159991 | 0.088 | 0.044 | 1 | 4 | Pdzd11 | 0.044 | Young |
| 0.083728 | 0.637374 | 0.088 | 0.045 | 1 | 4 | AU020206 | 0.043 | Young |
| 0.083906 | 0.475913 | 0.112 | 0.062 | 1 | 4 | Zfp329 | 0.05 | Young |
| 0.083969 | 0.696793 | 0.162 | 0.103 | 1 | 4 | Usp33 | 0.059 | Young |
| 0.083993 | 2.60668 | 0.025 | 0.007 | 1 | 4 | Apold1 | 0.018 | Young |
| 0.084267 | 0.433916 | 0.438 | 0.333 | 1 | 4 | Nckap1l | 0.105 | Young |
| 0.08428 | 1.475169 | 0.025 | 0.007 | 1 | 4 | Gm14305 | 0.018 | Young |
| 0.084337 | 0.513437 | 0.1 | 0.053 | 1 | 4 | Ciapin1 | 0.047 | Young |
| 0.08435 | 0.392002 | 0.275 | 0.198 | 1 | 4 | Mapk6 | 0.077 | Young |
| 0.084507 | 0.156493 | 0.4 | 0.29 | 1 | 4 | Etv5 | 0.11 | Young |
| 0.084603 | 0.037324 | 0.238 | 0.153 | 1 | 4 | Dr1 | 0.085 | Young |
| 0.085145 | 0.483724 | 0.25 | 0.173 | 1 | 4 | Dennd4c | 0.077 | Young |
| 0.085216 | 0.906644 | 0.075 | 0.037 | 1 | 4 | Tpm3-rs7 | 0.038 | Young |
| 0.085435 | 1.233291 | 0.025 | 0.007 | 1 | 4 | Gm28187 | 0.018 | Young |
| 0.085435 | 1.042799 | 0.025 | 0.007 | 1 | 4 | Mfsd6l | 0.018 | Young |
| 0.085551 | 0.852766 | 0.125 | 0.074 | 1 | 4 | Zdhhc5 | 0.051 | Young |
| 0.085573 | 0.509552 | 0.112 | 0.063 | 1 | 4 | Heatr6 | 0.049 | Young |
| 0.0856 | 0.920217 | 0.125 | 0.074 | 1 | 4 | Ndufv1 | 0.051 | Young |
| 0.085927 | 0.404773 | 0.5 | 0.405 | 1 | 4 | Sqstm1 | 0.095 | Young |
| 0.086167 | 0.353716 | 0.075 | 0.036 | 1 | 4 | Dnajc12 | 0.039 | Young |
| 0.086254 | 0.341116 | 0.512 | 0.423 | 1 | 4 | Rab11b | 0.089 | Young |
| 0.086433 | 1.962453 | 0.038 | 0.014 | 1 | 4 | Mr1 | 0.024 | Young |
| 0.086603 | 1.045463 | 0.025 | 0.007 | 1 | 4 | Tnfrsf22 | 0.018 | Young |
| 0.086603 | 0.946094 | 0.025 | 0.007 | 1 | 4 | Wdr35 | 0.018 | Young |
| 0.0868 | 0.507682 | 0.338 | 0.25 | 1 | 4 | Pkm | 0.088 | Young |
| 0.086836 | 0.344021 | 0.475 | 0.368 | 1 | 4 | Ptpra | 0.107 | Young |
| 0.08685 | 1.346743 | 0.05 | 0.021 | 1 | 4 | Zfp275 | 0.029 | Young |
| 0.086867 | 1.424296 | 0.038 | 0.014 | 1 | 4 | Cd300ld3 | 0.024 | Young |
| 0.086941 | 0.153906 | 0.95 | 0.917 | 1 | 4 | Rpl13 | 0.033 | Young |
| 0.087488 | 0.41136 | 0.025 | 0.007 | 1 | 4 | Sirt6 | 0.018 | Young |
| 0.0877 | 0.233426 | 0.662 | 0.556 | 1 | 4 | Ubl3 | 0.106 | Young |
| 0.087803 | 0.18603 | 0.812 | 0.659 | 1 | 4 | Ccl2 | 0.153 | Young |
| 0.088025 | 1.144917 | 0.062 | 0.029 | 1 | 4 | Efemp2 | 0.033 | Young |
| 0.088336 | 0.964998 | 0.062 | 0.029 | 1 | 4 | P2ry13 | 0.033 | Young |
| 0.088361 | 0.307727 | 0.625 | 0.522 | 1 | 4 | Eef1g | 0.103 | Young |
| 0.08838 | 0.017555 | 0.025 | 0.007 | 1 | 4 | Cygb | 0.018 | Young |
| 0.088444 | 0.797563 | 0.125 | 0.073 | 1 | 4 | Gnpda2 | 0.052 | Young |
| 0.088819 | 0.546113 | 0.1 | 0.053 | 1 | 4 | Camk2g | 0.047 | Young |
| 0.088855 | 0.505155 | 0.2 | 0.132 | 1 | 4 | Tnfaip1 | 0.068 | Young |

| | | | | | | | | |
|---|---|---|---|---|---|---|---|---|
| 0.088878 | 0.267345 | 0.675 | 0.572 | 1 | 4 | Tmcc1 | 0.103 | Young |
| 0.088938 | 0.472586 | 0.175 | 0.112 | 1 | 4 | Gps1 | 0.063 | Young |
| 0.089058 | 1.392914 | 0.075 | 0.037 | 1 | 4 | Nid2 | 0.038 | Young |
| 0.089064 | 0.623659 | 0.15 | 0.092 | 1 | 4 | Ipo8 | 0.058 | Young |
| 0.089494 | 0.945916 | 0.1 | 0.055 | 1 | 4 | Metap1d | 0.045 | Young |
| 0.089731 | 1.236953 | 0.038 | 0.014 | 1 | 4 | Ptpn13 | 0.024 | Young |
| 0.089738 | 1.024526 | 0.05 | 0.021 | 1 | 4 | Fam84b | 0.029 | Young |
| 0.089864 | 0.430794 | 0.325 | 0.233 | 1 | 4 | Gm6377 | 0.092 | Young |
| 0.090036 | 0.518222 | 0.1 | 0.054 | 1 | 4 | Arhgef5 | 0.046 | Young |
| 0.090223 | 0.666892 | 0.112 | 0.064 | 1 | 4 | Pop7 | 0.048 | Young |
| 0.090428 | 0.648274 | 0.188 | 0.125 | 1 | 4 | Bcor | 0.063 | Young |
| 0.090578 | 0.080534 | 0.062 | 0.028 | 1 | 4 | Cdc25a | 0.034 | Young |
| 0.091242 | 0.641938 | 0.1 | 0.055 | 1 | 4 | Slc25a16 | 0.045 | Young |
| 0.091286 | 0.735671 | 0.225 | 0.158 | 1 | 4 | Zc3h12a | 0.067 | Young |
| 0.091814 | 0.822348 | 0.2 | 0.14 | 1 | 4 | Strn4 | 0.06 | Young |
| 0.09183 | 0.870051 | 0.088 | 0.046 | 1 | 4 | Abhd17c | 0.042 | Young |
| 0.091884 | 0.383754 | 0.438 | 0.344 | 1 | 4 | Ube2r2 | 0.094 | Young |
| 0.092141 | 0.736207 | 0.05 | 0.021 | 1 | 4 | Hspbap1 | 0.029 | Young |
| 0.092257 | 0.883871 | 0.125 | 0.073 | 1 | 4 | Hsd17b10 | 0.052 | Young |
| 0.092328 | 0.920798 | 0.05 | 0.021 | 1 | 4 | Rab39 | 0.029 | Young |
| 0.09238 | 0.458116 | 0.362 | 0.266 | 1 | 4 | Cebpg | 0.096 | Young |
| 0.092516 | 0.552614 | 0.05 | 0.021 | 1 | 4 | Inpp5e | 0.029 | Young |
| 0.092849 | 0.354205 | 0.35 | 0.257 | 1 | 4 | Vasp | 0.093 | Young |
| 0.092854 | 0.608785 | 0.662 | 0.657 | 1 | 4 | Sdc4 | 0.005 | Young |
| 0.092899 | 0.745708 | 0.038 | 0.014 | 1 | 4 | Pkmyt1 | 0.024 | Young |
| 0.093244 | 0.434918 | 0.325 | 0.242 | 1 | 4 | Ubn1 | 0.083 | Young |
| 0.093477 | 0.564521 | 0.262 | 0.191 | 1 | 4 | Mylip | 0.071 | Young |
| 0.093584 | 0.773129 | 0.112 | 0.065 | 1 | 4 | Cog6 | 0.047 | Young |
| 0.093636 | 0.351575 | 0.15 | 0.092 | 1 | 4 | Lsm3 | 0.058 | Young |
| 0.093781 | 0.532662 | 0.088 | 0.045 | 1 | 4 | Nol6 | 0.043 | Young |
| 0.093979 | 0.446288 | 0.238 | 0.167 | 1 | 4 | Ndfip2 | 0.071 | Young |
| 0.094035 | 0.450463 | 0.3 | 0.223 | 1 | 4 | 1810058I2 | 0.077 | Young |
| 0.094052 | 0.462235 | 0.038 | 0.014 | 1 | 4 | Qprt | 0.024 | Young |
| 0.094071 | 0.551409 | 0.388 | 0.3 | 1 | 4 | Tmem55b | 0.088 | Young |
| 0.095061 | 0.536653 | 0.062 | 0.029 | 1 | 4 | Cdk7 | 0.033 | Young |
| 0.095476 | 0.937697 | 0.088 | 0.047 | 1 | 4 | Gzf1 | 0.041 | Young |
| 0.095675 | 0.94236 | 0.15 | 0.096 | 1 | 4 | Ndufs8 | 0.054 | Young |
| 0.095826 | 0.297471 | 0.388 | 0.292 | 1 | 4 | Acap2 | 0.096 | Young |
| 0.096579 | 0.27083 | 0.475 | 0.379 | 1 | 4 | Ube2b | 0.096 | Young |
| 0.096598 | 0.299721 | 0.562 | 0.451 | 1 | 4 | Cxcl16 | 0.111 | Young |
| 0.097054 | 0.368698 | 0.262 | 0.182 | 1 | 4 | Vps41 | 0.08 | Young |
| 0.097325 | 0.299269 | 0.475 | 0.377 | 1 | 4 | Ptpre | 0.098 | Young |
| 0.097433 | 0.6785 | 0.1 | 0.055 | 1 | 4 | 3110062M | 0.045 | Young |
| 0.097448 | 0.507237 | 0.175 | 0.114 | 1 | 4 | Usp37 | 0.061 | Young |
| 0.097629 | 0.419595 | 0.288 | 0.209 | 1 | 4 | Dyrk2 | 0.079 | Young |

| | | | | | | | | |
|---|---|---|---|---|---|---|---|---|
| 0.097706 | 0.517001 | 0.162 | 0.104 | 1 | 4 | Dpp7 | 0.058 | Young |
| 0.097787 | 0.582079 | 0.088 | 0.046 | 1 | 4 | Tsta3 | 0.042 | Young |
| 0.09884 | 1.056722 | 0.1 | 0.057 | 1 | 4 | Riok2 | 0.043 | Young |
| 0.098933 | 0.520556 | 0.15 | 0.093 | 1 | 4 | Pccb | 0.057 | Young |
| 0.09897 | 0.418748 | 0.488 | 0.395 | 1 | 4 | Sec14l1 | 0.093 | Young |
| 0.099101 | 0.172264 | 0.963 | 0.955 | 1 | 4 | Rps14 | 0.008 | Young |
| 0.099293 | 0.210239 | 0.988 | 0.979 | 1 | 4 | Fos | 0.009 | Young |
| 0.09942 | 0.896968 | 0.062 | 0.03 | 1 | 4 | Thap1 | 0.032 | Young |
| 0.09962 | 0.379983 | 0.537 | 0.434 | 1 | 4 | Hnrnpab | 0.103 | Young |
| 0.099623 | 0.343301 | 0.112 | 0.064 | 1 | 4 | Mtmr1 | 0.048 | Young |
| 0.099715 | 0.881947 | 0.162 | 0.107 | 1 | 4 | Zmym2 | 0.055 | Young |
| 0.099754 | 0.240812 | 0.2 | 0.129 | 1 | 4 | Fbxo3 | 0.071 | Young |
| 0.100421 | 0.123053 | 0.988 | 0.989 | 1 | 4 | mt-Nd1 | -0.001 | Young |
| 0.100898 | 0.940981 | 0.162 | 0.108 | 1 | 4 | Vav3 | 0.054 | Young |
| 0.100949 | 0.854062 | 0.088 | 0.046 | 1 | 4 | 6030458C1 | 0.042 | Young |
| 0.101227 | 0.276567 | 0.312 | 0.227 | 1 | 4 | Gspt1 | 0.085 | Young |
| 0.101346 | 0.236112 | 0.275 | 0.192 | 1 | 4 | Ppp2r2a | 0.083 | Young |
| 0.101586 | 0.325192 | 0.438 | 0.348 | 1 | 4 | Tpm4 | 0.09 | Young |
| 0.101736 | 0.4536 | 0.075 | 0.038 | 1 | 4 | Tmed8 | 0.037 | Young |
| 0.101855 | 0.387532 | 0.15 | 0.093 | 1 | 4 | Nploc4 | 0.057 | Young |
| 0.102429 | 1.035637 | 0.05 | 0.022 | 1 | 4 | Ndst1 | 0.028 | Young |
| 0.102908 | 0.639589 | 0.238 | 0.17 | 1 | 4 | Tgif1 | 0.068 | Young |
| 0.102974 | 0.263087 | 0.2 | 0.128 | 1 | 4 | Chordc1 | 0.072 | Young |
| 0.103047 | 0.193322 | 0.488 | 0.367 | 1 | 4 | Tax1bp1 | 0.121 | Young |
| 0.103055 | 0.294012 | 0.088 | 0.046 | 1 | 4 | Fahd1 | 0.042 | Young |
| 0.103209 | 0.451224 | 0.25 | 0.178 | 1 | 4 | Aoah | 0.072 | Young |
| 0.103352 | 0.715533 | 0.125 | 0.075 | 1 | 4 | Klhl18 | 0.05 | Young |
| 0.103405 | 0.768597 | 0.075 | 0.038 | 1 | 4 | Tradd | 0.037 | Young |
| 0.103454 | 0.543721 | 0.262 | 0.19 | 1 | 4 | Psma1 | 0.072 | Young |
| 0.103501 | 0.199232 | 0.85 | 0.77 | 1 | 4 | Rpl24 | 0.08 | Young |
| 0.10356 | 0.953635 | 0.075 | 0.038 | 1 | 4 | Ammecr1 | 0.037 | Young |
| 0.103605 | 0.222896 | 0.887 | 0.842 | 1 | 4 | Rpl18a | 0.045 | Young |
| 0.103686 | 0.16586 | 0.738 | 0.704 | 1 | 4 | Ctsz | 0.034 | Young |
| 0.103795 | 0.797196 | 0.1 | 0.057 | 1 | 4 | Rab11fip2 | 0.043 | Young |
| 0.104248 | 0.247902 | 0.15 | 0.092 | 1 | 4 | Iars | 0.058 | Young |
| 0.104478 | 0.514188 | 0.375 | 0.292 | 1 | 4 | Etf1 | 0.083 | Young |
| 0.104892 | 1.067318 | 0.075 | 0.039 | 1 | 4 | Cd200r4 | 0.036 | Young |
| 0.105126 | 0.422997 | 0.238 | 0.165 | 1 | 4 | Slc25a28 | 0.073 | Young |
| 0.105256 | 0.567206 | 0.162 | 0.105 | 1 | 4 | Apmap | 0.057 | Young |
| 0.105291 | 0.366519 | 0.275 | 0.194 | 1 | 4 | Ubr2 | 0.081 | Young |
| 0.105299 | 1.379694 | 0.038 | 0.014 | 1 | 4 | Gm41307 | 0.024 | Young |
| 0.105454 | 0.635691 | 0.162 | 0.105 | 1 | 4 | Mybbp1a | 0.057 | Young |
| 0.105797 | 1.105414 | 0.038 | 0.014 | 1 | 4 | Ntmt1 | 0.024 | Young |
| 0.105841 | 0.625079 | 0.062 | 0.03 | 1 | 4 | Pcid2 | 0.032 | Young |
| 0.106263 | 1.174195 | 0.062 | 0.03 | 1 | 4 | Tfpi | 0.032 | Young |

| | | | | | | | | |
|---|---|---|---|---|---|---|---|---|
| 0.10637 | 0.163961 | 0.975 | 0.987 | 1 | 4 | Cd74 | -0.012 | Young |
| 0.106992 | 0.359468 | 0.1 | 0.055 | 1 | 4 | Ranbp3 | 0.045 | Young |
| 0.10705 | 1.040151 | 0.038 | 0.014 | 1 | 4 | Sesn2 | 0.024 | Young |
| 0.107339 | 0.836163 | 0.1 | 0.057 | 1 | 4 | Arhgef7 | 0.043 | Young |
| 0.107449 | 0.659755 | 0.138 | 0.085 | 1 | 4 | Jmjd6 | 0.053 | Young |
| 0.107585 | 0.323116 | 0.15 | 0.093 | 1 | 4 | Hdac5 | 0.057 | Young |
| 0.107758 | 0.505302 | 0.138 | 0.084 | 1 | 4 | Gm38399 | 0.054 | Young |
| 0.107782 | 0.549932 | 0.262 | 0.188 | 1 | 4 | Psmb9 | 0.074 | Young |
| 0.107797 | 0.59957 | 0.275 | 0.209 | 1 | 4 | Ubxn4 | 0.066 | Young |
| 0.108061 | 0.845105 | 0.038 | 0.014 | 1 | 4 | Nle1 | 0.024 | Young |
| 0.10811 | 0.638557 | 0.1 | 0.056 | 1 | 4 | Brcc3 | 0.044 | Young |
| 0.108171 | 0.357641 | 0.05 | 0.022 | 1 | 4 | Eva1b | 0.028 | Young |
| 0.108367 | 1.887388 | 0.025 | 0.008 | 1 | 4 | Pigb | 0.017 | Young |
| 0.109032 | 0.582769 | 0.125 | 0.075 | 1 | 4 | Lipe | 0.05 | Young |
| 0.109135 | 0.253884 | 0.225 | 0.152 | 1 | 4 | Aimp1 | 0.073 | Young |
| 0.10916 | 0.302227 | 0.362 | 0.271 | 1 | 4 | Slc12a7 | 0.091 | Young |
| 0.109387 | 1.292488 | 0.025 | 0.008 | 1 | 4 | Ubtd2 | 0.017 | Young |
| 0.109395 | 0.068899 | 0.338 | 0.238 | 1 | 4 | Smim7 | 0.1 | Young |
| 0.109518 | 0.370294 | 0.225 | 0.156 | 1 | 4 | Eif4e3 | 0.069 | Young |
| 0.109592 | 0.170653 | 0.613 | 0.505 | 1 | 4 | Rnase4 | 0.108 | Young |
| 0.109811 | 0.269467 | 0.062 | 0.03 | 1 | 4 | Toe1 | 0.032 | Young |
| 0.110071 | 0.946719 | 0.025 | 0.008 | 1 | 4 | Creb3l3 | 0.017 | Young |
| 0.110145 | 0.347744 | 0.275 | 0.198 | 1 | 4 | Rwdd1 | 0.077 | Young |
| 0.110252 | 0.291924 | 0.537 | 0.447 | 1 | 4 | Laptm4a | 0.09 | Young |
| 0.110363 | 0.184343 | 0.038 | 0.014 | 1 | 4 | Gtf2h2 | 0.024 | Young |
| 0.110414 | 1.370639 | 0.025 | 0.008 | 1 | 4 | Ccdc173 | 0.017 | Young |
| 0.110702 | 0.258791 | 0.225 | 0.153 | 1 | 4 | Atp2c1 | 0.072 | Young |
| 0.110758 | 1.162428 | 0.025 | 0.008 | 1 | 4 | Gm29107 | 0.017 | Young |
| 0.110944 | 0.452907 | 0.262 | 0.192 | 1 | 4 | Mfsd14a | 0.07 | Young |
| 0.111102 | 0.5156 | 0.438 | 0.36 | 1 | 4 | Srgap2 | 0.078 | Young |
| 0.111103 | 0.637647 | 0.025 | 0.008 | 1 | 4 | Gm17259 | 0.017 | Young |
| 0.111109 | 1.182408 | 0.05 | 0.022 | 1 | 4 | Gstcd | 0.028 | Young |
| 0.111449 | 0.718133 | 0.025 | 0.008 | 1 | 4 | Gm10036 | 0.017 | Young |
| 0.111655 | 0.098514 | 0.062 | 0.03 | 1 | 4 | Macrod1 | 0.032 | Young |
| 0.11173 | 0.23132 | 0.662 | 0.595 | 1 | 4 | Rab14 | 0.067 | Young |
| 0.111744 | 1.205762 | 0.05 | 0.022 | 1 | 4 | Creld1 | 0.028 | Young |
| 0.111796 | 0.944763 | 0.025 | 0.008 | 1 | 4 | Taco1os | 0.017 | Young |
| 0.111796 | 0.47318 | 0.025 | 0.008 | 1 | 4 | Mettl22 | 0.017 | Young |
| 0.111898 | 0.132258 | 0.462 | 0.346 | 1 | 4 | Slco2b1 | 0.116 | Young |
| 0.112132 | 0.588383 | 0.362 | 0.314 | 1 | 4 | H2afj | 0.048 | Young |
| 0.112143 | 0.093674 | 0.025 | 0.008 | 1 | 4 | Sec16b | 0.017 | Young |
| 0.112492 | 0.56091 | 0.025 | 0.008 | 1 | 4 | Cdo1 | 0.017 | Young |
| 0.112765 | 0.584551 | 0.1 | 0.057 | 1 | 4 | Fem1a | 0.043 | Young |
| 0.112841 | 0.352764 | 0.025 | 0.008 | 1 | 4 | Cryba4 | 0.017 | Young |
| 0.112879 | 0.705821 | 0.15 | 0.097 | 1 | 4 | Atrnl1 | 0.053 | Young |

| | | | | | | | | |
|---|---|---|---|---|---|---|---|---|
| 0.112979 | 1.005271 | 0.088 | 0.049 | 1 | 4 | Etfdh | 0.039 | Young |
| 0.113191 | 0.687278 | 0.025 | 0.008 | 1 | 4 | Zfat | 0.017 | Young |
| 0.113205 | 0.174383 | 0.175 | 0.111 | 1 | 4 | Idh3b | 0.064 | Young |
| 0.113237 | 1.296956 | 0.05 | 0.022 | 1 | 4 | B230354K1 | 0.028 | Young |
| 0.113335 | 0.810178 | 0.062 | 0.03 | 1 | 4 | Haus3 | 0.032 | Young |
| 0.113932 | 0.35499 | 0.3 | 0.223 | 1 | 4 | N4bp2l2 | 0.077 | Young |
| 0.114824 | 0.905293 | 0.125 | 0.078 | 1 | 4 | Mxd1 | 0.047 | Young |
| 0.115066 | 0.569703 | 0.125 | 0.076 | 1 | 4 | Ctc1 | 0.049 | Young |
| 0.115077 | 0.314946 | 0.175 | 0.113 | 1 | 4 | Sec13 | 0.062 | Young |
| 0.115108 | 0.91031 | 0.112 | 0.067 | 1 | 4 | Tmem104 | 0.045 | Young |
| 0.115369 | 0.291419 | 0.25 | 0.174 | 1 | 4 | Helz | 0.076 | Young |
| 0.115431 | 0.685172 | 0.188 | 0.128 | 1 | 4 | Fam50a | 0.06 | Young |
| 0.115622 | 0.59299 | 0.088 | 0.047 | 1 | 4 | Mettl1 | 0.041 | Young |
| 0.115756 | 0.363015 | 0.175 | 0.115 | 1 | 4 | Kdm6a | 0.06 | Young |
| 0.11588 | 0.565424 | 0.138 | 0.084 | 1 | 4 | Tmem243 | 0.054 | Young |
| 0.116216 | 0.67571 | 0.188 | 0.128 | 1 | 4 | Nipsnap3b | 0.06 | Young |
| 0.116221 | 0.464595 | 0.188 | 0.125 | 1 | 4 | Wdfy4 | 0.063 | Young |
| 0.116239 | 0.115651 | 0.075 | 0.038 | 1 | 4 | Ier5l | 0.037 | Young |
| 0.116271 | 0.919152 | 0.05 | 0.022 | 1 | 4 | Tspyl3 | 0.028 | Young |
| 0.11658 | 0.622185 | 0.075 | 0.039 | 1 | 4 | Dnajb2 | 0.036 | Young |
| 0.116818 | 1.117548 | 0.088 | 0.049 | 1 | 4 | Ppm1m | 0.039 | Young |
| 0.1177 | 1.050385 | 0.15 | 0.1 | 1 | 4 | Pknox1 | 0.05 | Young |
| 0.11772 | 0.203424 | 0.662 | 0.563 | 1 | 4 | Ddx6 | 0.099 | Young |
| 0.117932 | 0.424328 | 0.225 | 0.159 | 1 | 4 | Dnajb11 | 0.066 | Young |
| 0.118131 | 0.544183 | 0.2 | 0.14 | 1 | 4 | Sqor | 0.06 | Young |
| 0.118346 | 0.513608 | 0.325 | 0.253 | 1 | 4 | Ttyh3 | 0.072 | Young |
| 0.118453 | 0.455766 | 0.075 | 0.039 | 1 | 4 | Leng1 | 0.036 | Young |
| 0.118476 | 0.666933 | 0.05 | 0.022 | 1 | 4 | Rabep2 | 0.028 | Young |
| 0.118549 | 0.587263 | 0.1 | 0.057 | 1 | 4 | Gins4 | 0.043 | Young |
| 0.118718 | 0.334931 | 0.412 | 0.334 | 1 | 4 | Tet3 | 0.078 | Young |
| 0.118796 | 0.490441 | 0.075 | 0.039 | 1 | 4 | 2410004B1 | 0.036 | Young |
| 0.118828 | 0.359879 | 0.162 | 0.104 | 1 | 4 | Ubfd1 | 0.058 | Young |
| 0.118958 | 0.541046 | 0.15 | 0.096 | 1 | 4 | Impa1 | 0.054 | Young |
| 0.119195 | 0.553739 | 0.112 | 0.067 | 1 | 4 | Tomm40 | 0.045 | Young |
| 0.119422 | 0.381853 | 0.088 | 0.047 | 1 | 4 | Fbxw5 | 0.041 | Young |
| 0.119613 | 0.226481 | 0.525 | 0.412 | 1 | 4 | Top1 | 0.113 | Young |
| 0.119881 | 1.476871 | 0.038 | 0.015 | 1 | 4 | Dnmt3b | 0.023 | Young |
| 0.119946 | 0.161985 | 0.162 | 0.102 | 1 | 4 | Cul4a | 0.06 | Young |
| 0.120243 | 0.729782 | 0.1 | 0.057 | 1 | 4 | Bcl2l12 | 0.043 | Young |
| 0.120344 | 0.431255 | 0.175 | 0.117 | 1 | 4 | Srek1ip1 | 0.058 | Young |
| 0.120604 | 1.035155 | 0.088 | 0.049 | 1 | 4 | D430042O( | 0.039 | Young |
| 0.120663 | 1.314979 | 0.062 | 0.031 | 1 | 4 | Dock4 | 0.031 | Young |
| 0.120892 | 0.44163 | 0.188 | 0.126 | 1 | 4 | Tslp | 0.062 | Young |
| 0.121006 | 0.506233 | 0.088 | 0.048 | 1 | 4 | Myh10 | 0.04 | Young |
| 0.121044 | 0.36738 | 0.075 | 0.039 | 1 | 4 | Btbd2 | 0.036 | Young |

| | | | | | | | | |
|---|---|---|---|---|---|---|---|---|
| 0.121438 | 0.173736 | 0.275 | 0.196 | 1 | 4 | Krtcap2 | 0.079 | Young |
| 0.121783 | 1.466467 | 0.038 | 0.015 | 1 | 4 | Btd | 0.023 | Young |
| 0.121849 | 0.25845 | 0.675 | 0.581 | 1 | 4 | Igf1 | 0.094 | Young |
| 0.121912 | 0.442992 | 0.175 | 0.115 | 1 | 4 | Preb | 0.06 | Young |
| 0.122218 | 0.63876 | 0.262 | 0.2 | 1 | 4 | Immt | 0.062 | Young |
| 0.122282 | 0.662746 | 0.1 | 0.057 | 1 | 4 | Mpp5 | 0.043 | Young |
| 0.122429 | 0.563304 | 0.15 | 0.097 | 1 | 4 | Faf2 | 0.053 | Young |
| 0.122606 | 0.563551 | 0.275 | 0.2 | 1 | 4 | Zfp710 | 0.075 | Young |
| 0.122679 | 0.087396 | 1 | 1 | 1 | 4 | Rps29 | 0 | Young |
| 0.122729 | 0.335151 | 0.225 | 0.155 | 1 | 4 | Smndc1 | 0.07 | Young |
| 0.122772 | 0.3319 | 0.362 | 0.279 | 1 | 4 | Ost4 | 0.083 | Young |
| 0.122783 | 2.501503 | 0.013 | 0.003 | 1 | 4 | Vasn | 0.01 | Young |
| 0.122783 | 2.499368 | 0.013 | 0.003 | 1 | 4 | Cdk5r1 | 0.01 | Young |
| 0.122783 | 2.207392 | 0.013 | 0.003 | 1 | 4 | Tnfrsf4 | 0.01 | Young |
| 0.122783 | 2.178727 | 0.013 | 0.003 | 1 | 4 | Dcdc2b | 0.01 | Young |
| 0.122783 | 2.159955 | 0.013 | 0.003 | 1 | 4 | Cd5l | 0.01 | Young |
| 0.122783 | 2.128653 | 0.013 | 0.003 | 1 | 4 | Abca8a | 0.01 | Young |
| 0.122783 | 1.962955 | 0.013 | 0.003 | 1 | 4 | Rufy4 | 0.01 | Young |
| 0.122783 | 1.829534 | 0.013 | 0.003 | 1 | 4 | Acot1 | 0.01 | Young |
| 0.122783 | 1.811016 | 0.013 | 0.003 | 1 | 4 | Acer2 | 0.01 | Young |
| 0.123167 | 1.080879 | 0.088 | 0.049 | 1 | 4 | Snip1 | 0.039 | Young |
| 0.123175 | 0.299009 | 0.562 | 0.47 | 1 | 4 | Hmgb1 | 0.092 | Young |
| 0.123321 | 0.294139 | 0.438 | 0.335 | 1 | 4 | Cggbp1 | 0.103 | Young |
| 0.123408 | 2.141931 | 0.013 | 0.003 | 1 | 4 | Smpd5 | 0.01 | Young |
| 0.123408 | 2.048554 | 0.013 | 0.003 | 1 | 4 | Adtrp | 0.01 | Young |
| 0.123408 | 1.988571 | 0.013 | 0.003 | 1 | 4 | Kif1a | 0.01 | Young |
| 0.123408 | 1.962077 | 0.013 | 0.003 | 1 | 4 | Slfn4 | 0.01 | Young |
| 0.123408 | 1.832742 | 0.013 | 0.003 | 1 | 4 | Lims2 | 0.01 | Young |
| 0.123408 | 1.758919 | 0.013 | 0.003 | 1 | 4 | Catsper2 | 0.01 | Young |
| 0.123408 | 1.544953 | 0.013 | 0.003 | 1 | 4 | Il18rap | 0.01 | Young |
| 0.123408 | 1.458389 | 0.013 | 0.003 | 1 | 4 | Gm8113 | 0.01 | Young |
| 0.123408 | 1.334281 | 0.013 | 0.003 | 1 | 4 | Gm14419 | 0.01 | Young |
| 0.123408 | 1.330401 | 0.013 | 0.003 | 1 | 4 | 4930550C1 | 0.01 | Young |
| 0.123408 | 1.209083 | 0.013 | 0.003 | 1 | 4 | Yes1 | 0.01 | Young |
| 0.123408 | 1.127635 | 0.013 | 0.003 | 1 | 4 | Acta2 | 0.01 | Young |
| 0.123408 | 0.914585 | 0.013 | 0.003 | 1 | 4 | Ung | 0.01 | Young |
| 0.123503 | 0.351485 | 0.075 | 0.039 | 1 | 4 | Cpt2 | 0.036 | Young |
| 0.123682 | 0.314795 | 0.162 | 0.105 | 1 | 4 | Actr10 | 0.057 | Young |
| 0.123818 | 0.295259 | 0.362 | 0.281 | 1 | 4 | Hdlbp | 0.081 | Young |
| 0.124035 | 1.838553 | 0.013 | 0.003 | 1 | 4 | Igsf10 | 0.01 | Young |
| 0.124035 | 1.689996 | 0.013 | 0.003 | 1 | 4 | Rspo1 | 0.01 | Young |
| 0.124035 | 1.568082 | 0.013 | 0.003 | 1 | 4 | Hist1h2ac | 0.01 | Young |
| 0.124035 | 1.534883 | 0.013 | 0.003 | 1 | 4 | Ccdc80 | 0.01 | Young |
| 0.124035 | 1.475617 | 0.013 | 0.003 | 1 | 4 | Gm20498 | 0.01 | Young |
| 0.124035 | 1.310477 | 0.013 | 0.003 | 1 | 4 | 2810457G( | 0.01 | Young |

| | | | | | | | | |
|---|---|---|---|---|---|---|---|---|
| 0.124035 | 1.115485 | 0.013 | 0.003 | 1 | 4 | AV099323 | 0.01 | Young |
| 0.124035 | 1.105396 | 0.013 | 0.003 | 1 | 4 | Sirpb1a | 0.01 | Young |
| 0.124035 | 1.069165 | 0.013 | 0.003 | 1 | 4 | Rnf144a | 0.01 | Young |
| 0.124035 | 1.048528 | 0.013 | 0.003 | 1 | 4 | Gm10320 | 0.01 | Young |
| 0.124035 | 1.014536 | 0.013 | 0.003 | 1 | 4 | Serpinb2 | 0.01 | Young |
| 0.124035 | 0.954344 | 0.013 | 0.003 | 1 | 4 | Galnt12 | 0.01 | Young |
| 0.124035 | 0.951403 | 0.013 | 0.003 | 1 | 4 | Ldhb | 0.01 | Young |
| 0.124035 | 0.774841 | 0.013 | 0.003 | 1 | 4 | Tmem229a | 0.01 | Young |
| 0.124035 | 0.545408 | 0.013 | 0.003 | 1 | 4 | Zfp960 | 0.01 | Young |
| 0.124125 | 1.008133 | 0.038 | 0.015 | 1 | 4 | Arl15 | 0.023 | Young |
| 0.124168 | 0.059801 | 0.1 | 0.056 | 1 | 4 | Slc25a20 | 0.044 | Young |
| 0.124236 | 0.244356 | 0.312 | 0.227 | 1 | 4 | Tpp2 | 0.085 | Young |
| 0.124475 | 0.599589 | 0.162 | 0.108 | 1 | 4 | Erlin2 | 0.054 | Young |
| 0.124541 | 0.984161 | 0.038 | 0.015 | 1 | 4 | Tmem107 | 0.023 | Young |
| 0.124664 | 1.326818 | 0.013 | 0.003 | 1 | 4 | Gm15972 | 0.01 | Young |
| 0.124664 | 1.210334 | 0.013 | 0.003 | 1 | 4 | B230398E0 | 0.01 | Young |
| 0.124664 | 0.908761 | 0.013 | 0.003 | 1 | 4 | Slc22a21 | 0.01 | Young |
| 0.124664 | 0.675528 | 0.013 | 0.003 | 1 | 4 | Cib3 | 0.01 | Young |
| 0.124664 | 0.664274 | 0.013 | 0.003 | 1 | 4 | Gm47389 | 0.01 | Young |
| 0.124664 | 0.632571 | 0.013 | 0.003 | 1 | 4 | Scn4b | 0.01 | Young |
| 0.124664 | 0.628985 | 0.013 | 0.003 | 1 | 4 | Frmd8os | 0.01 | Young |
| 0.124664 | 0.600374 | 0.013 | 0.003 | 1 | 4 | Pde3a | 0.01 | Young |
| 0.124664 | 0.597282 | 0.013 | 0.003 | 1 | 4 | Ica1l | 0.01 | Young |
| 0.124664 | 0.516973 | 0.013 | 0.003 | 1 | 4 | Parvg | 0.01 | Young |
| 0.124664 | 0.410142 | 0.013 | 0.003 | 1 | 4 | Egfl8 | 0.01 | Young |
| 0.124664 | 0.298689 | 0.013 | 0.003 | 1 | 4 | CT025521.1 | 0.01 | Young |
| 0.124664 | 0.041287 | 0.013 | 0.003 | 1 | 4 | Mastl | 0.01 | Young |
| 0.125099 | 0.956382 | 0.038 | 0.015 | 1 | 4 | Ecsit | 0.023 | Young |
| 0.125102 | 0.899714 | 0.112 | 0.069 | 1 | 4 | Zfp560 | 0.043 | Young |
| 0.125296 | 0.723738 | 0.013 | 0.003 | 1 | 4 | Nme3 | 0.01 | Young |
| 0.125296 | 0.555711 | 0.013 | 0.003 | 1 | 4 | Cdc20 | 0.01 | Young |
| 0.125296 | 0.494976 | 0.013 | 0.003 | 1 | 4 | Tmem182 | 0.01 | Young |
| 0.125296 | 0.36094 | 0.013 | 0.003 | 1 | 4 | Ldlrad1 | 0.01 | Young |
| 0.125296 | 0.323899 | 0.013 | 0.003 | 1 | 4 | Lgi4 | 0.01 | Young |
| 0.125296 | 0.251475 | 0.013 | 0.003 | 1 | 4 | Cyp2e1 | 0.01 | Young |
| 0.125296 | 0.224506 | 0.013 | 0.003 | 1 | 4 | Cetn4 | 0.01 | Young |
| 0.125296 | 0.106405 | 0.013 | 0.003 | 1 | 4 | 9330104G0 | 0.01 | Young |
| 0.125296 | 0.055089 | 0.013 | 0.003 | 1 | 4 | Gm6904 | 0.01 | Young |
| 0.125325 | 0.890536 | 0.088 | 0.048 | 1 | 4 | Man2b2 | 0.04 | Young |
| 0.125384 | 0.284385 | 0.375 | 0.292 | 1 | 4 | Cuta | 0.083 | Young |
| 0.125442 | 0.582449 | 0.138 | 0.087 | 1 | 4 | Plk3 | 0.051 | Young |
| 0.125998 | 0.271507 | 0.338 | 0.254 | 1 | 4 | Tnrc6a | 0.084 | Young |
| 0.126027 | 0.350753 | 0.4 | 0.314 | 1 | 4 | Uqcr10 | 0.086 | Young |
| 0.126036 | 0.387952 | 0.4 | 0.322 | 1 | 4 | Plekho1 | 0.078 | Young |
| 0.126053 | 0.341371 | 0.362 | 0.271 | 1 | 4 | Kpna4 | 0.091 | Young |

| | | | | | | | | |
|---|---|---|---|---|---|---|---|---|
| 0.126066 | 0.741665 | 0.1 | 0.057 | 1 | 4 | Gdpd1 | 0.043 | Young |
| 0.1261 | 0.524819 | 0.188 | 0.13 | 1 | 4 | Cenpx | 0.058 | Young |
| 0.126138 | 0.487088 | 0.088 | 0.048 | 1 | 4 | Nme4 | 0.04 | Young |
| 0.126239 | 0.4195 | 0.125 | 0.076 | 1 | 4 | Psmd10 | 0.049 | Young |
| 0.126401 | 0.416411 | 0.312 | 0.24 | 1 | 4 | Copb2 | 0.072 | Young |
| 0.12642 | 0.159021 | 0.912 | 0.841 | 1 | 4 | H3f3a | 0.071 | Young |
| 0.126507 | 0.113097 | 0.35 | 0.252 | 1 | 4 | Ctdnep1 | 0.098 | Young |
| 0.126727 | 0.433639 | 0.112 | 0.067 | 1 | 4 | Zfp266 | 0.045 | Young |
| 0.126783 | 0.814527 | 0.038 | 0.015 | 1 | 4 | Fam131a | 0.023 | Young |
| 0.126806 | 0.645326 | 0.1 | 0.058 | 1 | 4 | Cnrip1 | 0.042 | Young |
| 0.126873 | 0.345559 | 0.225 | 0.153 | 1 | 4 | Aco1 | 0.072 | Young |
| 0.127285 | 0.188134 | 0.725 | 0.653 | 1 | 4 | Gdi2 | 0.072 | Young |
| 0.127348 | 0.877592 | 0.038 | 0.015 | 1 | 4 | Sec22c | 0.023 | Young |
| 0.127921 | 0.226874 | 0.262 | 0.186 | 1 | 4 | Elk4 | 0.076 | Young |
| 0.128178 | 0.20304 | 0.112 | 0.066 | 1 | 4 | Adgre4 | 0.046 | Young |
| 0.128642 | 0.555674 | 0.212 | 0.15 | 1 | 4 | Cir1 | 0.062 | Young |
| 0.128702 | 0.528291 | 0.112 | 0.067 | 1 | 4 | Tceanc2 | 0.045 | Young |
| 0.12898 | 0.577709 | 0.175 | 0.119 | 1 | 4 | Maged1 | 0.056 | Young |
| 0.129058 | 0.350796 | 0.162 | 0.106 | 1 | 4 | Ahsa1 | 0.056 | Young |
| 0.129078 | 0.771913 | 0.138 | 0.09 | 1 | 4 | Sin3a | 0.048 | Young |
| 0.129091 | 0.635562 | 0.2 | 0.142 | 1 | 4 | Asph | 0.058 | Young |
| 0.129096 | 0.955564 | 0.188 | 0.134 | 1 | 4 | Aff1 | 0.054 | Young |
| 0.129485 | 0.388231 | 0.188 | 0.127 | 1 | 4 | Pqbp1 | 0.061 | Young |
| 0.129556 | 0.47697 | 0.112 | 0.067 | 1 | 4 | Ankrd49 | 0.045 | Young |
| 0.129711 | 0.37471 | 0.438 | 0.362 | 1 | 4 | Rock1 | 0.076 | Young |
| 0.129729 | 0.76838 | 0.075 | 0.04 | 1 | 4 | Ppp2r1b | 0.035 | Young |
| 0.130024 | 0.829861 | 0.1 | 0.059 | 1 | 4 | Cgrrf1 | 0.041 | Young |
| 0.130108 | 0.951494 | 0.162 | 0.108 | 1 | 4 | Ccl4 | 0.054 | Young |
| 0.130328 | 1.301989 | 0.112 | 0.07 | 1 | 4 | Tifab | 0.042 | Young |
| 0.130633 | 0.365954 | 0.238 | 0.169 | 1 | 4 | Stap1 | 0.069 | Young |
| 0.131216 | 0.917584 | 0.062 | 0.032 | 1 | 4 | Bcdin3d | 0.03 | Young |
| 0.131332 | 0.246695 | 0.1 | 0.057 | 1 | 4 | Ccnc | 0.043 | Young |
| 0.131831 | 1.251989 | 0.062 | 0.032 | 1 | 4 | Sec61a2 | 0.03 | Young |
| 0.132105 | 0.905396 | 0.188 | 0.133 | 1 | 4 | Mrpl57 | 0.055 | Young |
| 0.13307 | 0.32412 | 0.312 | 0.236 | 1 | 4 | Alox5 | 0.076 | Young |
| 0.133561 | 0.209085 | 0.85 | 0.776 | 1 | 4 | Rpl6 | 0.074 | Young |
| 0.133625 | 0.490156 | 0.138 | 0.088 | 1 | 4 | Ddx23 | 0.05 | Young |
| 0.134122 | 0.322998 | 0.625 | 0.574 | 1 | 4 | Vcam1 | 0.051 | Young |
| 0.134686 | 0.238215 | 0.562 | 0.486 | 1 | 4 | Atpif1 | 0.076 | Young |
| 0.134696 | 0.083191 | 0.075 | 0.04 | 1 | 4 | Adck5 | 0.035 | Young |
| 0.134834 | 0.620304 | 0.125 | 0.078 | 1 | 4 | Galnt4 | 0.047 | Young |
| 0.135147 | 0.505062 | 0.05 | 0.023 | 1 | 4 | Sparc | 0.027 | Young |
| 0.135267 | 0.465905 | 0.2 | 0.142 | 1 | 4 | Rnaseh2c | 0.058 | Young |
| 0.135464 | 0.564441 | 0.15 | 0.099 | 1 | 4 | Oas2 | 0.051 | Young |
| 0.135647 | 0.555075 | 0.088 | 0.049 | 1 | 4 | Ubr7 | 0.039 | Young |

| | | | | | | | | |
|---|---|---|---|---|---|---|---|---|
| 0.135677 | 1.643007 | 0.025 | 0.009 | 1 | 4 | Gm30025 | 0.016 | Young |
| 0.136068 | 1.629249 | 0.025 | 0.009 | 1 | 4 | Porcn | 0.016 | Young |
| 0.136068 | 1.339501 | 0.025 | 0.009 | 1 | 4 | Hykk | 0.016 | Young |
| 0.13626 | 0.171571 | 0.138 | 0.086 | 1 | 4 | Gars | 0.052 | Young |
| 0.136273 | 0.481772 | 0.188 | 0.13 | 1 | 4 | Ubr3 | 0.058 | Young |
| 0.136388 | 1.907398 | 0.038 | 0.016 | 1 | 4 | Zfp408 | 0.022 | Young |
| 0.136418 | 0.49651 | 0.162 | 0.108 | 1 | 4 | Atxn10 | 0.054 | Young |
| 0.136459 | 1.35666 | 0.025 | 0.009 | 1 | 4 | Hist1h1d | 0.016 | Young |
| 0.13657 | 0.573095 | 0.075 | 0.04 | 1 | 4 | Slc35g1 | 0.035 | Young |
| 0.136759 | 0.664185 | 0.075 | 0.04 | 1 | 4 | Nob1 | 0.035 | Young |
| 0.136787 | 0.518732 | 0.2 | 0.14 | 1 | 4 | Sertad2 | 0.06 | Young |
| 0.136851 | 1.331743 | 0.025 | 0.009 | 1 | 4 | Mks1 | 0.016 | Young |
| 0.136887 | 0.64442 | 0.175 | 0.12 | 1 | 4 | Polr2c | 0.055 | Young |
| 0.137017 | 0.728461 | 0.275 | 0.21 | 1 | 4 | Ssfa2 | 0.065 | Young |
| 0.137066 | 1.307782 | 0.05 | 0.024 | 1 | 4 | Tnfsf13b | 0.026 | Young |
| 0.137531 | 0.631964 | 0.088 | 0.049 | 1 | 4 | Zxdc | 0.039 | Young |
| 0.137638 | 1.274606 | 0.025 | 0.009 | 1 | 4 | Lgalsl | 0.016 | Young |
| 0.137638 | 1.25179 | 0.025 | 0.009 | 1 | 4 | Pcbp3 | 0.016 | Young |
| 0.13786 | 1.565933 | 0.038 | 0.016 | 1 | 4 | Taf1c | 0.022 | Young |
| 0.138033 | 0.910101 | 0.025 | 0.009 | 1 | 4 | Aqp1 | 0.016 | Young |
| 0.138036 | 0.756768 | 0.075 | 0.041 | 1 | 4 | Prkrip1 | 0.034 | Young |
| 0.138357 | 0.718913 | 0.1 | 0.059 | 1 | 4 | Fbxo7 | 0.041 | Young |
| 0.138429 | 1.077618 | 0.025 | 0.009 | 1 | 4 | Dhodh | 0.016 | Young |
| 0.138457 | 0.204317 | 0.238 | 0.167 | 1 | 4 | Tbc1d5 | 0.071 | Young |
| 0.138826 | 0.96373 | 0.025 | 0.009 | 1 | 4 | Acad10 | 0.016 | Young |
| 0.139046 | 1.217538 | 0.038 | 0.016 | 1 | 4 | Mical2 | 0.022 | Young |
| 0.139099 | 0.384048 | 0.438 | 0.354 | 1 | 4 | Tfe3 | 0.084 | Young |
| 0.139363 | 0.833168 | 0.075 | 0.041 | 1 | 4 | Clstn1 | 0.034 | Young |
| 0.139622 | 0.972712 | 0.025 | 0.009 | 1 | 4 | Trim59 | 0.016 | Young |
| 0.139622 | 0.733899 | 0.025 | 0.009 | 1 | 4 | Zfp503 | 0.016 | Young |
| 0.139705 | 0.492401 | 0.125 | 0.078 | 1 | 4 | Rnf168 | 0.047 | Young |
| 0.139968 | 0.974974 | 0.088 | 0.051 | 1 | 4 | Srprb | 0.037 | Young |
| 0.140021 | 0.848525 | 0.025 | 0.009 | 1 | 4 | Gm48066 | 0.016 | Young |
| 0.140033 | 0.456617 | 0.062 | 0.032 | 1 | 4 | Edrf1 | 0.03 | Young |
| 0.140239 | 1.268463 | 0.038 | 0.016 | 1 | 4 | Slc7a4 | 0.022 | Young |
| 0.140932 | 0.879638 | 0.062 | 0.032 | 1 | 4 | Ndufaf7 | 0.03 | Young |
| 0.140989 | 1.353579 | 0.075 | 0.042 | 1 | 4 | Tagap | 0.033 | Young |
| 0.141152 | 0.411525 | 0.188 | 0.129 | 1 | 4 | Pip5k1a | 0.059 | Young |
| 0.141224 | 0.4798 | 0.025 | 0.009 | 1 | 4 | Adamtsl4 | 0.016 | Young |
| 0.141224 | 0.055231 | 0.025 | 0.009 | 1 | 4 | Gsdme | 0.016 | Young |
| 0.141299 | 0.255908 | 0.525 | 0.451 | 1 | 4 | Cnbp | 0.074 | Young |
| 0.141304 | 0.635449 | 0.338 | 0.27 | 1 | 4 | Cd209a | 0.068 | Young |
| 0.141329 | 0.198112 | 0.262 | 0.191 | 1 | 4 | Tmem173 | 0.071 | Young |
| 0.141371 | 1.086449 | 0.075 | 0.042 | 1 | 4 | Ttc28 | 0.033 | Young |
| 0.141508 | 0.450283 | 0.3 | 0.231 | 1 | 4 | Snx20 | 0.069 | Young |

| | | | | | | | | |
|---|---|---|---|---|---|---|---|---|
| 0.141629 | 0.812641 | 0.125 | 0.08 | 1 | 4 | Wdr20 | 0.045 | Young |
| 0.141803 | 0.778549 | 0.088 | 0.05 | 1 | 4 | Rnf34 | 0.038 | Young |
| 0.141968 | 0.213116 | 0.288 | 0.21 | 1 | 4 | Eif3b | 0.078 | Young |
| 0.142091 | 0.442631 | 0.162 | 0.108 | 1 | 4 | Slc37a2 | 0.054 | Young |
| 0.142304 | 0.596991 | 0.15 | 0.1 | 1 | 4 | Zbtb11 | 0.05 | Young |
| 0.142486 | 0.985404 | 0.05 | 0.024 | 1 | 4 | Zfp770 | 0.026 | Young |
| 0.142526 | 0.354224 | 0.15 | 0.097 | 1 | 4 | Prrc1 | 0.053 | Young |
| 0.142529 | 0.105701 | 0.95 | 0.859 | 1 | 4 | Nfkbiz | 0.091 | Young |
| 0.142952 | 0.548552 | 0.2 | 0.144 | 1 | 4 | Nhlrc3 | 0.056 | Young |
| 0.143317 | 0.450235 | 0.362 | 0.295 | 1 | 4 | Tceal9 | 0.067 | Young |
| 0.143375 | 0.062352 | 0.25 | 0.174 | 1 | 4 | Slc25a11 | 0.076 | Young |
| 0.143484 | 0.380384 | 0.675 | 0.588 | 1 | 4 | Gmfg | 0.087 | Young |
| 0.143565 | 0.661227 | 0.088 | 0.05 | 1 | 4 | Emc4 | 0.038 | Young |
| 0.143648 | 0.046709 | 0.238 | 0.164 | 1 | 4 | Cops4 | 0.074 | Young |
| 0.143687 | 0.178418 | 0.075 | 0.04 | 1 | 4 | Donson | 0.035 | Young |
| 0.143713 | 0.05506 | 0.225 | 0.154 | 1 | 4 | Pcna | 0.071 | Young |
| 0.143739 | 0.99846 | 0.062 | 0.032 | 1 | 4 | Dhcr7 | 0.03 | Young |
| 0.143741 | 0.64817 | 0.05 | 0.024 | 1 | 4 | Cbr4 | 0.026 | Young |
| 0.144064 | 0.355698 | 0.2 | 0.14 | 1 | 4 | Errfi1 | 0.06 | Young |
| 0.144523 | 0.334306 | 0.112 | 0.069 | 1 | 4 | Vps53 | 0.043 | Young |
| 0.144612 | 0.772759 | 0.062 | 0.032 | 1 | 4 | Mettl26 | 0.03 | Young |
| 0.144641 | 0.251776 | 0.637 | 0.544 | 1 | 4 | Csnk1a1 | 0.093 | Young |
| 0.14475 | 0.67023 | 0.05 | 0.024 | 1 | 4 | A830010M | 0.026 | Young |
| 0.144912 | 0.259281 | 0.088 | 0.049 | 1 | 4 | Alkbh8 | 0.039 | Young |
| 0.145167 | 0.39902 | 0.338 | 0.266 | 1 | 4 | Ptpa | 0.072 | Young |
| 0.145272 | 0.081426 | 0.062 | 0.032 | 1 | 4 | Ppp1r12b | 0.03 | Young |
| 0.145311 | 0.508062 | 0.138 | 0.089 | 1 | 4 | Arl13b | 0.049 | Young |
| 0.145455 | 0.428817 | 0.075 | 0.041 | 1 | 4 | Rffl | 0.034 | Young |
| 0.14571 | 0.198605 | 0.2 | 0.136 | 1 | 4 | Eif4a3 | 0.064 | Young |
| 0.145973 | 0.586352 | 0.162 | 0.108 | 1 | 4 | Plscr3 | 0.054 | Young |
| 0.145997 | 0.205281 | 0.2 | 0.136 | 1 | 4 | Actr1a | 0.064 | Young |
| 0.146018 | 0.151695 | 0.038 | 0.016 | 1 | 4 | Gm26853 | 0.022 | Young |
| 0.146055 | 0.276498 | 0.738 | 0.663 | 1 | 4 | Rpl10 | 0.075 | Young |
| 0.146909 | 0.094643 | 0.988 | 0.978 | 1 | 4 | Rps21 | 0.01 | Young |
| 0.146946 | 0.301345 | 0.038 | 0.016 | 1 | 4 | 1810044D( | 0.022 | Young |
| 0.147125 | 0.4729 | 0.262 | 0.198 | 1 | 4 | Gabarapl1 | 0.064 | Young |
| 0.147308 | 0.367316 | 0.212 | 0.152 | 1 | 4 | Dnajc15 | 0.06 | Young |
| 0.147824 | 0.691703 | 0.188 | 0.134 | 1 | 4 | Rcor1 | 0.054 | Young |
| 0.148047 | 0.154614 | 0.425 | 0.312 | 1 | 4 | Ndufa7 | 0.113 | Young |
| 0.148101 | 0.173293 | 0.362 | 0.277 | 1 | 4 | Arhgef2 | 0.085 | Young |
| 0.148721 | 0.685508 | 0.225 | 0.167 | 1 | 4 | Iscu | 0.058 | Young |
| 0.14873 | 0.753824 | 0.112 | 0.07 | 1 | 4 | Ptpn23 | 0.042 | Young |
| 0.149086 | 0.431154 | 0.312 | 0.238 | 1 | 4 | Exoc5 | 0.074 | Young |
| 0.14928 | 0.477406 | 0.425 | 0.365 | 1 | 4 | Far1 | 0.06 | Young |
| 0.14935 | 0.446755 | 0.2 | 0.141 | 1 | 4 | Ubqln1 | 0.059 | Young |

| | | | | | | | | |
|---|---|---|---|---|---|---|---|---|
| 0.149369 | 0.930724 | 0.075 | 0.042 | 1 | 4 | Cstf1 | 0.033 | Young |
| 0.149501 | 0.407914 | 0.262 | 0.192 | 1 | 4 | Arhgap19 | 0.07 | Young |
| 0.149597 | 0.609856 | 0.175 | 0.121 | 1 | 4 | Ccdc34 | 0.054 | Young |
| 0.149721 | 0.505656 | 0.15 | 0.099 | 1 | 4 | Carhsp1 | 0.051 | Young |
| 0.149967 | 0.724121 | 0.075 | 0.042 | 1 | 4 | Cd300lg | 0.033 | Young |
| 0.150281 | 0.381046 | 0.188 | 0.128 | 1 | 4 | Wdr41 | 0.06 | Young |
| 0.150516 | 0.528049 | 0.188 | 0.129 | 1 | 4 | Trim26 | 0.059 | Young |
| 0.150643 | 0.292491 | 0.188 | 0.128 | 1 | 4 | Hipk2 | 0.06 | Young |
| 0.150845 | 0.120066 | 0.3 | 0.219 | 1 | 4 | Ciita | 0.081 | Young |
| 0.151285 | 0.63103 | 0.062 | 0.032 | 1 | 4 | Stambpl1 | 0.03 | Young |
| 0.151329 | 0.601008 | 0.112 | 0.07 | 1 | 4 | Poldip2 | 0.042 | Young |
| 0.151358 | 0.954457 | 0.088 | 0.051 | 1 | 4 | Smpdl3b | 0.037 | Young |
| 0.151446 | 0.276532 | 0.05 | 0.024 | 1 | 4 | Adam22 | 0.026 | Young |
| 0.151465 | 0.524104 | 0.125 | 0.078 | 1 | 4 | Ern1 | 0.047 | Young |
| 0.151708 | 0.245998 | 0.05 | 0.024 | 1 | 4 | Zfp874a | 0.026 | Young |
| 0.152165 | 0.260546 | 0.438 | 0.354 | 1 | 4 | Arrb2 | 0.084 | Young |
| 0.152378 | 1.092773 | 0.075 | 0.042 | 1 | 4 | Slc25a44 | 0.033 | Young |
| 0.152423 | 0.153581 | 0.787 | 0.695 | 1 | 4 | Rpl10a | 0.092 | Young |
| 0.152539 | 0.406175 | 0.225 | 0.161 | 1 | 4 | Rras | 0.064 | Young |
| 0.152742 | 0.191101 | 0.1 | 0.059 | 1 | 4 | Mrpl55 | 0.041 | Young |
| 0.153322 | 0.677994 | 0.175 | 0.123 | 1 | 4 | Jade1 | 0.052 | Young |
| 0.153409 | 0.139907 | 0.637 | 0.505 | 1 | 4 | Ccdc50 | 0.132 | Young |
| 0.153585 | 0.528285 | 0.188 | 0.13 | 1 | 4 | Prkar2b | 0.058 | Young |
| 0.153811 | 0.566175 | 0.2 | 0.142 | 1 | 4 | Gnl3 | 0.058 | Young |
| 0.154006 | 0.018064 | 0.212 | 0.144 | 1 | 4 | Eif4ebp1 | 0.068 | Young |
| 0.15402 | 0.256317 | 0.325 | 0.245 | 1 | 4 | Rab18 | 0.08 | Young |
| 0.15421 | 0.293511 | 0.362 | 0.279 | 1 | 4 | Tnpo1 | 0.083 | Young |
| 0.154539 | 0.387981 | 0.312 | 0.244 | 1 | 4 | Ccnt1 | 0.068 | Young |
| 0.15481 | 0.970764 | 0.062 | 0.033 | 1 | 4 | Sccpdh | 0.029 | Young |
| 0.15538 | 0.26501 | 0.325 | 0.252 | 1 | 4 | Fkbp15 | 0.073 | Young |
| 0.15557 | 0.131337 | 0.863 | 0.752 | 1 | 4 | Ucp2 | 0.111 | Young |
| 0.155586 | 0.353556 | 0.637 | 0.59 | 1 | 4 | Rpl7a | 0.047 | Young |
| 0.155825 | 0.452873 | 0.188 | 0.13 | 1 | 4 | Slf2 | 0.058 | Young |
| 0.15597 | 0.394152 | 0.262 | 0.196 | 1 | 4 | Was | 0.066 | Young |
| 0.156645 | 0.833724 | 0.188 | 0.136 | 1 | 4 | Dctn5 | 0.052 | Young |
| 0.156656 | 0.2557 | 0.275 | 0.204 | 1 | 4 | Fbl | 0.071 | Young |
| 0.156664 | 1.06599 | 0.05 | 0.024 | 1 | 4 | Crocc | 0.026 | Young |
| 0.156664 | 0.831292 | 0.05 | 0.024 | 1 | 4 | Oxnad1 | 0.026 | Young |
| 0.157462 | 0.839048 | 0.05 | 0.024 | 1 | 4 | Ankrd13d | 0.026 | Young |
| 0.157577 | 0.268969 | 0.2 | 0.14 | 1 | 4 | Zfp326 | 0.06 | Young |
| 0.157588 | 1.437926 | 0.038 | 0.016 | 1 | 4 | Fam13a | 0.022 | Young |
| 0.1578 | 0.500201 | 0.15 | 0.1 | 1 | 4 | Mrpl41 | 0.05 | Young |
| 0.158044 | 0.434882 | 0.35 | 0.271 | 1 | 4 | 2610001J0 | 0.079 | Young |
| 0.158428 | 1.128384 | 0.075 | 0.043 | 1 | 4 | Wipi1 | 0.032 | Young |
| 0.158533 | 0.325307 | 0.088 | 0.051 | 1 | 4 | Gm46224 | 0.037 | Young |

| | | | | | | | | |
|---|---|---|---|---|---|---|---|---|
| 0.158822 | 0.292031 | 0.138 | 0.089 | 1 | 4 | Kdm3b | 0.049 | Young |
| 0.158836 | 0.792219 | 0.125 | 0.082 | 1 | 4 | Rasgrp2 | 0.043 | Young |
| 0.15885 | 1.160855 | 0.088 | 0.053 | 1 | 4 | Cds1 | 0.035 | Young |
| 0.158923 | 0.604096 | 0.212 | 0.157 | 1 | 4 | Myd88 | 0.055 | Young |
| 0.15898 | 0.249128 | 0.525 | 0.445 | 1 | 4 | Bst2 | 0.08 | Young |
| 0.159168 | 0.535884 | 0.1 | 0.061 | 1 | 4 | Zfp938 | 0.039 | Young |
| 0.159412 | 1.167199 | 0.088 | 0.053 | 1 | 4 | Tmx4 | 0.035 | Young |
| 0.1596 | 0.224641 | 0.75 | 0.669 | 1 | 4 | Rps7 | 0.081 | Young |
| 0.160146 | 0.724625 | 0.05 | 0.024 | 1 | 4 | Zfp809 | 0.026 | Young |
| 0.160176 | 1.079719 | 0.038 | 0.016 | 1 | 4 | Ddx59 | 0.022 | Young |
| 0.160326 | 0.162816 | 0.45 | 0.353 | 1 | 4 | Eif4a2 | 0.097 | Young |
| 0.160502 | 1.193766 | 0.038 | 0.016 | 1 | 4 | Tada2a | 0.022 | Young |
| 0.160658 | 0.594828 | 0.112 | 0.07 | 1 | 4 | Kri1 | 0.042 | Young |
| 0.160716 | 0.376906 | 0.25 | 0.185 | 1 | 4 | Ndufa11 | 0.065 | Young |
| 0.160807 | 1.052236 | 0.088 | 0.052 | 1 | 4 | Nedd1 | 0.036 | Young |
| 0.160832 | 0.645032 | 0.062 | 0.033 | 1 | 4 | Pla2g12a | 0.029 | Young |
| 0.160906 | 0.25899 | 0.35 | 0.273 | 1 | 4 | Mapk1ip1l | 0.077 | Young |
| 0.160947 | 0.26874 | 0.212 | 0.151 | 1 | 4 | Slc45a4 | 0.061 | Young |
| 0.16118 | 0.529262 | 0.212 | 0.153 | 1 | 4 | Osbp | 0.059 | Young |
| 0.161283 | 0.593061 | 0.1 | 0.061 | 1 | 4 | Tpcn2 | 0.039 | Young |
| 0.161432 | 0.935719 | 0.125 | 0.083 | 1 | 4 | Gskip | 0.042 | Young |
| 0.161783 | 0.676928 | 0.188 | 0.132 | 1 | 4 | Rbm42 | 0.056 | Young |
| 0.162246 | 0.592555 | 0.062 | 0.033 | 1 | 4 | Hdac10 | 0.029 | Young |
| 0.162346 | 0.644477 | 0.125 | 0.081 | 1 | 4 | Engase | 0.044 | Young |
| 0.162671 | 1.988967 | 0.025 | 0.009 | 1 | 4 | Srd5a1 | 0.016 | Young |
| 0.162687 | 0.27639 | 0.388 | 0.316 | 1 | 4 | Dnajb14 | 0.072 | Young |
| 0.162812 | 0.480338 | 0.112 | 0.07 | 1 | 4 | Lztfl1 | 0.042 | Young |
| 0.163126 | 1.003977 | 0.038 | 0.016 | 1 | 4 | Mag | 0.022 | Young |
| 0.163126 | 0.913571 | 0.038 | 0.016 | 1 | 4 | Qtrt2 | 0.022 | Young |
| 0.163148 | 0.119331 | 0.262 | 0.188 | 1 | 4 | Coro1b | 0.074 | Young |
| 0.163458 | 0.511311 | 0.162 | 0.111 | 1 | 4 | Epc2 | 0.051 | Young |
| 0.163587 | 0.119953 | 0.9 | 0.859 | 1 | 4 | Unc93b1 | 0.041 | Young |
| 0.163616 | 0.239442 | 0.9 | 0.909 | 1 | 4 | Jun | -0.009 | Young |
| 0.163708 | 0.426894 | 0.375 | 0.304 | 1 | 4 | Cd300ld | 0.071 | Young |
| 0.163709 | 0.3111 | 0.275 | 0.208 | 1 | 4 | Scamp5 | 0.067 | Young |
| 0.163787 | 0.845075 | 0.038 | 0.016 | 1 | 4 | 1110004E0 | 0.022 | Young |
| 0.163957 | 0.51128 | 0.1 | 0.061 | 1 | 4 | Gm13212 | 0.039 | Young |
| 0.164026 | 0.384015 | 0.175 | 0.121 | 1 | 4 | Ufl1 | 0.054 | Young |
| 0.164027 | 0.189582 | 0.188 | 0.128 | 1 | 4 | Clip1 | 0.06 | Young |
| 0.164386 | 0.487339 | 0.062 | 0.033 | 1 | 4 | Gnl2 | 0.029 | Young |
| 0.164744 | 0.527201 | 0.2 | 0.139 | 1 | 4 | Clec2i | 0.061 | Young |
| 0.16499 | 0.633585 | 0.15 | 0.102 | 1 | 4 | Nf1 | 0.048 | Young |
| 0.165103 | 0.457374 | 0.062 | 0.033 | 1 | 4 | Med27 | 0.029 | Young |
| 0.165153 | 0.412142 | 0.188 | 0.132 | 1 | 4 | Cpeb2 | 0.056 | Young |
| 0.165616 | 0.224148 | 0.05 | 0.024 | 1 | 4 | Tlr8 | 0.026 | Young |

| | | | | | | | | |
|---|---|---|---|---|---|---|---|---|
| 0.165776 | 0.217661 | 0.338 | 0.256 | 1 | 4 | Mospd2 | 0.082 | Young |
| 0.165782 | 0.599775 | 0.038 | 0.016 | 1 | 4 | Wfdc18 | 0.022 | Young |
| 0.165855 | 0.407829 | 0.188 | 0.132 | 1 | 4 | Arhgap25 | 0.056 | Young |
| 0.165856 | 0.244069 | 0.775 | 0.693 | 1 | 4 | Fam46a | 0.082 | Young |
| 0.165897 | 1.325068 | 0.05 | 0.025 | 1 | 4 | Cxcr3 | 0.025 | Young |
| 0.166055 | 0.411881 | 0.188 | 0.133 | 1 | 4 | Rnf169 | 0.055 | Young |
| 0.166252 | 0.498306 | 0.112 | 0.07 | 1 | 4 | E2f4 | 0.042 | Young |
| 0.166452 | 0.536928 | 0.038 | 0.016 | 1 | 4 | Zfp523 | 0.022 | Young |
| 0.166452 | 0.479051 | 0.038 | 0.016 | 1 | 4 | Snrnp35 | 0.022 | Young |
| 0.166577 | 0.746164 | 0.15 | 0.101 | 1 | 4 | Tmem86a | 0.049 | Young |
| 0.166805 | 0.338726 | 0.175 | 0.121 | 1 | 4 | Ltbr | 0.054 | Young |
| 0.166918 | 0.594074 | 0.125 | 0.081 | 1 | 4 | Lyar | 0.044 | Young |
| 0.166997 | 1.478926 | 0.05 | 0.025 | 1 | 4 | Dcun1d4 | 0.025 | Young |
| 0.167227 | 0.848254 | 0.062 | 0.034 | 1 | 4 | Zc3h10 | 0.028 | Young |
| 0.167377 | 0.216174 | 0.212 | 0.149 | 1 | 4 | Vkorc1 | 0.063 | Young |
| 0.167541 | 0.147506 | 0.388 | 0.296 | 1 | 4 | Myl12a | 0.092 | Young |
| 0.167552 | 0.187197 | 0.362 | 0.277 | 1 | 4 | Fgd4 | 0.085 | Young |
| 0.167684 | 1.237029 | 0.1 | 0.063 | 1 | 4 | Traf7 | 0.037 | Young |
| 0.167776 | 0.398166 | 0.138 | 0.09 | 1 | 4 | Cds2 | 0.048 | Young |
| 0.167835 | 0.38892 | 0.25 | 0.189 | 1 | 4 | Slc39a7 | 0.061 | Young |
| 0.167872 | 0.52714 | 0.262 | 0.199 | 1 | 4 | Nab1 | 0.063 | Young |
| 0.167942 | 0.833969 | 0.025 | 0.009 | 1 | 4 | Paqr9 | 0.016 | Young |
| 0.168022 | 0.299564 | 0.275 | 0.208 | 1 | 4 | Csnk2a2 | 0.067 | Young |
| 0.16812 | 0.473361 | 0.112 | 0.07 | 1 | 4 | Spice1 | 0.042 | Young |
| 0.16813 | 0.160552 | 0.875 | 0.789 | 1 | 4 | Rpl18 | 0.086 | Young |
| 0.168146 | 0.825558 | 0.162 | 0.117 | 1 | 4 | As3mt | 0.045 | Young |
| 0.16815 | 0.289951 | 0.375 | 0.285 | 1 | 4 | Cox17 | 0.09 | Young |
| 0.168403 | 0.219085 | 0.05 | 0.024 | 1 | 4 | Tsen2 | 0.026 | Young |
| 0.16843 | 0.925272 | 0.062 | 0.034 | 1 | 4 | Zbtb17 | 0.028 | Young |
| 0.168656 | 1.232982 | 0.05 | 0.025 | 1 | 4 | Map3k10 | 0.025 | Young |
| 0.168884 | 0.327946 | 0.275 | 0.211 | 1 | 4 | Lsm4 | 0.064 | Young |
| 0.169194 | 0.406712 | 0.2 | 0.143 | 1 | 4 | Ufc1 | 0.057 | Young |
| 0.169323 | 0.192432 | 0.388 | 0.301 | 1 | 4 | Gna13 | 0.087 | Young |
| 0.169513 | 0.111529 | 0.15 | 0.099 | 1 | 4 | Strn | 0.051 | Young |
| 0.16952 | 0.426863 | 0.312 | 0.246 | 1 | 4 | Dusp22 | 0.066 | Young |
| 0.169536 | 0.425774 | 0.075 | 0.042 | 1 | 4 | Nubp2 | 0.033 | Young |
| 0.16954 | 0.640964 | 0.15 | 0.103 | 1 | 4 | Hk3 | 0.047 | Young |
| 0.169583 | 0.386988 | 0.25 | 0.188 | 1 | 4 | Lrrc8a | 0.062 | Young |
| 0.169662 | 0.217654 | 0.45 | 0.377 | 1 | 4 | Sf1 | 0.073 | Young |
| 0.169768 | 0.994426 | 0.05 | 0.025 | 1 | 4 | Rasgrp1 | 0.025 | Young |
| 0.169828 | 0.282427 | 0.038 | 0.016 | 1 | 4 | Tamm41 | 0.022 | Young |
| 0.170255 | 0.835592 | 0.075 | 0.043 | 1 | 4 | Fndc7 | 0.032 | Young |
| 0.170408 | 0.347558 | 0.075 | 0.042 | 1 | 4 | Gna15 | 0.033 | Young |
| 0.170625 | 0.568025 | 0.025 | 0.009 | 1 | 4 | Casz1 | 0.016 | Young |
| 0.170625 | 0.413609 | 0.025 | 0.009 | 1 | 4 | Cfap126 | 0.016 | Young |

| | | | | | | | | |
|---|---|---|---|---|---|---|---|---|
| 0.170823 | 0.362678 | 0.388 | 0.328 | 1 | 4 | Cab39 | 0.06 | Young |
| 0.170973 | 0.015886 | 0.388 | 0.29 | 1 | 4 | Brd4 | 0.098 | Young |
| 0.171002 | 0.393367 | 0.125 | 0.08 | 1 | 4 | Plau | 0.045 | Young |
| 0.171111 | 0.348569 | 0.325 | 0.26 | 1 | 4 | Phf21a | 0.065 | Young |
| 0.171337 | 0.065655 | 0.238 | 0.171 | 1 | 4 | Rab24 | 0.067 | Young |
| 0.171725 | 0.286825 | 0.188 | 0.13 | 1 | 4 | Zfp740 | 0.058 | Young |
| 0.171839 | 0.604803 | 0.112 | 0.072 | 1 | 4 | Rnf41 | 0.04 | Young |
| 0.172216 | 0.641155 | 0.075 | 0.043 | 1 | 4 | Nsun4 | 0.032 | Young |
| 0.172667 | 0.345116 | 0.5 | 0.428 | 1 | 4 | Skap2 | 0.072 | Young |
| 0.172995 | 0.219684 | 0.412 | 0.338 | 1 | 4 | Ralb | 0.074 | Young |
| 0.173176 | 0.455709 | 0.162 | 0.112 | 1 | 4 | Cltb | 0.05 | Young |
| 0.173515 | 0.373966 | 0.488 | 0.406 | 1 | 4 | Sec62 | 0.082 | Young |
| 0.173647 | 0.355417 | 0.338 | 0.267 | 1 | 4 | Hmox2 | 0.071 | Young |
| 0.173664 | 0.133087 | 0.388 | 0.3 | 1 | 4 | Msl1 | 0.088 | Young |
| 0.173702 | 0.512639 | 0.162 | 0.111 | 1 | 4 | Atf6b | 0.051 | Young |
| 0.173709 | 0.403302 | 0.125 | 0.081 | 1 | 4 | Fam96b | 0.044 | Young |
| 0.173736 | 0.445709 | 0.238 | 0.175 | 1 | 4 | Rnf111 | 0.063 | Young |
| 0.174018 | 0.421754 | 0.25 | 0.188 | 1 | 4 | Araf | 0.062 | Young |
| 0.17426 | 0.519458 | 0.112 | 0.072 | 1 | 4 | Bop1 | 0.04 | Young |
| 0.174275 | 0.975663 | 0.05 | 0.025 | 1 | 4 | Ccl25 | 0.025 | Young |
| 0.174434 | 0.600026 | 0.112 | 0.072 | 1 | 4 | Klhl20 | 0.04 | Young |
| 0.174585 | 0.054312 | 0.388 | 0.296 | 1 | 4 | Ccnd1 | 0.092 | Young |
| 0.174822 | 0.322786 | 0.075 | 0.042 | 1 | 4 | Cpeb3 | 0.033 | Young |
| 0.174925 | 0.068486 | 0.112 | 0.069 | 1 | 4 | Rrm2b | 0.043 | Young |
| 0.175033 | 0.243866 | 0.175 | 0.121 | 1 | 4 | Dnajc1 | 0.054 | Young |
| 0.175769 | 0.220636 | 0.488 | 0.389 | 1 | 4 | Cd2ap | 0.099 | Young |
| 0.175868 | 0.236152 | 0.2 | 0.14 | 1 | 4 | Nemf | 0.06 | Young |
| 0.17617 | 0.376119 | 0.238 | 0.178 | 1 | 4 | Trim28 | 0.06 | Young |
| 0.176249 | 0.058798 | 0.225 | 0.158 | 1 | 4 | Rps6kb1 | 0.067 | Young |
| 0.176615 | 1.017194 | 0.062 | 0.034 | 1 | 4 | Rmdn3 | 0.028 | Young |
| 0.176827 | 0.673047 | 0.288 | 0.235 | 1 | 4 | 1700017B0 | 0.053 | Young |
| 0.177394 | 0.728036 | 0.175 | 0.124 | 1 | 4 | Mlec | 0.051 | Young |
| 0.177561 | 0.202555 | 0.362 | 0.291 | 1 | 4 | Cox14 | 0.071 | Young |
| 0.177625 | 0.164168 | 0.338 | 0.251 | 1 | 4 | Ddrgk1 | 0.087 | Young |
| 0.177713 | 1.233155 | 0.075 | 0.044 | 1 | 4 | Flt3l | 0.031 | Young |
| 0.177777 | 0.291442 | 0.462 | 0.385 | 1 | 4 | Scarb2 | 0.077 | Young |
| 0.177868 | 0.232924 | 0.212 | 0.152 | 1 | 4 | Irs2 | 0.06 | Young |
| 0.17805 | 0.304135 | 0.162 | 0.112 | 1 | 4 | Nt5c | 0.05 | Young |
| 0.178106 | 0.372277 | 0.45 | 0.388 | 1 | 4 | Spi1 | 0.062 | Young |
| 0.178285 | 0.841683 | 0.062 | 0.034 | 1 | 4 | Lonrf1 | 0.028 | Young |
| 0.17858 | 0.526921 | 0.05 | 0.025 | 1 | 4 | Poll | 0.025 | Young |
| 0.178978 | 0.877593 | 0.112 | 0.071 | 1 | 4 | Ints9 | 0.041 | Young |
| 0.178996 | 0.349953 | 0.175 | 0.123 | 1 | 4 | Trp53 | 0.052 | Young |
| 0.178996 | 0.441171 | 0.15 | 0.101 | 1 | 4 | Ints6 | 0.049 | Young |
| 0.17916 | 0.567219 | 0.05 | 0.025 | 1 | 4 | Stk11ip | 0.025 | Young |

| | | | | | | | | |
|---|---|---|---|---|---|---|---|---|
| 0.179289 | 0.51066 | 0.15 | 0.103 | 1 | 4 | Mrps36 | 0.047 | Young |
| 0.179294 | 0.466405 | 0.062 | 0.034 | 1 | 4 | Fggy | 0.028 | Young |
| 0.179308 | 0.542037 | 0.125 | 0.082 | 1 | 4 | Zfp646 | 0.043 | Young |
| 0.179451 | 0.3479 | 0.05 | 0.025 | 1 | 4 | Aven | 0.025 | Young |
| 0.179768 | 0.179275 | 0.15 | 0.1 | 1 | 4 | Cep68 | 0.05 | Young |
| 0.180003 | 0.164599 | 0.35 | 0.267 | 1 | 4 | Dnajc5 | 0.083 | Young |
| 0.180034 | 0.267711 | 0.738 | 0.675 | 1 | 4 | Ctsd | 0.063 | Young |
| 0.18008 | 0.883236 | 0.138 | 0.096 | 1 | 4 | Ank | 0.042 | Young |
| 0.180467 | 0.342469 | 0.162 | 0.111 | 1 | 4 | Tmem250- | 0.051 | Young |
| 0.180539 | 0.983446 | 0.1 | 0.064 | 1 | 4 | Calcrl | 0.036 | Young |
| 0.180541 | 1.117583 | 0.038 | 0.017 | 1 | 4 | Tango6 | 0.021 | Young |
| 0.180851 | 0.639134 | 0.088 | 0.053 | 1 | 4 | 2810402E2 | 0.035 | Young |
| 0.181068 | 0.163062 | 0.238 | 0.172 | 1 | 4 | BC005624 | 0.066 | Young |
| 0.181369 | 1.016588 | 0.062 | 0.034 | 1 | 4 | Kctd3 | 0.028 | Young |
| 0.181748 | 0.329927 | 0.162 | 0.111 | 1 | 4 | Arhgap5 | 0.051 | Young |
| 0.181902 | 1.191541 | 0.1 | 0.065 | 1 | 4 | Crbn | 0.035 | Young |
| 0.181944 | 1.175004 | 0.038 | 0.017 | 1 | 4 | Emilin1 | 0.021 | Young |
| 0.182028 | 0.200437 | 0.462 | 0.371 | 1 | 4 | Ran | 0.091 | Young |
| 0.182081 | 0.459437 | 0.05 | 0.025 | 1 | 4 | 2310057M | 0.025 | Young |
| 0.182081 | 0.307473 | 0.05 | 0.025 | 1 | 4 | Fahd2a | 0.025 | Young |
| 0.182328 | 0.221932 | 0.325 | 0.252 | 1 | 4 | Zmynd11 | 0.073 | Young |
| 0.182339 | 0.680158 | 0.262 | 0.208 | 1 | 4 | Ophn1 | 0.054 | Young |
| 0.182617 | 0.412373 | 0.075 | 0.043 | 1 | 4 | Rhno1 | 0.032 | Young |
| 0.182649 | 1.122407 | 0.038 | 0.017 | 1 | 4 | Alms1 | 0.021 | Young |
| 0.182649 | 0.991312 | 0.038 | 0.017 | 1 | 4 | Trit1 | 0.021 | Young |
| 0.182889 | 0.558977 | 0.138 | 0.092 | 1 | 4 | Samd4 | 0.046 | Young |
| 0.183003 | 1.477611 | 0.038 | 0.017 | 1 | 4 | Gbp2b | 0.021 | Young |
| 0.183207 | 0.150357 | 0.725 | 0.667 | 1 | 4 | Srrm2 | 0.058 | Young |
| 0.183264 | 0.382397 | 0.375 | 0.31 | 1 | 4 | Mgat1 | 0.065 | Young |
| 0.18344 | 0.287539 | 0.15 | 0.101 | 1 | 4 | Kmo | 0.049 | Young |
| 0.18357 | 0.179781 | 0.588 | 0.499 | 1 | 4 | Rpl4 | 0.089 | Young |
| 0.183828 | 0.793112 | 0.075 | 0.043 | 1 | 4 | Zbed4 | 0.032 | Young |
| 0.184007 | 0.117443 | 0.375 | 0.285 | 1 | 4 | Ubxn1 | 0.09 | Young |
| 0.184108 | 0.144039 | 0.225 | 0.161 | 1 | 4 | P4ha1 | 0.064 | Young |
| 0.184374 | 0.318841 | 0.162 | 0.113 | 1 | 4 | Cd200r1 | 0.049 | Young |
| 0.184681 | 0.206193 | 0.537 | 0.449 | 1 | 4 | Ltc4s | 0.088 | Young |
| 0.185133 | 0.800572 | 0.038 | 0.017 | 1 | 4 | Prkar1b | 0.021 | Young |
| 0.185159 | 0.525152 | 0.188 | 0.136 | 1 | 4 | Uqcrc1 | 0.052 | Young |
| 0.185335 | 1.291703 | 0.05 | 0.026 | 1 | 4 | Zfp768 | 0.024 | Young |
| 0.185869 | 0.659774 | 0.088 | 0.052 | 1 | 4 | 5730455P1 | 0.036 | Young |
| 0.185986 | 0.296747 | 0.288 | 0.217 | 1 | 4 | Bcat2 | 0.071 | Young |
| 0.186186 | 1.140833 | 0.062 | 0.035 | 1 | 4 | Lrrc61 | 0.027 | Young |
| 0.18655 | 0.674485 | 0.125 | 0.084 | 1 | 4 | Aig1 | 0.041 | Young |
| 0.186584 | 0.130108 | 0.225 | 0.162 | 1 | 4 | Fbxo33 | 0.063 | Young |
| 0.186694 | 0.266484 | 0.125 | 0.081 | 1 | 4 | Scrib | 0.044 | Young |

| | | | | | | | | |
|---|---|---|---|---|---|---|---|---|
| 0.186839 | 0.400014 | 0.112 | 0.072 | 1 | 4 | Tmem214 | 0.04 | Young |
| 0.186922 | 0.634116 | 0.038 | 0.017 | 1 | 4 | Kat8 | 0.021 | Young |
| 0.187135 | 0.066203 | 0.738 | 0.646 | 1 | 4 | Pnrc1 | 0.092 | Young |
| 0.187145 | 0.165393 | 0.637 | 0.543 | 1 | 4 | Rpl31 | 0.094 | Young |
| 0.187281 | 0.649546 | 0.038 | 0.017 | 1 | 4 | Zfp609 | 0.021 | Young |
| 0.18738 | 0.609661 | 0.188 | 0.137 | 1 | 4 | Tcerg1 | 0.051 | Young |
| 0.187642 | 0.71068 | 0.175 | 0.128 | 1 | 4 | Upf1 | 0.047 | Young |
| 0.187958 | 0.671485 | 0.088 | 0.053 | 1 | 4 | Dvl3 | 0.035 | Young |
| 0.188011 | 2.309765 | 0.013 | 0.003 | 1 | 4 | Serpine2 | 0.01 | Young |
| 0.188011 | 2.205297 | 0.013 | 0.003 | 1 | 4 | Tmem72 | 0.01 | Young |
| 0.188011 | 2.139767 | 0.013 | 0.003 | 1 | 4 | Mybpc3 | 0.01 | Young |
| 0.188011 | 1.961858 | 0.013 | 0.003 | 1 | 4 | Fbxo44 | 0.01 | Young |
| 0.188011 | 1.879797 | 0.013 | 0.003 | 1 | 4 | Prdm16 | 0.01 | Young |
| 0.188011 | 1.859616 | 0.013 | 0.003 | 1 | 4 | Gm6277 | 0.01 | Young |
| 0.188115 | 0.490357 | 0.225 | 0.171 | 1 | 4 | Aup1 | 0.054 | Young |
| 0.188274 | 0.448384 | 0.15 | 0.103 | 1 | 4 | Cspp1 | 0.047 | Young |
| 0.188363 | 0.20308 | 0.212 | 0.153 | 1 | 4 | Camta2 | 0.059 | Young |
| 0.188594 | 0.962031 | 0.05 | 0.026 | 1 | 4 | Ssh3 | 0.024 | Young |
| 0.18867 | 0.182608 | 0.762 | 0.758 | 1 | 4 | mt-Nd5 | 0.004 | Young |
| 0.188799 | 2.674743 | 0.013 | 0.003 | 1 | 4 | Prg4 | 0.01 | Young |
| 0.188799 | 1.896585 | 0.013 | 0.003 | 1 | 4 | Itpr3 | 0.01 | Young |
| 0.188799 | 1.727671 | 0.013 | 0.003 | 1 | 4 | Mipol1 | 0.01 | Young |
| 0.188799 | 1.560932 | 0.013 | 0.003 | 1 | 4 | Pdxp | 0.01 | Young |
| 0.188799 | 1.198854 | 0.013 | 0.003 | 1 | 4 | Sox12 | 0.01 | Young |
| 0.188864 | 0.383631 | 0.212 | 0.157 | 1 | 4 | Gab2 | 0.055 | Young |
| 0.188899 | 0.550776 | 0.075 | 0.043 | 1 | 4 | Nfkbil1 | 0.032 | Young |
| 0.189088 | 0.415006 | 0.35 | 0.285 | 1 | 4 | Myo9b | 0.065 | Young |
| 0.189227 | 0.558357 | 0.138 | 0.094 | 1 | 4 | Mrpl4 | 0.044 | Young |
| 0.189548 | 0.409547 | 0.25 | 0.188 | 1 | 4 | 2-Mar | 0.062 | Young |
| 0.18959 | 1.697405 | 0.013 | 0.003 | 1 | 4 | Hist2h4 | 0.01 | Young |
| 0.18959 | 1.467696 | 0.013 | 0.003 | 1 | 4 | Grtp1 | 0.01 | Young |
| 0.18959 | 1.435304 | 0.013 | 0.003 | 1 | 4 | Dock3 | 0.01 | Young |
| 0.18959 | 1.3225 | 0.013 | 0.003 | 1 | 4 | Tmem170 | 0.01 | Young |
| 0.18959 | 1.280602 | 0.013 | 0.003 | 1 | 4 | Enkd1 | 0.01 | Young |
| 0.18959 | 1.260358 | 0.013 | 0.003 | 1 | 4 | Poln | 0.01 | Young |
| 0.18959 | 1.11403 | 0.013 | 0.003 | 1 | 4 | Gm10037 | 0.01 | Young |
| 0.18959 | 1.000447 | 0.013 | 0.003 | 1 | 4 | Plppr2 | 0.01 | Young |
| 0.18959 | 0.622033 | 0.013 | 0.003 | 1 | 4 | Plpp7 | 0.01 | Young |
| 0.189669 | 0.672747 | 0.188 | 0.137 | 1 | 4 | Myo1g | 0.051 | Young |
| 0.190383 | 1.14982 | 0.013 | 0.003 | 1 | 4 | B230217C1 | 0.01 | Young |
| 0.190383 | 1.084675 | 0.013 | 0.003 | 1 | 4 | Chaf1b | 0.01 | Young |
| 0.190383 | 1.049278 | 0.013 | 0.003 | 1 | 4 | Gm26759 | 0.01 | Young |
| 0.190383 | 0.972072 | 0.013 | 0.003 | 1 | 4 | Polh | 0.01 | Young |
| 0.190383 | 0.783782 | 0.013 | 0.003 | 1 | 4 | E430024P1 | 0.01 | Young |
| 0.190383 | 0.773143 | 0.013 | 0.003 | 1 | 4 | Car4 | 0.01 | Young |

| | | | | | | | | |
|---|---|---|---|---|---|---|---|---|
| 0.190383 | 0.452466 | 0.013 | 0.003 | 1 | 4 | D230017M | 0.01 | Young |
| 0.190383 | 0.346922 | 0.013 | 0.003 | 1 | 4 | Nbl1 | 0.01 | Young |
| 0.190899 | 0.631836 | 0.062 | 0.034 | 1 | 4 | Zfp429 | 0.028 | Young |
| 0.191179 | 1.11654 | 0.013 | 0.003 | 1 | 4 | 1810010D( | 0.01 | Young |
| 0.191179 | 1.064892 | 0.013 | 0.003 | 1 | 4 | Disp1 | 0.01 | Young |
| 0.191179 | 0.857092 | 0.013 | 0.003 | 1 | 4 | Aars2 | 0.01 | Young |
| 0.191179 | 0.705914 | 0.013 | 0.003 | 1 | 4 | Inca1 | 0.01 | Young |
| 0.191179 | 0.704712 | 0.013 | 0.003 | 1 | 4 | Pygm | 0.01 | Young |
| 0.191179 | 0.624955 | 0.013 | 0.003 | 1 | 4 | Cobll1 | 0.01 | Young |
| 0.191179 | 0.399601 | 0.013 | 0.003 | 1 | 4 | Gm26912 | 0.01 | Young |
| 0.191179 | 0.367046 | 0.013 | 0.003 | 1 | 4 | Sspn | 0.01 | Young |
| 0.191268 | 0.207825 | 0.038 | 0.017 | 1 | 4 | Eif2d | 0.021 | Young |
| 0.191607 | 0.1634 | 0.188 | 0.13 | 1 | 4 | Ccdc90b | 0.058 | Young |
| 0.191977 | 0.40345 | 0.013 | 0.003 | 1 | 4 | Galnt9 | 0.01 | Young |
| 0.191977 | 0.225663 | 0.013 | 0.003 | 1 | 4 | Gm34342 | 0.01 | Young |
| 0.191977 | 0.203196 | 0.013 | 0.003 | 1 | 4 | Ablim2 | 0.01 | Young |
| 0.191977 | 0.153874 | 0.013 | 0.003 | 1 | 4 | Zfp128 | 0.01 | Young |
| 0.191977 | 0.144106 | 0.013 | 0.003 | 1 | 4 | Gm20661 | 0.01 | Young |
| 0.191977 | 0.10028 | 0.013 | 0.003 | 1 | 4 | Stac3 | 0.01 | Young |
| 0.191977 | 0.08518 | 0.013 | 0.003 | 1 | 4 | Gm26809 | 0.01 | Young |
| 0.191977 | 0.024651 | 0.013 | 0.003 | 1 | 4 | Dnaaf2 | 0.01 | Young |
| 0.191977 | 0.012717 | 0.013 | 0.003 | 1 | 4 | Fgf9 | 0.01 | Young |
| 0.192127 | 0.202824 | 0.238 | 0.168 | 1 | 4 | Arl5a | 0.07 | Young |
| 0.192752 | 0.523014 | 0.188 | 0.137 | 1 | 4 | U2af1 | 0.051 | Young |
| 0.192776 | 0.330215 | 0.25 | 0.194 | 1 | 4 | Noc2l | 0.056 | Young |
| 0.19301 | 1.985466 | 0.025 | 0.01 | 1 | 4 | Sirpb1b | 0.015 | Young |
| 0.193308 | 0.299128 | 0.225 | 0.165 | 1 | 4 | Adprh | 0.06 | Young |
| 0.193392 | 0.348426 | 0.338 | 0.278 | 1 | 4 | Lrrc25 | 0.06 | Young |
| 0.193626 | 0.180484 | 0.6 | 0.493 | 1 | 4 | Fnbp1 | 0.107 | Young |
| 0.193951 | 1.105631 | 0.112 | 0.076 | 1 | 4 | Thap3 | 0.036 | Young |
| 0.193969 | 2.500825 | 0.025 | 0.01 | 1 | 4 | Map7d3 | 0.015 | Young |
| 0.194298 | 0.337 | 0.25 | 0.19 | 1 | 4 | Tbc1d2b | 0.06 | Young |
| 0.194351 | 0.729886 | 0.112 | 0.073 | 1 | 4 | Hsph1 | 0.039 | Young |
| 0.194931 | 1.573292 | 0.025 | 0.01 | 1 | 4 | Slc30a6 | 0.015 | Young |
| 0.194932 | 0.674813 | 0.05 | 0.026 | 1 | 4 | Pyurf | 0.024 | Young |
| 0.195384 | 0.556006 | 0.2 | 0.152 | 1 | 4 | Rftn1 | 0.048 | Young |
| 0.19539 | 0.656559 | 0.125 | 0.084 | 1 | 4 | Cd79b | 0.041 | Young |
| 0.195413 | 1.617607 | 0.025 | 0.01 | 1 | 4 | Abca2 | 0.015 | Young |
| 0.195501 | 0.446565 | 0.075 | 0.043 | 1 | 4 | Cnpy4 | 0.032 | Young |
| 0.195784 | 0.166314 | 0.088 | 0.052 | 1 | 4 | Zfp282 | 0.036 | Young |
| 0.196157 | 0.263484 | 0.05 | 0.026 | 1 | 4 | Tfdp2 | 0.024 | Young |
| 0.196304 | 0.040149 | 0.462 | 0.36 | 1 | 4 | Snrpb2 | 0.102 | Young |
| 0.196378 | 0.192332 | 0.225 | 0.161 | 1 | 4 | 2510039O1 | 0.064 | Young |
| 0.196536 | 0.789154 | 0.075 | 0.044 | 1 | 4 | Stx18 | 0.031 | Young |
| 0.196578 | 1.304503 | 0.05 | 0.026 | 1 | 4 | Cers4 | 0.024 | Young |

| | | | | | | | | |
|---|---|---|---|---|---|---|---|---|
| 0.196593 | 0.296361 | 0.225 | 0.165 | 1 | 4 | Anapc13 | 0.06 | Young |
| 0.196777 | 0.57989 | 0.138 | 0.096 | 1 | 4 | Ccdc186 | 0.042 | Young |
| 0.197322 | 0.895537 | 0.088 | 0.055 | 1 | 4 | Nat9 | 0.033 | Young |
| 0.197351 | 1.270238 | 0.025 | 0.01 | 1 | 4 | Pparg | 0.015 | Young |
| 0.197445 | 0.139585 | 0.112 | 0.071 | 1 | 4 | Mapk9 | 0.041 | Young |
| 0.197838 | 1.202563 | 0.025 | 0.01 | 1 | 4 | B4galt4 | 0.015 | Young |
| 0.198115 | 0.15895 | 0.35 | 0.262 | 1 | 4 | Golph3 | 0.088 | Young |
| 0.198489 | 0.836055 | 0.075 | 0.045 | 1 | 4 | Zfp830 | 0.03 | Young |
| 0.198789 | 0.151068 | 0.275 | 0.205 | 1 | 4 | Pank3 | 0.07 | Young |
| 0.19888 | 1.281352 | 0.038 | 0.018 | 1 | 4 | Cnst | 0.02 | Young |
| 0.199081 | 0.143902 | 0.175 | 0.123 | 1 | 4 | Zdhhc3 | 0.052 | Young |
| 0.199115 | 0.186834 | 0.388 | 0.31 | 1 | 4 | Cwc15 | 0.078 | Young |
| 0.199243 | 0.325768 | 0.05 | 0.026 | 1 | 4 | Zfp953 | 0.024 | Young |
| 0.199286 | 0.929014 | 0.162 | 0.119 | 1 | 4 | Jkamp | 0.043 | Young |
| 0.199303 | 1.134623 | 0.025 | 0.01 | 1 | 4 | Gm10762 | 0.015 | Young |
| 0.199317 | 0.355918 | 0.738 | 0.658 | 1 | 4 | Fcgr2b | 0.08 | Young |
| 0.199554 | 0.250033 | 0.05 | 0.026 | 1 | 4 | Dgcr14 | 0.024 | Young |
| 0.199794 | 0.994596 | 0.025 | 0.01 | 1 | 4 | Arap2 | 0.015 | Young |
| 0.199794 | 0.994045 | 0.025 | 0.01 | 1 | 4 | Cd28 | 0.015 | Young |
| 0.200215 | 0.715564 | 0.112 | 0.074 | 1 | 4 | Kpna6 | 0.038 | Young |
| 0.200234 | 0.268381 | 0.138 | 0.092 | 1 | 4 | Avl9 | 0.046 | Young |
| 0.200285 | 0.739794 | 0.025 | 0.01 | 1 | 4 | Nova1 | 0.015 | Young |
| 0.200731 | 0.434374 | 0.175 | 0.126 | 1 | 4 | Crat | 0.049 | Young |
| 0.200777 | 0.812032 | 0.025 | 0.01 | 1 | 4 | Pus7l | 0.015 | Young |
| 0.201007 | 0.241007 | 0.2 | 0.146 | 1 | 4 | Chd6 | 0.054 | Young |
| 0.201037 | 0.982388 | 0.1 | 0.065 | 1 | 4 | Mrpl27 | 0.035 | Young |
| 0.201084 | 0.691182 | 0.125 | 0.084 | 1 | 4 | Ppp2r3d | 0.041 | Young |
| 0.20121 | 0.274995 | 0.175 | 0.125 | 1 | 4 | Slc25a1 | 0.05 | Young |
| 0.20127 | 0.348694 | 0.025 | 0.01 | 1 | 4 | Tkfc | 0.015 | Young |
| 0.201359 | 0.257622 | 0.125 | 0.082 | 1 | 4 | Usp6nl | 0.043 | Young |
| 0.201855 | 0.318071 | 0.475 | 0.414 | 1 | 4 | Hacd4 | 0.061 | Young |
| 0.202202 | 0.072549 | 0.112 | 0.072 | 1 | 4 | Hat1 | 0.04 | Young |
| 0.202227 | 1.028748 | 0.038 | 0.018 | 1 | 4 | Ifitm5 | 0.02 | Young |
| 0.202434 | 0.200235 | 0.388 | 0.308 | 1 | 4 | Atp5c1 | 0.08 | Young |
| 0.202505 | 0.614433 | 0.088 | 0.054 | 1 | 4 | Xpo4 | 0.034 | Young |
| 0.202726 | 1.112979 | 0.05 | 0.026 | 1 | 4 | Zfp777 | 0.024 | Young |
| 0.202753 | 0.43907 | 0.025 | 0.01 | 1 | 4 | Zfp773 | 0.015 | Young |
| 0.202756 | 0.334389 | 0.138 | 0.093 | 1 | 4 | Nab2 | 0.045 | Young |
| 0.202802 | 0.655538 | 0.062 | 0.035 | 1 | 4 | Serpinb9 | 0.027 | Young |
| 0.20325 | 0.263879 | 0.025 | 0.01 | 1 | 4 | Plpp3 | 0.015 | Young |
| 0.203775 | 0.437625 | 0.075 | 0.044 | 1 | 4 | Mynn | 0.031 | Young |
| 0.204102 | 0.163023 | 0.275 | 0.205 | 1 | 4 | Hsd17b12 | 0.07 | Young |
| 0.204372 | 1.091224 | 0.075 | 0.045 | 1 | 4 | Gmpr2 | 0.03 | Young |
| 0.204615 | 0.378827 | 0.125 | 0.083 | 1 | 4 | Ralgapa1 | 0.042 | Young |
| 0.204713 | 0.516195 | 0.062 | 0.035 | 1 | 4 | Med19 | 0.027 | Young |

| | | | | | | | | |
|---|---|---|---|---|---|---|---|---|
| 0.204865 | 0.373844 | 0.175 | 0.126 | 1 | 4 | 4930523C0 | 0.049 | Young |
| 0.204872 | 0.403226 | 0.35 | 0.298 | 1 | 4 | D8Ertd738 | 0.052 | Young |
| 0.204927 | 0.215327 | 0.125 | 0.082 | 1 | 4 | Fam193b | 0.043 | Young |
| 0.205041 | 0.460716 | 0.15 | 0.105 | 1 | 4 | Zfp90 | 0.045 | Young |
| 0.20513 | 0.069441 | 0.1 | 0.063 | 1 | 4 | Bhlhe40 | 0.037 | Young |
| 0.205248 | 0.944666 | 0.088 | 0.055 | 1 | 4 | E4f1 | 0.033 | Young |
| 0.205664 | 0.202233 | 0.125 | 0.082 | 1 | 4 | Lpcat3 | 0.043 | Young |
| 0.205762 | 0.524226 | 0.125 | 0.084 | 1 | 4 | Mmadhc | 0.041 | Young |
| 0.205966 | 0.198232 | 0.1 | 0.063 | 1 | 4 | Hira | 0.037 | Young |
| 0.206247 | 0.05822 | 0.025 | 0.01 | 1 | 4 | Rpp38 | 0.015 | Young |
| 0.20685 | 0.212741 | 0.162 | 0.113 | 1 | 4 | Atmin | 0.049 | Young |
| 0.207212 | 0.819422 | 0.062 | 0.036 | 1 | 4 | Zmat3 | 0.026 | Young |
| 0.207515 | 0.66324 | 0.038 | 0.018 | 1 | 4 | Mios | 0.02 | Young |
| 0.208378 | 0.689782 | 0.05 | 0.026 | 1 | 4 | H2afx | 0.024 | Young |
| 0.209026 | 0.416049 | 0.15 | 0.105 | 1 | 4 | Pitpnb | 0.045 | Young |
| 0.209044 | 0.660855 | 0.038 | 0.018 | 1 | 4 | 1700008J0 | 0.02 | Young |
| 0.209129 | 0.417891 | 0.062 | 0.035 | 1 | 4 | Zfyve19 | 0.027 | Young |
| 0.209204 | 0.246875 | 0.738 | 0.724 | 1 | 4 | Cyba | 0.014 | Young |
| 0.209436 | 0.627435 | 0.212 | 0.165 | 1 | 4 | Klhl5 | 0.047 | Young |
| 0.209697 | 0.357531 | 0.138 | 0.094 | 1 | 4 | Akap8l | 0.044 | Young |
| 0.209929 | 0.060279 | 0.388 | 0.294 | 1 | 4 | Adam17 | 0.094 | Young |
| 0.210196 | 0.627302 | 0.038 | 0.018 | 1 | 4 | Fbxo32 | 0.02 | Young |
| 0.210354 | 0.512241 | 0.112 | 0.074 | 1 | 4 | Zbtb7b | 0.038 | Young |
| 0.210382 | 0.604654 | 0.1 | 0.065 | 1 | 4 | Kansl1l | 0.035 | Young |
| 0.210581 | 0.303139 | 0.038 | 0.018 | 1 | 4 | Gm20045 | 0.02 | Young |
| 0.210795 | 0.569209 | 0.162 | 0.115 | 1 | 4 | Dpagt1 | 0.047 | Young |
| 0.210824 | 0.09958 | 0.1 | 0.063 | 1 | 4 | Rasal2 | 0.037 | Young |
| 0.211173 | 0.650622 | 0.15 | 0.108 | 1 | 4 | Zfp992 | 0.042 | Young |
| 0.211302 | 0.267898 | 0.35 | 0.285 | 1 | 4 | Numa1 | 0.065 | Young |
| 0.211353 | 0.442079 | 0.038 | 0.018 | 1 | 4 | Harbi1 | 0.02 | Young |
| 0.211353 | 0.166585 | 0.038 | 0.018 | 1 | 4 | Zfp668 | 0.02 | Young |
| 0.211609 | 0.331611 | 0.2 | 0.149 | 1 | 4 | Slc48a1 | 0.051 | Young |
| 0.211616 | 0.535558 | 0.175 | 0.128 | 1 | 4 | Slc30a7 | 0.047 | Young |
| 0.21167 | 0.166536 | 0.225 | 0.165 | 1 | 4 | Camk2d | 0.06 | Young |
| 0.21196 | 0.723033 | 0.1 | 0.065 | 1 | 4 | Tmem161a | 0.035 | Young |
| 0.21232 | 0.417914 | 0.15 | 0.105 | 1 | 4 | Shprh | 0.045 | Young |
| 0.212409 | 0.451139 | 0.162 | 0.115 | 1 | 4 | 2310011J0 | 0.047 | Young |
| 0.212457 | 0.382741 | 0.138 | 0.094 | 1 | 4 | Dusp16 | 0.044 | Young |
| 0.212761 | 0.641691 | 0.112 | 0.075 | 1 | 4 | Chmp7 | 0.037 | Young |
| 0.212768 | 0.276238 | 0.062 | 0.035 | 1 | 4 | Cdk20 | 0.027 | Young |
| 0.213167 | 0.613032 | 0.075 | 0.045 | 1 | 4 | Med24 | 0.03 | Young |
| 0.213523 | 0.246904 | 0.288 | 0.223 | 1 | 4 | Pan3 | 0.065 | Young |
| 0.213615 | 0.136435 | 0.062 | 0.035 | 1 | 4 | Mllt1 | 0.027 | Young |
| 0.214181 | 0.20237 | 0.312 | 0.247 | 1 | 4 | Prpf4b | 0.065 | Young |
| 0.214273 | 0.113841 | 0.15 | 0.103 | 1 | 4 | Psmd12 | 0.047 | Young |

| | | | | | | | | |
|---|---|---|---|---|---|---|---|---|
| 0.214402 | 0.501233 | 0.138 | 0.096 | 1 | 4 | Foxk1 | 0.042 | Young |
| 0.214464 | 0.165252 | 0.388 | 0.318 | 1 | 4 | Polr1d | 0.07 | Young |
| 0.214493 | 0.285366 | 0.238 | 0.183 | 1 | 4 | Pcf11 | 0.055 | Young |
| 0.214641 | 1.290198 | 0.05 | 0.027 | 1 | 4 | Rrp9 | 0.023 | Young |
| 0.214791 | 0.336381 | 0.2 | 0.144 | 1 | 4 | Brox | 0.056 | Young |
| 0.215226 | 0.482797 | 0.2 | 0.148 | 1 | 4 | Hadha | 0.052 | Young |
| 0.215237 | 0.261306 | 0.225 | 0.17 | 1 | 4 | Gpalpp1 | 0.055 | Young |
| 0.215494 | 0.458986 | 0.312 | 0.26 | 1 | 4 | Crebrf | 0.052 | Young |
| 0.216523 | 0.375788 | 0.075 | 0.044 | 1 | 4 | Usp10 | 0.031 | Young |
| 0.216523 | 0.081604 | 0.075 | 0.044 | 1 | 4 | Dhx29 | 0.031 | Young |
| 0.216742 | 0.206663 | 0.05 | 0.026 | 1 | 4 | Tmem101 | 0.024 | Young |
| 0.216893 | 1.111454 | 0.05 | 0.027 | 1 | 4 | E330009J0 | 0.023 | Young |
| 0.217028 | 0.491906 | 0.212 | 0.162 | 1 | 4 | Gpat4 | 0.05 | Young |
| 0.217349 | 0.108598 | 0.238 | 0.177 | 1 | 4 | Bmt2 | 0.061 | Young |
| 0.217412 | 0.284625 | 0.312 | 0.255 | 1 | 4 | Os9 | 0.057 | Young |
| 0.217464 | 0.612802 | 0.075 | 0.045 | 1 | 4 | Alg9 | 0.03 | Young |
| 0.217819 | 0.363665 | 0.238 | 0.182 | 1 | 4 | Anapc5 | 0.056 | Young |
| 0.217925 | 0.636486 | 0.138 | 0.097 | 1 | 4 | Acsl1 | 0.041 | Young |
| 0.217962 | 0.348114 | 0.2 | 0.148 | 1 | 4 | Safb | 0.052 | Young |
| 0.218095 | 0.06471 | 0.4 | 0.318 | 1 | 4 | Ube2n | 0.082 | Young |
| 0.218169 | 0.070133 | 0.787 | 0.699 | 1 | 4 | Rpl14 | 0.088 | Young |
| 0.219006 | 0.444726 | 0.175 | 0.126 | 1 | 4 | Snap29 | 0.049 | Young |
| 0.219218 | 0.613827 | 0.162 | 0.117 | 1 | 4 | Iws1 | 0.045 | Young |
| 0.219385 | 0.208479 | 0.275 | 0.213 | 1 | 4 | AI314180 | 0.062 | Young |
| 0.219875 | 0.224643 | 0.325 | 0.264 | 1 | 4 | Pycard | 0.061 | Young |
| 0.219916 | 0.416064 | 0.138 | 0.095 | 1 | 4 | Dmtf1 | 0.043 | Young |
| 0.220126 | 0.375579 | 0.112 | 0.074 | 1 | 4 | Rcor3 | 0.038 | Young |
| 0.220512 | 0.166366 | 0.262 | 0.198 | 1 | 4 | Tmem9b | 0.064 | Young |
| 0.220549 | 0.999053 | 0.062 | 0.036 | 1 | 4 | Usp11 | 0.026 | Young |
| 0.220606 | 0.19956 | 0.462 | 0.383 | 1 | 4 | Pnisr | 0.079 | Young |
| 0.220621 | 0.305506 | 0.275 | 0.217 | 1 | 4 | Qdpr | 0.058 | Young |
| 0.220738 | 0.1689 | 0.35 | 0.281 | 1 | 4 | Neu1 | 0.069 | Young |
| 0.220965 | 0.06778 | 0.375 | 0.293 | 1 | 4 | Aff4 | 0.082 | Young |
| 0.221078 | 0.720416 | 0.138 | 0.097 | 1 | 4 | Acvrl1 | 0.041 | Young |
| 0.221113 | 0.157553 | 0.162 | 0.114 | 1 | 4 | Parp4 | 0.048 | Young |
| 0.221127 | 0.286854 | 0.312 | 0.25 | 1 | 4 | Dram2 | 0.062 | Young |
| 0.221634 | 0.650474 | 0.275 | 0.233 | 1 | 4 | Stat6 | 0.042 | Young |
| 0.22175 | 0.701726 | 0.125 | 0.085 | 1 | 4 | Nelfcd | 0.04 | Young |
| 0.222374 | 0.300797 | 0.212 | 0.159 | 1 | 4 | Ranbp1 | 0.053 | Young |
| 0.222398 | 0.435475 | 0.062 | 0.036 | 1 | 4 | Bnip1 | 0.026 | Young |
| 0.222432 | 0.907095 | 0.05 | 0.027 | 1 | 4 | Dusp12 | 0.023 | Young |
| 0.222454 | 0.253259 | 0.088 | 0.054 | 1 | 4 | Carm1 | 0.034 | Young |
| 0.222695 | 0.361541 | 0.162 | 0.115 | 1 | 4 | Stam2 | 0.047 | Young |
| 0.222769 | 0.266829 | 0.188 | 0.136 | 1 | 4 | Sh3gl1 | 0.052 | Young |
| 0.223091 | 0.862517 | 0.05 | 0.027 | 1 | 4 | Supt3 | 0.023 | Young |

| | | | | | | | | |
|---|---|---|---|---|---|---|---|---|
| 0.223404 | 1.042014 | 0.112 | 0.076 | 1 | 4 | Cwc25 | 0.036 | Young |
| 0.223745 | 0.425 | 0.238 | 0.186 | 1 | 4 | Kmt2d | 0.052 | Young |
| 0.223981 | 0.904598 | 0.062 | 0.036 | 1 | 4 | Timm8a1 | 0.026 | Young |
| 0.223991 | 0.183878 | 0.162 | 0.114 | 1 | 4 | Siglec1 | 0.048 | Young |
| 0.223999 | 0.08282 | 0.088 | 0.054 | 1 | 4 | Rbm4b | 0.034 | Young |
| 0.22441 | 0.406597 | 0.175 | 0.127 | 1 | 4 | Mast3 | 0.048 | Young |
| 0.224678 | 0.357143 | 0.075 | 0.045 | 1 | 4 | Gm20342 | 0.03 | Young |
| 0.224828 | 1.351274 | 0.038 | 0.018 | 1 | 4 | Zfp974 | 0.02 | Young |
| 0.225082 | 1.055253 | 0.062 | 0.037 | 1 | 4 | Alg8 | 0.025 | Young |
| 0.225256 | 0.058013 | 0.112 | 0.073 | 1 | 4 | Abcd3 | 0.039 | Young |
| 0.225889 | 0.262201 | 0.175 | 0.126 | 1 | 4 | Mrps9 | 0.049 | Young |
| 0.226023 | 1.069661 | 0.038 | 0.018 | 1 | 4 | Mto1 | 0.02 | Young |
| 0.226031 | 0.012733 | 0.088 | 0.054 | 1 | 4 | Adi1 | 0.034 | Young |
| 0.226399 | 2.127734 | 0.025 | 0.011 | 1 | 4 | Cep19 | 0.014 | Young |
| 0.226403 | 0.755589 | 0.05 | 0.027 | 1 | 4 | Mrrf | 0.023 | Young |
| 0.226692 | 0.141865 | 0.725 | 0.718 | 1 | 4 | Rpl36a | 0.007 | Young |
| 0.22672 | 0.16358 | 0.188 | 0.134 | 1 | 4 | Fbxl20 | 0.054 | Young |
| 0.227341 | 0.135363 | 0.7 | 0.65 | 1 | 4 | Calr | 0.05 | Young |
| 0.227356 | 0.088973 | 0.175 | 0.123 | 1 | 4 | Tmem170b | 0.052 | Young |
| 0.227531 | 0.429377 | 0.112 | 0.075 | 1 | 4 | Tapbpl | 0.037 | Young |
| 0.227542 | 0.118037 | 0.038 | 0.076 | 1 | 4 | Urod | -0.038 | Young |
| 0.227658 | 0.439309 | 0.475 | 0.424 | 1 | 4 | Dnaja1 | 0.051 | Young |
| 0.227676 | 0.207655 | 0.55 | 0.477 | 1 | 4 | Prrc2c | 0.073 | Young |
| 0.227934 | 0.155485 | 0.288 | 0.225 | 1 | 4 | 2810474O1 | 0.063 | Young |
| 0.227969 | 2.099718 | 0.025 | 0.011 | 1 | 4 | Catip | 0.014 | Young |
| 0.228068 | 0.133195 | 0.55 | 0.464 | 1 | 4 | 2610507B1 | 0.086 | Young |
| 0.228422 | 0.378344 | 0.225 | 0.175 | 1 | 4 | Pgd | 0.05 | Young |
| 0.228426 | 0.834999 | 0.038 | 0.018 | 1 | 4 | Msh3 | 0.02 | Young |
| 0.228461 | 0.430704 | 0.175 | 0.128 | 1 | 4 | Aip | 0.047 | Young |
| 0.228491 | 0.910549 | 0.075 | 0.046 | 1 | 4 | Tcp11l1 | 0.029 | Young |
| 0.228494 | 3.557858 | 0.025 | 0.011 | 1 | 4 | Cxcl9 | 0.014 | Young |
| 0.228513 | 0.136597 | 0.062 | 0.036 | 1 | 4 | Cep78 | 0.026 | Young |
| 0.22862 | 0.804116 | 0.1 | 0.067 | 1 | 4 | Stard3 | 0.033 | Young |
| 0.228806 | 0.113965 | 0.362 | 0.288 | 1 | 4 | Usp7 | 0.074 | Young |
| 0.228808 | 0.074846 | 0.062 | 0.036 | 1 | 4 | Wdr46 | 0.026 | Young |
| 0.228812 | 0.837756 | 0.112 | 0.078 | 1 | 4 | Opa3 | 0.034 | Young |
| 0.228828 | 0.780457 | 0.038 | 0.018 | 1 | 4 | Rnf123 | 0.02 | Young |
| 0.229231 | 0.792391 | 0.038 | 0.018 | 1 | 4 | 5430403G1 | 0.02 | Young |
| 0.229231 | 0.752544 | 0.038 | 0.018 | 1 | 4 | Plcg1 | 0.02 | Young |
| 0.229321 | 0.123128 | 0.2 | 0.144 | 1 | 4 | Slc28a2 | 0.056 | Young |
| 0.229438 | 0.196384 | 0.15 | 0.105 | 1 | 4 | Sptan1 | 0.045 | Young |
| 0.229508 | 0.18866 | 0.188 | 0.136 | 1 | 4 | Sh3pxd2a | 0.052 | Young |
| 0.229634 | 0.720624 | 0.038 | 0.018 | 1 | 4 | A230050P2 | 0.02 | Young |
| 0.229847 | 0.287321 | 0.65 | 0.596 | 1 | 4 | Gns | 0.054 | Young |
| 0.229872 | 0.235717 | 0.262 | 0.202 | 1 | 4 | Denr | 0.06 | Young |

| | | | | | | | | |
|---|---|---|---|---|---|---|---|---|
| 0.229992 | 0.446895 | 0.238 | 0.18 | 1 | 4 | Chchd1 | 0.058 | Young |
| 0.230045 | 0.212088 | 0.088 | 0.055 | 1 | 4 | Adnp2 | 0.033 | Young |
| 0.230074 | 1.198265 | 0.025 | 0.011 | 1 | 4 | Xrcc2 | 0.014 | Young |
| 0.230074 | 1.154159 | 0.025 | 0.011 | 1 | 4 | Zfp213 | 0.014 | Young |
| 0.230077 | 0.559533 | 0.062 | 0.036 | 1 | 4 | D17H6S53I | 0.026 | Young |
| 0.230245 | 0.343805 | 0.2 | 0.149 | 1 | 4 | Pdha1 | 0.051 | Young |
| 0.2305 | 0.346365 | 0.138 | 0.096 | 1 | 4 | Mpi | 0.042 | Young |
| 0.230826 | 0.467725 | 0.112 | 0.076 | 1 | 4 | Sh3glb2 | 0.036 | Young |
| 0.231132 | 1.104223 | 0.025 | 0.011 | 1 | 4 | Bcs1l | 0.014 | Young |
| 0.231132 | 1.089591 | 0.025 | 0.011 | 1 | 4 | Dffb | 0.014 | Young |
| 0.231132 | 1.066302 | 0.025 | 0.011 | 1 | 4 | Herc3 | 0.014 | Young |
| 0.231471 | 0.271983 | 0.375 | 0.31 | 1 | 4 | Map4k4 | 0.065 | Young |
| 0.231658 | 0.620246 | 0.038 | 0.018 | 1 | 4 | B230307C2 | 0.02 | Young |
| 0.231662 | 1.080428 | 0.025 | 0.011 | 1 | 4 | 3-Mar | 0.014 | Young |
| 0.231662 | 1.026334 | 0.025 | 0.011 | 1 | 4 | Eaf2 | 0.014 | Young |
| 0.231831 | 0.1142 | 0.975 | 0.953 | 1 | 4 | Cltc | 0.022 | Young |
| 0.23191 | 1.158003 | 0.125 | 0.09 | 1 | 4 | Rfc2 | 0.035 | Young |
| 0.231972 | 0.397455 | 0.25 | 0.2 | 1 | 4 | Slc25a39 | 0.05 | Young |
| 0.232152 | 0.263029 | 0.2 | 0.148 | 1 | 4 | Kbtbd2 | 0.052 | Young |
| 0.232402 | 0.149104 | 0.162 | 0.115 | 1 | 4 | Parp8 | 0.047 | Young |
| 0.232662 | 0.203775 | 0.188 | 0.138 | 1 | 4 | Dennd4b | 0.05 | Young |
| 0.232725 | 0.817444 | 0.025 | 0.011 | 1 | 4 | Gins2 | 0.014 | Young |
| 0.232725 | 0.731953 | 0.025 | 0.011 | 1 | 4 | Gm27003 | 0.014 | Young |
| 0.232776 | 0.11648 | 0.15 | 0.105 | 1 | 4 | Ppfia1 | 0.045 | Young |
| 0.232868 | 0.417692 | 0.25 | 0.197 | 1 | 4 | Zfp445 | 0.053 | Young |
| 0.23303 | 0.143287 | 0.112 | 0.074 | 1 | 4 | Ugdh | 0.038 | Young |
| 0.233352 | 0.91868 | 0.15 | 0.109 | 1 | 4 | Trabd | 0.041 | Young |
| 0.233614 | 0.298873 | 0.062 | 0.036 | 1 | 4 | Slc7a6os | 0.026 | Young |
| 0.233816 | 0.050696 | 0.2 | 0.144 | 1 | 4 | Mbd3 | 0.056 | Young |
| 0.233856 | 0.34286 | 0.225 | 0.171 | 1 | 4 | Med15 | 0.054 | Young |
| 0.233982 | 0.357803 | 0.075 | 0.045 | 1 | 4 | Ank2 | 0.03 | Young |
| 0.234086 | 0.30044 | 0.138 | 0.096 | 1 | 4 | Timm44 | 0.042 | Young |
| 0.234326 | 0.768899 | 0.025 | 0.011 | 1 | 4 | Adnp | 0.014 | Young |
| 0.234326 | 0.717566 | 0.025 | 0.011 | 1 | 4 | Cptp | 0.014 | Young |
| 0.234326 | 0.475984 | 0.025 | 0.011 | 1 | 4 | Sirt4 | 0.014 | Young |
| 0.234524 | 1.103539 | 0.062 | 0.037 | 1 | 4 | Igtp | 0.025 | Young |
| 0.234765 | 0.431871 | 0.138 | 0.096 | 1 | 4 | Taf13 | 0.042 | Young |
| 0.234861 | 0.724994 | 0.025 | 0.011 | 1 | 4 | Haus5 | 0.014 | Young |
| 0.23522 | 0.137493 | 0.275 | 0.207 | 1 | 4 | Rdx | 0.068 | Young |
| 0.235398 | 0.539776 | 0.025 | 0.011 | 1 | 4 | Tmem67 | 0.014 | Young |
| 0.235502 | 0.132623 | 0.2 | 0.146 | 1 | 4 | Mtch2 | 0.054 | Young |
| 0.235935 | 0.478444 | 0.025 | 0.011 | 1 | 4 | Tfb1m | 0.014 | Young |
| 0.236154 | 0.165018 | 0.038 | 0.018 | 1 | 4 | Plxna3 | 0.02 | Young |
| 0.236165 | 0.405058 | 0.462 | 0.37 | 1 | 4 | Coro1a | 0.092 | Young |
| 0.236389 | 0.385343 | 0.075 | 0.045 | 1 | 4 | Tars2 | 0.03 | Young |

| | | | | | | | | |
|---|---|---|---|---|---|---|---|---|
| 0.236566 | 0.487389 | 0.038 | 0.018 | 1 | 4 | 4930522L1 | 0.02 | Young |
| 0.236934 | 0.518723 | 0.075 | 0.046 | 1 | 4 | Trim23 | 0.029 | Young |
| 0.237374 | 0.088055 | 0.088 | 0.055 | 1 | 4 | Slc26a11 | 0.033 | Young |
| 0.237469 | 0.764537 | 0.075 | 0.046 | 1 | 4 | Spidr | 0.029 | Young |
| 0.237551 | 0.134526 | 0.025 | 0.011 | 1 | 4 | Apoo | 0.014 | Young |
| 0.237762 | 0.570824 | 0.1 | 0.067 | 1 | 4 | Hnmt | 0.033 | Young |
| 0.237791 | 1.004015 | 0.05 | 0.028 | 1 | 4 | Tpgs2 | 0.022 | Young |
| 0.238005 | 0.863847 | 0.162 | 0.124 | 1 | 4 | Nuak2 | 0.038 | Young |
| 0.238303 | 0.960703 | 0.05 | 0.028 | 1 | 4 | Usp46 | 0.022 | Young |
| 0.238357 | 0.205924 | 0.188 | 0.137 | 1 | 4 | Tgfbrap1 | 0.051 | Young |
| 0.238364 | 0.19598 | 0.088 | 0.055 | 1 | 4 | Alg14 | 0.033 | Young |
| 0.238716 | 0.184753 | 0.275 | 0.208 | 1 | 4 | Sltm | 0.067 | Young |
| 0.239295 | 0.696405 | 0.15 | 0.112 | 1 | 4 | Tpst2 | 0.038 | Young |
| 0.239337 | 0.949395 | 0.138 | 0.1 | 1 | 4 | Ten1 | 0.038 | Young |
| 0.239682 | 0.367228 | 0.15 | 0.107 | 1 | 4 | Kpna1 | 0.043 | Young |
| 0.239719 | 0.011469 | 0.025 | 0.011 | 1 | 4 | Zfp933 | 0.014 | Young |
| 0.240309 | 1.690023 | 0.05 | 0.028 | 1 | 4 | Glipr2 | 0.022 | Young |
| 0.240586 | 0.106624 | 0.138 | 0.094 | 1 | 4 | Pigz | 0.044 | Young |
| 0.240712 | 0.010197 | 0.038 | 0.018 | 1 | 4 | Exo5 | 0.02 | Young |
| 0.24104 | 0.376264 | 0.175 | 0.128 | 1 | 4 | Nap1l4 | 0.047 | Young |
| 0.24108 | 0.474548 | 0.2 | 0.155 | 1 | 4 | Golgb1 | 0.045 | Young |
| 0.241107 | 0.118858 | 0.738 | 0.691 | 1 | 4 | Ybx1 | 0.047 | Young |
| 0.241362 | 0.348092 | 0.2 | 0.15 | 1 | 4 | Slain2 | 0.05 | Young |
| 0.241374 | 0.190925 | 0.212 | 0.159 | 1 | 4 | Myo1c | 0.053 | Young |
| 0.241427 | 0.218599 | 0.175 | 0.127 | 1 | 4 | Fam114a2 | 0.048 | Young |
| 0.242095 | 0.634648 | 0.175 | 0.134 | 1 | 4 | Ahr | 0.041 | Young |
| 0.242178 | 0.311394 | 0.088 | 0.055 | 1 | 4 | Rpf2 | 0.033 | Young |
| 0.243327 | 0.259629 | 0.212 | 0.162 | 1 | 4 | Mknk1 | 0.05 | Young |
| 0.243949 | 0.379347 | 0.812 | 0.839 | 1 | 4 | Ccl8 | -0.027 | Young |
| 0.244322 | 0.147544 | 0.112 | 0.075 | 1 | 4 | Stx17 | 0.037 | Young |
| 0.244438 | 0.25111 | 0.1 | 0.066 | 1 | 4 | Trpv2 | 0.034 | Young |
| 0.244504 | 1.250597 | 0.038 | 0.019 | 1 | 4 | Mki67 | 0.019 | Young |
| 0.244728 | 0.436682 | 0.088 | 0.056 | 1 | 4 | Mpg | 0.032 | Young |
| 0.244895 | 0.680124 | 0.225 | 0.179 | 1 | 4 | Mepce | 0.046 | Young |
| 0.244949 | 0.239388 | 0.1 | 0.065 | 1 | 4 | Lix1l | 0.035 | Young |
| 0.245291 | 0.115637 | 0.2 | 0.149 | 1 | 4 | Bcl6 | 0.051 | Young |
| 0.24561 | 0.591561 | 0.062 | 0.037 | 1 | 4 | Efl1 | 0.025 | Young |
| 0.245757 | 0.656983 | 0.088 | 0.057 | 1 | 4 | Rtl8a | 0.031 | Young |
| 0.245879 | 0.495715 | 0.175 | 0.132 | 1 | 4 | Supt20 | 0.043 | Young |
| 0.245996 | 0.233572 | 0.537 | 0.442 | 1 | 4 | Tmed7 | 0.095 | Young |
| 0.246749 | 0.10839 | 0.075 | 0.045 | 1 | 4 | Plrg1 | 0.03 | Young |
| 0.246926 | 0.304113 | 0.138 | 0.096 | 1 | 4 | Pdgfb | 0.042 | Young |
| 0.247235 | 0.442351 | 0.088 | 0.056 | 1 | 4 | Lonp2 | 0.032 | Young |
| 0.247391 | 0.530327 | 0.1 | 0.067 | 1 | 4 | Tbc1d31 | 0.033 | Young |
| 0.247699 | 0.224411 | 0.375 | 0.314 | 1 | 4 | Abracl | 0.061 | Young |

| | | | | | | | | |
|---|---|---|---|---|---|---|---|---|
| 0.24798 | 0.511669 | 0.125 | 0.088 | 1 | 4 | Arfgef2 | 0.037 | Young |
| 0.248458 | 0.457551 | 0.388 | 0.326 | 1 | 4 | Zcchc11 | 0.062 | Young |
| 0.248553 | 0.662116 | 0.162 | 0.119 | 1 | 4 | Slc25a10 | 0.043 | Young |
| 0.248646 | 0.126535 | 0.15 | 0.105 | 1 | 4 | Lcor | 0.045 | Young |
| 0.248681 | 1.100632 | 0.038 | 0.019 | 1 | 4 | Nedd4l | 0.019 | Young |
| 0.248681 | 1.087775 | 0.038 | 0.019 | 1 | 4 | Atp5s | 0.019 | Young |
| 0.249359 | 0.618747 | 0.075 | 0.047 | 1 | 4 | Gm45902 | 0.028 | Young |
| 0.249523 | 0.95222 | 0.038 | 0.019 | 1 | 4 | Myom1 | 0.019 | Young |
| 0.249563 | 0.192416 | 0.125 | 0.084 | 1 | 4 | Blcap | 0.041 | Young |
| 0.249944 | 0.913704 | 0.038 | 0.019 | 1 | 4 | 6330418K( | 0.019 | Young |
| 0.250188 | 0.581075 | 0.112 | 0.078 | 1 | 4 | Cog8 | 0.034 | Young |
| 0.250789 | 0.908546 | 0.038 | 0.019 | 1 | 4 | Gm26518 | 0.019 | Young |
| 0.251635 | 0.945571 | 0.038 | 0.019 | 1 | 4 | Tbc1d22b | 0.019 | Young |
| 0.252118 | 0.281077 | 0.188 | 0.14 | 1 | 4 | Enox2 | 0.048 | Young |
| 0.252338 | 0.611782 | 0.25 | 0.21 | 1 | 4 | Rnf145 | 0.04 | Young |
| 0.25234 | 0.686066 | 0.15 | 0.111 | 1 | 4 | Nup153 | 0.039 | Young |
| 0.252484 | 0.777844 | 0.038 | 0.019 | 1 | 4 | Mfsd4b4 | 0.019 | Young |
| 0.252598 | 0.45381 | 0.238 | 0.192 | 1 | 4 | Ssr4 | 0.046 | Young |
| 0.253105 | 0.272135 | 0.212 | 0.162 | 1 | 4 | Fads1 | 0.05 | Young |
| 0.253305 | 0.227234 | 0.062 | 0.037 | 1 | 4 | Tlcd2 | 0.025 | Young |
| 0.253449 | 0.743652 | 0.062 | 0.038 | 1 | 4 | Tcta | 0.024 | Young |
| 0.254613 | 0.574709 | 0.038 | 0.019 | 1 | 4 | Cd209c | 0.019 | Young |
| 0.254981 | 0.648109 | 0.162 | 0.121 | 1 | 4 | Pld1 | 0.041 | Young |
| 0.255075 | 0.132337 | 0.3 | 0.232 | 1 | 4 | Ftl1-ps1 | 0.068 | Young |
| 0.255333 | 0.247973 | 0.275 | 0.213 | 1 | 4 | Nhsl2 | 0.062 | Young |
| 0.255343 | 0.785417 | 0.112 | 0.079 | 1 | 4 | Chchd5 | 0.033 | Young |
| 0.255674 | 0.147585 | 0.25 | 0.193 | 1 | 4 | Slamf7 | 0.057 | Young |
| 0.25582 | 0.186607 | 0.05 | 0.028 | 1 | 4 | B230118H( | 0.022 | Young |
| 0.256239 | 0.663986 | 0.062 | 0.038 | 1 | 4 | Zfp608 | 0.024 | Young |
| 0.25634 | 0.273695 | 0.112 | 0.076 | 1 | 4 | Prkab2 | 0.036 | Young |
| 0.25706 | 2.124937 | 0.013 | 0.004 | 1 | 4 | Gm45449 | 0.009 | Young |
| 0.25706 | 1.606387 | 0.013 | 0.004 | 1 | 4 | Rnls | 0.009 | Young |
| 0.25706 | 1.542389 | 0.013 | 0.004 | 1 | 4 | Gm9484 | 0.009 | Young |
| 0.257185 | 0.391225 | 0.038 | 0.019 | 1 | 4 | Sh3bp4 | 0.019 | Young |
| 0.257348 | 0.162715 | 0.9 | 0.831 | 1 | 4 | Rps13 | 0.069 | Young |
| 0.257456 | 0.221518 | 0.288 | 0.225 | 1 | 4 | Fam129a | 0.063 | Young |
| 0.257974 | 1.933071 | 0.013 | 0.004 | 1 | 4 | Gm45774 | 0.009 | Young |
| 0.257974 | 1.917889 | 0.013 | 0.004 | 1 | 4 | Gm33023 | 0.009 | Young |
| 0.257974 | 1.807196 | 0.013 | 0.004 | 1 | 4 | Gm42726 | 0.009 | Young |
| 0.257974 | 1.672793 | 0.013 | 0.004 | 1 | 4 | Ctla2a | 0.009 | Young |
| 0.257974 | 1.549863 | 0.013 | 0.004 | 1 | 4 | Gm26947 | 0.009 | Young |
| 0.257974 | 1.50212 | 0.013 | 0.004 | 1 | 4 | 1700020D( | 0.009 | Young |
| 0.257974 | 1.068433 | 0.013 | 0.004 | 1 | 4 | Pigl | 0.009 | Young |
| 0.258005 | 0.429372 | 0.212 | 0.166 | 1 | 4 | Syne2 | 0.046 | Young |
| 0.258047 | 0.38745 | 0.038 | 0.019 | 1 | 4 | 2900076A( | 0.019 | Young |

| | | | | | | | | |
|---|---|---|---|---|---|---|---|---|
| 0.258889 | 1.688004 | 0.013 | 0.004 | 1 | 4 | Akr1b7 | 0.009 | Young |
| 0.258889 | 1.2418 | 0.013 | 0.004 | 1 | 4 | Zfp28 | 0.009 | Young |
| 0.258889 | 1.184747 | 0.013 | 0.004 | 1 | 4 | A730063M | 0.009 | Young |
| 0.258889 | 0.844377 | 0.013 | 0.004 | 1 | 4 | Pim2 | 0.009 | Young |
| 0.258889 | 0.718928 | 0.013 | 0.004 | 1 | 4 | Dchs1 | 0.009 | Young |
| 0.258889 | 0.629565 | 0.013 | 0.004 | 1 | 4 | Gm15232 | 0.009 | Young |
| 0.25891 | 0.389641 | 0.038 | 0.019 | 1 | 4 | Sirt3 | 0.019 | Young |
| 0.259383 | 0.429258 | 0.288 | 0.234 | 1 | 4 | Diaph2 | 0.054 | Young |
| 0.259448 | 0.11358 | 0.55 | 0.507 | 1 | 4 | Reep5 | 0.043 | Young |
| 0.259581 | 0.284475 | 0.062 | 0.037 | 1 | 4 | Ric8b | 0.025 | Young |
| 0.259776 | 0.082555 | 0.038 | 0.019 | 1 | 4 | Pdk4 | 0.019 | Young |
| 0.259807 | 1.262991 | 0.013 | 0.004 | 1 | 4 | Olfr433 | 0.009 | Young |
| 0.259807 | 1.120255 | 0.013 | 0.004 | 1 | 4 | Zkscan4 | 0.009 | Young |
| 0.259807 | 1.1202 | 0.013 | 0.004 | 1 | 4 | Dnaja4 | 0.009 | Young |
| 0.259807 | 1.110939 | 0.013 | 0.004 | 1 | 4 | Lrrc32 | 0.009 | Young |
| 0.259807 | 1.00616 | 0.013 | 0.004 | 1 | 4 | Prps1l3 | 0.009 | Young |
| 0.259807 | 0.937126 | 0.013 | 0.004 | 1 | 4 | Pcolce2 | 0.009 | Young |
| 0.259807 | 0.80657 | 0.013 | 0.004 | 1 | 4 | Nup210l | 0.009 | Young |
| 0.259807 | 0.794322 | 0.013 | 0.004 | 1 | 4 | Pkp3 | 0.009 | Young |
| 0.259807 | 0.772271 | 0.013 | 0.004 | 1 | 4 | Slc16a14 | 0.009 | Young |
| 0.259807 | 0.707816 | 0.013 | 0.004 | 1 | 4 | Slc25a43 | 0.009 | Young |
| 0.259807 | 0.700637 | 0.013 | 0.004 | 1 | 4 | Prag1 | 0.009 | Young |
| 0.259807 | 0.660511 | 0.013 | 0.004 | 1 | 4 | BC055324 | 0.009 | Young |
| 0.259812 | 0.409617 | 0.388 | 0.32 | 1 | 4 | Ptp4a1 | 0.068 | Young |
| 0.259985 | 0.091367 | 0.112 | 0.076 | 1 | 4 | 2810004N2 | 0.036 | Young |
| 0.260003 | 0.270909 | 0.2 | 0.152 | 1 | 4 | Sike1 | 0.048 | Young |
| 0.2601 | 1.872413 | 0.025 | 0.011 | 1 | 4 | Eml6 | 0.014 | Young |
| 0.260168 | 0.179538 | 0.138 | 0.097 | 1 | 4 | Rbmxl1 | 0.041 | Young |
| 0.260209 | 0.389682 | 0.038 | 0.019 | 1 | 4 | Rtl6 | 0.019 | Young |
| 0.260686 | 0.2662 | 0.25 | 0.196 | 1 | 4 | Slc4a7 | 0.054 | Young |
| 0.260727 | 0.84605 | 0.013 | 0.004 | 1 | 4 | Myl2 | 0.009 | Young |
| 0.260727 | 0.835345 | 0.013 | 0.004 | 1 | 4 | Ppp1r13l | 0.009 | Young |
| 0.260727 | 0.803056 | 0.013 | 0.004 | 1 | 4 | Stard6 | 0.009 | Young |
| 0.260727 | 0.670281 | 0.013 | 0.004 | 1 | 4 | Trim62 | 0.009 | Young |
| 0.260727 | 0.648875 | 0.013 | 0.004 | 1 | 4 | Abraxas1 | 0.009 | Young |
| 0.260727 | 0.613587 | 0.013 | 0.004 | 1 | 4 | Rims3 | 0.009 | Young |
| 0.260727 | 0.397833 | 0.013 | 0.004 | 1 | 4 | Socs2 | 0.009 | Young |
| 0.260727 | 0.348627 | 0.013 | 0.004 | 1 | 4 | Kiss1r | 0.009 | Young |
| 0.260902 | 0.200994 | 0.1 | 0.066 | 1 | 4 | Tef | 0.034 | Young |
| 0.260937 | 0.615895 | 0.062 | 0.038 | 1 | 4 | Styx | 0.024 | Young |
| 0.260937 | 0.494783 | 0.062 | 0.038 | 1 | 4 | Mapkbp1 | 0.024 | Young |
| 0.26111 | 0.400604 | 0.075 | 0.047 | 1 | 4 | Papss2 | 0.028 | Young |
| 0.26111 | 0.288124 | 0.075 | 0.047 | 1 | 4 | Ttc17 | 0.028 | Young |
| 0.261237 | 0.047991 | 0.438 | 0.349 | 1 | 4 | Lamtor1 | 0.089 | Young |
| 0.261316 | 0.064448 | 0.138 | 0.096 | 1 | 4 | 1110038F1 | 0.042 | Young |

| | | | | | | | | |
|---|---|---|---|---|---|---|---|---|
| 0.261504 | 0.658422 | 0.125 | 0.088 | 1 | 4 | Xpa | 0.037 | Young |
| 0.26165 | 0.385231 | 0.013 | 0.004 | 1 | 4 | Ppp1r16b | 0.009 | Young |
| 0.26165 | 0.047707 | 0.013 | 0.004 | 1 | 4 | Hist1h4c | 0.009 | Young |
| 0.26165 | 0.02837 | 0.013 | 0.004 | 1 | 4 | Gfi1b | 0.009 | Young |
| 0.261914 | 0.314634 | 0.162 | 0.119 | 1 | 4 | Naa38 | 0.043 | Young |
| 0.262285 | 0.207045 | 0.425 | 0.361 | 1 | 4 | Ep300 | 0.064 | Young |
| 0.262427 | 0.651701 | 0.112 | 0.078 | 1 | 4 | Zfp277 | 0.034 | Young |
| 0.262474 | 0.003284 | 0.388 | 0.31 | 1 | 4 | Emc10 | 0.078 | Young |
| 0.262574 | 0.404195 | 0.013 | 0.004 | 1 | 4 | B230303O1 | 0.009 | Young |
| 0.262574 | 0.244795 | 0.013 | 0.004 | 1 | 4 | Fap | 0.009 | Young |
| 0.262574 | 0.117811 | 0.013 | 0.004 | 1 | 4 | Klc2 | 0.009 | Young |
| 0.262574 | 0.03326 | 0.013 | 0.004 | 1 | 4 | Fam171a1 | 0.009 | Young |
| 0.262617 | 0.666021 | 0.075 | 0.047 | 1 | 4 | Nop14 | 0.028 | Young |
| 0.262781 | 0.30321 | 0.25 | 0.2 | 1 | 4 | Mbtd1 | 0.05 | Young |
| 0.262861 | 0.372898 | 0.312 | 0.269 | 1 | 4 | Antxr2 | 0.043 | Young |
| 0.263257 | 0.11556 | 0.038 | 0.019 | 1 | 4 | Pfkm | 0.019 | Young |
| 0.263466 | 0.841118 | 0.062 | 0.038 | 1 | 4 | Golim4 | 0.024 | Young |
| 0.264419 | 0.493637 | 0.062 | 0.038 | 1 | 4 | Gabpb1 | 0.024 | Young |
| 0.264576 | 0.033118 | 0.225 | 0.169 | 1 | 4 | Cand1 | 0.056 | Young |
| 0.264759 | 0.062652 | 0.425 | 0.339 | 1 | 4 | Tsc22d1 | 0.086 | Young |
| 0.264953 | 0.590878 | 0.05 | 0.028 | 1 | 4 | Dcakd | 0.022 | Young |
| 0.265178 | 1.747319 | 0.025 | 0.011 | 1 | 4 | Pigw | 0.014 | Young |
| 0.265461 | 0.054166 | 0.537 | 0.457 | 1 | 4 | Fyb | 0.08 | Young |
| 0.265496 | 0.161703 | 0.662 | 0.556 | 1 | 4 | Akap13 | 0.106 | Young |
| 0.265637 | 0.275352 | 0.312 | 0.255 | 1 | 4 | Hip1 | 0.057 | Young |
| 0.265676 | 0.712239 | 0.088 | 0.058 | 1 | 4 | Sema4c | 0.03 | Young |
| 0.265682 | 0.562841 | 0.05 | 0.028 | 1 | 4 | Rnaseh1 | 0.022 | Young |
| 0.265806 | 0.607527 | 0.088 | 0.058 | 1 | 4 | Ppfibp1 | 0.03 | Young |
| 0.265858 | 0.453444 | 0.162 | 0.121 | 1 | 4 | D10Jhu81e | 0.041 | Young |
| 0.265915 | 0.250391 | 0.138 | 0.097 | 1 | 4 | Zbtb1 | 0.041 | Young |
| 0.266267 | 0.218122 | 0.15 | 0.109 | 1 | 4 | Spast | 0.041 | Young |
| 0.266522 | 0.291866 | 0.125 | 0.087 | 1 | 4 | Cep192 | 0.038 | Young |
| 0.266522 | 0.206452 | 0.125 | 0.087 | 1 | 4 | Zfp322a | 0.038 | Young |
| 0.266885 | 1.040263 | 0.025 | 0.011 | 1 | 4 | Dync2h1 | 0.014 | Young |
| 0.266885 | 0.859654 | 0.025 | 0.011 | 1 | 4 | Alkbh2 | 0.014 | Young |
| 0.267268 | 0.167013 | 0.125 | 0.086 | 1 | 4 | Acsl3 | 0.039 | Young |
| 0.267367 | 0.23771 | 0.212 | 0.163 | 1 | 4 | Cs | 0.049 | Young |
| 0.267456 | 1.233997 | 0.025 | 0.011 | 1 | 4 | Trip13 | 0.014 | Young |
| 0.267456 | 0.809421 | 0.025 | 0.011 | 1 | 4 | Dedd2 | 0.014 | Young |
| 0.267456 | 0.766709 | 0.025 | 0.011 | 1 | 4 | 9130221H1 | 0.014 | Young |
| 0.267456 | 0.6168 | 0.025 | 0.011 | 1 | 4 | Faap24 | 0.014 | Young |
| 0.26786 | 0.28809 | 0.4 | 0.323 | 1 | 4 | Psenen | 0.077 | Young |
| 0.267995 | 0.284776 | 0.075 | 0.047 | 1 | 4 | Ppat | 0.028 | Young |
| 0.268023 | 0.377579 | 0.1 | 0.067 | 1 | 4 | Psmd9 | 0.033 | Young |
| 0.268028 | 0.742864 | 0.025 | 0.011 | 1 | 4 | Bub1b | 0.014 | Young |

| | | | | | | | | |
|---|---|---|---|---|---|---|---|---|
| 0.268413 | 0.148262 | 0.188 | 0.14 | 1 | 4 | Gm37494 | 0.048 | Young |
| 0.268589 | 0.283154 | 0.125 | 0.088 | 1 | 4 | Trak2 | 0.037 | Young |
| 0.2686 | 0.680995 | 0.025 | 0.011 | 1 | 4 | AC139671. | 0.014 | Young |
| 0.268916 | 0.931505 | 0.175 | 0.139 | 1 | 4 | Thbd | 0.036 | Young |
| 0.269109 | 1.185233 | 0.05 | 0.029 | 1 | 4 | Ralgps1 | 0.021 | Young |
| 0.269173 | 0.496701 | 0.025 | 0.011 | 1 | 4 | Lcmt2 | 0.014 | Young |
| 0.269735 | 0.409916 | 0.075 | 0.047 | 1 | 4 | Pacs1 | 0.028 | Young |
| 0.26974 | 0.462966 | 0.1 | 0.067 | 1 | 4 | Dolpp1 | 0.033 | Young |
| 0.269748 | 0.707649 | 0.025 | 0.011 | 1 | 4 | L3mbtl2 | 0.014 | Young |
| 0.269749 | 0.210604 | 0.338 | 0.273 | 1 | 4 | Pxn | 0.065 | Young |
| 0.269884 | 0.220952 | 0.388 | 0.329 | 1 | 4 | Pacs2 | 0.059 | Young |
| 0.270478 | 0.393556 | 0.1 | 0.067 | 1 | 4 | Cmtm4 | 0.033 | Young |
| 0.270615 | 0.092389 | 0.2 | 0.15 | 1 | 4 | Ccny | 0.05 | Young |
| 0.270698 | 0.25622 | 0.325 | 0.262 | 1 | 4 | Aco2 | 0.063 | Young |
| 0.270705 | 0.165177 | 0.175 | 0.13 | 1 | 4 | Topors | 0.045 | Young |
| 0.271117 | 1.153184 | 0.05 | 0.029 | 1 | 4 | Atat1 | 0.021 | Young |
| 0.27116 | 0.256885 | 0.312 | 0.251 | 1 | 4 | Galc | 0.061 | Young |
| 0.27121 | 0.260914 | 0.125 | 0.086 | 1 | 4 | Zfp131 | 0.039 | Young |
| 0.271243 | 0.015426 | 0.2 | 0.148 | 1 | 4 | Tmem208 | 0.052 | Young |
| 0.271431 | 0.148986 | 0.225 | 0.173 | 1 | 4 | Cpeb4 | 0.052 | Young |
| 0.271475 | 0.07876 | 0.025 | 0.011 | 1 | 4 | Tmem143 | 0.014 | Young |
| 0.272053 | 0.313542 | 0.025 | 0.011 | 1 | 4 | C920006O1 | 0.014 | Young |
| 0.272156 | 0.219421 | 0.2 | 0.153 | 1 | 4 | Rtn1 | 0.047 | Young |
| 0.272191 | 0.101597 | 0.988 | 0.979 | 1 | 4 | Zfp36 | 0.009 | Young |
| 0.272231 | 0.67114 | 0.125 | 0.09 | 1 | 4 | Ppfia4 | 0.035 | Young |
| 0.2723 | 0.477484 | 0.138 | 0.099 | 1 | 4 | Pde7a | 0.039 | Young |
| 0.272448 | 0.130133 | 0.675 | 0.607 | 1 | 4 | Snx5 | 0.068 | Young |
| 0.272768 | 0.345147 | 0.1 | 0.069 | 1 | 4 | Plekhg2 | 0.031 | Young |
| 0.272782 | 0.59543 | 0.138 | 0.102 | 1 | 4 | Klhl6 | 0.036 | Young |
| 0.273277 | 0.153127 | 0.338 | 0.269 | 1 | 4 | 7-Mar | 0.069 | Young |
| 0.273427 | 0.206193 | 0.925 | 0.879 | 1 | 4 | Klf6 | 0.046 | Young |
| 0.273481 | 0.146814 | 0.225 | 0.173 | 1 | 4 | Srek1 | 0.052 | Young |
| 0.273597 | 0.194279 | 0.438 | 0.377 | 1 | 4 | Eif3c | 0.061 | Young |
| 0.27379 | 0.40669 | 0.05 | 0.028 | 1 | 4 | Dclre1b | 0.022 | Young |
| 0.27379 | 0.066564 | 0.025 | 0.011 | 1 | 4 | Mrps22 | 0.014 | Young |
| 0.274162 | 0.254367 | 0.05 | 0.028 | 1 | 4 | Dnajc18 | 0.022 | Young |
| 0.274212 | 0.898619 | 0.038 | 0.02 | 1 | 4 | Gm31718 | 0.018 | Young |
| 0.274539 | 0.107668 | 0.412 | 0.339 | 1 | 4 | Nacc2 | 0.073 | Young |
| 0.275047 | 0.089264 | 0.225 | 0.171 | 1 | 4 | Dhrs1 | 0.054 | Young |
| 0.27506 | 0.136772 | 0.088 | 0.057 | 1 | 4 | Eef1akmt1 | 0.031 | Young |
| 0.275099 | 0.91347 | 0.038 | 0.02 | 1 | 4 | Rbm12b1 | 0.018 | Young |
| 0.275099 | 0.896476 | 0.038 | 0.02 | 1 | 4 | AW209491 | 0.018 | Young |
| 0.275543 | 0.851004 | 0.038 | 0.02 | 1 | 4 | Tmtc4 | 0.018 | Young |
| 0.275557 | 0.156727 | 0.438 | 0.371 | 1 | 4 | Kdm7a | 0.067 | Young |
| 0.275918 | 0.006805 | 0.325 | 0.253 | 1 | 4 | Adam19 | 0.072 | Young |

| | | | | | | | | |
|---|---|---|---|---|---|---|---|---|
| 0.275987 | 0.880647 | 0.038 | 0.02 | 1 | 4 | Dynlt1f | 0.018 | Young |
| 0.276283 | 0.142574 | 0.188 | 0.138 | 1 | 4 | Lmbrd2 | 0.05 | Young |
| 0.276327 | 0.101499 | 0.45 | 0.377 | 1 | 4 | Arid1a | 0.073 | Young |
| 0.277582 | 0.170048 | 0.312 | 0.251 | 1 | 4 | Rab21 | 0.061 | Young |
| 0.277681 | 0.164812 | 0.062 | 0.038 | 1 | 4 | Tor4a | 0.024 | Young |
| 0.277868 | 0.153064 | 0.412 | 0.337 | 1 | 4 | Lrrc58 | 0.075 | Young |
| 0.277916 | 0.644203 | 0.075 | 0.048 | 1 | 4 | Tor2a | 0.027 | Young |
| 0.278311 | 0.234033 | 0.1 | 0.067 | 1 | 4 | Wdr77 | 0.033 | Young |
| 0.278665 | 0.660745 | 0.038 | 0.02 | 1 | 4 | Syne1 | 0.018 | Young |
| 0.278694 | 0.822958 | 0.05 | 0.029 | 1 | 4 | Zmat1 | 0.021 | Young |
| 0.278732 | 0.124699 | 0.288 | 0.229 | 1 | 4 | Slc25a36 | 0.059 | Young |
| 0.279171 | 0.354456 | 0.212 | 0.165 | 1 | 4 | Chd2 | 0.047 | Young |
| 0.279269 | 0.541751 | 0.088 | 0.059 | 1 | 4 | Mre11a | 0.029 | Young |
| 0.279454 | 0.165432 | 0.325 | 0.259 | 1 | 4 | Rap1gds1 | 0.066 | Young |
| 0.279712 | 0.059 | 0.3 | 0.233 | 1 | 4 | Prrc2b | 0.067 | Young |
| 0.279787 | 0.263723 | 0.3 | 0.237 | 1 | 4 | Cdk11b | 0.063 | Young |
| 0.280018 | 0.530089 | 0.075 | 0.047 | 1 | 4 | Tmcc2 | 0.028 | Young |
| 0.280109 | 0.205718 | 0.225 | 0.173 | 1 | 4 | Tmcc3 | 0.052 | Young |
| 0.280394 | 0.230738 | 0.55 | 0.47 | 1 | 4 | Msn | 0.08 | Young |
| 0.280455 | 0.108814 | 0.35 | 0.278 | 1 | 4 | Crk | 0.072 | Young |
| 0.28071 | 0.782268 | 0.075 | 0.048 | 1 | 4 | Eci1 | 0.027 | Young |
| 0.28108 | 0.466992 | 0.112 | 0.078 | 1 | 4 | Sord | 0.034 | Young |
| 0.2816 | 0.265522 | 0.388 | 0.328 | 1 | 4 | Etv1 | 0.06 | Young |
| 0.281727 | 0.120448 | 0.438 | 0.374 | 1 | 4 | Ski | 0.064 | Young |
| 0.282301 | 0.568457 | 0.088 | 0.058 | 1 | 4 | Ptpn7 | 0.03 | Young |
| 0.282499 | 0.475021 | 0.212 | 0.168 | 1 | 4 | Ap1g1 | 0.044 | Young |
| 0.283389 | 0.592832 | 0.088 | 0.058 | 1 | 4 | Rab10os | 0.03 | Young |
| 0.283492 | 0.220612 | 0.112 | 0.078 | 1 | 4 | Sec16a | 0.034 | Young |
| 0.284086 | 0.150342 | 0.863 | 0.816 | 1 | 4 | Cd36 | 0.047 | Young |
| 0.284128 | 0.294686 | 0.15 | 0.109 | 1 | 4 | Uimc1 | 0.041 | Young |
| 0.28429 | 0.062307 | 0.15 | 0.107 | 1 | 4 | Setd7 | 0.043 | Young |
| 0.284527 | 0.408688 | 0.038 | 0.02 | 1 | 4 | Gm13479 | 0.018 | Young |
| 0.285101 | 0.447006 | 0.212 | 0.167 | 1 | 4 | Ivns1abp | 0.045 | Young |
| 0.285163 | 0.630133 | 0.112 | 0.081 | 1 | 4 | Usp5 | 0.031 | Young |
| 0.285437 | 0.285095 | 0.038 | 0.02 | 1 | 4 | Dcbld2 | 0.018 | Young |
| 0.285496 | 0.471597 | 0.2 | 0.161 | 1 | 4 | Prelid1 | 0.039 | Young |
| 0.285703 | 0.505583 | 0.212 | 0.175 | 1 | 4 | Cyp4f18 | 0.037 | Young |
| 0.286004 | 1.028883 | 0.062 | 0.039 | 1 | 4 | Acot2 | 0.023 | Young |
| 0.286298 | 0.185115 | 0.1 | 0.067 | 1 | 4 | Smg9 | 0.033 | Young |
| 0.286511 | 0.306595 | 0.062 | 0.038 | 1 | 4 | Fgfr1op | 0.024 | Young |
| 0.287261 | 0.077151 | 0.038 | 0.02 | 1 | 4 | Lilra5 | 0.018 | Young |
| 0.287476 | 0.084544 | 0.2 | 0.151 | 1 | 4 | Ubald1 | 0.049 | Young |
| 0.287491 | 0.082641 | 0.462 | 0.381 | 1 | 4 | Myh9 | 0.081 | Young |
| 0.287513 | 0.494665 | 0.062 | 0.038 | 1 | 4 | Scd1 | 0.024 | Young |
| 0.28763 | 0.393238 | 0.088 | 0.058 | 1 | 4 | Dohh | 0.03 | Young |

| | | | | | | | | |
|---|---|---|---|---|---|---|---|---|
| 0.287721 | 1.490302 | 0.038 | 0.02 | 1 | 4 | Zfp959 | 0.018 | Young |
| 0.287755 | 0.831358 | 0.075 | 0.049 | 1 | 4 | Slc25a12 | 0.026 | Young |
| 0.288258 | 0.404149 | 0.112 | 0.079 | 1 | 4 | Parp12 | 0.033 | Young |
| 0.288398 | 0.156949 | 0.112 | 0.078 | 1 | 4 | Fam220a | 0.034 | Young |
| 0.288432 | 1.362928 | 0.062 | 0.04 | 1 | 4 | Tmem175 | 0.022 | Young |
| 0.288524 | 0.46327 | 0.262 | 0.217 | 1 | 4 | Edf1 | 0.045 | Young |
| 0.288621 | 0.509057 | 0.1 | 0.069 | 1 | 4 | Dlat | 0.031 | Young |
| 0.288646 | 0.27263 | 0.025 | 0.052 | 1 | 4 | Rab11fip1 | -0.027 | Young |
| 0.288851 | 0.37425 | 0.062 | 0.038 | 1 | 4 | Nek1 | 0.024 | Young |
| 0.288851 | 0.155563 | 0.062 | 0.038 | 1 | 4 | Leo1 | 0.024 | Young |
| 0.289024 | 0.352093 | 0.275 | 0.223 | 1 | 4 | Azin1 | 0.052 | Young |
| 0.289068 | 0.743293 | 0.1 | 0.07 | 1 | 4 | Farsb | 0.03 | Young |
| 0.289092 | 1.472131 | 0.062 | 0.04 | 1 | 4 | Ogfod2 | 0.022 | Young |
| 0.289181 | 0.955068 | 0.088 | 0.061 | 1 | 4 | Tmem181a | 0.027 | Young |
| 0.289273 | 0.302691 | 0.05 | 0.029 | 1 | 4 | E130307A1 | 0.021 | Young |
| 0.289316 | 0.137811 | 0.125 | 0.088 | 1 | 4 | Pik3c3 | 0.037 | Young |
| 0.289496 | 0.069172 | 0.175 | 0.128 | 1 | 4 | Ppie | 0.047 | Young |
| 0.289512 | 0.537048 | 0.1 | 0.069 | 1 | 4 | Tbc1d24 | 0.031 | Young |
| 0.289847 | 0.285999 | 0.125 | 0.089 | 1 | 4 | Agap3 | 0.036 | Young |
| 0.289858 | 0.253796 | 0.062 | 0.038 | 1 | 4 | Nmral1 | 0.024 | Young |
| 0.289938 | 0.212091 | 0.2 | 0.154 | 1 | 4 | Fbxw2 | 0.046 | Young |
| 0.290038 | 0.503819 | 0.05 | 0.029 | 1 | 4 | Slx1b | 0.021 | Young |
| 0.290074 | 0.217778 | 0.15 | 0.109 | 1 | 4 | Wars | 0.041 | Young |
| 0.290141 | 0.234247 | 0.275 | 0.219 | 1 | 4 | Psmc6 | 0.056 | Young |
| 0.290275 | 0.155458 | 0.388 | 0.321 | 1 | 4 | Tnks2 | 0.067 | Young |
| 0.290602 | 0.026199 | 0.438 | 0.347 | 1 | 4 | Manf | 0.091 | Young |
| 0.290833 | 0.19311 | 0.05 | 0.086 | 1 | 4 | Dph5 | -0.036 | Young |
| 0.291628 | 0.978477 | 0.075 | 0.049 | 1 | 4 | Pgm2 | 0.026 | Young |
| 0.291659 | 0.691118 | 0.062 | 0.039 | 1 | 4 | 10-Sep | 0.023 | Young |
| 0.291661 | 0.386721 | 0.138 | 0.101 | 1 | 4 | Zfp622 | 0.037 | Young |
| 0.291751 | 0.433633 | 0.188 | 0.149 | 1 | 4 | Psmd14 | 0.039 | Young |
| 0.291837 | 0.262757 | 0.212 | 0.167 | 1 | 4 | Kcnq1ot1 | 0.045 | Young |
| 0.291925 | 0.419401 | 0.075 | 0.048 | 1 | 4 | Pelp1 | 0.027 | Young |
| 0.292117 | 0.181566 | 0.6 | 0.536 | 1 | 4 | Cyth4 | 0.064 | Young |
| 0.292429 | 0.183978 | 0.362 | 0.295 | 1 | 4 | Mrps21 | 0.067 | Young |
| 0.292848 | 0.26039 | 0.175 | 0.13 | 1 | 4 | Foxo3 | 0.045 | Young |
| 0.29311 | 0.136377 | 0.162 | 0.119 | 1 | 4 | Rbl2 | 0.043 | Young |
| 0.293141 | 0.229255 | 0.15 | 0.11 | 1 | 4 | Babam2 | 0.04 | Young |
| 0.293159 | 0.312651 | 0.362 | 0.306 | 1 | 4 | Ptgs1 | 0.056 | Young |
| 0.293534 | 0.433178 | 0.262 | 0.215 | 1 | 4 | Szrd1 | 0.047 | Young |
| 0.293925 | 0.211835 | 0.662 | 0.613 | 1 | 4 | Hnrnpa3 | 0.049 | Young |
| 0.294009 | 0.530822 | 0.062 | 0.039 | 1 | 4 | Rhbdd2 | 0.023 | Young |
| 0.294379 | 0.325263 | 0.375 | 0.319 | 1 | 4 | Wdfy3 | 0.056 | Young |
| 0.294894 | 0.509712 | 0.1 | 0.069 | 1 | 4 | Ppp5c | 0.031 | Young |
| 0.294946 | 0.034126 | 0.162 | 0.119 | 1 | 4 | Washc5 | 0.043 | Young |

| | | | | | | | | |
|---|---|---|---|---|---|---|---|---|
| 0.294997 | 1.07305 | 0.038 | 0.02 | 1 | 4 | Ankrd39 | 0.018 | Young |
| 0.295312 | 0.04037 | 0.462 | 0.381 | 1 | 4 | Pcmtd1 | 0.081 | Young |
| 0.295333 | 0.156558 | 0.388 | 0.317 | 1 | 4 | Tsn | 0.071 | Young |
| 0.29617 | 0.110881 | 0.112 | 0.078 | 1 | 4 | Gm4951 | 0.034 | Young |
| 0.296198 | 0.215019 | 0.238 | 0.186 | 1 | 4 | Mgrn1 | 0.052 | Young |
| 0.296727 | 0.39386 | 0.125 | 0.09 | 1 | 4 | Med30 | 0.035 | Young |
| 0.297074 | 0.43485 | 0.112 | 0.08 | 1 | 4 | Creb5 | 0.032 | Young |
| 0.297194 | 0.245729 | 0.438 | 0.368 | 1 | 4 | Tmed5 | 0.07 | Young |
| 0.297443 | 0.250766 | 0.238 | 0.184 | 1 | 4 | Fbxl5 | 0.054 | Young |
| 0.297757 | 1.302176 | 0.038 | 0.02 | 1 | 4 | Iba57 | 0.018 | Young |
| 0.297757 | 0.852986 | 0.038 | 0.02 | 1 | 4 | Wbp1 | 0.018 | Young |
| 0.297807 | 0.34377 | 0.188 | 0.145 | 1 | 4 | Chmp1a | 0.043 | Young |
| 0.297985 | 0.344853 | 0.238 | 0.195 | 1 | 4 | Pik3cg | 0.043 | Young |
| 0.298265 | 0.015049 | 0.238 | 0.183 | 1 | 4 | Tspan4 | 0.055 | Young |
| 0.298288 | 0.239905 | 0.112 | 0.079 | 1 | 4 | Cdkn2aipnl | 0.033 | Young |
| 0.298502 | 0.251307 | 0.75 | 0.675 | 1 | 4 | Herpud1 | 0.075 | Young |
| 0.29864 | 0.072465 | 0.262 | 0.206 | 1 | 4 | Fam105a | 0.056 | Young |
| 0.298698 | 0.143618 | 0.8 | 0.739 | 1 | 4 | Fosb | 0.061 | Young |
| 0.299432 | 0.485034 | 0.2 | 0.155 | 1 | 4 | Uba5 | 0.045 | Young |
| 0.299454 | 0.819399 | 0.1 | 0.07 | 1 | 4 | Chid1 | 0.03 | Young |
| 0.299596 | 0.122824 | 0.725 | 0.69 | 1 | 4 | Pla2g2d | 0.035 | Young |
| 0.299727 | 0.025805 | 0.05 | 0.029 | 1 | 4 | Atg16l2 | 0.021 | Young |
| 0.299973 | 1.288725 | 0.05 | 0.03 | 1 | 4 | Apba3 | 0.02 | Young |
| 0.300121 | 0.514413 | 0.112 | 0.08 | 1 | 4 | Tesk1 | 0.032 | Young |
| 0.300194 | 1.146289 | 0.025 | 0.012 | 1 | 4 | Fam234b | 0.013 | Young |
| 0.300434 | 0.074643 | 0.162 | 0.121 | 1 | 4 | Synrg | 0.041 | Young |
| 0.300441 | 0.01073 | 0.188 | 0.141 | 1 | 4 | Trnau1ap | 0.047 | Young |
| 0.300442 | 0.294906 | 0.1 | 0.069 | 1 | 4 | Vrk3 | 0.031 | Young |
| 0.300488 | 0.40672 | 0.112 | 0.08 | 1 | 4 | Cryzl1 | 0.032 | Young |
| 0.300557 | 0.268976 | 0.3 | 0.242 | 1 | 4 | Usp24 | 0.058 | Young |
| 0.300568 | 0.098866 | 0.55 | 0.458 | 1 | 4 | Lnpep | 0.092 | Young |
| 0.300583 | 0.333939 | 0.088 | 0.059 | 1 | 4 | Hjurp | 0.029 | Young |
| 0.300584 | 0.487529 | 0.188 | 0.147 | 1 | 4 | Ube2m | 0.041 | Young |
| 0.300654 | 0.988055 | 0.05 | 0.03 | 1 | 4 | Inafm2 | 0.02 | Young |
| 0.30076 | 0.337431 | 0.125 | 0.09 | 1 | 4 | Tspan17 | 0.035 | Young |
| 0.300774 | 0.317766 | 0.125 | 0.09 | 1 | 4 | Gon4l | 0.035 | Young |
| 0.300797 | 1.055176 | 0.025 | 0.012 | 1 | 4 | Zfp799 | 0.013 | Young |
| 0.301429 | 0.388415 | 0.088 | 0.059 | 1 | 4 | Dip2c | 0.029 | Young |
| 0.302089 | 0.503298 | 0.088 | 0.059 | 1 | 4 | Dusp2 | 0.029 | Young |
| 0.302228 | 0.86134 | 0.075 | 0.049 | 1 | 4 | Mtmr4 | 0.026 | Young |
| 0.302268 | 0.081091 | 0.138 | 0.099 | 1 | 4 | Lage3 | 0.039 | Young |
| 0.302367 | 0.131133 | 0.537 | 0.46 | 1 | 4 | Trib1 | 0.077 | Young |
| 0.30261 | 0.86165 | 0.025 | 0.012 | 1 | 4 | C1qtnf6 | 0.013 | Young |
| 0.302862 | 0.813517 | 0.038 | 0.02 | 1 | 4 | Zfp619 | 0.018 | Young |
| 0.302862 | 0.499452 | 0.038 | 0.02 | 1 | 4 | Lsr | 0.018 | Young |

| | | | | | | | | |
|---|---|---|---|---|---|---|---|---|
| 0.30297 | 0.621844 | 0.1 | 0.07 | 1 | 4 | Ikbip | 0.03 | Young |
| 0.302993 | 0.799274 | 0.05 | 0.03 | 1 | 4 | Zfp874b | 0.02 | Young |
| 0.302993 | 0.615868 | 0.05 | 0.03 | 1 | 4 | Cep128 | 0.02 | Young |
| 0.303188 | 0.548029 | 0.05 | 0.03 | 1 | 4 | Lrch4 | 0.02 | Young |
| 0.303216 | 0.962161 | 0.025 | 0.012 | 1 | 4 | Cav1 | 0.013 | Young |
| 0.303243 | 0.179227 | 0.438 | 0.368 | 1 | 4 | Chmp4b | 0.07 | Young |
| 0.303329 | 0.671841 | 0.038 | 0.02 | 1 | 4 | Zmynd19 | 0.018 | Young |
| 0.303383 | 0.668538 | 0.05 | 0.03 | 1 | 4 | Cdkal1 | 0.02 | Young |
| 0.303563 | 0.616573 | 0.038 | 0.02 | 1 | 4 | Cd1d1 | 0.018 | Young |
| 0.303705 | 0.372708 | 0.175 | 0.134 | 1 | 4 | Vps50 | 0.041 | Young |
| 0.303726 | 0.185143 | 0.25 | 0.197 | 1 | 4 | Fam173a | 0.053 | Young |
| 0.303854 | 0.218225 | 0.175 | 0.133 | 1 | 4 | Nucb2 | 0.042 | Young |
| 0.303905 | 0.233249 | 0.462 | 0.396 | 1 | 4 | Bmp2k | 0.066 | Young |
| 0.304172 | 0.197295 | 0.3 | 0.243 | 1 | 4 | Cct2 | 0.057 | Young |
| 0.304264 | 0.622001 | 0.038 | 0.02 | 1 | 4 | Twnk | 0.018 | Young |
| 0.304328 | 1.050896 | 0.088 | 0.061 | 1 | 4 | Rmi1 | 0.027 | Young |
| 0.30443 | 0.749705 | 0.025 | 0.012 | 1 | 4 | Pter | 0.013 | Young |
| 0.30443 | 0.451481 | 0.025 | 0.012 | 1 | 4 | 8030462N1 | 0.013 | Young |
| 0.304617 | 0.32851 | 0.075 | 0.049 | 1 | 4 | Miga1 | 0.026 | Young |
| 0.304903 | 0.789628 | 0.062 | 0.04 | 1 | 4 | Zfp160 | 0.022 | Young |
| 0.304993 | 0.036563 | 0.262 | 0.203 | 1 | 4 | Tmem259 | 0.059 | Young |
| 0.305202 | 0.520157 | 0.038 | 0.02 | 1 | 4 | Golm1 | 0.018 | Young |
| 0.305202 | 0.18453 | 0.038 | 0.02 | 1 | 4 | Cgnl1 | 0.018 | Young |
| 0.305671 | 0.3898 | 0.038 | 0.02 | 1 | 4 | Pwwp2b | 0.018 | Young |
| 0.305709 | 0.417408 | 0.175 | 0.137 | 1 | 4 | Cdk19 | 0.038 | Young |
| 0.305734 | 0.195846 | 0.15 | 0.111 | 1 | 4 | Pds5b | 0.039 | Young |
| 0.305863 | 0.221433 | 0.162 | 0.121 | 1 | 4 | Srpk2 | 0.041 | Young |
| 0.306045 | 0.246355 | 0.3 | 0.246 | 1 | 4 | Npepps | 0.054 | Young |
| 0.306161 | 0.157049 | 0.25 | 0.197 | 1 | 4 | Atp6v1d | 0.053 | Young |
| 0.306229 | 0.831691 | 0.075 | 0.049 | 1 | 4 | Hmces | 0.026 | Young |
| 0.306257 | 0.611522 | 0.025 | 0.012 | 1 | 4 | Loxl3 | 0.013 | Young |
| 0.306868 | 0.542718 | 0.025 | 0.012 | 1 | 4 | Mtmr11 | 0.013 | Young |
| 0.306918 | 0.471013 | 0.125 | 0.092 | 1 | 4 | Zmym4 | 0.033 | Young |
| 0.306984 | 0.373334 | 0.2 | 0.158 | 1 | 4 | C1d | 0.042 | Young |
| 0.307259 | 0.501687 | 0.075 | 0.049 | 1 | 4 | Tmem140 | 0.026 | Young |
| 0.30765 | 0.648931 | 0.062 | 0.04 | 1 | 4 | Zfp729b | 0.022 | Young |
| 0.307665 | 1.074568 | 0.075 | 0.051 | 1 | 4 | Vps8 | 0.024 | Young |
| 0.307855 | 0.031141 | 0.238 | 0.184 | 1 | 4 | Nceh1 | 0.054 | Young |
| 0.307978 | 0.160369 | 0.2 | 0.155 | 1 | 4 | Klf11 | 0.045 | Young |
| 0.307983 | 0.701593 | 0.088 | 0.061 | 1 | 4 | Mfsd8 | 0.027 | Young |
| 0.308092 | 0.433665 | 0.025 | 0.012 | 1 | 4 | Lig4 | 0.013 | Young |
| 0.30843 | 0.169063 | 0.138 | 0.099 | 1 | 4 | Mplkip | 0.039 | Young |
| 0.308594 | 0.414244 | 0.175 | 0.138 | 1 | 4 | Chuk | 0.037 | Young |
| 0.308656 | 0.592479 | 0.088 | 0.06 | 1 | 4 | Dcaf10 | 0.028 | Young |
| 0.308724 | 0.486166 | 0.125 | 0.092 | 1 | 4 | Rexo1 | 0.033 | Young |

| | | | | | | | | |
|---|---|---|---|---|---|---|---|---|
| 0.308811 | 0.046231 | 0.212 | 0.162 | 1 | 4 | Ssu72 | 0.05 | Young |
| 0.309035 | 0.452597 | 0.088 | 0.059 | 1 | 4 | Nr2c1 | 0.029 | Young |
| 0.309287 | 0.498605 | 0.05 | 0.03 | 1 | 4 | Mad1l1 | 0.02 | Young |
| 0.309319 | 0.034382 | 0.025 | 0.012 | 1 | 4 | Ovca2 | 0.013 | Young |
| 0.309839 | 0.151174 | 0.488 | 0.415 | 1 | 4 | P4hb | 0.073 | Young |
| 0.31008 | 0.52945 | 0.05 | 0.03 | 1 | 4 | Rbm10 | 0.02 | Young |
| 0.310278 | 0.369922 | 0.05 | 0.03 | 1 | 4 | Trpm4 | 0.02 | Young |
| 0.310409 | 0.2481 | 0.2 | 0.157 | 1 | 4 | Cc2d1b | 0.043 | Young |
| 0.310422 | 0.406652 | 0.188 | 0.148 | 1 | 4 | Fam104a | 0.04 | Young |
| 0.310463 | 0.049048 | 1 | 0.998 | 1 | 4 | Itm2b | 0.002 | Young |
| 0.310696 | 0.090488 | 0.963 | 0.976 | 1 | 4 | Rplp2 | -0.013 | Young |
| 0.31076 | 0.570167 | 0.062 | 0.04 | 1 | 4 | Osgin2 | 0.022 | Young |
| 0.310817 | 0.151267 | 0.062 | 0.039 | 1 | 4 | Ogfod3 | 0.023 | Young |
| 0.311166 | 0.086952 | 0.062 | 0.039 | 1 | 4 | 9330160F1 | 0.023 | Young |
| 0.311331 | 0.066634 | 0.338 | 0.273 | 1 | 4 | Ndufv3 | 0.065 | Young |
| 0.311731 | 0.54816 | 0.1 | 0.071 | 1 | 4 | Snrpa1 | 0.029 | Young |
| 0.311826 | 1.663424 | 0.038 | 0.021 | 1 | 4 | Syt11 | 0.017 | Young |
| 0.311987 | 0.133621 | 0.625 | 0.553 | 1 | 4 | Ptprj | 0.072 | Young |
| 0.312045 | 0.976242 | 0.05 | 0.03 | 1 | 4 | Cks1b | 0.02 | Young |
| 0.312296 | 1.434272 | 0.038 | 0.021 | 1 | 4 | Bach2 | 0.017 | Young |
| 0.312305 | 0.065201 | 0.188 | 0.141 | 1 | 4 | Impad1 | 0.047 | Young |
| 0.312737 | 0.135593 | 0.15 | 0.111 | 1 | 4 | Spata2 | 0.039 | Young |
| 0.312768 | 0.234133 | 0.038 | 0.02 | 1 | 4 | Procr | 0.018 | Young |
| 0.313138 | 0.104889 | 0.975 | 0.985 | 1 | 4 | Rpl35a | -0.01 | Young |
| 0.313245 | 0.119128 | 0.038 | 0.02 | 1 | 4 | Zfp512b | 0.018 | Young |
| 0.313506 | 0.113629 | 0.55 | 0.468 | 1 | 4 | Pip4k2a | 0.082 | Young |
| 0.313626 | 1.425713 | 0.05 | 0.03 | 1 | 4 | Gm26881 | 0.02 | Young |
| 0.313662 | 0.10564 | 0.125 | 0.089 | 1 | 4 | Spic | 0.036 | Young |
| 0.313845 | 0.32505 | 0.162 | 0.123 | 1 | 4 | Casp8ap2 | 0.039 | Young |
| 0.313917 | 0.351006 | 0.088 | 0.059 | 1 | 4 | Snhg6 | 0.029 | Young |
| 0.313952 | 0.308975 | 0.1 | 0.07 | 1 | 4 | Elac1 | 0.03 | Young |
| 0.314365 | 0.453011 | 0.15 | 0.114 | 1 | 4 | Tlk1 | 0.036 | Young |
| 0.314613 | 0.312864 | 0.288 | 0.245 | 1 | 4 | Ppp1r10 | 0.043 | Young |
| 0.314844 | 0.394614 | 0.112 | 0.08 | 1 | 4 | Rnf144b | 0.032 | Young |
| 0.315368 | 0.249863 | 0.312 | 0.265 | 1 | 4 | Arid5a | 0.047 | Young |
| 0.315688 | 0.107059 | 0.15 | 0.11 | 1 | 4 | BC035044 | 0.04 | Young |
| 0.31583 | 0.330557 | 0.1 | 0.07 | 1 | 4 | Fas | 0.03 | Young |
| 0.31594 | 0.538322 | 0.112 | 0.082 | 1 | 4 | Slc30a9 | 0.03 | Young |
| 0.31599 | 0.451059 | 0.062 | 0.04 | 1 | 4 | Mgme1 | 0.022 | Young |
| 0.316252 | 0.138728 | 0.288 | 0.229 | 1 | 4 | Sipa1 | 0.059 | Young |
| 0.316279 | 0.099488 | 0.262 | 0.208 | 1 | 4 | P2rx7 | 0.054 | Young |
| 0.316908 | 0.948932 | 0.062 | 0.04 | 1 | 4 | Pag1 | 0.022 | Young |
| 0.317 | 0.335005 | 0.225 | 0.185 | 1 | 4 | Pgap2 | 0.04 | Young |
| 0.317356 | 0.701795 | 0.162 | 0.127 | 1 | 4 | Cdk12 | 0.035 | Young |
| 0.317971 | 0.139554 | 0.362 | 0.304 | 1 | 4 | H2afv | 0.058 | Young |

| | | | | | | | | |
|---|---|---|---|---|---|---|---|---|
| 0.318018 | 0.224274 | 0.35 | 0.3 | 1 | 4 | Zfp703 | 0.05 | Young |
| 0.318094 | 0.101923 | 0.362 | 0.3 | 1 | 4 | Cpsf6 | 0.062 | Young |
| 0.318105 | 0.233194 | 0.125 | 0.09 | 1 | 4 | 1110059G1 | 0.035 | Young |
| 0.318146 | 0.348443 | 0.212 | 0.17 | 1 | 4 | Zhx1 | 0.042 | Young |
| 0.318283 | 0.334427 | 0.275 | 0.231 | 1 | 4 | Msr1 | 0.044 | Young |
| 0.318399 | 0.748111 | 0.05 | 0.03 | 1 | 4 | Kbtbd7 | 0.02 | Young |
| 0.31861 | 0.173155 | 0.15 | 0.111 | 1 | 4 | Lactb2 | 0.039 | Young |
| 0.318633 | 0.126019 | 0.325 | 0.266 | 1 | 4 | Lamtor2 | 0.059 | Young |
| 0.319013 | 0.267076 | 0.112 | 0.08 | 1 | 4 | Thoc6 | 0.032 | Young |
| 0.319037 | 0.498786 | 0.138 | 0.104 | 1 | 4 | Brpf1 | 0.034 | Young |
| 0.319166 | 0.050245 | 0.65 | 0.567 | 1 | 4 | Btg1 | 0.083 | Young |
| 0.319283 | 0.532808 | 0.225 | 0.181 | 1 | 4 | Galk2 | 0.044 | Young |
| 0.3194 | 0.766712 | 0.05 | 0.03 | 1 | 4 | Vps37c | 0.02 | Young |
| 0.319413 | 0.67902 | 0.088 | 0.061 | 1 | 4 | Trp53bp2 | 0.027 | Young |
| 0.319709 | 0.038246 | 0.175 | 0.131 | 1 | 4 | Rapgef2 | 0.044 | Young |
| 0.319868 | 1.312856 | 0.038 | 0.021 | 1 | 4 | Rsl1 | 0.017 | Young |
| 0.32024 | 0.328188 | 0.262 | 0.213 | 1 | 4 | Plekha2 | 0.049 | Young |
| 0.320467 | 0.612186 | 0.075 | 0.05 | 1 | 4 | Pde12 | 0.025 | Young |
| 0.320485 | 0.484646 | 0.112 | 0.082 | 1 | 4 | Fkbp1b | 0.03 | Young |
| 0.321206 | 0.724842 | 0.05 | 0.03 | 1 | 4 | Ndufaf1 | 0.02 | Young |
| 0.321587 | 0.12974 | 0.312 | 0.26 | 1 | 4 | Zfp148 | 0.052 | Young |
| 0.321883 | 0.53475 | 0.1 | 0.071 | 1 | 4 | Snapc5 | 0.029 | Young |
| 0.322068 | 0.728018 | 0.088 | 0.059 | 1 | 4 | Heatr1 | 0.029 | Young |
| 0.322557 | 0.085954 | 0.5 | 0.41 | 1 | 4 | Atp6v0e | 0.09 | Young |
| 0.322817 | 0.625996 | 0.05 | 0.03 | 1 | 4 | Bet1l | 0.02 | Young |
| 0.322889 | 0.045256 | 0.262 | 0.209 | 1 | 4 | Dhrs7 | 0.053 | Young |
| 0.322956 | 0.261773 | 0.138 | 0.101 | 1 | 4 | Fxr2 | 0.037 | Young |
| 0.323743 | 0.031785 | 0.05 | 0.082 | 1 | 4 | Scyl2 | -0.032 | Young |
| 0.324282 | 0.543846 | 0.075 | 0.05 | 1 | 4 | D230025D1 | 0.025 | Young |
| 0.324465 | 0.23887 | 0.162 | 0.121 | 1 | 4 | Ccpg1 | 0.041 | Young |
| 0.324569 | 0.200074 | 0.088 | 0.059 | 1 | 4 | Yaf2 | 0.029 | Young |
| 0.324838 | 0.570434 | 0.05 | 0.03 | 1 | 4 | Lclat1 | 0.02 | Young |
| 0.324938 | 0.54971 | 0.088 | 0.061 | 1 | 4 | Med25 | 0.027 | Young |
| 0.325165 | 0.162198 | 0.125 | 0.092 | 1 | 4 | Pygb | 0.033 | Young |
| 0.325308 | 0.163172 | 0.4 | 0.339 | 1 | 4 | Cxcr4 | 0.061 | Young |
| 0.325308 | 1.20578 | 0.075 | 0.051 | 1 | 4 | Thoc5 | 0.024 | Young |
| 0.325391 | 0.995668 | 0.088 | 0.062 | 1 | 4 | Il1rl1 | 0.026 | Young |
| 0.325678 | 0.210395 | 0.275 | 0.229 | 1 | 4 | Psmb6 | 0.046 | Young |
| 0.326183 | 0.338022 | 0.112 | 0.081 | 1 | 4 | Zfp260 | 0.031 | Young |
| 0.326386 | 0.460921 | 0.175 | 0.139 | 1 | 4 | BC028528 | 0.036 | Young |
| 0.32647 | 0.325005 | 0.262 | 0.215 | 1 | 4 | Bod1l | 0.047 | Young |
| 0.326739 | 0.734687 | 0.1 | 0.07 | 1 | 4 | D6Wsu163 | 0.03 | Young |
| 0.327 | 3.143926 | 0.013 | 0.005 | 1 | 4 | Saa3 | 0.008 | Young |
| 0.327 | 1.9434 | 0.013 | 0.005 | 1 | 4 | Rnd1 | 0.008 | Young |
| 0.327 | 1.87977 | 0.013 | 0.005 | 1 | 4 | Epop | 0.008 | Young |

| | | | | | | | | |
|---|---|---|---|---|---|---|---|---|
| 0.327 | 1.69977 | 0.013 | 0.005 | 1 | 4 | 4930539J0 | 0.008 | Young |
| 0.327004 | 0.160572 | 0.3 | 0.238 | 1 | 4 | Pnn | 0.062 | Young |
| 0.327132 | 0.096665 | 0.138 | 0.101 | 1 | 4 | Gtf2h1 | 0.037 | Young |
| 0.327204 | 0.090698 | 0.1 | 0.069 | 1 | 4 | Tprkb | 0.031 | Young |
| 0.327345 | 0.051621 | 0.312 | 0.255 | 1 | 4 | Arhgef1 | 0.057 | Young |
| 0.327388 | 0.658975 | 0.175 | 0.138 | 1 | 4 | Gadd45g | 0.037 | Young |
| 0.32756 | 0.597732 | 0.038 | 0.021 | 1 | 4 | Mmgt2 | 0.017 | Young |
| 0.327571 | 0.142029 | 0.112 | 0.08 | 1 | 4 | Dnajc11 | 0.032 | Young |
| 0.328005 | 1.655023 | 0.013 | 0.005 | 1 | 4 | Gm45250 | 0.008 | Young |
| 0.328005 | 1.637084 | 0.013 | 0.005 | 1 | 4 | Cd200 | 0.008 | Young |
| 0.328005 | 1.52443 | 0.013 | 0.005 | 1 | 4 | Jam2 | 0.008 | Young |
| 0.328005 | 1.486109 | 0.013 | 0.005 | 1 | 4 | Tmc4 | 0.008 | Young |
| 0.328005 | 1.43922 | 0.013 | 0.005 | 1 | 4 | Iqcd | 0.008 | Young |
| 0.328005 | 1.231392 | 0.013 | 0.005 | 1 | 4 | Ccna2 | 0.008 | Young |
| 0.328958 | 0.214489 | 0.2 | 0.158 | 1 | 4 | Plpbp | 0.042 | Young |
| 0.329011 | 1.319296 | 0.013 | 0.005 | 1 | 4 | Glcci1 | 0.008 | Young |
| 0.329011 | 1.162656 | 0.013 | 0.005 | 1 | 4 | Gm16062 | 0.008 | Young |
| 0.329011 | 1.041047 | 0.013 | 0.005 | 1 | 4 | Gm15880 | 0.008 | Young |
| 0.329025 | 1.941851 | 0.025 | 0.013 | 1 | 4 | Tubg1 | 0.012 | Young |
| 0.329032 | 0.039358 | 0.45 | 0.372 | 1 | 4 | Atp5j | 0.078 | Young |
| 0.3293 | 0.17003 | 0.3 | 0.252 | 1 | 4 | Sbno1 | 0.048 | Young |
| 0.32965 | 1.806489 | 0.025 | 0.013 | 1 | 4 | Hmga1 | 0.012 | Young |
| 0.329722 | 0.351205 | 0.05 | 0.03 | 1 | 4 | Dgkq | 0.02 | Young |
| 0.329788 | 0.160145 | 0.125 | 0.091 | 1 | 4 | A630001G2 | 0.034 | Young |
| 0.33002 | 1.33055 | 0.013 | 0.005 | 1 | 4 | Eme2 | 0.008 | Young |
| 0.33002 | 1.242077 | 0.013 | 0.005 | 1 | 4 | Zfp791 | 0.008 | Young |
| 0.33002 | 1.17785 | 0.013 | 0.005 | 1 | 4 | Etohd2 | 0.008 | Young |
| 0.33002 | 1.109858 | 0.013 | 0.005 | 1 | 4 | Azin2 | 0.008 | Young |
| 0.33002 | 0.908307 | 0.013 | 0.005 | 1 | 4 | Ankrd28 | 0.008 | Young |
| 0.33002 | 0.908007 | 0.013 | 0.005 | 1 | 4 | 1200007C1 | 0.008 | Young |
| 0.33002 | 0.785615 | 0.013 | 0.005 | 1 | 4 | Thsd4 | 0.008 | Young |
| 0.33002 | 0.258415 | 0.013 | 0.005 | 1 | 4 | F5 | 0.008 | Young |
| 0.330034 | 0.447342 | 0.1 | 0.07 | 1 | 4 | Pfdn6 | 0.03 | Young |
| 0.330145 | 0.143152 | 0.175 | 0.133 | 1 | 4 | Commd6 | 0.042 | Young |
| 0.3309 | 0.279007 | 0.25 | 0.308 | 1 | 4 | Usmg5 | -0.058 | Young |
| 0.330927 | 0.27712 | 0.188 | 0.148 | 1 | 4 | D5Ertd579 | 0.04 | Young |
| 0.331031 | 1.168705 | 0.013 | 0.005 | 1 | 4 | Tcea2 | 0.008 | Young |
| 0.331031 | 0.672121 | 0.013 | 0.005 | 1 | 4 | Gm20513 | 0.008 | Young |
| 0.331031 | 0.532129 | 0.013 | 0.005 | 1 | 4 | 2610020C0 | 0.008 | Young |
| 0.331031 | 0.425518 | 0.013 | 0.005 | 1 | 4 | Dsn1 | 0.008 | Young |
| 0.331038 | 0.12522 | 0.575 | 0.528 | 1 | 4 | Nr3c1 | 0.047 | Young |
| 0.33128 | 0.094012 | 0.2 | 0.152 | 1 | 4 | Arid4b | 0.048 | Young |
| 0.331315 | 0.161258 | 0.938 | 0.891 | 1 | 4 | Rps4x | 0.047 | Young |
| 0.33151 | 0.132035 | 0.162 | 0.123 | 1 | 4 | Ncbp3 | 0.039 | Young |
| 0.331837 | 0.148516 | 0.025 | 0.05 | 1 | 4 | Agpat2 | -0.025 | Young |

| | | | | | | | | |
|---|---|---|---|---|---|---|---|---|
| 0.33191 | 1.377657 | 0.05 | 0.031 | 1 | 4 | AI504432 | 0.019 | Young |
| 0.331966 | 0.171032 | 0.1 | 0.07 | 1 | 4 | Ptgs2 | 0.03 | Young |
| 0.332043 | 0.907337 | 0.013 | 0.005 | 1 | 4 | Spp1 | 0.008 | Young |
| 0.332043 | 0.862486 | 0.013 | 0.005 | 1 | 4 | Spata24 | 0.008 | Young |
| 0.332043 | 0.672986 | 0.013 | 0.005 | 1 | 4 | Smim10l2a | 0.008 | Young |
| 0.332043 | 0.658354 | 0.013 | 0.005 | 1 | 4 | Fam149a | 0.008 | Young |
| 0.332043 | 0.656488 | 0.013 | 0.005 | 1 | 4 | Ccdc15 | 0.008 | Young |
| 0.332043 | 0.549853 | 0.013 | 0.005 | 1 | 4 | Ehd2 | 0.008 | Young |
| 0.332062 | 0.168771 | 0.425 | 0.362 | 1 | 4 | Il10rb | 0.063 | Young |
| 0.332157 | 1.545074 | 0.025 | 0.013 | 1 | 4 | Padi2 | 0.012 | Young |
| 0.332227 | 0.177808 | 0.1 | 0.07 | 1 | 4 | Scly | 0.03 | Young |
| 0.332487 | 0.405871 | 0.075 | 0.05 | 1 | 4 | Ascc2 | 0.025 | Young |
| 0.33274 | 0.221307 | 0.275 | 0.225 | 1 | 4 | Max | 0.05 | Young |
| 0.333058 | 0.403022 | 0.013 | 0.005 | 1 | 4 | Thbs2 | 0.008 | Young |
| 0.333058 | 0.367036 | 0.013 | 0.005 | 1 | 4 | Gm47230 | 0.008 | Young |
| 0.333058 | 0.239737 | 0.013 | 0.005 | 1 | 4 | Ppbp | 0.008 | Young |
| 0.333058 | 0.088748 | 0.013 | 0.005 | 1 | 4 | Gm5113 | 0.008 | Young |
| 0.333058 | 0.064369 | 0.013 | 0.005 | 1 | 4 | A930024E( | 0.008 | Young |
| 0.333322 | 0.784022 | 0.1 | 0.072 | 1 | 4 | Tmod1 | 0.028 | Young |
| 0.333416 | 1.138814 | 0.025 | 0.013 | 1 | 4 | Supv3l1 | 0.012 | Young |
| 0.333716 | 0.166596 | 0.138 | 0.101 | 1 | 4 | Sos2 | 0.037 | Young |
| 0.333871 | 0.528324 | 0.1 | 0.072 | 1 | 4 | Pcsk7 | 0.028 | Young |
| 0.333956 | 0.362913 | 0.162 | 0.127 | 1 | 4 | Dnttip2 | 0.035 | Young |
| 0.334075 | 0.331046 | 0.013 | 0.005 | 1 | 4 | Ube2v1 | 0.008 | Young |
| 0.334086 | 0.689084 | 0.088 | 0.061 | 1 | 4 | Mlycd | 0.027 | Young |
| 0.334241 | 0.223483 | 0.05 | 0.03 | 1 | 4 | Arrdc2 | 0.02 | Young |
| 0.334241 | 0.15304 | 0.05 | 0.03 | 1 | 4 | Ccdc58 | 0.02 | Young |
| 0.33434 | 0.115857 | 0.225 | 0.178 | 1 | 4 | Rab12 | 0.047 | Young |
| 0.334582 | 0.322284 | 0.2 | 0.161 | 1 | 4 | Cd40 | 0.039 | Young |
| 0.33467 | 0.338567 | 0.138 | 0.103 | 1 | 4 | Cd300lb | 0.035 | Young |
| 0.334677 | 1.094742 | 0.025 | 0.013 | 1 | 4 | Synj2 | 0.012 | Young |
| 0.334723 | 0.05521 | 0.125 | 0.09 | 1 | 4 | Mtmr14 | 0.035 | Young |
| 0.334879 | 0.41564 | 0.038 | 0.021 | 1 | 4 | Trim39 | 0.017 | Young |
| 0.335435 | 0.703773 | 0.062 | 0.041 | 1 | 4 | Tmc6 | 0.021 | Young |
| 0.335709 | 0.136854 | 0.125 | 0.091 | 1 | 4 | Fbrs | 0.034 | Young |
| 0.335863 | 0.094479 | 0.038 | 0.021 | 1 | 4 | Them4 | 0.017 | Young |
| 0.336007 | 0.48384 | 0.225 | 0.186 | 1 | 4 | Psmc5 | 0.039 | Young |
| 0.336301 | 0.126654 | 0.338 | 0.277 | 1 | 4 | Tmem189 | 0.061 | Young |
| 0.336308 | 0.212551 | 0.05 | 0.03 | 1 | 4 | Rhbdd3 | 0.02 | Young |
| 0.336525 | 0.498204 | 0.162 | 0.127 | 1 | 4 | H2-Q4 | 0.035 | Young |
| 0.336575 | 1.02591 | 0.025 | 0.013 | 1 | 4 | Zfp553 | 0.012 | Young |
| 0.336649 | 0.516108 | 0.15 | 0.116 | 1 | 4 | Ppm1b | 0.034 | Young |
| 0.336763 | 0.108007 | 0.138 | 0.1 | 1 | 4 | Ctr9 | 0.038 | Young |
| 0.336802 | 0.213917 | 0.3 | 0.249 | 1 | 4 | St6gal1 | 0.051 | Young |
| 0.336815 | 0.781925 | 0.05 | 0.031 | 1 | 4 | Zbtb16 | 0.019 | Young |

| | | | | | | | | |
|---|---|---|---|---|---|---|---|---|
| 0.336967 | 0.388699 | 0.1 | 0.07 | 1 | 4 | Pole3 | 0.03 | Young |
| 0.337191 | 0.244049 | 0.062 | 0.04 | 1 | 4 | Mri1 | 0.022 | Young |
| 0.337209 | 0.97721 | 0.025 | 0.013 | 1 | 4 | Chaf1a | 0.012 | Young |
| 0.337637 | 0.691255 | 0.05 | 0.031 | 1 | 4 | Smpd3 | 0.019 | Young |
| 0.337692 | 0.298452 | 0.188 | 0.144 | 1 | 4 | Slc16a10 | 0.044 | Young |
| 0.338031 | 0.219271 | 0.075 | 0.05 | 1 | 4 | Nhlrc2 | 0.025 | Young |
| 0.338033 | 0.078241 | 0.688 | 0.569 | 1 | 4 | Snrnp70 | 0.119 | Young |
| 0.338091 | 0.295795 | 0.162 | 0.125 | 1 | 4 | Gpatch2 | 0.037 | Young |
| 0.338479 | 0.726024 | 0.025 | 0.013 | 1 | 4 | Nipal3 | 0.012 | Young |
| 0.338548 | 0.053265 | 0.35 | 0.291 | 1 | 4 | Nedd8 | 0.059 | Young |
| 0.339015 | 0.262844 | 0.075 | 0.05 | 1 | 4 | Gtf2f2 | 0.025 | Young |
| 0.339038 | 0.234594 | 0.212 | 0.168 | 1 | 4 | Taf1 | 0.044 | Young |
| 0.339662 | 0.126872 | 0.288 | 0.232 | 1 | 4 | Hnrnpa1 | 0.056 | Young |
| 0.339722 | 0.197746 | 0.212 | 0.169 | 1 | 4 | Ociad1 | 0.043 | Young |
| 0.339753 | 0.625398 | 0.025 | 0.013 | 1 | 4 | Cad | 0.012 | Young |
| 0.34011 | 0.076911 | 0.138 | 0.101 | 1 | 4 | Sucla2 | 0.037 | Young |
| 0.340149 | 0.664208 | 0.075 | 0.051 | 1 | 4 | Uqcc1 | 0.024 | Young |
| 0.34081 | 0.078261 | 0.038 | 0.021 | 1 | 4 | Nrg2 | 0.017 | Young |
| 0.34085 | 0.586398 | 0.062 | 0.041 | 1 | 4 | Cdkn2aip | 0.021 | Young |
| 0.340853 | 0.268916 | 0.175 | 0.134 | 1 | 4 | Sf3b4 | 0.041 | Young |
| 0.341029 | 0.649765 | 0.025 | 0.013 | 1 | 4 | Rccd1 | 0.012 | Young |
| 0.341153 | 0.098233 | 0.112 | 0.081 | 1 | 4 | Sun1 | 0.031 | Young |
| 0.341595 | 0.393951 | 0.088 | 0.061 | 1 | 4 | Apeh | 0.027 | Young |
| 0.341762 | 0.045958 | 0.938 | 0.843 | 1 | 4 | Rpl9 | 0.095 | Young |
| 0.341814 | 0.465535 | 0.288 | 0.239 | 1 | 4 | Cmtm7 | 0.049 | Young |
| 0.34186 | 0.022588 | 0.438 | 0.358 | 1 | 4 | Leprot | 0.08 | Young |
| 0.342181 | 0.584591 | 0.05 | 0.031 | 1 | 4 | Hmgxb3 | 0.019 | Young |
| 0.34295 | 0.168819 | 0.025 | 0.013 | 1 | 4 | Tmem150b | 0.012 | Young |
| 0.34295 | 0.576182 | 0.025 | 0.013 | 1 | 4 | Fads3 | 0.012 | Young |
| 0.343062 | 0.119496 | 1 | 0.976 | 1 | 4 | Fau | 0.024 | Young |
| 0.343419 | 0.250954 | 0.125 | 0.092 | 1 | 4 | Ppp1r13b | 0.033 | Young |
| 0.343427 | 0.465738 | 0.05 | 0.031 | 1 | 4 | Phospho2 | 0.019 | Young |
| 0.343592 | 0.323057 | 0.025 | 0.013 | 1 | 4 | Atad5 | 0.012 | Young |
| 0.343744 | 0.420853 | 0.125 | 0.09 | 1 | 4 | 4833420G1 | 0.035 | Young |
| 0.343858 | 0.086288 | 0.388 | 0.313 | 1 | 4 | Kras | 0.075 | Young |
| 0.343952 | 0.284192 | 0.1 | 0.071 | 1 | 4 | Ppp6r2 | 0.029 | Young |
| 0.343965 | 0.105666 | 0.075 | 0.05 | 1 | 4 | Med31 | 0.025 | Young |
| 0.344208 | 0.197851 | 0.338 | 0.287 | 1 | 4 | Ppp3r1 | 0.051 | Young |
| 0.344231 | 0.158886 | 0.3 | 0.244 | 1 | 4 | Ubn2 | 0.056 | Young |
| 0.344867 | 0.131063 | 0.35 | 0.29 | 1 | 4 | Tprgl | 0.06 | Young |
| 0.34512 | 0.069274 | 0.488 | 0.424 | 1 | 4 | Cp | 0.064 | Young |
| 0.345175 | 0.782858 | 0.112 | 0.085 | 1 | 4 | Iars2 | 0.027 | Young |
| 0.345411 | 0.432754 | 0.075 | 0.051 | 1 | 4 | Uaca | 0.024 | Young |
| 0.345443 | 0.054622 | 0.188 | 0.144 | 1 | 4 | Churc1 | 0.044 | Young |
| 0.345509 | 0.685077 | 0.075 | 0.051 | 1 | 4 | Dcp1a | 0.024 | Young |

| | | | | | | | | |
|---|---|---|---|---|---|---|---|---|
| 0.345586 | 0.380039 | 0.1 | 0.072 | 1 | 4 | Letm1 | 0.028 | Young |
| 0.345694 | 0.040129 | 0.1 | 0.07 | 1 | 4 | Sgcb | 0.03 | Young |
| 0.345788 | 0.312916 | 0.15 | 0.114 | 1 | 4 | Syvn1 | 0.036 | Young |
| 0.345793 | 0.193927 | 0.225 | 0.178 | 1 | 4 | Zmym5 | 0.047 | Young |
| 0.346831 | 0.218891 | 0.1 | 0.07 | 1 | 4 | Tdg | 0.03 | Young |
| 0.347602 | 0.438771 | 0.05 | 0.031 | 1 | 4 | Akap1 | 0.019 | Young |
| 0.347606 | 0.516439 | 0.062 | 0.041 | 1 | 4 | Wfs1 | 0.021 | Young |
| 0.347657 | 0.192632 | 0.088 | 0.061 | 1 | 4 | Fam110b | 0.027 | Young |
| 0.348194 | 0.214175 | 0.2 | 0.159 | 1 | 4 | Acsl5 | 0.041 | Young |
| 0.348538 | 0.874191 | 0.075 | 0.052 | 1 | 4 | Ttc4 | 0.023 | Young |
| 0.348904 | 0.808589 | 0.138 | 0.107 | 1 | 4 | Ubac2 | 0.031 | Young |
| 0.349024 | 0.108776 | 0.175 | 0.223 | 1 | 4 | Ppp6c | -0.048 | Young |
| 0.349692 | 0.924518 | 0.038 | 0.022 | 1 | 4 | Rbm43 | 0.016 | Young |
| 0.349692 | 0.854485 | 0.038 | 0.022 | 1 | 4 | 1810024B0 | 0.016 | Young |
| 0.349692 | 0.775924 | 0.038 | 0.022 | 1 | 4 | Naif1 | 0.016 | Young |
| 0.349701 | 0.456911 | 0.05 | 0.031 | 1 | 4 | Ddx20 | 0.019 | Young |
| 0.35 | 0.370448 | 0.062 | 0.041 | 1 | 4 | Ercc5 | 0.021 | Young |
| 0.350513 | 0.917705 | 0.075 | 0.052 | 1 | 4 | Gcsh | 0.023 | Young |
| 0.350543 | 0.389175 | 0.05 | 0.031 | 1 | 4 | Ppih | 0.019 | Young |
| 0.350599 | 0.087096 | 0.338 | 0.275 | 1 | 4 | Morc3 | 0.063 | Young |
| 0.350617 | 0.242823 | 0.15 | 0.114 | 1 | 4 | Smurf2 | 0.036 | Young |
| 0.350822 | 0.008002 | 0.438 | 0.371 | 1 | 4 | Nipbl | 0.067 | Young |
| 0.351196 | 0.029949 | 0.262 | 0.208 | 1 | 4 | Eif3l | 0.054 | Young |
| 0.35125 | 0.334608 | 0.138 | 0.104 | 1 | 4 | Glrx2 | 0.034 | Young |
| 0.351306 | 0.275948 | 0.525 | 0.447 | 1 | 4 | Arf4 | 0.078 | Young |
| 0.351351 | 0.539652 | 0.138 | 0.107 | 1 | 4 | Fbxo38 | 0.031 | Young |
| 0.351357 | 0.245479 | 0.312 | 0.26 | 1 | 4 | Lamtor3 | 0.052 | Young |
| 0.35213 | 0.147579 | 0.188 | 0.146 | 1 | 4 | Rab1b | 0.042 | Young |
| 0.352414 | 0.044358 | 0.212 | 0.166 | 1 | 4 | Mta2 | 0.046 | Young |
| 0.352812 | 0.204415 | 0.112 | 0.082 | 1 | 4 | Plekhf2 | 0.03 | Young |
| 0.352812 | 0.190423 | 0.112 | 0.082 | 1 | 4 | Cops6 | 0.03 | Young |
| 0.353703 | 0.433083 | 0.062 | 0.041 | 1 | 4 | Hsf2 | 0.021 | Young |
| 0.353894 | 0.293546 | 0.25 | 0.208 | 1 | 4 | Anks1 | 0.042 | Young |
| 0.354348 | 0.136156 | 0.05 | 0.031 | 1 | 4 | Zfp961 | 0.019 | Young |
| 0.354666 | 0.198143 | 0.3 | 0.246 | 1 | 4 | Mat2b | 0.054 | Young |
| 0.354697 | 0.732253 | 0.038 | 0.022 | 1 | 4 | Nif3l1 | 0.016 | Young |
| 0.354697 | 0.597532 | 0.038 | 0.022 | 1 | 4 | Dmap1 | 0.016 | Young |
| 0.35498 | 0.753302 | 0.125 | 0.094 | 1 | 4 | Elf4 | 0.031 | Young |
| 0.3552 | 0.599586 | 0.038 | 0.022 | 1 | 4 | Cox18 | 0.016 | Young |
| 0.355313 | 0.077837 | 0.2 | 0.157 | 1 | 4 | Rgs19 | 0.043 | Young |
| 0.35553 | 0.055124 | 0.188 | 0.145 | 1 | 4 | Paox | 0.043 | Young |
| 0.355829 | 0.274463 | 0.2 | 0.161 | 1 | 4 | Zfp217 | 0.039 | Young |
| 0.356064 | 0.182058 | 0.162 | 0.126 | 1 | 4 | Pak1 | 0.036 | Young |
| 0.356483 | 0.644602 | 0.075 | 0.052 | 1 | 4 | Adprm | 0.023 | Young |
| 0.356715 | 0.14949 | 0.212 | 0.171 | 1 | 4 | Paf1 | 0.041 | Young |

| | | | | | | | | |
|---|---|---|---|---|---|---|---|---|
| 0.35673 | 0.186055 | 0.312 | 0.263 | 1 | 4 | Hivep2 | 0.049 | Young |
| 0.356917 | 0.012394 | 0.05 | 0.08 | 1 | 4 | Taf2 | -0.03 | Young |
| 0.357136 | 0.436292 | 0.075 | 0.051 | 1 | 4 | Zfp597 | 0.024 | Young |
| 0.357879 | 1.699821 | 0.038 | 0.022 | 1 | 4 | Gngt2 | 0.016 | Young |
| 0.357966 | 0.105121 | 0.225 | 0.179 | 1 | 4 | Stx16 | 0.046 | Young |
| 0.358133 | 0.201636 | 0.238 | 0.196 | 1 | 4 | 1500011K1 | 0.042 | Young |
| 0.358152 | 0.826336 | 0.075 | 0.052 | 1 | 4 | Utp15 | 0.023 | Young |
| 0.358546 | 0.12354 | 0.938 | 0.833 | 1 | 4 | Tgfbi | 0.105 | Young |
| 0.358949 | 0.019166 | 0.438 | 0.372 | 1 | 4 | Rnf19b | 0.066 | Young |
| 0.359 | 0.212086 | 0.375 | 0.327 | 1 | 4 | Ndufa13 | 0.048 | Young |
| 0.359046 | 0.045178 | 0.225 | 0.178 | 1 | 4 | Rtf1 | 0.047 | Young |
| 0.359241 | 0.457258 | 0.038 | 0.022 | 1 | 4 | Rmdn1 | 0.016 | Young |
| 0.359827 | 0.475442 | 0.075 | 0.051 | 1 | 4 | Pcnx3 | 0.024 | Young |
| 0.360067 | 0.31642 | 0.175 | 0.139 | 1 | 4 | Nudc | 0.036 | Young |
| 0.360103 | 0.119504 | 0.05 | 0.031 | 1 | 4 | C330018D2 | 0.019 | Young |
| 0.360389 | 0.287125 | 0.262 | 0.222 | 1 | 4 | Mtch1 | 0.04 | Young |
| 0.360679 | 0.281733 | 0.112 | 0.083 | 1 | 4 | Ssh1 | 0.029 | Young |
| 0.360977 | 0.215014 | 0.738 | 0.73 | 1 | 4 | Ehd4 | 0.008 | Young |
| 0.361718 | 0.108059 | 0.838 | 0.692 | 1 | 4 | Arhgdib | 0.146 | Young |
| 0.362691 | 0.064341 | 0.062 | 0.041 | 1 | 4 | Sssca1 | 0.021 | Young |
| 0.36274 | 0.792445 | 0.175 | 0.14 | 1 | 4 | Bbx | 0.035 | Young |
| 0.362819 | 0.048242 | 0.025 | 0.048 | 1 | 4 | Tmem18 | -0.023 | Young |
| 0.363271 | 0.351669 | 0.062 | 0.042 | 1 | 4 | Nmi | 0.02 | Young |
| 0.363466 | 0.228251 | 0.45 | 0.391 | 1 | 4 | Wasf2 | 0.059 | Young |
| 0.36362 | 0.365004 | 0.325 | 0.288 | 1 | 4 | Nsd1 | 0.037 | Young |
| 0.363647 | 0.475063 | 0.062 | 0.042 | 1 | 4 | Tinf2 | 0.02 | Young |
| 0.364182 | 0.375996 | 0.1 | 0.073 | 1 | 4 | Dnpep | 0.027 | Young |
| 0.364235 | 0.029357 | 0.075 | 0.051 | 1 | 4 | Srpk1 | 0.024 | Young |
| 0.364237 | 1.125758 | 0.1 | 0.075 | 1 | 4 | Slc15a2 | 0.025 | Young |
| 0.364317 | 0.206473 | 0.188 | 0.15 | 1 | 4 | Ndfip1 | 0.038 | Young |
| 0.364323 | 0.149764 | 0.175 | 0.135 | 1 | 4 | Brd8 | 0.04 | Young |
| 0.365145 | 0.487295 | 0.112 | 0.084 | 1 | 4 | Mfn2 | 0.028 | Young |
| 0.36524 | 0.086388 | 0.275 | 0.225 | 1 | 4 | Atp13a3 | 0.05 | Young |
| 0.365477 | 0.126546 | 0.262 | 0.216 | 1 | 4 | Bank1 | 0.046 | Young |
| 0.365794 | 0.238582 | 0.425 | 0.354 | 1 | 4 | Tiparp | 0.071 | Young |
| 0.36637 | 0.247465 | 0.1 | 0.072 | 1 | 4 | Kif3a | 0.028 | Young |
| 0.366807 | 0.709385 | 0.062 | 0.042 | 1 | 4 | Golga1 | 0.02 | Young |
| 0.367003 | 0.283037 | 0.088 | 0.062 | 1 | 4 | Timm29 | 0.026 | Young |
| 0.367003 | 0.186539 | 0.088 | 0.062 | 1 | 4 | Vps45 | 0.026 | Young |
| 0.367012 | 0.313365 | 0.125 | 0.094 | 1 | 4 | Rlf | 0.031 | Young |
| 0.367099 | 0.350964 | 0.225 | 0.189 | 1 | 4 | Arrdc3 | 0.036 | Young |
| 0.367399 | 0.693898 | 0.05 | 0.032 | 1 | 4 | Dimt1 | 0.018 | Young |
| 0.367485 | 0.360612 | 0.425 | 0.377 | 1 | 4 | Atp6v1e1 | 0.048 | Young |
| 0.367531 | 0.117794 | 0.062 | 0.095 | 1 | 4 | Ing5 | -0.033 | Young |
| 0.367602 | 1.416094 | 0.025 | 0.013 | 1 | 4 | Mir155hg | 0.012 | Young |

| | | | | | | | | |
|---|---|---|---|---|---|---|---|---|
| 0.367917 | 0.297506 | 0.325 | 0.281 | 1 | 4 | Hnrnpdl | 0.044 | Young |
| 0.367978 | 0.287879 | 0.075 | 0.051 | 1 | 4 | Limk1 | 0.024 | Young |
| 0.368257 | 1.394134 | 0.025 | 0.013 | 1 | 4 | B930036N1 | 0.012 | Young |
| 0.36869 | 0.44381 | 0.05 | 0.032 | 1 | 4 | Abhd14b | 0.018 | Young |
| 0.368906 | 0.412116 | 0.05 | 0.032 | 1 | 4 | Haus6 | 0.018 | Young |
| 0.369216 | 0.445098 | 0.1 | 0.074 | 1 | 4 | Rbsn | 0.026 | Young |
| 0.369305 | 0.350976 | 0.062 | 0.042 | 1 | 4 | Metrn | 0.02 | Young |
| 0.369428 | 0.157462 | 0.3 | 0.248 | 1 | 4 | Sh3kbp1 | 0.052 | Young |
| 0.36957 | 1.275459 | 0.025 | 0.013 | 1 | 4 | Zfp458 | 0.012 | Young |
| 0.369604 | 0.503779 | 0.125 | 0.094 | 1 | 4 | Tmem222 | 0.031 | Young |
| 0.369682 | 0.592514 | 0.138 | 0.11 | 1 | 4 | Casd1 | 0.028 | Young |
| 0.369798 | 0.108122 | 0.312 | 0.258 | 1 | 4 | Hspa4 | 0.054 | Young |
| 0.370019 | 0.256619 | 0.175 | 0.14 | 1 | 4 | Zbtb44 | 0.035 | Young |
| 0.370471 | 0.178111 | 0.288 | 0.246 | 1 | 4 | Akirin2 | 0.042 | Young |
| 0.370778 | 0.234038 | 0.275 | 0.227 | 1 | 4 | Samd4b | 0.048 | Young |
| 0.370975 | 0.160226 | 0.65 | 0.56 | 1 | 4 | Eef2 | 0.09 | Young |
| 0.371192 | 0.259717 | 0.312 | 0.274 | 1 | 4 | Tuba1c | 0.038 | Young |
| 0.371684 | 0.574426 | 0.075 | 0.052 | 1 | 4 | Rbm15b | 0.023 | Young |
| 0.37172 | 0.188231 | 0.388 | 0.339 | 1 | 4 | Herc1 | 0.049 | Young |
| 0.372643 | 0.513541 | 0.088 | 0.063 | 1 | 4 | Galk1 | 0.025 | Young |
| 0.372786 | 0.072685 | 0.075 | 0.051 | 1 | 4 | Nol4l | 0.024 | Young |
| 0.372864 | 1.078991 | 0.025 | 0.013 | 1 | 4 | Ak1 | 0.012 | Young |
| 0.372864 | 0.986975 | 0.025 | 0.013 | 1 | 4 | Vmac | 0.012 | Young |
| 0.373073 | 0.225836 | 0.188 | 0.151 | 1 | 4 | Tmem50b | 0.037 | Young |
| 0.373525 | 0.955433 | 0.025 | 0.013 | 1 | 4 | Layn | 0.012 | Young |
| 0.373644 | 0.910752 | 0.112 | 0.086 | 1 | 4 | Sumf1 | 0.026 | Young |
| 0.373871 | 0.054039 | 0.062 | 0.042 | 1 | 4 | Gapt | 0.02 | Young |
| 0.373986 | 0.6153 | 0.075 | 0.053 | 1 | 4 | Ltv1 | 0.022 | Young |
| 0.374017 | 0.775891 | 0.088 | 0.064 | 1 | 4 | Bola1 | 0.024 | Young |
| 0.374407 | 0.283223 | 0.088 | 0.061 | 1 | 4 | Auh | 0.027 | Young |
| 0.374462 | 0.712805 | 0.112 | 0.086 | 1 | 4 | Snx15 | 0.026 | Young |
| 0.374565 | 0.179914 | 0.088 | 0.062 | 1 | 4 | Grpel1 | 0.026 | Young |
| 0.37485 | 0.727105 | 0.025 | 0.013 | 1 | 4 | Rasgrf2 | 0.012 | Young |
| 0.375352 | 0.882575 | 0.038 | 0.022 | 1 | 4 | Iqcc | 0.016 | Young |
| 0.375468 | 0.584465 | 0.088 | 0.063 | 1 | 4 | Dpm2 | 0.025 | Young |
| 0.375513 | 1.016884 | 0.025 | 0.013 | 1 | 4 | Gm15853 | 0.012 | Young |
| 0.375513 | 0.621013 | 0.025 | 0.013 | 1 | 4 | Eml2 | 0.012 | Young |
| 0.375513 | 0.569074 | 0.025 | 0.013 | 1 | 4 | Xylt1 | 0.012 | Young |
| 0.375513 | 0.564743 | 0.025 | 0.013 | 1 | 4 | Gm29291 | 0.012 | Young |
| 0.375609 | 0.746804 | 0.038 | 0.022 | 1 | 4 | Ippk | 0.016 | Young |
| 0.375693 | 1.411013 | 0.05 | 0.033 | 1 | 4 | Gpr35 | 0.017 | Young |
| 0.375866 | 0.719565 | 0.038 | 0.022 | 1 | 4 | Hgf | 0.016 | Young |
| 0.375911 | 0.162218 | 0.375 | 0.323 | 1 | 4 | Lilr4b | 0.052 | Young |
| 0.375964 | 0.073116 | 0.425 | 0.374 | 1 | 4 | Ms4a6b | 0.051 | Young |
| 0.376039 | 0.561006 | 0.075 | 0.053 | 1 | 4 | Endog | 0.022 | Young |

| | | | | | | | | |
|---|---|---|---|---|---|---|---|---|
| 0.376104 | 0.454652 | 0.212 | 0.179 | 1 | 4 | Ankrd13c | 0.033 | Young |
| 0.37645 | 0.083672 | 0.3 | 0.246 | 1 | 4 | Cox7b | 0.054 | Young |
| 0.376724 | 0.077747 | 0.15 | 0.114 | 1 | 4 | Pdzd8 | 0.036 | Young |
| 0.37707 | 0.077553 | 0.112 | 0.082 | 1 | 4 | Zfp787 | 0.03 | Young |
| 0.377107 | 0.230187 | 0.088 | 0.062 | 1 | 4 | Hdgfl2 | 0.026 | Young |
| 0.377507 | 0.75729 | 0.025 | 0.013 | 1 | 4 | Ccdc171 | 0.012 | Young |
| 0.377524 | 0.302664 | 0.275 | 0.237 | 1 | 4 | Hexb | 0.038 | Young |
| 0.377993 | 0.166392 | 0.3 | 0.25 | 1 | 4 | Gtf2h5 | 0.05 | Young |
| 0.378173 | 0.627182 | 0.025 | 0.013 | 1 | 4 | Phc1 | 0.012 | Young |
| 0.378429 | 0.237619 | 0.388 | 0.337 | 1 | 4 | Taldo1 | 0.051 | Young |
| 0.378622 | 0.641698 | 0.088 | 0.063 | 1 | 4 | Brf1 | 0.025 | Young |
| 0.378682 | 0.244215 | 0.45 | 0.394 | 1 | 4 | Arhgap18 | 0.056 | Young |
| 0.378755 | 0.257074 | 0.55 | 0.467 | 1 | 4 | P2rx4 | 0.083 | Young |
| 0.378837 | 0.094033 | 0.738 | 0.704 | 1 | 4 | Rps2 | 0.034 | Young |
| 0.378837 | 0.371998 | 0.25 | 0.213 | 1 | 4 | Col4a3bp | 0.037 | Young |
| 0.37884 | 0.612113 | 0.025 | 0.013 | 1 | 4 | Jade3 | 0.012 | Young |
| 0.37884 | 0.528249 | 0.025 | 0.013 | 1 | 4 | Wdr19 | 0.012 | Young |
| 0.379507 | 0.537811 | 0.025 | 0.013 | 1 | 4 | Lmtk2 | 0.012 | Young |
| 0.379507 | 0.526252 | 0.025 | 0.013 | 1 | 4 | Asb10 | 0.012 | Young |
| 0.379688 | 0.59196 | 0.062 | 0.042 | 1 | 4 | Mrpl46 | 0.02 | Young |
| 0.37987 | 0.049042 | 0.112 | 0.082 | 1 | 4 | Rprd1b | 0.03 | Young |
| 0.380329 | 0.090766 | 0.312 | 0.252 | 1 | 4 | Yy1 | 0.06 | Young |
| 0.380773 | 0.728782 | 0.038 | 0.022 | 1 | 4 | Deaf1 | 0.016 | Young |
| 0.380773 | 0.56613 | 0.038 | 0.022 | 1 | 4 | Kif19a | 0.016 | Young |
| 0.381514 | 0.170687 | 0.025 | 0.013 | 1 | 4 | Ptch1 | 0.012 | Young |
| 0.381685 | 0.209549 | 0.15 | 0.117 | 1 | 4 | Slc25a45 | 0.033 | Young |
| 0.381694 | 0.367245 | 0.088 | 0.063 | 1 | 4 | Apobr | 0.025 | Young |
| 0.381757 | 0.744824 | 0.05 | 0.032 | 1 | 4 | 1110032A( | 0.018 | Young |
| 0.381767 | 0.161281 | 0.412 | 0.374 | 1 | 4 | Gng2 | 0.038 | Young |
| 0.382185 | 0.484935 | 0.025 | 0.013 | 1 | 4 | Zfp605 | 0.012 | Young |
| 0.382296 | 0.305534 | 0.212 | 0.168 | 1 | 4 | Ube2q2 | 0.044 | Young |
| 0.382719 | 0.577964 | 0.062 | 0.042 | 1 | 4 | Pdk1 | 0.02 | Young |
| 0.382735 | 0.187874 | 0.125 | 0.094 | 1 | 4 | Nfx1 | 0.031 | Young |
| 0.382761 | 0.803378 | 0.138 | 0.105 | 1 | 4 | Rnf19a | 0.033 | Young |
| 0.38285 | 0.454758 | 0.038 | 0.022 | 1 | 4 | 6330562C2 | 0.016 | Young |
| 0.382856 | 0.156561 | 0.025 | 0.013 | 1 | 4 | Gm4673 | 0.012 | Young |
| 0.383192 | 0.049705 | 0.025 | 0.013 | 1 | 4 | 2700049A( | 0.012 | Young |
| 0.383371 | 0.567612 | 0.038 | 0.022 | 1 | 4 | 9030025P2 | 0.016 | Young |
| 0.383511 | 0.258538 | 0.112 | 0.084 | 1 | 4 | Tmem68 | 0.028 | Young |
| 0.383882 | 0.235076 | 0.212 | 0.171 | 1 | 4 | Lsm14a | 0.041 | Young |
| 0.384176 | 0.280353 | 0.262 | 0.219 | 1 | 4 | Cxcl12 | 0.043 | Young |
| 0.384728 | 0.078174 | 0.162 | 0.125 | 1 | 4 | Dnajc2 | 0.037 | Young |
| 0.384747 | 1.217688 | 0.05 | 0.033 | 1 | 4 | Rabepk | 0.017 | Young |
| 0.385419 | 0.061964 | 1 | 1 | 1 | 4 | mt-Atp6 | 0 | Young |
| 0.385637 | 0.311203 | 0.1 | 0.073 | 1 | 4 | Eif2ak3 | 0.027 | Young |

| | | | | | | | | |
|---|---|---|---|---|---|---|---|---|
| 0.386235 | 0.190332 | 0.1 | 0.073 | 1 | 4 | G2e3 | 0.027 | Young |
| 0.387221 | 0.428014 | 0.088 | 0.063 | 1 | 4 | Fgd3 | 0.025 | Young |
| 0.38728 | 0.585845 | 0.125 | 0.099 | 1 | 4 | Gabpa | 0.026 | Young |
| 0.387288 | 0.131486 | 0.2 | 0.161 | 1 | 4 | Cuedc2 | 0.039 | Young |
| 0.387468 | 0.245655 | 0.088 | 0.063 | 1 | 4 | Cep162 | 0.025 | Young |
| 0.387536 | 0.648346 | 0.162 | 0.136 | 1 | 4 | Mrps24 | 0.026 | Young |
| 0.387655 | 0.481825 | 0.112 | 0.085 | 1 | 4 | Kifap3 | 0.027 | Young |
| 0.387688 | 0.428284 | 0.1 | 0.072 | 1 | 4 | Mthfr | 0.028 | Young |
| 0.387747 | 0.811547 | 0.075 | 0.054 | 1 | 4 | Fastk | 0.021 | Young |
| 0.38787 | 0.116901 | 0.438 | 0.378 | 1 | 4 | Ier3ip1 | 0.06 | Young |
| 0.388086 | 0.209656 | 0.25 | 0.204 | 1 | 4 | Arnt | 0.046 | Young |
| 0.388242 | 0.556292 | 0.3 | 0.264 | 1 | 4 | Polr2a | 0.036 | Young |
| 0.388264 | 0.137016 | 0.362 | 0.318 | 1 | 4 | Sesn1 | 0.044 | Young |
| 0.388686 | 0.21908 | 0.238 | 0.194 | 1 | 4 | Zfp644 | 0.044 | Young |
| 0.389125 | 0.200303 | 0.038 | 0.022 | 1 | 4 | Cdpf1 | 0.016 | Young |
| 0.389558 | 0.064084 | 0.438 | 0.366 | 1 | 4 | Sypl | 0.072 | Young |
| 0.389643 | 0.74568 | 0.112 | 0.087 | 1 | 4 | Cxcl1 | 0.025 | Young |
| 0.389733 | 0.406133 | 0.025 | 0.046 | 1 | 4 | Akr1b8 | -0.021 | Young |
| 0.3898 | 0.063987 | 0.1 | 0.072 | 1 | 4 | Gpd2 | 0.028 | Young |
| 0.389996 | 0.26034 | 0.488 | 0.432 | 1 | 4 | Cd68 | 0.056 | Young |
| 0.390047 | 0.750892 | 0.075 | 0.053 | 1 | 4 | Bloc1s6 | 0.022 | Young |
| 0.390113 | 0.296267 | 0.062 | 0.042 | 1 | 4 | Alg6 | 0.02 | Young |
| 0.39057 | 0.610652 | 0.075 | 0.053 | 1 | 4 | Maip1 | 0.022 | Young |
| 0.390982 | 0.058934 | 0.038 | 0.063 | 1 | 4 | Cinp | -0.025 | Young |
| 0.391488 | 0.09815 | 0.762 | 0.632 | 1 | 4 | Hnrnpu | 0.13 | Young |
| 0.391508 | 0.131315 | 0.088 | 0.063 | 1 | 4 | Rad51b | 0.025 | Young |
| 0.392136 | 0.185047 | 0.212 | 0.172 | 1 | 4 | Ptov1 | 0.04 | Young |
| 0.392368 | 0.241031 | 0.537 | 0.472 | 1 | 4 | Ets2 | 0.065 | Young |
| 0.392733 | 0.518078 | 0.088 | 0.064 | 1 | 4 | Dzip3 | 0.024 | Young |
| 0.393079 | 0.865961 | 0.088 | 0.065 | 1 | 4 | Daam1 | 0.023 | Young |
| 0.393172 | 0.078983 | 0.425 | 0.35 | 1 | 4 | Prex1 | 0.075 | Young |
| 0.393184 | 0.12321 | 0.4 | 0.358 | 1 | 4 | Tmem123 | 0.042 | Young |
| 0.39333 | 0.235729 | 0.188 | 0.151 | 1 | 4 | Znhit1 | 0.037 | Young |
| 0.393564 | 0.569424 | 0.062 | 0.043 | 1 | 4 | Abhd18 | 0.019 | Young |
| 0.393872 | 0.024929 | 0.038 | 0.022 | 1 | 4 | Dus2 | 0.016 | Young |
| 0.393887 | 0.072188 | 0.775 | 0.733 | 1 | 4 | Nrros | 0.042 | Young |
| 0.394478 | 0.174344 | 0.15 | 0.117 | 1 | 4 | Rnf5 | 0.033 | Young |
| 0.394647 | 0.084334 | 0.138 | 0.105 | 1 | 4 | Zc3h14 | 0.033 | Young |
| 0.39541 | 0.118435 | 0.2 | 0.161 | 1 | 4 | Cacul1 | 0.039 | Young |
| 0.395424 | 0.247474 | 0.05 | 0.032 | 1 | 4 | Kti12 | 0.018 | Young |
| 0.395539 | 0.159209 | 0.325 | 0.279 | 1 | 4 | Ik | 0.046 | Young |
| 0.395561 | 0.226231 | 0.188 | 0.15 | 1 | 4 | Siva1 | 0.038 | Young |
| 0.395647 | 0.451103 | 0.138 | 0.109 | 1 | 4 | Ostm1 | 0.029 | Young |
| 0.395651 | 0.054769 | 0.138 | 0.103 | 1 | 4 | Gm8186 | 0.035 | Young |
| 0.395683 | 0.304506 | 0.1 | 0.139 | 1 | 4 | Fh1 | -0.039 | Young |

| | | | | | | | | |
|---|---|---|---|---|---|---|---|---|
| 0.395684 | 0.201489 | 0.262 | 0.223 | 1 | 4 | Rela | 0.039 | Young |
| 0.395801 | 0.224437 | 0.225 | 0.186 | 1 | 4 | Slc25a4 | 0.039 | Young |
| 0.396075 | 2.225872 | 0.013 | 0.005 | 1 | 4 | Spry1 | 0.008 | Young |
| 0.396075 | 2.196658 | 0.013 | 0.005 | 1 | 4 | Haspin | 0.008 | Young |
| 0.396075 | 1.849562 | 0.013 | 0.005 | 1 | 4 | Zfp354c | 0.008 | Young |
| 0.396075 | 1.559051 | 0.013 | 0.005 | 1 | 4 | G0s2 | 0.008 | Young |
| 0.396075 | 1.546686 | 0.013 | 0.005 | 1 | 4 | Hrh2 | 0.008 | Young |
| 0.396192 | 0.194191 | 0.1 | 0.073 | 1 | 4 | Fbxo28 | 0.027 | Young |
| 0.396216 | 0.006567 | 0.212 | 0.266 | 1 | 4 | Slc29a3 | -0.054 | Young |
| 0.396706 | 0.862468 | 0.075 | 0.053 | 1 | 4 | Gpr137 | 0.022 | Young |
| 0.396801 | 0.029879 | 0.162 | 0.126 | 1 | 4 | Rnf166 | 0.036 | Young |
| 0.397144 | 1.515173 | 0.013 | 0.005 | 1 | 4 | AC152827. | 0.008 | Young |
| 0.397144 | 1.44718 | 0.013 | 0.005 | 1 | 4 | 1500002C1 | 0.008 | Young |
| 0.397144 | 1.316158 | 0.013 | 0.005 | 1 | 4 | Thrsp | 0.008 | Young |
| 0.398116 | 0.60276 | 0.075 | 0.053 | 1 | 4 | Cryzl2 | 0.022 | Young |
| 0.398157 | 0.061844 | 0.275 | 0.231 | 1 | 4 | Ccnt2 | 0.044 | Young |
| 0.398178 | 0.046727 | 0.875 | 0.816 | 1 | 4 | Mt1 | 0.059 | Young |
| 0.398255 | 0.086154 | 0.075 | 0.052 | 1 | 4 | Cbfa2t2 | 0.023 | Young |
| 0.398362 | 0.718594 | 0.112 | 0.086 | 1 | 4 | Akap7 | 0.026 | Young |
| 0.398447 | 0.39636 | 0.162 | 0.132 | 1 | 4 | Vrk1 | 0.03 | Young |
| 0.398626 | 0.354314 | 0.175 | 0.143 | 1 | 4 | Isy1 | 0.032 | Young |
| 0.39897 | 0.005605 | 0.075 | 0.052 | 1 | 4 | Prmt5 | 0.023 | Young |
| 0.398975 | 0.00512 | 0.2 | 0.159 | 1 | 4 | Txndc9 | 0.041 | Young |
| 0.399286 | 1.276196 | 0.013 | 0.005 | 1 | 4 | Ift122 | 0.008 | Young |
| 0.399286 | 1.060058 | 0.013 | 0.005 | 1 | 4 | Tal1 | 0.008 | Young |
| 0.399286 | 0.933673 | 0.013 | 0.005 | 1 | 4 | Gm38079 | 0.008 | Young |
| 0.399592 | 0.050239 | 1 | 0.999 | 1 | 4 | Tmsb4x | 0.001 | Young |
| 0.399595 | 0.315542 | 0.5 | 0.441 | 1 | 4 | Ap2s1 | 0.059 | Young |
| 0.399805 | 0.314519 | 0.112 | 0.086 | 1 | 4 | Usp53 | 0.026 | Young |
| 0.399812 | 0.170442 | 0.3 | 0.253 | 1 | 4 | Dynll2 | 0.047 | Young |
| 0.399942 | 0.072505 | 0.238 | 0.19 | 1 | 4 | Ndufb6 | 0.048 | Young |
| 0.400003 | 0.479352 | 0.125 | 0.097 | 1 | 4 | Asxl1 | 0.028 | Young |
| 0.400083 | 0.798217 | 0.112 | 0.088 | 1 | 4 | Maff | 0.024 | Young |
| 0.400359 | 1.020223 | 0.013 | 0.005 | 1 | 4 | B3gntl1 | 0.008 | Young |
| 0.400359 | 0.96412 | 0.013 | 0.005 | 1 | 4 | Mppe1 | 0.008 | Young |
| 0.400359 | 0.83905 | 0.013 | 0.005 | 1 | 4 | Cldn5 | 0.008 | Young |
| 0.400359 | 0.792126 | 0.013 | 0.005 | 1 | 4 | Srgap3 | 0.008 | Young |
| 0.400359 | 0.710594 | 0.013 | 0.005 | 1 | 4 | Armcx6 | 0.008 | Young |
| 0.400359 | 0.637638 | 0.013 | 0.005 | 1 | 4 | Ndc80 | 0.008 | Young |
| 0.400359 | 0.572152 | 0.013 | 0.005 | 1 | 4 | Tnfsf14 | 0.008 | Young |
| 0.400359 | 0.304699 | 0.013 | 0.005 | 1 | 4 | Jmjd4 | 0.008 | Young |
| 0.400359 | 0.071647 | 0.013 | 0.005 | 1 | 4 | Six5 | 0.008 | Young |
| 0.400538 | 0.249254 | 0.112 | 0.085 | 1 | 4 | Creb3 | 0.027 | Young |
| 0.400681 | 0.487633 | 0.188 | 0.155 | 1 | 4 | Sys1 | 0.033 | Young |
| 0.400813 | 0.326475 | 0.125 | 0.096 | 1 | 4 | Plcl2 | 0.029 | Young |

| | | | | | | | | |
|---|---|---|---|---|---|---|---|---|
| 0.401434 | 0.814597 | 0.013 | 0.005 | 1 | 4 | Gm16861 | 0.008 | Young |
| 0.401434 | 0.094471 | 0.013 | 0.005 | 1 | 4 | Dnph1 | 0.008 | Young |
| 0.40147 | 0.352999 | 0.15 | 0.12 | 1 | 4 | Npm3 | 0.03 | Young |
| 0.401789 | 0.510885 | 0.2 | 0.174 | 1 | 4 | Snd1 | 0.026 | Young |
| 0.401909 | 0.27805 | 0.25 | 0.211 | 1 | 4 | Dnajc8 | 0.039 | Young |
| 0.401932 | 0.203389 | 0.138 | 0.107 | 1 | 4 | Dnmt1 | 0.031 | Young |
| 0.401992 | 1.445525 | 0.025 | 0.014 | 1 | 4 | Kdelc1 | 0.011 | Young |
| 0.402336 | 0.129451 | 0.35 | 0.302 | 1 | 4 | Add1 | 0.048 | Young |
| 0.40247 | 0.076142 | 0.088 | 0.063 | 1 | 4 | Ap1m1 | 0.025 | Young |
| 0.402511 | 0.340781 | 0.013 | 0.005 | 1 | 4 | Jcad | 0.008 | Young |
| 0.402511 | 0.185943 | 0.013 | 0.005 | 1 | 4 | Fndc10 | 0.008 | Young |
| 0.402548 | 0.049369 | 0.162 | 0.126 | 1 | 4 | Dapk3 | 0.036 | Young |
| 0.402752 | 0.308472 | 0.2 | 0.161 | 1 | 4 | Trim56 | 0.039 | Young |
| 0.403547 | 0.363855 | 0.112 | 0.085 | 1 | 4 | Pgap1 | 0.027 | Young |
| 0.40359 | 0.061697 | 0.013 | 0.005 | 1 | 4 | Htra1 | 0.008 | Young |
| 0.404114 | 0.034962 | 0.088 | 0.063 | 1 | 4 | Gramd1b | 0.025 | Young |
| 0.404435 | 0.231944 | 0.075 | 0.053 | 1 | 4 | Jrkl | 0.022 | Young |
| 0.404516 | 1.262659 | 0.038 | 0.023 | 1 | 4 | Mapre3 | 0.015 | Young |
| 0.404671 | 0.058362 | 0.013 | 0.005 | 1 | 4 | Atp9a | 0.008 | Young |
| 0.404857 | 0.398463 | 0.075 | 0.053 | 1 | 4 | Btbd3 | 0.022 | Young |
| 0.404881 | 0.133232 | 0.312 | 0.262 | 1 | 4 | Tnpo3 | 0.05 | Young |
| 0.404911 | 0.080797 | 0.762 | 0.723 | 1 | 4 | Rplp0 | 0.039 | Young |
| 0.405087 | 0.303459 | 0.238 | 0.197 | 1 | 4 | Suz12 | 0.041 | Young |
| 0.405274 | 0.262678 | 0.275 | 0.236 | 1 | 4 | Ccar1 | 0.039 | Young |
| 0.406386 | 0.49848 | 0.05 | 0.033 | 1 | 4 | Gfer | 0.017 | Young |
| 0.406578 | 0.080042 | 0.475 | 0.417 | 1 | 4 | Syk | 0.058 | Young |
| 0.406643 | 0.582209 | 0.038 | 0.023 | 1 | 4 | Aldh4a1 | 0.015 | Young |
| 0.406904 | 0.236864 | 0.588 | 0.567 | 1 | 4 | Zbtb20 | 0.021 | Young |
| 0.407284 | 0.784095 | 0.05 | 0.033 | 1 | 4 | Mypopos | 0.017 | Young |
| 0.407351 | 0.267408 | 0.125 | 0.096 | 1 | 4 | Filip1l | 0.029 | Young |
| 0.407778 | 0.030861 | 0.388 | 0.331 | 1 | 4 | Ankrd11 | 0.057 | Young |
| 0.408039 | 0.23302 | 0.175 | 0.14 | 1 | 4 | Phlda1 | 0.035 | Young |
| 0.408173 | 0.171264 | 0.175 | 0.141 | 1 | 4 | Tia1 | 0.034 | Young |
| 0.408243 | 0.581444 | 0.038 | 0.023 | 1 | 4 | Gin1 | 0.015 | Young |
| 0.408405 | 0.068453 | 0.138 | 0.106 | 1 | 4 | Itgb1bp1 | 0.032 | Young |
| 0.408902 | 0.316799 | 0.3 | 0.263 | 1 | 4 | Rlim | 0.037 | Young |
| 0.410059 | 0.944552 | 0.062 | 0.044 | 1 | 4 | Plxna4os1 | 0.018 | Young |
| 0.410162 | 1.02203 | 0.025 | 0.014 | 1 | 4 | Zfp229 | 0.011 | Young |
| 0.410244 | 0.845716 | 0.075 | 0.055 | 1 | 4 | Inpp5a | 0.02 | Young |
| 0.410382 | 0.602091 | 0.038 | 0.023 | 1 | 4 | Sort1 | 0.015 | Young |
| 0.410396 | 0.604196 | 0.2 | 0.167 | 1 | 4 | R3hdm1 | 0.033 | Young |
| 0.410821 | 0.255592 | 0.15 | 0.119 | 1 | 4 | AI413582 | 0.031 | Young |
| 0.410847 | 1.016367 | 0.025 | 0.014 | 1 | 4 | Als2cl | 0.011 | Young |
| 0.411 | 0.585587 | 0.112 | 0.086 | 1 | 4 | Erg28 | 0.026 | Young |
| 0.411079 | 0.571634 | 0.062 | 0.043 | 1 | 4 | Gm26782 | 0.019 | Young |

| | | | | | | | | |
|---|---|---|---|---|---|---|---|---|
| 0.411081 | 0.045627 | 0.225 | 0.183 | 1 | 4 | Kansl1 | 0.042 | Young |
| 0.41165 | 0.129354 | 0.5 | 0.443 | 1 | 4 | Blvrb | 0.057 | Young |
| 0.411861 | 0.161049 | 0.3 | 0.252 | 1 | 4 | Nsd3 | 0.048 | Young |
| 0.411936 | 0.136504 | 0.5 | 0.45 | 1 | 4 | Tgfb1 | 0.05 | Young |
| 0.412026 | 0.292023 | 0.075 | 0.053 | 1 | 4 | Prr12 | 0.022 | Young |
| 0.412168 | 0.179984 | 0.35 | 0.306 | 1 | 4 | Ttc3 | 0.044 | Young |
| 0.412596 | 0.45657 | 0.162 | 0.131 | 1 | 4 | Mitf | 0.031 | Young |
| 0.412954 | 0.053373 | 0.262 | 0.316 | 1 | 4 | Ppig | -0.054 | Young |
| 0.413106 | 0.114686 | 0.55 | 0.498 | 1 | 4 | Clic1 | 0.052 | Young |
| 0.413112 | 0.303154 | 0.225 | 0.191 | 1 | 4 | Tnfrsf13b | 0.034 | Young |
| 0.413468 | 0.490207 | 0.075 | 0.054 | 1 | 4 | Aasdhppt | 0.021 | Young |
| 0.413595 | 0.672549 | 0.025 | 0.014 | 1 | 4 | Tnnt1 | 0.011 | Young |
| 0.413631 | 0.039659 | 0.15 | 0.116 | 1 | 4 | Slc44a1 | 0.034 | Young |
| 0.413957 | 0.120267 | 0.238 | 0.196 | 1 | 4 | Rufy3 | 0.042 | Young |
| 0.414207 | 0.22147 | 0.262 | 0.223 | 1 | 4 | Phf20l1 | 0.039 | Young |
| 0.414257 | 0.547128 | 0.062 | 0.043 | 1 | 4 | Dnttip1 | 0.019 | Young |
| 0.414284 | 0.70883 | 0.025 | 0.014 | 1 | 4 | Rpl9-ps6 | 0.011 | Young |
| 0.414514 | 0.443027 | 0.05 | 0.033 | 1 | 4 | Zfp64 | 0.017 | Young |
| 0.414675 | 0.097172 | 0.162 | 0.202 | 1 | 4 | Akirin1 | -0.04 | Young |
| 0.414679 | 0.27133 | 0.038 | 0.023 | 1 | 4 | Etl4 | 0.015 | Young |
| 0.414915 | 0.116444 | 0.075 | 0.053 | 1 | 4 | Atp11a | 0.022 | Young |
| 0.415223 | 0.42943 | 0.125 | 0.097 | 1 | 4 | Tob1 | 0.028 | Young |
| 0.415709 | 0.45463 | 0.162 | 0.132 | 1 | 4 | Zc3h4 | 0.03 | Young |
| 0.415932 | 0.197023 | 0.9 | 0.904 | 1 | 4 | Mcl1 | -0.004 | Young |
| 0.416298 | 0.184315 | 0.038 | 0.023 | 1 | 4 | Pomt2 | 0.015 | Young |
| 0.416838 | 0.130145 | 0.038 | 0.023 | 1 | 4 | Ift80 | 0.015 | Young |
| 0.417635 | 0.148802 | 0.338 | 0.293 | 1 | 4 | Ppp1r21 | 0.045 | Young |
| 0.417737 | 0.227193 | 0.025 | 0.014 | 1 | 4 | Ipp | 0.011 | Young |
| 0.418187 | 0.310632 | 0.15 | 0.121 | 1 | 4 | Cask | 0.029 | Young |
| 0.418365 | 0.105705 | 0.35 | 0.304 | 1 | 4 | Ankrd44 | 0.046 | Young |
| 0.418461 | 0.182704 | 0.038 | 0.023 | 1 | 4 | Meis3 | 0.015 | Young |
| 0.418925 | 0.006238 | 0.613 | 0.534 | 1 | 4 | Atp6ap2 | 0.079 | Young |
| 0.419452 | 0.468166 | 0.062 | 0.043 | 1 | 4 | Ust | 0.019 | Young |
| 0.419457 | 0.145643 | 0.238 | 0.201 | 1 | 4 | Agps | 0.037 | Young |
| 0.419992 | 0.188284 | 0.537 | 0.476 | 1 | 4 | Pomp | 0.061 | Young |
| 0.420088 | 0.067074 | 0.038 | 0.023 | 1 | 4 | Coq3 | 0.015 | Young |
| 0.420171 | 0.209214 | 0.875 | 0.874 | 1 | 4 | Ubc | 0.001 | Young |
| 0.420531 | 0.053543 | 0.062 | 0.043 | 1 | 4 | Tnni3 | 0.019 | Young |
| 0.420549 | 0.456561 | 0.138 | 0.111 | 1 | 4 | Klhl28 | 0.027 | Young |
| 0.421063 | 0.143823 | 0.175 | 0.14 | 1 | 4 | BC003331 | 0.035 | Young |
| 0.421107 | 0.261335 | 0.112 | 0.086 | 1 | 4 | Coro2b | 0.026 | Young |
| 0.421364 | 0.671461 | 0.05 | 0.034 | 1 | 4 | Rpap2 | 0.016 | Young |
| 0.421414 | 0.397906 | 0.075 | 0.054 | 1 | 4 | Sema4b | 0.021 | Young |
| 0.422085 | 0.134685 | 0.112 | 0.086 | 1 | 4 | Prrg1 | 0.026 | Young |
| 0.422608 | 0.474634 | 0.125 | 0.098 | 1 | 4 | Ebpl | 0.027 | Young |

| | | | | | | | | |
|---|---|---|---|---|---|---|---|---|
| 0.422623 | 0.290724 | 0.262 | 0.214 | 1 | 4 | Skp1a | 0.048 | Young |
| 0.422737 | 0.001903 | 0.05 | 0.033 | 1 | 4 | Polr3e | 0.017 | Young |
| 0.423563 | 0.34323 | 0.338 | 0.285 | 1 | 4 | Lmbrd1 | 0.053 | Young |
| 0.42363 | 0.225005 | 0.212 | 0.179 | 1 | 4 | Hcfc1r1 | 0.033 | Young |
| 0.42387 | 0.018652 | 0.525 | 0.453 | 1 | 4 | Mapkapk2 | 0.072 | Young |
| 0.423967 | 0.340061 | 0.15 | 0.12 | 1 | 4 | Trim33 | 0.03 | Young |
| 0.424756 | 0.739233 | 0.062 | 0.044 | 1 | 4 | Dhrs4 | 0.018 | Young |
| 0.424917 | 0.473245 | 0.138 | 0.111 | 1 | 4 | Gemin7 | 0.027 | Young |
| 0.425055 | 0.285659 | 0.075 | 0.054 | 1 | 4 | Rngtt | 0.021 | Young |
| 0.425102 | 0.105164 | 0.288 | 0.237 | 1 | 4 | Ppp4r3b | 0.051 | Young |
| 0.425972 | 0.453885 | 0.138 | 0.111 | 1 | 4 | Tmem251 | 0.027 | Young |
| 0.426136 | 0.704556 | 0.162 | 0.136 | 1 | 4 | Rab43 | 0.026 | Young |
| 0.426378 | 0.157191 | 0.775 | 0.713 | 1 | 4 | Rps6 | 0.062 | Young |
| 0.426382 | 0.650516 | 0.05 | 0.034 | 1 | 4 | Aimp2 | 0.016 | Young |
| 0.426387 | 0.309123 | 0.225 | 0.194 | 1 | 4 | Paics | 0.031 | Young |
| 0.426394 | 0.239158 | 0.112 | 0.086 | 1 | 4 | Uchl3 | 0.026 | Young |
| 0.4266 | 0.164813 | 0.588 | 0.556 | 1 | 4 | Ndufa4 | 0.032 | Young |
| 0.426707 | 0.277259 | 0.062 | 0.043 | 1 | 4 | Zdhhc23 | 0.019 | Young |
| 0.426986 | 0.086081 | 0.925 | 0.875 | 1 | 4 | Rpl17 | 0.05 | Young |
| 0.427112 | 0.420635 | 0.062 | 0.043 | 1 | 4 | C1ra | 0.019 | Young |
| 0.427574 | 0.170183 | 0.225 | 0.189 | 1 | 4 | Gm10073 | 0.036 | Young |
| 0.427811 | 0.531714 | 0.088 | 0.065 | 1 | 4 | Yod1 | 0.023 | Young |
| 0.428126 | 0.355796 | 0.062 | 0.043 | 1 | 4 | Wdsub1 | 0.019 | Young |
| 0.428347 | 0.313196 | 0.075 | 0.054 | 1 | 4 | Pycr2 | 0.021 | Young |
| 0.428371 | 0.167369 | 0.162 | 0.127 | 1 | 4 | Itpripl1 | 0.035 | Young |
| 0.428513 | 0.07016 | 0.575 | 0.51 | 1 | 4 | Eif4a1 | 0.065 | Young |
| 0.428979 | 0.677438 | 0.075 | 0.055 | 1 | 4 | Hmbs | 0.02 | Young |
| 0.429264 | 0.227255 | 0.125 | 0.096 | 1 | 4 | Brd9 | 0.029 | Young |
| 0.43003 | 0.81122 | 0.038 | 0.024 | 1 | 4 | Mipep | 0.014 | Young |
| 0.430105 | 0.059815 | 0.35 | 0.295 | 1 | 4 | Cd52 | 0.055 | Young |
| 0.43076 | 0.132058 | 0.262 | 0.226 | 1 | 4 | Rilpl2 | 0.036 | Young |
| 0.431257 | 0.086942 | 0.112 | 0.086 | 1 | 4 | Zfp871 | 0.026 | Young |
| 0.431673 | 0.039795 | 0.262 | 0.216 | 1 | 4 | 9030624J0 | 0.046 | Young |
| 0.432123 | 0.124084 | 0.162 | 0.129 | 1 | 4 | Trappc5 | 0.033 | Young |
| 0.433726 | 0.352431 | 0.25 | 0.21 | 1 | 4 | Cisd2 | 0.04 | Young |
| 0.433877 | 0.050215 | 0.238 | 0.198 | 1 | 4 | Mid1ip1 | 0.04 | Young |
| 0.433953 | 0.102252 | 0.475 | 0.423 | 1 | 4 | Ptger4 | 0.052 | Young |
| 0.434162 | 0.013671 | 0.375 | 0.312 | 1 | 4 | Lgals3bp | 0.063 | Young |
| 0.434332 | 0.150203 | 0.362 | 0.306 | 1 | 4 | Sec11c | 0.056 | Young |
| 0.434391 | 0.615254 | 0.038 | 0.024 | 1 | 4 | Tefm | 0.014 | Young |
| 0.434408 | 0.303447 | 0.175 | 0.144 | 1 | 4 | Samd8 | 0.031 | Young |
| 0.435322 | 0.161791 | 0.1 | 0.134 | 1 | 4 | Mrps15 | -0.034 | Young |
| 0.435485 | 0.580518 | 0.038 | 0.024 | 1 | 4 | L3mbtl3 | 0.014 | Young |
| 0.435755 | 0.08769 | 0.175 | 0.139 | 1 | 4 | Rsl1d1 | 0.036 | Young |
| 0.435789 | 0.299265 | 0.212 | 0.175 | 1 | 4 | Zfp292 | 0.037 | Young |

| | | | | | | | | |
|---|---|---|---|---|---|---|---|---|
| 0.435891 | 1.028375 | 0.05 | 0.034 | 1 | 4 | C87436 | 0.016 | Young |
| 0.436156 | 0.015868 | 0.125 | 0.096 | 1 | 4 | Myo18a | 0.029 | Young |
| 0.436368 | 0.512113 | 0.138 | 0.113 | 1 | 4 | Tnpo2 | 0.025 | Young |
| 0.436449 | 0.174404 | 0.212 | 0.175 | 1 | 4 | Gprc5c | 0.037 | Young |
| 0.436783 | 0.169425 | 0.175 | 0.142 | 1 | 4 | Nin | 0.033 | Young |
| 0.437787 | 0.12731 | 0.275 | 0.231 | 1 | 4 | Arhgap45 | 0.044 | Young |
| 0.437852 | 0.069741 | 0.2 | 0.164 | 1 | 4 | Wdr37 | 0.036 | Young |
| 0.438014 | 0.221957 | 0.275 | 0.246 | 1 | 4 | Retreg2 | 0.029 | Young |
| 0.438161 | 0.403284 | 0.075 | 0.055 | 1 | 4 | Nadk2 | 0.02 | Young |
| 0.438342 | 0.271952 | 0.288 | 0.248 | 1 | 4 | Tubb2a | 0.04 | Young |
| 0.438399 | 0.072825 | 0.875 | 0.881 | 1 | 4 | Zfp36l2 | -0.006 | Young |
| 0.438426 | 0.092895 | 0.438 | 0.377 | 1 | 4 | Slc38a2 | 0.061 | Young |
| 0.438549 | 0.158575 | 0.062 | 0.043 | 1 | 4 | Cdc14a | 0.019 | Young |
| 0.43873 | 0.700867 | 0.075 | 0.055 | 1 | 4 | Pgam5 | 0.02 | Young |
| 0.439084 | 0.071372 | 0.138 | 0.107 | 1 | 4 | Erap1 | 0.031 | Young |
| 0.439568 | 0.098344 | 0.138 | 0.108 | 1 | 4 | Stau1 | 0.03 | Young |
| 0.439897 | 0.021517 | 0.013 | 0.027 | 1 | 4 | Ccdc191 | -0.014 | Young |
| 0.440174 | 0.131563 | 0.075 | 0.054 | 1 | 4 | Cars2 | 0.021 | Young |
| 0.440428 | 0.141102 | 0.013 | 0.027 | 1 | 4 | Gja1 | -0.014 | Young |
| 0.440577 | 0.171687 | 0.225 | 0.19 | 1 | 4 | Vcpip1 | 0.035 | Young |
| 0.440744 | 1.172942 | 0.075 | 0.056 | 1 | 4 | Vopp1 | 0.019 | Young |
| 0.441335 | 1.364583 | 0.025 | 0.014 | 1 | 4 | Sorl1 | 0.011 | Young |
| 0.441337 | 0.092262 | 0.25 | 0.205 | 1 | 4 | Zbtb4 | 0.045 | Young |
| 0.441396 | 0.409309 | 0.062 | 0.044 | 1 | 4 | Myef2 | 0.018 | Young |
| 0.44168 | 0.035264 | 0.488 | 0.417 | 1 | 4 | Atp6v1f | 0.071 | Young |
| 0.441773 | 0.539444 | 0.2 | 0.171 | 1 | 4 | Bax | 0.029 | Young |
| 0.441865 | 0.384686 | 0.075 | 0.055 | 1 | 4 | 1600002K0 | 0.02 | Young |
| 0.442084 | 0.180519 | 0.038 | 0.024 | 1 | 4 | Gpr162 | 0.014 | Young |
| 0.442262 | 0.107483 | 0.062 | 0.043 | 1 | 4 | Eogt | 0.019 | Young |
| 0.442679 | 0.327124 | 0.125 | 0.099 | 1 | 4 | Il17ra | 0.026 | Young |
| 0.442719 | 0.497399 | 0.1 | 0.078 | 1 | 4 | Reps1 | 0.022 | Young |
| 0.442807 | 0.231824 | 0.062 | 0.088 | 1 | 4 | Golga5 | -0.026 | Young |
| 0.442938 | 0.028324 | 0.588 | 0.511 | 1 | 4 | Dock2 | 0.077 | Young |
| 0.443189 | 0.343639 | 0.038 | 0.024 | 1 | 4 | 2410131K1 | 0.014 | Young |
| 0.443307 | 0.388071 | 0.188 | 0.157 | 1 | 4 | Cnot3 | 0.031 | Young |
| 0.444073 | 0.336305 | 0.112 | 0.088 | 1 | 4 | Scaf4 | 0.024 | Young |
| 0.444426 | 0.152572 | 0.412 | 0.37 | 1 | 4 | Arf3 | 0.042 | Young |
| 0.444474 | 0.036022 | 0.138 | 0.107 | 1 | 4 | Fam193a | 0.031 | Young |
| 0.444568 | 0.210015 | 0.462 | 0.431 | 1 | 4 | Minos1 | 0.031 | Young |
| 0.444573 | 0.079773 | 0.65 | 0.592 | 1 | 4 | 2010107E0 | 0.058 | Young |
| 0.445443 | 0.079629 | 0.637 | 0.574 | 1 | 4 | Son | 0.063 | Young |
| 0.446325 | 0.277221 | 0.112 | 0.088 | 1 | 4 | Trim34a | 0.024 | Young |
| 0.446381 | 0.212024 | 0.125 | 0.097 | 1 | 4 | Slc11a2 | 0.028 | Young |
| 0.446509 | 0.242921 | 0.088 | 0.065 | 1 | 4 | Slc4a1ap | 0.023 | Young |
| 0.44663 | 0.002315 | 0.188 | 0.151 | 1 | 4 | Nsun2 | 0.037 | Young |

| | | | | | | | | |
|---|---|---|---|---|---|---|---|---|
| 0.446955 | 1.072468 | 0.025 | 0.014 | 1 | 4 | Txnl4b | 0.011 | Young |
| 0.447455 | 0.271575 | 0.075 | 0.055 | 1 | 4 | Ccdc117 | 0.02 | Young |
| 0.447602 | 0.084307 | 0.537 | 0.462 | 1 | 4 | Nktr | 0.075 | Young |
| 0.44766 | 0.82513 | 0.025 | 0.014 | 1 | 4 | Purg | 0.011 | Young |
| 0.447921 | 0.105145 | 0.212 | 0.177 | 1 | 4 | Rsbn1l | 0.035 | Young |
| 0.448352 | 0.065884 | 0.25 | 0.206 | 1 | 4 | Arpp19 | 0.044 | Young |
| 0.449073 | 0.797807 | 0.025 | 0.014 | 1 | 4 | Gm16124 | 0.011 | Young |
| 0.44957 | 0.428295 | 0.15 | 0.122 | 1 | 4 | Cpd | 0.028 | Young |
| 0.449691 | 0.196175 | 0.488 | 0.445 | 1 | 4 | Nfkb1 | 0.043 | Young |
| 0.450061 | 0.540802 | 0.062 | 0.044 | 1 | 4 | Aatf | 0.018 | Young |
| 0.450447 | 0.169555 | 0.138 | 0.109 | 1 | 4 | Cmas | 0.029 | Young |
| 0.451484 | 0.923601 | 0.125 | 0.102 | 1 | 4 | Ppm1g | 0.023 | Young |
| 0.451598 | 0.014959 | 0.088 | 0.065 | 1 | 4 | Gpn1 | 0.023 | Young |
| 0.451712 | 0.028582 | 0.238 | 0.196 | 1 | 4 | Tnrc6c | 0.042 | Young |
| 0.451846 | 0.126557 | 0.162 | 0.13 | 1 | 4 | Tmem230 | 0.032 | Young |
| 0.452617 | 0.497404 | 0.05 | 0.034 | 1 | 4 | 2610524H( | 0.016 | Young |
| 0.452627 | 0.032661 | 0.288 | 0.24 | 1 | 4 | Spcs3 | 0.048 | Young |
| 0.452689 | 0.343318 | 0.1 | 0.077 | 1 | 4 | Atxn7l3 | 0.023 | Young |
| 0.452953 | 0.743712 | 0.088 | 0.068 | 1 | 4 | Slc35e1 | 0.02 | Young |
| 0.452971 | 0.230951 | 0.062 | 0.044 | 1 | 4 | Scnm1 | 0.018 | Young |
| 0.453134 | 0.116194 | 0.35 | 0.306 | 1 | 4 | Ifnar1 | 0.044 | Young |
| 0.45332 | 0.637529 | 0.075 | 0.056 | 1 | 4 | Acvr1 | 0.019 | Young |
| 0.453326 | 0.658795 | 0.025 | 0.014 | 1 | 4 | Mapkapk5 | 0.011 | Young |
| 0.453326 | 0.484428 | 0.025 | 0.014 | 1 | 4 | Map2k6 | 0.011 | Young |
| 0.454474 | 0.189433 | 0.3 | 0.256 | 1 | 4 | Fxr1 | 0.044 | Young |
| 0.454598 | 0.852118 | 0.05 | 0.035 | 1 | 4 | Gbp5 | 0.015 | Young |
| 0.454749 | 0.54081 | 0.025 | 0.014 | 1 | 4 | Zfp688 | 0.011 | Young |
| 0.454969 | 0.49873 | 0.05 | 0.034 | 1 | 4 | Trp53rka | 0.016 | Young |
| 0.454969 | 0.451993 | 0.05 | 0.034 | 1 | 4 | Wdr70 | 0.016 | Young |
| 0.455454 | 0.134066 | 0.462 | 0.419 | 1 | 4 | Lyn | 0.043 | Young |
| 0.456105 | 0.017345 | 1 | 0.994 | 1 | 4 | Serinc3 | 0.006 | Young |
| 0.456123 | 0.14298 | 0.125 | 0.099 | 1 | 4 | Phf5a | 0.026 | Young |
| 0.456266 | 0.913286 | 0.038 | 0.024 | 1 | 4 | Plxna4 | 0.014 | Young |
| 0.456461 | 0.452108 | 0.15 | 0.124 | 1 | 4 | Dyrk1a | 0.026 | Young |
| 0.456821 | 0.893082 | 0.038 | 0.024 | 1 | 4 | Hist2h2be | 0.014 | Young |
| 0.456903 | 0.249709 | 0.25 | 0.212 | 1 | 4 | Snrpc | 0.038 | Young |
| 0.45717 | 0.290297 | 0.125 | 0.098 | 1 | 4 | Mfsd12 | 0.027 | Young |
| 0.457185 | 0.12172 | 0.662 | 0.626 | 1 | 4 | H2-DMa | 0.036 | Young |
| 0.457602 | 0.18778 | 0.025 | 0.014 | 1 | 4 | Kctd7 | 0.011 | Young |
| 0.457689 | 0.000999 | 0.112 | 0.086 | 1 | 4 | Slc35a2 | 0.026 | Young |
| 0.45788 | 0.403158 | 0.1 | 0.078 | 1 | 4 | Tnfrsf25 | 0.022 | Young |
| 0.458172 | 0.256626 | 0.088 | 0.066 | 1 | 4 | Cdr2 | 0.022 | Young |
| 0.458396 | 0.154577 | 0.338 | 0.294 | 1 | 4 | Tubb6 | 0.044 | Young |
| 0.458423 | 0.176655 | 0.425 | 0.383 | 1 | 4 | Ppp1r9b | 0.042 | Young |
| 0.458523 | 0.07263 | 0.138 | 0.108 | 1 | 4 | Prkag1 | 0.03 | Young |

| | | | | | | | | |
|---|---|---|---|---|---|---|---|---|
| 0.459692 | 0.449447 | 0.05 | 0.034 | 1 | 4 | Papolg | 0.016 | Young |
| 0.459748 | 0.160979 | 0.025 | 0.014 | 1 | 4 | Slfn9 | 0.011 | Young |
| 0.460146 | 0.03756 | 0.125 | 0.097 | 1 | 4 | Ticam2 | 0.028 | Young |
| 0.460166 | 0.433014 | 0.05 | 0.034 | 1 | 4 | Cdc42bpg | 0.016 | Young |
| 0.460177 | 0.132995 | 0.275 | 0.242 | 1 | 4 | Entpd1 | 0.033 | Young |
| 0.460408 | 0.012862 | 0.275 | 0.223 | 1 | 4 | Tmsb10 | 0.052 | Young |
| 0.460726 | 0.518981 | 0.112 | 0.09 | 1 | 4 | Ppp1r37 | 0.022 | Young |
| 0.461693 | 0.536807 | 0.088 | 0.067 | 1 | 4 | Wdr43 | 0.021 | Young |
| 0.461837 | 0.235603 | 0.738 | 0.691 | 1 | 4 | Pim1 | 0.047 | Young |
| 0.462064 | 0.209223 | 0.05 | 0.034 | 1 | 4 | Ints10 | 0.016 | Young |
| 0.462397 | 0.545418 | 0.038 | 0.024 | 1 | 4 | Wdr73 | 0.014 | Young |
| 0.462546 | 0.721601 | 0.088 | 0.069 | 1 | 4 | Mtif2 | 0.019 | Young |
| 0.462957 | 0.45715 | 0.038 | 0.024 | 1 | 4 | P2ry14 | 0.014 | Young |
| 0.463254 | 0.069218 | 0.225 | 0.184 | 1 | 4 | Fmnl1 | 0.041 | Young |
| 0.463276 | 1.933181 | 0.013 | 0.006 | 1 | 4 | Isoc2a | 0.007 | Young |
| 0.463517 | 0.555653 | 0.038 | 0.024 | 1 | 4 | Lag3 | 0.014 | Young |
| 0.463652 | 0.167321 | 0.112 | 0.088 | 1 | 4 | Tvp23b | 0.024 | Young |
| 0.464077 | 0.687709 | 0.038 | 0.024 | 1 | 4 | Mtrf1l | 0.014 | Young |
| 0.464077 | 0.384836 | 0.038 | 0.024 | 1 | 4 | Ulk3 | 0.014 | Young |
| 0.464386 | 1.576987 | 0.013 | 0.006 | 1 | 4 | Prodh | 0.007 | Young |
| 0.464386 | 1.438033 | 0.013 | 0.006 | 1 | 4 | Abhd3 | 0.007 | Young |
| 0.464386 | 1.367139 | 0.013 | 0.006 | 1 | 4 | Dpep1 | 0.007 | Young |
| 0.464386 | 1.002409 | 0.013 | 0.006 | 1 | 4 | Zfp647 | 0.007 | Young |
| 0.464827 | 0.724747 | 0.062 | 0.046 | 1 | 4 | Prmt3 | 0.016 | Young |
| 0.465199 | 0.450757 | 0.038 | 0.024 | 1 | 4 | Zfp568 | 0.014 | Young |
| 0.465498 | 1.229607 | 0.013 | 0.006 | 1 | 4 | Usp2 | 0.007 | Young |
| 0.465498 | 0.917085 | 0.013 | 0.006 | 1 | 4 | Ccnb2 | 0.007 | Young |
| 0.465689 | 0.391473 | 0.112 | 0.088 | 1 | 4 | Homer1 | 0.024 | Young |
| 0.465873 | 0.278567 | 0.05 | 0.034 | 1 | 4 | Zkscan8 | 0.016 | Young |
| 0.465873 | 0.257269 | 0.05 | 0.034 | 1 | 4 | Zdhhc13 | 0.016 | Young |
| 0.466098 | 0.216123 | 0.188 | 0.157 | 1 | 4 | Eef2k | 0.031 | Young |
| 0.466343 | 0.604602 | 0.062 | 0.045 | 1 | 4 | Rptor | 0.017 | Young |
| 0.466611 | 1.210134 | 0.013 | 0.006 | 1 | 4 | Ifi44 | 0.007 | Young |
| 0.466611 | 0.962693 | 0.013 | 0.006 | 1 | 4 | Oas1c | 0.007 | Young |
| 0.466611 | 0.749363 | 0.013 | 0.006 | 1 | 4 | Gm12355 | 0.007 | Young |
| 0.466791 | 0.20779 | 0.425 | 0.368 | 1 | 4 | Lmo2 | 0.057 | Young |
| 0.467726 | 0.974959 | 0.013 | 0.006 | 1 | 4 | Nadsyn1 | 0.007 | Young |
| 0.467726 | 0.798329 | 0.013 | 0.006 | 1 | 4 | Rangrf | 0.007 | Young |
| 0.467726 | 0.727823 | 0.013 | 0.006 | 1 | 4 | Jmjd7 | 0.007 | Young |
| 0.467726 | 0.627815 | 0.013 | 0.006 | 1 | 4 | Tjp3 | 0.007 | Young |
| 0.468435 | 0.486191 | 0.062 | 0.045 | 1 | 4 | Slc39a14 | 0.017 | Young |
| 0.468663 | 0.357262 | 0.15 | 0.121 | 1 | 4 | 4833439L1 | 0.029 | Young |
| 0.468729 | 0.246049 | 0.425 | 0.387 | 1 | 4 | Il10ra | 0.038 | Young |
| 0.46879 | 0.100861 | 0.975 | 0.945 | 1 | 4 | Rps20 | 0.03 | Young |
| 0.468843 | 0.845332 | 0.013 | 0.006 | 1 | 4 | 6720489N1 | 0.007 | Young |

| | | | | | | | | |
|---|---|---|---|---|---|---|---|---|
| 0.468843 | 0.751521 | 0.013 | 0.006 | 1 | 4 | Serpina3f | 0.007 | Young |
| 0.468843 | 0.714247 | 0.013 | 0.006 | 1 | 4 | Chchd10 | 0.007 | Young |
| 0.468843 | 0.589417 | 0.013 | 0.006 | 1 | 4 | Gm29336 | 0.007 | Young |
| 0.468843 | 0.439718 | 0.013 | 0.006 | 1 | 4 | Phf11a | 0.007 | Young |
| 0.468843 | 0.434986 | 0.013 | 0.006 | 1 | 4 | Armcx2 | 0.007 | Young |
| 0.468919 | 0.435589 | 0.112 | 0.088 | 1 | 4 | Tssc4 | 0.024 | Young |
| 0.469067 | 0.114041 | 0.613 | 0.562 | 1 | 4 | Rnasel | 0.051 | Young |
| 0.469138 | 0.268459 | 0.038 | 0.024 | 1 | 4 | Mlst8 | 0.014 | Young |
| 0.469931 | 0.166019 | 0.275 | 0.235 | 1 | 4 | Tle4 | 0.04 | Young |
| 0.46996 | 0.654613 | 0.013 | 0.006 | 1 | 4 | Ubxn11 | 0.007 | Young |
| 0.46996 | 0.407503 | 0.013 | 0.006 | 1 | 4 | Igfbp5 | 0.007 | Young |
| 0.46996 | 0.219572 | 0.013 | 0.006 | 1 | 4 | Gpr19 | 0.007 | Young |
| 0.47003 | 0.201938 | 0.275 | 0.24 | 1 | 4 | Dhx15 | 0.035 | Young |
| 0.470179 | 0.04516 | 0.05 | 0.034 | 1 | 4 | Cnnm2 | 0.016 | Young |
| 0.470454 | 0.319002 | 0.212 | 0.181 | 1 | 4 | Msrb1 | 0.031 | Young |
| 0.470685 | 0.064384 | 0.3 | 0.258 | 1 | 4 | Glmp | 0.042 | Young |
| 0.470757 | 0.038073 | 0.537 | 0.472 | 1 | 4 | Sppl2a | 0.065 | Young |
| 0.470832 | 0.076139 | 0.038 | 0.024 | 1 | 4 | Bcar3 | 0.014 | Young |
| 0.47108 | 0.29497 | 0.013 | 0.006 | 1 | 4 | Unc13b | 0.007 | Young |
| 0.47108 | 0.052527 | 0.013 | 0.006 | 1 | 4 | Abca5 | 0.007 | Young |
| 0.471174 | 0.178802 | 0.162 | 0.134 | 1 | 4 | Utp11 | 0.028 | Young |
| 0.471397 | 0.187987 | 0.038 | 0.024 | 1 | 4 | Zeb1 | 0.014 | Young |
| 0.471659 | 0.187432 | 0.062 | 0.045 | 1 | 4 | Narfl | 0.017 | Young |
| 0.471912 | 0.270403 | 0.125 | 0.099 | 1 | 4 | Lrrfip2 | 0.026 | Young |
| 0.472173 | 0.719164 | 0.075 | 0.057 | 1 | 4 | Sugp1 | 0.018 | Young |
| 0.472227 | 0.197907 | 0.3 | 0.26 | 1 | 4 | Dmxl1 | 0.04 | Young |
| 0.472646 | 0.174534 | 0.3 | 0.262 | 1 | 4 | Csf2rb | 0.038 | Young |
| 0.472712 | 0.117659 | 0.6 | 0.585 | 1 | 4 | Cox6a1 | 0.015 | Young |
| 0.472726 | 0.205017 | 0.25 | 0.209 | 1 | 4 | Npl | 0.041 | Young |
| 0.472806 | 0.391406 | 0.1 | 0.078 | 1 | 4 | Yeats2 | 0.022 | Young |
| 0.473289 | 0.902177 | 0.075 | 0.057 | 1 | 4 | Pbxip1 | 0.018 | Young |
| 0.473709 | 2.289391 | 0.025 | 0.015 | 1 | 4 | Coq6 | 0.01 | Young |
| 0.474065 | 0.053677 | 0.05 | 0.071 | 1 | 4 | Traf5 | -0.021 | Young |
| 0.474797 | 0.033033 | 0.038 | 0.024 | 1 | 4 | Slc25a15 | 0.014 | Young |
| 0.474852 | 0.033975 | 0.075 | 0.055 | 1 | 4 | Lsm10 | 0.02 | Young |
| 0.475394 | 0.46183 | 0.05 | 0.035 | 1 | 4 | F8a | 0.015 | Young |
| 0.475394 | 0.460287 | 0.05 | 0.035 | 1 | 4 | Rrs1 | 0.015 | Young |
| 0.475422 | 0.59895 | 0.088 | 0.068 | 1 | 4 | Ipo9 | 0.02 | Young |
| 0.475533 | 0.095899 | 0.175 | 0.144 | 1 | 4 | Mien1 | 0.031 | Young |
| 0.475743 | 0.002919 | 0.2 | 0.165 | 1 | 4 | Cacybp | 0.035 | Young |
| 0.475853 | 1.159815 | 0.025 | 0.015 | 1 | 4 | Zfp790 | 0.01 | Young |
| 0.475873 | 0.874889 | 0.05 | 0.035 | 1 | 4 | Zc3h6 | 0.015 | Young |
| 0.476352 | 0.326819 | 0.05 | 0.035 | 1 | 4 | Eif2ak4 | 0.015 | Young |
| 0.476852 | 0.414162 | 0.062 | 0.045 | 1 | 4 | Pdik1l | 0.017 | Young |
| 0.477181 | 1.083875 | 0.05 | 0.036 | 1 | 4 | Srbd1 | 0.014 | Young |

| | | | | | | | | |
|---|---|---|---|---|---|---|---|---|
| 0.477327 | 0.165797 | 0.375 | 0.331 | 1 | 4 | Rassf1 | 0.044 | Young |
| 0.477453 | 0.046753 | 0.125 | 0.099 | 1 | 4 | Pik3r4 | 0.026 | Young |
| 0.477593 | 0.226777 | 0.088 | 0.067 | 1 | 4 | Zmat5 | 0.021 | Young |
| 0.478439 | 0.202361 | 0.138 | 0.109 | 1 | 4 | Grap | 0.029 | Young |
| 0.47897 | 0.238158 | 0.1 | 0.077 | 1 | 4 | Sntb2 | 0.023 | Young |
| 0.479068 | 0.310682 | 0.075 | 0.056 | 1 | 4 | Cntln | 0.019 | Young |
| 0.479231 | 0.428918 | 0.05 | 0.035 | 1 | 4 | Ube2e2 | 0.015 | Young |
| 0.47925 | 1.188628 | 0.038 | 0.025 | 1 | 4 | Pcolce | 0.013 | Young |
| 0.479477 | 0.011957 | 0.112 | 0.144 | 1 | 4 | Snx12 | -0.032 | Young |
| 0.479712 | 0.128925 | 0.525 | 0.491 | 1 | 4 | Scp2 | 0.034 | Young |
| 0.48082 | 0.117331 | 0.188 | 0.155 | 1 | 4 | Ddx21 | 0.033 | Young |
| 0.480993 | 0.269984 | 0.075 | 0.056 | 1 | 4 | Nt5c3 | 0.019 | Young |
| 0.481711 | 0.54522 | 0.075 | 0.057 | 1 | 4 | Rusc1 | 0.018 | Young |
| 0.482671 | 0.47523 | 0.075 | 0.057 | 1 | 4 | Wdr18 | 0.018 | Young |
| 0.482793 | 0.25318 | 0.062 | 0.045 | 1 | 4 | Rapgef1 | 0.017 | Young |
| 0.482856 | 0.186913 | 0.125 | 0.1 | 1 | 4 | 9930021J0 | 0.025 | Young |
| 0.482877 | 0.31048 | 0.112 | 0.089 | 1 | 4 | Mtx1 | 0.023 | Young |
| 0.483205 | 0.738849 | 0.038 | 0.025 | 1 | 4 | Fasn | 0.013 | Young |
| 0.483369 | 0.193823 | 0.2 | 0.17 | 1 | 4 | Wdr89 | 0.03 | Young |
| 0.483761 | 1.041971 | 0.025 | 0.015 | 1 | 4 | Fads6 | 0.01 | Young |
| 0.483761 | 0.905015 | 0.025 | 0.015 | 1 | 4 | Mad2l1 | 0.01 | Young |
| 0.483761 | 0.096777 | 0.075 | 0.1 | 1 | 4 | Ilf2 | -0.025 | Young |
| 0.483866 | 0.141048 | 0.25 | 0.213 | 1 | 4 | Ube2s | 0.037 | Young |
| 0.483964 | 0.344576 | 0.088 | 0.067 | 1 | 4 | Itgav | 0.021 | Young |
| 0.484236 | 0.170683 | 0.412 | 0.368 | 1 | 4 | Nfic | 0.044 | Young |
| 0.485206 | 0.903349 | 0.025 | 0.015 | 1 | 4 | Mms22l | 0.01 | Young |
| 0.485206 | 0.862226 | 0.025 | 0.015 | 1 | 4 | Coil | 0.01 | Young |
| 0.485241 | 0.127544 | 0.075 | 0.056 | 1 | 4 | Tfdp1 | 0.019 | Young |
| 0.485242 | 0.109265 | 0.375 | 0.328 | 1 | 4 | Psd3 | 0.047 | Young |
| 0.485314 | 0.137833 | 0.4 | 0.372 | 1 | 4 | Osbpl9 | 0.028 | Young |
| 0.48593 | 0.731508 | 0.025 | 0.015 | 1 | 4 | Abcb6 | 0.01 | Young |
| 0.486265 | 0.073473 | 0.175 | 0.144 | 1 | 4 | Csnk1g2 | 0.031 | Young |
| 0.486454 | 0.133457 | 0.088 | 0.067 | 1 | 4 | Rnf170 | 0.021 | Young |
| 0.486654 | 0.877307 | 0.025 | 0.015 | 1 | 4 | 4933433G1 | 0.01 | Young |
| 0.486654 | 0.719467 | 0.025 | 0.015 | 1 | 4 | Fitm2 | 0.01 | Young |
| 0.486795 | 0.286087 | 0.2 | 0.171 | 1 | 4 | Smarca2 | 0.029 | Young |
| 0.48739 | 0.148934 | 0.1 | 0.078 | 1 | 4 | Cisd1 | 0.022 | Young |
| 0.48748 | 0.274958 | 0.238 | 0.202 | 1 | 4 | Dhx40 | 0.036 | Young |
| 0.487772 | 0.523962 | 0.075 | 0.057 | 1 | 4 | Urm1 | 0.018 | Young |
| 0.487903 | 0.22619 | 0.125 | 0.1 | 1 | 4 | Nop58 | 0.025 | Young |
| 0.488317 | 0.525078 | 0.038 | 0.025 | 1 | 4 | Immp2l | 0.013 | Young |
| 0.489089 | 0.063693 | 0.338 | 0.285 | 1 | 4 | Lasp1 | 0.053 | Young |
| 0.489382 | 0.29421 | 0.05 | 0.035 | 1 | 4 | Mrpl22 | 0.015 | Young |
| 0.489445 | 0.143378 | 0.212 | 0.18 | 1 | 4 | Ccnl2 | 0.032 | Young |
| 0.489556 | 1.079325 | 0.025 | 0.015 | 1 | 4 | Clec3b | 0.01 | Young |

| | | | | | | | | |
|---|---|---|---|---|---|---|---|---|
| 0.489556 | 0.521868 | 0.025 | 0.015 | 1 | 4 | Plcb2 | 0.01 | Young |
| 0.490086 | 0.086703 | 0.112 | 0.089 | 1 | 4 | Map2k4 | 0.023 | Young |
| 0.490205 | 0.0236 | 0.288 | 0.247 | 1 | 4 | Fam117b | 0.041 | Young |
| 0.490422 | 0.156391 | 0.138 | 0.111 | 1 | 4 | Stxbp6 | 0.027 | Young |
| 0.490462 | 0.244008 | 0.262 | 0.226 | 1 | 4 | Wbp2 | 0.036 | Young |
| 0.490599 | 0.95498 | 0.038 | 0.025 | 1 | 4 | H2-DMb2 | 0.013 | Young |
| 0.491011 | 0.719412 | 0.025 | 0.015 | 1 | 4 | Zswim7 | 0.01 | Young |
| 0.491155 | 0.004324 | 0.1 | 0.077 | 1 | 4 | Phf12 | 0.023 | Young |
| 0.49141 | 0.062412 | 0.9 | 0.864 | 1 | 4 | Irf2bp2 | 0.036 | Young |
| 0.491739 | 0.445048 | 0.025 | 0.015 | 1 | 4 | Spata7 | 0.01 | Young |
| 0.492084 | 0.327516 | 0.212 | 0.183 | 1 | 4 | Baiap2 | 0.029 | Young |
| 0.492228 | 0.334426 | 0.025 | 0.042 | 1 | 4 | Gtpbp8 | -0.017 | Young |
| 0.493198 | 0.311357 | 0.025 | 0.015 | 1 | 4 | 2700081O1 | 0.01 | Young |
| 0.493318 | 0.047428 | 0.2 | 0.167 | 1 | 4 | Rnpepl1 | 0.033 | Young |
| 0.493486 | 0.095061 | 0.475 | 0.431 | 1 | 4 | Galnt1 | 0.044 | Young |
| 0.493502 | 0.211035 | 0.062 | 0.045 | 1 | 4 | Zbtb14 | 0.017 | Young |
| 0.493938 | 0.20724 | 0.212 | 0.183 | 1 | 4 | Mnt | 0.029 | Young |
| 0.494178 | 0.251398 | 0.188 | 0.157 | 1 | 4 | Ncoa1 | 0.031 | Young |
| 0.494466 | 0.740688 | 0.05 | 0.036 | 1 | 4 | Ifitm1 | 0.014 | Young |
| 0.494511 | 0.238901 | 0.325 | 0.289 | 1 | 4 | Mxd4 | 0.036 | Young |
| 0.494536 | 0.002755 | 0.225 | 0.185 | 1 | 4 | Cnpy2 | 0.04 | Young |
| 0.494658 | 0.31393 | 0.025 | 0.015 | 1 | 4 | Ttc39c | 0.01 | Young |
| 0.495389 | 0.182078 | 0.025 | 0.015 | 1 | 4 | Rbm4 | 0.01 | Young |
| 0.495628 | 0.322315 | 0.075 | 0.057 | 1 | 4 | Ints12 | 0.018 | Young |
| 0.495864 | 0.234852 | 0.1 | 0.079 | 1 | 4 | Ap5m1 | 0.021 | Young |
| 0.49604 | 0.155507 | 0.3 | 0.258 | 1 | 4 | Cdv3 | 0.042 | Young |
| 0.496637 | 0.088037 | 0.088 | 0.114 | 1 | 4 | Zfp385a | -0.026 | Young |
| 0.496795 | 0.096673 | 0.787 | 0.772 | 1 | 4 | Capza2 | 0.015 | Young |
| 0.497379 | 0.342344 | 0.05 | 0.036 | 1 | 4 | Zfp511 | 0.014 | Young |
| 0.49767 | 0.119825 | 0.112 | 0.089 | 1 | 4 | Insig2 | 0.023 | Young |
| 0.498147 | 0.051212 | 0.412 | 0.366 | 1 | 4 | Cmklr1 | 0.046 | Young |
| 0.498296 | 0.363676 | 0.225 | 0.285 | 1 | 4 | Bmp2 | -0.06 | Young |
| 0.498319 | 0.11003 | 0.025 | 0.015 | 1 | 4 | Coq8a | 0.01 | Young |
| 0.498628 | 0.255189 | 0.038 | 0.025 | 1 | 4 | Ccdc66 | 0.013 | Young |
| 0.498673 | 0.075432 | 0.1 | 0.078 | 1 | 4 | Tsfm | 0.022 | Young |
| 0.499141 | 0.227494 | 0.075 | 0.057 | 1 | 4 | Zfp219 | 0.018 | Young |
| 0.499172 | 0.565238 | 0.212 | 0.184 | 1 | 4 | Pim3 | 0.028 | Young |
| 0.499204 | 0.218617 | 0.038 | 0.025 | 1 | 4 | Ap4e1 | 0.013 | Young |
| 0.499209 | 0.399927 | 0.088 | 0.068 | 1 | 4 | Pibf1 | 0.02 | Young |
| 0.499814 | 0.519267 | 0.05 | 0.036 | 1 | 4 | Pef1 | 0.014 | Young |
| 0.499879 | 0.182689 | 0.188 | 0.157 | 1 | 4 | Tomm34 | 0.031 | Young |
| 0.500212 | 0.109128 | 0.112 | 0.089 | 1 | 4 | 1700037H0 | 0.023 | Young |
| 0.500466 | 0.604112 | 0.125 | 0.103 | 1 | 4 | D730003I1 | 0.022 | Young |
| 0.500824 | 0.078213 | 0.388 | 0.333 | 1 | 4 | Atp2a2 | 0.055 | Young |
| 0.500931 | 0.036129 | 0.15 | 0.123 | 1 | 4 | Ube4b | 0.027 | Young |

| | | | | | | | | |
|---|---|---|---|---|---|---|---|---|
| 0.50137 | 0.175666 | 0.088 | 0.068 | 1 | 4 | Psmg3 | 0.02 | Young |
| 0.501962 | 0.612431 | 0.162 | 0.141 | 1 | 4 | Ide | 0.021 | Young |
| 0.50209 | 0.03284 | 0.038 | 0.025 | 1 | 4 | Ankrd26 | 0.013 | Young |
| 0.502093 | 0.091699 | 0.35 | 0.306 | 1 | 4 | Parp14 | 0.044 | Young |
| 0.503321 | 0.179909 | 0.138 | 0.113 | 1 | 4 | Cyp20a1 | 0.025 | Young |
| 0.503654 | 0.429136 | 0.275 | 0.246 | 1 | 4 | Rbm47 | 0.029 | Young |
| 0.503755 | 0.032062 | 0.575 | 0.514 | 1 | 4 | Eid1 | 0.061 | Young |
| 0.503806 | 0.366337 | 0.062 | 0.046 | 1 | 4 | Usp50 | 0.016 | Young |
| 0.503927 | 0.326298 | 0.1 | 0.08 | 1 | 4 | Cnnm3 | 0.02 | Young |
| 0.504248 | 0.71238 | 0.075 | 0.058 | 1 | 4 | Ptpn22 | 0.017 | Young |
| 0.504323 | 0.289037 | 0.125 | 0.157 | 1 | 4 | Traf1 | -0.032 | Young |
| 0.504431 | 0.408573 | 0.1 | 0.08 | 1 | 4 | Taf3 | 0.02 | Young |
| 0.504714 | 0.201749 | 0.038 | 0.057 | 1 | 4 | Cep290 | -0.019 | Young |
| 0.50484 | 0.044796 | 0.3 | 0.262 | 1 | 4 | Slc25a37 | 0.038 | Young |
| 0.504869 | 0.001931 | 0.175 | 0.143 | 1 | 4 | Mut | 0.032 | Young |
| 0.504986 | 0.121363 | 0.038 | 0.025 | 1 | 4 | Leng9 | 0.013 | Young |
| 0.505024 | 0.18198 | 0.075 | 0.057 | 1 | 4 | Tmem268 | 0.018 | Young |
| 0.505228 | 0.403895 | 0.075 | 0.057 | 1 | 4 | Inafm1 | 0.018 | Young |
| 0.505257 | 0.279194 | 0.388 | 0.34 | 1 | 4 | Ndufb10 | 0.048 | Young |
| 0.506208 | 0.282427 | 0.15 | 0.125 | 1 | 4 | Pqlc1 | 0.025 | Young |
| 0.506659 | 0.343001 | 0.162 | 0.136 | 1 | 4 | Ifi47 | 0.026 | Young |
| 0.507575 | 0.165162 | 0.2 | 0.171 | 1 | 4 | Tsg101 | 0.029 | Young |
| 0.508794 | 0.406464 | 0.15 | 0.123 | 1 | 4 | Apopt1 | 0.027 | Young |
| 0.509991 | 0.034348 | 0.6 | 0.562 | 1 | 4 | Tpd52 | 0.038 | Young |
| 0.51017 | 0.197926 | 0.45 | 0.424 | 1 | 4 | Tra2a | 0.026 | Young |
| 0.510476 | 0.82378 | 0.1 | 0.08 | 1 | 4 | Actn1 | 0.02 | Young |
| 0.510759 | 0.212462 | 0.225 | 0.199 | 1 | 4 | Trip11 | 0.026 | Young |
| 0.510824 | 0.191235 | 0.138 | 0.113 | 1 | 4 | Sft2d1 | 0.025 | Young |
| 0.510886 | 0.123012 | 0.188 | 0.158 | 1 | 4 | C1qbp | 0.03 | Young |
| 0.510902 | 0.14488 | 0.388 | 0.349 | 1 | 4 | Scaf11 | 0.039 | Young |
| 0.511313 | 0.264842 | 0.188 | 0.16 | 1 | 4 | Ktn1 | 0.028 | Young |
| 0.511586 | 0.164117 | 0.05 | 0.036 | 1 | 4 | Ppcdc | 0.014 | Young |
| 0.512012 | 0.327232 | 0.112 | 0.092 | 1 | 4 | Akt1s1 | 0.02 | Young |
| 0.512063 | 0.661685 | 0.075 | 0.058 | 1 | 4 | Gsto1 | 0.017 | Young |
| 0.512099 | 0.0367 | 0.088 | 0.067 | 1 | 4 | Ubxn7 | 0.021 | Young |
| 0.512252 | 0.002134 | 0.138 | 0.111 | 1 | 4 | Rnf14 | 0.027 | Young |
| 0.512267 | 0.099651 | 0.388 | 0.348 | 1 | 4 | Zfp91 | 0.04 | Young |
| 0.512502 | 0.177137 | 0.062 | 0.046 | 1 | 4 | Hsf1 | 0.016 | Young |
| 0.512792 | 0.271468 | 0.15 | 0.125 | 1 | 4 | Pgrmc2 | 0.025 | Young |
| 0.513126 | 0.389463 | 0.112 | 0.092 | 1 | 4 | Hs1bp3 | 0.02 | Young |
| 0.51314 | 0.650011 | 0.038 | 0.026 | 1 | 4 | Rubcnl | 0.012 | Young |
| 0.51314 | 0.730981 | 0.038 | 0.026 | 1 | 4 | Lix1 | 0.012 | Young |
| 0.513706 | 0.13177 | 0.088 | 0.068 | 1 | 4 | Tbpl1 | 0.02 | Young |
| 0.514183 | 0.082909 | 0.312 | 0.274 | 1 | 4 | Vamp3 | 0.038 | Young |
| 0.514343 | 0.055753 | 0.188 | 0.157 | 1 | 4 | Znfx1 | 0.031 | Young |

| | | | | | | | | |
|---|---|---|---|---|---|---|---|---|
| 0.515053 | 0.784763 | 0.05 | 0.036 | 1 | 4 | St3gal2 | 0.014 | Young |
| 0.515382 | 0.126689 | 0.588 | 0.57 | 1 | 4 | Jak1 | 0.018 | Young |
| 0.515545 | 0.5099 | 0.05 | 0.036 | 1 | 4 | Mib2 | 0.014 | Young |
| 0.516454 | 0.045623 | 0.262 | 0.229 | 1 | 4 | Ppt1 | 0.033 | Young |
| 0.516528 | 0.428183 | 0.05 | 0.036 | 1 | 4 | Scap | 0.014 | Young |
| 0.517115 | 0.176082 | 0.088 | 0.069 | 1 | 4 | Mafk | 0.019 | Young |
| 0.517275 | 0.264769 | 0.112 | 0.092 | 1 | 4 | Vps52 | 0.02 | Young |
| 0.517407 | 0.342411 | 0.1 | 0.08 | 1 | 4 | Dcaf15 | 0.02 | Young |
| 0.517516 | 1.039327 | 0.025 | 0.016 | 1 | 4 | Tmem150a | 0.009 | Young |
| 0.517527 | 0.064186 | 0.05 | 0.036 | 1 | 4 | Pex11b | 0.014 | Young |
| 0.518004 | 0.311709 | 0.162 | 0.136 | 1 | 4 | Spire1 | 0.026 | Young |
| 0.518078 | 0.621419 | 0.075 | 0.059 | 1 | 4 | Smim12 | 0.016 | Young |
| 0.518247 | 0.354624 | 0.125 | 0.104 | 1 | 4 | Ebna1bp2 | 0.021 | Young |
| 0.51825 | 0.757647 | 0.025 | 0.016 | 1 | 4 | Pigf | 0.009 | Young |
| 0.518985 | 0.848039 | 0.025 | 0.016 | 1 | 4 | Il18bp | 0.009 | Young |
| 0.51915 | 0.430746 | 0.075 | 0.058 | 1 | 4 | Top3b | 0.017 | Young |
| 0.519344 | 0.064138 | 0.675 | 0.622 | 1 | 4 | Tapbp | 0.053 | Young |
| 0.519518 | 0.386968 | 0.038 | 0.026 | 1 | 4 | Dph6 | 0.012 | Young |
| 0.51972 | 0.931796 | 0.025 | 0.016 | 1 | 4 | Kctd9 | 0.009 | Young |
| 0.520456 | 0.875865 | 0.025 | 0.016 | 1 | 4 | B3galt6 | 0.009 | Young |
| 0.520471 | 0.545161 | 0.05 | 0.036 | 1 | 4 | Dennd2c | 0.014 | Young |
| 0.520471 | 0.440443 | 0.05 | 0.036 | 1 | 4 | Prelid2 | 0.014 | Young |
| 0.520558 | 0.753707 | 0.188 | 0.168 | 1 | 4 | Dot1l | 0.02 | Young |
| 0.520683 | 0.05104 | 0.162 | 0.134 | 1 | 4 | Gripap1 | 0.028 | Young |
| 0.520694 | 0.056528 | 0.312 | 0.273 | 1 | 4 | Pds5a | 0.039 | Young |
| 0.520919 | 0.483959 | 0.238 | 0.209 | 1 | 4 | AW112010 | 0.029 | Young |
| 0.520949 | 0.34588 | 0.175 | 0.152 | 1 | 4 | Taf1d | 0.023 | Young |
| 0.520971 | 0.204003 | 0.125 | 0.102 | 1 | 4 | Cops3 | 0.023 | Young |
| 0.521036 | 0.096516 | 0.088 | 0.068 | 1 | 4 | Stx2 | 0.02 | Young |
| 0.521193 | 0.969793 | 0.025 | 0.016 | 1 | 4 | Slc22a18 | 0.009 | Young |
| 0.521261 | 0.455432 | 0.1 | 0.079 | 1 | 4 | Atg4b | 0.021 | Young |
| 0.521288 | 0.006439 | 0.112 | 0.09 | 1 | 4 | Abcd4 | 0.022 | Young |
| 0.521409 | 0.077679 | 0.225 | 0.194 | 1 | 4 | Cnih4 | 0.031 | Young |
| 0.52193 | 0.861098 | 0.025 | 0.016 | 1 | 4 | Cox11 | 0.009 | Young |
| 0.521937 | 0.72436 | 0.1 | 0.082 | 1 | 4 | Rad54l2 | 0.018 | Young |
| 0.522507 | 0.204465 | 0.15 | 0.126 | 1 | 4 | Mllt6 | 0.024 | Young |
| 0.522609 | 0.06432 | 0.075 | 0.057 | 1 | 4 | Cep57 | 0.018 | Young |
| 0.522892 | 0.487206 | 0.025 | 0.04 | 1 | 4 | Tbrg4 | -0.015 | Young |
| 0.5229 | 0.585974 | 0.112 | 0.091 | 1 | 4 | Eif2b5 | 0.021 | Young |
| 0.523406 | 0.771169 | 0.025 | 0.016 | 1 | 4 | Polr3h | 0.009 | Young |
| 0.523523 | 0.064665 | 0.613 | 0.547 | 1 | 4 | Ndufa2 | 0.066 | Young |
| 0.523558 | 0.261105 | 0.562 | 0.522 | 1 | 4 | Asah1 | 0.04 | Young |
| 0.524182 | 0.345563 | 0.038 | 0.026 | 1 | 4 | Nudcd1 | 0.012 | Young |
| 0.524399 | 0.354291 | 0.088 | 0.07 | 1 | 4 | Ufd1 | 0.018 | Young |
| 0.524645 | 0.017374 | 0.2 | 0.167 | 1 | 4 | Sugt1 | 0.033 | Young |

| | | | | | | | | |
|---|---|---|---|---|---|---|---|---|
| 0.524838 | 0.119655 | 0.112 | 0.091 | 1 | 4 | Adap1 | 0.021 | Young |
| 0.524926 | 0.503786 | 0.05 | 0.036 | 1 | 4 | Prune1 | 0.014 | Young |
| 0.525367 | 0.23553 | 0.05 | 0.069 | 1 | 4 | Slc30a4 | -0.019 | Young |
| 0.525435 | 0.28484 | 0.062 | 0.047 | 1 | 4 | Sec22a | 0.015 | Young |
| 0.525463 | 0.356662 | 0.112 | 0.092 | 1 | 4 | Qrich1 | 0.02 | Young |
| 0.526041 | 0.040916 | 0.188 | 0.156 | 1 | 4 | Gtpbp4 | 0.032 | Young |
| 0.526308 | 1.123499 | 0.038 | 0.026 | 1 | 4 | Tmub1 | 0.012 | Young |
| 0.526587 | 0.081093 | 0.875 | 0.856 | 1 | 4 | Ddx5 | 0.019 | Young |
| 0.527104 | 0.58636 | 0.025 | 0.016 | 1 | 4 | Cep76 | 0.009 | Young |
| 0.52736 | 0.04433 | 0.787 | 0.825 | 1 | 4 | Hspa5 | -0.038 | Young |
| 0.527907 | 0.253815 | 0.05 | 0.036 | 1 | 4 | Nsun3 | 0.014 | Young |
| 0.528052 | 1.715277 | 0.013 | 0.007 | 1 | 4 | Bend3 | 0.006 | Young |
| 0.528052 | 1.401007 | 0.013 | 0.007 | 1 | 4 | Nptxr | 0.006 | Young |
| 0.528281 | 0.024932 | 0.038 | 0.026 | 1 | 4 | Cmc4 | 0.012 | Young |
| 0.528626 | 1.01364 | 0.038 | 0.026 | 1 | 4 | Fcgr4 | 0.012 | Young |
| 0.529034 | 0.953632 | 0.088 | 0.071 | 1 | 4 | Rpp21 | 0.017 | Young |
| 0.529077 | 0.325093 | 0.075 | 0.058 | 1 | 4 | Ano10 | 0.017 | Young |
| 0.529187 | 1.398867 | 0.013 | 0.007 | 1 | 4 | Lrrc1 | 0.006 | Young |
| 0.529187 | 1.076316 | 0.013 | 0.007 | 1 | 4 | Gtf2ird1 | 0.006 | Young |
| 0.529211 | 0.151448 | 0.275 | 0.241 | 1 | 4 | Ralgds | 0.034 | Young |
| 0.5294 | 0.092567 | 0.05 | 0.036 | 1 | 4 | Man2c1 | 0.014 | Young |
| 0.529417 | 0.678139 | 0.112 | 0.093 | 1 | 4 | Bicral | 0.019 | Young |
| 0.529436 | 0.685131 | 0.112 | 0.096 | 1 | 4 | Akap11 | 0.016 | Young |
| 0.529727 | 0.208616 | 0.325 | 0.289 | 1 | 4 | Gbp7 | 0.036 | Young |
| 0.530073 | 0.378116 | 0.025 | 0.016 | 1 | 4 | Tctn3 | 0.009 | Young |
| 0.530073 | 0.396768 | 0.025 | 0.016 | 1 | 4 | Twsg1 | 0.009 | Young |
| 0.530324 | 1.365813 | 0.013 | 0.007 | 1 | 4 | Lrig3 | 0.006 | Young |
| 0.530324 | 1.11356 | 0.013 | 0.007 | 1 | 4 | N4bp3 | 0.006 | Young |
| 0.530324 | 0.988779 | 0.013 | 0.007 | 1 | 4 | Zfr2 | 0.006 | Young |
| 0.530324 | 0.963758 | 0.013 | 0.007 | 1 | 4 | Zscan18 | 0.006 | Young |
| 0.53054 | 0.142492 | 0.112 | 0.09 | 1 | 4 | Ube4a | 0.022 | Young |
| 0.531166 | 0.107923 | 0.062 | 0.047 | 1 | 4 | Dhx33 | 0.015 | Young |
| 0.531462 | 0.918695 | 0.013 | 0.007 | 1 | 4 | Lepr | 0.006 | Young |
| 0.532079 | 0.261368 | 0.2 | 0.171 | 1 | 4 | Btbd1 | 0.029 | Young |
| 0.532264 | 0.00428 | 0.188 | 0.157 | 1 | 4 | Rnf6 | 0.031 | Young |
| 0.532305 | 0.214707 | 0.025 | 0.016 | 1 | 4 | Tmem246 | 0.009 | Young |
| 0.532561 | 0.261743 | 0.225 | 0.198 | 1 | 4 | Cul1 | 0.027 | Young |
| 0.532601 | 0.799216 | 0.013 | 0.007 | 1 | 4 | Blm | 0.006 | Young |
| 0.532601 | 0.76671 | 0.013 | 0.007 | 1 | 4 | Rapgef5 | 0.006 | Young |
| 0.532601 | 0.729206 | 0.013 | 0.007 | 1 | 4 | Rftn2 | 0.006 | Young |
| 0.532912 | 0.303595 | 0.225 | 0.198 | 1 | 4 | Exoc3 | 0.027 | Young |
| 0.532924 | 0.123015 | 0.262 | 0.225 | 1 | 4 | Fkbp8 | 0.037 | Young |
| 0.533278 | 0.810291 | 0.038 | 0.026 | 1 | 4 | Gmnn | 0.012 | Young |
| 0.533672 | 0.6122 | 0.15 | 0.132 | 1 | 4 | Cdc123 | 0.018 | Young |
| 0.533742 | 0.821137 | 0.013 | 0.007 | 1 | 4 | Hps6 | 0.006 | Young |

| | | | | | | | | |
|---|---|---|---|---|---|---|---|---|
| 0.533742 | 0.354918 | 0.013 | 0.007 | 1 | 4 | Nutf2 | 0.006 | Young |
| 0.533742 | 0.348149 | 0.013 | 0.007 | 1 | 4 | Gm26520 | 0.006 | Young |
| 0.533795 | 0.174931 | 0.025 | 0.016 | 1 | 4 | Cox7a1 | 0.009 | Young |
| 0.534132 | 0.202457 | 0.112 | 0.09 | 1 | 4 | Zfp984 | 0.022 | Young |
| 0.53421 | 0.592852 | 0.05 | 0.037 | 1 | 4 | Pik3r2 | 0.013 | Young |
| 0.534276 | 0.181287 | 0.075 | 0.058 | 1 | 4 | Rabggta | 0.017 | Young |
| 0.534707 | 0.642835 | 0.05 | 0.037 | 1 | 4 | Nudt2 | 0.013 | Young |
| 0.534884 | 0.609074 | 0.013 | 0.007 | 1 | 4 | Ankrd24 | 0.006 | Young |
| 0.534884 | 0.538977 | 0.013 | 0.007 | 1 | 4 | Gm26542 | 0.006 | Young |
| 0.534884 | 0.498945 | 0.013 | 0.007 | 1 | 4 | Pmm1 | 0.006 | Young |
| 0.534884 | 0.288966 | 0.013 | 0.007 | 1 | 4 | Gemin8 | 0.006 | Young |
| 0.534884 | 0.143733 | 0.013 | 0.007 | 1 | 4 | Pkd1l2 | 0.006 | Young |
| 0.53489 | 0.34271 | 0.338 | 0.317 | 1 | 4 | Nono | 0.021 | Young |
| 0.53492 | 0.097751 | 0.475 | 0.421 | 1 | 4 | Bnip2 | 0.054 | Young |
| 0.535079 | 0.175804 | 0.475 | 0.439 | 1 | 4 | Nucks1 | 0.036 | Young |
| 0.535444 | 0.105448 | 0.388 | 0.345 | 1 | 4 | Sult1a1 | 0.043 | Young |
| 0.535597 | 0.100252 | 0.062 | 0.047 | 1 | 4 | Pola1 | 0.015 | Young |
| 0.535894 | 0.129142 | 0.088 | 0.069 | 1 | 4 | Idua | 0.019 | Young |
| 0.536027 | 0.573781 | 0.013 | 0.007 | 1 | 4 | 1700003F1 | 0.006 | Young |
| 0.536027 | 0.532055 | 0.013 | 0.007 | 1 | 4 | Cxcl3 | 0.006 | Young |
| 0.536027 | 0.28309 | 0.013 | 0.007 | 1 | 4 | Dtwd2 | 0.006 | Young |
| 0.536027 | 0.184944 | 0.013 | 0.007 | 1 | 4 | Ildr1 | 0.006 | Young |
| 0.536638 | 0.606343 | 0.038 | 0.055 | 1 | 4 | Gtf3a | -0.017 | Young |
| 0.537087 | 0.287157 | 0.075 | 0.058 | 1 | 4 | Bcas3 | 0.017 | Young |
| 0.537567 | 0.398562 | 0.15 | 0.128 | 1 | 4 | Tm9sf4 | 0.022 | Young |
| 0.53763 | 0.206093 | 0.125 | 0.103 | 1 | 4 | Tfec | 0.022 | Young |
| 0.538614 | 0.2131 | 0.1 | 0.08 | 1 | 4 | Ice1 | 0.02 | Young |
| 0.538888 | 0.271402 | 0.088 | 0.069 | 1 | 4 | Gm5547 | 0.019 | Young |
| 0.539018 | 0.492929 | 0.125 | 0.103 | 1 | 4 | Scaf8 | 0.022 | Young |
| 0.539038 | 0.222004 | 0.2 | 0.171 | 1 | 4 | Vbp1 | 0.029 | Young |
| 0.539118 | 0.340729 | 0.162 | 0.138 | 1 | 4 | Tbc1d23 | 0.024 | Young |
| 0.539186 | 0.45276 | 0.05 | 0.037 | 1 | 4 | Shmt2 | 0.013 | Young |
| 0.539528 | 0.146803 | 0.162 | 0.138 | 1 | 4 | Vps18 | 0.024 | Young |
| 0.539736 | 0.122826 | 0.3 | 0.269 | 1 | 4 | Camk1d | 0.031 | Young |
| 0.539905 | 0.154969 | 0.075 | 0.058 | 1 | 4 | Gab3 | 0.017 | Young |
| 0.541219 | 0.303437 | 0.162 | 0.142 | 1 | 4 | Basp1 | 0.02 | Young |
| 0.541725 | 0.157888 | 0.188 | 0.162 | 1 | 4 | Zcrb1 | 0.026 | Young |
| 0.541751 | 0.283222 | 0.15 | 0.125 | 1 | 4 | Nudt16 | 0.025 | Young |
| 0.54183 | 0.089976 | 0.275 | 0.244 | 1 | 4 | Tank | 0.031 | Young |
| 0.54362 | 0.270008 | 0.15 | 0.129 | 1 | 4 | Pdcd11 | 0.021 | Young |
| 0.544098 | 0.045702 | 0.138 | 0.113 | 1 | 4 | Slc17a5 | 0.025 | Young |
| 0.544998 | 0.573428 | 0.038 | 0.026 | 1 | 4 | Nbas | 0.012 | Young |
| 0.545824 | 0.675441 | 0.088 | 0.072 | 1 | 4 | Thap11 | 0.016 | Young |
| 0.546481 | 0.097356 | 0.15 | 0.126 | 1 | 4 | Ric8a | 0.024 | Young |
| 0.546694 | 0.421455 | 0.05 | 0.037 | 1 | 4 | Aldh6a1 | 0.013 | Young |

| | | | | | | | | |
|---|---|---|---|---|---|---|---|---|
| 0.546767 | 0.498988 | 0.038 | 0.026 | 1 | 4 | Pcgf6 | 0.012 | Young |
| 0.546767 | 0.255812 | 0.038 | 0.026 | 1 | 4 | 0610009O2 | 0.012 | Young |
| 0.546826 | 0.184441 | 0.238 | 0.204 | 1 | 4 | Ube2g2 | 0.034 | Young |
| 0.546944 | 0.010979 | 0.138 | 0.113 | 1 | 4 | Ngly1 | 0.025 | Young |
| 0.5474 | 0.3041 | 0.4 | 0.36 | 1 | 4 | Chp2 | 0.04 | Young |
| 0.54771 | 0.562894 | 0.038 | 0.054 | 1 | 4 | Fam98a | -0.016 | Young |
| 0.548571 | 0.190567 | 0.062 | 0.047 | 1 | 4 | Rrp7a | 0.015 | Young |
| 0.549109 | 0.279746 | 0.1 | 0.081 | 1 | 4 | Irf2bp1 | 0.019 | Young |
| 0.549759 | 0.267104 | 0.05 | 0.069 | 1 | 4 | Slc25a30 | -0.019 | Young |
| 0.549861 | 0.20974 | 0.375 | 0.334 | 1 | 4 | Nras | 0.041 | Young |
| 0.549872 | 0.344837 | 0.125 | 0.104 | 1 | 4 | Fnip2 | 0.021 | Young |
| 0.549908 | 0.570394 | 0.062 | 0.048 | 1 | 4 | Kat6b | 0.014 | Young |
| 0.550793 | 0.221974 | 0.075 | 0.059 | 1 | 4 | Actr8 | 0.016 | Young |
| 0.550905 | 0.117179 | 0.038 | 0.026 | 1 | 4 | Grwd1 | 0.012 | Young |
| 0.551025 | 0.114897 | 0.125 | 0.103 | 1 | 4 | Rars | 0.022 | Young |
| 0.551163 | 0.012771 | 0.238 | 0.204 | 1 | 4 | Rb1 | 0.034 | Young |
| 0.551198 | 0.24385 | 0.075 | 0.059 | 1 | 4 | Map3k3 | 0.016 | Young |
| 0.551223 | 0.182614 | 0.05 | 0.037 | 1 | 4 | Chkb | 0.013 | Young |
| 0.551728 | 0.173232 | 0.05 | 0.037 | 1 | 4 | Nt5dc2 | 0.013 | Young |
| 0.551868 | 0.01011 | 0.362 | 0.311 | 1 | 4 | Nfe2l1 | 0.051 | Young |
| 0.552088 | 0.696272 | 0.088 | 0.072 | 1 | 4 | Klhdc4 | 0.016 | Young |
| 0.552126 | 0.143949 | 0.112 | 0.092 | 1 | 4 | Cyp4f13 | 0.02 | Young |
| 0.552165 | 0.033905 | 0.138 | 0.114 | 1 | 4 | Ptgr2 | 0.024 | Young |
| 0.552179 | 0.215804 | 0.188 | 0.159 | 1 | 4 | Setd5 | 0.029 | Young |
| 0.552605 | 0.114072 | 0.1 | 0.08 | 1 | 4 | Epm2aip1 | 0.02 | Young |
| 0.552684 | 0.163304 | 0.038 | 0.026 | 1 | 4 | 1110019D1 | 0.012 | Young |
| 0.552833 | 0.531098 | 0.112 | 0.093 | 1 | 4 | Serhl | 0.019 | Young |
| 0.552966 | 0.157282 | 0.038 | 0.053 | 1 | 4 | B9d2 | -0.015 | Young |
| 0.553308 | 0.06426 | 0.412 | 0.375 | 1 | 4 | Ewsr1 | 0.037 | Young |
| 0.5536 | 0.0859 | 0.288 | 0.244 | 1 | 4 | Cnot1 | 0.044 | Young |
| 0.553654 | 0.041669 | 0.588 | 0.548 | 1 | 4 | Atp5g2 | 0.04 | Young |
| 0.55477 | 0.361752 | 0.075 | 0.059 | 1 | 4 | Sfxn2 | 0.016 | Young |
| 0.554881 | 0.446282 | 0.025 | 0.039 | 1 | 4 | Ints5.1 | -0.014 | Young |
| 0.555256 | 0.364098 | 0.188 | 0.161 | 1 | 4 | Vti1b | 0.027 | Young |
| 0.555601 | 0.103895 | 0.412 | 0.387 | 1 | 4 | Tardbp | 0.025 | Young |
| 0.555654 | 0.043546 | 0.038 | 0.026 | 1 | 4 | Zfp948 | 0.012 | Young |
| 0.555714 | 0.417958 | 0.088 | 0.069 | 1 | 4 | Armc10 | 0.019 | Young |
| 0.555715 | 0.369155 | 0.175 | 0.153 | 1 | 4 | Hgsnat | 0.022 | Young |
| 0.556417 | 0.196167 | 0.112 | 0.092 | 1 | 4 | Bahd1 | 0.02 | Young |
| 0.55649 | 0.746037 | 0.075 | 0.06 | 1 | 4 | Prpf31 | 0.015 | Young |
| 0.556572 | 0.8768 | 0.138 | 0.119 | 1 | 4 | Gdf15 | 0.019 | Young |
| 0.556688 | 0.122421 | 0.4 | 0.356 | 1 | 4 | Spcs2 | 0.044 | Young |
| 0.556936 | 0.837967 | 0.025 | 0.016 | 1 | 4 | Sh2b2 | 0.009 | Young |
| 0.557222 | 0.050708 | 0.088 | 0.069 | 1 | 4 | Triap1 | 0.019 | Young |
| 0.55744 | 0.07599 | 0.038 | 0.026 | 1 | 4 | Asb6 | 0.012 | Young |

| | | | | | | | | |
|---|---|---|---|---|---|---|---|---|
| 0.557486 | 0.403239 | 0.062 | 0.048 | 1 | 4 | Snta1 | 0.014 | Young |
| 0.557684 | 0.854259 | 0.025 | 0.016 | 1 | 4 | Cmtr2 | 0.009 | Young |
| 0.557945 | 0.384924 | 0.2 | 0.175 | 1 | 4 | Map2k3 | 0.025 | Young |
| 0.558305 | 0.106235 | 0.05 | 0.037 | 1 | 4 | Aptx | 0.013 | Young |
| 0.558511 | 0.297921 | 0.125 | 0.105 | 1 | 4 | Rab3d | 0.02 | Young |
| 0.558606 | 0.842154 | 0.1 | 0.085 | 1 | 4 | Katnbl1 | 0.015 | Young |
| 0.559028 | 0.202355 | 0.3 | 0.268 | 1 | 4 | Cat | 0.032 | Young |
| 0.559032 | 0.094382 | 0.112 | 0.092 | 1 | 4 | Gpr155 | 0.02 | Young |
| 0.559227 | 0.010202 | 0.175 | 0.148 | 1 | 4 | Lgals8 | 0.027 | Young |
| 0.559787 | 0.500693 | 0.1 | 0.081 | 1 | 4 | Dtymk | 0.019 | Young |
| 0.559835 | 0.040906 | 0.25 | 0.217 | 1 | 4 | Sav1 | 0.033 | Young |
| 0.559911 | 0.643146 | 0.05 | 0.038 | 1 | 4 | Bbc3 | 0.012 | Young |
| 0.560104 | 0.190178 | 0.375 | 0.366 | 1 | 4 | Prkacb | 0.009 | Young |
| 0.560534 | 0.545734 | 0.062 | 0.049 | 1 | 4 | Acot8 | 0.013 | Young |
| 0.561214 | 0.48505 | 0.188 | 0.167 | 1 | 4 | Lats2 | 0.021 | Young |
| 0.561533 | 0.906098 | 0.038 | 0.027 | 1 | 4 | Spryd3 | 0.011 | Young |
| 0.561866 | 0.269377 | 0.125 | 0.105 | 1 | 4 | Bex3 | 0.02 | Young |
| 0.562215 | 0.008036 | 0.038 | 0.026 | 1 | 4 | Zbed3 | 0.012 | Young |
| 0.562222 | 0.508241 | 0.088 | 0.071 | 1 | 4 | Timm22 | 0.017 | Young |
| 0.562673 | 0.003323 | 0.175 | 0.203 | 1 | 4 | Sec11a | -0.028 | Young |
| 0.562765 | 0.575192 | 0.062 | 0.049 | 1 | 4 | Brms1 | 0.013 | Young |
| 0.563004 | 0.033749 | 0.212 | 0.184 | 1 | 4 | Nfkbie | 0.028 | Young |
| 0.563069 | 0.197568 | 0.112 | 0.093 | 1 | 4 | Usp1 | 0.019 | Young |
| 0.563134 | 0.120003 | 0.1 | 0.081 | 1 | 4 | Zbtb22 | 0.019 | Young |
| 0.563345 | 0.539953 | 0.088 | 0.071 | 1 | 4 | Slc30a1 | 0.017 | Young |
| 0.563396 | 0.10771 | 0.338 | 0.304 | 1 | 4 | Paip2 | 0.034 | Young |
| 0.563898 | 0.138832 | 0.4 | 0.37 | 1 | 4 | H2afy | 0.03 | Young |
| 0.564112 | 0.55184 | 0.075 | 0.059 | 1 | 4 | Ctps2 | 0.016 | Young |
| 0.564468 | 1.042995 | 0.075 | 0.061 | 1 | 4 | Mitd1 | 0.014 | Young |
| 0.564524 | 0.059208 | 0.125 | 0.103 | 1 | 4 | Rsrc1 | 0.022 | Young |
| 0.564586 | 0.100459 | 0.162 | 0.198 | 1 | 4 | Tspyl1 | -0.036 | Young |
| 0.564928 | 0.299772 | 0.075 | 0.059 | 1 | 4 | Apex1 | 0.016 | Young |
| 0.56519 | 0.419264 | 0.025 | 0.016 | 1 | 4 | Tmem86b | 0.009 | Young |
| 0.56519 | 0.40846 | 0.025 | 0.016 | 1 | 4 | Znrd1as | 0.009 | Young |
| 0.56519 | 0.340353 | 0.025 | 0.016 | 1 | 4 | Paqr4 | 0.009 | Young |
| 0.565244 | 0.02959 | 0.013 | 0.022 | 1 | 4 | Zfp810 | -0.009 | Young |
| 0.565358 | 0.153032 | 0.262 | 0.241 | 1 | 4 | Skil | 0.021 | Young |
| 0.56579 | 0.011408 | 0.125 | 0.103 | 1 | 4 | Ctdspl2 | 0.022 | Young |
| 0.566115 | 0.225026 | 0.35 | 0.33 | 1 | 4 | Mrpl33 | 0.02 | Young |
| 0.566425 | 0.182506 | 0.188 | 0.162 | 1 | 4 | Taf6l | 0.026 | Young |
| 0.566866 | 0.692461 | 0.038 | 0.027 | 1 | 4 | Wrb | 0.011 | Young |
| 0.567492 | 0.969848 | 0.05 | 0.038 | 1 | 4 | Usp28 | 0.012 | Young |
| 0.567576 | 0.015936 | 0.138 | 0.115 | 1 | 4 | Zfp422 | 0.023 | Young |
| 0.567885 | 0.156572 | 0.175 | 0.148 | 1 | 4 | Kdm2b | 0.027 | Young |
| 0.567979 | 0.225162 | 0.088 | 0.07 | 1 | 4 | Gopc | 0.018 | Young |

| | | | | | | | | |
|---|---|---|---|---|---|---|---|---|
| 0.567996 | 1.343607 | 0.05 | 0.038 | 1 | 4 | Tes | 0.012 | Young |
| 0.568037 | 0.338146 | 0.075 | 0.06 | 1 | 4 | Slc39a3 | 0.015 | Young |
| 0.568263 | 0.201629 | 0.15 | 0.128 | 1 | 4 | Clpx | 0.022 | Young |
| 0.568783 | 0.244011 | 0.225 | 0.201 | 1 | 4 | Ogfrl1 | 0.024 | Young |
| 0.56901 | 0.228816 | 0.025 | 0.038 | 1 | 4 | Ppm1d | -0.013 | Young |
| 0.56945 | 0.281676 | 0.238 | 0.214 | 1 | 4 | Ppm1a | 0.024 | Young |
| 0.569717 | 0.18709 | 0.025 | 0.016 | 1 | 4 | Fancc | 0.009 | Young |
| 0.569788 | 0.100955 | 0.175 | 0.148 | 1 | 4 | Mbtps1 | 0.027 | Young |
| 0.570054 | 0.205508 | 0.112 | 0.093 | 1 | 4 | Thumpd1 | 0.019 | Young |
| 0.570473 | 0.170249 | 0.025 | 0.016 | 1 | 4 | Gstz1 | 0.009 | Young |
| 0.570896 | 0.13177 | 0.212 | 0.186 | 1 | 4 | Jtb | 0.026 | Young |
| 0.571818 | 0.285116 | 0.05 | 0.038 | 1 | 4 | Ripk2 | 0.012 | Young |
| 0.571887 | 0.004022 | 0.188 | 0.161 | 1 | 4 | Timm10b | 0.027 | Young |
| 0.573046 | 0.105656 | 0.25 | 0.223 | 1 | 4 | Eif3d | 0.027 | Young |
| 0.573343 | 0.278202 | 0.075 | 0.06 | 1 | 4 | Enpp4 | 0.015 | Young |
| 0.573566 | 0.120439 | 0.325 | 0.303 | 1 | 4 | Iqgap2 | 0.022 | Young |
| 0.574612 | 0.446999 | 0.038 | 0.027 | 1 | 4 | Lppos | 0.011 | Young |
| 0.575576 | 0.029065 | 0.212 | 0.184 | 1 | 4 | C1galt1 | 0.028 | Young |
| 0.576155 | 0.20129 | 0.05 | 0.038 | 1 | 4 | Rft1 | 0.012 | Young |
| 0.576155 | 0.201397 | 0.05 | 0.038 | 1 | 4 | Ankrd54 | 0.012 | Young |
| 0.576935 | 0.080826 | 0.738 | 0.698 | 1 | 4 | Ddx3x | 0.04 | Young |
| 0.577011 | 0.335164 | 0.1 | 0.083 | 1 | 4 | Gm47283 | 0.017 | Young |
| 0.577228 | 0.371376 | 0.1 | 0.082 | 1 | 4 | Dtnbp1 | 0.018 | Young |
| 0.577424 | 0.342661 | 0.162 | 0.141 | 1 | 4 | Mettl9 | 0.021 | Young |
| 0.577974 | 0.05577 | 1 | 0.998 | 1 | 4 | mt-Cytb | 0.002 | Young |
| 0.578145 | 0.026769 | 0.15 | 0.128 | 1 | 4 | Epb41l4aos | 0.022 | Young |
| 0.578221 | 0.088338 | 0.15 | 0.127 | 1 | 4 | Rhoc | 0.023 | Young |
| 0.578702 | 0.13056 | 0.188 | 0.161 | 1 | 4 | Gtpbp2 | 0.027 | Young |
| 0.57934 | 0.082201 | 0.025 | 0.038 | 1 | 4 | Pde8a | -0.013 | Young |
| 0.579645 | 0.246405 | 0.15 | 0.13 | 1 | 4 | Gpx3 | 0.02 | Young |
| 0.579676 | 0.376834 | 0.075 | 0.061 | 1 | 4 | Spaca6 | 0.014 | Young |
| 0.579696 | 0.420021 | 0.112 | 0.096 | 1 | 4 | Ssbp1 | 0.016 | Young |
| 0.579739 | 0.09865 | 0.175 | 0.149 | 1 | 4 | Samm50 | 0.026 | Young |
| 0.579746 | 0.081795 | 0.162 | 0.138 | 1 | 4 | Atad2b | 0.024 | Young |
| 0.580085 | 0.413843 | 0.075 | 0.061 | 1 | 4 | Nudt19 | 0.014 | Young |
| 0.580616 | 0.094302 | 0.262 | 0.228 | 1 | 4 | Becn1 | 0.034 | Young |
| 0.581861 | 0.257206 | 0.188 | 0.167 | 1 | 4 | Got1 | 0.021 | Young |
| 0.582378 | 0.03845 | 0.125 | 0.103 | 1 | 4 | Pex19 | 0.022 | Young |
| 0.582525 | 0.023009 | 0.138 | 0.115 | 1 | 4 | Nudt9 | 0.023 | Young |
| 0.582559 | 0.099666 | 0.05 | 0.038 | 1 | 4 | Alg11 | 0.012 | Young |
| 0.582897 | 0.281992 | 0.262 | 0.24 | 1 | 4 | Mvp | 0.022 | Young |
| 0.583569 | 0.378805 | 0.075 | 0.061 | 1 | 4 | Mea1 | 0.014 | Young |
| 0.58361 | 0.149938 | 0.075 | 0.06 | 1 | 4 | Rev1 | 0.015 | Young |
| 0.583917 | 0.024971 | 0.475 | 0.431 | 1 | 4 | Rab11fip5 | 0.044 | Young |
| 0.584488 | 0.04732 | 0.1 | 0.081 | 1 | 4 | Chst12 | 0.019 | Young |

| | | | | | | | | |
|---|---|---|---|---|---|---|---|---|
| 0.584511 | 0.209234 | 0.362 | 0.335 | 1 | 4 | Crebbp | 0.027 | Young |
| 0.584518 | 0.064731 | 0.088 | 0.07 | 1 | 4 | Trmt2a | 0.018 | Young |
| 0.584618 | 0.020573 | 0.312 | 0.279 | 1 | 4 | Erbin | 0.033 | Young |
| 0.584754 | 0.029192 | 0.275 | 0.24 | 1 | 4 | Zfand3 | 0.035 | Young |
| 0.585043 | 0.624976 | 0.1 | 0.084 | 1 | 4 | Zadh2 | 0.016 | Young |
| 0.585701 | 0.19186 | 0.125 | 0.106 | 1 | 4 | Clec4b1 | 0.019 | Young |
| 0.586146 | 0.127127 | 0.562 | 0.558 | 1 | 4 | Plekho2 | 0.004 | Young |
| 0.586651 | 0.506285 | 0.075 | 0.061 | 1 | 4 | Arhgef11 | 0.014 | Young |
| 0.586693 | 0.160498 | 0.162 | 0.14 | 1 | 4 | Wwc2 | 0.022 | Young |
| 0.586992 | 0.065291 | 0.688 | 0.621 | 1 | 4 | Luc7l2 | 0.067 | Young |
| 0.587536 | 0.501038 | 0.138 | 0.121 | 1 | 4 | Supt6 | 0.017 | Young |
| 0.587679 | 0.052035 | 0.15 | 0.125 | 1 | 4 | Cdk4 | 0.025 | Young |
| 0.587741 | 0.213864 | 0.075 | 0.06 | 1 | 4 | Zfp84 | 0.015 | Young |
| 0.588438 | 0.090621 | 0.488 | 0.42 | 1 | 4 | Cmtm6 | 0.068 | Young |
| 0.588676 | 0.433111 | 0.062 | 0.049 | 1 | 4 | Eci2 | 0.013 | Young |
| 0.588689 | 0.059942 | 0.2 | 0.166 | 1 | 4 | Nek9 | 0.034 | Young |
| 0.589255 | 0.141984 | 0.175 | 0.148 | 1 | 4 | Erlin1 | 0.027 | Young |
| 0.589774 | 0.10962 | 0.2 | 0.169 | 1 | 4 | Nfatc1 | 0.031 | Young |
| 0.590086 | 0.918065 | 0.025 | 0.017 | 1 | 4 | Dpy19l3 | 0.008 | Young |
| 0.590134 | 1.462149 | 0.013 | 0.007 | 1 | 4 | Cdh5 | 0.006 | Young |
| 0.590202 | 0.09328 | 0.45 | 0.422 | 1 | 4 | Gapvd1 | 0.028 | Young |
| 0.590841 | 0.843406 | 0.025 | 0.017 | 1 | 4 | E130308A1 | 0.008 | Young |
| 0.591282 | 1.223731 | 0.013 | 0.007 | 1 | 4 | Lhfp | 0.006 | Young |
| 0.591282 | 1.122255 | 0.013 | 0.007 | 1 | 4 | Bag2 | 0.006 | Young |
| 0.591282 | 1.04269 | 0.013 | 0.007 | 1 | 4 | Ccdc166 | 0.006 | Young |
| 0.591677 | 0.435892 | 0.088 | 0.072 | 1 | 4 | Coq8b | 0.016 | Young |
| 0.59178 | 0.16664 | 0.262 | 0.235 | 1 | 4 | Nfat5 | 0.027 | Young |
| 0.592076 | 0.1472 | 0.175 | 0.152 | 1 | 4 | Hcfc1 | 0.023 | Young |
| 0.592098 | 0.132464 | 0.1 | 0.082 | 1 | 4 | Alas1 | 0.018 | Young |
| 0.592352 | 0.774051 | 0.025 | 0.017 | 1 | 4 | Dnal4 | 0.008 | Young |
| 0.592373 | 0.06282 | 0.838 | 0.825 | 1 | 4 | Sdcbp | 0.013 | Young |
| 0.592431 | 1.057969 | 0.013 | 0.007 | 1 | 4 | Slc39a12 | 0.006 | Young |
| 0.592431 | 0.967368 | 0.013 | 0.007 | 1 | 4 | Arvcf | 0.006 | Young |
| 0.592431 | 0.906467 | 0.013 | 0.007 | 1 | 4 | Brf2 | 0.006 | Young |
| 0.59268 | 0.036081 | 0.55 | 0.518 | 1 | 4 | Mef2a | 0.032 | Young |
| 0.592698 | 0.705897 | 0.05 | 0.039 | 1 | 4 | Lncpint | 0.011 | Young |
| 0.592837 | 0.417374 | 0.075 | 0.061 | 1 | 4 | Slc12a2 | 0.014 | Young |
| 0.593041 | 0.268264 | 0.112 | 0.095 | 1 | 4 | Tango2 | 0.017 | Young |
| 0.593251 | 0.388099 | 0.075 | 0.061 | 1 | 4 | Cdyl | 0.014 | Young |
| 0.593354 | 0.061565 | 0.1 | 0.082 | 1 | 4 | Dbt | 0.018 | Young |
| 0.593367 | 0.30981 | 0.15 | 0.13 | 1 | 4 | Eif4e2 | 0.02 | Young |
| 0.593523 | 0.656454 | 0.062 | 0.049 | 1 | 4 | Mrps34 | 0.013 | Young |
| 0.59358 | 1.186899 | 0.013 | 0.007 | 1 | 4 | Gm30122 | 0.006 | Young |
| 0.59358 | 1.136481 | 0.013 | 0.007 | 1 | 4 | Tulp3 | 0.006 | Young |
| 0.59358 | 0.943515 | 0.013 | 0.007 | 1 | 4 | Anxa9 | 0.006 | Young |

| | | | | | | | | |
|---|---|---|---|---|---|---|---|---|
| 0.59358 | 0.877023 | 0.013 | 0.007 | 1 | 4 | Rnd2 | 0.006 | Young |
| 0.59358 | 0.866926 | 0.013 | 0.007 | 1 | 4 | Gm26536 | 0.006 | Young |
| 0.593702 | 0.698549 | 0.112 | 0.098 | 1 | 4 | Mrpl11 | 0.014 | Young |
| 0.594379 | 0.139218 | 0.075 | 0.06 | 1 | 4 | Tnfrsf17 | 0.015 | Young |
| 0.59455 | 0.092486 | 0.3 | 0.275 | 1 | 4 | Rb1cc1 | 0.025 | Young |
| 0.594623 | 0.688653 | 0.025 | 0.017 | 1 | 4 | Dnah2 | 0.008 | Young |
| 0.594731 | 0.792712 | 0.013 | 0.007 | 1 | 4 | B3gnt9 | 0.006 | Young |
| 0.594731 | 0.777159 | 0.013 | 0.007 | 1 | 4 | Hist3h2ba | 0.006 | Young |
| 0.595572 | 0.33968 | 0.312 | 0.302 | 1 | 4 | N4bp1 | 0.01 | Young |
| 0.595835 | 0.22652 | 0.112 | 0.094 | 1 | 4 | Atl2 | 0.018 | Young |
| 0.595883 | 0.811905 | 0.013 | 0.007 | 1 | 4 | AL591952.: | 0.006 | Young |
| 0.595883 | 0.612203 | 0.013 | 0.007 | 1 | 4 | Gm48529 | 0.006 | Young |
| 0.595883 | 0.352099 | 0.013 | 0.007 | 1 | 4 | Fam109b | 0.006 | Young |
| 0.595883 | 0.163097 | 0.013 | 0.007 | 1 | 4 | Gm527 | 0.006 | Young |
| 0.596209 | 0.308134 | 0.162 | 0.144 | 1 | 4 | Txlna | 0.018 | Young |
| 0.596522 | 0.160601 | 0.362 | 0.348 | 1 | 4 | Xiap | 0.014 | Young |
| 0.596607 | 0.063103 | 0.112 | 0.094 | 1 | 4 | S100pbp | 0.018 | Young |
| 0.59673 | 0.194149 | 0.088 | 0.109 | 1 | 4 | Lemd2 | -0.021 | Young |
| 0.596898 | 0.807598 | 0.025 | 0.017 | 1 | 4 | Mx1 | 0.008 | Young |
| 0.597036 | 0.591199 | 0.013 | 0.007 | 1 | 4 | Gm26756 | 0.006 | Young |
| 0.597036 | 0.412212 | 0.013 | 0.007 | 1 | 4 | Gm30978 | 0.006 | Young |
| 0.597036 | 0.348132 | 0.013 | 0.007 | 1 | 4 | Cenph | 0.006 | Young |
| 0.597036 | 0.145588 | 0.013 | 0.007 | 1 | 4 | 3110009E1 | 0.006 | Young |
| 0.598183 | 0.300849 | 0.162 | 0.144 | 1 | 4 | Pdap1 | 0.018 | Young |
| 0.59819 | 0.332001 | 0.013 | 0.007 | 1 | 4 | Mocs3 | 0.006 | Young |
| 0.59819 | 0.506109 | 0.013 | 0.007 | 1 | 4 | AC113595. | 0.006 | Young |
| 0.598307 | 0.474635 | 0.075 | 0.061 | 1 | 4 | Xpnpep1 | 0.014 | Young |
| 0.598862 | 0.462102 | 0.1 | 0.085 | 1 | 4 | Adh5 | 0.015 | Young |
| 0.599176 | 0.683107 | 0.025 | 0.017 | 1 | 4 | Crtc1 | 0.008 | Young |
| 0.599238 | 0.000874 | 0.188 | 0.161 | 1 | 4 | Rab5b | 0.027 | Young |
| 0.599345 | 0.459016 | 0.013 | 0.007 | 1 | 4 | Gm32856 | 0.006 | Young |
| 0.599345 | 0.032735 | 0.013 | 0.007 | 1 | 4 | E2f6 | 0.006 | Young |
| 0.599345 | 0.028358 | 0.013 | 0.007 | 1 | 4 | Tmem201 | 0.006 | Young |
| 0.599345 | 0.015249 | 0.013 | 0.007 | 1 | 4 | Gm43569 | 0.006 | Young |
| 0.599625 | 0.249976 | 0.05 | 0.038 | 1 | 4 | Rufy2 | 0.012 | Young |
| 0.599997 | 0.102657 | 0.025 | 0.037 | 1 | 4 | Gna11 | -0.012 | Young |
| 0.600131 | 0.112004 | 0.2 | 0.173 | 1 | 4 | Kdm1a | 0.027 | Young |
| 0.600396 | 0.230113 | 0.125 | 0.107 | 1 | 4 | Kpna3 | 0.018 | Young |
| 0.600423 | 0.03125 | 0.275 | 0.244 | 1 | 4 | Sbds | 0.031 | Young |
| 0.60047 | 0.210943 | 0.138 | 0.117 | 1 | 4 | Gid8 | 0.021 | Young |
| 0.600614 | 0.468393 | 0.062 | 0.05 | 1 | 4 | Stoml1 | 0.012 | Young |
| 0.600881 | 0.620838 | 0.05 | 0.039 | 1 | 4 | Srm | 0.011 | Young |
| 0.600929 | 0.122812 | 0.3 | 0.285 | 1 | 4 | Trim41 | 0.015 | Young |
| 0.601064 | 0.20441 | 0.025 | 0.037 | 1 | 4 | Rac3 | -0.012 | Young |
| 0.601802 | 0.010153 | 0.062 | 0.08 | 1 | 4 | Bin3 | -0.018 | Young |

| | | | | | | | | |
|---|---|---|---|---|---|---|---|---|
| 0.60264 | 0.152834 | 0.212 | 0.19 | 1 | 4 | Ndufaf8 | 0.022 | Young |
| 0.602855 | 0.339236 | 0.188 | 0.167 | 1 | 4 | Smu1 | 0.021 | Young |
| 0.602982 | 0.477246 | 0.025 | 0.017 | 1 | 4 | Xndc1 | 0.008 | Young |
| 0.602982 | 0.307224 | 0.025 | 0.017 | 1 | 4 | Cenpo | 0.008 | Young |
| 0.603237 | 0.039101 | 0.162 | 0.139 | 1 | 4 | Stxbp5 | 0.023 | Young |
| 0.603494 | 0.821526 | 0.1 | 0.085 | 1 | 4 | Prelid3b | 0.015 | Young |
| 0.603638 | 0.359173 | 0.038 | 0.028 | 1 | 4 | Smo | 0.01 | Young |
| 0.604187 | 0.110671 | 0.062 | 0.049 | 1 | 4 | Rpap3 | 0.013 | Young |
| 0.604801 | 0.100001 | 0.05 | 0.038 | 1 | 4 | Zfp367 | 0.012 | Young |
| 0.604822 | 0.015714 | 0.15 | 0.127 | 1 | 4 | Sf3a3 | 0.023 | Young |
| 0.604848 | 0.34979 | 0.038 | 0.028 | 1 | 4 | Bphl | 0.01 | Young |
| 0.605248 | 0.005318 | 0.2 | 0.175 | 1 | 4 | Pdcl3 | 0.025 | Young |
| 0.605434 | 0.515891 | 0.112 | 0.096 | 1 | 4 | Micu2 | 0.016 | Young |
| 0.606504 | 0.005201 | 0.637 | 0.595 | 1 | 4 | Pld4 | 0.042 | Young |
| 0.606617 | 0.163942 | 0.225 | 0.202 | 1 | 4 | 9-Sep | 0.023 | Young |
| 0.606887 | 0.034391 | 0.1 | 0.082 | 1 | 4 | Faap20 | 0.018 | Young |
| 0.607009 | 0.042832 | 0.212 | 0.189 | 1 | 4 | Xrn2 | 0.023 | Young |
| 0.607643 | 0.104104 | 0.25 | 0.226 | 1 | 4 | Ndufb4 | 0.024 | Young |
| 0.607713 | 0.996133 | 0.05 | 0.04 | 1 | 4 | AU022252 | 0.01 | Young |
| 0.607866 | 0.092846 | 0.112 | 0.094 | 1 | 4 | Tbck | 0.018 | Young |
| 0.60787 | 0.029697 | 0.25 | 0.222 | 1 | 4 | Brd7 | 0.028 | Young |
| 0.608329 | 0.088339 | 0.025 | 0.017 | 1 | 4 | Unc13d | 0.008 | Young |
| 0.608612 | 0.281017 | 0.175 | 0.156 | 1 | 4 | Uhrf1bp1l | 0.019 | Young |
| 0.608988 | 1.010922 | 0.038 | 0.028 | 1 | 4 | Kif21b | 0.01 | Young |
| 0.609116 | 0.328896 | 0.05 | 0.039 | 1 | 4 | Pam16 | 0.011 | Young |
| 0.61168 | 0.000255 | 0.35 | 0.31 | 1 | 4 | Trim8 | 0.04 | Young |
| 0.611699 | 0.382446 | 0.05 | 0.039 | 1 | 4 | Fam13b | 0.011 | Young |
| 0.611941 | 0.02885 | 0.212 | 0.185 | 1 | 4 | Tcf20 | 0.027 | Young |
| 0.611984 | 0.338193 | 0.112 | 0.094 | 1 | 4 | Egr3 | 0.018 | Young |
| 0.611993 | 0.080499 | 0.075 | 0.061 | 1 | 4 | 2210016L2 | 0.014 | Young |
| 0.612149 | 0.234099 | 0.2 | 0.181 | 1 | 4 | Gng12 | 0.019 | Young |
| 0.612584 | 0.06293 | 0.512 | 0.47 | 1 | 4 | Sbno2 | 0.042 | Young |
| 0.61284 | 0.010684 | 0.288 | 0.305 | 1 | 4 | Rsu1 | -0.017 | Young |
| 0.612899 | 0.705156 | 0.038 | 0.028 | 1 | 4 | Apool | 0.01 | Young |
| 0.613077 | 0.104041 | 0.575 | 0.524 | 1 | 4 | Gnaq | 0.051 | Young |
| 0.613252 | 0.141032 | 0.075 | 0.061 | 1 | 4 | Cyth2 | 0.014 | Young |
| 0.613272 | 0.112648 | 0.25 | 0.281 | 1 | 4 | Soat1 | -0.031 | Young |
| 0.613614 | 0.329542 | 0.112 | 0.098 | 1 | 4 | Fam98b | 0.014 | Young |
| 0.613873 | 0.358367 | 0.062 | 0.05 | 1 | 4 | Anks3 | 0.012 | Young |
| 0.614144 | 0.262211 | 0.138 | 0.161 | 1 | 4 | Lin7c | -0.023 | Young |
| 0.614565 | 0.10789 | 0.275 | 0.244 | 1 | 4 | Mfsd11 | 0.031 | Young |
| 0.614569 | 0.032314 | 0.038 | 0.028 | 1 | 4 | Hmgxb4 | 0.01 | Young |
| 0.615324 | 0.44591 | 0.05 | 0.039 | 1 | 4 | Cdkn2c | 0.011 | Young |
| 0.616663 | 0.076656 | 0.438 | 0.422 | 1 | 4 | Rnf149 | 0.016 | Young |
| 0.61681 | 0.126519 | 0.3 | 0.277 | 1 | 4 | Eif1b | 0.023 | Young |

| | | | | | | | | |
|---|---|---|---|---|---|---|---|---|
| 0.618403 | 0.405185 | 0.15 | 0.136 | 1 | 4 | Pdcd2l | 0.014 | Young |
| 0.618514 | 0.196293 | 0.162 | 0.142 | 1 | 4 | Ssbp4 | 0.02 | Young |
| 0.61901 | 1.32702 | 0.025 | 0.018 | 1 | 4 | Armc3 | 0.007 | Young |
| 0.619685 | 0.094053 | 0.3 | 0.265 | 1 | 4 | Uap1l1 | 0.035 | Young |
| 0.619999 | 0.314039 | 0.05 | 0.039 | 1 | 4 | Cnot11 | 0.011 | Young |
| 0.620379 | 0.036171 | 0.288 | 0.258 | 1 | 4 | Rragc | 0.03 | Young |
| 0.621286 | 1.17424 | 0.025 | 0.018 | 1 | 4 | Msl3l2 | 0.007 | Young |
| 0.621699 | 0.059552 | 0.15 | 0.13 | 1 | 4 | Gmps | 0.02 | Young |
| 0.622392 | 0.121626 | 0.3 | 0.265 | 1 | 4 | Dctn1 | 0.035 | Young |
| 0.62263 | 0.317921 | 0.062 | 0.05 | 1 | 4 | Zfp68 | 0.012 | Young |
| 0.622831 | 0.196565 | 0.15 | 0.131 | 1 | 4 | Uap1 | 0.019 | Young |
| 0.623603 | 0.136379 | 0.25 | 0.225 | 1 | 4 | Pdcd6 | 0.025 | Young |
| 0.623965 | 0.501088 | 0.112 | 0.097 | 1 | 4 | Myo1e | 0.015 | Young |
| 0.624573 | 0.04044 | 0.438 | 0.389 | 1 | 4 | Dynll1 | 0.049 | Young |
| 0.624597 | 0.413654 | 0.175 | 0.161 | 1 | 4 | Psme2b | 0.014 | Young |
| 0.625259 | 0.732937 | 0.062 | 0.051 | 1 | 4 | Trim36 | 0.011 | Young |
| 0.626777 | 0.815358 | 0.075 | 0.063 | 1 | 4 | Ppp1r8 | 0.012 | Young |
| 0.627738 | 0.49398 | 0.038 | 0.028 | 1 | 4 | Timmdc1 | 0.01 | Young |
| 0.627964 | 0.070231 | 0.075 | 0.061 | 1 | 4 | Orc3 | 0.014 | Young |
| 0.628036 | 0.170252 | 0.162 | 0.142 | 1 | 4 | Mob1b | 0.02 | Young |
| 0.628347 | 0.581334 | 0.038 | 0.028 | 1 | 4 | Itpr1 | 0.01 | Young |
| 0.628517 | 0.776145 | 0.025 | 0.018 | 1 | 4 | Pcdhga12 | 0.007 | Young |
| 0.62902 | 0.137318 | 0.075 | 0.061 | 1 | 4 | Bloc1s2 | 0.014 | Young |
| 0.630115 | 0.158197 | 0.15 | 0.132 | 1 | 4 | Ipmk | 0.018 | Young |
| 0.630259 | 0.09272 | 0.088 | 0.072 | 1 | 4 | Isca2 | 0.016 | Young |
| 0.631129 | 0.063585 | 0.175 | 0.153 | 1 | 4 | Ccdc115 | 0.022 | Young |
| 0.63119 | 0.804687 | 0.025 | 0.018 | 1 | 4 | Chd3os | 0.007 | Young |
| 0.631616 | 0.09434 | 0.013 | 0.02 | 1 | 4 | Bbs9 | -0.007 | Young |
| 0.632559 | 0.160868 | 0.212 | 0.193 | 1 | 4 | Bud31 | 0.019 | Young |
| 0.63257 | 0.064434 | 0.1 | 0.083 | 1 | 4 | Upf3a | 0.017 | Young |
| 0.63272 | 0.644226 | 0.025 | 0.018 | 1 | 4 | Lmnb2 | 0.007 | Young |
| 0.633228 | 0.231383 | 0.038 | 0.028 | 1 | 4 | Pck2 | 0.01 | Young |
| 0.633244 | 0.036232 | 0.787 | 0.717 | 1 | 4 | Akr1a1 | 0.07 | Young |
| 0.633803 | 0.766357 | 0.013 | 0.02 | 1 | 4 | Asb13 | -0.007 | Young |
| 0.633863 | 0.254284 | 0.3 | 0.279 | 1 | 4 | Ypel5 | 0.021 | Young |
| 0.634905 | 0.546065 | 0.062 | 0.051 | 1 | 4 | Anapc15 | 0.011 | Young |
| 0.635018 | 0.622868 | 0.025 | 0.018 | 1 | 4 | Nars2 | 0.007 | Young |
| 0.635062 | 0.295927 | 0.038 | 0.028 | 1 | 4 | Lysmd1 | 0.01 | Young |
| 0.635078 | 0.084467 | 0.2 | 0.175 | 1 | 4 | Phactr2 | 0.025 | Young |
| 0.635158 | 0.387146 | 0.075 | 0.063 | 1 | 4 | Coq5 | 0.012 | Young |
| 0.635598 | 0.078367 | 0.1 | 0.084 | 1 | 4 | Anapc4 | 0.016 | Young |
| 0.635851 | 1.106564 | 0.038 | 0.029 | 1 | 4 | Rnf219 | 0.009 | Young |
| 0.63597 | 0.658627 | 0.112 | 0.099 | 1 | 4 | Arid3a | 0.013 | Young |
| 0.636147 | 0.39841 | 0.05 | 0.04 | 1 | 4 | Cwc27 | 0.01 | Young |
| 0.636404 | 0.12805 | 0.125 | 0.107 | 1 | 4 | Ccng1 | 0.018 | Young |

| | | | | | | | | |
|---|---|---|---|---|---|---|---|---|
| 0.636574 | 0.816281 | 0.05 | 0.04 | 1 | 4 | Pilra | 0.01 | Young |
| 0.637319 | 0.58863 | 0.025 | 0.018 | 1 | 4 | Tmco4 | 0.007 | Young |
| 0.63755 | 0.045259 | 0.325 | 0.29 | 1 | 4 | Mbnl2 | 0.035 | Young |
| 0.637664 | 0.072873 | 0.662 | 0.626 | 1 | 4 | Gpx4 | 0.036 | Young |
| 0.637695 | 0.441221 | 0.025 | 0.036 | 1 | 4 | Comtd1 | -0.011 | Young |
| 0.63772 | 0.002908 | 0.55 | 0.494 | 1 | 4 | Serpinb8 | 0.056 | Young |
| 0.637995 | 0.068606 | 0.15 | 0.131 | 1 | 4 | Tnks | 0.019 | Young |
| 0.638087 | 0.603012 | 0.025 | 0.018 | 1 | 4 | Wdr53 | 0.007 | Young |
| 0.638087 | 0.158921 | 0.025 | 0.018 | 1 | 4 | 2010320M | 0.007 | Young |
| 0.638127 | 0.112454 | 0.112 | 0.096 | 1 | 4 | Hypk | 0.016 | Young |
| 0.638276 | 0.818967 | 0.038 | 0.029 | 1 | 4 | Repin1 | 0.009 | Young |
| 0.638448 | 0.321535 | 0.188 | 0.169 | 1 | 4 | Smap1 | 0.019 | Young |
| 0.638694 | 0.271272 | 0.025 | 0.035 | 1 | 4 | Fam122a | -0.01 | Young |
| 0.639129 | 0.362795 | 0.175 | 0.159 | 1 | 4 | Slc16a7 | 0.016 | Young |
| 0.63914 | 0.089668 | 0.8 | 0.803 | 1 | 4 | Rpl3 | -0.003 | Young |
| 0.639433 | 0.368187 | 0.15 | 0.132 | 1 | 4 | Zc3h18 | 0.018 | Young |
| 0.639623 | 0.405968 | 0.025 | 0.018 | 1 | 4 | Ccdc137 | 0.007 | Young |
| 0.639623 | 0.434027 | 0.025 | 0.018 | 1 | 4 | Tuba4a | 0.007 | Young |
| 0.639708 | 0.61849 | 0.075 | 0.062 | 1 | 4 | Plscr4 | 0.013 | Young |
| 0.640098 | 0.924027 | 0.038 | 0.029 | 1 | 4 | Tdrd3 | 0.009 | Young |
| 0.640315 | 0.234393 | 0.188 | 0.172 | 1 | 4 | Tgs1 | 0.016 | Young |
| 0.640963 | 0.12404 | 0.088 | 0.103 | 1 | 4 | Cd82 | -0.015 | Young |
| 0.641834 | 0.029151 | 0.963 | 0.912 | 1 | 4 | Npc2 | 0.051 | Young |
| 0.641931 | 0.339864 | 0.025 | 0.018 | 1 | 4 | Zfp949 | 0.007 | Young |
| 0.642 | 0.635032 | 0.125 | 0.111 | 1 | 4 | Rnf103 | 0.014 | Young |
| 0.642476 | 0.328464 | 0.062 | 0.051 | 1 | 4 | Eef2kmt | 0.011 | Young |
| 0.642949 | 0.204167 | 0.05 | 0.04 | 1 | 4 | 2310061I0 | 0.01 | Young |
| 0.642949 | 0.610732 | 0.05 | 0.04 | 1 | 4 | Mid1 | 0.01 | Young |
| 0.642949 | 0.245496 | 0.05 | 0.04 | 1 | 4 | Cipc | 0.01 | Young |
| 0.644086 | 0.173168 | 0.887 | 0.924 | 1 | 4 | Neat1 | -0.037 | Young |
| 0.644225 | 0.32859 | 0.1 | 0.086 | 1 | 4 | C2cd5 | 0.014 | Young |
| 0.644451 | 0.011102 | 0.438 | 0.399 | 1 | 4 | Cd164 | 0.039 | Young |
| 0.644639 | 0.062999 | 0.65 | 0.634 | 1 | 4 | Atp5e | 0.016 | Young |
| 0.645014 | 0.027331 | 0.025 | 0.018 | 1 | 4 | Zfp60 | 0.007 | Young |
| 0.645165 | 0.061861 | 0.462 | 0.431 | 1 | 4 | Eif5a | 0.031 | Young |
| 0.645211 | 0.126807 | 0.138 | 0.119 | 1 | 4 | Ccm2 | 0.019 | Young |
| 0.64547 | 0.073993 | 0.125 | 0.148 | 1 | 4 | Acaa1a | -0.023 | Young |
| 0.646624 | 0.04013 | 0.05 | 0.04 | 1 | 4 | Snx25 | 0.01 | Young |
| 0.647067 | 0.060714 | 0.125 | 0.148 | 1 | 4 | Stard7 | -0.023 | Young |
| 0.647218 | 0.130896 | 0.262 | 0.239 | 1 | 4 | Prkcsh | 0.023 | Young |
| 0.647543 | 0.062048 | 0.6 | 0.595 | 1 | 4 | Clec4a1 | 0.005 | Young |
| 0.647726 | 0.055749 | 0.1 | 0.085 | 1 | 4 | Unc119 | 0.015 | Young |
| 0.649237 | 0.552113 | 0.038 | 0.029 | 1 | 4 | Zkscan17 | 0.009 | Young |
| 0.649323 | 0.08236 | 0.288 | 0.26 | 1 | 4 | Arhgef10l | 0.028 | Young |
| 0.649425 | 1.245473 | 0.013 | 0.008 | 1 | 4 | Cryab | 0.005 | Young |

| | | | | | | | | |
|---|---|---|---|---|---|---|---|---|
| 0.649425 | 1.190433 | 0.013 | 0.008 | 1 | 4 | Gm15655 | 0.005 | Young |
| 0.649425 | 1.099146 | 0.013 | 0.008 | 1 | 4 | Apbb1 | 0.005 | Young |
| 0.6502 | 0.02755 | 0.112 | 0.096 | 1 | 4 | Commd10 | 0.016 | Young |
| 0.650282 | 0.232451 | 0.1 | 0.086 | 1 | 4 | CAAA0111[illegible] | 0.014 | Young |
| 0.650304 | 0.15685 | 0.1 | 0.085 | 1 | 4 | Fam53c | 0.015 | Young |
| 0.650531 | 0.182339 | 0.15 | 0.132 | 1 | 4 | Smpd1 | 0.018 | Young |
| 0.650575 | 1.420972 | 0.013 | 0.008 | 1 | 4 | C920021L1 | 0.005 | Young |
| 0.650575 | 1.386705 | 0.013 | 0.008 | 1 | 4 | Dpp4 | 0.005 | Young |
| 0.650575 | 1.183161 | 0.013 | 0.008 | 1 | 4 | Abhd8 | 0.005 | Young |
| 0.650575 | 1.038042 | 0.013 | 0.008 | 1 | 4 | Zfp58 | 0.005 | Young |
| 0.650575 | 0.998446 | 0.013 | 0.008 | 1 | 4 | Hells | 0.005 | Young |
| 0.651039 | 0.064839 | 0.212 | 0.191 | 1 | 4 | Cdc5l | 0.021 | Young |
| 0.651071 | 0.68833 | 0.038 | 0.029 | 1 | 4 | St7 | 0.009 | Young |
| 0.651335 | 0.194382 | 0.175 | 0.154 | 1 | 4 | Uba6 | 0.021 | Young |
| 0.651683 | 0.653826 | 0.038 | 0.029 | 1 | 4 | Rnf135 | 0.009 | Young |
| 0.651726 | 1.502591 | 0.013 | 0.008 | 1 | 4 | Ehhadh | 0.005 | Young |
| 0.651726 | 0.749822 | 0.013 | 0.008 | 1 | 4 | Klrk1 | 0.005 | Young |
| 0.652029 | 0.382544 | 0.288 | 0.276 | 1 | 4 | Secisbp2l | 0.012 | Young |
| 0.65206 | 0.172982 | 0.238 | 0.212 | 1 | 4 | Pip5k1c | 0.026 | Young |
| 0.652439 | 0.107343 | 0.112 | 0.097 | 1 | 4 | Cfap43 | 0.015 | Young |
| 0.652701 | 0.203003 | 0.15 | 0.133 | 1 | 4 | Ptk2 | 0.017 | Young |
| 0.652878 | 0.711779 | 0.013 | 0.008 | 1 | 4 | Icam2 | 0.005 | Young |
| 0.652878 | 0.696135 | 0.013 | 0.008 | 1 | 4 | Tarbp1 | 0.005 | Young |
| 0.652878 | 0.645045 | 0.013 | 0.008 | 1 | 4 | Tctn1 | 0.005 | Young |
| 0.652961 | 0.531175 | 0.112 | 0.097 | 1 | 4 | Nkiras2 | 0.015 | Young |
| 0.653032 | 0.22661 | 0.312 | 0.283 | 1 | 4 | B4galt1 | 0.029 | Young |
| 0.653124 | 0.41308 | 0.138 | 0.125 | 1 | 4 | Tmem205 | 0.013 | Young |
| 0.653871 | 0.2544 | 0.175 | 0.155 | 1 | 4 | Atg16l1 | 0.02 | Young |
| 0.65403 | 0.760979 | 0.013 | 0.008 | 1 | 4 | Rexo5 | 0.005 | Young |
| 0.65403 | 0.307155 | 0.013 | 0.008 | 1 | 4 | Aldh1b1 | 0.005 | Young |
| 0.654525 | 0.020971 | 0.025 | 0.035 | 1 | 4 | Slc25a22 | -0.01 | Young |
| 0.654588 | 0.022074 | 0.088 | 0.073 | 1 | 4 | Med20 | 0.015 | Young |
| 0.655046 | 0.049179 | 0.362 | 0.328 | 1 | 4 | Eif5 | 0.034 | Young |
| 0.655184 | 0.580583 | 0.013 | 0.008 | 1 | 4 | Rwdd2a | 0.005 | Young |
| 0.655184 | 0.409364 | 0.013 | 0.008 | 1 | 4 | Negr1 | 0.005 | Young |
| 0.655641 | 0.057758 | 0.312 | 0.284 | 1 | 4 | Tmem109 | 0.028 | Young |
| 0.655825 | 0.315713 | 0.138 | 0.121 | 1 | 4 | Rap2a | 0.017 | Young |
| 0.656248 | 0.066601 | 0.275 | 0.245 | 1 | 4 | Ppib | 0.03 | Young |
| 0.656338 | 0.362105 | 0.013 | 0.008 | 1 | 4 | Gm10642 | 0.005 | Young |
| 0.656397 | 0.334642 | 0.05 | 0.04 | 1 | 4 | Zfp839 | 0.01 | Young |
| 0.656442 | 0.052788 | 0.013 | 0.02 | 1 | 4 | Gstm5 | -0.007 | Young |
| 0.657226 | 0.033627 | 0.088 | 0.074 | 1 | 4 | Btbd10 | 0.014 | Young |
| 0.657448 | 0.075789 | 0.462 | 0.431 | 1 | 4 | Sf3b2 | 0.031 | Young |
| 0.657493 | 0.255229 | 0.013 | 0.008 | 1 | 4 | Gm28151 | 0.005 | Young |
| 0.657493 | 0.217303 | 0.013 | 0.008 | 1 | 4 | Slc26a6 | 0.005 | Young |

| | | | | | | | | |
|---|---|---|---|---|---|---|---|---|
| 0.657922 | 0.004895 | 0.062 | 0.051 | 1 | 4 | Pgghg | 0.011 | Young |
| 0.657952 | 0.967544 | 0.075 | 0.065 | 1 | 4 | Dgcr8 | 0.01 | Young |
| 0.658016 | 0.114699 | 0.088 | 0.074 | 1 | 4 | Idh3a | 0.014 | Young |
| 0.658063 | 0.165579 | 0.1 | 0.085 | 1 | 4 | Bcap29 | 0.015 | Young |
| 0.65855 | 0.259644 | 0.088 | 0.073 | 1 | 4 | Actr1b | 0.015 | Young |
| 0.658649 | 0.201849 | 0.013 | 0.008 | 1 | 4 | Cdc7 | 0.005 | Young |
| 0.658649 | 0.165819 | 0.013 | 0.008 | 1 | 4 | AW011738 | 0.005 | Young |
| 0.658649 | 0.038566 | 0.013 | 0.008 | 1 | 4 | Gm16845 | 0.005 | Young |
| 0.658981 | 0.366979 | 0.138 | 0.125 | 1 | 4 | Tmem106c | 0.013 | Young |
| 0.659393 | 0.3668 | 0.138 | 0.123 | 1 | 4 | Sdf2l1 | 0.015 | Young |
| 0.660963 | 0.15934 | 0.013 | 0.008 | 1 | 4 | Zfp870 | 0.005 | Young |
| 0.66097 | 0.193692 | 0.138 | 0.123 | 1 | 4 | Trmt1 | 0.015 | Young |
| 0.661397 | 0.246416 | 0.062 | 0.051 | 1 | 4 | Rrp15 | 0.011 | Young |
| 0.662093 | 0.112674 | 0.412 | 0.385 | 1 | 4 | Mindy2 | 0.027 | Young |
| 0.662107 | 0.213641 | 0.112 | 0.098 | 1 | 4 | Kmt5a | 0.014 | Young |
| 0.662597 | 1.28393 | 0.038 | 0.03 | 1 | 4 | Gstp3 | 0.008 | Young |
| 0.662614 | 0.189023 | 0.2 | 0.183 | 1 | 4 | Amdhd2 | 0.017 | Young |
| 0.662737 | 0.186282 | 0.038 | 0.029 | 1 | 4 | Slc7a13 | 0.009 | Young |
| 0.662865 | 0.345241 | 0.162 | 0.144 | 1 | 4 | Mrpl42 | 0.018 | Young |
| 0.662914 | 0.237963 | 0.288 | 0.264 | 1 | 4 | Psme2 | 0.024 | Young |
| 0.663161 | 0.088907 | 0.088 | 0.074 | 1 | 4 | Pla2g4a | 0.014 | Young |
| 0.663204 | 0.08434 | 0.2 | 0.175 | 1 | 4 | Trappc6b | 0.025 | Young |
| 0.663353 | 0.028623 | 0.488 | 0.486 | 1 | 4 | Tnfaip8 | 0.002 | Young |
| 0.663893 | 0.385222 | 0.062 | 0.052 | 1 | 4 | Sart3 | 0.01 | Young |
| 0.663932 | 0.057117 | 0.1 | 0.086 | 1 | 4 | Abi2 | 0.014 | Young |
| 0.664496 | 0.494724 | 0.238 | 0.225 | 1 | 4 | Dock8 | 0.013 | Young |
| 0.664518 | 0.33689 | 0.125 | 0.11 | 1 | 4 | Pdcd4 | 0.015 | Young |
| 0.665494 | 0.022051 | 0.25 | 0.278 | 1 | 4 | Bcap31 | -0.028 | Young |
| 0.665669 | 0.139803 | 0.138 | 0.121 | 1 | 4 | Washc1 | 0.017 | Young |
| 0.666465 | 0.02755 | 0.238 | 0.209 | 1 | 4 | Hsd17b4 | 0.029 | Young |
| 0.666518 | 0.194866 | 0.212 | 0.187 | 1 | 4 | Uvrag | 0.025 | Young |
| 0.667088 | 0.386559 | 0.162 | 0.146 | 1 | 4 | Tor1aip2 | 0.016 | Young |
| 0.667315 | 0.617771 | 0.025 | 0.018 | 1 | 4 | Ehbp1 | 0.007 | Young |
| 0.667458 | 0.174386 | 0.05 | 0.04 | 1 | 4 | Lin37 | 0.01 | Young |
| 0.667854 | 0.07588 | 0.15 | 0.131 | 1 | 4 | Trappc8 | 0.019 | Young |
| 0.668284 | 0.009054 | 0.15 | 0.175 | 1 | 4 | Sec22b | -0.025 | Young |
| 0.66923 | 0.136987 | 0.112 | 0.097 | 1 | 4 | Irgm1 | 0.015 | Young |
| 0.669287 | 0.120707 | 0.238 | 0.217 | 1 | 4 | Ssrp1 | 0.021 | Young |
| 0.669468 | 0.137801 | 0.162 | 0.145 | 1 | 4 | Fem1b | 0.017 | Young |
| 0.670545 | 0.749695 | 0.038 | 0.03 | 1 | 4 | Surf2 | 0.008 | Young |
| 0.670853 | 0.010931 | 0.1 | 0.116 | 1 | 4 | Ifi209 | -0.016 | Young |
| 0.671394 | 0.132283 | 0.512 | 0.493 | 1 | 4 | Rpl22l1 | 0.019 | Young |
| 0.671416 | 0.306466 | 0.05 | 0.063 | 1 | 4 | Rassf8 | -0.013 | Young |
| 0.67164 | 0.016981 | 0.088 | 0.104 | 1 | 4 | Zfp655 | -0.016 | Young |
| 0.671701 | 0.196582 | 0.112 | 0.097 | 1 | 4 | Proser1 | 0.015 | Young |

| | | | | | | | | |
|---|---|---|---|---|---|---|---|---|
| 0.671844 | 0.232269 | 0.238 | 0.223 | 1 | 4 | Tomm5 | 0.015 | Young |
| 0.671861 | 0.195269 | 0.175 | 0.158 | 1 | 4 | Zc3h15 | 0.017 | Young |
| 0.672006 | 0.125964 | 0.038 | 0.029 | 1 | 4 | Thbs1 | 0.009 | Young |
| 0.672029 | 0.010881 | 0.325 | 0.299 | 1 | 4 | 2-Mar | 0.026 | Young |
| 0.672381 | 0.001006 | 0.2 | 0.229 | 1 | 4 | Tmem87b | -0.029 | Young |
| 0.672384 | 0.1367 | 0.025 | 0.034 | 1 | 4 | Ppan | -0.009 | Young |
| 0.672433 | 0.397045 | 0.062 | 0.078 | 1 | 4 | Immp1l | -0.016 | Young |
| 0.673325 | 0.174151 | 0.325 | 0.308 | 1 | 4 | Smchd1 | 0.017 | Young |
| 0.673493 | 0.056445 | 0.225 | 0.201 | 1 | 4 | Tnfsf12 | 0.024 | Young |
| 0.673842 | 0.09244 | 0.25 | 0.225 | 1 | 4 | Atp6v1c1 | 0.025 | Young |
| 0.674216 | 0.335121 | 0.175 | 0.159 | 1 | 4 | Gdap10 | 0.016 | Young |
| 0.674228 | 0.125993 | 0.338 | 0.368 | 1 | 4 | Pik3r1 | -0.03 | Young |
| 0.674252 | 0.477723 | 0.025 | 0.018 | 1 | 4 | Olfr1167 | 0.007 | Young |
| 0.674524 | 0.137756 | 0.162 | 0.144 | 1 | 4 | Snf8 | 0.018 | Young |
| 0.674704 | 0.17309 | 0.025 | 0.034 | 1 | 4 | Zfp182 | -0.009 | Young |
| 0.675082 | 0.059463 | 0.425 | 0.383 | 1 | 4 | Ppp1ca | 0.042 | Young |
| 0.675202 | 0.272957 | 0.112 | 0.099 | 1 | 4 | Trappc1 | 0.013 | Young |
| 0.675526 | 0.823424 | 0.05 | 0.041 | 1 | 4 | Gm45606 | 0.009 | Young |
| 0.675852 | 0.043658 | 0.288 | 0.267 | 1 | 4 | Zranb2 | 0.021 | Young |
| 0.676176 | 0.152836 | 0.038 | 0.049 | 1 | 4 | Slc10a6 | -0.011 | Young |
| 0.677042 | 0.023674 | 0.238 | 0.211 | 1 | 4 | Eif2s3x | 0.027 | Young |
| 0.678506 | 0.012278 | 0.013 | 0.019 | 1 | 4 | Slc46a1 | -0.006 | Young |
| 0.679236 | 0.176523 | 0.412 | 0.447 | 1 | 4 | Ednrb | -0.035 | Young |
| 0.679635 | 0.041136 | 0.375 | 0.356 | 1 | 4 | Cd48 | 0.019 | Young |
| 0.679667 | 0.003696 | 0.025 | 0.018 | 1 | 4 | Setd6 | 0.007 | Young |
| 0.67982 | 0.655088 | 0.05 | 0.042 | 1 | 4 | Dcun1d3 | 0.008 | Young |
| 0.680305 | 0.215472 | 0.062 | 0.052 | 1 | 4 | Eri3 | 0.01 | Young |
| 0.680372 | 0.033443 | 0.162 | 0.146 | 1 | 4 | Ncaph2 | 0.016 | Young |
| 0.680376 | 0.71144 | 0.038 | 0.03 | 1 | 4 | Fam26f | 0.008 | Young |
| 0.680515 | 0.38883 | 0.025 | 0.034 | 1 | 4 | Gpatch1 | -0.009 | Young |
| 0.680609 | 0.056788 | 0.238 | 0.213 | 1 | 4 | Sh3pxd2b | 0.025 | Young |
| 0.681167 | 0.340624 | 0.075 | 0.064 | 1 | 4 | Zfp994 | 0.011 | Young |
| 0.681217 | 0.44074 | 0.125 | 0.113 | 1 | 4 | Sap18 | 0.012 | Young |
| 0.681355 | 0.026047 | 0.162 | 0.144 | 1 | 4 | Flot2 | 0.018 | Young |
| 0.681942 | 0.115519 | 0.075 | 0.063 | 1 | 4 | Dnajc30 | 0.012 | Young |
| 0.682465 | 0.857357 | 0.038 | 0.03 | 1 | 4 | Dlg4 | 0.008 | Young |
| 0.682504 | 0.057823 | 0.138 | 0.121 | 1 | 4 | Cyc1 | 0.017 | Young |
| 0.683002 | 0.250794 | 0.088 | 0.075 | 1 | 4 | Ruvbl1 | 0.013 | Young |
| 0.683157 | 0.705953 | 0.013 | 0.019 | 1 | 4 | Pcgf1 | -0.006 | Young |
| 0.68331 | 0.264185 | 0.075 | 0.064 | 1 | 4 | Nolc1 | 0.011 | Young |
| 0.683698 | 0.510865 | 0.112 | 0.103 | 1 | 4 | Clec5a | 0.009 | Young |
| 0.68469 | 0.070426 | 0.438 | 0.405 | 1 | 4 | Ssh2 | 0.033 | Young |
| 0.686437 | 0.008885 | 0.4 | 0.357 | 1 | 4 | Snrpe | 0.043 | Young |
| 0.687497 | 0.12305 | 0.338 | 0.314 | 1 | 4 | Zfand6 | 0.024 | Young |
| 0.687579 | 0.014891 | 0.575 | 0.518 | 1 | 4 | Slc43a2 | 0.057 | Young |

| | | | | | | | | |
|---|---|---|---|---|---|---|---|---|
| 0.687864 | 0.484034 | 0.062 | 0.053 | 1 | 4 | Pias3 | 0.009 | Young |
| 0.687948 | 0.191616 | 0.138 | 0.121 | 1 | 4 | Fam126b | 0.017 | Young |
| 0.689548 | 0.264749 | 0.025 | 0.034 | 1 | 4 | 4930555A( | -0.009 | Young |
| 0.690225 | 0.236331 | 0.05 | 0.061 | 1 | 4 | Arl2 | -0.011 | Young |
| 0.690263 | 0.196191 | 0.038 | 0.03 | 1 | 4 | Trim32 | 0.008 | Young |
| 0.690431 | 0.089071 | 0.125 | 0.109 | 1 | 4 | Ppp1r11 | 0.016 | Young |
| 0.691681 | 0.062925 | 0.138 | 0.122 | 1 | 4 | Socs4 | 0.016 | Young |
| 0.692103 | 0.358789 | 0.062 | 0.053 | 1 | 4 | Upf2 | 0.009 | Young |
| 0.692449 | 0.130589 | 0.325 | 0.306 | 1 | 4 | Rsrc2 | 0.019 | Young |
| 0.692733 | 0.058319 | 0.525 | 0.483 | 1 | 4 | Tacc1 | 0.042 | Young |
| 0.692754 | 0.208729 | 0.4 | 0.385 | 1 | 4 | Cebpd | 0.015 | Young |
| 0.692928 | 1.177831 | 0.025 | 0.019 | 1 | 4 | Cep152 | 0.006 | Young |
| 0.692968 | 0.033915 | 0.138 | 0.122 | 1 | 4 | Cd180 | 0.016 | Young |
| 0.693729 | 0.270481 | 0.1 | 0.121 | 1 | 4 | Usf1 | -0.021 | Young |
| 0.69407 | 0.102738 | 0.238 | 0.219 | 1 | 4 | Snx10 | 0.019 | Young |
| 0.694362 | 0.150787 | 0.225 | 0.202 | 1 | 4 | Ctsf | 0.023 | Young |
| 0.694422 | 0.002504 | 0.2 | 0.181 | 1 | 4 | Mtmr6 | 0.019 | Young |
| 0.694605 | 0.193189 | 0.038 | 0.03 | 1 | 4 | Ctdspl | 0.008 | Young |
| 0.695475 | 0.176817 | 0.088 | 0.076 | 1 | 4 | Nip7 | 0.012 | Young |
| 0.695752 | 0.085568 | 0.4 | 0.374 | 1 | 4 | Cox7a2 | 0.026 | Young |
| 0.695848 | 0.034848 | 0.038 | 0.03 | 1 | 4 | Tfcp2 | 0.008 | Young |
| 0.69588 | 0.698277 | 0.062 | 0.053 | 1 | 4 | Abhd5 | 0.009 | Young |
| 0.696772 | 0.923873 | 0.025 | 0.019 | 1 | 4 | Wdr76 | 0.006 | Young |
| 0.697039 | 0.044563 | 0.288 | 0.262 | 1 | 4 | Mrpl28 | 0.026 | Young |
| 0.697787 | 0.214419 | 0.125 | 0.113 | 1 | 4 | Slc39a10 | 0.012 | Young |
| 0.697801 | 0.043555 | 0.05 | 0.041 | 1 | 4 | Bin2 | 0.009 | Young |
| 0.697907 | 0.027106 | 0.288 | 0.312 | 1 | 4 | Itgb2 | -0.024 | Young |
| 0.698024 | 0.011378 | 0.325 | 0.302 | 1 | 4 | Nxf1 | 0.023 | Young |
| 0.698119 | 0.084354 | 0.312 | 0.282 | 1 | 4 | Tcea1 | 0.03 | Young |
| 0.699018 | 0.408469 | 0.025 | 0.034 | 1 | 4 | Akip1 | -0.009 | Young |
| 0.699053 | 0.076711 | 0.4 | 0.362 | 1 | 4 | Txn1 | 0.038 | Young |
| 0.699277 | 0.05772 | 0.438 | 0.403 | 1 | 4 | Larp4b | 0.035 | Young |
| 0.69947 | 0.184732 | 0.088 | 0.076 | 1 | 4 | Aars | 0.012 | Young |
| 0.699581 | 0.130278 | 0.375 | 0.355 | 1 | 4 | Stx7 | 0.02 | Young |
| 0.699999 | 0.129223 | 0.212 | 0.199 | 1 | 4 | Notch1 | 0.013 | Young |
| 0.700394 | 0.482411 | 0.05 | 0.042 | 1 | 4 | Plekhn1 | 0.008 | Young |
| 0.701159 | 0.098227 | 0.388 | 0.364 | 1 | 4 | Mapk14 | 0.024 | Young |
| 0.701394 | 1.109681 | 0.025 | 0.019 | 1 | 4 | Phykpl | 0.006 | Young |
| 0.701517 | 0.025239 | 0.388 | 0.354 | 1 | 4 | Pebp1 | 0.034 | Young |
| 0.702261 | 0.105749 | 0.25 | 0.227 | 1 | 4 | Irf5 | 0.023 | Young |
| 0.702276 | 0.406621 | 0.038 | 0.047 | 1 | 4 | Mrps11 | -0.009 | Young |
| 0.702348 | 0.170116 | 0.062 | 0.074 | 1 | 4 | Llgl1 | -0.012 | Young |
| 0.702509 | 0.193991 | 0.112 | 0.1 | 1 | 4 | Sra1 | 0.012 | Young |
| 0.702609 | 0.046737 | 0.05 | 0.041 | 1 | 4 | Slx4ip | 0.009 | Young |
| 0.70303 | 0.196169 | 0.188 | 0.175 | 1 | 4 | Mctp1 | 0.013 | Young |

| | | | | | | | | |
|---|---|---|---|---|---|---|---|---|
| 0.703048 | 0.3368 | 0.05 | 0.042 | 1 | 4 | Ap3m2 | 0.008 | Young |
| 0.7031 | 0.005274 | 0.013 | 0.018 | 1 | 4 | Wnt11 | -0.005 | Young |
| 0.70371 | 0.473522 | 0.025 | 0.019 | 1 | 4 | Lmnb1 | 0.006 | Young |
| 0.704057 | 0.231433 | 0.175 | 0.158 | 1 | 4 | Rnf114 | 0.017 | Young |
| 0.704456 | 0.086285 | 0.188 | 0.171 | 1 | 4 | Ddx46 | 0.017 | Young |
| 0.704519 | 0.647783 | 0.05 | 0.042 | 1 | 4 | Zfp358 | 0.008 | Young |
| 0.704556 | 0.038412 | 0.388 | 0.35 | 1 | 4 | Hnrnph2 | 0.038 | Young |
| 0.704611 | 0.408192 | 0.088 | 0.075 | 1 | 4 | Ado | 0.013 | Young |
| 0.704897 | 0.104245 | 0.1 | 0.088 | 1 | 4 | Cyp4f16 | 0.012 | Young |
| 0.705207 | 0.0038 | 0.2 | 0.182 | 1 | 4 | Abhd17b | 0.018 | Young |
| 0.705255 | 0.793237 | 0.025 | 0.019 | 1 | 4 | 3222401L1 | 0.006 | Young |
| 0.705567 | 0.025658 | 0.25 | 0.227 | 1 | 4 | Gdi1 | 0.023 | Young |
| 0.705584 | 0.031946 | 0.162 | 0.146 | 1 | 4 | Rspry1 | 0.016 | Young |
| 0.705929 | 1.414295 | 0.013 | 0.009 | 1 | 4 | Klhdc1 | 0.004 | Young |
| 0.705929 | 1.410499 | 0.013 | 0.009 | 1 | 4 | Fgfbp3 | 0.004 | Young |
| 0.705929 | 1.09049 | 0.013 | 0.009 | 1 | 4 | Neil2 | 0.004 | Young |
| 0.705929 | 0.984868 | 0.013 | 0.009 | 1 | 4 | Map10 | 0.004 | Young |
| 0.705977 | 0.226971 | 0.162 | 0.145 | 1 | 4 | Ldb1 | 0.017 | Young |
| 0.706075 | 0.579098 | 0.062 | 0.054 | 1 | 4 | Lztr1 | 0.008 | Young |
| 0.706238 | 0.336525 | 0.05 | 0.042 | 1 | 4 | Npat | 0.008 | Young |
| 0.707074 | 1.172632 | 0.013 | 0.009 | 1 | 4 | 0610040B1 | 0.004 | Young |
| 0.707074 | 1.112377 | 0.013 | 0.009 | 1 | 4 | B3gnt5 | 0.004 | Young |
| 0.707074 | 0.798153 | 0.013 | 0.009 | 1 | 4 | Fchsd1 | 0.004 | Young |
| 0.707074 | 0.798139 | 0.013 | 0.009 | 1 | 4 | Slc8a1 | 0.004 | Young |
| 0.707093 | 0.149085 | 0.013 | 0.018 | 1 | 4 | Cdkn1c | -0.005 | Young |
| 0.707438 | 0.461639 | 0.075 | 0.065 | 1 | 4 | Malsu1 | 0.01 | Young |
| 0.707632 | 0.182513 | 0.088 | 0.075 | 1 | 4 | Dera | 0.013 | Young |
| 0.707808 | 0.091806 | 0.138 | 0.123 | 1 | 4 | Slc38a10 | 0.015 | Young |
| 0.707893 | 0.583786 | 0.013 | 0.018 | 1 | 4 | Ydjc | -0.005 | Young |
| 0.707893 | 0.411984 | 0.013 | 0.018 | 1 | 4 | Ankrd52 | -0.005 | Young |
| 0.707903 | 0.195292 | 0.138 | 0.124 | 1 | 4 | Znrf1 | 0.014 | Young |
| 0.708219 | 0.760871 | 0.013 | 0.009 | 1 | 4 | 6230400D1 | 0.004 | Young |
| 0.708298 | 0.675708 | 0.038 | 0.03 | 1 | 4 | Mrpl44 | 0.008 | Young |
| 0.708653 | 0.111887 | 0.25 | 0.229 | 1 | 4 | Ythdf3 | 0.021 | Young |
| 0.708785 | 0.03863 | 0.362 | 0.337 | 1 | 4 | Arhgef12 | 0.025 | Young |
| 0.708917 | 0.395243 | 0.038 | 0.03 | 1 | 4 | Usp35 | 0.008 | Young |
| 0.709124 | 0.427467 | 0.025 | 0.019 | 1 | 4 | Mkks | 0.006 | Young |
| 0.709124 | 0.361154 | 0.025 | 0.019 | 1 | 4 | Bbs7 | 0.006 | Young |
| 0.709365 | 0.759173 | 0.013 | 0.009 | 1 | 4 | Pdss1 | 0.004 | Young |
| 0.709365 | 0.58681 | 0.013 | 0.009 | 1 | 4 | Aldh1l1 | 0.004 | Young |
| 0.709493 | 1.26307 | 0.013 | 0.018 | 1 | 4 | Tceal1 | -0.005 | Young |
| 0.709551 | 0.223592 | 0.25 | 0.235 | 1 | 4 | Fis1 | 0.015 | Young |
| 0.709579 | 0.037283 | 0.575 | 0.561 | 1 | 4 | Ifngr1 | 0.014 | Young |
| 0.709632 | 0.086162 | 0.062 | 0.053 | 1 | 4 | Nudt14 | 0.009 | Young |
| 0.709898 | 0.071097 | 0.025 | 0.019 | 1 | 4 | Trim14 | 0.006 | Young |

| | | | | | | | | |
|---|---|---|---|---|---|---|---|---|
| 0.710512 | 0.80439 | 0.013 | 0.009 | 1 | 4 | Zkscan5 | 0.004 | Young |
| 0.710512 | 0.641475 | 0.013 | 0.009 | 1 | 4 | Asap2 | 0.004 | Young |
| 0.711448 | 0.161206 | 0.025 | 0.019 | 1 | 4 | Serf1 | 0.006 | Young |
| 0.711448 | 0.046444 | 0.025 | 0.019 | 1 | 4 | Rdh13 | 0.006 | Young |
| 0.711659 | 0.629781 | 0.013 | 0.009 | 1 | 4 | Nxpe3 | 0.004 | Young |
| 0.711659 | 0.285007 | 0.013 | 0.009 | 1 | 4 | Nudt15 | 0.004 | Young |
| 0.711723 | 0.022858 | 0.975 | 0.958 | 1 | 4 | Laptm5 | 0.017 | Young |
| 0.711982 | 0.048448 | 0.025 | 0.033 | 1 | 4 | Cdk5rap2 | -0.008 | Young |
| 0.712224 | 0.207876 | 0.025 | 0.019 | 1 | 4 | Nudt22 | 0.006 | Young |
| 0.712224 | 0.152494 | 0.025 | 0.019 | 1 | 4 | Lbx2 | 0.006 | Young |
| 0.712612 | 0.152863 | 0.025 | 0.019 | 1 | 4 | Mars2 | 0.006 | Young |
| 0.712807 | 0.59124 | 0.013 | 0.009 | 1 | 4 | Gm35363 | 0.004 | Young |
| 0.712833 | 0.458763 | 0.125 | 0.116 | 1 | 4 | Mrpl54 | 0.009 | Young |
| 0.71287 | 0.160026 | 0.038 | 0.047 | 1 | 4 | Gtpbp1 | -0.009 | Young |
| 0.712982 | 0.512264 | 0.05 | 0.042 | 1 | 4 | Tmem126a | 0.008 | Young |
| 0.713 | 0.166053 | 0.025 | 0.019 | 1 | 4 | Spata5 | 0.006 | Young |
| 0.713375 | 0.167755 | 0.038 | 0.047 | 1 | 4 | Pus10 | -0.009 | Young |
| 0.713956 | 0.496909 | 0.013 | 0.009 | 1 | 4 | Rflnb | 0.004 | Young |
| 0.71474 | 0.157318 | 0.188 | 0.175 | 1 | 4 | St3gal6 | 0.013 | Young |
| 0.715121 | 0.223316 | 0.038 | 0.03 | 1 | 4 | Mthfsd | 0.008 | Young |
| 0.715413 | 0.097623 | 0.288 | 0.273 | 1 | 4 | Cct4 | 0.015 | Young |
| 0.715743 | 0.221462 | 0.038 | 0.03 | 1 | 4 | Fpgs | 0.008 | Young |
| 0.715809 | 0.054374 | 0.1 | 0.088 | 1 | 4 | Spns1 | 0.012 | Young |
| 0.716255 | 0.223507 | 0.013 | 0.009 | 1 | 4 | Klhl3 | 0.004 | Young |
| 0.716426 | 0.395372 | 0.212 | 0.2 | 1 | 4 | 4931406C0 | 0.012 | Young |
| 0.716431 | 0.728558 | 0.05 | 0.042 | 1 | 4 | Agk | 0.008 | Young |
| 0.717369 | 0.209604 | 0.162 | 0.15 | 1 | 4 | Ddx54 | 0.012 | Young |
| 0.717441 | 0.246171 | 0.15 | 0.139 | 1 | 4 | Herc2 | 0.011 | Young |
| 0.717844 | 0.113286 | 0.325 | 0.292 | 1 | 4 | Mia3 | 0.033 | Young |
| 0.71803 | 0.23901 | 0.038 | 0.046 | 1 | 4 | Arl14ep | -0.008 | Young |
| 0.71823 | 0.079992 | 0.038 | 0.03 | 1 | 4 | Tatdn1 | 0.008 | Young |
| 0.718329 | 0.166635 | 0.125 | 0.111 | 1 | 4 | Fut11 | 0.014 | Young |
| 0.71985 | 0.002962 | 0.05 | 0.042 | 1 | 4 | Hace1 | 0.008 | Young |
| 0.719933 | 0.790598 | 0.038 | 0.031 | 1 | 4 | Adck1 | 0.007 | Young |
| 0.719946 | 0.280635 | 0.112 | 0.101 | 1 | 4 | Cmip | 0.011 | Young |
| 0.720605 | 0.195828 | 0.525 | 0.505 | 1 | 4 | Chd4 | 0.02 | Young |
| 0.721407 | 0.167642 | 0.075 | 0.086 | 1 | 4 | Ech1 | -0.011 | Young |
| 0.721577 | 0.193736 | 0.025 | 0.032 | 1 | 4 | Dna2 | -0.007 | Young |
| 0.721684 | 0.091608 | 0.212 | 0.193 | 1 | 4 | Furin | 0.019 | Young |
| 0.721848 | 0.032685 | 0.462 | 0.449 | 1 | 4 | Rin2 | 0.013 | Young |
| 0.721967 | 0.09611 | 0.038 | 0.03 | 1 | 4 | Cul7 | 0.008 | Young |
| 0.722013 | 0.351414 | 0.05 | 0.042 | 1 | 4 | Mon1b | 0.008 | Young |
| 0.722797 | 0.118862 | 0.062 | 0.053 | 1 | 4 | Bag4 | 0.009 | Young |
| 0.723082 | 0.116178 | 0.825 | 0.791 | 1 | 4 | Serf2 | 0.034 | Young |
| 0.723791 | 0.105151 | 0.15 | 0.135 | 1 | 4 | Ttc39b | 0.015 | Young |

| | | | | | | | | |
|---|---|---|---|---|---|---|---|---|
| 0.724116 | 0.305412 | 0.075 | 0.066 | 1 | 4 | 6720427I0 | 0.009 | Young |
| 0.724304 | 0.064202 | 0.1 | 0.115 | 1 | 4 | Phf1 | -0.015 | Young |
| 0.724462 | 0.009704 | 0.038 | 0.03 | 1 | 4 | Dgkh | 0.008 | Young |
| 0.724759 | 0.073131 | 0.138 | 0.155 | 1 | 4 | Sp110 | -0.017 | Young |
| 0.724956 | 0.419524 | 0.062 | 0.054 | 1 | 4 | Snapc3 | 0.008 | Young |
| 0.726229 | 0.005936 | 0.45 | 0.408 | 1 | 4 | Eif5b | 0.042 | Young |
| 0.72633 | 0.254022 | 0.112 | 0.101 | 1 | 4 | Adam33 | 0.011 | Young |
| 0.72681 | 0.27471 | 0.05 | 0.042 | 1 | 4 | Glipr1 | 0.008 | Young |
| 0.727088 | 0.269605 | 0.062 | 0.053 | 1 | 4 | Rnf214 | 0.009 | Young |
| 0.727204 | 0.067752 | 0.175 | 0.16 | 1 | 4 | Foxj2 | 0.015 | Young |
| 0.728436 | 0.101816 | 0.212 | 0.237 | 1 | 4 | Smc3 | -0.025 | Young |
| 0.728585 | 0.48802 | 0.038 | 0.031 | 1 | 4 | Erich1 | 0.007 | Young |
| 0.729256 | 0.185075 | 0.275 | 0.265 | 1 | 4 | Pdxk | 0.01 | Young |
| 0.72948 | 0.297546 | 0.05 | 0.042 | 1 | 4 | Pls3 | 0.008 | Young |
| 0.729523 | 0.954246 | 0.025 | 0.02 | 1 | 4 | Slco3a1 | 0.005 | Young |
| 0.729859 | 0.097641 | 0.238 | 0.217 | 1 | 4 | Rab35 | 0.021 | Young |
| 0.730105 | 0.19642 | 0.162 | 0.149 | 1 | 4 | Tubgcp5 | 0.013 | Young |
| 0.73125 | 0.080396 | 0.013 | 0.018 | 1 | 4 | Aurkb | -0.005 | Young |
| 0.732303 | 0.49273 | 0.038 | 0.031 | 1 | 4 | Usp18 | 0.007 | Young |
| 0.732607 | 0.062304 | 0.1 | 0.089 | 1 | 4 | Guca1a | 0.011 | Young |
| 0.73279 | 0.001931 | 0.575 | 0.592 | 1 | 4 | Tmed10 | -0.017 | Young |
| 0.732823 | 0.040561 | 0.062 | 0.053 | 1 | 4 | Smarcd1 | 0.009 | Young |
| 0.73311 | 0.210951 | 0.388 | 0.371 | 1 | 4 | Bclaf1 | 0.017 | Young |
| 0.733222 | 0.228454 | 0.05 | 0.042 | 1 | 4 | Rdm1 | 0.008 | Young |
| 0.733933 | 0.061483 | 0.075 | 0.088 | 1 | 4 | Sgk1 | -0.013 | Young |
| 0.734544 | 0.425656 | 0.013 | 0.018 | 1 | 4 | Ogg1 | -0.005 | Young |
| 0.734639 | 0.275266 | 0.038 | 0.047 | 1 | 4 | Timm21 | -0.009 | Young |
| 0.735122 | 0.054131 | 0.075 | 0.065 | 1 | 4 | Taf4 | 0.01 | Young |
| 0.735386 | 0.17465 | 0.162 | 0.148 | 1 | 4 | Rnf216 | 0.014 | Young |
| 0.736725 | 0.062291 | 0.125 | 0.111 | 1 | 4 | Plekhm1 | 0.014 | Young |
| 0.737379 | 0.160125 | 0.1 | 0.09 | 1 | 4 | Zfp120 | 0.01 | Young |
| 0.737449 | 0.024026 | 0.2 | 0.184 | 1 | 4 | Marf1 | 0.016 | Young |
| 0.737751 | 0.022566 | 0.088 | 0.076 | 1 | 4 | Med14 | 0.012 | Young |
| 0.738537 | 0.083715 | 0.062 | 0.073 | 1 | 4 | Hikeshi | -0.011 | Young |
| 0.738964 | 0.040979 | 0.65 | 0.613 | 1 | 4 | Arpc3 | 0.037 | Young |
| 0.740415 | 0.417455 | 0.062 | 0.073 | 1 | 4 | Ubqln4 | -0.011 | Young |
| 0.740433 | 0.069972 | 0.112 | 0.101 | 1 | 4 | Rps6ka4 | 0.011 | Young |
| 0.741627 | 0.277618 | 0.038 | 0.031 | 1 | 4 | Dbp | 0.007 | Young |
| 0.743206 | 0.109165 | 0.175 | 0.159 | 1 | 4 | Ifi27 | 0.016 | Young |
| 0.743364 | 0.018381 | 0.2 | 0.213 | 1 | 4 | Dnajc7 | -0.013 | Young |
| 0.743497 | 0.353045 | 0.038 | 0.031 | 1 | 4 | Zfp143 | 0.007 | Young |
| 0.743671 | 0.552934 | 0.062 | 0.055 | 1 | 4 | Rfx7 | 0.007 | Young |
| 0.743951 | 0.213104 | 0.05 | 0.042 | 1 | 4 | Klhl22 | 0.008 | Young |
| 0.744432 | 0.258281 | 0.025 | 0.032 | 1 | 4 | Mrpl47 | -0.007 | Young |
| 0.744516 | 0.287797 | 0.075 | 0.066 | 1 | 4 | Fer | 0.009 | Young |

| | | | | | | | | |
|---|---|---|---|---|---|---|---|---|
| 0.744744 | 0.555082 | 0.038 | 0.031 | 1 | 4 | Tecpr2 | 0.007 | Young |
| 0.744754 | 0.047205 | 0.45 | 0.399 | 1 | 4 | Csrnp1 | 0.051 | Young |
| 0.744976 | 0.117027 | 0.025 | 0.02 | 1 | 4 | Etfbkmt | 0.005 | Young |
| 0.745278 | 0.014 | 0.162 | 0.182 | 1 | 4 | Pabpn1 | -0.02 | Young |
| 0.745565 | 0.014355 | 0.05 | 0.042 | 1 | 4 | Jpt2 | 0.008 | Young |
| 0.745752 | 0.145176 | 0.025 | 0.02 | 1 | 4 | Zfp995 | 0.005 | Young |
| 0.745752 | 0.015492 | 0.025 | 0.02 | 1 | 4 | Qtrt1 | 0.005 | Young |
| 0.745936 | 0.07272 | 0.162 | 0.176 | 1 | 4 | Lpcat1 | -0.014 | Young |
| 0.746139 | 0.041362 | 0.325 | 0.311 | 1 | 4 | Gmfb | 0.014 | Young |
| 0.746168 | 0.792532 | 0.038 | 0.032 | 1 | 4 | Rin3 | 0.006 | Young |
| 0.746482 | 0.016988 | 0.45 | 0.432 | 1 | 4 | Glg1 | 0.018 | Young |
| 0.746643 | 0.543226 | 0.062 | 0.055 | 1 | 4 | Rpp14 | 0.007 | Young |
| 0.746665 | 0.251787 | 0.1 | 0.091 | 1 | 4 | Gpaa1 | 0.009 | Young |
| 0.748079 | 0.110734 | 0.025 | 0.02 | 1 | 4 | 2310040G2 | 0.005 | Young |
| 0.748464 | 0.150413 | 0.4 | 0.376 | 1 | 4 | Ncf1 | 0.024 | Young |
| 0.748636 | 0.304216 | 0.262 | 0.256 | 1 | 4 | Vdac3 | 0.006 | Young |
| 0.748878 | 0.094307 | 0.15 | 0.165 | 1 | 4 | Ncoa6 | -0.015 | Young |
| 0.7489 | 0.064372 | 0.125 | 0.113 | 1 | 4 | Rnf220 | 0.012 | Young |
| 0.749114 | 0.08042 | 0.038 | 0.031 | 1 | 4 | Ofd1 | 0.007 | Young |
| 0.749633 | 0.063621 | 0.025 | 0.02 | 1 | 4 | Taf9b | 0.005 | Young |
| 0.75102 | 0.557496 | 0.05 | 0.043 | 1 | 4 | Pym1 | 0.007 | Young |
| 0.751905 | 0.091839 | 0.2 | 0.186 | 1 | 4 | Tm2d2 | 0.014 | Young |
| 0.752003 | 0.053713 | 0.713 | 0.716 | 1 | 4 | Ifitm2 | -0.003 | Young |
| 0.752719 | 0.114518 | 0.1 | 0.09 | 1 | 4 | Fhod1 | 0.01 | Young |
| 0.75337 | 0.45068 | 0.075 | 0.067 | 1 | 4 | Traf6 | 0.008 | Young |
| 0.753492 | 0.04663 | 0.038 | 0.031 | 1 | 4 | Diexf | 0.007 | Young |
| 0.7535 | 0.007026 | 0.088 | 0.077 | 1 | 4 | Oxa1l | 0.011 | Young |
| 0.754728 | 0.176475 | 0.238 | 0.229 | 1 | 4 | Snw1 | 0.009 | Young |
| 0.754979 | 0.199455 | 0.138 | 0.123 | 1 | 4 | Nipsnap2 | 0.015 | Young |
| 0.756078 | 0.849327 | 0.038 | 0.032 | 1 | 4 | Primpol | 0.006 | Young |
| 0.7564 | 0.577143 | 0.05 | 0.043 | 1 | 4 | Haus2 | 0.007 | Young |
| 0.757979 | 0.049051 | 0.088 | 0.099 | 1 | 4 | Atg12 | -0.011 | Young |
| 0.758009 | 0.235457 | 0.112 | 0.103 | 1 | 4 | Ppp1r35 | 0.009 | Young |
| 0.758039 | 0.373414 | 0.062 | 0.055 | 1 | 4 | Fam213a | 0.007 | Young |
| 0.759067 | 0.169466 | 0.05 | 0.043 | 1 | 4 | Ints13 | 0.007 | Young |
| 0.759067 | 0.030814 | 0.05 | 0.043 | 1 | 4 | Mthfd2l | 0.007 | Young |
| 0.759184 | 0.345382 | 0.038 | 0.032 | 1 | 4 | Rab38 | 0.006 | Young |
| 0.759856 | 0.02143 | 0.013 | 0.017 | 1 | 4 | Ndrg1 | -0.004 | Young |
| 0.760074 | 0.598313 | 0.025 | 0.032 | 1 | 4 | Cxcl10 | -0.007 | Young |
| 0.760704 | 0.00208 | 0.013 | 0.017 | 1 | 4 | Smim3 | -0.004 | Young |
| 0.760846 | 1.022451 | 0.013 | 0.009 | 1 | 4 | Zfp566 | 0.004 | Young |
| 0.760846 | 0.871612 | 0.013 | 0.009 | 1 | 4 | Tasp1 | 0.004 | Young |
| 0.761074 | 0.081622 | 0.338 | 0.323 | 1 | 4 | Usp25 | 0.015 | Young |
| 0.76114 | 0.070983 | 0.3 | 0.278 | 1 | 4 | Ifngr2 | 0.022 | Young |
| 0.761236 | 0.831839 | 0.088 | 0.082 | 1 | 4 | Ampd3 | 0.006 | Young |

| | | | | | | | | |
|---|---|---|---|---|---|---|---|---|
| 0.761279 | 0.169135 | 0.075 | 0.067 | 1 | 4 | Pgm1 | 0.008 | Young |
| 0.761636 | 0.145272 | 0.325 | 0.35 | 1 | 4 | Tmbim1 | -0.025 | Young |
| 0.76198 | 0.996069 | 0.013 | 0.009 | 1 | 4 | Tox2 | 0.004 | Young |
| 0.76198 | 0.890694 | 0.013 | 0.009 | 1 | 4 | Gm47232 | 0.004 | Young |
| 0.76205 | 0.029026 | 0.262 | 0.24 | 1 | 4 | Dpy19l1 | 0.022 | Young |
| 0.762237 | 0.05484 | 0.1 | 0.09 | 1 | 4 | Atp6v0a2 | 0.01 | Young |
| 0.762395 | 0.015968 | 0.262 | 0.25 | 1 | 4 | Ddx60 | 0.012 | Young |
| 0.763114 | 1.088232 | 0.013 | 0.009 | 1 | 4 | AA467197 | 0.004 | Young |
| 0.764099 | 0.500512 | 0.013 | 0.017 | 1 | 4 | Mtrf1 | -0.004 | Young |
| 0.764099 | 0.464963 | 0.013 | 0.017 | 1 | 4 | 2810454H( | -0.004 | Young |
| 0.764099 | 0.2633 | 0.013 | 0.017 | 1 | 4 | Nepro | -0.004 | Young |
| 0.764249 | 0.720004 | 0.013 | 0.009 | 1 | 4 | Ugt1a6a | 0.004 | Young |
| 0.764249 | 0.656727 | 0.013 | 0.009 | 1 | 4 | Arhgap22 | 0.004 | Young |
| 0.764943 | 0.323014 | 0.05 | 0.043 | 1 | 4 | Asb3 | 0.007 | Young |
| 0.764949 | 0.648277 | 0.013 | 0.017 | 1 | 4 | Rhobtb2 | -0.004 | Young |
| 0.764985 | 0.011461 | 0.05 | 0.043 | 1 | 4 | Drg2 | 0.007 | Young |
| 0.765385 | 0.4949 | 0.013 | 0.009 | 1 | 4 | Fam102a | 0.004 | Young |
| 0.765478 | 0.331348 | 0.05 | 0.043 | 1 | 4 | Cnot9 | 0.007 | Young |
| 0.76565 | 0.132801 | 0.062 | 0.055 | 1 | 4 | Hyal2 | 0.007 | Young |
| 0.765713 | 0.354324 | 0.138 | 0.154 | 1 | 4 | Polr2g | -0.016 | Young |
| 0.766521 | 0.529805 | 0.013 | 0.009 | 1 | 4 | Pecam1 | 0.004 | Young |
| 0.766521 | 0.403982 | 0.013 | 0.009 | 1 | 4 | Samd10 | 0.004 | Young |
| 0.766521 | 0.06653 | 0.013 | 0.009 | 1 | 4 | Zfyve9 | 0.004 | Young |
| 0.767088 | 0.708637 | 0.025 | 0.02 | 1 | 4 | Dcaf4 | 0.005 | Young |
| 0.76755 | 0.157466 | 0.112 | 0.101 | 1 | 4 | Naip2 | 0.011 | Young |
| 0.767658 | 0.510972 | 0.013 | 0.009 | 1 | 4 | Zkscan7 | 0.004 | Young |
| 0.767724 | 0.118465 | 0.125 | 0.115 | 1 | 4 | Zufsp | 0.01 | Young |
| 0.767985 | 0.04437 | 0.488 | 0.472 | 1 | 4 | Ctdsp2 | 0.016 | Young |
| 0.768795 | 0.197246 | 0.013 | 0.009 | 1 | 4 | Ube2t | 0.004 | Young |
| 0.768795 | 0.039946 | 0.013 | 0.009 | 1 | 4 | Gm13830 | 0.004 | Young |
| 0.768795 | 0.026254 | 0.013 | 0.009 | 1 | 4 | AC163018. | 0.004 | Young |
| 0.76882 | 0.45062 | 0.075 | 0.067 | 1 | 4 | Fam222b | 0.008 | Young |
| 0.770944 | 0.823753 | 0.025 | 0.02 | 1 | 4 | Zfp52 | 0.005 | Young |
| 0.771459 | 0.303817 | 0.088 | 0.08 | 1 | 4 | Abcb1b | 0.008 | Young |
| 0.771525 | 0.30775 | 0.125 | 0.116 | 1 | 4 | Wasl | 0.009 | Young |
| 0.772033 | 1.050398 | 0.1 | 0.091 | 1 | 4 | Iigp1 | 0.009 | Young |
| 0.772385 | 0.019638 | 0.075 | 0.086 | 1 | 4 | Irf3 | -0.011 | Young |
| 0.772979 | 0.474976 | 0.05 | 0.043 | 1 | 4 | Usb1 | 0.007 | Young |
| 0.773261 | 0.505385 | 0.025 | 0.02 | 1 | 4 | Polm | 0.005 | Young |
| 0.773403 | 0.088624 | 0.325 | 0.31 | 1 | 4 | Klhl24 | 0.015 | Young |
| 0.773829 | 0.337604 | 0.062 | 0.073 | 1 | 4 | Sertad3 | -0.011 | Young |
| 0.773883 | 0.343589 | 0.388 | 0.416 | 1 | 4 | Ptpn1 | -0.028 | Young |
| 0.773985 | 0.069189 | 0.2 | 0.188 | 1 | 4 | Lrrk1 | 0.012 | Young |
| 0.775192 | 0.058836 | 0.275 | 0.25 | 1 | 4 | Akt1 | 0.025 | Young |
| 0.775246 | 0.350955 | 0.225 | 0.211 | 1 | 4 | Hist1h1c | 0.014 | Young |

| | | | | | | | | |
|---|---|---|---|---|---|---|---|---|
| 0.776109 | 0.084113 | 0.188 | 0.172 | 1 | 4 | Chd8 | 0.016 | Young |
| 0.776231 | 0.205221 | 0.312 | 0.327 | 1 | 4 | Ppp1cb | -0.015 | Young |
| 0.776425 | 0.049957 | 0.425 | 0.403 | 1 | 4 | Usf2 | 0.022 | Young |
| 0.777121 | 0.243872 | 0.088 | 0.08 | 1 | 4 | Ecd | 0.008 | Young |
| 0.777126 | 0.294248 | 0.025 | 0.02 | 1 | 4 | Fam212a | 0.005 | Young |
| 0.777126 | 0.18808 | 0.025 | 0.02 | 1 | 4 | Eif5a2 | 0.005 | Young |
| 0.777359 | 0.484579 | 0.112 | 0.125 | 1 | 4 | Pigs | -0.013 | Young |
| 0.777496 | 0.280509 | 0.025 | 0.031 | 1 | 4 | Mtg1 | -0.006 | Young |
| 0.778737 | 0.152875 | 0.175 | 0.163 | 1 | 4 | Sun2 | 0.012 | Young |
| 0.778851 | 0.072052 | 0.3 | 0.279 | 1 | 4 | Tspan31 | 0.021 | Young |
| 0.778979 | 0.128141 | 0.7 | 0.645 | 1 | 4 | Ccnl1 | 0.055 | Young |
| 0.778989 | 0.501885 | 0.038 | 0.032 | 1 | 4 | Cdc42ep4 | 0.006 | Young |
| 0.779426 | 0.358848 | 0.05 | 0.043 | 1 | 4 | Hes6 | 0.007 | Young |
| 0.779498 | 0.080458 | 0.15 | 0.167 | 1 | 4 | Ctso | -0.017 | Young |
| 0.779537 | 0.153462 | 0.075 | 0.067 | 1 | 4 | Iffo1 | 0.008 | Young |
| 0.779581 | 0.230872 | 0.175 | 0.162 | 1 | 4 | Ncoa2 | 0.013 | Young |
| 0.779711 | 0.325362 | 0.088 | 0.08 | 1 | 4 | Rora | 0.008 | Young |
| 0.780154 | 0.026332 | 0.138 | 0.127 | 1 | 4 | Susd3 | 0.011 | Young |
| 0.780674 | 0.020047 | 0.112 | 0.103 | 1 | 4 | Gse1 | 0.009 | Young |
| 0.780996 | 0.08731 | 0.025 | 0.02 | 1 | 4 | Gm20604 | 0.005 | Young |
| 0.781205 | 0.194786 | 0.262 | 0.252 | 1 | 4 | Tmx1 | 0.01 | Young |
| 0.781327 | 0.496794 | 0.025 | 0.031 | 1 | 4 | Pank4 | -0.006 | Young |
| 0.781351 | 0.240372 | 0.4 | 0.375 | 1 | 4 | Phip | 0.025 | Young |
| 0.781625 | 0.026941 | 0.212 | 0.197 | 1 | 4 | Ppp1r2 | 0.015 | Young |
| 0.781872 | 0.021694 | 0.062 | 0.072 | 1 | 4 | Cela1 | -0.01 | Young |
| 0.781984 | 0.26159 | 0.075 | 0.068 | 1 | 4 | Krt80 | 0.007 | Young |
| 0.782713 | 0.715349 | 0.038 | 0.032 | 1 | 4 | Zfp629 | 0.006 | Young |
| 0.782843 | 0.068121 | 0.112 | 0.103 | 1 | 4 | Tmem147 | 0.009 | Young |
| 0.783255 | 0.14488 | 0.125 | 0.138 | 1 | 4 | Mysm1 | -0.013 | Young |
| 0.783804 | 0.079592 | 0.15 | 0.167 | 1 | 4 | Gatad1 | -0.017 | Young |
| 0.783951 | 0.483255 | 0.062 | 0.055 | 1 | 4 | Tirap | 0.007 | Young |
| 0.784217 | 0.037854 | 0.162 | 0.179 | 1 | 4 | Ehmt2 | -0.017 | Young |
| 0.784418 | 0.349189 | 0.088 | 0.08 | 1 | 4 | Pigyl | 0.008 | Young |
| 0.784629 | 0.246414 | 0.3 | 0.289 | 1 | 4 | Gm16286 | 0.011 | Young |
| 0.784855 | 0.103289 | 0.338 | 0.318 | 1 | 4 | Wapl | 0.02 | Young |
| 0.785132 | 0.301735 | 0.15 | 0.141 | 1 | 4 | Evl | 0.009 | Young |
| 0.78577 | 0.120614 | 0.275 | 0.251 | 1 | 4 | Dennd6a | 0.024 | Young |
| 0.786805 | 0.009963 | 0.062 | 0.071 | 1 | 4 | Mex3d | -0.009 | Young |
| 0.78683 | 0.139366 | 0.162 | 0.153 | 1 | 4 | Hdac1 | 0.009 | Young |
| 0.78705 | 0.288042 | 0.1 | 0.092 | 1 | 4 | Hes1 | 0.008 | Young |
| 0.787587 | 0.637851 | 0.05 | 0.059 | 1 | 4 | Kdm4c | -0.009 | Young |
| 0.787985 | 0.161353 | 0.275 | 0.292 | 1 | 4 | Mertk | -0.017 | Young |
| 0.788199 | 0.090162 | 0.062 | 0.055 | 1 | 4 | Cables1 | 0.007 | Young |
| 0.788252 | 0.004464 | 0.2 | 0.188 | 1 | 4 | Il13ra1 | 0.012 | Young |
| 0.788686 | 0.563169 | 0.1 | 0.094 | 1 | 4 | Man1b1 | 0.006 | Young |

| | | | | | | | | |
|---|---|---|---|---|---|---|---|---|
| 0.789106 | 0.164791 | 0.125 | 0.117 | 1 | 4 | Lrba | 0.008 | Young |
| 0.789247 | 0.009336 | 0.125 | 0.114 | 1 | 4 | Elovl1 | 0.011 | Young |
| 0.789416 | 0.116987 | 0.075 | 0.067 | 1 | 4 | Arhgef10 | 0.008 | Young |
| 0.789828 | 0.018605 | 0.388 | 0.345 | 1 | 4 | Wipf1 | 0.043 | Young |
| 0.790142 | 0.092452 | 0.45 | 0.418 | 1 | 4 | Ccr5 | 0.032 | Young |
| 0.790511 | 0.057107 | 0.175 | 0.196 | 1 | 4 | Trim12c | -0.021 | Young |
| 0.790734 | 0.192928 | 0.075 | 0.068 | 1 | 4 | Rbbp5 | 0.007 | Young |
| 0.791536 | 0.129513 | 0.013 | 0.016 | 1 | 4 | Eid2b | -0.003 | Young |
| 0.79157 | 0.040353 | 0.062 | 0.055 | 1 | 4 | Nxt2 | 0.007 | Young |
| 0.791578 | 0.24464 | 0.125 | 0.112 | 1 | 4 | Clcn7 | 0.013 | Young |
| 0.791585 | 0.641839 | 0.075 | 0.069 | 1 | 4 | Jmy | 0.006 | Young |
| 0.792056 | 0.188282 | 0.075 | 0.067 | 1 | 4 | Sirt1 | 0.008 | Young |
| 0.792866 | 0.086058 | 0.15 | 0.165 | 1 | 4 | Ufm1 | -0.015 | Young |
| 0.793148 | 0.566319 | 0.1 | 0.096 | 1 | 4 | Hspa4l | 0.004 | Young |
| 0.793284 | 0.309134 | 0.013 | 0.016 | 1 | 4 | Dcp1b | -0.003 | Young |
| 0.793284 | 0.162023 | 0.013 | 0.016 | 1 | 4 | D130040H: | -0.003 | Young |
| 0.793939 | 0.319881 | 0.138 | 0.128 | 1 | 4 | Lysmd3 | 0.01 | Young |
| 0.794424 | 0.101449 | 0.175 | 0.166 | 1 | 4 | Taf10 | 0.009 | Young |
| 0.794796 | 0.342182 | 0.038 | 0.044 | 1 | 4 | Cep85l | -0.006 | Young |
| 0.794964 | 0.089987 | 0.338 | 0.321 | 1 | 4 | Shtn1 | 0.017 | Young |
| 0.795067 | 0.264376 | 0.05 | 0.043 | 1 | 4 | Sprtn | 0.007 | Young |
| 0.795331 | 0.139898 | 0.212 | 0.204 | 1 | 4 | Atxn7 | 0.008 | Young |
| 0.795493 | 0.012212 | 0.212 | 0.198 | 1 | 4 | Thoc2 | 0.014 | Young |
| 0.795789 | 0.270341 | 0.038 | 0.032 | 1 | 4 | Znhit2 | 0.006 | Young |
| 0.797104 | 0.076867 | 0.05 | 0.058 | 1 | 4 | Slc9a8 | -0.008 | Young |
| 0.7972 | 0.170542 | 0.062 | 0.056 | 1 | 4 | Mrpl58 | 0.006 | Young |
| 0.797219 | 0.013479 | 0.225 | 0.211 | 1 | 4 | Ctbp1 | 0.014 | Young |
| 0.797582 | 0.666022 | 0.075 | 0.07 | 1 | 4 | Isg20l2 | 0.005 | Young |
| 0.797672 | 0.216841 | 0.2 | 0.215 | 1 | 4 | Atp13a2 | -0.015 | Young |
| 0.797753 | 0.092894 | 0.075 | 0.068 | 1 | 4 | Smarcal1 | 0.007 | Young |
| 0.798602 | 0.036978 | 0.025 | 0.03 | 1 | 4 | Nek4 | -0.005 | Young |
| 0.798641 | 0.339402 | 0.062 | 0.056 | 1 | 4 | Spg11 | 0.006 | Young |
| 0.799252 | 0.055505 | 0.025 | 0.03 | 1 | 4 | Asb1 | -0.005 | Young |
| 0.79942 | 0.678874 | 0.025 | 0.021 | 1 | 4 | Timeless | 0.004 | Young |
| 0.800658 | 0.09854 | 0.338 | 0.323 | 1 | 4 | Ccr1 | 0.015 | Young |
| 0.800671 | 0.193556 | 0.05 | 0.058 | 1 | 4 | Tmem80 | -0.008 | Young |
| 0.800957 | 0.724626 | 0.025 | 0.021 | 1 | 4 | Arfip2 | 0.004 | Young |
| 0.801049 | 0.069112 | 0.075 | 0.068 | 1 | 4 | Snx16 | 0.007 | Young |
| 0.801201 | 0.137266 | 0.025 | 0.03 | 1 | 4 | Zbtb11os1 | -0.005 | Young |
| 0.801475 | 0.068881 | 0.362 | 0.362 | 1 | 4 | Rab2a | 0 | Young |
| 0.801798 | 0.178516 | 0.05 | 0.044 | 1 | 4 | Mtap | 0.006 | Young |
| 0.802883 | 0.629439 | 0.038 | 0.033 | 1 | 4 | Prdm10 | 0.005 | Young |
| 0.803158 | 0.228634 | 0.175 | 0.167 | 1 | 4 | Eny2 | 0.008 | Young |
| 0.803597 | 0.28509 | 0.175 | 0.165 | 1 | 4 | Tlr4 | 0.01 | Young |
| 0.803987 | 0.039754 | 0.212 | 0.201 | 1 | 4 | Esd | 0.011 | Young |

| | | | | | | | | |
|---|---|---|---|---|---|---|---|---|
| 0.80416 | 0.41803 | 0.05 | 0.045 | 1 | 4 | Ccdc163 | 0.005 | Young |
| 0.804491 | 0.132395 | 0.05 | 0.044 | 1 | 4 | Acad9 | 0.006 | Young |
| 0.804538 | 0.496055 | 0.038 | 0.032 | 1 | 4 | Ptcd3 | 0.006 | Young |
| 0.804578 | 0.019009 | 0.537 | 0.491 | 1 | 4 | Tgoln1 | 0.046 | Young |
| 0.805102 | 0.556101 | 0.025 | 0.03 | 1 | 4 | Net1 | -0.005 | Young |
| 0.805569 | 0.07555 | 0.05 | 0.044 | 1 | 4 | Mllt11 | 0.006 | Young |
| 0.805572 | 0.833751 | 0.025 | 0.021 | 1 | 4 | Trdmt1 | 0.004 | Young |
| 0.805844 | 0.258461 | 0.138 | 0.131 | 1 | 4 | Chd1 | 0.007 | Young |
| 0.806548 | 0.091517 | 0.225 | 0.217 | 1 | 4 | Fbxl3 | 0.008 | Young |
| 0.807194 | 0.408416 | 0.175 | 0.187 | 1 | 4 | Aprt | -0.012 | Young |
| 0.807818 | 0.36723 | 0.062 | 0.057 | 1 | 4 | Arl10 | 0.005 | Young |
| 0.807883 | 0.376846 | 0.025 | 0.021 | 1 | 4 | Ccdc77 | 0.004 | Young |
| 0.80795 | 0.088969 | 0.138 | 0.149 | 1 | 4 | Paip1 | -0.011 | Young |
| 0.807966 | 0.050263 | 0.25 | 0.233 | 1 | 4 | Tifa | 0.017 | Young |
| 0.808268 | 0.204043 | 0.025 | 0.021 | 1 | 4 | Cyp2ab1 | 0.004 | Young |
| 0.808653 | 0.378968 | 0.025 | 0.021 | 1 | 4 | Zdhhc1 | 0.004 | Young |
| 0.808887 | 0.211996 | 0.162 | 0.153 | 1 | 4 | Psmb2 | 0.009 | Young |
| 0.809244 | 0.295466 | 0.062 | 0.057 | 1 | 4 | Git1 | 0.005 | Young |
| 0.809963 | 0.133481 | 0.05 | 0.057 | 1 | 4 | Khk | -0.007 | Young |
| 0.810273 | 0.292739 | 0.112 | 0.124 | 1 | 4 | Eps15l1 | -0.012 | Young |
| 0.810469 | 0.160284 | 0.2 | 0.193 | 1 | 4 | Trim47 | 0.007 | Young |
| 0.810511 | 0.025426 | 0.088 | 0.08 | 1 | 4 | Ttc13 | 0.008 | Young |
| 0.81081 | 0.00374 | 0.262 | 0.248 | 1 | 4 | Gxylt1 | 0.014 | Young |
| 0.810874 | 1.522991 | 0.013 | 0.01 | 1 | 4 | Fancm | 0.003 | Young |
| 0.810874 | 1.045926 | 0.013 | 0.01 | 1 | 4 | Cdkl3 | 0.003 | Young |
| 0.810874 | 0.994337 | 0.013 | 0.01 | 1 | 4 | Eya4 | 0.003 | Young |
| 0.810874 | 0.839675 | 0.013 | 0.01 | 1 | 4 | Usp27x | 0.003 | Young |
| 0.811279 | 0.070359 | 0.088 | 0.079 | 1 | 4 | Tmub2 | 0.009 | Young |
| 0.811462 | 0.299453 | 0.088 | 0.082 | 1 | 4 | Otud7b | 0.006 | Young |
| 0.812018 | 0.107048 | 0.088 | 0.095 | 1 | 4 | Trim24 | -0.007 | Young |
| 0.812044 | 0.026232 | 0.05 | 0.044 | 1 | 4 | Ift46 | 0.006 | Young |
| 0.812123 | 0.15427 | 0.025 | 0.021 | 1 | 4 | Wee1 | 0.004 | Young |
| 0.812751 | 0.018239 | 0.725 | 0.684 | 1 | 4 | Prdx1 | 0.041 | Young |
| 0.81292 | 0.553636 | 0.025 | 0.03 | 1 | 4 | Fbxo31 | -0.005 | Young |
| 0.813012 | 0.034945 | 0.025 | 0.03 | 1 | 4 | Trim7 | -0.005 | Young |
| 0.813111 | 0.697669 | 0.013 | 0.01 | 1 | 4 | Gpt | 0.003 | Young |
| 0.813189 | 1.236288 | 0.088 | 0.084 | 1 | 4 | Gbp2 | 0.004 | Young |
| 0.813478 | 0.048844 | 0.238 | 0.224 | 1 | 4 | Cnot6 | 0.014 | Young |
| 0.813539 | 0.259735 | 0.05 | 0.045 | 1 | 4 | Isca1 | 0.005 | Young |
| 0.814229 | 0.69506 | 0.013 | 0.01 | 1 | 4 | Pclaf | 0.003 | Young |
| 0.814229 | 0.570479 | 0.013 | 0.01 | 1 | 4 | D11Wsu47 | 0.003 | Young |
| 0.81438 | 0.201706 | 0.075 | 0.069 | 1 | 4 | Heatr3 | 0.006 | Young |
| 0.814388 | 0.062244 | 0.275 | 0.256 | 1 | 4 | Birc6 | 0.019 | Young |
| 0.814418 | 0.14253 | 0.112 | 0.105 | 1 | 4 | Grsf1 | 0.007 | Young |
| 0.815348 | 0.434273 | 0.013 | 0.01 | 1 | 4 | Ggact | 0.003 | Young |

| | | | | | | | | |
|---|---|---|---|---|---|---|---|---|
| 0.816278 | 0.186499 | 0.088 | 0.08 | 1 | 4 | Nasp | 0.008 | Young |
| 0.816468 | 0.68595 | 0.013 | 0.01 | 1 | 4 | 9330020H( | 0.003 | Young |
| 0.816468 | 0.587217 | 0.013 | 0.01 | 1 | 4 | Cmtr1 | 0.003 | Young |
| 0.816468 | 0.579119 | 0.013 | 0.01 | 1 | 4 | Acat2 | 0.003 | Young |
| 0.816468 | 0.364934 | 0.013 | 0.01 | 1 | 4 | C330027C( | 0.003 | Young |
| 0.81664 | 0.149259 | 0.1 | 0.093 | 1 | 4 | Parg | 0.007 | Young |
| 0.817307 | 0.132314 | 0.275 | 0.265 | 1 | 4 | Emilin2 | 0.01 | Young |
| 0.817588 | 0.456682 | 0.013 | 0.01 | 1 | 4 | BC052040 | 0.003 | Young |
| 0.817588 | 0.37894 | 0.013 | 0.01 | 1 | 4 | Nthl1 | 0.003 | Young |
| 0.817588 | 0.242474 | 0.013 | 0.01 | 1 | 4 | Tmem154 | 0.003 | Young |
| 0.817592 | 0.566174 | 0.038 | 0.044 | 1 | 4 | Mir17hg | -0.006 | Young |
| 0.817649 | 0.158823 | 0.05 | 0.057 | 1 | 4 | Rnf138 | -0.007 | Young |
| 0.818105 | 0.470898 | 0.05 | 0.045 | 1 | 4 | Bysl | 0.005 | Young |
| 0.818162 | 0.204003 | 0.138 | 0.128 | 1 | 4 | Cntrl | 0.01 | Young |
| 0.818648 | 0.062371 | 0.05 | 0.057 | 1 | 4 | Klf16 | -0.007 | Young |
| 0.818657 | 0.173386 | 0.2 | 0.196 | 1 | 4 | Cebpz | 0.004 | Young |
| 0.818708 | 0.348141 | 0.013 | 0.01 | 1 | 4 | Arl6 | 0.003 | Young |
| 0.818802 | 0.010056 | 0.225 | 0.235 | 1 | 4 | Acbd5 | -0.01 | Young |
| 0.819282 | 0.252007 | 0.112 | 0.125 | 1 | 4 | Pptc7 | -0.013 | Young |
| 0.819538 | 0.026397 | 0.1 | 0.11 | 1 | 4 | Golga3 | -0.01 | Young |
| 0.82026 | 0.003527 | 0.325 | 0.345 | 1 | 4 | Ldha | -0.02 | Young |
| 0.820316 | 0.047464 | 0.35 | 0.341 | 1 | 4 | Washc4 | 0.009 | Young |
| 0.820949 | 0.777018 | 0.088 | 0.097 | 1 | 4 | Prps2 | -0.009 | Young |
| 0.820949 | 0.086537 | 0.013 | 0.01 | 1 | 4 | Ptprm | 0.003 | Young |
| 0.820975 | 0.100624 | 0.15 | 0.14 | 1 | 4 | Ccnd3 | 0.01 | Young |
| 0.821125 | 0.020349 | 0.013 | 0.016 | 1 | 4 | Ccdc84 | -0.003 | Young |
| 0.822059 | 0.007288 | 0.938 | 0.925 | 1 | 4 | Eif1 | 0.013 | Young |
| 0.823242 | 0.103134 | 0.275 | 0.261 | 1 | 4 | Pkn1 | 0.014 | Young |
| 0.823484 | 0.185264 | 0.05 | 0.045 | 1 | 4 | Armc7 | 0.005 | Young |
| 0.824077 | 0.229338 | 0.062 | 0.057 | 1 | 4 | Ppwd1 | 0.005 | Young |
| 0.824373 | 0.055079 | 0.575 | 0.589 | 1 | 4 | Abca9 | -0.014 | Young |
| 0.824789 | 0.200783 | 0.1 | 0.092 | 1 | 4 | Vps26b | 0.008 | Young |
| 0.824807 | 0.252456 | 0.075 | 0.085 | 1 | 4 | Kmt5c | -0.01 | Young |
| 0.825036 | 0.260226 | 0.062 | 0.057 | 1 | 4 | Mmgt1 | 0.005 | Young |
| 0.825055 | 0.007228 | 0.075 | 0.083 | 1 | 4 | Vamp5 | -0.008 | Young |
| 0.825626 | 0.580741 | 0.013 | 0.016 | 1 | 4 | Nup43 | -0.003 | Young |
| 0.825626 | 0.510854 | 0.013 | 0.016 | 1 | 4 | Ttc30b | -0.003 | Young |
| 0.825626 | 0.417349 | 0.013 | 0.016 | 1 | 4 | Gm37305 | -0.003 | Young |
| 0.826188 | 0.186274 | 0.238 | 0.254 | 1 | 4 | Rab5a | -0.016 | Young |
| 0.826527 | 0.982705 | 0.013 | 0.016 | 1 | 4 | Dtwd1 | -0.003 | Young |
| 0.826527 | 0.816439 | 0.013 | 0.016 | 1 | 4 | Prpf40b | -0.003 | Young |
| 0.826527 | 0.675176 | 0.013 | 0.016 | 1 | 4 | Ybey | -0.003 | Young |
| 0.826681 | 0.141776 | 0.138 | 0.13 | 1 | 4 | Ahcyl1 | 0.008 | Young |
| 0.826686 | 0.325627 | 0.112 | 0.121 | 1 | 4 | Pcbd2 | -0.009 | Young |
| 0.82688 | 0.387865 | 0.112 | 0.108 | 1 | 4 | Tor1a | 0.004 | Young |

| | | | | | | | | |
|---|---|---|---|---|---|---|---|---|
| 0.826989 | 0.191845 | 0.075 | 0.069 | 1 | 4 | Larp1b | 0.006 | Young |
| 0.827314 | 0.045166 | 0.125 | 0.136 | 1 | 4 | Cse1l | -0.011 | Young |
| 0.827354 | 0.288809 | 0.138 | 0.146 | 1 | 4 | Odc1 | -0.008 | Young |
| 0.828066 | 0.814068 | 0.025 | 0.022 | 1 | 4 | Telo2 | 0.003 | Young |
| 0.82937 | 0.166062 | 0.188 | 0.178 | 1 | 4 | Kdm5a | 0.01 | Young |
| 0.830024 | 0.057305 | 0.275 | 0.262 | 1 | 4 | Fxyd5 | 0.013 | Young |
| 0.830205 | 0.160721 | 0.125 | 0.119 | 1 | 4 | Smarcad1 | 0.006 | Young |
| 0.830267 | 0.174743 | 0.038 | 0.033 | 1 | 4 | Katnal1 | 0.005 | Young |
| 0.830407 | 0.085636 | 0.088 | 0.08 | 1 | 4 | Casp3 | 0.008 | Young |
| 0.830579 | 0.026324 | 0.038 | 0.043 | 1 | 4 | 1300002E1 | -0.005 | Young |
| 0.830808 | 0.14957 | 0.112 | 0.125 | 1 | 4 | Csad | -0.013 | Young |
| 0.830852 | 0.341845 | 0.025 | 0.03 | 1 | 4 | Large1 | -0.005 | Young |
| 0.831442 | 0.306403 | 0.162 | 0.161 | 1 | 4 | 2410015M | 0.001 | Young |
| 0.83167 | 0.187677 | 0.138 | 0.15 | 1 | 4 | Gtf3c2 | -0.012 | Young |
| 0.831901 | 0.064079 | 0.55 | 0.544 | 1 | 4 | Rpl15 | 0.006 | Young |
| 0.831922 | 0.115223 | 0.112 | 0.105 | 1 | 4 | Fbrsl1 | 0.007 | Young |
| 0.832015 | 0.133371 | 0.138 | 0.132 | 1 | 4 | Cirbp | 0.006 | Young |
| 0.832493 | 0.077209 | 0.188 | 0.175 | 1 | 4 | Fes | 0.013 | Young |
| 0.833211 | 0.667066 | 0.075 | 0.071 | 1 | 4 | Terf2ip | 0.004 | Young |
| 0.833245 | 0.033058 | 0.162 | 0.173 | 1 | 4 | Nudt16l1 | -0.011 | Young |
| 0.833849 | 0.029866 | 0.35 | 0.337 | 1 | 4 | Gatad2b | 0.013 | Young |
| 0.8341 | 0.116759 | 0.125 | 0.118 | 1 | 4 | Naa20 | 0.007 | Young |
| 0.835004 | 0.033803 | 0.338 | 0.307 | 1 | 4 | Hnrnpul2 | 0.031 | Young |
| 0.835803 | 0.515984 | 0.062 | 0.059 | 1 | 4 | Vti1a | 0.003 | Young |
| 0.837246 | 0.486504 | 0.025 | 0.022 | 1 | 4 | Ighmbp2 | 0.003 | Young |
| 0.837485 | 0.360408 | 0.05 | 0.045 | 1 | 4 | Smox | 0.005 | Young |
| 0.83778 | 0.030089 | 0.15 | 0.141 | 1 | 4 | Zfp800 | 0.009 | Young |
| 0.838284 | 0.124058 | 0.312 | 0.304 | 1 | 4 | B230219D2 | 0.008 | Young |
| 0.838486 | 0.196023 | 0.062 | 0.057 | 1 | 4 | Mphosph1( | 0.005 | Young |
| 0.838651 | 0.057304 | 0.15 | 0.144 | 1 | 4 | Smim13 | 0.006 | Young |
| 0.839008 | 0.191012 | 0.1 | 0.094 | 1 | 4 | Wdr91 | 0.006 | Young |
| 0.839092 | 0.023355 | 0.775 | 0.743 | 1 | 4 | Slc25a3 | 0.032 | Young |
| 0.839334 | 0.472062 | 0.125 | 0.119 | 1 | 4 | Adrm1 | 0.006 | Young |
| 0.839446 | 0.047026 | 0.088 | 0.08 | 1 | 4 | Nbdy | 0.008 | Young |
| 0.839545 | 0.225522 | 0.025 | 0.022 | 1 | 4 | Kbtbd4 | 0.003 | Young |
| 0.839868 | 0.103901 | 0.225 | 0.213 | 1 | 4 | Pik3ca | 0.012 | Young |
| 0.840311 | 1.037482 | 0.025 | 0.022 | 1 | 4 | Zfp472 | 0.003 | Young |
| 0.840311 | 0.776593 | 0.025 | 0.022 | 1 | 4 | Tmem186 | 0.003 | Young |
| 0.841115 | 0.034609 | 0.262 | 0.257 | 1 | 4 | Hnrnpd | 0.005 | Young |
| 0.841519 | 0.200932 | 0.1 | 0.092 | 1 | 4 | Btaf1 | 0.008 | Young |
| 0.842006 | 0.171486 | 0.038 | 0.034 | 1 | 4 | Eef1akmt2 | 0.004 | Young |
| 0.842531 | 0.299567 | 0.25 | 0.252 | 1 | 4 | Cbx3 | -0.002 | Young |
| 0.843225 | 0.114723 | 0.05 | 0.055 | 1 | 4 | Tmem161b | -0.005 | Young |
| 0.843606 | 0.193924 | 0.138 | 0.15 | 1 | 4 | Usp14 | -0.012 | Young |
| 0.843848 | 0.043447 | 0.088 | 0.096 | 1 | 4 | Ddx52 | -0.008 | Young |

| | | | | | | | | |
|---|---|---|---|---|---|---|---|---|
| 0.843852 | 0.043609 | 0.025 | 0.029 | 1 | 4 | Capn15 | -0.004 | Young |
| 0.843883 | 0.032 | 0.288 | 0.273 | 1 | 4 | Arhgap15 | 0.015 | Young |
| 0.843998 | 0.113094 | 0.05 | 0.045 | 1 | 4 | Rnf157 | 0.005 | Young |
| 0.844353 | 0.003224 | 0.438 | 0.419 | 1 | 4 | Nfam1 | 0.019 | Young |
| 0.844857 | 0.657489 | 0.05 | 0.057 | 1 | 4 | Calcoco1 | -0.007 | Young |
| 0.844911 | 0.290374 | 0.138 | 0.131 | 1 | 4 | Anapc16 | 0.007 | Young |
| 0.844914 | 0.279092 | 0.025 | 0.022 | 1 | 4 | Hspg2 | 0.003 | Young |
| 0.844914 | 0.129118 | 0.025 | 0.022 | 1 | 4 | Hectd2 | 0.003 | Young |
| 0.845325 | 0.094726 | 0.212 | 0.202 | 1 | 4 | Mydgf | 0.01 | Young |
| 0.845546 | 0.137802 | 0.05 | 0.045 | 1 | 4 | Srcap | 0.005 | Young |
| 0.845711 | 0.124699 | 0.062 | 0.057 | 1 | 4 | Xab2 | 0.005 | Young |
| 0.846356 | 0.223952 | 0.038 | 0.034 | 1 | 4 | Def6 | 0.004 | Young |
| 0.846465 | 0.275971 | 0.5 | 0.488 | 1 | 4 | Gm26532 | 0.012 | Young |
| 0.846689 | 0.246479 | 0.038 | 0.043 | 1 | 4 | Abr | -0.005 | Young |
| 0.847034 | 0.094445 | 0.212 | 0.202 | 1 | 4 | Bub3 | 0.01 | Young |
| 0.847257 | 0.153995 | 0.075 | 0.069 | 1 | 4 | Kin | 0.006 | Young |
| 0.84733 | 0.074868 | 0.138 | 0.146 | 1 | 4 | Oxr1 | -0.008 | Young |
| 0.847512 | 0.408553 | 0.525 | 0.576 | 1 | 4 | Hsp90aa1 | -0.051 | Young |
| 0.847619 | 0.005456 | 0.438 | 0.404 | 1 | 4 | Cep170 | 0.034 | Young |
| 0.847863 | 0.011538 | 0.55 | 0.56 | 1 | 4 | Cox6c | -0.01 | Young |
| 0.848582 | 0.162261 | 0.262 | 0.256 | 1 | 4 | Akt2 | 0.006 | Young |
| 0.848798 | 0.089202 | 0.088 | 0.08 | 1 | 4 | Dcaf17 | 0.008 | Young |
| 0.849204 | 0.143214 | 0.075 | 0.07 | 1 | 4 | Dopey2 | 0.005 | Young |
| 0.850361 | 0.279992 | 0.138 | 0.135 | 1 | 4 | Pcyox1 | 0.003 | Young |
| 0.85039 | 0.020941 | 0.05 | 0.045 | 1 | 4 | Hic2 | 0.005 | Young |
| 0.850463 | 0.056314 | 0.138 | 0.129 | 1 | 4 | Fundc1 | 0.009 | Young |
| 0.851544 | 0.586103 | 0.062 | 0.059 | 1 | 4 | Matk | 0.003 | Young |
| 0.851679 | 0.173179 | 0.138 | 0.132 | 1 | 4 | Emc6 | 0.006 | Young |
| 0.851877 | 0.048713 | 0.138 | 0.131 | 1 | 4 | Siah2 | 0.007 | Young |
| 0.851919 | 0.3181 | 0.025 | 0.029 | 1 | 4 | Slc25a40 | -0.004 | Young |
| 0.853744 | 0.083916 | 0.138 | 0.145 | 1 | 4 | Relb | -0.007 | Young |
| 0.853825 | 0.086184 | 0.038 | 0.034 | 1 | 4 | Zfp113 | 0.004 | Young |
| 0.854391 | 0.144997 | 0.062 | 0.069 | 1 | 4 | Ddx41 | -0.007 | Young |
| 0.854482 | 0.531769 | 0.038 | 0.034 | 1 | 4 | Gnl3l | 0.004 | Young |
| 0.855009 | 0.075929 | 0.225 | 0.215 | 1 | 4 | Psmd1 | 0.01 | Young |
| 0.85537 | 0.288383 | 0.062 | 0.059 | 1 | 4 | Kif13a | 0.003 | Young |
| 0.855409 | 0.583727 | 0.038 | 0.034 | 1 | 4 | Tex30 | 0.004 | Young |
| 0.855804 | 0.145322 | 0.013 | 0.015 | 1 | 4 | Zfp580 | -0.002 | Young |
| 0.855804 | 0.123889 | 0.013 | 0.015 | 1 | 4 | Mblac1 | -0.002 | Young |
| 0.85673 | 0.2007 | 0.013 | 0.015 | 1 | 4 | Mvb12b | -0.002 | Young |
| 0.857451 | 0.014078 | 0.875 | 0.856 | 1 | 4 | Rpl8 | 0.019 | Young |
| 0.857894 | 0.007401 | 0.338 | 0.323 | 1 | 4 | Ttc14 | 0.015 | Young |
| 0.858363 | 0.15768 | 0.262 | 0.256 | 1 | 4 | Dclre1c | 0.006 | Young |
| 0.858583 | 0.37917 | 0.013 | 0.015 | 1 | 4 | F10 | -0.002 | Young |
| 0.858583 | 0.260943 | 0.013 | 0.015 | 1 | 4 | Rita1 | -0.002 | Young |

| | | | | | | | | |
|---|---|---|---|---|---|---|---|---|
| 0.859267 | 0.098961 | 0.088 | 0.082 | 1 | 4 | Igip | 0.006 | Young |
| 0.85951 | 0.596732 | 0.013 | 0.015 | 1 | 4 | Sap130 | -0.002 | Young |
| 0.85951 | 0.535512 | 0.013 | 0.015 | 1 | 4 | Eid2 | -0.002 | Young |
| 0.859534 | 1.144951 | 0.013 | 0.011 | 1 | 4 | Smtn | 0.002 | Young |
| 0.859534 | 1.012927 | 0.013 | 0.011 | 1 | 4 | Hhat | 0.002 | Young |
| 0.859612 | 0.422261 | 0.088 | 0.084 | 1 | 4 | Atic | 0.004 | Young |
| 0.860256 | 0.352537 | 0.162 | 0.155 | 1 | 4 | Slc30a5 | 0.007 | Young |
| 0.860633 | 0.609898 | 0.013 | 0.011 | 1 | 4 | Tnfrsf23 | 0.002 | Young |
| 0.861143 | 0.270626 | 0.062 | 0.058 | 1 | 4 | Sdhaf2 | 0.004 | Young |
| 0.861183 | 0.278227 | 0.312 | 0.302 | 1 | 4 | Ghitm | 0.01 | Young |
| 0.861472 | 0.143503 | 0.05 | 0.054 | 1 | 4 | Tanc1 | -0.004 | Young |
| 0.861593 | 0.079423 | 0.188 | 0.176 | 1 | 4 | Wbp11 | 0.012 | Young |
| 0.861593 | 0.254889 | 0.062 | 0.059 | 1 | 4 | Fam173b | 0.003 | Young |
| 0.861732 | 0.770449 | 0.013 | 0.011 | 1 | 4 | Nedd4 | 0.002 | Young |
| 0.862231 | 0.266011 | 0.1 | 0.109 | 1 | 4 | Ctns | -0.009 | Young |
| 0.862331 | 0.029592 | 0.1 | 0.094 | 1 | 4 | Arfrp1 | 0.006 | Young |
| 0.862747 | 0.169125 | 0.075 | 0.069 | 1 | 4 | Naglu | 0.006 | Young |
| 0.862832 | 0.273466 | 0.013 | 0.011 | 1 | 4 | Mboat1 | 0.002 | Young |
| 0.862944 | 0.859717 | 0.025 | 0.022 | 1 | 4 | Phtf1 | 0.003 | Young |
| 0.863799 | 0.146664 | 0.312 | 0.303 | 1 | 4 | Swap70 | 0.009 | Young |
| 0.86387 | 0.565893 | 0.075 | 0.072 | 1 | 4 | Atxn1l | 0.003 | Young |
| 0.863932 | 0.763735 | 0.013 | 0.011 | 1 | 4 | Gm12166 | 0.002 | Young |
| 0.863932 | 0.568677 | 0.013 | 0.011 | 1 | 4 | Phf19 | 0.002 | Young |
| 0.864975 | 0.022712 | 0.112 | 0.119 | 1 | 4 | Plcb3 | -0.007 | Young |
| 0.865032 | 0.263177 | 0.013 | 0.011 | 1 | 4 | Stox2 | 0.002 | Young |
| 0.866044 | 0.159195 | 0.15 | 0.146 | 1 | 4 | Mogs | 0.004 | Young |
| 0.866132 | 0.110769 | 0.013 | 0.011 | 1 | 4 | Gbp9 | 0.002 | Young |
| 0.866132 | 0.05829 | 0.013 | 0.011 | 1 | 4 | Mmp12 | 0.002 | Young |
| 0.8666 | 0.160562 | 0.138 | 0.146 | 1 | 4 | Ammecr1l | -0.008 | Young |
| 0.866671 | 0.095025 | 0.125 | 0.119 | 1 | 4 | Arhgap26 | 0.006 | Young |
| 0.867233 | 0.250435 | 0.013 | 0.011 | 1 | 4 | Tbc1d7 | 0.002 | Young |
| 0.867233 | 0.026195 | 0.013 | 0.011 | 1 | 4 | Mgll | 0.002 | Young |
| 0.867408 | 0.067989 | 0.088 | 0.082 | 1 | 4 | Mras | 0.006 | Young |
| 0.867712 | 0.231208 | 0.112 | 0.12 | 1 | 4 | Pafah1b2 | -0.008 | Young |
| 0.867993 | 0.746488 | 0.038 | 0.035 | 1 | 4 | Dmac2 | 0.003 | Young |
| 0.868269 | 0.506597 | 0.025 | 0.022 | 1 | 4 | Maml3 | 0.003 | Young |
| 0.868269 | 0.40802 | 0.025 | 0.022 | 1 | 4 | Phf7 | 0.003 | Young |
| 0.868596 | 0.099126 | 0.138 | 0.144 | 1 | 4 | Pcgf5 | -0.006 | Young |
| 0.868776 | 0.048188 | 0.05 | 0.046 | 1 | 4 | Gnpat | 0.004 | Young |
| 0.86917 | 0.498423 | 0.112 | 0.111 | 1 | 4 | Lrrc47 | 0.001 | Young |
| 0.869434 | 0.097417 | 0.013 | 0.011 | 1 | 4 | Gm31597 | 0.002 | Young |
| 0.869434 | 0.087734 | 0.013 | 0.011 | 1 | 4 | Magi3 | 0.002 | Young |
| 0.869836 | 0.663291 | 0.038 | 0.035 | 1 | 4 | Pex1 | 0.003 | Young |
| 0.872118 | 0.393207 | 0.038 | 0.034 | 1 | 4 | Gm11808 | 0.004 | Young |
| 0.872285 | 0.12247 | 0.362 | 0.372 | 1 | 4 | Hook3 | -0.01 | Young |

| | | | | | | | | |
|---|---|---|---|---|---|---|---|---|
| 0.872844 | 0.142808 | 0.025 | 0.028 | 1 | 4 | Mettl5 | -0.003 | Young |
| 0.874598 | 0.005235 | 0.038 | 0.034 | 1 | 4 | Atg13 | 0.004 | Young |
| 0.874818 | 0.509097 | 0.062 | 0.06 | 1 | 4 | Slc27a4 | 0.002 | Young |
| 0.87486 | 0.041295 | 0.088 | 0.095 | 1 | 4 | Scarb1 | -0.007 | Young |
| 0.875219 | 0.100045 | 0.038 | 0.034 | 1 | 4 | Zxdb | 0.004 | Young |
| 0.875904 | 0.00598 | 0.138 | 0.128 | 1 | 4 | Abl2 | 0.01 | Young |
| 0.876092 | 0.185565 | 0.125 | 0.121 | 1 | 4 | Senp2 | 0.004 | Young |
| 0.876205 | 0.075722 | 0.238 | 0.242 | 1 | 4 | Ddx24 | -0.004 | Young |
| 0.876227 | 0.159846 | 0.062 | 0.059 | 1 | 4 | Ppp2r3c | 0.003 | Young |
| 0.876262 | 0.348253 | 0.025 | 0.028 | 1 | 4 | Sema4d | -0.003 | Young |
| 0.876648 | 0.110385 | 0.025 | 0.022 | 1 | 4 | Ltb4r1 | 0.003 | Young |
| 0.876899 | 0.008587 | 0.088 | 0.083 | 1 | 4 | Ctbs | 0.005 | Young |
| 0.87692 | 0.316885 | 0.075 | 0.071 | 1 | 4 | Diablo | 0.004 | Young |
| 0.87752 | 0.191727 | 0.088 | 0.084 | 1 | 4 | Nfkbib | 0.004 | Young |
| 0.877523 | 0.041161 | 0.075 | 0.07 | 1 | 4 | Nr1d2 | 0.005 | Young |
| 0.877647 | 0.050229 | 0.3 | 0.279 | 1 | 4 | Wdr45b | 0.021 | Young |
| 0.877796 | 0.062796 | 0.162 | 0.174 | 1 | 4 | Fto | -0.012 | Young |
| 0.878174 | 0.382381 | 0.162 | 0.161 | 1 | 4 | Plgrkt | 0.001 | Young |
| 0.878188 | 0.022337 | 0.05 | 0.046 | 1 | 4 | Sclt1 | 0.004 | Young |
| 0.878221 | 0.098176 | 0.388 | 0.371 | 1 | 4 | Fuca1 | 0.017 | Young |
| 0.878424 | 0.313063 | 0.05 | 0.047 | 1 | 4 | Slamf6 | 0.003 | Young |
| 0.879413 | 0.434584 | 0.112 | 0.109 | 1 | 4 | Cdk17 | 0.003 | Young |
| 0.879921 | 0.08354 | 0.7 | 0.729 | 1 | 4 | Atp5l | -0.029 | Young |
| 0.879941 | 0.086699 | 0.175 | 0.171 | 1 | 4 | Dcun1d5 | 0.004 | Young |
| 0.88016 | 0.321123 | 0.188 | 0.181 | 1 | 4 | Zzef1 | 0.007 | Young |
| 0.880166 | 0.320925 | 0.075 | 0.082 | 1 | 4 | Timm10 | -0.007 | Young |
| 0.880185 | 0.275281 | 0.062 | 0.059 | 1 | 4 | Itga5 | 0.003 | Young |
| 0.88034 | 0.218168 | 0.05 | 0.053 | 1 | 4 | Aar2 | -0.003 | Young |
| 0.881135 | 0.117302 | 0.412 | 0.426 | 1 | 4 | Abi1 | -0.014 | Young |
| 0.882573 | 0.025186 | 0.45 | 0.436 | 1 | 4 | 5031439G( | 0.014 | Young |
| 0.882873 | 0.029159 | 0.188 | 0.193 | 1 | 4 | Orai3 | -0.005 | Young |
| 0.883172 | 0.016147 | 0.112 | 0.107 | 1 | 4 | Lrif1 | 0.005 | Young |
| 0.88373 | 0.244364 | 0.525 | 0.57 | 1 | 4 | Hnrnph1 | -0.045 | Young |
| 0.883789 | 0.666418 | 0.025 | 0.028 | 1 | 4 | 2700097O( | -0.003 | Young |
| 0.884311 | 0.312933 | 0.05 | 0.047 | 1 | 4 | Fam129b | 0.003 | Young |
| 0.88501 | 0.015502 | 0.462 | 0.443 | 1 | 4 | Tmem59 | 0.019 | Young |
| 0.885361 | 0.013552 | 0.112 | 0.107 | 1 | 4 | Polr2i | 0.005 | Young |
| 0.885531 | 0.437692 | 0.075 | 0.072 | 1 | 4 | Nfyb | 0.003 | Young |
| 0.885941 | 0.042256 | 0.25 | 0.267 | 1 | 4 | Cenpb | -0.017 | Young |
| 0.886235 | 0.075067 | 0.112 | 0.106 | 1 | 4 | Akr1e1 | 0.006 | Young |
| 0.886609 | 0.031519 | 0.762 | 0.738 | 1 | 4 | Gm10076 | 0.024 | Young |
| 0.886722 | 0.278479 | 0.05 | 0.047 | 1 | 4 | Ypel2 | 0.003 | Young |
| 0.88699 | 0.129908 | 0.05 | 0.047 | 1 | 4 | Gfm2 | 0.003 | Young |
| 0.887115 | 0.135043 | 0.112 | 0.118 | 1 | 4 | Lrpprc | -0.006 | Young |
| 0.887509 | 0.095681 | 0.25 | 0.239 | 1 | 4 | Setd1b | 0.011 | Young |

| | | | | | | | | |
|---|---|---|---|---|---|---|---|---|
| 0.888098 | 0.21976 | 0.1 | 0.108 | 1 | 4 | Nudt4 | -0.008 | Young |
| 0.888806 | 0.011719 | 0.613 | 0.602 | 1 | 4 | Erp29 | 0.011 | Young |
| 0.889007 | 0.0382 | 0.7 | 0.69 | 1 | 4 | Ube2d3 | 0.01 | Young |
| 0.889311 | 0.315011 | 0.013 | 0.014 | 1 | 4 | Mtg2 | -0.001 | Young |
| 0.889575 | 0.94162 | 0.025 | 0.023 | 1 | 4 | Ccp110 | 0.002 | Young |
| 0.889796 | 0.137832 | 0.15 | 0.145 | 1 | 4 | Pa2g4 | 0.005 | Young |
| 0.889873 | 0.081589 | 0.275 | 0.261 | 1 | 4 | Psmb8 | 0.014 | Young |
| 0.890834 | 0.003751 | 0.025 | 0.028 | 1 | 4 | 9930104L0 | -0.003 | Young |
| 0.890891 | 0.599179 | 0.05 | 0.047 | 1 | 4 | Rnase6 | 0.003 | Young |
| 0.891216 | 0.46747 | 0.013 | 0.014 | 1 | 4 | Syngr1 | -0.001 | Young |
| 0.891654 | 0.477122 | 0.075 | 0.071 | 1 | 4 | Zfp362 | 0.004 | Young |
| 0.89168 | 0.063266 | 0.362 | 0.357 | 1 | 4 | Psme1 | 0.005 | Young |
| 0.892049 | 0.207781 | 0.238 | 0.235 | 1 | 4 | Efcab14 | 0.003 | Young |
| 0.892094 | 0.240201 | 0.075 | 0.071 | 1 | 4 | Hras | 0.004 | Young |
| 0.892238 | 0.181307 | 0.138 | 0.136 | 1 | 4 | Fam103a1 | 0.002 | Young |
| 0.892615 | 0.177027 | 0.038 | 0.035 | 1 | 4 | Zfp866 | 0.003 | Young |
| 0.892653 | 0.012255 | 0.188 | 0.199 | 1 | 4 | Peak1 | -0.011 | Young |
| 0.892818 | 0.211357 | 0.038 | 0.041 | 1 | 4 | Klra3 | -0.003 | Young |
| 0.892966 | 0.121442 | 0.075 | 0.072 | 1 | 4 | Usp36 | 0.003 | Young |
| 0.893121 | 0.364188 | 0.013 | 0.014 | 1 | 4 | Gm17586 | -0.001 | Young |
| 0.893121 | 0.513116 | 0.013 | 0.014 | 1 | 4 | Klhl42 | -0.001 | Young |
| 0.89401 | 0.162958 | 0.088 | 0.084 | 1 | 4 | Unkl | 0.004 | Young |
| 0.894074 | 0.52842 | 0.013 | 0.014 | 1 | 4 | Tti1 | -0.001 | Young |
| 0.895116 | 0.091169 | 0.038 | 0.041 | 1 | 4 | Tmem185a | -0.003 | Young |
| 0.895194 | 0.081986 | 0.075 | 0.07 | 1 | 4 | Rinl | 0.005 | Young |
| 0.895989 | 0.349279 | 0.075 | 0.073 | 1 | 4 | 2610002M | 0.002 | Young |
| 0.896265 | 0.169926 | 0.038 | 0.041 | 1 | 4 | Ift172 | -0.003 | Young |
| 0.896365 | 0.510585 | 0.025 | 0.023 | 1 | 4 | Gm26669 | 0.002 | Young |
| 0.896932 | 0.087803 | 0.15 | 0.147 | 1 | 4 | Mpp6 | 0.003 | Young |
| 0.896936 | 0.15634 | 0.038 | 0.035 | 1 | 4 | Paxx | 0.003 | Young |
| 0.897733 | 0.759805 | 0.062 | 0.061 | 1 | 4 | Rce1 | 0.001 | Young |
| 0.898414 | 0.001256 | 0.95 | 0.939 | 1 | 4 | Rpl7 | 0.011 | Young |
| 0.898788 | 0.136881 | 0.038 | 0.035 | 1 | 4 | Pde6d | 0.003 | Young |
| 0.898878 | 0.173816 | 0.062 | 0.059 | 1 | 4 | Selenoi | 0.003 | Young |
| 0.899158 | 0.254311 | 0.025 | 0.028 | 1 | 4 | Itgb3bp | -0.003 | Young |
| 0.899714 | 0.299302 | 0.038 | 0.041 | 1 | 4 | Tbccd1 | -0.003 | Young |
| 0.899766 | 0.101978 | 0.075 | 0.08 | 1 | 4 | Ascc1 | -0.005 | Young |
| 0.900141 | 0.446139 | 0.025 | 0.023 | 1 | 4 | Mtor | 0.002 | Young |
| 0.900429 | 0.031783 | 0.075 | 0.072 | 1 | 4 | Bad | 0.003 | Young |
| 0.900547 | 0.265055 | 0.025 | 0.028 | 1 | 4 | 4632415L0 | -0.003 | Young |
| 0.900548 | 0.438908 | 0.05 | 0.053 | 1 | 4 | Ccdc32 | -0.003 | Young |
| 0.900602 | 0.083931 | 0.075 | 0.08 | 1 | 4 | Tm7sf3 | -0.005 | Young |
| 0.900869 | 0.053229 | 0.15 | 0.145 | 1 | 4 | Txndc15 | 0.005 | Young |
| 0.900889 | 0.021364 | 0.175 | 0.165 | 1 | 4 | Inppl1 | 0.01 | Young |
| 0.90136 | 0.268919 | 0.1 | 0.098 | 1 | 4 | Mta1 | 0.002 | Young |

| | | | | | | | | |
|---|---|---|---|---|---|---|---|---|
| 0.901652 | 0.664015 | 0.025 | 0.023 | 1 | 4 | P2ry10b | 0.002 | Young |
| 0.902246 | 0.052775 | 0.088 | 0.084 | 1 | 4 | Lyst | 0.004 | Young |
| 0.902557 | 0.008752 | 0.988 | 0.976 | 1 | 4 | Rps3a1 | 0.012 | Young |
| 0.902631 | 0.504572 | 0.05 | 0.055 | 1 | 4 | Hsdl2 | -0.005 | Young |
| 0.903222 | 0.419195 | 0.062 | 0.067 | 1 | 4 | Tufm | -0.005 | Young |
| 0.903388 | 0.02821 | 0.188 | 0.183 | 1 | 4 | Dennd1b | 0.005 | Young |
| 0.903626 | 0.393304 | 0.038 | 0.036 | 1 | 4 | Trim21 | 0.002 | Young |
| 0.904091 | 0.113526 | 0.05 | 0.053 | 1 | 4 | Cd274 | -0.003 | Young |
| 0.905431 | 0.239251 | 0.025 | 0.023 | 1 | 4 | Lnx2 | 0.002 | Young |
| 0.905629 | 0.058667 | 0.2 | 0.195 | 1 | 4 | Heca | 0.005 | Young |
| 0.906629 | 0.370343 | 0.05 | 0.047 | 1 | 4 | Hint3 | 0.003 | Young |
| 0.906894 | 0.854731 | 0.013 | 0.011 | 1 | 4 | Gnb5 | 0.002 | Young |
| 0.906943 | 0.199314 | 0.025 | 0.023 | 1 | 4 | Tfip11 | 0.002 | Young |
| 0.907419 | 0.061577 | 0.175 | 0.181 | 1 | 4 | Chp1 | -0.006 | Young |
| 0.9077 | 0.375191 | 0.025 | 0.023 | 1 | 4 | Tmem29 | 0.002 | Young |
| 0.907972 | 0.525092 | 0.013 | 0.011 | 1 | 4 | Napb | 0.002 | Young |
| 0.908244 | 0.015136 | 0.062 | 0.067 | 1 | 4 | Mtmr2 | -0.005 | Young |
| 0.90905 | 0.747477 | 0.013 | 0.011 | 1 | 4 | Slc9a4 | 0.002 | Young |
| 0.90905 | 0.359795 | 0.013 | 0.011 | 1 | 4 | Opn3 | 0.002 | Young |
| 0.909212 | 0.112508 | 0.025 | 0.023 | 1 | 4 | Dbf4 | 0.002 | Young |
| 0.909254 | 0.131508 | 0.125 | 0.121 | 1 | 4 | Stk40 | 0.004 | Young |
| 0.909557 | 0.110602 | 0.15 | 0.152 | 1 | 4 | Lpar1 | -0.002 | Young |
| 0.909693 | 0.054865 | 0.038 | 0.04 | 1 | 4 | Zfp672 | -0.002 | Young |
| 0.909756 | 0.385379 | 0.038 | 0.036 | 1 | 4 | Nup160 | 0.002 | Young |
| 0.910985 | 0.086388 | 0.062 | 0.067 | 1 | 4 | Tstd2 | -0.005 | Young |
| 0.911171 | 0.014673 | 0.05 | 0.047 | 1 | 4 | Card6 | 0.003 | Young |
| 0.911206 | 0.523578 | 0.013 | 0.011 | 1 | 4 | Zfp438 | 0.002 | Young |
| 0.911206 | 0.372974 | 0.013 | 0.011 | 1 | 4 | Wdr34 | 0.002 | Young |
| 0.911561 | 0.063094 | 0.088 | 0.084 | 1 | 4 | Sipa1l3 | 0.004 | Young |
| 0.91221 | 0.244052 | 0.038 | 0.036 | 1 | 4 | Tbc1d32 | 0.002 | Young |
| 0.912326 | 0.331345 | 0.162 | 0.157 | 1 | 4 | Dap | 0.005 | Young |
| 0.912334 | 0.155044 | 0.562 | 0.537 | 1 | 4 | Tpr | 0.025 | Young |
| 0.912387 | 0.381267 | 0.075 | 0.071 | 1 | 4 | Nit1 | 0.004 | Young |
| 0.912824 | 0.310729 | 0.038 | 0.036 | 1 | 4 | Ptpn21 | 0.002 | Young |
| 0.913928 | 0.023023 | 0.188 | 0.182 | 1 | 4 | Nr4a2 | 0.006 | Young |
| 0.915521 | 0.089466 | 0.013 | 0.011 | 1 | 4 | Zcwpw1 | 0.002 | Young |
| 0.915521 | 0.050224 | 0.013 | 0.011 | 1 | 4 | Erbb3 | 0.002 | Young |
| 0.915693 | 0.064247 | 0.088 | 0.092 | 1 | 4 | Sil1 | -0.004 | Young |
| 0.915733 | 0.340994 | 0.062 | 0.061 | 1 | 4 | Cep63 | 0.001 | Young |
| 0.916013 | 0.149914 | 0.062 | 0.067 | 1 | 4 | Prpsap1 | -0.005 | Young |
| 0.9166 | 0.080348 | 0.013 | 0.011 | 1 | 4 | Pop1 | 0.002 | Young |
| 0.916771 | 0.585735 | 0.188 | 0.181 | 1 | 4 | Taz | 0.007 | Young |
| 0.917053 | 0.031287 | 0.05 | 0.047 | 1 | 4 | Cebpzos | 0.003 | Young |
| 0.917134 | 0.058276 | 0.15 | 0.158 | 1 | 4 | Adipor2 | -0.008 | Young |
| 0.917358 | 0.143463 | 0.088 | 0.085 | 1 | 4 | Trappc3 | 0.003 | Young |

| | | | | | | | | |
|---|---|---|---|---|---|---|---|---|
| 0.917406 | 0.05373 | 0.162 | 0.159 | 1 | 4 | Blvra | 0.003 | Young |
| 0.917694 | 0.072172 | 0.088 | 0.084 | 1 | 4 | Slc46a3 | 0.004 | Young |
| 0.918042 | 0.036331 | 0.2 | 0.208 | 1 | 4 | Csnk1g3 | -0.008 | Young |
| 0.918428 | 0.160731 | 0.062 | 0.059 | 1 | 4 | Rnf40 | 0.003 | Young |
| 0.91897 | 0.070087 | 0.112 | 0.119 | 1 | 4 | Eif3g | -0.007 | Young |
| 0.919332 | 0.061749 | 0.112 | 0.119 | 1 | 4 | Oasl2 | -0.007 | Young |
| 0.920178 | 0.046956 | 0.288 | 0.283 | 1 | 4 | Anp32e | 0.005 | Young |
| 0.920313 | 0.22309 | 0.05 | 0.053 | 1 | 4 | Haus8 | -0.003 | Young |
| 0.920915 | 0.041692 | 0.112 | 0.109 | 1 | 4 | Prkaca | 0.003 | Young |
| 0.921231 | 0.123982 | 0.025 | 0.027 | 1 | 4 | Mrpl39 | -0.002 | Young |
| 0.92142 | 0.017643 | 0.038 | 0.036 | 1 | 4 | Cdt1 | 0.002 | Young |
| 0.921592 | 0.567486 | 0.025 | 0.024 | 1 | 4 | Jpx | 0.001 | Young |
| 0.922349 | 0.068209 | 0.637 | 0.594 | 1 | 4 | Prkcd | 0.043 | Young |
| 0.922386 | 0.396313 | 0.075 | 0.073 | 1 | 4 | Pmm2 | 0.002 | Young |
| 0.922486 | 0.061793 | 0.013 | 0.014 | 1 | 4 | Adat1 | -0.001 | Young |
| 0.922486 | 0.043533 | 0.013 | 0.014 | 1 | 4 | 1110051M | -0.001 | Young |
| 0.922851 | 0.201803 | 0.05 | 0.053 | 1 | 4 | Gon7 | -0.003 | Young |
| 0.922994 | 0.022021 | 0.062 | 0.066 | 1 | 4 | Slc9a6 | -0.004 | Young |
| 0.923533 | 3.83E-05 | 0.138 | 0.141 | 1 | 4 | Ubp1 | -0.003 | Young |
| 0.923951 | 0.046507 | 0.05 | 0.053 | 1 | 4 | Pou5f2 | -0.003 | Young |
| 0.924048 | 0.333628 | 0.025 | 0.027 | 1 | 4 | Bri3bp | -0.002 | Young |
| 0.92463 | 0.025512 | 0.325 | 0.326 | 1 | 4 | Stx4a | -0.001 | Young |
| 0.924815 | 0.820159 | 0.038 | 0.036 | 1 | 4 | Lyrm9 | 0.002 | Young |
| 0.925544 | 0.234132 | 0.05 | 0.048 | 1 | 4 | Ikbkap | 0.002 | Young |
| 0.926077 | 0.1591 | 0.05 | 0.048 | 1 | 4 | Sugp2 | 0.002 | Young |
| 0.926807 | 0.116159 | 0.062 | 0.065 | 1 | 4 | Zfp467 | -0.003 | Young |
| 0.926829 | 0.620792 | 0.025 | 0.024 | 1 | 4 | Rps6kb2 | 0.001 | Young |
| 0.926866 | 0.466126 | 0.025 | 0.027 | 1 | 4 | Nsdhl | -0.002 | Young |
| 0.926913 | 0.024263 | 0.088 | 0.092 | 1 | 4 | Arl6ip4 | -0.004 | Young |
| 0.927165 | 0.016996 | 0.188 | 0.186 | 1 | 4 | Igbp1 | 0.002 | Young |
| 0.927378 | 0.270119 | 0.013 | 0.014 | 1 | 4 | Armcx1 | -0.001 | Young |
| 0.927392 | 0.160776 | 0.125 | 0.12 | 1 | 4 | Snapc1 | 0.005 | Young |
| 0.927414 | 0.193388 | 0.075 | 0.073 | 1 | 4 | Nsmce2 | 0.002 | Young |
| 0.928088 | 0.136428 | 0.1 | 0.098 | 1 | 4 | Stim1 | 0.002 | Young |
| 0.92818 | 0.096649 | 0.038 | 0.036 | 1 | 4 | Mterf4 | 0.002 | Young |
| 0.928326 | 0.539824 | 0.025 | 0.024 | 1 | 4 | Cluap1 | 0.001 | Young |
| 0.928357 | 0.214108 | 0.013 | 0.014 | 1 | 4 | Thg1l | -0.001 | Young |
| 0.928738 | 0.156643 | 0.05 | 0.048 | 1 | 4 | Mrpl50 | 0.002 | Young |
| 0.929444 | 0.632172 | 0.038 | 0.04 | 1 | 4 | Zc3hc1 | -0.002 | Young |
| 0.9295 | 0.32514 | 0.062 | 0.061 | 1 | 4 | Abhd2 | 0.001 | Young |
| 0.9298 | 0.10343 | 0.1 | 0.097 | 1 | 4 | Zswim8 | 0.003 | Young |
| 0.929803 | 0.16346 | 0.05 | 0.048 | 1 | 4 | Dusp18 | 0.002 | Young |
| 0.930262 | 0.089123 | 0.088 | 0.092 | 1 | 4 | Ppip5k2 | -0.004 | Young |
| 0.93073 | 0.087587 | 0.162 | 0.161 | 1 | 4 | Synj2bp | 0.001 | Young |
| 0.930942 | 0.426441 | 0.075 | 0.08 | 1 | 4 | Phka2 | -0.005 | Young |

| | | | | | | | | |
|---|---|---|---|---|---|---|---|---|
| 0.930974 | 0.248195 | 0.112 | 0.117 | 1 | 4 | Fubp3 | -0.005 | Young |
| 0.931293 | 0.818493 | 0.013 | 0.014 | 1 | 4 | Strbp | -0.001 | Young |
| 0.932132 | 0.33655 | 0.038 | 0.036 | 1 | 4 | Saal1 | 0.002 | Young |
| 0.93239 | 0.11327 | 0.112 | 0.117 | 1 | 4 | Rangap1 | -0.005 | Young |
| 0.932465 | 0.117587 | 0.05 | 0.048 | 1 | 4 | Nubp1 | 0.002 | Young |
| 0.932756 | 0.486605 | 0.05 | 0.053 | 1 | 4 | Rcan3 | -0.003 | Young |
| 0.93351 | 0.113851 | 0.125 | 0.125 | 1 | 4 | Rnh1 | 0 | Young |
| 0.933697 | 0.109448 | 0.062 | 0.061 | 1 | 4 | Kctd6 | 0.001 | Young |
| 0.933911 | 0.164748 | 0.162 | 0.16 | 1 | 4 | Ezh1 | 0.002 | Young |
| 0.934572 | 0.235596 | 0.038 | 0.036 | 1 | 4 | Slc12a4 | 0.002 | Young |
| 0.935094 | 0.14398 | 0.162 | 0.159 | 1 | 4 | Dnase1l1 | 0.003 | Young |
| 0.935352 | 0.068985 | 0.5 | 0.511 | 1 | 4 | Tmem30a | -0.011 | Young |
| 0.935701 | 0.123824 | 0.05 | 0.053 | 1 | 4 | Nelfb | -0.003 | Young |
| 0.936194 | 0.060748 | 0.05 | 0.048 | 1 | 4 | Lig1 | 0.002 | Young |
| 0.936787 | 0.051105 | 0.112 | 0.118 | 1 | 4 | Uxs1 | -0.006 | Young |
| 0.937041 | 0.095361 | 0.062 | 0.061 | 1 | 4 | Timm9 | 0.001 | Young |
| 0.937312 | 0.10781 | 0.025 | 0.024 | 1 | 4 | Fut4 | 0.001 | Young |
| 0.937425 | 0.114125 | 0.075 | 0.078 | 1 | 4 | Cass4 | -0.003 | Young |
| 0.937749 | 0.478495 | 0.075 | 0.074 | 1 | 4 | Zbtb37 | 0.001 | Young |
| 0.938147 | 0.673899 | 0.025 | 0.027 | 1 | 4 | D3Ertd254 | -0.002 | Young |
| 0.938233 | 0.286434 | 0.125 | 0.129 | 1 | 4 | Txndc11 | -0.004 | Young |
| 0.938436 | 0.064437 | 0.112 | 0.109 | 1 | 4 | Smim4 | 0.003 | Young |
| 0.938448 | 0.018434 | 0.125 | 0.123 | 1 | 4 | Spr | 0.002 | Young |
| 0.938621 | 0.038677 | 0.112 | 0.117 | 1 | 4 | St3gal4 | -0.005 | Young |
| 0.938811 | 0.033652 | 0.025 | 0.024 | 1 | 4 | Mthfsl | 0.001 | Young |
| 0.938892 | 0.003054 | 0.162 | 0.163 | 1 | 4 | Fam168a | -0.001 | Young |
| 0.939284 | 0.032968 | 0.225 | 0.217 | 1 | 4 | Cbx1 | 0.008 | Young |
| 0.939454 | 0.123853 | 0.038 | 0.036 | 1 | 4 | Cul4b | 0.002 | Young |
| 0.939461 | 0.021279 | 0.75 | 0.765 | 1 | 4 | Aif1 | -0.015 | Young |
| 0.939669 | 0.075864 | 0.075 | 0.073 | 1 | 4 | Pilrb2 | 0.002 | Young |
| 0.939756 | 0.216783 | 0.038 | 0.04 | 1 | 4 | Rbmx2 | -0.002 | Young |
| 0.940133 | 0.415901 | 0.088 | 0.092 | 1 | 4 | Oxsr1 | -0.004 | Young |
| 0.940291 | 0.023403 | 0.025 | 0.026 | 1 | 4 | Gcfc2 | -0.001 | Young |
| 0.940304 | 0.171094 | 0.05 | 0.053 | 1 | 4 | Nbn | -0.003 | Young |
| 0.941601 | 0.167174 | 0.062 | 0.065 | 1 | 4 | Cops7a | -0.003 | Young |
| 0.942776 | 0.04596 | 0.062 | 0.061 | 1 | 4 | Phkb | 0.001 | Young |
| 0.944358 | 0.088886 | 0.1 | 0.098 | 1 | 4 | Polr2b | 0.002 | Young |
| 0.94556 | 0.153318 | 0.038 | 0.036 | 1 | 4 | Triobp | 0.002 | Young |
| 0.946002 | 0.107033 | 0.025 | 0.026 | 1 | 4 | Iqce | -0.001 | Young |
| 0.946018 | 0.005595 | 0.262 | 0.254 | 1 | 4 | Arrb1 | 0.008 | Young |
| 0.946645 | 0.24112 | 0.162 | 0.163 | 1 | 4 | Mpc1 | -0.001 | Young |
| 0.947153 | 0.053557 | 0.375 | 0.359 | 1 | 4 | 9530068E0 | 0.016 | Young |
| 0.947683 | 0.239452 | 0.05 | 0.052 | 1 | 4 | Zkscan1 | -0.002 | Young |
| 0.948447 | 0.059214 | 0.312 | 0.339 | 1 | 4 | Tcf25 | -0.027 | Young |
| 0.948693 | 0.029955 | 0.525 | 0.518 | 1 | 4 | Epsti1 | 0.007 | Young |

| | | | | | | | | |
|---|---|---|---|---|---|---|---|---|
| 0.949365 | 0.125291 | 0.225 | 0.238 | 1 | 4 | Pon2 | -0.013 | Young |
| 0.94985 | 1.112623 | 0.013 | 0.012 | 1 | 4 | Armc6 | 0.001 | Young |
| 0.949978 | 0.461017 | 0.075 | 0.078 | 1 | 4 | Gmeb1 | -0.003 | Young |
| 0.950288 | 0.250173 | 0.025 | 0.026 | 1 | 4 | 3110002H1 | -0.001 | Young |
| 0.950288 | 0.227084 | 0.025 | 0.026 | 1 | 4 | Car13 | -0.001 | Young |
| 0.950904 | 0.829938 | 0.013 | 0.012 | 1 | 4 | Socs1 | 0.001 | Young |
| 0.950904 | 0.80564 | 0.013 | 0.012 | 1 | 4 | Rtn2 | 0.001 | Young |
| 0.950904 | 0.683786 | 0.013 | 0.012 | 1 | 4 | Pex11g | 0.001 | Young |
| 0.951655 | 0.008897 | 0.588 | 0.601 | 1 | 4 | Snx3 | -0.013 | Young |
| 0.951846 | 0.347507 | 0.062 | 0.062 | 1 | 4 | Surf1 | 0 | Young |
| 0.952083 | 0.091603 | 0.162 | 0.171 | 1 | 4 | N4bp2l1 | -0.009 | Young |
| 0.953065 | 0.075084 | 0.225 | 0.225 | 1 | 4 | Krit1 | 0 | Young |
| 0.953613 | 0.144181 | 0.088 | 0.086 | 1 | 4 | Riok1 | 0.002 | Young |
| 0.95386 | 0.345561 | 0.025 | 0.026 | 1 | 4 | Pkdcc | -0.001 | Young |
| 0.954068 | 0.609543 | 0.013 | 0.012 | 1 | 4 | Zfp607b | 0.001 | Young |
| 0.954068 | 0.57105 | 0.013 | 0.012 | 1 | 4 | Spryd4 | 0.001 | Young |
| 0.954172 | 0.244274 | 0.062 | 0.063 | 1 | 4 | Gm2000 | -0.001 | Young |
| 0.954231 | 0.232477 | 0.075 | 0.078 | 1 | 4 | Pkd1 | -0.003 | Young |
| 0.954603 | 0.071124 | 0.05 | 0.049 | 1 | 4 | Dgka | 0.001 | Young |
| 0.954747 | 0.011974 | 0.688 | 0.665 | 1 | 4 | Ptprc | 0.023 | Young |
| 0.954793 | 0.071312 | 0.262 | 0.265 | 1 | 4 | Itga9 | -0.003 | Young |
| 0.955082 | 0.065667 | 0.362 | 0.369 | 1 | 4 | Diaph1 | -0.007 | Young |
| 0.955123 | 0.386965 | 0.013 | 0.012 | 1 | 4 | Nlrp1b | 0.001 | Young |
| 0.955123 | 0.255811 | 0.013 | 0.012 | 1 | 4 | Casp12 | 0.001 | Young |
| 0.955183 | 0.128795 | 0.075 | 0.074 | 1 | 4 | Osbpl2 | 0.001 | Young |
| 0.955571 | 0.043492 | 0.212 | 0.219 | 1 | 4 | Renbp | -0.007 | Young |
| 0.956178 | 0.203433 | 0.013 | 0.012 | 1 | 4 | Zfp189 | 0.001 | Young |
| 0.956284 | 0.299826 | 0.1 | 0.101 | 1 | 4 | Arsa | -0.001 | Young |
| 0.956763 | 0.02033 | 0.637 | 0.654 | 1 | 4 | Ctsl | -0.017 | Young |
| 0.957093 | 0.050264 | 0.062 | 0.061 | 1 | 4 | Bcl2 | 0.001 | Young |
| 0.957276 | 0.059041 | 0.55 | 0.559 | 1 | 4 | Serp1 | -0.009 | Young |
| 0.957375 | 0.058249 | 0.975 | 0.962 | 1 | 4 | F13a1 | 0.013 | Young |
| 0.957433 | 0.162729 | 0.025 | 0.026 | 1 | 4 | Mettl13 | -0.001 | Young |
| 0.957513 | 0.284901 | 0.05 | 0.049 | 1 | 4 | Nlrc5 | 0.001 | Young |
| 0.957928 | 0.043425 | 0.038 | 0.039 | 1 | 4 | Zfp658 | -0.001 | Young |
| 0.958148 | 0.36995 | 0.025 | 0.026 | 1 | 4 | Gcdh | -0.001 | Young |
| 0.958288 | 0.061713 | 0.013 | 0.012 | 1 | 4 | Ptar1 | 0.001 | Young |
| 0.958834 | 0.482243 | 0.025 | 0.024 | 1 | 4 | Pnkp | 0.001 | Young |
| 0.959685 | 0.188133 | 0.088 | 0.09 | 1 | 4 | Scaf1 | -0.002 | Young |
| 0.961007 | 0.502364 | 0.025 | 0.026 | 1 | 4 | Cog1 | -0.001 | Young |
| 0.96185 | 0.087753 | 0.438 | 0.433 | 1 | 4 | Ndufc2 | 0.005 | Young |
| 0.961994 | 0.128738 | 0.088 | 0.088 | 1 | 4 | Coasy | 0 | Young |
| 0.962073 | 0.237403 | 0.038 | 0.039 | 1 | 4 | Enpp1 | -0.001 | Young |
| 0.962224 | 0.103906 | 0.212 | 0.213 | 1 | 4 | Borcs6 | -0.001 | Young |
| 0.962306 | 0.417442 | 0.013 | 0.013 | 1 | 4 | Zfp442 | 0 | Young |

| | | | | | | | | |
|---|---|---|---|---|---|---|---|---|
| 0.962306 | 0.181845 | 0.013 | 0.013 | 1 | 4 | Ppp1r3e | 0 | Young |
| 0.962657 | 0.005601 | 0.175 | 0.178 | 1 | 4 | Gak | -0.003 | Young |
| 0.962743 | 0.152698 | 0.1 | 0.104 | 1 | 4 | Afdn | -0.004 | Young |
| 0.96331 | 0.266118 | 0.013 | 0.013 | 1 | 4 | Adat2 | 0 | Young |
| 0.96331 | 0.241326 | 0.013 | 0.013 | 1 | 4 | Adck2 | 0 | Young |
| 0.96331 | 0.115962 | 0.013 | 0.013 | 1 | 4 | Cyb5rl | 0 | Young |
| 0.963335 | 0.159382 | 0.112 | 0.111 | 1 | 4 | Atg101 | 0.001 | Young |
| 0.963968 | 0.027039 | 0.875 | 0.864 | 1 | 4 | Rpl19 | 0.011 | Young |
| 0.964315 | 0.284096 | 0.013 | 0.013 | 1 | 4 | Mmaa | 0 | Young |
| 0.964315 | 0.244356 | 0.013 | 0.013 | 1 | 4 | Setd4 | 0 | Young |
| 0.964315 | 0.240231 | 0.013 | 0.013 | 1 | 4 | Hsf4 | 0 | Young |
| 0.964502 | 0.060815 | 0.162 | 0.168 | 1 | 4 | Trim35 | -0.006 | Young |
| 0.965216 | 0.172193 | 0.2 | 0.198 | 1 | 4 | Adgre5 | 0.002 | Young |
| 0.966152 | 0.027996 | 0.4 | 0.389 | 1 | 4 | Tsc22d4 | 0.011 | Young |
| 0.966849 | 0.124103 | 0.038 | 0.038 | 1 | 4 | 1600010M | 0 | Young |
| 0.967651 | 0.017149 | 0.088 | 0.088 | 1 | 4 | Togaram1 | 0 | Young |
| 0.968945 | 0.032486 | 0.162 | 0.16 | 1 | 4 | Rab22a | 0.002 | Young |
| 0.969568 | 0.413504 | 0.088 | 0.088 | 1 | 4 | 2310001H1 | 0 | Young |
| 0.969579 | 0.080224 | 0.062 | 0.064 | 1 | 4 | Chmp6 | -0.002 | Young |
| 0.970365 | 0.166867 | 0.038 | 0.039 | 1 | 4 | Usp30 | -0.001 | Young |
| 0.970365 | 0.661453 | 0.038 | 0.039 | 1 | 4 | BC002059 | -0.001 | Young |
| 0.97205 | 0.290478 | 0.438 | 0.433 | 1 | 4 | Arhgef6 | 0.005 | Young |
| 0.972875 | 0.134582 | 0.075 | 0.075 | 1 | 4 | Lcmt1 | 0 | Young |
| 0.972987 | 0.025391 | 0.038 | 0.037 | 1 | 4 | Piga | 0.001 | Young |
| 0.973033 | 0.455792 | 0.175 | 0.18 | 1 | 4 | Mtm1 | -0.005 | Young |
| 0.973661 | 0.034139 | 0.025 | 0.024 | 1 | 4 | Zfp944 | 0.001 | Young |
| 0.974295 | 0.271284 | 0.025 | 0.026 | 1 | 4 | Prnp | -0.001 | Young |
| 0.974751 | 0.105758 | 0.075 | 0.073 | 1 | 4 | Abcf3 | 0.002 | Young |
| 0.974948 | 0.008748 | 0.075 | 0.074 | 1 | 4 | Scarf1 | 0.001 | Young |
| 0.975019 | 0.081902 | 0.025 | 0.026 | 1 | 4 | Nme6 | -0.001 | Young |
| 0.975316 | 0.099719 | 0.238 | 0.238 | 1 | 4 | Tpi1 | 0 | Young |
| 0.975661 | 0.067921 | 0.062 | 0.064 | 1 | 4 | Atrn | -0.002 | Young |
| 0.976797 | 0.052358 | 0.088 | 0.086 | 1 | 4 | Ppme1 | 0.002 | Young |
| 0.977077 | 0.047739 | 0.188 | 0.183 | 1 | 4 | Blmh | 0.005 | Young |
| 0.977725 | 0.037279 | 0.112 | 0.112 | 1 | 4 | Mprip | 0 | Young |
| 0.977915 | 0.249725 | 0.025 | 0.026 | 1 | 4 | D2hgdh | -0.001 | Young |
| 0.977983 | 0.347509 | 0.075 | 0.078 | 1 | 4 | Ggps1 | -0.003 | Young |
| 0.978148 | 0.06222 | 0.55 | 0.534 | 1 | 4 | Atf4 | 0.016 | Young |
| 0.978157 | 0.042303 | 0.2 | 0.197 | 1 | 4 | Pcm1 | 0.003 | Young |
| 0.978557 | 0.062775 | 0.088 | 0.086 | 1 | 4 | Zfp330 | 0.002 | Young |
| 0.978744 | 0.225298 | 0.05 | 0.05 | 1 | 4 | Mfng | 0 | Young |
| 0.97935 | 0.139788 | 0.15 | 0.155 | 1 | 4 | Zbtb18 | -0.005 | Young |
| 0.979526 | 0.415083 | 0.038 | 0.038 | 1 | 4 | Rmnd5b | 0 | Young |
| 0.979986 | 0.071714 | 0.038 | 0.038 | 1 | 4 | Smyd3 | 0 | Young |
| 0.980201 | 0.038122 | 0.05 | 0.049 | 1 | 4 | Atad3a | 0.001 | Young |

| | | | | | | | | |
|---|---|---|---|---|---|---|---|---|
| 0.980956 | 0.003645 | 0.05 | 0.051 | 1 | 4 | Setd1a | -0.001 | Young |
| 0.981535 | 0.512592 | 0.025 | 0.026 | 1 | 4 | Ldlrad3 | -0.001 | Young |
| 0.981987 | 0.121015 | 0.075 | 0.075 | 1 | 4 | Faf1 | 0 | Young |
| 0.981999 | 0.075731 | 0.05 | 0.051 | 1 | 4 | Cpsf1 | -0.001 | Young |
| 0.982259 | 0.349275 | 0.025 | 0.026 | 1 | 4 | BC024978 | -0.001 | Young |
| 0.982555 | 0.398635 | 0.075 | 0.076 | 1 | 4 | Sfmbt1 | -0.001 | Young |
| 0.98262 | 0.396241 | 0.075 | 0.077 | 1 | 4 | H2-Ke6 | -0.002 | Young |
| 0.982777 | 0.573048 | 0.025 | 0.025 | 1 | 4 | Tmed1 | 0 | Young |
| 0.982777 | 0.343537 | 0.025 | 0.025 | 1 | 4 | Mrgbp | 0 | Young |
| 0.982985 | 0.13116 | 0.075 | 0.076 | 1 | 4 | Ralgapb | -0.001 | Young |
| 0.984409 | 0.260963 | 0.062 | 0.063 | 1 | 4 | Zcchc8 | -0.001 | Young |
| 0.985155 | 0.469225 | 0.025 | 0.026 | 1 | 4 | Bcl2a1a | -0.001 | Young |
| 0.985216 | 0.267517 | 0.05 | 0.051 | 1 | 4 | Cox19 | -0.001 | Young |
| 0.986373 | 0.121693 | 0.288 | 0.281 | 1 | 4 | Sumo1 | 0.007 | Young |
| 0.987127 | 0.131183 | 0.062 | 0.064 | 1 | 4 | Ginm1 | -0.002 | Young |
| 0.987327 | 0.425403 | 0.025 | 0.026 | 1 | 4 | Blzf1 | -0.001 | Young |
| 0.988606 | 0.453097 | 0.1 | 0.105 | 1 | 4 | Tubgcp4 | -0.005 | Young |
| 0.988775 | 0.531579 | 0.025 | 0.026 | 1 | 4 | Zkscan14 | -0.001 | Young |
| 0.989297 | 0.069483 | 0.288 | 0.285 | 1 | 4 | Ggnbp2 | 0.003 | Young |
| 0.989372 | 0.3323 | 0.025 | 0.025 | 1 | 4 | Prickle3 | 0 | Young |
| 0.989406 | 0.360154 | 0.075 | 0.076 | 1 | 4 | Mfsd10 | -0.001 | Young |
| 0.989499 | 0.686306 | 0.025 | 0.026 | 1 | 4 | Scimp | -0.001 | Young |
| 0.990271 | 0.010009 | 0.075 | 0.076 | 1 | 4 | Ifi35 | -0.001 | Young |
| 0.990838 | 0.364118 | 0.025 | 0.025 | 1 | 4 | Ears2 | 0 | Young |
| 0.991021 | 0.44583 | 0.075 | 0.078 | 1 | 4 | Ints3 | -0.003 | Young |
| 0.991571 | 0.290104 | 0.025 | 0.025 | 1 | 4 | Gtpbp3 | 0 | Young |
| 0.99176 | 1.724041 | 0.013 | 0.013 | 1 | 4 | Hdhd5 | 0 | Young |
| 0.991858 | 0.256684 | 0.1 | 0.101 | 1 | 4 | Trim27 | -0.001 | Young |
| 0.99246 | 0.547508 | 0.088 | 0.092 | 1 | 4 | Elp5 | -0.004 | Young |
| 0.992481 | 0.032097 | 0.262 | 0.267 | 1 | 4 | Cdkn1b | -0.005 | Young |
| 0.99249 | 0.338871 | 0.075 | 0.077 | 1 | 4 | Pnkd | -0.002 | Young |
| 0.992535 | 0.00631 | 0.138 | 0.137 | 1 | 4 | Rara | 0.001 | Young |
| 0.992881 | 0.083754 | 0.25 | 0.256 | 1 | 4 | Tomm22 | -0.006 | Young |
| 0.993073 | 0.086334 | 0.138 | 0.139 | 1 | 4 | 5031425E2 | -0.001 | Young |
| 0.99312 | 0.681707 | 0.025 | 0.026 | 1 | 4 | E130102H2 | -0.001 | Young |
| 0.99382 | 0.67525 | 0.013 | 0.013 | 1 | 4 | Lipt2 | 0 | Young |
| 0.99382 | 0.618523 | 0.013 | 0.013 | 1 | 4 | Tgm2 | 0 | Young |
| 0.99382 | 0.439132 | 0.013 | 0.013 | 1 | 4 | Bbs2 | 0 | Young |
| 0.993941 | 0.235439 | 0.1 | 0.101 | 1 | 4 | Dpy30 | -0.001 | Young |
| 0.99485 | 0.556944 | 0.013 | 0.013 | 1 | 4 | Gk | 0 | Young |
| 0.99485 | 0.537223 | 0.013 | 0.013 | 1 | 4 | Ifi208 | 0 | Young |
| 0.994888 | 0.062788 | 0.1 | 0.101 | 1 | 4 | Chpt1 | -0.001 | Young |
| 0.995169 | 0.213972 | 0.112 | 0.115 | 1 | 4 | Hist1h2bc | -0.003 | Young |
| 0.995293 | 0.705686 | 0.025 | 0.026 | 1 | 4 | Shb | -0.001 | Young |
| 0.99559 | 0.318241 | 0.05 | 0.051 | 1 | 4 | Zfp407 | -0.001 | Young |

| | | | | | | | | |
|---|---|---|---|---|---|---|---|---|
| 0.99588 | 0.325533 | 0.013 | 0.013 | 1 | 4 | E430018J2: | 0 | Young |
| 0.995994 | 0.171704 | 0.088 | 0.09 | 1 | 4 | Ccdc127 | -0.002 | Young |
| 0.996054 | 0.108124 | 0.188 | 0.186 | 1 | 4 | Tap1 | 0.002 | Young |
| 0.996086 | 0.112441 | 0.05 | 0.051 | 1 | 4 | Gm48065 | -0.001 | Young |
| 0.996547 | 0.023924 | 0.35 | 0.364 | 1 | 4 | G3bp1 | -0.014 | Young |
| 0.996628 | 0.551876 | 0.05 | 0.051 | 1 | 4 | H6pd | -0.001 | Young |
| 0.99691 | 0.583941 | 0.013 | 0.013 | 1 | 4 | Fam92a | 0 | Young |
| 0.99691 | 0.555406 | 0.013 | 0.013 | 1 | 4 | Tut1 | 0 | Young |
| 0.99691 | 0.485934 | 0.013 | 0.013 | 1 | 4 | Gpsm2 | 0 | Young |
| 0.99691 | 0.427197 | 0.013 | 0.013 | 1 | 4 | Ctla2b | 0 | Young |
| 0.99691 | 0.397906 | 0.013 | 0.013 | 1 | 4 | A430033K( | 0 | Young |
| 0.998377 | 0.003463 | 0.45 | 0.457 | 1 | 4 | Trim25 | -0.007 | Young |
| 0.999634 | 0.12548 | 0.025 | 0.025 | 1 | 4 | Mcm7 | 0 | Young |
| 5.44E-41 | 0.569367 | 1 | 0.98 | 1.69E-36 | 5 | Cd74 | 0.02 | Old |
| 5.25E-35 | 1.032352 | 0.916 | 0.52 | 1.63E-30 | 5 | Xist | 0.396 | Old |
| 4.84E-25 | 1.073162 | 0.803 | 0.584 | 1.50E-20 | 5 | C4b | 0.219 | Old |
| 8.25E-22 | 1.039604 | 0.832 | 0.584 | 2.56E-17 | 5 | Vcam1 | 0.248 | Old |
| 1.26E-18 | 0.443361 | 0.993 | 0.947 | 3.90E-14 | 5 | H2-Eb1 | 0.046 | Old |
| 1.70E-17 | 0.307688 | 1 | 1 | 5.29E-13 | 5 | Cst3 | 0 | Old |
| 4.16E-14 | 0.871529 | 0.785 | 0.66 | 1.29E-09 | 5 | AC149090. | 0.125 | Old |
| 4.74E-14 | 0.35317 | 1 | 0.972 | 1.47E-09 | 5 | H2-Aa | 0.028 | Old |
| 8.67E-14 | 0.346498 | 0.998 | 1 | 2.69E-09 | 5 | Ftl1 | -0.002 | Old |
| 4.60E-13 | 0.357806 | 0.995 | 0.959 | 1.43E-08 | 5 | H2-Ab1 | 0.036 | Old |
| 3.00E-12 | 0.329066 | 0.993 | 0.982 | 9.33E-08 | 5 | H2-D1 | 0.011 | Old |
| 1.00E-11 | 1.051768 | 0.657 | 0.447 | 3.12E-07 | 5 | Ccl8 | 0.21 | Old |
| 1.35E-11 | 0.478065 | 0.973 | 0.901 | 4.19E-07 | 5 | Clec10a | 0.072 | Old |
| 1.77E-11 | 0.34977 | 0.99 | 0.995 | 5.49E-07 | 5 | Hspa8 | -0.005 | Old |
| 3.12E-11 | 0.272624 | 1 | 0.997 | 9.68E-07 | 5 | C1qb | 0.003 | Old |
| 7.59E-11 | 0.353756 | 0.976 | 0.942 | 2.36E-06 | 5 | Trf | 0.034 | Old |
| 1.23E-10 | 0.832912 | 0.593 | 0.416 | 3.82E-06 | 5 | Slc11a1 | 0.177 | Old |
| 1.64E-10 | 0.60715 | 0.798 | 0.662 | 5.08E-06 | 5 | Cd68 | 0.136 | Old |
| 2.63E-10 | 1.378082 | 0.234 | 0.079 | 8.16E-06 | 5 | C6 | 0.155 | Old |
| 3.23E-10 | 0.341189 | 0.975 | 0.924 | 1.00E-05 | 5 | Cd81 | 0.051 | Old |
| 5.02E-10 | 0.649621 | 0.783 | 0.64 | 1.56E-05 | 5 | Pla2g2d | 0.143 | Old |
| 7.43E-10 | 2.975167 | 0.12 | 0.013 | 2.31E-05 | 5 | Gdf3 | 0.107 | Old |
| 7.63E-10 | 0.470076 | 0.912 | 0.853 | 2.37E-05 | 5 | Ly6e | 0.059 | Old |
| 1.25E-09 | 0.868328 | 0.414 | 0.231 | 3.88E-05 | 5 | Mylk | 0.183 | Old |
| 1.53E-09 | 0.261106 | 1 | 0.995 | 4.76E-05 | 5 | Ctsb | 0.005 | Old |
| 1.82E-09 | 2.449998 | 0.143 | 0.028 | 5.66E-05 | 5 | C130026I2 | 0.115 | Old |
| 8.86E-09 | 0.640942 | 0.734 | 0.635 | 0.000275 | 5 | Cd86 | 0.099 | Old |
| 1.81E-08 | 2.161069 | 0.141 | 0.033 | 0.000562 | 5 | Clec1a | 0.108 | Old |
| 1.88E-08 | 0.618372 | 0.579 | 0.404 | 0.000582 | 5 | Cxcl12 | 0.175 | Old |
| 1.99E-08 | 0.424069 | 0.801 | 0.668 | 0.000619 | 5 | Lilr4b | 0.133 | Old |
| 2.24E-08 | 0.361689 | 0.953 | 0.886 | 0.000696 | 5 | C5ar1 | 0.067 | Old |
| 2.37E-08 | 0.769171 | 0.416 | 0.246 | 0.000735 | 5 | Gpx3 | 0.17 | Old |

| | | | | | | | | |
|---|---|---|---|---|---|---|---|---|
| 3.13E-08 | 0.395133 | 0.825 | 0.695 | 0.000973 | 5 | Mmp9 | 0.13 | Old |
| 4.03E-08 | 0.481778 | 0.79 | 0.65 | 0.001251 | 5 | Ninj1 | 0.14 | Old |
| 5.36E-08 | 1.347841 | 0.29 | 0.15 | 0.001664 | 5 | Slc7a13 | 0.14 | Old |
| 5.56E-08 | 0.682655 | 0.652 | 0.475 | 0.001728 | 5 | Cd209f | 0.177 | Old |
| 9.26E-08 | 0.618835 | 0.542 | 0.373 | 0.002876 | 5 | Clec4b1 | 0.169 | Old |
| 1.37E-07 | 0.651724 | 0.554 | 0.386 | 0.004243 | 5 | Ciita | 0.168 | Old |
| 1.41E-07 | 1.922157 | 0.101 | 0.015 | 0.004381 | 5 | Cacna1a | 0.086 | Old |
| 1.42E-07 | 1.040737 | 0.308 | 0.162 | 0.004415 | 5 | Dusp8 | 0.146 | Old |
| 1.63E-07 | 0.909282 | 0.517 | 0.371 | 0.00506 | 5 | Tfrc | 0.146 | Old |
| 1.75E-07 | 0.328817 | 0.946 | 0.921 | 0.005425 | 5 | Btg2 | 0.025 | Old |
| 3.30E-07 | 0.194945 | 0.975 | 0.97 | 0.01025 | 5 | Tyrobp | 0.005 | Old |
| 4.59E-07 | 0.861474 | 0.382 | 0.239 | 0.014262 | 5 | Ly96 | 0.143 | Old |
| 4.61E-07 | 0.378078 | 0.884 | 0.794 | 0.014314 | 5 | Cd163 | 0.09 | Old |
| 5.65E-07 | 0.534317 | 0.847 | 0.782 | 0.017554 | 5 | Hspa1a | 0.065 | Old |
| 7.40E-07 | 0.668589 | 0.428 | 0.274 | 0.022993 | 5 | Emb | 0.154 | Old |
| 8.24E-07 | 0.298771 | 0.949 | 0.86 | 0.025594 | 5 | Mgl2 | 0.089 | Old |
| 8.47E-07 | 0.596331 | 0.554 | 0.406 | 0.026313 | 5 | Lilra5 | 0.148 | Old |
| 9.69E-07 | 0.402569 | 0.808 | 0.703 | 0.030098 | 5 | Creg1 | 0.105 | Old |
| 1.20E-06 | 0.297366 | 0.948 | 0.883 | 0.037317 | 5 | Tmem176a | 0.065 | Old |
| 1.23E-06 | 2.068654 | 0.091 | 0.015 | 0.038195 | 5 | Paqr9 | 0.076 | Old |
| 1.30E-06 | 1.822273 | 0.133 | 0.041 | 0.040218 | 5 | Il18r1 | 0.092 | Old |
| 1.63E-06 | 1.905536 | 0.116 | 0.03 | 0.050675 | 5 | Spon1 | 0.086 | Old |
| 1.89E-06 | 0.68937 | 0.675 | 0.586 | 0.058601 | 5 | Ccl24 | 0.089 | Old |
| 1.95E-06 | 0.729974 | 0.367 | 0.223 | 0.060538 | 5 | Dot1l | 0.144 | Old |
| 2.51E-06 | 0.179734 | 0.998 | 1 | 0.078064 | 5 | Lyz2 | -0.002 | Old |
| 2.67E-06 | 0.349004 | 0.906 | 0.815 | 0.083052 | 5 | H2-DMa | 0.091 | Old |
| 3.07E-06 | 0.304836 | 0.87 | 0.784 | 0.095446 | 5 | Vmp1 | 0.086 | Old |
| 3.11E-06 | 0.437884 | 0.66 | 0.53 | 0.096644 | 5 | P2ry6 | 0.13 | Old |
| 4.53E-06 | 0.574148 | 0.559 | 0.429 | 0.140643 | 5 | Cd209g | 0.13 | Old |
| 5.93E-06 | 0.623447 | 0.48 | 0.332 | 0.183998 | 5 | Tank | 0.148 | Old |
| 6.44E-06 | 1.412927 | 0.114 | 0.033 | 0.199877 | 5 | Igkc | 0.081 | Old |
| 6.68E-06 | 0.373655 | 0.976 | 0.952 | 0.207544 | 5 | Ubc | 0.024 | Old |
| 6.88E-06 | 0.604133 | 0.458 | 0.31 | 0.213606 | 5 | Slc15a2 | 0.148 | Old |
| 7.44E-06 | 1.402429 | 0.175 | 0.076 | 0.231056 | 5 | Cd55 | 0.099 | Old |
| 8.52E-06 | 0.318772 | 0.818 | 0.708 | 0.264559 | 5 | Kdm6b | 0.11 | Old |
| 9.20E-06 | 0.498625 | 0.545 | 0.401 | 0.285588 | 5 | Slamf7 | 0.144 | Old |
| 9.29E-06 | 0.329828 | 0.879 | 0.777 | 0.288348 | 5 | H2-DMb1 | 0.102 | Old |
| 1.29E-05 | 0.898275 | 0.254 | 0.14 | 0.401019 | 5 | Slamf8 | 0.114 | Old |
| 1.41E-05 | 0.255735 | 0.894 | 0.805 | 0.439274 | 5 | Pim1 | 0.089 | Old |
| 1.48E-05 | 0.18229 | 0.997 | 0.982 | 0.459576 | 5 | Csf1r | 0.015 | Old |
| 1.49E-05 | 0.484594 | 0.623 | 0.5 | 0.464175 | 5 | Ndel1 | 0.123 | Old |
| 1.68E-05 | 0.574322 | 0.544 | 0.429 | 0.520224 | 5 | Ftl1-ps1 | 0.115 | Old |
| 1.76E-05 | 0.761189 | 0.313 | 0.195 | 0.54708 | 5 | Rcn3 | 0.118 | Old |
| 1.92E-05 | 0.488525 | 0.801 | 0.746 | 0.596352 | 5 | Hspa1b | 0.055 | Old |
| 2.04E-05 | 0.299179 | 0.923 | 0.871 | 0.632536 | 5 | Ifitm3 | 0.052 | Old |

| | | | | | | | | |
|---|---|---|---|---|---|---|---|---|
| 2.18E-05 | 0.764005 | 0.448 | 0.325 | 0.677852 | 5 | Cd209d | 0.123 | Old |
| 2.48E-05 | 0.440635 | 0.791 | 0.746 | 0.769773 | 5 | Cxcl2 | 0.045 | Old |
| 4.21E-05 | 0.622242 | 0.402 | 0.289 | 1 | 5 | Plbd2 | 0.113 | Old |
| 4.63E-05 | 0.362761 | 0.763 | 0.703 | 1 | 5 | Pirb | 0.06 | Old |
| 4.72E-05 | 0.693965 | 0.345 | 0.226 | 1 | 5 | AW112010 | 0.119 | Old |
| 4.85E-05 | 2.778448 | 0.047 | 0.003 | 1 | 5 | Peak1os | 0.044 | Old |
| 5.44E-05 | 0.224996 | 0.973 | 0.947 | 1 | 5 | Tmem176b | 0.026 | Old |
| 5.50E-05 | 0.267376 | 0.946 | 0.924 | 1 | 5 | Nfkbiz | 0.022 | Old |
| 8.54E-05 | 0.463956 | 0.672 | 0.553 | 1 | 5 | Ccr5 | 0.119 | Old |
| 9.13E-05 | 0.379119 | 0.502 | 0.373 | 1 | 5 | Ddost | 0.129 | Old |
| 0.000117 | 0.518391 | 0.424 | 0.307 | 1 | 5 | Gstm1 | 0.117 | Old |
| 0.000126 | 0.472642 | 0.497 | 0.365 | 1 | 5 | Lgals3bp | 0.132 | Old |
| 0.000136 | 0.150061 | 1 | 1 | 1 | 5 | mt-Nd2 | 0 | Old |
| 0.00015 | 0.218174 | 0.928 | 0.878 | 1 | 5 | Ehd4 | 0.05 | Old |
| 0.000152 | 0.415765 | 0.436 | 0.302 | 1 | 5 | Mt2 | 0.134 | Old |
| 0.000155 | 2.728336 | 0.052 | 0.008 | 1 | 5 | Cdh6 | 0.044 | Old |
| 0.000163 | 3.173029 | 0.035 | 0 | 1 | 5 | Srpx | 0.035 | Old |
| 0.000164 | 0.663935 | 0.298 | 0.19 | 1 | 5 | BC028528 | 0.108 | Old |
| 0.000167 | 2.241775 | 0.052 | 0.008 | 1 | 5 | Ankrd33b | 0.044 | Old |
| 0.000187 | 0.335513 | 0.729 | 0.635 | 1 | 5 | Spi1 | 0.094 | Old |
| 0.000209 | 0.616874 | 0.138 | 0.063 | 1 | 5 | Nlrc5 | 0.075 | Old |
| 0.000209 | 0.294885 | 0.87 | 0.794 | 1 | 5 | Mat2a | 0.076 | Old |
| 0.000219 | 0.778504 | 0.298 | 0.201 | 1 | 5 | Il2rg | 0.097 | Old |
| 0.00022 | 0.256419 | 0.966 | 0.924 | 1 | 5 | Iqgap1 | 0.042 | Old |
| 0.000227 | 0.473989 | 0.463 | 0.348 | 1 | 5 | Mknk1 | 0.115 | Old |
| 0.000228 | 0.264993 | 0.906 | 0.83 | 1 | 5 | Calr | 0.076 | Old |
| 0.000228 | 0.375089 | 0.872 | 0.807 | 1 | 5 | Ubb | 0.065 | Old |
| 0.000245 | 0.799663 | 0.29 | 0.188 | 1 | 5 | Tmod1 | 0.102 | Old |
| 0.000245 | 1.065888 | 0.145 | 0.071 | 1 | 5 | Rusc2 | 0.074 | Old |
| 0.000246 | 0.14438 | 0.993 | 0.985 | 1 | 5 | Fcer1g | 0.008 | Old |
| 0.000273 | 0.728409 | 0.271 | 0.173 | 1 | 5 | Phf11b | 0.098 | Old |
| 0.000284 | 0.662466 | 0.332 | 0.231 | 1 | 5 | A4galt | 0.101 | Old |
| 0.000289 | 0.370832 | 0.67 | 0.566 | 1 | 5 | Csf2ra | 0.104 | Old |
| 0.000291 | 1.436215 | 0.103 | 0.041 | 1 | 5 | Gng11 | 0.062 | Old |
| 0.000301 | 2.011701 | 0.059 | 0.013 | 1 | 5 | Bmpr1a | 0.046 | Old |
| 0.000332 | 0.586685 | 0.357 | 0.254 | 1 | 5 | Vamp2 | 0.103 | Old |
| 0.000344 | 0.33105 | 0.798 | 0.701 | 1 | 5 | Srgn | 0.097 | Old |
| 0.000346 | 0.337813 | 0.603 | 0.5 | 1 | 5 | Bank1 | 0.103 | Old |
| 0.00035 | 1.35814 | 0.13 | 0.061 | 1 | 5 | Syne1 | 0.069 | Old |
| 0.000382 | 0.236998 | 0.808 | 0.751 | 1 | 5 | Ifi27l2a | 0.057 | Old |
| 0.000416 | 0.341195 | 0.513 | 0.398 | 1 | 5 | Saraf | 0.115 | Old |
| 0.000419 | 0.130225 | 1 | 1 | 1 | 5 | Selenop | 0 | Old |
| 0.000436 | 1.359789 | 0.079 | 0.025 | 1 | 5 | Csprs | 0.054 | Old |
| 0.000438 | 0.818857 | 0.145 | 0.071 | 1 | 5 | Tnfrsf17 | 0.074 | Old |
| 0.000445 | 0.797025 | 0.215 | 0.129 | 1 | 5 | Agmo | 0.086 | Old |

| | | | | | | | | |
|---|---|---|---|---|---|---|---|---|
| 0.000461 | 0.579979 | 0.335 | 0.231 | 1 | 5 | Myh10 | 0.104 | Old |
| 0.000493 | 2.807707 | 0.03 | 0 | 1 | 5 | Kif1a | 0.03 | Old |
| 0.000505 | 0.415611 | 0.495 | 0.401 | 1 | 5 | Itgb2 | 0.094 | Old |
| 0.000564 | 0.584703 | 0.276 | 0.183 | 1 | 5 | Pigs | 0.093 | Old |
| 0.000586 | 0.872318 | 0.168 | 0.091 | 1 | 5 | Batf | 0.077 | Old |
| 0.000587 | 0.308149 | 0.522 | 0.406 | 1 | 5 | Ptgs1 | 0.116 | Old |
| 0.000596 | 0.982189 | 0.114 | 0.051 | 1 | 5 | Camsap1 | 0.063 | Old |
| 0.000608 | 1.11173 | 0.093 | 0.036 | 1 | 5 | Enho | 0.057 | Old |
| 0.000611 | 0.20291 | 0.953 | 0.919 | 1 | 5 | Marcksl1 | 0.034 | Old |
| 0.000645 | 0.445582 | 0.391 | 0.282 | 1 | 5 | Gm6377 | 0.109 | Old |
| 0.000666 | 0.297403 | 0.8 | 0.744 | 1 | 5 | Pnrc1 | 0.056 | Old |
| 0.000667 | 0.330618 | 0.955 | 0.942 | 1 | 5 | Neat1 | 0.013 | Old |
| 0.000681 | 0.914027 | 0.165 | 0.091 | 1 | 5 | Hvcn1 | 0.074 | Old |
| 0.000724 | 0.20765 | 0.904 | 0.845 | 1 | 5 | Fcgr3 | 0.059 | Old |
| 0.000752 | 0.528691 | 0.247 | 0.155 | 1 | 5 | Gm48065 | 0.092 | Old |
| 0.000753 | 0.266477 | 0.662 | 0.561 | 1 | 5 | Tgoln1 | 0.101 | Old |
| 0.000761 | 0.38742 | 0.394 | 0.282 | 1 | 5 | Ddx24 | 0.112 | Old |
| 0.000816 | 0.441447 | 0.352 | 0.249 | 1 | 5 | Siglec1 | 0.103 | Old |
| 0.000818 | 1.695738 | 0.074 | 0.025 | 1 | 5 | Atg9a | 0.049 | Old |
| 0.000819 | 0.68815 | 0.256 | 0.175 | 1 | 5 | Tmem205 | 0.081 | Old |
| 0.000872 | 0.653494 | 0.239 | 0.15 | 1 | 5 | Hk3 | 0.089 | Old |
| 0.000873 | 0.647398 | 0.337 | 0.244 | 1 | 5 | Pdcd2l | 0.093 | Old |
| 0.000898 | 0.686411 | 0.249 | 0.162 | 1 | 5 | Fh1 | 0.087 | Old |
| 0.000907 | 0.333661 | 0.628 | 0.528 | 1 | 5 | Cp | 0.1 | Old |
| 0.000917 | 0.78628 | 0.128 | 0.061 | 1 | 5 | Isg15 | 0.067 | Old |
| 0.000929 | 1.791257 | 0.066 | 0.02 | 1 | 5 | Tanc1 | 0.046 | Old |
| 0.000943 | 0.175981 | 0.886 | 0.881 | 1 | 5 | Ifitm2 | 0.005 | Old |
| 0.000944 | 0.375565 | 0.512 | 0.401 | 1 | 5 | Ppp1r10 | 0.111 | Old |
| 0.000944 | 0.492218 | 0.46 | 0.371 | 1 | 5 | Id2 | 0.089 | Old |
| 0.000981 | 0.837406 | 0.15 | 0.081 | 1 | 5 | Unc5a | 0.069 | Old |
| 0.001052 | 0.561191 | 0.322 | 0.226 | 1 | 5 | Zfp950 | 0.096 | Old |
| 0.001143 | 0.157032 | 0.98 | 0.959 | 1 | 5 | Ddx5 | 0.021 | Old |
| 0.001166 | 0.617351 | 0.414 | 0.325 | 1 | 5 | Midn | 0.089 | Old |
| 0.001166 | 0.595264 | 0.227 | 0.142 | 1 | 5 | Ssr2 | 0.085 | Old |
| 0.001208 | 0.697336 | 0.172 | 0.099 | 1 | 5 | 1600012H( | 0.073 | Old |
| 0.001227 | 0.490228 | 0.394 | 0.299 | 1 | 5 | Bcl6 | 0.095 | Old |
| 0.00126 | 2.532931 | 0.032 | 0.003 | 1 | 5 | Sox7 | 0.029 | Old |
| 0.001304 | 0.470774 | 0.189 | 0.109 | 1 | 5 | Scimp | 0.08 | Old |
| 0.001341 | 2.025363 | 0.032 | 0.003 | 1 | 5 | Hoxb7 | 0.029 | Old |
| 0.001364 | 0.275795 | 0.646 | 0.51 | 1 | 5 | Lbh | 0.136 | Old |
| 0.001371 | 0.293815 | 0.779 | 0.734 | 1 | 5 | Clec4a3 | 0.045 | Old |
| 0.001388 | 0.260217 | 0.771 | 0.711 | 1 | 5 | Tmem106a | 0.06 | Old |
| 0.001389 | 0.272847 | 0.709 | 0.642 | 1 | 5 | Aldh2 | 0.067 | Old |
| 0.001409 | 0.783991 | 0.113 | 0.053 | 1 | 5 | Dpt | 0.06 | Old |
| 0.001433 | 0.432561 | 0.286 | 0.195 | 1 | 5 | Tmem219 | 0.091 | Old |

| | | | | | | | | |
|---|---|---|---|---|---|---|---|---|
| 0.001453 | 0.463573 | 0.342 | 0.249 | 1 | 5 | Acvrl1 | 0.093 | Old |
| 0.001492 | 2.580648 | 0.025 | 0 | 1 | 5 | AC168977. | 0.025 | Old |
| 0.001492 | 1.124384 | 0.136 | 0.074 | 1 | 5 | Gdpd1 | 0.062 | Old |
| 0.001548 | 0.288641 | 0.704 | 0.624 | 1 | 5 | Litaf | 0.08 | Old |
| 0.001581 | 0.657013 | 0.232 | 0.157 | 1 | 5 | Ppm1h | 0.075 | Old |
| 0.001793 | 0.59514 | 0.217 | 0.137 | 1 | 5 | Rasgrp2 | 0.08 | Old |
| 0.001818 | 0.426101 | 0.338 | 0.241 | 1 | 5 | Krtcap2 | 0.097 | Old |
| 0.001819 | 0.859191 | 0.15 | 0.084 | 1 | 5 | Tpst1 | 0.066 | Old |
| 0.001824 | 0.651479 | 0.234 | 0.152 | 1 | 5 | Tmem88 | 0.082 | Old |
| 0.001919 | 0.356398 | 0.614 | 0.546 | 1 | 5 | Spag9 | 0.068 | Old |
| 0.002028 | 0.96696 | 0.146 | 0.084 | 1 | 5 | Tnfrsf12a | 0.062 | Old |
| 0.002043 | 0.538773 | 0.308 | 0.221 | 1 | 5 | Pgrmc1 | 0.087 | Old |
| 0.002049 | 0.188903 | 0.97 | 0.987 | 1 | 5 | Zfp36 | -0.017 | Old |
| 0.002149 | 0.227718 | 0.806 | 0.734 | 1 | 5 | Man2b1 | 0.072 | Old |
| 0.002159 | 0.486757 | 0.291 | 0.203 | 1 | 5 | Ralgds | 0.088 | Old |
| 0.002186 | 0.435933 | 0.259 | 0.173 | 1 | 5 | B3galnt1 | 0.086 | Old |
| 0.002186 | 0.761256 | 0.394 | 0.315 | 1 | 5 | Cd209a | 0.079 | Old |
| 0.00219 | 0.188756 | 0.911 | 0.865 | 1 | 5 | Rpl10 | 0.046 | Old |
| 0.00222 | 1.183448 | 0.098 | 0.046 | 1 | 5 | Gphn | 0.052 | Old |
| 0.002275 | 0.863148 | 0.12 | 0.061 | 1 | 5 | Ndst1 | 0.059 | Old |
| 0.002289 | 0.33378 | 0.567 | 0.447 | 1 | 5 | Stap1 | 0.12 | Old |
| 0.002315 | 1.165828 | 0.088 | 0.038 | 1 | 5 | Scarf1 | 0.05 | Old |
| 0.00244 | 0.212945 | 0.855 | 0.815 | 1 | 5 | Ctsz | 0.04 | Old |
| 0.002471 | 0.592993 | 0.246 | 0.168 | 1 | 5 | Fam219a | 0.078 | Old |
| 0.002494 | 0.262482 | 0.522 | 0.426 | 1 | 5 | Wac | 0.096 | Old |
| 0.00255 | 0.601881 | 0.16 | 0.094 | 1 | 5 | Mtx1 | 0.066 | Old |
| 0.002563 | 0.552015 | 0.227 | 0.155 | 1 | 5 | Zfp512 | 0.072 | Old |
| 0.002583 | 0.361419 | 0.515 | 0.411 | 1 | 5 | H2-M3 | 0.104 | Old |
| 0.002599 | 0.543122 | 0.19 | 0.117 | 1 | 5 | Phf10 | 0.073 | Old |
| 0.00263 | 0.521143 | 0.355 | 0.279 | 1 | 5 | Card19 | 0.076 | Old |
| 0.002664 | 0.612263 | 0.177 | 0.107 | 1 | 5 | Slc41a2 | 0.07 | Old |
| 0.002743 | 1.07679 | 0.14 | 0.081 | 1 | 5 | Ppid | 0.059 | Old |
| 0.002772 | 4.292524 | 0.034 | 0.005 | 1 | 5 | Rnase2a | 0.029 | Old |
| 0.002801 | 0.415127 | 0.613 | 0.543 | 1 | 5 | Clic4 | 0.07 | Old |
| 0.002804 | 0.796746 | 0.125 | 0.066 | 1 | 5 | Mark4 | 0.059 | Old |
| 0.002886 | 0.491852 | 0.323 | 0.241 | 1 | 5 | Pim3 | 0.082 | Old |
| 0.002939 | 0.684984 | 0.483 | 0.398 | 1 | 5 | Gm26532 | 0.085 | Old |
| 0.002977 | 1.294623 | 0.067 | 0.025 | 1 | 5 | Gtf2h4 | 0.042 | Old |
| 0.003109 | 0.38526 | 0.52 | 0.442 | 1 | 5 | Ppib | 0.078 | Old |
| 0.003118 | 0.219264 | 0.808 | 0.751 | 1 | 5 | Plekho2 | 0.057 | Old |
| 0.003248 | 0.207149 | 0.47 | 0.373 | 1 | 5 | Ms4a6d | 0.097 | Old |
| 0.003287 | 0.653067 | 0.291 | 0.213 | 1 | 5 | Tspan32 | 0.078 | Old |
| 0.003305 | 0.262556 | 0.396 | 0.297 | 1 | 5 | Nfkbie | 0.099 | Old |
| 0.003338 | 0.577163 | 0.19 | 0.119 | 1 | 5 | Rbm14 | 0.071 | Old |
| 0.003434 | 1.069034 | 0.103 | 0.051 | 1 | 5 | Negr1 | 0.052 | Old |

| | | | | | | | | |
|---|---|---|---|---|---|---|---|---|
| 0.00347 | 0.311768 | 0.505 | 0.411 | 1 | 5 | Etf1 | 0.094 | Old |
| 0.003521 | 0.312403 | 0.375 | 0.287 | 1 | 5 | Pmaip1 | 0.088 | Old |
| 0.003611 | 0.178014 | 0.83 | 0.772 | 1 | 5 | Eif4a1 | 0.058 | Old |
| 0.003623 | 0.46915 | 0.315 | 0.231 | 1 | 5 | Cd209b | 0.084 | Old |
| 0.003626 | 0.126934 | 0.933 | 0.901 | 1 | 5 | Hexa | 0.032 | Old |
| 0.003663 | 0.555303 | 0.165 | 0.099 | 1 | 5 | C3 | 0.066 | Old |
| 0.003901 | 0.212847 | 0.83 | 0.766 | 1 | 5 | Ifi207 | 0.064 | Old |
| 0.003966 | 1.148439 | 0.088 | 0.041 | 1 | 5 | Lrch4 | 0.047 | Old |
| 0.004115 | 0.357593 | 0.451 | 0.368 | 1 | 5 | Tnfrsf1a | 0.083 | Old |
| 0.004136 | 1.837988 | 0.032 | 0.005 | 1 | 5 | Pdlim3 | 0.027 | Old |
| 0.004232 | 1.24613 | 0.062 | 0.023 | 1 | 5 | Akr1c12 | 0.039 | Old |
| 0.004239 | 0.462646 | 0.295 | 0.211 | 1 | 5 | Ly9 | 0.084 | Old |
| 0.004257 | 1.623412 | 0.032 | 0.005 | 1 | 5 | Cnbd2 | 0.027 | Old |
| 0.004262 | 0.221396 | 0.593 | 0.485 | 1 | 5 | Reps2 | 0.108 | Old |
| 0.00431 | 0.642821 | 0.242 | 0.17 | 1 | 5 | Eif3g | 0.072 | Old |
| 0.004339 | 0.829382 | 0.146 | 0.086 | 1 | 5 | Aspa | 0.06 | Old |
| 0.004414 | 0.586028 | 0.237 | 0.162 | 1 | 5 | Pdxdc1 | 0.075 | Old |
| 0.004446 | 1.04552 | 0.066 | 0.025 | 1 | 5 | A230050P2 | 0.041 | Old |
| 0.004485 | 0.462735 | 0.264 | 0.188 | 1 | 5 | Bicd2 | 0.076 | Old |
| 0.004572 | 1.202457 | 0.106 | 0.056 | 1 | 5 | Acp5 | 0.05 | Old |
| 0.004592 | 0.174988 | 0.556 | 0.457 | 1 | 5 | Sash1 | 0.099 | Old |
| 0.004608 | 0.324058 | 0.485 | 0.393 | 1 | 5 | Coq10b | 0.092 | Old |
| 0.004734 | 0.45762 | 0.273 | 0.19 | 1 | 5 | Ldlr | 0.083 | Old |
| 0.004777 | 0.40078 | 0.323 | 0.246 | 1 | 5 | Napa | 0.077 | Old |
| 0.004787 | 2.282013 | 0.04 | 0.01 | 1 | 5 | 1110002J0 | 0.03 | Old |
| 0.004805 | 0.438585 | 0.421 | 0.353 | 1 | 5 | Nbeal1 | 0.068 | Old |
| 0.0049 | 0.374832 | 0.579 | 0.513 | 1 | 5 | Tcn2 | 0.066 | Old |
| 0.004913 | 0.48762 | 0.263 | 0.188 | 1 | 5 | Cenpx | 0.075 | Old |
| 0.004927 | 0.475592 | 0.278 | 0.198 | 1 | 5 | Tmem9b | 0.08 | Old |
| 0.00499 | 0.130587 | 0.938 | 0.901 | 1 | 5 | Hspa5 | 0.037 | Old |
| 0.00517 | 0.2203 | 0.544 | 0.452 | 1 | 5 | Fgl2 | 0.092 | Old |
| 0.005214 | 0.187406 | 0.641 | 0.566 | 1 | 5 | Cd38 | 0.075 | Old |
| 0.005255 | 0.183504 | 0.515 | 0.414 | 1 | 5 | Tspan3 | 0.101 | Old |
| 0.00528 | 0.292398 | 0.621 | 0.546 | 1 | 5 | Taok3 | 0.075 | Old |
| 0.005396 | 0.517708 | 0.241 | 0.173 | 1 | 5 | Cnp | 0.068 | Old |
| 0.005435 | 1.293736 | 0.064 | 0.025 | 1 | 5 | Lin37 | 0.039 | Old |
| 0.005543 | 0.163711 | 0.84 | 0.772 | 1 | 5 | Clec2d | 0.068 | Old |
| 0.005561 | 0.341128 | 0.375 | 0.279 | 1 | 5 | Trpv4 | 0.096 | Old |
| 0.005581 | 1.761525 | 0.035 | 0.008 | 1 | 5 | A530040E1 | 0.027 | Old |
| 0.005597 | 0.928508 | 0.116 | 0.063 | 1 | 5 | Coro2b | 0.053 | Old |
| 0.005641 | 1.055673 | 0.114 | 0.063 | 1 | 5 | Rnf157 | 0.051 | Old |
| 0.005674 | 1.986202 | 0.025 | 0.003 | 1 | 5 | Zfp213 | 0.022 | Old |
| 0.005738 | 0.426443 | 0.352 | 0.277 | 1 | 5 | Gpcpd1 | 0.075 | Old |
| 0.005764 | 0.394303 | 0.318 | 0.241 | 1 | 5 | Nfya | 0.077 | Old |
| 0.005793 | 0.354306 | 0.556 | 0.492 | 1 | 5 | Hnrnph2 | 0.064 | Old |

| | | | | | | | | |
|---|---|---|---|---|---|---|---|---|
| 0.005871 | 1.704089 | 0.03 | 0.005 | 1 | 5 | Cabp4 | 0.025 | Old |
| 0.005873 | 0.830828 | 0.096 | 0.048 | 1 | 5 | Phf13 | 0.048 | Old |
| 0.005888 | 0.313597 | 0.621 | 0.523 | 1 | 5 | Cd48 | 0.098 | Old |
| 0.005901 | 0.21664 | 0.747 | 0.645 | 1 | 5 | Dnaja1 | 0.102 | Old |
| 0.005989 | 0.457408 | 0.316 | 0.234 | 1 | 5 | Il16 | 0.082 | Old |
| 0.006006 | 1.702497 | 0.03 | 0.005 | 1 | 5 | Fzd8 | 0.025 | Old |
| 0.006039 | 0.234776 | 0.579 | 0.513 | 1 | 5 | P4hb | 0.066 | Old |
| 0.006102 | 0.819788 | 0.185 | 0.122 | 1 | 5 | Rrad | 0.063 | Old |
| 0.0062 | 0.086035 | 1 | 1 | 1 | 5 | C1qc | 0 | Old |
| 0.006244 | 0.134194 | 0.87 | 0.807 | 1 | 5 | Nr4a1 | 0.063 | Old |
| 0.006267 | 0.641839 | 0.157 | 0.096 | 1 | 5 | Sgce | 0.061 | Old |
| 0.006272 | 1.164861 | 0.091 | 0.046 | 1 | 5 | Gstp1 | 0.045 | Old |
| 0.006346 | 0.190673 | 0.389 | 0.307 | 1 | 5 | Slc40a1 | 0.082 | Old |
| 0.006493 | 0.23495 | 0.695 | 0.632 | 1 | 5 | Tra2a | 0.063 | Old |
| 0.006503 | 0.161416 | 0.948 | 0.906 | 1 | 5 | Cd83 | 0.042 | Old |
| 0.006504 | 0.341061 | 0.352 | 0.272 | 1 | 5 | Cic | 0.08 | Old |
| 0.006506 | 0.302615 | 0.449 | 0.368 | 1 | 5 | Ppp1r15b | 0.081 | Old |
| 0.006513 | 0.144189 | 0.961 | 0.931 | 1 | 5 | Timp2 | 0.03 | Old |
| 0.006549 | 0.378739 | 0.34 | 0.256 | 1 | 5 | Ifi27 | 0.084 | Old |
| 0.006615 | 0.586899 | 0.194 | 0.129 | 1 | 5 | Edem2 | 0.065 | Old |
| 0.006735 | 0.279522 | 0.648 | 0.571 | 1 | 5 | Pnisr | 0.077 | Old |
| 0.006759 | 0.714602 | 0.12 | 0.066 | 1 | 5 | Dtx3 | 0.054 | Old |
| 0.006765 | 0.210873 | 0.774 | 0.718 | 1 | 5 | H2-T23 | 0.056 | Old |
| 0.006771 | 0.19099 | 0.875 | 0.82 | 1 | 5 | Rps18 | 0.055 | Old |
| 0.006784 | 1.126703 | 0.056 | 0.02 | 1 | 5 | Macrod1 | 0.036 | Old |
| 0.007114 | 0.190181 | 0.924 | 0.886 | 1 | 5 | Cd63 | 0.038 | Old |
| 0.007168 | 0.428347 | 0.473 | 0.396 | 1 | 5 | Pld3 | 0.077 | Old |
| 0.00722 | 0.339231 | 0.497 | 0.414 | 1 | 5 | Egr2 | 0.083 | Old |
| 0.007313 | 0.508828 | 0.194 | 0.127 | 1 | 5 | Ttpal | 0.067 | Old |
| 0.007318 | 0.69711 | 0.163 | 0.104 | 1 | 5 | Pdgfa | 0.059 | Old |
| 0.007341 | 0.339021 | 0.226 | 0.155 | 1 | 5 | Sec24a | 0.071 | Old |
| 0.007341 | 0.86956 | 0.141 | 0.086 | 1 | 5 | Cers4 | 0.055 | Old |
| 0.007427 | 0.522799 | 0.281 | 0.213 | 1 | 5 | Rheb | 0.068 | Old |
| 0.007559 | 1.142751 | 0.089 | 0.046 | 1 | 5 | Klhl7 | 0.043 | Old |
| 0.007754 | 0.431318 | 0.313 | 0.234 | 1 | 5 | Adgre4 | 0.079 | Old |
| 0.007766 | 0.173872 | 0.894 | 0.817 | 1 | 5 | C3ar1 | 0.077 | Old |
| 0.007802 | 0.405088 | 0.298 | 0.218 | 1 | 5 | Ephx1 | 0.08 | Old |
| 0.007932 | 0.60612 | 0.231 | 0.168 | 1 | 5 | Ticam2 | 0.063 | Old |
| 0.008073 | 0.372477 | 0.379 | 0.299 | 1 | 5 | Micu1 | 0.08 | Old |
| 0.008239 | 0.364443 | 0.492 | 0.414 | 1 | 5 | Atxn2l | 0.078 | Old |
| 0.008349 | 1.380579 | 0.024 | 0.003 | 1 | 5 | Zscan20 | 0.021 | Old |
| 0.008401 | 1.652446 | 0.029 | 0.005 | 1 | 5 | Plxnb1 | 0.024 | Old |
| 0.008446 | 0.7133 | 0.098 | 0.051 | 1 | 5 | Ppan | 0.047 | Old |
| 0.008532 | 0.210978 | 0.684 | 0.594 | 1 | 5 | Itsn1 | 0.09 | Old |
| 0.008574 | 0.286434 | 0.419 | 0.33 | 1 | 5 | Sec61a1 | 0.089 | Old |

| | | | | | | | | |
|---|---|---|---|---|---|---|---|---|
| 0.008642 | 0.260072 | 0.547 | 0.482 | 1 | 5 | Tubb2a | 0.065 | Old |
| 0.00879 | 0.513344 | 0.253 | 0.185 | 1 | 5 | Tnfaip1 | 0.068 | Old |
| 0.008799 | 0.15861 | 0.884 | 0.815 | 1 | 5 | Cfp | 0.069 | Old |
| 0.008872 | 0.169983 | 0.401 | 0.307 | 1 | 5 | Hs6st1 | 0.094 | Old |
| 0.008945 | 1.280774 | 0.054 | 0.02 | 1 | 5 | Cpeb1 | 0.034 | Old |
| 0.009281 | 0.533649 | 0.236 | 0.173 | 1 | 5 | Cds2 | 0.063 | Old |
| 0.009306 | 0.492563 | 0.138 | 0.084 | 1 | 5 | Idi1 | 0.054 | Old |
| 0.009407 | 0.114445 | 0.981 | 0.952 | 1 | 5 | Hsp90b1 | 0.029 | Old |
| 0.009426 | 0.260919 | 0.83 | 0.815 | 1 | 5 | Ier3 | 0.015 | Old |
| 0.009476 | 0.25451 | 0.406 | 0.325 | 1 | 5 | Maoa | 0.081 | Old |
| 0.009537 | 0.146156 | 0.694 | 0.579 | 1 | 5 | Plxnb2 | 0.115 | Old |
| 0.009569 | 0.2876 | 0.343 | 0.264 | 1 | 5 | Anapc5 | 0.079 | Old |
| 0.009576 | 0.19126 | 0.631 | 0.525 | 1 | 5 | Slamf9 | 0.106 | Old |
| 0.009615 | 0.42908 | 0.165 | 0.107 | 1 | 5 | Ctps2 | 0.058 | Old |
| 0.009693 | 2.233267 | 0.017 | 0 | 1 | 5 | Poli | 0.017 | Old |
| 0.009693 | 2.163116 | 0.017 | 0 | 1 | 5 | Lrrc29 | 0.017 | Old |
| 0.009693 | 2.003596 | 0.017 | 0 | 1 | 5 | Fzd1 | 0.017 | Old |
| 0.009697 | 0.441067 | 0.407 | 0.343 | 1 | 5 | Sptbn1 | 0.064 | Old |
| 0.009736 | 0.883482 | 0.271 | 0.206 | 1 | 5 | Gdf15 | 0.065 | Old |
| 0.009936 | 0.369649 | 0.333 | 0.254 | 1 | 5 | Sertad1 | 0.079 | Old |
| 0.009948 | 0.185686 | 0.542 | 0.439 | 1 | 5 | D8Ertd738 | 0.103 | Old |
| 0.009957 | 0.713259 | 0.15 | 0.096 | 1 | 5 | Kif3a | 0.054 | Old |
| 0.010048 | 0.297246 | 0.456 | 0.376 | 1 | 5 | Ilk | 0.08 | Old |
| 0.010134 | 0.658409 | 0.153 | 0.096 | 1 | 5 | Trp53inp2 | 0.057 | Old |
| 0.010156 | 0.458019 | 0.17 | 0.109 | 1 | 5 | Dpp3 | 0.061 | Old |
| 0.010197 | 0.341357 | 0.335 | 0.264 | 1 | 5 | Zcchc7 | 0.071 | Old |
| 0.010209 | 0.663535 | 0.17 | 0.112 | 1 | 5 | Celf4 | 0.058 | Old |
| 0.010273 | 0.217554 | 0.478 | 0.376 | 1 | 5 | Srsf1 | 0.102 | Old |
| 0.010285 | 0.549676 | 0.157 | 0.099 | 1 | 5 | Ints1 | 0.058 | Old |
| 0.010338 | 0.245044 | 0.579 | 0.5 | 1 | 5 | Dnajc3 | 0.079 | Old |
| 0.010374 | 0.496929 | 0.239 | 0.173 | 1 | 5 | Prdx4 | 0.066 | Old |
| 0.010396 | 0.016904 | 0.557 | 0.614 | 1 | 5 | Retnla | -0.057 | Old |
| 0.010468 | 0.289654 | 0.463 | 0.371 | 1 | 5 | Rasa4 | 0.092 | Old |
| 0.010614 | 0.372551 | 0.305 | 0.226 | 1 | 5 | Plxnc1 | 0.079 | Old |
| 0.010677 | 0.137585 | 0.444 | 0.348 | 1 | 5 | Snx29 | 0.096 | Old |
| 0.010793 | 0.242567 | 0.702 | 0.642 | 1 | 5 | Atf4 | 0.06 | Old |
| 0.011216 | 0.365099 | 0.717 | 0.675 | 1 | 5 | Ccl7 | 0.042 | Old |
| 0.011221 | 0.755187 | 0.138 | 0.086 | 1 | 5 | Ttc32 | 0.052 | Old |
| 0.011304 | 0.491388 | 0.138 | 0.084 | 1 | 5 | Nelfe | 0.054 | Old |
| 0.011466 | 0.343035 | 0.483 | 0.411 | 1 | 5 | Sh3pxd2b | 0.072 | Old |
| 0.011504 | 0.752388 | 0.131 | 0.081 | 1 | 5 | Hic2 | 0.05 | Old |
| 0.011542 | 0.467918 | 0.29 | 0.221 | 1 | 5 | Ang | 0.069 | Old |
| 0.011589 | 0.738213 | 0.094 | 0.051 | 1 | 5 | D16Ertd47 | 0.043 | Old |
| 0.011635 | 0.160582 | 0.714 | 0.629 | 1 | 5 | Blvrb | 0.085 | Old |
| 0.011638 | 0.617303 | 0.032 | 0.008 | 1 | 5 | Tedc2 | 0.024 | Old |

| | | | | | | | | |
|---|---|---|---|---|---|---|---|---|
| 0.011648 | 0.442082 | 0.247 | 0.18 | 1 | 5 | Ppp1r11 | 0.067 | Old |
| 0.011652 | 0.209635 | 0.603 | 0.508 | 1 | 5 | Nckap1l | 0.095 | Old |
| 0.011689 | 1.967573 | 0.022 | 0.003 | 1 | 5 | Eef1akmt4 | 0.019 | Old |
| 0.011692 | 0.661105 | 0.195 | 0.14 | 1 | 5 | Tbrg1 | 0.055 | Old |
| 0.011726 | 2.004059 | 0.022 | 0.003 | 1 | 5 | Smoc2 | 0.019 | Old |
| 0.011758 | 0.927023 | 0.081 | 0.041 | 1 | 5 | Decr2 | 0.04 | Old |
| 0.011799 | 0.478748 | 0.219 | 0.155 | 1 | 5 | Usp37 | 0.064 | Old |
| 0.011814 | 0.433461 | 0.286 | 0.218 | 1 | 5 | Tlr4 | 0.068 | Old |
| 0.011839 | 0.764549 | 0.108 | 0.061 | 1 | 5 | Mgp | 0.047 | Old |
| 0.011913 | 1.570448 | 0.027 | 0.005 | 1 | 5 | Gm17586 | 0.022 | Old |
| 0.011947 | 1.656202 | 0.027 | 0.005 | 1 | 5 | 1700109H( | 0.022 | Old |
| 0.011949 | 0.626623 | 0.16 | 0.104 | 1 | 5 | Necap1 | 0.056 | Old |
| 0.012045 | 0.387772 | 0.327 | 0.251 | 1 | 5 | Cyp27a1 | 0.076 | Old |
| 0.01209 | 0.235502 | 0.505 | 0.414 | 1 | 5 | Ankrd17 | 0.091 | Old |
| 0.012093 | 0.623659 | 0.146 | 0.094 | 1 | 5 | Grpel1 | 0.052 | Old |
| 0.012122 | 0.237853 | 0.62 | 0.548 | 1 | 5 | Colgalt1 | 0.072 | Old |
| 0.012141 | 0.495135 | 0.256 | 0.19 | 1 | 5 | Xdh | 0.066 | Old |
| 0.012216 | 1.110296 | 0.027 | 0.005 | 1 | 5 | Syde1 | 0.022 | Old |
| 0.012238 | 0.157397 | 0.854 | 0.835 | 1 | 5 | Fcrls | 0.019 | Old |
| 0.012325 | 0.358621 | 0.187 | 0.124 | 1 | 5 | Carhsp1 | 0.063 | Old |
| 0.012624 | 0.549255 | 0.184 | 0.124 | 1 | 5 | Agap3 | 0.06 | Old |
| 0.012853 | 0.127481 | 0.924 | 0.886 | 1 | 5 | Aplp2 | 0.038 | Old |
| 0.012912 | 0.464047 | 0.316 | 0.246 | 1 | 5 | Retreg1 | 0.07 | Old |
| 0.012964 | 0.307091 | 0.665 | 0.619 | 1 | 5 | P2rx4 | 0.046 | Old |
| 0.013273 | 0.427398 | 0.308 | 0.239 | 1 | 5 | Zrsr2 | 0.069 | Old |
| 0.013306 | 0.187182 | 0.747 | 0.647 | 1 | 5 | Emp3 | 0.1 | Old |
| 0.013338 | 0.721786 | 0.056 | 0.023 | 1 | 5 | Eif2ak4 | 0.033 | Old |
| 0.013388 | 0.74283 | 0.089 | 0.048 | 1 | 5 | Zfp687 | 0.041 | Old |
| 0.013501 | 1.183904 | 0.155 | 0.104 | 1 | 5 | Ptgs2 | 0.051 | Old |
| 0.013562 | 0.209334 | 0.441 | 0.368 | 1 | 5 | Srrm1 | 0.073 | Old |
| 0.013605 | 1.236842 | 0.044 | 0.015 | 1 | 5 | Oasl1 | 0.029 | Old |
| 0.013634 | 1.175791 | 0.044 | 0.015 | 1 | 5 | Pyroxd2 | 0.029 | Old |
| 0.013755 | 0.689329 | 0.128 | 0.079 | 1 | 5 | Arhgap10 | 0.049 | Old |
| 0.013811 | 0.803333 | 0.106 | 0.061 | 1 | 5 | Flnb | 0.045 | Old |
| 0.013837 | 0.170795 | 0.961 | 0.919 | 1 | 5 | Adgre1 | 0.042 | Old |
| 0.013886 | 0.523698 | 0.141 | 0.089 | 1 | 5 | Stxbp1 | 0.052 | Old |
| 0.01403 | 0.251138 | 0.576 | 0.49 | 1 | 5 | Sdf4 | 0.086 | Old |
| 0.014146 | 0.871304 | 0.099 | 0.056 | 1 | 5 | Arfgap3 | 0.043 | Old |
| 0.014185 | 1.976339 | 0.015 | 0 | 1 | 5 | Gm43614 | 0.015 | Old |
| 0.014185 | 1.946407 | 0.015 | 0 | 1 | 5 | Gm28187 | 0.015 | Old |
| 0.014185 | 1.886939 | 0.015 | 0 | 1 | 5 | Timp4 | 0.015 | Old |
| 0.014242 | 0.762615 | 0.035 | 0.01 | 1 | 5 | Gm15675 | 0.025 | Old |
| 0.014249 | 0.312912 | 0.33 | 0.261 | 1 | 5 | Srebf2 | 0.069 | Old |
| 0.014273 | 0.45128 | 0.246 | 0.183 | 1 | 5 | Arrdc4 | 0.063 | Old |
| 0.014274 | 0.258599 | 0.939 | 0.888 | 1 | 5 | Ccl6 | 0.051 | Old |

| | | | | | | | | |
|---|---|---|---|---|---|---|---|---|
| 0.014387 | 0.52659 | 0.178 | 0.122 | 1 | 5 | Itpk1 | 0.056 | Old |
| 0.014494 | 0.525226 | 0.175 | 0.119 | 1 | 5 | Diablo | 0.056 | Old |
| 0.014622 | 0.202721 | 0.921 | 0.898 | 1 | 5 | Rrbp1 | 0.023 | Old |
| 0.014718 | 0.295505 | 0.556 | 0.49 | 1 | 5 | Dnajb1 | 0.066 | Old |
| 0.01475 | 0.554401 | 0.19 | 0.132 | 1 | 5 | Gsdmd | 0.058 | Old |
| 0.014755 | 0.295719 | 0.439 | 0.365 | 1 | 5 | BC017643 | 0.074 | Old |
| 0.014772 | 0.37233 | 0.15 | 0.096 | 1 | 5 | Taf3 | 0.054 | Old |
| 0.014785 | 0.550844 | 0.178 | 0.124 | 1 | 5 | Adrm1 | 0.054 | Old |
| 0.015326 | 0.59012 | 0.113 | 0.066 | 1 | 5 | Pigz | 0.047 | Old |
| 0.015329 | 0.529623 | 0.158 | 0.107 | 1 | 5 | Tbc1d22a | 0.051 | Old |
| 0.015437 | 1.194001 | 0.054 | 0.023 | 1 | 5 | Dnajc27 | 0.031 | Old |
| 0.015476 | 1.530906 | 0.03 | 0.008 | 1 | 5 | Atrip | 0.022 | Old |
| 0.015607 | 0.286749 | 0.47 | 0.396 | 1 | 5 | Cpsf6 | 0.074 | Old |
| 0.015705 | 0.718244 | 0.173 | 0.122 | 1 | 5 | Tars | 0.051 | Old |
| 0.015707 | 0.501612 | 0.263 | 0.201 | 1 | 5 | Deptor | 0.062 | Old |
| 0.015755 | 0.32493 | 0.48 | 0.409 | 1 | 5 | Cdk2ap2 | 0.071 | Old |
| 0.015959 | 0.006233 | 0.88 | 0.794 | 1 | 5 | Hsp90aa1 | 0.086 | Old |
| 0.015971 | 0.150288 | 0.776 | 0.652 | 1 | 5 | Plbd1 | 0.124 | Old |
| 0.016026 | 0.445417 | 0.263 | 0.201 | 1 | 5 | Tbc1d9b | 0.062 | Old |
| 0.016028 | 1.465318 | 0.03 | 0.008 | 1 | 5 | Zfp37 | 0.022 | Old |
| 0.016077 | 0.827301 | 0.145 | 0.096 | 1 | 5 | Vps39 | 0.049 | Old |
| 0.016388 | 0.986193 | 0.057 | 0.025 | 1 | 5 | Etl4 | 0.032 | Old |
| 0.016405 | 0.283538 | 0.446 | 0.36 | 1 | 5 | Rilpl2 | 0.086 | Old |
| 0.016415 | 1.163508 | 0.054 | 0.023 | 1 | 5 | Fads6 | 0.031 | Old |
| 0.016427 | 1.782209 | 0.025 | 0.005 | 1 | 5 | Tsacc | 0.02 | Old |
| 0.016496 | 0.189726 | 0.781 | 0.734 | 1 | 5 | Clec12a | 0.047 | Old |
| 0.016659 | 0.193551 | 0.438 | 0.355 | 1 | 5 | Cdk6 | 0.083 | Old |
| 0.016687 | 0.327146 | 0.332 | 0.256 | 1 | 5 | Lars2 | 0.076 | Old |
| 0.016763 | 0.183136 | 0.652 | 0.594 | 1 | 5 | Stat3 | 0.058 | Old |
| 0.016809 | 2.069698 | 0.02 | 0.003 | 1 | 5 | Gm26826 | 0.017 | Old |
| 0.016848 | 0.443473 | 0.291 | 0.226 | 1 | 5 | Vrk2 | 0.065 | Old |
| 0.016862 | 1.829241 | 0.02 | 0.003 | 1 | 5 | Acy3 | 0.017 | Old |
| 0.016941 | 0.997182 | 0.064 | 0.03 | 1 | 5 | Polr3d | 0.034 | Old |
| 0.017008 | 0.332182 | 0.316 | 0.249 | 1 | 5 | Asap1 | 0.067 | Old |
| 0.017022 | 0.239009 | 0.515 | 0.431 | 1 | 5 | Pik3cd | 0.084 | Old |
| 0.017047 | 0.732421 | 0.14 | 0.091 | 1 | 5 | Tufm | 0.049 | Old |
| 0.017048 | 0.526814 | 0.093 | 0.051 | 1 | 5 | Ube2e2 | 0.042 | Old |
| 0.017073 | 1.508595 | 0.025 | 0.005 | 1 | 5 | Cdk5rap1 | 0.02 | Old |
| 0.017084 | 0.17385 | 0.609 | 0.538 | 1 | 5 | Sec61g | 0.071 | Old |
| 0.0171 | 0.262666 | 0.429 | 0.358 | 1 | 5 | Tmsb10 | 0.071 | Old |
| 0.017514 | 0.202579 | 0.36 | 0.287 | 1 | 5 | Nceh1 | 0.073 | Old |
| 0.017557 | 0.248098 | 0.409 | 0.332 | 1 | 5 | 7-Mar | 0.077 | Old |
| 0.01767 | 0.066203 | 1 | 0.995 | 1 | 5 | B2m | 0.005 | Old |
| 0.017687 | 0.221074 | 0.52 | 0.439 | 1 | 5 | Ost4 | 0.081 | Old |
| 0.017706 | 0.408456 | 0.256 | 0.193 | 1 | 5 | Clip1 | 0.063 | Old |

| | | | | | | | | |
|---|---|---|---|---|---|---|---|---|
| 0.01771 | 0.47075 | 0.17 | 0.114 | 1 | 5 | Cdk16 | 0.056 | Old |
| 0.017788 | 1.303538 | 0.042 | 0.015 | 1 | 5 | Sncg | 0.027 | Old |
| 0.017806 | 0.717426 | 0.125 | 0.079 | 1 | 5 | Vps45 | 0.046 | Old |
| 0.018097 | 0.535284 | 0.121 | 0.074 | 1 | 5 | Zfp398 | 0.047 | Old |
| 0.018139 | 0.67236 | 0.123 | 0.076 | 1 | 5 | Senp3 | 0.047 | Old |
| 0.018169 | 0.464675 | 0.217 | 0.157 | 1 | 5 | Abcb1b | 0.06 | Old |
| 0.018297 | 0.502333 | 0.184 | 0.127 | 1 | 5 | Opa3 | 0.057 | Old |
| 0.018316 | 0.92762 | 0.081 | 0.043 | 1 | 5 | Mrpl40 | 0.038 | Old |
| 0.018328 | 0.29473 | 0.549 | 0.475 | 1 | 5 | Map1lc3b | 0.074 | Old |
| 0.018407 | 0.273083 | 0.411 | 0.33 | 1 | 5 | Azin1 | 0.081 | Old |
| 0.018501 | 0.356158 | 0.29 | 0.218 | 1 | 5 | Nrd1 | 0.072 | Old |
| 0.018639 | 0.713895 | 0.042 | 0.015 | 1 | 5 | Ndufaf1 | 0.027 | Old |
| 0.018819 | 0.731588 | 0.094 | 0.053 | 1 | 5 | Mrps30 | 0.041 | Old |
| 0.018871 | 1.031623 | 0.067 | 0.033 | 1 | 5 | Osbpl7 | 0.034 | Old |
| 0.018879 | 0.284016 | 0.833 | 0.807 | 1 | 5 | Ccl2 | 0.026 | Old |
| 0.018898 | 0.88979 | 0.106 | 0.063 | 1 | 5 | St3gal3 | 0.043 | Old |
| 0.019053 | 0.16053 | 0.788 | 0.716 | 1 | 5 | Tnfaip3 | 0.072 | Old |
| 0.019319 | 0.543531 | 0.256 | 0.195 | 1 | 5 | Cdr2 | 0.061 | Old |
| 0.019366 | 0.188555 | 0.579 | 0.482 | 1 | 5 | Prr13 | 0.097 | Old |
| 0.019608 | 1.381539 | 0.052 | 0.023 | 1 | 5 | Epas1 | 0.029 | Old |
| 0.019665 | 0.270464 | 0.236 | 0.173 | 1 | 5 | Psmd3 | 0.063 | Old |
| 0.019712 | 1.37037 | 0.049 | 0.02 | 1 | 5 | Gpx7 | 0.029 | Old |
| 0.019731 | 0.451361 | 0.123 | 0.076 | 1 | 5 | Donson | 0.047 | Old |
| 0.019773 | 0.269144 | 0.337 | 0.264 | 1 | 5 | H2-Q4 | 0.073 | Old |
| 0.019963 | 0.462914 | 0.187 | 0.129 | 1 | 5 | Rnf10 | 0.058 | Old |
| 0.019995 | 0.24354 | 0.544 | 0.475 | 1 | 5 | Hsd17b11 | 0.069 | Old |
| 0.02033 | 0.293002 | 0.443 | 0.371 | 1 | 5 | Mvp | 0.072 | Old |
| 0.020507 | 0.39495 | 0.205 | 0.147 | 1 | 5 | Irak2 | 0.058 | Old |
| 0.020642 | 0.923914 | 0.045 | 0.018 | 1 | 5 | Mkks | 0.027 | Old |
| 0.020645 | 0.678224 | 0.143 | 0.096 | 1 | 5 | Lonp2 | 0.047 | Old |
| 0.020832 | 2.133956 | 0.013 | 0 | 1 | 5 | Prom2 | 0.013 | Old |
| 0.020832 | 1.702175 | 0.013 | 0 | 1 | 5 | 2310047D( | 0.013 | Old |
| 0.020832 | 1.684752 | 0.013 | 0 | 1 | 5 | 1700030J2 | 0.013 | Old |
| 0.020832 | 1.570695 | 0.013 | 0 | 1 | 5 | Rnf144a | 0.013 | Old |
| 0.021166 | 0.236923 | 0.67 | 0.647 | 1 | 5 | Lipa | 0.023 | Old |
| 0.021307 | 0.585939 | 0.145 | 0.094 | 1 | 5 | Arhgap4 | 0.051 | Old |
| 0.021331 | 0.288179 | 0.35 | 0.279 | 1 | 5 | Slc23a2 | 0.071 | Old |
| 0.021397 | 0.644742 | 0.14 | 0.091 | 1 | 5 | Snx14 | 0.049 | Old |
| 0.021488 | 0.292962 | 0.343 | 0.277 | 1 | 5 | Madd | 0.066 | Old |
| 0.021532 | 0.718585 | 0.116 | 0.074 | 1 | 5 | Scly | 0.042 | Old |
| 0.021606 | 0.699071 | 0.091 | 0.051 | 1 | 5 | Lig1 | 0.04 | Old |
| 0.021621 | 0.327039 | 0.15 | 0.096 | 1 | 5 | Kifap3 | 0.054 | Old |
| 0.021746 | 0.714199 | 0.106 | 0.063 | 1 | 5 | Adcy3 | 0.043 | Old |
| 0.022086 | 0.48482 | 0.244 | 0.185 | 1 | 5 | Relb | 0.059 | Old |
| 0.022105 | 0.396006 | 0.342 | 0.272 | 1 | 5 | Dnase1l3 | 0.07 | Old |

| | | | | | | | | |
|---|---|---|---|---|---|---|---|---|
| 0.022137 | 0.18691 | 0.806 | 0.739 | 1 | 5 | Nfkbid | 0.067 | Old |
| 0.022943 | 0.982646 | 0.086 | 0.048 | 1 | 5 | Zfp182 | 0.038 | Old |
| 0.023001 | 0.453048 | 0.133 | 0.086 | 1 | 5 | Slc25a25 | 0.047 | Old |
| 0.023039 | 0.168498 | 0.645 | 0.543 | 1 | 5 | Lmo2 | 0.102 | Old |
| 0.023158 | 0.245328 | 0.623 | 0.52 | 1 | 5 | Sf1 | 0.103 | Old |
| 0.02325 | 1.365214 | 0.04 | 0.015 | 1 | 5 | Gm31243 | 0.025 | Old |
| 0.023767 | 1.031595 | 0.062 | 0.03 | 1 | 5 | Cd200r4 | 0.032 | Old |
| 0.023988 | 1.726887 | 0.024 | 0.005 | 1 | 5 | Olfr1033 | 0.019 | Old |
| 0.024078 | 0.547949 | 0.152 | 0.102 | 1 | 5 | Taf7 | 0.05 | Old |
| 0.024383 | 0.169948 | 0.891 | 0.812 | 1 | 5 | Tagln2 | 0.079 | Old |
| 0.024383 | 0.333952 | 0.286 | 0.228 | 1 | 5 | Cpeb4 | 0.058 | Old |
| 0.024436 | 1.740243 | 0.019 | 0.003 | 1 | 5 | Lrrc32 | 0.016 | Old |
| 0.024448 | 0.31344 | 0.379 | 0.312 | 1 | 5 | Impact | 0.067 | Old |
| 0.024505 | 0.25447 | 0.357 | 0.287 | 1 | 5 | Errfi1 | 0.07 | Old |
| 0.024512 | 1.627593 | 0.019 | 0.003 | 1 | 5 | 1810055G( | 0.016 | Old |
| 0.024742 | 1.685112 | 0.019 | 0.003 | 1 | 5 | Tgfbr3 | 0.016 | Old |
| 0.024807 | 0.271692 | 0.162 | 0.109 | 1 | 5 | Fnip2 | 0.053 | Old |
| 0.02513 | 0.760098 | 0.089 | 0.051 | 1 | 5 | Ccdc106 | 0.038 | Old |
| 0.025208 | 1.478942 | 0.019 | 0.003 | 1 | 5 | Art2b | 0.016 | Old |
| 0.025208 | 1.390265 | 0.019 | 0.003 | 1 | 5 | Palm | 0.016 | Old |
| 0.025235 | 1.355621 | 0.057 | 0.028 | 1 | 5 | Plxna3 | 0.029 | Old |
| 0.025295 | 0.550823 | 0.093 | 0.053 | 1 | 5 | Wdyhv1 | 0.04 | Old |
| 0.025428 | 0.815866 | 0.103 | 0.063 | 1 | 5 | Pskh1 | 0.04 | Old |
| 0.02553 | 0.374372 | 0.162 | 0.109 | 1 | 5 | Trappc3 | 0.053 | Old |
| 0.025683 | 0.398365 | 0.458 | 0.404 | 1 | 5 | Lfng | 0.054 | Old |
| 0.025957 | 0.395679 | 0.263 | 0.201 | 1 | 5 | Ifitm6 | 0.062 | Old |
| 0.025974 | 0.439248 | 0.244 | 0.185 | 1 | 5 | Dnajb4 | 0.059 | Old |
| 0.025997 | 0.472413 | 0.118 | 0.074 | 1 | 5 | Ocel1 | 0.044 | Old |
| 0.026094 | 0.972991 | 0.044 | 0.018 | 1 | 5 | Tle2 | 0.026 | Old |
| 0.026096 | 0.390847 | 0.2 | 0.142 | 1 | 5 | Zcchc14 | 0.058 | Old |
| 0.026124 | 0.707169 | 0.111 | 0.069 | 1 | 5 | Arhgef5 | 0.042 | Old |
| 0.026153 | 0.642913 | 0.079 | 0.043 | 1 | 5 | Ltbp3 | 0.036 | Old |
| 0.026358 | 0.637448 | 0.126 | 0.081 | 1 | 5 | Mtif2 | 0.045 | Old |
| 0.026506 | 0.181543 | 0.374 | 0.294 | 1 | 5 | Hadhb | 0.08 | Old |
| 0.026546 | 0.132608 | 0.663 | 0.584 | 1 | 5 | Etv3 | 0.079 | Old |
| 0.026637 | 0.652698 | 0.101 | 0.061 | 1 | 5 | Actn1 | 0.04 | Old |
| 0.026637 | 0.536524 | 0.101 | 0.061 | 1 | 5 | Cd209e | 0.04 | Old |
| 0.026728 | 0.517156 | 0.146 | 0.099 | 1 | 5 | Zswim4 | 0.047 | Old |
| 0.026782 | 0.321389 | 0.37 | 0.302 | 1 | 5 | Tnrc6c | 0.068 | Old |
| 0.026955 | 0.403977 | 0.21 | 0.155 | 1 | 5 | Lrrfip1 | 0.055 | Old |
| 0.027042 | 0.32805 | 0.342 | 0.269 | 1 | 5 | Dusp16 | 0.073 | Old |
| 0.027046 | 0.278785 | 0.594 | 0.53 | 1 | 5 | Casp4 | 0.064 | Old |
| 0.027323 | 0.192923 | 0.901 | 0.863 | 1 | 5 | Cd14 | 0.038 | Old |
| 0.027345 | 0.315183 | 0.65 | 0.604 | 1 | 5 | Il1b | 0.046 | Old |
| 0.027387 | 0.249543 | 0.512 | 0.429 | 1 | 5 | Spcs2 | 0.083 | Old |

| | | | | | | | | |
|---|---|---|---|---|---|---|---|---|
| 0.027612 | 0.40698 | 0.226 | 0.17 | 1 | 5 | Kmo | 0.056 | Old |
| 0.027748 | 1.379876 | 0.077 | 0.043 | 1 | 5 | Bambi | 0.034 | Old |
| 0.027766 | 0.624572 | 0.136 | 0.091 | 1 | 5 | Fgd3 | 0.045 | Old |
| 0.027884 | 0.412574 | 0.207 | 0.152 | 1 | 5 | Impdh2 | 0.055 | Old |
| 0.027953 | 0.314698 | 0.172 | 0.119 | 1 | 5 | Flvcr1 | 0.053 | Old |
| 0.028077 | 0.150464 | 0.667 | 0.604 | 1 | 5 | Csrnp1 | 0.063 | Old |
| 0.028108 | 0.388444 | 0.244 | 0.188 | 1 | 5 | Atf6b | 0.056 | Old |
| 0.028346 | 0.098924 | 0.904 | 0.825 | 1 | 5 | Clec4a1 | 0.079 | Old |
| 0.02846 | 0.924376 | 0.047 | 0.02 | 1 | 5 | Zmynd19 | 0.027 | Old |
| 0.028466 | 0.419453 | 0.286 | 0.234 | 1 | 5 | Yipf3 | 0.052 | Old |
| 0.028773 | 0.858537 | 0.12 | 0.079 | 1 | 5 | St8sia6 | 0.041 | Old |
| 0.028802 | 0.815221 | 0.081 | 0.046 | 1 | 5 | Lclat1 | 0.035 | Old |
| 0.028875 | 0.419434 | 0.184 | 0.132 | 1 | 5 | Mrps12 | 0.052 | Old |
| 0.028897 | 0.189289 | 0.48 | 0.414 | 1 | 5 | Abracl | 0.066 | Old |
| 0.029 | 0.878226 | 0.131 | 0.086 | 1 | 5 | Cxcl10 | 0.045 | Old |
| 0.029118 | 0.537272 | 0.091 | 0.053 | 1 | 5 | Leng9 | 0.038 | Old |
| 0.029147 | 0.359596 | 0.311 | 0.251 | 1 | 5 | Oxct1 | 0.06 | Old |
| 0.029216 | 0.334108 | 0.222 | 0.165 | 1 | 5 | Mau2 | 0.057 | Old |
| 0.029288 | 0.812293 | 0.126 | 0.084 | 1 | 5 | Tmem37 | 0.042 | Old |
| 0.029439 | 0.748244 | 0.047 | 0.02 | 1 | 5 | Papss2 | 0.027 | Old |
| 0.029447 | 0.763139 | 0.077 | 0.043 | 1 | 5 | Scap | 0.034 | Old |
| 0.029525 | 0.197302 | 0.753 | 0.718 | 1 | 5 | Laptm4a | 0.035 | Old |
| 0.0296 | 0.176646 | 0.722 | 0.655 | 1 | 5 | Jpt1 | 0.067 | Old |
| 0.029655 | 0.396086 | 0.202 | 0.147 | 1 | 5 | Ube4a | 0.055 | Old |
| 0.02975 | 0.427574 | 0.18 | 0.129 | 1 | 5 | Slc39a9 | 0.051 | Old |
| 0.029838 | 0.236126 | 0.224 | 0.162 | 1 | 5 | Bcat2 | 0.062 | Old |
| 0.029922 | 1.179319 | 0.035 | 0.013 | 1 | 5 | Plpp3 | 0.022 | Old |
| 0.02995 | 0.792189 | 0.054 | 0.025 | 1 | 5 | Tnfsf13os | 0.029 | Old |
| 0.030048 | 0.912041 | 0.054 | 0.025 | 1 | 5 | Vill | 0.029 | Old |
| 0.030126 | 1.687709 | 0.027 | 0.008 | 1 | 5 | Dph7 | 0.019 | Old |
| 0.030171 | 0.270597 | 0.382 | 0.307 | 1 | 5 | Oat | 0.075 | Old |
| 0.030175 | 0.34822 | 0.258 | 0.195 | 1 | 5 | Dyrk1a | 0.063 | Old |
| 0.030204 | 0.197897 | 0.628 | 0.553 | 1 | 5 | St3gal6 | 0.075 | Old |
| 0.030301 | 1.059016 | 0.064 | 0.033 | 1 | 5 | Slc10a6 | 0.031 | Old |
| 0.030485 | 0.96008 | 0.035 | 0.013 | 1 | 5 | 1110019D1 | 0.022 | Old |
| 0.030582 | 0.118172 | 0.463 | 0.378 | 1 | 5 | Hip1 | 0.085 | Old |
| 0.030587 | 0.236329 | 0.303 | 0.239 | 1 | 5 | Sun2 | 0.064 | Old |
| 0.030729 | 2.138862 | 0.012 | 0 | 1 | 5 | Gm10603 | 0.012 | Old |
| 0.030729 | 1.906973 | 0.012 | 0 | 1 | 5 | Rpusd2 | 0.012 | Old |
| 0.030729 | 1.770075 | 0.012 | 0 | 1 | 5 | Gm26536 | 0.012 | Old |
| 0.030729 | 1.740982 | 0.012 | 0 | 1 | 5 | Pou5f1 | 0.012 | Old |
| 0.030729 | 1.549712 | 0.012 | 0 | 1 | 5 | Tcf15 | 0.012 | Old |
| 0.030729 | 1.548078 | 0.012 | 0 | 1 | 5 | Adm | 0.012 | Old |
| 0.030729 | 1.494365 | 0.012 | 0 | 1 | 5 | Gm10053 | 0.012 | Old |
| 0.030729 | 1.488282 | 0.012 | 0 | 1 | 5 | A630001O1 | 0.012 | Old |

| | | | | | | | | |
|---|---|---|---|---|---|---|---|---|
| 0.030729 | 1.450243 | 0.012 | 0 | 1 | 5 | Proca1 | 0.012 | Old |
| 0.030743 | 0.299193 | 0.375 | 0.315 | 1 | 5 | Rab18 | 0.06 | Old |
| 0.030754 | 0.283314 | 0.254 | 0.195 | 1 | 5 | Oas2 | 0.059 | Old |
| 0.030861 | 1.195561 | 0.027 | 0.008 | 1 | 5 | Cxcl9 | 0.019 | Old |
| 0.030881 | 0.273644 | 0.816 | 0.792 | 1 | 5 | Atf3 | 0.024 | Old |
| 0.030934 | 0.215337 | 0.609 | 0.523 | 1 | 5 | Irf1 | 0.086 | Old |
| 0.030955 | 0.601282 | 0.114 | 0.074 | 1 | 5 | Ryk | 0.04 | Old |
| 0.031073 | 0.725234 | 0.103 | 0.063 | 1 | 5 | Zfp771 | 0.04 | Old |
| 0.031122 | 0.355719 | 0.36 | 0.297 | 1 | 5 | Calu | 0.063 | Old |
| 0.031153 | 0.418269 | 0.155 | 0.107 | 1 | 5 | Hjurp | 0.048 | Old |
| 0.0312 | 0.290389 | 0.303 | 0.246 | 1 | 5 | Rnf4 | 0.057 | Old |
| 0.03148 | 0.178262 | 0.616 | 0.558 | 1 | 5 | Sec62 | 0.058 | Old |
| 0.031514 | 0.810165 | 0.071 | 0.038 | 1 | 5 | Pank4 | 0.033 | Old |
| 0.031539 | 0.368057 | 0.305 | 0.244 | 1 | 5 | Ahsa1 | 0.061 | Old |
| 0.031733 | 0.101104 | 0.37 | 0.305 | 1 | 5 | Jup | 0.065 | Old |
| 0.032077 | 0.307032 | 0.337 | 0.272 | 1 | 5 | Chic2 | 0.065 | Old |
| 0.032182 | 0.082112 | 0.382 | 0.299 | 1 | 5 | Dnajb9 | 0.083 | Old |
| 0.032247 | 0.57649 | 0.061 | 0.03 | 1 | 5 | Gcdh | 0.031 | Old |
| 0.032302 | 0.14314 | 0.877 | 0.825 | 1 | 5 | Ube2d3 | 0.052 | Old |
| 0.032607 | 0.217793 | 0.48 | 0.411 | 1 | 5 | Glmp | 0.069 | Old |
| 0.032644 | 0.734879 | 0.071 | 0.038 | 1 | 5 | Zfp938 | 0.033 | Old |
| 0.032661 | 0.040807 | 1 | 1 | 1 | 5 | mt-Atp6 | 0 | Old |
| 0.032857 | 0.728262 | 0.099 | 0.061 | 1 | 5 | Foxo4 | 0.038 | Old |
| 0.032959 | 0.191968 | 0.731 | 0.678 | 1 | 5 | Cd93 | 0.053 | Old |
| 0.033054 | 0.349044 | 0.274 | 0.211 | 1 | 5 | Por | 0.063 | Old |
| 0.033145 | 0.456333 | 0.168 | 0.119 | 1 | 5 | Cyp4f13 | 0.049 | Old |
| 0.033556 | 0.141993 | 0.596 | 0.515 | 1 | 5 | Epn1 | 0.081 | Old |
| 0.033577 | 0.838999 | 0.067 | 0.036 | 1 | 5 | D17H6S53I | 0.031 | Old |
| 0.033731 | 0.089033 | 0.357 | 0.277 | 1 | 5 | Anapc11 | 0.08 | Old |
| 0.034104 | 0.595239 | 0.114 | 0.074 | 1 | 5 | Tada3 | 0.04 | Old |
| 0.0346 | 1.349112 | 0.022 | 0.005 | 1 | 5 | Mansc1 | 0.017 | Old |
| 0.034624 | 0.173459 | 0.756 | 0.728 | 1 | 5 | Trps1 | 0.028 | Old |
| 0.034784 | 1.343516 | 0.022 | 0.005 | 1 | 5 | Dph2 | 0.017 | Old |
| 0.034912 | 0.528377 | 0.209 | 0.155 | 1 | 5 | Lrrc8c | 0.054 | Old |
| 0.03497 | 1.105181 | 0.022 | 0.005 | 1 | 5 | Icam2 | 0.017 | Old |
| 0.034972 | 0.287494 | 0.424 | 0.371 | 1 | 5 | Siglece | 0.053 | Old |
| 0.034983 | 0.261217 | 0.418 | 0.35 | 1 | 5 | Nop53 | 0.068 | Old |
| 0.035011 | 1.194416 | 0.03 | 0.01 | 1 | 5 | Hmga1 | 0.02 | Old |
| 0.035043 | 0.137652 | 0.82 | 0.792 | 1 | 5 | Rock2 | 0.028 | Old |
| 0.035066 | 0.510764 | 0.386 | 0.327 | 1 | 5 | Lgals3 | 0.059 | Old |
| 0.035337 | 0.544212 | 0.14 | 0.096 | 1 | 5 | Bad | 0.044 | Old |
| 0.035502 | 0.516838 | 0.064 | 0.033 | 1 | 5 | Fut4 | 0.031 | Old |
| 0.035585 | 1.847457 | 0.017 | 0.003 | 1 | 5 | Igha | 0.014 | Old |
| 0.035791 | 0.244903 | 0.268 | 0.203 | 1 | 5 | B4galt5 | 0.065 | Old |
| 0.035813 | 0.491249 | 0.099 | 0.061 | 1 | 5 | Snx11 | 0.038 | Old |

| | | | | | | | | |
|---|---|---|---|---|---|---|---|---|
| 0.035892 | 0.672819 | 0.042 | 0.018 | 1 | 5 | Kcnj2 | 0.024 | Old |
| 0.035916 | 1.595363 | 0.017 | 0.003 | 1 | 5 | Zbtb7c | 0.014 | Old |
| 0.036 | 0.267108 | 0.226 | 0.168 | 1 | 5 | Mgst1 | 0.058 | Old |
| 0.036053 | 0.568749 | 0.093 | 0.056 | 1 | 5 | Mfn1 | 0.037 | Old |
| 0.036072 | 0.228726 | 0.581 | 0.513 | 1 | 5 | Pura | 0.068 | Old |
| 0.036246 | 0.182711 | 0.71 | 0.637 | 1 | 5 | Rgl1 | 0.073 | Old |
| 0.036473 | 1.393507 | 0.017 | 0.003 | 1 | 5 | Gm7879 | 0.014 | Old |
| 0.036473 | 1.260781 | 0.017 | 0.003 | 1 | 5 | Hsf4 | 0.014 | Old |
| 0.036473 | 1.251353 | 0.017 | 0.003 | 1 | 5 | Ndor1 | 0.014 | Old |
| 0.036746 | 1.045703 | 0.045 | 0.02 | 1 | 5 | Kti12 | 0.025 | Old |
| 0.03677 | 0.2161 | 0.471 | 0.406 | 1 | 5 | Tmbim1 | 0.065 | Old |
| 0.036823 | 0.677707 | 0.13 | 0.089 | 1 | 5 | Klhl20 | 0.041 | Old |
| 0.036909 | 0.232861 | 0.295 | 0.231 | 1 | 5 | Plod3 | 0.064 | Old |
| 0.037354 | 0.643544 | 0.104 | 0.066 | 1 | 5 | Hagh | 0.038 | Old |
| 0.037513 | 0.384329 | 0.221 | 0.17 | 1 | 5 | Pilrb1 | 0.051 | Old |
| 0.037796 | 0.782898 | 0.114 | 0.076 | 1 | 5 | Dap3 | 0.038 | Old |
| 0.037918 | 0.118028 | 0.774 | 0.67 | 1 | 5 | Pdia6 | 0.104 | Old |
| 0.037952 | 0.538016 | 0.189 | 0.14 | 1 | 5 | Krt80 | 0.049 | Old |
| 0.038072 | 0.173746 | 0.672 | 0.604 | 1 | 5 | Rabac1 | 0.068 | Old |
| 0.038279 | 0.225107 | 0.313 | 0.246 | 1 | 5 | Clptm1 | 0.067 | Old |
| 0.038311 | 0.582274 | 0.189 | 0.142 | 1 | 5 | Pspc1 | 0.047 | Old |
| 0.038397 | 0.529672 | 0.12 | 0.079 | 1 | 5 | Ttc19 | 0.041 | Old |
| 0.038669 | 0.200153 | 0.722 | 0.657 | 1 | 5 | Mapkapk2 | 0.065 | Old |
| 0.0387 | 0.440397 | 0.202 | 0.15 | 1 | 5 | Siah2 | 0.052 | Old |
| 0.038939 | 0.725937 | 0.079 | 0.046 | 1 | 5 | Plekhg1 | 0.033 | Old |
| 0.038978 | 1.293188 | 0.034 | 0.013 | 1 | 5 | Zfr2 | 0.021 | Old |
| 0.038982 | 0.41913 | 0.285 | 0.231 | 1 | 5 | Tlnrd1 | 0.054 | Old |
| 0.039037 | 0.296142 | 0.298 | 0.241 | 1 | 5 | Glb1 | 0.057 | Old |
| 0.039128 | 0.509343 | 0.099 | 0.061 | 1 | 5 | Neurl1a | 0.038 | Old |
| 0.039289 | 0.185887 | 0.759 | 0.698 | 1 | 5 | Apobec3 | 0.061 | Old |
| 0.039405 | 0.15674 | 0.588 | 0.508 | 1 | 5 | Man1c1 | 0.08 | Old |
| 0.039416 | 0.829305 | 0.099 | 0.063 | 1 | 5 | Etfdh | 0.036 | Old |
| 0.039511 | 0.811724 | 0.066 | 0.036 | 1 | 5 | Zc2hc1a | 0.03 | Old |
| 0.039603 | 0.23503 | 0.367 | 0.302 | 1 | 5 | Tcea1 | 0.065 | Old |
| 0.0397 | 0.287393 | 0.424 | 0.365 | 1 | 5 | Bcl2a1b | 0.059 | Old |
| 0.040193 | 0.577344 | 0.069 | 0.038 | 1 | 5 | Gstz1 | 0.031 | Old |
| 0.040206 | 0.751524 | 0.052 | 0.025 | 1 | 5 | Cryz | 0.027 | Old |
| 0.040338 | 0.029846 | 0.997 | 0.99 | 1 | 5 | H2-K1 | 0.007 | Old |
| 0.04053 | 0.352906 | 0.167 | 0.119 | 1 | 5 | Naxe | 0.048 | Old |
| 0.040625 | 1.789358 | 0.025 | 0.008 | 1 | 5 | Adam8 | 0.017 | Old |
| 0.040668 | 0.863532 | 0.056 | 0.028 | 1 | 5 | Barx2 | 0.028 | Old |
| 0.040757 | 0.76691 | 0.066 | 0.036 | 1 | 5 | Fbxl15 | 0.03 | Old |
| 0.040784 | 0.425832 | 0.125 | 0.084 | 1 | 5 | Sc5d | 0.041 | Old |
| 0.040795 | 0.553756 | 0.158 | 0.112 | 1 | 5 | Vta1 | 0.046 | Old |
| 0.040919 | 0.705302 | 0.056 | 0.028 | 1 | 5 | Cdc42bpa | 0.028 | Old |

| | | | | | | | | |
|---|---|---|---|---|---|---|---|---|
| 0.041085 | 0.571006 | 0.076 | 0.043 | 1 | 5 | Tmem35b | 0.033 | Old |
| 0.041136 | 0.218503 | 0.598 | 0.541 | 1 | 5 | Metrnl | 0.057 | Old |
| 0.041173 | 0.292624 | 0.199 | 0.147 | 1 | 5 | Ccdc59 | 0.052 | Old |
| 0.041427 | 1.24693 | 0.037 | 0.015 | 1 | 5 | Zfp667 | 0.022 | Old |
| 0.041572 | 0.224421 | 0.32 | 0.251 | 1 | 5 | Txnrd1 | 0.069 | Old |
| 0.041617 | 0.22445 | 0.549 | 0.48 | 1 | 5 | Srsf6 | 0.069 | Old |
| 0.041999 | 0.689288 | 0.084 | 0.051 | 1 | 5 | Prkar1b | 0.033 | Old |
| 0.042004 | 0.269508 | 0.237 | 0.18 | 1 | 5 | Abce1 | 0.057 | Old |
| 0.042093 | 1.321194 | 0.025 | 0.008 | 1 | 5 | Ptpn13 | 0.017 | Old |
| 0.042292 | 1.192453 | 0.025 | 0.008 | 1 | 5 | Hells | 0.017 | Old |
| 0.042394 | 0.900433 | 0.093 | 0.058 | 1 | 5 | Sorl1 | 0.035 | Old |
| 0.042438 | 0.538289 | 0.15 | 0.107 | 1 | 5 | Msmo1 | 0.043 | Old |
| 0.04244 | 0.162189 | 0.426 | 0.348 | 1 | 5 | Fkbp2 | 0.078 | Old |
| 0.042523 | 0.400006 | 0.204 | 0.157 | 1 | 5 | Pfdn1 | 0.047 | Old |
| 0.042532 | 0.512319 | 0.079 | 0.046 | 1 | 5 | Casp2 | 0.033 | Old |
| 0.042753 | 0.291232 | 0.421 | 0.353 | 1 | 5 | Yy1 | 0.068 | Old |
| 0.04295 | 0.585256 | 0.155 | 0.112 | 1 | 5 | Cdk2 | 0.043 | Old |
| 0.043097 | 1.14548 | 0.025 | 0.008 | 1 | 5 | Zfp202 | 0.017 | Old |
| 0.043199 | 0.45768 | 0.202 | 0.152 | 1 | 5 | Smarcb1 | 0.05 | Old |
| 0.043592 | 0.141644 | 0.598 | 0.523 | 1 | 5 | Axl | 0.075 | Old |
| 0.043616 | 0.271245 | 0.328 | 0.266 | 1 | 5 | P3h2 | 0.062 | Old |
| 0.043856 | 0.913905 | 0.037 | 0.015 | 1 | 5 | Gm29107 | 0.022 | Old |
| 0.044075 | 0.430354 | 0.192 | 0.14 | 1 | 5 | Fgd6 | 0.052 | Old |
| 0.044203 | 0.234979 | 0.426 | 0.365 | 1 | 5 | Os9 | 0.061 | Old |
| 0.044757 | 0.119657 | 0.551 | 0.472 | 1 | 5 | Adap2 | 0.079 | Old |
| 0.044863 | 0.961322 | 0.067 | 0.038 | 1 | 5 | Smug1 | 0.029 | Old |
| 0.045051 | 0.730823 | 0.116 | 0.079 | 1 | 5 | Cyp4v3 | 0.037 | Old |
| 0.045077 | 0.40321 | 0.276 | 0.228 | 1 | 5 | Wwp2 | 0.048 | Old |
| 0.045099 | 0.48967 | 0.242 | 0.195 | 1 | 5 | Palld | 0.047 | Old |
| 0.045192 | 0.579626 | 0.099 | 0.063 | 1 | 5 | Ppp2r5d | 0.036 | Old |
| 0.045274 | 0.166412 | 0.547 | 0.477 | 1 | 5 | Ywhag | 0.07 | Old |
| 0.045302 | 0.426349 | 0.229 | 0.178 | 1 | 5 | Oaz2 | 0.051 | Old |
| 0.045385 | 0.305874 | 0.241 | 0.188 | 1 | 5 | Jmjd6 | 0.053 | Old |
| 0.045449 | 0.485428 | 0.152 | 0.109 | 1 | 5 | Atn1 | 0.043 | Old |
| 0.045591 | 1.888515 | 0.01 | 0 | 1 | 5 | Gm8113 | 0.01 | Old |
| 0.045591 | 1.721426 | 0.01 | 0 | 1 | 5 | Ccdc39 | 0.01 | Old |
| 0.045591 | 1.672206 | 0.01 | 0 | 1 | 5 | Llgl2 | 0.01 | Old |
| 0.045591 | 1.623356 | 0.01 | 0 | 1 | 5 | Gm15834 | 0.01 | Old |
| 0.045591 | 1.602564 | 0.01 | 0 | 1 | 5 | Hic1 | 0.01 | Old |
| 0.045591 | 1.569698 | 0.01 | 0 | 1 | 5 | Apbb2 | 0.01 | Old |
| 0.045591 | 1.559417 | 0.01 | 0 | 1 | 5 | AC122413. | 0.01 | Old |
| 0.045591 | 1.544546 | 0.01 | 0 | 1 | 5 | Akr1c19 | 0.01 | Old |
| 0.045591 | 1.507702 | 0.01 | 0 | 1 | 5 | Ablim3 | 0.01 | Old |
| 0.045591 | 1.405387 | 0.01 | 0 | 1 | 5 | Adam4 | 0.01 | Old |
| 0.045591 | 1.378845 | 0.01 | 0 | 1 | 5 | Gm26912 | 0.01 | Old |

| | | | | | | | | |
|---|---|---|---|---|---|---|---|---|
| 0.045591 | 1.374974 | 0.01 | 0 | 1 | 5 | Gm47392 | 0.01 | Old |
| 0.045591 | 1.361517 | 0.01 | 0 | 1 | 5 | Mapk12 | 0.01 | Old |
| 0.045591 | 1.320579 | 0.01 | 0 | 1 | 5 | Aldh1l1 | 0.01 | Old |
| 0.045591 | 1.304087 | 0.01 | 0 | 1 | 5 | Kcnd1 | 0.01 | Old |
| 0.045591 | 1.288214 | 0.01 | 0 | 1 | 5 | Gm37584 | 0.01 | Old |
| 0.045591 | 1.191151 | 0.01 | 0 | 1 | 5 | Spag5 | 0.01 | Old |
| 0.0456 | 1.095637 | 0.04 | 0.018 | 1 | 5 | Mllt3 | 0.022 | Old |
| 0.045627 | 0.726518 | 0.2 | 0.16 | 1 | 5 | Lsm8 | 0.04 | Old |
| 0.04572 | 0.792122 | 0.074 | 0.043 | 1 | 5 | Ppih | 0.031 | Old |
| 0.046148 | 0.624777 | 0.157 | 0.114 | 1 | 5 | Fibp | 0.043 | Old |
| 0.04616 | 0.400233 | 0.17 | 0.124 | 1 | 5 | Usp3 | 0.046 | Old |
| 0.046398 | 0.894978 | 0.086 | 0.053 | 1 | 5 | 5730455P1 | 0.033 | Old |
| 0.046479 | 0.517484 | 0.114 | 0.076 | 1 | 5 | Stard4 | 0.038 | Old |
| 0.046485 | 0.854858 | 0.064 | 0.036 | 1 | 5 | Usp42 | 0.028 | Old |
| 0.046573 | 0.539092 | 0.192 | 0.147 | 1 | 5 | Gpr108 | 0.045 | Old |
| 0.046641 | 0.245706 | 0.449 | 0.388 | 1 | 5 | Aoah | 0.061 | Old |
| 0.046976 | 0.110633 | 0.914 | 0.876 | 1 | 5 | Ctsa | 0.038 | Old |
| 0.047364 | 0.102813 | 0.965 | 0.962 | 1 | 5 | Mcl1 | 0.003 | Old |
| 0.047461 | 0.463436 | 0.184 | 0.14 | 1 | 5 | Pcbd2 | 0.044 | Old |
| 0.047507 | 0.349897 | 0.231 | 0.178 | 1 | 5 | Ube2m | 0.053 | Old |
| 0.047792 | 0.146495 | 0.766 | 0.713 | 1 | 5 | Rpl4 | 0.053 | Old |
| 0.047897 | 0.204197 | 0.515 | 0.444 | 1 | 5 | Cers2 | 0.071 | Old |
| 0.048119 | 0.556171 | 0.114 | 0.076 | 1 | 5 | Arl10 | 0.038 | Old |
| 0.048449 | 0.971675 | 0.076 | 0.046 | 1 | 5 | Tmem185a | 0.03 | Old |
| 0.048456 | 0.330902 | 0.21 | 0.16 | 1 | 5 | Ltbr | 0.05 | Old |
| 0.048484 | 0.335291 | 0.328 | 0.272 | 1 | 5 | Rap2b | 0.056 | Old |
| 0.048551 | 0.105664 | 0.993 | 0.985 | 1 | 5 | Actg1 | 0.008 | Old |
| 0.048652 | 0.48987 | 0.177 | 0.132 | 1 | 5 | Rac3 | 0.045 | Old |
| 0.048714 | 0.303917 | 0.281 | 0.226 | 1 | 5 | Srrt | 0.055 | Old |
| 0.048902 | 0.191262 | 0.221 | 0.165 | 1 | 5 | Egr3 | 0.056 | Old |
| 0.048902 | 0.253539 | 0.207 | 0.155 | 1 | 5 | Wdr82 | 0.052 | Old |
| 0.048933 | 1.243786 | 0.02 | 0.005 | 1 | 5 | Psd | 0.015 | Old |
| 0.048966 | 0.281913 | 0.244 | 0.19 | 1 | 5 | 2610037D0 | 0.054 | Old |
| 0.049005 | 0.592361 | 0.241 | 0.193 | 1 | 5 | Nr4a3 | 0.048 | Old |
| 0.049033 | 0.364631 | 0.254 | 0.206 | 1 | 5 | Tmem51 | 0.048 | Old |
| 0.049053 | 0.620838 | 0.071 | 0.041 | 1 | 5 | Pomgnt1 | 0.03 | Old |
| 0.049062 | 0.598858 | 0.054 | 0.028 | 1 | 5 | Sec61a2 | 0.026 | Old |
| 0.049069 | 0.157104 | 0.232 | 0.175 | 1 | 5 | Zfp217 | 0.057 | Old |
| 0.049118 | 0.547893 | 0.099 | 0.063 | 1 | 5 | Gla | 0.036 | Old |
| 0.049186 | 0.878051 | 0.047 | 0.023 | 1 | 5 | Optn | 0.024 | Old |
| 0.049268 | 0.994772 | 0.029 | 0.01 | 1 | 5 | Creb3l1 | 0.019 | Old |
| 0.049342 | 0.618613 | 0.081 | 0.048 | 1 | 5 | Gabpb1 | 0.033 | Old |
| 0.049543 | 0.169848 | 0.616 | 0.548 | 1 | 5 | Ctnnb1 | 0.068 | Old |
| 0.049836 | 0.822459 | 0.051 | 0.025 | 1 | 5 | Zfp111 | 0.026 | Old |
| 0.050102 | 0.177632 | 0.163 | 0.114 | 1 | 5 | Nup214 | 0.049 | Old |

| | | | | | | | | |
|---|---|---|---|---|---|---|---|---|
| 0.050114 | 0.451655 | 0.184 | 0.137 | 1 | 5 | Snrnp48 | 0.047 | Old |
| 0.050148 | 0.743955 | 0.051 | 0.025 | 1 | 5 | Zfp747 | 0.026 | Old |
| 0.050544 | 0.863249 | 0.029 | 0.01 | 1 | 5 | Ddx28 | 0.019 | Old |
| 0.050725 | 0.146246 | 0.53 | 0.462 | 1 | 5 | Trir | 0.068 | Old |
| 0.050976 | 0.821635 | 0.029 | 0.01 | 1 | 5 | Agbl3 | 0.019 | Old |
| 0.051008 | 0.163416 | 0.207 | 0.152 | 1 | 5 | Plk3 | 0.055 | Old |
| 0.051209 | 0.29285 | 0.227 | 0.178 | 1 | 5 | Snd1 | 0.049 | Old |
| 0.051258 | 0.254805 | 0.742 | 0.721 | 1 | 5 | Gadd45b | 0.021 | Old |
| 0.051294 | 0.221248 | 0.224 | 0.175 | 1 | 5 | Smg7 | 0.049 | Old |
| 0.051339 | 0.549674 | 0.067 | 0.038 | 1 | 5 | Cpt2 | 0.029 | Old |
| 0.051346 | 0.199118 | 0.488 | 0.421 | 1 | 5 | Iqgap2 | 0.067 | Old |
| 0.051561 | 0.257009 | 0.291 | 0.234 | 1 | 5 | Siglech | 0.057 | Old |
| 0.051626 | 0.72394 | 0.064 | 0.036 | 1 | 5 | Syp | 0.028 | Old |
| 0.0519 | 0.489829 | 0.12 | 0.081 | 1 | 5 | Urod | 0.039 | Old |
| 0.05193 | 1.727316 | 0.015 | 0.003 | 1 | 5 | Haspin | 0.012 | Old |
| 0.052014 | 0.498715 | 0.187 | 0.142 | 1 | 5 | As3mt | 0.045 | Old |
| 0.05214 | 0.197766 | 0.69 | 0.617 | 1 | 5 | Hacd4 | 0.073 | Old |
| 0.052248 | 1.680095 | 0.015 | 0.003 | 1 | 5 | Jcad | 0.012 | Old |
| 0.052248 | 1.558239 | 0.015 | 0.003 | 1 | 5 | Adgra3 | 0.012 | Old |
| 0.052248 | 1.538983 | 0.015 | 0.003 | 1 | 5 | Rbp7 | 0.012 | Old |
| 0.0523 | 0.432949 | 0.126 | 0.086 | 1 | 5 | Letmd1 | 0.04 | Old |
| 0.052319 | 0.149405 | 0.702 | 0.635 | 1 | 5 | Ptpre | 0.067 | Old |
| 0.052496 | 0.550184 | 0.096 | 0.061 | 1 | 5 | Kdm4c | 0.035 | Old |
| 0.052605 | 0.330858 | 0.138 | 0.096 | 1 | 5 | Fam98c | 0.042 | Old |
| 0.052629 | 0.178431 | 0.253 | 0.195 | 1 | 5 | Csf3r | 0.058 | Old |
| 0.05292 | 0.127861 | 0.929 | 0.919 | 1 | 5 | Rpl36 | 0.01 | Old |
| 0.053129 | 1.235176 | 0.032 | 0.013 | 1 | 5 | Dus2 | 0.019 | Old |
| 0.053311 | 0.215861 | 0.145 | 0.102 | 1 | 5 | Rgs3 | 0.043 | Old |
| 0.053331 | 0.646447 | 0.082 | 0.051 | 1 | 5 | Fbxl12 | 0.031 | Old |
| 0.053373 | 0.961566 | 0.015 | 0.003 | 1 | 5 | Fam198b | 0.012 | Old |
| 0.053443 | 0.043523 | 0.497 | 0.414 | 1 | 5 | Dclre1c | 0.083 | Old |
| 0.053485 | 0.630067 | 0.057 | 0.03 | 1 | 5 | Endog | 0.027 | Old |
| 0.053626 | 0.188679 | 0.36 | 0.294 | 1 | 5 | Mfsd14b | 0.066 | Old |
| 0.05373 | 0.370983 | 0.222 | 0.175 | 1 | 5 | Trappc5 | 0.047 | Old |
| 0.053956 | 0.152643 | 0.798 | 0.754 | 1 | 5 | Mpeg1 | 0.044 | Old |
| 0.054046 | 0.318078 | 0.184 | 0.14 | 1 | 5 | Tuba1a | 0.044 | Old |
| 0.054248 | 0.543273 | 0.12 | 0.081 | 1 | 5 | Tyw1 | 0.039 | Old |
| 0.054399 | 0.38887 | 0.219 | 0.173 | 1 | 5 | Fam50a | 0.046 | Old |
| 0.054688 | 0.222828 | 0.707 | 0.66 | 1 | 5 | Plin2 | 0.047 | Old |
| 0.054711 | 0.345898 | 0.153 | 0.109 | 1 | 5 | Mta1 | 0.044 | Old |
| 0.054754 | 0.134055 | 0.396 | 0.335 | 1 | 5 | Tap2 | 0.061 | Old |
| 0.055037 | 0.40153 | 0.261 | 0.211 | 1 | 5 | Mrps14 | 0.05 | Old |
| 0.055109 | 0.373753 | 0.241 | 0.19 | 1 | 5 | Jade2 | 0.051 | Old |
| 0.055176 | 0.380221 | 0.125 | 0.086 | 1 | 5 | Usp5 | 0.039 | Old |
| 0.055204 | 0.36514 | 0.212 | 0.16 | 1 | 5 | Cycs | 0.052 | Old |

| | | | | | | | | |
|---|---|---|---|---|---|---|---|---|
| 0.055208 | 0.163849 | 0.641 | 0.584 | 1 | 5 | Tbxas1 | 0.057 | Old |
| 0.055227 | 0.170014 | 0.544 | 0.495 | 1 | 5 | Zbtb7a | 0.049 | Old |
| 0.055595 | 0.747903 | 0.079 | 0.048 | 1 | 5 | Aatf | 0.031 | Old |
| 0.055949 | 0.118441 | 0.7 | 0.627 | 1 | 5 | Nktr | 0.073 | Old |
| 0.05608 | 0.35639 | 0.246 | 0.195 | 1 | 5 | Bcl2 | 0.051 | Old |
| 0.056108 | 0.464788 | 0.098 | 0.063 | 1 | 5 | Gins4 | 0.035 | Old |
| 0.056453 | 0.1859 | 0.285 | 0.228 | 1 | 5 | Tox4 | 0.057 | Old |
| 0.056521 | 0.437513 | 0.226 | 0.18 | 1 | 5 | Pld1 | 0.046 | Old |
| 0.056732 | 0.232382 | 0.285 | 0.234 | 1 | 5 | Rsbn1l | 0.051 | Old |
| 0.057114 | 0.399307 | 0.194 | 0.147 | 1 | 5 | Ppm1g | 0.047 | Old |
| 0.057218 | 1.128989 | 0.077 | 0.048 | 1 | 5 | Mphosph1 | 0.029 | Old |
| 0.057321 | 0.491337 | 0.116 | 0.079 | 1 | 5 | Ms4a14 | 0.037 | Old |
| 0.057404 | 0.201046 | 0.47 | 0.411 | 1 | 5 | Cmtm3 | 0.059 | Old |
| 0.057559 | 0.192816 | 0.466 | 0.398 | 1 | 5 | Tns1 | 0.068 | Old |
| 0.057572 | 1.229139 | 0.039 | 0.018 | 1 | 5 | Ift140 | 0.021 | Old |
| 0.057572 | 1.204416 | 0.039 | 0.018 | 1 | 5 | Btbd19 | 0.021 | Old |
| 0.057844 | 1.282689 | 0.024 | 0.008 | 1 | 5 | Bora | 0.016 | Old |
| 0.057899 | 0.108255 | 0.715 | 0.66 | 1 | 5 | Rap1a | 0.055 | Old |
| 0.058111 | 1.396226 | 0.024 | 0.008 | 1 | 5 | Grhpr | 0.016 | Old |
| 0.058326 | 0.467401 | 0.116 | 0.079 | 1 | 5 | Spred2 | 0.037 | Old |
| 0.058637 | 0.630066 | 0.094 | 0.061 | 1 | 5 | Nsun6 | 0.033 | Old |
| 0.058673 | 0.563528 | 0.172 | 0.129 | 1 | 5 | Rtp4 | 0.043 | Old |
| 0.058688 | 0.916152 | 0.042 | 0.02 | 1 | 5 | 0610040J0 | 0.022 | Old |
| 0.058728 | 0.444984 | 0.13 | 0.091 | 1 | 5 | Golph3l | 0.039 | Old |
| 0.059143 | 0.876309 | 0.049 | 0.025 | 1 | 5 | Nr1h3 | 0.024 | Old |
| 0.059272 | 0.135228 | 0.896 | 0.848 | 1 | 5 | Celf2 | 0.048 | Old |
| 0.059475 | 0.240712 | 0.229 | 0.175 | 1 | 5 | Mien1 | 0.054 | Old |
| 0.059914 | 0.373367 | 0.256 | 0.211 | 1 | 5 | Bex3 | 0.045 | Old |
| 0.060057 | 0.316048 | 0.295 | 0.246 | 1 | 5 | Zmiz2 | 0.049 | Old |
| 0.060259 | 0.203583 | 0.551 | 0.48 | 1 | 5 | Ptges3 | 0.071 | Old |
| 0.060305 | 0.31702 | 0.301 | 0.256 | 1 | 5 | Htt | 0.045 | Old |
| 0.060408 | 0.903657 | 0.069 | 0.041 | 1 | 5 | Thrb | 0.028 | Old |
| 0.060519 | 0.686201 | 0.059 | 0.033 | 1 | 5 | Hist3h2ba | 0.026 | Old |
| 0.060567 | 0.564745 | 0.088 | 0.056 | 1 | 5 | Sdccag8 | 0.032 | Old |
| 0.060567 | 0.384714 | 0.16 | 0.117 | 1 | 5 | Mrpl11 | 0.043 | Old |
| 0.060584 | 0.177652 | 0.3 | 0.241 | 1 | 5 | Tap1 | 0.059 | Old |
| 0.060771 | 0.878526 | 0.042 | 0.02 | 1 | 5 | Ogg1 | 0.022 | Old |
| 0.06098 | 0.267528 | 0.236 | 0.183 | 1 | 5 | Aip | 0.053 | Old |
| 0.061009 | 0.245537 | 0.887 | 0.873 | 1 | 5 | Mt1 | 0.014 | Old |
| 0.061128 | 0.183415 | 0.692 | 0.637 | 1 | 5 | Ywhah | 0.055 | Old |
| 0.061137 | 0.122898 | 0.519 | 0.442 | 1 | 5 | Slc29a1 | 0.077 | Old |
| 0.061175 | 0.372308 | 0.185 | 0.14 | 1 | 5 | Zbtb11 | 0.045 | Old |
| 0.061241 | 0.821202 | 0.039 | 0.018 | 1 | 5 | Itfg2 | 0.021 | Old |
| 0.061579 | 0.860625 | 0.077 | 0.048 | 1 | 5 | Cwc27 | 0.029 | Old |
| 0.061618 | 0.409905 | 0.091 | 0.058 | 1 | 5 | Mlx | 0.033 | Old |

| | | | | | | | | |
|---|---|---|---|---|---|---|---|---|
| 0.061689 | 0.24323 | 0.313 | 0.256 | 1 | 5 | Nhlrc3 | 0.057 | Old |
| 0.061954 | 0.220714 | 0.219 | 0.17 | 1 | 5 | Txndc11 | 0.049 | Old |
| 0.062036 | 0.136397 | 0.566 | 0.518 | 1 | 5 | Cd302 | 0.048 | Old |
| 0.062395 | 0.261656 | 0.133 | 0.094 | 1 | 5 | Man2b2 | 0.039 | Old |
| 0.06244 | 0.622061 | 0.091 | 0.058 | 1 | 5 | Cavin1 | 0.033 | Old |
| 0.062619 | 0.303012 | 0.342 | 0.287 | 1 | 5 | Clptm1l | 0.055 | Old |
| 0.062686 | 0.719048 | 0.059 | 0.033 | 1 | 5 | Wfs1 | 0.026 | Old |
| 0.062915 | 0.845006 | 0.049 | 0.025 | 1 | 5 | Gm36839 | 0.024 | Old |
| 0.062963 | 0.359605 | 0.189 | 0.142 | 1 | 5 | AW549877 | 0.047 | Old |
| 0.063172 | 0.382344 | 0.167 | 0.124 | 1 | 5 | Tob1 | 0.043 | Old |
| 0.063202 | 0.900467 | 0.049 | 0.025 | 1 | 5 | Zfp932 | 0.024 | Old |
| 0.063273 | 0.249009 | 0.635 | 0.609 | 1 | 5 | Rel | 0.026 | Old |
| 0.063306 | 0.621837 | 0.096 | 0.063 | 1 | 5 | Dvl3 | 0.033 | Old |
| 0.06335 | 0.230738 | 0.347 | 0.297 | 1 | 5 | Mctp1 | 0.05 | Old |
| 0.063446 | 1.655976 | 0.027 | 0.01 | 1 | 5 | Zcchc3 | 0.017 | Old |
| 0.063689 | 0.720404 | 0.062 | 0.036 | 1 | 5 | Ier5l | 0.026 | Old |
| 0.063777 | 0.71924 | 0.066 | 0.038 | 1 | 5 | Gm26669 | 0.028 | Old |
| 0.063854 | 0.371283 | 0.251 | 0.203 | 1 | 5 | Psen1 | 0.048 | Old |
| 0.063947 | 0.893147 | 0.052 | 0.028 | 1 | 5 | Tmem184c | 0.024 | Old |
| 0.063971 | 0.171421 | 0.093 | 0.058 | 1 | 5 | Isyna1 | 0.035 | Old |
| 0.063993 | 0.780518 | 0.052 | 0.028 | 1 | 5 | Tjap1 | 0.024 | Old |
| 0.064077 | 0.189856 | 0.65 | 0.591 | 1 | 5 | Epsti1 | 0.059 | Old |
| 0.06425 | 0.488898 | 0.125 | 0.086 | 1 | 5 | Cfap97 | 0.039 | Old |
| 0.064327 | 0.248813 | 0.522 | 0.464 | 1 | 5 | Manf | 0.058 | Old |
| 0.064653 | 1.212907 | 0.027 | 0.01 | 1 | 5 | Serping1 | 0.017 | Old |
| 0.064758 | 0.222946 | 0.271 | 0.218 | 1 | 5 | Egln2 | 0.053 | Old |
| 0.064995 | 0.149967 | 0.365 | 0.312 | 1 | 5 | Smpdl3b | 0.053 | Old |
| 0.065046 | 0.225149 | 0.79 | 0.751 | 1 | 5 | Zfand5 | 0.039 | Old |
| 0.065047 | 0.177137 | 0.285 | 0.223 | 1 | 5 | Myd88 | 0.062 | Old |
| 0.065203 | 0.376899 | 0.266 | 0.221 | 1 | 5 | Kdelr2 | 0.045 | Old |
| 0.065322 | 0.586971 | 0.128 | 0.091 | 1 | 5 | BC005561 | 0.037 | Old |
| 0.065366 | 0.257573 | 0.246 | 0.198 | 1 | 5 | Aagab | 0.048 | Old |
| 0.065761 | 0.363777 | 0.239 | 0.193 | 1 | 5 | Thra | 0.046 | Old |
| 0.065802 | 0.128164 | 0.377 | 0.315 | 1 | 5 | Chp2 | 0.062 | Old |
| 0.065849 | 0.391205 | 0.056 | 0.03 | 1 | 5 | Letm2 | 0.026 | Old |
| 0.065889 | 0.356713 | 0.308 | 0.269 | 1 | 5 | Crtc3 | 0.039 | Old |
| 0.065934 | 0.538096 | 0.049 | 0.025 | 1 | 5 | Map7d3 | 0.024 | Old |
| 0.066534 | 0.322146 | 0.189 | 0.145 | 1 | 5 | Smad2 | 0.044 | Old |
| 0.06655 | 0.51706 | 0.076 | 0.046 | 1 | 5 | Zfp511 | 0.03 | Old |
| 0.066567 | 0.966485 | 0.027 | 0.01 | 1 | 5 | Afg1l | 0.017 | Old |
| 0.066671 | 0.156504 | 0.877 | 0.855 | 1 | 5 | Fcgr2b | 0.022 | Old |
| 0.066676 | 0.765269 | 0.074 | 0.046 | 1 | 5 | Alg2 | 0.028 | Old |
| 0.066801 | 0.232238 | 0.364 | 0.317 | 1 | 5 | Igf1r | 0.047 | Old |
| 0.066801 | 0.651325 | 0.052 | 0.028 | 1 | 5 | Ap4e1 | 0.024 | Old |
| 0.066898 | 0.670985 | 0.052 | 0.028 | 1 | 5 | Foxp4 | 0.024 | Old |

| | | | | | | | | |
|---|---|---|---|---|---|---|---|---|
| 0.066984 | 1.213144 | 0.027 | 0.01 | 1 | 5 | Rnf24 | 0.017 | Old |
| 0.06704 | 0.221333 | 0.596 | 0.551 | 1 | 5 | Notch2 | 0.045 | Old |
| 0.067125 | 0.455588 | 0.143 | 0.104 | 1 | 5 | Ybx3 | 0.039 | Old |
| 0.067241 | 0.871973 | 0.074 | 0.046 | 1 | 5 | Usp53 | 0.028 | Old |
| 0.067653 | 0.133947 | 0.293 | 0.236 | 1 | 5 | Paf1 | 0.057 | Old |
| 0.067721 | 0.593314 | 0.189 | 0.15 | 1 | 5 | Snrnp27 | 0.039 | Old |
| 0.067899 | 0.060044 | 0.987 | 0.942 | 1 | 5 | Hsp90ab1 | 0.045 | Old |
| 0.068189 | 0.565987 | 0.088 | 0.056 | 1 | 5 | Stk16 | 0.032 | Old |
| 0.06826 | 1.679473 | 0.019 | 0.005 | 1 | 5 | Hunk | 0.014 | Old |
| 0.068274 | 0.32356 | 0.182 | 0.14 | 1 | 5 | 1810022K0 | 0.042 | Old |
| 0.068313 | 0.485286 | 0.081 | 0.051 | 1 | 5 | Efemp2 | 0.03 | Old |
| 0.068435 | 1.553986 | 0.019 | 0.005 | 1 | 5 | Tlr11 | 0.014 | Old |
| 0.068634 | 0.017525 | 0.626 | 0.571 | 1 | 5 | Fosl2 | 0.055 | Old |
| 0.068672 | 0.413613 | 0.195 | 0.152 | 1 | 5 | Trak1 | 0.043 | Old |
| 0.068781 | 0.388132 | 0.153 | 0.114 | 1 | 5 | Sestd1 | 0.039 | Old |
| 0.068869 | 0.692826 | 0.056 | 0.03 | 1 | 5 | Htr1b | 0.026 | Old |
| 0.068997 | 0.29205 | 0.418 | 0.36 | 1 | 5 | H2-Q7 | 0.058 | Old |
| 0.069022 | 0.603081 | 0.108 | 0.074 | 1 | 5 | Ets1 | 0.034 | Old |
| 0.069097 | 0.717974 | 0.027 | 0.01 | 1 | 5 | Plppr2 | 0.017 | Old |
| 0.069222 | 0.186766 | 0.355 | 0.297 | 1 | 5 | AI314180 | 0.058 | Old |
| 0.069243 | 0.319654 | 0.266 | 0.221 | 1 | 5 | Eno1 | 0.045 | Old |
| 0.069312 | 0.282068 | 0.138 | 0.099 | 1 | 5 | Mxra7 | 0.039 | Old |
| 0.069314 | 1.272909 | 0.019 | 0.005 | 1 | 5 | Zfp438 | 0.014 | Old |
| 0.069515 | 0.509682 | 0.125 | 0.089 | 1 | 5 | Cep85 | 0.036 | Old |
| 0.06982 | 0.260573 | 0.081 | 0.051 | 1 | 5 | Cyb561d2 | 0.03 | Old |
| 0.069846 | 1.124845 | 0.019 | 0.005 | 1 | 5 | Foxd2 | 0.014 | Old |
| 0.069846 | 1.110366 | 0.019 | 0.005 | 1 | 5 | Tcerg1l | 0.014 | Old |
| 0.070005 | 1.260376 | 0.03 | 0.013 | 1 | 5 | Cdyl2 | 0.017 | Old |
| 0.070023 | 0.14852 | 0.584 | 0.513 | 1 | 5 | St8sia4 | 0.071 | Old |
| 0.070118 | 0.323743 | 0.315 | 0.266 | 1 | 5 | Xpo6 | 0.049 | Old |
| 0.070602 | 0.355695 | 0.273 | 0.226 | 1 | 5 | Foxn2 | 0.047 | Old |
| 0.070657 | 0.307498 | 0.234 | 0.185 | 1 | 5 | Cyc1 | 0.049 | Old |
| 0.070788 | 0.267441 | 0.327 | 0.277 | 1 | 5 | Heatr5a | 0.05 | Old |
| 0.070857 | 0.479512 | 0.214 | 0.17 | 1 | 5 | Id3 | 0.044 | Old |
| 0.071066 | 0.125849 | 0.704 | 0.652 | 1 | 5 | Eps8 | 0.052 | Old |
| 0.071111 | 0.164345 | 0.2 | 0.152 | 1 | 5 | Tex264 | 0.048 | Old |
| 0.071282 | 0.629624 | 0.019 | 0.005 | 1 | 5 | Syn1 | 0.014 | Old |
| 0.07132 | 0.307106 | 0.29 | 0.241 | 1 | 5 | Zranb1 | 0.049 | Old |
| 0.071363 | 0.992851 | 0.03 | 0.013 | 1 | 5 | Ap4b1 | 0.017 | Old |
| 0.071507 | 0.433683 | 0.273 | 0.221 | 1 | 5 | Isy1 | 0.052 | Old |
| 0.072086 | 0.293225 | 0.507 | 0.467 | 1 | 5 | Itpkb | 0.04 | Old |
| 0.072111 | 0.314551 | 0.261 | 0.216 | 1 | 5 | BC031181 | 0.045 | Old |
| 0.072142 | 0.272146 | 0.189 | 0.142 | 1 | 5 | Sar1b | 0.047 | Old |
| 0.072275 | 0.27114 | 0.138 | 0.099 | 1 | 5 | Lin54 | 0.039 | Old |
| 0.072309 | 0.317004 | 0.204 | 0.157 | 1 | 5 | Lpar1 | 0.047 | Old |

| | | | | | | | | |
|---|---|---|---|---|---|---|---|---|
| 0.072458 | 0.130264 | 0.816 | 0.764 | 1 | 5 | Ubl3 | 0.052 | Old |
| 0.072722 | 0.223229 | 0.163 | 0.122 | 1 | 5 | Ebpl | 0.041 | Old |
| 0.072742 | 1.647303 | 0.03 | 0.013 | 1 | 5 | Rflnb | 0.017 | Old |
| 0.072795 | 0.635702 | 0.123 | 0.089 | 1 | 5 | Rab11fip2 | 0.034 | Old |
| 0.073529 | 0.459326 | 0.093 | 0.061 | 1 | 5 | Yaf2 | 0.032 | Old |
| 0.073838 | 0.259231 | 0.145 | 0.102 | 1 | 5 | Mettl23 | 0.043 | Old |
| 0.074227 | 1.342133 | 0.04 | 0.02 | 1 | 5 | Gnb4 | 0.02 | Old |
| 0.07435 | 0.474154 | 0.114 | 0.079 | 1 | 5 | Scrib | 0.035 | Old |
| 0.074359 | 0.193048 | 0.424 | 0.371 | 1 | 5 | Ccnt1 | 0.053 | Old |
| 0.074441 | 0.320455 | 0.308 | 0.259 | 1 | 5 | Lrpap1 | 0.049 | Old |
| 0.07465 | 0.213358 | 0.259 | 0.206 | 1 | 5 | Dlgap4 | 0.053 | Old |
| 0.07498 | 0.21183 | 0.17 | 0.124 | 1 | 5 | Dld | 0.046 | Old |
| 0.075372 | 1.156636 | 0.034 | 0.015 | 1 | 5 | Adamtsl4 | 0.019 | Old |
| 0.075542 | 1.017259 | 0.044 | 0.023 | 1 | 5 | Gm10791 | 0.021 | Old |
| 0.075585 | 0.01273 | 0.283 | 0.226 | 1 | 5 | Hsph1 | 0.057 | Old |
| 0.075642 | 0.791507 | 0.034 | 0.015 | 1 | 5 | Thop1 | 0.019 | Old |
| 0.07584 | 0.307411 | 0.074 | 0.046 | 1 | 5 | Pdzd11 | 0.028 | Old |
| 0.075937 | 0.711733 | 0.082 | 0.053 | 1 | 5 | Lyrm9 | 0.029 | Old |
| 0.075974 | 1.76965 | 0.013 | 0.003 | 1 | 5 | Cttn | 0.01 | Old |
| 0.075974 | 1.460655 | 0.013 | 0.003 | 1 | 5 | Cyp2ab1 | 0.01 | Old |
| 0.075974 | 1.4472 | 0.013 | 0.003 | 1 | 5 | Islr | 0.01 | Old |
| 0.076371 | 0.073678 | 1 | 1 | 1 | 5 | Malat1 | 0 | Old |
| 0.07679 | 1.129168 | 0.04 | 0.02 | 1 | 5 | Mgll | 0.02 | Old |
| 0.076893 | 1.219234 | 0.013 | 0.003 | 1 | 5 | Sbsn | 0.01 | Old |
| 0.077015 | 0.283912 | 0.279 | 0.226 | 1 | 5 | Maff | 0.053 | Old |
| 0.077046 | 0.220528 | 0.529 | 0.487 | 1 | 5 | Cox7a2 | 0.042 | Old |
| 0.077124 | 1.167594 | 0.013 | 0.003 | 1 | 5 | Mmp23 | 0.01 | Old |
| 0.077124 | 1.100372 | 0.013 | 0.003 | 1 | 5 | Wdr78 | 0.01 | Old |
| 0.077223 | 0.329126 | 0.118 | 0.081 | 1 | 5 | Cpsf4 | 0.037 | Old |
| 0.077282 | 0.816474 | 0.157 | 0.119 | 1 | 5 | Cxcl13 | 0.038 | Old |
| 0.077295 | 0.122341 | 0.362 | 0.305 | 1 | 5 | Psma4 | 0.057 | Old |
| 0.077312 | 0.80145 | 0.037 | 0.018 | 1 | 5 | Slc6a8 | 0.019 | Old |
| 0.077588 | 0.914261 | 0.013 | 0.003 | 1 | 5 | 2200002D( | 0.01 | Old |
| 0.077665 | 0.11931 | 0.212 | 0.165 | 1 | 5 | Eif4e2 | 0.047 | Old |
| 0.077808 | 0.185064 | 0.421 | 0.368 | 1 | 5 | Pdlim5 | 0.053 | Old |
| 0.078271 | 0.868546 | 0.047 | 0.025 | 1 | 5 | Rnf26 | 0.022 | Old |
| 0.078852 | 0.12241 | 0.355 | 0.299 | 1 | 5 | Ppp2r2a | 0.056 | Old |
| 0.079344 | 0.231894 | 0.187 | 0.142 | 1 | 5 | 2210016F1 | 0.045 | Old |
| 0.079376 | 0.411784 | 0.2 | 0.16 | 1 | 5 | Cklf | 0.04 | Old |
| 0.079491 | 0.347093 | 0.285 | 0.241 | 1 | 5 | Atp6v1h | 0.044 | Old |
| 0.079615 | 0.175462 | 0.793 | 0.754 | 1 | 5 | Ptprj | 0.039 | Old |
| 0.079753 | 0.130995 | 0.623 | 0.571 | 1 | 5 | Cndp2 | 0.052 | Old |
| 0.07979 | 0.205097 | 0.384 | 0.335 | 1 | 5 | Tbc1d5 | 0.049 | Old |
| 0.079809 | 0.398373 | 0.158 | 0.119 | 1 | 5 | Sacm1l | 0.039 | Old |
| 0.079959 | 0.701747 | 0.064 | 0.038 | 1 | 5 | Eif2b3 | 0.026 | Old |

| | | | | | | | | |
|---|---|---|---|---|---|---|---|---|
| 0.080387 | 0.240388 | 0.103 | 0.069 | 1 | 5 | Rnf31 | 0.034 | Old |
| 0.080499 | 0.299679 | 0.184 | 0.142 | 1 | 5 | Aph1c | 0.042 | Old |
| 0.0805 | 0.720145 | 0.054 | 0.03 | 1 | 5 | Lzic | 0.024 | Old |
| 0.080513 | 0.341637 | 0.077 | 0.048 | 1 | 5 | Acad9 | 0.029 | Old |
| 0.080643 | 0.294928 | 0.121 | 0.086 | 1 | 5 | Srxn1 | 0.035 | Old |
| 0.0807 | 0.258081 | 0.219 | 0.173 | 1 | 5 | Mitf | 0.046 | Old |
| 0.080725 | 1.149621 | 0.022 | 0.008 | 1 | 5 | Dpyd | 0.014 | Old |
| 0.080832 | 0.438891 | 0.054 | 0.03 | 1 | 5 | Cipc | 0.024 | Old |
| 0.080899 | 0.784517 | 0.04 | 0.02 | 1 | 5 | Znhit2 | 0.02 | Old |
| 0.080911 | 0.16802 | 0.527 | 0.477 | 1 | 5 | Brd2 | 0.05 | Old |
| 0.081443 | 0.641077 | 0.086 | 0.056 | 1 | 5 | Ifitm1 | 0.03 | Old |
| 0.081462 | 0.301825 | 0.071 | 0.043 | 1 | 5 | Mterf4 | 0.028 | Old |
| 0.081534 | 0.325951 | 0.057 | 0.033 | 1 | 5 | Zcchc4 | 0.024 | Old |
| 0.081662 | 0.428066 | 0.093 | 0.061 | 1 | 5 | Tspyl2 | 0.032 | Old |
| 0.081827 | 0.663324 | 0.044 | 0.023 | 1 | 5 | Map2k6 | 0.021 | Old |
| 0.081847 | 0.229453 | 0.155 | 0.114 | 1 | 5 | Pibf1 | 0.041 | Old |
| 0.081941 | 0.31606 | 0.067 | 0.041 | 1 | 5 | Parp6 | 0.026 | Old |
| 0.082078 | 0.223976 | 0.18 | 0.135 | 1 | 5 | Cib2 | 0.045 | Old |
| 0.082125 | 0.441903 | 0.121 | 0.086 | 1 | 5 | Sh2b1 | 0.035 | Old |
| 0.082279 | 0.204102 | 0.234 | 0.188 | 1 | 5 | Fam213b | 0.046 | Old |
| 0.082365 | 0.747122 | 0.022 | 0.008 | 1 | 5 | Prodh | 0.014 | Old |
| 0.082568 | 0.64918 | 0.072 | 0.046 | 1 | 5 | Asb13 | 0.026 | Old |
| 0.082733 | 0.93774 | 0.022 | 0.008 | 1 | 5 | Creg2 | 0.014 | Old |
| 0.082831 | 0.163612 | 0.301 | 0.254 | 1 | 5 | Clock | 0.047 | Old |
| 0.082848 | 0.256319 | 0.348 | 0.305 | 1 | 5 | Fam234a | 0.043 | Old |
| 0.082871 | 0.2122 | 0.401 | 0.345 | 1 | 5 | Cuta | 0.056 | Old |
| 0.082918 | 0.988651 | 0.022 | 0.008 | 1 | 5 | Slc9a4 | 0.014 | Old |
| 0.082918 | 0.793708 | 0.022 | 0.008 | 1 | 5 | Pi16 | 0.014 | Old |
| 0.082925 | 0.627481 | 0.061 | 0.036 | 1 | 5 | Pcolce | 0.025 | Old |
| 0.08305 | 0.432551 | 0.189 | 0.15 | 1 | 5 | Irf3 | 0.039 | Old |
| 0.083145 | 0.312946 | 0.089 | 0.058 | 1 | 5 | Mtmr4 | 0.031 | Old |
| 0.083537 | 0.734559 | 0.076 | 0.048 | 1 | 5 | Slf1 | 0.028 | Old |
| 0.083885 | 0.484191 | 0.109 | 0.076 | 1 | 5 | Dnajb2 | 0.033 | Old |
| 0.083903 | 0.280756 | 0.337 | 0.289 | 1 | 5 | Aup1 | 0.048 | Old |
| 0.083906 | 0.515977 | 0.094 | 0.063 | 1 | 5 | Zfhx2 | 0.031 | Old |
| 0.083927 | 0.259442 | 0.2 | 0.157 | 1 | 5 | Cetn3 | 0.043 | Old |
| 0.083965 | 0.129332 | 0.63 | 0.561 | 1 | 5 | Tmem59 | 0.069 | Old |
| 0.084146 | 0.217476 | 0.313 | 0.261 | 1 | 5 | Myo1c | 0.052 | Old |
| 0.084755 | 0.569326 | 0.079 | 0.051 | 1 | 5 | Gimap6 | 0.028 | Old |
| 0.084938 | 0.198742 | 0.253 | 0.203 | 1 | 5 | Crim1 | 0.05 | Old |
| 0.085015 | 0.403601 | 0.184 | 0.145 | 1 | 5 | Glrx | 0.039 | Old |
| 0.085274 | 0.151985 | 0.732 | 0.685 | 1 | 5 | Jmjd1c | 0.047 | Old |
| 0.085283 | 0.814068 | 0.069 | 0.043 | 1 | 5 | Ttc4 | 0.026 | Old |
| 0.085838 | 0.325429 | 0.086 | 0.056 | 1 | 5 | Dtx2 | 0.03 | Old |
| 0.086032 | 0.308464 | 0.209 | 0.162 | 1 | 5 | Pml | 0.047 | Old |

| | | | | | | | | |
|---|---|---|---|---|---|---|---|---|
| 0.08639 | 0.495963 | 0.15 | 0.114 | 1 | 5 | Sec23ip | 0.036 | Old |
| 0.086404 | 0.312834 | 0.04 | 0.02 | 1 | 5 | Dtd1 | 0.02 | Old |
| 0.086421 | 0.676357 | 0.044 | 0.023 | 1 | 5 | Zfp948 | 0.021 | Old |
| 0.086509 | 0.560687 | 0.197 | 0.16 | 1 | 5 | Clec2i | 0.037 | Old |
| 0.087055 | 0.397795 | 0.209 | 0.168 | 1 | 5 | Pih1d1 | 0.041 | Old |
| 0.087478 | 0.142094 | 0.387 | 0.33 | 1 | 5 | Cct7 | 0.057 | Old |
| 0.087593 | 0.29158 | 0.209 | 0.162 | 1 | 5 | Dhx30 | 0.047 | Old |
| 0.087691 | 0.144709 | 0.082 | 0.053 | 1 | 5 | Hspb1 | 0.029 | Old |
| 0.088064 | 0.102967 | 0.611 | 0.566 | 1 | 5 | Gpi1 | 0.045 | Old |
| 0.088086 | 0.497857 | 0.099 | 0.069 | 1 | 5 | Zfp410 | 0.03 | Old |
| 0.088222 | 0.151908 | 0.285 | 0.231 | 1 | 5 | Pcyox1 | 0.054 | Old |
| 0.088422 | 0.252473 | 0.431 | 0.371 | 1 | 5 | Limd1 | 0.06 | Old |
| 0.089046 | 0.339538 | 0.178 | 0.14 | 1 | 5 | Prkab2 | 0.038 | Old |
| 0.089059 | 1.044423 | 0.025 | 0.01 | 1 | 5 | Tnks1bp1 | 0.015 | Old |
| 0.08917 | 0.283345 | 0.411 | 0.373 | 1 | 5 | N4bp1 | 0.038 | Old |
| 0.089419 | 0.805606 | 0.025 | 0.01 | 1 | 5 | Zfp820 | 0.015 | Old |
| 0.089493 | 0.36106 | 0.109 | 0.076 | 1 | 5 | Hikeshi | 0.033 | Old |
| 0.08958 | 0.127129 | 0.409 | 0.345 | 1 | 5 | Kmt2c | 0.064 | Old |
| 0.089589 | 0.207799 | 0.47 | 0.411 | 1 | 5 | Ifnar1 | 0.059 | Old |
| 0.089734 | 0.638545 | 0.108 | 0.076 | 1 | 5 | Crebl2 | 0.032 | Old |
| 0.08981 | 0.078982 | 0.729 | 0.657 | 1 | 5 | Top1 | 0.072 | Old |
| 0.089911 | 0.334035 | 0.187 | 0.15 | 1 | 5 | Erf | 0.037 | Old |
| 0.089962 | 1.168615 | 0.025 | 0.01 | 1 | 5 | Igfbp7 | 0.015 | Old |
| 0.090063 | 0.316882 | 0.379 | 0.34 | 1 | 5 | Crlf2 | 0.039 | Old |
| 0.090117 | 0.465507 | 0.143 | 0.107 | 1 | 5 | Hlcs | 0.036 | Old |
| 0.090133 | 0.253033 | 0.399 | 0.35 | 1 | 5 | Slc12a7 | 0.049 | Old |
| 0.090322 | 0.49063 | 0.125 | 0.091 | 1 | 5 | Syap1 | 0.034 | Old |
| 0.090326 | 0.790929 | 0.025 | 0.01 | 1 | 5 | Pop1 | 0.015 | Old |
| 0.090478 | 0.331884 | 0.316 | 0.277 | 1 | 5 | Acsl5 | 0.039 | Old |
| 0.090508 | 1.115329 | 0.025 | 0.01 | 1 | 5 | Ercc8 | 0.015 | Old |
| 0.090677 | 0.283116 | 0.17 | 0.129 | 1 | 5 | Aig1 | 0.041 | Old |
| 0.090795 | 0.174504 | 0.559 | 0.505 | 1 | 5 | Dnmt3a | 0.054 | Old |
| 0.090802 | 0.096398 | 0.443 | 0.383 | 1 | 5 | Entpd1 | 0.06 | Old |
| 0.090873 | 0.905328 | 0.025 | 0.01 | 1 | 5 | Rem1 | 0.015 | Old |
| 0.091239 | 0.892346 | 0.025 | 0.01 | 1 | 5 | Kcnq1 | 0.015 | Old |
| 0.091468 | 0.431468 | 0.178 | 0.14 | 1 | 5 | Mfsd10 | 0.038 | Old |
| 0.091595 | 0.417151 | 0.133 | 0.099 | 1 | 5 | Trpm2 | 0.034 | Old |
| 0.092474 | 0.39887 | 0.072 | 0.046 | 1 | 5 | Zbtb17 | 0.026 | Old |
| 0.09262 | 0.585027 | 0.062 | 0.038 | 1 | 5 | Rab34 | 0.024 | Old |
| 0.092918 | 0.189018 | 0.364 | 0.312 | 1 | 5 | Rybp | 0.052 | Old |
| 0.093295 | 0.68772 | 0.113 | 0.081 | 1 | 5 | 4930455G( | 0.032 | Old |
| 0.093414 | 0.160993 | 0.519 | 0.457 | 1 | 5 | Eef1d | 0.062 | Old |
| 0.093421 | 0.290323 | 0.232 | 0.19 | 1 | 5 | Ddx54 | 0.042 | Old |
| 0.093485 | 0.103809 | 0.519 | 0.457 | 1 | 5 | Scaf11 | 0.062 | Old |
| 0.093515 | 0.812829 | 0.056 | 0.033 | 1 | 5 | Nipa1 | 0.023 | Old |

| | | | | | | | | |
|---|---|---|---|---|---|---|---|---|
| 0.093542 | 0.044399 | 0.611 | 0.533 | 1 | 5 | Arf4 | 0.078 | Old |
| 0.093603 | 0.239867 | 0.202 | 0.157 | 1 | 5 | Zfp131 | 0.045 | Old |
| 0.093685 | 0.850392 | 0.052 | 0.03 | 1 | 5 | Park2 | 0.022 | Old |
| 0.093795 | 0.125325 | 0.453 | 0.383 | 1 | 5 | Rab6a | 0.07 | Old |
| 0.093886 | 0.404661 | 0.258 | 0.221 | 1 | 5 | Tmem5 | 0.037 | Old |
| 0.093937 | 0.934513 | 0.052 | 0.03 | 1 | 5 | Dsel | 0.022 | Old |
| 0.094075 | 0.534091 | 0.135 | 0.102 | 1 | 5 | Rev1 | 0.033 | Old |
| 0.094148 | 0.319872 | 0.091 | 0.061 | 1 | 5 | Pde8a | 0.03 | Old |
| 0.094289 | 0.66616 | 0.059 | 0.036 | 1 | 5 | Tbcc | 0.023 | Old |
| 0.094319 | 1.15084 | 0.029 | 0.013 | 1 | 5 | Cep89 | 0.016 | Old |
| 0.094694 | 0.64773 | 0.077 | 0.051 | 1 | 5 | Ttyh2 | 0.026 | Old |
| 0.094804 | 0.178743 | 0.468 | 0.393 | 1 | 5 | Antxr2 | 0.075 | Old |
| 0.095147 | 0.519333 | 0.069 | 0.043 | 1 | 5 | Qsox2 | 0.026 | Old |
| 0.09522 | 0.144237 | 0.433 | 0.371 | 1 | 5 | Ist1 | 0.062 | Old |
| 0.095257 | 0.533054 | 0.072 | 0.046 | 1 | 5 | Ldlrad3 | 0.026 | Old |
| 0.095419 | 0.137416 | 0.778 | 0.723 | 1 | 5 | Vsir | 0.055 | Old |
| 0.095463 | 0.02022 | 0.375 | 0.312 | 1 | 5 | Sema6d | 0.063 | Old |
| 0.095565 | 0.368344 | 0.076 | 0.048 | 1 | 5 | Eif2b2 | 0.028 | Old |
| 0.095575 | 0.542487 | 0.066 | 0.041 | 1 | 5 | Slc37a3 | 0.025 | Old |
| 0.09572 | 0.914454 | 0.052 | 0.03 | 1 | 5 | Mrps31 | 0.022 | Old |
| 0.095731 | 0.799531 | 0.029 | 0.013 | 1 | 5 | Uros | 0.016 | Old |
| 0.095917 | 0.173244 | 0.412 | 0.36 | 1 | 5 | Tmed3 | 0.052 | Old |
| 0.096264 | 0.783063 | 0.029 | 0.013 | 1 | 5 | Shb | 0.016 | Old |
| 0.096603 | 1.226428 | 0.017 | 0.005 | 1 | 5 | Efna2 | 0.012 | Old |
| 0.096603 | 0.159117 | 0.488 | 0.424 | 1 | 5 | Tubb6 | 0.064 | Old |
| 0.096759 | 0.258014 | 0.345 | 0.302 | 1 | 5 | Setd2 | 0.043 | Old |
| 0.097085 | 1.488718 | 0.017 | 0.005 | 1 | 5 | Col15a1 | 0.012 | Old |
| 0.097107 | 0.266193 | 0.325 | 0.282 | 1 | 5 | Vps13a | 0.043 | Old |
| 0.097159 | 0.85877 | 0.029 | 0.013 | 1 | 5 | 1110035H1 | 0.016 | Old |
| 0.097311 | 0.356718 | 0.118 | 0.084 | 1 | 5 | Man2c1 | 0.034 | Old |
| 0.097519 | 0.718841 | 0.029 | 0.013 | 1 | 5 | Msh5 | 0.016 | Old |
| 0.09757 | 1.188488 | 0.017 | 0.005 | 1 | 5 | Ndrg2 | 0.012 | Old |
| 0.097767 | 0.423798 | 0.093 | 0.063 | 1 | 5 | Taf12 | 0.03 | Old |
| 0.098204 | 0.225081 | 0.227 | 0.185 | 1 | 5 | Ldah | 0.042 | Old |
| 0.098312 | 0.384244 | 0.126 | 0.091 | 1 | 5 | Egfr | 0.035 | Old |
| 0.098544 | 1.107685 | 0.017 | 0.005 | 1 | 5 | Anxa9 | 0.012 | Old |
| 0.098789 | 1.267016 | 0.017 | 0.005 | 1 | 5 | Plac8 | 0.012 | Old |
| 0.098789 | 1.007788 | 0.017 | 0.005 | 1 | 5 | Top1mt | 0.012 | Old |
| 0.098839 | 0.792913 | 0.052 | 0.03 | 1 | 5 | Eps8l1 | 0.022 | Old |
| 0.098854 | 0.307668 | 0.323 | 0.272 | 1 | 5 | Sec11a | 0.051 | Old |
| 0.099035 | 1.186807 | 0.017 | 0.005 | 1 | 5 | Fbxo44 | 0.012 | Old |
| 0.099281 | 1.250798 | 0.017 | 0.005 | 1 | 5 | Nes | 0.012 | Old |
| 0.099281 | 1.0346 | 0.017 | 0.005 | 1 | 5 | Tnfsf10 | 0.012 | Old |
| 0.099527 | 1.105712 | 0.017 | 0.005 | 1 | 5 | Frk | 0.012 | Old |
| 0.099527 | 1.024197 | 0.017 | 0.005 | 1 | 5 | Macrod2 | 0.012 | Old |

| | | | | | | | | |
|---|---|---|---|---|---|---|---|---|
| 0.099657 | 0.239794 | 0.391 | 0.34 | 1 | 5 | Tcp1 | 0.051 | Old |
| 0.099956 | 0.539925 | 0.377 | 0.345 | 1 | 5 | Gda | 0.032 | Old |
| 0.099962 | 0.112363 | 0.813 | 0.741 | 1 | 5 | Hnrnpu | 0.072 | Old |
| 0.100021 | 0.624275 | 0.017 | 0.005 | 1 | 5 | Plppr3 | 0.012 | Old |
| 0.100081 | 0.363631 | 0.104 | 0.074 | 1 | 5 | Traf6 | 0.03 | Old |
| 0.10021 | 0.62342 | 0.074 | 0.048 | 1 | 5 | Col4a1 | 0.026 | Old |
| 0.100643 | 0.877415 | 0.032 | 0.015 | 1 | 5 | Timp3 | 0.017 | Old |
| 0.100804 | 0.551787 | 0.029 | 0.013 | 1 | 5 | Ispd | 0.016 | Old |
| 0.100817 | 0.828889 | 0.032 | 0.015 | 1 | 5 | Apex2 | 0.017 | Old |
| 0.100829 | 0.182536 | 0.549 | 0.505 | 1 | 5 | Ewsr1 | 0.044 | Old |
| 0.100921 | 0.321997 | 0.189 | 0.152 | 1 | 5 | Nop58 | 0.037 | Old |
| 0.100955 | 0.813378 | 0.045 | 0.025 | 1 | 5 | Tti1 | 0.02 | Old |
| 0.100974 | 0.476231 | 0.101 | 0.071 | 1 | 5 | Fam98a | 0.03 | Old |
| 0.101174 | 0.281161 | 0.029 | 0.013 | 1 | 5 | L3mbtl2 | 0.016 | Old |
| 0.101249 | 0.938424 | 0.035 | 0.018 | 1 | 5 | Zgpat | 0.017 | Old |
| 0.101273 | 0.021287 | 1 | 0.997 | 1 | 5 | mt-Cytb | 0.003 | Old |
| 0.101487 | 0.124702 | 0.992 | 0.982 | 1 | 5 | H3f3b | 0.01 | Old |
| 0.1015 | 0.535027 | 0.113 | 0.081 | 1 | 5 | Tmx4 | 0.032 | Old |
| 0.101514 | 0.909217 | 0.032 | 0.015 | 1 | 5 | Ppif | 0.017 | Old |
| 0.101616 | 0.215977 | 0.562 | 0.51 | 1 | 5 | Pias1 | 0.052 | Old |
| 0.101798 | 0.075228 | 0.278 | 0.226 | 1 | 5 | Ctbp2 | 0.052 | Old |
| 0.10184 | 0.268966 | 0.481 | 0.452 | 1 | 5 | Rnf19b | 0.029 | Old |
| 0.101913 | 0.255639 | 0.375 | 0.325 | 1 | 5 | Bcl3 | 0.05 | Old |
| 0.101966 | 0.820481 | 0.045 | 0.025 | 1 | 5 | Kptn | 0.02 | Old |
| 0.101985 | 0.136387 | 0.175 | 0.135 | 1 | 5 | Gm46224 | 0.04 | Old |
| 0.101994 | 0.698718 | 0.042 | 0.023 | 1 | 5 | Gm16638 | 0.019 | Old |
| 0.102335 | 0.500869 | 0.13 | 0.096 | 1 | 5 | Dusp23 | 0.034 | Old |
| 0.102567 | 0.659971 | 0.032 | 0.015 | 1 | 5 | Dennd5b | 0.017 | Old |
| 0.102653 | 0.768125 | 0.067 | 0.043 | 1 | 5 | Med9 | 0.024 | Old |
| 0.102768 | 0.158799 | 0.244 | 0.201 | 1 | 5 | Nup88 | 0.043 | Old |
| 0.102832 | 0.508031 | 0.081 | 0.053 | 1 | 5 | Klhl13 | 0.028 | Old |
| 0.103096 | 0.530479 | 0.032 | 0.015 | 1 | 5 | Ficd | 0.017 | Old |
| 0.103235 | 0.323854 | 0.2 | 0.16 | 1 | 5 | Eri1 | 0.04 | Old |
| 0.103312 | 0.337528 | 0.163 | 0.127 | 1 | 5 | Epor | 0.036 | Old |
| 0.103649 | 0.130157 | 0.517 | 0.467 | 1 | 5 | Slfn5 | 0.05 | Old |
| 0.104199 | 0.201614 | 0.153 | 0.114 | 1 | 5 | Armc5 | 0.039 | Old |
| 0.10443 | 0.493297 | 0.146 | 0.112 | 1 | 5 | Atp8b4 | 0.034 | Old |
| 0.10469 | 0.163493 | 0.109 | 0.079 | 1 | 5 | Agpat2 | 0.03 | Old |
| 0.10483 | 0.367296 | 0.29 | 0.249 | 1 | 5 | Mink1 | 0.041 | Old |
| 0.104988 | 0.480237 | 0.143 | 0.109 | 1 | 5 | Fkbp9 | 0.034 | Old |
| 0.105494 | 0.557694 | 0.045 | 0.025 | 1 | 5 | Tmem206 | 0.02 | Old |
| 0.105733 | 0.131583 | 0.684 | 0.635 | 1 | 5 | Gnas | 0.049 | Old |
| 0.106029 | 0.564455 | 0.049 | 0.028 | 1 | 5 | Fads3 | 0.021 | Old |
| 0.106241 | 0.76799 | 0.045 | 0.025 | 1 | 5 | Ankrd52 | 0.02 | Old |
| 0.106542 | 0.805683 | 0.072 | 0.048 | 1 | 5 | Pym1 | 0.024 | Old |

| | | | | | | | | |
|---|---|---|---|---|---|---|---|---|
| 0.106681 | 0.129691 | 0.34 | 0.282 | 1 | 5 | Tulp4 | 0.058 | Old |
| 0.106686 | 0.61795 | 0.123 | 0.091 | 1 | 5 | Scyl1 | 0.032 | Old |
| 0.106734 | 0.471541 | 0.14 | 0.107 | 1 | 5 | Ptk2 | 0.033 | Old |
| 0.107093 | 0.113342 | 0.813 | 0.741 | 1 | 5 | Cyth4 | 0.072 | Old |
| 0.107438 | 0.316711 | 0.276 | 0.236 | 1 | 5 | Abcc5 | 0.04 | Old |
| 0.107455 | 0.374292 | 0.104 | 0.074 | 1 | 5 | Alg8 | 0.03 | Old |
| 0.107533 | 0.248934 | 0.406 | 0.365 | 1 | 5 | Lima1 | 0.041 | Old |
| 0.10769 | 0.539176 | 0.071 | 0.046 | 1 | 5 | Cyp2j6 | 0.025 | Old |
| 0.107757 | 0.234414 | 0.209 | 0.168 | 1 | 5 | Nt5c | 0.041 | Old |
| 0.107779 | 0.217313 | 0.394 | 0.35 | 1 | 5 | Psmd11 | 0.044 | Old |
| 0.107831 | 0.094165 | 0.099 | 0.069 | 1 | 5 | Rsad2 | 0.03 | Old |
| 0.108198 | 0.482016 | 0.155 | 0.122 | 1 | 5 | Map3k3 | 0.033 | Old |
| 0.108375 | 0.614231 | 0.079 | 0.053 | 1 | 5 | Tpm3-rs7 | 0.026 | Old |
| 0.108387 | 0.47839 | 0.086 | 0.058 | 1 | 5 | Agfg2 | 0.028 | Old |
| 0.108508 | 0.463237 | 0.045 | 0.025 | 1 | 5 | Abhd14a | 0.02 | Old |
| 0.108539 | 0.174921 | 0.569 | 0.538 | 1 | 5 | Kitl | 0.031 | Old |
| 0.10868 | 0.322209 | 0.17 | 0.135 | 1 | 5 | Iah1 | 0.035 | Old |
| 0.108759 | 0.201857 | 0.273 | 0.226 | 1 | 5 | Plagl2 | 0.047 | Old |
| 0.109314 | 1.560828 | 0.02 | 0.008 | 1 | 5 | Ggnbp1 | 0.012 | Old |
| 0.109609 | 0.199631 | 0.269 | 0.226 | 1 | 5 | Fez2 | 0.043 | Old |
| 0.109653 | 0.202812 | 0.392 | 0.353 | 1 | 5 | Phc3 | 0.039 | Old |
| 0.109682 | 0.401989 | 0.141 | 0.107 | 1 | 5 | Hcfc2 | 0.034 | Old |
| 0.110014 | 0.22811 | 0.343 | 0.297 | 1 | 5 | Ctsf | 0.046 | Old |
| 0.110029 | 1.479205 | 0.02 | 0.008 | 1 | 5 | Taf4b | 0.012 | Old |
| 0.110332 | 0.344988 | 0.219 | 0.178 | 1 | 5 | Hspa4l | 0.041 | Old |
| 0.110509 | 1.335845 | 0.02 | 0.008 | 1 | 5 | Insl6 | 0.012 | Old |
| 0.110544 | 0.360685 | 0.232 | 0.195 | 1 | 5 | Dnajc15 | 0.037 | Old |
| 0.110709 | 0.215821 | 0.165 | 0.127 | 1 | 5 | Zfp622 | 0.038 | Old |
| 0.110862 | 0.127477 | 0.32 | 0.269 | 1 | 5 | Stip1 | 0.051 | Old |
| 0.111012 | 0.125058 | 0.081 | 0.053 | 1 | 5 | Nt5dc3 | 0.028 | Old |
| 0.111114 | 0.257636 | 0.157 | 0.119 | 1 | 5 | Creb3 | 0.038 | Old |
| 0.11117 | 0.34716 | 0.162 | 0.124 | 1 | 5 | Slc35c2 | 0.038 | Old |
| 0.111491 | 1.555767 | 0.012 | 0.003 | 1 | 5 | Gm3435 | 0.009 | Old |
| 0.111491 | 1.528949 | 0.012 | 0.003 | 1 | 5 | Osbpl5 | 0.009 | Old |
| 0.111665 | 0.668574 | 0.084 | 0.058 | 1 | 5 | Rnf8 | 0.026 | Old |
| 0.111713 | 0.134522 | 0.337 | 0.282 | 1 | 5 | Snx13 | 0.055 | Old |
| 0.111714 | 1.115212 | 0.02 | 0.008 | 1 | 5 | Zfp58 | 0.012 | Old |
| 0.111821 | 1.264568 | 0.012 | 0.003 | 1 | 5 | Hsd17b14 | 0.009 | Old |
| 0.111821 | 1.137399 | 0.012 | 0.003 | 1 | 5 | Cep170b | 0.009 | Old |
| 0.111821 | 1.121297 | 0.012 | 0.003 | 1 | 5 | Sfxn4 | 0.009 | Old |
| 0.111919 | 0.468296 | 0.162 | 0.127 | 1 | 5 | Bcl2a1d | 0.035 | Old |
| 0.112152 | 1.345354 | 0.012 | 0.003 | 1 | 5 | Zscan18 | 0.009 | Old |
| 0.112152 | 1.089445 | 0.012 | 0.003 | 1 | 5 | Cfb | 0.009 | Old |
| 0.112207 | 0.118207 | 0.658 | 0.602 | 1 | 5 | Dusp3 | 0.056 | Old |
| 0.112213 | 0.127296 | 0.424 | 0.365 | 1 | 5 | Tnfsf9 | 0.059 | Old |

| | | | | | | | | |
|---|---|---|---|---|---|---|---|---|
| 0.112358 | 0.523324 | 0.125 | 0.094 | 1 | 5 | Hnmt | 0.031 | Old |
| 0.112442 | 1.208754 | 0.02 | 0.008 | 1 | 5 | Hspa12a | 0.012 | Old |
| 0.112484 | 1.298871 | 0.012 | 0.003 | 1 | 5 | Hspb6 | 0.009 | Old |
| 0.112484 | 1.11913 | 0.012 | 0.003 | 1 | 5 | A830052D1 | 0.009 | Old |
| 0.112484 | 1.024479 | 0.012 | 0.003 | 1 | 5 | Zfp128 | 0.009 | Old |
| 0.112484 | 0.45813 | 0.194 | 0.16 | 1 | 5 | Arid2 | 0.034 | Old |
| 0.112594 | 0.329142 | 0.222 | 0.183 | 1 | 5 | Baiap2 | 0.039 | Old |
| 0.112673 | 0.041192 | 0.7 | 0.624 | 1 | 5 | Ifrd1 | 0.076 | Old |
| 0.112685 | 0.184379 | 0.229 | 0.188 | 1 | 5 | 4930402H2 | 0.041 | Old |
| 0.112842 | 0.100261 | 1 | 0.997 | 1 | 5 | Ctsc | 0.003 | Old |
| 0.112906 | 0.285588 | 0.367 | 0.322 | 1 | 5 | Csnk1d | 0.045 | Old |
| 0.11293 | 1.023451 | 0.02 | 0.008 | 1 | 5 | Adamts5 | 0.012 | Old |
| 0.11293 | 1.012387 | 0.02 | 0.008 | 1 | 5 | Pomgnt2 | 0.012 | Old |
| 0.113008 | 0.832803 | 0.054 | 0.033 | 1 | 5 | Zfp637 | 0.021 | Old |
| 0.113142 | 0.459866 | 0.067 | 0.043 | 1 | 5 | Gstm5 | 0.024 | Old |
| 0.11315 | 1.0837 | 0.012 | 0.003 | 1 | 5 | Nudt6 | 0.009 | Old |
| 0.113172 | 0.328316 | 0.165 | 0.129 | 1 | 5 | Ppp5c | 0.036 | Old |
| 0.113174 | 1.070016 | 0.02 | 0.008 | 1 | 5 | Tgfa | 0.012 | Old |
| 0.113421 | 0.092448 | 0.204 | 0.16 | 1 | 5 | Syne2 | 0.044 | Old |
| 0.113484 | 0.711243 | 0.012 | 0.003 | 1 | 5 | Dact1 | 0.009 | Old |
| 0.113497 | 0.277349 | 0.278 | 0.236 | 1 | 5 | Plekhm2 | 0.042 | Old |
| 0.113664 | 1.011498 | 0.02 | 0.008 | 1 | 5 | Sphk1 | 0.012 | Old |
| 0.113676 | 0.488955 | 0.114 | 0.084 | 1 | 5 | Slc39a13 | 0.03 | Old |
| 0.113819 | 0.779384 | 0.012 | 0.003 | 1 | 5 | 4833413G1 | 0.009 | Old |
| 0.113819 | 0.629775 | 0.012 | 0.003 | 1 | 5 | Tmem240 | 0.009 | Old |
| 0.113886 | 0.720961 | 0.084 | 0.058 | 1 | 5 | Pxmp4 | 0.026 | Old |
| 0.113908 | 0.153306 | 0.468 | 0.416 | 1 | 5 | Clcn5 | 0.052 | Old |
| 0.113961 | 0.537965 | 0.101 | 0.074 | 1 | 5 | 2410002F2 | 0.027 | Old |
| 0.114013 | 0.23523 | 0.207 | 0.165 | 1 | 5 | Mrps15 | 0.042 | Old |
| 0.114214 | 0.283056 | 0.138 | 0.104 | 1 | 5 | Eya3 | 0.034 | Old |
| 0.114261 | 0.15255 | 0.409 | 0.353 | 1 | 5 | Tlr7 | 0.056 | Old |
| 0.114468 | 0.646902 | 0.057 | 0.036 | 1 | 5 | Clip2 | 0.021 | Old |
| 0.114497 | 0.120231 | 0.246 | 0.201 | 1 | 5 | Tsg101 | 0.045 | Old |
| 0.11465 | 0.77306 | 0.02 | 0.008 | 1 | 5 | Jag1 | 0.012 | Old |
| 0.114762 | 0.227647 | 0.222 | 0.183 | 1 | 5 | Slc30a7 | 0.039 | Old |
| 0.114935 | 0.124299 | 0.224 | 0.183 | 1 | 5 | Slc20a1 | 0.041 | Old |
| 0.115145 | 0.265578 | 0.561 | 0.503 | 1 | 5 | AC160336. | 0.058 | Old |
| 0.115232 | 0.065465 | 0.655 | 0.576 | 1 | 5 | Slc9a9 | 0.079 | Old |
| 0.115239 | 0.243324 | 0.121 | 0.089 | 1 | 5 | Ankrd27 | 0.032 | Old |
| 0.115742 | 0.356124 | 0.298 | 0.259 | 1 | 5 | Mfsd14a | 0.039 | Old |
| 0.116382 | 0.24031 | 0.195 | 0.155 | 1 | 5 | Elovl1 | 0.04 | Old |
| 0.116497 | 0.136655 | 0.278 | 0.228 | 1 | 5 | Ssbp4 | 0.05 | Old |
| 0.1165 | 0.532225 | 0.064 | 0.041 | 1 | 5 | Recql5 | 0.023 | Old |
| 0.116583 | 0.406962 | 0.106 | 0.076 | 1 | 5 | Avpi1 | 0.03 | Old |
| 0.116706 | 0.181838 | 0.236 | 0.188 | 1 | 5 | Gsk3a | 0.048 | Old |

| | | | | | | | | |
|---|---|---|---|---|---|---|---|---|
| 0.116966 | 0.195917 | 0.279 | 0.234 | 1 | 5 | Rufy3 | 0.045 | Old |
| 0.117052 | 0.188114 | 0.113 | 0.081 | 1 | 5 | Srprb | 0.032 | Old |
| 0.117067 | 0.416164 | 0.094 | 0.066 | 1 | 5 | Dnmbp | 0.028 | Old |
| 0.117095 | 0.229459 | 0.254 | 0.216 | 1 | 5 | Pisd | 0.038 | Old |
| 0.117268 | 0.034518 | 0.507 | 0.439 | 1 | 5 | Txn1 | 0.068 | Old |
| 0.117436 | 0.103039 | 0.641 | 0.609 | 1 | 5 | Tspo | 0.032 | Old |
| 0.117456 | 0.235551 | 0.128 | 0.094 | 1 | 5 | Lman2l | 0.034 | Old |
| 0.117565 | 0.500219 | 0.138 | 0.107 | 1 | 5 | Ap5m1 | 0.031 | Old |
| 0.117701 | 0.242262 | 0.256 | 0.213 | 1 | 5 | Rab22a | 0.043 | Old |
| 0.117902 | 0.597475 | 0.084 | 0.058 | 1 | 5 | Atat1 | 0.026 | Old |
| 0.117989 | 1.13439 | 0.024 | 0.01 | 1 | 5 | 3-Mar | 0.014 | Old |
| 0.118118 | 0.101428 | 0.86 | 0.817 | 1 | 5 | Gm10076 | 0.043 | Old |
| 0.118237 | 0.468176 | 0.079 | 0.053 | 1 | 5 | Nudt2 | 0.026 | Old |
| 0.118452 | 1.173356 | 0.024 | 0.01 | 1 | 5 | Kpna2 | 0.014 | Old |
| 0.118548 | 0.357896 | 0.338 | 0.299 | 1 | 5 | Kdm5a | 0.039 | Old |
| 0.118578 | 0.307899 | 0.152 | 0.117 | 1 | 5 | Rmi1 | 0.035 | Old |
| 0.118748 | 0.16648 | 0.591 | 0.556 | 1 | 5 | Eif5a | 0.035 | Old |
| 0.119001 | 0.300289 | 0.12 | 0.089 | 1 | 5 | Slc25a12 | 0.031 | Old |
| 0.119629 | 0.320057 | 0.123 | 0.091 | 1 | 5 | Rbmx | 0.032 | Old |
| 0.120317 | 0.684958 | 0.024 | 0.01 | 1 | 5 | Chac2 | 0.014 | Old |
| 0.120339 | 0.15738 | 0.207 | 0.168 | 1 | 5 | Naga | 0.039 | Old |
| 0.120699 | 1.130428 | 0.04 | 0.023 | 1 | 5 | Slx4 | 0.017 | Old |
| 0.120781 | 0.144549 | 0.34 | 0.289 | 1 | 5 | Prkx | 0.051 | Old |
| 0.121007 | 0.245322 | 0.082 | 0.056 | 1 | 5 | Limk1 | 0.026 | Old |
| 0.121067 | 0.636398 | 0.057 | 0.036 | 1 | 5 | Nbas | 0.021 | Old |
| 0.121125 | 0.367161 | 0.094 | 0.066 | 1 | 5 | Sp4 | 0.028 | Old |
| 0.121162 | 0.468635 | 0.194 | 0.16 | 1 | 5 | Senp1 | 0.034 | Old |
| 0.121197 | 0.281573 | 0.591 | 0.558 | 1 | 5 | Peli1 | 0.033 | Old |
| 0.121533 | 0.272472 | 0.311 | 0.266 | 1 | 5 | Timm10b | 0.045 | Old |
| 0.122444 | 1.288957 | 0.024 | 0.01 | 1 | 5 | Itgax | 0.014 | Old |
| 0.122485 | 0.290938 | 0.418 | 0.381 | 1 | 5 | Atp13a3 | 0.037 | Old |
| 0.122676 | 0.225497 | 0.574 | 0.515 | 1 | 5 | Arhgap17 | 0.059 | Old |
| 0.122682 | 0.923699 | 0.024 | 0.01 | 1 | 5 | Npr1 | 0.014 | Old |
| 0.122704 | 0.219559 | 0.194 | 0.15 | 1 | 5 | Twistnb | 0.044 | Old |
| 0.122774 | 0.457296 | 0.088 | 0.061 | 1 | 5 | Slc25a19 | 0.027 | Old |
| 0.12292 | 0.708419 | 0.024 | 0.01 | 1 | 5 | Zfp668 | 0.014 | Old |
| 0.123325 | 0.24404 | 0.231 | 0.19 | 1 | 5 | Chd1 | 0.041 | Old |
| 0.123645 | 0.126769 | 0.902 | 0.883 | 1 | 5 | Cyba | 0.019 | Old |
| 0.123652 | 0.480614 | 0.069 | 0.046 | 1 | 5 | Srrd | 0.023 | Old |
| 0.123808 | 1.207772 | 0.03 | 0.015 | 1 | 5 | Spef1 | 0.015 | Old |
| 0.123878 | 0.612182 | 0.024 | 0.01 | 1 | 5 | Rims3 | 0.014 | Old |
| 0.124215 | 0.918451 | 0.074 | 0.051 | 1 | 5 | Zbtb46 | 0.023 | Old |
| 0.124241 | 1.027262 | 0.04 | 0.023 | 1 | 5 | Abcb4 | 0.017 | Old |
| 0.124464 | 0.702155 | 0.051 | 0.03 | 1 | 5 | Tgds | 0.021 | Old |
| 0.124593 | 0.440363 | 0.076 | 0.051 | 1 | 5 | Fgr | 0.025 | Old |

| | | | | | | | | |
|---|---|---|---|---|---|---|---|---|
| 0.124845 | 0.425806 | 0.168 | 0.135 | 1 | 5 | Trim27 | 0.033 | Old |
| 0.125076 | 0.479042 | 0.126 | 0.096 | 1 | 5 | Pigq | 0.03 | Old |
| 0.125128 | 0.214801 | 0.215 | 0.175 | 1 | 5 | Ubald1 | 0.04 | Old |
| 0.125349 | 0.134413 | 0.498 | 0.442 | 1 | 5 | Fgd4 | 0.056 | Old |
| 0.12539 | 0.399973 | 0.2 | 0.165 | 1 | 5 | Plgrkt | 0.035 | Old |
| 0.125476 | 0.133885 | 0.261 | 0.213 | 1 | 5 | Slc50a1 | 0.048 | Old |
| 0.125586 | 0.619151 | 0.051 | 0.03 | 1 | 5 | Zfp319 | 0.021 | Old |
| 0.125658 | 0.103469 | 0.88 | 0.815 | 1 | 5 | Tm6sf1 | 0.065 | Old |
| 0.125701 | 0.433969 | 0.135 | 0.104 | 1 | 5 | Mrps7 | 0.031 | Old |
| 0.125867 | 0.270846 | 0.173 | 0.135 | 1 | 5 | Ppil4 | 0.038 | Old |
| 0.12626 | 0.600682 | 0.066 | 0.043 | 1 | 5 | Med6 | 0.023 | Old |
| 0.126439 | 0.348117 | 0.256 | 0.216 | 1 | 5 | C1qtnf1 | 0.04 | Old |
| 0.126676 | 1.065407 | 0.044 | 0.025 | 1 | 5 | Gm20045 | 0.019 | Old |
| 0.126683 | 0.607657 | 0.066 | 0.043 | 1 | 5 | Zscan21 | 0.023 | Old |
| 0.127298 | 0.155265 | 0.421 | 0.371 | 1 | 5 | Zyx | 0.05 | Old |
| 0.127516 | 0.158058 | 0.064 | 0.041 | 1 | 5 | Src | 0.023 | Old |
| 0.127579 | 0.102412 | 0.481 | 0.421 | 1 | 5 | Acer3 | 0.06 | Old |
| 0.127607 | 0.181927 | 0.468 | 0.421 | 1 | 5 | Nup98 | 0.047 | Old |
| 0.12783 | 0.547718 | 0.069 | 0.046 | 1 | 5 | 2310061I0 | 0.023 | Old |
| 0.127877 | 1.215125 | 0.034 | 0.018 | 1 | 5 | Zdhhc24 | 0.016 | Old |
| 0.127901 | 0.814378 | 0.027 | 0.013 | 1 | 5 | Cacna1f | 0.014 | Old |
| 0.127956 | 0.230701 | 0.276 | 0.234 | 1 | 5 | Baz2a | 0.042 | Old |
| 0.127984 | 0.119973 | 0.675 | 0.624 | 1 | 5 | Tra2b | 0.051 | Old |
| 0.127986 | 0.094351 | 0.547 | 0.497 | 1 | 5 | Ctnnd1 | 0.05 | Old |
| 0.128003 | 0.388779 | 0.18 | 0.145 | 1 | 5 | Mboat7 | 0.035 | Old |
| 0.128074 | 0.079553 | 0.492 | 0.424 | 1 | 5 | Eif1a | 0.068 | Old |
| 0.128415 | 0.197515 | 0.481 | 0.439 | 1 | 5 | Hivep2 | 0.042 | Old |
| 0.128604 | 0.158127 | 0.146 | 0.112 | 1 | 5 | Wsb2 | 0.034 | Old |
| 0.128834 | 0.14697 | 0.458 | 0.406 | 1 | 5 | Ppt1 | 0.052 | Old |
| 0.128835 | 0.54085 | 0.051 | 0.03 | 1 | 5 | 4933433G1 | 0.021 | Old |
| 0.128949 | 0.237331 | 0.327 | 0.282 | 1 | 5 | Ext1 | 0.045 | Old |
| 0.12909 | 0.266054 | 0.146 | 0.112 | 1 | 5 | Kpna6 | 0.034 | Old |
| 0.129274 | 0.088689 | 0.818 | 0.754 | 1 | 5 | Cox7c | 0.064 | Old |
| 0.12941 | 0.287325 | 0.099 | 0.071 | 1 | 5 | Wdsub1 | 0.028 | Old |
| 0.129583 | 0.217566 | 0.109 | 0.079 | 1 | 5 | Fcf1 | 0.03 | Old |
| 0.129704 | 0.904726 | 0.034 | 0.018 | 1 | 5 | Pcgf1 | 0.016 | Old |
| 0.129704 | 0.859575 | 0.04 | 0.023 | 1 | 5 | Zfp316 | 0.017 | Old |
| 0.129889 | 0.875464 | 0.04 | 0.023 | 1 | 5 | Svip | 0.017 | Old |
| 0.129931 | 0.215784 | 0.357 | 0.31 | 1 | 5 | Ndufs7 | 0.047 | Old |
| 0.130102 | 0.389209 | 0.298 | 0.256 | 1 | 5 | Azi2 | 0.042 | Old |
| 0.130163 | 0.145873 | 0.397 | 0.355 | 1 | 5 | Ndufa1 | 0.042 | Old |
| 0.130765 | 0.101897 | 0.685 | 0.619 | 1 | 5 | Sgpl1 | 0.066 | Old |
| 0.131067 | 0.25238 | 0.061 | 0.038 | 1 | 5 | Mapk1ip1 | 0.023 | Old |
| 0.13113 | 0.504209 | 0.072 | 0.048 | 1 | 5 | Capn15 | 0.024 | Old |
| 0.131138 | 1.049632 | 0.034 | 0.018 | 1 | 5 | Det1 | 0.016 | Old |

| | | | | | | | | |
|---|---|---|---|---|---|---|---|---|
| 0.131379 | 0.70773 | 0.04 | 0.023 | 1 | 5 | Cmss1 | 0.017 | Old |
| 0.131905 | 0.199487 | 0.396 | 0.35 | 1 | 5 | Usp24 | 0.046 | Old |
| 0.131952 | 0.1622 | 0.215 | 0.178 | 1 | 5 | Senp2 | 0.037 | Old |
| 0.13206 | 0.760731 | 0.027 | 0.013 | 1 | 5 | Tnfrsf26 | 0.014 | Old |
| 0.132317 | 0.738784 | 0.04 | 0.023 | 1 | 5 | Cd300ld2 | 0.017 | Old |
| 0.132317 | 0.614802 | 0.04 | 0.023 | 1 | 5 | Gm15859 | 0.017 | Old |
| 0.132637 | 0.481116 | 0.114 | 0.086 | 1 | 5 | Cdc16 | 0.028 | Old |
| 0.13267 | 0.76018 | 0.104 | 0.079 | 1 | 5 | Plekhn1 | 0.025 | Old |
| 0.132945 | 0.363252 | 0.133 | 0.102 | 1 | 5 | Lpcat3 | 0.031 | Old |
| 0.133532 | 0.34597 | 0.148 | 0.117 | 1 | 5 | Fam96b | 0.031 | Old |
| 0.133876 | 0.128799 | 0.205 | 0.165 | 1 | 5 | Eng | 0.04 | Old |
| 0.133942 | 0.246955 | 0.221 | 0.183 | 1 | 5 | Ube2d1 | 0.038 | Old |
| 0.134198 | 0.170869 | 0.774 | 0.728 | 1 | 5 | Lst1 | 0.046 | Old |
| 0.134256 | 0.397517 | 0.152 | 0.119 | 1 | 5 | Faf2 | 0.033 | Old |
| 0.134549 | 0.041372 | 1 | 1 | 1 | 5 | C1qa | 0 | Old |
| 0.134657 | 0.270887 | 0.332 | 0.289 | 1 | 5 | Tmcc3 | 0.043 | Old |
| 0.134758 | 0.081645 | 0.16 | 0.124 | 1 | 5 | Tesk1 | 0.036 | Old |
| 0.134904 | 0.166753 | 0.493 | 0.437 | 1 | 5 | Mbnl2 | 0.056 | Old |
| 0.135022 | 0.709738 | 0.03 | 0.015 | 1 | 5 | Map4k1 | 0.015 | Old |
| 0.135344 | 0.335991 | 0.084 | 0.058 | 1 | 5 | Pgm1 | 0.026 | Old |
| 0.135511 | 0.172826 | 0.439 | 0.388 | 1 | 5 | Slc15a3 | 0.051 | Old |
| 0.135673 | 0.14247 | 0.739 | 0.726 | 1 | 5 | Tgfb1 | 0.013 | Old |
| 0.135692 | 0.434507 | 0.03 | 0.015 | 1 | 5 | Rpl9-ps6 | 0.015 | Old |
| 0.135927 | 0.263641 | 0.212 | 0.175 | 1 | 5 | Spast | 0.037 | Old |
| 0.135984 | 0.554153 | 0.081 | 0.056 | 1 | 5 | Mthfsl | 0.025 | Old |
| 0.136169 | 0.307515 | 0.076 | 0.051 | 1 | 5 | Pgghg | 0.025 | Old |
| 0.136191 | 0.17602 | 0.224 | 0.183 | 1 | 5 | Dscr3 | 0.041 | Old |
| 0.136202 | 0.230796 | 0.229 | 0.19 | 1 | 5 | Tcf7l2 | 0.039 | Old |
| 0.136365 | 0.574518 | 0.044 | 0.025 | 1 | 5 | Mrpl49 | 0.019 | Old |
| 0.136614 | 0.113455 | 0.689 | 0.624 | 1 | 5 | Dock2 | 0.065 | Old |
| 0.136742 | 0.563101 | 0.091 | 0.066 | 1 | 5 | Phf6 | 0.025 | Old |
| 0.136788 | 0.418206 | 0.034 | 0.018 | 1 | 5 | Fam222a | 0.016 | Old |
| 0.136818 | 1.49143 | 0.015 | 0.005 | 1 | 5 | Stab2 | 0.01 | Old |
| 0.137053 | 0.222194 | 0.121 | 0.089 | 1 | 5 | Gsr | 0.032 | Old |
| 0.137287 | 0.192031 | 0.182 | 0.145 | 1 | 5 | Cfdp1 | 0.037 | Old |
| 0.137827 | 0.554824 | 0.138 | 0.109 | 1 | 5 | Ern1 | 0.029 | Old |
| 0.137837 | 0.434763 | 0.131 | 0.102 | 1 | 5 | Nvl | 0.029 | Old |
| 0.137945 | 0.764752 | 0.03 | 0.015 | 1 | 5 | Endov | 0.015 | Old |
| 0.138105 | 0.437556 | 0.074 | 0.051 | 1 | 5 | Ss18l2 | 0.023 | Old |
| 0.138162 | 0.299758 | 0.209 | 0.173 | 1 | 5 | Txndc5 | 0.036 | Old |
| 0.138248 | 0.152187 | 0.222 | 0.183 | 1 | 5 | Tmem229b | 0.039 | Old |
| 0.138371 | 0.708901 | 0.059 | 0.038 | 1 | 5 | Fancg | 0.021 | Old |
| 0.138479 | 1.186506 | 0.015 | 0.005 | 1 | 5 | Slc16a12 | 0.01 | Old |
| 0.138479 | 1.17374 | 0.015 | 0.005 | 1 | 5 | En2 | 0.01 | Old |
| 0.138481 | 0.141233 | 0.458 | 0.414 | 1 | 5 | Mapk1 | 0.044 | Old |

| | | | | | | | | |
|---|---|---|---|---|---|---|---|---|
| 0.138504 | 0.084316 | 0.921 | 0.888 | 1 | 5 | Ptprc | 0.033 | Old |
| 0.138628 | 0.376752 | 0.062 | 0.041 | 1 | 5 | Sh3d19 | 0.021 | Old |
| 0.138643 | 0.172906 | 0.253 | 0.208 | 1 | 5 | Oasl2 | 0.045 | Old |
| 0.138789 | 0.211273 | 0.037 | 0.02 | 1 | 5 | Dhodh | 0.017 | Old |
| 0.138813 | 1.168381 | 0.015 | 0.005 | 1 | 5 | Ghr | 0.01 | Old |
| 0.138842 | 0.164689 | 0.281 | 0.236 | 1 | 5 | Arih2 | 0.045 | Old |
| 0.13901 | 0.553456 | 0.059 | 0.038 | 1 | 5 | Atp6v0d2 | 0.021 | Old |
| 0.139147 | 1.03357 | 0.015 | 0.005 | 1 | 5 | Gm5113 | 0.01 | Old |
| 0.139183 | 0.274789 | 0.204 | 0.165 | 1 | 5 | Nf1 | 0.039 | Old |
| 0.139326 | 0.074704 | 0.396 | 0.335 | 1 | 5 | Inpp4a | 0.061 | Old |
| 0.139358 | 0.336496 | 0.034 | 0.018 | 1 | 5 | Pcdhga12 | 0.016 | Old |
| 0.139437 | 0.47753 | 0.145 | 0.117 | 1 | 5 | Mafk | 0.028 | Old |
| 0.139482 | 1.097002 | 0.015 | 0.005 | 1 | 5 | Gspt2 | 0.01 | Old |
| 0.139482 | 1.030674 | 0.015 | 0.005 | 1 | 5 | Nsl1 | 0.01 | Old |
| 0.139535 | 0.124435 | 0.579 | 0.528 | 1 | 5 | Rpl27 | 0.051 | Old |
| 0.139608 | 0.48202 | 0.037 | 0.02 | 1 | 5 | Ada | 0.017 | Old |
| 0.139788 | 0.28976 | 0.355 | 0.327 | 1 | 5 | Rhoc | 0.028 | Old |
| 0.139818 | 1.160043 | 0.015 | 0.005 | 1 | 5 | Mlph | 0.01 | Old |
| 0.139863 | 0.346575 | 0.345 | 0.307 | 1 | 5 | Arl5b | 0.038 | Old |
| 0.140084 | 0.30433 | 0.089 | 0.063 | 1 | 5 | C1ra | 0.026 | Old |
| 0.140284 | 0.057692 | 0.088 | 0.061 | 1 | 5 | Zkscan8 | 0.027 | Old |
| 0.140533 | 0.325215 | 0.168 | 0.135 | 1 | 5 | Mrpl34 | 0.033 | Old |
| 0.141007 | 0.100874 | 0.239 | 0.198 | 1 | 5 | Slc48a1 | 0.041 | Old |
| 0.141244 | 0.283873 | 0.229 | 0.188 | 1 | 5 | Dync1li1 | 0.041 | Old |
| 0.141331 | 0.141088 | 0.589 | 0.548 | 1 | 5 | Rab3il1 | 0.041 | Old |
| 0.141506 | 0.782027 | 0.015 | 0.005 | 1 | 5 | Fam149a | 0.01 | Old |
| 0.141506 | 0.454277 | 0.015 | 0.005 | 1 | 5 | Serpina3f | 0.01 | Old |
| 0.141734 | 0.11898 | 0.832 | 0.82 | 1 | 5 | Igf1 | 0.012 | Old |
| 0.141846 | 0.112807 | 0.015 | 0.005 | 1 | 5 | 1700056N1 | 0.01 | Old |
| 0.14217 | 0.132668 | 0.818 | 0.787 | 1 | 5 | Prdx1 | 0.031 | Old |
| 0.142186 | 0.598878 | 0.015 | 0.005 | 1 | 5 | Chic1 | 0.01 | Old |
| 0.14243 | 0.430017 | 0.167 | 0.135 | 1 | 5 | 5-Mar | 0.032 | Old |
| 0.142522 | 0.185939 | 0.29 | 0.244 | 1 | 5 | Slc31a2 | 0.046 | Old |
| 0.143124 | 0.33488 | 0.128 | 0.096 | 1 | 5 | Ppp1r37 | 0.032 | Old |
| 0.143801 | 0.302578 | 0.192 | 0.157 | 1 | 5 | Tnpo2 | 0.035 | Old |
| 0.143871 | 0.426785 | 0.14 | 0.109 | 1 | 5 | Ado | 0.031 | Old |
| 0.144031 | 0.05532 | 0.311 | 0.264 | 1 | 5 | Ebp | 0.047 | Old |
| 0.144176 | 0.136672 | 0.247 | 0.203 | 1 | 5 | Pak1 | 0.044 | Old |
| 0.144667 | 0.287358 | 0.189 | 0.155 | 1 | 5 | Dhx38 | 0.034 | Old |
| 0.144902 | 0.229946 | 0.167 | 0.132 | 1 | 5 | Ppfia1 | 0.035 | Old |
| 0.144941 | 0.109039 | 0.429 | 0.378 | 1 | 5 | Mertk | 0.051 | Old |
| 0.145033 | 0.255553 | 0.077 | 0.053 | 1 | 5 | Snx25 | 0.024 | Old |
| 0.145107 | 1.095071 | 0.045 | 0.028 | 1 | 5 | Odf2l | 0.017 | Old |
| 0.145529 | 0.490753 | 0.098 | 0.071 | 1 | 5 | Scai | 0.027 | Old |
| 0.145729 | 0.370982 | 0.056 | 0.036 | 1 | 5 | Inpp5e | 0.02 | Old |

| | | | | | | | | |
|---|---|---|---|---|---|---|---|---|
| 0.145931 | 0.235839 | 0.081 | 0.056 | 1 | 5 | Mrpl58 | 0.025 | Old |
| 0.146064 | 0.13125 | 0.332 | 0.284 | 1 | 5 | Il3ra | 0.048 | Old |
| 0.146137 | 0.066798 | 0.618 | 0.548 | 1 | 5 | Plekho1 | 0.07 | Old |
| 0.146182 | 0.16284 | 0.19 | 0.155 | 1 | 5 | Ccpg1 | 0.035 | Old |
| 0.146213 | 0.284682 | 0.281 | 0.249 | 1 | 5 | Mepce | 0.032 | Old |
| 0.146255 | 0.489693 | 0.082 | 0.058 | 1 | 5 | Plppr4 | 0.024 | Old |
| 0.146787 | 0.044057 | 0.502 | 0.444 | 1 | 5 | Diaph1 | 0.058 | Old |
| 0.147111 | 0.515712 | 0.059 | 0.038 | 1 | 5 | 5730409E0 | 0.021 | Old |
| 0.147692 | 0.469245 | 0.17 | 0.14 | 1 | 5 | Samd1 | 0.03 | Old |
| 0.1477 | 0.161376 | 0.347 | 0.289 | 1 | 5 | Ttc7 | 0.058 | Old |
| 0.147856 | 0.393929 | 0.081 | 0.056 | 1 | 5 | Tom1l2 | 0.025 | Old |
| 0.147986 | 1.157421 | 0.121 | 0.091 | 1 | 5 | Il1rn | 0.03 | Old |
| 0.148148 | 0.46196 | 0.205 | 0.175 | 1 | 5 | Rbmxl1 | 0.03 | Old |
| 0.14816 | 0.139617 | 0.305 | 0.259 | 1 | 5 | Cd37 | 0.046 | Old |
| 0.148224 | 0.460089 | 0.086 | 0.061 | 1 | 5 | Prkce | 0.025 | Old |
| 0.148391 | 0.312503 | 0.168 | 0.135 | 1 | 5 | Syvn1 | 0.033 | Old |
| 0.14847 | 0.067646 | 0.29 | 0.241 | 1 | 5 | Elk4 | 0.049 | Old |
| 0.148509 | 0.168764 | 0.254 | 0.216 | 1 | 5 | Ano6 | 0.038 | Old |
| 0.149179 | 0.522614 | 0.056 | 0.036 | 1 | 5 | Rsl1 | 0.02 | Old |
| 0.149353 | 0.524854 | 0.056 | 0.036 | 1 | 5 | Prelid2 | 0.02 | Old |
| 0.149669 | 0.134409 | 0.375 | 0.335 | 1 | 5 | Pabpn1 | 0.04 | Old |
| 0.149774 | 0.179278 | 0.367 | 0.33 | 1 | 5 | Marf1 | 0.037 | Old |
| 0.150237 | 0.277896 | 0.121 | 0.091 | 1 | 5 | Smg9 | 0.03 | Old |
| 0.150273 | 0.18733 | 0.261 | 0.221 | 1 | 5 | Raf1 | 0.04 | Old |
| 0.150395 | 0.060421 | 0.966 | 0.97 | 1 | 5 | Lamp1 | -0.004 | Old |
| 0.150679 | 0.329474 | 0.125 | 0.094 | 1 | 5 | Bloc1s4 | 0.031 | Old |
| 0.150798 | 0.138401 | 0.342 | 0.289 | 1 | 5 | Dnajc7 | 0.053 | Old |
| 0.15091 | 0.243666 | 0.059 | 0.038 | 1 | 5 | C87436 | 0.021 | Old |
| 0.151032 | 0.335425 | 0.168 | 0.137 | 1 | 5 | Tm7sf3 | 0.031 | Old |
| 0.151356 | 1.359995 | 0.019 | 0.008 | 1 | 5 | Arap2 | 0.011 | Old |
| 0.151526 | 0.243674 | 0.167 | 0.135 | 1 | 5 | Tmem214 | 0.032 | Old |
| 0.151991 | 1.257196 | 0.019 | 0.008 | 1 | 5 | Gm16174 | 0.011 | Old |
| 0.152126 | 0.379581 | 0.182 | 0.15 | 1 | 5 | Cd80 | 0.032 | Old |
| 0.152202 | 0.154421 | 0.438 | 0.388 | 1 | 5 | Ttyh3 | 0.05 | Old |
| 0.152309 | 1.41257 | 0.019 | 0.008 | 1 | 5 | Ifi44 | 0.011 | Old |
| 0.152361 | 0.584767 | 0.076 | 0.053 | 1 | 5 | 1110034G2 | 0.023 | Old |
| 0.152499 | 0.325062 | 0.067 | 0.046 | 1 | 5 | Yae1d1 | 0.021 | Old |
| 0.152812 | 0.277901 | 0.226 | 0.188 | 1 | 5 | Tmem170b | 0.038 | Old |
| 0.152836 | 0.569709 | 0.049 | 0.03 | 1 | 5 | Cep70 | 0.019 | Old |
| 0.153 | 0.944051 | 0.045 | 0.028 | 1 | 5 | Gm44067 | 0.017 | Old |
| 0.153103 | 0.048474 | 0.773 | 0.69 | 1 | 5 | Ctsd | 0.083 | Old |
| 0.153248 | 0.170866 | 0.094 | 0.069 | 1 | 5 | Rgl3 | 0.025 | Old |
| 0.153267 | 1.081138 | 0.019 | 0.008 | 1 | 5 | Tmem171 | 0.011 | Old |
| 0.153618 | 0.094125 | 0.133 | 0.102 | 1 | 5 | Fam213a | 0.031 | Old |
| 0.153909 | 1.011037 | 0.019 | 0.008 | 1 | 5 | Cyp2j9 | 0.011 | Old |

| | | | | | | | | |
|---|---|---|---|---|---|---|---|---|
| 0.154012 | 0.147126 | 0.818 | 0.779 | 1 | 5 | Zcchc6 | 0.039 | Old |
| 0.154086 | 0.407136 | 0.111 | 0.084 | 1 | 5 | Cdc27 | 0.027 | Old |
| 0.154286 | 0.14882 | 0.34 | 0.299 | 1 | 5 | Gmip | 0.041 | Old |
| 0.154737 | 0.552059 | 0.093 | 0.069 | 1 | 5 | Acot8 | 0.024 | Old |
| 0.15474 | 0.415724 | 0.152 | 0.122 | 1 | 5 | Gt(ROSA)2( | 0.03 | Old |
| 0.154874 | 0.772262 | 0.019 | 0.008 | 1 | 5 | Medag | 0.011 | Old |
| 0.155105 | 0.128501 | 0.253 | 0.211 | 1 | 5 | Cdc37l1 | 0.042 | Old |
| 0.155147 | 0.277139 | 0.071 | 0.048 | 1 | 5 | Polr3e | 0.023 | Old |
| 0.155212 | 0.239789 | 0.291 | 0.261 | 1 | 5 | Ube2e1 | 0.03 | Old |
| 0.15552 | 0.81889 | 0.019 | 0.008 | 1 | 5 | Csrnp2 | 0.011 | Old |
| 0.155693 | 0.292706 | 0.106 | 0.079 | 1 | 5 | 0610010K1 | 0.027 | Old |
| 0.155876 | 0.359378 | 0.152 | 0.119 | 1 | 5 | 5031425F1 | 0.033 | Old |
| 0.156287 | 0.29292 | 0.153 | 0.122 | 1 | 5 | Tfeb | 0.031 | Old |
| 0.156435 | 0.048146 | 0.451 | 0.404 | 1 | 5 | Parp14 | 0.047 | Old |
| 0.156609 | 0.23881 | 0.226 | 0.188 | 1 | 5 | Htra2 | 0.038 | Old |
| 0.156614 | 0.496827 | 0.194 | 0.162 | 1 | 5 | Swt1 | 0.032 | Old |
| 0.156819 | 0.89518 | 0.019 | 0.008 | 1 | 5 | A430106G1 | 0.011 | Old |
| 0.156819 | 0.854331 | 0.019 | 0.008 | 1 | 5 | Ajuba | 0.011 | Old |
| 0.156994 | 0.804524 | 0.042 | 0.025 | 1 | 5 | Cav1 | 0.017 | Old |
| 0.157145 | 0.629947 | 0.019 | 0.008 | 1 | 5 | Ric3 | 0.011 | Old |
| 0.157191 | 0.57902 | 0.079 | 0.056 | 1 | 5 | Golga1 | 0.023 | Old |
| 0.157337 | 0.287004 | 0.125 | 0.096 | 1 | 5 | Rrp7a | 0.029 | Old |
| 0.157702 | 0.28874 | 0.125 | 0.096 | 1 | 5 | Trmt2a | 0.029 | Old |
| 0.157788 | 0.283333 | 0.162 | 0.129 | 1 | 5 | Mfap1b | 0.033 | Old |
| 0.158075 | 0.134562 | 0.232 | 0.188 | 1 | 5 | Nuak2 | 0.044 | Old |
| 0.15829 | 0.652085 | 0.019 | 0.008 | 1 | 5 | Dqx1 | 0.011 | Old |
| 0.158544 | 0.234156 | 0.269 | 0.231 | 1 | 5 | Smarca4 | 0.038 | Old |
| 0.158576 | 0.204442 | 0.205 | 0.17 | 1 | 5 | Svil | 0.035 | Old |
| 0.158979 | 0.163179 | 0.318 | 0.274 | 1 | 5 | Pole4 | 0.044 | Old |
| 0.159012 | 0.103883 | 0.113 | 0.084 | 1 | 5 | Pfkl | 0.029 | Old |
| 0.159156 | 0.517417 | 0.052 | 0.033 | 1 | 5 | Pced1a | 0.019 | Old |
| 0.159175 | 0.327025 | 0.146 | 0.117 | 1 | 5 | Usp38 | 0.029 | Old |
| 0.159374 | 0.496322 | 0.049 | 0.03 | 1 | 5 | Gstt3 | 0.019 | Old |
| 0.159509 | 0.640492 | 0.042 | 0.025 | 1 | 5 | Tmem51os | 0.017 | Old |
| 0.15962 | 0.051381 | 0.059 | 0.038 | 1 | 5 | B230118H( | 0.021 | Old |
| 0.159779 | 0.291244 | 0.241 | 0.206 | 1 | 5 | BC004004 | 0.035 | Old |
| 0.160102 | 0.366687 | 0.019 | 0.008 | 1 | 5 | Zswim1 | 0.011 | Old |
| 0.160204 | 0.148199 | 0.359 | 0.315 | 1 | 5 | Akirin2 | 0.044 | Old |
| 0.160315 | 0.077587 | 0.579 | 0.536 | 1 | 5 | Atp6v1a | 0.043 | Old |
| 0.160595 | 0.448389 | 0.135 | 0.107 | 1 | 5 | Mavs | 0.028 | Old |
| 0.160746 | 0.567726 | 0.049 | 0.03 | 1 | 5 | Prickle3 | 0.019 | Old |
| 0.160779 | 0.873671 | 0.042 | 0.025 | 1 | 5 | BC030499 | 0.017 | Old |
| 0.160922 | 0.168916 | 0.387 | 0.34 | 1 | 5 | Snw1 | 0.047 | Old |
| 0.161207 | 1.294798 | 0.025 | 0.013 | 1 | 5 | Ehhadh | 0.012 | Old |
| 0.161611 | 0.146976 | 0.46 | 0.398 | 1 | 5 | Mob3c | 0.062 | Old |

| | | | | | | | | |
|---|---|---|---|---|---|---|---|---|
| 0.162092 | 0.175454 | 0.418 | 0.383 | 1 | 5 | Tmem258 | 0.035 | Old |
| 0.16215 | 0.192274 | 0.14 | 0.109 | 1 | 5 | Dcaf11 | 0.031 | Old |
| 0.162183 | 0.122955 | 0.377 | 0.338 | 1 | 5 | Prrc2b | 0.039 | Old |
| 0.162395 | 0.465377 | 0.091 | 0.066 | 1 | 5 | Cxxc1 | 0.025 | Old |
| 0.162456 | 0.27946 | 0.306 | 0.269 | 1 | 5 | Zc3h12a | 0.037 | Old |
| 0.162483 | 0.595113 | 0.042 | 0.025 | 1 | 5 | Tenm4 | 0.017 | Old |
| 0.162546 | 0.246309 | 0.116 | 0.089 | 1 | 5 | Gtf2f1 | 0.027 | Old |
| 0.162897 | 0.36977 | 0.093 | 0.069 | 1 | 5 | Cog4 | 0.024 | Old |
| 0.16304 | 0.239325 | 0.136 | 0.107 | 1 | 5 | Slc25a33 | 0.029 | Old |
| 0.163085 | 0.122638 | 0.278 | 0.234 | 1 | 5 | Trappc2l | 0.044 | Old |
| 0.16311 | 0.271195 | 0.172 | 0.14 | 1 | 5 | Sra1 | 0.032 | Old |
| 0.163172 | 1.006919 | 0.022 | 0.01 | 1 | 5 | AC151836. | 0.012 | Old |
| 0.163223 | 0.746224 | 0.039 | 0.023 | 1 | 5 | Tubgcp6 | 0.016 | Old |
| 0.163493 | 0.072815 | 0.63 | 0.574 | 1 | 5 | Slfn2 | 0.056 | Old |
| 0.163734 | 1.076474 | 0.025 | 0.013 | 1 | 5 | 2010007H( | 0.012 | Old |
| 0.163753 | 0.518251 | 0.089 | 0.066 | 1 | 5 | Dip2a | 0.023 | Old |
| 0.163955 | 0.109896 | 0.17 | 0.135 | 1 | 5 | Atxn7l3 | 0.035 | Old |
| 0.164017 | 1.231804 | 0.025 | 0.013 | 1 | 5 | A430033K( | 0.012 | Old |
| 0.164017 | 1.123303 | 0.025 | 0.013 | 1 | 5 | Kdr | 0.012 | Old |
| 0.164089 | 0.785229 | 0.022 | 0.01 | 1 | 5 | Wdr62 | 0.012 | Old |
| 0.164201 | 0.525924 | 0.042 | 0.025 | 1 | 5 | Simc1 | 0.017 | Old |
| 0.164224 | 1.207701 | 0.01 | 0.003 | 1 | 5 | B3gnt2 | 0.007 | Old |
| 0.164224 | 1.135418 | 0.01 | 0.003 | 1 | 5 | Tmem182 | 0.007 | Old |
| 0.164224 | 1.058778 | 0.01 | 0.003 | 1 | 5 | Atp9a | 0.007 | Old |
| 0.164224 | 0.966353 | 0.01 | 0.003 | 1 | 5 | Gm45890 | 0.007 | Old |
| 0.164505 | 0.102498 | 0.052 | 0.033 | 1 | 5 | Enpp1 | 0.019 | Old |
| 0.164647 | 0.208828 | 0.222 | 0.19 | 1 | 5 | Slc7a7 | 0.032 | Old |
| 0.164697 | 0.399141 | 0.631 | 0.622 | 1 | 5 | Ccl12 | 0.009 | Old |
| 0.1647 | 1.388982 | 0.01 | 0.003 | 1 | 5 | Kbtbd8 | 0.007 | Old |
| 0.1647 | 1.186314 | 0.01 | 0.003 | 1 | 5 | Gm26606 | 0.007 | Old |
| 0.1647 | 1.18011 | 0.01 | 0.003 | 1 | 5 | Cntnap1 | 0.007 | Old |
| 0.1647 | 1.176386 | 0.01 | 0.003 | 1 | 5 | Mfsd6l | 0.007 | Old |
| 0.1647 | 1.139025 | 0.01 | 0.003 | 1 | 5 | Rtkn | 0.007 | Old |
| 0.1647 | 1.114635 | 0.01 | 0.003 | 1 | 5 | Gpd1 | 0.007 | Old |
| 0.1647 | 1.082477 | 0.01 | 0.003 | 1 | 5 | Camk2n2 | 0.007 | Old |
| 0.1647 | 1.079773 | 0.01 | 0.003 | 1 | 5 | Gm10269 | 0.007 | Old |
| 0.164716 | 1.216555 | 0.032 | 0.018 | 1 | 5 | Caprin2 | 0.014 | Old |
| 0.16501 | 0.854759 | 0.022 | 0.01 | 1 | 5 | Tmtc2 | 0.012 | Old |
| 0.165177 | 1.037303 | 0.01 | 0.003 | 1 | 5 | Nxpe4 | 0.007 | Old |
| 0.165177 | 0.928245 | 0.01 | 0.003 | 1 | 5 | Tnxb | 0.007 | Old |
| 0.165177 | 0.833272 | 0.01 | 0.003 | 1 | 5 | Ankrd9 | 0.007 | Old |
| 0.165203 | 0.263321 | 0.269 | 0.239 | 1 | 5 | Rab24 | 0.03 | Old |
| 0.165251 | 0.38808 | 0.064 | 0.043 | 1 | 5 | Tgm1 | 0.021 | Old |
| 0.165262 | 0.122857 | 0.589 | 0.528 | 1 | 5 | Fubp1 | 0.061 | Old |
| 0.165388 | 0.195651 | 0.182 | 0.147 | 1 | 5 | Hdac9 | 0.035 | Old |

| | | | | | | | | |
|---|---|---|---|---|---|---|---|---|
| 0.165476 | 0.635909 | 0.039 | 0.023 | 1 | 5 | Gm16973 | 0.016 | Old |
| 0.165558 | 0.075684 | 0.332 | 0.282 | 1 | 5 | Edf1 | 0.05 | Old |
| 0.165655 | 1.157513 | 0.01 | 0.003 | 1 | 5 | Il4 | 0.007 | Old |
| 0.165655 | 1.044796 | 0.01 | 0.003 | 1 | 5 | Mxra8 | 0.007 | Old |
| 0.165655 | 1.013195 | 0.01 | 0.003 | 1 | 5 | Cyb5d1 | 0.007 | Old |
| 0.165939 | 0.3023 | 0.155 | 0.122 | 1 | 5 | Med8 | 0.033 | Old |
| 0.166047 | 0.304907 | 0.116 | 0.089 | 1 | 5 | Med29 | 0.027 | Old |
| 0.166133 | 1.040758 | 0.01 | 0.003 | 1 | 5 | Prelp | 0.007 | Old |
| 0.166219 | 0.948672 | 0.032 | 0.018 | 1 | 5 | 3110009E1 | 0.014 | Old |
| 0.166244 | 0.68155 | 0.022 | 0.01 | 1 | 5 | Zfp951 | 0.012 | Old |
| 0.166613 | 1.028634 | 0.01 | 0.003 | 1 | 5 | Arc | 0.007 | Old |
| 0.166613 | 0.774949 | 0.01 | 0.003 | 1 | 5 | 4930539E0 | 0.007 | Old |
| 0.166613 | 0.749036 | 0.01 | 0.003 | 1 | 5 | Mettl4 | 0.007 | Old |
| 0.16666 | 0.26853 | 0.16 | 0.127 | 1 | 5 | Rfwd3 | 0.033 | Old |
| 0.166764 | 0.648314 | 0.057 | 0.038 | 1 | 5 | Arvcf | 0.019 | Old |
| 0.166764 | 0.627815 | 0.057 | 0.038 | 1 | 5 | Zc3h10 | 0.019 | Old |
| 0.166913 | 0.203791 | 0.253 | 0.213 | 1 | 5 | Rcn2 | 0.04 | Old |
| 0.167094 | 0.666802 | 0.01 | 0.003 | 1 | 5 | Mnd1 | 0.007 | Old |
| 0.167094 | 0.633932 | 0.01 | 0.003 | 1 | 5 | Zfp993 | 0.007 | Old |
| 0.167094 | 0.522274 | 0.01 | 0.003 | 1 | 5 | Hbb-bs | 0.007 | Old |
| 0.167094 | 0.435445 | 0.01 | 0.003 | 1 | 5 | 1700013D2 | 0.007 | Old |
| 0.167117 | 0.086518 | 0.466 | 0.411 | 1 | 5 | Hif1a | 0.055 | Old |
| 0.167174 | 0.641172 | 0.022 | 0.01 | 1 | 5 | Klc2 | 0.012 | Old |
| 0.167174 | 0.233172 | 0.022 | 0.01 | 1 | 5 | Rab23 | 0.012 | Old |
| 0.167287 | 0.37186 | 0.103 | 0.076 | 1 | 5 | Phf11d | 0.027 | Old |
| 0.167461 | 0.240436 | 0.222 | 0.185 | 1 | 5 | Vti1b | 0.037 | Old |
| 0.167468 | 0.105352 | 0.197 | 0.16 | 1 | 5 | Rab43 | 0.037 | Old |
| 0.167724 | 0.954909 | 0.025 | 0.013 | 1 | 5 | Abca14 | 0.012 | Old |
| 0.16778 | 0.11629 | 0.197 | 0.16 | 1 | 5 | Mmd | 0.037 | Old |
| 0.167862 | 0.060325 | 0.79 | 0.726 | 1 | 5 | Cd300ld | 0.064 | Old |
| 0.168012 | 0.85449 | 0.025 | 0.013 | 1 | 5 | Smarcd3 | 0.012 | Old |
| 0.168012 | 0.814418 | 0.025 | 0.013 | 1 | 5 | Mrps22 | 0.012 | Old |
| 0.168372 | 0.156819 | 0.2 | 0.16 | 1 | 5 | Pld2 | 0.04 | Old |
| 0.168467 | 0.798259 | 0.035 | 0.02 | 1 | 5 | Zfp512b | 0.015 | Old |
| 0.168588 | 0.909231 | 0.025 | 0.013 | 1 | 5 | Gm26522 | 0.012 | Old |
| 0.168618 | 0.367536 | 0.215 | 0.185 | 1 | 5 | Spin1 | 0.03 | Old |
| 0.168661 | 0.403988 | 0.152 | 0.122 | 1 | 5 | Rnpc3 | 0.03 | Old |
| 0.168816 | 0.803194 | 0.029 | 0.015 | 1 | 5 | Lrmda | 0.014 | Old |
| 0.169035 | 0.05955 | 0.197 | 0.16 | 1 | 5 | Eif6 | 0.037 | Old |
| 0.16959 | 0.491603 | 0.098 | 0.074 | 1 | 5 | Dalrd3 | 0.024 | Old |
| 0.170196 | 0.359912 | 0.207 | 0.175 | 1 | 5 | Imp3 | 0.032 | Old |
| 0.170302 | 0.39437 | 0.022 | 0.01 | 1 | 5 | Gm20513 | 0.012 | Old |
| 0.170323 | 0.0738 | 0.276 | 0.231 | 1 | 5 | Hexim1 | 0.045 | Old |
| 0.170399 | 0.201978 | 0.045 | 0.028 | 1 | 5 | Rnf123 | 0.017 | Old |
| 0.170441 | 0.463655 | 0.081 | 0.058 | 1 | 5 | Jagn1 | 0.023 | Old |

| | | | | | | | | |
|---|---|---|---|---|---|---|---|---|
| 0.170713 | 0.752602 | 0.029 | 0.015 | 1 | 5 | Bag2 | 0.014 | Old |
| 0.17091 | 0.236723 | 0.025 | 0.013 | 1 | 5 | Rgcc | 0.012 | Old |
| 0.17119 | 0.138885 | 0.261 | 0.218 | 1 | 5 | Ascc3 | 0.043 | Old |
| 0.171472 | 0.44679 | 0.072 | 0.051 | 1 | 5 | Adam22 | 0.021 | Old |
| 0.171622 | 0.083655 | 0.874 | 0.807 | 1 | 5 | Arhgdia | 0.067 | Old |
| 0.171622 | 0.125588 | 0.463 | 0.406 | 1 | 5 | Coro1b | 0.057 | Old |
| 0.17174 | 0.281362 | 0.343 | 0.302 | 1 | 5 | Trim47 | 0.041 | Old |
| 0.171804 | 0.604879 | 0.029 | 0.015 | 1 | 5 | Mex3b | 0.014 | Old |
| 0.172352 | 0.502246 | 0.035 | 0.02 | 1 | 5 | Taf1c | 0.015 | Old |
| 0.172373 | 0.197857 | 0.025 | 0.013 | 1 | 5 | Pced1b | 0.012 | Old |
| 0.172597 | 0.250452 | 0.035 | 0.02 | 1 | 5 | Slc25a53 | 0.015 | Old |
| 0.172607 | 0.400616 | 0.039 | 0.023 | 1 | 5 | Casp12 | 0.016 | Old |
| 0.17296 | 0.898753 | 0.025 | 0.013 | 1 | 5 | Slc22a18 | 0.012 | Old |
| 0.172987 | 0.201634 | 0.16 | 0.127 | 1 | 5 | Cd79b | 0.033 | Old |
| 0.173252 | 0.188656 | 0.261 | 0.223 | 1 | 5 | Dido1 | 0.038 | Old |
| 0.173438 | 0.045616 | 0.431 | 0.383 | 1 | 5 | Myo1f | 0.048 | Old |
| 0.173451 | 0.400398 | 0.029 | 0.015 | 1 | 5 | Pdia5 | 0.014 | Old |
| 0.173842 | 0.522714 | 0.072 | 0.051 | 1 | 5 | Acrbp | 0.021 | Old |
| 0.174058 | 0.231288 | 0.091 | 0.066 | 1 | 5 | Klhdc10 | 0.025 | Old |
| 0.174107 | 0.104644 | 0.214 | 0.175 | 1 | 5 | Selenom | 0.039 | Old |
| 0.174392 | 0.178632 | 0.172 | 0.137 | 1 | 5 | Arsa | 0.035 | Old |
| 0.174566 | 0.307321 | 0.035 | 0.02 | 1 | 5 | Tbc1d30 | 0.015 | Old |
| 0.174733 | 0.642232 | 0.025 | 0.013 | 1 | 5 | Rtl6 | 0.012 | Old |
| 0.174813 | 0.422653 | 0.035 | 0.02 | 1 | 5 | Aldh4a1 | 0.015 | Old |
| 0.175086 | 0.331032 | 0.081 | 0.058 | 1 | 5 | Nod1 | 0.023 | Old |
| 0.175187 | 0.359896 | 0.167 | 0.137 | 1 | 5 | Ipo9 | 0.03 | Old |
| 0.175387 | 0.438232 | 0.029 | 0.015 | 1 | 5 | P3h1 | 0.014 | Old |
| 0.1754 | 0.294906 | 0.145 | 0.114 | 1 | 5 | Rnf14 | 0.031 | Old |
| 0.175493 | 0.139823 | 0.177 | 0.145 | 1 | 5 | Rnf139 | 0.032 | Old |
| 0.175739 | 0.408092 | 0.12 | 0.094 | 1 | 5 | 2310022A1 | 0.026 | Old |
| 0.175943 | 0.442167 | 0.029 | 0.015 | 1 | 5 | Gpm6b | 0.014 | Old |
| 0.17602 | 0.121126 | 0.407 | 0.358 | 1 | 5 | Rb1cc1 | 0.049 | Old |
| 0.17628 | 0.112145 | 0.091 | 0.066 | 1 | 5 | Carm1 | 0.025 | Old |
| 0.176384 | 0.470595 | 0.081 | 0.058 | 1 | 5 | Rtn4rl1 | 0.023 | Old |
| 0.176443 | 0.424999 | 0.084 | 0.061 | 1 | 5 | Prdm4 | 0.023 | Old |
| 0.176519 | 0.464726 | 0.025 | 0.013 | 1 | 5 | Hcar2 | 0.012 | Old |
| 0.176643 | 0.208036 | 0.123 | 0.094 | 1 | 5 | Dnajc16 | 0.029 | Old |
| 0.176694 | 0.138928 | 0.242 | 0.203 | 1 | 5 | A630001G2 | 0.039 | Old |
| 0.177004 | 0.218773 | 0.281 | 0.236 | 1 | 5 | Tug1 | 0.045 | Old |
| 0.177035 | 0.175596 | 0.298 | 0.259 | 1 | 5 | Rnpepl1 | 0.039 | Old |
| 0.177213 | 0.53569 | 0.057 | 0.038 | 1 | 5 | Arhgef10 | 0.019 | Old |
| 0.17729 | 0.308104 | 0.032 | 0.018 | 1 | 5 | Gm20300 | 0.014 | Old |
| 0.177547 | 0.281687 | 0.261 | 0.226 | 1 | 5 | Ssbp3 | 0.035 | Old |
| 0.17787 | 0.328548 | 0.088 | 0.063 | 1 | 5 | Kri1 | 0.025 | Old |
| 0.177985 | 0.369036 | 0.234 | 0.206 | 1 | 5 | Pygb | 0.028 | Old |

| | | | | | | | | |
|---|---|---|---|---|---|---|---|---|
| 0.178524 | 0.279144 | 0.039 | 0.023 | 1 | 5 | Asf1b | 0.016 | Old |
| 0.178843 | 0.449197 | 0.184 | 0.157 | 1 | 5 | Pafah1b2 | 0.027 | Old |
| 0.17916 | 0.136871 | 0.313 | 0.264 | 1 | 5 | Tomm34 | 0.049 | Old |
| 0.179358 | 0.068385 | 0.042 | 0.025 | 1 | 5 | 1600020E0 | 0.017 | Old |
| 0.179436 | 0.277717 | 0.096 | 0.071 | 1 | 5 | Ndufaf4 | 0.025 | Old |
| 0.179815 | 0.490825 | 0.098 | 0.074 | 1 | 5 | Gfm1 | 0.024 | Old |
| 0.180127 | 0.384833 | 0.136 | 0.109 | 1 | 5 | Mbtps2 | 0.027 | Old |
| 0.180129 | 0.103054 | 0.18 | 0.145 | 1 | 5 | Rps6ka4 | 0.035 | Old |
| 0.180279 | 0.534233 | 0.077 | 0.056 | 1 | 5 | Spats2 | 0.021 | Old |
| 0.180528 | 0.228405 | 0.266 | 0.234 | 1 | 5 | Cdk12 | 0.032 | Old |
| 0.180857 | 0.694864 | 0.051 | 0.033 | 1 | 5 | Fam131a | 0.018 | Old |
| 0.180873 | 0.213478 | 0.214 | 0.18 | 1 | 5 | Crat | 0.034 | Old |
| 0.180876 | 0.208883 | 0.195 | 0.16 | 1 | 5 | Arid3a | 0.035 | Old |
| 0.181002 | 0.602466 | 0.071 | 0.051 | 1 | 5 | St3gal2 | 0.02 | Old |
| 0.18151 | 0.116722 | 0.933 | 0.898 | 1 | 5 | Rsrp1 | 0.035 | Old |
| 0.181805 | 0.110257 | 0.293 | 0.254 | 1 | 5 | Vezf1 | 0.039 | Old |
| 0.182036 | 0.262933 | 0.173 | 0.142 | 1 | 5 | Zfp397 | 0.031 | Old |
| 0.182086 | 0.218639 | 0.333 | 0.299 | 1 | 5 | Ergic3 | 0.034 | Old |
| 0.182763 | 0.084771 | 0.949 | 0.916 | 1 | 5 | Cbr2 | 0.033 | Old |
| 0.182823 | 0.462625 | 0.077 | 0.056 | 1 | 5 | Dynlt1f | 0.021 | Old |
| 0.18283 | 0.266543 | 0.212 | 0.18 | 1 | 5 | Ube2j2 | 0.032 | Old |
| 0.183069 | 0.02677 | 0.241 | 0.203 | 1 | 5 | Bmt2 | 0.038 | Old |
| 0.183143 | 0.138116 | 0.172 | 0.137 | 1 | 5 | B4galt3 | 0.035 | Old |
| 0.18315 | 0.259978 | 0.113 | 0.086 | 1 | 5 | Mapkapk3 | 0.027 | Old |
| 0.183541 | 0.261295 | 0.209 | 0.175 | 1 | 5 | Scn1b | 0.034 | Old |
| 0.183753 | 0.270518 | 0.136 | 0.109 | 1 | 5 | Lrch1 | 0.027 | Old |
| 0.184416 | 0.469375 | 0.066 | 0.046 | 1 | 5 | Mto1 | 0.02 | Old |
| 0.184765 | 0.177791 | 0.273 | 0.239 | 1 | 5 | Rnf169 | 0.034 | Old |
| 0.184772 | 0.341414 | 0.143 | 0.114 | 1 | 5 | Wdr91 | 0.029 | Old |
| 0.18514 | 0.321241 | 0.17 | 0.142 | 1 | 5 | Kdelc2 | 0.028 | Old |
| 0.185298 | 0.076761 | 0.621 | 0.574 | 1 | 5 | Tet3 | 0.047 | Old |
| 0.185392 | 0.123175 | 0.253 | 0.216 | 1 | 5 | Tmem165 | 0.037 | Old |
| 0.185501 | 0.145466 | 0.069 | 0.048 | 1 | 5 | Disc1 | 0.021 | Old |
| 0.185732 | 0.135119 | 0.426 | 0.383 | 1 | 5 | Hfe | 0.043 | Old |
| 0.185808 | 0.11673 | 0.433 | 0.378 | 1 | 5 | Washc2 | 0.055 | Old |
| 0.185814 | 0.144309 | 0.13 | 0.102 | 1 | 5 | Ddx56 | 0.028 | Old |
| 0.186649 | 0.188674 | 0.229 | 0.193 | 1 | 5 | Rbbp8 | 0.036 | Old |
| 0.186687 | 0.13026 | 0.574 | 0.52 | 1 | 5 | Rp2 | 0.054 | Old |
| 0.186751 | 0.156931 | 0.239 | 0.198 | 1 | 5 | Tbc1d2b | 0.041 | Old |
| 0.186976 | 0.712394 | 0.047 | 0.03 | 1 | 5 | Ptcd1 | 0.017 | Old |
| 0.18721 | 0.146211 | 0.249 | 0.211 | 1 | 5 | Fam53b | 0.038 | Old |
| 0.18787 | 0.29663 | 0.126 | 0.099 | 1 | 5 | Sntb2 | 0.027 | Old |
| 0.188256 | 0.509539 | 0.088 | 0.066 | 1 | 5 | Myo1e | 0.022 | Old |
| 0.188659 | 0.170719 | 0.29 | 0.249 | 1 | 5 | Ctcf | 0.041 | Old |
| 0.188692 | 0.272306 | 0.101 | 0.076 | 1 | 5 | Cul9 | 0.025 | Old |

| | | | | | | | | |
|---|---|---|---|---|---|---|---|---|
| 0.189057 | 0.162327 | 0.465 | 0.442 | 1 | 5 | Tmed9 | 0.023 | Old |
| 0.189432 | 0.004786 | 0.226 | 0.188 | 1 | 5 | Ctso | 0.038 | Old |
| 0.190106 | 0.308162 | 0.14 | 0.109 | 1 | 5 | Zfp598 | 0.031 | Old |
| 0.191432 | 0.163837 | 0.305 | 0.264 | 1 | 5 | Capn2 | 0.041 | Old |
| 0.191433 | 0.447042 | 0.074 | 0.053 | 1 | 5 | Ano8 | 0.021 | Old |
| 0.191482 | 0.179662 | 0.485 | 0.447 | 1 | 5 | Klf7 | 0.038 | Old |
| 0.191507 | 0.103823 | 0.209 | 0.173 | 1 | 5 | Smim13 | 0.036 | Old |
| 0.192104 | 0.305271 | 0.141 | 0.114 | 1 | 5 | Zfp330 | 0.027 | Old |
| 0.192253 | 0.038147 | 0.614 | 0.558 | 1 | 5 | Tmed7 | 0.056 | Old |
| 0.192389 | 0.304923 | 0.215 | 0.188 | 1 | 5 | Nsun2 | 0.027 | Old |
| 0.192494 | 0.947282 | 0.04 | 0.025 | 1 | 5 | Tuba4a | 0.015 | Old |
| 0.192516 | 0.480481 | 0.074 | 0.053 | 1 | 5 | Rmnd1 | 0.021 | Old |
| 0.192521 | 0.432031 | 0.094 | 0.071 | 1 | 5 | Rnf121 | 0.023 | Old |
| 0.19286 | 0.316482 | 0.072 | 0.051 | 1 | 5 | Nubp2 | 0.021 | Old |
| 0.19309 | 0.169641 | 0.279 | 0.249 | 1 | 5 | Inppl1 | 0.03 | Old |
| 0.193125 | 0.163656 | 0.418 | 0.371 | 1 | 5 | Arhgap31 | 0.047 | Old |
| 0.193522 | 0.521512 | 0.079 | 0.058 | 1 | 5 | Fndc4 | 0.021 | Old |
| 0.193762 | 1.298732 | 0.013 | 0.005 | 1 | 5 | Plpp7 | 0.008 | Old |
| 0.193762 | 1.175487 | 0.013 | 0.005 | 1 | 5 | Med18 | 0.008 | Old |
| 0.193762 | 0.987841 | 0.013 | 0.005 | 1 | 5 | Gpr21 | 0.008 | Old |
| 0.193938 | 0.128212 | 0.202 | 0.165 | 1 | 5 | Tmem230 | 0.037 | Old |
| 0.193954 | 0.75604 | 0.044 | 0.028 | 1 | 5 | 4930555A( | 0.016 | Old |
| 0.194213 | 1.290762 | 0.013 | 0.005 | 1 | 5 | Ptprb | 0.008 | Old |
| 0.194213 | 1.033643 | 0.013 | 0.005 | 1 | 5 | Rnd1 | 0.008 | Old |
| 0.194213 | 0.98643 | 0.013 | 0.005 | 1 | 5 | Pcdhga9 | 0.008 | Old |
| 0.194213 | 0.967453 | 0.013 | 0.005 | 1 | 5 | Eml2 | 0.008 | Old |
| 0.194551 | 0.063579 | 0.85 | 0.797 | 1 | 5 | Itm2c | 0.053 | Old |
| 0.194746 | 0.37545 | 0.175 | 0.147 | 1 | 5 | Nae1 | 0.028 | Old |
| 0.194928 | 0.117735 | 0.448 | 0.406 | 1 | 5 | Fndc3a | 0.042 | Old |
| 0.195097 | 0.867143 | 0.037 | 0.023 | 1 | 5 | Frs3 | 0.014 | Old |
| 0.195179 | 0.229074 | 0.14 | 0.112 | 1 | 5 | Jak2 | 0.028 | Old |
| 0.1954 | 0.263211 | 0.089 | 0.066 | 1 | 5 | Pcid2 | 0.023 | Old |
| 0.195477 | 0.13319 | 0.177 | 0.145 | 1 | 5 | Nudt9 | 0.032 | Old |
| 0.19557 | 0.969409 | 0.013 | 0.005 | 1 | 5 | Gm45894 | 0.008 | Old |
| 0.195653 | 0.603239 | 0.059 | 0.041 | 1 | 5 | Cidec | 0.018 | Old |
| 0.195658 | 0.178547 | 0.29 | 0.251 | 1 | 5 | Mtmr3 | 0.039 | Old |
| 0.195804 | 0.587969 | 0.088 | 0.066 | 1 | 5 | Kif21b | 0.022 | Old |
| 0.195873 | 0.20469 | 0.21 | 0.175 | 1 | 5 | Cbx6 | 0.035 | Old |
| 0.196024 | 1.005613 | 0.013 | 0.005 | 1 | 5 | Kcnt2 | 0.008 | Old |
| 0.196024 | 0.832879 | 0.013 | 0.005 | 1 | 5 | Sfn | 0.008 | Old |
| 0.196034 | 0.46609 | 0.113 | 0.089 | 1 | 5 | Sun1 | 0.024 | Old |
| 0.196066 | 0.148858 | 0.18 | 0.147 | 1 | 5 | Cyb5r1 | 0.033 | Old |
| 0.196178 | 0.048066 | 0.421 | 0.376 | 1 | 5 | Swi5 | 0.045 | Old |
| 0.196624 | 0.499388 | 0.116 | 0.091 | 1 | 5 | Angptl4 | 0.025 | Old |
| 0.196934 | 1.132556 | 0.013 | 0.005 | 1 | 5 | Pecam1 | 0.008 | Old |

| | | | | | | | | |
|---|---|---|---|---|---|---|---|---|
| 0.196934 | 0.879913 | 0.013 | 0.005 | 1 | 5 | Pole2 | 0.008 | Old |
| 0.19739 | 0.809449 | 0.013 | 0.005 | 1 | 5 | 4931428F0 | 0.008 | Old |
| 0.19739 | 0.706022 | 0.013 | 0.005 | 1 | 5 | 9330020H( | 0.008 | Old |
| 0.197462 | 0.817464 | 0.04 | 0.025 | 1 | 5 | Mrpl47 | 0.015 | Old |
| 0.197564 | 0.291089 | 0.051 | 0.033 | 1 | 5 | Pla2g12a | 0.018 | Old |
| 0.197847 | 0.707988 | 0.013 | 0.005 | 1 | 5 | Otud3 | 0.008 | Old |
| 0.197847 | 0.790938 | 0.013 | 0.005 | 1 | 5 | Gm26759 | 0.008 | Old |
| 0.197847 | 0.750296 | 0.013 | 0.005 | 1 | 5 | 4833445I0 | 0.008 | Old |
| 0.198616 | 0.399396 | 0.094 | 0.071 | 1 | 5 | Terf1 | 0.023 | Old |
| 0.198616 | 0.464099 | 0.082 | 0.061 | 1 | 5 | Aamdc | 0.021 | Old |
| 0.198646 | 0.313288 | 0.077 | 0.056 | 1 | 5 | Hdhd2 | 0.021 | Old |
| 0.198734 | 0.208063 | 0.114 | 0.089 | 1 | 5 | Manbal | 0.025 | Old |
| 0.198825 | 0.141367 | 0.682 | 0.642 | 1 | 5 | Sema4a | 0.04 | Old |
| 0.19899 | 0.501267 | 0.044 | 0.028 | 1 | 5 | Csrp2 | 0.016 | Old |
| 0.199085 | 0.343684 | 0.15 | 0.122 | 1 | 5 | Ciz1 | 0.028 | Old |
| 0.199223 | 0.475866 | 0.013 | 0.005 | 1 | 5 | C1s1 | 0.008 | Old |
| 0.199322 | 0.076233 | 0.261 | 0.221 | 1 | 5 | Sdhd | 0.04 | Old |
| 0.199461 | 0.320625 | 0.162 | 0.132 | 1 | 5 | Pmpcb | 0.03 | Old |
| 0.199524 | 0.263096 | 0.178 | 0.147 | 1 | 5 | Ccnh | 0.031 | Old |
| 0.199683 | 0.602638 | 0.013 | 0.005 | 1 | 5 | Spdl1 | 0.008 | Old |
| 0.199683 | 0.353564 | 0.013 | 0.005 | 1 | 5 | Trpt1 | 0.008 | Old |
| 0.199683 | 0.352136 | 0.013 | 0.005 | 1 | 5 | Amot | 0.008 | Old |
| 0.199973 | 0.355127 | 0.106 | 0.081 | 1 | 5 | Susd1 | 0.025 | Old |
| 0.199994 | 0.064722 | 0.912 | 0.863 | 1 | 5 | Arpc1b | 0.049 | Old |
| 0.200143 | 0.610926 | 0.013 | 0.005 | 1 | 5 | Gm12355 | 0.008 | Old |
| 0.200267 | 0.250324 | 0.224 | 0.19 | 1 | 5 | Zfp871 | 0.034 | Old |
| 0.200283 | 0.039635 | 0.766 | 0.713 | 1 | 5 | Atpif1 | 0.053 | Old |
| 0.200597 | 0.201885 | 0.217 | 0.183 | 1 | 5 | Dazap1 | 0.034 | Old |
| 0.200933 | 0.283996 | 0.276 | 0.239 | 1 | 5 | Dusp2 | 0.037 | Old |
| 0.201023 | 0.171642 | 0.131 | 0.104 | 1 | 5 | Scfd1 | 0.027 | Old |
| 0.201058 | 0.152837 | 0.409 | 0.365 | 1 | 5 | Dock7 | 0.044 | Old |
| 0.201067 | 0.07577 | 0.013 | 0.005 | 1 | 5 | Magi1 | 0.008 | Old |
| 0.20136 | 0.320144 | 0.168 | 0.142 | 1 | 5 | Dpp7 | 0.026 | Old |
| 0.201464 | 0.287229 | 0.178 | 0.152 | 1 | 5 | Aldh7a1 | 0.026 | Old |
| 0.201503 | 0.685175 | 0.04 | 0.025 | 1 | 5 | Rttn | 0.015 | Old |
| 0.201682 | 0.021367 | 0.715 | 0.652 | 1 | 5 | Zbtb20 | 0.063 | Old |
| 0.201752 | 0.237184 | 0.184 | 0.155 | 1 | 5 | Dhrs7b | 0.029 | Old |
| 0.201824 | 0.428616 | 0.111 | 0.086 | 1 | 5 | Asb7 | 0.025 | Old |
| 0.201889 | 0.124064 | 0.498 | 0.467 | 1 | 5 | Stx4a | 0.031 | Old |
| 0.201955 | 0.314498 | 0.123 | 0.096 | 1 | 5 | Zmynd15 | 0.027 | Old |
| 0.202268 | 0.697289 | 0.04 | 0.025 | 1 | 5 | Tcaf1 | 0.015 | Old |
| 0.202356 | 0.048119 | 0.259 | 0.221 | 1 | 5 | Slc25a45 | 0.038 | Old |
| 0.202364 | 0.071252 | 0.404 | 0.358 | 1 | 5 | Sgk3 | 0.046 | Old |
| 0.202748 | 0.083348 | 0.305 | 0.264 | 1 | 5 | Mdh2 | 0.041 | Old |
| 0.202755 | 0.562822 | 0.056 | 0.038 | 1 | 5 | 1700096K1 | 0.018 | Old |

| | | | | | | | | |
|---|---|---|---|---|---|---|---|---|
| 0.202881 | 0.404697 | 0.093 | 0.071 | 1 | 5 | Rhbdf1 | 0.022 | Old |
| 0.203165 | 0.508661 | 0.079 | 0.058 | 1 | 5 | Pcyt1b | 0.021 | Old |
| 0.203288 | 0.109197 | 0.626 | 0.581 | 1 | 5 | Wsb1 | 0.045 | Old |
| 0.203387 | 0.195044 | 0.234 | 0.203 | 1 | 5 | Ufsp2 | 0.031 | Old |
| 0.203506 | 0.109258 | 0.426 | 0.393 | 1 | 5 | Mcub | 0.033 | Old |
| 0.20409 | 0.215938 | 0.109 | 0.084 | 1 | 5 | Ero1lb | 0.025 | Old |
| 0.20434 | 0.089103 | 0.668 | 0.622 | 1 | 5 | Ptpra | 0.046 | Old |
| 0.204453 | 0.527075 | 0.086 | 0.066 | 1 | 5 | Cox19 | 0.02 | Old |
| 0.204851 | 0.44855 | 0.104 | 0.081 | 1 | 5 | Rbms2 | 0.023 | Old |
| 0.20487 | 0.334666 | 0.219 | 0.19 | 1 | 5 | Engase | 0.029 | Old |
| 0.204939 | 0.032066 | 0.355 | 0.307 | 1 | 5 | Fam49a | 0.048 | Old |
| 0.205025 | 0.133745 | 0.379 | 0.335 | 1 | 5 | Efcab14 | 0.044 | Old |
| 0.205477 | 0.099423 | 0.766 | 0.734 | 1 | 5 | Tmed10 | 0.032 | Old |
| 0.205553 | 0.933106 | 0.066 | 0.048 | 1 | 5 | Rhbdd3 | 0.018 | Old |
| 0.205679 | 0.262082 | 0.116 | 0.089 | 1 | 5 | Haghl | 0.027 | Old |
| 0.205715 | 0.116379 | 0.379 | 0.322 | 1 | 5 | Snrpg | 0.057 | Old |
| 0.205766 | 0.159892 | 0.269 | 0.231 | 1 | 5 | Mtr | 0.038 | Old |
| 0.205772 | 0.222743 | 0.85 | 0.843 | 1 | 5 | Fosb | 0.007 | Old |
| 0.206301 | 0.62921 | 0.067 | 0.048 | 1 | 5 | Ccdc32 | 0.019 | Old |
| 0.206374 | 0.305357 | 0.106 | 0.081 | 1 | 5 | Arl4a | 0.025 | Old |
| 0.206468 | 0.33456 | 0.131 | 0.107 | 1 | 5 | Ttc9c | 0.024 | Old |
| 0.20649 | 0.111879 | 0.423 | 0.383 | 1 | 5 | Tmod3 | 0.04 | Old |
| 0.206704 | 0.041688 | 0.657 | 0.609 | 1 | 5 | Atp6ap2 | 0.048 | Old |
| 0.20677 | 0.192393 | 0.202 | 0.168 | 1 | 5 | Abhd17b | 0.034 | Old |
| 0.206776 | 0.162642 | 0.066 | 0.046 | 1 | 5 | Prkrip1 | 0.02 | Old |
| 0.206835 | 0.073852 | 0.397 | 0.363 | 1 | 5 | Ncstn | 0.034 | Old |
| 0.206992 | 0.079947 | 0.327 | 0.284 | 1 | 5 | Ppm1a | 0.043 | Old |
| 0.207368 | 0.429853 | 0.172 | 0.145 | 1 | 5 | Ngdn | 0.027 | Old |
| 0.207528 | 0.268462 | 0.091 | 0.069 | 1 | 5 | Metrn | 0.022 | Old |
| 0.207659 | 0.094784 | 0.564 | 0.528 | 1 | 5 | Tram1 | 0.036 | Old |
| 0.208138 | 0.46014 | 0.059 | 0.041 | 1 | 5 | Utp23 | 0.018 | Old |
| 0.208487 | 0.566863 | 0.076 | 0.056 | 1 | 5 | Atg4d | 0.02 | Old |
| 0.208683 | 0.393369 | 0.037 | 0.023 | 1 | 5 | Cad | 0.014 | Old |
| 0.208742 | 0.256017 | 0.165 | 0.135 | 1 | 5 | Tsc1 | 0.03 | Old |
| 0.208803 | 1.170071 | 0.017 | 0.008 | 1 | 5 | Cacna2d4 | 0.009 | Old |
| 0.209243 | 0.529368 | 0.04 | 0.025 | 1 | 5 | Dhrs11 | 0.015 | Old |
| 0.20934 | 0.210691 | 0.082 | 0.061 | 1 | 5 | Unc45a | 0.021 | Old |
| 0.210029 | 0.26598 | 0.04 | 0.025 | 1 | 5 | Coq4 | 0.015 | Old |
| 0.210262 | 0.237036 | 0.168 | 0.137 | 1 | 5 | Chchd7 | 0.031 | Old |
| 0.21032 | 0.813428 | 0.034 | 0.02 | 1 | 5 | Ptch1 | 0.014 | Old |
| 0.210439 | 0.279503 | 0.099 | 0.076 | 1 | 5 | Hook2 | 0.023 | Old |
| 0.210609 | 0.821777 | 0.034 | 0.02 | 1 | 5 | Oxnad1 | 0.014 | Old |
| 0.210776 | 0.086974 | 0.471 | 0.421 | 1 | 5 | Pkn1 | 0.05 | Old |
| 0.2109 | 1.011437 | 0.017 | 0.008 | 1 | 5 | Tagap1 | 0.009 | Old |
| 0.2109 | 0.899208 | 0.017 | 0.008 | 1 | 5 | Pard6a | 0.009 | Old |

| | | | | | | | | |
|---|---|---|---|---|---|---|---|---|
| 0.211701 | 0.583235 | 0.037 | 0.023 | 1 | 5 | Gm26870 | 0.014 | Old |
| 0.211845 | 0.394815 | 0.096 | 0.074 | 1 | 5 | 6720427I0 | 0.022 | Old |
| 0.212146 | 0.139335 | 0.135 | 0.107 | 1 | 5 | Gclm | 0.028 | Old |
| 0.212165 | 0.796489 | 0.017 | 0.008 | 1 | 5 | Aebp1 | 0.009 | Old |
| 0.21228 | 0.095212 | 0.251 | 0.213 | 1 | 5 | Nin | 0.038 | Old |
| 0.21228 | 0.24372 | 0.387 | 0.358 | 1 | 5 | Rlim | 0.029 | Old |
| 0.212529 | 0.247986 | 0.037 | 0.023 | 1 | 5 | AC162302. | 0.014 | Old |
| 0.212539 | 0.382303 | 0.067 | 0.048 | 1 | 5 | Spn | 0.019 | Old |
| 0.212588 | 0.944247 | 0.017 | 0.008 | 1 | 5 | Id1 | 0.009 | Old |
| 0.212588 | 0.846138 | 0.017 | 0.008 | 1 | 5 | Cx3cl1 | 0.009 | Old |
| 0.212588 | 0.721647 | 0.017 | 0.008 | 1 | 5 | Txk | 0.009 | Old |
| 0.21272 | 0.31344 | 0.16 | 0.132 | 1 | 5 | Wdr61 | 0.028 | Old |
| 0.212814 | 0.190633 | 0.428 | 0.398 | 1 | 5 | Kras | 0.03 | Old |
| 0.212869 | 0.398991 | 0.157 | 0.129 | 1 | 5 | B3glct | 0.028 | Old |
| 0.212928 | 0.404326 | 0.04 | 0.025 | 1 | 5 | 9930104L0 | 0.015 | Old |
| 0.212978 | 0.255973 | 0.221 | 0.193 | 1 | 5 | Dnmt1 | 0.028 | Old |
| 0.213012 | 0.876867 | 0.017 | 0.008 | 1 | 5 | Mical1 | 0.009 | Old |
| 0.213193 | 0.311063 | 0.04 | 0.025 | 1 | 5 | 3110040N1 | 0.015 | Old |
| 0.213214 | 1.077221 | 0.024 | 0.013 | 1 | 5 | Gm44777 | 0.011 | Old |
| 0.213217 | 0.333689 | 0.125 | 0.099 | 1 | 5 | Rcan3 | 0.026 | Old |
| 0.213263 | 0.639808 | 0.094 | 0.074 | 1 | 5 | Rfx1 | 0.02 | Old |
| 0.213436 | 0.76026 | 0.017 | 0.008 | 1 | 5 | Mpzl3 | 0.009 | Old |
| 0.213458 | 0.408862 | 0.04 | 0.025 | 1 | 5 | Depdc7 | 0.015 | Old |
| 0.213861 | 0.982468 | 0.017 | 0.008 | 1 | 5 | Tyw3 | 0.009 | Old |
| 0.214033 | 0.410772 | 0.108 | 0.084 | 1 | 5 | Smad3 | 0.024 | Old |
| 0.214103 | 0.090247 | 0.581 | 0.497 | 1 | 5 | Shisa5 | 0.084 | Old |
| 0.214145 | 0.136125 | 0.439 | 0.396 | 1 | 5 | Creb5 | 0.043 | Old |
| 0.214434 | 0.451305 | 0.131 | 0.107 | 1 | 5 | Ell2 | 0.024 | Old |
| 0.214454 | 0.056549 | 0.739 | 0.695 | 1 | 5 | Selenok | 0.044 | Old |
| 0.214508 | 0.13907 | 0.418 | 0.368 | 1 | 5 | Pdia4 | 0.05 | Old |
| 0.214627 | 0.476288 | 0.064 | 0.046 | 1 | 5 | Ttl | 0.018 | Old |
| 0.214788 | 0.148273 | 0.18 | 0.15 | 1 | 5 | Crtc2 | 0.03 | Old |
| 0.214928 | 0.244905 | 0.096 | 0.074 | 1 | 5 | Chkb | 0.022 | Old |
| 0.214971 | 0.671105 | 0.034 | 0.02 | 1 | 5 | Hectd2 | 0.014 | Old |
| 0.215373 | 0.082688 | 0.35 | 0.307 | 1 | 5 | Csf2rb | 0.043 | Old |
| 0.215724 | 0.118292 | 0.448 | 0.416 | 1 | 5 | Arhgef1 | 0.032 | Old |
| 0.2158 | 0.142139 | 0.268 | 0.228 | 1 | 5 | C1galt1 | 0.04 | Old |
| 0.215909 | 0.879431 | 0.03 | 0.018 | 1 | 5 | Tox2 | 0.012 | Old |
| 0.215994 | 0.386826 | 0.017 | 0.008 | 1 | 5 | Gm49368 | 0.009 | Old |
| 0.216004 | 0.276336 | 0.143 | 0.117 | 1 | 5 | Dda1 | 0.026 | Old |
| 0.216023 | 0.328956 | 0.116 | 0.091 | 1 | 5 | Atg7 | 0.025 | Old |
| 0.216333 | 0.610883 | 0.049 | 0.033 | 1 | 5 | Hexdc | 0.016 | Old |
| 0.216531 | 0.664016 | 0.03 | 0.018 | 1 | 5 | 6330562C2 | 0.012 | Old |
| 0.216719 | 0.257382 | 0.082 | 0.061 | 1 | 5 | Ikbkap | 0.021 | Old |
| 0.216752 | 0.363899 | 0.128 | 0.104 | 1 | 5 | Mir142hg | 0.024 | Old |

| | | | | | | | | |
|---|---|---|---|---|---|---|---|---|
| 0.216875 | 0.55551 | 0.103 | 0.081 | 1 | 5 | Tmco3 | 0.022 | Old |
| 0.217093 | 0.321162 | 0.082 | 0.061 | 1 | 5 | Myg1 | 0.021 | Old |
| 0.21711 | 0.957943 | 0.024 | 0.013 | 1 | 5 | C030006K1 | 0.011 | Old |
| 0.217111 | 0.617279 | 0.061 | 0.043 | 1 | 5 | Cnot11 | 0.018 | Old |
| 0.217126 | 0.511926 | 0.108 | 0.086 | 1 | 5 | Nom1 | 0.022 | Old |
| 0.217263 | 0.456787 | 0.064 | 0.046 | 1 | 5 | 3222401L1 | 0.018 | Old |
| 0.217293 | 0.374624 | 0.052 | 0.036 | 1 | 5 | Rrp9 | 0.016 | Old |
| 0.217528 | 0.518407 | 0.052 | 0.036 | 1 | 5 | Mthfd1 | 0.016 | Old |
| 0.217711 | 0.580328 | 0.017 | 0.008 | 1 | 5 | 1810034E1 | 0.009 | Old |
| 0.217711 | 0.435373 | 0.017 | 0.008 | 1 | 5 | Ccl11 | 0.009 | Old |
| 0.218048 | 0.319751 | 0.093 | 0.071 | 1 | 5 | Dhx58 | 0.022 | Old |
| 0.218155 | 0.212889 | 0.231 | 0.201 | 1 | 5 | 1810013L2 | 0.03 | Old |
| 0.218204 | 0.266003 | 0.288 | 0.259 | 1 | 5 | Gtf2h5 | 0.029 | Old |
| 0.218236 | 0.162203 | 0.153 | 0.124 | 1 | 5 | Ginm1 | 0.029 | Old |
| 0.218507 | 0.475211 | 0.034 | 0.02 | 1 | 5 | Gm17745 | 0.014 | Old |
| 0.218578 | 0.185987 | 0.123 | 0.096 | 1 | 5 | Exosc8 | 0.027 | Old |
| 0.218667 | 0.122572 | 0.626 | 0.581 | 1 | 5 | Syk | 0.045 | Old |
| 0.218763 | 0.582248 | 0.049 | 0.033 | 1 | 5 | Inpp5b | 0.016 | Old |
| 0.219236 | 0.276341 | 0.079 | 0.058 | 1 | 5 | Haus4 | 0.021 | Old |
| 0.219252 | 0.170318 | 0.099 | 0.076 | 1 | 5 | Tmem203 | 0.023 | Old |
| 0.219554 | 0.270814 | 0.071 | 0.051 | 1 | 5 | Dcps | 0.02 | Old |
| 0.220079 | 0.197735 | 0.347 | 0.32 | 1 | 5 | Tmem8 | 0.027 | Old |
| 0.220299 | 0.066732 | 0.197 | 0.165 | 1 | 5 | Pttg1 | 0.032 | Old |
| 0.220336 | 1.028808 | 0.024 | 0.013 | 1 | 5 | Tpmt | 0.011 | Old |
| 0.220701 | 0.203686 | 0.219 | 0.188 | 1 | 5 | Tcf20 | 0.031 | Old |
| 0.220788 | 0.254473 | 0.204 | 0.173 | 1 | 5 | Fam160b1 | 0.031 | Old |
| 0.220919 | 0.440703 | 0.03 | 0.018 | 1 | 5 | Efnb1 | 0.012 | Old |
| 0.221103 | 0.668378 | 0.027 | 0.015 | 1 | 5 | Mmaa | 0.012 | Old |
| 0.221128 | 0.207307 | 0.189 | 0.157 | 1 | 5 | Csrp1 | 0.032 | Old |
| 0.221419 | 0.753871 | 0.024 | 0.013 | 1 | 5 | Cptp | 0.011 | Old |
| 0.22144 | 0.754746 | 0.027 | 0.015 | 1 | 5 | Gpr162 | 0.012 | Old |
| 0.221493 | 0.252388 | 0.226 | 0.195 | 1 | 5 | Cpne2 | 0.031 | Old |
| 0.221691 | 0.274127 | 0.148 | 0.122 | 1 | 5 | Mrpl12 | 0.026 | Old |
| 0.221868 | 0.620666 | 0.03 | 0.018 | 1 | 5 | Gca | 0.012 | Old |
| 0.222322 | 0.290901 | 0.103 | 0.079 | 1 | 5 | Josd2 | 0.024 | Old |
| 0.222535 | 0.469858 | 0.111 | 0.089 | 1 | 5 | Esr1 | 0.022 | Old |
| 0.222735 | 0.130216 | 0.231 | 0.198 | 1 | 5 | Csnk1g2 | 0.033 | Old |
| 0.222806 | 0.165914 | 0.16 | 0.129 | 1 | 5 | Adamtsl5 | 0.031 | Old |
| 0.223196 | 0.054984 | 0.64 | 0.594 | 1 | 5 | Tsc22d1 | 0.046 | Old |
| 0.223233 | 0.68734 | 0.024 | 0.013 | 1 | 5 | Ddx59 | 0.011 | Old |
| 0.223296 | 0.174936 | 0.527 | 0.497 | 1 | 5 | Aes | 0.03 | Old |
| 0.223312 | 0.34096 | 0.116 | 0.094 | 1 | 5 | Ndufv1 | 0.022 | Old |
| 0.223566 | 0.298102 | 0.281 | 0.251 | 1 | 5 | Atxn7l1 | 0.03 | Old |
| 0.223678 | 0.139066 | 0.444 | 0.404 | 1 | 5 | Polr2a | 0.04 | Old |
| 0.22387 | 0.452573 | 0.072 | 0.053 | 1 | 5 | Zfp275 | 0.019 | Old |

| | | | | | | | | |
|---|---|---|---|---|---|---|---|---|
| 0.223873 | 0.430419 | 0.14 | 0.117 | 1 | 5 | Tmem71 | 0.023 | Old |
| 0.223909 | 0.184462 | 0.409 | 0.376 | 1 | 5 | Rassf1 | 0.033 | Old |
| 0.224029 | 0.400533 | 0.02 | 0.01 | 1 | 5 | Enpp2 | 0.01 | Old |
| 0.224041 | 0.142459 | 0.217 | 0.18 | 1 | 5 | Abl1 | 0.037 | Old |
| 0.224269 | 0.159517 | 0.247 | 0.211 | 1 | 5 | Prpsap2 | 0.036 | Old |
| 0.224403 | 0.052458 | 0.254 | 0.216 | 1 | 5 | Ahcyl1 | 0.038 | Old |
| 0.224435 | 0.210088 | 0.121 | 0.096 | 1 | 5 | Traf3 | 0.025 | Old |
| 0.224509 | 0.389007 | 0.184 | 0.157 | 1 | 5 | Plekhm3 | 0.027 | Old |
| 0.224653 | 0.112591 | 0.394 | 0.355 | 1 | 5 | Dpm3 | 0.039 | Old |
| 0.224757 | 0.215931 | 0.131 | 0.107 | 1 | 5 | Prpf38a | 0.024 | Old |
| 0.225173 | 0.353091 | 0.135 | 0.109 | 1 | 5 | Cox16 | 0.026 | Old |
| 0.225341 | 0.384696 | 0.125 | 0.102 | 1 | 5 | Rbbp5 | 0.023 | Old |
| 0.225411 | 0.432224 | 0.069 | 0.051 | 1 | 5 | Slc35d2 | 0.018 | Old |
| 0.225423 | 0.162376 | 0.024 | 0.013 | 1 | 5 | Mcat | 0.011 | Old |
| 0.225505 | 0.653714 | 0.027 | 0.015 | 1 | 5 | Lum | 0.012 | Old |
| 0.225621 | 0.40452 | 0.02 | 0.01 | 1 | 5 | Cep97 | 0.01 | Old |
| 0.225868 | 0.146197 | 0.126 | 0.102 | 1 | 5 | Usf3 | 0.024 | Old |
| 0.226156 | 0.826412 | 0.024 | 0.013 | 1 | 5 | Cit | 0.011 | Old |
| 0.226156 | 0.591087 | 0.024 | 0.013 | 1 | 5 | Apoo | 0.011 | Old |
| 0.226171 | 0.50902 | 0.049 | 0.033 | 1 | 5 | Prdm10 | 0.016 | Old |
| 0.226349 | 0.439158 | 0.084 | 0.063 | 1 | 5 | Clmp | 0.021 | Old |
| 0.226654 | 0.292248 | 0.03 | 0.018 | 1 | 5 | Dnmt3b | 0.012 | Old |
| 0.226852 | 0.178832 | 0.138 | 0.112 | 1 | 5 | Brwd1 | 0.026 | Old |
| 0.226904 | 0.23344 | 0.108 | 0.084 | 1 | 5 | Mcc | 0.024 | Old |
| 0.227494 | 0.577246 | 0.061 | 0.043 | 1 | 5 | Cul7 | 0.018 | Old |
| 0.227961 | 0.334407 | 0.072 | 0.053 | 1 | 5 | Maip1 | 0.019 | Old |
| 0.22799 | 0.438266 | 0.094 | 0.074 | 1 | 5 | Fbxl6 | 0.02 | Old |
| 0.22828 | 0.186077 | 0.244 | 0.216 | 1 | 5 | Srf | 0.028 | Old |
| 0.228297 | 0.171444 | 0.37 | 0.33 | 1 | 5 | Agtrap | 0.04 | Old |
| 0.228541 | 0.114185 | 0.476 | 0.437 | 1 | 5 | Myl12a | 0.039 | Old |
| 0.22884 | 0.059174 | 0.732 | 0.665 | 1 | 5 | Nfe2l2 | 0.067 | Old |
| 0.228934 | 0.503158 | 0.027 | 0.015 | 1 | 5 | Nhej1 | 0.012 | Old |
| 0.229177 | 0.269094 | 0.231 | 0.201 | 1 | 5 | Bmyc | 0.03 | Old |
| 0.229554 | 0.141864 | 0.549 | 0.513 | 1 | 5 | Dnase2a | 0.036 | Old |
| 0.229969 | 0.685172 | 0.054 | 0.038 | 1 | 5 | Laptm4b | 0.016 | Old |
| 0.230149 | 0.338649 | 0.172 | 0.145 | 1 | 5 | Pias4 | 0.027 | Old |
| 0.230161 | 0.254287 | 0.143 | 0.117 | 1 | 5 | Nolc1 | 0.026 | Old |
| 0.23023 | 0.122869 | 0.301 | 0.266 | 1 | 5 | Lrp10 | 0.035 | Old |
| 0.230254 | 0.608574 | 0.045 | 0.03 | 1 | 5 | Tmem150b | 0.015 | Old |
| 0.230447 | 0.424157 | 0.049 | 0.033 | 1 | 5 | Intu | 0.016 | Old |
| 0.23075 | 0.428226 | 0.098 | 0.076 | 1 | 5 | Dlg4 | 0.022 | Old |
| 0.230789 | 0.134737 | 0.308 | 0.274 | 1 | 5 | Arf6 | 0.034 | Old |
| 0.23096 | 0.20472 | 0.306 | 0.274 | 1 | 5 | Cd180 | 0.032 | Old |
| 0.231121 | 0.184831 | 0.295 | 0.266 | 1 | 5 | Usp4 | 0.029 | Old |
| 0.231293 | 0.187335 | 0.497 | 0.462 | 1 | 5 | Gpr65 | 0.035 | Old |

| | | | | | | | | |
|---|---|---|---|---|---|---|---|---|
| 0.231716 | 0.390393 | 0.049 | 0.033 | 1 | 5 | Diexf | 0.016 | Old |
| 0.231774 | 0.089836 | 0.079 | 0.058 | 1 | 5 | Tbl3 | 0.021 | Old |
| 0.232707 | 0.118978 | 0.224 | 0.19 | 1 | 5 | Uba52 | 0.034 | Old |
| 0.232887 | 0.712394 | 0.045 | 0.03 | 1 | 5 | Areg | 0.015 | Old |
| 0.232989 | 0.139556 | 0.076 | 0.056 | 1 | 5 | Hacd2 | 0.02 | Old |
| 0.233309 | 0.10462 | 0.466 | 0.429 | 1 | 5 | Psmb8 | 0.037 | Old |
| 0.233393 | 0.244211 | 0.2 | 0.173 | 1 | 5 | Rfc2 | 0.027 | Old |
| 0.233563 | 0.054927 | 0.478 | 0.442 | 1 | 5 | Serpinf1 | 0.036 | Old |
| 0.23382 | 0.074257 | 0.189 | 0.16 | 1 | 5 | Dgcr2 | 0.029 | Old |
| 0.234939 | 0.303801 | 0.076 | 0.056 | 1 | 5 | Gm35154 | 0.02 | Old |
| 0.235215 | 0.161593 | 0.416 | 0.383 | 1 | 5 | Srsf7 | 0.033 | Old |
| 0.235295 | 0.230781 | 0.175 | 0.147 | 1 | 5 | 9930021J0 | 0.028 | Old |
| 0.235362 | 0.024921 | 0.79 | 0.754 | 1 | 5 | Asah1 | 0.036 | Old |
| 0.235424 | 0.183122 | 0.242 | 0.211 | 1 | 5 | Mtch2 | 0.031 | Old |
| 0.235775 | 0.12639 | 0.259 | 0.228 | 1 | 5 | Nudc | 0.031 | Old |
| 0.235943 | 0.312047 | 0.18 | 0.155 | 1 | 5 | Bag6 | 0.025 | Old |
| 0.236137 | 0.67155 | 0.042 | 0.028 | 1 | 5 | Taf5 | 0.014 | Old |
| 0.236826 | 0.095637 | 0.569 | 0.53 | 1 | 5 | Sbno2 | 0.039 | Old |
| 0.236887 | 0.073912 | 0.325 | 0.287 | 1 | 5 | Pdcd10 | 0.038 | Old |
| 0.237077 | 0.078335 | 0.158 | 0.129 | 1 | 5 | Plekhf2 | 0.029 | Old |
| 0.237332 | 0.046974 | 0.825 | 0.779 | 1 | 5 | Ms4a7 | 0.046 | Old |
| 0.23737 | 0.303606 | 0.236 | 0.211 | 1 | 5 | Snx24 | 0.025 | Old |
| 0.238188 | 0.278085 | 0.113 | 0.089 | 1 | 5 | AL590144.: | 0.024 | Old |
| 0.238277 | 0.127749 | 0.337 | 0.299 | 1 | 5 | Copb1 | 0.038 | Old |
| 0.238461 | 0.254379 | 0.153 | 0.127 | 1 | 5 | Prpf19 | 0.026 | Old |
| 0.238754 | 0.350103 | 0.045 | 0.03 | 1 | 5 | Gstt2 | 0.015 | Old |
| 0.238777 | 0.17801 | 0.178 | 0.15 | 1 | 5 | Plcl2 | 0.028 | Old |
| 0.23917 | 0.108763 | 0.613 | 0.576 | 1 | 5 | Psd3 | 0.037 | Old |
| 0.239181 | 0.067238 | 0.109 | 0.086 | 1 | 5 | Casp9 | 0.023 | Old |
| 0.239272 | 0.471371 | 0.118 | 0.096 | 1 | 5 | Mrpl16 | 0.022 | Old |
| 0.239316 | 0.035938 | 0.029 | 0.043 | 1 | 5 | Pcbp4 | -0.014 | Old |
| 0.239357 | 0.153294 | 0.416 | 0.378 | 1 | 5 | Bcap31 | 0.038 | Old |
| 0.23941 | 0.036484 | 0.564 | 0.513 | 1 | 5 | Selplg | 0.051 | Old |
| 0.239617 | 0.072728 | 0.49 | 0.434 | 1 | 5 | Ndufa13 | 0.056 | Old |
| 0.239983 | 0.229448 | 0.372 | 0.332 | 1 | 5 | Ccl3 | 0.04 | Old |
| 0.240275 | 0.137143 | 0.224 | 0.193 | 1 | 5 | Mesd | 0.031 | Old |
| 0.240578 | 0.19368 | 0.123 | 0.099 | 1 | 5 | 1190002N: | 0.024 | Old |
| 0.240887 | 0.528184 | 0.042 | 0.028 | 1 | 5 | Tle6 | 0.014 | Old |
| 0.241054 | 0.259129 | 0.217 | 0.19 | 1 | 5 | Ipo7 | 0.027 | Old |
| 0.241138 | 0.41019 | 0.069 | 0.051 | 1 | 5 | Noc4l | 0.018 | Old |
| 0.241141 | 0.298361 | 0.18 | 0.155 | 1 | 5 | Atg2a | 0.025 | Old |
| 0.241168 | 0.53419 | 0.042 | 0.028 | 1 | 5 | P3h3 | 0.014 | Old |
| 0.241804 | 0.350681 | 0.227 | 0.198 | 1 | 5 | Maea | 0.029 | Old |
| 0.241926 | 0.605125 | 0.054 | 0.038 | 1 | 5 | Eif2b1 | 0.016 | Old |
| 0.242273 | 0.246649 | 0.256 | 0.231 | 1 | 5 | Klhl18 | 0.025 | Old |

| | | | | | | | | |
|---|---|---|---|---|---|---|---|---|
| 0.242536 | 0.135691 | 0.071 | 0.051 | 1 | 5 | Spryd7 | 0.02 | Old |
| 0.242539 | 0.417327 | 0.076 | 0.056 | 1 | 5 | Dusp10 | 0.02 | Old |
| 0.242579 | 0.04951 | 0.16 | 0.129 | 1 | 5 | Sec24d | 0.031 | Old |
| 0.242728 | 0.074024 | 0.15 | 0.122 | 1 | 5 | Timm8b | 0.028 | Old |
| 0.242805 | 0.09733 | 0.492 | 0.439 | 1 | 5 | Mcfd2 | 0.053 | Old |
| 0.243028 | 0.142668 | 0.138 | 0.112 | 1 | 5 | Bcl9 | 0.026 | Old |
| 0.243145 | 0.432686 | 0.042 | 0.028 | 1 | 5 | Amd2 | 0.014 | Old |
| 0.243943 | 0.35538 | 0.136 | 0.112 | 1 | 5 | Per3 | 0.024 | Old |
| 0.244258 | 0.027101 | 0.333 | 0.292 | 1 | 5 | Arid5a | 0.041 | Old |
| 0.24454 | 0.222516 | 0.091 | 0.069 | 1 | 5 | Osbpl2 | 0.022 | Old |
| 0.245944 | 0.492978 | 0.066 | 0.048 | 1 | 5 | Pcca | 0.018 | Old |
| 0.246448 | 0.154567 | 0.101 | 0.079 | 1 | 5 | 5430427O1 | 0.022 | Old |
| 0.246672 | 0.487956 | 0.057 | 0.041 | 1 | 5 | Comtd1 | 0.016 | Old |
| 0.246729 | 0.282724 | 0.172 | 0.145 | 1 | 5 | Gcn1l1 | 0.027 | Old |
| 0.247373 | 0.136371 | 0.566 | 0.523 | 1 | 5 | Git2 | 0.043 | Old |
| 0.2474 | 0.54123 | 0.062 | 0.046 | 1 | 5 | Thap4 | 0.016 | Old |
| 0.247433 | 0.449009 | 0.054 | 0.038 | 1 | 5 | Fam151b | 0.016 | Old |
| 0.247859 | 0.487904 | 0.096 | 0.076 | 1 | 5 | Rbm12b2 | 0.02 | Old |
| 0.248463 | 1.353092 | 0.032 | 0.02 | 1 | 5 | Ehf | 0.012 | Old |
| 0.248601 | 0.385056 | 0.234 | 0.206 | 1 | 5 | Igsf9 | 0.028 | Old |
| 0.248763 | 0.14938 | 0.259 | 0.226 | 1 | 5 | Tmem248 | 0.033 | Old |
| 0.248855 | 0.426306 | 0.042 | 0.028 | 1 | 5 | Blm | 0.014 | Old |
| 0.249212 | 0.091833 | 0.352 | 0.315 | 1 | 5 | Rb1 | 0.037 | Old |
| 0.249273 | 0.273196 | 0.146 | 0.122 | 1 | 5 | Thap3 | 0.024 | Old |
| 0.249279 | 0.028309 | 0.556 | 0.505 | 1 | 5 | Prkcb | 0.051 | Old |
| 0.249443 | 0.159625 | 0.311 | 0.274 | 1 | 5 | Slc39a7 | 0.037 | Old |
| 0.249746 | 0.102399 | 0.333 | 0.307 | 1 | 5 | Nudt3 | 0.026 | Old |
| 0.250243 | 0.101825 | 0.13 | 0.104 | 1 | 5 | Dgat1 | 0.026 | Old |
| 0.250613 | 0.109107 | 0.148 | 0.122 | 1 | 5 | Foxj3 | 0.026 | Old |
| 0.250735 | 0.360512 | 0.088 | 0.069 | 1 | 5 | Lsm1 | 0.019 | Old |
| 0.250888 | 0.149981 | 0.414 | 0.381 | 1 | 5 | Stard9 | 0.033 | Old |
| 0.25097 | 0.154256 | 0.241 | 0.208 | 1 | 5 | Lamtor5 | 0.033 | Old |
| 0.251329 | 0.19211 | 0.185 | 0.157 | 1 | 5 | Ptdss1 | 0.028 | Old |
| 0.251497 | 0.726913 | 0.051 | 0.036 | 1 | 5 | Alms1 | 0.015 | Old |
| 0.251562 | 0.32558 | 0.103 | 0.081 | 1 | 5 | Taf5l | 0.022 | Old |
| 0.251628 | 0.379234 | 0.111 | 0.089 | 1 | 5 | Vti1a | 0.022 | Old |
| 0.251849 | 0.595316 | 0.039 | 0.025 | 1 | 5 | 6330403L0 | 0.014 | Old |
| 0.251922 | 0.068055 | 0.815 | 0.769 | 1 | 5 | Dhrs3 | 0.046 | Old |
| 0.252179 | 0.638565 | 0.076 | 0.058 | 1 | 5 | Tex30 | 0.018 | Old |
| 0.252213 | 0.243683 | 0.114 | 0.091 | 1 | 5 | Snrpa1 | 0.023 | Old |
| 0.252316 | 0.337484 | 0.082 | 0.063 | 1 | 5 | Fyn | 0.019 | Old |
| 0.252352 | 0.623964 | 0.062 | 0.046 | 1 | 5 | Commd9 | 0.016 | Old |
| 0.252722 | 0.165911 | 0.069 | 0.051 | 1 | 5 | Cd274 | 0.018 | Old |
| 0.252723 | 0.083503 | 0.364 | 0.33 | 1 | 5 | Sh3bp1 | 0.034 | Old |
| 0.252762 | 0.334483 | 0.081 | 0.061 | 1 | 5 | Pin1 | 0.02 | Old |

| | | | | | | | | |
|---|---|---|---|---|---|---|---|---|
| 0.252895 | 0.185997 | 0.106 | 0.084 | 1 | 5 | Exosc5 | 0.022 | Old |
| 0.253282 | 0.328018 | 0.337 | 0.312 | 1 | 5 | Abcc1 | 0.025 | Old |
| 0.253574 | 0.116338 | 0.281 | 0.246 | 1 | 5 | 2810403A( | 0.035 | Old |
| 0.254018 | 0.562538 | 0.062 | 0.046 | 1 | 5 | Lrrc61 | 0.016 | Old |
| 0.254026 | 0.221908 | 0.116 | 0.094 | 1 | 5 | Srcap | 0.022 | Old |
| 0.254134 | 0.059589 | 0.189 | 0.157 | 1 | 5 | Rnf146 | 0.032 | Old |
| 0.254182 | 0.129843 | 0.246 | 0.211 | 1 | 5 | Abca3 | 0.035 | Old |
| 0.254349 | 0.132162 | 0.293 | 0.256 | 1 | 5 | Vps18 | 0.037 | Old |
| 0.254359 | 0.533069 | 0.035 | 0.023 | 1 | 5 | Tbl2 | 0.012 | Old |
| 0.254702 | 0.22341 | 0.244 | 0.216 | 1 | 5 | Map3k2 | 0.028 | Old |
| 0.255189 | 0.056531 | 0.221 | 0.188 | 1 | 5 | Tmem243 | 0.033 | Old |
| 0.255805 | 0.21626 | 0.039 | 0.025 | 1 | 5 | Tchp | 0.014 | Old |
| 0.255963 | 0.104144 | 0.214 | 0.18 | 1 | 5 | Utp14b | 0.034 | Old |
| 0.256418 | 0.395164 | 0.039 | 0.025 | 1 | 5 | Abcb6 | 0.014 | Old |
| 0.257053 | 0.044225 | 0.712 | 0.673 | 1 | 5 | Cflar | 0.039 | Old |
| 0.257438 | 0.141137 | 0.231 | 0.203 | 1 | 5 | Smpd1 | 0.028 | Old |
| 0.257723 | 0.064839 | 0.591 | 0.548 | 1 | 5 | Leng8 | 0.043 | Old |
| 0.258189 | 0.198039 | 0.285 | 0.254 | 1 | 5 | Nsf | 0.031 | Old |
| 0.258445 | 0.102853 | 0.221 | 0.19 | 1 | 5 | Ncln | 0.031 | Old |
| 0.258484 | 0.018574 | 0.145 | 0.117 | 1 | 5 | Rarg | 0.028 | Old |
| 0.259032 | 0.127541 | 0.126 | 0.102 | 1 | 5 | Dtnbp1 | 0.024 | Old |
| 0.259197 | 0.890725 | 0.032 | 0.02 | 1 | 5 | Acot1 | 0.012 | Old |
| 0.259206 | 0.256592 | 0.247 | 0.216 | 1 | 5 | Rabgef1 | 0.031 | Old |
| 0.259359 | 0.13498 | 0.116 | 0.094 | 1 | 5 | Elof1 | 0.022 | Old |
| 0.259616 | 0.179293 | 0.264 | 0.234 | 1 | 5 | Eif4e3 | 0.03 | Old |
| 0.259685 | 0.108273 | 0.527 | 0.467 | 1 | 5 | Gpr132 | 0.06 | Old |
| 0.259805 | 0.088956 | 0.039 | 0.025 | 1 | 5 | Hyal1 | 0.014 | Old |
| 0.259818 | 0.092711 | 0.638 | 0.581 | 1 | 5 | Skap2 | 0.057 | Old |
| 0.25987 | 0.246341 | 0.168 | 0.145 | 1 | 5 | Zbtb37 | 0.023 | Old |
| 0.26015 | 1.422579 | 0.035 | 0.023 | 1 | 5 | Ly6a | 0.012 | Old |
| 0.260261 | 0.203058 | 0.212 | 0.185 | 1 | 5 | Trim12a | 0.027 | Old |
| 0.260274 | 0.328528 | 0.062 | 0.046 | 1 | 5 | Lmf1 | 0.016 | Old |
| 0.260607 | 0.095265 | 0.33 | 0.297 | 1 | 5 | Eif3m | 0.033 | Old |
| 0.260785 | 0.326531 | 0.082 | 0.063 | 1 | 5 | Ccdc85b | 0.019 | Old |
| 0.260903 | 0.897333 | 0.032 | 0.02 | 1 | 5 | Gm35363 | 0.012 | Old |
| 0.260966 | 0.421286 | 0.034 | 0.048 | 1 | 5 | Steap4 | -0.014 | Old |
| 0.26107 | 0.292894 | 0.103 | 0.081 | 1 | 5 | Asnsd1 | 0.022 | Old |
| 0.261094 | 0.087543 | 0.753 | 0.726 | 1 | 5 | Gns | 0.027 | Old |
| 0.261803 | 0.581439 | 0.059 | 0.043 | 1 | 5 | Pnpo | 0.016 | Old |
| 0.261849 | 0.098986 | 0.522 | 0.475 | 1 | 5 | Blnk | 0.047 | Old |
| 0.261888 | 0.431392 | 0.088 | 0.069 | 1 | 5 | Nedd1 | 0.019 | Old |
| 0.262273 | 0.702558 | 0.032 | 0.02 | 1 | 5 | Hirip3 | 0.012 | Old |
| 0.262552 | 0.05491 | 0.611 | 0.581 | 1 | 5 | Gapvd1 | 0.03 | Old |
| 0.262945 | 0.210946 | 0.066 | 0.048 | 1 | 5 | Tsr1 | 0.018 | Old |
| 0.263156 | 0.072399 | 0.722 | 0.695 | 1 | 5 | Tpp1 | 0.027 | Old |

| | | | | | | | | |
|---|---|---|---|---|---|---|---|---|
| 0.26352 | 0.276396 | 0.069 | 0.051 | 1 | 5 | Hspb11 | 0.018 | Old |
| 0.263593 | 0.302389 | 0.177 | 0.152 | 1 | 5 | CAAA0111‹ | 0.025 | Old |
| 0.263597 | 0.325817 | 0.13 | 0.107 | 1 | 5 | Rhbdf2 | 0.023 | Old |
| 0.263953 | 0.120954 | 0.463 | 0.421 | 1 | 5 | Ube2n | 0.042 | Old |
| 0.264467 | 0.271386 | 0.094 | 0.074 | 1 | 5 | Dkc1 | 0.02 | Old |
| 0.264485 | 0.15286 | 0.33 | 0.297 | 1 | 5 | Akt2 | 0.033 | Old |
| 0.264643 | 0.138263 | 0.089 | 0.069 | 1 | 5 | Zfp943 | 0.02 | Old |
| 0.264862 | 0.148963 | 0.298 | 0.266 | 1 | 5 | Idh3g | 0.032 | Old |
| 0.265045 | 0.361163 | 0.035 | 0.023 | 1 | 5 | Fdxr | 0.012 | Old |
| 0.265069 | 0.409953 | 0.059 | 0.043 | 1 | 5 | 1700021F0 | 0.016 | Old |
| 0.265096 | 0.090554 | 0.099 | 0.079 | 1 | 5 | Taf1b | 0.02 | Old |
| 0.265586 | 0.243583 | 0.285 | 0.264 | 1 | 5 | Selenon | 0.021 | Old |
| 0.265907 | 0.194799 | 0.258 | 0.228 | 1 | 5 | Ak2 | 0.03 | Old |
| 0.26594 | 0.236841 | 0.091 | 0.071 | 1 | 5 | Dhx34 | 0.02 | Old |
| 0.266031 | 0.374258 | 0.035 | 0.023 | 1 | 5 | Fchsd1 | 0.012 | Old |
| 0.266053 | 0.019682 | 0.495 | 0.462 | 1 | 5 | Map7d1 | 0.033 | Old |
| 0.266179 | 0.169479 | 0.465 | 0.434 | 1 | 5 | Ppp1r21 | 0.031 | Old |
| 0.266359 | 0.211527 | 0.276 | 0.251 | 1 | 5 | Dennd1c | 0.025 | Old |
| 0.267129 | 0.317582 | 0.062 | 0.046 | 1 | 5 | BC002059 | 0.016 | Old |
| 0.267399 | 0.462218 | 0.116 | 0.096 | 1 | 5 | Zbtb16 | 0.02 | Old |
| 0.268432 | 0.174527 | 0.126 | 0.104 | 1 | 5 | Arhgef11 | 0.022 | Old |
| 0.268527 | 0.237296 | 0.096 | 0.076 | 1 | 5 | Ncbp1 | 0.02 | Old |
| 0.268536 | 0.082145 | 0.258 | 0.221 | 1 | 5 | Cdk13 | 0.037 | Old |
| 0.269006 | 0.411516 | 0.035 | 0.023 | 1 | 5 | Nkrf | 0.012 | Old |
| 0.269633 | 0.145095 | 0.404 | 0.365 | 1 | 5 | Top2b | 0.039 | Old |
| 0.270078 | 0.009927 | 0.114 | 0.091 | 1 | 5 | Ppp1r13b | 0.023 | Old |
| 0.270137 | 0.520213 | 0.108 | 0.089 | 1 | 5 | Syt11 | 0.019 | Old |
| 0.270316 | 0.347358 | 0.189 | 0.165 | 1 | 5 | Mras | 0.024 | Old |
| 0.27078 | 0.085246 | 0.453 | 0.414 | 1 | 5 | Larp1 | 0.039 | Old |
| 0.270878 | 0.133661 | 0.224 | 0.198 | 1 | 5 | Ndufb2 | 0.026 | Old |
| 0.271265 | 0.224058 | 0.088 | 0.069 | 1 | 5 | Tpk1 | 0.019 | Old |
| 0.271267 | 0.234869 | 0.231 | 0.201 | 1 | 5 | Ppfibp2 | 0.03 | Old |
| 0.271621 | 0.186226 | 0.066 | 0.048 | 1 | 5 | B3gnt8 | 0.018 | Old |
| 0.271651 | 0.587352 | 0.032 | 0.02 | 1 | 5 | Gba2 | 0.012 | Old |
| 0.27185 | 0.288095 | 0.062 | 0.046 | 1 | 5 | Zc3hc1 | 0.016 | Old |
| 0.271898 | 0.325078 | 0.088 | 0.069 | 1 | 5 | Lsg1 | 0.019 | Old |
| 0.271935 | 0.413789 | 0.128 | 0.107 | 1 | 5 | Mdc1 | 0.021 | Old |
| 0.272083 | 0.241984 | 0.207 | 0.183 | 1 | 5 | Traf1 | 0.024 | Old |
| 0.272124 | 0.195703 | 0.175 | 0.147 | 1 | 5 | Mrpl4 | 0.028 | Old |
| 0.272327 | 0.404197 | 0.059 | 0.043 | 1 | 5 | Ranbp6 | 0.016 | Old |
| 0.272455 | 0.450337 | 0.059 | 0.043 | 1 | 5 | Eif5a2 | 0.016 | Old |
| 0.272601 | 0.219795 | 0.062 | 0.046 | 1 | 5 | Lix1 | 0.016 | Old |
| 0.27306 | 0.497463 | 0.032 | 0.02 | 1 | 5 | Gstt1 | 0.012 | Old |
| 0.27306 | 0.451546 | 0.032 | 0.02 | 1 | 5 | Gtf3c5 | 0.012 | Old |
| 0.273326 | 0.02637 | 1 | 1 | 1 | 5 | Itm2b | 0 | Old |

| | | | | | | | | |
|---|---|---|---|---|---|---|---|---|
| 0.273607 | 0.232767 | 0.244 | 0.218 | 1 | 5 | Cbx7 | 0.026 | Old |
| 0.273609 | 0.488135 | 0.114 | 0.094 | 1 | 5 | Gbe1 | 0.02 | Old |
| 0.274251 | 0.152432 | 0.104 | 0.084 | 1 | 5 | Tusc2 | 0.02 | Old |
| 0.274304 | 0.052644 | 0.293 | 0.251 | 1 | 5 | 4933434E2 | 0.042 | Old |
| 0.274658 | 1.255977 | 0.012 | 0.005 | 1 | 5 | Rarres2 | 0.007 | Old |
| 0.274658 | 1.027745 | 0.012 | 0.005 | 1 | 5 | Fam177a | 0.007 | Old |
| 0.274974 | 0.511911 | 0.071 | 0.053 | 1 | 5 | Arel1 | 0.018 | Old |
| 0.275224 | 0.172894 | 0.082 | 0.063 | 1 | 5 | Kantr | 0.019 | Old |
| 0.275267 | 1.086984 | 0.012 | 0.005 | 1 | 5 | Ccdc88c | 0.007 | Old |
| 0.27534 | 0.126057 | 0.38 | 0.353 | 1 | 5 | Ssr4 | 0.027 | Old |
| 0.275401 | 0.1017 | 0.259 | 0.228 | 1 | 5 | Gigyf2 | 0.031 | Old |
| 0.275484 | 0.273935 | 0.239 | 0.213 | 1 | 5 | Npepl1 | 0.026 | Old |
| 0.275576 | 0.550886 | 0.029 | 0.018 | 1 | 5 | Ripply3 | 0.011 | Old |
| 0.275678 | 0.058617 | 0.412 | 0.371 | 1 | 5 | Mgea5 | 0.041 | Old |
| 0.275747 | 0.214509 | 0.209 | 0.18 | 1 | 5 | Fxr2 | 0.029 | Old |
| 0.275795 | 0.199326 | 0.158 | 0.132 | 1 | 5 | Cyhr1 | 0.026 | Old |
| 0.275828 | 0.690354 | 0.056 | 0.041 | 1 | 5 | Thap1 | 0.015 | Old |
| 0.275876 | 0.954233 | 0.012 | 0.005 | 1 | 5 | Col27a1 | 0.007 | Old |
| 0.275876 | 0.915745 | 0.012 | 0.005 | 1 | 5 | Htra3 | 0.007 | Old |
| 0.275876 | 0.84823 | 0.012 | 0.005 | 1 | 5 | Mtus2 | 0.007 | Old |
| 0.275976 | 0.238718 | 0.067 | 0.051 | 1 | 5 | Tmem18 | 0.016 | Old |
| 0.276059 | 0.171788 | 0.322 | 0.294 | 1 | 5 | Fam126a | 0.028 | Old |
| 0.276113 | 0.133832 | 0.603 | 0.546 | 1 | 5 | Arid1a | 0.057 | Old |
| 0.276486 | 0.989832 | 0.012 | 0.005 | 1 | 5 | Car11 | 0.007 | Old |
| 0.276486 | 0.851206 | 0.012 | 0.005 | 1 | 5 | Tonsl | 0.007 | Old |
| 0.276604 | 0.19791 | 0.032 | 0.02 | 1 | 5 | Bckdhb | 0.012 | Old |
| 0.276631 | 0.08315 | 0.279 | 0.246 | 1 | 5 | Cnpy3 | 0.033 | Old |
| 0.276764 | 0.225534 | 0.082 | 0.063 | 1 | 5 | Zbp1 | 0.019 | Old |
| 0.277083 | 0.795121 | 0.029 | 0.018 | 1 | 5 | Cks1b | 0.011 | Old |
| 0.277097 | 0.717873 | 0.012 | 0.005 | 1 | 5 | 2810021J2 | 0.007 | Old |
| 0.277097 | 0.687641 | 0.012 | 0.005 | 1 | 5 | Synm | 0.007 | Old |
| 0.27725 | 0.181515 | 0.051 | 0.036 | 1 | 5 | Tcea3 | 0.015 | Old |
| 0.277529 | 0.931144 | 0.025 | 0.015 | 1 | 5 | Gm21781 | 0.01 | Old |
| 0.277843 | 0.134559 | 0.175 | 0.15 | 1 | 5 | Lrch3 | 0.025 | Old |
| 0.277939 | 0.314544 | 0.109 | 0.089 | 1 | 5 | Ikzf2 | 0.02 | Old |
| 0.278236 | 0.230472 | 0.157 | 0.132 | 1 | 5 | Med12 | 0.025 | Old |
| 0.278322 | 0.537002 | 0.012 | 0.005 | 1 | 5 | 4930430E1 | 0.007 | Old |
| 0.278322 | 0.379223 | 0.012 | 0.005 | 1 | 5 | Nkapl | 0.007 | Old |
| 0.278362 | 0.312261 | 0.172 | 0.15 | 1 | 5 | Pde7a | 0.022 | Old |
| 0.278464 | 0.083533 | 0.286 | 0.251 | 1 | 5 | Tm2d2 | 0.035 | Old |
| 0.278936 | 0.7054 | 0.012 | 0.005 | 1 | 5 | Fam120c | 0.007 | Old |
| 0.278936 | 0.691456 | 0.012 | 0.005 | 1 | 5 | Sorbs1 | 0.007 | Old |
| 0.278936 | 0.607254 | 0.012 | 0.005 | 1 | 5 | Gm11713 | 0.007 | Old |
| 0.278936 | 0.291414 | 0.012 | 0.005 | 1 | 5 | Gm45844 | 0.007 | Old |
| 0.27904 | 0.229463 | 0.091 | 0.071 | 1 | 5 | Apobr | 0.02 | Old |

| | | | | | | | | |
|---|---|---|---|---|---|---|---|---|
| 0.279041 | 0.3725 | 0.056 | 0.041 | 1 | 5 | Nup188 | 0.015 | Old |
| 0.279267 | 0.179392 | 0.638 | 0.617 | 1 | 5 | Tlr2 | 0.021 | Old |
| 0.279318 | 0.161341 | 0.226 | 0.201 | 1 | 5 | Znfx1 | 0.025 | Old |
| 0.279347 | 0.095936 | 0.722 | 0.703 | 1 | 5 | Slc3a2 | 0.019 | Old |
| 0.279551 | 0.542473 | 0.012 | 0.005 | 1 | 5 | Mrm3 | 0.007 | Old |
| 0.279551 | 0.449623 | 0.012 | 0.005 | 1 | 5 | Gm44851 | 0.007 | Old |
| 0.279551 | 0.3666 | 0.012 | 0.005 | 1 | 5 | Uprt | 0.007 | Old |
| 0.279551 | 0.348707 | 0.012 | 0.005 | 1 | 5 | Gm34934 | 0.007 | Old |
| 0.279637 | 0.087243 | 0.153 | 0.127 | 1 | 5 | Uba7 | 0.026 | Old |
| 0.280027 | 0.279117 | 0.106 | 0.086 | 1 | 5 | Rbm15b | 0.02 | Old |
| 0.280208 | 0.257289 | 0.071 | 0.053 | 1 | 5 | Cblb | 0.018 | Old |
| 0.280438 | 0.154098 | 0.088 | 0.069 | 1 | 5 | Nr2c1 | 0.019 | Old |
| 0.280638 | 0.078134 | 0.323 | 0.282 | 1 | 5 | Araf | 0.041 | Old |
| 0.281189 | 0.205244 | 0.13 | 0.107 | 1 | 5 | Ing3 | 0.023 | Old |
| 0.281709 | 0.323164 | 0.178 | 0.155 | 1 | 5 | Drg1 | 0.023 | Old |
| 0.282019 | 0.220349 | 0.012 | 0.005 | 1 | 5 | Gvin1 | 0.007 | Old |
| 0.282026 | 0.154462 | 0.152 | 0.127 | 1 | 5 | Tom1 | 0.025 | Old |
| 0.282253 | 0.966864 | 0.022 | 0.013 | 1 | 5 | Car13 | 0.009 | Old |
| 0.282253 | 0.946344 | 0.022 | 0.013 | 1 | 5 | Qrsl1 | 0.009 | Old |
| 0.282344 | 0.20407 | 0.212 | 0.185 | 1 | 5 | Dmtf1 | 0.027 | Old |
| 0.282361 | 0.357147 | 0.076 | 0.058 | 1 | 5 | Zfp839 | 0.018 | Old |
| 0.282419 | 0.083092 | 0.574 | 0.528 | 1 | 5 | Nlrp3 | 0.046 | Old |
| 0.28251 | 0.037403 | 0.237 | 0.206 | 1 | 5 | Gpbp1l1 | 0.031 | Old |
| 0.282704 | 0.336953 | 0.101 | 0.081 | 1 | 5 | Bpnt1 | 0.02 | Old |
| 0.282779 | 1.265037 | 0.015 | 0.008 | 1 | 5 | Trmt44 | 0.007 | Old |
| 0.282998 | 0.62564 | 0.069 | 0.053 | 1 | 5 | Primpol | 0.016 | Old |
| 0.283166 | 0.339448 | 0.029 | 0.018 | 1 | 5 | Zhx2 | 0.011 | Old |
| 0.283558 | 0.142463 | 0.522 | 0.495 | 1 | 5 | Kdm2a | 0.027 | Old |
| 0.283809 | 0.259048 | 0.067 | 0.051 | 1 | 5 | Serpinb9 | 0.016 | Old |
| 0.284079 | 0.052598 | 0.623 | 0.556 | 1 | 5 | Basp1 | 0.067 | Old |
| 0.284135 | 0.200244 | 0.394 | 0.358 | 1 | 5 | Cul3 | 0.036 | Old |
| 0.284244 | 0.202199 | 0.131 | 0.109 | 1 | 5 | Arid3b | 0.022 | Old |
| 0.284378 | 0.16062 | 0.478 | 0.447 | 1 | 5 | Lman2 | 0.031 | Old |
| 0.284482 | 0.113091 | 0.195 | 0.168 | 1 | 5 | Wdr33 | 0.027 | Old |
| 0.284785 | 0.253226 | 0.096 | 0.076 | 1 | 5 | Fam222b | 0.02 | Old |
| 0.284852 | 0.739538 | 0.025 | 0.015 | 1 | 5 | Pdk2 | 0.01 | Old |
| 0.285171 | 0.342921 | 0.118 | 0.096 | 1 | 5 | Pqlc3 | 0.022 | Old |
| 0.285395 | 0.046564 | 0.338 | 0.297 | 1 | 5 | Tmem259 | 0.041 | Old |
| 0.28551 | 0.158084 | 0.483 | 0.48 | 1 | 5 | Smarca5 | 0.003 | Old |
| 0.285605 | 0.096416 | 0.428 | 0.396 | 1 | 5 | Mrpl23 | 0.032 | Old |
| 0.28567 | 0.156989 | 0.141 | 0.117 | 1 | 5 | Lnpk | 0.024 | Old |
| 0.285708 | 0.238496 | 0.135 | 0.112 | 1 | 5 | Bmi1 | 0.023 | Old |
| 0.285788 | 0.850903 | 0.022 | 0.013 | 1 | 5 | Tmem218 | 0.009 | Old |
| 0.285788 | 0.721532 | 0.022 | 0.013 | 1 | 5 | Gm17259 | 0.009 | Old |
| 0.285791 | 0.166044 | 0.067 | 0.051 | 1 | 5 | Mkrn2 | 0.016 | Old |

| | | | | | | | | |
|---|---|---|---|---|---|---|---|---|
| 0.285855 | 0.296461 | 0.029 | 0.018 | 1 | 5 | Herc3 | 0.011 | Old |
| 0.28586 | 0.427289 | 0.076 | 0.058 | 1 | 5 | Tor4a | 0.018 | Old |
| 0.28656 | 1.073349 | 0.015 | 0.008 | 1 | 5 | Tmem41a | 0.007 | Old |
| 0.286708 | 0.183174 | 0.182 | 0.157 | 1 | 5 | Sfswap | 0.025 | Old |
| 0.286758 | 0.492335 | 0.064 | 0.048 | 1 | 5 | Polr3a | 0.016 | Old |
| 0.287121 | 0.717284 | 0.022 | 0.013 | 1 | 5 | Lysmd1 | 0.009 | Old |
| 0.287121 | 0.641776 | 0.022 | 0.013 | 1 | 5 | N4bp3 | 0.009 | Old |
| 0.287647 | 0.748601 | 0.015 | 0.008 | 1 | 5 | Junos | 0.007 | Old |
| 0.288062 | 0.038841 | 0.953 | 0.939 | 1 | 5 | Irf2bp2 | 0.014 | Old |
| 0.288284 | 0.185108 | 0.047 | 0.033 | 1 | 5 | Immp2l | 0.014 | Old |
| 0.288345 | 0.387265 | 0.084 | 0.066 | 1 | 5 | Nfrkb | 0.018 | Old |
| 0.288458 | 0.656411 | 0.022 | 0.013 | 1 | 5 | Rccd1 | 0.009 | Old |
| 0.288635 | 0.191566 | 0.14 | 0.117 | 1 | 5 | Tmem104 | 0.023 | Old |
| 0.288738 | 0.101098 | 0.222 | 0.19 | 1 | 5 | Unc50 | 0.032 | Old |
| 0.2888 | 0.316524 | 0.16 | 0.137 | 1 | 5 | Dopey1 | 0.023 | Old |
| 0.288901 | 0.714032 | 0.019 | 0.01 | 1 | 5 | Ldlrad4 | 0.009 | Old |
| 0.288947 | 0.097718 | 0.209 | 0.18 | 1 | 5 | Ubxn6 | 0.029 | Old |
| 0.289049 | 0.316579 | 0.19 | 0.168 | 1 | 5 | Eps15l1 | 0.022 | Old |
| 0.289137 | 0.296211 | 0.145 | 0.124 | 1 | 5 | Cog5 | 0.021 | Old |
| 0.28978 | 0.161372 | 0.044 | 0.03 | 1 | 5 | Gcfc2 | 0.014 | Old |
| 0.289799 | 0.482543 | 0.022 | 0.013 | 1 | 5 | Sgsm3 | 0.009 | Old |
| 0.289857 | 0.13532 | 0.377 | 0.35 | 1 | 5 | 4932438A1 | 0.027 | Old |
| 0.28988 | 0.792414 | 0.019 | 0.01 | 1 | 5 | Gm30411 | 0.009 | Old |
| 0.290162 | 0.418517 | 0.106 | 0.086 | 1 | 5 | Nelfb | 0.02 | Old |
| 0.290247 | 0.389008 | 0.044 | 0.03 | 1 | 5 | Bud13 | 0.014 | Old |
| 0.290247 | 0.598569 | 0.022 | 0.013 | 1 | 5 | Gm26935 | 0.009 | Old |
| 0.290247 | 0.581821 | 0.022 | 0.013 | 1 | 5 | Jaml | 0.009 | Old |
| 0.290258 | 0.142931 | 0.121 | 0.099 | 1 | 5 | Rnf170 | 0.022 | Old |
| 0.290376 | 0.432496 | 0.015 | 0.008 | 1 | 5 | Zfp772 | 0.007 | Old |
| 0.290559 | 0.239553 | 0.044 | 0.03 | 1 | 5 | Gm49359 | 0.014 | Old |
| 0.290611 | 0.072671 | 0.18 | 0.152 | 1 | 5 | Ehmt1 | 0.028 | Old |
| 0.29086 | 0.698835 | 0.019 | 0.01 | 1 | 5 | D11Wsu47 | 0.009 | Old |
| 0.290924 | 0.436638 | 0.015 | 0.008 | 1 | 5 | Zkscan5 | 0.007 | Old |
| 0.291054 | 0.251811 | 0.025 | 0.015 | 1 | 5 | 1110051M | 0.01 | Old |
| 0.291145 | 0.649124 | 0.022 | 0.013 | 1 | 5 | Gpr19 | 0.009 | Old |
| 0.291171 | 0.063199 | 0.49 | 0.447 | 1 | 5 | Nrp2 | 0.043 | Old |
| 0.291203 | 0.499757 | 0.04 | 0.028 | 1 | 5 | Mettl13 | 0.012 | Old |
| 0.291305 | 0.159305 | 0.104 | 0.084 | 1 | 5 | Dhx8 | 0.02 | Old |
| 0.291594 | 0.500401 | 0.022 | 0.013 | 1 | 5 | Eid2 | 0.009 | Old |
| 0.291595 | 0.147387 | 0.266 | 0.239 | 1 | 5 | Kif1c | 0.027 | Old |
| 0.291736 | 0.173102 | 0.224 | 0.201 | 1 | 5 | Polr2g | 0.023 | Old |
| 0.291844 | 1.098518 | 0.019 | 0.01 | 1 | 5 | Xlr4c | 0.009 | Old |
| 0.291844 | 0.777909 | 0.019 | 0.01 | 1 | 5 | Gbp2b | 0.009 | Old |
| 0.291844 | 0.666276 | 0.019 | 0.01 | 1 | 5 | Ube2t | 0.009 | Old |
| 0.291887 | 0.284634 | 0.025 | 0.015 | 1 | 5 | A530032D1 | 0.01 | Old |

| | | | | | | | | |
|---|---|---|---|---|---|---|---|---|
| 0.292572 | 0.093904 | 0.015 | 0.008 | 1 | 5 | Dnaja4 | 0.007 | Old |
| 0.292696 | 0.129496 | 0.155 | 0.132 | 1 | 5 | Aldh3a2 | 0.023 | Old |
| 0.292829 | 0.538299 | 0.019 | 0.01 | 1 | 5 | Slc9a3r2 | 0.009 | Old |
| 0.292829 | 0.533472 | 0.019 | 0.01 | 1 | 5 | Gpihbp1 | 0.009 | Old |
| 0.292829 | 0.447127 | 0.019 | 0.01 | 1 | 5 | Zfp775 | 0.009 | Old |
| 0.292983 | 0.148953 | 0.495 | 0.457 | 1 | 5 | Srsf11 | 0.038 | Old |
| 0.293308 | 0.341476 | 0.064 | 0.048 | 1 | 5 | Bud23 | 0.016 | Old |
| 0.293534 | 0.17431 | 0.118 | 0.094 | 1 | 5 | Rnf34 | 0.024 | Old |
| 0.293629 | 0.264385 | 0.18 | 0.155 | 1 | 5 | Axin1 | 0.025 | Old |
| 0.293817 | 0.42025 | 0.019 | 0.01 | 1 | 5 | Tmem138 | 0.009 | Old |
| 0.293848 | 0.110223 | 0.374 | 0.343 | 1 | 5 | Ankrd12 | 0.031 | Old |
| 0.293963 | 0.116116 | 0.369 | 0.332 | 1 | 5 | Lilrb4a | 0.037 | Old |
| 0.293998 | 0.517474 | 0.086 | 0.069 | 1 | 5 | Fam114a1 | 0.017 | Old |
| 0.294226 | 0.47061 | 0.015 | 0.008 | 1 | 5 | Gne | 0.007 | Old |
| 0.294301 | 0.362141 | 0.022 | 0.013 | 1 | 5 | Plcxd2 | 0.009 | Old |
| 0.294311 | 0.601613 | 0.019 | 0.01 | 1 | 5 | Myo10 | 0.009 | Old |
| 0.294409 | 0.297434 | 0.146 | 0.124 | 1 | 5 | E2f4 | 0.022 | Old |
| 0.294415 | 0.220792 | 0.253 | 0.228 | 1 | 5 | Tor1b | 0.025 | Old |
| 0.294699 | 0.419777 | 0.116 | 0.096 | 1 | 5 | Nxt2 | 0.02 | Old |
| 0.294778 | 0.144026 | 0.256 | 0.228 | 1 | 5 | Gpr107 | 0.028 | Old |
| 0.294876 | 0.039674 | 0.167 | 0.142 | 1 | 5 | Mrpl54 | 0.025 | Old |
| 0.295047 | 0.196568 | 0.125 | 0.104 | 1 | 5 | Nacc1 | 0.021 | Old |
| 0.295085 | 0.030508 | 0.461 | 0.404 | 1 | 5 | Chmp2a | 0.057 | Old |
| 0.295322 | 0.601945 | 0.061 | 0.046 | 1 | 5 | Dnajc25 | 0.015 | Old |
| 0.295389 | 0.289789 | 0.155 | 0.135 | 1 | 5 | Bahd1 | 0.02 | Old |
| 0.295601 | 0.055455 | 0.128 | 0.104 | 1 | 5 | Ccdc107 | 0.024 | Old |
| 0.295636 | 0.300233 | 0.114 | 0.094 | 1 | 5 | Scarb1 | 0.02 | Old |
| 0.296019 | 0.383985 | 0.086 | 0.069 | 1 | 5 | Pmvk | 0.017 | Old |
| 0.296114 | 0.41685 | 0.022 | 0.013 | 1 | 5 | Opn3 | 0.009 | Old |
| 0.296114 | 0.259814 | 0.022 | 0.013 | 1 | 5 | Cpt1c | 0.009 | Old |
| 0.29637 | 0.683576 | 0.037 | 0.025 | 1 | 5 | Pdss2 | 0.012 | Old |
| 0.296695 | 0.031363 | 0.894 | 0.868 | 1 | 5 | Rps7 | 0.026 | Old |
| 0.296802 | 0.491157 | 0.197 | 0.175 | 1 | 5 | Mex3c | 0.022 | Old |
| 0.296923 | 0.151978 | 0.025 | 0.015 | 1 | 5 | Hs3st1 | 0.01 | Old |
| 0.297024 | 0.993083 | 0.022 | 0.013 | 1 | 5 | Rgs5 | 0.009 | Old |
| 0.297291 | 0.332278 | 0.019 | 0.01 | 1 | 5 | Pex26 | 0.009 | Old |
| 0.297578 | 0.026555 | 0.502 | 0.452 | 1 | 5 | Smdt1 | 0.05 | Old |
| 0.297639 | 0.113281 | 0.126 | 0.104 | 1 | 5 | 3110062M | 0.022 | Old |
| 0.298066 | 0.129997 | 0.215 | 0.185 | 1 | 5 | Myo1g | 0.03 | Old |
| 0.298105 | 0.163449 | 0.301 | 0.272 | 1 | 5 | Sec63 | 0.029 | Old |
| 0.298289 | 0.287725 | 0.019 | 0.01 | 1 | 5 | Il4i1 | 0.009 | Old |
| 0.298291 | 0.116863 | 0.204 | 0.178 | 1 | 5 | Cmip | 0.026 | Old |
| 0.298309 | 0.242973 | 0.071 | 0.053 | 1 | 5 | Col4a2 | 0.018 | Old |
| 0.298557 | 0.074666 | 0.187 | 0.16 | 1 | 5 | Cd200r1 | 0.027 | Old |
| 0.298716 | 0.083818 | 0.495 | 0.459 | 1 | 5 | Socs3 | 0.036 | Old |

| | | | | | | | | |
|---|---|---|---|---|---|---|---|---|
| 0.299053 | 0.293803 | 0.052 | 0.038 | 1 | 5 | Rad9a | 0.014 | Old |
| 0.299163 | 0.412913 | 0.15 | 0.129 | 1 | 5 | Gbp3 | 0.021 | Old |
| 0.299589 | 0.413745 | 0.069 | 0.053 | 1 | 5 | Kctd18 | 0.016 | Old |
| 0.299735 | 0.373363 | 0.04 | 0.028 | 1 | 5 | Fbxw9 | 0.012 | Old |
| 0.299817 | 0.489574 | 0.037 | 0.025 | 1 | 5 | Rab3a | 0.012 | Old |
| 0.299953 | 0.150756 | 0.283 | 0.259 | 1 | 5 | Dlst | 0.024 | Old |
| 0.30018 | 0.287636 | 0.114 | 0.094 | 1 | 5 | Dpm2 | 0.02 | Old |
| 0.300262 | 0.178496 | 0.056 | 0.041 | 1 | 5 | Cnnm2 | 0.015 | Old |
| 0.300454 | 0.055592 | 0.249 | 0.218 | 1 | 5 | Yipf5 | 0.031 | Old |
| 0.300793 | 0.282938 | 0.019 | 0.01 | 1 | 5 | 1700030K0 | 0.009 | Old |
| 0.301561 | 0.000948 | 0.195 | 0.168 | 1 | 5 | Ndufs8 | 0.027 | Old |
| 0.301594 | 0.100248 | 0.217 | 0.188 | 1 | 5 | Tmem260 | 0.029 | Old |
| 0.301723 | 0.153093 | 0.067 | 0.051 | 1 | 5 | Mtfr1 | 0.016 | Old |
| 0.302089 | 0.139725 | 0.237 | 0.208 | 1 | 5 | Luzp1 | 0.029 | Old |
| 0.302257 | 0.148199 | 0.247 | 0.218 | 1 | 5 | Fam104a | 0.029 | Old |
| 0.302567 | 0.104587 | 0.17 | 0.145 | 1 | 5 | Kansl3 | 0.025 | Old |
| 0.302846 | 0.105962 | 0.273 | 0.249 | 1 | 5 | Trio | 0.024 | Old |
| 0.303056 | 0.065925 | 0.044 | 0.03 | 1 | 5 | Mus81 | 0.014 | Old |
| 0.303136 | 0.44069 | 0.069 | 0.053 | 1 | 5 | Cela1 | 0.016 | Old |
| 0.303268 | 0.20885 | 0.114 | 0.094 | 1 | 5 | Fbxl17 | 0.02 | Old |
| 0.303362 | 0.053967 | 0.404 | 0.365 | 1 | 5 | Sec11c | 0.039 | Old |
| 0.303625 | 0.200912 | 0.121 | 0.099 | 1 | 5 | Htatip2 | 0.022 | Old |
| 0.304989 | 0.280648 | 0.271 | 0.251 | 1 | 5 | Rbm33 | 0.02 | Old |
| 0.305189 | 0.257643 | 0.052 | 0.038 | 1 | 5 | Tysnd1 | 0.014 | Old |
| 0.305306 | 0.143907 | 0.254 | 0.228 | 1 | 5 | Hcfc1 | 0.026 | Old |
| 0.305404 | 0.317447 | 0.04 | 0.028 | 1 | 5 | Tmtc4 | 0.012 | Old |
| 0.305577 | 0.214419 | 0.173 | 0.15 | 1 | 5 | Lrrc41 | 0.023 | Old |
| 0.305592 | 0.183455 | 0.111 | 0.091 | 1 | 5 | Immp1l | 0.02 | Old |
| 0.306629 | 0.095027 | 0.104 | 0.084 | 1 | 5 | Yif1a | 0.02 | Old |
| 0.307029 | 0.039749 | 0.263 | 0.228 | 1 | 5 | Ube2f | 0.035 | Old |
| 0.30726 | 0.136218 | 0.359 | 0.33 | 1 | 5 | Spcs3 | 0.029 | Old |
| 0.307312 | 0.094928 | 0.296 | 0.261 | 1 | 5 | Cdk14 | 0.035 | Old |
| 0.307312 | 0.10152 | 0.481 | 0.437 | 1 | 5 | Magt1 | 0.044 | Old |
| 0.30775 | 0.243574 | 0.209 | 0.185 | 1 | 5 | Pilrb2 | 0.024 | Old |
| 0.307924 | 0.292867 | 0.116 | 0.096 | 1 | 5 | Nos1ap | 0.02 | Old |
| 0.308235 | 0.023255 | 0.103 | 0.084 | 1 | 5 | Dglucy | 0.019 | Old |
| 0.308552 | 0.451261 | 0.037 | 0.025 | 1 | 5 | Zfp52 | 0.012 | Old |
| 0.30876 | 0.02366 | 0.345 | 0.302 | 1 | 5 | Szrd1 | 0.043 | Old |
| 0.308776 | 0.115793 | 0.601 | 0.576 | 1 | 5 | Cep170 | 0.025 | Old |
| 0.309623 | 0.253147 | 0.052 | 0.038 | 1 | 5 | Dimt1 | 0.014 | Old |
| 0.309701 | 0.39916 | 0.088 | 0.071 | 1 | 5 | Nmi | 0.017 | Old |
| 0.310016 | 0.295864 | 0.173 | 0.15 | 1 | 5 | Afg3l1 | 0.023 | Old |
| 0.310464 | 0.380979 | 0.04 | 0.028 | 1 | 5 | Ttc8 | 0.012 | Old |
| 0.310498 | 0.097567 | 0.259 | 0.226 | 1 | 5 | Vat1 | 0.033 | Old |
| 0.310737 | 0.322891 | 0.114 | 0.096 | 1 | 5 | Triap1 | 0.018 | Old |

| | | | | | | | | |
|---|---|---|---|---|---|---|---|---|
| 0.311738 | 0.441352 | 0.037 | 0.025 | 1 | 5 | Nudt22 | 0.012 | Old |
| 0.311984 | 0.003902 | 0.923 | 0.876 | 1 | 5 | Rpl11 | 0.047 | Old |
| 0.312426 | 0.288265 | 0.173 | 0.152 | 1 | 5 | Plekhm1 | 0.021 | Old |
| 0.312448 | 0.459157 | 0.037 | 0.025 | 1 | 5 | Psph | 0.012 | Old |
| 0.312491 | 0.181633 | 0.323 | 0.297 | 1 | 5 | Pcyt1a | 0.026 | Old |
| 0.31252 | 0.358659 | 0.086 | 0.069 | 1 | 5 | Tes | 0.017 | Old |
| 0.312719 | 0.015483 | 0.17 | 0.145 | 1 | 5 | Pgrmc2 | 0.025 | Old |
| 0.312804 | 0.344968 | 0.037 | 0.025 | 1 | 5 | Pus3 | 0.012 | Old |
| 0.312823 | 0.257883 | 0.103 | 0.084 | 1 | 5 | Reep4 | 0.019 | Old |
| 0.312837 | 0.220513 | 0.123 | 0.102 | 1 | 5 | Med23 | 0.021 | Old |
| 0.312983 | 0.0638 | 0.17 | 0.142 | 1 | 5 | Ccdc124 | 0.028 | Old |
| 0.313054 | 0.003118 | 0.242 | 0.211 | 1 | 5 | Dctn3 | 0.031 | Old |
| 0.313305 | 0.017088 | 0.305 | 0.269 | 1 | 5 | Fnbp1l | 0.036 | Old |
| 0.313432 | 0.038875 | 0.449 | 0.416 | 1 | 5 | Tnpo3 | 0.033 | Old |
| 0.313454 | 0.117664 | 0.195 | 0.168 | 1 | 5 | Upf1 | 0.027 | Old |
| 0.31391 | 0.3484 | 0.049 | 0.036 | 1 | 5 | Cavin2 | 0.013 | Old |
| 0.314146 | 0.206412 | 0.118 | 0.096 | 1 | 5 | Iqsec2 | 0.022 | Old |
| 0.31448 | 0.198957 | 0.136 | 0.117 | 1 | 5 | Fkbp5 | 0.019 | Old |
| 0.314839 | 0.49818 | 0.049 | 0.036 | 1 | 5 | AI839979 | 0.013 | Old |
| 0.314945 | 0.448815 | 0.061 | 0.046 | 1 | 5 | Abcg3 | 0.015 | Old |
| 0.315915 | 0.071547 | 0.428 | 0.396 | 1 | 5 | Cox14 | 0.032 | Old |
| 0.31606 | 0.666676 | 0.034 | 0.023 | 1 | 5 | Kctd9 | 0.011 | Old |
| 0.316083 | 0.002425 | 0.33 | 0.294 | 1 | 5 | Ndufs5 | 0.036 | Old |
| 0.316427 | 0.161732 | 0.396 | 0.373 | 1 | 5 | Ggnbp2 | 0.023 | Old |
| 0.316499 | 0.186821 | 0.209 | 0.185 | 1 | 5 | Zwint | 0.024 | Old |
| 0.31691 | 0.341931 | 0.082 | 0.066 | 1 | 5 | Pak4 | 0.016 | Old |
| 0.316982 | 0.081837 | 0.71 | 0.703 | 1 | 5 | Birc3 | 0.007 | Old |
| 0.317069 | 0.232749 | 0.12 | 0.099 | 1 | 5 | Mkl1 | 0.021 | Old |
| 0.317943 | 0.791501 | 0.034 | 0.023 | 1 | 5 | Ptprm | 0.011 | Old |
| 0.317943 | 0.606967 | 0.034 | 0.023 | 1 | 5 | Nynrin | 0.011 | Old |
| 0.318417 | 0.181153 | 0.414 | 0.381 | 1 | 5 | Phlda1 | 0.033 | Old |
| 0.319325 | 0.206605 | 0.146 | 0.124 | 1 | 5 | Dcun1d1 | 0.022 | Old |
| 0.319354 | 0.311149 | 0.141 | 0.122 | 1 | 5 | Clns1a | 0.019 | Old |
| 0.319405 | 0.024819 | 0.185 | 0.157 | 1 | 5 | Nab2 | 0.028 | Old |
| 0.319478 | 0.005597 | 0.623 | 0.574 | 1 | 5 | Tfe3 | 0.049 | Old |
| 0.319957 | 0.106548 | 0.199 | 0.17 | 1 | 5 | Spen | 0.029 | Old |
| 0.320133 | 0.576324 | 0.045 | 0.033 | 1 | 5 | Rps6kb2 | 0.012 | Old |
| 0.320306 | 0.002628 | 0.141 | 0.117 | 1 | 5 | Mrps6 | 0.024 | Old |
| 0.320491 | 0.232037 | 0.141 | 0.119 | 1 | 5 | Med21 | 0.022 | Old |
| 0.32077 | 0.05529 | 0.507 | 0.477 | 1 | 5 | Ankrd44 | 0.03 | Old |
| 0.320971 | 0.534638 | 0.034 | 0.023 | 1 | 5 | AC163032. | 0.011 | Old |
| 0.321326 | 0.065497 | 0.465 | 0.434 | 1 | 5 | Hnrnpdl | 0.031 | Old |
| 0.321525 | 0.233395 | 0.079 | 0.063 | 1 | 5 | Nt5dc1 | 0.016 | Old |
| 0.321639 | 0.020645 | 0.116 | 0.096 | 1 | 5 | Tbc1d10a | 0.02 | Old |
| 0.321931 | 0.014994 | 0.242 | 0.218 | 1 | 5 | Sptlc1 | 0.024 | Old |

| | | | | | | | | |
|---|---|---|---|---|---|---|---|---|
| 0.321955 | 0.164596 | 0.155 | 0.132 | 1 | 5 | Naip2 | 0.023 | Old |
| 0.322032 | 0.176077 | 0.173 | 0.15 | 1 | 5 | Foxk1 | 0.023 | Old |
| 0.32221 | 0.031577 | 0.498 | 0.449 | 1 | 5 | Lbr | 0.049 | Old |
| 0.322227 | 0.065283 | 0.182 | 0.157 | 1 | 5 | Desi2 | 0.025 | Old |
| 0.322341 | 0.187058 | 0.418 | 0.391 | 1 | 5 | U2surp | 0.027 | Old |
| 0.322765 | 0.179041 | 0.081 | 0.063 | 1 | 5 | Dhx33 | 0.018 | Old |
| 0.322987 | 0.181809 | 0.143 | 0.122 | 1 | 5 | Tmem120a | 0.021 | Old |
| 0.322996 | 0.245488 | 0.113 | 0.094 | 1 | 5 | Mtmr1 | 0.019 | Old |
| 0.323008 | 0.053913 | 0.32 | 0.289 | 1 | 5 | Itga9 | 0.031 | Old |
| 0.323024 | 0.122848 | 0.131 | 0.109 | 1 | 5 | Sesn3 | 0.022 | Old |
| 0.323042 | 0.051185 | 0.51 | 0.464 | 1 | 5 | Itsn2 | 0.046 | Old |
| 0.323255 | 0.380408 | 0.034 | 0.023 | 1 | 5 | Ippk | 0.011 | Old |
| 0.323329 | 0.073003 | 0.429 | 0.386 | 1 | 5 | Ppp2r5a | 0.043 | Old |
| 0.323414 | 0.021497 | 0.274 | 0.244 | 1 | 5 | Plekhg5 | 0.03 | Old |
| 0.323966 | 0.301374 | 0.118 | 0.099 | 1 | 5 | Cdk5rap3 | 0.019 | Old |
| 0.324592 | 0.017253 | 0.231 | 0.201 | 1 | 5 | Tmem50b | 0.03 | Old |
| 0.32464 | 0.826701 | 0.03 | 0.02 | 1 | 5 | Cbr3 | 0.01 | Old |
| 0.325359 | 0.054729 | 0.234 | 0.206 | 1 | 5 | Mtmr12 | 0.028 | Old |
| 0.325396 | 0.098183 | 0.34 | 0.315 | 1 | 5 | Nfatc1 | 0.025 | Old |
| 0.325452 | 0.084618 | 0.402 | 0.365 | 1 | 5 | Slc35b2 | 0.037 | Old |
| 0.325645 | 0.123776 | 0.348 | 0.317 | 1 | 5 | Pycard | 0.031 | Old |
| 0.326233 | 0.534898 | 0.079 | 0.063 | 1 | 5 | Kif19a | 0.016 | Old |
| 0.326498 | 0.146986 | 0.162 | 0.137 | 1 | 5 | Slc38a7 | 0.025 | Old |
| 0.3267 | 0.567943 | 0.034 | 0.023 | 1 | 5 | Carmil1 | 0.011 | Old |
| 0.326738 | 0.073661 | 0.194 | 0.168 | 1 | 5 | Ric8a | 0.026 | Old |
| 0.327331 | 0.280483 | 0.18 | 0.16 | 1 | 5 | Rnf103 | 0.02 | Old |
| 0.328005 | 0.175819 | 0.163 | 0.142 | 1 | 5 | Nasp | 0.021 | Old |
| 0.328325 | 0.04345 | 0.38 | 0.343 | 1 | 5 | Kat2b | 0.037 | Old |
| 0.328602 | 0.006297 | 0.101 | 0.081 | 1 | 5 | Vps26b | 0.02 | Old |
| 0.328648 | 0.188014 | 0.18 | 0.157 | 1 | 5 | Dok1 | 0.023 | Old |
| 0.329054 | 0.193853 | 0.057 | 0.043 | 1 | 5 | Gpatch11 | 0.014 | Old |
| 0.329218 | 0.273153 | 0.249 | 0.226 | 1 | 5 | Tnfaip2 | 0.023 | Old |
| 0.329434 | 0.298525 | 0.152 | 0.132 | 1 | 5 | Cep192 | 0.02 | Old |
| 0.329579 | 0.110116 | 0.51 | 0.467 | 1 | 5 | Znrf2 | 0.043 | Old |
| 0.329636 | 0.396868 | 0.045 | 0.033 | 1 | 5 | Ccdc66 | 0.012 | Old |
| 0.329723 | 0.024517 | 0.579 | 0.536 | 1 | 5 | Fam174a | 0.043 | Old |
| 0.32982 | 0.083262 | 0.387 | 0.35 | 1 | 5 | Cox5b | 0.037 | Old |
| 0.330188 | 0.082071 | 0.286 | 0.254 | 1 | 5 | Ccdc12 | 0.032 | Old |
| 0.330786 | 0.03173 | 0.061 | 0.046 | 1 | 5 | Ankrd26 | 0.015 | Old |
| 0.330788 | 0.319455 | 0.088 | 0.071 | 1 | 5 | Bola3 | 0.017 | Old |
| 0.330866 | 0.099867 | 0.234 | 0.203 | 1 | 5 | Dctn2 | 0.031 | Old |
| 0.331017 | 0.08973 | 0.212 | 0.188 | 1 | 5 | 0610030E2 | 0.024 | Old |
| 0.331119 | 0.319244 | 0.057 | 0.043 | 1 | 5 | Qpctl | 0.014 | Old |
| 0.33151 | 0.285041 | 0.082 | 0.066 | 1 | 5 | Pi4k2b | 0.016 | Old |
| 0.331544 | 0.592866 | 0.03 | 0.02 | 1 | 5 | Gm4631 | 0.01 | Old |

| | | | | | | | | |
|---|---|---|---|---|---|---|---|---|
| 0.331613 | 0.223361 | 0.246 | 0.221 | 1 | 5 | Mast3 | 0.025 | Old |
| 0.331761 | 0.202999 | 0.14 | 0.119 | 1 | 5 | Hk1 | 0.021 | Old |
| 0.33188 | 0.413397 | 0.082 | 0.066 | 1 | 5 | Fam160b2 | 0.016 | Old |
| 0.331961 | 0.016104 | 0.726 | 0.695 | 1 | 5 | Abhd12 | 0.031 | Old |
| 0.332236 | 0.273856 | 0.074 | 0.058 | 1 | 5 | Clasrp | 0.016 | Old |
| 0.332466 | 0.078836 | 0.345 | 0.312 | 1 | 5 | Adam9 | 0.033 | Old |
| 0.332972 | 0.033833 | 0.197 | 0.168 | 1 | 5 | Aak1 | 0.029 | Old |
| 0.333209 | 0.083032 | 0.135 | 0.114 | 1 | 5 | Stx8 | 0.021 | Old |
| 0.333243 | 0.238519 | 0.168 | 0.147 | 1 | 5 | Vps4a | 0.021 | Old |
| 0.333315 | 0.142221 | 0.116 | 0.096 | 1 | 5 | Traf7 | 0.02 | Old |
| 0.333621 | 0.269401 | 0.045 | 0.033 | 1 | 5 | Ift27 | 0.012 | Old |
| 0.333672 | 0.218055 | 0.3 | 0.272 | 1 | 5 | Fcgr4 | 0.028 | Old |
| 0.333918 | 0.062853 | 0.535 | 0.497 | 1 | 5 | Uqcrq | 0.038 | Old |
| 0.33404 | 0.704668 | 0.042 | 0.03 | 1 | 5 | Cep78 | 0.012 | Old |
| 0.334139 | 0.021021 | 0.401 | 0.363 | 1 | 5 | Kpna4 | 0.038 | Old |
| 0.334186 | 0.017314 | 0.431 | 0.391 | 1 | 5 | Ctdnep1 | 0.04 | Old |
| 0.334288 | 0.322104 | 0.045 | 0.033 | 1 | 5 | Trim30c | 0.012 | Old |
| 0.334564 | 0.052181 | 0.47 | 0.442 | 1 | 5 | Raly | 0.028 | Old |
| 0.334921 | 0.16468 | 0.214 | 0.19 | 1 | 5 | Setd7 | 0.024 | Old |
| 0.335121 | 0.144842 | 0.165 | 0.145 | 1 | 5 | Edc4 | 0.02 | Old |
| 0.335268 | 0.458697 | 0.054 | 0.041 | 1 | 5 | Slc2a9 | 0.013 | Old |
| 0.33531 | 0.163872 | 0.114 | 0.094 | 1 | 5 | Cisd1 | 0.02 | Old |
| 0.335729 | 0.114493 | 0.111 | 0.091 | 1 | 5 | Mrps18a | 0.02 | Old |
| 0.335775 | 0.581516 | 0.042 | 0.03 | 1 | 5 | Klhl36 | 0.012 | Old |
| 0.335809 | 0.203816 | 0.618 | 0.64 | 1 | 5 | Trib1 | -0.022 | Old |
| 0.33589 | 0.442511 | 0.084 | 0.069 | 1 | 5 | Mettl21a | 0.015 | Old |
| 0.336062 | 0.547764 | 0.03 | 0.02 | 1 | 5 | Msl3l2 | 0.01 | Old |
| 0.33626 | 0.029721 | 0.537 | 0.49 | 1 | 5 | Mdfic | 0.047 | Old |
| 0.336475 | 0.337004 | 0.03 | 0.02 | 1 | 5 | Mrps27 | 0.01 | Old |
| 0.336496 | 0.438027 | 0.054 | 0.041 | 1 | 5 | Zfp142 | 0.013 | Old |
| 0.336712 | 0.052883 | 0.194 | 0.17 | 1 | 5 | Stat2 | 0.024 | Old |
| 0.337018 | 0.177296 | 0.374 | 0.35 | 1 | 5 | Armc8 | 0.024 | Old |
| 0.337066 | 0.207856 | 0.066 | 0.051 | 1 | 5 | Ulk3 | 0.015 | Old |
| 0.337728 | 0.195006 | 0.19 | 0.168 | 1 | 5 | Xpot | 0.022 | Old |
| 0.337774 | 0.357892 | 0.143 | 0.124 | 1 | 5 | Mrpl18 | 0.019 | Old |
| 0.338783 | 0.237232 | 0.197 | 0.173 | 1 | 5 | Sf3a2 | 0.024 | Old |
| 0.33941 | 0.27603 | 0.096 | 0.079 | 1 | 5 | Naglu | 0.017 | Old |
| 0.339562 | 0.09828 | 0.098 | 0.079 | 1 | 5 | C030034L1 | 0.019 | Old |
| 0.339612 | 0.524518 | 0.042 | 0.03 | 1 | 5 | Rasgrf2 | 0.012 | Old |
| 0.339788 | 0.521357 | 0.03 | 0.02 | 1 | 5 | Ccdc69 | 0.01 | Old |
| 0.339811 | 0.253374 | 0.098 | 0.081 | 1 | 5 | Ddx49 | 0.017 | Old |
| 0.340222 | 0.018372 | 0.178 | 0.15 | 1 | 5 | Casp3 | 0.028 | Old |
| 0.340329 | 0.134596 | 0.088 | 0.071 | 1 | 5 | Il18 | 0.017 | Old |
| 0.34056 | 0.091503 | 0.204 | 0.178 | 1 | 5 | Lats1 | 0.026 | Old |
| 0.340577 | 0.00894 | 0.298 | 0.261 | 1 | 5 | Smim11 | 0.037 | Old |

| | | | | | | | | |
|---|---|---|---|---|---|---|---|---|
| 0.340714 | 0.149784 | 0.165 | 0.142 | 1 | 5 | Tmem41b | 0.023 | Old |
| 0.340901 | 0.375252 | 0.071 | 0.056 | 1 | 5 | Anapc7 | 0.015 | Old |
| 0.3412 | 0.079187 | 0.3 | 0.272 | 1 | 5 | Slfn8 | 0.028 | Old |
| 0.341436 | 0.383911 | 0.054 | 0.041 | 1 | 5 | Pde4d | 0.013 | Old |
| 0.34151 | 0.107694 | 0.49 | 0.459 | 1 | 5 | Pros1 | 0.031 | Old |
| 0.341701 | 0.204919 | 0.306 | 0.284 | 1 | 5 | Cisd2 | 0.022 | Old |
| 0.341717 | 0.530283 | 0.042 | 0.03 | 1 | 5 | Sdhaf3 | 0.012 | Old |
| 0.341884 | 0.033841 | 0.357 | 0.317 | 1 | 5 | Myof | 0.04 | Old |
| 0.341928 | 0.063138 | 0.116 | 0.096 | 1 | 5 | Coq5 | 0.02 | Old |
| 0.342035 | 0.226533 | 0.116 | 0.096 | 1 | 5 | Tmem55a | 0.02 | Old |
| 0.342056 | 0.280849 | 0.054 | 0.041 | 1 | 5 | Hmgn5 | 0.013 | Old |
| 0.342099 | 0.302132 | 0.062 | 0.048 | 1 | 5 | Cep162 | 0.014 | Old |
| 0.342168 | 0.246023 | 0.057 | 0.043 | 1 | 5 | Slc37a4 | 0.014 | Old |
| 0.342242 | 0.63307 | 0.027 | 0.018 | 1 | 5 | Gemin8 | 0.009 | Old |
| 0.342263 | 0.056101 | 0.399 | 0.355 | 1 | 5 | Cnot1 | 0.044 | Old |
| 0.342391 | 0.55089 | 0.067 | 0.053 | 1 | 5 | Hmgxb3 | 0.014 | Old |
| 0.342564 | 0.092304 | 0.261 | 0.228 | 1 | 5 | Mogs | 0.033 | Old |
| 0.342685 | 0.591245 | 0.027 | 0.018 | 1 | 5 | Pms1 | 0.009 | Old |
| 0.343007 | 0.079006 | 0.3 | 0.274 | 1 | 5 | Rnf114 | 0.026 | Old |
| 0.343008 | 0.09312 | 0.3 | 0.272 | 1 | 5 | 2310035C2 | 0.028 | Old |
| 0.34304 | 0.032285 | 0.463 | 0.421 | 1 | 5 | Arglu1 | 0.042 | Old |
| 0.343133 | 0.045507 | 0.217 | 0.19 | 1 | 5 | Stk24 | 0.027 | Old |
| 0.343193 | 0.069326 | 0.131 | 0.109 | 1 | 5 | Ikzf5 | 0.022 | Old |
| 0.343614 | 0.018524 | 0.343 | 0.307 | 1 | 5 | Lrmp | 0.036 | Old |
| 0.344233 | 0.21018 | 0.168 | 0.147 | 1 | 5 | Spata2 | 0.021 | Old |
| 0.344611 | 0.384423 | 0.088 | 0.071 | 1 | 5 | Patz1 | 0.017 | Old |
| 0.345466 | 0.320574 | 0.14 | 0.122 | 1 | 5 | Mum1 | 0.018 | Old |
| 0.345519 | 0.132765 | 0.177 | 0.155 | 1 | 5 | Ube3c | 0.022 | Old |
| 0.345524 | 0.118346 | 0.367 | 0.338 | 1 | 5 | Fem1c | 0.029 | Old |
| 0.345793 | 0.23933 | 0.254 | 0.234 | 1 | 5 | Ddx46 | 0.02 | Old |
| 0.346034 | 0.026821 | 1 | 1 | 1 | 5 | mt-Nd4 | 0 | Old |
| 0.346352 | 0.225668 | 0.121 | 0.102 | 1 | 5 | Vps9d1 | 0.019 | Old |
| 0.346421 | 0.281151 | 0.054 | 0.041 | 1 | 5 | Zkscan17 | 0.013 | Old |
| 0.346476 | 0.194589 | 0.03 | 0.02 | 1 | 5 | Lnx2 | 0.01 | Old |
| 0.346691 | 0.536706 | 0.027 | 0.018 | 1 | 5 | Clec3b | 0.009 | Old |
| 0.346748 | 0.266406 | 0.263 | 0.244 | 1 | 5 | Gnl3 | 0.019 | Old |
| 0.346938 | 0.059993 | 0.153 | 0.129 | 1 | 5 | Gga3 | 0.024 | Old |
| 0.34695 | 0.057048 | 0.423 | 0.386 | 1 | 5 | Mdh1 | 0.037 | Old |
| 0.347085 | 0.075644 | 0.456 | 0.416 | 1 | 5 | Polr1d | 0.04 | Old |
| 0.347675 | 0.099502 | 0.054 | 0.041 | 1 | 5 | Tmem115 | 0.013 | Old |
| 0.348186 | 0.185278 | 0.088 | 0.071 | 1 | 5 | Gabbr1 | 0.017 | Old |
| 0.34853 | 0.265925 | 0.192 | 0.173 | 1 | 5 | Polr2c | 0.019 | Old |
| 0.348543 | 0.364573 | 0.111 | 0.094 | 1 | 5 | Tmem97 | 0.017 | Old |
| 0.348771 | 0.293893 | 0.072 | 0.058 | 1 | 5 | Umps | 0.014 | Old |
| 0.348938 | 0.107049 | 0.229 | 0.206 | 1 | 5 | Plin3 | 0.023 | Old |

| | | | | | | | | |
|---|---|---|---|---|---|---|---|---|
| 0.349378 | 0.241163 | 0.027 | 0.018 | 1 | 5 | Car5b | 0.009 | Old |
| 0.349428 | 0.051242 | 0.03 | 0.02 | 1 | 5 | Stk11ip | 0.01 | Old |
| 0.349634 | 0.212402 | 0.215 | 0.19 | 1 | 5 | Rras | 0.025 | Old |
| 0.349638 | 0.395506 | 0.103 | 0.086 | 1 | 5 | Psd4 | 0.017 | Old |
| 0.349695 | 0.277234 | 0.093 | 0.076 | 1 | 5 | Thoc5 | 0.017 | Old |
| 0.350079 | 0.54688 | 0.039 | 0.028 | 1 | 5 | P2ry14 | 0.011 | Old |
| 0.350219 | 0.171746 | 0.042 | 0.03 | 1 | 5 | Plcg1 | 0.012 | Old |
| 0.350366 | 0.188809 | 0.13 | 0.112 | 1 | 5 | Anapc4 | 0.018 | Old |
| 0.350631 | 0.283681 | 0.066 | 0.051 | 1 | 5 | Myom1 | 0.015 | Old |
| 0.350884 | 0.288675 | 0.141 | 0.124 | 1 | 5 | Commd1 | 0.017 | Old |
| 0.351614 | 0.059843 | 0.325 | 0.294 | 1 | 5 | Taf6l | 0.031 | Old |
| 0.351813 | 0.096945 | 0.17 | 0.147 | 1 | 5 | Ppp4r1 | 0.023 | Old |
| 0.352105 | 0.117798 | 0.342 | 0.315 | 1 | 5 | Gpbp1 | 0.027 | Old |
| 0.352728 | 0.152625 | 0.165 | 0.142 | 1 | 5 | Nsmce2 | 0.023 | Old |
| 0.35283 | 0.207799 | 0.114 | 0.096 | 1 | 5 | Grcc10 | 0.018 | Old |
| 0.353533 | 0.158281 | 0.384 | 0.348 | 1 | 5 | Mid1ip1 | 0.036 | Old |
| 0.353663 | 0.529226 | 0.059 | 0.046 | 1 | 5 | Zbtb5 | 0.013 | Old |
| 0.353758 | 0.798486 | 0.024 | 0.015 | 1 | 5 | Zfp287 | 0.009 | Old |
| 0.353923 | 0.183799 | 0.229 | 0.208 | 1 | 5 | Rara | 0.021 | Old |
| 0.354027 | 0.292467 | 0.14 | 0.122 | 1 | 5 | C2cd3 | 0.018 | Old |
| 0.354203 | 0.110312 | 0.362 | 0.33 | 1 | 5 | Nfat5 | 0.032 | Old |
| 0.354702 | 0.084198 | 0.005 | 0.01 | 1 | 5 | Zfp72 | -0.005 | Old |
| 0.3548 | 0.101018 | 0.194 | 0.17 | 1 | 5 | Txndc12 | 0.024 | Old |
| 0.354913 | 0.029824 | 0.488 | 0.444 | 1 | 5 | Sypl | 0.044 | Old |
| 0.354953 | 0.245423 | 0.062 | 0.048 | 1 | 5 | 4930503L1 | 0.014 | Old |
| 0.355159 | 0.160167 | 0.197 | 0.175 | 1 | 5 | Copg1 | 0.022 | Old |
| 0.355245 | 0.241911 | 0.027 | 0.018 | 1 | 5 | Naprt | 0.009 | Old |
| 0.355394 | 0.187788 | 0.269 | 0.251 | 1 | 5 | Tpst2 | 0.018 | Old |
| 0.355463 | 0.058993 | 0.125 | 0.104 | 1 | 5 | Stoml2 | 0.021 | Old |
| 0.355552 | 0.078023 | 0.848 | 0.794 | 1 | 5 | Hnrnpa3 | 0.054 | Old |
| 0.355792 | 0.182709 | 0.12 | 0.102 | 1 | 5 | Upf2 | 0.018 | Old |
| 0.355904 | 0.148438 | 0.244 | 0.218 | 1 | 5 | Uap1 | 0.026 | Old |
| 0.356025 | 0.109477 | 0.411 | 0.398 | 1 | 5 | Ptpa | 0.013 | Old |
| 0.356244 | 0.042231 | 0.303 | 0.272 | 1 | 5 | Chordc1 | 0.031 | Old |
| 0.356273 | 0.337567 | 0.113 | 0.094 | 1 | 5 | Myc | 0.019 | Old |
| 0.356533 | 0.117386 | 0.082 | 0.066 | 1 | 5 | Brcc3 | 0.016 | Old |
| 0.356542 | 0.105816 | 0.194 | 0.168 | 1 | 5 | Phf1 | 0.026 | Old |
| 0.356586 | 1.020771 | 0.064 | 0.051 | 1 | 5 | Hes1 | 0.013 | Old |
| 0.356834 | 0.114039 | 0.404 | 0.381 | 1 | 5 | Phf3 | 0.023 | Old |
| 0.35685 | 0.109091 | 0.054 | 0.041 | 1 | 5 | Zfp84 | 0.013 | Old |
| 0.356867 | 0.02756 | 0.079 | 0.063 | 1 | 5 | Ptcd2 | 0.016 | Old |
| 0.357479 | 0.050399 | 0.269 | 0.241 | 1 | 5 | Ssu72 | 0.028 | Old |
| 0.357646 | 0.046721 | 0.086 | 0.069 | 1 | 5 | Gtf2e2 | 0.017 | Old |
| 0.357647 | 0.377738 | 0.024 | 0.015 | 1 | 5 | Tmem67 | 0.009 | Old |
| 0.357842 | 0.109438 | 0.264 | 0.234 | 1 | 5 | Nfix | 0.03 | Old |

| | | | | | | | | |
|---|---|---|---|---|---|---|---|---|
| 0.357905 | 0.175385 | 0.099 | 0.081 | 1 | 5 | Ring1 | 0.018 | Old |
| 0.357934 | 0.346267 | 0.094 | 0.079 | 1 | 5 | Sbf1 | 0.015 | Old |
| 0.358107 | 0.099721 | 0.168 | 0.15 | 1 | 5 | Sae1 | 0.018 | Old |
| 0.358613 | 0.078362 | 0.18 | 0.157 | 1 | 5 | Micu2 | 0.023 | Old |
| 0.358623 | 0.610945 | 0.024 | 0.015 | 1 | 5 | Ndnf | 0.009 | Old |
| 0.358859 | 0.039182 | 0.554 | 0.515 | 1 | 5 | Swap70 | 0.039 | Old |
| 0.359657 | 0.13105 | 0.439 | 0.419 | 1 | 5 | Zmynd8 | 0.02 | Old |
| 0.359703 | 0.377492 | 0.072 | 0.058 | 1 | 5 | Ttc38 | 0.014 | Old |
| 0.35974 | 0.088204 | 0.638 | 0.589 | 1 | 5 | Fli1 | 0.049 | Old |
| 0.359941 | 0.119478 | 0.086 | 0.069 | 1 | 5 | Elp3 | 0.017 | Old |
| 0.360025 | 0.146225 | 0.101 | 0.084 | 1 | 5 | Agl | 0.017 | Old |
| 0.360091 | 0.658069 | 0.024 | 0.015 | 1 | 5 | Rdh5 | 0.009 | Old |
| 0.360091 | 0.555918 | 0.024 | 0.015 | 1 | 5 | Thg1l | 0.009 | Old |
| 0.360308 | 0.272496 | 0.096 | 0.079 | 1 | 5 | Slc10a7 | 0.017 | Old |
| 0.360749 | 0.378279 | 0.098 | 0.081 | 1 | 5 | Lsm2 | 0.017 | Old |
| 0.360941 | 0.688032 | 0.047 | 0.036 | 1 | 5 | Adck2 | 0.011 | Old |
| 0.36139 | 0.088518 | 0.227 | 0.195 | 1 | 5 | Spire1 | 0.032 | Old |
| 0.362438 | 0.120395 | 0.157 | 0.135 | 1 | 5 | Trappc6a | 0.022 | Old |
| 0.362449 | 0.16451 | 0.412 | 0.381 | 1 | 5 | 8-Sep | 0.031 | Old |
| 0.362546 | 0.423293 | 0.024 | 0.015 | 1 | 5 | Bcs1l | 0.009 | Old |
| 0.362673 | 0.153238 | 0.158 | 0.137 | 1 | 5 | Thoc6 | 0.021 | Old |
| 0.362734 | 0.338186 | 0.247 | 0.228 | 1 | 5 | Cpeb2 | 0.019 | Old |
| 0.363178 | 0.038252 | 0.209 | 0.185 | 1 | 5 | Cstf3 | 0.024 | Old |
| 0.363262 | 0.189332 | 0.051 | 0.038 | 1 | 5 | Gm20703 | 0.013 | Old |
| 0.363515 | 0.031299 | 0.387 | 0.358 | 1 | 5 | Cast | 0.029 | Old |
| 0.363649 | 0.086267 | 0.084 | 0.069 | 1 | 5 | Eogt | 0.015 | Old |
| 0.36394 | 0.038644 | 0.352 | 0.315 | 1 | 5 | Asph | 0.037 | Old |
| 0.364276 | 0.867693 | 0.02 | 0.013 | 1 | 5 | 4732440D( | 0.007 | Old |
| 0.364547 | 0.234747 | 0.121 | 0.102 | 1 | 5 | Gps2 | 0.019 | Old |
| 0.364837 | 0.074534 | 0.407 | 0.376 | 1 | 5 | Fkbp15 | 0.031 | Old |
| 0.36506 | 0.349829 | 0.081 | 0.066 | 1 | 5 | Ttc37 | 0.015 | Old |
| 0.365082 | 0.06318 | 0.162 | 0.14 | 1 | 5 | Lrba | 0.022 | Old |
| 0.36537 | 0.11479 | 0.215 | 0.193 | 1 | 5 | Ndufv2 | 0.022 | Old |
| 0.365929 | 0.26494 | 0.051 | 0.038 | 1 | 5 | Xndc1 | 0.013 | Old |
| 0.366 | 0.056979 | 0.024 | 0.015 | 1 | 5 | Unc13b | 0.009 | Old |
| 0.366111 | 0.127153 | 0.116 | 0.099 | 1 | 5 | Gart | 0.017 | Old |
| 0.366417 | 0.982535 | 0.02 | 0.013 | 1 | 5 | Spry1 | 0.007 | Old |
| 0.367385 | 0.056717 | 0.322 | 0.292 | 1 | 5 | Zfp704 | 0.03 | Old |
| 0.367623 | 0.347903 | 0.101 | 0.084 | 1 | 5 | D6Wsu163 | 0.017 | Old |
| 0.367883 | 0.05589 | 0.512 | 0.477 | 1 | 5 | Acin1 | 0.035 | Old |
| 0.367972 | 0.272203 | 0.081 | 0.066 | 1 | 5 | Tecpr2 | 0.015 | Old |
| 0.367983 | 0.48749 | 0.024 | 0.015 | 1 | 5 | Elovl6 | 0.009 | Old |
| 0.368231 | 0.149976 | 0.224 | 0.206 | 1 | 5 | Cldnd1 | 0.018 | Old |
| 0.368258 | 0.042106 | 0.98 | 0.962 | 1 | 5 | Lcp1 | 0.018 | Old |
| 0.368476 | 0.182333 | 0.19 | 0.168 | 1 | 5 | Fbrsl1 | 0.022 | Old |

| | | | | | | | | |
|---|---|---|---|---|---|---|---|---|
| 0.368647 | 0.294631 | 0.089 | 0.074 | 1 | 5 | Rtl8b | 0.015 | Old |
| 0.36876 | 0.370904 | 0.14 | 0.122 | 1 | 5 | Gpr137b | 0.018 | Old |
| 0.36905 | 0.040927 | 0.197 | 0.175 | 1 | 5 | Anapc2 | 0.022 | Old |
| 0.36931 | 0.347846 | 0.059 | 0.046 | 1 | 5 | Nubp1 | 0.013 | Old |
| 0.369799 | 0.077812 | 0.182 | 0.16 | 1 | 5 | Cdkn2aipnl | 0.022 | Old |
| 0.369953 | 0.04682 | 0.338 | 0.302 | 1 | 5 | Dusp11 | 0.036 | Old |
| 0.370637 | 0.285427 | 0.071 | 0.056 | 1 | 5 | Alkbh8 | 0.015 | Old |
| 0.370751 | 0.317672 | 0.114 | 0.099 | 1 | 5 | Zfyve27 | 0.015 | Old |
| 0.371201 | 0.237116 | 0.064 | 0.051 | 1 | 5 | Zfand2b | 0.013 | Old |
| 0.371654 | 0.300407 | 0.047 | 0.036 | 1 | 5 | Bco2 | 0.011 | Old |
| 0.371681 | 0.106757 | 0.163 | 0.142 | 1 | 5 | Homer1 | 0.021 | Old |
| 0.371747 | 0.202167 | 0.062 | 0.048 | 1 | 5 | Prkd2 | 0.014 | Old |
| 0.371806 | 0.529571 | 0.02 | 0.013 | 1 | 5 | Tbc1d25 | 0.007 | Old |
| 0.371861 | 0.2509 | 0.113 | 0.094 | 1 | 5 | Zfp277 | 0.019 | Old |
| 0.37205 | 0.295281 | 0.126 | 0.109 | 1 | 5 | Cpsf1 | 0.017 | Old |
| 0.372315 | 0.128142 | 0.219 | 0.198 | 1 | 5 | Banf1 | 0.021 | Old |
| 0.372317 | 0.060673 | 0.056 | 0.071 | 1 | 5 | Ddx31 | -0.015 | Old |
| 0.372347 | 0.552254 | 0.02 | 0.013 | 1 | 5 | Pigo | 0.007 | Old |
| 0.372889 | 0.803884 | 0.02 | 0.013 | 1 | 5 | Pla2g6 | 0.007 | Old |
| 0.373432 | 0.308083 | 0.02 | 0.013 | 1 | 5 | Ptger2 | 0.007 | Old |
| 0.373783 | 0.052553 | 0.071 | 0.056 | 1 | 5 | Gm15417 | 0.015 | Old |
| 0.373975 | 0.43574 | 0.02 | 0.013 | 1 | 5 | Gm28707 | 0.007 | Old |
| 0.373996 | 0.227903 | 0.039 | 0.028 | 1 | 5 | Zfyve9 | 0.011 | Old |
| 0.374203 | 0.082708 | 0.399 | 0.371 | 1 | 5 | Prdx2 | 0.028 | Old |
| 0.374518 | 0.569106 | 0.02 | 0.013 | 1 | 5 | Rpgrip1l | 0.007 | Old |
| 0.374772 | 0.146767 | 0.039 | 0.028 | 1 | 5 | Trim39 | 0.011 | Old |
| 0.374846 | 0.12832 | 0.308 | 0.282 | 1 | 5 | Flii | 0.026 | Old |
| 0.375083 | 1.008028 | 0.017 | 0.01 | 1 | 5 | Zfp446 | 0.007 | Old |
| 0.375115 | 0.109445 | 0.136 | 0.114 | 1 | 5 | Mpnd | 0.022 | Old |
| 0.375311 | 0.12319 | 0.195 | 0.178 | 1 | 5 | Zdhhc3 | 0.017 | Old |
| 0.375502 | 0.234876 | 0.047 | 0.036 | 1 | 5 | Lmln | 0.011 | Old |
| 0.375626 | 0.014843 | 0.751 | 0.716 | 1 | 5 | Prkcd | 0.035 | Old |
| 0.3757 | 0.132852 | 0.051 | 0.038 | 1 | 5 | Ccdc58 | 0.013 | Old |
| 0.376132 | 0.206536 | 0.067 | 0.053 | 1 | 5 | Ppp2r5b | 0.014 | Old |
| 0.376282 | 1.176216 | 0.017 | 0.01 | 1 | 5 | Tmem150a | 0.007 | Old |
| 0.376882 | 0.923945 | 0.017 | 0.01 | 1 | 5 | Mvd | 0.007 | Old |
| 0.377483 | 0.86964 | 0.017 | 0.01 | 1 | 5 | Abca5 | 0.007 | Old |
| 0.377486 | 0.445343 | 0.069 | 0.056 | 1 | 5 | Map4k5 | 0.013 | Old |
| 0.377726 | 0.281176 | 0.103 | 0.086 | 1 | 5 | Zfp991 | 0.017 | Old |
| 0.378085 | 1.62284 | 0.017 | 0.01 | 1 | 5 | Cxcl3 | 0.007 | Old |
| 0.378208 | 0.60183 | 0.066 | 0.053 | 1 | 5 | Taf6 | 0.013 | Old |
| 0.378423 | 0.107536 | 0.051 | 0.038 | 1 | 5 | Poll | 0.013 | Old |
| 0.378728 | 0.090013 | 0.116 | 0.099 | 1 | 5 | Phkb | 0.017 | Old |
| 0.378884 | 0.429972 | 0.02 | 0.013 | 1 | 5 | Ccl27a | 0.007 | Old |
| 0.378923 | 0.064962 | 0.108 | 0.089 | 1 | 5 | Mrpl37 | 0.019 | Old |

| | | | | | | | | |
|---|---|---|---|---|---|---|---|---|
| 0.378955 | 0.082694 | 0.502 | 0.457 | 1 | 5 | Ythdc1 | 0.045 | Old |
| 0.379289 | 0.82827 | 0.017 | 0.01 | 1 | 5 | Gm10762 | 0.007 | Old |
| 0.379432 | 0.211788 | 0.02 | 0.013 | 1 | 5 | Pcdhb22 | 0.007 | Old |
| 0.379631 | 0.062565 | 0.726 | 0.68 | 1 | 5 | Rab1a | 0.046 | Old |
| 0.379892 | 0.722012 | 0.017 | 0.01 | 1 | 5 | Rabl2 | 0.007 | Old |
| 0.379959 | 0.049058 | 0.412 | 0.393 | 1 | 5 | Arih1 | 0.019 | Old |
| 0.379964 | 0.037048 | 0.035 | 0.025 | 1 | 5 | Eno3 | 0.01 | Old |
| 0.379971 | 0.172079 | 0.067 | 0.053 | 1 | 5 | Tfb2m | 0.014 | Old |
| 0.38013 | 0.133213 | 0.143 | 0.124 | 1 | 5 | Cnnm3 | 0.019 | Old |
| 0.380132 | 0.145778 | 0.051 | 0.038 | 1 | 5 | Pgm3 | 0.013 | Old |
| 0.380613 | 0.280185 | 0.296 | 0.272 | 1 | 5 | Samd4 | 0.024 | Old |
| 0.380924 | 0.15539 | 0.271 | 0.249 | 1 | 5 | Gaa | 0.022 | Old |
| 0.381058 | 0.139859 | 0.072 | 0.058 | 1 | 5 | Mettl6 | 0.014 | Old |
| 0.381101 | 0.511502 | 0.017 | 0.01 | 1 | 5 | Dchs1 | 0.007 | Old |
| 0.381127 | 0.284494 | 0.056 | 0.043 | 1 | 5 | Pdik1l | 0.013 | Old |
| 0.381132 | 0.041193 | 0.141 | 0.119 | 1 | 5 | Rbm15 | 0.022 | Old |
| 0.381806 | 0.172692 | 0.219 | 0.198 | 1 | 5 | Sdccag3 | 0.021 | Old |
| 0.381855 | 0.079112 | 0.259 | 0.239 | 1 | 5 | Polb | 0.02 | Old |
| 0.381896 | 0.359429 | 0.232 | 0.216 | 1 | 5 | Arl5c | 0.016 | Old |
| 0.382523 | 0.040979 | 0.145 | 0.122 | 1 | 5 | Zfp664 | 0.023 | Old |
| 0.382581 | 0.010993 | 0.039 | 0.028 | 1 | 5 | Ftsj1 | 0.011 | Old |
| 0.382672 | 0.073181 | 0.517 | 0.48 | 1 | 5 | Ywhaq | 0.037 | Old |
| 0.382777 | 0.249271 | 0.072 | 0.058 | 1 | 5 | Tfdp1 | 0.014 | Old |
| 0.382777 | 0.212535 | 0.072 | 0.058 | 1 | 5 | Wwox | 0.014 | Old |
| 0.382931 | 0.126871 | 0.185 | 0.162 | 1 | 5 | Suclg1 | 0.023 | Old |
| 0.382949 | 0.066285 | 0.227 | 0.203 | 1 | 5 | Gpr146 | 0.024 | Old |
| 0.383246 | 0.356347 | 0.035 | 0.025 | 1 | 5 | Zbtb8a | 0.01 | Old |
| 0.383524 | 0.694662 | 0.017 | 0.01 | 1 | 5 | Azin2 | 0.007 | Old |
| 0.383524 | 0.051286 | 0.017 | 0.01 | 1 | 5 | Otub2 | 0.007 | Old |
| 0.383564 | 1.099352 | 0.013 | 0.008 | 1 | 5 | Adgrf5 | 0.005 | Old |
| 0.383855 | 0.205236 | 0.244 | 0.223 | 1 | 5 | Cdip1 | 0.021 | Old |
| 0.383904 | 0.038809 | 0.274 | 0.246 | 1 | 5 | Sp110 | 0.028 | Old |
| 0.383987 | 0.926136 | 0.01 | 0.005 | 1 | 5 | Tbx6 | 0.005 | Old |
| 0.383987 | 0.919725 | 0.01 | 0.005 | 1 | 5 | Egfl8 | 0.005 | Old |
| 0.383987 | 0.739029 | 0.01 | 0.005 | 1 | 5 | Hist1h4h | 0.005 | Old |
| 0.384096 | 0.018109 | 0.264 | 0.239 | 1 | 5 | BC035044 | 0.025 | Old |
| 0.384132 | 0.543327 | 0.017 | 0.01 | 1 | 5 | Coq8a | 0.007 | Old |
| 0.384372 | 0.275349 | 0.123 | 0.107 | 1 | 5 | Prr14l | 0.016 | Old |
| 0.3845 | 0.191652 | 0.072 | 0.058 | 1 | 5 | Loxl3 | 0.014 | Old |
| 0.38474 | 0.304972 | 0.017 | 0.01 | 1 | 5 | Heg1 | 0.007 | Old |
| 0.384789 | 0.933349 | 0.01 | 0.005 | 1 | 5 | Tmem178b | 0.005 | Old |
| 0.384789 | 0.890987 | 0.01 | 0.005 | 1 | 5 | Vwa7 | 0.005 | Old |
| 0.384789 | 0.706238 | 0.01 | 0.005 | 1 | 5 | Fhad1 | 0.005 | Old |
| 0.385348 | 0.541696 | 0.017 | 0.01 | 1 | 5 | Col16a1 | 0.007 | Old |
| 0.38537 | 0.101571 | 0.079 | 0.063 | 1 | 5 | Chchd4 | 0.016 | Old |

| | | | | | | | | |
|---|---|---|---|---|---|---|---|---|
| 0.385463 | 0.333929 | 0.061 | 0.048 | 1 | 5 | Cdt1 | 0.013 | Old |
| 0.385592 | 1.075857 | 0.01 | 0.005 | 1 | 5 | Aqp7 | 0.005 | Old |
| 0.385592 | 0.996766 | 0.01 | 0.005 | 1 | 5 | Tedc1 | 0.005 | Old |
| 0.385592 | 0.943247 | 0.01 | 0.005 | 1 | 5 | Ascl2 | 0.005 | Old |
| 0.385625 | 0.200246 | 0.067 | 0.053 | 1 | 5 | Hyal2 | 0.014 | Old |
| 0.385758 | 0.656312 | 0.032 | 0.023 | 1 | 5 | Oas3 | 0.009 | Old |
| 0.386305 | 0.816297 | 0.013 | 0.008 | 1 | 5 | Zfp692 | 0.005 | Old |
| 0.386305 | 0.805947 | 0.013 | 0.008 | 1 | 5 | 1700020D( | 0.005 | Old |
| 0.386305 | 0.757615 | 0.013 | 0.008 | 1 | 5 | Acer2 | 0.005 | Old |
| 0.386305 | 0.637665 | 0.013 | 0.008 | 1 | 5 | Gm2415 | 0.005 | Old |
| 0.38638 | 0.127339 | 0.167 | 0.145 | 1 | 5 | Alkbh1 | 0.022 | Old |
| 0.386396 | 0.928778 | 0.01 | 0.005 | 1 | 5 | Gm26698 | 0.005 | Old |
| 0.386396 | 0.828216 | 0.01 | 0.005 | 1 | 5 | Slc39a12 | 0.005 | Old |
| 0.386396 | 0.817622 | 0.01 | 0.005 | 1 | 5 | Nudt8.1 | 0.005 | Old |
| 0.386627 | 0.609377 | 0.032 | 0.023 | 1 | 5 | Mb21d2 | 0.009 | Old |
| 0.386706 | 0.038697 | 0.279 | 0.256 | 1 | 5 | Gm34084 | 0.023 | Old |
| 0.386993 | 0.838007 | 0.013 | 0.008 | 1 | 5 | Bbs5 | 0.005 | Old |
| 0.387084 | 0.106797 | 0.306 | 0.279 | 1 | 5 | Nus1 | 0.027 | Old |
| 0.387177 | 0.51095 | 0.017 | 0.01 | 1 | 5 | Ttll5 | 0.007 | Old |
| 0.3872 | 0.831856 | 0.01 | 0.005 | 1 | 5 | Zfp493 | 0.005 | Old |
| 0.3872 | 0.830452 | 0.01 | 0.005 | 1 | 5 | Slc25a27 | 0.005 | Old |
| 0.387277 | 0.098097 | 0.268 | 0.241 | 1 | 5 | Aprt | 0.027 | Old |
| 0.387445 | 0.279787 | 0.044 | 0.033 | 1 | 5 | Cpne8 | 0.011 | Old |
| 0.387654 | 0.112001 | 0.576 | 0.543 | 1 | 5 | Ralbp1 | 0.033 | Old |
| 0.387681 | 0.679063 | 0.013 | 0.008 | 1 | 5 | Gpt | 0.005 | Old |
| 0.388006 | 0.811486 | 0.01 | 0.005 | 1 | 5 | AL591952.: | 0.005 | Old |
| 0.388006 | 0.662841 | 0.01 | 0.005 | 1 | 5 | Slc16a14 | 0.005 | Old |
| 0.388006 | 0.59572 | 0.01 | 0.005 | 1 | 5 | Lonrf3 | 0.005 | Old |
| 0.388135 | 0.149273 | 0.077 | 0.063 | 1 | 5 | Jpt2 | 0.014 | Old |
| 0.388233 | 0.478637 | 0.074 | 0.061 | 1 | 5 | Plcb2 | 0.013 | Old |
| 0.388267 | 0.236212 | 0.047 | 0.036 | 1 | 5 | Zfp944 | 0.011 | Old |
| 0.388293 | 0.206405 | 0.111 | 0.094 | 1 | 5 | Dcaf1 | 0.017 | Old |
| 0.388813 | 0.652911 | 0.01 | 0.005 | 1 | 5 | Wdr5b | 0.005 | Old |
| 0.388813 | 0.544401 | 0.01 | 0.005 | 1 | 5 | Scn3b | 0.005 | Old |
| 0.388813 | 0.461922 | 0.01 | 0.005 | 1 | 5 | Fbxl22 | 0.005 | Old |
| 0.38901 | 0.057196 | 0.241 | 0.216 | 1 | 5 | Tdrd7 | 0.025 | Old |
| 0.389059 | 0.703223 | 0.013 | 0.008 | 1 | 5 | Lipt2 | 0.005 | Old |
| 0.389079 | 0.16279 | 0.106 | 0.089 | 1 | 5 | Mrto4 | 0.017 | Old |
| 0.38951 | 0.046318 | 0.113 | 0.094 | 1 | 5 | Sh3tc1 | 0.019 | Old |
| 0.389542 | 0.16378 | 0.114 | 0.096 | 1 | 5 | Tmem63b | 0.018 | Old |
| 0.389602 | 0.152904 | 0.278 | 0.256 | 1 | 5 | Inpp5k | 0.022 | Old |
| 0.389621 | 0.533992 | 0.01 | 0.005 | 1 | 5 | Kifc2 | 0.005 | Old |
| 0.389621 | 0.474958 | 0.01 | 0.005 | 1 | 5 | 5330429C( | 0.005 | Old |
| 0.389749 | 0.658019 | 0.013 | 0.008 | 1 | 5 | Ebf1 | 0.005 | Old |
| 0.389749 | 0.620242 | 0.013 | 0.008 | 1 | 5 | A730063M | 0.005 | Old |

| | | | | | | | | |
|---|---|---|---|---|---|---|---|---|
| 0.390121 | 0.221078 | 0.108 | 0.091 | 1 | 5 | Slc25a30 | 0.017 | Old |
| 0.39044 | 0.629496 | 0.013 | 0.008 | 1 | 5 | Unc5b | 0.005 | Old |
| 0.39044 | 0.570582 | 0.013 | 0.008 | 1 | 5 | Slc25a43 | 0.005 | Old |
| 0.391195 | 0.219381 | 0.121 | 0.104 | 1 | 5 | Iffo2 | 0.017 | Old |
| 0.39124 | 0.481218 | 0.01 | 0.005 | 1 | 5 | Gprc5b | 0.005 | Old |
| 0.391363 | 0.195632 | 0.155 | 0.135 | 1 | 5 | Zbtb21 | 0.02 | Old |
| 0.391594 | 0.201514 | 0.069 | 0.056 | 1 | 5 | Dxo | 0.013 | Old |
| 0.391741 | 0.101214 | 0.362 | 0.335 | 1 | 5 | Fes | 0.027 | Old |
| 0.39177 | 0.327399 | 0.125 | 0.109 | 1 | 5 | Incenp | 0.016 | Old |
| 0.391824 | 0.445088 | 0.013 | 0.008 | 1 | 5 | Lca5 | 0.005 | Old |
| 0.391824 | 0.391325 | 0.013 | 0.008 | 1 | 5 | Dtna | 0.005 | Old |
| 0.392051 | 0.239919 | 0.01 | 0.005 | 1 | 5 | Pm20d1 | 0.005 | Old |
| 0.392051 | 0.190244 | 0.01 | 0.005 | 1 | 5 | Rilpl1 | 0.005 | Old |
| 0.392051 | 0.162448 | 0.01 | 0.005 | 1 | 5 | Nat8f1 | 0.005 | Old |
| 0.392051 | 0.138047 | 0.01 | 0.005 | 1 | 5 | Zfp850 | 0.005 | Old |
| 0.392051 | 0.077825 | 0.01 | 0.005 | 1 | 5 | Elmo3 | 0.005 | Old |
| 0.392189 | 0.152586 | 0.205 | 0.185 | 1 | 5 | Zswim8 | 0.02 | Old |
| 0.392258 | 0.172008 | 0.14 | 0.122 | 1 | 5 | Dpagt1 | 0.018 | Old |
| 0.3923 | 0.459684 | 0.032 | 0.023 | 1 | 5 | A930007I1 | 0.009 | Old |
| 0.392592 | 0.026987 | 0.221 | 0.195 | 1 | 5 | Slc25a51 | 0.026 | Old |
| 0.392654 | 0.003641 | 0.402 | 0.371 | 1 | 5 | Pan3 | 0.031 | Old |
| 0.392831 | 0.118723 | 0.178 | 0.155 | 1 | 5 | Zfp384 | 0.023 | Old |
| 0.393211 | 0.256364 | 0.013 | 0.008 | 1 | 5 | Gbp8 | 0.005 | Old |
| 0.393211 | 0.142509 | 0.013 | 0.008 | 1 | 5 | Cd300lg | 0.005 | Old |
| 0.393675 | 0.363015 | 0.01 | 0.005 | 1 | 5 | Pde8b | 0.005 | Old |
| 0.393675 | 0.065354 | 0.01 | 0.005 | 1 | 5 | Cc2d2a | 0.005 | Old |
| 0.393675 | 0.052251 | 0.01 | 0.005 | 1 | 5 | Lefty1 | 0.005 | Old |
| 0.393906 | 0.328208 | 0.013 | 0.008 | 1 | 5 | Bik | 0.005 | Old |
| 0.394377 | 0.040239 | 0.178 | 0.157 | 1 | 5 | Mbd6 | 0.021 | Old |
| 0.394417 | 0.197061 | 0.052 | 0.041 | 1 | 5 | Dcun1d4 | 0.011 | Old |
| 0.394461 | 0.004669 | 0.146 | 0.168 | 1 | 5 | Gclc | -0.022 | Old |
| 0.394959 | 0.090448 | 0.067 | 0.053 | 1 | 5 | Acsf2 | 0.014 | Old |
| 0.395103 | 0.274236 | 0.052 | 0.041 | 1 | 5 | Actr6 | 0.011 | Old |
| 0.395229 | 0.119306 | 0.279 | 0.256 | 1 | 5 | Slc25a11 | 0.023 | Old |
| 0.395297 | 0.350072 | 0.013 | 0.008 | 1 | 5 | Hyls1 | 0.005 | Old |
| 0.395338 | 0.212172 | 0.118 | 0.102 | 1 | 5 | Hsd17b10 | 0.016 | Old |
| 0.395829 | 0.052441 | 0.444 | 0.411 | 1 | 5 | Ranbp2 | 0.033 | Old |
| 0.395994 | 0.276951 | 0.013 | 0.008 | 1 | 5 | Arl16 | 0.005 | Old |
| 0.395994 | 0.150266 | 0.013 | 0.008 | 1 | 5 | Gstp2 | 0.005 | Old |
| 0.396416 | 0.377183 | 0.074 | 0.061 | 1 | 5 | Slc25a32 | 0.013 | Old |
| 0.396552 | 0.112937 | 0.165 | 0.142 | 1 | 5 | Ppm1k | 0.023 | Old |
| 0.396692 | 0.238069 | 0.013 | 0.008 | 1 | 5 | Gm20655 | 0.005 | Old |
| 0.39756 | 0.0368 | 0.505 | 0.464 | 1 | 5 | Aamp | 0.041 | Old |
| 0.397722 | 0.15058 | 0.072 | 0.058 | 1 | 5 | Cnot9 | 0.014 | Old |
| 0.398023 | 0.400374 | 0.032 | 0.023 | 1 | 5 | Aaas | 0.009 | Old |

| | | | | | | | | |
|---|---|---|---|---|---|---|---|---|
| 0.398138 | 0.329393 | 0.077 | 0.063 | 1 | 5 | Cdc42ep2 | 0.014 | Old |
| 0.398283 | 0.135216 | 0.123 | 0.104 | 1 | 5 | Mocs2 | 0.019 | Old |
| 0.398588 | 0.076095 | 0.586 | 0.553 | 1 | 5 | Naaa | 0.033 | Old |
| 0.398866 | 0.182508 | 0.121 | 0.104 | 1 | 5 | Kif1bp | 0.017 | Old |
| 0.398969 | 0.178838 | 0.342 | 0.32 | 1 | 5 | Igsf6 | 0.022 | Old |
| 0.398973 | 0.558335 | 0.04 | 0.03 | 1 | 5 | Ripk3 | 0.01 | Old |
| 0.399088 | 0.294273 | 0.146 | 0.129 | 1 | 5 | 1110038F1 | 0.017 | Old |
| 0.399336 | 0.019594 | 1 | 1 | 1 | 5 | mt-Co1 | 0 | Old |
| 0.399641 | 0.111978 | 0.776 | 0.756 | 1 | 5 | Csde1 | 0.02 | Old |
| 0.40002 | 0.69719 | 0.029 | 0.02 | 1 | 5 | Ttll1 | 0.009 | Old |
| 0.401049 | 0.241073 | 0.082 | 0.069 | 1 | 5 | Drg2 | 0.013 | Old |
| 0.401267 | 0.059573 | 0.951 | 0.929 | 1 | 5 | Calm1 | 0.022 | Old |
| 0.401308 | 0.246364 | 0.044 | 0.033 | 1 | 5 | Aifm2 | 0.011 | Old |
| 0.401654 | 0.357481 | 0.052 | 0.041 | 1 | 5 | Ssh3 | 0.011 | Old |
| 0.402489 | 0.420231 | 0.037 | 0.028 | 1 | 5 | Wars2 | 0.009 | Old |
| 0.403109 | 0.102645 | 0.545 | 0.52 | 1 | 5 | Trip12 | 0.025 | Old |
| 0.403213 | 0.083816 | 0.552 | 0.523 | 1 | 5 | Arl8a | 0.029 | Old |
| 0.403317 | 0.061161 | 0.49 | 0.462 | 1 | 5 | Hsbp1 | 0.028 | Old |
| 0.403697 | 0.134735 | 0.108 | 0.091 | 1 | 5 | Ube2l6 | 0.017 | Old |
| 0.403797 | 0.26772 | 0.032 | 0.023 | 1 | 5 | Adipoq | 0.009 | Old |
| 0.403797 | 0.160352 | 0.032 | 0.023 | 1 | 5 | A830010M | 0.009 | Old |
| 0.404084 | 0.354384 | 0.052 | 0.041 | 1 | 5 | Pold1 | 0.011 | Old |
| 0.40489 | 0.18556 | 0.091 | 0.076 | 1 | 5 | Ccs | 0.015 | Old |
| 0.404953 | 0.236979 | 0.072 | 0.058 | 1 | 5 | Pdlim2 | 0.014 | Old |
| 0.405095 | 0.160116 | 0.126 | 0.112 | 1 | 5 | Malt1 | 0.014 | Old |
| 0.405261 | 0.076403 | 0.114 | 0.096 | 1 | 5 | Dctpp1 | 0.018 | Old |
| 0.405362 | 0.071055 | 0.438 | 0.411 | 1 | 5 | Med13l | 0.027 | Old |
| 0.405404 | 0.519282 | 0.049 | 0.038 | 1 | 5 | Mageh1 | 0.011 | Old |
| 0.405883 | 0.475187 | 0.057 | 0.046 | 1 | 5 | Pls3 | 0.011 | Old |
| 0.405979 | 0.036387 | 0.357 | 0.327 | 1 | 5 | Idh1 | 0.03 | Old |
| 0.406004 | 0.250926 | 0.126 | 0.112 | 1 | 5 | Rin3 | 0.014 | Old |
| 0.406465 | 0.266605 | 0.04 | 0.03 | 1 | 5 | Gnpnat1 | 0.01 | Old |
| 0.406631 | 0.195773 | 0.044 | 0.033 | 1 | 5 | A130010J1 | 0.011 | Old |
| 0.406712 | 0.230533 | 0.077 | 0.063 | 1 | 5 | Larp1b | 0.014 | Old |
| 0.406775 | 0.364485 | 0.155 | 0.14 | 1 | 5 | Tbcel | 0.015 | Old |
| 0.406862 | 0.254423 | 0.04 | 0.03 | 1 | 5 | Ttc27 | 0.01 | Old |
| 0.406875 | 0.208396 | 0.096 | 0.081 | 1 | 5 | Ppp2cb | 0.015 | Old |
| 0.406946 | 0.448686 | 0.076 | 0.063 | 1 | 5 | Atp8b2 | 0.013 | Old |
| 0.407706 | 0.002081 | 0.2 | 0.175 | 1 | 5 | Ei24 | 0.025 | Old |
| 0.407717 | 0.130628 | 0.316 | 0.292 | 1 | 5 | D1Ertd622 | 0.024 | Old |
| 0.407823 | 0.17746 | 0.032 | 0.023 | 1 | 5 | Amotl2 | 0.009 | Old |
| 0.408074 | 0.129062 | 0.148 | 0.129 | 1 | 5 | Scaf8 | 0.019 | Old |
| 0.408146 | 0.274749 | 0.069 | 0.056 | 1 | 5 | Cradd | 0.013 | Old |
| 0.40821 | 0.125453 | 0.239 | 0.221 | 1 | 5 | Gabarapl2 | 0.018 | Old |
| 0.408449 | 0.185492 | 0.113 | 0.096 | 1 | 5 | Gnptg | 0.017 | Old |

| | | | | | | | | |
|---|---|---|---|---|---|---|---|---|
| 0.408822 | 0.170336 | 0.072 | 0.058 | 1 | 5 | Mrpl1 | 0.014 | Old |
| 0.409408 | 0.053342 | 0.997 | 0.98 | 1 | 5 | Eif1 | 0.017 | Old |
| 0.409467 | 0.079917 | 0.138 | 0.122 | 1 | 5 | Ints3 | 0.016 | Old |
| 0.409532 | 0.028145 | 0.5 | 0.47 | 1 | 5 | Scand1 | 0.03 | Old |
| 0.410007 | 0.091005 | 0.094 | 0.079 | 1 | 5 | Hilpda | 0.015 | Old |
| 0.410172 | 0.061541 | 0.476 | 0.442 | 1 | 5 | Ddi2 | 0.034 | Old |
| 0.410216 | 0.074367 | 0.157 | 0.137 | 1 | 5 | Cog7 | 0.02 | Old |
| 0.410458 | 0.095517 | 0.044 | 0.033 | 1 | 5 | Deaf1 | 0.011 | Old |
| 0.41069 | 0.053605 | 0.03 | 0.041 | 1 | 5 | Adat1 | -0.011 | Old |
| 0.41149 | 0.073128 | 0.86 | 0.886 | 1 | 5 | Txnip | -0.026 | Old |
| 0.412026 | 0.065098 | 0.737 | 0.688 | 1 | 5 | Srsf3 | 0.049 | Old |
| 0.412215 | 0.121982 | 0.195 | 0.173 | 1 | 5 | Rnh1 | 0.022 | Old |
| 0.412659 | 0.025592 | 0.929 | 0.906 | 1 | 5 | Serf2 | 0.023 | Old |
| 0.412851 | 0.127812 | 0.165 | 0.145 | 1 | 5 | Pogz | 0.02 | Old |
| 0.412908 | 0.32799 | 0.167 | 0.152 | 1 | 5 | Dars | 0.015 | Old |
| 0.412967 | 0.13157 | 0.525 | 0.523 | 1 | 5 | Mapk14 | 0.002 | Old |
| 0.413068 | 0.05402 | 0.077 | 0.063 | 1 | 5 | Pex16 | 0.014 | Old |
| 0.413243 | 0.174403 | 0.266 | 0.246 | 1 | 5 | Cpsf2 | 0.02 | Old |
| 0.413271 | 0.371209 | 0.029 | 0.02 | 1 | 5 | Cyb5d2 | 0.009 | Old |
| 0.413571 | 0.292092 | 0.057 | 0.046 | 1 | 5 | Mars | 0.011 | Old |
| 0.413942 | 0.065922 | 0.184 | 0.162 | 1 | 5 | Ccdc71l | 0.022 | Old |
| 0.414078 | 0.114795 | 0.588 | 0.576 | 1 | 5 | Gm2a | 0.012 | Old |
| 0.414281 | 0.529383 | 0.202 | 0.185 | 1 | 5 | Hbegf | 0.017 | Old |
| 0.414475 | 0.124199 | 0.247 | 0.223 | 1 | 5 | Vps13b | 0.024 | Old |
| 0.414666 | 0.127624 | 0.072 | 0.058 | 1 | 5 | Sharpin | 0.014 | Old |
| 0.414706 | 0.550316 | 0.029 | 0.02 | 1 | 5 | Nepro | 0.009 | Old |
| 0.415416 | 0.047059 | 0.369 | 0.343 | 1 | 5 | Setd1b | 0.026 | Old |
| 0.415646 | 0.27949 | 0.04 | 0.03 | 1 | 5 | Tsen2 | 0.01 | Old |
| 0.415654 | 0.576018 | 0.101 | 0.089 | 1 | 5 | Rnf144b | 0.012 | Old |
| 0.415664 | 0.232639 | 0.081 | 0.066 | 1 | 5 | Gm45716 | 0.015 | Old |
| 0.415664 | 0.454741 | 0.029 | 0.02 | 1 | 5 | Crym | 0.009 | Old |
| 0.415762 | 0.189035 | 0.074 | 0.061 | 1 | 5 | Ezh2 | 0.013 | Old |
| 0.416291 | 0.014853 | 0.189 | 0.17 | 1 | 5 | Abhd17a | 0.019 | Old |
| 0.416375 | 0.061478 | 0.177 | 0.155 | 1 | 5 | Nfu1 | 0.022 | Old |
| 0.416624 | 0.216916 | 0.029 | 0.02 | 1 | 5 | Med26 | 0.009 | Old |
| 0.416624 | 0.091486 | 0.022 | 0.03 | 1 | 5 | Sirt3 | -0.008 | Old |
| 0.417187 | 0.052694 | 0.487 | 0.442 | 1 | 5 | Morf4l2 | 0.045 | Old |
| 0.417228 | 0.048414 | 0.313 | 0.294 | 1 | 5 | Tpi1 | 0.019 | Old |
| 0.417284 | 0.10911 | 0.094 | 0.079 | 1 | 5 | Pparg | 0.015 | Old |
| 0.417536 | 0.184422 | 0.143 | 0.124 | 1 | 5 | Mtf1 | 0.019 | Old |
| 0.417683 | 0.168101 | 0.094 | 0.079 | 1 | 5 | Mrps28 | 0.015 | Old |
| 0.417938 | 0.546884 | 0.062 | 0.051 | 1 | 5 | Mri1 | 0.011 | Old |
| 0.417995 | 0.212133 | 0.074 | 0.061 | 1 | 5 | Dnajc9 | 0.013 | Old |
| 0.418066 | 0.520302 | 0.029 | 0.02 | 1 | 5 | Amacr | 0.009 | Old |
| 0.418066 | 0.319639 | 0.029 | 0.02 | 1 | 5 | Oas1g | 0.009 | Old |

| | | | | | | | | |
|---|---|---|---|---|---|---|---|---|
| 0.41855 | 0.178566 | 0.246 | 0.223 | 1 | 5 | Cd300lb | 0.023 | Old |
| 0.418754 | 0.10624 | 0.222 | 0.201 | 1 | 5 | Lsm12 | 0.021 | Old |
| 0.419028 | 0.086408 | 0.409 | 0.373 | 1 | 5 | Pdxk | 0.036 | Old |
| 0.419029 | 0.369381 | 0.029 | 0.02 | 1 | 5 | Ppp1r18os | 0.009 | Old |
| 0.419206 | 0.363285 | 0.049 | 0.038 | 1 | 5 | 6330418K0 | 0.011 | Old |
| 0.419368 | 0.060085 | 0.185 | 0.165 | 1 | 5 | Gemin7 | 0.02 | Old |
| 0.419482 | 0.045627 | 0.431 | 0.421 | 1 | 5 | Zfp395 | 0.01 | Old |
| 0.419511 | 0.143572 | 0.029 | 0.02 | 1 | 5 | Ms4a4a | 0.009 | Old |
| 0.419676 | 0.151363 | 0.04 | 0.03 | 1 | 5 | Gemin5 | 0.01 | Old |
| 0.419712 | 0.058646 | 0.428 | 0.409 | 1 | 5 | Sipa1 | 0.019 | Old |
| 0.419839 | 0.427823 | 0.071 | 0.058 | 1 | 5 | Ttbk2 | 0.013 | Old |
| 0.419926 | 0.012592 | 0.492 | 0.439 | 1 | 5 | Atp5c1 | 0.053 | Old |
| 0.420068 | 0.206071 | 0.217 | 0.203 | 1 | 5 | Amdhd2 | 0.014 | Old |
| 0.420076 | 0.533325 | 0.067 | 0.056 | 1 | 5 | Eef1akmt2 | 0.011 | Old |
| 0.420899 | 0.170791 | 0.15 | 0.132 | 1 | 5 | Zfp422 | 0.018 | Old |
| 0.421042 | 0.277779 | 0.049 | 0.038 | 1 | 5 | Cmpk2 | 0.011 | Old |
| 0.421416 | 0.071358 | 0.103 | 0.086 | 1 | 5 | Cbll1 | 0.017 | Old |
| 0.4217 | 0.164002 | 0.04 | 0.03 | 1 | 5 | Pnpla1 | 0.01 | Old |
| 0.421891 | 0.234207 | 0.293 | 0.277 | 1 | 5 | Rad23a | 0.016 | Old |
| 0.422228 | 0.020655 | 0.12 | 0.102 | 1 | 5 | Parp11 | 0.018 | Old |
| 0.422333 | 0.350511 | 0.093 | 0.079 | 1 | 5 | Snx16 | 0.014 | Old |
| 0.422464 | 0.058411 | 0.153 | 0.135 | 1 | 5 | Patl1 | 0.018 | Old |
| 0.422635 | 0.1289 | 0.217 | 0.195 | 1 | 5 | Cryl1 | 0.022 | Old |
| 0.42283 | 0.003348 | 0.173 | 0.15 | 1 | 5 | Sil1 | 0.023 | Old |
| 0.423161 | 0.230259 | 0.045 | 0.036 | 1 | 5 | Catip | 0.009 | Old |
| 0.423285 | 0.147367 | 0.148 | 0.129 | 1 | 5 | Atg4b | 0.019 | Old |
| 0.423378 | 0.214189 | 0.029 | 0.02 | 1 | 5 | Hrh2 | 0.009 | Old |
| 0.423378 | 0.133265 | 0.029 | 0.02 | 1 | 5 | 1700008J0 | 0.009 | Old |
| 0.423538 | 0.021267 | 0.68 | 0.652 | 1 | 5 | Chd4 | 0.028 | Old |
| 0.423714 | 0.711046 | 0.025 | 0.018 | 1 | 5 | Agbl5 | 0.007 | Old |
| 0.423717 | 0.177109 | 0.098 | 0.084 | 1 | 5 | Parp10 | 0.014 | Old |
| 0.423948 | 0.364948 | 0.076 | 0.063 | 1 | 5 | Parp3 | 0.013 | Old |
| 0.424149 | 0.325499 | 0.066 | 0.053 | 1 | 5 | Phospho2 | 0.013 | Old |
| 0.424348 | 0.455444 | 0.114 | 0.102 | 1 | 5 | Tpcn2 | 0.012 | Old |
| 0.424864 | 0.376114 | 0.054 | 0.043 | 1 | 5 | Endod1 | 0.011 | Old |
| 0.425009 | 0.368864 | 0.079 | 0.066 | 1 | 5 | Triobp | 0.013 | Old |
| 0.425297 | 0.220158 | 0.098 | 0.084 | 1 | 5 | Zfp639 | 0.014 | Old |
| 0.425599 | 0.024502 | 0.239 | 0.211 | 1 | 5 | Hlx | 0.028 | Old |
| 0.425873 | 0.147529 | 0.116 | 0.102 | 1 | 5 | Brf1 | 0.014 | Old |
| 0.426158 | 0.086881 | 0.143 | 0.124 | 1 | 5 | Rnf181 | 0.019 | Old |
| 0.426461 | 0.240076 | 0.091 | 0.076 | 1 | 5 | Tmtc3 | 0.015 | Old |
| 0.426477 | 0.001899 | 0.333 | 0.312 | 1 | 5 | Scamp5 | 0.021 | Old |
| 0.426579 | 0.194915 | 0.04 | 0.03 | 1 | 5 | Ptdss2 | 0.01 | Old |
| 0.426602 | 0.192916 | 0.045 | 0.036 | 1 | 5 | Nipal3 | 0.009 | Old |
| 0.426982 | 0.048891 | 0.483 | 0.462 | 1 | 5 | Tmem109 | 0.021 | Old |

| | | | | | | | | |
|---|---|---|---|---|---|---|---|---|
| 0.427079 | 0.076762 | 0.173 | 0.152 | 1 | 5 | Tpbgl | 0.021 | Old |
| 0.427362 | 0.121757 | 0.16 | 0.142 | 1 | 5 | Babam1 | 0.018 | Old |
| 0.428374 | 0.541457 | 0.025 | 0.018 | 1 | 5 | Cald1 | 0.007 | Old |
| 0.428635 | 0.067181 | 0.212 | 0.193 | 1 | 5 | H2-T22 | 0.019 | Old |
| 0.42959 | 0.152558 | 0.066 | 0.053 | 1 | 5 | Mllt11 | 0.013 | Old |
| 0.429609 | 0.309414 | 0.126 | 0.112 | 1 | 5 | Kin | 0.014 | Old |
| 0.429739 | 0.304394 | 0.037 | 0.028 | 1 | 5 | Dmac2 | 0.009 | Old |
| 0.430732 | 0.023367 | 0.249 | 0.226 | 1 | 5 | Sec24b | 0.023 | Old |
| 0.4311 | 0.09071 | 0.261 | 0.244 | 1 | 5 | Ints6 | 0.017 | Old |
| 0.431317 | 0.062043 | 0.301 | 0.274 | 1 | 5 | Psmb3 | 0.027 | Old |
| 0.431393 | 0.021526 | 0.278 | 0.249 | 1 | 5 | Psmb7 | 0.029 | Old |
| 0.431879 | 0.410923 | 0.037 | 0.028 | 1 | 5 | Qtrt1 | 0.009 | Old |
| 0.432019 | 0.313934 | 0.025 | 0.018 | 1 | 5 | Cwf19l1 | 0.007 | Old |
| 0.432542 | 0.267804 | 0.025 | 0.018 | 1 | 5 | Klhdc1 | 0.007 | Old |
| 0.433337 | 0.169597 | 0.091 | 0.076 | 1 | 5 | Polr2h | 0.015 | Old |
| 0.433534 | 0.026583 | 0.387 | 0.358 | 1 | 5 | Hspa4 | 0.029 | Old |
| 0.433595 | 0.072509 | 0.185 | 0.165 | 1 | 5 | Ubfd1 | 0.02 | Old |
| 0.433601 | 0.061964 | 0.387 | 0.35 | 1 | 5 | Gm26740 | 0.037 | Old |
| 0.433722 | 0.097539 | 0.316 | 0.299 | 1 | 5 | Ddx50 | 0.017 | Old |
| 0.433849 | 0.644412 | 0.025 | 0.018 | 1 | 5 | Jchain | 0.007 | Old |
| 0.433876 | 0.105858 | 0.104 | 0.089 | 1 | 5 | Mea1 | 0.015 | Old |
| 0.434634 | 0.344791 | 0.025 | 0.018 | 1 | 5 | Tmem220 | 0.007 | Old |
| 0.434788 | 0.139566 | 0.32 | 0.297 | 1 | 5 | Poldip3 | 0.023 | Old |
| 0.435113 | 0.273947 | 0.071 | 0.058 | 1 | 5 | Zfp597 | 0.013 | Old |
| 0.435786 | 0.191976 | 0.12 | 0.104 | 1 | 5 | Snrnp40 | 0.016 | Old |
| 0.435819 | 0.001919 | 0.266 | 0.236 | 1 | 5 | Anapc13 | 0.03 | Old |
| 0.435903 | 0.064684 | 0.305 | 0.269 | 1 | 5 | Taz | 0.036 | Old |
| 0.436207 | 0.455544 | 0.025 | 0.018 | 1 | 5 | Klhl33 | 0.007 | Old |
| 0.436409 | 0.025115 | 0.891 | 0.86 | 1 | 5 | Sh3bgrl3 | 0.031 | Old |
| 0.437882 | 0.102922 | 0.421 | 0.401 | 1 | 5 | Plxnd1 | 0.02 | Old |
| 0.43831 | 0.428139 | 0.025 | 0.018 | 1 | 5 | Lrrc20 | 0.007 | Old |
| 0.438589 | 0.06278 | 0.365 | 0.345 | 1 | 5 | Stt3a | 0.02 | Old |
| 0.439211 | 0.039157 | 0.318 | 0.297 | 1 | 5 | Slc8b1 | 0.021 | Old |
| 0.439701 | 0.235222 | 0.109 | 0.094 | 1 | 5 | Timm22 | 0.015 | Old |
| 0.43975 | 0.376014 | 0.045 | 0.036 | 1 | 5 | Sh3bp4 | 0.009 | Old |
| 0.439966 | 0.300168 | 0.062 | 0.051 | 1 | 5 | Iqcb1 | 0.011 | Old |
| 0.440122 | 0.28698 | 0.054 | 0.043 | 1 | 5 | Snrnp35 | 0.011 | Old |
| 0.440536 | 0.056251 | 0.175 | 0.155 | 1 | 5 | Cmtm4 | 0.02 | Old |
| 0.441211 | 0.024474 | 0.14 | 0.122 | 1 | 5 | Trip4 | 0.018 | Old |
| 0.441475 | 0.25806 | 0.025 | 0.018 | 1 | 5 | Sac3d1 | 0.007 | Old |
| 0.441558 | 0.321884 | 0.054 | 0.043 | 1 | 5 | Atg13 | 0.011 | Old |
| 0.441562 | 0.204652 | 0.17 | 0.155 | 1 | 5 | Abcd2 | 0.015 | Old |
| 0.441801 | 0.11811 | 0.037 | 0.028 | 1 | 5 | 5730480H( | 0.009 | Old |
| 0.442248 | 0.175227 | 0.123 | 0.109 | 1 | 5 | Xaf1 | 0.014 | Old |
| 0.442255 | 0.230429 | 0.079 | 0.066 | 1 | 5 | Ndc1 | 0.013 | Old |

| | | | | | | | | |
|---|---|---|---|---|---|---|---|---|
| 0.4423 | 0.088057 | 0.145 | 0.127 | 1 | 5 | Slc12a2 | 0.018 | Old |
| 0.442638 | 0.132285 | 0.12 | 0.104 | 1 | 5 | Cd2bp2 | 0.016 | Old |
| 0.443233 | 0.195673 | 0.253 | 0.239 | 1 | 5 | Vps28 | 0.014 | Old |
| 0.444011 | 8.62E-05 | 0.332 | 0.302 | 1 | 5 | Cr1l | 0.03 | Old |
| 0.444534 | 0.083361 | 0.182 | 0.16 | 1 | 5 | Supt4a | 0.022 | Old |
| 0.444731 | 0.060884 | 0.926 | 0.909 | 1 | 5 | Rps17 | 0.017 | Old |
| 0.444887 | 0.112472 | 0.337 | 0.325 | 1 | 5 | Api5 | 0.012 | Old |
| 0.445215 | 0.0505 | 0.244 | 0.223 | 1 | 5 | Kdm5c | 0.021 | Old |
| 0.445481 | 0.005911 | 0.291 | 0.264 | 1 | 5 | Syf2 | 0.027 | Old |
| 0.445718 | 0.012269 | 0.037 | 0.028 | 1 | 5 | Ncapd2 | 0.009 | Old |
| 0.445749 | 0.152028 | 0.178 | 0.16 | 1 | 5 | Ccdc6 | 0.018 | Old |
| 0.446101 | 0.05422 | 0.049 | 0.038 | 1 | 5 | Abhd14b | 0.011 | Old |
| 0.446193 | 0.096495 | 0.096 | 0.081 | 1 | 5 | Zer1 | 0.015 | Old |
| 0.4465 | 0.153839 | 0.062 | 0.051 | 1 | 5 | Pde6d | 0.011 | Old |
| 0.447029 | 0.173446 | 0.037 | 0.028 | 1 | 5 | Ubiad1 | 0.009 | Old |
| 0.447156 | 0.176385 | 0.226 | 0.211 | 1 | 5 | Tbc1d10b | 0.015 | Old |
| 0.447555 | 0.043669 | 0.222 | 0.201 | 1 | 5 | Sdf2l1 | 0.021 | Old |
| 0.447641 | 0.17427 | 0.079 | 0.066 | 1 | 5 | Bod1 | 0.013 | Old |
| 0.447687 | 0.177418 | 0.054 | 0.043 | 1 | 5 | St7 | 0.011 | Old |
| 0.448151 | 0.118075 | 0.441 | 0.424 | 1 | 5 | C5ar2 | 0.017 | Old |
| 0.44827 | 0.0226 | 0.542 | 0.497 | 1 | 5 | Snx6 | 0.045 | Old |
| 0.44838 | 0.143257 | 0.045 | 0.036 | 1 | 5 | Card6 | 0.009 | Old |
| 0.44842 | 0.067415 | 0.145 | 0.127 | 1 | 5 | Pkd2 | 0.018 | Old |
| 0.448699 | 0.708397 | 0.022 | 0.015 | 1 | 5 | Lekr1 | 0.007 | Old |
| 0.449444 | 0.18733 | 0.079 | 0.066 | 1 | 5 | Nrg2 | 0.013 | Old |
| 0.449847 | 0.394279 | 0.034 | 0.025 | 1 | 5 | Slc16a3 | 0.009 | Old |
| 0.45012 | 0.32425 | 0.103 | 0.089 | 1 | 5 | Thoc3 | 0.014 | Old |
| 0.450278 | 0.070024 | 0.535 | 0.513 | 1 | 5 | Hook3 | 0.022 | Old |
| 0.450907 | 0.041623 | 0.296 | 0.266 | 1 | 5 | Gprc5c | 0.03 | Old |
| 0.451113 | 0.081993 | 0.212 | 0.193 | 1 | 5 | Tor3a | 0.019 | Old |
| 0.451553 | 0.120099 | 0.079 | 0.066 | 1 | 5 | Mfsd8 | 0.013 | Old |
| 0.451687 | 0.394661 | 0.034 | 0.025 | 1 | 5 | Slc22a15 | 0.009 | Old |
| 0.452043 | 0.174757 | 0.059 | 0.048 | 1 | 5 | Cog2 | 0.011 | Old |
| 0.452196 | 0.366094 | 0.239 | 0.226 | 1 | 5 | Noct | 0.013 | Old |
| 0.452303 | 0.439759 | 0.077 | 0.066 | 1 | 5 | Nostrin | 0.011 | Old |
| 0.452809 | 0.167294 | 0.345 | 0.315 | 1 | 5 | Sf3b3 | 0.03 | Old |
| 0.452863 | 0.062043 | 0.104 | 0.124 | 1 | 5 | Dip2c | -0.02 | Old |
| 0.453213 | 0.0628 | 0.184 | 0.165 | 1 | 5 | Kif2a | 0.019 | Old |
| 0.453297 | 0.622388 | 0.022 | 0.015 | 1 | 5 | Dnajb13 | 0.007 | Old |
| 0.453874 | 0.452413 | 0.022 | 0.015 | 1 | 5 | 9230111E0 | 0.007 | Old |
| 0.454388 | 0.304334 | 0.099 | 0.086 | 1 | 5 | Rinl | 0.013 | Old |
| 0.454451 | 0.51746 | 0.022 | 0.015 | 1 | 5 | Dna2 | 0.007 | Old |
| 0.454451 | 0.473425 | 0.022 | 0.015 | 1 | 5 | Pex12 | 0.007 | Old |
| 0.454456 | 0.11853 | 0.273 | 0.254 | 1 | 5 | Senp6 | 0.019 | Old |
| 0.454511 | 0.048355 | 0.21 | 0.188 | 1 | 5 | Btaf1 | 0.022 | Old |

| | | | | | | | | |
|---|---|---|---|---|---|---|---|---|
| 0.454616 | 0.148894 | 0.21 | 0.193 | 1 | 5 | Gramd1a | 0.017 | Old |
| 0.454617 | 0.147458 | 0.062 | 0.051 | 1 | 5 | Irf7 | 0.011 | Old |
| 0.454859 | 0.112763 | 0.158 | 0.14 | 1 | 5 | Atp11b | 0.018 | Old |
| 0.455029 | 0.555861 | 0.022 | 0.015 | 1 | 5 | Rfesd | 0.007 | Old |
| 0.4554 | 0.050069 | 0.106 | 0.124 | 1 | 5 | Tppp | -0.018 | Old |
| 0.455607 | 0.13087 | 0.022 | 0.015 | 1 | 5 | Zfp12 | 0.007 | Old |
| 0.456764 | 0.197939 | 0.022 | 0.015 | 1 | 5 | Agrn | 0.007 | Old |
| 0.456768 | 0.330449 | 0.034 | 0.025 | 1 | 5 | Dlg3 | 0.009 | Old |
| 0.456879 | 0.284481 | 0.067 | 0.056 | 1 | 5 | Tns4 | 0.011 | Old |
| 0.456902 | 0.14989 | 0.318 | 0.299 | 1 | 5 | Mcmbp | 0.019 | Old |
| 0.457343 | 0.112423 | 0.022 | 0.015 | 1 | 5 | Gm13205 | 0.007 | Old |
| 0.457513 | 0.025906 | 0.221 | 0.195 | 1 | 5 | Mtm1 | 0.026 | Old |
| 0.457948 | 0.120357 | 0.448 | 0.426 | 1 | 5 | Ppp1cb | 0.022 | Old |
| 0.458295 | 0.024432 | 0.242 | 0.223 | 1 | 5 | Chuk | 0.019 | Old |
| 0.459031 | 0.173632 | 0.173 | 0.157 | 1 | 5 | Supt6 | 0.016 | Old |
| 0.460157 | 0.193109 | 0.269 | 0.254 | 1 | 5 | Zzef1 | 0.015 | Old |
| 0.460827 | 0.026327 | 0.022 | 0.015 | 1 | 5 | Ppcs | 0.007 | Old |
| 0.461098 | 0.139673 | 0.359 | 0.332 | 1 | 5 | Pcnp | 0.027 | Old |
| 0.461579 | 0.182579 | 0.133 | 0.117 | 1 | 5 | Prrc1 | 0.016 | Old |
| 0.461991 | 0.243285 | 0.022 | 0.015 | 1 | 5 | Dppa3 | 0.007 | Old |
| 0.461999 | 0.302943 | 0.081 | 0.069 | 1 | 5 | Unk | 0.012 | Old |
| 0.462194 | 0.093054 | 0.608 | 0.591 | 1 | 5 | Dync1h1 | 0.017 | Old |
| 0.462347 | 0.296893 | 0.034 | 0.025 | 1 | 5 | Qpct | 0.009 | Old |
| 0.462986 | 0.197506 | 0.14 | 0.124 | 1 | 5 | Vegfb | 0.016 | Old |
| 0.463101 | 0.003644 | 0.079 | 0.066 | 1 | 5 | Gna15 | 0.013 | Old |
| 0.46313 | 0.040291 | 0.736 | 0.744 | 1 | 5 | Tgfbr2 | -0.008 | Old |
| 0.463157 | 0.129467 | 0.022 | 0.015 | 1 | 5 | Unc119b | 0.007 | Old |
| 0.463372 | 0.022666 | 0.283 | 0.254 | 1 | 5 | Nek9 | 0.029 | Old |
| 0.464215 | 0.099266 | 0.153 | 0.137 | 1 | 5 | Slc35a2 | 0.016 | Old |
| 0.46441 | 0.345456 | 0.051 | 0.041 | 1 | 5 | Dexi | 0.01 | Old |
| 0.464723 | 0.74564 | 0.019 | 0.013 | 1 | 5 | Trmu | 0.006 | Old |
| 0.464909 | 0.348399 | 0.096 | 0.084 | 1 | 5 | Arl3 | 0.012 | Old |
| 0.465151 | 0.317998 | 0.034 | 0.025 | 1 | 5 | 9130221H1 | 0.009 | Old |
| 0.465336 | 0.354297 | 0.072 | 0.061 | 1 | 5 | Zdhhc8 | 0.011 | Old |
| 0.465513 | 0.226539 | 0.232 | 0.223 | 1 | 5 | Rictor | 0.009 | Old |
| 0.465744 | 0.371379 | 0.051 | 0.041 | 1 | 5 | Slc12a5 | 0.01 | Old |
| 0.465963 | 0.024048 | 0.21 | 0.188 | 1 | 5 | Mysm1 | 0.022 | Old |
| 0.466085 | 0.145104 | 0.062 | 0.051 | 1 | 5 | Sufu | 0.011 | Old |
| 0.466419 | 0.004378 | 0.138 | 0.122 | 1 | 5 | Pla2g4a | 0.016 | Old |
| 0.466421 | 0.236565 | 0.059 | 0.048 | 1 | 5 | Cdc6 | 0.011 | Old |
| 0.466552 | 0.050644 | 0.126 | 0.109 | 1 | 5 | Tlcd2 | 0.017 | Old |
| 0.466634 | 0.382198 | 0.069 | 0.058 | 1 | 5 | Stk35 | 0.011 | Old |
| 0.46672 | 0.041081 | 0.5 | 0.48 | 1 | 5 | Ndufa3 | 0.02 | Old |
| 0.466734 | 0.097202 | 0.487 | 0.464 | 1 | 5 | Trpc4ap | 0.023 | Old |
| 0.466734 | 0.094217 | 0.054 | 0.043 | 1 | 5 | Ubxn8 | 0.011 | Old |

| | | | | | | | | |
|---|---|---|---|---|---|---|---|---|
| 0.466737 | 0.036625 | 0.343 | 0.317 | 1 | 5 | Mbtd1 | 0.026 | Old |
| 0.466867 | 0.050204 | 0.764 | 0.734 | 1 | 5 | Tpd52 | 0.03 | Old |
| 0.467058 | 0.065885 | 0.17 | 0.152 | 1 | 5 | Aurkaip1 | 0.018 | Old |
| 0.467252 | 0.223302 | 0.022 | 0.015 | 1 | 5 | Faap24 | 0.007 | Old |
| 0.467479 | 0.214659 | 0.241 | 0.223 | 1 | 5 | Lats2 | 0.018 | Old |
| 0.467507 | 0.012662 | 0.19 | 0.173 | 1 | 5 | Acadvl | 0.017 | Old |
| 0.467514 | 0.030771 | 0.781 | 0.731 | 1 | 5 | Rab7b | 0.05 | Old |
| 0.467643 | 0.049761 | 0.177 | 0.157 | 1 | 5 | Proser1 | 0.02 | Old |
| 0.467644 | 0.345299 | 0.077 | 0.066 | 1 | 5 | Memo1 | 0.011 | Old |
| 0.467705 | 0.003435 | 0.079 | 0.066 | 1 | 5 | Fdx1 | 0.013 | Old |
| 0.468347 | 0.021838 | 0.045 | 0.036 | 1 | 5 | Ints5.1 | 0.009 | Old |
| 0.468356 | 0.11552 | 0.237 | 0.216 | 1 | 5 | Phf14 | 0.021 | Old |
| 0.46886 | 0.064756 | 0.152 | 0.135 | 1 | 5 | Hmg20b | 0.017 | Old |
| 0.469513 | 0.00223 | 0.981 | 0.982 | 1 | 5 | Rpl37 | -0.001 | Old |
| 0.469759 | 0.121767 | 0.067 | 0.056 | 1 | 5 | Lin9 | 0.011 | Old |
| 0.469827 | 0.806701 | 0.019 | 0.013 | 1 | 5 | Cacnb1 | 0.006 | Old |
| 0.469842 | 0.028357 | 0.032 | 0.041 | 1 | 5 | Cdkn2a | -0.009 | Old |
| 0.469931 | 0.029981 | 0.116 | 0.102 | 1 | 5 | Ermp1 | 0.014 | Old |
| 0.470016 | 0.072081 | 0.071 | 0.058 | 1 | 5 | Wdcp | 0.013 | Old |
| 0.470645 | 0.101207 | 0.288 | 0.269 | 1 | 5 | Mllt10 | 0.019 | Old |
| 0.470717 | 0.320158 | 0.051 | 0.041 | 1 | 5 | Rnaseh1 | 0.01 | Old |
| 0.470787 | 0.079755 | 0.027 | 0.036 | 1 | 5 | Txnl4b | -0.009 | Old |
| 0.471067 | 0.088947 | 0.426 | 0.404 | 1 | 5 | Arhgef12 | 0.022 | Old |
| 0.471108 | 0.680722 | 0.019 | 0.013 | 1 | 5 | Igfbp5 | 0.006 | Old |
| 0.471484 | 0.279204 | 0.051 | 0.041 | 1 | 5 | Cmc4 | 0.01 | Old |
| 0.471591 | 0.468706 | 0.039 | 0.03 | 1 | 5 | Gem | 0.009 | Old |
| 0.471807 | 0.430544 | 0.03 | 0.023 | 1 | 5 | Dtnb | 0.007 | Old |
| 0.472029 | 0.421748 | 0.039 | 0.03 | 1 | 5 | Uevld | 0.009 | Old |
| 0.472303 | 0.769349 | 0.03 | 0.023 | 1 | 5 | Trim6 | 0.007 | Old |
| 0.472456 | 0.273596 | 0.152 | 0.137 | 1 | 5 | Guca1a | 0.015 | Old |
| 0.473033 | 0.452393 | 0.019 | 0.013 | 1 | 5 | Acy1 | 0.006 | Old |
| 0.473344 | 0.562415 | 0.039 | 0.03 | 1 | 5 | Vsig10 | 0.009 | Old |
| 0.473605 | 0.091393 | 0.879 | 0.878 | 1 | 5 | Srsf5 | 0.001 | Old |
| 0.474286 | 0.414407 | 0.084 | 0.074 | 1 | 5 | Dnaaf3 | 0.01 | Old |
| 0.474318 | 0.626833 | 0.019 | 0.013 | 1 | 5 | Rbp1 | 0.006 | Old |
| 0.474432 | 0.167593 | 0.067 | 0.056 | 1 | 5 | Pdp1 | 0.011 | Old |
| 0.474962 | 0.879463 | 0.019 | 0.013 | 1 | 5 | A330069E1 | 0.006 | Old |
| 0.475236 | 0.579942 | 0.047 | 0.038 | 1 | 5 | Prelid3a | 0.009 | Old |
| 0.475415 | 0.034361 | 0.293 | 0.274 | 1 | 5 | Tgs1 | 0.019 | Old |
| 0.475788 | 0.33406 | 0.03 | 0.023 | 1 | 5 | Tsen54 | 0.007 | Old |
| 0.475864 | 0.180467 | 0.5 | 0.482 | 1 | 5 | Ccl4 | 0.018 | Old |
| 0.476432 | 0.513938 | 0.047 | 0.038 | 1 | 5 | Dhx37 | 0.009 | Old |
| 0.476439 | 0.113974 | 0.214 | 0.195 | 1 | 5 | Aldh3b1 | 0.019 | Old |
| 0.476787 | 0.094738 | 0.219 | 0.198 | 1 | 5 | Insig1 | 0.021 | Old |
| 0.476918 | 0.111428 | 0.32 | 0.299 | 1 | 5 | Zc3hav1 | 0.021 | Old |

| | | | | | | | | |
|---|---|---|---|---|---|---|---|---|
| 0.477081 | 0.28715 | 0.039 | 0.03 | 1 | 5 | Cnst | 0.009 | Old |
| 0.477336 | 0.146806 | 0.337 | 0.317 | 1 | 5 | Atp2c1 | 0.02 | Old |
| 0.478187 | 0.671893 | 0.019 | 0.013 | 1 | 5 | Gas8 | 0.006 | Old |
| 0.478187 | 0.394075 | 0.019 | 0.013 | 1 | 5 | Mpzl1 | 0.006 | Old |
| 0.47825 | 0.023834 | 0.402 | 0.381 | 1 | 5 | Samd4b | 0.021 | Old |
| 0.478434 | 0.055298 | 0.944 | 0.906 | 1 | 5 | Egr1 | 0.038 | Old |
| 0.478833 | 0.438666 | 0.019 | 0.013 | 1 | 5 | Mttp | 0.006 | Old |
| 0.479127 | 0.115843 | 0.106 | 0.091 | 1 | 5 | E2f5 | 0.015 | Old |
| 0.47948 | 0.371462 | 0.019 | 0.013 | 1 | 5 | Eya4 | 0.006 | Old |
| 0.47948 | 0.30758 | 0.019 | 0.013 | 1 | 5 | Rftn2 | 0.006 | Old |
| 0.479549 | 0.10029 | 0.108 | 0.094 | 1 | 5 | Poglut1 | 0.014 | Old |
| 0.479744 | 0.029212 | 0.135 | 0.157 | 1 | 5 | Tpra1 | -0.022 | Old |
| 0.479925 | 0.15493 | 0.076 | 0.063 | 1 | 5 | Arl11 | 0.013 | Old |
| 0.480024 | 0.174338 | 0.01 | 0.015 | 1 | 5 | Pik3ip1 | -0.005 | Old |
| 0.480128 | 0.342507 | 0.019 | 0.013 | 1 | 5 | Gm13822 | 0.006 | Old |
| 0.480691 | 0.061807 | 0.547 | 0.51 | 1 | 5 | Bclaf1 | 0.037 | Old |
| 0.480775 | 0.219212 | 0.019 | 0.013 | 1 | 5 | Zfp707 | 0.006 | Old |
| 0.481424 | 0.23107 | 0.019 | 0.013 | 1 | 5 | 6720489N1 | 0.006 | Old |
| 0.481559 | 0.648328 | 0.052 | 0.043 | 1 | 5 | Gm26601 | 0.009 | Old |
| 0.481694 | 0.26723 | 0.077 | 0.066 | 1 | 5 | Nop9 | 0.011 | Old |
| 0.482753 | 0.02713 | 0.897 | 0.878 | 1 | 5 | Cd53 | 0.019 | Old |
| 0.482836 | 0.172731 | 0.261 | 0.246 | 1 | 5 | Phb2 | 0.015 | Old |
| 0.483064 | 0.055923 | 0.428 | 0.393 | 1 | 5 | Arrb1 | 0.035 | Old |
| 0.483266 | 0.024895 | 0.763 | 0.716 | 1 | 5 | Nrp1 | 0.047 | Old |
| 0.483372 | 0.05554 | 0.019 | 0.013 | 1 | 5 | Cbarp | 0.006 | Old |
| 0.483573 | 0.073661 | 0.421 | 0.401 | 1 | 5 | Apobec1 | 0.02 | Old |
| 0.483655 | 0.03585 | 0.776 | 0.751 | 1 | 5 | Tapbp | 0.025 | Old |
| 0.484579 | 0.05604 | 0.251 | 0.228 | 1 | 5 | Sqor | 0.023 | Old |
| 0.485121 | 0.251824 | 0.056 | 0.046 | 1 | 5 | Kctd11 | 0.01 | Old |
| 0.485221 | 0.222949 | 0.086 | 0.074 | 1 | 5 | Slc33a1 | 0.012 | Old |
| 0.485572 | 0.069498 | 0.185 | 0.17 | 1 | 5 | Calhm2 | 0.015 | Old |
| 0.485671 | 0.065426 | 0.249 | 0.223 | 1 | 5 | Ccdc82 | 0.026 | Old |
| 0.486617 | 0.076236 | 0.487 | 0.472 | 1 | 5 | Rpl13a | 0.015 | Old |
| 0.486759 | 0.021265 | 0.052 | 0.063 | 1 | 5 | Acvr2b | -0.011 | Old |
| 0.487724 | 0.233744 | 0.056 | 0.046 | 1 | 5 | Ercc4 | 0.01 | Old |
| 0.487802 | 0.281715 | 0.069 | 0.058 | 1 | 5 | Nrde2 | 0.011 | Old |
| 0.48789 | 0.017664 | 0.084 | 0.071 | 1 | 5 | Psmd5 | 0.013 | Old |
| 0.487933 | 0.169757 | 0.019 | 0.013 | 1 | 5 | 0610040B1 | 0.006 | Old |
| 0.488156 | 0.237255 | 0.229 | 0.216 | 1 | 5 | Lrrc59 | 0.013 | Old |
| 0.488395 | 0.974142 | 0.027 | 0.02 | 1 | 5 | Znrf3 | 0.007 | Old |
| 0.488418 | 0.14254 | 0.146 | 0.132 | 1 | 5 | Ptpn22 | 0.014 | Old |
| 0.488842 | 0.179936 | 0.056 | 0.046 | 1 | 5 | E130309D0 | 0.01 | Old |
| 0.489095 | 0.040696 | 0.187 | 0.168 | 1 | 5 | Cnrip1 | 0.019 | Old |
| 0.489232 | 0.051637 | 0.593 | 0.541 | 1 | 5 | Ppp1ca | 0.052 | Old |
| 0.489644 | 0.061378 | 0.788 | 0.751 | 1 | 5 | Rab14 | 0.037 | Old |

| | | | | | | | | |
|---|---|---|---|---|---|---|---|---|
| 0.490125 | 0.038174 | 0.163 | 0.147 | 1 | 5 | Crtap | 0.016 | Old |
| 0.490375 | 0.017066 | 0.03 | 0.023 | 1 | 5 | Lancl2 | 0.007 | Old |
| 0.490469 | 0.044902 | 0.042 | 0.033 | 1 | 5 | Prr7 | 0.009 | Old |
| 0.490644 | 0.893445 | 0.015 | 0.01 | 1 | 5 | Mical3 | 0.005 | Old |
| 0.490724 | 0.127978 | 0.165 | 0.147 | 1 | 5 | Wdr20 | 0.018 | Old |
| 0.490851 | 0.040929 | 0.343 | 0.315 | 1 | 5 | Stx16 | 0.028 | Old |
| 0.49137 | 0.917907 | 0.015 | 0.01 | 1 | 5 | C1qtnf6 | 0.005 | Old |
| 0.49137 | 0.785706 | 0.015 | 0.01 | 1 | 5 | Nipsnap1 | 0.005 | Old |
| 0.491955 | 0.036813 | 0.313 | 0.287 | 1 | 5 | Mfsd11 | 0.026 | Old |
| 0.492172 | 0.057231 | 0.086 | 0.074 | 1 | 5 | Surf1 | 0.012 | Old |
| 0.492274 | 0.11541 | 0.059 | 0.048 | 1 | 5 | Fen1 | 0.011 | Old |
| 0.492422 | 0.585708 | 0.044 | 0.036 | 1 | 5 | Gm49339 | 0.008 | Old |
| 0.49247 | 0.08649 | 0.051 | 0.041 | 1 | 5 | Sergef | 0.01 | Old |
| 0.492649 | 0.048919 | 0.517 | 0.485 | 1 | 5 | Lrrc25 | 0.032 | Old |
| 0.49279 | 0.035326 | 0.084 | 0.071 | 1 | 5 | Fbxo9 | 0.013 | Old |
| 0.492862 | 0.221407 | 0.039 | 0.03 | 1 | 5 | Ccdc181 | 0.009 | Old |
| 0.493086 | 0.226411 | 0.109 | 0.096 | 1 | 5 | Fkbp1b | 0.013 | Old |
| 0.493148 | 0.04578 | 0.303 | 0.274 | 1 | 5 | Rp9 | 0.029 | Old |
| 0.49355 | 0.053636 | 0.059 | 0.048 | 1 | 5 | Plekha8 | 0.011 | Old |
| 0.49419 | 0.444637 | 0.071 | 0.061 | 1 | 5 | Cog1 | 0.01 | Old |
| 0.494281 | 0.654647 | 0.015 | 0.01 | 1 | 5 | Creb3l3 | 0.005 | Old |
| 0.494475 | 0.09562 | 0.108 | 0.094 | 1 | 5 | Ubtd1 | 0.014 | Old |
| 0.494626 | 0.104557 | 0.52 | 0.5 | 1 | 5 | Nufip2 | 0.02 | Old |
| 0.495163 | 0.3784 | 0.061 | 0.051 | 1 | 5 | Rft1 | 0.01 | Old |
| 0.495527 | 0.161961 | 0.109 | 0.096 | 1 | 5 | Unc119 | 0.013 | Old |
| 0.495642 | 0.23526 | 0.234 | 0.221 | 1 | 5 | Usp47 | 0.013 | Old |
| 0.495656 | 0.105515 | 0.396 | 0.373 | 1 | 5 | Zfp148 | 0.023 | Old |
| 0.495739 | 0.952235 | 0.015 | 0.01 | 1 | 5 | Ndufaf6 | 0.005 | Old |
| 0.495816 | 0.058663 | 0.604 | 0.561 | 1 | 5 | Ubap2l | 0.043 | Old |
| 0.496033 | 0.050897 | 0.184 | 0.168 | 1 | 5 | Pik3r5 | 0.016 | Old |
| 0.496148 | 0.074717 | 0.704 | 0.655 | 1 | 5 | Wdr1 | 0.049 | Old |
| 0.49698 | 0.063556 | 0.219 | 0.203 | 1 | 5 | Sos2 | 0.016 | Old |
| 0.496998 | 0.018554 | 0.264 | 0.244 | 1 | 5 | Mecp2 | 0.02 | Old |
| 0.497085 | 0.551726 | 0.057 | 0.048 | 1 | 5 | Zfp444 | 0.009 | Old |
| 0.4972 | 0.576843 | 0.015 | 0.01 | 1 | 5 | Napb | 0.005 | Old |
| 0.497306 | 0.133781 | 0.135 | 0.119 | 1 | 5 | Vgll4 | 0.016 | Old |
| 0.497311 | 0.078806 | 0.833 | 0.815 | 1 | 5 | Ccnl1 | 0.018 | Old |
| 0.498213 | 0.082698 | 0.263 | 0.241 | 1 | 5 | Plpbp | 0.022 | Old |
| 0.49848 | 0.226111 | 0.121 | 0.109 | 1 | 5 | Sart3 | 0.012 | Old |
| 0.498531 | 0.196085 | 0.077 | 0.066 | 1 | 5 | Gm20696 | 0.011 | Old |
| 0.498663 | 0.433943 | 0.015 | 0.01 | 1 | 5 | Dffb | 0.005 | Old |
| 0.498663 | 0.161269 | 0.015 | 0.01 | 1 | 5 | Rtn2 | 0.005 | Old |
| 0.498768 | 0.012357 | 0.498 | 0.467 | 1 | 5 | Akr1b3 | 0.031 | Old |
| 0.498819 | 0.472962 | 0.101 | 0.091 | 1 | 5 | Tubgcp3 | 0.01 | Old |
| 0.499108 | 0.235094 | 0.246 | 0.226 | 1 | 5 | Sowahc | 0.02 | Old |

| | | | | | | | | |
|---|---|---|---|---|---|---|---|---|
| 0.499274 | 0.032383 | 0.96 | 0.916 | 1 | 5 | Wwp1 | 0.044 | Old |
| 0.500003 | 0.30021 | 0.093 | 0.081 | 1 | 5 | Cars | 0.012 | Old |
| 0.500129 | 0.420838 | 0.015 | 0.01 | 1 | 5 | Atp2a3 | 0.005 | Old |
| 0.500129 | 0.146372 | 0.015 | 0.01 | 1 | 5 | Wdr74 | 0.005 | Old |
| 0.50026 | 0.58301 | 0.027 | 0.02 | 1 | 5 | AW146154 | 0.007 | Old |
| 0.50084 | 0.249235 | 0.15 | 0.137 | 1 | 5 | Rsrc1 | 0.013 | Old |
| 0.500862 | 0.459516 | 0.015 | 0.01 | 1 | 5 | Pkd1l2 | 0.005 | Old |
| 0.500862 | 0.396635 | 0.015 | 0.01 | 1 | 5 | Ninl | 0.005 | Old |
| 0.500862 | 0.390974 | 0.015 | 0.01 | 1 | 5 | 1700052K1 | 0.005 | Old |
| 0.501292 | 0.434299 | 0.044 | 0.036 | 1 | 5 | Crocc | 0.008 | Old |
| 0.501596 | 0.312089 | 0.015 | 0.01 | 1 | 5 | Gins3 | 0.005 | Old |
| 0.501596 | 0.19595 | 0.015 | 0.01 | 1 | 5 | Mfsd13a | 0.005 | Old |
| 0.501897 | 0.540063 | 0.062 | 0.053 | 1 | 5 | Rhno1 | 0.009 | Old |
| 0.501991 | 0.213756 | 0.215 | 0.201 | 1 | 5 | Pnpla7 | 0.014 | Old |
| 0.502008 | 0.078305 | 0.189 | 0.173 | 1 | 5 | Lsm3 | 0.016 | Old |
| 0.502111 | 0.50166 | 0.035 | 0.028 | 1 | 5 | Ears2 | 0.007 | Old |
| 0.502331 | 0.291084 | 0.015 | 0.01 | 1 | 5 | Them6 | 0.005 | Old |
| 0.502647 | 0.069093 | 0.12 | 0.104 | 1 | 5 | Orc4 | 0.016 | Old |
| 0.502717 | 0.154882 | 0.145 | 0.129 | 1 | 5 | Polg | 0.016 | Old |
| 0.5029 | 0.060169 | 0.059 | 0.048 | 1 | 5 | Hist1h4i | 0.011 | Old |
| 0.503523 | 0.364892 | 0.027 | 0.02 | 1 | 5 | Tnni3 | 0.007 | Old |
| 0.503562 | 0.067314 | 0.205 | 0.185 | 1 | 5 | Pofut1 | 0.02 | Old |
| 0.503628 | 0.093408 | 0.199 | 0.185 | 1 | 5 | Bag3 | 0.014 | Old |
| 0.503802 | 0.137313 | 0.015 | 0.01 | 1 | 5 | Tesk2 | 0.005 | Old |
| 0.503875 | 0.198829 | 0.082 | 0.071 | 1 | 5 | Zfp358 | 0.011 | Old |
| 0.504068 | 0.187364 | 0.027 | 0.02 | 1 | 5 | Nck2 | 0.007 | Old |
| 0.504078 | 0.099498 | 0.086 | 0.074 | 1 | 5 | Atg16l2 | 0.012 | Old |
| 0.504538 | 0.101857 | 0.015 | 0.01 | 1 | 5 | Usp27x | 0.005 | Old |
| 0.504538 | 0.071511 | 0.015 | 0.01 | 1 | 5 | Pusl1 | 0.005 | Old |
| 0.504794 | 0.083881 | 0.047 | 0.038 | 1 | 5 | Ms4a4c | 0.009 | Old |
| 0.504842 | 0.128633 | 0.239 | 0.223 | 1 | 5 | Xpo7 | 0.016 | Old |
| 0.504893 | 0.083497 | 0.126 | 0.112 | 1 | 5 | Erh | 0.014 | Old |
| 0.50528 | 0.174641 | 0.135 | 0.119 | 1 | 5 | Tmem192 | 0.016 | Old |
| 0.50531 | 0.010046 | 0.355 | 0.34 | 1 | 5 | Tcf12 | 0.015 | Old |
| 0.505722 | 0.189473 | 0.118 | 0.104 | 1 | 5 | Med7 | 0.014 | Old |
| 0.505925 | 0.062339 | 0.187 | 0.17 | 1 | 5 | Tceal8 | 0.017 | Old |
| 0.50625 | 0.458687 | 0.027 | 0.02 | 1 | 5 | Gm26624 | 0.007 | Old |
| 0.506372 | 0.223133 | 0.167 | 0.152 | 1 | 5 | Ifi211 | 0.015 | Old |
| 0.506751 | 0.206983 | 0.015 | 0.01 | 1 | 5 | Tnfrsf14 | 0.005 | Old |
| 0.506852 | 0.428632 | 0.035 | 0.028 | 1 | 5 | Riox1 | 0.007 | Old |
| 0.506852 | 0.363381 | 0.035 | 0.028 | 1 | 5 | Amd1 | 0.007 | Old |
| 0.507328 | 0.620653 | 0.035 | 0.028 | 1 | 5 | Fam216a | 0.007 | Old |
| 0.507453 | 0.159694 | 0.064 | 0.053 | 1 | 5 | Slc18b1 | 0.011 | Old |
| 0.507552 | 0.086332 | 0.109 | 0.096 | 1 | 5 | E4f1 | 0.013 | Old |
| 0.508636 | 0.108165 | 0.338 | 0.322 | 1 | 5 | Nab1 | 0.016 | Old |

| | | | | | | | | |
|---|---|---|---|---|---|---|---|---|
| 0.508863 | 0.132889 | 0.152 | 0.137 | 1 | 5 | Ltn1 | 0.015 | Old |
| 0.508954 | 0.21103 | 0.116 | 0.104 | 1 | 5 | AI504432 | 0.012 | Old |
| 0.509713 | 0.063905 | 0.82 | 0.794 | 1 | 5 | Luc7l2 | 0.026 | Old |
| 0.509713 | 0.029142 | 0.335 | 0.315 | 1 | 5 | Map3k11 | 0.02 | Old |
| 0.509834 | 0.041359 | 0.271 | 0.249 | 1 | 5 | Tmem127 | 0.022 | Old |
| 0.510185 | 0.318577 | 0.035 | 0.028 | 1 | 5 | Fam136a | 0.007 | Old |
| 0.51049 | 0.108726 | 0.173 | 0.157 | 1 | 5 | Hdac2 | 0.016 | Old |
| 0.510505 | 0.03703 | 0.308 | 0.287 | 1 | 5 | Ndufa5 | 0.021 | Old |
| 0.510629 | 0.30176 | 0.027 | 0.02 | 1 | 5 | Rtn4ip1 | 0.007 | Old |
| 0.510711 | 0.074874 | 0.101 | 0.089 | 1 | 5 | Wdr77 | 0.012 | Old |
| 0.511124 | 0.075384 | 0.33 | 0.307 | 1 | 5 | Atraid | 0.023 | Old |
| 0.51142 | 0.034829 | 0.335 | 0.31 | 1 | 5 | Ppp4c | 0.025 | Old |
| 0.51196 | 0.295851 | 0.044 | 0.036 | 1 | 5 | Tfpt | 0.008 | Old |
| 0.511986 | 0.218132 | 0.126 | 0.114 | 1 | 5 | Tmem159 | 0.012 | Old |
| 0.512082 | 0.098746 | 0.178 | 0.162 | 1 | 5 | Ilkap | 0.016 | Old |
| 0.512572 | 0.305062 | 0.035 | 0.028 | 1 | 5 | Zfp53 | 0.007 | Old |
| 0.512572 | 0.300269 | 0.035 | 0.028 | 1 | 5 | Cenpt | 0.007 | Old |
| 0.512605 | 0.15344 | 0.146 | 0.132 | 1 | 5 | Ube2a | 0.014 | Old |
| 0.512812 | 0.022353 | 0.199 | 0.183 | 1 | 5 | Slc36a1 | 0.016 | Old |
| 0.513051 | 0.389173 | 0.035 | 0.028 | 1 | 5 | Jmjd8 | 0.007 | Old |
| 0.51309 | 1.100407 | 0.012 | 0.008 | 1 | 5 | Gimap1 | 0.004 | Old |
| 0.51309 | 0.784595 | 0.012 | 0.008 | 1 | 5 | Hyal3 | 0.004 | Old |
| 0.51309 | 0.647244 | 0.012 | 0.008 | 1 | 5 | Tmem120b | 0.004 | Old |
| 0.513937 | 0.831095 | 0.012 | 0.008 | 1 | 5 | D330050G2 | 0.004 | Old |
| 0.514642 | 0.032093 | 0.815 | 0.794 | 1 | 5 | Eef2 | 0.021 | Old |
| 0.514784 | 1.034689 | 0.012 | 0.008 | 1 | 5 | Gm16845 | 0.004 | Old |
| 0.515632 | 0.729428 | 0.012 | 0.008 | 1 | 5 | Mybpc3 | 0.004 | Old |
| 0.516097 | 0.174251 | 0.18 | 0.165 | 1 | 5 | Bet1 | 0.015 | Old |
| 0.516476 | 0.188946 | 0.044 | 0.036 | 1 | 5 | Tti2 | 0.008 | Old |
| 0.516481 | 0.958243 | 0.012 | 0.008 | 1 | 5 | Ugt1a7c | 0.004 | Old |
| 0.516481 | 0.756666 | 0.012 | 0.008 | 1 | 5 | Rasip1 | 0.004 | Old |
| 0.516653 | 0.008075 | 0.269 | 0.249 | 1 | 5 | Gdap10 | 0.02 | Old |
| 0.516684 | 0.105074 | 0.027 | 0.02 | 1 | 5 | Mfsd4a | 0.007 | Old |
| 0.516684 | 0.044875 | 0.027 | 0.02 | 1 | 5 | Slc5a6 | 0.007 | Old |
| 0.516852 | 0.194092 | 0.056 | 0.046 | 1 | 5 | Hpse | 0.01 | Old |
| 0.516972 | 0.036272 | 0.906 | 0.86 | 1 | 5 | Rps3 | 0.046 | Old |
| 0.517236 | 0.028322 | 0.027 | 0.02 | 1 | 5 | Acad10 | 0.007 | Old |
| 0.51733 | 0.749897 | 0.012 | 0.008 | 1 | 5 | Nek8 | 0.004 | Old |
| 0.517334 | 0.141046 | 0.061 | 0.051 | 1 | 5 | Unc13d | 0.01 | Old |
| 0.517365 | 0.041385 | 0.035 | 0.028 | 1 | 5 | Zbtb24 | 0.007 | Old |
| 0.51787 | 0.041772 | 0.776 | 0.739 | 1 | 5 | Bri3 | 0.037 | Old |
| 0.517941 | 0.127627 | 0.066 | 0.056 | 1 | 5 | Ubac1 | 0.01 | Old |
| 0.51818 | 0.674106 | 0.012 | 0.008 | 1 | 5 | 1700113A1 | 0.004 | Old |
| 0.51818 | 0.082951 | 0.008 | 0.013 | 1 | 5 | Aars2 | -0.005 | Old |
| 0.518602 | 0.026992 | 0.561 | 0.528 | 1 | 5 | Fermt3 | 0.033 | Old |

| | | | | | | | | |
|---|---|---|---|---|---|---|---|---|
| 0.51864 | 0.199383 | 0.089 | 0.104 | 1 | 5 | Zfp322a | -0.015 | Old |
| 0.518807 | 0.174834 | 0.035 | 0.028 | 1 | 5 | A930033H1 | 0.007 | Old |
| 0.518848 | 0.088684 | 0.157 | 0.142 | 1 | 5 | Htatsf1 | 0.015 | Old |
| 0.519031 | 0.527271 | 0.012 | 0.008 | 1 | 5 | Pter | 0.004 | Old |
| 0.519031 | 0.515456 | 0.012 | 0.008 | 1 | 5 | Cgnl1 | 0.004 | Old |
| 0.519254 | 0.008403 | 0.298 | 0.274 | 1 | 5 | Rnf6 | 0.024 | Old |
| 0.519383 | 0.058843 | 0.209 | 0.195 | 1 | 5 | Cers5 | 0.014 | Old |
| 0.519448 | 0.074896 | 0.027 | 0.02 | 1 | 5 | Cyb5rl | 0.007 | Old |
| 0.519882 | 0.701858 | 0.012 | 0.008 | 1 | 5 | Flt1 | 0.004 | Old |
| 0.519882 | 0.596566 | 0.012 | 0.008 | 1 | 5 | Fmo2 | 0.004 | Old |
| 0.519882 | 0.524422 | 0.012 | 0.008 | 1 | 5 | Gbp6 | 0.004 | Old |
| 0.520199 | 0.149044 | 0.091 | 0.079 | 1 | 5 | AC165953. | 0.012 | Old |
| 0.520579 | 0.349692 | 0.044 | 0.036 | 1 | 5 | Zhx3 | 0.008 | Old |
| 0.520735 | 0.560906 | 0.012 | 0.008 | 1 | 5 | Cox6a2 | 0.004 | Old |
| 0.520735 | 0.447375 | 0.012 | 0.008 | 1 | 5 | Gm33973 | 0.004 | Old |
| 0.520735 | 0.393215 | 0.012 | 0.008 | 1 | 5 | Nid1 | 0.004 | Old |
| 0.520832 | 0.130706 | 0.061 | 0.051 | 1 | 5 | Me2 | 0.01 | Old |
| 0.520852 | 0.066286 | 0.158 | 0.142 | 1 | 5 | Oxsr1 | 0.016 | Old |
| 0.520886 | 0.088431 | 0.173 | 0.155 | 1 | 5 | Hpf1 | 0.018 | Old |
| 0.521327 | 0.483985 | 0.049 | 0.041 | 1 | 5 | C2cd2 | 0.008 | Old |
| 0.521458 | 0.035987 | 0.507 | 0.477 | 1 | 5 | Mkln1 | 0.03 | Old |
| 0.521584 | 0.149086 | 0.024 | 0.03 | 1 | 5 | Rnf122 | -0.006 | Old |
| 0.521588 | 0.595134 | 0.012 | 0.008 | 1 | 5 | Fbln1 | 0.004 | Old |
| 0.521588 | 0.237438 | 0.012 | 0.008 | 1 | 5 | Gm15964 | 0.004 | Old |
| 0.522217 | 0.682296 | 0.054 | 0.046 | 1 | 5 | Dclre1a | 0.008 | Old |
| 0.522421 | 0.308268 | 0.035 | 0.028 | 1 | 5 | Blzf1 | 0.007 | Old |
| 0.522441 | 0.349448 | 0.012 | 0.008 | 1 | 5 | Tm7sf2 | 0.004 | Old |
| 0.522544 | 0.187676 | 0.145 | 0.132 | 1 | 5 | Washc5 | 0.013 | Old |
| 0.522574 | 0.076993 | 0.104 | 0.091 | 1 | 5 | Acot9 | 0.013 | Old |
| 0.522784 | 0.286292 | 0.143 | 0.129 | 1 | 5 | Lrrfip2 | 0.014 | Old |
| 0.523294 | 0.312867 | 0.167 | 0.155 | 1 | 5 | Braf | 0.012 | Old |
| 0.523296 | 0.394466 | 0.012 | 0.008 | 1 | 5 | Jmjd7 | 0.004 | Old |
| 0.523296 | 0.39168 | 0.012 | 0.008 | 1 | 5 | Upb1 | 0.004 | Old |
| 0.523296 | 0.373975 | 0.012 | 0.008 | 1 | 5 | B230369F2 | 0.004 | Old |
| 0.523296 | 0.305484 | 0.012 | 0.008 | 1 | 5 | Lpar2 | 0.004 | Old |
| 0.523296 | 0.147675 | 0.012 | 0.008 | 1 | 5 | Ulk4 | 0.004 | Old |
| 0.523905 | 0.097612 | 0.168 | 0.155 | 1 | 5 | Nfx1 | 0.013 | Old |
| 0.523955 | 0.140252 | 0.241 | 0.223 | 1 | 5 | Foxred2 | 0.018 | Old |
| 0.524326 | 0.037856 | 0.074 | 0.063 | 1 | 5 | Prmt7 | 0.011 | Old |
| 0.524423 | 0.245943 | 0.088 | 0.076 | 1 | 5 | Sap30bp | 0.012 | Old |
| 0.524538 | 0.192475 | 0.337 | 0.33 | 1 | 5 | Ubald2 | 0.007 | Old |
| 0.52456 | 0.03453 | 0.731 | 0.69 | 1 | 5 | Clic1 | 0.041 | Old |
| 0.524817 | 0.355676 | 0.076 | 0.066 | 1 | 5 | Slc25a44 | 0.01 | Old |
| 0.525007 | 0.209244 | 0.012 | 0.008 | 1 | 5 | Gm12184 | 0.004 | Old |
| 0.525202 | 0.447748 | 0.077 | 0.069 | 1 | 5 | Mynn | 0.008 | Old |

| | | | | | | | | |
|---|---|---|---|---|---|---|---|---|
| 0.525583 | 0.49036 | 0.057 | 0.048 | 1 | 5 | Extl2 | 0.009 | Old |
| 0.525634 | 0.438916 | 0.067 | 0.058 | 1 | 5 | Bet1l | 0.009 | Old |
| 0.525863 | 0.203228 | 0.012 | 0.008 | 1 | 5 | Vnn3 | 0.004 | Old |
| 0.525863 | 0.180061 | 0.012 | 0.008 | 1 | 5 | Rexo5 | 0.004 | Old |
| 0.525863 | 0.179551 | 0.012 | 0.008 | 1 | 5 | Gm6133 | 0.004 | Old |
| 0.525946 | 0.271851 | 0.071 | 0.061 | 1 | 5 | Bcl2a1a | 0.01 | Old |
| 0.526583 | 0.023299 | 0.628 | 0.569 | 1 | 5 | Atp6v1b2 | 0.059 | Old |
| 0.526772 | 0.511747 | 0.024 | 0.018 | 1 | 5 | Gstm4 | 0.006 | Old |
| 0.527135 | 0.478754 | 0.062 | 0.053 | 1 | 5 | Gmcl1 | 0.009 | Old |
| 0.527206 | 0.269162 | 0.222 | 0.211 | 1 | 5 | Rnf11 | 0.011 | Old |
| 0.527247 | 0.106918 | 0.293 | 0.277 | 1 | 5 | Dennd4b | 0.016 | Old |
| 0.527324 | 0.201186 | 0.106 | 0.094 | 1 | 5 | Mill2 | 0.012 | Old |
| 0.527436 | 0.023473 | 0.936 | 0.942 | 1 | 5 | Ctsh | -0.006 | Old |
| 0.527947 | 0.465365 | 0.032 | 0.025 | 1 | 5 | Spata5 | 0.007 | Old |
| 0.528437 | 0.098093 | 0.012 | 0.008 | 1 | 5 | 2700046G( | 0.004 | Old |
| 0.528514 | 0.380738 | 0.04 | 0.033 | 1 | 5 | 2700049A( | 0.007 | Old |
| 0.528955 | 0.16423 | 0.066 | 0.056 | 1 | 5 | Gm5086 | 0.01 | Old |
| 0.528998 | 0.130508 | 0.052 | 0.043 | 1 | 5 | Ptpn4 | 0.009 | Old |
| 0.529156 | 0.498233 | 0.024 | 0.018 | 1 | 5 | Zfp280b | 0.006 | Old |
| 0.529709 | 0.116815 | 0.044 | 0.036 | 1 | 5 | Bend6 | 0.008 | Old |
| 0.529717 | 0.095721 | 0.135 | 0.122 | 1 | 5 | Vamp7 | 0.013 | Old |
| 0.529752 | 0.510584 | 0.024 | 0.018 | 1 | 5 | B3galt4 | 0.006 | Old |
| 0.529932 | 0.209647 | 0.035 | 0.028 | 1 | 5 | Rida | 0.007 | Old |
| 0.529983 | 0.256007 | 0.108 | 0.096 | 1 | 5 | Gtdc1 | 0.012 | Old |
| 0.530068 | 0.010619 | 0.305 | 0.282 | 1 | 5 | Cxcl1 | 0.023 | Old |
| 0.53023 | 0.008835 | 0.603 | 0.561 | 1 | 5 | Lacc1 | 0.042 | Old |
| 0.530361 | 0.147598 | 0.126 | 0.114 | 1 | 5 | Fbxo7 | 0.012 | Old |
| 0.530905 | 0.099177 | 0.035 | 0.028 | 1 | 5 | 2410131K1 | 0.007 | Old |
| 0.531456 | 0.203719 | 0.03 | 0.038 | 1 | 5 | Pfas | -0.008 | Old |
| 0.531483 | 0.134603 | 0.182 | 0.168 | 1 | 5 | Casp1 | 0.014 | Old |
| 0.531544 | 0.543842 | 0.024 | 0.018 | 1 | 5 | Gm29367 | 0.006 | Old |
| 0.531655 | 0.051826 | 0.21 | 0.236 | 1 | 5 | Adprh | -0.026 | Old |
| 0.532007 | 0.111562 | 0.29 | 0.274 | 1 | 5 | Mgat2 | 0.016 | Old |
| 0.53212 | 0.326021 | 0.135 | 0.124 | 1 | 5 | Med25 | 0.011 | Old |
| 0.532147 | 0.022797 | 0.64 | 0.612 | 1 | 5 | Neurl3 | 0.028 | Old |
| 0.532179 | 0.017573 | 0.517 | 0.492 | 1 | 5 | Psenen | 0.025 | Old |
| 0.532401 | 0.486715 | 0.054 | 0.046 | 1 | 5 | Ggcx | 0.008 | Old |
| 0.532627 | 0.231973 | 0.118 | 0.107 | 1 | 5 | Mrpl10 | 0.011 | Old |
| 0.532685 | 0.06123 | 0.288 | 0.272 | 1 | 5 | Exoc6b | 0.016 | Old |
| 0.533458 | 0.055488 | 0.433 | 0.419 | 1 | 5 | Cwc15 | 0.014 | Old |
| 0.533597 | 0.181352 | 0.106 | 0.094 | 1 | 5 | Hsdl2 | 0.012 | Old |
| 0.533606 | 0.009286 | 0.131 | 0.117 | 1 | 5 | Ubl7 | 0.014 | Old |
| 0.533609 | 0.035027 | 0.066 | 0.056 | 1 | 5 | Tmem175 | 0.01 | Old |
| 0.53383 | 0.021661 | 0.035 | 0.028 | 1 | 5 | Armcx2 | 0.007 | Old |
| 0.534044 | 0.079607 | 0.071 | 0.061 | 1 | 5 | Smpd4 | 0.01 | Old |

| | | | | | | | | |
|---|---|---|---|---|---|---|---|---|
| 0.534475 | 0.253443 | 0.076 | 0.066 | 1 | 5 | Edem3 | 0.01 | Old |
| 0.534508 | 0.132835 | 0.079 | 0.069 | 1 | 5 | Fbxo4 | 0.01 | Old |
| 0.534538 | 0.11341 | 0.024 | 0.018 | 1 | 5 | Btnl2 | 0.006 | Old |
| 0.535099 | 0.558727 | 0.032 | 0.025 | 1 | 5 | Zfp141 | 0.007 | Old |
| 0.535136 | 0.081594 | 0.057 | 0.048 | 1 | 5 | Mettl5 | 0.009 | Old |
| 0.535342 | 0.336429 | 0.049 | 0.041 | 1 | 5 | Traf3ip1 | 0.008 | Old |
| 0.535494 | 0.119228 | 0.121 | 0.109 | 1 | 5 | Mfn2 | 0.012 | Old |
| 0.536017 | 0.028552 | 0.598 | 0.556 | 1 | 5 | Serpinb8 | 0.042 | Old |
| 0.536138 | 0.291423 | 0.162 | 0.152 | 1 | 5 | Tpm1 | 0.01 | Old |
| 0.536223 | 0.14131 | 0.052 | 0.043 | 1 | 5 | Gatc | 0.009 | Old |
| 0.536238 | 0.032773 | 0.217 | 0.201 | 1 | 5 | Rab6b | 0.016 | Old |
| 0.536793 | 0.14768 | 0.185 | 0.17 | 1 | 5 | Aph1b | 0.015 | Old |
| 0.536898 | 0.157552 | 0.157 | 0.142 | 1 | 5 | Chm | 0.015 | Old |
| 0.537153 | 0.268164 | 0.04 | 0.033 | 1 | 5 | Gm46430 | 0.007 | Old |
| 0.537173 | 0.156065 | 0.328 | 0.315 | 1 | 5 | Il1rl1 | 0.013 | Old |
| 0.53753 | 0.217359 | 0.178 | 0.165 | 1 | 5 | Cabin1 | 0.013 | Old |
| 0.53754 | 0.32541 | 0.024 | 0.018 | 1 | 5 | Phf24 | 0.006 | Old |
| 0.537665 | 0.369887 | 0.032 | 0.025 | 1 | 5 | Surf6 | 0.007 | Old |
| 0.538693 | 0.412793 | 0.032 | 0.025 | 1 | 5 | Gm6563 | 0.007 | Old |
| 0.538767 | 0.00195 | 0.8 | 0.782 | 1 | 5 | Snrnp70 | 0.018 | Old |
| 0.539808 | 0.302729 | 0.062 | 0.053 | 1 | 5 | Erich1 | 0.009 | Old |
| 0.539948 | 0.339056 | 0.024 | 0.018 | 1 | 5 | 5830416I1 | 0.006 | Old |
| 0.540009 | 0.030081 | 0.463 | 0.437 | 1 | 5 | Metap2 | 0.026 | Old |
| 0.540764 | 0.244746 | 0.153 | 0.142 | 1 | 5 | Stam2 | 0.011 | Old |
| 0.541268 | 0.33727 | 0.032 | 0.025 | 1 | 5 | Glmn | 0.007 | Old |
| 0.541796 | 0.079075 | 0.375 | 0.358 | 1 | 5 | Mapk6 | 0.017 | Old |
| 0.542002 | 0.255001 | 0.094 | 0.084 | 1 | 5 | Med4 | 0.01 | Old |
| 0.542214 | 0.113126 | 0.237 | 0.223 | 1 | 5 | Mrpl20 | 0.014 | Old |
| 0.542545 | 0.078954 | 0.229 | 0.216 | 1 | 5 | Ndufa12 | 0.013 | Old |
| 0.543172 | 0.028784 | 0.269 | 0.251 | 1 | 5 | Evl | 0.018 | Old |
| 0.543632 | 0.042089 | 0.146 | 0.132 | 1 | 5 | Chchd3 | 0.014 | Old |
| 0.543709 | 0.131522 | 0.089 | 0.079 | 1 | 5 | Sass6 | 0.01 | Old |
| 0.543831 | 0.02859 | 0.32 | 0.302 | 1 | 5 | Ostc | 0.018 | Old |
| 0.54451 | 0.276847 | 0.049 | 0.041 | 1 | 5 | Emc9 | 0.008 | Old |
| 0.54475 | 0.025504 | 0.451 | 0.434 | 1 | 5 | Rbm5 | 0.017 | Old |
| 0.54478 | 0.138002 | 0.024 | 0.018 | 1 | 5 | Timmdc1 | 0.006 | Old |
| 0.544891 | 0.075465 | 0.094 | 0.084 | 1 | 5 | Eci2 | 0.01 | Old |
| 0.544929 | 0.221621 | 0.049 | 0.041 | 1 | 5 | Dnajc4 | 0.008 | Old |
| 0.545163 | 0.006584 | 0.246 | 0.223 | 1 | 5 | Blvra | 0.023 | Old |
| 0.545642 | 0.075432 | 0.226 | 0.213 | 1 | 5 | Filip1l | 0.013 | Old |
| 0.545843 | 0.037223 | 0.221 | 0.208 | 1 | 5 | Chmp1a | 0.013 | Old |
| 0.546401 | 0.021726 | 0.375 | 0.36 | 1 | 5 | Prkcsh | 0.015 | Old |
| 0.546523 | 0.155398 | 0.089 | 0.079 | 1 | 5 | Adnp2 | 0.01 | Old |
| 0.546721 | 0.01649 | 0.18 | 0.165 | 1 | 5 | Rbm12 | 0.015 | Old |
| 0.546878 | 0.188741 | 0.152 | 0.14 | 1 | 5 | Luc7l | 0.012 | Old |

| | | | | | | | | |
|---|---|---|---|---|---|---|---|---|
| 0.54722 | 0.181168 | 0.079 | 0.069 | 1 | 5 | Fmc1 | 0.01 | Old |
| 0.547244 | 0.336872 | 0.04 | 0.033 | 1 | 5 | Mfsd7a | 0.007 | Old |
| 0.547685 | 0.508747 | 0.045 | 0.038 | 1 | 5 | Ogfod2 | 0.007 | Old |
| 0.548148 | 0.147429 | 0.071 | 0.061 | 1 | 5 | Alad | 0.01 | Old |
| 0.549044 | 0.097397 | 0.29 | 0.272 | 1 | 5 | Mga | 0.018 | Old |
| 0.549099 | 0.075447 | 0.163 | 0.15 | 1 | 5 | Commd4 | 0.013 | Old |
| 0.549167 | 0.030797 | 0.168 | 0.15 | 1 | 5 | Kbtbd11 | 0.018 | Old |
| 0.549526 | 0.383054 | 0.064 | 0.056 | 1 | 5 | Gm20707 | 0.008 | Old |
| 0.549965 | 0.105187 | 0.049 | 0.041 | 1 | 5 | Gm26881 | 0.008 | Old |
| 0.550153 | 0.167359 | 0.086 | 0.076 | 1 | 5 | Oas1a | 0.01 | Old |
| 0.550441 | 1.076042 | 0.02 | 0.015 | 1 | 5 | Chst1 | 0.005 | Old |
| 0.550602 | 0.169972 | 0.089 | 0.079 | 1 | 5 | Ticam1 | 0.01 | Old |
| 0.550678 | 0.076504 | 0.029 | 0.036 | 1 | 5 | C030034I2 | -0.007 | Old |
| 0.550883 | 0.059619 | 0.305 | 0.284 | 1 | 5 | Soat1 | 0.021 | Old |
| 0.551069 | 0.241942 | 0.146 | 0.137 | 1 | 5 | Ppp1r35 | 0.009 | Old |
| 0.551448 | 0.017767 | 0.894 | 0.883 | 1 | 5 | Rpl12 | 0.011 | Old |
| 0.551546 | 0.495686 | 0.064 | 0.056 | 1 | 5 | Parn | 0.008 | Old |
| 0.551651 | 0.165081 | 0.283 | 0.272 | 1 | 5 | Slc15a4 | 0.011 | Old |
| 0.551962 | 0.22973 | 0.101 | 0.091 | 1 | 5 | Mex3d | 0.01 | Old |
| 0.552669 | 0.052517 | 0.032 | 0.025 | 1 | 5 | C1rl | 0.007 | Old |
| 0.552707 | 0.081349 | 0.263 | 0.292 | 1 | 5 | Adgre5 | -0.029 | Old |
| 0.552865 | 0.161044 | 0.054 | 0.046 | 1 | 5 | Fas | 0.008 | Old |
| 0.553054 | 0.073476 | 0.157 | 0.142 | 1 | 5 | Rlf | 0.015 | Old |
| 0.553068 | 0.759736 | 0.02 | 0.015 | 1 | 5 | Cavin3 | 0.005 | Old |
| 0.553192 | 0.174685 | 0.086 | 0.076 | 1 | 5 | Slc9a6 | 0.01 | Old |
| 0.55322 | 0.221419 | 0.131 | 0.119 | 1 | 5 | Prkch | 0.012 | Old |
| 0.553583 | 0.021159 | 0.343 | 0.322 | 1 | 5 | Mydgf | 0.021 | Old |
| 0.553726 | 0.83526 | 0.02 | 0.015 | 1 | 5 | Gdf9 | 0.005 | Old |
| 0.554211 | 0.429617 | 0.045 | 0.038 | 1 | 5 | Phpt1 | 0.007 | Old |
| 0.554453 | 0.048487 | 0.239 | 0.223 | 1 | 5 | Ap1g1 | 0.016 | Old |
| 0.554457 | 0.349493 | 0.059 | 0.051 | 1 | 5 | Pam16 | 0.008 | Old |
| 0.554648 | 0.242139 | 0.045 | 0.038 | 1 | 5 | Ublcp1 | 0.007 | Old |
| 0.554907 | 0.030822 | 0.067 | 0.079 | 1 | 5 | Cops7b | -0.012 | Old |
| 0.555203 | 0.05319 | 0.343 | 0.32 | 1 | 5 | Cdc37 | 0.023 | Old |
| 0.555277 | 0.049992 | 0.032 | 0.025 | 1 | 5 | Atp5s | 0.007 | Old |
| 0.555445 | 0.372831 | 0.101 | 0.089 | 1 | 5 | Mmp13 | 0.012 | Old |
| 0.555487 | 0.125332 | 0.266 | 0.254 | 1 | 5 | Stk40 | 0.012 | Old |
| 0.555521 | 0.406729 | 0.045 | 0.038 | 1 | 5 | BC017158 | 0.007 | Old |
| 0.555702 | 0.640963 | 0.02 | 0.015 | 1 | 5 | Cbwd1 | 0.005 | Old |
| 0.556131 | 0.161963 | 0.194 | 0.18 | 1 | 5 | Pip5k1a | 0.014 | Old |
| 0.556656 | 0.086029 | 0.015 | 0.02 | 1 | 5 | Mrm1 | -0.005 | Old |
| 0.556664 | 0.032951 | 0.377 | 0.353 | 1 | 5 | Rmnd5a | 0.024 | Old |
| 0.557023 | 0.025159 | 0.374 | 0.35 | 1 | 5 | Trmt112 | 0.024 | Old |
| 0.557138 | 0.228358 | 0.042 | 0.051 | 1 | 5 | Zmat3 | -0.009 | Old |
| 0.557367 | 0.162312 | 0.032 | 0.025 | 1 | 5 | Zfp35 | 0.007 | Old |

| | | | | | | | | |
|---|---|---|---|---|---|---|---|---|
| 0.557896 | 0.022585 | 0.877 | 0.853 | 1 | 5 | Ddx3x | 0.024 | Old |
| 0.558669 | 0.158152 | 0.175 | 0.165 | 1 | 5 | Ppp2r5e | 0.01 | Old |
| 0.558832 | 0.075642 | 0.357 | 0.34 | 1 | 5 | 2810474O1 | 0.017 | Old |
| 0.559003 | 0.540355 | 0.02 | 0.015 | 1 | 5 | BC051226 | 0.005 | Old |
| 0.559459 | 0.402686 | 0.045 | 0.038 | 1 | 5 | Prnp | 0.007 | Old |
| 0.5596 | 0.004399 | 0.564 | 0.548 | 1 | 5 | Tmem256 | 0.016 | Old |
| 0.559605 | 0.068203 | 0.416 | 0.404 | 1 | 5 | Casp8 | 0.012 | Old |
| 0.559804 | 0.147779 | 0.254 | 0.239 | 1 | 5 | Coro7 | 0.015 | Old |
| 0.560322 | 0.424091 | 0.185 | 0.178 | 1 | 5 | Dcn | 0.007 | Old |
| 0.560418 | 0.333359 | 0.037 | 0.03 | 1 | 5 | Nudt1 | 0.007 | Old |
| 0.560989 | 0.190619 | 0.02 | 0.015 | 1 | 5 | Gm14296 | 0.005 | Old |
| 0.561118 | 0.045968 | 0.084 | 0.074 | 1 | 5 | Gprasp1 | 0.01 | Old |
| 0.562377 | 0.03603 | 0.148 | 0.135 | 1 | 5 | Dcxr | 0.013 | Old |
| 0.562555 | 0.099294 | 0.04 | 0.033 | 1 | 5 | Exosc1 | 0.007 | Old |
| 0.562593 | 0.175658 | 0.114 | 0.104 | 1 | 5 | Pde4a | 0.01 | Old |
| 0.563298 | 0.037705 | 0.15 | 0.135 | 1 | 5 | Fam193b | 0.015 | Old |
| 0.563853 | 0.014487 | 0.152 | 0.137 | 1 | 5 | Smox | 0.015 | Old |
| 0.563916 | 0.08867 | 0.402 | 0.378 | 1 | 5 | Vma21 | 0.024 | Old |
| 0.563991 | 0.078649 | 0.727 | 0.711 | 1 | 5 | Lifr | 0.016 | Old |
| 0.564305 | 0.240289 | 0.02 | 0.015 | 1 | 5 | Zfp566 | 0.005 | Old |
| 0.564969 | 0.238053 | 0.02 | 0.015 | 1 | 5 | Shf | 0.005 | Old |
| 0.565163 | 0.185517 | 0.054 | 0.046 | 1 | 5 | Osbpl3 | 0.008 | Old |
| 0.565402 | 0.083762 | 0.322 | 0.302 | 1 | 5 | Camkk2 | 0.02 | Old |
| 0.565593 | 0.163199 | 0.192 | 0.178 | 1 | 5 | Pknox1 | 0.014 | Old |
| 0.565634 | 0.332294 | 0.02 | 0.015 | 1 | 5 | 1110002L0 | 0.005 | Old |
| 0.565735 | 0.073431 | 0.278 | 0.264 | 1 | 5 | Psmb5 | 0.014 | Old |
| 0.566299 | 0.40909 | 0.02 | 0.015 | 1 | 5 | Zfp963 | 0.005 | Old |
| 0.566971 | 0.03165 | 0.232 | 0.216 | 1 | 5 | Hmgcl | 0.016 | Old |
| 0.56756 | 0.01745 | 0.158 | 0.142 | 1 | 5 | Utrn | 0.016 | Old |
| 0.567627 | 0.22181 | 0.145 | 0.135 | 1 | 5 | Zfp110 | 0.01 | Old |
| 0.56763 | 0.547688 | 0.02 | 0.015 | 1 | 5 | Dzip1 | 0.005 | Old |
| 0.567648 | 0.501902 | 0.029 | 0.023 | 1 | 5 | Armc2 | 0.006 | Old |
| 0.568296 | 0.283108 | 0.02 | 0.015 | 1 | 5 | Flrt2 | 0.005 | Old |
| 0.568478 | 0.201105 | 0.167 | 0.155 | 1 | 5 | Galnt10 | 0.012 | Old |
| 0.568655 | 0.004002 | 0.057 | 0.048 | 1 | 5 | Fam241a | 0.009 | Old |
| 0.568944 | 0.058211 | 0.333 | 0.312 | 1 | 5 | Cept1 | 0.021 | Old |
| 0.569123 | 0.023647 | 0.141 | 0.157 | 1 | 5 | Slc30a1 | -0.016 | Old |
| 0.56914 | 0.29286 | 0.059 | 0.051 | 1 | 5 | Lppos | 0.008 | Old |
| 0.569318 | 0.204468 | 0.029 | 0.023 | 1 | 5 | Trp53i11 | 0.006 | Old |
| 0.569516 | 0.330791 | 0.082 | 0.074 | 1 | 5 | Gm37233 | 0.008 | Old |
| 0.569892 | 0.475632 | 0.084 | 0.076 | 1 | 5 | Hinfp | 0.008 | Old |
| 0.569934 | 0.059843 | 0.322 | 0.305 | 1 | 5 | Ccnl2 | 0.017 | Old |
| 0.570025 | 0.029156 | 0.52 | 0.487 | 1 | 5 | Zfand6 | 0.033 | Old |
| 0.570145 | 0.109133 | 0.094 | 0.084 | 1 | 5 | Nop14 | 0.01 | Old |
| 0.570171 | 0.195911 | 0.037 | 0.03 | 1 | 5 | Cenpo | 0.007 | Old |

| | | | | | | | | |
|---|---|---|---|---|---|---|---|---|
| 0.570175 | 0.072829 | 0.473 | 0.462 | 1 | 5 | Dynll1 | 0.011 | Old |
| 0.570448 | 0.230032 | 0.064 | 0.056 | 1 | 5 | Fabp5 | 0.008 | Old |
| 0.570654 | 0.212657 | 0.072 | 0.063 | 1 | 5 | Zbed4 | 0.009 | Old |
| 0.570831 | 0.016522 | 0.143 | 0.129 | 1 | 5 | Ivd | 0.014 | Old |
| 0.571633 | 0.157244 | 0.02 | 0.015 | 1 | 5 | Fgf11 | 0.005 | Old |
| 0.571823 | 0.00427 | 0.357 | 0.332 | 1 | 5 | Lman1 | 0.025 | Old |
| 0.571834 | 0.547487 | 0.069 | 0.061 | 1 | 5 | Fbxl18 | 0.008 | Old |
| 0.572018 | 4.15E-05 | 0.153 | 0.137 | 1 | 5 | Mosmo | 0.016 | Old |
| 0.572318 | 0.170636 | 0.118 | 0.107 | 1 | 5 | Prkaa1 | 0.011 | Old |
| 0.572608 | 0.046142 | 0.527 | 0.497 | 1 | 5 | Atp6v0a1 | 0.03 | Old |
| 0.572666 | 0.385201 | 0.029 | 0.023 | 1 | 5 | Gatsl3 | 0.006 | Old |
| 0.572684 | 0.230567 | 0.045 | 0.038 | 1 | 5 | Usp30 | 0.007 | Old |
| 0.572774 | 0.098818 | 0.136 | 0.124 | 1 | 5 | Siae | 0.012 | Old |
| 0.572951 | 0.181174 | 0.094 | 0.084 | 1 | 5 | Tmem209 | 0.01 | Old |
| 0.572971 | 0.265251 | 0.02 | 0.015 | 1 | 5 | Nr3c2 | 0.005 | Old |
| 0.572971 | 0.098133 | 0.02 | 0.015 | 1 | 5 | Zfp652os | 0.005 | Old |
| 0.57306 | 0.000392 | 0.453 | 0.434 | 1 | 5 | Nadk | 0.019 | Old |
| 0.573164 | 0.10716 | 0.2 | 0.185 | 1 | 5 | Kcnk13 | 0.015 | Old |
| 0.573167 | 0.114084 | 0.103 | 0.091 | 1 | 5 | Anapc10 | 0.012 | Old |
| 0.573209 | 0.390711 | 0.051 | 0.043 | 1 | 5 | Ppp1r12b | 0.008 | Old |
| 0.573461 | 0.049413 | 0.076 | 0.066 | 1 | 5 | Snupn | 0.01 | Old |
| 0.573603 | 0.405096 | 0.037 | 0.03 | 1 | 5 | Padi2 | 0.007 | Old |
| 0.573668 | 0.485039 | 0.022 | 0.028 | 1 | 5 | Ankzf1 | -0.006 | Old |
| 0.573785 | 0.371863 | 0.029 | 0.023 | 1 | 5 | Trim7 | 0.006 | Old |
| 0.573853 | 0.073682 | 0.172 | 0.16 | 1 | 5 | Spg20 | 0.012 | Old |
| 0.574635 | 0.136356 | 0.126 | 0.114 | 1 | 5 | 1700123O2 | 0.012 | Old |
| 0.574642 | 0.180601 | 0.152 | 0.142 | 1 | 5 | Eftud2 | 0.01 | Old |
| 0.574962 | 0.089068 | 0.067 | 0.058 | 1 | 5 | Clpp | 0.009 | Old |
| 0.57594 | 0.0161 | 0.259 | 0.244 | 1 | 5 | Cdc26 | 0.015 | Old |
| 0.576584 | 0.267666 | 0.029 | 0.023 | 1 | 5 | Gm340 | 0.006 | Old |
| 0.576801 | 0.289262 | 0.096 | 0.086 | 1 | 5 | Nr1d2 | 0.01 | Old |
| 0.576867 | 0.008661 | 0.424 | 0.401 | 1 | 5 | Rapgef6 | 0.023 | Old |
| 0.576991 | 0.151096 | 0.02 | 0.015 | 1 | 5 | Bgn | 0.005 | Old |
| 0.577347 | 0.110474 | 0.067 | 0.058 | 1 | 5 | Nop16 | 0.009 | Old |
| 0.578061 | 0.072067 | 0.136 | 0.124 | 1 | 5 | Stk25 | 0.012 | Old |
| 0.578254 | 0.283221 | 0.135 | 0.124 | 1 | 5 | Hsdl1 | 0.011 | Old |
| 0.578516 | 0.015596 | 0.685 | 0.665 | 1 | 5 | Btg1 | 0.02 | Old |
| 0.57885 | 0.117929 | 0.258 | 0.244 | 1 | 5 | Emsy | 0.014 | Old |
| 0.579191 | 0.549135 | 0.047 | 0.041 | 1 | 5 | Galnt12 | 0.006 | Old |
| 0.579721 | 0.300158 | 0.109 | 0.099 | 1 | 5 | Akr1e1 | 0.01 | Old |
| 0.580207 | 0.197149 | 0.089 | 0.079 | 1 | 5 | Tfip11 | 0.01 | Old |
| 0.580766 | 0.070407 | 0.133 | 0.122 | 1 | 5 | Sash3 | 0.011 | Old |
| 0.580825 | 0.05873 | 0.184 | 0.17 | 1 | 5 | Akr1b10 | 0.014 | Old |
| 0.581281 | 0.101927 | 0.195 | 0.183 | 1 | 5 | Srp54a | 0.012 | Old |
| 0.581553 | 0.064706 | 0.141 | 0.127 | 1 | 5 | Ctnnbip1 | 0.014 | Old |

| | | | | | | | | |
|---|---|---|---|---|---|---|---|---|
| 0.58156 | 0.216933 | 0.086 | 0.076 | 1 | 5 | Exosc2 | 0.01 | Old |
| 0.581607 | 0.123952 | 0.163 | 0.15 | 1 | 5 | Cdc42bpb | 0.013 | Old |
| 0.581881 | 0.102746 | 0.886 | 0.86 | 1 | 5 | Fus | 0.026 | Old |
| 0.582266 | 0.299141 | 0.143 | 0.135 | 1 | 5 | Anpep | 0.008 | Old |
| 0.58259 | 0.063018 | 0.182 | 0.168 | 1 | 5 | Tef | 0.014 | Old |
| 0.582809 | 0.065966 | 0.261 | 0.244 | 1 | 5 | Hipk2 | 0.017 | Old |
| 0.583193 | 0.282725 | 0.226 | 0.221 | 1 | 5 | Fbrs | 0.005 | Old |
| 0.583211 | 0.080641 | 0.391 | 0.378 | 1 | 5 | Tnrc6a | 0.013 | Old |
| 0.583426 | 0.617024 | 0.042 | 0.036 | 1 | 5 | Fndc7 | 0.006 | Old |
| 0.583544 | 0.077419 | 0.146 | 0.135 | 1 | 5 | Tmem19 | 0.011 | Old |
| 0.583583 | 0.122981 | 0.135 | 0.124 | 1 | 5 | Prpf6 | 0.011 | Old |
| 0.583713 | 0.147787 | 0.232 | 0.223 | 1 | 5 | Vps13d | 0.009 | Old |
| 0.584093 | 0.107696 | 0.091 | 0.081 | 1 | 5 | Skiv2l2 | 0.01 | Old |
| 0.584264 | 0.055997 | 0.045 | 0.038 | 1 | 5 | Rnf219 | 0.007 | Old |
| 0.584438 | 0.278524 | 0.162 | 0.152 | 1 | 5 | Tubgcp4 | 0.01 | Old |
| 0.584701 | 0.113297 | 0.194 | 0.18 | 1 | 5 | Srpk2 | 0.014 | Old |
| 0.584816 | 0.465714 | 0.042 | 0.036 | 1 | 5 | Tab1 | 0.006 | Old |
| 0.585422 | 0.141542 | 0.069 | 0.061 | 1 | 5 | Dusp12 | 0.008 | Old |
| 0.585445 | 0.180561 | 0.037 | 0.03 | 1 | 5 | Ccdc77 | 0.007 | Old |
| 0.586028 | 0.199554 | 0.148 | 0.14 | 1 | 5 | Umad1 | 0.008 | Old |
| 0.586474 | 0.14088 | 0.123 | 0.114 | 1 | 5 | Slc1a5 | 0.009 | Old |
| 0.586924 | 0.066881 | 0.387 | 0.376 | 1 | 5 | Mapk1ip1l | 0.011 | Old |
| 0.587277 | 0.073491 | 0.295 | 0.282 | 1 | 5 | Vps29 | 0.013 | Old |
| 0.58783 | 0.141823 | 0.29 | 0.272 | 1 | 5 | Mospd2 | 0.018 | Old |
| 0.588298 | 0.0977 | 0.045 | 0.038 | 1 | 5 | Nsdhl | 0.007 | Old |
| 0.588494 | 0.053538 | 0.054 | 0.046 | 1 | 5 | Fra10ac1 | 0.008 | Old |
| 0.589051 | 0.915592 | 0.017 | 0.013 | 1 | 5 | Nupl2 | 0.004 | Old |
| 0.589364 | 0.063314 | 0.116 | 0.104 | 1 | 5 | Cmc1 | 0.012 | Old |
| 0.589659 | 0.123651 | 0.439 | 0.437 | 1 | 5 | Arhgap15 | 0.002 | Old |
| 0.589792 | 0.732366 | 0.017 | 0.013 | 1 | 5 | Ybey | 0.004 | Old |
| 0.590266 | 0.153122 | 0.077 | 0.069 | 1 | 5 | Frs2 | 0.008 | Old |
| 0.591292 | 0.081486 | 0.29 | 0.272 | 1 | 5 | Ube2g2 | 0.018 | Old |
| 0.592391 | 0.039973 | 0.33 | 0.31 | 1 | 5 | Anks1 | 0.02 | Old |
| 0.593122 | 0.12312 | 0.051 | 0.043 | 1 | 5 | Gpr180 | 0.008 | Old |
| 0.593359 | 0.019822 | 0.19 | 0.175 | 1 | 5 | Vps36 | 0.015 | Old |
| 0.593748 | 0.415007 | 0.034 | 0.028 | 1 | 5 | Cenpv | 0.006 | Old |
| 0.593748 | 0.243989 | 0.034 | 0.028 | 1 | 5 | Map3k15 | 0.006 | Old |
| 0.594061 | 0.176751 | 0.157 | 0.147 | 1 | 5 | Ttc5 | 0.01 | Old |
| 0.5944 | 0.065889 | 0.037 | 0.03 | 1 | 5 | Eefsec | 0.007 | Old |
| 0.5944 | 0.019641 | 0.037 | 0.03 | 1 | 5 | Coq9 | 0.007 | Old |
| 0.594478 | 0.246805 | 0.059 | 0.051 | 1 | 5 | Eva1b | 0.008 | Old |
| 0.594596 | 0.076218 | 0.064 | 0.056 | 1 | 5 | Rhbdd2 | 0.008 | Old |
| 0.594864 | 0.034927 | 0.084 | 0.074 | 1 | 5 | Anln | 0.01 | Old |
| 0.594995 | 0.22574 | 0.017 | 0.013 | 1 | 5 | Tspoap1 | 0.004 | Old |
| 0.595268 | 0.130613 | 0.182 | 0.17 | 1 | 5 | Ino80e | 0.012 | Old |

| | | | | | | | | |
|---|---|---|---|---|---|---|---|---|
| 0.595472 | 0.102697 | 0.306 | 0.287 | 1 | 5 | Khsrp | 0.019 | Old |
| 0.595732 | 0.03564 | 0.438 | 0.419 | 1 | 5 | Eif4g1 | 0.019 | Old |
| 0.595749 | 0.037229 | 0.232 | 0.216 | 1 | 5 | Fbxo33 | 0.016 | Old |
| 0.596031 | 0.268768 | 0.069 | 0.061 | 1 | 5 | Steap3 | 0.008 | Old |
| 0.596031 | 0.2747 | 0.069 | 0.061 | 1 | 5 | Ino80b | 0.008 | Old |
| 0.596408 | 0.83929 | 0.025 | 0.02 | 1 | 5 | Klra3 | 0.005 | Old |
| 0.596466 | 0.108941 | 0.042 | 0.036 | 1 | 5 | Ccdc28a | 0.006 | Old |
| 0.597171 | 0.155878 | 0.177 | 0.168 | 1 | 5 | Clasp1 | 0.009 | Old |
| 0.597231 | 0.532857 | 0.017 | 0.013 | 1 | 5 | Polr1e | 0.004 | Old |
| 0.598513 | 0.019146 | 0.323 | 0.302 | 1 | 5 | Tmem173 | 0.021 | Old |
| 0.59869 | 0.269076 | 0.093 | 0.084 | 1 | 5 | Pias3 | 0.009 | Old |
| 0.598693 | 0.001968 | 0.31 | 0.332 | 1 | 5 | Phf20l1 | -0.022 | Old |
| 0.598724 | 0.305039 | 0.017 | 0.013 | 1 | 5 | Gm47232 | 0.004 | Old |
| 0.599434 | 0.56129 | 0.025 | 0.02 | 1 | 5 | 2010320M | 0.005 | Old |
| 0.599471 | 0.029154 | 0.013 | 0.018 | 1 | 5 | Cldn5 | -0.005 | Old |
| 0.599592 | 0.08462 | 0.121 | 0.109 | 1 | 5 | Fam219b | 0.012 | Old |
| 0.599709 | 0.11751 | 0.069 | 0.061 | 1 | 5 | Zfp758 | 0.008 | Old |
| 0.599959 | 0.04701 | 0.027 | 0.033 | 1 | 5 | Porcn | -0.006 | Old |
| 0.600037 | 0.190924 | 0.234 | 0.223 | 1 | 5 | Erlin1 | 0.011 | Old |
| 0.600089 | 0.029235 | 0.02 | 0.025 | 1 | 5 | Gm42722 | -0.005 | Old |
| 0.6005 | 0.057265 | 0.027 | 0.033 | 1 | 5 | Syne3 | -0.006 | Old |
| 0.600966 | 0.267624 | 0.017 | 0.013 | 1 | 5 | Fam161a | 0.004 | Old |
| 0.601073 | 0.411333 | 0.034 | 0.028 | 1 | 5 | Nif3l1 | 0.006 | Old |
| 0.601306 | 0.005466 | 0.189 | 0.178 | 1 | 5 | Gpalpp1 | 0.011 | Old |
| 0.60164 | 0.20169 | 0.098 | 0.089 | 1 | 5 | Josd1 | 0.009 | Old |
| 0.601715 | 0.092187 | 0.013 | 0.018 | 1 | 5 | B3gntl1 | -0.005 | Old |
| 0.60186 | 0.581845 | 0.025 | 0.02 | 1 | 5 | Gas1 | 0.005 | Old |
| 0.601908 | 0.077838 | 0.187 | 0.178 | 1 | 5 | Rnf167 | 0.009 | Old |
| 0.602065 | 0.40884 | 0.076 | 0.069 | 1 | 5 | Tmem242 | 0.007 | Old |
| 0.60212 | 0.034991 | 0.549 | 0.508 | 1 | 5 | Tmco1 | 0.041 | Old |
| 0.602202 | 0.199299 | 0.047 | 0.056 | 1 | 5 | Gpn2 | -0.009 | Old |
| 0.602324 | 0.072886 | 0.256 | 0.241 | 1 | 5 | Prpf8 | 0.015 | Old |
| 0.602549 | 0.151724 | 0.244 | 0.231 | 1 | 5 | Dcun1d5 | 0.013 | Old |
| 0.602648 | 0.470381 | 0.034 | 0.028 | 1 | 5 | Grwd1 | 0.006 | Old |
| 0.603001 | 0.087048 | 0.184 | 0.17 | 1 | 5 | Ifih1 | 0.014 | Old |
| 0.603711 | 0.176904 | 0.263 | 0.251 | 1 | 5 | Adam33 | 0.012 | Old |
| 0.603977 | 0.330354 | 0.042 | 0.036 | 1 | 5 | Tcta | 0.006 | Old |
| 0.604019 | 0.0186 | 0.281 | 0.261 | 1 | 5 | Cenpb | 0.02 | Old |
| 0.604054 | 0.017674 | 0.212 | 0.198 | 1 | 5 | Dhdh | 0.014 | Old |
| 0.604394 | 0.06842 | 0.056 | 0.063 | 1 | 5 | Gstp3 | -0.007 | Old |
| 0.604437 | 0.004507 | 0.204 | 0.223 | 1 | 5 | Rnaset2a | -0.019 | Old |
| 0.604922 | 0.101561 | 0.064 | 0.056 | 1 | 5 | Kat14 | 0.008 | Old |
| 0.605004 | 0.077527 | 0.355 | 0.343 | 1 | 5 | Rer1 | 0.012 | Old |
| 0.605389 | 0.172816 | 0.042 | 0.036 | 1 | 5 | F8a | 0.006 | Old |
| 0.605507 | 0.415134 | 0.025 | 0.02 | 1 | 5 | Pms2 | 0.005 | Old |

| | | | | | | | | |
|---|---|---|---|---|---|---|---|---|
| 0.605903 | 0.107482 | 0.082 | 0.074 | 1 | 5 | Mib2 | 0.008 | Old |
| 0.606116 | 0.488896 | 0.025 | 0.02 | 1 | 5 | Tprn | 0.005 | Old |
| 0.606227 | 0.235421 | 0.056 | 0.048 | 1 | 5 | Prkdc | 0.008 | Old |
| 0.606396 | 0.212037 | 0.093 | 0.084 | 1 | 5 | Otud7b | 0.009 | Old |
| 0.606725 | 0.397141 | 0.025 | 0.02 | 1 | 5 | Fgfrl1 | 0.005 | Old |
| 0.606729 | 0.078221 | 0.552 | 0.536 | 1 | 5 | Tm9sf2 | 0.016 | Old |
| 0.606858 | 0.108866 | 0.034 | 0.028 | 1 | 5 | Pigb | 0.006 | Old |
| 0.60759 | 0.167636 | 0.13 | 0.119 | 1 | 5 | Iqsec1 | 0.011 | Old |
| 0.607838 | 0.060896 | 0.069 | 0.061 | 1 | 5 | E130307A1 | 0.008 | Old |
| 0.607944 | 0.308006 | 0.025 | 0.02 | 1 | 5 | Txnrd3 | 0.005 | Old |
| 0.608439 | 0.275274 | 0.034 | 0.028 | 1 | 5 | Sort1 | 0.006 | Old |
| 0.608554 | 0.348227 | 0.025 | 0.02 | 1 | 5 | Hs3st3a1 | 0.005 | Old |
| 0.608554 | 0.211876 | 0.025 | 0.02 | 1 | 5 | Pde2a | 0.005 | Old |
| 0.608595 | 0.089887 | 0.232 | 0.218 | 1 | 5 | Gab2 | 0.014 | Old |
| 0.608664 | 0.026027 | 0.146 | 0.135 | 1 | 5 | Itgb1bp1 | 0.011 | Old |
| 0.610064 | 0.146614 | 0.069 | 0.061 | 1 | 5 | Tada2b | 0.008 | Old |
| 0.610365 | 0.00986 | 0.362 | 0.335 | 1 | 5 | Clasp2 | 0.027 | Old |
| 0.610494 | 0.43557 | 0.101 | 0.094 | 1 | 5 | Gan | 0.007 | Old |
| 0.610692 | 0.165312 | 0.074 | 0.066 | 1 | 5 | Carnmt1 | 0.008 | Old |
| 0.610996 | 0.304037 | 0.025 | 0.02 | 1 | 5 | Aplf | 0.005 | Old |
| 0.611629 | 0.053218 | 0.177 | 0.168 | 1 | 5 | Insig2 | 0.009 | Old |
| 0.612129 | 0.011032 | 0.226 | 0.213 | 1 | 5 | Pdcl3 | 0.013 | Old |
| 0.612586 | 0.055208 | 0.072 | 0.063 | 1 | 5 | Mycbp | 0.009 | Old |
| 0.612715 | 0.170921 | 0.199 | 0.188 | 1 | 5 | Tlk1 | 0.011 | Old |
| 0.613424 | 0.124592 | 0.042 | 0.036 | 1 | 5 | B4galt7 | 0.006 | Old |
| 0.613443 | 0.27174 | 0.025 | 0.02 | 1 | 5 | Adnp | 0.005 | Old |
| 0.613668 | 0.367855 | 0.066 | 0.058 | 1 | 5 | Gm12840 | 0.008 | Old |
| 0.613699 | 0.140669 | 0.29 | 0.282 | 1 | 5 | Rbfa | 0.008 | Old |
| 0.613905 | 0.085479 | 0.742 | 0.713 | 1 | 5 | Frmd4b | 0.029 | Old |
| 0.613951 | 0.223311 | 0.138 | 0.132 | 1 | 5 | Banp | 0.006 | Old |
| 0.614275 | 0.083705 | 0.141 | 0.129 | 1 | 5 | Ptpn9 | 0.012 | Old |
| 0.614912 | 0.008809 | 0.232 | 0.218 | 1 | 5 | Bbx | 0.014 | Old |
| 0.614979 | 0.141284 | 0.202 | 0.188 | 1 | 5 | Ankle2 | 0.014 | Old |
| 0.615281 | 0.05749 | 0.025 | 0.02 | 1 | 5 | Casz1 | 0.005 | Old |
| 0.615353 | 0.010853 | 0.039 | 0.046 | 1 | 5 | Ahdc1 | -0.007 | Old |
| 0.615428 | 0.00216 | 0.256 | 0.239 | 1 | 5 | Slc25a4 | 0.017 | Old |
| 0.615438 | 0.135207 | 0.047 | 0.041 | 1 | 5 | Ighd | 0.006 | Old |
| 0.616219 | 0.270346 | 0.061 | 0.053 | 1 | 5 | Spsb3 | 0.008 | Old |
| 0.616333 | 0.153734 | 0.13 | 0.119 | 1 | 5 | Mapkap1 | 0.011 | Old |
| 0.616671 | 0.039592 | 0.104 | 0.094 | 1 | 5 | Gpt2 | 0.01 | Old |
| 0.617469 | 0.135216 | 0.212 | 0.201 | 1 | 5 | Tk2 | 0.011 | Old |
| 0.61749 | 0.600769 | 0.03 | 0.025 | 1 | 5 | Stx3 | 0.005 | Old |
| 0.617493 | 0.404936 | 0.081 | 0.074 | 1 | 5 | Plrg1 | 0.007 | Old |
| 0.617878 | 0.091159 | 0.456 | 0.434 | 1 | 5 | Hdlbp | 0.022 | Old |
| 0.618577 | 0.283864 | 0.047 | 0.041 | 1 | 5 | 2310057M | 0.006 | Old |

| | | | | | | | | |
|---|---|---|---|---|---|---|---|---|
| 0.618616 | 0.09151 | 0.056 | 0.048 | 1 | 5 | Rmnd5b | 0.008 | Old |
| 0.619168 | 0.057842 | 0.202 | 0.216 | 1 | 5 | Pbx1 | -0.014 | Old |
| 0.61924 | 0.024582 | 0.184 | 0.173 | 1 | 5 | Soga1 | 0.011 | Old |
| 0.619549 | 0.06673 | 0.158 | 0.147 | 1 | 5 | Stim1 | 0.011 | Old |
| 0.619881 | 0.014957 | 0.101 | 0.091 | 1 | 5 | Magohb | 0.01 | Old |
| 0.619925 | 0.118107 | 0.047 | 0.041 | 1 | 5 | Mrpl39 | 0.006 | Old |
| 0.620236 | 0.17504 | 0.13 | 0.119 | 1 | 5 | Gatsl2 | 0.011 | Old |
| 0.620418 | 0.30591 | 0.071 | 0.063 | 1 | 5 | Tmem70 | 0.008 | Old |
| 0.620759 | 0.065844 | 0.271 | 0.259 | 1 | 5 | Gng12 | 0.012 | Old |
| 0.620891 | 0.192321 | 0.079 | 0.071 | 1 | 5 | Papolg | 0.008 | Old |
| 0.621836 | 0.066273 | 0.237 | 0.221 | 1 | 5 | Ago4 | 0.016 | Old |
| 0.62216 | 0.072025 | 0.229 | 0.218 | 1 | 5 | Rsl1d1 | 0.011 | Old |
| 0.622179 | 0.230982 | 0.061 | 0.053 | 1 | 5 | Mir155hg | 0.008 | Old |
| 0.622187 | 0.067968 | 0.133 | 0.122 | 1 | 5 | Prkci | 0.011 | Old |
| 0.622221 | 0.201076 | 0.074 | 0.066 | 1 | 5 | Slc35e4 | 0.008 | Old |
| 0.622886 | 0.11676 | 0.17 | 0.16 | 1 | 5 | Rrm2b | 0.01 | Old |
| 0.623273 | 0.032276 | 0.025 | 0.02 | 1 | 5 | 8030462N1 | 0.005 | Old |
| 0.623491 | 0.01038 | 0.253 | 0.239 | 1 | 5 | Ddx42 | 0.014 | Old |
| 0.624377 | 0.096485 | 0.178 | 0.168 | 1 | 5 | C1d | 0.01 | Old |
| 0.624666 | 0.070391 | 0.12 | 0.109 | 1 | 5 | Hrh1 | 0.011 | Old |
| 0.624904 | 0.091869 | 0.16 | 0.15 | 1 | 5 | Inpp5f | 0.01 | Old |
| 0.624954 | 0.114866 | 0.221 | 0.211 | 1 | 5 | Eif4a3 | 0.01 | Old |
| 0.625425 | 0.149555 | 0.109 | 0.099 | 1 | 5 | Rprd1b | 0.01 | Old |
| 0.625477 | 0.228532 | 0.094 | 0.086 | 1 | 5 | Slc13a3 | 0.008 | Old |
| 0.625714 | 0.006739 | 0.298 | 0.277 | 1 | 5 | Uso1 | 0.021 | Old |
| 0.626472 | 0.280348 | 0.057 | 0.051 | 1 | 5 | Cd300ld3 | 0.006 | Old |
| 0.626664 | 0.186424 | 0.066 | 0.058 | 1 | 5 | Tipin | 0.008 | Old |
| 0.62703 | 0.254056 | 0.079 | 0.071 | 1 | 5 | Ikbke | 0.008 | Old |
| 0.627587 | 0.097167 | 0.047 | 0.041 | 1 | 5 | Maged2 | 0.006 | Old |
| 0.627605 | 0.07567 | 0.192 | 0.183 | 1 | 5 | Ergic2 | 0.009 | Old |
| 0.627722 | 0.056348 | 0.025 | 0.03 | 1 | 5 | Fdxacb1 | -0.005 | Old |
| 0.628144 | 0.029481 | 0.224 | 0.208 | 1 | 5 | Jtb | 0.016 | Old |
| 0.628316 | 0.371769 | 0.057 | 0.051 | 1 | 5 | Carf | 0.006 | Old |
| 0.628866 | 0.026281 | 0.103 | 0.094 | 1 | 5 | Nfyb | 0.009 | Old |
| 0.629958 | 0.331072 | 0.057 | 0.051 | 1 | 5 | Gm45606 | 0.006 | Old |
| 0.630151 | 0.143712 | 0.195 | 0.185 | 1 | 5 | Zfp281 | 0.01 | Old |
| 0.631293 | 0.184072 | 0.074 | 0.066 | 1 | 5 | Cadm1 | 0.008 | Old |
| 0.631532 | 0.002175 | 0.056 | 0.048 | 1 | 5 | Mul1 | 0.008 | Old |
| 0.632477 | 0.735241 | 0.013 | 0.01 | 1 | 5 | Gm4924 | 0.003 | Old |
| 0.633126 | 0.116701 | 0.451 | 0.431 | 1 | 5 | Cln8 | 0.02 | Old |
| 0.633306 | 0.398485 | 0.062 | 0.056 | 1 | 5 | Abhd17c | 0.006 | Old |
| 0.633331 | 0.588448 | 0.013 | 0.01 | 1 | 5 | Tjp3 | 0.003 | Old |
| 0.633393 | 0.046409 | 0.168 | 0.157 | 1 | 5 | Trim65 | 0.011 | Old |
| 0.634058 | 0.306745 | 0.052 | 0.046 | 1 | 5 | Vwa8 | 0.006 | Old |
| 0.634058 | 0.063088 | 0.052 | 0.046 | 1 | 5 | Tdrd3 | 0.006 | Old |

| | | | | | | | | |
|---|---|---|---|---|---|---|---|---|
| 0.634077 | 0.044022 | 0.52 | 0.505 | 1 | 5 | Tpm4 | 0.015 | Old |
| 0.634167 | 0.016394 | 0.185 | 0.203 | 1 | 5 | Mrpl17 | -0.018 | Old |
| 0.634718 | 0.694373 | 0.022 | 0.018 | 1 | 5 | Mphosph9 | 0.004 | Old |
| 0.635136 | 0.088004 | 0.066 | 0.058 | 1 | 5 | Ppip5k1 | 0.008 | Old |
| 0.635897 | 0.519965 | 0.013 | 0.01 | 1 | 5 | Pdzd4 | 0.003 | Old |
| 0.636887 | 0.23546 | 0.039 | 0.033 | 1 | 5 | Dusp18 | 0.006 | Old |
| 0.6372 | 0.132191 | 0.392 | 0.381 | 1 | 5 | Cpne3 | 0.011 | Old |
| 0.63761 | 0.459114 | 0.013 | 0.01 | 1 | 5 | Neurl1b | 0.003 | Old |
| 0.63761 | 0.436603 | 0.013 | 0.01 | 1 | 5 | Plxnb3 | 0.003 | Old |
| 0.637704 | 0.112557 | 0.153 | 0.145 | 1 | 5 | Tmem64 | 0.008 | Old |
| 0.638467 | 0.500427 | 0.013 | 0.01 | 1 | 5 | Zfp324 | 0.003 | Old |
| 0.638467 | 0.334289 | 0.013 | 0.01 | 1 | 5 | Tnfsf14 | 0.003 | Old |
| 0.638577 | 0.017497 | 0.914 | 0.858 | 1 | 5 | Picalm | 0.056 | Old |
| 0.639325 | 0.5188 | 0.013 | 0.01 | 1 | 5 | C130050O1 | 0.003 | Old |
| 0.63939 | 0.243624 | 0.066 | 0.058 | 1 | 5 | Men1 | 0.008 | Old |
| 0.639777 | 0.138787 | 0.066 | 0.058 | 1 | 5 | Rrp8 | 0.008 | Old |
| 0.639781 | 0.025577 | 0.025 | 0.03 | 1 | 5 | Aldh18a1 | -0.005 | Old |
| 0.640032 | 0.261106 | 0.062 | 0.056 | 1 | 5 | Zfp367 | 0.006 | Old |
| 0.640049 | 0.652344 | 0.022 | 0.018 | 1 | 5 | Adh1 | 0.004 | Old |
| 0.640098 | 0.063881 | 0.374 | 0.363 | 1 | 5 | Vasp | 0.011 | Old |
| 0.640183 | 0.295334 | 0.013 | 0.01 | 1 | 5 | Spa17 | 0.003 | Old |
| 0.640183 | 0.280043 | 0.013 | 0.01 | 1 | 5 | Ccdc17 | 0.003 | Old |
| 0.640344 | 0.196056 | 0.032 | 0.038 | 1 | 5 | Dpysl3 | -0.006 | Old |
| 0.640546 | 0.281106 | 0.03 | 0.025 | 1 | 5 | Aacs | 0.005 | Old |
| 0.640597 | 0.29539 | 0.044 | 0.038 | 1 | 5 | Pus7 | 0.006 | Old |
| 0.640947 | 0.096643 | 0.14 | 0.129 | 1 | 5 | Gm4951 | 0.011 | Old |
| 0.64168 | 0.343971 | 0.03 | 0.025 | 1 | 5 | Nicn1 | 0.005 | Old |
| 0.641901 | 0.195114 | 0.013 | 0.01 | 1 | 5 | Bbs1 | 0.003 | Old |
| 0.642722 | 0.429133 | 0.022 | 0.018 | 1 | 5 | Fam46c | 0.004 | Old |
| 0.64276 | 0.17547 | 0.16 | 0.15 | 1 | 5 | Phyh | 0.01 | Old |
| 0.643157 | 0.209704 | 0.057 | 0.051 | 1 | 5 | Adprhl2 | 0.006 | Old |
| 0.643238 | 0.005756 | 0.128 | 0.117 | 1 | 5 | Cyp51 | 0.011 | Old |
| 0.643505 | 0.01827 | 0.362 | 0.345 | 1 | 5 | Gpr160 | 0.017 | Old |
| 0.643621 | 0.250898 | 0.013 | 0.01 | 1 | 5 | D130017N( | 0.003 | Old |
| 0.643621 | 0.236899 | 0.013 | 0.01 | 1 | 5 | Cerkl | 0.003 | Old |
| 0.643621 | 0.104365 | 0.013 | 0.01 | 1 | 5 | Wdr19 | 0.003 | Old |
| 0.643621 | 0.060945 | 0.013 | 0.01 | 1 | 5 | Gm10521 | 0.003 | Old |
| 0.643667 | 0.011254 | 0.253 | 0.236 | 1 | 5 | Smndc1 | 0.017 | Old |
| 0.643854 | 0.178474 | 0.076 | 0.069 | 1 | 5 | Kctd3 | 0.007 | Old |
| 0.64395 | 0.017851 | 0.03 | 0.025 | 1 | 5 | Tmlhe | 0.005 | Old |
| 0.644351 | 0.101752 | 0.264 | 0.256 | 1 | 5 | Trp53 | 0.008 | Old |
| 0.644401 | 0.242197 | 0.057 | 0.051 | 1 | 5 | Ofd1 | 0.006 | Old |
| 0.644481 | 0.139359 | 0.013 | 0.01 | 1 | 5 | Phf21b | 0.003 | Old |
| 0.644712 | 0.183448 | 0.082 | 0.076 | 1 | 5 | Slc7a1 | 0.006 | Old |
| 0.6454 | 0.439506 | 0.022 | 0.018 | 1 | 5 | Ccdc51 | 0.004 | Old |

| | | | | | | | | |
|---|---|---|---|---|---|---|---|---|
| 0.6454 | 0.247742 | 0.022 | 0.018 | 1 | 5 | Zfp799 | 0.004 | Old |
| 0.645655 | 0.055619 | 0.268 | 0.254 | 1 | 5 | Tmem33 | 0.014 | Old |
| 0.64607 | 0.395451 | 0.022 | 0.018 | 1 | 5 | Ss18l1 | 0.004 | Old |
| 0.646077 | 0.102582 | 0.177 | 0.168 | 1 | 5 | Adgrg6 | 0.009 | Old |
| 0.646179 | 0.063301 | 0.052 | 0.046 | 1 | 5 | Rrnad1 | 0.006 | Old |
| 0.646204 | 0.028661 | 0.013 | 0.01 | 1 | 5 | Tmem267 | 0.003 | Old |
| 0.646204 | 0.020802 | 0.013 | 0.01 | 1 | 5 | Gm45250 | 0.003 | Old |
| 0.646242 | 0.201642 | 0.177 | 0.17 | 1 | 5 | Pex2 | 0.007 | Old |
| 0.64674 | 0.489441 | 0.022 | 0.018 | 1 | 5 | Gm48066 | 0.004 | Old |
| 0.646879 | 0.066979 | 0.296 | 0.284 | 1 | 5 | Rgs19 | 0.012 | Old |
| 0.647045 | 0.314205 | 0.072 | 0.066 | 1 | 5 | Clcn6 | 0.006 | Old |
| 0.647069 | 0.002819 | 0.247 | 0.234 | 1 | 5 | Mlec | 0.013 | Old |
| 0.647097 | 0.255977 | 0.049 | 0.043 | 1 | 5 | Zfp719 | 0.006 | Old |
| 0.647367 | 0.053804 | 0.495 | 0.48 | 1 | 5 | Comt | 0.015 | Old |
| 0.648539 | 0.316873 | 0.054 | 0.048 | 1 | 5 | Cox10 | 0.006 | Old |
| 0.648624 | 0.003199 | 0.177 | 0.19 | 1 | 5 | Bcl7c | -0.013 | Old |
| 0.648719 | 0.114228 | 0.205 | 0.195 | 1 | 5 | Ranbp9 | 0.01 | Old |
| 0.648792 | 0.119624 | 0.013 | 0.01 | 1 | 5 | Tulp3 | 0.003 | Old |
| 0.648921 | 0.063394 | 0.241 | 0.226 | 1 | 5 | Ankrd40 | 0.015 | Old |
| 0.649037 | 0.261692 | 0.012 | 0.015 | 1 | 5 | Ryr1 | -0.003 | Old |
| 0.64907 | 0.158567 | 0.03 | 0.025 | 1 | 5 | Pmm1 | 0.005 | Old |
| 0.649542 | 0.054788 | 0.219 | 0.213 | 1 | 5 | Ptpn2 | 0.006 | Old |
| 0.649809 | 0.289828 | 0.086 | 0.079 | 1 | 5 | Tlr6 | 0.007 | Old |
| 0.650423 | 0.07582 | 0.088 | 0.079 | 1 | 5 | Cmc2 | 0.009 | Old |
| 0.650464 | 0.088325 | 0.039 | 0.033 | 1 | 5 | Glb1l | 0.006 | Old |
| 0.650466 | 0.162458 | 0.081 | 0.074 | 1 | 5 | Tmem198b | 0.007 | Old |
| 0.650507 | 0.024407 | 0.152 | 0.165 | 1 | 5 | Rasgrp3 | -0.013 | Old |
| 0.650661 | 0.270598 | 0.067 | 0.061 | 1 | 5 | Ralgapa2 | 0.006 | Old |
| 0.650768 | 0.444164 | 0.022 | 0.018 | 1 | 5 | Fxn | 0.004 | Old |
| 0.650768 | 0.153723 | 0.022 | 0.018 | 1 | 5 | Abcb8 | 0.004 | Old |
| 0.650947 | 0.006247 | 0.761 | 0.761 | 1 | 5 | Tacc1 | 0 | Old |
| 0.65118 | 0.054513 | 0.079 | 0.089 | 1 | 5 | Pi4kb | -0.01 | Old |
| 0.651359 | 0.043773 | 0.949 | 0.97 | 1 | 5 | Fos | -0.021 | Old |
| 0.65144 | 0.207134 | 0.022 | 0.018 | 1 | 5 | Sirpb1b | 0.004 | Old |
| 0.6515 | 0.04241 | 0.379 | 0.358 | 1 | 5 | Gpsm3 | 0.021 | Old |
| 0.651883 | 0.009779 | 0.214 | 0.203 | 1 | 5 | Clk3 | 0.011 | Old |
| 0.652717 | 0.012609 | 0.493 | 0.467 | 1 | 5 | Elf1 | 0.026 | Old |
| 0.653073 | 0.630072 | 0.035 | 0.03 | 1 | 5 | Rfc4 | 0.005 | Old |
| 0.653192 | 0.094913 | 0.012 | 0.015 | 1 | 5 | Lysmd4 | -0.003 | Old |
| 0.653394 | 0.005316 | 0.066 | 0.058 | 1 | 5 | Rpusd4 | 0.008 | Old |
| 0.653582 | 0.233039 | 0.086 | 0.079 | 1 | 5 | Ints13 | 0.007 | Old |
| 0.653618 | 0.041905 | 0.182 | 0.173 | 1 | 5 | Prkaca | 0.009 | Old |
| 0.65362 | 0.009549 | 0.2 | 0.188 | 1 | 5 | Stxbp5 | 0.012 | Old |
| 0.654054 | 0.140181 | 0.2 | 0.195 | 1 | 5 | Slc35a3 | 0.005 | Old |
| 0.654383 | 0.075627 | 0.057 | 0.051 | 1 | 5 | Fam122a | 0.006 | Old |

| | | | | | | | | |
|---|---|---|---|---|---|---|---|---|
| 0.654492 | 0.072947 | 0.402 | 0.401 | 1 | 5 | Gm21188 | 0.001 | Old |
| 0.654509 | 0.32686 | 0.049 | 0.043 | 1 | 5 | Phtf1 | 0.006 | Old |
| 0.654593 | 0.32808 | 0.032 | 0.038 | 1 | 5 | Sparcl1 | -0.006 | Old |
| 0.654806 | 0.22057 | 0.044 | 0.038 | 1 | 5 | Lbx2 | 0.006 | Old |
| 0.65548 | 0.005377 | 0.022 | 0.018 | 1 | 5 | Sirt6 | 0.004 | Old |
| 0.656565 | 0.247442 | 0.082 | 0.076 | 1 | 5 | Ttll3 | 0.006 | Old |
| 0.656829 | 0.05976 | 0.022 | 0.018 | 1 | 5 | Camk2b | 0.004 | Old |
| 0.657068 | 0.109215 | 0.03 | 0.025 | 1 | 5 | Dpy19l3 | 0.005 | Old |
| 0.657068 | 0.002175 | 0.03 | 0.025 | 1 | 5 | 2010013B2 | 0.005 | Old |
| 0.657315 | 0.118361 | 0.076 | 0.069 | 1 | 5 | Fzd5 | 0.007 | Old |
| 0.657646 | 0.039054 | 0.453 | 0.429 | 1 | 5 | Trim41 | 0.024 | Old |
| 0.657972 | 0.016032 | 0.098 | 0.089 | 1 | 5 | Nfs1 | 0.009 | Old |
| 0.658179 | 0.001652 | 0.022 | 0.018 | 1 | 5 | Kctd21 | 0.004 | Old |
| 0.658612 | 0.062106 | 0.199 | 0.19 | 1 | 5 | Acp1 | 0.009 | Old |
| 0.658887 | 0.327373 | 0.035 | 0.03 | 1 | 5 | Zfp961 | 0.005 | Old |
| 0.658979 | 0.249448 | 0.057 | 0.051 | 1 | 5 | Zfp809 | 0.006 | Old |
| 0.659095 | 0.043737 | 0.212 | 0.198 | 1 | 5 | Actr1a | 0.014 | Old |
| 0.659403 | 0.126016 | 0.251 | 0.239 | 1 | 5 | Acsl1 | 0.012 | Old |
| 0.659416 | 0.304834 | 0.035 | 0.03 | 1 | 5 | Parvg | 0.005 | Old |
| 0.659936 | 0.03777 | 0.106 | 0.117 | 1 | 5 | Cherp | -0.011 | Old |
| 0.660377 | 0.061984 | 0.697 | 0.69 | 1 | 5 | Xbp1 | 0.007 | Old |
| 0.660408 | 0.03713 | 0.507 | 0.495 | 1 | 5 | Macf1 | 0.012 | Old |
| 0.660901 | 0.114282 | 0.061 | 0.053 | 1 | 5 | Vcl | 0.008 | Old |
| 0.660969 | 0.022829 | 0.35 | 0.33 | 1 | 5 | Sar1a | 0.02 | Old |
| 0.66119 | 0.019771 | 0.153 | 0.145 | 1 | 5 | Pex19 | 0.008 | Old |
| 0.661553 | 0.0054 | 0.077 | 0.086 | 1 | 5 | Slc7a6os | -0.009 | Old |
| 0.662095 | 0.090795 | 0.037 | 0.043 | 1 | 5 | Pold2 | -0.006 | Old |
| 0.6624 | 0.080513 | 0.094 | 0.104 | 1 | 5 | Thumpd1 | -0.01 | Old |
| 0.662785 | 0.059024 | 0.295 | 0.277 | 1 | 5 | Tbc1d23 | 0.018 | Old |
| 0.662936 | 0.035971 | 0.731 | 0.708 | 1 | 5 | Runx1 | 0.023 | Old |
| 0.663421 | 0.065752 | 0.303 | 0.289 | 1 | 5 | Epb41 | 0.014 | Old |
| 0.663604 | 0.157226 | 0.062 | 0.056 | 1 | 5 | 1810011H1 | 0.006 | Old |
| 0.664221 | 0.042669 | 0.074 | 0.066 | 1 | 5 | Bcl2l13 | 0.008 | Old |
| 0.66438 | 0.035227 | 0.071 | 0.063 | 1 | 5 | Agk | 0.008 | Old |
| 0.665218 | 0.787694 | 0.027 | 0.023 | 1 | 5 | Dpep2 | 0.004 | Old |
| 0.665596 | 0.157396 | 0.104 | 0.096 | 1 | 5 | Crybg1 | 0.008 | Old |
| 0.665791 | 0.087127 | 0.042 | 0.048 | 1 | 5 | Ctdspl | -0.006 | Old |
| 0.666179 | 0.011757 | 0.168 | 0.157 | 1 | 5 | Scaf1 | 0.011 | Old |
| 0.666257 | 0.001675 | 0.03 | 0.025 | 1 | 5 | Aven | 0.005 | Old |
| 0.666264 | 0.097389 | 0.044 | 0.038 | 1 | 5 | Msto1 | 0.006 | Old |
| 0.666278 | 0.099417 | 0.071 | 0.063 | 1 | 5 | Flt3l | 0.008 | Old |
| 0.666497 | 0.107345 | 0.165 | 0.157 | 1 | 5 | Hccs | 0.008 | Old |
| 0.666548 | 0.418863 | 0.04 | 0.036 | 1 | 5 | Galm | 0.004 | Old |
| 0.666658 | 0.08704 | 0.071 | 0.063 | 1 | 5 | Exoc8 | 0.008 | Old |
| 0.666865 | 0.19495 | 0.138 | 0.129 | 1 | 5 | R3hcc1l | 0.009 | Old |

| | | | | | | | | |
|---|---|---|---|---|---|---|---|---|
| 0.667143 | 0.039122 | 0.328 | 0.353 | 1 | 5 | Lpp | -0.025 | Old |
| 0.667181 | 0.007996 | 0.042 | 0.048 | 1 | 5 | Ddx20 | -0.006 | Old |
| 0.668623 | 0.127217 | 0.091 | 0.084 | 1 | 5 | Extl3 | 0.007 | Old |
| 0.669134 | 0.005116 | 0.308 | 0.292 | 1 | 5 | Bub3 | 0.016 | Old |
| 0.66932 | 0.108906 | 0.071 | 0.063 | 1 | 5 | Rras2 | 0.008 | Old |
| 0.669501 | 0.278398 | 0.108 | 0.102 | 1 | 5 | BC003965 | 0.006 | Old |
| 0.66951 | 0.163902 | 0.035 | 0.03 | 1 | 5 | Cdpf1 | 0.005 | Old |
| 0.669908 | 0.002632 | 0.061 | 0.053 | 1 | 5 | Paxx | 0.008 | Old |
| 0.669984 | 0.04409 | 0.152 | 0.14 | 1 | 5 | Pgap1 | 0.012 | Old |
| 0.670504 | 0.029215 | 0.258 | 0.246 | 1 | 5 | Cpq | 0.012 | Old |
| 0.671026 | 0.444391 | 0.04 | 0.036 | 1 | 5 | Nmnat1 | 0.004 | Old |
| 0.67106 | 0.00077 | 0.022 | 0.018 | 1 | 5 | Ovca2 | 0.004 | Old |
| 0.671883 | 0.199391 | 0.03 | 0.036 | 1 | 5 | Dennd2a | -0.006 | Old |
| 0.672303 | 0.035797 | 0.081 | 0.074 | 1 | 5 | Depdc5 | 0.007 | Old |
| 0.6724 | 0.156931 | 0.205 | 0.198 | 1 | 5 | Chsy1 | 0.007 | Old |
| 0.672574 | 0.052734 | 0.291 | 0.279 | 1 | 5 | Zkscan3 | 0.012 | Old |
| 0.672608 | 0.215388 | 0.072 | 0.066 | 1 | 5 | 1600010M | 0.006 | Old |
| 0.672748 | 0.106386 | 0.071 | 0.063 | 1 | 5 | Ppa2 | 0.008 | Old |
| 0.673191 | 0.051984 | 0.557 | 0.546 | 1 | 5 | Gsk3b | 0.011 | Old |
| 0.673508 | 0.059934 | 0.03 | 0.036 | 1 | 5 | Ift43 | -0.006 | Old |
| 0.673876 | 0.001323 | 0.313 | 0.294 | 1 | 5 | Arhgap12 | 0.019 | Old |
| 0.674581 | 0.272783 | 0.133 | 0.124 | 1 | 5 | Gm13212 | 0.009 | Old |
| 0.674629 | 0.1066 | 0.165 | 0.155 | 1 | 5 | Casd1 | 0.01 | Old |
| 0.674846 | 0.004952 | 0.035 | 0.03 | 1 | 5 | Zbtb12 | 0.005 | Old |
| 0.675055 | 0.108034 | 0.067 | 0.061 | 1 | 5 | Narfl | 0.006 | Old |
| 0.675578 | 0.15463 | 0.049 | 0.043 | 1 | 5 | Adgrl2 | 0.006 | Old |
| 0.67572 | 0.037726 | 0.315 | 0.302 | 1 | 5 | Kcmf1 | 0.013 | Old |
| 0.675776 | 0.11393 | 0.096 | 0.089 | 1 | 5 | Eri3 | 0.007 | Old |
| 0.676692 | 0.143943 | 0.145 | 0.137 | 1 | 5 | Tax1bp3 | 0.008 | Old |
| 0.676764 | 0.110659 | 0.03 | 0.036 | 1 | 5 | Sacs | -0.006 | Old |
| 0.677521 | 0.204747 | 0.035 | 0.03 | 1 | 5 | Als2cl | 0.005 | Old |
| 0.678542 | 0.320665 | 0.079 | 0.074 | 1 | 5 | Gm2000 | 0.005 | Old |
| 0.678577 | 0.660479 | 0.01 | 0.008 | 1 | 5 | Ropn1l | 0.002 | Old |
| 0.679993 | 0.038245 | 0.047 | 0.053 | 1 | 5 | Ppp2r3a | -0.006 | Old |
| 0.680605 | 0.879828 | 0.01 | 0.008 | 1 | 5 | Dynlt1c | 0.002 | Old |
| 0.680605 | 0.796427 | 0.01 | 0.008 | 1 | 5 | Krba1 | 0.002 | Old |
| 0.680605 | 0.54992 | 0.01 | 0.008 | 1 | 5 | Dennd6b | 0.002 | Old |
| 0.680781 | 0.092368 | 0.109 | 0.102 | 1 | 5 | Snhg12 | 0.007 | Old |
| 0.681125 | 0.212728 | 0.162 | 0.155 | 1 | 5 | Asxl1 | 0.007 | Old |
| 0.681214 | 0.245141 | 0.059 | 0.053 | 1 | 5 | Entpd5 | 0.006 | Old |
| 0.681435 | 0.056045 | 0.173 | 0.162 | 1 | 5 | Wbp4 | 0.011 | Old |
| 0.68144 | 0.125909 | 0.114 | 0.107 | 1 | 5 | Trak2 | 0.007 | Old |
| 0.681619 | 0.680967 | 0.01 | 0.008 | 1 | 5 | Ppp1r3d | 0.002 | Old |
| 0.681619 | 0.674833 | 0.01 | 0.008 | 1 | 5 | AC125149. | 0.002 | Old |
| 0.682345 | 0.045146 | 0.035 | 0.03 | 1 | 5 | Alg3 | 0.005 | Old |

| | | | | | | | | |
|---|---|---|---|---|---|---|---|---|
| 0.682356 | 0.017983 | 0.234 | 0.223 | 1 | 5 | Wdr37 | 0.011 | Old |
| 0.683252 | 0.329527 | 0.114 | 0.109 | 1 | 5 | Pnkd | 0.005 | Old |
| 0.68336 | 0.548036 | 0.019 | 0.015 | 1 | 5 | Prim1 | 0.004 | Old |
| 0.68365 | 0.75876 | 0.01 | 0.008 | 1 | 5 | Gbgt1 | 0.002 | Old |
| 0.68365 | 0.63616 | 0.01 | 0.008 | 1 | 5 | Fgd5 | 0.002 | Old |
| 0.68365 | 0.628487 | 0.01 | 0.008 | 1 | 5 | Plekha5 | 0.002 | Old |
| 0.68365 | 0.403457 | 0.01 | 0.008 | 1 | 5 | Adgrl1 | 0.002 | Old |
| 0.683745 | 0.097618 | 0.077 | 0.071 | 1 | 5 | Btbd3 | 0.006 | Old |
| 0.683802 | 0.096217 | 0.145 | 0.157 | 1 | 5 | Asgr2 | -0.012 | Old |
| 0.683998 | 0.112162 | 0.195 | 0.188 | 1 | 5 | Map3k14 | 0.007 | Old |
| 0.684116 | 0.125308 | 0.072 | 0.066 | 1 | 5 | Lrrc40 | 0.006 | Old |
| 0.684493 | 0.174189 | 0.035 | 0.03 | 1 | 5 | Vwf | 0.005 | Old |
| 0.684667 | 0.297176 | 0.01 | 0.008 | 1 | 5 | Rad54l | 0.002 | Old |
| 0.684669 | 0.31431 | 0.108 | 0.102 | 1 | 5 | 1700025G( | 0.006 | Old |
| 0.685211 | 0.034517 | 0.077 | 0.071 | 1 | 5 | Mrs2 | 0.006 | Old |
| 0.685314 | 0.04883 | 0.076 | 0.069 | 1 | 5 | Dolk | 0.007 | Old |
| 0.685472 | 0.035281 | 0.104 | 0.114 | 1 | 5 | Nkiras2 | -0.01 | Old |
| 0.68568 | 0.019365 | 0.059 | 0.066 | 1 | 5 | Atg10 | -0.007 | Old |
| 0.685683 | 0.348967 | 0.01 | 0.008 | 1 | 5 | Gnai1 | 0.002 | Old |
| 0.685683 | 0.334735 | 0.01 | 0.008 | 1 | 5 | Tmppe | 0.002 | Old |
| 0.68579 | 0.034084 | 0.088 | 0.081 | 1 | 5 | Zfp729a | 0.007 | Old |
| 0.685815 | 0.065564 | 0.241 | 0.226 | 1 | 5 | Sugt1 | 0.015 | Old |
| 0.686075 | 0.169891 | 0.101 | 0.094 | 1 | 5 | Pprc1 | 0.007 | Old |
| 0.686085 | 0.455739 | 0.045 | 0.041 | 1 | 5 | Dgkq | 0.004 | Old |
| 0.686379 | 0.035424 | 0.813 | 0.764 | 1 | 5 | Tmcc1 | 0.049 | Old |
| 0.686391 | 0.035701 | 0.71 | 0.673 | 1 | 5 | Kdm7a | 0.037 | Old |
| 0.686701 | 0.310946 | 0.01 | 0.008 | 1 | 5 | Gm6277 | 0.002 | Old |
| 0.687097 | 0.000875 | 0.968 | 0.967 | 1 | 5 | Unc93b1 | 0.001 | Old |
| 0.687513 | 0.022523 | 0.062 | 0.056 | 1 | 5 | Zfp729b | 0.006 | Old |
| 0.687719 | 0.326924 | 0.01 | 0.008 | 1 | 5 | Pcbp3 | 0.002 | Old |
| 0.687719 | 0.221382 | 0.01 | 0.008 | 1 | 5 | B3gnt9 | 0.002 | Old |
| 0.687719 | 0.163331 | 0.01 | 0.008 | 1 | 5 | Kyat3 | 0.002 | Old |
| 0.688049 | 0.229132 | 0.04 | 0.036 | 1 | 5 | Slc19a1 | 0.004 | Old |
| 0.688656 | 0.007826 | 0.219 | 0.213 | 1 | 5 | Rspry1 | 0.006 | Old |
| 0.688738 | 0.133759 | 0.01 | 0.008 | 1 | 5 | Nnat | 0.002 | Old |
| 0.688791 | 0.218423 | 0.024 | 0.028 | 1 | 5 | Traf3ip3 | -0.004 | Old |
| 0.689055 | 0.280147 | 0.04 | 0.036 | 1 | 5 | Dcbld2 | 0.004 | Old |
| 0.689702 | 0.007137 | 0.178 | 0.193 | 1 | 5 | Zfyve16 | -0.015 | Old |
| 0.689757 | 0.004534 | 0.01 | 0.008 | 1 | 5 | Marveld2 | 0.002 | Old |
| 0.690036 | 0.067562 | 0.354 | 0.345 | 1 | 5 | Furin | 0.009 | Old |
| 0.690353 | 0.475076 | 0.045 | 0.041 | 1 | 5 | Cd28 | 0.004 | Old |
| 0.690392 | 0.02928 | 0.426 | 0.414 | 1 | 5 | Cox17 | 0.012 | Old |
| 0.690444 | 0.050862 | 0.966 | 0.942 | 1 | 5 | Klf6 | 0.024 | Old |
| 0.690599 | 0.046778 | 0.512 | 0.505 | 1 | 5 | Cdc42se2 | 0.007 | Old |
| 0.690641 | 0.118517 | 0.027 | 0.023 | 1 | 5 | Gm26982 | 0.004 | Old |

| | | | | | | | | |
|---|---|---|---|---|---|---|---|---|
| 0.69072 | 0.13736 | 0.077 | 0.071 | 1 | 5 | Trappc2 | 0.006 | Old |
| 0.690776 | 0.472119 | 0.01 | 0.008 | 1 | 5 | Olfr373 | 0.002 | Old |
| 0.690797 | 0.39176 | 0.019 | 0.015 | 1 | 5 | 4931414P1 | 0.004 | Old |
| 0.690797 | 0.388992 | 0.019 | 0.015 | 1 | 5 | Prdm9 | 0.004 | Old |
| 0.691222 | 0.001637 | 0.185 | 0.175 | 1 | 5 | Aimp1 | 0.01 | Old |
| 0.691289 | 0.020419 | 0.253 | 0.277 | 1 | 5 | Pcmt1 | -0.024 | Old |
| 0.691302 | 0.476939 | 0.045 | 0.041 | 1 | 5 | Dcun1d2 | 0.004 | Old |
| 0.691722 | 0.187517 | 0.184 | 0.178 | 1 | 5 | Trappc8 | 0.006 | Old |
| 0.691875 | 0.171457 | 0.027 | 0.023 | 1 | 5 | Wdr24 | 0.004 | Old |
| 0.69279 | 0.115987 | 0.114 | 0.107 | 1 | 5 | Tapbpl | 0.007 | Old |
| 0.692897 | 0.036331 | 0.094 | 0.104 | 1 | 5 | Atrn | -0.01 | Old |
| 0.693215 | 0.519423 | 0.032 | 0.028 | 1 | 5 | Brat1 | 0.004 | Old |
| 0.693256 | 0.015072 | 0.328 | 0.315 | 1 | 5 | Anxa4 | 0.013 | Old |
| 0.693417 | 0.053435 | 0.226 | 0.221 | 1 | 5 | Mob1b | 0.005 | Old |
| 0.693548 | 0.134467 | 0.069 | 0.063 | 1 | 5 | Tbp | 0.006 | Old |
| 0.693891 | 0.123162 | 0.547 | 0.546 | 1 | 5 | Psme1 | 0.001 | Old |
| 0.694632 | 0.066572 | 0.163 | 0.157 | 1 | 5 | Psmc2 | 0.006 | Old |
| 0.695159 | 0.022875 | 0.155 | 0.145 | 1 | 5 | Polr2i | 0.01 | Old |
| 0.695274 | 0.244367 | 0.019 | 0.015 | 1 | 5 | Dnajc17 | 0.004 | Old |
| 0.695447 | 0.432408 | 0.051 | 0.046 | 1 | 5 | Trim14 | 0.005 | Old |
| 0.69561 | 0.16009 | 0.04 | 0.036 | 1 | 5 | Jade3 | 0.004 | Old |
| 0.695642 | 0.080595 | 0.091 | 0.084 | 1 | 5 | Gipc1 | 0.007 | Old |
| 0.69574 | 0.038195 | 0.111 | 0.104 | 1 | 5 | Gpatch2l | 0.007 | Old |
| 0.695779 | 0.034588 | 0.143 | 0.135 | 1 | 5 | Stx17 | 0.008 | Old |
| 0.695832 | 0.002912 | 0.31 | 0.302 | 1 | 5 | Agps | 0.008 | Old |
| 0.696251 | 0.050584 | 0.327 | 0.325 | 1 | 5 | Stat6 | 0.002 | Old |
| 0.696749 | 0.178819 | 0.017 | 0.02 | 1 | 5 | Tpm2 | -0.003 | Old |
| 0.697436 | 0.286632 | 0.074 | 0.069 | 1 | 5 | Sppl2b | 0.005 | Old |
| 0.697563 | 0.042518 | 0.497 | 0.47 | 1 | 5 | Hprt | 0.027 | Old |
| 0.698204 | 0.274824 | 0.017 | 0.02 | 1 | 5 | Zfp30 | -0.003 | Old |
| 0.698374 | 0.032178 | 0.093 | 0.102 | 1 | 5 | Ppp1r8 | -0.009 | Old |
| 0.698415 | 0.127827 | 0.113 | 0.107 | 1 | 5 | Gm43848 | 0.006 | Old |
| 0.698465 | 0.195888 | 0.088 | 0.081 | 1 | 5 | Traf5 | 0.007 | Old |
| 0.698878 | 0.373109 | 0.029 | 0.033 | 1 | 5 | Fpr1 | -0.004 | Old |
| 0.699694 | 0.334663 | 0.074 | 0.069 | 1 | 5 | Zmym1 | 0.005 | Old |
| 0.699695 | 0.01032 | 0.17 | 0.162 | 1 | 5 | Shprh | 0.008 | Old |
| 0.700826 | 0.28913 | 0.045 | 0.041 | 1 | 5 | Zfp764 | 0.004 | Old |
| 0.700826 | 0.079085 | 0.128 | 0.14 | 1 | 5 | Crebzf | -0.012 | Old |
| 0.701001 | 0.020344 | 0.133 | 0.124 | 1 | 5 | Gle1 | 0.009 | Old |
| 0.701155 | 0.13687 | 0.106 | 0.099 | 1 | 5 | Usp6nl | 0.007 | Old |
| 0.701212 | 0.024064 | 0.419 | 0.406 | 1 | 5 | Ensa | 0.013 | Old |
| 0.701423 | 0.04867 | 0.18 | 0.173 | 1 | 5 | Cltb | 0.007 | Old |
| 0.701592 | 0.051933 | 0.503 | 0.48 | 1 | 5 | Ralb | 0.023 | Old |
| 0.701716 | 0.176099 | 0.032 | 0.028 | 1 | 5 | Nptxr | 0.004 | Old |
| 0.701844 | 0.038904 | 0.017 | 0.02 | 1 | 5 | Nod2 | -0.003 | Old |

| | | | | | | | | |
|---|---|---|---|---|---|---|---|---|
| 0.702534 | 0.012273 | 0.126 | 0.137 | 1 | 5 | Zadh2 | -0.011 | Old |
| 0.702793 | 0.121218 | 0.175 | 0.168 | 1 | 5 | Naa35 | 0.007 | Old |
| 0.703161 | 0.012019 | 0.276 | 0.266 | 1 | 5 | Slc28a2 | 0.01 | Old |
| 0.703365 | 0.148979 | 0.217 | 0.208 | 1 | 5 | Bcor | 0.009 | Old |
| 0.704275 | 0.208544 | 0.059 | 0.053 | 1 | 5 | Coq10a | 0.006 | Old |
| 0.704495 | 0.082779 | 0.109 | 0.102 | 1 | 5 | Prmt3 | 0.007 | Old |
| 0.704561 | 0.02404 | 0.907 | 0.896 | 1 | 5 | Rps6 | 0.011 | Old |
| 0.704565 | 0.131465 | 0.096 | 0.089 | 1 | 5 | Phlpp2 | 0.007 | Old |
| 0.704655 | 0.067161 | 0.298 | 0.289 | 1 | 5 | Erp44 | 0.009 | Old |
| 0.705258 | 0.213714 | 0.079 | 0.074 | 1 | 5 | Tmcc2 | 0.005 | Old |
| 0.705291 | 0.214991 | 0.138 | 0.132 | 1 | 5 | Pogk | 0.006 | Old |
| 0.70541 | 0.22168 | 0.051 | 0.046 | 1 | 5 | Tram2 | 0.005 | Old |
| 0.705631 | 0.075915 | 0.01 | 0.013 | 1 | 5 | Ptges3l | -0.003 | Old |
| 0.705824 | 0.152776 | 0.125 | 0.119 | 1 | 5 | Nkap | 0.006 | Old |
| 0.706192 | 0.251437 | 0.086 | 0.081 | 1 | 5 | Cryzl2 | 0.005 | Old |
| 0.706773 | 0.320306 | 0.051 | 0.046 | 1 | 5 | Mrgbp | 0.005 | Old |
| 0.707441 | 0.325615 | 0.061 | 0.056 | 1 | 5 | Slc20a2 | 0.005 | Old |
| 0.707872 | 0.230129 | 0.077 | 0.086 | 1 | 5 | Ddx27 | -0.009 | Old |
| 0.708076 | 0.183016 | 0.059 | 0.053 | 1 | 5 | Mast2 | 0.006 | Old |
| 0.708133 | 0.001005 | 0.109 | 0.102 | 1 | 5 | Ppard | 0.007 | Old |
| 0.708141 | 0.068333 | 0.963 | 0.926 | 1 | 5 | Crip1 | 0.037 | Old |
| 0.708415 | 0.159046 | 0.017 | 0.02 | 1 | 5 | Tubd1 | -0.003 | Old |
| 0.708425 | 0.488513 | 0.01 | 0.013 | 1 | 5 | Gm48551 | -0.003 | Old |
| 0.708958 | 0.043586 | 0.045 | 0.041 | 1 | 5 | Oplah | 0.004 | Old |
| 0.709203 | 0.344831 | 0.056 | 0.051 | 1 | 5 | Il1rap | 0.005 | Old |
| 0.709251 | 0.039678 | 0.145 | 0.137 | 1 | 5 | Prmt2 | 0.008 | Old |
| 0.709358 | 0.112173 | 0.01 | 0.013 | 1 | 5 | Cercam | -0.003 | Old |
| 0.710093 | 0.00203 | 0.431 | 0.411 | 1 | 5 | Aff4 | 0.02 | Old |
| 0.710254 | 0.029933 | 0.029 | 0.033 | 1 | 5 | Lrfn1 | -0.004 | Old |
| 0.710314 | 0.082563 | 0.04 | 0.036 | 1 | 5 | Osgin1 | 0.004 | Old |
| 0.710447 | 0.021454 | 0.158 | 0.15 | 1 | 5 | Atp11c | 0.008 | Old |
| 0.711224 | 0.040528 | 0.01 | 0.013 | 1 | 5 | Tgtp2 | -0.003 | Old |
| 0.711284 | 0.031978 | 0.279 | 0.272 | 1 | 5 | Safb | 0.007 | Old |
| 0.712014 | 0.295764 | 0.071 | 0.066 | 1 | 5 | Mad1l1 | 0.005 | Old |
| 0.712233 | 0.365929 | 0.051 | 0.046 | 1 | 5 | Ogfod3 | 0.005 | Old |
| 0.712469 | 0.049926 | 0.239 | 0.254 | 1 | 5 | Cyp4f18 | -0.015 | Old |
| 0.712706 | 0.036531 | 0.589 | 0.591 | 1 | 5 | Lpl | -0.002 | Old |
| 0.713459 | 0.101969 | 0.069 | 0.063 | 1 | 5 | Atm | 0.006 | Old |
| 0.713738 | 0.035269 | 0.488 | 0.487 | 1 | 5 | Crk | 0.001 | Old |
| 0.714216 | 0.124983 | 0.125 | 0.117 | 1 | 5 | Nup133 | 0.008 | Old |
| 0.714379 | 0.139377 | 0.148 | 0.14 | 1 | 5 | Chst12 | 0.008 | Old |
| 0.714654 | 0.02823 | 0.394 | 0.388 | 1 | 5 | Ppp6r3 | 0.006 | Old |
| 0.714961 | 0.236446 | 0.01 | 0.013 | 1 | 5 | Ctif | -0.003 | Old |
| 0.71514 | 0.042167 | 0.33 | 0.32 | 1 | 5 | Sri | 0.01 | Old |
| 0.715853 | 0.009435 | 0.596 | 0.624 | 1 | 5 | Tmed2 | -0.028 | Old |

| | | | | | | | | |
|---|---|---|---|---|---|---|---|---|
| 0.716364 | 0.263175 | 0.01 | 0.013 | 1 | 5 | Ccdc63 | -0.003 | Old |
| 0.716642 | 0.088271 | 0.045 | 0.041 | 1 | 5 | Fastkd2 | 0.004 | Old |
| 0.716723 | 0.042361 | 0.205 | 0.198 | 1 | 5 | Mlxip | 0.007 | Old |
| 0.716843 | 0.186963 | 0.088 | 0.081 | 1 | 5 | Thyn1 | 0.007 | Old |
| 0.71711 | 0.05936 | 0.032 | 0.028 | 1 | 5 | Anks6 | 0.004 | Old |
| 0.717123 | 0.135041 | 0.045 | 0.041 | 1 | 5 | Fdft1 | 0.004 | Old |
| 0.717242 | 0.011096 | 0.221 | 0.208 | 1 | 5 | Rabl6 | 0.013 | Old |
| 0.717806 | 0.093396 | 0.082 | 0.076 | 1 | 5 | Mtmr9 | 0.006 | Old |
| 0.717856 | 0.116812 | 0.096 | 0.089 | 1 | 5 | Ms4a8a | 0.007 | Old |
| 0.717898 | 0.1012 | 0.088 | 0.081 | 1 | 5 | Dhx57 | 0.007 | Old |
| 0.718164 | 0.207244 | 0.051 | 0.046 | 1 | 5 | Sema6b | 0.005 | Old |
| 0.719095 | 0.030202 | 0.059 | 0.053 | 1 | 5 | Bclaf3 | 0.006 | Old |
| 0.719974 | 0.039003 | 0.032 | 0.028 | 1 | 5 | Rasal3 | 0.004 | Old |
| 0.720762 | 0.012489 | 0.249 | 0.241 | 1 | 5 | Snrnp200 | 0.008 | Old |
| 0.722076 | 0.093441 | 0.354 | 0.343 | 1 | 5 | Srpr | 0.011 | Old |
| 0.722225 | 0.097618 | 0.052 | 0.058 | 1 | 5 | Wdr46 | -0.006 | Old |
| 0.722234 | 0.155609 | 0.074 | 0.069 | 1 | 5 | Mcm3ap | 0.005 | Old |
| 0.722268 | 0.047697 | 0.032 | 0.028 | 1 | 5 | Zfp41 | 0.004 | Old |
| 0.722391 | 0.001971 | 0.121 | 0.114 | 1 | 5 | Pdcl | 0.007 | Old |
| 0.722436 | 0.130839 | 0.306 | 0.302 | 1 | 5 | Ankhd1 | 0.004 | Old |
| 0.722583 | 0.432568 | 0.024 | 0.02 | 1 | 5 | Oas1c | 0.004 | Old |
| 0.722841 | 0.095868 | 0.152 | 0.142 | 1 | 5 | Slc46a3 | 0.01 | Old |
| 0.723224 | 0.000716 | 0.135 | 0.145 | 1 | 5 | Trappc9 | -0.01 | Old |
| 0.723756 | 0.036669 | 0.057 | 0.063 | 1 | 5 | A530017D2 | -0.006 | Old |
| 0.724827 | 0.051634 | 1 | 1 | 1 | 5 | Psap | 0 | Old |
| 0.725029 | 0.078057 | 0.051 | 0.046 | 1 | 5 | Zfp768 | 0.005 | Old |
| 0.725325 | 0.132897 | 0.056 | 0.051 | 1 | 5 | Asb6 | 0.005 | Old |
| 0.725391 | 0.241849 | 0.061 | 0.056 | 1 | 5 | Smg8 | 0.005 | Old |
| 0.725549 | 0.17303 | 0.084 | 0.079 | 1 | 5 | Med31 | 0.005 | Old |
| 0.725714 | 0.056594 | 0.032 | 0.028 | 1 | 5 | Mmp19 | 0.004 | Old |
| 0.725754 | 0.10363 | 0.242 | 0.239 | 1 | 5 | Samd9l | 0.003 | Old |
| 0.725782 | 0.023016 | 0.077 | 0.084 | 1 | 5 | Prmt5 | -0.007 | Old |
| 0.726055 | 0.030355 | 0.397 | 0.393 | 1 | 5 | Limd2 | 0.004 | Old |
| 0.726564 | 0.047245 | 0.466 | 0.467 | 1 | 5 | Rela | -0.001 | Old |
| 0.726581 | 0.067671 | 0.093 | 0.102 | 1 | 5 | Slc35b3 | -0.009 | Old |
| 0.72679 | 0.274419 | 0.042 | 0.038 | 1 | 5 | 2700081O1 | 0.004 | Old |
| 0.726956 | 0.404867 | 0.076 | 0.071 | 1 | 5 | B9d2 | 0.005 | Old |
| 0.727279 | 0.274401 | 0.037 | 0.033 | 1 | 5 | Habp4 | 0.004 | Old |
| 0.727345 | 0.325282 | 0.066 | 0.061 | 1 | 5 | Ccdc28b | 0.005 | Old |
| 0.728376 | 0.019902 | 0.529 | 0.528 | 1 | 5 | Rock1 | 0.001 | Old |
| 0.728396 | 0.016296 | 0.045 | 0.051 | 1 | 5 | Eid3 | -0.006 | Old |
| 0.729168 | 0.283268 | 0.061 | 0.056 | 1 | 5 | Cep135 | 0.005 | Old |
| 0.729921 | 0.017888 | 0.064 | 0.058 | 1 | 5 | Slc35e2 | 0.006 | Old |
| 0.731192 | 0.0469 | 0.069 | 0.063 | 1 | 5 | Rad51d | 0.006 | Old |
| 0.732206 | 0.05913 | 0.108 | 0.102 | 1 | 5 | Fbxo30 | 0.006 | Old |

| | | | | | | | | |
|---|---|---|---|---|---|---|---|---|
| 0.732816 | 0.090446 | 0.234 | 0.228 | 1 | 5 | Hnrnph3 | 0.006 | Old |
| 0.733223 | 0.049569 | 0.253 | 0.244 | 1 | 5 | Taf10 | 0.009 | Old |
| 0.733959 | 0.135491 | 0.024 | 0.02 | 1 | 5 | Eral1 | 0.004 | Old |
| 0.734765 | 0.018974 | 0.034 | 0.038 | 1 | 5 | Abcc4 | -0.004 | Old |
| 0.735187 | 0.149028 | 0.098 | 0.107 | 1 | 5 | Las1l | -0.009 | Old |
| 0.735642 | 0.150369 | 0.076 | 0.071 | 1 | 5 | Wdr18 | 0.005 | Old |
| 0.735647 | 0.04784 | 0.182 | 0.173 | 1 | 5 | Tmem128 | 0.009 | Old |
| 0.735667 | 0.028581 | 0.146 | 0.155 | 1 | 5 | Tm2d3 | -0.009 | Old |
| 0.737625 | 0.21765 | 0.104 | 0.099 | 1 | 5 | Rnf215 | 0.005 | Old |
| 0.73816 | 0.008763 | 0.177 | 0.165 | 1 | 5 | Zfp992 | 0.012 | Old |
| 0.738633 | 0.545313 | 0.015 | 0.013 | 1 | 5 | Tm4sf5 | 0.002 | Old |
| 0.738652 | 0.032432 | 0.152 | 0.142 | 1 | 5 | Vezt | 0.01 | Old |
| 0.738677 | 0.24311 | 0.066 | 0.061 | 1 | 5 | Cep128 | 0.005 | Old |
| 0.738836 | 0.125254 | 0.264 | 0.256 | 1 | 5 | Casc3 | 0.008 | Old |
| 0.738911 | 0.02615 | 0.067 | 0.061 | 1 | 5 | Aar2 | 0.006 | Old |
| 0.739595 | 0.024673 | 0.242 | 0.234 | 1 | 5 | 2-Mar | 0.008 | Old |
| 0.739856 | 0.119272 | 0.202 | 0.195 | 1 | 5 | Rnaseh2c | 0.007 | Old |
| 0.740237 | 0.201425 | 0.204 | 0.198 | 1 | 5 | Dlg1 | 0.006 | Old |
| 0.740663 | 0.006985 | 0.554 | 0.551 | 1 | 5 | Rab11fip5 | 0.003 | Old |
| 0.741152 | 0.332555 | 0.015 | 0.013 | 1 | 5 | Tmem107 | 0.002 | Old |
| 0.741223 | 0.113466 | 0.114 | 0.107 | 1 | 5 | Lamc1 | 0.007 | Old |
| 0.741354 | 0.265993 | 0.042 | 0.038 | 1 | 5 | Ttc7b | 0.004 | Old |
| 0.742265 | 0.093414 | 0.064 | 0.058 | 1 | 5 | Zfp629 | 0.006 | Old |
| 0.742361 | 0.05662 | 0.042 | 0.038 | 1 | 5 | Syt3 | 0.004 | Old |
| 0.742399 | 0.007872 | 0.975 | 0.985 | 1 | 5 | Junb | -0.01 | Old |
| 0.742452 | 0.074573 | 0.056 | 0.051 | 1 | 5 | Kdm1b | 0.005 | Old |
| 0.742833 | 0.414653 | 0.015 | 0.013 | 1 | 5 | Osbpl10 | 0.002 | Old |
| 0.743343 | 0.105948 | 0.099 | 0.094 | 1 | 5 | Cog6 | 0.005 | Old |
| 0.743675 | 0.351529 | 0.081 | 0.076 | 1 | 5 | Rasgrp1 | 0.005 | Old |
| 0.744183 | 0.000737 | 0.397 | 0.386 | 1 | 5 | Arhgap9 | 0.011 | Old |
| 0.746199 | 0.170133 | 0.015 | 0.013 | 1 | 5 | Tmem53 | 0.002 | Old |
| 0.746556 | 0.06615 | 0.327 | 0.322 | 1 | 5 | Atxn2 | 0.005 | Old |
| 0.746582 | 0.010971 | 0.747 | 0.761 | 1 | 5 | Foxp1 | -0.014 | Old |
| 0.74662 | 0.380923 | 0.015 | 0.013 | 1 | 5 | Ltbp4 | 0.002 | Old |
| 0.746807 | 0.059485 | 0.064 | 0.058 | 1 | 5 | Pop4 | 0.006 | Old |
| 0.747639 | 0.232784 | 0.047 | 0.043 | 1 | 5 | Ccdc167 | 0.004 | Old |
| 0.747655 | 0.04994 | 0.113 | 0.107 | 1 | 5 | Erg28 | 0.006 | Old |
| 0.747884 | 0.203219 | 0.015 | 0.013 | 1 | 5 | Slc13a2 | 0.002 | Old |
| 0.747884 | 0.086694 | 0.015 | 0.013 | 1 | 5 | C130013H( | 0.002 | Old |
| 0.748503 | 0.030158 | 0.141 | 0.135 | 1 | 5 | Zdhhc17 | 0.006 | Old |
| 0.748633 | 0.009869 | 0.261 | 0.254 | 1 | 5 | Rasa3 | 0.007 | Old |
| 0.748727 | 0.339077 | 0.015 | 0.013 | 1 | 5 | Zfp28 | 0.002 | Old |
| 0.748727 | 0.17838 | 0.015 | 0.013 | 1 | 5 | A630072M | 0.002 | Old |
| 0.749121 | 0.099993 | 0.241 | 0.234 | 1 | 5 | Zmym5 | 0.007 | Old |
| 0.749224 | 0.208099 | 0.029 | 0.025 | 1 | 5 | Gm9493 | 0.004 | Old |

| | | | | | | | | |
|---|---|---|---|---|---|---|---|---|
| 0.749836 | 0.293241 | 0.029 | 0.025 | 1 | 5 | Efcab2 | 0.004 | Old |
| 0.749964 | 0.00967 | 0.227 | 0.218 | 1 | 5 | Myo7a | 0.009 | Old |
| 0.750414 | 0.14497 | 0.015 | 0.013 | 1 | 5 | Fam189b | 0.002 | Old |
| 0.750414 | 0.097339 | 0.015 | 0.013 | 1 | 5 | Mocs3 | 0.002 | Old |
| 0.75082 | 0.013599 | 0.172 | 0.165 | 1 | 5 | Fam107b | 0.007 | Old |
| 0.751452 | 0.127217 | 0.039 | 0.043 | 1 | 5 | Tmem129 | -0.004 | Old |
| 0.752139 | 0.00838 | 0.155 | 0.147 | 1 | 5 | Mmadhc | 0.008 | Old |
| 0.752665 | 0.07163 | 0.488 | 0.482 | 1 | 5 | Aldoa | 0.006 | Old |
| 0.7529 | 0.261382 | 0.081 | 0.076 | 1 | 5 | Plscr4 | 0.005 | Old |
| 0.753425 | 0.058906 | 0.234 | 0.228 | 1 | 5 | Hdac1 | 0.006 | Old |
| 0.75344 | 0.239159 | 0.077 | 0.074 | 1 | 5 | Itpkc | 0.003 | Old |
| 0.753507 | 0.218793 | 0.029 | 0.025 | 1 | 5 | Smyd2 | 0.004 | Old |
| 0.753567 | 0.068087 | 0.18 | 0.173 | 1 | 5 | Cyth3 | 0.007 | Old |
| 0.753618 | 0.029116 | 0.219 | 0.211 | 1 | 5 | Abl2 | 0.008 | Old |
| 0.753747 | 0.17347 | 0.168 | 0.165 | 1 | 5 | Plxna1 | 0.003 | Old |
| 0.753921 | 0.146289 | 0.015 | 0.018 | 1 | 5 | Mvk | -0.003 | Old |
| 0.75395 | 0.009294 | 0.096 | 0.104 | 1 | 5 | Ap4s1 | -0.008 | Old |
| 0.753983 | 0.130875 | 0.042 | 0.038 | 1 | 5 | Zfp770 | 0.004 | Old |
| 0.754119 | 0.351409 | 0.029 | 0.025 | 1 | 5 | Nectin1 | 0.004 | Old |
| 0.754795 | 0.212856 | 0.052 | 0.048 | 1 | 5 | Map3k10 | 0.004 | Old |
| 0.755092 | 0.021049 | 0.278 | 0.264 | 1 | 5 | Txnl1 | 0.014 | Old |
| 0.755543 | 0.04404 | 0.219 | 0.211 | 1 | 5 | Pdap1 | 0.008 | Old |
| 0.756357 | 0.098323 | 0.126 | 0.122 | 1 | 5 | Ccnk | 0.004 | Old |
| 0.756406 | 0.006949 | 0.328 | 0.31 | 1 | 5 | Smc1a | 0.018 | Old |
| 0.757045 | 0.080533 | 0.056 | 0.051 | 1 | 5 | Plscr1 | 0.005 | Old |
| 0.757746 | 0.01029 | 0.328 | 0.34 | 1 | 5 | Emc7 | -0.012 | Old |
| 0.758604 | 0.014431 | 0.19 | 0.183 | 1 | 5 | Opa1 | 0.007 | Old |
| 0.758607 | 0.011396 | 0.34 | 0.327 | 1 | 5 | Ino80d | 0.013 | Old |
| 0.75901 | 0.124555 | 0.197 | 0.193 | 1 | 5 | 3110043O2 | 0.004 | Old |
| 0.759355 | 0.219871 | 0.052 | 0.048 | 1 | 5 | Ip6k2 | 0.004 | Old |
| 0.759465 | 0.021426 | 0.359 | 0.348 | 1 | 5 | Rpl36al | 0.011 | Old |
| 0.759811 | 0.218436 | 0.052 | 0.048 | 1 | 5 | Nprl2 | 0.004 | Old |
| 0.760258 | 0.01872 | 0.015 | 0.018 | 1 | 5 | Morn2 | -0.003 | Old |
| 0.760354 | 0.22447 | 0.088 | 0.084 | 1 | 5 | Retsat | 0.004 | Old |
| 0.760953 | 0.167184 | 0.052 | 0.048 | 1 | 5 | Gmppb | 0.004 | Old |
| 0.761051 | 0.084888 | 0.015 | 0.018 | 1 | 5 | 3300005D0 | -0.003 | Old |
| 0.761669 | 0.348822 | 0.034 | 0.03 | 1 | 5 | Btd | 0.004 | Old |
| 0.761986 | 0.059874 | 0.155 | 0.165 | 1 | 5 | Qrich1 | -0.01 | Old |
| 0.762094 | 0.019705 | 0.029 | 0.025 | 1 | 5 | Hspa1l | 0.004 | Old |
| 0.762649 | 0.217806 | 0.108 | 0.104 | 1 | 5 | Acot2 | 0.004 | Old |
| 0.762875 | 0.09291 | 0.057 | 0.053 | 1 | 5 | Ankrd13d | 0.004 | Old |
| 0.763081 | 0.048185 | 0.259 | 0.246 | 1 | 5 | Sod2 | 0.013 | Old |
| 0.763263 | 0.039342 | 0.14 | 0.15 | 1 | 5 | Tango2 | -0.01 | Old |
| 0.763429 | 0.084209 | 0.076 | 0.071 | 1 | 5 | Pwp1 | 0.005 | Old |
| 0.764227 | 0.199878 | 0.015 | 0.018 | 1 | 5 | Bend3 | -0.003 | Old |

| | | | | | | | | |
|---|---|---|---|---|---|---|---|---|
| 0.764901 | 0.014266 | 0.143 | 0.137 | 1 | 5 | Med10 | 0.006 | Old |
| 0.765152 | 0.329383 | 0.039 | 0.043 | 1 | 5 | Prkca | -0.004 | Old |
| 0.765535 | 0.145726 | 0.027 | 0.03 | 1 | 5 | Rmdn1 | -0.003 | Old |
| 0.765816 | 0.380807 | 0.015 | 0.018 | 1 | 5 | Alkbh2 | -0.003 | Old |
| 0.76602 | 0.166359 | 0.153 | 0.15 | 1 | 5 | Tubb4b | 0.003 | Old |
| 0.766123 | 0.09095 | 0.084 | 0.079 | 1 | 5 | Emc4 | 0.005 | Old |
| 0.767779 | 0.426114 | 0.02 | 0.018 | 1 | 5 | 2510017J1 | 0.002 | Old |
| 0.768526 | 0.006501 | 0.707 | 0.683 | 1 | 5 | Ehd1 | 0.024 | Old |
| 0.76854 | 0.111774 | 0.062 | 0.058 | 1 | 5 | Slc46a1 | 0.004 | Old |
| 0.768907 | 0.081422 | 0.135 | 0.129 | 1 | 5 | Entpd7 | 0.006 | Old |
| 0.769198 | 0.078266 | 0.239 | 0.234 | 1 | 5 | Fam114a2 | 0.005 | Old |
| 0.769409 | 0.125822 | 0.185 | 0.18 | 1 | 5 | Nfkbib | 0.005 | Old |
| 0.769721 | 0.139663 | 0.16 | 0.155 | 1 | 5 | Nrf1 | 0.005 | Old |
| 0.76981 | 0.026232 | 0.123 | 0.117 | 1 | 5 | Ankrd10 | 0.006 | Old |
| 0.770459 | 0.326779 | 0.039 | 0.036 | 1 | 5 | Nlrx1 | 0.003 | Old |
| 0.770656 | 0.135422 | 0.116 | 0.112 | 1 | 5 | AY036118 | 0.004 | Old |
| 0.771033 | 0.162106 | 0.091 | 0.086 | 1 | 5 | Zfyve1 | 0.005 | Old |
| 0.771387 | 0.086107 | 0.153 | 0.147 | 1 | 5 | Parp2 | 0.006 | Old |
| 0.771437 | 0.817097 | 0.02 | 0.018 | 1 | 5 | Trim59 | 0.002 | Old |
| 0.771775 | 0.257485 | 0.042 | 0.038 | 1 | 5 | Pdk4 | 0.004 | Old |
| 0.772169 | 0.496124 | 0.02 | 0.018 | 1 | 5 | Trmo | 0.002 | Old |
| 0.772445 | 0.179708 | 0.261 | 0.261 | 1 | 5 | Abcd1 | 0 | Old |
| 0.772545 | 0.014181 | 0.114 | 0.109 | 1 | 5 | Ubxn7 | 0.005 | Old |
| 0.772837 | 0.02462 | 0.195 | 0.188 | 1 | 5 | Idnk | 0.007 | Old |
| 0.773099 | 0.183111 | 0.077 | 0.074 | 1 | 5 | Adi1 | 0.003 | Old |
| 0.773163 | 0.179797 | 0.044 | 0.041 | 1 | 5 | Micall1 | 0.003 | Old |
| 0.773403 | 0.047127 | 0.076 | 0.071 | 1 | 5 | Mir17hg | 0.005 | Old |
| 0.773501 | 0.308784 | 0.098 | 0.094 | 1 | 5 | Ap4m1 | 0.004 | Old |
| 0.773578 | 0.082085 | 0.034 | 0.03 | 1 | 5 | Prkcg | 0.004 | Old |
| 0.774402 | 0.003289 | 0.133 | 0.127 | 1 | 5 | Fundc2 | 0.006 | Old |
| 0.7751 | 0.305077 | 0.02 | 0.018 | 1 | 5 | Gcat | 0.002 | Old |
| 0.775473 | 0.009932 | 0.645 | 0.627 | 1 | 5 | Sumo2 | 0.018 | Old |
| 0.776021 | 0.125472 | 0.103 | 0.099 | 1 | 5 | Decr1 | 0.004 | Old |
| 0.776458 | 0.068301 | 0.044 | 0.048 | 1 | 5 | Alg13 | -0.004 | Old |
| 0.776567 | 0.255292 | 0.02 | 0.018 | 1 | 5 | Phf7 | 0.002 | Old |
| 0.776585 | 0.049815 | 0.167 | 0.16 | 1 | 5 | Crkl | 0.007 | Old |
| 0.778135 | 0.125001 | 0.052 | 0.048 | 1 | 5 | Uxt | 0.004 | Old |
| 0.778154 | 0.164076 | 0.044 | 0.041 | 1 | 5 | Apbb1 | 0.003 | Old |
| 0.778192 | 0.172996 | 0.19 | 0.188 | 1 | 5 | Slirp | 0.002 | Old |
| 0.778372 | 0.003944 | 0.756 | 0.734 | 1 | 5 | Plek | 0.022 | Old |
| 0.778798 | 0.071445 | 0.091 | 0.086 | 1 | 5 | Metap1d | 0.005 | Old |
| 0.779186 | 0.058851 | 0.19 | 0.183 | 1 | 5 | Magoh | 0.007 | Old |
| 0.780085 | 0.030173 | 0.434 | 0.424 | 1 | 5 | Tbc1d14 | 0.01 | Old |
| 0.780407 | 0.08086 | 0.034 | 0.03 | 1 | 5 | Ddr2 | 0.004 | Old |
| 0.780639 | 0.023625 | 0.281 | 0.277 | 1 | 5 | Snrpd1 | 0.004 | Old |

| | | | | | | | | |
|---|---|---|---|---|---|---|---|---|
| 0.781455 | 0.08182 | 0.008 | 0.01 | 1 | 5 | Ubox5 | -0.002 | Old |
| 0.781455 | 0.03943 | 0.008 | 0.01 | 1 | 5 | Adamts4 | -0.002 | Old |
| 0.781877 | 0.149571 | 0.086 | 0.081 | 1 | 5 | Cep63 | 0.005 | Old |
| 0.782047 | 0.115874 | 0.135 | 0.145 | 1 | 5 | AI413582 | -0.01 | Old |
| 0.782058 | 0.091559 | 0.086 | 0.081 | 1 | 5 | Ccnc | 0.005 | Old |
| 0.782492 | 0.039373 | 0.231 | 0.226 | 1 | 5 | Sdhb | 0.005 | Old |
| 0.782518 | 0.095531 | 0.008 | 0.01 | 1 | 5 | Pkia | -0.002 | Old |
| 0.782518 | 0.08398 | 0.008 | 0.01 | 1 | 5 | Asap2 | -0.002 | Old |
| 0.782518 | 0.010031 | 0.008 | 0.01 | 1 | 5 | Fam102a | -0.002 | Old |
| 0.783177 | 0.492126 | 0.02 | 0.018 | 1 | 5 | Slc16a9 | 0.002 | Old |
| 0.783177 | 0.151596 | 0.02 | 0.018 | 1 | 5 | Ahrr | 0.002 | Old |
| 0.783581 | 0.116982 | 0.008 | 0.01 | 1 | 5 | Gpr137c | -0.002 | Old |
| 0.783581 | 0.110996 | 0.008 | 0.01 | 1 | 5 | C330027C( | -0.002 | Old |
| 0.783581 | 0.008685 | 0.008 | 0.01 | 1 | 5 | Zik1 | -0.002 | Old |
| 0.783581 | 0.003721 | 0.008 | 0.01 | 1 | 5 | Dnph1 | -0.002 | Old |
| 0.783659 | 0.024422 | 0.136 | 0.129 | 1 | 5 | Idua | 0.007 | Old |
| 0.783678 | 0.056437 | 0.232 | 0.249 | 1 | 5 | Arhgap5 | -0.017 | Old |
| 0.783912 | 0.022033 | 0.02 | 0.018 | 1 | 5 | Hykk | 0.002 | Old |
| 0.784029 | 0.555823 | 0.025 | 0.023 | 1 | 5 | Zfp995 | 0.002 | Old |
| 0.784398 | 0.198173 | 0.084 | 0.091 | 1 | 5 | Usb1 | -0.007 | Old |
| 0.784644 | 0.168037 | 0.008 | 0.01 | 1 | 5 | Padi3 | -0.002 | Old |
| 0.784644 | 0.066067 | 0.008 | 0.01 | 1 | 5 | Catsperg1 | -0.002 | Old |
| 0.784974 | 0.038864 | 0.072 | 0.069 | 1 | 5 | N6amt1 | 0.003 | Old |
| 0.785166 | 0.063365 | 0.138 | 0.132 | 1 | 5 | Tfec | 0.006 | Old |
| 0.785224 | 0.05705 | 0.148 | 0.142 | 1 | 5 | Esrra | 0.006 | Old |
| 0.78559 | 0.097808 | 0.02 | 0.023 | 1 | 5 | Gpr85 | -0.003 | Old |
| 0.785692 | 0.146809 | 0.057 | 0.053 | 1 | 5 | Dubr | 0.004 | Old |
| 0.78612 | 0.084467 | 0.02 | 0.018 | 1 | 5 | Gm13373 | 0.002 | Old |
| 0.786326 | 0.021489 | 0.271 | 0.282 | 1 | 5 | Ssrp1 | -0.011 | Old |
| 0.786337 | 0.028309 | 0.184 | 0.178 | 1 | 5 | Pou2f1 | 0.006 | Old |
| 0.786372 | 0.12724 | 0.113 | 0.109 | 1 | 5 | H2-Q6 | 0.004 | Old |
| 0.787047 | 0.015737 | 0.306 | 0.297 | 1 | 5 | Tpcn1 | 0.009 | Old |
| 0.787049 | 0.110418 | 0.21 | 0.208 | 1 | 5 | Gabpb2 | 0.002 | Old |
| 0.787522 | 0.083652 | 0.081 | 0.076 | 1 | 5 | Ap3m2 | 0.005 | Old |
| 0.788172 | 0.267887 | 0.093 | 0.089 | 1 | 5 | Atp11a | 0.004 | Old |
| 0.788515 | 0.039255 | 0.108 | 0.102 | 1 | 5 | Gucd1 | 0.006 | Old |
| 0.788967 | 0.060777 | 0.034 | 0.03 | 1 | 5 | Asb10 | 0.004 | Old |
| 0.789408 | 0.111056 | 0.184 | 0.18 | 1 | 5 | Tmem147 | 0.004 | Old |
| 0.789802 | 0.065894 | 0.02 | 0.018 | 1 | 5 | D130040H: | 0.002 | Old |
| 0.790499 | 0.067668 | 0.02 | 0.023 | 1 | 5 | Pus7l | -0.003 | Old |
| 0.790936 | 0.145287 | 0.098 | 0.094 | 1 | 5 | Wiz | 0.004 | Old |
| 0.791424 | 0.034484 | 0.327 | 0.315 | 1 | 5 | Ndufb4 | 0.012 | Old |
| 0.791688 | 0.074238 | 0.118 | 0.112 | 1 | 5 | Odf2 | 0.006 | Old |
| 0.791855 | 0.007373 | 0.271 | 0.279 | 1 | 5 | Chd8 | -0.008 | Old |
| 0.792674 | 0.013001 | 0.402 | 0.429 | 1 | 5 | Pcmtd1 | -0.027 | Old |

| | | | | | | | | |
|---|---|---|---|---|---|---|---|---|
| 0.79349 | 0.033531 | 0.02 | 0.018 | 1 | 5 | Gm4707 | 0.002 | Old |
| 0.793588 | 0.146409 | 0.135 | 0.132 | 1 | 5 | Map3k4 | 0.003 | Old |
| 0.794011 | 0.238233 | 0.02 | 0.023 | 1 | 5 | Psat1 | -0.003 | Old |
| 0.794046 | 0.187708 | 0.141 | 0.137 | 1 | 5 | Ric1 | 0.004 | Old |
| 0.794228 | 0.005503 | 0.02 | 0.018 | 1 | 5 | Trmt61a | 0.002 | Old |
| 0.794547 | 0.376172 | 0.025 | 0.023 | 1 | 5 | Ugt1a6a | 0.002 | Old |
| 0.794686 | 0.077535 | 0.044 | 0.041 | 1 | 5 | Mon1a | 0.003 | Old |
| 0.794725 | 0.145839 | 0.062 | 0.058 | 1 | 5 | Rrn3 | 0.004 | Old |
| 0.795324 | 0.014036 | 0.168 | 0.157 | 1 | 5 | Zdhhc21 | 0.011 | Old |
| 0.795816 | 0.073001 | 0.072 | 0.069 | 1 | 5 | Lxn | 0.003 | Old |
| 0.796327 | 0.051725 | 0.204 | 0.195 | 1 | 5 | Acsl3 | 0.009 | Old |
| 0.796523 | 0.188901 | 0.025 | 0.023 | 1 | 5 | Paqr4 | 0.002 | Old |
| 0.796582 | 0.031438 | 0.067 | 0.063 | 1 | 5 | Cactin | 0.004 | Old |
| 0.797112 | 0.050156 | 0.074 | 0.079 | 1 | 5 | Fggy | -0.005 | Old |
| 0.797437 | 0.225062 | 0.059 | 0.056 | 1 | 5 | Msh6 | 0.003 | Old |
| 0.797605 | 0.15568 | 0.067 | 0.063 | 1 | 5 | Nufip1 | 0.004 | Old |
| 0.797643 | 0.08723 | 0.047 | 0.051 | 1 | 5 | Rbm19 | -0.004 | Old |
| 0.797792 | 0.622027 | 0.012 | 0.01 | 1 | 5 | Zfp870 | 0.002 | Old |
| 0.797841 | 0.368541 | 0.025 | 0.023 | 1 | 5 | Gm15564 | 0.002 | Old |
| 0.798031 | 0.173049 | 0.081 | 0.076 | 1 | 5 | Cpped1 | 0.005 | Old |
| 0.798198 | 0.055412 | 0.251 | 0.266 | 1 | 5 | Stt3b | -0.015 | Old |
| 0.798218 | 0.056425 | 0.067 | 0.063 | 1 | 5 | Rbbp9 | 0.004 | Old |
| 0.798702 | 0.543562 | 0.03 | 0.028 | 1 | 5 | Ccdc171 | 0.002 | Old |
| 0.799413 | 0.007853 | 0.99 | 0.995 | 1 | 5 | Dab2 | -0.005 | Old |
| 0.800476 | 0.277457 | 0.054 | 0.051 | 1 | 5 | Fpgs | 0.003 | Old |
| 0.800663 | 0.052555 | 0.325 | 0.33 | 1 | 5 | Dennd1b | -0.005 | Old |
| 0.800696 | 0.751822 | 0.012 | 0.01 | 1 | 5 | Etohd2 | 0.002 | Old |
| 0.800696 | 0.587517 | 0.012 | 0.01 | 1 | 5 | Stbd1 | 0.002 | Old |
| 0.800696 | 0.460506 | 0.012 | 0.01 | 1 | 5 | Prss23 | 0.002 | Old |
| 0.801139 | 0.201579 | 0.025 | 0.023 | 1 | 5 | Gm10605 | 0.002 | Old |
| 0.80141 | 0.219268 | 0.084 | 0.081 | 1 | 5 | 2210016L2 | 0.003 | Old |
| 0.802041 | 0.06228 | 0.057 | 0.053 | 1 | 5 | 2900089D1 | 0.004 | Old |
| 0.803744 | 0.055269 | 0.173 | 0.168 | 1 | 5 | Trappc1 | 0.005 | Old |
| 0.804441 | 0.056299 | 0.025 | 0.023 | 1 | 5 | 4933406I1 | 0.002 | Old |
| 0.804571 | 0.52876 | 0.012 | 0.01 | 1 | 5 | 4930522L1 | 0.002 | Old |
| 0.804571 | 0.369369 | 0.012 | 0.01 | 1 | 5 | Tm4sf1 | 0.002 | Old |
| 0.805341 | 0.105173 | 0.082 | 0.079 | 1 | 5 | Zfp593 | 0.003 | Old |
| 0.806016 | 0.038181 | 0.182 | 0.175 | 1 | 5 | Fam3a | 0.007 | Old |
| 0.806063 | 0.039138 | 0.657 | 0.629 | 1 | 5 | Hpgds | 0.028 | Old |
| 0.808452 | 0.289558 | 0.012 | 0.01 | 1 | 5 | Msrb3 | 0.002 | Old |
| 0.808452 | 0.225598 | 0.012 | 0.01 | 1 | 5 | Plin1 | 0.002 | Old |
| 0.808452 | 0.158941 | 0.012 | 0.01 | 1 | 5 | Itpr1 | 0.002 | Old |
| 0.808452 | 0.090735 | 0.012 | 0.01 | 1 | 5 | Pcx | 0.002 | Old |
| 0.808564 | 0.022629 | 0.116 | 0.112 | 1 | 5 | Dph3 | 0.004 | Old |
| 0.808944 | 0.472139 | 0.03 | 0.028 | 1 | 5 | Cbx8 | 0.002 | Old |

| | | | | | | | | |
|---|---|---|---|---|---|---|---|---|
| 0.809231 | 0.043734 | 0.025 | 0.028 | 1 | 5 | Exog | -0.003 | Old |
| 0.809399 | 0.032 | 0.414 | 0.409 | 1 | 5 | Elavl1 | 0.005 | Old |
| 0.809422 | 0.21057 | 0.012 | 0.01 | 1 | 5 | Tm9sf1 | 0.002 | Old |
| 0.810231 | 0.211981 | 0.114 | 0.112 | 1 | 5 | Ttc1 | 0.002 | Old |
| 0.810274 | 0.076857 | 0.259 | 0.256 | 1 | 5 | Fam172a | 0.003 | Old |
| 0.810371 | 0.086244 | 0.145 | 0.152 | 1 | 5 | Emc8 | -0.007 | Old |
| 0.810394 | 0.282564 | 0.012 | 0.01 | 1 | 5 | 1700086PC | 0.002 | Old |
| 0.810394 | 0.253226 | 0.012 | 0.01 | 1 | 5 | Lbx1 | 0.002 | Old |
| 0.810394 | 0.205989 | 0.012 | 0.01 | 1 | 5 | Ctps | 0.002 | Old |
| 0.810781 | 0.087883 | 0.251 | 0.249 | 1 | 5 | Mpp6 | 0.002 | Old |
| 0.811006 | 0.088524 | 0.116 | 0.124 | 1 | 5 | Ambra1 | -0.008 | Old |
| 0.811055 | 0.082738 | 0.025 | 0.023 | 1 | 5 | AW209491 | 0.002 | Old |
| 0.811061 | 0.050548 | 0.305 | 0.299 | 1 | 5 | Snrpc | 0.006 | Old |
| 0.811415 | 0.079466 | 0.143 | 0.137 | 1 | 5 | Mfsd12 | 0.006 | Old |
| 0.811574 | 0.028443 | 0.103 | 0.099 | 1 | 5 | Oma1 | 0.004 | Old |
| 0.811752 | 0.053238 | 0.118 | 0.114 | 1 | 5 | Fkrp | 0.004 | Old |
| 0.812038 | 0.140054 | 0.064 | 0.061 | 1 | 5 | Ints2 | 0.003 | Old |
| 0.812066 | 0.000899 | 0.941 | 0.901 | 1 | 5 | Pltp | 0.04 | Old |
| 0.812263 | 0.163981 | 0.059 | 0.056 | 1 | 5 | Ap5z1 | 0.003 | Old |
| 0.812337 | 0.08368 | 0.012 | 0.01 | 1 | 5 | Rwdd2a | 0.002 | Old |
| 0.812566 | 0.337478 | 0.03 | 0.028 | 1 | 5 | 1810024BC | 0.002 | Old |
| 0.813465 | 0.244347 | 0.074 | 0.071 | 1 | 5 | Lias | 0.003 | Old |
| 0.813895 | 0.021283 | 0.111 | 0.107 | 1 | 5 | Fbxw7 | 0.004 | Old |
| 0.813911 | 0.000921 | 0.495 | 0.47 | 1 | 5 | Rsrc2 | 0.025 | Old |
| 0.814147 | 0.075918 | 0.278 | 0.269 | 1 | 5 | Derl2 | 0.009 | Old |
| 0.814149 | 0.04088 | 0.101 | 0.096 | 1 | 5 | Trmt6 | 0.005 | Old |
| 0.81417 | 0.011234 | 0.744 | 0.741 | 1 | 5 | Pou2f2 | 0.003 | Old |
| 0.814916 | 0.006765 | 0.057 | 0.053 | 1 | 5 | Prune1 | 0.004 | Old |
| 0.815254 | 0.094766 | 0.012 | 0.01 | 1 | 5 | Ifi205 | 0.002 | Old |
| 0.816699 | 0.019528 | 0.983 | 0.972 | 1 | 5 | Zeb2 | 0.011 | Old |
| 0.81687 | 0.217126 | 0.158 | 0.157 | 1 | 5 | Exoc2 | 0.001 | Old |
| 0.817719 | 0.034859 | 0.204 | 0.211 | 1 | 5 | Irs2 | -0.007 | Old |
| 0.818329 | 0.065461 | 0.103 | 0.099 | 1 | 5 | Alg1 | 0.004 | Old |
| 0.818649 | 0.338814 | 0.035 | 0.033 | 1 | 5 | Ccdc84 | 0.002 | Old |
| 0.818726 | 0.051565 | 0.232 | 0.223 | 1 | 5 | Scamp3 | 0.009 | Old |
| 0.818886 | 0.059403 | 0.042 | 0.046 | 1 | 5 | Smyd5 | -0.004 | Old |
| 0.819102 | 0.070062 | 0.271 | 0.282 | 1 | 5 | Rabep1 | -0.011 | Old |
| 0.819209 | 0.177495 | 0.035 | 0.033 | 1 | 5 | Bphl | 0.002 | Old |
| 0.819823 | 0.122135 | 0.03 | 0.028 | 1 | 5 | Nav2 | 0.002 | Old |
| 0.820608 | 0.021417 | 0.219 | 0.231 | 1 | 5 | Rnft1 | -0.012 | Old |
| 0.82094 | 0.167217 | 0.013 | 0.015 | 1 | 5 | Ccdc173 | -0.002 | Old |
| 0.821005 | 0.194867 | 0.054 | 0.051 | 1 | 5 | BC049352 | 0.003 | Old |
| 0.821263 | 0.05531 | 0.098 | 0.094 | 1 | 5 | Isoc1 | 0.004 | Old |
| 0.823335 | 0.091676 | 0.148 | 0.145 | 1 | 5 | Pkp4 | 0.003 | Old |
| 0.823536 | 0.008388 | 0.013 | 0.015 | 1 | 5 | Tnni2 | -0.002 | Old |

| | | | | | | | | |
|---|---|---|---|---|---|---|---|---|
| 0.824767 | 0.097501 | 0.104 | 0.102 | 1 | 5 | Xpo4 | 0.002 | Old |
| 0.825363 | 0.066181 | 0.076 | 0.071 | 1 | 5 | Coro2a | 0.005 | Old |
| 0.825754 | 0.11373 | 0.072 | 0.076 | 1 | 5 | Dhx29 | -0.004 | Old |
| 0.826285 | 0.145599 | 0.049 | 0.046 | 1 | 5 | BC024978 | 0.003 | Old |
| 0.826433 | 0.130791 | 0.268 | 0.266 | 1 | 5 | Btbd7 | 0.002 | Old |
| 0.826696 | 0.04841 | 0.197 | 0.193 | 1 | 5 | Map3k5 | 0.004 | Old |
| 0.826976 | 0.067928 | 0.072 | 0.069 | 1 | 5 | Msantd2 | 0.003 | Old |
| 0.827459 | 0.013802 | 0.214 | 0.223 | 1 | 5 | Sh3gl1 | -0.009 | Old |
| 0.827468 | 0.141701 | 0.086 | 0.081 | 1 | 5 | Tspan13 | 0.005 | Old |
| 0.827469 | 0.113967 | 0.135 | 0.132 | 1 | 5 | Mtf2 | 0.003 | Old |
| 0.827867 | 0.047351 | 0.013 | 0.015 | 1 | 5 | Nlrp1a | -0.002 | Old |
| 0.828169 | 0.09134 | 0.131 | 0.14 | 1 | 5 | Ints9 | -0.009 | Old |
| 0.829318 | 0.079191 | 0.067 | 0.071 | 1 | 5 | Gpr89 | -0.004 | Old |
| 0.829523 | 0.123698 | 0.03 | 0.028 | 1 | 5 | Smim1 | 0.002 | Old |
| 0.829567 | 0.065676 | 0.165 | 0.162 | 1 | 5 | Srsf4 | 0.003 | Old |
| 0.829635 | 0.13763 | 0.108 | 0.104 | 1 | 5 | 2010315B0 | 0.004 | Old |
| 0.829923 | 0.050983 | 0.079 | 0.076 | 1 | 5 | Knop1 | 0.003 | Old |
| 0.83008 | 0.050167 | 0.49 | 0.482 | 1 | 5 | Ppp1r12c | 0.008 | Old |
| 0.830168 | 0.322356 | 0.042 | 0.046 | 1 | 5 | Twsg1 | -0.004 | Old |
| 0.830232 | 0.112709 | 0.059 | 0.056 | 1 | 5 | Utp4 | 0.003 | Old |
| 0.831191 | 0.016163 | 0.062 | 0.066 | 1 | 5 | Tmc6 | -0.004 | Old |
| 0.831312 | 0.13669 | 0.116 | 0.114 | 1 | 5 | Faim | 0.002 | Old |
| 0.831335 | 0.160981 | 0.013 | 0.015 | 1 | 5 | Polrmt | -0.002 | Old |
| 0.831414 | 0.168047 | 0.069 | 0.066 | 1 | 5 | Cep44 | 0.003 | Old |
| 0.832124 | 0.229 | 0.035 | 0.033 | 1 | 5 | Lrrc49 | 0.002 | Old |
| 0.832203 | 0.207243 | 0.013 | 0.015 | 1 | 5 | Phf19 | -0.002 | Old |
| 0.832429 | 0.173123 | 0.059 | 0.056 | 1 | 5 | Pde12 | 0.003 | Old |
| 0.83335 | 0.287441 | 0.086 | 0.084 | 1 | 5 | Ptgr1 | 0.002 | Old |
| 0.833475 | 0.011455 | 0.131 | 0.137 | 1 | 5 | Cyth2 | -0.006 | Old |
| 0.834807 | 0.403454 | 0.013 | 0.015 | 1 | 5 | Efemp1 | -0.002 | Old |
| 0.83541 | 0.320438 | 0.017 | 0.015 | 1 | 5 | Zbtb25 | 0.002 | Old |
| 0.836112 | 0.071894 | 0.04 | 0.038 | 1 | 5 | Qtrt2 | 0.002 | Old |
| 0.836142 | 0.060111 | 0.322 | 0.317 | 1 | 5 | Vav1 | 0.005 | Old |
| 0.836234 | 0.005152 | 0.077 | 0.081 | 1 | 5 | Ap2a1 | -0.004 | Old |
| 0.836585 | 0.123746 | 0.074 | 0.079 | 1 | 5 | Ralgps1 | -0.005 | Old |
| 0.837466 | 0.244059 | 0.045 | 0.043 | 1 | 5 | Rtca | 0.002 | Old |
| 0.837851 | 0.520918 | 0.017 | 0.015 | 1 | 5 | 1700037C1 | 0.002 | Old |
| 0.837851 | 0.268937 | 0.017 | 0.015 | 1 | 5 | Fam92a | 0.002 | Old |
| 0.837851 | 0.179666 | 0.017 | 0.015 | 1 | 5 | Adal | 0.002 | Old |
| 0.838665 | 0.262053 | 0.017 | 0.015 | 1 | 5 | Sox12 | 0.002 | Old |
| 0.838665 | 0.182576 | 0.017 | 0.015 | 1 | 5 | Cdc42ep5 | 0.002 | Old |
| 0.839222 | 0.026877 | 0.342 | 0.335 | 1 | 5 | Map3k8 | 0.007 | Old |
| 0.83925 | 0.011961 | 0.03 | 0.028 | 1 | 5 | Urb2 | 0.002 | Old |
| 0.839257 | 0.067617 | 0.12 | 0.114 | 1 | 5 | Smarcal1 | 0.006 | Old |
| 0.839479 | 0.154449 | 0.017 | 0.015 | 1 | 5 | Cd22 | 0.002 | Old |

| | | | | | | | | |
|---|---|---|---|---|---|---|---|---|
| 0.841107 | 0.202613 | 0.017 | 0.015 | 1 | 5 | Ube2v1 | 0.002 | Old |
| 0.841788 | 0.033351 | 0.177 | 0.175 | 1 | 5 | Ndufs1 | 0.002 | Old |
| 0.841889 | 0.040205 | 0.418 | 0.419 | 1 | 5 | Ppig | -0.001 | Old |
| 0.841922 | 0.241968 | 0.017 | 0.015 | 1 | 5 | Siah1b | 0.002 | Old |
| 0.842042 | 0.063504 | 0.322 | 0.327 | 1 | 5 | Batf3 | -0.005 | Old |
| 0.842736 | 0.074548 | 0.017 | 0.015 | 1 | 5 | Crip2 | 0.002 | Old |
| 0.842839 | 0.273683 | 0.056 | 0.053 | 1 | 5 | Trp53i13 | 0.003 | Old |
| 0.843001 | 0.051254 | 0.113 | 0.109 | 1 | 5 | Pcsk7 | 0.004 | Old |
| 0.843116 | 0.359896 | 0.03 | 0.033 | 1 | 5 | Mtif3 | -0.003 | Old |
| 0.843906 | 0.085068 | 0.064 | 0.061 | 1 | 5 | Mettl3 | 0.003 | Old |
| 0.844519 | 0.007001 | 0.035 | 0.033 | 1 | 5 | Bri3bp | 0.002 | Old |
| 0.84482 | 0.153423 | 0.04 | 0.038 | 1 | 5 | Mpst | 0.002 | Old |
| 0.845445 | 0.033004 | 0.155 | 0.152 | 1 | 5 | Cyp20a1 | 0.003 | Old |
| 0.84626 | 0.073215 | 0.173 | 0.17 | 1 | 5 | Fam133b | 0.003 | Old |
| 0.846323 | 0.023088 | 0.101 | 0.096 | 1 | 5 | Nit1 | 0.005 | Old |
| 0.84646 | 0.038679 | 0.136 | 0.132 | 1 | 5 | Pfdn4 | 0.004 | Old |
| 0.846861 | 0.128605 | 0.074 | 0.079 | 1 | 5 | Acvr1b | -0.005 | Old |
| 0.847031 | 0.125697 | 0.116 | 0.114 | 1 | 5 | Farsb | 0.002 | Old |
| 0.847866 | 0.413571 | 0.022 | 0.02 | 1 | 5 | Ndufaf5 | 0.002 | Old |
| 0.84858 | 0.2931 | 0.022 | 0.02 | 1 | 5 | Mamld1 | 0.002 | Old |
| 0.849628 | 0.283951 | 0.056 | 0.053 | 1 | 5 | Cdc42ep4 | 0.003 | Old |
| 0.850076 | 0.044844 | 0.017 | 0.015 | 1 | 5 | Fndc10 | 0.002 | Old |
| 0.850265 | 0.034203 | 0.184 | 0.19 | 1 | 5 | Cln6 | -0.006 | Old |
| 0.850429 | 0.228973 | 0.045 | 0.043 | 1 | 5 | C1qtnf12 | 0.002 | Old |
| 0.851478 | 0.018488 | 0.261 | 0.254 | 1 | 5 | Plau | 0.007 | Old |
| 0.851564 | 0.071471 | 0.019 | 0.02 | 1 | 5 | Scd2 | -0.001 | Old |
| 0.851564 | 0.062542 | 0.019 | 0.02 | 1 | 5 | Ehd2 | -0.001 | Old |
| 0.851802 | 0.051889 | 0.148 | 0.145 | 1 | 5 | Acadsb | 0.003 | Old |
| 0.852055 | 0.089089 | 0.177 | 0.173 | 1 | 5 | Maml1 | 0.004 | Old |
| 0.852771 | 0.29515 | 0.052 | 0.051 | 1 | 5 | Haus6 | 0.001 | Old |
| 0.853097 | 0.463567 | 0.027 | 0.025 | 1 | 5 | Phf11a | 0.002 | Old |
| 0.853342 | 0.088241 | 0.017 | 0.015 | 1 | 5 | Tarsl2 | 0.002 | Old |
| 0.853581 | 0.376077 | 0.022 | 0.02 | 1 | 5 | Snapc4 | 0.002 | Old |
| 0.853816 | 0.706796 | 0.019 | 0.02 | 1 | 5 | Serpine1 | -0.001 | Old |
| 0.853892 | 0.092967 | 0.057 | 0.061 | 1 | 5 | Xpnpep1 | -0.004 | Old |
| 0.854419 | 0.095488 | 0.207 | 0.206 | 1 | 5 | Gatad1 | 0.001 | Old |
| 0.854605 | 0.165419 | 0.04 | 0.038 | 1 | 5 | Ccp110 | 0.002 | Old |
| 0.855016 | 0.089132 | 0.212 | 0.211 | 1 | 5 | Zfp385a | 0.001 | Old |
| 0.855664 | 0.083388 | 0.04 | 0.038 | 1 | 5 | Ercc1 | 0.002 | Old |
| 0.856442 | 0.317194 | 0.022 | 0.02 | 1 | 5 | Gm14286 | 0.002 | Old |
| 0.856724 | 0.098343 | 0.04 | 0.038 | 1 | 5 | Zfp846 | 0.002 | Old |
| 0.856827 | 0.232285 | 0.064 | 0.069 | 1 | 5 | Wrb | -0.005 | Old |
| 0.856882 | 0.130476 | 0.056 | 0.053 | 1 | 5 | Mrps18b | 0.003 | Old |
| 0.858376 | 0.010908 | 0.308 | 0.312 | 1 | 5 | Cnih4 | -0.004 | Old |
| 0.858588 | 0.480468 | 0.022 | 0.02 | 1 | 5 | Gm26510 | 0.002 | Old |

| | | | | | | | | |
|---|---|---|---|---|---|---|---|---|
| 0.858588 | 0.334112 | 0.022 | 0.02 | 1 | 5 | Lins1 | 0.002 | Old |
| 0.859342 | 0.084945 | 0.187 | 0.185 | 1 | 5 | Clcn7 | 0.002 | Old |
| 0.859401 | 0.260671 | 0.061 | 0.058 | 1 | 5 | Sgf29 | 0.003 | Old |
| 0.859785 | 0.013105 | 0.079 | 0.084 | 1 | 5 | Mpg | -0.005 | Old |
| 0.860054 | 0.078655 | 0.061 | 0.058 | 1 | 5 | Nxpe3 | 0.003 | Old |
| 0.860441 | 0.00551 | 0.088 | 0.091 | 1 | 5 | D430042O( | -0.003 | Old |
| 0.861279 | 0.078922 | 0.214 | 0.211 | 1 | 5 | Pdcd4 | 0.003 | Old |
| 0.862418 | 0.454282 | 0.062 | 0.061 | 1 | 5 | Srbd1 | 0.001 | Old |
| 0.862885 | 0.161978 | 0.022 | 0.02 | 1 | 5 | AC163018. | 0.002 | Old |
| 0.863396 | 0.031994 | 0.305 | 0.31 | 1 | 5 | Tmx1 | -0.005 | Old |
| 0.863601 | 0.293509 | 0.022 | 0.02 | 1 | 5 | Armc6 | 0.002 | Old |
| 0.863908 | 0.013217 | 0.103 | 0.099 | 1 | 5 | Khk | 0.004 | Old |
| 0.864286 | 0.02868 | 0.214 | 0.206 | 1 | 5 | Oser1 | 0.008 | Old |
| 0.865349 | 0.000249 | 0.099 | 0.096 | 1 | 5 | Mak16 | 0.003 | Old |
| 0.865757 | 0.034372 | 0.237 | 0.244 | 1 | 5 | Fgfr1 | -0.007 | Old |
| 0.865967 | 0.038277 | 0.093 | 0.089 | 1 | 5 | Atic | 0.004 | Old |
| 0.866017 | 0.045615 | 0.492 | 0.495 | 1 | 5 | Klf13 | -0.003 | Old |
| 0.866421 | 0.041209 | 0.056 | 0.053 | 1 | 5 | Trip10 | 0.003 | Old |
| 0.866468 | 0.172061 | 0.022 | 0.02 | 1 | 5 | Phlpp1 | 0.002 | Old |
| 0.86681 | 0.094796 | 0.061 | 0.058 | 1 | 5 | Ubap2 | 0.003 | Old |
| 0.86707 | 0.074095 | 0.094 | 0.099 | 1 | 5 | Exosc9 | -0.005 | Old |
| 0.867185 | 0.174802 | 0.022 | 0.02 | 1 | 5 | Ap5b1 | 0.002 | Old |
| 0.868026 | 0.085462 | 0.194 | 0.193 | 1 | 5 | Tpd52l2 | 0.001 | Old |
| 0.868183 | 0.028753 | 0.12 | 0.117 | 1 | 5 | Coq8b | 0.003 | Old |
| 0.868461 | 0.034887 | 0.2 | 0.198 | 1 | 5 | Golgb1 | 0.002 | Old |
| 0.86915 | 0.20632 | 0.056 | 0.053 | 1 | 5 | Drosha | 0.003 | Old |
| 0.869337 | 0.0147 | 0.022 | 0.02 | 1 | 5 | Haus1 | 0.002 | Old |
| 0.869977 | 0.096263 | 0.194 | 0.193 | 1 | 5 | Psmd14 | 0.001 | Old |
| 0.870536 | 0.080612 | 0.254 | 0.259 | 1 | 5 | Sertad2 | -0.005 | Old |
| 0.870586 | 0.073281 | 0.224 | 0.234 | 1 | 5 | Srd5a3 | -0.01 | Old |
| 0.872052 | 0.034966 | 0.158 | 0.165 | 1 | 5 | Tatdn2 | -0.007 | Old |
| 0.872649 | 0.286154 | 0.12 | 0.119 | 1 | 5 | Dph5 | 0.001 | Old |
| 0.872789 | 0.269656 | 0.153 | 0.155 | 1 | 5 | Amn1 | -0.002 | Old |
| 0.872862 | 0.058358 | 0.029 | 0.03 | 1 | 5 | Pvt1 | -0.001 | Old |
| 0.873122 | 0.047649 | 0.389 | 0.391 | 1 | 5 | Rreb1 | -0.002 | Old |
| 0.873357 | 0.114657 | 0.061 | 0.058 | 1 | 5 | Efl1 | 0.003 | Old |
| 0.873635 | 0.004012 | 0.066 | 0.063 | 1 | 5 | Uqcc1 | 0.003 | Old |
| 0.875953 | 0.049395 | 0.279 | 0.282 | 1 | 5 | Eprs | -0.003 | Old |
| 0.876166 | 0.029644 | 0.061 | 0.063 | 1 | 5 | Zfp160 | -0.002 | Old |
| 0.877091 | 0.076968 | 0.034 | 0.036 | 1 | 5 | Msrb2 | -0.002 | Old |
| 0.877321 | 0.024474 | 0.364 | 0.358 | 1 | 5 | Tpp2 | 0.006 | Old |
| 0.878197 | 0.006906 | 0.795 | 0.759 | 1 | 5 | Atp5a1 | 0.036 | Old |
| 0.87856 | 0.016277 | 0.071 | 0.069 | 1 | 5 | Akt3 | 0.002 | Old |
| 0.8795 | 0.137161 | 0.125 | 0.122 | 1 | 5 | Uchl3 | 0.003 | Old |
| 0.87958 | 0.003926 | 0.027 | 0.025 | 1 | 5 | Trmt10b | 0.002 | Old |

| | | | | | | | | |
|---|---|---|---|---|---|---|---|---|
| 0.879834 | 0.008108 | 0.239 | 0.244 | 1 | 5 | Dnajb11 | -0.005 | Old |
| 0.881425 | 0.052469 | 0.202 | 0.198 | 1 | 5 | Atxn1 | 0.004 | Old |
| 0.881674 | 0.131585 | 0.182 | 0.183 | 1 | 5 | Sirt7 | -0.001 | Old |
| 0.88168 | 0.058891 | 0.18 | 0.178 | 1 | 5 | Mrps36 | 0.002 | Old |
| 0.882225 | 0.076475 | 0.032 | 0.03 | 1 | 5 | Mlst8 | 0.002 | Old |
| 0.882228 | 0.08389 | 0.219 | 0.218 | 1 | 5 | Chd6 | 0.001 | Old |
| 0.882349 | 0.029174 | 0.458 | 0.454 | 1 | 5 | Adam19 | 0.004 | Old |
| 0.882419 | 0.007682 | 0.082 | 0.086 | 1 | 5 | Snhg6 | -0.004 | Old |
| 0.882686 | 0.109825 | 0.029 | 0.03 | 1 | 5 | Gm31718 | -0.001 | Old |
| 0.882861 | 0.005775 | 0.185 | 0.183 | 1 | 5 | Erap1 | 0.002 | Old |
| 0.883217 | 0.004259 | 0.845 | 0.853 | 1 | 5 | Snx5 | -0.008 | Old |
| 0.883316 | 0.053666 | 0.066 | 0.063 | 1 | 5 | Zfp760 | 0.003 | Old |
| 0.883615 | 0.115308 | 0.051 | 0.053 | 1 | 5 | Surf2 | -0.002 | Old |
| 0.883727 | 0.223626 | 0.037 | 0.036 | 1 | 5 | Taf9b | 0.001 | Old |
| 0.884125 | 0.187777 | 0.057 | 0.056 | 1 | 5 | Gm16536 | 0.001 | Old |
| 0.884893 | 0.077742 | 0.13 | 0.129 | 1 | 5 | Mis12 | 0.001 | Old |
| 0.885145 | 0.026982 | 0.029 | 0.03 | 1 | 5 | Snap47 | -0.001 | Old |
| 0.885412 | 0.147416 | 0.027 | 0.025 | 1 | 5 | Nmb | 0.002 | Old |
| 0.885796 | 0.042734 | 0.032 | 0.03 | 1 | 5 | Mapkapk5 | 0.002 | Old |
| 0.886094 | 0.087096 | 0.032 | 0.03 | 1 | 5 | Kcng2 | 0.002 | Old |
| 0.886305 | 0.035858 | 0.268 | 0.274 | 1 | 5 | Sft2d1 | -0.006 | Old |
| 0.886392 | 0.109991 | 0.032 | 0.03 | 1 | 5 | 1810032O( | 0.002 | Old |
| 0.886915 | 0.019492 | 0.061 | 0.058 | 1 | 5 | Gnl2 | 0.003 | Old |
| 0.887836 | 0.165539 | 0.051 | 0.053 | 1 | 5 | Mettl1 | -0.002 | Old |
| 0.888837 | 0.071367 | 0.145 | 0.142 | 1 | 5 | Ahsa2 | 0.003 | Old |
| 0.889046 | 0.021743 | 0.805 | 0.81 | 1 | 5 | Gltp | -0.005 | Old |
| 0.889186 | 0.018331 | 0.167 | 0.173 | 1 | 5 | Trnt1 | -0.006 | Old |
| 0.890066 | 0.082998 | 0.184 | 0.183 | 1 | 5 | Klhl6 | 0.001 | Old |
| 0.89017 | 0.008401 | 0.217 | 0.216 | 1 | 5 | Cul5 | 0.001 | Old |
| 0.890577 | 0.074668 | 0.067 | 0.066 | 1 | 5 | Fbxl19 | 0.001 | Old |
| 0.891784 | 0.017327 | 1 | 0.997 | 1 | 5 | Serinc3 | 0.003 | Old |
| 0.892109 | 0.140315 | 0.062 | 0.061 | 1 | 5 | Cdk5rap2 | 0.001 | Old |
| 0.892349 | 0.044424 | 0.032 | 0.03 | 1 | 5 | Slc22a17 | 0.002 | Old |
| 0.892493 | 0.031664 | 0.668 | 0.665 | 1 | 5 | Rnase4 | 0.003 | Old |
| 0.892586 | 0.013709 | 0.17 | 0.175 | 1 | 5 | Man1b1 | -0.005 | Old |
| 0.893066 | 0.187391 | 0.042 | 0.041 | 1 | 5 | Nt5m | 0.001 | Old |
| 0.893125 | 0.228048 | 0.088 | 0.086 | 1 | 5 | Pck2 | 0.002 | Old |
| 0.893148 | 0.168723 | 0.037 | 0.036 | 1 | 5 | 9030025P2 | 0.001 | Old |
| 0.893368 | 0.075362 | 0.077 | 0.076 | 1 | 5 | Med22 | 0.001 | Old |
| 0.89355 | 0.003188 | 0.054 | 0.056 | 1 | 5 | Ncdn | -0.002 | Old |
| 0.895022 | 0.086032 | 0.061 | 0.058 | 1 | 5 | Nsun3 | 0.003 | Old |
| 0.896002 | 0.0202 | 0.076 | 0.074 | 1 | 5 | Usp10 | 0.002 | Old |
| 0.89775 | 0.097641 | 0.082 | 0.081 | 1 | 5 | Mov10 | 0.001 | Old |
| 0.897968 | 0.01271 | 0.274 | 0.269 | 1 | 5 | Dguok | 0.005 | Old |
| 0.898685 | 0.032458 | 0.424 | 0.437 | 1 | 5 | Vdac2 | -0.013 | Old |

| | | | | | | | | |
|---|---|---|---|---|---|---|---|---|
| 0.898988 | 0.012312 | 0.566 | 0.586 | 1 | 5 | Fxyd5 | -0.02 | Old |
| 0.899001 | 0.093832 | 0.047 | 0.046 | 1 | 5 | Pcgf6 | 0.001 | Old |
| 0.89994 | 0.120607 | 0.066 | 0.069 | 1 | 5 | Riox2 | -0.003 | Old |
| 0.900807 | 0.083017 | 0.114 | 0.114 | 1 | 5 | Mdn1 | 0 | Old |
| 0.900855 | 0.053005 | 0.125 | 0.129 | 1 | 5 | Apeh | -0.004 | Old |
| 0.901056 | 0.073732 | 0.365 | 0.368 | 1 | 5 | Anxa7 | -0.003 | Old |
| 0.901539 | 0.001573 | 0.081 | 0.084 | 1 | 5 | Eml3 | -0.003 | Old |
| 0.901623 | 0.020731 | 0.012 | 0.013 | 1 | 5 | Stard10 | -0.001 | Old |
| 0.90232 | 0.097273 | 0.057 | 0.056 | 1 | 5 | Foxred1 | 0.001 | Old |
| 0.902397 | 0.178434 | 0.052 | 0.051 | 1 | 5 | Elac2 | 0.001 | Old |
| 0.902496 | 0.018022 | 0.098 | 0.096 | 1 | 5 | Acvr1 | 0.002 | Old |
| 0.902573 | 0.06171 | 0.012 | 0.013 | 1 | 5 | Pdxp | -0.001 | Old |
| 0.902573 | 0.01587 | 0.012 | 0.013 | 1 | 5 | Layn | -0.001 | Old |
| 0.902666 | 0.029577 | 0.167 | 0.175 | 1 | 5 | Pkd1 | -0.008 | Old |
| 0.902822 | 0.026446 | 0.131 | 0.132 | 1 | 5 | Lrpprc | -0.001 | Old |
| 0.903524 | 0.04529 | 0.012 | 0.013 | 1 | 5 | Sgtb | -0.001 | Old |
| 0.903933 | 0.015805 | 0.295 | 0.287 | 1 | 5 | Eif3l | 0.008 | Old |
| 0.903939 | 0.047715 | 0.047 | 0.046 | 1 | 5 | Rcl1 | 0.001 | Old |
| 0.903973 | 0.105205 | 0.113 | 0.112 | 1 | 5 | Nosip | 0.001 | Old |
| 0.904265 | 0.011391 | 0.465 | 0.47 | 1 | 5 | Atp6v0d1 | -0.005 | Old |
| 0.904474 | 0.090893 | 0.012 | 0.013 | 1 | 5 | Mocos | -0.001 | Old |
| 0.904679 | 0.064561 | 0.076 | 0.079 | 1 | 5 | Nt5dc2 | -0.003 | Old |
| 0.904801 | 0.165484 | 0.042 | 0.041 | 1 | 5 | Ifi208 | 0.001 | Old |
| 0.905425 | 0.300909 | 0.012 | 0.013 | 1 | 5 | Lgals4 | -0.001 | Old |
| 0.905808 | 0.066924 | 0.357 | 0.355 | 1 | 5 | Slk | 0.002 | Old |
| 0.905873 | 0.261178 | 0.013 | 0.013 | 1 | 5 | Tmem238 | 0 | Old |
| 0.905948 | 0.138702 | 0.29 | 0.302 | 1 | 5 | Pdgfb | -0.012 | Old |
| 0.906111 | 0.765211 | 0.019 | 0.018 | 1 | 5 | Serpinh1 | 0.001 | Old |
| 0.906769 | 0.091762 | 0.049 | 0.051 | 1 | 5 | Rab39 | -0.002 | Old |
| 0.906787 | 0.369421 | 0.013 | 0.013 | 1 | 5 | Xrcc5 | 0 | Old |
| 0.906937 | 0.007591 | 0.419 | 0.429 | 1 | 5 | Icam1 | -0.01 | Old |
| 0.90715 | 0.153509 | 0.093 | 0.096 | 1 | 5 | Zfp157 | -0.003 | Old |
| 0.907327 | 0.153929 | 0.012 | 0.013 | 1 | 5 | Zfp59 | -0.001 | Old |
| 0.907327 | 0.140877 | 0.012 | 0.013 | 1 | 5 | Tmem69 | -0.001 | Old |
| 0.907769 | 0.048155 | 0.101 | 0.104 | 1 | 5 | Abcd4 | -0.003 | Old |
| 0.908111 | 0.027211 | 0.724 | 0.744 | 1 | 5 | Cbl | -0.02 | Old |
| 0.908278 | 0.214123 | 0.012 | 0.013 | 1 | 5 | Gm49085 | -0.001 | Old |
| 0.908434 | 0.003231 | 0.727 | 0.728 | 1 | 5 | Tob2 | -0.001 | Old |
| 0.908616 | 0.404216 | 0.013 | 0.013 | 1 | 5 | Gm10602 | 0 | Old |
| 0.908853 | 0.055275 | 0.162 | 0.162 | 1 | 5 | Trmt1l | 0 | Old |
| 0.909013 | 0.010493 | 0.236 | 0.236 | 1 | 5 | Mprip | 0 | Old |
| 0.909295 | 0.108679 | 0.091 | 0.094 | 1 | 5 | Lrrc57 | -0.003 | Old |
| 0.909458 | 0.037973 | 0.017 | 0.018 | 1 | 5 | Dgat2 | -0.001 | Old |
| 0.90953 | 0.478812 | 0.013 | 0.013 | 1 | 5 | Gm47389 | 0 | Old |
| 0.90953 | 0.449163 | 0.013 | 0.013 | 1 | 5 | Neil2 | 0 | Old |

| | | | | | | | | |
|---|---|---|---|---|---|---|---|---|
| 0.910029 | 0.023823 | 0.133 | 0.132 | 1 | 5 | Ccdc97 | 0.001 | Old |
| 0.910445 | 0.368921 | 0.013 | 0.013 | 1 | 5 | Fah | 0 | Old |
| 0.910445 | 0.193688 | 0.013 | 0.013 | 1 | 5 | Ly6c1 | 0 | Old |
| 0.910918 | 0.086426 | 0.071 | 0.074 | 1 | 5 | Zxdb | -0.003 | Old |
| 0.91136 | 0.18995 | 0.013 | 0.013 | 1 | 5 | Timeless | 0 | Old |
| 0.911933 | 0.122107 | 0.088 | 0.086 | 1 | 5 | Alg9 | 0.002 | Old |
| 0.912118 | 0.07919 | 0.086 | 0.089 | 1 | 5 | Pafah1b3 | -0.003 | Old |
| 0.912274 | 0.081713 | 0.013 | 0.013 | 1 | 5 | Gpr35 | 0 | Old |
| 0.913189 | 0.032436 | 0.013 | 0.013 | 1 | 5 | Rps6ka2 | 0 | Old |
| 0.913466 | 0.101141 | 0.017 | 0.018 | 1 | 5 | Thnsl1 | -0.001 | Old |
| 0.913891 | 0.005469 | 0.448 | 0.437 | 1 | 5 | Sirt2 | 0.011 | Old |
| 0.914854 | 0.096062 | 0.128 | 0.132 | 1 | 5 | Naa10 | -0.004 | Old |
| 0.914928 | 0.065624 | 0.049 | 0.051 | 1 | 5 | 2610021A0 | -0.002 | Old |
| 0.91502 | 0.27209 | 0.013 | 0.013 | 1 | 5 | Dnaaf5 | 0 | Old |
| 0.91502 | 0.22334 | 0.013 | 0.013 | 1 | 5 | Mtrf1 | 0 | Old |
| 0.91502 | 0.132165 | 0.013 | 0.013 | 1 | 5 | Ubtd2 | 0 | Old |
| 0.915021 | 0.116316 | 0.062 | 0.061 | 1 | 5 | Bin2 | 0.001 | Old |
| 0.915311 | 0.062638 | 0.047 | 0.046 | 1 | 5 | Spout1 | 0.001 | Old |
| 0.915462 | 0.526203 | 0.019 | 0.018 | 1 | 5 | Gm9796 | 0.001 | Old |
| 0.915491 | 0.05966 | 0.098 | 0.096 | 1 | 5 | Ell | 0.002 | Old |
| 0.915935 | 0.140945 | 0.013 | 0.013 | 1 | 5 | Zfp677 | 0 | Old |
| 0.915935 | 0.002713 | 0.013 | 0.013 | 1 | 5 | 4930404N1 | 0 | Old |
| 0.916067 | 0.016439 | 0.197 | 0.195 | 1 | 5 | Sh3bp2 | 0.002 | Old |
| 0.916674 | 0.015708 | 0.017 | 0.018 | 1 | 5 | Magi3 | -0.001 | Old |
| 0.917476 | 0.009763 | 0.017 | 0.018 | 1 | 5 | Ankrd24 | -0.001 | Old |
| 0.917802 | 0.298978 | 0.019 | 0.018 | 1 | 5 | Ccdc138 | 0.001 | Old |
| 0.918233 | 0.053695 | 0.082 | 0.081 | 1 | 5 | Supt3 | 0.001 | Old |
| 0.918559 | 0.179114 | 0.044 | 0.046 | 1 | 5 | Ipo4 | -0.002 | Old |
| 0.919363 | 0.217319 | 0.019 | 0.018 | 1 | 5 | Slc7a5 | 0.001 | Old |
| 0.919597 | 0.012332 | 0.013 | 0.013 | 1 | 5 | Dync2h1 | 0 | Old |
| 0.920525 | 0.059299 | 0.234 | 0.231 | 1 | 5 | Ddx21 | 0.003 | Old |
| 0.920807 | 0.029557 | 0.165 | 0.162 | 1 | 5 | Kdm3b | 0.003 | Old |
| 0.921088 | 0.218566 | 0.044 | 0.046 | 1 | 5 | Polr1a | -0.002 | Old |
| 0.921168 | 0.082099 | 0.246 | 0.246 | 1 | 5 | Nmt1 | 0 | Old |
| 0.921185 | 0.33069 | 0.024 | 0.023 | 1 | 5 | Zbtb48 | 0.001 | Old |
| 0.921953 | 0.048696 | 0.062 | 0.061 | 1 | 5 | Rap1gap2 | 0.001 | Old |
| 0.922265 | 0.306405 | 0.044 | 0.043 | 1 | 5 | Kif3c | 0.001 | Old |
| 0.922291 | 0.22648 | 0.017 | 0.018 | 1 | 5 | Nr6a1 | -0.001 | Old |
| 0.922603 | 0.024388 | 0.062 | 0.061 | 1 | 5 | Pik3r2 | 0.001 | Old |
| 0.922644 | 0.026101 | 0.109 | 0.109 | 1 | 5 | Gmeb1 | 0 | Old |
| 0.922912 | 0.05933 | 0.121 | 0.124 | 1 | 5 | Ola1 | -0.003 | Old |
| 0.923954 | 0.321067 | 0.024 | 0.023 | 1 | 5 | Prr5l | 0.001 | Old |
| 0.924319 | 0.445362 | 0.039 | 0.038 | 1 | 5 | Aasdh | 0.001 | Old |
| 0.925376 | 0.031268 | 0.064 | 0.066 | 1 | 5 | Rdm1 | -0.002 | Old |
| 0.925489 | 0.052178 | 0.15 | 0.15 | 1 | 5 | Tbc1d31 | 0 | Old |

| | | | | | | | | |
|---|---|---|---|---|---|---|---|---|
| 0.925624 | 0.059125 | 0.054 | 0.056 | 1 | 5 | Ammecr1 | -0.002 | Old |
| 0.925866 | 0.033461 | 0.364 | 0.363 | 1 | 5 | Grk3 | 0.001 | Old |
| 0.925917 | 0.070226 | 0.12 | 0.119 | 1 | 5 | Bin3 | 0.001 | Old |
| 0.92666 | 0.03801 | 0.074 | 0.076 | 1 | 5 | Gtpbp6 | -0.002 | Old |
| 0.92689 | 0.249854 | 0.022 | 0.023 | 1 | 5 | Slc2a3 | -0.001 | Old |
| 0.92689 | 0.205138 | 0.022 | 0.023 | 1 | 5 | Setd6 | -0.001 | Old |
| 0.92689 | 0.043165 | 0.022 | 0.023 | 1 | 5 | Hspg2 | -0.001 | Old |
| 0.927818 | 0.221863 | 0.113 | 0.114 | 1 | 5 | Ankrd46 | -0.001 | Old |
| 0.927949 | 0.013213 | 0.019 | 0.018 | 1 | 5 | Rfc5 | 0.001 | Old |
| 0.928636 | 0.026539 | 0.394 | 0.396 | 1 | 5 | Dyrk2 | -0.002 | Old |
| 0.928715 | 0.336273 | 0.017 | 0.018 | 1 | 5 | Pafah2 | -0.001 | Old |
| 0.928715 | 0.267749 | 0.017 | 0.018 | 1 | 5 | Gm12166 | -0.001 | Old |
| 0.929181 | 0.196732 | 0.034 | 0.033 | 1 | 5 | Tgif2 | 0.001 | Old |
| 0.929935 | 0.04043 | 0.311 | 0.305 | 1 | 5 | Puf60 | 0.006 | Old |
| 0.930352 | 0.374866 | 0.049 | 0.048 | 1 | 5 | Rpgrip1 | 0.001 | Old |
| 0.930958 | 0.052789 | 0.279 | 0.279 | 1 | 5 | Bcl7b | 0 | Old |
| 0.931817 | 0.126574 | 0.093 | 0.091 | 1 | 5 | Rfxap | 0.002 | Old |
| 0.931846 | 0.058611 | 0.022 | 0.023 | 1 | 5 | Rabl3 | -0.001 | Old |
| 0.931855 | 0.005761 | 0.019 | 0.018 | 1 | 5 | Stard13 | 0.001 | Old |
| 0.931989 | 0.056168 | 0.347 | 0.353 | 1 | 5 | Tsen34 | -0.006 | Old |
| 0.931996 | 0.068363 | 0.311 | 0.315 | 1 | 5 | Lamtor3 | -0.004 | Old |
| 0.932975 | 0.074542 | 0.054 | 0.056 | 1 | 5 | Fbxo45 | -0.002 | Old |
| 0.933719 | 0.03063 | 0.288 | 0.292 | 1 | 5 | Tle3 | -0.004 | Old |
| 0.93378 | 0.212307 | 0.074 | 0.074 | 1 | 5 | Mettl14 | 0 | Old |
| 0.934141 | 0.02895 | 0.027 | 0.028 | 1 | 5 | Abhd11 | -0.001 | Old |
| 0.934141 | 0.009959 | 0.027 | 0.028 | 1 | 5 | Hes7 | -0.001 | Old |
| 0.934346 | 0.005287 | 0.024 | 0.023 | 1 | 5 | Fadd | 0.001 | Old |
| 0.935009 | 0.105247 | 0.066 | 0.069 | 1 | 5 | Ppm1f | -0.003 | Old |
| 0.935292 | 0.102268 | 0.029 | 0.028 | 1 | 5 | Lmnb2 | 0.001 | Old |
| 0.935789 | 0.081729 | 0.047 | 0.048 | 1 | 5 | Pwwp2b | -0.001 | Old |
| 0.936528 | 0.094155 | 0.3 | 0.305 | 1 | 5 | Cnot6 | -0.005 | Old |
| 0.936719 | 0.079262 | 0.094 | 0.096 | 1 | 5 | Pctp | -0.002 | Old |
| 0.937588 | 0.095389 | 0.155 | 0.155 | 1 | 5 | Srp68 | 0 | Old |
| 0.937623 | 0.277586 | 0.044 | 0.043 | 1 | 5 | Wbp1 | 0.001 | Old |
| 0.9379 | 0.180873 | 0.039 | 0.038 | 1 | 5 | Wfdc18 | 0.001 | Old |
| 0.938222 | 0.318366 | 0.022 | 0.023 | 1 | 5 | Gm12689 | -0.001 | Old |
| 0.938372 | 0.080484 | 0.049 | 0.048 | 1 | 5 | Nit2 | 0.001 | Old |
| 0.938419 | 0.099658 | 0.114 | 0.114 | 1 | 5 | Pigu | 0 | Old |
| 0.938765 | 0.147306 | 0.099 | 0.099 | 1 | 5 | Fam214a | 0 | Old |
| 0.939064 | 0.085836 | 0.347 | 0.355 | 1 | 5 | Pcf11 | -0.008 | Old |
| 0.939069 | 0.256101 | 0.034 | 0.033 | 1 | 5 | Diras2 | 0.001 | Old |
| 0.939093 | 0.100552 | 0.167 | 0.168 | 1 | 5 | Ppp2r3d | -0.001 | Old |
| 0.9392 | 0.281252 | 0.024 | 0.023 | 1 | 5 | Gm45871 | 0.001 | Old |
| 0.93927 | 0.09407 | 0.027 | 0.028 | 1 | 5 | Noa1 | -0.001 | Old |
| 0.939582 | 0.18179 | 0.136 | 0.142 | 1 | 5 | Slc25a46 | -0.006 | Old |

| | | | | | | | | |
|---|---|---|---|---|---|---|---|---|
| 0.940225 | 0.029631 | 0.032 | 0.033 | 1 | 5 | 4632404H1 | -0.001 | Old |
| 0.940287 | 0.072285 | 0.042 | 0.043 | 1 | 5 | Them4 | -0.001 | Old |
| 0.940587 | 0.013834 | 0.024 | 0.023 | 1 | 5 | Ccdc94 | 0.001 | Old |
| 0.941195 | 0.19455 | 0.027 | 0.028 | 1 | 5 | Zdhhc1 | -0.001 | Old |
| 0.941397 | 0.182478 | 0.034 | 0.033 | 1 | 5 | Strbp | 0.001 | Old |
| 0.941589 | 0.064127 | 0.126 | 0.129 | 1 | 5 | Gmppa | -0.003 | Old |
| 0.941766 | 0.327018 | 0.022 | 0.023 | 1 | 5 | Ptchd1 | -0.001 | Old |
| 0.941974 | 0.061379 | 0.024 | 0.023 | 1 | 5 | Il17rd | 0.001 | Old |
| 0.941979 | 0.115741 | 0.034 | 0.033 | 1 | 5 | Gm13431 | 0.001 | Old |
| 0.943068 | 0.017726 | 0.205 | 0.203 | 1 | 5 | Maged1 | 0.002 | Old |
| 0.943143 | 0.43614 | 0.034 | 0.033 | 1 | 5 | Ear2 | 0.001 | Old |
| 0.943343 | 0.035444 | 0.916 | 0.896 | 1 | 5 | Anxa5 | 0.02 | Old |
| 0.943515 | 0.176282 | 0.054 | 0.053 | 1 | 5 | Mrps10 | 0.001 | Old |
| 0.943882 | 0.005398 | 0.039 | 0.038 | 1 | 5 | Kbtbd4 | 0.001 | Old |
| 0.943964 | 0.192132 | 0.049 | 0.048 | 1 | 5 | Smim3 | 0.001 | Old |
| 0.94443 | 0.104118 | 0.029 | 0.028 | 1 | 5 | Chek2 | 0.001 | Old |
| 0.944493 | 0.002894 | 0.064 | 0.063 | 1 | 5 | Fam20c | 0.001 | Old |
| 0.944526 | 0.065315 | 0.059 | 0.058 | 1 | 5 | Mthfsd | 0.001 | Old |
| 0.945045 | 0.299849 | 0.027 | 0.028 | 1 | 5 | Aste1 | -0.001 | Old |
| 0.945314 | 0.006808 | 0.278 | 0.279 | 1 | 5 | Hck | -0.001 | Old |
| 0.945424 | 0.050789 | 0.049 | 0.048 | 1 | 5 | Alyref2 | 0.001 | Old |
| 0.945439 | 0.055726 | 0.374 | 0.373 | 1 | 5 | Chd9 | 0.001 | Old |
| 0.945744 | 0.057544 | 0.194 | 0.193 | 1 | 5 | Prr14 | 0.001 | Old |
| 0.945838 | 0.14326 | 0.032 | 0.033 | 1 | 5 | Vmac | -0.001 | Old |
| 0.946254 | 0.030036 | 0.512 | 0.505 | 1 | 5 | Bptf | 0.007 | Old |
| 0.946526 | 0.088265 | 0.111 | 0.114 | 1 | 5 | Ero1l | -0.003 | Old |
| 0.946952 | 0.078277 | 0.029 | 0.028 | 1 | 5 | Bbs9 | 0.001 | Old |
| 0.947292 | 0.001596 | 0.037 | 0.038 | 1 | 5 | Clybl | -0.001 | Old |
| 0.948467 | 0.002389 | 0.343 | 0.345 | 1 | 5 | Ubxn4 | -0.002 | Old |
| 0.948526 | 0.240933 | 0.076 | 0.079 | 1 | 5 | Gna11 | -0.003 | Old |
| 0.94877 | 0.31444 | 0.084 | 0.086 | 1 | 5 | Il1a | -0.002 | Old |
| 0.948857 | 0.511599 | 0.022 | 0.023 | 1 | 5 | 1110004E0 | -0.001 | Old |
| 0.949867 | 0.124113 | 0.039 | 0.038 | 1 | 5 | Pigf | 0.001 | Old |
| 0.950163 | 0.065944 | 0.113 | 0.112 | 1 | 5 | Glipr1 | 0.001 | Old |
| 0.950425 | 0.009787 | 0.173 | 0.173 | 1 | 5 | Suco | 0 | Old |
| 0.951997 | 0.128711 | 0.029 | 0.028 | 1 | 5 | D3Ertd254 | 0.001 | Old |
| 0.952061 | 0.15547 | 0.074 | 0.076 | 1 | 5 | Gm26782 | -0.002 | Old |
| 0.952635 | 0.023225 | 0.786 | 0.774 | 1 | 5 | Fnbp1 | 0.012 | Old |
| 0.952904 | 0.171566 | 0.434 | 0.439 | 1 | 5 | Fabp4 | -0.005 | Old |
| 0.953015 | 0.154411 | 0.136 | 0.137 | 1 | 5 | Brpf1 | -0.001 | Old |
| 0.953173 | 0.073533 | 0.131 | 0.132 | 1 | 5 | D10Wsu10 | -0.001 | Old |
| 0.953532 | 0.001618 | 0.263 | 0.266 | 1 | 5 | Herpud2 | -0.003 | Old |
| 0.954521 | 0.002673 | 0.212 | 0.216 | 1 | 5 | Pigp | -0.004 | Old |
| 0.954566 | 0.070461 | 0.13 | 0.132 | 1 | 5 | Cwc25 | -0.002 | Old |
| 0.955046 | 0.003649 | 0.916 | 0.914 | 1 | 5 | Sdc3 | 0.002 | Old |

| | | | | | | | | |
|---|---|---|---|---|---|---|---|---|
| 0.9562 | 0.024019 | 0.283 | 0.287 | 1 | 5 | Arpc1a | -0.004 | Old |
| 0.956261 | 0.203897 | 0.162 | 0.162 | 1 | 5 | Ecm1 | 0 | Old |
| 0.95644 | 0.056544 | 0.278 | 0.284 | 1 | 5 | Appl1 | -0.006 | Old |
| 0.956816 | 0.011339 | 0.081 | 0.081 | 1 | 5 | Btbd2 | 0 | Old |
| 0.956954 | 0.094575 | 0.923 | 0.919 | 1 | 5 | Jun | 0.004 | Old |
| 0.957257 | 0.005746 | 0.215 | 0.213 | 1 | 5 | Lrrc8a | 0.002 | Old |
| 0.95838 | 0.015234 | 0.153 | 0.152 | 1 | 5 | Ing5 | 0.001 | Old |
| 0.959544 | 0.037399 | 0.088 | 0.089 | 1 | 5 | Atr | -0.001 | Old |
| 0.962326 | 0.055342 | 0.054 | 0.053 | 1 | 5 | Ptcd3 | 0.001 | Old |
| 0.962935 | 0.006495 | 0.582 | 0.581 | 1 | 5 | Evi2a | 0.001 | Old |
| 0.963316 | 0.168644 | 0.047 | 0.048 | 1 | 5 | B4galt4 | -0.001 | Old |
| 0.963507 | 0.048783 | 0.194 | 0.193 | 1 | 5 | Lrrc8d | 0.001 | Old |
| 0.966998 | 0.024068 | 0.232 | 0.239 | 1 | 5 | Sec22b | -0.007 | Old |
| 0.968832 | 0.014103 | 0.269 | 0.269 | 1 | 5 | Ncf4 | 0 | Old |
| 0.970552 | 0.04315 | 0.084 | 0.084 | 1 | 5 | Cwc22 | 0 | Old |
| 0.970622 | 0.031085 | 0.064 | 0.063 | 1 | 5 | Pola1 | 0.001 | Old |
| 0.970681 | 0.055884 | 0.367 | 0.368 | 1 | 5 | Gbp7 | -0.001 | Old |
| 0.971138 | 0.009732 | 0.094 | 0.094 | 1 | 5 | Gm38399 | 0 | Old |
| 0.972114 | 0.350778 | 0.037 | 0.038 | 1 | 5 | Trmt10c | -0.001 | Old |
| 0.973161 | 0.010199 | 0.242 | 0.241 | 1 | 5 | Polr2f | 0.001 | Old |
| 0.973206 | 0.002024 | 0.948 | 0.947 | 1 | 5 | Cybb | 0.001 | Old |
| 0.97642 | 0.161018 | 0.072 | 0.074 | 1 | 5 | Ppcdc | -0.002 | Old |
| 0.976817 | 0.136582 | 0.099 | 0.102 | 1 | 5 | Slc41a3 | -0.003 | Old |
| 0.977748 | 0.142427 | 0.057 | 0.058 | 1 | 5 | Mospd1 | -0.001 | Old |
| 0.977871 | 0.028582 | 0.209 | 0.208 | 1 | 5 | Usp14 | 0.001 | Old |
| 0.977871 | 0.116222 | 0.135 | 0.137 | 1 | 5 | Wdr48 | -0.002 | Old |
| 0.978536 | 0.074705 | 0.067 | 0.069 | 1 | 5 | Cc2d1a | -0.002 | Old |
| 0.979452 | 0.048591 | 0.779 | 0.777 | 1 | 5 | Folr2 | 0.002 | Old |
| 0.979534 | 0.398952 | 0.084 | 0.086 | 1 | 5 | Helb | -0.002 | Old |
| 0.979813 | 0.122586 | 0.061 | 0.061 | 1 | 5 | Gga2 | 0 | Old |
| 0.980111 | 0.096119 | 0.096 | 0.096 | 1 | 5 | Strip1 | 0 | Old |
| 0.980239 | 0.031911 | 0.759 | 0.754 | 1 | 5 | Rassf4 | 0.005 | Old |
| 0.981372 | 0.036887 | 0.785 | 0.774 | 1 | 5 | Clk1 | 0.011 | Old |
| 0.981576 | 0.073928 | 0.141 | 0.142 | 1 | 5 | Atl2 | -0.001 | Old |
| 0.981919 | 0.024939 | 0.242 | 0.246 | 1 | 5 | Appbp2 | -0.004 | Old |
| 0.982513 | 0.005337 | 0.17 | 0.17 | 1 | 5 | Chfr | 0 | Old |
| 0.983901 | 0.03901 | 0.104 | 0.104 | 1 | 5 | Nup205 | 0 | Old |
| 0.983949 | 0.170406 | 0.184 | 0.19 | 1 | 5 | Chd7 | -0.006 | Old |
| 0.984364 | 0.065889 | 0.276 | 0.282 | 1 | 5 | Slc16a10 | -0.006 | Old |
| 0.984663 | 0.055767 | 0.045 | 0.046 | 1 | 5 | Gstcd | -0.001 | Old |
| 0.985191 | 0.096426 | 0.082 | 0.084 | 1 | 5 | Micu3 | -0.002 | Old |
| 0.98537 | 0.101552 | 0.04 | 0.041 | 1 | 5 | Rcc1l | -0.001 | Old |
| 0.985416 | 0.117217 | 0.066 | 0.066 | 1 | 5 | Zfp68 | 0 | Old |
| 0.985474 | 0.015079 | 0.192 | 0.193 | 1 | 5 | Ilf3 | -0.001 | Old |
| 0.986941 | 0.062439 | 0.173 | 0.175 | 1 | 5 | Snf8 | -0.002 | Old |

| | | | | | | | | |
|---|---|---|---|---|---|---|---|---|
| 0.986945 | 0.003122 | 0.076 | 0.076 | 1 | 5 | Sec23a | 0 | Old |
| 0.986945 | 0.01241 | 0.076 | 0.076 | 1 | 5 | Gon7 | 0 | Old |
| 0.987092 | 0.025966 | 0.256 | 0.256 | 1 | 5 | Ntan1 | 0 | Old |
| 0.987434 | 0.074596 | 0.088 | 0.089 | 1 | 5 | Nol4l | -0.001 | Old |
| 0.988406 | 0.069295 | 0.059 | 0.058 | 1 | 5 | Ppox | 0.001 | Old |
| 0.988408 | 0.105179 | 0.157 | 0.16 | 1 | 5 | Fbxo38 | -0.003 | Old |
| 0.988457 | 0.12211 | 0.02 | 0.02 | 1 | 5 | Nsun5 | 0 | Old |
| 0.98853 | 0.010232 | 0.333 | 0.335 | 1 | 5 | Zranb2 | -0.002 | Old |
| 0.988768 | 0.004664 | 0.071 | 0.071 | 1 | 5 | Guf1 | 0 | Old |
| 0.990488 | 0.015623 | 0.066 | 0.066 | 1 | 5 | Rnf135 | 0 | Old |
| 0.990703 | 0.05623 | 0.076 | 0.076 | 1 | 5 | Fbxl4 | 0 | Old |
| 0.991652 | 0.126977 | 0.025 | 0.025 | 1 | 5 | Acat2 | 0 | Old |
| 0.991749 | 0.218979 | 0.03 | 0.03 | 1 | 5 | Praf2 | 0 | Old |
| 0.991756 | 0.160891 | 0.066 | 0.066 | 1 | 5 | Zfyve21 | 0 | Old |
| 0.992141 | 0.019502 | 0.01 | 0.01 | 1 | 5 | Gm12258 | 0 | Old |
| 0.992382 | 0.061404 | 0.461 | 0.459 | 1 | 5 | Hspa9 | 0.002 | Old |
| 0.992708 | 0.04425 | 0.045 | 0.046 | 1 | 5 | Meis3 | -0.001 | Old |
| 0.992709 | 0.648027 | 0.015 | 0.015 | 1 | 5 | Gmpr | 0 | Old |
| 0.992852 | 0.041043 | 0.071 | 0.071 | 1 | 5 | G6pc3 | 0 | Old |
| 0.993276 | 0.012946 | 0.121 | 0.122 | 1 | 5 | Lmf2 | -0.001 | Old |
| 0.993714 | 0.201164 | 0.045 | 0.046 | 1 | 5 | Dis3 | -0.001 | Old |
| 0.994217 | 0.009908 | 0.045 | 0.046 | 1 | 5 | Arfip2 | -0.001 | Old |
| 0.994237 | 0.129945 | 0.01 | 0.01 | 1 | 5 | Gm14548 | 0 | Old |
| 0.994237 | 0.059929 | 0.01 | 0.01 | 1 | 5 | Cd27 | 0 | Old |
| 0.994425 | 0.295066 | 0.015 | 0.015 | 1 | 5 | Tmem177 | 0 | Old |
| 0.994449 | 0.298301 | 0.069 | 0.071 | 1 | 5 | Pnpla6 | -0.002 | Old |
| 0.995159 | 0.328899 | 0.02 | 0.02 | 1 | 5 | Ipp | 0 | Old |
| 0.995282 | 0.367117 | 0.015 | 0.015 | 1 | 5 | Gtf2h3 | 0 | Old |
| 0.995285 | 0.172941 | 0.01 | 0.01 | 1 | 5 | Iqgap3 | 0 | Old |
| 0.995285 | 0.091742 | 0.01 | 0.01 | 1 | 5 | Gm49249 | 0 | Old |
| 0.995285 | 0.050647 | 0.01 | 0.01 | 1 | 5 | Aldh1b1 | 0 | Old |
| 0.99538 | 0.021348 | 0.088 | 0.089 | 1 | 5 | Tmem167b | -0.001 | Old |
| 0.99565 | 0.032349 | 0.207 | 0.211 | 1 | 5 | Gm36161 | -0.004 | Old |
| 0.995745 | 0.043072 | 0.035 | 0.036 | 1 | 5 | Rrp12 | -0.001 | Old |
| 0.996119 | 0.052771 | 0.088 | 0.089 | 1 | 5 | Hectd3 | -0.001 | Old |
| 0.996332 | 0.161029 | 0.01 | 0.01 | 1 | 5 | Ccl5 | 0 | Old |
| 0.996768 | 0.018908 | 0.182 | 0.185 | 1 | 5 | Imp4 | -0.003 | Old |
| 0.996998 | 0.10393 | 0.015 | 0.015 | 1 | 5 | Tnfrsf23 | 0 | Old |
| 0.99738 | 0.122029 | 0.01 | 0.01 | 1 | 5 | Klhl11 | 0 | Old |
| 0.997499 | 0.012621 | 0.377 | 0.381 | 1 | 5 | Lmbrd1 | -0.004 | Old |
| 0.997671 | 0.053626 | 0.111 | 0.112 | 1 | 5 | Fam45a | -0.001 | Old |
| 0.997856 | 0.260806 | 0.015 | 0.015 | 1 | 5 | Ccdc141 | 0 | Old |
| 0.997856 | 0.172326 | 0.015 | 0.015 | 1 | 5 | Wdr34 | 0 | Old |
| 0.997856 | 0.172011 | 0.015 | 0.015 | 1 | 5 | 9130008F2 | 0 | Old |
| 0.997856 | 0.090477 | 0.015 | 0.015 | 1 | 5 | Phlda3 | 0 | Old |

| | | | | | | | | |
|---|---|---|---|---|---|---|---|---|
| 0.997861 | 0.131873 | 0.03 | 0.03 | 1 | 5 | Lmbr1 | 0 | Old |
| 0.998138 | 0.492174 | 0.02 | 0.02 | 1 | 5 | D3Ertd751 | 0 | Old |
| 0.998241 | 0.053542 | 0.077 | 0.079 | 1 | 5 | Mrpl55 | -0.002 | Old |
| 0.998428 | 0.740309 | 0.01 | 0.01 | 1 | 5 | Fastkd3 | 0 | Old |
| 0.998428 | 0.289645 | 0.01 | 0.01 | 1 | 5 | Zfp971 | 0 | Old |
| 0.998713 | 0.24303 | 0.015 | 0.015 | 1 | 5 | Gorab | 0 | Old |
| 0.998857 | 0.228252 | 0.056 | 0.056 | 1 | 5 | AI597479 | 0 | Old |
| 0.998883 | 0.21937 | 0.02 | 0.02 | 1 | 5 | Qprt | 0 | Old |
| 0.998883 | 0.012291 | 0.02 | 0.02 | 1 | 5 | Il27ra | 0 | Old |
| 0.99904 | 0.014538 | 0.081 | 0.081 | 1 | 5 | Tsfm | 0 | Old |
| 0.999476 | 0.44065 | 0.01 | 0.01 | 1 | 5 | Hist2h4 | 0 | Old |
| 0.999476 | 0.280617 | 0.01 | 0.01 | 1 | 5 | Zfp784 | 0 | Old |
| 0.999476 | 0.228626 | 0.01 | 0.01 | 1 | 5 | Gm9774 | 0 | Old |
| 0.999545 | 0.020213 | 0.838 | 0.838 | 1 | 5 | Pld4 | 0 | Old |
| 0.999626 | 0.032063 | 0.086 | 0.086 | 1 | 5 | Dbt | 0 | Old |
| 4.40E-29 | 3.449494 | 0.264 | 0.027 | 1.37E-24 | 5 | Ddx3y | 0.237 | Young |
| 2.02E-26 | 2.589342 | 0.358 | 0.089 | 6.29E-22 | 5 | Cx3cr1 | 0.269 | Young |
| 1.60E-21 | 0.564842 | 0.985 | 0.981 | 4.98E-17 | 5 | Lgmn | 0.004 | Young |
| 7.89E-20 | 0.379425 | 1 | 0.992 | 2.45E-15 | 5 | Rps9 | 0.008 | Young |
| 1.46E-18 | 0.364895 | 0.997 | 0.998 | 4.52E-14 | 5 | Rpl37a | -0.001 | Young |
| 2.09E-18 | 0.358011 | 0.995 | 0.99 | 6.49E-14 | 5 | Rplp2 | 0.005 | Young |
| 4.42E-13 | 0.946128 | 0.66 | 0.483 | 1.37E-08 | 5 | Tsc22d3 | 0.177 | Young |
| 5.16E-12 | 0.232482 | 1 | 1 | 1.60E-07 | 5 | Rps29 | 0 | Young |
| 6.17E-12 | 1.811573 | 0.299 | 0.128 | 1.91E-07 | 5 | Ccr1 | 0.171 | Young |
| 1.69E-10 | 0.762056 | 0.774 | 0.699 | 5.23E-06 | 5 | Abca1 | 0.075 | Young |
| 1.98E-10 | 2.232824 | 0.157 | 0.042 | 6.16E-06 | 5 | Lpcat2 | 0.115 | Young |
| 5.94E-10 | 0.309551 | 0.99 | 0.988 | 1.85E-05 | 5 | Rps28 | 0.002 | Young |
| 9.38E-10 | 0.276005 | 0.98 | 0.981 | 2.91E-05 | 5 | Rps26 | -0.001 | Young |
| 2.11E-09 | 0.2291 | 0.997 | 1 | 6.56E-05 | 5 | Tmsb4x | -0.003 | Young |
| 2.29E-09 | 2.234873 | 0.14 | 0.037 | 7.12E-05 | 5 | Mtus1 | 0.103 | Young |
| 3.10E-09 | 0.340165 | 0.99 | 0.99 | 9.63E-05 | 5 | Rps27a | 0 | Young |
| 3.62E-09 | 0.201604 | 0.997 | 0.995 | 0.000112 | 5 | Tpt1 | 0.002 | Young |
| 8.61E-09 | 0.214513 | 0.997 | 0.995 | 0.000267 | 5 | Rpl41 | 0.002 | Young |
| 1.45E-08 | 2.477023 | 0.081 | 0.01 | 0.000451 | 5 | Eif2s3y | 0.071 | Young |
| 1.80E-08 | 2.14506 | 0.112 | 0.025 | 0.000558 | 5 | Tln2 | 0.087 | Young |
| 5.71E-08 | 0.254853 | 0.97 | 0.987 | 0.001772 | 5 | Rpsa | -0.017 | Young |
| 6.16E-08 | 0.241505 | 0.987 | 0.988 | 0.001911 | 5 | Rpl38 | -0.001 | Young |
| 6.23E-08 | 2.433311 | 0.109 | 0.027 | 0.001936 | 5 | Colec12 | 0.082 | Young |
| 6.89E-08 | 0.458763 | 1 | 1 | 0.002139 | 5 | Gm42418 | 0 | Young |
| 7.39E-08 | 0.914252 | 0.327 | 0.177 | 0.002294 | 5 | Lsp1 | 0.15 | Young |
| 1.41E-07 | 0.396096 | 0.84 | 0.806 | 0.004391 | 5 | Arpc5 | 0.034 | Young |
| 2.80E-07 | 0.321243 | 0.954 | 0.938 | 0.00869 | 5 | Ahnak | 0.016 | Young |
| 3.48E-07 | 1.247654 | 0.185 | 0.077 | 0.010816 | 5 | Emp1 | 0.108 | Young |
| 3.66E-07 | 0.557489 | 0.637 | 0.508 | 0.011357 | 5 | Cd33 | 0.129 | Young |
| 3.74E-07 | 0.256279 | 0.997 | 0.995 | 0.011611 | 5 | Rpl35a | 0.002 | Young |

| | | | | | | | | |
|---|---|---|---|---|---|---|---|---|
| 3.83E-07 | 0.196268 | 0.997 | 0.992 | 0.011893 | 5 | Rps21 | 0.005 | Young |
| 7.12E-07 | 1.496438 | 0.18 | 0.077 | 0.022106 | 5 | Arsb | 0.103 | Young |
| 7.75E-07 | 0.541428 | 0.766 | 0.689 | 0.024071 | 5 | Fam46a | 0.077 | Young |
| 1.24E-06 | 0.650126 | 0.553 | 0.436 | 0.038396 | 5 | Rab31 | 0.117 | Young |
| 1.95E-06 | 0.253689 | 0.992 | 0.993 | 0.060441 | 5 | Rps14 | -0.001 | Young |
| 2.55E-06 | 0.177109 | 1 | 0.993 | 0.079072 | 5 | Rplp1 | 0.007 | Young |
| 2.94E-06 | 0.336975 | 0.919 | 0.912 | 0.091297 | 5 | BC005537 | 0.007 | Young |
| 3.60E-06 | 1.250447 | 0.142 | 0.056 | 0.111781 | 5 | Tagap | 0.086 | Young |
| 4.04E-06 | 0.247883 | 0.97 | 0.958 | 0.125516 | 5 | Rpl23a | 0.012 | Young |
| 4.32E-06 | 1.294643 | 0.251 | 0.146 | 0.134209 | 5 | Trem2 | 0.105 | Young |
| 4.43E-06 | 0.916339 | 0.34 | 0.217 | 0.137455 | 5 | S100a6 | 0.123 | Young |
| 4.73E-06 | 0.335807 | 0.848 | 0.813 | 0.147016 | 5 | Gdi2 | 0.035 | Young |
| 4.88E-06 | 0.295996 | 0.99 | 0.975 | 0.151493 | 5 | Maf | 0.015 | Young |
| 5.13E-06 | 0.344703 | 0.883 | 0.865 | 0.159213 | 5 | Cox4i1 | 0.018 | Young |
| 7.50E-06 | 0.711955 | 0.449 | 0.323 | 0.233007 | 5 | Dusp6 | 0.126 | Young |
| 8.37E-06 | 0.624573 | 0.553 | 0.431 | 0.259781 | 5 | Cd44 | 0.122 | Young |
| 9.91E-06 | 0.709339 | 0.464 | 0.348 | 0.307663 | 5 | Cttnbp2nl | 0.116 | Young |
| 1.04E-05 | 0.448252 | 0.924 | 0.906 | 0.322768 | 5 | Mafb | 0.018 | Young |
| 1.13E-05 | 0.348537 | 0.792 | 0.695 | 0.350718 | 5 | Snx2 | 0.097 | Young |
| 1.18E-05 | 1.433827 | 0.165 | 0.077 | 0.367944 | 5 | S100a4 | 0.088 | Young |
| 1.20E-05 | 1.701069 | 0.058 | 0.01 | 0.372772 | 5 | Kdm5d | 0.048 | Young |
| 1.35E-05 | 0.960349 | 0.307 | 0.197 | 0.420585 | 5 | Rbbp7 | 0.11 | Young |
| 1.65E-05 | 0.285675 | 0.954 | 0.951 | 0.511101 | 5 | Rpl22 | 0.003 | Young |
| 1.65E-05 | 0.261388 | 0.919 | 0.902 | 0.512875 | 5 | Rpl3 | 0.017 | Young |
| 1.70E-05 | 0.304919 | 0.921 | 0.923 | 0.529042 | 5 | Dazap2 | -0.002 | Young |
| 1.76E-05 | 0.673619 | 0.612 | 0.522 | 0.546258 | 5 | Ssh2 | 0.09 | Young |
| 1.92E-05 | 0.220847 | 0.995 | 0.993 | 0.595492 | 5 | Rps8 | 0.002 | Young |
| 2.01E-05 | 2.672448 | 0.056 | 0.01 | 0.622984 | 5 | Mmp14 | 0.046 | Young |
| 2.29E-05 | 0.219954 | 0.997 | 0.997 | 0.712273 | 5 | Marcks | 0 | Young |
| 3.06E-05 | 0.198244 | 0.995 | 0.997 | 0.950202 | 5 | Fau | -0.002 | Young |
| 3.23E-05 | 0.215162 | 0.957 | 0.949 | 1 | 5 | Rpl8 | 0.008 | Young |
| 3.71E-05 | 0.261034 | 0.967 | 0.953 | 1 | 5 | Rps10 | 0.014 | Young |
| 4.42E-05 | 1.149881 | 0.226 | 0.133 | 1 | 5 | Cpd | 0.093 | Young |
| 5.55E-05 | 1.153527 | 0.14 | 0.062 | 1 | 5 | Cfd | 0.078 | Young |
| 6.05E-05 | 0.240438 | 0.982 | 0.985 | 1 | 5 | Ptma | -0.003 | Young |
| 6.56E-05 | 0.476521 | 0.685 | 0.608 | 1 | 5 | Rbm3 | 0.077 | Young |
| 6.68E-05 | 0.185395 | 0.995 | 0.995 | 1 | 5 | Rps24 | 0 | Young |
| 6.73E-05 | 0.446067 | 0.617 | 0.545 | 1 | 5 | Arpc4 | 0.072 | Young |
| 7.80E-05 | 0.20025 | 1 | 0.998 | 1 | 5 | Fth1 | 0.002 | Young |
| 8.03E-05 | 0.932758 | 0.302 | 0.202 | 1 | 5 | Lmo4 | 0.1 | Young |
| 8.50E-05 | 0.780166 | 0.401 | 0.301 | 1 | 5 | Elk3 | 0.1 | Young |
| 8.55E-05 | 0.563168 | 0.409 | 0.29 | 1 | 5 | Etfb | 0.119 | Young |
| 8.58E-05 | 0.188534 | 0.992 | 0.993 | 1 | 5 | Rps12 | -0.001 | Young |
| 0.000132 | 1.87201 | 0.112 | 0.049 | 1 | 5 | Pmepa1 | 0.063 | Young |
| 0.000134 | 0.313355 | 0.876 | 0.84 | 1 | 5 | Qk | 0.036 | Young |

| | | | | | | | | |
|---|---|---|---|---|---|---|---|---|
| 0.000138 | 0.269385 | 0.911 | 0.897 | 1 | 5 | Igfbp4 | 0.014 | Young |
| 0.00015 | 0.325639 | 0.827 | 0.805 | 1 | 5 | Tmem50a | 0.022 | Young |
| 0.000161 | 0.265953 | 0.985 | 0.975 | 1 | 5 | Kctd12 | 0.01 | Young |
| 0.000165 | 0.520324 | 0.528 | 0.424 | 1 | 5 | Gpr34 | 0.104 | Young |
| 0.000167 | 0.776697 | 0.315 | 0.212 | 1 | 5 | Anxa2 | 0.103 | Young |
| 0.000192 | 2.389026 | 0.033 | 0.003 | 1 | 5 | Tmem119 | 0.03 | Young |
| 0.000202 | 0.979578 | 0.198 | 0.116 | 1 | 5 | Parp12 | 0.082 | Young |
| 0.000216 | 1.196364 | 0.114 | 0.051 | 1 | 5 | Slc25a16 | 0.063 | Young |
| 0.000216 | 0.175125 | 0.98 | 0.97 | 1 | 5 | Rpl7 | 0.01 | Young |
| 0.000217 | 2.405486 | 0.023 | 0 | 1 | 5 | Abi3 | 0.023 | Young |
| 0.000229 | 0.784448 | 0.302 | 0.209 | 1 | 5 | Pdlim4 | 0.093 | Young |
| 0.000244 | 1.39075 | 0.094 | 0.037 | 1 | 5 | Mrvi1 | 0.057 | Young |
| 0.000246 | 0.376903 | 0.612 | 0.513 | 1 | 5 | Nsa2 | 0.099 | Young |
| 0.000253 | 0.276522 | 0.665 | 0.569 | 1 | 5 | Pepd | 0.096 | Young |
| 0.000257 | 0.899385 | 0.236 | 0.15 | 1 | 5 | Pdlim1 | 0.086 | Young |
| 0.000259 | 0.254416 | 0.871 | 0.896 | 1 | 5 | Capza2 | -0.025 | Young |
| 0.000307 | 0.260512 | 0.977 | 0.97 | 1 | 5 | Zfp36l1 | 0.007 | Young |
| 0.000309 | 2.118135 | 0.036 | 0.005 | 1 | 5 | Spats2l | 0.031 | Young |
| 0.000311 | 1.04089 | 0.069 | 0.022 | 1 | 5 | Runx3 | 0.047 | Young |
| 0.00034 | 0.354342 | 0.746 | 0.645 | 1 | 5 | Hpgd | 0.101 | Young |
| 0.000371 | 0.735352 | 0.35 | 0.256 | 1 | 5 | Anxa3 | 0.094 | Young |
| 0.000386 | 0.271298 | 0.843 | 0.859 | 1 | 5 | Chchd2 | -0.016 | Young |
| 0.00039 | 0.627962 | 0.47 | 0.387 | 1 | 5 | Acat1 | 0.083 | Young |
| 0.0004 | 0.17188 | 0.985 | 0.992 | 1 | 5 | Cltc | -0.007 | Young |
| 0.0004 | 0.702068 | 0.365 | 0.268 | 1 | 5 | Ccr2 | 0.097 | Young |
| 0.00048 | 1.53708 | 0.048 | 0.012 | 1 | 5 | Pald1 | 0.036 | Young |
| 0.000486 | 1.775098 | 0.048 | 0.012 | 1 | 5 | Zscan12 | 0.036 | Young |
| 0.000502 | 2.154119 | 0.025 | 0.002 | 1 | 5 | Pck1 | 0.023 | Young |
| 0.000512 | 0.297097 | 0.708 | 0.672 | 1 | 5 | Atp5g2 | 0.036 | Young |
| 0.000529 | 0.954433 | 0.287 | 0.2 | 1 | 5 | Hist1h1c | 0.087 | Young |
| 0.000533 | 1.697845 | 0.086 | 0.035 | 1 | 5 | Pbxip1 | 0.051 | Young |
| 0.000547 | 0.156617 | 0.985 | 0.988 | 1 | 5 | Rps3a1 | -0.003 | Young |
| 0.000576 | 0.176251 | 0.99 | 0.983 | 1 | 5 | Rpl27a | 0.007 | Young |
| 0.000612 | 0.240415 | 0.904 | 0.894 | 1 | 5 | Rpl24 | 0.01 | Young |
| 0.000626 | 0.410558 | 0.708 | 0.648 | 1 | 5 | Gatm | 0.06 | Young |
| 0.000629 | 1.064163 | 0.079 | 0.03 | 1 | 5 | Plaur | 0.049 | Young |
| 0.000632 | 0.274081 | 0.799 | 0.731 | 1 | 5 | Zfhx3 | 0.068 | Young |
| 0.000644 | 1.095975 | 0.091 | 0.039 | 1 | 5 | Gypc | 0.052 | Young |
| 0.000653 | 0.92522 | 0.183 | 0.108 | 1 | 5 | Pon3 | 0.075 | Young |
| 0.000655 | 2.305891 | 0.033 | 0.005 | 1 | 5 | Sox4 | 0.028 | Young |
| 0.000691 | 0.173459 | 0.977 | 0.975 | 1 | 5 | Rps16 | 0.002 | Young |
| 0.000694 | 0.388207 | 0.665 | 0.598 | 1 | 5 | Rassf2 | 0.067 | Young |
| 0.000706 | 1.798759 | 0.056 | 0.017 | 1 | 5 | Gtpbp3 | 0.039 | Young |
| 0.000727 | 0.707703 | 0.142 | 0.074 | 1 | 5 | Pggt1b | 0.068 | Young |
| 0.000768 | 1.083972 | 0.213 | 0.133 | 1 | 5 | Pfkfb3 | 0.08 | Young |

| | | | | | | | | |
|---|---|---|---|---|---|---|---|---|
| 0.000802 | 0.527528 | 0.378 | 0.281 | 1 | 5 | S100a10 | 0.097 | Young |
| 0.000841 | 0.717324 | 0.228 | 0.145 | 1 | 5 | Nub1 | 0.083 | Young |
| 0.000905 | 1.961808 | 0.028 | 0.003 | 1 | 5 | Des | 0.025 | Young |
| 0.000915 | 0.334324 | 0.754 | 0.695 | 1 | 5 | Npm1 | 0.059 | Young |
| 0.000947 | 0.362672 | 0.683 | 0.631 | 1 | 5 | Eid1 | 0.052 | Young |
| 0.000957 | 0.369742 | 0.832 | 0.791 | 1 | 5 | Uqcrh | 0.041 | Young |
| 0.00098 | 0.167302 | 0.939 | 0.938 | 1 | 5 | Rack1 | 0.001 | Young |
| 0.000998 | 0.871296 | 0.17 | 0.099 | 1 | 5 | Tppp3 | 0.071 | Young |
| 0.001059 | 1.718287 | 0.114 | 0.059 | 1 | 5 | Cd4 | 0.055 | Young |
| 0.001059 | 0.28916 | 0.784 | 0.766 | 1 | 5 | Arf5 | 0.018 | Young |
| 0.001071 | 0.255802 | 0.799 | 0.778 | 1 | 5 | Cyfip1 | 0.021 | Young |
| 0.001081 | 0.258337 | 0.843 | 0.832 | 1 | 5 | Pcbp2 | 0.011 | Young |
| 0.001104 | 0.246944 | 0.799 | 0.774 | 1 | 5 | Cotl1 | 0.025 | Young |
| 0.001175 | 0.643356 | 0.36 | 0.264 | 1 | 5 | Smc3 | 0.096 | Young |
| 0.001227 | 0.541342 | 0.365 | 0.269 | 1 | 5 | 9030624J0 | 0.096 | Young |
| 0.001282 | 0.323003 | 0.731 | 0.704 | 1 | 5 | Scamp2 | 0.027 | Young |
| 0.001306 | 1.010934 | 0.188 | 0.118 | 1 | 5 | Polr2b | 0.07 | Young |
| 0.001363 | 0.530004 | 0.292 | 0.207 | 1 | 5 | Arrdc3 | 0.085 | Young |
| 0.001366 | 0.324004 | 0.713 | 0.677 | 1 | 5 | Tab2 | 0.036 | Young |
| 0.001369 | 0.237188 | 0.838 | 0.8 | 1 | 5 | Ywhaz | 0.038 | Young |
| 0.001389 | 0.790828 | 0.244 | 0.165 | 1 | 5 | Rab32 | 0.079 | Young |
| 0.0014 | 0.558813 | 0.36 | 0.273 | 1 | 5 | Atp6v1c1 | 0.087 | Young |
| 0.00141 | 0.800321 | 0.053 | 0.017 | 1 | 5 | 4930430F0 | 0.036 | Young |
| 0.001411 | 1.12798 | 0.15 | 0.086 | 1 | 5 | Zfp518a | 0.064 | Young |
| 0.001413 | 0.544325 | 0.355 | 0.269 | 1 | 5 | Ppp1r14b | 0.086 | Young |
| 0.00149 | 0.371981 | 0.599 | 0.561 | 1 | 5 | Atp5g3 | 0.038 | Young |
| 0.00151 | 0.091475 | 1 | 0.997 | 1 | 5 | Eef1a1 | 0.003 | Young |
| 0.001615 | 0.820313 | 0.208 | 0.133 | 1 | 5 | Tmem86a | 0.075 | Young |
| 0.001628 | 0.572333 | 0.266 | 0.182 | 1 | 5 | Hist3h2a | 0.084 | Young |
| 0.001673 | 0.363932 | 0.683 | 0.621 | 1 | 5 | Gm26917 | 0.062 | Young |
| 0.001676 | 0.540324 | 0.442 | 0.355 | 1 | 5 | Epb41l2 | 0.087 | Young |
| 0.001743 | 0.482761 | 0.373 | 0.285 | 1 | 5 | R3hdm2 | 0.088 | Young |
| 0.001821 | 2.20082 | 0.033 | 0.007 | 1 | 5 | F11r | 0.026 | Young |
| 0.001844 | 0.390256 | 0.551 | 0.48 | 1 | 5 | Nedd8 | 0.071 | Young |
| 0.001916 | 0.484711 | 0.31 | 0.221 | 1 | 5 | Smarce1 | 0.089 | Young |
| 0.00194 | 0.284433 | 0.863 | 0.837 | 1 | 5 | mt-Nd4l | 0.026 | Young |
| 0.001956 | 2.068922 | 0.025 | 0.003 | 1 | 5 | Spata13 | 0.022 | Young |
| 0.00198 | 2.145819 | 0.025 | 0.003 | 1 | 5 | G0s2 | 0.022 | Young |
| 0.002035 | 0.177231 | 0.99 | 0.978 | 1 | 5 | Rps20 | 0.012 | Young |
| 0.002066 | 0.4407 | 0.538 | 0.456 | 1 | 5 | Rin2 | 0.082 | Young |
| 0.002237 | 0.840732 | 0.18 | 0.113 | 1 | 5 | Ttc28 | 0.067 | Young |
| 0.002259 | 0.284897 | 0.736 | 0.715 | 1 | 5 | Serbp1 | 0.021 | Young |
| 0.002266 | 0.261632 | 0.701 | 0.635 | 1 | 5 | Gnaq | 0.066 | Young |
| 0.002349 | 0.509557 | 0.129 | 0.071 | 1 | 5 | Sgk1 | 0.058 | Young |
| 0.002541 | 2.562852 | 0.02 | 0.002 | 1 | 5 | Olfml3 | 0.018 | Young |

| | | | | | | | | |
|---|---|---|---|---|---|---|---|---|
| 0.002563 | 0.325849 | 0.69 | 0.638 | 1 | 5 | Lgals1 | 0.052 | Young |
| 0.00257 | 0.636864 | 0.246 | 0.167 | 1 | 5 | Prpf18 | 0.079 | Young |
| 0.002576 | 0.97594 | 0.129 | 0.072 | 1 | 5 | Dock4 | 0.057 | Young |
| 0.002576 | 2.282539 | 0.02 | 0.002 | 1 | 5 | 9130019O2 | 0.018 | Young |
| 0.002609 | 0.401544 | 0.569 | 0.502 | 1 | 5 | Ap2s1 | 0.067 | Young |
| 0.00262 | 0.173372 | 0.906 | 0.919 | 1 | 5 | Ptp4a2 | -0.013 | Young |
| 0.002632 | 0.427231 | 0.68 | 0.652 | 1 | 5 | Klf2 | 0.028 | Young |
| 0.002635 | 0.912593 | 0.129 | 0.072 | 1 | 5 | Trp53inp1 | 0.057 | Young |
| 0.002694 | 0.731394 | 0.249 | 0.177 | 1 | 5 | Nr2c2 | 0.072 | Young |
| 0.002695 | 0.220172 | 0.957 | 0.958 | 1 | 5 | Ctss | -0.001 | Young |
| 0.002709 | 0.489304 | 0.35 | 0.268 | 1 | 5 | Immt | 0.082 | Young |
| 0.002901 | 0.307511 | 0.906 | 0.909 | 1 | 5 | Tgfbi | -0.003 | Young |
| 0.002931 | 1.156566 | 0.107 | 0.057 | 1 | 5 | Gm38843 | 0.05 | Young |
| 0.002954 | 1.006657 | 0.109 | 0.057 | 1 | 5 | Sparc | 0.052 | Young |
| 0.003038 | 0.643061 | 0.142 | 0.082 | 1 | 5 | Mrpl9 | 0.06 | Young |
| 0.003108 | 0.721386 | 0.132 | 0.074 | 1 | 5 | Lztr1 | 0.058 | Young |
| 0.003156 | 0.21689 | 0.916 | 0.912 | 1 | 5 | Glul | 0.004 | Young |
| 0.003214 | 0.263772 | 0.751 | 0.714 | 1 | 5 | Itgb1 | 0.037 | Young |
| 0.003277 | 0.422529 | 0.594 | 0.507 | 1 | 5 | Nucks1 | 0.087 | Young |
| 0.003329 | 0.731689 | 0.165 | 0.103 | 1 | 5 | Rap2a | 0.062 | Young |
| 0.003348 | 0.304556 | 0.594 | 0.519 | 1 | 5 | Eif5b | 0.075 | Young |
| 0.00337 | 0.865015 | 0.124 | 0.069 | 1 | 5 | Snrnp25 | 0.055 | Young |
| 0.003382 | 0.209576 | 0.871 | 0.899 | 1 | 5 | Rplp0 | -0.028 | Young |
| 0.003416 | 0.203588 | 0.893 | 0.894 | 1 | 5 | Capzb | -0.001 | Young |
| 0.00342 | 0.485499 | 0.241 | 0.163 | 1 | 5 | Gnl1 | 0.078 | Young |
| 0.003487 | 0.507379 | 0.437 | 0.357 | 1 | 5 | Smim10l1 | 0.08 | Young |
| 0.003512 | 1.813166 | 0.043 | 0.013 | 1 | 5 | Uty | 0.03 | Young |
| 0.003599 | 0.783594 | 0.099 | 0.051 | 1 | 5 | Rprd1a | 0.048 | Young |
| 0.00366 | 0.639101 | 0.475 | 0.407 | 1 | 5 | Arl4c | 0.068 | Young |
| 0.003674 | 0.846684 | 0.15 | 0.091 | 1 | 5 | Amz2 | 0.059 | Young |
| 0.003795 | 1.561878 | 0.03 | 0.007 | 1 | 5 | Dact3 | 0.023 | Young |
| 0.003813 | 0.209412 | 0.805 | 0.8 | 1 | 5 | Naca | 0.005 | Young |
| 0.003839 | 0.924334 | 0.127 | 0.072 | 1 | 5 | Prps1 | 0.055 | Young |
| 0.003913 | 0.441355 | 0.467 | 0.377 | 1 | 5 | Sesn1 | 0.09 | Young |
| 0.003978 | 1.058612 | 0.086 | 0.042 | 1 | 5 | AU022252 | 0.044 | Young |
| 0.004026 | 1.298257 | 0.046 | 0.015 | 1 | 5 | Gngt2 | 0.031 | Young |
| 0.004144 | 0.456005 | 0.464 | 0.394 | 1 | 5 | Adipor1 | 0.07 | Young |
| 0.004176 | 0.524463 | 0.266 | 0.194 | 1 | 5 | Tbc1d9 | 0.072 | Young |
| 0.004259 | 1.856063 | 0.023 | 0.003 | 1 | 5 | Havcr2 | 0.02 | Young |
| 0.004292 | 2.307118 | 0.023 | 0.003 | 1 | 5 | Dock9 | 0.02 | Young |
| 0.004319 | 0.223416 | 0.815 | 0.798 | 1 | 5 | Sh3glb1 | 0.017 | Young |
| 0.004384 | 0.145699 | 0.98 | 0.992 | 1 | 5 | Rpl23 | -0.012 | Young |
| 0.004498 | 0.411089 | 0.505 | 0.439 | 1 | 5 | Srp9 | 0.066 | Young |
| 0.004624 | 0.968215 | 0.079 | 0.037 | 1 | 5 | Zdhhc13 | 0.042 | Young |
| 0.004723 | 0.778919 | 0.152 | 0.094 | 1 | 5 | Auh | 0.058 | Young |

| | | | | | | | | |
|---|---|---|---|---|---|---|---|---|
| 0.004745 | 0.963362 | 0.051 | 0.019 | 1 | 5 | Acta2 | 0.032 | Young |
| 0.004776 | 0.241381 | 0.815 | 0.795 | 1 | 5 | Vim | 0.02 | Young |
| 0.004798 | 0.372759 | 0.269 | 0.194 | 1 | 5 | Ednrb | 0.075 | Young |
| 0.004825 | 0.701808 | 0.137 | 0.081 | 1 | 5 | Ufd1 | 0.056 | Young |
| 0.004834 | 0.746072 | 0.18 | 0.12 | 1 | 5 | Rsl24d1 | 0.06 | Young |
| 0.004866 | 0.49845 | 0.152 | 0.096 | 1 | 5 | Fn1 | 0.056 | Young |
| 0.004993 | 0.300884 | 0.52 | 0.431 | 1 | 5 | Snrpf | 0.089 | Young |
| 0.005011 | 0.408184 | 0.373 | 0.293 | 1 | 5 | Rbm8a | 0.08 | Young |
| 0.005042 | 0.833276 | 0.16 | 0.101 | 1 | 5 | Fpr2 | 0.059 | Young |
| 0.005059 | 0.553462 | 0.378 | 0.298 | 1 | 5 | Elmo1 | 0.08 | Young |
| 0.005149 | 0.243198 | 0.838 | 0.8 | 1 | 5 | Rbms1 | 0.038 | Young |
| 0.005206 | 0.524151 | 0.429 | 0.357 | 1 | 5 | Smim15 | 0.072 | Young |
| 0.005306 | 0.666656 | 0.152 | 0.093 | 1 | 5 | Neurl4 | 0.059 | Young |
| 0.005321 | 0.26357 | 0.708 | 0.662 | 1 | 5 | Akap13 | 0.046 | Young |
| 0.005451 | 0.420779 | 0.513 | 0.439 | 1 | 5 | Sdha | 0.074 | Young |
| 0.005561 | 0.352925 | 0.477 | 0.387 | 1 | 5 | S100a13 | 0.09 | Young |
| 0.005576 | 0.470767 | 0.34 | 0.261 | 1 | 5 | Smarcd2 | 0.079 | Young |
| 0.005642 | 0.399997 | 0.365 | 0.279 | 1 | 5 | Rnf44 | 0.086 | Young |
| 0.005682 | 0.624622 | 0.201 | 0.136 | 1 | 5 | Zyg11b | 0.065 | Young |
| 0.005692 | 2.452742 | 0.018 | 0.002 | 1 | 5 | Pigl | 0.016 | Young |
| 0.005692 | 1.787442 | 0.018 | 0.002 | 1 | 5 | Gm41307 | 0.016 | Young |
| 0.005712 | 0.151362 | 0.975 | 0.973 | 1 | 5 | F13a1 | 0.002 | Young |
| 0.005758 | 1.23225 | 0.119 | 0.071 | 1 | 5 | Itga6 | 0.048 | Young |
| 0.005769 | 1.918 | 0.018 | 0.002 | 1 | 5 | Rgs7bp | 0.016 | Young |
| 0.00577 | 0.698028 | 0.117 | 0.066 | 1 | 5 | Tfam | 0.051 | Young |
| 0.00589 | 0.168965 | 0.949 | 0.946 | 1 | 5 | Gas6 | 0.003 | Young |
| 0.005956 | 2.047091 | 0.013 | 0 | 1 | 5 | Colq | 0.013 | Young |
| 0.005956 | 1.431425 | 0.013 | 0 | 1 | 5 | Atp2b4 | 0.013 | Young |
| 0.006052 | 0.451448 | 0.36 | 0.285 | 1 | 5 | Atp13a2 | 0.075 | Young |
| 0.006077 | 0.587546 | 0.102 | 0.054 | 1 | 5 | Gzf1 | 0.048 | Young |
| 0.006159 | 0.389735 | 0.528 | 0.492 | 1 | 5 | Eif3c | 0.036 | Young |
| 0.006335 | 0.763835 | 0.109 | 0.061 | 1 | 5 | Gfer | 0.048 | Young |
| 0.006379 | 0.400545 | 0.543 | 0.493 | 1 | 5 | Hdgf | 0.05 | Young |
| 0.006497 | 0.171856 | 0.972 | 0.961 | 1 | 5 | Rpl18a | 0.011 | Young |
| 0.006505 | 0.658217 | 0.155 | 0.096 | 1 | 5 | Rars | 0.059 | Young |
| 0.006573 | 0.437785 | 0.449 | 0.372 | 1 | 5 | Dmac1 | 0.077 | Young |
| 0.006782 | 0.483348 | 0.353 | 0.278 | 1 | 5 | Tspan14 | 0.075 | Young |
| 0.006822 | 0.570168 | 0.36 | 0.296 | 1 | 5 | Timm13 | 0.064 | Young |
| 0.006875 | 0.681013 | 0.162 | 0.104 | 1 | 5 | Sipa1l2 | 0.058 | Young |
| 0.007056 | 0.636955 | 0.173 | 0.113 | 1 | 5 | Mfsd6 | 0.06 | Young |
| 0.007073 | 1.325949 | 0.043 | 0.015 | 1 | 5 | Tstd3 | 0.028 | Young |
| 0.00712 | 0.290551 | 0.868 | 0.825 | 1 | 5 | Stk17b | 0.043 | Young |
| 0.007238 | 1.463546 | 0.063 | 0.029 | 1 | 5 | Smpd2 | 0.034 | Young |
| 0.007259 | 1.506963 | 0.043 | 0.015 | 1 | 5 | Col3a1 | 0.028 | Young |
| 0.007558 | 0.586135 | 0.282 | 0.219 | 1 | 5 | 1700017B0 | 0.063 | Young |

| | | | | | | | | |
|---|---|---|---|---|---|---|---|---|
| 0.007582 | 0.577994 | 0.277 | 0.207 | 1 | 5 | Rxra | 0.07 | Young |
| 0.007732 | 0.66753 | 0.15 | 0.093 | 1 | 5 | Kif13a | 0.057 | Young |
| 0.007733 | 1.376894 | 0.051 | 0.02 | 1 | 5 | Col1a2 | 0.031 | Young |
| 0.007784 | 1.095218 | 0.051 | 0.02 | 1 | 5 | Cdc42ep3 | 0.031 | Young |
| 0.007784 | 1.410454 | 0.028 | 0.007 | 1 | 5 | Lyrm1 | 0.021 | Young |
| 0.007812 | 0.505199 | 0.302 | 0.232 | 1 | 5 | Max | 0.07 | Young |
| 0.007813 | 0.122632 | 0.967 | 0.985 | 1 | 5 | Rps15 | -0.018 | Young |
| 0.007816 | 0.833806 | 0.152 | 0.098 | 1 | 5 | Ptp4a3 | 0.054 | Young |
| 0.00791 | 0.800745 | 0.165 | 0.109 | 1 | 5 | Wars | 0.056 | Young |
| 0.007958 | 0.178591 | 0.827 | 0.796 | 1 | 5 | Sat1 | 0.031 | Young |
| 0.008003 | 0.367886 | 0.589 | 0.535 | 1 | 5 | Ddx39b | 0.054 | Young |
| 0.008183 | 0.937269 | 0.104 | 0.059 | 1 | 5 | Smc2 | 0.045 | Young |
| 0.008317 | 1.059082 | 0.053 | 0.022 | 1 | 5 | Plp2 | 0.031 | Young |
| 0.008318 | 0.731397 | 0.16 | 0.104 | 1 | 5 | Plk2 | 0.056 | Young |
| 0.008318 | 0.278711 | 0.734 | 0.702 | 1 | 5 | Dbi | 0.032 | Young |
| 0.008399 | 0.803282 | 0.173 | 0.116 | 1 | 5 | Hacd3 | 0.057 | Young |
| 0.008424 | 0.890525 | 0.071 | 0.034 | 1 | 5 | Btf3l4 | 0.037 | Young |
| 0.008427 | 0.457337 | 0.376 | 0.303 | 1 | 5 | Peak1 | 0.073 | Young |
| 0.0085 | 0.756701 | 0.145 | 0.091 | 1 | 5 | Wdr5 | 0.054 | Young |
| 0.008753 | 0.171064 | 0.921 | 0.934 | 1 | 5 | Ucp2 | -0.013 | Young |
| 0.008823 | 0.708433 | 0.216 | 0.158 | 1 | 5 | Wdr11 | 0.058 | Young |
| 0.009096 | 0.652209 | 0.132 | 0.081 | 1 | 5 | Epb41l1 | 0.051 | Young |
| 0.009153 | 1.612589 | 0.02 | 0.003 | 1 | 5 | Zfp956 | 0.017 | Young |
| 0.009201 | 1.476936 | 0.03 | 0.008 | 1 | 5 | Zfp688 | 0.022 | Young |
| 0.009227 | 1.027482 | 0.094 | 0.052 | 1 | 5 | Gpn3 | 0.042 | Young |
| 0.009485 | 0.304266 | 0.794 | 0.79 | 1 | 5 | Tuba1b | 0.004 | Young |
| 0.00964 | 0.520014 | 0.211 | 0.146 | 1 | 5 | Smc5 | 0.065 | Young |
| 0.009734 | 0.535359 | 0.249 | 0.182 | 1 | 5 | Rps12-ps3 | 0.067 | Young |
| 0.00991 | 0.434685 | 0.449 | 0.384 | 1 | 5 | Tprgl | 0.065 | Young |
| 0.00992 | 0.602865 | 0.221 | 0.157 | 1 | 5 | Gcnt1 | 0.064 | Young |
| 0.009957 | 0.196257 | 0.855 | 0.842 | 1 | 5 | Ptbp3 | 0.013 | Young |
| 0.010219 | 0.294429 | 0.558 | 0.497 | 1 | 5 | Prkacb | 0.061 | Young |
| 0.010264 | 0.560067 | 0.152 | 0.096 | 1 | 5 | Aga | 0.056 | Young |
| 0.010508 | 1.037305 | 0.091 | 0.051 | 1 | 5 | AU020206 | 0.04 | Young |
| 0.010509 | 1.24384 | 0.069 | 0.034 | 1 | 5 | Ggta1 | 0.035 | Young |
| 0.01051 | 0.681704 | 0.272 | 0.21 | 1 | 5 | P2ry12 | 0.062 | Young |
| 0.010579 | 1.05876 | 0.074 | 0.037 | 1 | 5 | Mpi | 0.037 | Young |
| 0.010586 | 1.261777 | 0.033 | 0.01 | 1 | 5 | B930036N1 | 0.023 | Young |
| 0.011047 | 0.561241 | 0.284 | 0.214 | 1 | 5 | Hltf | 0.07 | Young |
| 0.011122 | 0.29698 | 0.703 | 0.684 | 1 | 5 | Cox7a2l | 0.019 | Young |
| 0.011157 | 0.517333 | 0.348 | 0.281 | 1 | 5 | Fam105a | 0.067 | Young |
| 0.011264 | 0.378613 | 0.282 | 0.212 | 1 | 5 | Dock1 | 0.07 | Young |
| 0.011372 | 0.362131 | 0.53 | 0.463 | 1 | 5 | Ppp2r5c | 0.067 | Young |
| 0.011426 | 0.470795 | 0.282 | 0.215 | 1 | 5 | Col4a3bp | 0.067 | Young |
| 0.011502 | 0.374531 | 0.431 | 0.367 | 1 | 5 | Mcrip1 | 0.064 | Young |

| | | | | | | | | |
|---|---|---|---|---|---|---|---|---|
| 0.011548 | 0.539497 | 0.254 | 0.189 | 1 | 5 | Ing4 | 0.065 | Young |
| 0.011723 | 0.974469 | 0.036 | 0.012 | 1 | 5 | Dennd2d | 0.024 | Young |
| 0.011833 | 0.219424 | 0.797 | 0.842 | 1 | 5 | Ddx17 | -0.045 | Young |
| 0.012059 | 0.749868 | 0.157 | 0.104 | 1 | 5 | Zcchc17 | 0.053 | Young |
| 0.012135 | 0.450472 | 0.254 | 0.187 | 1 | 5 | Pgap2 | 0.067 | Young |
| 0.012156 | 0.463302 | 0.261 | 0.194 | 1 | 5 | Eapp | 0.067 | Young |
| 0.012165 | 0.819952 | 0.089 | 0.049 | 1 | 5 | Tsta3 | 0.04 | Young |
| 0.01222 | 0.679666 | 0.094 | 0.052 | 1 | 5 | Eif1ad | 0.042 | Young |
| 0.012232 | 0.533418 | 0.317 | 0.251 | 1 | 5 | Atp8a1 | 0.066 | Young |
| 0.012299 | 1.324024 | 0.056 | 0.025 | 1 | 5 | Fig4 | 0.031 | Young |
| 0.012554 | 0.498518 | 0.234 | 0.172 | 1 | 5 | Echs1 | 0.062 | Young |
| 0.01268 | 1.863691 | 0.023 | 0.005 | 1 | 5 | Ifit2 | 0.018 | Young |
| 0.012719 | 1.089204 | 0.058 | 0.027 | 1 | 5 | Pygl | 0.031 | Young |
| 0.012804 | 1.499807 | 0.043 | 0.017 | 1 | 5 | Rapgef5 | 0.026 | Young |
| 0.012806 | 0.490548 | 0.185 | 0.125 | 1 | 5 | Ccdc90b | 0.06 | Young |
| 0.012816 | 0.510719 | 0.355 | 0.291 | 1 | 5 | Ctbp1 | 0.064 | Young |
| 0.012842 | 1.746074 | 0.015 | 0.002 | 1 | 5 | Bbs12 | 0.013 | Young |
| 0.012848 | 0.372937 | 0.457 | 0.397 | 1 | 5 | Arhgap45 | 0.06 | Young |
| 0.012871 | 0.630964 | 0.195 | 0.138 | 1 | 5 | Smarcad1 | 0.057 | Young |
| 0.012898 | 1.842435 | 0.015 | 0.002 | 1 | 5 | Zfp651 | 0.013 | Young |
| 0.012929 | 0.769102 | 0.038 | 0.013 | 1 | 5 | Etfbkmt | 0.025 | Young |
| 0.013096 | 0.409658 | 0.315 | 0.247 | 1 | 5 | Dync1li2 | 0.068 | Young |
| 0.013127 | 1.40986 | 0.015 | 0.002 | 1 | 5 | Gm13830 | 0.013 | Young |
| 0.013177 | 0.643722 | 0.15 | 0.098 | 1 | 5 | Sms | 0.052 | Young |
| 0.013185 | 1.469986 | 0.015 | 0.002 | 1 | 5 | Arhgap29 | 0.013 | Young |
| 0.01334 | 0.733363 | 0.15 | 0.099 | 1 | 5 | Dhdds | 0.051 | Young |
| 0.013423 | 0.807371 | 0.107 | 0.064 | 1 | 5 | Wdtc1 | 0.043 | Young |
| 0.013458 | 0.625808 | 0.145 | 0.094 | 1 | 5 | Mrpl41 | 0.051 | Young |
| 0.01347 | 1.09574 | 0.053 | 0.024 | 1 | 5 | Supv3l1 | 0.029 | Young |
| 0.013696 | 0.207246 | 0.865 | 0.875 | 1 | 5 | Rbpj | -0.01 | Young |
| 0.013865 | 0.155868 | 0.881 | 0.896 | 1 | 5 | Gnb2 | -0.015 | Young |
| 0.013947 | 0.63962 | 0.145 | 0.094 | 1 | 5 | Pank1 | 0.051 | Young |
| 0.013956 | 2.029079 | 0.01 | 0 | 1 | 5 | Sult1e1 | 0.01 | Young |
| 0.013956 | 1.832504 | 0.01 | 0 | 1 | 5 | Hoxa7 | 0.01 | Young |
| 0.013956 | 1.820948 | 0.01 | 0 | 1 | 5 | Prtn3 | 0.01 | Young |
| 0.013956 | 1.70176 | 0.01 | 0 | 1 | 5 | Hist1h2ae | 0.01 | Young |
| 0.013956 | 1.44342 | 0.01 | 0 | 1 | 5 | Mucl1 | 0.01 | Young |
| 0.013956 | 1.419719 | 0.01 | 0 | 1 | 5 | Micalcl | 0.01 | Young |
| 0.013956 | 1.396515 | 0.01 | 0 | 1 | 5 | Rnf113a1 | 0.01 | Young |
| 0.013956 | 1.323286 | 0.01 | 0 | 1 | 5 | Ahcy | 0.01 | Young |
| 0.013956 | 1.28643 | 0.01 | 0 | 1 | 5 | Plxdc1 | 0.01 | Young |
| 0.013956 | 1.174532 | 0.01 | 0 | 1 | 5 | Efna1 | 0.01 | Young |
| 0.013996 | 0.404839 | 0.409 | 0.342 | 1 | 5 | Hnrnpa1 | 0.067 | Young |
| 0.014013 | 0.86272 | 0.119 | 0.074 | 1 | 5 | Tctex1d2 | 0.045 | Young |
| 0.014223 | 1.058344 | 0.081 | 0.044 | 1 | 5 | Card9 | 0.037 | Young |

| | | | | | | | | |
|---|---|---|---|---|---|---|---|---|
| 0.014304 | 0.640409 | 0.152 | 0.099 | 1 | 5 | Vps53 | 0.053 | Young |
| 0.014466 | 0.285045 | 0.645 | 0.577 | 1 | 5 | Sec61b | 0.068 | Young |
| 0.014492 | 0.461535 | 0.365 | 0.3 | 1 | 5 | Camk2d | 0.065 | Young |
| 0.014499 | 0.184197 | 0.843 | 0.855 | 1 | 5 | Actr2 | -0.012 | Young |
| 0.014515 | 0.197987 | 0.751 | 0.744 | 1 | 5 | Tln1 | 0.007 | Young |
| 0.01465 | 0.400997 | 0.388 | 0.32 | 1 | 5 | Commd8 | 0.068 | Young |
| 0.014791 | 1.433604 | 0.069 | 0.035 | 1 | 5 | Ramp1 | 0.034 | Young |
| 0.014869 | 0.194119 | 0.459 | 0.37 | 1 | 5 | Adrb2 | 0.089 | Young |
| 0.014911 | 0.442994 | 0.256 | 0.189 | 1 | 5 | Prpf39 | 0.067 | Young |
| 0.015044 | 0.150903 | 0.98 | 0.98 | 1 | 5 | Rpl39 | 0 | Young |
| 0.015186 | 0.891586 | 0.079 | 0.042 | 1 | 5 | Lrrc14 | 0.037 | Young |
| 0.015324 | 1.091864 | 0.074 | 0.039 | 1 | 5 | Inpp4b | 0.035 | Young |
| 0.015373 | 0.343455 | 0.388 | 0.315 | 1 | 5 | Cbx1 | 0.073 | Young |
| 0.015427 | 0.850885 | 0.135 | 0.088 | 1 | 5 | Timd4 | 0.047 | Young |
| 0.015524 | 0.937272 | 0.076 | 0.04 | 1 | 5 | Bloc1s5 | 0.036 | Young |
| 0.015571 | 0.685746 | 0.079 | 0.042 | 1 | 5 | Alg6 | 0.037 | Young |
| 0.015718 | 0.399973 | 0.482 | 0.431 | 1 | 5 | Zcchc24 | 0.051 | Young |
| 0.015945 | 0.769016 | 0.142 | 0.094 | 1 | 5 | Lrrcc1 | 0.048 | Young |
| 0.015972 | 0.232264 | 0.827 | 0.823 | 1 | 5 | Eef1b2 | 0.004 | Young |
| 0.016091 | 0.489203 | 0.188 | 0.131 | 1 | 5 | Wwc2 | 0.057 | Young |
| 0.016131 | 0.3861 | 0.348 | 0.278 | 1 | 5 | Zmat2 | 0.07 | Young |
| 0.01623 | 0.302155 | 0.64 | 0.599 | 1 | 5 | Scpep1 | 0.041 | Young |
| 0.016255 | 0.731325 | 0.145 | 0.096 | 1 | 5 | Il17ra | 0.049 | Young |
| 0.016497 | 0.384036 | 0.434 | 0.386 | 1 | 5 | Cmpk1 | 0.048 | Young |
| 0.01676 | 0.57165 | 0.282 | 0.215 | 1 | 5 | Tspyl1 | 0.067 | Young |
| 0.016788 | 0.230976 | 0.827 | 0.788 | 1 | 5 | Ptafr | 0.039 | Young |
| 0.016891 | 1.61185 | 0.028 | 0.008 | 1 | 5 | Gramd4 | 0.02 | Young |
| 0.016974 | 0.876361 | 0.099 | 0.059 | 1 | 5 | Bcas3 | 0.04 | Young |
| 0.01715 | 0.204118 | 0.883 | 0.875 | 1 | 5 | Rpl10a | 0.008 | Young |
| 0.017164 | 0.185644 | 0.939 | 0.941 | 1 | 5 | Rpl18 | -0.002 | Young |
| 0.017189 | 0.473032 | 0.266 | 0.202 | 1 | 5 | Taok2 | 0.064 | Young |
| 0.017345 | 0.676914 | 0.185 | 0.131 | 1 | 5 | Naxd | 0.054 | Young |
| 0.017474 | 0.718964 | 0.107 | 0.064 | 1 | 5 | Coasy | 0.043 | Young |
| 0.017768 | 0.133261 | 0.734 | 0.682 | 1 | 5 | Wdr26 | 0.052 | Young |
| 0.018035 | 0.753658 | 0.074 | 0.039 | 1 | 5 | Rbm10 | 0.035 | Young |
| 0.018102 | 0.52576 | 0.208 | 0.152 | 1 | 5 | Mplkip | 0.056 | Young |
| 0.018309 | 0.450643 | 0.355 | 0.291 | 1 | 5 | Ints6l | 0.064 | Young |
| 0.018327 | 0.179678 | 0.906 | 0.921 | 1 | 5 | mt-Nd5 | -0.015 | Young |
| 0.018329 | 0.82403 | 0.086 | 0.049 | 1 | 5 | Nrm | 0.037 | Young |
| 0.018415 | 0.292089 | 0.602 | 0.564 | 1 | 5 | Psmb4 | 0.038 | Young |
| 0.018434 | 0.643493 | 0.028 | 0.008 | 1 | 5 | Dmwd | 0.02 | Young |
| 0.018447 | 2.982642 | 0.048 | 0.022 | 1 | 5 | Cyr61 | 0.026 | Young |
| 0.018489 | 0.319373 | 0.579 | 0.529 | 1 | 5 | Slc38a2 | 0.05 | Young |
| 0.018583 | 0.365556 | 0.195 | 0.136 | 1 | 5 | Plec | 0.059 | Young |
| 0.019163 | 0.531522 | 0.226 | 0.165 | 1 | 5 | Brd8 | 0.061 | Young |

| | | | | | | | | |
|---|---|---|---|---|---|---|---|---|
| 0.019267 | 0.475068 | 0.109 | 0.066 | 1 | 5 | Zfp626 | 0.043 | Young |
| 0.019408 | 1.09308 | 0.079 | 0.044 | 1 | 5 | Xlr | 0.035 | Young |
| 0.019462 | 0.501503 | 0.292 | 0.226 | 1 | 5 | Borcs6 | 0.066 | Young |
| 0.01949 | 0.376659 | 0.467 | 0.411 | 1 | 5 | Cmah | 0.056 | Young |
| 0.019593 | 0.125624 | 0.84 | 0.827 | 1 | 5 | Ywhae | 0.013 | Young |
| 0.019688 | 0.362022 | 0.52 | 0.483 | 1 | 5 | H2afy | 0.037 | Young |
| 0.019707 | 1.057891 | 0.03 | 0.01 | 1 | 5 | Zfp251 | 0.02 | Young |
| 0.019721 | 0.968005 | 0.081 | 0.045 | 1 | 5 | Rnf180 | 0.036 | Young |
| 0.019857 | 1.062824 | 0.061 | 0.03 | 1 | 5 | Afdn | 0.031 | Young |
| 0.01986 | 0.231016 | 0.734 | 0.721 | 1 | 5 | Myo5a | 0.013 | Young |
| 0.019938 | 0.177743 | 0.888 | 0.907 | 1 | 5 | Stab1 | -0.019 | Young |
| 0.019985 | 0.156546 | 0.916 | 0.912 | 1 | 5 | Pabpc1 | 0.004 | Young |
| 0.019985 | 1.397022 | 0.018 | 0.003 | 1 | 5 | Glipr2 | 0.015 | Young |
| 0.019985 | 1.096294 | 0.018 | 0.003 | 1 | 5 | Dhcr24 | 0.015 | Young |
| 0.020003 | 0.241314 | 0.556 | 0.465 | 1 | 5 | Srp14 | 0.091 | Young |
| 0.020196 | 0.212909 | 0.624 | 0.581 | 1 | 5 | Cops9 | 0.043 | Young |
| 0.020293 | 1.26623 | 0.048 | 0.022 | 1 | 5 | Fut10 | 0.026 | Young |
| 0.020323 | 0.433272 | 0.223 | 0.163 | 1 | 5 | Lsm7 | 0.06 | Young |
| 0.020521 | 1.601847 | 0.033 | 0.012 | 1 | 5 | Spata7 | 0.021 | Young |
| 0.02053 | 0.846672 | 0.109 | 0.067 | 1 | 5 | Cytip | 0.042 | Young |
| 0.020569 | 0.726726 | 0.165 | 0.116 | 1 | 5 | Gpatch2 | 0.049 | Young |
| 0.020871 | 0.11838 | 0.957 | 0.948 | 1 | 5 | Rpl5 | 0.009 | Young |
| 0.020938 | 0.219138 | 0.622 | 0.577 | 1 | 5 | Mfsd1 | 0.045 | Young |
| 0.021086 | 0.205541 | 0.916 | 0.943 | 1 | 5 | Myl6 | -0.027 | Young |
| 0.021106 | 0.707192 | 0.132 | 0.086 | 1 | 5 | Oard1 | 0.046 | Young |
| 0.021147 | 0.104958 | 0.604 | 0.557 | 1 | 5 | Klf4 | 0.047 | Young |
| 0.021169 | 0.864214 | 0.099 | 0.059 | 1 | 5 | Acad8 | 0.04 | Young |
| 0.02123 | 0.682091 | 0.175 | 0.126 | 1 | 5 | Anxa1 | 0.049 | Young |
| 0.021262 | 0.763742 | 0.142 | 0.096 | 1 | 5 | Gfra2 | 0.046 | Young |
| 0.021401 | 1.355125 | 0.048 | 0.022 | 1 | 5 | Itgb3bp | 0.026 | Young |
| 0.021402 | 0.912741 | 0.033 | 0.012 | 1 | 5 | Ift88 | 0.021 | Young |
| 0.021846 | 0.549877 | 0.282 | 0.226 | 1 | 5 | Sh3pxd2a | 0.056 | Young |
| 0.021952 | 0.715819 | 0.137 | 0.091 | 1 | 5 | Cpox | 0.046 | Young |
| 0.021972 | 0.878869 | 0.036 | 0.013 | 1 | 5 | Gm826 | 0.023 | Young |
| 0.021978 | 0.767392 | 0.203 | 0.153 | 1 | 5 | Slc12a9 | 0.05 | Young |
| 0.02206 | 0.692732 | 0.185 | 0.133 | 1 | 5 | Emilin2 | 0.052 | Young |
| 0.022176 | 1.051959 | 0.038 | 0.015 | 1 | 5 | Snx33 | 0.023 | Young |
| 0.022325 | 1.064633 | 0.038 | 0.015 | 1 | 5 | Fn3krp | 0.023 | Young |
| 0.022574 | 0.92225 | 0.096 | 0.059 | 1 | 5 | Hmbs | 0.037 | Young |
| 0.022674 | 0.11575 | 0.924 | 0.97 | 1 | 5 | Arpc2 | -0.046 | Young |
| 0.022678 | 0.9793 | 0.038 | 0.015 | 1 | 5 | Kdelc1 | 0.023 | Young |
| 0.022681 | 0.901974 | 0.051 | 0.024 | 1 | 5 | Cry1 | 0.027 | Young |
| 0.022808 | 1.069134 | 0.041 | 0.017 | 1 | 5 | Inpp1 | 0.024 | Young |
| 0.022826 | 1.025726 | 0.048 | 0.022 | 1 | 5 | Tmem186 | 0.026 | Young |
| 0.023106 | 0.393654 | 0.449 | 0.386 | 1 | 5 | H2afv | 0.063 | Young |

| | | | | | | | | |
|---|---|---|---|---|---|---|---|---|
| 0.023206 | 0.445457 | 0.188 | 0.133 | 1 | 5 | Ipo11 | 0.055 | Young |
| 0.023296 | 0.900287 | 0.043 | 0.019 | 1 | 5 | Prmt6 | 0.024 | Young |
| 0.023559 | 0.791339 | 0.195 | 0.15 | 1 | 5 | Mrpl36 | 0.045 | Young |
| 0.023839 | 0.546153 | 0.206 | 0.15 | 1 | 5 | Nudt4 | 0.056 | Young |
| 0.023963 | 0.12844 | 0.977 | 0.973 | 1 | 5 | Cdc42 | 0.004 | Young |
| 0.024044 | 0.710066 | 0.129 | 0.086 | 1 | 5 | Mtmr14 | 0.043 | Young |
| 0.024377 | 0.839484 | 0.112 | 0.071 | 1 | 5 | Paxip1 | 0.041 | Young |
| 0.024634 | 0.404601 | 0.459 | 0.402 | 1 | 5 | 1700020I1 | 0.057 | Young |
| 0.024803 | 1.689686 | 0.02 | 0.005 | 1 | 5 | Ggact | 0.015 | Young |
| 0.024862 | 0.454842 | 0.297 | 0.241 | 1 | 5 | Nxpe5 | 0.056 | Young |
| 0.024874 | 0.165089 | 0.985 | 0.98 | 1 | 5 | Rpl30 | 0.005 | Young |
| 0.024955 | 0.446474 | 0.206 | 0.148 | 1 | 5 | Snx12 | 0.058 | Young |
| 0.025021 | 0.496099 | 0.368 | 0.308 | 1 | 5 | Sh3bp5 | 0.06 | Young |
| 0.025046 | 1.429729 | 0.02 | 0.005 | 1 | 5 | Adig | 0.015 | Young |
| 0.025065 | 0.294613 | 0.566 | 0.524 | 1 | 5 | Lamtor4 | 0.042 | Young |
| 0.025107 | 0.546667 | 0.208 | 0.155 | 1 | 5 | Tbk1 | 0.053 | Young |
| 0.025179 | 0.3171 | 0.371 | 0.3 | 1 | 5 | Psmd4 | 0.071 | Young |
| 0.025262 | 0.5874 | 0.272 | 0.215 | 1 | 5 | Coa5 | 0.057 | Young |
| 0.025286 | 0.907999 | 0.086 | 0.051 | 1 | 5 | Cbfa2t2 | 0.035 | Young |
| 0.025372 | 1.658672 | 0.02 | 0.005 | 1 | 5 | Gm26802 | 0.015 | Young |
| 0.025454 | 1.243122 | 0.02 | 0.005 | 1 | 5 | Rab4b | 0.015 | Young |
| 0.025525 | 1.028178 | 0.063 | 0.034 | 1 | 5 | Trim37 | 0.029 | Young |
| 0.025537 | 1.354707 | 0.02 | 0.005 | 1 | 5 | Zfp354c | 0.015 | Young |
| 0.025537 | 1.310002 | 0.02 | 0.005 | 1 | 5 | Slc2a6 | 0.015 | Young |
| 0.025869 | 0.948521 | 0.02 | 0.005 | 1 | 5 | Gm26756 | 0.015 | Young |
| 0.026104 | 0.407403 | 0.317 | 0.256 | 1 | 5 | Pik3ca | 0.061 | Young |
| 0.026221 | 0.301193 | 0.612 | 0.549 | 1 | 5 | Snrpb | 0.063 | Young |
| 0.026239 | 0.244223 | 0.766 | 0.749 | 1 | 5 | Ms4a6c | 0.017 | Young |
| 0.026247 | 0.319317 | 0.518 | 0.461 | 1 | 5 | Ube2l3 | 0.057 | Young |
| 0.026407 | 0.232856 | 0.127 | 0.081 | 1 | 5 | Srebf1 | 0.046 | Young |
| 0.02646 | 0.776081 | 0.086 | 0.051 | 1 | 5 | Rdh11 | 0.035 | Young |
| 0.026745 | 0.43434 | 0.299 | 0.244 | 1 | 5 | Pde3b | 0.055 | Young |
| 0.026885 | 0.671271 | 0.117 | 0.076 | 1 | 5 | Dgcr8 | 0.041 | Young |
| 0.026948 | 0.125814 | 0.957 | 0.953 | 1 | 5 | Ppia | 0.004 | Young |
| 0.027165 | 0.468073 | 0.264 | 0.209 | 1 | 5 | Glrx3 | 0.055 | Young |
| 0.027205 | 0.393424 | 0.36 | 0.298 | 1 | 5 | Klhl24 | 0.062 | Young |
| 0.027256 | 0.095244 | 0.997 | 0.997 | 1 | 5 | Laptm5 | 0 | Young |
| 0.027368 | 0.715611 | 0.221 | 0.17 | 1 | 5 | Ntpcr | 0.051 | Young |
| 0.027433 | 0.43978 | 0.264 | 0.202 | 1 | 5 | Fyttd1 | 0.062 | Young |
| 0.027468 | 0.570105 | 0.119 | 0.077 | 1 | 5 | Alcam | 0.042 | Young |
| 0.027561 | 0.164817 | 0.919 | 0.914 | 1 | 5 | Mef2c | 0.005 | Young |
| 0.027639 | 0.441355 | 0.315 | 0.253 | 1 | 5 | Ncoa1 | 0.062 | Young |
| 0.028037 | 0.605778 | 0.099 | 0.061 | 1 | 5 | Scaper | 0.038 | Young |
| 0.028159 | 0.566667 | 0.251 | 0.2 | 1 | 5 | Bdp1 | 0.051 | Young |
| 0.028303 | 0.209847 | 0.878 | 0.82 | 1 | 5 | Aif1 | 0.058 | Young |

| | | | | | | | | |
|---|---|---|---|---|---|---|---|---|
| 0.028414 | 0.63062 | 0.175 | 0.128 | 1 | 5 | Ezr | 0.047 | Young |
| 0.028591 | 0.486041 | 0.241 | 0.185 | 1 | 5 | Tmem131 | 0.056 | Young |
| 0.028729 | 0.425863 | 0.36 | 0.305 | 1 | 5 | Llph | 0.055 | Young |
| 0.02886 | 0.580167 | 0.256 | 0.205 | 1 | 5 | Hcst | 0.051 | Young |
| 0.028924 | 0.468913 | 0.19 | 0.138 | 1 | 5 | Ncbp2 | 0.052 | Young |
| 0.028954 | 0.897154 | 0.122 | 0.082 | 1 | 5 | Zbtb18 | 0.04 | Young |
| 0.02902 | 1.685446 | 0.023 | 0.007 | 1 | 5 | Wdr35 | 0.016 | Young |
| 0.029055 | 1.833484 | 0.013 | 0.002 | 1 | 5 | Ccdc166 | 0.011 | Young |
| 0.029055 | 1.402234 | 0.013 | 0.002 | 1 | 5 | Tpx2 | 0.011 | Young |
| 0.029073 | 0.430585 | 0.279 | 0.217 | 1 | 5 | Fam76b | 0.062 | Young |
| 0.02918 | 1.795673 | 0.013 | 0.002 | 1 | 5 | Arg1 | 0.011 | Young |
| 0.02918 | 1.435036 | 0.013 | 0.002 | 1 | 5 | Ccnf | 0.011 | Young |
| 0.02918 | 1.249137 | 0.013 | 0.002 | 1 | 5 | Zfyve28 | 0.011 | Young |
| 0.029297 | 0.355178 | 0.404 | 0.337 | 1 | 5 | Supt16 | 0.067 | Young |
| 0.029305 | 1.901036 | 0.013 | 0.002 | 1 | 5 | Kif18a | 0.011 | Young |
| 0.029305 | 1.650751 | 0.013 | 0.002 | 1 | 5 | Nrep | 0.011 | Young |
| 0.029305 | 1.489103 | 0.013 | 0.002 | 1 | 5 | 4930532G1 | 0.011 | Young |
| 0.029361 | 1.404291 | 0.023 | 0.007 | 1 | 5 | Fignl1 | 0.016 | Young |
| 0.029431 | 1.182183 | 0.013 | 0.002 | 1 | 5 | Afmid | 0.011 | Young |
| 0.029509 | 0.589342 | 0.249 | 0.197 | 1 | 5 | Snapin | 0.052 | Young |
| 0.029557 | 0.793022 | 0.091 | 0.056 | 1 | 5 | Dnaja3 | 0.035 | Young |
| 0.029557 | 1.347371 | 0.013 | 0.002 | 1 | 5 | Gm47079 | 0.011 | Young |
| 0.029727 | 0.568545 | 0.17 | 0.123 | 1 | 5 | Mfap1a | 0.047 | Young |
| 0.029958 | 0.327428 | 0.503 | 0.436 | 1 | 5 | Tceal9 | 0.067 | Young |
| 0.029996 | 0.332467 | 0.34 | 0.278 | 1 | 5 | Ap3d1 | 0.062 | Young |
| 0.030165 | 0.22556 | 0.797 | 0.783 | 1 | 5 | Mtss1 | 0.014 | Young |
| 0.030406 | 0.953164 | 0.023 | 0.007 | 1 | 5 | Platr17 | 0.016 | Young |
| 0.030454 | 1.044456 | 0.074 | 0.042 | 1 | 5 | Bend4 | 0.032 | Young |
| 0.030516 | 0.12787 | 0.982 | 0.981 | 1 | 5 | Rps4x | 0.001 | Young |
| 0.030589 | 0.247058 | 0.599 | 0.547 | 1 | 5 | Set | 0.052 | Young |
| 0.031101 | 0.227572 | 0.627 | 0.599 | 1 | 5 | Glg1 | 0.028 | Young |
| 0.031265 | 0.45593 | 0.124 | 0.081 | 1 | 5 | Slc35e3 | 0.043 | Young |
| 0.031295 | 0.185551 | 0.766 | 0.751 | 1 | 5 | Ccdc50 | 0.015 | Young |
| 0.031458 | 0.520065 | 0.213 | 0.158 | 1 | 5 | Pdcd7 | 0.055 | Young |
| 0.031504 | 0.732379 | 0.102 | 0.064 | 1 | 5 | Map3k20 | 0.038 | Young |
| 0.031993 | 0.42975 | 0.259 | 0.204 | 1 | 5 | Rabgap1 | 0.055 | Young |
| 0.032042 | 0.63833 | 0.061 | 0.032 | 1 | 5 | Nprl3 | 0.029 | Young |
| 0.03207 | 0.592929 | 0.142 | 0.098 | 1 | 5 | Git1 | 0.044 | Young |
| 0.032258 | 0.225558 | 0.784 | 0.811 | 1 | 5 | Atp5l | -0.027 | Young |
| 0.032316 | 1.098681 | 0.051 | 0.025 | 1 | 5 | 4632415L0 | 0.026 | Young |
| 0.03234 | 0.524753 | 0.249 | 0.197 | 1 | 5 | Yipf4 | 0.052 | Young |
| 0.032346 | 0.416572 | 0.175 | 0.123 | 1 | 5 | Cdk17 | 0.052 | Young |
| 0.032413 | 0.372806 | 0.261 | 0.205 | 1 | 5 | Slc30a5 | 0.056 | Young |
| 0.032489 | 0.203515 | 0.807 | 0.786 | 1 | 5 | Arpc3 | 0.021 | Young |
| 0.032512 | 1.225801 | 0.063 | 0.035 | 1 | 5 | Sema4c | 0.028 | Young |

| | | | | | | | | |
|---|---|---|---|---|---|---|---|---|
| 0.032546 | 0.145154 | 0.843 | 0.832 | 1 | 5 | Mob1a | 0.011 | Young |
| 0.032617 | 0.437444 | 0.376 | 0.327 | 1 | 5 | Bag1 | 0.049 | Young |
| 0.032795 | 0.554526 | 0.307 | 0.259 | 1 | 5 | Tomm5 | 0.048 | Young |
| 0.032836 | 1.243829 | 0.051 | 0.025 | 1 | 5 | Fam26f | 0.026 | Young |
| 0.032905 | 0.307332 | 0.487 | 0.419 | 1 | 5 | Add1 | 0.068 | Young |
| 0.033042 | 0.704786 | 0.119 | 0.079 | 1 | 5 | Psmd9 | 0.04 | Young |
| 0.033066 | 1.199144 | 0.025 | 0.008 | 1 | 5 | Il1rl2 | 0.017 | Young |
| 0.033079 | 0.29788 | 0.475 | 0.418 | 1 | 5 | Map4k4 | 0.057 | Young |
| 0.033131 | 0.473781 | 0.345 | 0.295 | 1 | 5 | 2510039O1 | 0.05 | Young |
| 0.033135 | 1.126911 | 0.048 | 0.024 | 1 | 5 | Rfx2 | 0.024 | Young |
| 0.033196 | 0.813046 | 0.102 | 0.064 | 1 | 5 | Dennd2c | 0.038 | Young |
| 0.033257 | 0.376333 | 0.292 | 0.229 | 1 | 5 | Gas2l3 | 0.063 | Young |
| 0.033476 | 0.946736 | 0.109 | 0.072 | 1 | 5 | Gsto1 | 0.037 | Young |
| 0.033537 | 0.201079 | 0.744 | 0.756 | 1 | 5 | Btf3 | -0.012 | Young |
| 0.033556 | 0.615291 | 0.112 | 0.072 | 1 | 5 | Rpl27-ps3 | 0.04 | Young |
| 0.033777 | 0.933655 | 0.053 | 0.027 | 1 | 5 | 1600014C1 | 0.026 | Young |
| 0.03382 | 0.469337 | 0.183 | 0.133 | 1 | 5 | Zfp984 | 0.05 | Young |
| 0.033845 | 0.264983 | 0.518 | 0.46 | 1 | 5 | Ncf1 | 0.058 | Young |
| 0.034325 | 0.351063 | 0.348 | 0.286 | 1 | 5 | Dhx40 | 0.062 | Young |
| 0.034335 | 0.78115 | 0.096 | 0.061 | 1 | 5 | Hsf2 | 0.035 | Young |
| 0.0345 | 1.085824 | 0.051 | 0.025 | 1 | 5 | Mzt2 | 0.026 | Young |
| 0.034513 | 0.615384 | 0.132 | 0.089 | 1 | 5 | Ormdl1 | 0.043 | Young |
| 0.034562 | 0.253383 | 0.645 | 0.618 | 1 | 5 | Rnf13 | 0.027 | Young |
| 0.034583 | 0.340435 | 0.467 | 0.412 | 1 | 5 | Psmd2 | 0.055 | Young |
| 0.034788 | 0.759521 | 0.117 | 0.077 | 1 | 5 | Bckdha | 0.04 | Young |
| 0.034806 | 0.443328 | 0.294 | 0.237 | 1 | 5 | Ddx60 | 0.057 | Young |
| 0.034928 | 0.859591 | 0.051 | 0.025 | 1 | 5 | Me1 | 0.026 | Young |
| 0.034962 | 0.392745 | 0.246 | 0.189 | 1 | 5 | Sidt2 | 0.057 | Young |
| 0.035281 | 1.282365 | 0.028 | 0.01 | 1 | 5 | Mamdc2 | 0.018 | Young |
| 0.035304 | 0.272787 | 0.558 | 0.507 | 1 | 5 | Xiap | 0.051 | Young |
| 0.035522 | 1.036557 | 0.046 | 0.022 | 1 | 5 | Tmed8 | 0.024 | Young |
| 0.035588 | 0.902123 | 0.046 | 0.022 | 1 | 5 | R3hcc1 | 0.024 | Young |
| 0.035618 | 1.057156 | 0.063 | 0.035 | 1 | 5 | Stradb | 0.028 | Young |
| 0.03563 | 0.765478 | 0.074 | 0.042 | 1 | 5 | Zbtb6 | 0.032 | Young |
| 0.03568 | 0.176551 | 0.865 | 0.816 | 1 | 5 | Cfh | 0.049 | Young |
| 0.035719 | 0.199345 | 0.764 | 0.766 | 1 | 5 | Fam49b | -0.002 | Young |
| 0.035723 | 0.836982 | 0.028 | 0.01 | 1 | 5 | Hps6 | 0.018 | Young |
| 0.035959 | 0.498009 | 0.162 | 0.116 | 1 | 5 | Dcaf17 | 0.046 | Young |
| 0.03611 | 1.115846 | 0.03 | 0.012 | 1 | 5 | Sap18b | 0.018 | Young |
| 0.036111 | 1.237881 | 0.033 | 0.013 | 1 | 5 | Zkscan7 | 0.02 | Young |
| 0.036192 | 1.183644 | 0.033 | 0.013 | 1 | 5 | Ift57 | 0.02 | Young |
| 0.036195 | 1.278683 | 0.03 | 0.012 | 1 | 5 | Hopx | 0.018 | Young |
| 0.036212 | 0.841064 | 0.107 | 0.069 | 1 | 5 | Xpa | 0.038 | Young |
| 0.036277 | 0.453461 | 0.305 | 0.253 | 1 | 5 | Zc3h13 | 0.052 | Young |
| 0.036365 | 0.432202 | 0.206 | 0.152 | 1 | 5 | Hmg20a | 0.054 | Young |

| | | | | | | | | |
|---|---|---|---|---|---|---|---|---|
| 0.03657 | 0.329123 | 0.523 | 0.463 | 1 | 5 | Wipf1 | 0.06 | Young |
| 0.036599 | 1.375555 | 0.033 | 0.013 | 1 | 5 | Szt2 | 0.02 | Young |
| 0.036626 | 0.260841 | 0.536 | 0.478 | 1 | 5 | Trim25 | 0.058 | Young |
| 0.036763 | 1.375189 | 0.033 | 0.013 | 1 | 5 | Enpep | 0.02 | Young |
| 0.036804 | 0.239495 | 0.706 | 0.687 | 1 | 5 | Pdcd6ip | 0.019 | Young |
| 0.03691 | 0.163653 | 0.799 | 0.793 | 1 | 5 | Vamp8 | 0.006 | Young |
| 0.036928 | 1.149977 | 0.033 | 0.013 | 1 | 5 | Neil1 | 0.02 | Young |
| 0.036938 | 0.428938 | 0.223 | 0.17 | 1 | 5 | Tmem65 | 0.053 | Young |
| 0.036971 | 0.370303 | 0.332 | 0.266 | 1 | 5 | Nudcd3 | 0.066 | Young |
| 0.037054 | 0.363624 | 0.429 | 0.38 | 1 | 5 | Nme1 | 0.049 | Young |
| 0.037072 | 0.541737 | 0.228 | 0.175 | 1 | 5 | Csgalnact2 | 0.053 | Young |
| 0.037075 | 0.493924 | 0.236 | 0.187 | 1 | 5 | Gm47283 | 0.049 | Young |
| 0.037108 | 0.396546 | 0.536 | 0.508 | 1 | 5 | Eif4h | 0.028 | Young |
| 0.037415 | 0.510862 | 0.15 | 0.106 | 1 | 5 | Atad2 | 0.044 | Young |
| 0.037484 | 0.286803 | 0.447 | 0.392 | 1 | 5 | Mrpl52 | 0.055 | Young |
| 0.037562 | 1.030859 | 0.091 | 0.057 | 1 | 5 | Ripk2 | 0.034 | Young |
| 0.037635 | 0.742252 | 0.041 | 0.019 | 1 | 5 | Cep112 | 0.022 | Young |
| 0.037844 | 1.029188 | 0.033 | 0.013 | 1 | 5 | Fbxo32 | 0.02 | Young |
| 0.037932 | 0.875305 | 0.041 | 0.019 | 1 | 5 | Bcdin3d | 0.022 | Young |
| 0.038298 | 0.931564 | 0.036 | 0.015 | 1 | 5 | Gng7 | 0.021 | Young |
| 0.038423 | 0.458836 | 0.226 | 0.175 | 1 | 5 | Wtip | 0.051 | Young |
| 0.038431 | 0.503254 | 0.216 | 0.163 | 1 | 5 | Zc3h4 | 0.053 | Young |
| 0.038558 | 0.620198 | 0.102 | 0.066 | 1 | 5 | Fam207a | 0.036 | Young |
| 0.038683 | 0.696138 | 0.094 | 0.059 | 1 | 5 | Nid2 | 0.035 | Young |
| 0.0387 | 0.275446 | 0.579 | 0.522 | 1 | 5 | Rnf141 | 0.057 | Young |
| 0.038737 | 0.490899 | 0.089 | 0.054 | 1 | 5 | Cnnm4 | 0.035 | Young |
| 0.038866 | 0.897817 | 0.033 | 0.013 | 1 | 5 | Zfp60 | 0.02 | Young |
| 0.038954 | 0.339248 | 0.31 | 0.253 | 1 | 5 | Alyref | 0.057 | Young |
| 0.039176 | 0.776574 | 0.117 | 0.079 | 1 | 5 | Pgp | 0.038 | Young |
| 0.039201 | 0.832499 | 0.036 | 0.015 | 1 | 5 | Yars2 | 0.021 | Young |
| 0.039296 | 0.730064 | 0.041 | 0.019 | 1 | 5 | Coa4 | 0.022 | Young |
| 0.039502 | 0.45548 | 0.289 | 0.236 | 1 | 5 | Ndufb6 | 0.053 | Young |
| 0.039586 | 0.574288 | 0.142 | 0.099 | 1 | 5 | Cdc14b | 0.043 | Young |
| 0.039697 | 0.720287 | 0.038 | 0.017 | 1 | 5 | Coq3 | 0.021 | Young |
| 0.039918 | 0.676508 | 0.084 | 0.051 | 1 | 5 | Zbtb41 | 0.033 | Young |
| 0.040132 | 0.432246 | 0.442 | 0.401 | 1 | 5 | Stub1 | 0.041 | Young |
| 0.040179 | 0.271496 | 0.548 | 0.497 | 1 | 5 | Irf2 | 0.051 | Young |
| 0.040222 | 0.851733 | 0.081 | 0.049 | 1 | 5 | Rars2 | 0.032 | Young |
| 0.040457 | 0.543485 | 0.183 | 0.136 | 1 | 5 | Lipo3 | 0.047 | Young |
| 0.040514 | 0.281662 | 0.49 | 0.426 | 1 | 5 | Msr1 | 0.064 | Young |
| 0.040543 | 0.754427 | 0.041 | 0.019 | 1 | 5 | E330009J0 | 0.022 | Young |
| 0.040785 | 0.468418 | 0.305 | 0.251 | 1 | 5 | Dynlt3 | 0.054 | Young |
| 0.040949 | 3.365542 | 0.015 | 0.003 | 1 | 5 | Saa3 | 0.012 | Young |
| 0.041158 | 0.482619 | 0.259 | 0.207 | 1 | 5 | Tomm70a | 0.052 | Young |
| 0.041239 | 1.776864 | 0.015 | 0.003 | 1 | 5 | Tmem17 | 0.012 | Young |

| | | | | | | | | |
|---|---|---|---|---|---|---|---|---|
| 0.041385 | 1.14181 | 0.015 | 0.003 | 1 | 5 | Cyfip2 | 0.012 | Young |
| 0.041532 | 1.676415 | 0.015 | 0.003 | 1 | 5 | Ccdc18 | 0.012 | Young |
| 0.041543 | 0.4656 | 0.404 | 0.347 | 1 | 5 | Smim14 | 0.057 | Young |
| 0.041546 | 0.613777 | 0.162 | 0.12 | 1 | 5 | Uba3 | 0.042 | Young |
| 0.041752 | 0.470055 | 0.203 | 0.155 | 1 | 5 | Piezo1 | 0.048 | Young |
| 0.041775 | 0.723629 | 0.109 | 0.072 | 1 | 5 | Arl6ip4 | 0.037 | Young |
| 0.04179 | 0.547266 | 0.066 | 0.037 | 1 | 5 | Ndufaf2 | 0.029 | Young |
| 0.041826 | 1.865114 | 0.015 | 0.003 | 1 | 5 | Socs2 | 0.012 | Young |
| 0.041826 | 1.29212 | 0.015 | 0.003 | 1 | 5 | Wdr66 | 0.012 | Young |
| 0.042419 | 0.896282 | 0.015 | 0.003 | 1 | 5 | Pdzd2 | 0.012 | Young |
| 0.042478 | 0.144289 | 0.822 | 0.801 | 1 | 5 | Akr1a1 | 0.021 | Young |
| 0.042602 | 0.445303 | 0.32 | 0.266 | 1 | 5 | Pon2 | 0.054 | Young |
| 0.042958 | 0.271189 | 0.495 | 0.438 | 1 | 5 | Cmtm7 | 0.057 | Young |
| 0.043079 | 0.546119 | 0.244 | 0.192 | 1 | 5 | Slc38a9 | 0.052 | Young |
| 0.043089 | 0.165773 | 0.83 | 0.825 | 1 | 5 | Rpl7a | 0.005 | Young |
| 0.043562 | 0.680815 | 0.117 | 0.079 | 1 | 5 | Nhlrc2 | 0.038 | Young |
| 0.043617 | 0.326974 | 0.264 | 0.204 | 1 | 5 | Tbc1d4 | 0.06 | Young |
| 0.043941 | 0.145531 | 0.845 | 0.867 | 1 | 5 | Ybx1 | -0.022 | Young |
| 0.044337 | 0.38562 | 0.619 | 0.586 | 1 | 5 | Bach1 | 0.033 | Young |
| 0.044517 | 0.281826 | 0.772 | 0.744 | 1 | 5 | Ier2 | 0.028 | Young |
| 0.044584 | 1.042261 | 0.058 | 0.032 | 1 | 5 | Hgf | 0.026 | Young |
| 0.044629 | 0.239274 | 0.525 | 0.468 | 1 | 5 | Ttc3 | 0.057 | Young |
| 0.044713 | 0.249714 | 0.503 | 0.443 | 1 | 5 | Snrpb2 | 0.06 | Young |
| 0.0448 | 0.334397 | 0.406 | 0.355 | 1 | 5 | Edem1 | 0.051 | Young |
| 0.044847 | 0.567141 | 0.203 | 0.157 | 1 | 5 | Nsd2 | 0.046 | Young |
| 0.045097 | 0.52979 | 0.099 | 0.064 | 1 | 5 | Slc26a11 | 0.035 | Young |
| 0.045198 | 0.114563 | 0.967 | 0.981 | 1 | 5 | Cfl1 | -0.014 | Young |
| 0.045352 | 0.950763 | 0.056 | 0.03 | 1 | 5 | Noc3l | 0.026 | Young |
| 0.045484 | 0.498493 | 0.129 | 0.089 | 1 | 5 | Zscan29 | 0.04 | Young |
| 0.045493 | 0.677582 | 0.079 | 0.047 | 1 | 5 | Rab2b | 0.032 | Young |
| 0.045835 | 0.197318 | 0.678 | 0.65 | 1 | 5 | Brk1 | 0.028 | Young |
| 0.04585 | 0.656991 | 0.074 | 0.044 | 1 | 5 | Ints7 | 0.03 | Young |
| 0.045976 | 0.429643 | 0.353 | 0.301 | 1 | 5 | Fam91a1 | 0.052 | Young |
| 0.046231 | 0.579772 | 0.145 | 0.103 | 1 | 5 | Nckipsd | 0.042 | Young |
| 0.04669 | 0.67854 | 0.102 | 0.067 | 1 | 5 | Mgme1 | 0.035 | Young |
| 0.046934 | 0.564042 | 0.223 | 0.172 | 1 | 5 | Zfp326 | 0.051 | Young |
| 0.046956 | 0.277645 | 0.264 | 0.215 | 1 | 5 | Baz1a | 0.049 | Young |
| 0.047566 | 0.312746 | 0.467 | 0.419 | 1 | 5 | Rad21 | 0.048 | Young |
| 0.047798 | 0.825412 | 0.053 | 0.029 | 1 | 5 | Gm45902 | 0.024 | Young |
| 0.047802 | 0.69117 | 0.155 | 0.114 | 1 | 5 | Cib1 | 0.041 | Young |
| 0.047814 | 0.339722 | 0.14 | 0.098 | 1 | 5 | Usp36 | 0.042 | Young |
| 0.04784 | 0.444679 | 0.213 | 0.165 | 1 | 5 | Ccdc115 | 0.048 | Young |
| 0.048014 | 0.327767 | 0.485 | 0.444 | 1 | 5 | 9530068E0 | 0.041 | Young |
| 0.048025 | 0.172072 | 0.675 | 0.63 | 1 | 5 | Tpr | 0.045 | Young |
| 0.04843 | 0.577335 | 0.168 | 0.125 | 1 | 5 | Slc25a10 | 0.043 | Young |

| | | | | | | | | |
|---|---|---|---|---|---|---|---|---|
| 0.049002 | 0.425515 | 0.239 | 0.185 | 1 | 5 | Rcor1 | 0.054 | Young |
| 0.049254 | 0.73141 | 0.107 | 0.071 | 1 | 5 | Rpa2 | 0.036 | Young |
| 0.049294 | 1.426807 | 0.018 | 0.005 | 1 | 5 | 9130023H2 | 0.013 | Young |
| 0.049413 | 0.304967 | 0.581 | 0.564 | 1 | 5 | Cox5a | 0.017 | Young |
| 0.049421 | 0.763044 | 0.081 | 0.051 | 1 | 5 | Cdk7 | 0.03 | Young |
| 0.049751 | 1.238463 | 0.018 | 0.005 | 1 | 5 | Gm3336 | 0.013 | Young |
| 0.049783 | 0.390552 | 0.398 | 0.345 | 1 | 5 | Emc10 | 0.053 | Young |
| 0.049917 | 0.301306 | 0.457 | 0.397 | 1 | 5 | Stag2 | 0.06 | Young |
| 0.049942 | 0.340702 | 0.497 | 0.461 | 1 | 5 | Cysltr1 | 0.036 | Young |
| 0.050348 | 0.178016 | 0.799 | 0.779 | 1 | 5 | Arl6ip1 | 0.02 | Young |
| 0.050366 | 1.29078 | 0.018 | 0.005 | 1 | 5 | Gm26901 | 0.013 | Young |
| 0.050675 | 1.40472 | 0.018 | 0.005 | 1 | 5 | Ift122 | 0.013 | Young |
| 0.050677 | 0.677794 | 0.102 | 0.067 | 1 | 5 | Sema4d | 0.035 | Young |
| 0.050739 | 0.466054 | 0.14 | 0.098 | 1 | 5 | Ddx52 | 0.042 | Young |
| 0.050829 | 0.646092 | 0.16 | 0.12 | 1 | 5 | Ssbp1 | 0.04 | Young |
| 0.050831 | 1.052298 | 0.018 | 0.005 | 1 | 5 | Gm14326 | 0.013 | Young |
| 0.051223 | 0.827789 | 0.084 | 0.052 | 1 | 5 | Ubxn2b | 0.032 | Young |
| 0.051244 | 0.239969 | 0.693 | 0.695 | 1 | 5 | Pip4k2a | -0.002 | Young |
| 0.05139 | 0.579748 | 0.132 | 0.093 | 1 | 5 | Rapgef1 | 0.039 | Young |
| 0.051457 | 0.958098 | 0.018 | 0.005 | 1 | 5 | Gm14698 | 0.013 | Young |
| 0.051723 | 0.347304 | 0.299 | 0.242 | 1 | 5 | Zfc3h1 | 0.057 | Young |
| 0.051735 | 0.276166 | 0.614 | 0.567 | 1 | 5 | Ms4a6b | 0.047 | Young |
| 0.05194 | 0.182269 | 0.741 | 0.702 | 1 | 5 | Cxcl16 | 0.039 | Young |
| 0.052087 | 0.992428 | 0.048 | 0.025 | 1 | 5 | Mocs1 | 0.023 | Young |
| 0.05222 | 0.097643 | 0.985 | 0.99 | 1 | 5 | Rps23 | -0.005 | Young |
| 0.052543 | 0.168275 | 0.546 | 0.488 | 1 | 5 | Zfp703 | 0.058 | Young |
| 0.052551 | 0.483607 | 0.129 | 0.089 | 1 | 5 | Zfp467 | 0.04 | Young |
| 0.0526 | 0.132084 | 0.97 | 0.955 | 1 | 5 | Rpl19 | 0.015 | Young |
| 0.052841 | 0.354973 | 0.332 | 0.271 | 1 | 5 | Mff | 0.061 | Young |
| 0.052847 | 0.595171 | 0.094 | 0.061 | 1 | 5 | Cacna1d | 0.033 | Young |
| 0.052866 | 0.167998 | 0.675 | 0.64 | 1 | 5 | Arap1 | 0.035 | Young |
| 0.052968 | 0.850066 | 0.046 | 0.024 | 1 | 5 | Gys1 | 0.022 | Young |
| 0.052988 | 0.447348 | 0.416 | 0.364 | 1 | 5 | Itga4 | 0.052 | Young |
| 0.053261 | 0.212665 | 0.779 | 0.739 | 1 | 5 | Hnrnpa0 | 0.04 | Young |
| 0.053284 | 0.8747 | 0.063 | 0.037 | 1 | 5 | Trim23 | 0.026 | Young |
| 0.053359 | 0.212724 | 0.678 | 0.645 | 1 | 5 | Ppp1r18 | 0.033 | Young |
| 0.053522 | 0.438008 | 0.325 | 0.273 | 1 | 5 | Bcas2 | 0.052 | Young |
| 0.05362 | 0.420173 | 0.297 | 0.244 | 1 | 5 | Smad1 | 0.053 | Young |
| 0.053627 | 0.289632 | 0.5 | 0.465 | 1 | 5 | Eif3e | 0.035 | Young |
| 0.053649 | 0.576097 | 0.14 | 0.101 | 1 | 5 | Qars | 0.039 | Young |
| 0.053938 | 0.957359 | 0.076 | 0.047 | 1 | 5 | 2510009E0 | 0.029 | Young |
| 0.053948 | 0.31964 | 0.383 | 0.322 | 1 | 5 | Morc3 | 0.061 | Young |
| 0.054091 | 0.500368 | 0.152 | 0.111 | 1 | 5 | Slc30a9 | 0.041 | Young |
| 0.054104 | 0.453206 | 0.294 | 0.249 | 1 | 5 | 9-Sep | 0.045 | Young |
| 0.054998 | 0.250288 | 0.51 | 0.449 | 1 | 5 | Ifngr2 | 0.061 | Young |

| | | | | | | | | |
|---|---|---|---|---|---|---|---|---|
| 0.055013 | 0.116964 | 0.886 | 0.894 | 1 | 5 | Rps2 | -0.008 | Young |
| 0.055014 | 0.216413 | 0.642 | 0.616 | 1 | 5 | Fkbp1a | 0.026 | Young |
| 0.055101 | 0.438938 | 0.188 | 0.141 | 1 | 5 | Rpl10-ps3 | 0.047 | Young |
| 0.055204 | 0.112239 | 0.855 | 0.879 | 1 | 5 | Rpl14 | -0.024 | Young |
| 0.055242 | 0.588686 | 0.657 | 0.635 | 1 | 5 | Hmox1 | 0.022 | Young |
| 0.055521 | 1.38604 | 0.02 | 0.007 | 1 | 5 | Gm16104 | 0.013 | Young |
| 0.055619 | 0.272324 | 0.358 | 0.295 | 1 | 5 | Gga1 | 0.063 | Young |
| 0.055766 | 0.815916 | 0.063 | 0.037 | 1 | 5 | Pdpr | 0.026 | Young |
| 0.055823 | 0.237637 | 0.434 | 0.379 | 1 | 5 | Ncoa3 | 0.055 | Young |
| 0.055845 | 0.297339 | 0.472 | 0.439 | 1 | 5 | Spop | 0.033 | Young |
| 0.056157 | 0.586234 | 0.089 | 0.057 | 1 | 5 | Oxsm | 0.032 | Young |
| 0.056303 | 0.155515 | 0.853 | 0.86 | 1 | 5 | Hnrnpf | -0.007 | Young |
| 0.056391 | 0.468669 | 0.226 | 0.177 | 1 | 5 | Lysmd3 | 0.049 | Young |
| 0.056446 | 1.147461 | 0.02 | 0.007 | 1 | 5 | Icos | 0.013 | Young |
| 0.056571 | 0.271078 | 0.556 | 0.515 | 1 | 5 | Prex1 | 0.041 | Young |
| 0.056748 | 0.558006 | 0.112 | 0.076 | 1 | 5 | Galk1 | 0.036 | Young |
| 0.056838 | 0.153275 | 0.353 | 0.291 | 1 | 5 | Ppp4r3b | 0.062 | Young |
| 0.056914 | 1.168265 | 0.02 | 0.007 | 1 | 5 | Gm26947 | 0.013 | Young |
| 0.056957 | 1.054425 | 0.058 | 0.034 | 1 | 5 | Tmem132a | 0.024 | Young |
| 0.057074 | 0.929165 | 0.076 | 0.047 | 1 | 5 | Col14a1 | 0.029 | Young |
| 0.057227 | 0.951876 | 0.02 | 0.007 | 1 | 5 | Gm48604 | 0.013 | Young |
| 0.057431 | 0.240887 | 0.576 | 0.547 | 1 | 5 | Ppp1r15a | 0.029 | Young |
| 0.057662 | 0.581253 | 0.124 | 0.088 | 1 | 5 | Arhgap6 | 0.036 | Young |
| 0.05774 | 0.492038 | 0.099 | 0.066 | 1 | 5 | Ercc6l2 | 0.033 | Young |
| 0.057908 | 0.227129 | 0.769 | 0.744 | 1 | 5 | G3bp2 | 0.025 | Young |
| 0.058148 | 0.1026 | 0.881 | 0.884 | 1 | 5 | Pid1 | -0.003 | Young |
| 0.058173 | 0.987668 | 0.038 | 0.019 | 1 | 5 | Tubg1 | 0.019 | Young |
| 0.058175 | 0.707984 | 0.02 | 0.007 | 1 | 5 | Hist1h4d | 0.013 | Young |
| 0.058287 | 0.621756 | 0.107 | 0.072 | 1 | 5 | Lix1l | 0.035 | Young |
| 0.058335 | 0.822001 | 0.02 | 0.007 | 1 | 5 | Rbms3 | 0.013 | Young |
| 0.058507 | 1.636818 | 0.023 | 0.008 | 1 | 5 | Ifit3b | 0.015 | Young |
| 0.058587 | 0.4664 | 0.157 | 0.116 | 1 | 5 | Nr2f6 | 0.041 | Young |
| 0.058655 | 0.40309 | 0.02 | 0.007 | 1 | 5 | Xlr3b | 0.013 | Young |
| 0.058656 | 0.493452 | 0.142 | 0.103 | 1 | 5 | 2310033P0 | 0.039 | Young |
| 0.058749 | 0.248841 | 0.563 | 0.513 | 1 | 5 | Kif5b | 0.05 | Young |
| 0.058759 | 0.633063 | 0.145 | 0.106 | 1 | 5 | Gopc | 0.039 | Young |
| 0.058804 | 1.6083 | 0.023 | 0.008 | 1 | 5 | P2ry10b | 0.015 | Young |
| 0.058906 | 0.22111 | 0.612 | 0.581 | 1 | 5 | Dpysl2 | 0.031 | Young |
| 0.059185 | 0.21701 | 0.52 | 0.466 | 1 | 5 | Rnpep | 0.054 | Young |
| 0.059191 | 0.594786 | 0.104 | 0.071 | 1 | 5 | Stxbp6 | 0.033 | Young |
| 0.059214 | 0.458241 | 0.193 | 0.15 | 1 | 5 | Gatd1 | 0.043 | Young |
| 0.059263 | 0.51149 | 0.066 | 0.039 | 1 | 5 | Ap5s1 | 0.027 | Young |
| 0.059334 | 0.336236 | 0.302 | 0.242 | 1 | 5 | Wdfy4 | 0.06 | Young |
| 0.059382 | 1.087765 | 0.038 | 0.019 | 1 | 5 | Fancc | 0.019 | Young |
| 0.059419 | 0.643187 | 0.089 | 0.057 | 1 | 5 | Sec22a | 0.032 | Young |

| | | | | | | | | |
|---|---|---|---|---|---|---|---|---|
| 0.059426 | 0.112403 | 0.939 | 0.944 | 1 | 5 | Hnrnpa2b1 | -0.005 | Young |
| 0.059495 | 0.113272 | 0.975 | 0.978 | 1 | 5 | Atp2b1 | -0.003 | Young |
| 0.059565 | 0.625473 | 0.135 | 0.098 | 1 | 5 | Zfp236 | 0.037 | Young |
| 0.059619 | 0.947362 | 0.056 | 0.032 | 1 | 5 | Ccl25 | 0.024 | Young |
| 0.059715 | 0.966394 | 0.038 | 0.019 | 1 | 5 | Pstpip2 | 0.019 | Young |
| 0.059827 | 0.587508 | 0.076 | 0.047 | 1 | 5 | Tbcd | 0.029 | Young |
| 0.059854 | 1.185012 | 0.023 | 0.008 | 1 | 5 | Tns2 | 0.015 | Young |
| 0.06005 | 0.905569 | 0.038 | 0.019 | 1 | 5 | Zfp952 | 0.019 | Young |
| 0.060146 | 1.356005 | 0.033 | 0.015 | 1 | 5 | Tldc1 | 0.018 | Young |
| 0.060157 | 1.306271 | 0.023 | 0.008 | 1 | 5 | Gins1 | 0.015 | Young |
| 0.060308 | 0.886097 | 0.023 | 0.008 | 1 | 5 | Cfap45 | 0.015 | Young |
| 0.060329 | 0.525021 | 0.089 | 0.057 | 1 | 5 | Eif2b4 | 0.032 | Young |
| 0.060386 | 0.751224 | 0.038 | 0.019 | 1 | 5 | Ctu1 | 0.019 | Young |
| 0.060597 | 0.270407 | 0.449 | 0.404 | 1 | 5 | Creb1 | 0.045 | Young |
| 0.060613 | 1.068865 | 0.023 | 0.008 | 1 | 5 | Gm42670 | 0.015 | Young |
| 0.060616 | 0.236566 | 0.091 | 0.059 | 1 | 5 | Lonrf1 | 0.032 | Young |
| 0.060747 | 1.464735 | 0.028 | 0.012 | 1 | 5 | Zfp553 | 0.016 | Young |
| 0.060757 | 0.299653 | 0.261 | 0.207 | 1 | 5 | Ireb2 | 0.054 | Young |
| 0.060777 | 1.377919 | 0.025 | 0.01 | 1 | 5 | Zbtb49 | 0.015 | Young |
| 0.060975 | 1.18051 | 0.036 | 0.017 | 1 | 5 | Zfp748 | 0.019 | Young |
| 0.061045 | 0.779082 | 0.107 | 0.072 | 1 | 5 | Cntln | 0.035 | Young |
| 0.061113 | 0.210558 | 0.404 | 0.335 | 1 | 5 | Tnfrsf11a | 0.069 | Young |
| 0.061125 | 1.089485 | 0.033 | 0.015 | 1 | 5 | Zfp1 | 0.018 | Young |
| 0.0612 | 0.460235 | 0.155 | 0.113 | 1 | 5 | Mrps23 | 0.042 | Young |
| 0.06128 | 0.296453 | 0.183 | 0.136 | 1 | 5 | Mbd1 | 0.047 | Young |
| 0.061288 | 0.313693 | 0.325 | 0.269 | 1 | 5 | Ppp3cb | 0.056 | Young |
| 0.061416 | 0.267097 | 0.51 | 0.449 | 1 | 5 | Dnajc13 | 0.061 | Young |
| 0.061685 | 0.799105 | 0.036 | 0.017 | 1 | 5 | Gtf2ird1 | 0.019 | Young |
| 0.061991 | 1.147715 | 0.033 | 0.015 | 1 | 5 | Enc1 | 0.018 | Young |
| 0.062181 | 1.054078 | 0.071 | 0.044 | 1 | 5 | Ampd2 | 0.027 | Young |
| 0.062363 | 0.851972 | 0.058 | 0.034 | 1 | 5 | Serf1 | 0.024 | Young |
| 0.06241 | 0.603537 | 0.16 | 0.121 | 1 | 5 | Plxdc2 | 0.039 | Young |
| 0.062524 | 1.038708 | 0.028 | 0.012 | 1 | 5 | BC055324 | 0.016 | Young |
| 0.062524 | 0.993188 | 0.028 | 0.012 | 1 | 5 | Kctd17 | 0.016 | Young |
| 0.062663 | 0.973458 | 0.025 | 0.01 | 1 | 5 | Aurkb | 0.015 | Young |
| 0.06283 | 0.304198 | 0.594 | 0.547 | 1 | 5 | Tardbp | 0.047 | Young |
| 0.062936 | 0.42852 | 0.211 | 0.165 | 1 | 5 | Fam118a | 0.046 | Young |
| 0.062964 | 0.555631 | 0.19 | 0.146 | 1 | 5 | Kmt2b | 0.044 | Young |
| 0.063108 | 0.530352 | 0.223 | 0.178 | 1 | 5 | Mllt6 | 0.045 | Young |
| 0.063119 | 0.803307 | 0.081 | 0.052 | 1 | 5 | Zfp830 | 0.029 | Young |
| 0.063168 | 0.668961 | 0.074 | 0.045 | 1 | 5 | Fbxo21 | 0.029 | Young |
| 0.063259 | 0.260151 | 0.183 | 0.136 | 1 | 5 | Abhd4 | 0.047 | Young |
| 0.063406 | 0.731856 | 0.071 | 0.044 | 1 | 5 | Cdo1 | 0.027 | Young |
| 0.063463 | 0.468419 | 0.157 | 0.116 | 1 | 5 | Tbck | 0.041 | Young |
| 0.063487 | 0.763298 | 0.102 | 0.069 | 1 | 5 | Abcg1 | 0.033 | Young |

| | | | | | | | | |
|---|---|---|---|---|---|---|---|---|
| 0.063488 | 0.325499 | 0.264 | 0.214 | 1 | 5 | Mrpl14 | 0.05 | Young |
| 0.063568 | 0.533568 | 0.266 | 0.219 | 1 | 5 | Trpm7 | 0.047 | Young |
| 0.063744 | 0.311352 | 0.376 | 0.322 | 1 | 5 | Pdcd5 | 0.054 | Young |
| 0.063804 | 0.070499 | 0.962 | 0.941 | 1 | 5 | Rps5 | 0.021 | Young |
| 0.063825 | 0.581198 | 0.058 | 0.034 | 1 | 5 | Pgm2 | 0.024 | Young |
| 0.063846 | 1.025805 | 0.03 | 0.013 | 1 | 5 | Polg2 | 0.017 | Young |
| 0.064152 | 0.25233 | 0.231 | 0.178 | 1 | 5 | Cse1l | 0.053 | Young |
| 0.064213 | 0.301963 | 0.594 | 0.567 | 1 | 5 | Hnrnpm | 0.027 | Young |
| 0.064383 | 0.835003 | 0.03 | 0.013 | 1 | 5 | Zfp763 | 0.017 | Young |
| 0.064642 | 0.565573 | 0.147 | 0.108 | 1 | 5 | Cdk10 | 0.039 | Young |
| 0.064653 | 0.984524 | 0.03 | 0.013 | 1 | 5 | Rab4a | 0.017 | Young |
| 0.0647 | 0.052754 | 0.904 | 0.896 | 1 | 5 | Zfp36l2 | 0.008 | Young |
| 0.064998 | 0.292953 | 0.503 | 0.465 | 1 | 5 | Nras | 0.038 | Young |
| 0.065057 | 0.696498 | 0.028 | 0.012 | 1 | 5 | Hdhd5 | 0.016 | Young |
| 0.065221 | 0.577588 | 0.058 | 0.034 | 1 | 5 | Pick1 | 0.024 | Young |
| 0.065343 | 0.83926 | 0.028 | 0.012 | 1 | 5 | Spc25 | 0.016 | Young |
| 0.065431 | 0.405921 | 0.487 | 0.456 | 1 | 5 | Fam102b | 0.031 | Young |
| 0.065617 | 0.527189 | 0.165 | 0.126 | 1 | 5 | Mapk8 | 0.039 | Young |
| 0.065741 | 0.388791 | 0.213 | 0.168 | 1 | 5 | Anapc16 | 0.045 | Young |
| 0.065957 | 1.952768 | 0.01 | 0.002 | 1 | 5 | Apoc1 | 0.008 | Young |
| 0.065957 | 1.890259 | 0.01 | 0.002 | 1 | 5 | Frat2 | 0.008 | Young |
| 0.065957 | 1.616161 | 0.01 | 0.002 | 1 | 5 | Bloc1s6os | 0.008 | Young |
| 0.065957 | 1.468138 | 0.01 | 0.002 | 1 | 5 | Ablim2 | 0.008 | Young |
| 0.065957 | 1.458877 | 0.01 | 0.002 | 1 | 5 | Nme2 | 0.008 | Young |
| 0.065957 | 0.996028 | 0.01 | 0.002 | 1 | 5 | Gm14221 | 0.008 | Young |
| 0.06623 | 1.60764 | 0.01 | 0.002 | 1 | 5 | Fam187b | 0.008 | Young |
| 0.06623 | 1.507227 | 0.01 | 0.002 | 1 | 5 | Fam71e1 | 0.008 | Young |
| 0.06623 | 1.330788 | 0.01 | 0.002 | 1 | 5 | Gm4285 | 0.008 | Young |
| 0.06623 | 1.145065 | 0.01 | 0.002 | 1 | 5 | Dclk1 | 0.008 | Young |
| 0.06623 | 0.932179 | 0.01 | 0.002 | 1 | 5 | Gm15448 | 0.008 | Young |
| 0.066309 | 0.198525 | 0.678 | 0.653 | 1 | 5 | Pafah1b1 | 0.025 | Young |
| 0.066504 | 1.689955 | 0.01 | 0.002 | 1 | 5 | Gm6297 | 0.008 | Young |
| 0.066504 | 1.668705 | 0.01 | 0.002 | 1 | 5 | Epb41l3 | 0.008 | Young |
| 0.066504 | 1.527018 | 0.01 | 0.002 | 1 | 5 | Igfbp3 | 0.008 | Young |
| 0.066504 | 1.310074 | 0.01 | 0.002 | 1 | 5 | Vegfc | 0.008 | Young |
| 0.066504 | 1.210706 | 0.01 | 0.002 | 1 | 5 | Reep6 | 0.008 | Young |
| 0.066504 | 1.035202 | 0.01 | 0.002 | 1 | 5 | Creb3l4 | 0.008 | Young |
| 0.066504 | 1.018246 | 0.01 | 0.002 | 1 | 5 | Tinagl1 | 0.008 | Young |
| 0.066529 | 0.34682 | 0.256 | 0.209 | 1 | 5 | Ap3b1 | 0.047 | Young |
| 0.066779 | 1.236091 | 0.01 | 0.002 | 1 | 5 | Gramd1c | 0.008 | Young |
| 0.066779 | 1.162963 | 0.01 | 0.002 | 1 | 5 | Knl1 | 0.008 | Young |
| 0.066779 | 1.084802 | 0.01 | 0.002 | 1 | 5 | Gm12454 | 0.008 | Young |
| 0.066779 | 1.030384 | 0.01 | 0.002 | 1 | 5 | Prkcq | 0.008 | Young |
| 0.066779 | 1.005713 | 0.01 | 0.002 | 1 | 5 | Sh3d21 | 0.008 | Young |
| 0.066779 | 0.859659 | 0.01 | 0.002 | 1 | 5 | Lhfpl2 | 0.008 | Young |

| | | | | | | | | |
|---|---|---|---|---|---|---|---|---|
| 0.066779 | 0.856489 | 0.01 | 0.002 | 1 | 5 | Nim1k | 0.008 | Young |
| 0.066779 | 0.841394 | 0.01 | 0.002 | 1 | 5 | Gm7334 | 0.008 | Young |
| 0.066823 | 0.738829 | 0.056 | 0.032 | 1 | 5 | Zc3h6 | 0.024 | Young |
| 0.066918 | 0.624954 | 0.155 | 0.116 | 1 | 5 | Exoc7 | 0.039 | Young |
| 0.067045 | 0.426645 | 0.025 | 0.01 | 1 | 5 | Gm29562 | 0.015 | Young |
| 0.067055 | 0.949811 | 0.01 | 0.002 | 1 | 5 | Gm9889 | 0.008 | Young |
| 0.067055 | 0.695287 | 0.01 | 0.002 | 1 | 5 | Ptgis | 0.008 | Young |
| 0.067055 | 0.622418 | 0.01 | 0.002 | 1 | 5 | Ankef1 | 0.008 | Young |
| 0.067055 | 0.610186 | 0.01 | 0.002 | 1 | 5 | Pacsin3 | 0.008 | Young |
| 0.067448 | 0.38695 | 0.152 | 0.111 | 1 | 5 | Usp40 | 0.041 | Young |
| 0.067791 | 0.369295 | 0.317 | 0.271 | 1 | 5 | Kmt2d | 0.046 | Young |
| 0.067798 | 0.336616 | 0.492 | 0.458 | 1 | 5 | Ypel3 | 0.034 | Young |
| 0.067951 | 0.41396 | 0.234 | 0.189 | 1 | 5 | Slc44a1 | 0.045 | Young |
| 0.068299 | 0.416941 | 0.34 | 0.293 | 1 | 5 | Rnf213 | 0.047 | Young |
| 0.068415 | 0.513237 | 0.241 | 0.197 | 1 | 5 | Zzz3 | 0.044 | Young |
| 0.068462 | 0.512111 | 0.086 | 0.056 | 1 | 5 | Ccdc163 | 0.03 | Young |
| 0.068519 | 0.388565 | 0.284 | 0.234 | 1 | 5 | Suz12 | 0.05 | Young |
| 0.068872 | 0.633261 | 0.251 | 0.205 | 1 | 5 | 8-Mar | 0.046 | Young |
| 0.068944 | 0.083794 | 0.939 | 0.941 | 1 | 5 | Rap1b | -0.002 | Young |
| 0.068997 | 0.392263 | 0.234 | 0.184 | 1 | 5 | Glrx5 | 0.05 | Young |
| 0.069018 | 0.202524 | 0.459 | 0.391 | 1 | 5 | Ocrl | 0.068 | Young |
| 0.069169 | 0.352541 | 0.5 | 0.461 | 1 | 5 | Scarb2 | 0.039 | Young |
| 0.069658 | 0.265263 | 0.604 | 0.564 | 1 | 5 | Minos1 | 0.04 | Young |
| 0.070176 | 0.462654 | 0.234 | 0.19 | 1 | 5 | Brd3 | 0.044 | Young |
| 0.070344 | 0.136503 | 0.736 | 0.739 | 1 | 5 | Glud1 | -0.003 | Young |
| 0.070517 | 0.377165 | 0.409 | 0.359 | 1 | 5 | Mgat1 | 0.05 | Young |
| 0.070621 | 0.223141 | 0.338 | 0.278 | 1 | 5 | Klc1 | 0.06 | Young |
| 0.070757 | 0.306665 | 0.256 | 0.207 | 1 | 5 | Wdr89 | 0.049 | Young |
| 0.070904 | 0.578929 | 0.132 | 0.094 | 1 | 5 | Nudt5 | 0.038 | Young |
| 0.071151 | 0.628191 | 0.137 | 0.101 | 1 | 5 | Abraxas2 | 0.036 | Young |
| 0.071611 | 0.229169 | 0.68 | 0.685 | 1 | 5 | Sub1 | -0.005 | Young |
| 0.071616 | 0.37487 | 0.239 | 0.19 | 1 | 5 | Tgfbrap1 | 0.049 | Young |
| 0.07182 | 0.557483 | 0.091 | 0.061 | 1 | 5 | Cdc14a | 0.03 | Young |
| 0.072093 | 0.540981 | 0.071 | 0.044 | 1 | 5 | Bnip1 | 0.027 | Young |
| 0.072186 | 0.363166 | 0.348 | 0.295 | 1 | 5 | Snrpd2 | 0.053 | Young |
| 0.072533 | 0.469491 | 0.155 | 0.113 | 1 | 5 | Bnip3 | 0.042 | Young |
| 0.072621 | 0.26792 | 0.497 | 0.46 | 1 | 5 | Arf3 | 0.037 | Young |
| 0.072732 | 0.789824 | 0.066 | 0.04 | 1 | 5 | Pyroxd1 | 0.026 | Young |
| 0.072778 | 0.440395 | 0.289 | 0.246 | 1 | 5 | Esd | 0.043 | Young |
| 0.072785 | 0.41301 | 0.228 | 0.18 | 1 | 5 | Psme3 | 0.048 | Young |
| 0.072977 | 0.466112 | 0.053 | 0.03 | 1 | 5 | Slc16a1 | 0.023 | Young |
| 0.073217 | 0.379153 | 0.249 | 0.197 | 1 | 5 | Pts | 0.052 | Young |
| 0.073268 | 0.403457 | 0.259 | 0.212 | 1 | 5 | Trim35 | 0.047 | Young |
| 0.07354 | 0.537651 | 0.079 | 0.051 | 1 | 5 | Clk2 | 0.028 | Young |
| 0.073719 | 0.571774 | 0.234 | 0.187 | 1 | 5 | Fam43a | 0.047 | Young |

| | | | | | | | | |
|---|---|---|---|---|---|---|---|---|
| 0.073844 | 0.520699 | 0.053 | 0.03 | 1 | 5 | Trim2 | 0.023 | Young |
| 0.074034 | 0.875569 | 0.048 | 0.027 | 1 | 5 | Tnk2 | 0.021 | Young |
| 0.074584 | 0.31753 | 0.495 | 0.471 | 1 | 5 | Ssb | 0.024 | Young |
| 0.074825 | 0.817951 | 0.091 | 0.062 | 1 | 5 | Mrps26 | 0.029 | Young |
| 0.074876 | 1.045868 | 0.081 | 0.054 | 1 | 5 | Rmdn3 | 0.027 | Young |
| 0.074901 | 0.370837 | 0.315 | 0.268 | 1 | 5 | St6galnac4 | 0.047 | Young |
| 0.075345 | 0.495005 | 0.14 | 0.104 | 1 | 5 | Ptk2b | 0.036 | Young |
| 0.075726 | 0.442839 | 0.165 | 0.125 | 1 | 5 | Ccdc22 | 0.04 | Young |
| 0.075821 | 0.479827 | 0.218 | 0.177 | 1 | 5 | Rab3gap1 | 0.041 | Young |
| 0.0761 | 0.331235 | 0.348 | 0.291 | 1 | 5 | Tnfrsf13b | 0.057 | Young |
| 0.076105 | 0.299591 | 0.378 | 0.33 | 1 | 5 | Gtf2a1 | 0.048 | Young |
| 0.076136 | 0.453008 | 0.335 | 0.279 | 1 | 5 | Maz | 0.056 | Young |
| 0.076368 | 0.258051 | 0.363 | 0.308 | 1 | 5 | Fbl | 0.055 | Young |
| 0.076473 | 0.440967 | 0.173 | 0.133 | 1 | 5 | Tssc4 | 0.04 | Young |
| 0.076518 | 0.938935 | 0.063 | 0.039 | 1 | 5 | Arhgap23 | 0.024 | Young |
| 0.076614 | 0.485636 | 0.132 | 0.096 | 1 | 5 | Sgcb | 0.036 | Young |
| 0.076718 | 0.387513 | 0.203 | 0.158 | 1 | 5 | Vps37a | 0.045 | Young |
| 0.07688 | 0.423774 | 0.117 | 0.082 | 1 | 5 | Ascc2 | 0.035 | Young |
| 0.076944 | 0.841727 | 0.071 | 0.045 | 1 | 5 | Dus3l | 0.026 | Young |
| 0.077053 | 0.24829 | 0.533 | 0.485 | 1 | 5 | Eif4ebp2 | 0.048 | Young |
| 0.077065 | 0.391683 | 0.104 | 0.071 | 1 | 5 | Megf8 | 0.033 | Young |
| 0.077191 | 0.254596 | 0.398 | 0.343 | 1 | 5 | Myo9b | 0.055 | Young |
| 0.077315 | 0.346707 | 0.299 | 0.247 | 1 | 5 | Rasgef1b | 0.052 | Young |
| 0.077954 | 0.211899 | 0.68 | 0.641 | 1 | 5 | Ndufa2 | 0.039 | Young |
| 0.078688 | 0.475166 | 0.178 | 0.136 | 1 | 5 | Ebna1bp2 | 0.042 | Young |
| 0.078851 | 0.99811 | 0.046 | 0.025 | 1 | 5 | Rbm45 | 0.021 | Young |
| 0.078901 | 0.232964 | 0.345 | 0.288 | 1 | 5 | Apaf1 | 0.057 | Young |
| 0.078996 | 0.27477 | 0.36 | 0.306 | 1 | 5 | Arrdc1 | 0.054 | Young |
| 0.079156 | 0.331304 | 0.34 | 0.288 | 1 | 5 | Diaph2 | 0.052 | Young |
| 0.079257 | 0.184551 | 0.662 | 0.638 | 1 | 5 | 5031439G( | 0.024 | Young |
| 0.0794 | 0.340458 | 0.32 | 0.274 | 1 | 5 | Srsf10 | 0.046 | Young |
| 0.079414 | 0.437633 | 0.439 | 0.399 | 1 | 5 | Ndufb10 | 0.04 | Young |
| 0.079999 | 0.436272 | 0.152 | 0.114 | 1 | 5 | Ccdc34 | 0.038 | Young |
| 0.080538 | 0.294439 | 0.117 | 0.081 | 1 | 5 | Lrwd1 | 0.036 | Young |
| 0.080814 | 0.215941 | 0.401 | 0.36 | 1 | 5 | Ptpn12 | 0.041 | Young |
| 0.08084 | 0.425211 | 0.287 | 0.241 | 1 | 5 | Agpat3 | 0.046 | Young |
| 0.081079 | 0.361019 | 0.145 | 0.104 | 1 | 5 | Ift52 | 0.041 | Young |
| 0.081082 | 0.309184 | 0.373 | 0.337 | 1 | 5 | Lpin2 | 0.036 | Young |
| 0.081255 | 0.564184 | 0.246 | 0.207 | 1 | 5 | Psmc5 | 0.039 | Young |
| 0.081406 | 0.089643 | 0.964 | 0.961 | 1 | 5 | Rps19 | 0.003 | Young |
| 0.08141 | 0.389286 | 0.211 | 0.168 | 1 | 5 | Ash2l | 0.043 | Young |
| 0.08147 | 0.181434 | 0.703 | 0.675 | 1 | 5 | Celf1 | 0.028 | Young |
| 0.081706 | 0.341257 | 0.14 | 0.103 | 1 | 5 | Rbm34 | 0.037 | Young |
| 0.081986 | 0.016909 | 0.231 | 0.185 | 1 | 5 | Rnd3 | 0.046 | Young |
| 0.082255 | 0.415313 | 0.241 | 0.197 | 1 | 5 | Foxj2 | 0.044 | Young |

| | | | | | | | | |
|---|---|---|---|---|---|---|---|---|
| 0.083081 | 0.577686 | 0.109 | 0.077 | 1 | 5 | Kat2a | 0.032 | Young |
| 0.083107 | 0.335163 | 0.261 | 0.212 | 1 | 5 | Nfatc3 | 0.049 | Young |
| 0.083442 | 0.221002 | 0.338 | 0.291 | 1 | 5 | Mtmr6 | 0.047 | Young |
| 0.083601 | 0.301116 | 0.142 | 0.106 | 1 | 5 | Ppp1r7 | 0.036 | Young |
| 0.08364 | 0.395152 | 0.124 | 0.089 | 1 | 5 | Psmd10 | 0.035 | Young |
| 0.083676 | 0.580384 | 0.168 | 0.128 | 1 | 5 | Cbx5 | 0.04 | Young |
| 0.083798 | 0.344621 | 0.355 | 0.298 | 1 | 5 | Apc | 0.057 | Young |
| 0.084181 | 0.166215 | 0.736 | 0.722 | 1 | 5 | Nisch | 0.014 | Young |
| 0.084219 | 0.267041 | 0.497 | 0.453 | 1 | 5 | Psmb1 | 0.044 | Young |
| 0.084324 | 0.42686 | 0.165 | 0.126 | 1 | 5 | Terf2 | 0.039 | Young |
| 0.08441 | 0.309711 | 0.142 | 0.104 | 1 | 5 | Tmem11 | 0.038 | Young |
| 0.084513 | 0.510072 | 0.076 | 0.049 | 1 | 5 | 2610008E1 | 0.027 | Young |
| 0.084516 | 0.379736 | 0.216 | 0.17 | 1 | 5 | Shoc2 | 0.046 | Young |
| 0.084634 | 0.445478 | 0.145 | 0.108 | 1 | 5 | Isca2 | 0.037 | Young |
| 0.084653 | 0.218947 | 0.523 | 0.492 | 1 | 5 | Dek | 0.031 | Young |
| 0.084865 | 0.844834 | 0.043 | 0.024 | 1 | 5 | Top2a | 0.019 | Young |
| 0.085549 | 0.788463 | 0.043 | 0.024 | 1 | 5 | Gm47675 | 0.019 | Young |
| 0.085588 | 0.556694 | 0.089 | 0.061 | 1 | 5 | Spice1 | 0.028 | Young |
| 0.085596 | 0.457033 | 0.15 | 0.111 | 1 | 5 | Pex3 | 0.039 | Young |
| 0.085845 | 0.232163 | 0.647 | 0.641 | 1 | 5 | Ctdsp2 | 0.006 | Young |
| 0.085938 | 0.034987 | 0.165 | 0.214 | 1 | 5 | Vav2 | -0.049 | Young |
| 0.086006 | 0.79185 | 0.079 | 0.052 | 1 | 5 | Naip6 | 0.027 | Young |
| 0.086469 | 0.94287 | 0.041 | 0.022 | 1 | 5 | Polr3h | 0.019 | Young |
| 0.086513 | 0.885932 | 0.043 | 0.024 | 1 | 5 | Rsad1 | 0.019 | Young |
| 0.086619 | 0.320995 | 0.48 | 0.461 | 1 | 5 | Hnrnpul2 | 0.019 | Young |
| 0.086652 | 0.518403 | 0.043 | 0.024 | 1 | 5 | 2010111I0 | 0.019 | Young |
| 0.08684 | 1.308844 | 0.013 | 0.003 | 1 | 5 | Tlcd1 | 0.01 | Young |
| 0.08684 | 0.941491 | 0.013 | 0.003 | 1 | 5 | Gm45151 | 0.01 | Young |
| 0.086849 | 0.375552 | 0.338 | 0.293 | 1 | 5 | Aebp2 | 0.045 | Young |
| 0.086884 | 0.507146 | 0.061 | 0.037 | 1 | 5 | Tmem110 | 0.024 | Young |
| 0.086899 | 1.411204 | 0.038 | 0.02 | 1 | 5 | Vars2 | 0.018 | Young |
| 0.086938 | 0.176306 | 0.68 | 0.657 | 1 | 5 | Nap1l1 | 0.023 | Young |
| 0.087129 | 1.149088 | 0.013 | 0.003 | 1 | 5 | Gm10501 | 0.01 | Young |
| 0.087129 | 1.014131 | 0.013 | 0.003 | 1 | 5 | Cdca7l | 0.01 | Young |
| 0.087374 | 0.440683 | 0.19 | 0.152 | 1 | 5 | Wdr81 | 0.038 | Young |
| 0.087419 | 1.246954 | 0.013 | 0.003 | 1 | 5 | Enkd1 | 0.01 | Young |
| 0.087419 | 1.121547 | 0.013 | 0.003 | 1 | 5 | Lpcat4 | 0.01 | Young |
| 0.087419 | 1.107408 | 0.013 | 0.003 | 1 | 5 | Ttc30b | 0.01 | Young |
| 0.087526 | 0.561258 | 0.185 | 0.148 | 1 | 5 | Fzr1 | 0.037 | Young |
| 0.087617 | 0.309298 | 0.68 | 0.69 | 1 | 5 | Atp5h | -0.01 | Young |
| 0.087903 | 0.422406 | 0.16 | 0.123 | 1 | 5 | Polr2j | 0.037 | Young |
| 0.088001 | 0.636828 | 0.013 | 0.003 | 1 | 5 | Gm16740 | 0.01 | Young |
| 0.088108 | 0.348755 | 0.18 | 0.138 | 1 | 5 | Nploc4 | 0.042 | Young |
| 0.088198 | 1.130269 | 0.041 | 0.022 | 1 | 5 | Apold1 | 0.019 | Young |
| 0.088437 | 0.315073 | 0.404 | 0.364 | 1 | 5 | Pkig | 0.04 | Young |

| | | | | | | | | |
|---|---|---|---|---|---|---|---|---|
| 0.088489 | 0.843471 | 0.041 | 0.022 | 1 | 5 | Gpatch3 | 0.019 | Young |
| 0.088685 | 0.537232 | 0.165 | 0.128 | 1 | 5 | Scoc | 0.037 | Young |
| 0.08878 | 0.766987 | 0.041 | 0.022 | 1 | 5 | B3galt6 | 0.019 | Young |
| 0.08879 | 0.286067 | 0.132 | 0.096 | 1 | 5 | Lactb | 0.036 | Young |
| 0.08888 | 0.830631 | 0.013 | 0.003 | 1 | 5 | Gsta3 | 0.01 | Young |
| 0.088957 | 0.339884 | 0.188 | 0.143 | 1 | 5 | Dnajc2 | 0.045 | Young |
| 0.089365 | 0.254894 | 0.409 | 0.364 | 1 | 5 | Huwe1 | 0.045 | Young |
| 0.089774 | 0.581374 | 0.152 | 0.116 | 1 | 5 | Pdrg1 | 0.036 | Young |
| 0.089848 | 0.592631 | 0.104 | 0.074 | 1 | 5 | Sfxn2 | 0.03 | Young |
| 0.089884 | 0.46526 | 0.178 | 0.138 | 1 | 5 | Seh1l | 0.04 | Young |
| 0.089886 | 0.814564 | 0.079 | 0.052 | 1 | 5 | Rfx3 | 0.027 | Young |
| 0.089935 | 0.217433 | 0.637 | 0.591 | 1 | 5 | Synj1 | 0.046 | Young |
| 0.090006 | 0.155538 | 0.297 | 0.244 | 1 | 5 | Kcnq1ot1 | 0.053 | Young |
| 0.090065 | 0.965442 | 0.038 | 0.02 | 1 | 5 | Gmnn | 0.018 | Young |
| 0.090153 | 0.73771 | 0.056 | 0.034 | 1 | 5 | Dcp1b | 0.022 | Young |
| 0.090198 | 0.674813 | 0.069 | 0.044 | 1 | 5 | Slc39a11 | 0.025 | Young |
| 0.090335 | 0.259501 | 0.482 | 0.449 | 1 | 5 | Nipa2 | 0.033 | Young |
| 0.090494 | 1.066129 | 0.036 | 0.019 | 1 | 5 | Zswim9 | 0.017 | Young |
| 0.090502 | 0.602859 | 0.117 | 0.084 | 1 | 5 | Abca7 | 0.033 | Young |
| 0.090642 | 0.680927 | 0.069 | 0.044 | 1 | 5 | 9330160F1 | 0.025 | Young |
| 0.090941 | 0.399374 | 0.218 | 0.175 | 1 | 5 | Klhdc3 | 0.043 | Young |
| 0.091555 | 0.429492 | 0.231 | 0.194 | 1 | 5 | Mfap3 | 0.037 | Young |
| 0.091736 | 0.179306 | 0.612 | 0.572 | 1 | 5 | Pum2 | 0.04 | Young |
| 0.091908 | 0.426752 | 0.19 | 0.15 | 1 | 5 | Plaa | 0.04 | Young |
| 0.091937 | 1.162145 | 0.036 | 0.019 | 1 | 5 | Cby1 | 0.017 | Young |
| 0.092023 | 0.623731 | 0.119 | 0.088 | 1 | 5 | 6430548M | 0.031 | Young |
| 0.092188 | 0.441886 | 0.251 | 0.215 | 1 | 5 | Akna | 0.036 | Young |
| 0.092448 | 0.647691 | 0.104 | 0.074 | 1 | 5 | Nln | 0.03 | Young |
| 0.092576 | 0.850807 | 0.094 | 0.066 | 1 | 5 | Zfp524 | 0.028 | Young |
| 0.09276 | 0.264173 | 0.381 | 0.337 | 1 | 5 | Ccni | 0.044 | Young |
| 0.092785 | 0.412797 | 0.081 | 0.054 | 1 | 5 | Timm9 | 0.027 | Young |
| 0.093248 | 0.450194 | 0.041 | 0.022 | 1 | 5 | Senp8 | 0.019 | Young |
| 0.09326 | 0.444688 | 0.239 | 0.199 | 1 | 5 | Elf4 | 0.04 | Young |
| 0.093563 | 1.027135 | 0.036 | 0.019 | 1 | 5 | Ildr2 | 0.017 | Young |
| 0.093931 | 0.383331 | 0.211 | 0.168 | 1 | 5 | Cir1 | 0.043 | Young |
| 0.093993 | 0.306817 | 0.19 | 0.15 | 1 | 5 | Hadh | 0.04 | Young |
| 0.093998 | 0.381223 | 0.289 | 0.244 | 1 | 5 | Emc3 | 0.045 | Young |
| 0.094125 | 0.412808 | 0.188 | 0.148 | 1 | 5 | Gsap | 0.04 | Young |
| 0.094413 | 0.405769 | 0.109 | 0.077 | 1 | 5 | Nat6 | 0.032 | Young |
| 0.094549 | 0.98012 | 0.036 | 0.019 | 1 | 5 | Cenpq | 0.017 | Young |
| 0.094829 | 0.184707 | 0.292 | 0.237 | 1 | 5 | Evi5 | 0.055 | Young |
| 0.094907 | 0.767228 | 0.038 | 0.02 | 1 | 5 | Ank2 | 0.018 | Young |
| 0.095063 | 0.211467 | 0.665 | 0.678 | 1 | 5 | Hmgb1 | -0.013 | Young |
| 0.095165 | 0.399091 | 0.249 | 0.209 | 1 | 5 | Utp11 | 0.04 | Young |
| 0.095226 | 0.339068 | 0.038 | 0.02 | 1 | 5 | Kbtbd3 | 0.018 | Young |

| | | | | | | | | |
|---|---|---|---|---|---|---|---|---|
| 0.095315 | 0.468046 | 0.211 | 0.17 | 1 | 5 | Snx1 | 0.041 | Young |
| 0.095349 | 0.636635 | 0.079 | 0.052 | 1 | 5 | Mier2 | 0.027 | Young |
| 0.095573 | 0.412219 | 0.261 | 0.222 | 1 | 5 | Mdm2 | 0.039 | Young |
| 0.095939 | 0.33311 | 0.31 | 0.263 | 1 | 5 | Gspt1 | 0.047 | Young |
| 0.096003 | 0.472012 | 0.208 | 0.167 | 1 | 5 | Marveld1 | 0.041 | Young |
| 0.096444 | 0.105171 | 0.629 | 0.582 | 1 | 5 | Rhob | 0.047 | Young |
| 0.096467 | 0.195338 | 0.655 | 0.623 | 1 | 5 | Tmem30a | 0.032 | Young |
| 0.096605 | 0.353113 | 0.165 | 0.125 | 1 | 5 | Lipe | 0.04 | Young |
| 0.096678 | 0.403684 | 0.132 | 0.098 | 1 | 5 | Daxx | 0.034 | Young |
| 0.096796 | 0.351026 | 0.287 | 0.242 | 1 | 5 | Vps13c | 0.045 | Young |
| 0.097139 | 0.396771 | 0.353 | 0.315 | 1 | 5 | Rassf3 | 0.038 | Young |
| 0.097257 | 0.872617 | 0.051 | 0.03 | 1 | 5 | 1500015A0 | 0.021 | Young |
| 0.097264 | 0.291267 | 0.325 | 0.276 | 1 | 5 | Insr | 0.049 | Young |
| 0.097498 | 0.545102 | 0.104 | 0.074 | 1 | 5 | Ganc | 0.03 | Young |
| 0.097534 | 1.270811 | 0.015 | 0.005 | 1 | 5 | Zfp607b | 0.01 | Young |
| 0.097649 | 0.440586 | 0.038 | 0.02 | 1 | 5 | Mtrr | 0.018 | Young |
| 0.097693 | 0.224778 | 0.175 | 0.135 | 1 | 5 | Dnajc10 | 0.04 | Young |
| 0.097727 | 0.545957 | 0.203 | 0.163 | 1 | 5 | Scd1 | 0.04 | Young |
| 0.098067 | 0.689182 | 0.036 | 0.019 | 1 | 5 | Lyrm4 | 0.017 | Young |
| 0.098242 | 0.385892 | 0.381 | 0.342 | 1 | 5 | Ophn1 | 0.039 | Young |
| 0.098408 | 0.643008 | 0.036 | 0.019 | 1 | 5 | Zfp619 | 0.017 | Young |
| 0.098432 | 0.177787 | 0.797 | 0.754 | 1 | 5 | Mef2a | 0.043 | Young |
| 0.098494 | 0.286117 | 0.368 | 0.328 | 1 | 5 | Tle4 | 0.04 | Young |
| 0.098786 | 0.548168 | 0.218 | 0.182 | 1 | 5 | Acaa2 | 0.036 | Young |
| 0.098854 | 0.328531 | 0.244 | 0.199 | 1 | 5 | Zfp451 | 0.045 | Young |
| 0.099135 | 0.506677 | 0.244 | 0.2 | 1 | 5 | Dapp1 | 0.044 | Young |
| 0.099153 | 0.618769 | 0.053 | 0.032 | 1 | 5 | Gtf2h2 | 0.021 | Young |
| 0.099792 | 0.918292 | 0.015 | 0.005 | 1 | 5 | Fermt2 | 0.01 | Young |
| 0.099821 | 0.143179 | 0.662 | 0.64 | 1 | 5 | Rpn1 | 0.022 | Young |
| 0.100029 | 0.507351 | 0.185 | 0.15 | 1 | 5 | Epb41l4aos | 0.035 | Young |
| 0.100078 | 0.990852 | 0.015 | 0.005 | 1 | 5 | Lyplal1 | 0.01 | Young |
| 0.100078 | 0.869464 | 0.015 | 0.005 | 1 | 5 | Homez | 0.01 | Young |
| 0.100364 | 0.755642 | 0.015 | 0.005 | 1 | 5 | Oaf | 0.01 | Young |
| 0.100393 | 0.496187 | 0.221 | 0.184 | 1 | 5 | Emc6 | 0.037 | Young |
| 0.10047 | 0.561213 | 0.036 | 0.019 | 1 | 5 | Sars2 | 0.017 | Young |
| 0.100525 | 0.662106 | 0.099 | 0.071 | 1 | 5 | Tnfrsf21 | 0.028 | Young |
| 0.100572 | 0.302539 | 0.376 | 0.338 | 1 | 5 | Slc25a36 | 0.038 | Young |
| 0.100683 | 0.327237 | 0.376 | 0.338 | 1 | 5 | Snx30 | 0.038 | Young |
| 0.100781 | 0.413075 | 0.305 | 0.264 | 1 | 5 | Prkar2a | 0.041 | Young |
| 0.101085 | 0.944138 | 0.19 | 0.153 | 1 | 5 | Hgsnat | 0.037 | Young |
| 0.101225 | 0.716249 | 0.015 | 0.005 | 1 | 5 | P4ha2 | 0.01 | Young |
| 0.101373 | 0.319688 | 0.434 | 0.382 | 1 | 5 | Ndufc1 | 0.052 | Young |
| 0.101452 | 0.394472 | 0.119 | 0.086 | 1 | 5 | Cxxc5 | 0.033 | Young |
| 0.101616 | 0.218072 | 0.515 | 0.488 | 1 | 5 | Zdhhc20 | 0.027 | Young |
| 0.10162 | 0.639963 | 0.099 | 0.071 | 1 | 5 | Abhd6 | 0.028 | Young |

| | | | | | | | | |
|---|---|---|---|---|---|---|---|---|
| 0.101744 | 0.235458 | 0.338 | 0.293 | 1 | 5 | Uhrf2 | 0.045 | Young |
| 0.101803 | 0.218009 | 0.015 | 0.005 | 1 | 5 | Gm8013 | 0.01 | Young |
| 0.101832 | 0.17816 | 0.718 | 0.707 | 1 | 5 | Msn | 0.011 | Young |
| 0.101883 | 0.501386 | 0.033 | 0.017 | 1 | 5 | Nup37 | 0.016 | Young |
| 0.102093 | 0.523268 | 0.015 | 0.005 | 1 | 5 | Dus4l | 0.01 | Young |
| 0.102437 | 0.294547 | 0.398 | 0.357 | 1 | 5 | Higd2a | 0.041 | Young |
| 0.102515 | 0.271337 | 0.386 | 0.34 | 1 | 5 | Ccar1 | 0.046 | Young |
| 0.102549 | 0.415281 | 0.079 | 0.052 | 1 | 5 | 10-Sep | 0.027 | Young |
| 0.102567 | 0.354088 | 0.036 | 0.019 | 1 | 5 | Tbc1d8b | 0.017 | Young |
| 0.102581 | 0.329915 | 0.348 | 0.306 | 1 | 5 | Retreg2 | 0.042 | Young |
| 0.102885 | 0.227145 | 0.556 | 0.527 | 1 | 5 | Bnip2 | 0.029 | Young |
| 0.102926 | 0.352053 | 0.183 | 0.143 | 1 | 5 | Naa20 | 0.04 | Young |
| 0.103132 | 1.23353 | 0.046 | 0.027 | 1 | 5 | 1810043G( | 0.019 | Young |
| 0.103167 | 0.642269 | 0.033 | 0.017 | 1 | 5 | Gm26520 | 0.016 | Young |
| 0.103437 | 0.134049 | 0.901 | 0.923 | 1 | 5 | Ly86 | -0.022 | Young |
| 0.103512 | 1.456621 | 0.018 | 0.007 | 1 | 5 | Col6a3 | 0.011 | Young |
| 0.103895 | 0.450382 | 0.208 | 0.172 | 1 | 5 | Ubac2 | 0.036 | Young |
| 0.103931 | 0.34243 | 0.206 | 0.165 | 1 | 5 | Thoc7 | 0.041 | Young |
| 0.104044 | 1.096913 | 0.018 | 0.007 | 1 | 5 | Serpinb1a | 0.011 | Young |
| 0.104385 | 0.641087 | 0.084 | 0.057 | 1 | 5 | Aldh6a1 | 0.027 | Young |
| 0.104675 | 0.494785 | 0.109 | 0.079 | 1 | 5 | Tlr5 | 0.03 | Young |
| 0.10478 | 0.219091 | 0.307 | 0.256 | 1 | 5 | Rps27rt | 0.051 | Young |
| 0.1051 | 0.462373 | 0.145 | 0.109 | 1 | 5 | Ate1 | 0.036 | Young |
| 0.105115 | 1.180994 | 0.018 | 0.007 | 1 | 5 | Napsa | 0.011 | Young |
| 0.105115 | 1.108418 | 0.018 | 0.007 | 1 | 5 | Ttc30a1 | 0.011 | Young |
| 0.105168 | 0.627951 | 0.084 | 0.057 | 1 | 5 | Abcg2 | 0.027 | Young |
| 0.105185 | 0.410317 | 0.211 | 0.17 | 1 | 5 | Gcc2 | 0.041 | Young |
| 0.105384 | 1.065653 | 0.018 | 0.007 | 1 | 5 | Rad9b | 0.011 | Young |
| 0.10544 | 0.625542 | 0.18 | 0.146 | 1 | 5 | Trnau1ap | 0.034 | Young |
| 0.105505 | 0.539947 | 0.084 | 0.057 | 1 | 5 | Rpp14 | 0.027 | Young |
| 0.105548 | 0.424334 | 0.246 | 0.205 | 1 | 5 | Paics | 0.041 | Young |
| 0.105597 | 0.213304 | 0.447 | 0.392 | 1 | 5 | Ubtf | 0.055 | Young |
| 0.105654 | 1.095546 | 0.018 | 0.007 | 1 | 5 | Pxmp2 | 0.011 | Young |
| 0.105668 | 0.351549 | 0.051 | 0.03 | 1 | 5 | Zcchc10 | 0.021 | Young |
| 0.106195 | 0.643543 | 0.018 | 0.007 | 1 | 5 | Gm5141 | 0.011 | Young |
| 0.106226 | 0.487535 | 0.03 | 0.015 | 1 | 5 | Lrrc8b | 0.015 | Young |
| 0.106343 | 1.102399 | 0.025 | 0.012 | 1 | 5 | Taf1a | 0.013 | Young |
| 0.106423 | 1.180872 | 0.02 | 0.008 | 1 | 5 | Fancm | 0.012 | Young |
| 0.10672 | 1.525172 | 0.023 | 0.01 | 1 | 5 | Vsig4 | 0.013 | Young |
| 0.106782 | 1.307267 | 0.025 | 0.012 | 1 | 5 | Smad7 | 0.013 | Young |
| 0.106782 | 0.977813 | 0.025 | 0.012 | 1 | 5 | Cyp2e1 | 0.013 | Young |
| 0.106806 | 0.402567 | 0.119 | 0.088 | 1 | 5 | Gm28875 | 0.031 | Young |
| 0.10683 | 0.576105 | 0.081 | 0.056 | 1 | 5 | Nudt14 | 0.025 | Young |
| 0.106913 | 0.218476 | 0.556 | 0.529 | 1 | 5 | Nptn | 0.027 | Young |
| 0.106973 | 0.294375 | 0.201 | 0.158 | 1 | 5 | Ncbp3 | 0.043 | Young |

| | | | | | | | | |
|---|---|---|---|---|---|---|---|---|
| 0.107002 | 1.155839 | 0.025 | 0.012 | 1 | 5 | A430046D1 | 0.013 | Young |
| 0.107011 | 0.839382 | 0.018 | 0.007 | 1 | 5 | Ccdc102a | 0.011 | Young |
| 0.107091 | 0.399559 | 0.16 | 0.125 | 1 | 5 | 3830406C1 | 0.035 | Young |
| 0.107362 | 0.321976 | 0.124 | 0.091 | 1 | 5 | Ecd | 0.033 | Young |
| 0.107427 | 0.899838 | 0.02 | 0.008 | 1 | 5 | Vcpkmt | 0.012 | Young |
| 0.107557 | 0.676719 | 0.018 | 0.007 | 1 | 5 | Plpp1 | 0.011 | Young |
| 0.107679 | 1.069763 | 0.02 | 0.008 | 1 | 5 | Gm19325 | 0.012 | Young |
| 0.107739 | 0.39931 | 0.17 | 0.133 | 1 | 5 | Meaf6 | 0.037 | Young |
| 0.107743 | 0.642408 | 0.127 | 0.096 | 1 | 5 | Ccng2 | 0.031 | Young |
| 0.107779 | 0.368523 | 0.388 | 0.357 | 1 | 5 | Nrip1 | 0.031 | Young |
| 0.107852 | 0.330353 | 0.327 | 0.29 | 1 | 5 | Mtch1 | 0.037 | Young |
| 0.107931 | 0.938546 | 0.02 | 0.008 | 1 | 5 | Sirpb1c | 0.012 | Young |
| 0.108106 | 0.595326 | 0.018 | 0.007 | 1 | 5 | Ddx11 | 0.011 | Young |
| 0.108146 | 0.895543 | 0.028 | 0.013 | 1 | 5 | Adat2 | 0.015 | Young |
| 0.108567 | 0.52739 | 0.028 | 0.013 | 1 | 5 | Atp1a2 | 0.015 | Young |
| 0.108567 | 0.478402 | 0.028 | 0.013 | 1 | 5 | Ckap2l | 0.015 | Young |
| 0.108601 | 1.075315 | 0.023 | 0.01 | 1 | 5 | Stxbp4 | 0.013 | Young |
| 0.108861 | 0.228945 | 0.596 | 0.542 | 1 | 5 | Zfp706 | 0.054 | Young |
| 0.109044 | 0.607128 | 0.157 | 0.126 | 1 | 5 | Sgsh | 0.031 | Young |
| 0.109076 | 1.122363 | 0.023 | 0.01 | 1 | 5 | Fxyd1 | 0.013 | Young |
| 0.10922 | 0.854282 | 0.025 | 0.012 | 1 | 5 | Ttc12 | 0.013 | Young |
| 0.109382 | 0.680143 | 0.107 | 0.077 | 1 | 5 | Tmem140 | 0.03 | Young |
| 0.109401 | 0.566768 | 0.124 | 0.093 | 1 | 5 | Phkg2 | 0.031 | Young |
| 0.10942 | 0.291914 | 0.03 | 0.015 | 1 | 5 | Fgfbp3 | 0.015 | Young |
| 0.109457 | 0.662417 | 0.02 | 0.008 | 1 | 5 | Atp23 | 0.012 | Young |
| 0.109552 | 0.972452 | 0.023 | 0.01 | 1 | 5 | Nlrc3 | 0.013 | Young |
| 0.109713 | 0.426118 | 0.168 | 0.131 | 1 | 5 | Mrps25 | 0.037 | Young |
| 0.109857 | 0.478105 | 0.157 | 0.121 | 1 | 5 | Mpdu1 | 0.036 | Young |
| 0.109903 | 0.677306 | 0.061 | 0.039 | 1 | 5 | Xrcc6 | 0.022 | Young |
| 0.110029 | 0.77999 | 0.023 | 0.01 | 1 | 5 | 6-Sep | 0.013 | Young |
| 0.110065 | 0.798858 | 0.086 | 0.061 | 1 | 5 | Plpp6 | 0.025 | Young |
| 0.11023 | 0.455632 | 0.388 | 0.354 | 1 | 5 | Jdp2 | 0.034 | Young |
| 0.110269 | 0.952875 | 0.023 | 0.01 | 1 | 5 | Cd226 | 0.013 | Young |
| 0.110465 | 0.385119 | 0.051 | 0.03 | 1 | 5 | Zmym6 | 0.021 | Young |
| 0.110483 | 0.629198 | 0.02 | 0.008 | 1 | 5 | Trmt5 | 0.012 | Young |
| 0.110525 | 0.302429 | 0.185 | 0.143 | 1 | 5 | Psmc4 | 0.042 | Young |
| 0.110551 | 0.446956 | 0.063 | 0.04 | 1 | 5 | Nob1 | 0.023 | Young |
| 0.110701 | 0.012434 | 0.231 | 0.283 | 1 | 5 | 1810058I24 | -0.052 | Young |
| 0.111141 | 0.802529 | 0.046 | 0.027 | 1 | 5 | Zfp617 | 0.019 | Young |
| 0.111474 | 0.544473 | 0.025 | 0.012 | 1 | 5 | Trmt12 | 0.013 | Young |
| 0.111632 | 0.443281 | 0.15 | 0.116 | 1 | 5 | Rfx7 | 0.034 | Young |
| 0.11193 | 0.541302 | 0.025 | 0.012 | 1 | 5 | Rapsn | 0.013 | Young |
| 0.111981 | 0.299684 | 0.028 | 0.013 | 1 | 5 | Klf8 | 0.015 | Young |
| 0.112074 | 0.116389 | 0.876 | 0.855 | 1 | 5 | Rpl36a | 0.021 | Young |
| 0.112248 | 0.363399 | 0.168 | 0.131 | 1 | 5 | Tcp11l2 | 0.037 | Young |

| | | | | | | | | |
|---|---|---|---|---|---|---|---|---|
| 0.112388 | 0.380156 | 0.327 | 0.286 | 1 | 5 | Pgd | 0.041 | Young |
| 0.113168 | 0.172179 | 0.569 | 0.539 | 1 | 5 | Itpripl2 | 0.03 | Young |
| 0.113272 | 0.130526 | 0.774 | 0.771 | 1 | 5 | Coro1a | 0.003 | Young |
| 0.113815 | 0.644785 | 0.195 | 0.158 | 1 | 5 | Mxi1 | 0.037 | Young |
| 0.113966 | 0.151516 | 0.726 | 0.732 | 1 | 5 | Ap1b1 | -0.006 | Young |
| 0.113993 | 0.191357 | 0.421 | 0.369 | 1 | 5 | Kat6a | 0.052 | Young |
| 0.114017 | 0.702613 | 0.079 | 0.054 | 1 | 5 | Zfp180 | 0.025 | Young |
| 0.114328 | 0.509432 | 0.18 | 0.143 | 1 | 5 | Ctbs | 0.037 | Young |
| 0.11514 | 0.67139 | 0.076 | 0.052 | 1 | 5 | Tlr3 | 0.024 | Young |
| 0.11533 | 0.390418 | 0.129 | 0.096 | 1 | 5 | Sigmar1 | 0.033 | Young |
| 0.115721 | 0.393247 | 0.32 | 0.278 | 1 | 5 | Usp48 | 0.042 | Young |
| 0.115806 | 0.494179 | 0.137 | 0.106 | 1 | 5 | Thbd | 0.031 | Young |
| 0.116138 | 0.35923 | 0.084 | 0.057 | 1 | 5 | Fbxo25 | 0.027 | Young |
| 0.116365 | 0.282669 | 0.34 | 0.296 | 1 | 5 | Fbxl3 | 0.044 | Young |
| 0.116399 | 0.350044 | 0.325 | 0.285 | 1 | 5 | Acly | 0.04 | Young |
| 0.116547 | 0.293809 | 0.259 | 0.219 | 1 | 5 | Rbm27 | 0.04 | Young |
| 0.116697 | 0.692464 | 0.043 | 0.025 | 1 | 5 | Zbed3 | 0.018 | Young |
| 0.116918 | 0.566548 | 0.119 | 0.089 | 1 | 5 | Zfp24 | 0.03 | Young |
| 0.117534 | 0.300152 | 0.52 | 0.473 | 1 | 5 | Vps4b | 0.047 | Young |
| 0.11776 | 0.671215 | 0.046 | 0.027 | 1 | 5 | Gm26778 | 0.019 | Young |
| 0.118001 | 0.512817 | 0.074 | 0.049 | 1 | 5 | Acvr2a | 0.025 | Young |
| 0.118242 | 0.307854 | 0.292 | 0.246 | 1 | 5 | Rtn1 | 0.046 | Young |
| 0.118359 | 0.410736 | 0.368 | 0.33 | 1 | 5 | Bmpr2 | 0.038 | Young |
| 0.118453 | 0.566943 | 0.185 | 0.148 | 1 | 5 | Cramp1l | 0.037 | Young |
| 0.118735 | 0.494662 | 0.109 | 0.079 | 1 | 5 | Zfp26 | 0.03 | Young |
| 0.118893 | 0.227087 | 0.424 | 0.375 | 1 | 5 | Sin3b | 0.049 | Young |
| 0.119418 | 0.225538 | 0.307 | 0.261 | 1 | 5 | Becn1 | 0.046 | Young |
| 0.11944 | 0.481831 | 0.165 | 0.128 | 1 | 5 | Ssbp2 | 0.037 | Young |
| 0.119627 | 0.287444 | 0.414 | 0.367 | 1 | 5 | Acsl4 | 0.047 | Young |
| 0.119675 | 0.114747 | 0.931 | 0.968 | 1 | 5 | Pfn1 | -0.037 | Young |
| 0.119903 | 0.393822 | 0.294 | 0.254 | 1 | 5 | 1500011K1 | 0.04 | Young |
| 0.120024 | 0.127165 | 0.701 | 0.643 | 1 | 5 | Stard8 | 0.058 | Young |
| 0.12028 | 0.328808 | 0.091 | 0.064 | 1 | 5 | Sclt1 | 0.027 | Young |
| 0.120464 | 0.394647 | 0.406 | 0.365 | 1 | 5 | Cebpd | 0.041 | Young |
| 0.120566 | 0.21367 | 0.129 | 0.096 | 1 | 5 | Chmp7 | 0.033 | Young |
| 0.121011 | 0.511193 | 0.046 | 0.027 | 1 | 5 | Ssx2ip | 0.019 | Young |
| 0.121135 | 0.812159 | 0.056 | 0.035 | 1 | 5 | Tapt1 | 0.021 | Young |
| 0.121153 | 0.190975 | 0.213 | 0.172 | 1 | 5 | Arhgap25 | 0.041 | Young |
| 0.121279 | 0.528672 | 0.221 | 0.185 | 1 | 5 | Dapk1 | 0.036 | Young |
| 0.121754 | 0.586202 | 0.056 | 0.035 | 1 | 5 | Panx1 | 0.021 | Young |
| 0.121772 | 0.615787 | 0.069 | 0.045 | 1 | 5 | Tab3 | 0.024 | Young |
| 0.122041 | 0.464666 | 0.099 | 0.072 | 1 | 5 | Urm1 | 0.027 | Young |
| 0.122151 | 0.277663 | 0.437 | 0.394 | 1 | 5 | Dnajc5 | 0.043 | Young |
| 0.122339 | 0.615301 | 0.043 | 0.025 | 1 | 5 | Lilra6 | 0.018 | Young |
| 0.122419 | 0.491264 | 0.132 | 0.099 | 1 | 5 | Kansl2 | 0.033 | Young |

| | | | | | | | | |
|---|---|---|---|---|---|---|---|---|
| 0.12245 | 0.242291 | 0.365 | 0.327 | 1 | 5 | Usp15 | 0.038 | Young |
| 0.122532 | 0.755119 | 0.056 | 0.035 | 1 | 5 | Bysl | 0.021 | Young |
| 0.122557 | 0.331675 | 0.14 | 0.106 | 1 | 5 | Sp2 | 0.034 | Young |
| 0.122631 | 0.426032 | 0.223 | 0.189 | 1 | 5 | Arl6ip5 | 0.034 | Young |
| 0.122659 | 0.201329 | 0.563 | 0.529 | 1 | 5 | Prkar1a | 0.034 | Young |
| 0.123141 | 0.846027 | 0.081 | 0.057 | 1 | 5 | Coq7 | 0.024 | Young |
| 0.123239 | 0.461819 | 0.043 | 0.025 | 1 | 5 | Tagln | 0.018 | Young |
| 0.123489 | 0.500068 | 0.109 | 0.081 | 1 | 5 | Slc36a4 | 0.028 | Young |
| 0.123544 | 1.027534 | 0.041 | 0.024 | 1 | 5 | 5430403G1 | 0.017 | Young |
| 0.123893 | 0.403499 | 0.203 | 0.163 | 1 | 5 | Psmg4 | 0.04 | Young |
| 0.123955 | 0.363404 | 0.198 | 0.16 | 1 | 5 | Zcrb1 | 0.038 | Young |
| 0.124079 | 0.286219 | 0.292 | 0.251 | 1 | 5 | Bcl2l11 | 0.041 | Young |
| 0.124232 | 0.192192 | 0.629 | 0.596 | 1 | 5 | Gng2 | 0.033 | Young |
| 0.124421 | 0.257358 | 0.391 | 0.359 | 1 | 5 | Tmem14c | 0.032 | Young |
| 0.124658 | 0.269174 | 0.325 | 0.281 | 1 | 5 | Psmd8 | 0.044 | Young |
| 0.124688 | 0.507305 | 0.188 | 0.153 | 1 | 5 | Nsmce1 | 0.035 | Young |
| 0.125103 | 0.321227 | 0.213 | 0.175 | 1 | 5 | Lsm5 | 0.038 | Young |
| 0.125302 | 0.219295 | 0.589 | 0.566 | 1 | 5 | Ubl5 | 0.023 | Young |
| 0.125646 | 0.595141 | 0.15 | 0.118 | 1 | 5 | Mrpl43 | 0.032 | Young |
| 0.125654 | 0.303788 | 0.244 | 0.205 | 1 | 5 | Ubr3 | 0.039 | Young |
| 0.125673 | 0.40139 | 0.157 | 0.123 | 1 | 5 | Fer | 0.034 | Young |
| 0.125744 | 1.047081 | 0.038 | 0.022 | 1 | 5 | Rab38 | 0.016 | Young |
| 0.126134 | 0.317795 | 0.206 | 0.163 | 1 | 5 | Zmpste24 | 0.043 | Young |
| 0.126143 | 0.165637 | 0.274 | 0.237 | 1 | 5 | Fcna | 0.037 | Young |
| 0.126898 | 0.653925 | 0.053 | 0.034 | 1 | 5 | Coil | 0.019 | Young |
| 0.127065 | 0.277688 | 0.419 | 0.387 | 1 | 5 | Anp32e | 0.032 | Young |
| 0.127232 | 0.428283 | 0.198 | 0.165 | 1 | 5 | Ckap5 | 0.033 | Young |
| 0.127356 | 0.26365 | 0.157 | 0.121 | 1 | 5 | Ptgr2 | 0.036 | Young |
| 0.127365 | 0.319689 | 0.292 | 0.253 | 1 | 5 | Psmd1 | 0.039 | Young |
| 0.127499 | 0.703562 | 0.066 | 0.044 | 1 | 5 | Wdr12 | 0.022 | Young |
| 0.128154 | 0.377917 | 0.173 | 0.14 | 1 | 5 | Zpr1 | 0.033 | Young |
| 0.128244 | 0.368382 | 0.33 | 0.296 | 1 | 5 | Uhmk1 | 0.034 | Young |
| 0.128346 | 0.255832 | 0.287 | 0.239 | 1 | 5 | Mta2 | 0.048 | Young |
| 0.128386 | 0.575439 | 0.074 | 0.051 | 1 | 5 | Pam | 0.023 | Young |
| 0.128481 | 0.340362 | 0.168 | 0.13 | 1 | 5 | Sap30l | 0.038 | Young |
| 0.128513 | 0.242476 | 0.305 | 0.261 | 1 | 5 | Ythdf1 | 0.044 | Young |
| 0.12858 | 0.274187 | 0.17 | 0.133 | 1 | 5 | Usf1 | 0.037 | Young |
| 0.128724 | 0.30491 | 0.264 | 0.227 | 1 | 5 | Epc2 | 0.037 | Young |
| 0.128774 | 0.429845 | 0.282 | 0.249 | 1 | 5 | Tmf1 | 0.033 | Young |
| 0.129106 | 0.894046 | 0.038 | 0.022 | 1 | 5 | Katnb1 | 0.016 | Young |
| 0.129223 | 0.687495 | 0.074 | 0.051 | 1 | 5 | Atg2b | 0.023 | Young |
| 0.129325 | 0.518868 | 0.239 | 0.202 | 1 | 5 | Rwdd1 | 0.037 | Young |
| 0.129514 | 0.492467 | 0.145 | 0.113 | 1 | 5 | Gpaa1 | 0.032 | Young |
| 0.129677 | 0.328723 | 0.201 | 0.163 | 1 | 5 | Prkd3 | 0.038 | Young |
| 0.129765 | 0.228071 | 0.236 | 0.195 | 1 | 5 | Slain2 | 0.041 | Young |

| | | | | | | | | |
|---|---|---|---|---|---|---|---|---|
| 0.129831 | 0.199389 | 0.378 | 0.332 | 1 | 5 | Ago3 | 0.046 | Young |
| 0.129954 | 0.25866 | 0.439 | 0.412 | 1 | 5 | Spcs1 | 0.027 | Young |
| 0.13002 | 0.682746 | 0.041 | 0.024 | 1 | 5 | Dcaf4 | 0.017 | Young |
| 0.130573 | 0.573 | 0.142 | 0.111 | 1 | 5 | Gtf3c6 | 0.031 | Young |
| 0.130975 | 0.581325 | 0.102 | 0.074 | 1 | 5 | Mief1 | 0.028 | Young |
| 0.130978 | 0.464417 | 0.198 | 0.162 | 1 | 5 | Ppie | 0.036 | Young |
| 0.130982 | 0.402672 | 0.147 | 0.114 | 1 | 5 | Acbd6 | 0.033 | Young |
| 0.130985 | 0.514332 | 0.124 | 0.094 | 1 | 5 | Agpat1 | 0.03 | Young |
| 0.131013 | 0.298173 | 0.279 | 0.236 | 1 | 5 | Ubr5 | 0.043 | Young |
| 0.131048 | 0.203993 | 0.652 | 0.633 | 1 | 5 | Selenof | 0.019 | Young |
| 0.131341 | 0.489422 | 0.14 | 0.108 | 1 | 5 | Ccdc174 | 0.032 | Young |
| 0.131436 | 0.13424 | 0.261 | 0.209 | 1 | 5 | Gm37494 | 0.052 | Young |
| 0.131513 | 0.446998 | 0.056 | 0.035 | 1 | 5 | Paqr7 | 0.021 | Young |
| 0.131518 | 0.212957 | 0.629 | 0.63 | 1 | 5 | Inpp5d | -0.001 | Young |
| 0.131557 | 0.33559 | 0.178 | 0.14 | 1 | 5 | Arl13b | 0.038 | Young |
| 0.131582 | 0.546058 | 0.041 | 0.024 | 1 | 5 | Zc4h2 | 0.017 | Young |
| 0.13159 | 0.613004 | 0.104 | 0.077 | 1 | 5 | Nbdy | 0.027 | Young |
| 0.131811 | 0.553961 | 0.099 | 0.072 | 1 | 5 | Ttc13 | 0.027 | Young |
| 0.131938 | 0.247465 | 0.107 | 0.077 | 1 | 5 | Plekha3 | 0.03 | Young |
| 0.132381 | 0.218735 | 0.396 | 0.35 | 1 | 5 | Jarid2 | 0.046 | Young |
| 0.132508 | 0.171854 | 0.619 | 0.574 | 1 | 5 | Galnt1 | 0.045 | Young |
| 0.13281 | 0.336133 | 0.355 | 0.318 | 1 | 5 | Rbm26 | 0.037 | Young |
| 0.132831 | 0.155098 | 0.69 | 0.667 | 1 | 5 | Ncl | 0.023 | Young |
| 0.133365 | 0.303593 | 0.137 | 0.104 | 1 | 5 | Lrig2 | 0.033 | Young |
| 0.133757 | 0.262014 | 0.279 | 0.237 | 1 | 5 | Med15 | 0.042 | Young |
| 0.134059 | 0.255816 | 0.64 | 0.64 | 1 | 5 | Reep3 | 0 | Young |
| 0.134075 | 0.231925 | 0.261 | 0.219 | 1 | 5 | Mnt | 0.042 | Young |
| 0.134334 | 0.531187 | 0.132 | 0.101 | 1 | 5 | Mtx2 | 0.031 | Young |
| 0.13445 | 0.238892 | 0.228 | 0.189 | 1 | 5 | Smad5 | 0.039 | Young |
| 0.134719 | 0.381697 | 0.246 | 0.205 | 1 | 5 | Nap1l4 | 0.041 | Young |
| 0.134771 | 0.325114 | 0.203 | 0.167 | 1 | 5 | Sec23b | 0.036 | Young |
| 0.134795 | 0.286547 | 0.5 | 0.465 | 1 | 5 | Bin1 | 0.035 | Young |
| 0.134944 | 0.347392 | 0.223 | 0.185 | 1 | 5 | Ube2j1 | 0.038 | Young |
| 0.134979 | 0.332506 | 0.086 | 0.061 | 1 | 5 | Caap1 | 0.025 | Young |
| 0.135504 | 0.260838 | 0.421 | 0.384 | 1 | 5 | Fam120a | 0.037 | Young |
| 0.135605 | 0.250258 | 0.185 | 0.145 | 1 | 5 | Golga3 | 0.04 | Young |
| 0.135837 | 0.537673 | 0.063 | 0.042 | 1 | 5 | Dnajb5 | 0.021 | Young |
| 0.135996 | 0.300627 | 0.053 | 0.034 | 1 | 5 | Accs | 0.019 | Young |
| 0.136155 | 0.130563 | 0.708 | 0.714 | 1 | 5 | Reep5 | -0.006 | Young |
| 0.136776 | 0.650779 | 0.051 | 0.032 | 1 | 5 | Fdps | 0.019 | Young |
| 0.13688 | 1.009873 | 0.036 | 0.02 | 1 | 5 | Dusp19 | 0.016 | Young |
| 0.13698 | 0.292285 | 0.444 | 0.399 | 1 | 5 | Dip2b | 0.045 | Young |
| 0.137058 | 0.125015 | 0.766 | 0.742 | 1 | 5 | H2afz | 0.024 | Young |
| 0.137353 | 0.578577 | 0.086 | 0.062 | 1 | 5 | Wdr7 | 0.024 | Young |
| 0.137726 | 0.456583 | 0.147 | 0.114 | 1 | 5 | Cops7a | 0.033 | Young |

| | | | | | | | | |
|---|---|---|---|---|---|---|---|---|
| 0.137809 | 0.656101 | 0.099 | 0.074 | 1 | 5 | Mgst3 | 0.025 | Young |
| 0.137967 | 0.952997 | 0.036 | 0.02 | 1 | 5 | Coq6 | 0.016 | Young |
| 0.138197 | 0.495659 | 0.137 | 0.106 | 1 | 5 | Ddx1 | 0.031 | Young |
| 0.138404 | 1.085509 | 0.036 | 0.02 | 1 | 5 | 2810454H( | 0.016 | Young |
| 0.138554 | 0.316458 | 0.338 | 0.3 | 1 | 5 | Fam117b | 0.038 | Young |
| 0.138577 | 0.166013 | 0.056 | 0.035 | 1 | 5 | Gm42031 | 0.021 | Young |
| 0.138587 | 0.394358 | 0.094 | 0.067 | 1 | 5 | Ccdc112 | 0.027 | Young |
| 0.138921 | 0.426152 | 0.117 | 0.088 | 1 | 5 | Fbxo6 | 0.029 | Young |
| 0.139648 | 0.634889 | 0.051 | 0.032 | 1 | 5 | 2900093K2 | 0.019 | Young |
| 0.140003 | 0.079447 | 0.033 | 0.054 | 1 | 5 | Mettl7a1 | -0.021 | Young |
| 0.140117 | 0.306033 | 0.444 | 0.426 | 1 | 5 | Cct8 | 0.018 | Young |
| 0.140502 | 0.473976 | 0.142 | 0.113 | 1 | 5 | Ftsj3 | 0.029 | Young |
| 0.140723 | 0.318414 | 0.155 | 0.121 | 1 | 5 | Mb21d1 | 0.034 | Young |
| 0.140902 | 0.324112 | 0.411 | 0.386 | 1 | 5 | Efr3a | 0.025 | Young |
| 0.141848 | 0.395226 | 0.137 | 0.106 | 1 | 5 | Med30 | 0.031 | Young |
| 0.141892 | 0.669006 | 0.096 | 0.071 | 1 | 5 | Ints14 | 0.025 | Young |
| 0.142159 | 0.554569 | 0.061 | 0.04 | 1 | 5 | Thrsp | 0.021 | Young |
| 0.142184 | 0.458049 | 0.129 | 0.099 | 1 | 5 | Smurf1 | 0.03 | Young |
| 0.142258 | 0.192632 | 0.571 | 0.53 | 1 | 5 | Capns1 | 0.041 | Young |
| 0.142348 | 0.347763 | 0.223 | 0.189 | 1 | 5 | Psmd6 | 0.034 | Young |
| 0.142494 | 0.458913 | 0.114 | 0.086 | 1 | 5 | Fbxo8 | 0.028 | Young |
| 0.142581 | 0.249351 | 0.464 | 0.421 | 1 | 5 | Dennd4a | 0.043 | Young |
| 0.142645 | 0.295333 | 0.185 | 0.15 | 1 | 5 | Fryl | 0.035 | Young |
| 0.142703 | 0.890806 | 0.048 | 0.03 | 1 | 5 | H2-T24 | 0.018 | Young |
| 0.142945 | 0.336036 | 0.081 | 0.057 | 1 | 5 | Zmat1 | 0.024 | Young |
| 0.143851 | 0.280008 | 0.157 | 0.123 | 1 | 5 | Ube2g1 | 0.034 | Young |
| 0.143865 | 0.647788 | 0.069 | 0.047 | 1 | 5 | Slc25a23 | 0.022 | Young |
| 0.144035 | 0.189329 | 0.541 | 0.5 | 1 | 5 | Skil | 0.041 | Young |
| 0.144204 | 0.454199 | 0.19 | 0.155 | 1 | 5 | Utp3 | 0.035 | Young |
| 0.144251 | 0.419867 | 0.287 | 0.258 | 1 | 5 | Brox | 0.029 | Young |
| 0.14473 | 0.420419 | 0.216 | 0.182 | 1 | 5 | Pxk | 0.034 | Young |
| 0.144989 | 0.689237 | 0.094 | 0.069 | 1 | 5 | Spidr | 0.025 | Young |
| 0.145511 | 0.629557 | 0.061 | 0.04 | 1 | 5 | Zfp568 | 0.021 | Young |
| 0.145842 | 0.367589 | 0.173 | 0.138 | 1 | 5 | Tmem63a | 0.035 | Young |
| 0.146132 | 0.152226 | 0.195 | 0.157 | 1 | 5 | Smc4 | 0.038 | Young |
| 0.146255 | 0.227555 | 0.368 | 0.333 | 1 | 5 | Psme4 | 0.035 | Young |
| 0.14659 | 0.221372 | 0.279 | 0.236 | 1 | 5 | Pigk | 0.043 | Young |
| 0.146631 | 0.373533 | 0.145 | 0.113 | 1 | 5 | Iffo1 | 0.032 | Young |
| 0.146758 | 0.572777 | 0.099 | 0.072 | 1 | 5 | Ankrd16 | 0.027 | Young |
| 0.14676 | 0.417421 | 0.231 | 0.194 | 1 | 5 | Dennd4c | 0.037 | Young |
| 0.146812 | 0.301431 | 0.122 | 0.093 | 1 | 5 | Hebp1 | 0.029 | Young |
| 0.146855 | 0.484914 | 0.096 | 0.071 | 1 | 5 | Trim36 | 0.025 | Young |
| 0.147025 | 0.421089 | 0.071 | 0.049 | 1 | 5 | Gapt | 0.022 | Young |
| 0.147155 | 0.470779 | 0.081 | 0.057 | 1 | 5 | Arl14ep | 0.024 | Young |
| 0.147183 | 0.211201 | 0.546 | 0.508 | 1 | 5 | Eif4b | 0.038 | Young |

| | | | | | | | | |
|---|---|---|---|---|---|---|---|---|
| 0.147407 | 0.145894 | 0.551 | 0.502 | 1 | 5 | Ube2b | 0.049 | Young |
| 0.147457 | 0.214533 | 0.277 | 0.236 | 1 | 5 | Slc16a6 | 0.041 | Young |
| 0.14784 | 0.734941 | 0.033 | 0.019 | 1 | 5 | Twnk | 0.014 | Young |
| 0.14797 | 0.514226 | 0.071 | 0.049 | 1 | 5 | Zfp65 | 0.022 | Young |
| 0.148019 | 0.414666 | 0.155 | 0.123 | 1 | 5 | Agtpbp1 | 0.032 | Young |
| 0.148231 | 0.936638 | 0.081 | 0.059 | 1 | 5 | Orc2 | 0.022 | Young |
| 0.148758 | 0.443302 | 0.152 | 0.123 | 1 | 5 | Slc25a1 | 0.029 | Young |
| 0.148825 | 0.706292 | 0.046 | 0.029 | 1 | 5 | Alkbh7 | 0.017 | Young |
| 0.149204 | 0.376237 | 0.036 | 0.02 | 1 | 5 | Ciart | 0.016 | Young |
| 0.149368 | 0.515493 | 0.155 | 0.125 | 1 | 5 | Phax | 0.03 | Young |
| 0.14942 | 0.128233 | 0.713 | 0.704 | 1 | 5 | Rpl29 | 0.009 | Young |
| 0.149593 | 1.087022 | 0.03 | 0.017 | 1 | 5 | Bhlhb9 | 0.013 | Young |
| 0.149657 | 0.586921 | 0.15 | 0.12 | 1 | 5 | Rrm1 | 0.03 | Young |
| 0.149748 | 0.237781 | 0.124 | 0.094 | 1 | 5 | Nf2 | 0.03 | Young |
| 0.149874 | 0.445269 | 0.071 | 0.049 | 1 | 5 | Nxt1 | 0.022 | Young |
| 0.149909 | 0.592776 | 0.15 | 0.121 | 1 | 5 | Riok1 | 0.029 | Young |
| 0.150169 | 0.093404 | 0.957 | 0.975 | 1 | 5 | Mbnl1 | -0.018 | Young |
| 0.150397 | 0.503944 | 0.089 | 0.064 | 1 | 5 | Nol6 | 0.025 | Young |
| 0.150485 | 0.301222 | 0.335 | 0.293 | 1 | 5 | Cux1 | 0.042 | Young |
| 0.150754 | 0.372701 | 0.256 | 0.222 | 1 | 5 | Zfp292 | 0.034 | Young |
| 0.1511 | 0.354626 | 0.122 | 0.091 | 1 | 5 | Rngtt | 0.031 | Young |
| 0.151109 | 1.193378 | 0.03 | 0.017 | 1 | 5 | Cnr2 | 0.013 | Young |
| 0.151314 | 0.520737 | 0.071 | 0.049 | 1 | 5 | Mcm2 | 0.022 | Young |
| 0.151521 | 0.364356 | 0.061 | 0.04 | 1 | 5 | Tmem39b | 0.021 | Young |
| 0.151526 | 0.491802 | 0.173 | 0.141 | 1 | 5 | Ifi47 | 0.032 | Young |
| 0.151613 | 0.53895 | 0.157 | 0.126 | 1 | 5 | Smpdl3a | 0.031 | Young |
| 0.151714 | 0.329499 | 0.203 | 0.167 | 1 | 5 | Icosl | 0.036 | Young |
| 0.151783 | 0.511458 | 0.048 | 0.03 | 1 | 5 | Ncaph | 0.018 | Young |
| 0.151947 | 0.506584 | 0.033 | 0.019 | 1 | 5 | Gm26853 | 0.014 | Young |
| 0.152052 | 0.596013 | 0.046 | 0.029 | 1 | 5 | Zfp709 | 0.017 | Young |
| 0.15228 | 0.592818 | 0.114 | 0.088 | 1 | 5 | Rpa1 | 0.026 | Young |
| 0.152371 | 0.426977 | 0.091 | 0.066 | 1 | 5 | Mthfd2l | 0.025 | Young |
| 0.152713 | 0.462748 | 0.294 | 0.266 | 1 | 5 | Acbd3 | 0.028 | Young |
| 0.152829 | 0.384292 | 0.266 | 0.232 | 1 | 5 | Smarcc1 | 0.034 | Young |
| 0.153065 | 0.271157 | 0.449 | 0.411 | 1 | 5 | Fgfr1op2 | 0.038 | Young |
| 0.153168 | 0.313009 | 0.112 | 0.084 | 1 | 5 | Nudcd2 | 0.028 | Young |
| 0.153334 | 0.710985 | 0.066 | 0.045 | 1 | 5 | Fmnl3 | 0.021 | Young |
| 0.153417 | 0.126297 | 0.036 | 0.02 | 1 | 5 | Tefm | 0.016 | Young |
| 0.153624 | 0.262768 | 0.426 | 0.391 | 1 | 5 | Dram2 | 0.035 | Young |
| 0.15367 | 0.808728 | 0.066 | 0.045 | 1 | 5 | Haus3 | 0.021 | Young |
| 0.153712 | 0.261213 | 0.084 | 0.059 | 1 | 5 | Polr3b | 0.025 | Young |
| 0.153901 | 0.162678 | 0.5 | 0.46 | 1 | 5 | Hnrnpul1 | 0.04 | Young |
| 0.154066 | 0.155932 | 0.355 | 0.298 | 1 | 5 | Slbp | 0.057 | Young |
| 0.154092 | 0.106057 | 0.602 | 0.566 | 1 | 5 | Tmed5 | 0.036 | Young |
| 0.154382 | 0.148714 | 0.787 | 0.796 | 1 | 5 | Wfdc17 | -0.009 | Young |

| | | | | | | | | |
|---|---|---|---|---|---|---|---|---|
| 0.154548 | 0.447685 | 0.307 | 0.279 | 1 | 5 | Cd72 | 0.028 | Young |
| 0.154631 | 0.399416 | 0.17 | 0.138 | 1 | 5 | Map3k7 | 0.032 | Young |
| 0.154913 | 0.323523 | 0.112 | 0.084 | 1 | 5 | Tor2a | 0.028 | Young |
| 0.155355 | 0.139224 | 0.17 | 0.133 | 1 | 5 | Atp13a1 | 0.037 | Young |
| 0.155468 | 0.868646 | 0.03 | 0.017 | 1 | 5 | Gins2 | 0.013 | Young |
| 0.155579 | 0.308543 | 0.251 | 0.21 | 1 | 5 | Taf1 | 0.041 | Young |
| 0.155727 | 0.816479 | 0.03 | 0.017 | 1 | 5 | Fastkd5 | 0.013 | Young |
| 0.155899 | 0.398737 | 0.213 | 0.177 | 1 | 5 | Rubcn | 0.036 | Young |
| 0.156052 | 0.378927 | 0.218 | 0.185 | 1 | 5 | Kxd1 | 0.033 | Young |
| 0.156117 | 0.716517 | 0.102 | 0.077 | 1 | 5 | Gpkow | 0.025 | Young |
| 0.156162 | 0.229442 | 0.739 | 0.717 | 1 | 5 | Gmfg | 0.022 | Young |
| 0.156379 | 0.636113 | 0.066 | 0.045 | 1 | 5 | Oraov1 | 0.021 | Young |
| 0.156824 | 0.429362 | 0.096 | 0.071 | 1 | 5 | Msh2 | 0.025 | Young |
| 0.156862 | 0.157909 | 0.211 | 0.175 | 1 | 5 | Ccnd2 | 0.036 | Young |
| 0.157255 | 0.249239 | 0.437 | 0.409 | 1 | 5 | Erbin | 0.028 | Young |
| 0.157369 | 0.295541 | 0.188 | 0.152 | 1 | 5 | Actr10 | 0.036 | Young |
| 0.157377 | 0.291005 | 0.251 | 0.214 | 1 | 5 | Txndc16 | 0.037 | Young |
| 0.157525 | 0.438586 | 0.122 | 0.094 | 1 | 5 | G2e3 | 0.028 | Young |
| 0.158017 | 0.306085 | 0.302 | 0.263 | 1 | 5 | Elf2 | 0.039 | Young |
| 0.158069 | 0.350857 | 0.467 | 0.438 | 1 | 5 | Etnk1 | 0.029 | Young |
| 0.158339 | 0.568084 | 0.03 | 0.017 | 1 | 5 | Top3a | 0.013 | Young |
| 0.158339 | 0.498875 | 0.03 | 0.017 | 1 | 5 | Rcc1 | 0.013 | Young |
| 0.15904 | 0.369349 | 0.396 | 0.374 | 1 | 5 | Prune2 | 0.022 | Young |
| 0.159083 | 0.262872 | 0.492 | 0.475 | 1 | 5 | Mier1 | 0.017 | Young |
| 0.1595 | 0.284247 | 0.188 | 0.153 | 1 | 5 | Fam98b | 0.035 | Young |
| 0.159699 | 0.280209 | 0.251 | 0.212 | 1 | 5 | Dtx3l | 0.039 | Young |
| 0.159759 | 0.160527 | 0.604 | 0.581 | 1 | 5 | Myh9 | 0.023 | Young |
| 0.159816 | 1.041511 | 0.028 | 0.015 | 1 | 5 | Mad2l1 | 0.013 | Young |
| 0.159817 | 0.810122 | 0.056 | 0.037 | 1 | 5 | Urb1 | 0.019 | Young |
| 0.160195 | 0.070713 | 1 | 1 | 1 | 5 | mt-Co3 | 0 | Young |
| 0.160199 | 0.248458 | 0.497 | 0.466 | 1 | 5 | Hmgb2 | 0.031 | Young |
| 0.160567 | 0.382477 | 0.046 | 0.029 | 1 | 5 | 2-Sep | 0.017 | Young |
| 0.160928 | 0.851465 | 0.028 | 0.015 | 1 | 5 | Tmem144 | 0.013 | Young |
| 0.16106 | 0.347511 | 0.147 | 0.116 | 1 | 5 | 2310001H1 | 0.031 | Young |
| 0.161577 | 0.18173 | 0.602 | 0.599 | 1 | 5 | Ube2d2a | 0.003 | Young |
| 0.161766 | 0.849827 | 0.028 | 0.015 | 1 | 5 | Elp6 | 0.013 | Young |
| 0.162046 | 0.891379 | 0.028 | 0.015 | 1 | 5 | Tigar | 0.013 | Young |
| 0.162218 | 0.412954 | 0.109 | 0.082 | 1 | 5 | Smim12 | 0.027 | Young |
| 0.162262 | 0.870822 | 0.061 | 0.042 | 1 | 5 | Pdk1 | 0.019 | Young |
| 0.162451 | 0.161984 | 0.119 | 0.089 | 1 | 5 | Nsmce3 | 0.03 | Young |
| 0.162605 | 0.606364 | 0.066 | 0.045 | 1 | 5 | Tomm40l | 0.021 | Young |
| 0.162889 | 0.795886 | 0.028 | 0.015 | 1 | 5 | Gm15880 | 0.013 | Young |
| 0.162923 | 0.917901 | 0.043 | 0.027 | 1 | 5 | Dlc1 | 0.016 | Young |
| 0.162942 | 0.376751 | 0.34 | 0.313 | 1 | 5 | Ubn1 | 0.027 | Young |
| 0.16301 | 0.277526 | 0.365 | 0.33 | 1 | 5 | Crebrf | 0.035 | Young |

| | | | | | | | | |
|---|---|---|---|---|---|---|---|---|
| 0.163103 | 0.708391 | 0.071 | 0.051 | 1 | 5 | 2610524H( | 0.02 | Young |
| 0.163522 | 0.543193 | 0.129 | 0.101 | 1 | 5 | Clec5a | 0.028 | Young |
| 0.163677 | 0.248202 | 0.31 | 0.268 | 1 | 5 | Mgrn1 | 0.042 | Young |
| 0.163799 | 0.497723 | 0.183 | 0.15 | 1 | 5 | Msi2 | 0.033 | Young |
| 0.163973 | 0.156439 | 0.538 | 0.525 | 1 | 5 | Rab11a | 0.013 | Young |
| 0.164743 | 0.244773 | 0.383 | 0.347 | 1 | 5 | Ddrgk1 | 0.036 | Young |
| 0.164977 | 0.45168 | 0.14 | 0.111 | 1 | 5 | Sphk2 | 0.029 | Young |
| 0.165571 | 0.640371 | 0.056 | 0.037 | 1 | 5 | B230307C2 | 0.019 | Young |
| 0.16596 | 0.735579 | 0.056 | 0.037 | 1 | 5 | Usp35 | 0.019 | Young |
| 0.166171 | 0.979778 | 0.041 | 0.025 | 1 | 5 | Fam212a | 0.016 | Young |
| 0.16642 | 0.158226 | 0.69 | 0.714 | 1 | 5 | Eef1g | -0.024 | Young |
| 0.166473 | 0.234777 | 0.345 | 0.308 | 1 | 5 | Ndufs2 | 0.037 | Young |
| 0.166837 | 0.13461 | 0.827 | 0.828 | 1 | 5 | Eps15 | -0.001 | Young |
| 0.16684 | 0.518298 | 0.107 | 0.081 | 1 | 5 | Crbn | 0.026 | Young |
| 0.16684 | 0.398673 | 0.107 | 0.081 | 1 | 5 | Abcf2 | 0.026 | Young |
| 0.166908 | 0.366202 | 0.168 | 0.136 | 1 | 5 | Ddx10 | 0.032 | Young |
| 0.167296 | 0.873199 | 0.053 | 0.035 | 1 | 5 | Ccdc61 | 0.018 | Young |
| 0.1673 | 0.194058 | 0.495 | 0.468 | 1 | 5 | Lamtor1 | 0.027 | Young |
| 0.167891 | 0.588642 | 0.074 | 0.052 | 1 | 5 | Gramd1b | 0.022 | Young |
| 0.168012 | 0.347291 | 0.353 | 0.32 | 1 | 5 | Was | 0.033 | Young |
| 0.168024 | 0.856171 | 0.041 | 0.025 | 1 | 5 | Tmub1 | 0.016 | Young |
| 0.168052 | 0.931176 | 0.025 | 0.013 | 1 | 5 | Bub1b | 0.012 | Young |
| 0.168356 | 0.660947 | 0.025 | 0.013 | 1 | 5 | Gm11772 | 0.012 | Young |
| 0.168444 | 0.305784 | 0.226 | 0.19 | 1 | 5 | Ufc1 | 0.036 | Young |
| 0.16847 | 0.184691 | 0.058 | 0.039 | 1 | 5 | Tbrg4 | 0.019 | Young |
| 0.168493 | 0.261819 | 0.216 | 0.178 | 1 | 5 | Anxa11 | 0.038 | Young |
| 0.168582 | 0.354522 | 0.206 | 0.172 | 1 | 5 | Cnot10 | 0.034 | Young |
| 0.168791 | 0.318348 | 0.261 | 0.227 | 1 | 5 | Golga7 | 0.034 | Young |
| 0.169098 | 0.290658 | 0.388 | 0.365 | 1 | 5 | Acbd5 | 0.023 | Young |
| 0.169373 | 0.348959 | 0.127 | 0.099 | 1 | 5 | Sdhaf2 | 0.028 | Young |
| 0.169554 | 0.181902 | 0.35 | 0.303 | 1 | 5 | Eif3j1 | 0.047 | Young |
| 0.16963 | 0.284503 | 0.292 | 0.254 | 1 | 5 | Arhgef10l | 0.038 | Young |
| 0.169879 | 0.935568 | 0.025 | 0.013 | 1 | 5 | Trim45 | 0.012 | Young |
| 0.170362 | 0.840909 | 0.041 | 0.025 | 1 | 5 | Cenpl | 0.016 | Young |
| 0.171104 | 0.359857 | 0.096 | 0.071 | 1 | 5 | AI987944 | 0.025 | Young |
| 0.171105 | 0.668156 | 0.025 | 0.013 | 1 | 5 | Prag1 | 0.012 | Young |
| 0.171461 | 0.229097 | 0.244 | 0.205 | 1 | 5 | Man1a2 | 0.039 | Young |
| 0.171844 | 0.282056 | 0.31 | 0.271 | 1 | 5 | Uqcrc2 | 0.039 | Young |
| 0.17196 | 0.192214 | 0.124 | 0.094 | 1 | 5 | Bcl2l12 | 0.03 | Young |
| 0.172016 | 0.314842 | 0.485 | 0.461 | 1 | 5 | Mapre2 | 0.024 | Young |
| 0.17202 | 0.268896 | 0.307 | 0.268 | 1 | 5 | Rap2c | 0.039 | Young |
| 0.172338 | 0.790812 | 0.025 | 0.013 | 1 | 5 | Gm13091 | 0.012 | Young |
| 0.172487 | 0.696651 | 0.041 | 0.025 | 1 | 5 | Def8 | 0.016 | Young |
| 0.173024 | 0.369499 | 0.198 | 0.167 | 1 | 5 | Paxbp1 | 0.031 | Young |
| 0.173086 | 0.422044 | 0.107 | 0.081 | 1 | 5 | Rbpms | 0.026 | Young |

| | | | | | | | | |
|---|---|---|---|---|---|---|---|---|
| 0.173578 | 0.634289 | 0.025 | 0.013 | 1 | 5 | Cfap126 | 0.012 | Young |
| 0.173592 | 0.410914 | 0.053 | 0.035 | 1 | 5 | Msh3 | 0.018 | Young |
| 0.173627 | 0.300844 | 0.305 | 0.264 | 1 | 5 | Lsm14a | 0.041 | Young |
| 0.174018 | 0.647471 | 0.099 | 0.076 | 1 | 5 | Fbxo42 | 0.023 | Young |
| 0.174451 | 0.926249 | 0.023 | 0.012 | 1 | 5 | Trim62 | 0.011 | Young |
| 0.174527 | 0.219315 | 0.459 | 0.424 | 1 | 5 | Map3k1 | 0.035 | Young |
| 0.174647 | 0.491017 | 0.152 | 0.125 | 1 | 5 | Nabp2 | 0.027 | Young |
| 0.174893 | 0.526792 | 0.119 | 0.094 | 1 | 5 | Ap1ar | 0.025 | Young |
| 0.175017 | 0.528254 | 0.14 | 0.113 | 1 | 5 | Pold3 | 0.027 | Young |
| 0.175136 | 0.379717 | 0.025 | 0.013 | 1 | 5 | Magee1 | 0.012 | Young |
| 0.175531 | 0.42704 | 0.244 | 0.209 | 1 | 5 | Zhx1 | 0.035 | Young |
| 0.175688 | 0.241877 | 0.345 | 0.305 | 1 | 5 | Ripk1 | 0.04 | Young |
| 0.175763 | 0.471967 | 0.025 | 0.013 | 1 | 5 | Zcwpw1 | 0.012 | Young |
| 0.176026 | 0.343017 | 0.19 | 0.16 | 1 | 5 | Mob3b | 0.03 | Young |
| 0.17607 | 0.094376 | 0.843 | 0.801 | 1 | 5 | Ppp3ca | 0.042 | Young |
| 0.176281 | 0.550587 | 0.081 | 0.059 | 1 | 5 | Mrpl22 | 0.022 | Young |
| 0.177112 | 0.27975 | 0.365 | 0.337 | 1 | 5 | Mkrn1 | 0.028 | Young |
| 0.177126 | 0.257386 | 0.168 | 0.136 | 1 | 5 | Suds3 | 0.032 | Young |
| 0.177205 | 0.977306 | 0.01 | 0.003 | 1 | 5 | 6230400D1 | 0.007 | Young |
| 0.177451 | 0.177385 | 0.376 | 0.333 | 1 | 5 | Smarcc2 | 0.043 | Young |
| 0.177593 | 0.390511 | 0.195 | 0.163 | 1 | 5 | Irak3 | 0.032 | Young |
| 0.177652 | 0.353156 | 0.025 | 0.013 | 1 | 5 | Phldb3 | 0.012 | Young |
| 0.177749 | 0.955539 | 0.01 | 0.003 | 1 | 5 | Card11 | 0.007 | Young |
| 0.178046 | 0.335399 | 0.066 | 0.045 | 1 | 5 | Acd | 0.021 | Young |
| 0.17823 | 0.578313 | 0.193 | 0.168 | 1 | 5 | Rangap1 | 0.025 | Young |
| 0.178294 | 1.692334 | 0.01 | 0.003 | 1 | 5 | Map10 | 0.007 | Young |
| 0.178294 | 1.42843 | 0.01 | 0.003 | 1 | 5 | 2010310C0 | 0.007 | Young |
| 0.178294 | 1.168814 | 0.01 | 0.003 | 1 | 5 | Hist1h4c | 0.007 | Young |
| 0.178294 | 1.044425 | 0.01 | 0.003 | 1 | 5 | Rsph9 | 0.007 | Young |
| 0.178294 | 0.967912 | 0.01 | 0.003 | 1 | 5 | Adamts6 | 0.007 | Young |
| 0.178294 | 0.786426 | 0.01 | 0.003 | 1 | 5 | Gm17655 | 0.007 | Young |
| 0.178441 | 0.318001 | 0.284 | 0.247 | 1 | 5 | Maf1 | 0.037 | Young |
| 0.178737 | 0.39694 | 0.041 | 0.025 | 1 | 5 | Scrn3 | 0.016 | Young |
| 0.17884 | 1.11167 | 0.01 | 0.003 | 1 | 5 | 6530409C1 | 0.007 | Young |
| 0.17884 | 0.983535 | 0.01 | 0.003 | 1 | 5 | Ltb4r1 | 0.007 | Young |
| 0.17931 | 0.815587 | 0.038 | 0.024 | 1 | 5 | Nlrp1b | 0.014 | Young |
| 0.179317 | 0.453394 | 0.211 | 0.18 | 1 | 5 | Ykt6 | 0.031 | Young |
| 0.179388 | 1.034386 | 0.01 | 0.003 | 1 | 5 | 9330104G0 | 0.007 | Young |
| 0.179388 | 0.97507 | 0.01 | 0.003 | 1 | 5 | Zfp300 | 0.007 | Young |
| 0.179388 | 0.759127 | 0.01 | 0.003 | 1 | 5 | Actg2 | 0.007 | Young |
| 0.179516 | 0.244054 | 0.365 | 0.327 | 1 | 5 | Tet2 | 0.038 | Young |
| 0.179672 | 0.135318 | 0.599 | 0.564 | 1 | 5 | Tsc22d4 | 0.035 | Young |
| 0.179784 | 0.373911 | 0.109 | 0.082 | 1 | 5 | Abhd2 | 0.027 | Young |
| 0.179937 | 0.865859 | 0.01 | 0.003 | 1 | 5 | Gm48882 | 0.007 | Young |
| 0.179937 | 0.836568 | 0.01 | 0.003 | 1 | 5 | Fam171a1 | 0.007 | Young |

| | | | | | | | | |
|---|---|---|---|---|---|---|---|---|
| 0.180033 | 0.437445 | 0.063 | 0.044 | 1 | 5 | Tmem241 | 0.019 | Young |
| 0.180189 | 0.442913 | 0.124 | 0.098 | 1 | 5 | Ckap4 | 0.026 | Young |
| 0.180444 | 0.32888 | 0.178 | 0.145 | 1 | 5 | Ranbp10 | 0.033 | Young |
| 0.180487 | 0.684857 | 0.01 | 0.003 | 1 | 5 | Kdm8 | 0.007 | Young |
| 0.180487 | 0.624698 | 0.01 | 0.003 | 1 | 5 | E430018J2 | 0.007 | Young |
| 0.180487 | 0.578595 | 0.01 | 0.003 | 1 | 5 | Tnfrsf13c | 0.007 | Young |
| 0.180554 | 0.287937 | 0.332 | 0.298 | 1 | 5 | Alkbh5 | 0.034 | Young |
| 0.180748 | 0.640423 | 0.076 | 0.056 | 1 | 5 | Jmy | 0.02 | Young |
| 0.180861 | 0.660872 | 0.084 | 0.062 | 1 | 5 | Usp11 | 0.022 | Young |
| 0.180987 | 0.37631 | 0.185 | 0.153 | 1 | 5 | Stx6 | 0.032 | Young |
| 0.181038 | 0.426778 | 0.01 | 0.003 | 1 | 5 | Flt3 | 0.007 | Young |
| 0.181091 | 0.22021 | 0.193 | 0.155 | 1 | 5 | Herc6 | 0.038 | Young |
| 0.181103 | 0.320784 | 0.198 | 0.163 | 1 | 5 | Ccnd3 | 0.035 | Young |
| 0.181441 | 0.200128 | 0.137 | 0.108 | 1 | 5 | Nde1 | 0.029 | Young |
| 0.181581 | 0.31183 | 0.195 | 0.162 | 1 | 5 | Pik3r6 | 0.033 | Young |
| 0.181775 | 0.456937 | 0.15 | 0.125 | 1 | 5 | Smg6 | 0.025 | Young |
| 0.181897 | 0.538025 | 0.221 | 0.189 | 1 | 5 | Gadd45g | 0.032 | Young |
| 0.182013 | 0.609572 | 0.051 | 0.034 | 1 | 5 | Smyd4 | 0.017 | Young |
| 0.182017 | 0.278619 | 0.381 | 0.34 | 1 | 5 | Pde4b | 0.041 | Young |
| 0.182305 | 0.216249 | 0.411 | 0.369 | 1 | 5 | Sbno1 | 0.042 | Young |
| 0.182342 | 0.501154 | 0.117 | 0.091 | 1 | 5 | Fem1a | 0.026 | Young |
| 0.182643 | 0.301335 | 0.386 | 0.348 | 1 | 5 | Cdkn1b | 0.038 | Young |
| 0.18272 | 0.140044 | 0.457 | 0.428 | 1 | 5 | Adcy7 | 0.029 | Young |
| 0.182841 | 0.026227 | 0.464 | 0.522 | 1 | 5 | Fam168b | -0.058 | Young |
| 0.182957 | 0.221452 | 0.074 | 0.052 | 1 | 5 | Zfp658 | 0.022 | Young |
| 0.183366 | 0.106164 | 0.721 | 0.692 | 1 | 5 | Ncf2 | 0.029 | Young |
| 0.183454 | 0.529262 | 0.079 | 0.057 | 1 | 5 | Invs | 0.022 | Young |
| 0.183469 | 0.60466 | 0.069 | 0.049 | 1 | 5 | Nol9 | 0.02 | Young |
| 0.183529 | 0.296258 | 0.228 | 0.194 | 1 | 5 | Tia1 | 0.034 | Young |
| 0.183888 | 0.076125 | 0.023 | 0.012 | 1 | 5 | Camk2n1 | 0.011 | Young |
| 0.183892 | 0.616336 | 0.038 | 0.024 | 1 | 5 | Izumo4 | 0.014 | Young |
| 0.184363 | 0.811532 | 0.066 | 0.047 | 1 | 5 | Pex11b | 0.019 | Young |
| 0.184504 | 0.341969 | 0.206 | 0.168 | 1 | 5 | Zfp592 | 0.038 | Young |
| 0.184679 | 0.549022 | 0.079 | 0.057 | 1 | 5 | Ldlrap1 | 0.022 | Young |
| 0.184735 | 0.695307 | 0.02 | 0.01 | 1 | 5 | Gm11131 | 0.01 | Young |
| 0.184735 | 0.659679 | 0.02 | 0.01 | 1 | 5 | Nle1 | 0.01 | Young |
| 0.184813 | 0.56545 | 0.084 | 0.062 | 1 | 5 | 2510002D2 | 0.022 | Young |
| 0.184865 | 0.408193 | 0.112 | 0.086 | 1 | 5 | Pum3 | 0.026 | Young |
| 0.185197 | 1.048796 | 0.018 | 0.008 | 1 | 5 | Xk | 0.01 | Young |
| 0.185289 | 0.314643 | 0.206 | 0.173 | 1 | 5 | Ccndbp1 | 0.033 | Young |
| 0.185431 | 0.338703 | 0.122 | 0.096 | 1 | 5 | Usp1 | 0.026 | Young |
| 0.185439 | 0.688792 | 0.038 | 0.024 | 1 | 5 | Zfp87 | 0.014 | Young |
| 0.185577 | 0.400702 | 0.084 | 0.062 | 1 | 5 | Selenoi | 0.022 | Young |
| 0.185612 | 0.234527 | 0.023 | 0.012 | 1 | 5 | Gale | 0.011 | Young |
| 0.185694 | 0.368943 | 0.109 | 0.084 | 1 | 5 | Lancl1 | 0.025 | Young |

| | | | | | | | | |
|---|---|---|---|---|---|---|---|---|
| 0.185842 | 0.655396 | 0.02 | 0.01 | 1 | 5 | Zfp449 | 0.01 | Young |
| 0.18589 | 0.213185 | 0.701 | 0.673 | 1 | 5 | Pak2 | 0.028 | Young |
| 0.186034 | 0.983601 | 0.048 | 0.032 | 1 | 5 | Fam84b | 0.016 | Young |
| 0.186371 | 0.692898 | 0.096 | 0.074 | 1 | 5 | Nol12 | 0.022 | Young |
| 0.186451 | 0.366364 | 0.431 | 0.396 | 1 | 5 | Stard3nl | 0.035 | Young |
| 0.186816 | 0.335817 | 0.34 | 0.306 | 1 | 5 | Vdac3 | 0.034 | Young |
| 0.186824 | 0.476431 | 0.132 | 0.106 | 1 | 5 | Mzt1 | 0.026 | Young |
| 0.186868 | 0.212353 | 0.635 | 0.616 | 1 | 5 | Lrp1 | 0.019 | Young |
| 0.186941 | 0.276675 | 0.183 | 0.148 | 1 | 5 | St3gal4 | 0.035 | Young |
| 0.187011 | 0.847461 | 0.056 | 0.039 | 1 | 5 | Smyd3 | 0.017 | Young |
| 0.187063 | 0.449583 | 0.282 | 0.251 | 1 | 5 | Igbp1 | 0.031 | Young |
| 0.187196 | 0.861396 | 0.018 | 0.008 | 1 | 5 | Gm527 | 0.01 | Young |
| 0.187271 | 1.081742 | 0.013 | 0.005 | 1 | 5 | Ttc23 | 0.008 | Young |
| 0.187326 | 0.716708 | 0.02 | 0.01 | 1 | 5 | Zranb3 | 0.01 | Young |
| 0.187415 | 0.278349 | 0.254 | 0.215 | 1 | 5 | Flcn | 0.039 | Young |
| 0.187597 | 0.868128 | 0.018 | 0.008 | 1 | 5 | Myl2 | 0.01 | Young |
| 0.187614 | 0.305273 | 0.135 | 0.106 | 1 | 5 | Cdadc1 | 0.029 | Young |
| 0.187772 | 0.310224 | 0.18 | 0.15 | 1 | 5 | mt-Nd6 | 0.03 | Young |
| 0.187914 | 0.1751 | 0.335 | 0.293 | 1 | 5 | Ppp6r1 | 0.042 | Young |
| 0.188192 | 0.287794 | 0.35 | 0.32 | 1 | 5 | Snx20 | 0.03 | Young |
| 0.188239 | 0.349746 | 0.251 | 0.222 | 1 | 5 | Smap1 | 0.029 | Young |
| 0.188254 | 1.273274 | 0.013 | 0.005 | 1 | 5 | Hoxc8 | 0.008 | Young |
| 0.188254 | 1.224766 | 0.013 | 0.005 | 1 | 5 | Tjp1 | 0.008 | Young |
| 0.188254 | 1.108222 | 0.013 | 0.005 | 1 | 5 | Zfp869 | 0.008 | Young |
| 0.188392 | 0.379177 | 0.071 | 0.051 | 1 | 5 | Mbd4 | 0.02 | Young |
| 0.188419 | 0.422564 | 0.124 | 0.098 | 1 | 5 | Rad54l2 | 0.026 | Young |
| 0.188747 | 0.947472 | 0.013 | 0.005 | 1 | 5 | Uhrf1 | 0.008 | Young |
| 0.188806 | 0.995866 | 0.018 | 0.008 | 1 | 5 | Gstm2 | 0.01 | Young |
| 0.188806 | 0.953325 | 0.018 | 0.008 | 1 | 5 | Cep72 | 0.01 | Young |
| 0.188836 | 0.615234 | 0.058 | 0.04 | 1 | 5 | 3110082I1 | 0.018 | Young |
| 0.189235 | 0.258226 | 0.091 | 0.067 | 1 | 5 | Rrp36 | 0.024 | Young |
| 0.189241 | 1.166477 | 0.013 | 0.005 | 1 | 5 | AC152827. | 0.008 | Young |
| 0.189241 | 1.050859 | 0.013 | 0.005 | 1 | 5 | Cenph | 0.008 | Young |
| 0.189241 | 0.808586 | 0.013 | 0.005 | 1 | 5 | Inf2 | 0.008 | Young |
| 0.189419 | 0.250505 | 0.624 | 0.645 | 1 | 5 | 2010107E0 | -0.021 | Young |
| 0.189498 | 0.289235 | 0.117 | 0.089 | 1 | 5 | Kdm5b | 0.028 | Young |
| 0.189547 | 0.489807 | 0.086 | 0.064 | 1 | 5 | Xxylt1 | 0.022 | Young |
| 0.189736 | 0.736585 | 0.013 | 0.005 | 1 | 5 | Vps25 | 0.008 | Young |
| 0.18974 | 0.967475 | 0.015 | 0.007 | 1 | 5 | AC113595. | 0.008 | Young |
| 0.189756 | 0.787203 | 0.036 | 0.022 | 1 | 5 | Cluap1 | 0.014 | Young |
| 0.189771 | 0.49262 | 0.061 | 0.042 | 1 | 5 | Amigo1 | 0.019 | Young |
| 0.189804 | 0.2443 | 0.16 | 0.126 | 1 | 5 | Iars | 0.034 | Young |
| 0.189943 | 0.443378 | 0.02 | 0.01 | 1 | 5 | Col1a1 | 0.01 | Young |
| 0.19002 | 0.775282 | 0.018 | 0.008 | 1 | 5 | Mms22l | 0.01 | Young |
| 0.19002 | 0.766475 | 0.018 | 0.008 | 1 | 5 | 5430416N0 | 0.01 | Young |

| | | | | | | | | |
|---|---|---|---|---|---|---|---|---|
| 0.190232 | 0.897412 | 0.013 | 0.005 | 1 | 5 | Snhg10 | 0.008 | Young |
| 0.190232 | 0.731722 | 0.013 | 0.005 | 1 | 5 | Foxm1 | 0.008 | Young |
| 0.190278 | 0.41552 | 0.279 | 0.241 | 1 | 5 | Ezh1 | 0.038 | Young |
| 0.19032 | 0.284826 | 0.02 | 0.01 | 1 | 5 | Glis2 | 0.01 | Young |
| 0.190381 | 0.282517 | 0.112 | 0.086 | 1 | 5 | Zc3h7b | 0.026 | Young |
| 0.190471 | 0.485851 | 0.084 | 0.062 | 1 | 5 | Narf | 0.022 | Young |
| 0.190729 | 0.877346 | 0.013 | 0.005 | 1 | 5 | Elavl4 | 0.008 | Young |
| 0.190729 | 0.777755 | 0.013 | 0.005 | 1 | 5 | Zfp583 | 0.008 | Young |
| 0.190729 | 0.629606 | 0.013 | 0.005 | 1 | 5 | Zc3hav1l | 0.008 | Young |
| 0.190833 | 0.743769 | 0.018 | 0.008 | 1 | 5 | Oxld1 | 0.01 | Young |
| 0.191022 | 0.418565 | 0.234 | 0.204 | 1 | 5 | Hspa14 | 0.03 | Young |
| 0.19107 | 0.452357 | 0.069 | 0.049 | 1 | 5 | Rusc1 | 0.02 | Young |
| 0.191201 | 0.436568 | 0.195 | 0.168 | 1 | 5 | Erlin2 | 0.027 | Young |
| 0.191226 | 0.666446 | 0.013 | 0.005 | 1 | 5 | Gas2 | 0.008 | Young |
| 0.191226 | 0.603687 | 0.013 | 0.005 | 1 | 5 | Hist4h4 | 0.008 | Young |
| 0.191226 | 0.444399 | 0.013 | 0.005 | 1 | 5 | Cuedc1 | 0.008 | Young |
| 0.19152 | 0.825151 | 0.015 | 0.007 | 1 | 5 | Xylt1 | 0.008 | Young |
| 0.191551 | 0.394921 | 0.162 | 0.135 | 1 | 5 | Manea | 0.027 | Young |
| 0.19171 | 0.237689 | 0.081 | 0.059 | 1 | 5 | Exosc7 | 0.022 | Young |
| 0.191725 | 0.769137 | 0.013 | 0.005 | 1 | 5 | Cetn4 | 0.008 | Young |
| 0.191725 | 0.525251 | 0.013 | 0.005 | 1 | 5 | Jmjd4 | 0.008 | Young |
| 0.191725 | 0.3415 | 0.013 | 0.005 | 1 | 5 | Fam221a | 0.008 | Young |
| 0.191823 | 0.477588 | 0.061 | 0.042 | 1 | 5 | Zfp429 | 0.019 | Young |
| 0.191967 | 0.963404 | 0.015 | 0.007 | 1 | 5 | Gm43672 | 0.008 | Young |
| 0.192018 | 0.716374 | 0.048 | 0.032 | 1 | 5 | Trit1 | 0.016 | Young |
| 0.192225 | 0.888952 | 0.013 | 0.005 | 1 | 5 | Gm17092 | 0.008 | Young |
| 0.192225 | 0.493225 | 0.013 | 0.005 | 1 | 5 | Ankrd28 | 0.008 | Young |
| 0.192225 | 0.485918 | 0.013 | 0.005 | 1 | 5 | Zfp947 | 0.008 | Young |
| 0.192225 | 0.393409 | 0.013 | 0.005 | 1 | 5 | Smim10l2a | 0.008 | Young |
| 0.192365 | 0.338346 | 0.137 | 0.109 | 1 | 5 | Nmt2 | 0.028 | Young |
| 0.192468 | 0.302382 | 0.371 | 0.343 | 1 | 5 | Gdi1 | 0.028 | Young |
| 0.192493 | 0.708049 | 0.036 | 0.022 | 1 | 5 | Klhl42 | 0.014 | Young |
| 0.192505 | 0.267458 | 0.129 | 0.101 | 1 | 5 | H2-Ke6 | 0.028 | Young |
| 0.192828 | 0.50342 | 0.155 | 0.126 | 1 | 5 | Topbp1 | 0.029 | Young |
| 0.192906 | 0.403287 | 0.094 | 0.071 | 1 | 5 | Fasn | 0.023 | Young |
| 0.192951 | 0.76921 | 0.048 | 0.032 | 1 | 5 | Zfp386 | 0.016 | Young |
| 0.193227 | 0.389314 | 0.013 | 0.005 | 1 | 5 | Arhgef40 | 0.008 | Young |
| 0.193227 | 0.367948 | 0.013 | 0.005 | 1 | 5 | Tsku | 0.008 | Young |
| 0.193583 | 0.26166 | 0.081 | 0.059 | 1 | 5 | Eif2ak3 | 0.022 | Young |
| 0.193729 | 0.217898 | 0.013 | 0.005 | 1 | 5 | Armcx1 | 0.008 | Young |
| 0.193762 | 0.703365 | 0.015 | 0.007 | 1 | 5 | 9130230L2 | 0.008 | Young |
| 0.193762 | 0.573197 | 0.015 | 0.007 | 1 | 5 | Psmc3ip | 0.008 | Young |
| 0.193833 | 0.595593 | 0.074 | 0.054 | 1 | 5 | Rgp1 | 0.02 | Young |
| 0.194067 | 0.57526 | 0.094 | 0.071 | 1 | 5 | Stx11 | 0.023 | Young |
| 0.194099 | 0.232186 | 0.061 | 0.042 | 1 | 5 | Abhd18 | 0.019 | Young |

| | | | | | | | | |
|---|---|---|---|---|---|---|---|---|
| 0.194213 | 0.576778 | 0.015 | 0.007 | 1 | 5 | Chaf1b | 0.008 | Young |
| 0.194368 | 0.203065 | 0.322 | 0.283 | 1 | 5 | Cep350 | 0.039 | Young |
| 0.194448 | 0.140786 | 0.246 | 0.209 | 1 | 5 | B3gat3 | 0.037 | Young |
| 0.194517 | 0.233333 | 0.16 | 0.128 | 1 | 5 | Rps6kc1 | 0.032 | Young |
| 0.194592 | 0.509808 | 0.048 | 0.032 | 1 | 5 | 4933427D1 | 0.016 | Young |
| 0.195168 | 0.271088 | 0.282 | 0.246 | 1 | 5 | Stk10 | 0.036 | Young |
| 0.195538 | 0.952062 | 0.036 | 0.022 | 1 | 5 | Isg20 | 0.014 | Young |
| 0.19556 | 0.158187 | 0.665 | 0.657 | 1 | 5 | Eif3f | 0.008 | Young |
| 0.195658 | 0.200326 | 0.431 | 0.399 | 1 | 5 | 2-Mar | 0.032 | Young |
| 0.195662 | 0.240739 | 0.464 | 0.441 | 1 | 5 | Necap2 | 0.023 | Young |
| 0.195784 | 0.525314 | 0.058 | 0.04 | 1 | 5 | Iba57 | 0.018 | Young |
| 0.195905 | 0.441607 | 0.162 | 0.136 | 1 | 5 | Bcap29 | 0.026 | Young |
| 0.196196 | 0.425903 | 0.107 | 0.082 | 1 | 5 | Champ1 | 0.025 | Young |
| 0.196244 | 0.258862 | 0.414 | 0.38 | 1 | 5 | Hipk1 | 0.034 | Young |
| 0.196723 | 0.514994 | 0.069 | 0.049 | 1 | 5 | Chd3os | 0.02 | Young |
| 0.196815 | 0.25585 | 0.061 | 0.042 | 1 | 5 | Pnkp | 0.019 | Young |
| 0.197006 | 0.136991 | 0.119 | 0.093 | 1 | 5 | Slc38a1 | 0.026 | Young |
| 0.197066 | 0.630663 | 0.104 | 0.081 | 1 | 5 | Fbxw8 | 0.023 | Young |
| 0.197273 | 0.276362 | 0.289 | 0.253 | 1 | 5 | Ndufab1 | 0.036 | Young |
| 0.197493 | 0.619537 | 0.036 | 0.022 | 1 | 5 | Olfr920 | 0.014 | Young |
| 0.197493 | 0.245399 | 0.236 | 0.202 | 1 | 5 | Slc38a6 | 0.034 | Young |
| 0.197551 | 0.240258 | 0.457 | 0.416 | 1 | 5 | Rap1gds1 | 0.041 | Young |
| 0.19776 | 0.089815 | 0.937 | 0.946 | 1 | 5 | H3f3a | -0.009 | Young |
| 0.197827 | 0.258407 | 0.122 | 0.094 | 1 | 5 | Trap1 | 0.028 | Young |
| 0.19799 | 0.353935 | 0.129 | 0.103 | 1 | 5 | Mgat5 | 0.026 | Young |
| 0.198286 | 0.357602 | 0.104 | 0.079 | 1 | 5 | AI606181 | 0.025 | Young |
| 0.19883 | 0.315481 | 0.079 | 0.057 | 1 | 5 | Cdkn2c | 0.022 | Young |
| 0.198911 | 0.558069 | 0.127 | 0.101 | 1 | 5 | Irgm2 | 0.026 | Young |
| 0.199469 | 0.181327 | 0.5 | 0.478 | 1 | 5 | Ptpro | 0.022 | Young |
| 0.199565 | 0.142255 | 0.538 | 0.498 | 1 | 5 | Hmgn1 | 0.04 | Young |
| 0.199689 | 0.270477 | 0.19 | 0.155 | 1 | 5 | Bms1 | 0.035 | Young |
| 0.199775 | 0.370858 | 0.091 | 0.069 | 1 | 5 | Dcakd | 0.022 | Young |
| 0.200072 | 0.238461 | 0.234 | 0.2 | 1 | 5 | Eif1ax | 0.034 | Young |
| 0.200085 | 0.494117 | 0.058 | 0.04 | 1 | 5 | Lrrc42 | 0.018 | Young |
| 0.200136 | 0.268689 | 0.32 | 0.285 | 1 | 5 | Tmem164 | 0.035 | Young |
| 0.200198 | 0.737394 | 0.071 | 0.052 | 1 | 5 | Numb | 0.019 | Young |
| 0.200561 | 0.193177 | 0.576 | 0.556 | 1 | 5 | Dnajb6 | 0.02 | Young |
| 0.200825 | 0.386133 | 0.114 | 0.089 | 1 | 5 | Uri1 | 0.025 | Young |
| 0.20097 | 0.329409 | 0.094 | 0.071 | 1 | 5 | Top3b | 0.023 | Young |
| 0.20151 | 0.267649 | 0.239 | 0.205 | 1 | 5 | Zufsp | 0.034 | Young |
| 0.201519 | 0.189764 | 0.551 | 0.512 | 1 | 5 | Slc12a6 | 0.039 | Young |
| 0.201562 | 0.147423 | 0.673 | 0.652 | 1 | 5 | Pomp | 0.021 | Young |
| 0.201626 | 0.304201 | 0.442 | 0.418 | 1 | 5 | Ccdc88a | 0.024 | Young |
| 0.201732 | 0.455689 | 0.036 | 0.022 | 1 | 5 | Rbm4 | 0.014 | Young |
| 0.202184 | 0.357913 | 0.317 | 0.279 | 1 | 5 | Setd3 | 0.038 | Young |

| | | | | | | | | |
|---|---|---|---|---|---|---|---|---|
| 0.202261 | 0.436503 | 0.058 | 0.04 | 1 | 5 | Snhg9 | 0.018 | Young |
| 0.202423 | 0.350455 | 0.099 | 0.076 | 1 | 5 | Ddx55 | 0.023 | Young |
| 0.202555 | 0.907812 | 0.033 | 0.02 | 1 | 5 | Erbb3 | 0.013 | Young |
| 0.202577 | 0.319758 | 0.236 | 0.204 | 1 | 5 | Ndfip1 | 0.032 | Young |
| 0.202741 | 0.640903 | 0.071 | 0.052 | 1 | 5 | Znhit6 | 0.019 | Young |
| 0.203181 | 0.476737 | 0.089 | 0.067 | 1 | 5 | Utp20 | 0.022 | Young |
| 0.203186 | 0.425604 | 0.066 | 0.047 | 1 | 5 | Orc5 | 0.019 | Young |
| 0.203206 | 0.158515 | 0.782 | 0.793 | 1 | 5 | Rpl15 | -0.011 | Young |
| 0.203498 | 0.556712 | 0.135 | 0.109 | 1 | 5 | Mcm6 | 0.026 | Young |
| 0.203708 | 0.261995 | 0.069 | 0.049 | 1 | 5 | Fam60a | 0.02 | Young |
| 0.203911 | 0.274545 | 0.069 | 0.049 | 1 | 5 | Mthfd2 | 0.02 | Young |
| 0.204103 | 0.346715 | 0.147 | 0.118 | 1 | 5 | Nfyc | 0.029 | Young |
| 0.204252 | 0.252927 | 0.431 | 0.399 | 1 | 5 | Thrap3 | 0.032 | Young |
| 0.20437 | 0.231505 | 0.18 | 0.148 | 1 | 5 | Commd6 | 0.032 | Young |
| 0.204646 | 0.137075 | 0.543 | 0.493 | 1 | 5 | Rab2a | 0.05 | Young |
| 0.204667 | 0.288355 | 0.109 | 0.084 | 1 | 5 | Galnt4 | 0.025 | Young |
| 0.205075 | 0.30527 | 0.165 | 0.135 | 1 | 5 | Washc3 | 0.03 | Young |
| 0.20511 | 0.70966 | 0.071 | 0.052 | 1 | 5 | Slc39a6 | 0.019 | Young |
| 0.205526 | 0.200137 | 0.284 | 0.249 | 1 | 5 | Fto | 0.035 | Young |
| 0.205635 | 0.151292 | 0.388 | 0.348 | 1 | 5 | Snx27 | 0.04 | Young |
| 0.205763 | 0.093932 | 0.919 | 0.934 | 1 | 5 | Wnk1 | -0.015 | Young |
| 0.205867 | 0.537564 | 0.066 | 0.047 | 1 | 5 | Tsr2 | 0.019 | Young |
| 0.205875 | 0.251982 | 0.421 | 0.394 | 1 | 5 | Gnai3 | 0.027 | Young |
| 0.206348 | 0.243359 | 0.069 | 0.049 | 1 | 5 | Smo | 0.02 | Young |
| 0.206436 | 0.892771 | 0.033 | 0.02 | 1 | 5 | Sap130 | 0.013 | Young |
| 0.206539 | 0.122317 | 0.612 | 0.598 | 1 | 5 | Vwa5a | 0.014 | Young |
| 0.206546 | 0.492775 | 0.089 | 0.067 | 1 | 5 | Prep | 0.022 | Young |
| 0.206585 | 0.524873 | 0.094 | 0.072 | 1 | 5 | Tex2 | 0.022 | Young |
| 0.206603 | 0.185293 | 0.16 | 0.13 | 1 | 5 | Ube3b | 0.03 | Young |
| 0.207038 | 0.746127 | 0.033 | 0.02 | 1 | 5 | A130077B1 | 0.013 | Young |
| 0.207094 | 0.571548 | 0.063 | 0.045 | 1 | 5 | Gtpbp8 | 0.018 | Young |
| 0.207534 | 0.2447 | 0.52 | 0.505 | 1 | 5 | Plekhb2 | 0.015 | Young |
| 0.207739 | 0.411488 | 0.129 | 0.104 | 1 | 5 | Sdad1 | 0.025 | Young |
| 0.207925 | 0.366887 | 0.157 | 0.13 | 1 | 5 | Zdhhc7 | 0.027 | Young |
| 0.208155 | 0.522029 | 0.063 | 0.045 | 1 | 5 | Nat10 | 0.018 | Young |
| 0.20868 | 0.415423 | 0.152 | 0.123 | 1 | 5 | Abcd3 | 0.029 | Young |
| 0.208807 | 0.167559 | 0.949 | 0.944 | 1 | 5 | Rps13 | 0.005 | Young |
| 0.209007 | 0.188863 | 0.269 | 0.236 | 1 | 5 | Arid4b | 0.033 | Young |
| 0.209116 | 0.347987 | 0.058 | 0.04 | 1 | 5 | Repin1 | 0.018 | Young |
| 0.209194 | 0.18017 | 0.518 | 0.481 | 1 | 5 | Luc7l3 | 0.037 | Young |
| 0.209458 | 0.614919 | 0.033 | 0.02 | 1 | 5 | Gm16124 | 0.013 | Young |
| 0.209689 | 0.276941 | 0.36 | 0.328 | 1 | 5 | Map2k2 | 0.032 | Young |
| 0.209835 | 0.361938 | 0.074 | 0.054 | 1 | 5 | Med27 | 0.02 | Young |
| 0.210067 | 0.582269 | 0.033 | 0.02 | 1 | 5 | Zfand1 | 0.013 | Young |
| 0.210091 | 0.453904 | 0.109 | 0.086 | 1 | 5 | Ccdc93 | 0.023 | Young |

| | | | | | | | | |
|---|---|---|---|---|---|---|---|---|
| 0.210371 | 0.743915 | 0.033 | 0.02 | 1 | 5 | Tyms | 0.013 | Young |
| 0.210606 | 0.264662 | 0.104 | 0.081 | 1 | 5 | Cep290 | 0.023 | Young |
| 0.210718 | 0.529547 | 0.063 | 0.045 | 1 | 5 | Gpr157 | 0.018 | Young |
| 0.210769 | 0.325206 | 0.069 | 0.049 | 1 | 5 | Irgq | 0.02 | Young |
| 0.210942 | 0.047419 | 0.041 | 0.059 | 1 | 5 | 2700038G2 | -0.018 | Young |
| 0.211328 | 0.129066 | 0.873 | 0.852 | 1 | 5 | Slc6a6 | 0.021 | Young |
| 0.211344 | 0.658547 | 0.053 | 0.037 | 1 | 5 | Zfp931 | 0.016 | Young |
| 0.211475 | 0.630016 | 0.043 | 0.029 | 1 | 5 | Dyrk1b | 0.014 | Young |
| 0.211626 | 0.558099 | 0.086 | 0.066 | 1 | 5 | Gm5547 | 0.02 | Young |
| 0.21171 | 0.279235 | 0.122 | 0.096 | 1 | 5 | Zbtb34 | 0.026 | Young |
| 0.211761 | 0.032965 | 0.058 | 0.081 | 1 | 5 | Nmral1 | -0.023 | Young |
| 0.21177 | 0.225747 | 0.437 | 0.412 | 1 | 5 | Wapl | 0.025 | Young |
| 0.211865 | 0.142577 | 0.779 | 0.769 | 1 | 5 | Man1a | 0.01 | Young |
| 0.211944 | 0.178596 | 0.122 | 0.096 | 1 | 5 | Rpa3 | 0.026 | Young |
| 0.212336 | 0.697851 | 0.071 | 0.052 | 1 | 5 | Ercc5 | 0.019 | Young |
| 0.21261 | 0.018463 | 0.14 | 0.175 | 1 | 5 | Aldh16a1 | -0.035 | Young |
| 0.212655 | 0.425021 | 0.157 | 0.13 | 1 | 5 | Fam118b | 0.027 | Young |
| 0.212957 | 0.405616 | 0.292 | 0.266 | 1 | 5 | Clec4n | 0.026 | Young |
| 0.212975 | 0.183776 | 0.665 | 0.64 | 1 | 5 | Sec14l1 | 0.025 | Young |
| 0.213143 | 0.352344 | 0.208 | 0.177 | 1 | 5 | Klhl5 | 0.031 | Young |
| 0.213736 | 0.534506 | 0.063 | 0.045 | 1 | 5 | Usp49 | 0.018 | Young |
| 0.213798 | 0.196073 | 0.259 | 0.221 | 1 | 5 | Zfp800 | 0.038 | Young |
| 0.213846 | 0.450789 | 0.14 | 0.114 | 1 | 5 | Dhx16 | 0.026 | Young |
| 0.213988 | 0.399874 | 0.239 | 0.21 | 1 | 5 | Fkbp3 | 0.029 | Young |
| 0.214178 | 0.560235 | 0.203 | 0.175 | 1 | 5 | Dag1 | 0.028 | Young |
| 0.214196 | 0.31602 | 0.16 | 0.131 | 1 | 5 | Utp14a | 0.029 | Young |
| 0.214287 | 0.43272 | 0.046 | 0.03 | 1 | 5 | Scfd2 | 0.016 | Young |
| 0.214287 | 0.258857 | 0.414 | 0.384 | 1 | 5 | Etv5 | 0.03 | Young |
| 0.21442 | 0.29244 | 0.135 | 0.108 | 1 | 5 | Gnpda2 | 0.027 | Young |
| 0.21457 | 0.203134 | 0.231 | 0.195 | 1 | 5 | Stom | 0.036 | Young |
| 0.214575 | 0.276229 | 0.152 | 0.123 | 1 | 5 | Cdk2ap1 | 0.029 | Young |
| 0.214768 | 0.287538 | 0.325 | 0.291 | 1 | 5 | Helz | 0.034 | Young |
| 0.215113 | 0.219344 | 0.307 | 0.269 | 1 | 5 | Dbnl | 0.038 | Young |
| 0.215206 | 0.261347 | 0.14 | 0.111 | 1 | 5 | Tmem68 | 0.029 | Young |
| 0.215401 | 0.209425 | 0.239 | 0.205 | 1 | 5 | Ktn1 | 0.034 | Young |
| 0.21585 | 0.132209 | 0.033 | 0.051 | 1 | 5 | Il18bp | -0.018 | Young |
| 0.216104 | 0.201457 | 0.51 | 0.473 | 1 | 5 | Taldo1 | 0.037 | Young |
| 0.216201 | 0.205909 | 0.497 | 0.468 | 1 | 5 | Selenos | 0.029 | Young |
| 0.216267 | 0.261128 | 0.312 | 0.281 | 1 | 5 | Exoc3 | 0.031 | Young |
| 0.216806 | 0.115025 | 0.706 | 0.705 | 1 | 5 | Pfdn5 | 0.001 | Young |
| 0.217258 | 0.284046 | 0.302 | 0.271 | 1 | 5 | Zfx | 0.031 | Young |
| 0.21731 | 0.251612 | 0.518 | 0.5 | 1 | 5 | Nfxl1 | 0.018 | Young |
| 0.217425 | 0.578036 | 0.061 | 0.044 | 1 | 5 | Fscn1 | 0.017 | Young |
| 0.217872 | 0.586014 | 0.061 | 0.044 | 1 | 5 | Pik3cb | 0.017 | Young |
| 0.217974 | 0.310843 | 0.119 | 0.094 | 1 | 5 | Mis18a | 0.025 | Young |

| | | | | | | | | |
|---|---|---|---|---|---|---|---|---|
| 0.218692 | 0.404672 | 0.096 | 0.074 | 1 | 5 | Tmem199 | 0.022 | Young |
| 0.218989 | 0.171353 | 0.332 | 0.295 | 1 | 5 | Plekhj1 | 0.037 | Young |
| 0.219231 | 0.606522 | 0.076 | 0.057 | 1 | 5 | Tubgcp2 | 0.019 | Young |
| 0.219657 | 0.332725 | 0.066 | 0.047 | 1 | 5 | Dhx35 | 0.019 | Young |
| 0.220061 | 0.009905 | 0.058 | 0.04 | 1 | 5 | Sgsm2 | 0.018 | Young |
| 0.220142 | 0.273786 | 0.452 | 0.412 | 1 | 5 | Eif3h | 0.04 | Young |
| 0.220797 | 0.636348 | 0.061 | 0.044 | 1 | 5 | Rbm12b1 | 0.017 | Young |
| 0.221224 | 0.034638 | 0.058 | 0.04 | 1 | 5 | E2f3 | 0.018 | Young |
| 0.221294 | 0.369191 | 0.249 | 0.222 | 1 | 5 | Orai3 | 0.027 | Young |
| 0.22132 | 0.345303 | 0.211 | 0.18 | 1 | 5 | Dpy19l4 | 0.031 | Young |
| 0.221674 | 0.266142 | 0.264 | 0.232 | 1 | 5 | F630028O1 | 0.032 | Young |
| 0.221994 | 0.42452 | 0.086 | 0.066 | 1 | 5 | Pag1 | 0.02 | Young |
| 0.222016 | 0.261683 | 0.533 | 0.507 | 1 | 5 | Cnot6l | 0.026 | Young |
| 0.222112 | 0.440459 | 0.15 | 0.125 | 1 | 5 | Lap3 | 0.025 | Young |
| 0.222267 | 0.430312 | 0.195 | 0.168 | 1 | 5 | Mrpl51 | 0.027 | Young |
| 0.22231 | 0.511851 | 0.063 | 0.045 | 1 | 5 | Mtx3 | 0.018 | Young |
| 0.222344 | 0.271026 | 0.325 | 0.286 | 1 | 5 | Dcaf12 | 0.039 | Young |
| 0.22242 | 0.576642 | 0.132 | 0.108 | 1 | 5 | Rcan1 | 0.024 | Young |
| 0.222473 | 0.519739 | 0.03 | 0.019 | 1 | 5 | 2700062C0 | 0.011 | Young |
| 0.222604 | 0.323161 | 0.223 | 0.19 | 1 | 5 | Cnot8 | 0.033 | Young |
| 0.222942 | 0.158059 | 0.619 | 0.603 | 1 | 5 | Klf3 | 0.016 | Young |
| 0.22297 | 0.36721 | 0.112 | 0.088 | 1 | 5 | Llgl1 | 0.024 | Young |
| 0.223061 | 0.286648 | 0.335 | 0.31 | 1 | 5 | Hps3 | 0.025 | Young |
| 0.223235 | 0.33237 | 0.081 | 0.061 | 1 | 5 | Prrg1 | 0.02 | Young |
| 0.223466 | 0.552011 | 0.03 | 0.019 | 1 | 5 | Megf9 | 0.011 | Young |
| 0.223561 | 0.463195 | 0.396 | 0.389 | 1 | 5 | Prdx5 | 0.007 | Young |
| 0.223933 | 0.39259 | 0.236 | 0.21 | 1 | 5 | Yeats4 | 0.026 | Young |
| 0.224129 | 0.504109 | 0.03 | 0.019 | 1 | 5 | 2810001G2 | 0.011 | Young |
| 0.224361 | 0.334575 | 0.071 | 0.052 | 1 | 5 | Zfp369 | 0.019 | Young |
| 0.224667 | 0.201934 | 0.33 | 0.296 | 1 | 5 | Skp1a | 0.034 | Young |
| 0.224889 | 0.403581 | 0.198 | 0.17 | 1 | 5 | Snrk | 0.028 | Young |
| 0.225128 | 0.677629 | 0.03 | 0.019 | 1 | 5 | Ska2 | 0.011 | Young |
| 0.225128 | 0.606565 | 0.03 | 0.019 | 1 | 5 | Klra2 | 0.011 | Young |
| 0.225307 | 0.209426 | 0.074 | 0.054 | 1 | 5 | Traf2 | 0.02 | Young |
| 0.22544 | 0.199629 | 0.332 | 0.303 | 1 | 5 | Noc2l | 0.029 | Young |
| 0.225824 | 0.308841 | 0.129 | 0.104 | 1 | 5 | Fyco1 | 0.025 | Young |
| 0.225836 | 0.100442 | 0.929 | 0.939 | 1 | 5 | Rps25 | -0.01 | Young |
| 0.225901 | 0.257705 | 0.424 | 0.394 | 1 | 5 | Fmnl1 | 0.03 | Young |
| 0.225983 | 0.363247 | 0.234 | 0.207 | 1 | 5 | Mospd3 | 0.027 | Young |
| 0.226549 | 0.109047 | 0.68 | 0.66 | 1 | 5 | Irf8 | 0.02 | Young |
| 0.226678 | 0.298006 | 0.137 | 0.111 | 1 | 5 | Dnpep | 0.026 | Young |
| 0.226799 | 0.553005 | 0.03 | 0.019 | 1 | 5 | Igfbp6 | 0.011 | Young |
| 0.227438 | 0.410377 | 0.102 | 0.079 | 1 | 5 | Nelfcd | 0.023 | Young |
| 0.227652 | 0.337162 | 0.061 | 0.044 | 1 | 5 | Gm43936 | 0.017 | Young |
| 0.227885 | 0.31061 | 0.249 | 0.219 | 1 | 5 | Dpp8 | 0.03 | Young |

| | | | | | | | | |
|---|---|---|---|---|---|---|---|---|
| 0.227913 | 0.301438 | 0.216 | 0.18 | 1 | 5 | Kctd2 | 0.036 | Young |
| 0.228295 | 0.352155 | 0.124 | 0.099 | 1 | 5 | 4930523C0 | 0.025 | Young |
| 0.228581 | 0.382433 | 0.18 | 0.157 | 1 | 5 | Polr3gl | 0.023 | Young |
| 0.22907 | 0.268133 | 0.19 | 0.157 | 1 | 5 | Fam192a | 0.033 | Young |
| 0.229071 | 0.184739 | 0.063 | 0.045 | 1 | 5 | Zfp653 | 0.018 | Young |
| 0.229073 | 0.485353 | 0.069 | 0.051 | 1 | 5 | Iqce | 0.018 | Young |
| 0.229101 | 0.162071 | 0.68 | 0.7 | 1 | 5 | Ywhab | -0.02 | Young |
| 0.229242 | 0.182631 | 0.596 | 0.567 | 1 | 5 | Ndufa6 | 0.029 | Young |
| 0.229274 | 0.664866 | 0.061 | 0.044 | 1 | 5 | Tma16 | 0.017 | Young |
| 0.22973 | 0.58206 | 0.109 | 0.088 | 1 | 5 | Dopey2 | 0.021 | Young |
| 0.23023 | 0.259745 | 0.071 | 0.052 | 1 | 5 | Sprtn | 0.019 | Young |
| 0.230288 | 0.533074 | 0.091 | 0.071 | 1 | 5 | Atad3a | 0.02 | Young |
| 0.231169 | 0.346978 | 0.168 | 0.14 | 1 | 5 | Maml2 | 0.028 | Young |
| 0.231231 | 0.765871 | 0.041 | 0.027 | 1 | 5 | Nedd4l | 0.014 | Young |
| 0.231864 | 0.310829 | 0.03 | 0.019 | 1 | 5 | Klhl8 | 0.011 | Young |
| 0.231954 | 0.454289 | 0.084 | 0.064 | 1 | 5 | Cep57 | 0.02 | Young |
| 0.23227 | 0.204688 | 0.365 | 0.332 | 1 | 5 | Eif2s3x | 0.033 | Young |
| 0.232345 | 0.836664 | 0.028 | 0.017 | 1 | 5 | Tnfaip8l1 | 0.011 | Young |
| 0.232361 | 0.5392 | 0.069 | 0.051 | 1 | 5 | H2afx | 0.018 | Young |
| 0.232402 | 0.225203 | 0.203 | 0.17 | 1 | 5 | Cntrl | 0.033 | Young |
| 0.232606 | 0.580425 | 0.058 | 0.042 | 1 | 5 | Borcs5 | 0.016 | Young |
| 0.232797 | 0.081664 | 0.86 | 0.859 | 1 | 5 | Hnrnpk | 0.001 | Young |
| 0.23284 | 0.359173 | 0.34 | 0.318 | 1 | 5 | Ralgps2 | 0.022 | Young |
| 0.233097 | 0.312615 | 0.015 | 0.027 | 1 | 5 | Bcr | -0.012 | Young |
| 0.233209 | 0.107411 | 0.201 | 0.234 | 1 | 5 | Cbx4 | -0.033 | Young |
| 0.233425 | 0.400867 | 0.165 | 0.14 | 1 | 5 | Rab29 | 0.025 | Young |
| 0.233442 | 0.125308 | 0.746 | 0.746 | 1 | 5 | Ssr3 | 0 | Young |
| 0.233663 | 0.261024 | 0.089 | 0.067 | 1 | 5 | Vhl | 0.022 | Young |
| 0.233684 | 0.283843 | 0.104 | 0.081 | 1 | 5 | Pigyl | 0.023 | Young |
| 0.233942 | 0.307777 | 0.292 | 0.261 | 1 | 5 | Car3 | 0.031 | Young |
| 0.234323 | 0.23758 | 0.18 | 0.148 | 1 | 5 | Ahctf1 | 0.032 | Young |
| 0.234358 | 0.245943 | 0.178 | 0.148 | 1 | 5 | Eif4enif1 | 0.03 | Young |
| 0.234633 | 0.33092 | 0.152 | 0.125 | 1 | 5 | Sdhaf4 | 0.027 | Young |
| 0.234849 | 0.971543 | 0.028 | 0.017 | 1 | 5 | Ccdc15 | 0.011 | Young |
| 0.235002 | 0.301082 | 0.251 | 0.224 | 1 | 5 | Cs | 0.027 | Young |
| 0.235083 | 0.187137 | 0.244 | 0.209 | 1 | 5 | Bbip1 | 0.035 | Young |
| 0.235456 | 0.104039 | 0.36 | 0.325 | 1 | 5 | H1f0 | 0.035 | Young |
| 0.235851 | 0.385442 | 0.104 | 0.081 | 1 | 5 | Crls1 | 0.023 | Young |
| 0.235928 | 0.194393 | 0.036 | 0.052 | 1 | 5 | Zfand2a | -0.016 | Young |
| 0.236525 | 0.368737 | 0.135 | 0.109 | 1 | 5 | Foxo1 | 0.026 | Young |
| 0.237069 | 0.558424 | 0.041 | 0.027 | 1 | 5 | Dynlt1a | 0.014 | Young |
| 0.237376 | 0.54504 | 0.135 | 0.111 | 1 | 5 | Vps33a | 0.024 | Young |
| 0.237614 | 0.416262 | 0.099 | 0.077 | 1 | 5 | Ipo13 | 0.022 | Young |
| 0.237701 | 0.271351 | 0.03 | 0.019 | 1 | 5 | Cox18 | 0.011 | Young |
| 0.237912 | 0.128182 | 0.18 | 0.148 | 1 | 5 | Plcb3 | 0.032 | Young |

| | | | | | | | | |
|---|---|---|---|---|---|---|---|---|
| 0.237946 | 0.05305 | 1 | 0.998 | 1 | 5 | Mrc1 | 0.002 | Young |
| 0.238514 | 0.153234 | 0.223 | 0.189 | 1 | 5 | Rab3gap2 | 0.034 | Young |
| 0.238696 | 0.127192 | 0.495 | 0.461 | 1 | 5 | Caprin1 | 0.034 | Young |
| 0.23872 | 0.442137 | 0.091 | 0.071 | 1 | 5 | Gbp5 | 0.02 | Young |
| 0.238788 | 0.622945 | 0.114 | 0.094 | 1 | 5 | Lrp12 | 0.02 | Young |
| 0.238789 | 0.141731 | 0.635 | 0.636 | 1 | 5 | B4galt6 | -0.001 | Young |
| 0.239003 | 0.163573 | 0.429 | 0.396 | 1 | 5 | Clint1 | 0.033 | Young |
| 0.239375 | 0.44912 | 0.051 | 0.035 | 1 | 5 | Emilin1 | 0.016 | Young |
| 0.239461 | 0.246582 | 0.381 | 0.362 | 1 | 5 | R3hdm4 | 0.019 | Young |
| 0.239495 | 0.220893 | 0.226 | 0.197 | 1 | 5 | Akap9 | 0.029 | Young |
| 0.23955 | 0.499483 | 0.028 | 0.017 | 1 | 5 | L2hgdh | 0.011 | Young |
| 0.239562 | 0.76447 | 0.048 | 0.034 | 1 | 5 | Plk4 | 0.014 | Young |
| 0.239674 | 0.400865 | 0.239 | 0.21 | 1 | 5 | Stk11 | 0.029 | Young |
| 0.239832 | 0.70282 | 0.048 | 0.034 | 1 | 5 | Fam13a | 0.014 | Young |
| 0.239892 | 0.350757 | 0.061 | 0.044 | 1 | 5 | Katnal1 | 0.017 | Young |
| 0.239969 | 0.193239 | 0.345 | 0.311 | 1 | 5 | Stag1 | 0.034 | Young |
| 0.240584 | 0.720347 | 0.038 | 0.025 | 1 | 5 | Tsr3 | 0.013 | Young |
| 0.240621 | 0.364454 | 0.041 | 0.027 | 1 | 5 | 2610035D1 | 0.014 | Young |
| 0.240644 | 0.199336 | 0.096 | 0.074 | 1 | 5 | Prkag2 | 0.022 | Young |
| 0.240826 | 0.503543 | 0.132 | 0.109 | 1 | 5 | Phtf2 | 0.023 | Young |
| 0.240919 | 0.579867 | 0.041 | 0.027 | 1 | 5 | Rrs1 | 0.014 | Young |
| 0.241009 | 0.650041 | 0.028 | 0.017 | 1 | 5 | Zfp105 | 0.011 | Young |
| 0.241875 | 0.068196 | 0.332 | 0.285 | 1 | 5 | Mdm4 | 0.047 | Young |
| 0.241967 | 0.534693 | 0.132 | 0.109 | 1 | 5 | Smim4 | 0.023 | Young |
| 0.241993 | 0.305442 | 0.467 | 0.453 | 1 | 5 | Cdc42se1 | 0.014 | Young |
| 0.242082 | 0.219133 | 0.14 | 0.111 | 1 | 5 | Psip1 | 0.029 | Young |
| 0.242212 | 0.024935 | 0.071 | 0.094 | 1 | 5 | Flywch1 | -0.023 | Young |
| 0.242298 | 0.278324 | 0.061 | 0.044 | 1 | 5 | Ints10 | 0.017 | Young |
| 0.24271 | 0.370902 | 0.041 | 0.027 | 1 | 5 | Gm26518 | 0.014 | Young |
| 0.242771 | 0.542773 | 0.17 | 0.15 | 1 | 5 | Eed | 0.02 | Young |
| 0.242806 | 0.303297 | 0.076 | 0.057 | 1 | 5 | Psmg2 | 0.019 | Young |
| 0.242881 | 0.178911 | 0.284 | 0.253 | 1 | 5 | Crybg3 | 0.031 | Young |
| 0.243265 | 0.417057 | 0.061 | 0.044 | 1 | 5 | Nek1 | 0.017 | Young |
| 0.243512 | 0.235866 | 0.383 | 0.357 | 1 | 5 | Ulk2 | 0.026 | Young |
| 0.243546 | 0.472507 | 0.058 | 0.042 | 1 | 5 | Retn | 0.016 | Young |
| 0.243738 | 0.431157 | 0.074 | 0.056 | 1 | 5 | Ogfod1 | 0.018 | Young |
| 0.243948 | 0.298122 | 0.028 | 0.017 | 1 | 5 | Cdk20 | 0.011 | Young |
| 0.244115 | 0.16798 | 0.266 | 0.231 | 1 | 5 | Lpar6 | 0.035 | Young |
| 0.244478 | 0.533137 | 0.107 | 0.086 | 1 | 5 | Zfp560 | 0.021 | Young |
| 0.245025 | 0.704995 | 0.063 | 0.047 | 1 | 5 | Map4k2 | 0.016 | Young |
| 0.245026 | 0.227228 | 0.325 | 0.288 | 1 | 5 | Rps6ka1 | 0.037 | Young |
| 0.245056 | 0.39614 | 0.028 | 0.017 | 1 | 5 | Ecsit | 0.011 | Young |
| 0.24522 | 0.681507 | 0.038 | 0.025 | 1 | 5 | Amer1 | 0.013 | Young |
| 0.245496 | 0.672252 | 0.074 | 0.056 | 1 | 5 | Snip1 | 0.018 | Young |
| 0.245568 | 0.253962 | 0.188 | 0.158 | 1 | 5 | Spr | 0.03 | Young |

| | | | | | | | | |
|---|---|---|---|---|---|---|---|---|
| 0.245778 | 0.227641 | 0.119 | 0.094 | 1 | 5 | Pex14 | 0.025 | Young |
| 0.246208 | 0.307231 | 0.244 | 0.212 | 1 | 5 | Usp22 | 0.032 | Young |
| 0.246538 | 0.464578 | 0.069 | 0.051 | 1 | 5 | Zc3h12d | 0.018 | Young |
| 0.246847 | 0.204291 | 0.175 | 0.146 | 1 | 5 | Cops5 | 0.029 | Young |
| 0.247091 | 0.665754 | 0.038 | 0.025 | 1 | 5 | Fam117a | 0.013 | Young |
| 0.247154 | 0.052042 | 0.216 | 0.18 | 1 | 5 | Rbl2 | 0.036 | Young |
| 0.247436 | 0.313182 | 0.383 | 0.369 | 1 | 5 | Srsf9 | 0.014 | Young |
| 0.247602 | 0.223477 | 0.112 | 0.088 | 1 | 5 | Ift46 | 0.024 | Young |
| 0.247747 | 0.206825 | 0.35 | 0.315 | 1 | 5 | Npc1 | 0.035 | Young |
| 0.247971 | 0.204161 | 0.368 | 0.35 | 1 | 5 | Rdx | 0.018 | Young |
| 0.248023 | 0.176862 | 0.411 | 0.379 | 1 | 5 | Sf3b5 | 0.032 | Young |
| 0.248052 | 0.287034 | 0.305 | 0.264 | 1 | 5 | Stat1 | 0.041 | Young |
| 0.248146 | 0.252964 | 0.221 | 0.189 | 1 | 5 | Pa2g4 | 0.032 | Young |
| 0.248309 | 0.360242 | 0.198 | 0.17 | 1 | 5 | Lars | 0.028 | Young |
| 0.248338 | 0.541947 | 0.066 | 0.049 | 1 | 5 | Tfcp2 | 0.017 | Young |
| 0.248768 | 0.275132 | 0.195 | 0.167 | 1 | 5 | Arhgap39 | 0.028 | Young |
| 0.248781 | 0.102039 | 0.31 | 0.345 | 1 | 5 | Bmp2 | -0.035 | Young |
| 0.248783 | 0.099177 | 0.711 | 0.697 | 1 | 5 | Ddx6 | 0.014 | Young |
| 0.248955 | 0.778971 | 0.025 | 0.015 | 1 | 5 | Bcar3 | 0.01 | Young |
| 0.248973 | 0.400818 | 0.038 | 0.025 | 1 | 5 | Fam217b | 0.013 | Young |
| 0.248973 | 0.560866 | 0.038 | 0.025 | 1 | 5 | Tal1 | 0.013 | Young |
| 0.249188 | 0.140424 | 0.558 | 0.537 | 1 | 5 | Atp5j2 | 0.021 | Young |
| 0.249693 | 0.442525 | 0.084 | 0.064 | 1 | 5 | Slc27a4 | 0.02 | Young |
| 0.249714 | 0.692433 | 0.056 | 0.04 | 1 | 5 | Cdkn2d | 0.016 | Young |
| 0.250136 | 0.646972 | 0.025 | 0.015 | 1 | 5 | Slc25a14 | 0.01 | Young |
| 0.250136 | 0.543189 | 0.025 | 0.015 | 1 | 5 | Vsig10l | 0.01 | Young |
| 0.250245 | 0.331795 | 0.162 | 0.138 | 1 | 5 | Hspbp1 | 0.024 | Young |
| 0.250605 | 0.070264 | 0.485 | 0.461 | 1 | 5 | Gsn | 0.024 | Young |
| 0.250925 | 0.604711 | 0.025 | 0.015 | 1 | 5 | Rbak | 0.01 | Young |
| 0.250969 | 0.351701 | 0.223 | 0.194 | 1 | 5 | Cstf2t | 0.029 | Young |
| 0.251002 | 0.422438 | 0.094 | 0.074 | 1 | 5 | Usp50 | 0.02 | Young |
| 0.251223 | 0.084059 | 0.315 | 0.276 | 1 | 5 | Gripap1 | 0.039 | Young |
| 0.251321 | 0.586716 | 0.025 | 0.015 | 1 | 5 | Dnm1 | 0.01 | Young |
| 0.251375 | 0.282459 | 0.282 | 0.249 | 1 | 5 | Lgals9 | 0.033 | Young |
| 0.251542 | 0.396624 | 0.058 | 0.042 | 1 | 5 | Abhd5 | 0.016 | Young |
| 0.251716 | 0.737907 | 0.025 | 0.015 | 1 | 5 | Zfp229 | 0.01 | Young |
| 0.252272 | 0.165074 | 0.386 | 0.348 | 1 | 5 | Bola2 | 0.038 | Young |
| 0.252682 | 0.783382 | 0.071 | 0.054 | 1 | 5 | Apex1 | 0.017 | Young |
| 0.252906 | 0.435648 | 0.025 | 0.015 | 1 | 5 | Ppp1r3e | 0.01 | Young |
| 0.252939 | 0.20996 | 0.622 | 0.623 | 1 | 5 | Ctsl | -0.001 | Young |
| 0.252968 | 0.261286 | 0.114 | 0.091 | 1 | 5 | Desi1 | 0.023 | Young |
| 0.253059 | 0.067323 | 0.216 | 0.18 | 1 | 5 | AI467606 | 0.036 | Young |
| 0.253095 | 0.41846 | 0.218 | 0.192 | 1 | 5 | Cracr2b | 0.026 | Young |
| 0.253304 | 0.599347 | 0.025 | 0.015 | 1 | 5 | Rhebl1 | 0.01 | Young |
| 0.253636 | 0.332424 | 0.107 | 0.084 | 1 | 5 | Atp6v0a2 | 0.023 | Young |

| | | | | | | | | |
|---|---|---|---|---|---|---|---|---|
| 0.253693 | 0.077835 | 0.863 | 0.869 | 1 | 5 | App | -0.006 | Young |
| 0.2541 | 0.557997 | 0.025 | 0.015 | 1 | 5 | Gemin2 | 0.01 | Young |
| 0.254532 | 0.138125 | 0.449 | 0.418 | 1 | 5 | Vapa | 0.031 | Young |
| 0.255297 | 0.457309 | 0.025 | 0.015 | 1 | 5 | Zfp408 | 0.01 | Young |
| 0.255495 | 0.183148 | 0.594 | 0.569 | 1 | 5 | Rtn3 | 0.025 | Young |
| 0.255739 | 0.318439 | 0.127 | 0.104 | 1 | 5 | Smcr8 | 0.023 | Young |
| 0.255898 | 0.649392 | 0.025 | 0.015 | 1 | 5 | Il15ra | 0.01 | Young |
| 0.256126 | 0.394358 | 0.244 | 0.217 | 1 | 5 | Cript | 0.027 | Young |
| 0.256264 | 0.253424 | 0.277 | 0.247 | 1 | 5 | Lsm6 | 0.03 | Young |
| 0.256298 | 0.232537 | 0.574 | 0.569 | 1 | 5 | Lrp6 | 0.005 | Young |
| 0.256499 | 0.4079 | 0.025 | 0.015 | 1 | 5 | Lig4 | 0.01 | Young |
| 0.257219 | 0.418526 | 0.102 | 0.081 | 1 | 5 | 2310022B0 | 0.021 | Young |
| 0.257386 | 0.382201 | 0.127 | 0.104 | 1 | 5 | Cfap20 | 0.023 | Young |
| 0.257542 | 0.020827 | 0.736 | 0.763 | 1 | 5 | Slco2b1 | -0.027 | Young |
| 0.257545 | 0.504637 | 0.17 | 0.15 | 1 | 5 | Napg | 0.02 | Young |
| 0.257865 | 0.965535 | 0.046 | 0.032 | 1 | 5 | Cutc | 0.014 | Young |
| 0.258431 | 0.505954 | 0.071 | 0.054 | 1 | 5 | Apool | 0.017 | Young |
| 0.258532 | 0.375145 | 0.038 | 0.025 | 1 | 5 | Mdm1 | 0.013 | Young |
| 0.258679 | 0.788686 | 0.069 | 0.052 | 1 | 5 | Prmt9 | 0.017 | Young |
| 0.258847 | 0.717399 | 0.036 | 0.024 | 1 | 5 | Lamb2 | 0.012 | Young |
| 0.258948 | 0.746255 | 0.053 | 0.039 | 1 | 5 | Fbxw17 | 0.014 | Young |
| 0.259129 | 0.106423 | 0.83 | 0.788 | 1 | 5 | Rac1 | 0.042 | Young |
| 0.259273 | 0.466076 | 0.063 | 0.047 | 1 | 5 | C330018D2 | 0.016 | Young |
| 0.259276 | 0.254687 | 0.185 | 0.155 | 1 | 5 | Tmem57 | 0.03 | Young |
| 0.259943 | 0.3205 | 0.289 | 0.258 | 1 | 5 | Btk | 0.031 | Young |
| 0.259965 | 0.149446 | 0.048 | 0.034 | 1 | 5 | Gm20604 | 0.014 | Young |
| 0.26015 | 0.419374 | 0.038 | 0.025 | 1 | 5 | Mrpl44 | 0.013 | Young |
| 0.260202 | 0.544256 | 0.046 | 0.032 | 1 | 5 | Armc3 | 0.014 | Young |
| 0.260397 | 0.237999 | 0.289 | 0.261 | 1 | 5 | Gpr183 | 0.028 | Young |
| 0.260745 | 0.26695 | 0.198 | 0.168 | 1 | 5 | Snrpa | 0.03 | Young |
| 0.260891 | 0.460322 | 0.086 | 0.067 | 1 | 5 | B3galnt2 | 0.019 | Young |
| 0.261264 | 0.351238 | 0.294 | 0.266 | 1 | 5 | Arhgap19 | 0.028 | Young |
| 0.261364 | 0.235529 | 0.398 | 0.384 | 1 | 5 | Prpf4b | 0.014 | Young |
| 0.261398 | 0.352424 | 0.104 | 0.082 | 1 | 5 | Abhd13 | 0.022 | Young |
| 0.26176 | 0.14845 | 0.67 | 0.7 | 1 | 5 | Adam15 | -0.03 | Young |
| 0.261848 | 0.245752 | 0.437 | 0.406 | 1 | 5 | Nop10 | 0.031 | Young |
| 0.261991 | 0.712121 | 0.063 | 0.047 | 1 | 5 | Ndufaf7 | 0.016 | Young |
| 0.262128 | 0.232358 | 0.14 | 0.114 | 1 | 5 | Blcap | 0.026 | Young |
| 0.26252 | 0.086058 | 0.409 | 0.377 | 1 | 5 | Lyve1 | 0.032 | Young |
| 0.262743 | 0.107115 | 0.662 | 0.648 | 1 | 5 | Atp5f1 | 0.014 | Young |
| 0.262881 | 0.010292 | 0.145 | 0.118 | 1 | 5 | Sp140 | 0.027 | Young |
| 0.263346 | 0.455796 | 0.023 | 0.013 | 1 | 5 | Shmt1 | 0.01 | Young |
| 0.264031 | 0.818208 | 0.046 | 0.032 | 1 | 5 | Mettl17 | 0.014 | Young |
| 0.264113 | 0.558295 | 0.132 | 0.109 | 1 | 5 | Agap1 | 0.023 | Young |
| 0.26421 | 0.834763 | 0.023 | 0.013 | 1 | 5 | B130006D0 | 0.01 | Young |

| | | | | | | | | |
|---|---|---|---|---|---|---|---|---|
| 0.264231 | 0.518056 | 0.036 | 0.024 | 1 | 5 | Nme6 | 0.012 | Young |
| 0.264485 | 0.347809 | 0.094 | 0.074 | 1 | 5 | Ric8b | 0.02 | Young |
| 0.264665 | 0.170627 | 0.081 | 0.062 | 1 | 5 | Inip | 0.019 | Young |
| 0.26492 | 0.628771 | 0.046 | 0.032 | 1 | 5 | Setdb2 | 0.014 | Young |
| 0.265135 | 0.180264 | 0.338 | 0.303 | 1 | 5 | Ube2z | 0.035 | Young |
| 0.265155 | 0.110647 | 0.201 | 0.168 | 1 | 5 | Ergic1 | 0.033 | Young |
| 0.265473 | 0.35737 | 0.079 | 0.061 | 1 | 5 | Chn2 | 0.018 | Young |
| 0.265509 | 0.717214 | 0.023 | 0.013 | 1 | 5 | Zfp518b | 0.01 | Young |
| 0.265666 | 0.558144 | 0.084 | 0.066 | 1 | 5 | Pcyt2 | 0.018 | Young |
| 0.265927 | 0.203925 | 0.617 | 0.635 | 1 | 5 | Sulf2 | -0.018 | Young |
| 0.266159 | 0.142203 | 0.183 | 0.153 | 1 | 5 | Gabpa | 0.03 | Young |
| 0.266378 | 0.666358 | 0.023 | 0.013 | 1 | 5 | Gadd45a | 0.01 | Young |
| 0.26643 | 0.337203 | 0.277 | 0.247 | 1 | 5 | Vcpip1 | 0.03 | Young |
| 0.266565 | 0.16219 | 0.221 | 0.192 | 1 | 5 | Fbxw2 | 0.029 | Young |
| 0.266575 | 0.401957 | 0.074 | 0.056 | 1 | 5 | Ankmy2 | 0.018 | Young |
| 0.266603 | 0.200603 | 0.325 | 0.293 | 1 | 5 | Ivns1abp | 0.032 | Young |
| 0.266665 | 0.368654 | 0.15 | 0.125 | 1 | 5 | Zdhhc18 | 0.025 | Young |
| 0.26675 | 0.206731 | 0.513 | 0.485 | 1 | 5 | Wdfy3 | 0.028 | Young |
| 0.266792 | 0.322719 | 0.203 | 0.177 | 1 | 5 | Wdr41 | 0.026 | Young |
| 0.266813 | 0.809075 | 0.023 | 0.013 | 1 | 5 | Wdr60 | 0.01 | Young |
| 0.267202 | 0.12201 | 0.297 | 0.263 | 1 | 5 | Cxcr4 | 0.034 | Young |
| 0.267612 | 0.409433 | 0.551 | 0.534 | 1 | 5 | Hexb | 0.017 | Young |
| 0.268061 | 0.151824 | 0.124 | 0.101 | 1 | 5 | Hmces | 0.023 | Young |
| 0.268133 | 0.183043 | 0.482 | 0.444 | 1 | 5 | Ndufb7 | 0.038 | Young |
| 0.268501 | 0.500326 | 0.094 | 0.074 | 1 | 5 | Sesn2 | 0.02 | Young |
| 0.268559 | 0.417419 | 0.023 | 0.013 | 1 | 5 | Mmab | 0.01 | Young |
| 0.269067 | 0.201139 | 0.048 | 0.034 | 1 | 5 | Natd1 | 0.014 | Young |
| 0.269141 | 0.329708 | 0.17 | 0.143 | 1 | 5 | Sympk | 0.027 | Young |
| 0.26917 | 0.331171 | 0.335 | 0.31 | 1 | 5 | Tmem168 | 0.025 | Young |
| 0.269348 | 0.529552 | 0.036 | 0.024 | 1 | 5 | Rrp1b | 0.012 | Young |
| 0.269435 | 0.615303 | 0.023 | 0.013 | 1 | 5 | Nmrk1 | 0.01 | Young |
| 0.270026 | 0.191303 | 0.157 | 0.131 | 1 | 5 | Ptpn7 | 0.026 | Young |
| 0.270312 | 0.425293 | 0.023 | 0.013 | 1 | 5 | Lrrc1 | 0.01 | Young |
| 0.27034 | 0.013227 | 0.038 | 0.054 | 1 | 5 | Lrsam1 | -0.016 | Young |
| 0.270583 | 0.235263 | 0.325 | 0.298 | 1 | 5 | Dctn4 | 0.027 | Young |
| 0.270602 | 0.26422 | 0.066 | 0.049 | 1 | 5 | Pigg | 0.017 | Young |
| 0.270795 | 0.106542 | 0.074 | 0.056 | 1 | 5 | Polr3f | 0.018 | Young |
| 0.271025 | 0.496462 | 0.063 | 0.047 | 1 | 5 | Slx4ip | 0.016 | Young |
| 0.271641 | 0.261957 | 0.251 | 0.219 | 1 | 5 | Mbd3 | 0.032 | Young |
| 0.271996 | 0.045684 | 0.467 | 0.544 | 1 | 5 | Far1 | -0.077 | Young |
| 0.272191 | 0.341406 | 0.056 | 0.04 | 1 | 5 | Ccdc71 | 0.016 | Young |
| 0.272556 | 0.080684 | 0.81 | 0.791 | 1 | 5 | Cd47 | 0.019 | Young |
| 0.272645 | 0.243649 | 0.269 | 0.241 | 1 | 5 | Ss18 | 0.028 | Young |
| 0.272652 | 0.101839 | 0.472 | 0.418 | 1 | 5 | Ssr1 | 0.054 | Young |
| 0.273159 | 0.243511 | 0.124 | 0.101 | 1 | 5 | 1700037H( | 0.023 | Young |

| | | | | | | | | |
|---|---|---|---|---|---|---|---|---|
| 0.273693 | 0.026362 | 0.137 | 0.167 | 1 | 5 | Nlrc4 | -0.03 | Young |
| 0.273744 | 0.669591 | 0.053 | 0.039 | 1 | 5 | Nup107 | 0.014 | Young |
| 0.273918 | 0.299724 | 0.236 | 0.21 | 1 | 5 | Irak4 | 0.026 | Young |
| 0.274153 | 0.172589 | 0.36 | 0.325 | 1 | 5 | Rassf5 | 0.035 | Young |
| 0.274395 | 0.268838 | 0.244 | 0.219 | 1 | 5 | Vkorc1 | 0.025 | Young |
| 0.274901 | 0.324338 | 0.145 | 0.121 | 1 | 5 | Rufy1 | 0.024 | Young |
| 0.275183 | 0.150988 | 0.32 | 0.279 | 1 | 5 | Rtf2 | 0.041 | Young |
| 0.275349 | 0.266728 | 0.137 | 0.113 | 1 | 5 | Pygo2 | 0.024 | Young |
| 0.275554 | 0.297825 | 0.086 | 0.067 | 1 | 5 | Abtb1 | 0.019 | Young |
| 0.275578 | 0.193386 | 0.036 | 0.024 | 1 | 5 | Naif1 | 0.012 | Young |
| 0.275619 | 0.331152 | 0.023 | 0.013 | 1 | 5 | Gm37305 | 0.01 | Young |
| 0.275719 | 0.361713 | 0.091 | 0.072 | 1 | 5 | Hras | 0.019 | Young |
| 0.276061 | 0.142818 | 0.675 | 0.652 | 1 | 5 | Uqcr11 | 0.023 | Young |
| 0.276108 | 0.332715 | 0.124 | 0.103 | 1 | 5 | Pilra | 0.021 | Young |
| 0.276352 | 0.337576 | 0.386 | 0.365 | 1 | 5 | Zfp652 | 0.021 | Young |
| 0.276393 | 0.310231 | 0.063 | 0.047 | 1 | 5 | Fuom | 0.016 | Young |
| 0.276874 | 0.280677 | 0.086 | 0.067 | 1 | 5 | Inpp5a | 0.019 | Young |
| 0.277397 | 0.30174 | 0.124 | 0.103 | 1 | 5 | Zbtb22 | 0.021 | Young |
| 0.277561 | 0.382941 | 0.084 | 0.066 | 1 | 5 | Ppp1r3b | 0.018 | Young |
| 0.277955 | 0.470069 | 0.053 | 0.039 | 1 | 5 | Casp7 | 0.014 | Young |
| 0.277971 | 0.735839 | 0.051 | 0.037 | 1 | 5 | Rbm48 | 0.014 | Young |
| 0.278082 | 0.584076 | 0.043 | 0.03 | 1 | 5 | Mettl8 | 0.013 | Young |
| 0.27831 | 0.261224 | 0.178 | 0.15 | 1 | 5 | Ech1 | 0.028 | Young |
| 0.278451 | 0.388057 | 0.16 | 0.135 | 1 | 5 | Armc1 | 0.025 | Young |
| 0.278779 | 0.496372 | 0.122 | 0.101 | 1 | 5 | Rgs1 | 0.021 | Young |
| 0.278803 | 0.100681 | 0.812 | 0.805 | 1 | 5 | Slc25a3 | 0.007 | Young |
| 0.27889 | 0.344945 | 0.14 | 0.116 | 1 | 5 | Ankra2 | 0.024 | Young |
| 0.278912 | 0.122987 | 0.066 | 0.049 | 1 | 5 | Hist2h2be | 0.017 | Young |
| 0.279004 | 0.322996 | 0.206 | 0.178 | 1 | 5 | Gab1 | 0.028 | Young |
| 0.279258 | 0.232535 | 0.409 | 0.377 | 1 | 5 | Rragc | 0.032 | Young |
| 0.279482 | 0.243135 | 0.404 | 0.37 | 1 | 5 | Csk | 0.034 | Young |
| 0.280042 | 0.377103 | 0.046 | 0.032 | 1 | 5 | Gm10138 | 0.014 | Young |
| 0.280111 | 0.436579 | 0.147 | 0.126 | 1 | 5 | 4833420G1 | 0.021 | Young |
| 0.280193 | 0.962502 | 0.02 | 0.012 | 1 | 5 | Gm12195 | 0.008 | Young |
| 0.280198 | 0.134425 | 0.287 | 0.254 | 1 | 5 | Nabp1 | 0.033 | Young |
| 0.281359 | 0.202393 | 0.162 | 0.138 | 1 | 5 | Manba | 0.024 | Young |
| 0.281474 | 0.014991 | 0.883 | 0.906 | 1 | 5 | Ier5 | -0.023 | Young |
| 0.281801 | 0.295885 | 0.216 | 0.187 | 1 | 5 | Zbtb38 | 0.029 | Young |
| 0.28206 | 0.66265 | 0.033 | 0.022 | 1 | 5 | Acot7 | 0.011 | Young |
| 0.282109 | 0.318997 | 0.185 | 0.16 | 1 | 5 | Cops6 | 0.025 | Young |
| 0.282112 | 0.690864 | 0.02 | 0.012 | 1 | 5 | Cenpe | 0.008 | Young |
| 0.282117 | 0.355145 | 0.132 | 0.111 | 1 | 5 | Nol11 | 0.021 | Young |
| 0.282593 | 0.991907 | 0.02 | 0.012 | 1 | 5 | Gm38394 | 0.008 | Young |
| 0.28282 | 0.457901 | 0.109 | 0.089 | 1 | 5 | Tex10 | 0.02 | Young |
| 0.283216 | 0.31873 | 0.165 | 0.14 | 1 | 5 | Mxd1 | 0.025 | Young |

| | | | | | | | | |
|---|---|---|---|---|---|---|---|---|
| 0.283783 | 0.50472 | 0.051 | 0.037 | 1 | 5 | Nup35 | 0.014 | Young |
| 0.2839 | 0.538126 | 0.033 | 0.022 | 1 | 5 | Stk39 | 0.011 | Young |
| 0.285006 | 0.337863 | 0.147 | 0.125 | 1 | 5 | Glt8d1 | 0.022 | Young |
| 0.285008 | 0.562181 | 0.02 | 0.012 | 1 | 5 | Synj2 | 0.008 | Young |
| 0.285008 | 0.499099 | 0.02 | 0.012 | 1 | 5 | 5830432E0 | 0.008 | Young |
| 0.285198 | 0.281713 | 0.358 | 0.338 | 1 | 5 | Pum1 | 0.02 | Young |
| 0.285749 | 0.683492 | 0.033 | 0.022 | 1 | 5 | Mx1 | 0.011 | Young |
| 0.285966 | 0.348307 | 0.168 | 0.143 | 1 | 5 | Hypk | 0.025 | Young |
| 0.28612 | 0.458914 | 0.033 | 0.022 | 1 | 5 | Mthfd1l | 0.011 | Young |
| 0.286251 | 0.168244 | 0.046 | 0.032 | 1 | 5 | Fam129b | 0.014 | Young |
| 0.28638 | 0.252986 | 0.201 | 0.173 | 1 | 5 | Tyk2 | 0.028 | Young |
| 0.286388 | 0.154611 | 0.528 | 0.508 | 1 | 5 | Mxd4 | 0.02 | Young |
| 0.286462 | 0.351294 | 0.086 | 0.067 | 1 | 5 | Tspan5 | 0.019 | Young |
| 0.286548 | 0.258227 | 0.393 | 0.357 | 1 | 5 | Ppp3r1 | 0.036 | Young |
| 0.286654 | 0.368906 | 0.135 | 0.113 | 1 | 5 | Abcc3 | 0.022 | Young |
| 0.286867 | 0.302498 | 0.447 | 0.418 | 1 | 5 | Rgs2 | 0.029 | Young |
| 0.287431 | 0.330438 | 0.221 | 0.194 | 1 | 5 | Keap1 | 0.027 | Young |
| 0.287852 | 0.431832 | 0.175 | 0.153 | 1 | 5 | Nuak1 | 0.022 | Young |
| 0.288088 | 0.455487 | 0.122 | 0.103 | 1 | 5 | Polr3c | 0.019 | Young |
| 0.288122 | 0.271371 | 0.327 | 0.295 | 1 | 5 | Frmd6 | 0.032 | Young |
| 0.288352 | 0.490253 | 0.033 | 0.022 | 1 | 5 | Socs1 | 0.011 | Young |
| 0.288371 | 0.232487 | 0.449 | 0.423 | 1 | 5 | Fnip1 | 0.026 | Young |
| 0.288961 | 0.592607 | 0.081 | 0.064 | 1 | 5 | Asb3 | 0.017 | Young |
| 0.289526 | 0.331627 | 0.061 | 0.045 | 1 | 5 | Hmgxb4 | 0.016 | Young |
| 0.290441 | 0.217238 | 0.234 | 0.205 | 1 | 5 | Rbm22 | 0.029 | Young |
| 0.290595 | 0.52322 | 0.033 | 0.022 | 1 | 5 | Pecr | 0.011 | Young |
| 0.290873 | 0.559037 | 0.112 | 0.093 | 1 | 5 | Mrpl15 | 0.019 | Young |
| 0.291352 | 0.262981 | 0.02 | 0.012 | 1 | 5 | Myl9 | 0.008 | Young |
| 0.291371 | 0.432058 | 0.074 | 0.057 | 1 | 5 | Wdr75 | 0.017 | Young |
| 0.291695 | 0.444716 | 0.061 | 0.045 | 1 | 5 | Maml3 | 0.016 | Young |
| 0.291871 | 0.173402 | 0.721 | 0.71 | 1 | 5 | Serp1 | 0.011 | Young |
| 0.291882 | 0.167226 | 0.157 | 0.131 | 1 | 5 | Uchl5 | 0.026 | Young |
| 0.29201 | 0.419918 | 0.102 | 0.081 | 1 | 5 | Svbp | 0.021 | Young |
| 0.292143 | 0.208406 | 0.345 | 0.316 | 1 | 5 | Lsm4 | 0.029 | Young |
| 0.292655 | 0.206085 | 0.221 | 0.189 | 1 | 5 | Rnf20 | 0.032 | Young |
| 0.292707 | 0.704619 | 0.079 | 0.062 | 1 | 5 | Ddx19b | 0.017 | Young |
| 0.292897 | 0.442954 | 0.19 | 0.17 | 1 | 5 | Rhoh | 0.02 | Young |
| 0.293174 | 0.226013 | 0.165 | 0.138 | 1 | 5 | Apmap | 0.027 | Young |
| 0.293353 | 0.246952 | 0.129 | 0.108 | 1 | 5 | Faf1 | 0.021 | Young |
| 0.293641 | 0.200564 | 0.284 | 0.247 | 1 | 5 | Eif2a | 0.037 | Young |
| 0.293695 | 0.124916 | 0.589 | 0.566 | 1 | 5 | Mpp1 | 0.023 | Young |
| 0.2937 | 0.285999 | 0.135 | 0.113 | 1 | 5 | Vps37b | 0.022 | Young |
| 0.293731 | 0.098522 | 0.665 | 0.663 | 1 | 5 | Sod1 | 0.002 | Young |
| 0.294393 | 0.161963 | 0.541 | 0.524 | 1 | 5 | Tnrc18 | 0.017 | Young |
| 0.294818 | 0.221662 | 0.015 | 0.025 | 1 | 5 | Rad18 | -0.01 | Young |

| | | | | | | | | |
|---|---|---|---|---|---|---|---|---|
| 0.294818 | 0.034335 | 0.015 | 0.025 | 1 | 5 | Tspan7 | -0.01 | Young |
| 0.294947 | 0.164832 | 0.114 | 0.093 | 1 | 5 | Chchd5 | 0.021 | Young |
| 0.295242 | 0.201549 | 0.061 | 0.045 | 1 | 5 | Ccar2 | 0.016 | Young |
| 0.295357 | 0.434716 | 0.107 | 0.088 | 1 | 5 | Gtf3a | 0.019 | Young |
| 0.295708 | 0.299752 | 0.129 | 0.108 | 1 | 5 | Pik3c3 | 0.021 | Young |
| 0.295766 | 0.463891 | 0.058 | 0.044 | 1 | 5 | Fry | 0.014 | Young |
| 0.295853 | 0.238266 | 0.447 | 0.429 | 1 | 5 | Ncoa4 | 0.018 | Young |
| 0.296084 | 0.274404 | 0.17 | 0.145 | 1 | 5 | Wdr13 | 0.025 | Young |
| 0.296336 | 0.230782 | 0.266 | 0.236 | 1 | 5 | Atp6v1d | 0.03 | Young |
| 0.296487 | 0.291557 | 0.091 | 0.072 | 1 | 5 | Heatr3 | 0.019 | Young |
| 0.296588 | 0.884875 | 0.03 | 0.02 | 1 | 5 | Ehbp1 | 0.01 | Young |
| 0.296621 | 0.120928 | 0.607 | 0.571 | 1 | 5 | Rab10 | 0.036 | Young |
| 0.296723 | 0.229534 | 0.424 | 0.391 | 1 | 5 | Nfe2l1 | 0.033 | Young |
| 0.297258 | 0.365254 | 0.069 | 0.052 | 1 | 5 | Usp28 | 0.017 | Young |
| 0.297267 | 0.315816 | 0.201 | 0.177 | 1 | 5 | Fut11 | 0.024 | Young |
| 0.297447 | 0.129651 | 0.216 | 0.185 | 1 | 5 | Trim33 | 0.031 | Young |
| 0.29759 | 0.195485 | 0.107 | 0.086 | 1 | 5 | Gadd45gip | 0.021 | Young |
| 0.298173 | 0.172635 | 0.102 | 0.081 | 1 | 5 | Zfp329 | 0.021 | Young |
| 0.298289 | 0.557811 | 0.058 | 0.044 | 1 | 5 | Pdk3 | 0.014 | Young |
| 0.29876 | 0.276167 | 0.277 | 0.251 | 1 | 5 | R3hdm1 | 0.026 | Young |
| 0.298884 | 0.592123 | 0.018 | 0.01 | 1 | 5 | Hhat | 0.008 | Young |
| 0.299143 | 0.220358 | 0.188 | 0.16 | 1 | 5 | Grsf1 | 0.028 | Young |
| 0.299155 | 0.270896 | 0.221 | 0.194 | 1 | 5 | Cep120 | 0.027 | Young |
| 0.299507 | 0.181294 | 0.147 | 0.123 | 1 | 5 | Galnt7 | 0.024 | Young |
| 0.299958 | 0.633987 | 0.018 | 0.01 | 1 | 5 | Mettl18 | 0.008 | Young |
| 0.300119 | 0.095192 | 0.604 | 0.593 | 1 | 5 | Abi1 | 0.011 | Young |
| 0.300143 | 0.274793 | 0.076 | 0.059 | 1 | 5 | Borcs7 | 0.017 | Young |
| 0.300235 | 0.168035 | 0.185 | 0.16 | 1 | 5 | Cmas | 0.025 | Young |
| 0.300285 | 0.42158 | 0.066 | 0.051 | 1 | 5 | 3110001I2 | 0.015 | Young |
| 0.300497 | 0.711152 | 0.018 | 0.01 | 1 | 5 | Zfp976 | 0.008 | Young |
| 0.300497 | 0.668799 | 0.018 | 0.01 | 1 | 5 | Gm39469 | 0.008 | Young |
| 0.300681 | 0.271812 | 0.302 | 0.274 | 1 | 5 | Uggt1 | 0.028 | Young |
| 0.300743 | 0.076942 | 0.345 | 0.303 | 1 | 5 | Pfdn2 | 0.042 | Young |
| 0.300821 | 0.064642 | 0.363 | 0.424 | 1 | 5 | Lamtor2 | -0.061 | Young |
| 0.300908 | 0.182333 | 0.444 | 0.414 | 1 | 5 | Tnpo1 | 0.03 | Young |
| 0.301036 | 0.609226 | 0.018 | 0.01 | 1 | 5 | Lrrk2 | 0.008 | Young |
| 0.301099 | 0.402364 | 0.051 | 0.037 | 1 | 5 | Wdr6 | 0.014 | Young |
| 0.301505 | 0.332456 | 0.129 | 0.108 | 1 | 5 | Timm44 | 0.021 | Young |
| 0.301575 | 0.501832 | 0.018 | 0.01 | 1 | 5 | Bicc1 | 0.008 | Young |
| 0.301625 | 0.19013 | 0.216 | 0.187 | 1 | 5 | Sfxn3 | 0.029 | Young |
| 0.301734 | 0.427988 | 0.132 | 0.111 | 1 | 5 | Mcee | 0.021 | Young |
| 0.302115 | 0.534128 | 0.018 | 0.01 | 1 | 5 | Spry2 | 0.008 | Young |
| 0.302388 | 0.420757 | 0.071 | 0.056 | 1 | 5 | Nbn | 0.015 | Young |
| 0.303034 | 0.152321 | 0.586 | 0.586 | 1 | 5 | Khdrbs1 | 0 | Young |
| 0.303536 | 0.313397 | 0.132 | 0.111 | 1 | 5 | Tlr13 | 0.021 | Young |

| | | | | | | | | |
|---|---|---|---|---|---|---|---|---|
| 0.303578 | 0.146387 | 0.396 | 0.372 | 1 | 5 | Nenf | 0.024 | Young |
| 0.30374 | 0.469232 | 0.018 | 0.01 | 1 | 5 | Gm14305 | 0.008 | Young |
| 0.303779 | 0.507146 | 0.091 | 0.074 | 1 | 5 | Ttc33 | 0.017 | Young |
| 0.303951 | 0.845486 | 0.038 | 0.027 | 1 | 5 | Zfp825 | 0.011 | Young |
| 0.304282 | 0.348084 | 0.018 | 0.01 | 1 | 5 | Zfp7 | 0.008 | Young |
| 0.304712 | 0.283994 | 0.096 | 0.077 | 1 | 5 | Tubb2b | 0.019 | Young |
| 0.30475 | 0.397934 | 0.241 | 0.221 | 1 | 5 | Galk2 | 0.02 | Young |
| 0.305021 | 0.012221 | 0.246 | 0.288 | 1 | 5 | Plekhg3 | -0.042 | Young |
| 0.305071 | 0.44415 | 0.086 | 0.069 | 1 | 5 | Epn2 | 0.017 | Young |
| 0.30536 | 0.29426 | 0.069 | 0.052 | 1 | 5 | Zbtb45 | 0.017 | Young |
| 0.305366 | 0.610948 | 0.03 | 0.02 | 1 | 5 | Ak1 | 0.01 | Young |
| 0.30537 | 0.359285 | 0.018 | 0.01 | 1 | 5 | Alkbh4 | 0.008 | Young |
| 0.305374 | 0.5151 | 0.058 | 0.044 | 1 | 5 | Map2k3os | 0.014 | Young |
| 0.30544 | 0.234242 | 0.487 | 0.468 | 1 | 5 | Kdelr1 | 0.019 | Young |
| 0.305566 | 0.322808 | 0.081 | 0.064 | 1 | 5 | Ppa1 | 0.017 | Young |
| 0.305931 | 0.401143 | 0.175 | 0.153 | 1 | 5 | Sars | 0.022 | Young |
| 0.306172 | 0.593577 | 0.03 | 0.02 | 1 | 5 | Abca6 | 0.01 | Young |
| 0.306243 | 0.471928 | 0.071 | 0.056 | 1 | 5 | Sssca1 | 0.015 | Young |
| 0.306317 | 0.474534 | 0.041 | 0.029 | 1 | 5 | Tacc2 | 0.012 | Young |
| 0.30646 | 0.261756 | 0.018 | 0.01 | 1 | 5 | Zfp605 | 0.008 | Young |
| 0.306558 | 0.337547 | 0.14 | 0.118 | 1 | 5 | Zmat5 | 0.022 | Young |
| 0.306564 | 0.446547 | 0.216 | 0.195 | 1 | 5 | Tmem106c | 0.021 | Young |
| 0.306597 | 0.221471 | 0.365 | 0.345 | 1 | 5 | Zfp638 | 0.02 | Young |
| 0.307006 | 0.151343 | 0.018 | 0.01 | 1 | 5 | Aff3 | 0.008 | Young |
| 0.307529 | 0.536662 | 0.084 | 0.067 | 1 | 5 | Bbc3 | 0.017 | Young |
| 0.307538 | 0.335086 | 0.051 | 0.037 | 1 | 5 | Cdc42bpg | 0.014 | Young |
| 0.307712 | 0.320634 | 0.119 | 0.099 | 1 | 5 | Casp6 | 0.02 | Young |
| 0.307828 | 0.199869 | 0.096 | 0.077 | 1 | 5 | Ap3s2 | 0.019 | Young |
| 0.307908 | 0.12114 | 0.503 | 0.476 | 1 | 5 | Capza1 | 0.027 | Young |
| 0.307939 | 0.091434 | 0.096 | 0.077 | 1 | 5 | Ppm1m | 0.019 | Young |
| 0.308049 | 0.437447 | 0.041 | 0.029 | 1 | 5 | Brca2 | 0.012 | Young |
| 0.30824 | 0.200285 | 0.241 | 0.214 | 1 | 5 | Pfkfb4 | 0.027 | Young |
| 0.308577 | 0.451468 | 0.099 | 0.081 | 1 | 5 | Aasdhppt | 0.018 | Young |
| 0.308841 | 0.263275 | 0.246 | 0.221 | 1 | 5 | Stxbp2 | 0.025 | Young |
| 0.309006 | 0.564181 | 0.03 | 0.02 | 1 | 5 | Tlr12 | 0.01 | Young |
| 0.309076 | 0.272193 | 0.444 | 0.428 | 1 | 5 | Numa1 | 0.016 | Young |
| 0.309356 | 0.797571 | 0.046 | 0.034 | 1 | 5 | Dbr1 | 0.012 | Young |
| 0.310007 | 0.350174 | 0.086 | 0.069 | 1 | 5 | Txnrd2 | 0.017 | Young |
| 0.310097 | 0.090896 | 0.348 | 0.389 | 1 | 5 | Akt1 | -0.041 | Young |
| 0.310248 | 0.349488 | 0.19 | 0.168 | 1 | 5 | Tmem135 | 0.022 | Young |
| 0.310362 | 0.308261 | 0.086 | 0.069 | 1 | 5 | Slc9a1 | 0.017 | Young |
| 0.311333 | 0.505487 | 0.096 | 0.079 | 1 | 5 | Ankrd49 | 0.017 | Young |
| 0.311516 | 0.271958 | 0.269 | 0.241 | 1 | 5 | Sde2 | 0.028 | Young |
| 0.311569 | 0.046981 | 0.789 | 0.776 | 1 | 5 | Eif3a | 0.013 | Young |
| 0.311727 | 0.258905 | 0.228 | 0.204 | 1 | 5 | Gtf2i | 0.024 | Young |

| | | | | | | | | |
|---|---|---|---|---|---|---|---|---|
| 0.311919 | 0.333049 | 0.112 | 0.091 | 1 | 5 | Rnf185 | 0.021 | Young |
| 0.312038 | 0.49151 | 0.168 | 0.152 | 1 | 5 | Wdr36 | 0.016 | Young |
| 0.31209 | 0.104705 | 0.411 | 0.374 | 1 | 5 | Ap1s2 | 0.037 | Young |
| 0.313026 | 0.183533 | 0.574 | 0.571 | 1 | 5 | Hnrnpc | 0.003 | Young |
| 0.313054 | 0.347786 | 0.287 | 0.269 | 1 | 5 | Bcl2l1 | 0.018 | Young |
| 0.313524 | 0.210356 | 0.371 | 0.352 | 1 | 5 | Dynlrb1 | 0.019 | Young |
| 0.313539 | 0.615355 | 0.056 | 0.042 | 1 | 5 | Zfp954 | 0.014 | Young |
| 0.313681 | 0.405186 | 0.147 | 0.126 | 1 | 5 | Nedd9 | 0.021 | Young |
| 0.314101 | 0.765765 | 0.015 | 0.008 | 1 | 5 | B430306N( | 0.007 | Young |
| 0.314157 | 0.24868 | 0.259 | 0.234 | 1 | 5 | Blmh | 0.025 | Young |
| 0.314322 | 0.245067 | 0.208 | 0.182 | 1 | 5 | Fam20b | 0.026 | Young |
| 0.314351 | 0.815464 | 0.038 | 0.027 | 1 | 5 | Zfp933 | 0.011 | Young |
| 0.314449 | 0.233692 | 0.122 | 0.101 | 1 | 5 | 2810004N2 | 0.021 | Young |
| 0.314893 | 0.398321 | 0.221 | 0.2 | 1 | 5 | Arhgap1 | 0.021 | Young |
| 0.31524 | 0.296575 | 0.066 | 0.051 | 1 | 5 | Gmeb2 | 0.015 | Young |
| 0.315307 | 0.849324 | 0.015 | 0.008 | 1 | 5 | Ccl22 | 0.007 | Young |
| 0.315307 | 0.747174 | 0.015 | 0.008 | 1 | 5 | Grasp | 0.007 | Young |
| 0.315395 | 0.207441 | 0.256 | 0.229 | 1 | 5 | Kdm1a | 0.027 | Young |
| 0.31542 | 0.208316 | 0.132 | 0.109 | 1 | 5 | Med11 | 0.023 | Young |
| 0.315911 | 1.125861 | 0.015 | 0.008 | 1 | 5 | Pkn3 | 0.007 | Young |
| 0.315911 | 0.79583 | 0.015 | 0.008 | 1 | 5 | 1700006J1 | 0.007 | Young |
| 0.315911 | 0.658894 | 0.015 | 0.008 | 1 | 5 | Gm17089 | 0.007 | Young |
| 0.315932 | 0.380732 | 0.048 | 0.035 | 1 | 5 | Sag | 0.013 | Young |
| 0.316623 | 0.566069 | 0.076 | 0.061 | 1 | 5 | Terf2ip | 0.015 | Young |
| 0.317122 | 1.008667 | 0.015 | 0.008 | 1 | 5 | Paqr8 | 0.007 | Young |
| 0.317122 | 0.896519 | 0.015 | 0.008 | 1 | 5 | Adcy6 | 0.007 | Young |
| 0.317122 | 0.731441 | 0.015 | 0.008 | 1 | 5 | Car9 | 0.007 | Young |
| 0.317122 | 0.6828 | 0.015 | 0.008 | 1 | 5 | Gm45733 | 0.007 | Young |
| 0.317205 | 0.17509 | 0.152 | 0.128 | 1 | 5 | Uxs1 | 0.024 | Young |
| 0.317543 | 0.339922 | 0.188 | 0.167 | 1 | 5 | Cep68 | 0.021 | Young |
| 0.317915 | 0.271111 | 0.335 | 0.311 | 1 | 5 | Dennd6a | 0.024 | Young |
| 0.317939 | 0.334875 | 0.183 | 0.162 | 1 | 5 | Hibadh | 0.021 | Young |
| 0.317961 | 0.834483 | 0.028 | 0.019 | 1 | 5 | Synpo | 0.009 | Young |
| 0.318162 | 0.236765 | 0.272 | 0.242 | 1 | 5 | Slf2 | 0.03 | Young |
| 0.318381 | 0.346314 | 0.109 | 0.091 | 1 | 5 | Lcmt1 | 0.018 | Young |
| 0.318825 | 0.69922 | 0.028 | 0.019 | 1 | 5 | Tctn3 | 0.009 | Young |
| 0.318887 | 0.264468 | 0.147 | 0.125 | 1 | 5 | Fbxo28 | 0.022 | Young |
| 0.318905 | 0.004481 | 0.279 | 0.311 | 1 | 5 | Dck | -0.032 | Young |
| 0.318943 | 1.062422 | 0.015 | 0.008 | 1 | 5 | Gm2814 | 0.007 | Young |
| 0.318943 | 0.649142 | 0.015 | 0.008 | 1 | 5 | Lockd | 0.007 | Young |
| 0.318976 | 0.136129 | 0.563 | 0.527 | 1 | 5 | Rhog | 0.036 | Young |
| 0.319267 | 0.132771 | 0.03 | 0.02 | 1 | 5 | Dock6 | 0.01 | Young |
| 0.319542 | 0.176873 | 0.185 | 0.158 | 1 | 5 | Abhd16a | 0.027 | Young |
| 0.319552 | 0.499736 | 0.015 | 0.008 | 1 | 5 | Cenps | 0.007 | Young |
| 0.31958 | 0.091102 | 0.058 | 0.044 | 1 | 5 | Cd9 | 0.014 | Young |

| | | | | | | | | |
|---|---|---|---|---|---|---|---|---|
| 0.32017 | 0.249026 | 0.251 | 0.222 | 1 | 5 | Parp1 | 0.029 | Young |
| 0.320191 | 0.219898 | 0.497 | 0.481 | 1 | 5 | Dnajb14 | 0.016 | Young |
| 0.320199 | 0.1302 | 0.35 | 0.316 | 1 | 5 | Psmb10 | 0.034 | Young |
| 0.320415 | 0.264738 | 0.36 | 0.337 | 1 | 5 | Nol7 | 0.023 | Young |
| 0.320604 | 0.415378 | 0.127 | 0.106 | 1 | 5 | Caml | 0.021 | Young |
| 0.320705 | 0.113044 | 0.14 | 0.116 | 1 | 5 | Fbxo22 | 0.024 | Young |
| 0.320772 | 0.251979 | 0.015 | 0.008 | 1 | 5 | Cdk1 | 0.007 | Young |
| 0.320782 | 0.262955 | 0.048 | 0.035 | 1 | 5 | Zfp874a | 0.013 | Young |
| 0.320804 | 0.091108 | 0.571 | 0.547 | 1 | 5 | Ptpn6 | 0.024 | Young |
| 0.321383 | 0.386426 | 0.015 | 0.008 | 1 | 5 | 9130011E1 | 0.007 | Young |
| 0.321759 | 0.138628 | 0.574 | 0.554 | 1 | 5 | Usf2 | 0.02 | Young |
| 0.321784 | 0.58574 | 0.147 | 0.13 | 1 | 5 | Brip1os | 0.017 | Young |
| 0.321856 | 0.258033 | 0.066 | 0.051 | 1 | 5 | Stambpl1 | 0.015 | Young |
| 0.321944 | 0.641136 | 0.069 | 0.054 | 1 | 5 | Dus1l | 0.015 | Young |
| 0.321994 | 0.283021 | 0.015 | 0.008 | 1 | 5 | Rad51c | 0.007 | Young |
| 0.322163 | 0.184043 | 0.429 | 0.409 | 1 | 5 | Hp1bp3 | 0.02 | Young |
| 0.322317 | 0.244546 | 0.16 | 0.136 | 1 | 5 | Bccip | 0.024 | Young |
| 0.322442 | 0.188545 | 0.629 | 0.655 | 1 | 5 | Spred1 | -0.026 | Young |
| 0.322602 | 0.379115 | 0.173 | 0.15 | 1 | 5 | Cenpa | 0.023 | Young |
| 0.322603 | 0.24712 | 0.246 | 0.221 | 1 | 5 | Paip1 | 0.025 | Young |
| 0.322607 | 0.27202 | 0.015 | 0.008 | 1 | 5 | Lgalsl | 0.007 | Young |
| 0.322733 | 0.832895 | 0.028 | 0.019 | 1 | 5 | Tfb1m | 0.009 | Young |
| 0.322858 | 0.241682 | 0.041 | 0.029 | 1 | 5 | Fahd2a | 0.012 | Young |
| 0.322865 | 0.256011 | 0.261 | 0.237 | 1 | 5 | Pi4k2a | 0.024 | Young |
| 0.322978 | 0.132239 | 0.838 | 0.865 | 1 | 5 | Rpl35 | -0.027 | Young |
| 0.32322 | 0.422637 | 0.015 | 0.008 | 1 | 5 | Garnl3 | 0.007 | Young |
| 0.32322 | 0.192292 | 0.015 | 0.008 | 1 | 5 | Zscan22 | 0.007 | Young |
| 0.32353 | 0.184391 | 0.424 | 0.409 | 1 | 5 | Tsn | 0.015 | Young |
| 0.324045 | 0.151813 | 0.487 | 0.453 | 1 | 5 | Ehbp1l1 | 0.034 | Young |
| 0.324202 | 0.232177 | 0.363 | 0.34 | 1 | 5 | Ankrd13a | 0.023 | Young |
| 0.324266 | 0.241951 | 0.246 | 0.221 | 1 | 5 | Frg1 | 0.025 | Young |
| 0.324291 | 0.431783 | 0.127 | 0.108 | 1 | 5 | Slc17a5 | 0.019 | Young |
| 0.324325 | 0.393263 | 0.246 | 0.227 | 1 | 5 | Gm10073 | 0.019 | Young |
| 0.324413 | 0.894668 | 0.053 | 0.04 | 1 | 5 | Mrrf | 0.013 | Young |
| 0.324488 | 0.562377 | 0.084 | 0.067 | 1 | 5 | Mon1b | 0.017 | Young |
| 0.324649 | 0.34396 | 0.145 | 0.125 | 1 | 5 | Ddx19a | 0.02 | Young |
| 0.324917 | 0.308837 | 0.137 | 0.116 | 1 | 5 | Rad50 | 0.021 | Young |
| 0.32506 | 0.608559 | 0.046 | 0.034 | 1 | 5 | Asah2 | 0.012 | Young |
| 0.325354 | 0.538728 | 0.038 | 0.027 | 1 | 5 | Rdh13 | 0.011 | Young |
| 0.325497 | 0.341511 | 0.063 | 0.049 | 1 | 5 | Bid | 0.014 | Young |
| 0.325532 | 0.106571 | 0.698 | 0.714 | 1 | 5 | Arf1 | -0.016 | Young |
| 0.32609 | 0.442944 | 0.119 | 0.101 | 1 | 5 | Tceanc2 | 0.018 | Young |
| 0.32722 | 0.076633 | 0.612 | 0.604 | 1 | 5 | Mapre1 | 0.008 | Young |
| 0.327564 | 0.040038 | 0.49 | 0.534 | 1 | 5 | Washc4 | -0.044 | Young |
| 0.327795 | 0.029879 | 0.558 | 0.591 | 1 | 5 | Wasf2 | -0.033 | Young |

| | | | | | | | | |
|---|---|---|---|---|---|---|---|---|
| 0.327832 | 0.372012 | 0.094 | 0.076 | 1 | 5 | Alkbh6 | 0.018 | Young |
| 0.328006 | 0.121497 | 0.685 | 0.707 | 1 | 5 | Sf3b1 | -0.022 | Young |
| 0.328044 | 0.223181 | 0.051 | 0.037 | 1 | 5 | Nradd | 0.014 | Young |
| 0.328064 | 0.203791 | 0.188 | 0.163 | 1 | 5 | Plxna4 | 0.025 | Young |
| 0.328146 | 0.305568 | 0.343 | 0.318 | 1 | 5 | Cbx3 | 0.025 | Young |
| 0.32831 | 0.438956 | 0.086 | 0.069 | 1 | 5 | Homer3 | 0.017 | Young |
| 0.328432 | 0.496751 | 0.028 | 0.019 | 1 | 5 | Tacc3 | 0.009 | Young |
| 0.328703 | 0.459382 | 0.038 | 0.027 | 1 | 5 | Ppargc1b | 0.011 | Young |
| 0.329237 | 0.428385 | 0.071 | 0.056 | 1 | 5 | Entpd6 | 0.015 | Young |
| 0.3293 | 0.292135 | 0.277 | 0.254 | 1 | 5 | Sppl3 | 0.023 | Young |
| 0.32945 | 0.635408 | 0.038 | 0.027 | 1 | 5 | Plekhh3 | 0.011 | Young |
| 0.329571 | 0.246799 | 0.109 | 0.089 | 1 | 5 | Cog8 | 0.02 | Young |
| 0.329605 | 0.451889 | 0.107 | 0.089 | 1 | 5 | Chid1 | 0.018 | Young |
| 0.329648 | 0.190244 | 0.48 | 0.466 | 1 | 5 | Zcchc11 | 0.014 | Young |
| 0.329705 | 0.91547 | 0.112 | 0.093 | 1 | 5 | Lyz1 | 0.019 | Young |
| 0.329922 | 0.158809 | 0.198 | 0.172 | 1 | 5 | Chrac1 | 0.026 | Young |
| 0.330168 | 0.938231 | 0.036 | 0.025 | 1 | 5 | 2810006K2 | 0.011 | Young |
| 0.330199 | 0.575358 | 0.038 | 0.027 | 1 | 5 | Colca2 | 0.011 | Young |
| 0.3306 | 0.191904 | 0.47 | 0.451 | 1 | 5 | Adam17 | 0.019 | Young |
| 0.330616 | 0.138354 | 0.322 | 0.295 | 1 | 5 | Cdk11b | 0.027 | Young |
| 0.330863 | 0.283196 | 0.071 | 0.056 | 1 | 5 | 1110032A0 | 0.015 | Young |
| 0.330948 | 0.59672 | 0.038 | 0.027 | 1 | 5 | Mettl2 | 0.011 | Young |
| 0.331085 | 0.405565 | 0.028 | 0.019 | 1 | 5 | Rbfox2 | 0.009 | Young |
| 0.331168 | 0.250758 | 0.289 | 0.269 | 1 | 5 | Uhrf1bp1l | 0.02 | Young |
| 0.331504 | 0.256907 | 0.223 | 0.197 | 1 | 5 | Rnps1 | 0.026 | Young |
| 0.331574 | 0.29076 | 0.259 | 0.237 | 1 | 5 | Nbr1 | 0.022 | Young |
| 0.33173 | 1.561866 | 0.013 | 0.007 | 1 | 5 | Clec2g | 0.006 | Young |
| 0.33173 | 0.994815 | 0.013 | 0.007 | 1 | 5 | 4930579G2 | 0.006 | Young |
| 0.332047 | 0.173133 | 0.454 | 0.443 | 1 | 5 | Cebpg | 0.011 | Young |
| 0.332194 | 0.086336 | 0.165 | 0.194 | 1 | 5 | Pitpnb | -0.029 | Young |
| 0.332419 | 0.973101 | 0.013 | 0.007 | 1 | 5 | 2700012I2 | 0.006 | Young |
| 0.332419 | 0.920095 | 0.013 | 0.007 | 1 | 5 | Gm27003 | 0.006 | Young |
| 0.332419 | 0.78664 | 0.013 | 0.007 | 1 | 5 | Parva | 0.006 | Young |
| 0.332419 | 0.72244 | 0.013 | 0.007 | 1 | 5 | Fam78b | 0.006 | Young |
| 0.332419 | 0.683584 | 0.013 | 0.007 | 1 | 5 | Ubxn11 | 0.006 | Young |
| 0.332455 | 0.148278 | 0.548 | 0.525 | 1 | 5 | Cmklr1 | 0.023 | Young |
| 0.332481 | 0.327362 | 0.175 | 0.153 | 1 | 5 | 4833439L1 | 0.022 | Young |
| 0.33311 | 0.809285 | 0.013 | 0.007 | 1 | 5 | Ar | 0.006 | Young |
| 0.33311 | 0.751374 | 0.013 | 0.007 | 1 | 5 | Nadsyn1 | 0.006 | Young |
| 0.33311 | 0.728978 | 0.013 | 0.007 | 1 | 5 | BC048403 | 0.006 | Young |
| 0.333272 | 0.76331 | 0.036 | 0.025 | 1 | 5 | Cep41 | 0.011 | Young |
| 0.333359 | 0.486807 | 0.046 | 0.034 | 1 | 5 | Pomt2 | 0.012 | Young |
| 0.333458 | 0.226903 | 0.246 | 0.222 | 1 | 5 | Trappc6b | 0.024 | Young |
| 0.33347 | 0.201597 | 0.302 | 0.281 | 1 | 5 | Fbxl5 | 0.021 | Young |
| 0.3335 | 0.428381 | 0.061 | 0.047 | 1 | 5 | Fahd1 | 0.014 | Young |

| | | | | | | | | |
|---|---|---|---|---|---|---|---|---|
| 0.333802 | 1.009408 | 0.013 | 0.007 | 1 | 5 | Cdkl2 | 0.006 | Young |
| 0.333802 | 0.624255 | 0.013 | 0.007 | 1 | 5 | Cxcl14 | 0.006 | Young |
| 0.33405 | 0.053933 | 0.023 | 0.034 | 1 | 5 | AU041133 | -0.011 | Young |
| 0.334054 | 0.497993 | 0.119 | 0.101 | 1 | 5 | Klhdc2 | 0.018 | Young |
| 0.334109 | 0.300048 | 0.135 | 0.114 | 1 | 5 | Fam199x | 0.021 | Young |
| 0.334197 | 0.339947 | 0.028 | 0.019 | 1 | 5 | Chchd6 | 0.009 | Young |
| 0.334216 | 0.338084 | 0.294 | 0.281 | 1 | 5 | Tmem160 | 0.013 | Young |
| 0.334494 | 0.894695 | 0.013 | 0.007 | 1 | 5 | Nbl1 | 0.006 | Young |
| 0.334726 | 0.323071 | 0.246 | 0.226 | 1 | 5 | Prdm2 | 0.02 | Young |
| 0.334727 | 0.38571 | 0.046 | 0.034 | 1 | 5 | Ift80 | 0.012 | Young |
| 0.334782 | 0.195865 | 0.411 | 0.389 | 1 | 5 | Ghitm | 0.022 | Young |
| 0.335188 | 0.797569 | 0.013 | 0.007 | 1 | 5 | Lep | 0.006 | Young |
| 0.335188 | 0.61925 | 0.013 | 0.007 | 1 | 5 | 6430573P0 | 0.006 | Young |
| 0.335188 | 0.575934 | 0.013 | 0.007 | 1 | 5 | Arv1 | 0.006 | Young |
| 0.335204 | 0.223464 | 0.157 | 0.135 | 1 | 5 | Slc4a2 | 0.022 | Young |
| 0.33533 | 0.281076 | 0.297 | 0.271 | 1 | 5 | Gabarapl1 | 0.026 | Young |
| 0.335459 | 0.118292 | 0.563 | 0.515 | 1 | 5 | Sft2d2 | 0.048 | Young |
| 0.335783 | 0.146242 | 0.208 | 0.18 | 1 | 5 | Mrps17 | 0.028 | Young |
| 0.335872 | 0.055347 | 0.942 | 0.934 | 1 | 5 | Rpl6 | 0.008 | Young |
| 0.335882 | 0.604926 | 0.013 | 0.007 | 1 | 5 | 2810457G0 | 0.006 | Young |
| 0.335882 | 0.557221 | 0.013 | 0.007 | 1 | 5 | Rgs12 | 0.006 | Young |
| 0.335882 | 0.51229 | 0.013 | 0.007 | 1 | 5 | Usp2 | 0.006 | Young |
| 0.336548 | 0.436349 | 0.089 | 0.072 | 1 | 5 | Heatr1 | 0.017 | Young |
| 0.336628 | 0.419931 | 0.053 | 0.04 | 1 | 5 | Klhl15 | 0.013 | Young |
| 0.336737 | 0.244646 | 0.406 | 0.394 | 1 | 5 | Pebp1 | 0.012 | Young |
| 0.337171 | 0.222887 | 0.104 | 0.086 | 1 | 5 | Gpr137 | 0.018 | Young |
| 0.337177 | 0.71863 | 0.036 | 0.025 | 1 | 5 | Gdpd5 | 0.011 | Young |
| 0.337219 | 0.445527 | 0.086 | 0.069 | 1 | 5 | Ddx41 | 0.017 | Young |
| 0.337262 | 0.199293 | 0.053 | 0.04 | 1 | 5 | Zfp143 | 0.013 | Young |
| 0.337273 | 0.589461 | 0.013 | 0.007 | 1 | 5 | Pomc | 0.006 | Young |
| 0.337273 | 0.271662 | 0.013 | 0.007 | 1 | 5 | Rangrf | 0.006 | Young |
| 0.337512 | 0.200876 | 0.216 | 0.189 | 1 | 5 | Clcc1 | 0.027 | Young |
| 0.337678 | 0.614893 | 0.066 | 0.052 | 1 | 5 | AW554918 | 0.014 | Young |
| 0.337971 | 0.519391 | 0.013 | 0.007 | 1 | 5 | Timm23 | 0.006 | Young |
| 0.337971 | 0.268549 | 0.013 | 0.007 | 1 | 5 | Morc4 | 0.006 | Young |
| 0.338043 | 0.333013 | 0.069 | 0.054 | 1 | 5 | Mcu | 0.015 | Young |
| 0.338474 | 0.180875 | 0.345 | 0.338 | 1 | 5 | Smim7 | 0.007 | Young |
| 0.338508 | 0.524085 | 0.046 | 0.034 | 1 | 5 | Dedd2 | 0.012 | Young |
| 0.338669 | 0.321061 | 0.013 | 0.007 | 1 | 5 | Zg16 | 0.006 | Young |
| 0.338669 | 0.295548 | 0.013 | 0.007 | 1 | 5 | Fkbpl | 0.006 | Young |
| 0.338669 | 0.240349 | 0.013 | 0.007 | 1 | 5 | C920006O1 | 0.006 | Young |
| 0.33879 | 0.19481 | 0.241 | 0.219 | 1 | 5 | Uqcrc1 | 0.022 | Young |
| 0.339152 | 0.373962 | 0.183 | 0.163 | 1 | 5 | Mcoln1 | 0.02 | Young |
| 0.339368 | 0.31591 | 0.013 | 0.007 | 1 | 5 | Cdh24 | 0.006 | Young |
| 0.339406 | 0.20158 | 0.145 | 0.121 | 1 | 5 | Kars | 0.024 | Young |

| | | | | | | | | |
|---|---|---|---|---|---|---|---|---|
| 0.339499 | 0.323524 | 0.084 | 0.067 | 1 | 5 | Kctd13 | 0.017 | Young |
| 0.339994 | 0.172492 | 0.048 | 0.035 | 1 | 5 | Ccdc130 | 0.013 | Young |
| 0.340067 | 0.217547 | 0.013 | 0.007 | 1 | 5 | Timp1 | 0.006 | Young |
| 0.340076 | 0.119196 | 0.536 | 0.524 | 1 | 5 | Bmp2k | 0.012 | Young |
| 0.340148 | 0.376404 | 0.135 | 0.114 | 1 | 5 | Tecpr1 | 0.021 | Young |
| 0.340209 | 0.597861 | 0.058 | 0.045 | 1 | 5 | Zfp866 | 0.013 | Young |
| 0.340357 | 0.789872 | 0.025 | 0.017 | 1 | 5 | Calml4 | 0.008 | Young |
| 0.340367 | 0.390113 | 0.414 | 0.402 | 1 | 5 | Il4ra | 0.012 | Young |
| 0.340474 | 0.833126 | 0.043 | 0.032 | 1 | 5 | Chst15 | 0.011 | Young |
| 0.340502 | 0.156888 | 0.312 | 0.286 | 1 | 5 | Riok3 | 0.026 | Young |
| 0.340884 | 0.336266 | 0.157 | 0.138 | 1 | 5 | Arhgap26 | 0.019 | Young |
| 0.341057 | 0.029441 | 0.843 | 0.83 | 1 | 5 | Efhd2 | 0.013 | Young |
| 0.34122 | 0.098092 | 0.454 | 0.414 | 1 | 5 | Prpf40a | 0.04 | Young |
| 0.341777 | 0.961025 | 0.025 | 0.017 | 1 | 5 | Ccdc137 | 0.008 | Young |
| 0.341836 | 0.317948 | 0.071 | 0.056 | 1 | 5 | Rbm38 | 0.015 | Young |
| 0.3424 | 0.539774 | 0.061 | 0.047 | 1 | 5 | Mcm5 | 0.014 | Young |
| 0.342405 | 0.308037 | 0.091 | 0.074 | 1 | 5 | Coa6 | 0.017 | Young |
| 0.342426 | 0.284894 | 0.104 | 0.086 | 1 | 5 | Rwdd4a | 0.018 | Young |
| 0.343359 | 0.372048 | 0.109 | 0.093 | 1 | 5 | Fiz1 | 0.016 | Young |
| 0.34337 | 0.299804 | 0.299 | 0.281 | 1 | 5 | Acadm | 0.018 | Young |
| 0.343467 | 0.139816 | 0.183 | 0.158 | 1 | 5 | Tifab | 0.025 | Young |
| 0.343676 | 0.849409 | 0.025 | 0.017 | 1 | 5 | Bhlhe40 | 0.008 | Young |
| 0.343806 | 0.342949 | 0.069 | 0.054 | 1 | 5 | Cdca4 | 0.015 | Young |
| 0.343883 | 0.61573 | 0.036 | 0.025 | 1 | 5 | Prpf40b | 0.011 | Young |
| 0.344164 | 0.152947 | 0.873 | 0.906 | 1 | 5 | Sem1 | -0.033 | Young |
| 0.344401 | 0.396684 | 0.223 | 0.205 | 1 | 5 | Slc37a2 | 0.018 | Young |
| 0.344678 | 0.685637 | 0.036 | 0.025 | 1 | 5 | Lrrc28 | 0.011 | Young |
| 0.344762 | 0.084103 | 0.114 | 0.094 | 1 | 5 | Stx2 | 0.02 | Young |
| 0.344867 | 0.12622 | 0.533 | 0.503 | 1 | 5 | Rbx1 | 0.03 | Young |
| 0.345008 | 1.501097 | 0.01 | 0.005 | 1 | 5 | Zfp93 | 0.005 | Young |
| 0.345008 | 1.41577 | 0.01 | 0.005 | 1 | 5 | Rab42 | 0.005 | Young |
| 0.345582 | 0.756973 | 0.025 | 0.017 | 1 | 5 | Rbks | 0.008 | Young |
| 0.34581 | 1.5389 | 0.01 | 0.005 | 1 | 5 | Lrg1 | 0.005 | Young |
| 0.34581 | 1.395844 | 0.01 | 0.005 | 1 | 5 | Kcnj10 | 0.005 | Young |
| 0.346071 | 0.192796 | 0.069 | 0.054 | 1 | 5 | Stk38l | 0.015 | Young |
| 0.346312 | 0.217729 | 0.114 | 0.094 | 1 | 5 | Mkl2 | 0.02 | Young |
| 0.346538 | 0.727514 | 0.025 | 0.017 | 1 | 5 | Armc9 | 0.008 | Young |
| 0.346597 | 0.292567 | 0.162 | 0.14 | 1 | 5 | Parl | 0.022 | Young |
| 0.346978 | 0.248755 | 0.114 | 0.094 | 1 | 5 | Cars2 | 0.02 | Young |
| 0.347016 | 0.628574 | 0.025 | 0.017 | 1 | 5 | Impa2 | 0.008 | Young |
| 0.347016 | 0.615702 | 0.025 | 0.017 | 1 | 5 | Srr | 0.008 | Young |
| 0.347416 | 0.999518 | 0.01 | 0.005 | 1 | 5 | C920021L1 | 0.005 | Young |
| 0.347416 | 0.716504 | 0.01 | 0.005 | 1 | 5 | Ncs1 | 0.005 | Young |
| 0.347416 | 0.55487 | 0.01 | 0.005 | 1 | 5 | Clec9a | 0.005 | Young |
| 0.347485 | 0.283864 | 0.135 | 0.114 | 1 | 5 | Crot | 0.021 | Young |

| | | | | | | | | |
|---|---|---|---|---|---|---|---|---|
| 0.347495 | 0.723394 | 0.025 | 0.017 | 1 | 5 | Amigo3 | 0.008 | Young |
| 0.347623 | 0.040387 | 0.112 | 0.091 | 1 | 5 | Tarbp2 | 0.021 | Young |
| 0.347739 | 0.09501 | 0.02 | 0.03 | 1 | 5 | Slfn1 | -0.01 | Young |
| 0.347974 | 0.416955 | 0.025 | 0.017 | 1 | 5 | Arsi | 0.008 | Young |
| 0.348221 | 0.891283 | 0.01 | 0.005 | 1 | 5 | 4-Sep | 0.005 | Young |
| 0.348221 | 0.885343 | 0.01 | 0.005 | 1 | 5 | Arl6 | 0.005 | Young |
| 0.348221 | 0.811284 | 0.01 | 0.005 | 1 | 5 | Fam71a | 0.005 | Young |
| 0.34825 | 0.341413 | 0.096 | 0.079 | 1 | 5 | Samsn1 | 0.017 | Young |
| 0.348268 | 0.414101 | 0.036 | 0.025 | 1 | 5 | Rhobtb2 | 0.011 | Young |
| 0.34863 | 0.377284 | 0.274 | 0.253 | 1 | 5 | Npl | 0.021 | Young |
| 0.348736 | 0.136154 | 0.558 | 0.547 | 1 | 5 | Sf3b2 | 0.011 | Young |
| 0.348934 | 0.491118 | 0.025 | 0.017 | 1 | 5 | Tarbp1 | 0.008 | Young |
| 0.348956 | 0.22845 | 0.046 | 0.034 | 1 | 5 | Miga2 | 0.012 | Young |
| 0.349021 | 0.348093 | 0.061 | 0.047 | 1 | 5 | Kbtbd7 | 0.014 | Young |
| 0.349027 | 0.799581 | 0.01 | 0.005 | 1 | 5 | Palmd | 0.005 | Young |
| 0.349027 | 0.665245 | 0.01 | 0.005 | 1 | 5 | 2610301B2 | 0.005 | Young |
| 0.349027 | 0.516632 | 0.01 | 0.005 | 1 | 5 | Thsd1 | 0.005 | Young |
| 0.349047 | 0.194832 | 0.419 | 0.402 | 1 | 5 | Tnfsf12 | 0.017 | Young |
| 0.349428 | 0.070691 | 0.02 | 0.03 | 1 | 5 | Bpgm | -0.01 | Young |
| 0.349818 | 0.358919 | 0.091 | 0.076 | 1 | 5 | Zcchc8 | 0.015 | Young |
| 0.349834 | 0.773579 | 0.01 | 0.005 | 1 | 5 | Gm15853 | 0.005 | Young |
| 0.349834 | 0.743202 | 0.01 | 0.005 | 1 | 5 | Matn2 | 0.005 | Young |
| 0.349834 | 0.653175 | 0.01 | 0.005 | 1 | 5 | Tmem216 | 0.005 | Young |
| 0.349834 | 0.614777 | 0.01 | 0.005 | 1 | 5 | Gm15545 | 0.005 | Young |
| 0.349834 | 0.525075 | 0.01 | 0.005 | 1 | 5 | Stard6 | 0.005 | Young |
| 0.350034 | 0.253857 | 0.19 | 0.17 | 1 | 5 | Psmd12 | 0.02 | Young |
| 0.350105 | 0.097613 | 0.218 | 0.189 | 1 | 5 | Etv6 | 0.029 | Young |
| 0.350158 | 0.459365 | 0.109 | 0.091 | 1 | 5 | Pja1 | 0.018 | Young |
| 0.350642 | 0.527141 | 0.01 | 0.005 | 1 | 5 | Rax | 0.005 | Young |
| 0.350674 | 0.468189 | 0.036 | 0.025 | 1 | 5 | Arl15 | 0.011 | Young |
| 0.350878 | 0.56848 | 0.043 | 0.032 | 1 | 5 | Zfp46 | 0.011 | Young |
| 0.351009 | 0.989088 | 0.041 | 0.03 | 1 | 5 | Mturn | 0.011 | Young |
| 0.351145 | 0.207563 | 0.096 | 0.079 | 1 | 5 | Rfx5 | 0.017 | Young |
| 0.351347 | 0.117374 | 0.497 | 0.458 | 1 | 5 | Tbca | 0.039 | Young |
| 0.351452 | 0.406081 | 0.01 | 0.005 | 1 | 5 | Gm31597 | 0.005 | Young |
| 0.351452 | 0.488915 | 0.01 | 0.005 | 1 | 5 | Cenpw | 0.005 | Young |
| 0.351452 | 0.476684 | 0.01 | 0.005 | 1 | 5 | Gsg1 | 0.005 | Young |
| 0.351452 | 0.470358 | 0.01 | 0.005 | 1 | 5 | D930028M | 0.005 | Young |
| 0.351452 | 0.432172 | 0.01 | 0.005 | 1 | 5 | Man2c1os | 0.005 | Young |
| 0.351452 | 0.396149 | 0.01 | 0.005 | 1 | 5 | Tmem72 | 0.005 | Young |
| 0.351452 | 0.395343 | 0.01 | 0.005 | 1 | 5 | Cdca7 | 0.005 | Young |
| 0.351452 | 0.362273 | 0.01 | 0.005 | 1 | 5 | Slc19a2 | 0.005 | Young |
| 0.351452 | 0.198808 | 0.01 | 0.005 | 1 | 5 | Cenpi | 0.005 | Young |
| 0.351512 | 0.254144 | 0.201 | 0.177 | 1 | 5 | S100pbp | 0.024 | Young |
| 0.352265 | 0.218691 | 0.482 | 0.471 | 1 | 5 | Snrpe | 0.011 | Young |

| | | | | | | | | |
|---|---|---|---|---|---|---|---|---|
| 0.353003 | 0.280613 | 0.343 | 0.328 | 1 | 5 | Golga4 | 0.015 | Young |
| 0.353607 | 0.183431 | 0.261 | 0.234 | 1 | 5 | Dpy19l1 | 0.027 | Young |
| 0.353888 | 0.042086 | 0.01 | 0.005 | 1 | 5 | Zfp691 | 0.005 | Young |
| 0.354014 | 0.188816 | 0.34 | 0.313 | 1 | 5 | Usp19 | 0.027 | Young |
| 0.354702 | 0.155537 | 0.01 | 0.005 | 1 | 5 | Cystm1 | 0.005 | Young |
| 0.354809 | 0.528253 | 0.081 | 0.066 | 1 | 5 | Trim21 | 0.015 | Young |
| 0.354842 | 0.41051 | 0.122 | 0.104 | 1 | 5 | Ascc1 | 0.018 | Young |
| 0.354979 | 0.14961 | 0.241 | 0.214 | 1 | 5 | Vamp4 | 0.027 | Young |
| 0.355213 | 0.236495 | 0.025 | 0.017 | 1 | 5 | Akap17b | 0.008 | Young |
| 0.355228 | 0.263642 | 0.069 | 0.054 | 1 | 5 | Nsun4 | 0.015 | Young |
| 0.355795 | 0.465923 | 0.104 | 0.088 | 1 | 5 | Rnf19a | 0.016 | Young |
| 0.355925 | 0.349235 | 0.036 | 0.025 | 1 | 5 | Shq1 | 0.011 | Young |
| 0.355969 | 0.421614 | 0.099 | 0.082 | 1 | 5 | Timm17b | 0.017 | Young |
| 0.355974 | 0.204778 | 0.195 | 0.172 | 1 | 5 | Ten1 | 0.023 | Young |
| 0.356163 | 0.457096 | 0.066 | 0.052 | 1 | 5 | Zfp810 | 0.014 | Young |
| 0.356586 | 0.07675 | 0.114 | 0.096 | 1 | 5 | Prkra | 0.018 | Young |
| 0.356672 | 0.30258 | 0.025 | 0.017 | 1 | 5 | Trip13 | 0.008 | Young |
| 0.356849 | 0.378472 | 0.127 | 0.109 | 1 | 5 | Slc4a1ap | 0.018 | Young |
| 0.357082 | 0.262674 | 0.201 | 0.178 | 1 | 5 | Rbm28 | 0.023 | Young |
| 0.357241 | 0.126932 | 0.503 | 0.487 | 1 | 5 | Klhl9 | 0.016 | Young |
| 0.357391 | 0.284271 | 0.089 | 0.072 | 1 | 5 | Kctd5 | 0.017 | Young |
| 0.357463 | 0.29901 | 0.147 | 0.126 | 1 | 5 | Aars | 0.021 | Young |
| 0.357514 | 0.068001 | 0.114 | 0.094 | 1 | 5 | Evi2 | 0.02 | Young |
| 0.357626 | 0.125978 | 0.079 | 0.062 | 1 | 5 | Yod1 | 0.017 | Young |
| 0.357733 | 0.377419 | 0.16 | 0.141 | 1 | 5 | Letm1 | 0.019 | Young |
| 0.357753 | 0.159996 | 0.221 | 0.194 | 1 | 5 | Myo18a | 0.027 | Young |
| 0.357754 | 0.291081 | 0.17 | 0.15 | 1 | 5 | Nipsnap2 | 0.02 | Young |
| 0.357802 | 0.567805 | 0.043 | 0.032 | 1 | 5 | Telo2 | 0.011 | Young |
| 0.357858 | 0.46389 | 0.051 | 0.039 | 1 | 5 | Rabepk | 0.012 | Young |
| 0.357932 | 0.182878 | 0.406 | 0.379 | 1 | 5 | Snx18 | 0.027 | Young |
| 0.358104 | 0.496998 | 0.058 | 0.045 | 1 | 5 | Armt1 | 0.013 | Young |
| 0.358184 | 0.325749 | 0.165 | 0.143 | 1 | 5 | Papss1 | 0.022 | Young |
| 0.35875 | 0.212624 | 0.218 | 0.19 | 1 | 5 | Sf3a3 | 0.028 | Young |
| 0.358772 | 0.166509 | 0.036 | 0.025 | 1 | 5 | Ice2 | 0.011 | Young |
| 0.359045 | 0.431484 | 0.058 | 0.045 | 1 | 5 | 2810402E2 | 0.013 | Young |
| 0.359503 | 0.100766 | 0.723 | 0.731 | 1 | 5 | Pitpna | -0.008 | Young |
| 0.360128 | 0.14615 | 0.371 | 0.348 | 1 | 5 | Mib1 | 0.023 | Young |
| 0.360218 | 0.189866 | 0.368 | 0.34 | 1 | 5 | Ccnt2 | 0.028 | Young |
| 0.360276 | 0.226601 | 0.193 | 0.17 | 1 | 5 | Phf2 | 0.023 | Young |
| 0.360351 | 0.162267 | 0.513 | 0.503 | 1 | 5 | Drap1 | 0.01 | Young |
| 0.360405 | 0.194894 | 0.036 | 0.025 | 1 | 5 | Hotairm1 | 0.011 | Young |
| 0.360437 | 0.021718 | 1 | 1 | 1 | 5 | mt-Co2 | 0 | Young |
| 0.36095 | 0.245453 | 0.162 | 0.143 | 1 | 5 | Glo1 | 0.019 | Young |
| 0.361025 | 0.109118 | 0.65 | 0.63 | 1 | 5 | Nfkb1 | 0.02 | Young |
| 0.361224 | 0.18106 | 0.036 | 0.025 | 1 | 5 | Zkscan6 | 0.011 | Young |

| | | | | | | | | |
|---|---|---|---|---|---|---|---|---|
| 0.361269 | 0.156517 | 0.447 | 0.423 | 1 | 5 | Smchd1 | 0.024 | Young |
| 0.361283 | 0.259047 | 0.53 | 0.54 | 1 | 5 | Aftph | -0.01 | Young |
| 0.361289 | 0.147807 | 0.226 | 0.2 | 1 | 5 | Lgals8 | 0.026 | Young |
| 0.361414 | 0.317274 | 0.109 | 0.091 | 1 | 5 | Lrp5 | 0.018 | Young |
| 0.361459 | 0.286067 | 0.081 | 0.066 | 1 | 5 | Zdhhc16 | 0.015 | Young |
| 0.361629 | 0.360976 | 0.094 | 0.077 | 1 | 5 | Kat5 | 0.017 | Young |
| 0.361655 | 0.40682 | 0.18 | 0.162 | 1 | 5 | Ibtk | 0.018 | Young |
| 0.36167 | 0.516414 | 0.071 | 0.057 | 1 | 5 | Slc39a14 | 0.014 | Young |
| 0.361954 | 0.130232 | 0.147 | 0.125 | 1 | 5 | Zfp646 | 0.022 | Young |
| 0.362074 | 0.828501 | 0.033 | 0.024 | 1 | 5 | 1110012L1 | 0.009 | Young |
| 0.3621 | 0.206277 | 0.563 | 0.557 | 1 | 5 | Syngr2 | 0.006 | Young |
| 0.362377 | 0.296594 | 0.094 | 0.077 | 1 | 5 | Tomm6 | 0.017 | Young |
| 0.362675 | 0.597726 | 0.063 | 0.051 | 1 | 5 | Cpeb3 | 0.012 | Young |
| 0.362982 | 0.171113 | 0.201 | 0.177 | 1 | 5 | Nmd3 | 0.024 | Young |
| 0.363163 | 0.183085 | 0.076 | 0.061 | 1 | 5 | Trmt2b | 0.015 | Young |
| 0.363452 | 0.337932 | 0.277 | 0.258 | 1 | 5 | Tecr | 0.019 | Young |
| 0.363575 | 0.197525 | 0.137 | 0.116 | 1 | 5 | Avl9 | 0.021 | Young |
| 0.363832 | 0.275049 | 0.353 | 0.337 | 1 | 5 | Emc1 | 0.016 | Young |
| 0.364031 | 0.128427 | 0.386 | 0.357 | 1 | 5 | Sel1l | 0.029 | Young |
| 0.364132 | 0.30907 | 0.119 | 0.101 | 1 | 5 | Slc9a8 | 0.018 | Young |
| 0.364757 | 0.347704 | 0.142 | 0.123 | 1 | 5 | Fam208b | 0.019 | Young |
| 0.365044 | 0.309691 | 0.058 | 0.045 | 1 | 5 | Edrf1 | 0.013 | Young |
| 0.365053 | 0.185494 | 0.47 | 0.468 | 1 | 5 | Syncrip | 0.002 | Young |
| 0.365101 | 0.414111 | 0.074 | 0.059 | 1 | 5 | Rnf113a2 | 0.015 | Young |
| 0.365226 | 0.179407 | 0.35 | 0.323 | 1 | 5 | Uvrag | 0.027 | Young |
| 0.365342 | 0.685351 | 0.023 | 0.015 | 1 | 5 | Smco4 | 0.008 | Young |
| 0.36547 | 0.323334 | 0.15 | 0.128 | 1 | 5 | Wasl | 0.022 | Young |
| 0.365555 | 0.394058 | 0.043 | 0.032 | 1 | 5 | Zfp930 | 0.011 | Young |
| 0.365819 | 0.118423 | 0.079 | 0.062 | 1 | 5 | Stambp | 0.017 | Young |
| 0.365908 | 0.688791 | 0.033 | 0.024 | 1 | 5 | Acp6 | 0.009 | Young |
| 0.366104 | 0.238003 | 0.213 | 0.189 | 1 | 5 | Txlna | 0.024 | Young |
| 0.366126 | 0.278199 | 0.145 | 0.126 | 1 | 5 | Prorsd1 | 0.019 | Young |
| 0.366232 | 0.0274 | 0.074 | 0.059 | 1 | 5 | Vipas39 | 0.015 | Young |
| 0.366298 | 0.237157 | 0.129 | 0.109 | 1 | 5 | Sirt1 | 0.02 | Young |
| 0.366384 | 0.965974 | 0.023 | 0.015 | 1 | 5 | Asb4 | 0.008 | Young |
| 0.366591 | 0.332078 | 0.091 | 0.076 | 1 | 5 | Mecr | 0.015 | Young |
| 0.366731 | 0.354726 | 0.16 | 0.14 | 1 | 5 | Ostm1 | 0.02 | Young |
| 0.366746 | 0.137364 | 0.165 | 0.141 | 1 | 5 | Phf12 | 0.024 | Young |
| 0.366817 | 0.26718 | 0.168 | 0.143 | 1 | 5 | Tlr1 | 0.025 | Young |
| 0.367485 | 0.158611 | 0.114 | 0.096 | 1 | 5 | Ttf1 | 0.018 | Young |
| 0.367487 | 0.060467 | 0.015 | 0.024 | 1 | 5 | Skint3 | -0.009 | Young |
| 0.367533 | 0.309106 | 0.086 | 0.071 | 1 | 5 | Hectd4 | 0.015 | Young |
| 0.36762 | 0.746792 | 0.033 | 0.024 | 1 | 5 | Tnfrsf22 | 0.009 | Young |
| 0.367864 | 0.262707 | 0.289 | 0.268 | 1 | 5 | Ldb1 | 0.021 | Young |
| 0.36789 | 0.390801 | 0.094 | 0.077 | 1 | 5 | Fbxw5 | 0.017 | Young |

| | | | | | | | | |
|---|---|---|---|---|---|---|---|---|
| 0.368007 | 0.162355 | 0.749 | 0.727 | 1 | 5 | Nr3c1 | 0.022 | Young |
| 0.368051 | 0.221233 | 0.051 | 0.039 | 1 | 5 | Elk1 | 0.012 | Young |
| 0.368173 | 0.270861 | 0.107 | 0.089 | 1 | 5 | 2610002M | 0.018 | Young |
| 0.368226 | 0.316588 | 0.058 | 0.045 | 1 | 5 | Lsm11 | 0.013 | Young |
| 0.368572 | 0.468233 | 0.127 | 0.111 | 1 | 5 | Eif2ak1 | 0.016 | Young |
| 0.368628 | 0.290366 | 0.157 | 0.136 | 1 | 5 | Hdac3 | 0.021 | Young |
| 0.368674 | 0.292047 | 0.401 | 0.387 | 1 | 5 | Eif1b | 0.014 | Young |
| 0.368788 | 0.88042 | 0.041 | 0.03 | 1 | 5 | 1700066M | 0.011 | Young |
| 0.369156 | 0.18432 | 0.401 | 0.387 | 1 | 5 | Mrps33 | 0.014 | Young |
| 0.369363 | 0.342956 | 0.254 | 0.232 | 1 | 5 | Asxl2 | 0.022 | Young |
| 0.369469 | 0.022579 | 0.104 | 0.125 | 1 | 5 | Igtp | -0.021 | Young |
| 0.369616 | 0.227498 | 0.053 | 0.04 | 1 | 5 | Crcp | 0.013 | Young |
| 0.369624 | 0.330901 | 0.119 | 0.101 | 1 | 5 | Sfi1 | 0.018 | Young |
| 0.370609 | 0.343264 | 0.15 | 0.131 | 1 | 5 | Yeats2 | 0.019 | Young |
| 0.370635 | 0.273274 | 0.096 | 0.081 | 1 | 5 | Ppm1d | 0.015 | Young |
| 0.370653 | 0.205967 | 0.343 | 0.316 | 1 | 5 | Psmd13 | 0.027 | Young |
| 0.37067 | 0.166166 | 0.241 | 0.214 | 1 | 5 | Hbp1 | 0.027 | Young |
| 0.370821 | 0.016099 | 0.327 | 0.369 | 1 | 5 | Med28 | -0.042 | Young |
| 0.371028 | 0.096687 | 0.241 | 0.21 | 1 | 5 | Ubqln1 | 0.031 | Young |
| 0.371097 | 0.805112 | 0.023 | 0.015 | 1 | 5 | Suox | 0.008 | Young |
| 0.371488 | 0.480653 | 0.051 | 0.039 | 1 | 5 | Gin1 | 0.012 | Young |
| 0.371538 | 0.195552 | 0.254 | 0.231 | 1 | 5 | Ptpn11 | 0.023 | Young |
| 0.371622 | 0.585876 | 0.023 | 0.015 | 1 | 5 | Cd276 | 0.008 | Young |
| 0.371631 | 0.366609 | 0.096 | 0.081 | 1 | 5 | Tmem161b | 0.015 | Young |
| 0.371725 | 0.255781 | 0.201 | 0.18 | 1 | 5 | Slc35a5 | 0.021 | Young |
| 0.372092 | 0.053768 | 0.619 | 0.598 | 1 | 5 | Rab7 | 0.021 | Young |
| 0.373009 | 0.235408 | 0.31 | 0.288 | 1 | 5 | Tifa | 0.022 | Young |
| 0.373085 | 0.131017 | 0.579 | 0.571 | 1 | 5 | Ap2m1 | 0.008 | Young |
| 0.37324 | 0.124698 | 0.683 | 0.694 | 1 | 5 | Ckb | -0.011 | Young |
| 0.373278 | 0.182971 | 0.15 | 0.13 | 1 | 5 | Med14 | 0.02 | Young |
| 0.373297 | 0.365924 | 0.102 | 0.086 | 1 | 5 | Nup93 | 0.016 | Young |
| 0.373665 | 0.063039 | 0.053 | 0.04 | 1 | 5 | Slc12a4 | 0.013 | Young |
| 0.373731 | 0.718207 | 0.023 | 0.015 | 1 | 5 | Tut1 | 0.008 | Young |
| 0.374606 | 0.269913 | 0.162 | 0.14 | 1 | 5 | Thap12 | 0.022 | Young |
| 0.375005 | 0.514847 | 0.102 | 0.086 | 1 | 5 | Cds1 | 0.016 | Young |
| 0.3751 | 0.133457 | 0.505 | 0.478 | 1 | 5 | Elob | 0.027 | Young |
| 0.375317 | 0.484127 | 0.084 | 0.069 | 1 | 5 | Fgfr1op | 0.015 | Young |
| 0.375317 | 0.681706 | 0.023 | 0.015 | 1 | 5 | Epm2a | 0.008 | Young |
| 0.375317 | 0.649258 | 0.023 | 0.015 | 1 | 5 | Akap12 | 0.008 | Young |
| 0.375317 | 0.581268 | 0.023 | 0.015 | 1 | 5 | Eid2b | 0.008 | Young |
| 0.375562 | 0.309057 | 0.104 | 0.088 | 1 | 5 | Tnfsf13b | 0.016 | Young |
| 0.375681 | 0.080403 | 0.127 | 0.106 | 1 | 5 | Dcaf15 | 0.021 | Young |
| 0.376509 | 0.204283 | 0.086 | 0.071 | 1 | 5 | Sfxn1 | 0.015 | Young |
| 0.376691 | 0.245068 | 0.033 | 0.024 | 1 | 5 | Hepacam2 | 0.009 | Young |
| 0.376793 | 0.354426 | 0.066 | 0.052 | 1 | 5 | Med19 | 0.014 | Young |

| | | | | | | | | |
|---|---|---|---|---|---|---|---|---|
| 0.37681 | 0.29237 | 0.043 | 0.032 | 1 | 5 | Slc25a15 | 0.011 | Young |
| 0.376907 | 0.701167 | 0.023 | 0.015 | 1 | 5 | Smtn | 0.008 | Young |
| 0.376907 | 0.586432 | 0.023 | 0.015 | 1 | 5 | Cysltr2 | 0.008 | Young |
| 0.376937 | 0.090548 | 0.82 | 0.815 | 1 | 5 | Prcp | 0.005 | Young |
| 0.377061 | 0.420397 | 0.114 | 0.098 | 1 | 5 | Akt1s1 | 0.016 | Young |
| 0.377091 | 0.465997 | 0.089 | 0.074 | 1 | 5 | Pelp1 | 0.015 | Young |
| 0.377204 | 0.323916 | 0.274 | 0.251 | 1 | 5 | Arid1b | 0.023 | Young |
| 0.377438 | 0.466256 | 0.023 | 0.015 | 1 | 5 | Zfp867 | 0.008 | Young |
| 0.377505 | 0.176614 | 0.173 | 0.15 | 1 | 5 | Rbm18 | 0.023 | Young |
| 0.377657 | 0.289755 | 0.114 | 0.096 | 1 | 5 | Pigc | 0.018 | Young |
| 0.378203 | 0.358727 | 0.175 | 0.158 | 1 | 5 | Higd1a | 0.017 | Young |
| 0.378331 | 0.195137 | 0.381 | 0.357 | 1 | 5 | Klf10 | 0.024 | Young |
| 0.378501 | 0.689817 | 0.023 | 0.015 | 1 | 5 | Abcc10 | 0.008 | Young |
| 0.37855 | 0.279459 | 0.084 | 0.069 | 1 | 5 | Zfp414 | 0.015 | Young |
| 0.378739 | 0.081893 | 0.124 | 0.104 | 1 | 5 | Ddah2 | 0.02 | Young |
| 0.379354 | 0.034783 | 0.368 | 0.412 | 1 | 5 | Kmt2a | -0.044 | Young |
| 0.379441 | 0.298228 | 0.086 | 0.071 | 1 | 5 | Mrpl45 | 0.015 | Young |
| 0.379467 | 0.08202 | 0.043 | 0.032 | 1 | 5 | Fam234b | 0.011 | Young |
| 0.379642 | 0.075462 | 0.997 | 0.998 | 1 | 5 | Pf4 | -0.001 | Young |
| 0.379661 | 0.438664 | 0.096 | 0.081 | 1 | 5 | Tirap | 0.015 | Young |
| 0.3801 | 0.547072 | 0.023 | 0.015 | 1 | 5 | Cd320 | 0.008 | Young |
| 0.380183 | 0.225542 | 0.033 | 0.024 | 1 | 5 | Cldn34c1 | 0.009 | Young |
| 0.380378 | 0.220513 | 0.327 | 0.298 | 1 | 5 | Arhgef3 | 0.029 | Young |
| 0.38055 | 0.389797 | 0.234 | 0.215 | 1 | 5 | Mylip | 0.019 | Young |
| 0.380622 | 0.236581 | 0.107 | 0.089 | 1 | 5 | Golt1b | 0.018 | Young |
| 0.380903 | 0.102953 | 0.216 | 0.246 | 1 | 5 | Ccm2 | -0.03 | Young |
| 0.381058 | 0.052671 | 0.089 | 0.108 | 1 | 5 | Vps37c | -0.019 | Young |
| 0.381168 | 0.483769 | 0.023 | 0.015 | 1 | 5 | Amz1 | 0.008 | Young |
| 0.38166 | 0.316254 | 0.274 | 0.256 | 1 | 5 | Hmbox1 | 0.018 | Young |
| 0.381787 | 0.395581 | 0.15 | 0.133 | 1 | 5 | Pcgf3 | 0.017 | Young |
| 0.381954 | 0.256618 | 0.315 | 0.293 | 1 | 5 | Zfp445 | 0.022 | Young |
| 0.382241 | 0.230998 | 0.53 | 0.512 | 1 | 5 | Ptger4 | 0.018 | Young |
| 0.382456 | 0.177499 | 0.327 | 0.301 | 1 | 5 | Osgep | 0.026 | Young |
| 0.382637 | 0.312581 | 0.201 | 0.18 | 1 | 5 | Rcbtb2 | 0.021 | Young |
| 0.382672 | 0.259672 | 0.094 | 0.077 | 1 | 5 | Elmod2 | 0.017 | Young |
| 0.382773 | 0.235408 | 0.023 | 0.015 | 1 | 5 | Capn10 | 0.008 | Young |
| 0.382814 | 0.242331 | 0.033 | 0.024 | 1 | 5 | Mpp7 | 0.009 | Young |
| 0.382963 | 0.330383 | 0.112 | 0.094 | 1 | 5 | Adap1 | 0.018 | Young |
| 0.383651 | 0.073455 | 0.807 | 0.79 | 1 | 5 | Il6ra | 0.017 | Young |
| 0.38386 | 0.266969 | 0.206 | 0.182 | 1 | 5 | 2410015M | 0.024 | Young |
| 0.384222 | 0.181615 | 0.074 | 0.091 | 1 | 5 | Cinp | -0.017 | Young |
| 0.384815 | 0.200699 | 0.043 | 0.032 | 1 | 5 | Olfr1167 | 0.011 | Young |
| 0.384888 | 0.06937 | 0.119 | 0.099 | 1 | 5 | Trappc11 | 0.02 | Young |
| 0.38496 | 0.368038 | 0.18 | 0.162 | 1 | 5 | Siva1 | 0.018 | Young |
| 0.385629 | 0.678292 | 0.053 | 0.042 | 1 | 5 | Trp53rka | 0.011 | Young |

| | | | | | | | | |
|---|---|---|---|---|---|---|---|---|
| 0.385721 | 0.09543 | 0.678 | 0.672 | 1 | 5 | Tnfaip8 | 0.006 | Young |
| 0.386704 | 0.832584 | 0.046 | 0.035 | 1 | 5 | Tmem38a | 0.011 | Young |
| 0.386856 | 0.283901 | 0.071 | 0.057 | 1 | 5 | Pira2 | 0.014 | Young |
| 0.387041 | 0.130614 | 0.33 | 0.31 | 1 | 5 | Ccdc47 | 0.02 | Young |
| 0.387395 | 0.180215 | 0.36 | 0.328 | 1 | 5 | Rc3h1 | 0.032 | Young |
| 0.387464 | 0.118454 | 0.548 | 0.537 | 1 | 5 | Cstb | 0.011 | Young |
| 0.387594 | 0.219775 | 0.195 | 0.173 | 1 | 5 | Cirbp | 0.022 | Young |
| 0.387668 | 0.042949 | 0.033 | 0.024 | 1 | 5 | Tmem191c | 0.009 | Young |
| 0.387908 | 0.308915 | 0.041 | 0.03 | 1 | 5 | Etaa1 | 0.011 | Young |
| 0.388551 | 0.125689 | 0.19 | 0.215 | 1 | 5 | Trim30d | -0.025 | Young |
| 0.388573 | 0.206858 | 0.173 | 0.153 | 1 | 5 | Acox1 | 0.02 | Young |
| 0.388984 | 0.022712 | 0.036 | 0.047 | 1 | 5 | Gm47754 | -0.011 | Young |
| 0.389079 | 0.328684 | 0.109 | 0.093 | 1 | 5 | Camk2g | 0.016 | Young |
| 0.38909 | 0.581652 | 0.099 | 0.084 | 1 | 5 | Arhgap21 | 0.015 | Young |
| 0.389099 | 0.044994 | 0.081 | 0.098 | 1 | 5 | Thoc1 | -0.017 | Young |
| 0.389412 | 0.051184 | 0.249 | 0.286 | 1 | 5 | BC005624 | -0.037 | Young |
| 0.389463 | 0.420992 | 0.104 | 0.089 | 1 | 5 | Atrnl1 | 0.015 | Young |
| 0.389695 | 0.257666 | 0.076 | 0.062 | 1 | 5 | Ubl4a | 0.014 | Young |
| 0.39043 | 0.141015 | 0.005 | 0.01 | 1 | 5 | Lims2 | -0.005 | Young |
| 0.39064 | 0.087946 | 0.703 | 0.689 | 1 | 5 | Abca9 | 0.014 | Young |
| 0.391175 | 0.627043 | 0.03 | 0.022 | 1 | 5 | Pigw | 0.008 | Young |
| 0.391238 | 0.140011 | 0.102 | 0.084 | 1 | 5 | Tstd2 | 0.018 | Young |
| 0.391566 | 0.10007 | 0.114 | 0.096 | 1 | 5 | Dcaf10 | 0.018 | Young |
| 0.391668 | 0.348835 | 0.056 | 0.044 | 1 | 5 | Gatb | 0.012 | Young |
| 0.3921 | 0.610309 | 0.03 | 0.022 | 1 | 5 | Sec22c | 0.008 | Young |
| 0.392612 | 0.156816 | 0.388 | 0.367 | 1 | 5 | Use1 | 0.021 | Young |
| 0.392862 | 0.32766 | 0.005 | 0.01 | 1 | 5 | Tnfrsf4 | -0.005 | Young |
| 0.393167 | 0.350486 | 0.145 | 0.126 | 1 | 5 | Vac14 | 0.019 | Young |
| 0.39336 | 0.219979 | 0.056 | 0.044 | 1 | 5 | Zfp715 | 0.012 | Young |
| 0.393368 | 0.246828 | 0.211 | 0.189 | 1 | 5 | Ank | 0.022 | Young |
| 0.39352 | 0.112738 | 0.378 | 0.35 | 1 | 5 | Osbpl8 | 0.028 | Young |
| 0.393577 | 0.243784 | 0.175 | 0.155 | 1 | 5 | Ipmk | 0.02 | Young |
| 0.393625 | 0.479536 | 0.074 | 0.061 | 1 | 5 | Slc25a22 | 0.013 | Young |
| 0.393675 | 0.159703 | 0.005 | 0.01 | 1 | 5 | Thnsl2 | -0.005 | Young |
| 0.394078 | 0.328189 | 0.142 | 0.126 | 1 | 5 | Ulk1 | 0.016 | Young |
| 0.394223 | 0.458419 | 0.094 | 0.079 | 1 | 5 | Miga1 | 0.015 | Young |
| 0.394724 | 0.142723 | 0.142 | 0.167 | 1 | 5 | Ubap1 | -0.025 | Young |
| 0.395265 | 0.531202 | 0.053 | 0.042 | 1 | 5 | Psmg1 | 0.011 | Young |
| 0.395349 | 0.554303 | 0.03 | 0.022 | 1 | 5 | Trdmt1 | 0.008 | Young |
| 0.395505 | 0.251561 | 0.198 | 0.178 | 1 | 5 | Ilvbl | 0.02 | Young |
| 0.396338 | 0.155016 | 0.287 | 0.261 | 1 | 5 | Renbp | 0.026 | Young |
| 0.396855 | 0.443317 | 0.137 | 0.121 | 1 | 5 | Cav2 | 0.016 | Young |
| 0.396859 | 0.264036 | 0.063 | 0.051 | 1 | 5 | Ints4 | 0.012 | Young |
| 0.397051 | 0.172422 | 0.14 | 0.121 | 1 | 5 | Uspl1 | 0.019 | Young |
| 0.397233 | 0.322861 | 0.132 | 0.114 | 1 | 5 | Ice1 | 0.018 | Young |

| | | | | | | | | |
|---|---|---|---|---|---|---|---|---|
| 0.397468 | 0.346682 | 0.297 | 0.286 | 1 | 5 | Dcaf7 | 0.011 | Young |
| 0.397899 | 0.331486 | 0.102 | 0.086 | 1 | 5 | Ormdl3 | 0.016 | Young |
| 0.398326 | 0.249008 | 0.213 | 0.194 | 1 | 5 | Chp1 | 0.019 | Young |
| 0.39833 | 0.172865 | 0.112 | 0.094 | 1 | 5 | Dlat | 0.018 | Young |
| 0.398798 | 0.298671 | 0.091 | 0.076 | 1 | 5 | Aspscr1 | 0.015 | Young |
| 0.399201 | 0.277646 | 0.193 | 0.173 | 1 | 5 | Gon4l | 0.02 | Young |
| 0.399202 | 0.6688 | 0.02 | 0.013 | 1 | 5 | Shpk | 0.007 | Young |
| 0.399264 | 0.201369 | 0.345 | 0.328 | 1 | 5 | Sec31a | 0.017 | Young |
| 0.399486 | 0.247879 | 0.056 | 0.044 | 1 | 5 | Tbccd1 | 0.012 | Young |
| 0.399785 | 0.775318 | 0.02 | 0.013 | 1 | 5 | Polr3g | 0.007 | Young |
| 0.399785 | 0.716811 | 0.02 | 0.013 | 1 | 5 | Lrp4 | 0.007 | Young |
| 0.400092 | 0.194863 | 0.198 | 0.177 | 1 | 5 | Spata6 | 0.021 | Young |
| 0.400539 | 0.275165 | 0.084 | 0.069 | 1 | 5 | Gfm2 | 0.015 | Young |
| 0.40154 | 0.52972 | 0.137 | 0.123 | 1 | 5 | Ipo8 | 0.014 | Young |
| 0.401583 | 0.316954 | 0.102 | 0.086 | 1 | 5 | Mpp5 | 0.016 | Young |
| 0.401639 | 0.295441 | 0.112 | 0.133 | 1 | 5 | Exosc10 | -0.021 | Young |
| 0.401855 | 0.323434 | 0.223 | 0.202 | 1 | 5 | Supt20 | 0.021 | Young |
| 0.401921 | 0.56692 | 0.084 | 0.071 | 1 | 5 | Plxna4os1 | 0.013 | Young |
| 0.402066 | 0.714473 | 0.046 | 0.035 | 1 | 5 | Nudt19 | 0.011 | Young |
| 0.402554 | 0.153485 | 0.305 | 0.279 | 1 | 5 | Brd7 | 0.026 | Young |
| 0.402587 | 0.48851 | 0.053 | 0.042 | 1 | 5 | Rab13 | 0.011 | Young |
| 0.40284 | 0.142797 | 0.525 | 0.525 | 1 | 5 | Mapk3 | 0 | Young |
| 0.402952 | 0.265814 | 0.206 | 0.187 | 1 | 5 | Zdhhc14 | 0.019 | Young |
| 0.403292 | 0.717524 | 0.02 | 0.013 | 1 | 5 | Sfxn5 | 0.007 | Young |
| 0.403292 | 0.57785 | 0.02 | 0.013 | 1 | 5 | Map3k12 | 0.007 | Young |
| 0.403414 | 0.168487 | 0.071 | 0.057 | 1 | 5 | Pop7 | 0.014 | Young |
| 0.403701 | 0.172466 | 0.416 | 0.394 | 1 | 5 | Gm16286 | 0.022 | Young |
| 0.403779 | 0.49039 | 0.03 | 0.022 | 1 | 5 | Sbk1 | 0.008 | Young |
| 0.404251 | 0.280453 | 0.03 | 0.022 | 1 | 5 | Hspbap1 | 0.008 | Young |
| 0.404345 | 0.553283 | 0.079 | 0.066 | 1 | 5 | Itpa | 0.013 | Young |
| 0.404412 | 0.615736 | 0.038 | 0.029 | 1 | 5 | Tspan33 | 0.009 | Young |
| 0.404536 | 0.034281 | 0.061 | 0.076 | 1 | 5 | Zdhhc23 | -0.015 | Young |
| 0.405379 | 0.121751 | 0.041 | 0.03 | 1 | 5 | Nol10 | 0.011 | Young |
| 0.405578 | 0.119845 | 0.183 | 0.162 | 1 | 5 | Cops2 | 0.021 | Young |
| 0.405815 | 0.150012 | 0.142 | 0.123 | 1 | 5 | Kctd10 | 0.019 | Young |
| 0.405828 | 0.321337 | 0.35 | 0.342 | 1 | 5 | Gxylt1 | 0.008 | Young |
| 0.405976 | 0.460005 | 0.061 | 0.049 | 1 | 5 | Ttll4 | 0.012 | Young |
| 0.406082 | 0.177887 | 0.34 | 0.315 | 1 | 5 | Larp4 | 0.025 | Young |
| 0.40614 | 0.179293 | 0.03 | 0.022 | 1 | 5 | Arhgap35 | 0.008 | Young |
| 0.40647 | 0.196867 | 0.343 | 0.322 | 1 | 5 | Vps26a | 0.021 | Young |
| 0.406655 | 0.207511 | 0.124 | 0.106 | 1 | 5 | Taf2 | 0.018 | Young |
| 0.406979 | 0.245857 | 0.137 | 0.118 | 1 | 5 | Zfp318 | 0.019 | Young |
| 0.407052 | 0.220674 | 0.056 | 0.044 | 1 | 5 | Aen | 0.012 | Young |
| 0.407111 | 0.14271 | 0.193 | 0.172 | 1 | 5 | Tslp | 0.021 | Young |
| 0.407446 | 0.165343 | 0.069 | 0.056 | 1 | 5 | Pabpc4 | 0.013 | Young |

| | | | | | | | | |
|---|---|---|---|---|---|---|---|---|
| 0.407618 | 0.212077 | 0.492 | 0.47 | 1 | 5 | Lmna | 0.022 | Young |
| 0.407668 | 0.174396 | 0.327 | 0.298 | 1 | 5 | Zdhhc9 | 0.029 | Young |
| 0.40772 | 0.257907 | 0.236 | 0.217 | 1 | 5 | Oxr1 | 0.019 | Young |
| 0.407768 | 0.480491 | 0.038 | 0.029 | 1 | 5 | Nfib | 0.009 | Young |
| 0.407792 | 0.050404 | 0.168 | 0.145 | 1 | 5 | Kdm3a | 0.023 | Young |
| 0.407909 | 0.077283 | 0.19 | 0.167 | 1 | 5 | Cops8 | 0.023 | Young |
| 0.408348 | 0.201065 | 0.371 | 0.354 | 1 | 5 | Pnpla8 | 0.017 | Young |
| 0.408412 | 0.314468 | 0.124 | 0.106 | 1 | 5 | Mapk8ip3 | 0.018 | Young |
| 0.408556 | 0.050063 | 0.193 | 0.221 | 1 | 5 | Ufl1 | -0.028 | Young |
| 0.408614 | 0.244795 | 0.137 | 0.12 | 1 | 5 | Tmem222 | 0.017 | Young |
| 0.408782 | 0.332186 | 0.061 | 0.049 | 1 | 5 | Cdk5 | 0.012 | Young |
| 0.409028 | 0.442652 | 0.124 | 0.109 | 1 | 5 | Pign | 0.015 | Young |
| 0.409175 | 0.220577 | 0.249 | 0.231 | 1 | 5 | 11-Sep | 0.018 | Young |
| 0.409177 | 0.416871 | 0.02 | 0.013 | 1 | 5 | D830025C( | 0.007 | Young |
| 0.409177 | 0.414886 | 0.02 | 0.013 | 1 | 5 | Zfp458 | 0.007 | Young |
| 0.409356 | 0.412978 | 0.15 | 0.135 | 1 | 5 | Bak1 | 0.015 | Young |
| 0.409874 | 0.161574 | 0.761 | 0.771 | 1 | 5 | Gng5 | -0.01 | Young |
| 0.409904 | 0.286169 | 0.218 | 0.199 | 1 | 5 | Setdb1 | 0.019 | Young |
| 0.409918 | 0.247746 | 0.201 | 0.182 | 1 | 5 | Hnrnpll | 0.019 | Young |
| 0.410105 | 0.179971 | 0.122 | 0.104 | 1 | 5 | Pigm | 0.018 | Young |
| 0.410355 | 0.301907 | 0.094 | 0.079 | 1 | 5 | Tmem181a | 0.015 | Young |
| 0.41036 | 0.182961 | 0.02 | 0.013 | 1 | 5 | Gm35339 | 0.007 | Young |
| 0.410491 | 0.17316 | 0.34 | 0.322 | 1 | 5 | S1pr1 | 0.018 | Young |
| 0.410823 | 0.047141 | 0.947 | 0.978 | 1 | 5 | Oaz1 | -0.031 | Young |
| 0.410871 | 0.071764 | 0.195 | 0.17 | 1 | 5 | Nagpa | 0.025 | Young |
| 0.410953 | 0.329831 | 0.02 | 0.013 | 1 | 5 | Zfp119b | 0.007 | Young |
| 0.41107 | 0.325555 | 0.218 | 0.2 | 1 | 5 | Rbck1 | 0.018 | Young |
| 0.411434 | 0.238288 | 0.061 | 0.049 | 1 | 5 | Nup160 | 0.012 | Young |
| 0.411546 | 0.14733 | 0.02 | 0.013 | 1 | 5 | Tmem81 | 0.007 | Young |
| 0.411873 | 0.038601 | 0.434 | 0.475 | 1 | 5 | Slc29a3 | -0.041 | Young |
| 0.411975 | 0.128142 | 0.711 | 0.719 | 1 | 5 | Vcp | -0.008 | Young |
| 0.412209 | 0.200402 | 0.195 | 0.173 | 1 | 5 | Ppfia4 | 0.022 | Young |
| 0.412481 | 0.11678 | 0.525 | 0.502 | 1 | 5 | U2af2 | 0.023 | Young |
| 0.412733 | 0.192286 | 0.02 | 0.013 | 1 | 5 | Gm48653 | 0.007 | Young |
| 0.412764 | 0.262917 | 0.046 | 0.035 | 1 | 5 | Mpv17l | 0.011 | Young |
| 0.413097 | 0.002089 | 0.053 | 0.067 | 1 | 5 | D930016D( | -0.014 | Young |
| 0.413647 | 0.176799 | 0.327 | 0.301 | 1 | 5 | Hnrnpr | 0.026 | Young |
| 0.413762 | 0.395471 | 0.061 | 0.049 | 1 | 5 | Lmbr1l | 0.012 | Young |
| 0.413935 | 0.321263 | 0.162 | 0.146 | 1 | 5 | Sap18 | 0.016 | Young |
| 0.414042 | 0.07905 | 0.246 | 0.219 | 1 | 5 | Zfp263 | 0.027 | Young |
| 0.414306 | 0.093151 | 0.576 | 0.562 | 1 | 5 | Arhgef6 | 0.014 | Young |
| 0.41453 | 0.480854 | 0.038 | 0.029 | 1 | 5 | Tfap4 | 0.009 | Young |
| 0.414685 | 0.340794 | 0.147 | 0.128 | 1 | 5 | Phf5a | 0.019 | Young |
| 0.415047 | 0.225618 | 0.193 | 0.17 | 1 | 5 | Tlk2 | 0.023 | Young |
| 0.415402 | 0.001934 | 0.056 | 0.044 | 1 | 5 | Mmgt2 | 0.012 | Young |

| | | | | | | | | |
|---|---|---|---|---|---|---|---|---|
| 0.415471 | 0.371812 | 0.066 | 0.054 | 1 | 5 | Alpk1 | 0.012 | Young |
| 0.41571 | 0.037613 | 0.02 | 0.013 | 1 | 5 | Wdr25 | 0.007 | Young |
| 0.415919 | 0.39498 | 0.137 | 0.12 | 1 | 5 | Tmem250- | 0.017 | Young |
| 0.416022 | 0.26702 | 0.261 | 0.241 | 1 | 5 | Ing2 | 0.02 | Young |
| 0.416927 | 0.48704 | 0.089 | 0.076 | 1 | 5 | Cep295 | 0.013 | Young |
| 0.417397 | 0.30934 | 0.188 | 0.168 | 1 | 5 | Trrap | 0.02 | Young |
| 0.41753 | 0.159305 | 0.157 | 0.182 | 1 | 5 | Atad2b | -0.025 | Young |
| 0.417804 | 0.569308 | 0.114 | 0.101 | 1 | 5 | Nudt13 | 0.013 | Young |
| 0.41791 | 0.212565 | 0.145 | 0.128 | 1 | 5 | Pot1a | 0.017 | Young |
| 0.417961 | 0.320916 | 0.165 | 0.148 | 1 | 5 | Tmub2 | 0.017 | Young |
| 0.418091 | 0.484141 | 0.058 | 0.047 | 1 | 5 | Mrps35 | 0.011 | Young |
| 0.418215 | 0.020392 | 0.173 | 0.197 | 1 | 5 | Dusp7 | -0.024 | Young |
| 0.418223 | 0.212593 | 0.14 | 0.121 | 1 | 5 | Hint2 | 0.019 | Young |
| 0.418595 | 0.146518 | 0.213 | 0.194 | 1 | 5 | Aqr | 0.019 | Young |
| 0.418636 | 0.250426 | 0.137 | 0.12 | 1 | 5 | Kif23 | 0.017 | Young |
| 0.418691 | 0.202231 | 0.299 | 0.276 | 1 | 5 | Uba2 | 0.023 | Young |
| 0.418789 | 1.041052 | 0.036 | 0.027 | 1 | 5 | Gm30329 | 0.009 | Young |
| 0.418987 | 0.652967 | 0.043 | 0.034 | 1 | 5 | Cep57l1 | 0.009 | Young |
| 0.419682 | 0.060257 | 0.142 | 0.123 | 1 | 5 | Tbpl1 | 0.019 | Young |
| 0.41969 | 0.339547 | 0.084 | 0.071 | 1 | 5 | Slc35g1 | 0.013 | Young |
| 0.419876 | 0.004045 | 0.365 | 0.397 | 1 | 5 | Idh2 | -0.032 | Young |
| 0.420023 | 0.197433 | 0.368 | 0.347 | 1 | 5 | Thoc2 | 0.021 | Young |
| 0.420138 | 0.292883 | 0.155 | 0.136 | 1 | 5 | 0610037L1 | 0.019 | Young |
| 0.420376 | 0.149148 | 0.34 | 0.315 | 1 | 5 | Ubr2 | 0.025 | Young |
| 0.4205 | 0.432164 | 0.038 | 0.029 | 1 | 5 | Crtc1 | 0.009 | Young |
| 0.42094 | 0.220078 | 0.203 | 0.184 | 1 | 5 | Pptc7 | 0.019 | Young |
| 0.420987 | 0.333754 | 0.244 | 0.227 | 1 | 5 | Ccny | 0.017 | Young |
| 0.421252 | 0.278214 | 0.249 | 0.232 | 1 | 5 | Rchy1 | 0.017 | Young |
| 0.421301 | 0.714334 | 0.028 | 0.02 | 1 | 5 | Cep95 | 0.008 | Young |
| 0.421303 | 0.08643 | 0.754 | 0.763 | 1 | 5 | Ptms | -0.009 | Young |
| 0.421467 | 0.316635 | 0.061 | 0.049 | 1 | 5 | Med24 | 0.012 | Young |
| 0.421719 | 0.143002 | 0.33 | 0.303 | 1 | 5 | Csnk2a1 | 0.027 | Young |
| 0.422063 | 0.265647 | 0.129 | 0.111 | 1 | 5 | Mrpl32 | 0.018 | Young |
| 0.422171 | 0.254424 | 0.079 | 0.066 | 1 | 5 | Cnpy4 | 0.013 | Young |
| 0.422298 | 0.242012 | 0.129 | 0.111 | 1 | 5 | Wipi2 | 0.018 | Young |
| 0.422488 | 0.023385 | 0.079 | 0.064 | 1 | 5 | Leo1 | 0.015 | Young |
| 0.423007 | 0.118846 | 0.193 | 0.168 | 1 | 5 | Ube4b | 0.025 | Young |
| 0.423203 | 0.048865 | 0.15 | 0.128 | 1 | 5 | Oxa1l | 0.022 | Young |
| 0.423595 | 0.263223 | 0.269 | 0.253 | 1 | 5 | Got1 | 0.016 | Young |
| 0.423609 | 0.408707 | 0.086 | 0.072 | 1 | 5 | Nat9 | 0.014 | Young |
| 0.423654 | 0.111033 | 0.619 | 0.621 | 1 | 5 | Atp6v0e | -0.002 | Young |
| 0.423676 | 0.397865 | 0.147 | 0.131 | 1 | 5 | Aggf1 | 0.016 | Young |
| 0.424004 | 0.481829 | 0.066 | 0.054 | 1 | 5 | Zmym3 | 0.012 | Young |
| 0.42434 | 0.247058 | 0.099 | 0.084 | 1 | 5 | Thap11 | 0.015 | Young |
| 0.424452 | 0.323277 | 0.124 | 0.109 | 1 | 5 | Timm50 | 0.015 | Young |

| | | | | | | | | |
|---|---|---|---|---|---|---|---|---|
| 0.424542 | 0.123935 | 0.03 | 0.04 | 1 | 5 | Slc25a40 | -0.01 | Young |
| 0.424661 | 0.117412 | 0.642 | 0.653 | 1 | 5 | Cd52 | -0.011 | Young |
| 0.424974 | 0.172256 | 0.602 | 0.601 | 1 | 5 | Il10ra | 0.001 | Young |
| 0.425391 | 0.697926 | 0.043 | 0.034 | 1 | 5 | Bcorl1 | 0.009 | Young |
| 0.425484 | 0.400162 | 0.165 | 0.148 | 1 | 5 | Vrk1 | 0.017 | Young |
| 0.425561 | 0.274111 | 0.145 | 0.128 | 1 | 5 | Rxrb | 0.017 | Young |
| 0.425578 | 0.262761 | 0.168 | 0.152 | 1 | 5 | Otud4 | 0.016 | Young |
| 0.425846 | 0.279152 | 0.028 | 0.02 | 1 | 5 | Fam135a | 0.008 | Young |
| 0.425922 | 0.005348 | 0.102 | 0.118 | 1 | 5 | Rab28 | -0.016 | Young |
| 0.426242 | 0.011804 | 0.65 | 0.684 | 1 | 5 | Serpinb6a | -0.034 | Young |
| 0.426643 | 0.232233 | 0.241 | 0.221 | 1 | 5 | Tubgcp5 | 0.02 | Young |
| 0.427104 | 0.275905 | 0.19 | 0.173 | 1 | 5 | Usp12 | 0.017 | Young |
| 0.427124 | 0.876087 | 0.018 | 0.012 | 1 | 5 | Tmem14a | 0.006 | Young |
| 0.42742 | 0.068016 | 0.135 | 0.157 | 1 | 5 | Mrps16 | -0.022 | Young |
| 0.427431 | 0.341885 | 0.091 | 0.077 | 1 | 5 | Sdhaf1 | 0.014 | Young |
| 0.427487 | 0.083267 | 0.596 | 0.599 | 1 | 5 | Gna12 | -0.003 | Young |
| 0.427508 | 0.038115 | 0.173 | 0.197 | 1 | 5 | Cdc123 | -0.024 | Young |
| 0.427596 | 0.162828 | 0.269 | 0.246 | 1 | 5 | BC037034 | 0.023 | Young |
| 0.427659 | 0.630085 | 0.036 | 0.027 | 1 | 5 | Nop2 | 0.009 | Young |
| 0.427734 | 0.230015 | 0.36 | 0.352 | 1 | 5 | Cdv3 | 0.008 | Young |
| 0.428213 | 0.142882 | 0.251 | 0.229 | 1 | 5 | Spopl | 0.022 | Young |
| 0.428497 | 0.087386 | 0.084 | 0.069 | 1 | 5 | Styx | 0.015 | Young |
| 0.4294 | 0.977559 | 0.028 | 0.02 | 1 | 5 | Ttf2 | 0.008 | Young |
| 0.4294 | 0.461907 | 0.028 | 0.02 | 1 | 5 | Zfp523 | 0.008 | Young |
| 0.429719 | 0.513217 | 0.018 | 0.012 | 1 | 5 | Ppp1r3f | 0.006 | Young |
| 0.429805 | 0.033617 | 0.056 | 0.069 | 1 | 5 | Dym | -0.013 | Young |
| 0.431152 | 0.086404 | 0.259 | 0.29 | 1 | 5 | Cyth1 | -0.031 | Young |
| 0.431697 | 0.205935 | 0.099 | 0.116 | 1 | 5 | Pofut2 | -0.017 | Young |
| 0.431723 | 0.15059 | 0.431 | 0.426 | 1 | 5 | H2afj | 0.005 | Young |
| 0.43218 | 0.175492 | 0.376 | 0.352 | 1 | 5 | Rab5a | 0.024 | Young |
| 0.432322 | 0.652841 | 0.018 | 0.012 | 1 | 5 | Bcam | 0.006 | Young |
| 0.432457 | 0.323071 | 0.089 | 0.074 | 1 | 5 | Zfp738 | 0.015 | Young |
| 0.432459 | 0.334215 | 0.086 | 0.072 | 1 | 5 | Ncapg2 | 0.014 | Young |
| 0.43276 | 0.25869 | 0.147 | 0.13 | 1 | 5 | Nrbf2 | 0.017 | Young |
| 0.432778 | 0.28958 | 0.074 | 0.061 | 1 | 5 | Cep83 | 0.013 | Young |
| 0.432974 | 0.360526 | 0.018 | 0.012 | 1 | 5 | Gm15987 | 0.006 | Young |
| 0.433069 | 0.34242 | 0.043 | 0.034 | 1 | 5 | Cd3eap | 0.009 | Young |
| 0.433353 | 0.109496 | 0.338 | 0.316 | 1 | 5 | Pnn | 0.022 | Young |
| 0.433482 | 0.398012 | 0.036 | 0.027 | 1 | 5 | Gm42715 | 0.009 | Young |
| 0.433483 | 0.461856 | 0.028 | 0.02 | 1 | 5 | Ercc6 | 0.008 | Young |
| 0.433496 | 0.295755 | 0.251 | 0.237 | 1 | 5 | Qdpr | 0.014 | Young |
| 0.433617 | 0.255253 | 0.17 | 0.155 | 1 | 5 | Arid5b | 0.015 | Young |
| 0.433727 | 0.140511 | 0.525 | 0.522 | 1 | 5 | Lrrc58 | 0.003 | Young |
| 0.43396 | 0.039503 | 0.03 | 0.04 | 1 | 5 | Mki67 | -0.01 | Young |
| 0.434123 | 0.348951 | 0.117 | 0.103 | 1 | 5 | Pmf1 | 0.014 | Young |

| | | | | | | | | |
|---|---|---|---|---|---|---|---|---|
| 0.434237 | 0.019468 | 0.431 | 0.473 | 1 | 5 | Rbm47 | -0.042 | Young |
| 0.434316 | 0.076868 | 0.157 | 0.136 | 1 | 5 | Chpf2 | 0.021 | Young |
| 0.434402 | 0.457023 | 0.089 | 0.076 | 1 | 5 | Zbtb2 | 0.013 | Young |
| 0.434558 | 0.171504 | 0.335 | 0.311 | 1 | 5 | Emg1 | 0.024 | Young |
| 0.434863 | 0.379538 | 0.155 | 0.138 | 1 | 5 | Kif13b | 0.017 | Young |
| 0.435015 | 0.291613 | 0.058 | 0.047 | 1 | 5 | Wrnip1 | 0.011 | Young |
| 0.435154 | 0.217273 | 0.241 | 0.221 | 1 | 5 | Atf1 | 0.02 | Young |
| 0.435608 | 0.047223 | 0.972 | 0.973 | 1 | 5 | Rpl17 | -0.001 | Young |
| 0.435612 | 0.21421 | 0.195 | 0.178 | 1 | 5 | Mtfr1l | 0.017 | Young |
| 0.436009 | 0.174976 | 0.411 | 0.397 | 1 | 5 | Mknk2 | 0.014 | Young |
| 0.436244 | 0.229185 | 0.018 | 0.012 | 1 | 5 | Wdpcp | 0.006 | Young |
| 0.436316 | 0.434375 | 0.063 | 0.052 | 1 | 5 | Ust | 0.011 | Young |
| 0.43662 | 0.380242 | 0.086 | 0.074 | 1 | 5 | Eef1e1 | 0.012 | Young |
| 0.437209 | 0.012233 | 0.142 | 0.121 | 1 | 5 | Mdp1 | 0.021 | Young |
| 0.437556 | 0.461649 | 0.018 | 0.012 | 1 | 5 | Map1b | 0.006 | Young |
| 0.437621 | 0.388938 | 0.084 | 0.071 | 1 | 5 | Fbxo34 | 0.013 | Young |
| 0.437641 | 0.05332 | 0.348 | 0.387 | 1 | 5 | Cerk | -0.039 | Young |
| 0.437993 | 0.241556 | 0.036 | 0.027 | 1 | 5 | Cep131 | 0.009 | Young |
| 0.438079 | 0.322292 | 0.109 | 0.094 | 1 | 5 | Rhbdd1 | 0.015 | Young |
| 0.438524 | 0.218678 | 0.137 | 0.12 | 1 | 5 | BC029722 | 0.017 | Young |
| 0.439061 | 0.236894 | 0.36 | 0.352 | 1 | 5 | Snx17 | 0.008 | Young |
| 0.439158 | 0.448027 | 0.15 | 0.135 | 1 | 5 | Erlec1 | 0.015 | Young |
| 0.439453 | 0.280205 | 0.325 | 0.303 | 1 | 5 | Lpxn | 0.022 | Young |
| 0.439528 | 0.416575 | 0.018 | 0.012 | 1 | 5 | Spata2l | 0.006 | Young |
| 0.43973 | 0.33938 | 0.135 | 0.12 | 1 | 5 | Cass4 | 0.015 | Young |
| 0.439805 | 0.383149 | 0.036 | 0.027 | 1 | 5 | E130311K1 | 0.009 | Young |
| 0.440051 | 0.139883 | 0.066 | 0.054 | 1 | 5 | Fmn1 | 0.012 | Young |
| 0.440187 | 0.255024 | 0.018 | 0.012 | 1 | 5 | Slc22a5 | 0.006 | Young |
| 0.44041 | 0.098658 | 0.563 | 0.544 | 1 | 5 | Tnfrsf1b | 0.019 | Young |
| 0.440537 | 0.242957 | 0.081 | 0.067 | 1 | 5 | Sertad3 | 0.014 | Young |
| 0.440856 | 0.042467 | 0.226 | 0.202 | 1 | 5 | Rest | 0.024 | Young |
| 0.44088 | 0.233616 | 0.071 | 0.059 | 1 | 5 | Gtf2f2 | 0.012 | Young |
| 0.440946 | 0.171344 | 0.018 | 0.025 | 1 | 5 | Sardh | -0.007 | Young |
| 0.441044 | 0.217447 | 0.066 | 0.054 | 1 | 5 | Mrps11 | 0.012 | Young |
| 0.441237 | 0.023718 | 0.132 | 0.153 | 1 | 5 | Elovl5 | -0.021 | Young |
| 0.44155 | 0.204782 | 0.497 | 0.487 | 1 | 5 | Camk1 | 0.01 | Young |
| 0.441874 | 0.075523 | 0.094 | 0.109 | 1 | 5 | Rora | -0.015 | Young |
| 0.441918 | 0.016073 | 0.15 | 0.168 | 1 | 5 | Ccz1 | -0.018 | Young |
| 0.442187 | 0.175322 | 0.124 | 0.146 | 1 | 5 | Senp5 | -0.022 | Young |
| 0.442234 | 0.159939 | 0.028 | 0.02 | 1 | 5 | Fam206a | 0.008 | Young |
| 0.442476 | 0.274907 | 0.071 | 0.059 | 1 | 5 | Trub2 | 0.012 | Young |
| 0.442497 | 0.421795 | 0.051 | 0.04 | 1 | 5 | Mipep | 0.011 | Young |
| 0.442827 | 0.259731 | 0.018 | 0.012 | 1 | 5 | Ptgfrn | 0.006 | Young |
| 0.442827 | 0.195666 | 0.018 | 0.012 | 1 | 5 | Arsg | 0.006 | Young |
| 0.443112 | 0.161409 | 0.378 | 0.365 | 1 | 5 | Wdr45b | 0.013 | Young |

| | | | | | | | | |
|---|---|---|---|---|---|---|---|---|
| 0.443134 | 0.108166 | 0.404 | 0.391 | 1 | 5 | Pik3cg | 0.013 | Young |
| 0.443292 | 0.170387 | 0.043 | 0.034 | 1 | 5 | Bach2 | 0.009 | Young |
| 0.443335 | 0.267551 | 0.292 | 0.278 | 1 | 5 | Vdac1 | 0.014 | Young |
| 0.443367 | 0.285323 | 0.089 | 0.076 | 1 | 5 | Jrkl | 0.013 | Young |
| 0.443753 | 0.077993 | 0.089 | 0.074 | 1 | 5 | B4gat1 | 0.015 | Young |
| 0.444116 | 0.284509 | 0.043 | 0.034 | 1 | 5 | Ulbp1 | 0.009 | Young |
| 0.444149 | 0.752099 | 0.053 | 0.044 | 1 | 5 | Fam110a | 0.009 | Young |
| 0.444367 | 0.108979 | 0.294 | 0.271 | 1 | 5 | Rtf1 | 0.023 | Young |
| 0.444394 | 0.305332 | 0.071 | 0.059 | 1 | 5 | Dcun1d3 | 0.012 | Young |
| 0.444752 | 0.100925 | 0.68 | 0.699 | 1 | 5 | Gpx4 | -0.019 | Young |
| 0.445242 | 0.195001 | 0.096 | 0.116 | 1 | 5 | Socs5 | -0.02 | Young |
| 0.445353 | 0.221751 | 0.386 | 0.377 | 1 | 5 | Sp1 | 0.009 | Young |
| 0.445526 | 0.277218 | 0.218 | 0.202 | 1 | 5 | Pwwp2a | 0.016 | Young |
| 0.445623 | 0.188866 | 0.096 | 0.082 | 1 | 5 | Myef2 | 0.014 | Young |
| 0.446089 | 0.389483 | 0.063 | 0.052 | 1 | 5 | Tmem94 | 0.011 | Young |
| 0.446143 | 0.267332 | 0.165 | 0.148 | 1 | 5 | Sfmbt1 | 0.017 | Young |
| 0.446639 | 0.091511 | 0.056 | 0.069 | 1 | 5 | Fam13b | -0.013 | Young |
| 0.446681 | 0.247816 | 0.18 | 0.165 | 1 | 5 | Gtf2h1 | 0.015 | Young |
| 0.447658 | 0.025294 | 0.142 | 0.125 | 1 | 5 | Mettl16 | 0.017 | Young |
| 0.447844 | 0.240545 | 0.018 | 0.025 | 1 | 5 | AW011738 | -0.007 | Young |
| 0.448012 | 0.217808 | 0.036 | 0.027 | 1 | 5 | Cdc25a | 0.009 | Young |
| 0.448107 | 0.216294 | 0.14 | 0.123 | 1 | 5 | Topors | 0.017 | Young |
| 0.448207 | 0.250189 | 0.18 | 0.162 | 1 | 5 | Ctr9 | 0.018 | Young |
| 0.448249 | 0.464865 | 0.104 | 0.091 | 1 | 5 | Pacs1 | 0.013 | Young |
| 0.448755 | 0.058606 | 0.137 | 0.157 | 1 | 5 | Tmem245 | -0.02 | Young |
| 0.448908 | 0.101826 | 0.505 | 0.481 | 1 | 5 | Papola | 0.024 | Young |
| 0.449046 | 0.078737 | 0.926 | 0.971 | 1 | 5 | Cd36 | -0.045 | Young |
| 0.449661 | 0.0891 | 0.731 | 0.749 | 1 | 5 | Gas7 | -0.018 | Young |
| 0.44967 | 0.147803 | 0.34 | 0.318 | 1 | 5 | Ggt5 | 0.022 | Young |
| 0.449983 | 0.662653 | 0.041 | 0.032 | 1 | 5 | Rbm41 | 0.009 | Young |
| 0.450013 | 0.0516 | 0.294 | 0.325 | 1 | 5 | Psmb9 | -0.031 | Young |
| 0.450106 | 0.183491 | 0.069 | 0.056 | 1 | 5 | Sephs2 | 0.013 | Young |
| 0.450307 | 0.080863 | 0.668 | 0.663 | 1 | 5 | Tma7 | 0.005 | Young |
| 0.450386 | 0.084484 | 0.764 | 0.783 | 1 | 5 | Samhd1 | -0.019 | Young |
| 0.45051 | 0.409124 | 0.063 | 0.052 | 1 | 5 | Gtf2e1 | 0.011 | Young |
| 0.45068 | 0.255436 | 0.14 | 0.123 | 1 | 5 | Hspa13 | 0.017 | Young |
| 0.450697 | 0.215657 | 0.213 | 0.194 | 1 | 5 | Atg5 | 0.019 | Young |
| 0.450769 | 0.183725 | 0.416 | 0.423 | 1 | 5 | Gna13 | -0.007 | Young |
| 0.4508 | 0.33373 | 0.081 | 0.069 | 1 | 5 | Ripor1 | 0.012 | Young |
| 0.450814 | 0.110517 | 0.053 | 0.042 | 1 | 5 | Nt5c3b | 0.011 | Young |
| 0.45093 | 0.34692 | 0.173 | 0.16 | 1 | 5 | Hk2 | 0.013 | Young |
| 0.451497 | 0.193132 | 0.297 | 0.279 | 1 | 5 | Ppp6c | 0.018 | Young |
| 0.451672 | 0.137964 | 0.185 | 0.162 | 1 | 5 | Ppt2 | 0.023 | Young |
| 0.451687 | 0.14847 | 0.036 | 0.027 | 1 | 5 | Pigh | 0.009 | Young |
| 0.451776 | 0.284532 | 0.15 | 0.131 | 1 | 5 | Trim34a | 0.019 | Young |

| | | | | | | | | |
|---|---|---|---|---|---|---|---|---|
| 0.451794 | 0.008897 | 0.239 | 0.261 | 1 | 5 | Dgkz | -0.022 | Young |
| 0.451897 | 0.126152 | 0.266 | 0.291 | 1 | 5 | Crem | -0.025 | Young |
| 0.452412 | 0.163808 | 0.175 | 0.158 | 1 | 5 | Stard7 | 0.017 | Young |
| 0.452516 | 0.128807 | 0.107 | 0.091 | 1 | 5 | Slc41a1 | 0.016 | Young |
| 0.452993 | 0.039157 | 0.254 | 0.281 | 1 | 5 | Mgat4b | -0.027 | Young |
| 0.453155 | 0.257313 | 0.122 | 0.108 | 1 | 5 | E2f1 | 0.014 | Young |
| 0.453404 | 0.13668 | 0.066 | 0.054 | 1 | 5 | Trmt13 | 0.012 | Young |
| 0.453413 | 0.159654 | 0.563 | 0.551 | 1 | 5 | Bzw1 | 0.012 | Young |
| 0.453497 | 0.071799 | 0.838 | 0.837 | 1 | 5 | Ccl9 | 0.001 | Young |
| 0.453686 | 0.165661 | 0.107 | 0.126 | 1 | 5 | Mapk9 | -0.019 | Young |
| 0.453798 | 0.541398 | 0.048 | 0.039 | 1 | 5 | Poc5 | 0.009 | Young |
| 0.454092 | 0.151857 | 0.208 | 0.187 | 1 | 5 | Mbtps1 | 0.021 | Young |
| 0.454351 | 0.145384 | 0.165 | 0.146 | 1 | 5 | Cul4a | 0.019 | Young |
| 0.454622 | 0.178481 | 0.437 | 0.412 | 1 | 5 | Ik | 0.025 | Young |
| 0.454916 | 0.188656 | 0.036 | 0.027 | 1 | 5 | Hsd17b7 | 0.009 | Young |
| 0.455168 | 0.196919 | 0.074 | 0.061 | 1 | 5 | Ciapin1 | 0.013 | Young |
| 0.456557 | 0.253279 | 0.332 | 0.315 | 1 | 5 | Alox5 | 0.017 | Young |
| 0.45707 | 0.040066 | 0.251 | 0.226 | 1 | 5 | Fbxo11 | 0.025 | Young |
| 0.45716 | 0.063455 | 0.462 | 0.431 | 1 | 5 | Mndal | 0.031 | Young |
| 0.457351 | 0.755623 | 0.025 | 0.019 | 1 | 5 | Ppbp | 0.006 | Young |
| 0.457351 | 0.653061 | 0.025 | 0.019 | 1 | 5 | Dtwd1 | 0.006 | Young |
| 0.457376 | 0.182021 | 0.089 | 0.076 | 1 | 5 | Stx18 | 0.013 | Young |
| 0.457885 | 0.180489 | 0.201 | 0.184 | 1 | 5 | Mybbp1a | 0.017 | Young |
| 0.458014 | 0.25763 | 0.411 | 0.392 | 1 | 5 | Usp8 | 0.019 | Young |
| 0.458142 | 0.147206 | 0.173 | 0.153 | 1 | 5 | Dpy30 | 0.02 | Young |
| 0.458367 | 0.069882 | 0.201 | 0.177 | 1 | 5 | Atmin | 0.024 | Young |
| 0.458595 | 0.018112 | 0.14 | 0.162 | 1 | 5 | Timm17a | -0.022 | Young |
| 0.458623 | 0.127589 | 0.036 | 0.027 | 1 | 5 | Zfp959 | 0.009 | Young |
| 0.458889 | 0.034367 | 0.249 | 0.276 | 1 | 5 | Fam173a | -0.027 | Young |
| 0.458897 | 0.128968 | 0.538 | 0.517 | 1 | 5 | Pten | 0.021 | Young |
| 0.4589 | 0.053692 | 0.244 | 0.278 | 1 | 5 | Rbbp4 | -0.034 | Young |
| 0.459008 | 0.52395 | 0.025 | 0.019 | 1 | 5 | Hdac6 | 0.006 | Young |
| 0.459126 | 0.342944 | 0.152 | 0.136 | 1 | 5 | Arpc5l | 0.016 | Young |
| 0.459199 | 0.03628 | 0.388 | 0.426 | 1 | 5 | Safb2 | -0.038 | Young |
| 0.459317 | 0.164762 | 0.071 | 0.086 | 1 | 5 | Cdkn2aip | -0.015 | Young |
| 0.45949 | 0.320719 | 0.127 | 0.111 | 1 | 5 | Ppip5k2 | 0.016 | Young |
| 0.459632 | 0.490199 | 0.239 | 0.229 | 1 | 5 | Paox | 0.01 | Young |
| 0.459679 | 0.032012 | 0.162 | 0.141 | 1 | 5 | Eif2s1 | 0.021 | Young |
| 0.459833 | 0.119851 | 0.244 | 0.224 | 1 | 5 | Pex6 | 0.02 | Young |
| 0.459957 | 0.124829 | 0.137 | 0.157 | 1 | 5 | Il10 | -0.02 | Young |
| 0.460114 | 0.592036 | 0.025 | 0.019 | 1 | 5 | Klhl25 | 0.006 | Young |
| 0.460138 | 0.339786 | 0.137 | 0.121 | 1 | 5 | Cog3 | 0.016 | Young |
| 0.460284 | 0.527468 | 0.041 | 0.032 | 1 | 5 | Tbc1d16 | 0.009 | Young |
| 0.460473 | 0.229597 | 0.393 | 0.375 | 1 | 5 | Tgif1 | 0.018 | Young |
| 0.460568 | 0.37592 | 0.071 | 0.059 | 1 | 5 | Pstk | 0.012 | Young |

| | | | | | | | | |
|---|---|---|---|---|---|---|---|---|
| 0.460614 | 0.119436 | 0.48 | 0.461 | 1 | 5 | Arhgef2 | 0.019 | Young |
| 0.460668 | 0.440017 | 0.025 | 0.019 | 1 | 5 | Setd4 | 0.006 | Young |
| 0.460673 | 0.555965 | 0.086 | 0.074 | 1 | 5 | Ints12 | 0.012 | Young |
| 0.460859 | 0.562186 | 0.033 | 0.025 | 1 | 5 | Pkib | 0.008 | Young |
| 0.461222 | 0.454476 | 0.025 | 0.019 | 1 | 5 | BC052040 | 0.006 | Young |
| 0.461324 | 0.127371 | 0.157 | 0.138 | 1 | 5 | Tbc1d1 | 0.019 | Young |
| 0.46149 | 0.112877 | 0.789 | 0.81 | 1 | 5 | Ighm | -0.021 | Young |
| 0.461518 | 0.271461 | 0.203 | 0.187 | 1 | 5 | Mgat4a | 0.016 | Young |
| 0.461617 | 0.270537 | 0.043 | 0.034 | 1 | 5 | Tmco6 | 0.009 | Young |
| 0.462211 | 0.078207 | 0.264 | 0.288 | 1 | 5 | 0610012G( | -0.024 | Young |
| 0.462245 | 0.021636 | 0.736 | 0.786 | 1 | 5 | Atp5e | -0.05 | Young |
| 0.462331 | 0.443269 | 0.025 | 0.019 | 1 | 5 | Ceacam1 | 0.006 | Young |
| 0.462447 | 0.533878 | 0.041 | 0.032 | 1 | 5 | Zfp608 | 0.009 | Young |
| 0.462787 | 0.852104 | 0.033 | 0.025 | 1 | 5 | 1110065P2 | 0.008 | Young |
| 0.462908 | 0.146019 | 0.15 | 0.133 | 1 | 5 | Ruvbl1 | 0.017 | Young |
| 0.463198 | 0.129398 | 0.256 | 0.234 | 1 | 5 | Ubr4 | 0.022 | Young |
| 0.463376 | 0.192709 | 0.079 | 0.066 | 1 | 5 | Tars2 | 0.013 | Young |
| 0.464494 | 0.93329 | 0.015 | 0.01 | 1 | 5 | Frg2f1 | 0.005 | Young |
| 0.464703 | 0.051109 | 0.124 | 0.143 | 1 | 5 | Adh5 | -0.019 | Young |
| 0.465157 | 0.160915 | 0.119 | 0.103 | 1 | 5 | Clstn1 | 0.016 | Young |
| 0.465489 | 0.179929 | 0.274 | 0.247 | 1 | 5 | Mcts1 | 0.027 | Young |
| 0.465668 | 0.474002 | 0.025 | 0.019 | 1 | 5 | Fbxl8 | 0.006 | Young |
| 0.465722 | 0.309248 | 0.089 | 0.076 | 1 | 5 | Haus8 | 0.013 | Young |
| 0.465764 | 0.121561 | 0.345 | 0.328 | 1 | 5 | Cebpa | 0.017 | Young |
| 0.465932 | 0.234909 | 0.084 | 0.071 | 1 | 5 | Arhgap22 | 0.013 | Young |
| 0.465961 | 0.94677 | 0.015 | 0.01 | 1 | 5 | Dok4 | 0.005 | Young |
| 0.465961 | 0.839574 | 0.015 | 0.01 | 1 | 5 | Srd5a1 | 0.005 | Young |
| 0.465961 | 0.711656 | 0.015 | 0.01 | 1 | 5 | Taco1os | 0.005 | Young |
| 0.466226 | 0.384226 | 0.025 | 0.019 | 1 | 5 | Trim68 | 0.006 | Young |
| 0.466604 | 0.098274 | 0.393 | 0.36 | 1 | 5 | Slc25a37 | 0.033 | Young |
| 0.466783 | 0.482225 | 0.025 | 0.019 | 1 | 5 | Znrd1as | 0.006 | Young |
| 0.466783 | 0.478001 | 0.025 | 0.019 | 1 | 5 | Osgepl1 | 0.006 | Young |
| 0.466791 | 0.006149 | 0.074 | 0.088 | 1 | 5 | Zfp654 | -0.014 | Young |
| 0.467118 | 0.301337 | 0.104 | 0.091 | 1 | 5 | Gtf3c3 | 0.013 | Young |
| 0.467391 | 0.006836 | 0.246 | 0.266 | 1 | 5 | Tspan4 | -0.02 | Young |
| 0.46743 | 0.483856 | 0.015 | 0.01 | 1 | 5 | Fam120ao: | 0.005 | Young |
| 0.467628 | 0.634933 | 0.033 | 0.025 | 1 | 5 | Nup43 | 0.008 | Young |
| 0.467792 | 0.322916 | 0.19 | 0.175 | 1 | 5 | Fblim1 | 0.015 | Young |
| 0.468073 | 0.369995 | 0.129 | 0.114 | 1 | 5 | Glrx2 | 0.015 | Young |
| 0.468166 | 0.943359 | 0.015 | 0.01 | 1 | 5 | Mettl15 | 0.005 | Young |
| 0.468329 | 0.155442 | 0.261 | 0.239 | 1 | 5 | Rnf2 | 0.022 | Young |
| 0.468812 | 0.181353 | 0.127 | 0.111 | 1 | 5 | Hax1 | 0.016 | Young |
| 0.468902 | 0.816985 | 0.015 | 0.01 | 1 | 5 | Zfp169 | 0.005 | Young |
| 0.469078 | 0.596009 | 0.061 | 0.051 | 1 | 5 | Wdr47 | 0.01 | Young |
| 0.469086 | 0.349307 | 0.033 | 0.025 | 1 | 5 | Kdm4b | 0.008 | Young |

| | | | | | | | | |
|---|---|---|---|---|---|---|---|---|
| 0.469162 | 0.373326 | 0.053 | 0.044 | 1 | 5 | Rpp30 | 0.009 | Young |
| 0.469515 | 0.124921 | 0.086 | 0.072 | 1 | 5 | Ctu2 | 0.014 | Young |
| 0.469639 | 0.509097 | 0.015 | 0.01 | 1 | 5 | Ildr1 | 0.005 | Young |
| 0.46968 | 0.063114 | 0.061 | 0.074 | 1 | 5 | Lncpint | -0.013 | Young |
| 0.469747 | 0.086692 | 0.142 | 0.125 | 1 | 5 | Phrf1 | 0.017 | Young |
| 0.469841 | 0.229831 | 0.437 | 0.419 | 1 | 5 | Slc7a8 | 0.018 | Young |
| 0.469929 | 0.322202 | 0.198 | 0.178 | 1 | 5 | Ifi209 | 0.02 | Young |
| 0.470075 | 0.22548 | 0.056 | 0.045 | 1 | 5 | Tsix | 0.011 | Young |
| 0.470281 | 0.142955 | 0.358 | 0.34 | 1 | 5 | Lypla2 | 0.018 | Young |
| 0.470377 | 0.367264 | 0.015 | 0.01 | 1 | 5 | Mmachc | 0.005 | Young |
| 0.470389 | 0.446536 | 0.069 | 0.057 | 1 | 5 | Rundc1 | 0.012 | Young |
| 0.4704 | 0.293158 | 0.15 | 0.135 | 1 | 5 | Dirc2 | 0.015 | Young |
| 0.470713 | 0.265965 | 0.206 | 0.192 | 1 | 5 | Coa3 | 0.014 | Young |
| 0.470719 | 0.202659 | 0.14 | 0.123 | 1 | 5 | Lemd3 | 0.017 | Young |
| 0.47074 | 0.210185 | 0.107 | 0.093 | 1 | 5 | Taf13 | 0.014 | Young |
| 0.470987 | 0.229476 | 0.137 | 0.12 | 1 | 5 | Ccdc134 | 0.017 | Young |
| 0.471115 | 0.438004 | 0.015 | 0.01 | 1 | 5 | Gm31410 | 0.005 | Young |
| 0.471115 | 0.403884 | 0.015 | 0.01 | 1 | 5 | Rcn1 | 0.005 | Young |
| 0.471115 | 0.397904 | 0.015 | 0.01 | 1 | 5 | Il2 | 0.005 | Young |
| 0.471115 | 0.155437 | 0.015 | 0.01 | 1 | 5 | Rpp38 | 0.005 | Young |
| 0.471231 | 0.106354 | 0.254 | 0.234 | 1 | 5 | Polr2e | 0.02 | Young |
| 0.471344 | 0.286173 | 0.114 | 0.099 | 1 | 5 | Tmem60 | 0.015 | Young |
| 0.471714 | 0.182569 | 0.084 | 0.071 | 1 | 5 | Isca1 | 0.013 | Young |
| 0.471854 | 0.399567 | 0.015 | 0.01 | 1 | 5 | Ydjc | 0.005 | Young |
| 0.472314 | 0.180228 | 0.211 | 0.19 | 1 | 5 | Scamp1 | 0.021 | Young |
| 0.472536 | 0.046999 | 0.34 | 0.308 | 1 | 5 | Plekha2 | 0.032 | Young |
| 0.472985 | 0.130631 | 0.482 | 0.466 | 1 | 5 | Txn2 | 0.016 | Young |
| 0.473334 | 0.274102 | 0.015 | 0.01 | 1 | 5 | Ddit4 | 0.005 | Young |
| 0.473804 | 0.262965 | 0.056 | 0.045 | 1 | 5 | Nek4 | 0.011 | Young |
| 0.473844 | 0.179181 | 0.546 | 0.557 | 1 | 5 | Dync1i2 | -0.011 | Young |
| 0.473982 | 0.492147 | 0.074 | 0.062 | 1 | 5 | Atpaf1 | 0.012 | Young |
| 0.474074 | 0.641055 | 0.015 | 0.01 | 1 | 5 | Itih5 | 0.005 | Young |
| 0.474074 | 0.385331 | 0.015 | 0.01 | 1 | 5 | Fcho1 | 0.005 | Young |
| 0.474074 | 0.35222 | 0.015 | 0.01 | 1 | 5 | 4930518I1! | 0.005 | Young |
| 0.47443 | 0.127163 | 0.553 | 0.557 | 1 | 5 | Crebbp | -0.004 | Young |
| 0.474816 | 0.82163 | 0.015 | 0.01 | 1 | 5 | Gm19951 | 0.005 | Young |
| 0.474816 | 0.258799 | 0.015 | 0.01 | 1 | 5 | Rita1 | 0.005 | Young |
| 0.475347 | 0.177598 | 0.254 | 0.229 | 1 | 5 | Wipf2 | 0.025 | Young |
| 0.475419 | 0.169747 | 0.211 | 0.192 | 1 | 5 | Pnrc2 | 0.019 | Young |
| 0.475431 | 0.578017 | 0.033 | 0.025 | 1 | 5 | 9430015G1 | 0.008 | Young |
| 0.475558 | 0.245538 | 0.015 | 0.01 | 1 | 5 | Pard6g | 0.005 | Young |
| 0.475558 | 0.225081 | 0.015 | 0.01 | 1 | 5 | Rnls | 0.005 | Young |
| 0.475558 | 0.199286 | 0.015 | 0.01 | 1 | 5 | Ptar1 | 0.005 | Young |
| 0.475735 | 0.013628 | 0.703 | 0.668 | 1 | 5 | Arhgap18 | 0.035 | Young |
| 0.476301 | 0.230684 | 0.015 | 0.01 | 1 | 5 | AU022793 | 0.005 | Young |

| | | | | | | | | |
|---|---|---|---|---|---|---|---|---|
| 0.476623 | 0.200713 | 0.114 | 0.099 | 1 | 5 | Cetn2 | 0.015 | Young |
| 0.476929 | 0.342964 | 0.297 | 0.283 | 1 | 5 | Pcm1 | 0.014 | Young |
| 0.47728 | 0.32516 | 0.061 | 0.051 | 1 | 5 | Spsb2 | 0.01 | Young |
| 0.47734 | 0.094416 | 0.787 | 0.79 | 1 | 5 | Pea15a | -0.003 | Young |
| 0.477455 | 0.403634 | 0.16 | 0.145 | 1 | 5 | Pqlc1 | 0.015 | Young |
| 0.477788 | 0.145594 | 0.015 | 0.01 | 1 | 5 | Gm34342 | 0.005 | Young |
| 0.478022 | 0.020417 | 0.162 | 0.143 | 1 | 5 | Ipo5 | 0.019 | Young |
| 0.478142 | 0.239396 | 0.404 | 0.394 | 1 | 5 | Cct4 | 0.01 | Young |
| 0.478211 | 0.166054 | 0.393 | 0.374 | 1 | 5 | Ids | 0.019 | Young |
| 0.478433 | 0.026285 | 0.234 | 0.259 | 1 | 5 | Fam32a | -0.025 | Young |
| 0.478504 | 0.173676 | 0.048 | 0.039 | 1 | 5 | Tatdn3 | 0.009 | Young |
| 0.478694 | 0.115261 | 0.299 | 0.278 | 1 | 5 | Nsfl1c | 0.021 | Young |
| 0.47874 | 0.241957 | 0.173 | 0.157 | 1 | 5 | Vps54 | 0.016 | Young |
| 0.47897 | 0.102017 | 0.063 | 0.076 | 1 | 5 | Agpat4 | -0.013 | Young |
| 0.479018 | 0.366302 | 0.058 | 0.049 | 1 | 5 | Fam160a2 | 0.009 | Young |
| 0.479397 | 0.380401 | 0.135 | 0.121 | 1 | 5 | Ccdc9 | 0.014 | Young |
| 0.479503 | 0.375759 | 0.157 | 0.145 | 1 | 5 | Klhl28 | 0.012 | Young |
| 0.479638 | 0.462826 | 0.071 | 0.061 | 1 | 5 | Rassf8 | 0.01 | Young |
| 0.479765 | 0.061558 | 0.858 | 0.909 | 1 | 5 | Gabarap | -0.051 | Young |
| 0.480024 | 0.069946 | 0.015 | 0.01 | 1 | 5 | Grk4 | 0.005 | Young |
| 0.480415 | 0.234137 | 0.119 | 0.104 | 1 | 5 | Epm2aip1 | 0.015 | Young |
| 0.480728 | 0.044644 | 0.282 | 0.259 | 1 | 5 | U2af1 | 0.023 | Young |
| 0.480741 | 0.081249 | 0.244 | 0.273 | 1 | 5 | Parp8 | -0.029 | Young |
| 0.481081 | 0.119089 | 0.063 | 0.052 | 1 | 5 | Rint1 | 0.011 | Young |
| 0.481112 | 0.01964 | 0.289 | 0.31 | 1 | 5 | Cul1 | -0.021 | Young |
| 0.481395 | 0.272406 | 0.165 | 0.15 | 1 | 5 | Parp4 | 0.015 | Young |
| 0.482073 | 0.097515 | 0.013 | 0.019 | 1 | 5 | Krtcap3 | -0.006 | Young |
| 0.482073 | 0.016271 | 0.013 | 0.019 | 1 | 5 | Vps51 | -0.006 | Young |
| 0.482108 | 0.026462 | 0.952 | 0.963 | 1 | 5 | Rps11 | -0.011 | Young |
| 0.482317 | 0.132591 | 0.033 | 0.025 | 1 | 5 | Dnajc28 | 0.008 | Young |
| 0.482526 | 0.317667 | 0.076 | 0.064 | 1 | 5 | Snhg20 | 0.012 | Young |
| 0.482917 | 0.392223 | 0.066 | 0.056 | 1 | 5 | Wrap73 | 0.01 | Young |
| 0.483241 | 0.014137 | 0.117 | 0.135 | 1 | 5 | Sos1 | -0.018 | Young |
| 0.483367 | 0.108921 | 0.647 | 0.653 | 1 | 5 | Fcho2 | -0.006 | Young |
| 0.483537 | 0.284929 | 0.094 | 0.081 | 1 | 5 | Gyg | 0.013 | Young |
| 0.483567 | 0.105336 | 0.089 | 0.076 | 1 | 5 | Setd1a | 0.013 | Young |
| 0.483586 | 0.831658 | 0.051 | 0.042 | 1 | 5 | Rnaseh2b | 0.009 | Young |
| 0.483681 | 0.322571 | 0.094 | 0.082 | 1 | 5 | Serinc5 | 0.012 | Young |
| 0.483883 | 0.230667 | 0.317 | 0.303 | 1 | 5 | Pgk1 | 0.014 | Young |
| 0.48389 | 0.069 | 0.109 | 0.094 | 1 | 5 | Ptrhd1 | 0.015 | Young |
| 0.483914 | 0.178311 | 0.104 | 0.091 | 1 | 5 | Hps1 | 0.013 | Young |
| 0.484022 | 0.124573 | 0.013 | 0.019 | 1 | 5 | Ahi1 | -0.006 | Young |
| 0.484149 | 0.433704 | 0.058 | 0.049 | 1 | 5 | Pcyox1l | 0.009 | Young |
| 0.484229 | 0.093347 | 0.452 | 0.428 | 1 | 5 | Il13ra1 | 0.024 | Young |
| 0.484422 | 0.814704 | 0.03 | 0.024 | 1 | 5 | Slc37a1 | 0.006 | Young |

| | | | | | | | | |
|---|---|---|---|---|---|---|---|---|
| 0.484607 | 0.248595 | 0.234 | 0.219 | 1 | 5 | Gpat4 | 0.015 | Young |
| 0.484898 | 0.141614 | 0.033 | 0.042 | 1 | 5 | Clec4e | -0.009 | Young |
| 0.484934 | 0.106566 | 0.619 | 0.62 | 1 | 5 | Atp6v1f | -0.001 | Young |
| 0.485496 | 0.264097 | 0.096 | 0.084 | 1 | 5 | Faap100 | 0.012 | Young |
| 0.485559 | 0.282889 | 0.061 | 0.051 | 1 | 5 | Timm8a1 | 0.01 | Young |
| 0.485723 | 0.050552 | 0.244 | 0.268 | 1 | 5 | Ncoa7 | -0.024 | Young |
| 0.486072 | 0.25968 | 0.373 | 0.36 | 1 | 5 | Ndufb5 | 0.013 | Young |
| 0.486125 | 0.232998 | 0.089 | 0.076 | 1 | 5 | Tmem80 | 0.013 | Young |
| 0.486162 | 0.384865 | 0.152 | 0.14 | 1 | 5 | Gpr155 | 0.012 | Young |
| 0.48658 | 0.229921 | 0.203 | 0.189 | 1 | 5 | Uba5 | 0.014 | Young |
| 0.486609 | 0.016038 | 0.033 | 0.042 | 1 | 5 | Tpgs2 | -0.009 | Young |
| 0.486921 | 0.107706 | 0.287 | 0.268 | 1 | 5 | Phactr2 | 0.019 | Young |
| 0.487685 | 0.116681 | 0.452 | 0.495 | 1 | 5 | Kif1b | -0.043 | Young |
| 0.48779 | 0.132005 | 0.282 | 0.31 | 1 | 5 | Cdc5l | -0.028 | Young |
| 0.488116 | 0.018957 | 0.266 | 0.242 | 1 | 5 | Bax | 0.024 | Young |
| 0.488375 | 0.285552 | 0.079 | 0.067 | 1 | 5 | Mcm4 | 0.012 | Young |
| 0.488501 | 0.042328 | 0.071 | 0.084 | 1 | 5 | Stoml1 | -0.013 | Young |
| 0.488647 | 0.552219 | 0.046 | 0.037 | 1 | 5 | Pcnx4 | 0.009 | Young |
| 0.488768 | 0.226054 | 0.236 | 0.222 | 1 | 5 | Pcmtd2 | 0.014 | Young |
| 0.489312 | 0.26773 | 0.269 | 0.256 | 1 | 5 | Srp72 | 0.013 | Young |
| 0.489818 | 0.116627 | 0.386 | 0.37 | 1 | 5 | Tmx3 | 0.016 | Young |
| 0.490004 | 0.082565 | 0.365 | 0.389 | 1 | 5 | Ap2b1 | -0.024 | Young |
| 0.49055 | 0.065264 | 0.013 | 0.019 | 1 | 5 | Aldh1a2 | -0.006 | Young |
| 0.490908 | 0.001311 | 0.462 | 0.495 | 1 | 5 | Pla2g15 | -0.033 | Young |
| 0.491295 | 0.224452 | 0.155 | 0.14 | 1 | 5 | Nlk | 0.015 | Young |
| 0.491456 | 0.024971 | 0.063 | 0.076 | 1 | 5 | Cep85l | -0.013 | Young |
| 0.491748 | 0.070956 | 0.033 | 0.025 | 1 | 5 | Zfp420 | 0.008 | Young |
| 0.491965 | 0.155296 | 0.041 | 0.032 | 1 | 5 | Eif2d | 0.009 | Young |
| 0.492002 | 0.346726 | 0.046 | 0.037 | 1 | 5 | Zfp934 | 0.009 | Young |
| 0.492029 | 0.492138 | 0.117 | 0.104 | 1 | 5 | Ino80c | 0.013 | Young |
| 0.492032 | 0.285389 | 0.074 | 0.062 | 1 | 5 | Cdc23 | 0.012 | Young |
| 0.492152 | 0.927368 | 0.03 | 0.024 | 1 | 5 | Gm36266 | 0.006 | Young |
| 0.492597 | 0.480548 | 0.053 | 0.044 | 1 | 5 | Phactr1 | 0.009 | Young |
| 0.492689 | 0.041027 | 0.091 | 0.077 | 1 | 5 | Armc10 | 0.014 | Young |
| 0.492878 | 0.175527 | 0.393 | 0.379 | 1 | 5 | Pigt | 0.014 | Young |
| 0.493204 | 0.249863 | 0.122 | 0.108 | 1 | 5 | Sh3glb2 | 0.014 | Young |
| 0.493375 | 0.29509 | 0.053 | 0.044 | 1 | 5 | Rabep2 | 0.009 | Young |
| 0.493443 | 0.638577 | 0.023 | 0.017 | 1 | 5 | 1110046J0 | 0.006 | Young |
| 0.493564 | 0.362731 | 0.124 | 0.111 | 1 | 5 | Atxn1l | 0.013 | Young |
| 0.493669 | 0.127427 | 0.734 | 0.736 | 1 | 5 | Mrfap1 | -0.002 | Young |
| 0.493967 | 0.053492 | 0.18 | 0.158 | 1 | 5 | Elp2 | 0.022 | Young |
| 0.494224 | 0.644233 | 0.03 | 0.024 | 1 | 5 | Nnt | 0.006 | Young |
| 0.494241 | 0.189566 | 0.122 | 0.106 | 1 | 5 | Dera | 0.016 | Young |
| 0.494441 | 0.011799 | 0.023 | 0.03 | 1 | 5 | Sirt4 | -0.007 | Young |
| 0.49459 | 0.057831 | 0.137 | 0.121 | 1 | 5 | Snap29 | 0.016 | Young |

| | | | | | | | | |
|---|---|---|---|---|---|---|---|---|
| 0.494649 | 0.206386 | 0.307 | 0.291 | 1 | 5 | Mbd2 | 0.016 | Young |
| 0.495169 | 0.197662 | 0.424 | 0.416 | 1 | 5 | Tmem55b | 0.008 | Young |
| 0.495454 | 0.051497 | 0.17 | 0.189 | 1 | 5 | Adar | -0.019 | Young |
| 0.495661 | 0.053963 | 0.477 | 0.446 | 1 | 5 | Tmem123 | 0.031 | Young |
| 0.496468 | 0.877144 | 0.023 | 0.017 | 1 | 5 | Procr | 0.006 | Young |
| 0.496767 | 0.15431 | 0.437 | 0.416 | 1 | 5 | Usp25 | 0.021 | Young |
| 0.496821 | 0.752648 | 0.03 | 0.024 | 1 | 5 | Homer2 | 0.006 | Young |
| 0.496821 | 0.675319 | 0.03 | 0.024 | 1 | 5 | Atad5 | 0.006 | Young |
| 0.496929 | 0.314158 | 0.107 | 0.094 | 1 | 5 | Foxk2 | 0.013 | Young |
| 0.497032 | 0.313757 | 0.178 | 0.162 | 1 | 5 | Ccdc186 | 0.016 | Young |
| 0.497058 | 0.533139 | 0.046 | 0.037 | 1 | 5 | Tdp1 | 0.009 | Young |
| 0.497075 | 0.48601 | 0.023 | 0.017 | 1 | 5 | Hdac8 | 0.006 | Young |
| 0.497342 | 0.123325 | 0.254 | 0.279 | 1 | 5 | Rapgef2 | -0.025 | Young |
| 0.497419 | 0.205527 | 0.195 | 0.217 | 1 | 5 | Slc5a3 | -0.022 | Young |
| 0.498013 | 0.122715 | 0.023 | 0.03 | 1 | 5 | Meis1 | -0.007 | Young |
| 0.49804 | 0.016038 | 0.188 | 0.167 | 1 | 5 | Zfp280d | 0.021 | Young |
| 0.498098 | 0.151305 | 0.17 | 0.152 | 1 | 5 | Jkamp | 0.018 | Young |
| 0.498196 | 0.20736 | 0.228 | 0.209 | 1 | 5 | Znhit1 | 0.019 | Young |
| 0.49824 | 0.348794 | 0.033 | 0.042 | 1 | 5 | Rab11fip1 | -0.009 | Young |
| 0.498288 | 0.521404 | 0.023 | 0.017 | 1 | 5 | 1110008L1 | 0.006 | Young |
| 0.498474 | 0.242886 | 0.175 | 0.158 | 1 | 5 | Fam120b | 0.017 | Young |
| 0.498663 | 0.296473 | 0.322 | 0.323 | 1 | 5 | Brd1 | -0.001 | Young |
| 0.49871 | 0.409414 | 0.119 | 0.106 | 1 | 5 | Pbdc1 | 0.013 | Young |
| 0.498853 | 0.002135 | 0.487 | 0.519 | 1 | 5 | Ube3a | -0.032 | Young |
| 0.498904 | 0.442652 | 0.03 | 0.024 | 1 | 5 | Ccdc73 | 0.006 | Young |
| 0.498907 | 0.231615 | 0.47 | 0.46 | 1 | 5 | Nxf1 | 0.01 | Young |
| 0.498946 | 0.195298 | 0.231 | 0.217 | 1 | 5 | Fam76a | 0.014 | Young |
| 0.498995 | 0.020418 | 0.089 | 0.104 | 1 | 5 | Slc30a4 | -0.015 | Young |
| 0.49903 | 0.108851 | 0.52 | 0.503 | 1 | 5 | Eif5 | 0.017 | Young |
| 0.49926 | 0.143132 | 0.14 | 0.123 | 1 | 5 | Sumf1 | 0.017 | Young |
| 0.499327 | 0.377223 | 0.226 | 0.214 | 1 | 5 | Eif2ak2 | 0.012 | Young |
| 0.499595 | 0.197349 | 0.421 | 0.418 | 1 | 5 | Kpnb1 | 0.003 | Young |
| 0.499668 | 0.0319 | 0.949 | 0.976 | 1 | 5 | Gpx1 | -0.027 | Young |
| 0.499754 | 0.084237 | 0.414 | 0.396 | 1 | 5 | Baz1b | 0.018 | Young |
| 0.499912 | 0.263666 | 0.208 | 0.19 | 1 | 5 | Vps11 | 0.018 | Young |
| 0.500317 | 0.272345 | 0.327 | 0.313 | 1 | 5 | Rbm6 | 0.014 | Young |
| 0.50032 | 0.255096 | 0.307 | 0.301 | 1 | 5 | Eif3b | 0.006 | Young |
| 0.500573 | 0.043783 | 0.447 | 0.473 | 1 | 5 | Degs1 | -0.026 | Young |
| 0.50072 | 0.48299 | 0.023 | 0.017 | 1 | 5 | Fkbp14 | 0.006 | Young |
| 0.50072 | 0.385821 | 0.023 | 0.017 | 1 | 5 | H2-DMb2 | 0.006 | Young |
| 0.500732 | 0.149409 | 0.162 | 0.146 | 1 | 5 | Golga2 | 0.016 | Young |
| 0.500836 | 0.486274 | 0.102 | 0.091 | 1 | 5 | Timm29 | 0.011 | Young |
| 0.500934 | 0.178164 | 0.17 | 0.153 | 1 | 5 | Csnk1g1 | 0.017 | Young |
| 0.500936 | 0.226461 | 0.482 | 0.476 | 1 | 5 | Ptpn1 | 0.006 | Young |
| 0.501329 | 0.527155 | 0.023 | 0.017 | 1 | 5 | Nckap5l | 0.006 | Young |

| | | | | | | | | |
|---|---|---|---|---|---|---|---|---|
| 0.501488 | 0.202599 | 0.135 | 0.118 | 1 | 5 | Nelfa | 0.017 | Young |
| 0.502153 | 0.89587 | 0.013 | 0.008 | 1 | 5 | Rnd2 | 0.005 | Young |
| 0.502223 | 0.406418 | 0.091 | 0.081 | 1 | 5 | Zfp407 | 0.01 | Young |
| 0.502252 | 0.130696 | 0.548 | 0.545 | 1 | 5 | Fam129a | 0.003 | Young |
| 0.502568 | 0.086562 | 0.137 | 0.121 | 1 | 5 | Esco1 | 0.016 | Young |
| 0.502678 | 0.599987 | 0.081 | 0.071 | 1 | 5 | Dnlz | 0.01 | Young |
| 0.502784 | 0.164 | 0.099 | 0.086 | 1 | 5 | Mrps5 | 0.013 | Young |
| 0.502821 | 0.468288 | 0.069 | 0.059 | 1 | 5 | Tdrkh | 0.01 | Young |
| 0.502895 | 0.102845 | 0.091 | 0.079 | 1 | 5 | Zkscan1 | 0.012 | Young |
| 0.502912 | 0.181373 | 0.376 | 0.359 | 1 | 5 | Naa50 | 0.017 | Young |
| 0.50299 | 0.652162 | 0.013 | 0.008 | 1 | 5 | Rab26os | 0.005 | Young |
| 0.503071 | 0.150632 | 0.117 | 0.103 | 1 | 5 | Rab33b | 0.014 | Young |
| 0.503095 | 0.072475 | 0.048 | 0.039 | 1 | 5 | Prdm11 | 0.009 | Young |
| 0.503427 | 0.140535 | 0.241 | 0.222 | 1 | 5 | Klf9 | 0.019 | Young |
| 0.503558 | 0.178808 | 0.147 | 0.131 | 1 | 5 | Vars | 0.016 | Young |
| 0.503822 | 0.037252 | 0.061 | 0.051 | 1 | 5 | Otud1 | 0.01 | Young |
| 0.504127 | 0.245571 | 0.261 | 0.244 | 1 | 5 | Csnk1g3 | 0.017 | Young |
| 0.504162 | 0.293306 | 0.211 | 0.195 | 1 | 5 | Aida | 0.016 | Young |
| 0.504288 | 0.063564 | 0.373 | 0.406 | 1 | 5 | Rab21 | -0.033 | Young |
| 0.504396 | 0.064128 | 0.099 | 0.113 | 1 | 5 | Dvl2 | -0.014 | Young |
| 0.504802 | 0.141692 | 0.302 | 0.283 | 1 | 5 | Bscl2 | 0.019 | Young |
| 0.505505 | 0.576242 | 0.013 | 0.008 | 1 | 5 | Zfp773 | 0.005 | Young |
| 0.505705 | 0.481571 | 0.03 | 0.024 | 1 | 5 | Tcaim | 0.006 | Young |
| 0.505877 | 0.193138 | 0.396 | 0.382 | 1 | 5 | Eloc | 0.014 | Young |
| 0.50623 | 0.623745 | 0.03 | 0.024 | 1 | 5 | Aplnr | 0.006 | Young |
| 0.506336 | 0.226497 | 0.038 | 0.03 | 1 | 5 | Stn1 | 0.008 | Young |
| 0.506345 | 0.943126 | 0.013 | 0.008 | 1 | 5 | Gm42595 | 0.005 | Young |
| 0.506345 | 0.7048 | 0.013 | 0.008 | 1 | 5 | Tmem202 | 0.005 | Young |
| 0.506472 | 0.272861 | 0.183 | 0.17 | 1 | 5 | Trappc10 | 0.013 | Young |
| 0.506569 | 0.075667 | 0.071 | 0.061 | 1 | 5 | Gm20342 | 0.01 | Young |
| 0.506686 | 0.412084 | 0.185 | 0.177 | 1 | 5 | Psen2 | 0.008 | Young |
| 0.506828 | 0.326306 | 0.023 | 0.017 | 1 | 5 | Tha1 | 0.006 | Young |
| 0.507186 | 0.978272 | 0.013 | 0.008 | 1 | 5 | Gm26714 | 0.005 | Young |
| 0.507186 | 0.812593 | 0.013 | 0.008 | 1 | 5 | Hp | 0.005 | Young |
| 0.507186 | 0.792027 | 0.013 | 0.008 | 1 | 5 | Stac3 | 0.005 | Young |
| 0.50749 | 0.070455 | 0.01 | 0.015 | 1 | 5 | 1700056E2 | -0.005 | Young |
| 0.507635 | 0.236385 | 0.18 | 0.167 | 1 | 5 | Tmem251 | 0.013 | Young |
| 0.507745 | 0.389891 | 0.074 | 0.062 | 1 | 5 | Dhcr7 | 0.012 | Young |
| 0.507757 | 0.033252 | 0.168 | 0.19 | 1 | 5 | Nup153 | -0.022 | Young |
| 0.507841 | 0.00439 | 0.056 | 0.067 | 1 | 5 | Ch25h | -0.011 | Young |
| 0.50819 | 0.204018 | 0.091 | 0.079 | 1 | 5 | Rbl1 | 0.012 | Young |
| 0.508229 | 0.016449 | 0.01 | 0.015 | 1 | 5 | Gm16062 | -0.005 | Young |
| 0.508478 | 0.266547 | 0.297 | 0.285 | 1 | 5 | Dhx9 | 0.012 | Young |
| 0.508506 | 0.284084 | 0.129 | 0.116 | 1 | 5 | Dzip3 | 0.013 | Young |
| 0.508811 | 0.064915 | 0.069 | 0.081 | 1 | 5 | Zfp579 | -0.012 | Young |

| | | | | | | | | |
|---|---|---|---|---|---|---|---|---|
| 0.50886 | 0.483607 | 0.03 | 0.024 | 1 | 5 | Pex11g | 0.006 | Young |
| 0.508869 | 0.675856 | 0.013 | 0.008 | 1 | 5 | Eri2 | 0.005 | Young |
| 0.5093 | 0.157667 | 0.19 | 0.173 | 1 | 5 | Ndufa9 | 0.017 | Young |
| 0.509678 | 0.237596 | 0.261 | 0.244 | 1 | 5 | Lpcat1 | 0.017 | Young |
| 0.509712 | 0.608489 | 0.013 | 0.008 | 1 | 5 | Gm16867 | 0.005 | Young |
| 0.509712 | 0.484088 | 0.013 | 0.008 | 1 | 5 | Mdga1 | 0.005 | Young |
| 0.510056 | 0.220242 | 0.317 | 0.303 | 1 | 5 | Irf9 | 0.014 | Young |
| 0.510219 | 0.158788 | 0.178 | 0.158 | 1 | 5 | Tor1a | 0.02 | Young |
| 0.510511 | 0.318659 | 0.023 | 0.017 | 1 | 5 | Fuk | 0.006 | Young |
| 0.510577 | 0.092462 | 0.081 | 0.069 | 1 | 5 | Ruvbl2 | 0.012 | Young |
| 0.511102 | 0.292377 | 0.046 | 0.037 | 1 | 5 | Fzd7 | 0.009 | Young |
| 0.511133 | 0.381469 | 0.104 | 0.093 | 1 | 5 | Arhgef7 | 0.011 | Young |
| 0.511317 | 0.305554 | 0.076 | 0.066 | 1 | 5 | Mvb12b | 0.01 | Young |
| 0.51136 | 0.0515 | 0.761 | 0.758 | 1 | 5 | Gnb1 | 0.003 | Young |
| 0.5114 | 0.655168 | 0.013 | 0.008 | 1 | 5 | Bmp1 | 0.005 | Young |
| 0.5114 | 0.649747 | 0.013 | 0.008 | 1 | 5 | Aldoc | 0.005 | Young |
| 0.5114 | 0.419919 | 0.013 | 0.008 | 1 | 5 | Zfp455 | 0.005 | Young |
| 0.5114 | 0.357806 | 0.013 | 0.008 | 1 | 5 | Sdc1 | 0.005 | Young |
| 0.51151 | 0.286809 | 0.17 | 0.155 | 1 | 5 | Mcm3 | 0.015 | Young |
| 0.511777 | 0.159161 | 0.421 | 0.412 | 1 | 5 | Nsd1 | 0.009 | Young |
| 0.511783 | 0.072894 | 0.162 | 0.146 | 1 | 5 | Dgkh | 0.016 | Young |
| 0.511803 | 0.005901 | 0.358 | 0.379 | 1 | 5 | Rgs18 | -0.021 | Young |
| 0.512094 | 0.091457 | 0.028 | 0.035 | 1 | 5 | Zfp40 | -0.007 | Young |
| 0.512229 | 0.222013 | 0.404 | 0.406 | 1 | 5 | B4galt1 | -0.002 | Young |
| 0.512245 | 0.621251 | 0.013 | 0.008 | 1 | 5 | Nlrp10 | 0.005 | Young |
| 0.512245 | 0.355991 | 0.013 | 0.008 | 1 | 5 | 4732471J0 | 0.005 | Young |
| 0.512364 | 0.227294 | 0.292 | 0.278 | 1 | 5 | Cebpz | 0.014 | Young |
| 0.512559 | 0.26899 | 0.155 | 0.14 | 1 | 5 | Cul2 | 0.015 | Young |
| 0.513016 | 0.158378 | 0.254 | 0.236 | 1 | 5 | Ide | 0.018 | Young |
| 0.513021 | 0.199999 | 0.119 | 0.104 | 1 | 5 | Trim3 | 0.015 | Young |
| 0.51309 | 0.544095 | 0.013 | 0.008 | 1 | 5 | Ppm1l | 0.005 | Young |
| 0.51309 | 0.376462 | 0.013 | 0.008 | 1 | 5 | Dmpk | 0.005 | Young |
| 0.51309 | 0.31721 | 0.013 | 0.008 | 1 | 5 | Ccdc80 | 0.005 | Young |
| 0.51309 | 0.312208 | 0.013 | 0.008 | 1 | 5 | Mss51 | 0.005 | Young |
| 0.513418 | 0.051228 | 0.193 | 0.21 | 1 | 5 | Zfp90 | -0.017 | Young |
| 0.513529 | 0.259767 | 0.195 | 0.18 | 1 | 5 | Cab39l | 0.015 | Young |
| 0.513663 | 0.183107 | 0.343 | 0.338 | 1 | 5 | Copb2 | 0.005 | Young |
| 0.513822 | 0.042876 | 0.124 | 0.138 | 1 | 5 | Matk | -0.014 | Young |
| 0.513937 | 0.276382 | 0.013 | 0.008 | 1 | 5 | Sh2d1b1 | 0.005 | Young |
| 0.514056 | 0.029704 | 0.472 | 0.449 | 1 | 5 | Gatad2b | 0.023 | Young |
| 0.514325 | 0.147526 | 0.038 | 0.03 | 1 | 5 | Tada2a | 0.008 | Young |
| 0.514353 | 0.131502 | 0.269 | 0.254 | 1 | 5 | Yme1l1 | 0.015 | Young |
| 0.514514 | 0.117268 | 0.244 | 0.226 | 1 | 5 | Slc14a1 | 0.018 | Young |
| 0.514695 | 0.185051 | 0.178 | 0.162 | 1 | 5 | Prdx6 | 0.016 | Young |
| 0.514739 | 0.182716 | 0.297 | 0.286 | 1 | 5 | Pank3 | 0.011 | Young |

| | | | | | | | | |
|---|---|---|---|---|---|---|---|---|
| 0.514784 | 0.411202 | 0.013 | 0.008 | 1 | 5 | Dnhd1 | 0.005 | Young |
| 0.514784 | 0.269494 | 0.013 | 0.008 | 1 | 5 | Pidd1 | 0.005 | Young |
| 0.515131 | 0.091152 | 0.802 | 0.798 | 1 | 5 | Pmp22 | 0.004 | Young |
| 0.515201 | 0.207506 | 0.03 | 0.024 | 1 | 5 | Slc25a26 | 0.006 | Young |
| 0.515399 | 0.105796 | 0.046 | 0.037 | 1 | 5 | Mtg1 | 0.009 | Young |
| 0.515443 | 0.00178 | 0.023 | 0.017 | 1 | 5 | Zfp873 | 0.006 | Young |
| 0.515632 | 0.328258 | 0.013 | 0.008 | 1 | 5 | Gm17435 | 0.005 | Young |
| 0.515632 | 0.143716 | 0.013 | 0.008 | 1 | 5 | Fancf | 0.005 | Young |
| 0.515746 | 0.153255 | 0.368 | 0.355 | 1 | 5 | Tns3 | 0.013 | Young |
| 0.515804 | 0.352002 | 0.081 | 0.071 | 1 | 5 | Pole3 | 0.01 | Young |
| 0.516261 | 0.186731 | 0.114 | 0.099 | 1 | 5 | Hdgfl2 | 0.015 | Young |
| 0.516271 | 0.173505 | 0.244 | 0.229 | 1 | 5 | Vps41 | 0.015 | Young |
| 0.516273 | 0.069972 | 0.081 | 0.094 | 1 | 5 | Ttc17 | -0.013 | Young |
| 0.516419 | 0.19807 | 0.165 | 0.15 | 1 | 5 | Faap20 | 0.015 | Young |
| 0.516481 | 0.279563 | 0.013 | 0.008 | 1 | 5 | Zfp937 | 0.005 | Young |
| 0.516793 | 0.21402 | 0.03 | 0.024 | 1 | 5 | Mtfmt | 0.006 | Young |
| 0.516793 | 0.02018 | 0.03 | 0.024 | 1 | 5 | Eml6 | 0.006 | Young |
| 0.516945 | 0.356935 | 0.114 | 0.101 | 1 | 5 | Cln5 | 0.013 | Young |
| 0.517241 | 0.424615 | 0.051 | 0.042 | 1 | 5 | Nfatc2ip | 0.009 | Young |
| 0.51733 | 0.139625 | 0.013 | 0.008 | 1 | 5 | Pex10 | 0.005 | Young |
| 0.51733 | 0.060824 | 0.013 | 0.008 | 1 | 5 | Smpd5 | 0.005 | Young |
| 0.51751 | 0.020936 | 0.231 | 0.251 | 1 | 5 | Txndc9 | -0.02 | Young |
| 0.517534 | 0.124272 | 0.358 | 0.33 | 1 | 5 | Zc3h7a | 0.028 | Young |
| 0.517661 | 0.018196 | 0.335 | 0.362 | 1 | 5 | Cyb5a | -0.027 | Young |
| 0.518128 | 0.532932 | 0.079 | 0.069 | 1 | 5 | Bag5 | 0.01 | Young |
| 0.51818 | 0.118772 | 0.013 | 0.008 | 1 | 5 | Zfp273 | 0.005 | Young |
| 0.518313 | 0.358955 | 0.03 | 0.039 | 1 | 5 | L3mbtl3 | -0.009 | Young |
| 0.518387 | 0.034294 | 0.03 | 0.024 | 1 | 5 | 2610044O1 | 0.006 | Young |
| 0.518624 | 0.201219 | 0.063 | 0.054 | 1 | 5 | Hars2 | 0.009 | Young |
| 0.51886 | 0.16503 | 0.411 | 0.397 | 1 | 5 | Pds5a | 0.014 | Young |
| 0.519054 | 0.102715 | 0.038 | 0.03 | 1 | 5 | Mtmr11 | 0.008 | Young |
| 0.519667 | 0.284248 | 0.142 | 0.13 | 1 | 5 | Snapc1 | 0.012 | Young |
| 0.519687 | 0.485691 | 0.056 | 0.047 | 1 | 5 | Hddc2 | 0.009 | Young |
| 0.519696 | 0.462402 | 0.061 | 0.052 | 1 | 5 | Smim8 | 0.009 | Young |
| 0.519757 | 0.144404 | 0.551 | 0.556 | 1 | 5 | Myl12b | -0.005 | Young |
| 0.519882 | 0.128299 | 0.013 | 0.008 | 1 | 5 | Col6a2 | 0.005 | Young |
| 0.520001 | 0.025628 | 0.251 | 0.273 | 1 | 5 | Emd | -0.022 | Young |
| 0.520099 | 0.269842 | 0.051 | 0.042 | 1 | 5 | Tmem29 | 0.009 | Young |
| 0.520186 | 0.063513 | 0.168 | 0.152 | 1 | 5 | Nup62 | 0.016 | Young |
| 0.520631 | 0.180407 | 0.632 | 0.646 | 1 | 5 | Ccnd1 | -0.014 | Young |
| 0.520886 | 0.544295 | 0.036 | 0.029 | 1 | 5 | Zc3h8 | 0.007 | Young |
| 0.520958 | 0.003494 | 0.19 | 0.212 | 1 | 5 | Grk6 | -0.022 | Young |
| 0.520982 | 0.142354 | 0.497 | 0.493 | 1 | 5 | Fam111a | 0.004 | Young |
| 0.521412 | 0.037727 | 0.67 | 0.657 | 1 | 5 | 2610507B1 | 0.013 | Young |
| 0.52243 | 0.206641 | 0.261 | 0.244 | 1 | 5 | Dhrs7 | 0.017 | Young |

| | | | | | | | | |
|---|---|---|---|---|---|---|---|---|
| 0.522434 | 0.214178 | 0.142 | 0.128 | 1 | 5 | Vps52 | 0.014 | Young |
| 0.522618 | 0.164023 | 0.127 | 0.113 | 1 | 5 | Med20 | 0.014 | Young |
| 0.522767 | 0.223596 | 0.294 | 0.285 | 1 | 5 | Rala | 0.009 | Young |
| 0.522911 | 0.208284 | 0.122 | 0.109 | 1 | 5 | Tada1 | 0.013 | Young |
| 0.522921 | 0.172685 | 0.256 | 0.242 | 1 | 5 | Cyb5r3 | 0.014 | Young |
| 0.522942 | 0.010072 | 0.241 | 0.259 | 1 | 5 | Rhoq | -0.018 | Young |
| 0.523048 | 0.538888 | 0.079 | 0.069 | 1 | 5 | Etfrf1 | 0.01 | Young |
| 0.523296 | 0.013037 | 0.008 | 0.012 | 1 | 5 | Magi2 | -0.004 | Young |
| 0.52333 | 0.368632 | 0.02 | 0.027 | 1 | 5 | Ifrd2 | -0.007 | Young |
| 0.523905 | 0.085653 | 0.038 | 0.047 | 1 | 5 | Hoxb4 | -0.009 | Young |
| 0.523925 | 0.099181 | 0.066 | 0.056 | 1 | 5 | Dolpp1 | 0.01 | Young |
| 0.524002 | 0.120507 | 0.266 | 0.293 | 1 | 5 | Spty2d1 | -0.027 | Young |
| 0.524082 | 0.161562 | 0.312 | 0.296 | 1 | 5 | Prelid1 | 0.016 | Young |
| 0.524374 | 0.163839 | 0.378 | 0.369 | 1 | 5 | Rasa1 | 0.009 | Young |
| 0.524704 | 0.245171 | 0.135 | 0.121 | 1 | 5 | D730003I1 | 0.014 | Young |
| 0.525393 | 0.241203 | 0.152 | 0.138 | 1 | 5 | Fam53c | 0.014 | Young |
| 0.525465 | 0.199151 | 0.373 | 0.359 | 1 | 5 | Etv1 | 0.014 | Young |
| 0.52584 | 0.194977 | 0.051 | 0.042 | 1 | 5 | Acad12 | 0.009 | Young |
| 0.525863 | 0.007819 | 0.008 | 0.012 | 1 | 5 | Jag2 | -0.004 | Young |
| 0.526392 | 0.276208 | 0.043 | 0.035 | 1 | 5 | Bcl7a | 0.008 | Young |
| 0.526435 | 0.110321 | 0.591 | 0.593 | 1 | 5 | Lair1 | -0.002 | Young |
| 0.526752 | 0.089529 | 0.183 | 0.204 | 1 | 5 | Pstpip1 | -0.021 | Young |
| 0.526815 | 0.185026 | 0.193 | 0.177 | 1 | 5 | Stau1 | 0.016 | Young |
| 0.52682 | 0.293221 | 0.091 | 0.081 | 1 | 5 | Fam173b | 0.01 | Young |
| 0.527188 | 0.057782 | 0.284 | 0.315 | 1 | 5 | Mfsd5 | -0.031 | Young |
| 0.527341 | 0.008812 | 0.464 | 0.497 | 1 | 5 | Neu1 | -0.033 | Young |
| 0.527579 | 0.062355 | 0.008 | 0.012 | 1 | 5 | Wnt11 | -0.004 | Young |
| 0.527716 | 0.014163 | 0.256 | 0.273 | 1 | 5 | Ehmt2 | -0.017 | Young |
| 0.527817 | 0.002991 | 0.084 | 0.096 | 1 | 5 | Mbd5 | -0.012 | Young |
| 0.528414 | 0.406091 | 0.251 | 0.241 | 1 | 5 | Chchd1 | 0.01 | Young |
| 0.528833 | 0.059279 | 0.65 | 0.66 | 1 | 5 | Gls | -0.01 | Young |
| 0.528864 | 0.517154 | 0.056 | 0.047 | 1 | 5 | Zdhhc4 | 0.009 | Young |
| 0.529178 | 0.686098 | 0.02 | 0.015 | 1 | 5 | Hgh1 | 0.005 | Young |
| 0.529348 | 0.027973 | 0.393 | 0.429 | 1 | 5 | Mrpl33 | -0.036 | Young |
| 0.529372 | 0.239241 | 0.071 | 0.061 | 1 | 5 | Eci1 | 0.01 | Young |
| 0.529436 | 0.285838 | 0.086 | 0.076 | 1 | 5 | Gosr1 | 0.01 | Young |
| 0.529446 | 0.173735 | 0.49 | 0.49 | 1 | 5 | Atp6v1e1 | 0 | Young |
| 0.529446 | 0.076661 | 0.028 | 0.035 | 1 | 5 | Wdr92 | -0.007 | Young |
| 0.529712 | 0.29267 | 0.094 | 0.082 | 1 | 5 | Nip7 | 0.012 | Young |
| 0.529832 | 0.109109 | 0.216 | 0.199 | 1 | 5 | Katna1 | 0.017 | Young |
| 0.530157 | 0.420499 | 0.008 | 0.012 | 1 | 5 | Hist1h2be | -0.004 | Young |
| 0.530177 | 0.21457 | 0.256 | 0.242 | 1 | 5 | Arl5a | 0.014 | Young |
| 0.53023 | 0.482132 | 0.036 | 0.029 | 1 | 5 | Cenpj | 0.007 | Young |
| 0.530264 | 0.012881 | 0.863 | 0.892 | 1 | 5 | Ap2a2 | -0.029 | Young |
| 0.53032 | 0.218381 | 0.251 | 0.237 | 1 | 5 | Cd300c2 | 0.014 | Young |

| | | | | | | | | |
|---|---|---|---|---|---|---|---|---|
| 0.530373 | 0.261324 | 0.063 | 0.054 | 1 | 5 | Fpgt | 0.009 | Young |
| 0.530515 | 0.361669 | 0.091 | 0.081 | 1 | 5 | Btbd10 | 0.01 | Young |
| 0.530552 | 0.168967 | 0.211 | 0.195 | 1 | 5 | Fam103a1 | 0.016 | Young |
| 0.530574 | 0.4633 | 0.129 | 0.12 | 1 | 5 | Metap1 | 0.009 | Young |
| 0.53059 | 0.060427 | 0.074 | 0.062 | 1 | 5 | Dbf4 | 0.012 | Young |
| 0.530648 | 0.177203 | 0.147 | 0.133 | 1 | 5 | Larp7 | 0.014 | Young |
| 0.530659 | 0.235616 | 0.124 | 0.111 | 1 | 5 | Slc35e1 | 0.013 | Young |
| 0.530712 | 0.08589 | 0.053 | 0.044 | 1 | 5 | Dph6 | 0.009 | Young |
| 0.530863 | 0.245101 | 0.043 | 0.035 | 1 | 5 | I830077J02 | 0.008 | Young |
| 0.531 | 0.088851 | 0.043 | 0.052 | 1 | 5 | B3gnt3 | -0.009 | Young |
| 0.531011 | 0.479928 | 0.112 | 0.101 | 1 | 5 | P2ry13 | 0.011 | Young |
| 0.531018 | 0.027818 | 0.008 | 0.012 | 1 | 5 | Glb1l2 | -0.004 | Young |
| 0.531536 | 0.151555 | 0.307 | 0.286 | 1 | 5 | Birc2 | 0.021 | Young |
| 0.531694 | 0.100263 | 0.279 | 0.261 | 1 | 5 | Psmb2 | 0.018 | Young |
| 0.531942 | 0.039116 | 0.046 | 0.056 | 1 | 5 | Espn | -0.01 | Young |
| 0.531948 | 0.115869 | 0.492 | 0.48 | 1 | 5 | Lims1 | 0.012 | Young |
| 0.532059 | 0.201636 | 0.241 | 0.226 | 1 | 5 | Itpripl1 | 0.015 | Young |
| 0.532703 | 0.47563 | 0.036 | 0.029 | 1 | 5 | Wdr53 | 0.007 | Young |
| 0.532703 | 0.347737 | 0.036 | 0.029 | 1 | 5 | 3110002H1 | 0.007 | Young |
| 0.533006 | 0.342057 | 0.099 | 0.088 | 1 | 5 | Chmp6 | 0.011 | Young |
| 0.53305 | 0.151328 | 0.175 | 0.16 | 1 | 5 | Irf4 | 0.015 | Young |
| 0.533246 | 0.041257 | 0.353 | 0.372 | 1 | 5 | Psmc3 | -0.019 | Young |
| 0.53384 | 0.001432 | 0.365 | 0.375 | 1 | 5 | Golph3 | -0.01 | Young |
| 0.533898 | 0.182096 | 0.277 | 0.268 | 1 | 5 | Ubash3b | 0.009 | Young |
| 0.534197 | 0.019939 | 0.32 | 0.338 | 1 | 5 | Sav1 | -0.018 | Young |
| 0.534301 | 0.201678 | 0.15 | 0.135 | 1 | 5 | Spg7 | 0.015 | Young |
| 0.534358 | 0.093897 | 0.246 | 0.231 | 1 | 5 | Slc27a1 | 0.015 | Young |
| 0.534957 | 0.218272 | 0.117 | 0.104 | 1 | 5 | Icmt | 0.013 | Young |
| 0.535218 | 0.290727 | 0.14 | 0.128 | 1 | 5 | Sipa1l3 | 0.012 | Young |
| 0.535594 | 0.004591 | 0.84 | 0.864 | 1 | 5 | Sdc4 | -0.024 | Young |
| 0.535739 | 0.419833 | 0.114 | 0.103 | 1 | 5 | Ranbp3 | 0.011 | Young |
| 0.535816 | 0.50258 | 0.02 | 0.015 | 1 | 5 | Nup210 | 0.005 | Young |
| 0.535994 | 0.093474 | 0.391 | 0.365 | 1 | 5 | Cox7b | 0.026 | Young |
| 0.536057 | 0.018426 | 0.228 | 0.251 | 1 | 5 | Taf1d | -0.023 | Young |
| 0.536082 | 0.280624 | 0.104 | 0.093 | 1 | 5 | Tprkb | 0.011 | Young |
| 0.536197 | 0.226197 | 0.398 | 0.384 | 1 | 5 | Eea1 | 0.014 | Young |
| 0.536344 | 0.336369 | 0.056 | 0.047 | 1 | 5 | Tbc1d32 | 0.009 | Young |
| 0.536482 | 0.570629 | 0.02 | 0.015 | 1 | 5 | Nars2 | 0.005 | Young |
| 0.536484 | 0.171976 | 0.063 | 0.054 | 1 | 5 | Aptx | 0.009 | Young |
| 0.536719 | 0.23123 | 0.307 | 0.305 | 1 | 5 | Hcfc1r1 | 0.002 | Young |
| 0.536869 | 0.055735 | 0.165 | 0.148 | 1 | 5 | Slc25a17 | 0.017 | Young |
| 0.536898 | 0.292964 | 0.157 | 0.146 | 1 | 5 | Pi4ka | 0.011 | Young |
| 0.537058 | 0.395433 | 0.089 | 0.079 | 1 | 5 | Afap1l1 | 0.01 | Young |
| 0.537149 | 0.21249 | 0.02 | 0.015 | 1 | 5 | Mterf1a | 0.005 | Young |
| 0.537803 | 0.118263 | 0.477 | 0.468 | 1 | 5 | Man2a1 | 0.009 | Young |

| | | | | | | | | |
|---|---|---|---|---|---|---|---|---|
| 0.537815 | 0.060537 | 0.464 | 0.453 | 1 | 5 | Cd300a | 0.011 | Young |
| 0.537913 | 0.378386 | 0.132 | 0.12 | 1 | 5 | Ralgapa1 | 0.012 | Young |
| 0.538056 | 0.095314 | 0.086 | 0.099 | 1 | 5 | Pcnx3 | -0.013 | Young |
| 0.538082 | 0.25735 | 0.223 | 0.214 | 1 | 5 | Commd3 | 0.009 | Young |
| 0.538148 | 0.185498 | 0.129 | 0.116 | 1 | 5 | Rabggtb | 0.013 | Young |
| 0.538265 | 0.463639 | 0.048 | 0.04 | 1 | 5 | Ghdc | 0.008 | Young |
| 0.538483 | 0.320527 | 0.02 | 0.015 | 1 | 5 | Tmc4 | 0.005 | Young |
| 0.53855 | 0.132063 | 0.505 | 0.505 | 1 | 5 | Sumo3 | 0 | Young |
| 0.538628 | 0.237024 | 0.299 | 0.281 | 1 | 5 | Cpsf7 | 0.018 | Young |
| 0.538684 | 0.325898 | 0.102 | 0.091 | 1 | 5 | Togaram1 | 0.011 | Young |
| 0.539031 | 0.174346 | 0.122 | 0.108 | 1 | 5 | Pycr2 | 0.014 | Young |
| 0.539151 | 0.354318 | 0.104 | 0.093 | 1 | 5 | Akr7a5 | 0.011 | Young |
| 0.539195 | 0.001809 | 0.081 | 0.093 | 1 | 5 | Anapc15 | -0.012 | Young |
| 0.539273 | 0.229195 | 0.287 | 0.276 | 1 | 5 | Cnppd1 | 0.011 | Young |
| 0.539289 | 0.053616 | 0.091 | 0.103 | 1 | 5 | Mrpl19 | -0.012 | Young |
| 0.539411 | 0.19558 | 0.043 | 0.035 | 1 | 5 | Bzw2 | 0.008 | Young |
| 0.53942 | 0.18173 | 0.127 | 0.114 | 1 | 5 | A430005L1 | 0.013 | Young |
| 0.539501 | 0.223483 | 0.249 | 0.234 | 1 | 5 | Ap1s1 | 0.015 | Young |
| 0.539819 | 0.512537 | 0.02 | 0.015 | 1 | 5 | Phldb1 | 0.005 | Young |
| 0.539819 | 0.340155 | 0.02 | 0.015 | 1 | 5 | Ttc39c | 0.005 | Young |
| 0.540019 | 0.270047 | 0.112 | 0.101 | 1 | 5 | Rpf1 | 0.011 | Young |
| 0.540247 | 0.05839 | 0.239 | 0.221 | 1 | 5 | Kidins220 | 0.018 | Young |
| 0.540318 | 0.45612 | 0.142 | 0.131 | 1 | 5 | Kank2 | 0.011 | Young |
| 0.540332 | 0.395124 | 0.096 | 0.086 | 1 | 5 | Rnf214 | 0.01 | Young |
| 0.540371 | 0.224007 | 0.216 | 0.2 | 1 | 5 | Bckdk | 0.016 | Young |
| 0.540419 | 0.602692 | 0.028 | 0.022 | 1 | 5 | Spryd4 | 0.006 | Young |
| 0.540487 | 0.700248 | 0.02 | 0.015 | 1 | 5 | Pdlim7 | 0.005 | Young |
| 0.540841 | 0.073938 | 0.312 | 0.328 | 1 | 5 | Krcc1 | -0.016 | Young |
| 0.540893 | 0.207653 | 0.124 | 0.111 | 1 | 5 | Rbm4b | 0.013 | Young |
| 0.540898 | 0.257029 | 0.114 | 0.101 | 1 | 5 | Ago1 | 0.013 | Young |
| 0.54094 | 0.180692 | 0.132 | 0.118 | 1 | 5 | Rnf40 | 0.014 | Young |
| 0.541156 | 0.405791 | 0.02 | 0.015 | 1 | 5 | 2900060B1 | 0.005 | Young |
| 0.541197 | 0.02506 | 0.099 | 0.113 | 1 | 5 | Pold4 | -0.014 | Young |
| 0.54126 | 0.402916 | 0.147 | 0.136 | 1 | 5 | Ddx23 | 0.011 | Young |
| 0.54127 | 0.513858 | 0.053 | 0.045 | 1 | 5 | Galt | 0.008 | Young |
| 0.541403 | 0.182597 | 0.089 | 0.077 | 1 | 5 | Ltv1 | 0.012 | Young |
| 0.541409 | 0.039722 | 0.201 | 0.217 | 1 | 5 | Elmo2 | -0.016 | Young |
| 0.541826 | 0.387132 | 0.02 | 0.015 | 1 | 5 | Nphp1 | 0.005 | Young |
| 0.541889 | 0.071361 | 0.051 | 0.061 | 1 | 5 | Actr5 | -0.01 | Young |
| 0.542125 | 0.213782 | 0.043 | 0.035 | 1 | 5 | Krt10 | 0.008 | Young |
| 0.542542 | 0.49073 | 0.117 | 0.108 | 1 | 5 | Utp6 | 0.009 | Young |
| 0.542665 | 0.148171 | 0.102 | 0.089 | 1 | 5 | Hace1 | 0.013 | Young |
| 0.542838 | 0.050753 | 0.051 | 0.042 | 1 | 5 | Zfp74 | 0.009 | Young |
| 0.543124 | 0.240694 | 0.066 | 0.057 | 1 | 5 | Tmem126a | 0.009 | Young |
| 0.543138 | 0.077574 | 0.17 | 0.153 | 1 | 5 | Ilf2 | 0.017 | Young |

| | | | | | | | | |
|---|---|---|---|---|---|---|---|---|
| 0.543139 | 0.201897 | 0.107 | 0.096 | 1 | 5 | Upf3b | 0.011 | Young |
| 0.543166 | 0.408494 | 0.02 | 0.015 | 1 | 5 | Dnal1 | 0.005 | Young |
| 0.543299 | 0.196509 | 0.246 | 0.234 | 1 | 5 | Synj2bp | 0.012 | Young |
| 0.543342 | 0.27759 | 0.188 | 0.175 | 1 | 5 | AU040320 | 0.013 | Young |
| 0.543483 | 0.091541 | 0.363 | 0.394 | 1 | 5 | Rad23b | -0.031 | Young |
| 0.54349 | 0.125565 | 0.208 | 0.192 | 1 | 5 | 1810030O( | 0.016 | Young |
| 0.543814 | 0.389725 | 0.028 | 0.022 | 1 | 5 | Satb1 | 0.006 | Young |
| 0.543836 | 0.405642 | 0.02 | 0.015 | 1 | 5 | Itpr3 | 0.005 | Young |
| 0.544016 | 0.459261 | 0.398 | 0.423 | 1 | 5 | Ltc4s | -0.025 | Young |
| 0.544105 | 0.180141 | 0.178 | 0.162 | 1 | 5 | Zc3h14 | 0.016 | Young |
| 0.544107 | 0.204584 | 0.294 | 0.276 | 1 | 5 | 1110008F1 | 0.018 | Young |
| 0.544299 | 0.362191 | 0.107 | 0.096 | 1 | 5 | Qsox1 | 0.011 | Young |
| 0.544643 | 0.209311 | 0.228 | 0.212 | 1 | 5 | Slc45a4 | 0.016 | Young |
| 0.545179 | 0.362979 | 0.02 | 0.015 | 1 | 5 | Patj | 0.005 | Young |
| 0.545179 | 0.322564 | 0.02 | 0.015 | 1 | 5 | Gm28151 | 0.005 | Young |
| 0.54537 | 0.076275 | 0.942 | 0.936 | 1 | 5 | Rpl34 | 0.006 | Young |
| 0.545513 | 0.198694 | 0.31 | 0.291 | 1 | 5 | Csnk2b | 0.019 | Young |
| 0.545747 | 0.07324 | 0.122 | 0.108 | 1 | 5 | Rtl8a | 0.014 | Young |
| 0.545811 | 0.086271 | 0.429 | 0.406 | 1 | 5 | Anp32b | 0.023 | Young |
| 0.545851 | 0.252355 | 0.02 | 0.015 | 1 | 5 | Hist2h3c1 | 0.005 | Young |
| 0.545964 | 0.126156 | 0.109 | 0.096 | 1 | 5 | Tbc1d24 | 0.013 | Young |
| 0.545996 | 0.460572 | 0.063 | 0.054 | 1 | 5 | Mllt1 | 0.009 | Young |
| 0.546052 | 0.197023 | 0.251 | 0.232 | 1 | 5 | Hipk3 | 0.019 | Young |
| 0.546087 | 0.16182 | 0.614 | 0.645 | 1 | 5 | Slc25a5 | -0.031 | Young |
| 0.546266 | 0.19876 | 0.056 | 0.047 | 1 | 5 | Mrps2 | 0.009 | Young |
| 0.546362 | 0.07831 | 0.249 | 0.268 | 1 | 5 | Slc25a28 | -0.019 | Young |
| 0.546523 | 0.270279 | 0.02 | 0.015 | 1 | 5 | Pfkfb2 | 0.005 | Young |
| 0.546916 | 0.177893 | 0.305 | 0.295 | 1 | 5 | Vav3 | 0.01 | Young |
| 0.547082 | 0.028449 | 0.977 | 0.985 | 1 | 5 | Grn | -0.008 | Young |
| 0.547083 | 0.292124 | 0.234 | 0.222 | 1 | 5 | Egln1 | 0.012 | Young |
| 0.54716 | 0.505857 | 0.036 | 0.029 | 1 | 5 | Cygb | 0.007 | Young |
| 0.547186 | 0.410103 | 0.061 | 0.052 | 1 | 5 | Mmgt1 | 0.009 | Young |
| 0.547196 | 0.372666 | 0.02 | 0.015 | 1 | 5 | Hist1h2ap | 0.005 | Young |
| 0.547218 | 0.22738 | 0.028 | 0.022 | 1 | 5 | Nat2 | 0.006 | Young |
| 0.547255 | 0.280662 | 0.317 | 0.313 | 1 | 5 | Rnf145 | 0.004 | Young |
| 0.547285 | 0.105972 | 0.264 | 0.247 | 1 | 5 | Cyld | 0.017 | Young |
| 0.547715 | 0.125961 | 0.254 | 0.236 | 1 | 5 | Ubp1 | 0.018 | Young |
| 0.54774 | 0.049672 | 0.19 | 0.172 | 1 | 5 | Ampd3 | 0.018 | Young |
| 0.547876 | 0.315102 | 0.066 | 0.057 | 1 | 5 | Eef2kmt | 0.009 | Young |
| 0.548247 | 0.206959 | 0.063 | 0.054 | 1 | 5 | Mgmt | 0.009 | Young |
| 0.548781 | 0.588002 | 0.053 | 0.045 | 1 | 5 | Akip1 | 0.008 | Young |
| 0.548934 | 0.435633 | 0.109 | 0.099 | 1 | 5 | Dhx32 | 0.01 | Young |
| 0.548975 | 0.385275 | 0.132 | 0.12 | 1 | 5 | Rabggta | 0.012 | Young |
| 0.549474 | 0.041815 | 0.246 | 0.271 | 1 | 5 | Gba | -0.025 | Young |
| 0.549494 | 0.22217 | 0.028 | 0.022 | 1 | 5 | Erc1 | 0.006 | Young |

| | | | | | | | | |
|---|---|---|---|---|---|---|---|---|
| 0.549556 | 0.047425 | 0.921 | 0.963 | 1 | 5 | Gnai2 | -0.042 | Young |
| 0.549634 | 0.108596 | 0.018 | 0.024 | 1 | 5 | Cd24a | -0.006 | Young |
| 0.549672 | 0.329994 | 0.036 | 0.029 | 1 | 5 | Sh3bp5l | 0.007 | Young |
| 0.549823 | 0.394227 | 0.058 | 0.051 | 1 | 5 | Naa16 | 0.007 | Young |
| 0.549874 | 0.112401 | 0.332 | 0.364 | 1 | 5 | Kcnk6 | -0.032 | Young |
| 0.549941 | 0.057942 | 0.185 | 0.2 | 1 | 5 | Znrd1 | -0.015 | Young |
| 0.550219 | 0.252313 | 0.18 | 0.167 | 1 | 5 | Rftn1 | 0.013 | Young |
| 0.550522 | 0.145612 | 0.599 | 0.589 | 1 | 5 | 1810037I1 | 0.01 | Young |
| 0.550565 | 0.244681 | 0.02 | 0.015 | 1 | 5 | Gm42726 | 0.005 | Young |
| 0.550724 | 0.243557 | 0.266 | 0.254 | 1 | 5 | Mark3 | 0.012 | Young |
| 0.550902 | 0.15796 | 0.071 | 0.061 | 1 | 5 | Clpb | 0.01 | Young |
| 0.550929 | 0.149767 | 0.074 | 0.086 | 1 | 5 | Gcc1 | -0.012 | Young |
| 0.551179 | 0.248005 | 0.066 | 0.057 | 1 | 5 | Dnajc30 | 0.009 | Young |
| 0.551219 | 0.223664 | 0.043 | 0.035 | 1 | 5 | Platr25 | 0.008 | Young |
| 0.551714 | 0.216091 | 0.335 | 0.318 | 1 | 5 | Osbpl11 | 0.017 | Young |
| 0.551734 | 0.147126 | 0.117 | 0.103 | 1 | 5 | Irf2bp1 | 0.014 | Young |
| 0.552415 | 0.005412 | 0.584 | 0.609 | 1 | 5 | Dennd5a | -0.025 | Young |
| 0.552592 | 0.062545 | 0.02 | 0.015 | 1 | 5 | Zbtb42 | 0.005 | Young |
| 0.552694 | 0.222088 | 0.036 | 0.029 | 1 | 5 | Zfp456 | 0.007 | Young |
| 0.553021 | 0.085702 | 0.566 | 0.566 | 1 | 5 | Susd6 | 0 | Young |
| 0.553037 | 0.383936 | 0.19 | 0.185 | 1 | 5 | Kdm4a | 0.005 | Young |
| 0.553184 | 0.046739 | 0.299 | 0.274 | 1 | 5 | Dr1 | 0.025 | Young |
| 0.553788 | 0.118717 | 0.203 | 0.222 | 1 | 5 | Ift20 | -0.019 | Young |
| 0.553898 | 0.224074 | 0.018 | 0.024 | 1 | 5 | 4930481A1 | -0.006 | Young |
| 0.553936 | 0.928152 | 0.01 | 0.007 | 1 | 5 | Tmem82 | 0.003 | Young |
| 0.553936 | 0.748669 | 0.01 | 0.007 | 1 | 5 | B230303O1 | 0.003 | Young |
| 0.553975 | 0.007172 | 0.203 | 0.224 | 1 | 5 | Ammecr1l | -0.021 | Young |
| 0.554208 | 0.138633 | 0.036 | 0.029 | 1 | 5 | Mccc2 | 0.007 | Young |
| 0.55472 | 0.159839 | 0.376 | 0.362 | 1 | 5 | Sdf2 | 0.014 | Young |
| 0.554722 | 0.207507 | 0.213 | 0.199 | 1 | 5 | Cdk4 | 0.014 | Young |
| 0.554919 | 0.714338 | 0.01 | 0.007 | 1 | 5 | Xrcc2 | 0.003 | Young |
| 0.554919 | 0.092158 | 0.01 | 0.007 | 1 | 5 | Gpr68 | 0.003 | Young |
| 0.554989 | 0.120927 | 0.381 | 0.367 | 1 | 5 | Dmxl1 | 0.014 | Young |
| 0.555091 | 0.30871 | 0.079 | 0.069 | 1 | 5 | Strada | 0.01 | Young |
| 0.555219 | 0.145295 | 0.036 | 0.029 | 1 | 5 | Mcph1 | 0.007 | Young |
| 0.555902 | 0.671021 | 0.01 | 0.007 | 1 | 5 | 1200007C1 | 0.003 | Young |
| 0.555902 | 0.576613 | 0.01 | 0.007 | 1 | 5 | Tmem246 | 0.003 | Young |
| 0.555902 | 0.449297 | 0.01 | 0.007 | 1 | 5 | Gpsm2 | 0.003 | Young |
| 0.555902 | 0.406702 | 0.01 | 0.007 | 1 | 5 | Gm28086 | 0.003 | Young |
| 0.555902 | 0.374797 | 0.01 | 0.007 | 1 | 5 | Ccdc14 | 0.003 | Young |
| 0.556034 | 0.013099 | 0.033 | 0.04 | 1 | 5 | Zfp788 | -0.007 | Young |
| 0.55636 | 0.13673 | 0.201 | 0.187 | 1 | 5 | Psmd7 | 0.014 | Young |
| 0.55644 | 0.053339 | 0.058 | 0.069 | 1 | 5 | Gtpbp10 | -0.011 | Young |
| 0.556887 | 0.64319 | 0.01 | 0.007 | 1 | 5 | Gm11491 | 0.003 | Young |
| 0.556887 | 0.575851 | 0.01 | 0.007 | 1 | 5 | Net1 | 0.003 | Young |

| | | | | | | | | |
|---|---|---|---|---|---|---|---|---|
| 0.556887 | 0.448073 | 0.01 | 0.007 | 1 | 5 | Lrrc75a | 0.003 | Young |
| 0.556887 | 0.418443 | 0.01 | 0.007 | 1 | 5 | Dtwd2 | 0.003 | Young |
| 0.556887 | 0.311595 | 0.01 | 0.007 | 1 | 5 | Tmem91 | 0.003 | Young |
| 0.556901 | 0.808608 | 0.046 | 0.039 | 1 | 5 | Tspyl3 | 0.007 | Young |
| 0.557448 | 0.03496 | 0.084 | 0.096 | 1 | 5 | Skp2 | -0.012 | Young |
| 0.557872 | 0.691317 | 0.01 | 0.007 | 1 | 5 | Tkfc | 0.003 | Young |
| 0.557872 | 0.464743 | 0.01 | 0.007 | 1 | 5 | Slc7a6 | 0.003 | Young |
| 0.557899 | 0.168137 | 0.31 | 0.293 | 1 | 5 | 6-Mar | 0.017 | Young |
| 0.558045 | 0.042223 | 0.388 | 0.421 | 1 | 5 | Wbp2 | -0.033 | Young |
| 0.558065 | 0.002504 | 0.117 | 0.131 | 1 | 5 | Stard3 | -0.014 | Young |
| 0.558359 | 0.058439 | 0.033 | 0.04 | 1 | 5 | Supt7l | -0.007 | Young |
| 0.558436 | 0.183249 | 0.048 | 0.04 | 1 | 5 | Nme7 | 0.008 | Young |
| 0.558584 | 0.237178 | 0.119 | 0.108 | 1 | 5 | Polr3k | 0.011 | Young |
| 0.559262 | 0.114995 | 0.586 | 0.591 | 1 | 5 | Prrc2a | -0.005 | Young |
| 0.559262 | 0.102973 | 0.419 | 0.406 | 1 | 5 | Ldha | 0.013 | Young |
| 0.559404 | 0.088729 | 0.183 | 0.2 | 1 | 5 | Wdr44 | -0.017 | Young |
| 0.559406 | 0.064556 | 0.345 | 0.367 | 1 | 5 | Aco2 | -0.022 | Young |
| 0.559624 | 0.022613 | 0.114 | 0.128 | 1 | 5 | Relt | -0.014 | Young |
| 0.559777 | 0.116741 | 0.137 | 0.125 | 1 | 5 | Sucla2 | 0.012 | Young |
| 0.559836 | 0.446636 | 0.053 | 0.045 | 1 | 5 | Mrpl46 | 0.008 | Young |
| 0.559845 | 0.27418 | 0.01 | 0.007 | 1 | 5 | Zfp39 | 0.003 | Young |
| 0.559845 | 0.193261 | 0.01 | 0.007 | 1 | 5 | Slit1 | 0.003 | Young |
| 0.559863 | 0.206967 | 0.348 | 0.345 | 1 | 5 | Snrpd3 | 0.003 | Young |
| 0.560108 | 0.357017 | 0.084 | 0.074 | 1 | 5 | Ctdp1 | 0.01 | Young |
| 0.560297 | 0.110566 | 0.312 | 0.295 | 1 | 5 | Ssfa2 | 0.017 | Young |
| 0.560442 | 0.180203 | 0.099 | 0.089 | 1 | 5 | Zfp513 | 0.01 | Young |
| 0.56051 | 0.103322 | 0.025 | 0.032 | 1 | 5 | Znhit3 | -0.007 | Young |
| 0.560833 | 0.246872 | 0.01 | 0.007 | 1 | 5 | Gm43796 | 0.003 | Young |
| 0.560833 | 0.184536 | 0.01 | 0.007 | 1 | 5 | Pclaf | 0.003 | Young |
| 0.56095 | 0.246545 | 0.109 | 0.098 | 1 | 5 | Usp45 | 0.011 | Young |
| 0.561286 | 0.15641 | 0.058 | 0.069 | 1 | 5 | Ypel2 | -0.011 | Young |
| 0.561496 | 0.363116 | 0.089 | 0.079 | 1 | 5 | Naf1 | 0.01 | Young |
| 0.561718 | 0.05663 | 0.898 | 0.911 | 1 | 5 | Lamp2 | -0.013 | Young |
| 0.561766 | 0.145897 | 0.19 | 0.178 | 1 | 5 | Mrpl30 | 0.012 | Young |
| 0.561822 | 0.235871 | 0.01 | 0.007 | 1 | 5 | Tube1 | 0.003 | Young |
| 0.561822 | 0.146729 | 0.01 | 0.007 | 1 | 5 | Gm21988 | 0.003 | Young |
| 0.561822 | 0.133822 | 0.01 | 0.007 | 1 | 5 | Ino80dos | 0.003 | Young |
| 0.561996 | 0.32011 | 0.152 | 0.143 | 1 | 5 | Pcnt | 0.009 | Young |
| 0.562084 | 0.240622 | 0.033 | 0.027 | 1 | 5 | Tnnt1 | 0.006 | Young |
| 0.562425 | 0.289612 | 0.102 | 0.091 | 1 | 5 | Gpn1 | 0.011 | Young |
| 0.562474 | 0.451416 | 0.018 | 0.024 | 1 | 5 | AC139671. | -0.006 | Young |
| 0.562602 | 0.012474 | 0.066 | 0.076 | 1 | 5 | Cebpzos | -0.01 | Young |
| 0.562647 | 0.250643 | 0.183 | 0.17 | 1 | 5 | Gatad2a | 0.013 | Young |
| 0.562685 | 0.124273 | 0.218 | 0.204 | 1 | 5 | Herc2 | 0.014 | Young |
| 0.562999 | 0.403058 | 0.066 | 0.057 | 1 | 5 | Utp18 | 0.009 | Young |

| | | | | | | | | |
|---|---|---|---|---|---|---|---|---|
| 0.563058 | 0.085596 | 0.398 | 0.439 | 1 | 5 | Txndc17 | -0.041 | Young |
| 0.563185 | 0.178976 | 0.464 | 0.453 | 1 | 5 | Tax1bp1 | 0.011 | Young |
| 0.563431 | 0.317738 | 0.185 | 0.172 | 1 | 5 | Map1s | 0.013 | Young |
| 0.563865 | 0.367801 | 0.117 | 0.106 | 1 | 5 | Abcf3 | 0.011 | Young |
| 0.564042 | 0.002542 | 0.157 | 0.17 | 1 | 5 | Naa38 | -0.013 | Young |
| 0.564219 | 0.16343 | 0.129 | 0.114 | 1 | 5 | Zfp260 | 0.015 | Young |
| 0.564376 | 0.076839 | 0.937 | 0.943 | 1 | 5 | Rps15a | -0.006 | Young |
| 0.564404 | 0.233779 | 0.18 | 0.168 | 1 | 5 | Pdcd11 | 0.012 | Young |
| 0.56449 | 0.15107 | 0.464 | 0.448 | 1 | 5 | Itch | 0.016 | Young |
| 0.564506 | 0.094994 | 0.82 | 0.827 | 1 | 5 | Rnf130 | -0.007 | Young |
| 0.564793 | 0.00716 | 0.01 | 0.007 | 1 | 5 | Zfp712 | 0.003 | Young |
| 0.565081 | 0.037171 | 0.457 | 0.468 | 1 | 5 | Amfr | -0.011 | Young |
| 0.565281 | 0.309525 | 0.051 | 0.061 | 1 | 5 | 2310009B1 | -0.01 | Young |
| 0.565418 | 0.42433 | 0.041 | 0.034 | 1 | 5 | Fbxo46 | 0.007 | Young |
| 0.565612 | 0.003891 | 0.038 | 0.045 | 1 | 5 | Slc17a9 | -0.007 | Young |
| 0.565634 | 0.477279 | 0.041 | 0.049 | 1 | 5 | F830016B0 | -0.008 | Young |
| 0.56581 | 0.135851 | 0.315 | 0.3 | 1 | 5 | Snx4 | 0.015 | Young |
| 0.5659 | 0.351089 | 0.102 | 0.091 | 1 | 5 | Klhl21 | 0.011 | Young |
| 0.565933 | 0.203539 | 0.244 | 0.224 | 1 | 5 | Lin7c | 0.02 | Young |
| 0.565981 | 0.212331 | 0.079 | 0.069 | 1 | 5 | Utp15 | 0.01 | Young |
| 0.566821 | 0.534004 | 0.033 | 0.027 | 1 | 5 | Gtf2ird2 | 0.006 | Young |
| 0.566888 | 0.082141 | 0.388 | 0.382 | 1 | 5 | B230219D2 | 0.006 | Young |
| 0.566969 | 0.099421 | 0.272 | 0.253 | 1 | 5 | C330007P0 | 0.019 | Young |
| 0.566988 | 0.204775 | 0.17 | 0.155 | 1 | 5 | Hmgcs1 | 0.015 | Young |
| 0.567275 | 0.164675 | 0.185 | 0.17 | 1 | 5 | Hmgn2 | 0.015 | Young |
| 0.567312 | 0.140734 | 0.358 | 0.345 | 1 | 5 | Rere | 0.013 | Young |
| 0.567399 | 0.027762 | 0.226 | 0.246 | 1 | 5 | Ube2q2 | -0.02 | Young |
| 0.567831 | 0.244036 | 0.081 | 0.071 | 1 | 5 | Alg14 | 0.01 | Young |
| 0.567861 | 0.225133 | 0.129 | 0.118 | 1 | 5 | Nifk | 0.011 | Young |
| 0.567952 | 0.10985 | 0.335 | 0.322 | 1 | 5 | Galc | 0.013 | Young |
| 0.567998 | 0.096335 | 0.726 | 0.739 | 1 | 5 | Rgs10 | -0.013 | Young |
| 0.568 | 0.178005 | 0.223 | 0.212 | 1 | 5 | Gtpbp4 | 0.011 | Young |
| 0.568251 | 0.052134 | 0.147 | 0.131 | 1 | 5 | Esf1 | 0.016 | Young |
| 0.568404 | 0.292532 | 0.033 | 0.027 | 1 | 5 | Wdr73 | 0.006 | Young |
| 0.568411 | 0.222268 | 0.056 | 0.047 | 1 | 5 | Gdpd3 | 0.009 | Young |
| 0.568926 | 0.10696 | 0.292 | 0.318 | 1 | 5 | Tbl1x | -0.026 | Young |
| 0.569375 | 0.117185 | 0.338 | 0.327 | 1 | 5 | Olfm1 | 0.011 | Young |
| 0.569443 | 0.159091 | 0.416 | 0.396 | 1 | 5 | Coro1c | 0.02 | Young |
| 0.569642 | 0.204617 | 0.431 | 0.433 | 1 | 5 | Nars | -0.002 | Young |
| 0.56995 | 0.004511 | 0.693 | 0.724 | 1 | 5 | Jak1 | -0.031 | Young |
| 0.570023 | 0.026403 | 0.058 | 0.067 | 1 | 5 | 2300009A0 | -0.009 | Young |
| 0.570125 | 0.248904 | 0.086 | 0.076 | 1 | 5 | B4galnt1 | 0.01 | Young |
| 0.570147 | 0.112128 | 0.129 | 0.146 | 1 | 5 | Btbd9 | -0.017 | Young |
| 0.570324 | 0.22352 | 0.089 | 0.103 | 1 | 5 | Rnase6 | -0.014 | Young |
| 0.570684 | 0.173076 | 0.315 | 0.31 | 1 | 5 | Pkn2 | 0.005 | Young |

| | | | | | | | | |
|---|---|---|---|---|---|---|---|---|
| 0.570729 | 0.380211 | 0.081 | 0.072 | 1 | 5 | Dnajc24 | 0.009 | Young |
| 0.571383 | 0.034859 | 0.18 | 0.195 | 1 | 5 | Kif3b | -0.015 | Young |
| 0.571874 | 0.427497 | 0.109 | 0.099 | 1 | 5 | Hsf1 | 0.01 | Young |
| 0.572153 | 0.358712 | 0.076 | 0.067 | 1 | 5 | 6030458C1 | 0.009 | Young |
| 0.5724 | 0.092594 | 0.279 | 0.263 | 1 | 5 | Phc2 | 0.016 | Young |
| 0.572483 | 0.033075 | 0.056 | 0.047 | 1 | 5 | Rnf25 | 0.009 | Young |
| 0.572661 | 0.345024 | 0.053 | 0.045 | 1 | 5 | B230354K1 | 0.008 | Young |
| 0.572692 | 0.244686 | 0.066 | 0.057 | 1 | 5 | Usp46 | 0.009 | Young |
| 0.572914 | 0.027308 | 0.353 | 0.379 | 1 | 5 | Chtop | -0.026 | Young |
| 0.573114 | 0.822841 | 0.025 | 0.02 | 1 | 5 | Cmtr1 | 0.005 | Young |
| 0.573382 | 0.145309 | 0.437 | 0.431 | 1 | 5 | Arcn1 | 0.006 | Young |
| 0.573461 | 0.064252 | 0.066 | 0.076 | 1 | 5 | Zbtb33 | -0.01 | Young |
| 0.573507 | 0.150526 | 0.315 | 0.305 | 1 | 5 | Cbfb | 0.01 | Young |
| 0.57357 | 0.139658 | 0.546 | 0.557 | 1 | 5 | Atp5k | -0.011 | Young |
| 0.573696 | 0.500683 | 0.033 | 0.027 | 1 | 5 | Zkscan14 | 0.006 | Young |
| 0.573781 | 0.353728 | 0.137 | 0.125 | 1 | 5 | Aim2 | 0.012 | Young |
| 0.573787 | 0.236604 | 0.193 | 0.182 | 1 | 5 | Susd3 | 0.011 | Young |
| 0.573879 | 0.021114 | 0.797 | 0.857 | 1 | 5 | Nrros | -0.06 | Young |
| 0.574103 | 0.090541 | 0.081 | 0.071 | 1 | 5 | Xrcc4 | 0.01 | Young |
| 0.5742 | 0.270442 | 0.096 | 0.086 | 1 | 5 | Mtpap | 0.01 | Young |
| 0.574736 | 0.117691 | 0.084 | 0.096 | 1 | 5 | Zbtb14 | -0.012 | Young |
| 0.574747 | 0.009675 | 0.442 | 0.478 | 1 | 5 | Trim8 | -0.036 | Young |
| 0.574894 | 0.096264 | 0.195 | 0.178 | 1 | 5 | Sf3a1 | 0.017 | Young |
| 0.574942 | 0.408814 | 0.076 | 0.067 | 1 | 5 | Prrg2 | 0.009 | Young |
| 0.575431 | 0.027649 | 0.249 | 0.264 | 1 | 5 | Ccrl2 | -0.015 | Young |
| 0.575498 | 0.078228 | 0.693 | 0.7 | 1 | 5 | Arrb2 | -0.007 | Young |
| 0.576151 | 0.788214 | 0.025 | 0.02 | 1 | 5 | Spred3 | 0.005 | Young |
| 0.576246 | 0.183099 | 0.063 | 0.074 | 1 | 5 | Usp31 | -0.011 | Young |
| 0.57632 | 0.078627 | 0.015 | 0.02 | 1 | 5 | Tasp1 | -0.005 | Young |
| 0.576351 | 0.306774 | 0.033 | 0.027 | 1 | 5 | Racgap1 | 0.006 | Young |
| 0.576511 | 0.773541 | 0.018 | 0.013 | 1 | 5 | Chaf1a | 0.005 | Young |
| 0.576847 | 0.138045 | 0.152 | 0.14 | 1 | 5 | Senp7 | 0.012 | Young |
| 0.577108 | 0.273 | 0.147 | 0.136 | 1 | 5 | Itgav | 0.011 | Young |
| 0.577244 | 0.377558 | 0.053 | 0.045 | 1 | 5 | Pomt1 | 0.008 | Young |
| 0.577245 | 0.872739 | 0.018 | 0.013 | 1 | 5 | 5033421B0 | 0.005 | Young |
| 0.577544 | 0.088179 | 0.086 | 0.076 | 1 | 5 | Cryzl1 | 0.01 | Young |
| 0.577944 | 0.095757 | 0.368 | 0.396 | 1 | 5 | Trip11 | -0.028 | Young |
| 0.578605 | 0.187886 | 0.213 | 0.197 | 1 | 5 | Arfgap2 | 0.016 | Young |
| 0.578715 | 0.846965 | 0.018 | 0.013 | 1 | 5 | Pim2 | 0.005 | Young |
| 0.578853 | 0.178569 | 0.266 | 0.295 | 1 | 5 | P4ha1 | -0.029 | Young |
| 0.578943 | 0.267207 | 0.206 | 0.195 | 1 | 5 | Emc2 | 0.011 | Young |
| 0.579044 | 0.206121 | 0.142 | 0.16 | 1 | 5 | Yrdc | -0.018 | Young |
| 0.57945 | 0.453107 | 0.018 | 0.013 | 1 | 5 | Gm26542 | 0.005 | Young |
| 0.579804 | 0.527876 | 0.025 | 0.02 | 1 | 5 | Hemk1 | 0.005 | Young |
| 0.579913 | 0.023321 | 0.244 | 0.259 | 1 | 5 | Rbm42 | -0.015 | Young |

| | | | | | | | | |
|---|---|---|---|---|---|---|---|---|
| 0.579925 | 0.048189 | 0.076 | 0.088 | 1 | 5 | Polk | -0.012 | Young |
| 0.579955 | 0.11888 | 0.495 | 0.545 | 1 | 5 | Trim30a | -0.05 | Young |
| 0.580186 | 0.813862 | 0.018 | 0.013 | 1 | 5 | Syngr1 | 0.005 | Young |
| 0.580236 | 0.117749 | 0.388 | 0.367 | 1 | 5 | Ppp2r1a | 0.021 | Young |
| 0.580344 | 0.163488 | 0.206 | 0.19 | 1 | 5 | Atxn3 | 0.016 | Young |
| 0.580414 | 0.441241 | 0.025 | 0.02 | 1 | 5 | Dusp28 | 0.005 | Young |
| 0.580547 | 0.106506 | 0.404 | 0.404 | 1 | 5 | Osbpl9 | 0 | Young |
| 0.580559 | 0.189886 | 0.117 | 0.104 | 1 | 5 | Scmh1 | 0.013 | Young |
| 0.580861 | 0.2046 | 0.061 | 0.052 | 1 | 5 | Ksr1 | 0.009 | Young |
| 0.5809 | 0.106588 | 0.371 | 0.359 | 1 | 5 | Pikfyve | 0.012 | Young |
| 0.580932 | 0.08386 | 0.193 | 0.21 | 1 | 5 | Ythdf2 | -0.017 | Young |
| 0.581144 | 0.383724 | 0.033 | 0.027 | 1 | 5 | Trim32 | 0.006 | Young |
| 0.581368 | 0.003533 | 0.437 | 0.46 | 1 | 5 | Strn3 | -0.023 | Young |
| 0.581489 | 0.165833 | 0.046 | 0.054 | 1 | 5 | Txlng | -0.008 | Young |
| 0.581563 | 0.117837 | 0.096 | 0.086 | 1 | 5 | Unkl | 0.01 | Young |
| 0.581659 | 0.534166 | 0.018 | 0.013 | 1 | 5 | Zswim3 | 0.005 | Young |
| 0.5817 | 0.2739 | 0.132 | 0.121 | 1 | 5 | Zfp120 | 0.011 | Young |
| 0.581903 | 0.080477 | 0.396 | 0.419 | 1 | 5 | Ikzf1 | -0.023 | Young |
| 0.582062 | 0.123514 | 0.449 | 0.448 | 1 | 5 | Dynll2 | 0.001 | Young |
| 0.582396 | 0.393573 | 0.018 | 0.013 | 1 | 5 | Gm30122 | 0.005 | Young |
| 0.582478 | 0.358389 | 0.033 | 0.027 | 1 | 5 | Tepsin | 0.006 | Young |
| 0.583872 | 0.349705 | 0.018 | 0.013 | 1 | 5 | Bbs4 | 0.005 | Young |
| 0.583933 | 0.193649 | 0.109 | 0.098 | 1 | 5 | Psmg3 | 0.011 | Young |
| 0.583973 | 0.261441 | 0.315 | 0.305 | 1 | 5 | Mcur1 | 0.01 | Young |
| 0.584051 | 0.087417 | 0.652 | 0.657 | 1 | 5 | Rab5c | -0.005 | Young |
| 0.584107 | 0.167506 | 0.345 | 0.386 | 1 | 5 | Tnf | -0.041 | Young |
| 0.584203 | 0.356097 | 0.046 | 0.039 | 1 | 5 | Orc6 | 0.007 | Young |
| 0.584204 | 0.114379 | 0.655 | 0.687 | 1 | 5 | Hcls1 | -0.032 | Young |
| 0.584243 | 0.386078 | 0.157 | 0.145 | 1 | 5 | Fbxo18 | 0.012 | Young |
| 0.584337 | 0.298329 | 0.119 | 0.109 | 1 | 5 | Fbxl14 | 0.01 | Young |
| 0.584426 | 0.03081 | 0.683 | 0.67 | 1 | 5 | Tm9sf3 | 0.013 | Young |
| 0.584611 | 0.742441 | 0.018 | 0.013 | 1 | 5 | Slc39a8 | 0.005 | Young |
| 0.584611 | 0.53682 | 0.018 | 0.013 | 1 | 5 | Zfp235 | 0.005 | Young |
| 0.584793 | 0.182992 | 0.094 | 0.084 | 1 | 5 | Rtl5 | 0.01 | Young |
| 0.585301 | 0.304208 | 0.081 | 0.072 | 1 | 5 | Zfp942 | 0.009 | Young |
| 0.585304 | 0.526717 | 0.025 | 0.02 | 1 | 5 | Slamf6 | 0.005 | Young |
| 0.585508 | 0.142766 | 0.086 | 0.076 | 1 | 5 | Rfng | 0.01 | Young |
| 0.585603 | 0.202411 | 0.246 | 0.237 | 1 | 5 | Nemf | 0.009 | Young |
| 0.585611 | 0.434334 | 0.025 | 0.02 | 1 | 5 | Gm4673 | 0.005 | Young |
| 0.585753 | 0.199963 | 0.091 | 0.104 | 1 | 5 | Ino80 | -0.013 | Young |
| 0.586017 | 0.046899 | 0.046 | 0.054 | 1 | 5 | Acap3 | -0.008 | Young |
| 0.586089 | 0.324282 | 0.018 | 0.013 | 1 | 5 | Dbndd2 | 0.005 | Young |
| 0.586152 | 0.04541 | 0.023 | 0.029 | 1 | 5 | Ntmt1 | -0.006 | Young |
| 0.586157 | 0.024704 | 0.5 | 0.535 | 1 | 5 | Dgkd | -0.035 | Young |
| 0.586425 | 0.366489 | 0.015 | 0.02 | 1 | 5 | Brdt | -0.005 | Young |

| | | | | | | | | |
|---|---|---|---|---|---|---|---|---|
| 0.58653 | 0.163251 | 0.025 | 0.02 | 1 | 5 | Tshz2 | 0.005 | Young |
| 0.587001 | 0.121426 | 0.259 | 0.241 | 1 | 5 | Med1 | 0.018 | Young |
| 0.587523 | 0.347728 | 0.213 | 0.205 | 1 | 5 | Rnf115 | 0.008 | Young |
| 0.58754 | 0.069121 | 0.061 | 0.071 | 1 | 5 | Urgcp | -0.01 | Young |
| 0.587757 | 0.683826 | 0.025 | 0.02 | 1 | 5 | Cox7a1 | 0.005 | Young |
| 0.587801 | 0.025643 | 0.629 | 0.598 | 1 | 5 | St13 | 0.031 | Young |
| 0.58831 | 0.370022 | 0.018 | 0.013 | 1 | 5 | AC129328. | 0.005 | Young |
| 0.588533 | 0.198208 | 0.122 | 0.111 | 1 | 5 | Thap2 | 0.011 | Young |
| 0.588637 | 0.119026 | 0.033 | 0.027 | 1 | 5 | Slc16a13 | 0.006 | Young |
| 0.588976 | 0.417071 | 0.122 | 0.113 | 1 | 5 | Fam8a1 | 0.009 | Young |
| 0.588979 | 0.024106 | 0.023 | 0.029 | 1 | 5 | Sh2b2 | -0.006 | Young |
| 0.589051 | 0.393421 | 0.018 | 0.013 | 1 | 5 | Cox11 | 0.005 | Young |
| 0.589051 | 0.242315 | 0.018 | 0.013 | 1 | 5 | Clspn | 0.005 | Young |
| 0.589158 | 0.241934 | 0.076 | 0.067 | 1 | 5 | Alg11 | 0.009 | Young |
| 0.589297 | 0.047001 | 0.47 | 0.503 | 1 | 5 | Zfp207 | -0.033 | Young |
| 0.589324 | 0.381536 | 0.051 | 0.044 | 1 | 5 | Zfp595 | 0.007 | Young |
| 0.589599 | 0.400664 | 0.025 | 0.02 | 1 | 5 | Mks1 | 0.005 | Young |
| 0.589599 | 0.098972 | 0.025 | 0.02 | 1 | 5 | Mettl25 | 0.005 | Young |
| 0.589792 | 0.332995 | 0.018 | 0.013 | 1 | 5 | Guca1b | 0.005 | Young |
| 0.590112 | 0.260455 | 0.023 | 0.029 | 1 | 5 | Tmem42 | -0.006 | Young |
| 0.590318 | 0.342947 | 0.099 | 0.089 | 1 | 5 | Ddx39 | 0.01 | Young |
| 0.590322 | 0.16427 | 0.383 | 0.38 | 1 | 5 | Rnf7 | 0.003 | Young |
| 0.590323 | 0.222836 | 0.33 | 0.332 | 1 | 5 | Nfatc2 | -0.002 | Young |
| 0.590541 | 0.039663 | 0.145 | 0.158 | 1 | 5 | Cd151 | -0.013 | Young |
| 0.59078 | 0.005397 | 0.089 | 0.077 | 1 | 5 | Ercc3 | 0.012 | Young |
| 0.591444 | 0.417714 | 0.025 | 0.02 | 1 | 5 | Itgal | 0.005 | Young |
| 0.591444 | 0.39054 | 0.025 | 0.02 | 1 | 5 | Ptpdc1 | 0.005 | Young |
| 0.591838 | 0.0838 | 0.117 | 0.104 | 1 | 5 | Slc25a20 | 0.013 | Young |
| 0.592019 | 0.178102 | 0.018 | 0.013 | 1 | 5 | Thumpd2 | 0.005 | Young |
| 0.592105 | 0.074213 | 0.183 | 0.199 | 1 | 5 | Slc35a4 | -0.016 | Young |
| 0.592441 | 0.117074 | 0.155 | 0.141 | 1 | 5 | Hs1bp3 | 0.014 | Young |
| 0.592609 | 0.052675 | 0.127 | 0.114 | 1 | 5 | Ston2 | 0.013 | Young |
| 0.592735 | 0.138642 | 0.107 | 0.096 | 1 | 5 | Actr8 | 0.011 | Young |
| 0.592748 | 0.070789 | 0.424 | 0.406 | 1 | 5 | St6gal1 | 0.018 | Young |
| 0.592949 | 0.132568 | 0.023 | 0.029 | 1 | 5 | Fam167b | -0.006 | Young |
| 0.593225 | 0.18004 | 0.076 | 0.067 | 1 | 5 | Nup155 | 0.009 | Young |
| 0.594297 | 0.040876 | 0.119 | 0.108 | 1 | 5 | Rnf138 | 0.011 | Young |
| 0.594859 | 0.499089 | 0.074 | 0.066 | 1 | 5 | Aktip | 0.008 | Young |
| 0.594964 | 0.10766 | 0.165 | 0.152 | 1 | 5 | Akr1c13 | 0.013 | Young |
| 0.595193 | 0.060914 | 0.041 | 0.034 | 1 | 5 | Gpsm1 | 0.007 | Young |
| 0.595218 | 0.011203 | 0.36 | 0.387 | 1 | 5 | Psme2 | -0.027 | Young |
| 0.595284 | 0.151864 | 0.091 | 0.081 | 1 | 5 | Calcoco1 | 0.01 | Young |
| 0.595337 | 0.242664 | 0.203 | 0.194 | 1 | 5 | Txndc15 | 0.009 | Young |
| 0.595343 | 0.284623 | 0.15 | 0.14 | 1 | 5 | Prps2 | 0.01 | Young |
| 0.595358 | 0.029486 | 0.358 | 0.377 | 1 | 5 | Dst | -0.019 | Young |

| | | | | | | | | |
|---|---|---|---|---|---|---|---|---|
| 0.595679 | 0.07527 | 0.14 | 0.128 | 1 | 5 | Arfrp1 | 0.012 | Young |
| 0.59574 | 0.080422 | 0.018 | 0.013 | 1 | 5 | Zc3h11a | 0.005 | Young |
| 0.596157 | 0.12587 | 0.223 | 0.209 | 1 | 5 | Msl3 | 0.014 | Young |
| 0.596168 | 0.144005 | 0.119 | 0.108 | 1 | 5 | Rabgap1l | 0.011 | Young |
| 0.596342 | 0.143315 | 0.147 | 0.16 | 1 | 5 | Ctc1 | -0.013 | Young |
| 0.596482 | 0.152821 | 0.14 | 0.128 | 1 | 5 | Cops3 | 0.012 | Young |
| 0.596554 | 0.088147 | 0.175 | 0.16 | 1 | 5 | Agpat5 | 0.015 | Young |
| 0.59675 | 0.264941 | 0.155 | 0.14 | 1 | 5 | Med16 | 0.015 | Young |
| 0.596768 | 0.066527 | 0.307 | 0.295 | 1 | 5 | Atp5g1 | 0.012 | Young |
| 0.597141 | 0.029677 | 0.764 | 0.788 | 1 | 5 | Ogt | -0.024 | Young |
| 0.597188 | 0.401279 | 0.19 | 0.184 | 1 | 5 | Fam96a | 0.006 | Young |
| 0.597403 | 0.505366 | 0.061 | 0.054 | 1 | 5 | Ick | 0.007 | Young |
| 0.597541 | 0.032005 | 0.279 | 0.298 | 1 | 5 | Exoc4 | -0.019 | Young |
| 0.597791 | 0.060486 | 0.193 | 0.21 | 1 | 5 | Pank2 | -0.017 | Young |
| 0.597913 | 0.172074 | 0.261 | 0.249 | 1 | 5 | Xrn2 | 0.012 | Young |
| 0.598287 | 0.067498 | 0.266 | 0.291 | 1 | 5 | Chmp3 | -0.025 | Young |
| 0.598318 | 0.116887 | 0.569 | 0.556 | 1 | 5 | Ankrd11 | 0.013 | Young |
| 0.598579 | 0.012146 | 0.206 | 0.224 | 1 | 5 | Atp9b | -0.018 | Young |
| 0.598964 | 0.26976 | 0.089 | 0.079 | 1 | 5 | Pcnx | 0.01 | Young |
| 0.599344 | 0.035104 | 0.289 | 0.269 | 1 | 5 | 1110004F1 | 0.02 | Young |
| 0.599521 | 0.158389 | 0.071 | 0.081 | 1 | 5 | Papd7 | -0.01 | Young |
| 0.599531 | 0.008752 | 0.091 | 0.081 | 1 | 5 | Tusc3 | 0.01 | Young |
| 0.59955 | 0.212657 | 0.112 | 0.103 | 1 | 5 | Zfp362 | 0.009 | Young |
| 0.599558 | 0.073513 | 0.056 | 0.064 | 1 | 5 | Mblac2 | -0.008 | Young |
| 0.599566 | 0.171249 | 0.223 | 0.207 | 1 | 5 | Abcf1 | 0.016 | Young |
| 0.5999 | 0.133226 | 0.063 | 0.072 | 1 | 5 | Gm5617 | -0.009 | Young |
| 0.6 | 0.114887 | 0.269 | 0.251 | 1 | 5 | Trim12c | 0.018 | Young |
| 0.600123 | 0.367302 | 0.051 | 0.044 | 1 | 5 | Polr2d | 0.007 | Young |
| 0.600439 | 0.116584 | 0.282 | 0.264 | 1 | 5 | Iscu | 0.018 | Young |
| 0.6007 | 0.154608 | 0.183 | 0.17 | 1 | 5 | Pds5b | 0.013 | Young |
| 0.601407 | 0.150248 | 0.256 | 0.239 | 1 | 5 | Ncoa6 | 0.017 | Young |
| 0.601715 | 0.10838 | 0.013 | 0.017 | 1 | 5 | Wwtr1 | -0.004 | Young |
| 0.601778 | 0.093026 | 0.081 | 0.093 | 1 | 5 | Nupr1 | -0.012 | Young |
| 0.60184 | 0.064526 | 0.051 | 0.059 | 1 | 5 | Coprs | -0.008 | Young |
| 0.601923 | 0.01216 | 0.168 | 0.184 | 1 | 5 | Nucb2 | -0.016 | Young |
| 0.602057 | 0.37313 | 0.063 | 0.056 | 1 | 5 | Sft2d3 | 0.007 | Young |
| 0.602805 | 0.079956 | 0.371 | 0.352 | 1 | 5 | Pik3c2a | 0.019 | Young |
| 0.603067 | 0.133407 | 0.127 | 0.116 | 1 | 5 | Ccser2 | 0.011 | Young |
| 0.603254 | 0.061244 | 0.769 | 0.81 | 1 | 5 | Rpl31 | -0.041 | Young |
| 0.603341 | 0.769721 | 0.03 | 0.025 | 1 | 5 | Tyw5 | 0.005 | Young |
| 0.603524 | 0.38336 | 0.165 | 0.155 | 1 | 5 | Tgfbr1 | 0.01 | Young |
| 0.603889 | 0.111572 | 0.543 | 0.544 | 1 | 5 | Tcf25 | -0.001 | Young |
| 0.604099 | 0.044206 | 0.201 | 0.217 | 1 | 5 | Rps6kb1 | -0.016 | Young |
| 0.604278 | 0.041091 | 0.19 | 0.177 | 1 | 5 | Ube2w | 0.013 | Young |
| 0.604562 | 0.439209 | 0.056 | 0.049 | 1 | 5 | Zfp81 | 0.007 | Young |

| | | | | | | | | |
|---|---|---|---|---|---|---|---|---|
| 0.604779 | 0.026434 | 0.363 | 0.374 | 1 | 5 | Zmynd11 | -0.011 | Young |
| 0.604874 | 0.1504 | 0.391 | 0.386 | 1 | 5 | Adam10 | 0.005 | Young |
| 0.605028 | 1.065047 | 0.03 | 0.025 | 1 | 5 | Thbs1 | 0.005 | Young |
| 0.605906 | 0.061213 | 0.16 | 0.178 | 1 | 5 | Trove2 | -0.018 | Young |
| 0.60614 | 0.054121 | 0.119 | 0.108 | 1 | 5 | Inafm1 | 0.011 | Young |
| 0.606154 | 0.499241 | 0.03 | 0.025 | 1 | 5 | Slc52a2 | 0.005 | Young |
| 0.606213 | 0.08082 | 0.013 | 0.017 | 1 | 5 | Slc2a1 | -0.004 | Young |
| 0.606345 | 0.120556 | 0.063 | 0.056 | 1 | 5 | Zfp946 | 0.007 | Young |
| 0.606384 | 0.09497 | 0.053 | 0.045 | 1 | 5 | Dtd2 | 0.008 | Young |
| 0.606471 | 0.168118 | 0.079 | 0.069 | 1 | 5 | Asna1 | 0.01 | Young |
| 0.606675 | 0.218139 | 0.188 | 0.175 | 1 | 5 | Chpt1 | 0.013 | Young |
| 0.606751 | 0.167963 | 0.076 | 0.067 | 1 | 5 | Rnaseh2a | 0.009 | Young |
| 0.606965 | 0.020898 | 0.013 | 0.017 | 1 | 5 | Rbp4 | -0.004 | Young |
| 0.607145 | 0.520949 | 0.053 | 0.047 | 1 | 5 | Zw10 | 0.006 | Young |
| 0.607208 | 0.15298 | 0.315 | 0.306 | 1 | 5 | Pja2 | 0.009 | Young |
| 0.607388 | 0.13995 | 0.223 | 0.212 | 1 | 5 | Flot2 | 0.011 | Young |
| 0.607535 | 0.134936 | 0.185 | 0.172 | 1 | 5 | Mms19 | 0.013 | Young |
| 0.607673 | 0.277791 | 0.056 | 0.049 | 1 | 5 | Zfp317 | 0.007 | Young |
| 0.607712 | 0.436741 | 0.066 | 0.059 | 1 | 5 | Wipi1 | 0.007 | Young |
| 0.608178 | 0.139553 | 0.269 | 0.259 | 1 | 5 | Rcsd1 | 0.01 | Young |
| 0.608552 | 0.188838 | 0.127 | 0.116 | 1 | 5 | Zbtb8os | 0.011 | Young |
| 0.608553 | 0.086041 | 0.14 | 0.13 | 1 | 5 | Tvp23b | 0.01 | Young |
| 0.609076 | 0.20521 | 0.299 | 0.288 | 1 | 5 | Dnajc8 | 0.011 | Young |
| 0.609087 | 0.198343 | 0.114 | 0.104 | 1 | 5 | St7l | 0.01 | Young |
| 0.609148 | 0.043992 | 0.071 | 0.081 | 1 | 5 | Spg11 | -0.01 | Young |
| 0.609537 | 0.633781 | 0.03 | 0.025 | 1 | 5 | AI480526 | 0.005 | Young |
| 0.609572 | 0.256557 | 0.096 | 0.088 | 1 | 5 | Prpf31 | 0.008 | Young |
| 0.610546 | 0.074633 | 0.18 | 0.197 | 1 | 5 | Ufm1 | -0.017 | Young |
| 0.610595 | 0.118319 | 0.157 | 0.143 | 1 | 5 | Hs2st1 | 0.014 | Young |
| 0.610727 | 0.014464 | 0.013 | 0.017 | 1 | 5 | Kif20b | -0.004 | Young |
| 0.611259 | 0.042471 | 0.147 | 0.158 | 1 | 5 | Vrk3 | -0.011 | Young |
| 0.611391 | 0.11249 | 0.211 | 0.231 | 1 | 5 | Tiprl | -0.02 | Young |
| 0.611481 | 0.100216 | 0.013 | 0.017 | 1 | 5 | Gm16287 | -0.004 | Young |
| 0.611523 | 0.064541 | 0.102 | 0.091 | 1 | 5 | Prpf4 | 0.011 | Young |
| 0.611618 | 0.040411 | 0.442 | 0.418 | 1 | 5 | Tsc22d2 | 0.024 | Young |
| 0.611972 | 0.077076 | 0.147 | 0.133 | 1 | 5 | Uimc1 | 0.014 | Young |
| 0.611998 | 0.127749 | 0.211 | 0.195 | 1 | 5 | Fem1b | 0.016 | Young |
| 0.612033 | 0.085025 | 0.635 | 0.631 | 1 | 5 | Ppp2ca | 0.004 | Young |
| 0.612048 | 0.167748 | 0.251 | 0.244 | 1 | 5 | Trappc4 | 0.007 | Young |
| 0.612158 | 0.145108 | 0.685 | 0.692 | 1 | 5 | Bst2 | -0.007 | Young |
| 0.612235 | 0.08239 | 0.013 | 0.017 | 1 | 5 | Cyp2r1 | -0.004 | Young |
| 0.612249 | 0.444143 | 0.056 | 0.049 | 1 | 5 | Tiam1 | 0.007 | Young |
| 0.612399 | 0.205773 | 0.109 | 0.099 | 1 | 5 | Daam1 | 0.01 | Young |
| 0.61246 | 0.232344 | 0.091 | 0.082 | 1 | 5 | Ppme1 | 0.009 | Young |
| 0.612761 | 0.45092 | 0.051 | 0.044 | 1 | 5 | Fance | 0.007 | Young |

| | | | | | | | | |
|---|---|---|---|---|---|---|---|---|
| 0.612941 | 0.006982 | 0.571 | 0.596 | 1 | 5 | Sppl2a | -0.025 | Young |
| 0.613189 | 0.196091 | 0.178 | 0.165 | 1 | 5 | Tmem184b | 0.013 | Young |
| 0.614068 | 0.42002 | 0.061 | 0.054 | 1 | 5 | Fam35a | 0.007 | Young |
| 0.614333 | 0.416805 | 0.081 | 0.074 | 1 | 5 | Dusp5 | 0.007 | Young |
| 0.614612 | 0.10444 | 0.069 | 0.061 | 1 | 5 | Dclre1b | 0.008 | Young |
| 0.615022 | 0.165886 | 0.241 | 0.231 | 1 | 5 | Mrpl48 | 0.01 | Young |
| 0.61513 | 0.19145 | 0.345 | 0.333 | 1 | 5 | Arpp19 | 0.012 | Young |
| 0.615272 | 0.22696 | 0.17 | 0.16 | 1 | 5 | Sec16a | 0.01 | Young |
| 0.615321 | 0.381236 | 0.043 | 0.037 | 1 | 5 | Nphp3 | 0.006 | Young |
| 0.615551 | 0.772833 | 0.023 | 0.019 | 1 | 5 | Gm11696 | 0.004 | Young |
| 0.615789 | 0.176943 | 0.038 | 0.032 | 1 | 5 | Gm42047 | 0.006 | Young |
| 0.615894 | 0.056704 | 0.02 | 0.025 | 1 | 5 | Rpl36a-ps1 | -0.005 | Young |
| 0.616215 | 0.113728 | 0.058 | 0.051 | 1 | 5 | Adprm | 0.007 | Young |
| 0.616319 | 0.506465 | 0.114 | 0.106 | 1 | 5 | Qser1 | 0.008 | Young |
| 0.616364 | 0.109516 | 0.094 | 0.104 | 1 | 5 | Hps4 | -0.01 | Young |
| 0.616423 | 0.331135 | 0.056 | 0.049 | 1 | 5 | Zfyve19 | 0.007 | Young |
| 0.616819 | 0.017843 | 0.266 | 0.281 | 1 | 5 | Ppp1r2 | -0.015 | Young |
| 0.617464 | 0.217131 | 0.03 | 0.025 | 1 | 5 | Zfp953 | 0.005 | Young |
| 0.617697 | 0.387091 | 0.043 | 0.037 | 1 | 5 | Recql | 0.006 | Young |
| 0.617744 | 0.037217 | 0.942 | 0.936 | 1 | 5 | Sirpa | 0.006 | Young |
| 0.617955 | 0.115747 | 0.505 | 0.503 | 1 | 5 | Vps35 | 0.002 | Young |
| 0.618096 | 0.455917 | 0.056 | 0.049 | 1 | 5 | Mtor | 0.007 | Young |
| 0.618412 | 0.16009 | 0.124 | 0.114 | 1 | 5 | Rab40c | 0.01 | Young |
| 0.618542 | 0.014227 | 0.096 | 0.086 | 1 | 5 | Dis3l2 | 0.01 | Young |
| 0.619095 | 0.036949 | 0.36 | 0.345 | 1 | 5 | Psmb6 | 0.015 | Young |
| 0.619223 | 0.214365 | 0.236 | 0.227 | 1 | 5 | Tmem208 | 0.009 | Young |
| 0.619281 | 0.239347 | 0.358 | 0.354 | 1 | 5 | Stk38 | 0.004 | Young |
| 0.619563 | 0.10471 | 0.221 | 0.204 | 1 | 5 | Nudt21 | 0.017 | Young |
| 0.619577 | 0.328242 | 0.114 | 0.106 | 1 | 5 | AU019823 | 0.008 | Young |
| 0.619579 | 0.003306 | 0.02 | 0.025 | 1 | 5 | Gm13479 | -0.005 | Young |
| 0.619622 | 0.334875 | 0.218 | 0.209 | 1 | 5 | Churc1 | 0.009 | Young |
| 0.619985 | 0.284293 | 0.109 | 0.099 | 1 | 5 | Peli2 | 0.01 | Young |
| 0.620076 | 0.423663 | 0.043 | 0.037 | 1 | 5 | Slc44a2 | 0.006 | Young |
| 0.620168 | 0.069922 | 0.264 | 0.251 | 1 | 5 | Dcp2 | 0.013 | Young |
| 0.620245 | 0.110679 | 0.508 | 0.503 | 1 | 5 | Snap23 | 0.005 | Young |
| 0.621334 | 0.063688 | 0.135 | 0.148 | 1 | 5 | Hbs1l | -0.013 | Young |
| 0.621339 | 0.140705 | 0.213 | 0.202 | 1 | 5 | Tmem131l | 0.011 | Young |
| 0.621393 | 0.240799 | 0.089 | 0.081 | 1 | 5 | Hira | 0.008 | Young |
| 0.621677 | 0.2608 | 0.19 | 0.178 | 1 | 5 | Selenbp1 | 0.012 | Young |
| 0.621958 | 0.303804 | 0.312 | 0.308 | 1 | 5 | Atg3 | 0.004 | Young |
| 0.622174 | 0.006339 | 0.495 | 0.52 | 1 | 5 | Camk1d | -0.025 | Young |
| 0.622186 | 0.224288 | 0.091 | 0.082 | 1 | 5 | Xrcc1 | 0.009 | Young |
| 0.622191 | 0.29444 | 0.069 | 0.061 | 1 | 5 | Zfp280c | 0.008 | Young |
| 0.622291 | 0.040524 | 0.353 | 0.333 | 1 | 5 | Chmp5 | 0.02 | Young |
| 0.622444 | 0.079551 | 0.091 | 0.103 | 1 | 5 | Cenpc1 | -0.012 | Young |

| | | | | | | | | |
|---|---|---|---|---|---|---|---|---|
| 0.622584 | 0.343688 | 0.03 | 0.025 | 1 | 5 | Zfp212 | 0.005 | Young |
| 0.622897 | 0.174873 | 0.239 | 0.224 | 1 | 5 | 5031425E2 | 0.015 | Young |
| 0.623066 | 0.199648 | 0.226 | 0.214 | 1 | 5 | Tmem106b | 0.012 | Young |
| 0.623154 | 0.39377 | 0.03 | 0.025 | 1 | 5 | Rab11fip3 | 0.005 | Young |
| 0.623286 | 0.096563 | 0.086 | 0.077 | 1 | 5 | Mphosph6 | 0.009 | Young |
| 0.624067 | 0.01364 | 0.145 | 0.13 | 1 | 5 | Sord | 0.015 | Young |
| 0.624443 | 0.035288 | 0.152 | 0.167 | 1 | 5 | Flot1 | -0.015 | Young |
| 0.624944 | 0.226373 | 0.353 | 0.35 | 1 | 5 | Ogfrl1 | 0.003 | Young |
| 0.625001 | 0.027832 | 0.081 | 0.089 | 1 | 5 | Bcl9l | -0.008 | Young |
| 0.625067 | 0.137422 | 0.454 | 0.448 | 1 | 5 | Sumo1 | 0.006 | Young |
| 0.625685 | 0.374313 | 0.058 | 0.051 | 1 | 5 | Upp2 | 0.007 | Young |
| 0.626031 | 0.045222 | 0.287 | 0.308 | 1 | 5 | Dnajc19 | -0.021 | Young |
| 0.626099 | 0.253794 | 0.137 | 0.128 | 1 | 5 | Eaf1 | 0.009 | Young |
| 0.626114 | 0.135306 | 0.178 | 0.16 | 1 | 5 | Atxn10 | 0.018 | Young |
| 0.626232 | 0.828733 | 0.048 | 0.042 | 1 | 5 | Gm20186 | 0.006 | Young |
| 0.626239 | 0.205639 | 0.368 | 0.365 | 1 | 5 | Ubxn1 | 0.003 | Young |
| 0.626414 | 0.009784 | 0.043 | 0.051 | 1 | 5 | Vps33b | -0.008 | Young |
| 0.626493 | 0.219319 | 0.091 | 0.082 | 1 | 5 | Bola1 | 0.009 | Young |
| 0.626759 | 0.207993 | 0.043 | 0.037 | 1 | 5 | Wdr83 | 0.006 | Young |
| 0.626865 | 0.083339 | 0.107 | 0.098 | 1 | 5 | Commd10 | 0.009 | Young |
| 0.626934 | 0.114807 | 0.234 | 0.215 | 1 | 5 | Arsk | 0.019 | Young |
| 0.627198 | 0.266007 | 0.142 | 0.133 | 1 | 5 | Hps5 | 0.009 | Young |
| 0.627284 | 0.131424 | 0.051 | 0.044 | 1 | 5 | Zfp777 | 0.007 | Young |
| 0.62745 | 0.007852 | 0.266 | 0.281 | 1 | 5 | Xpr1 | -0.015 | Young |
| 0.627608 | 0.204921 | 0.168 | 0.158 | 1 | 5 | St6galnac6 | 0.01 | Young |
| 0.627813 | 0.007516 | 0.503 | 0.534 | 1 | 5 | Stxbp3 | -0.031 | Young |
| 0.627816 | 0.078942 | 0.193 | 0.178 | 1 | 5 | Chmp2b | 0.015 | Young |
| 0.628169 | 0.230438 | 0.124 | 0.114 | 1 | 5 | Vkorc1l1 | 0.01 | Young |
| 0.628674 | 0.225732 | 0.043 | 0.037 | 1 | 5 | Fam53a | 0.006 | Young |
| 0.628743 | 0.618454 | 0.023 | 0.019 | 1 | 5 | Dhfr | 0.004 | Young |
| 0.628818 | 0.1798 | 0.091 | 0.082 | 1 | 5 | Ppil3 | 0.009 | Young |
| 0.629099 | 0.077685 | 0.368 | 0.391 | 1 | 5 | Trafd1 | -0.023 | Young |
| 0.629123 | 0.01731 | 0.188 | 0.205 | 1 | 5 | Rsbn1 | -0.017 | Young |
| 0.629405 | 0.448056 | 0.023 | 0.019 | 1 | 5 | Stox2 | 0.004 | Young |
| 0.629451 | 0.582304 | 0.02 | 0.025 | 1 | 5 | Gja1 | -0.005 | Young |
| 0.629767 | 0.12206 | 0.675 | 0.675 | 1 | 5 | Cox6c | 0 | Young |
| 0.629973 | 0.030434 | 0.312 | 0.333 | 1 | 5 | Fkbp8 | -0.021 | Young |
| 0.630012 | 0.628902 | 0.03 | 0.025 | 1 | 5 | Cish | 0.005 | Young |
| 0.630052 | 0.143446 | 0.206 | 0.194 | 1 | 5 | Zc3h18 | 0.012 | Young |
| 0.630201 | 0.043473 | 0.109 | 0.121 | 1 | 5 | Pitpnm1 | -0.012 | Young |
| 0.630239 | 0.148759 | 0.307 | 0.291 | 1 | 5 | Zfr | 0.016 | Young |
| 0.630388 | 0.052833 | 0.142 | 0.155 | 1 | 5 | Ccdc88b | -0.013 | Young |
| 0.630407 | 0.017306 | 0.089 | 0.079 | 1 | 5 | Nemp1 | 0.01 | Young |
| 0.630886 | 0.855252 | 0.015 | 0.012 | 1 | 5 | Slc43a3 | 0.003 | Young |
| 0.631076 | 0.111872 | 0.069 | 0.077 | 1 | 5 | Alkbh3 | -0.008 | Young |

| | | | | | | | | |
|---|---|---|---|---|---|---|---|---|
| 0.631173 | 0.20284 | 0.254 | 0.249 | 1 | 5 | Ndfip2 | 0.005 | Young |
| 0.631238 | 0.11272 | 0.528 | 0.544 | 1 | 5 | 7-Sep | -0.016 | Young |
| 0.631552 | 0.483039 | 0.043 | 0.037 | 1 | 5 | H2-Ob | 0.006 | Young |
| 0.631615 | 0.140566 | 0.327 | 0.316 | 1 | 5 | Ube2k | 0.011 | Young |
| 0.631668 | 0.051315 | 0.297 | 0.285 | 1 | 5 | Ncoa2 | 0.012 | Young |
| 0.63174 | 0.280887 | 0.094 | 0.086 | 1 | 5 | Golim4 | 0.008 | Young |
| 0.631788 | 0.158881 | 0.345 | 0.335 | 1 | 5 | Cct2 | 0.01 | Young |
| 0.632336 | 0.063459 | 0.18 | 0.165 | 1 | 5 | Rnaset2b | 0.015 | Young |
| 0.632394 | 0.049031 | 0.056 | 0.049 | 1 | 5 | Hscb | 0.007 | Young |
| 0.632532 | 0.194636 | 0.127 | 0.118 | 1 | 5 | Fut8 | 0.009 | Young |
| 0.632556 | 0.127229 | 0.574 | 0.582 | 1 | 5 | Cmtm6 | -0.008 | Young |
| 0.632606 | 0.264948 | 0.14 | 0.13 | 1 | 5 | Yipf6 | 0.01 | Young |
| 0.633071 | 0.032074 | 0.091 | 0.101 | 1 | 5 | Riok2 | -0.01 | Young |
| 0.633242 | 0.032307 | 0.919 | 0.965 | 1 | 5 | Rtn4 | -0.046 | Young |
| 0.633246 | 0.305756 | 0.048 | 0.042 | 1 | 5 | Cdan1 | 0.006 | Young |
| 0.633331 | 0.10681 | 0.571 | 0.569 | 1 | 5 | Foxn3 | 0.002 | Young |
| 0.633339 | 0.028738 | 0.206 | 0.189 | 1 | 5 | Rnf38 | 0.017 | Young |
| 0.633347 | 0.320193 | 0.036 | 0.03 | 1 | 5 | Il27 | 0.006 | Young |
| 0.633348 | 0.64563 | 0.015 | 0.012 | 1 | 5 | Pianp | 0.003 | Young |
| 0.633348 | 0.587576 | 0.015 | 0.012 | 1 | 5 | Abcb9 | 0.003 | Young |
| 0.633348 | 0.469224 | 0.015 | 0.012 | 1 | 5 | Gm13465 | 0.003 | Young |
| 0.633457 | 0.211859 | 0.028 | 0.034 | 1 | 5 | Egfl7 | -0.006 | Young |
| 0.633697 | 0.107618 | 0.528 | 0.532 | 1 | 5 | Selenot | -0.004 | Young |
| 0.633843 | 0.037417 | 0.076 | 0.067 | 1 | 5 | Mterf3 | 0.009 | Young |
| 0.634017 | 0.065831 | 0.069 | 0.061 | 1 | 5 | Fam58b | 0.008 | Young |
| 0.634053 | 0.266533 | 0.023 | 0.019 | 1 | 5 | Chpf | 0.004 | Young |
| 0.634178 | 0.174247 | 0.284 | 0.273 | 1 | 5 | Setx | 0.011 | Young |
| 0.634412 | 0.278409 | 0.02 | 0.025 | 1 | 5 | Wdr4 | -0.005 | Young |
| 0.634544 | 0.026959 | 0.216 | 0.231 | 1 | 5 | Cacul1 | -0.015 | Young |
| 0.634918 | 0.00688 | 0.251 | 0.263 | 1 | 5 | Vbp1 | -0.012 | Young |
| 0.634929 | 0.205323 | 0.259 | 0.249 | 1 | 5 | Plod1 | 0.01 | Young |
| 0.635117 | 0.082582 | 0.845 | 0.877 | 1 | 5 | Tcf4 | -0.032 | Young |
| 0.635383 | 0.306555 | 0.023 | 0.019 | 1 | 5 | Acbd4 | 0.004 | Young |
| 0.635386 | 0.329881 | 0.061 | 0.054 | 1 | 5 | Rhoj | 0.007 | Young |
| 0.6354 | 0.090512 | 0.226 | 0.215 | 1 | 5 | Lat2 | 0.011 | Young |
| 0.635426 | 0.290102 | 0.104 | 0.096 | 1 | 5 | Mthfs | 0.008 | Young |
| 0.635523 | 0.103376 | 0.376 | 0.359 | 1 | 5 | Fndc3b | 0.017 | Young |
| 0.63562 | 0.134901 | 0.223 | 0.212 | 1 | 5 | Cyb5r4 | 0.011 | Young |
| 0.635624 | 0.143956 | 0.459 | 0.448 | 1 | 5 | Cab39 | 0.011 | Young |
| 0.635815 | 0.468645 | 0.015 | 0.012 | 1 | 5 | D130020LC | 0.003 | Young |
| 0.636202 | 0.201011 | 0.056 | 0.049 | 1 | 5 | Ube2o | 0.007 | Young |
| 0.636306 | 0.193608 | 0.147 | 0.136 | 1 | 5 | Gbf1 | 0.011 | Young |
| 0.63634 | 0.130897 | 0.063 | 0.056 | 1 | 5 | Fastk | 0.007 | Young |
| 0.636562 | 0.02298 | 0.048 | 0.056 | 1 | 5 | Dbp | -0.008 | Young |
| 0.636601 | 0.211685 | 0.061 | 0.054 | 1 | 5 | Jpx | 0.007 | Young |

| | | | | | | | | |
|---|---|---|---|---|---|---|---|---|
| 0.636638 | 0.555279 | 0.015 | 0.012 | 1 | 5 | Hacl1 | 0.003 | Young |
| 0.636976 | 0.375021 | 0.076 | 0.069 | 1 | 5 | Zfp780b | 0.007 | Young |
| 0.636989 | 0.004047 | 0.175 | 0.187 | 1 | 5 | Mob3a | -0.012 | Young |
| 0.637242 | 0.080458 | 0.975 | 0.973 | 1 | 5 | Dusp1 | 0.002 | Young |
| 0.637461 | 0.332424 | 0.015 | 0.012 | 1 | 5 | Bcl6b | 0.003 | Young |
| 0.638287 | 0.313803 | 0.036 | 0.042 | 1 | 5 | Rpap2 | -0.006 | Young |
| 0.638297 | 0.248615 | 0.028 | 0.034 | 1 | 5 | Arl4d | -0.006 | Young |
| 0.638447 | 0.08845 | 0.102 | 0.093 | 1 | 5 | Cnksr3 | 0.009 | Young |
| 0.638597 | 0.027505 | 0.053 | 0.061 | 1 | 5 | 2700097O( | -0.008 | Young |
| 0.63899 | 0.00229 | 0.398 | 0.384 | 1 | 5 | Itgb5 | 0.014 | Young |
| 0.639308 | 0.044499 | 0.069 | 0.077 | 1 | 5 | Sugp1 | -0.008 | Young |
| 0.639347 | 0.034604 | 0.282 | 0.3 | 1 | 5 | Tmbim4 | -0.018 | Young |
| 0.639442 | 0.27231 | 0.061 | 0.054 | 1 | 5 | Xlr4b | 0.007 | Young |
| 0.639442 | 0.238057 | 0.061 | 0.054 | 1 | 5 | Fbf1 | 0.007 | Young |
| 0.639496 | 0.154501 | 0.449 | 0.444 | 1 | 5 | Wdfy2 | 0.005 | Young |
| 0.63972 | 0.011953 | 0.051 | 0.044 | 1 | 5 | Pomk | 0.007 | Young |
| 0.639735 | 0.088524 | 0.127 | 0.14 | 1 | 5 | Ccdc127 | -0.013 | Young |
| 0.639789 | 0.19586 | 0.221 | 0.209 | 1 | 5 | Jade1 | 0.012 | Young |
| 0.639801 | 0.011503 | 0.358 | 0.328 | 1 | 5 | Hnrnpd | 0.03 | Young |
| 0.639818 | 0.214266 | 0.213 | 0.204 | 1 | 5 | Tm2d1 | 0.009 | Young |
| 0.639894 | 0.150724 | 0.14 | 0.128 | 1 | 5 | Cdk8 | 0.012 | Young |
| 0.639934 | 0.452765 | 0.015 | 0.012 | 1 | 5 | Asrgl1 | 0.003 | Young |
| 0.639937 | 0.15069 | 0.381 | 0.384 | 1 | 5 | Papd4 | -0.003 | Young |
| 0.640164 | 0.0822 | 0.058 | 0.066 | 1 | 5 | Cstf1 | -0.008 | Young |
| 0.640582 | 0.247875 | 0.109 | 0.101 | 1 | 5 | Rnf168 | 0.008 | Young |
| 0.640717 | 0.242688 | 0.023 | 0.019 | 1 | 5 | Zfp472 | 0.004 | Young |
| 0.640717 | 0.186651 | 0.023 | 0.019 | 1 | 5 | Tctn1 | 0.004 | Young |
| 0.640925 | 0.173627 | 0.223 | 0.212 | 1 | 5 | Sarnp | 0.011 | Young |
| 0.641042 | 0.034992 | 0.01 | 0.013 | 1 | 5 | Mlkl | -0.003 | Young |
| 0.641069 | 0.010254 | 0.168 | 0.184 | 1 | 5 | Zfp62 | -0.016 | Young |
| 0.641385 | 0.219343 | 0.023 | 0.019 | 1 | 5 | Mblac1 | 0.004 | Young |
| 0.641385 | 0.176416 | 0.023 | 0.019 | 1 | 5 | Tnfaip8l3 | 0.004 | Young |
| 0.641585 | 0.319596 | 0.015 | 0.012 | 1 | 5 | Slc26a6 | 0.003 | Young |
| 0.641684 | 0.267185 | 0.135 | 0.125 | 1 | 5 | Prkar2b | 0.01 | Young |
| 0.642031 | 0.088169 | 0.15 | 0.14 | 1 | 5 | Fnbp4 | 0.01 | Young |
| 0.642146 | 0.056777 | 0.056 | 0.049 | 1 | 5 | Aimp2 | 0.007 | Young |
| 0.642172 | 0.145158 | 0.216 | 0.204 | 1 | 5 | Zbtb44 | 0.012 | Young |
| 0.642412 | 0.646209 | 0.015 | 0.012 | 1 | 5 | Ccsap | 0.003 | Young |
| 0.642527 | 0.029052 | 0.142 | 0.153 | 1 | 5 | Rpe | -0.011 | Young |
| 0.642867 | 0.081484 | 0.642 | 0.643 | 1 | 5 | Anp32a | -0.001 | Young |
| 0.643529 | 0.017343 | 0.396 | 0.414 | 1 | 5 | mt-Atp8 | -0.018 | Young |
| 0.643791 | 0.309642 | 0.053 | 0.047 | 1 | 5 | Tmem221 | 0.006 | Young |
| 0.643931 | 0.059381 | 0.35 | 0.374 | 1 | 5 | Vamp3 | -0.024 | Young |
| 0.644065 | 0.215855 | 0.015 | 0.012 | 1 | 5 | Trim13 | 0.003 | Young |
| 0.644233 | 0.39418 | 0.028 | 0.034 | 1 | 5 | Zfp606 | -0.006 | Young |

| | | | | | | | | |
|---|---|---|---|---|---|---|---|---|
| 0.644274 | 0.045818 | 0.056 | 0.049 | 1 | 5 | Ppdpf | 0.007 | Young |
| 0.644301 | 0.053727 | 0.124 | 0.113 | 1 | 5 | Mrpl21 | 0.011 | Young |
| 0.64464 | 0.161784 | 0.089 | 0.081 | 1 | 5 | Klhl2 | 0.008 | Young |
| 0.645229 | 0.173284 | 0.119 | 0.109 | 1 | 5 | Pfdn6 | 0.01 | Young |
| 0.645331 | 0.095961 | 0.241 | 0.226 | 1 | 5 | Slc16a7 | 0.015 | Young |
| 0.645334 | 0.003573 | 0.272 | 0.288 | 1 | 5 | Armcx3 | -0.016 | Young |
| 0.6454 | 0.165735 | 0.023 | 0.019 | 1 | 5 | Srfbp1 | 0.004 | Young |
| 0.645638 | 0.111043 | 0.272 | 0.291 | 1 | 5 | Stat5b | -0.019 | Young |
| 0.645721 | 0.120576 | 0.015 | 0.012 | 1 | 5 | Cd46 | 0.003 | Young |
| 0.645721 | 0.116085 | 0.015 | 0.012 | 1 | 5 | Dagla | 0.003 | Young |
| 0.645721 | 0.01719 | 0.015 | 0.012 | 1 | 5 | L1cam | 0.003 | Young |
| 0.646679 | 0.270196 | 0.089 | 0.081 | 1 | 5 | Ppp2r3c | 0.008 | Young |
| 0.646697 | 0.008768 | 0.571 | 0.604 | 1 | 5 | Matr3 | -0.033 | Young |
| 0.646747 | 0.163692 | 0.147 | 0.138 | 1 | 5 | Lrrc47 | 0.009 | Young |
| 0.646991 | 0.117305 | 0.043 | 0.037 | 1 | 5 | Ppat | 0.006 | Young |
| 0.647191 | 0.260976 | 0.218 | 0.21 | 1 | 5 | Ddhd2 | 0.008 | Young |
| 0.647251 | 0.266093 | 0.226 | 0.222 | 1 | 5 | Ankrd13c | 0.004 | Young |
| 0.647374 | 0.399161 | 0.048 | 0.042 | 1 | 5 | Chd1l | 0.006 | Young |
| 0.647411 | 0.025963 | 0.023 | 0.019 | 1 | 5 | Nectin2 | 0.004 | Young |
| 0.647531 | 0.231504 | 0.162 | 0.153 | 1 | 5 | Gps1 | 0.009 | Young |
| 0.647929 | 0.240111 | 0.01 | 0.013 | 1 | 5 | Sh3rf1 | -0.003 | Young |
| 0.64798 | 0.077847 | 0.371 | 0.372 | 1 | 5 | Ythdf3 | -0.001 | Young |
| 0.648082 | 0.004676 | 0.018 | 0.022 | 1 | 5 | Zfp958 | -0.004 | Young |
| 0.648207 | 0.19929 | 0.015 | 0.012 | 1 | 5 | Col6a1 | 0.003 | Young |
| 0.648907 | 0.031238 | 0.228 | 0.214 | 1 | 5 | Fmr1 | 0.014 | Young |
| 0.649028 | 0.054767 | 0.069 | 0.061 | 1 | 5 | Klhl22 | 0.008 | Young |
| 0.649037 | 0.131275 | 0.015 | 0.012 | 1 | 5 | Wee1 | 0.003 | Young |
| 0.649313 | 0.138772 | 0.477 | 0.466 | 1 | 5 | Eif2s2 | 0.011 | Young |
| 0.650026 | 0.009982 | 0.063 | 0.071 | 1 | 5 | Gar1 | -0.008 | Young |
| 0.650082 | 0.062166 | 0.482 | 0.466 | 1 | 5 | Ppp1cc | 0.016 | Young |
| 0.6504 | 0.0645 | 0.096 | 0.088 | 1 | 5 | Rexo4 | 0.008 | Young |
| 0.650582 | 0.295414 | 0.048 | 0.042 | 1 | 5 | Lmnb1 | 0.006 | Young |
| 0.650713 | 0.047579 | 0.16 | 0.148 | 1 | 5 | Akap10 | 0.012 | Young |
| 0.65093 | 0.192982 | 0.269 | 0.264 | 1 | 5 | Cdc40 | 0.005 | Young |
| 0.651105 | 0.0992 | 0.635 | 0.66 | 1 | 5 | Prrc2c | -0.025 | Young |
| 0.651201 | 0.011113 | 0.467 | 0.48 | 1 | 5 | Dok2 | -0.013 | Young |
| 0.651219 | 0.10031 | 0.277 | 0.269 | 1 | 5 | Zfp516 | 0.008 | Young |
| 0.651313 | 0.227108 | 0.376 | 0.384 | 1 | 5 | Ndufb8 | -0.008 | Young |
| 0.651383 | 0.280084 | 0.01 | 0.013 | 1 | 5 | Plcl1 | -0.003 | Young |
| 0.651598 | 0.191572 | 0.178 | 0.167 | 1 | 5 | Cnot2 | 0.011 | Young |
| 0.651778 | 0.004026 | 0.637 | 0.643 | 1 | 5 | Cited2 | -0.006 | Young |
| 0.651941 | 0.122134 | 0.053 | 0.061 | 1 | 5 | Dgcr6 | -0.008 | Young |
| 0.652015 | 0.055255 | 0.398 | 0.391 | 1 | 5 | Zbtb4 | 0.007 | Young |
| 0.652357 | 0.319147 | 0.096 | 0.089 | 1 | 5 | Rasal2 | 0.007 | Young |
| 0.652499 | 0.277299 | 0.036 | 0.03 | 1 | 5 | Elmod3 | 0.006 | Young |

| | | | | | | | | |
|---|---|---|---|---|---|---|---|---|
| 0.652525 | 0.017626 | 0.193 | 0.205 | 1 | 5 | Socs4 | -0.012 | Young |
| 0.652561 | 0.063584 | 0.292 | 0.286 | 1 | 5 | Rgl2 | 0.006 | Young |
| 0.652658 | 0.057371 | 0.213 | 0.2 | 1 | 5 | Nr4a2 | 0.013 | Young |
| 0.652661 | 0.001994 | 0.063 | 0.071 | 1 | 5 | Mfge8 | -0.008 | Young |
| 0.653044 | 0.079915 | 0.391 | 0.424 | 1 | 5 | Arid4a | -0.033 | Young |
| 0.653156 | 0.002737 | 0.231 | 0.246 | 1 | 5 | Rsf1 | -0.015 | Young |
| 0.653328 | 0.358057 | 0.117 | 0.109 | 1 | 5 | Tnip1 | 0.008 | Young |
| 0.65334 | 0.336213 | 0.058 | 0.052 | 1 | 5 | Gpank1 | 0.006 | Young |
| 0.65357 | 0.235518 | 0.036 | 0.03 | 1 | 5 | Ccnyl1 | 0.006 | Young |
| 0.653765 | 0.128882 | 0.076 | 0.069 | 1 | 5 | Naa25 | 0.007 | Young |
| 0.653808 | 0.045302 | 0.35 | 0.364 | 1 | 5 | Eif3d | -0.014 | Young |
| 0.654256 | 0.118786 | 0.048 | 0.042 | 1 | 5 | Slco3a1 | 0.006 | Young |
| 0.654459 | 0.279974 | 0.053 | 0.047 | 1 | 5 | Orai2 | 0.006 | Young |
| 0.65452 | 0.038835 | 0.168 | 0.155 | 1 | 5 | Lypla1 | 0.013 | Young |
| 0.654676 | 0.126647 | 0.096 | 0.088 | 1 | 5 | Uckl1 | 0.008 | Young |
| 0.655004 | 0.123425 | 0.292 | 0.279 | 1 | 5 | Agfg1 | 0.013 | Young |
| 0.655423 | 0.103984 | 0.069 | 0.061 | 1 | 5 | Exd2 | 0.008 | Young |
| 0.65548 | 0.074933 | 0.018 | 0.022 | 1 | 5 | D930030I0 | -0.004 | Young |
| 0.655636 | 0.075292 | 0.048 | 0.042 | 1 | 5 | Bloc1s3 | 0.006 | Young |
| 0.656014 | 0.058175 | 0.127 | 0.14 | 1 | 5 | Pex5 | -0.013 | Young |
| 0.656634 | 0.142322 | 0.234 | 0.224 | 1 | 5 | Esyt1 | 0.01 | Young |
| 0.656814 | 0.073146 | 0.079 | 0.071 | 1 | 5 | Sema4b | 0.008 | Young |
| 0.656827 | 0.213087 | 0.124 | 0.116 | 1 | 5 | Ncoa5 | 0.008 | Young |
| 0.656983 | 0.259448 | 0.096 | 0.088 | 1 | 5 | Nt5c3 | 0.008 | Young |
| 0.657401 | 0.377422 | 0.109 | 0.103 | 1 | 5 | Zfpl1 | 0.006 | Young |
| 0.657967 | 0.130256 | 0.497 | 0.492 | 1 | 5 | Pik3r1 | 0.005 | Young |
| 0.658011 | 0.046674 | 0.371 | 0.352 | 1 | 5 | Ube2s | 0.019 | Young |
| 0.65805 | 0.019791 | 0.168 | 0.18 | 1 | 5 | Uqcc2 | -0.012 | Young |
| 0.658321 | 0.10406 | 0.228 | 0.212 | 1 | 5 | Strn4 | 0.016 | Young |
| 0.658669 | 0.169473 | 0.266 | 0.256 | 1 | 5 | Nudt16l1 | 0.01 | Young |
| 0.659148 | 0.040941 | 0.145 | 0.157 | 1 | 5 | Gm8186 | -0.012 | Young |
| 0.659216 | 0.10921 | 0.14 | 0.128 | 1 | 5 | Ubxn2a | 0.012 | Young |
| 0.659569 | 0.01484 | 0.211 | 0.226 | 1 | 5 | Iws1 | -0.015 | Young |
| 0.659571 | 0.010538 | 0.038 | 0.044 | 1 | 5 | Pycrl | -0.006 | Young |
| 0.659627 | 0.004736 | 0.228 | 0.244 | 1 | 5 | Sec24c | -0.016 | Young |
| 0.660116 | 0.041118 | 0.246 | 0.231 | 1 | 5 | Rbm17 | 0.015 | Young |
| 0.660171 | 0.02061 | 1 | 1 | 1 | 5 | mt-Nd1 | 0 | Young |
| 0.661491 | 0.070821 | 0.404 | 0.419 | 1 | 5 | P2rx7 | -0.015 | Young |
| 0.66216 | 0.105381 | 0.124 | 0.135 | 1 | 5 | Pop5 | -0.011 | Young |
| 0.662233 | 0.079269 | 0.518 | 0.53 | 1 | 5 | Tkt | -0.012 | Young |
| 0.662347 | 0.211646 | 0.272 | 0.259 | 1 | 5 | Enox2 | 0.013 | Young |
| 0.66259 | 0.09868 | 0.091 | 0.082 | 1 | 5 | Apba1 | 0.009 | Young |
| 0.663012 | 0.203408 | 0.041 | 0.035 | 1 | 5 | Piga | 0.006 | Young |
| 0.663012 | 0.144935 | 0.218 | 0.207 | 1 | 5 | Anapc1 | 0.011 | Young |
| 0.663283 | 0.014449 | 0.515 | 0.505 | 1 | 5 | Lcp2 | 0.01 | Young |

| | | | | | | | | |
|---|---|---|---|---|---|---|---|---|
| 0.66338 | 0.073726 | 0.025 | 0.03 | 1 | 5 | Cd209c | -0.005 | Young |
| 0.66338 | 0.00809 | 0.025 | 0.03 | 1 | 5 | Ift74 | -0.005 | Young |
| 0.663465 | 0.061856 | 0.584 | 0.586 | 1 | 5 | Gusb | -0.002 | Young |
| 0.663478 | 0.157714 | 0.048 | 0.042 | 1 | 5 | D030056L2 | 0.006 | Young |
| 0.663832 | 0.024477 | 0.17 | 0.158 | 1 | 5 | Acox3 | 0.012 | Young |
| 0.664758 | 0.085882 | 0.046 | 0.052 | 1 | 5 | Ubr7 | -0.006 | Young |
| 0.664914 | 0.544188 | 0.028 | 0.024 | 1 | 5 | Mfsd3 | 0.004 | Young |
| 0.665121 | 0.528894 | 0.102 | 0.096 | 1 | 5 | Afg3l2 | 0.006 | Young |
| 0.665288 | 0.196989 | 0.091 | 0.084 | 1 | 5 | Fam78a | 0.007 | Young |
| 0.665522 | 0.494576 | 0.028 | 0.024 | 1 | 5 | Mr1 | 0.004 | Young |
| 0.665665 | 0.011224 | 0.165 | 0.178 | 1 | 5 | Cd82 | -0.013 | Young |
| 0.665717 | 0.244691 | 0.145 | 0.135 | 1 | 5 | Scaf4 | 0.01 | Young |
| 0.665914 | 0.310797 | 0.069 | 0.062 | 1 | 5 | Zfp64 | 0.007 | Young |
| 0.666334 | 0.12239 | 0.135 | 0.126 | 1 | 5 | Gm11808 | 0.009 | Young |
| 0.666473 | 0.220536 | 0.076 | 0.069 | 1 | 5 | Orc3 | 0.007 | Young |
| 0.666545 | 0.447059 | 0.046 | 0.04 | 1 | 5 | Heatr5b | 0.006 | Young |
| 0.666688 | 0.102972 | 0.14 | 0.13 | 1 | 5 | Iars2 | 0.01 | Young |
| 0.666747 | 0.157781 | 0.119 | 0.109 | 1 | 5 | Osgin2 | 0.01 | Young |
| 0.66759 | 0.160119 | 0.173 | 0.189 | 1 | 5 | Tsnax | -0.016 | Young |
| 0.667747 | 0.03768 | 0.239 | 0.226 | 1 | 5 | Uba6 | 0.013 | Young |
| 0.667957 | 0.259357 | 0.028 | 0.024 | 1 | 5 | Tmem143 | 0.004 | Young |
| 0.668424 | 0.11161 | 0.19 | 0.205 | 1 | 5 | Idh3b | -0.015 | Young |
| 0.668437 | 0.309863 | 0.122 | 0.114 | 1 | 5 | Igip | 0.008 | Young |
| 0.668471 | 0.292868 | 0.137 | 0.13 | 1 | 5 | Ubr1 | 0.007 | Young |
| 0.668546 | 0.040454 | 0.086 | 0.077 | 1 | 5 | Saysd1 | 0.009 | Young |
| 0.668562 | 0.17663 | 0.025 | 0.03 | 1 | 5 | Gm5431 | -0.005 | Young |
| 0.66875 | 0.102316 | 0.132 | 0.121 | 1 | 5 | Zfp787 | 0.011 | Young |
| 0.668861 | 0.138295 | 0.183 | 0.173 | 1 | 5 | Cd99l2 | 0.01 | Young |
| 0.66888 | 0.057553 | 0.368 | 0.389 | 1 | 5 | Twf2 | -0.021 | Young |
| 0.668901 | 0.238892 | 0.038 | 0.044 | 1 | 5 | Hdac7 | -0.006 | Young |
| 0.668915 | 0.161536 | 0.046 | 0.04 | 1 | 5 | Selenoo | 0.006 | Young |
| 0.668988 | 0.005934 | 0.447 | 0.46 | 1 | 5 | Mef2d | -0.013 | Young |
| 0.669011 | 0.056334 | 0.325 | 0.337 | 1 | 5 | N4bp2l2 | -0.012 | Young |
| 0.669138 | 0.33803 | 0.025 | 0.03 | 1 | 5 | Gm15446 | -0.005 | Young |
| 0.669176 | 0.208583 | 0.028 | 0.024 | 1 | 5 | Gm26890 | 0.004 | Young |
| 0.669415 | 0.112466 | 0.292 | 0.285 | 1 | 5 | Polr2k | 0.007 | Young |
| 0.669567 | 0.152582 | 0.094 | 0.086 | 1 | 5 | Smim24 | 0.008 | Young |
| 0.669668 | 0.173755 | 0.15 | 0.14 | 1 | 5 | Fech | 0.01 | Young |
| 0.669722 | 0.217509 | 0.109 | 0.101 | 1 | 5 | Mrpl27 | 0.008 | Young |
| 0.669842 | 0.222653 | 0.223 | 0.217 | 1 | 5 | Dnttip2 | 0.006 | Young |
| 0.669848 | 0.234072 | 0.142 | 0.135 | 1 | 5 | Impa1 | 0.007 | Young |
| 0.669905 | 0.133445 | 0.18 | 0.172 | 1 | 5 | Pdha1 | 0.008 | Young |
| 0.66996 | 0.210293 | 0.147 | 0.14 | 1 | 5 | Brd9 | 0.007 | Young |
| 0.669972 | 0.236973 | 0.114 | 0.108 | 1 | 5 | Rexo1 | 0.006 | Young |
| 0.670423 | 0.344789 | 0.079 | 0.072 | 1 | 5 | Pigv | 0.007 | Young |

| | | | | | | | | |
|---|---|---|---|---|---|---|---|---|
| 0.670793 | 0.244523 | 0.188 | 0.178 | 1 | 5 | Slu7 | 0.01 | Young |
| 0.671099 | 0.016029 | 0.195 | 0.207 | 1 | 5 | Mrps24 | -0.012 | Young |
| 0.671551 | 0.029581 | 0.053 | 0.061 | 1 | 5 | AC127341. | -0.008 | Young |
| 0.672028 | 0.050732 | 0.046 | 0.052 | 1 | 5 | Lin52 | -0.006 | Young |
| 0.672052 | 0.050449 | 0.234 | 0.246 | 1 | 5 | Rexo2 | -0.012 | Young |
| 0.672077 | 0.425663 | 0.041 | 0.035 | 1 | 5 | Tmco4 | 0.006 | Young |
| 0.672204 | 0.07779 | 0.185 | 0.2 | 1 | 5 | Ppp1r9a | -0.015 | Young |
| 0.672568 | 0.059458 | 0.482 | 0.473 | 1 | 5 | Herc1 | 0.009 | Young |
| 0.672864 | 0.155523 | 0.292 | 0.279 | 1 | 5 | Ggh | 0.013 | Young |
| 0.672866 | 0.185626 | 0.157 | 0.146 | 1 | 5 | Clpx | 0.011 | Young |
| 0.672886 | 0.322782 | 0.071 | 0.064 | 1 | 5 | Wdr70 | 0.007 | Young |
| 0.673285 | 0.262458 | 0.16 | 0.153 | 1 | 5 | Cep250 | 0.007 | Young |
| 0.674147 | 0.027361 | 0.048 | 0.042 | 1 | 5 | Nudt7 | 0.006 | Young |
| 0.674193 | 0.126235 | 0.211 | 0.204 | 1 | 5 | Gch1 | 0.007 | Young |
| 0.674248 | 0.058808 | 0.419 | 0.441 | 1 | 5 | Rsu1 | -0.022 | Young |
| 0.674334 | 0.120428 | 0.157 | 0.146 | 1 | 5 | Rif1 | 0.011 | Young |
| 0.674646 | 0.371149 | 0.109 | 0.103 | 1 | 5 | Fam69a | 0.006 | Young |
| 0.674727 | 0.11784 | 0.414 | 0.401 | 1 | 5 | Nucb1 | 0.013 | Young |
| 0.674757 | 0.103774 | 0.081 | 0.074 | 1 | 5 | Msra | 0.007 | Young |
| 0.675035 | 0.124258 | 0.368 | 0.404 | 1 | 5 | Rab20 | -0.036 | Young |
| 0.675071 | 0.525864 | 0.02 | 0.017 | 1 | 5 | Snx32 | 0.003 | Young |
| 0.675282 | 0.137217 | 0.028 | 0.024 | 1 | 5 | S100a16 | 0.004 | Young |
| 0.675618 | 0.01733 | 0.244 | 0.259 | 1 | 5 | Sik1 | -0.015 | Young |
| 0.675873 | 0.13987 | 0.038 | 0.044 | 1 | 5 | 1300002E1 | -0.006 | Young |
| 0.676151 | 0.369712 | 0.107 | 0.099 | 1 | 5 | C1galt1c1 | 0.008 | Young |
| 0.676275 | 0.059432 | 0.67 | 0.684 | 1 | 5 | Cox6b1 | -0.014 | Young |
| 0.676505 | 0.20316 | 0.137 | 0.13 | 1 | 5 | Mrpl2 | 0.007 | Young |
| 0.676636 | 0.213556 | 0.127 | 0.12 | 1 | 5 | Ttll12 | 0.007 | Young |
| 0.676836 | 0.106174 | 0.038 | 0.044 | 1 | 5 | Flad1 | -0.006 | Young |
| 0.676992 | 0.055739 | 0.109 | 0.118 | 1 | 5 | Ak3 | -0.009 | Young |
| 0.677127 | 0.098077 | 0.482 | 0.483 | 1 | 5 | Wls | -0.001 | Young |
| 0.677199 | 0.105231 | 0.234 | 0.258 | 1 | 5 | Akirin1 | -0.024 | Young |
| 0.677227 | 0.726521 | 0.02 | 0.017 | 1 | 5 | C2 | 0.003 | Young |
| 0.677347 | 0.470298 | 0.076 | 0.071 | 1 | 5 | Stmn1 | 0.005 | Young |
| 0.677557 | 0.183223 | 0.183 | 0.175 | 1 | 5 | Dicer1 | 0.008 | Young |
| 0.677567 | 0.212074 | 0.089 | 0.081 | 1 | 5 | Eipr1 | 0.008 | Young |
| 0.677597 | 0.220914 | 0.198 | 0.19 | 1 | 5 | Rfc1 | 0.008 | Young |
| 0.677662 | 0.102872 | 0.348 | 0.345 | 1 | 5 | Ago2 | 0.003 | Young |
| 0.67776 | 0.126029 | 0.089 | 0.098 | 1 | 5 | Timm10 | -0.009 | Young |
| 0.677813 | 0.206787 | 0.025 | 0.03 | 1 | 5 | Kyat1 | -0.005 | Young |
| 0.677896 | 0.062877 | 0.916 | 0.911 | 1 | 5 | Cebpb | 0.005 | Young |
| 0.677901 | 0.069583 | 0.464 | 0.455 | 1 | 5 | Irf5 | 0.009 | Young |
| 0.677966 | 0.06383 | 0.195 | 0.212 | 1 | 5 | Cyb5b | -0.017 | Young |
| 0.678301 | 0.152774 | 0.439 | 0.428 | 1 | 5 | Fchsd2 | 0.011 | Young |
| 0.678965 | 0.046496 | 0.353 | 0.372 | 1 | 5 | Cct3 | -0.019 | Young |

| | | | | | | | | |
|---|---|---|---|---|---|---|---|---|
| 0.679054 | 0.17299 | 0.228 | 0.217 | 1 | 5 | Aaed1 | 0.011 | Young |
| 0.679133 | 0.309018 | 0.033 | 0.029 | 1 | 5 | Ankrd39 | 0.004 | Young |
| 0.679295 | 0.180094 | 0.084 | 0.076 | 1 | 5 | Nectin4 | 0.008 | Young |
| 0.679486 | 0.297015 | 0.102 | 0.094 | 1 | 5 | Klf16 | 0.008 | Young |
| 0.679643 | 0.080527 | 0.14 | 0.15 | 1 | 5 | Pigx | -0.01 | Young |
| 0.679662 | 0.201176 | 0.236 | 0.231 | 1 | 5 | Lpin1 | 0.005 | Young |
| 0.679695 | 0.590471 | 0.033 | 0.029 | 1 | 5 | Gm29291 | 0.004 | Young |
| 0.679859 | 0.426081 | 0.046 | 0.04 | 1 | 5 | Creld1 | 0.006 | Young |
| 0.680005 | 0.074013 | 0.084 | 0.093 | 1 | 5 | Ano10 | -0.009 | Young |
| 0.680432 | 0.287139 | 0.107 | 0.099 | 1 | 5 | Exosc3 | 0.008 | Young |
| 0.680521 | 0.125853 | 0.543 | 0.549 | 1 | 5 | Ski | -0.006 | Young |
| 0.68078 | 0.226699 | 0.089 | 0.082 | 1 | 5 | Smn1 | 0.007 | Young |
| 0.680827 | 0.431703 | 0.02 | 0.017 | 1 | 5 | Atp6v0c | 0.003 | Young |
| 0.680843 | 0.003455 | 0.368 | 0.396 | 1 | 5 | Tmem234 | -0.028 | Young |
| 0.680939 | 0.267728 | 0.355 | 0.36 | 1 | 5 | Glyr1 | -0.005 | Young |
| 0.681022 | 0.291062 | 0.317 | 0.318 | 1 | 5 | Cdipt | -0.001 | Young |
| 0.681652 | 0.074396 | 0.132 | 0.123 | 1 | 5 | Spns1 | 0.009 | Young |
| 0.681746 | 0.057364 | 0.107 | 0.114 | 1 | 5 | Eif2b5 | -0.007 | Young |
| 0.68177 | 0.238626 | 0.046 | 0.04 | 1 | 5 | Zeb1 | 0.006 | Young |
| 0.68181 | 0.23374 | 0.063 | 0.057 | 1 | 5 | Cdkal1 | 0.006 | Young |
| 0.681827 | 0.354836 | 0.145 | 0.138 | 1 | 5 | Bicral | 0.007 | Young |
| 0.681865 | 0.17056 | 0.129 | 0.121 | 1 | 5 | Ssh1 | 0.008 | Young |
| 0.682203 | 0.005279 | 0.036 | 0.03 | 1 | 5 | Cep152 | 0.006 | Young |
| 0.682268 | 0.142937 | 0.02 | 0.017 | 1 | 5 | Cdiptos | 0.003 | Young |
| 0.682352 | 0.07171 | 0.142 | 0.152 | 1 | 5 | Mrps9 | -0.01 | Young |
| 0.683175 | 0.113701 | 0.223 | 0.217 | 1 | 5 | Fbxl20 | 0.006 | Young |
| 0.683235 | 0.304965 | 0.063 | 0.057 | 1 | 5 | Tsen15 | 0.006 | Young |
| 0.683253 | 0.007142 | 0.985 | 0.983 | 1 | 5 | Rpl13 | 0.002 | Young |
| 0.683487 | 0.474203 | 0.091 | 0.084 | 1 | 5 | Gbp2 | 0.007 | Young |
| 0.683535 | 0.229672 | 0.315 | 0.316 | 1 | 5 | Rraga | -0.001 | Young |
| 0.683731 | 0.392697 | 0.041 | 0.035 | 1 | 5 | Polm | 0.006 | Young |
| 0.683974 | 0.16898 | 0.228 | 0.253 | 1 | 5 | Frmd4a | -0.025 | Young |
| 0.684506 | 0.192617 | 0.081 | 0.074 | 1 | 5 | Ermard | 0.007 | Young |
| 0.684673 | 0.189785 | 0.464 | 0.483 | 1 | 5 | Hspe1 | -0.019 | Young |
| 0.684723 | 0.203104 | 0.107 | 0.099 | 1 | 5 | Trim11 | 0.008 | Young |
| 0.685177 | 0.398225 | 0.043 | 0.039 | 1 | 5 | Jak3 | 0.004 | Young |
| 0.685186 | 0.196007 | 0.162 | 0.153 | 1 | 5 | Rnf166 | 0.009 | Young |
| 0.68586 | 0.276104 | 0.188 | 0.182 | 1 | 5 | Npm3 | 0.006 | Young |
| 0.685878 | 0.537766 | 0.02 | 0.017 | 1 | 5 | Gm11423 | 0.003 | Young |
| 0.685878 | 0.451468 | 0.02 | 0.017 | 1 | 5 | Farp2 | 0.003 | Young |
| 0.685878 | 0.207713 | 0.02 | 0.017 | 1 | 5 | Nemp2 | 0.003 | Young |
| 0.685879 | 0.438677 | 0.033 | 0.029 | 1 | 5 | Hibch | 0.004 | Young |
| 0.685879 | 0.410207 | 0.033 | 0.029 | 1 | 5 | Mcoln2 | 0.004 | Young |
| 0.685927 | 0.034838 | 0.34 | 0.325 | 1 | 5 | Sh2b3 | 0.015 | Young |
| 0.685994 | 0.527106 | 0.061 | 0.056 | 1 | 5 | Lig3 | 0.005 | Young |

| | | | | | | | | |
|---|---|---|---|---|---|---|---|---|
| 0.685994 | 0.127043 | 0.424 | 0.431 | 1 | 5 | Snu13 | -0.007 | Young |
| 0.686075 | 0.282944 | 0.046 | 0.04 | 1 | 5 | Tmem86b | 0.006 | Young |
| 0.686172 | 0.168163 | 0.051 | 0.045 | 1 | 5 | 0610009O2 | 0.006 | Young |
| 0.686601 | 0.155971 | 0.02 | 0.017 | 1 | 5 | Inmt | 0.003 | Young |
| 0.686628 | 0.044759 | 0.173 | 0.184 | 1 | 5 | BC003331 | -0.011 | Young |
| 0.687109 | 0.069009 | 0.14 | 0.13 | 1 | 5 | Alas1 | 0.01 | Young |
| 0.687245 | 0.139039 | 0.051 | 0.057 | 1 | 5 | Pnpt1 | -0.006 | Young |
| 0.687324 | 0.228107 | 0.241 | 0.259 | 1 | 5 | D5Ertd579 | -0.018 | Young |
| 0.687326 | 0.271332 | 0.109 | 0.101 | 1 | 5 | Arl2 | 0.008 | Young |
| 0.688448 | 0.175417 | 0.284 | 0.276 | 1 | 5 | Cox20 | 0.008 | Young |
| 0.688508 | 0.020223 | 0.65 | 0.677 | 1 | 5 | Cnbp | -0.027 | Young |
| 0.688771 | 0.188001 | 0.02 | 0.017 | 1 | 5 | 1700019L1 | 0.003 | Young |
| 0.688873 | 0.206302 | 0.297 | 0.293 | 1 | 5 | St3gal1 | 0.004 | Young |
| 0.68935 | 0.121892 | 0.063 | 0.057 | 1 | 5 | Npat | 0.006 | Young |
| 0.689442 | 0.023403 | 0.142 | 0.131 | 1 | 5 | Got2 | 0.011 | Young |
| 0.689495 | 0.252828 | 0.02 | 0.017 | 1 | 5 | Nmnat3 | 0.003 | Young |
| 0.689495 | 0.15278 | 0.02 | 0.017 | 1 | 5 | Knstrn | 0.003 | Young |
| 0.690219 | 0.262758 | 0.02 | 0.017 | 1 | 5 | Gm15247 | 0.003 | Young |
| 0.690267 | 0.041295 | 0.827 | 0.86 | 1 | 5 | Alox5ap | -0.033 | Young |
| 0.690391 | 0.100382 | 0.381 | 0.364 | 1 | 5 | Dnaja2 | 0.017 | Young |
| 0.690776 | 0.106095 | 0.008 | 0.01 | 1 | 5 | Dock5 | -0.002 | Young |
| 0.690955 | 0.286874 | 0.033 | 0.029 | 1 | 5 | Dgke | 0.004 | Young |
| 0.691424 | 0.103035 | 0.231 | 0.247 | 1 | 5 | Mpc1 | -0.016 | Young |
| 0.691524 | 0.131953 | 0.17 | 0.16 | 1 | 5 | Chmp1b | 0.01 | Young |
| 0.691633 | 0.179061 | 0.381 | 0.367 | 1 | 5 | Smagp | 0.014 | Young |
| 0.691797 | 0.68754 | 0.008 | 0.01 | 1 | 5 | 2900026A0 | -0.002 | Young |
| 0.691797 | 0.225284 | 0.008 | 0.01 | 1 | 5 | Phgdh | -0.002 | Young |
| 0.691797 | 0.074417 | 0.008 | 0.01 | 1 | 5 | Gm40773 | -0.002 | Young |
| 0.691873 | 0.201874 | 0.254 | 0.244 | 1 | 5 | Etfa | 0.01 | Young |
| 0.691998 | 0.148882 | 0.525 | 0.512 | 1 | 5 | Ash1l | 0.013 | Young |
| 0.692143 | 0.128749 | 0.051 | 0.057 | 1 | 5 | Cables2 | -0.006 | Young |
| 0.692201 | 0.173234 | 0.244 | 0.237 | 1 | 5 | Polr2m | 0.007 | Young |
| 0.692222 | 0.009812 | 0.132 | 0.143 | 1 | 5 | Usp33 | -0.011 | Young |
| 0.692502 | 0.143432 | 0.145 | 0.135 | 1 | 5 | Appl2 | 0.01 | Young |
| 0.692644 | 0.615406 | 0.013 | 0.01 | 1 | 5 | Cdca8 | 0.003 | Young |
| 0.692818 | 0.007601 | 0.008 | 0.01 | 1 | 5 | Capn5 | -0.002 | Young |
| 0.693839 | 0.347212 | 0.008 | 0.01 | 1 | 5 | Gm20219 | -0.002 | Young |
| 0.693839 | 0.251129 | 0.008 | 0.01 | 1 | 5 | Aatk | -0.002 | Young |
| 0.693839 | 0.209434 | 0.008 | 0.01 | 1 | 5 | Cd34 | -0.002 | Young |
| 0.693839 | 0.064511 | 0.008 | 0.01 | 1 | 5 | Gm37800 | -0.002 | Young |
| 0.693848 | 0.083222 | 0.056 | 0.062 | 1 | 5 | Snx19 | -0.006 | Young |
| 0.694174 | 0.109482 | 0.409 | 0.399 | 1 | 5 | Csnk1e | 0.01 | Young |
| 0.694337 | 0.140704 | 0.142 | 0.155 | 1 | 5 | Dapk3 | -0.013 | Young |
| 0.694541 | 0.075696 | 0.67 | 0.673 | 1 | 5 | Dstn | -0.003 | Young |
| 0.69457 | 0.203998 | 0.02 | 0.017 | 1 | 5 | BC043934 | 0.003 | Young |

| | | | | | | | | |
|---|---|---|---|---|---|---|---|---|
| 0.694585 | 0.011289 | 0.094 | 0.086 | 1 | 5 | Gnpat | 0.008 | Young |
| 0.694605 | 0.018985 | 0.421 | 0.439 | 1 | 5 | Atp5d | -0.018 | Young |
| 0.694682 | 0.056594 | 0.586 | 0.589 | 1 | 5 | Mtpn | -0.003 | Young |
| 0.694861 | 0.405563 | 0.008 | 0.01 | 1 | 5 | Neo1 | -0.002 | Young |
| 0.695314 | 0.041975 | 0.36 | 0.38 | 1 | 5 | Ddb1 | -0.02 | Young |
| 0.695337 | 0.041122 | 0.289 | 0.298 | 1 | 5 | Phf23 | -0.009 | Young |
| 0.695398 | 0.145669 | 0.124 | 0.116 | 1 | 5 | 0610010F0 | 0.008 | Young |
| 0.695896 | 0.021181 | 0.127 | 0.12 | 1 | 5 | Ankrd50 | 0.007 | Young |
| 0.696054 | 0.195232 | 0.137 | 0.126 | 1 | 5 | Odc1 | 0.011 | Young |
| 0.696749 | 0.179355 | 0.02 | 0.017 | 1 | 5 | Adap2os | 0.003 | Young |
| 0.696809 | 0.108357 | 0.563 | 0.554 | 1 | 5 | Eif4a2 | 0.009 | Young |
| 0.696929 | 0.072587 | 0.69 | 0.715 | 1 | 5 | Ifngr1 | -0.025 | Young |
| 0.697272 | 0.560295 | 0.013 | 0.01 | 1 | 5 | Amotl1 | 0.003 | Young |
| 0.697318 | 0.036767 | 0.292 | 0.278 | 1 | 5 | Setd5 | 0.014 | Young |
| 0.69742 | 0.459121 | 0.089 | 0.082 | 1 | 5 | Rffl | 0.007 | Young |
| 0.697507 | 0.002285 | 0.041 | 0.035 | 1 | 5 | Glcci1 | 0.006 | Young |
| 0.697979 | 0.058242 | 0.102 | 0.111 | 1 | 5 | Ifi35 | -0.009 | Young |
| 0.697988 | 0.287923 | 0.157 | 0.15 | 1 | 5 | Socs6 | 0.007 | Young |
| 0.698387 | 0.041821 | 0.048 | 0.054 | 1 | 5 | Dgcr14 | -0.006 | Young |
| 0.698388 | 0.563611 | 0.038 | 0.034 | 1 | 5 | Ncapd3 | 0.004 | Young |
| 0.698734 | 0.301124 | 0.066 | 0.061 | 1 | 5 | Rps19bp1 | 0.005 | Young |
| 0.698792 | 0.0263 | 0.673 | 0.699 | 1 | 5 | Myadm | -0.026 | Young |
| 0.698931 | 0.070178 | 0.02 | 0.017 | 1 | 5 | Fhit | 0.003 | Young |
| 0.699053 | 0.044137 | 0.046 | 0.04 | 1 | 5 | Phykpl | 0.006 | Young |
| 0.699126 | 0.431333 | 0.013 | 0.01 | 1 | 5 | Vegfa | 0.003 | Young |
| 0.699155 | 0.239461 | 0.099 | 0.093 | 1 | 5 | Lsm10 | 0.006 | Young |
| 0.699655 | 0.288996 | 0.114 | 0.108 | 1 | 5 | Stk19 | 0.006 | Young |
| 0.70004 | 0.100634 | 0.094 | 0.086 | 1 | 5 | Lonp1 | 0.008 | Young |
| 0.700258 | 0.019689 | 0.142 | 0.13 | 1 | 5 | Cpsf3 | 0.012 | Young |
| 0.700509 | 0.061761 | 0.114 | 0.106 | 1 | 5 | Gamt | 0.008 | Young |
| 0.700982 | 0.472539 | 0.013 | 0.01 | 1 | 5 | Cep295nl | 0.003 | Young |
| 0.700982 | 0.314596 | 0.013 | 0.01 | 1 | 5 | Gm10382 | 0.003 | Young |
| 0.701148 | 0.134585 | 0.033 | 0.029 | 1 | 5 | Mapkbp1 | 0.004 | Young |
| 0.701588 | 0.122508 | 0.038 | 0.044 | 1 | 5 | Fbxo31 | -0.006 | Young |
| 0.701614 | 0.229101 | 0.294 | 0.295 | 1 | 5 | Rtcb | -0.001 | Young |
| 0.70165 | 0.107038 | 0.404 | 0.396 | 1 | 5 | Mia3 | 0.008 | Young |
| 0.701762 | 0.15694 | 0.058 | 0.052 | 1 | 5 | Syce2 | 0.006 | Young |
| 0.701887 | 0.043032 | 0.307 | 0.32 | 1 | 5 | Tbcb | -0.013 | Young |
| 0.701911 | 0.353845 | 0.013 | 0.01 | 1 | 5 | Kif9 | 0.003 | Young |
| 0.701911 | 0.306549 | 0.013 | 0.01 | 1 | 5 | Prps1l3 | 0.003 | Young |
| 0.702183 | 0.10029 | 0.157 | 0.148 | 1 | 5 | Hyou1 | 0.009 | Young |
| 0.702234 | 0.238766 | 0.168 | 0.162 | 1 | 5 | Socs7 | 0.006 | Young |
| 0.702284 | 0.378476 | 0.109 | 0.104 | 1 | 5 | Tigd2 | 0.005 | Young |
| 0.702476 | 0.497486 | 0.076 | 0.071 | 1 | 5 | Cry2 | 0.005 | Young |
| 0.702695 | 0.027194 | 0.036 | 0.04 | 1 | 5 | Pde1b | -0.004 | Young |

| | | | | | | | | |
|---|---|---|---|---|---|---|---|---|
| 0.702741 | 0.533755 | 0.074 | 0.069 | 1 | 5 | Ctnnbl1 | 0.005 | Young |
| 0.70276 | 0.505643 | 0.056 | 0.051 | 1 | 5 | Il6 | 0.005 | Young |
| 0.702841 | 0.716864 | 0.013 | 0.01 | 1 | 5 | Hist2h3c2 | 0.003 | Young |
| 0.702841 | 0.359435 | 0.013 | 0.01 | 1 | 5 | Xpo5 | 0.003 | Young |
| 0.702924 | 0.12071 | 0.404 | 0.409 | 1 | 5 | Sf3b6 | -0.005 | Young |
| 0.703117 | 0.090464 | 0.467 | 0.463 | 1 | 5 | Dnm2 | 0.004 | Young |
| 0.703263 | 0.027374 | 0.058 | 0.052 | 1 | 5 | Haus7 | 0.006 | Young |
| 0.703303 | 0.01734 | 0.02 | 0.017 | 1 | 5 | Ggct | 0.003 | Young |
| 0.703739 | 0.068867 | 0.17 | 0.158 | 1 | 5 | Phf8 | 0.012 | Young |
| 0.70377 | 0.175515 | 0.013 | 0.01 | 1 | 5 | Samd10 | 0.003 | Young |
| 0.703922 | 0.431567 | 0.038 | 0.034 | 1 | 5 | Evi5l | 0.004 | Young |
| 0.704203 | 0.155224 | 0.142 | 0.133 | 1 | 5 | Phactr4 | 0.009 | Young |
| 0.704313 | 0.219331 | 0.061 | 0.056 | 1 | 5 | Dgka | 0.005 | Young |
| 0.704461 | 0.279224 | 0.084 | 0.077 | 1 | 5 | Ccdc43 | 0.007 | Young |
| 0.704506 | 0.255314 | 0.056 | 0.051 | 1 | 5 | Mical2 | 0.005 | Young |
| 0.704563 | 0.155778 | 0.114 | 0.106 | 1 | 5 | Rpp25l | 0.008 | Young |
| 0.704674 | 0.324998 | 0.071 | 0.066 | 1 | 5 | Batf2 | 0.005 | Young |
| 0.7047 | 0.384336 | 0.013 | 0.01 | 1 | 5 | Map3k6 | 0.003 | Young |
| 0.7047 | 0.317891 | 0.013 | 0.01 | 1 | 5 | 4930524J0 | 0.003 | Young |
| 0.7047 | 0.291837 | 0.013 | 0.01 | 1 | 5 | Noxo1 | 0.003 | Young |
| 0.70504 | 0.129563 | 0.604 | 0.606 | 1 | 5 | Rnasel | -0.002 | Young |
| 0.705166 | 0.022007 | 0.079 | 0.086 | 1 | 5 | Ngrn | -0.007 | Young |
| 0.705223 | 0.047729 | 0.604 | 0.641 | 1 | 5 | Ifi204 | -0.037 | Young |
| 0.70542 | 0.182882 | 0.175 | 0.167 | 1 | 5 | Arl6ip6 | 0.008 | Young |
| 0.705631 | 0.145211 | 0.013 | 0.01 | 1 | 5 | Myo1b | 0.003 | Young |
| 0.705951 | 0.079012 | 0.335 | 0.322 | 1 | 5 | Birc6 | 0.013 | Young |
| 0.706441 | 0.077273 | 0.272 | 0.285 | 1 | 5 | Pip5k1c | -0.013 | Young |
| 0.706514 | 0.066821 | 0.208 | 0.2 | 1 | 5 | Dstyk | 0.008 | Young |
| 0.706562 | 0.097822 | 0.013 | 0.01 | 1 | 5 | Cdkl3 | 0.003 | Young |
| 0.706757 | 0.000937 | 0.048 | 0.054 | 1 | 5 | Psmf1 | -0.006 | Young |
| 0.706868 | 0.097114 | 0.492 | 0.492 | 1 | 5 | Crlf3 | 0 | Young |
| 0.707091 | 0.460556 | 0.038 | 0.034 | 1 | 5 | Thap7 | 0.004 | Young |
| 0.707252 | 0.300857 | 0.18 | 0.173 | 1 | 5 | Arntl | 0.007 | Young |
| 0.707263 | 0.064332 | 0.015 | 0.019 | 1 | 5 | Pradc1 | -0.004 | Young |
| 0.707458 | 0.212398 | 0.094 | 0.088 | 1 | 5 | Stat5a | 0.006 | Young |
| 0.707493 | 0.287828 | 0.013 | 0.01 | 1 | 5 | Sec16b | 0.003 | Young |
| 0.707562 | 0.072973 | 0.426 | 0.438 | 1 | 5 | Smap2 | -0.012 | Young |
| 0.707739 | 0.240368 | 0.109 | 0.103 | 1 | 5 | Crnkl1 | 0.006 | Young |
| 0.708121 | 0.054541 | 0.865 | 0.899 | 1 | 5 | Fcgrt | -0.034 | Young |
| 0.708216 | 0.015852 | 0.548 | 0.539 | 1 | 5 | Cdkn1a | 0.009 | Young |
| 0.708425 | 0.431786 | 0.013 | 0.01 | 1 | 5 | Pcdhgb4 | 0.003 | Young |
| 0.708767 | 0.222433 | 0.015 | 0.019 | 1 | 5 | Cryab | -0.004 | Young |
| 0.708849 | 0.061831 | 0.063 | 0.057 | 1 | 5 | Shmt2 | 0.006 | Young |
| 0.708883 | 0.010428 | 0.066 | 0.072 | 1 | 5 | Sugp2 | -0.006 | Young |
| 0.708951 | 0.10978 | 0.14 | 0.131 | 1 | 5 | Golga5 | 0.009 | Young |

| | | | | | | | | |
|---|---|---|---|---|---|---|---|---|
| 0.709041 | 0.071499 | 0.084 | 0.091 | 1 | 5 | 1600002K0 | -0.007 | Young |
| 0.709106 | 0.053637 | 0.081 | 0.074 | 1 | 5 | Ppfibp1 | 0.007 | Young |
| 0.709358 | 0.116852 | 0.013 | 0.01 | 1 | 5 | Ccdc114 | 0.003 | Young |
| 0.709358 | 0.096363 | 0.013 | 0.01 | 1 | 5 | Mast4 | 0.003 | Young |
| 0.709358 | 0.06047 | 0.013 | 0.01 | 1 | 5 | Atp1a3 | 0.003 | Young |
| 0.710049 | 0.556725 | 0.025 | 0.022 | 1 | 5 | Sema6a | 0.003 | Young |
| 0.710337 | 0.113539 | 0.391 | 0.38 | 1 | 5 | Tbl1xr1 | 0.011 | Young |
| 0.710481 | 0.117735 | 0.152 | 0.141 | 1 | 5 | 1110059G1 | 0.011 | Young |
| 0.710579 | 0.132887 | 0.132 | 0.125 | 1 | 5 | Atg12 | 0.007 | Young |
| 0.711304 | 0.046895 | 0.249 | 0.263 | 1 | 5 | Chd3 | -0.014 | Young |
| 0.711593 | 0.097874 | 0.086 | 0.079 | 1 | 5 | Prkab1 | 0.007 | Young |
| 0.711668 | 0.048391 | 0.584 | 0.577 | 1 | 5 | Rnf150 | 0.007 | Young |
| 0.711669 | 0.305013 | 0.063 | 0.071 | 1 | 5 | Rpia | -0.008 | Young |
| 0.71189 | 0.091963 | 0.68 | 0.682 | 1 | 5 | Scp2 | -0.002 | Young |
| 0.712128 | 0.146556 | 0.109 | 0.101 | 1 | 5 | Pou5f2 | 0.008 | Young |
| 0.712384 | 0.166866 | 0.038 | 0.034 | 1 | 5 | Zfp574 | 0.004 | Young |
| 0.712531 | 0.124924 | 0.015 | 0.019 | 1 | 5 | Haus5 | -0.004 | Young |
| 0.712614 | 0.107856 | 0.185 | 0.173 | 1 | 5 | Lta4h | 0.012 | Young |
| 0.712832 | 0.25198 | 0.109 | 0.103 | 1 | 5 | Tnfrsf25 | 0.006 | Young |
| 0.712911 | 0.069898 | 0.581 | 0.577 | 1 | 5 | Taok1 | 0.004 | Young |
| 0.713065 | 0.141407 | 0.444 | 0.439 | 1 | 5 | Ube2r2 | 0.005 | Young |
| 0.713225 | 0.360734 | 0.102 | 0.094 | 1 | 5 | Naip5 | 0.008 | Young |
| 0.713297 | 0.368587 | 0.025 | 0.022 | 1 | 5 | Coa7 | 0.003 | Young |
| 0.713372 | 0.083633 | 0.416 | 0.438 | 1 | 5 | Twf1 | -0.022 | Young |
| 0.713532 | 0.00076 | 0.429 | 0.449 | 1 | 5 | Gmfb | -0.02 | Young |
| 0.713679 | 0.005213 | 0.033 | 0.029 | 1 | 5 | Bivm | 0.004 | Young |
| 0.713856 | 0.068807 | 0.175 | 0.185 | 1 | 5 | Nipsnap3b | -0.01 | Young |
| 0.714724 | 0.10061 | 0.274 | 0.261 | 1 | 5 | Ogfr | 0.013 | Young |
| 0.71476 | 0.15517 | 0.211 | 0.2 | 1 | 5 | Gfpt1 | 0.011 | Young |
| 0.714804 | 0.120514 | 0.211 | 0.202 | 1 | 5 | Kctd20 | 0.009 | Young |
| 0.714832 | 0.124133 | 0.135 | 0.128 | 1 | 5 | Zbtb1 | 0.007 | Young |
| 0.714968 | 0.137627 | 0.269 | 0.261 | 1 | 5 | Tollip | 0.008 | Young |
| 0.714991 | 0.254809 | 0.127 | 0.121 | 1 | 5 | Smim20 | 0.006 | Young |
| 0.715034 | 0.023381 | 0.046 | 0.04 | 1 | 5 | Dnal4 | 0.006 | Young |
| 0.71548 | 0.166675 | 0.023 | 0.027 | 1 | 5 | Wdr76 | -0.004 | Young |
| 0.715493 | 0.038342 | 0.264 | 0.273 | 1 | 5 | Fip1l1 | -0.009 | Young |
| 0.715535 | 0.056177 | 0.081 | 0.074 | 1 | 5 | Bloc1s1 | 0.007 | Young |
| 0.715677 | 0.219279 | 0.043 | 0.049 | 1 | 5 | Trub1 | -0.006 | Young |
| 0.715956 | 0.162177 | 0.063 | 0.071 | 1 | 5 | Mapre3 | -0.008 | Young |
| 0.716062 | 0.190201 | 0.061 | 0.056 | 1 | 5 | Brms1l | 0.005 | Young |
| 0.716364 | 0.091625 | 0.536 | 0.557 | 1 | 5 | Grina | -0.021 | Young |
| 0.716415 | 0.126666 | 0.142 | 0.133 | 1 | 5 | Calcrl | 0.009 | Young |
| 0.71654 | 0.162567 | 0.142 | 0.155 | 1 | 5 | Elp5 | -0.013 | Young |
| 0.716803 | 0.39617 | 0.043 | 0.039 | 1 | 5 | Rbm43 | 0.004 | Young |
| 0.716843 | 0.284311 | 0.089 | 0.082 | 1 | 5 | Tuft1 | 0.007 | Young |

| | | | | | | | | |
|---|---|---|---|---|---|---|---|---|
| 0.71714 | 0.134051 | 0.416 | 0.412 | 1 | 5 | Bnip3l | 0.004 | Young |
| 0.717147 | 0.011163 | 0.472 | 0.487 | 1 | 5 | Fcgr1 | -0.015 | Young |
| 0.71753 | 0.161246 | 0.135 | 0.145 | 1 | 5 | Fam220a | -0.01 | Young |
| 0.717546 | 0.175697 | 0.089 | 0.082 | 1 | 5 | Bop1 | 0.007 | Young |
| 0.717672 | 0.029736 | 0.183 | 0.192 | 1 | 5 | Trim24 | -0.009 | Young |
| 0.717853 | 0.315718 | 0.025 | 0.022 | 1 | 5 | Abca2 | 0.003 | Young |
| 0.717892 | 0.171231 | 0.363 | 0.355 | 1 | 5 | Med13 | 0.008 | Young |
| 0.718468 | 0.246412 | 0.076 | 0.071 | 1 | 5 | Edc3 | 0.005 | Young |
| 0.718946 | 0.018144 | 0.599 | 0.611 | 1 | 5 | Ostf1 | -0.012 | Young |
| 0.718976 | 0.231644 | 0.152 | 0.145 | 1 | 5 | Plekhg2 | 0.007 | Young |
| 0.719224 | 0.291611 | 0.076 | 0.071 | 1 | 5 | Nqo2 | 0.005 | Young |
| 0.719406 | 0.105323 | 0.541 | 0.564 | 1 | 5 | Sgpp1 | -0.023 | Young |
| 0.719437 | 0.631094 | 0.03 | 0.027 | 1 | 5 | Ddx51 | 0.003 | Young |
| 0.71964 | 0.383267 | 0.048 | 0.044 | 1 | 5 | Cyb561d1 | 0.004 | Young |
| 0.719735 | 0.000938 | 0.091 | 0.084 | 1 | 5 | Pip4k2c | 0.007 | Young |
| 0.719741 | 0.106252 | 0.266 | 0.278 | 1 | 5 | Add3 | -0.012 | Young |
| 0.720461 | 0.235335 | 0.025 | 0.022 | 1 | 5 | Hus1 | 0.003 | Young |
| 0.720475 | 0.078609 | 0.528 | 0.564 | 1 | 5 | Sbf2 | -0.036 | Young |
| 0.720489 | 0.134703 | 0.048 | 0.054 | 1 | 5 | Rdh10 | -0.006 | Young |
| 0.721012 | 0.082484 | 0.563 | 0.594 | 1 | 5 | Hnrnpab | -0.031 | Young |
| 0.721462 | 0.167569 | 0.239 | 0.229 | 1 | 5 | Mettl9 | 0.01 | Young |
| 0.721641 | 0.046708 | 0.135 | 0.143 | 1 | 5 | Hmgcr | -0.008 | Young |
| 0.721682 | 0.051127 | 0.332 | 0.352 | 1 | 5 | Sp3 | -0.02 | Young |
| 0.721711 | 0.000134 | 0.157 | 0.168 | 1 | 5 | Dok3 | -0.011 | Young |
| 0.721922 | 0.350722 | 0.063 | 0.059 | 1 | 5 | Wdfy1 | 0.004 | Young |
| 0.72212 | 0.007394 | 0.571 | 0.566 | 1 | 5 | Rpl22l1 | 0.005 | Young |
| 0.722213 | 0.071433 | 0.038 | 0.034 | 1 | 5 | Dnajc18 | 0.004 | Young |
| 0.722367 | 0.063506 | 0.487 | 0.49 | 1 | 5 | Romo1 | -0.003 | Young |
| 0.722479 | 0.117863 | 0.038 | 0.034 | 1 | 5 | Sumf2 | 0.004 | Young |
| 0.722565 | 0.198425 | 0.119 | 0.113 | 1 | 5 | Pmpca | 0.006 | Young |
| 0.722806 | 0.201998 | 0.061 | 0.056 | 1 | 5 | Prpf3 | 0.005 | Young |
| 0.722843 | 0.049625 | 0.127 | 0.135 | 1 | 5 | Gtf3c1 | -0.008 | Young |
| 0.72298 | 0.093881 | 0.127 | 0.12 | 1 | 5 | Hat1 | 0.007 | Young |
| 0.723143 | 0.079985 | 0.386 | 0.406 | 1 | 5 | Tmem189 | -0.02 | Young |
| 0.723342 | 0.042529 | 0.239 | 0.247 | 1 | 5 | Pbx3 | -0.008 | Young |
| 0.723367 | 0.070383 | 0.201 | 0.209 | 1 | 5 | Ap3m1 | -0.008 | Young |
| 0.723389 | 0.003658 | 0.041 | 0.045 | 1 | 5 | Nudt18 | -0.004 | Young |
| 0.723545 | 0.189555 | 0.038 | 0.034 | 1 | 5 | Trmt11 | 0.004 | Young |
| 0.723622 | 0.182579 | 0.096 | 0.106 | 1 | 5 | Tsc2 | -0.01 | Young |
| 0.723725 | 0.106015 | 0.025 | 0.022 | 1 | 5 | Zbtb26 | 0.003 | Young |
| 0.723922 | 0.002169 | 0.241 | 0.249 | 1 | 5 | Slc35f6 | -0.008 | Young |
| 0.724225 | 0.016897 | 0.145 | 0.153 | 1 | 5 | Slc25a24 | -0.008 | Young |
| 0.724383 | 0.211691 | 0.162 | 0.157 | 1 | 5 | Strn | 0.005 | Young |
| 0.725263 | 0.075671 | 0.122 | 0.131 | 1 | 5 | Adk | -0.009 | Young |
| 0.725697 | 0.141717 | 0.18 | 0.173 | 1 | 5 | Cd40 | 0.007 | Young |

| | | | | | | | | |
|---|---|---|---|---|---|---|---|---|
| 0.726088 | 0.069719 | 0.779 | 0.79 | 1 | 5 | Son | -0.011 | Young |
| 0.726153 | 0.00841 | 0.102 | 0.094 | 1 | 5 | Msantd4 | 0.008 | Young |
| 0.72629 | 0.166797 | 0.043 | 0.039 | 1 | 5 | Fam210b | 0.004 | Young |
| 0.726458 | 0.408594 | 0.063 | 0.059 | 1 | 5 | Atg4c | 0.004 | Young |
| 0.726559 | 0.579587 | 0.03 | 0.027 | 1 | 5 | Cxcr3 | 0.003 | Young |
| 0.727254 | 0.078713 | 0.234 | 0.253 | 1 | 5 | Nhsl2 | -0.019 | Young |
| 0.727641 | 0.010887 | 0.157 | 0.163 | 1 | 5 | Tmed4 | -0.006 | Young |
| 0.72765 | 0.148109 | 0.282 | 0.305 | 1 | 5 | Tmpo | -0.023 | Young |
| 0.727901 | 0.205693 | 0.36 | 0.362 | 1 | 5 | Rcc2 | -0.002 | Young |
| 0.728292 | 0.349119 | 0.043 | 0.039 | 1 | 5 | Dis3l | 0.004 | Young |
| 0.728325 | 0.115542 | 0.602 | 0.625 | 1 | 5 | Rps27l | -0.023 | Young |
| 0.728367 | 0.020575 | 0.226 | 0.232 | 1 | 5 | Synrg | -0.006 | Young |
| 0.728793 | 0.424634 | 0.043 | 0.039 | 1 | 5 | Ccdc91 | 0.004 | Young |
| 0.728799 | 0.023718 | 0.287 | 0.303 | 1 | 5 | Psmc6 | -0.016 | Young |
| 0.729253 | 0.021911 | 0.414 | 0.397 | 1 | 5 | Gnpda1 | 0.017 | Young |
| 0.729693 | 0.420765 | 0.074 | 0.069 | 1 | 5 | Zfp746 | 0.005 | Young |
| 0.729863 | 0.10365 | 0.401 | 0.386 | 1 | 5 | Actn4 | 0.015 | Young |
| 0.729937 | 0.013772 | 0.02 | 0.024 | 1 | 5 | Tfdp2 | -0.004 | Young |
| 0.729954 | 0.119408 | 0.096 | 0.089 | 1 | 5 | Klhdc4 | 0.007 | Young |
| 0.730159 | 0.026263 | 0.447 | 0.475 | 1 | 5 | Fxyd2 | -0.028 | Young |
| 0.730188 | 0.011914 | 0.216 | 0.227 | 1 | 5 | Rpl7l1 | -0.011 | Young |
| 0.730269 | 0.109212 | 0.025 | 0.022 | 1 | 5 | Armcx5 | 0.003 | Young |
| 0.730305 | 0.09436 | 0.221 | 0.237 | 1 | 5 | Ylpm1 | -0.016 | Young |
| 0.730576 | 0.255101 | 0.132 | 0.126 | 1 | 5 | Tmx2 | 0.006 | Young |
| 0.730678 | 0.118368 | 0.274 | 0.266 | 1 | 5 | Map2k7 | 0.008 | Young |
| 0.730782 | 0.109329 | 0.069 | 0.076 | 1 | 5 | Gramd3 | -0.007 | Young |
| 0.731137 | 0.330026 | 0.086 | 0.081 | 1 | 5 | Naa40 | 0.005 | Young |
| 0.731935 | 0.019603 | 0.269 | 0.278 | 1 | 5 | Gorasp2 | -0.009 | Young |
| 0.732095 | 0.211934 | 0.041 | 0.045 | 1 | 5 | Gde1 | -0.004 | Young |
| 0.732953 | 0.278886 | 0.114 | 0.108 | 1 | 5 | Rps6ka5 | 0.006 | Young |
| 0.732955 | 0.178742 | 0.269 | 0.266 | 1 | 5 | Srp19 | 0.003 | Young |
| 0.733059 | 0.110055 | 0.084 | 0.077 | 1 | 5 | Tmem9 | 0.007 | Young |
| 0.733774 | 0.125349 | 0.028 | 0.032 | 1 | 5 | Tbc1d19 | -0.004 | Young |
| 0.733941 | 0.157973 | 0.482 | 0.476 | 1 | 5 | Bcl10 | 0.006 | Young |
| 0.734028 | 0.010989 | 0.058 | 0.064 | 1 | 5 | Pgam5 | -0.006 | Young |
| 0.734205 | 0.122369 | 0.025 | 0.022 | 1 | 5 | Fam109b | 0.003 | Young |
| 0.734205 | 0.026706 | 0.025 | 0.022 | 1 | 5 | Cntrob | 0.003 | Young |
| 0.734792 | 0.061421 | 0.079 | 0.072 | 1 | 5 | Epg5 | 0.007 | Young |
| 0.734931 | 0.056686 | 0.107 | 0.114 | 1 | 5 | Ciao1 | -0.007 | Young |
| 0.735083 | 0.227756 | 0.32 | 0.318 | 1 | 5 | Rab35 | 0.002 | Young |
| 0.735209 | 0.012779 | 0.201 | 0.185 | 1 | 5 | Tcerg1 | 0.016 | Young |
| 0.735248 | 0.273236 | 0.099 | 0.093 | 1 | 5 | Vamp5 | 0.006 | Young |
| 0.735416 | 0.19442 | 0.099 | 0.093 | 1 | 5 | Kctd6 | 0.006 | Young |
| 0.735548 | 0.183572 | 0.074 | 0.081 | 1 | 5 | Gm44751 | -0.007 | Young |
| 0.735666 | 0.09358 | 0.48 | 0.481 | 1 | 5 | Shtn1 | -0.001 | Young |

| | | | | | | | | |
|---|---|---|---|---|---|---|---|---|
| 0.735805 | 0.109666 | 0.127 | 0.135 | 1 | 5 | Pias2 | -0.008 | Young |
| 0.736169 | 0.088973 | 0.525 | 0.532 | 1 | 5 | Mindy2 | -0.007 | Young |
| 0.736383 | 0.096579 | 0.112 | 0.121 | 1 | 5 | Alg5 | -0.009 | Young |
| 0.736633 | 0.068672 | 0.325 | 0.311 | 1 | 5 | Pitpnc1 | 0.014 | Young |
| 0.736681 | 0.198165 | 0.145 | 0.14 | 1 | 5 | Fubp3 | 0.005 | Young |
| 0.736888 | 0.13789 | 0.464 | 0.46 | 1 | 5 | Uqcr10 | 0.004 | Young |
| 0.737473 | 0.081847 | 0.398 | 0.397 | 1 | 5 | Rps6ka3 | 0.001 | Young |
| 0.737604 | 0.489273 | 0.066 | 0.061 | 1 | 5 | Arrdc2 | 0.005 | Young |
| 0.73766 | 0.215658 | 0.206 | 0.199 | 1 | 5 | Tcof1 | 0.007 | Young |
| 0.737741 | 0.148613 | 0.155 | 0.146 | 1 | 5 | Ndrg3 | 0.009 | Young |
| 0.737929 | 0.024988 | 0.178 | 0.189 | 1 | 5 | Slc11a2 | -0.011 | Young |
| 0.738048 | 0.394476 | 0.063 | 0.059 | 1 | 5 | Sephs1 | 0.004 | Young |
| 0.738108 | 0.133662 | 0.048 | 0.044 | 1 | 5 | Mtg2 | 0.004 | Young |
| 0.73827 | 0.118327 | 0.236 | 0.229 | 1 | 5 | Cnpy2 | 0.007 | Young |
| 0.738308 | 0.016179 | 0.117 | 0.125 | 1 | 5 | Prr12 | -0.008 | Young |
| 0.738597 | 0.137736 | 0.251 | 0.266 | 1 | 5 | Atp7a | -0.015 | Young |
| 0.738736 | 0.065163 | 0.195 | 0.184 | 1 | 5 | Zmym2 | 0.011 | Young |
| 0.738743 | 0.025322 | 0.718 | 0.715 | 1 | 5 | Csnk1a1 | 0.003 | Young |
| 0.738841 | 0.091554 | 0.122 | 0.114 | 1 | 5 | Ap1m1 | 0.008 | Young |
| 0.739294 | 0.337516 | 0.063 | 0.059 | 1 | 5 | 2310039H( | 0.004 | Young |
| 0.739749 | 0.10749 | 0.211 | 0.224 | 1 | 5 | Ubqln2 | -0.013 | Young |
| 0.739773 | 0.048634 | 0.074 | 0.079 | 1 | 5 | Mpv17l2 | -0.005 | Young |
| 0.740428 | 0.06642 | 0.18 | 0.19 | 1 | 5 | Nr1h2 | -0.01 | Young |
| 0.740828 | 0.327307 | 0.234 | 0.231 | 1 | 5 | Ganab | 0.003 | Young |
| 0.740837 | 0.237515 | 0.041 | 0.045 | 1 | 5 | Tmed1 | -0.004 | Young |
| 0.740874 | 0.376091 | 0.03 | 0.027 | 1 | 5 | Polr1b | 0.003 | Young |
| 0.740895 | 0.050754 | 0.107 | 0.099 | 1 | 5 | 9430038I0: | 0.008 | Young |
| 0.741351 | 0.12431 | 0.02 | 0.024 | 1 | 5 | Alg12 | -0.004 | Young |
| 0.741435 | 0.100663 | 0.107 | 0.116 | 1 | 5 | Heatr6 | -0.009 | Young |
| 0.741467 | 0.182188 | 0.071 | 0.066 | 1 | 5 | Lyrm2 | 0.005 | Young |
| 0.741739 | 0.060917 | 0.033 | 0.037 | 1 | 5 | Tmem38b | -0.004 | Young |
| 0.741994 | 0.152705 | 0.058 | 0.054 | 1 | 5 | Prim2 | 0.004 | Young |
| 0.742176 | 0.069529 | 0.152 | 0.162 | 1 | 5 | Usp32 | -0.01 | Young |
| 0.742433 | 0.106823 | 0.142 | 0.153 | 1 | 5 | Pfkp | -0.011 | Young |
| 0.742654 | 0.04623 | 0.893 | 0.921 | 1 | 5 | Canx | -0.028 | Young |
| 0.742698 | 0.129551 | 0.02 | 0.024 | 1 | 5 | Gsdme | -0.004 | Young |
| 0.7428 | 0.188637 | 0.228 | 0.219 | 1 | 5 | Ncaph2 | 0.009 | Young |
| 0.742868 | 0.160531 | 0.19 | 0.185 | 1 | 5 | Slc31a1 | 0.005 | Young |
| 0.74287 | 0.112124 | 0.33 | 0.323 | 1 | 5 | 4931406C( | 0.007 | Young |
| 0.742871 | 0.229489 | 0.018 | 0.015 | 1 | 5 | Zfp51 | 0.003 | Young |
| 0.742989 | 0.178084 | 0.266 | 0.266 | 1 | 5 | Pdcd6 | 0 | Young |
| 0.74316 | 0.027702 | 0.114 | 0.121 | 1 | 5 | Rai1 | -0.007 | Young |
| 0.743466 | 0.069549 | 0.579 | 0.572 | 1 | 5 | Acp2 | 0.007 | Young |
| 0.744051 | 0.082572 | 0.246 | 0.237 | 1 | 5 | Fam3c | 0.009 | Young |
| 0.744259 | 0.07352 | 0.145 | 0.155 | 1 | 5 | Prkag1 | -0.01 | Young |

| | | | | | | | | |
|---|---|---|---|---|---|---|---|---|
| 0.744289 | 0.065243 | 0.063 | 0.069 | 1 | 5 | Ccdc126 | -0.006 | Young |
| 0.74438 | 0.174911 | 0.152 | 0.146 | 1 | 5 | Dcaf13 | 0.006 | Young |
| 0.744392 | 0.060233 | 0.145 | 0.136 | 1 | 5 | Abcb7 | 0.009 | Young |
| 0.744446 | 0.431756 | 0.018 | 0.015 | 1 | 5 | Rad52 | 0.003 | Young |
| 0.744561 | 0.1285 | 0.066 | 0.061 | 1 | 5 | Tmem126b | 0.005 | Young |
| 0.744628 | 0.27105 | 0.071 | 0.066 | 1 | 5 | Pus10 | 0.005 | Young |
| 0.744883 | 0.053878 | 0.043 | 0.039 | 1 | 5 | Mlh1 | 0.004 | Young |
| 0.744908 | 0.004431 | 0.223 | 0.234 | 1 | 5 | Eif4ebp1 | -0.011 | Young |
| 0.745047 | 0.102579 | 0.68 | 0.707 | 1 | 5 | Ndufa4 | -0.027 | Young |
| 0.745275 | 0.061372 | 0.393 | 0.429 | 1 | 5 | Psma3 | -0.036 | Young |
| 0.745293 | 0.506147 | 0.051 | 0.047 | 1 | 5 | Tusc1 | 0.004 | Young |
| 0.745625 | 0.423804 | 0.036 | 0.032 | 1 | 5 | Atp6v1g2 | 0.004 | Young |
| 0.745784 | 0.141614 | 0.234 | 0.222 | 1 | 5 | Ndufa8 | 0.012 | Young |
| 0.745814 | 0.15271 | 0.071 | 0.066 | 1 | 5 | Ddhd1 | 0.005 | Young |
| 0.746025 | 0.069334 | 0.066 | 0.072 | 1 | 5 | Pkdcc | -0.006 | Young |
| 0.746156 | 0.306831 | 0.053 | 0.049 | 1 | 5 | Fancl | 0.004 | Young |
| 0.746397 | 0.097906 | 0.043 | 0.039 | 1 | 5 | Zfp628 | 0.004 | Young |
| 0.746462 | 0.101177 | 0.431 | 0.424 | 1 | 5 | Ebi3 | 0.007 | Young |
| 0.746574 | 0.174688 | 0.096 | 0.091 | 1 | 5 | Alg10b | 0.005 | Young |
| 0.746742 | 0.231787 | 0.02 | 0.024 | 1 | 5 | E2f6 | -0.004 | Young |
| 0.746803 | 0.214214 | 0.071 | 0.066 | 1 | 5 | Rpf2 | 0.005 | Young |
| 0.746902 | 0.183342 | 0.043 | 0.039 | 1 | 5 | Zc3h12c | 0.004 | Young |
| 0.747151 | 0.155546 | 0.358 | 0.357 | 1 | 5 | Arfgef1 | 0.001 | Young |
| 0.747198 | 0.164323 | 0.269 | 0.264 | 1 | 5 | Bsdc1 | 0.005 | Young |
| 0.747259 | 0.127662 | 0.198 | 0.19 | 1 | 5 | Cask | 0.008 | Young |
| 0.74728 | 0.441573 | 0.041 | 0.037 | 1 | 5 | Cox6b2 | 0.004 | Young |
| 0.747281 | 0.221617 | 0.398 | 0.399 | 1 | 5 | Ogdh | -0.001 | Young |
| 0.747373 | 0.122756 | 0.155 | 0.165 | 1 | 5 | Sptan1 | -0.01 | Young |
| 0.747602 | 0.062145 | 0.124 | 0.116 | 1 | 5 | Akap7 | 0.008 | Young |
| 0.747844 | 0.236282 | 0.086 | 0.081 | 1 | 5 | Ikbkg | 0.005 | Young |
| 0.74803 | 0.265442 | 0.063 | 0.059 | 1 | 5 | Timm21 | 0.004 | Young |
| 0.748041 | 0.014142 | 0.244 | 0.232 | 1 | 5 | Strap | 0.012 | Young |
| 0.74865 | 0.302368 | 0.206 | 0.205 | 1 | 5 | Tec | 0.001 | Young |
| 0.748666 | 0.245208 | 0.03 | 0.027 | 1 | 5 | Fitm2 | 0.003 | Young |
| 0.748842 | 0.043156 | 0.264 | 0.273 | 1 | 5 | Myo9a | -0.009 | Young |
| 0.748904 | 0.332324 | 0.084 | 0.079 | 1 | 5 | Arl2bp | 0.005 | Young |
| 0.748959 | 0.054001 | 0.173 | 0.163 | 1 | 5 | Pvr | 0.01 | Young |
| 0.749376 | 0.055282 | 0.071 | 0.066 | 1 | 5 | Zbtb43 | 0.005 | Young |
| 0.749566 | 0.184117 | 0.373 | 0.369 | 1 | 5 | Msl2 | 0.004 | Young |
| 0.749637 | 0.125351 | 0.195 | 0.192 | 1 | 5 | Nt5c2 | 0.003 | Young |
| 0.750005 | 0.093623 | 0.15 | 0.158 | 1 | 5 | Wdr3 | -0.008 | Young |
| 0.750053 | 0.121625 | 0.076 | 0.071 | 1 | 5 | Gm27010 | 0.005 | Young |
| 0.750167 | 0.037941 | 0.03 | 0.027 | 1 | 5 | Gpatch4 | 0.003 | Young |
| 0.75019 | 0.119356 | 0.31 | 0.31 | 1 | 5 | Fis1 | 0 | Young |
| 0.750468 | 0.052369 | 0.03 | 0.027 | 1 | 5 | Zfp146 | 0.003 | Young |

| | | | | | | | | |
|---|---|---|---|---|---|---|---|---|
| 0.75072 | 0.102119 | 0.246 | 0.237 | 1 | 5 | Kdm6a | 0.009 | Young |
| 0.750758 | 0.897661 | 0.018 | 0.015 | 1 | 5 | Lif | 0.003 | Young |
| 0.750767 | 0.089781 | 0.142 | 0.135 | 1 | 5 | Apopt1 | 0.007 | Young |
| 0.750772 | 0.18677 | 0.053 | 0.059 | 1 | 5 | Prr5 | -0.006 | Young |
| 0.750883 | 0.022115 | 0.104 | 0.109 | 1 | 5 | Tspan17 | -0.005 | Young |
| 0.751107 | 0.157411 | 0.058 | 0.054 | 1 | 5 | Eepd1 | 0.004 | Young |
| 0.751314 | 0.131577 | 0.046 | 0.051 | 1 | 5 | Nubpl | -0.005 | Young |
| 0.751462 | 0.141161 | 0.048 | 0.044 | 1 | 5 | Cacfd1 | 0.004 | Young |
| 0.751548 | 0.03201 | 0.018 | 0.015 | 1 | 5 | Usp54 | 0.003 | Young |
| 0.751729 | 0.004823 | 0.277 | 0.263 | 1 | 5 | Gtf3c2 | 0.014 | Young |
| 0.751829 | 0.132363 | 0.32 | 0.313 | 1 | 5 | Stx12 | 0.007 | Young |
| 0.752014 | 0.078147 | 0.769 | 0.827 | 1 | 5 | Pcbp1 | -0.058 | Young |
| 0.752275 | 0.147266 | 0.226 | 0.221 | 1 | 5 | Zscan26 | 0.005 | Young |
| 0.752374 | 0.23509 | 0.236 | 0.234 | 1 | 5 | Rfk | 0.002 | Young |
| 0.75249 | 0.197831 | 0.127 | 0.121 | 1 | 5 | Wrn | 0.006 | Young |
| 0.752493 | 0.034874 | 0.571 | 0.564 | 1 | 5 | Phip | 0.007 | Young |
| 0.752764 | 0.166078 | 0.069 | 0.064 | 1 | 5 | Ifitm5 | 0.005 | Young |
| 0.753057 | 0.063722 | 0.294 | 0.286 | 1 | 5 | Ypel5 | 0.008 | Young |
| 0.753153 | 0.273112 | 0.122 | 0.116 | 1 | 5 | Minpp1 | 0.006 | Young |
| 0.753233 | 0.172564 | 0.19 | 0.202 | 1 | 5 | Itfg1 | -0.012 | Young |
| 0.753376 | 0.400898 | 0.048 | 0.044 | 1 | 5 | Mtrf1l | 0.004 | Young |
| 0.753524 | 0.166867 | 0.226 | 0.219 | 1 | 5 | Lpgat1 | 0.007 | Young |
| 0.753721 | 0.005849 | 0.051 | 0.056 | 1 | 5 | Zfp672 | -0.005 | Young |
| 0.753791 | 0.110092 | 0.013 | 0.015 | 1 | 5 | Sdr42e1 | -0.002 | Young |
| 0.753827 | 0.074852 | 0.635 | 0.648 | 1 | 5 | Hint1 | -0.013 | Young |
| 0.754291 | 0.220597 | 0.063 | 0.059 | 1 | 5 | Mccc1 | 0.004 | Young |
| 0.754422 | 0.098141 | 0.14 | 0.131 | 1 | 5 | Plcg2 | 0.009 | Young |
| 0.754636 | 0.107721 | 0.013 | 0.015 | 1 | 5 | Lzts3 | -0.002 | Young |
| 0.754636 | 0.038852 | 0.013 | 0.015 | 1 | 5 | 1500002C1 | -0.002 | Young |
| 0.754869 | 0.096367 | 0.246 | 0.237 | 1 | 5 | Foxo3 | 0.009 | Young |
| 0.754873 | 0.007472 | 0.137 | 0.145 | 1 | 5 | Otud6b | -0.008 | Young |
| 0.755084 | 0.023622 | 0.551 | 0.571 | 1 | 5 | Nfam1 | -0.02 | Young |
| 0.755252 | 0.113017 | 0.477 | 0.49 | 1 | 5 | Tiparp | -0.013 | Young |
| 0.755481 | 0.0424 | 0.013 | 0.015 | 1 | 5 | 2210408I2 | -0.002 | Young |
| 0.756327 | 0.329771 | 0.013 | 0.015 | 1 | 5 | Gm10642 | -0.002 | Young |
| 0.756346 | 0.087334 | 0.165 | 0.157 | 1 | 5 | S100a11 | 0.008 | Young |
| 0.756387 | 0.011904 | 0.074 | 0.067 | 1 | 5 | 9130401M | 0.007 | Young |
| 0.756483 | 0.089087 | 0.03 | 0.027 | 1 | 5 | Dnah2 | 0.003 | Young |
| 0.756484 | 0.16371 | 0.251 | 0.247 | 1 | 5 | Mpc2 | 0.004 | Young |
| 0.756576 | 0.175225 | 0.076 | 0.071 | 1 | 5 | Map2k5 | 0.005 | Young |
| 0.756719 | 0.050619 | 0.69 | 0.689 | 1 | 5 | Tomm7 | 0.001 | Young |
| 0.757325 | 0.02382 | 0.378 | 0.396 | 1 | 5 | Dennd1a | -0.018 | Young |
| 0.757459 | 0.010977 | 0.373 | 0.389 | 1 | 5 | Psma6 | -0.016 | Young |
| 0.757806 | 0.080841 | 0.081 | 0.076 | 1 | 5 | Nr2c2ap | 0.005 | Young |
| 0.757835 | 0.000801 | 0.731 | 0.763 | 1 | 5 | Hnrnph1 | -0.032 | Young |

| | | | | | | | | |
|---|---|---|---|---|---|---|---|---|
| 0.758171 | 0.218292 | 0.162 | 0.157 | 1 | 5 | Cnep1r1 | 0.005 | Young |
| 0.75862 | 0.013034 | 0.135 | 0.126 | 1 | 5 | Trappc12 | 0.009 | Young |
| 0.759096 | 0.004344 | 0.241 | 0.251 | 1 | 5 | Psmc1 | -0.01 | Young |
| 0.759209 | 0.039462 | 0.157 | 0.148 | 1 | 5 | Atg101 | 0.009 | Young |
| 0.760065 | 0.300415 | 0.036 | 0.032 | 1 | 5 | Zswim7 | 0.004 | Young |
| 0.760097 | 0.037523 | 0.272 | 0.283 | 1 | 5 | Nfia | -0.011 | Young |
| 0.760461 | 0.436628 | 0.061 | 0.057 | 1 | 5 | Rrp15 | 0.004 | Young |
| 0.760692 | 0.221845 | 0.058 | 0.054 | 1 | 5 | Zc3h3 | 0.004 | Young |
| 0.760796 | 0.107851 | 0.629 | 0.66 | 1 | 5 | Lnpep | -0.031 | Young |
| 0.760872 | 0.048376 | 0.096 | 0.089 | 1 | 5 | Ppwd1 | 0.007 | Young |
| 0.760907 | 0.251139 | 0.117 | 0.113 | 1 | 5 | Polr1c | 0.004 | Young |
| 0.761047 | 0.079376 | 0.043 | 0.047 | 1 | 5 | Hint3 | -0.004 | Young |
| 0.761447 | 0.094682 | 0.18 | 0.17 | 1 | 5 | Ctdspl2 | 0.01 | Young |
| 0.761704 | 0.141537 | 0.033 | 0.037 | 1 | 5 | Dut | -0.004 | Young |
| 0.761737 | 0.1372 | 0.036 | 0.032 | 1 | 5 | Fntb | 0.004 | Young |
| 0.762299 | 0.09917 | 0.294 | 0.285 | 1 | 5 | Ociad1 | 0.009 | Young |
| 0.76241 | 0.237991 | 0.218 | 0.214 | 1 | 5 | Atf2 | 0.004 | Young |
| 0.76247 | 0.034081 | 0.046 | 0.051 | 1 | 5 | Ercc2 | -0.005 | Young |
| 0.762674 | 0.133041 | 0.086 | 0.081 | 1 | 5 | Dvl1 | 0.005 | Young |
| 0.762801 | 0.146269 | 0.091 | 0.086 | 1 | 5 | Mre11a | 0.005 | Young |
| 0.76287 | 0.191318 | 0.16 | 0.153 | 1 | 5 | Znrf1 | 0.007 | Young |
| 0.762878 | 0.21187 | 0.071 | 0.067 | 1 | 5 | Ddt | 0.004 | Young |
| 0.763034 | 0.051489 | 0.114 | 0.121 | 1 | 5 | Mrpl38 | -0.007 | Young |
| 0.763118 | 0.264796 | 0.038 | 0.042 | 1 | 5 | Peg10 | -0.004 | Young |
| 0.763323 | 0.012581 | 0.025 | 0.029 | 1 | 5 | Slc22a23 | -0.004 | Young |
| 0.76333 | 0.10225 | 0.033 | 0.037 | 1 | 5 | Isoc2b | -0.004 | Young |
| 0.763707 | 0.325595 | 0.094 | 0.089 | 1 | 5 | Dhrs4 | 0.005 | Young |
| 0.763811 | 0.17652 | 0.353 | 0.36 | 1 | 5 | Park7 | -0.007 | Young |
| 0.763868 | 0.127424 | 0.185 | 0.175 | 1 | 5 | Ddx18 | 0.01 | Young |
| 0.764012 | 0.065393 | 0.165 | 0.158 | 1 | 5 | Arhgap11a | 0.007 | Young |
| 0.764243 | 0.107503 | 0.391 | 0.392 | 1 | 5 | Smg1 | -0.001 | Young |
| 0.76471 | 0.0576 | 0.53 | 0.562 | 1 | 5 | Cd164 | -0.032 | Young |
| 0.764824 | 0.083524 | 0.371 | 0.364 | 1 | 5 | Psma2 | 0.007 | Young |
| 0.765498 | 0.024078 | 0.058 | 0.054 | 1 | 5 | 1500011B0 | 0.004 | Young |
| 0.765849 | 0.028626 | 0.096 | 0.103 | 1 | 5 | Krr1 | -0.007 | Young |
| 0.76598 | 0.023672 | 0.195 | 0.207 | 1 | 5 | Ndufs3 | -0.012 | Young |
| 0.766001 | 0.046606 | 0.056 | 0.061 | 1 | 5 | Atf5 | -0.005 | Young |
| 0.766094 | 0.042597 | 0.089 | 0.082 | 1 | 5 | Adck5 | 0.007 | Young |
| 0.766503 | 0.206936 | 0.142 | 0.138 | 1 | 5 | Rnf41 | 0.004 | Young |
| 0.766639 | 0.06147 | 0.277 | 0.293 | 1 | 5 | Bud31 | -0.016 | Young |
| 0.766765 | 0.189813 | 0.046 | 0.042 | 1 | 5 | Abhd10 | 0.004 | Young |
| 0.767255 | 0.303229 | 0.096 | 0.093 | 1 | 5 | Cep164 | 0.003 | Young |
| 0.767281 | 0.387478 | 0.061 | 0.057 | 1 | 5 | Rubcnl | 0.004 | Young |
| 0.76745 | 0.013107 | 0.122 | 0.114 | 1 | 5 | Mphosph8 | 0.008 | Young |
| 0.76755 | 0.206353 | 0.15 | 0.145 | 1 | 5 | Smg5 | 0.005 | Young |

| | | | | | | | | |
|---|---|---|---|---|---|---|---|---|
| 0.76763 | 0.234668 | 0.025 | 0.029 | 1 | 5 | Akr1b8 | -0.004 | Young |
| 0.767675 | 0.22105 | 0.114 | 0.109 | 1 | 5 | Stard5 | 0.005 | Young |
| 0.767892 | 0.118244 | 0.183 | 0.192 | 1 | 5 | Nfil3 | -0.009 | Young |
| 0.767952 | 0.018356 | 0.383 | 0.404 | 1 | 5 | Fxr1 | -0.021 | Young |
| 0.768134 | 0.083349 | 0.102 | 0.096 | 1 | 5 | Isg20l2 | 0.006 | Young |
| 0.768246 | 0.221293 | 0.025 | 0.029 | 1 | 5 | Zfp974 | -0.004 | Young |
| 0.76869 | 0.114153 | 0.061 | 0.066 | 1 | 5 | Ptpn21 | -0.005 | Young |
| 0.768853 | 0.052397 | 0.348 | 0.332 | 1 | 5 | Ppp4r2 | 0.016 | Young |
| 0.769006 | 0.173082 | 0.102 | 0.108 | 1 | 5 | Camsap2 | -0.006 | Young |
| 0.769478 | 0.079745 | 0.025 | 0.029 | 1 | 5 | Mcrip2 | -0.004 | Young |
| 0.769795 | 0.752548 | 0.01 | 0.008 | 1 | 5 | Dpep1 | 0.002 | Young |
| 0.769987 | 0.16859 | 0.109 | 0.103 | 1 | 5 | Fmo5 | 0.006 | Young |
| 0.770058 | 0.218398 | 0.211 | 0.207 | 1 | 5 | Esyt2 | 0.004 | Young |
| 0.770185 | 0.149979 | 0.107 | 0.101 | 1 | 5 | Mier3 | 0.006 | Young |
| 0.770853 | 0.756604 | 0.01 | 0.008 | 1 | 5 | Bbs2 | 0.002 | Young |
| 0.771002 | 0.030605 | 0.607 | 0.589 | 1 | 5 | Serinc1 | 0.018 | Young |
| 0.771062 | 0.20021 | 0.31 | 0.318 | 1 | 5 | Ndufv3 | -0.008 | Young |
| 0.771064 | 0.217498 | 0.193 | 0.192 | 1 | 5 | Msrb1 | 0.001 | Young |
| 0.771775 | 0.089075 | 0.038 | 0.042 | 1 | 5 | Gm3448 | -0.004 | Young |
| 0.771818 | 0.040684 | 0.5 | 0.48 | 1 | 5 | Dock8 | 0.02 | Young |
| 0.772 | 0.011945 | 0.091 | 0.086 | 1 | 5 | N4bp2 | 0.005 | Young |
| 0.772115 | 0.059634 | 0.195 | 0.205 | 1 | 5 | Rit1 | -0.01 | Young |
| 0.772374 | 0.112561 | 0.16 | 0.153 | 1 | 5 | C2cd5 | 0.007 | Young |
| 0.772502 | 0.052603 | 0.168 | 0.16 | 1 | 5 | Adss | 0.008 | Young |
| 0.772671 | 0.185811 | 0.094 | 0.099 | 1 | 5 | Exoc6 | -0.005 | Young |
| 0.772707 | 0.014189 | 0.67 | 0.69 | 1 | 5 | Atp6v1g1 | -0.02 | Young |
| 0.772779 | 0.165823 | 0.041 | 0.037 | 1 | 5 | Als2 | 0.004 | Young |
| 0.772853 | 0.181891 | 0.086 | 0.081 | 1 | 5 | Pdhx | 0.005 | Young |
| 0.772943 | 0.141027 | 0.058 | 0.054 | 1 | 5 | Rpap1 | 0.004 | Young |
| 0.77297 | 0.412984 | 0.01 | 0.008 | 1 | 5 | Spag1 | 0.002 | Young |
| 0.773252 | 0.120798 | 0.18 | 0.177 | 1 | 5 | Kmt5a | 0.003 | Young |
| 0.773589 | 0.053356 | 0.063 | 0.059 | 1 | 5 | Slc10a3 | 0.004 | Young |
| 0.77366 | 0.479839 | 0.046 | 0.042 | 1 | 5 | Mtap | 0.004 | Young |
| 0.774029 | 0.441138 | 0.01 | 0.008 | 1 | 5 | Fgf9 | 0.002 | Young |
| 0.774029 | 0.418757 | 0.01 | 0.008 | 1 | 5 | Bbs10 | 0.002 | Young |
| 0.774029 | 0.32067 | 0.01 | 0.008 | 1 | 5 | Iqcd | 0.002 | Young |
| 0.774144 | 0.427849 | 0.051 | 0.047 | 1 | 5 | Mbip | 0.004 | Young |
| 0.774837 | 0.156303 | 0.038 | 0.042 | 1 | 5 | 2310040G2 | -0.004 | Young |
| 0.775139 | 0.079627 | 0.553 | 0.576 | 1 | 5 | Atp5b | -0.023 | Young |
| 0.775148 | 0.015182 | 0.261 | 0.271 | 1 | 5 | Ttc39b | -0.01 | Young |
| 0.775379 | 0.037407 | 0.048 | 0.052 | 1 | 5 | Slx1b | -0.004 | Young |
| 0.775415 | 0.028132 | 0.485 | 0.502 | 1 | 5 | Nfic | -0.017 | Young |
| 0.775569 | 0.08763 | 0.431 | 0.451 | 1 | 5 | Snx8 | -0.02 | Young |
| 0.775727 | 0.05035 | 0.404 | 0.419 | 1 | 5 | Aldh9a1 | -0.015 | Young |
| 0.775814 | 0.02932 | 0.305 | 0.293 | 1 | 5 | Uap1l1 | 0.012 | Young |

| | | | | | | | | |
|---|---|---|---|---|---|---|---|---|
| 0.77589 | 0.194965 | 0.19 | 0.184 | 1 | 5 | Pcif1 | 0.006 | Young |
| 0.776085 | 0.000987 | 0.175 | 0.167 | 1 | 5 | Itpr2 | 0.008 | Young |
| 0.776149 | 0.46654 | 0.01 | 0.008 | 1 | 5 | Gm45449 | 0.002 | Young |
| 0.776149 | 0.217557 | 0.01 | 0.008 | 1 | 5 | Cped1 | 0.002 | Young |
| 0.776271 | 0.044743 | 0.036 | 0.032 | 1 | 5 | Arpin | 0.004 | Young |
| 0.776297 | 0.071575 | 0.053 | 0.049 | 1 | 5 | Abt1 | 0.004 | Young |
| 0.776406 | 0.021352 | 0.47 | 0.481 | 1 | 5 | Leprot | -0.011 | Young |
| 0.776424 | 0.165372 | 0.312 | 0.313 | 1 | 5 | Tcirg1 | -0.001 | Young |
| 0.776678 | 0.244059 | 0.091 | 0.088 | 1 | 5 | H6pd | 0.003 | Young |
| 0.776869 | 0.023965 | 0.84 | 0.875 | 1 | 5 | Atp6v0b | -0.035 | Young |
| 0.777019 | 0.221642 | 0.069 | 0.064 | 1 | 5 | Pef1 | 0.005 | Young |
| 0.77702 | 0.1243 | 0.452 | 0.451 | 1 | 5 | Mrps21 | 0.001 | Young |
| 0.777087 | 0.201766 | 0.079 | 0.074 | 1 | 5 | Smarcd1 | 0.005 | Young |
| 0.777194 | 0.195505 | 0.023 | 0.02 | 1 | 5 | Abhd8 | 0.003 | Young |
| 0.777194 | 0.060555 | 0.023 | 0.02 | 1 | 5 | C730034F0 | 0.003 | Young |
| 0.777209 | 0.527716 | 0.01 | 0.008 | 1 | 5 | 9930012K1 | 0.002 | Young |
| 0.777209 | 0.353779 | 0.01 | 0.008 | 1 | 5 | Zfp503 | 0.002 | Young |
| 0.777209 | 0.256768 | 0.01 | 0.008 | 1 | 5 | Gm40645 | 0.002 | Young |
| 0.777209 | 0.210642 | 0.01 | 0.008 | 1 | 5 | Fcor | 0.002 | Young |
| 0.777953 | 0.003598 | 0.036 | 0.032 | 1 | 5 | Sqle | 0.004 | Young |
| 0.778011 | 0.033909 | 0.497 | 0.524 | 1 | 5 | Ttc14 | -0.027 | Young |
| 0.778012 | 0.526695 | 0.254 | 0.246 | 1 | 5 | Chka | 0.008 | Young |
| 0.778035 | 0.061956 | 0.198 | 0.189 | 1 | 5 | Tm9sf4 | 0.009 | Young |
| 0.778115 | 0.134797 | 0.447 | 0.458 | 1 | 5 | Tuba1c | -0.011 | Young |
| 0.778121 | 0.154187 | 0.135 | 0.141 | 1 | 5 | Chst7 | -0.006 | Young |
| 0.778195 | 0.235415 | 0.185 | 0.182 | 1 | 5 | Atad1 | 0.003 | Young |
| 0.778224 | 0.192436 | 0.173 | 0.167 | 1 | 5 | Dnajc14 | 0.006 | Young |
| 0.778238 | 0.121749 | 0.069 | 0.064 | 1 | 5 | Plpp2 | 0.005 | Young |
| 0.77827 | 0.278717 | 0.01 | 0.008 | 1 | 5 | Gm48529 | 0.002 | Young |
| 0.778425 | 0.371897 | 0.056 | 0.052 | 1 | 5 | Pik3ap1 | 0.004 | Young |
| 0.778442 | 0.026063 | 0.094 | 0.099 | 1 | 5 | Kat6b | -0.005 | Young |
| 0.778888 | 0.244799 | 0.203 | 0.202 | 1 | 5 | Il11ra1 | 0.001 | Young |
| 0.779276 | 0.007481 | 0.147 | 0.155 | 1 | 5 | Creb3l2 | -0.008 | Young |
| 0.779331 | 0.291214 | 0.01 | 0.008 | 1 | 5 | Ercc6l | 0.002 | Young |
| 0.779331 | 0.184328 | 0.01 | 0.008 | 1 | 5 | Pars2 | 0.002 | Young |
| 0.779331 | 0.173858 | 0.01 | 0.008 | 1 | 5 | Nek3 | 0.002 | Young |
| 0.779331 | 0.069307 | 0.01 | 0.008 | 1 | 5 | Fam57a | 0.002 | Young |
| 0.779587 | 0.202547 | 0.041 | 0.037 | 1 | 5 | Def6 | 0.004 | Young |
| 0.779828 | 0.063124 | 0.183 | 0.19 | 1 | 5 | Preb | -0.007 | Young |
| 0.779969 | 0.067799 | 0.058 | 0.054 | 1 | 5 | Leng1 | 0.004 | Young |
| 0.780099 | 0.589643 | 0.028 | 0.025 | 1 | 5 | Asb2 | 0.003 | Young |
| 0.780393 | 0.404139 | 0.01 | 0.008 | 1 | 5 | Eme2 | 0.002 | Young |
| 0.780393 | 0.192105 | 0.01 | 0.008 | 1 | 5 | Rpgr | 0.002 | Young |
| 0.780636 | 0.076026 | 0.041 | 0.037 | 1 | 5 | Slc30a6 | 0.004 | Young |
| 0.780689 | 0.18154 | 0.023 | 0.02 | 1 | 5 | Tbc1d7 | 0.003 | Young |

| | | | | | | | | |
|---|---|---|---|---|---|---|---|---|
| 0.78088 | 0.069162 | 0.259 | 0.249 | 1 | 5 | Chtf8 | 0.01 | Young |
| 0.780976 | 0.049298 | 0.099 | 0.104 | 1 | 5 | Csf2rb2 | -0.005 | Young |
| 0.781188 | 0.028996 | 0.137 | 0.13 | 1 | 5 | Phb | 0.007 | Young |
| 0.781455 | 0.184386 | 0.01 | 0.008 | 1 | 5 | Slc7a4 | 0.002 | Young |
| 0.781455 | 0.059439 | 0.01 | 0.008 | 1 | 5 | Gm7967 | 0.002 | Young |
| 0.781455 | 0.042911 | 0.01 | 0.008 | 1 | 5 | Adora2b | 0.002 | Young |
| 0.781455 | 0.03361 | 0.01 | 0.008 | 1 | 5 | Dctd | 0.002 | Young |
| 0.781485 | 0.361393 | 0.038 | 0.042 | 1 | 5 | Pgap3 | -0.004 | Young |
| 0.781643 | 0.302365 | 0.051 | 0.047 | 1 | 5 | Rbmx2 | 0.004 | Young |
| 0.781777 | 0.022859 | 0.188 | 0.177 | 1 | 5 | Ing1 | 0.011 | Young |
| 0.781853 | 0.028486 | 0.596 | 0.586 | 1 | 5 | Rab11b | 0.01 | Young |
| 0.781897 | 0.164315 | 0.069 | 0.064 | 1 | 5 | Gtf3c4 | 0.005 | Young |
| 0.782212 | 0.075529 | 0.272 | 0.264 | 1 | 5 | Eif3i | 0.008 | Young |
| 0.782256 | 0.214202 | 0.076 | 0.072 | 1 | 5 | Cstf2 | 0.004 | Young |
| 0.782304 | 0.124269 | 0.069 | 0.064 | 1 | 5 | Ccdc86 | 0.005 | Young |
| 0.78245 | 0.05056 | 0.076 | 0.071 | 1 | 5 | Pan2 | 0.005 | Young |
| 0.782518 | 0.011703 | 0.01 | 0.008 | 1 | 5 | Actr3b | 0.002 | Young |
| 0.782687 | 0.163392 | 0.03 | 0.034 | 1 | 5 | Gm8369 | -0.004 | Young |
| 0.782788 | 0.216395 | 0.023 | 0.02 | 1 | 5 | Zfp790 | 0.003 | Young |
| 0.782796 | 0.002389 | 0.188 | 0.197 | 1 | 5 | Ptprs | -0.009 | Young |
| 0.783556 | 0.080581 | 0.279 | 0.269 | 1 | 5 | Mrpl28 | 0.01 | Young |
| 0.783793 | 0.086278 | 0.074 | 0.069 | 1 | 5 | Pot1b | 0.005 | Young |
| 0.784306 | 0.01784 | 0.602 | 0.616 | 1 | 5 | Mycbp2 | -0.014 | Young |
| 0.78431 | 0.043028 | 0.041 | 0.037 | 1 | 5 | Osbpl1a | 0.004 | Young |
| 0.78462 | 0.029733 | 0.317 | 0.311 | 1 | 5 | Map2k3 | 0.006 | Young |
| 0.784644 | 0.166094 | 0.01 | 0.008 | 1 | 5 | Neu3 | 0.002 | Young |
| 0.784644 | 0.035927 | 0.01 | 0.008 | 1 | 5 | Gm17025 | 0.002 | Young |
| 0.784967 | 0.14193 | 0.066 | 0.062 | 1 | 5 | Acyp1 | 0.004 | Young |
| 0.785142 | 0.215877 | 0.056 | 0.052 | 1 | 5 | Cdkn1c | 0.004 | Young |
| 0.785163 | 0.039527 | 0.657 | 0.687 | 1 | 5 | Clec4a2 | -0.03 | Young |
| 0.785507 | 0.122146 | 0.322 | 0.328 | 1 | 5 | Pgls | -0.006 | Young |
| 0.78559 | 0.115312 | 0.023 | 0.02 | 1 | 5 | Tspyl4 | 0.003 | Young |
| 0.785803 | 0.235478 | 0.168 | 0.162 | 1 | 5 | Ncor2 | 0.006 | Young |
| 0.785909 | 0.204864 | 0.234 | 0.231 | 1 | 5 | Pqbp1 | 0.003 | Young |
| 0.785916 | 0.087681 | 0.274 | 0.266 | 1 | 5 | Cop1 | 0.008 | Young |
| 0.785953 | 0.162053 | 0.155 | 0.167 | 1 | 5 | Kpna3 | -0.012 | Young |
| 0.786183 | 0.106935 | 0.274 | 0.271 | 1 | 5 | Leprotl1 | 0.003 | Young |
| 0.786346 | 0.168265 | 0.157 | 0.168 | 1 | 5 | Mut | -0.011 | Young |
| 0.786417 | 0.170229 | 0.094 | 0.089 | 1 | 5 | Elac1 | 0.005 | Young |
| 0.786502 | 0.190116 | 0.173 | 0.168 | 1 | 5 | Trmt1 | 0.005 | Young |
| 0.786649 | 0.39656 | 0.033 | 0.03 | 1 | 5 | Lrrc45 | 0.003 | Young |
| 0.786672 | 0.099578 | 0.102 | 0.098 | 1 | 5 | Tsga10 | 0.004 | Young |
| 0.786799 | 0.181567 | 0.124 | 0.118 | 1 | 5 | Mtmr2 | 0.006 | Young |
| 0.78706 | 0.015168 | 0.893 | 0.928 | 1 | 5 | Rpl21 | -0.035 | Young |
| 0.787151 | 0.266234 | 0.086 | 0.082 | 1 | 5 | Kcnab2 | 0.004 | Young |

| | | | | | | | | |
|---|---|---|---|---|---|---|---|---|
| 0.787377 | 0.06727 | 0.335 | 0.327 | 1 | 5 | Gigyf1 | 0.008 | Young |
| 0.78764 | 0.137335 | 0.467 | 0.481 | 1 | 5 | Ran | -0.014 | Young |
| 0.787675 | 0.175173 | 0.058 | 0.054 | 1 | 5 | Pdcd2 | 0.004 | Young |
| 0.787757 | 0.021781 | 0.206 | 0.195 | 1 | 5 | Cln3 | 0.011 | Young |
| 0.787819 | 0.095457 | 0.223 | 0.236 | 1 | 5 | Pgm2l1 | -0.013 | Young |
| 0.788044 | 0.033697 | 0.556 | 0.588 | 1 | 5 | Ppp1r9b | -0.032 | Young |
| 0.788296 | 0.10556 | 0.114 | 0.109 | 1 | 5 | Pmm2 | 0.005 | Young |
| 0.788317 | 0.064325 | 0.817 | 0.845 | 1 | 5 | Ptpn18 | -0.028 | Young |
| 0.788892 | 0.101571 | 0.069 | 0.074 | 1 | 5 | Rev3l | -0.005 | Young |
| 0.789161 | 0.170089 | 0.119 | 0.114 | 1 | 5 | Thumpd3 | 0.005 | Young |
| 0.789195 | 0.055859 | 0.848 | 0.891 | 1 | 5 | Calm2 | -0.043 | Young |
| 0.789636 | 0.027075 | 0.096 | 0.103 | 1 | 5 | Prelid3b | -0.007 | Young |
| 0.78985 | 0.173496 | 0.142 | 0.152 | 1 | 5 | Fam214b | -0.01 | Young |
| 0.789941 | 0.222956 | 0.14 | 0.135 | 1 | 5 | Fam204a | 0.005 | Young |
| 0.79071 | 0.06996 | 0.34 | 0.338 | 1 | 5 | Eif4e | 0.002 | Young |
| 0.790738 | 0.048507 | 0.332 | 0.35 | 1 | 5 | Sbds | -0.018 | Young |
| 0.790984 | 0.134508 | 0.053 | 0.057 | 1 | 5 | Sepsecs | -0.004 | Young |
| 0.790991 | 0.149843 | 0.335 | 0.36 | 1 | 5 | Nacc2 | -0.025 | Young |
| 0.791388 | 0.103716 | 0.282 | 0.298 | 1 | 5 | Pgam1 | -0.016 | Young |
| 0.791391 | 0.154355 | 0.294 | 0.313 | 1 | 5 | Pnp | -0.019 | Young |
| 0.791425 | 0.359599 | 0.107 | 0.103 | 1 | 5 | Mlycd | 0.004 | Young |
| 0.791697 | 0.163789 | 0.147 | 0.14 | 1 | 5 | Abi2 | 0.007 | Young |
| 0.792004 | 0.326617 | 0.124 | 0.121 | 1 | 5 | Kat7 | 0.003 | Young |
| 0.792152 | 0.235514 | 0.061 | 0.057 | 1 | 5 | Hes6 | 0.004 | Young |
| 0.792176 | 0.14109 | 0.168 | 0.175 | 1 | 5 | Gm49336 | -0.007 | Young |
| 0.792202 | 0.068775 | 0.442 | 0.436 | 1 | 5 | Uba1 | 0.006 | Young |
| 0.79245 | 0.271598 | 0.157 | 0.157 | 1 | 5 | Pccb | 0 | Young |
| 0.792457 | 0.159197 | 0.157 | 0.155 | 1 | 5 | Lcor | 0.002 | Young |
| 0.79273 | 0.394956 | 0.028 | 0.025 | 1 | 5 | Mrm2 | 0.003 | Young |
| 0.79274 | 0.135202 | 0.294 | 0.293 | 1 | 5 | Hsd17b4 | 0.001 | Young |
| 0.792748 | 0.269607 | 0.272 | 0.276 | 1 | 5 | Slc38a10 | -0.004 | Young |
| 0.792785 | 0.068188 | 0.32 | 0.327 | 1 | 5 | Atf7 | -0.007 | Young |
| 0.793363 | 0.351663 | 0.028 | 0.025 | 1 | 5 | Phka1 | 0.003 | Young |
| 0.79346 | 0.075134 | 0.411 | 0.414 | 1 | 5 | Rac2 | -0.003 | Young |
| 0.79371 | 0.097265 | 0.155 | 0.146 | 1 | 5 | Cfl2 | 0.009 | Young |
| 0.79378 | 0.174429 | 0.173 | 0.167 | 1 | 5 | Kpna1 | 0.006 | Young |
| 0.793914 | 0.036455 | 0.391 | 0.386 | 1 | 5 | Ndufb9 | 0.005 | Young |
| 0.79414 | 0.036387 | 0.147 | 0.153 | 1 | 5 | Pes1 | -0.006 | Young |
| 0.794228 | 0.076793 | 0.018 | 0.02 | 1 | 5 | Aqp1 | -0.002 | Young |
| 0.794388 | 0.182001 | 0.043 | 0.047 | 1 | 5 | Ppp1r16a | -0.004 | Young |
| 0.79456 | 0.164349 | 0.467 | 0.481 | 1 | 5 | Sptssa | -0.014 | Young |
| 0.79467 | 0.189967 | 0.17 | 0.165 | 1 | 5 | Tnip2 | 0.005 | Young |
| 0.794887 | 0.020531 | 0.35 | 0.357 | 1 | 5 | Slc4a7 | -0.007 | Young |
| 0.794916 | 0.31839 | 0.036 | 0.039 | 1 | 5 | Cox15 | -0.003 | Young |
| 0.795268 | 0.016352 | 0.157 | 0.148 | 1 | 5 | Sin3a | 0.009 | Young |

| | | | | | | | | |
|---|---|---|---|---|---|---|---|---|
| 0.795325 | 0.147033 | 0.543 | 0.566 | 1 | 5 | Atxn7l3b | -0.023 | Young |
| 0.795418 | 0.014092 | 0.373 | 0.394 | 1 | 5 | Rtraf | -0.021 | Young |
| 0.795679 | 0.094064 | 0.157 | 0.152 | 1 | 5 | G6pdx | 0.005 | Young |
| 0.795984 | 0.031521 | 0.662 | 0.719 | 1 | 5 | Tubb5 | -0.057 | Young |
| 0.796247 | 0.098651 | 0.056 | 0.061 | 1 | 5 | Ccdc122 | -0.005 | Young |
| 0.796286 | 0.286541 | 0.198 | 0.199 | 1 | 5 | Map2k1 | -0.001 | Young |
| 0.796443 | 0.060349 | 0.018 | 0.02 | 1 | 5 | Gm12216 | -0.002 | Young |
| 0.797392 | 0.152475 | 0.061 | 0.066 | 1 | 5 | Cbr1 | -0.005 | Young |
| 0.797551 | 0.142754 | 0.266 | 0.264 | 1 | 5 | Uqcrfs1 | 0.002 | Young |
| 0.79817 | 0.102616 | 0.518 | 0.544 | 1 | 5 | Kmt2e | -0.026 | Young |
| 0.798336 | 0.231693 | 0.112 | 0.118 | 1 | 5 | Rab3d | -0.006 | Young |
| 0.798461 | 0.139733 | 0.081 | 0.086 | 1 | 5 | Dnttip1 | -0.005 | Young |
| 0.798562 | 0.265514 | 0.033 | 0.03 | 1 | 5 | Slfn9 | 0.003 | Young |
| 0.798622 | 0.010265 | 0.992 | 0.997 | 1 | 5 | Jund | -0.005 | Young |
| 0.798627 | 0.009561 | 0.069 | 0.064 | 1 | 5 | Ppp2r1b | 0.005 | Young |
| 0.79866 | 0.316897 | 0.018 | 0.02 | 1 | 5 | Mtcp1 | -0.002 | Young |
| 0.799283 | 0.094059 | 0.53 | 0.535 | 1 | 5 | Pbrm1 | -0.005 | Young |
| 0.799536 | 0.03261 | 0.076 | 0.081 | 1 | 5 | Zfp266 | -0.005 | Young |
| 0.7997 | 0.036492 | 0.028 | 0.025 | 1 | 5 | Slc25a13 | 0.003 | Young |
| 0.799702 | 0.087421 | 0.165 | 0.172 | 1 | 5 | Ccng1 | -0.007 | Young |
| 0.799775 | 0.02513 | 0.391 | 0.404 | 1 | 5 | Supt5 | -0.013 | Young |
| 0.799973 | 0.099443 | 0.157 | 0.155 | 1 | 5 | Cspp1 | 0.002 | Young |
| 0.800004 | 0.078698 | 0.051 | 0.047 | 1 | 5 | Nedd4 | 0.004 | Young |
| 0.800499 | 0.048393 | 0.096 | 0.091 | 1 | 5 | Nme4 | 0.005 | Young |
| 0.800506 | 0.038446 | 0.203 | 0.21 | 1 | 5 | Aco1 | -0.007 | Young |
| 0.802273 | 0.511441 | 0.038 | 0.035 | 1 | 5 | Bahcc1 | 0.003 | Young |
| 0.802325 | 0.031114 | 0.5 | 0.498 | 1 | 5 | Paip2 | 0.002 | Young |
| 0.802366 | 0.191386 | 0.051 | 0.047 | 1 | 5 | Sco2 | 0.004 | Young |
| 0.802383 | 0.033035 | 0.137 | 0.145 | 1 | 5 | Akap11 | -0.008 | Young |
| 0.802513 | 0.021329 | 0.089 | 0.094 | 1 | 5 | Shkbp1 | -0.005 | Young |
| 0.802736 | 0.013008 | 0.041 | 0.044 | 1 | 5 | Btg3 | -0.003 | Young |
| 0.80299 | 0.094409 | 0.147 | 0.157 | 1 | 5 | Suclg2 | -0.01 | Young |
| 0.803316 | 0.134225 | 0.437 | 0.436 | 1 | 5 | Notch1 | 0.001 | Young |
| 0.80337 | 0.149383 | 0.142 | 0.136 | 1 | 5 | Mcrs1 | 0.006 | Young |
| 0.803403 | 0.153459 | 0.046 | 0.042 | 1 | 5 | Zbtb40 | 0.004 | Young |
| 0.803461 | 0.131744 | 0.114 | 0.12 | 1 | 5 | Katnbl1 | -0.006 | Young |
| 0.803509 | 0.01755 | 0.028 | 0.025 | 1 | 5 | Numbl | 0.003 | Young |
| 0.803569 | 0.089937 | 0.206 | 0.215 | 1 | 5 | Zfp740 | -0.009 | Young |
| 0.803785 | 0.133538 | 0.34 | 0.357 | 1 | 5 | 2610001J0 | -0.017 | Young |
| 0.80398 | 0.186197 | 0.14 | 0.146 | 1 | 5 | Washc1 | -0.006 | Young |
| 0.805105 | 0.132645 | 0.404 | 0.421 | 1 | 5 | Ndufb11 | -0.017 | Young |
| 0.805136 | 0.141999 | 0.211 | 0.215 | 1 | 5 | Yipf1 | -0.004 | Young |
| 0.805416 | 0.043878 | 0.028 | 0.025 | 1 | 5 | Fam109a | 0.003 | Young |
| 0.805557 | 0.304614 | 0.033 | 0.03 | 1 | 5 | Lmo1 | 0.003 | Young |
| 0.805671 | 0.229006 | 0.046 | 0.049 | 1 | 5 | Ccdc191 | -0.003 | Young |

| | | | | | | | | |
|---|---|---|---|---|---|---|---|---|
| 0.80576 | 0.012516 | 0.041 | 0.044 | 1 | 5 | Pola2 | -0.003 | Young |
| 0.80578 | 0.285191 | 0.086 | 0.082 | 1 | 5 | Ccdc152 | 0.004 | Young |
| 0.805795 | 0.142483 | 0.234 | 0.229 | 1 | 5 | Tnfaip8l2 | 0.005 | Young |
| 0.805905 | 0.043701 | 0.069 | 0.072 | 1 | 5 | Xpc | -0.003 | Young |
| 0.806084 | 0.026818 | 0.129 | 0.136 | 1 | 5 | Zdhhc5 | -0.007 | Young |
| 0.806141 | 0.254677 | 0.033 | 0.03 | 1 | 5 | Mcm7 | 0.003 | Young |
| 0.806615 | 0.369746 | 0.038 | 0.035 | 1 | 5 | Lag3 | 0.003 | Young |
| 0.80679 | 0.091419 | 0.17 | 0.167 | 1 | 5 | Capg | 0.003 | Young |
| 0.807094 | 0.293436 | 0.069 | 0.072 | 1 | 5 | Galns | -0.003 | Young |
| 0.80735 | 0.034721 | 0.299 | 0.31 | 1 | 5 | Snx10 | -0.011 | Young |
| 0.807843 | 0.30528 | 0.107 | 0.103 | 1 | 5 | Snapc3 | 0.004 | Young |
| 0.80815 | 0.117778 | 0.332 | 0.337 | 1 | 5 | Zfand3 | -0.005 | Young |
| 0.808199 | 0.266417 | 0.104 | 0.101 | 1 | 5 | Actl6a | 0.003 | Young |
| 0.808332 | 0.034232 | 0.058 | 0.062 | 1 | 5 | Bag4 | -0.004 | Young |
| 0.808852 | 0.747386 | 0.015 | 0.013 | 1 | 5 | Cryba4 | 0.002 | Young |
| 0.809058 | 0.03417 | 0.741 | 0.774 | 1 | 5 | Grk2 | -0.033 | Young |
| 0.809332 | 0.026276 | 0.109 | 0.114 | 1 | 5 | Srpk1 | -0.005 | Young |
| 0.809451 | 0.04435 | 0.056 | 0.052 | 1 | 5 | Acaca | 0.004 | Young |
| 0.809714 | 0.678683 | 0.015 | 0.013 | 1 | 5 | 9530082P2 | 0.002 | Young |
| 0.81015 | 0.006442 | 0.071 | 0.076 | 1 | 5 | Gnl3l | -0.005 | Young |
| 0.810329 | 0.007426 | 0.315 | 0.316 | 1 | 5 | Ptbp1 | -0.001 | Young |
| 0.81055 | 0.292889 | 0.226 | 0.222 | 1 | 5 | N4bp2l1 | 0.004 | Young |
| 0.810576 | 0.016395 | 0.627 | 0.635 | 1 | 5 | Pla2g7 | -0.008 | Young |
| 0.810633 | 0.154165 | 0.178 | 0.173 | 1 | 5 | Tle1 | 0.005 | Young |
| 0.810723 | 0.383764 | 0.117 | 0.114 | 1 | 5 | Scyl2 | 0.003 | Young |
| 0.810807 | 0.04748 | 0.041 | 0.044 | 1 | 5 | Abcb10 | -0.003 | Young |
| 0.810814 | 0.176987 | 0.033 | 0.03 | 1 | 5 | Rab3ip | 0.003 | Young |
| 0.810894 | 0.429096 | 0.109 | 0.106 | 1 | 5 | Ugdh | 0.003 | Young |
| 0.810905 | 0.001961 | 0.396 | 0.389 | 1 | 5 | Phf21a | 0.007 | Young |
| 0.81096 | 0.02241 | 0.048 | 0.052 | 1 | 5 | Kank3 | -0.004 | Young |
| 0.811081 | 0.262363 | 0.053 | 0.051 | 1 | 5 | Borcs8 | 0.002 | Young |
| 0.811514 | 0.325268 | 0.036 | 0.039 | 1 | 5 | Sccpdh | -0.003 | Young |
| 0.811518 | 0.000961 | 0.348 | 0.357 | 1 | 5 | Dhx15 | -0.009 | Young |
| 0.811728 | 0.21564 | 0.147 | 0.145 | 1 | 5 | Sart1 | 0.002 | Young |
| 0.811781 | 0.11009 | 0.289 | 0.286 | 1 | 5 | Rab8a | 0.003 | Young |
| 0.812337 | 0.020811 | 0.01 | 0.012 | 1 | 5 | Lrrc56 | -0.002 | Young |
| 0.812554 | 0.036771 | 0.376 | 0.386 | 1 | 5 | Eif4g3 | -0.01 | Young |
| 0.812845 | 0.051231 | 0.074 | 0.077 | 1 | 5 | Slc35a1 | -0.003 | Young |
| 0.812868 | 0.350208 | 0.165 | 0.165 | 1 | 5 | Mrpl13 | 0 | Young |
| 0.812876 | 0.015474 | 0.137 | 0.143 | 1 | 5 | Asl | -0.006 | Young |
| 0.813309 | 0.377844 | 0.01 | 0.012 | 1 | 5 | Pcgf2 | -0.002 | Young |
| 0.813492 | 0.116159 | 0.051 | 0.047 | 1 | 5 | Ctla2b | 0.004 | Young |
| 0.813648 | 0.045992 | 0.086 | 0.091 | 1 | 5 | Sik3 | -0.005 | Young |
| 0.813749 | 0.074232 | 0.343 | 0.333 | 1 | 5 | Tomm22 | 0.01 | Young |
| 0.813764 | 0.379466 | 0.043 | 0.04 | 1 | 5 | D2hgdh | 0.003 | Young |

| | | | | | | | | |
|---|---|---|---|---|---|---|---|---|
| 0.813808 | 0.008334 | 0.322 | 0.328 | 1 | 5 | Gtpbp2 | -0.006 | Young |
| 0.813827 | 0.136964 | 0.086 | 0.091 | 1 | 5 | Zfp994 | -0.005 | Young |
| 0.813838 | 0.102803 | 0.041 | 0.044 | 1 | 5 | Suv39h1 | -0.003 | Young |
| 0.813899 | 0.085494 | 0.137 | 0.143 | 1 | 5 | Yif1b | -0.006 | Young |
| 0.814013 | 0.097913 | 0.561 | 0.569 | 1 | 5 | Zfp91 | -0.008 | Young |
| 0.814027 | 0.282505 | 0.015 | 0.013 | 1 | 5 | Lcmt2 | 0.002 | Young |
| 0.814027 | 0.18378 | 0.015 | 0.013 | 1 | 5 | Gm32856 | 0.002 | Young |
| 0.814323 | 0.044706 | 0.033 | 0.03 | 1 | 5 | Bbs7 | 0.003 | Young |
| 0.814468 | 0.011167 | 0.855 | 0.88 | 1 | 5 | Rhoa | -0.025 | Young |
| 0.814493 | 0.015297 | 0.213 | 0.224 | 1 | 5 | Mrpl42 | -0.011 | Young |
| 0.814772 | 0.247974 | 0.038 | 0.035 | 1 | 5 | Zfp335 | 0.003 | Young |
| 0.814803 | 0.208143 | 0.185 | 0.184 | 1 | 5 | Ankib1 | 0.001 | Young |
| 0.814886 | 0.150514 | 0.223 | 0.221 | 1 | 5 | Cops4 | 0.002 | Young |
| 0.814922 | 0.054552 | 0.137 | 0.143 | 1 | 5 | Gpd1l | -0.006 | Young |
| 0.815254 | 0.171847 | 0.01 | 0.012 | 1 | 5 | Fhl1 | -0.002 | Young |
| 0.815278 | 0.084303 | 0.566 | 0.588 | 1 | 5 | 2900097C1 | -0.022 | Young |
| 0.815533 | 0.046394 | 0.216 | 0.204 | 1 | 5 | Ppil2 | 0.012 | Young |
| 0.815754 | 0.17339 | 0.015 | 0.013 | 1 | 5 | Wrap53 | 0.002 | Young |
| 0.815785 | 0.0463 | 0.056 | 0.052 | 1 | 5 | Zfp609 | 0.004 | Young |
| 0.815828 | 0.322785 | 0.109 | 0.106 | 1 | 5 | Rbsn | 0.003 | Young |
| 0.816145 | 0.135233 | 0.145 | 0.148 | 1 | 5 | Capn7 | -0.003 | Young |
| 0.816152 | 0.065797 | 0.086 | 0.091 | 1 | 5 | Gcsh | -0.005 | Young |
| 0.816226 | 0.08125 | 0.01 | 0.012 | 1 | 5 | Gpr150 | -0.002 | Young |
| 0.816227 | 0.149237 | 0.36 | 0.375 | 1 | 5 | Acap2 | -0.015 | Young |
| 0.816618 | 0.137566 | 0.015 | 0.013 | 1 | 5 | Exo5 | 0.002 | Young |
| 0.817008 | 0.294885 | 0.048 | 0.052 | 1 | 5 | Hdac10 | -0.004 | Young |
| 0.817119 | 0.05612 | 0.132 | 0.136 | 1 | 5 | Tmem183a | -0.004 | Young |
| 0.817482 | 0.561215 | 0.015 | 0.013 | 1 | 5 | Fsd2 | 0.002 | Young |
| 0.817746 | 0.078025 | 0.236 | 0.231 | 1 | 5 | Eml4 | 0.005 | Young |
| 0.817983 | 0.0927 | 0.299 | 0.316 | 1 | 5 | Clk4 | -0.017 | Young |
| 0.818306 | 0.046346 | 0.619 | 0.645 | 1 | 5 | Rbm25 | -0.026 | Young |
| 0.818415 | 0.182625 | 0.084 | 0.089 | 1 | 5 | Acot13 | -0.005 | Young |
| 0.818427 | 0.10301 | 0.297 | 0.298 | 1 | 5 | Dnase1l1 | -0.001 | Young |
| 0.8185 | 0.214656 | 0.036 | 0.039 | 1 | 5 | Apba3 | -0.003 | Young |
| 0.818505 | 0.082454 | 0.201 | 0.192 | 1 | 5 | Tbc1d13 | 0.009 | Young |
| 0.819005 | 0.085231 | 0.074 | 0.077 | 1 | 5 | Ifi206 | -0.003 | Young |
| 0.819122 | 0.073332 | 0.135 | 0.128 | 1 | 5 | Spaca6 | 0.007 | Young |
| 0.819146 | 0.068171 | 0.244 | 0.237 | 1 | 5 | Xpo1 | 0.007 | Young |
| 0.819287 | 0.034485 | 0.315 | 0.325 | 1 | 5 | Eloa | -0.01 | Young |
| 0.819879 | 0.101844 | 0.221 | 0.229 | 1 | 5 | Cc2d1b | -0.008 | Young |
| 0.819885 | 0.063596 | 0.279 | 0.271 | 1 | 5 | Smc6 | 0.008 | Young |
| 0.819946 | 0.114478 | 0.096 | 0.101 | 1 | 5 | Glce | -0.005 | Young |
| 0.819961 | 0.206933 | 0.292 | 0.286 | 1 | 5 | Mat2b | 0.006 | Young |
| 0.820041 | 0.074499 | 0.416 | 0.441 | 1 | 5 | Cdk9 | -0.025 | Young |
| 0.820075 | 0.095683 | 0.015 | 0.013 | 1 | 5 | Mfsd4b4 | 0.002 | Young |

| | | | | | | | | |
|---|---|---|---|---|---|---|---|---|
| 0.820075 | 0.05087 | 0.015 | 0.013 | 1 | 5 | Diaph3 | 0.002 | Young |
| 0.820189 | 0.045254 | 0.112 | 0.106 | 1 | 5 | Hif1an | 0.006 | Young |
| 0.820336 | 0.04008 | 0.023 | 0.025 | 1 | 5 | Zbtb11os1 | -0.002 | Young |
| 0.820944 | 0.250474 | 0.157 | 0.155 | 1 | 5 | Tmem43 | 0.002 | Young |
| 0.821 | 0.143394 | 0.023 | 0.025 | 1 | 5 | Gm20632 | -0.002 | Young |
| 0.821005 | 0.069355 | 0.051 | 0.054 | 1 | 5 | Rfc3 | -0.003 | Young |
| 0.821065 | 0.367788 | 0.086 | 0.084 | 1 | 5 | Zdhhc6 | 0.002 | Young |
| 0.821307 | 0.230184 | 0.17 | 0.167 | 1 | 5 | Smurf2 | 0.003 | Young |
| 0.82157 | 0.047939 | 0.058 | 0.062 | 1 | 5 | C2cd2l | -0.004 | Young |
| 0.821696 | 0.061108 | 0.32 | 0.313 | 1 | 5 | Trim28 | 0.007 | Young |
| 0.82173 | 0.391549 | 0.036 | 0.039 | 1 | 5 | Mios | -0.003 | Young |
| 0.821805 | 0.035128 | 0.015 | 0.013 | 1 | 5 | Palb2 | 0.002 | Young |
| 0.82181 | 0.158754 | 0.081 | 0.077 | 1 | 5 | Prcc | 0.004 | Young |
| 0.822039 | 0.023862 | 0.145 | 0.148 | 1 | 5 | Mroh1 | -0.003 | Young |
| 0.822067 | 0.117022 | 0.157 | 0.165 | 1 | 5 | Mpv17 | -0.008 | Young |
| 0.822075 | 0.196754 | 0.383 | 0.392 | 1 | 5 | Stk4 | -0.009 | Young |
| 0.822404 | 0.088375 | 0.038 | 0.035 | 1 | 5 | Nbeal2 | 0.003 | Young |
| 0.822859 | 0.380047 | 0.058 | 0.056 | 1 | 5 | Pphln1 | 0.002 | Young |
| 0.822902 | 0.013652 | 0.127 | 0.133 | 1 | 5 | Zfp945 | -0.006 | Young |
| 0.823044 | 0.522241 | 0.01 | 0.012 | 1 | 5 | Gm26825 | -0.002 | Young |
| 0.823345 | 0.110916 | 0.168 | 0.16 | 1 | 5 | Slc39a10 | 0.008 | Young |
| 0.823458 | 0.117849 | 0.028 | 0.03 | 1 | 5 | Mad2l2 | -0.002 | Young |
| 0.823785 | 0.023952 | 0.259 | 0.266 | 1 | 5 | Ube2e3 | -0.007 | Young |
| 0.8239 | 0.0714 | 0.325 | 0.327 | 1 | 5 | Sgms1 | -0.002 | Young |
| 0.824019 | 0.406903 | 0.01 | 0.012 | 1 | 5 | D230017M | -0.002 | Young |
| 0.824064 | 0.105886 | 0.028 | 0.03 | 1 | 5 | Gpr84 | -0.002 | Young |
| 0.824719 | 0.20245 | 0.132 | 0.13 | 1 | 5 | Ggps1 | 0.002 | Young |
| 0.824767 | 0.081381 | 0.107 | 0.101 | 1 | 5 | Atl3 | 0.006 | Young |
| 0.824857 | 0.024166 | 0.198 | 0.202 | 1 | 5 | Dnajc1 | -0.004 | Young |
| 0.825044 | 0.071919 | 0.259 | 0.266 | 1 | 5 | Psme2b | -0.007 | Young |
| 0.825097 | 0.153844 | 0.119 | 0.114 | 1 | 5 | Dcaf5 | 0.005 | Young |
| 0.825513 | 0.276771 | 0.058 | 0.056 | 1 | 5 | Clp1 | 0.002 | Young |
| 0.825614 | 0.138665 | 0.122 | 0.118 | 1 | 5 | Tbc1d8 | 0.004 | Young |
| 0.82598 | 0.111219 | 0.076 | 0.081 | 1 | 5 | Ints11 | -0.005 | Young |
| 0.826061 | 0.054848 | 0.236 | 0.239 | 1 | 5 | Sike1 | -0.003 | Young |
| 0.826205 | 0.052315 | 0.173 | 0.165 | 1 | 5 | Lrif1 | 0.008 | Young |
| 0.826351 | 0.082338 | 0.967 | 0.983 | 1 | 5 | mt-Nd3 | -0.016 | Young |
| 0.826359 | 0.103765 | 0.076 | 0.081 | 1 | 5 | Rpain | -0.005 | Young |
| 0.827055 | 0.261488 | 0.084 | 0.081 | 1 | 5 | Gtpbp1 | 0.003 | Young |
| 0.827212 | 0.183744 | 0.094 | 0.089 | 1 | 5 | Cd1d1 | 0.005 | Young |
| 0.827488 | 0.015369 | 0.046 | 0.049 | 1 | 5 | Sall2 | -0.003 | Young |
| 0.827622 | 0.491884 | 0.02 | 0.019 | 1 | 5 | Arhgef18 | 0.001 | Young |
| 0.827717 | 0.070265 | 0.294 | 0.301 | 1 | 5 | Pdpk1 | -0.007 | Young |
| 0.828036 | 0.108759 | 0.541 | 0.545 | 1 | 5 | Gapdh | -0.004 | Young |
| 0.828075 | 0.049908 | 0.112 | 0.106 | 1 | 5 | Slc35b1 | 0.006 | Young |

| | | | | | | | | |
|---|---|---|---|---|---|---|---|---|
| 0.828183 | 0.1152 | 0.505 | 0.519 | 1 | 5 | Usp9x | -0.014 | Young |
| 0.828717 | 0.287555 | 0.124 | 0.123 | 1 | 5 | Acads | 0.001 | Young |
| 0.829413 | 0.049567 | 0.317 | 0.306 | 1 | 5 | Arnt | 0.011 | Young |
| 0.829546 | 0.070876 | 0.041 | 0.044 | 1 | 5 | Dmap1 | -0.003 | Young |
| 0.829607 | 0.103061 | 0.185 | 0.184 | 1 | 5 | Fam126b | 0.001 | Young |
| 0.829626 | 0.031848 | 0.259 | 0.269 | 1 | 5 | Cacybp | -0.01 | Young |
| 0.829938 | 0.027984 | 0.741 | 0.778 | 1 | 5 | Snx3 | -0.037 | Young |
| 0.830242 | 0.20186 | 0.244 | 0.244 | 1 | 5 | Slmap | 0 | Young |
| 0.830321 | 0.128273 | 0.157 | 0.153 | 1 | 5 | Naa60 | 0.004 | Young |
| 0.830616 | 0.215947 | 0.046 | 0.049 | 1 | 5 | Pwp2 | -0.003 | Young |
| 0.830841 | 0.208332 | 0.076 | 0.072 | 1 | 5 | Bmf | 0.004 | Young |
| 0.830891 | 0.192204 | 0.119 | 0.126 | 1 | 5 | Dnajc21 | -0.007 | Young |
| 0.831125 | 0.059008 | 0.195 | 0.202 | 1 | 5 | Sec13 | -0.007 | Young |
| 0.831385 | 0.088753 | 0.231 | 0.222 | 1 | 5 | Epc1 | 0.009 | Young |
| 0.831636 | 0.045126 | 0.129 | 0.133 | 1 | 5 | Mta3 | -0.004 | Young |
| 0.831719 | 0.014422 | 0.637 | 0.658 | 1 | 5 | Raph1 | -0.021 | Young |
| 0.831959 | 0.089772 | 0.104 | 0.108 | 1 | 5 | Nup85 | -0.004 | Young |
| 0.832152 | 0.160284 | 0.14 | 0.136 | 1 | 5 | Smim19 | 0.004 | Young |
| 0.832424 | 0.137426 | 0.114 | 0.111 | 1 | 5 | Zfp219 | 0.003 | Young |
| 0.832772 | 0.097285 | 0.213 | 0.217 | 1 | 5 | Ahcyl2 | -0.004 | Young |
| 0.832773 | 0.036766 | 0.063 | 0.067 | 1 | 5 | Uck1 | -0.004 | Young |
| 0.832964 | 0.114076 | 0.348 | 0.343 | 1 | 5 | Atxn7 | 0.005 | Young |
| 0.833187 | 0.118478 | 0.063 | 0.067 | 1 | 5 | Yars | -0.004 | Young |
| 0.833398 | 0.073263 | 0.398 | 0.404 | 1 | 5 | Sdhc | -0.006 | Young |
| 0.834513 | 0.000713 | 0.744 | 0.778 | 1 | 5 | Grb2 | -0.034 | Young |
| 0.834635 | 0.278894 | 0.147 | 0.143 | 1 | 5 | Hhex | 0.004 | Young |
| 0.834643 | 0.136816 | 0.119 | 0.116 | 1 | 5 | Hdac4 | 0.003 | Young |
| 0.834874 | 0.009286 | 0.972 | 0.983 | 1 | 5 | Rps27 | -0.011 | Young |
| 0.835034 | 0.060205 | 0.251 | 0.258 | 1 | 5 | Plscr3 | -0.007 | Young |
| 0.835084 | 0.03077 | 0.056 | 0.052 | 1 | 5 | Tatdn1 | 0.004 | Young |
| 0.835975 | 0.153393 | 0.127 | 0.123 | 1 | 5 | Kctd12b | 0.004 | Young |
| 0.836036 | 0.058923 | 0.162 | 0.168 | 1 | 5 | Igsf8 | -0.006 | Young |
| 0.836436 | 0.12204 | 0.066 | 0.062 | 1 | 5 | Ptrh2 | 0.004 | Young |
| 0.836448 | 0.026416 | 0.345 | 0.33 | 1 | 5 | Dcaf8 | 0.015 | Young |
| 0.836514 | 0.008276 | 0.256 | 0.266 | 1 | 5 | Adipor2 | -0.01 | Young |
| 0.836613 | 0.163296 | 0.132 | 0.128 | 1 | 5 | Actr1b | 0.004 | Young |
| 0.83692 | 0.10872 | 1 | 0.997 | 1 | 5 | Apoe | 0.003 | Young |
| 0.836959 | 0.028326 | 0.142 | 0.136 | 1 | 5 | Arfip1 | 0.006 | Young |
| 0.837351 | 0.02861 | 0.472 | 0.46 | 1 | 5 | Cat | 0.012 | Young |
| 0.837367 | 0.057647 | 0.043 | 0.04 | 1 | 5 | Zfp113 | 0.003 | Young |
| 0.837416 | 0.053567 | 0.071 | 0.067 | 1 | 5 | Nomo1 | 0.004 | Young |
| 0.837592 | 0.121372 | 0.208 | 0.221 | 1 | 5 | Trim26 | -0.013 | Young |
| 0.837932 | 0.268172 | 0.18 | 0.19 | 1 | 5 | Tor1aip2 | -0.01 | Young |
| 0.837932 | 0.480912 | 0.081 | 0.079 | 1 | 5 | Smim26 | 0.002 | Young |
| 0.838078 | 0.172974 | 0.02 | 0.019 | 1 | 5 | Ogn | 0.001 | Young |

| | | | | | | | | |
|---|---|---|---|---|---|---|---|---|
| 0.838133 | 0.055454 | 0.533 | 0.562 | 1 | 5 | Nipbl | -0.029 | Young |
| 0.83859 | 0.001238 | 0.66 | 0.662 | 1 | 5 | Atp1a1 | -0.002 | Young |
| 0.83941 | 0.091833 | 0.429 | 0.423 | 1 | 5 | Lasp1 | 0.006 | Young |
| 0.839454 | 0.048143 | 0.096 | 0.101 | 1 | 5 | Rptor | -0.005 | Young |
| 0.839574 | 0.285368 | 0.02 | 0.019 | 1 | 5 | Tgm2 | 0.001 | Young |
| 0.8396 | 0.104618 | 0.239 | 0.234 | 1 | 5 | Ndufaf8 | 0.005 | Young |
| 0.839609 | 0.05192 | 0.147 | 0.145 | 1 | 5 | Iigp1 | 0.002 | Young |
| 0.840323 | 0.290493 | 0.02 | 0.019 | 1 | 5 | 1810044D( | 0.001 | Young |
| 0.840353 | 0.078466 | 0.299 | 0.293 | 1 | 5 | Mrpl24 | 0.006 | Young |
| 0.840467 | 0.090532 | 0.028 | 0.03 | 1 | 5 | E130308A1 | -0.002 | Young |
| 0.840896 | 0.093766 | 0.218 | 0.212 | 1 | 5 | Stx5a | 0.006 | Young |
| 0.841076 | 0.660259 | 0.03 | 0.029 | 1 | 5 | Lhpp | 0.001 | Young |
| 0.841249 | 0.368801 | 0.117 | 0.114 | 1 | 5 | Pxdc1 | 0.003 | Young |
| 0.841317 | 0.097811 | 0.048 | 0.045 | 1 | 5 | Gpatch1 | 0.003 | Young |
| 0.841617 | 0.158916 | 0.099 | 0.096 | 1 | 5 | Mthfr | 0.003 | Young |
| 0.84195 | 0.037796 | 0.338 | 0.335 | 1 | 5 | Dusp22 | 0.003 | Young |
| 0.842338 | 0.020967 | 0.525 | 0.539 | 1 | 5 | Sult1a1 | -0.014 | Young |
| 0.843119 | 0.012341 | 0.19 | 0.185 | 1 | 5 | Rnf220 | 0.005 | Young |
| 0.843174 | 0.013265 | 0.079 | 0.074 | 1 | 5 | Ece1 | 0.005 | Young |
| 0.843189 | 0.12374 | 0.274 | 0.264 | 1 | 5 | Dpf2 | 0.01 | Young |
| 0.843317 | 0.104503 | 0.02 | 0.019 | 1 | 5 | Snhg18 | 0.001 | Young |
| 0.843317 | 0.003089 | 0.02 | 0.019 | 1 | 5 | Purg | 0.001 | Young |
| 0.843349 | 0.50769 | 0.051 | 0.049 | 1 | 5 | Lzts2 | 0.002 | Young |
| 0.843406 | 0.148476 | 0.282 | 0.281 | 1 | 5 | Fads1 | 0.001 | Young |
| 0.843434 | 0.169415 | 0.16 | 0.158 | 1 | 5 | Virma | 0.002 | Young |
| 0.84346 | 0.004884 | 0.195 | 0.194 | 1 | 5 | Chml | 0.001 | Young |
| 0.843499 | 0.616813 | 0.041 | 0.039 | 1 | 5 | Snn | 0.002 | Young |
| 0.843618 | 0.267078 | 0.201 | 0.199 | 1 | 5 | Lrrk1 | 0.002 | Young |
| 0.843775 | 0.137894 | 0.48 | 0.497 | 1 | 5 | Ugcg | -0.017 | Young |
| 0.843907 | 0.033758 | 0.071 | 0.067 | 1 | 5 | Cfap36 | 0.004 | Young |
| 0.844366 | 0.112447 | 0.015 | 0.017 | 1 | 5 | Gm38251 | -0.002 | Young |
| 0.844519 | 0.097437 | 0.033 | 0.035 | 1 | 5 | Cbr4 | -0.002 | Young |
| 0.844536 | 0.079736 | 0.099 | 0.094 | 1 | 5 | 2410089E( | 0.005 | Young |
| 0.844677 | 0.122079 | 0.18 | 0.189 | 1 | 5 | Mif4gd | -0.009 | Young |
| 0.844815 | 0.084486 | 0.02 | 0.019 | 1 | 5 | Elp4 | 0.001 | Young |
| 0.844847 | 0.014134 | 0.094 | 0.089 | 1 | 5 | Scnm1 | 0.005 | Young |
| 0.844977 | 0.102703 | 0.178 | 0.185 | 1 | 5 | Tbc1d17 | -0.007 | Young |
| 0.845059 | 0.160444 | 0.289 | 0.288 | 1 | 5 | Pbx2 | 0.001 | Young |
| 0.845175 | 0.102973 | 0.201 | 0.195 | 1 | 5 | Pcgf5 | 0.006 | Young |
| 0.845484 | 0.005803 | 0.223 | 0.227 | 1 | 5 | Zeb2os | -0.004 | Young |
| 0.845565 | 0.130141 | 0.02 | 0.019 | 1 | 5 | Cep76 | 0.001 | Young |
| 0.845641 | 0.088549 | 0.056 | 0.059 | 1 | 5 | Zfp821 | -0.003 | Young |
| 0.845871 | 0.277849 | 0.096 | 0.093 | 1 | 5 | Ccdc117 | 0.003 | Young |
| 0.845909 | 0.048269 | 0.343 | 0.343 | 1 | 5 | Exoc5 | 0 | Young |
| 0.845967 | 0.210476 | 0.107 | 0.103 | 1 | 5 | Pak1ip1 | 0.004 | Young |

| | | | | | | | | |
|---|---|---|---|---|---|---|---|---|
| 0.845967 | 0.16084 | 0.107 | 0.103 | 1 | 5 | Ralgapb | 0.004 | Young |
| 0.845987 | 0.292771 | 0.025 | 0.024 | 1 | 5 | Zfp975 | 0.001 | Young |
| 0.845997 | 0.092027 | 0.015 | 0.017 | 1 | 5 | Csf1 | -0.002 | Young |
| 0.846042 | 0.059792 | 0.65 | 0.71 | 1 | 5 | Slc43a2 | -0.06 | Young |
| 0.846082 | 0.043593 | 0.056 | 0.059 | 1 | 5 | Dcaf6 | -0.003 | Young |
| 0.846317 | 0.071076 | 0.066 | 0.062 | 1 | 5 | Wdr45 | 0.004 | Young |
| 0.846382 | 0.155933 | 0.109 | 0.114 | 1 | 5 | Taf4 | -0.005 | Young |
| 0.846383 | 0.037733 | 0.203 | 0.21 | 1 | 5 | Impad1 | -0.007 | Young |
| 0.846715 | 0.114971 | 0.305 | 0.313 | 1 | 5 | Smarca2 | -0.008 | Young |
| 0.847 | 0.282494 | 0.063 | 0.061 | 1 | 5 | Exosc4 | 0.002 | Young |
| 0.847018 | 0.086808 | 0.188 | 0.197 | 1 | 5 | Khnyn | -0.009 | Young |
| 0.847161 | 0.021715 | 0.043 | 0.04 | 1 | 5 | Snx21 | 0.003 | Young |
| 0.847244 | 0.442399 | 0.112 | 0.111 | 1 | 5 | Sp3os | 0.001 | Young |
| 0.847613 | 0.285068 | 0.051 | 0.049 | 1 | 5 | Usp18 | 0.002 | Young |
| 0.847799 | 0.038508 | 0.32 | 0.306 | 1 | 5 | Tmem87b | 0.014 | Young |
| 0.847855 | 0.265903 | 0.063 | 0.061 | 1 | 5 | Fam149b | 0.002 | Young |
| 0.847993 | 0.409998 | 0.025 | 0.024 | 1 | 5 | Wdr55 | 0.001 | Young |
| 0.848093 | 0.049681 | 0.086 | 0.089 | 1 | 5 | Bloc1s6 | -0.003 | Young |
| 0.848152 | 0.061762 | 0.048 | 0.045 | 1 | 5 | Zfp276 | 0.003 | Young |
| 0.848329 | 0.067726 | 0.49 | 0.487 | 1 | 5 | Cap1 | 0.003 | Young |
| 0.848974 | 0.050212 | 0.33 | 0.332 | 1 | 5 | Anxa6 | -0.002 | Young |
| 0.848986 | 0.022561 | 0.307 | 0.318 | 1 | 5 | Ap3s1 | -0.011 | Young |
| 0.849318 | 0.080746 | 0.147 | 0.153 | 1 | 5 | Fnta | -0.006 | Young |
| 0.849441 | 0.02293 | 0.32 | 0.327 | 1 | 5 | Taf15 | -0.007 | Young |
| 0.850097 | 0.090896 | 0.208 | 0.204 | 1 | 5 | Samd8 | 0.004 | Young |
| 0.850488 | 0.004218 | 0.464 | 0.481 | 1 | 5 | Uqcrb | -0.017 | Young |
| 0.850978 | 0.361414 | 0.168 | 0.167 | 1 | 5 | Nhp2 | 0.001 | Young |
| 0.851035 | 0.154355 | 0.129 | 0.126 | 1 | 5 | Uvssa | 0.003 | Young |
| 0.851069 | 0.132281 | 0.409 | 0.441 | 1 | 5 | Usp7 | -0.032 | Young |
| 0.851191 | 0.000559 | 0.525 | 0.522 | 1 | 5 | Sp100 | 0.003 | Young |
| 0.851416 | 0.032559 | 0.929 | 0.929 | 1 | 5 | Sdcbp | 0 | Young |
| 0.851503 | 0.349317 | 0.074 | 0.072 | 1 | 5 | Trp53bp1 | 0.002 | Young |
| 0.851575 | 0.097782 | 0.15 | 0.155 | 1 | 5 | Mrps18c | -0.005 | Young |
| 0.851579 | 0.638147 | 0.036 | 0.034 | 1 | 5 | Ifit1 | 0.002 | Young |
| 0.851649 | 0.123933 | 0.15 | 0.146 | 1 | 5 | Ripor2 | 0.004 | Young |
| 0.85171 | 0.103496 | 0.058 | 0.056 | 1 | 5 | Pramef8 | 0.002 | Young |
| 0.851754 | 0.05093 | 0.104 | 0.101 | 1 | 5 | Fam110b | 0.003 | Young |
| 0.851928 | 0.135057 | 0.165 | 0.173 | 1 | 5 | Tmem2 | -0.008 | Young |
| 0.85221 | 0.137879 | 0.096 | 0.093 | 1 | 5 | Nsmf | 0.003 | Young |
| 0.852525 | 0.190256 | 0.015 | 0.017 | 1 | 5 | Zfp607a | -0.002 | Young |
| 0.852549 | 0.044891 | 0.14 | 0.145 | 1 | 5 | Phka2 | -0.005 | Young |
| 0.852788 | 0.063414 | 0.122 | 0.118 | 1 | 5 | Cdc34 | 0.004 | Young |
| 0.852811 | 0.234658 | 0.114 | 0.12 | 1 | 5 | Fhod1 | -0.006 | Young |
| 0.852993 | 0.390483 | 0.036 | 0.034 | 1 | 5 | Rad1 | 0.002 | Young |
| 0.853017 | 0.040081 | 0.038 | 0.04 | 1 | 5 | Saal1 | -0.002 | Young |

| | | | | | | | | |
|---|---|---|---|---|---|---|---|---|
| 0.853215 | 0.116126 | 0.317 | 0.32 | 1 | 5 | Mob4 | -0.003 | Young |
| 0.853293 | 0.010705 | 0.48 | 0.5 | 1 | 5 | Tomm20 | -0.02 | Young |
| 0.853347 | 0.049112 | 0.025 | 0.024 | 1 | 5 | Iqcc | 0.001 | Young |
| 0.854075 | 0.014921 | 0.312 | 0.306 | 1 | 5 | Zfp644 | 0.006 | Young |
| 0.854124 | 0.126682 | 0.033 | 0.035 | 1 | 5 | Toe1 | -0.002 | Young |
| 0.854379 | 0.123234 | 0.107 | 0.103 | 1 | 5 | Tdg | 0.004 | Young |
| 0.854495 | 0.367778 | 0.03 | 0.029 | 1 | 5 | 2900076A0 | 0.001 | Young |
| 0.854744 | 0.071706 | 0.218 | 0.226 | 1 | 5 | Milr1 | -0.008 | Young |
| 0.855135 | 0.414636 | 0.038 | 0.04 | 1 | 5 | Tcim | -0.002 | Young |
| 0.855242 | 0.163527 | 0.254 | 0.251 | 1 | 5 | Rab1b | 0.003 | Young |
| 0.855715 | 0.199744 | 0.228 | 0.229 | 1 | 5 | Zcchc2 | -0.001 | Young |
| 0.855784 | 0.076178 | 0.439 | 0.438 | 1 | 5 | H13 | 0.001 | Young |
| 0.855793 | 0.132483 | 0.015 | 0.017 | 1 | 5 | Zgrf1 | -0.002 | Young |
| 0.856005 | 0.015499 | 0.65 | 0.648 | 1 | 5 | Srsf2 | 0.002 | Young |
| 0.85628 | 0.12817 | 0.162 | 0.172 | 1 | 5 | Akap8 | -0.01 | Young |
| 0.856466 | 0.009641 | 0.048 | 0.045 | 1 | 5 | Gorasp1 | 0.003 | Young |
| 0.856694 | 0.021645 | 0.338 | 0.335 | 1 | 5 | Sh3kbp1 | 0.003 | Young |
| 0.856697 | 0.131173 | 0.025 | 0.024 | 1 | 5 | BC147527 | 0.001 | Young |
| 0.856804 | 0.004378 | 0.122 | 0.126 | 1 | 5 | Gpd2 | -0.004 | Young |
| 0.856914 | 0.061199 | 0.132 | 0.126 | 1 | 5 | Lage3 | 0.006 | Young |
| 0.856922 | 0.048708 | 0.152 | 0.157 | 1 | 5 | Rad51b | -0.005 | Young |
| 0.857046 | 0.189409 | 0.201 | 0.199 | 1 | 5 | Ppp2r2d | 0.002 | Young |
| 0.857537 | 0.006941 | 0.495 | 0.498 | 1 | 5 | Ptp4a1 | -0.003 | Young |
| 0.857592 | 0.251504 | 0.122 | 0.12 | 1 | 5 | Usp21 | 0.002 | Young |
| 0.857821 | 0.002049 | 0.543 | 0.561 | 1 | 5 | Slc39a1 | -0.018 | Young |
| 0.857947 | 0.017192 | 0.109 | 0.104 | 1 | 5 | Mcm9 | 0.005 | Young |
| 0.858016 | 0.108817 | 0.15 | 0.146 | 1 | 5 | Gtf2b | 0.004 | Young |
| 0.858032 | 0.080077 | 0.223 | 0.231 | 1 | 5 | Kbtbd2 | -0.008 | Young |
| 0.858038 | 0.143609 | 0.025 | 0.024 | 1 | 5 | Acsf3 | 0.001 | Young |
| 0.858175 | 0.309239 | 0.046 | 0.044 | 1 | 5 | Cks2 | 0.002 | Young |
| 0.858368 | 0.36694 | 0.036 | 0.034 | 1 | 5 | Gemin6 | 0.002 | Young |
| 0.858368 | 0.304835 | 0.036 | 0.034 | 1 | 5 | Nkiras1 | 0.002 | Young |
| 0.858393 | 0.217564 | 0.272 | 0.279 | 1 | 5 | Kmt5b | -0.007 | Young |
| 0.858473 | 0.111583 | 0.414 | 0.411 | 1 | 5 | Pxn | 0.003 | Young |
| 0.858763 | 0.051825 | 0.353 | 0.365 | 1 | 5 | Pacsin2 | -0.012 | Young |
| 0.858926 | 0.011716 | 0.312 | 0.3 | 1 | 5 | Spg21 | 0.012 | Young |
| 0.859129 | 0.019304 | 0.086 | 0.082 | 1 | 5 | Nadk2 | 0.004 | Young |
| 0.859862 | 0.154357 | 0.234 | 0.237 | 1 | 5 | Mif | -0.003 | Young |
| 0.86007 | 0.045879 | 0.094 | 0.098 | 1 | 5 | Poldip2 | -0.004 | Young |
| 0.860288 | 0.128829 | 0.287 | 0.295 | 1 | 5 | Man2a2 | -0.008 | Young |
| 0.860468 | 0.056117 | 0.17 | 0.165 | 1 | 5 | Ctns | 0.005 | Young |
| 0.860491 | 0.043072 | 0.157 | 0.162 | 1 | 5 | D10Jhu81e | -0.005 | Young |
| 0.860709 | 0.166209 | 0.155 | 0.152 | 1 | 5 | Mvb12a | 0.003 | Young |
| 0.861151 | 0.085934 | 0.206 | 0.215 | 1 | 5 | Eny2 | -0.009 | Young |
| 0.861453 | 0.025711 | 0.142 | 0.146 | 1 | 5 | Hist1h2bc | -0.004 | Young |

| | | | | | | | | |
|---|---|---|---|---|---|---|---|---|
| 0.861475 | 0.076201 | 0.102 | 0.098 | 1 | 5 | Rcor3 | 0.004 | Young |
| 0.861644 | 0.195507 | 0.228 | 0.242 | 1 | 5 | Irf2bpl | -0.014 | Young |
| 0.861695 | 0.096778 | 0.114 | 0.111 | 1 | 5 | Upf3a | 0.003 | Young |
| 0.861725 | 0.039298 | 0.114 | 0.118 | 1 | 5 | Brwd3 | -0.004 | Young |
| 0.861876 | 0.06632 | 0.208 | 0.205 | 1 | 5 | Sipa1l1 | 0.003 | Young |
| 0.86188 | 0.046088 | 0.398 | 0.399 | 1 | 5 | Zfp710 | -0.001 | Young |
| 0.862096 | 0.101135 | 0.066 | 0.069 | 1 | 5 | Pou6f1 | -0.003 | Young |
| 0.862284 | 0.000931 | 0.201 | 0.197 | 1 | 5 | Mtmr10 | 0.004 | Young |
| 0.862671 | 0.011051 | 0.404 | 0.419 | 1 | 5 | Nsd3 | -0.015 | Young |
| 0.862795 | 0.303262 | 0.086 | 0.091 | 1 | 5 | Tmem161a | -0.005 | Young |
| 0.863198 | 0.202704 | 0.089 | 0.086 | 1 | 5 | Morc2a | 0.003 | Young |
| 0.863405 | 0.174271 | 0.025 | 0.024 | 1 | 5 | Pkmyt1 | 0.001 | Young |
| 0.863652 | 0.021475 | 0.211 | 0.205 | 1 | 5 | Prmt1 | 0.006 | Young |
| 0.863746 | 0.009865 | 0.396 | 0.412 | 1 | 5 | 1-Mar | -0.016 | Young |
| 0.863967 | 0.093165 | 0.109 | 0.104 | 1 | 5 | Dnajb12 | 0.005 | Young |
| 0.864324 | 0.05509 | 0.079 | 0.082 | 1 | 5 | 4930453N2 | -0.003 | Young |
| 0.865096 | 0.15477 | 0.058 | 0.056 | 1 | 5 | 0610009B2 | 0.002 | Young |
| 0.865209 | 0.113975 | 0.038 | 0.04 | 1 | 5 | Zfp426 | -0.002 | Young |
| 0.865422 | 0.024423 | 0.119 | 0.114 | 1 | 5 | Wdr43 | 0.005 | Young |
| 0.865487 | 0.094751 | 0.241 | 0.241 | 1 | 5 | Cdc73 | 0 | Young |
| 0.86565 | 0.097494 | 0.185 | 0.178 | 1 | 5 | Rhot1 | 0.007 | Young |
| 0.866123 | 0.275689 | 0.051 | 0.054 | 1 | 5 | Mapk7 | -0.003 | Young |
| 0.86617 | 0.073642 | 0.206 | 0.214 | 1 | 5 | Stk3 | -0.008 | Young |
| 0.866467 | 0.170953 | 0.137 | 0.135 | 1 | 5 | Arfgef2 | 0.002 | Young |
| 0.866817 | 0.032261 | 0.429 | 0.438 | 1 | 5 | Nono | -0.009 | Young |
| 0.867492 | 0.061921 | 0.299 | 0.298 | 1 | 5 | Ube2h | 0.001 | Young |
| 0.86753 | 0.225225 | 0.195 | 0.195 | 1 | 5 | Sf3b4 | 0 | Young |
| 0.867605 | 0.075984 | 0.129 | 0.135 | 1 | 5 | Cnot7 | -0.006 | Young |
| 0.86773 | 0.138748 | 0.53 | 0.557 | 1 | 5 | Flna | -0.027 | Young |
| 0.867846 | 0.205734 | 0.086 | 0.084 | 1 | 5 | Tbkbp1 | 0.002 | Young |
| 0.868067 | 0.038631 | 0.157 | 0.162 | 1 | 5 | Gars | -0.005 | Young |
| 0.868449 | 0.000271 | 0.058 | 0.056 | 1 | 5 | Rfxank | 0.002 | Young |
| 0.868652 | 0.090113 | 0.119 | 0.116 | 1 | 5 | Secisbp2 | 0.003 | Young |
| 0.868694 | 0.046303 | 0.452 | 0.456 | 1 | 5 | Dock10 | -0.004 | Young |
| 0.869007 | 0.129331 | 0.52 | 0.559 | 1 | 5 | Ncor1 | -0.039 | Young |
| 0.869186 | 0.080237 | 0.353 | 0.343 | 1 | 5 | Mfhas1 | 0.01 | Young |
| 0.869657 | 0.105308 | 0.282 | 0.298 | 1 | 5 | Nrbp1 | -0.016 | Young |
| 0.869776 | 0.11217 | 0.175 | 0.17 | 1 | 5 | Hdac5 | 0.005 | Young |
| 0.870236 | 0.113037 | 0.429 | 0.441 | 1 | 5 | Tspan31 | -0.012 | Young |
| 0.870936 | 0.125976 | 0.14 | 0.145 | 1 | 5 | Ptpn23 | -0.005 | Young |
| 0.871022 | 0.036063 | 0.03 | 0.029 | 1 | 5 | Gk5 | 0.001 | Young |
| 0.871052 | 0.065519 | 0.041 | 0.039 | 1 | 5 | Kiz | 0.002 | Young |
| 0.871268 | 0.148045 | 0.104 | 0.099 | 1 | 5 | Tbce | 0.005 | Young |
| 0.871445 | 0.074058 | 0.175 | 0.172 | 1 | 5 | Ormdl2 | 0.003 | Young |
| 0.871635 | 0.101615 | 0.03 | 0.029 | 1 | 5 | Trmt10a | 0.001 | Young |

| | | | | | | | | |
|---|---|---|---|---|---|---|---|---|
| 0.871911 | 0.060985 | 0.457 | 0.47 | 1 | 5 | Atf6 | -0.013 | Young |
| 0.871933 | 0.089974 | 0.193 | 0.197 | 1 | 5 | Lcorl | -0.004 | Young |
| 0.872149 | 0.112259 | 0.251 | 0.244 | 1 | 5 | Fkbp4 | 0.007 | Young |
| 0.872207 | 0.054114 | 0.02 | 0.022 | 1 | 5 | Atp1b1 | -0.002 | Young |
| 0.87225 | 0.156126 | 0.096 | 0.094 | 1 | 5 | Mrpl35 | 0.002 | Young |
| 0.872609 | 0.087095 | 0.416 | 0.443 | 1 | 5 | Irak1 | -0.027 | Young |
| 0.872862 | 0.016688 | 0.03 | 0.029 | 1 | 5 | Ftx | 0.001 | Young |
| 0.872873 | 0.069149 | 0.185 | 0.18 | 1 | 5 | 2310011J0 | 0.005 | Young |
| 0.872925 | 0.138468 | 0.02 | 0.022 | 1 | 5 | Rmdn2 | -0.002 | Young |
| 0.873431 | 0.089493 | 0.322 | 0.315 | 1 | 5 | Smad4 | 0.007 | Young |
| 0.873608 | 0.047028 | 0.124 | 0.128 | 1 | 5 | AI837181 | -0.004 | Young |
| 0.873754 | 0.008801 | 0.254 | 0.259 | 1 | 5 | Pink1 | -0.005 | Young |
| 0.873887 | 0.111478 | 0.279 | 0.285 | 1 | 5 | Denr | -0.006 | Young |
| 0.873895 | 0.118823 | 0.076 | 0.079 | 1 | 5 | Dohh | -0.003 | Young |
| 0.874073 | 0.033454 | 0.954 | 0.941 | 1 | 5 | Rpl26 | 0.013 | Young |
| 0.874497 | 0.024261 | 0.36 | 0.352 | 1 | 5 | Copz1 | 0.008 | Young |
| 0.874772 | 0.271371 | 0.294 | 0.3 | 1 | 5 | Sptlc2 | -0.006 | Young |
| 0.874818 | 0.09296 | 0.036 | 0.034 | 1 | 5 | Smpd3 | 0.002 | Young |
| 0.874919 | 0.069348 | 0.299 | 0.311 | 1 | 5 | Rex1bd | -0.012 | Young |
| 0.874928 | 0.143138 | 0.089 | 0.086 | 1 | 5 | Rufy2 | 0.003 | Young |
| 0.875056 | 0.16217 | 0.206 | 0.217 | 1 | 5 | Pin4 | -0.011 | Young |
| 0.875135 | 0.129963 | 0.439 | 0.449 | 1 | 5 | Hnrnpl | -0.01 | Young |
| 0.875316 | 0.081072 | 0.03 | 0.029 | 1 | 5 | Tlr8 | 0.001 | Young |
| 0.875447 | 0.016797 | 0.614 | 0.621 | 1 | 5 | Tor1aip1 | -0.007 | Young |
| 0.875524 | 0.031027 | 0.053 | 0.056 | 1 | 5 | Dffa | -0.003 | Young |
| 0.875629 | 0.055409 | 0.221 | 0.231 | 1 | 5 | Dhx36 | -0.01 | Young |
| 0.87593 | 0.133322 | 0.03 | 0.029 | 1 | 5 | Snapc2 | 0.001 | Young |
| 0.87608 | 0.224353 | 0.183 | 0.184 | 1 | 5 | Gmps | -0.001 | Young |
| 0.876081 | 0.118796 | 0.401 | 0.406 | 1 | 5 | Snx9 | -0.005 | Young |
| 0.876135 | 0.101553 | 0.152 | 0.148 | 1 | 5 | Fhl3 | 0.004 | Young |
| 0.876161 | 0.106635 | 0.518 | 0.515 | 1 | 5 | Zmiz1 | 0.003 | Young |
| 0.876264 | 0.273843 | 0.175 | 0.177 | 1 | 5 | Ddx58 | -0.002 | Young |
| 0.876686 | 0.133244 | 0.132 | 0.13 | 1 | 5 | Pla2g16 | 0.002 | Young |
| 0.876796 | 0.162338 | 0.353 | 0.348 | 1 | 5 | Ndufs6 | 0.005 | Young |
| 0.876944 | 0.063687 | 0.492 | 0.513 | 1 | 5 | Ifi203 | -0.021 | Young |
| 0.877022 | 0.020959 | 0.107 | 0.104 | 1 | 5 | Gid4 | 0.003 | Young |
| 0.877175 | 0.134165 | 0.294 | 0.291 | 1 | 5 | Ppp4r3a | 0.003 | Young |
| 0.877235 | 0.338408 | 0.02 | 0.022 | 1 | 5 | Fastkd1 | -0.002 | Young |
| 0.877235 | 0.197728 | 0.02 | 0.022 | 1 | 5 | Gm43569 | -0.002 | Young |
| 0.877575 | 0.234222 | 0.056 | 0.054 | 1 | 5 | Rnf126 | 0.002 | Young |
| 0.877708 | 0.045873 | 0.109 | 0.104 | 1 | 5 | Lsm14b | 0.005 | Young |
| 0.877725 | 0.024526 | 0.066 | 0.069 | 1 | 5 | Gnptab | -0.003 | Young |
| 0.877739 | 0.0824 | 0.074 | 0.071 | 1 | 5 | Eef1akmt1 | 0.003 | Young |
| 0.87792 | 0.038403 | 0.173 | 0.168 | 1 | 5 | Sgta | 0.005 | Young |
| 0.878164 | 0.105929 | 0.51 | 0.53 | 1 | 5 | Per1 | -0.02 | Young |

| | | | | | | | | |
|---|---|---|---|---|---|---|---|---|
| 0.878343 | 0.09849 | 0.221 | 0.219 | 1 | 5 | Gas2l1 | 0.002 | Young |
| 0.878386 | 0.181504 | 0.03 | 0.029 | 1 | 5 | Camk2a | 0.001 | Young |
| 0.878573 | 0.181145 | 0.119 | 0.118 | 1 | 5 | Nsrp1 | 0.001 | Young |
| 0.878612 | 0.081735 | 0.244 | 0.251 | 1 | 5 | Otub1 | -0.007 | Young |
| 0.878616 | 0.020761 | 0.198 | 0.2 | 1 | 5 | Pdzd8 | -0.002 | Young |
| 0.878985 | 0.298631 | 0.137 | 0.138 | 1 | 5 | Rpp21 | -0.001 | Young |
| 0.879365 | 0.067493 | 0.036 | 0.034 | 1 | 5 | Gss | 0.002 | Young |
| 0.87946 | 0.233691 | 0.16 | 0.168 | 1 | 5 | Trim5 | -0.008 | Young |
| 0.879596 | 0.011402 | 0.086 | 0.089 | 1 | 5 | Rgs14 | -0.003 | Young |
| 0.879734 | 0.123391 | 0.234 | 0.229 | 1 | 5 | Ugp2 | 0.005 | Young |
| 0.879818 | 0.049103 | 0.287 | 0.288 | 1 | 5 | Prdm1 | -0.001 | Young |
| 0.879877 | 0.212267 | 0.226 | 0.224 | 1 | 5 | Ethe1 | 0.002 | Young |
| 0.880205 | 0.315032 | 0.145 | 0.145 | 1 | 5 | Ndst2 | 0 | Young |
| 0.88049 | 0.031234 | 0.043 | 0.045 | 1 | 5 | Lsr | -0.002 | Young |
| 0.881083 | 0.148482 | 0.193 | 0.189 | 1 | 5 | Tbc1d20 | 0.004 | Young |
| 0.881402 | 0.056102 | 0.64 | 0.663 | 1 | 5 | Atox1 | -0.023 | Young |
| 0.881495 | 0.045817 | 0.878 | 0.865 | 1 | 5 | Clta | 0.013 | Young |
| 0.881688 | 0.166013 | 0.122 | 0.12 | 1 | 5 | Dhps | 0.002 | Young |
| 0.881747 | 0.026991 | 0.135 | 0.128 | 1 | 5 | Slc35f5 | 0.007 | Young |
| 0.881769 | 0.025239 | 0.302 | 0.295 | 1 | 5 | Chd2 | 0.007 | Young |
| 0.88209 | 0.020948 | 0.226 | 0.224 | 1 | 5 | Smim27 | 0.002 | Young |
| 0.882357 | 0.198663 | 0.051 | 0.049 | 1 | 5 | Rpap3 | 0.002 | Young |
| 0.882413 | 0.036138 | 0.538 | 0.529 | 1 | 5 | Prpf38b | 0.009 | Young |
| 0.882505 | 0.12325 | 0.462 | 0.485 | 1 | 5 | Eif3k | -0.023 | Young |
| 0.88271 | 0.094259 | 0.203 | 0.199 | 1 | 5 | Acadl | 0.004 | Young |
| 0.882741 | 0.056807 | 0.503 | 0.525 | 1 | 5 | Fuca1 | -0.022 | Young |
| 0.882752 | 0.085449 | 0.046 | 0.044 | 1 | 5 | Tbc1d22b | 0.002 | Young |
| 0.882793 | 0.15209 | 0.231 | 0.234 | 1 | 5 | Ythdc2 | -0.003 | Young |
| 0.88304 | 0.047686 | 0.315 | 0.311 | 1 | 5 | Krit1 | 0.004 | Young |
| 0.883363 | 0.04237 | 0.761 | 0.779 | 1 | 5 | Sh3bgrl | -0.018 | Young |
| 0.883461 | 0.078296 | 0.16 | 0.155 | 1 | 5 | Exoc1 | 0.005 | Young |
| 0.884001 | 0.198888 | 0.112 | 0.109 | 1 | 5 | Dnajc11 | 0.003 | Young |
| 0.884145 | 0.048115 | 0.137 | 0.14 | 1 | 5 | Commd2 | -0.003 | Young |
| 0.884356 | 0.235519 | 0.041 | 0.039 | 1 | 5 | Spryd3 | 0.002 | Young |
| 0.884418 | 0.203435 | 0.495 | 0.515 | 1 | 5 | Srgap2 | -0.02 | Young |
| 0.88449 | 0.184492 | 0.193 | 0.199 | 1 | 5 | Tnip3 | -0.006 | Young |
| 0.884913 | 0.136835 | 0.487 | 0.492 | 1 | 5 | Stx7 | -0.005 | Young |
| 0.884918 | 0.048212 | 0.365 | 0.364 | 1 | 5 | Cpt1a | 0.001 | Young |
| 0.885003 | 0.086097 | 0.122 | 0.125 | 1 | 5 | Rae1 | -0.003 | Young |
| 0.885152 | 0.036611 | 0.421 | 0.421 | 1 | 5 | Mlf2 | 0 | Young |
| 0.885338 | 0.055622 | 0.241 | 0.244 | 1 | 5 | Wbp1l | -0.003 | Young |
| 0.885557 | 0.104719 | 0.066 | 0.069 | 1 | 5 | Usp39 | -0.003 | Young |
| 0.885864 | 0.484913 | 0.02 | 0.022 | 1 | 5 | Nlgn2 | -0.002 | Young |
| 0.886279 | 0.0912 | 0.081 | 0.084 | 1 | 5 | Ssna1 | -0.003 | Young |
| 0.886384 | 0.002184 | 0.107 | 0.109 | 1 | 5 | Fam162a | -0.002 | Young |

| | | | | | | | | |
|---|---|---|---|---|---|---|---|---|
| 0.886747 | 0.043235 | 0.079 | 0.076 | 1 | 5 | Gm7694 | 0.003 | Young |
| 0.887191 | 0.167444 | 0.213 | 0.214 | 1 | 5 | Cwf19l2 | -0.001 | Young |
| 0.887424 | 0.199319 | 0.099 | 0.098 | 1 | 5 | Tmem87a | 0.001 | Young |
| 0.88774 | 0.048936 | 0.792 | 0.83 | 1 | 5 | Pdia3 | -0.038 | Young |
| 0.888286 | 0.119429 | 0.046 | 0.044 | 1 | 5 | Mypopos | 0.002 | Young |
| 0.888333 | 0.06919 | 0.18 | 0.185 | 1 | 5 | Babam2 | -0.005 | Young |
| 0.888464 | 0.100738 | 0.127 | 0.125 | 1 | 5 | Trp53bp2 | 0.002 | Young |
| 0.888524 | 0.054776 | 0.056 | 0.054 | 1 | 5 | Trpm4 | 0.002 | Young |
| 0.88853 | 0.184223 | 0.467 | 0.481 | 1 | 5 | Msl1 | -0.014 | Young |
| 0.888563 | 0.098165 | 0.193 | 0.189 | 1 | 5 | Plekha1 | 0.004 | Young |
| 0.888702 | 0.057902 | 0.18 | 0.175 | 1 | 5 | Ptbp2 | 0.005 | Young |
| 0.888705 | 0.11844 | 0.076 | 0.074 | 1 | 5 | Zxdc | 0.002 | Young |
| 0.889045 | 0.104544 | 0.226 | 0.226 | 1 | 5 | Ndufa10 | 0 | Young |
| 0.889163 | 0.096018 | 0.076 | 0.079 | 1 | 5 | Arfgap1 | -0.003 | Young |
| 0.889218 | 0.058647 | 0.5 | 0.495 | 1 | 5 | Surf4 | 0.005 | Young |
| 0.88922 | 0.058119 | 0.292 | 0.301 | 1 | 5 | Ep400 | -0.009 | Young |
| 0.889282 | 0.812361 | 0.013 | 0.012 | 1 | 5 | Rasl2-9 | 0.001 | Young |
| 0.889397 | 0.082776 | 0.48 | 0.492 | 1 | 5 | Pttg1ip | -0.012 | Young |
| 0.88968 | 0.153696 | 0.114 | 0.111 | 1 | 5 | Nup50 | 0.003 | Young |
| 0.889898 | 0.109512 | 0.091 | 0.094 | 1 | 5 | Fopnl | -0.003 | Young |
| 0.890048 | 0.3621 | 0.043 | 0.045 | 1 | 5 | Fam122b | -0.002 | Young |
| 0.89023 | 0.562039 | 0.013 | 0.012 | 1 | 5 | Zfp383 | 0.001 | Young |
| 0.890284 | 0.082172 | 0.109 | 0.111 | 1 | 5 | Arap3 | -0.002 | Young |
| 0.890416 | 0.03407 | 0.094 | 0.098 | 1 | 5 | Serhl | -0.004 | Young |
| 0.890628 | 0.141747 | 0.129 | 0.126 | 1 | 5 | Nans | 0.003 | Young |
| 0.890858 | 0.084147 | 0.061 | 0.059 | 1 | 5 | Txnl4a | 0.002 | Young |
| 0.890934 | 0.056018 | 0.728 | 0.764 | 1 | 5 | Cox6a1 | -0.036 | Young |
| 0.890945 | 0.068237 | 0.109 | 0.111 | 1 | 5 | Brpf3 | -0.002 | Young |
| 0.891154 | 0.021145 | 0.277 | 0.274 | 1 | 5 | Fam168a | 0.003 | Young |
| 0.891765 | 0.185549 | 0.168 | 0.175 | 1 | 5 | Il21r | -0.007 | Young |
| 0.891852 | 0.799817 | 0.018 | 0.017 | 1 | 5 | Nova1 | 0.001 | Young |
| 0.89186 | 0.127301 | 0.277 | 0.285 | 1 | 5 | Rnf187 | -0.008 | Young |
| 0.891945 | 0.029938 | 0.409 | 0.414 | 1 | 5 | Dctn1 | -0.005 | Young |
| 0.892412 | 0.071147 | 0.137 | 0.141 | 1 | 5 | Mid1 | -0.004 | Young |
| 0.892475 | 0.024536 | 0.201 | 0.205 | 1 | 5 | Clcn3 | -0.004 | Young |
| 0.892549 | 0.116859 | 0.025 | 0.027 | 1 | 5 | Proser3 | -0.002 | Young |
| 0.893077 | 0.819706 | 0.013 | 0.012 | 1 | 5 | C230037L1 | 0.001 | Young |
| 0.893077 | 0.333245 | 0.013 | 0.012 | 1 | 5 | Fam185a | 0.001 | Young |
| 0.893077 | 0.27807 | 0.013 | 0.012 | 1 | 5 | Gm39459 | 0.001 | Young |
| 0.893086 | 0.332501 | 0.074 | 0.072 | 1 | 5 | Tcp11l1 | 0.002 | Young |
| 0.893199 | 0.191721 | 0.025 | 0.027 | 1 | 5 | Uhrf1bp1 | -0.002 | Young |
| 0.893474 | 0.115183 | 0.335 | 0.342 | 1 | 5 | Hsd17b12 | -0.007 | Young |
| 0.89351 | 0.121616 | 0.266 | 0.263 | 1 | 5 | Naa15 | 0.003 | Young |
| 0.893935 | 0.03001 | 0.094 | 0.098 | 1 | 5 | Snta1 | -0.004 | Young |
| 0.894026 | 0.165991 | 0.013 | 0.012 | 1 | 5 | Swsap1 | 0.001 | Young |

| | | | | | | | | |
|---|---|---|---|---|---|---|---|---|
| 0.894085 | 0.165602 | 0.071 | 0.074 | 1 | 5 | Adssl1 | -0.003 | Young |
| 0.894677 | 0.082155 | 0.14 | 0.141 | 1 | 5 | Pex13 | -0.001 | Young |
| 0.894687 | 0.230822 | 0.132 | 0.138 | 1 | 5 | Gskip | -0.006 | Young |
| 0.89479 | 0.203835 | 0.048 | 0.051 | 1 | 5 | Fkbp7 | -0.003 | Young |
| 0.894803 | 0.163929 | 0.058 | 0.061 | 1 | 5 | Aifm1 | -0.003 | Young |
| 0.894975 | 0.194237 | 0.013 | 0.012 | 1 | 5 | Cdc25b | 0.001 | Young |
| 0.895026 | 0.058428 | 0.381 | 0.387 | 1 | 5 | Secisbp2l | -0.006 | Young |
| 0.895683 | 0.087756 | 0.071 | 0.074 | 1 | 5 | Rad17 | -0.003 | Young |
| 0.895836 | 0.053287 | 0.056 | 0.054 | 1 | 5 | Zbtb10 | 0.002 | Young |
| 0.895836 | 0.018469 | 0.056 | 0.054 | 1 | 5 | Ndufaf3 | 0.002 | Young |
| 0.896315 | 0.141096 | 0.335 | 0.33 | 1 | 5 | Ctdsp1 | 0.005 | Young |
| 0.896365 | 0.063397 | 0.294 | 0.305 | 1 | 5 | Vapb | -0.011 | Young |
| 0.896649 | 0.562446 | 0.018 | 0.017 | 1 | 5 | Cobll1 | 0.001 | Young |
| 0.897182 | 0.033718 | 0.97 | 0.944 | 1 | 5 | Npc2 | 0.026 | Young |
| 0.89735 | 0.191433 | 0.173 | 0.173 | 1 | 5 | Map1lc3a | 0 | Young |
| 0.897665 | 0.160739 | 0.056 | 0.054 | 1 | 5 | 2410004B1 | 0.002 | Young |
| 0.897823 | 0.128471 | 0.013 | 0.012 | 1 | 5 | Ndrg1 | 0.001 | Young |
| 0.898134 | 0.13185 | 0.251 | 0.249 | 1 | 5 | Atf7ip | 0.002 | Young |
| 0.898173 | 0.066 | 0.076 | 0.074 | 1 | 5 | Nol8 | 0.002 | Young |
| 0.898296 | 0.02375 | 0.104 | 0.106 | 1 | 5 | Fktn | -0.002 | Young |
| 0.898733 | 0.067618 | 0.178 | 0.182 | 1 | 5 | Acss1 | -0.004 | Young |
| 0.898735 | 0.024035 | 0.383 | 0.397 | 1 | 5 | Ifnar2 | -0.014 | Young |
| 0.898751 | 0.167921 | 0.251 | 0.258 | 1 | 5 | Rnf216 | -0.007 | Young |
| 0.898773 | 0.17681 | 0.013 | 0.012 | 1 | 5 | Klhl3 | 0.001 | Young |
| 0.898773 | 0.095954 | 0.013 | 0.012 | 1 | 5 | Cdh5 | 0.001 | Young |
| 0.898773 | 0.030578 | 0.013 | 0.012 | 1 | 5 | Mfsd9 | 0.001 | Young |
| 0.899068 | 0.102552 | 0.081 | 0.079 | 1 | 5 | Slc35c1 | 0.002 | Young |
| 0.899504 | 0.022842 | 0.693 | 0.692 | 1 | 5 | Rab8b | 0.001 | Young |
| 0.899723 | 0.464881 | 0.013 | 0.012 | 1 | 5 | Gnb1l | 0.001 | Young |
| 0.899723 | 0.16957 | 0.013 | 0.012 | 1 | 5 | Cmtr2 | 0.001 | Young |
| 0.899723 | 0.017525 | 0.013 | 0.012 | 1 | 5 | Abraxas1 | 0.001 | Young |
| 0.899874 | 0.006892 | 0.046 | 0.044 | 1 | 5 | Zdhhc12 | 0.002 | Young |
| 0.900033 | 0.040341 | 0.168 | 0.165 | 1 | 5 | Ext2 | 0.003 | Young |
| 0.90041 | 0.217527 | 0.053 | 0.056 | 1 | 5 | Klc4 | -0.003 | Young |
| 0.900464 | 0.161249 | 0.14 | 0.136 | 1 | 5 | Ddx47 | 0.004 | Young |
| 0.90049 | 0.055503 | 0.629 | 0.652 | 1 | 5 | M6pr | -0.023 | Young |
| 0.900549 | 0.122933 | 0.071 | 0.069 | 1 | 5 | Grk5 | 0.002 | Young |
| 0.900916 | 0.05122 | 0.279 | 0.278 | 1 | 5 | Fbxw4 | 0.001 | Young |
| 0.901169 | 0.079878 | 0.383 | 0.387 | 1 | 5 | Hectd1 | -0.004 | Young |
| 0.901461 | 0.057843 | 0.439 | 0.451 | 1 | 5 | Dad1 | -0.012 | Young |
| 0.901562 | 0.194951 | 0.079 | 0.077 | 1 | 5 | Gmpr2 | 0.002 | Young |
| 0.901623 | 0.11156 | 0.013 | 0.012 | 1 | 5 | Pdss1 | 0.001 | Young |
| 0.901674 | 0.430084 | 0.137 | 0.14 | 1 | 5 | Rgmb | -0.003 | Young |
| 0.901914 | 0.125301 | 0.376 | 0.379 | 1 | 5 | Aph1a | -0.003 | Young |
| 0.902491 | 0.115427 | 0.147 | 0.153 | 1 | 5 | Reps1 | -0.006 | Young |

| | | | | | | | | |
|---|---|---|---|---|---|---|---|---|
| 0.902859 | 0.153287 | 0.523 | 0.549 | 1 | 5 | Ep300 | -0.026 | Young |
| 0.90286 | 0.073885 | 0.231 | 0.229 | 1 | 5 | Pip4k2b | 0.002 | Young |
| 0.90305 | 0.254906 | 0.018 | 0.017 | 1 | 5 | Slc8a1 | 0.001 | Young |
| 0.903405 | 0.017821 | 0.046 | 0.044 | 1 | 5 | Pno1 | 0.002 | Young |
| 0.903408 | 0.030671 | 0.145 | 0.15 | 1 | 5 | Kifc3 | -0.005 | Young |
| 0.903476 | 0.001035 | 0.216 | 0.21 | 1 | 5 | C1qbp | 0.006 | Young |
| 0.903524 | 0.132928 | 0.013 | 0.012 | 1 | 5 | Abhd3 | 0.001 | Young |
| 0.903851 | 0.444694 | 0.018 | 0.017 | 1 | 5 | Cep19 | 0.001 | Young |
| 0.903955 | 0.045096 | 0.447 | 0.456 | 1 | 5 | Specc1l | -0.009 | Young |
| 0.904025 | 0.048272 | 0.497 | 0.507 | 1 | 5 | Psma7 | -0.01 | Young |
| 0.904066 | 0.164986 | 0.254 | 0.258 | 1 | 5 | Polr2l | -0.004 | Young |
| 0.90422 | 0.132709 | 0.259 | 0.258 | 1 | 5 | Trim56 | 0.001 | Young |
| 0.904222 | 0.050174 | 0.142 | 0.138 | 1 | 5 | Espl1 | 0.004 | Young |
| 0.904283 | 0.199465 | 0.074 | 0.072 | 1 | 5 | Mrps34 | 0.002 | Young |
| 0.904301 | 0.292104 | 0.053 | 0.056 | 1 | 5 | Asf1a | -0.003 | Young |
| 0.904554 | 0.054267 | 0.551 | 0.559 | 1 | 5 | 9930111J2 | -0.008 | Young |
| 0.9046 | 0.001251 | 0.683 | 0.695 | 1 | 5 | Purb | -0.012 | Young |
| 0.904609 | 0.032132 | 0.411 | 0.419 | 1 | 5 | Ubn2 | -0.008 | Young |
| 0.904652 | 0.432599 | 0.018 | 0.017 | 1 | 5 | Sh3bgrl2 | 0.001 | Young |
| 0.904656 | 0.098626 | 0.132 | 0.13 | 1 | 5 | Rnmt | 0.002 | Young |
| 0.904883 | 0.265624 | 0.071 | 0.074 | 1 | 5 | Tbc1d12 | -0.003 | Young |
| 0.905046 | 0.036573 | 0.731 | 0.788 | 1 | 5 | Sfpq | -0.057 | Young |
| 0.905323 | 0.11208 | 0.041 | 0.042 | 1 | 5 | Ctla2a | -0.001 | Young |
| 0.905441 | 0.22442 | 0.332 | 0.343 | 1 | 5 | Nampt | -0.011 | Young |
| 0.905803 | 0.374512 | 0.094 | 0.093 | 1 | 5 | Ctnnd2 | 0.001 | Young |
| 0.90581 | 0.018888 | 0.051 | 0.049 | 1 | 5 | Eaf2 | 0.002 | Young |
| 0.905824 | 0.002604 | 0.855 | 0.877 | 1 | 5 | Rbm39 | -0.022 | Young |
| 0.90583 | 0.316005 | 0.079 | 0.077 | 1 | 5 | Zfp865 | 0.002 | Young |
| 0.905929 | 0.120566 | 0.201 | 0.205 | 1 | 5 | Xylt2 | -0.004 | Young |
| 0.905959 | 0.114636 | 0.18 | 0.185 | 1 | 5 | Capn1 | -0.005 | Young |
| 0.906254 | 0.305449 | 0.018 | 0.017 | 1 | 5 | Rilp | 0.001 | Young |
| 0.906631 | 0.160446 | 0.051 | 0.052 | 1 | 5 | Thada | -0.001 | Young |
| 0.90687 | 0.050333 | 0.404 | 0.401 | 1 | 5 | Tex261 | 0.003 | Young |
| 0.907013 | 0.025128 | 0.299 | 0.306 | 1 | 5 | Wbp11 | -0.007 | Young |
| 0.907029 | 0.116228 | 0.036 | 0.037 | 1 | 5 | Nudcd1 | -0.001 | Young |
| 0.907192 | 0.013723 | 0.183 | 0.175 | 1 | 5 | Casp8ap2 | 0.008 | Young |
| 0.907455 | 0.193598 | 0.018 | 0.017 | 1 | 5 | Zfp623 | 0.001 | Young |
| 0.907774 | 0.026603 | 0.477 | 0.488 | 1 | 5 | Cggbp1 | -0.011 | Young |
| 0.907929 | 0.224292 | 0.142 | 0.141 | 1 | 5 | Parg | 0.001 | Young |
| 0.908199 | 0.010619 | 0.272 | 0.271 | 1 | 5 | Rab12 | 0.001 | Young |
| 0.908533 | 0.043982 | 0.508 | 0.503 | 1 | 5 | Fam208a | 0.005 | Young |
| 0.908657 | 0.295425 | 0.018 | 0.017 | 1 | 5 | Nthl1 | 0.001 | Young |
| 0.908657 | 0.279365 | 0.018 | 0.017 | 1 | 5 | Gm15706 | 0.001 | Young |
| 0.908783 | 0.068572 | 0.244 | 0.241 | 1 | 5 | Sla | 0.003 | Young |
| 0.909124 | 0.026526 | 0.129 | 0.131 | 1 | 5 | Trabd | -0.002 | Young |

| | | | | | | | | |
|---|---|---|---|---|---|---|---|---|
| 0.909305 | 0.019312 | 0.213 | 0.219 | 1 | 5 | Slc25a39 | -0.006 | Young |
| 0.909339 | 0.296371 | 0.038 | 0.037 | 1 | 5 | Gstk1 | 0.001 | Young |
| 0.909458 | 0.273828 | 0.018 | 0.017 | 1 | 5 | Wdr90 | 0.001 | Young |
| 0.909885 | 0.054284 | 0.373 | 0.375 | 1 | 5 | D17Wsu92 | -0.002 | Young |
| 0.910023 | 0.006552 | 0.041 | 0.042 | 1 | 5 | Golm1 | -0.001 | Young |
| 0.910128 | 0.102736 | 0.066 | 0.064 | 1 | 5 | Kremen1 | 0.002 | Young |
| 0.91026 | 0.20507 | 0.018 | 0.017 | 1 | 5 | Isoc2a | 0.001 | Young |
| 0.91026 | 0.168708 | 0.018 | 0.017 | 1 | 5 | Xkr8 | 0.001 | Young |
| 0.910417 | 0.144113 | 0.168 | 0.167 | 1 | 5 | Nsmaf | 0.001 | Young |
| 0.910473 | 0.034109 | 0.046 | 0.044 | 1 | 5 | Limk2 | 0.002 | Young |
| 0.910514 | 0.396467 | 0.053 | 0.052 | 1 | 5 | Snx7 | 0.001 | Young |
| 0.910683 | 0.112877 | 0.14 | 0.138 | 1 | 5 | Shc1 | 0.002 | Young |
| 0.910711 | 0.132669 | 0.15 | 0.153 | 1 | 5 | Pex7 | -0.003 | Young |
| 0.910717 | 0.127059 | 0.279 | 0.288 | 1 | 5 | Slc9a3r1 | -0.009 | Young |
| 0.910967 | 0.065163 | 0.931 | 0.926 | 1 | 5 | Rpl9 | 0.005 | Young |
| 0.911016 | 0.003657 | 0.878 | 0.899 | 1 | 5 | Tpm3 | -0.021 | Young |
| 0.911061 | 0.191052 | 0.018 | 0.017 | 1 | 5 | Slc25a42 | 0.001 | Young |
| 0.911061 | 0.088602 | 0.018 | 0.017 | 1 | 5 | Gnb5 | 0.001 | Young |
| 0.911112 | 0.033797 | 0.147 | 0.143 | 1 | 5 | Naa30 | 0.004 | Young |
| 0.911234 | 0.003695 | 0.327 | 0.33 | 1 | 5 | Kansl1 | -0.003 | Young |
| 0.911521 | 0.155396 | 0.109 | 0.113 | 1 | 5 | Gse1 | -0.004 | Young |
| 0.911591 | 0.14075 | 0.041 | 0.042 | 1 | 5 | Pex11a | -0.001 | Young |
| 0.911903 | 0.085514 | 0.221 | 0.219 | 1 | 5 | Rprd2 | 0.002 | Young |
| 0.912039 | 0.767245 | 0.023 | 0.022 | 1 | 5 | Hist1h1e | 0.001 | Young |
| 0.91214 | 0.072022 | 0.102 | 0.099 | 1 | 5 | Pgs1 | 0.003 | Young |
| 0.912382 | 0.047087 | 0.096 | 0.094 | 1 | 5 | Tinf2 | 0.002 | Young |
| 0.912536 | 0.196926 | 0.19 | 0.187 | 1 | 5 | Vps50 | 0.003 | Young |
| 0.912729 | 0.228874 | 0.058 | 0.057 | 1 | 5 | Gm17018 | 0.001 | Young |
| 0.912792 | 0.024371 | 0.581 | 0.608 | 1 | 5 | Tanc2 | -0.027 | Young |
| 0.9128 | 0.178969 | 0.069 | 0.071 | 1 | 5 | Per2 | -0.002 | Young |
| 0.913111 | 0.056238 | 0.807 | 0.842 | 1 | 5 | Arhgdib | -0.035 | Young |
| 0.91324 | 0.024101 | 0.162 | 0.165 | 1 | 5 | Akap8l | -0.003 | Young |
| 0.913452 | 0.285495 | 0.023 | 0.022 | 1 | 5 | Kif5c | 0.001 | Young |
| 0.91373 | 0.030337 | 0.51 | 0.515 | 1 | 5 | Cd84 | -0.005 | Young |
| 0.913837 | 0.184715 | 0.081 | 0.079 | 1 | 5 | Anks3 | 0.002 | Young |
| 0.914241 | 0.208572 | 0.107 | 0.106 | 1 | 5 | Stam | 0.001 | Young |
| 0.91429 | 0.19657 | 0.028 | 0.027 | 1 | 5 | Fignl2 | 0.001 | Young |
| 0.914344 | 0.130588 | 0.246 | 0.244 | 1 | 5 | Zc3h15 | 0.002 | Young |
| 0.914448 | 0.05388 | 0.17 | 0.165 | 1 | 5 | Pnpla2 | 0.005 | Young |
| 0.914862 | 0.057447 | 0.244 | 0.254 | 1 | 5 | Cant1 | -0.01 | Young |
| 0.91502 | 0.131963 | 0.013 | 0.013 | 1 | 5 | Zfp189 | 0 | Young |
| 0.91502 | 0.439241 | 0.043 | 0.045 | 1 | 5 | Akap1 | -0.002 | Young |
| 0.915101 | 0.074394 | 0.32 | 0.313 | 1 | 5 | Dock11 | 0.007 | Young |
| 0.915255 | 0.104846 | 0.317 | 0.32 | 1 | 5 | Usp34 | -0.003 | Young |
| 0.915372 | 0.184894 | 0.036 | 0.037 | 1 | 5 | Bhlhe41 | -0.001 | Young |

| | | | | | | | | |
|---|---|---|---|---|---|---|---|---|
| 0.915411 | 0.171195 | 0.058 | 0.057 | 1 | 5 | Atpaf2 | 0.001 | Young |
| 0.91552 | 0.062964 | 0.642 | 0.65 | 1 | 5 | Arhgap30 | -0.008 | Young |
| 0.915564 | 0.203196 | 0.178 | 0.182 | 1 | 5 | Mrpl57 | -0.004 | Young |
| 0.915572 | 0.134772 | 0.023 | 0.022 | 1 | 5 | Gbp9 | 0.001 | Young |
| 0.915872 | 0.122019 | 0.018 | 0.017 | 1 | 5 | Xlr4a | 0.001 | Young |
| 0.915872 | 0.057437 | 0.018 | 0.017 | 1 | 5 | Zbtb39 | 0.001 | Young |
| 0.916054 | 0.197811 | 0.221 | 0.232 | 1 | 5 | Trim44 | -0.011 | Young |
| 0.916279 | 0.160818 | 0.023 | 0.022 | 1 | 5 | Cyp2d22 | 0.001 | Young |
| 0.916501 | 0.276233 | 0.071 | 0.074 | 1 | 5 | Fmnl2 | -0.003 | Young |
| 0.916607 | 0.132206 | 0.231 | 0.232 | 1 | 5 | Csad | -0.001 | Young |
| 0.917476 | 0.054348 | 0.018 | 0.017 | 1 | 5 | Zfp119a | 0.001 | Young |
| 0.917488 | 0.085337 | 0.028 | 0.027 | 1 | 5 | Ptges2 | 0.001 | Young |
| 0.918313 | 0.020362 | 0.546 | 0.554 | 1 | 5 | Rell1 | -0.008 | Young |
| 0.918586 | 0.069933 | 0.185 | 0.189 | 1 | 5 | Skiv2l | -0.004 | Young |
| 0.918776 | 0.093753 | 0.046 | 0.047 | 1 | 5 | Mettl26 | -0.001 | Young |
| 0.919129 | 0.060124 | 0.556 | 0.569 | 1 | 5 | Ndufc2 | -0.013 | Young |
| 0.919271 | 0.376478 | 0.046 | 0.047 | 1 | 5 | Cul4b | -0.001 | Young |
| 0.919557 | 0.1083 | 0.355 | 0.374 | 1 | 5 | Parvb | -0.019 | Young |
| 0.919572 | 0.185491 | 0.033 | 0.032 | 1 | 5 | Usp20 | 0.001 | Young |
| 0.919597 | 0.328909 | 0.013 | 0.013 | 1 | 5 | St18 | 0 | Young |
| 0.9198 | 0.086884 | 0.109 | 0.111 | 1 | 5 | Glod4 | -0.002 | Young |
| 0.919814 | 0.136814 | 0.023 | 0.022 | 1 | 5 | Atg4a | 0.001 | Young |
| 0.920083 | 0.24448 | 0.119 | 0.123 | 1 | 5 | Ifi213 | -0.004 | Young |
| 0.920162 | 0.126433 | 0.033 | 0.032 | 1 | 5 | Nr1d1 | 0.001 | Young |
| 0.920369 | 0.056217 | 0.117 | 0.116 | 1 | 5 | Tomm40 | 0.001 | Young |
| 0.920382 | 0.201024 | 0.036 | 0.037 | 1 | 5 | Ttc21b | -0.001 | Young |
| 0.92043 | 0.130285 | 0.335 | 0.342 | 1 | 5 | Ube2i | -0.007 | Young |
| 0.920479 | 0.1958 | 0.041 | 0.042 | 1 | 5 | Tmem101 | -0.001 | Young |
| 0.920522 | 0.302618 | 0.023 | 0.022 | 1 | 5 | Zfp949 | 0.001 | Young |
| 0.920582 | 0.055887 | 0.515 | 0.513 | 1 | 5 | Fgd2 | 0.002 | Young |
| 0.920785 | 0.295591 | 0.053 | 0.052 | 1 | 5 | Paip2b | 0.001 | Young |
| 0.920839 | 0.068707 | 0.15 | 0.153 | 1 | 5 | Bcl2l2 | -0.003 | Young |
| 0.921171 | 0.062857 | 0.17 | 0.175 | 1 | 5 | 4931406P1 | -0.005 | Young |
| 0.921219 | 0.084124 | 0.137 | 0.135 | 1 | 5 | Specc1 | 0.002 | Young |
| 0.921462 | 0.183508 | 0.109 | 0.111 | 1 | 5 | Siah1a | -0.002 | Young |
| 0.921502 | 0.128002 | 0.102 | 0.099 | 1 | 5 | Ubqln4 | 0.003 | Young |
| 0.921506 | 0.159137 | 0.378 | 0.369 | 1 | 5 | Tial1 | 0.009 | Young |
| 0.921519 | 0.233221 | 0.061 | 0.062 | 1 | 5 | Mag | -0.001 | Young |
| 0.921816 | 0.139041 | 0.178 | 0.178 | 1 | 5 | Cers6 | 0 | Young |
| 0.921856 | 0.241165 | 0.198 | 0.199 | 1 | 5 | S1pr2 | -0.001 | Young |
| 0.921923 | 0.940004 | 0.03 | 0.032 | 1 | 5 | Ifit3 | -0.002 | Young |
| 0.92225 | 0.034019 | 0.244 | 0.236 | 1 | 5 | Rogdi | 0.008 | Young |
| 0.92229 | 0.048619 | 0.079 | 0.081 | 1 | 5 | Mob2 | -0.002 | Young |
| 0.922331 | 0.307021 | 0.036 | 0.037 | 1 | 5 | Tceal1 | -0.001 | Young |
| 0.922644 | 0.008007 | 0.023 | 0.022 | 1 | 5 | Pinx1 | 0.001 | Young |

| | | | | | | | | |
|---|---|---|---|---|---|---|---|---|
| 0.922738 | 0.179332 | 0.046 | 0.047 | 1 | 5 | Uaca | -0.001 | Young |
| 0.922788 | 0.046826 | 0.34 | 0.338 | 1 | 5 | Heca | 0.002 | Young |
| 0.922806 | 0.0549 | 0.31 | 0.311 | 1 | 5 | Kdm2b | -0.001 | Young |
| 0.92315 | 0.100601 | 0.122 | 0.12 | 1 | 5 | Zfyve26 | 0.002 | Young |
| 0.923204 | 0.07858 | 0.64 | 0.633 | 1 | 5 | Ets2 | 0.007 | Young |
| 0.923433 | 0.080372 | 0.203 | 0.207 | 1 | 5 | Pom121 | -0.004 | Young |
| 0.923434 | 0.056065 | 0.071 | 0.069 | 1 | 5 | Map4k3 | 0.002 | Young |
| 0.924177 | 0.151304 | 0.013 | 0.013 | 1 | 5 | Txlnb | 0 | Young |
| 0.924289 | 0.229899 | 0.033 | 0.032 | 1 | 5 | Fuz | 0.001 | Young |
| 0.92453 | 0.366434 | 0.028 | 0.027 | 1 | 5 | Wdhd1 | 0.001 | Young |
| 0.924536 | 0.013184 | 0.051 | 0.052 | 1 | 5 | Pyurf | -0.001 | Young |
| 0.92477 | 0.089707 | 0.383 | 0.391 | 1 | 5 | Baz2b | -0.008 | Young |
| 0.92508 | 0.094784 | 0.056 | 0.057 | 1 | 5 | Adck1 | -0.001 | Young |
| 0.925241 | 0.142702 | 0.208 | 0.205 | 1 | 5 | Ube2v2 | 0.003 | Young |
| 0.925288 | 0.342407 | 0.069 | 0.067 | 1 | 5 | Cep104 | 0.002 | Young |
| 0.925389 | 0.077115 | 0.119 | 0.121 | 1 | 5 | Dmxl2 | -0.002 | Young |
| 0.92548 | 0.113644 | 0.119 | 0.118 | 1 | 5 | Rcbtb1 | 0.001 | Young |
| 0.925522 | 0.026091 | 0.299 | 0.301 | 1 | 5 | Phf20 | -0.002 | Young |
| 0.92557 | 0.158956 | 0.058 | 0.061 | 1 | 5 | D230025D: | -0.003 | Young |
| 0.925953 | 0.194838 | 0.036 | 0.037 | 1 | 5 | Vamp1 | -0.001 | Young |
| 0.925999 | 0.005874 | 0.315 | 0.32 | 1 | 5 | Lyl1 | -0.005 | Young |
| 0.92601 | 0.458337 | 0.013 | 0.013 | 1 | 5 | St3gal5 | 0 | Young |
| 0.926014 | 0.048089 | 0.099 | 0.101 | 1 | 5 | Ebag9 | -0.002 | Young |
| 0.926056 | 0.050801 | 0.429 | 0.443 | 1 | 5 | Atp2a2 | -0.014 | Young |
| 0.926084 | 0.188709 | 0.053 | 0.056 | 1 | 5 | Mlh3 | -0.003 | Young |
| 0.926297 | 0.031253 | 0.528 | 0.542 | 1 | 5 | Calm3 | -0.014 | Young |
| 0.926303 | 0.130271 | 0.043 | 0.042 | 1 | 5 | Stau2 | 0.001 | Young |
| 0.926451 | 0.20361 | 0.028 | 0.027 | 1 | 5 | Parp16 | 0.001 | Young |
| 0.92665 | 0.013617 | 0.665 | 0.662 | 1 | 5 | Lyn | 0.003 | Young |
| 0.926657 | 0.034024 | 0.34 | 0.333 | 1 | 5 | Rnf111 | 0.007 | Young |
| 0.926663 | 0.033714 | 0.254 | 0.259 | 1 | 5 | Rbm7 | -0.005 | Young |
| 0.926955 | 0.447924 | 0.048 | 0.047 | 1 | 5 | Large1 | 0.001 | Young |
| 0.92697 | 0.022196 | 0.764 | 0.769 | 1 | 5 | Dse | -0.005 | Young |
| 0.927119 | 0.226248 | 0.157 | 0.163 | 1 | 5 | Papd5 | -0.006 | Young |
| 0.927828 | 0.196653 | 0.033 | 0.032 | 1 | 5 | Rabif | 0.001 | Young |
| 0.927844 | 0.05664 | 0.091 | 0.094 | 1 | 5 | Pitrm1 | -0.003 | Young |
| 0.927849 | 0.12536 | 0.089 | 0.091 | 1 | 5 | Sorbs3 | -0.002 | Young |
| 0.927937 | 0.073861 | 0.069 | 0.071 | 1 | 5 | Cdyl | -0.002 | Young |
| 0.928211 | 0.388704 | 0.03 | 0.032 | 1 | 5 | Zfp868 | -0.002 | Young |
| 0.928744 | 0.001279 | 0.865 | 0.911 | 1 | 5 | Morf4l1 | -0.046 | Young |
| 0.928766 | 0.020624 | 0.264 | 0.254 | 1 | 5 | Ranbp1 | 0.01 | Young |
| 0.929045 | 0.009589 | 0.396 | 0.397 | 1 | 5 | Hpcal1 | -0.001 | Young |
| 0.92907 | 0.154776 | 0.124 | 0.126 | 1 | 5 | Get4 | -0.002 | Young |
| 0.929287 | 0.067987 | 0.355 | 0.347 | 1 | 5 | Rc3h2 | 0.008 | Young |
| 0.930008 | 0.017042 | 0.871 | 0.902 | 1 | 5 | Actr3 | -0.031 | Young |

| | | | | | | | | |
|---|---|---|---|---|---|---|---|---|
| 0.930121 | 0.07446 | 0.221 | 0.219 | 1 | 5 | Guk1 | 0.002 | Young |
| 0.930425 | 0.007757 | 0.041 | 0.042 | 1 | 5 | Zfp874b | -0.001 | Young |
| 0.930453 | 0.026254 | 0.305 | 0.313 | 1 | 5 | Rhobtb1 | -0.008 | Young |
| 0.930487 | 0.036892 | 0.409 | 0.426 | 1 | 5 | Nek7 | -0.017 | Young |
| 0.930606 | 0.07813 | 0.272 | 0.278 | 1 | 5 | Cope | -0.006 | Young |
| 0.930671 | 0.044743 | 0.904 | 0.931 | 1 | 5 | Nfkbia | -0.027 | Young |
| 0.930852 | 0.016326 | 0.221 | 0.226 | 1 | 5 | Fbxw11 | -0.005 | Young |
| 0.931063 | 0.095154 | 0.439 | 0.449 | 1 | 5 | Daglb | -0.01 | Young |
| 0.931069 | 0.055488 | 0.053 | 0.052 | 1 | 5 | Neurl2 | 0.001 | Young |
| 0.931073 | 0.20802 | 0.058 | 0.057 | 1 | 5 | Scyl3 | 0.001 | Young |
| 0.931131 | 0.091166 | 0.246 | 0.242 | 1 | 5 | Csnk2a2 | 0.004 | Young |
| 0.931365 | 0.2543 | 0.063 | 0.062 | 1 | 5 | Ints8 | 0.001 | Young |
| 0.931385 | 0.202666 | 0.112 | 0.111 | 1 | 5 | Kdsr | 0.001 | Young |
| 0.931476 | 0.115828 | 0.188 | 0.187 | 1 | 5 | Gtf2a2 | 0.001 | Young |
| 0.931511 | 0.394192 | 0.013 | 0.013 | 1 | 5 | Mars2 | 0 | Young |
| 0.931644 | 0.175354 | 0.096 | 0.099 | 1 | 5 | Asb8 | -0.003 | Young |
| 0.931803 | 0.049455 | 0.444 | 0.439 | 1 | 5 | Cct5 | 0.005 | Young |
| 0.931855 | 0.051224 | 0.018 | 0.019 | 1 | 5 | Tmem62 | -0.001 | Young |
| 0.931934 | 0.087237 | 0.444 | 0.463 | 1 | 5 | Derl1 | -0.019 | Young |
| 0.93218 | 0.010696 | 0.297 | 0.305 | 1 | 5 | Arf2 | -0.008 | Young |
| 0.932259 | 0.019267 | 0.203 | 0.199 | 1 | 5 | Trpv2 | 0.004 | Young |
| 0.932273 | 0.004701 | 0.107 | 0.108 | 1 | 5 | Dtymk | -0.001 | Young |
| 0.932372 | 0.235696 | 0.127 | 0.125 | 1 | 5 | Frmd8 | 0.002 | Young |
| 0.932487 | 0.027908 | 0.647 | 0.677 | 1 | 5 | Rpn2 | -0.03 | Young |
| 0.932637 | 0.003647 | 0.018 | 0.019 | 1 | 5 | 4833418N( | -0.001 | Young |
| 0.933063 | 0.041892 | 0.439 | 0.455 | 1 | 5 | Ankfy1 | -0.016 | Young |
| 0.933111 | 0.082485 | 0.574 | 0.589 | 1 | 5 | Itgam | -0.015 | Young |
| 0.933165 | 0.184602 | 0.249 | 0.261 | 1 | 5 | Tmem141 | -0.012 | Young |
| 0.933334 | 0.102983 | 0.048 | 0.047 | 1 | 5 | 1110008P1 | 0.001 | Young |
| 0.933338 | 0.072559 | 0.368 | 0.374 | 1 | 5 | Elmsan1 | -0.006 | Young |
| 0.933418 | 0.031807 | 0.018 | 0.019 | 1 | 5 | Gm38079 | -0.001 | Young |
| 0.933418 | 0.262507 | 0.018 | 0.019 | 1 | 5 | Zfp442 | -0.001 | Young |
| 0.933501 | 0.080155 | 0.061 | 0.059 | 1 | 5 | Gmds | 0.002 | Young |
| 0.933545 | 0.048202 | 0.066 | 0.067 | 1 | 5 | Rdh14 | -0.001 | Young |
| 0.933694 | 0.068426 | 0.378 | 0.389 | 1 | 5 | Orai1 | -0.011 | Young |
| 0.934085 | 0.106822 | 0.256 | 0.258 | 1 | 5 | Dctn5 | -0.002 | Young |
| 0.934103 | 0.045657 | 0.109 | 0.111 | 1 | 5 | Pik3r4 | -0.002 | Young |
| 0.934141 | 0.044556 | 0.51 | 0.505 | 1 | 5 | Chmp4b | 0.005 | Young |
| 0.9342 | 0.042045 | 0.018 | 0.019 | 1 | 5 | Tamm41 | -0.001 | Young |
| 0.934346 | 0.127361 | 0.023 | 0.024 | 1 | 5 | Gm7160 | -0.001 | Young |
| 0.934676 | 0.209519 | 0.19 | 0.187 | 1 | 5 | Cdk19 | 0.003 | Young |
| 0.934749 | 0.030736 | 0.718 | 0.737 | 1 | 5 | Atp6ap1 | -0.019 | Young |
| 0.93518 | 1.940986 | 0.013 | 0.013 | 1 | 5 | Gbp4 | 0 | Young |
| 0.93519 | 0.040487 | 0.168 | 0.167 | 1 | 5 | Adcy9 | 0.001 | Young |
| 0.935373 | 0.085402 | 0.129 | 0.131 | 1 | 5 | Copg2 | -0.002 | Young |

| | | | | | | | | |
|---|---|---|---|---|---|---|---|---|
| 0.935692 | 0.02453 | 0.152 | 0.148 | 1 | 5 | Frrs1 | 0.004 | Young |
| 0.935708 | 0.001549 | 0.256 | 0.259 | 1 | 5 | Ahr | -0.003 | Young |
| 0.935763 | 0.302971 | 0.018 | 0.019 | 1 | 5 | Rsph3b | -0.001 | Young |
| 0.936141 | 0.051178 | 0.043 | 0.042 | 1 | 5 | Asb1 | 0.001 | Young |
| 0.936204 | 0.050348 | 0.269 | 0.269 | 1 | 5 | Lmbrd2 | 0 | Young |
| 0.936413 | 0.015884 | 0.094 | 0.094 | 1 | 5 | Sik2 | 0 | Young |
| 0.936444 | 0.044105 | 0.168 | 0.167 | 1 | 5 | Parp9 | 0.001 | Young |
| 0.936455 | 0.080138 | 0.624 | 0.653 | 1 | 5 | Mtdh | -0.029 | Young |
| 0.936609 | 0.17825 | 0.102 | 0.099 | 1 | 5 | Pde4dip | 0.003 | Young |
| 0.936864 | 0.221579 | 0.178 | 0.177 | 1 | 5 | Retreg3 | 0.001 | Young |
| 0.936917 | 0.00902 | 0.223 | 0.226 | 1 | 5 | Otulin | -0.003 | Young |
| 0.93697 | 0.063097 | 0.071 | 0.072 | 1 | 5 | Vps8 | -0.001 | Young |
| 0.937119 | 0.0088 | 0.023 | 0.024 | 1 | 5 | AI427809 | -0.001 | Young |
| 0.937182 | 0.091909 | 0.269 | 0.273 | 1 | 5 | Btbd1 | -0.004 | Young |
| 0.937378 | 0.201688 | 0.175 | 0.173 | 1 | 5 | Nudt16 | 0.002 | Young |
| 0.937685 | 0.114731 | 0.112 | 0.111 | 1 | 5 | Impdh1 | 0.001 | Young |
| 0.937862 | 0.004544 | 0.033 | 0.032 | 1 | 5 | Zfp580 | 0.001 | Young |
| 0.937927 | 0.225758 | 0.038 | 0.037 | 1 | 5 | Tnfsfm13 | 0.001 | Young |
| 0.938108 | 0.280144 | 0.018 | 0.019 | 1 | 5 | Hspa2 | -0.001 | Young |
| 0.938108 | 0.261399 | 0.018 | 0.019 | 1 | 5 | Gk | -0.001 | Young |
| 0.938108 | 0.261156 | 0.018 | 0.019 | 1 | 5 | H2-Q10 | -0.001 | Young |
| 0.938258 | 0.065512 | 0.063 | 0.062 | 1 | 5 | Fam210a | 0.001 | Young |
| 0.938376 | 0.198082 | 0.14 | 0.138 | 1 | 5 | Dram1 | 0.002 | Young |
| 0.93841 | 0.091778 | 0.299 | 0.295 | 1 | 5 | Aff1 | 0.004 | Young |
| 0.938566 | 0.209392 | 0.066 | 0.067 | 1 | 5 | Prr3 | -0.001 | Young |
| 0.938657 | 0.014838 | 0.114 | 0.116 | 1 | 5 | Dtx4 | -0.002 | Young |
| 0.938735 | 0.218892 | 0.048 | 0.047 | 1 | 5 | Tmem237 | 0.001 | Young |
| 0.938916 | 0.038348 | 0.094 | 0.094 | 1 | 5 | Bloc1s2 | 0 | Young |
| 0.939049 | 0.216319 | 0.19 | 0.192 | 1 | 5 | Ndufs4 | -0.002 | Young |
| 0.939089 | 0.044991 | 0.061 | 0.062 | 1 | 5 | Enoph1 | -0.001 | Young |
| 0.939154 | 0.028893 | 0.244 | 0.246 | 1 | 5 | Hadha | -0.002 | Young |
| 0.939264 | 0.00641 | 0.272 | 0.271 | 1 | 5 | Clcn4 | 0.001 | Young |
| 0.939342 | 0.079136 | 0.53 | 0.569 | 1 | 5 | Atp1b3 | -0.039 | Young |
| 0.939498 | 0.079609 | 0.411 | 0.421 | 1 | 5 | Rbbp6 | -0.01 | Young |
| 0.93999 | 0.202422 | 0.147 | 0.146 | 1 | 5 | Ngly1 | 0.001 | Young |
| 0.94015 | 0.063078 | 0.492 | 0.493 | 1 | 5 | Atp5j | -0.001 | Young |
| 0.940259 | 0.110017 | 0.099 | 0.101 | 1 | 5 | Tfpi | -0.002 | Young |
| 0.940587 | 0.249339 | 0.023 | 0.024 | 1 | 5 | Zfat | -0.001 | Young |
| 0.940734 | 0.450608 | 0.041 | 0.04 | 1 | 5 | Slc25a38 | 0.001 | Young |
| 0.941221 | 0.093206 | 0.089 | 0.089 | 1 | 5 | Hpn | 0 | Young |
| 0.941597 | 0.088809 | 0.074 | 0.072 | 1 | 5 | 1110059E2 | 0.002 | Young |
| 0.941676 | 0.128808 | 0.274 | 0.285 | 1 | 5 | Tmem167 | -0.011 | Young |
| 0.942059 | 0.077089 | 0.391 | 0.392 | 1 | 5 | Wdr83os | -0.001 | Young |
| 0.942244 | 0.047938 | 0.406 | 0.416 | 1 | 5 | Pacs2 | -0.01 | Young |
| 0.942319 | 0.102566 | 0.069 | 0.067 | 1 | 5 | Mrpl50 | 0.002 | Young |

| | | | | | | | | |
|---|---|---|---|---|---|---|---|---|
| 0.942595 | 0.084238 | 0.079 | 0.077 | 1 | 5 | Nek6 | 0.002 | Young |
| 0.94275 | 0.098487 | 0.094 | 0.093 | 1 | 5 | Gab3 | 0.001 | Young |
| 0.942801 | 0.092072 | 0.018 | 0.019 | 1 | 5 | C130074G1 | -0.001 | Young |
| 0.942855 | 0.024968 | 0.168 | 0.165 | 1 | 5 | Dpp9 | 0.003 | Young |
| 0.942863 | 0.14128 | 0.277 | 0.281 | 1 | 5 | Gpatch8 | -0.004 | Young |
| 0.943179 | 0.003373 | 0.033 | 0.032 | 1 | 5 | Trim30b | 0.001 | Young |
| 0.94326 | 0.101746 | 0.043 | 0.044 | 1 | 5 | Inafm2 | -0.001 | Young |
| 0.943261 | 0.066598 | 0.294 | 0.293 | 1 | 5 | Cnn2 | 0.001 | Young |
| 0.943287 | 0.058416 | 0.244 | 0.249 | 1 | 5 | Bfar | -0.005 | Young |
| 0.943361 | 0.164753 | 0.023 | 0.024 | 1 | 5 | Clec16a | -0.001 | Young |
| 0.943365 | 0.15457 | 0.109 | 0.109 | 1 | 5 | Trappc13 | 0 | Young |
| 0.943427 | 0.140734 | 0.127 | 0.126 | 1 | 5 | Prpsap1 | 0.001 | Young |
| 0.943608 | 0.19273 | 0.066 | 0.066 | 1 | 5 | Haus2 | 0 | Young |
| 0.943702 | 0.190781 | 0.264 | 0.274 | 1 | 5 | Psma5 | -0.01 | Young |
| 0.943982 | 0.109238 | 0.069 | 0.067 | 1 | 5 | Vopp1 | 0.002 | Young |
| 0.943986 | 0.002714 | 0.038 | 0.037 | 1 | 5 | Sirt5 | 0.001 | Young |
| 0.944265 | 0.217294 | 0.193 | 0.197 | 1 | 5 | S100a1 | -0.004 | Young |
| 0.9443 | 0.092346 | 0.213 | 0.212 | 1 | 5 | Fam193a | 0.001 | Young |
| 0.944581 | 0.167872 | 0.279 | 0.279 | 1 | 5 | Gak | 0 | Young |
| 0.944612 | 0.096885 | 0.249 | 0.251 | 1 | 5 | Camta2 | -0.002 | Young |
| 0.944643 | 0.007608 | 0.053 | 0.052 | 1 | 5 | Gm15445 | 0.001 | Young |
| 0.944885 | 0.096996 | 0.119 | 0.12 | 1 | 5 | Dpm1 | -0.001 | Young |
| 0.944903 | 0.046156 | 0.426 | 0.433 | 1 | 5 | Brd4 | -0.007 | Young |
| 0.945092 | 0.219853 | 0.165 | 0.167 | 1 | 5 | Osm | -0.002 | Young |
| 0.945154 | 0.080536 | 0.533 | 0.544 | 1 | 5 | Larp4b | -0.011 | Young |
| 0.945194 | 0.019156 | 0.152 | 0.152 | 1 | 5 | Hars | 0 | Young |
| 0.946164 | 0.111027 | 0.327 | 0.325 | 1 | 5 | Ip6k1 | 0.002 | Young |
| 0.946321 | 0.065837 | 0.028 | 0.029 | 1 | 5 | Zrsr1 | -0.001 | Young |
| 0.94637 | 0.057078 | 0.104 | 0.103 | 1 | 5 | Lyst | 0.001 | Young |
| 0.946831 | 0.208711 | 0.023 | 0.024 | 1 | 5 | Rpusd1 | -0.001 | Young |
| 0.947502 | 0.104715 | 0.363 | 0.367 | 1 | 5 | Apbb1ip | -0.004 | Young |
| 0.947642 | 0.038766 | 0.444 | 0.444 | 1 | 5 | Mia2 | 0 | Young |
| 0.947706 | 0.037174 | 0.091 | 0.093 | 1 | 5 | Malsu1 | -0.002 | Young |
| 0.947753 | 0.024549 | 0.096 | 0.094 | 1 | 5 | Pus1 | 0.002 | Young |
| 0.948042 | 0.226372 | 0.129 | 0.13 | 1 | 5 | Tbc1d15 | -0.001 | Young |
| 0.948384 | 0.068376 | 0.033 | 0.034 | 1 | 5 | Ighmbp2 | -0.001 | Young |
| 0.948568 | 0.217941 | 0.213 | 0.221 | 1 | 5 | Galnt11 | -0.008 | Young |
| 0.948687 | 0.091449 | 0.269 | 0.271 | 1 | 5 | Nfkb2 | -0.002 | Young |
| 0.948992 | 0.346555 | 0.046 | 0.045 | 1 | 5 | Acad11 | 0.001 | Young |
| 0.949315 | 0.02731 | 0.277 | 0.276 | 1 | 5 | Sltm | 0.001 | Young |
| 0.949369 | 0.083792 | 0.16 | 0.157 | 1 | 5 | Map2k4 | 0.003 | Young |
| 0.94941 | 0.189808 | 0.228 | 0.229 | 1 | 5 | Commd7 | -0.001 | Young |
| 0.949607 | 0.253745 | 0.023 | 0.024 | 1 | 5 | Kcnn4 | -0.001 | Young |
| 0.949839 | 0.108061 | 0.15 | 0.153 | 1 | 5 | Rnf5 | -0.003 | Young |
| 0.949893 | 0.042664 | 0.533 | 0.532 | 1 | 5 | Nav1 | 0.001 | Young |

| | | | | | | | | |
|---|---|---|---|---|---|---|---|---|
| 0.950219 | 0.221413 | 0.061 | 0.061 | 1 | 5 | Adamts10 | 0 | Young |
| 0.950301 | 0.189619 | 0.023 | 0.024 | 1 | 5 | Wdr59 | -0.001 | Young |
| 0.950524 | 0.162593 | 0.107 | 0.109 | 1 | 5 | Mitd1 | -0.002 | Young |
| 0.950627 | 0.208989 | 0.216 | 0.222 | 1 | 5 | Slc26a2 | -0.006 | Young |
| 0.950686 | 0.105474 | 0.421 | 0.439 | 1 | 5 | Zfp106 | -0.018 | Young |
| 0.950717 | 0.11754 | 0.152 | 0.15 | 1 | 5 | Tcf3 | 0.002 | Young |
| 0.950996 | 0.227796 | 0.023 | 0.024 | 1 | 5 | Klhl26 | -0.001 | Young |
| 0.951043 | 0.024526 | 0.142 | 0.145 | 1 | 5 | Ddit3 | -0.003 | Young |
| 0.951136 | 0.130347 | 0.416 | 0.418 | 1 | 5 | Ppp1r12a | -0.002 | Young |
| 0.95126 | 0.036389 | 0.685 | 0.719 | 1 | 5 | Bsg | -0.034 | Young |
| 0.952384 | 0.40107 | 0.023 | 0.024 | 1 | 5 | Btbd6 | -0.001 | Young |
| 0.952434 | 0.08533 | 0.703 | 0.702 | 1 | 5 | Cox8a | 0.001 | Young |
| 0.952448 | 0.176198 | 0.419 | 0.444 | 1 | 5 | Usmg5 | -0.025 | Young |
| 0.952452 | 0.024139 | 0.193 | 0.194 | 1 | 5 | Dnm1l | -0.001 | Young |
| 0.952644 | 0.281952 | 0.051 | 0.052 | 1 | 5 | Yipf2 | -0.001 | Young |
| 0.952907 | 0.019356 | 0.251 | 0.251 | 1 | 5 | Cuedc2 | 0 | Young |
| 0.952943 | 0.076859 | 0.475 | 0.48 | 1 | 5 | Tep1 | -0.005 | Young |
| 0.95325 | 0.01806 | 0.165 | 0.163 | 1 | 5 | Mindy3 | 0.002 | Young |
| 0.953415 | 0.015385 | 0.363 | 0.364 | 1 | 5 | Npepps | -0.001 | Young |
| 0.953449 | 0.098022 | 0.18 | 0.177 | 1 | 5 | Spic | 0.003 | Young |
| 0.953748 | 0.132575 | 0.081 | 0.081 | 1 | 5 | Btrc | 0 | Young |
| 0.95379 | 0.274782 | 0.129 | 0.133 | 1 | 5 | Sap30 | -0.004 | Young |
| 0.953852 | 0.034614 | 0.371 | 0.369 | 1 | 5 | Pet100 | 0.002 | Young |
| 0.953935 | 0.185866 | 0.19 | 0.194 | 1 | 5 | Lemd2 | -0.004 | Young |
| 0.953987 | 0.224973 | 0.094 | 0.096 | 1 | 5 | Cgrrf1 | -0.002 | Young |
| 0.954346 | 0.084828 | 0.091 | 0.091 | 1 | 5 | Slc2a8 | 0 | Young |
| 0.954521 | 0.081351 | 0.028 | 0.029 | 1 | 5 | Pex1 | -0.001 | Young |
| 0.954739 | 0.128014 | 0.152 | 0.155 | 1 | 5 | Creld2 | -0.003 | Young |
| 0.954884 | 0.183978 | 0.122 | 0.126 | 1 | 5 | Rasa2 | -0.004 | Young |
| 0.954984 | 0.187165 | 0.114 | 0.114 | 1 | 5 | Slc35b4 | 0 | Young |
| 0.955041 | 0.052716 | 0.124 | 0.123 | 1 | 5 | Xab2 | 0.001 | Young |
| 0.955273 | 0.055251 | 0.102 | 0.103 | 1 | 5 | Idh3a | -0.001 | Young |
| 0.955399 | 0.075362 | 0.109 | 0.108 | 1 | 5 | Ap1g2 | 0.001 | Young |
| 0.955588 | 0.212665 | 0.053 | 0.054 | 1 | 5 | Taf8 | -0.001 | Young |
| 0.955662 | 0.043359 | 0.079 | 0.077 | 1 | 5 | Pithd1 | 0.002 | Young |
| 0.955703 | 0.0232 | 0.505 | 0.529 | 1 | 5 | Cd2ap | -0.024 | Young |
| 0.956623 | 0.281829 | 0.036 | 0.035 | 1 | 5 | Harbi1 | 0.001 | Young |
| 0.956663 | 0.124857 | 0.254 | 0.251 | 1 | 5 | Smu1 | 0.003 | Young |
| 0.957045 | 0.395302 | 0.028 | 0.029 | 1 | 5 | Dars2 | -0.001 | Young |
| 0.957166 | 0.264014 | 0.094 | 0.094 | 1 | 5 | Dcp1a | 0 | Young |
| 0.957756 | 0.632875 | 0.036 | 0.035 | 1 | 5 | Zfp346 | 0.001 | Young |
| 0.957764 | 0.064188 | 0.302 | 0.306 | 1 | 5 | Spag7 | -0.004 | Young |
| 0.958462 | 0.277309 | 0.066 | 0.067 | 1 | 5 | Tpgs1 | -0.001 | Young |
| 0.958594 | 0.089369 | 0.129 | 0.131 | 1 | 5 | Bicra | -0.002 | Young |
| 0.95897 | 0.319687 | 0.071 | 0.071 | 1 | 5 | Camta1 | 0 | Young |

| | | | | | | | | |
|---|---|---|---|---|---|---|---|---|
| 0.959598 | 0.291287 | 0.137 | 0.143 | 1 | 5 | Hivep3 | -0.006 | Young |
| 0.95992 | 0.044152 | 0.343 | 0.345 | 1 | 5 | Rasgrp4 | -0.002 | Young |
| 0.959938 | 0.111473 | 0.162 | 0.162 | 1 | 5 | Nagk | 0 | Young |
| 0.960049 | 0.096115 | 0.071 | 0.072 | 1 | 5 | Gdap2 | -0.001 | Young |
| 0.960066 | 0.090214 | 0.066 | 0.066 | 1 | 5 | Trip6 | 0 | Young |
| 0.96023 | 0.012166 | 1 | 1 | 1 | 5 | Actb | 0 | Young |
| 0.960245 | 0.029351 | 0.457 | 0.463 | 1 | 5 | Copa | -0.006 | Young |
| 0.960311 | 0.022391 | 0.784 | 0.79 | 1 | 5 | Erp29 | -0.006 | Young |
| 0.960716 | 0.005966 | 0.731 | 0.753 | 1 | 5 | Fyb | -0.022 | Young |
| 0.961634 | 0.120162 | 0.213 | 0.221 | 1 | 5 | Atg16l1 | -0.008 | Young |
| 0.961702 | 0.11722 | 0.216 | 0.221 | 1 | 5 | Hivep1 | -0.005 | Young |
| 0.96203 | 0.116015 | 0.127 | 0.131 | 1 | 5 | Nop56 | -0.004 | Young |
| 0.962112 | 0.15384 | 0.234 | 0.236 | 1 | 5 | Nsmce4a | -0.002 | Young |
| 0.962119 | 0.094276 | 0.074 | 0.072 | 1 | 5 | Atg14 | 0.002 | Young |
| 0.962952 | 0.111131 | 0.033 | 0.034 | 1 | 5 | E130102H2 | -0.001 | Young |
| 0.963119 | 0.175004 | 0.269 | 0.273 | 1 | 5 | Cnih1 | -0.004 | Young |
| 0.963334 | 0.141263 | 0.114 | 0.116 | 1 | 5 | Snapc5 | -0.002 | Young |
| 0.963535 | 0.430709 | 0.033 | 0.034 | 1 | 5 | Tango6 | -0.001 | Young |
| 0.964392 | 0.085456 | 0.168 | 0.17 | 1 | 5 | Fuca2 | -0.002 | Young |
| 0.964423 | 0.162829 | 0.056 | 0.057 | 1 | 5 | Ankrd54 | -0.001 | Young |
| 0.964701 | 0.367321 | 0.033 | 0.034 | 1 | 5 | Ppil1 | -0.001 | Young |
| 0.964846 | 0.029757 | 0.457 | 0.468 | 1 | 5 | Ctnna1 | -0.011 | Young |
| 0.964927 | 0.370846 | 0.046 | 0.047 | 1 | 5 | Adrb1 | -0.001 | Young |
| 0.965245 | 0.038076 | 0.112 | 0.111 | 1 | 5 | Dctn6 | 0.001 | Young |
| 0.965403 | 0.152033 | 0.218 | 0.221 | 1 | 5 | Gosr2 | -0.003 | Young |
| 0.965641 | 0.111821 | 0.132 | 0.133 | 1 | 5 | Tshz1 | -0.001 | Young |
| 0.965882 | 0.148271 | 0.35 | 0.362 | 1 | 5 | Ube2q1 | -0.012 | Young |
| 0.965922 | 0.237858 | 0.046 | 0.047 | 1 | 5 | 1190007I0 | -0.001 | Young |
| 0.966171 | 0.082803 | 0.206 | 0.205 | 1 | 5 | Pdhb | 0.001 | Young |
| 0.966373 | 0.001533 | 0.119 | 0.116 | 1 | 5 | Brap | 0.003 | Young |
| 0.966435 | 0.235694 | 0.061 | 0.061 | 1 | 5 | Tmem268 | 0 | Young |
| 0.966761 | 0.008068 | 0.074 | 0.074 | 1 | 5 | Lztfl1 | 0 | Young |
| 0.966775 | 0.109402 | 0.386 | 0.402 | 1 | 5 | Arl8b | -0.016 | Young |
| 0.96707 | 0.011011 | 0.091 | 0.091 | 1 | 5 | Tdp2 | 0 | Young |
| 0.967136 | 0.049355 | 0.203 | 0.205 | 1 | 5 | Cfap43 | -0.002 | Young |
| 0.967671 | 0.040422 | 0.053 | 0.054 | 1 | 5 | Grpel2 | -0.001 | Young |
| 0.967714 | 0.071882 | 0.261 | 0.263 | 1 | 5 | Dhrs1 | -0.002 | Young |
| 0.967823 | 0.342757 | 0.048 | 0.049 | 1 | 5 | Ift172 | -0.001 | Young |
| 0.967905 | 0.182761 | 0.14 | 0.14 | 1 | 5 | Il15 | 0 | Young |
| 0.968145 | 0.066311 | 0.168 | 0.17 | 1 | 5 | Tnks | -0.002 | Young |
| 0.968364 | 0.079473 | 0.183 | 0.184 | 1 | 5 | Srek1ip1 | -0.001 | Young |
| 0.968442 | 0.083497 | 0.251 | 0.253 | 1 | 5 | Samm50 | -0.002 | Young |
| 0.968708 | 0.214544 | 0.246 | 0.253 | 1 | 5 | Srek1 | -0.007 | Young |
| 0.968816 | 0.044897 | 0.518 | 0.524 | 1 | 5 | G3bp1 | -0.006 | Young |
| 0.968911 | 0.023465 | 0.865 | 0.88 | 1 | 5 | Eif4g2 | -0.015 | Young |

| | | | | | | | | |
|---|---|---|---|---|---|---|---|---|
| 0.968924 | 0.030819 | 0.145 | 0.146 | 1 | 5 | Zcchc9 | -0.001 | Young |
| 0.968925 | 0.153182 | 0.079 | 0.079 | 1 | 5 | Kif16b | 0 | Young |
| 0.968932 | 0.027218 | 0.107 | 0.106 | 1 | 5 | Rab9 | 0.001 | Young |
| 0.969068 | 0.189977 | 0.135 | 0.14 | 1 | 5 | Bst1 | -0.005 | Young |
| 0.969191 | 0.127064 | 0.109 | 0.108 | 1 | 5 | Selenoh | 0.001 | Young |
| 0.969387 | 0.023086 | 0.896 | 0.933 | 1 | 5 | Tmbim6 | -0.037 | Young |
| 0.969461 | 0.143095 | 0.089 | 0.091 | 1 | 5 | Coq2 | -0.002 | Young |
| 0.969528 | 0.331224 | 0.107 | 0.109 | 1 | 5 | Mnat1 | -0.002 | Young |
| 0.969639 | 0.083602 | 0.096 | 0.096 | 1 | 5 | Snx15 | 0 | Young |
| 0.969735 | 0.025195 | 0.302 | 0.298 | 1 | 5 | Themis2 | 0.004 | Young |
| 0.970224 | 0.073944 | 0.353 | 0.36 | 1 | 5 | Bod1l | -0.007 | Young |
| 0.970432 | 0.106859 | 0.188 | 0.192 | 1 | 5 | Sys1 | -0.004 | Young |
| 0.970439 | 0.006662 | 0.239 | 0.237 | 1 | 5 | AB124611 | 0.002 | Young |
| 0.970491 | 0.204027 | 0.152 | 0.155 | 1 | 5 | Zmym4 | -0.003 | Young |
| 0.9706 | 0.080193 | 0.312 | 0.325 | 1 | 5 | Psma1 | -0.013 | Young |
| 0.971054 | 0.123776 | 0.152 | 0.152 | 1 | 5 | Lactb2 | 0 | Young |
| 0.971106 | 0.02907 | 0.076 | 0.077 | 1 | 5 | Xpnpep3 | -0.001 | Young |
| 0.971236 | 0.090725 | 0.096 | 0.096 | 1 | 5 | Ppp6r2 | 0 | Young |
| 0.971303 | 0.115075 | 0.129 | 0.131 | 1 | 5 | Irgm1 | -0.002 | Young |
| 0.971564 | 0.156158 | 0.596 | 0.626 | 1 | 5 | Sqstm1 | -0.03 | Young |
| 0.971933 | 0.22533 | 0.264 | 0.271 | 1 | 5 | Map4 | -0.007 | Young |
| 0.972025 | 0.190569 | 0.051 | 0.051 | 1 | 5 | Spindoc | 0 | Young |
| 0.972283 | 0.237127 | 0.033 | 0.034 | 1 | 5 | Phc1 | -0.001 | Young |
| 0.972959 | 0.285884 | 0.025 | 0.025 | 1 | 5 | Acyp2 | 0 | Young |
| 0.973024 | 0.132603 | 0.043 | 0.044 | 1 | 5 | Slc35d1 | -0.001 | Young |
| 0.973393 | 0.000615 | 0.089 | 0.088 | 1 | 5 | Vps72 | 0.001 | Young |
| 0.973449 | 0.302476 | 0.033 | 0.034 | 1 | 5 | Acss2 | -0.001 | Young |
| 0.973603 | 0.036138 | 0.241 | 0.241 | 1 | 5 | Cnot4 | 0 | Young |
| 0.97367 | 0.136345 | 0.041 | 0.04 | 1 | 5 | Ipcef1 | 0.001 | Young |
| 0.973676 | 0.000354 | 0.683 | 0.715 | 1 | 5 | Herpud1 | -0.032 | Young |
| 0.973714 | 0.133013 | 0.056 | 0.056 | 1 | 5 | Miip | 0 | Young |
| 0.973995 | 0.16733 | 0.086 | 0.088 | 1 | 5 | Rhot2 | -0.002 | Young |
| 0.97402 | 0.016234 | 0.914 | 0.933 | 1 | 5 | Rpl32 | -0.019 | Young |
| 0.974029 | 0.298518 | 0.03 | 0.03 | 1 | 5 | Ddb2 | 0 | Young |
| 0.974181 | 0.168779 | 0.053 | 0.054 | 1 | 5 | Pdp2 | -0.001 | Young |
| 0.97443 | 0.00267 | 0.206 | 0.202 | 1 | 5 | Acaa1a | 0.004 | Young |
| 0.974565 | 0.060074 | 0.043 | 0.044 | 1 | 5 | Echdc1 | -0.001 | Young |
| 0.975381 | 0.32111 | 0.104 | 0.108 | 1 | 5 | Abr | -0.004 | Young |
| 0.97562 | 0.097765 | 0.048 | 0.049 | 1 | 5 | Prdm15 | -0.001 | Young |
| 0.975743 | 0.244809 | 0.261 | 0.271 | 1 | 5 | Eef2k | -0.01 | Young |
| 0.975767 | 0.047548 | 0.063 | 0.064 | 1 | 5 | Farsa | -0.001 | Young |
| 0.976456 | 0.001012 | 0.119 | 0.12 | 1 | 5 | Pgpep1 | -0.001 | Young |
| 0.976767 | 0.07721 | 0.576 | 0.579 | 1 | 5 | Rnf149 | -0.003 | Young |
| 0.976768 | 0.129178 | 0.081 | 0.081 | 1 | 5 | Brms1 | 0 | Young |
| 0.976915 | 0.349625 | 0.074 | 0.076 | 1 | 5 | 4921524J1 | -0.002 | Young |

| | | | | | | | | |
|---|---|---|---|---|---|---|---|---|
| 0.97715 | 0.101083 | 0.274 | 0.276 | 1 | 5 | Xrn1 | -0.002 | Young |
| 0.977337 | 0.027158 | 0.462 | 0.444 | 1 | 5 | Hmox2 | 0.018 | Young |
| 0.977371 | 0.147243 | 0.056 | 0.056 | 1 | 5 | Kat8 | 0 | Young |
| 0.977402 | 0.021622 | 0.213 | 0.207 | 1 | 5 | Grap | 0.006 | Young |
| 0.977998 | 0.117253 | 0.14 | 0.14 | 1 | 5 | Kmt5c | 0 | Young |
| 0.978075 | 0.142627 | 0.287 | 0.296 | 1 | 5 | Mark2 | -0.009 | Young |
| 0.978339 | 0.283033 | 0.063 | 0.064 | 1 | 5 | Tradd | -0.001 | Young |
| 0.978367 | 0.076857 | 0.053 | 0.054 | 1 | 5 | Srm | -0.001 | Young |
| 0.97839 | 0.17425 | 0.198 | 0.205 | 1 | 5 | Tfg | -0.007 | Young |
| 0.978522 | 0.004243 | 0.249 | 0.246 | 1 | 5 | Cand1 | 0.003 | Young |
| 0.978644 | 0.084952 | 0.173 | 0.173 | 1 | 5 | Itga5 | 0 | Young |
| 0.978768 | 0.062754 | 0.063 | 0.064 | 1 | 5 | Dnajc12 | -0.001 | Young |
| 0.97926 | 0.031955 | 0.84 | 0.865 | 1 | 5 | Srrm2 | -0.025 | Young |
| 0.979282 | 0.192881 | 0.038 | 0.039 | 1 | 5 | Enpp4 | -0.001 | Young |
| 0.979436 | 0.024448 | 0.132 | 0.131 | 1 | 5 | Nup54 | 0.001 | Young |
| 0.979594 | 0.034627 | 0.061 | 0.061 | 1 | 5 | Gm29340 | 0 | Young |
| 0.979624 | 0.166604 | 0.112 | 0.113 | 1 | 5 | Tmem185b | -0.001 | Young |
| 0.980113 | 0.029001 | 0.056 | 0.056 | 1 | 5 | Tjp2 | 0 | Young |
| 0.980256 | 0.203768 | 0.416 | 0.436 | 1 | 5 | Sfr1 | -0.02 | Young |
| 0.980266 | 0.383345 | 0.02 | 0.02 | 1 | 5 | Sco1 | 0 | Young |
| 0.980408 | 0.015701 | 0.14 | 0.14 | 1 | 5 | Adpgk | 0 | Young |
| 0.980482 | 0.115488 | 0.175 | 0.178 | 1 | 5 | Pdgfc | -0.003 | Young |
| 0.980617 | 0.096295 | 0.112 | 0.113 | 1 | 5 | Bap1 | -0.001 | Young |
| 0.981365 | 0.084271 | 0.246 | 0.253 | 1 | 5 | Usp16 | -0.007 | Young |
| 0.982093 | 0.094755 | 0.231 | 0.234 | 1 | 5 | Arl1 | -0.003 | Young |
| 0.982156 | 0.087466 | 0.452 | 0.46 | 1 | 5 | Tnrc6b | -0.008 | Young |
| 0.982531 | 0.015378 | 0.104 | 0.104 | 1 | 5 | Brix1 | 0 | Young |
| 0.983008 | 0.223664 | 0.081 | 0.082 | 1 | 5 | Ak6 | -0.001 | Young |
| 0.983097 | 0.355165 | 0.038 | 0.039 | 1 | 5 | Mad2l1bp | -0.001 | Young |
| 0.983221 | 0.042087 | 0.599 | 0.611 | 1 | 5 | Atrx | -0.012 | Young |
| 0.983406 | 0.230492 | 0.046 | 0.045 | 1 | 5 | Pfkm | 0.001 | Young |
| 0.98343 | 0.124994 | 0.193 | 0.195 | 1 | 5 | Mrpl3 | -0.002 | Young |
| 0.983497 | 0.096655 | 0.31 | 0.311 | 1 | 5 | Pcna | -0.001 | Young |
| 0.984117 | 0.086274 | 0.188 | 0.187 | 1 | 5 | Klf11 | 0.001 | Young |
| 0.984217 | 0.071247 | 0.198 | 0.199 | 1 | 5 | Stim2 | -0.001 | Young |
| 0.984239 | 0.106621 | 0.099 | 0.099 | 1 | 5 | Tmem39a | 0 | Young |
| 0.984281 | 0.132662 | 0.244 | 0.249 | 1 | 5 | Cnot3 | -0.005 | Young |
| 0.984416 | 0.158507 | 0.03 | 0.03 | 1 | 5 | Fbxo10 | 0 | Young |
| 0.984452 | 0.022914 | 0.221 | 0.219 | 1 | 5 | Mindy1 | 0.002 | Young |
| 0.984517 | 0.110723 | 0.19 | 0.192 | 1 | 5 | Osbp | -0.002 | Young |
| 0.984535 | 0.537825 | 0.033 | 0.034 | 1 | 5 | Prepl | -0.001 | Young |
| 0.984649 | 0.17392 | 0.216 | 0.219 | 1 | 5 | Kansl1l | -0.003 | Young |
| 0.984707 | 0.132532 | 0.084 | 0.084 | 1 | 5 | Zfp282 | 0 | Young |
| 0.984725 | 0.045847 | 0.15 | 0.153 | 1 | 5 | Cables1 | -0.003 | Young |
| 0.984839 | 0.117264 | 0.058 | 0.059 | 1 | 5 | Dedd | -0.001 | Young |

| | | | | | | | | |
|---|---|---|---|---|---|---|---|---|
| 0.984963 | 0.0839 | 0.165 | 0.165 | 1 | 5 | Mbp | 0 | Young |
| 0.98499 | 0.226841 | 0.015 | 0.015 | 1 | 5 | Gm28557 | 0 | Young |
| 0.98508 | 0.142515 | 0.14 | 0.14 | 1 | 5 | Rab10os | 0 | Young |
| 0.9858 | 0.126391 | 0.206 | 0.21 | 1 | 5 | Fbxo3 | -0.004 | Young |
| 0.985848 | 0.360079 | 0.015 | 0.015 | 1 | 5 | L3hypdh | 0 | Young |
| 0.985891 | 0.161981 | 0.051 | 0.051 | 1 | 5 | Pqlc2 | 0 | Young |
| 0.986013 | 0.092698 | 0.079 | 0.079 | 1 | 5 | Cisd3 | 0 | Young |
| 0.986266 | 0.139593 | 0.183 | 0.184 | 1 | 5 | Taf11 | -0.001 | Young |
| 0.986346 | 0.392011 | 0.048 | 0.049 | 1 | 5 | Chst14 | -0.001 | Young |
| 0.986358 | 0.087336 | 0.104 | 0.106 | 1 | 5 | Lyar | -0.002 | Young |
| 0.986369 | 0.096518 | 0.051 | 0.051 | 1 | 5 | Fars2 | 0 | Young |
| 0.986705 | 0.205865 | 0.015 | 0.015 | 1 | 5 | Plin4 | 0 | Young |
| 0.987563 | 0.323575 | 0.015 | 0.015 | 1 | 5 | Cnn3 | 0 | Young |
| 0.987563 | 0.151639 | 0.015 | 0.015 | 1 | 5 | Lss | 0 | Young |
| 0.987736 | 0.036928 | 0.921 | 0.944 | 1 | 5 | Rpl28 | -0.023 | Young |
| 0.987804 | 0.105169 | 0.036 | 0.035 | 1 | 5 | Plpp5 | 0.001 | Young |
| 0.987899 | 0.249812 | 0.129 | 0.131 | 1 | 5 | Zbtb7b | -0.002 | Young |
| 0.98795 | 0.367218 | 0.01 | 0.01 | 1 | 5 | Zfp341 | 0 | Young |
| 0.988115 | 0.075914 | 0.401 | 0.402 | 1 | 5 | Pkm | -0.001 | Young |
| 0.988136 | 0.422852 | 0.053 | 0.054 | 1 | 5 | Apip | -0.001 | Young |
| 0.988214 | 0.159938 | 0.487 | 0.497 | 1 | 5 | Hspd1 | -0.01 | Young |
| 0.988536 | 0.083596 | 0.442 | 0.449 | 1 | 5 | Ikbkb | -0.007 | Young |
| 0.989202 | 0.017506 | 0.02 | 0.02 | 1 | 5 | Gm4070 | 0 | Young |
| 0.989389 | 0.169984 | 0.305 | 0.318 | 1 | 5 | Ptov1 | -0.013 | Young |
| 0.98951 | 0.124313 | 0.079 | 0.079 | 1 | 5 | Nfkbil1 | 0 | Young |
| 0.989522 | 0.132946 | 0.096 | 0.096 | 1 | 5 | Ccdc25 | 0 | Young |
| 0.989649 | 0.186382 | 0.025 | 0.025 | 1 | 5 | Ift81 | 0 | Young |
| 0.989691 | 0.126528 | 0.046 | 0.045 | 1 | 5 | Ift22 | 0.001 | Young |
| 0.989784 | 0.072757 | 0.462 | 0.483 | 1 | 5 | Ndufa7 | -0.021 | Young |
| 0.989916 | 0.003851 | 0.581 | 0.589 | 1 | 5 | Selenow | -0.008 | Young |
| 0.990045 | 0.220155 | 0.01 | 0.01 | 1 | 5 | Pcdhgc4 | 0 | Young |
| 0.990136 | 0.22596 | 0.015 | 0.015 | 1 | 5 | Tceanc | 0 | Young |
| 0.9902 | 0.016982 | 0.135 | 0.135 | 1 | 5 | Poc1b | 0 | Young |
| 0.990211 | 0.011014 | 0.457 | 0.465 | 1 | 5 | Ndufb3 | -0.008 | Young |
| 0.990317 | 0.32832 | 0.025 | 0.025 | 1 | 5 | Focad | 0 | Young |
| 0.990948 | 0.102736 | 0.046 | 0.045 | 1 | 5 | Zfp101 | 0.001 | Young |
| 0.991093 | 0.298738 | 0.01 | 0.01 | 1 | 5 | Plag1 | 0 | Young |
| 0.991093 | 0.20406 | 0.01 | 0.01 | 1 | 5 | Tmem263 | 0 | Young |
| 0.991093 | 0.195597 | 0.01 | 0.01 | 1 | 5 | 2510046G1 | 0 | Young |
| 0.991297 | 0.017182 | 0.292 | 0.293 | 1 | 5 | Rab5b | -0.001 | Young |
| 0.991379 | 0.158865 | 0.256 | 0.259 | 1 | 5 | Wtap | -0.003 | Young |
| 0.991436 | 0.132242 | 0.02 | 0.02 | 1 | 5 | Trim16 | 0 | Young |
| 0.991571 | 0.081167 | 0.201 | 0.202 | 1 | 5 | Otud5 | -0.001 | Young |
| 0.992043 | 0.289727 | 0.152 | 0.155 | 1 | 5 | Cyp4f16 | -0.003 | Young |
| 0.992093 | 0.111119 | 0.429 | 0.446 | 1 | 5 | Tnks2 | -0.017 | Young |

| | | | | | | | | |
|---|---|---|---|---|---|---|---|---|
| 0.992141 | 0.422392 | 0.01 | 0.01 | 1 | 5 | Brf2 | 0 | Young |
| 0.992141 | 0.111519 | 0.01 | 0.01 | 1 | 5 | Gm30025 | 0 | Young |
| 0.992503 | 0.104434 | 0.48 | 0.503 | 1 | 5 | Ier3ip1 | -0.023 | Young |
| 0.99253 | 0.1321 | 0.234 | 0.234 | 1 | 5 | Dap | 0 | Young |
| 0.992999 | 0.093725 | 0.335 | 0.342 | 1 | 5 | Rrp1 | -0.007 | Young |
| 0.993189 | 0.347527 | 0.01 | 0.01 | 1 | 5 | Gm10851 | 0 | Young |
| 0.993189 | 0.104564 | 0.01 | 0.01 | 1 | 5 | Fstl1 | 0 | Young |
| 0.993189 | 0.046302 | 0.01 | 0.01 | 1 | 5 | Adarb1 | 0 | Young |
| 0.993222 | 0.003956 | 0.378 | 0.377 | 1 | 5 | Il6st | 0.001 | Young |
| 0.993752 | 0.007515 | 0.102 | 0.101 | 1 | 5 | Rce1 | 0.001 | Young |
| 0.993787 | 0.120284 | 0.426 | 0.444 | 1 | 5 | Mon2 | -0.018 | Young |
| 0.993954 | 0.135069 | 0.183 | 0.185 | 1 | 5 | Uqcc3 | -0.002 | Young |
| 0.99406 | 0.150474 | 0.33 | 0.34 | 1 | 5 | Atp5o.1 | -0.01 | Young |
| 0.994075 | 0.132173 | 0.173 | 0.173 | 1 | 5 | Gid8 | 0 | Young |
| 0.994204 | 0.209712 | 0.069 | 0.069 | 1 | 5 | Copz2 | 0 | Young |
| 0.994449 | 0.033561 | 0.589 | 0.594 | 1 | 5 | Mafg | -0.005 | Young |
| 0.9945 | 0.030216 | 0.051 | 0.051 | 1 | 5 | Ppp3cc | 0 | Young |
| 0.994638 | 0.152299 | 0.063 | 0.064 | 1 | 5 | Armc7 | -0.001 | Young |
| 0.994928 | 0.315781 | 0.071 | 0.072 | 1 | 5 | Mfng | -0.001 | Young |
| 0.995094 | 0.020742 | 0.165 | 0.167 | 1 | 5 | Gkap1 | -0.002 | Young |
| 0.995282 | 0.163881 | 0.015 | 0.015 | 1 | 5 | Tmem201 | 0 | Young |
| 0.995285 | 0.249676 | 0.01 | 0.01 | 1 | 5 | Gm16486 | 0 | Young |
| 0.995285 | 0.174537 | 0.01 | 0.01 | 1 | 5 | Helq | 0 | Young |
| 0.995285 | 0.151472 | 0.01 | 0.01 | 1 | 5 | Gm49207 | 0 | Young |
| 0.995522 | 0.14683 | 0.206 | 0.21 | 1 | 5 | Fundc1 | -0.004 | Young |
| 0.996187 | 0.020734 | 0.178 | 0.178 | 1 | 5 | Ppm1b | 0 | Young |
| 0.996433 | 0.287483 | 0.058 | 0.059 | 1 | 5 | Whamm | -0.001 | Young |
| 0.996513 | 0.049723 | 0.223 | 0.224 | 1 | 5 | Angel2 | -0.001 | Young |
| 0.996592 | 0.065108 | 0.16 | 0.162 | 1 | 5 | Prdx3 | -0.002 | Young |
| 0.997055 | 0.122599 | 0.32 | 0.33 | 1 | 5 | Herc4 | -0.01 | Young |
| 0.997173 | 0.522538 | 0.043 | 0.044 | 1 | 5 | Clec4d | -0.001 | Young |
| 0.997592 | 0.159009 | 0.16 | 0.163 | 1 | 5 | Zfp655 | -0.003 | Young |
| 0.997634 | 0.105831 | 0.091 | 0.091 | 1 | 5 | Slc39a3 | 0 | Young |
| 0.997723 | 0.07816 | 0.099 | 0.099 | 1 | 5 | Vps16 | 0 | Young |
| 0.997754 | 0.068514 | 0.071 | 0.071 | 1 | 5 | Cluh | 0 | Young |
| 0.997856 | 0.041481 | 0.015 | 0.015 | 1 | 5 | Map3k9 | 0 | Young |
| 0.998243 | 0.012221 | 0.67 | 0.672 | 1 | 5 | Il10rb | -0.002 | Young |
| 0.999144 | 0.099418 | 0.104 | 0.104 | 1 | 5 | Ikbip | 0 | Young |
| 0.999375 | 0.159151 | 0.221 | 0.222 | 1 | 5 | Nck1 | -0.001 | Young |
| 0.999638 | 0.123374 | 0.241 | 0.244 | 1 | 5 | Ndufa11 | -0.003 | Young |
| 0.999757 | 0.002055 | 0.236 | 0.239 | 1 | 5 | Cbfa2t3 | -0.003 | Young |
| 0.999784 | 0.007954 | 0.063 | 0.062 | 1 | 5 | Adsl | 0.001 | Young |
| 2.80E-15 | 4.517819 | 0.143 | 0.015 | 8.68E-11 | 6 | Igkc | 0.128 | Old |
| 6.18E-10 | 4.104167 | 0.083 | 0.007 | 1.92E-05 | 6 | Iglc2 | 0.076 | Old |
| 4.95E-09 | 0.564859 | 0.9 | 0.849 | 0.000154 | 6 | Hsp90b1 | 0.051 | Old |

| | | | | | | | | |
|---|---|---|---|---|---|---|---|---|
| 5.67E-09 | 3.007926 | 0.071 | 0.005 | 0.000176 | 6 | Cd79a | 0.066 | Old |
| 7.08E-09 | 4.943124 | 0.066 | 0.003 | 0.00022 | 6 | Mzb1 | 0.063 | Old |
| 1.28E-08 | 0.245844 | 1 | 0.998 | 0.000397 | 6 | mt-Co1 | 0.002 | Old |
| 4.56E-08 | 3.687557 | 0.06 | 0.003 | 0.001418 | 6 | Ly6d | 0.057 | Old |
| 5.55E-08 | 4.585887 | 0.054 | 0.002 | 0.001723 | 6 | Ctla4 | 0.052 | Old |
| 6.30E-08 | 3.952223 | 0.08 | 0.012 | 0.001956 | 6 | Ly6a | 0.068 | Old |
| 8.42E-08 | 3.070056 | 0.063 | 0.005 | 0.002615 | 6 | Gimap1 | 0.058 | Old |
| 2.20E-07 | 0.265061 | 1 | 1 | 0.006831 | 6 | mt-Co3 | 0 | Old |
| 4.32E-07 | 0.445306 | 0.96 | 0.935 | 0.01341 | 6 | B2m | 0.025 | Old |
| 4.58E-07 | 3.941775 | 0.057 | 0.005 | 0.014213 | 6 | Iglc3 | 0.052 | Old |
| 7.91E-07 | 3.33818 | 0.071 | 0.012 | 0.024557 | 6 | Cd79b | 0.059 | Old |
| 9.31E-07 | 1.656724 | 0.3 | 0.177 | 0.028899 | 6 | Ly6e | 0.123 | Old |
| 1.14E-06 | 3.077347 | 0.054 | 0.005 | 0.035376 | 6 | Trp53i11 | 0.049 | Old |
| 1.61E-06 | 3.681013 | 0.049 | 0.003 | 0.050053 | 6 | Iglc1 | 0.046 | Old |
| 2.01E-06 | 4.779022 | 0.043 | 0.002 | 0.062426 | 6 | Gimap3 | 0.041 | Old |
| 2.10E-06 | 3.766875 | 0.043 | 0.002 | 0.065206 | 6 | Ms4a1 | 0.041 | Old |
| 2.67E-06 | 1.824103 | 0.123 | 0.042 | 0.083011 | 6 | Gsr | 0.081 | Old |
| 2.68E-06 | 3.931028 | 0.051 | 0.005 | 0.083254 | 6 | Ptprcap | 0.046 | Old |
| 4.08E-06 | 2.963217 | 0.046 | 0.003 | 0.126543 | 6 | Pou2af1 | 0.043 | Old |
| 4.10E-06 | 3.642907 | 0.066 | 0.012 | 0.127356 | 6 | Jchain | 0.054 | Old |
| 4.25E-06 | 2.979151 | 0.066 | 0.012 | 0.131897 | 6 | Ets1 | 0.054 | Old |
| 4.64E-06 | 0.258111 | 0.997 | 0.997 | 0.144206 | 6 | mt-Atp6 | 0 | Old |
| 4.94E-06 | 4.463903 | 0.04 | 0.002 | 0.153387 | 6 | Gimap4 | 0.038 | Old |
| 4.99E-06 | 3.529213 | 0.04 | 0.002 | 0.155059 | 6 | Derl3 | 0.038 | Old |
| 5.32E-06 | 0.354486 | 0.983 | 0.96 | 0.165147 | 6 | mt-Cytb | 0.023 | Old |
| 7.84E-06 | 0.319138 | 0.897 | 0.868 | 0.243357 | 6 | mt-Nd3 | 0.029 | Old |
| 8.01E-06 | 2.63804 | 0.06 | 0.01 | 0.248613 | 6 | Dennd5b | 0.05 | Old |
| 9.73E-06 | 3.246881 | 0.043 | 0.003 | 0.302076 | 6 | Cd55 | 0.04 | Old |
| 1.05E-05 | 0.685837 | 0.58 | 0.46 | 0.325134 | 6 | Ccnd2 | 0.12 | Old |
| 1.29E-05 | 2.448703 | 0.037 | 0.002 | 0.401977 | 6 | Gstt2 | 0.035 | Old |
| 1.41E-05 | 0.464673 | 0.843 | 0.751 | 0.439133 | 6 | Srsf5 | 0.092 | Old |
| 1.61E-05 | 2.850414 | 0.074 | 0.018 | 0.49853 | 6 | C130026I2 | 0.056 | Old |
| 1.73E-05 | 2.246056 | 0.071 | 0.017 | 0.536804 | 6 | Cxcl13 | 0.054 | Old |
| 1.83E-05 | 2.226538 | 0.057 | 0.01 | 0.568959 | 6 | A530032D1 | 0.047 | Old |
| 2.41E-05 | 0.134626 | 0.989 | 0.998 | 0.748999 | 6 | Actb | -0.009 | Old |
| 3.29E-05 | 3.773376 | 0.029 | 0 | 1 | 6 | Blk | 0.029 | Old |
| 3.66E-05 | 2.009478 | 0.091 | 0.03 | 1 | 6 | Prmt7 | 0.061 | Old |
| 4.83E-05 | 1.391949 | 0.22 | 0.125 | 1 | 6 | Ckap4 | 0.095 | Old |
| 5.25E-05 | 0.384536 | 0.909 | 0.895 | 1 | 6 | H3f3b | 0.014 | Old |
| 5.54E-05 | 3.08676 | 0.037 | 0.003 | 1 | 6 | Phgdh | 0.034 | Old |
| 6.04E-05 | 2.465629 | 0.037 | 0.003 | 1 | 6 | Ikzf3 | 0.034 | Old |
| 6.11E-05 | 0.792881 | 0.223 | 0.124 | 1 | 6 | Dhx36 | 0.099 | Old |
| 7.07E-05 | 0.205174 | 0.917 | 0.946 | 1 | 6 | Cfl1 | -0.029 | Old |
| 7.26E-05 | 1.753715 | 0.223 | 0.13 | 1 | 6 | AW112010 | 0.093 | Old |
| 7.35E-05 | 2.026429 | 0.223 | 0.13 | 1 | 6 | Txndc5 | 0.093 | Old |

| | | | | | | | | |
|---|---|---|---|---|---|---|---|---|
| 7.56E-05 | 3.942831 | 0.031 | 0.002 | 1 | 6 | Gimap5 | 0.029 | Old |
| 8.24E-05 | 5.149801 | 0.026 | 0 | 1 | 6 | Iglv1 | 0.026 | Old |
| 8.24E-05 | 3.880276 | 0.026 | 0 | 1 | 6 | Gm30211 | 0.026 | Old |
| 8.24E-05 | 3.47261 | 0.026 | 0 | 1 | 6 | Faah | 0.026 | Old |
| 8.24E-05 | 2.939049 | 0.026 | 0 | 1 | 6 | Il5ra | 0.026 | Old |
| 8.24E-05 | 2.635541 | 0.026 | 0 | 1 | 6 | Hook1 | 0.026 | Old |
| 0.000101 | 1.276004 | 0.297 | 0.194 | 1 | 6 | Hist1h1c | 0.103 | Old |
| 0.000132 | 0.659338 | 0.483 | 0.373 | 1 | 6 | Coro2a | 0.11 | Old |
| 0.000138 | 1.983858 | 0.06 | 0.015 | 1 | 6 | Ankrd28 | 0.045 | Old |
| 0.00014 | 2.357103 | 0.034 | 0.003 | 1 | 6 | Bhlhe41 | 0.031 | Old |
| 0.00014 | 2.408385 | 0.034 | 0.003 | 1 | 6 | Lcmt2 | 0.031 | Old |
| 0.00015 | 3.180589 | 0.046 | 0.008 | 1 | 6 | Tbc1d10c | 0.038 | Old |
| 0.000155 | 1.175679 | 0.094 | 0.035 | 1 | 6 | Gipc1 | 0.059 | Old |
| 0.00019 | 2.884055 | 0.029 | 0.002 | 1 | 6 | Capn10 | 0.027 | Old |
| 0.00019 | 2.816467 | 0.029 | 0.002 | 1 | 6 | Zfp961 | 0.027 | Old |
| 0.000191 | 0.801704 | 0.374 | 0.271 | 1 | 6 | Psmb9 | 0.103 | Old |
| 0.000194 | 2.521204 | 0.029 | 0.002 | 1 | 6 | Cacna1e | 0.027 | Old |
| 0.000203 | 2.996696 | 0.037 | 0.005 | 1 | 6 | Eaf2 | 0.032 | Old |
| 0.000207 | 6.302168 | 0.023 | 0 | 1 | 6 | Gata3 | 0.023 | Old |
| 0.000207 | 4.752632 | 0.023 | 0 | 1 | 6 | Thy1 | 0.023 | Old |
| 0.000207 | 3.88495 | 0.023 | 0 | 1 | 6 | Gimap7 | 0.023 | Old |
| 0.000207 | 3.737253 | 0.023 | 0 | 1 | 6 | Gimap9 | 0.023 | Old |
| 0.000207 | 2.75817 | 0.023 | 0 | 1 | 6 | Snn | 0.023 | Old |
| 0.000343 | 4.06733 | 0.031 | 0.003 | 1 | 6 | Slpi | 0.028 | Old |
| 0.000352 | 0.197274 | 0.991 | 0.975 | 1 | 6 | mt-Nd1 | 0.016 | Old |
| 0.000451 | 1.621715 | 0.046 | 0.01 | 1 | 6 | Pick1 | 0.036 | Old |
| 0.000456 | 1.861439 | 0.1 | 0.043 | 1 | 6 | Fam214a | 0.057 | Old |
| 0.000465 | 3.852796 | 0.026 | 0.002 | 1 | 6 | Tnfrsf25 | 0.024 | Old |
| 0.00047 | 2.693454 | 0.026 | 0.002 | 1 | 6 | Cd28 | 0.024 | Old |
| 0.000484 | 2.004672 | 0.026 | 0.002 | 1 | 6 | Atp1b1 | 0.024 | Old |
| 0.000487 | 2.530455 | 0.026 | 0.002 | 1 | 6 | Gimap8 | 0.024 | Old |
| 0.0005 | 0.243631 | 0.917 | 0.918 | 1 | 6 | Pfn1 | -0.001 | Old |
| 0.000523 | 4.479766 | 0.02 | 0 | 1 | 6 | Bcl11b | 0.02 | Old |
| 0.000523 | 4.001874 | 0.02 | 0 | 1 | 6 | Trbc1 | 0.02 | Old |
| 0.000523 | 3.407237 | 0.02 | 0 | 1 | 6 | A230050P2 | 0.02 | Old |
| 0.000523 | 3.091767 | 0.02 | 0 | 1 | 6 | Prg2 | 0.02 | Old |
| 0.000523 | 2.445937 | 0.02 | 0 | 1 | 6 | Zfp40 | 0.02 | Old |
| 0.000548 | 0.798642 | 0.149 | 0.077 | 1 | 6 | Xpot | 0.072 | Old |
| 0.000569 | 2.175444 | 0.054 | 0.015 | 1 | 6 | Smpd2 | 0.039 | Old |
| 0.000646 | 0.754919 | 0.343 | 0.241 | 1 | 6 | Gorasp2 | 0.102 | Old |
| 0.000758 | 0.414605 | 0.671 | 0.592 | 1 | 6 | Tuba1b | 0.079 | Old |
| 0.000774 | 4.216773 | 0.029 | 0.003 | 1 | 6 | Il2ra | 0.026 | Old |
| 0.00082 | 1.523303 | 0.04 | 0.008 | 1 | 6 | Gimap6 | 0.032 | Old |
| 0.000924 | 0.565254 | 0.449 | 0.333 | 1 | 6 | Hspa9 | 0.116 | Old |
| 0.000935 | 0.554389 | 0.54 | 0.462 | 1 | 6 | Ran | 0.078 | Old |

| | | | | | | | | |
|---|---|---|---|---|---|---|---|---|
| 0.000946 | 0.272127 | 0.966 | 0.965 | 1 | 6 | H2-K1 | 0.001 | Old |
| 0.000947 | 0.664538 | 0.134 | 0.069 | 1 | 6 | Hsf1 | 0.065 | Old |
| 0.00104 | 0.250967 | 0.854 | 0.836 | 1 | 6 | Actr2 | 0.018 | Old |
| 0.001066 | 1.875957 | 0.071 | 0.027 | 1 | 6 | Sh3bp5 | 0.044 | Old |
| 0.001083 | 0.145959 | 1 | 1 | 1 | 6 | mt-Co2 | 0 | Old |
| 0.001089 | 1.935596 | 0.063 | 0.022 | 1 | 6 | Plac8 | 0.041 | Old |
| 0.001128 | 2.398613 | 0.031 | 0.005 | 1 | 6 | 2900060B1 | 0.026 | Old |
| 0.001135 | 1.264792 | 0.183 | 0.109 | 1 | 6 | Atp2a3 | 0.074 | Old |
| 0.001158 | 0.707044 | 0.114 | 0.055 | 1 | 6 | Dcaf10 | 0.059 | Old |
| 0.001159 | 0.950365 | 0.303 | 0.217 | 1 | 6 | Pdia4 | 0.086 | Old |
| 0.001168 | 2.574923 | 0.023 | 0.002 | 1 | 6 | Smco4 | 0.021 | Old |
| 0.001176 | 1.441538 | 0.031 | 0.005 | 1 | 6 | F8a | 0.026 | Old |
| 0.00118 | 2.610319 | 0.023 | 0.002 | 1 | 6 | Cped1 | 0.021 | Old |
| 0.00119 | 0.591475 | 0.283 | 0.194 | 1 | 6 | Nfatc3 | 0.089 | Old |
| 0.001192 | 3.002159 | 0.023 | 0.002 | 1 | 6 | Il2rb | 0.021 | Old |
| 0.001192 | 1.84258 | 0.023 | 0.002 | 1 | 6 | Slain1 | 0.021 | Old |
| 0.001205 | 1.497578 | 0.023 | 0.002 | 1 | 6 | Cd19 | 0.021 | Old |
| 0.001249 | 0.869341 | 0.16 | 0.09 | 1 | 6 | Agpat5 | 0.07 | Old |
| 0.00133 | 4.776111 | 0.017 | 0 | 1 | 6 | Cd3g | 0.017 | Old |
| 0.00133 | 3.671286 | 0.017 | 0 | 1 | 6 | Fcmr | 0.017 | Old |
| 0.00133 | 3.386591 | 0.017 | 0 | 1 | 6 | Tnfsf10 | 0.017 | Old |
| 0.00133 | 3.318559 | 0.017 | 0 | 1 | 6 | Lck | 0.017 | Old |
| 0.00133 | 2.904336 | 0.017 | 0 | 1 | 6 | Gm15675 | 0.017 | Old |
| 0.00133 | 2.739974 | 0.017 | 0 | 1 | 6 | Sh3bgrl2 | 0.017 | Old |
| 0.00133 | 2.710713 | 0.017 | 0 | 1 | 6 | 1810046K0 | 0.017 | Old |
| 0.00133 | 2.477701 | 0.017 | 0 | 1 | 6 | Alkbh4 | 0.017 | Old |
| 0.00133 | 2.392231 | 0.017 | 0 | 1 | 6 | 2010007H0 | 0.017 | Old |
| 0.00133 | 2.388236 | 0.017 | 0 | 1 | 6 | Zfp507 | 0.017 | Old |
| 0.00133 | 2.070093 | 0.017 | 0 | 1 | 6 | Krt222 | 0.017 | Old |
| 0.00133 | 2.043802 | 0.017 | 0 | 1 | 6 | Pla2g6 | 0.017 | Old |
| 0.00133 | 1.921478 | 0.017 | 0 | 1 | 6 | Zfp963 | 0.017 | Old |
| 0.00133 | 1.440654 | 0.017 | 0 | 1 | 6 | Bhlha15 | 0.017 | Old |
| 0.001443 | 3.099661 | 0.089 | 0.04 | 1 | 6 | Cd69 | 0.049 | Old |
| 0.001468 | 0.878707 | 0.209 | 0.134 | 1 | 6 | Creld2 | 0.075 | Old |
| 0.001494 | 0.29202 | 0.803 | 0.773 | 1 | 6 | Hnrnpa3 | 0.03 | Old |
| 0.001594 | 0.800441 | 0.131 | 0.069 | 1 | 6 | Card11 | 0.062 | Old |
| 0.001742 | 1.171667 | 0.08 | 0.033 | 1 | 6 | AC163354. | 0.047 | Old |
| 0.001833 | 1.29406 | 0.077 | 0.032 | 1 | 6 | Zfp746 | 0.045 | Old |
| 0.001877 | 0.275775 | 0.806 | 0.773 | 1 | 6 | Fam49b | 0.033 | Old |
| 0.001926 | 1.929579 | 0.026 | 0.003 | 1 | 6 | Ficd | 0.023 | Old |
| 0.001944 | 2.327626 | 0.026 | 0.003 | 1 | 6 | Zcwpw1 | 0.023 | Old |
| 0.001974 | 1.153131 | 0.071 | 0.028 | 1 | 6 | Pdzd11 | 0.043 | Old |
| 0.002005 | 0.911795 | 0.026 | 0.003 | 1 | 6 | Lax1 | 0.023 | Old |
| 0.00208 | 1.936922 | 0.06 | 0.022 | 1 | 6 | Rnf123 | 0.038 | Old |
| 0.002137 | 1.319474 | 0.103 | 0.05 | 1 | 6 | Ccl8 | 0.053 | Old |

| | | | | | | | | |
|---|---|---|---|---|---|---|---|---|
| 0.002155 | 1.140491 | 0.066 | 0.025 | 1 | 6 | Mkrn2 | 0.041 | Old |
| 0.002155 | 1.563845 | 0.06 | 0.022 | 1 | 6 | E2f3 | 0.038 | Old |
| 0.002228 | 1.729773 | 0.054 | 0.018 | 1 | 6 | Bloc1s5 | 0.036 | Old |
| 0.00228 | 0.24442 | 0.829 | 0.784 | 1 | 6 | Samhd1 | 0.045 | Old |
| 0.002384 | 1.311872 | 0.086 | 0.038 | 1 | 6 | Prkdc | 0.048 | Old |
| 0.002413 | 1.150843 | 0.106 | 0.054 | 1 | 6 | Txndc12 | 0.052 | Old |
| 0.002599 | 0.56491 | 0.151 | 0.087 | 1 | 6 | Atg4d | 0.064 | Old |
| 0.00268 | 1.684797 | 0.029 | 0.005 | 1 | 6 | Ccdc137 | 0.024 | Old |
| 0.002756 | 1.318803 | 0.029 | 0.005 | 1 | 6 | AC125149. | 0.024 | Old |
| 0.002809 | 0.395892 | 0.614 | 0.548 | 1 | 6 | Rpn1 | 0.066 | Old |
| 0.002911 | 3.105792 | 0.02 | 0.002 | 1 | 6 | AC168977. | 0.018 | Old |
| 0.00294 | 1.619121 | 0.02 | 0.002 | 1 | 6 | Oosp1 | 0.018 | Old |
| 0.002958 | 0.663101 | 0.274 | 0.191 | 1 | 6 | Zfp292 | 0.083 | Old |
| 0.00296 | 1.04705 | 0.114 | 0.06 | 1 | 6 | Ppp2r3c | 0.054 | Old |
| 0.002968 | 1.150358 | 0.074 | 0.032 | 1 | 6 | Zfp513 | 0.042 | Old |
| 0.002985 | 1.596159 | 0.02 | 0.002 | 1 | 6 | Fkbp11 | 0.018 | Old |
| 0.003092 | 0.829947 | 0.177 | 0.112 | 1 | 6 | Txndc11 | 0.065 | Old |
| 0.00318 | 0.760698 | 0.283 | 0.206 | 1 | 6 | Slc25a24 | 0.077 | Old |
| 0.003261 | 1.411091 | 0.069 | 0.028 | 1 | 6 | Mrpl22 | 0.041 | Old |
| 0.003411 | 3.764256 | 0.014 | 0 | 1 | 6 | Sh2d2a | 0.014 | Old |
| 0.003411 | 3.661215 | 0.014 | 0 | 1 | 6 | Itk | 0.014 | Old |
| 0.003411 | 2.729423 | 0.014 | 0 | 1 | 6 | Nid1 | 0.014 | Old |
| 0.003411 | 2.212791 | 0.014 | 0 | 1 | 6 | Gm34215 | 0.014 | Old |
| 0.003411 | 2.149009 | 0.014 | 0 | 1 | 6 | Fam169b | 0.014 | Old |
| 0.003411 | 2.105498 | 0.014 | 0 | 1 | 6 | Gm3336 | 0.014 | Old |
| 0.003411 | 1.561701 | 0.014 | 0 | 1 | 6 | Pls1 | 0.014 | Old |
| 0.003447 | 1.079554 | 0.086 | 0.04 | 1 | 6 | Med23 | 0.046 | Old |
| 0.003626 | 2.106868 | 0.034 | 0.008 | 1 | 6 | Mgst1 | 0.026 | Old |
| 0.003657 | 1.121385 | 0.103 | 0.054 | 1 | 6 | Ifi47 | 0.049 | Old |
| 0.003739 | 0.55149 | 0.451 | 0.365 | 1 | 6 | Srsf1 | 0.086 | Old |
| 0.00375 | 1.036053 | 0.109 | 0.057 | 1 | 6 | Irgm1 | 0.052 | Old |
| 0.003871 | 0.313687 | 0.634 | 0.55 | 1 | 6 | 11-Sep | 0.084 | Old |
| 0.003874 | 1.292028 | 0.114 | 0.062 | 1 | 6 | D10Wsu10 | 0.052 | Old |
| 0.003925 | 0.39586 | 0.337 | 0.244 | 1 | 6 | Tcirg1 | 0.093 | Old |
| 0.003955 | 1.847733 | 0.106 | 0.057 | 1 | 6 | Trp53inp1 | 0.049 | Old |
| 0.003989 | 1.626068 | 0.057 | 0.022 | 1 | 6 | Arl14ep | 0.035 | Old |
| 0.004008 | 2.078929 | 0.06 | 0.023 | 1 | 6 | Atxn1 | 0.037 | Old |
| 0.004121 | 2.527563 | 0.04 | 0.012 | 1 | 6 | Tespa1 | 0.028 | Old |
| 0.004237 | 1.063422 | 0.077 | 0.035 | 1 | 6 | Zc3hc1 | 0.042 | Old |
| 0.00424 | 1.200772 | 0.08 | 0.037 | 1 | 6 | Strbp | 0.043 | Old |
| 0.004326 | 1.584652 | 0.04 | 0.012 | 1 | 6 | Chst11 | 0.028 | Old |
| 0.004416 | 4.578104 | 0.023 | 0.003 | 1 | 6 | Cxcr6 | 0.02 | Old |
| 0.004459 | 1.206775 | 0.04 | 0.012 | 1 | 6 | Csprs | 0.028 | Old |
| 0.004485 | 1.508781 | 0.043 | 0.013 | 1 | 6 | Mrpl47 | 0.03 | Old |
| 0.004512 | 0.381669 | 0.306 | 0.221 | 1 | 6 | Usp16 | 0.085 | Old |

| | | | | | | | | |
|---|---|---|---|---|---|---|---|---|
| 0.004533 | 2.455507 | 0.023 | 0.003 | 1 | 6 | Mterf1a | 0.02 | Old |
| 0.004553 | 2.280726 | 0.023 | 0.003 | 1 | 6 | Enpp1 | 0.02 | Old |
| 0.004582 | 0.92182 | 0.251 | 0.179 | 1 | 6 | Cd37 | 0.072 | Old |
| 0.004648 | 0.193136 | 0.906 | 0.885 | 1 | 6 | Hnrnpa2b1 | 0.021 | Old |
| 0.004783 | 0.947661 | 0.054 | 0.02 | 1 | 6 | Vegfa | 0.034 | Old |
| 0.005145 | 0.74177 | 0.163 | 0.1 | 1 | 6 | Tars | 0.063 | Old |
| 0.005251 | 1.569105 | 0.217 | 0.149 | 1 | 6 | Odc1 | 0.068 | Old |
| 0.005379 | 0.784443 | 0.186 | 0.12 | 1 | 6 | Pum3 | 0.066 | Old |
| 0.005714 | 0.294595 | 0.774 | 0.699 | 1 | 6 | Zfp36l2 | 0.075 | Old |
| 0.005844 | 1.40769 | 0.086 | 0.042 | 1 | 6 | Dtx3 | 0.044 | Old |
| 0.005848 | 0.274613 | 0.763 | 0.721 | 1 | 6 | Atp5f1 | 0.042 | Old |
| 0.005967 | 0.7699 | 0.16 | 0.1 | 1 | 6 | Rpl7l1 | 0.06 | Old |
| 0.006134 | 0.978286 | 0.12 | 0.069 | 1 | 6 | Tceanc2 | 0.051 | Old |
| 0.006134 | 0.273794 | 0.026 | 0.005 | 1 | 6 | Ly6c2 | 0.021 | Old |
| 0.006215 | 1.131541 | 0.129 | 0.077 | 1 | 6 | Rpia | 0.052 | Old |
| 0.006406 | 0.75631 | 0.309 | 0.231 | 1 | 6 | Pnn | 0.078 | Old |
| 0.006684 | 0.548451 | 0.217 | 0.147 | 1 | 6 | Ppie | 0.07 | Old |
| 0.006692 | 0.837996 | 0.094 | 0.048 | 1 | 6 | Rrn3 | 0.046 | Old |
| 0.00675 | 0.380661 | 0.8 | 0.844 | 1 | 6 | Naaa | -0.044 | Old |
| 0.006778 | 2.511724 | 0.029 | 0.007 | 1 | 6 | Irgm2 | 0.022 | Old |
| 0.00683 | 0.869047 | 0.074 | 0.035 | 1 | 6 | Dclre1a | 0.039 | Old |
| 0.006838 | 0.90516 | 0.077 | 0.037 | 1 | 6 | Entpd6 | 0.04 | Old |
| 0.007002 | 0.757244 | 0.137 | 0.082 | 1 | 6 | Commd2 | 0.055 | Old |
| 0.007013 | 2.146167 | 0.054 | 0.022 | 1 | 6 | AA467197 | 0.032 | Old |
| 0.007053 | 0.53526 | 0.563 | 0.492 | 1 | 6 | Fubp1 | 0.071 | Old |
| 0.007153 | 0.862884 | 0.114 | 0.064 | 1 | 6 | Gart | 0.05 | Old |
| 0.007193 | 2.041038 | 0.017 | 0.002 | 1 | 6 | Igfbp7 | 0.015 | Old |
| 0.007218 | 0.651712 | 0.234 | 0.166 | 1 | 6 | Stat1 | 0.068 | Old |
| 0.007265 | 2.493845 | 0.017 | 0.002 | 1 | 6 | Jakmip1 | 0.015 | Old |
| 0.007274 | 1.094779 | 0.029 | 0.007 | 1 | 6 | Yes1 | 0.022 | Old |
| 0.007338 | 1.623333 | 0.017 | 0.002 | 1 | 6 | Dusp4 | 0.015 | Old |
| 0.007406 | 2.749487 | 0.031 | 0.008 | 1 | 6 | Ablim1 | 0.023 | Old |
| 0.007429 | 1.184303 | 0.029 | 0.007 | 1 | 6 | Hip1r | 0.022 | Old |
| 0.007552 | 1.47691 | 0.071 | 0.033 | 1 | 6 | Actr5 | 0.038 | Old |
| 0.007603 | 2.350919 | 0.031 | 0.008 | 1 | 6 | Igf2r | 0.023 | Old |
| 0.007721 | 0.801321 | 0.149 | 0.092 | 1 | 6 | Crebzf | 0.057 | Old |
| 0.007752 | 0.58572 | 0.266 | 0.191 | 1 | 6 | Nudcd3 | 0.075 | Old |
| 0.007759 | 0.584992 | 0.306 | 0.231 | 1 | 6 | Aldh9a1 | 0.075 | Old |
| 0.007764 | 1.131261 | 0.051 | 0.02 | 1 | 6 | Akr1e1 | 0.031 | Old |
| 0.007798 | 0.754957 | 0.243 | 0.169 | 1 | 6 | Phc3 | 0.074 | Old |
| 0.008 | 0.150612 | 0.929 | 0.933 | 1 | 6 | Cdc42 | -0.004 | Old |
| 0.008041 | 1.167917 | 0.123 | 0.072 | 1 | 6 | Eif2b2 | 0.051 | Old |
| 0.00811 | 1.346448 | 0.049 | 0.018 | 1 | 6 | Pecam1 | 0.031 | Old |
| 0.008253 | 1.768503 | 0.046 | 0.017 | 1 | 6 | Lzic | 0.029 | Old |
| 0.008307 | 1.698968 | 0.034 | 0.01 | 1 | 6 | Abcb8 | 0.024 | Old |

| | | | | | | | | |
|---|---|---|---|---|---|---|---|---|
| 0.00833 | 1.485249 | 0.094 | 0.05 | 1 | 6 | Dcn | 0.044 | Old |
| 0.008376 | 1.319509 | 0.066 | 0.03 | 1 | 6 | Cmah | 0.036 | Old |
| 0.0084 | 1.20249 | 0.114 | 0.065 | 1 | 6 | Zbp1 | 0.049 | Old |
| 0.008402 | 1.331899 | 0.046 | 0.017 | 1 | 6 | Fcrla | 0.029 | Old |
| 0.008499 | 0.644054 | 0.289 | 0.214 | 1 | 6 | Psmd14 | 0.075 | Old |
| 0.008551 | 0.808037 | 0.083 | 0.042 | 1 | 6 | Pnkp | 0.041 | Old |
| 0.008597 | 1.049 | 0.069 | 0.032 | 1 | 6 | Tlr4 | 0.037 | Old |
| 0.008599 | 0.507894 | 0.414 | 0.333 | 1 | 6 | Pgam1 | 0.081 | Old |
| 0.008659 | 0.95768 | 0.04 | 0.013 | 1 | 6 | Med26 | 0.027 | Old |
| 0.008745 | 0.817102 | 0.137 | 0.084 | 1 | 6 | Grpel1 | 0.053 | Old |
| 0.008769 | 1.132663 | 0.08 | 0.04 | 1 | 6 | Elp3 | 0.04 | Old |
| 0.008796 | 1.902223 | 0.074 | 0.037 | 1 | 6 | Ralgps2 | 0.037 | Old |
| 0.008863 | 4.174062 | 0.011 | 0 | 1 | 6 | Klrg1 | 0.011 | Old |
| 0.008863 | 3.994439 | 0.011 | 0 | 1 | 6 | Skap1 | 0.011 | Old |
| 0.008863 | 3.987086 | 0.011 | 0 | 1 | 6 | Trac | 0.011 | Old |
| 0.008863 | 3.239605 | 0.011 | 0 | 1 | 6 | Trbc2 | 0.011 | Old |
| 0.008863 | 3.045517 | 0.011 | 0 | 1 | 6 | Sugct | 0.011 | Old |
| 0.008863 | 2.720817 | 0.011 | 0 | 1 | 6 | Igkv3-2 | 0.011 | Old |
| 0.008863 | 2.571612 | 0.011 | 0 | 1 | 6 | Bcl2l15 | 0.011 | Old |
| 0.008863 | 2.533757 | 0.011 | 0 | 1 | 6 | Tnik | 0.011 | Old |
| 0.008863 | 2.493859 | 0.011 | 0 | 1 | 6 | Krt8 | 0.011 | Old |
| 0.008863 | 2.262191 | 0.011 | 0 | 1 | 6 | Wnt10a | 0.011 | Old |
| 0.008863 | 2.208345 | 0.011 | 0 | 1 | 6 | Chrnb1 | 0.011 | Old |
| 0.008863 | 2.040755 | 0.011 | 0 | 1 | 6 | Tst | 0.011 | Old |
| 0.008863 | 2.035219 | 0.011 | 0 | 1 | 6 | Nutf2 | 0.011 | Old |
| 0.008863 | 2.02091 | 0.011 | 0 | 1 | 6 | Il9r | 0.011 | Old |
| 0.008863 | 2.012001 | 0.011 | 0 | 1 | 6 | Spc24 | 0.011 | Old |
| 0.008863 | 1.997628 | 0.011 | 0 | 1 | 6 | Ahnak2 | 0.011 | Old |
| 0.008863 | 1.955887 | 0.011 | 0 | 1 | 6 | Egln3 | 0.011 | Old |
| 0.008863 | 1.77511 | 0.011 | 0 | 1 | 6 | Pik3c2b | 0.011 | Old |
| 0.008863 | 1.732113 | 0.011 | 0 | 1 | 6 | Rtl6 | 0.011 | Old |
| 0.008863 | 1.715645 | 0.011 | 0 | 1 | 6 | Hoxa7 | 0.011 | Old |
| 0.008863 | 1.651659 | 0.011 | 0 | 1 | 6 | AC133103. | 0.011 | Old |
| 0.008863 | 1.62437 | 0.011 | 0 | 1 | 6 | Slc6a9 | 0.011 | Old |
| 0.008863 | 1.599334 | 0.011 | 0 | 1 | 6 | Spag1 | 0.011 | Old |
| 0.008863 | 1.5345 | 0.011 | 0 | 1 | 6 | H3f3aos | 0.011 | Old |
| 0.008863 | 1.365993 | 0.011 | 0 | 1 | 6 | Snx32 | 0.011 | Old |
| 0.008863 | 1.31461 | 0.011 | 0 | 1 | 6 | Plekhh3 | 0.011 | Old |
| 0.008863 | 1.202793 | 0.011 | 0 | 1 | 6 | Tmem150a | 0.011 | Old |
| 0.008863 | 1.044422 | 0.011 | 0 | 1 | 6 | Aurka | 0.011 | Old |
| 0.008863 | 0.941533 | 0.011 | 0 | 1 | 6 | Iglv3 | 0.011 | Old |
| 0.008924 | 0.436304 | 0.189 | 0.122 | 1 | 6 | Flvcr1 | 0.067 | Old |
| 0.008991 | 0.496714 | 0.28 | 0.206 | 1 | 6 | Rbbp4 | 0.074 | Old |
| 0.009091 | 0.854367 | 0.069 | 0.032 | 1 | 6 | Mtg1 | 0.037 | Old |
| 0.009255 | 0.425043 | 0.42 | 0.331 | 1 | 6 | Copa | 0.089 | Old |

| | | | | | | | | |
|---|---|---|---|---|---|---|---|---|
| 0.009448 | 1.173388 | 0.08 | 0.04 | 1 | 6 | Wrnip1 | 0.04 | Old |
| 0.009639 | 0.823239 | 0.129 | 0.079 | 1 | 6 | Pprc1 | 0.05 | Old |
| 0.009671 | 0.631892 | 0.174 | 0.115 | 1 | 6 | Tatdn2 | 0.059 | Old |
| 0.00974 | 0.971428 | 0.066 | 0.03 | 1 | 6 | Paxx | 0.036 | Old |
| 0.009787 | 0.268302 | 0.734 | 0.689 | 1 | 6 | Prkcd | 0.045 | Old |
| 0.009959 | 0.735902 | 0.231 | 0.164 | 1 | 6 | Xist | 0.067 | Old |
| 0.00996 | 0.408378 | 0.403 | 0.321 | 1 | 6 | Tnrc18 | 0.082 | Old |
| 0.010129 | 0.884576 | 0.126 | 0.075 | 1 | 6 | Ifi27 | 0.051 | Old |
| 0.010158 | 0.999613 | 0.063 | 0.028 | 1 | 6 | Zw10 | 0.035 | Old |
| 0.010172 | 0.811145 | 0.174 | 0.115 | 1 | 6 | Mrpl13 | 0.059 | Old |
| 0.010217 | 0.363207 | 0.48 | 0.4 | 1 | 6 | Zfp207 | 0.08 | Old |
| 0.010291 | 0.682776 | 0.146 | 0.09 | 1 | 6 | Wsb2 | 0.056 | Old |
| 0.010341 | 0.514572 | 0.14 | 0.085 | 1 | 6 | E2f4 | 0.055 | Old |
| 0.010342 | 0.518072 | 0.42 | 0.349 | 1 | 6 | Psma2 | 0.071 | Old |
| 0.010391 | 0.779295 | 0.209 | 0.144 | 1 | 6 | H1f0 | 0.065 | Old |
| 0.010508 | 0.821187 | 0.08 | 0.04 | 1 | 6 | Ncoa5 | 0.04 | Old |
| 0.01055 | 2.096725 | 0.02 | 0.003 | 1 | 6 | Xlr3b | 0.017 | Old |
| 0.010684 | 2.317589 | 0.02 | 0.003 | 1 | 6 | Gm36839 | 0.017 | Old |
| 0.010688 | 0.64471 | 0.226 | 0.161 | 1 | 6 | Rchy1 | 0.065 | Old |
| 0.010729 | 1.857166 | 0.02 | 0.003 | 1 | 6 | Gpr55 | 0.017 | Old |
| 0.010772 | 0.809087 | 0.126 | 0.075 | 1 | 6 | Ppa2 | 0.051 | Old |
| 0.010819 | 2.035111 | 0.02 | 0.003 | 1 | 6 | Mss51 | 0.017 | Old |
| 0.010819 | 1.616216 | 0.02 | 0.003 | 1 | 6 | Phtf1os | 0.017 | Old |
| 0.010864 | 1.444048 | 0.02 | 0.003 | 1 | 6 | Coro2b | 0.017 | Old |
| 0.011014 | 0.801439 | 0.117 | 0.07 | 1 | 6 | Ccdc107 | 0.047 | Old |
| 0.01102 | 0.456878 | 0.289 | 0.212 | 1 | 6 | Slbp | 0.077 | Old |
| 0.011321 | 0.598032 | 0.411 | 0.336 | 1 | 6 | Il2rg | 0.075 | Old |
| 0.011499 | 1.533974 | 0.071 | 0.035 | 1 | 6 | Zc3h7b | 0.036 | Old |
| 0.011531 | 0.815556 | 0.137 | 0.085 | 1 | 6 | Uchl3 | 0.052 | Old |
| 0.011592 | 0.920648 | 0.123 | 0.074 | 1 | 6 | Stk25 | 0.049 | Old |
| 0.01161 | 0.693066 | 0.154 | 0.099 | 1 | 6 | Senp1 | 0.055 | Old |
| 0.011878 | 0.798809 | 0.203 | 0.14 | 1 | 6 | Ube2j1 | 0.063 | Old |
| 0.011932 | 2.023619 | 0.051 | 0.022 | 1 | 6 | Spint2 | 0.029 | Old |
| 0.011944 | 0.754948 | 0.06 | 0.027 | 1 | 6 | Ankrd39 | 0.033 | Old |
| 0.011979 | 0.715625 | 0.143 | 0.089 | 1 | 6 | Parg | 0.054 | Old |
| 0.011987 | 1.276396 | 0.083 | 0.043 | 1 | 6 | AU020206 | 0.04 | Old |
| 0.012282 | 0.785795 | 0.294 | 0.227 | 1 | 6 | Cd164 | 0.067 | Old |
| 0.01238 | 0.702433 | 0.143 | 0.09 | 1 | 6 | Prps1 | 0.053 | Old |
| 0.012482 | 1.13134 | 0.054 | 0.023 | 1 | 6 | 1500011B( | 0.031 | Old |
| 0.012688 | 0.325675 | 0.62 | 0.569 | 1 | 6 | Atp5g3 | 0.051 | Old |
| 0.012698 | 0.864471 | 0.203 | 0.145 | 1 | 6 | Tle4 | 0.058 | Old |
| 0.012742 | 0.034643 | 0.066 | 0.115 | 1 | 6 | Brwd1 | -0.049 | Old |
| 0.012785 | 0.976759 | 0.054 | 0.023 | 1 | 6 | Atg10 | 0.031 | Old |
| 0.012881 | 1.135062 | 0.051 | 0.022 | 1 | 6 | Tmem80 | 0.029 | Old |
| 0.012907 | 0.615784 | 0.183 | 0.124 | 1 | 6 | Mrpl51 | 0.059 | Old |

| | | | | | | | | |
|---|---|---|---|---|---|---|---|---|
| 0.013025 | 0.902526 | 0.117 | 0.07 | 1 | 6 | Ddx1 | 0.047 | Old |
| 0.01324 | 1.7215 | 0.023 | 0.005 | 1 | 6 | Zfp951 | 0.018 | Old |
| 0.01324 | 1.480259 | 0.023 | 0.005 | 1 | 6 | Fam160a2 | 0.018 | Old |
| 0.013258 | 0.392148 | 0.549 | 0.473 | 1 | 6 | Nucks1 | 0.076 | Old |
| 0.013289 | 1.797393 | 0.023 | 0.005 | 1 | 6 | Rap1gap | 0.018 | Old |
| 0.013307 | 0.538079 | 0.237 | 0.171 | 1 | 6 | Prpf39 | 0.066 | Old |
| 0.013328 | 1.115308 | 0.097 | 0.055 | 1 | 6 | Gm14548 | 0.042 | Old |
| 0.01333 | 0.565099 | 0.394 | 0.324 | 1 | 6 | Gsk3b | 0.07 | Old |
| 0.013388 | 1.502278 | 0.023 | 0.005 | 1 | 6 | Dpt | 0.018 | Old |
| 0.013766 | 0.228356 | 0.854 | 0.898 | 1 | 6 | Actr3 | -0.044 | Old |
| 0.013856 | 0.563324 | 0.166 | 0.11 | 1 | 6 | Ptbp2 | 0.056 | Old |
| 0.013908 | 1.583726 | 0.046 | 0.018 | 1 | 6 | Tdp1 | 0.028 | Old |
| 0.014195 | 0.285955 | 0.783 | 0.742 | 1 | 6 | Cd44 | 0.041 | Old |
| 0.014196 | 0.838077 | 0.066 | 0.032 | 1 | 6 | Slc16a6 | 0.034 | Old |
| 0.014209 | 1.073523 | 0.051 | 0.022 | 1 | 6 | Gm20045 | 0.029 | Old |
| 0.014249 | 0.180961 | 0.957 | 0.953 | 1 | 6 | Myl6 | 0.004 | Old |
| 0.014305 | 1.10199 | 0.066 | 0.032 | 1 | 6 | Drosha | 0.034 | Old |
| 0.014394 | 0.814153 | 0.071 | 0.035 | 1 | 6 | Tinf2 | 0.036 | Old |
| 0.014826 | 1.554355 | 0.046 | 0.018 | 1 | 6 | Fam189b | 0.028 | Old |
| 0.014861 | 1.219163 | 0.046 | 0.018 | 1 | 6 | Atg4a | 0.028 | Old |
| 0.015047 | 1.431254 | 0.026 | 0.007 | 1 | 6 | Eral1 | 0.019 | Old |
| 0.015052 | 1.011748 | 0.086 | 0.047 | 1 | 6 | Ints7 | 0.039 | Old |
| 0.015098 | 1.862357 | 0.026 | 0.007 | 1 | 6 | Nr1d1 | 0.019 | Old |
| 0.015144 | 1.32905 | 0.046 | 0.018 | 1 | 6 | Fam102a | 0.028 | Old |
| 0.015194 | 0.665465 | 0.049 | 0.02 | 1 | 6 | Miga1 | 0.029 | Old |
| 0.015216 | 1.268127 | 0.046 | 0.018 | 1 | 6 | Hirip3 | 0.028 | Old |
| 0.015216 | 0.677427 | 0.217 | 0.154 | 1 | 6 | Dicer1 | 0.063 | Old |
| 0.015221 | 0.833815 | 0.12 | 0.074 | 1 | 6 | Plcl2 | 0.046 | Old |
| 0.015301 | 0.638852 | 0.16 | 0.105 | 1 | 6 | Tcf3 | 0.055 | Old |
| 0.015309 | 1.415018 | 0.043 | 0.017 | 1 | 6 | Pml | 0.026 | Old |
| 0.015314 | 1.089993 | 0.083 | 0.045 | 1 | 6 | Abhd4 | 0.038 | Old |
| 0.015359 | 1.700639 | 0.046 | 0.018 | 1 | 6 | Rab27a | 0.028 | Old |
| 0.015359 | 1.1264 | 0.046 | 0.018 | 1 | 6 | Cdc25a | 0.028 | Old |
| 0.015461 | 2.737599 | 0.037 | 0.013 | 1 | 6 | Ltb | 0.024 | Old |
| 0.015521 | 0.506748 | 0.454 | 0.405 | 1 | 6 | Manf | 0.049 | Old |
| 0.015534 | 1.230751 | 0.043 | 0.017 | 1 | 6 | Cd300lf | 0.026 | Old |
| 0.015572 | 0.690211 | 0.14 | 0.089 | 1 | 6 | Vta1 | 0.051 | Old |
| 0.015649 | 0.67415 | 0.36 | 0.286 | 1 | 6 | Ubash3b | 0.074 | Old |
| 0.015691 | 0.546196 | 0.211 | 0.154 | 1 | 6 | Hyou1 | 0.057 | Old |
| 0.015747 | 2.774853 | 0.029 | 0.008 | 1 | 6 | Birc5 | 0.021 | Old |
| 0.015747 | 1.790118 | 0.029 | 0.008 | 1 | 6 | Zfp119a | 0.021 | Old |
| 0.015796 | 1.830614 | 0.029 | 0.008 | 1 | 6 | Poll | 0.021 | Old |
| 0.016014 | 0.325163 | 0.36 | 0.279 | 1 | 6 | Zfand6 | 0.081 | Old |
| 0.016044 | 1.58156 | 0.037 | 0.013 | 1 | 6 | Zfp653 | 0.024 | Old |
| 0.016044 | 1.706229 | 0.029 | 0.008 | 1 | 6 | Cep57l1 | 0.021 | Old |

| | | | | | | | | |
|---|---|---|---|---|---|---|---|---|
| 0.016129 | 1.726488 | 0.037 | 0.013 | 1 | 6 | Slc7a5 | 0.024 | Old |
| 0.016383 | 0.722569 | 0.103 | 0.06 | 1 | 6 | Pggt1b | 0.043 | Old |
| 0.016392 | 0.565932 | 0.146 | 0.094 | 1 | 6 | Camk2d | 0.052 | Old |
| 0.016477 | 0.961921 | 0.034 | 0.012 | 1 | 6 | Cacna1s | 0.022 | Old |
| 0.01649 | 1.617612 | 0.031 | 0.01 | 1 | 6 | 2810001G2 | 0.021 | Old |
| 0.016748 | 0.8878 | 0.229 | 0.169 | 1 | 6 | Man1a2 | 0.06 | Old |
| 0.016892 | 0.471849 | 0.337 | 0.266 | 1 | 6 | Casp8 | 0.071 | Old |
| 0.01691 | 0.587563 | 0.437 | 0.378 | 1 | 6 | Lgals1 | 0.059 | Old |
| 0.016914 | 0.870583 | 0.097 | 0.055 | 1 | 6 | Gnptg | 0.042 | Old |
| 0.016973 | 0.551717 | 0.089 | 0.048 | 1 | 6 | Psmg3 | 0.041 | Old |
| 0.017019 | 0.767006 | 0.029 | 0.008 | 1 | 6 | Atp23 | 0.021 | Old |
| 0.017143 | 0.60717 | 0.074 | 0.038 | 1 | 6 | Ruvbl2 | 0.036 | Old |
| 0.01722 | 0.275556 | 0.454 | 0.38 | 1 | 6 | Ep300 | 0.074 | Old |
| 0.017315 | 1.161261 | 0.037 | 0.013 | 1 | 6 | Xndc1 | 0.024 | Old |
| 0.017322 | 0.705742 | 0.074 | 0.038 | 1 | 6 | Maip1 | 0.036 | Old |
| 0.017439 | 0.862246 | 0.06 | 0.028 | 1 | 6 | Zfp280c | 0.032 | Old |
| 0.017457 | 0.434757 | 0.191 | 0.134 | 1 | 6 | Hdac9 | 0.057 | Old |
| 0.017608 | 0.607892 | 0.109 | 0.065 | 1 | 6 | Mettl1 | 0.044 | Old |
| 0.017625 | 0.435272 | 0.611 | 0.565 | 1 | 6 | Hif1a | 0.046 | Old |
| 0.017654 | 0.170188 | 0.986 | 0.982 | 1 | 6 | mt-Nd4 | 0.004 | Old |
| 0.017742 | 0.452802 | 0.417 | 0.348 | 1 | 6 | Psmd4 | 0.069 | Old |
| 0.017755 | 0.38269 | 0.234 | 0.166 | 1 | 6 | Epc1 | 0.068 | Old |
| 0.017931 | 0.693887 | 0.123 | 0.075 | 1 | 6 | Hdac7 | 0.048 | Old |
| 0.017978 | 0.806512 | 0.171 | 0.119 | 1 | 6 | Mpnd | 0.052 | Old |
| 0.017999 | 1.840268 | 0.014 | 0.002 | 1 | 6 | Vcan | 0.012 | Old |
| 0.018022 | 1.011907 | 0.089 | 0.05 | 1 | 6 | Adk | 0.039 | Old |
| 0.018068 | 1.025259 | 0.08 | 0.043 | 1 | 6 | Manea | 0.037 | Old |
| 0.018107 | 0.462134 | 0.229 | 0.166 | 1 | 6 | Phf11b | 0.063 | Old |
| 0.018174 | 1.885603 | 0.014 | 0.002 | 1 | 6 | Dnm3 | 0.012 | Old |
| 0.018174 | 1.594711 | 0.014 | 0.002 | 1 | 6 | Gstm2 | 0.012 | Old |
| 0.018174 | 1.588148 | 0.014 | 0.002 | 1 | 6 | Cdh5 | 0.012 | Old |
| 0.018262 | 1.865131 | 0.014 | 0.002 | 1 | 6 | Chst3 | 0.012 | Old |
| 0.018262 | 1.817045 | 0.014 | 0.002 | 1 | 6 | Adgrl2 | 0.012 | Old |
| 0.018262 | 1.761737 | 0.014 | 0.002 | 1 | 6 | Eml5 | 0.012 | Old |
| 0.018262 | 1.381517 | 0.014 | 0.002 | 1 | 6 | Zfyve9 | 0.012 | Old |
| 0.018279 | 1.834872 | 0.069 | 0.035 | 1 | 6 | Zfp369 | 0.034 | Old |
| 0.018317 | 0.438132 | 0.351 | 0.281 | 1 | 6 | Clptm1l | 0.07 | Old |
| 0.01835 | 1.505043 | 0.014 | 0.002 | 1 | 6 | Agbl3 | 0.012 | Old |
| 0.01835 | 1.456936 | 0.014 | 0.002 | 1 | 6 | Rab23 | 0.012 | Old |
| 0.01835 | 1.104798 | 0.014 | 0.002 | 1 | 6 | Rapgef3 | 0.012 | Old |
| 0.018439 | 1.652298 | 0.014 | 0.002 | 1 | 6 | Pax5 | 0.012 | Old |
| 0.018439 | 1.280005 | 0.014 | 0.002 | 1 | 6 | Gpr174 | 0.012 | Old |
| 0.018439 | 0.752163 | 0.014 | 0.002 | 1 | 6 | Ighv11-2 | 0.012 | Old |
| 0.018439 | 0.62028 | 0.014 | 0.002 | 1 | 6 | St14 | 0.012 | Old |
| 0.018439 | 0.300229 | 0.014 | 0.002 | 1 | 6 | Loxl3 | 0.012 | Old |

| | | | | | | | | |
|---|---|---|---|---|---|---|---|---|
| 0.01844 | 1.030116 | 0.117 | 0.074 | 1 | 6 | Tmem154 | 0.043 | Old |
| 0.018462 | 0.686492 | 0.134 | 0.085 | 1 | 6 | Rbm34 | 0.049 | Old |
| 0.018629 | 0.536119 | 0.437 | 0.37 | 1 | 6 | Surf4 | 0.067 | Old |
| 0.018753 | 0.611436 | 0.191 | 0.134 | 1 | 6 | Tmem127 | 0.057 | Old |
| 0.018842 | 0.522783 | 0.497 | 0.458 | 1 | 6 | Ifi27l2a | 0.039 | Old |
| 0.018974 | 0.481252 | 0.294 | 0.227 | 1 | 6 | Zranb2 | 0.067 | Old |
| 0.019152 | 1.106237 | 0.069 | 0.035 | 1 | 6 | Mfsd8 | 0.034 | Old |
| 0.019227 | 0.299466 | 0.637 | 0.595 | 1 | 6 | Uqcr11 | 0.042 | Old |
| 0.019321 | 0.898402 | 0.137 | 0.09 | 1 | 6 | Srp68 | 0.047 | Old |
| 0.019335 | 0.492938 | 0.203 | 0.144 | 1 | 6 | Tcof1 | 0.059 | Old |
| 0.019398 | 0.86557 | 0.206 | 0.147 | 1 | 6 | Osbp | 0.059 | Old |
| 0.01942 | 0.590255 | 0.229 | 0.169 | 1 | 6 | Asxl2 | 0.06 | Old |
| 0.019443 | 0.466028 | 0.343 | 0.271 | 1 | 6 | Alyref | 0.072 | Old |
| 0.019514 | 0.582935 | 0.057 | 0.027 | 1 | 6 | Rmi1 | 0.03 | Old |
| 0.019532 | 0.944125 | 0.069 | 0.035 | 1 | 6 | Spindoc | 0.034 | Old |
| 0.01986 | 0.411102 | 0.277 | 0.216 | 1 | 6 | Ap3s1 | 0.061 | Old |
| 0.01988 | 1.234772 | 0.054 | 0.025 | 1 | 6 | Flad1 | 0.029 | Old |
| 0.020407 | 0.342273 | 0.354 | 0.284 | 1 | 6 | Ppp4c | 0.07 | Old |
| 0.020411 | 0.379578 | 0.426 | 0.356 | 1 | 6 | Magt1 | 0.07 | Old |
| 0.020439 | 1.007025 | 0.066 | 0.033 | 1 | 6 | Fam78b | 0.033 | Old |
| 0.020727 | 0.780542 | 0.077 | 0.042 | 1 | 6 | Ppil3 | 0.035 | Old |
| 0.020737 | 0.497424 | 0.111 | 0.067 | 1 | 6 | Fdps | 0.044 | Old |
| 0.020925 | 0.984894 | 0.066 | 0.033 | 1 | 6 | Atg13 | 0.033 | Old |
| 0.021077 | 1.082207 | 0.066 | 0.033 | 1 | 6 | Exosc7 | 0.033 | Old |
| 0.021321 | 1.212891 | 0.051 | 0.023 | 1 | 6 | Pex1 | 0.028 | Old |
| 0.021322 | 0.552892 | 0.146 | 0.095 | 1 | 6 | Ilvbl | 0.051 | Old |
| 0.021469 | 0.644189 | 0.129 | 0.082 | 1 | 6 | Pno1 | 0.047 | Old |
| 0.021734 | 0.614433 | 0.169 | 0.114 | 1 | 6 | Sirt7 | 0.055 | Old |
| 0.021746 | 0.635884 | 0.317 | 0.254 | 1 | 6 | Ccnt2 | 0.063 | Old |
| 0.021839 | 0.412376 | 0.517 | 0.436 | 1 | 6 | Hnrnpl | 0.081 | Old |
| 0.022057 | 0.472614 | 0.22 | 0.157 | 1 | 6 | Hs6st1 | 0.063 | Old |
| 0.022113 | 0.339907 | 0.471 | 0.401 | 1 | 6 | Tor1aip1 | 0.07 | Old |
| 0.022132 | 1.520562 | 0.049 | 0.022 | 1 | 6 | Igtp | 0.027 | Old |
| 0.022365 | 0.702422 | 0.066 | 0.033 | 1 | 6 | Cdca4 | 0.033 | Old |
| 0.022737 | 1.052608 | 0.074 | 0.04 | 1 | 6 | Gnpnat1 | 0.034 | Old |
| 0.02292 | 0.940227 | 0.051 | 0.023 | 1 | 6 | Rfxank | 0.028 | Old |
| 0.02318 | 0.852463 | 0.146 | 0.099 | 1 | 6 | Mrpl38 | 0.047 | Old |
| 0.023263 | 1.24871 | 0.046 | 0.02 | 1 | 6 | Tmem101 | 0.026 | Old |
| 0.023307 | 0.530486 | 0.129 | 0.082 | 1 | 6 | Eef1e1 | 0.047 | Old |
| 0.023658 | 0.539235 | 0.197 | 0.142 | 1 | 6 | Eif2s1 | 0.055 | Old |
| 0.023864 | 0.520721 | 0.18 | 0.125 | 1 | 6 | Tbc1d23 | 0.055 | Old |
| 0.024019 | 0.232479 | 0.297 | 0.221 | 1 | 6 | Puf60 | 0.076 | Old |
| 0.024419 | 0.917909 | 0.1 | 0.06 | 1 | 6 | Hcfc2 | 0.04 | Old |
| 0.024461 | 1.104327 | 0.046 | 0.02 | 1 | 6 | Rbm12b2 | 0.026 | Old |
| 0.0245 | 0.724178 | 0.049 | 0.022 | 1 | 6 | Pinx1 | 0.027 | Old |

| | | | | | | | | |
|---|---|---|---|---|---|---|---|---|
| 0.024633 | 1.767863 | 0.017 | 0.003 | 1 | 6 | Atp1a2 | 0.014 | Old |
| 0.024663 | 0.27611 | 0.649 | 0.604 | 1 | 6 | Plekho2 | 0.045 | Old |
| 0.024834 | 0.593419 | 0.36 | 0.293 | 1 | 6 | Rexo2 | 0.067 | Old |
| 0.024932 | 2.785688 | 0.017 | 0.003 | 1 | 6 | Snhg18 | 0.014 | Old |
| 0.024932 | 1.364915 | 0.017 | 0.003 | 1 | 6 | Tbc1d30 | 0.014 | Old |
| 0.025163 | 0.47104 | 0.32 | 0.251 | 1 | 6 | Tap2 | 0.069 | Old |
| 0.025335 | 0.584044 | 0.017 | 0.003 | 1 | 6 | Gm3604 | 0.014 | Old |
| 0.025337 | 0.619348 | 0.063 | 0.032 | 1 | 6 | Isg15 | 0.031 | Old |
| 0.025355 | 0.546922 | 0.091 | 0.054 | 1 | 6 | Gm45902 | 0.037 | Old |
| 0.025539 | 0.752827 | 0.017 | 0.003 | 1 | 6 | B3gntl1 | 0.014 | Old |
| 0.025612 | 0.504211 | 0.486 | 0.426 | 1 | 6 | Srsf6 | 0.06 | Old |
| 0.025707 | 0.508278 | 0.38 | 0.321 | 1 | 6 | 1700020I1 | 0.059 | Old |
| 0.025744 | 0.933922 | 0.017 | 0.003 | 1 | 6 | Disp1 | 0.014 | Old |
| 0.025832 | 0.303816 | 0.491 | 0.415 | 1 | 6 | Anxa7 | 0.076 | Old |
| 0.025848 | 0.734326 | 0.143 | 0.095 | 1 | 6 | Polr3k | 0.048 | Old |
| 0.025902 | 0.501427 | 0.103 | 0.062 | 1 | 6 | Pex16 | 0.041 | Old |
| 0.026124 | 0.686712 | 0.14 | 0.094 | 1 | 6 | Pycr2 | 0.046 | Old |
| 0.026233 | 0.596064 | 0.117 | 0.074 | 1 | 6 | Smg6 | 0.043 | Old |
| 0.026374 | 0.363539 | 0.429 | 0.366 | 1 | 6 | Ywhaq | 0.063 | Old |
| 0.026554 | 1.174528 | 0.043 | 0.018 | 1 | 6 | Sft2d3 | 0.025 | Old |
| 0.026668 | 0.690211 | 0.157 | 0.107 | 1 | 6 | Snrnp40 | 0.05 | Old |
| 0.026925 | 0.5684 | 0.174 | 0.12 | 1 | 6 | Clasp1 | 0.054 | Old |
| 0.026972 | 0.837695 | 0.043 | 0.018 | 1 | 6 | Apba3 | 0.025 | Old |
| 0.027024 | 0.491812 | 0.194 | 0.139 | 1 | 6 | Suclg1 | 0.055 | Old |
| 0.027147 | 0.643312 | 0.071 | 0.038 | 1 | 6 | Ddx58 | 0.033 | Old |
| 0.027338 | 0.738274 | 0.12 | 0.077 | 1 | 6 | Mrps18a | 0.043 | Old |
| 0.027479 | 0.8984 | 0.057 | 0.028 | 1 | 6 | Amd2 | 0.029 | Old |
| 0.027647 | 0.620985 | 0.12 | 0.077 | 1 | 6 | Lap3 | 0.043 | Old |
| 0.027673 | 0.707759 | 0.089 | 0.052 | 1 | 6 | Fbxo6 | 0.037 | Old |
| 0.027898 | 1.287929 | 0.069 | 0.037 | 1 | 6 | Asb3 | 0.032 | Old |
| 0.028049 | 1.485881 | 0.057 | 0.028 | 1 | 6 | Gbp4 | 0.029 | Old |
| 0.028419 | 0.284022 | 0.709 | 0.666 | 1 | 6 | Srsf3 | 0.043 | Old |
| 0.028634 | 0.320913 | 0.386 | 0.324 | 1 | 6 | Rbbp6 | 0.062 | Old |
| 0.028684 | 0.859027 | 0.057 | 0.028 | 1 | 6 | Scyl3 | 0.029 | Old |
| 0.02869 | 2.177152 | 0.02 | 0.005 | 1 | 6 | Cd200 | 0.015 | Old |
| 0.028709 | 1.175049 | 0.037 | 0.015 | 1 | 6 | Gkap1 | 0.022 | Old |
| 0.028893 | 1.550732 | 0.02 | 0.005 | 1 | 6 | Zrsr1 | 0.015 | Old |
| 0.028934 | 0.284136 | 0.686 | 0.625 | 1 | 6 | Pdia3 | 0.061 | Old |
| 0.02906 | 2.181057 | 0.034 | 0.013 | 1 | 6 | Otud7b | 0.021 | Old |
| 0.029065 | 0.487366 | 0.283 | 0.221 | 1 | 6 | Atp8b4 | 0.062 | Old |
| 0.02908 | 1.709612 | 0.051 | 0.025 | 1 | 6 | Eif2b1 | 0.026 | Old |
| 0.029098 | 2.301829 | 0.02 | 0.005 | 1 | 6 | Dusp14 | 0.015 | Old |
| 0.029099 | 0.461274 | 0.454 | 0.4 | 1 | 6 | Ikzf1 | 0.054 | Old |
| 0.029404 | 1.100576 | 0.037 | 0.015 | 1 | 6 | Mtg2 | 0.022 | Old |
| 0.029406 | 1.456787 | 0.02 | 0.005 | 1 | 6 | Zfp169 | 0.015 | Old |

| | | | | | | | | |
|---|---|---|---|---|---|---|---|---|
| 0.029457 | 0.663018 | 0.103 | 0.064 | 1 | 6 | Tmem161b | 0.039 | Old |
| 0.029573 | 1.359312 | 0.034 | 0.013 | 1 | 6 | Gbp8 | 0.021 | Old |
| 0.029614 | 1.422982 | 0.02 | 0.005 | 1 | 6 | Plxna3 | 0.015 | Old |
| 0.029647 | 1.740067 | 0.034 | 0.013 | 1 | 6 | Chchd10 | 0.021 | Old |
| 0.029686 | 0.888558 | 0.037 | 0.015 | 1 | 6 | Tepsin | 0.022 | Old |
| 0.030033 | 0.947607 | 0.02 | 0.005 | 1 | 6 | Zkscan7 | 0.015 | Old |
| 0.030062 | 0.139654 | 0.974 | 0.985 | 1 | 6 | mt-Nd2 | -0.011 | Old |
| 0.030066 | 0.826818 | 0.094 | 0.057 | 1 | 6 | Ptcd2 | 0.037 | Old |
| 0.030184 | 1.040833 | 0.037 | 0.015 | 1 | 6 | Grpel2 | 0.022 | Old |
| 0.030363 | 0.502059 | 0.371 | 0.311 | 1 | 6 | Emg1 | 0.06 | Old |
| 0.030471 | 1.497472 | 0.034 | 0.013 | 1 | 6 | 1-Sep | 0.021 | Old |
| 0.030477 | 0.801699 | 0.12 | 0.079 | 1 | 6 | Ccdc69 | 0.041 | Old |
| 0.030526 | 0.4155 | 0.12 | 0.077 | 1 | 6 | Gng12 | 0.043 | Old |
| 0.03069 | 0.745322 | 0.037 | 0.015 | 1 | 6 | Fpgs | 0.022 | Old |
| 0.030852 | 1.500505 | 0.034 | 0.013 | 1 | 6 | Pced1b | 0.021 | Old |
| 0.030897 | 1.042804 | 0.066 | 0.035 | 1 | 6 | Fam149b | 0.031 | Old |
| 0.030929 | 0.983159 | 0.034 | 0.013 | 1 | 6 | Nmnat3 | 0.021 | Old |
| 0.030958 | 1.241453 | 0.109 | 0.069 | 1 | 6 | Bcl2 | 0.04 | Old |
| 0.031009 | 0.395547 | 0.149 | 0.1 | 1 | 6 | Rnf139 | 0.049 | Old |
| 0.03111 | 0.760634 | 0.094 | 0.057 | 1 | 6 | Abhd17c | 0.037 | Old |
| 0.031136 | 0.229179 | 0.8 | 0.789 | 1 | 6 | Cd53 | 0.011 | Old |
| 0.031284 | 1.659011 | 0.023 | 0.007 | 1 | 6 | Xlr4a | 0.016 | Old |
| 0.03136 | 0.168107 | 0.649 | 0.594 | 1 | 6 | Arpc4 | 0.055 | Old |
| 0.031383 | 1.675553 | 0.023 | 0.007 | 1 | 6 | Timp3 | 0.016 | Old |
| 0.031462 | 1.294156 | 0.031 | 0.012 | 1 | 6 | Aldh7a1 | 0.019 | Old |
| 0.031484 | 1.748839 | 0.023 | 0.007 | 1 | 6 | Krt7 | 0.016 | Old |
| 0.031484 | 1.375913 | 0.023 | 0.007 | 1 | 6 | Armc3 | 0.016 | Old |
| 0.031584 | 1.668499 | 0.023 | 0.007 | 1 | 6 | Serping1 | 0.016 | Old |
| 0.031658 | 2.162843 | 0.029 | 0.01 | 1 | 6 | Xcr1 | 0.019 | Old |
| 0.031694 | 0.433538 | 0.057 | 0.028 | 1 | 6 | Snrnp35 | 0.029 | Old |
| 0.031752 | 0.068606 | 0.954 | 0.995 | 1 | 6 | Psap | -0.041 | Old |
| 0.031794 | 1.77873 | 0.026 | 0.008 | 1 | 6 | Prnp | 0.018 | Old |
| 0.03181 | 1.236973 | 0.051 | 0.025 | 1 | 6 | Gm42047 | 0.026 | Old |
| 0.031862 | 1.049393 | 0.034 | 0.013 | 1 | 6 | Etaa1 | 0.021 | Old |
| 0.03196 | 0.392151 | 0.1 | 0.06 | 1 | 6 | Gtpbp1 | 0.04 | Old |
| 0.031982 | 1.774295 | 0.026 | 0.008 | 1 | 6 | Inpp1 | 0.018 | Old |
| 0.031988 | 1.48893 | 0.023 | 0.007 | 1 | 6 | Tulp3 | 0.016 | Old |
| 0.032068 | 0.791223 | 0.066 | 0.035 | 1 | 6 | Tfdp2 | 0.031 | Old |
| 0.032192 | 1.057474 | 0.023 | 0.007 | 1 | 6 | Ada | 0.016 | Old |
| 0.032238 | 0.541533 | 0.16 | 0.112 | 1 | 6 | Pex6 | 0.048 | Old |
| 0.032284 | 0.329396 | 0.517 | 0.45 | 1 | 6 | Psmd13 | 0.067 | Old |
| 0.032306 | 1.145154 | 0.051 | 0.025 | 1 | 6 | Taf6 | 0.026 | Old |
| 0.032647 | 1.669153 | 0.026 | 0.008 | 1 | 6 | Pclaf | 0.018 | Old |
| 0.032743 | 1.485674 | 0.026 | 0.008 | 1 | 6 | Cep95 | 0.018 | Old |
| 0.032743 | 1.234604 | 0.026 | 0.008 | 1 | 6 | Slc16a13 | 0.018 | Old |

| | | | | | | | | |
|---|---|---|---|---|---|---|---|---|
| 0.032809 | 0.701892 | 0.051 | 0.025 | 1 | 6 | Rpap3 | 0.026 | Old |
| 0.032839 | 0.936274 | 0.026 | 0.008 | 1 | 6 | Smug1 | 0.018 | Old |
| 0.032901 | 0.399978 | 0.029 | 0.01 | 1 | 6 | Trpm2 | 0.019 | Old |
| 0.032935 | 0.404641 | 0.026 | 0.008 | 1 | 6 | Gm8113 | 0.018 | Old |
| 0.033109 | 0.301877 | 0.577 | 0.525 | 1 | 6 | M6pr | 0.052 | Old |
| 0.033255 | 0.796019 | 0.051 | 0.025 | 1 | 6 | Gatb | 0.026 | Old |
| 0.03343 | 0.712857 | 0.126 | 0.084 | 1 | 6 | Dkc1 | 0.042 | Old |
| 0.033466 | 1.017648 | 0.074 | 0.042 | 1 | 6 | Itpr3 | 0.032 | Old |
| 0.033533 | 1.31227 | 0.049 | 0.023 | 1 | 6 | Zfp994 | 0.026 | Old |
| 0.033731 | 0.497201 | 0.303 | 0.241 | 1 | 6 | Nrd1 | 0.062 | Old |
| 0.033945 | 0.321654 | 0.651 | 0.605 | 1 | 6 | Arl6ip1 | 0.046 | Old |
| 0.03402 | 0.776944 | 0.134 | 0.09 | 1 | 6 | Gm5547 | 0.044 | Old |
| 0.034181 | 0.768776 | 0.086 | 0.05 | 1 | 6 | Tnfrsf13c | 0.036 | Old |
| 0.034332 | 1.03611 | 0.106 | 0.067 | 1 | 6 | Fam120b | 0.039 | Old |
| 0.034359 | 0.348168 | 0.614 | 0.58 | 1 | 6 | Egr1 | 0.034 | Old |
| 0.034615 | 1.299641 | 0.06 | 0.032 | 1 | 6 | Zfp146 | 0.028 | Old |
| 0.034877 | 0.421191 | 0.086 | 0.05 | 1 | 6 | Lztr1 | 0.036 | Old |
| 0.034902 | 0.520196 | 0.154 | 0.105 | 1 | 6 | Ncbp2 | 0.049 | Old |
| 0.035966 | 1.164619 | 0.06 | 0.032 | 1 | 6 | Sec24d | 0.028 | Old |
| 0.035995 | 0.554687 | 0.317 | 0.253 | 1 | 6 | Mgea5 | 0.064 | Old |
| 0.036197 | 0.19866 | 0.4 | 0.326 | 1 | 6 | Snrpd3 | 0.074 | Old |
| 0.036202 | 0.48329 | 0.151 | 0.104 | 1 | 6 | Phf12 | 0.047 | Old |
| 0.036266 | 0.977743 | 0.094 | 0.059 | 1 | 6 | Dag1 | 0.035 | Old |
| 0.03628 | 1.076518 | 0.06 | 0.032 | 1 | 6 | Adnp2 | 0.028 | Old |
| 0.036344 | 0.438542 | 0.129 | 0.085 | 1 | 6 | Ppp2r3d | 0.044 | Old |
| 0.036621 | 0.720446 | 0.203 | 0.152 | 1 | 6 | Lman1 | 0.051 | Old |
| 0.036628 | 0.2618 | 0.349 | 0.286 | 1 | 6 | Akt2 | 0.063 | Old |
| 0.036756 | 0.127338 | 0.963 | 0.96 | 1 | 6 | Ptma | 0.003 | Old |
| 0.036871 | 1.060152 | 0.086 | 0.052 | 1 | 6 | Mrps18b | 0.034 | Old |
| 0.036962 | 0.659412 | 0.111 | 0.072 | 1 | 6 | Hhex | 0.039 | Old |
| 0.0371 | 0.706899 | 0.083 | 0.048 | 1 | 6 | Slc33a1 | 0.035 | Old |
| 0.037223 | 1.48905 | 0.057 | 0.03 | 1 | 6 | Pwp1 | 0.027 | Old |
| 0.038017 | 0.323314 | 0.14 | 0.095 | 1 | 6 | Cul2 | 0.045 | Old |
| 0.038114 | 0.598947 | 0.194 | 0.142 | 1 | 6 | Cpeb4 | 0.052 | Old |
| 0.038132 | 1.160686 | 0.046 | 0.022 | 1 | 6 | Gem | 0.024 | Old |
| 0.038188 | 0.456088 | 0.106 | 0.067 | 1 | 6 | Pnkd | 0.039 | Old |
| 0.038316 | 0.778371 | 0.08 | 0.047 | 1 | 6 | Ddx55 | 0.033 | Old |
| 0.038428 | 0.379227 | 0.134 | 0.09 | 1 | 6 | Lrrc41 | 0.044 | Old |
| 0.038439 | 1.141033 | 0.046 | 0.022 | 1 | 6 | A530040E1 | 0.024 | Old |
| 0.038628 | 1.140202 | 0.057 | 0.03 | 1 | 6 | Qtrt1 | 0.027 | Old |
| 0.038698 | 0.355332 | 0.649 | 0.627 | 1 | 6 | Pdia6 | 0.022 | Old |
| 0.038774 | 0.370014 | 0.374 | 0.314 | 1 | 6 | Vapa | 0.06 | Old |
| 0.038953 | 0.697878 | 0.126 | 0.085 | 1 | 6 | Acadvl | 0.041 | Old |
| 0.03917 | 0.345606 | 0.449 | 0.401 | 1 | 6 | Trir | 0.048 | Old |
| 0.039235 | 0.276635 | 0.711 | 0.651 | 1 | 6 | Serbp1 | 0.06 | Old |

| | | | | | | | | |
|---|---|---|---|---|---|---|---|---|
| 0.03945 | 0.620997 | 0.534 | 0.477 | 1 | 6 | Fgl2 | 0.057 | Old |
| 0.039475 | 1.220699 | 0.06 | 0.032 | 1 | 6 | Mrgbp | 0.028 | Old |
| 0.039612 | 0.423686 | 0.323 | 0.264 | 1 | 6 | Leprotl1 | 0.059 | Old |
| 0.039768 | 1.491454 | 0.043 | 0.02 | 1 | 6 | Impa2 | 0.023 | Old |
| 0.039788 | 0.335944 | 0.131 | 0.087 | 1 | 6 | Tubb4b | 0.044 | Old |
| 0.03993 | 0.225352 | 0.457 | 0.39 | 1 | 6 | Pigt | 0.067 | Old |
| 0.040015 | 1.27194 | 0.043 | 0.02 | 1 | 6 | 0610009B2 | 0.023 | Old |
| 0.040503 | 0.651373 | 0.157 | 0.11 | 1 | 6 | Dido1 | 0.047 | Old |
| 0.040584 | 0.491956 | 0.123 | 0.08 | 1 | 6 | Vac14 | 0.043 | Old |
| 0.040604 | 0.85763 | 0.157 | 0.115 | 1 | 6 | Ccdc82 | 0.042 | Old |
| 0.040849 | 1.06942 | 0.043 | 0.02 | 1 | 6 | Usp46 | 0.023 | Old |
| 0.04093 | 0.425663 | 0.346 | 0.281 | 1 | 6 | Uhmk1 | 0.065 | Old |
| 0.041186 | 1.196953 | 0.043 | 0.02 | 1 | 6 | Tmem86b | 0.023 | Old |
| 0.04127 | 0.639722 | 0.077 | 0.045 | 1 | 6 | Wdr75 | 0.032 | Old |
| 0.041459 | 2.279277 | 0.04 | 0.018 | 1 | 6 | Tasp1 | 0.022 | Old |
| 0.041496 | 0.383706 | 0.203 | 0.151 | 1 | 6 | Inpp5k | 0.052 | Old |
| 0.041526 | 1.246074 | 0.043 | 0.02 | 1 | 6 | Nup107 | 0.023 | Old |
| 0.041538 | 0.803258 | 0.14 | 0.097 | 1 | 6 | Tnfrsf18 | 0.043 | Old |
| 0.041548 | 2.178273 | 0.04 | 0.018 | 1 | 6 | Wipi1 | 0.022 | Old |
| 0.041865 | 0.901275 | 0.057 | 0.03 | 1 | 6 | Dusp12 | 0.027 | Old |
| 0.042234 | 0.541533 | 0.22 | 0.169 | 1 | 6 | Mgat2 | 0.051 | Old |
| 0.042386 | 0.560625 | 0.043 | 0.02 | 1 | 6 | Pank1 | 0.023 | Old |
| 0.042426 | 0.299151 | 0.494 | 0.423 | 1 | 6 | P4hb | 0.071 | Old |
| 0.042477 | 0.599562 | 0.114 | 0.075 | 1 | 6 | Tcp11l1 | 0.039 | Old |
| 0.042716 | 0.426574 | 0.226 | 0.171 | 1 | 6 | Taz | 0.055 | Old |
| 0.042724 | 0.668763 | 0.069 | 0.038 | 1 | 6 | Ogfod1 | 0.031 | Old |
| 0.042935 | 0.488832 | 0.209 | 0.156 | 1 | 6 | Slc50a1 | 0.053 | Old |
| 0.043235 | 0.345614 | 0.406 | 0.336 | 1 | 6 | Tmco1 | 0.07 | Old |
| 0.043364 | 0.51752 | 0.069 | 0.038 | 1 | 6 | Bloc1s4 | 0.031 | Old |
| 0.04372 | 1.499977 | 0.04 | 0.018 | 1 | 6 | Gm46430 | 0.022 | Old |
| 0.043766 | 0.269121 | 0.58 | 0.543 | 1 | 6 | Nme1 | 0.037 | Old |
| 0.043883 | 0.944798 | 0.043 | 0.02 | 1 | 6 | Rpgrip1 | 0.023 | Old |
| 0.044064 | 1.113095 | 0.091 | 0.059 | 1 | 6 | Tjp2 | 0.032 | Old |
| 0.044157 | 0.476797 | 0.149 | 0.104 | 1 | 6 | Ifi35 | 0.045 | Old |
| 0.044176 | 0.582438 | 0.097 | 0.062 | 1 | 6 | 2610002M | 0.035 | Old |
| 0.044216 | 0.192908 | 0.303 | 0.237 | 1 | 6 | Copb2 | 0.066 | Old |
| 0.044471 | 0.20534 | 0.454 | 0.378 | 1 | 6 | Tcp1 | 0.076 | Old |
| 0.04482 | 0.473299 | 0.066 | 0.037 | 1 | 6 | Exosc2 | 0.029 | Old |
| 0.044896 | 0.322845 | 0.503 | 0.435 | 1 | 6 | Twf2 | 0.068 | Old |
| 0.045148 | 2.054565 | 0.011 | 0.002 | 1 | 6 | Cobll1 | 0.009 | Old |
| 0.045148 | 2.000023 | 0.011 | 0.002 | 1 | 6 | Flrt2 | 0.009 | Old |
| 0.045148 | 1.432296 | 0.011 | 0.002 | 1 | 6 | Sox5 | 0.009 | Old |
| 0.045148 | 1.390884 | 0.011 | 0.002 | 1 | 6 | Wdr54 | 0.009 | Old |
| 0.045196 | 0.297435 | 0.34 | 0.271 | 1 | 6 | Supt5 | 0.069 | Old |
| 0.045217 | 0.478626 | 0.329 | 0.274 | 1 | 6 | Maz | 0.055 | Old |

| | | | | | | | | |
|---|---|---|---|---|---|---|---|---|
| 0.04522 | 1.291416 | 0.04 | 0.018 | 1 | 6 | Cenpt | 0.022 | Old |
| 0.045358 | 1.466066 | 0.011 | 0.002 | 1 | 6 | 4931414P1 | 0.009 | Old |
| 0.045358 | 0.905906 | 0.011 | 0.002 | 1 | 6 | Zscan20 | 0.009 | Old |
| 0.04557 | 1.973192 | 0.011 | 0.002 | 1 | 6 | Hsd11b1 | 0.009 | Old |
| 0.04557 | 1.964699 | 0.011 | 0.002 | 1 | 6 | Cspg5 | 0.009 | Old |
| 0.04557 | 1.737089 | 0.011 | 0.002 | 1 | 6 | Slc22a23 | 0.009 | Old |
| 0.04557 | 1.712545 | 0.011 | 0.002 | 1 | 6 | Fbln1 | 0.009 | Old |
| 0.04557 | 1.251088 | 0.011 | 0.002 | 1 | 6 | 1700048O2 | 0.009 | Old |
| 0.04557 | 1.23175 | 0.011 | 0.002 | 1 | 6 | Gm7160 | 0.009 | Old |
| 0.045616 | 0.210244 | 0.991 | 0.995 | 1 | 6 | Malat1 | -0.004 | Old |
| 0.045782 | 1.713927 | 0.011 | 0.002 | 1 | 6 | Igkv14-126 | 0.009 | Old |
| 0.045782 | 1.658197 | 0.011 | 0.002 | 1 | 6 | Prr5 | 0.009 | Old |
| 0.045782 | 1.289464 | 0.011 | 0.002 | 1 | 6 | Ighg3 | 0.009 | Old |
| 0.045782 | 1.06477 | 0.011 | 0.002 | 1 | 6 | Rab19 | 0.009 | Old |
| 0.045782 | 0.676217 | 0.011 | 0.002 | 1 | 6 | Ttc41 | 0.009 | Old |
| 0.045814 | 0.909732 | 0.063 | 0.035 | 1 | 6 | Espl1 | 0.028 | Old |
| 0.045995 | 1.019833 | 0.011 | 0.002 | 1 | 6 | Zfp784 | 0.009 | Old |
| 0.045995 | 0.97956 | 0.011 | 0.002 | 1 | 6 | Lmbr1 | 0.009 | Old |
| 0.045995 | 0.898242 | 0.011 | 0.002 | 1 | 6 | Zfp229 | 0.009 | Old |
| 0.045995 | 0.691789 | 0.011 | 0.002 | 1 | 6 | Mtmr11 | 0.009 | Old |
| 0.046039 | 0.644936 | 0.177 | 0.132 | 1 | 6 | Cdk5rap3 | 0.045 | Old |
| 0.046378 | 0.760223 | 0.071 | 0.042 | 1 | 6 | Usp35 | 0.029 | Old |
| 0.04671 | 1.04043 | 0.063 | 0.035 | 1 | 6 | Trit1 | 0.028 | Old |
| 0.046833 | 0.611029 | 0.054 | 0.028 | 1 | 6 | Alad | 0.026 | Old |
| 0.046915 | 0.573042 | 0.126 | 0.084 | 1 | 6 | Gmppa | 0.042 | Old |
| 0.047299 | 0.446411 | 0.1 | 0.064 | 1 | 6 | Slc30a1 | 0.036 | Old |
| 0.047511 | 0.298239 | 0.22 | 0.164 | 1 | 6 | Fndc3b | 0.056 | Old |
| 0.047668 | 0.288562 | 0.24 | 0.186 | 1 | 6 | Polr2f | 0.054 | Old |
| 0.047724 | 0.76262 | 0.209 | 0.161 | 1 | 6 | Dip2b | 0.048 | Old |
| 0.048275 | 0.900379 | 0.109 | 0.072 | 1 | 6 | Trap1 | 0.037 | Old |
| 0.048546 | 0.856687 | 0.063 | 0.035 | 1 | 6 | Patz1 | 0.028 | Old |
| 0.048574 | 0.758606 | 0.089 | 0.055 | 1 | 6 | Tmem60 | 0.034 | Old |
| 0.04894 | 1.519985 | 0.06 | 0.033 | 1 | 6 | Psd3 | 0.027 | Old |
| 0.049009 | 0.953308 | 0.037 | 0.017 | 1 | 6 | Abca2 | 0.02 | Old |
| 0.049883 | 0.565537 | 0.063 | 0.035 | 1 | 6 | Fam122b | 0.028 | Old |
| 0.049925 | 0.225768 | 0.606 | 0.55 | 1 | 6 | Shisa5 | 0.056 | Old |
| 0.050212 | 0.256036 | 0.569 | 0.497 | 1 | 6 | Selenot | 0.072 | Old |
| 0.050281 | 0.687203 | 0.211 | 0.162 | 1 | 6 | Phf8 | 0.049 | Old |
| 0.050335 | 0.66136 | 0.12 | 0.082 | 1 | 6 | Tubgcp4 | 0.038 | Old |
| 0.050603 | 0.80902 | 0.063 | 0.035 | 1 | 6 | Cdkal1 | 0.028 | Old |
| 0.051007 | 0.875689 | 0.063 | 0.035 | 1 | 6 | Zfp830 | 0.028 | Old |
| 0.051133 | 0.924589 | 0.131 | 0.09 | 1 | 6 | Tmem41b | 0.041 | Old |
| 0.051142 | 0.947281 | 0.06 | 0.033 | 1 | 6 | Dgcr6 | 0.027 | Old |
| 0.051475 | 0.59564 | 0.051 | 0.027 | 1 | 6 | Plpp5 | 0.024 | Old |
| 0.051499 | 0.071526 | 0.037 | 0.017 | 1 | 6 | Tbcc | 0.02 | Old |

| | | | | | | | | |
|---|---|---|---|---|---|---|---|---|
| 0.051724 | 0.382043 | 0.243 | 0.187 | 1 | 6 | Ube2j2 | 0.056 | Old |
| 0.051813 | 0.479127 | 0.134 | 0.092 | 1 | 6 | Jkamp | 0.042 | Old |
| 0.05196 | 0.692924 | 0.151 | 0.11 | 1 | 6 | Parp4 | 0.041 | Old |
| 0.052077 | 0.544379 | 0.109 | 0.072 | 1 | 6 | Mapk9 | 0.037 | Old |
| 0.052109 | 1.032708 | 0.034 | 0.015 | 1 | 6 | Sars2 | 0.019 | Old |
| 0.052167 | 1.004104 | 0.083 | 0.052 | 1 | 6 | Arhgap21 | 0.031 | Old |
| 0.052389 | 0.250122 | 0.037 | 0.017 | 1 | 6 | Tdrkh | 0.02 | Old |
| 0.052412 | 0.843973 | 0.16 | 0.117 | 1 | 6 | Unc50 | 0.043 | Old |
| 0.052441 | 0.410614 | 0.383 | 0.326 | 1 | 6 | Bak1 | 0.057 | Old |
| 0.052831 | 0.44928 | 0.254 | 0.199 | 1 | 6 | Cacybp | 0.055 | Old |
| 0.052906 | 0.479638 | 0.186 | 0.135 | 1 | 6 | Tor1aip2 | 0.051 | Old |
| 0.053061 | 0.576935 | 0.374 | 0.321 | 1 | 6 | Strn3 | 0.053 | Old |
| 0.053153 | 0.550522 | 0.151 | 0.109 | 1 | 6 | Amdhd2 | 0.042 | Old |
| 0.053167 | 0.896601 | 0.034 | 0.015 | 1 | 6 | Kctd14 | 0.019 | Old |
| 0.053418 | 0.41104 | 0.117 | 0.079 | 1 | 6 | Copg2 | 0.038 | Old |
| 0.053522 | 0.686218 | 0.137 | 0.097 | 1 | 6 | Keap1 | 0.04 | Old |
| 0.053526 | 0.691582 | 0.189 | 0.142 | 1 | 6 | Nabp2 | 0.047 | Old |
| 0.053537 | 0.740244 | 0.129 | 0.089 | 1 | 6 | Dpm1 | 0.04 | Old |
| 0.05389 | 0.329091 | 0.466 | 0.411 | 1 | 6 | Ranbp2 | 0.055 | Old |
| 0.053974 | 0.549304 | 0.231 | 0.181 | 1 | 6 | Lin7c | 0.05 | Old |
| 0.054061 | 0.492266 | 0.069 | 0.04 | 1 | 6 | Trim34a | 0.029 | Old |
| 0.05427 | 0.414299 | 0.057 | 0.032 | 1 | 6 | Iqce | 0.025 | Old |
| 0.054326 | 0.431146 | 0.471 | 0.421 | 1 | 6 | Csk | 0.05 | Old |
| 0.05457 | 1.303167 | 0.031 | 0.013 | 1 | 6 | Mios | 0.018 | Old |
| 0.054575 | 0.609895 | 0.154 | 0.112 | 1 | 6 | Atg16l1 | 0.042 | Old |
| 0.05471 | 0.521561 | 0.069 | 0.04 | 1 | 6 | Scfd2 | 0.029 | Old |
| 0.055029 | 0.686993 | 0.106 | 0.07 | 1 | 6 | Ddx56 | 0.036 | Old |
| 0.055414 | 0.278327 | 0.274 | 0.217 | 1 | 6 | Snd1 | 0.057 | Old |
| 0.055886 | 0.302385 | 0.671 | 0.624 | 1 | 6 | Hnrnpab | 0.047 | Old |
| 0.055926 | 2.467479 | 0.046 | 0.023 | 1 | 6 | C3 | 0.023 | Old |
| 0.056452 | 0.985815 | 0.057 | 0.032 | 1 | 6 | Wdr81 | 0.025 | Old |
| 0.056648 | 1.951231 | 0.029 | 0.012 | 1 | 6 | Actn1 | 0.017 | Old |
| 0.056731 | 1.927772 | 0.014 | 0.003 | 1 | 6 | Mirt1 | 0.011 | Old |
| 0.056759 | 0.457711 | 0.217 | 0.274 | 1 | 6 | Cxcl2 | -0.057 | Old |
| 0.056925 | 0.662628 | 0.151 | 0.109 | 1 | 6 | Ccdc22 | 0.042 | Old |
| 0.057163 | 2.543294 | 0.014 | 0.003 | 1 | 6 | Rflnb | 0.011 | Old |
| 0.057163 | 1.848271 | 0.014 | 0.003 | 1 | 6 | Zfp12 | 0.011 | Old |
| 0.057163 | 1.658268 | 0.014 | 0.003 | 1 | 6 | Apod | 0.011 | Old |
| 0.057256 | 0.786923 | 0.103 | 0.069 | 1 | 6 | Mrps30 | 0.034 | Old |
| 0.057292 | 0.32553 | 0.323 | 0.266 | 1 | 6 | Fkbp2 | 0.057 | Old |
| 0.05738 | 2.904119 | 0.014 | 0.003 | 1 | 6 | Ly6c1 | 0.011 | Old |
| 0.05738 | 1.97043 | 0.014 | 0.003 | 1 | 6 | Zhx2 | 0.011 | Old |
| 0.05738 | 1.910757 | 0.014 | 0.003 | 1 | 6 | Bub1b | 0.011 | Old |
| 0.05738 | 1.611005 | 0.014 | 0.003 | 1 | 6 | Zfp975 | 0.011 | Old |
| 0.057565 | 0.614006 | 0.031 | 0.013 | 1 | 6 | Hvcn1 | 0.018 | Old |

| | | | | | | | | |
|---|---|---|---|---|---|---|---|---|
| 0.057584 | 0.287329 | 0.174 | 0.129 | 1 | 6 | Ylpm1 | 0.045 | Old |
| 0.057597 | 2.222508 | 0.014 | 0.003 | 1 | 6 | Cystm1 | 0.011 | Old |
| 0.057597 | 1.102567 | 0.014 | 0.003 | 1 | 6 | Arl16 | 0.011 | Old |
| 0.057611 | 1.724005 | 0.046 | 0.023 | 1 | 6 | Gm26870 | 0.023 | Old |
| 0.057626 | 2.003637 | 0.029 | 0.012 | 1 | 6 | Bicd1 | 0.017 | Old |
| 0.057816 | 1.895949 | 0.014 | 0.003 | 1 | 6 | Ankrd55 | 0.011 | Old |
| 0.057816 | 1.3871 | 0.014 | 0.003 | 1 | 6 | Gm17552 | 0.011 | Old |
| 0.057816 | 0.301499 | 0.617 | 0.589 | 1 | 6 | Colgalt1 | 0.028 | Old |
| 0.057908 | 1.162399 | 0.029 | 0.012 | 1 | 6 | Dhrs11 | 0.017 | Old |
| 0.05792 | 0.496343 | 0.131 | 0.092 | 1 | 6 | Ap1m1 | 0.039 | Old |
| 0.057982 | 0.513804 | 0.357 | 0.301 | 1 | 6 | Psmc3 | 0.056 | Old |
| 0.057983 | 0.794954 | 0.123 | 0.085 | 1 | 6 | Nifk | 0.038 | Old |
| 0.058 | 0.296641 | 0.049 | 0.025 | 1 | 6 | Nfatc2ip | 0.024 | Old |
| 0.058035 | 1.938929 | 0.014 | 0.003 | 1 | 6 | Coq8a | 0.011 | Old |
| 0.058035 | 1.429307 | 0.014 | 0.003 | 1 | 6 | Olfr920 | 0.011 | Old |
| 0.058035 | 1.293931 | 0.014 | 0.003 | 1 | 6 | Rab30 | 0.011 | Old |
| 0.058035 | 0.848869 | 0.014 | 0.003 | 1 | 6 | Etl4 | 0.011 | Old |
| 0.058192 | 0.356212 | 0.254 | 0.196 | 1 | 6 | Yipf3 | 0.058 | Old |
| 0.058307 | 0.676809 | 0.066 | 0.038 | 1 | 6 | Prpf4 | 0.028 | Old |
| 0.058332 | 0.456867 | 0.346 | 0.289 | 1 | 6 | Polr2k | 0.057 | Old |
| 0.058389 | 0.593123 | 0.1 | 0.065 | 1 | 6 | Vps45 | 0.035 | Old |
| 0.058475 | 0.88835 | 0.014 | 0.003 | 1 | 6 | Mcts2 | 0.011 | Old |
| 0.058475 | 0.55048 | 0.014 | 0.003 | 1 | 6 | Prlr | 0.011 | Old |
| 0.058727 | 0.676089 | 0.246 | 0.194 | 1 | 6 | Pde7a | 0.052 | Old |
| 0.0588 | 0.875513 | 0.08 | 0.05 | 1 | 6 | Tmem106c | 0.03 | Old |
| 0.058802 | 0.846812 | 0.057 | 0.032 | 1 | 6 | Znhit2 | 0.025 | Old |
| 0.058897 | 0.687681 | 0.154 | 0.112 | 1 | 6 | Cdk2 | 0.042 | Old |
| 0.058979 | 0.524741 | 0.206 | 0.159 | 1 | 6 | Cpsf7 | 0.047 | Old |
| 0.059047 | 1.527043 | 0.029 | 0.012 | 1 | 6 | Ccne1 | 0.017 | Old |
| 0.059185 | 0.600444 | 0.057 | 0.032 | 1 | 6 | Fam136a | 0.025 | Old |
| 0.05923 | 0.649521 | 0.306 | 0.258 | 1 | 6 | 6-Mar | 0.048 | Old |
| 0.059337 | 0.476715 | 0.091 | 0.059 | 1 | 6 | Prdm4 | 0.032 | Old |
| 0.059375 | 0.378042 | 0.374 | 0.318 | 1 | 6 | Lfng | 0.056 | Old |
| 0.059558 | 0.783413 | 0.046 | 0.023 | 1 | 6 | Ccdc163 | 0.023 | Old |
| 0.059624 | 1.150145 | 0.029 | 0.012 | 1 | 6 | Bod1 | 0.017 | Old |
| 0.059789 | 0.884195 | 0.083 | 0.052 | 1 | 6 | Fam53b | 0.031 | Old |
| 0.059837 | 0.316034 | 0.323 | 0.263 | 1 | 6 | Myo5a | 0.06 | Old |
| 0.060135 | 1.361632 | 0.043 | 0.022 | 1 | 6 | Zfp157 | 0.021 | Old |
| 0.060279 | 0.651794 | 0.089 | 0.057 | 1 | 6 | P2ry10b | 0.032 | Old |
| 0.060352 | 0.387203 | 0.029 | 0.012 | 1 | 6 | Mcat | 0.017 | Old |
| 0.060466 | 0.329997 | 0.26 | 0.207 | 1 | 6 | Fam32a | 0.053 | Old |
| 0.060544 | 0.701901 | 0.057 | 0.032 | 1 | 6 | Epg5 | 0.025 | Old |
| 0.060818 | 0.126859 | 0.14 | 0.097 | 1 | 6 | Pigm | 0.043 | Old |
| 0.060844 | 0.351169 | 0.166 | 0.122 | 1 | 6 | Tsen34 | 0.044 | Old |
| 0.060874 | 0.474957 | 0.166 | 0.12 | 1 | 6 | Rpe | 0.046 | Old |

| | | | | | | | | |
|---|---|---|---|---|---|---|---|---|
| 0.060941 | 0.32451 | 0.157 | 0.114 | 1 | 6 | Fnbp4 | 0.043 | Old |
| 0.061199 | 0.240122 | 0.114 | 0.077 | 1 | 6 | Zscan26 | 0.037 | Old |
| 0.06136 | 0.643501 | 0.089 | 0.057 | 1 | 6 | Kdm5d | 0.032 | Old |
| 0.061421 | 0.697899 | 0.12 | 0.084 | 1 | 6 | Haus8 | 0.036 | Old |
| 0.061532 | 0.684987 | 0.109 | 0.074 | 1 | 6 | Nop9 | 0.035 | Old |
| 0.061588 | 0.945254 | 0.18 | 0.142 | 1 | 6 | Casp8ap2 | 0.038 | Old |
| 0.061589 | 1.072028 | 0.117 | 0.082 | 1 | 6 | Ppt2 | 0.035 | Old |
| 0.061704 | 0.36542 | 0.231 | 0.179 | 1 | 6 | Hspa14 | 0.052 | Old |
| 0.061745 | 1.027855 | 0.063 | 0.037 | 1 | 6 | Sgf29 | 0.026 | Old |
| 0.061853 | 0.888836 | 0.094 | 0.062 | 1 | 6 | Hmces | 0.032 | Old |
| 0.06191 | 0.914958 | 0.054 | 0.03 | 1 | 6 | Ankrd54 | 0.024 | Old |
| 0.061956 | 0.356615 | 0.137 | 0.095 | 1 | 6 | Ruvbl1 | 0.042 | Old |
| 0.061987 | 1.32035 | 0.043 | 0.022 | 1 | 6 | Nop2 | 0.021 | Old |
| 0.062113 | 1.010051 | 0.026 | 0.01 | 1 | 6 | Zfyve19 | 0.016 | Old |
| 0.06217 | 0.741976 | 0.197 | 0.156 | 1 | 6 | Usp12 | 0.041 | Old |
| 0.062273 | 1.623899 | 0.026 | 0.01 | 1 | 6 | Neil1 | 0.016 | Old |
| 0.062273 | 0.910694 | 0.026 | 0.01 | 1 | 6 | Gale | 0.016 | Old |
| 0.062374 | 1.832659 | 0.017 | 0.005 | 1 | 6 | Serpina3g | 0.012 | Old |
| 0.062424 | 0.421584 | 0.12 | 0.084 | 1 | 6 | Tmem185k | 0.036 | Old |
| 0.062501 | 0.512846 | 0.223 | 0.174 | 1 | 6 | Commd7 | 0.049 | Old |
| 0.062787 | 2.045704 | 0.017 | 0.005 | 1 | 6 | Dtnb | 0.012 | Old |
| 0.062787 | 1.048915 | 0.017 | 0.005 | 1 | 6 | Xlr4c | 0.012 | Old |
| 0.062914 | 0.579739 | 0.026 | 0.01 | 1 | 6 | Nck2 | 0.016 | Old |
| 0.062994 | 1.143966 | 0.017 | 0.005 | 1 | 6 | Sco1 | 0.012 | Old |
| 0.063011 | 0.38434 | 0.22 | 0.169 | 1 | 6 | Atp8a1 | 0.051 | Old |
| 0.063246 | 0.566885 | 0.16 | 0.117 | 1 | 6 | Upf3b | 0.043 | Old |
| 0.06341 | 1.34715 | 0.017 | 0.005 | 1 | 6 | Asrgl1 | 0.012 | Old |
| 0.06341 | 1.110684 | 0.017 | 0.005 | 1 | 6 | Mmp23 | 0.012 | Old |
| 0.063553 | 1.542058 | 0.023 | 0.008 | 1 | 6 | Pdss1 | 0.015 | Old |
| 0.063553 | 0.602844 | 0.023 | 0.008 | 1 | 6 | Tnfsf8 | 0.015 | Old |
| 0.063605 | 0.042793 | 0.189 | 0.14 | 1 | 6 | Acp1 | 0.049 | Old |
| 0.06362 | 0.503506 | 0.123 | 0.085 | 1 | 6 | Nelfcd | 0.038 | Old |
| 0.063673 | 1.614185 | 0.02 | 0.007 | 1 | 6 | Nudt15 | 0.013 | Old |
| 0.063727 | 1.300589 | 0.023 | 0.008 | 1 | 6 | Cpeb3 | 0.015 | Old |
| 0.063828 | 1.126203 | 0.017 | 0.005 | 1 | 6 | Pomt1 | 0.012 | Old |
| 0.063902 | 1.304825 | 0.023 | 0.008 | 1 | 6 | Mcoln2 | 0.015 | Old |
| 0.063978 | 0.393078 | 0.169 | 0.125 | 1 | 6 | Trappc8 | 0.044 | Old |
| 0.064078 | 0.081228 | 0.874 | 0.896 | 1 | 6 | Capzb | -0.022 | Old |
| 0.064248 | 0.728888 | 0.017 | 0.005 | 1 | 6 | Trmt44 | 0.012 | Old |
| 0.064345 | 0.645191 | 0.106 | 0.072 | 1 | 6 | Dhps | 0.034 | Old |
| 0.064346 | 1.235554 | 0.051 | 0.028 | 1 | 6 | Acot2 | 0.023 | Old |
| 0.064428 | 1.029142 | 0.023 | 0.008 | 1 | 6 | Htr7 | 0.015 | Old |
| 0.064435 | 1.990174 | 0.02 | 0.007 | 1 | 6 | Nmb | 0.013 | Old |
| 0.064435 | 1.197771 | 0.02 | 0.007 | 1 | 6 | Tmem138 | 0.013 | Old |
| 0.06454 | 0.41544 | 0.026 | 0.01 | 1 | 6 | Setbp1 | 0.016 | Old |

| | | | | | | | | |
|---|---|---|---|---|---|---|---|---|
| 0.064605 | 1.343179 | 0.023 | 0.008 | 1 | 6 | Hspb1 | 0.015 | Old |
| 0.064626 | 1.367748 | 0.02 | 0.007 | 1 | 6 | Sirt5 | 0.013 | Old |
| 0.064654 | 0.797813 | 0.086 | 0.055 | 1 | 6 | Zfp398 | 0.031 | Old |
| 0.064671 | 0.774385 | 0.017 | 0.005 | 1 | 6 | Kif3c | 0.012 | Old |
| 0.064818 | 1.196453 | 0.02 | 0.007 | 1 | 6 | Gm49339 | 0.013 | Old |
| 0.064891 | 0.538686 | 0.249 | 0.202 | 1 | 6 | Mfhas1 | 0.047 | Old |
| 0.064931 | 0.622525 | 0.126 | 0.089 | 1 | 6 | Rxra | 0.037 | Old |
| 0.065026 | 0.876648 | 0.086 | 0.055 | 1 | 6 | Lias | 0.031 | Old |
| 0.065136 | 0.706305 | 0.023 | 0.008 | 1 | 6 | Cyp2d22 | 0.015 | Old |
| 0.065161 | 0.332882 | 0.306 | 0.251 | 1 | 6 | AI467606 | 0.055 | Old |
| 0.065204 | 0.996487 | 0.02 | 0.007 | 1 | 6 | Gas8 | 0.013 | Old |
| 0.065397 | 1.192087 | 0.02 | 0.007 | 1 | 6 | Hist1h2ap | 0.013 | Old |
| 0.065397 | 1.081296 | 0.02 | 0.007 | 1 | 6 | Uhrf1bp1 | 0.013 | Old |
| 0.065422 | 0.381375 | 0.331 | 0.276 | 1 | 6 | Aco2 | 0.055 | Old |
| 0.06547 | 1.012051 | 0.054 | 0.03 | 1 | 6 | Dhx33 | 0.024 | Old |
| 0.065474 | 0.658844 | 0.3 | 0.251 | 1 | 6 | Man1a | 0.049 | Old |
| 0.065492 | 0.671291 | 0.023 | 0.008 | 1 | 6 | Tmem140 | 0.015 | Old |
| 0.0656 | 0.774909 | 0.063 | 0.037 | 1 | 6 | Brms1l | 0.026 | Old |
| 0.065708 | 0.854889 | 0.043 | 0.022 | 1 | 6 | Ubxn2b | 0.021 | Old |
| 0.066299 | 0.750966 | 0.137 | 0.1 | 1 | 6 | Drg1 | 0.037 | Old |
| 0.066371 | 0.902786 | 0.02 | 0.007 | 1 | 6 | Hid1 | 0.013 | Old |
| 0.066374 | 0.123329 | 0.889 | 0.883 | 1 | 6 | Ywhaz | 0.006 | Old |
| 0.066729 | 0.629992 | 0.071 | 0.043 | 1 | 6 | Ccdc28b | 0.028 | Old |
| 0.067159 | 0.526658 | 0.02 | 0.007 | 1 | 6 | Ifi213 | 0.013 | Old |
| 0.067266 | 0.438143 | 0.22 | 0.169 | 1 | 6 | Kcnq1ot1 | 0.051 | Old |
| 0.067279 | 0.208089 | 0.611 | 0.565 | 1 | 6 | Cops9 | 0.046 | Old |
| 0.067357 | 0.040011 | 0.02 | 0.007 | 1 | 6 | Zfp870 | 0.013 | Old |
| 0.067669 | 0.980486 | 0.051 | 0.028 | 1 | 6 | Adprhl2 | 0.023 | Old |
| 0.068039 | 0.518998 | 0.183 | 0.137 | 1 | 6 | Aldh3b1 | 0.046 | Old |
| 0.06805 | 0.249947 | 0.603 | 0.557 | 1 | 6 | Atp5k | 0.046 | Old |
| 0.068485 | 0.248406 | 0.714 | 0.766 | 1 | 6 | Sdc3 | -0.052 | Old |
| 0.068969 | 0.697518 | 0.043 | 0.022 | 1 | 6 | Isg20 | 0.021 | Old |
| 0.069268 | 1.229865 | 0.051 | 0.028 | 1 | 6 | Arrdc3 | 0.023 | Old |
| 0.069289 | 0.656734 | 0.134 | 0.097 | 1 | 6 | Cdk17 | 0.037 | Old |
| 0.069313 | 0.412618 | 0.12 | 0.084 | 1 | 6 | Cnot10 | 0.036 | Old |
| 0.069387 | 0.182896 | 0.303 | 0.246 | 1 | 6 | Usp15 | 0.057 | Old |
| 0.069504 | 0.125068 | 0.08 | 0.05 | 1 | 6 | Rev1 | 0.03 | Old |
| 0.069741 | 1.257841 | 0.04 | 0.02 | 1 | 6 | Rhbdd3 | 0.02 | Old |
| 0.069853 | 0.016015 | 0.231 | 0.176 | 1 | 6 | Atf1 | 0.055 | Old |
| 0.070152 | 0.468361 | 0.129 | 0.09 | 1 | 6 | Tomm40 | 0.039 | Old |
| 0.070209 | 0.604545 | 0.131 | 0.094 | 1 | 6 | Ankrd10 | 0.037 | Old |
| 0.070403 | 1.743264 | 0.037 | 0.018 | 1 | 6 | Zfp617 | 0.019 | Old |
| 0.070413 | 0.655949 | 0.04 | 0.02 | 1 | 6 | Casp7 | 0.02 | Old |
| 0.070492 | 0.340598 | 0.151 | 0.109 | 1 | 6 | Pop5 | 0.042 | Old |
| 0.070694 | 0.75252 | 0.08 | 0.05 | 1 | 6 | Cables2 | 0.03 | Old |

| | | | | | | | | |
|---|---|---|---|---|---|---|---|---|
| 0.0708 | 0.74323 | 0.063 | 0.037 | 1 | 6 | Gmpr2 | 0.026 | Old |
| 0.071298 | 0.718351 | 0.203 | 0.161 | 1 | 6 | 9030624J0 | 0.042 | Old |
| 0.071448 | 0.21901 | 0.706 | 0.679 | 1 | 6 | H2-T23 | 0.027 | Old |
| 0.071613 | 0.132158 | 0.857 | 0.886 | 1 | 6 | Efhd2 | -0.029 | Old |
| 0.071614 | 0.426835 | 0.166 | 0.124 | 1 | 6 | Papd5 | 0.042 | Old |
| 0.071691 | 0.196181 | 0.731 | 0.702 | 1 | 6 | Ywhae | 0.029 | Old |
| 0.071802 | 0.246522 | 0.171 | 0.127 | 1 | 6 | Nr1h2 | 0.044 | Old |
| 0.071893 | 0.845562 | 0.143 | 0.105 | 1 | 6 | Lrpprc | 0.038 | Old |
| 0.072283 | 0.658713 | 0.16 | 0.12 | 1 | 6 | Syvn1 | 0.04 | Old |
| 0.072395 | 0.873896 | 0.06 | 0.035 | 1 | 6 | Ly9 | 0.025 | Old |
| 0.072424 | 0.150184 | 0.486 | 0.413 | 1 | 6 | Prrc2a | 0.073 | Old |
| 0.072834 | 0.309683 | 0.357 | 0.299 | 1 | 6 | Cstb | 0.058 | Old |
| 0.072838 | 0.475486 | 0.129 | 0.092 | 1 | 6 | Elk3 | 0.037 | Old |
| 0.073398 | 0.456875 | 0.309 | 0.259 | 1 | 6 | Sypl | 0.05 | Old |
| 0.073597 | 0.384022 | 0.106 | 0.072 | 1 | 6 | Josd1 | 0.034 | Old |
| 0.073889 | 0.650509 | 0.086 | 0.055 | 1 | 6 | Cog7 | 0.031 | Old |
| 0.074046 | 0.243703 | 0.7 | 0.649 | 1 | 6 | Hnrnph1 | 0.051 | Old |
| 0.074113 | 0.528102 | 0.126 | 0.089 | 1 | 6 | Kin | 0.037 | Old |
| 0.074434 | 0.611955 | 0.117 | 0.082 | 1 | 6 | Ppme1 | 0.035 | Old |
| 0.074651 | 0.355622 | 0.32 | 0.264 | 1 | 6 | Tecr | 0.056 | Old |
| 0.074713 | 1.111331 | 0.049 | 0.027 | 1 | 6 | Zfp160 | 0.022 | Old |
| 0.07478 | 0.653391 | 0.251 | 0.211 | 1 | 6 | Birc2 | 0.04 | Old |
| 0.075176 | 0.551117 | 0.06 | 0.035 | 1 | 6 | Dus3l | 0.025 | Old |
| 0.075306 | 1.020709 | 0.037 | 0.018 | 1 | 6 | Ppp1r3b | 0.019 | Old |
| 0.075311 | 0.453304 | 0.12 | 0.084 | 1 | 6 | Crkl | 0.036 | Old |
| 0.075601 | 0.949077 | 0.049 | 0.027 | 1 | 6 | Znhit6 | 0.022 | Old |
| 0.075659 | 0.616763 | 0.117 | 0.082 | 1 | 6 | Iars2 | 0.035 | Old |
| 0.075828 | 3.733395 | 0.046 | 0.025 | 1 | 6 | Ikzf2 | 0.021 | Old |
| 0.075836 | 0.243577 | 0.286 | 0.236 | 1 | 6 | Ndufa8 | 0.05 | Old |
| 0.076656 | 0.371281 | 0.06 | 0.035 | 1 | 6 | Ints4 | 0.025 | Old |
| 0.076687 | 0.328962 | 0.24 | 0.187 | 1 | 6 | Ndufb2 | 0.053 | Old |
| 0.076895 | 0.195089 | 0.64 | 0.607 | 1 | 6 | Psmb4 | 0.033 | Old |
| 0.076958 | 0.015244 | 0.066 | 0.1 | 1 | 6 | Sorl1 | -0.034 | Old |
| 0.076969 | 1.085581 | 0.203 | 0.162 | 1 | 6 | Itgav | 0.041 | Old |
| 0.077087 | 0.339722 | 0.66 | 0.63 | 1 | 6 | Snrnp70 | 0.03 | Old |
| 0.077102 | 0.663828 | 0.037 | 0.018 | 1 | 6 | Cd200r4 | 0.019 | Old |
| 0.077232 | 0.123209 | 0.063 | 0.037 | 1 | 6 | Zfp821 | 0.026 | Old |
| 0.077336 | 0.792539 | 0.049 | 0.027 | 1 | 6 | Rdm1 | 0.022 | Old |
| 0.077635 | 0.810272 | 0.057 | 0.033 | 1 | 6 | Rhot2 | 0.024 | Old |
| 0.077658 | 0.448167 | 0.366 | 0.324 | 1 | 6 | Luc7l3 | 0.042 | Old |
| 0.077982 | 0.705792 | 0.094 | 0.064 | 1 | 6 | Pus1 | 0.03 | Old |
| 0.078213 | 0.354393 | 0.126 | 0.089 | 1 | 6 | Ndufv1 | 0.037 | Old |
| 0.0783 | 0.579277 | 0.134 | 0.099 | 1 | 6 | Xdh | 0.035 | Old |
| 0.07871 | 0.76147 | 0.057 | 0.033 | 1 | 6 | Metap1d | 0.024 | Old |
| 0.078753 | 0.554374 | 0.18 | 0.142 | 1 | 6 | Trim30d | 0.038 | Old |

| | | | | | | | | |
|---|---|---|---|---|---|---|---|---|
| 0.078789 | 0.523528 | 0.437 | 0.398 | 1 | 6 | H2-Q7 | 0.039 | Old |
| 0.07888 | 1.293916 | 0.034 | 0.017 | 1 | 6 | Ighmbp2 | 0.017 | Old |
| 0.078934 | 0.328289 | 0.251 | 0.202 | 1 | 6 | Rgs19 | 0.049 | Old |
| 0.07907 | 0.994913 | 0.057 | 0.033 | 1 | 6 | Dusp7 | 0.024 | Old |
| 0.079379 | 0.347287 | 0.083 | 0.054 | 1 | 6 | Jpt2 | 0.029 | Old |
| 0.079411 | 0.380462 | 0.203 | 0.156 | 1 | 6 | Mrpl42 | 0.047 | Old |
| 0.079495 | 0.586036 | 0.457 | 0.426 | 1 | 6 | Apobec3 | 0.031 | Old |
| 0.079587 | 0.247143 | 0.754 | 0.734 | 1 | 6 | mt-Nd5 | 0.02 | Old |
| 0.079614 | 0.452203 | 0.077 | 0.048 | 1 | 6 | Mast4 | 0.029 | Old |
| 0.079832 | 1.090039 | 0.071 | 0.045 | 1 | 6 | Msantd4 | 0.026 | Old |
| 0.080138 | 0.438972 | 0.271 | 0.229 | 1 | 6 | H2-Q6 | 0.042 | Old |
| 0.080415 | 0.204858 | 0.571 | 0.508 | 1 | 6 | Top1 | 0.063 | Old |
| 0.080583 | 0.41997 | 0.323 | 0.274 | 1 | 6 | Apaf1 | 0.049 | Old |
| 0.080771 | 0.43654 | 0.203 | 0.157 | 1 | 6 | Fbxw2 | 0.046 | Old |
| 0.080899 | 0.307897 | 0.126 | 0.089 | 1 | 6 | Gpr155 | 0.037 | Old |
| 0.080971 | 0.720272 | 0.066 | 0.04 | 1 | 6 | Fam210b | 0.026 | Old |
| 0.08113 | 1.150768 | 0.063 | 0.038 | 1 | 6 | Fmo5 | 0.025 | Old |
| 0.081134 | 0.339389 | 0.483 | 0.433 | 1 | 6 | Eif4a2 | 0.05 | Old |
| 0.081243 | 0.552361 | 0.049 | 0.027 | 1 | 6 | Ercc2 | 0.022 | Old |
| 0.082122 | 0.341702 | 0.171 | 0.129 | 1 | 6 | Pspc1 | 0.042 | Old |
| 0.082382 | 0.485962 | 0.109 | 0.075 | 1 | 6 | Tmem71 | 0.034 | Old |
| 0.08244 | 0.451315 | 0.251 | 0.206 | 1 | 6 | Fbxl3 | 0.045 | Old |
| 0.082706 | 0.136591 | 0.823 | 0.799 | 1 | 6 | Chchd2 | 0.024 | Old |
| 0.082724 | 0.507288 | 0.294 | 0.244 | 1 | 6 | Slc1a5 | 0.05 | Old |
| 0.083215 | 0.332238 | 0.374 | 0.321 | 1 | 6 | Tcea1 | 0.053 | Old |
| 0.083287 | 0.259666 | 0.386 | 0.323 | 1 | 6 | Cers2 | 0.063 | Old |
| 0.083602 | 1.108529 | 0.063 | 0.038 | 1 | 6 | Pym1 | 0.025 | Old |
| 0.083841 | 0.550097 | 0.363 | 0.309 | 1 | 6 | Nrip1 | 0.054 | Old |
| 0.084326 | 0.813208 | 0.034 | 0.017 | 1 | 6 | Miga2 | 0.017 | Old |
| 0.084487 | 0.140828 | 0.817 | 0.804 | 1 | 6 | Hnrnpf | 0.013 | Old |
| 0.084615 | 0.732557 | 0.237 | 0.192 | 1 | 6 | Nus1 | 0.045 | Old |
| 0.085037 | 0.346063 | 0.314 | 0.261 | 1 | 6 | Ddb1 | 0.053 | Old |
| 0.085155 | 0.751543 | 0.057 | 0.033 | 1 | 6 | Pla2g12a | 0.024 | Old |
| 0.085175 | 0.172142 | 0.529 | 0.47 | 1 | 6 | Hipk1 | 0.059 | Old |
| 0.085177 | 0.540777 | 0.034 | 0.017 | 1 | 6 | Fbxl8 | 0.017 | Old |
| 0.085523 | 0.508331 | 0.234 | 0.194 | 1 | 6 | Glrx | 0.04 | Old |
| 0.085908 | 0.626545 | 0.209 | 0.166 | 1 | 6 | Ide | 0.043 | Old |
| 0.086369 | 0.709441 | 0.123 | 0.089 | 1 | 6 | Ddhd2 | 0.034 | Old |
| 0.086529 | 0.404368 | 0.18 | 0.139 | 1 | 6 | Tmem147 | 0.041 | Old |
| 0.086591 | 0.430223 | 0.189 | 0.145 | 1 | 6 | Ddx18 | 0.044 | Old |
| 0.086621 | 0.412108 | 0.123 | 0.087 | 1 | 6 | Nsrp1 | 0.036 | Old |
| 0.086676 | 0.601081 | 0.046 | 0.025 | 1 | 6 | Tfpt | 0.021 | Old |
| 0.086882 | 0.5413 | 0.12 | 0.085 | 1 | 6 | Smim4 | 0.035 | Old |
| 0.086917 | 0.312456 | 0.074 | 0.047 | 1 | 6 | Hivep3 | 0.027 | Old |
| 0.087095 | 0.607086 | 0.594 | 0.562 | 1 | 6 | Xbp1 | 0.032 | Old |

| | | | | | | | | |
|---|---|---|---|---|---|---|---|---|
| 0.087256 | 0.550956 | 0.169 | 0.129 | 1 | 6 | Thoc6 | 0.04 | Old |
| 0.087556 | 0.222419 | 0.474 | 0.426 | 1 | 6 | Copz1 | 0.048 | Old |
| 0.08756 | 0.367167 | 0.046 | 0.025 | 1 | 6 | Gm36161 | 0.021 | Old |
| 0.087749 | 0.308783 | 0.271 | 0.217 | 1 | 6 | Zcchc7 | 0.054 | Old |
| 0.087919 | 0.179441 | 0.743 | 0.722 | 1 | 6 | Atp5a1 | 0.021 | Old |
| 0.088134 | 1.058233 | 0.06 | 0.037 | 1 | 6 | Dnaja3 | 0.023 | Old |
| 0.08833 | 1.185138 | 0.031 | 0.015 | 1 | 6 | Ildr1 | 0.016 | Old |
| 0.088355 | 0.310885 | 0.117 | 0.084 | 1 | 6 | Anapc2 | 0.033 | Old |
| 0.08868 | 1.182766 | 0.051 | 0.03 | 1 | 6 | Ppp1r13b | 0.021 | Old |
| 0.089049 | 0.796145 | 0.046 | 0.025 | 1 | 6 | Gde1 | 0.021 | Old |
| 0.089152 | 0.382929 | 0.137 | 0.099 | 1 | 6 | Man1b1 | 0.038 | Old |
| 0.089225 | 0.750288 | 0.094 | 0.064 | 1 | 6 | Cpox | 0.03 | Old |
| 0.089693 | 0.310693 | 0.406 | 0.368 | 1 | 6 | Eif4g1 | 0.038 | Old |
| 0.089718 | 0.313408 | 0.291 | 0.241 | 1 | 6 | Wdr45b | 0.05 | Old |
| 0.090099 | 0.814036 | 0.097 | 0.067 | 1 | 6 | Fubp3 | 0.03 | Old |
| 0.090654 | 0.593953 | 0.094 | 0.064 | 1 | 6 | Slco3a1 | 0.03 | Old |
| 0.09081 | 0.140532 | 0.117 | 0.082 | 1 | 6 | Prpsap2 | 0.035 | Old |
| 0.090942 | 1.037848 | 0.031 | 0.015 | 1 | 6 | Rtn4rl1 | 0.016 | Old |
| 0.091065 | 0.281023 | 0.094 | 0.064 | 1 | 6 | Golim4 | 0.03 | Old |
| 0.091123 | 0.460409 | 0.7 | 0.689 | 1 | 6 | Serinc3 | 0.011 | Old |
| 0.091285 | 1.344801 | 0.043 | 0.023 | 1 | 6 | Ttc7b | 0.02 | Old |
| 0.091296 | 0.164753 | 0.831 | 0.821 | 1 | 6 | Tagln2 | 0.01 | Old |
| 0.091458 | 0.208991 | 0.217 | 0.169 | 1 | 6 | Relt | 0.048 | Old |
| 0.092083 | 0.421664 | 0.146 | 0.109 | 1 | 6 | Prkag1 | 0.037 | Old |
| 0.092271 | 0.95207 | 0.031 | 0.015 | 1 | 6 | Fbf1 | 0.016 | Old |
| 0.092271 | 0.642903 | 0.031 | 0.015 | 1 | 6 | Spaca9 | 0.016 | Old |
| 0.092277 | 1.26368 | 0.06 | 0.037 | 1 | 6 | Smyd5 | 0.023 | Old |
| 0.092654 | 0.720252 | 0.031 | 0.015 | 1 | 6 | Bloc1s3 | 0.016 | Old |
| 0.092669 | 0.224389 | 0.491 | 0.443 | 1 | 6 | Txn2 | 0.048 | Old |
| 0.092803 | 0.265432 | 0.371 | 0.326 | 1 | 6 | Lta4h | 0.045 | Old |
| 0.093323 | 0.480695 | 0.063 | 0.038 | 1 | 6 | Gpr180 | 0.025 | Old |
| 0.093369 | 0.470913 | 0.103 | 0.072 | 1 | 6 | Brap | 0.031 | Old |
| 0.093616 | 0.69829 | 0.031 | 0.015 | 1 | 6 | Nmrk1 | 0.016 | Old |
| 0.093801 | 1.188278 | 0.051 | 0.03 | 1 | 6 | Ano8 | 0.021 | Old |
| 0.093835 | 1.952333 | 0.029 | 0.013 | 1 | 6 | Parvg | 0.016 | Old |
| 0.094215 | 0.917564 | 0.1 | 0.07 | 1 | 6 | Bambi | 0.03 | Old |
| 0.094653 | 1.285362 | 0.029 | 0.013 | 1 | 6 | Zfp868 | 0.016 | Old |
| 0.095064 | 1.564449 | 0.029 | 0.013 | 1 | 6 | Pus7l | 0.016 | Old |
| 0.09507 | 0.408497 | 0.406 | 0.355 | 1 | 6 | Elf1 | 0.051 | Old |
| 0.095172 | 0.334848 | 0.031 | 0.015 | 1 | 6 | Bcdin3d | 0.016 | Old |
| 0.09545 | 0.229864 | 0.649 | 0.619 | 1 | 6 | Gda | 0.03 | Old |
| 0.095468 | 0.762352 | 0.043 | 0.023 | 1 | 6 | Eml2 | 0.02 | Old |
| 0.09567 | 0.222388 | 0.614 | 0.542 | 1 | 6 | Myl12b | 0.072 | Old |
| 0.095718 | 0.318548 | 0.371 | 0.323 | 1 | 6 | Polr2e | 0.048 | Old |
| 0.095726 | 0.449461 | 0.077 | 0.05 | 1 | 6 | Dhx29 | 0.027 | Old |

| | | | | | | | | |
|---|---|---|---|---|---|---|---|---|
| 0.095899 | 0.496658 | 0.166 | 0.127 | 1 | 6 | Snx12 | 0.039 | Old |
| 0.095957 | 0.422373 | 0.031 | 0.015 | 1 | 6 | Inpp5e | 0.016 | Old |
| 0.095957 | 0.161374 | 0.031 | 0.015 | 1 | 6 | Lyrm4 | 0.016 | Old |
| 0.095996 | 0.335278 | 0.1 | 0.069 | 1 | 6 | Qsox1 | 0.031 | Old |
| 0.096212 | 0.644382 | 0.114 | 0.082 | 1 | 6 | Scaf4 | 0.032 | Old |
| 0.096532 | 0.655681 | 0.089 | 0.06 | 1 | 6 | Mief1 | 0.029 | Old |
| 0.096555 | 0.906399 | 0.131 | 0.099 | 1 | 6 | Zfp384 | 0.032 | Old |
| 0.097126 | 0.206163 | 0.171 | 0.132 | 1 | 6 | Prdx3 | 0.039 | Old |
| 0.09714 | 0.78253 | 0.131 | 0.097 | 1 | 6 | Uba5 | 0.034 | Old |
| 0.097147 | 0.358446 | 0.071 | 0.045 | 1 | 6 | Slfn8 | 0.026 | Old |
| 0.09727 | 0.07381 | 0.92 | 0.968 | 1 | 6 | Gpx1 | -0.048 | Old |
| 0.097572 | 0.345195 | 0.386 | 0.334 | 1 | 6 | Arglu1 | 0.052 | Old |
| 0.097906 | 0.349704 | 0.351 | 0.306 | 1 | 6 | Ubxn4 | 0.045 | Old |
| 0.098194 | 1.113603 | 0.029 | 0.013 | 1 | 6 | Coq4 | 0.016 | Old |
| 0.098507 | 0.131918 | 0.26 | 0.207 | 1 | 6 | C1qbp | 0.053 | Old |
| 0.09855 | 0.328877 | 0.177 | 0.137 | 1 | 6 | Acadl | 0.04 | Old |
| 0.098639 | 0.553645 | 0.069 | 0.043 | 1 | 6 | Chmp7 | 0.026 | Old |
| 0.099042 | 0.81684 | 0.029 | 0.013 | 1 | 6 | Zfp692 | 0.016 | Old |
| 0.09929 | 0.682983 | 0.14 | 0.105 | 1 | 6 | Ddx19a | 0.035 | Old |
| 0.099308 | 0.893507 | 0.051 | 0.03 | 1 | 6 | Cmss1 | 0.021 | Old |
| 0.099379 | 0.632331 | 0.117 | 0.084 | 1 | 6 | Rprd1b | 0.033 | Old |
| 0.099459 | 0.476138 | 0.22 | 0.174 | 1 | 6 | Gsk3a | 0.046 | Old |
| 0.099787 | 0.196317 | 0.651 | 0.634 | 1 | 6 | Set | 0.017 | Old |
| 0.099896 | 0.591699 | 0.029 | 0.013 | 1 | 6 | Aste1 | 0.016 | Old |
| 0.099997 | 0.490654 | 0.286 | 0.237 | 1 | 6 | Mcts1 | 0.049 | Old |
| 0.100213 | 0.997332 | 0.077 | 0.052 | 1 | 6 | Flcn | 0.025 | Old |
| 0.100224 | 0.613164 | 0.166 | 0.127 | 1 | 6 | Tiprl | 0.039 | Old |
| 0.100225 | 0.861395 | 0.051 | 0.03 | 1 | 6 | Dtx2 | 0.021 | Old |
| 0.100324 | 0.835426 | 0.06 | 0.037 | 1 | 6 | Tbl3 | 0.023 | Old |
| 0.100372 | 0.29095 | 0.811 | 0.834 | 1 | 6 | Neat1 | -0.023 | Old |
| 0.100714 | 0.566429 | 0.1 | 0.07 | 1 | 6 | Gpn1 | 0.03 | Old |
| 0.100747 | 0.659897 | 0.06 | 0.037 | 1 | 6 | Pon3 | 0.023 | Old |
| 0.100757 | 0.55917 | 0.029 | 0.013 | 1 | 6 | Homer3 | 0.016 | Old |
| 0.100863 | 0.237154 | 0.46 | 0.418 | 1 | 6 | Ddx39b | 0.042 | Old |
| 0.100953 | 0.484459 | 0.146 | 0.109 | 1 | 6 | Vps9d1 | 0.037 | Old |
| 0.100972 | 0.420133 | 0.029 | 0.013 | 1 | 6 | Rnf135 | 0.016 | Old |
| 0.101064 | 0.611056 | 0.146 | 0.11 | 1 | 6 | Psip1 | 0.036 | Old |
| 0.101405 | 0.677109 | 0.029 | 0.013 | 1 | 6 | 2610044O1 | 0.016 | Old |
| 0.101811 | 0.914731 | 0.12 | 0.089 | 1 | 6 | Gpr68 | 0.031 | Old |
| 0.10184 | 0.242754 | 0.029 | 0.013 | 1 | 6 | Cbx6 | 0.016 | Old |
| 0.101903 | 0.714734 | 0.08 | 0.054 | 1 | 6 | 1700017B0 | 0.026 | Old |
| 0.10209 | 1.501509 | 0.026 | 0.012 | 1 | 6 | Slc26a6 | 0.014 | Old |
| 0.102322 | 2.256133 | 0.026 | 0.012 | 1 | 6 | Ebf1 | 0.014 | Old |
| 0.102458 | 0.312578 | 0.097 | 0.067 | 1 | 6 | Alas1 | 0.03 | Old |
| 0.102628 | 0.159472 | 0.14 | 0.184 | 1 | 6 | Cish | -0.044 | Old |

| | | | | | | | | |
|---|---|---|---|---|---|---|---|---|
| 0.102785 | 1.939395 | 0.026 | 0.012 | 1 | 6 | Cers4 | 0.014 | Old |
| 0.103138 | 0.896547 | 0.074 | 0.048 | 1 | 6 | Zfp646 | 0.026 | Old |
| 0.103387 | 0.348407 | 0.291 | 0.242 | 1 | 6 | Psmd1 | 0.049 | Old |
| 0.103536 | 0.790235 | 0.04 | 0.022 | 1 | 6 | Tfip11 | 0.018 | Old |
| 0.104423 | 0.399927 | 0.254 | 0.204 | 1 | 6 | Golga4 | 0.05 | Old |
| 0.104587 | 0.523137 | 0.097 | 0.067 | 1 | 6 | Cyp4f13 | 0.03 | Old |
| 0.1047 | 0.642414 | 0.18 | 0.14 | 1 | 6 | Aagab | 0.04 | Old |
| 0.104808 | 0.702087 | 0.04 | 0.022 | 1 | 6 | Zswim7 | 0.018 | Old |
| 0.105016 | 0.525357 | 0.16 | 0.122 | 1 | 6 | Slc35a4 | 0.038 | Old |
| 0.105357 | 0.659934 | 0.04 | 0.022 | 1 | 6 | Mul1 | 0.018 | Old |
| 0.105509 | 0.714263 | 0.066 | 0.042 | 1 | 6 | Edem3 | 0.024 | Old |
| 0.105509 | 0.5176 | 0.066 | 0.042 | 1 | 6 | Nr2c2ap | 0.024 | Old |
| 0.10584 | 1.094768 | 0.026 | 0.012 | 1 | 6 | Ftx | 0.014 | Old |
| 0.106078 | 1.286108 | 0.026 | 0.012 | 1 | 6 | Rarb | 0.014 | Old |
| 0.106078 | 0.993713 | 0.026 | 0.012 | 1 | 6 | Nipsnap1 | 0.014 | Old |
| 0.106078 | 0.674694 | 0.026 | 0.012 | 1 | 6 | Rfx2 | 0.014 | Old |
| 0.106118 | 0.443333 | 0.154 | 0.117 | 1 | 6 | Rarg | 0.037 | Old |
| 0.106397 | 0.206274 | 0.511 | 0.467 | 1 | 6 | Matr3 | 0.044 | Old |
| 0.106588 | 0.375951 | 0.343 | 0.299 | 1 | 6 | Mcrip1 | 0.044 | Old |
| 0.106705 | 0.198462 | 0.3 | 0.247 | 1 | 6 | Psmc6 | 0.053 | Old |
| 0.107546 | 0.4099 | 0.12 | 0.087 | 1 | 6 | Anapc15 | 0.033 | Old |
| 0.107561 | 0.570803 | 0.22 | 0.181 | 1 | 6 | Ss18 | 0.039 | Old |
| 0.107727 | 0.527184 | 0.151 | 0.115 | 1 | 6 | Dync1li2 | 0.036 | Old |
| 0.107765 | 0.915884 | 0.077 | 0.052 | 1 | 6 | Rasgrp1 | 0.025 | Old |
| 0.108062 | 0.45887 | 0.094 | 0.065 | 1 | 6 | Lyrm2 | 0.029 | Old |
| 0.108146 | 0.664104 | 0.091 | 0.064 | 1 | 6 | Aars | 0.027 | Old |
| 0.108298 | 1.170738 | 0.037 | 0.02 | 1 | 6 | Rundc1 | 0.017 | Old |
| 0.108581 | 0.753613 | 0.003 | 0.013 | 1 | 6 | Zfp629 | -0.01 | Old |
| 0.108668 | 0.877826 | 0.117 | 0.087 | 1 | 6 | Psmd9 | 0.03 | Old |
| 0.108871 | 0.856025 | 0.077 | 0.052 | 1 | 6 | Eef2kmt | 0.025 | Old |
| 0.108882 | 1.347472 | 0.054 | 0.033 | 1 | 6 | Myo1e | 0.021 | Old |
| 0.108966 | 0.428406 | 0.026 | 0.012 | 1 | 6 | Zfp35 | 0.014 | Old |
| 0.10902 | 0.325811 | 0.42 | 0.376 | 1 | 6 | Apbb1ip | 0.044 | Old |
| 0.109093 | 0.294287 | 0.057 | 0.035 | 1 | 6 | Zbed4 | 0.022 | Old |
| 0.109105 | 0.528257 | 0.1 | 0.07 | 1 | 6 | Phkb | 0.03 | Old |
| 0.109139 | 0.328718 | 0.126 | 0.092 | 1 | 6 | Agtrap | 0.034 | Old |
| 0.109174 | 1.154787 | 0.063 | 0.04 | 1 | 6 | Akt1s1 | 0.023 | Old |
| 0.109183 | 0.578009 | 0.186 | 0.149 | 1 | 6 | Phf2 | 0.037 | Old |
| 0.109402 | 0.646292 | 0.057 | 0.035 | 1 | 6 | Ccdc86 | 0.022 | Old |
| 0.109612 | 0.364151 | 0.229 | 0.182 | 1 | 6 | Kdm6a | 0.047 | Old |
| 0.109643 | 0.427607 | 0.04 | 0.022 | 1 | 6 | Oas1a | 0.018 | Old |
| 0.109698 | 0.236374 | 0.026 | 0.012 | 1 | 6 | Tbc1d24 | 0.014 | Old |
| 0.109993 | 0.399778 | 0.274 | 0.227 | 1 | 6 | Kdm5a | 0.047 | Old |
| 0.110591 | 0.570853 | 0.071 | 0.047 | 1 | 6 | Mtpap | 0.024 | Old |
| 0.110682 | 0.013632 | 0.263 | 0.316 | 1 | 6 | Dnmt3a | -0.053 | Old |

| | | | | | | | | |
|---|---|---|---|---|---|---|---|---|
| 0.110691 | 0.333105 | 0.254 | 0.207 | 1 | 6 | Samm50 | 0.047 | Old |
| 0.110799 | 0.653331 | 0.057 | 0.035 | 1 | 6 | Kat14 | 0.022 | Old |
| 0.110852 | 1.117702 | 0.037 | 0.02 | 1 | 6 | Rasgrp3 | 0.017 | Old |
| 0.110867 | 0.529454 | 0.071 | 0.047 | 1 | 6 | Psmg2 | 0.024 | Old |
| 0.11156 | 0.403294 | 0.066 | 0.042 | 1 | 6 | 1110065P2 | 0.024 | Old |
| 0.111653 | 0.407868 | 0.049 | 0.028 | 1 | 6 | Rrnad1 | 0.021 | Old |
| 0.111701 | 0.293839 | 0.243 | 0.201 | 1 | 6 | Setx | 0.042 | Old |
| 0.111904 | 0.724 | 0.1 | 0.072 | 1 | 6 | Klhl20 | 0.028 | Old |
| 0.111917 | 0.131215 | 0.309 | 0.256 | 1 | 6 | Kmt2c | 0.053 | Old |
| 0.112071 | 0.477399 | 0.066 | 0.042 | 1 | 6 | Unc13d | 0.024 | Old |
| 0.112112 | 1.568155 | 0.023 | 0.01 | 1 | 6 | Zfat | 0.013 | Old |
| 0.11228 | 0.403324 | 0.403 | 0.366 | 1 | 6 | Spcs2 | 0.037 | Old |
| 0.112825 | 0.248425 | 0.369 | 0.319 | 1 | 6 | Ap2b1 | 0.05 | Old |
| 0.112854 | 0.500747 | 0.129 | 0.095 | 1 | 6 | Dennd4c | 0.034 | Old |
| 0.112911 | 1.690974 | 0.023 | 0.01 | 1 | 6 | Cavin2 | 0.013 | Old |
| 0.112911 | 1.161023 | 0.023 | 0.01 | 1 | 6 | Mapk13 | 0.013 | Old |
| 0.112911 | 1.047468 | 0.023 | 0.01 | 1 | 6 | Glb1l | 0.013 | Old |
| 0.113182 | 0.44307 | 0.106 | 0.075 | 1 | 6 | Skiv2l2 | 0.031 | Old |
| 0.113274 | 0.156903 | 0.477 | 0.423 | 1 | 6 | Psmd2 | 0.054 | Old |
| 0.11357 | 0.102042 | 0.094 | 0.065 | 1 | 6 | Borcs5 | 0.029 | Old |
| 0.113984 | 0.971263 | 0.023 | 0.01 | 1 | 6 | Zbtb25 | 0.013 | Old |
| 0.114201 | 0.412406 | 0.414 | 0.38 | 1 | 6 | Vdac2 | 0.034 | Old |
| 0.114304 | 0.335173 | 0.094 | 0.065 | 1 | 6 | Rpa1 | 0.029 | Old |
| 0.114324 | 0.226537 | 0.223 | 0.179 | 1 | 6 | 4931406C0 | 0.044 | Old |
| 0.114465 | 0.776967 | 0.037 | 0.02 | 1 | 6 | Eogt | 0.017 | Old |
| 0.114523 | 1.196523 | 0.023 | 0.01 | 1 | 6 | Rccd1 | 0.013 | Old |
| 0.114809 | 0.146926 | 0.177 | 0.137 | 1 | 6 | Mospd2 | 0.04 | Old |
| 0.11508 | 0.083365 | 0.1 | 0.07 | 1 | 6 | Fam98a | 0.03 | Old |
| 0.115224 | 1.146183 | 0.06 | 0.038 | 1 | 6 | Zfp942 | 0.022 | Old |
| 0.115487 | 0.461566 | 0.077 | 0.052 | 1 | 6 | Zfp654 | 0.025 | Old |
| 0.11569 | 0.50973 | 0.037 | 0.02 | 1 | 6 | Pigh | 0.017 | Old |
| 0.116044 | 0.351527 | 0.057 | 0.035 | 1 | 6 | Abhd11 | 0.022 | Old |
| 0.116154 | 0.974519 | 0.023 | 0.01 | 1 | 6 | Lamb3 | 0.013 | Old |
| 0.116317 | 0.169458 | 0.297 | 0.244 | 1 | 6 | Uso1 | 0.053 | Old |
| 0.11659 | 0.661837 | 0.18 | 0.142 | 1 | 6 | Pcm1 | 0.038 | Old |
| 0.116778 | 0.404812 | 0.263 | 0.219 | 1 | 6 | Txnl1 | 0.044 | Old |
| 0.117119 | 0.630927 | 0.083 | 0.057 | 1 | 6 | Rfx1 | 0.026 | Old |
| 0.117228 | 0.414106 | 0.369 | 0.323 | 1 | 6 | Erbin | 0.046 | Old |
| 0.117257 | 0.112067 | 0.049 | 0.028 | 1 | 6 | Gm20707 | 0.021 | Old |
| 0.117689 | 0.241448 | 0.134 | 0.1 | 1 | 6 | Inppl1 | 0.034 | Old |
| 0.117894 | 0.327906 | 0.114 | 0.082 | 1 | 6 | Adar | 0.032 | Old |
| 0.118681 | 0.222926 | 0.56 | 0.517 | 1 | 6 | Atp5j | 0.043 | Old |
| 0.118847 | 1.024156 | 0.06 | 0.038 | 1 | 6 | Neurl2 | 0.022 | Old |
| 0.119424 | 0.453157 | 0.183 | 0.144 | 1 | 6 | Polr2c | 0.039 | Old |
| 0.119516 | 0.974163 | 0.054 | 0.033 | 1 | 6 | Naglu | 0.021 | Old |

| | | | | | | | | |
|---|---|---|---|---|---|---|---|---|
| 0.119797 | 0.289087 | 0.211 | 0.169 | 1 | 6 | Sec23b | 0.042 | Old |
| 0.119835 | 0.33147 | 0.309 | 0.261 | 1 | 6 | Eprs | 0.048 | Old |
| 0.119851 | 0.297082 | 0.254 | 0.207 | 1 | 6 | Capn1 | 0.047 | Old |
| 0.120026 | 1.146092 | 0.054 | 0.033 | 1 | 6 | Bag5 | 0.021 | Old |
| 0.120084 | 0.421144 | 0.32 | 0.273 | 1 | 6 | Krit1 | 0.047 | Old |
| 0.120224 | 0.964276 | 0.109 | 0.08 | 1 | 6 | Tmem65 | 0.029 | Old |
| 0.120261 | 1.235939 | 0.02 | 0.008 | 1 | 6 | Zfp58 | 0.012 | Old |
| 0.120399 | 0.346332 | 0.383 | 0.333 | 1 | 6 | Leng8 | 0.05 | Old |
| 0.120425 | 0.272486 | 0.4 | 0.349 | 1 | 6 | Ndufs6 | 0.051 | Old |
| 0.120854 | 0.208741 | 0.337 | 0.281 | 1 | 6 | Tm9sf2 | 0.056 | Old |
| 0.121165 | 0.683134 | 0.083 | 0.057 | 1 | 6 | Serpinb1a | 0.026 | Old |
| 0.121448 | 0.196352 | 0.446 | 0.406 | 1 | 6 | Psme2 | 0.04 | Old |
| 0.121482 | 0.036441 | 0.057 | 0.035 | 1 | 6 | Galt | 0.022 | Old |
| 0.121674 | 0.736736 | 0.089 | 0.062 | 1 | 6 | Tubgcp3 | 0.027 | Old |
| 0.121728 | 0.062161 | 0.746 | 0.697 | 1 | 6 | Eif3a | 0.049 | Old |
| 0.121987 | 0.694428 | 0.083 | 0.057 | 1 | 6 | H2afx | 0.026 | Old |
| 0.12199 | 0.202205 | 0.197 | 0.154 | 1 | 6 | Tmem33 | 0.043 | Old |
| 0.122089 | 0.993098 | 0.02 | 0.008 | 1 | 6 | 4930512J1 | 0.012 | Old |
| 0.122153 | 0.697601 | 0.063 | 0.04 | 1 | 6 | Terf2ip | 0.023 | Old |
| 0.122494 | 1.021076 | 0.091 | 0.065 | 1 | 6 | Pibf1 | 0.026 | Old |
| 0.122671 | 0.122317 | 0.689 | 0.664 | 1 | 6 | Hnrnpa0 | 0.025 | Old |
| 0.122729 | 0.323286 | 0.343 | 0.293 | 1 | 6 | Sulf2 | 0.05 | Old |
| 0.123011 | 0.644521 | 0.02 | 0.008 | 1 | 6 | Ncs1 | 0.012 | Old |
| 0.123125 | 0.26289 | 0.357 | 0.309 | 1 | 6 | Ostc | 0.048 | Old |
| 0.123629 | 0.942937 | 0.02 | 0.008 | 1 | 6 | Cptp | 0.012 | Old |
| 0.123629 | 0.769405 | 0.02 | 0.008 | 1 | 6 | Cwf19l1 | 0.012 | Old |
| 0.124561 | 0.52459 | 0.02 | 0.008 | 1 | 6 | Zfp280b | 0.012 | Old |
| 0.124589 | 0.200862 | 0.171 | 0.13 | 1 | 6 | Atl2 | 0.041 | Old |
| 0.124613 | 0.40791 | 0.089 | 0.062 | 1 | 6 | Ttc1 | 0.027 | Old |
| 0.124856 | 0.505148 | 0.046 | 0.027 | 1 | 6 | Mettl3 | 0.019 | Old |
| 0.124866 | 1.0222 | 0.054 | 0.033 | 1 | 6 | Rhof | 0.021 | Old |
| 0.125042 | 0.659274 | 0.054 | 0.033 | 1 | 6 | Rnaseh1 | 0.021 | Old |
| 0.125165 | 0.280762 | 0.343 | 0.291 | 1 | 6 | Sec31a | 0.052 | Old |
| 0.125422 | 0.017785 | 0.994 | 1 | 1 | 6 | Tmsb4x | -0.006 | Old |
| 0.125496 | 0.262939 | 0.163 | 0.124 | 1 | 6 | Fcf1 | 0.039 | Old |
| 0.12565 | 0.313866 | 0.103 | 0.072 | 1 | 6 | Ecd | 0.031 | Old |
| 0.125905 | 0.769465 | 0.034 | 0.018 | 1 | 6 | Usp20 | 0.016 | Old |
| 0.125996 | 0.160685 | 0.169 | 0.13 | 1 | 6 | Naa35 | 0.039 | Old |
| 0.126013 | 1.700726 | 0.017 | 0.007 | 1 | 6 | Fam185a | 0.01 | Old |
| 0.126013 | 1.384762 | 0.017 | 0.007 | 1 | 6 | Tmem186 | 0.01 | Old |
| 0.126027 | 1.164275 | 0.051 | 0.032 | 1 | 6 | Cdk5rap2 | 0.019 | Old |
| 0.126357 | 1.436105 | 0.017 | 0.007 | 1 | 6 | Gpc1 | 0.01 | Old |
| 0.126407 | 0.358792 | 0.146 | 0.112 | 1 | 6 | Mtmr4 | 0.034 | Old |
| 0.126527 | 0.523779 | 0.06 | 0.038 | 1 | 6 | Ublcp1 | 0.022 | Old |
| 0.126671 | 0.330042 | 0.22 | 0.181 | 1 | 6 | Agps | 0.039 | Old |

| | | | | | | | | |
|---|---|---|---|---|---|---|---|---|
| 0.126702 | 1.646717 | 0.017 | 0.007 | 1 | 6 | Dennd6b | 0.01 | Old |
| 0.12719 | 0.192032 | 0.437 | 0.368 | 1 | 6 | Tsn | 0.069 | Old |
| 0.12722 | 0.25554 | 0.406 | 0.355 | 1 | 6 | Lypla2 | 0.051 | Old |
| 0.127631 | 0.527188 | 0.149 | 0.115 | 1 | 6 | Ubfd1 | 0.034 | Old |
| 0.127971 | 0.339072 | 0.034 | 0.018 | 1 | 6 | Mettl17 | 0.016 | Old |
| 0.128089 | 1.689494 | 0.017 | 0.007 | 1 | 6 | Clec3b | 0.01 | Old |
| 0.128167 | 0.65195 | 0.171 | 0.139 | 1 | 6 | Ing4 | 0.032 | Old |
| 0.12832 | 0.55634 | 0.214 | 0.181 | 1 | 6 | Bloc1s2 | 0.033 | Old |
| 0.128438 | 1.522964 | 0.017 | 0.007 | 1 | 6 | Mgll | 0.01 | Old |
| 0.128438 | 1.337669 | 0.017 | 0.007 | 1 | 6 | AW011738 | 0.01 | Old |
| 0.128479 | 2.857696 | 0.011 | 0.003 | 1 | 6 | Ctla2a | 0.008 | Old |
| 0.128479 | 1.134316 | 0.011 | 0.003 | 1 | 6 | Mamld1 | 0.008 | Old |
| 0.128621 | 0.310876 | 0.077 | 0.052 | 1 | 6 | Naa25 | 0.025 | Old |
| 0.128666 | 0.455263 | 0.034 | 0.018 | 1 | 6 | Lmln | 0.016 | Old |
| 0.128709 | 0.795887 | 0.089 | 0.062 | 1 | 6 | Eml3 | 0.027 | Old |
| 0.128787 | 1.13741 | 0.017 | 0.007 | 1 | 6 | Gm42726 | 0.01 | Old |
| 0.129056 | 0.82234 | 0.086 | 0.06 | 1 | 6 | Exosc9 | 0.026 | Old |
| 0.129132 | 0.303244 | 0.343 | 0.299 | 1 | 6 | Cyld | 0.044 | Old |
| 0.129193 | 0.660401 | 0.043 | 0.025 | 1 | 6 | Pcyox1l | 0.018 | Old |
| 0.129381 | 2.4463 | 0.011 | 0.003 | 1 | 6 | Sorbs1 | 0.008 | Old |
| 0.129381 | 2.078003 | 0.011 | 0.003 | 1 | 6 | Cd4 | 0.008 | Old |
| 0.129381 | 1.789019 | 0.011 | 0.003 | 1 | 6 | Klhdc8b | 0.008 | Old |
| 0.129381 | 1.724285 | 0.011 | 0.003 | 1 | 6 | Gm39469 | 0.008 | Old |
| 0.129381 | 1.598497 | 0.011 | 0.003 | 1 | 6 | Bbs2 | 0.008 | Old |
| 0.129381 | 1.51336 | 0.011 | 0.003 | 1 | 6 | Gm7967 | 0.008 | Old |
| 0.129381 | 1.509119 | 0.011 | 0.003 | 1 | 6 | Kank2 | 0.008 | Old |
| 0.129381 | 1.191242 | 0.011 | 0.003 | 1 | 6 | Zfp869 | 0.008 | Old |
| 0.129381 | 1.064724 | 0.011 | 0.003 | 1 | 6 | Adamts5 | 0.008 | Old |
| 0.129381 | 1.054328 | 0.011 | 0.003 | 1 | 6 | Asb4 | 0.008 | Old |
| 0.129423 | 0.053035 | 0.057 | 0.085 | 1 | 6 | Zc3h12c | -0.028 | Old |
| 0.129488 | 0.443335 | 0.017 | 0.007 | 1 | 6 | Erc1 | 0.01 | Old |
| 0.129538 | 0.631987 | 0.369 | 0.336 | 1 | 6 | Spcs1 | 0.033 | Old |
| 0.12962 | 0.66004 | 0.109 | 0.08 | 1 | 6 | Atg12 | 0.029 | Old |
| 0.129794 | 0.495441 | 0.117 | 0.087 | 1 | 6 | Pfkfb4 | 0.03 | Old |
| 0.129834 | 2.147515 | 0.011 | 0.003 | 1 | 6 | Inmt | 0.008 | Old |
| 0.129834 | 1.54326 | 0.011 | 0.003 | 1 | 6 | Serpina3f | 0.008 | Old |
| 0.129834 | 1.448074 | 0.011 | 0.003 | 1 | 6 | Hist1h1b | 0.008 | Old |
| 0.129834 | 1.396005 | 0.011 | 0.003 | 1 | 6 | Tjp3 | 0.008 | Old |
| 0.129834 | 1.28977 | 0.011 | 0.003 | 1 | 6 | Chil5 | 0.008 | Old |
| 0.129834 | 1.165124 | 0.011 | 0.003 | 1 | 6 | 2200002D( | 0.008 | Old |
| 0.12984 | 0.326334 | 0.017 | 0.007 | 1 | 6 | Zfp764 | 0.01 | Old |
| 0.129849 | 0.398569 | 0.126 | 0.094 | 1 | 6 | Samd1 | 0.032 | Old |
| 0.130027 | 0.597241 | 0.089 | 0.062 | 1 | 6 | Cstf2 | 0.027 | Old |
| 0.130192 | 0.575226 | 0.017 | 0.007 | 1 | 6 | Dmwd | 0.01 | Old |
| 0.130192 | 0.400586 | 0.017 | 0.007 | 1 | 6 | Slamf6 | 0.01 | Old |

| | | | | | | | | |
|---|---|---|---|---|---|---|---|---|
| 0.130251 | 0.294996 | 0.426 | 0.38 | 1 | 6 | Aamp | 0.046 | Old |
| 0.130265 | 0.40078 | 0.349 | 0.301 | 1 | 6 | Ncoa3 | 0.048 | Old |
| 0.130287 | 1.863533 | 0.011 | 0.003 | 1 | 6 | Asf1b | 0.008 | Old |
| 0.130287 | 1.80664 | 0.011 | 0.003 | 1 | 6 | Gcsam | 0.008 | Old |
| 0.130287 | 1.715623 | 0.011 | 0.003 | 1 | 6 | Dock9 | 0.008 | Old |
| 0.130287 | 1.281377 | 0.011 | 0.003 | 1 | 6 | Nfib | 0.008 | Old |
| 0.130287 | 1.113357 | 0.011 | 0.003 | 1 | 6 | Ankrd9 | 0.008 | Old |
| 0.130287 | 1.036111 | 0.011 | 0.003 | 1 | 6 | Slc43a3 | 0.008 | Old |
| 0.130287 | 0.833434 | 0.011 | 0.003 | 1 | 6 | Ndor1 | 0.008 | Old |
| 0.130545 | 0.411829 | 0.017 | 0.007 | 1 | 6 | Ndst1 | 0.01 | Old |
| 0.13057 | 0.336876 | 0.18 | 0.142 | 1 | 6 | Srgap3 | 0.038 | Old |
| 0.130628 | 0.260352 | 0.549 | 0.522 | 1 | 6 | Pkm | 0.027 | Old |
| 0.130677 | 0.861149 | 0.151 | 0.119 | 1 | 6 | Cdc34 | 0.032 | Old |
| 0.130692 | 0.781143 | 0.097 | 0.07 | 1 | 6 | Tspan17 | 0.027 | Old |
| 0.130743 | 1.373918 | 0.011 | 0.003 | 1 | 6 | Adam4 | 0.008 | Old |
| 0.130743 | 1.08286 | 0.011 | 0.003 | 1 | 6 | Gm49101 | 0.008 | Old |
| 0.130743 | 0.979581 | 0.011 | 0.003 | 1 | 6 | Nme2 | 0.008 | Old |
| 0.130743 | 0.737448 | 0.011 | 0.003 | 1 | 6 | Rpp40 | 0.008 | Old |
| 0.130743 | 0.690261 | 0.011 | 0.003 | 1 | 6 | Xk | 0.008 | Old |
| 0.130743 | 0.683851 | 0.011 | 0.003 | 1 | 6 | Zfp251 | 0.008 | Old |
| 0.130743 | 0.513286 | 0.011 | 0.003 | 1 | 6 | 2900026A( | 0.008 | Old |
| 0.130743 | 0.480568 | 0.011 | 0.003 | 1 | 6 | Zfp473 | 0.008 | Old |
| 0.131085 | 0.704677 | 0.08 | 0.055 | 1 | 6 | Fam173b | 0.025 | Old |
| 0.131183 | 1.100907 | 0.014 | 0.005 | 1 | 6 | Angptl6 | 0.009 | Old |
| 0.131199 | 0.992334 | 0.011 | 0.003 | 1 | 6 | Tyw3 | 0.008 | Old |
| 0.131199 | 0.84629 | 0.011 | 0.003 | 1 | 6 | Gm12216 | 0.008 | Old |
| 0.131199 | 0.824532 | 0.011 | 0.003 | 1 | 6 | Bbs9 | 0.008 | Old |
| 0.131199 | 0.561909 | 0.011 | 0.003 | 1 | 6 | Ttyh2 | 0.008 | Old |
| 0.131199 | 0.542362 | 0.011 | 0.003 | 1 | 6 | Zfp623 | 0.008 | Old |
| 0.131288 | 0.274729 | 0.437 | 0.395 | 1 | 6 | Lman2 | 0.042 | Old |
| 0.13158 | 2.118342 | 0.014 | 0.005 | 1 | 6 | Kif5c | 0.009 | Old |
| 0.13158 | 1.098058 | 0.014 | 0.005 | 1 | 6 | Wnt4 | 0.009 | Old |
| 0.13158 | 0.987566 | 0.014 | 0.005 | 1 | 6 | Fancm | 0.009 | Old |
| 0.131606 | 0.491101 | 0.074 | 0.05 | 1 | 6 | Tradd | 0.024 | Old |
| 0.131608 | 0.622619 | 0.017 | 0.007 | 1 | 6 | Homez | 0.01 | Old |
| 0.131608 | 0.43846 | 0.017 | 0.007 | 1 | 6 | Zfp81 | 0.01 | Old |
| 0.131815 | 0.158573 | 0.3 | 0.251 | 1 | 6 | Tmem259 | 0.049 | Old |
| 0.131867 | 0.156121 | 0.214 | 0.171 | 1 | 6 | Gba | 0.043 | Old |
| 0.131978 | 1.840376 | 0.014 | 0.005 | 1 | 6 | Hes7 | 0.009 | Old |
| 0.132319 | 0.636313 | 0.086 | 0.06 | 1 | 6 | Tfam | 0.026 | Old |
| 0.132376 | 1.022045 | 0.014 | 0.005 | 1 | 6 | Tnfrsf17 | 0.009 | Old |
| 0.132406 | 0.304221 | 0.303 | 0.258 | 1 | 6 | Ppp6r1 | 0.045 | Old |
| 0.132462 | 0.793383 | 0.086 | 0.06 | 1 | 6 | Tmem260 | 0.026 | Old |
| 0.132493 | 0.363987 | 0.131 | 0.099 | 1 | 6 | 4930455G( | 0.032 | Old |
| 0.132601 | 0.064086 | 0.129 | 0.095 | 1 | 6 | Elp2 | 0.034 | Old |

| | | | | | | | | |
|---|---|---|---|---|---|---|---|---|
| 0.132657 | 0.147191 | 0.034 | 0.018 | 1 | 6 | 2700062C0 | 0.016 | Old |
| 0.13304 | 0.323121 | 0.269 | 0.229 | 1 | 6 | Eif1ax | 0.04 | Old |
| 0.133328 | 0.738563 | 0.066 | 0.043 | 1 | 6 | Hsd17b7 | 0.023 | Old |
| 0.133509 | 0.163562 | 0.686 | 0.645 | 1 | 6 | Tapbp | 0.041 | Old |
| 0.133577 | 0.695996 | 0.014 | 0.005 | 1 | 6 | Atp9a | 0.009 | Old |
| 0.133577 | 0.690119 | 0.014 | 0.005 | 1 | 6 | Cdk1 | 0.009 | Old |
| 0.133701 | 0.467978 | 0.111 | 0.082 | 1 | 6 | Mtx1 | 0.029 | Old |
| 0.133784 | 0.141213 | 0.174 | 0.135 | 1 | 6 | Fam208b | 0.039 | Old |
| 0.13384 | 0.091777 | 0.403 | 0.348 | 1 | 6 | Sp100 | 0.055 | Old |
| 0.134383 | 0.346698 | 0.014 | 0.005 | 1 | 6 | Rbak | 0.009 | Old |
| 0.134694 | 1.033211 | 0.031 | 0.017 | 1 | 6 | Gk5 | 0.014 | Old |
| 0.134808 | 0.643739 | 0.043 | 0.025 | 1 | 6 | Cutc | 0.018 | Old |
| 0.134946 | 1.598277 | 0.031 | 0.017 | 1 | 6 | Hist2h2be | 0.014 | Old |
| 0.135004 | 0.72581 | 0.151 | 0.119 | 1 | 6 | Ufl1 | 0.032 | Old |
| 0.135059 | 0.358707 | 0.166 | 0.127 | 1 | 6 | Pcif1 | 0.039 | Old |
| 0.135192 | 0.283136 | 0.014 | 0.005 | 1 | 6 | Efcab2 | 0.009 | Old |
| 0.135238 | 0.23249 | 0.466 | 0.425 | 1 | 6 | Rps6ka3 | 0.041 | Old |
| 0.135266 | 0.122486 | 0.109 | 0.079 | 1 | 6 | Prpf19 | 0.03 | Old |
| 0.135399 | 0.322673 | 0.223 | 0.184 | 1 | 6 | Ociad1 | 0.039 | Old |
| 0.135784 | 0.000887 | 0.46 | 0.53 | 1 | 6 | Pum2 | -0.07 | Old |
| 0.135785 | 0.491405 | 0.154 | 0.12 | 1 | 6 | Mpdu1 | 0.034 | Old |
| 0.13587 | 0.451419 | 0.149 | 0.114 | 1 | 6 | Ppp1r16b | 0.035 | Old |
| 0.13607 | 0.619636 | 0.16 | 0.127 | 1 | 6 | Rasa1 | 0.033 | Old |
| 0.13632 | 0.135978 | 0.1 | 0.07 | 1 | 6 | Thoc5 | 0.03 | Old |
| 0.13633 | 0.328424 | 0.106 | 0.077 | 1 | 6 | Mtx2 | 0.029 | Old |
| 0.136462 | 0.879358 | 0.031 | 0.017 | 1 | 6 | Dtd1 | 0.014 | Old |
| 0.136544 | 0.652415 | 0.151 | 0.119 | 1 | 6 | Vps50 | 0.032 | Old |
| 0.136587 | 0.920649 | 0.049 | 0.03 | 1 | 6 | Gm30978 | 0.019 | Old |
| 0.13808 | 0.696955 | 0.34 | 0.309 | 1 | 6 | Slc3a2 | 0.031 | Old |
| 0.138083 | 0.25016 | 0.257 | 0.214 | 1 | 6 | Fam91a1 | 0.043 | Old |
| 0.138137 | 0.549722 | 0.1 | 0.074 | 1 | 6 | Ms4a6d | 0.026 | Old |
| 0.138525 | 0.209784 | 0.58 | 0.547 | 1 | 6 | Khdrbs1 | 0.033 | Old |
| 0.138627 | 0.286289 | 0.149 | 0.114 | 1 | 6 | Pak1ip1 | 0.035 | Old |
| 0.138762 | 0.734453 | 0.031 | 0.017 | 1 | 6 | Tysnd1 | 0.014 | Old |
| 0.138917 | 0.893416 | 0.057 | 0.037 | 1 | 6 | Fen1 | 0.02 | Old |
| 0.139019 | 0.707085 | 0.031 | 0.017 | 1 | 6 | B3galt6 | 0.014 | Old |
| 0.139112 | 0.379262 | 0.089 | 0.062 | 1 | 6 | Lipo3 | 0.027 | Old |
| 0.139183 | 1.272478 | 0.049 | 0.03 | 1 | 6 | Alg1 | 0.019 | Old |
| 0.139232 | 0.38203 | 0.251 | 0.207 | 1 | 6 | Noc2l | 0.044 | Old |
| 0.139277 | 0.707091 | 0.031 | 0.017 | 1 | 6 | Zbtb26 | 0.014 | Old |
| 0.139638 | 0.171305 | 0.269 | 0.222 | 1 | 6 | Ndufaf8 | 0.047 | Old |
| 0.139738 | 0.014372 | 0.043 | 0.025 | 1 | 6 | Zfp276 | 0.018 | Old |
| 0.139793 | 0.573282 | 0.031 | 0.017 | 1 | 6 | Zgpat | 0.014 | Old |
| 0.140351 | 0.090466 | 0.274 | 0.226 | 1 | 6 | Otub1 | 0.048 | Old |
| 0.14044 | 0.373967 | 0.1 | 0.072 | 1 | 6 | Ldlrap1 | 0.028 | Old |

| | | | | | | | | |
|---|---|---|---|---|---|---|---|---|
| 0.140548 | 1.186554 | 0.04 | 0.023 | 1 | 6 | Acp5 | 0.017 | Old |
| 0.141 | 0.916016 | 0.04 | 0.023 | 1 | 6 | Fam210a | 0.017 | Old |
| 0.141395 | 0.246365 | 0.417 | 0.361 | 1 | 6 | Rab11a | 0.056 | Old |
| 0.141401 | 0.339397 | 0.246 | 0.204 | 1 | 6 | Ccr1 | 0.042 | Old |
| 0.141409 | 0.680576 | 0.049 | 0.03 | 1 | 6 | Chpf | 0.019 | Old |
| 0.141905 | 0.293147 | 0.283 | 0.239 | 1 | 6 | Acsl4 | 0.044 | Old |
| 0.141911 | 0.581861 | 0.197 | 0.161 | 1 | 6 | Aqr | 0.036 | Old |
| 0.142125 | 0.3347 | 0.186 | 0.149 | 1 | 6 | Exoc2 | 0.037 | Old |
| 0.142225 | 1.156241 | 0.049 | 0.03 | 1 | 6 | Mrpl44 | 0.019 | Old |
| 0.142275 | 0.560107 | 0.091 | 0.065 | 1 | 6 | Stard5 | 0.026 | Old |
| 0.142307 | 0.510181 | 0.111 | 0.084 | 1 | 6 | Ftsj3 | 0.027 | Old |
| 0.142466 | 0.734173 | 0.146 | 0.115 | 1 | 6 | BC037034 | 0.031 | Old |
| 0.142491 | 0.520476 | 0.117 | 0.087 | 1 | 6 | Nlk | 0.03 | Old |
| 0.142592 | 0.484328 | 0.194 | 0.161 | 1 | 6 | Frmd4b | 0.033 | Old |
| 0.143048 | 0.521093 | 0.04 | 0.023 | 1 | 6 | Cipc | 0.017 | Old |
| 0.143186 | 0.247521 | 0.051 | 0.032 | 1 | 6 | Zer1 | 0.019 | Old |
| 0.143823 | 0.480924 | 0.066 | 0.043 | 1 | 6 | Zfp954 | 0.023 | Old |
| 0.144077 | 0.866709 | 0.054 | 0.035 | 1 | 6 | Ripk3 | 0.019 | Old |
| 0.144269 | 0.273401 | 0.186 | 0.149 | 1 | 6 | Slc30a5 | 0.037 | Old |
| 0.144681 | 0.140782 | 0.737 | 0.756 | 1 | 6 | Gnas | -0.019 | Old |
| 0.145228 | 0.521342 | 0.126 | 0.095 | 1 | 6 | Wdr91 | 0.031 | Old |
| 0.145306 | 0.206885 | 0.031 | 0.017 | 1 | 6 | Wdr6 | 0.014 | Old |
| 0.145873 | 0.368012 | 0.406 | 0.363 | 1 | 6 | Birc6 | 0.043 | Old |
| 0.14609 | 1.339186 | 0.029 | 0.015 | 1 | 6 | Taf1a | 0.014 | Old |
| 0.146309 | 0.472661 | 0.126 | 0.095 | 1 | 6 | Cpsf3 | 0.031 | Old |
| 0.147116 | 0.548293 | 0.129 | 0.099 | 1 | 6 | Lats1 | 0.03 | Old |
| 0.148484 | 0.595885 | 0.111 | 0.084 | 1 | 6 | Mir17hg | 0.027 | Old |
| 0.148891 | 0.344404 | 0.203 | 0.166 | 1 | 6 | Atp2c1 | 0.037 | Old |
| 0.149008 | 0.325352 | 0.423 | 0.383 | 1 | 6 | Hspd1 | 0.04 | Old |
| 0.149363 | 0.32361 | 0.094 | 0.067 | 1 | 6 | Srek1ip1 | 0.027 | Old |
| 0.14968 | 0.435926 | 0.277 | 0.242 | 1 | 6 | Cop1 | 0.035 | Old |
| 0.149777 | 1.083699 | 0.029 | 0.015 | 1 | 6 | Cox15 | 0.014 | Old |
| 0.14984 | 0.213104 | 0.074 | 0.104 | 1 | 6 | Cln3 | -0.03 | Old |
| 0.149948 | 0.404194 | 0.049 | 0.03 | 1 | 6 | Zfp346 | 0.019 | Old |
| 0.150022 | 0.203015 | 0.323 | 0.279 | 1 | 6 | Polr2a | 0.044 | Old |
| 0.150243 | 0.012665 | 0.077 | 0.107 | 1 | 6 | Il7r | -0.03 | Old |
| 0.150266 | 0.525148 | 0.111 | 0.084 | 1 | 6 | Bop1 | 0.027 | Old |
| 0.150544 | 0.169807 | 0.157 | 0.12 | 1 | 6 | Sf3a1 | 0.037 | Old |
| 0.150812 | 0.564873 | 0.089 | 0.064 | 1 | 6 | Tmco3 | 0.025 | Old |
| 0.151214 | 0.895927 | 0.029 | 0.015 | 1 | 6 | Sec61a2 | 0.014 | Old |
| 0.151503 | 0.826405 | 0.029 | 0.015 | 1 | 6 | Aldh18a1 | 0.014 | Old |
| 0.151747 | 0.913528 | 0.106 | 0.079 | 1 | 6 | Cabin1 | 0.027 | Old |
| 0.152009 | 0.144776 | 0.806 | 0.788 | 1 | 6 | Hnrnpk | 0.018 | Old |
| 0.152073 | 0.773952 | 0.091 | 0.067 | 1 | 6 | Specc1 | 0.024 | Old |
| 0.152161 | 0.501615 | 0.331 | 0.286 | 1 | 6 | Prpf4b | 0.045 | Old |

| | | | | | | | | |
|---|---|---|---|---|---|---|---|---|
| 0.152662 | 0.708935 | 0.029 | 0.015 | 1 | 6 | Chchd6 | 0.014 | Old |
| 0.152746 | 0.216501 | 0.663 | 0.624 | 1 | 6 | Cnbp | 0.039 | Old |
| 0.152952 | 0.967408 | 0.029 | 0.015 | 1 | 6 | A630072M | 0.014 | Old |
| 0.152989 | 0.49984 | 0.114 | 0.085 | 1 | 6 | Sdhaf4 | 0.029 | Old |
| 0.153572 | 0.236341 | 0.086 | 0.06 | 1 | 6 | Ago1 | 0.026 | Old |
| 0.153715 | 0.970191 | 0.066 | 0.045 | 1 | 6 | Armc5 | 0.021 | Old |
| 0.153734 | 0.694868 | 0.134 | 0.104 | 1 | 6 | Slc8b1 | 0.03 | Old |
| 0.153986 | 0.451533 | 0.166 | 0.134 | 1 | 6 | Rgs1 | 0.032 | Old |
| 0.154087 | 0.917875 | 0.103 | 0.077 | 1 | 6 | Stam2 | 0.026 | Old |
| 0.154226 | 0.322168 | 0.414 | 0.376 | 1 | 6 | Git2 | 0.038 | Old |
| 0.154448 | 0.16693 | 0.057 | 0.037 | 1 | 6 | Alkbh7 | 0.02 | Old |
| 0.154625 | 0.139769 | 0.417 | 0.366 | 1 | 6 | Nadk | 0.051 | Old |
| 0.154706 | 0.395426 | 0.029 | 0.015 | 1 | 6 | Taf9b | 0.014 | Old |
| 0.154798 | 0.409491 | 0.246 | 0.207 | 1 | 6 | Zfr | 0.039 | Old |
| 0.154868 | 0.28748 | 0.489 | 0.457 | 1 | 6 | Ncor1 | 0.032 | Old |
| 0.154986 | 0.293023 | 0.071 | 0.048 | 1 | 6 | Rnf121 | 0.023 | Old |
| 0.155341 | 0.254767 | 0.54 | 0.487 | 1 | 6 | Xiap | 0.053 | Old |
| 0.155347 | 0.388679 | 0.094 | 0.069 | 1 | 6 | Emc8 | 0.025 | Old |
| 0.155429 | 0.255545 | 0.634 | 0.624 | 1 | 6 | Snrpe | 0.01 | Old |
| 0.155473 | 0.832247 | 0.12 | 0.092 | 1 | 6 | Nfyb | 0.028 | Old |
| 0.155709 | 0.650196 | 0.063 | 0.042 | 1 | 6 | AC165953. | 0.021 | Old |
| 0.155834 | 0.470667 | 0.209 | 0.169 | 1 | 6 | Nup153 | 0.04 | Old |
| 0.155848 | 0.270745 | 0.171 | 0.137 | 1 | 6 | Gabpb2 | 0.034 | Old |
| 0.155856 | 0.296426 | 0.18 | 0.144 | 1 | 6 | Tbrg1 | 0.036 | Old |
| 0.156341 | 0.233823 | 0.214 | 0.176 | 1 | 6 | Psmd7 | 0.038 | Old |
| 0.156474 | 0.539884 | 0.029 | 0.015 | 1 | 6 | Itgb3bp | 0.014 | Old |
| 0.156686 | 0.348188 | 0.303 | 0.259 | 1 | 6 | Mrpl28 | 0.044 | Old |
| 0.156796 | 0.490954 | 0.089 | 0.064 | 1 | 6 | Topors | 0.025 | Old |
| 0.157078 | 0.29279 | 0.249 | 0.209 | 1 | 6 | Tnfrsf21 | 0.04 | Old |
| 0.157273 | 0.586355 | 0.111 | 0.084 | 1 | 6 | Slc39a9 | 0.027 | Old |
| 0.157294 | 0.066871 | 0.406 | 0.463 | 1 | 6 | Snx5 | -0.057 | Old |
| 0.157627 | 0.225924 | 0.443 | 0.393 | 1 | 6 | Ptprj | 0.05 | Old |
| 0.15779 | 0.125422 | 0.803 | 0.858 | 1 | 6 | Olfm1 | -0.055 | Old |
| 0.158163 | 0.138742 | 0.469 | 0.411 | 1 | 6 | Rtn3 | 0.058 | Old |
| 0.158349 | 0.320566 | 0.323 | 0.279 | 1 | 6 | Slc12a6 | 0.044 | Old |
| 0.158446 | 0.257868 | 0.063 | 0.089 | 1 | 6 | 4930402H2 | -0.026 | Old |
| 0.158561 | 0.633617 | 0.089 | 0.064 | 1 | 6 | Sharpin | 0.025 | Old |
| 0.158708 | 0.217896 | 0.166 | 0.129 | 1 | 6 | Mrps12 | 0.037 | Old |
| 0.158728 | 0.577548 | 0.069 | 0.047 | 1 | 6 | Fbxo9 | 0.022 | Old |
| 0.158758 | 0.326789 | 0.054 | 0.035 | 1 | 6 | Haus3 | 0.019 | Old |
| 0.159042 | 0.812172 | 0.037 | 0.022 | 1 | 6 | Cwc27 | 0.015 | Old |
| 0.159671 | 0.36759 | 0.077 | 0.054 | 1 | 6 | D730003I1 | 0.023 | Old |
| 0.160014 | 0.36166 | 0.054 | 0.035 | 1 | 6 | Gmnn | 0.019 | Old |
| 0.160017 | 0.929091 | 0.111 | 0.085 | 1 | 6 | Sfi1 | 0.026 | Old |
| 0.16007 | 0.405008 | 0.203 | 0.167 | 1 | 6 | BC004004 | 0.036 | Old |

| | | | | | | | | |
|---|---|---|---|---|---|---|---|---|
| 0.160332 | 1.053351 | 0.037 | 0.022 | 1 | 6 | Polr1b | 0.015 | Old |
| 0.160369 | 0.476571 | 0.134 | 0.104 | 1 | 6 | Vhl | 0.03 | Old |
| 0.160536 | 0.843571 | 0.06 | 0.04 | 1 | 6 | Fkbp5 | 0.02 | Old |
| 0.160911 | 1.044545 | 0.051 | 0.033 | 1 | 6 | Tmem110 | 0.018 | Old |
| 0.160932 | 0.213061 | 0.257 | 0.214 | 1 | 6 | Hmgn1 | 0.043 | Old |
| 0.161103 | 0.216289 | 0.171 | 0.135 | 1 | 6 | Tmem39a | 0.036 | Old |
| 0.161109 | 0.445942 | 0.177 | 0.144 | 1 | 6 | Clcn3 | 0.033 | Old |
| 0.161114 | 0.130201 | 0.071 | 0.048 | 1 | 6 | Uba7 | 0.023 | Old |
| 0.161208 | 0.399733 | 0.354 | 0.311 | 1 | 6 | Sf3b3 | 0.043 | Old |
| 0.161598 | 0.207181 | 0.114 | 0.085 | 1 | 6 | Naxe | 0.029 | Old |
| 0.16206 | 0.623286 | 0.106 | 0.08 | 1 | 6 | Exoc1 | 0.026 | Old |
| 0.162293 | 0.429539 | 0.24 | 0.202 | 1 | 6 | Ero1lb | 0.038 | Old |
| 0.162383 | 0.391046 | 0.317 | 0.269 | 1 | 6 | Senp6 | 0.048 | Old |
| 0.162646 | 0.874056 | 0.051 | 0.033 | 1 | 6 | Clcn6 | 0.018 | Old |
| 0.16302 | 0.363097 | 0.131 | 0.102 | 1 | 6 | Fam220a | 0.029 | Old |
| 0.163094 | 0.65158 | 0.103 | 0.077 | 1 | 6 | Ddx49 | 0.026 | Old |
| 0.163185 | 0.677263 | 0.054 | 0.035 | 1 | 6 | Snx19 | 0.019 | Old |
| 0.163188 | 1.147416 | 0.046 | 0.028 | 1 | 6 | Rab40c | 0.018 | Old |
| 0.163488 | 0.332702 | 0.329 | 0.289 | 1 | 6 | Tmem14c | 0.04 | Old |
| 0.163689 | 0.594891 | 0.143 | 0.114 | 1 | 6 | Rgs14 | 0.029 | Old |
| 0.163717 | 0.175673 | 0.077 | 0.054 | 1 | 6 | Ubac1 | 0.023 | Old |
| 0.163762 | 0.112157 | 0.171 | 0.209 | 1 | 6 | Spty2d1 | -0.038 | Old |
| 0.163982 | 0.211506 | 0.2 | 0.161 | 1 | 6 | Slc39a7 | 0.039 | Old |
| 0.164184 | 0.449152 | 0.123 | 0.094 | 1 | 6 | Clcc1 | 0.029 | Old |
| 0.164278 | 0.333503 | 0.257 | 0.217 | 1 | 6 | Vamp2 | 0.04 | Old |
| 0.164613 | 0.500184 | 0.074 | 0.052 | 1 | 6 | Tmem29 | 0.022 | Old |
| 0.164832 | 0.459445 | 0.294 | 0.258 | 1 | 6 | Gm26740 | 0.036 | Old |
| 0.164885 | 0.399757 | 0.074 | 0.052 | 1 | 6 | Atm | 0.022 | Old |
| 0.165 | 0.494125 | 0.12 | 0.092 | 1 | 6 | Smurf2 | 0.028 | Old |
| 0.165084 | 0.446468 | 0.571 | 0.543 | 1 | 6 | Gm26917 | 0.028 | Old |
| 0.165452 | 0.27432 | 0.231 | 0.189 | 1 | 6 | Chd9 | 0.042 | Old |
| 0.165551 | 0.519279 | 0.097 | 0.072 | 1 | 6 | Atg9a | 0.025 | Old |
| 0.165759 | 0.183671 | 0.329 | 0.278 | 1 | 6 | Mpc2 | 0.051 | Old |
| 0.165972 | 0.230938 | 0.174 | 0.139 | 1 | 6 | Itgb1bp1 | 0.035 | Old |
| 0.166134 | 0.724302 | 0.071 | 0.05 | 1 | 6 | Mtor | 0.021 | Old |
| 0.166246 | 0.450652 | 0.046 | 0.028 | 1 | 6 | Twsg1 | 0.018 | Old |
| 0.166658 | 0.185299 | 0.629 | 0.584 | 1 | 6 | Psme1 | 0.045 | Old |
| 0.166958 | 0.30827 | 0.046 | 0.028 | 1 | 6 | Fcgr4 | 0.018 | Old |
| 0.16703 | 0.187162 | 0.331 | 0.288 | 1 | 6 | 1700025G( | 0.043 | Old |
| 0.167059 | 0.31733 | 0.2 | 0.164 | 1 | 6 | Rbm28 | 0.036 | Old |
| 0.167297 | 0.652288 | 0.126 | 0.099 | 1 | 6 | Pds5b | 0.027 | Old |
| 0.167458 | 0.434993 | 0.097 | 0.072 | 1 | 6 | C1galt1c1 | 0.025 | Old |
| 0.167657 | 0.282677 | 0.214 | 0.177 | 1 | 6 | Tor1a | 0.037 | Old |
| 0.167939 | 0.798385 | 0.026 | 0.013 | 1 | 6 | Gm36445 | 0.013 | Old |
| 0.168351 | 0.265801 | 0.237 | 0.196 | 1 | 6 | Rrp1 | 0.041 | Old |

| | | | | | | | | |
|---|---|---|---|---|---|---|---|---|
| 0.168354 | 0.4436 | 0.083 | 0.059 | 1 | 6 | Kpna6 | 0.024 | Old |
| 0.168505 | 0.397658 | 0.037 | 0.022 | 1 | 6 | Snupn | 0.015 | Old |
| 0.168519 | 0.412368 | 0.28 | 0.241 | 1 | 6 | Pabpn1 | 0.039 | Old |
| 0.168749 | 1.01444 | 0.077 | 0.055 | 1 | 6 | Pbx1 | 0.022 | Old |
| 0.169129 | 0.101995 | 0.197 | 0.156 | 1 | 6 | Mvb12a | 0.041 | Old |
| 0.169262 | 1.420276 | 0.026 | 0.013 | 1 | 6 | Il18r1 | 0.013 | Old |
| 0.169608 | 1.27917 | 0.034 | 0.02 | 1 | 6 | Zfp760 | 0.014 | Old |
| 0.169713 | 0.169875 | 0.76 | 0.763 | 1 | 6 | Nap1l1 | -0.003 | Old |
| 0.169859 | 0.624864 | 0.089 | 0.065 | 1 | 6 | Epb41l1 | 0.024 | Old |
| 0.169876 | 0.073822 | 0.126 | 0.095 | 1 | 6 | Slc31a1 | 0.031 | Old |
| 0.17067 | 0.640231 | 0.066 | 0.045 | 1 | 6 | Fra10ac1 | 0.021 | Old |
| 0.170749 | 0.113063 | 0.171 | 0.135 | 1 | 6 | Vps41 | 0.036 | Old |
| 0.170853 | 0.722268 | 0.051 | 0.033 | 1 | 6 | Lag3 | 0.018 | Old |
| 0.170853 | 0.563148 | 0.051 | 0.033 | 1 | 6 | Bcl9l | 0.018 | Old |
| 0.170886 | 0.925417 | 0.043 | 0.027 | 1 | 6 | Isoc2b | 0.016 | Old |
| 0.171035 | 0.245823 | 0.046 | 0.028 | 1 | 6 | Gcdh | 0.018 | Old |
| 0.171174 | 0.50265 | 0.114 | 0.087 | 1 | 6 | Rwdd4a | 0.027 | Old |
| 0.171497 | 0.330569 | 0.391 | 0.37 | 1 | 6 | Palld | 0.021 | Old |
| 0.172285 | 0.759848 | 0.111 | 0.085 | 1 | 6 | Naa20 | 0.026 | Old |
| 0.172406 | 0.276376 | 0.069 | 0.047 | 1 | 6 | Osbpl2 | 0.022 | Old |
| 0.172657 | 0.239155 | 0.28 | 0.242 | 1 | 6 | Bud31 | 0.038 | Old |
| 0.173001 | 1.268884 | 0.034 | 0.02 | 1 | 6 | Pim2 | 0.014 | Old |
| 0.173015 | 0.109604 | 0.046 | 0.069 | 1 | 6 | Plekha3 | -0.023 | Old |
| 0.173045 | 0.694121 | 0.08 | 0.057 | 1 | 6 | Scd1 | 0.023 | Old |
| 0.173201 | 0.457921 | 0.149 | 0.117 | 1 | 6 | Esco1 | 0.032 | Old |
| 0.17358 | 0.631006 | 0.051 | 0.033 | 1 | 6 | Gmds | 0.018 | Old |
| 0.174425 | 0.161656 | 0.626 | 0.595 | 1 | 6 | Tpm4 | 0.031 | Old |
| 0.175003 | 1.071247 | 0.086 | 0.064 | 1 | 6 | Noc4l | 0.022 | Old |
| 0.175162 | 0.585469 | 0.043 | 0.027 | 1 | 6 | Gm26518 | 0.016 | Old |
| 0.175554 | 0.345218 | 0.06 | 0.04 | 1 | 6 | Mettl5 | 0.02 | Old |
| 0.175679 | 0.165711 | 0.046 | 0.028 | 1 | 6 | Alg2 | 0.018 | Old |
| 0.175774 | 0.425662 | 0.08 | 0.057 | 1 | 6 | 2510002D2 | 0.023 | Old |
| 0.175798 | 0.830692 | 0.043 | 0.027 | 1 | 6 | Cadm3 | 0.016 | Old |
| 0.175866 | 0.928944 | 0.034 | 0.02 | 1 | 6 | Jpx | 0.014 | Old |
| 0.175866 | 0.727579 | 0.034 | 0.02 | 1 | 6 | Asb6 | 0.014 | Old |
| 0.176407 | 0.469912 | 0.083 | 0.06 | 1 | 6 | Trappc3 | 0.023 | Old |
| 0.176684 | 0.223638 | 0.394 | 0.336 | 1 | 6 | Lrrc58 | 0.058 | Old |
| 0.176885 | 0.09707 | 0.674 | 0.64 | 1 | 6 | Son | 0.034 | Old |
| 0.176979 | 0.686594 | 0.1 | 0.075 | 1 | 6 | Mphosph6 | 0.025 | Old |
| 0.177035 | 0.449269 | 0.051 | 0.033 | 1 | 6 | Bcar3 | 0.018 | Old |
| 0.177184 | 0.134181 | 0.269 | 0.224 | 1 | 6 | Phf14 | 0.045 | Old |
| 0.17728 | 0.133603 | 0.589 | 0.542 | 1 | 6 | Ndufb7 | 0.047 | Old |
| 0.177301 | 0.285105 | 0.066 | 0.045 | 1 | 6 | Ikbkg | 0.021 | Old |
| 0.177331 | 0.362193 | 0.34 | 0.301 | 1 | 6 | Eif3b | 0.039 | Old |
| 0.177361 | 0.12771 | 0.1 | 0.074 | 1 | 6 | Lmbrd2 | 0.026 | Old |

| | | | | | | | | |
|---|---|---|---|---|---|---|---|---|
| 0.177459 | 0.419161 | 0.043 | 0.027 | 1 | 6 | Polr3e | 0.016 | Old |
| 0.177532 | 0.570533 | 0.237 | 0.202 | 1 | 6 | Ell2 | 0.035 | Old |
| 0.177811 | 0.635149 | 0.066 | 0.045 | 1 | 6 | Unkl | 0.021 | Old |
| 0.178159 | 0.504589 | 0.123 | 0.094 | 1 | 6 | Ccdc25 | 0.029 | Old |
| 0.178186 | 0.380341 | 0.06 | 0.085 | 1 | 6 | E2f1 | -0.025 | Old |
| 0.178186 | 0.287401 | 0.1 | 0.074 | 1 | 6 | Sdc1 | 0.026 | Old |
| 0.178188 | 0.184844 | 0.329 | 0.284 | 1 | 6 | Mtch1 | 0.045 | Old |
| 0.178479 | 0.090175 | 0.143 | 0.174 | 1 | 6 | Sfswap | -0.031 | Old |
| 0.178744 | 0.628465 | 0.043 | 0.027 | 1 | 6 | Supt7l | 0.016 | Old |
| 0.179352 | 0.669617 | 0.034 | 0.02 | 1 | 6 | R3hcc1 | 0.014 | Old |
| 0.179843 | 0.187829 | 0.126 | 0.097 | 1 | 6 | Farsb | 0.029 | Old |
| 0.17988 | 0.073709 | 0.303 | 0.258 | 1 | 6 | Stat6 | 0.045 | Old |
| 0.18007 | 0.39503 | 0.051 | 0.033 | 1 | 6 | Btrc | 0.018 | Old |
| 0.180073 | 0.400905 | 0.211 | 0.179 | 1 | 6 | Abcf1 | 0.032 | Old |
| 0.180089 | 0.381899 | 0.137 | 0.107 | 1 | 6 | Lcor | 0.03 | Old |
| 0.180203 | 0.57344 | 0.089 | 0.065 | 1 | 6 | Tpst1 | 0.024 | Old |
| 0.180575 | 1.212425 | 0.023 | 0.012 | 1 | 6 | Chst1 | 0.011 | Old |
| 0.180674 | 0.264239 | 0.22 | 0.184 | 1 | 6 | Chfr | 0.036 | Old |
| 0.180815 | 0.53344 | 0.043 | 0.027 | 1 | 6 | Nrg2 | 0.016 | Old |
| 0.1809 | 0.104249 | 0.086 | 0.062 | 1 | 6 | Arid3a | 0.024 | Old |
| 0.18112 | 0.579355 | 0.083 | 0.06 | 1 | 6 | Nlrc5 | 0.023 | Old |
| 0.181277 | 1.239603 | 0.049 | 0.032 | 1 | 6 | Vps37c | 0.017 | Old |
| 0.181316 | 0.973811 | 0.023 | 0.012 | 1 | 6 | Kyat1 | 0.011 | Old |
| 0.181325 | 0.533436 | 0.14 | 0.112 | 1 | 6 | Rnf157 | 0.028 | Old |
| 0.181685 | 0.517362 | 0.1 | 0.075 | 1 | 6 | Aen | 0.025 | Old |
| 0.181763 | 0.191471 | 0.649 | 0.622 | 1 | 6 | Srp9 | 0.027 | Old |
| 0.181859 | 0.003291 | 0.22 | 0.266 | 1 | 6 | Rap1gds1 | -0.046 | Old |
| 0.181903 | 0.062914 | 0.017 | 0.032 | 1 | 6 | Pard6a | -0.015 | Old |
| 0.182051 | 0.650571 | 0.131 | 0.104 | 1 | 6 | Ric1 | 0.027 | Old |
| 0.182274 | 0.703217 | 0.057 | 0.038 | 1 | 6 | Rdh14 | 0.019 | Old |
| 0.182313 | 0.222304 | 0.326 | 0.281 | 1 | 6 | Osgep | 0.045 | Old |
| 0.182398 | 0.164523 | 0.743 | 0.744 | 1 | 6 | 2010107E0 | -0.001 | Old |
| 0.182498 | 0.903144 | 0.057 | 0.038 | 1 | 6 | Rabepk | 0.019 | Old |
| 0.182589 | 0.266888 | 0.083 | 0.06 | 1 | 6 | Dolpp1 | 0.023 | Old |
| 0.182641 | 0.190237 | 0.043 | 0.027 | 1 | 6 | Serinc5 | 0.016 | Old |
| 0.182737 | 0.414387 | 0.157 | 0.127 | 1 | 6 | Cog5 | 0.03 | Old |
| 0.182769 | 0.674251 | 0.074 | 0.054 | 1 | 6 | 2310039H0 | 0.02 | Old |
| 0.182773 | 0.460256 | 0.083 | 0.06 | 1 | 6 | Wdr5 | 0.023 | Old |
| 0.182853 | 0.366121 | 0.274 | 0.236 | 1 | 6 | Ski | 0.038 | Old |
| 0.182859 | 0.667225 | 0.211 | 0.182 | 1 | 6 | Trappc6b | 0.029 | Old |
| 0.183026 | 0.335128 | 0.171 | 0.139 | 1 | 6 | Gpr108 | 0.032 | Old |
| 0.183137 | 0.389232 | 0.191 | 0.157 | 1 | 6 | Nrp1 | 0.034 | Old |
| 0.18316 | 0.180206 | 0.78 | 0.776 | 1 | 6 | Ube2d3 | 0.004 | Old |
| 0.18318 | 1.746323 | 0.023 | 0.012 | 1 | 6 | Atp6v1g2 | 0.011 | Old |
| 0.18318 | 1.135715 | 0.023 | 0.012 | 1 | 6 | Ddah2 | 0.011 | Old |

| | | | | | | | | |
|---|---|---|---|---|---|---|---|---|
| 0.183245 | 0.7013 | 0.14 | 0.114 | 1 | 6 | Pfdn4 | 0.026 | Old |
| 0.183468 | 0.773349 | 0.049 | 0.032 | 1 | 6 | Nat10 | 0.017 | Old |
| 0.183996 | 0.725662 | 0.063 | 0.043 | 1 | 6 | Arl2 | 0.02 | Old |
| 0.184049 | 0.987032 | 0.094 | 0.07 | 1 | 6 | Lpar6 | 0.024 | Old |
| 0.184145 | 0.305157 | 0.36 | 0.328 | 1 | 6 | Cct3 | 0.032 | Old |
| 0.184377 | 0.398534 | 0.034 | 0.02 | 1 | 6 | 1810043G( | 0.014 | Old |
| 0.185065 | 0.509345 | 0.257 | 0.226 | 1 | 6 | U2af1 | 0.031 | Old |
| 0.185079 | 0.429016 | 0.071 | 0.05 | 1 | 6 | Tapbpl | 0.021 | Old |
| 0.185277 | 0.297636 | 0.123 | 0.094 | 1 | 6 | Atad2 | 0.029 | Old |
| 0.185834 | 0.076477 | 0.563 | 0.517 | 1 | 6 | Cox17 | 0.046 | Old |
| 0.186009 | 0.288043 | 0.151 | 0.12 | 1 | 6 | Mdn1 | 0.031 | Old |
| 0.18607 | 0.444575 | 0.043 | 0.027 | 1 | 6 | Tmem156 | 0.016 | Old |
| 0.186087 | 0.23017 | 0.071 | 0.05 | 1 | 6 | Mki67 | 0.021 | Old |
| 0.186191 | 0.482936 | 0.023 | 0.012 | 1 | 6 | Commd9 | 0.011 | Old |
| 0.186206 | 0.621334 | 0.103 | 0.079 | 1 | 6 | Arhgef12 | 0.024 | Old |
| 0.186287 | 0.058081 | 0.111 | 0.144 | 1 | 6 | Tlk2 | -0.033 | Old |
| 0.186521 | 0.203801 | 0.377 | 0.331 | 1 | 6 | Slc39a1 | 0.046 | Old |
| 0.186643 | 0.113073 | 0.391 | 0.344 | 1 | 6 | Atp6v1e1 | 0.047 | Old |
| 0.186787 | 0.319982 | 0.234 | 0.197 | 1 | 6 | Vps28 | 0.037 | Old |
| 0.187099 | 0.430096 | 0.071 | 0.05 | 1 | 6 | Fam53a | 0.021 | Old |
| 0.187381 | 0.158342 | 0.111 | 0.084 | 1 | 6 | Mkl2 | 0.027 | Old |
| 0.187463 | 0.558898 | 0.163 | 0.132 | 1 | 6 | Rsl24d1 | 0.031 | Old |
| 0.1876 | 0.214607 | 0.697 | 0.667 | 1 | 6 | Grb2 | 0.03 | Old |
| 0.187602 | 0.423002 | 0.126 | 0.097 | 1 | 6 | Med10 | 0.029 | Old |
| 0.187635 | 0.740279 | 0.097 | 0.074 | 1 | 6 | Prkch | 0.023 | Old |
| 0.18798 | 0.257907 | 0.491 | 0.467 | 1 | 6 | Dok2 | 0.024 | Old |
| 0.188154 | 0.698888 | 0.097 | 0.074 | 1 | 6 | Cd2 | 0.023 | Old |
| 0.188471 | 0.095796 | 0.043 | 0.027 | 1 | 6 | Slc38a6 | 0.016 | Old |
| 0.188491 | 0.229029 | 0.091 | 0.067 | 1 | 6 | Tmem251 | 0.024 | Old |
| 0.188629 | 0.891234 | 0.069 | 0.048 | 1 | 6 | Utp15 | 0.021 | Old |
| 0.188819 | 0.348061 | 0.274 | 0.236 | 1 | 6 | Mark2 | 0.038 | Old |
| 0.189031 | 0.409144 | 0.223 | 0.184 | 1 | 6 | Cntrl | 0.039 | Old |
| 0.189118 | 0.269624 | 0.417 | 0.381 | 1 | 6 | Etf1 | 0.036 | Old |
| 0.189239 | 0.013207 | 0.023 | 0.012 | 1 | 6 | Tnk2 | 0.011 | Old |
| 0.189623 | 0.107613 | 0.023 | 0.012 | 1 | 6 | Sprtn | 0.011 | Old |
| 0.189962 | 0.484539 | 0.123 | 0.095 | 1 | 6 | Cnot7 | 0.028 | Old |
| 0.190226 | 0.007774 | 0.417 | 0.465 | 1 | 6 | Dnajb14 | -0.048 | Old |
| 0.190245 | 0.263885 | 0.294 | 0.263 | 1 | 6 | Bst2 | 0.031 | Old |
| 0.190286 | 0.266106 | 0.151 | 0.124 | 1 | 6 | Med8 | 0.027 | Old |
| 0.190683 | 0.061132 | 0.146 | 0.177 | 1 | 6 | Maf1 | -0.031 | Old |
| 0.190683 | 0.276665 | 0.057 | 0.038 | 1 | 6 | Wiz | 0.019 | Old |
| 0.190683 | 0.262275 | 0.057 | 0.038 | 1 | 6 | Zfp64 | 0.019 | Old |
| 0.190827 | 1.17331 | 0.031 | 0.018 | 1 | 6 | Fbxw17 | 0.013 | Old |
| 0.191449 | 0.263765 | 0.057 | 0.08 | 1 | 6 | Mfng | -0.023 | Old |
| 0.191498 | 0.361082 | 0.237 | 0.202 | 1 | 6 | Sike1 | 0.035 | Old |

| | | | | | | | | |
|---|---|---|---|---|---|---|---|---|
| 0.191831 | 0.40282 | 0.077 | 0.055 | 1 | 6 | Prr14l | 0.022 | Old |
| 0.191918 | 0.384892 | 0.243 | 0.209 | 1 | 6 | Kcnn4 | 0.034 | Old |
| 0.192108 | 0.82626 | 0.031 | 0.018 | 1 | 6 | Gpr82 | 0.013 | Old |
| 0.192555 | 0.242237 | 0.189 | 0.154 | 1 | 6 | Chml | 0.035 | Old |
| 0.193246 | 0.293933 | 0.423 | 0.388 | 1 | 6 | Raly | 0.035 | Old |
| 0.193768 | 0.463372 | 0.286 | 0.246 | 1 | 6 | Hnrnpr | 0.04 | Old |
| 0.193865 | 0.6322 | 0.106 | 0.08 | 1 | 6 | Senp5 | 0.026 | Old |
| 0.193927 | 0.207623 | 0.523 | 0.482 | 1 | 6 | Cap1 | 0.041 | Old |
| 0.194114 | 0.780889 | 0.066 | 0.047 | 1 | 6 | Psat1 | 0.019 | Old |
| 0.194249 | 0.68441 | 0.04 | 0.025 | 1 | 6 | Wrap73 | 0.015 | Old |
| 0.194281 | 0.356721 | 0.12 | 0.092 | 1 | 6 | Cwf19l2 | 0.028 | Old |
| 0.194289 | 0.409076 | 0.183 | 0.151 | 1 | 6 | Parp14 | 0.032 | Old |
| 0.194649 | 0.143815 | 0.354 | 0.314 | 1 | 6 | Psma1 | 0.04 | Old |
| 0.194697 | 0.185342 | 0.16 | 0.127 | 1 | 6 | Ppfibp1 | 0.033 | Old |
| 0.194835 | 0.140951 | 0.543 | 0.5 | 1 | 6 | Prkar1a | 0.043 | Old |
| 0.194946 | 0.474217 | 0.114 | 0.089 | 1 | 6 | B3galnt2 | 0.025 | Old |
| 0.19501 | 0.012691 | 0.246 | 0.286 | 1 | 6 | Lrmp | -0.04 | Old |
| 0.195015 | 1.082041 | 0.031 | 0.018 | 1 | 6 | Abhd10 | 0.013 | Old |
| 0.195054 | 0.152194 | 0.171 | 0.139 | 1 | 6 | Dpf2 | 0.032 | Old |
| 0.195384 | 0.598406 | 0.04 | 0.025 | 1 | 6 | Abhd18 | 0.015 | Old |
| 0.195654 | 0.078507 | 0.071 | 0.05 | 1 | 6 | Plcb3 | 0.021 | Old |
| 0.195899 | 0.332674 | 0.154 | 0.124 | 1 | 6 | Sec22b | 0.03 | Old |
| 0.195991 | 1.105096 | 0.031 | 0.018 | 1 | 6 | Uevld | 0.013 | Old |
| 0.19713 | 0.093526 | 0.011 | 0.023 | 1 | 6 | Fmnl3 | -0.012 | Old |
| 0.19725 | 0.584941 | 0.186 | 0.156 | 1 | 6 | Camta2 | 0.03 | Old |
| 0.197379 | 0.279985 | 0.354 | 0.319 | 1 | 6 | Ppp1r14b | 0.035 | Old |
| 0.197405 | 0.398453 | 0.097 | 0.074 | 1 | 6 | Prdx4 | 0.023 | Old |
| 0.197546 | 0.240334 | 0.151 | 0.122 | 1 | 6 | Eif4enif1 | 0.029 | Old |
| 0.197626 | 1.115341 | 0.031 | 0.018 | 1 | 6 | Abhd14b | 0.013 | Old |
| 0.197801 | 0.226684 | 0.049 | 0.032 | 1 | 6 | B3gnt5 | 0.017 | Old |
| 0.197864 | 0.340297 | 0.083 | 0.06 | 1 | 6 | Slc7a7 | 0.023 | Old |
| 0.198282 | 1.041268 | 0.031 | 0.018 | 1 | 6 | Top3a | 0.013 | Old |
| 0.198318 | 0.763143 | 0.049 | 0.032 | 1 | 6 | Ppif | 0.017 | Old |
| 0.198495 | 0.516957 | 0.063 | 0.043 | 1 | 6 | Hddc2 | 0.02 | Old |
| 0.19858 | 0.520121 | 0.143 | 0.115 | 1 | 6 | Lyl1 | 0.028 | Old |
| 0.19868 | 0.623475 | 0.109 | 0.084 | 1 | 6 | Ergic1 | 0.025 | Old |
| 0.198992 | 0.340164 | 0.154 | 0.124 | 1 | 6 | Hk1 | 0.03 | Old |
| 0.199053 | 1.699322 | 0.02 | 0.01 | 1 | 6 | Gtpbp3 | 0.01 | Old |
| 0.1991 | 0.566733 | 0.109 | 0.085 | 1 | 6 | Snx29 | 0.024 | Old |
| 0.199304 | 0.681845 | 0.094 | 0.072 | 1 | 6 | Tmem198b | 0.022 | Old |
| 0.199407 | 0.284964 | 0.514 | 0.475 | 1 | 6 | Ube2d2a | 0.039 | Old |
| 0.199472 | 0.331567 | 0.214 | 0.179 | 1 | 6 | Rasa3 | 0.035 | Old |
| 0.19967 | 0.328315 | 0.191 | 0.154 | 1 | 6 | Mob1b | 0.037 | Old |
| 0.199753 | 0.312396 | 0.289 | 0.251 | 1 | 6 | Bax | 0.038 | Old |
| 0.199929 | 0.419617 | 0.269 | 0.239 | 1 | 6 | Vps36 | 0.03 | Old |

| | | | | | | | | |
|---|---|---|---|---|---|---|---|---|
| 0.199931 | 0.75847 | 0.031 | 0.018 | 1 | 6 | Lmnb2 | 0.013 | Old |
| 0.200336 | 1.043574 | 0.02 | 0.01 | 1 | 6 | Zbtb32 | 0.01 | Old |
| 0.20072 | 0.870236 | 0.109 | 0.085 | 1 | 6 | Nbeal2 | 0.024 | Old |
| 0.200736 | 0.646413 | 0.069 | 0.048 | 1 | 6 | Mpg | 0.021 | Old |
| 0.200749 | 0.567464 | 0.06 | 0.042 | 1 | 6 | Stambp | 0.018 | Old |
| 0.200849 | 0.046647 | 0.294 | 0.339 | 1 | 6 | Hipk2 | -0.045 | Old |
| 0.201016 | 0.710866 | 0.129 | 0.104 | 1 | 6 | Idh3a | 0.025 | Old |
| 0.201422 | 0.603429 | 0.109 | 0.084 | 1 | 6 | Cep250 | 0.025 | Old |
| 0.201499 | 0.54118 | 0.054 | 0.037 | 1 | 6 | Idua | 0.017 | Old |
| 0.201509 | 0.524435 | 0.103 | 0.079 | 1 | 6 | Mettl16 | 0.024 | Old |
| 0.20159 | 0.511526 | 0.031 | 0.018 | 1 | 6 | Fam198b | 0.013 | Old |
| 0.20159 | 0.488749 | 0.031 | 0.018 | 1 | 6 | Cenpo | 0.013 | Old |
| 0.201621 | 0.741108 | 0.091 | 0.069 | 1 | 6 | Traf3ip3 | 0.022 | Old |
| 0.201626 | 1.403297 | 0.02 | 0.01 | 1 | 6 | Suv39h2 | 0.01 | Old |
| 0.202057 | 1.542961 | 0.02 | 0.01 | 1 | 6 | Txnl4b | 0.01 | Old |
| 0.202057 | 1.433025 | 0.02 | 0.01 | 1 | 6 | Pms1 | 0.01 | Old |
| 0.202113 | 0.563781 | 0.134 | 0.107 | 1 | 6 | Med21 | 0.027 | Old |
| 0.202204 | 0.316184 | 0.449 | 0.406 | 1 | 6 | Fgfr1op2 | 0.043 | Old |
| 0.202477 | 0.339109 | 0.546 | 0.518 | 1 | 6 | Nktr | 0.028 | Old |
| 0.202541 | 0.204886 | 0.146 | 0.115 | 1 | 6 | Dpp3 | 0.031 | Old |
| 0.202669 | 0.740257 | 0.066 | 0.047 | 1 | 6 | 2310061I0 | 0.019 | Old |
| 0.202745 | 0.212722 | 0.18 | 0.145 | 1 | 6 | Ppp2r5e | 0.035 | Old |
| 0.202921 | 0.90752 | 0.02 | 0.01 | 1 | 6 | Hgh1 | 0.01 | Old |
| 0.202921 | 0.765769 | 0.02 | 0.01 | 1 | 6 | Pou5f1 | 0.01 | Old |
| 0.203051 | 0.18052 | 0.089 | 0.065 | 1 | 6 | Plagl2 | 0.024 | Old |
| 0.203354 | 0.534594 | 0.02 | 0.01 | 1 | 6 | Cenpe | 0.01 | Old |
| 0.203546 | 0.217308 | 0.049 | 0.032 | 1 | 6 | Adprm | 0.017 | Old |
| 0.203658 | 0.403482 | 0.206 | 0.172 | 1 | 6 | Dynll2 | 0.034 | Old |
| 0.203788 | 0.608923 | 0.02 | 0.01 | 1 | 6 | Zfp472 | 0.01 | Old |
| 0.203833 | 0.650683 | 0.123 | 0.097 | 1 | 6 | Enox2 | 0.026 | Old |
| 0.203971 | 0.22231 | 0.411 | 0.373 | 1 | 6 | Clec4a3 | 0.038 | Old |
| 0.204458 | 0.446123 | 0.143 | 0.115 | 1 | 6 | Txndc16 | 0.028 | Old |
| 0.2046 | 0.615113 | 0.031 | 0.018 | 1 | 6 | Mettl2 | 0.013 | Old |
| 0.204637 | 0.106683 | 0.04 | 0.025 | 1 | 6 | Fzd5 | 0.015 | Old |
| 0.205183 | 0.672057 | 0.046 | 0.03 | 1 | 6 | Calcrl | 0.016 | Old |
| 0.205504 | 0.184748 | 0.549 | 0.512 | 1 | 6 | Cmpk1 | 0.037 | Old |
| 0.205825 | 0.630448 | 0.18 | 0.154 | 1 | 6 | Dennd6a | 0.026 | Old |
| 0.205945 | 0.868439 | 0.077 | 0.057 | 1 | 6 | Slc25a16 | 0.02 | Old |
| 0.205994 | 0.233107 | 0.203 | 0.166 | 1 | 6 | Eml4 | 0.037 | Old |
| 0.206149 | 0.572272 | 0.103 | 0.079 | 1 | 6 | Ap5m1 | 0.024 | Old |
| 0.206325 | 0.364611 | 0.16 | 0.129 | 1 | 6 | Riok1 | 0.031 | Old |
| 0.206405 | 0.287538 | 0.02 | 0.01 | 1 | 6 | Gm10320 | 0.01 | Old |
| 0.206844 | 0.341831 | 0.02 | 0.01 | 1 | 6 | Tti1 | 0.01 | Old |
| 0.207246 | 0.176957 | 0.129 | 0.1 | 1 | 6 | Pafah1b2 | 0.029 | Old |
| 0.207791 | 0.769131 | 0.051 | 0.035 | 1 | 6 | Sacs | 0.016 | Old |

| | | | | | | | | |
|---|---|---|---|---|---|---|---|---|
| 0.207986 | 0.501752 | 0.22 | 0.189 | 1 | 6 | Prrc2b | 0.031 | Old |
| 0.208021 | 0.012182 | 0.243 | 0.202 | 1 | 6 | Otulin | 0.041 | Old |
| 0.208025 | 0.325334 | 0.129 | 0.159 | 1 | 6 | Ifitm1 | -0.03 | Old |
| 0.208338 | 0.417679 | 0.026 | 0.042 | 1 | 6 | BC024978 | -0.016 | Old |
| 0.208409 | 0.392578 | 0.086 | 0.064 | 1 | 6 | Bap1 | 0.022 | Old |
| 0.209065 | 0.289192 | 0.326 | 0.289 | 1 | 6 | Mia3 | 0.037 | Old |
| 0.209404 | 0.296451 | 0.086 | 0.064 | 1 | 6 | Cwc22 | 0.022 | Old |
| 0.209405 | 0.287984 | 0.103 | 0.079 | 1 | 6 | Pgp | 0.024 | Old |
| 0.209437 | 0.557798 | 0.14 | 0.114 | 1 | 6 | Fam193a | 0.026 | Old |
| 0.209994 | 0.55884 | 0.054 | 0.037 | 1 | 6 | Pnpt1 | 0.017 | Old |
| 0.210026 | 0.121594 | 0.549 | 0.507 | 1 | 6 | Myl12a | 0.042 | Old |
| 0.210329 | 0.403596 | 0.134 | 0.107 | 1 | 6 | Exosc3 | 0.027 | Old |
| 0.210371 | 0.2403 | 0.38 | 0.344 | 1 | 6 | Elmo2 | 0.036 | Old |
| 0.210893 | 0.232104 | 0.529 | 0.51 | 1 | 6 | Ptpn18 | 0.019 | Old |
| 0.211002 | 0.246219 | 0.511 | 0.487 | 1 | 6 | Tram1 | 0.024 | Old |
| 0.21106 | 0.14759 | 0.109 | 0.084 | 1 | 6 | Zfp236 | 0.025 | Old |
| 0.211919 | 0.724689 | 0.094 | 0.072 | 1 | 6 | Nudt9 | 0.022 | Old |
| 0.211968 | 0.758542 | 0.109 | 0.085 | 1 | 6 | Rtl8a | 0.024 | Old |
| 0.212348 | 0.250246 | 0.629 | 0.6 | 1 | 6 | Jpt1 | 0.029 | Old |
| 0.213172 | 0.229186 | 0.094 | 0.07 | 1 | 6 | Pex13 | 0.024 | Old |
| 0.213175 | 0.271279 | 0.251 | 0.212 | 1 | 6 | B4galt1 | 0.039 | Old |
| 0.213608 | 0.270777 | 0.12 | 0.094 | 1 | 6 | Zmat5 | 0.026 | Old |
| 0.213699 | 1.01846 | 0.083 | 0.062 | 1 | 6 | Tm7sf3 | 0.021 | Old |
| 0.214255 | 0.499195 | 0.063 | 0.045 | 1 | 6 | Slc20a2 | 0.018 | Old |
| 0.214612 | 0.63492 | 0.046 | 0.03 | 1 | 6 | Cyb561d1 | 0.016 | Old |
| 0.214723 | 1.022778 | 0.029 | 0.017 | 1 | 6 | Gm17089 | 0.012 | Old |
| 0.214989 | 0.016049 | 0.206 | 0.247 | 1 | 6 | Emp1 | -0.041 | Old |
| 0.215037 | 0.654224 | 0.037 | 0.023 | 1 | 6 | Cd3eap | 0.014 | Old |
| 0.215332 | 1.577526 | 0.003 | 0.01 | 1 | 6 | Pram1 | -0.007 | Old |
| 0.215798 | 0.096215 | 0.457 | 0.5 | 1 | 6 | Rac2 | -0.043 | Old |
| 0.215806 | 0.405482 | 0.263 | 0.231 | 1 | 6 | Lamtor3 | 0.032 | Old |
| 0.216001 | 0.086352 | 0.226 | 0.268 | 1 | 6 | 6-Sep | -0.042 | Old |
| 0.216426 | 0.256298 | 0.131 | 0.164 | 1 | 6 | Fmr1 | -0.033 | Old |
| 0.216493 | 0.502288 | 0.129 | 0.102 | 1 | 6 | Milr1 | 0.027 | Old |
| 0.216553 | 1.659608 | 0.029 | 0.017 | 1 | 6 | Tlr3 | 0.012 | Old |
| 0.216923 | 0.545963 | 0.054 | 0.037 | 1 | 6 | Cep57 | 0.017 | Old |
| 0.217114 | 0.040665 | 0.166 | 0.132 | 1 | 6 | Tlk1 | 0.034 | Old |
| 0.21725 | 0.516749 | 0.131 | 0.105 | 1 | 6 | Arid2 | 0.026 | Old |
| 0.217258 | 0.241203 | 0.48 | 0.448 | 1 | 6 | Txn1 | 0.032 | Old |
| 0.217921 | 0.6484 | 0.12 | 0.094 | 1 | 6 | Taf5l | 0.026 | Old |
| 0.218025 | 0.838633 | 0.029 | 0.017 | 1 | 6 | Pgap3 | 0.012 | Old |
| 0.218073 | 2.095855 | 0.017 | 0.008 | 1 | 6 | Gm4631 | 0.009 | Old |
| 0.218503 | 0.334091 | 0.171 | 0.144 | 1 | 6 | Utp18 | 0.027 | Old |
| 0.218657 | 0.31477 | 0.126 | 0.099 | 1 | 6 | Hdac5 | 0.027 | Old |
| 0.218764 | 1.027958 | 0.029 | 0.017 | 1 | 6 | Abhd6 | 0.012 | Old |

| | | | | | | | | |
|---|---|---|---|---|---|---|---|---|
| 0.218764 | 0.451155 | 0.029 | 0.017 | 1 | 6 | Zfp810 | 0.012 | Old |
| 0.218848 | 0.909861 | 0.037 | 0.023 | 1 | 6 | Polg2 | 0.014 | Old |
| 0.219286 | 0.641015 | 0.106 | 0.084 | 1 | 6 | Ttc5 | 0.022 | Old |
| 0.219443 | 0.408371 | 0.046 | 0.03 | 1 | 6 | Coq9 | 0.016 | Old |
| 0.219836 | 0.073054 | 0.091 | 0.069 | 1 | 6 | Arsa | 0.022 | Old |
| 0.219875 | 0.649713 | 0.029 | 0.017 | 1 | 6 | Slc22a21 | 0.012 | Old |
| 0.21996 | 0.246013 | 0.4 | 0.361 | 1 | 6 | Psma6 | 0.039 | Old |
| 0.22006 | 1.498136 | 0.017 | 0.008 | 1 | 6 | Msl3l2 | 0.009 | Old |
| 0.22006 | 1.189276 | 0.017 | 0.008 | 1 | 6 | Zbtb12 | 0.009 | Old |
| 0.220125 | 0.914879 | 0.057 | 0.04 | 1 | 6 | Pias3 | 0.017 | Old |
| 0.220647 | 0.466788 | 0.2 | 0.169 | 1 | 6 | Srek1 | 0.031 | Old |
| 0.220775 | 0.390208 | 0.117 | 0.092 | 1 | 6 | Retreg3 | 0.025 | Old |
| 0.220859 | 0.39297 | 0.18 | 0.149 | 1 | 6 | Tuba4a | 0.031 | Old |
| 0.220991 | 0.465212 | 0.029 | 0.017 | 1 | 6 | Fam118a | 0.012 | Old |
| 0.220991 | 0.347722 | 0.029 | 0.017 | 1 | 6 | Gprin3 | 0.012 | Old |
| 0.221047 | 0.236344 | 0.029 | 0.045 | 1 | 6 | Slf1 | -0.016 | Old |
| 0.221059 | 1.017015 | 0.017 | 0.008 | 1 | 6 | Lysmd1 | 0.009 | Old |
| 0.221617 | 0.384511 | 0.086 | 0.064 | 1 | 6 | Zfp282 | 0.022 | Old |
| 0.221737 | 0.718289 | 0.029 | 0.017 | 1 | 6 | Arfip2 | 0.012 | Old |
| 0.222006 | 0.306753 | 0.214 | 0.181 | 1 | 6 | Ilf3 | 0.033 | Old |
| 0.22206 | 0.602354 | 0.037 | 0.023 | 1 | 6 | Lrrc14 | 0.014 | Old |
| 0.22206 | 0.235577 | 0.037 | 0.023 | 1 | 6 | Zbtb40 | 0.014 | Old |
| 0.222134 | 0.439014 | 0.077 | 0.057 | 1 | 6 | Adora2b | 0.02 | Old |
| 0.222161 | 0.875835 | 0.057 | 0.04 | 1 | 6 | Cbll1 | 0.017 | Old |
| 0.222323 | 0.258179 | 0.286 | 0.246 | 1 | 6 | Sys1 | 0.04 | Old |
| 0.222384 | 0.45011 | 0.037 | 0.023 | 1 | 6 | Triap1 | 0.014 | Old |
| 0.222435 | 0.401759 | 0.106 | 0.082 | 1 | 6 | Eipr1 | 0.024 | Old |
| 0.222562 | 1.618937 | 0.017 | 0.008 | 1 | 6 | Pi16 | 0.009 | Old |
| 0.222562 | 1.002217 | 0.017 | 0.008 | 1 | 6 | D3Ertd254 | 0.009 | Old |
| 0.222625 | 0.361323 | 0.246 | 0.216 | 1 | 6 | Cyb5b | 0.03 | Old |
| 0.222873 | 0.405508 | 0.306 | 0.268 | 1 | 6 | Calu | 0.038 | Old |
| 0.223065 | 1.216889 | 0.017 | 0.008 | 1 | 6 | Zfp956 | 0.009 | Old |
| 0.223569 | 0.867254 | 0.017 | 0.008 | 1 | 6 | Ska2 | 0.009 | Old |
| 0.22359 | 0.151804 | 0.394 | 0.346 | 1 | 6 | Cct7 | 0.048 | Old |
| 0.223775 | 0.447751 | 0.114 | 0.089 | 1 | 6 | Tfrc | 0.025 | Old |
| 0.224057 | 0.127878 | 0.666 | 0.644 | 1 | 6 | Wdr1 | 0.022 | Old |
| 0.224074 | 0.760395 | 0.017 | 0.008 | 1 | 6 | Pfkfb2 | 0.009 | Old |
| 0.224329 | 0.781009 | 0.037 | 0.023 | 1 | 6 | Zxdc | 0.014 | Old |
| 0.224531 | 1.217152 | 0.043 | 0.028 | 1 | 6 | Aven | 0.015 | Old |
| 0.224531 | 1.011842 | 0.043 | 0.028 | 1 | 6 | Arhgap27 | 0.015 | Old |
| 0.224655 | 0.493364 | 0.037 | 0.023 | 1 | 6 | Edc3 | 0.014 | Old |
| 0.224722 | 0.351399 | 0.194 | 0.162 | 1 | 6 | Rnmt | 0.032 | Old |
| 0.224983 | 0.613899 | 0.057 | 0.04 | 1 | 6 | Nr2c1 | 0.017 | Old |
| 0.225068 | 0.591718 | 0.069 | 0.05 | 1 | 6 | Uxt | 0.019 | Old |
| 0.225085 | 1.060136 | 0.017 | 0.008 | 1 | 6 | Pms2 | 0.009 | Old |

| | | | | | | | | |
|---|---|---|---|---|---|---|---|---|
| 0.225085 | 0.317296 | 0.017 | 0.008 | 1 | 6 | Them6 | 0.009 | Old |
| 0.225116 | 0.084192 | 0.029 | 0.017 | 1 | 6 | Cd101 | 0.012 | Old |
| 0.225178 | 0.398649 | 0.169 | 0.14 | 1 | 6 | Rreb1 | 0.029 | Old |
| 0.225585 | 0.314176 | 0.091 | 0.069 | 1 | 6 | Tbck | 0.022 | Old |
| 0.225819 | 0.891171 | 0.063 | 0.045 | 1 | 6 | Vps39 | 0.018 | Old |
| 0.225872 | 1.794892 | 0.029 | 0.017 | 1 | 6 | Zfand2a | 0.012 | Old |
| 0.2261 | 0.972529 | 0.017 | 0.008 | 1 | 6 | Gnal | 0.009 | Old |
| 0.2261 | 0.707318 | 0.017 | 0.008 | 1 | 6 | Hist1h4c | 0.009 | Old |
| 0.226287 | 0.01761 | 0.037 | 0.023 | 1 | 6 | Clip2 | 0.014 | Old |
| 0.226484 | 0.307426 | 0.251 | 0.219 | 1 | 6 | Sdf2l1 | 0.032 | Old |
| 0.226772 | 0.513945 | 0.123 | 0.099 | 1 | 6 | Ngly1 | 0.024 | Old |
| 0.226923 | 0.347847 | 0.377 | 0.348 | 1 | 6 | Cpsf6 | 0.029 | Old |
| 0.227112 | 0.465333 | 0.111 | 0.087 | 1 | 6 | Tpst2 | 0.024 | Old |
| 0.227118 | 0.336089 | 0.017 | 0.008 | 1 | 6 | Dtl | 0.009 | Old |
| 0.227271 | 0.265177 | 0.037 | 0.023 | 1 | 6 | Fam122a | 0.014 | Old |
| 0.227628 | 0.198947 | 0.017 | 0.008 | 1 | 6 | Zscan12 | 0.009 | Old |
| 0.227628 | 0.177801 | 0.017 | 0.008 | 1 | 6 | Fbxw9 | 0.009 | Old |
| 0.227959 | 1.374457 | 0.034 | 0.022 | 1 | 6 | Slc25a40 | 0.012 | Old |
| 0.228015 | 0.438485 | 0.117 | 0.094 | 1 | 6 | Snapc1 | 0.023 | Old |
| 0.228029 | 0.464796 | 0.017 | 0.03 | 1 | 6 | Lonrf1 | -0.013 | Old |
| 0.228139 | 0.496055 | 0.017 | 0.008 | 1 | 6 | Dhodh | 0.009 | Old |
| 0.228689 | 0.151185 | 0.331 | 0.291 | 1 | 6 | Pura | 0.04 | Old |
| 0.229163 | 0.263117 | 0.051 | 0.035 | 1 | 6 | Slc37a2 | 0.016 | Old |
| 0.229163 | 0.293946 | 0.017 | 0.008 | 1 | 6 | Slc16a1 | 0.009 | Old |
| 0.229241 | 0.430989 | 0.106 | 0.082 | 1 | 6 | Dnmt1 | 0.024 | Old |
| 0.229452 | 1.049127 | 0.049 | 0.033 | 1 | 6 | Mast2 | 0.016 | Old |
| 0.229462 | 0.233025 | 0.594 | 0.569 | 1 | 6 | Cd24a | 0.025 | Old |
| 0.229958 | 0.643117 | 0.111 | 0.089 | 1 | 6 | Nsmce1 | 0.022 | Old |
| 0.229994 | 0.500109 | 0.051 | 0.035 | 1 | 6 | Hk3 | 0.016 | Old |
| 0.230054 | 0.217481 | 0.226 | 0.191 | 1 | 6 | Kpna1 | 0.035 | Old |
| 0.230124 | 0.224232 | 0.363 | 0.314 | 1 | 6 | Metap2 | 0.049 | Old |
| 0.230697 | 0.20725 | 0.494 | 0.453 | 1 | 6 | Prr13 | 0.041 | Old |
| 0.231108 | 0.082517 | 0.08 | 0.059 | 1 | 6 | Pja1 | 0.021 | Old |
| 0.231454 | 0.567014 | 0.097 | 0.075 | 1 | 6 | Pip4k2c | 0.022 | Old |
| 0.23153 | 0.182417 | 0.703 | 0.697 | 1 | 6 | Luc7l2 | 0.006 | Old |
| 0.231914 | 0.017927 | 0.077 | 0.057 | 1 | 6 | Dnajc11 | 0.02 | Old |
| 0.232016 | 0.927957 | 0.063 | 0.045 | 1 | 6 | Timm29 | 0.018 | Old |
| 0.232027 | 0.041714 | 0.051 | 0.072 | 1 | 6 | Usp40 | -0.021 | Old |
| 0.232041 | 0.359512 | 0.089 | 0.067 | 1 | 6 | Zfp511 | 0.022 | Old |
| 0.232206 | 0.411978 | 0.226 | 0.194 | 1 | 6 | Psme3 | 0.032 | Old |
| 0.232246 | 0.603315 | 0.283 | 0.256 | 1 | 6 | Sel1l | 0.027 | Old |
| 0.232494 | 0.27204 | 0.191 | 0.161 | 1 | 6 | Hmbox1 | 0.03 | Old |
| 0.232672 | 0.936763 | 0.126 | 0.102 | 1 | 6 | C2cd5 | 0.024 | Old |
| 0.23273 | 0.476368 | 0.043 | 0.028 | 1 | 6 | Galns | 0.015 | Old |
| 0.232979 | 0.521409 | 0.123 | 0.099 | 1 | 6 | Cramp1l | 0.024 | Old |

| | | | | | | | | |
|---|---|---|---|---|---|---|---|---|
| 0.233044 | 0.239031 | 0.109 | 0.084 | 1 | 6 | Cd274 | 0.025 | Old |
| 0.233089 | 0.715672 | 0.12 | 0.097 | 1 | 6 | Sfxn1 | 0.023 | Old |
| 0.233132 | 0.199745 | 0.214 | 0.184 | 1 | 6 | Mprip | 0.03 | Old |
| 0.233149 | 0.438128 | 0.103 | 0.08 | 1 | 6 | Sik3 | 0.023 | Old |
| 0.233489 | 0.206486 | 0.526 | 0.507 | 1 | 6 | Psmb8 | 0.019 | Old |
| 0.234323 | 0.833608 | 0.069 | 0.05 | 1 | 6 | Ppwd1 | 0.019 | Old |
| 0.234378 | 0.435418 | 0.171 | 0.145 | 1 | 6 | Preb | 0.026 | Old |
| 0.234782 | 0.492974 | 0.091 | 0.07 | 1 | 6 | Asb8 | 0.021 | Old |
| 0.23489 | 0.42375 | 0.043 | 0.028 | 1 | 6 | Xylt2 | 0.015 | Old |
| 0.235086 | 1.724444 | 0.026 | 0.015 | 1 | 6 | Gpx3 | 0.011 | Old |
| 0.235168 | 0.353404 | 0.111 | 0.087 | 1 | 6 | Zbtb38 | 0.024 | Old |
| 0.235374 | 0.172099 | 0.383 | 0.338 | 1 | 6 | Cggbp1 | 0.045 | Old |
| 0.235403 | 0.374511 | 0.186 | 0.157 | 1 | 6 | Hps5 | 0.029 | Old |
| 0.236105 | 0.348476 | 0.231 | 0.197 | 1 | 6 | Tnrc6c | 0.034 | Old |
| 0.236208 | 0.562138 | 0.08 | 0.06 | 1 | 6 | Fbxl14 | 0.02 | Old |
| 0.236212 | 0.271854 | 0.474 | 0.441 | 1 | 6 | Selenos | 0.033 | Old |
| 0.236363 | 0.149405 | 0.083 | 0.062 | 1 | 6 | Eif2ak3 | 0.021 | Old |
| 0.236613 | 1.830859 | 0.034 | 0.022 | 1 | 6 | Asb1 | 0.012 | Old |
| 0.238312 | 0.564798 | 0.043 | 0.028 | 1 | 6 | Noc3l | 0.015 | Old |
| 0.238382 | 1.289584 | 0.026 | 0.015 | 1 | 6 | Ulbp1 | 0.011 | Old |
| 0.238383 | 0.261782 | 0.12 | 0.095 | 1 | 6 | Tssc4 | 0.025 | Old |
| 0.238608 | 0.145337 | 0.343 | 0.301 | 1 | 6 | Tmed9 | 0.042 | Old |
| 0.238757 | 0.402746 | 0.42 | 0.393 | 1 | 6 | Plin2 | 0.027 | Old |
| 0.238843 | 0.248815 | 0.063 | 0.045 | 1 | 6 | Zfp626 | 0.018 | Old |
| 0.238939 | 0.425844 | 0.043 | 0.028 | 1 | 6 | Cmc4 | 0.015 | Old |
| 0.239211 | 1.347547 | 0.026 | 0.015 | 1 | 6 | Zfp930 | 0.011 | Old |
| 0.239636 | 0.191973 | 0.249 | 0.212 | 1 | 6 | Lpgat1 | 0.037 | Old |
| 0.240311 | 0.8488 | 0.066 | 0.048 | 1 | 6 | Tada3 | 0.018 | Old |
| 0.240509 | 0.745487 | 0.043 | 0.028 | 1 | 6 | Zkscan1 | 0.015 | Old |
| 0.240627 | 0.053996 | 0.134 | 0.107 | 1 | 6 | Map4k2 | 0.027 | Old |
| 0.240633 | 0.25062 | 0.1 | 0.077 | 1 | 6 | Tsc1 | 0.023 | Old |
| 0.240674 | 0.076828 | 0.206 | 0.172 | 1 | 6 | Sipa1 | 0.034 | Old |
| 0.240785 | 0.19285 | 0.54 | 0.512 | 1 | 6 | Srebf2 | 0.028 | Old |
| 0.240875 | 1.176782 | 0.026 | 0.015 | 1 | 6 | Akap17b | 0.011 | Old |
| 0.241203 | 0.615948 | 0.034 | 0.022 | 1 | 6 | Ndc1 | 0.012 | Old |
| 0.241203 | 0.301106 | 0.034 | 0.022 | 1 | 6 | Mrpl1 | 0.012 | Old |
| 0.241421 | 2.234403 | 0.014 | 0.007 | 1 | 6 | Scml4 | 0.007 | Old |
| 0.241559 | 0.35497 | 0.034 | 0.022 | 1 | 6 | Tmtc4 | 0.012 | Old |
| 0.241597 | 0.581232 | 0.049 | 0.033 | 1 | 6 | Cd209g | 0.016 | Old |
| 0.242272 | 0.757291 | 0.034 | 0.022 | 1 | 6 | Acap3 | 0.012 | Old |
| 0.24244 | 0.855087 | 0.077 | 0.059 | 1 | 6 | Terf2 | 0.018 | Old |
| 0.242599 | 1.383675 | 0.014 | 0.007 | 1 | 6 | Pglyrp1 | 0.007 | Old |
| 0.242599 | 0.981781 | 0.014 | 0.007 | 1 | 6 | Gga2 | 0.007 | Old |
| 0.242817 | 0.152706 | 0.131 | 0.105 | 1 | 6 | Scarb1 | 0.026 | Old |
| 0.243099 | 0.469071 | 0.134 | 0.109 | 1 | 6 | Armc1 | 0.025 | Old |

| | | | | | | | | |
|---|---|---|---|---|---|---|---|---|
| 0.243189 | 1.205361 | 0.014 | 0.007 | 1 | 6 | P2rx5 | 0.007 | Old |
| 0.243268 | 0.487842 | 0.057 | 0.04 | 1 | 6 | Prkab1 | 0.017 | Old |
| 0.243277 | 0.543197 | 0.106 | 0.084 | 1 | 6 | Adpgk | 0.022 | Old |
| 0.243343 | 0.321443 | 0.034 | 0.022 | 1 | 6 | Nup188 | 0.012 | Old |
| 0.243422 | 0.058594 | 0.154 | 0.189 | 1 | 6 | Tifa | -0.035 | Old |
| 0.24378 | 0.981788 | 0.014 | 0.007 | 1 | 6 | Rab3a | 0.007 | Old |
| 0.24378 | 0.834626 | 0.014 | 0.007 | 1 | 6 | Cd200r3 | 0.007 | Old |
| 0.24378 | 0.807659 | 0.014 | 0.007 | 1 | 6 | AU041133 | 0.007 | Old |
| 0.24378 | 0.737657 | 0.014 | 0.007 | 1 | 6 | Zfp677 | 0.007 | Old |
| 0.24378 | 0.481948 | 0.014 | 0.007 | 1 | 6 | Myo10 | 0.007 | Old |
| 0.244112 | 0.285735 | 0.34 | 0.303 | 1 | 6 | Cdipt | 0.037 | Old |
| 0.244338 | 0.012332 | 0.446 | 0.513 | 1 | 6 | Myo1g | -0.067 | Old |
| 0.244373 | 0.388932 | 0.014 | 0.007 | 1 | 6 | Gpt2 | 0.007 | Old |
| 0.244761 | 0.238417 | 0.151 | 0.124 | 1 | 6 | Sdad1 | 0.027 | Old |
| 0.244777 | 0.926991 | 0.034 | 0.022 | 1 | 6 | AI480526 | 0.012 | Old |
| 0.244852 | 0.890473 | 0.049 | 0.033 | 1 | 6 | Shmt2 | 0.016 | Old |
| 0.244966 | 0.983972 | 0.014 | 0.007 | 1 | 6 | Dhrs13 | 0.007 | Old |
| 0.244966 | 0.622924 | 0.014 | 0.007 | 1 | 6 | Gm10762 | 0.007 | Old |
| 0.245422 | 0.396059 | 0.131 | 0.105 | 1 | 6 | Ppa1 | 0.026 | Old |
| 0.24544 | 0.315103 | 0.206 | 0.172 | 1 | 6 | Spag7 | 0.034 | Old |
| 0.245463 | 0.265699 | 0.543 | 0.5 | 1 | 6 | Srsf2 | 0.043 | Old |
| 0.245561 | 0.711539 | 0.014 | 0.007 | 1 | 6 | Cdc42bpg | 0.007 | Old |
| 0.245561 | 0.572202 | 0.014 | 0.007 | 1 | 6 | Wdr34 | 0.007 | Old |
| 0.245561 | 0.469296 | 0.014 | 0.007 | 1 | 6 | Psmc3ip | 0.007 | Old |
| 0.245561 | 0.238102 | 0.014 | 0.007 | 1 | 6 | 6330562C2 | 0.007 | Old |
| 0.245753 | 0.379578 | 0.097 | 0.075 | 1 | 6 | Smim26 | 0.022 | Old |
| 0.246074 | 0.441334 | 0.283 | 0.253 | 1 | 6 | Exoc5 | 0.03 | Old |
| 0.24608 | 0.369606 | 0.069 | 0.05 | 1 | 6 | Ube2o | 0.019 | Old |
| 0.246156 | 0.530067 | 0.014 | 0.007 | 1 | 6 | Gm14325 | 0.007 | Old |
| 0.246159 | 0.148121 | 0.663 | 0.637 | 1 | 6 | Ssr3 | 0.026 | Old |
| 0.246527 | 0.106754 | 0.246 | 0.207 | 1 | 6 | Snw1 | 0.039 | Old |
| 0.246578 | 0.252518 | 0.034 | 0.022 | 1 | 6 | Gtf2h4 | 0.012 | Old |
| 0.246578 | 0.270577 | 0.069 | 0.05 | 1 | 6 | Mtmr2 | 0.019 | Old |
| 0.24658 | 0.325006 | 0.246 | 0.211 | 1 | 6 | Wbp2 | 0.035 | Old |
| 0.246753 | 0.395005 | 0.014 | 0.007 | 1 | 6 | Eno3 | 0.007 | Old |
| 0.246753 | 0.15718 | 0.014 | 0.007 | 1 | 6 | Rtel1 | 0.007 | Old |
| 0.24735 | 1.041019 | 0.014 | 0.007 | 1 | 6 | Abhd8 | 0.007 | Old |
| 0.24735 | 0.584518 | 0.014 | 0.007 | 1 | 6 | Zfp458 | 0.007 | Old |
| 0.24735 | 0.399465 | 0.014 | 0.007 | 1 | 6 | Hist1h1e | 0.007 | Old |
| 0.24735 | 0.392596 | 0.014 | 0.007 | 1 | 6 | Magi3 | 0.007 | Old |
| 0.247633 | 0.371332 | 0.191 | 0.159 | 1 | 6 | Bccip | 0.032 | Old |
| 0.247698 | 0.446939 | 0.123 | 0.097 | 1 | 6 | Slc25a28 | 0.026 | Old |
| 0.248331 | 0.497928 | 0.237 | 0.209 | 1 | 6 | Mat2b | 0.028 | Old |
| 0.248548 | 0.310598 | 0.014 | 0.007 | 1 | 6 | Chrna5 | 0.007 | Old |
| 0.248548 | 0.191553 | 0.014 | 0.007 | 1 | 6 | Fmn1 | 0.007 | Old |

| | | | | | | | | |
|---|---|---|---|---|---|---|---|---|
| 0.248737 | 0.386642 | 0.049 | 0.033 | 1 | 6 | Thap7 | 0.016 | Old |
| 0.248919 | 0.079876 | 0.149 | 0.12 | 1 | 6 | Akap8 | 0.029 | Old |
| 0.249149 | 0.435587 | 0.014 | 0.007 | 1 | 6 | Lrp8 | 0.007 | Old |
| 0.249401 | 0.018908 | 0.154 | 0.124 | 1 | 6 | 1700123O2 | 0.03 | Old |
| 0.249589 | 0.077063 | 0.057 | 0.04 | 1 | 6 | Aimp2 | 0.017 | Old |
| 0.249745 | 0.200042 | 0.026 | 0.015 | 1 | 6 | Zmym6 | 0.011 | Old |
| 0.249867 | 0.478741 | 0.149 | 0.12 | 1 | 6 | Snx30 | 0.029 | Old |
| 0.24995 | 0.161792 | 0.097 | 0.075 | 1 | 6 | Tnfrsf11a | 0.022 | Old |
| 0.250243 | 0.496836 | 0.049 | 0.033 | 1 | 6 | Timmdc1 | 0.016 | Old |
| 0.250729 | 0.24753 | 0.36 | 0.329 | 1 | 6 | Arf6 | 0.031 | Old |
| 0.250811 | 0.004526 | 0.743 | 0.769 | 1 | 6 | Atp2b1 | -0.026 | Old |
| 0.250824 | 0.176178 | 0.251 | 0.216 | 1 | 6 | Mdm4 | 0.035 | Old |
| 0.251362 | 0.334503 | 0.151 | 0.125 | 1 | 6 | Slc12a9 | 0.026 | Old |
| 0.251683 | 0.033086 | 0.18 | 0.212 | 1 | 6 | Rraga | -0.032 | Old |
| 0.251872 | 0.376876 | 0.054 | 0.038 | 1 | 6 | Clasrp | 0.016 | Old |
| 0.252233 | 0.082671 | 0.123 | 0.097 | 1 | 6 | 2210016F1 | 0.026 | Old |
| 0.252752 | 0.182467 | 0.026 | 0.015 | 1 | 6 | Slc22a18 | 0.011 | Old |
| 0.253185 | 0.181505 | 0.063 | 0.045 | 1 | 6 | Hes6 | 0.018 | Old |
| 0.253231 | 0.457869 | 0.209 | 0.179 | 1 | 6 | Ube2g2 | 0.03 | Old |
| 0.253276 | 0.863326 | 0.04 | 0.027 | 1 | 6 | Champ1 | 0.013 | Old |
| 0.253471 | 0.778541 | 0.091 | 0.072 | 1 | 6 | Apeh | 0.019 | Old |
| 0.253504 | 0.419154 | 0.289 | 0.259 | 1 | 6 | Zfp638 | 0.03 | Old |
| 0.253551 | 1.205241 | 0.046 | 0.032 | 1 | 6 | Tmem185a | 0.014 | Old |
| 0.253578 | 0.309188 | 0.049 | 0.033 | 1 | 6 | Eva1b | 0.016 | Old |
| 0.254392 | 0.362196 | 0.129 | 0.104 | 1 | 6 | Scaf8 | 0.025 | Old |
| 0.254596 | 0.775602 | 0.08 | 0.062 | 1 | 6 | Cstf2t | 0.018 | Old |
| 0.254734 | 0.466228 | 0.174 | 0.149 | 1 | 6 | Med30 | 0.025 | Old |
| 0.255265 | 0.525688 | 0.123 | 0.099 | 1 | 6 | Tmem68 | 0.024 | Old |
| 0.255299 | 0.322083 | 0.16 | 0.13 | 1 | 6 | Ormdl2 | 0.03 | Old |
| 0.25539 | 0.357919 | 0.06 | 0.043 | 1 | 6 | Tmem97 | 0.017 | Old |
| 0.255876 | 0.475154 | 0.129 | 0.105 | 1 | 6 | Bzw2 | 0.024 | Old |
| 0.256018 | 0.175427 | 0.137 | 0.11 | 1 | 6 | Tnpo2 | 0.027 | Old |
| 0.256221 | 0.295817 | 0.546 | 0.528 | 1 | 6 | Celf1 | 0.018 | Old |
| 0.256333 | 0.286166 | 0.1 | 0.079 | 1 | 6 | Uck2 | 0.021 | Old |
| 0.256506 | 0.120395 | 0.074 | 0.055 | 1 | 6 | Ptger2 | 0.019 | Old |
| 0.256509 | 0.258138 | 0.226 | 0.192 | 1 | 6 | Ldb1 | 0.034 | Old |
| 0.256586 | 0.538695 | 0.077 | 0.059 | 1 | 6 | Gm19951 | 0.018 | Old |
| 0.256886 | 0.406382 | 0.086 | 0.065 | 1 | 6 | Nsmce3 | 0.021 | Old |
| 0.257107 | 0.337362 | 0.211 | 0.182 | 1 | 6 | Sh2b3 | 0.029 | Old |
| 0.257151 | 0.153538 | 0.331 | 0.299 | 1 | 6 | Hpcal1 | 0.032 | Old |
| 0.257354 | 0.294715 | 0.314 | 0.278 | 1 | 6 | Syf2 | 0.036 | Old |
| 0.25768 | 0.650589 | 0.04 | 0.027 | 1 | 6 | Mex3d | 0.013 | Old |
| 0.257896 | 0.154124 | 0.437 | 0.4 | 1 | 6 | Spcs3 | 0.037 | Old |
| 0.257905 | 0.335409 | 0.169 | 0.14 | 1 | 6 | Vps26a | 0.029 | Old |
| 0.258205 | 0.050501 | 0.471 | 0.51 | 1 | 6 | Mrfap1 | -0.039 | Old |

| | | | | | | | | |
|---|---|---|---|---|---|---|---|---|
| 0.258515 | 0.657763 | 0.077 | 0.059 | 1 | 6 | Cbx7 | 0.018 | Old |
| 0.258542 | 0.175711 | 0.397 | 0.375 | 1 | 6 | Tbxas1 | 0.022 | Old |
| 0.258639 | 0.182857 | 0.654 | 0.634 | 1 | 6 | Akr1a1 | 0.02 | Old |
| 0.258785 | 0.323453 | 0.054 | 0.038 | 1 | 6 | Ccdc58 | 0.016 | Old |
| 0.259072 | 0.165848 | 0.543 | 0.492 | 1 | 6 | G3bp1 | 0.051 | Old |
| 0.259143 | 0.893003 | 0.074 | 0.057 | 1 | 6 | Zfyve26 | 0.017 | Old |
| 0.259272 | 0.941636 | 0.066 | 0.048 | 1 | 6 | Gmppb | 0.018 | Old |
| 0.259365 | 0.162902 | 0.126 | 0.1 | 1 | 6 | Yipf6 | 0.026 | Old |
| 0.259371 | 0.11881 | 0.274 | 0.236 | 1 | 6 | Papd4 | 0.038 | Old |
| 0.259509 | 0.513362 | 0.06 | 0.043 | 1 | 6 | Gpank1 | 0.017 | Old |
| 0.259972 | 0.053908 | 0.123 | 0.099 | 1 | 6 | Gbp2 | 0.024 | Old |
| 0.26018 | 0.98839 | 0.046 | 0.032 | 1 | 6 | Lpcat4 | 0.014 | Old |
| 0.260241 | 0.372716 | 0.206 | 0.177 | 1 | 6 | Ogfr | 0.029 | Old |
| 0.260314 | 0.217768 | 0.014 | 0.025 | 1 | 6 | Fam213a | -0.011 | Old |
| 0.260324 | 0.101398 | 0.14 | 0.114 | 1 | 6 | Mta3 | 0.026 | Old |
| 0.260898 | 0.096409 | 0.229 | 0.266 | 1 | 6 | Csnk1g3 | -0.037 | Old |
| 0.261602 | 0.047581 | 0.14 | 0.114 | 1 | 6 | Fam204a | 0.026 | Old |
| 0.261647 | 0.587751 | 0.066 | 0.048 | 1 | 6 | Mdc1 | 0.018 | Old |
| 0.261744 | 0.133564 | 0.686 | 0.676 | 1 | 6 | Ywhab | 0.01 | Old |
| 0.262072 | 0.506651 | 0.094 | 0.074 | 1 | 6 | Mum1 | 0.02 | Old |
| 0.26214 | 0.933805 | 0.011 | 0.005 | 1 | 6 | AC129328. | 0.006 | Old |
| 0.262558 | 0.7753 | 0.06 | 0.043 | 1 | 6 | Fdx1 | 0.017 | Old |
| 0.262566 | 0.332445 | 0.137 | 0.114 | 1 | 6 | Rhob | 0.023 | Old |
| 0.262677 | 0.345084 | 0.18 | 0.154 | 1 | 6 | Slc25a37 | 0.026 | Old |
| 0.262743 | 0.212307 | 0.569 | 0.56 | 1 | 6 | Tmem30a | 0.009 | Old |
| 0.262752 | 0.180439 | 0.26 | 0.222 | 1 | 6 | Kansl1 | 0.038 | Old |
| 0.262846 | 1.42088 | 0.011 | 0.005 | 1 | 6 | Gm15706 | 0.006 | Old |
| 0.262849 | 0.650483 | 0.103 | 0.082 | 1 | 6 | 2310001H1 | 0.021 | Old |
| 0.263211 | 0.272424 | 0.163 | 0.135 | 1 | 6 | Tmem106b | 0.028 | Old |
| 0.263401 | 0.279969 | 0.337 | 0.311 | 1 | 6 | Mfsd14a | 0.026 | Old |
| 0.263553 | 0.965312 | 0.011 | 0.005 | 1 | 6 | Mppe1 | 0.006 | Old |
| 0.263553 | 0.92819 | 0.011 | 0.005 | 1 | 6 | Zdhhc24 | 0.006 | Old |
| 0.264211 | 0.108149 | 0.231 | 0.202 | 1 | 6 | Dcp2 | 0.029 | Old |
| 0.264309 | 0.193818 | 0.383 | 0.348 | 1 | 6 | Ost4 | 0.035 | Old |
| 0.264987 | 0.125395 | 0.091 | 0.07 | 1 | 6 | Tmem181a | 0.021 | Old |
| 0.265558 | 0.308357 | 0.291 | 0.261 | 1 | 6 | Mlf2 | 0.03 | Old |
| 0.265683 | 1.020147 | 0.011 | 0.005 | 1 | 6 | Ptpn21 | 0.006 | Old |
| 0.265683 | 0.792331 | 0.011 | 0.005 | 1 | 6 | Nupl2 | 0.006 | Old |
| 0.265683 | 0.761565 | 0.011 | 0.005 | 1 | 6 | Tsacc | 0.006 | Old |
| 0.265683 | 0.691515 | 0.011 | 0.005 | 1 | 6 | Epha2 | 0.006 | Old |
| 0.265683 | 0.682616 | 0.011 | 0.005 | 1 | 6 | Calml4 | 0.006 | Old |
| 0.265788 | 0.557252 | 0.014 | 0.025 | 1 | 6 | Zc3h11a | -0.011 | Old |
| 0.265913 | 0.340253 | 0.06 | 0.043 | 1 | 6 | Zfp866 | 0.017 | Old |
| 0.266395 | 1.039616 | 0.011 | 0.005 | 1 | 6 | 4632415L0 | 0.006 | Old |
| 0.266395 | 0.799391 | 0.011 | 0.005 | 1 | 6 | Gdf3 | 0.006 | Old |

| | | | | | | | | |
|---|---|---|---|---|---|---|---|---|
| 0.266395 | 0.792318 | 0.011 | 0.005 | 1 | 6 | Fblim1 | 0.006 | Old |
| 0.266395 | 0.758091 | 0.011 | 0.005 | 1 | 6 | Gm42595 | 0.006 | Old |
| 0.266395 | 0.684459 | 0.011 | 0.005 | 1 | 6 | Dnph1 | 0.006 | Old |
| 0.266395 | 0.657493 | 0.011 | 0.005 | 1 | 6 | Gm49207 | 0.006 | Old |
| 0.266395 | 0.595112 | 0.011 | 0.005 | 1 | 6 | Ubtd2 | 0.006 | Old |
| 0.266439 | 0.029226 | 0.049 | 0.033 | 1 | 6 | Emc9 | 0.016 | Old |
| 0.266543 | 0.105772 | 0.32 | 0.284 | 1 | 6 | Safb2 | 0.036 | Old |
| 0.266723 | 0.189857 | 0.274 | 0.246 | 1 | 6 | Stt3a | 0.028 | Old |
| 0.266911 | 0.103648 | 0.08 | 0.06 | 1 | 6 | Srm | 0.02 | Old |
| 0.267109 | 1.056969 | 0.011 | 0.005 | 1 | 6 | Rhoj | 0.006 | Old |
| 0.267109 | 0.482017 | 0.011 | 0.005 | 1 | 6 | Gm45238 | 0.006 | Old |
| 0.267109 | 0.394083 | 0.011 | 0.005 | 1 | 6 | Stxbp4 | 0.006 | Old |
| 0.267445 | 0.044 | 0.123 | 0.152 | 1 | 6 | Arhgef3 | -0.029 | Old |
| 0.267602 | 0.559901 | 0.06 | 0.043 | 1 | 6 | Dmac2 | 0.017 | Old |
| 0.267833 | 0.731541 | 0.031 | 0.02 | 1 | 6 | Mlh3 | 0.011 | Old |
| 0.267954 | 0.229604 | 0.126 | 0.102 | 1 | 6 | Grcc10 | 0.024 | Old |
| 0.267989 | 0.2322 | 0.36 | 0.321 | 1 | 6 | Mapk3 | 0.039 | Old |
| 0.268076 | 0.199063 | 0.171 | 0.142 | 1 | 6 | Iws1 | 0.029 | Old |
| 0.268104 | 0.44819 | 0.16 | 0.134 | 1 | 6 | Afg3l1 | 0.026 | Old |
| 0.268112 | 0.1055 | 0.243 | 0.209 | 1 | 6 | Taf1 | 0.034 | Old |
| 0.268303 | 0.215116 | 0.094 | 0.074 | 1 | 6 | Tarbp2 | 0.02 | Old |
| 0.268348 | 0.186126 | 0.16 | 0.132 | 1 | 6 | Fgfr1op | 0.028 | Old |
| 0.26854 | 0.778466 | 0.011 | 0.005 | 1 | 6 | Aplf | 0.006 | Old |
| 0.26854 | 0.559303 | 0.011 | 0.005 | 1 | 6 | Exog | 0.006 | Old |
| 0.268552 | 0.59328 | 0.046 | 0.032 | 1 | 6 | Gm26782 | 0.014 | Old |
| 0.268571 | 0.470044 | 0.006 | 0.013 | 1 | 6 | Tmc8 | -0.007 | Old |
| 0.268643 | 0.284981 | 0.317 | 0.286 | 1 | 6 | Lrrc8c | 0.031 | Old |
| 0.268713 | 0.653871 | 0.083 | 0.065 | 1 | 6 | Camsap1 | 0.018 | Old |
| 0.268857 | 0.447421 | 0.091 | 0.07 | 1 | 6 | Ccser2 | 0.021 | Old |
| 0.268954 | 1.374918 | 0.023 | 0.013 | 1 | 6 | Tigar | 0.01 | Old |
| 0.269069 | 0.503498 | 0.191 | 0.167 | 1 | 6 | Rnf20 | 0.024 | Old |
| 0.269258 | 0.317617 | 0.011 | 0.005 | 1 | 6 | Rac3 | 0.006 | Old |
| 0.269258 | 0.031491 | 0.011 | 0.005 | 1 | 6 | Naip1 | 0.006 | Old |
| 0.269864 | 0.864513 | 0.06 | 0.043 | 1 | 6 | Recql5 | 0.017 | Old |
| 0.269909 | 1.010939 | 0.023 | 0.013 | 1 | 6 | Fchsd1 | 0.01 | Old |
| 0.269977 | 0.161563 | 0.011 | 0.005 | 1 | 6 | Xkr8 | 0.006 | Old |
| 0.270174 | 0.401264 | 0.143 | 0.117 | 1 | 6 | Zyg11b | 0.026 | Old |
| 0.270359 | 0.166374 | 0.8 | 0.798 | 1 | 6 | Rhoa | 0.002 | Old |
| 0.270378 | 0.178248 | 0.006 | 0.013 | 1 | 6 | Klc2 | -0.007 | Old |
| 0.270521 | 0.277761 | 0.306 | 0.274 | 1 | 6 | Lsm12 | 0.032 | Old |
| 0.270698 | 0.168288 | 0.011 | 0.005 | 1 | 6 | Hmga1b | 0.006 | Old |
| 0.271032 | 1.019726 | 0.031 | 0.02 | 1 | 6 | A430078G: | 0.011 | Old |
| 0.271046 | 0.408495 | 0.077 | 0.059 | 1 | 6 | Actr8 | 0.018 | Old |
| 0.271413 | 0.957751 | 0.051 | 0.037 | 1 | 6 | Cstf1 | 0.014 | Old |
| 0.271603 | 0.455015 | 0.177 | 0.151 | 1 | 6 | Rabl6 | 0.026 | Old |

| | | | | | | | | |
|---|---|---|---|---|---|---|---|---|
| 0.271827 | 0.68242 | 0.023 | 0.013 | 1 | 6 | Abcc10 | 0.01 | Old |
| 0.271827 | 0.550715 | 0.023 | 0.013 | 1 | 6 | Mkks | 0.01 | Old |
| 0.271833 | 0.251983 | 0.12 | 0.097 | 1 | 6 | Calhm2 | 0.023 | Old |
| 0.271998 | 0.253165 | 0.306 | 0.274 | 1 | 6 | Ube2s | 0.032 | Old |
| 0.272261 | 0.10198 | 0.246 | 0.211 | 1 | 6 | Tspan31 | 0.035 | Old |
| 0.272566 | 0.23212 | 0.254 | 0.222 | 1 | 6 | Strap | 0.032 | Old |
| 0.272768 | 0.190711 | 0.643 | 0.607 | 1 | 6 | Pak2 | 0.036 | Old |
| 0.272806 | 1.128403 | 0.046 | 0.032 | 1 | 6 | Dexi | 0.014 | Old |
| 0.272933 | 0.355203 | 0.28 | 0.247 | 1 | 6 | Rad23a | 0.033 | Old |
| 0.272953 | 0.718535 | 0.051 | 0.037 | 1 | 6 | Paip2b | 0.014 | Old |
| 0.273303 | 0.402139 | 0.494 | 0.468 | 1 | 6 | Edem1 | 0.026 | Old |
| 0.273406 | 0.400116 | 0.134 | 0.11 | 1 | 6 | Ap4s1 | 0.024 | Old |
| 0.273449 | 0.774776 | 0.031 | 0.02 | 1 | 6 | Lrrc1 | 0.011 | Old |
| 0.273571 | 0.857482 | 0.051 | 0.037 | 1 | 6 | Fcho1 | 0.014 | Old |
| 0.27363 | 0.743877 | 0.069 | 0.052 | 1 | 6 | Ccdc134 | 0.017 | Old |
| 0.273716 | 0.127943 | 0.874 | 0.88 | 1 | 6 | Arf5 | -0.006 | Old |
| 0.273754 | 0.757272 | 0.023 | 0.013 | 1 | 6 | Btd | 0.01 | Old |
| 0.273908 | 0.231033 | 0.054 | 0.038 | 1 | 6 | Fez2 | 0.016 | Old |
| 0.274001 | 0.22694 | 0.537 | 0.502 | 1 | 6 | Naga | 0.035 | Old |
| 0.274257 | 0.465437 | 0.031 | 0.02 | 1 | 6 | Tbp | 0.011 | Old |
| 0.274514 | 0.162298 | 0.166 | 0.137 | 1 | 6 | Lgals9 | 0.029 | Old |
| 0.274929 | 0.059288 | 0.071 | 0.094 | 1 | 6 | Ash2l | -0.023 | Old |
| 0.275319 | 0.295008 | 0.094 | 0.074 | 1 | 6 | Golga3 | 0.02 | Old |
| 0.275474 | 0.92567 | 0.031 | 0.02 | 1 | 6 | Mical3 | 0.011 | Old |
| 0.275582 | 0.28215 | 0.543 | 0.522 | 1 | 6 | Rpn2 | 0.021 | Old |
| 0.275636 | 0.245146 | 0.509 | 0.483 | 1 | 6 | Txnip | 0.026 | Old |
| 0.27569 | 0.28751 | 0.177 | 0.149 | 1 | 6 | Ehmt1 | 0.028 | Old |
| 0.275728 | 0.507964 | 0.163 | 0.142 | 1 | 6 | Rgl2 | 0.021 | Old |
| 0.275733 | 0.031764 | 0.24 | 0.271 | 1 | 6 | Chp1 | -0.031 | Old |
| 0.275897 | 0.28739 | 0.163 | 0.135 | 1 | 6 | Cnpy2 | 0.028 | Old |
| 0.275945 | 0.026249 | 0.171 | 0.14 | 1 | 6 | Atxn10 | 0.031 | Old |
| 0.276167 | 0.414344 | 0.1 | 0.079 | 1 | 6 | Ppp1r8 | 0.021 | Old |
| 0.276171 | 0.301192 | 0.349 | 0.324 | 1 | 6 | Stk24 | 0.025 | Old |
| 0.276176 | 0.58642 | 0.023 | 0.013 | 1 | 6 | Spryd4 | 0.01 | Old |
| 0.276432 | 0.206175 | 0.28 | 0.249 | 1 | 6 | Nudt21 | 0.031 | Old |
| 0.276693 | 0.337057 | 0.031 | 0.02 | 1 | 6 | Agpat3 | 0.011 | Old |
| 0.276899 | 0.157187 | 0.04 | 0.027 | 1 | 6 | Ccdc167 | 0.013 | Old |
| 0.276974 | 0.133725 | 0.16 | 0.191 | 1 | 6 | Rbm22 | -0.031 | Old |
| 0.277107 | 0.334885 | 0.046 | 0.032 | 1 | 6 | Zbtb41 | 0.014 | Old |
| 0.277263 | 0.077969 | 0.229 | 0.196 | 1 | 6 | Snx17 | 0.033 | Old |
| 0.277309 | 0.374124 | 0.289 | 0.263 | 1 | 6 | Klf10 | 0.026 | Old |
| 0.277494 | 0.270771 | 0.334 | 0.298 | 1 | 6 | Ubtf | 0.036 | Old |
| 0.277636 | 0.588333 | 0.023 | 0.013 | 1 | 6 | S1pr4 | 0.01 | Old |
| 0.27788 | 0.088493 | 0.429 | 0.478 | 1 | 6 | Sgpl1 | -0.049 | Old |
| 0.277982 | 0.264585 | 0.449 | 0.423 | 1 | 6 | Itgb2 | 0.026 | Old |

| | | | | | | | | |
|---|---|---|---|---|---|---|---|---|
| 0.277989 | 0.189939 | 0.217 | 0.184 | 1 | 6 | Kdelr2 | 0.033 | Old |
| 0.277996 | 0.625047 | 0.08 | 0.062 | 1 | 6 | Dvl1 | 0.018 | Old |
| 0.278124 | 0.514533 | 0.023 | 0.013 | 1 | 6 | Magee1 | 0.01 | Old |
| 0.278295 | 0.61922 | 0.074 | 0.057 | 1 | 6 | Fdft1 | 0.017 | Old |
| 0.278471 | 0.284311 | 0.46 | 0.436 | 1 | 6 | Dnajc3 | 0.024 | Old |
| 0.278773 | 0.393894 | 0.046 | 0.032 | 1 | 6 | Zbtb14 | 0.014 | Old |
| 0.279101 | 0.172051 | 0.023 | 0.013 | 1 | 6 | Krtcap3 | 0.01 | Old |
| 0.279284 | 0.36224 | 0.106 | 0.084 | 1 | 6 | Dnajc25 | 0.022 | Old |
| 0.27929 | 0.576369 | 0.109 | 0.087 | 1 | 6 | Prpf31 | 0.022 | Old |
| 0.279535 | 0.333339 | 0.209 | 0.177 | 1 | 6 | Wtap | 0.032 | Old |
| 0.279545 | 0.210045 | 0.189 | 0.161 | 1 | 6 | Babam2 | 0.028 | Old |
| 0.279591 | 0.426567 | 0.023 | 0.013 | 1 | 6 | Gm12166 | 0.01 | Old |
| 0.279953 | 0.017162 | 0.28 | 0.309 | 1 | 6 | Glyr1 | -0.029 | Old |
| 0.280572 | 0.547337 | 0.023 | 0.013 | 1 | 6 | Hus1 | 0.01 | Old |
| 0.280821 | 0.141297 | 0.511 | 0.468 | 1 | 6 | Ewsr1 | 0.043 | Old |
| 0.281231 | 0.318684 | 0.126 | 0.102 | 1 | 6 | Lage3 | 0.024 | Old |
| 0.28124 | 0.83556 | 0.071 | 0.055 | 1 | 6 | Nt5dc3 | 0.016 | Old |
| 0.281613 | 0.280235 | 0.22 | 0.194 | 1 | 6 | Mtch2 | 0.026 | Old |
| 0.281616 | 0.311642 | 0.186 | 0.159 | 1 | 6 | Pih1d1 | 0.027 | Old |
| 0.282006 | 0.311812 | 0.186 | 0.161 | 1 | 6 | Rfk | 0.025 | Old |
| 0.282228 | 0.771488 | 0.069 | 0.052 | 1 | 6 | Inip | 0.017 | Old |
| 0.282503 | 0.423186 | 0.169 | 0.142 | 1 | 6 | Plgrkt | 0.027 | Old |
| 0.282759 | 0.063154 | 0.434 | 0.393 | 1 | 6 | Tspan13 | 0.041 | Old |
| 0.282781 | 0.070895 | 0.16 | 0.187 | 1 | 6 | Clk3 | -0.027 | Old |
| 0.282858 | 0.341504 | 0.403 | 0.375 | 1 | 6 | Sgk3 | 0.028 | Old |
| 0.282954 | 0.263832 | 0.157 | 0.134 | 1 | 6 | Derl2 | 0.023 | Old |
| 0.282962 | 0.381266 | 0.091 | 0.072 | 1 | 6 | Lonp1 | 0.019 | Old |
| 0.283069 | 0.088586 | 0.089 | 0.069 | 1 | 6 | Rnf126 | 0.02 | Old |
| 0.28343 | 0.287344 | 0.214 | 0.187 | 1 | 6 | Smarcc2 | 0.027 | Old |
| 0.283501 | 0.349368 | 0.08 | 0.062 | 1 | 6 | Coasy | 0.018 | Old |
| 0.28381 | 0.220809 | 0.226 | 0.196 | 1 | 6 | Ifi209 | 0.03 | Old |
| 0.283813 | 0.137706 | 0.046 | 0.032 | 1 | 6 | Heatr5b | 0.014 | Old |
| 0.284232 | 0.120251 | 0.229 | 0.197 | 1 | 6 | Ggh | 0.032 | Old |
| 0.284511 | 0.190683 | 0.597 | 0.577 | 1 | 6 | Rap1a | 0.02 | Old |
| 0.284526 | 0.140681 | 0.357 | 0.319 | 1 | 6 | Ctbp1 | 0.038 | Old |
| 0.284673 | 0.159546 | 0.086 | 0.067 | 1 | 6 | A430005L1 | 0.019 | Old |
| 0.284996 | 0.175893 | 0.68 | 0.679 | 1 | 6 | Ctsz | 0.001 | Old |
| 0.285653 | 1.158513 | 0.037 | 0.025 | 1 | 6 | Nt5m | 0.012 | Old |
| 0.285792 | 0.315611 | 0.097 | 0.077 | 1 | 6 | Ss18l2 | 0.02 | Old |
| 0.286599 | 0.816176 | 0.037 | 0.025 | 1 | 6 | Sufu | 0.012 | Old |
| 0.286906 | 0.322371 | 0.183 | 0.159 | 1 | 6 | Uimc1 | 0.024 | Old |
| 0.287851 | 0.318461 | 0.223 | 0.194 | 1 | 6 | Srf | 0.029 | Old |
| 0.288063 | 0.669631 | 0.186 | 0.159 | 1 | 6 | Fabp4 | 0.027 | Old |
| 0.288073 | 0.187374 | 0.094 | 0.074 | 1 | 6 | Grsf1 | 0.02 | Old |
| 0.288873 | 0.233981 | 0.109 | 0.134 | 1 | 6 | Ccl7 | -0.025 | Old |

| | | | | | | | | |
|---|---|---|---|---|---|---|---|---|
| 0.289093 | 0.206845 | 0.697 | 0.701 | 1 | 6 | Gnb1 | -0.004 | Old |
| 0.289297 | 0.47147 | 0.1 | 0.08 | 1 | 6 | Cyp20a1 | 0.02 | Old |
| 0.289419 | 0.240838 | 0.24 | 0.211 | 1 | 6 | Cope | 0.029 | Old |
| 0.289469 | 0.26946 | 0.237 | 0.207 | 1 | 6 | Bub3 | 0.03 | Old |
| 0.290215 | 0.612113 | 0.037 | 0.025 | 1 | 6 | Stxbp1 | 0.012 | Old |
| 0.290434 | 0.148788 | 0.094 | 0.074 | 1 | 6 | Heatr3 | 0.02 | Old |
| 0.290803 | 0.031183 | 0.109 | 0.132 | 1 | 6 | Zmpste24 | -0.023 | Old |
| 0.291713 | 0.404118 | 0.057 | 0.042 | 1 | 6 | Slx4ip | 0.015 | Old |
| 0.291747 | 0.764486 | 0.037 | 0.025 | 1 | 6 | Kctd12b | 0.012 | Old |
| 0.291747 | 0.452219 | 0.037 | 0.025 | 1 | 6 | Cdk7 | 0.012 | Old |
| 0.291908 | 0.25822 | 0.051 | 0.037 | 1 | 6 | Eif2d | 0.014 | Old |
| 0.292065 | 0.231893 | 0.186 | 0.157 | 1 | 6 | Snrpa | 0.029 | Old |
| 0.292632 | 0.490603 | 0.169 | 0.144 | 1 | 6 | Dhrs7 | 0.025 | Old |
| 0.293018 | 0.576453 | 0.146 | 0.125 | 1 | 6 | Msi2 | 0.021 | Old |
| 0.293391 | 0.457717 | 0.16 | 0.137 | 1 | 6 | Cyp4f16 | 0.023 | Old |
| 0.293791 | 0.301329 | 0.263 | 0.237 | 1 | 6 | Srp19 | 0.026 | Old |
| 0.293795 | 0.272827 | 0.443 | 0.406 | 1 | 6 | Cdc42se2 | 0.037 | Old |
| 0.293898 | 0.030071 | 0.077 | 0.059 | 1 | 6 | Wdr18 | 0.018 | Old |
| 0.293944 | 0.119358 | 0.171 | 0.201 | 1 | 6 | Arl5a | -0.03 | Old |
| 0.29397 | 1.235167 | 0.029 | 0.018 | 1 | 6 | Ffar4 | 0.011 | Old |
| 0.294082 | 0.757281 | 0.043 | 0.03 | 1 | 6 | Mrpl39 | 0.013 | Old |
| 0.294116 | 0.543801 | 0.071 | 0.055 | 1 | 6 | Gfm1 | 0.016 | Old |
| 0.294826 | 0.579245 | 0.037 | 0.025 | 1 | 6 | Cdkn2c | 0.012 | Old |
| 0.294826 | 0.414731 | 0.037 | 0.025 | 1 | 6 | Creld1 | 0.012 | Old |
| 0.295084 | 0.293822 | 0.057 | 0.042 | 1 | 6 | Metrn | 0.015 | Old |
| 0.29547 | 0.111885 | 0.406 | 0.365 | 1 | 6 | Prdx2 | 0.041 | Old |
| 0.295747 | 0.876696 | 0.029 | 0.018 | 1 | 6 | Mmp25 | 0.011 | Old |
| 0.295987 | 0.309229 | 0.037 | 0.025 | 1 | 6 | Zfp386 | 0.012 | Old |
| 0.296157 | 0.263788 | 0.169 | 0.14 | 1 | 6 | Wdr82 | 0.029 | Old |
| 0.296307 | 0.143386 | 0.697 | 0.679 | 1 | 6 | H13 | 0.018 | Old |
| 0.296328 | 0.189043 | 0.023 | 0.035 | 1 | 6 | Rhobtb2 | -0.012 | Old |
| 0.296374 | 0.556304 | 0.037 | 0.025 | 1 | 6 | Aldh4a1 | 0.012 | Old |
| 0.296414 | 0.544061 | 0.234 | 0.209 | 1 | 6 | Man2a1 | 0.025 | Old |
| 0.296426 | 0.427405 | 0.18 | 0.154 | 1 | 6 | Fnbp1l | 0.026 | Old |
| 0.296452 | 0.234287 | 0.403 | 0.36 | 1 | 6 | Sumo1 | 0.043 | Old |
| 0.296511 | 0.134179 | 0.457 | 0.425 | 1 | 6 | Ier3ip1 | 0.032 | Old |
| 0.296701 | 0.226049 | 0.137 | 0.112 | 1 | 6 | Mcrs1 | 0.025 | Old |
| 0.296964 | 0.158024 | 0.177 | 0.147 | 1 | 6 | Dap3 | 0.03 | Old |
| 0.29715 | 0.375915 | 0.037 | 0.025 | 1 | 6 | Ift22 | 0.012 | Old |
| 0.297363 | 0.293234 | 0.391 | 0.363 | 1 | 6 | Anxa1 | 0.028 | Old |
| 0.297555 | 0.167814 | 0.086 | 0.067 | 1 | 6 | Tfdp1 | 0.019 | Old |
| 0.297585 | 0.276053 | 0.354 | 0.323 | 1 | 6 | 2-Mar | 0.031 | Old |
| 0.298057 | 0.006307 | 0.52 | 0.569 | 1 | 6 | Dek | -0.049 | Old |
| 0.298292 | 0.225968 | 0.369 | 0.336 | 1 | 6 | Rhog | 0.033 | Old |
| 0.29858 | 0.266672 | 0.117 | 0.095 | 1 | 6 | Scfd1 | 0.022 | Old |

| | | | | | | | | |
|---|---|---|---|---|---|---|---|---|
| 0.298696 | 0.267657 | 0.149 | 0.124 | 1 | 6 | Ttc9c | 0.025 | Old |
| 0.298991 | 0.78819 | 0.054 | 0.04 | 1 | 6 | Trmt2b | 0.014 | Old |
| 0.299096 | 0.365142 | 0.037 | 0.025 | 1 | 6 | Tmem132a | 0.012 | Old |
| 0.299192 | 0.401806 | 0.129 | 0.105 | 1 | 6 | Lrch3 | 0.024 | Old |
| 0.299303 | 0.10979 | 0.149 | 0.174 | 1 | 6 | Atf6b | -0.025 | Old |
| 0.299375 | 0.167325 | 0.009 | 0.017 | 1 | 6 | Traf4 | -0.008 | Old |
| 0.299403 | 0.217343 | 0.051 | 0.037 | 1 | 6 | Zfp182 | 0.014 | Old |
| 0.29945 | 0.810642 | 0.02 | 0.012 | 1 | 6 | Armc6 | 0.008 | Old |
| 0.299824 | 0.536774 | 0.086 | 0.067 | 1 | 6 | Zfpl1 | 0.019 | Old |
| 0.299964 | 0.397177 | 0.131 | 0.109 | 1 | 6 | Smad1 | 0.022 | Old |
| 0.299998 | 1.37143 | 0.02 | 0.012 | 1 | 6 | Mov10 | 0.008 | Old |
| 0.300025 | 0.61354 | 0.106 | 0.087 | 1 | 6 | Smg9 | 0.019 | Old |
| 0.300034 | 0.24257 | 0.057 | 0.042 | 1 | 6 | Tmem161a | 0.015 | Old |
| 0.300261 | 0.600166 | 0.054 | 0.04 | 1 | 6 | Tex2 | 0.014 | Old |
| 0.300625 | 0.142263 | 0.083 | 0.064 | 1 | 6 | Thoc1 | 0.019 | Old |
| 0.300719 | 0.115984 | 0.051 | 0.037 | 1 | 6 | Zfp938 | 0.014 | Old |
| 0.300819 | 0.470383 | 0.063 | 0.047 | 1 | 6 | Prkd2 | 0.016 | Old |
| 0.300936 | 0.43308 | 0.171 | 0.147 | 1 | 6 | Txndc15 | 0.024 | Old |
| 0.300942 | 0.403934 | 0.126 | 0.104 | 1 | 6 | St3gal3 | 0.022 | Old |
| 0.301442 | 0.044235 | 0.037 | 0.025 | 1 | 6 | Pced1a | 0.012 | Old |
| 0.301661 | 0.327951 | 0.177 | 0.152 | 1 | 6 | Fkbp3 | 0.025 | Old |
| 0.301981 | 0.164336 | 0.063 | 0.082 | 1 | 6 | Cntln | -0.019 | Old |
| 0.302059 | 0.205349 | 0.371 | 0.343 | 1 | 6 | Stard3nl | 0.028 | Old |
| 0.302143 | 0.213545 | 0.246 | 0.214 | 1 | 6 | Arl6ip5 | 0.032 | Old |
| 0.302156 | 0.344361 | 0.089 | 0.07 | 1 | 6 | Nup85 | 0.019 | Old |
| 0.302196 | 0.218059 | 0.303 | 0.276 | 1 | 6 | Sla | 0.027 | Old |
| 0.302198 | 1.425533 | 0.02 | 0.012 | 1 | 6 | Tmem72 | 0.008 | Old |
| 0.302219 | 0.01577 | 0.331 | 0.371 | 1 | 6 | Atxn7l3b | -0.04 | Old |
| 0.302227 | 0.093211 | 0.037 | 0.025 | 1 | 6 | Ap4b1 | 0.012 | Old |
| 0.302405 | 0.552339 | 0.089 | 0.07 | 1 | 6 | Sirt1 | 0.019 | Old |
| 0.302478 | 0.693976 | 0.029 | 0.018 | 1 | 6 | Sema4b | 0.011 | Old |
| 0.303189 | 0.209149 | 0.157 | 0.134 | 1 | 6 | Actl6a | 0.023 | Old |
| 0.303441 | 0.846631 | 0.043 | 0.03 | 1 | 6 | Psph | 0.013 | Old |
| 0.303502 | 0.108915 | 0.7 | 0.692 | 1 | 6 | Sdcbp | 0.008 | Old |
| 0.303584 | 0.350859 | 0.214 | 0.187 | 1 | 6 | C1d | 0.027 | Old |
| 0.304075 | 0.854591 | 0.049 | 0.035 | 1 | 6 | Lrrc57 | 0.014 | Old |
| 0.304094 | 0.730927 | 0.054 | 0.04 | 1 | 6 | Myc | 0.014 | Old |
| 0.304169 | 0.40356 | 0.043 | 0.03 | 1 | 6 | Msh6 | 0.013 | Old |
| 0.304169 | 0.086423 | 0.043 | 0.03 | 1 | 6 | Oasl2 | 0.013 | Old |
| 0.304289 | 0.827901 | 0.029 | 0.018 | 1 | 6 | Tcta | 0.011 | Old |
| 0.304512 | 0.417049 | 0.026 | 0.038 | 1 | 6 | Taf4b | -0.012 | Old |
| 0.304864 | 0.487251 | 0.071 | 0.055 | 1 | 6 | Scyl1 | 0.016 | Old |
| 0.304977 | 0.179633 | 0.137 | 0.115 | 1 | 6 | Tnfaip2 | 0.022 | Old |
| 0.305518 | 1.006144 | 0.02 | 0.012 | 1 | 6 | Ppef2 | 0.008 | Old |
| 0.305518 | 0.550579 | 0.02 | 0.012 | 1 | 6 | Disc1 | 0.008 | Old |

| | | | | | | | | |
|---|---|---|---|---|---|---|---|---|
| 0.305653 | 0.759442 | 0.029 | 0.018 | 1 | 6 | Zfp953 | 0.011 | Old |
| 0.305909 | 0.259498 | 0.089 | 0.07 | 1 | 6 | Anapc4 | 0.019 | Old |
| 0.305948 | 0.10718 | 0.069 | 0.052 | 1 | 6 | Git1 | 0.017 | Old |
| 0.305994 | 0.767555 | 0.043 | 0.03 | 1 | 6 | Leng9 | 0.013 | Old |
| 0.306073 | 0.72594 | 0.02 | 0.012 | 1 | 6 | Serf1 | 0.008 | Old |
| 0.306119 | 0.186983 | 0.111 | 0.089 | 1 | 6 | Golga2 | 0.022 | Old |
| 0.306427 | 0.053095 | 0.677 | 0.644 | 1 | 6 | Wdr26 | 0.033 | Old |
| 0.306633 | 0.009481 | 0.443 | 0.483 | 1 | 6 | Arf3 | -0.04 | Old |
| 0.307085 | 0.077546 | 0.117 | 0.094 | 1 | 6 | Pkp3 | 0.023 | Old |
| 0.307092 | 0.568959 | 0.043 | 0.03 | 1 | 6 | Pold2 | 0.013 | Old |
| 0.307187 | 1.283821 | 0.02 | 0.012 | 1 | 6 | Mylk | 0.008 | Old |
| 0.307187 | 1.133585 | 0.02 | 0.012 | 1 | 6 | Slc37a3 | 0.008 | Old |
| 0.307187 | 0.778519 | 0.02 | 0.012 | 1 | 6 | Gvin1 | 0.008 | Old |
| 0.307187 | 0.514507 | 0.02 | 0.012 | 1 | 6 | Dolk | 0.008 | Old |
| 0.307188 | 0.185158 | 0.309 | 0.276 | 1 | 6 | Srpr | 0.033 | Old |
| 0.30737 | 0.428567 | 0.149 | 0.125 | 1 | 6 | Smc5 | 0.024 | Old |
| 0.307464 | 0.36485 | 0.1 | 0.08 | 1 | 6 | Msra | 0.02 | Old |
| 0.307724 | 0.36494 | 0.251 | 0.224 | 1 | 6 | Cmas | 0.027 | Old |
| 0.307724 | 0.2532 | 0.254 | 0.222 | 1 | 6 | Ebp | 0.032 | Old |
| 0.307942 | 0.527397 | 0.071 | 0.055 | 1 | 6 | Pcsk7 | 0.016 | Old |
| 0.308087 | 0.128936 | 0.554 | 0.542 | 1 | 6 | Arf4 | 0.012 | Old |
| 0.308283 | 0.580623 | 0.054 | 0.04 | 1 | 6 | 2010315B0 | 0.014 | Old |
| 0.308392 | 0.737123 | 0.029 | 0.018 | 1 | 6 | Vamp1 | 0.011 | Old |
| 0.308524 | 0.372665 | 0.143 | 0.12 | 1 | 6 | Oxsr1 | 0.023 | Old |
| 0.308861 | 0.299452 | 0.02 | 0.012 | 1 | 6 | Hist1h4d | 0.008 | Old |
| 0.30893 | 0.317241 | 0.206 | 0.179 | 1 | 6 | Hdac1 | 0.027 | Old |
| 0.309073 | 0.327906 | 0.194 | 0.169 | 1 | 6 | Ergic2 | 0.025 | Old |
| 0.30972 | 0.086779 | 0.7 | 0.716 | 1 | 6 | Calr | -0.016 | Old |
| 0.309768 | 0.173695 | 0.029 | 0.018 | 1 | 6 | Nme6 | 0.011 | Old |
| 0.309802 | 0.221864 | 0.157 | 0.132 | 1 | 6 | Rnaseh2c | 0.025 | Old |
| 0.309995 | 0.430676 | 0.157 | 0.132 | 1 | 6 | Neu1 | 0.025 | Old |
| 0.310172 | 0.284759 | 0.149 | 0.124 | 1 | 6 | Lmnb1 | 0.025 | Old |
| 0.310228 | 0.411693 | 0.029 | 0.018 | 1 | 6 | Incenp | 0.011 | Old |
| 0.310433 | 0.30188 | 0.166 | 0.142 | 1 | 6 | Rufy3 | 0.024 | Old |
| 0.310542 | 0.421938 | 0.02 | 0.012 | 1 | 6 | Cnst | 0.008 | Old |
| 0.310965 | 0.377386 | 0.111 | 0.09 | 1 | 6 | Gcc2 | 0.021 | Old |
| 0.311104 | 0.447844 | 0.02 | 0.012 | 1 | 6 | Upp2 | 0.008 | Old |
| 0.311104 | 0.142997 | 0.02 | 0.012 | 1 | 6 | Thbs3 | 0.008 | Old |
| 0.311421 | 0.178844 | 0.246 | 0.209 | 1 | 6 | Tomm70a | 0.037 | Old |
| 0.311666 | 0.38379 | 0.02 | 0.012 | 1 | 6 | Cyb5rl | 0.008 | Old |
| 0.311763 | 0.078847 | 0.486 | 0.452 | 1 | 6 | Lat2 | 0.034 | Old |
| 0.312006 | 0.233446 | 0.069 | 0.052 | 1 | 6 | Iffo2 | 0.017 | Old |
| 0.312081 | 0.502688 | 0.317 | 0.279 | 1 | 6 | AC149090. | 0.038 | Old |
| 0.312108 | 0.113273 | 0.137 | 0.112 | 1 | 6 | Gapt | 0.025 | Old |
| 0.312229 | 0.408013 | 0.02 | 0.012 | 1 | 6 | 1110051M | 0.008 | Old |

| | | | | | | | | |
|---|---|---|---|---|---|---|---|---|
| 0.312238 | 0.133773 | 0.131 | 0.109 | 1 | 6 | Commd1 | 0.022 | Old |
| 0.312395 | 0.214926 | 0.08 | 0.062 | 1 | 6 | Nol6 | 0.018 | Old |
| 0.312403 | 0.068014 | 0.417 | 0.376 | 1 | 6 | Hnrnpul2 | 0.041 | Old |
| 0.312609 | 0.189397 | 0.417 | 0.385 | 1 | 6 | Serinc1 | 0.032 | Old |
| 0.312793 | 0.238668 | 0.02 | 0.012 | 1 | 6 | Tnks1bp1 | 0.008 | Old |
| 0.312837 | 0.638659 | 0.054 | 0.04 | 1 | 6 | Snx11 | 0.014 | Old |
| 0.312851 | 0.051138 | 0.166 | 0.194 | 1 | 6 | Ssbp3 | -0.028 | Old |
| 0.312946 | 0.737638 | 0.066 | 0.05 | 1 | 6 | Papolg | 0.016 | Old |
| 0.313474 | 0.389425 | 0.217 | 0.192 | 1 | 6 | Ccng1 | 0.025 | Old |
| 0.313839 | 0.569241 | 0.22 | 0.199 | 1 | 6 | Gpbp1l1 | 0.021 | Old |
| 0.313855 | 0.028567 | 0.44 | 0.472 | 1 | 6 | Tet2 | -0.032 | Old |
| 0.314482 | 0.046288 | 0.111 | 0.135 | 1 | 6 | Pttg1 | -0.024 | Old |
| 0.314554 | 0.629526 | 0.06 | 0.045 | 1 | 6 | Mcu | 0.015 | Old |
| 0.314847 | 0.174739 | 0.029 | 0.018 | 1 | 6 | Sdhaf3 | 0.011 | Old |
| 0.315507 | 0.364359 | 0.14 | 0.119 | 1 | 6 | Dhrs1 | 0.021 | Old |
| 0.315893 | 0.260376 | 0.206 | 0.182 | 1 | 6 | Vps29 | 0.024 | Old |
| 0.316102 | 0.117485 | 0.117 | 0.095 | 1 | 6 | Wdr61 | 0.022 | Old |
| 0.316213 | 0.332211 | 0.189 | 0.164 | 1 | 6 | 4933434E2 | 0.025 | Old |
| 0.316242 | 0.113137 | 0.029 | 0.018 | 1 | 6 | Trpv4 | 0.011 | Old |
| 0.316274 | 0.640899 | 0.034 | 0.023 | 1 | 6 | Mospd1 | 0.011 | Old |
| 0.316482 | 0.150848 | 0.206 | 0.177 | 1 | 6 | Exosc5 | 0.029 | Old |
| 0.316512 | 0.209598 | 0.066 | 0.05 | 1 | 6 | Ppcdc | 0.016 | Old |
| 0.316674 | 0.022114 | 0.126 | 0.102 | 1 | 6 | Hint3 | 0.024 | Old |
| 0.317008 | 0.164652 | 0.691 | 0.699 | 1 | 6 | Slc25a5 | -0.008 | Old |
| 0.317111 | 0.022636 | 0.063 | 0.047 | 1 | 6 | Pgam5 | 0.016 | Old |
| 0.31747 | 0.523455 | 0.043 | 0.03 | 1 | 6 | D230025D: | 0.013 | Old |
| 0.317728 | 0.215712 | 0.177 | 0.151 | 1 | 6 | Nucb2 | 0.026 | Old |
| 0.317851 | 0.749352 | 0.069 | 0.054 | 1 | 6 | Slc39a3 | 0.015 | Old |
| 0.318305 | 0.438593 | 0.066 | 0.05 | 1 | 6 | Guf1 | 0.016 | Old |
| 0.318377 | 0.708477 | 0.034 | 0.023 | 1 | 6 | Ntmt1 | 0.011 | Old |
| 0.318495 | 0.053918 | 0.043 | 0.059 | 1 | 6 | Zfp367 | -0.016 | Old |
| 0.318664 | 0.970504 | 0.04 | 0.028 | 1 | 6 | Stoml1 | 0.012 | Old |
| 0.318798 | 1.17488 | 0.034 | 0.023 | 1 | 6 | 2-Sep | 0.011 | Old |
| 0.318798 | 0.389103 | 0.034 | 0.023 | 1 | 6 | Dcaf6 | 0.011 | Old |
| 0.318949 | 0.177141 | 0.46 | 0.435 | 1 | 6 | St13 | 0.025 | Old |
| 0.31947 | 0.256374 | 0.294 | 0.266 | 1 | 6 | Psmb5 | 0.028 | Old |
| 0.319476 | 0.204371 | 0.24 | 0.212 | 1 | 6 | Polr2l | 0.028 | Old |
| 0.32025 | 0.175343 | 0.094 | 0.075 | 1 | 6 | Cks1b | 0.019 | Old |
| 0.321131 | 0.080729 | 0.074 | 0.057 | 1 | 6 | Txnrd2 | 0.017 | Old |
| 0.322113 | 0.272881 | 0.049 | 0.035 | 1 | 6 | Tmem18 | 0.014 | Old |
| 0.322269 | 0.572603 | 0.011 | 0.02 | 1 | 6 | Acsf3 | -0.009 | Old |
| 0.32258 | 0.278776 | 0.234 | 0.206 | 1 | 6 | Tmbim4 | 0.028 | Old |
| 0.32304 | 0.748842 | 0.071 | 0.057 | 1 | 6 | Timm22 | 0.014 | Old |
| 0.323119 | 0.260727 | 0.066 | 0.05 | 1 | 6 | Ppp6r2 | 0.016 | Old |
| 0.323156 | 0.197029 | 0.346 | 0.309 | 1 | 6 | Psmd11 | 0.037 | Old |

| | | | | | | | | |
|---|---|---|---|---|---|---|---|---|
| 0.32345 | 0.216399 | 0.597 | 0.597 | 1 | 6 | Prdx1 | 0 | Old |
| 0.323459 | 0.357456 | 0.034 | 0.023 | 1 | 6 | Phykpl | 0.011 | Old |
| 0.323632 | 0.307172 | 0.24 | 0.211 | 1 | 6 | Slfn5 | 0.029 | Old |
| 0.323888 | 0.398017 | 0.011 | 0.02 | 1 | 6 | Bend3 | -0.009 | Old |
| 0.323951 | 0.412738 | 0.146 | 0.124 | 1 | 6 | Hr | 0.022 | Old |
| 0.324004 | 0.405898 | 0.077 | 0.097 | 1 | 6 | Areg | -0.02 | Old |
| 0.32428 | 0.225993 | 0.371 | 0.343 | 1 | 6 | Gfra2 | 0.028 | Old |
| 0.32433 | 0.235599 | 0.046 | 0.062 | 1 | 6 | Cog6 | -0.016 | Old |
| 0.324496 | 0.27266 | 0.429 | 0.4 | 1 | 6 | Ogdh | 0.029 | Old |
| 0.325081 | 0.159243 | 0.106 | 0.085 | 1 | 6 | Sin3a | 0.021 | Old |
| 0.325174 | 1.346007 | 0.046 | 0.033 | 1 | 6 | Parp10 | 0.013 | Old |
| 0.32531 | 0.45345 | 0.171 | 0.149 | 1 | 6 | Lypla1 | 0.022 | Old |
| 0.325462 | 0.264653 | 0.314 | 0.286 | 1 | 6 | Metrnl | 0.028 | Old |
| 0.32602 | 0.531541 | 0.034 | 0.023 | 1 | 6 | Ip6k2 | 0.011 | Old |
| 0.326091 | 0.109554 | 0.054 | 0.04 | 1 | 6 | Elovl6 | 0.014 | Old |
| 0.326824 | 0.304718 | 0.091 | 0.074 | 1 | 6 | Nfyc | 0.017 | Old |
| 0.326877 | 0.803424 | 0.034 | 0.023 | 1 | 6 | Fxn | 0.011 | Old |
| 0.326912 | 0.280462 | 0.066 | 0.05 | 1 | 6 | Ipo13 | 0.016 | Old |
| 0.327113 | 0.285472 | 0.1 | 0.08 | 1 | 6 | Upf2 | 0.02 | Old |
| 0.327249 | 0.678617 | 0.08 | 0.064 | 1 | 6 | Pcyt2 | 0.016 | Old |
| 0.327342 | 0.39343 | 0.14 | 0.119 | 1 | 6 | Shc1 | 0.021 | Old |
| 0.328078 | 0.154709 | 0.111 | 0.135 | 1 | 6 | Lztfl1 | -0.024 | Old |
| 0.328159 | 0.604767 | 0.069 | 0.054 | 1 | 6 | Rcbtb2 | 0.015 | Old |
| 0.328183 | 0.188957 | 0.214 | 0.186 | 1 | 6 | Emsy | 0.028 | Old |
| 0.328523 | 0.189766 | 0.189 | 0.166 | 1 | 6 | Med15 | 0.023 | Old |
| 0.328549 | 0.114839 | 0.177 | 0.152 | 1 | 6 | Slc44a1 | 0.025 | Old |
| 0.328593 | 0.276999 | 0.249 | 0.222 | 1 | 6 | Tmem164 | 0.027 | Old |
| 0.328594 | 0.566174 | 0.034 | 0.023 | 1 | 6 | Polr3h | 0.011 | Old |
| 0.328617 | 0.462559 | 0.103 | 0.085 | 1 | 6 | Gmeb1 | 0.018 | Old |
| 0.328641 | 0.358301 | 0.14 | 0.117 | 1 | 6 | Mmadhc | 0.023 | Old |
| 0.328705 | 0.091142 | 0.791 | 0.763 | 1 | 6 | Tmbim6 | 0.028 | Old |
| 0.328896 | 0.202461 | 0.066 | 0.05 | 1 | 6 | Zfp329 | 0.016 | Old |
| 0.329038 | 0.221735 | 0.209 | 0.181 | 1 | 6 | Nans | 0.028 | Old |
| 0.329122 | 0.584451 | 0.12 | 0.1 | 1 | 6 | Cdc42ep3 | 0.02 | Old |
| 0.329334 | 0.207507 | 0.089 | 0.07 | 1 | 6 | Ascc2 | 0.019 | Old |
| 0.329388 | 0.02576 | 0.4 | 0.438 | 1 | 6 | Fcho2 | -0.038 | Old |
| 0.32995 | 0.263655 | 0.337 | 0.309 | 1 | 6 | Cnot6 | 0.028 | Old |
| 0.330228 | 0.105257 | 0.403 | 0.446 | 1 | 6 | Rgs2 | -0.043 | Old |
| 0.33034 | 0.62024 | 0.057 | 0.043 | 1 | 6 | 1190002N1 | 0.014 | Old |
| 0.330463 | 0.11593 | 0.189 | 0.164 | 1 | 6 | Cnot3 | 0.025 | Old |
| 0.330625 | 0.397631 | 0.063 | 0.048 | 1 | 6 | Alkbh6 | 0.015 | Old |
| 0.331151 | 0.144581 | 0.263 | 0.236 | 1 | 6 | Rnf44 | 0.027 | Old |
| 0.331404 | 0.715244 | 0.109 | 0.092 | 1 | 6 | Trim36 | 0.017 | Old |
| 0.331522 | 0.192137 | 0.314 | 0.276 | 1 | 6 | Taf15 | 0.038 | Old |
| 0.331889 | 0.275862 | 0.163 | 0.14 | 1 | 6 | Chmp1b | 0.023 | Old |

| | | | | | | | | |
|---|---|---|---|---|---|---|---|---|
| 0.332127 | 0.185566 | 0.171 | 0.147 | 1 | 6 | Hars | 0.024 | Old |
| 0.332195 | 0.019909 | 0.243 | 0.268 | 1 | 6 | Eny2 | -0.025 | Old |
| 0.332503 | 0.83809 | 0.063 | 0.048 | 1 | 6 | Stmn1 | 0.015 | Old |
| 0.332527 | 0.791036 | 0.046 | 0.033 | 1 | 6 | Ormdl3 | 0.013 | Old |
| 0.332793 | 0.033033 | 0.063 | 0.08 | 1 | 6 | Setd1a | -0.017 | Old |
| 0.332921 | 0.274158 | 0.2 | 0.176 | 1 | 6 | H2-T22 | 0.024 | Old |
| 0.332945 | 0.317679 | 0.229 | 0.204 | 1 | 6 | Mta2 | 0.025 | Old |
| 0.333174 | 0.198449 | 0.137 | 0.115 | 1 | 6 | Zbtb7b | 0.022 | Old |
| 0.333197 | 0.425165 | 0.14 | 0.12 | 1 | 6 | Dcaf13 | 0.02 | Old |
| 0.333457 | 0.071687 | 0.129 | 0.107 | 1 | 6 | Cpne2 | 0.022 | Old |
| 0.33357 | 0.685206 | 0.051 | 0.038 | 1 | 6 | Eif2ak4 | 0.013 | Old |
| 0.333663 | 0.202351 | 0.229 | 0.201 | 1 | 6 | Rnpepl1 | 0.028 | Old |
| 0.333919 | 0.665665 | 0.051 | 0.038 | 1 | 6 | Sec22c | 0.013 | Old |
| 0.333919 | 0.539222 | 0.051 | 0.038 | 1 | 6 | Slc37a4 | 0.013 | Old |
| 0.334567 | 0.116821 | 0.129 | 0.107 | 1 | 6 | Nop16 | 0.022 | Old |
| 0.334652 | 0.187684 | 0.034 | 0.023 | 1 | 6 | Dnajc18 | 0.011 | Old |
| 0.33506 | 0.146954 | 0.311 | 0.276 | 1 | 6 | Ivns1abp | 0.035 | Old |
| 0.335088 | 0.350917 | 0.034 | 0.023 | 1 | 6 | Tprn | 0.011 | Old |
| 0.335095 | 0.042182 | 0.729 | 0.839 | 1 | 6 | Mgl2 | -0.11 | Old |
| 0.335126 | 0.530086 | 0.046 | 0.033 | 1 | 6 | Fdxacb1 | 0.013 | Old |
| 0.335396 | 0.094914 | 0.04 | 0.028 | 1 | 6 | Mlst8 | 0.012 | Old |
| 0.335701 | 0.131725 | 0.389 | 0.36 | 1 | 6 | Ctnnb1 | 0.029 | Old |
| 0.335955 | 0.467456 | 0.069 | 0.054 | 1 | 6 | Socs5 | 0.015 | Old |
| 0.336354 | 1.013205 | 0.026 | 0.017 | 1 | 6 | Trip6 | 0.009 | Old |
| 0.336502 | 0.008828 | 0.089 | 0.07 | 1 | 6 | Isca2 | 0.019 | Old |
| 0.336732 | 0.05574 | 0.203 | 0.232 | 1 | 6 | Rapgef6 | -0.029 | Old |
| 0.337101 | 0.336393 | 0.14 | 0.119 | 1 | 6 | Ebpl | 0.021 | Old |
| 0.337319 | 0.072593 | 0.486 | 0.523 | 1 | 6 | Slc38a1 | -0.037 | Old |
| 0.337324 | 0.033397 | 0.966 | 0.983 | 1 | 6 | Rpl38 | -0.017 | Old |
| 0.337563 | 1.321576 | 0.017 | 0.01 | 1 | 6 | Tyms | 0.007 | Old |
| 0.337663 | 0.288339 | 0.134 | 0.114 | 1 | 6 | Sgta | 0.02 | Old |
| 0.3378 | 0.264946 | 0.129 | 0.107 | 1 | 6 | Trim26 | 0.022 | Old |
| 0.337925 | 0.781393 | 0.046 | 0.033 | 1 | 6 | Lig1 | 0.013 | Old |
| 0.338117 | 0.294892 | 0.189 | 0.164 | 1 | 6 | Gak | 0.025 | Old |
| 0.338384 | 0.39717 | 0.103 | 0.084 | 1 | 6 | Rapgef1 | 0.019 | Old |
| 0.33842 | 0.129495 | 0.18 | 0.154 | 1 | 6 | Ccdc47 | 0.026 | Old |
| 0.338844 | 1.457302 | 0.017 | 0.01 | 1 | 6 | Bcr | 0.007 | Old |
| 0.339253 | 0.002374 | 0.071 | 0.055 | 1 | 6 | Crcp | 0.016 | Old |
| 0.339387 | 0.094532 | 0.1 | 0.08 | 1 | 6 | Naa40 | 0.02 | Old |
| 0.339448 | 4.093902 | 0.291 | 0.279 | 1 | 6 | Ighm | 0.012 | Old |
| 0.339498 | 0.278314 | 0.08 | 0.064 | 1 | 6 | Nfs1 | 0.016 | Old |
| 0.339576 | 0.275383 | 0.3 | 0.279 | 1 | 6 | Snrpd1 | 0.021 | Old |
| 0.339713 | 0.364502 | 0.066 | 0.05 | 1 | 6 | Trmt13 | 0.016 | Old |
| 0.339714 | 0.029764 | 0.14 | 0.115 | 1 | 6 | Mta1 | 0.025 | Old |
| 0.340435 | 0.061885 | 0.274 | 0.241 | 1 | 6 | Rictor | 0.033 | Old |

| | | | | | | | | |
|---|---|---|---|---|---|---|---|---|
| 0.34086 | 0.276277 | 0.129 | 0.109 | 1 | 6 | Rars | 0.02 | Old |
| 0.341202 | 0.242303 | 0.337 | 0.309 | 1 | 6 | Hectd1 | 0.028 | Old |
| 0.341414 | 1.281589 | 0.017 | 0.01 | 1 | 6 | Muc1 | 0.007 | Old |
| 0.341519 | 0.308378 | 0.126 | 0.105 | 1 | 6 | Ddx39 | 0.021 | Old |
| 0.341869 | 0.253374 | 0.046 | 0.033 | 1 | 6 | Cox10 | 0.013 | Old |
| 0.342059 | 0.849947 | 0.017 | 0.01 | 1 | 6 | Ropn1l | 0.007 | Old |
| 0.342059 | 0.65287 | 0.017 | 0.01 | 1 | 6 | Gm44067 | 0.007 | Old |
| 0.342104 | 0.530842 | 0.034 | 0.023 | 1 | 6 | Tango6 | 0.011 | Old |
| 0.342284 | 0.384229 | 0.057 | 0.043 | 1 | 6 | Arfgap3 | 0.014 | Old |
| 0.342354 | 0.805923 | 0.137 | 0.119 | 1 | 6 | Gbf1 | 0.018 | Old |
| 0.342704 | 0.830797 | 0.017 | 0.01 | 1 | 6 | Acap1 | 0.007 | Old |
| 0.342704 | 0.700282 | 0.017 | 0.01 | 1 | 6 | Asb2 | 0.007 | Old |
| 0.343664 | 0.411866 | 0.163 | 0.139 | 1 | 6 | Tsnax | 0.024 | Old |
| 0.343847 | 0.086226 | 0.26 | 0.231 | 1 | 6 | Sdhc | 0.029 | Old |
| 0.344137 | 0.275205 | 0.126 | 0.105 | 1 | 6 | Faim | 0.021 | Old |
| 0.34422 | 0.274697 | 0.214 | 0.194 | 1 | 6 | Dctn4 | 0.02 | Old |
| 0.344536 | 0.245821 | 0.143 | 0.12 | 1 | 6 | Aldoc | 0.023 | Old |
| 0.34456 | 0.138324 | 0.026 | 0.017 | 1 | 6 | Smyd4 | 0.009 | Old |
| 0.345034 | 0.696489 | 0.094 | 0.079 | 1 | 6 | BC029722 | 0.015 | Old |
| 0.345293 | 1.273872 | 0.017 | 0.01 | 1 | 6 | Cdo1 | 0.007 | Old |
| 0.345293 | 0.800795 | 0.017 | 0.01 | 1 | 6 | Tmem86a | 0.007 | Old |
| 0.345595 | 0.511999 | 0.026 | 0.017 | 1 | 6 | Arl6 | 0.009 | Old |
| 0.34562 | 0.120965 | 0.14 | 0.117 | 1 | 6 | Timm17a | 0.023 | Old |
| 0.345622 | 0.260698 | 0.163 | 0.14 | 1 | 6 | Atf7 | 0.023 | Old |
| 0.345627 | 0.003225 | 0.143 | 0.119 | 1 | 6 | Arl6ip4 | 0.024 | Old |
| 0.346077 | 0.460356 | 0.083 | 0.067 | 1 | 6 | Srcap | 0.016 | Old |
| 0.346441 | 0.514234 | 0.203 | 0.181 | 1 | 6 | Scamp3 | 0.022 | Old |
| 0.346721 | 0.123955 | 0.68 | 0.701 | 1 | 6 | Pld4 | -0.021 | Old |
| 0.347133 | 0.436122 | 0.063 | 0.048 | 1 | 6 | Ifi204 | 0.015 | Old |
| 0.347405 | 0.207681 | 0.671 | 0.661 | 1 | 6 | mt-Nd4l | 0.01 | Old |
| 0.347843 | 0.189901 | 0.103 | 0.125 | 1 | 6 | Mapkapk3 | -0.022 | Old |
| 0.348131 | 0.040964 | 0.017 | 0.027 | 1 | 6 | Ice2 | -0.01 | Old |
| 0.348668 | 0.199403 | 0.449 | 0.42 | 1 | 6 | Pnisr | 0.029 | Old |
| 0.348697 | 0.398909 | 0.046 | 0.033 | 1 | 6 | Wdr7 | 0.013 | Old |
| 0.349067 | 0.400346 | 0.049 | 0.064 | 1 | 6 | Qser1 | -0.015 | Old |
| 0.349138 | 0.544309 | 0.051 | 0.038 | 1 | 6 | Ano10 | 0.013 | Old |
| 0.349138 | 0.230896 | 0.051 | 0.038 | 1 | 6 | Cisd3 | 0.013 | Old |
| 0.349277 | 0.030552 | 0.211 | 0.186 | 1 | 6 | Cirbp | 0.025 | Old |
| 0.349798 | 0.292093 | 0.1 | 0.082 | 1 | 6 | Pam16 | 0.018 | Old |
| 0.350298 | 0.56415 | 0.031 | 0.022 | 1 | 6 | Mad2l1 | 0.009 | Old |
| 0.350768 | 0.13923 | 0.177 | 0.152 | 1 | 6 | Ebna1bp2 | 0.025 | Old |
| 0.351044 | 0.567306 | 0.017 | 0.027 | 1 | 6 | Tsen15 | -0.01 | Old |
| 0.351194 | 0.188794 | 0.266 | 0.234 | 1 | 6 | 4932438A1 | 0.032 | Old |
| 0.351294 | 0.388102 | 0.14 | 0.12 | 1 | 6 | Ftl1-ps1 | 0.02 | Old |
| 0.351315 | 0.178476 | 0.231 | 0.206 | 1 | 6 | Ppdpf | 0.025 | Old |

| | | | | | | | | |
|---|---|---|---|---|---|---|---|---|
| 0.351376 | 0.237911 | 0.046 | 0.033 | 1 | 6 | Dvl2 | 0.013 | Old |
| 0.351698 | 1.07918 | 0.031 | 0.022 | 1 | 6 | Zbtb10 | 0.009 | Old |
| 0.35182 | 0.069083 | 0.017 | 0.01 | 1 | 6 | Gm16094 | 0.007 | Old |
| 0.351869 | 0.09655 | 0.103 | 0.084 | 1 | 6 | Rab3il1 | 0.019 | Old |
| 0.352047 | 0.1495 | 0.166 | 0.142 | 1 | 6 | Adrm1 | 0.024 | Old |
| 0.352326 | 0.481439 | 0.08 | 0.064 | 1 | 6 | Gps2 | 0.016 | Old |
| 0.353046 | 0.290372 | 0.263 | 0.236 | 1 | 6 | Anxa6 | 0.027 | Old |
| 0.353105 | 0.06277 | 0.149 | 0.125 | 1 | 6 | Slc38a10 | 0.024 | Old |
| 0.353126 | 1.184087 | 0.043 | 0.032 | 1 | 6 | Setdb2 | 0.011 | Old |
| 0.353625 | 0.213236 | 0.02 | 0.03 | 1 | 6 | Zfp719 | -0.01 | Old |
| 0.353743 | 0.001164 | 0.317 | 0.349 | 1 | 6 | Crybg1 | -0.032 | Old |
| 0.354389 | 0.493995 | 0.106 | 0.087 | 1 | 6 | Med16 | 0.019 | Old |
| 0.354978 | 1.078894 | 0.031 | 0.022 | 1 | 6 | Ulk3 | 0.009 | Old |
| 0.355073 | 0.130459 | 0.263 | 0.236 | 1 | 6 | Alkbh5 | 0.027 | Old |
| 0.355683 | 0.771962 | 0.031 | 0.022 | 1 | 6 | Gm15228 | 0.009 | Old |
| 0.355766 | 0.121152 | 0.389 | 0.351 | 1 | 6 | U2af2 | 0.038 | Old |
| 0.355864 | 0.161484 | 0.234 | 0.207 | 1 | 6 | Hexim1 | 0.027 | Old |
| 0.355891 | 0.249744 | 0.306 | 0.284 | 1 | 6 | Fkbp8 | 0.022 | Old |
| 0.355988 | 0.137309 | 0.463 | 0.43 | 1 | 6 | Rnf141 | 0.033 | Old |
| 0.356062 | 0.57239 | 0.06 | 0.047 | 1 | 6 | Stx18 | 0.013 | Old |
| 0.356738 | 0.137681 | 0.291 | 0.261 | 1 | 6 | Rsu1 | 0.03 | Old |
| 0.357018 | 0.050356 | 0.551 | 0.503 | 1 | 6 | Zcchc6 | 0.048 | Old |
| 0.357277 | 0.251818 | 0.426 | 0.395 | 1 | 6 | Paip2 | 0.031 | Old |
| 0.357425 | 0.139347 | 0.154 | 0.13 | 1 | 6 | Mrpl17 | 0.024 | Old |
| 0.35751 | 0.128497 | 0.197 | 0.171 | 1 | 6 | Sars | 0.026 | Old |
| 0.357606 | 0.307067 | 0.117 | 0.097 | 1 | 6 | Cep83 | 0.02 | Old |
| 0.358115 | 0.853834 | 0.071 | 0.057 | 1 | 6 | Cyth3 | 0.014 | Old |
| 0.358123 | 0.203644 | 0.023 | 0.033 | 1 | 6 | Rrs1 | -0.01 | Old |
| 0.358296 | 0.1031 | 0.074 | 0.059 | 1 | 6 | Atad3a | 0.015 | Old |
| 0.35857 | 0.188497 | 0.369 | 0.333 | 1 | 6 | Rlim | 0.036 | Old |
| 0.358974 | 0.179505 | 0.283 | 0.254 | 1 | 6 | Khsrp | 0.029 | Old |
| 0.359412 | 0.131408 | 0.534 | 0.512 | 1 | 6 | Psma3 | 0.022 | Old |
| 0.359594 | 0.185052 | 0.094 | 0.114 | 1 | 6 | Rad50 | -0.02 | Old |
| 0.35974 | 0.669207 | 0.097 | 0.079 | 1 | 6 | Sun1 | 0.018 | Old |
| 0.360008 | 0.079177 | 0.006 | 0.012 | 1 | 6 | Msrb2 | -0.006 | Old |
| 0.360095 | 0.672491 | 0.02 | 0.03 | 1 | 6 | Galnt6 | -0.01 | Old |
| 0.36017 | 0.789448 | 0.031 | 0.022 | 1 | 6 | Dffa | 0.009 | Old |
| 0.360595 | 0.816761 | 0.066 | 0.052 | 1 | 6 | Rab11fip2 | 0.014 | Old |
| 0.360644 | 0.595474 | 0.031 | 0.022 | 1 | 6 | Rabl3 | 0.009 | Old |
| 0.360761 | 1.387102 | 0.114 | 0.099 | 1 | 6 | Rabgap1l | 0.015 | Old |
| 0.360776 | 0.072182 | 0.006 | 0.012 | 1 | 6 | Hexdc | -0.006 | Old |
| 0.361546 | 0.219469 | 0.006 | 0.012 | 1 | 6 | Gm47232 | -0.006 | Old |
| 0.361594 | 0.64207 | 0.031 | 0.022 | 1 | 6 | Limk2 | 0.009 | Old |
| 0.361875 | 0.119751 | 0.18 | 0.157 | 1 | 6 | Mindy3 | 0.023 | Old |
| 0.36207 | 0.208019 | 0.031 | 0.022 | 1 | 6 | Slc9a4 | 0.009 | Old |

| | | | | | | | | |
|---|---|---|---|---|---|---|---|---|
| 0.362138 | 0.303093 | 0.151 | 0.132 | 1 | 6 | Tnip2 | 0.019 | Old |
| 0.362261 | 0.084141 | 0.2 | 0.172 | 1 | 6 | Nsmce4a | 0.028 | Old |
| 0.362272 | 0.11035 | 0.1 | 0.082 | 1 | 6 | Atp13a2 | 0.018 | Old |
| 0.362317 | 0.150584 | 0.006 | 0.012 | 1 | 6 | Slc40a1 | -0.006 | Old |
| 0.362521 | 0.66803 | 0.083 | 0.067 | 1 | 6 | Rpp25l | 0.016 | Old |
| 0.362546 | 0.896854 | 0.031 | 0.022 | 1 | 6 | Dock5 | 0.009 | Old |
| 0.36278 | 0.675142 | 0.06 | 0.047 | 1 | 6 | Rrp9 | 0.013 | Old |
| 0.362802 | 0.820998 | 0.037 | 0.027 | 1 | 6 | Cd99l2 | 0.01 | Old |
| 0.363088 | 0.244245 | 0.006 | 0.012 | 1 | 6 | Maml2 | -0.006 | Old |
| 0.363088 | 0.096568 | 0.006 | 0.012 | 1 | 6 | Trem1 | -0.006 | Old |
| 0.363192 | 0.385989 | 0.077 | 0.062 | 1 | 6 | Jade1 | 0.015 | Old |
| 0.363356 | 0.215277 | 0.229 | 0.266 | 1 | 6 | Numa1 | -0.037 | Old |
| 0.363795 | 0.177975 | 0.137 | 0.115 | 1 | 6 | Lrpap1 | 0.022 | Old |
| 0.364003 | 0.201177 | 0.454 | 0.431 | 1 | 6 | Ssr4 | 0.023 | Old |
| 0.364107 | 0.59545 | 0.037 | 0.027 | 1 | 6 | Pola1 | 0.01 | Old |
| 0.364178 | 0.307175 | 0.091 | 0.112 | 1 | 6 | Armc10 | -0.021 | Old |
| 0.364449 | 0.294568 | 0.194 | 0.172 | 1 | 6 | Nsun2 | 0.022 | Old |
| 0.364454 | 0.585385 | 0.031 | 0.022 | 1 | 6 | Plat | 0.009 | Old |
| 0.36453 | 0.163781 | 0.174 | 0.149 | 1 | 6 | Rcn2 | 0.025 | Old |
| 0.364634 | 1.791628 | 0.006 | 0.012 | 1 | 6 | Rbpms2 | -0.006 | Old |
| 0.364634 | 0.430299 | 0.006 | 0.012 | 1 | 6 | Wdr5b | -0.006 | Old |
| 0.364634 | 0.042992 | 0.006 | 0.012 | 1 | 6 | Zbed3 | -0.006 | Old |
| 0.364652 | 0.520923 | 0.094 | 0.079 | 1 | 6 | Phyh | 0.015 | Old |
| 0.364718 | 0.183944 | 0.069 | 0.087 | 1 | 6 | Fmnl2 | -0.018 | Old |
| 0.364917 | 0.134802 | 0.029 | 0.04 | 1 | 6 | Klhl7 | -0.011 | Old |
| 0.365153 | 0.128825 | 0.063 | 0.048 | 1 | 6 | Zfp422 | 0.015 | Old |
| 0.365441 | 0.180643 | 0.034 | 0.047 | 1 | 6 | Lrrc40 | -0.013 | Old |
| 0.365551 | 0.340186 | 0.094 | 0.077 | 1 | 6 | Minpp1 | 0.017 | Old |
| 0.365975 | 0.3671 | 0.111 | 0.094 | 1 | 6 | Slc4a2 | 0.017 | Old |
| 0.366091 | 0.072898 | 0.891 | 0.911 | 1 | 6 | Ddx5 | -0.02 | Old |
| 0.366095 | 0.37767 | 0.094 | 0.077 | 1 | 6 | Mid1ip1 | 0.017 | Old |
| 0.366412 | 0.463524 | 0.223 | 0.202 | 1 | 6 | Dnajc15 | 0.021 | Old |
| 0.366508 | 0.522285 | 0.037 | 0.027 | 1 | 6 | Tti2 | 0.01 | Old |
| 0.366779 | 0.057889 | 0.171 | 0.147 | 1 | 6 | Chd6 | 0.024 | Old |
| 0.366951 | 0.949154 | 0.054 | 0.042 | 1 | 6 | Wdr70 | 0.012 | Old |
| 0.367154 | 0.161554 | 0.063 | 0.048 | 1 | 6 | Bcl2l13 | 0.015 | Old |
| 0.367306 | 0.153129 | 0.166 | 0.144 | 1 | 6 | B3gat3 | 0.022 | Old |
| 0.367548 | 0.652592 | 0.043 | 0.032 | 1 | 6 | Trim7 | 0.011 | Old |
| 0.367639 | 0.175761 | 0.074 | 0.092 | 1 | 6 | Tsfm | -0.018 | Old |
| 0.367954 | 0.649811 | 0.043 | 0.032 | 1 | 6 | Cd38 | 0.011 | Old |
| 0.368211 | 1.062934 | 0.06 | 0.047 | 1 | 6 | Drg2 | 0.013 | Old |
| 0.368221 | 1.524908 | 0.023 | 0.015 | 1 | 6 | 2310057M | 0.008 | Old |
| 0.368313 | 0.151205 | 0.511 | 0.478 | 1 | 6 | Dync1h1 | 0.033 | Old |
| 0.368385 | 0.251225 | 0.377 | 0.351 | 1 | 6 | Stk4 | 0.026 | Old |
| 0.368623 | 0.432399 | 0.237 | 0.216 | 1 | 6 | Frg1 | 0.021 | Old |

| | | | | | | | | |
|---|---|---|---|---|---|---|---|---|
| 0.368634 | 0.033823 | 0.16 | 0.137 | 1 | 6 | Selenoi | 0.023 | Old |
| 0.369359 | 0.048033 | 0.111 | 0.092 | 1 | 6 | Pak4 | 0.019 | Old |
| 0.369376 | 0.020845 | 0.029 | 0.04 | 1 | 6 | Ints8 | -0.011 | Old |
| 0.369732 | 0.175152 | 0.031 | 0.022 | 1 | 6 | Gm26853 | 0.009 | Old |
| 0.370714 | 0.014307 | 0.303 | 0.269 | 1 | 6 | Pole4 | 0.034 | Old |
| 0.370805 | 0.201226 | 0.266 | 0.241 | 1 | 6 | Pim3 | 0.025 | Old |
| 0.370942 | 0.18099 | 0.114 | 0.095 | 1 | 6 | Nudt5 | 0.019 | Old |
| 0.371353 | 0.587024 | 0.049 | 0.037 | 1 | 6 | 2410004B1 | 0.012 | Old |
| 0.371407 | 0.246363 | 0.2 | 0.174 | 1 | 6 | Grina | 0.026 | Old |
| 0.371509 | 0.047363 | 0.143 | 0.169 | 1 | 6 | Eif2s3y | -0.026 | Old |
| 0.371645 | 0.8546 | 0.023 | 0.015 | 1 | 6 | Tmtc3 | 0.008 | Old |
| 0.372056 | 0.099579 | 0.031 | 0.043 | 1 | 6 | Extl2 | -0.012 | Old |
| 0.372127 | 0.312258 | 0.16 | 0.139 | 1 | 6 | Arpc5l | 0.021 | Old |
| 0.372596 | 0.035891 | 0.106 | 0.087 | 1 | 6 | Lgals3bp | 0.019 | Old |
| 0.372983 | 0.227609 | 0.089 | 0.072 | 1 | 6 | Ccnh | 0.017 | Old |
| 0.373193 | 0.144149 | 0.323 | 0.293 | 1 | 6 | Ppp6r3 | 0.03 | Old |
| 0.373289 | 0.412618 | 0.109 | 0.092 | 1 | 6 | Nvl | 0.017 | Old |
| 0.373459 | 0.107797 | 0.471 | 0.435 | 1 | 6 | Fermt3 | 0.036 | Old |
| 0.373835 | 0.287956 | 0.137 | 0.119 | 1 | 6 | Mrpl2 | 0.018 | Old |
| 0.374035 | 0.108653 | 0.329 | 0.299 | 1 | 6 | Vav3 | 0.03 | Old |
| 0.374176 | 0.099539 | 0.123 | 0.102 | 1 | 6 | Poglut1 | 0.021 | Old |
| 0.374249 | 0.16987 | 0.514 | 0.505 | 1 | 6 | Sf1 | 0.009 | Old |
| 0.374451 | 0.146017 | 0.294 | 0.268 | 1 | 6 | Sec11a | 0.026 | Old |
| 0.3748 | 0.173228 | 0.171 | 0.199 | 1 | 6 | Evl | -0.028 | Old |
| 0.374992 | 0.535138 | 0.094 | 0.079 | 1 | 6 | Atrnl1 | 0.015 | Old |
| 0.375096 | 0.495618 | 0.037 | 0.027 | 1 | 6 | Ggct | 0.01 | Old |
| 0.375118 | 0.258989 | 0.351 | 0.324 | 1 | 6 | Nisch | 0.027 | Old |
| 0.375138 | 0.52132 | 0.094 | 0.077 | 1 | 6 | Cyfip2 | 0.017 | Old |
| 0.375138 | 0.172888 | 0.094 | 0.077 | 1 | 6 | Ubl7 | 0.017 | Old |
| 0.375275 | 0.043746 | 0.417 | 0.446 | 1 | 6 | Tcf4 | -0.029 | Old |
| 0.375367 | 0.310366 | 0.091 | 0.075 | 1 | 6 | Insig2 | 0.016 | Old |
| 0.375573 | 0.452551 | 0.066 | 0.052 | 1 | 6 | Ift46 | 0.014 | Old |
| 0.375646 | 0.618358 | 0.091 | 0.075 | 1 | 6 | Exosc10 | 0.016 | Old |
| 0.375761 | 0.594653 | 0.129 | 0.11 | 1 | 6 | Shprh | 0.019 | Old |
| 0.375959 | 0.039505 | 0.174 | 0.199 | 1 | 6 | Slc25a20 | -0.025 | Old |
| 0.376339 | 0.112786 | 0.766 | 0.766 | 1 | 6 | Ptms | 0 | Old |
| 0.376399 | 0.7188 | 0.063 | 0.05 | 1 | 6 | Plekha5 | 0.013 | Old |
| 0.376451 | 0.597438 | 0.06 | 0.047 | 1 | 6 | St7l | 0.013 | Old |
| 0.376451 | 0.477688 | 0.06 | 0.047 | 1 | 6 | Zfp595 | 0.013 | Old |
| 0.37651 | 0.223308 | 0.163 | 0.14 | 1 | 6 | Pank2 | 0.023 | Old |
| 0.37654 | 0.403526 | 0.043 | 0.032 | 1 | 6 | Sephs2 | 0.011 | Old |
| 0.376812 | 0.22772 | 0.497 | 0.473 | 1 | 6 | Dnajb6 | 0.024 | Old |
| 0.376819 | 0.959009 | 0.023 | 0.015 | 1 | 6 | Cep89 | 0.008 | Old |
| 0.377681 | 0.0316 | 0.231 | 0.258 | 1 | 6 | Psmc5 | -0.027 | Old |
| 0.377957 | 0.363618 | 0.123 | 0.105 | 1 | 6 | 9130401M | 0.018 | Old |

| | | | | | | | | |
|---|---|---|---|---|---|---|---|---|
| 0.377972 | 0.086691 | 0.817 | 0.809 | 1 | 6 | Eif4g2 | 0.008 | Old |
| 0.378019 | 0.508991 | 0.094 | 0.079 | 1 | 6 | Patl1 | 0.015 | Old |
| 0.378367 | 0.92875 | 0.046 | 0.035 | 1 | 6 | Tram2 | 0.011 | Old |
| 0.378426 | 0.569304 | 0.183 | 0.162 | 1 | 6 | Zranb1 | 0.021 | Old |
| 0.378554 | 0.906599 | 0.023 | 0.015 | 1 | 6 | Zfp668 | 0.008 | Old |
| 0.379133 | 0.722509 | 0.023 | 0.015 | 1 | 6 | BC017158 | 0.008 | Old |
| 0.379619 | 0.431977 | 0.054 | 0.042 | 1 | 6 | Prr7 | 0.012 | Old |
| 0.379713 | 1.270756 | 0.023 | 0.015 | 1 | 6 | Gadd45a | 0.008 | Old |
| 0.379713 | 0.788021 | 0.023 | 0.015 | 1 | 6 | Wdr24 | 0.008 | Old |
| 0.379713 | 0.572943 | 0.023 | 0.015 | 1 | 6 | Zfp60 | 0.008 | Old |
| 0.379791 | 0.102206 | 0.143 | 0.12 | 1 | 6 | Dpy30 | 0.023 | Old |
| 0.379869 | 0.116789 | 0.76 | 0.766 | 1 | 6 | Ptbp3 | -0.006 | Old |
| 0.379874 | 0.52666 | 0.066 | 0.052 | 1 | 6 | Abcd4 | 0.014 | Old |
| 0.379892 | 0.186434 | 0.157 | 0.137 | 1 | 6 | Usp45 | 0.02 | Old |
| 0.381228 | 0.39559 | 0.237 | 0.216 | 1 | 6 | Psmc1 | 0.021 | Old |
| 0.381253 | 0.006296 | 0.146 | 0.124 | 1 | 6 | Kcnab2 | 0.022 | Old |
| 0.38145 | 0.575955 | 0.054 | 0.042 | 1 | 6 | Tmem94 | 0.012 | Old |
| 0.381455 | 0.729203 | 0.023 | 0.015 | 1 | 6 | Nle1 | 0.008 | Old |
| 0.381753 | 0.220963 | 0.503 | 0.487 | 1 | 6 | H2-DMb2 | 0.016 | Old |
| 0.381873 | 0.203262 | 0.289 | 0.259 | 1 | 6 | Eif1b | 0.03 | Old |
| 0.381916 | 0.56607 | 0.043 | 0.032 | 1 | 6 | 2610008E1 | 0.011 | Old |
| 0.382428 | 0.195967 | 0.731 | 0.726 | 1 | 6 | Lbh | 0.005 | Old |
| 0.382551 | 0.544485 | 0.054 | 0.042 | 1 | 6 | Lig3 | 0.012 | Old |
| 0.382551 | 0.506896 | 0.054 | 0.042 | 1 | 6 | Tatdn3 | 0.012 | Old |
| 0.38262 | 0.171359 | 0.023 | 0.015 | 1 | 6 | Dus2 | 0.008 | Old |
| 0.382827 | 0.50477 | 0.134 | 0.115 | 1 | 6 | Wdr44 | 0.019 | Old |
| 0.38287 | 0.274358 | 0.066 | 0.052 | 1 | 6 | Foxp4 | 0.014 | Old |
| 0.38304 | 2.285421 | 0.014 | 0.008 | 1 | 6 | Itgb3 | 0.006 | Old |
| 0.383203 | 0.578625 | 0.023 | 0.015 | 1 | 6 | Gm15564 | 0.008 | Old |
| 0.383203 | 0.558997 | 0.023 | 0.015 | 1 | 6 | Wdr90 | 0.008 | Old |
| 0.383708 | 0.201405 | 0.049 | 0.037 | 1 | 6 | Asb7 | 0.012 | Old |
| 0.383731 | 0.176074 | 0.277 | 0.254 | 1 | 6 | Copb1 | 0.023 | Old |
| 0.383771 | 0.024706 | 0.26 | 0.231 | 1 | 6 | Trim12c | 0.029 | Old |
| 0.384259 | 0.004759 | 0.503 | 0.535 | 1 | 6 | Rock1 | -0.032 | Old |
| 0.384343 | 0.030653 | 0.177 | 0.154 | 1 | 6 | Commd3 | 0.023 | Old |
| 0.384371 | 0.337924 | 0.023 | 0.015 | 1 | 6 | Nrxn2 | 0.008 | Old |
| 0.384703 | 0.391112 | 0.106 | 0.09 | 1 | 6 | Ift52 | 0.016 | Old |
| 0.384751 | 0.464892 | 0.077 | 0.062 | 1 | 6 | Tsr1 | 0.015 | Old |
| 0.384922 | 0.114526 | 0.711 | 0.709 | 1 | 6 | Ywhah | 0.002 | Old |
| 0.384943 | 0.243928 | 0.091 | 0.075 | 1 | 6 | Las1l | 0.016 | Old |
| 0.385039 | 0.133173 | 0.029 | 0.04 | 1 | 6 | Alg13 | -0.011 | Old |
| 0.385085 | 0.380636 | 0.14 | 0.122 | 1 | 6 | Bckdk | 0.018 | Old |
| 0.385316 | 0.822466 | 0.014 | 0.008 | 1 | 6 | Exo5 | 0.006 | Old |
| 0.385506 | 0.138481 | 0.06 | 0.047 | 1 | 6 | Tmem268 | 0.013 | Old |
| 0.385541 | 0.636454 | 0.023 | 0.015 | 1 | 6 | Zfp46 | 0.008 | Old |

| | | | | | | | | |
|---|---|---|---|---|---|---|---|---|
| 0.385541 | 0.362624 | 0.023 | 0.015 | 1 | 6 | Nemp2 | 0.008 | Old |
| 0.385555 | 0.211152 | 0.263 | 0.236 | 1 | 6 | Ppp4r2 | 0.027 | Old |
| 0.385848 | 0.029436 | 0.286 | 0.254 | 1 | 6 | 1110004F1 | 0.032 | Old |
| 0.386237 | 0.067689 | 0.054 | 0.042 | 1 | 6 | Golga1 | 0.012 | Old |
| 0.386633 | 0.46466 | 0.1 | 0.084 | 1 | 6 | Dcps | 0.016 | Old |
| 0.386839 | 2.014802 | 0.014 | 0.008 | 1 | 6 | Ms4a4b | 0.006 | Old |
| 0.386839 | 0.718934 | 0.014 | 0.008 | 1 | 6 | Gm20604 | 0.006 | Old |
| 0.386999 | 0.100188 | 0.511 | 0.478 | 1 | 6 | Adam15 | 0.033 | Old |
| 0.387382 | 0.348283 | 0.2 | 0.177 | 1 | 6 | Strn4 | 0.023 | Old |
| 0.387602 | 0.892049 | 0.014 | 0.008 | 1 | 6 | Kyat3 | 0.006 | Old |
| 0.387602 | 0.657417 | 0.014 | 0.008 | 1 | 6 | Zbtb8a | 0.006 | Old |
| 0.388047 | 0.328562 | 0.46 | 0.44 | 1 | 6 | Ankrd11 | 0.02 | Old |
| 0.388365 | 1.224773 | 0.014 | 0.008 | 1 | 6 | Nrg1 | 0.006 | Old |
| 0.388365 | 0.776865 | 0.014 | 0.008 | 1 | 6 | Rilp | 0.006 | Old |
| 0.388365 | 0.741242 | 0.014 | 0.008 | 1 | 6 | Col3a1 | 0.006 | Old |
| 0.388365 | 0.641157 | 0.014 | 0.008 | 1 | 6 | C730034F0 | 0.006 | Old |
| 0.388365 | 0.513165 | 0.014 | 0.008 | 1 | 6 | Amotl2 | 0.006 | Old |
| 0.388387 | 0.164879 | 0.043 | 0.032 | 1 | 6 | Irgq | 0.011 | Old |
| 0.388472 | 0.307563 | 0.126 | 0.107 | 1 | 6 | Rnf181 | 0.019 | Old |
| 0.388475 | 0.265522 | 0.023 | 0.015 | 1 | 6 | Dennd2d | 0.008 | Old |
| 0.388616 | 0.234587 | 0.24 | 0.216 | 1 | 6 | H2-M3 | 0.024 | Old |
| 0.388707 | 0.112169 | 0.171 | 0.149 | 1 | 6 | Dars | 0.022 | Old |
| 0.388823 | 0.413064 | 0.086 | 0.105 | 1 | 6 | Pik3cb | -0.019 | Old |
| 0.389064 | 0.147961 | 0.023 | 0.015 | 1 | 6 | Pla1a | 0.008 | Old |
| 0.389129 | 0.565563 | 0.014 | 0.008 | 1 | 6 | Dnaaf5 | 0.006 | Old |
| 0.389129 | 0.407094 | 0.014 | 0.008 | 1 | 6 | Farp2 | 0.006 | Old |
| 0.389129 | 0.346365 | 0.014 | 0.008 | 1 | 6 | Adal | 0.006 | Old |
| 0.389413 | 0.060663 | 0.437 | 0.477 | 1 | 6 | Ppp2r5a | -0.04 | Old |
| 0.389625 | 0.274932 | 0.326 | 0.301 | 1 | 6 | Ankrd13a | 0.025 | Old |
| 0.389652 | 1.30182 | 0.034 | 0.025 | 1 | 6 | Tbccd1 | 0.009 | Old |
| 0.389653 | 0.577675 | 0.023 | 0.015 | 1 | 6 | L2hgdh | 0.008 | Old |
| 0.389729 | 0.304707 | 0.06 | 0.047 | 1 | 6 | Pgm3 | 0.013 | Old |
| 0.389895 | 0.750264 | 0.014 | 0.008 | 1 | 6 | Dyrk3 | 0.006 | Old |
| 0.389916 | 0.130791 | 0.066 | 0.052 | 1 | 6 | 1600012H0 | 0.014 | Old |
| 0.390086 | 0.242917 | 0.111 | 0.094 | 1 | 6 | Mrpl10 | 0.017 | Old |
| 0.390143 | 0.152785 | 0.426 | 0.383 | 1 | 6 | Stxbp3 | 0.043 | Old |
| 0.390438 | 0.312593 | 0.151 | 0.132 | 1 | 6 | 3-Sep | 0.019 | Old |
| 0.390609 | 0.219043 | 0.194 | 0.174 | 1 | 6 | Nampt | 0.02 | Old |
| 0.390781 | 0.140957 | 0.191 | 0.171 | 1 | 6 | Gars | 0.02 | Old |
| 0.391023 | 0.102243 | 0.183 | 0.159 | 1 | 6 | Bcor | 0.024 | Old |
| 0.391164 | 0.091529 | 0.223 | 0.197 | 1 | 6 | Srrt | 0.026 | Old |
| 0.391328 | 0.066996 | 0.08 | 0.097 | 1 | 6 | Naa30 | -0.017 | Old |
| 0.391428 | 0.309258 | 0.014 | 0.008 | 1 | 6 | Ccdc157 | 0.006 | Old |
| 0.391551 | 0.573244 | 0.009 | 0.015 | 1 | 6 | 1110035H1 | -0.006 | Old |
| 0.391697 | 0.019644 | 0.083 | 0.067 | 1 | 6 | Hira | 0.016 | Old |

| | | | | | | | | |
|---|---|---|---|---|---|---|---|---|
| 0.391796 | 0.942662 | 0.057 | 0.045 | 1 | 6 | Nufip1 | 0.012 | Old |
| 0.3919 | 0.268002 | 0.077 | 0.062 | 1 | 6 | Il15 | 0.015 | Old |
| 0.392037 | 0.235938 | 0.429 | 0.401 | 1 | 6 | Ago2 | 0.028 | Old |
| 0.392108 | 0.22512 | 0.097 | 0.08 | 1 | 6 | Impdh1 | 0.017 | Old |
| 0.392126 | 0.24204 | 0.14 | 0.12 | 1 | 6 | Ubqln2 | 0.02 | Old |
| 0.392196 | 0.872034 | 0.014 | 0.008 | 1 | 6 | Zfp580 | 0.006 | Old |
| 0.392196 | 0.229643 | 0.014 | 0.008 | 1 | 6 | Gm15448 | 0.006 | Old |
| 0.392196 | 0.127307 | 0.014 | 0.008 | 1 | 6 | Agrn | 0.006 | Old |
| 0.392196 | 0.125656 | 0.014 | 0.008 | 1 | 6 | Hes1 | 0.006 | Old |
| 0.392221 | 0.132034 | 0.123 | 0.104 | 1 | 6 | Ykt6 | 0.019 | Old |
| 0.392296 | 0.497823 | 0.074 | 0.06 | 1 | 6 | Unc45a | 0.014 | Old |
| 0.392355 | 0.493828 | 0.077 | 0.064 | 1 | 6 | Gpn3 | 0.013 | Old |
| 0.392423 | 0.224831 | 0.089 | 0.072 | 1 | 6 | Zfp524 | 0.017 | Old |
| 0.392916 | 0.972555 | 0.029 | 0.02 | 1 | 6 | Herc3 | 0.009 | Old |
| 0.392965 | 0.568283 | 0.014 | 0.008 | 1 | 6 | Trim16 | 0.006 | Old |
| 0.392965 | 0.395945 | 0.014 | 0.008 | 1 | 6 | Tsix | 0.006 | Old |
| 0.393079 | 0.377041 | 0.174 | 0.154 | 1 | 6 | Mtfr1l | 0.02 | Old |
| 0.393126 | 0.168297 | 0.14 | 0.119 | 1 | 6 | Tufm | 0.021 | Old |
| 0.393393 | 0.124214 | 0.126 | 0.107 | 1 | 6 | Mtmr1 | 0.019 | Old |
| 0.393533 | 0.235755 | 0.049 | 0.037 | 1 | 6 | Wdr55 | 0.012 | Old |
| 0.393539 | 0.315265 | 0.146 | 0.125 | 1 | 6 | Tceal9 | 0.021 | Old |
| 0.393735 | 0.311237 | 0.014 | 0.008 | 1 | 6 | 2900076A( | 0.006 | Old |
| 0.393981 | 0.504536 | 0.06 | 0.047 | 1 | 6 | Dip2a | 0.013 | Old |
| 0.394091 | 0.16878 | 0.091 | 0.075 | 1 | 6 | Ceacam1 | 0.016 | Old |
| 0.394506 | 0.174624 | 0.014 | 0.008 | 1 | 6 | Tsen2 | 0.006 | Old |
| 0.394645 | 0.679029 | 0.063 | 0.05 | 1 | 6 | Mynn | 0.013 | Old |
| 0.395119 | 0.044294 | 0.049 | 0.037 | 1 | 6 | Slc25a32 | 0.012 | Old |
| 0.396217 | 0.345557 | 0.111 | 0.095 | 1 | 6 | Uspl1 | 0.016 | Old |
| 0.397235 | 0.031057 | 0.054 | 0.042 | 1 | 6 | Gstp1 | 0.012 | Old |
| 0.397418 | 0.261585 | 0.214 | 0.196 | 1 | 6 | Map3k11 | 0.018 | Old |
| 0.397579 | 0.021889 | 0.134 | 0.154 | 1 | 6 | Socs7 | -0.02 | Old |
| 0.398036 | 0.39747 | 0.157 | 0.137 | 1 | 6 | Trabd | 0.02 | Old |
| 0.398168 | 0.103675 | 0.337 | 0.308 | 1 | 6 | Leprot | 0.029 | Old |
| 0.398287 | 0.153337 | 0.486 | 0.46 | 1 | 6 | Bclaf1 | 0.026 | Old |
| 0.399508 | 0.177038 | 0.397 | 0.375 | 1 | 6 | Lrrfip1 | 0.022 | Old |
| 0.400046 | 0.311596 | 0.149 | 0.129 | 1 | 6 | Smg5 | 0.02 | Old |
| 0.400077 | 0.892706 | 0.034 | 0.025 | 1 | 6 | Zkscan14 | 0.009 | Old |
| 0.400104 | 0.548906 | 0.1 | 0.084 | 1 | 6 | Nsmf | 0.016 | Old |
| 0.400266 | 0.549231 | 0.029 | 0.02 | 1 | 6 | Plcg1 | 0.009 | Old |
| 0.400521 | 0.170949 | 0.28 | 0.259 | 1 | 6 | Thoc7 | 0.021 | Old |
| 0.401517 | 0.101438 | 0.086 | 0.104 | 1 | 6 | Banp | -0.018 | Old |
| 0.402357 | 0.284281 | 0.16 | 0.14 | 1 | 6 | Rbfa | 0.02 | Old |
| 0.4028 | 0.119182 | 0.614 | 0.595 | 1 | 6 | Hnrnpm | 0.019 | Old |
| 0.402819 | 0.187183 | 0.731 | 0.749 | 1 | 6 | Srrm2 | -0.018 | Old |
| 0.40291 | 0.351394 | 0.029 | 0.02 | 1 | 6 | 1500015A( | 0.009 | Old |

| | | | | | | | | |
|---|---|---|---|---|---|---|---|---|
| 0.40304 | 0.13532 | 0.209 | 0.184 | 1 | 6 | H2-Q4 | 0.025 | Old |
| 0.403141 | 0.432175 | 0.206 | 0.186 | 1 | 6 | Paxbp1 | 0.02 | Old |
| 0.403442 | 1.111931 | 0.046 | 0.035 | 1 | 6 | Mccc1 | 0.011 | Old |
| 0.403442 | 0.181301 | 0.046 | 0.035 | 1 | 6 | Aasdh | 0.011 | Old |
| 0.403451 | 0.359298 | 0.309 | 0.289 | 1 | 6 | Arhgef2 | 0.02 | Old |
| 0.403542 | 0.052107 | 0.103 | 0.085 | 1 | 6 | Ppid | 0.018 | Old |
| 0.403622 | 0.112857 | 0.189 | 0.166 | 1 | 6 | Tnks | 0.023 | Old |
| 0.403646 | 0.274115 | 0.057 | 0.045 | 1 | 6 | Dnajc24 | 0.012 | Old |
| 0.403946 | 0.41591 | 0.077 | 0.064 | 1 | 6 | Thyn1 | 0.013 | Old |
| 0.404283 | 0.219853 | 0.1 | 0.084 | 1 | 6 | Foxred2 | 0.016 | Old |
| 0.404422 | 0.216898 | 0.2 | 0.177 | 1 | 6 | Ddhd1 | 0.023 | Old |
| 0.404702 | 0.255114 | 0.283 | 0.254 | 1 | 6 | Acsl5 | 0.029 | Old |
| 0.405161 | 0.332635 | 0.16 | 0.14 | 1 | 6 | Ipo5 | 0.02 | Old |
| 0.405355 | 0.659116 | 0.094 | 0.079 | 1 | 6 | Rbm15 | 0.015 | Old |
| 0.405625 | 0.757674 | 0.083 | 0.069 | 1 | 6 | Caap1 | 0.014 | Old |
| 0.405647 | 0.406839 | 0.04 | 0.03 | 1 | 6 | Dars2 | 0.01 | Old |
| 0.405834 | 0.843074 | 0.034 | 0.025 | 1 | 6 | Dbr1 | 0.009 | Old |
| 0.406014 | 0.038441 | 0.12 | 0.14 | 1 | 6 | Agtpbp1 | -0.02 | Old |
| 0.406091 | 0.767938 | 0.04 | 0.03 | 1 | 6 | Stradb | 0.01 | Old |
| 0.406356 | 0.983776 | 0.1 | 0.085 | 1 | 6 | Irf8 | 0.015 | Old |
| 0.406523 | 0.630592 | 0.1 | 0.084 | 1 | 6 | Mrpl55 | 0.016 | Old |
| 0.406535 | 0.837343 | 0.04 | 0.03 | 1 | 6 | Fam213b | 0.01 | Old |
| 0.406895 | 0.031883 | 0.909 | 0.933 | 1 | 6 | Calm1 | -0.024 | Old |
| 0.406962 | 0.050563 | 0.126 | 0.105 | 1 | 6 | Cdk16 | 0.021 | Old |
| 0.40715 | 0.471193 | 0.057 | 0.045 | 1 | 6 | Tubgcp2 | 0.012 | Old |
| 0.407447 | 0.275652 | 0.3 | 0.274 | 1 | 6 | Sdf2 | 0.026 | Old |
| 0.407607 | 0.202598 | 0.34 | 0.309 | 1 | 6 | Sltm | 0.031 | Old |
| 0.40767 | 0.116998 | 0.117 | 0.099 | 1 | 6 | Med12 | 0.018 | Old |
| 0.407993 | 0.5139 | 0.046 | 0.035 | 1 | 6 | Uck1 | 0.011 | Old |
| 0.408379 | 0.143654 | 0.569 | 0.55 | 1 | 6 | Prrc2c | 0.019 | Old |
| 0.408388 | 0.156342 | 0.203 | 0.179 | 1 | 6 | Golgb1 | 0.024 | Old |
| 0.40859 | 0.047268 | 0.046 | 0.059 | 1 | 6 | Syne3 | -0.013 | Old |
| 0.409158 | 0.024981 | 0.089 | 0.072 | 1 | 6 | Fbxo34 | 0.017 | Old |
| 0.40919 | 0.200974 | 0.149 | 0.127 | 1 | 6 | Crat | 0.022 | Old |
| 0.409215 | 0.548002 | 0.034 | 0.025 | 1 | 6 | Mccc2 | 0.009 | Old |
| 0.409308 | 0.571822 | 0.083 | 0.069 | 1 | 6 | Jak3 | 0.014 | Old |
| 0.409661 | 0.002724 | 0.563 | 0.604 | 1 | 6 | Sbno2 | -0.041 | Old |
| 0.409699 | 0.542673 | 0.034 | 0.025 | 1 | 6 | Mettl13 | 0.009 | Old |
| 0.409836 | 0.576215 | 0.029 | 0.02 | 1 | 6 | Mypopos | 0.009 | Old |
| 0.410205 | 0.094872 | 0.38 | 0.421 | 1 | 6 | Stk38 | -0.041 | Old |
| 0.410409 | 0.000106 | 0.26 | 0.231 | 1 | 6 | Smc6 | 0.029 | Old |
| 0.410547 | 0.019241 | 0.217 | 0.241 | 1 | 6 | Rab31 | -0.024 | Old |
| 0.410568 | 0.008085 | 0.197 | 0.219 | 1 | 6 | Trio | -0.022 | Old |
| 0.41081 | 0.357563 | 0.114 | 0.097 | 1 | 6 | Dpp9 | 0.017 | Old |
| 0.41089 | 0.216549 | 0.614 | 0.584 | 1 | 6 | Slc38a2 | 0.03 | Old |

| | | | | | | | | |
|---|---|---|---|---|---|---|---|---|
| 0.411312 | 0.299369 | 0.011 | 0.018 | 1 | 6 | Brca2 | -0.007 | Old |
| 0.411367 | 0.41922 | 0.194 | 0.172 | 1 | 6 | Gtpbp2 | 0.022 | Old |
| 0.411792 | 0.064338 | 0.794 | 0.818 | 1 | 6 | Cox7c | -0.024 | Old |
| 0.412126 | 0.49205 | 0.034 | 0.025 | 1 | 6 | Mtx3 | 0.009 | Old |
| 0.412195 | 0.256774 | 0.089 | 0.074 | 1 | 6 | Ndufaf4 | 0.015 | Old |
| 0.412366 | 0.219696 | 0.183 | 0.162 | 1 | 6 | Synj2bp | 0.021 | Old |
| 0.412867 | 0.077367 | 0.114 | 0.134 | 1 | 6 | Vps37a | -0.02 | Old |
| 0.413181 | 0.044281 | 0.354 | 0.319 | 1 | 6 | Arhgef1 | 0.035 | Old |
| 0.413238 | 0.237826 | 0.063 | 0.05 | 1 | 6 | Zfp787 | 0.013 | Old |
| 0.413586 | 0.819519 | 0.034 | 0.025 | 1 | 6 | E130308A1 | 0.009 | Old |
| 0.414073 | 0.635614 | 0.034 | 0.025 | 1 | 6 | Ermard | 0.009 | Old |
| 0.414073 | 0.345878 | 0.034 | 0.025 | 1 | 6 | B930036N1 | 0.009 | Old |
| 0.414127 | 0.385218 | 0.04 | 0.03 | 1 | 6 | Tma16 | 0.01 | Old |
| 0.414247 | 0.621198 | 0.046 | 0.035 | 1 | 6 | Dut | 0.011 | Old |
| 0.414557 | 0.463157 | 0.011 | 0.018 | 1 | 6 | Fndc4 | -0.007 | Old |
| 0.414557 | 0.364008 | 0.011 | 0.018 | 1 | 6 | Adam23 | -0.007 | Old |
| 0.415085 | 0.531531 | 0.046 | 0.035 | 1 | 6 | Pde1b | 0.011 | Old |
| 0.41518 | 0.602636 | 0.083 | 0.069 | 1 | 6 | Pcnx | 0.014 | Old |
| 0.415801 | 0.541525 | 0.083 | 0.069 | 1 | 6 | C2cd3 | 0.014 | Old |
| 0.416281 | 0.015393 | 0.189 | 0.167 | 1 | 6 | Dopey1 | 0.022 | Old |
| 0.416706 | 1.076766 | 0.069 | 0.057 | 1 | 6 | Il21r | 0.012 | Old |
| 0.416916 | 0.145603 | 0.191 | 0.171 | 1 | 6 | Dld | 0.02 | Old |
| 0.417324 | 0.235174 | 0.194 | 0.176 | 1 | 6 | Acbd3 | 0.018 | Old |
| 0.417601 | 0.29813 | 0.106 | 0.09 | 1 | 6 | Rpf2 | 0.016 | Old |
| 0.417795 | 0.438634 | 0.103 | 0.087 | 1 | 6 | Prpsap1 | 0.016 | Old |
| 0.41817 | 0.330779 | 0.269 | 0.247 | 1 | 6 | Rps6ka1 | 0.022 | Old |
| 0.418594 | 0.178127 | 0.08 | 0.097 | 1 | 6 | Dnajb12 | -0.017 | Old |
| 0.418716 | 0.004932 | 0.109 | 0.127 | 1 | 6 | Topbp1 | -0.018 | Old |
| 0.418724 | 0.298339 | 0.091 | 0.075 | 1 | 6 | Mphosph1 | 0.016 | Old |
| 0.418966 | 0.269669 | 0.034 | 0.025 | 1 | 6 | Cpt2 | 0.009 | Old |
| 0.419007 | 0.784867 | 0.083 | 0.07 | 1 | 6 | Trmt2a | 0.013 | Old |
| 0.419125 | 0.194979 | 0.011 | 0.018 | 1 | 6 | Tmed1 | -0.007 | Old |
| 0.419289 | 0.18202 | 0.2 | 0.177 | 1 | 6 | Twf1 | 0.023 | Old |
| 0.419991 | 0.042923 | 0.157 | 0.135 | 1 | 6 | Trim41 | 0.022 | Old |
| 0.420255 | 0.230264 | 0.077 | 0.064 | 1 | 6 | Gpr160 | 0.013 | Old |
| 0.420483 | 0.223344 | 0.486 | 0.465 | 1 | 6 | Ndufa1 | 0.021 | Old |
| 0.420765 | 0.31162 | 0.163 | 0.144 | 1 | 6 | Pgrmc1 | 0.019 | Old |
| 0.420896 | 0.167446 | 0.04 | 0.03 | 1 | 6 | Usp21 | 0.01 | Old |
| 0.421055 | 0.150773 | 0.417 | 0.39 | 1 | 6 | Hnrnpdl | 0.027 | Old |
| 0.421263 | 0.318463 | 0.146 | 0.127 | 1 | 6 | Opa1 | 0.019 | Old |
| 0.421334 | 0.182483 | 0.163 | 0.144 | 1 | 6 | Atic | 0.019 | Old |
| 0.421823 | 0.392029 | 0.12 | 0.105 | 1 | 6 | Aldh3a2 | 0.015 | Old |
| 0.422271 | 0.643894 | 0.046 | 0.059 | 1 | 6 | Sclt1 | -0.013 | Old |
| 0.422497 | 0.011953 | 0.069 | 0.055 | 1 | 6 | Ttc32 | 0.014 | Old |
| 0.42325 | 0.186117 | 0.109 | 0.092 | 1 | 6 | Snrnp25 | 0.017 | Old |

| | | | | | | | | |
|---|---|---|---|---|---|---|---|---|
| 0.423281 | 0.218222 | 0.029 | 0.038 | 1 | 6 | Ap5z1 | -0.009 | Old |
| 0.423622 | 0.623473 | 0.04 | 0.03 | 1 | 6 | Ranbp6 | 0.01 | Old |
| 0.423722 | 0.718249 | 0.02 | 0.013 | 1 | 6 | Fn3krp | 0.007 | Old |
| 0.423858 | 0.251027 | 0.174 | 0.154 | 1 | 6 | Nup88 | 0.02 | Old |
| 0.423926 | 0.249565 | 0.329 | 0.308 | 1 | 6 | Atp6v0d1 | 0.021 | Old |
| 0.424077 | 0.403289 | 0.04 | 0.03 | 1 | 6 | Ecsit | 0.01 | Old |
| 0.425646 | 0.325219 | 0.046 | 0.035 | 1 | 6 | Whamm | 0.011 | Old |
| 0.425701 | 0.88316 | 0.02 | 0.013 | 1 | 6 | Aars2 | 0.007 | Old |
| 0.425878 | 0.281698 | 0.2 | 0.182 | 1 | 6 | Eea1 | 0.018 | Old |
| 0.425963 | 0.202475 | 0.486 | 0.468 | 1 | 6 | Eif5b | 0.018 | Old |
| 0.426022 | 0.893221 | 0.049 | 0.038 | 1 | 6 | Nol9 | 0.011 | Old |
| 0.426156 | 0.055828 | 0.186 | 0.164 | 1 | 6 | Seh1l | 0.022 | Old |
| 0.426211 | 0.015499 | 0.157 | 0.137 | 1 | 6 | Trnau1ap | 0.02 | Old |
| 0.426362 | 0.671928 | 0.02 | 0.013 | 1 | 6 | Mtrf1 | 0.007 | Old |
| 0.426647 | 0.134273 | 0.823 | 0.863 | 1 | 6 | Arhgdib | -0.04 | Old |
| 0.426958 | 0.580826 | 0.071 | 0.059 | 1 | 6 | Dbt | 0.012 | Old |
| 0.426969 | 0.457529 | 0.151 | 0.134 | 1 | 6 | Rab3d | 0.017 | Old |
| 0.4276 | 0.779314 | 0.014 | 0.022 | 1 | 6 | Zfp770 | -0.008 | Old |
| 0.427736 | 0.336291 | 0.071 | 0.087 | 1 | 6 | Zdhhc18 | -0.016 | Old |
| 0.427858 | 0.050985 | 0.023 | 0.032 | 1 | 6 | Hpgds | -0.009 | Old |
| 0.427998 | 0.900995 | 0.069 | 0.057 | 1 | 6 | Krr1 | 0.012 | Old |
| 0.428348 | 1.078128 | 0.02 | 0.013 | 1 | 6 | Kcnip3 | 0.007 | Old |
| 0.428348 | 0.593042 | 0.02 | 0.013 | 1 | 6 | Tfcp2 | 0.007 | Old |
| 0.428369 | 0.077871 | 0.109 | 0.092 | 1 | 6 | Slc23a2 | 0.017 | Old |
| 0.428717 | 0.127569 | 0.123 | 0.142 | 1 | 6 | Zfp318 | -0.019 | Old |
| 0.428814 | 0.226796 | 0.057 | 0.07 | 1 | 6 | Pop4 | -0.013 | Old |
| 0.428905 | 0.250346 | 0.017 | 0.025 | 1 | 6 | 1110002J0 | -0.008 | Old |
| 0.428972 | 0.024006 | 0.703 | 0.732 | 1 | 6 | Gltp | -0.029 | Old |
| 0.428988 | 0.673703 | 0.08 | 0.067 | 1 | 6 | Limk1 | 0.013 | Old |
| 0.429012 | 0.49676 | 0.02 | 0.013 | 1 | 6 | Zfp566 | 0.007 | Old |
| 0.42902 | 0.743814 | 0.154 | 0.139 | 1 | 6 | Nfatc1 | 0.015 | Old |
| 0.42906 | 0.433856 | 0.063 | 0.05 | 1 | 6 | Clk2 | 0.013 | Old |
| 0.429295 | 0.474707 | 0.131 | 0.115 | 1 | 6 | Med14 | 0.016 | Old |
| 0.429676 | 1.008112 | 0.02 | 0.013 | 1 | 6 | Bbs4 | 0.007 | Old |
| 0.429676 | 0.608764 | 0.02 | 0.013 | 1 | 6 | Iqcc | 0.007 | Old |
| 0.429676 | 0.12215 | 0.02 | 0.013 | 1 | 6 | Cryz | 0.007 | Old |
| 0.429686 | 0.003643 | 0.051 | 0.04 | 1 | 6 | Btbd2 | 0.011 | Old |
| 0.430167 | 0.077329 | 0.449 | 0.423 | 1 | 6 | Notch2 | 0.026 | Old |
| 0.43034 | 0.429984 | 0.02 | 0.013 | 1 | 6 | Gm31718 | 0.007 | Old |
| 0.430632 | 0.017864 | 0.514 | 0.564 | 1 | 6 | Nfam1 | -0.05 | Old |
| 0.430848 | 0.240763 | 0.2 | 0.179 | 1 | 6 | Angel2 | 0.021 | Old |
| 0.431187 | 0.559467 | 0.117 | 0.1 | 1 | 6 | Mphosph8 | 0.017 | Old |
| 0.431847 | 0.021751 | 0.029 | 0.038 | 1 | 6 | Nbas | -0.009 | Old |
| 0.43188 | 0.007296 | 0.577 | 0.545 | 1 | 6 | Arf1 | 0.032 | Old |
| 0.432121 | 0.085491 | 0.12 | 0.102 | 1 | 6 | Rabgap1 | 0.018 | Old |

| | | | | | | | | |
|---|---|---|---|---|---|---|---|---|
| 0.432127 | 0.135784 | 0.194 | 0.172 | 1 | 6 | Cd36 | 0.022 | Old |
| 0.432573 | 0.415675 | 0.197 | 0.176 | 1 | 6 | Atp6v1d | 0.021 | Old |
| 0.432842 | 0.077492 | 0.283 | 0.253 | 1 | 6 | Mrpl54 | 0.03 | Old |
| 0.432849 | 0.25129 | 0.089 | 0.074 | 1 | 6 | Camsap2 | 0.015 | Old |
| 0.433214 | 0.344953 | 0.06 | 0.048 | 1 | 6 | AI837181 | 0.012 | Old |
| 0.433347 | 0.040416 | 0.046 | 0.035 | 1 | 6 | Clpb | 0.011 | Old |
| 0.433457 | 0.649953 | 0.1 | 0.087 | 1 | 6 | Naxd | 0.013 | Old |
| 0.433617 | 1.716782 | 0.026 | 0.018 | 1 | 6 | Sell | 0.008 | Old |
| 0.433719 | 0.255979 | 0.714 | 0.756 | 1 | 6 | Fos | -0.042 | Old |
| 0.433834 | 0.191487 | 0.2 | 0.177 | 1 | 6 | 2610001J0 | 0.023 | Old |
| 0.434047 | 0.572108 | 0.143 | 0.13 | 1 | 6 | Rab3gap1 | 0.013 | Old |
| 0.43434 | 0.239275 | 0.02 | 0.013 | 1 | 6 | Bivm | 0.007 | Old |
| 0.434395 | 0.250155 | 0.16 | 0.142 | 1 | 6 | Sumf1 | 0.018 | Old |
| 0.434606 | 0.12866 | 0.714 | 0.722 | 1 | 6 | Serp1 | -0.008 | Old |
| 0.434723 | 0.131629 | 0.471 | 0.446 | 1 | 6 | Rab11b | 0.025 | Old |
| 0.434894 | 0.170219 | 0.086 | 0.072 | 1 | 6 | Faap100 | 0.014 | Old |
| 0.435174 | 0.201477 | 0.034 | 0.045 | 1 | 6 | Socs2 | -0.011 | Old |
| 0.435276 | 0.317184 | 0.386 | 0.368 | 1 | 6 | 7-Mar | 0.018 | Old |
| 0.435302 | 0.721909 | 0.143 | 0.129 | 1 | 6 | Supt20 | 0.014 | Old |
| 0.435313 | 0.300687 | 0.209 | 0.189 | 1 | 6 | Srp54a | 0.02 | Old |
| 0.435541 | 0.151103 | 0.611 | 0.627 | 1 | 6 | Ap2a2 | -0.016 | Old |
| 0.435605 | 0.186378 | 0.177 | 0.157 | 1 | 6 | Pja2 | 0.02 | Old |
| 0.435657 | 1.253049 | 0.011 | 0.007 | 1 | 6 | Gm10645 | 0.004 | Old |
| 0.435678 | 0.258598 | 0.02 | 0.013 | 1 | 6 | Ssx2ip | 0.007 | Old |
| 0.436005 | 0.171264 | 0.197 | 0.174 | 1 | 6 | 2510039O1 | 0.023 | Old |
| 0.436089 | 1.529497 | 0.031 | 0.023 | 1 | 6 | Mtrf1l | 0.008 | Old |
| 0.436351 | 0.241425 | 0.106 | 0.09 | 1 | 6 | Snhg6 | 0.016 | Old |
| 0.436571 | 0.127315 | 0.257 | 0.232 | 1 | 6 | Ncf4 | 0.025 | Old |
| 0.436571 | 1.213236 | 0.011 | 0.007 | 1 | 6 | Senp8 | 0.004 | Old |
| 0.436683 | 0.609935 | 0.02 | 0.013 | 1 | 6 | Fam216a | 0.007 | Old |
| 0.437227 | 0.051668 | 0.046 | 0.035 | 1 | 6 | Zfp687 | 0.011 | Old |
| 0.43725 | 0.539494 | 0.066 | 0.054 | 1 | 6 | Fdxr | 0.012 | Old |
| 0.437487 | 1.380414 | 0.011 | 0.007 | 1 | 6 | Adgrl1 | 0.004 | Old |
| 0.437487 | 1.201901 | 0.011 | 0.007 | 1 | 6 | Gpr157 | 0.004 | Old |
| 0.437807 | 0.123982 | 0.18 | 0.201 | 1 | 6 | Gm47283 | -0.021 | Old |
| 0.437964 | 0.355408 | 0.294 | 0.279 | 1 | 6 | Ap1s2 | 0.015 | Old |
| 0.437996 | 0.352927 | 0.131 | 0.114 | 1 | 6 | Klhl28 | 0.017 | Old |
| 0.438403 | 1.719318 | 0.011 | 0.007 | 1 | 6 | Ifit1 | 0.004 | Old |
| 0.438403 | 1.384195 | 0.011 | 0.007 | 1 | 6 | Tlr12 | 0.004 | Old |
| 0.438403 | 0.810043 | 0.011 | 0.007 | 1 | 6 | Nap1l2 | 0.004 | Old |
| 0.438424 | 1.260329 | 0.037 | 0.028 | 1 | 6 | Haus7 | 0.009 | Old |
| 0.439222 | 0.461456 | 0.129 | 0.112 | 1 | 6 | Hibadh | 0.017 | Old |
| 0.439296 | 0.168669 | 0.26 | 0.234 | 1 | 6 | Ptpn2 | 0.026 | Old |
| 0.43932 | 1.355797 | 0.011 | 0.007 | 1 | 6 | Zfp606 | 0.004 | Old |
| 0.43932 | 1.020336 | 0.011 | 0.007 | 1 | 6 | G630030J0 | 0.004 | Old |

| | | | | | | | | |
|---|---|---|---|---|---|---|---|---|
| 0.43932 | 0.794772 | 0.011 | 0.007 | 1 | 6 | Ube2t | 0.004 | Old |
| 0.440239 | 1.137677 | 0.011 | 0.007 | 1 | 6 | Tagap1 | 0.004 | Old |
| 0.440239 | 0.968142 | 0.011 | 0.007 | 1 | 6 | Klk8 | 0.004 | Old |
| 0.440239 | 0.748383 | 0.011 | 0.007 | 1 | 6 | Gm26935 | 0.004 | Old |
| 0.440239 | 0.68634 | 0.011 | 0.007 | 1 | 6 | Smtn | 0.004 | Old |
| 0.440239 | 0.659197 | 0.011 | 0.007 | 1 | 6 | Clec2g | 0.004 | Old |
| 0.440239 | 0.495991 | 0.011 | 0.007 | 1 | 6 | Bcs1l | 0.004 | Old |
| 0.440239 | 0.470299 | 0.011 | 0.007 | 1 | 6 | AC139347. | 0.004 | Old |
| 0.440535 | 0.303038 | 0.511 | 0.512 | 1 | 6 | Sec11c | -0.001 | Old |
| 0.441158 | 1.404672 | 0.011 | 0.007 | 1 | 6 | Stk26 | 0.004 | Old |
| 0.441158 | 0.641563 | 0.011 | 0.007 | 1 | 6 | Prdm9 | 0.004 | Old |
| 0.441158 | 0.551199 | 0.011 | 0.007 | 1 | 6 | Bbs7 | 0.004 | Old |
| 0.441746 | 0.489023 | 0.157 | 0.14 | 1 | 6 | Sdccag8 | 0.017 | Old |
| 0.442041 | 0.013795 | 0.634 | 0.614 | 1 | 6 | Sec61g | 0.02 | Old |
| 0.442079 | 0.852051 | 0.011 | 0.007 | 1 | 6 | Gemin8 | 0.004 | Old |
| 0.442079 | 0.801153 | 0.011 | 0.007 | 1 | 6 | Nup37 | 0.004 | Old |
| 0.442079 | 0.377114 | 0.011 | 0.007 | 1 | 6 | Intu | 0.004 | Old |
| 0.442079 | 0.374491 | 0.011 | 0.007 | 1 | 6 | Ctif | 0.004 | Old |
| 0.442079 | 0.055888 | 0.011 | 0.007 | 1 | 6 | Klhdc1 | 0.004 | Old |
| 0.442396 | 0.062209 | 0.24 | 0.259 | 1 | 6 | Crem | -0.019 | Old |
| 0.442888 | 0.066983 | 0.731 | 0.734 | 1 | 6 | Erp29 | -0.003 | Old |
| 0.443 | 0.341855 | 0.011 | 0.007 | 1 | 6 | Tmem218 | 0.004 | Old |
| 0.443117 | 0.002486 | 0.229 | 0.254 | 1 | 6 | Rbbp7 | -0.025 | Old |
| 0.443155 | 0.08947 | 0.246 | 0.268 | 1 | 6 | Peli1 | -0.022 | Old |
| 0.443515 | 0.240838 | 0.129 | 0.112 | 1 | 6 | Khnyn | 0.017 | Old |
| 0.443721 | 0.478216 | 0.066 | 0.054 | 1 | 6 | Eef1akmt2 | 0.012 | Old |
| 0.443923 | 0.446337 | 0.011 | 0.007 | 1 | 6 | Gnb1l | 0.004 | Old |
| 0.443923 | 0.216083 | 0.011 | 0.007 | 1 | 6 | Lgals4 | 0.004 | Old |
| 0.444259 | 0.364929 | 0.077 | 0.064 | 1 | 6 | Srprb | 0.013 | Old |
| 0.444474 | 0.308095 | 0.18 | 0.161 | 1 | 6 | Nfu1 | 0.019 | Old |
| 0.444483 | 0.749253 | 0.031 | 0.023 | 1 | 6 | Arl11 | 0.008 | Old |
| 0.444552 | 0.033235 | 0.237 | 0.211 | 1 | 6 | Dock7 | 0.026 | Old |
| 0.444685 | 0.226717 | 0.229 | 0.207 | 1 | 6 | Tcp11l2 | 0.022 | Old |
| 0.444847 | 0.438776 | 0.011 | 0.007 | 1 | 6 | Dpy19l3 | 0.004 | Old |
| 0.444847 | 0.300672 | 0.011 | 0.007 | 1 | 6 | Kif22 | 0.004 | Old |
| 0.444847 | 0.190648 | 0.011 | 0.007 | 1 | 6 | Zfp428 | 0.004 | Old |
| 0.444851 | 0.214159 | 0.286 | 0.264 | 1 | 6 | Ndufa5 | 0.022 | Old |
| 0.445058 | 0.063587 | 0.171 | 0.191 | 1 | 6 | 1500011K1 | -0.02 | Old |
| 0.445596 | 0.26385 | 0.077 | 0.064 | 1 | 6 | Slc45a4 | 0.013 | Old |
| 0.445643 | 1.311989 | 0.037 | 0.028 | 1 | 6 | Osbpl3 | 0.009 | Old |
| 0.445772 | 0.119531 | 0.011 | 0.007 | 1 | 6 | Gorasp1 | 0.004 | Old |
| 0.445892 | 1.044348 | 0.026 | 0.018 | 1 | 6 | Aldh6a1 | 0.008 | Old |
| 0.445936 | 0.128973 | 0.637 | 0.632 | 1 | 6 | St8sia4 | 0.005 | Old |
| 0.446219 | 0.212916 | 0.291 | 0.319 | 1 | 6 | Siglecg | -0.028 | Old |
| 0.446252 | 0.543459 | 0.066 | 0.054 | 1 | 6 | Snx25 | 0.012 | Old |

| | | | | | | | | |
|---|---|---|---|---|---|---|---|---|
| 0.446266 | 0.531397 | 0.077 | 0.064 | 1 | 6 | Rngtt | 0.013 | Old |
| 0.446295 | 0.228555 | 0.189 | 0.216 | 1 | 6 | Nemf | -0.027 | Old |
| 0.446362 | 0.311979 | 0.06 | 0.048 | 1 | 6 | Cog8 | 0.012 | Old |
| 0.446698 | 0.04935 | 0.011 | 0.007 | 1 | 6 | Dok4 | 0.004 | Old |
| 0.446977 | 0.1665 | 0.109 | 0.094 | 1 | 6 | Chchd5 | 0.015 | Old |
| 0.447071 | 0.411772 | 0.026 | 0.018 | 1 | 6 | Arhgap32 | 0.008 | Old |
| 0.447124 | 0.597459 | 0.031 | 0.023 | 1 | 6 | Acrbp | 0.008 | Old |
| 0.44715 | 0.446165 | 0.054 | 0.043 | 1 | 6 | Arel1 | 0.011 | Old |
| 0.447548 | 0.093877 | 0.974 | 0.983 | 1 | 6 | Junb | -0.009 | Old |
| 0.447829 | 0.266824 | 0.209 | 0.189 | 1 | 6 | Trp53 | 0.02 | Old |
| 0.448007 | 0.255678 | 0.023 | 0.032 | 1 | 6 | Mthfsd | -0.009 | Old |
| 0.448102 | 0.250651 | 0.074 | 0.062 | 1 | 6 | Ttc33 | 0.012 | Old |
| 0.448183 | 0.566849 | 0.031 | 0.023 | 1 | 6 | Akip1 | 0.008 | Old |
| 0.448201 | 0.017435 | 0.166 | 0.145 | 1 | 6 | Tpm1 | 0.021 | Old |
| 0.448832 | 0.115222 | 0.357 | 0.324 | 1 | 6 | Tomm5 | 0.033 | Old |
| 0.448843 | 0.541162 | 0.026 | 0.018 | 1 | 6 | Mbnl3 | 0.008 | Old |
| 0.448843 | 0.404191 | 0.026 | 0.018 | 1 | 6 | P2ry14 | 0.008 | Old |
| 0.449312 | 0.416245 | 0.063 | 0.052 | 1 | 6 | Mthfd1 | 0.011 | Old |
| 0.449363 | 0.163538 | 0.171 | 0.152 | 1 | 6 | Mex3c | 0.019 | Old |
| 0.449589 | 0.02735 | 0.123 | 0.105 | 1 | 6 | Slc9a6 | 0.018 | Old |
| 0.449597 | 0.284326 | 0.129 | 0.11 | 1 | 6 | Gtf2h1 | 0.019 | Old |
| 0.449815 | 0.028648 | 0.937 | 0.972 | 1 | 6 | H2-D1 | -0.035 | Old |
| 0.450027 | 0.329798 | 0.026 | 0.018 | 1 | 6 | Trmt10c | 0.008 | Old |
| 0.450306 | 0.605781 | 0.031 | 0.023 | 1 | 6 | Ercc8 | 0.008 | Old |
| 0.450306 | 0.434709 | 0.031 | 0.023 | 1 | 6 | 4933427D1 | 0.008 | Old |
| 0.45037 | 0.089017 | 0.211 | 0.186 | 1 | 6 | Parp1 | 0.025 | Old |
| 0.450619 | 0.339029 | 0.026 | 0.018 | 1 | 6 | Ndufaf1 | 0.008 | Old |
| 0.450758 | 0.21453 | 0.403 | 0.378 | 1 | 6 | Ensa | 0.025 | Old |
| 0.450928 | 0.148957 | 0.151 | 0.132 | 1 | 6 | Asxl1 | 0.019 | Old |
| 0.451169 | 0.079003 | 0.177 | 0.156 | 1 | 6 | Tmem170b | 0.021 | Old |
| 0.451279 | 0.15633 | 0.134 | 0.117 | 1 | 6 | Rnf220 | 0.017 | Old |
| 0.451703 | 0.237088 | 0.066 | 0.054 | 1 | 6 | Spata5 | 0.012 | Old |
| 0.451806 | 0.294236 | 0.026 | 0.018 | 1 | 6 | Nek6 | 0.008 | Old |
| 0.451858 | 0.212271 | 0.074 | 0.062 | 1 | 6 | Fam60a | 0.012 | Old |
| 0.451953 | 1.08697 | 0.037 | 0.028 | 1 | 6 | 2610035D1 | 0.009 | Old |
| 0.452102 | 0.304736 | 0.08 | 0.067 | 1 | 6 | Lrwd1 | 0.013 | Old |
| 0.452399 | 0.395948 | 0.026 | 0.018 | 1 | 6 | Ccdc66 | 0.008 | Old |
| 0.452478 | 0.182968 | 0.234 | 0.217 | 1 | 6 | Rgs18 | 0.017 | Old |
| 0.452532 | 0.322561 | 0.069 | 0.057 | 1 | 6 | Arhgap10 | 0.012 | Old |
| 0.452566 | 0.095981 | 0.129 | 0.112 | 1 | 6 | Ube2w | 0.017 | Old |
| 0.452739 | 0.016722 | 0.38 | 0.408 | 1 | 6 | Ghitm | -0.028 | Old |
| 0.452752 | 0.669526 | 0.054 | 0.043 | 1 | 6 | Pelp1 | 0.011 | Old |
| 0.45293 | 0.271222 | 0.423 | 0.42 | 1 | 6 | Adam8 | 0.003 | Old |
| 0.453094 | 0.372621 | 0.08 | 0.067 | 1 | 6 | Mrps34 | 0.013 | Old |
| 0.453588 | 0.48067 | 0.026 | 0.018 | 1 | 6 | Als2 | 0.008 | Old |

| | | | | | | | | |
|---|---|---|---|---|---|---|---|---|
| 0.453588 | 0.305273 | 0.026 | 0.018 | 1 | 6 | Pld1 | 0.008 | Old |
| 0.453712 | 0.033987 | 0.066 | 0.054 | 1 | 6 | Cep295 | 0.012 | Old |
| 0.453804 | 0.164576 | 0.049 | 0.038 | 1 | 6 | Rubcnl | 0.011 | Old |
| 0.453849 | 0.500479 | 0.043 | 0.033 | 1 | 6 | Nrde2 | 0.01 | Old |
| 0.453905 | 1.060297 | 0.037 | 0.028 | 1 | 6 | Tmco4 | 0.009 | Old |
| 0.453957 | 0.501068 | 0.054 | 0.043 | 1 | 6 | Alkbh3 | 0.011 | Old |
| 0.454017 | 0.507718 | 0.049 | 0.038 | 1 | 6 | Tmem150b | 0.011 | Old |
| 0.454034 | 0.944583 | 0.031 | 0.023 | 1 | 6 | Ccl5 | 0.008 | Old |
| 0.454095 | 0.070584 | 0.166 | 0.145 | 1 | 6 | Wdfy3 | 0.021 | Old |
| 0.454185 | 0.031674 | 0.1 | 0.115 | 1 | 6 | Trak1 | -0.015 | Old |
| 0.454394 | 0.430838 | 0.037 | 0.028 | 1 | 6 | Armc7 | 0.009 | Old |
| 0.454779 | 0.182578 | 0.026 | 0.018 | 1 | 6 | Haus1 | 0.008 | Old |
| 0.454868 | 0.181115 | 0.049 | 0.038 | 1 | 6 | Ttf2 | 0.011 | Old |
| 0.455382 | 0.072226 | 0.709 | 0.729 | 1 | 6 | Nfkbid | -0.02 | Old |
| 0.455436 | 0.487127 | 0.091 | 0.079 | 1 | 6 | Mpv17l2 | 0.012 | Old |
| 0.455862 | 0.349981 | 0.037 | 0.028 | 1 | 6 | 3110040N1 | 0.009 | Old |
| 0.456081 | 0.262928 | 0.143 | 0.127 | 1 | 6 | Cir1 | 0.016 | Old |
| 0.456119 | 0.479183 | 0.043 | 0.033 | 1 | 6 | Eci1 | 0.01 | Old |
| 0.456165 | 0.213002 | 0.28 | 0.254 | 1 | 6 | Setd3 | 0.026 | Old |
| 0.456331 | 0.082513 | 0.071 | 0.059 | 1 | 6 | Akap7 | 0.012 | Old |
| 0.456569 | 0.155853 | 0.026 | 0.018 | 1 | 6 | Pycrl | 0.008 | Old |
| 0.456708 | 0.766665 | 0.031 | 0.023 | 1 | 6 | Nxpe3 | 0.008 | Old |
| 0.456761 | 0.030398 | 0.634 | 0.682 | 1 | 6 | Eps8 | -0.048 | Old |
| 0.456933 | 0.353083 | 0.143 | 0.125 | 1 | 6 | Qrich1 | 0.018 | Old |
| 0.457127 | 0.374933 | 0.131 | 0.115 | 1 | 6 | Arl3 | 0.016 | Old |
| 0.457298 | 0.205033 | 0.151 | 0.134 | 1 | 6 | Cdk4 | 0.017 | Old |
| 0.457418 | 0.202951 | 0.194 | 0.174 | 1 | 6 | Smu1 | 0.02 | Old |
| 0.457482 | 0.317876 | 0.1 | 0.085 | 1 | 6 | Desi1 | 0.015 | Old |
| 0.457764 | 0.037251 | 0.026 | 0.018 | 1 | 6 | Eml6 | 0.008 | Old |
| 0.457823 | 0.299991 | 0.037 | 0.028 | 1 | 6 | Cmtm4 | 0.009 | Old |
| 0.458316 | 0.804378 | 0.031 | 0.023 | 1 | 6 | Cbarp | 0.008 | Old |
| 0.458388 | 0.178557 | 0.06 | 0.048 | 1 | 6 | Hmgn5 | 0.012 | Old |
| 0.458784 | 0.006002 | 0.583 | 0.612 | 1 | 6 | Macf1 | -0.029 | Old |
| 0.458816 | 0.06463 | 0.206 | 0.181 | 1 | 6 | Pdap1 | 0.025 | Old |
| 0.458846 | 0.094532 | 0.066 | 0.054 | 1 | 6 | Fbxl12 | 0.012 | Old |
| 0.458935 | 0.001715 | 0.077 | 0.092 | 1 | 6 | Orc4 | -0.015 | Old |
| 0.459064 | 0.024057 | 0.146 | 0.166 | 1 | 6 | Syne2 | -0.02 | Old |
| 0.459947 | 0.153324 | 0.106 | 0.09 | 1 | 6 | Fbxw4 | 0.016 | Old |
| 0.460402 | 0.209182 | 0.163 | 0.144 | 1 | 6 | Usp38 | 0.019 | Old |
| 0.460928 | 0.285156 | 0.14 | 0.124 | 1 | 6 | Cops8 | 0.016 | Old |
| 0.46117 | 0.015094 | 0.151 | 0.132 | 1 | 6 | Gm21188 | 0.019 | Old |
| 0.461335 | 0.055867 | 0.966 | 0.99 | 1 | 6 | Crip1 | -0.024 | Old |
| 0.461483 | 0.1258 | 0.077 | 0.064 | 1 | 6 | Xrcc1 | 0.013 | Old |
| 0.461579 | 0.105415 | 0.146 | 0.164 | 1 | 6 | Zbtb46 | -0.018 | Old |
| 0.461595 | 0.735829 | 0.043 | 0.033 | 1 | 6 | Gm6563 | 0.01 | Old |

| | | | | | | | | |
|---|---|---|---|---|---|---|---|---|
| 0.46163 | 0.347187 | 0.177 | 0.161 | 1 | 6 | Clock | 0.016 | Old |
| 0.461664 | 0.055097 | 0.06 | 0.048 | 1 | 6 | Top3b | 0.012 | Old |
| 0.461722 | 0.065917 | 0.163 | 0.144 | 1 | 6 | Dcun1d1 | 0.019 | Old |
| 0.461769 | 0.496556 | 0.077 | 0.065 | 1 | 6 | Stxbp5 | 0.012 | Old |
| 0.462053 | 0.560578 | 0.043 | 0.033 | 1 | 6 | Gm11772 | 0.01 | Old |
| 0.462077 | 0.298706 | 0.08 | 0.067 | 1 | 6 | Strip1 | 0.013 | Old |
| 0.462197 | 0.15663 | 0.743 | 0.739 | 1 | 6 | Hspa5 | 0.004 | Old |
| 0.462204 | 0.223369 | 0.163 | 0.142 | 1 | 6 | Hip1 | 0.021 | Old |
| 0.462215 | 0.269636 | 0.103 | 0.089 | 1 | 6 | Mfap3 | 0.014 | Old |
| 0.462338 | 0.06024 | 0.157 | 0.177 | 1 | 6 | Sdccag3 | -0.02 | Old |
| 0.462853 | 0.002979 | 0.054 | 0.043 | 1 | 6 | Bag4 | 0.011 | Old |
| 0.463152 | 0.249642 | 0.137 | 0.122 | 1 | 6 | Rab29 | 0.015 | Old |
| 0.463227 | 0.362392 | 0.077 | 0.092 | 1 | 6 | Tbc1d4 | -0.015 | Old |
| 0.463329 | 0.57166 | 0.089 | 0.075 | 1 | 6 | Snhg12 | 0.014 | Old |
| 0.463347 | 0.230498 | 0.094 | 0.08 | 1 | 6 | Arhgap12 | 0.014 | Old |
| 0.463382 | 0.202741 | 0.36 | 0.343 | 1 | 6 | Pbxip1 | 0.017 | Old |
| 0.46358 | 0.158674 | 0.337 | 0.314 | 1 | 6 | Adam10 | 0.023 | Old |
| 0.463697 | 0.318953 | 0.117 | 0.1 | 1 | 6 | Auh | 0.017 | Old |
| 0.46385 | 0.133196 | 0.32 | 0.293 | 1 | 6 | Adss | 0.027 | Old |
| 0.463865 | 0.254773 | 0.191 | 0.176 | 1 | 6 | Snx8 | 0.015 | Old |
| 0.463931 | 0.257605 | 0.2 | 0.181 | 1 | 6 | Gtf3c2 | 0.019 | Old |
| 0.464088 | 0.056213 | 0.251 | 0.222 | 1 | 6 | Elf4 | 0.029 | Old |
| 0.464241 | 0.050629 | 0.031 | 0.023 | 1 | 6 | Rad9a | 0.008 | Old |
| 0.464279 | 0.163341 | 0.229 | 0.207 | 1 | 6 | Cpsf2 | 0.022 | Old |
| 0.464341 | 0.451333 | 0.106 | 0.092 | 1 | 6 | Nbr1 | 0.014 | Old |
| 0.464796 | 0.078495 | 0.214 | 0.194 | 1 | 6 | Gdi1 | 0.02 | Old |
| 0.464875 | 0.172714 | 0.274 | 0.256 | 1 | 6 | Chic2 | 0.018 | Old |
| 0.465084 | 0.003957 | 0.18 | 0.157 | 1 | 6 | Parl | 0.023 | Old |
| 0.465089 | 0.145438 | 0.534 | 0.513 | 1 | 6 | Hnrnpc | 0.021 | Old |
| 0.465241 | 0.136471 | 0.309 | 0.288 | 1 | 6 | Sar1a | 0.021 | Old |
| 0.465266 | 0.427482 | 0.043 | 0.033 | 1 | 6 | Ier5l | 0.01 | Old |
| 0.465323 | 0.146447 | 0.031 | 0.023 | 1 | 6 | Fastkd2 | 0.008 | Old |
| 0.465962 | 0.048523 | 0.111 | 0.127 | 1 | 6 | Rbm4b | -0.016 | Old |
| 0.46653 | 0.543101 | 0.077 | 0.065 | 1 | 6 | Cryzl1 | 0.012 | Old |
| 0.467073 | 0.220339 | 0.097 | 0.082 | 1 | 6 | Fopnl | 0.015 | Old |
| 0.467235 | 0.055594 | 0.303 | 0.329 | 1 | 6 | Hopx | -0.026 | Old |
| 0.467388 | 0.060366 | 0.183 | 0.162 | 1 | 6 | Ndufa9 | 0.021 | Old |
| 0.467861 | 0.081588 | 0.4 | 0.431 | 1 | 6 | Selenof | -0.031 | Old |
| 0.46798 | 0.212233 | 0.503 | 0.492 | 1 | 6 | Tmem123 | 0.011 | Old |
| 0.468491 | 0.112519 | 0.043 | 0.033 | 1 | 6 | Mrvi1 | 0.01 | Old |
| 0.468696 | 0.101308 | 0.414 | 0.443 | 1 | 6 | Inpp5d | -0.029 | Old |
| 0.469013 | 0.489906 | 0.117 | 0.102 | 1 | 6 | Adap1 | 0.015 | Old |
| 0.469026 | 0.305187 | 0.066 | 0.054 | 1 | 6 | Ormdl1 | 0.012 | Old |
| 0.469841 | 0.15102 | 0.443 | 0.483 | 1 | 6 | Zbtb20 | -0.04 | Old |
| 0.470088 | 0.244473 | 0.271 | 0.246 | 1 | 6 | Chuk | 0.025 | Old |

| | | | | | | | | |
|---|---|---|---|---|---|---|---|---|
| 0.470208 | 0.065003 | 0.031 | 0.023 | 1 | 6 | Kctd9 | 0.008 | Old |
| 0.470553 | 0.345221 | 0.074 | 0.062 | 1 | 6 | Osbpl11 | 0.012 | Old |
| 0.470764 | 0.050242 | 0.351 | 0.323 | 1 | 6 | Snx9 | 0.028 | Old |
| 0.470802 | 0.144326 | 0.043 | 0.033 | 1 | 6 | Inafm2 | 0.01 | Old |
| 0.471096 | 0.062365 | 0.22 | 0.246 | 1 | 6 | Ak2 | -0.026 | Old |
| 0.471778 | 0.065284 | 0.391 | 0.415 | 1 | 6 | Plec | -0.024 | Old |
| 0.472033 | 0.496712 | 0.103 | 0.089 | 1 | 6 | Tmem19 | 0.014 | Old |
| 0.472183 | 0.362986 | 0.037 | 0.028 | 1 | 6 | Gm20300 | 0.009 | Old |
| 0.472417 | 0.204806 | 0.291 | 0.271 | 1 | 6 | Ggnbp2 | 0.02 | Old |
| 0.472667 | 0.040917 | 0.054 | 0.043 | 1 | 6 | Dgcr8 | 0.011 | Old |
| 0.473119 | 0.047309 | 0.043 | 0.033 | 1 | 6 | Cmtr1 | 0.01 | Old |
| 0.473129 | 0.608838 | 0.211 | 0.197 | 1 | 6 | Cxcr4 | 0.014 | Old |
| 0.473156 | 0.154853 | 0.114 | 0.1 | 1 | 6 | Xab2 | 0.014 | Old |
| 0.47348 | 0.026879 | 0.031 | 0.023 | 1 | 6 | Vcpkmt | 0.008 | Old |
| 0.473807 | 0.160113 | 0.3 | 0.283 | 1 | 6 | Bcl2l11 | 0.017 | Old |
| 0.474536 | 0.088904 | 0.203 | 0.182 | 1 | 6 | Ankhd1 | 0.021 | Old |
| 0.474684 | 0.003484 | 0.037 | 0.028 | 1 | 6 | Vipas39 | 0.009 | Old |
| 0.474718 | 0.9876 | 0.04 | 0.032 | 1 | 6 | Cep70 | 0.008 | Old |
| 0.474871 | 0.732775 | 0.057 | 0.047 | 1 | 6 | Papd7 | 0.01 | Old |
| 0.474947 | 0.388463 | 0.377 | 0.348 | 1 | 6 | Hmgb2 | 0.029 | Old |
| 0.474978 | 0.196852 | 0.043 | 0.033 | 1 | 6 | Aar2 | 0.01 | Old |
| 0.475953 | 0.052764 | 0.569 | 0.599 | 1 | 6 | Emb | -0.03 | Old |
| 0.475958 | 0.193908 | 0.226 | 0.207 | 1 | 6 | Mafb | 0.019 | Old |
| 0.47598 | 0.223498 | 0.086 | 0.072 | 1 | 6 | St3gal2 | 0.014 | Old |
| 0.476012 | 0.479585 | 0.063 | 0.052 | 1 | 6 | Ksr1 | 0.011 | Old |
| 0.476028 | 0.402075 | 0.251 | 0.232 | 1 | 6 | Abce1 | 0.019 | Old |
| 0.476071 | 0.335598 | 0.057 | 0.047 | 1 | 6 | Thap11 | 0.01 | Old |
| 0.476132 | 0.378708 | 0.077 | 0.065 | 1 | 6 | Anapc10 | 0.012 | Old |
| 0.476633 | 0.775852 | 0.04 | 0.032 | 1 | 6 | Flywch1 | 0.008 | Old |
| 0.476777 | 0.329843 | 0.063 | 0.052 | 1 | 6 | Parn | 0.011 | Old |
| 0.477167 | 0.341328 | 0.077 | 0.065 | 1 | 6 | Tomm6 | 0.012 | Old |
| 0.477392 | 0.208152 | 0.337 | 0.309 | 1 | 6 | Gm16286 | 0.028 | Old |
| 0.477697 | 0.794567 | 0.051 | 0.042 | 1 | 6 | Tmem214 | 0.009 | Old |
| 0.478745 | 1.412273 | 0.017 | 0.012 | 1 | 6 | 4732440D( | 0.005 | Old |
| 0.478924 | 0.342555 | 0.586 | 0.59 | 1 | 6 | Dab2 | -0.004 | Old |
| 0.478959 | 0.174146 | 0.223 | 0.199 | 1 | 6 | Plekha2 | 0.024 | Old |
| 0.479141 | 0.062505 | 0.08 | 0.067 | 1 | 6 | Gpr89 | 0.013 | Old |
| 0.4793 | 0.204797 | 0.083 | 0.07 | 1 | 6 | Parp3 | 0.013 | Old |
| 0.479339 | 0.206126 | 0.391 | 0.365 | 1 | 6 | Snrpd2 | 0.026 | Old |
| 0.479504 | 0.032617 | 0.049 | 0.038 | 1 | 6 | Fbxo7 | 0.011 | Old |
| 0.479869 | 0.319162 | 0.309 | 0.293 | 1 | 6 | Cnot1 | 0.016 | Old |
| 0.480051 | 0.01577 | 0.169 | 0.149 | 1 | 6 | Impa1 | 0.02 | Old |
| 0.480166 | 0.227396 | 0.117 | 0.102 | 1 | 6 | Ino80e | 0.015 | Old |
| 0.480196 | 0.041003 | 0.094 | 0.11 | 1 | 6 | Med11 | -0.016 | Old |
| 0.481081 | 0.161968 | 0.18 | 0.164 | 1 | 6 | Ncoa2 | 0.016 | Old |

| | | | | | | | | |
|---|---|---|---|---|---|---|---|---|
| 0.481382 | 0.315349 | 0.166 | 0.151 | 1 | 6 | Zfp950 | 0.015 | Old |
| 0.481475 | 0.005713 | 0.517 | 0.564 | 1 | 6 | Ldlr | -0.047 | Old |
| 0.481598 | 0.196676 | 0.26 | 0.236 | 1 | 6 | Dnajb11 | 0.024 | Old |
| 0.481742 | 0.150071 | 0.177 | 0.199 | 1 | 6 | Rnf114 | -0.022 | Old |
| 0.482196 | 0.13523 | 0.074 | 0.087 | 1 | 6 | Plekhm1 | -0.013 | Old |
| 0.482444 | 0.046344 | 0.686 | 0.679 | 1 | 6 | Ddx3x | 0.007 | Old |
| 0.482557 | 0.293861 | 0.143 | 0.127 | 1 | 6 | Brpf1 | 0.016 | Old |
| 0.482823 | 0.422193 | 0.091 | 0.079 | 1 | 6 | Def6 | 0.012 | Old |
| 0.483139 | 0.331826 | 0.214 | 0.197 | 1 | 6 | Atp6v1c1 | 0.017 | Old |
| 0.483371 | 0.257921 | 0.046 | 0.057 | 1 | 6 | Rcl1 | -0.011 | Old |
| 0.483572 | 0.354662 | 0.174 | 0.156 | 1 | 6 | Ilkap | 0.018 | Old |
| 0.483575 | 0.085144 | 0.186 | 0.166 | 1 | 6 | Pip4k2b | 0.02 | Old |
| 0.483692 | 0.062459 | 0.563 | 0.574 | 1 | 6 | Qk | -0.011 | Old |
| 0.484064 | 0.810724 | 0.017 | 0.012 | 1 | 6 | Gm26982 | 0.005 | Old |
| 0.484064 | 0.753481 | 0.017 | 0.012 | 1 | 6 | Ppcs | 0.005 | Old |
| 0.484117 | 0.248901 | 0.057 | 0.047 | 1 | 6 | Pcid2 | 0.01 | Old |
| 0.484246 | 0.112259 | 0.069 | 0.082 | 1 | 6 | Ticam1 | -0.013 | Old |
| 0.484629 | 0.749738 | 0.006 | 0.01 | 1 | 6 | Setd4 | -0.004 | Old |
| 0.484694 | 0.131878 | 0.103 | 0.089 | 1 | 6 | Ambra1 | 0.014 | Old |
| 0.484864 | 0.063263 | 0.16 | 0.142 | 1 | 6 | Plaa | 0.018 | Old |
| 0.485032 | 0.001561 | 0.28 | 0.296 | 1 | 6 | Man2a2 | -0.016 | Old |
| 0.485178 | 0.474411 | 0.134 | 0.12 | 1 | 6 | Abcb7 | 0.014 | Old |
| 0.485508 | 0.481562 | 0.12 | 0.107 | 1 | 6 | Ncln | 0.013 | Old |
| 0.485547 | 0.195046 | 0.34 | 0.324 | 1 | 6 | Ppp1r15b | 0.016 | Old |
| 0.485548 | 0.052487 | 0.337 | 0.366 | 1 | 6 | Kdm2b | -0.029 | Old |
| 0.485589 | 0.489282 | 0.017 | 0.012 | 1 | 6 | Kpna2 | 0.005 | Old |
| 0.485599 | 0.923277 | 0.006 | 0.01 | 1 | 6 | Slfn1 | -0.004 | Old |
| 0.485867 | 0.193548 | 0.091 | 0.079 | 1 | 6 | Tmem203 | 0.012 | Old |
| 0.48623 | 0.139085 | 0.137 | 0.12 | 1 | 6 | Ppil4 | 0.017 | Old |
| 0.486353 | 0.508388 | 0.017 | 0.012 | 1 | 6 | Shank1 | 0.005 | Old |
| 0.486353 | 0.45714 | 0.017 | 0.012 | 1 | 6 | Gm42670 | 0.005 | Old |
| 0.48657 | 0.503432 | 0.006 | 0.01 | 1 | 6 | 4933423P2 | -0.004 | Old |
| 0.48657 | 0.059822 | 0.006 | 0.01 | 1 | 6 | Diaph3 | -0.004 | Old |
| 0.486759 | 0.794493 | 0.04 | 0.032 | 1 | 6 | Ngrn | 0.008 | Old |
| 0.48695 | 0.311718 | 0.057 | 0.047 | 1 | 6 | Ttll12 | 0.01 | Old |
| 0.487461 | 0.124291 | 0.394 | 0.365 | 1 | 6 | Mapk1 | 0.029 | Old |
| 0.487488 | 0.11692 | 0.094 | 0.08 | 1 | 6 | Arid5b | 0.014 | Old |
| 0.48757 | 0.050042 | 0.094 | 0.079 | 1 | 6 | Uba3 | 0.015 | Old |
| 0.487882 | 0.137543 | 0.017 | 0.012 | 1 | 6 | Tslp | 0.005 | Old |
| 0.487912 | 0.361388 | 0.211 | 0.194 | 1 | 6 | Ing1 | 0.017 | Old |
| 0.488017 | 0.04226 | 0.223 | 0.244 | 1 | 6 | Rbck1 | -0.021 | Old |
| 0.488165 | 0.116869 | 0.1 | 0.087 | 1 | 6 | Slc27a1 | 0.013 | Old |
| 0.488184 | 0.08308 | 0.594 | 0.584 | 1 | 6 | Chd4 | 0.01 | Old |
| 0.488285 | 0.391441 | 0.146 | 0.132 | 1 | 6 | Supt6 | 0.014 | Old |
| 0.488639 | 0.055291 | 0.106 | 0.092 | 1 | 6 | Ankra2 | 0.014 | Old |

| | | | | | | | | |
|---|---|---|---|---|---|---|---|---|
| 0.488648 | 0.409195 | 0.017 | 0.012 | 1 | 6 | Tcaim | 0.005 | Old |
| 0.488966 | 0.095052 | 0.497 | 0.472 | 1 | 6 | Evi2a | 0.025 | Old |
| 0.489166 | 0.226427 | 0.071 | 0.084 | 1 | 6 | Dnajb2 | -0.013 | Old |
| 0.48917 | 0.301817 | 0.317 | 0.288 | 1 | 6 | Fyn | 0.029 | Old |
| 0.489267 | 0.232989 | 0.037 | 0.047 | 1 | 6 | Rgp1 | -0.01 | Old |
| 0.489415 | 0.215386 | 0.017 | 0.012 | 1 | 6 | Cbwd1 | 0.005 | Old |
| 0.489481 | 0.108632 | 0.14 | 0.122 | 1 | 6 | Slc25a51 | 0.018 | Old |
| 0.48949 | 1.017007 | 0.006 | 0.01 | 1 | 6 | Sox4 | -0.004 | Old |
| 0.48974 | 0.071696 | 0.437 | 0.416 | 1 | 6 | Baz1a | 0.021 | Old |
| 0.490182 | 0.243669 | 0.017 | 0.012 | 1 | 6 | Tcea2 | 0.005 | Old |
| 0.490406 | 0.052964 | 0.123 | 0.107 | 1 | 6 | Mrpl41 | 0.016 | Old |
| 0.490461 | 0.04652 | 0.049 | 0.059 | 1 | 6 | Rabggta | -0.01 | Old |
| 0.491152 | 0.13479 | 0.069 | 0.057 | 1 | 6 | Yars | 0.012 | Old |
| 0.491421 | 0.301438 | 0.057 | 0.047 | 1 | 6 | Abcc4 | 0.01 | Old |
| 0.491561 | 0.123653 | 0.369 | 0.346 | 1 | 6 | Sec61a1 | 0.023 | Old |
| 0.491614 | 0.661172 | 0.089 | 0.077 | 1 | 6 | Pknox1 | 0.012 | Old |
| 0.491639 | 0.037205 | 0.183 | 0.201 | 1 | 6 | Ndnf | -0.018 | Old |
| 0.491717 | 0.682632 | 0.017 | 0.012 | 1 | 6 | Trim2 | 0.005 | Old |
| 0.491717 | 0.376946 | 0.017 | 0.012 | 1 | 6 | Gm16867 | 0.005 | Old |
| 0.49175 | 0.072697 | 0.263 | 0.281 | 1 | 6 | Mkln1 | -0.018 | Old |
| 0.491854 | 0.310683 | 0.08 | 0.069 | 1 | 6 | Prr3 | 0.011 | Old |
| 0.492188 | 0.082249 | 0.051 | 0.042 | 1 | 6 | Cd72 | 0.009 | Old |
| 0.492236 | 0.12406 | 0.057 | 0.047 | 1 | 6 | Rab6b | 0.01 | Old |
| 0.492345 | 0.0836 | 0.169 | 0.151 | 1 | 6 | Cops6 | 0.018 | Old |
| 0.492486 | 0.097592 | 0.017 | 0.012 | 1 | 6 | Dhfr | 0.005 | Old |
| 0.492689 | 0.289319 | 0.163 | 0.145 | 1 | 6 | Smad2 | 0.018 | Old |
| 0.492736 | 0.712687 | 0.034 | 0.027 | 1 | 6 | Trmt11 | 0.007 | Old |
| 0.492776 | 0.141708 | 0.103 | 0.089 | 1 | 6 | Rnf31 | 0.014 | Old |
| 0.492884 | 0.025707 | 0.357 | 0.38 | 1 | 6 | Lsm6 | -0.023 | Old |
| 0.492906 | 1.659601 | 0.023 | 0.017 | 1 | 6 | Rora | 0.006 | Old |
| 0.493169 | 0.597487 | 0.046 | 0.037 | 1 | 6 | Dlg4 | 0.009 | Old |
| 0.493256 | 0.079787 | 0.017 | 0.012 | 1 | 6 | Creb3l1 | 0.005 | Old |
| 0.493326 | 0.285229 | 0.163 | 0.147 | 1 | 6 | Ola1 | 0.016 | Old |
| 0.493562 | 0.880464 | 0.023 | 0.017 | 1 | 6 | Dffb | 0.006 | Old |
| 0.493804 | 0.238871 | 0.083 | 0.07 | 1 | 6 | Atr | 0.013 | Old |
| 0.493821 | 0.215542 | 0.257 | 0.239 | 1 | 6 | Zfp148 | 0.018 | Old |
| 0.493996 | 0.218606 | 0.214 | 0.197 | 1 | 6 | Cd81 | 0.017 | Old |
| 0.494026 | 0.106136 | 0.017 | 0.012 | 1 | 6 | Them4 | 0.005 | Old |
| 0.494082 | 0.515741 | 0.046 | 0.037 | 1 | 6 | Arhgef10 | 0.009 | Old |
| 0.494365 | 0.219503 | 0.106 | 0.092 | 1 | 6 | Cep192 | 0.014 | Old |
| 0.49466 | 0.895265 | 0.029 | 0.022 | 1 | 6 | Alg12 | 0.007 | Old |
| 0.49472 | 0.275501 | 0.186 | 0.169 | 1 | 6 | Gm49336 | 0.017 | Old |
| 0.49476 | 0.278456 | 0.146 | 0.13 | 1 | 6 | Gps1 | 0.016 | Old |
| 0.494875 | 1.155187 | 0.023 | 0.017 | 1 | 6 | Haus5 | 0.006 | Old |
| 0.495007 | 0.430545 | 0.231 | 0.214 | 1 | 6 | Ncaph2 | 0.017 | Old |

| | | | | | | | | |
|---|---|---|---|---|---|---|---|---|
| 0.495166 | 0.233416 | 0.306 | 0.293 | 1 | 6 | Slc20a1 | 0.013 | Old |
| 0.495673 | 0.096223 | 0.109 | 0.094 | 1 | 6 | Itpkc | 0.015 | Old |
| 0.495806 | 0.15711 | 0.334 | 0.311 | 1 | 6 | Gna13 | 0.023 | Old |
| 0.49603 | 0.274088 | 0.3 | 0.284 | 1 | 6 | Pdcd6 | 0.016 | Old |
| 0.496341 | 0.317247 | 0.017 | 0.012 | 1 | 6 | Gm26890 | 0.005 | Old |
| 0.496405 | 0.26769 | 0.183 | 0.204 | 1 | 6 | Tmem64 | -0.021 | Old |
| 0.496495 | 0.412695 | 0.051 | 0.042 | 1 | 6 | Nol4l | 0.009 | Old |
| 0.496855 | 0.248046 | 0.169 | 0.152 | 1 | 6 | Acaa2 | 0.017 | Old |
| 0.497187 | 0.306892 | 0.089 | 0.077 | 1 | 6 | Mrpl37 | 0.012 | Old |
| 0.497332 | 0.444548 | 0.091 | 0.079 | 1 | 6 | Sh3gl1 | 0.012 | Old |
| 0.497579 | 0.570001 | 0.029 | 0.022 | 1 | 6 | Ftsj1 | 0.007 | Old |
| 0.497981 | 0.675035 | 0.04 | 0.032 | 1 | 6 | Dnaaf3 | 0.008 | Old |
| 0.498568 | 0.463002 | 0.034 | 0.027 | 1 | 6 | Caml | 0.007 | Old |
| 0.498846 | 0.257894 | 0.049 | 0.06 | 1 | 6 | Rdh11 | -0.011 | Old |
| 0.499192 | 0.005031 | 0.831 | 0.898 | 1 | 6 | Tmsb10 | -0.067 | Old |
| 0.49924 | 0.022703 | 0.5 | 0.52 | 1 | 6 | Rab8b | -0.02 | Old |
| 0.499598 | 0.110572 | 0.123 | 0.107 | 1 | 6 | Ltv1 | 0.016 | Old |
| 0.500147 | 0.831358 | 0.023 | 0.017 | 1 | 6 | A130010J1 | 0.006 | Old |
| 0.500379 | 0.014353 | 0.08 | 0.067 | 1 | 6 | Pgm1 | 0.013 | Old |
| 0.500388 | 0.228791 | 0.051 | 0.042 | 1 | 6 | Memo1 | 0.009 | Old |
| 0.50044 | 0.349376 | 0.126 | 0.11 | 1 | 6 | Mtf2 | 0.016 | Old |
| 0.500506 | 0.556081 | 0.029 | 0.022 | 1 | 6 | Exoc8 | 0.007 | Old |
| 0.500563 | 0.166139 | 0.046 | 0.057 | 1 | 6 | Scly | -0.011 | Old |
| 0.500784 | 0.100708 | 0.109 | 0.094 | 1 | 6 | Manba | 0.015 | Old |
| 0.500959 | 0.334323 | 0.046 | 0.037 | 1 | 6 | Alg9 | 0.009 | Old |
| 0.50102 | 0.131144 | 0.509 | 0.503 | 1 | 6 | Atp5b | 0.006 | Old |
| 0.501288 | 0.101394 | 0.18 | 0.162 | 1 | 6 | Trim12a | 0.018 | Old |
| 0.50147 | 0.503722 | 0.023 | 0.017 | 1 | 6 | Katnal1 | 0.006 | Old |
| 0.502121 | 0.026687 | 0.46 | 0.478 | 1 | 6 | Cep170 | -0.018 | Old |
| 0.502163 | 0.153342 | 0.686 | 0.714 | 1 | 6 | Myadm | -0.028 | Old |
| 0.502569 | 0.167293 | 0.2 | 0.182 | 1 | 6 | Adssl1 | 0.018 | Old |
| 0.502633 | 0.405879 | 0.146 | 0.129 | 1 | 6 | Zc3h14 | 0.017 | Old |
| 0.50283 | 0.211521 | 0.306 | 0.284 | 1 | 6 | Tpi1 | 0.022 | Old |
| 0.502832 | 1.437367 | 0.034 | 0.027 | 1 | 6 | Il10 | 0.007 | Old |
| 0.502855 | 0.973827 | 0.029 | 0.022 | 1 | 6 | Slc41a2 | 0.007 | Old |
| 0.503105 | 0.173534 | 0.557 | 0.565 | 1 | 6 | Cd9 | -0.008 | Old |
| 0.503198 | 0.063922 | 0.171 | 0.152 | 1 | 6 | Pigp | 0.019 | Old |
| 0.503366 | 0.425305 | 0.034 | 0.027 | 1 | 6 | Cmtm8 | 0.007 | Old |
| 0.503396 | 0.524633 | 0.04 | 0.032 | 1 | 6 | Kat2a | 0.008 | Old |
| 0.503427 | 0.166053 | 0.051 | 0.042 | 1 | 6 | Tpgs2 | 0.009 | Old |
| 0.503458 | 0.423508 | 0.023 | 0.017 | 1 | 6 | Yars2 | 0.006 | Old |
| 0.504025 | 0.517618 | 0.029 | 0.037 | 1 | 6 | Pgap1 | -0.008 | Old |
| 0.505138 | 0.319744 | 0.183 | 0.166 | 1 | 6 | Anapc16 | 0.017 | Old |
| 0.505308 | 0.167797 | 0.189 | 0.169 | 1 | 6 | Siah2 | 0.02 | Old |
| 0.505449 | 0.386871 | 0.023 | 0.017 | 1 | 6 | Fuz | 0.006 | Old |

| | | | | | | | | |
|---|---|---|---|---|---|---|---|---|
| 0.505497 | 0.200079 | 0.151 | 0.135 | 1 | 6 | Sp140 | 0.016 | Old |
| 0.505507 | 0.760741 | 0.034 | 0.027 | 1 | 6 | Orc5 | 0.007 | Old |
| 0.505732 | 0.038886 | 0.031 | 0.04 | 1 | 6 | Zfp429 | -0.009 | Old |
| 0.505762 | 0.148594 | 0.129 | 0.114 | 1 | 6 | AI413582 | 0.015 | Old |
| 0.505831 | 0.297799 | 0.16 | 0.142 | 1 | 6 | Larp1b | 0.018 | Old |
| 0.506075 | 0.099311 | 0.263 | 0.242 | 1 | 6 | Tmem51 | 0.021 | Old |
| 0.506208 | 0.09318 | 0.149 | 0.132 | 1 | 6 | Sf3b4 | 0.017 | Old |
| 0.506367 | 0.195989 | 0.026 | 0.033 | 1 | 6 | Rft1 | -0.007 | Old |
| 0.506389 | 0.152527 | 0.029 | 0.022 | 1 | 6 | Clec16a | 0.007 | Old |
| 0.506597 | 0.284087 | 0.146 | 0.129 | 1 | 6 | Ubxn2a | 0.017 | Old |
| 0.506642 | 0.012265 | 0.2 | 0.179 | 1 | 6 | Rabep1 | 0.021 | Old |
| 0.50667 | 0.535402 | 0.06 | 0.072 | 1 | 6 | Hacd3 | -0.012 | Old |
| 0.507361 | 0.00247 | 0.409 | 0.383 | 1 | 6 | Rnf13 | 0.026 | Old |
| 0.507368 | 0.667006 | 0.131 | 0.119 | 1 | 6 | Zbtb1 | 0.012 | Old |
| 0.507423 | 0.196515 | 0.034 | 0.043 | 1 | 6 | Accs | -0.009 | Old |
| 0.50751 | 0.235193 | 0.249 | 0.232 | 1 | 6 | Tubb2a | 0.017 | Old |
| 0.508024 | 0.100121 | 0.651 | 0.649 | 1 | 6 | Cyth4 | 0.002 | Old |
| 0.508111 | 0.463449 | 0.023 | 0.017 | 1 | 6 | Polr3g | 0.006 | Old |
| 0.508222 | 0.168353 | 0.051 | 0.042 | 1 | 6 | Fam45a | 0.009 | Old |
| 0.50835 | 0.277766 | 0.046 | 0.037 | 1 | 6 | Mthfd1l | 0.009 | Old |
| 0.508597 | 0.094961 | 0.106 | 0.09 | 1 | 6 | Abhd16a | 0.016 | Old |
| 0.508632 | 0.086487 | 0.94 | 0.967 | 1 | 6 | Ahnak | -0.027 | Old |
| 0.509013 | 0.043857 | 0.471 | 0.44 | 1 | 6 | Hdgf | 0.031 | Old |
| 0.509135 | 0.323995 | 0.151 | 0.135 | 1 | 6 | Ing5 | 0.016 | Old |
| 0.509264 | 0.88422 | 0.034 | 0.027 | 1 | 6 | Trim23 | 0.007 | Old |
| 0.509344 | 0.557879 | 0.029 | 0.022 | 1 | 6 | Dok3 | 0.007 | Old |
| 0.509344 | 0.205683 | 0.029 | 0.022 | 1 | 6 | Mus81 | 0.007 | Old |
| 0.509502 | 0.067384 | 0.077 | 0.065 | 1 | 6 | Ercc5 | 0.012 | Old |
| 0.509748 | 0.022978 | 0.009 | 0.013 | 1 | 6 | Gm28043 | -0.004 | Old |
| 0.509936 | 0.467705 | 0.029 | 0.022 | 1 | 6 | Skp2 | 0.007 | Old |
| 0.509936 | 0.134566 | 0.029 | 0.022 | 1 | 6 | Pigg | 0.007 | Old |
| 0.510064 | 0.370475 | 0.026 | 0.033 | 1 | 6 | Etnk2 | -0.007 | Old |
| 0.510341 | 0.119152 | 0.034 | 0.027 | 1 | 6 | Telo2 | 0.007 | Old |
| 0.510445 | 0.194793 | 0.377 | 0.355 | 1 | 6 | Map2k2 | 0.022 | Old |
| 0.511121 | 0.671478 | 0.029 | 0.022 | 1 | 6 | N4bp3 | 0.007 | Old |
| 0.51137 | 0.421355 | 0.166 | 0.152 | 1 | 6 | Dmxl1 | 0.014 | Old |
| 0.511418 | 0.808574 | 0.034 | 0.027 | 1 | 6 | Cd151 | 0.007 | Old |
| 0.51143 | 0.05611 | 0.134 | 0.151 | 1 | 6 | Slc16a3 | -0.017 | Old |
| 0.511451 | 0.806083 | 0.009 | 0.013 | 1 | 6 | Abhd14a | -0.004 | Old |
| 0.511536 | 0.58506 | 0.091 | 0.08 | 1 | 6 | Creb3l2 | 0.011 | Old |
| 0.511714 | 0.330338 | 0.029 | 0.022 | 1 | 6 | Zfp74 | 0.007 | Old |
| 0.511826 | 0.385376 | 0.04 | 0.032 | 1 | 6 | Galnt4 | 0.008 | Old |
| 0.512303 | 0.388486 | 0.009 | 0.013 | 1 | 6 | Ldlrad1 | -0.004 | Old |
| 0.512303 | 0.280081 | 0.009 | 0.013 | 1 | 6 | Atp2b4 | -0.004 | Old |
| 0.512449 | 0.455443 | 0.151 | 0.139 | 1 | 6 | Csnk1g1 | 0.012 | Old |

| | | | | | | | | |
|---|---|---|---|---|---|---|---|---|
| 0.512823 | 0.209458 | 0.04 | 0.032 | 1 | 6 | Kctd3 | 0.008 | Old |
| 0.512902 | 0.390415 | 0.029 | 0.022 | 1 | 6 | Zkscan8 | 0.007 | Old |
| 0.513008 | 0.129086 | 0.24 | 0.264 | 1 | 6 | Ubn1 | -0.024 | Old |
| 0.513473 | 0.120881 | 0.18 | 0.162 | 1 | 6 | Mrps17 | 0.018 | Old |
| 0.51357 | 0.459153 | 0.066 | 0.055 | 1 | 6 | Nadk2 | 0.011 | Old |
| 0.513812 | 0.974124 | 0.054 | 0.045 | 1 | 6 | Piga | 0.009 | Old |
| 0.51382 | 0.078112 | 0.04 | 0.032 | 1 | 6 | Fance | 0.008 | Old |
| 0.514011 | 0.428059 | 0.009 | 0.013 | 1 | 6 | Tarsl2 | -0.004 | Old |
| 0.514011 | 0.351239 | 0.009 | 0.013 | 1 | 6 | Gm9856 | -0.004 | Old |
| 0.514128 | 0.148132 | 0.023 | 0.017 | 1 | 6 | Rnf26 | 0.006 | Old |
| 0.514164 | 0.141104 | 0.106 | 0.09 | 1 | 6 | Nmi | 0.016 | Old |
| 0.514525 | 0.00663 | 0.294 | 0.313 | 1 | 6 | Kras | -0.019 | Old |
| 0.514585 | 0.112347 | 0.357 | 0.336 | 1 | 6 | Mapk6 | 0.021 | Old |
| 0.514739 | 0.284688 | 0.111 | 0.127 | 1 | 6 | Stk3 | -0.016 | Old |
| 0.515032 | 0.14899 | 0.1 | 0.087 | 1 | 6 | Sh3glb2 | 0.013 | Old |
| 0.515252 | 0.063028 | 0.103 | 0.089 | 1 | 6 | Gid8 | 0.014 | Old |
| 0.515326 | 0.664386 | 0.066 | 0.057 | 1 | 6 | Ankmy2 | 0.009 | Old |
| 0.515361 | 0.253231 | 0.186 | 0.172 | 1 | 6 | Cdk19 | 0.014 | Old |
| 0.515976 | 0.013929 | 0.394 | 0.365 | 1 | 6 | Ap2s1 | 0.029 | Old |
| 0.516196 | 0.092391 | 0.657 | 0.627 | 1 | 6 | Ddx17 | 0.03 | Old |
| 0.516366 | 0.198551 | 0.654 | 0.657 | 1 | 6 | Csnk1a1 | -0.003 | Old |
| 0.516823 | 0.344578 | 0.034 | 0.027 | 1 | 6 | Iqsec2 | 0.007 | Old |
| 0.517038 | 0.505658 | 0.109 | 0.097 | 1 | 6 | Rrp7a | 0.012 | Old |
| 0.517261 | 0.099997 | 0.317 | 0.296 | 1 | 6 | Tmem189 | 0.021 | Old |
| 0.517284 | 0.366081 | 0.049 | 0.04 | 1 | 6 | Nat9 | 0.009 | Old |
| 0.517359 | 0.427398 | 0.083 | 0.099 | 1 | 6 | Cadm1 | -0.016 | Old |
| 0.517399 | 0.510776 | 0.074 | 0.064 | 1 | 6 | Pphln1 | 0.01 | Old |
| 0.517552 | 0.305283 | 0.283 | 0.268 | 1 | 6 | Emc7 | 0.015 | Old |
| 0.517844 | 0.223325 | 0.066 | 0.055 | 1 | 6 | Nudt13 | 0.011 | Old |
| 0.517875 | 0.068628 | 0.231 | 0.211 | 1 | 6 | Tmpo | 0.02 | Old |
| 0.518136 | 0.015178 | 0.046 | 0.037 | 1 | 6 | Zfyve1 | 0.009 | Old |
| 0.518371 | 0.310276 | 0.246 | 0.229 | 1 | 6 | Csnk2a1 | 0.017 | Old |
| 0.518628 | 0.213775 | 0.209 | 0.191 | 1 | 6 | Snx4 | 0.018 | Old |
| 0.518696 | 0.365209 | 0.146 | 0.132 | 1 | 6 | Cdc14a | 0.014 | Old |
| 0.51886 | 0.243438 | 0.029 | 0.022 | 1 | 6 | Map2k3os | 0.007 | Old |
| 0.519245 | 0.036126 | 0.591 | 0.62 | 1 | 6 | Ppp2ca | -0.029 | Old |
| 0.520056 | 0.00039 | 0.02 | 0.027 | 1 | 6 | Cnr2 | -0.007 | Old |
| 0.520207 | 0.267023 | 0.183 | 0.169 | 1 | 6 | Pcna | 0.014 | Old |
| 0.520332 | 0.108556 | 0.149 | 0.132 | 1 | 6 | Gpr107 | 0.017 | Old |
| 0.52057 | 0.428128 | 0.023 | 0.03 | 1 | 6 | Dpp7 | -0.007 | Old |
| 0.520671 | 0.192716 | 0.017 | 0.023 | 1 | 6 | P2ry1 | -0.006 | Old |
| 0.521151 | 0.379531 | 0.177 | 0.164 | 1 | 6 | Emc2 | 0.013 | Old |
| 0.521168 | 0.77189 | 0.034 | 0.027 | 1 | 6 | Alg11 | 0.007 | Old |
| 0.521261 | 0.057541 | 0.303 | 0.323 | 1 | 6 | Cbx3 | -0.02 | Old |
| 0.521316 | 0.049402 | 0.3 | 0.321 | 1 | 6 | Map7d1 | -0.021 | Old |

| | | | | | | | | |
|---|---|---|---|---|---|---|---|---|
| 0.52145 | 0.017411 | 0.674 | 0.722 | 1 | 6 | Pip4k2a | -0.048 | Old |
| 0.521802 | 0.392537 | 0.077 | 0.09 | 1 | 6 | Ctsl | -0.013 | Old |
| 0.52185 | 0.344405 | 0.174 | 0.159 | 1 | 6 | Slmap | 0.015 | Old |
| 0.521969 | 0.200871 | 0.106 | 0.092 | 1 | 6 | Fam199x | 0.014 | Old |
| 0.522081 | 0.273914 | 0.389 | 0.375 | 1 | 6 | Dhx15 | 0.014 | Old |
| 0.522086 | 0.319145 | 0.069 | 0.059 | 1 | 6 | Tmem63a | 0.01 | Old |
| 0.522248 | 0.06895 | 0.066 | 0.077 | 1 | 6 | Zfp397 | -0.011 | Old |
| 0.522272 | 0.165014 | 0.651 | 0.639 | 1 | 6 | Clic1 | 0.012 | Old |
| 0.522899 | 0.356601 | 0.2 | 0.189 | 1 | 6 | Cse1l | 0.011 | Old |
| 0.523164 | 0.097772 | 0.054 | 0.065 | 1 | 6 | Cfap97 | -0.011 | Old |
| 0.523167 | 0.547055 | 0.049 | 0.04 | 1 | 6 | Wwox | 0.009 | Old |
| 0.523334 | 0.327892 | 0.011 | 0.017 | 1 | 6 | Sap18b | -0.006 | Old |
| 0.523453 | 0.129856 | 0.063 | 0.052 | 1 | 6 | B4galt4 | 0.011 | Old |
| 0.52365 | 0.671987 | 0.054 | 0.045 | 1 | 6 | Snhg9 | 0.009 | Old |
| 0.524047 | 0.205291 | 0.406 | 0.378 | 1 | 6 | Rad23b | 0.028 | Old |
| 0.524295 | 0.149998 | 0.18 | 0.164 | 1 | 6 | Tor1b | 0.016 | Old |
| 0.524524 | 0.048631 | 0.017 | 0.023 | 1 | 6 | Pdgfa | -0.006 | Old |
| 0.52485 | 0.005596 | 0.586 | 0.604 | 1 | 6 | Lamp2 | -0.018 | Old |
| 0.524912 | 0.094982 | 0.257 | 0.278 | 1 | 6 | Vdac3 | -0.021 | Old |
| 0.525168 | 0.264053 | 0.017 | 0.023 | 1 | 6 | Fam149a | -0.006 | Old |
| 0.52562 | 0.582532 | 0.097 | 0.087 | 1 | 6 | Cdk10 | 0.01 | Old |
| 0.525658 | 0.111206 | 0.183 | 0.166 | 1 | 6 | Slirp | 0.017 | Old |
| 0.525672 | 0.331294 | 0.014 | 0.02 | 1 | 6 | Zfp141 | -0.006 | Old |
| 0.525803 | 0.005326 | 0.066 | 0.077 | 1 | 6 | Nom1 | -0.011 | Old |
| 0.526375 | 0.186878 | 0.04 | 0.032 | 1 | 6 | Rint1 | 0.008 | Old |
| 0.527336 | 0.122307 | 0.489 | 0.472 | 1 | 6 | Dpysl2 | 0.017 | Old |
| 0.527767 | 0.08812 | 0.014 | 0.02 | 1 | 6 | 1700096K1 | -0.006 | Old |
| 0.528318 | 0.026219 | 0.297 | 0.328 | 1 | 6 | Becn1 | -0.031 | Old |
| 0.52834 | 0.194032 | 0.72 | 0.707 | 1 | 6 | Sfpq | 0.013 | Old |
| 0.528356 | 0.090088 | 0.663 | 0.639 | 1 | 6 | Grk2 | 0.024 | Old |
| 0.528427 | 0.10351 | 0.174 | 0.159 | 1 | 6 | Ndfip2 | 0.015 | Old |
| 0.528456 | 0.609007 | 0.069 | 0.082 | 1 | 6 | Inf2 | -0.013 | Old |
| 0.528737 | 0.171125 | 0.171 | 0.156 | 1 | 6 | Atg101 | 0.015 | Old |
| 0.528788 | 0.123023 | 0.2 | 0.186 | 1 | 6 | Mif4gd | 0.014 | Old |
| 0.528801 | 0.362763 | 0.177 | 0.157 | 1 | 6 | Ankib1 | 0.02 | Old |
| 0.528908 | 0.445678 | 0.029 | 0.037 | 1 | 6 | Ccdc32 | -0.008 | Old |
| 0.52954 | 0.379849 | 0.049 | 0.04 | 1 | 6 | Wdr45 | 0.009 | Old |
| 0.529633 | 0.212666 | 0.023 | 0.03 | 1 | 6 | Ercc6l | -0.007 | Old |
| 0.529648 | 0.23002 | 0.086 | 0.074 | 1 | 6 | Washc5 | 0.012 | Old |
| 0.52977 | 0.24594 | 0.297 | 0.281 | 1 | 6 | Cebpz | 0.016 | Old |
| 0.52993 | 0.022363 | 0.146 | 0.13 | 1 | 6 | Pnpla2 | 0.016 | Old |
| 0.529952 | 0.003427 | 0.483 | 0.505 | 1 | 6 | Larp4b | -0.022 | Old |
| 0.530238 | 0.149549 | 0.491 | 0.477 | 1 | 6 | Irf5 | 0.014 | Old |
| 0.530366 | 0.174644 | 0.454 | 0.44 | 1 | 6 | Cnot6l | 0.014 | Old |
| 0.530539 | 0.284128 | 0.189 | 0.174 | 1 | 6 | Med1 | 0.015 | Old |

| | | | | | | | | |
|---|---|---|---|---|---|---|---|---|
| 0.530583 | 0.263673 | 0.274 | 0.259 | 1 | 6 | Mcmbp | 0.015 | Old |
| 0.53091 | 0.157479 | 0.323 | 0.306 | 1 | 6 | Mif | 0.017 | Old |
| 0.530948 | 0.083974 | 0.389 | 0.373 | 1 | 6 | Ddi2 | 0.016 | Old |
| 0.531268 | 0.063413 | 0.014 | 0.02 | 1 | 6 | Ccdc84 | -0.006 | Old |
| 0.531325 | 0.400892 | 0.3 | 0.289 | 1 | 6 | Galnt1 | 0.011 | Old |
| 0.531803 | 0.205741 | 0.109 | 0.095 | 1 | 6 | Nhlrc2 | 0.014 | Old |
| 0.532468 | 0.535418 | 0.12 | 0.107 | 1 | 6 | Rcsd1 | 0.013 | Old |
| 0.53315 | 0.197384 | 0.043 | 0.052 | 1 | 6 | Cep85 | -0.009 | Old |
| 0.535128 | 0.181151 | 0.063 | 0.054 | 1 | 6 | Uty | 0.009 | Old |
| 0.53535 | 0.054421 | 0.103 | 0.09 | 1 | 6 | 2810004N2 | 0.013 | Old |
| 0.535565 | 0.249614 | 0.18 | 0.162 | 1 | 6 | B4galt3 | 0.018 | Old |
| 0.536079 | 0.170427 | 0.157 | 0.142 | 1 | 6 | Rab22a | 0.015 | Old |
| 0.536164 | 0.170313 | 0.16 | 0.144 | 1 | 6 | Vars | 0.016 | Old |
| 0.536198 | 0.380361 | 0.046 | 0.055 | 1 | 6 | Frs2 | -0.009 | Old |
| 0.536221 | 0.177901 | 0.149 | 0.135 | 1 | 6 | Snx1 | 0.014 | Old |
| 0.536966 | 0.066362 | 0.074 | 0.064 | 1 | 6 | Nudt2 | 0.01 | Old |
| 0.537745 | 0.271803 | 0.109 | 0.097 | 1 | 6 | Rusc1 | 0.012 | Old |
| 0.537888 | 0.408184 | 0.057 | 0.048 | 1 | 6 | Rnf40 | 0.009 | Old |
| 0.538299 | 0.330145 | 0.16 | 0.147 | 1 | 6 | Maea | 0.013 | Old |
| 0.538356 | 0.559314 | 0.063 | 0.054 | 1 | 6 | Mrps11 | 0.009 | Old |
| 0.538707 | 0.08634 | 0.606 | 0.649 | 1 | 6 | Dnaja1 | -0.043 | Old |
| 0.539071 | 0.127921 | 0.463 | 0.448 | 1 | 6 | Zfp706 | 0.015 | Old |
| 0.539117 | 0.265652 | 0.171 | 0.159 | 1 | 6 | BC035044 | 0.012 | Old |
| 0.539143 | 0.309235 | 0.106 | 0.094 | 1 | 6 | Flt3l | 0.012 | Old |
| 0.539143 | 0.058499 | 0.46 | 0.446 | 1 | 6 | Mdh2 | 0.014 | Old |
| 0.539204 | 0.294285 | 0.074 | 0.064 | 1 | 6 | Dalrd3 | 0.01 | Old |
| 0.53924 | 0.457094 | 0.054 | 0.045 | 1 | 6 | Wdhd1 | 0.009 | Old |
| 0.539646 | 0.290842 | 0.1 | 0.114 | 1 | 6 | Tmem245 | -0.014 | Old |
| 0.539785 | 0.130348 | 0.969 | 0.985 | 1 | 6 | Jund | -0.016 | Old |
| 0.539822 | 0.849315 | 0.037 | 0.03 | 1 | 6 | Mfsd4a | 0.007 | Old |
| 0.540141 | 0.155253 | 0.249 | 0.229 | 1 | 6 | Rcc2 | 0.02 | Old |
| 0.540805 | 0.336731 | 0.077 | 0.067 | 1 | 6 | Prmt5 | 0.01 | Old |
| 0.54119 | 0.371547 | 0.106 | 0.094 | 1 | 6 | Gtf2i | 0.012 | Old |
| 0.541355 | 0.124456 | 0.103 | 0.117 | 1 | 6 | Pcnt | -0.014 | Old |
| 0.541537 | 0.253492 | 0.077 | 0.067 | 1 | 6 | Tex264 | 0.01 | Old |
| 0.541998 | 0.496326 | 0.063 | 0.054 | 1 | 6 | Rgcc | 0.009 | Old |
| 0.542344 | 0.06683 | 0.091 | 0.104 | 1 | 6 | Vps52 | -0.013 | Old |
| 0.542446 | 0.545079 | 0.071 | 0.062 | 1 | 6 | Wdfy1 | 0.009 | Old |
| 0.542549 | 0.113031 | 0.046 | 0.055 | 1 | 6 | Rrp36 | -0.009 | Old |
| 0.542889 | 0.007627 | 0.206 | 0.184 | 1 | 6 | Brd8 | 0.022 | Old |
| 0.542945 | 0.16134 | 0.366 | 0.338 | 1 | 6 | Bcap31 | 0.028 | Old |
| 0.54332 | 0.012006 | 0.074 | 0.064 | 1 | 6 | Mmgt1 | 0.01 | Old |
| 0.543557 | 0.299019 | 0.106 | 0.094 | 1 | 6 | Trim24 | 0.012 | Old |
| 0.543665 | 0.77193 | 0.031 | 0.025 | 1 | 6 | Gm26669 | 0.006 | Old |
| 0.544522 | 0.100037 | 0.774 | 0.809 | 1 | 6 | Capza2 | -0.035 | Old |

| | | | | | | | | |
|---|---|---|---|---|---|---|---|---|
| 0.544615 | 0.561733 | 0.051 | 0.043 | 1 | 6 | Vdr | 0.008 | Old |
| 0.544649 | 0.200849 | 0.137 | 0.124 | 1 | 6 | Mrpl3 | 0.013 | Old |
| 0.544915 | 0.052026 | 0.22 | 0.202 | 1 | 6 | Rspry1 | 0.018 | Old |
| 0.545061 | 0.077178 | 0.677 | 0.684 | 1 | 6 | Hnrnpu | -0.007 | Old |
| 0.545109 | 0.732631 | 0.037 | 0.03 | 1 | 6 | Zfp944 | 0.007 | Old |
| 0.545973 | 0.95284 | 0.031 | 0.025 | 1 | 6 | Ccdc181 | 0.006 | Old |
| 0.546106 | 0.361871 | 0.229 | 0.212 | 1 | 6 | R3hdm1 | 0.017 | Old |
| 0.54647 | 0.094152 | 0.194 | 0.176 | 1 | 6 | Wbp1l | 0.018 | Old |
| 0.546688 | 0.176607 | 0.051 | 0.062 | 1 | 6 | Sash1 | -0.011 | Old |
| 0.547006 | 1.418887 | 0.014 | 0.01 | 1 | 6 | Glis2 | 0.004 | Old |
| 0.547006 | 0.977916 | 0.014 | 0.01 | 1 | 6 | Zfp873 | 0.004 | Old |
| 0.547121 | 0.483313 | 0.054 | 0.045 | 1 | 6 | Spice1 | 0.009 | Old |
| 0.547755 | 0.299303 | 0.126 | 0.112 | 1 | 6 | Pbdc1 | 0.014 | Old |
| 0.547903 | 0.101015 | 0.191 | 0.172 | 1 | 6 | Tagap | 0.019 | Old |
| 0.547965 | 0.308372 | 0.049 | 0.04 | 1 | 6 | Trim39 | 0.009 | Old |
| 0.548505 | 0.174832 | 0.171 | 0.156 | 1 | 6 | Tmem9b | 0.015 | Old |
| 0.548616 | 0.33068 | 0.037 | 0.045 | 1 | 6 | Pnpla6 | -0.008 | Old |
| 0.548772 | 0.934364 | 0.014 | 0.01 | 1 | 6 | Npl | 0.004 | Old |
| 0.548772 | 0.837341 | 0.014 | 0.01 | 1 | 6 | B3galt2 | 0.004 | Old |
| 0.548944 | 0.056871 | 0.091 | 0.08 | 1 | 6 | Cenpc1 | 0.011 | Old |
| 0.549358 | 0.595822 | 0.037 | 0.03 | 1 | 6 | Trim14 | 0.007 | Old |
| 0.54971 | 0.002082 | 0.749 | 0.764 | 1 | 6 | Csde1 | -0.015 | Old |
| 0.550104 | 0.086034 | 0.737 | 0.756 | 1 | 6 | Canx | -0.019 | Old |
| 0.55054 | 1.483445 | 0.014 | 0.01 | 1 | 6 | Gm10642 | 0.004 | Old |
| 0.550566 | 0.133424 | 0.114 | 0.102 | 1 | 6 | Prelid2 | 0.012 | Old |
| 0.550642 | 0.431763 | 0.043 | 0.052 | 1 | 6 | Hmgxb4 | -0.009 | Old |
| 0.550741 | 0.2856 | 0.14 | 0.156 | 1 | 6 | Pcyt1a | -0.016 | Old |
| 0.550756 | 0.041138 | 0.049 | 0.04 | 1 | 6 | Nudt18 | 0.009 | Old |
| 0.550962 | 0.355559 | 0.086 | 0.075 | 1 | 6 | Faf2 | 0.011 | Old |
| 0.55099 | 0.010744 | 0.054 | 0.064 | 1 | 6 | Msh2 | -0.01 | Old |
| 0.551275 | 0.076119 | 0.489 | 0.53 | 1 | 6 | Synj1 | -0.041 | Old |
| 0.551425 | 0.594971 | 0.014 | 0.01 | 1 | 6 | Rai14 | 0.004 | Old |
| 0.551629 | 0.411091 | 0.083 | 0.072 | 1 | 6 | Ap3s2 | 0.011 | Old |
| 0.551844 | 0.736919 | 0.077 | 0.067 | 1 | 6 | Rrm2b | 0.01 | Old |
| 0.552028 | 1.000449 | 0.051 | 0.043 | 1 | 6 | Iqsec1 | 0.008 | Old |
| 0.552111 | 0.028224 | 0.691 | 0.661 | 1 | 6 | Cox6a1 | 0.03 | Old |
| 0.552152 | 0.180625 | 0.451 | 0.457 | 1 | 6 | Lifr | -0.006 | Old |
| 0.552166 | 0.399538 | 0.083 | 0.072 | 1 | 6 | Crls1 | 0.011 | Old |
| 0.552213 | 0.401232 | 0.077 | 0.067 | 1 | 6 | Brip1os | 0.01 | Old |
| 0.552569 | 0.420283 | 0.343 | 0.331 | 1 | 6 | Raph1 | 0.012 | Old |
| 0.552634 | 0.225612 | 0.043 | 0.035 | 1 | 6 | Ptcd3 | 0.008 | Old |
| 0.5529 | 0.706334 | 0.026 | 0.02 | 1 | 6 | Fam109b | 0.006 | Old |
| 0.552927 | 0.962407 | 0.031 | 0.025 | 1 | 6 | Ralgps1 | 0.006 | Old |
| 0.553131 | 0.325122 | 0.043 | 0.035 | 1 | 6 | Tstd2 | 0.008 | Old |
| 0.553131 | 0.013403 | 0.043 | 0.035 | 1 | 6 | Rfc3 | 0.008 | Old |

| | | | | | | | | |
|---|---|---|---|---|---|---|---|---|
| 0.553156 | 0.376592 | 0.051 | 0.043 | 1 | 6 | Clec4g | 0.008 | Old |
| 0.553197 | 0.688402 | 0.014 | 0.01 | 1 | 6 | Nectin4 | 0.004 | Old |
| 0.553197 | 0.683449 | 0.014 | 0.01 | 1 | 6 | Gcfc2 | 0.004 | Old |
| 0.553197 | 0.55688 | 0.014 | 0.01 | 1 | 6 | Aldh1b1 | 0.004 | Old |
| 0.553217 | 0.670747 | 0.057 | 0.048 | 1 | 6 | Sdhaf1 | 0.009 | Old |
| 0.553508 | 0.697261 | 0.031 | 0.025 | 1 | 6 | Polr3a | 0.006 | Old |
| 0.553542 | 0.040918 | 0.32 | 0.349 | 1 | 6 | Csnk2b | -0.029 | Old |
| 0.553607 | 0.8057 | 0.051 | 0.043 | 1 | 6 | Jade2 | 0.008 | Old |
| 0.553874 | 0.193039 | 0.311 | 0.294 | 1 | 6 | Ece1 | 0.017 | Old |
| 0.553916 | 0.116086 | 0.186 | 0.169 | 1 | 6 | Ubr3 | 0.017 | Old |
| 0.554014 | 0.173078 | 0.089 | 0.077 | 1 | 6 | Ocel1 | 0.012 | Old |
| 0.554049 | 0.328634 | 0.211 | 0.197 | 1 | 6 | Cdc73 | 0.014 | Old |
| 0.554084 | 1.100029 | 0.014 | 0.01 | 1 | 6 | Pcbp3 | 0.004 | Old |
| 0.554245 | 0.047095 | 0.069 | 0.059 | 1 | 6 | Orc3 | 0.01 | Old |
| 0.554271 | 0.055558 | 0.491 | 0.457 | 1 | 6 | Pik3cd | 0.034 | Old |
| 0.554359 | 0.235613 | 0.166 | 0.151 | 1 | 6 | Klhdc3 | 0.015 | Old |
| 0.554409 | 0.136606 | 0.343 | 0.323 | 1 | 6 | Tap1 | 0.02 | Old |
| 0.554727 | 0.04751 | 0.251 | 0.232 | 1 | 6 | Ap3b1 | 0.019 | Old |
| 0.554877 | 1.375935 | 0.02 | 0.015 | 1 | 6 | Mmp14 | 0.005 | Old |
| 0.554972 | 0.045315 | 0.014 | 0.01 | 1 | 6 | Gm12355 | 0.004 | Old |
| 0.554978 | 0.296581 | 0.186 | 0.172 | 1 | 6 | Zfx | 0.014 | Old |
| 0.554981 | 0.67845 | 0.066 | 0.057 | 1 | 6 | Fbxw5 | 0.009 | Old |
| 0.555063 | 0.084886 | 0.503 | 0.493 | 1 | 6 | Atp6ap1 | 0.01 | Old |
| 0.555808 | 0.002332 | 0.277 | 0.299 | 1 | 6 | Adipor1 | -0.022 | Old |
| 0.555832 | 0.285019 | 0.071 | 0.062 | 1 | 6 | G2e3 | 0.009 | Old |
| 0.55586 | 1.28437 | 0.014 | 0.01 | 1 | 6 | Cp | 0.004 | Old |
| 0.55586 | 0.661131 | 0.014 | 0.01 | 1 | 6 | Tm7sf2 | 0.004 | Old |
| 0.556302 | 0.048172 | 0.086 | 0.097 | 1 | 6 | Rfwd3 | -0.011 | Old |
| 0.55675 | 0.400073 | 0.014 | 0.01 | 1 | 6 | Rad51c | 0.004 | Old |
| 0.55675 | 0.300515 | 0.014 | 0.01 | 1 | 6 | Klhl25 | 0.004 | Old |
| 0.557094 | 1.356015 | 0.02 | 0.015 | 1 | 6 | Prdm11 | 0.005 | Old |
| 0.557101 | 0.068979 | 0.266 | 0.291 | 1 | 6 | Gtpbp4 | -0.025 | Old |
| 0.557111 | 0.035373 | 0.043 | 0.035 | 1 | 6 | Tab3 | 0.008 | Old |
| 0.557286 | 0.192879 | 0.1 | 0.089 | 1 | 6 | Ppip5k2 | 0.011 | Old |
| 0.55764 | 0.314858 | 0.014 | 0.01 | 1 | 6 | Slc1a4 | 0.004 | Old |
| 0.557904 | 0.358493 | 0.037 | 0.03 | 1 | 6 | Mfsd13a | 0.007 | Old |
| 0.557944 | 0.276614 | 0.046 | 0.055 | 1 | 6 | Tdrd3 | -0.009 | Old |
| 0.55817 | 0.371789 | 0.031 | 0.025 | 1 | 6 | Alg14 | 0.006 | Old |
| 0.55853 | 0.913395 | 0.014 | 0.01 | 1 | 6 | Tmem170 | 0.004 | Old |
| 0.55853 | 0.321168 | 0.014 | 0.01 | 1 | 6 | Palm | 0.004 | Old |
| 0.55853 | 0.274086 | 0.014 | 0.01 | 1 | 6 | Il4i1 | 0.004 | Old |
| 0.55853 | 0.186092 | 0.014 | 0.01 | 1 | 6 | Gm14305 | 0.004 | Old |
| 0.55853 | 0.151877 | 0.014 | 0.01 | 1 | 6 | Jmjd8 | 0.004 | Old |
| 0.558656 | 0.1455 | 0.126 | 0.112 | 1 | 6 | Zc3h4 | 0.014 | Old |
| 0.55894 | 0.04691 | 0.626 | 0.63 | 1 | 6 | Tgfb1 | -0.004 | Old |

| | | | | | | | | |
|---|---|---|---|---|---|---|---|---|
| 0.559094 | 0.142993 | 0.586 | 0.585 | 1 | 6 | Ccnd1 | 0.001 | Old |
| 0.559318 | 0.098021 | 0.506 | 0.475 | 1 | 6 | Clec2d | 0.031 | Old |
| 0.559369 | 0.990775 | 0.026 | 0.02 | 1 | 6 | Zswim1 | 0.006 | Old |
| 0.559422 | 0.356984 | 0.014 | 0.01 | 1 | 6 | Cep78 | 0.004 | Old |
| 0.559761 | 0.288756 | 0.12 | 0.134 | 1 | 6 | Ppfia1 | -0.014 | Old |
| 0.56005 | 0.176644 | 0.037 | 0.03 | 1 | 6 | 2810006K2 | 0.007 | Old |
| 0.560121 | 0.279823 | 0.151 | 0.137 | 1 | 6 | Slc35e1 | 0.014 | Old |
| 0.560754 | 0.16731 | 0.077 | 0.067 | 1 | 6 | Prep | 0.01 | Old |
| 0.56086 | 0.018621 | 0.323 | 0.313 | 1 | 6 | Sh3bp1 | 0.01 | Old |
| 0.56115 | 0.064389 | 0.037 | 0.045 | 1 | 6 | Zbtb17 | -0.008 | Old |
| 0.561186 | 0.874636 | 0.046 | 0.038 | 1 | 6 | Steap3 | 0.008 | Old |
| 0.561424 | 0.135866 | 0.763 | 0.794 | 1 | 6 | Arhgdia | -0.031 | Old |
| 0.56168 | 0.566806 | 0.031 | 0.025 | 1 | 6 | Pde8a | 0.006 | Old |
| 0.56168 | 0.307554 | 0.031 | 0.025 | 1 | 6 | Lancl2 | 0.006 | Old |
| 0.561927 | 0.06282 | 0.251 | 0.231 | 1 | 6 | Lars2 | 0.02 | Old |
| 0.561959 | 0.168938 | 0.309 | 0.291 | 1 | 6 | Mbd2 | 0.018 | Old |
| 0.562246 | 0.062194 | 0.077 | 0.067 | 1 | 6 | Zbtb22 | 0.01 | Old |
| 0.562284 | 1.080665 | 0.02 | 0.015 | 1 | 6 | Fcer2a | 0.005 | Old |
| 0.562603 | 0.10959 | 0.074 | 0.064 | 1 | 6 | Exoc7 | 0.01 | Old |
| 0.563132 | 0.43859 | 0.109 | 0.097 | 1 | 6 | Tbce | 0.012 | Old |
| 0.56327 | 0.009762 | 0.091 | 0.104 | 1 | 6 | Ttc19 | -0.013 | Old |
| 0.563598 | 0.077634 | 0.4 | 0.366 | 1 | 6 | Fis1 | 0.034 | Old |
| 0.563638 | 0.497375 | 0.123 | 0.112 | 1 | 6 | Cbx5 | 0.011 | Old |
| 0.563692 | 0.29519 | 0.12 | 0.109 | 1 | 6 | Lrif1 | 0.011 | Old |
| 0.563816 | 0.463654 | 0.037 | 0.03 | 1 | 6 | E330009J0 | 0.007 | Old |
| 0.564025 | 0.10022 | 0.031 | 0.025 | 1 | 6 | Zfp574 | 0.006 | Old |
| 0.564036 | 0.068123 | 0.197 | 0.216 | 1 | 6 | Atf7ip | -0.019 | Old |
| 0.564108 | 0.022485 | 0.343 | 0.358 | 1 | 6 | Ndufs2 | -0.015 | Old |
| 0.564359 | 0.010924 | 0.383 | 0.408 | 1 | 6 | Kdm2a | -0.025 | Old |
| 0.564569 | 0.417153 | 0.194 | 0.181 | 1 | 6 | Glipr1 | 0.013 | Old |
| 0.564612 | 0.873557 | 0.031 | 0.025 | 1 | 6 | Mpzl3 | 0.006 | Old |
| 0.5652 | 0.264689 | 0.031 | 0.025 | 1 | 6 | Inpp5b | 0.006 | Old |
| 0.5654 | 0.158057 | 0.117 | 0.132 | 1 | 6 | Uri1 | -0.015 | Old |
| 0.566375 | 0.394496 | 0.031 | 0.025 | 1 | 6 | Mgp | 0.006 | Old |
| 0.566418 | 0.016931 | 0.334 | 0.348 | 1 | 6 | Dnm2 | -0.014 | Old |
| 0.566656 | 0.335819 | 0.071 | 0.062 | 1 | 6 | Psmd5 | 0.009 | Old |
| 0.566774 | 0.606723 | 0.051 | 0.043 | 1 | 6 | Zfp780b | 0.008 | Old |
| 0.56711 | 0.069171 | 0.037 | 0.045 | 1 | 6 | Spidr | -0.008 | Old |
| 0.567498 | 0.538757 | 0.02 | 0.015 | 1 | 6 | Fut4 | 0.005 | Old |
| 0.567594 | 0.348285 | 0.037 | 0.03 | 1 | 6 | Sephs1 | 0.007 | Old |
| 0.568141 | 0.188282 | 0.091 | 0.08 | 1 | 6 | Dctn6 | 0.011 | Old |
| 0.568201 | 0.180328 | 0.126 | 0.114 | 1 | 6 | Slc30a9 | 0.012 | Old |
| 0.568245 | 0.137988 | 0.02 | 0.015 | 1 | 6 | Rrp1b | 0.005 | Old |
| 0.568487 | 0.618454 | 0.026 | 0.02 | 1 | 6 | Sgsm3 | 0.006 | Old |
| 0.568562 | 0.612191 | 0.106 | 0.095 | 1 | 6 | Ino80 | 0.011 | Old |

| | | | | | | | | |
|---|---|---|---|---|---|---|---|---|
| 0.56897 | 0.266747 | 0.034 | 0.042 | 1 | 6 | Tmem175 | -0.008 | Old |
| 0.568992 | 0.401431 | 0.02 | 0.015 | 1 | 6 | Wtip | 0.005 | Old |
| 0.569399 | 0.483696 | 0.046 | 0.038 | 1 | 6 | Igf1 | 0.008 | Old |
| 0.569496 | 0.087059 | 0.063 | 0.054 | 1 | 6 | Mier2 | 0.009 | Old |
| 0.56967 | 0.01223 | 0.054 | 0.064 | 1 | 6 | Xxylt1 | -0.01 | Old |
| 0.56979 | 0.057919 | 0.26 | 0.239 | 1 | 6 | Akirin1 | 0.021 | Old |
| 0.569796 | 0.298279 | 0.143 | 0.13 | 1 | 6 | Zmym2 | 0.013 | Old |
| 0.569909 | 0.200834 | 0.031 | 0.025 | 1 | 6 | Gtf3c5 | 0.006 | Old |
| 0.570053 | 0.127701 | 0.18 | 0.164 | 1 | 6 | Agfg1 | 0.016 | Old |
| 0.570326 | 0.01034 | 0.397 | 0.425 | 1 | 6 | Capn2 | -0.028 | Old |
| 0.57096 | 0.121881 | 0.034 | 0.042 | 1 | 6 | Phldb1 | -0.008 | Old |
| 0.571089 | 0.326033 | 0.066 | 0.057 | 1 | 6 | Pcnx3 | 0.009 | Old |
| 0.571196 | 0.545532 | 0.083 | 0.074 | 1 | 6 | Rinl | 0.009 | Old |
| 0.57131 | 0.097314 | 0.38 | 0.365 | 1 | 6 | Usp25 | 0.015 | Old |
| 0.571441 | 0.275201 | 0.114 | 0.102 | 1 | 6 | Rasa2 | 0.012 | Old |
| 0.571733 | 0.244925 | 0.131 | 0.119 | 1 | 6 | Tmem222 | 0.012 | Old |
| 0.57202 | 0.342244 | 0.223 | 0.206 | 1 | 6 | Kmt5a | 0.017 | Old |
| 0.572091 | 0.24914 | 0.111 | 0.1 | 1 | 6 | Ube2l6 | 0.011 | Old |
| 0.572228 | 0.350611 | 0.231 | 0.219 | 1 | 6 | Sarnp | 0.012 | Old |
| 0.572416 | 0.377764 | 0.026 | 0.02 | 1 | 6 | Fhl3 | 0.006 | Old |
| 0.572746 | 0.180903 | 0.297 | 0.281 | 1 | 6 | Tnrc6a | 0.016 | Old |
| 0.572797 | 0.448481 | 0.046 | 0.038 | 1 | 6 | Terf1 | 0.008 | Old |
| 0.572863 | 0.21569 | 0.031 | 0.038 | 1 | 6 | Asf1a | -0.007 | Old |
| 0.573009 | 0.115776 | 0.346 | 0.363 | 1 | 6 | Ubxn1 | -0.017 | Old |
| 0.573072 | 0.206604 | 0.026 | 0.02 | 1 | 6 | Gm20703 | 0.006 | Old |
| 0.573072 | 0.102657 | 0.026 | 0.02 | 1 | 6 | Gm28151 | 0.006 | Old |
| 0.57363 | 0.085284 | 0.203 | 0.222 | 1 | 6 | Poldip3 | -0.019 | Old |
| 0.573891 | 0.030469 | 0.249 | 0.226 | 1 | 6 | Sp3 | 0.023 | Old |
| 0.573902 | 0.116072 | 0.031 | 0.038 | 1 | 6 | Asap2 | -0.007 | Old |
| 0.57398 | 0.08076 | 0.117 | 0.13 | 1 | 6 | Camk2g | -0.013 | Old |
| 0.574364 | 0.130904 | 0.26 | 0.241 | 1 | 6 | Tmem219 | 0.019 | Old |
| 0.574944 | 0.108792 | 0.177 | 0.159 | 1 | 6 | Znhit1 | 0.018 | Old |
| 0.574986 | 0.081151 | 0.02 | 0.015 | 1 | 6 | Tmem267 | 0.005 | Old |
| 0.575034 | 0.14278 | 0.417 | 0.401 | 1 | 6 | Ist1 | 0.016 | Old |
| 0.575078 | 0.127287 | 0.12 | 0.109 | 1 | 6 | Vps54 | 0.011 | Old |
| 0.575481 | 0.526629 | 0.051 | 0.043 | 1 | 6 | AL590144.: | 0.008 | Old |
| 0.575614 | 0.211178 | 0.129 | 0.117 | 1 | 6 | Usp1 | 0.012 | Old |
| 0.575732 | 0.027963 | 0.92 | 0.943 | 1 | 6 | Iqgap1 | -0.023 | Old |
| 0.575737 | 0.185585 | 0.02 | 0.015 | 1 | 6 | Gm36266 | 0.005 | Old |
| 0.575796 | 0.348111 | 0.1 | 0.089 | 1 | 6 | Gga3 | 0.011 | Old |
| 0.576358 | 0.40127 | 0.026 | 0.02 | 1 | 6 | Timp1 | 0.006 | Old |
| 0.576358 | 0.007703 | 0.026 | 0.02 | 1 | 6 | Pkn3 | 0.006 | Old |
| 0.576447 | 0.570087 | 0.083 | 0.074 | 1 | 6 | Pex3 | 0.009 | Old |
| 0.576764 | 0.573635 | 0.04 | 0.033 | 1 | 6 | Hrh1 | 0.007 | Old |
| 0.57678 | 0.023312 | 0.089 | 0.079 | 1 | 6 | Adora2a | 0.01 | Old |

| | | | | | | | | |
|---|---|---|---|---|---|---|---|---|
| 0.577237 | 0.737871 | 0.074 | 0.065 | 1 | 6 | Rps19bp1 | 0.009 | Old |
| 0.577255 | 0.125355 | 0.143 | 0.129 | 1 | 6 | Rnf146 | 0.014 | Old |
| 0.577364 | 0.072676 | 0.446 | 0.416 | 1 | 6 | Dad1 | 0.03 | Old |
| 0.5776 | 0.096501 | 0.257 | 0.239 | 1 | 6 | Rbm47 | 0.018 | Old |
| 0.577616 | 0.090402 | 0.111 | 0.1 | 1 | 6 | Cisd1 | 0.011 | Old |
| 0.577825 | 0.072471 | 0.217 | 0.201 | 1 | 6 | Rtl5 | 0.016 | Old |
| 0.577994 | 0.045155 | 0.02 | 0.015 | 1 | 6 | Sag | 0.005 | Old |
| 0.578273 | 0.006267 | 0.637 | 0.672 | 1 | 6 | Tubb5 | -0.035 | Old |
| 0.578458 | 0.065312 | 0.169 | 0.154 | 1 | 6 | Xpo7 | 0.015 | Old |
| 0.57887 | 0.211009 | 0.214 | 0.197 | 1 | 6 | Hmox2 | 0.017 | Old |
| 0.578988 | 0.425768 | 0.08 | 0.07 | 1 | 6 | Mfsd6 | 0.01 | Old |
| 0.579111 | 0.307838 | 0.234 | 0.219 | 1 | 6 | Fip1l1 | 0.015 | Old |
| 0.579252 | 0.230871 | 0.126 | 0.114 | 1 | 6 | Vps4a | 0.012 | Old |
| 0.579442 | 0.051859 | 0.789 | 0.829 | 1 | 6 | Tpm3 | -0.04 | Old |
| 0.579498 | 0.205976 | 0.286 | 0.266 | 1 | 6 | Chmp5 | 0.02 | Old |
| 0.579565 | 0.410549 | 0.089 | 0.1 | 1 | 6 | Serpinb6a | -0.011 | Old |
| 0.579593 | 0.074235 | 0.734 | 0.751 | 1 | 6 | Hsp90aa1 | -0.017 | Old |
| 0.579629 | 0.147495 | 0.363 | 0.348 | 1 | 6 | Mapk14 | 0.015 | Old |
| 0.580111 | 0.503714 | 0.046 | 0.038 | 1 | 6 | Sergef | 0.008 | Old |
| 0.58028 | 0.124095 | 0.089 | 0.102 | 1 | 6 | Arap2 | -0.013 | Old |
| 0.5812 | 0.062236 | 0.169 | 0.187 | 1 | 6 | Casp3 | -0.018 | Old |
| 0.581854 | 0.115622 | 0.32 | 0.301 | 1 | 6 | Cd84 | 0.019 | Old |
| 0.58205 | 0.286233 | 0.223 | 0.212 | 1 | 6 | Stt3b | 0.011 | Old |
| 0.582279 | 0.018899 | 0.037 | 0.03 | 1 | 6 | Tecpr2 | 0.007 | Old |
| 0.58229 | 0.413317 | 0.066 | 0.057 | 1 | 6 | Med24 | 0.009 | Old |
| 0.582299 | 0.305394 | 0.186 | 0.174 | 1 | 6 | Pigk | 0.012 | Old |
| 0.58236 | 0.199867 | 0.031 | 0.025 | 1 | 6 | Stbd1 | 0.006 | Old |
| 0.582559 | 0.155494 | 0.046 | 0.038 | 1 | 6 | Capn15 | 0.008 | Old |
| 0.58282 | 0.117896 | 0.114 | 0.127 | 1 | 6 | Pgs1 | -0.013 | Old |
| 0.582946 | 0.288933 | 0.194 | 0.182 | 1 | 6 | Mrps16 | 0.012 | Old |
| 0.583276 | 0.411467 | 0.114 | 0.104 | 1 | 6 | Sirpb1b | 0.01 | Old |
| 0.583313 | 0.404379 | 0.066 | 0.057 | 1 | 6 | Rhno1 | 0.009 | Old |
| 0.583451 | 0.133556 | 0.28 | 0.263 | 1 | 6 | Rtcb | 0.017 | Old |
| 0.58346 | 0.161885 | 0.371 | 0.355 | 1 | 6 | Eif4g3 | 0.016 | Old |
| 0.583493 | 0.002679 | 0.223 | 0.237 | 1 | 6 | Elk4 | -0.014 | Old |
| 0.583586 | 0.050912 | 0.389 | 0.376 | 1 | 6 | Ndufb8 | 0.013 | Old |
| 0.58377 | 0.35468 | 0.074 | 0.065 | 1 | 6 | Htatip2 | 0.009 | Old |
| 0.583883 | 0.221046 | 0.137 | 0.152 | 1 | 6 | Kidins220 | -0.015 | Old |
| 0.584144 | 0.092885 | 0.169 | 0.154 | 1 | 6 | Acox3 | 0.015 | Old |
| 0.584578 | 0.043008 | 0.091 | 0.104 | 1 | 6 | Oxr1 | -0.013 | Old |
| 0.584613 | 0.203303 | 0.171 | 0.156 | 1 | 6 | Dph3 | 0.015 | Old |
| 0.58472 | 0.155605 | 0.229 | 0.216 | 1 | 6 | Paics | 0.013 | Old |
| 0.584926 | 0.555205 | 0.037 | 0.045 | 1 | 6 | Ripor2 | -0.008 | Old |
| 0.585008 | 0.323832 | 0.103 | 0.092 | 1 | 6 | Brix1 | 0.011 | Old |
| 0.585018 | 0.179268 | 0.029 | 0.035 | 1 | 6 | Zfp444 | -0.006 | Old |

| | | | | | | | | |
|---|---|---|---|---|---|---|---|---|
| 0.585087 | 0.450812 | 0.054 | 0.047 | 1 | 6 | Tmed8 | 0.007 | Old |
| 0.58509 | 0.549077 | 0.043 | 0.052 | 1 | 6 | 3110082I1 | -0.009 | Old |
| 0.585502 | 0.163673 | 0.046 | 0.038 | 1 | 6 | Rcc1l | 0.008 | Old |
| 0.585587 | 0.015551 | 0.351 | 0.365 | 1 | 6 | Dusp11 | -0.014 | Old |
| 0.585617 | 0.436467 | 0.089 | 0.079 | 1 | 6 | Arid3b | 0.01 | Old |
| 0.585617 | 0.114834 | 0.089 | 0.079 | 1 | 6 | Selenom | 0.01 | Old |
| 0.585679 | 0.123978 | 0.183 | 0.201 | 1 | 6 | Tgs1 | -0.018 | Old |
| 0.585908 | 0.071515 | 0.209 | 0.224 | 1 | 6 | Mga | -0.015 | Old |
| 0.586084 | 0.181351 | 0.063 | 0.072 | 1 | 6 | Pde4a | -0.009 | Old |
| 0.587282 | 0.175096 | 0.146 | 0.161 | 1 | 6 | Kansl3 | -0.015 | Old |
| 0.587547 | 0.14921 | 0.52 | 0.502 | 1 | 6 | 7-Sep | 0.018 | Old |
| 0.587652 | 0.118615 | 0.094 | 0.084 | 1 | 6 | Sesn2 | 0.01 | Old |
| 0.588237 | 0.048807 | 0.066 | 0.057 | 1 | 6 | Klhdc2 | 0.009 | Old |
| 0.588302 | 0.292866 | 0.143 | 0.132 | 1 | 6 | Vgll4 | 0.011 | Old |
| 0.588306 | 0.129377 | 0.429 | 0.457 | 1 | 6 | Clint1 | -0.028 | Old |
| 0.588823 | 0.137878 | 0.046 | 0.054 | 1 | 6 | Mpv17 | -0.008 | Old |
| 0.588931 | 0.199665 | 0.06 | 0.052 | 1 | 6 | Sugp2 | 0.008 | Old |
| 0.589069 | 0.585651 | 0.069 | 0.06 | 1 | 6 | Ndrg1 | 0.009 | Old |
| 0.589737 | 0.049159 | 0.174 | 0.189 | 1 | 6 | Ppt1 | -0.015 | Old |
| 0.590268 | 0.123417 | 0.129 | 0.115 | 1 | 6 | Wdr43 | 0.014 | Old |
| 0.590272 | 1.311221 | 0.034 | 0.028 | 1 | 6 | Tcf7l2 | 0.006 | Old |
| 0.590396 | 0.144262 | 0.194 | 0.174 | 1 | 6 | Ehmt2 | 0.02 | Old |
| 0.590489 | 0.621604 | 0.054 | 0.047 | 1 | 6 | Bex3 | 0.007 | Old |
| 0.590702 | 0.367641 | 0.109 | 0.097 | 1 | 6 | Heg1 | 0.012 | Old |
| 0.590752 | 0.197761 | 0.246 | 0.229 | 1 | 6 | Ttc3 | 0.017 | Old |
| 0.590944 | 0.162048 | 0.2 | 0.187 | 1 | 6 | Plbd2 | 0.013 | Old |
| 0.590948 | 0.167221 | 0.089 | 0.079 | 1 | 6 | Nudt19 | 0.01 | Old |
| 0.591228 | 0.806878 | 0.049 | 0.042 | 1 | 6 | Il27ra | 0.007 | Old |
| 0.591416 | 0.23991 | 0.123 | 0.112 | 1 | 6 | Slc25a1 | 0.011 | Old |
| 0.592196 | 0.008535 | 0.089 | 0.079 | 1 | 6 | Dhdh | 0.01 | Old |
| 0.592277 | 0.086878 | 0.474 | 0.467 | 1 | 6 | Ifitm6 | 0.007 | Old |
| 0.592345 | 0.013646 | 0.529 | 0.548 | 1 | 6 | 5031439G( | -0.019 | Old |
| 0.592548 | 1.080998 | 0.034 | 0.028 | 1 | 6 | 4921524J1 | 0.006 | Old |
| 0.593055 | 0.16948 | 0.126 | 0.115 | 1 | 6 | Fbxw7 | 0.011 | Old |
| 0.593437 | 0.298599 | 0.151 | 0.139 | 1 | 6 | Nsd2 | 0.012 | Old |
| 0.593733 | 0.112957 | 0.586 | 0.572 | 1 | 6 | Ezr | 0.014 | Old |
| 0.593771 | 0.3119 | 0.174 | 0.162 | 1 | 6 | Exoc3 | 0.012 | Old |
| 0.593957 | 0.088906 | 0.126 | 0.139 | 1 | 6 | Fasn | -0.013 | Old |
| 0.59455 | 0.364587 | 0.106 | 0.095 | 1 | 6 | Dtx3l | 0.011 | Old |
| 0.594553 | 0.061264 | 0.24 | 0.258 | 1 | 6 | Ttyh3 | -0.018 | Old |
| 0.594604 | 0.262342 | 0.04 | 0.033 | 1 | 6 | Slc52a2 | 0.007 | Old |
| 0.59486 | 0.255973 | 0.274 | 0.259 | 1 | 6 | Polr2m | 0.015 | Old |
| 0.594927 | 0.007493 | 0.029 | 0.035 | 1 | 6 | Alg3 | -0.006 | Old |
| 0.595048 | 0.088932 | 0.049 | 0.057 | 1 | 6 | Slc25a13 | -0.008 | Old |
| 0.595096 | 0.222192 | 0.063 | 0.054 | 1 | 6 | Elac1 | 0.009 | Old |

| | | | | | | | | |
|---|---|---|---|---|---|---|---|---|
| 0.59574 | 0.39771 | 0.063 | 0.055 | 1 | 6 | Trp53bp2 | 0.008 | Old |
| 0.59577 | 0.170909 | 0.063 | 0.072 | 1 | 6 | Tcn2 | -0.009 | Old |
| 0.59587 | 0.034237 | 0.143 | 0.156 | 1 | 6 | Rnf10 | -0.013 | Old |
| 0.596046 | 0.251903 | 0.32 | 0.304 | 1 | 6 | Zdhhc20 | 0.016 | Old |
| 0.596135 | 0.254902 | 0.06 | 0.069 | 1 | 6 | Senp3 | -0.009 | Old |
| 0.596582 | 0.501189 | 0.063 | 0.055 | 1 | 6 | Dtnbp1 | 0.008 | Old |
| 0.59666 | 0.330498 | 0.089 | 0.079 | 1 | 6 | Rit1 | 0.01 | Old |
| 0.596819 | 0.037298 | 0.051 | 0.043 | 1 | 6 | Ttc4 | 0.008 | Old |
| 0.597111 | 0.025644 | 0.46 | 0.435 | 1 | 6 | Dbnl | 0.025 | Old |
| 0.597326 | 0.094869 | 0.063 | 0.072 | 1 | 6 | Hjurp | -0.009 | Old |
| 0.597436 | 0.115953 | 0.186 | 0.172 | 1 | 6 | Nck1 | 0.014 | Old |
| 0.597546 | 0.09379 | 0.194 | 0.182 | 1 | 6 | Pik3cg | 0.012 | Old |
| 0.597645 | 0.109835 | 0.491 | 0.507 | 1 | 6 | Atp5j2 | -0.016 | Old |
| 0.59818 | 0.508814 | 0.054 | 0.047 | 1 | 6 | B230307C2 | 0.007 | Old |
| 0.598273 | 0.102309 | 0.706 | 0.702 | 1 | 6 | Rrbp1 | 0.004 | Old |
| 0.59883 | 0.236889 | 0.08 | 0.09 | 1 | 6 | Ripk2 | -0.01 | Old |
| 0.598833 | 0.166073 | 0.046 | 0.038 | 1 | 6 | Trub2 | 0.008 | Old |
| 0.598925 | 0.104578 | 0.149 | 0.162 | 1 | 6 | Ocrl | -0.013 | Old |
| 0.59929 | 0.059632 | 0.454 | 0.477 | 1 | 6 | Mia2 | -0.023 | Old |
| 0.599342 | 0.005358 | 0.051 | 0.06 | 1 | 6 | Mtif2 | -0.009 | Old |
| 0.599546 | 0.154077 | 0.091 | 0.102 | 1 | 6 | Tbk1 | -0.011 | Old |
| 0.600915 | 0.044444 | 0.271 | 0.254 | 1 | 6 | Dnajc19 | 0.017 | Old |
| 0.60107 | 0.084328 | 0.4 | 0.44 | 1 | 6 | Aebp2 | -0.04 | Old |
| 0.60123 | 0.189223 | 0.06 | 0.052 | 1 | 6 | Plod1 | 0.008 | Old |
| 0.601644 | 0.103555 | 0.223 | 0.237 | 1 | 6 | Ddx46 | -0.014 | Old |
| 0.601922 | 0.06521 | 0.114 | 0.102 | 1 | 6 | Mknk1 | 0.012 | Old |
| 0.602554 | 0.249398 | 0.04 | 0.033 | 1 | 6 | Phospho2 | 0.007 | Old |
| 0.602843 | 0.949597 | 0.034 | 0.028 | 1 | 6 | C030034I2 | 0.006 | Old |
| 0.603475 | 0.156188 | 0.606 | 0.595 | 1 | 6 | Tmed10 | 0.011 | Old |
| 0.603475 | 0.360819 | 0.089 | 0.079 | 1 | 6 | Ccdc127 | 0.01 | Old |
| 0.603617 | 0.142921 | 0.04 | 0.033 | 1 | 6 | Endog | 0.007 | Old |
| 0.604013 | 0.054843 | 0.123 | 0.109 | 1 | 6 | Mut | 0.014 | Old |
| 0.604416 | 0.851501 | 0.04 | 0.033 | 1 | 6 | Cog2 | 0.007 | Old |
| 0.604444 | 0.131159 | 0.097 | 0.089 | 1 | 6 | Scap | 0.008 | Old |
| 0.604612 | 0.164271 | 0.337 | 0.324 | 1 | 6 | Dnajc7 | 0.013 | Old |
| 0.604845 | 0.32093 | 0.134 | 0.124 | 1 | 6 | Cspp1 | 0.01 | Old |
| 0.605111 | 0.079281 | 0.163 | 0.177 | 1 | 6 | Vps13c | -0.014 | Old |
| 0.605129 | 0.181592 | 0.086 | 0.095 | 1 | 6 | Acsl1 | -0.009 | Old |
| 0.605163 | 0.247836 | 0.209 | 0.194 | 1 | 6 | Wdr37 | 0.015 | Old |
| 0.605203 | 0.18914 | 0.066 | 0.057 | 1 | 6 | Tipin | 0.009 | Old |
| 0.605432 | 0.105997 | 0.046 | 0.054 | 1 | 6 | Ptk2 | -0.008 | Old |
| 0.605519 | 0.05593 | 0.154 | 0.167 | 1 | 6 | Plpbp | -0.013 | Old |
| 0.605691 | 0.110717 | 0.257 | 0.276 | 1 | 6 | Nucb1 | -0.019 | Old |
| 0.605737 | 0.178503 | 0.234 | 0.219 | 1 | 6 | Homer1 | 0.015 | Old |
| 0.605874 | 0.045658 | 0.12 | 0.107 | 1 | 6 | Cdkn2aipnl | 0.013 | Old |

| | | | | | | | | |
|---|---|---|---|---|---|---|---|---|
| 0.605917 | 0.258942 | 0.054 | 0.047 | 1 | 6 | Bloc1s1 | 0.007 | Old |
| 0.606136 | 0.027619 | 0.231 | 0.249 | 1 | 6 | Kdm4a | -0.018 | Old |
| 0.606173 | 0.001925 | 0.197 | 0.179 | 1 | 6 | Lrrc59 | 0.018 | Old |
| 0.606263 | 0.114618 | 0.026 | 0.032 | 1 | 6 | Aurkb | -0.006 | Old |
| 0.606298 | 0.339156 | 0.246 | 0.232 | 1 | 6 | Abhd17b | 0.014 | Old |
| 0.606419 | 0.122334 | 0.106 | 0.095 | 1 | 6 | Smad5 | 0.011 | Old |
| 0.606506 | 0.387293 | 0.097 | 0.087 | 1 | 6 | Klhl2 | 0.01 | Old |
| 0.606657 | 0.20639 | 0.069 | 0.06 | 1 | 6 | Wasl | 0.009 | Old |
| 0.606798 | 0.135232 | 0.46 | 0.452 | 1 | 6 | Bsg | 0.008 | Old |
| 0.606869 | 0.490001 | 0.034 | 0.028 | 1 | 6 | Coprs | 0.006 | Old |
| 0.60687 | 0.255916 | 0.054 | 0.064 | 1 | 6 | Mterf3 | -0.01 | Old |
| 0.607221 | 0.198878 | 0.023 | 0.028 | 1 | 6 | Haus4 | -0.005 | Old |
| 0.607221 | 0.111924 | 0.023 | 0.028 | 1 | 6 | Ctdspl | -0.005 | Old |
| 0.607445 | 0.293382 | 0.034 | 0.028 | 1 | 6 | Zfp934 | 0.006 | Old |
| 0.607505 | 0.281599 | 0.194 | 0.179 | 1 | 6 | Gigyf2 | 0.015 | Old |
| 0.607675 | 0.145547 | 0.069 | 0.06 | 1 | 6 | Coq5 | 0.009 | Old |
| 0.607711 | 0.319959 | 0.043 | 0.05 | 1 | 6 | Atg16l2 | -0.007 | Old |
| 0.607841 | 0.03012 | 0.023 | 0.028 | 1 | 6 | Trp53rka | -0.005 | Old |
| 0.608216 | 1.267966 | 0.029 | 0.023 | 1 | 6 | Tom1l2 | 0.006 | Old |
| 0.608486 | 0.348732 | 0.131 | 0.122 | 1 | 6 | Selenoh | 0.009 | Old |
| 0.608658 | 0.361181 | 0.054 | 0.047 | 1 | 6 | Zfp771 | 0.007 | Old |
| 0.609018 | 0.163605 | 0.14 | 0.127 | 1 | 6 | Cenpb | 0.013 | Old |
| 0.609567 | 0.269204 | 0.16 | 0.147 | 1 | 6 | Hivep1 | 0.013 | Old |
| 0.60978 | 0.030926 | 0.389 | 0.37 | 1 | 6 | Amfr | 0.019 | Old |
| 0.609838 | 0.014485 | 0.254 | 0.268 | 1 | 6 | Cmtm3 | -0.014 | Old |
| 0.609926 | 0.059742 | 0.071 | 0.062 | 1 | 6 | Reep4 | 0.009 | Old |
| 0.610113 | 0.670898 | 0.029 | 0.023 | 1 | 6 | Coa4 | 0.006 | Old |
| 0.610288 | 0.048513 | 0.083 | 0.074 | 1 | 6 | Gpatch2 | 0.009 | Old |
| 0.610406 | 0.09407 | 0.283 | 0.268 | 1 | 6 | Gmfb | 0.015 | Old |
| 0.610555 | 0.114388 | 0.214 | 0.199 | 1 | 6 | Map2k1 | 0.015 | Old |
| 0.610615 | 0.084452 | 0.751 | 0.796 | 1 | 6 | Ier2 | -0.045 | Old |
| 0.61066 | 0.840765 | 0.043 | 0.037 | 1 | 6 | Dgkh | 0.006 | Old |
| 0.610715 | 0.074863 | 0.103 | 0.092 | 1 | 6 | Gtf3a | 0.011 | Old |
| 0.610789 | 0.524701 | 0.103 | 0.094 | 1 | 6 | Trim33 | 0.009 | Old |
| 0.610802 | 0.055693 | 0.02 | 0.025 | 1 | 6 | Lyve1 | -0.005 | Old |
| 0.61092 | 0.079485 | 0.117 | 0.129 | 1 | 6 | Slc25a30 | -0.012 | Old |
| 0.611112 | 0.082826 | 0.166 | 0.152 | 1 | 6 | Ube2d1 | 0.014 | Old |
| 0.611222 | 0.097845 | 0.306 | 0.298 | 1 | 6 | Klf7 | 0.008 | Old |
| 0.611321 | 0.081624 | 0.194 | 0.179 | 1 | 6 | Tspan14 | 0.015 | Old |
| 0.611379 | 0.737466 | 0.029 | 0.023 | 1 | 6 | Eefsec | 0.006 | Old |
| 0.612373 | 0.090809 | 0.229 | 0.212 | 1 | 6 | Acbd5 | 0.017 | Old |
| 0.612692 | 0.042451 | 0.04 | 0.033 | 1 | 6 | Cenpq | 0.007 | Old |
| 0.612993 | 0.185979 | 0.063 | 0.072 | 1 | 6 | Trp53bp1 | -0.009 | Old |
| 0.613218 | 0.284437 | 0.034 | 0.028 | 1 | 6 | Mrps31 | 0.006 | Old |
| 0.613581 | 0.069328 | 0.289 | 0.301 | 1 | 6 | Prdx6 | -0.012 | Old |

| | | | | | | | | |
|---|---|---|---|---|---|---|---|---|
| 0.613693 | 0.15761 | 0.274 | 0.261 | 1 | 6 | Erlin1 | 0.013 | Old |
| 0.613811 | 0.250104 | 0.103 | 0.094 | 1 | 6 | Gtf2f2 | 0.009 | Old |
| 0.614528 | 0.282455 | 0.049 | 0.042 | 1 | 6 | Card9 | 0.007 | Old |
| 0.61455 | 0.716738 | 0.029 | 0.023 | 1 | 6 | Sort1 | 0.006 | Old |
| 0.615027 | 0.201448 | 0.069 | 0.06 | 1 | 6 | Orc2 | 0.009 | Old |
| 0.61529 | 0.032711 | 0.963 | 0.97 | 1 | 6 | Rpl8 | -0.007 | Old |
| 0.615598 | 0.200895 | 0.146 | 0.134 | 1 | 6 | Epb41 | 0.012 | Old |
| 0.615861 | 0.102569 | 0.449 | 0.463 | 1 | 6 | Nipbl | -0.014 | Old |
| 0.615916 | 0.133208 | 0.023 | 0.028 | 1 | 6 | Setd6 | -0.005 | Old |
| 0.616316 | 0.133839 | 0.046 | 0.038 | 1 | 6 | Zfp26 | 0.008 | Old |
| 0.616659 | 0.121991 | 0.134 | 0.122 | 1 | 6 | Plod3 | 0.012 | Old |
| 0.616832 | 0.233828 | 0.34 | 0.321 | 1 | 6 | Pdcd4 | 0.019 | Old |
| 0.616929 | 0.182765 | 0.569 | 0.574 | 1 | 6 | Far1 | -0.005 | Old |
| 0.617092 | 0.623665 | 0.029 | 0.023 | 1 | 6 | Kbtbd3 | 0.006 | Old |
| 0.61737 | 0.22784 | 0.154 | 0.142 | 1 | 6 | Cdc123 | 0.012 | Old |
| 0.617854 | 0.403807 | 0.034 | 0.028 | 1 | 6 | Zfp758 | 0.006 | Old |
| 0.618088 | 0.590701 | 0.02 | 0.025 | 1 | 6 | Pomt2 | -0.005 | Old |
| 0.618188 | 0.493695 | 0.103 | 0.094 | 1 | 6 | 2310033P0 | 0.009 | Old |
| 0.618365 | 0.405462 | 0.029 | 0.023 | 1 | 6 | Zfp971 | 0.006 | Old |
| 0.619044 | 0.070834 | 0.231 | 0.214 | 1 | 6 | Dcaf7 | 0.017 | Old |
| 0.6195 | 0.07921 | 0.423 | 0.401 | 1 | 6 | Itsn2 | 0.022 | Old |
| 0.619612 | 0.254762 | 0.143 | 0.156 | 1 | 6 | Itpr2 | -0.013 | Old |
| 0.619714 | 0.04714 | 0.174 | 0.189 | 1 | 6 | Atxn2 | -0.015 | Old |
| 0.62003 | 0.039258 | 0.146 | 0.159 | 1 | 6 | Gsto1 | -0.013 | Old |
| 0.620062 | 0.071174 | 0.597 | 0.572 | 1 | 6 | Rbm25 | 0.025 | Old |
| 0.620081 | 0.309225 | 0.02 | 0.025 | 1 | 6 | Trub1 | -0.005 | Old |
| 0.620178 | 0.122377 | 0.034 | 0.028 | 1 | 6 | Mgme1 | 0.006 | Old |
| 0.620234 | 0.101729 | 0.234 | 0.249 | 1 | 6 | Smarca4 | -0.015 | Old |
| 0.620276 | 0.345336 | 0.029 | 0.023 | 1 | 6 | Actr6 | 0.006 | Old |
| 0.620709 | 0.006536 | 0.22 | 0.234 | 1 | 6 | Sun2 | -0.014 | Old |
| 0.62133 | 0.022049 | 0.126 | 0.114 | 1 | 6 | Snrpa1 | 0.012 | Old |
| 0.621341 | 0.18315 | 0.034 | 0.028 | 1 | 6 | Aig1 | 0.006 | Old |
| 0.62178 | 0.662891 | 0.051 | 0.045 | 1 | 6 | Sirpb1a | 0.006 | Old |
| 0.622064 | 0.157656 | 0.494 | 0.473 | 1 | 6 | Bnip2 | 0.021 | Old |
| 0.622241 | 0.077552 | 0.163 | 0.149 | 1 | 6 | Hmg20b | 0.014 | Old |
| 0.622301 | 0.062715 | 0.186 | 0.199 | 1 | 6 | Ctcf | -0.013 | Old |
| 0.622306 | 0.166356 | 0.12 | 0.109 | 1 | 6 | Glo1 | 0.011 | Old |
| 0.622541 | 0.002595 | 0.257 | 0.278 | 1 | 6 | Sema4d | -0.021 | Old |
| 0.622778 | 0.785378 | 0.043 | 0.037 | 1 | 6 | Sgsm2 | 0.006 | Old |
| 0.623302 | 0.004109 | 0.083 | 0.092 | 1 | 6 | Pias4 | -0.009 | Old |
| 0.623503 | 0.001074 | 0.103 | 0.114 | 1 | 6 | Zfp62 | -0.011 | Old |
| 0.623754 | 0.114255 | 0.194 | 0.181 | 1 | 6 | Gnl3 | 0.013 | Old |
| 0.623984 | 0.705155 | 0.057 | 0.05 | 1 | 6 | Bcl2l1 | 0.007 | Old |
| 0.624181 | 0.245537 | 0.14 | 0.129 | 1 | 6 | Smarcb1 | 0.011 | Old |
| 0.624254 | 0.19483 | 0.077 | 0.069 | 1 | 6 | Utp6 | 0.008 | Old |

| | | | | | | | | |
|---|---|---|---|---|---|---|---|---|
| 0.624333 | 0.889416 | 0.043 | 0.037 | 1 | 6 | Gabbr1 | 0.006 | Old |
| 0.624649 | 0.249412 | 0.123 | 0.112 | 1 | 6 | Tmem230 | 0.011 | Old |
| 0.625554 | 0.313557 | 0.111 | 0.102 | 1 | 6 | AI839979 | 0.009 | Old |
| 0.625716 | 0.187415 | 0.063 | 0.055 | 1 | 6 | Zfp991 | 0.008 | Old |
| 0.625798 | 0.049368 | 0.469 | 0.455 | 1 | 6 | Romo1 | 0.014 | Old |
| 0.625809 | 0.246045 | 0.18 | 0.166 | 1 | 6 | Cklf | 0.014 | Old |
| 0.625882 | 0.63866 | 0.023 | 0.018 | 1 | 6 | Hells | 0.005 | Old |
| 0.625931 | 0.347466 | 0.063 | 0.055 | 1 | 6 | Zdhhc6 | 0.008 | Old |
| 0.626 | 0.040952 | 0.263 | 0.269 | 1 | 6 | Mrpl23 | -0.006 | Old |
| 0.626025 | 0.468967 | 0.029 | 0.023 | 1 | 6 | Pdss2 | 0.006 | Old |
| 0.626559 | 0.081636 | 0.194 | 0.179 | 1 | 6 | Psmb2 | 0.015 | Old |
| 0.626712 | 0.366874 | 0.103 | 0.114 | 1 | 6 | Slc9a9 | -0.011 | Old |
| 0.627102 | 0.699844 | 0.066 | 0.059 | 1 | 6 | Gins4 | 0.007 | Old |
| 0.627149 | 0.176268 | 0.08 | 0.07 | 1 | 6 | Taf11 | 0.01 | Old |
| 0.627186 | 0.162825 | 0.094 | 0.085 | 1 | 6 | Borcs8 | 0.009 | Old |
| 0.627369 | 0.244699 | 0.194 | 0.184 | 1 | 6 | Dynlt3 | 0.01 | Old |
| 0.627549 | 0.068435 | 0.174 | 0.159 | 1 | 6 | Suco | 0.015 | Old |
| 0.628128 | 0.086148 | 0.123 | 0.11 | 1 | 6 | Lpin1 | 0.013 | Old |
| 0.62826 | 0.27268 | 0.16 | 0.176 | 1 | 6 | Rapgef2 | -0.016 | Old |
| 0.628269 | 0.009149 | 0.111 | 0.1 | 1 | 6 | Enoph1 | 0.011 | Old |
| 0.628512 | 0.154068 | 0.063 | 0.055 | 1 | 6 | Tmx2 | 0.008 | Old |
| 0.628587 | 0.078457 | 0.029 | 0.023 | 1 | 6 | Zc3h3 | 0.006 | Old |
| 0.628878 | 0.132921 | 0.554 | 0.56 | 1 | 6 | 9-Sep | -0.006 | Old |
| 0.628878 | 0.052728 | 0.163 | 0.147 | 1 | 6 | Mrpl18 | 0.016 | Old |
| 0.62918 | 0.065735 | 0.163 | 0.151 | 1 | 6 | Csrp1 | 0.012 | Old |
| 0.629236 | 0.136444 | 0.077 | 0.087 | 1 | 6 | Polr3gl | -0.01 | Old |
| 0.629266 | 0.61767 | 0.043 | 0.037 | 1 | 6 | Prkca | 0.006 | Old |
| 0.629392 | 0.374259 | 0.143 | 0.156 | 1 | 6 | Grap2 | -0.013 | Old |
| 0.629423 | 0.162597 | 0.437 | 0.426 | 1 | 6 | Fam120a | 0.011 | Old |
| 0.629872 | 0.515862 | 0.111 | 0.122 | 1 | 6 | Maf | -0.011 | Old |
| 0.629892 | 0.19591 | 0.263 | 0.251 | 1 | 6 | Mdfic | 0.012 | Old |
| 0.629984 | 1.475917 | 0.017 | 0.013 | 1 | 6 | 9130008F2 | 0.004 | Old |
| 0.630192 | 0.41004 | 0.017 | 0.022 | 1 | 6 | Ddx28 | -0.005 | Old |
| 0.630513 | 0.114295 | 0.143 | 0.156 | 1 | 6 | Hspa4l | -0.013 | Old |
| 0.630717 | 0.015283 | 0.786 | 0.803 | 1 | 6 | Pcbp2 | -0.017 | Old |
| 0.631064 | 0.149103 | 0.423 | 0.416 | 1 | 6 | Ywhag | 0.007 | Old |
| 0.631537 | 0.539128 | 0.143 | 0.135 | 1 | 6 | Gabpa | 0.008 | Old |
| 0.631754 | 1.011496 | 0.011 | 0.008 | 1 | 6 | Elp6 | 0.003 | Old |
| 0.631854 | 0.311079 | 0.243 | 0.234 | 1 | 6 | Ppp4r3a | 0.009 | Old |
| 0.632795 | 1.269901 | 0.011 | 0.008 | 1 | 6 | Fam120aos | 0.003 | Old |
| 0.632795 | 0.617791 | 0.011 | 0.008 | 1 | 6 | Spaca6 | 0.003 | Old |
| 0.63291 | 0.011309 | 0.054 | 0.062 | 1 | 6 | Polk | -0.008 | Old |
| 0.632934 | 0.015665 | 0.18 | 0.164 | 1 | 6 | Herc2 | 0.016 | Old |
| 0.633082 | 0.136471 | 0.029 | 0.023 | 1 | 6 | B4galt7 | 0.006 | Old |
| 0.63322 | 0.27636 | 0.169 | 0.157 | 1 | 6 | Cltb | 0.012 | Old |

| | | | | | | | | |
|---|---|---|---|---|---|---|---|---|
| 0.633728 | 0.028804 | 0.214 | 0.229 | 1 | 6 | Stxbp2 | -0.015 | Old |
| 0.633837 | 1.312668 | 0.011 | 0.008 | 1 | 6 | Adarb1 | 0.003 | Old |
| 0.633837 | 1.154852 | 0.011 | 0.008 | 1 | 6 | Mfsd4b4 | 0.003 | Old |
| 0.634169 | 0.756471 | 0.017 | 0.013 | 1 | 6 | Mtcp1 | 0.004 | Old |
| 0.634412 | 0.118104 | 0.334 | 0.358 | 1 | 6 | Nsd1 | -0.024 | Old |
| 0.634516 | 0.256899 | 0.023 | 0.018 | 1 | 6 | Mrps27 | 0.005 | Old |
| 0.634573 | 0.399908 | 0.08 | 0.072 | 1 | 6 | Ubqln4 | 0.008 | Old |
| 0.634573 | 1.432154 | 0.037 | 0.032 | 1 | 6 | Ercc6 | 0.005 | Old |
| 0.63488 | 0.896326 | 0.011 | 0.008 | 1 | 6 | Gorab | 0.003 | Old |
| 0.635401 | 0.051161 | 0.089 | 0.099 | 1 | 6 | Med7 | -0.01 | Old |
| 0.635609 | 0.140625 | 0.571 | 0.562 | 1 | 6 | Ccdc50 | 0.009 | Old |
| 0.635662 | 0.356374 | 0.091 | 0.082 | 1 | 6 | Kdm5b | 0.009 | Old |
| 0.635705 | 0.439838 | 0.057 | 0.05 | 1 | 6 | Gphn | 0.007 | Old |
| 0.635876 | 0.193366 | 0.111 | 0.102 | 1 | 6 | Ccnc | 0.009 | Old |
| 0.635923 | 1.039576 | 0.011 | 0.008 | 1 | 6 | Tmem81 | 0.003 | Old |
| 0.635923 | 0.8337 | 0.011 | 0.008 | 1 | 6 | Glcci1 | 0.003 | Old |
| 0.635923 | 0.530589 | 0.011 | 0.008 | 1 | 6 | 6330403L0 | 0.003 | Old |
| 0.635979 | 0.005767 | 0.223 | 0.209 | 1 | 6 | Smarca2 | 0.014 | Old |
| 0.636301 | 0.280061 | 0.029 | 0.023 | 1 | 6 | Vars2 | 0.006 | Old |
| 0.636462 | 0.044998 | 0.36 | 0.385 | 1 | 6 | Tpd52 | -0.025 | Old |
| 0.636544 | 0.151598 | 0.169 | 0.156 | 1 | 6 | Vav2 | 0.013 | Old |
| 0.636686 | 0.574242 | 0.017 | 0.013 | 1 | 6 | Kbtbd4 | 0.004 | Old |
| 0.636806 | 0.29497 | 0.24 | 0.226 | 1 | 6 | Rab35 | 0.014 | Old |
| 0.636946 | 0.053704 | 0.029 | 0.023 | 1 | 6 | Rfc5 | 0.006 | Old |
| 0.636967 | 0.621615 | 0.011 | 0.008 | 1 | 6 | Cdkn1c | 0.003 | Old |
| 0.636967 | 0.365341 | 0.011 | 0.008 | 1 | 6 | Acad10 | 0.003 | Old |
| 0.636967 | 0.251772 | 0.011 | 0.008 | 1 | 6 | Frg2f1 | 0.003 | Old |
| 0.636967 | 0.223186 | 0.011 | 0.008 | 1 | 6 | Il2 | 0.003 | Old |
| 0.636995 | 0.323825 | 0.249 | 0.237 | 1 | 6 | Tsc22d1 | 0.012 | Old |
| 0.637405 | 0.250803 | 0.017 | 0.022 | 1 | 6 | Sgcb | -0.005 | Old |
| 0.637738 | 0.11206 | 0.494 | 0.477 | 1 | 6 | Runx1 | 0.017 | Old |
| 0.638007 | 0.241082 | 0.057 | 0.065 | 1 | 6 | Sugp1 | -0.008 | Old |
| 0.638013 | 0.107546 | 0.423 | 0.406 | 1 | 6 | Rbx1 | 0.017 | Old |
| 0.638087 | 0.034606 | 0.669 | 0.689 | 1 | 6 | Pcbp1 | -0.02 | Old |
| 0.638095 | 1.05283 | 0.037 | 0.043 | 1 | 6 | Ttc28 | -0.006 | Old |
| 0.638236 | 0.423471 | 0.029 | 0.023 | 1 | 6 | Rhox8 | 0.006 | Old |
| 0.638427 | 0.17645 | 0.3 | 0.303 | 1 | 6 | Tsc22d2 | -0.003 | Old |
| 0.638821 | 0.152372 | 0.334 | 0.328 | 1 | 6 | Klhl24 | 0.006 | Old |
| 0.638952 | 0.315672 | 0.011 | 0.015 | 1 | 6 | Immp2l | -0.004 | Old |
| 0.639008 | 0.076975 | 0.671 | 0.711 | 1 | 6 | Tma7 | -0.04 | Old |
| 0.639058 | 0.930802 | 0.011 | 0.008 | 1 | 6 | Zfp408 | 0.003 | Old |
| 0.639058 | 0.78841 | 0.011 | 0.008 | 1 | 6 | Mmp12 | 0.003 | Old |
| 0.639058 | 0.314904 | 0.011 | 0.008 | 1 | 6 | Cmtr2 | 0.003 | Old |
| 0.639058 | 0.236329 | 0.011 | 0.008 | 1 | 6 | Spa17 | 0.003 | Old |
| 0.639058 | 0.030217 | 0.011 | 0.008 | 1 | 6 | 6230400D1 | 0.003 | Old |

| | | | | | | | | |
|---|---|---|---|---|---|---|---|---|
| 0.639117 | 0.147622 | 0.491 | 0.487 | 1 | 6 | Zfp106 | 0.004 | Old |
| 0.639207 | 0.557954 | 0.017 | 0.013 | 1 | 6 | Pex12 | 0.004 | Old |
| 0.639478 | 0.207105 | 0.114 | 0.105 | 1 | 6 | Tmem229b | 0.009 | Old |
| 0.639571 | 0.551523 | 0.08 | 0.072 | 1 | 6 | Slc9a1 | 0.008 | Old |
| 0.640048 | 0.201645 | 0.017 | 0.013 | 1 | 6 | Car9 | 0.004 | Old |
| 0.640049 | 0.29486 | 0.274 | 0.263 | 1 | 6 | Sp1 | 0.011 | Old |
| 0.640104 | 0.598118 | 0.011 | 0.008 | 1 | 6 | Tlr7 | 0.003 | Old |
| 0.640104 | 0.40489 | 0.011 | 0.008 | 1 | 6 | Insl6 | 0.003 | Old |
| 0.640104 | 0.07498 | 0.011 | 0.008 | 1 | 6 | Gab1 | 0.003 | Old |
| 0.640424 | 0.209945 | 0.071 | 0.064 | 1 | 6 | Nedd4l | 0.007 | Old |
| 0.640835 | 0.104591 | 0.063 | 0.055 | 1 | 6 | Tex30 | 0.008 | Old |
| 0.64089 | 0.976926 | 0.017 | 0.013 | 1 | 6 | Rad54l | 0.004 | Old |
| 0.641017 | 0.324687 | 0.043 | 0.037 | 1 | 6 | Gfod1 | 0.006 | Old |
| 0.641151 | 0.728479 | 0.011 | 0.008 | 1 | 6 | A430033K0 | 0.003 | Old |
| 0.641226 | 0.412278 | 0.014 | 0.018 | 1 | 6 | Mars2 | -0.004 | Old |
| 0.641226 | 0.120818 | 0.014 | 0.018 | 1 | 6 | Tmem120b | -0.004 | Old |
| 0.641496 | 0.035902 | 0.154 | 0.144 | 1 | 6 | Hdac2 | 0.01 | Old |
| 0.641587 | 0.847376 | 0.094 | 0.087 | 1 | 6 | Mcm6 | 0.007 | Old |
| 0.64166 | 0.083198 | 0.209 | 0.194 | 1 | 6 | Rtf2 | 0.015 | Old |
| 0.641663 | 0.151177 | 0.229 | 0.222 | 1 | 6 | Abca7 | 0.007 | Old |
| 0.641732 | 0.187344 | 0.017 | 0.013 | 1 | 6 | Bora | 0.004 | Old |
| 0.641808 | 0.189007 | 0.183 | 0.171 | 1 | 6 | Actr10 | 0.012 | Old |
| 0.641821 | 0.058828 | 0.506 | 0.53 | 1 | 6 | Tax1bp1 | -0.024 | Old |
| 0.641925 | 0.100036 | 0.137 | 0.127 | 1 | 6 | Ubald1 | 0.01 | Old |
| 0.642137 | 0.191418 | 0.111 | 0.102 | 1 | 6 | Hspa13 | 0.009 | Old |
| 0.642574 | 0.112379 | 0.017 | 0.013 | 1 | 6 | C1qtnf6 | 0.004 | Old |
| 0.642804 | 0.161029 | 0.014 | 0.018 | 1 | 6 | 3110009E1 | -0.004 | Old |
| 0.643248 | 0.414628 | 0.011 | 0.008 | 1 | 6 | Fastkd5 | 0.003 | Old |
| 0.643248 | 0.026009 | 0.011 | 0.008 | 1 | 6 | Usp54 | 0.003 | Old |
| 0.643417 | 1.019193 | 0.017 | 0.013 | 1 | 6 | Acyp2 | 0.004 | Old |
| 0.643417 | 0.646678 | 0.017 | 0.013 | 1 | 6 | Bcl2a1c | 0.004 | Old |
| 0.643417 | 0.321204 | 0.017 | 0.013 | 1 | 6 | Gm4707 | 0.004 | Old |
| 0.643417 | 0.28999 | 0.017 | 0.013 | 1 | 6 | Pkmyt1 | 0.004 | Old |
| 0.643417 | 0.270566 | 0.017 | 0.013 | 1 | 6 | Lrrc56 | 0.004 | Old |
| 0.643594 | 0.352046 | 0.014 | 0.018 | 1 | 6 | Thbd | -0.004 | Old |
| 0.643925 | 0.314059 | 0.023 | 0.018 | 1 | 6 | Tmem263 | 0.005 | Old |
| 0.643931 | 0.022839 | 0.086 | 0.077 | 1 | 6 | Naip6 | 0.009 | Old |
| 0.644261 | 0.58413 | 0.017 | 0.013 | 1 | 6 | Spib | 0.004 | Old |
| 0.644517 | 0.011219 | 0.054 | 0.062 | 1 | 6 | Zfp518a | -0.008 | Old |
| 0.644846 | 0.181975 | 0.211 | 0.197 | 1 | 6 | Rsl1d1 | 0.014 | Old |
| 0.64489 | 0.048869 | 0.16 | 0.145 | 1 | 6 | Hacd4 | 0.015 | Old |
| 0.645042 | 0.178887 | 0.066 | 0.059 | 1 | 6 | Frmd4a | 0.007 | Old |
| 0.645105 | 0.150063 | 0.017 | 0.013 | 1 | 6 | Mrps22 | 0.004 | Old |
| 0.645442 | 0.027756 | 0.794 | 0.828 | 1 | 6 | Anxa2 | -0.034 | Old |
| 0.645683 | 0.682219 | 0.009 | 0.012 | 1 | 6 | Btbd6 | -0.003 | Old |

| | | | | | | | | |
|---|---|---|---|---|---|---|---|---|
| 0.645862 | 0.065421 | 0.074 | 0.065 | 1 | 6 | Cfap36 | 0.009 | Old |
| 0.645945 | 0.104445 | 0.011 | 0.015 | 1 | 6 | Nradd | -0.004 | Old |
| 0.645949 | 0.238476 | 0.017 | 0.013 | 1 | 6 | Adat1 | 0.004 | Old |
| 0.645949 | 0.174687 | 0.017 | 0.013 | 1 | 6 | Prr15 | 0.004 | Old |
| 0.645965 | 0.193718 | 0.014 | 0.018 | 1 | 6 | Ick | -0.004 | Old |
| 0.64615 | 0.300822 | 0.057 | 0.05 | 1 | 6 | Fbxo8 | 0.007 | Old |
| 0.64657 | 0.145809 | 0.129 | 0.142 | 1 | 6 | Uhrf1bp1l | -0.013 | Old |
| 0.646606 | 0.197047 | 0.057 | 0.05 | 1 | 6 | Pold1 | 0.007 | Old |
| 0.64669 | 0.135107 | 0.069 | 0.077 | 1 | 6 | Zfp672 | -0.008 | Old |
| 0.646729 | 0.132892 | 0.24 | 0.224 | 1 | 6 | 2310035C2 | 0.016 | Old |
| 0.646821 | 0.447748 | 0.011 | 0.015 | 1 | 6 | Card6 | -0.004 | Old |
| 0.646821 | 0.43025 | 0.011 | 0.015 | 1 | 6 | Pigl | -0.004 | Old |
| 0.646821 | 0.1708 | 0.011 | 0.015 | 1 | 6 | Gm21781 | -0.004 | Old |
| 0.646832 | 0.020489 | 0.023 | 0.018 | 1 | 6 | St8sia1 | 0.005 | Old |
| 0.646923 | 0.037845 | 0.154 | 0.139 | 1 | 6 | Coa3 | 0.015 | Old |
| 0.64821 | 0.332528 | 0.134 | 0.125 | 1 | 6 | Eif2b5 | 0.009 | Old |
| 0.648334 | 0.053652 | 0.497 | 0.482 | 1 | 6 | Grk3 | 0.015 | Old |
| 0.648339 | 0.217726 | 0.014 | 0.018 | 1 | 6 | Hdhd5 | -0.004 | Old |
| 0.648897 | 0.022189 | 0.051 | 0.059 | 1 | 6 | Mocs2 | -0.008 | Old |
| 0.649081 | 0.323926 | 0.066 | 0.059 | 1 | 6 | Nudt14 | 0.007 | Old |
| 0.649115 | 0.314107 | 0.057 | 0.05 | 1 | 6 | Dhcr7 | 0.007 | Old |
| 0.649226 | 0.08147 | 0.154 | 0.142 | 1 | 6 | Atf2 | 0.012 | Old |
| 0.649331 | 0.049011 | 0.017 | 0.013 | 1 | 6 | Tk1 | 0.004 | Old |
| 0.649453 | 0.23268 | 0.011 | 0.015 | 1 | 6 | Paqr8 | -0.004 | Old |
| 0.649682 | 0.208458 | 0.137 | 0.129 | 1 | 6 | Dcaf15 | 0.008 | Old |
| 0.649775 | 0.098504 | 0.229 | 0.242 | 1 | 6 | Vps13a | -0.013 | Old |
| 0.650013 | 0.033935 | 0.654 | 0.709 | 1 | 6 | Bhlhe40 | -0.055 | Old |
| 0.650165 | 0.036777 | 0.18 | 0.197 | 1 | 6 | Chmp1a | -0.017 | Old |
| 0.650249 | 0.069973 | 0.471 | 0.505 | 1 | 6 | Selenow | -0.034 | Old |
| 0.650284 | 0.328291 | 0.137 | 0.129 | 1 | 6 | Cox16 | 0.008 | Old |
| 0.65113 | 0.075771 | 0.626 | 0.667 | 1 | 6 | Nptn | -0.041 | Old |
| 0.65121 | 0.009707 | 0.011 | 0.015 | 1 | 6 | Gm45051 | -0.004 | Old |
| 0.651291 | 0.393511 | 0.051 | 0.045 | 1 | 6 | Zmat3 | 0.006 | Old |
| 0.651325 | 0.007053 | 0.48 | 0.473 | 1 | 6 | Snrpg | 0.007 | Old |
| 0.65152 | 0.059938 | 0.76 | 0.789 | 1 | 6 | Rbm39 | -0.029 | Old |
| 0.651647 | 0.176083 | 0.103 | 0.114 | 1 | 6 | Flot1 | -0.011 | Old |
| 0.651677 | 0.036312 | 0.009 | 0.012 | 1 | 6 | Il15ra | -0.003 | Old |
| 0.651912 | 0.092224 | 0.157 | 0.144 | 1 | 6 | Micu2 | 0.013 | Old |
| 0.652068 | 0.012126 | 0.246 | 0.227 | 1 | 6 | Rdx | 0.019 | Old |
| 0.652089 | 0.266252 | 0.011 | 0.015 | 1 | 6 | Zfp946 | -0.004 | Old |
| 0.652582 | 0.256098 | 0.483 | 0.512 | 1 | 6 | Ear2 | -0.029 | Old |
| 0.652595 | 0.082367 | 0.043 | 0.037 | 1 | 6 | Gm9844 | 0.006 | Old |
| 0.652678 | 1.083869 | 0.009 | 0.012 | 1 | 6 | Nbl1 | -0.003 | Old |
| 0.652678 | 0.623173 | 0.009 | 0.012 | 1 | 6 | Itga9 | -0.003 | Old |
| 0.652678 | 0.1706 | 0.009 | 0.012 | 1 | 6 | Gm4285 | -0.003 | Old |

| | | | | | | | | |
|---|---|---|---|---|---|---|---|---|
| 0.652678 | 0.137431 | 0.009 | 0.012 | 1 | 6 | Cit | -0.003 | Old |
| 0.653 | 0.152342 | 0.057 | 0.05 | 1 | 6 | Mtap | 0.007 | Old |
| 0.653023 | 0.339529 | 0.157 | 0.145 | 1 | 6 | Pabpc4 | 0.012 | Old |
| 0.653038 | 0.065776 | 0.103 | 0.112 | 1 | 6 | Tfeb | -0.009 | Old |
| 0.653275 | 0.059285 | 0.146 | 0.135 | 1 | 6 | Gsap | 0.011 | Old |
| 0.65368 | 0.704782 | 0.009 | 0.012 | 1 | 6 | Gtf2ird2 | -0.003 | Old |
| 0.653816 | 0.16443 | 0.323 | 0.304 | 1 | 6 | Itgal | 0.019 | Old |
| 0.654011 | 0.232221 | 0.109 | 0.119 | 1 | 6 | Zfp592 | -0.01 | Old |
| 0.654129 | 0.046098 | 0.071 | 0.064 | 1 | 6 | Cav2 | 0.007 | Old |
| 0.65423 | 0.151736 | 0.369 | 0.353 | 1 | 6 | Lsm4 | 0.016 | Old |
| 0.654269 | 0.157765 | 0.097 | 0.089 | 1 | 6 | Fut8 | 0.008 | Old |
| 0.654682 | 0.192845 | 0.009 | 0.012 | 1 | 6 | Cd209e | -0.003 | Old |
| 0.654889 | 0.292738 | 0.069 | 0.077 | 1 | 6 | Rem1 | -0.008 | Old |
| 0.654909 | 0.00925 | 0.334 | 0.319 | 1 | 6 | Api5 | 0.015 | Old |
| 0.655045 | 0.000111 | 0.563 | 0.569 | 1 | 6 | Rpl13a | -0.006 | Old |
| 0.655422 | 0.132617 | 0.263 | 0.253 | 1 | 6 | Slamf8 | 0.01 | Old |
| 0.655543 | 0.038472 | 0.677 | 0.712 | 1 | 6 | Rsrp1 | -0.035 | Old |
| 0.655565 | 0.042177 | 0.166 | 0.179 | 1 | 6 | Ppfia4 | -0.013 | Old |
| 0.655667 | 0.040231 | 0.423 | 0.403 | 1 | 6 | Cdk2ap2 | 0.02 | Old |
| 0.655748 | 0.162703 | 0.057 | 0.05 | 1 | 6 | Poldip2 | 0.007 | Old |
| 0.656196 | 0.075015 | 0.209 | 0.192 | 1 | 6 | Nudc | 0.017 | Old |
| 0.656197 | 0.133692 | 0.137 | 0.125 | 1 | 6 | Dcaf11 | 0.012 | Old |
| 0.656296 | 0.394421 | 0.043 | 0.037 | 1 | 6 | Cdadc1 | 0.006 | Old |
| 0.656492 | 0.518587 | 0.011 | 0.015 | 1 | 6 | Mcrip2 | -0.004 | Old |
| 0.656665 | 0.177144 | 0.189 | 0.174 | 1 | 6 | Wipf2 | 0.015 | Old |
| 0.656689 | 0.442115 | 0.009 | 0.012 | 1 | 6 | Zfp949 | -0.003 | Old |
| 0.657305 | 0.073614 | 0.114 | 0.125 | 1 | 6 | Lipe | -0.011 | Old |
| 0.657355 | 0.04695 | 0.043 | 0.037 | 1 | 6 | Gstcd | 0.006 | Old |
| 0.657375 | 0.370796 | 0.011 | 0.015 | 1 | 6 | Rbbp9 | -0.004 | Old |
| 0.657832 | 0.000592 | 0.817 | 0.848 | 1 | 6 | Arpc5 | -0.031 | Old |
| 0.657885 | 0.35708 | 0.283 | 0.278 | 1 | 6 | Aprt | 0.005 | Old |
| 0.658108 | 0.935427 | 0.054 | 0.048 | 1 | 6 | Gabpb1 | 0.006 | Old |
| 0.658647 | 0.047543 | 0.609 | 0.614 | 1 | 6 | Ncl | -0.005 | Old |
| 0.658696 | 0.478169 | 0.06 | 0.054 | 1 | 6 | Prkrip1 | 0.006 | Old |
| 0.658871 | 0.022449 | 0.097 | 0.107 | 1 | 6 | Jdp2 | -0.01 | Old |
| 0.658945 | 0.391971 | 0.043 | 0.037 | 1 | 6 | Gbp9 | 0.006 | Old |
| 0.659004 | 0.055847 | 0.48 | 0.5 | 1 | 6 | Chmp4b | -0.02 | Old |
| 0.659012 | 0.103904 | 0.351 | 0.329 | 1 | 6 | Aftph | 0.022 | Old |
| 0.659156 | 0.236684 | 0.037 | 0.043 | 1 | 6 | Slc16a10 | -0.006 | Old |
| 0.659215 | 0.075259 | 0.209 | 0.221 | 1 | 6 | Bdp1 | -0.012 | Old |
| 0.659297 | 0.487858 | 0.04 | 0.047 | 1 | 6 | Mpp5 | -0.007 | Old |
| 0.659888 | 0.171681 | 0.171 | 0.161 | 1 | 6 | Kctd2 | 0.01 | Old |
| 0.659939 | 0.362354 | 0.046 | 0.04 | 1 | 6 | Bclaf3 | 0.006 | Old |
| 0.661091 | 0.272961 | 0.22 | 0.212 | 1 | 6 | Ptpn22 | 0.008 | Old |
| 0.661483 | 0.006019 | 0.603 | 0.602 | 1 | 6 | Pomp | 0.001 | Old |

| | | | | | | | | |
|---|---|---|---|---|---|---|---|---|
| 0.661774 | 0.142152 | 0.389 | 0.375 | 1 | 6 | Actn4 | 0.014 | Old |
| 0.661955 | 0.043532 | 0.149 | 0.137 | 1 | 6 | Smim3 | 0.012 | Old |
| 0.661994 | 0.580075 | 0.031 | 0.027 | 1 | 6 | Prorsd1 | 0.004 | Old |
| 0.662128 | 0.090334 | 0.103 | 0.114 | 1 | 6 | Rexo1 | -0.011 | Old |
| 0.662162 | 0.028344 | 0.614 | 0.619 | 1 | 6 | Cltc | -0.005 | Old |
| 0.662383 | 0.252716 | 0.269 | 0.258 | 1 | 6 | Sec24b | 0.011 | Old |
| 0.662639 | 0.117372 | 0.411 | 0.436 | 1 | 6 | Usmg5 | -0.025 | Old |
| 0.662639 | 0.338476 | 0.057 | 0.05 | 1 | 6 | Usp31 | 0.007 | Old |
| 0.662869 | 0.169712 | 0.049 | 0.055 | 1 | 6 | Cep290 | -0.006 | Old |
| 0.663898 | 0.08063 | 0.397 | 0.376 | 1 | 6 | Zfp91 | 0.021 | Old |
| 0.664028 | 0.214817 | 0.174 | 0.164 | 1 | 6 | Aim2 | 0.01 | Old |
| 0.664183 | 0.077044 | 0.094 | 0.102 | 1 | 6 | Gnl3l | -0.008 | Old |
| 0.664309 | 0.075237 | 0.589 | 0.59 | 1 | 6 | Atp6v1g1 | -0.001 | Old |
| 0.664442 | 0.002376 | 0.12 | 0.13 | 1 | 6 | Edem2 | -0.01 | Old |
| 0.664481 | 0.093102 | 0.069 | 0.062 | 1 | 6 | Mfn2 | 0.007 | Old |
| 0.665684 | 0.231068 | 0.137 | 0.127 | 1 | 6 | Atad1 | 0.01 | Old |
| 0.666422 | 0.094672 | 0.106 | 0.095 | 1 | 6 | Cluh | 0.011 | Old |
| 0.666535 | 0.041064 | 0.231 | 0.244 | 1 | 6 | Arpc1a | -0.013 | Old |
| 0.666674 | 0.102911 | 0.246 | 0.258 | 1 | 6 | Pcf11 | -0.012 | Old |
| 0.666961 | 0.028206 | 0.114 | 0.124 | 1 | 6 | Tmx4 | -0.01 | Old |
| 0.667071 | 0.307963 | 0.134 | 0.147 | 1 | 6 | Arhgap6 | -0.013 | Old |
| 0.667207 | 0.568936 | 0.06 | 0.054 | 1 | 6 | Erich1 | 0.006 | Old |
| 0.667541 | 0.250684 | 0.14 | 0.129 | 1 | 6 | Nop56 | 0.011 | Old |
| 0.667608 | 0.469533 | 0.08 | 0.074 | 1 | 6 | Tbc1d2b | 0.006 | Old |
| 0.668169 | 0.382262 | 0.034 | 0.04 | 1 | 6 | Selenoo | -0.006 | Old |
| 0.668191 | 0.62942 | 0.031 | 0.027 | 1 | 6 | Ttbk2 | 0.004 | Old |
| 0.668327 | 0.250042 | 0.089 | 0.08 | 1 | 6 | Atg4b | 0.009 | Old |
| 0.668338 | 0.012029 | 0.031 | 0.037 | 1 | 6 | Cryzl2 | -0.006 | Old |
| 0.668555 | 0.207168 | 0.06 | 0.054 | 1 | 6 | Mlycd | 0.006 | Old |
| 0.669171 | 0.133606 | 0.051 | 0.045 | 1 | 6 | Jrkl | 0.006 | Old |
| 0.669205 | 0.277183 | 0.123 | 0.114 | 1 | 6 | Mrto4 | 0.009 | Old |
| 0.669707 | 0.076606 | 0.786 | 0.809 | 1 | 6 | Celf2 | -0.023 | Old |
| 0.669915 | 0.24003 | 0.143 | 0.134 | 1 | 6 | Dhx16 | 0.009 | Old |
| 0.670454 | 0.767978 | 0.111 | 0.102 | 1 | 6 | Hax1 | 0.009 | Old |
| 0.671222 | 0.350584 | 0.046 | 0.04 | 1 | 6 | Pvt1 | 0.006 | Old |
| 0.671343 | 0.215575 | 0.137 | 0.145 | 1 | 6 | Sepsecs | -0.008 | Old |
| 0.671609 | 0.018061 | 0.206 | 0.217 | 1 | 6 | Card19 | -0.011 | Old |
| 0.671668 | 0.16332 | 0.169 | 0.156 | 1 | 6 | Chchd3 | 0.013 | Old |
| 0.67192 | 0.428282 | 0.031 | 0.027 | 1 | 6 | Tpcn2 | 0.004 | Old |
| 0.672131 | 0.030777 | 0.251 | 0.231 | 1 | 6 | Uhrf2 | 0.02 | Old |
| 0.673295 | 0.560623 | 0.14 | 0.13 | 1 | 6 | Insr | 0.01 | Old |
| 0.67351 | 0.024673 | 0.386 | 0.4 | 1 | 6 | Scamp2 | -0.014 | Old |
| 0.673982 | 0.222114 | 0.031 | 0.037 | 1 | 6 | Nfkbil1 | -0.006 | Old |
| 0.674269 | 0.23418 | 0.146 | 0.137 | 1 | 6 | Bin3 | 0.009 | Old |
| 0.674613 | 0.217678 | 0.08 | 0.072 | 1 | 6 | Ttc13 | 0.008 | Old |

| | | | | | | | | |
|---|---|---|---|---|---|---|---|---|
| 0.674918 | 0.374777 | 0.043 | 0.048 | 1 | 6 | Cdk6 | -0.005 | Old |
| 0.674927 | 0.001452 | 0.131 | 0.12 | 1 | 6 | BC003331 | 0.011 | Old |
| 0.674989 | 0.038789 | 0.271 | 0.283 | 1 | 6 | Lemd2 | -0.012 | Old |
| 0.675049 | 0.057338 | 0.423 | 0.45 | 1 | 6 | Ciita | -0.027 | Old |
| 0.676043 | 0.21373 | 0.057 | 0.05 | 1 | 6 | 4930453N2 | 0.007 | Old |
| 0.676275 | 0.252482 | 0.049 | 0.055 | 1 | 6 | Arhgef10l | -0.006 | Old |
| 0.676891 | 0.235884 | 0.046 | 0.04 | 1 | 6 | Usp42 | 0.006 | Old |
| 0.676911 | 0.241007 | 0.091 | 0.084 | 1 | 6 | Camk1 | 0.007 | Old |
| 0.677377 | 0.12514 | 0.031 | 0.037 | 1 | 6 | Dtd2 | -0.006 | Old |
| 0.6779 | 0.296194 | 0.083 | 0.075 | 1 | 6 | Cherp | 0.008 | Old |
| 0.678202 | 0.183407 | 0.446 | 0.438 | 1 | 6 | Cdc42se1 | 0.008 | Old |
| 0.678288 | 0.065454 | 0.206 | 0.192 | 1 | 6 | Rras2 | 0.014 | Old |
| 0.678809 | 0.071528 | 0.594 | 0.615 | 1 | 6 | Tacc1 | -0.021 | Old |
| 0.678862 | 0.03193 | 0.131 | 0.122 | 1 | 6 | Stard9 | 0.009 | Old |
| 0.678872 | 0.018577 | 0.12 | 0.13 | 1 | 6 | Nploc4 | -0.01 | Old |
| 0.678957 | 0.469547 | 0.046 | 0.04 | 1 | 6 | Rufy2 | 0.006 | Old |
| 0.679064 | 1.391094 | 0.026 | 0.022 | 1 | 6 | Dnajc27 | 0.004 | Old |
| 0.679304 | 0.464834 | 0.054 | 0.048 | 1 | 6 | Smpd4 | 0.006 | Old |
| 0.680113 | 0.381992 | 0.077 | 0.07 | 1 | 6 | Dera | 0.007 | Old |
| 0.680416 | 0.136913 | 0.083 | 0.075 | 1 | 6 | Gm5617 | 0.008 | Old |
| 0.680507 | 0.012888 | 0.643 | 0.692 | 1 | 6 | Cnn2 | -0.049 | Old |
| 0.680625 | 0.346444 | 0.137 | 0.13 | 1 | 6 | Gskip | 0.007 | Old |
| 0.681279 | 0.036873 | 0.031 | 0.027 | 1 | 6 | Mblac2 | 0.004 | Old |
| 0.681346 | 0.158168 | 0.031 | 0.037 | 1 | 6 | Gtpbp10 | -0.006 | Old |
| 0.681488 | 0.506986 | 0.04 | 0.035 | 1 | 6 | Plau | 0.005 | Old |
| 0.68155 | 0.073165 | 0.251 | 0.239 | 1 | 6 | Themis2 | 0.012 | Old |
| 0.681692 | 0.584503 | 0.086 | 0.079 | 1 | 6 | Vkorc1l1 | 0.007 | Old |
| 0.681849 | 0.016464 | 0.051 | 0.045 | 1 | 6 | Tusc2 | 0.006 | Old |
| 0.681915 | 0.315114 | 0.1 | 0.109 | 1 | 6 | Rptor | -0.009 | Old |
| 0.68225 | 0.119279 | 0.246 | 0.236 | 1 | 6 | Pfdn2 | 0.01 | Old |
| 0.682511 | 0.025555 | 0.129 | 0.139 | 1 | 6 | Hilpda | -0.01 | Old |
| 0.682551 | 0.106972 | 0.143 | 0.154 | 1 | 6 | Reps1 | -0.011 | Old |
| 0.682585 | 0.68117 | 0.094 | 0.087 | 1 | 6 | Abl2 | 0.007 | Old |
| 0.682867 | 0.115328 | 0.18 | 0.169 | 1 | 6 | Ccdc59 | 0.011 | Old |
| 0.683203 | 0.030638 | 0.029 | 0.033 | 1 | 6 | Ercc4 | -0.004 | Old |
| 0.683249 | 0.019543 | 0.049 | 0.055 | 1 | 6 | Slc35d1 | -0.006 | Old |
| 0.683372 | 0.189372 | 0.08 | 0.089 | 1 | 6 | Nol8 | -0.009 | Old |
| 0.683424 | 0.285706 | 0.094 | 0.104 | 1 | 6 | A630001G2 | -0.01 | Old |
| 0.683611 | 0.120234 | 0.169 | 0.177 | 1 | 6 | Braf | -0.008 | Old |
| 0.683615 | 0.176785 | 0.046 | 0.04 | 1 | 6 | Med22 | 0.006 | Old |
| 0.683729 | 0.487512 | 0.089 | 0.082 | 1 | 6 | Plk2 | 0.007 | Old |
| 0.684592 | 0.294152 | 0.166 | 0.159 | 1 | 6 | Fbxl5 | 0.007 | Old |
| 0.684619 | 0.23808 | 0.103 | 0.114 | 1 | 6 | Ralgapb | -0.011 | Old |
| 0.684725 | 0.227708 | 0.257 | 0.246 | 1 | 6 | Emc6 | 0.011 | Old |
| 0.684757 | 0.223636 | 0.097 | 0.09 | 1 | 6 | Sympk | 0.007 | Old |

| | | | | | | | | |
|---|---|---|---|---|---|---|---|---|
| 0.684769 | 0.941744 | 0.049 | 0.043 | 1 | 6 | Gtdc1 | 0.006 | Old |
| 0.684986 | 0.117986 | 0.211 | 0.219 | 1 | 6 | Ppp1r11 | -0.008 | Old |
| 0.685122 | 0.506812 | 0.063 | 0.07 | 1 | 6 | Tnni2 | -0.007 | Old |
| 0.685122 | 0.141359 | 0.063 | 0.07 | 1 | 6 | Nup133 | -0.007 | Old |
| 0.685301 | 0.041434 | 0.111 | 0.102 | 1 | 6 | Rbm14 | 0.009 | Old |
| 0.685437 | 0.023231 | 0.151 | 0.159 | 1 | 6 | Elovl1 | -0.008 | Old |
| 0.685584 | 0.257955 | 0.029 | 0.033 | 1 | 6 | D6Wsu163 | -0.004 | Old |
| 0.686191 | 0.075141 | 0.271 | 0.256 | 1 | 6 | Ddx24 | 0.015 | Old |
| 0.686837 | 0.324364 | 0.086 | 0.079 | 1 | 6 | 0610010F0 | 0.007 | Old |
| 0.686938 | 0.043601 | 0.12 | 0.11 | 1 | 6 | Srsf4 | 0.01 | Old |
| 0.687686 | 0.037659 | 0.426 | 0.436 | 1 | 6 | Cmtm6 | -0.01 | Old |
| 0.688092 | 0.059243 | 0.586 | 0.585 | 1 | 6 | Ppp1r18 | 0.001 | Old |
| 0.688496 | 0.080916 | 0.057 | 0.064 | 1 | 6 | Crip2 | -0.007 | Old |
| 0.688537 | 0.060132 | 0.163 | 0.172 | 1 | 6 | Tnf | -0.009 | Old |
| 0.688635 | 0.140372 | 0.34 | 0.358 | 1 | 6 | Emc10 | -0.018 | Old |
| 0.688742 | 0.01007 | 0.351 | 0.351 | 1 | 6 | Lims1 | 0 | Old |
| 0.688757 | 0.271333 | 0.186 | 0.179 | 1 | 6 | Dnajc1 | 0.007 | Old |
| 0.688806 | 0.035777 | 0.154 | 0.145 | 1 | 6 | Snx18 | 0.009 | Old |
| 0.689032 | 0.007763 | 0.111 | 0.102 | 1 | 6 | Cblb | 0.009 | Old |
| 0.689274 | 0.088237 | 0.263 | 0.254 | 1 | 6 | Iscu | 0.009 | Old |
| 0.68976 | 0.662228 | 0.029 | 0.033 | 1 | 6 | Fktn | -0.004 | Old |
| 0.689948 | 0.092062 | 0.094 | 0.085 | 1 | 6 | Usp50 | 0.009 | Old |
| 0.690256 | 0.099966 | 0.234 | 0.224 | 1 | 6 | Lmbrd1 | 0.01 | Old |
| 0.690375 | 0.022914 | 0.683 | 0.706 | 1 | 6 | Rac1 | -0.023 | Old |
| 0.69044 | 0.174078 | 0.36 | 0.356 | 1 | 6 | Psmb3 | 0.004 | Old |
| 0.690594 | 0.325262 | 0.06 | 0.069 | 1 | 6 | Rbm15b | -0.009 | Old |
| 0.690837 | 0.272128 | 0.026 | 0.022 | 1 | 6 | Zfhx2 | 0.004 | Old |
| 0.69095 | 0.46649 | 0.109 | 0.102 | 1 | 6 | Ppp1r35 | 0.007 | Old |
| 0.691306 | 0.064163 | 0.24 | 0.229 | 1 | 6 | Anapc11 | 0.011 | Old |
| 0.691352 | 0.057668 | 0.091 | 0.084 | 1 | 6 | Mea1 | 0.007 | Old |
| 0.69139 | 0.197597 | 0.134 | 0.127 | 1 | 6 | Clns1a | 0.007 | Old |
| 0.691683 | 0.085159 | 0.431 | 0.405 | 1 | 6 | Sdha | 0.026 | Old |
| 0.691891 | 0.225119 | 0.183 | 0.172 | 1 | 6 | Mbtps1 | 0.011 | Old |
| 0.692053 | 0.284802 | 0.226 | 0.217 | 1 | 6 | Dr1 | 0.009 | Old |
| 0.692161 | 0.106942 | 0.206 | 0.194 | 1 | 6 | Marf1 | 0.012 | Old |
| 0.692184 | 0.070411 | 0.331 | 0.358 | 1 | 6 | Rap2a | -0.027 | Old |
| 0.692235 | 0.464231 | 0.074 | 0.067 | 1 | 6 | Txnl4a | 0.007 | Old |
| 0.692353 | 0.290544 | 0.077 | 0.07 | 1 | 6 | Eci2 | 0.007 | Old |
| 0.692361 | 0.229195 | 0.089 | 0.082 | 1 | 6 | Metap1 | 0.007 | Old |
| 0.693564 | 0.11963 | 0.16 | 0.149 | 1 | 6 | Sf3a3 | 0.011 | Old |
| 0.693578 | 0.112433 | 0.054 | 0.048 | 1 | 6 | AI987944 | 0.006 | Old |
| 0.694008 | 0.479631 | 0.046 | 0.04 | 1 | 6 | Aifm2 | 0.006 | Old |
| 0.694057 | 0.060195 | 0.206 | 0.214 | 1 | 6 | Eif4e2 | -0.008 | Old |
| 0.694449 | 0.724455 | 0.031 | 0.037 | 1 | 6 | Ift57 | -0.006 | Old |
| 0.69478 | 0.33968 | 0.04 | 0.035 | 1 | 6 | Dusp23 | 0.005 | Old |

| | | | | | | | | |
|---|---|---|---|---|---|---|---|---|
| 0.695014 | 0.074805 | 0.309 | 0.319 | 1 | 6 | Bcl11a | -0.01 | Old |
| 0.695431 | 0.10476 | 0.554 | 0.567 | 1 | 6 | Acer3 | -0.013 | Old |
| 0.695436 | 0.01166 | 0.343 | 0.36 | 1 | 6 | Cd300a | -0.017 | Old |
| 0.695572 | 0.199489 | 0.046 | 0.04 | 1 | 6 | Qtrt2 | 0.006 | Old |
| 0.695641 | 0.331313 | 0.197 | 0.189 | 1 | 6 | B4galt5 | 0.008 | Old |
| 0.695862 | 0.133596 | 0.151 | 0.14 | 1 | 6 | Chka | 0.011 | Old |
| 0.695927 | 0.046343 | 0.083 | 0.09 | 1 | 6 | Trip4 | -0.007 | Old |
| 0.695976 | 0.128015 | 0.137 | 0.127 | 1 | 6 | Ppp2r2d | 0.01 | Old |
| 0.696404 | 0.150774 | 0.026 | 0.022 | 1 | 6 | Lrrc45 | 0.004 | Old |
| 0.696436 | 0.111122 | 0.231 | 0.239 | 1 | 6 | Uap1 | -0.008 | Old |
| 0.697005 | 0.275989 | 0.04 | 0.035 | 1 | 6 | Spryd3 | 0.005 | Old |
| 0.697101 | 0.20455 | 0.026 | 0.022 | 1 | 6 | Zbtb24 | 0.004 | Old |
| 0.697679 | 0.076963 | 0.211 | 0.201 | 1 | 6 | Glrx2 | 0.01 | Old |
| 0.697819 | 0.057965 | 0.18 | 0.167 | 1 | 6 | Ppfibp2 | 0.013 | Old |
| 0.697915 | 0.084083 | 0.374 | 0.36 | 1 | 6 | Prpf40a | 0.014 | Old |
| 0.69804 | 0.546742 | 0.06 | 0.055 | 1 | 6 | Tmem209 | 0.005 | Old |
| 0.698072 | 0.093582 | 0.877 | 0.965 | 1 | 6 | S100a6 | -0.088 | Old |
| 0.698162 | 0.116689 | 0.323 | 0.304 | 1 | 6 | Ifi203 | 0.019 | Old |
| 0.698219 | 0.064965 | 0.337 | 0.353 | 1 | 6 | Vrk1 | -0.016 | Old |
| 0.698523 | 0.044906 | 0.426 | 0.436 | 1 | 6 | Taok1 | -0.01 | Old |
| 0.698801 | 0.132631 | 0.146 | 0.135 | 1 | 6 | Tmem5 | 0.011 | Old |
| 0.699234 | 0.44726 | 0.134 | 0.129 | 1 | 6 | Smad7 | 0.005 | Old |
| 0.699553 | 0.142579 | 0.129 | 0.137 | 1 | 6 | Irf3 | -0.008 | Old |
| 0.699772 | 0.073514 | 0.211 | 0.222 | 1 | 6 | Rfx7 | -0.011 | Old |
| 0.700521 | 0.13394 | 0.04 | 0.045 | 1 | 6 | C87436 | -0.005 | Old |
| 0.700774 | 0.363115 | 0.026 | 0.03 | 1 | 6 | Mipep | -0.004 | Old |
| 0.701128 | 0.574853 | 0.057 | 0.052 | 1 | 6 | Rnaseh2b | 0.005 | Old |
| 0.701424 | 0.122852 | 0.326 | 0.338 | 1 | 6 | Mapre2 | -0.012 | Old |
| 0.701466 | 1.214097 | 0.02 | 0.017 | 1 | 6 | Krt10 | 0.003 | Old |
| 0.701572 | 0.125593 | 0.294 | 0.309 | 1 | 6 | Pkn1 | -0.015 | Old |
| 0.701791 | 0.373809 | 0.091 | 0.084 | 1 | 6 | Ggps1 | 0.007 | Old |
| 0.702247 | 0.364554 | 0.063 | 0.057 | 1 | 6 | Cdc16 | 0.006 | Old |
| 0.702483 | 0.210115 | 0.249 | 0.234 | 1 | 6 | Lrp10 | 0.015 | Old |
| 0.702698 | 0.049243 | 0.714 | 0.717 | 1 | 6 | Sf3b1 | -0.003 | Old |
| 0.703052 | 0.5332 | 0.02 | 0.017 | 1 | 6 | D3Ertd751 | 0.003 | Old |
| 0.703519 | 0.00419 | 0.123 | 0.114 | 1 | 6 | AW549877 | 0.009 | Old |
| 0.703846 | 0.622232 | 0.02 | 0.017 | 1 | 6 | Trim30c | 0.003 | Old |
| 0.703864 | 0.28501 | 0.214 | 0.229 | 1 | 6 | Fam107b | -0.015 | Old |
| 0.704224 | 0.14124 | 0.043 | 0.048 | 1 | 6 | Abcb1b | -0.005 | Old |
| 0.704571 | 0.461408 | 0.026 | 0.03 | 1 | 6 | Pam | -0.004 | Old |
| 0.704724 | 0.111823 | 0.043 | 0.048 | 1 | 6 | Mgat5 | -0.005 | Old |
| 0.70487 | 0.313272 | 0.186 | 0.177 | 1 | 6 | Zmym5 | 0.009 | Old |
| 0.704988 | 0.120521 | 0.271 | 0.281 | 1 | 6 | Jaml | -0.01 | Old |
| 0.705323 | 0.027489 | 0.431 | 0.436 | 1 | 6 | Caprin1 | -0.005 | Old |
| 0.70542 | 0.422768 | 0.049 | 0.043 | 1 | 6 | Fgd6 | 0.006 | Old |

| | | | | | | | | |
|---|---|---|---|---|---|---|---|---|
| 0.705485 | 0.29276 | 0.026 | 0.022 | 1 | 6 | Kdm4b | 0.004 | Old |
| 0.70564 | 0.021404 | 0.066 | 0.072 | 1 | 6 | Mrpl45 | -0.006 | Old |
| 0.705664 | 0.025244 | 0.414 | 0.421 | 1 | 6 | Itpkb | -0.007 | Old |
| 0.705886 | 0.013137 | 0.114 | 0.122 | 1 | 6 | Mrps23 | -0.008 | Old |
| 0.706164 | 0.070325 | 0.226 | 0.237 | 1 | 6 | Mrps15 | -0.011 | Old |
| 0.706228 | 0.569016 | 0.02 | 0.017 | 1 | 6 | Adck2 | 0.003 | Old |
| 0.70633 | 0.126196 | 0.297 | 0.288 | 1 | 6 | Ube2i | 0.009 | Old |
| 0.706389 | 0.241967 | 0.089 | 0.097 | 1 | 6 | Parp9 | -0.008 | Old |
| 0.706499 | 0.388899 | 0.077 | 0.07 | 1 | 6 | Mfap1b | 0.007 | Old |
| 0.706659 | 0.008905 | 0.437 | 0.457 | 1 | 6 | Ubap2l | -0.02 | Old |
| 0.706819 | 0.171088 | 0.123 | 0.134 | 1 | 6 | St8sia6 | -0.011 | Old |
| 0.707321 | 0.11477 | 0.269 | 0.253 | 1 | 6 | Lsm5 | 0.016 | Old |
| 0.707564 | 0.083036 | 0.117 | 0.109 | 1 | 6 | Rnpc3 | 0.008 | Old |
| 0.707805 | 0.610076 | 0.066 | 0.06 | 1 | 6 | Fiz1 | 0.006 | Old |
| 0.707954 | 0.329412 | 0.049 | 0.043 | 1 | 6 | Dusp6 | 0.006 | Old |
| 0.708204 | 0.104166 | 0.057 | 0.064 | 1 | 6 | Lxn | -0.007 | Old |
| 0.708373 | 0.354036 | 0.163 | 0.154 | 1 | 6 | Rnf11 | 0.009 | Old |
| 0.708373 | 0.270244 | 0.151 | 0.161 | 1 | 6 | C330007PC | -0.01 | Old |
| 0.708598 | 0.761603 | 0.057 | 0.052 | 1 | 6 | Rad51d | 0.005 | Old |
| 0.709008 | 0.097945 | 0.403 | 0.43 | 1 | 6 | Brd2 | -0.027 | Old |
| 0.70903 | 0.033656 | 0.714 | 0.724 | 1 | 6 | Akap13 | -0.01 | Old |
| 0.709268 | 1.215858 | 0.034 | 0.03 | 1 | 6 | Ddx31 | 0.004 | Old |
| 0.709837 | 0.133833 | 0.234 | 0.224 | 1 | 6 | Rbm26 | 0.01 | Old |
| 0.709868 | 0.226969 | 0.063 | 0.069 | 1 | 6 | Med4 | -0.006 | Old |
| 0.710016 | 0.203236 | 0.429 | 0.423 | 1 | 6 | Egr2 | 0.006 | Old |
| 0.710039 | 0.318409 | 0.097 | 0.09 | 1 | 6 | Zfp992 | 0.007 | Old |
| 0.710067 | 0.048804 | 0.897 | 0.921 | 1 | 6 | Gm10076 | -0.024 | Old |
| 0.71026 | 0.321292 | 0.471 | 0.468 | 1 | 6 | Herpud1 | 0.003 | Old |
| 0.710364 | 0.043808 | 0.04 | 0.045 | 1 | 6 | Mrpl50 | -0.005 | Old |
| 0.710483 | 0.330952 | 0.237 | 0.231 | 1 | 6 | Zfand3 | 0.006 | Old |
| 0.710507 | 0.252222 | 0.154 | 0.147 | 1 | 6 | Arl6ip6 | 0.007 | Old |
| 0.710959 | 0.043076 | 0.614 | 0.609 | 1 | 6 | Tm9sf3 | 0.005 | Old |
| 0.711237 | 0.016655 | 0.429 | 0.423 | 1 | 6 | Hnrnph2 | 0.006 | Old |
| 0.71126 | 0.40763 | 0.114 | 0.109 | 1 | 6 | Bahd1 | 0.005 | Old |
| 0.711272 | 0.123399 | 0.36 | 0.351 | 1 | 6 | Cuta | 0.009 | Old |
| 0.711317 | 0.143151 | 0.346 | 0.339 | 1 | 6 | Psmb10 | 0.007 | Old |
| 0.711639 | 0.332848 | 0.063 | 0.057 | 1 | 6 | Blcap | 0.006 | Old |
| 0.712525 | 0.219725 | 0.071 | 0.065 | 1 | 6 | Smn1 | 0.006 | Old |
| 0.71259 | 0.033089 | 0.134 | 0.125 | 1 | 6 | Itfg1 | 0.009 | Old |
| 0.713158 | 0.346652 | 0.209 | 0.194 | 1 | 6 | Rangap1 | 0.015 | Old |
| 0.713456 | 0.182031 | 0.203 | 0.194 | 1 | 6 | Arnt | 0.009 | Old |
| 0.713924 | 0.05357 | 0.08 | 0.087 | 1 | 6 | Rnf169 | -0.007 | Old |
| 0.713925 | 0.260982 | 0.051 | 0.057 | 1 | 6 | Arhgap4 | -0.006 | Old |
| 0.713966 | 0.049727 | 0.294 | 0.276 | 1 | 6 | Taf1d | 0.018 | Old |
| 0.714191 | 0.755524 | 0.02 | 0.017 | 1 | 6 | Ndufaf6 | 0.003 | Old |

| | | | | | | | | |
|---|---|---|---|---|---|---|---|---|
| 0.714191 | 0.652587 | 0.02 | 0.017 | 1 | 6 | Mtus1 | 0.003 | Old |
| 0.715059 | 0.107346 | 0.177 | 0.167 | 1 | 6 | 1810013L2 | 0.01 | Old |
| 0.71522 | 0.636635 | 0.12 | 0.114 | 1 | 6 | Rbl2 | 0.006 | Old |
| 0.715435 | 0.094074 | 0.054 | 0.048 | 1 | 6 | Cebpzos | 0.006 | Old |
| 0.715439 | 0.180347 | 0.103 | 0.095 | 1 | 6 | Zbtb4 | 0.008 | Old |
| 0.715676 | 0.114559 | 0.054 | 0.048 | 1 | 6 | Cars | 0.006 | Old |
| 0.716084 | 0.417185 | 0.049 | 0.043 | 1 | 6 | Mbip | 0.006 | Old |
| 0.716586 | 1.008542 | 0.02 | 0.017 | 1 | 6 | H6pd | 0.003 | Old |
| 0.717244 | 0.128878 | 0.409 | 0.405 | 1 | 6 | Ccdc12 | 0.004 | Old |
| 0.717625 | 0.085615 | 0.223 | 0.209 | 1 | 6 | Psme2b | 0.014 | Old |
| 0.71773 | 0.13557 | 0.069 | 0.062 | 1 | 6 | Ercc6l2 | 0.007 | Old |
| 0.717975 | 0.38461 | 0.057 | 0.052 | 1 | 6 | Tmem87a | 0.005 | Old |
| 0.718374 | 0.170389 | 0.063 | 0.057 | 1 | 6 | Crbn | 0.006 | Old |
| 0.71897 | 0.127889 | 0.086 | 0.092 | 1 | 6 | Blvrb | -0.006 | Old |
| 0.71897 | 0.092382 | 0.086 | 0.092 | 1 | 6 | Sos1 | -0.006 | Old |
| 0.719247 | 0.247673 | 0.034 | 0.03 | 1 | 6 | Agk | 0.004 | Old |
| 0.719456 | 0.193382 | 0.16 | 0.152 | 1 | 6 | Rogdi | 0.008 | Old |
| 0.719499 | 0.220242 | 0.123 | 0.115 | 1 | 6 | Rap2b | 0.008 | Old |
| 0.719779 | 0.156561 | 0.306 | 0.296 | 1 | 6 | Usp4 | 0.01 | Old |
| 0.719818 | 0.501973 | 0.106 | 0.099 | 1 | 6 | Mbd6 | 0.007 | Old |
| 0.71983 | 0.526555 | 0.026 | 0.03 | 1 | 6 | Tbc1d16 | -0.004 | Old |
| 0.719832 | 0.001695 | 0.214 | 0.227 | 1 | 6 | Mgrn1 | -0.013 | Old |
| 0.720158 | 0.135312 | 0.32 | 0.309 | 1 | 6 | Ndufb4 | 0.011 | Old |
| 0.720225 | 0.049124 | 0.023 | 0.027 | 1 | 6 | Tlr6 | -0.004 | Old |
| 0.720583 | 0.082641 | 0.02 | 0.017 | 1 | 6 | Zfyve21 | 0.003 | Old |
| 0.72077 | 0.112938 | 0.16 | 0.152 | 1 | 6 | Sigmar1 | 0.008 | Old |
| 0.720866 | 0.00065 | 0.8 | 0.823 | 1 | 6 | H2afy | -0.023 | Old |
| 0.72095 | 0.101668 | 0.077 | 0.07 | 1 | 6 | E130309D( | 0.007 | Old |
| 0.721222 | 0.076811 | 0.066 | 0.072 | 1 | 6 | Cdc42ep4 | -0.006 | Old |
| 0.721673 | 0.344935 | 0.034 | 0.03 | 1 | 6 | Calcoco1 | 0.004 | Old |
| 0.722679 | 0.169683 | 0.057 | 0.052 | 1 | 6 | Lsg1 | 0.005 | Old |
| 0.722815 | 0.161585 | 0.4 | 0.386 | 1 | 6 | Swi5 | 0.014 | Old |
| 0.72315 | 0.252004 | 0.057 | 0.052 | 1 | 6 | Gatsl2 | 0.005 | Old |
| 0.723319 | 0.327309 | 0.149 | 0.14 | 1 | 6 | Usp24 | 0.009 | Old |
| 0.723486 | 0.055503 | 0.131 | 0.122 | 1 | 6 | Zfp281 | 0.009 | Old |
| 0.723515 | 0.61571 | 0.029 | 0.025 | 1 | 6 | Pramef8 | 0.004 | Old |
| 0.72417 | 0.073934 | 0.143 | 0.134 | 1 | 6 | Nek9 | 0.009 | Old |
| 0.72418 | 1.274773 | 0.014 | 0.012 | 1 | 6 | Fosl1 | 0.002 | Old |
| 0.72418 | 0.960152 | 0.014 | 0.012 | 1 | 6 | Tspan2 | 0.002 | Old |
| 0.724324 | 0.146962 | 0.303 | 0.298 | 1 | 6 | Chn2 | 0.005 | Old |
| 0.724421 | 0.046768 | 0.483 | 0.458 | 1 | 6 | Cct2 | 0.025 | Old |
| 0.724736 | 0.281266 | 0.134 | 0.127 | 1 | 6 | Aaed1 | 0.007 | Old |
| 0.724904 | 0.250032 | 0.066 | 0.06 | 1 | 6 | Usp53 | 0.006 | Old |
| 0.725132 | 0.747264 | 0.014 | 0.012 | 1 | 6 | Zfp512b | 0.002 | Old |
| 0.725132 | 0.647493 | 0.014 | 0.012 | 1 | 6 | Rpgrip1l | 0.002 | Old |

| | | | | | | | | |
|---|---|---|---|---|---|---|---|---|
| 0.725313 | 0.208213 | 0.18 | 0.171 | 1 | 6 | Zdhhc21 | 0.009 | Old |
| 0.72581 | 0.14501 | 0.177 | 0.186 | 1 | 6 | Ptk2b | -0.009 | Old |
| 0.725895 | 0.185259 | 0.203 | 0.194 | 1 | 6 | Pqbp1 | 0.009 | Old |
| 0.725926 | 0.16023 | 0.034 | 0.03 | 1 | 6 | Zbtb11os1 | 0.004 | Old |
| 0.72618 | 0.38024 | 0.077 | 0.072 | 1 | 6 | Nol11 | 0.005 | Old |
| 0.72641 | 0.258881 | 0.109 | 0.119 | 1 | 6 | 2610037D( | -0.01 | Old |
| 0.726743 | 0.071102 | 0.091 | 0.085 | 1 | 6 | Nrm | 0.006 | Old |
| 0.726751 | 0.005043 | 0.209 | 0.216 | 1 | 6 | Ppp1r21 | -0.007 | Old |
| 0.726802 | 0.288975 | 0.297 | 0.296 | 1 | 6 | Sptbn1 | 0.001 | Old |
| 0.726937 | 0.211585 | 0.063 | 0.057 | 1 | 6 | Acd | 0.006 | Old |
| 0.726996 | 0.12526 | 0.243 | 0.234 | 1 | 6 | Klc1 | 0.009 | Old |
| 0.727088 | 0.381054 | 0.08 | 0.087 | 1 | 6 | Ssna1 | -0.007 | Old |
| 0.727569 | 0.003085 | 0.549 | 0.548 | 1 | 6 | Dusp3 | 0.001 | Old |
| 0.727675 | 0.110321 | 0.554 | 0.54 | 1 | 6 | Hcls1 | 0.014 | Old |
| 0.727867 | 0.538142 | 0.057 | 0.052 | 1 | 6 | Gmeb2 | 0.005 | Old |
| 0.727867 | 0.190258 | 0.057 | 0.052 | 1 | 6 | Nt5c3 | 0.005 | Old |
| 0.727929 | 0.988456 | 0.043 | 0.038 | 1 | 6 | Cd209f | 0.005 | Old |
| 0.72799 | 0.57682 | 0.014 | 0.012 | 1 | 6 | Slc9a7 | 0.002 | Old |
| 0.72799 | 0.421393 | 0.014 | 0.012 | 1 | 6 | Slc25a53 | 0.002 | Old |
| 0.728184 | 0.108852 | 0.5 | 0.532 | 1 | 6 | Snrpb | -0.032 | Old |
| 0.728287 | 0.173387 | 0.186 | 0.177 | 1 | 6 | Ndufs4 | 0.009 | Old |
| 0.72836 | 0.376675 | 0.034 | 0.03 | 1 | 6 | Nxt2 | 0.004 | Old |
| 0.728389 | 0.108546 | 0.151 | 0.161 | 1 | 6 | Zfp740 | -0.01 | Old |
| 0.728392 | 0.152986 | 0.403 | 0.403 | 1 | 6 | P2ry10 | 0 | Old |
| 0.728811 | 0.155354 | 0.057 | 0.052 | 1 | 6 | Lclat1 | 0.005 | Old |
| 0.728943 | 0.611954 | 0.014 | 0.012 | 1 | 6 | Gm15445 | 0.002 | Old |
| 0.728943 | 0.315297 | 0.014 | 0.012 | 1 | 6 | Mlkl | 0.002 | Old |
| 0.729311 | 0.211301 | 0.066 | 0.06 | 1 | 6 | Depdc5 | 0.006 | Old |
| 0.729321 | 0.38695 | 0.149 | 0.144 | 1 | 6 | Stat2 | 0.005 | Old |
| 0.729558 | 0.017147 | 0.397 | 0.383 | 1 | 6 | Acly | 0.014 | Old |
| 0.729559 | 0.423112 | 0.043 | 0.038 | 1 | 6 | Mrpl46 | 0.005 | Old |
| 0.72971 | 1.052798 | 0.023 | 0.027 | 1 | 6 | Izumo4 | -0.004 | Old |
| 0.729897 | 0.496837 | 0.014 | 0.012 | 1 | 6 | Dnase1l3 | 0.002 | Old |
| 0.730069 | 0.045238 | 0.126 | 0.134 | 1 | 6 | 2310011J0 | -0.008 | Old |
| 0.730226 | 0.172091 | 0.149 | 0.139 | 1 | 6 | Psmd12 | 0.01 | Old |
| 0.730404 | 0.014051 | 0.091 | 0.084 | 1 | 6 | Vps26b | 0.007 | Old |
| 0.73065 | 0.081524 | 0.086 | 0.079 | 1 | 6 | Aak1 | 0.007 | Old |
| 0.730852 | 0.603807 | 0.014 | 0.012 | 1 | 6 | Nectin2 | 0.002 | Old |
| 0.730852 | 0.148886 | 0.014 | 0.012 | 1 | 6 | Spc25 | 0.002 | Old |
| 0.731068 | 1.200224 | 0.023 | 0.027 | 1 | 6 | Ryk | -0.004 | Old |
| 0.731134 | 0.699632 | 0.1 | 0.094 | 1 | 6 | Rhot1 | 0.006 | Old |
| 0.731195 | 0.19428 | 0.231 | 0.227 | 1 | 6 | Tmem168 | 0.004 | Old |
| 0.731806 | 0.261129 | 0.014 | 0.012 | 1 | 6 | Aoah | 0.002 | Old |
| 0.73216 | 0.477396 | 0.029 | 0.025 | 1 | 6 | Pwp2 | 0.004 | Old |
| 0.732406 | 0.198229 | 0.086 | 0.079 | 1 | 6 | Rnf138 | 0.007 | Old |

| | | | | | | | | |
|---|---|---|---|---|---|---|---|---|
| 0.732485 | 0.042405 | 0.16 | 0.151 | 1 | 6 | Atp13a1 | 0.009 | Old |
| 0.732514 | 0.427413 | 0.02 | 0.023 | 1 | 6 | Gm47230 | -0.003 | Old |
| 0.732764 | 0.416656 | 0.089 | 0.084 | 1 | 6 | Dnajc16 | 0.005 | Old |
| 0.732802 | 0.311806 | 0.191 | 0.182 | 1 | 6 | Arhgap5 | 0.009 | Old |
| 0.732943 | 0.131122 | 0.04 | 0.035 | 1 | 6 | Vamp5 | 0.005 | Old |
| 0.733018 | 0.069636 | 0.769 | 0.856 | 1 | 6 | Pid1 | -0.087 | Old |
| 0.733053 | 0.334139 | 0.074 | 0.069 | 1 | 6 | Endod1 | 0.005 | Old |
| 0.733124 | 0.076409 | 0.397 | 0.385 | 1 | 6 | Snx10 | 0.012 | Old |
| 0.733164 | 0.00111 | 0.134 | 0.125 | 1 | 6 | Casp6 | 0.009 | Old |
| 0.733317 | 0.386489 | 0.574 | 0.59 | 1 | 6 | Il1rl1 | -0.016 | Old |
| 0.733367 | 0.254773 | 0.043 | 0.038 | 1 | 6 | Ino80b | 0.005 | Old |
| 0.733679 | 0.119611 | 0.06 | 0.055 | 1 | 6 | Synpo | 0.005 | Old |
| 0.733713 | 0.318351 | 0.209 | 0.206 | 1 | 6 | Tug1 | 0.003 | Old |
| 0.734208 | 0.002774 | 0.08 | 0.074 | 1 | 6 | Pck2 | 0.006 | Old |
| 0.734415 | 0.195951 | 0.38 | 0.371 | 1 | 6 | Ppp1r9b | 0.009 | Old |
| 0.734555 | 0.256531 | 0.077 | 0.084 | 1 | 6 | Actr1b | -0.007 | Old |
| 0.734617 | 0.21615 | 0.083 | 0.077 | 1 | 6 | Nr1d2 | 0.006 | Old |
| 0.734625 | 0.077561 | 0.603 | 0.59 | 1 | 6 | Gls | 0.013 | Old |
| 0.735068 | 0.041798 | 0.034 | 0.03 | 1 | 6 | Gpn2 | 0.004 | Old |
| 0.735188 | 0.282328 | 0.249 | 0.241 | 1 | 6 | Idi1 | 0.008 | Old |
| 0.736475 | 0.42162 | 0.094 | 0.087 | 1 | 6 | Usp33 | 0.007 | Old |
| 0.736868 | 0.328973 | 0.129 | 0.139 | 1 | 6 | Ndufs1 | -0.01 | Old |
| 0.737066 | 0.03184 | 0.229 | 0.239 | 1 | 6 | Tifab | -0.01 | Old |
| 0.737366 | 0.056188 | 0.129 | 0.12 | 1 | 6 | Lrrk2 | 0.009 | Old |
| 0.737498 | 0.396302 | 0.029 | 0.025 | 1 | 6 | Atrip | 0.004 | Old |
| 0.73764 | 0.13494 | 0.089 | 0.082 | 1 | 6 | Pias2 | 0.007 | Old |
| 0.737701 | 0.02505 | 0.534 | 0.547 | 1 | 6 | Tmed2 | -0.013 | Old |
| 0.737911 | 0.153841 | 0.163 | 0.174 | 1 | 6 | Map3k7 | -0.011 | Old |
| 0.737982 | 0.238624 | 0.083 | 0.077 | 1 | 6 | Naa60 | 0.006 | Old |
| 0.738109 | 0.016987 | 0.254 | 0.242 | 1 | 6 | Pdxk | 0.012 | Old |
| 0.738435 | 0.257767 | 0.123 | 0.117 | 1 | 6 | Plxna1 | 0.006 | Old |
| 0.738502 | 0.155369 | 0.014 | 0.012 | 1 | 6 | Armcx2 | 0.002 | Old |
| 0.738834 | 0.691819 | 0.029 | 0.025 | 1 | 6 | Galnt9 | 0.004 | Old |
| 0.738925 | 0.068726 | 0.091 | 0.099 | 1 | 6 | Hotairm1 | -0.008 | Old |
| 0.738957 | 0.111473 | 0.123 | 0.13 | 1 | 6 | Ascc1 | -0.007 | Old |
| 0.739016 | 0.155193 | 0.197 | 0.184 | 1 | 6 | Ndufs3 | 0.013 | Old |
| 0.739121 | 0.019153 | 0.36 | 0.341 | 1 | 6 | Mrps33 | 0.019 | Old |
| 0.739766 | 0.363139 | 0.083 | 0.077 | 1 | 6 | Slc10a7 | 0.006 | Old |
| 0.740223 | 0.092025 | 0.1 | 0.092 | 1 | 6 | Ate1 | 0.008 | Old |
| 0.740651 | 0.138655 | 0.057 | 0.052 | 1 | 6 | Pithd1 | 0.005 | Old |
| 0.74084 | 0.830132 | 0.029 | 0.025 | 1 | 6 | Vcam1 | 0.004 | Old |
| 0.741124 | 0.018387 | 0.197 | 0.207 | 1 | 6 | Btk | -0.01 | Old |
| 0.74168 | 0.220198 | 0.071 | 0.067 | 1 | 6 | D430042O( | 0.004 | Old |
| 0.741796 | 0.369849 | 0.117 | 0.125 | 1 | 6 | Trappc1 | -0.008 | Old |
| 0.742153 | 0.109304 | 0.26 | 0.269 | 1 | 6 | Rnf187 | -0.009 | Old |

| | | | | | | | | |
|---|---|---|---|---|---|---|---|---|
| 0.742178 | 0.214695 | 0.029 | 0.025 | 1 | 6 | 1600014C1 | 0.004 | Old |
| 0.743059 | 0.155131 | 0.171 | 0.179 | 1 | 6 | Foxo1 | -0.008 | Old |
| 0.743456 | 0.230484 | 0.171 | 0.166 | 1 | 6 | Tm2d1 | 0.005 | Old |
| 0.743524 | 0.396378 | 0.171 | 0.167 | 1 | 6 | Mtm1 | 0.004 | Old |
| 0.743536 | 0.041097 | 0.129 | 0.137 | 1 | 6 | Gbp7 | -0.008 | Old |
| 0.744164 | 0.196673 | 0.217 | 0.212 | 1 | 6 | Adipor2 | 0.005 | Old |
| 0.744165 | 0.17889 | 0.297 | 0.286 | 1 | 6 | Mrpl20 | 0.011 | Old |
| 0.744187 | 0.199431 | 0.029 | 0.025 | 1 | 6 | 5730409E0 | 0.004 | Old |
| 0.744266 | 0.04569 | 0.22 | 0.229 | 1 | 6 | Ypel5 | -0.009 | Old |
| 0.744299 | 0.470386 | 0.034 | 0.038 | 1 | 6 | Zkscan6 | -0.004 | Old |
| 0.744453 | 0.023374 | 0.057 | 0.052 | 1 | 6 | Alg8 | 0.005 | Old |
| 0.74481 | 0.180155 | 0.283 | 0.281 | 1 | 6 | Setd2 | 0.002 | Old |
| 0.744857 | 0.545081 | 0.029 | 0.025 | 1 | 6 | Sparc | 0.004 | Old |
| 0.745086 | 0.067809 | 0.237 | 0.251 | 1 | 6 | Pnp | -0.014 | Old |
| 0.745259 | 0.259884 | 0.103 | 0.097 | 1 | 6 | Slc35a2 | 0.006 | Old |
| 0.745582 | 0.050862 | 0.311 | 0.314 | 1 | 6 | Cbfb | -0.003 | Old |
| 0.746208 | 0.175121 | 0.063 | 0.069 | 1 | 6 | Abl1 | -0.006 | Old |
| 0.74622 | 0.187952 | 0.274 | 0.261 | 1 | 6 | Ube2h | 0.013 | Old |
| 0.746583 | 0.161432 | 0.066 | 0.06 | 1 | 6 | Mfsd12 | 0.006 | Old |
| 0.74685 | 0.218215 | 0.24 | 0.254 | 1 | 6 | Hsd17b12 | -0.014 | Old |
| 0.746873 | 0.012929 | 0.634 | 0.645 | 1 | 6 | Myh9 | -0.011 | Old |
| 0.747026 | 0.327146 | 0.077 | 0.072 | 1 | 6 | Tor2a | 0.005 | Old |
| 0.747574 | 0.408951 | 0.054 | 0.06 | 1 | 6 | Itga1 | -0.006 | Old |
| 0.747609 | 0.761707 | 0.017 | 0.02 | 1 | 6 | Mapkbp1 | -0.003 | Old |
| 0.747935 | 0.138574 | 0.137 | 0.145 | 1 | 6 | Plekhm2 | -0.008 | Old |
| 0.748065 | 0.126721 | 0.069 | 0.064 | 1 | 6 | Zfp24 | 0.005 | Old |
| 0.748534 | 0.397297 | 0.02 | 0.023 | 1 | 6 | Zfp319 | -0.003 | Old |
| 0.748669 | 0.490497 | 0.077 | 0.072 | 1 | 6 | Prps2 | 0.005 | Old |
| 0.748881 | 0.577835 | 0.029 | 0.025 | 1 | 6 | Agap2 | 0.004 | Old |
| 0.748881 | 0.190156 | 0.029 | 0.025 | 1 | 6 | Katnb1 | 0.004 | Old |
| 0.748919 | 0.243674 | 0.037 | 0.033 | 1 | 6 | Trmt61a | 0.004 | Old |
| 0.749074 | 0.044281 | 0.106 | 0.112 | 1 | 6 | Zfp664 | -0.006 | Old |
| 0.749078 | 0.201598 | 0.051 | 0.047 | 1 | 6 | Plk4 | 0.004 | Old |
| 0.749179 | 0.004618 | 0.38 | 0.388 | 1 | 6 | Acin1 | -0.008 | Old |
| 0.749252 | 0.068627 | 0.066 | 0.06 | 1 | 6 | Lrp12 | 0.006 | Old |
| 0.749269 | 0.207397 | 0.163 | 0.156 | 1 | 6 | Stx6 | 0.007 | Old |
| 0.749414 | 0.061243 | 0.609 | 0.622 | 1 | 6 | Selplg | -0.013 | Old |
| 0.749439 | 0.055473 | 0.28 | 0.294 | 1 | 6 | Dgkd | -0.014 | Old |
| 0.749874 | 0.148826 | 0.389 | 0.386 | 1 | 6 | Ap2m1 | 0.003 | Old |
| 0.74989 | 0.005651 | 0.203 | 0.212 | 1 | 6 | Cat | -0.009 | Old |
| 0.749896 | 0.249177 | 0.243 | 0.236 | 1 | 6 | Hprt | 0.007 | Old |
| 0.749962 | 0.023758 | 0.357 | 0.349 | 1 | 6 | Ralb | 0.008 | Old |
| 0.750134 | 0.034133 | 0.403 | 0.428 | 1 | 6 | Tmed7 | -0.025 | Old |
| 0.750185 | 0.093294 | 0.737 | 0.731 | 1 | 6 | Dock2 | 0.006 | Old |
| 0.750224 | 0.094705 | 0.029 | 0.025 | 1 | 6 | Dtx4 | 0.004 | Old |

| | | | | | | | | |
|---|---|---|---|---|---|---|---|---|
| 0.750442 | 0.070626 | 0.314 | 0.323 | 1 | 6 | Krtcap2 | -0.009 | Old |
| 0.750569 | 0.060847 | 0.034 | 0.038 | 1 | 6 | Ptdss2 | -0.004 | Old |
| 0.750595 | 0.005345 | 0.106 | 0.099 | 1 | 6 | Rae1 | 0.007 | Old |
| 0.750701 | 0.178621 | 0.406 | 0.425 | 1 | 6 | Gng2 | -0.019 | Old |
| 0.750821 | 0.29781 | 0.177 | 0.171 | 1 | 6 | Ptov1 | 0.006 | Old |
| 0.751111 | 0.078019 | 0.186 | 0.176 | 1 | 6 | Fam50a | 0.01 | Old |
| 0.75114 | 0.257735 | 0.034 | 0.038 | 1 | 6 | Pdhx | -0.004 | Old |
| 0.752076 | 0.064814 | 0.057 | 0.052 | 1 | 6 | Ptrh2 | 0.005 | Old |
| 0.752192 | 0.309963 | 0.094 | 0.089 | 1 | 6 | Ttpal | 0.005 | Old |
| 0.752912 | 0.134142 | 0.029 | 0.025 | 1 | 6 | Pus3 | 0.004 | Old |
| 0.753227 | 0.022586 | 0.18 | 0.169 | 1 | 6 | Trappc2l | 0.011 | Old |
| 0.753333 | 0.054706 | 0.111 | 0.119 | 1 | 6 | Txlna | -0.008 | Old |
| 0.753561 | 0.031525 | 0.36 | 0.338 | 1 | 6 | Eif3l | 0.022 | Old |
| 0.753811 | 0.638268 | 0.077 | 0.072 | 1 | 6 | Ccdc97 | 0.005 | Old |
| 0.753914 | 0.2376 | 0.046 | 0.042 | 1 | 6 | Kif16b | 0.004 | Old |
| 0.754581 | 0.0872 | 0.171 | 0.164 | 1 | 6 | Cdc26 | 0.007 | Old |
| 0.754683 | 0.108756 | 0.083 | 0.077 | 1 | 6 | Acaca | 0.006 | Old |
| 0.754704 | 1.093347 | 0.037 | 0.042 | 1 | 6 | Ttc39c | -0.005 | Old |
| 0.754872 | 0.018273 | 0.094 | 0.087 | 1 | 6 | Mesd | 0.007 | Old |
| 0.754883 | 0.058613 | 0.083 | 0.077 | 1 | 6 | Pik3c3 | 0.006 | Old |
| 0.754974 | 0.57186 | 0.046 | 0.042 | 1 | 6 | Slc35a1 | 0.004 | Old |
| 0.75503 | 0.13675 | 0.151 | 0.144 | 1 | 6 | Stx5a | 0.007 | Old |
| 0.755039 | 0.035532 | 0.263 | 0.251 | 1 | 6 | Smarcd2 | 0.012 | Old |
| 0.755059 | 0.278788 | 0.263 | 0.247 | 1 | 6 | Irs2 | 0.016 | Old |
| 0.755145 | 0.053033 | 0.306 | 0.293 | 1 | 6 | Cdk11b | 0.013 | Old |
| 0.755148 | 0.104795 | 0.203 | 0.216 | 1 | 6 | Cdc40 | -0.013 | Old |
| 0.755598 | 0.162566 | 0.22 | 0.229 | 1 | 6 | N4bp2l1 | -0.009 | Old |
| 0.755603 | 0.056422 | 0.029 | 0.025 | 1 | 6 | Zfp87 | 0.004 | Old |
| 0.755664 | 0.216366 | 0.077 | 0.072 | 1 | 6 | Uckl1 | 0.005 | Old |
| 0.755812 | 0.157123 | 0.269 | 0.256 | 1 | 6 | Bscl2 | 0.013 | Old |
| 0.755971 | 0.813443 | 0.037 | 0.033 | 1 | 6 | Mmp9 | 0.004 | Old |
| 0.756276 | 0.496695 | 0.029 | 0.025 | 1 | 6 | Cxcl12 | 0.004 | Old |
| 0.75629 | 0.034203 | 0.017 | 0.02 | 1 | 6 | Fzd4 | -0.003 | Old |
| 0.756767 | 0.009465 | 0.426 | 0.431 | 1 | 6 | Esyt1 | -0.005 | Old |
| 0.757737 | 0.40721 | 0.037 | 0.033 | 1 | 6 | Ppox | 0.004 | Old |
| 0.758209 | 0.058211 | 0.36 | 0.381 | 1 | 6 | Tuba1a | -0.021 | Old |
| 0.758297 | 0.58149 | 0.029 | 0.025 | 1 | 6 | Ppm1f | 0.004 | Old |
| 0.758527 | 0.170977 | 0.243 | 0.237 | 1 | 6 | Inpp4a | 0.006 | Old |
| 0.758688 | 0.459609 | 0.054 | 0.05 | 1 | 6 | Fbxo32 | 0.004 | Old |
| 0.758723 | 0.265812 | 0.086 | 0.08 | 1 | 6 | Slc25a25 | 0.006 | Old |
| 0.759059 | 0.208573 | 0.177 | 0.184 | 1 | 6 | Kif1b | -0.007 | Old |
| 0.759154 | 0.409647 | 0.023 | 0.02 | 1 | 6 | Armcx5 | 0.003 | Old |
| 0.759408 | 0.132467 | 0.16 | 0.167 | 1 | 6 | Fry | -0.007 | Old |
| 0.759434 | 0.435465 | 0.223 | 0.216 | 1 | 6 | Ubp1 | 0.007 | Old |
| 0.759611 | 0.021421 | 0.046 | 0.05 | 1 | 6 | Hlx | -0.004 | Old |

| | | | | | | | | |
|---|---|---|---|---|---|---|---|---|
| 0.760262 | 0.284897 | 0.114 | 0.107 | 1 | 6 | Hdac4 | 0.007 | Old |
| 0.76028 | 0.251411 | 0.046 | 0.042 | 1 | 6 | Fxyd2 | 0.004 | Old |
| 0.760793 | 0.241995 | 0.131 | 0.124 | 1 | 6 | Nf1 | 0.007 | Old |
| 0.760921 | 0.291707 | 0.069 | 0.064 | 1 | 6 | Ints11 | 0.005 | Old |
| 0.761127 | 0.431655 | 0.054 | 0.05 | 1 | 6 | Fam222b | 0.004 | Old |
| 0.761308 | 0.102362 | 0.043 | 0.038 | 1 | 6 | Mto1 | 0.005 | Old |
| 0.761381 | 0.160395 | 0.16 | 0.151 | 1 | 6 | D5Ertd579 | 0.009 | Old |
| 0.762114 | 0.102714 | 0.186 | 0.196 | 1 | 6 | Phf5a | -0.01 | Old |
| 0.762367 | 0.048176 | 0.437 | 0.425 | 1 | 6 | Mndal | 0.012 | Old |
| 0.762553 | 0.10098 | 0.283 | 0.274 | 1 | 6 | Aph1c | 0.009 | Old |
| 0.762629 | 0.044587 | 0.251 | 0.242 | 1 | 6 | Araf | 0.009 | Old |
| 0.762915 | 0.50004 | 0.023 | 0.02 | 1 | 6 | Bst1 | 0.003 | Old |
| 0.762916 | 0.319785 | 0.243 | 0.258 | 1 | 6 | Rara | -0.015 | Old |
| 0.763132 | 0.057624 | 0.046 | 0.05 | 1 | 6 | Fam43a | -0.004 | Old |
| 0.763229 | 0.260174 | 0.1 | 0.107 | 1 | 6 | Slc25a12 | -0.007 | Old |
| 0.763355 | 0.400509 | 0.091 | 0.085 | 1 | 6 | Pde12 | 0.006 | Old |
| 0.763633 | 0.176954 | 0.037 | 0.033 | 1 | 6 | Oas3 | 0.004 | Old |
| 0.763668 | 0.641049 | 0.023 | 0.02 | 1 | 6 | Prune2 | 0.003 | Old |
| 0.763715 | 0.058911 | 0.383 | 0.38 | 1 | 6 | Ash1l | 0.003 | Old |
| 0.763745 | 0.018578 | 0.24 | 0.249 | 1 | 6 | Abhd17a | -0.009 | Old |
| 0.763868 | 0.123675 | 0.451 | 0.448 | 1 | 6 | Prpf38b | 0.003 | Old |
| 0.764157 | 0.088551 | 0.363 | 0.363 | 1 | 6 | Arf2 | 0 | Old |
| 0.764232 | 0.060275 | 0.289 | 0.281 | 1 | 6 | Snx2 | 0.008 | Old |
| 0.764414 | 0.201263 | 0.174 | 0.166 | 1 | 6 | Yipf4 | 0.008 | Old |
| 0.765168 | 0.052116 | 0.294 | 0.281 | 1 | 6 | Pias1 | 0.013 | Old |
| 0.765175 | 0.618213 | 0.023 | 0.02 | 1 | 6 | Ifrd2 | 0.003 | Old |
| 0.765249 | 0.036843 | 0.063 | 0.069 | 1 | 6 | Nhlrc3 | -0.006 | Old |
| 0.765404 | 0.236471 | 0.037 | 0.033 | 1 | 6 | Taf8 | 0.004 | Old |
| 0.765433 | 1.213328 | 0.031 | 0.028 | 1 | 6 | Vill | 0.003 | Old |
| 0.76594 | 0.338677 | 0.091 | 0.099 | 1 | 6 | Ipo8 | -0.008 | Old |
| 0.766036 | 0.100791 | 0.077 | 0.084 | 1 | 6 | 6430548M | -0.007 | Old |
| 0.766072 | 1.640983 | 0.031 | 0.028 | 1 | 6 | Dst | 0.003 | Old |
| 0.766286 | 0.189129 | 0.466 | 0.45 | 1 | 6 | Pgk1 | 0.016 | Old |
| 0.766524 | 0.266141 | 0.246 | 0.239 | 1 | 6 | Brd7 | 0.007 | Old |
| 0.766661 | 0.567098 | 0.046 | 0.042 | 1 | 6 | Havcr2 | 0.004 | Old |
| 0.766988 | 0.965602 | 0.063 | 0.069 | 1 | 6 | Fyco1 | -0.006 | Old |
| 0.767152 | 0.113122 | 0.194 | 0.202 | 1 | 6 | Mff | -0.008 | Old |
| 0.767162 | 0.015959 | 0.046 | 0.05 | 1 | 6 | Rpusd4 | -0.004 | Old |
| 0.767242 | 0.093334 | 0.146 | 0.156 | 1 | 6 | Ppm1k | -0.01 | Old |
| 0.767352 | 0.972219 | 0.031 | 0.028 | 1 | 6 | Timd4 | 0.003 | Old |
| 0.767441 | 0.161604 | 0.214 | 0.227 | 1 | 6 | Cyb5a | -0.013 | Old |
| 0.767637 | 0.143151 | 0.231 | 0.224 | 1 | 6 | Ufc1 | 0.007 | Old |
| 0.767992 | 0.690923 | 0.031 | 0.028 | 1 | 6 | Rpap2 | 0.003 | Old |
| 0.768712 | 0.182294 | 0.357 | 0.36 | 1 | 6 | Mdh1 | -0.003 | Old |
| 0.768949 | 0.08496 | 0.24 | 0.227 | 1 | 6 | Pa2g4 | 0.013 | Old |

| | | | | | | | | |
|---|---|---|---|---|---|---|---|---|
| 0.769811 | 0.017591 | 0.251 | 0.264 | 1 | 6 | Snrpc | -0.013 | Old |
| 0.770008 | 0.242044 | 0.383 | 0.366 | 1 | 6 | Crlf3 | 0.017 | Old |
| 0.770143 | 0.119834 | 0.169 | 0.161 | 1 | 6 | Lars | 0.008 | Old |
| 0.770325 | 0.091744 | 0.66 | 0.674 | 1 | 6 | Jun | -0.014 | Old |
| 0.770391 | 0.190566 | 0.046 | 0.042 | 1 | 6 | Parp6 | 0.004 | Old |
| 0.77045 | 0.16206 | 0.131 | 0.139 | 1 | 6 | Dse | -0.008 | Old |
| 0.770533 | 0.457732 | 0.014 | 0.017 | 1 | 6 | Acpp | -0.003 | Old |
| 0.770865 | 0.063603 | 0.251 | 0.261 | 1 | 6 | Ndufab1 | -0.01 | Old |
| 0.771195 | 0.426556 | 0.031 | 0.028 | 1 | 6 | Mxra7 | 0.003 | Old |
| 0.771196 | 0.07873 | 0.277 | 0.288 | 1 | 6 | Gm6377 | -0.011 | Old |
| 0.77121 | 0.333565 | 0.023 | 0.02 | 1 | 6 | AW146154 | 0.003 | Old |
| 0.771299 | 0.506979 | 0.109 | 0.104 | 1 | 6 | Lrba | 0.005 | Old |
| 0.771304 | 0.375962 | 0.06 | 0.055 | 1 | 6 | Decr1 | 0.005 | Old |
| 0.771364 | 0.052172 | 0.203 | 0.212 | 1 | 6 | Pik3r6 | -0.009 | Old |
| 0.771403 | 0.342976 | 0.014 | 0.017 | 1 | 6 | Mfsd6l | -0.003 | Old |
| 0.771416 | 0.056205 | 0.28 | 0.271 | 1 | 6 | Erp44 | 0.009 | Old |
| 0.771479 | 0.072022 | 0.451 | 0.448 | 1 | 6 | Uqcr10 | 0.003 | Old |
| 0.771836 | 0.902172 | 0.031 | 0.028 | 1 | 6 | Nprl2 | 0.003 | Old |
| 0.772705 | 0.056424 | 0.094 | 0.1 | 1 | 6 | Otud6b | -0.006 | Old |
| 0.773129 | 0.074388 | 0.22 | 0.226 | 1 | 6 | Fryl | -0.006 | Old |
| 0.773476 | 0.630077 | 0.023 | 0.02 | 1 | 6 | Camk2n1 | 0.003 | Old |
| 0.774014 | 0.364926 | 0.014 | 0.017 | 1 | 6 | Mfsd3 | -0.003 | Old |
| 0.774059 | 0.030996 | 0.531 | 0.538 | 1 | 6 | Klf2 | -0.007 | Old |
| 0.774409 | 0.257378 | 0.231 | 0.247 | 1 | 6 | Prkcsh | -0.016 | Old |
| 0.774486 | 0.159727 | 0.077 | 0.072 | 1 | 6 | Fem1a | 0.005 | Old |
| 0.774532 | 0.042384 | 0.594 | 0.632 | 1 | 6 | Vcp | -0.038 | Old |
| 0.774597 | 0.361361 | 0.12 | 0.115 | 1 | 6 | Wdr48 | 0.005 | Old |
| 0.774658 | 0.175438 | 0.363 | 0.361 | 1 | 6 | Fkbp1a | 0.002 | Old |
| 0.774884 | 0.113521 | 0.014 | 0.017 | 1 | 6 | Mmab | -0.003 | Old |
| 0.774963 | 0.009097 | 0.891 | 0.901 | 1 | 6 | Sem1 | -0.01 | Old |
| 0.775276 | 0.058001 | 0.403 | 0.396 | 1 | 6 | Atf6 | 0.007 | Old |
| 0.776667 | 0.095359 | 0.286 | 0.294 | 1 | 6 | Chd3 | -0.008 | Old |
| 0.776798 | 0.342093 | 0.046 | 0.042 | 1 | 6 | Atpaf2 | 0.004 | Old |
| 0.776818 | 0.151742 | 0.343 | 0.361 | 1 | 6 | Tsc22d3 | -0.018 | Old |
| 0.777613 | 0.685794 | 0.031 | 0.028 | 1 | 6 | Toe1 | 0.003 | Old |
| 0.777637 | 0.345921 | 0.086 | 0.08 | 1 | 6 | Ypel2 | 0.006 | Old |
| 0.777669 | 0.131993 | 0.36 | 0.381 | 1 | 6 | Lamtor1 | -0.021 | Old |
| 0.778143 | 0.220644 | 0.2 | 0.192 | 1 | 6 | Pqlc3 | 0.008 | Old |
| 0.778267 | 0.37455 | 0.091 | 0.099 | 1 | 6 | Dennd1a | -0.008 | Old |
| 0.778387 | 0.111905 | 0.277 | 0.266 | 1 | 6 | Srsf10 | 0.011 | Old |
| 0.778737 | 0.062668 | 0.117 | 0.11 | 1 | 6 | Slu7 | 0.007 | Old |
| 0.778757 | 0.184249 | 0.309 | 0.304 | 1 | 6 | Rc3h1 | 0.005 | Old |
| 0.779243 | 0.104873 | 0.014 | 0.017 | 1 | 6 | Pcnx4 | -0.003 | Old |
| 0.779371 | 0.218857 | 0.06 | 0.065 | 1 | 6 | Itpa | -0.005 | Old |
| 0.780035 | 0.15183 | 0.211 | 0.202 | 1 | 6 | Ibtk | 0.009 | Old |

| | | | | | | | | |
|---|---|---|---|---|---|---|---|---|
| 0.780212 | 0.201198 | 0.06 | 0.055 | 1 | 6 | Arhgef11 | 0.005 | Old |
| 0.780783 | 0.194681 | 0.277 | 0.269 | 1 | 6 | Tmem109 | 0.008 | Old |
| 0.780909 | 0.458913 | 0.057 | 0.054 | 1 | 6 | Slc2a1 | 0.003 | Old |
| 0.780988 | 0.248137 | 0.014 | 0.017 | 1 | 6 | Tut1 | -0.003 | Old |
| 0.781552 | 0.045532 | 0.08 | 0.075 | 1 | 6 | Nt5c2 | 0.005 | Old |
| 0.781586 | 0.115493 | 0.086 | 0.09 | 1 | 6 | Xpo4 | -0.004 | Old |
| 0.781631 | 0.114436 | 0.069 | 0.074 | 1 | 6 | Ercc3 | -0.005 | Old |
| 0.781861 | 0.146279 | 0.014 | 0.017 | 1 | 6 | 2900093K2 | -0.003 | Old |
| 0.781892 | 0.194756 | 0.154 | 0.162 | 1 | 6 | Ppm1b | -0.008 | Old |
| 0.782115 | 0.526932 | 0.031 | 0.028 | 1 | 6 | Inpp4b | 0.003 | Old |
| 0.783563 | 0.167408 | 0.074 | 0.069 | 1 | 6 | Yod1 | 0.005 | Old |
| 0.783608 | 0.346644 | 0.014 | 0.017 | 1 | 6 | Txk | -0.003 | Old |
| 0.784082 | 0.059695 | 0.091 | 0.085 | 1 | 6 | Plscr1 | 0.006 | Old |
| 0.785322 | 0.05626 | 0.8 | 0.833 | 1 | 6 | Ptp4a2 | -0.033 | Old |
| 0.785364 | 0.070804 | 0.046 | 0.042 | 1 | 6 | Mars | 0.004 | Old |
| 0.785583 | 0.385408 | 0.094 | 0.102 | 1 | 6 | Lsm14b | -0.008 | Old |
| 0.785847 | 0.120863 | 0.171 | 0.164 | 1 | 6 | Gmps | 0.007 | Old |
| 0.785979 | 0.505898 | 0.031 | 0.028 | 1 | 6 | Phc1 | 0.003 | Old |
| 0.786131 | 0.287216 | 0.074 | 0.08 | 1 | 6 | Dcaf17 | -0.006 | Old |
| 0.786347 | 0.074788 | 0.174 | 0.166 | 1 | 6 | Mllt6 | 0.008 | Old |
| 0.786361 | 0.418309 | 0.031 | 0.035 | 1 | 6 | B4gat1 | -0.004 | Old |
| 0.78663 | 0.086574 | 0.311 | 0.324 | 1 | 6 | Srp72 | -0.013 | Old |
| 0.786633 | 0.022198 | 0.054 | 0.05 | 1 | 6 | Zbtb43 | 0.004 | Old |
| 0.78667 | 0.05663 | 0.231 | 0.222 | 1 | 6 | Fyb | 0.009 | Old |
| 0.786684 | 0.065438 | 0.226 | 0.234 | 1 | 6 | Smim15 | -0.008 | Old |
| 0.786919 | 0.061369 | 0.469 | 0.49 | 1 | 6 | Wnt11 | -0.021 | Old |
| 0.787126 | 0.126196 | 0.054 | 0.05 | 1 | 6 | Pdpr | 0.004 | Old |
| 0.787132 | 0.263906 | 0.1 | 0.094 | 1 | 6 | Htra2 | 0.006 | Old |
| 0.787195 | 0.153844 | 0.149 | 0.14 | 1 | 6 | Arfgap2 | 0.009 | Old |
| 0.787217 | 0.387147 | 0.04 | 0.043 | 1 | 6 | Irf2bp1 | -0.003 | Old |
| 0.787267 | 0.356203 | 0.083 | 0.079 | 1 | 6 | Pik3r4 | 0.004 | Old |
| 0.787268 | 0.426956 | 0.031 | 0.028 | 1 | 6 | Fam58b | 0.003 | Old |
| 0.787472 | 0.047311 | 0.071 | 0.067 | 1 | 6 | Rps6kc1 | 0.004 | Old |
| 0.78748 | 0.08457 | 0.12 | 0.112 | 1 | 6 | Mak16 | 0.008 | Old |
| 0.788401 | 0.058188 | 0.094 | 0.089 | 1 | 6 | Sec23a | 0.005 | Old |
| 0.788421 | 0.257839 | 0.046 | 0.05 | 1 | 6 | Camta1 | -0.004 | Old |
| 0.788493 | 0.01545 | 0.151 | 0.157 | 1 | 6 | Ogfrl1 | -0.006 | Old |
| 0.788501 | 0.052234 | 0.08 | 0.075 | 1 | 6 | Fabp5 | 0.005 | Old |
| 0.788704 | 0.456276 | 0.1 | 0.095 | 1 | 6 | Donson | 0.005 | Old |
| 0.78909 | 0.018766 | 0.254 | 0.251 | 1 | 6 | Sp110 | 0.003 | Old |
| 0.789848 | 0.259593 | 0.031 | 0.028 | 1 | 6 | Prim2 | 0.003 | Old |
| 0.790084 | 0.402006 | 0.049 | 0.054 | 1 | 6 | Rcor3 | -0.005 | Old |
| 0.790306 | 0.275184 | 0.077 | 0.084 | 1 | 6 | Gfer | -0.007 | Old |
| 0.790494 | 0.195136 | 0.031 | 0.028 | 1 | 6 | Bcorl1 | 0.003 | Old |
| 0.790563 | 0.146317 | 0.054 | 0.059 | 1 | 6 | Zdhhc16 | -0.005 | Old |

| | | | | | | | | |
|---|---|---|---|---|---|---|---|---|
| 0.790596 | 0.164667 | 0.191 | 0.186 | 1 | 6 | Ahr | 0.005 | Old |
| 0.790601 | 0.0491 | 0.043 | 0.047 | 1 | 6 | Zfp410 | -0.004 | Old |
| 0.790769 | 0.083794 | 0.42 | 0.443 | 1 | 6 | Nop10 | -0.023 | Old |
| 0.791073 | 0.21411 | 0.231 | 0.229 | 1 | 6 | Rassf3 | 0.002 | Old |
| 0.79109 | 0.269869 | 0.029 | 0.032 | 1 | 6 | Gm49359 | -0.003 | Old |
| 0.791332 | 0.054436 | 0.057 | 0.062 | 1 | 6 | Serhl | -0.005 | Old |
| 0.791482 | 0.608559 | 0.017 | 0.015 | 1 | 6 | 4930522L1 | 0.002 | Old |
| 0.791818 | 0.006187 | 0.454 | 0.443 | 1 | 6 | Stx7 | 0.011 | Old |
| 0.791874 | 0.403363 | 0.04 | 0.037 | 1 | 6 | Ubxn8 | 0.003 | Old |
| 0.79199 | 0.178138 | 0.24 | 0.229 | 1 | 6 | Yeats4 | 0.011 | Old |
| 0.792059 | 0.291616 | 0.054 | 0.05 | 1 | 6 | Bcl9 | 0.004 | Old |
| 0.792114 | 0.099656 | 0.503 | 0.5 | 1 | 6 | Sema4a | 0.003 | Old |
| 0.792571 | 0.048359 | 0.354 | 0.37 | 1 | 6 | Pds5a | -0.016 | Old |
| 0.792621 | 0.229454 | 0.069 | 0.074 | 1 | 6 | Mbd5 | -0.005 | Old |
| 0.792891 | 0.138673 | 0.174 | 0.169 | 1 | 6 | Znrd1 | 0.005 | Old |
| 0.792895 | 0.265841 | 0.083 | 0.079 | 1 | 6 | Smim20 | 0.004 | Old |
| 0.792948 | 0.056435 | 0.12 | 0.112 | 1 | 6 | Rai1 | 0.008 | Old |
| 0.792971 | 0.282699 | 0.126 | 0.119 | 1 | 6 | Ubap1 | 0.007 | Old |
| 0.793077 | 0.070359 | 0.031 | 0.028 | 1 | 6 | 0610009O2 | 0.003 | Old |
| 0.793827 | 0.021504 | 0.409 | 0.396 | 1 | 6 | Trip12 | 0.013 | Old |
| 0.794112 | 0.485067 | 0.017 | 0.015 | 1 | 6 | B3galt4 | 0.002 | Old |
| 0.794208 | 0.153298 | 0.011 | 0.013 | 1 | 6 | Zfp607a | -0.002 | Old |
| 0.794208 | 0.040461 | 0.011 | 0.013 | 1 | 6 | Proser3 | -0.002 | Old |
| 0.794592 | 0.112158 | 0.38 | 0.368 | 1 | 6 | Tomm22 | 0.012 | Old |
| 0.795016 | 0.494957 | 0.031 | 0.028 | 1 | 6 | Mpv17l | 0.003 | Old |
| 0.795296 | 0.108016 | 0.206 | 0.201 | 1 | 6 | Phf23 | 0.005 | Old |
| 0.795308 | 0.086706 | 0.04 | 0.037 | 1 | 6 | Mtfr1 | 0.003 | Old |
| 0.795513 | 0.115563 | 0.354 | 0.344 | 1 | 6 | Ptpa | 0.01 | Old |
| 0.795536 | 0.059111 | 0.029 | 0.032 | 1 | 6 | St3gal6 | -0.003 | Old |
| 0.795543 | 0.114668 | 0.746 | 0.731 | 1 | 6 | Fus | 0.015 | Old |
| 0.795866 | 1.216505 | 0.017 | 0.015 | 1 | 6 | Pik3ap1 | 0.002 | Old |
| 0.796251 | 0.21708 | 0.057 | 0.054 | 1 | 6 | Jagn1 | 0.003 | Old |
| 0.796534 | 0.089182 | 0.16 | 0.166 | 1 | 6 | Ifnar2 | -0.006 | Old |
| 0.796632 | 0.220281 | 0.466 | 0.48 | 1 | 6 | Rab10 | -0.014 | Old |
| 0.797146 | 0.03396 | 0.011 | 0.013 | 1 | 6 | Rita1 | -0.002 | Old |
| 0.797622 | 0.375108 | 0.017 | 0.015 | 1 | 6 | Txnrd3 | 0.002 | Old |
| 0.797909 | 0.083003 | 0.32 | 0.313 | 1 | 6 | Dnajc5 | 0.007 | Old |
| 0.79805 | 0.128728 | 0.557 | 0.574 | 1 | 6 | Ogt | -0.017 | Old |
| 0.79806 | 0.405503 | 0.106 | 0.102 | 1 | 6 | Mosmo | 0.004 | Old |
| 0.798126 | 0.183574 | 0.011 | 0.013 | 1 | 6 | H2-T24 | -0.002 | Old |
| 0.79813 | 0.048146 | 0.077 | 0.082 | 1 | 6 | Sp2 | -0.005 | Old |
| 0.798175 | 0.232678 | 0.057 | 0.054 | 1 | 6 | Cnot9 | 0.003 | Old |
| 0.798261 | 0.031146 | 0.046 | 0.042 | 1 | 6 | Cep162 | 0.004 | Old |
| 0.79828 | 0.151448 | 0.114 | 0.109 | 1 | 6 | Ado | 0.005 | Old |
| 0.798615 | 0.182876 | 0.186 | 0.179 | 1 | 6 | Sppl3 | 0.007 | Old |

| | | | | | | | | |
|---|---|---|---|---|---|---|---|---|
| 0.798656 | 0.031104 | 0.449 | 0.43 | 1 | 6 | Ube2l3 | 0.019 | Old |
| 0.799106 | 0.13036 | 0.011 | 0.013 | 1 | 6 | Clec1a | -0.002 | Old |
| 0.799318 | 0.122721 | 0.531 | 0.535 | 1 | 6 | Bzw1 | -0.004 | Old |
| 0.799463 | 0.087642 | 0.711 | 0.711 | 1 | 6 | Clk1 | 0 | Old |
| 0.79954 | 0.397577 | 0.077 | 0.074 | 1 | 6 | Dclre1b | 0.003 | Old |
| 0.799545 | 0.064768 | 0.031 | 0.028 | 1 | 6 | D2hgdh | 0.003 | Old |
| 0.799641 | 0.051269 | 0.097 | 0.092 | 1 | 6 | Pitpnm1 | 0.005 | Old |
| 0.799979 | 0.265224 | 0.294 | 0.283 | 1 | 6 | Ythdf3 | 0.011 | Old |
| 0.79999 | 0.219259 | 0.029 | 0.032 | 1 | 6 | Prpf3 | -0.003 | Old |
| 0.800087 | 0.262736 | 0.011 | 0.013 | 1 | 6 | Tmsb15b1 | -0.002 | Old |
| 0.800203 | 0.035735 | 0.089 | 0.085 | 1 | 6 | Pmm2 | 0.004 | Old |
| 0.800678 | 0.465841 | 0.074 | 0.07 | 1 | 6 | Smc2 | 0.004 | Old |
| 0.801003 | 0.058314 | 0.129 | 0.135 | 1 | 6 | Prkar2b | -0.006 | Old |
| 0.801372 | 0.188241 | 0.091 | 0.087 | 1 | 6 | Rnf168 | 0.004 | Old |
| 0.802015 | 0.228805 | 0.017 | 0.015 | 1 | 6 | C2cd2 | 0.002 | Old |
| 0.802049 | 0.631845 | 0.011 | 0.013 | 1 | 6 | Cenpj | -0.002 | Old |
| 0.802049 | 0.364034 | 0.011 | 0.013 | 1 | 6 | Gm340 | -0.002 | Old |
| 0.802222 | 0.238735 | 0.2 | 0.194 | 1 | 6 | Hcst | 0.006 | Old |
| 0.802258 | 0.363178 | 0.111 | 0.105 | 1 | 6 | Srebf1 | 0.006 | Old |
| 0.802437 | 0.021334 | 0.089 | 0.084 | 1 | 6 | Psmd10 | 0.005 | Old |
| 0.802459 | 0.146218 | 0.04 | 0.043 | 1 | 6 | Hars2 | -0.003 | Old |
| 0.802509 | 0.0137 | 0.126 | 0.119 | 1 | 6 | Ctbp2 | 0.007 | Old |
| 0.8027 | 0.931845 | 0.066 | 0.062 | 1 | 6 | Gnpda2 | 0.004 | Old |
| 0.802829 | 0.061114 | 0.283 | 0.288 | 1 | 6 | Ptpn12 | -0.005 | Old |
| 0.80294 | 0.232447 | 0.166 | 0.157 | 1 | 6 | Slc41a1 | 0.009 | Old |
| 0.803031 | 0.716019 | 0.011 | 0.013 | 1 | 6 | Prim1 | -0.002 | Old |
| 0.803031 | 0.331737 | 0.011 | 0.013 | 1 | 6 | Rmdn1 | -0.002 | Old |
| 0.803031 | 0.238373 | 0.011 | 0.013 | 1 | 6 | Mtif3 | -0.002 | Old |
| 0.803111 | 0.049506 | 0.106 | 0.1 | 1 | 6 | Chchd7 | 0.006 | Old |
| 0.803436 | 0.171611 | 0.054 | 0.05 | 1 | 6 | 1600002K0 | 0.004 | Old |
| 0.803774 | 0.145912 | 0.017 | 0.015 | 1 | 6 | Gm45894 | 0.002 | Old |
| 0.80391 | 0.350896 | 0.04 | 0.037 | 1 | 6 | 2010111I0 | 0.003 | Old |
| 0.804427 | 0.068844 | 0.054 | 0.05 | 1 | 6 | Ppan | 0.004 | Old |
| 0.804654 | 0.172217 | 0.017 | 0.015 | 1 | 6 | Dyrk1b | 0.002 | Old |
| 0.804833 | 0.044154 | 0.191 | 0.196 | 1 | 6 | Map2k7 | -0.005 | Old |
| 0.805073 | 0.495118 | 0.106 | 0.102 | 1 | 6 | Nrf1 | 0.004 | Old |
| 0.805174 | 0.152823 | 0.214 | 0.206 | 1 | 6 | Fto | 0.008 | Old |
| 0.805486 | 0.168781 | 0.134 | 0.142 | 1 | 6 | Ccz1 | -0.008 | Old |
| 0.805826 | 0.012789 | 0.317 | 0.324 | 1 | 6 | Glipr2 | -0.007 | Old |
| 0.805938 | 0.322826 | 0.197 | 0.191 | 1 | 6 | Dazap1 | 0.006 | Old |
| 0.806404 | 0.00125 | 0.097 | 0.092 | 1 | 6 | Utp20 | 0.005 | Old |
| 0.806415 | 0.193317 | 0.371 | 0.373 | 1 | 6 | Ppp1r12a | -0.002 | Old |
| 0.806445 | 0.025385 | 0.523 | 0.533 | 1 | 6 | Mtpn | -0.01 | Old |
| 0.806675 | 0.061238 | 0.031 | 0.028 | 1 | 6 | Itfg2 | 0.003 | Old |
| 0.806929 | 0.348387 | 0.08 | 0.077 | 1 | 6 | Brcc3 | 0.003 | Old |

| | | | | | | | | |
|---|---|---|---|---|---|---|---|---|
| 0.807328 | 0.531705 | 0.057 | 0.054 | 1 | 6 | Cdt1 | 0.003 | Old |
| 0.807804 | 0.183609 | 0.026 | 0.028 | 1 | 6 | Thop1 | -0.002 | Old |
| 0.807912 | 0.025268 | 0.14 | 0.134 | 1 | 6 | Lst1 | 0.006 | Old |
| 0.808176 | 0.141823 | 0.017 | 0.015 | 1 | 6 | Proser2 | 0.002 | Old |
| 0.808176 | 0.102296 | 0.017 | 0.015 | 1 | 6 | Tmc4 | 0.002 | Old |
| 0.808232 | 0.033143 | 0.14 | 0.134 | 1 | 6 | D10Jhu81e | 0.006 | Old |
| 0.80912 | 0.148543 | 0.214 | 0.209 | 1 | 6 | Zfp217 | 0.005 | Old |
| 0.809644 | 0.128234 | 0.103 | 0.107 | 1 | 6 | B9d2 | -0.004 | Old |
| 0.809933 | 0.23249 | 0.334 | 0.344 | 1 | 6 | Sbno1 | -0.01 | Old |
| 0.810424 | 0.21928 | 0.054 | 0.059 | 1 | 6 | Sh3pxd2a | -0.005 | Old |
| 0.810857 | 0.48318 | 0.026 | 0.023 | 1 | 6 | Gm41307 | 0.003 | Old |
| 0.811318 | 0.539208 | 0.06 | 0.057 | 1 | 6 | Gpatch11 | 0.003 | Old |
| 0.811454 | 0.06031 | 0.26 | 0.271 | 1 | 6 | R3hdm4 | -0.011 | Old |
| 0.811512 | 0.062541 | 0.206 | 0.209 | 1 | 6 | Kmt5b | -0.003 | Old |
| 0.811578 | 0.038893 | 0.646 | 0.667 | 1 | 6 | Itm2c | -0.021 | Old |
| 0.811673 | 0.610311 | 0.057 | 0.054 | 1 | 6 | Polr3b | 0.003 | Old |
| 0.811729 | 0.23846 | 0.08 | 0.085 | 1 | 6 | Atrn | -0.005 | Old |
| 0.811871 | 0.220589 | 0.031 | 0.028 | 1 | 6 | Polm | 0.003 | Old |
| 0.812156 | 0.204545 | 0.057 | 0.054 | 1 | 6 | Gtf3c3 | 0.003 | Old |
| 0.81227 | 0.448534 | 0.083 | 0.079 | 1 | 6 | Rnf215 | 0.004 | Old |
| 0.81227 | 0.015737 | 0.126 | 0.12 | 1 | 6 | Rassf8 | 0.006 | Old |
| 0.812536 | 0.573595 | 0.04 | 0.037 | 1 | 6 | Tmco6 | 0.003 | Old |
| 0.812668 | 0.178867 | 0.217 | 0.214 | 1 | 6 | Cd300c2 | 0.003 | Old |
| 0.813302 | 0.273702 | 0.077 | 0.074 | 1 | 6 | Ankrd16 | 0.003 | Old |
| 0.813479 | 0.40451 | 0.129 | 0.124 | 1 | 6 | Gm28875 | 0.005 | Old |
| 0.813511 | 0.572191 | 0.077 | 0.074 | 1 | 6 | Phlpp1 | 0.003 | Old |
| 0.813954 | 0.03861 | 0.24 | 0.244 | 1 | 6 | Fam49a | -0.004 | Old |
| 0.814195 | 0.163698 | 0.317 | 0.308 | 1 | 6 | Cul3 | 0.009 | Old |
| 0.814374 | 0.367203 | 0.037 | 0.04 | 1 | 6 | Cbfa2t2 | -0.003 | Old |
| 0.814432 | 0.001977 | 0.134 | 0.139 | 1 | 6 | Zcchc24 | -0.005 | Old |
| 0.814435 | 0.255237 | 0.026 | 0.023 | 1 | 6 | Apex2 | 0.003 | Old |
| 0.81479 | 0.405434 | 0.131 | 0.127 | 1 | 6 | Tmem183a | 0.004 | Old |
| 0.814892 | 0.022176 | 0.306 | 0.314 | 1 | 6 | Cebpg | -0.008 | Old |
| 0.815063 | 0.298429 | 0.163 | 0.161 | 1 | 6 | Pikfyve | 0.002 | Old |
| 0.815215 | 0.978382 | 0.026 | 0.028 | 1 | 6 | Chpt1 | -0.002 | Old |
| 0.815416 | 0.292394 | 0.04 | 0.037 | 1 | 6 | Fbxo31 | 0.003 | Old |
| 0.8163 | 0.651762 | 0.051 | 0.048 | 1 | 6 | Tecpr1 | 0.003 | Old |
| 0.816325 | 0.073995 | 0.351 | 0.343 | 1 | 6 | Herc1 | 0.008 | Old |
| 0.817432 | 0.234141 | 0.069 | 0.065 | 1 | 6 | Pigu | 0.004 | Old |
| 0.817532 | 0.084815 | 0.191 | 0.186 | 1 | 6 | Slain2 | 0.005 | Old |
| 0.817726 | 0.78669 | 0.031 | 0.028 | 1 | 6 | Fcna | 0.003 | Old |
| 0.817938 | 0.154105 | 0.251 | 0.247 | 1 | 6 | Acap2 | 0.004 | Old |
| 0.8187 | 0.514552 | 0.146 | 0.145 | 1 | 6 | Kif1c | 0.001 | Old |
| 0.818733 | 0.032873 | 0.08 | 0.075 | 1 | 6 | Fam219a | 0.005 | Old |
| 0.818838 | 0.833496 | 0.051 | 0.048 | 1 | 6 | Tdp2 | 0.003 | Old |

| | | | | | | | | |
|---|---|---|---|---|---|---|---|---|
| 0.819502 | 0.186534 | 0.111 | 0.107 | 1 | 6 | Plekhf2 | 0.004 | Old |
| 0.819606 | 0.076245 | 0.177 | 0.171 | 1 | 6 | Ugp2 | 0.006 | Old |
| 0.819721 | 0.084256 | 0.166 | 0.172 | 1 | 6 | Npc1 | -0.006 | Old |
| 0.819991 | 0.265288 | 0.009 | 0.01 | 1 | 6 | Adamts4 | -0.001 | Old |
| 0.820166 | 0.072797 | 0.026 | 0.023 | 1 | 6 | Cdca8 | 0.003 | Old |
| 0.820166 | 0.279003 | 0.026 | 0.023 | 1 | 6 | H2-Q10 | 0.003 | Old |
| 0.820625 | 0.101951 | 0.223 | 0.217 | 1 | 6 | Atad2b | 0.006 | Old |
| 0.820883 | 0.258054 | 0.026 | 0.023 | 1 | 6 | Zfp65 | 0.003 | Old |
| 0.82142 | 0.001218 | 0.08 | 0.075 | 1 | 6 | Dopey2 | 0.005 | Old |
| 0.821457 | 0.057737 | 0.109 | 0.104 | 1 | 6 | Mfsd10 | 0.005 | Old |
| 0.821793 | 0.035332 | 0.274 | 0.284 | 1 | 6 | Fam105a | -0.01 | Old |
| 0.821868 | 0.040865 | 0.086 | 0.09 | 1 | 6 | Pdrg1 | -0.004 | Old |
| 0.821885 | 0.242107 | 0.143 | 0.149 | 1 | 6 | Jmjd6 | -0.006 | Old |
| 0.821926 | 0.12769 | 0.226 | 0.219 | 1 | 6 | Ppp3cb | 0.007 | Old |
| 0.821967 | 0.273275 | 0.026 | 0.028 | 1 | 6 | Ankrd13b | -0.002 | Old |
| 0.821981 | 0.284302 | 0.069 | 0.074 | 1 | 6 | Zdhhc17 | -0.005 | Old |
| 0.822143 | 0.574102 | 0.046 | 0.05 | 1 | 6 | Nrep | -0.004 | Old |
| 0.822313 | 0.173162 | 0.034 | 0.032 | 1 | 6 | Lppos | 0.002 | Old |
| 0.822318 | 0.228513 | 0.026 | 0.023 | 1 | 6 | Urb1 | 0.003 | Old |
| 0.822366 | 0.093147 | 0.091 | 0.097 | 1 | 6 | Ulk1 | -0.006 | Old |
| 0.822453 | 0.014706 | 0.2 | 0.207 | 1 | 6 | Phc2 | -0.007 | Old |
| 0.822868 | 0.325053 | 0.226 | 0.224 | 1 | 6 | Fam208a | 0.002 | Old |
| 0.823137 | 0.100832 | 0.077 | 0.074 | 1 | 6 | Stk16 | 0.003 | Old |
| 0.823263 | 0.570033 | 0.04 | 0.043 | 1 | 6 | Zdhhc4 | -0.003 | Old |
| 0.823288 | 0.082355 | 0.057 | 0.054 | 1 | 6 | Tns4 | 0.003 | Old |
| 0.823406 | 0.366167 | 0.009 | 0.01 | 1 | 6 | Amacr | -0.001 | Old |
| 0.823644 | 0.303738 | 0.043 | 0.04 | 1 | 6 | Stk39 | 0.003 | Old |
| 0.823815 | 0.024068 | 0.177 | 0.169 | 1 | 6 | Ncoa7 | 0.008 | Old |
| 0.823905 | 0.010563 | 0.151 | 0.144 | 1 | 6 | Anks1 | 0.007 | Old |
| 0.824106 | 0.282193 | 0.171 | 0.169 | 1 | 6 | Zfc3h1 | 0.002 | Old |
| 0.824269 | 0.075614 | 0.089 | 0.084 | 1 | 6 | Urm1 | 0.005 | Old |
| 0.824389 | 0.598266 | 0.066 | 0.07 | 1 | 6 | Eaf1 | -0.004 | Old |
| 0.824545 | 0.039729 | 0.009 | 0.01 | 1 | 6 | Ccdc114 | -0.001 | Old |
| 0.824545 | 0.015247 | 0.009 | 0.01 | 1 | 6 | Il11 | -0.001 | Old |
| 0.824983 | 0.050496 | 0.226 | 0.216 | 1 | 6 | Prkce | 0.01 | Old |
| 0.825109 | 0.186974 | 0.24 | 0.234 | 1 | 6 | Cep350 | 0.006 | Old |
| 0.825184 | 0.048294 | 0.406 | 0.395 | 1 | 6 | 9930111J2 | 0.011 | Old |
| 0.8253 | 0.236061 | 0.066 | 0.062 | 1 | 6 | Cxxc1 | 0.004 | Old |
| 0.825684 | 0.370632 | 0.009 | 0.01 | 1 | 6 | Aoc2 | -0.001 | Old |
| 0.825684 | 0.344473 | 0.009 | 0.01 | 1 | 6 | Gm9938 | -0.001 | Old |
| 0.825684 | 0.059072 | 0.009 | 0.01 | 1 | 6 | Mettl22 | -0.001 | Old |
| 0.825691 | 0.067669 | 0.286 | 0.274 | 1 | 6 | Plekha1 | 0.012 | Old |
| 0.825726 | 0.102473 | 0.611 | 0.61 | 1 | 6 | Cd2ap | 0.001 | Old |
| 0.825842 | 0.023851 | 0.64 | 0.644 | 1 | 6 | Reep5 | -0.004 | Old |
| 0.825908 | 0.271795 | 0.026 | 0.023 | 1 | 6 | Zfp358 | 0.003 | Old |

| | | | | | | | | |
|---|---|---|---|---|---|---|---|---|
| 0.826416 | 0.229364 | 0.143 | 0.137 | 1 | 6 | Dnajb4 | 0.006 | Old |
| 0.826471 | 0.077142 | 0.077 | 0.082 | 1 | 6 | Mtr | -0.005 | Old |
| 0.826824 | 0.446857 | 0.009 | 0.01 | 1 | 6 | Adam33 | -0.001 | Old |
| 0.826824 | 0.129185 | 0.009 | 0.01 | 1 | 6 | Rab38 | -0.001 | Old |
| 0.826828 | 1.224486 | 0.011 | 0.01 | 1 | 6 | Gm28187 | 0.001 | Old |
| 0.826828 | 1.143375 | 0.011 | 0.01 | 1 | 6 | Kcnj2 | 0.001 | Old |
| 0.827904 | 0.22718 | 0.034 | 0.032 | 1 | 6 | Gemin6 | 0.002 | Old |
| 0.82791 | 1.409499 | 0.011 | 0.01 | 1 | 6 | 1700008J0 | 0.001 | Old |
| 0.827964 | 0.429885 | 0.009 | 0.01 | 1 | 6 | Gm43569 | -0.001 | Old |
| 0.827964 | 0.085707 | 0.009 | 0.01 | 1 | 6 | 1110019D1 | -0.001 | Old |
| 0.828525 | 0.291005 | 0.034 | 0.032 | 1 | 6 | Abcb10 | 0.002 | Old |
| 0.828651 | 0.429396 | 0.043 | 0.04 | 1 | 6 | Atg14 | 0.003 | Old |
| 0.828657 | 0.054268 | 0.297 | 0.288 | 1 | 6 | Baiap2 | 0.009 | Old |
| 0.828811 | 0.006453 | 0.083 | 0.087 | 1 | 6 | Taf7 | -0.004 | Old |
| 0.829105 | 0.379723 | 0.009 | 0.01 | 1 | 6 | Zfp707 | -0.001 | Old |
| 0.829105 | 0.234919 | 0.009 | 0.01 | 1 | 6 | Scrn2 | -0.001 | Old |
| 0.82919 | 0.135569 | 0.037 | 0.04 | 1 | 6 | Mon1b | -0.003 | Old |
| 0.829769 | 0.153764 | 0.034 | 0.032 | 1 | 6 | Rfesd | 0.002 | Old |
| 0.830236 | 0.144492 | 0.143 | 0.137 | 1 | 6 | Ccdc71l | 0.006 | Old |
| 0.83054 | 0.341065 | 0.051 | 0.048 | 1 | 6 | Rbm10 | 0.003 | Old |
| 0.8306 | 1.007913 | 0.043 | 0.04 | 1 | 6 | Nupr1 | 0.003 | Old |
| 0.830695 | 0.171406 | 0.12 | 0.125 | 1 | 6 | Ankrd40 | -0.005 | Old |
| 0.830911 | 0.079463 | 0.226 | 0.234 | 1 | 6 | Bcas2 | -0.008 | Old |
| 0.831093 | 0.140752 | 0.231 | 0.246 | 1 | 6 | Pilra | -0.015 | Old |
| 0.831109 | 0.235668 | 0.077 | 0.074 | 1 | 6 | Ppp2r5d | 0.003 | Old |
| 0.8312 | 0.29465 | 0.066 | 0.062 | 1 | 6 | Sap30 | 0.004 | Old |
| 0.831387 | 0.563344 | 0.009 | 0.01 | 1 | 6 | Faap24 | -0.001 | Old |
| 0.831636 | 0.388548 | 0.034 | 0.032 | 1 | 6 | Tmem237 | 0.002 | Old |
| 0.831714 | 0.324398 | 0.043 | 0.04 | 1 | 6 | Alkbh8 | 0.003 | Old |
| 0.831902 | 0.115581 | 0.094 | 0.09 | 1 | 6 | Ptpn4 | 0.004 | Old |
| 0.831949 | 0.113775 | 0.283 | 0.293 | 1 | 6 | Tulp4 | -0.01 | Old |
| 0.832038 | 0.054377 | 0.117 | 0.112 | 1 | 6 | Bms1 | 0.005 | Old |
| 0.832114 | 0.012783 | 0.331 | 0.334 | 1 | 6 | Papola | -0.003 | Old |
| 0.832242 | 1.182039 | 0.011 | 0.01 | 1 | 6 | Tuft1 | 0.001 | Old |
| 0.832242 | 0.752277 | 0.011 | 0.01 | 1 | 6 | Sfxn5 | 0.001 | Old |
| 0.832258 | 0.053637 | 0.034 | 0.032 | 1 | 6 | Tsr3 | 0.002 | Old |
| 0.832529 | 0.904377 | 0.009 | 0.01 | 1 | 6 | Dppa3 | -0.001 | Old |
| 0.83273 | 0.143597 | 0.317 | 0.308 | 1 | 6 | Cs | 0.009 | Old |
| 0.832883 | 0.219992 | 0.129 | 0.132 | 1 | 6 | Rnf5 | -0.003 | Old |
| 0.833124 | 0.097183 | 0.469 | 0.462 | 1 | 6 | Ssb | 0.007 | Old |
| 0.833326 | 1.159657 | 0.011 | 0.01 | 1 | 6 | Lamc1 | 0.001 | Old |
| 0.833326 | 0.672629 | 0.011 | 0.01 | 1 | 6 | Itga6 | 0.001 | Old |
| 0.833326 | 0.600629 | 0.011 | 0.01 | 1 | 6 | 1700066M | 0.001 | Old |
| 0.833326 | 0.447645 | 0.011 | 0.01 | 1 | 6 | Susd1 | 0.001 | Old |
| 0.833357 | 0.253223 | 0.306 | 0.296 | 1 | 6 | Cab39 | 0.01 | Old |

| | | | | | | | | |
|---|---|---|---|---|---|---|---|---|
| 0.83369 | 0.05204 | 0.814 | 0.833 | 1 | 6 | Cd47 | -0.019 | Old |
| 0.833817 | 0.010749 | 0.023 | 0.025 | 1 | 6 | Card10 | -0.002 | Old |
| 0.834189 | 0.063836 | 0.286 | 0.293 | 1 | 6 | Plekhm3 | -0.007 | Old |
| 0.834206 | 0.049944 | 0.914 | 0.967 | 1 | 6 | Ptprc | -0.053 | Old |
| 0.834264 | 0.363244 | 0.04 | 0.043 | 1 | 6 | Mettl14 | -0.003 | Old |
| 0.834333 | 0.072597 | 0.037 | 0.04 | 1 | 6 | Phf11d | -0.003 | Old |
| 0.83441 | 1.176479 | 0.011 | 0.01 | 1 | 6 | Copz2 | 0.001 | Old |
| 0.83441 | 0.420347 | 0.011 | 0.01 | 1 | 6 | Rab11fip3 | 0.001 | Old |
| 0.834521 | 0.009055 | 0.211 | 0.204 | 1 | 6 | Ktn1 | 0.007 | Old |
| 0.834537 | 0.075339 | 0.023 | 0.025 | 1 | 6 | B630019K0 | -0.002 | Old |
| 0.83465 | 0.09005 | 0.266 | 0.256 | 1 | 6 | Sidt2 | 0.01 | Old |
| 0.834905 | 0.240767 | 0.037 | 0.04 | 1 | 6 | Gm49368 | -0.003 | Old |
| 0.834923 | 0.170734 | 0.18 | 0.189 | 1 | 6 | Cdc37l1 | -0.009 | Old |
| 0.835166 | 0.056075 | 0.263 | 0.269 | 1 | 6 | BC017643 | -0.006 | Old |
| 0.835977 | 0.42086 | 0.023 | 0.025 | 1 | 6 | Dgke | -0.002 | Old |
| 0.836363 | 0.046239 | 0.106 | 0.102 | 1 | 6 | Sec23ip | 0.004 | Old |
| 0.83658 | 2.024213 | 0.011 | 0.01 | 1 | 6 | Klre1 | 0.001 | Old |
| 0.83658 | 0.327254 | 0.011 | 0.01 | 1 | 6 | D130040H1 | 0.001 | Old |
| 0.836944 | 0.329465 | 0.046 | 0.048 | 1 | 6 | Nfrkb | -0.002 | Old |
| 0.837209 | 0.023984 | 0.071 | 0.075 | 1 | 6 | Ubap2 | -0.004 | Old |
| 0.837223 | 0.142374 | 0.086 | 0.09 | 1 | 6 | Espn | -0.004 | Old |
| 0.837418 | 0.423519 | 0.023 | 0.025 | 1 | 6 | Pla2g2d | -0.002 | Old |
| 0.837418 | 0.010952 | 0.026 | 0.023 | 1 | 6 | Gm12159 | 0.003 | Old |
| 0.837665 | 0.547174 | 0.011 | 0.01 | 1 | 6 | Dsel | 0.001 | Old |
| 0.837779 | 0.157689 | 0.309 | 0.301 | 1 | 6 | Tyk2 | 0.008 | Old |
| 0.837864 | 0.041642 | 0.034 | 0.032 | 1 | 6 | Clybl | 0.002 | Old |
| 0.838003 | 0.135086 | 0.091 | 0.087 | 1 | 6 | Kat7 | 0.004 | Old |
| 0.83875 | 0.462129 | 0.011 | 0.01 | 1 | 6 | Coa7 | 0.001 | Old |
| 0.839007 | 0.116988 | 0.12 | 0.115 | 1 | 6 | Prdm1 | 0.005 | Old |
| 0.839432 | 0.099118 | 0.223 | 0.229 | 1 | 6 | Stag1 | -0.006 | Old |
| 0.839836 | 0.186852 | 0.011 | 0.01 | 1 | 6 | Afap1 | 0.001 | Old |
| 0.839836 | 0.107868 | 0.011 | 0.01 | 1 | 6 | Cenpm | 0.001 | Old |
| 0.839873 | 0.233742 | 0.129 | 0.135 | 1 | 6 | Tbc1d8 | -0.006 | Old |
| 0.840082 | 0.067559 | 0.043 | 0.04 | 1 | 6 | Adck1 | 0.003 | Old |
| 0.840281 | 0.293473 | 0.251 | 0.246 | 1 | 6 | Ankrd12 | 0.005 | Old |
| 0.840301 | 0.093609 | 0.023 | 0.025 | 1 | 6 | Enho | -0.002 | Old |
| 0.84043 | 0.005025 | 0.223 | 0.224 | 1 | 6 | Gnl1 | -0.001 | Old |
| 0.840949 | 0.014068 | 0.243 | 0.232 | 1 | 6 | Dctn3 | 0.011 | Old |
| 0.840997 | 0.581284 | 0.057 | 0.06 | 1 | 6 | Dcp1a | -0.003 | Old |
| 0.841022 | 0.549081 | 0.023 | 0.025 | 1 | 6 | Pde4d | -0.002 | Old |
| 0.841022 | 0.318044 | 0.023 | 0.025 | 1 | 6 | Msrb3 | -0.002 | Old |
| 0.841527 | 0.062841 | 0.257 | 0.246 | 1 | 6 | Sugt1 | 0.011 | Old |
| 0.84156 | 0.040464 | 0.06 | 0.057 | 1 | 6 | Sphk2 | 0.003 | Old |
| 0.841683 | 0.005877 | 0.306 | 0.309 | 1 | 6 | Uqcrc2 | -0.003 | Old |
| 0.842008 | 0.042947 | 0.011 | 0.01 | 1 | 6 | 5430403G1 | 0.001 | Old |

| | | | | | | | | |
|---|---|---|---|---|---|---|---|---|
| 0.84232 | 0.127338 | 0.194 | 0.192 | 1 | 6 | Rsbn1l | 0.002 | Old |
| 0.842372 | 0.075702 | 0.634 | 0.654 | 1 | 6 | Tomm7 | -0.02 | Old |
| 0.842519 | 0.165147 | 0.269 | 0.281 | 1 | 6 | Fam168a | -0.012 | Old |
| 0.842519 | 0.048686 | 0.166 | 0.169 | 1 | 6 | Lats2 | -0.003 | Old |
| 0.842583 | 0.116893 | 0.097 | 0.102 | 1 | 6 | Mcm3 | -0.005 | Old |
| 0.842846 | 0.183299 | 0.56 | 0.605 | 1 | 6 | Ldha | -0.045 | Old |
| 0.842942 | 0.130662 | 0.103 | 0.1 | 1 | 6 | Aggf1 | 0.003 | Old |
| 0.842955 | 0.129285 | 0.06 | 0.064 | 1 | 6 | Zfp945 | -0.004 | Old |
| 0.843035 | 0.05499 | 0.329 | 0.324 | 1 | 6 | Dnaja2 | 0.005 | Old |
| 0.843095 | 0.148233 | 0.011 | 0.01 | 1 | 6 | Sac3d1 | 0.001 | Old |
| 0.843095 | 0.071488 | 0.011 | 0.01 | 1 | 6 | Slc4a11 | 0.001 | Old |
| 0.84322 | 0.040962 | 0.343 | 0.333 | 1 | 6 | Unc119 | 0.01 | Old |
| 0.843489 | 0.236335 | 0.126 | 0.132 | 1 | 6 | Nubp1 | -0.006 | Old |
| 0.843595 | 0.028499 | 0.369 | 0.376 | 1 | 6 | Hfe | -0.007 | Old |
| 0.84397 | 0.056764 | 0.383 | 0.361 | 1 | 6 | Ube2k | 0.022 | Old |
| 0.844182 | 0.168203 | 0.011 | 0.01 | 1 | 6 | Gng11 | 0.001 | Old |
| 0.844868 | 0.090662 | 0.389 | 0.388 | 1 | 6 | Nsd3 | 0.001 | Old |
| 0.845309 | 0.492249 | 0.083 | 0.089 | 1 | 6 | Tbcel | -0.006 | Old |
| 0.845608 | 0.093719 | 0.086 | 0.09 | 1 | 6 | Tmtc2 | -0.004 | Old |
| 0.845836 | 0.075345 | 0.329 | 0.321 | 1 | 6 | Rer1 | 0.008 | Old |
| 0.846452 | 0.11128 | 0.263 | 0.273 | 1 | 6 | Cr1l | -0.01 | Old |
| 0.846531 | 0.454385 | 0.054 | 0.052 | 1 | 6 | Tgm2 | 0.002 | Old |
| 0.846779 | 0.02469 | 0.091 | 0.087 | 1 | 6 | Zbtb16 | 0.004 | Old |
| 0.846812 | 0.145645 | 0.291 | 0.299 | 1 | 6 | Mxd4 | -0.008 | Old |
| 0.847277 | 0.125138 | 0.146 | 0.151 | 1 | 6 | Psmc4 | -0.005 | Old |
| 0.847401 | 0.773633 | 0.029 | 0.027 | 1 | 6 | Serpinf1 | 0.002 | Old |
| 0.847799 | 0.194549 | 0.169 | 0.164 | 1 | 6 | Usp22 | 0.005 | Old |
| 0.848016 | 0.050395 | 0.094 | 0.09 | 1 | 6 | Gdap2 | 0.004 | Old |
| 0.848081 | 0.276293 | 0.029 | 0.027 | 1 | 6 | Cul9 | 0.002 | Old |
| 0.848109 | 0.193858 | 0.086 | 0.082 | 1 | 6 | Vti1a | 0.004 | Old |
| 0.84816 | 0.410131 | 0.071 | 0.077 | 1 | 6 | Ppp1r37 | -0.006 | Old |
| 0.848232 | 0.117956 | 0.211 | 0.214 | 1 | 6 | Irak4 | -0.003 | Old |
| 0.848233 | 0.067785 | 0.317 | 0.331 | 1 | 6 | Sri | -0.014 | Old |
| 0.84824 | 0.537541 | 0.023 | 0.025 | 1 | 6 | Satb1 | -0.002 | Old |
| 0.848305 | 0.198294 | 0.063 | 0.067 | 1 | 6 | Rab33b | -0.004 | Old |
| 0.848415 | 0.162686 | 0.14 | 0.135 | 1 | 6 | Napg | 0.005 | Old |
| 0.84885 | 0.085727 | 0.214 | 0.207 | 1 | 6 | BC005624 | 0.007 | Old |
| 0.848948 | 0.042552 | 0.046 | 0.048 | 1 | 6 | Ciao1 | -0.002 | Old |
| 0.849438 | 0.132876 | 0.151 | 0.157 | 1 | 6 | Tec | -0.006 | Old |
| 0.849708 | 0.092207 | 0.454 | 0.485 | 1 | 6 | Rpl36al | -0.031 | Old |
| 0.849913 | 0.450955 | 0.02 | 0.018 | 1 | 6 | Gm10053 | 0.002 | Old |
| 0.850118 | 0.334796 | 0.08 | 0.085 | 1 | 6 | Gbp5 | -0.005 | Old |
| 0.850124 | 0.478384 | 0.029 | 0.027 | 1 | 6 | Mrrf | 0.002 | Old |
| 0.850245 | 0.019953 | 0.169 | 0.174 | 1 | 6 | Naa38 | -0.005 | Old |
| 0.850604 | 0.034259 | 0.72 | 0.784 | 1 | 6 | Mob1a | -0.064 | Old |

| | | | | | | | | |
|---|---|---|---|---|---|---|---|---|
| 0.850667 | 0.146621 | 0.049 | 0.052 | 1 | 6 | Prr12 | -0.003 | Old |
| 0.85071 | 0.043323 | 0.229 | 0.227 | 1 | 6 | Rps12-ps3 | 0.002 | Old |
| 0.850805 | 1.029498 | 0.029 | 0.027 | 1 | 6 | Dhx34 | 0.002 | Old |
| 0.850865 | 0.163564 | 0.203 | 0.202 | 1 | 6 | Azi2 | 0.001 | Old |
| 0.850924 | 0.075147 | 0.106 | 0.102 | 1 | 6 | Cnnm4 | 0.004 | Old |
| 0.850978 | 0.295214 | 0.051 | 0.048 | 1 | 6 | Ahi1 | 0.003 | Old |
| 0.85145 | 0.184459 | 0.069 | 0.072 | 1 | 6 | Cep63 | -0.003 | Old |
| 0.851503 | 0.44838 | 0.054 | 0.052 | 1 | 6 | 2510009E0 | 0.002 | Old |
| 0.851561 | 0.065043 | 0.394 | 0.388 | 1 | 6 | Brd4 | 0.006 | Old |
| 0.851623 | 0.128869 | 0.329 | 0.326 | 1 | 6 | Vps4b | 0.003 | Old |
| 0.852001 | 0.594158 | 0.054 | 0.052 | 1 | 6 | 2410089E0 | 0.002 | Old |
| 0.852034 | 0.059868 | 0.2 | 0.192 | 1 | 6 | Meaf6 | 0.008 | Old |
| 0.852314 | 0.383142 | 0.037 | 0.035 | 1 | 6 | Nccrp1 | 0.002 | Old |
| 0.852359 | 0.750409 | 0.02 | 0.018 | 1 | 6 | Adora3 | 0.002 | Old |
| 0.852359 | 0.356213 | 0.02 | 0.018 | 1 | 6 | Pmepa1 | 0.002 | Old |
| 0.852613 | 0.209076 | 0.034 | 0.037 | 1 | 6 | Acvr1b | -0.003 | Old |
| 0.852766 | 0.07408 | 0.4 | 0.393 | 1 | 6 | Cdv3 | 0.007 | Old |
| 0.852803 | 0.172818 | 0.114 | 0.11 | 1 | 6 | Mast3 | 0.004 | Old |
| 0.852883 | 0.110042 | 0.171 | 0.169 | 1 | 6 | Psmd3 | 0.002 | Old |
| 0.85299 | 0.276308 | 0.069 | 0.065 | 1 | 6 | Lsm1 | 0.004 | Old |
| 0.853511 | 0.626861 | 0.037 | 0.035 | 1 | 6 | Hmgxb3 | 0.002 | Old |
| 0.854002 | 0.059026 | 0.206 | 0.199 | 1 | 6 | Ndufa12 | 0.007 | Old |
| 0.854034 | 0.235821 | 0.374 | 0.376 | 1 | 6 | Gm26532 | -0.002 | Old |
| 0.854329 | 0.262319 | 0.069 | 0.065 | 1 | 6 | Usp10 | 0.004 | Old |
| 0.854331 | 0.278882 | 0.111 | 0.107 | 1 | 6 | Ncoa4 | 0.004 | Old |
| 0.855776 | 0.022768 | 0.246 | 0.249 | 1 | 6 | Lsm14a | -0.003 | Old |
| 0.856203 | 0.037857 | 0.34 | 0.346 | 1 | 6 | Arl8b | -0.006 | Old |
| 0.856437 | 0.27057 | 0.02 | 0.018 | 1 | 6 | Blm | 0.002 | Old |
| 0.856488 | 0.167958 | 0.24 | 0.232 | 1 | 6 | Wbp11 | 0.008 | Old |
| 0.856616 | 0.077722 | 0.051 | 0.048 | 1 | 6 | Rcan1 | 0.003 | Old |
| 0.85693 | 1.005037 | 0.046 | 0.043 | 1 | 6 | 1700037H0 | 0.003 | Old |
| 0.857079 | 0.030451 | 0.826 | 0.838 | 1 | 6 | Slc25a3 | -0.012 | Old |
| 0.857103 | 0.559819 | 0.037 | 0.035 | 1 | 6 | Nudt1 | 0.002 | Old |
| 0.857103 | 0.043942 | 0.037 | 0.035 | 1 | 6 | Rmnd1 | 0.002 | Old |
| 0.85724 | 0.225803 | 0.163 | 0.172 | 1 | 6 | Tollip | -0.009 | Old |
| 0.857343 | 0.066855 | 0.469 | 0.47 | 1 | 6 | St3gal5 | -0.001 | Old |
| 0.857662 | 0.355085 | 0.02 | 0.018 | 1 | 6 | Ap5s1 | 0.002 | Old |
| 0.857808 | 0.109385 | 0.366 | 0.383 | 1 | 6 | Fmnl1 | -0.017 | Old |
| 0.858077 | 0.005272 | 0.231 | 0.224 | 1 | 6 | Vma21 | 0.007 | Old |
| 0.858242 | 0.21555 | 0.366 | 0.355 | 1 | 6 | Rbm5 | 0.011 | Old |
| 0.858287 | 0.010785 | 0.12 | 0.124 | 1 | 6 | Ahctf1 | -0.004 | Old |
| 0.858483 | 0.118299 | 0.089 | 0.09 | 1 | 6 | Trappc9 | -0.001 | Old |
| 0.859037 | 0.162334 | 0.097 | 0.102 | 1 | 6 | Wdr20 | -0.005 | Old |
| 0.859059 | 0.083424 | 0.46 | 0.455 | 1 | 6 | Nipa2 | 0.005 | Old |
| 0.859166 | 0.152572 | 0.123 | 0.127 | 1 | 6 | Mob3c | -0.004 | Old |

| | | | | | | | | |
|---|---|---|---|---|---|---|---|---|
| 0.859206 | 0.045793 | 0.34 | 0.331 | 1 | 6 | Yy1 | 0.009 | Old |
| 0.859323 | 0.001795 | 0.4 | 0.393 | 1 | 6 | Arid1a | 0.007 | Old |
| 0.859655 | 0.01015 | 0.126 | 0.122 | 1 | 6 | Pbx2 | 0.004 | Old |
| 0.859689 | 0.142855 | 0.254 | 0.251 | 1 | 6 | Herc4 | 0.003 | Old |
| 0.859699 | 0.069377 | 0.069 | 0.072 | 1 | 6 | Nop14 | -0.003 | Old |
| 0.859707 | 0.879529 | 0.02 | 0.022 | 1 | 6 | 2610021A( | -0.002 | Old |
| 0.859707 | 0.178625 | 0.02 | 0.022 | 1 | 6 | Gm4673 | -0.002 | Old |
| 0.859736 | 0.025796 | 0.203 | 0.194 | 1 | 6 | Rps6kb1 | 0.009 | Old |
| 0.859816 | 0.969476 | 0.023 | 0.025 | 1 | 6 | Enpp4 | -0.002 | Old |
| 0.85997 | 0.182141 | 0.054 | 0.052 | 1 | 6 | Tsga10 | 0.002 | Old |
| 0.860461 | 0.355443 | 0.129 | 0.134 | 1 | 6 | Nup54 | -0.005 | Old |
| 0.860485 | 0.275488 | 0.066 | 0.069 | 1 | 6 | Slc7a6os | -0.003 | Old |
| 0.86052 | 0.37907 | 0.02 | 0.018 | 1 | 6 | Dnal4 | 0.002 | Old |
| 0.860925 | 0.107158 | 0.814 | 0.866 | 1 | 6 | Lcp1 | -0.052 | Old |
| 0.861019 | 0.101392 | 0.183 | 0.179 | 1 | 6 | Slc35a3 | 0.004 | Old |
| 0.861191 | 0.035116 | 0.354 | 0.366 | 1 | 6 | Hspa4 | -0.012 | Old |
| 0.861337 | 0.022936 | 0.291 | 0.289 | 1 | 6 | Csnk1g2 | 0.002 | Old |
| 0.861337 | 0.631989 | 0.02 | 0.018 | 1 | 6 | Zfp825 | 0.002 | Old |
| 0.861337 | 0.554795 | 0.02 | 0.018 | 1 | 6 | Klhl22 | 0.002 | Old |
| 0.861396 | 0.058433 | 0.08 | 0.084 | 1 | 6 | St6galnac4 | -0.004 | Old |
| 0.861619 | 0.101899 | 0.257 | 0.269 | 1 | 6 | Xpr1 | -0.012 | Old |
| 0.862213 | 0.099775 | 0.066 | 0.062 | 1 | 6 | Ipcef1 | 0.004 | Old |
| 0.862693 | 0.291119 | 0.057 | 0.06 | 1 | 6 | Gk | -0.003 | Old |
| 0.862811 | 0.918891 | 0.02 | 0.022 | 1 | 6 | Purg | -0.002 | Old |
| 0.862971 | 0.082812 | 0.02 | 0.018 | 1 | 6 | Fam177a | 0.002 | Old |
| 0.863138 | 0.258915 | 0.251 | 0.256 | 1 | 6 | mt-Atp8 | -0.005 | Old |
| 0.863587 | 0.11479 | 0.02 | 0.022 | 1 | 6 | Atpaf1 | -0.002 | Old |
| 0.863877 | 0.599093 | 0.114 | 0.122 | 1 | 6 | Amn1 | -0.008 | Old |
| 0.864445 | 0.049218 | 0.231 | 0.236 | 1 | 6 | Siva1 | -0.005 | Old |
| 0.865117 | 0.25276 | 0.137 | 0.134 | 1 | 6 | Brd3 | 0.003 | Old |
| 0.865123 | 0.199155 | 0.109 | 0.112 | 1 | 6 | Tusc1 | -0.003 | Old |
| 0.86513 | 0.079918 | 0.029 | 0.027 | 1 | 6 | Riox1 | 0.002 | Old |
| 0.865921 | 0.111019 | 0.331 | 0.329 | 1 | 6 | Asph | 0.002 | Old |
| 0.866612 | 0.058513 | 0.117 | 0.114 | 1 | 6 | Sv2b | 0.003 | Old |
| 0.866635 | 0.259447 | 0.271 | 0.276 | 1 | 6 | Etnk1 | -0.005 | Old |
| 0.867072 | 0.394394 | 0.149 | 0.149 | 1 | 6 | Fam46c | 0 | Old |
| 0.86715 | 0.092008 | 0.066 | 0.069 | 1 | 6 | Snapc3 | -0.003 | Old |
| 0.867264 | 0.319504 | 0.146 | 0.142 | 1 | 6 | Sec24c | 0.004 | Old |
| 0.867367 | 0.001335 | 0.16 | 0.161 | 1 | 6 | Atp9b | -0.001 | Old |
| 0.86747 | 0.180175 | 0.02 | 0.022 | 1 | 6 | Rad18 | -0.002 | Old |
| 0.867541 | 0.010089 | 0.146 | 0.151 | 1 | 6 | Fbxo4 | -0.005 | Old |
| 0.867802 | 0.007603 | 0.457 | 0.452 | 1 | 6 | Srsf11 | 0.005 | Old |
| 0.867878 | 0.28228 | 0.02 | 0.018 | 1 | 6 | Tbx21 | 0.002 | Old |
| 0.868141 | 0.216665 | 0.097 | 0.094 | 1 | 6 | 2310022A1 | 0.003 | Old |
| 0.86915 | 0.068982 | 0.031 | 0.033 | 1 | 6 | Ogfod3 | -0.002 | Old |

| | | | | | | | | |
|---|---|---|---|---|---|---|---|---|
| 0.869231 | 0.320935 | 0.029 | 0.027 | 1 | 6 | Dis3 | 0.002 | Old |
| 0.869399 | 0.284474 | 0.034 | 0.037 | 1 | 6 | Qrsl1 | -0.003 | Old |
| 0.869958 | 0.062548 | 0.44 | 0.448 | 1 | 6 | Ptpn6 | -0.008 | Old |
| 0.870605 | 0.026391 | 0.417 | 0.413 | 1 | 6 | Plekhb2 | 0.004 | Old |
| 0.870623 | 0.204616 | 0.377 | 0.36 | 1 | 6 | Rab21 | 0.017 | Old |
| 0.870706 | 0.015154 | 0.183 | 0.182 | 1 | 6 | Shoc2 | 0.001 | Old |
| 0.871659 | 0.093211 | 0.229 | 0.231 | 1 | 6 | Rbm33 | -0.002 | Old |
| 0.871783 | 0.321063 | 0.243 | 0.241 | 1 | 6 | 2810403A0 | 0.002 | Old |
| 0.87211 | 0.190929 | 0.046 | 0.043 | 1 | 6 | Kptn | 0.003 | Old |
| 0.872381 | 0.101615 | 0.043 | 0.045 | 1 | 6 | Hagh | -0.002 | Old |
| 0.872463 | 0.261167 | 0.2 | 0.204 | 1 | 6 | Fundc1 | -0.004 | Old |
| 0.87284 | 0.179887 | 0.157 | 0.152 | 1 | 6 | Gabarapl1 | 0.005 | Old |
| 0.873088 | 0.01949 | 0.086 | 0.082 | 1 | 6 | Tbc1d17 | 0.004 | Old |
| 0.873126 | 0.064609 | 0.086 | 0.089 | 1 | 6 | Susd3 | -0.003 | Old |
| 0.873345 | 0.226879 | 0.369 | 0.383 | 1 | 6 | Pik3r1 | -0.014 | Old |
| 0.873366 | 0.343561 | 0.071 | 0.075 | 1 | 6 | Togaram1 | -0.004 | Old |
| 0.873547 | 0.253767 | 0.031 | 0.033 | 1 | 6 | Kctd6 | -0.002 | Old |
| 0.873603 | 0.174309 | 0.074 | 0.072 | 1 | 6 | Exoc6 | 0.002 | Old |
| 0.87366 | 0.341982 | 0.297 | 0.299 | 1 | 6 | Derl1 | -0.002 | Old |
| 0.873723 | 0.234198 | 0.043 | 0.04 | 1 | 6 | Cep104 | 0.003 | Old |
| 0.873906 | 0.748258 | 0.037 | 0.035 | 1 | 6 | Zfp142 | 0.002 | Old |
| 0.874016 | 0.037175 | 0.154 | 0.157 | 1 | 6 | Usp14 | -0.003 | Old |
| 0.874206 | 0.589921 | 0.034 | 0.037 | 1 | 6 | Gpr171 | -0.003 | Old |
| 0.874553 | 0.069069 | 0.303 | 0.299 | 1 | 6 | Rnh1 | 0.004 | Old |
| 0.87534 | 0.184136 | 0.097 | 0.1 | 1 | 6 | Cd2bp2 | -0.003 | Old |
| 0.875433 | 0.16279 | 0.031 | 0.033 | 1 | 6 | Zbtb34 | -0.002 | Old |
| 0.875433 | 0.045943 | 0.031 | 0.033 | 1 | 6 | Gstk1 | -0.002 | Old |
| 0.875754 | 0.090639 | 0.097 | 0.094 | 1 | 6 | Papss1 | 0.003 | Old |
| 0.875951 | 0.270434 | 0.057 | 0.06 | 1 | 6 | Bloc1s6 | -0.003 | Old |
| 0.875975 | 0.208174 | 0.04 | 0.042 | 1 | 6 | Zc3h12d | -0.002 | Old |
| 0.876274 | 0.091661 | 0.097 | 0.1 | 1 | 6 | Tbpl1 | -0.003 | Old |
| 0.876403 | 0.158381 | 0.074 | 0.079 | 1 | 6 | Scoc | -0.005 | Old |
| 0.877504 | 0.182868 | 0.077 | 0.08 | 1 | 6 | Polg | -0.003 | Old |
| 0.877514 | 0.174134 | 0.037 | 0.035 | 1 | 6 | 1600020E0 | 0.002 | Old |
| 0.877889 | 0.122197 | 0.157 | 0.151 | 1 | 6 | Wbp4 | 0.006 | Old |
| 0.877998 | 0.111238 | 0.411 | 0.4 | 1 | 6 | Sec62 | 0.011 | Old |
| 0.878191 | 0.039468 | 0.089 | 0.085 | 1 | 6 | Fam160b1 | 0.004 | Old |
| 0.878311 | 0.01434 | 0.1 | 0.097 | 1 | 6 | Slc41a3 | 0.003 | Old |
| 0.878336 | 0.034207 | 0.126 | 0.12 | 1 | 6 | Gpd2 | 0.006 | Old |
| 0.880261 | 0.098448 | 0.046 | 0.043 | 1 | 6 | L3mbtl3 | 0.003 | Old |
| 0.880295 | 0.120352 | 0.063 | 0.06 | 1 | 6 | Poc1b | 0.003 | Old |
| 0.880315 | 0.041179 | 0.654 | 0.657 | 1 | 6 | Cbfa2t3 | -0.003 | Old |
| 0.880773 | 0.124675 | 0.369 | 0.373 | 1 | 6 | Smap2 | -0.004 | Old |
| 0.880821 | 0.03346 | 0.363 | 0.349 | 1 | 6 | Msl1 | 0.014 | Old |
| 0.881476 | 0.165025 | 0.091 | 0.087 | 1 | 6 | Proser1 | 0.004 | Old |

| | | | | | | | | |
|---|---|---|---|---|---|---|---|---|
| 0.881497 | 0.011423 | 0.1 | 0.095 | 1 | 6 | Taf4 | 0.005 | Old |
| 0.881621 | 0.122084 | 0.043 | 0.045 | 1 | 6 | Pskh1 | -0.002 | Old |
| 0.881858 | 0.210338 | 0.134 | 0.13 | 1 | 6 | Ap1ar | 0.004 | Old |
| 0.882169 | 0.357276 | 0.06 | 0.064 | 1 | 6 | Gm45716 | -0.004 | Old |
| 0.882359 | 0.033567 | 0.14 | 0.135 | 1 | 6 | Trappc10 | 0.005 | Old |
| 0.882606 | 0.119309 | 0.263 | 0.258 | 1 | 6 | Map3k14 | 0.005 | Old |
| 0.882738 | 0.056807 | 0.066 | 0.069 | 1 | 6 | 2210016L2 | -0.003 | Old |
| 0.882931 | 0.100857 | 0.037 | 0.035 | 1 | 6 | Rnaset2b | 0.002 | Old |
| 0.88367 | 0.089097 | 0.32 | 0.309 | 1 | 6 | Tnrc6b | 0.011 | Old |
| 0.884186 | 0.016998 | 0.471 | 0.465 | 1 | 6 | Capg | 0.006 | Old |
| 0.884781 | 0.250539 | 0.197 | 0.192 | 1 | 6 | BC031181 | 0.005 | Old |
| 0.885026 | 0.673789 | 0.031 | 0.03 | 1 | 6 | Hmgn3 | 0.001 | Old |
| 0.885451 | 0.152912 | 0.097 | 0.099 | 1 | 6 | Exoc6b | -0.002 | Old |
| 0.885792 | 0.011771 | 0.231 | 0.231 | 1 | 6 | Hmgcs1 | 0 | Old |
| 0.88616 | 0.161708 | 0.337 | 0.341 | 1 | 6 | Mfsd14b | -0.004 | Old |
| 0.886177 | 0.436896 | 0.086 | 0.089 | 1 | 6 | Rce1 | -0.003 | Old |
| 0.886552 | 0.202986 | 0.111 | 0.117 | 1 | 6 | Clec4e | -0.006 | Old |
| 0.886742 | 0.341535 | 0.049 | 0.047 | 1 | 6 | 8-Sep | 0.002 | Old |
| 0.886949 | 0.031753 | 0.074 | 0.072 | 1 | 6 | AC127341. | 0.002 | Old |
| 0.887247 | 0.129741 | 0.131 | 0.135 | 1 | 6 | Narf | -0.004 | Old |
| 0.887337 | 0.073253 | 0.351 | 0.349 | 1 | 6 | Dpp4 | 0.002 | Old |
| 0.887466 | 0.026816 | 0.054 | 0.052 | 1 | 6 | Smcr8 | 0.002 | Old |
| 0.887774 | 0.116828 | 0.554 | 0.54 | 1 | 6 | 1810037I1 | 0.014 | Old |
| 0.887996 | 0.205721 | 0.137 | 0.135 | 1 | 6 | Sar1b | 0.002 | Old |
| 0.888011 | 0.181301 | 0.049 | 0.05 | 1 | 6 | Nudcd1 | -0.001 | Old |
| 0.888051 | 0.076926 | 0.114 | 0.117 | 1 | 6 | Ptgir | -0.003 | Old |
| 0.8889 | 0.079403 | 0.126 | 0.129 | 1 | 6 | Esrra | -0.003 | Old |
| 0.888956 | 0.051846 | 0.037 | 0.035 | 1 | 6 | Kansl1l | 0.002 | Old |
| 0.889486 | 0.036374 | 0.249 | 0.242 | 1 | 6 | Htt | 0.007 | Old |
| 0.88994 | 0.002026 | 0.077 | 0.08 | 1 | 6 | Bad | -0.003 | Old |
| 0.889982 | 0.159513 | 0.16 | 0.157 | 1 | 6 | Zzz3 | 0.003 | Old |
| 0.890545 | 0.500879 | 0.031 | 0.033 | 1 | 6 | Zfp839 | -0.002 | Old |
| 0.890559 | 0.018496 | 0.34 | 0.334 | 1 | 6 | Usp7 | 0.006 | Old |
| 0.890596 | 0.008832 | 0.157 | 0.152 | 1 | 6 | Aldh16a1 | 0.005 | Old |
| 0.890884 | 0.90896 | 0.031 | 0.03 | 1 | 6 | Edrf1 | 0.001 | Old |
| 0.89117 | 0.115845 | 0.18 | 0.176 | 1 | 6 | Scd2 | 0.004 | Old |
| 0.891556 | 0.05036 | 0.017 | 0.018 | 1 | 6 | E2f8 | -0.001 | Old |
| 0.891673 | 0.100431 | 0.029 | 0.03 | 1 | 6 | 2810402E2 | -0.001 | Old |
| 0.891806 | 0.711755 | 0.031 | 0.033 | 1 | 6 | Smad3 | -0.002 | Old |
| 0.892231 | 0.058541 | 0.051 | 0.054 | 1 | 6 | Rfng | -0.003 | Old |
| 0.892402 | 0.145568 | 0.017 | 0.018 | 1 | 6 | Ankrd33b | -0.001 | Old |
| 0.892597 | 0.339897 | 0.151 | 0.159 | 1 | 6 | Rab7b | -0.008 | Old |
| 0.892838 | 0.215676 | 0.031 | 0.03 | 1 | 6 | Zfp932 | 0.001 | Old |
| 0.892893 | 0.025844 | 0.04 | 0.042 | 1 | 6 | 2610524H( | -0.002 | Old |
| 0.892912 | 0.486856 | 0.086 | 0.089 | 1 | 6 | Mfap1a | -0.003 | Old |

| | | | | | | | | |
|---|---|---|---|---|---|---|---|---|
| 0.893066 | 0.310713 | 0.04 | 0.038 | 1 | 6 | Ncdn | 0.002 | Old |
| 0.893444 | 0.407309 | 0.091 | 0.09 | 1 | 6 | Gadd45gip | 0.001 | Old |
| 0.893674 | 0.153345 | 0.054 | 0.057 | 1 | 6 | Ddit3 | -0.003 | Old |
| 0.89371 | 1.742561 | 0.014 | 0.013 | 1 | 6 | Clstn1 | 0.001 | Old |
| 0.894249 | 0.199049 | 0.211 | 0.206 | 1 | 6 | Bmt2 | 0.005 | Old |
| 0.894328 | 0.171788 | 0.149 | 0.149 | 1 | 6 | Rap1gap2 | 0 | Old |
| 0.894794 | 0.561969 | 0.066 | 0.069 | 1 | 6 | Col4a2 | -0.003 | Old |
| 0.894938 | 0.318747 | 0.017 | 0.018 | 1 | 6 | Scamp5 | -0.001 | Old |
| 0.894987 | 0.099352 | 0.029 | 0.03 | 1 | 6 | Gm26778 | -0.001 | Old |
| 0.895019 | 0.2492 | 0.06 | 0.062 | 1 | 6 | Ubr7 | -0.002 | Old |
| 0.895365 | 0.073015 | 0.477 | 0.488 | 1 | 6 | Reep3 | -0.011 | Old |
| 0.895581 | 0.013466 | 0.314 | 0.319 | 1 | 6 | Nfat5 | -0.005 | Old |
| 0.895688 | 0.183255 | 0.086 | 0.089 | 1 | 6 | Phkg2 | -0.003 | Old |
| 0.895772 | 0.132873 | 0.129 | 0.127 | 1 | 6 | Prpf38a | 0.002 | Old |
| 0.89619 | 0.169488 | 0.063 | 0.06 | 1 | 6 | Glt8d1 | 0.003 | Old |
| 0.897014 | 0.026542 | 0.106 | 0.107 | 1 | 6 | Taf3 | -0.001 | Old |
| 0.897076 | 0.26259 | 0.086 | 0.089 | 1 | 6 | Gpatch2l | -0.003 | Old |
| 0.897076 | 0.242406 | 0.086 | 0.089 | 1 | 6 | Isyna1 | -0.003 | Old |
| 0.897337 | 0.010399 | 0.129 | 0.13 | 1 | 6 | 3110043O2 | -0.001 | Old |
| 0.897409 | 0.062806 | 0.449 | 0.47 | 1 | 6 | Parvb | -0.021 | Old |
| 0.897476 | 0.356834 | 0.017 | 0.018 | 1 | 6 | Dcakd | -0.001 | Old |
| 0.897476 | 0.194749 | 0.017 | 0.018 | 1 | 6 | Coq3 | -0.001 | Old |
| 0.897476 | 0.142941 | 0.017 | 0.018 | 1 | 6 | Mical2 | -0.001 | Old |
| 0.897586 | 0.530372 | 0.023 | 0.022 | 1 | 6 | E2f6 | 0.001 | Old |
| 0.89781 | 0.167141 | 0.077 | 0.075 | 1 | 6 | Alg10b | 0.002 | Old |
| 0.897822 | 0.083957 | 0.037 | 0.038 | 1 | 6 | Rasal3 | -0.001 | Old |
| 0.897874 | 0.15994 | 0.237 | 0.237 | 1 | 6 | Saraf | 0 | Old |
| 0.89795 | 0.150991 | 0.326 | 0.329 | 1 | 6 | 9530068E0 | -0.003 | Old |
| 0.898216 | 0.02144 | 0.774 | 0.799 | 1 | 6 | Flna | -0.025 | Old |
| 0.898536 | 0.613731 | 0.014 | 0.013 | 1 | 6 | Spp1 | 0.001 | Old |
| 0.898536 | 0.601497 | 0.014 | 0.013 | 1 | 6 | Atp5s | 0.001 | Old |
| 0.899074 | 0.201101 | 0.143 | 0.147 | 1 | 6 | Pes1 | -0.004 | Old |
| 0.899258 | 0.039722 | 0.051 | 0.054 | 1 | 6 | Ctdp1 | -0.003 | Old |
| 0.899652 | 0.07342 | 0.151 | 0.149 | 1 | 6 | Ppm1g | 0.002 | Old |
| 0.900862 | 0.369946 | 0.017 | 0.018 | 1 | 6 | Enah | -0.001 | Old |
| 0.900986 | 0.392167 | 0.049 | 0.047 | 1 | 6 | Slc25a38 | 0.002 | Old |
| 0.901088 | 0.302704 | 0.04 | 0.042 | 1 | 6 | Lrrc8b | -0.002 | Old |
| 0.901402 | 0.514677 | 0.023 | 0.022 | 1 | 6 | Il1r1 | 0.001 | Old |
| 0.901434 | 0.981572 | 0.014 | 0.013 | 1 | 6 | Zfp931 | 0.001 | Old |
| 0.901434 | 0.326821 | 0.014 | 0.013 | 1 | 6 | A530099J1 | 0.001 | Old |
| 0.901441 | 0.87104 | 0.126 | 0.125 | 1 | 6 | Dgat1 | 0.001 | Old |
| 0.901491 | 0.247782 | 0.057 | 0.055 | 1 | 6 | Slc44a2 | 0.002 | Old |
| 0.90155 | 0.475705 | 0.114 | 0.119 | 1 | 6 | Ncbp1 | -0.005 | Old |
| 0.901992 | 0.204013 | 0.286 | 0.289 | 1 | 6 | Phf20l1 | -0.003 | Old |
| 0.902092 | 0.171416 | 0.277 | 0.281 | 1 | 6 | U2surp | -0.004 | Old |

| | | | | | | | | |
|---|---|---|---|---|---|---|---|---|
| 0.902349 | 0.317563 | 0.163 | 0.169 | 1 | 6 | Zbtb11 | -0.006 | Old |
| 0.902362 | 0.172264 | 0.194 | 0.201 | 1 | 6 | Evi5 | -0.007 | Old |
| 0.902834 | 0.482987 | 0.049 | 0.047 | 1 | 6 | Nt5dc1 | 0.002 | Old |
| 0.902909 | 0.190266 | 0.071 | 0.07 | 1 | 6 | Gpkow | 0.001 | Old |
| 0.902947 | 0.854123 | 0.029 | 0.03 | 1 | 6 | Wdcp | -0.001 | Old |
| 0.902993 | 0.003654 | 0.106 | 0.102 | 1 | 6 | Map1s | 0.004 | Old |
| 0.903367 | 0.246607 | 0.014 | 0.013 | 1 | 6 | Zfp628 | 0.001 | Old |
| 0.903693 | 0.257147 | 0.023 | 0.022 | 1 | 6 | Lyrm1 | 0.001 | Old |
| 0.903891 | 0.148061 | 0.163 | 0.162 | 1 | 6 | Sesn3 | 0.001 | Old |
| 0.904333 | 0.447887 | 0.014 | 0.013 | 1 | 6 | Gm20513 | 0.001 | Old |
| 0.904333 | 0.236927 | 0.014 | 0.013 | 1 | 6 | Zfp7 | 0.001 | Old |
| 0.904456 | 0.239068 | 0.023 | 0.022 | 1 | 6 | Zfp747 | 0.001 | Old |
| 0.904684 | 0.005649 | 0.049 | 0.047 | 1 | 6 | Zfp414 | 0.002 | Old |
| 0.904713 | 0.201832 | 0.06 | 0.059 | 1 | 6 | Pla2g4a | 0.001 | Old |
| 0.90522 | 0.401594 | 0.023 | 0.022 | 1 | 6 | Zfp933 | 0.001 | Old |
| 0.9053 | 0.121573 | 0.014 | 0.013 | 1 | 6 | Zbtb39 | 0.001 | Old |
| 0.90533 | 0.09726 | 0.6 | 0.619 | 1 | 6 | Lamp1 | -0.019 | Old |
| 0.905602 | 0.512378 | 0.029 | 0.03 | 1 | 6 | E2f5 | -0.001 | Old |
| 0.905627 | 0.410825 | 0.043 | 0.042 | 1 | 6 | Zfp874a | 0.001 | Old |
| 0.905794 | 0.306742 | 0.06 | 0.062 | 1 | 6 | Spata13 | -0.002 | Old |
| 0.905984 | 0.019766 | 0.023 | 0.022 | 1 | 6 | Tlr13 | 0.001 | Old |
| 0.906267 | 0.533614 | 0.014 | 0.013 | 1 | 6 | S1pr1 | 0.001 | Old |
| 0.906267 | 0.106811 | 0.014 | 0.013 | 1 | 6 | Srr | 0.001 | Old |
| 0.907269 | 0.209424 | 0.137 | 0.134 | 1 | 6 | Pfdn6 | 0.003 | Old |
| 0.90754 | 0.20811 | 0.094 | 0.097 | 1 | 6 | Traf3 | -0.003 | Old |
| 0.90784 | 0.080152 | 0.031 | 0.03 | 1 | 6 | Atf5 | 0.001 | Old |
| 0.908384 | 0.009118 | 0.049 | 0.047 | 1 | 6 | Armt1 | 0.002 | Old |
| 0.908795 | 0.100457 | 0.22 | 0.219 | 1 | 6 | Morc3 | 0.001 | Old |
| 0.908835 | 0.354664 | 0.163 | 0.164 | 1 | 6 | Fkbp4 | -0.001 | Old |
| 0.909006 | 0.009535 | 0.046 | 0.047 | 1 | 6 | Akr1b10 | -0.001 | Old |
| 0.909006 | 0.087803 | 0.131 | 0.129 | 1 | 6 | Ostm1 | 0.002 | Old |
| 0.909064 | 0.012578 | 0.126 | 0.122 | 1 | 6 | Dnm1l | 0.004 | Old |
| 0.90912 | 0.052287 | 0.083 | 0.08 | 1 | 6 | Heatr1 | 0.003 | Old |
| 0.909143 | 0.004955 | 0.346 | 0.353 | 1 | 6 | Irf4 | -0.007 | Old |
| 0.909513 | 0.096465 | 0.183 | 0.176 | 1 | 6 | Cited2 | 0.007 | Old |
| 0.909525 | 0.001235 | 0.294 | 0.288 | 1 | 6 | Pcnp | 0.006 | Old |
| 0.909835 | 0.04885 | 0.331 | 0.334 | 1 | 6 | Lrrfip2 | -0.003 | Old |
| 0.910042 | 0.007656 | 0.129 | 0.127 | 1 | 6 | Erlin2 | 0.002 | Old |
| 0.910066 | 0.127802 | 0.054 | 0.052 | 1 | 6 | Sh2b1 | 0.002 | Old |
| 0.910135 | 0.360486 | 0.014 | 0.013 | 1 | 6 | Ube2c | 0.001 | Old |
| 0.910318 | 0.160612 | 0.186 | 0.187 | 1 | 6 | Setd5 | -0.001 | Old |
| 0.910446 | 0.005237 | 0.583 | 0.607 | 1 | 6 | Tra2b | -0.024 | Old |
| 0.910453 | 0.009493 | 0.031 | 0.03 | 1 | 6 | Eef1akmt4 | 0.001 | Old |
| 0.910501 | 0.4856 | 0.066 | 0.064 | 1 | 6 | Ms4a6b | 0.002 | Old |
| 0.910501 | 0.093437 | 0.066 | 0.064 | 1 | 6 | Ttc37 | 0.002 | Old |

| | | | | | | | | |
|---|---|---|---|---|---|---|---|---|
| 0.910569 | 0.115303 | 0.023 | 0.022 | 1 | 6 | Hoxb4 | 0.001 | Old |
| 0.910641 | 0.082147 | 0.286 | 0.289 | 1 | 6 | Riok3 | -0.003 | Old |
| 0.910745 | 0.001182 | 0.451 | 0.45 | 1 | 6 | Kmt2e | 0.001 | Old |
| 0.911103 | 0.109296 | 0.014 | 0.013 | 1 | 6 | Zfp974 | 0.001 | Old |
| 0.911103 | 0.061639 | 0.014 | 0.013 | 1 | 6 | Phf19 | 0.001 | Old |
| 0.911103 | 0.00569 | 0.014 | 0.013 | 1 | 6 | Wdr74 | 0.001 | Old |
| 0.911106 | 0.092875 | 0.031 | 0.03 | 1 | 6 | Blzf1 | 0.001 | Old |
| 0.911192 | 0.218452 | 0.357 | 0.358 | 1 | 6 | Dennd1b | -0.001 | Old |
| 0.911759 | 0.478817 | 0.031 | 0.03 | 1 | 6 | Nek1 | 0.001 | Old |
| 0.91199 | 0.093022 | 0.377 | 0.385 | 1 | 6 | Slc43a2 | -0.008 | Old |
| 0.912515 | 0.140009 | 0.111 | 0.109 | 1 | 6 | Galnt7 | 0.002 | Old |
| 0.913038 | 0.096933 | 0.014 | 0.013 | 1 | 6 | Zfp518b | 0.001 | Old |
| 0.913603 | 0.119343 | 0.18 | 0.177 | 1 | 6 | N4bp1 | 0.003 | Old |
| 0.913627 | 0.020483 | 0.023 | 0.022 | 1 | 6 | Lsm11 | 0.001 | Old |
| 0.913898 | 0.172437 | 0.263 | 0.261 | 1 | 6 | Trim28 | 0.002 | Old |
| 0.913952 | 0.069579 | 0.114 | 0.115 | 1 | 6 | Il1rn | -0.001 | Old |
| 0.914444 | 0.292452 | 0.134 | 0.137 | 1 | 6 | Fam20b | -0.003 | Old |
| 0.914702 | 0.133108 | 0.057 | 0.055 | 1 | 6 | Acyp1 | 0.002 | Old |
| 0.914922 | 0.139353 | 0.454 | 0.46 | 1 | 6 | Smchd1 | -0.006 | Old |
| 0.915121 | 0.017812 | 0.566 | 0.579 | 1 | 6 | Gpx4 | -0.013 | Old |
| 0.915224 | 0.611562 | 0.034 | 0.033 | 1 | 6 | Apool | 0.001 | Old |
| 0.915617 | 0.127337 | 0.714 | 0.729 | 1 | 6 | Pmaip1 | -0.015 | Old |
| 0.916198 | 0.062315 | 0.22 | 0.219 | 1 | 6 | Nfil3 | 0.001 | Old |
| 0.91647 | 0.202787 | 0.049 | 0.05 | 1 | 6 | Mamdc2 | -0.001 | Old |
| 0.916988 | 0.388971 | 0.031 | 0.03 | 1 | 6 | Yipf2 | 0.001 | Old |
| 0.917008 | 0.30757 | 0.111 | 0.109 | 1 | 6 | Pfkl | 0.002 | Old |
| 0.917027 | 0.135047 | 0.309 | 0.301 | 1 | 6 | Ppp4r3b | 0.008 | Old |
| 0.917036 | 0.139387 | 0.369 | 0.376 | 1 | 6 | Itch | -0.007 | Old |
| 0.91723 | 0.420728 | 0.226 | 0.226 | 1 | 6 | Baz2a | 0 | Old |
| 0.91723 | 0.006124 | 0.223 | 0.227 | 1 | 6 | Tppp3 | -0.004 | Old |
| 0.917392 | 0.267676 | 0.08 | 0.079 | 1 | 6 | Mtmr12 | 0.001 | Old |
| 0.917829 | 0.072678 | 0.537 | 0.538 | 1 | 6 | Eif2s2 | -0.001 | Old |
| 0.917905 | 0.246946 | 0.083 | 0.084 | 1 | 6 | Abcf2 | -0.001 | Old |
| 0.918296 | 0.116885 | 0.031 | 0.03 | 1 | 6 | 4833407H1 | 0.001 | Old |
| 0.918479 | 0.303383 | 0.014 | 0.015 | 1 | 6 | Crtc1 | -0.001 | Old |
| 0.918575 | 0.230258 | 0.069 | 0.07 | 1 | 6 | Dnttip1 | -0.001 | Old |
| 0.918846 | 0.048965 | 0.16 | 0.162 | 1 | 6 | Il3ra | -0.002 | Old |
| 0.919403 | 0.100078 | 0.106 | 0.104 | 1 | 6 | Vps53 | 0.002 | Old |
| 0.919615 | 0.225593 | 0.143 | 0.142 | 1 | 6 | Ccm2 | 0.001 | Old |
| 0.919663 | 0.107627 | 0.54 | 0.554 | 1 | 6 | Mapre1 | -0.014 | Old |
| 0.920157 | 0.128605 | 0.3 | 0.298 | 1 | 6 | Ndufs5 | 0.002 | Old |
| 0.920347 | 0.628168 | 0.014 | 0.015 | 1 | 6 | Chaf1b | -0.001 | Old |
| 0.921968 | 0.005913 | 0.171 | 0.172 | 1 | 6 | Tbc1d14 | -0.001 | Old |
| 0.922502 | 0.027354 | 0.163 | 0.161 | 1 | 6 | Dyrk1a | 0.002 | Old |
| 0.922812 | 0.477538 | 0.077 | 0.08 | 1 | 6 | Rhoh | -0.003 | Old |

| | | | | | | | | |
|---|---|---|---|---|---|---|---|---|
| 0.922827 | 0.346045 | 0.083 | 0.085 | 1 | 6 | Arl4a | -0.002 | Old |
| 0.92287 | 0.141128 | 0.171 | 0.167 | 1 | 6 | Pi4k2a | 0.004 | Old |
| 0.922875 | 0.417239 | 0.031 | 0.03 | 1 | 6 | Sppl2b | 0.001 | Old |
| 0.923342 | 0.085062 | 0.457 | 0.485 | 1 | 6 | Asap1 | -0.028 | Old |
| 0.923663 | 0.077739 | 0.134 | 0.132 | 1 | 6 | Exosc8 | 0.002 | Old |
| 0.924084 | 0.02428 | 0.014 | 0.015 | 1 | 6 | Sh3bp5l | -0.001 | Old |
| 0.924139 | 0.005791 | 0.323 | 0.324 | 1 | 6 | Lpl | -0.001 | Old |
| 0.924294 | 0.203121 | 0.114 | 0.117 | 1 | 6 | Nipsnap2 | -0.003 | Old |
| 0.924306 | 0.12363 | 0.037 | 0.038 | 1 | 6 | Trmt10a | -0.001 | Old |
| 0.924437 | 0.668556 | 0.026 | 0.025 | 1 | 6 | Dcp1b | 0.001 | Old |
| 0.924477 | 0.218159 | 0.237 | 0.247 | 1 | 6 | Dnajc2 | -0.01 | Old |
| 0.924956 | 0.307267 | 0.426 | 0.445 | 1 | 6 | Id2 | -0.019 | Old |
| 0.924976 | 0.032664 | 0.1 | 0.104 | 1 | 6 | Trim5 | -0.004 | Old |
| 0.925005 | 0.302402 | 0.14 | 0.139 | 1 | 6 | Fbxo3 | 0.001 | Old |
| 0.925018 | 0.112143 | 0.014 | 0.015 | 1 | 6 | Ddb2 | -0.001 | Old |
| 0.925108 | 0.076767 | 0.254 | 0.256 | 1 | 6 | Dctn1 | -0.002 | Old |
| 0.925289 | 0.064438 | 0.157 | 0.156 | 1 | 6 | Spred1 | 0.001 | Old |
| 0.925379 | 0.027573 | 0.3 | 0.289 | 1 | 6 | Wls | 0.011 | Old |
| 0.925439 | 0.01205 | 0.294 | 0.289 | 1 | 6 | Ccnl2 | 0.005 | Old |
| 0.925519 | 0.10578 | 0.126 | 0.129 | 1 | 6 | Nacc2 | -0.003 | Old |
| 0.925725 | 0.250363 | 0.106 | 0.109 | 1 | 6 | Mrpl9 | -0.003 | Old |
| 0.925907 | 0.086289 | 0.109 | 0.107 | 1 | 6 | Tmem141 | 0.002 | Old |
| 0.925969 | 0.013133 | 0.177 | 0.176 | 1 | 6 | Tmem238 | 0.001 | Old |
| 0.92623 | 0.016317 | 0.177 | 0.174 | 1 | 6 | St6gal1 | 0.003 | Old |
| 0.926271 | 0.15102 | 0.366 | 0.366 | 1 | 6 | Mbnl2 | 0 | Old |
| 0.926298 | 0.132213 | 0.071 | 0.074 | 1 | 6 | Mcm5 | -0.003 | Old |
| 0.926344 | 0.251326 | 0.043 | 0.042 | 1 | 6 | Slc10a3 | 0.001 | Old |
| 0.926346 | 0.060103 | 0.257 | 0.258 | 1 | 6 | Arid4b | -0.001 | Old |
| 0.926549 | 0.090065 | 0.62 | 0.637 | 1 | 6 | Lnpep | -0.017 | Old |
| 0.927076 | 0.873701 | 0.034 | 0.033 | 1 | 6 | B230354K1 | 0.001 | Old |
| 0.927679 | 0.25006 | 0.026 | 0.027 | 1 | 6 | Dbf4 | -0.001 | Old |
| 0.927844 | 0.130028 | 0.037 | 0.038 | 1 | 6 | Mettl26 | -0.001 | Old |
| 0.928035 | 0.046995 | 0.223 | 0.226 | 1 | 6 | Mrps24 | -0.003 | Old |
| 0.928083 | 0.330525 | 0.089 | 0.089 | 1 | 6 | Usp5 | 0 | Old |
| 0.928284 | 0.408675 | 0.151 | 0.156 | 1 | 6 | Fchsd2 | -0.005 | Old |
| 0.928757 | 0.28308 | 0.014 | 0.015 | 1 | 6 | AC163040. | -0.001 | Old |
| 0.928757 | 0.122741 | 0.014 | 0.015 | 1 | 6 | Dtwd2 | -0.001 | Old |
| 0.929326 | 0.269576 | 0.171 | 0.177 | 1 | 6 | Crtc2 | -0.006 | Old |
| 0.929692 | 0.155 | 0.014 | 0.015 | 1 | 6 | Lins1 | -0.001 | Old |
| 0.929692 | 0.072595 | 0.014 | 0.015 | 1 | 6 | Lix1l | -0.001 | Old |
| 0.92989 | 0.154579 | 0.386 | 0.398 | 1 | 6 | Ddost | -0.012 | Old |
| 0.929948 | 0.057159 | 0.783 | 0.794 | 1 | 6 | Pitpna | -0.011 | Old |
| 0.929995 | 0.134277 | 0.189 | 0.187 | 1 | 6 | L1cam | 0.002 | Old |
| 0.930135 | 0.106432 | 0.471 | 0.478 | 1 | 6 | Usf2 | -0.007 | Old |
| 0.930197 | 0.619114 | 0.034 | 0.033 | 1 | 6 | Hps6 | 0.001 | Old |

| | | | | | | | | |
|---|---|---|---|---|---|---|---|---|
| 0.93041 | 0.065995 | 0.411 | 0.411 | 1 | 6 | Timm13 | 0 | Old |
| 0.931115 | 0.105745 | 0.411 | 0.413 | 1 | 6 | Scpep1 | -0.002 | Old |
| 0.931523 | 0.239622 | 0.217 | 0.227 | 1 | 6 | Myo9a | -0.01 | Old |
| 0.931562 | 1.427493 | 0.014 | 0.015 | 1 | 6 | Rnf152 | -0.001 | Old |
| 0.931562 | 0.635778 | 0.014 | 0.015 | 1 | 6 | Lrrc3 | -0.001 | Old |
| 0.931562 | 0.353613 | 0.014 | 0.015 | 1 | 6 | Ifi205 | -0.001 | Old |
| 0.931562 | 0.330567 | 0.014 | 0.015 | 1 | 6 | Pusl1 | -0.001 | Old |
| 0.931562 | 0.269908 | 0.014 | 0.015 | 1 | 6 | Sh3bp4 | -0.001 | Old |
| 0.93159 | 0.187628 | 0.22 | 0.214 | 1 | 6 | Tmem248 | 0.006 | Old |
| 0.931845 | 0.065605 | 0.126 | 0.127 | 1 | 6 | Gtf3c1 | -0.001 | Old |
| 0.932407 | 0.176466 | 0.286 | 0.288 | 1 | 6 | Atxn7l1 | -0.002 | Old |
| 0.933115 | 0.136892 | 0.054 | 0.054 | 1 | 6 | Carm1 | 0 | Old |
| 0.933343 | 0.043336 | 0.069 | 0.067 | 1 | 6 | Phax | 0.002 | Old |
| 0.933433 | 0.572093 | 0.014 | 0.015 | 1 | 6 | D830025C( | -0.001 | Old |
| 0.933433 | 0.367989 | 0.014 | 0.015 | 1 | 6 | Arhgap35 | -0.001 | Old |
| 0.933445 | 0.693716 | 0.046 | 0.045 | 1 | 6 | Pef1 | 0.001 | Old |
| 0.933452 | 0.10841 | 0.349 | 0.339 | 1 | 6 | Eif2s3x | 0.01 | Old |
| 0.93356 | 0.072205 | 0.183 | 0.186 | 1 | 6 | Ncstn | -0.003 | Old |
| 0.9336 | 0.626199 | 0.049 | 0.05 | 1 | 6 | Atp11a | -0.001 | Old |
| 0.933716 | 0.14452 | 0.106 | 0.109 | 1 | 6 | Pofut2 | -0.003 | Old |
| 0.934014 | 0.283266 | 0.026 | 0.027 | 1 | 6 | Letm2 | -0.001 | Old |
| 0.934301 | 0.35558 | 0.034 | 0.035 | 1 | 6 | 1110034G2 | -0.001 | Old |
| 0.934352 | 0.047916 | 0.143 | 0.14 | 1 | 6 | Pbx3 | 0.003 | Old |
| 0.934718 | 0.25783 | 0.026 | 0.027 | 1 | 6 | Rtn4ip1 | -0.001 | Old |
| 0.934806 | 0.135134 | 0.143 | 0.145 | 1 | 6 | Ccdc88c | -0.002 | Old |
| 0.934925 | 0.138362 | 0.037 | 0.038 | 1 | 6 | Zfp959 | -0.001 | Old |
| 0.934951 | 0.137398 | 0.209 | 0.206 | 1 | 6 | Klrk1 | 0.003 | Old |
| 0.935177 | 0.271071 | 0.226 | 0.231 | 1 | 6 | Tsg101 | -0.005 | Old |
| 0.935194 | 0.225043 | 0.034 | 0.033 | 1 | 6 | 10-Sep | 0.001 | Old |
| 0.935547 | 0.115065 | 0.157 | 0.159 | 1 | 6 | Zkscan3 | -0.002 | Old |
| 0.936053 | 0.039244 | 0.346 | 0.343 | 1 | 6 | Tmem167 | 0.003 | Old |
| 0.936118 | 0.165994 | 0.123 | 0.125 | 1 | 6 | Cyb5r1 | -0.002 | Old |
| 0.936224 | 0.097384 | 0.383 | 0.396 | 1 | 6 | Adcy7 | -0.013 | Old |
| 0.936348 | 0.04415 | 0.814 | 0.893 | 1 | 6 | H2-DMb1 | -0.079 | Old |
| 0.936464 | 0.29387 | 0.117 | 0.12 | 1 | 6 | Ip6k1 | -0.003 | Old |
| 0.936724 | 0.230198 | 0.071 | 0.07 | 1 | 6 | Slc35c1 | 0.001 | Old |
| 0.937315 | 0.373926 | 0.129 | 0.127 | 1 | 6 | Upf1 | 0.002 | Old |
| 0.937355 | 0.128737 | 0.277 | 0.281 | 1 | 6 | Cd226 | -0.004 | Old |
| 0.937537 | 0.161626 | 0.063 | 0.062 | 1 | 6 | Slc7a8 | 0.001 | Old |
| 0.937831 | 0.169099 | 0.169 | 0.167 | 1 | 6 | Raf1 | 0.002 | Old |
| 0.937919 | 0.229242 | 0.083 | 0.082 | 1 | 6 | Slc35a5 | 0.001 | Old |
| 0.93808 | 0.41561 | 0.026 | 0.025 | 1 | 6 | Cradd | 0.001 | Old |
| 0.938092 | 0.178815 | 0.097 | 0.095 | 1 | 6 | Helb | 0.002 | Old |
| 0.938513 | 0.317084 | 0.066 | 0.067 | 1 | 6 | Heatr6 | -0.001 | Old |
| 0.93872 | 0.002854 | 0.28 | 0.269 | 1 | 6 | Ssrp1 | 0.011 | Old |

| | | | | | | | | |
|---|---|---|---|---|---|---|---|---|
| 0.938898 | 0.039815 | 0.094 | 0.092 | 1 | 6 | Lncpint | 0.002 | Old |
| 0.939217 | 0.184632 | 0.074 | 0.077 | 1 | 6 | Ssh1 | -0.003 | Old |
| 0.93992 | 0.119711 | 0.086 | 0.087 | 1 | 6 | Bid | -0.001 | Old |
| 0.940432 | 0.05688 | 0.549 | 0.554 | 1 | 6 | Tomm20 | -0.005 | Old |
| 0.94059 | 0.076754 | 0.423 | 0.418 | 1 | 6 | Wsb1 | 0.005 | Old |
| 0.941077 | 0.266044 | 0.157 | 0.157 | 1 | 6 | Echs1 | 0 | Old |
| 0.941116 | 0.08045 | 0.054 | 0.054 | 1 | 6 | Ccs | 0 | Old |
| 0.941989 | 0.078941 | 0.091 | 0.09 | 1 | 6 | Oard1 | 0.001 | Old |
| 0.942069 | 0.081783 | 0.034 | 0.033 | 1 | 6 | Scimp | 0.001 | Old |
| 0.942225 | 0.033313 | 0.071 | 0.07 | 1 | 6 | Morc2a | 0.001 | Old |
| 0.942364 | 0.115926 | 0.274 | 0.276 | 1 | 6 | Rc3h2 | -0.002 | Old |
| 0.94241 | 0.117401 | 0.077 | 0.075 | 1 | 6 | Sipa1l1 | 0.002 | Old |
| 0.942515 | 0.080001 | 0.137 | 0.139 | 1 | 6 | Taf13 | -0.002 | Old |
| 0.942615 | 0.362912 | 0.157 | 0.156 | 1 | 6 | Socs4 | 0.001 | Old |
| 0.943172 | 0.084009 | 0.351 | 0.353 | 1 | 6 | Nxf1 | -0.002 | Old |
| 0.943591 | 0.115537 | 0.189 | 0.191 | 1 | 6 | Ndufs8 | -0.002 | Old |
| 0.943645 | 0.232305 | 0.423 | 0.423 | 1 | 6 | Usp9x | 0 | Old |
| 0.943744 | 0.040985 | 0.409 | 0.416 | 1 | 6 | Sf3b6 | -0.007 | Old |
| 0.944092 | 0.331518 | 0.071 | 0.074 | 1 | 6 | Ammecr1 | -0.003 | Old |
| 0.944231 | 1.652124 | 0.017 | 0.017 | 1 | 6 | Agbl5 | 0 | Old |
| 0.944624 | 0.143344 | 0.129 | 0.13 | 1 | 6 | Abhd2 | -0.001 | Old |
| 0.944896 | 0.089513 | 0.229 | 0.234 | 1 | 6 | Gtf2a1 | -0.005 | Old |
| 0.945043 | 0.144139 | 0.254 | 0.259 | 1 | 6 | Psma5 | -0.005 | Old |
| 0.94537 | 0.100135 | 0.054 | 0.055 | 1 | 6 | Pot1a | -0.001 | Old |
| 0.945512 | 0.100339 | 0.286 | 0.276 | 1 | 6 | Ptbp1 | 0.01 | Old |
| 0.94564 | 0.025112 | 0.143 | 0.139 | 1 | 6 | Ccdc186 | 0.004 | Old |
| 0.945682 | 0.23504 | 0.074 | 0.075 | 1 | 6 | Avl9 | -0.001 | Old |
| 0.945723 | 0.091387 | 0.1 | 0.102 | 1 | 6 | Mapk8ip3 | -0.002 | Old |
| 0.945987 | 0.568178 | 0.026 | 0.025 | 1 | 6 | Gm20506 | 0.001 | Old |
| 0.945991 | 1.529871 | 0.026 | 0.027 | 1 | 6 | Trim21 | -0.001 | Old |
| 0.946048 | 0.047278 | 0.463 | 0.45 | 1 | 6 | Dock8 | 0.013 | Old |
| 0.94611 | 0.061879 | 0.074 | 0.075 | 1 | 6 | Cfap20 | -0.001 | Old |
| 0.94628 | 0.279179 | 0.091 | 0.094 | 1 | 6 | Matk | -0.003 | Old |
| 0.94648 | 0.068304 | 0.68 | 0.696 | 1 | 6 | Tkt | -0.016 | Old |
| 0.947142 | 0.053829 | 0.086 | 0.087 | 1 | 6 | Rnf41 | -0.001 | Old |
| 0.947387 | 0.194849 | 0.209 | 0.206 | 1 | 6 | Cnot4 | 0.003 | Old |
| 0.947436 | 0.122671 | 0.283 | 0.279 | 1 | 6 | Dhx9 | 0.004 | Old |
| 0.94772 | 0.196738 | 0.326 | 0.321 | 1 | 6 | Smg1 | 0.005 | Old |
| 0.948275 | 0.004583 | 0.34 | 0.338 | 1 | 6 | Slc4a7 | 0.002 | Old |
| 0.948461 | 0.066035 | 0.086 | 0.085 | 1 | 6 | Gnpat | 0.001 | Old |
| 0.948658 | 0.140676 | 0.131 | 0.13 | 1 | 6 | Usp48 | 0.001 | Old |
| 0.948809 | 0.112863 | 0.097 | 0.095 | 1 | 6 | Gm13212 | 0.002 | Old |
| 0.948863 | 0.101392 | 0.026 | 0.025 | 1 | 6 | Cyb5d2 | 0.001 | Old |
| 0.949202 | 0.120936 | 0.357 | 0.356 | 1 | 6 | Nras | 0.001 | Old |
| 0.94953 | 0.209889 | 0.251 | 0.256 | 1 | 6 | Limd2 | -0.005 | Old |

| | | | | | | | | |
|---|---|---|---|---|---|---|---|---|
| 0.949694 | 0.38316 | 0.066 | 0.065 | 1 | 6 | Tm2d3 | 0.001 | Old |
| 0.9497 | 0.469499 | 0.034 | 0.035 | 1 | 6 | Mtfmt | -0.001 | Old |
| 0.949752 | 0.383384 | 0.109 | 0.109 | 1 | 6 | Bola3 | 0 | Old |
| 0.949921 | 0.030422 | 0.043 | 0.042 | 1 | 6 | Ddx41 | 0.001 | Old |
| 0.949968 | 0.085858 | 0.174 | 0.177 | 1 | 6 | Rbm7 | -0.003 | Old |
| 0.950026 | 0.253049 | 0.069 | 0.07 | 1 | 6 | Atn1 | -0.001 | Old |
| 0.950302 | 0.014463 | 0.026 | 0.025 | 1 | 6 | Gtpbp8 | 0.001 | Old |
| 0.950371 | 0.787242 | 0.017 | 0.017 | 1 | 6 | Csf3r | 0 | Old |
| 0.950825 | 0.089396 | 0.034 | 0.033 | 1 | 6 | Slc7a6 | 0.001 | Old |
| 0.951057 | 0.069006 | 0.083 | 0.082 | 1 | 6 | Ints13 | 0.001 | Old |
| 0.951288 | 0.085717 | 0.306 | 0.308 | 1 | 6 | Elmo1 | -0.002 | Old |
| 0.951703 | 0.409795 | 0.037 | 0.037 | 1 | 6 | Gin1 | 0 | Old |
| 0.952109 | 0.038256 | 0.12 | 0.119 | 1 | 6 | Vps16 | 0.001 | Old |
| 0.952132 | 0.265248 | 0.16 | 0.161 | 1 | 6 | Cpq | -0.001 | Old |
| 0.952489 | 0.335957 | 0.063 | 0.062 | 1 | 6 | Cmc2 | 0.001 | Old |
| 0.95265 | 0.315022 | 0.24 | 0.237 | 1 | 6 | Pou2f2 | 0.003 | Old |
| 0.952681 | 0.298868 | 0.089 | 0.089 | 1 | 6 | Mindy1 | 0 | Old |
| 0.9527 | 0.1947 | 0.083 | 0.082 | 1 | 6 | Spin1 | 0.001 | Old |
| 0.952782 | 0.435638 | 0.034 | 0.035 | 1 | 6 | Prmt2 | -0.001 | Old |
| 0.952861 | 0.17076 | 0.063 | 0.064 | 1 | 6 | Afg3l2 | -0.001 | Old |
| 0.952911 | 0.108097 | 0.403 | 0.401 | 1 | 6 | Tiparp | 0.002 | Old |
| 0.953003 | 0.294117 | 0.017 | 0.017 | 1 | 6 | Rnf24 | 0 | Old |
| 0.953244 | 0.045888 | 0.226 | 0.222 | 1 | 6 | Rnf6 | 0.004 | Old |
| 0.953263 | 0.057067 | 0.111 | 0.114 | 1 | 6 | Acp2 | -0.003 | Old |
| 0.953612 | 0.031105 | 0.389 | 0.38 | 1 | 6 | Necap2 | 0.009 | Old |
| 0.953949 | 0.07294 | 0.263 | 0.251 | 1 | 6 | Rere | 0.012 | Old |
| 0.954375 | 0.111762 | 0.066 | 0.067 | 1 | 6 | Chpf2 | -0.001 | Old |
| 0.954522 | 0.076394 | 0.557 | 0.564 | 1 | 6 | Pdcd6ip | -0.007 | Old |
| 0.9551 | 0.655335 | 0.046 | 0.047 | 1 | 6 | Bank1 | -0.001 | Old |
| 0.955292 | 0.325371 | 0.054 | 0.055 | 1 | 6 | Gramd3 | -0.001 | Old |
| 0.955338 | 0.032181 | 0.026 | 0.025 | 1 | 6 | Hist3h2ba | 0.001 | Old |
| 0.955659 | 0.128272 | 0.2 | 0.197 | 1 | 6 | Mettl23 | 0.003 | Old |
| 0.955928 | 0.071303 | 0.226 | 0.222 | 1 | 6 | Trip11 | 0.004 | Old |
| 0.956094 | 0.258859 | 0.109 | 0.109 | 1 | 6 | Kmt5c | 0 | Old |
| 0.956513 | 0.539639 | 0.017 | 0.017 | 1 | 6 | Cacna1d | 0 | Old |
| 0.957126 | 0.210417 | 0.069 | 0.07 | 1 | 6 | Sertad3 | -0.001 | Old |
| 0.95739 | 0.371217 | 0.017 | 0.017 | 1 | 6 | Nuak1 | 0 | Old |
| 0.95739 | 0.262347 | 0.017 | 0.017 | 1 | 6 | Surf6 | 0 | Old |
| 0.95749 | 0.066514 | 0.1 | 0.1 | 1 | 6 | Trmt1l | 0 | Old |
| 0.957585 | 0.127465 | 0.246 | 0.246 | 1 | 6 | Skp1a | 0 | Old |
| 0.957692 | 0.067909 | 0.346 | 0.344 | 1 | 6 | Bmp2k | 0.002 | Old |
| 0.958182 | 0.002444 | 0.4 | 0.401 | 1 | 6 | Pebp1 | -0.001 | Old |
| 0.958268 | 0.209637 | 0.017 | 0.017 | 1 | 6 | Tmem206 | 0 | Old |
| 0.958284 | 0.146606 | 0.023 | 0.023 | 1 | 6 | Tmem229a | 0 | Old |
| 0.958533 | 0.006659 | 0.263 | 0.261 | 1 | 6 | Pcmtd1 | 0.002 | Old |

| | | | | | | | | |
|---|---|---|---|---|---|---|---|---|
| 0.95855 | 0.002545 | 0.074 | 0.075 | 1 | 6 | Hgsnat | -0.001 | Old |
| 0.959137 | 0.720058 | 0.054 | 0.054 | 1 | 6 | Mecr | 0 | Old |
| 0.959212 | 0.042544 | 0.477 | 0.467 | 1 | 6 | G3bp2 | 0.01 | Old |
| 0.959379 | 0.086583 | 0.354 | 0.353 | 1 | 6 | Irak1 | 0.001 | Old |
| 0.959504 | 0.012933 | 0.063 | 0.062 | 1 | 6 | Hace1 | 0.001 | Old |
| 0.959564 | 0.049508 | 0.474 | 0.488 | 1 | 6 | Shtn1 | -0.014 | Old |
| 0.959582 | 0.179921 | 0.08 | 0.079 | 1 | 6 | Med25 | 0.001 | Old |
| 0.959601 | 0.173149 | 0.134 | 0.134 | 1 | 6 | Arrdc4 | 0 | Old |
| 0.959786 | 0.032701 | 0.023 | 0.023 | 1 | 6 | Nit2 | 0 | Old |
| 0.960006 | 0.136084 | 0.074 | 0.074 | 1 | 6 | Gosr1 | 0 | Old |
| 0.960659 | 0.22747 | 0.08 | 0.08 | 1 | 6 | Zbtb37 | 0 | Old |
| 0.960902 | 0.351351 | 0.017 | 0.017 | 1 | 6 | Gatsl3 | 0 | Old |
| 0.960978 | 0.12837 | 0.186 | 0.186 | 1 | 6 | Lrp6 | 0 | Old |
| 0.96178 | 0.624689 | 0.017 | 0.017 | 1 | 6 | Tmem67 | 0 | Old |
| 0.96178 | 0.075182 | 0.017 | 0.017 | 1 | 6 | C330027C0 | 0 | Old |
| 0.961877 | 0.154689 | 0.109 | 0.11 | 1 | 6 | Tnfaip1 | -0.001 | Old |
| 0.962061 | 0.088444 | 0.151 | 0.151 | 1 | 6 | Katna1 | 0 | Old |
| 0.962161 | 0.083892 | 0.18 | 0.184 | 1 | 6 | D1Ertd622 | -0.004 | Old |
| 0.962772 | 0.016023 | 0.18 | 0.179 | 1 | 6 | Gdap10 | 0.001 | Old |
| 0.962945 | 0.037322 | 0.066 | 0.065 | 1 | 6 | Sesn1 | 0.001 | Old |
| 0.962952 | 0.087543 | 0.051 | 0.052 | 1 | 6 | Abcc5 | -0.001 | Old |
| 0.963693 | 0.164587 | 0.037 | 0.037 | 1 | 6 | Slc39a13 | 0 | Old |
| 0.964213 | 0.015937 | 0.546 | 0.538 | 1 | 6 | Ctsa | 0.008 | Old |
| 0.964414 | 0.122806 | 0.017 | 0.017 | 1 | 6 | Chek2 | 0 | Old |
| 0.964531 | 0.134173 | 0.237 | 0.234 | 1 | 6 | Ap3d1 | 0.003 | Old |
| 0.964543 | 0.101503 | 0.211 | 0.211 | 1 | 6 | Fkbp15 | 0 | Old |
| 0.964647 | 0.217678 | 0.223 | 0.227 | 1 | 6 | Zmynd11 | -0.004 | Old |
| 0.964868 | 0.167631 | 0.123 | 0.122 | 1 | 6 | Mplkip | 0.001 | Old |
| 0.964994 | 0.011186 | 0.257 | 0.256 | 1 | 6 | Atp6v1a | 0.001 | Old |
| 0.965399 | 0.014859 | 0.226 | 0.221 | 1 | 6 | Chd8 | 0.005 | Old |
| 0.965735 | 0.279738 | 0.031 | 0.032 | 1 | 6 | Top2a | -0.001 | Old |
| 0.965968 | 0.038974 | 0.457 | 0.465 | 1 | 6 | Capza1 | -0.008 | Old |
| 0.966778 | 0.131163 | 0.374 | 0.378 | 1 | 6 | Elavl1 | -0.004 | Old |
| 0.966824 | 0.071324 | 0.186 | 0.184 | 1 | 6 | Acaa1a | 0.002 | Old |
| 0.96701 | 0.506666 | 0.1 | 0.104 | 1 | 6 | Gt(ROSA)2 | -0.004 | Old |
| 0.967248 | 0.163412 | 0.063 | 0.064 | 1 | 6 | Scaper | -0.001 | Old |
| 0.967298 | 0.267774 | 0.023 | 0.023 | 1 | 6 | Adat2 | 0 | Old |
| 0.967481 | 0.097384 | 0.251 | 0.253 | 1 | 6 | Qdpr | -0.002 | Old |
| 0.967762 | 0.085849 | 0.011 | 0.012 | 1 | 6 | Zfp275 | -0.001 | Old |
| 0.968364 | 0.031865 | 0.714 | 0.717 | 1 | 6 | Rtn4 | -0.003 | Old |
| 0.968384 | 0.035427 | 0.329 | 0.333 | 1 | 6 | Clec4a2 | -0.004 | Old |
| 0.968455 | 0.08484 | 0.491 | 0.505 | 1 | 6 | Aldoa | -0.014 | Old |
| 0.968576 | 0.137869 | 0.546 | 0.543 | 1 | 6 | Arhgap30 | 0.003 | Old |
| 0.968871 | 0.032912 | 0.111 | 0.11 | 1 | 6 | Tpd52l2 | 0.001 | Old |
| 0.969313 | 0.00146 | 0.94 | 0.945 | 1 | 6 | Pim1 | -0.005 | Old |

| | | | | | | | | |
|---|---|---|---|---|---|---|---|---|
| 0.969381 | 0.130591 | 0.114 | 0.115 | 1 | 6 | Nelfe | -0.001 | Old |
| 0.96969 | 0.024724 | 0.469 | 0.468 | 1 | 6 | Psma4 | 0.001 | Old |
| 0.969875 | 0.068667 | 0.011 | 0.012 | 1 | 6 | Dbndd2 | -0.001 | Old |
| 0.969923 | 0.048628 | 0.311 | 0.313 | 1 | 6 | Clk4 | -0.002 | Old |
| 0.970291 | 0.752219 | 0.037 | 0.037 | 1 | 6 | Mcph1 | 0 | Old |
| 0.970457 | 0.072511 | 0.026 | 0.025 | 1 | 6 | Gm37233 | 0.001 | Old |
| 0.970653 | 0.087278 | 0.271 | 0.269 | 1 | 6 | Osbpl8 | 0.002 | Old |
| 0.97081 | 0.040061 | 0.329 | 0.324 | 1 | 6 | Creb1 | 0.005 | Old |
| 0.971 | 0.157677 | 0.074 | 0.075 | 1 | 6 | Pcgf3 | -0.001 | Old |
| 0.971055 | 0.641685 | 0.023 | 0.023 | 1 | 6 | Dnmbp | 0 | Old |
| 0.971074 | 0.201285 | 0.091 | 0.09 | 1 | 6 | Hmg20a | 0.001 | Old |
| 0.971178 | 0.067498 | 0.094 | 0.092 | 1 | 6 | Cdyl | 0.002 | Old |
| 0.9712 | 0.096408 | 0.18 | 0.177 | 1 | 6 | Anxa11 | 0.003 | Old |
| 0.971988 | 1.430422 | 0.011 | 0.012 | 1 | 6 | Uros | -0.001 | Old |
| 0.972007 | 0.063278 | 0.091 | 0.092 | 1 | 6 | Timm17b | -0.001 | Old |
| 0.97209 | 0.366094 | 0.037 | 0.037 | 1 | 6 | Pkp4 | 0 | Old |
| 0.972128 | 0.054348 | 0.211 | 0.207 | 1 | 6 | Churc1 | 0.004 | Old |
| 0.972474 | 0.062186 | 0.649 | 0.661 | 1 | 6 | Cox5a | -0.012 | Old |
| 0.972523 | 0.415811 | 0.034 | 0.035 | 1 | 6 | Mycl | -0.001 | Old |
| 0.972547 | 0.083648 | 0.066 | 0.065 | 1 | 6 | Ubtd1 | 0.001 | Old |
| 0.972763 | 0.106119 | 0.111 | 0.112 | 1 | 6 | Dpagt1 | -0.001 | Old |
| 0.972812 | 0.225775 | 0.169 | 0.166 | 1 | 6 | Myo18a | 0.003 | Old |
| 0.972902 | 0.080931 | 0.209 | 0.207 | 1 | 6 | Mob3a | 0.002 | Old |
| 0.972932 | 0.12224 | 0.134 | 0.135 | 1 | 6 | Pkig | -0.001 | Old |
| 0.973262 | 0.096979 | 0.489 | 0.505 | 1 | 6 | Mbtd1 | -0.016 | Old |
| 0.973432 | 0.252735 | 0.066 | 0.067 | 1 | 6 | Ralgapa2 | -0.001 | Old |
| 0.973784 | 0.577419 | 0.143 | 0.144 | 1 | 6 | Slc5a3 | -0.001 | Old |
| 0.973797 | 0.31671 | 0.109 | 0.11 | 1 | 6 | Ddx27 | -0.001 | Old |
| 0.974102 | 0.280998 | 0.011 | 0.012 | 1 | 6 | Pfkm | -0.001 | Old |
| 0.974222 | 0.033593 | 0.36 | 0.361 | 1 | 6 | Dcaf12 | -0.001 | Old |
| 0.97434 | 0.378465 | 0.066 | 0.067 | 1 | 6 | Wdyhv1 | -0.001 | Old |
| 0.97435 | 0.125416 | 0.317 | 0.324 | 1 | 6 | Thrap3 | -0.007 | Old |
| 0.974457 | 0.557786 | 0.049 | 0.048 | 1 | 6 | Grap | 0.001 | Old |
| 0.974545 | 0.021998 | 0.643 | 0.654 | 1 | 6 | Ndufa4 | -0.011 | Old |
| 0.974859 | 0.24137 | 0.057 | 0.057 | 1 | 6 | Gmcl1 | 0 | Old |
| 0.975158 | 0.890019 | 0.011 | 0.012 | 1 | 6 | Pou6f1 | -0.001 | Old |
| 0.975158 | 0.45363 | 0.011 | 0.012 | 1 | 6 | Ugt1a6a | -0.001 | Old |
| 0.975624 | 0.127142 | 0.229 | 0.224 | 1 | 6 | Smc3 | 0.005 | Old |
| 0.97569 | 0.398817 | 0.037 | 0.037 | 1 | 6 | Kctd18 | 0 | Old |
| 0.975702 | 0.341243 | 0.066 | 0.067 | 1 | 6 | Hook2 | -0.001 | Old |
| 0.976215 | 0.580852 | 0.011 | 0.012 | 1 | 6 | Cldn10 | -0.001 | Old |
| 0.976215 | 0.470161 | 0.011 | 0.012 | 1 | 6 | Tcf19 | -0.001 | Old |
| 0.976323 | 0.45328 | 0.057 | 0.057 | 1 | 6 | Sap30bp | 0 | Old |
| 0.97714 | 0.422904 | 0.054 | 0.055 | 1 | 6 | Zfp90 | -0.001 | Old |
| 0.977158 | 0.221954 | 0.169 | 0.169 | 1 | 6 | Lima1 | 0 | Old |

| | | | | | | | | |
|---|---|---|---|---|---|---|---|---|
| 0.977272 | 0.500695 | 0.011 | 0.012 | 1 | 6 | Cyp2r1 | -0.001 | Old |
| 0.977651 | 0.114144 | 0.163 | 0.161 | 1 | 6 | Dhx38 | 0.002 | Old |
| 0.978324 | 0.036149 | 0.214 | 0.211 | 1 | 6 | Ppp2r2a | 0.003 | Old |
| 0.978329 | 1.539754 | 0.011 | 0.012 | 1 | 6 | Lockd | -0.001 | Old |
| 0.978329 | 0.033334 | 0.011 | 0.012 | 1 | 6 | Mief2 | -0.001 | Old |
| 0.978432 | 0.063735 | 0.22 | 0.217 | 1 | 6 | Sirt2 | 0.003 | Old |
| 0.979483 | 0.027781 | 0.377 | 0.375 | 1 | 6 | Bag1 | 0.002 | Old |
| 0.979757 | 0.024212 | 0.569 | 0.569 | 1 | 6 | Hspe1 | 0 | Old |
| 0.9798 | 0.059067 | 0.14 | 0.139 | 1 | 6 | Rpp21 | 0.001 | Old |
| 0.980118 | 0.08551 | 0.151 | 0.154 | 1 | 6 | Ppil2 | -0.003 | Old |
| 0.980634 | 0.077819 | 0.583 | 0.607 | 1 | 6 | App | -0.024 | Old |
| 0.980724 | 0.074225 | 0.06 | 0.06 | 1 | 6 | Hdhd2 | 0 | Old |
| 0.980944 | 0.022493 | 0.251 | 0.251 | 1 | 6 | Uqcrc1 | 0 | Old |
| 0.9814 | 0.117078 | 0.146 | 0.144 | 1 | 6 | Trim27 | 0.002 | Old |
| 0.981894 | 0.162125 | 0.031 | 0.032 | 1 | 6 | Kif23 | -0.001 | Old |
| 0.982024 | 0.058916 | 0.454 | 0.458 | 1 | 6 | Ndufa3 | -0.004 | Old |
| 0.982089 | 0.176746 | 0.16 | 0.162 | 1 | 6 | 1810030O( | -0.002 | Old |
| 0.98237 | 0.056679 | 0.209 | 0.204 | 1 | 6 | Yme1l1 | 0.005 | Old |
| 0.9825 | 0.006374 | 0.1 | 0.1 | 1 | 6 | Clcn5 | 0 | Old |
| 0.983257 | 0.005935 | 0.129 | 0.127 | 1 | 6 | Erlec1 | 0.002 | Old |
| 0.983382 | 0.178769 | 0.263 | 0.261 | 1 | 6 | Rab43 | 0.002 | Old |
| 0.983435 | 0.047021 | 0.706 | 0.722 | 1 | 6 | Rap1b | -0.016 | Old |
| 0.983833 | 0.050657 | 0.626 | 0.649 | 1 | 6 | Itgb1 | -0.023 | Old |
| 0.983929 | 0.23902 | 0.206 | 0.207 | 1 | 6 | Tspan32 | -0.001 | Old |
| 0.984443 | 0.166942 | 0.3 | 0.304 | 1 | 6 | Arap1 | -0.004 | Old |
| 0.984523 | 0.087718 | 0.111 | 0.11 | 1 | 6 | Pex7 | 0.001 | Old |
| 0.984535 | 0.097966 | 0.217 | 0.219 | 1 | 6 | Slc25a4 | -0.002 | Old |
| 0.984774 | 0.080787 | 0.083 | 0.082 | 1 | 6 | Ankrd27 | 0.001 | Old |
| 0.984779 | 0.114534 | 0.063 | 0.062 | 1 | 6 | Emc4 | 0.001 | Old |
| 0.984793 | 0.065853 | 0.149 | 0.149 | 1 | 6 | Sash3 | 0 | Old |
| 0.98487 | 0.72899 | 0.034 | 0.035 | 1 | 6 | Unk | -0.001 | Old |
| 0.98501 | 0.182069 | 0.169 | 0.171 | 1 | 6 | Sptlc1 | -0.002 | Old |
| 0.985368 | 0.109551 | 0.266 | 0.266 | 1 | 6 | Mrps14 | 0 | Old |
| 0.985459 | 0.120769 | 0.323 | 0.321 | 1 | 6 | Lactb | 0.002 | Old |
| 0.985773 | 0.054513 | 0.031 | 0.032 | 1 | 6 | Ppp1r3d | -0.001 | Old |
| 0.98642 | 0.114773 | 0.031 | 0.032 | 1 | 6 | Dhx32 | -0.001 | Old |
| 0.987454 | 0.198395 | 0.137 | 0.134 | 1 | 6 | Pisd | 0.003 | Old |
| 0.987607 | 0.347563 | 0.146 | 0.149 | 1 | 6 | Retreg1 | -0.003 | Old |
| 0.987732 | 0.591772 | 0.086 | 0.087 | 1 | 6 | Abraxas2 | -0.001 | Old |
| 0.987987 | 0.306782 | 0.251 | 0.259 | 1 | 6 | Pkn2 | -0.008 | Old |
| 0.988064 | 0.214137 | 0.089 | 0.089 | 1 | 6 | Thumpd3 | 0 | Old |
| 0.988152 | 0.008444 | 0.18 | 0.181 | 1 | 6 | Gpr65 | -0.001 | Old |
| 0.988787 | 0.160775 | 0.24 | 0.244 | 1 | 6 | Immt | -0.004 | Old |
| 0.98901 | 0.039067 | 0.051 | 0.052 | 1 | 6 | Elk1 | -0.001 | Old |
| 0.989017 | 0.104914 | 0.066 | 0.065 | 1 | 6 | Nup93 | 0.001 | Old |

| | | | | | | | | |
|---|---|---|---|---|---|---|---|---|
| 0.989042 | 0.122991 | 0.26 | 0.258 | 1 | 6 | Ppp1cc | 0.002 | Old |
| 0.989291 | 0.149433 | 0.109 | 0.109 | 1 | 6 | Zfp330 | 0 | Old |
| 0.989315 | 0.012348 | 0.289 | 0.286 | 1 | 6 | Rasgrp4 | 0.003 | Old |
| 0.989358 | 0.110883 | 0.374 | 0.39 | 1 | 6 | Coro1c | -0.016 | Old |
| 0.989547 | 0.062759 | 0.546 | 0.57 | 1 | 6 | Coq10b | -0.024 | Old |
| 0.98957 | 0.270941 | 0.249 | 0.253 | 1 | 6 | Senp2 | -0.004 | Old |
| 0.989575 | 0.059537 | 0.317 | 0.321 | 1 | 6 | Ranbp1 | -0.004 | Old |
| 0.989766 | 0.181819 | 0.06 | 0.06 | 1 | 6 | Mtmr10 | 0 | Old |
| 0.989991 | 0.049406 | 0.057 | 0.057 | 1 | 6 | Elmod2 | 0 | Old |
| 0.990009 | 0.261239 | 0.063 | 0.064 | 1 | 6 | Ttc17 | -0.001 | Old |
| 0.990504 | 0.090389 | 0.094 | 0.094 | 1 | 6 | Carnmt1 | 0 | Old |
| 0.990799 | 0.270967 | 0.051 | 0.052 | 1 | 6 | Acsl3 | -0.001 | Old |
| 0.991746 | 0.112885 | 0.117 | 0.117 | 1 | 6 | Prkaca | 0 | Old |
| 0.992193 | 0.009293 | 0.463 | 0.455 | 1 | 6 | Ptges3 | 0.008 | Old |
| 0.992348 | 0.094684 | 0.414 | 0.423 | 1 | 6 | Tet3 | -0.009 | Old |
| 0.992613 | 0.08586 | 0.786 | 0.798 | 1 | 6 | BC005537 | -0.012 | Old |
| 0.992652 | 0.172766 | 0.111 | 0.112 | 1 | 6 | Mrpl11 | -0.001 | Old |
| 0.992886 | 0.389845 | 0.031 | 0.032 | 1 | 6 | Ptch1 | -0.001 | Old |
| 0.992929 | 0.097009 | 0.251 | 0.253 | 1 | 6 | Rtf1 | -0.002 | Old |
| 0.993061 | 0.21727 | 0.069 | 0.069 | 1 | 6 | mt-Nd6 | 0 | Old |
| 0.993371 | 0.047773 | 0.62 | 0.651 | 1 | 6 | Rassf4 | -0.031 | Old |
| 0.993887 | 0.068301 | 0.143 | 0.144 | 1 | 6 | Galk2 | -0.001 | Old |
| 0.993941 | 0.06719 | 0.02 | 0.02 | 1 | 6 | Trmt10b | 0 | Old |
| 0.993999 | 0.581577 | 0.069 | 0.07 | 1 | 6 | Il18rap | -0.001 | Old |
| 0.994405 | 0.170707 | 0.354 | 0.363 | 1 | 6 | Tnpo1 | -0.009 | Old |
| 0.994436 | 0.159877 | 0.191 | 0.192 | 1 | 6 | Vbp1 | -0.001 | Old |
| 0.994482 | 0.059827 | 0.106 | 0.105 | 1 | 6 | Ahsa2 | 0.001 | Old |
| 0.994896 | 0.458309 | 0.029 | 0.028 | 1 | 6 | Prune1 | 0.001 | Old |
| 0.994911 | 0.066269 | 0.297 | 0.299 | 1 | 6 | Dlst | -0.002 | Old |
| 0.994959 | 0.433471 | 0.071 | 0.072 | 1 | 6 | Serpinb8 | -0.001 | Old |
| 0.994961 | 0.241627 | 0.094 | 0.094 | 1 | 6 | Dpm2 | 0 | Old |
| 0.99569 | 0.23171 | 0.157 | 0.161 | 1 | 6 | Fam76b | -0.004 | Old |
| 0.995786 | 0.084067 | 0.049 | 0.048 | 1 | 6 | Lancl1 | 0.001 | Old |
| 0.995927 | 0.148504 | 0.309 | 0.309 | 1 | 6 | Srsf7 | 0 | Old |
| 0.995942 | 0.069602 | 0.289 | 0.284 | 1 | 6 | Ppp1r2 | 0.005 | Old |
| 0.9963 | 0.07256 | 0.143 | 0.144 | 1 | 6 | S100pbp | -0.001 | Old |
| 0.996313 | 0.173645 | 0.049 | 0.048 | 1 | 6 | Nol10 | 0.001 | Old |
| 0.996404 | 0.120298 | 0.137 | 0.139 | 1 | 6 | Nckipsd | -0.002 | Old |
| 0.996421 | 0.014233 | 0.383 | 0.388 | 1 | 6 | Gnaq | -0.005 | Old |
| 0.99646 | 0.355008 | 0.08 | 0.08 | 1 | 6 | Tmem205 | 0 | Old |
| 0.996653 | 0.236906 | 0.146 | 0.147 | 1 | 6 | Cfdp1 | -0.001 | Old |
| 0.99684 | 0.133379 | 0.049 | 0.048 | 1 | 6 | Cdc27 | 0.001 | Old |
| 0.997283 | 0.042312 | 0.249 | 0.247 | 1 | 6 | Setd1b | 0.002 | Old |
| 0.997728 | 0.213529 | 0.166 | 0.166 | 1 | 6 | Grk6 | 0 | Old |
| 0.99798 | 0.58583 | 0.04 | 0.04 | 1 | 6 | Stk11ip | 0 | Old |

| | | | | | | | | |
|---|---|---|---|---|---|---|---|---|
| 0.998196 | 0.156063 | 0.169 | 0.169 | 1 | 6 | Trim44 | 0 | Old |
| 0.998288 | 0.014185 | 0.237 | 0.236 | 1 | 6 | Pon2 | 0.001 | Old |
| 0.998372 | 0.219154 | 0.217 | 0.219 | 1 | 6 | Egln1 | -0.002 | Old |
| 0.998557 | 0.025719 | 0.04 | 0.04 | 1 | 6 | Mtmr9 | 0 | Old |
| 0.998751 | 0.132589 | 0.08 | 0.08 | 1 | 6 | Kdsr | 0 | Old |
| 0.998913 | 0.025048 | 0.157 | 0.156 | 1 | 6 | Ncoa1 | 0.001 | Old |
| 0.999031 | 0.102752 | 0.094 | 0.094 | 1 | 6 | Ctso | 0 | Old |
| 0.999096 | 0.323092 | 0.109 | 0.11 | 1 | 6 | Eef1akmt1 | -0.001 | Old |
| 0.999231 | 0.039018 | 0.414 | 0.411 | 1 | 6 | Mtss1 | 0.003 | Old |
| 0.999586 | 0.031451 | 0.523 | 0.533 | 1 | 6 | Hnrnpa1 | -0.01 | Old |
| 0.999596 | 0.318041 | 0.02 | 0.02 | 1 | 6 | 2700097O( | 0 | Old |
| 0.999596 | 0.129315 | 0.02 | 0.02 | 1 | 6 | Mpp7 | 0 | Old |
| 0.999596 | 0.022816 | 0.02 | 0.02 | 1 | 6 | Noa1 | 0 | Old |
| 0.99966 | 0.011446 | 0.029 | 0.028 | 1 | 6 | Large1 | 0.001 | Old |
| 0.999661 | 0.087119 | 0.126 | 0.127 | 1 | 6 | Tusc3 | -0.001 | Old |
| 6.14E-15 | 0.363522 | 0.99 | 0.969 | 1.91E-10 | 6 | Rps14 | 0.021 | Young |
| 6.22E-14 | 0.42796 | 0.975 | 0.929 | 1.93E-09 | 6 | Rpl18a | 0.046 | Young |
| 1.93E-13 | 0.348352 | 0.985 | 0.96 | 5.99E-09 | 6 | Rpl23 | 0.025 | Young |
| 3.64E-13 | 0.506374 | 0.915 | 0.866 | 1.13E-08 | 6 | Rpl11 | 0.049 | Young |
| 5.01E-13 | 0.370866 | 0.973 | 0.963 | 1.56E-08 | 6 | Rps16 | 0.01 | Young |
| 7.66E-13 | 0.356627 | 0.98 | 0.963 | 2.38E-08 | 6 | Rpl30 | 0.017 | Young |
| 9.41E-13 | 0.330176 | 0.973 | 0.949 | 2.92E-08 | 6 | Rps4x | 0.024 | Young |
| 1.11E-12 | 0.444948 | 0.941 | 0.88 | 3.44E-08 | 6 | Rps7 | 0.061 | Young |
| 1.76E-12 | 0.285908 | 0.993 | 0.98 | 5.46E-08 | 6 | Rps9 | 0.013 | Young |
| 1.84E-12 | 0.397 | 0.97 | 0.946 | 5.73E-08 | 6 | Rps13 | 0.024 | Young |
| 4.31E-12 | 0.904591 | 0.505 | 0.269 | 1.34E-07 | 6 | Hspa1a | 0.236 | Young |
| 9.26E-12 | 0.280571 | 0.998 | 0.963 | 2.87E-07 | 6 | Cd74 | 0.035 | Young |
| 1.71E-11 | 0.326245 | 0.982 | 0.943 | 5.30E-07 | 6 | Rpl27a | 0.039 | Young |
| 2.95E-11 | 0.315225 | 0.992 | 0.974 | 9.17E-07 | 6 | Rpl13 | 0.018 | Young |
| 3.43E-11 | 0.335985 | 0.975 | 0.943 | 1.07E-06 | 6 | Rps5 | 0.032 | Young |
| 3.44E-11 | 0.354197 | 0.982 | 0.949 | 1.07E-06 | 6 | Rps10 | 0.033 | Young |
| 4.24E-11 | 0.345174 | 0.97 | 0.937 | 1.32E-06 | 6 | Rpl17 | 0.033 | Young |
| 4.26E-10 | 0.406603 | 0.95 | 0.886 | 1.32E-05 | 6 | Rpl32 | 0.064 | Young |
| 6.86E-10 | 0.396167 | 0.935 | 0.797 | 2.13E-05 | 6 | Fcer1g | 0.138 | Young |
| 1.22E-09 | 0.373349 | 0.943 | 0.909 | 3.77E-05 | 6 | Rpl26 | 0.034 | Young |
| 2.36E-09 | 0.253344 | 0.992 | 0.986 | 7.32E-05 | 6 | Fau | 0.006 | Young |
| 4.47E-09 | 0.212888 | 0.997 | 0.98 | 0.000139 | 6 | Tpt1 | 0.017 | Young |
| 5.25E-09 | 0.3537 | 0.955 | 0.911 | 0.000163 | 6 | Rpl34 | 0.044 | Young |
| 7.97E-09 | 0.243374 | 0.99 | 0.983 | 0.000248 | 6 | Rplp1 | 0.007 | Young |
| 9.51E-09 | 0.645649 | 0.699 | 0.523 | 0.000295 | 6 | Nr4a3 | 0.176 | Young |
| 1.47E-08 | 0.341611 | 0.953 | 0.929 | 0.000455 | 6 | Rpl18 | 0.024 | Young |
| 2.03E-08 | 0.304169 | 0.96 | 0.926 | 0.00063 | 6 | Rps11 | 0.034 | Young |
| 2.41E-08 | 0.34783 | 0.945 | 0.889 | 0.000748 | 6 | Rps6 | 0.056 | Young |
| 2.98E-08 | 0.237872 | 0.992 | 0.977 | 0.000924 | 6 | Rps24 | 0.015 | Young |
| 3.76E-08 | 0.299504 | 0.941 | 0.917 | 0.001167 | 6 | Rps25 | 0.024 | Young |

| | | | | | | | | |
|---|---|---|---|---|---|---|---|---|
| 6.10E-08 | 0.322286 | 0.941 | 0.9 | 0.001894 | 6 | Rpl21 | 0.041 | Young |
| 6.78E-08 | 1.735033 | 0.207 | 0.077 | 0.002106 | 6 | Plxdc2 | 0.13 | Young |
| 7.82E-08 | 0.809082 | 0.446 | 0.274 | 0.002428 | 6 | Cd163 | 0.172 | Young |
| 9.96E-08 | 0.376564 | 0.92 | 0.763 | 0.003094 | 6 | Tnip3 | 0.157 | Young |
| 1.45E-07 | 0.322872 | 0.928 | 0.883 | 0.004503 | 6 | Rpl10a | 0.045 | Young |
| 1.74E-07 | 0.292545 | 0.955 | 0.917 | 0.005411 | 6 | Rpl9 | 0.038 | Young |
| 2.28E-07 | 0.229746 | 0.98 | 0.92 | 0.00709 | 6 | Cst3 | 0.06 | Young |
| 2.58E-07 | 0.24316 | 0.98 | 0.957 | 0.008012 | 6 | Rps23 | 0.023 | Young |
| 3.57E-07 | 0.223901 | 0.988 | 0.96 | 0.011079 | 6 | Rps27a | 0.028 | Young |
| 3.64E-07 | 0.481429 | 0.758 | 0.646 | 0.011308 | 6 | Trf | 0.112 | Young |
| 3.78E-07 | 0.393849 | 0.915 | 0.794 | 0.011744 | 6 | Il1b | 0.121 | Young |
| 4.32E-07 | 0.172236 | 0.987 | 0.986 | 0.013424 | 6 | Rps29 | 0.001 | Young |
| 7.00E-07 | 0.674003 | 0.485 | 0.323 | 0.02175 | 6 | Hspa1b | 0.162 | Young |
| 8.89E-07 | 0.248208 | 0.998 | 0.957 | 0.027597 | 6 | H2-Aa | 0.041 | Young |
| 9.15E-07 | 0.337705 | 0.915 | 0.783 | 0.028406 | 6 | Tyrobp | 0.132 | Young |
| 9.45E-07 | 0.307729 | 0.9 | 0.874 | 0.029359 | 6 | Rps3 | 0.026 | Young |
| 9.47E-07 | 0.342107 | 0.957 | 0.84 | 0.029422 | 6 | Cd83 | 0.117 | Young |
| 9.53E-07 | 2.681663 | 0.129 | 0.034 | 0.029591 | 6 | Hpgd | 0.095 | Young |
| 1.14E-06 | 0.297652 | 0.911 | 0.869 | 0.035285 | 6 | Rpl12 | 0.042 | Young |
| 1.18E-06 | 0.230647 | 0.973 | 0.943 | 0.036681 | 6 | Rps19 | 0.03 | Young |
| 1.19E-06 | 0.424864 | 0.821 | 0.717 | 0.036881 | 6 | Rel | 0.104 | Young |
| 1.44E-06 | 0.535143 | 0.801 | 0.706 | 0.044774 | 6 | Mt1 | 0.095 | Young |
| 1.47E-06 | 0.52049 | 0.747 | 0.651 | 0.045663 | 6 | Cdkn1a | 0.096 | Young |
| 1.54E-06 | 0.322032 | 0.9 | 0.843 | 0.047723 | 6 | Rps18 | 0.057 | Young |
| 1.71E-06 | 0.657792 | 0.629 | 0.503 | 0.053066 | 6 | Cd14 | 0.126 | Young |
| 1.90E-06 | 0.409462 | 0.828 | 0.706 | 0.05914 | 6 | Tmem176b | 0.122 | Young |
| 2.19E-06 | 0.295172 | 0.938 | 0.903 | 0.067939 | 6 | Rpl36a | 0.035 | Young |
| 2.37E-06 | 0.812449 | 0.446 | 0.303 | 0.073736 | 6 | Apoe | 0.143 | Young |
| 2.88E-06 | 0.193476 | 0.977 | 0.966 | 0.089411 | 6 | Rps12 | 0.011 | Young |
| 2.93E-06 | 0.502078 | 0.901 | 0.811 | 0.090911 | 6 | Lyz2 | 0.09 | Young |
| 3.53E-06 | 0.19626 | 0.99 | 0.983 | 0.109617 | 6 | Rps8 | 0.007 | Young |
| 3.72E-06 | 0.241772 | 0.977 | 0.957 | 0.115394 | 6 | Fth1 | 0.02 | Young |
| 3.99E-06 | 0.344021 | 0.911 | 0.829 | 0.124021 | 6 | Nfkbiz | 0.082 | Young |
| 4.91E-06 | 0.251424 | 0.948 | 0.909 | 0.152345 | 6 | Rpl28 | 0.039 | Young |
| 5.52E-06 | 0.336813 | 0.831 | 0.734 | 0.171316 | 6 | Plek | 0.097 | Young |
| 8.39E-06 | 0.333543 | 0.868 | 0.814 | 0.260686 | 6 | Rpl10 | 0.054 | Young |
| 8.50E-06 | 0.309328 | 0.858 | 0.749 | 0.263889 | 6 | Kdm6b | 0.109 | Young |
| 1.09E-05 | 0.607398 | 0.543 | 0.414 | 0.339177 | 6 | Skil | 0.129 | Young |
| 1.12E-05 | 0.169356 | 0.987 | 0.957 | 0.347358 | 6 | Rps15 | 0.03 | Young |
| 1.31E-05 | 0.226491 | 0.967 | 0.937 | 0.4062 | 6 | Rpl5 | 0.03 | Young |
| 1.38E-05 | 0.334299 | 0.878 | 0.783 | 0.428862 | 6 | Rpl15 | 0.095 | Young |
| 1.75E-05 | 0.204659 | 0.97 | 0.969 | 0.543778 | 6 | Rpl19 | 0.001 | Young |
| 1.78E-05 | 0.249494 | 0.95 | 0.914 | 0.552909 | 6 | Rps15a | 0.036 | Young |
| 1.98E-05 | 0.216331 | 0.972 | 0.934 | 0.613425 | 6 | Rplp0 | 0.038 | Young |
| 2.11E-05 | 0.214335 | 0.975 | 0.951 | 0.654059 | 6 | Rps20 | 0.024 | Young |

| | | | | | | | | |
|---|---|---|---|---|---|---|---|---|
| 2.42E-05 | 0.607603 | 0.535 | 0.406 | 0.750521 | 6 | Dot1l | 0.129 | Young |
| 2.74E-05 | 0.69346 | 0.338 | 0.214 | 0.850913 | 6 | Ptger4 | 0.124 | Young |
| 3.14E-05 | 0.2524 | 0.946 | 0.88 | 0.976253 | 6 | Ftl1 | 0.066 | Young |
| 3.94E-05 | 0.615384 | 0.453 | 0.334 | 1 | 6 | Dennd5a | 0.119 | Young |
| 4.26E-05 | 0.241415 | 0.923 | 0.891 | 1 | 6 | Srgn | 0.032 | Young |
| 4.30E-05 | 0.306704 | 0.878 | 0.769 | 1 | 6 | Wfdc17 | 0.109 | Young |
| 5.11E-05 | 0.314716 | 0.773 | 0.666 | 1 | 6 | Fcgr2b | 0.107 | Young |
| 5.15E-05 | 0.262766 | 0.91 | 0.863 | 1 | 6 | Rpl35 | 0.047 | Young |
| 5.34E-05 | 0.556402 | 0.502 | 0.363 | 1 | 6 | Bcl2a1d | 0.139 | Young |
| 5.35E-05 | 0.595794 | 0.535 | 0.391 | 1 | 6 | Egr3 | 0.144 | Young |
| 5.52E-05 | 0.190839 | 0.972 | 0.946 | 1 | 6 | Rpl22 | 0.026 | Young |
| 5.81E-05 | 0.196492 | 0.963 | 0.926 | 1 | 6 | Pabpc1 | 0.037 | Young |
| 6.65E-05 | 0.300719 | 0.796 | 0.703 | 1 | 6 | Purb | 0.093 | Young |
| 6.74E-05 | 0.325002 | 0.843 | 0.769 | 1 | 6 | Dennd4a | 0.074 | Young |
| 7.26E-05 | 0.589765 | 0.485 | 0.36 | 1 | 6 | Abca9 | 0.125 | Young |
| 8.20E-05 | 0.44238 | 0.709 | 0.591 | 1 | 6 | Marcksl1 | 0.118 | Young |
| 8.35E-05 | 0.631455 | 0.533 | 0.414 | 1 | 6 | Mt2 | 0.119 | Young |
| 8.41E-05 | 0.160744 | 0.995 | 0.971 | 1 | 6 | Rplp2 | 0.024 | Young |
| 9.24E-05 | 0.830591 | 0.209 | 0.111 | 1 | 6 | Sult1a1 | 0.098 | Young |
| 9.72E-05 | 0.539131 | 0.559 | 0.437 | 1 | 6 | Nfe2l2 | 0.122 | Young |
| 0.000101 | 0.160313 | 0.992 | 0.974 | 1 | 6 | Rpl35a | 0.018 | Young |
| 0.000113 | 0.199072 | 0.963 | 0.957 | 1 | 6 | Rps27 | 0.006 | Young |
| 0.000125 | 0.28524 | 0.783 | 0.677 | 1 | 6 | Gadd45b | 0.106 | Young |
| 0.000154 | 0.281817 | 0.943 | 0.843 | 1 | 6 | Marcks | 0.1 | Young |
| 0.00016 | 0.330657 | 0.714 | 0.586 | 1 | 6 | Eif3k | 0.128 | Young |
| 0.00018 | 0.217939 | 0.94 | 0.883 | 1 | 6 | Rpl36 | 0.057 | Young |
| 0.000184 | 0.294155 | 0.794 | 0.751 | 1 | 6 | Cox7a2l | 0.043 | Young |
| 0.000186 | 0.170834 | 0.978 | 0.954 | 1 | 6 | Rack1 | 0.024 | Young |
| 0.000188 | 0.340751 | 0.691 | 0.597 | 1 | 6 | Pnrc1 | 0.094 | Young |
| 0.000213 | 0.352799 | 0.754 | 0.66 | 1 | 6 | Kdm7a | 0.094 | Young |
| 0.000226 | 0.511592 | 0.296 | 0.191 | 1 | 6 | Mcemp1 | 0.105 | Young |
| 0.00026 | 0.332741 | 0.615 | 0.5 | 1 | 6 | Znrf2 | 0.115 | Young |
| 0.000261 | 0.190932 | 0.997 | 0.937 | 1 | 6 | H2-Eb1 | 0.06 | Young |
| 0.000279 | 0.153301 | 0.987 | 0.974 | 1 | 6 | Rps21 | 0.013 | Young |
| 0.000282 | 1.427091 | 0.192 | 0.106 | 1 | 6 | Lyz1 | 0.086 | Young |
| 0.000332 | 0.625137 | 0.375 | 0.271 | 1 | 6 | Rasgef1b | 0.104 | Young |
| 0.000333 | 0.375265 | 0.736 | 0.669 | 1 | 6 | Cox8a | 0.067 | Young |
| 0.000336 | 0.089548 | 1 | 1 | 1 | 6 | Gm42418 | 0 | Young |
| 0.000374 | 0.834459 | 0.279 | 0.177 | 1 | 6 | Dnajb1 | 0.102 | Young |
| 0.000404 | 0.686169 | 0.156 | 0.077 | 1 | 6 | Zfp395 | 0.079 | Young |
| 0.000415 | 1.072071 | 0.13 | 0.057 | 1 | 6 | Gm48065 | 0.073 | Young |
| 0.000432 | 0.196716 | 0.935 | 0.863 | 1 | 6 | Eef1b2 | 0.072 | Young |
| 0.000513 | 0.211948 | 0.936 | 0.854 | 1 | 6 | H3f3a | 0.082 | Young |
| 0.000524 | 0.291787 | 0.339 | 0.24 | 1 | 6 | Igsf9 | 0.099 | Young |
| 0.000542 | 0.258544 | 0.871 | 0.809 | 1 | 6 | Rpl7a | 0.062 | Young |

| | | | | | | | | |
|---|---|---|---|---|---|---|---|---|
| 0.00057 | 0.296785 | 0.669 | 0.569 | 1 | 6 | Ikbkb | 0.1 | Young |
| 0.00058 | 0.24144 | 0.784 | 0.703 | 1 | 6 | Fosb | 0.081 | Young |
| 0.000582 | 0.172447 | 0.977 | 0.951 | 1 | 6 | Rpl23a | 0.026 | Young |
| 0.000632 | 0.248404 | 0.784 | 0.677 | 1 | 6 | Cybb | 0.107 | Young |
| 0.000638 | 0.294291 | 0.727 | 0.64 | 1 | 6 | Spag9 | 0.087 | Young |
| 0.000693 | 0.468649 | 0.497 | 0.403 | 1 | 6 | Elmsan1 | 0.094 | Young |
| 0.000733 | 0.553419 | 0.408 | 0.303 | 1 | 6 | Dyrk2 | 0.105 | Young |
| 0.000741 | 0.265474 | 0.756 | 0.646 | 1 | 6 | Ptafr | 0.11 | Young |
| 0.000744 | 0.944526 | 0.231 | 0.146 | 1 | 6 | Zrsr2 | 0.085 | Young |
| 0.000748 | 0.461235 | 0.462 | 0.363 | 1 | 6 | Lmo4 | 0.099 | Young |
| 0.000753 | 0.262724 | 0.846 | 0.737 | 1 | 6 | Prcp | 0.109 | Young |
| 0.000754 | 0.681371 | 0.294 | 0.2 | 1 | 6 | Nlrp3 | 0.094 | Young |
| 0.000785 | 0.130505 | 0.987 | 0.966 | 1 | 6 | Rps3a1 | 0.021 | Young |
| 0.000787 | 0.991725 | 0.211 | 0.129 | 1 | 6 | Tnfrsf12a | 0.082 | Young |
| 0.000805 | 0.621264 | 0.316 | 0.214 | 1 | 6 | Cxcl16 | 0.102 | Young |
| 0.000809 | 0.346508 | 0.692 | 0.594 | 1 | 6 | Nsa2 | 0.098 | Young |
| 0.000821 | 0.145811 | 0.975 | 0.957 | 1 | 6 | Rpl39 | 0.018 | Young |
| 0.000881 | 0.51916 | 0.421 | 0.317 | 1 | 6 | Rnf150 | 0.104 | Young |
| 0.000899 | 0.363391 | 0.577 | 0.474 | 1 | 6 | Slamf7 | 0.103 | Young |
| 0.000931 | 0.330358 | 0.614 | 0.494 | 1 | 6 | Sowahc | 0.12 | Young |
| 0.00095 | 0.620303 | 0.344 | 0.251 | 1 | 6 | Arhgap31 | 0.093 | Young |
| 0.000958 | 0.175557 | 0.957 | 0.94 | 1 | 6 | Hspa8 | 0.017 | Young |
| 0.000959 | 0.620596 | 0.284 | 0.189 | 1 | 6 | Mefv | 0.095 | Young |
| 0.001025 | 0.640394 | 0.308 | 0.214 | 1 | 6 | Daglb | 0.094 | Young |
| 0.001045 | 2.505268 | 0.037 | 0.003 | 1 | 6 | Zkscan17 | 0.034 | Young |
| 0.001105 | 0.824477 | 0.201 | 0.12 | 1 | 6 | Mb21d1 | 0.081 | Young |
| 0.001137 | 0.313407 | 0.701 | 0.611 | 1 | 6 | Aplp2 | 0.09 | Young |
| 0.001201 | 0.35234 | 0.615 | 0.531 | 1 | 6 | Gpi1 | 0.084 | Young |
| 0.001238 | 0.375825 | 0.5 | 0.414 | 1 | 6 | Cyfip1 | 0.086 | Young |
| 0.001292 | 1.275439 | 0.072 | 0.023 | 1 | 6 | Pmp22 | 0.049 | Young |
| 0.001353 | 0.250339 | 0.759 | 0.689 | 1 | 6 | Gdi2 | 0.07 | Young |
| 0.001381 | 0.346325 | 0.632 | 0.543 | 1 | 6 | Gapdh | 0.089 | Young |
| 0.001383 | 0.637147 | 0.358 | 0.269 | 1 | 6 | Tmcc3 | 0.089 | Young |
| 0.001388 | 1.245095 | 0.072 | 0.023 | 1 | 6 | Ptgfrn | 0.049 | Young |
| 0.001478 | 0.380647 | 0.617 | 0.494 | 1 | 6 | Clec4b1 | 0.123 | Young |
| 0.001523 | 1.512017 | 0.052 | 0.011 | 1 | 6 | Eif5a2 | 0.041 | Young |
| 0.001527 | 0.170646 | 0.905 | 0.846 | 1 | 6 | H2afz | 0.059 | Young |
| 0.001535 | 0.797545 | 0.259 | 0.174 | 1 | 6 | Pf4 | 0.085 | Young |
| 0.001625 | 0.765966 | 0.301 | 0.22 | 1 | 6 | Csf1r | 0.081 | Young |
| 0.001676 | 1.064484 | 0.092 | 0.037 | 1 | 6 | Col27a1 | 0.055 | Young |
| 0.001677 | 0.709655 | 0.314 | 0.231 | 1 | 6 | Clec12a | 0.083 | Young |
| 0.001751 | 0.22372 | 0.858 | 0.806 | 1 | 6 | Sub1 | 0.052 | Young |
| 0.001756 | 0.185528 | 0.92 | 0.886 | 1 | 6 | Rps2 | 0.034 | Young |
| 0.001784 | 0.868676 | 0.192 | 0.117 | 1 | 6 | Fyttd1 | 0.075 | Young |
| 0.001808 | 0.245953 | 0.622 | 0.523 | 1 | 6 | Gmfg | 0.099 | Young |

| | | | | | | | | |
|---|---|---|---|---|---|---|---|---|
| 0.001875 | 0.438216 | 0.527 | 0.449 | 1 | 6 | Fam129a | 0.078 | Young |
| 0.001894 | 0.639256 | 0.288 | 0.2 | 1 | 6 | Svil | 0.088 | Young |
| 0.001932 | 0.389667 | 0.423 | 0.32 | 1 | 6 | Iqgap2 | 0.103 | Young |
| 0.001988 | 0.199042 | 0.893 | 0.834 | 1 | 6 | Cyba | 0.059 | Young |
| 0.002052 | 0.699662 | 0.253 | 0.169 | 1 | 6 | Ube2e3 | 0.084 | Young |
| 0.002122 | 0.271197 | 0.806 | 0.703 | 1 | 6 | Mrc1 | 0.103 | Young |
| 0.002138 | 0.176483 | 0.998 | 0.951 | 1 | 6 | H2-Ab1 | 0.047 | Young |
| 0.002191 | 0.613836 | 0.329 | 0.24 | 1 | 6 | Lipa | 0.089 | Young |
| 0.002339 | 0.138579 | 0.968 | 0.943 | 1 | 6 | Eif1 | 0.025 | Young |
| 0.002362 | 2.355589 | 0.038 | 0.006 | 1 | 6 | Fcgr1 | 0.032 | Young |
| 0.002446 | 0.268822 | 0.706 | 0.583 | 1 | 6 | Tnfsf9 | 0.123 | Young |
| 0.002688 | 0.550977 | 0.381 | 0.297 | 1 | 6 | Pttg1ip | 0.084 | Young |
| 0.002708 | 0.557125 | 0.217 | 0.134 | 1 | 6 | Parp8 | 0.083 | Young |
| 0.002783 | 0.812024 | 0.151 | 0.083 | 1 | 6 | Slc2a3 | 0.068 | Young |
| 0.002928 | 0.657544 | 0.227 | 0.149 | 1 | 6 | Vps37b | 0.078 | Young |
| 0.003046 | 0.505467 | 0.324 | 0.237 | 1 | 6 | Prkab2 | 0.087 | Young |
| 0.003106 | 0.380149 | 0.522 | 0.42 | 1 | 6 | Litaf | 0.102 | Young |
| 0.003209 | 0.164309 | 0.868 | 0.794 | 1 | 6 | Eef2 | 0.074 | Young |
| 0.003317 | 0.482356 | 0.411 | 0.323 | 1 | 6 | Fuca1 | 0.088 | Young |
| 0.003353 | 0.139543 | 0.973 | 0.943 | 1 | 6 | Rpl37 | 0.03 | Young |
| 0.003418 | 0.11709 | 0.714 | 0.606 | 1 | 6 | Fam46a | 0.108 | Young |
| 0.00342 | 0.461112 | 0.52 | 0.437 | 1 | 6 | Selenop | 0.083 | Young |
| 0.003504 | 0.844689 | 0.192 | 0.123 | 1 | 6 | Sec14l1 | 0.069 | Young |
| 0.003505 | 0.463103 | 0.488 | 0.403 | 1 | 6 | Ank | 0.085 | Young |
| 0.003559 | 0.984624 | 0.172 | 0.103 | 1 | 6 | Ccl22 | 0.069 | Young |
| 0.003634 | 0.207107 | 0.908 | 0.866 | 1 | 6 | Cox4i1 | 0.042 | Young |
| 0.003656 | 0.479669 | 0.341 | 0.254 | 1 | 6 | Elovl5 | 0.087 | Young |
| 0.003685 | 1.125161 | 0.102 | 0.049 | 1 | 6 | Flrt3 | 0.053 | Young |
| 0.003705 | 0.49522 | 0.452 | 0.366 | 1 | 6 | Icam1 | 0.086 | Young |
| 0.003715 | 0.223181 | 0.855 | 0.766 | 1 | 6 | Tmem176a | 0.089 | Young |
| 0.003879 | 0.156541 | 0.953 | 0.929 | 1 | 6 | Rpl3 | 0.024 | Young |
| 0.004001 | 1.877141 | 0.03 | 0.003 | 1 | 6 | Mx1 | 0.027 | Young |
| 0.004056 | 1.258975 | 0.097 | 0.046 | 1 | 6 | Sh3tc1 | 0.051 | Young |
| 0.004098 | 0.689463 | 0.145 | 0.083 | 1 | 6 | Tctex1d2 | 0.062 | Young |
| 0.004145 | 0.807609 | 0.094 | 0.043 | 1 | 6 | Ppard | 0.051 | Young |
| 0.004166 | 0.452198 | 0.548 | 0.466 | 1 | 6 | Krt80 | 0.082 | Young |
| 0.004292 | 0.235715 | 0.816 | 0.737 | 1 | 6 | Eif3f | 0.079 | Young |
| 0.004392 | 0.355171 | 0.517 | 0.437 | 1 | 6 | Use1 | 0.08 | Young |
| 0.004563 | 1.883346 | 0.035 | 0.006 | 1 | 6 | Trim47 | 0.029 | Young |
| 0.004742 | 1.068169 | 0.097 | 0.046 | 1 | 6 | Cd33 | 0.051 | Young |
| 0.004814 | 0.395124 | 0.803 | 0.714 | 1 | 6 | Retnla | 0.089 | Young |
| 0.004886 | 1.185194 | 0.125 | 0.069 | 1 | 6 | Gpr141 | 0.056 | Young |
| 0.004938 | 0.485293 | 0.428 | 0.351 | 1 | 6 | Fcgr3 | 0.077 | Young |
| 0.004948 | 0.661716 | 0.296 | 0.22 | 1 | 6 | Armc8 | 0.076 | Young |
| 0.00496 | 0.574598 | 0.363 | 0.283 | 1 | 6 | Gpr35 | 0.08 | Young |

| | | | | | | | | |
|---|---|---|---|---|---|---|---|---|
| 0.004997 | 0.187321 | 0.953 | 0.909 | 1 | 6 | Rpl24 | 0.044 | Young |
| 0.005033 | 0.263535 | 0.731 | 0.646 | 1 | 6 | Napsa | 0.085 | Young |
| 0.005156 | 0.188242 | 0.982 | 0.914 | 1 | 6 | Dusp1 | 0.068 | Young |
| 0.00518 | 0.497222 | 0.371 | 0.297 | 1 | 6 | Il13ra1 | 0.074 | Young |
| 0.005233 | 0.610095 | 0.288 | 0.214 | 1 | 6 | Dusp22 | 0.074 | Young |
| 0.005396 | 0.408709 | 0.406 | 0.309 | 1 | 6 | Sf3b5 | 0.097 | Young |
| 0.005427 | 0.317734 | 0.368 | 0.289 | 1 | 6 | Hepacam2 | 0.079 | Young |
| 0.005482 | 0.257407 | 0.545 | 0.44 | 1 | 6 | Got1 | 0.105 | Young |
| 0.005515 | 2.295551 | 0.022 | 0 | 1 | 6 | Deaf1 | 0.022 | Young |
| 0.005528 | 0.230955 | 0.518 | 0.411 | 1 | 6 | Nav1 | 0.107 | Young |
| 0.005563 | 0.32548 | 0.577 | 0.483 | 1 | 6 | Pde4b | 0.094 | Young |
| 0.005625 | 0.710071 | 0.197 | 0.129 | 1 | 6 | Il17ra | 0.068 | Young |
| 0.005677 | 0.288702 | 0.462 | 0.374 | 1 | 6 | Nckap1l | 0.088 | Young |
| 0.005876 | 1.285641 | 0.087 | 0.04 | 1 | 6 | Efnb2 | 0.047 | Young |
| 0.005922 | 0.355764 | 0.513 | 0.443 | 1 | 6 | Lrp1 | 0.07 | Young |
| 0.005933 | 0.282182 | 0.44 | 0.349 | 1 | 6 | Lgmn | 0.091 | Young |
| 0.005996 | 0.447961 | 0.092 | 0.043 | 1 | 6 | Svbp | 0.049 | Young |
| 0.006156 | 1.099175 | 0.129 | 0.074 | 1 | 6 | Suclg2 | 0.055 | Young |
| 0.006203 | 0.339561 | 0.513 | 0.437 | 1 | 6 | Mpp6 | 0.076 | Young |
| 0.00648 | 0.167269 | 0.923 | 0.851 | 1 | 6 | Btg1 | 0.072 | Young |
| 0.006534 | 0.304859 | 0.483 | 0.394 | 1 | 6 | Furin | 0.089 | Young |
| 0.006818 | 0.322905 | 0.538 | 0.466 | 1 | 6 | Ncf1 | 0.072 | Young |
| 0.006881 | 0.276856 | 0.545 | 0.443 | 1 | 6 | Arhgap18 | 0.102 | Young |
| 0.006946 | 0.187249 | 0.916 | 0.851 | 1 | 6 | Npc2 | 0.065 | Young |
| 0.006977 | 0.531759 | 0.177 | 0.111 | 1 | 6 | Coro7 | 0.066 | Young |
| 0.006981 | 0.359902 | 0.465 | 0.389 | 1 | 6 | Rtraf | 0.076 | Young |
| 0.007101 | 1.543271 | 0.077 | 0.034 | 1 | 6 | Tcea3 | 0.043 | Young |
| 0.00711 | 0.491701 | 0.405 | 0.326 | 1 | 6 | Nfkbie | 0.079 | Young |
| 0.007241 | 1.593828 | 0.065 | 0.026 | 1 | 6 | P2rx7 | 0.039 | Young |
| 0.007533 | 0.168073 | 0.846 | 0.754 | 1 | 6 | Rpl31 | 0.092 | Young |
| 0.007539 | 1.583846 | 0.052 | 0.017 | 1 | 6 | Sema6b | 0.035 | Young |
| 0.007706 | 0.382764 | 0.548 | 0.474 | 1 | 6 | Pacs2 | 0.074 | Young |
| 0.007823 | 1.167389 | 0.132 | 0.077 | 1 | 6 | Rrad | 0.055 | Young |
| 0.00791 | 0.291315 | 0.502 | 0.42 | 1 | 6 | Hexb | 0.082 | Young |
| 0.008082 | 0.244977 | 0.729 | 0.626 | 1 | 6 | Ctsc | 0.103 | Young |
| 0.008149 | 0.176566 | 0.635 | 0.566 | 1 | 6 | Atf3 | 0.069 | Young |
| 0.008153 | 0.873137 | 0.052 | 0.017 | 1 | 6 | Tbc1d12 | 0.035 | Young |
| 0.008355 | 2.39963 | 0.032 | 0.006 | 1 | 6 | Wars2 | 0.026 | Young |
| 0.008416 | 0.70709 | 0.304 | 0.24 | 1 | 6 | Cd80 | 0.064 | Young |
| 0.008416 | 0.947937 | 0.095 | 0.049 | 1 | 6 | Stat4 | 0.046 | Young |
| 0.008499 | 1.715338 | 0.032 | 0.006 | 1 | 6 | Dnajc12 | 0.026 | Young |
| 0.008831 | 0.814695 | 0.177 | 0.114 | 1 | 6 | Dclre1c | 0.063 | Young |
| 0.009033 | 0.580178 | 0.289 | 0.22 | 1 | 6 | Adam17 | 0.069 | Young |
| 0.009115 | 0.283368 | 0.368 | 0.28 | 1 | 6 | Txnrd1 | 0.088 | Young |
| 0.009272 | 1.699247 | 0.055 | 0.02 | 1 | 6 | Fbxl4 | 0.035 | Young |

| | | | | | | | | |
|---|---|---|---|---|---|---|---|---|
| 0.009289 | 1.432288 | 0.037 | 0.009 | 1 | 6 | Zc2hc1a | 0.028 | Young |
| 0.009374 | 0.217626 | 0.908 | 0.88 | 1 | 6 | Uqcrh | 0.028 | Young |
| 0.009387 | 0.324836 | 0.458 | 0.371 | 1 | 6 | Ehd1 | 0.087 | Young |
| 0.009544 | 1.343634 | 0.094 | 0.049 | 1 | 6 | Ints12 | 0.045 | Young |
| 0.009676 | 0.156745 | 0.933 | 0.909 | 1 | 6 | Rpl14 | 0.024 | Young |
| 0.009794 | 0.500573 | 0.396 | 0.326 | 1 | 6 | Arl4c | 0.07 | Young |
| 0.009898 | 0.126751 | 0.988 | 0.971 | 1 | 6 | Rps28 | 0.017 | Young |
| 0.010035 | 0.286611 | 0.545 | 0.471 | 1 | 6 | Ms4a6c | 0.074 | Young |
| 0.010246 | 0.218919 | 0.711 | 0.646 | 1 | 6 | Rpl29 | 0.065 | Young |
| 0.010266 | 0.498481 | 0.348 | 0.277 | 1 | 6 | Dhx40 | 0.071 | Young |
| 0.010337 | 0.339698 | 0.582 | 0.514 | 1 | 6 | Nup98 | 0.068 | Young |
| 0.01036 | 0.206146 | 0.746 | 0.663 | 1 | 6 | Clic4 | 0.083 | Young |
| 0.010365 | 0.186979 | 0.868 | 0.794 | 1 | 6 | Rpl4 | 0.074 | Young |
| 0.010453 | 0.469764 | 0.249 | 0.174 | 1 | 6 | Ptgs2 | 0.075 | Young |
| 0.010532 | 0.191838 | 0.604 | 0.503 | 1 | 6 | Dapk1 | 0.101 | Young |
| 0.010568 | 0.120107 | 0.953 | 0.943 | 1 | 6 | Rpl6 | 0.01 | Young |
| 0.010584 | 0.179683 | 0.913 | 0.857 | 1 | 6 | Rps17 | 0.056 | Young |
| 0.010691 | 0.305172 | 0.375 | 0.294 | 1 | 6 | Arid5a | 0.081 | Young |
| 0.010768 | 2.911308 | 0.018 | 0 | 1 | 6 | 2610301B2 | 0.018 | Young |
| 0.011066 | 0.429413 | 0.375 | 0.3 | 1 | 6 | Zfp516 | 0.075 | Young |
| 0.011088 | 0.491093 | 0.383 | 0.311 | 1 | 6 | Arl5c | 0.072 | Young |
| 0.01135 | 0.168204 | 0.537 | 0.457 | 1 | 6 | Tgoln1 | 0.08 | Young |
| 0.01152 | 0.121632 | 0.952 | 0.906 | 1 | 6 | Nfkbia | 0.046 | Young |
| 0.011555 | 0.296171 | 0.545 | 0.474 | 1 | 6 | Ier3 | 0.071 | Young |
| 0.011625 | 1.928183 | 0.03 | 0.006 | 1 | 6 | Ankrd26 | 0.024 | Young |
| 0.011661 | 0.114341 | 0.547 | 0.454 | 1 | 6 | Neurl3 | 0.093 | Young |
| 0.011757 | 0.336661 | 0.324 | 0.243 | 1 | 6 | Aif1 | 0.081 | Young |
| 0.011911 | 1.689729 | 0.035 | 0.009 | 1 | 6 | Dhrs9 | 0.026 | Young |
| 0.012111 | 0.583711 | 0.191 | 0.129 | 1 | 6 | Crim1 | 0.062 | Young |
| 0.012122 | 0.273396 | 0.418 | 0.334 | 1 | 6 | Tnfrsf1b | 0.084 | Young |
| 0.012154 | 0.2594 | 0.712 | 0.64 | 1 | 6 | Pfdn5 | 0.072 | Young |
| 0.012346 | 0.165263 | 0.793 | 0.706 | 1 | 6 | Glud1 | 0.087 | Young |
| 0.012521 | 0.23843 | 0.458 | 0.377 | 1 | 6 | Casp4 | 0.081 | Young |
| 0.012558 | 0.933255 | 0.082 | 0.04 | 1 | 6 | Cpd | 0.042 | Young |
| 0.01258 | 0.244676 | 0.666 | 0.58 | 1 | 6 | Ncf2 | 0.086 | Young |
| 0.012609 | 0.627646 | 0.328 | 0.257 | 1 | 6 | Rab6a | 0.071 | Young |
| 0.013139 | 0.471664 | 0.351 | 0.286 | 1 | 6 | Cyth1 | 0.065 | Young |
| 0.013143 | 0.412399 | 0.356 | 0.286 | 1 | 6 | Pan3 | 0.07 | Young |
| 0.013156 | 0.339879 | 0.462 | 0.386 | 1 | 6 | Ralbp1 | 0.076 | Young |
| 0.013244 | 0.508692 | 0.304 | 0.234 | 1 | 6 | Arrb2 | 0.07 | Young |
| 0.013398 | 0.420595 | 0.333 | 0.26 | 1 | 6 | Nfix | 0.073 | Young |
| 0.013449 | 0.510724 | 0.283 | 0.214 | 1 | 6 | Flt3 | 0.069 | Young |
| 0.013637 | 0.41922 | 0.445 | 0.369 | 1 | 6 | Klrb1b | 0.076 | Young |
| 0.01368 | 0.463827 | 0.308 | 0.243 | 1 | 6 | Trappc5 | 0.065 | Young |
| 0.01374 | 0.105031 | 0.985 | 0.969 | 1 | 6 | Rpl7 | 0.016 | Young |

| | | | | | | | | |
|---|---|---|---|---|---|---|---|---|
| 0.013805 | 0.164576 | 0.873 | 0.766 | 1 | 6 | Mpeg1 | 0.107 | Young |
| 0.014016 | 0.727914 | 0.107 | 0.06 | 1 | 6 | Zfyve27 | 0.047 | Young |
| 0.014153 | 0.213847 | 0.692 | 0.631 | 1 | 6 | Psma7 | 0.061 | Young |
| 0.014231 | 0.292669 | 0.423 | 0.357 | 1 | 6 | Cdk14 | 0.066 | Young |
| 0.014362 | 0.586147 | 0.268 | 0.206 | 1 | 6 | Bcl2a1a | 0.062 | Young |
| 0.014459 | 0.112259 | 0.465 | 0.394 | 1 | 6 | Per1 | 0.071 | Young |
| 0.014562 | 1.025936 | 0.069 | 0.031 | 1 | 6 | Rnase4 | 0.038 | Young |
| 0.014819 | 0.195053 | 0.851 | 0.746 | 1 | 6 | Ccl6 | 0.105 | Young |
| 0.014947 | 1.062159 | 0.084 | 0.043 | 1 | 6 | Cep85l | 0.041 | Young |
| 0.015094 | 2.523879 | 0.017 | 0 | 1 | 6 | Gm26510 | 0.017 | Young |
| 0.015094 | 2.279394 | 0.017 | 0 | 1 | 6 | Fam109a | 0.017 | Young |
| 0.015094 | 2.265408 | 0.017 | 0 | 1 | 6 | Epop | 0.017 | Young |
| 0.015137 | 0.38577 | 0.391 | 0.317 | 1 | 6 | 2810474O1 | 0.074 | Young |
| 0.015141 | 0.314495 | 0.411 | 0.34 | 1 | 6 | Baz2b | 0.071 | Young |
| 0.015292 | 0.351756 | 0.458 | 0.38 | 1 | 6 | Bcl2a1b | 0.078 | Young |
| 0.015386 | 0.208087 | 0.666 | 0.611 | 1 | 6 | Bach1 | 0.055 | Young |
| 0.015673 | 0.546744 | 0.202 | 0.143 | 1 | 6 | Dnajb9 | 0.059 | Young |
| 0.01585 | 2.1817 | 0.028 | 0.006 | 1 | 6 | Platr25 | 0.022 | Young |
| 0.015869 | 0.201545 | 0.786 | 0.74 | 1 | 6 | Cytip | 0.046 | Young |
| 0.015908 | 0.232161 | 0.734 | 0.651 | 1 | 6 | Ifitm2 | 0.083 | Young |
| 0.015915 | 0.431068 | 0.139 | 0.086 | 1 | 6 | Tle1 | 0.053 | Young |
| 0.015923 | 0.294144 | 0.201 | 0.14 | 1 | 6 | Opa3 | 0.061 | Young |
| 0.016087 | 1.01986 | 0.08 | 0.04 | 1 | 6 | Crebl2 | 0.04 | Young |
| 0.016143 | 0.840875 | 0.177 | 0.123 | 1 | 6 | Wwp1 | 0.054 | Young |
| 0.016192 | 0.721625 | 0.072 | 0.034 | 1 | 6 | Fancl | 0.038 | Young |
| 0.016269 | 0.391891 | 0.261 | 0.189 | 1 | 6 | Zbtb18 | 0.072 | Young |
| 0.016408 | 0.187176 | 0.886 | 0.771 | 1 | 6 | Alox5ap | 0.115 | Young |
| 0.016662 | 0.194891 | 0.712 | 0.674 | 1 | 6 | Tob2 | 0.038 | Young |
| 0.016766 | 0.934398 | 0.105 | 0.06 | 1 | 6 | Zfp560 | 0.045 | Young |
| 0.016769 | 0.694979 | 0.129 | 0.08 | 1 | 6 | Ccnk | 0.049 | Young |
| 0.017037 | 0.246424 | 0.465 | 0.394 | 1 | 6 | Zcchc11 | 0.071 | Young |
| 0.017244 | 0.643208 | 0.084 | 0.043 | 1 | 6 | 2300009A0 | 0.041 | Young |
| 0.017253 | 0.087875 | 0.329 | 0.254 | 1 | 6 | Glul | 0.075 | Young |
| 0.017338 | 0.3 | 0.445 | 0.366 | 1 | 6 | Upb1 | 0.079 | Young |
| 0.017434 | 0.137906 | 0.923 | 0.88 | 1 | 6 | Vim | 0.043 | Young |
| 0.017471 | 0.183239 | 0.749 | 0.671 | 1 | 6 | Ccdc88a | 0.078 | Young |
| 0.017594 | 0.448838 | 0.334 | 0.26 | 1 | 6 | Bcl3 | 0.074 | Young |
| 0.017641 | 0.476144 | 0.263 | 0.197 | 1 | 6 | Gch1 | 0.066 | Young |
| 0.01794 | 0.213302 | 0.701 | 0.634 | 1 | 6 | Vmp1 | 0.067 | Young |
| 0.018302 | 0.321717 | 0.36 | 0.286 | 1 | 6 | AB124611 | 0.074 | Young |
| 0.018421 | 0.495142 | 0.187 | 0.129 | 1 | 6 | Tmem8 | 0.058 | Young |
| 0.018558 | 0.2033 | 0.701 | 0.62 | 1 | 6 | Nfkb1 | 0.081 | Young |
| 0.018682 | 0.228326 | 0.425 | 0.36 | 1 | 6 | Hook3 | 0.065 | Young |
| 0.018857 | 0.533069 | 0.313 | 0.251 | 1 | 6 | Higd1a | 0.062 | Young |
| 0.018966 | 0.193558 | 0.635 | 0.56 | 1 | 6 | Tep1 | 0.075 | Young |

| | | | | | | | | |
|---|---|---|---|---|---|---|---|---|
| 0.019039 | 0.363236 | 0.346 | 0.277 | 1 | 6 | Gtf2h5 | 0.069 | Young |
| 0.019201 | 0.377224 | 0.253 | 0.183 | 1 | 6 | Pts | 0.07 | Young |
| 0.019273 | 0.383529 | 0.42 | 0.357 | 1 | 6 | H2afv | 0.063 | Young |
| 0.019353 | 0.195462 | 0.692 | 0.643 | 1 | 6 | Ndufa2 | 0.049 | Young |
| 0.019599 | 0.363935 | 0.152 | 0.097 | 1 | 6 | Fam103a1 | 0.055 | Young |
| 0.019611 | 0.428881 | 0.244 | 0.186 | 1 | 6 | Stom | 0.058 | Young |
| 0.019708 | 1.392117 | 0.042 | 0.014 | 1 | 6 | Ms4a7 | 0.028 | Young |
| 0.019817 | 0.359111 | 0.405 | 0.331 | 1 | 6 | Gpsm3 | 0.074 | Young |
| 0.019925 | 0.087761 | 0.853 | 0.763 | 1 | 6 | Tnfaip3 | 0.09 | Young |
| 0.019973 | 0.204088 | 0.503 | 0.431 | 1 | 6 | Trim25 | 0.072 | Young |
| 0.020063 | 0.437577 | 0.199 | 0.137 | 1 | 6 | Cd209c | 0.062 | Young |
| 0.020155 | 1.784903 | 0.022 | 0.003 | 1 | 6 | Afg1l | 0.019 | Young |
| 0.020273 | 0.94548 | 0.114 | 0.069 | 1 | 6 | Kcne3 | 0.045 | Young |
| 0.020409 | 1.559184 | 0.022 | 0.003 | 1 | 6 | Pomk | 0.019 | Young |
| 0.020524 | 0.731293 | 0.079 | 0.04 | 1 | 6 | Cd93 | 0.039 | Young |
| 0.020544 | 0.240657 | 0.681 | 0.62 | 1 | 6 | Ubb | 0.061 | Young |
| 0.020582 | 0.257705 | 0.363 | 0.289 | 1 | 6 | Med13 | 0.074 | Young |
| 0.020903 | 0.666953 | 0.169 | 0.114 | 1 | 6 | Chmp2b | 0.055 | Young |
| 0.021088 | 0.675152 | 0.062 | 0.029 | 1 | 6 | Rnf144b | 0.033 | Young |
| 0.021165 | 0.190009 | 0.871 | 0.831 | 1 | 6 | Fxyd5 | 0.04 | Young |
| 0.021317 | 0.309093 | 0.508 | 0.434 | 1 | 6 | Dpy19l1 | 0.074 | Young |
| 0.021528 | 0.237639 | 0.467 | 0.4 | 1 | 6 | Samd4b | 0.067 | Young |
| 0.021782 | 1.658271 | 0.027 | 0.006 | 1 | 6 | Tspan4 | 0.021 | Young |
| 0.021796 | 1.421954 | 0.102 | 0.06 | 1 | 6 | Gpr183 | 0.042 | Young |
| 0.021989 | 0.664157 | 0.074 | 0.037 | 1 | 6 | Daam1 | 0.037 | Young |
| 0.022011 | 0.204305 | 0.704 | 0.634 | 1 | 6 | Csf2ra | 0.07 | Young |
| 0.022055 | 0.269601 | 0.303 | 0.231 | 1 | 6 | Stx16 | 0.072 | Young |
| 0.022143 | 1.445279 | 0.027 | 0.006 | 1 | 6 | Vegfb | 0.021 | Young |
| 0.022172 | 0.205359 | 0.309 | 0.243 | 1 | 6 | Kmt2a | 0.066 | Young |
| 0.022637 | 0.205865 | 0.614 | 0.526 | 1 | 6 | Pirb | 0.088 | Young |
| 0.023138 | 1.64277 | 0.057 | 0.026 | 1 | 6 | Rbm41 | 0.031 | Young |
| 0.02316 | 0.220111 | 0.552 | 0.48 | 1 | 6 | Bri3 | 0.072 | Young |
| 0.023256 | 0.307198 | 0.498 | 0.446 | 1 | 6 | Midn | 0.052 | Young |
| 0.023397 | 0.092293 | 0.863 | 0.791 | 1 | 6 | Klf6 | 0.072 | Young |
| 0.023609 | 0.502259 | 0.286 | 0.226 | 1 | 6 | Uba52 | 0.06 | Young |
| 0.023876 | 0.129467 | 0.441 | 0.351 | 1 | 6 | Fn1 | 0.09 | Young |
| 0.02393 | 0.58356 | 0.261 | 0.2 | 1 | 6 | Washc2 | 0.061 | Young |
| 0.023943 | 0.533479 | 0.062 | 0.029 | 1 | 6 | Ikbkap | 0.033 | Young |
| 0.02401 | 0.821529 | 0.08 | 0.043 | 1 | 6 | Crot | 0.037 | Young |
| 0.024914 | 0.969723 | 0.08 | 0.043 | 1 | 6 | Sgsh | 0.037 | Young |
| 0.02515 | 0.237505 | 0.625 | 0.563 | 1 | 6 | Sat1 | 0.062 | Young |
| 0.025155 | 0.159922 | 0.931 | 0.863 | 1 | 6 | Cd52 | 0.068 | Young |
| 0.025225 | 0.144644 | 0.963 | 0.906 | 1 | 6 | Btg2 | 0.057 | Young |
| 0.025291 | 0.733995 | 0.102 | 0.06 | 1 | 6 | Dhrs7b | 0.042 | Young |
| 0.025457 | 0.089877 | 0.992 | 0.983 | 1 | 6 | Eef1a1 | 0.009 | Young |

| | | | | | | | | |
|---|---|---|---|---|---|---|---|---|
| 0.025627 | 0.638573 | 0.154 | 0.103 | 1 | 6 | Pfkfb3 | 0.051 | Young |
| 0.025728 | 0.26883 | 0.383 | 0.317 | 1 | 6 | Prkd3 | 0.066 | Young |
| 0.025786 | 0.444768 | 0.324 | 0.251 | 1 | 6 | Tlr2 | 0.073 | Young |
| 0.025881 | 0.526962 | 0.062 | 0.029 | 1 | 6 | Cgrrf1 | 0.033 | Young |
| 0.026015 | 0.456871 | 0.281 | 0.217 | 1 | 6 | Notch1 | 0.064 | Young |
| 0.026044 | 0.307227 | 0.201 | 0.137 | 1 | 6 | Heatr5a | 0.064 | Young |
| 0.026062 | 0.219396 | 0.202 | 0.146 | 1 | 6 | Ccng2 | 0.056 | Young |
| 0.02633 | 0.447918 | 0.186 | 0.131 | 1 | 6 | Dync1li1 | 0.055 | Young |
| 0.02642 | 0.11205 | 0.487 | 0.4 | 1 | 6 | Skint3 | 0.087 | Young |
| 0.026505 | 0.191077 | 0.564 | 0.477 | 1 | 6 | Fbl | 0.087 | Young |
| 0.026527 | 0.258975 | 0.323 | 0.257 | 1 | 6 | Samsn1 | 0.066 | Young |
| 0.026638 | 0.358239 | 0.11 | 0.066 | 1 | 6 | Blvra | 0.044 | Young |
| 0.026812 | 2.286203 | 0.035 | 0.011 | 1 | 6 | Itgb5 | 0.024 | Young |
| 0.026943 | 0.15127 | 0.226 | 0.169 | 1 | 6 | Rnpep | 0.057 | Young |
| 0.026995 | 2.119121 | 0.035 | 0.011 | 1 | 6 | Lrrc49 | 0.024 | Young |
| 0.027148 | 0.33752 | 0.355 | 0.286 | 1 | 6 | Zc3h12a | 0.069 | Young |
| 0.02721 | 0.254676 | 0.52 | 0.463 | 1 | 6 | Ctdsp2 | 0.057 | Young |
| 0.027415 | 0.335194 | 0.194 | 0.137 | 1 | 6 | Lsr | 0.057 | Young |
| 0.027431 | 0.43171 | 0.365 | 0.306 | 1 | 6 | Dnajc8 | 0.059 | Young |
| 0.027505 | 1.688434 | 0.02 | 0.003 | 1 | 6 | Fkbp7 | 0.017 | Young |
| 0.027519 | 0.56527 | 0.224 | 0.169 | 1 | 6 | Fam69a | 0.055 | Young |
| 0.027559 | 0.44666 | 0.249 | 0.191 | 1 | 6 | Banf1 | 0.058 | Young |
| 0.027584 | 0.160596 | 0.707 | 0.651 | 1 | 6 | Ankrd44 | 0.056 | Young |
| 0.027898 | 0.668762 | 0.157 | 0.109 | 1 | 6 | Fbrs | 0.048 | Young |
| 0.027954 | 0.429265 | 0.246 | 0.186 | 1 | 6 | Sh3pxd2b | 0.06 | Young |
| 0.028379 | 0.4509 | 0.151 | 0.1 | 1 | 6 | Trim56 | 0.051 | Young |
| 0.02853 | 0.361839 | 0.321 | 0.254 | 1 | 6 | Nudt3 | 0.067 | Young |
| 0.028626 | 0.77981 | 0.246 | 0.194 | 1 | 6 | Slc15a4 | 0.052 | Young |
| 0.028663 | 0.642971 | 0.154 | 0.106 | 1 | 6 | Igsf6 | 0.048 | Young |
| 0.028681 | 0.565702 | 0.139 | 0.091 | 1 | 6 | Tbc1d1 | 0.048 | Young |
| 0.028708 | 0.110904 | 0.714 | 0.629 | 1 | 6 | Eif3e | 0.085 | Young |
| 0.028777 | 1.348224 | 0.124 | 0.083 | 1 | 6 | Grk5 | 0.041 | Young |
| 0.029027 | 1.688113 | 0.07 | 0.037 | 1 | 6 | Tbc1d9b | 0.033 | Young |
| 0.029355 | 2.055876 | 0.025 | 0.006 | 1 | 6 | Hdac10 | 0.019 | Young |
| 0.029684 | 0.706303 | 0.06 | 0.029 | 1 | 6 | Pcyox1 | 0.031 | Young |
| 0.029832 | 0.792958 | 0.09 | 0.051 | 1 | 6 | Tns1 | 0.039 | Young |
| 0.029835 | 1.82522 | 0.025 | 0.006 | 1 | 6 | Ptcd1 | 0.019 | Young |
| 0.029916 | 2.045165 | 0.025 | 0.006 | 1 | 6 | Gm33973 | 0.019 | Young |
| 0.029924 | 2.102575 | 0.013 | 0 | 1 | 6 | Stac3 | 0.013 | Young |
| 0.029924 | 1.904702 | 0.013 | 0 | 1 | 6 | Tmem42 | 0.013 | Young |
| 0.029924 | 1.873784 | 0.013 | 0 | 1 | 6 | Rusc2 | 0.013 | Young |
| 0.029924 | 1.841 | 0.013 | 0 | 1 | 6 | Cd300c | 0.013 | Young |
| 0.030024 | 0.421824 | 0.115 | 0.071 | 1 | 6 | Hat1 | 0.044 | Young |
| 0.030209 | 0.701838 | 0.147 | 0.1 | 1 | 6 | Iah1 | 0.047 | Young |
| 0.030406 | 0.336001 | 0.311 | 0.246 | 1 | 6 | Slc48a1 | 0.065 | Young |

| | | | | | | | | |
|---|---|---|---|---|---|---|---|---|
| 0.030438 | 0.164366 | 0.921 | 0.834 | 1 | 6 | H2-DMa | 0.087 | Young |
| 0.030546 | 0.452909 | 0.182 | 0.129 | 1 | 6 | Cacul1 | 0.053 | Young |
| 0.030623 | 0.441463 | 0.298 | 0.237 | 1 | 6 | Dusp16 | 0.061 | Young |
| 0.030631 | 0.432547 | 0.157 | 0.106 | 1 | 6 | Arhgef18 | 0.051 | Young |
| 0.030748 | 0.214235 | 0.346 | 0.28 | 1 | 6 | Nr3c1 | 0.066 | Young |
| 0.030819 | 0.924946 | 0.149 | 0.103 | 1 | 6 | Ticam2 | 0.046 | Young |
| 0.031352 | 0.438935 | 0.276 | 0.22 | 1 | 6 | Park7 | 0.056 | Young |
| 0.031382 | 0.119077 | 0.952 | 0.914 | 1 | 6 | Mcl1 | 0.038 | Young |
| 0.031443 | 0.894366 | 0.043 | 0.017 | 1 | 6 | Alox5 | 0.026 | Young |
| 0.031647 | 0.642274 | 0.1 | 0.06 | 1 | 6 | Josd2 | 0.04 | Young |
| 0.031649 | 0.171686 | 0.806 | 0.769 | 1 | 6 | Atp5h | 0.037 | Young |
| 0.031675 | 0.265986 | 0.552 | 0.509 | 1 | 6 | Atp1b3 | 0.043 | Young |
| 0.031678 | 1.35692 | 0.077 | 0.043 | 1 | 6 | Wdr36 | 0.034 | Young |
| 0.031726 | 0.762392 | 0.139 | 0.094 | 1 | 6 | Cd82 | 0.045 | Young |
| 0.031883 | 0.66898 | 0.092 | 0.054 | 1 | 6 | Rev3l | 0.038 | Young |
| 0.032224 | 0.382267 | 0.309 | 0.246 | 1 | 6 | Borcs6 | 0.063 | Young |
| 0.032562 | 0.224853 | 0.512 | 0.446 | 1 | 6 | 4930523C0 | 0.066 | Young |
| 0.032732 | 0.814907 | 0.079 | 0.043 | 1 | 6 | Fndc7 | 0.036 | Young |
| 0.032735 | 0.233173 | 0.686 | 0.591 | 1 | 6 | Nr4a2 | 0.095 | Young |
| 0.03281 | 0.528263 | 0.169 | 0.117 | 1 | 6 | Xylt1 | 0.052 | Young |
| 0.033768 | 0.404646 | 0.346 | 0.294 | 1 | 6 | Ptpn11 | 0.052 | Young |
| 0.033836 | 0.377612 | 0.246 | 0.189 | 1 | 6 | Il4ra | 0.057 | Young |
| 0.033889 | 1.236257 | 0.047 | 0.02 | 1 | 6 | Wfdc18 | 0.027 | Young |
| 0.033982 | 1.103053 | 0.122 | 0.08 | 1 | 6 | Kdm3a | 0.042 | Young |
| 0.034005 | 0.471909 | 0.115 | 0.071 | 1 | 6 | 4931406P1 | 0.044 | Young |
| 0.034012 | 0.330784 | 0.47 | 0.417 | 1 | 6 | Smarca5 | 0.053 | Young |
| 0.034253 | 0.13636 | 0.885 | 0.849 | 1 | 6 | Arpc1b | 0.036 | Young |
| 0.034273 | 0.34486 | 0.445 | 0.389 | 1 | 6 | Ndufb5 | 0.056 | Young |
| 0.034276 | 0.595006 | 0.264 | 0.206 | 1 | 6 | H2-Ob | 0.058 | Young |
| 0.034296 | 0.822389 | 0.097 | 0.057 | 1 | 6 | Gla | 0.04 | Young |
| 0.034389 | 0.36869 | 0.373 | 0.326 | 1 | 6 | Lamtor2 | 0.047 | Young |
| 0.034404 | 1.253706 | 0.038 | 0.014 | 1 | 6 | Rnd1 | 0.024 | Young |
| 0.034634 | 0.122745 | 0.338 | 0.266 | 1 | 6 | Lcp2 | 0.072 | Young |
| 0.034987 | 0.238112 | 0.515 | 0.454 | 1 | 6 | Ctnna1 | 0.061 | Young |
| 0.035207 | 0.795189 | 0.13 | 0.086 | 1 | 6 | Slc15a2 | 0.044 | Young |
| 0.035229 | 0.768294 | 0.171 | 0.123 | 1 | 6 | Cd40 | 0.048 | Young |
| 0.035254 | 0.180213 | 0.786 | 0.749 | 1 | 6 | Stk17b | 0.037 | Young |
| 0.035309 | 0.740201 | 0.164 | 0.117 | 1 | 6 | Slc35f6 | 0.047 | Young |
| 0.035344 | 0.354127 | 0.276 | 0.217 | 1 | 6 | Emc1 | 0.059 | Young |
| 0.035372 | 0.988271 | 0.059 | 0.029 | 1 | 6 | Smagp | 0.03 | Young |
| 0.035383 | 0.410346 | 0.097 | 0.057 | 1 | 6 | Stx2 | 0.04 | Young |
| 0.035563 | 0.859571 | 0.038 | 0.014 | 1 | 6 | Nxpe5 | 0.024 | Young |
| 0.035621 | 0.253164 | 0.293 | 0.231 | 1 | 6 | 1810058I24 | 0.062 | Young |
| 0.035681 | 0.613437 | 0.189 | 0.137 | 1 | 6 | Cd300ld | 0.052 | Young |
| 0.035747 | 0.34537 | 0.079 | 0.043 | 1 | 6 | Clec9a | 0.036 | Young |

| | | | | | | | | |
|---|---|---|---|---|---|---|---|---|
| 0.035755 | 0.041652 | 0.995 | 0.983 | 1 | 6 | Rpsa | 0.012 | Young |
| 0.03587 | 1.414726 | 0.059 | 0.029 | 1 | 6 | Gnptab | 0.03 | Young |
| 0.035883 | 0.427672 | 0.286 | 0.229 | 1 | 6 | Dmac1 | 0.057 | Young |
| 0.036029 | 0.36137 | 0.104 | 0.063 | 1 | 6 | Pianp | 0.041 | Young |
| 0.03608 | 1.092561 | 0.038 | 0.014 | 1 | 6 | Nkrf | 0.024 | Young |
| 0.036089 | 0.622743 | 0.11 | 0.069 | 1 | 6 | Tspan5 | 0.041 | Young |
| 0.036271 | 0.915777 | 0.149 | 0.106 | 1 | 6 | Pgap2 | 0.043 | Young |
| 0.036287 | 0.86537 | 0.112 | 0.071 | 1 | 6 | Tmem11 | 0.041 | Young |
| 0.036354 | 0.300051 | 0.398 | 0.34 | 1 | 6 | Bcl10 | 0.058 | Young |
| 0.036924 | 0.86851 | 0.077 | 0.043 | 1 | 6 | Tubgcp5 | 0.034 | Young |
| 0.037214 | 0.256612 | 0.58 | 0.514 | 1 | 6 | Man2b1 | 0.066 | Young |
| 0.037292 | 1.372654 | 0.033 | 0.011 | 1 | 6 | Rttn | 0.022 | Young |
| 0.037311 | 0.138922 | 0.793 | 0.743 | 1 | 6 | Morf4l1 | 0.05 | Young |
| 0.037484 | 0.393311 | 0.431 | 0.386 | 1 | 6 | Ndufa7 | 0.045 | Young |
| 0.037545 | 0.342987 | 0.164 | 0.114 | 1 | 6 | Lin54 | 0.05 | Young |
| 0.03755 | 0.133384 | 0.793 | 0.717 | 1 | 6 | Csrnp1 | 0.076 | Young |
| 0.037772 | 0.573845 | 0.11 | 0.069 | 1 | 6 | Dock4 | 0.041 | Young |
| 0.03806 | 0.854888 | 0.132 | 0.089 | 1 | 6 | Ecm1 | 0.043 | Young |
| 0.038291 | 0.461132 | 0.286 | 0.231 | 1 | 6 | Mafg | 0.055 | Young |
| 0.03836 | 0.450277 | 0.36 | 0.32 | 1 | 6 | 1-Mar | 0.04 | Young |
| 0.03854 | 0.205271 | 0.453 | 0.4 | 1 | 6 | Ube2b | 0.053 | Young |
| 0.038782 | 1.798506 | 0.018 | 0.003 | 1 | 6 | Patj | 0.015 | Young |
| 0.038957 | 0.69432 | 0.157 | 0.111 | 1 | 6 | Soga1 | 0.046 | Young |
| 0.039225 | 0.998956 | 0.042 | 0.017 | 1 | 6 | Dram1 | 0.025 | Young |
| 0.039606 | 0.691216 | 0.042 | 0.017 | 1 | 6 | Tnfsf13b | 0.025 | Young |
| 0.039871 | 0.438439 | 0.154 | 0.106 | 1 | 6 | P2ry6 | 0.048 | Young |
| 0.039951 | 0.140809 | 0.656 | 0.606 | 1 | 6 | Foxp1 | 0.05 | Young |
| 0.040185 | 1.528143 | 0.028 | 0.009 | 1 | 6 | BC002059 | 0.019 | Young |
| 0.040445 | 0.187601 | 0.687 | 0.617 | 1 | 6 | Ifrd1 | 0.07 | Young |
| 0.040485 | 0.099743 | 0.323 | 0.257 | 1 | 6 | Ccl2 | 0.066 | Young |
| 0.040623 | 1.79661 | 0.023 | 0.006 | 1 | 6 | Gm43672 | 0.017 | Young |
| 0.040764 | 1.205239 | 0.028 | 0.009 | 1 | 6 | Sipa1l2 | 0.019 | Young |
| 0.040818 | 0.644766 | 0.102 | 0.063 | 1 | 6 | Gas2l1 | 0.039 | Young |
| 0.040839 | 0.395777 | 0.281 | 0.223 | 1 | 6 | Slc27a3 | 0.058 | Young |
| 0.040944 | 0.674052 | 0.122 | 0.08 | 1 | 6 | Med20 | 0.042 | Young |
| 0.041036 | 0.309403 | 0.395 | 0.326 | 1 | 6 | Cd209d | 0.069 | Young |
| 0.041131 | 0.346875 | 0.192 | 0.14 | 1 | 6 | Spire1 | 0.052 | Young |
| 0.041166 | 1.263939 | 0.023 | 0.006 | 1 | 6 | Scrn3 | 0.017 | Young |
| 0.041261 | 0.194187 | 0.324 | 0.26 | 1 | 6 | Gfpt1 | 0.064 | Young |
| 0.041318 | 0.788762 | 0.065 | 0.034 | 1 | 6 | Klrb1a | 0.031 | Young |
| 0.041419 | 0.85526 | 0.162 | 0.117 | 1 | 6 | Mir155hg | 0.045 | Young |
| 0.041687 | 0.101035 | 0.438 | 0.369 | 1 | 6 | Ssh2 | 0.069 | Young |
| 0.041831 | 0.539199 | 0.12 | 0.08 | 1 | 6 | Mgat4a | 0.04 | Young |
| 0.042204 | 0.141628 | 0.87 | 0.743 | 1 | 6 | Cd209a | 0.127 | Young |
| 0.042387 | 2.025684 | 0.012 | 0 | 1 | 6 | Nphp1 | 0.012 | Young |

| | | | | | | | | |
|---|---|---|---|---|---|---|---|---|
| 0.042387 | 2.001342 | 0.012 | 0 | 1 | 6 | Prelid3a | 0.012 | Young |
| 0.042387 | 1.794568 | 0.012 | 0 | 1 | 6 | Capsl | 0.012 | Young |
| 0.042387 | 1.756498 | 0.012 | 0 | 1 | 6 | Gm26759 | 0.012 | Young |
| 0.042387 | 1.727802 | 0.012 | 0 | 1 | 6 | 2810405F1 | 0.012 | Young |
| 0.042387 | 1.699175 | 0.012 | 0 | 1 | 6 | Ggcx | 0.012 | Young |
| 0.042387 | 1.330943 | 0.012 | 0 | 1 | 6 | Akr1c18 | 0.012 | Young |
| 0.042559 | 0.245164 | 0.423 | 0.351 | 1 | 6 | Rassf1 | 0.072 | Young |
| 0.042578 | 0.600028 | 0.102 | 0.063 | 1 | 6 | Fbxo18 | 0.039 | Young |
| 0.042921 | 0.469567 | 0.222 | 0.171 | 1 | 6 | Fbxo11 | 0.051 | Young |
| 0.043438 | 1.188149 | 0.037 | 0.014 | 1 | 6 | F11r | 0.023 | Young |
| 0.043737 | 0.768837 | 0.157 | 0.111 | 1 | 6 | Deptor | 0.046 | Young |
| 0.043962 | 0.558613 | 0.229 | 0.177 | 1 | 6 | Apc | 0.052 | Young |
| 0.044297 | 0.779631 | 0.09 | 0.054 | 1 | 6 | Rnaseh2a | 0.036 | Young |
| 0.044366 | 0.280628 | 0.288 | 0.237 | 1 | 6 | Tpp2 | 0.051 | Young |
| 0.044424 | 0.170896 | 0.398 | 0.34 | 1 | 6 | Tmcc1 | 0.058 | Young |
| 0.044432 | 0.204985 | 0.661 | 0.6 | 1 | 6 | Rpl27 | 0.061 | Young |
| 0.044482 | 0.734434 | 0.065 | 0.034 | 1 | 6 | Tigd2 | 0.031 | Young |
| 0.044563 | 0.268999 | 0.159 | 0.111 | 1 | 6 | Atxn3 | 0.048 | Young |
| 0.044962 | 0.563484 | 0.105 | 0.066 | 1 | 6 | Ptgs1 | 0.039 | Young |
| 0.04498 | 0.137288 | 0.44 | 0.36 | 1 | 6 | Ramp1 | 0.08 | Young |
| 0.044992 | 0.288536 | 0.61 | 0.546 | 1 | 6 | Atf4 | 0.064 | Young |
| 0.04509 | 0.5024 | 0.184 | 0.137 | 1 | 6 | Pdpk1 | 0.047 | Young |
| 0.045125 | 0.910507 | 0.075 | 0.043 | 1 | 6 | Cdkn2aip | 0.032 | Young |
| 0.045222 | 0.540666 | 0.161 | 0.114 | 1 | 6 | Tnfsf12 | 0.047 | Young |
| 0.045347 | 0.116858 | 0.677 | 0.629 | 1 | 6 | Birc3 | 0.048 | Young |
| 0.045503 | 0.245277 | 0.373 | 0.303 | 1 | 6 | Mknk2 | 0.07 | Young |
| 0.045526 | 0.616815 | 0.045 | 0.02 | 1 | 6 | Cmklr1 | 0.025 | Young |
| 0.045543 | 1.124447 | 0.057 | 0.029 | 1 | 6 | Xkrx | 0.028 | Young |
| 0.046095 | 0.982755 | 0.079 | 0.046 | 1 | 6 | Cnnm3 | 0.033 | Young |
| 0.046264 | 0.403148 | 0.169 | 0.123 | 1 | 6 | St6galnac6 | 0.046 | Young |
| 0.046486 | 0.928092 | 0.145 | 0.103 | 1 | 6 | Samd8 | 0.042 | Young |
| 0.046599 | 0.604225 | 0.097 | 0.06 | 1 | 6 | Zswim4 | 0.037 | Young |
| 0.046664 | 0.504104 | 0.054 | 0.026 | 1 | 6 | Fars2 | 0.028 | Young |
| 0.046753 | 0.33113 | 0.284 | 0.231 | 1 | 6 | Arl8a | 0.053 | Young |
| 0.046796 | 0.380823 | 0.231 | 0.177 | 1 | 6 | Hpf1 | 0.054 | Young |
| 0.04692 | 0.655202 | 0.201 | 0.154 | 1 | 6 | Noct | 0.047 | Young |
| 0.046967 | 0.275069 | 0.177 | 0.129 | 1 | 6 | Pkib | 0.048 | Young |
| 0.047363 | 0.645086 | 0.214 | 0.166 | 1 | 6 | Mras | 0.048 | Young |
| 0.047439 | 0.611079 | 0.217 | 0.169 | 1 | 6 | C1qc | 0.048 | Young |
| 0.047531 | 0.343757 | 0.349 | 0.294 | 1 | 6 | Ccrl2 | 0.055 | Young |
| 0.047651 | 0.4235 | 0.258 | 0.209 | 1 | 6 | Il10rb | 0.049 | Young |
| 0.047733 | 0.261823 | 0.604 | 0.534 | 1 | 6 | Socs3 | 0.07 | Young |
| 0.047745 | 0.734844 | 0.095 | 0.06 | 1 | 6 | Ap2a1 | 0.035 | Young |
| 0.047799 | 0.62087 | 0.079 | 0.046 | 1 | 6 | Ubl4a | 0.033 | Young |
| 0.047868 | 0.895429 | 0.092 | 0.057 | 1 | 6 | Kantr | 0.035 | Young |

| | | | | | | | | |
|---|---|---|---|---|---|---|---|---|
| 0.048004 | 0.578314 | 0.159 | 0.114 | 1 | 6 | Otud4 | 0.045 | Young |
| 0.04837 | 0.450108 | 0.082 | 0.049 | 1 | 6 | Gpr137 | 0.033 | Young |
| 0.048506 | 0.212884 | 0.386 | 0.331 | 1 | 6 | Plaur | 0.055 | Young |
| 0.048555 | 0.299613 | 0.421 | 0.374 | 1 | 6 | Tmem256 | 0.047 | Young |
| 0.048605 | 0.729329 | 0.085 | 0.051 | 1 | 6 | Nf2 | 0.034 | Young |
| 0.048636 | 0.612393 | 0.124 | 0.083 | 1 | 6 | Apobr | 0.041 | Young |
| 0.048752 | 0.555266 | 0.268 | 0.214 | 1 | 6 | Bfar | 0.054 | Young |
| 0.049003 | 0.740439 | 0.06 | 0.031 | 1 | 6 | Apip | 0.029 | Young |
| 0.049057 | 0.320683 | 0.204 | 0.151 | 1 | 6 | Tbc1d5 | 0.053 | Young |
| 0.049145 | 0.397085 | 0.187 | 0.14 | 1 | 6 | Map1lc3a | 0.047 | Young |
| 0.049233 | 1.012869 | 0.048 | 0.023 | 1 | 6 | Gm27010 | 0.025 | Young |
| 0.04945 | 0.368845 | 0.236 | 0.183 | 1 | 6 | Rab32 | 0.053 | Young |
| 0.049737 | 0.35198 | 0.087 | 0.051 | 1 | 6 | Tango2 | 0.036 | Young |
| 0.049754 | 0.774896 | 0.186 | 0.143 | 1 | 6 | Nfkbib | 0.043 | Young |
| 0.049897 | 0.22748 | 0.654 | 0.594 | 1 | 6 | Gng5 | 0.06 | Young |
| 0.049945 | 0.889759 | 0.112 | 0.074 | 1 | 6 | Abr | 0.038 | Young |
| 0.049967 | 0.324148 | 0.263 | 0.209 | 1 | 6 | Fbrsl1 | 0.054 | Young |
| 0.049971 | 1.397536 | 0.032 | 0.011 | 1 | 6 | Ift172 | 0.021 | Young |
| 0.050599 | 0.183323 | 0.599 | 0.537 | 1 | 6 | Sfr1 | 0.062 | Young |
| 0.050721 | 0.099079 | 0.654 | 0.577 | 1 | 6 | Trps1 | 0.077 | Young |
| 0.050733 | 0.277695 | 0.246 | 0.191 | 1 | 6 | Stx3 | 0.055 | Young |
| 0.05085 | 0.358608 | 0.478 | 0.429 | 1 | 6 | Stap1 | 0.049 | Young |
| 0.050889 | 0.144307 | 0.687 | 0.594 | 1 | 6 | Slc6a6 | 0.093 | Young |
| 0.051142 | 0.15669 | 0.268 | 0.211 | 1 | 6 | Slc15a3 | 0.057 | Young |
| 0.051171 | 0.265358 | 0.463 | 0.397 | 1 | 6 | Slfn2 | 0.066 | Young |
| 0.051371 | 0.269829 | 0.281 | 0.223 | 1 | 6 | Soat1 | 0.058 | Young |
| 0.051517 | 0.50081 | 0.154 | 0.111 | 1 | 6 | Zfp871 | 0.043 | Young |
| 0.051625 | 0.238767 | 0.314 | 0.257 | 1 | 6 | Rp9 | 0.057 | Young |
| 0.052257 | 0.478805 | 0.079 | 0.046 | 1 | 6 | Rmnd5b | 0.033 | Young |
| 0.052363 | 0.667487 | 0.119 | 0.08 | 1 | 6 | Fam118b | 0.039 | Young |
| 0.052364 | 1.53 | 0.043 | 0.02 | 1 | 6 | Svip | 0.023 | Young |
| 0.052413 | 0.547766 | 0.1 | 0.063 | 1 | 6 | P3h2 | 0.037 | Young |
| 0.052431 | 0.355183 | 0.107 | 0.069 | 1 | 6 | Pitpnc1 | 0.038 | Young |
| 0.052511 | 0.443949 | 0.365 | 0.317 | 1 | 6 | Bnip3l | 0.048 | Young |
| 0.052555 | 0.786782 | 0.157 | 0.114 | 1 | 6 | Kri1 | 0.043 | Young |
| 0.052896 | 0.518134 | 0.181 | 0.134 | 1 | 6 | Adgre4 | 0.047 | Young |
| 0.053157 | 0.821538 | 0.157 | 0.117 | 1 | 6 | Fam162a | 0.04 | Young |
| 0.053197 | 1.22164 | 0.074 | 0.043 | 1 | 6 | Atxn1l | 0.031 | Young |
| 0.053425 | 0.987404 | 0.052 | 0.026 | 1 | 6 | Ccdc112 | 0.026 | Young |
| 0.053685 | 0.923318 | 0.027 | 0.009 | 1 | 6 | Cdr2 | 0.018 | Young |
| 0.053687 | 0.832688 | 0.052 | 0.026 | 1 | 6 | Npat | 0.026 | Young |
| 0.053773 | 2.121047 | 0.017 | 0.003 | 1 | 6 | Hyal1 | 0.014 | Young |
| 0.053773 | 1.524318 | 0.017 | 0.003 | 1 | 6 | Adamts6 | 0.014 | Young |
| 0.054103 | 2.214397 | 0.017 | 0.003 | 1 | 6 | Dqx1 | 0.014 | Young |
| 0.054156 | 0.504277 | 0.105 | 0.069 | 1 | 6 | Gm13373 | 0.036 | Young |

| | | | | | | | | |
|---|---|---|---|---|---|---|---|---|
| 0.054245 | 0.3625 | 0.344 | 0.297 | 1 | 6 | B230219D2 | 0.047 | Young |
| 0.054268 | 1.939908 | 0.017 | 0.003 | 1 | 6 | Zc3hav1l | 0.014 | Young |
| 0.054268 | 1.668026 | 0.017 | 0.003 | 1 | 6 | Ldlrad4 | 0.014 | Young |
| 0.05452 | 0.477373 | 0.104 | 0.066 | 1 | 6 | G6pc3 | 0.038 | Young |
| 0.054768 | 1.306871 | 0.017 | 0.003 | 1 | 6 | 1810055G0 | 0.014 | Young |
| 0.054883 | 0.065862 | 0.881 | 0.803 | 1 | 6 | Rbpj | 0.078 | Young |
| 0.055025 | 0.392433 | 0.294 | 0.249 | 1 | 6 | Fes | 0.045 | Young |
| 0.05544 | 0.663036 | 0.017 | 0.003 | 1 | 6 | Epas1 | 0.014 | Young |
| 0.055599 | 0.391186 | 0.171 | 0.126 | 1 | 6 | Eif4a3 | 0.045 | Young |
| 0.055632 | 0.268484 | 0.216 | 0.163 | 1 | 6 | Ccdc124 | 0.053 | Young |
| 0.055645 | 1.073056 | 0.043 | 0.02 | 1 | 6 | Usp28 | 0.023 | Young |
| 0.055846 | 0.135151 | 0.171 | 0.123 | 1 | 6 | Galnt11 | 0.048 | Young |
| 0.055862 | 1.5583 | 0.022 | 0.006 | 1 | 6 | Polrmt | 0.016 | Young |
| 0.055881 | 0.167326 | 0.52 | 0.443 | 1 | 6 | Mapkapk2 | 0.077 | Young |
| 0.055891 | 0.199555 | 0.605 | 0.566 | 1 | 6 | Atrx | 0.039 | Young |
| 0.055933 | 0.242101 | 0.293 | 0.237 | 1 | 6 | Csnk1e | 0.056 | Young |
| 0.056151 | 0.043097 | 0.259 | 0.203 | 1 | 6 | B4galnt1 | 0.056 | Young |
| 0.056186 | 1.606065 | 0.035 | 0.014 | 1 | 6 | Ddx51 | 0.021 | Young |
| 0.056385 | 0.359813 | 0.249 | 0.203 | 1 | 6 | Cand1 | 0.046 | Young |
| 0.056555 | 0.490912 | 0.247 | 0.197 | 1 | 6 | Bckdhb | 0.05 | Young |
| 0.056682 | 0.217342 | 0.624 | 0.583 | 1 | 6 | Hint1 | 0.041 | Young |
| 0.056762 | 0.336434 | 0.306 | 0.249 | 1 | 6 | Ddr1 | 0.057 | Young |
| 0.056834 | 0.329591 | 0.326 | 0.269 | 1 | 6 | Mbp | 0.057 | Young |
| 0.057039 | 1.170378 | 0.022 | 0.006 | 1 | 6 | Dusp19 | 0.016 | Young |
| 0.05735 | 0.747861 | 0.08 | 0.049 | 1 | 6 | Gclm | 0.031 | Young |
| 0.057659 | 0.243628 | 0.239 | 0.186 | 1 | 6 | Sde2 | 0.053 | Young |
| 0.057661 | 0.249914 | 0.321 | 0.271 | 1 | 6 | Ube2f | 0.05 | Young |
| 0.057722 | 0.393941 | 0.338 | 0.274 | 1 | 6 | Dusp2 | 0.064 | Young |
| 0.058062 | 0.765471 | 0.043 | 0.02 | 1 | 6 | Ahdc1 | 0.023 | Young |
| 0.05859 | 0.100993 | 0.967 | 0.926 | 1 | 6 | Hsp90ab1 | 0.041 | Young |
| 0.058682 | 0.919599 | 0.065 | 0.037 | 1 | 6 | Mcm7 | 0.028 | Young |
| 0.058688 | 0.38679 | 0.328 | 0.277 | 1 | 6 | Hp1bp3 | 0.051 | Young |
| 0.059001 | 0.106688 | 0.908 | 0.791 | 1 | 6 | Ccl9 | 0.117 | Young |
| 0.059102 | 0.376112 | 0.244 | 0.197 | 1 | 6 | Appl1 | 0.047 | Young |
| 0.059289 | 0.918421 | 0.08 | 0.049 | 1 | 6 | Rnf214 | 0.031 | Young |
| 0.059372 | 0.123264 | 0.96 | 0.911 | 1 | 6 | Lsp1 | 0.049 | Young |
| 0.059539 | 0.471007 | 0.099 | 0.063 | 1 | 6 | Fam53c | 0.036 | Young |
| 0.059867 | 0.355099 | 0.186 | 0.14 | 1 | 6 | Tspan33 | 0.046 | Young |
| 0.060379 | 2.501554 | 0.01 | 0 | 1 | 6 | Gata6 | 0.01 | Young |
| 0.060379 | 2.195943 | 0.01 | 0 | 1 | 6 | Zfp39 | 0.01 | Young |
| 0.060379 | 1.986589 | 0.01 | 0 | 1 | 6 | Doc2g | 0.01 | Young |
| 0.060379 | 1.913539 | 0.01 | 0 | 1 | 6 | 9530036M | 0.01 | Young |
| 0.060379 | 1.829371 | 0.01 | 0 | 1 | 6 | Gm34923 | 0.01 | Young |
| 0.060379 | 1.640104 | 0.01 | 0 | 1 | 6 | Fbxo5 | 0.01 | Young |
| 0.060379 | 1.41152 | 0.01 | 0 | 1 | 6 | Nr6a1 | 0.01 | Young |

| | | | | | | | | |
|---|---|---|---|---|---|---|---|---|
| 0.060379 | 1.309753 | 0.01 | 0 | 1 | 6 | Fam110b | 0.01 | Young |
| 0.060379 | 1.215602 | 0.01 | 0 | 1 | 6 | Cdkl2 | 0.01 | Young |
| 0.060379 | 1.181447 | 0.01 | 0 | 1 | 6 | Eid2b | 0.01 | Young |
| 0.060379 | 1.080243 | 0.01 | 0 | 1 | 6 | Gm47815 | 0.01 | Young |
| 0.060434 | 0.72908 | 0.119 | 0.08 | 1 | 6 | Sbf2 | 0.039 | Young |
| 0.06074 | 1.531263 | 0.05 | 0.026 | 1 | 6 | Plrg1 | 0.024 | Young |
| 0.060807 | 0.340771 | 0.276 | 0.22 | 1 | 6 | Dhcr24 | 0.056 | Young |
| 0.061352 | 0.089084 | 0.659 | 0.594 | 1 | 6 | Atp6v0e | 0.065 | Young |
| 0.06155 | 0.228621 | 0.465 | 0.411 | 1 | 6 | Arl5b | 0.054 | Young |
| 0.061583 | 0.056776 | 0.263 | 0.206 | 1 | 6 | Slc39a10 | 0.057 | Young |
| 0.061651 | 0.379979 | 0.189 | 0.143 | 1 | 6 | Lrrc25 | 0.046 | Young |
| 0.061667 | 0.279877 | 0.137 | 0.097 | 1 | 6 | Bend4 | 0.04 | Young |
| 0.061802 | 0.523607 | 0.102 | 0.066 | 1 | 6 | Pla2g7 | 0.036 | Young |
| 0.061924 | 0.124757 | 0.543 | 0.477 | 1 | 6 | Zc3hav1 | 0.066 | Young |
| 0.06197 | 0.197666 | 0.186 | 0.14 | 1 | 6 | Ap1g1 | 0.046 | Young |
| 0.062149 | 0.248837 | 0.258 | 0.203 | 1 | 6 | Adam19 | 0.055 | Young |
| 0.062229 | 0.481387 | 0.207 | 0.16 | 1 | 6 | Bmpr2 | 0.047 | Young |
| 0.062491 | 0.314612 | 0.147 | 0.106 | 1 | 6 | Stim1 | 0.041 | Young |
| 0.06316 | 1.787461 | 0.03 | 0.011 | 1 | 6 | Zscan21 | 0.019 | Young |
| 0.063189 | 0.394076 | 0.105 | 0.069 | 1 | 6 | Mrps26 | 0.036 | Young |
| 0.063495 | 0.448203 | 0.216 | 0.171 | 1 | 6 | Fam173a | 0.045 | Young |
| 0.063795 | 0.256039 | 0.467 | 0.423 | 1 | 6 | Ndufb9 | 0.044 | Young |
| 0.063869 | 0.424207 | 0.144 | 0.103 | 1 | 6 | Asl | 0.041 | Young |
| 0.06451 | 1.290118 | 0.03 | 0.011 | 1 | 6 | Coq6 | 0.019 | Young |
| 0.064572 | 0.934536 | 0.065 | 0.037 | 1 | 6 | Zfp639 | 0.028 | Young |
| 0.064587 | 0.274679 | 0.291 | 0.234 | 1 | 6 | Cul1 | 0.057 | Young |
| 0.065205 | 0.771954 | 0.065 | 0.037 | 1 | 6 | Rars2 | 0.028 | Young |
| 0.065269 | 0.737671 | 0.097 | 0.063 | 1 | 6 | Rnf122 | 0.034 | Young |
| 0.065285 | 1.094772 | 0.075 | 0.046 | 1 | 6 | Map2k5 | 0.029 | Young |
| 0.065382 | 0.390506 | 0.214 | 0.169 | 1 | 6 | Klhl6 | 0.045 | Young |
| 0.065473 | 0.717364 | 0.149 | 0.109 | 1 | 6 | Ube4b | 0.04 | Young |
| 0.065584 | 0.742357 | 0.11 | 0.074 | 1 | 6 | Igfbp4 | 0.036 | Young |
| 0.065607 | 0.022087 | 0.03 | 0.011 | 1 | 6 | Ippk | 0.019 | Young |
| 0.065732 | 0.321978 | 0.249 | 0.197 | 1 | 6 | Rragc | 0.052 | Young |
| 0.065822 | 0.360971 | 0.582 | 0.54 | 1 | 6 | Klf4 | 0.042 | Young |
| 0.066244 | 0.51062 | 0.184 | 0.14 | 1 | 6 | Ifi207 | 0.044 | Young |
| 0.066804 | 0.29647 | 0.124 | 0.083 | 1 | 6 | Cxcl10 | 0.041 | Young |
| 0.066881 | 0.153597 | 0.565 | 0.5 | 1 | 6 | Tle3 | 0.065 | Young |
| 0.067008 | 0.569737 | 0.08 | 0.049 | 1 | 6 | Dpy19l4 | 0.031 | Young |
| 0.067557 | 0.325636 | 0.219 | 0.169 | 1 | 6 | Klf9 | 0.05 | Young |
| 0.067726 | 0.152152 | 0.574 | 0.52 | 1 | 6 | Brk1 | 0.054 | Young |
| 0.067831 | 0.20079 | 0.446 | 0.389 | 1 | 6 | Tsc22d4 | 0.057 | Young |
| 0.068084 | 0.312282 | 0.239 | 0.186 | 1 | 6 | Zmynd8 | 0.053 | Young |
| 0.068477 | 1.703436 | 0.042 | 0.02 | 1 | 6 | Aptx | 0.022 | Young |
| 0.068503 | 0.53043 | 0.171 | 0.129 | 1 | 6 | Stard8 | 0.042 | Young |

| | | | | | | | | |
|---|---|---|---|---|---|---|---|---|
| 0.068652 | 0.798744 | 0.122 | 0.086 | 1 | 6 | Dtymk | 0.036 | Young |
| 0.06868 | 0.368875 | 0.273 | 0.223 | 1 | 6 | Rgs10 | 0.05 | Young |
| 0.068766 | 0.602279 | 0.097 | 0.063 | 1 | 6 | Fam214b | 0.034 | Young |
| 0.068878 | 0.039757 | 0.109 | 0.071 | 1 | 6 | 1110059G1 | 0.038 | Young |
| 0.069748 | 1.093021 | 0.057 | 0.031 | 1 | 6 | Rnf113a2 | 0.026 | Young |
| 0.070012 | 0.240925 | 0.376 | 0.323 | 1 | 6 | Tmem234 | 0.053 | Young |
| 0.070057 | 0.243868 | 0.498 | 0.437 | 1 | 6 | Lamtor4 | 0.061 | Young |
| 0.070415 | 0.65002 | 0.132 | 0.094 | 1 | 6 | Pink1 | 0.038 | Young |
| 0.070829 | 0.931801 | 0.082 | 0.051 | 1 | 6 | Fam207a | 0.031 | Young |
| 0.070864 | 0.49149 | 0.134 | 0.094 | 1 | 6 | Ccl17 | 0.04 | Young |
| 0.07138 | 0.145461 | 0.42 | 0.357 | 1 | 6 | Ddx3y | 0.063 | Young |
| 0.071406 | 0.260052 | 0.152 | 0.109 | 1 | 6 | Akt3 | 0.043 | Young |
| 0.071693 | 0.558091 | 0.156 | 0.114 | 1 | 6 | Ago3 | 0.042 | Young |
| 0.071869 | 0.919199 | 0.079 | 0.049 | 1 | 6 | Il1r2 | 0.03 | Young |
| 0.071899 | 1.313673 | 0.025 | 0.009 | 1 | 6 | 2700049A0 | 0.016 | Young |
| 0.071998 | 0.224562 | 0.401 | 0.346 | 1 | 6 | St3gal4 | 0.055 | Young |
| 0.072017 | 0.331658 | 0.261 | 0.214 | 1 | 6 | Ripk1 | 0.047 | Young |
| 0.072071 | 0.285656 | 0.254 | 0.209 | 1 | 6 | Nbeal1 | 0.045 | Young |
| 0.072343 | 0.304596 | 0.318 | 0.269 | 1 | 6 | Ccar1 | 0.049 | Young |
| 0.072433 | 0.44522 | 0.124 | 0.086 | 1 | 6 | Rpl27-ps3 | 0.038 | Young |
| 0.072456 | 0.066504 | 0.513 | 0.451 | 1 | 6 | Rnf130 | 0.062 | Young |
| 0.072466 | 0.485407 | 0.162 | 0.12 | 1 | 6 | Inpp5f | 0.042 | Young |
| 0.072665 | 0.161846 | 0.376 | 0.32 | 1 | 6 | Crebbp | 0.056 | Young |
| 0.072725 | 1.200178 | 0.025 | 0.009 | 1 | 6 | Gcnt1 | 0.016 | Young |
| 0.072874 | 1.954036 | 0.033 | 0.014 | 1 | 6 | Gm39121 | 0.019 | Young |
| 0.073089 | 0.170763 | 0.559 | 0.494 | 1 | 6 | Tbca | 0.065 | Young |
| 0.07328 | 0.618344 | 0.082 | 0.051 | 1 | 6 | Haao | 0.031 | Young |
| 0.07367 | 0.183156 | 0.482 | 0.411 | 1 | 6 | Csf2rb | 0.071 | Young |
| 0.073758 | 0.096534 | 0.714 | 0.649 | 1 | 6 | Clec10a | 0.065 | Young |
| 0.073766 | 0.228167 | 0.309 | 0.254 | 1 | 6 | Fam102b | 0.055 | Young |
| 0.073976 | 0.118795 | 0.808 | 0.746 | 1 | 6 | Atox1 | 0.062 | Young |
| 0.0741 | 0.310291 | 0.166 | 0.123 | 1 | 6 | Batf | 0.043 | Young |
| 0.074166 | 0.248791 | 0.182 | 0.134 | 1 | 6 | Kif3b | 0.048 | Young |
| 0.074351 | 0.802459 | 0.082 | 0.051 | 1 | 6 | Abtb1 | 0.031 | Young |
| 0.074781 | 0.185522 | 0.336 | 0.277 | 1 | 6 | Cux1 | 0.059 | Young |
| 0.075009 | 0.28121 | 0.443 | 0.4 | 1 | 6 | Atp6ap2 | 0.043 | Young |
| 0.075078 | 0.210522 | 0.483 | 0.44 | 1 | 6 | Pgls | 0.043 | Young |
| 0.07509 | 0.610456 | 0.145 | 0.106 | 1 | 6 | Pcmtd2 | 0.039 | Young |
| 0.075527 | 0.608992 | 0.069 | 0.04 | 1 | 6 | Ephx1 | 0.029 | Young |
| 0.07564 | 1.798479 | 0.015 | 0.003 | 1 | 6 | Tbc1d2 | 0.012 | Young |
| 0.075669 | 0.122264 | 0.518 | 0.457 | 1 | 6 | Fcgrt | 0.061 | Young |
| 0.075869 | 1.684659 | 0.015 | 0.003 | 1 | 6 | Ift88 | 0.012 | Young |
| 0.075869 | 1.380341 | 0.015 | 0.003 | 1 | 6 | Gm10269 | 0.012 | Young |
| 0.07604 | 1.001347 | 0.033 | 0.014 | 1 | 6 | Prepl | 0.019 | Young |
| 0.076139 | 0.382804 | 0.11 | 0.074 | 1 | 6 | Nkiras2 | 0.036 | Young |

| | | | | | | | | |
|---|---|---|---|---|---|---|---|---|
| 0.076541 | 0.642262 | 0.085 | 0.054 | 1 | 6 | Syne1 | 0.031 | Young |
| 0.076559 | 1.160092 | 0.015 | 0.003 | 1 | 6 | Clec4d | 0.012 | Young |
| 0.076627 | 0.618033 | 0.129 | 0.091 | 1 | 6 | Khk | 0.038 | Young |
| 0.076737 | 1.478163 | 0.02 | 0.006 | 1 | 6 | Zfyve28 | 0.014 | Young |
| 0.076778 | 0.326716 | 0.229 | 0.183 | 1 | 6 | Gabarapl2 | 0.046 | Young |
| 0.076923 | 0.689887 | 0.033 | 0.014 | 1 | 6 | Nim1k | 0.019 | Young |
| 0.077022 | 1.292202 | 0.015 | 0.003 | 1 | 6 | Macrod2 | 0.012 | Young |
| 0.077022 | 1.111003 | 0.015 | 0.003 | 1 | 6 | Grk4 | 0.012 | Young |
| 0.07704 | 0.103907 | 0.841 | 0.751 | 1 | 6 | Ctsh | 0.09 | Young |
| 0.077105 | 0.130492 | 0.672 | 0.62 | 1 | 6 | Eif5a | 0.052 | Young |
| 0.077254 | 0.877003 | 0.015 | 0.003 | 1 | 6 | Ackr1 | 0.012 | Young |
| 0.077327 | 1.469051 | 0.02 | 0.006 | 1 | 6 | Zfp523 | 0.014 | Young |
| 0.077487 | 0.656066 | 0.015 | 0.003 | 1 | 6 | Hes2 | 0.012 | Young |
| 0.077511 | 0.101801 | 0.045 | 0.074 | 1 | 6 | Bcl2l12 | -0.029 | Young |
| 0.077524 | 1.364896 | 0.02 | 0.006 | 1 | 6 | Aaas | 0.014 | Young |
| 0.07778 | 1.038642 | 0.064 | 0.037 | 1 | 6 | Col4a1 | 0.027 | Young |
| 0.07778 | 0.05175 | 0.043 | 0.071 | 1 | 6 | Tyw1 | -0.028 | Young |
| 0.07792 | 1.292881 | 0.02 | 0.006 | 1 | 6 | Cep41 | 0.014 | Young |
| 0.078073 | 0.254626 | 0.53 | 0.48 | 1 | 6 | Fem1c | 0.05 | Young |
| 0.078174 | 0.194087 | 0.629 | 0.597 | 1 | 6 | Stat3 | 0.032 | Young |
| 0.078318 | 1.222525 | 0.02 | 0.006 | 1 | 6 | Slc25a23 | 0.014 | Young |
| 0.078517 | 0.813915 | 0.02 | 0.006 | 1 | 6 | Gm16553 | 0.014 | Young |
| 0.079151 | 0.362794 | 0.192 | 0.149 | 1 | 6 | Asnsd1 | 0.043 | Young |
| 0.079151 | 0.250301 | 0.194 | 0.146 | 1 | 6 | Rest | 0.048 | Young |
| 0.079482 | 0.119913 | 0.719 | 0.637 | 1 | 6 | Slamf9 | 0.082 | Young |
| 0.07963 | 0.732297 | 0.134 | 0.097 | 1 | 6 | Dcstamp | 0.037 | Young |
| 0.079911 | 0.496996 | 0.125 | 0.089 | 1 | 6 | Ext2 | 0.036 | Young |
| 0.080103 | 0.182358 | 0.579 | 0.52 | 1 | 6 | Arhgef6 | 0.059 | Young |
| 0.080177 | 0.322484 | 0.229 | 0.186 | 1 | 6 | Eloa | 0.043 | Young |
| 0.080178 | 0.176111 | 0.079 | 0.049 | 1 | 6 | Lmtk2 | 0.03 | Young |
| 0.080262 | 0.15049 | 0.144 | 0.103 | 1 | 6 | Rin2 | 0.041 | Young |
| 0.080625 | 1.090683 | 0.07 | 0.043 | 1 | 6 | Cd300lb | 0.027 | Young |
| 0.08073 | 0.305204 | 0.089 | 0.057 | 1 | 6 | Pxmp4 | 0.032 | Young |
| 0.080865 | 1.154339 | 0.08 | 0.051 | 1 | 6 | Nrbf2 | 0.029 | Young |
| 0.081029 | 0.121608 | 0.154 | 0.111 | 1 | 6 | Bcl7b | 0.043 | Young |
| 0.081173 | 0.085597 | 0.901 | 0.846 | 1 | 6 | Gnai2 | 0.055 | Young |
| 0.081295 | 0.06675 | 0.557 | 0.477 | 1 | 6 | Vasp | 0.08 | Young |
| 0.081339 | 0.817214 | 0.064 | 0.037 | 1 | 6 | Mib2 | 0.027 | Young |
| 0.081429 | 0.454871 | 0.222 | 0.18 | 1 | 6 | Icosl | 0.042 | Young |
| 0.081467 | 0.277265 | 0.323 | 0.28 | 1 | 6 | Ipo7 | 0.043 | Young |
| 0.081709 | 0.957095 | 0.048 | 0.026 | 1 | 6 | Cenpl | 0.022 | Young |
| 0.081927 | 0.228363 | 0.373 | 0.317 | 1 | 6 | Ube2r2 | 0.056 | Young |
| 0.082123 | 0.292934 | 0.446 | 0.406 | 1 | 6 | Oxct1 | 0.04 | Young |
| 0.082288 | 0.470914 | 0.099 | 0.066 | 1 | 6 | Pin1 | 0.033 | Young |
| 0.0825 | 0.379545 | 0.371 | 0.331 | 1 | 6 | Cflar | 0.04 | Young |

| | | | | | | | | |
|---|---|---|---|---|---|---|---|---|
| 0.082541 | 0.161404 | 0.416 | 0.363 | 1 | 6 | Vsir | 0.053 | Young |
| 0.083 | 0.329527 | 0.403 | 0.36 | 1 | 6 | Ptpra | 0.043 | Young |
| 0.083091 | 0.688196 | 0.099 | 0.066 | 1 | 6 | Pex5 | 0.033 | Young |
| 0.08326 | 0.687244 | 0.077 | 0.049 | 1 | 6 | Tgfbrap1 | 0.028 | Young |
| 0.083263 | 0.769137 | 0.161 | 0.123 | 1 | 6 | P4ha1 | 0.038 | Young |
| 0.083355 | 0.185335 | 0.184 | 0.137 | 1 | 6 | Pank3 | 0.047 | Young |
| 0.083361 | 0.335813 | 0.227 | 0.18 | 1 | 6 | Specc1l | 0.047 | Young |
| 0.08351 | 0.341086 | 0.274 | 0.226 | 1 | 6 | Zzef1 | 0.048 | Young |
| 0.083644 | 0.360474 | 0.403 | 0.346 | 1 | 6 | Arsb | 0.057 | Young |
| 0.083886 | 0.496155 | 0.127 | 0.091 | 1 | 6 | Mis18a | 0.036 | Young |
| 0.08406 | 0.488916 | 0.134 | 0.097 | 1 | 6 | Sec24a | 0.037 | Young |
| 0.084079 | 0.167049 | 0.659 | 0.58 | 1 | 6 | Fgfr1 | 0.079 | Young |
| 0.084331 | 0.355782 | 0.266 | 0.217 | 1 | 6 | Cox20 | 0.049 | Young |
| 0.084459 | 0.313612 | 0.226 | 0.18 | 1 | 6 | Mark3 | 0.046 | Young |
| 0.084716 | 0.352469 | 0.249 | 0.203 | 1 | 6 | Bcl6 | 0.046 | Young |
| 0.08481 | 0.217467 | 0.351 | 0.3 | 1 | 6 | Cnih4 | 0.051 | Young |
| 0.084941 | 0.233104 | 0.545 | 0.486 | 1 | 6 | Creg1 | 0.059 | Young |
| 0.085365 | 0.344428 | 0.187 | 0.146 | 1 | 6 | Nr2c2 | 0.041 | Young |
| 0.085903 | 0.15636 | 0.313 | 0.263 | 1 | 6 | Ppp2r1a | 0.05 | Young |
| 0.085998 | 0.082897 | 0.952 | 0.9 | 1 | 6 | Kctd12 | 0.052 | Young |
| 0.086052 | 0.283033 | 0.095 | 0.063 | 1 | 6 | AU019823 | 0.032 | Young |
| 0.086643 | 0.362723 | 0.289 | 0.243 | 1 | 6 | Zfhx3 | 0.046 | Young |
| 0.086645 | 1.163478 | 0.055 | 0.031 | 1 | 6 | Car3 | 0.024 | Young |
| 0.086764 | 0.24733 | 0.147 | 0.106 | 1 | 6 | Rab12 | 0.041 | Young |
| 0.086936 | 0.125686 | 0.888 | 0.837 | 1 | 6 | Ubc | 0.051 | Young |
| 0.087043 | 0.081074 | 0.696 | 0.609 | 1 | 6 | Grn | 0.087 | Young |
| 0.087094 | 0.821612 | 0.037 | 0.017 | 1 | 6 | 1810024B0 | 0.02 | Young |
| 0.087288 | 0.412741 | 0.028 | 0.011 | 1 | 6 | Plcb1 | 0.017 | Young |
| 0.087381 | 0.57303 | 0.171 | 0.134 | 1 | 6 | Mrpl34 | 0.037 | Young |
| 0.087568 | 0.379905 | 0.236 | 0.191 | 1 | 6 | Csnk2a2 | 0.045 | Young |
| 0.087846 | 0.411329 | 0.321 | 0.277 | 1 | 6 | Ebi3 | 0.044 | Young |
| 0.088079 | 0.625081 | 0.241 | 0.206 | 1 | 6 | Dnase2a | 0.035 | Young |
| 0.088116 | 0.149189 | 0.256 | 0.206 | 1 | 6 | Prkacb | 0.05 | Young |
| 0.088339 | 0.451482 | 0.037 | 0.017 | 1 | 6 | Gm16536 | 0.02 | Young |
| 0.088442 | 0.276288 | 0.064 | 0.037 | 1 | 6 | 2310022B0 | 0.027 | Young |
| 0.088702 | 0.566805 | 0.227 | 0.186 | 1 | 6 | Supt4a | 0.041 | Young |
| 0.088768 | 0.433745 | 0.318 | 0.274 | 1 | 6 | Smpdl3a | 0.044 | Young |
| 0.088913 | 0.179202 | 0.401 | 0.351 | 1 | 6 | Fgd2 | 0.05 | Young |
| 0.089049 | 1.432047 | 0.04 | 0.02 | 1 | 6 | Slc25a33 | 0.02 | Young |
| 0.089228 | 0.282303 | 0.502 | 0.46 | 1 | 6 | Cox7b | 0.042 | Young |
| 0.090122 | 0.080904 | 0.747 | 0.689 | 1 | 6 | Alcam | 0.058 | Young |
| 0.090443 | 0.427649 | 0.283 | 0.24 | 1 | 6 | Ndufa10 | 0.043 | Young |
| 0.090753 | 0.418176 | 0.015 | 0.031 | 1 | 6 | Ppp3cc | -0.016 | Young |
| 0.091024 | 0.235897 | 0.355 | 0.314 | 1 | 6 | Akt1 | 0.041 | Young |
| 0.091696 | 0.161522 | 0.415 | 0.349 | 1 | 6 | Eif1a | 0.066 | Young |

| | | | | | | | | |
|---|---|---|---|---|---|---|---|---|
| 0.091974 | 0.12852 | 0.458 | 0.389 | 1 | 6 | Srsf9 | 0.069 | Young |
| 0.092089 | 0.72546 | 0.04 | 0.02 | 1 | 6 | Ccdc61 | 0.02 | Young |
| 0.092634 | 0.651288 | 0.104 | 0.071 | 1 | 6 | Nagk | 0.033 | Young |
| 0.093079 | 0.550341 | 0.191 | 0.151 | 1 | 6 | Tmx1 | 0.04 | Young |
| 0.093125 | 0.477348 | 0.261 | 0.22 | 1 | 6 | Smarcc1 | 0.041 | Young |
| 0.093297 | 1.975476 | 0.023 | 0.009 | 1 | 6 | Paqr4 | 0.014 | Young |
| 0.093519 | 0.122842 | 0.284 | 0.237 | 1 | 6 | Ep400 | 0.047 | Young |
| 0.0938 | 0.14861 | 0.485 | 0.429 | 1 | 6 | Jarid2 | 0.056 | Young |
| 0.09416 | 0.102832 | 0.488 | 0.426 | 1 | 6 | Arhgap9 | 0.062 | Young |
| 0.094199 | 0.168994 | 0.711 | 0.68 | 1 | 6 | Dbi | 0.031 | Young |
| 0.094387 | 0.314833 | 0.363 | 0.317 | 1 | 6 | Sptssa | 0.046 | Young |
| 0.09449 | 0.211467 | 0.457 | 0.411 | 1 | 6 | Ptp4a1 | 0.046 | Young |
| 0.094714 | 0.62254 | 0.08 | 0.051 | 1 | 6 | Nedd4 | 0.029 | Young |
| 0.094978 | 1.699019 | 0.023 | 0.009 | 1 | 6 | Slc19a2 | 0.014 | Young |
| 0.095351 | 0.32246 | 0.385 | 0.331 | 1 | 6 | Ifnar1 | 0.054 | Young |
| 0.095402 | 1.411074 | 0.023 | 0.009 | 1 | 6 | Gpatch3 | 0.014 | Young |
| 0.095482 | 0.200744 | 0.087 | 0.057 | 1 | 6 | Nkap | 0.03 | Young |
| 0.095625 | 0.662025 | 0.052 | 0.029 | 1 | 6 | Abca6 | 0.023 | Young |
| 0.095982 | 0.238525 | 0.176 | 0.134 | 1 | 6 | Chsy1 | 0.042 | Young |
| 0.09599 | 0.142192 | 0.314 | 0.26 | 1 | 6 | Rab5a | 0.054 | Young |
| 0.096155 | 0.482399 | 0.212 | 0.169 | 1 | 6 | Eif4e3 | 0.043 | Young |
| 0.096225 | 0.656372 | 0.117 | 0.083 | 1 | 6 | Rbm18 | 0.034 | Young |
| 0.096431 | 0.127221 | 0.657 | 0.591 | 1 | 6 | Mycbp2 | 0.066 | Young |
| 0.096495 | 0.773454 | 0.059 | 0.034 | 1 | 6 | Gbe1 | 0.025 | Young |
| 0.096499 | 1.021099 | 0.032 | 0.014 | 1 | 6 | Ehf | 0.018 | Young |
| 0.096683 | 1.106923 | 0.023 | 0.009 | 1 | 6 | Fbxo46 | 0.014 | Young |
| 0.096912 | 0.340392 | 0.329 | 0.286 | 1 | 6 | Crlf2 | 0.043 | Young |
| 0.097197 | 0.673494 | 0.104 | 0.071 | 1 | 6 | Mthfs | 0.033 | Young |
| 0.097531 | 0.181755 | 0.559 | 0.506 | 1 | 6 | Prkar2a | 0.053 | Young |
| 0.097836 | 0.222886 | 0.473 | 0.431 | 1 | 6 | Cct4 | 0.042 | Young |
| 0.097964 | 0.953487 | 0.065 | 0.04 | 1 | 6 | Neurl1b | 0.025 | Young |
| 0.09802 | 0.220244 | 0.386 | 0.343 | 1 | 6 | Bptf | 0.043 | Young |
| 0.098142 | 0.036419 | 0.197 | 0.154 | 1 | 6 | Xpo6 | 0.043 | Young |
| 0.098347 | 0.401198 | 0.169 | 0.131 | 1 | 6 | S100a1 | 0.038 | Young |
| 0.098362 | 0.634935 | 0.115 | 0.083 | 1 | 6 | Ranbp10 | 0.032 | Young |
| 0.098436 | 0.146878 | 0.147 | 0.109 | 1 | 6 | Ilf2 | 0.038 | Young |
| 0.098821 | 0.486712 | 0.283 | 0.24 | 1 | 6 | Tnfrsf1a | 0.043 | Young |
| 0.099112 | 1.191477 | 0.043 | 0.023 | 1 | 6 | Abcd2 | 0.02 | Young |
| 0.099112 | 0.808959 | 0.043 | 0.023 | 1 | 6 | Ltc4s | 0.02 | Young |
| 0.099147 | 0.243541 | 0.142 | 0.103 | 1 | 6 | Pofut1 | 0.039 | Young |
| 0.0992 | 0.208131 | 0.592 | 0.549 | 1 | 6 | Mrpl33 | 0.043 | Young |
| 0.099489 | 0.498517 | 0.202 | 0.163 | 1 | 6 | Padi2 | 0.039 | Young |
| 0.099507 | 0.451332 | 0.023 | 0.009 | 1 | 6 | Socs1 | 0.014 | Young |
| 0.099517 | 0.648682 | 0.109 | 0.077 | 1 | 6 | Zfp655 | 0.032 | Young |
| 0.099617 | 0.933108 | 0.032 | 0.014 | 1 | 6 | Hacd1 | 0.018 | Young |

| | | | | | | | | |
|---|---|---|---|---|---|---|---|---|
| 0.099642 | 0.807785 | 0.092 | 0.063 | 1 | 6 | Alpk1 | 0.029 | Young |
| 0.099709 | 0.556991 | 0.072 | 0.046 | 1 | 6 | Cnn3 | 0.026 | Young |
| 0.099913 | 0.316143 | 0.426 | 0.391 | 1 | 6 | Arid4a | 0.035 | Young |
| 0.100007 | 0.471355 | 0.095 | 0.066 | 1 | 6 | Atg7 | 0.029 | Young |
| 0.100345 | 0.310418 | 0.189 | 0.149 | 1 | 6 | Sptlc2 | 0.04 | Young |
| 0.100375 | 0.800103 | 0.072 | 0.046 | 1 | 6 | Tsc2 | 0.026 | Young |
| 0.100468 | 0.390355 | 0.264 | 0.22 | 1 | 6 | Hcfc1r1 | 0.044 | Young |
| 0.10056 | 0.162676 | 0.779 | 0.746 | 1 | 6 | Rps27l | 0.033 | Young |
| 0.100737 | 0.500426 | 0.032 | 0.014 | 1 | 6 | Fbxo21 | 0.018 | Young |
| 0.100808 | 0.63067 | 0.14 | 0.106 | 1 | 6 | Hmgcl | 0.034 | Young |
| 0.100861 | 0.174739 | 0.734 | 0.683 | 1 | 6 | Tln1 | 0.051 | Young |
| 0.100918 | 0.751264 | 0.119 | 0.086 | 1 | 6 | D930030I0 | 0.033 | Young |
| 0.10122 | 0.03997 | 0.473 | 0.414 | 1 | 6 | Ltb4r1 | 0.059 | Young |
| 0.101319 | 0.631282 | 0.043 | 0.023 | 1 | 6 | Knstrn | 0.02 | Young |
| 0.101363 | 0.097951 | 0.878 | 0.809 | 1 | 6 | Naca | 0.069 | Young |
| 0.101382 | 0.397065 | 0.192 | 0.151 | 1 | 6 | Vamp4 | 0.041 | Young |
| 0.101638 | 0.135877 | 0.467 | 0.411 | 1 | 6 | Mef2c | 0.056 | Young |
| 0.102395 | 0.766778 | 0.072 | 0.046 | 1 | 6 | Siglece | 0.026 | Young |
| 0.102506 | 0.133917 | 0.86 | 0.849 | 1 | 6 | Itm2b | 0.011 | Young |
| 0.102677 | 1.865063 | 0.018 | 0.006 | 1 | 6 | 4933433G1 | 0.012 | Young |
| 0.102682 | 0.116797 | 0.891 | 0.863 | 1 | 6 | Serf2 | 0.028 | Young |
| 0.102758 | 0.597783 | 0.043 | 0.023 | 1 | 6 | Cdc23 | 0.02 | Young |
| 0.103047 | 0.034794 | 0.187 | 0.143 | 1 | 6 | Rtn1 | 0.044 | Young |
| 0.103123 | 0.238862 | 0.386 | 0.346 | 1 | 6 | Smim14 | 0.04 | Young |
| 0.10371 | 1.790295 | 0.018 | 0.006 | 1 | 6 | Vipr1 | 0.012 | Young |
| 0.103942 | 0.325513 | 0.368 | 0.323 | 1 | 6 | Tubb6 | 0.045 | Young |
| 0.103969 | 1.455857 | 0.018 | 0.006 | 1 | 6 | Atxn7l2 | 0.012 | Young |
| 0.104238 | 0.329424 | 0.246 | 0.197 | 1 | 6 | Msrb1 | 0.049 | Young |
| 0.104521 | 0.775869 | 0.065 | 0.04 | 1 | 6 | Mitd1 | 0.025 | Young |
| 0.10461 | 0.170846 | 0.147 | 0.109 | 1 | 6 | Ctnnd1 | 0.038 | Young |
| 0.10493 | 0.629022 | 0.082 | 0.054 | 1 | 6 | Adcy9 | 0.028 | Young |
| 0.105273 | 2.774591 | 0.018 | 0.006 | 1 | 6 | Rsad2 | 0.012 | Young |
| 0.105533 | 0.850744 | 0.047 | 0.026 | 1 | 6 | Pkd2 | 0.021 | Young |
| 0.105953 | 0.184753 | 0.147 | 0.109 | 1 | 6 | Zwint | 0.038 | Young |
| 0.105966 | 0.072008 | 0.935 | 0.849 | 1 | 6 | Zeb2 | 0.086 | Young |
| 0.106027 | 1.822488 | 0.013 | 0.003 | 1 | 6 | Gm17435 | 0.01 | Young |
| 0.106027 | 1.686386 | 0.013 | 0.003 | 1 | 6 | Hs3st3b1 | 0.01 | Young |
| 0.10608 | 0.212265 | 0.339 | 0.283 | 1 | 6 | Slc35b2 | 0.056 | Young |
| 0.106318 | 0.257699 | 0.559 | 0.517 | 1 | 6 | H2afj | 0.042 | Young |
| 0.106343 | 1.992963 | 0.013 | 0.003 | 1 | 6 | Siglech | 0.01 | Young |
| 0.106343 | 1.826196 | 0.013 | 0.003 | 1 | 6 | Marveld2 | 0.01 | Young |
| 0.106343 | 1.587724 | 0.013 | 0.003 | 1 | 6 | Thumpd2 | 0.01 | Young |
| 0.106343 | 1.527509 | 0.013 | 0.003 | 1 | 6 | Samd4 | 0.01 | Young |
| 0.106343 | 1.467135 | 0.013 | 0.003 | 1 | 6 | 3300005D0 | 0.01 | Young |
| 0.106343 | 1.365296 | 0.013 | 0.003 | 1 | 6 | Dcbld2 | 0.01 | Young |

| | | | | | | | | |
|---|---|---|---|---|---|---|---|---|
| 0.106591 | 0.906868 | 0.018 | 0.006 | 1 | 6 | Rnf219 | 0.012 | Young |
| 0.106591 | 0.851478 | 0.018 | 0.006 | 1 | 6 | Odf2l | 0.012 | Young |
| 0.10662 | 0.549136 | 0.122 | 0.089 | 1 | 6 | Orai3 | 0.033 | Young |
| 0.106661 | 1.865605 | 0.013 | 0.003 | 1 | 6 | Gm44777 | 0.01 | Young |
| 0.106979 | 1.851935 | 0.013 | 0.003 | 1 | 6 | Focad | 0.01 | Young |
| 0.107194 | 0.262307 | 0.281 | 0.237 | 1 | 6 | Pvr | 0.044 | Young |
| 0.10739 | 1.089553 | 0.035 | 0.017 | 1 | 6 | Slco2b1 | 0.018 | Young |
| 0.107938 | 0.999966 | 0.013 | 0.003 | 1 | 6 | Otud3 | 0.01 | Young |
| 0.108049 | 0.129792 | 0.818 | 0.757 | 1 | 6 | Etv3 | 0.061 | Young |
| 0.108504 | 0.398517 | 0.179 | 0.14 | 1 | 6 | Tmem184b | 0.039 | Young |
| 0.108513 | 0.394774 | 0.398 | 0.354 | 1 | 6 | Ttc7 | 0.044 | Young |
| 0.108564 | 0.589231 | 0.047 | 0.026 | 1 | 6 | Kit | 0.021 | Young |
| 0.10863 | 0.908895 | 0.05 | 0.029 | 1 | 6 | Ndufaf5 | 0.021 | Young |
| 0.108703 | 1.58619 | 0.027 | 0.011 | 1 | 6 | Tmem69 | 0.016 | Young |
| 0.108715 | 0.215148 | 0.222 | 0.183 | 1 | 6 | Btbd1 | 0.039 | Young |
| 0.108887 | 0.307146 | 0.14 | 0.106 | 1 | 6 | Usp37 | 0.034 | Young |
| 0.108888 | 0.901427 | 0.035 | 0.017 | 1 | 6 | Klc4 | 0.018 | Young |
| 0.10923 | 0.157819 | 0.237 | 0.197 | 1 | 6 | Smad4 | 0.04 | Young |
| 0.109454 | 1.041872 | 0.035 | 0.017 | 1 | 6 | Fuom | 0.018 | Young |
| 0.109576 | 1.139078 | 0.067 | 0.043 | 1 | 6 | Ipo9 | 0.024 | Young |
| 0.109617 | 0.210593 | 0.48 | 0.431 | 1 | 6 | Eif4h | 0.049 | Young |
| 0.109643 | 0.241018 | 0.022 | 0.04 | 1 | 6 | Ydjc | -0.018 | Young |
| 0.109742 | 0.58904 | 0.124 | 0.091 | 1 | 6 | Erf | 0.033 | Young |
| 0.110028 | 0.253869 | 0.405 | 0.354 | 1 | 6 | Taf10 | 0.051 | Young |
| 0.110383 | 0.585552 | 0.109 | 0.077 | 1 | 6 | Acadsb | 0.032 | Young |
| 0.110871 | 0.168488 | 0.383 | 0.326 | 1 | 6 | Itpripl2 | 0.057 | Young |
| 0.110887 | 0.502607 | 0.286 | 0.249 | 1 | 6 | Psme4 | 0.037 | Young |
| 0.110891 | 0.885995 | 0.054 | 0.031 | 1 | 6 | Ccl12 | 0.023 | Young |
| 0.110896 | 1.333557 | 0.027 | 0.011 | 1 | 6 | Acat2 | 0.016 | Young |
| 0.111249 | 0.420291 | 0.217 | 0.18 | 1 | 6 | Ubr4 | 0.037 | Young |
| 0.111357 | 0.524383 | 0.035 | 0.017 | 1 | 6 | Nepro | 0.018 | Young |
| 0.111498 | 0.284308 | 0.047 | 0.026 | 1 | 6 | Cul4b | 0.021 | Young |
| 0.111598 | 1.295558 | 0.038 | 0.02 | 1 | 6 | Xrcc4 | 0.018 | Young |
| 0.111599 | 0.469301 | 0.075 | 0.049 | 1 | 6 | Cdc42bpb | 0.026 | Young |
| 0.112429 | 0.225732 | 0.485 | 0.429 | 1 | 6 | Eif3h | 0.056 | Young |
| 0.112438 | 0.22077 | 0.177 | 0.137 | 1 | 6 | Filip1l | 0.04 | Young |
| 0.112453 | 1.1868 | 0.027 | 0.011 | 1 | 6 | Slc2a9 | 0.016 | Young |
| 0.112585 | 0.982408 | 0.087 | 0.06 | 1 | 6 | Slc36a1 | 0.027 | Young |
| 0.112697 | 1.025271 | 0.038 | 0.02 | 1 | 6 | Rhoc | 0.018 | Young |
| 0.113327 | 0.181268 | 0.44 | 0.391 | 1 | 6 | Rab2a | 0.049 | Young |
| 0.113393 | 0.544405 | 0.095 | 0.066 | 1 | 6 | Akap10 | 0.029 | Young |
| 0.114003 | 0.127639 | 0.301 | 0.254 | 1 | 6 | Cic | 0.047 | Young |
| 0.114114 | 0.796112 | 0.129 | 0.097 | 1 | 6 | Vopp1 | 0.032 | Young |
| 0.114361 | 0.967243 | 0.038 | 0.02 | 1 | 6 | Zfp68 | 0.018 | Young |
| 0.114406 | 0.084265 | 0.425 | 0.366 | 1 | 6 | Hnrnpd | 0.059 | Young |

| | | | | | | | | |
|---|---|---|---|---|---|---|---|---|
| 0.114758 | 0.129521 | 0.747 | 0.651 | 1 | 6 | Lgals3 | 0.096 | Young |
| 0.114918 | 0.033359 | 0.692 | 0.626 | 1 | 6 | Slk | 0.066 | Young |
| 0.115111 | 0.107 | 0.95 | 0.911 | 1 | 6 | Mbnl1 | 0.039 | Young |
| 0.115198 | 0.58723 | 0.164 | 0.129 | 1 | 6 | Bet1 | 0.035 | Young |
| 0.115214 | 0.380637 | 0.145 | 0.109 | 1 | 6 | Cln6 | 0.036 | Young |
| 0.115257 | 0.386415 | 0.236 | 0.2 | 1 | 6 | Ufm1 | 0.036 | Young |
| 0.115451 | 0.572299 | 0.082 | 0.054 | 1 | 6 | Thap3 | 0.028 | Young |
| 0.115966 | 0.273391 | 0.216 | 0.177 | 1 | 6 | Tpp1 | 0.039 | Young |
| 0.116068 | 0.107061 | 0.747 | 0.697 | 1 | 6 | Atp1a1 | 0.05 | Young |
| 0.116171 | 0.143804 | 0.423 | 0.374 | 1 | 6 | Akr1b3 | 0.049 | Young |
| 0.116173 | 0.22626 | 0.443 | 0.394 | 1 | 6 | Rock2 | 0.049 | Young |
| 0.116483 | 0.18727 | 0.396 | 0.349 | 1 | 6 | Ccnd3 | 0.047 | Young |
| 0.11674 | 0.087311 | 0.836 | 0.751 | 1 | 6 | Ckb | 0.085 | Young |
| 0.116798 | 0.228623 | 0.381 | 0.331 | 1 | 6 | Eloc | 0.05 | Young |
| 0.116803 | 0.096179 | 0.863 | 0.817 | 1 | 6 | Btf3 | 0.046 | Young |
| 0.117317 | 0.740353 | 0.06 | 0.037 | 1 | 6 | Lrrc20 | 0.023 | Young |
| 0.117378 | 0.513114 | 0.217 | 0.18 | 1 | 6 | Npepl1 | 0.037 | Young |
| 0.117919 | 0.358895 | 0.027 | 0.011 | 1 | 6 | Gm42722 | 0.016 | Young |
| 0.118029 | 0.362938 | 0.416 | 0.371 | 1 | 6 | Ubn2 | 0.045 | Young |
| 0.118425 | 0.161902 | 0.43 | 0.377 | 1 | 6 | Tgif1 | 0.053 | Young |
| 0.118527 | 0.367069 | 0.057 | 0.034 | 1 | 6 | Arfgap1 | 0.023 | Young |
| 0.118743 | 1.174287 | 0.042 | 0.023 | 1 | 6 | Plxdc1 | 0.019 | Young |
| 0.118818 | 0.182887 | 0.202 | 0.16 | 1 | 6 | Itga5 | 0.042 | Young |
| 0.118963 | 0.340366 | 0.206 | 0.166 | 1 | 6 | Tspyl1 | 0.04 | Young |
| 0.119029 | 0.199822 | 0.38 | 0.331 | 1 | 6 | Fxr1 | 0.049 | Young |
| 0.119084 | 0.673851 | 0.038 | 0.02 | 1 | 6 | Prdm10 | 0.018 | Young |
| 0.1192 | 0.501044 | 0.095 | 0.066 | 1 | 6 | Kdm4c | 0.029 | Young |
| 0.119663 | 0.479249 | 0.117 | 0.086 | 1 | 6 | Igsf8 | 0.031 | Young |
| 0.119678 | 0.19654 | 0.176 | 0.137 | 1 | 6 | Bicd2 | 0.039 | Young |
| 0.119749 | 0.402408 | 0.145 | 0.111 | 1 | 6 | Slc30a4 | 0.034 | Young |
| 0.119934 | 0.261224 | 0.151 | 0.117 | 1 | 6 | Dlgap4 | 0.034 | Young |
| 0.120158 | 0.049217 | 0.925 | 0.797 | 1 | 6 | Plbd1 | 0.128 | Young |
| 0.120168 | 0.143789 | 0.232 | 0.189 | 1 | 6 | Ttc39b | 0.043 | Young |
| 0.120186 | 0.725607 | 0.067 | 0.043 | 1 | 6 | Spryd7 | 0.024 | Young |
| 0.120455 | 0.461876 | 0.05 | 0.029 | 1 | 6 | Mfge8 | 0.021 | Young |
| 0.121299 | 0.022541 | 0.676 | 0.64 | 1 | 6 | Itga4 | 0.036 | Young |
| 0.121612 | 0.486431 | 0.177 | 0.143 | 1 | 6 | Lyar | 0.034 | Young |
| 0.121629 | 1.51577 | 0.03 | 0.014 | 1 | 6 | Tada2b | 0.016 | Young |
| 0.121929 | 0.103491 | 0.472 | 0.411 | 1 | 6 | Syncrip | 0.061 | Young |
| 0.122074 | 1.453044 | 0.03 | 0.014 | 1 | 6 | Adi1 | 0.016 | Young |
| 0.122077 | 0.171025 | 0.186 | 0.146 | 1 | 6 | Zfp652 | 0.04 | Young |
| 0.122253 | 0.140056 | 0.682 | 0.646 | 1 | 6 | Ctsb | 0.036 | Young |
| 0.122632 | 0.932882 | 0.042 | 0.023 | 1 | 6 | Mex3b | 0.019 | Young |
| 0.12282 | 0.855341 | 0.042 | 0.023 | 1 | 6 | Slc17a5 | 0.019 | Young |
| 0.123393 | 0.247624 | 0.413 | 0.366 | 1 | 6 | Higd2a | 0.047 | Young |

| | | | | | | | | |
|---|---|---|---|---|---|---|---|---|
| 0.123995 | 0.642677 | 0.067 | 0.043 | 1 | 6 | Rfx3 | 0.024 | Young |
| 0.124769 | 1.190341 | 0.03 | 0.014 | 1 | 6 | Wdr76 | 0.016 | Young |
| 0.124893 | 2.290073 | 0.022 | 0.009 | 1 | 6 | 0610040B1 | 0.013 | Young |
| 0.124944 | 0.282296 | 0.388 | 0.343 | 1 | 6 | Chd7 | 0.045 | Young |
| 0.125352 | 0.082611 | 0.858 | 0.743 | 1 | 6 | Ifitm3 | 0.115 | Young |
| 0.125355 | 0.1693 | 0.336 | 0.289 | 1 | 6 | Psmb6 | 0.047 | Young |
| 0.12545 | 0.592614 | 0.03 | 0.014 | 1 | 6 | Zfp456 | 0.016 | Young |
| 0.125557 | 0.140917 | 0.53 | 0.483 | 1 | 6 | Cast | 0.047 | Young |
| 0.125731 | 0.406024 | 0.144 | 0.109 | 1 | 6 | Ddx52 | 0.035 | Young |
| 0.12598 | 1.684884 | 0.022 | 0.009 | 1 | 6 | Adck5 | 0.013 | Young |
| 0.126233 | 0.666746 | 0.09 | 0.063 | 1 | 6 | Il1rl2 | 0.027 | Young |
| 0.12631 | 0.2667 | 0.47 | 0.42 | 1 | 6 | Pea15a | 0.05 | Young |
| 0.126438 | 0.451568 | 0.08 | 0.054 | 1 | 6 | Fam3c | 0.026 | Young |
| 0.126792 | 0.243091 | 0.179 | 0.143 | 1 | 6 | Mecp2 | 0.036 | Young |
| 0.1268 | 1.403852 | 0.022 | 0.009 | 1 | 6 | 4930481A1 | 0.013 | Young |
| 0.127005 | 0.338906 | 0.246 | 0.206 | 1 | 6 | Mctp1 | 0.04 | Young |
| 0.127025 | 0.100796 | 0.244 | 0.2 | 1 | 6 | Klrd1 | 0.044 | Young |
| 0.127067 | 0.437138 | 0.06 | 0.037 | 1 | 6 | Sestd1 | 0.023 | Young |
| 0.127227 | 0.750084 | 0.06 | 0.037 | 1 | 6 | Acvr1 | 0.023 | Young |
| 0.127329 | 0.581224 | 0.129 | 0.097 | 1 | 6 | Fgd4 | 0.032 | Young |
| 0.127378 | 0.215682 | 0.413 | 0.369 | 1 | 6 | Asah1 | 0.044 | Young |
| 0.127409 | 0.310667 | 0.271 | 0.229 | 1 | 6 | Vcl | 0.042 | Young |
| 0.127624 | 1.415128 | 0.022 | 0.009 | 1 | 6 | Pls3 | 0.013 | Young |
| 0.127899 | 1.267366 | 0.022 | 0.009 | 1 | 6 | Ppp2r3a | 0.013 | Young |
| 0.127899 | 1.178908 | 0.022 | 0.009 | 1 | 6 | Mtbp | 0.013 | Young |
| 0.12809 | 0.136546 | 0.338 | 0.286 | 1 | 6 | Tfe3 | 0.052 | Young |
| 0.1281 | 0.530152 | 0.104 | 0.074 | 1 | 6 | Smurf1 | 0.03 | Young |
| 0.128104 | 0.199174 | 0.535 | 0.503 | 1 | 6 | Cox7a2 | 0.032 | Young |
| 0.128613 | 0.444527 | 0.231 | 0.191 | 1 | 6 | Elp5 | 0.04 | Young |
| 0.128744 | 0.121829 | 0.619 | 0.557 | 1 | 6 | Tnfaip8 | 0.062 | Young |
| 0.129144 | 0.003363 | 0.505 | 0.449 | 1 | 6 | Scp2 | 0.056 | Young |
| 0.129284 | 1.092372 | 0.022 | 0.009 | 1 | 6 | Map3k9 | 0.013 | Young |
| 0.129402 | 0.558507 | 0.045 | 0.026 | 1 | 6 | 3110002H1 | 0.019 | Young |
| 0.129892 | 0.409362 | 0.135 | 0.103 | 1 | 6 | Atl3 | 0.032 | Young |
| 0.129924 | 0.209113 | 0.182 | 0.143 | 1 | 6 | Elf2 | 0.039 | Young |
| 0.129937 | 0.195979 | 0.463 | 0.429 | 1 | 6 | Cct8 | 0.034 | Young |
| 0.130091 | 0.222768 | 0.204 | 0.166 | 1 | 6 | Btbd7 | 0.038 | Young |
| 0.130121 | 1.153776 | 0.022 | 0.009 | 1 | 6 | Pfas | 0.013 | Young |
| 0.130262 | 0.100937 | 0.731 | 0.686 | 1 | 6 | Arpc3 | 0.045 | Young |
| 0.130267 | 0.0063 | 0.58 | 0.514 | 1 | 6 | Ifngr1 | 0.066 | Young |
| 0.130388 | 0.210613 | 0.433 | 0.391 | 1 | 6 | Zbtb7a | 0.042 | Young |
| 0.130946 | 1.441857 | 0.033 | 0.017 | 1 | 6 | Gpr84 | 0.016 | Young |
| 0.131057 | 0.238637 | 0.493 | 0.469 | 1 | 6 | Ndufb10 | 0.024 | Young |
| 0.131131 | 0.241454 | 0.216 | 0.177 | 1 | 6 | Lrrk1 | 0.039 | Young |
| 0.131676 | 0.061272 | 0.545 | 0.491 | 1 | 6 | Ahcyl2 | 0.054 | Young |

| | | | | | | | | |
|---|---|---|---|---|---|---|---|---|
| 0.132089 | 0.372857 | 0.064 | 0.04 | 1 | 6 | Itsn1 | 0.024 | Young |
| 0.132116 | 0.218187 | 0.003 | 0.011 | 1 | 6 | Spon1 | -0.008 | Young |
| 0.132116 | 0.163372 | 0.003 | 0.011 | 1 | 6 | Cygb | -0.008 | Young |
| 0.132241 | 0.136371 | 0.828 | 0.766 | 1 | 6 | Picalm | 0.062 | Young |
| 0.132255 | 0.270397 | 0.06 | 0.037 | 1 | 6 | Aacs | 0.023 | Young |
| 0.132622 | 0.580635 | 0.045 | 0.026 | 1 | 6 | Cluap1 | 0.019 | Young |
| 0.132968 | 0.359371 | 0.303 | 0.263 | 1 | 6 | Ppp1r10 | 0.04 | Young |
| 0.133149 | 0.280439 | 0.273 | 0.231 | 1 | 6 | Fam104a | 0.042 | Young |
| 0.133415 | 0.311315 | 0.169 | 0.134 | 1 | 6 | Trrap | 0.035 | Young |
| 0.13445 | 0.495473 | 0.112 | 0.083 | 1 | 6 | Nmt2 | 0.029 | Young |
| 0.134539 | 1.209473 | 0.059 | 0.037 | 1 | 6 | Gas2l3 | 0.022 | Young |
| 0.134619 | 0.347555 | 0.1 | 0.071 | 1 | 6 | Ciapin1 | 0.029 | Young |
| 0.134697 | 0.505891 | 0.236 | 0.2 | 1 | 6 | Basp1 | 0.036 | Young |
| 0.134716 | 0.121206 | 0.263 | 0.214 | 1 | 6 | Rbm27 | 0.049 | Young |
| 0.134754 | 1.01833 | 0.033 | 0.017 | 1 | 6 | F3 | 0.016 | Young |
| 0.134783 | 0.177999 | 0.338 | 0.303 | 1 | 6 | Mpc1 | 0.035 | Young |
| 0.134964 | 0.138381 | 0.074 | 0.049 | 1 | 6 | Idh2 | 0.025 | Young |
| 0.135075 | 0.31456 | 0.11 | 0.08 | 1 | 6 | Slc36a4 | 0.03 | Young |
| 0.135138 | 0.166913 | 0.358 | 0.311 | 1 | 6 | Myo9b | 0.047 | Young |
| 0.135301 | 0.149624 | 0.276 | 0.229 | 1 | 6 | Cd86 | 0.047 | Young |
| 0.135723 | 0.78912 | 0.059 | 0.037 | 1 | 6 | Wbp1 | 0.022 | Young |
| 0.135891 | 0.966665 | 0.033 | 0.017 | 1 | 6 | Alms1 | 0.016 | Young |
| 0.135916 | 0.490087 | 0.145 | 0.114 | 1 | 6 | Mms19 | 0.031 | Young |
| 0.136122 | 0.66815 | 0.109 | 0.08 | 1 | 6 | Gm46224 | 0.029 | Young |
| 0.136254 | 0.059495 | 0.187 | 0.149 | 1 | 6 | Rab5b | 0.038 | Young |
| 0.136616 | 0.105035 | 0.502 | 0.454 | 1 | 6 | Atp5c1 | 0.048 | Young |
| 0.136628 | 0.464401 | 0.08 | 0.054 | 1 | 6 | A530064D( | 0.026 | Young |
| 0.136879 | 0.604785 | 0.045 | 0.026 | 1 | 6 | E130307A1 | 0.019 | Young |
| 0.137167 | 0.108916 | 0.052 | 0.077 | 1 | 6 | Slc11a2 | -0.025 | Young |
| 0.137373 | 0.527949 | 0.164 | 0.129 | 1 | 6 | Ppm1h | 0.035 | Young |
| 0.137701 | 0.427512 | 0.074 | 0.049 | 1 | 6 | Gm34084 | 0.025 | Young |
| 0.138013 | 0.307457 | 0.356 | 0.314 | 1 | 6 | D17Wsu92 | 0.042 | Young |
| 0.138186 | 0.95442 | 0.033 | 0.017 | 1 | 6 | Zfp568 | 0.016 | Young |
| 0.138274 | 0.743765 | 0.052 | 0.031 | 1 | 6 | Acot13 | 0.021 | Young |
| 0.138291 | 0.161784 | 0.179 | 0.14 | 1 | 6 | Hint2 | 0.039 | Young |
| 0.138417 | 1.025625 | 0.033 | 0.017 | 1 | 6 | Hdac6 | 0.016 | Young |
| 0.138648 | 0.69473 | 0.033 | 0.017 | 1 | 6 | Nsun3 | 0.016 | Young |
| 0.13981 | 0.967752 | 0.033 | 0.017 | 1 | 6 | Bmp2 | 0.016 | Young |
| 0.139883 | 0.771981 | 0.145 | 0.114 | 1 | 6 | Ezh2 | 0.031 | Young |
| 0.140276 | 0.44201 | 0.033 | 0.017 | 1 | 6 | Gpr34 | 0.016 | Young |
| 0.14073 | 1.760526 | 0.017 | 0.006 | 1 | 6 | Nipa1 | 0.011 | Young |
| 0.140757 | 0.119923 | 0.458 | 0.411 | 1 | 6 | Tmem59 | 0.047 | Young |
| 0.141073 | 1.620489 | 0.017 | 0.006 | 1 | 6 | Naif1 | 0.011 | Young |
| 0.141088 | 0.951268 | 0.072 | 0.049 | 1 | 6 | Sbk1 | 0.023 | Young |
| 0.1411 | 1.731014 | 0.025 | 0.011 | 1 | 6 | Wdr53 | 0.014 | Young |

| | | | | | | | | |
|---|---|---|---|---|---|---|---|---|
| 0.14113 | 0.276815 | 0.385 | 0.343 | 1 | 6 | Hsd17b11 | 0.042 | Young |
| 0.14127 | 0.244715 | 0.226 | 0.186 | 1 | 6 | Tmem88 | 0.04 | Young |
| 0.141619 | 0.529381 | 0.099 | 0.071 | 1 | 6 | Eya3 | 0.028 | Young |
| 0.141868 | 0.88209 | 0.062 | 0.04 | 1 | 6 | Kank3 | 0.022 | Young |
| 0.141958 | 0.07296 | 0.935 | 0.869 | 1 | 6 | Gm2a | 0.066 | Young |
| 0.142155 | 0.984405 | 0.033 | 0.017 | 1 | 6 | Trem2 | 0.016 | Young |
| 0.142155 | 0.486379 | 0.033 | 0.017 | 1 | 6 | Zfp609 | 0.016 | Young |
| 0.142196 | 0.612646 | 0.154 | 0.12 | 1 | 6 | Gm20186 | 0.034 | Young |
| 0.142196 | 0.250786 | 0.114 | 0.083 | 1 | 6 | Gcnt2 | 0.031 | Young |
| 0.142206 | 0.422432 | 0.132 | 0.1 | 1 | 6 | AY036118 | 0.032 | Young |
| 0.14252 | 0.102453 | 0.221 | 0.177 | 1 | 6 | Golga7 | 0.044 | Young |
| 0.14258 | 0.171437 | 0.37 | 0.326 | 1 | 6 | Ankrd17 | 0.044 | Young |
| 0.142703 | 0.111629 | 0.059 | 0.086 | 1 | 6 | Mrpl19 | -0.027 | Young |
| 0.142795 | 1.370711 | 0.017 | 0.006 | 1 | 6 | Nphp3 | 0.011 | Young |
| 0.142891 | 0.075494 | 0.587 | 0.543 | 1 | 6 | Ppp1r15a | 0.044 | Young |
| 0.143055 | 0.101166 | 0.318 | 0.266 | 1 | 6 | Pdcd10 | 0.052 | Young |
| 0.143142 | 1.110663 | 0.017 | 0.006 | 1 | 6 | Agmo | 0.011 | Young |
| 0.143145 | 1.19198 | 0.072 | 0.049 | 1 | 6 | Ccl24 | 0.023 | Young |
| 0.143251 | 0.131399 | 0.564 | 0.509 | 1 | 6 | Itpr1 | 0.055 | Young |
| 0.143355 | 0.674977 | 0.082 | 0.057 | 1 | 6 | AI606181 | 0.025 | Young |
| 0.14372 | 0.217812 | 0.037 | 0.02 | 1 | 6 | Kynu | 0.017 | Young |
| 0.143836 | 0.206466 | 0.403 | 0.363 | 1 | 6 | Srrm1 | 0.04 | Young |
| 0.143836 | 1.079523 | 0.017 | 0.006 | 1 | 6 | Nmnat1 | 0.011 | Young |
| 0.143836 | 1.062914 | 0.017 | 0.006 | 1 | 6 | Cdiptos | 0.011 | Young |
| 0.143859 | 0.208827 | 0.296 | 0.254 | 1 | 6 | Chd2 | 0.042 | Young |
| 0.143932 | 0.29194 | 0.065 | 0.043 | 1 | 6 | Isoc1 | 0.022 | Young |
| 0.144185 | 1.572474 | 0.017 | 0.006 | 1 | 6 | Adm | 0.011 | Young |
| 0.144192 | 0.686208 | 0.095 | 0.069 | 1 | 6 | Slc9a8 | 0.026 | Young |
| 0.144332 | 0.958228 | 0.107 | 0.08 | 1 | 6 | Phka2 | 0.027 | Young |
| 0.144376 | 0.214261 | 0.595 | 0.566 | 1 | 6 | Ubl5 | 0.029 | Young |
| 0.144441 | 0.524526 | 0.122 | 0.091 | 1 | 6 | Cds1 | 0.031 | Young |
| 0.144564 | 0.291455 | 0.381 | 0.343 | 1 | 6 | Tbc1d9 | 0.038 | Young |
| 0.144883 | 0.84214 | 0.017 | 0.006 | 1 | 6 | Cfap45 | 0.011 | Young |
| 0.145585 | 0.146009 | 0.017 | 0.006 | 1 | 6 | Bcam | 0.011 | Young |
| 0.145585 | 0.098811 | 0.017 | 0.006 | 1 | 6 | Gsg1 | 0.011 | Young |
| 0.146032 | 0.670622 | 0.075 | 0.051 | 1 | 6 | Gm9530 | 0.024 | Young |
| 0.14622 | 0.834824 | 0.037 | 0.02 | 1 | 6 | Zfp467 | 0.017 | Young |
| 0.146355 | 0.663146 | 0.062 | 0.04 | 1 | 6 | Mzt2 | 0.022 | Young |
| 0.147054 | 0.276557 | 0.102 | 0.074 | 1 | 6 | Akap8l | 0.028 | Young |
| 0.147097 | 1.514977 | 0.04 | 0.023 | 1 | 6 | Smpdl3b | 0.017 | Young |
| 0.147103 | 0.323973 | 0.216 | 0.177 | 1 | 6 | Sertad1 | 0.039 | Young |
| 0.147248 | 0.651897 | 0.114 | 0.086 | 1 | 6 | Ssbp1 | 0.028 | Young |
| 0.147768 | 0.552069 | 0.114 | 0.086 | 1 | 6 | Ssbp2 | 0.028 | Young |
| 0.148211 | 0.098027 | 0.635 | 0.597 | 1 | 6 | Plxnd1 | 0.038 | Young |
| 0.148304 | 0.693456 | 0.125 | 0.097 | 1 | 6 | Rsrc1 | 0.028 | Young |

| | | | | | | | | |
|---|---|---|---|---|---|---|---|---|
| 0.149301 | 1.096295 | 0.04 | 0.023 | 1 | 6 | Gys1 | 0.017 | Young |
| 0.149408 | 0.054979 | 0.326 | 0.277 | 1 | 6 | Mrpl24 | 0.049 | Young |
| 0.14953 | 1.686561 | 0.012 | 0.003 | 1 | 6 | 4931440F1 | 0.009 | Young |
| 0.14953 | 1.650769 | 0.012 | 0.003 | 1 | 6 | Hgf | 0.009 | Young |
| 0.149799 | 0.974582 | 0.115 | 0.089 | 1 | 6 | Wdr33 | 0.026 | Young |
| 0.149834 | 0.856293 | 0.079 | 0.054 | 1 | 6 | Cln5 | 0.025 | Young |
| 0.149969 | 1.458316 | 0.012 | 0.003 | 1 | 6 | Gm15050 | 0.009 | Young |
| 0.149969 | 1.305708 | 0.012 | 0.003 | 1 | 6 | Ccl27a | 0.009 | Young |
| 0.150008 | 0.101365 | 0.766 | 0.706 | 1 | 6 | Atp5g2 | 0.06 | Young |
| 0.150162 | 0.126448 | 0.644 | 0.603 | 1 | 6 | Clta | 0.041 | Young |
| 0.150409 | 1.930144 | 0.012 | 0.003 | 1 | 6 | 9130011E1 | 0.009 | Young |
| 0.150409 | 1.39295 | 0.012 | 0.003 | 1 | 6 | Teddm3 | 0.009 | Young |
| 0.150635 | 0.800872 | 0.04 | 0.023 | 1 | 6 | Suox | 0.017 | Young |
| 0.150849 | 1.007961 | 0.012 | 0.003 | 1 | 6 | Ctu1 | 0.009 | Young |
| 0.150851 | 0.545613 | 0.037 | 0.02 | 1 | 6 | Pdk1 | 0.017 | Young |
| 0.151224 | 0.291013 | 0.515 | 0.471 | 1 | 6 | Avpi1 | 0.044 | Young |
| 0.151291 | 1.336538 | 0.012 | 0.003 | 1 | 6 | Flt1 | 0.009 | Young |
| 0.151291 | 1.064223 | 0.012 | 0.003 | 1 | 6 | Usp2 | 0.009 | Young |
| 0.151662 | 0.069944 | 0.704 | 0.671 | 1 | 6 | Tmem50a | 0.033 | Young |
| 0.151733 | 0.987747 | 0.012 | 0.003 | 1 | 6 | Phf11c | 0.009 | Young |
| 0.151733 | 0.928602 | 0.012 | 0.003 | 1 | 6 | Poli | 0.009 | Young |
| 0.151733 | 0.877191 | 0.012 | 0.003 | 1 | 6 | AC162302. | 0.009 | Young |
| 0.151733 | 0.860533 | 0.012 | 0.003 | 1 | 6 | Pecr | 0.009 | Young |
| 0.151733 | 0.844459 | 0.012 | 0.003 | 1 | 6 | Serpinb6b | 0.009 | Young |
| 0.151754 | 1.011313 | 0.04 | 0.023 | 1 | 6 | Peli2 | 0.017 | Young |
| 0.152163 | 0.386037 | 0.151 | 0.117 | 1 | 6 | Plk3 | 0.034 | Young |
| 0.152177 | 0.833351 | 0.012 | 0.003 | 1 | 6 | Pcbp4 | 0.009 | Young |
| 0.152177 | 0.77244 | 0.012 | 0.003 | 1 | 6 | Gm26826 | 0.009 | Young |
| 0.152177 | 0.549777 | 0.012 | 0.003 | 1 | 6 | Nptxr | 0.009 | Young |
| 0.15237 | 1.123144 | 0.082 | 0.057 | 1 | 6 | Pld3 | 0.025 | Young |
| 0.15266 | 0.037135 | 0.565 | 0.517 | 1 | 6 | Rab14 | 0.048 | Young |
| 0.153104 | 0.592617 | 0.072 | 0.049 | 1 | 6 | Pou5f2 | 0.023 | Young |
| 0.153123 | 0.12852 | 0.117 | 0.154 | 1 | 6 | Hbs1l | -0.037 | Young |
| 0.153682 | 0.305006 | 0.037 | 0.02 | 1 | 6 | Gamt | 0.017 | Young |
| 0.153854 | 0.143429 | 0.569 | 0.531 | 1 | 6 | Pafah1b1 | 0.038 | Young |
| 0.153982 | 0.295855 | 0.127 | 0.097 | 1 | 6 | Pde3b | 0.03 | Young |
| 0.153994 | 0.064283 | 0.087 | 0.06 | 1 | 6 | Phf13 | 0.027 | Young |
| 0.154011 | 0.81798 | 0.04 | 0.023 | 1 | 6 | Tamm41 | 0.017 | Young |
| 0.154207 | 0.209827 | 0.211 | 0.174 | 1 | 6 | Rftn1 | 0.037 | Young |
| 0.154655 | 0.035405 | 0.809 | 0.723 | 1 | 6 | S100a4 | 0.086 | Young |
| 0.1552 | 0.286831 | 0.038 | 0.06 | 1 | 6 | Msto1 | -0.022 | Young |
| 0.155239 | 0.08987 | 0.547 | 0.5 | 1 | 6 | Capns1 | 0.047 | Young |
| 0.155341 | 0.546285 | 0.065 | 0.043 | 1 | 6 | Ifih1 | 0.022 | Young |
| 0.155363 | 0.365457 | 0.159 | 0.126 | 1 | 6 | Rsbn1 | 0.033 | Young |
| 0.155661 | 0.224731 | 0.263 | 0.223 | 1 | 6 | Akna | 0.04 | Young |

| | | | | | | | | |
|---|---|---|---|---|---|---|---|---|
| 0.155837 | 0.871418 | 0.12 | 0.091 | 1 | 6 | Ccl3 | 0.029 | Young |
| 0.155852 | 0.293625 | 0.134 | 0.103 | 1 | 6 | Rxrb | 0.031 | Young |
| 0.156063 | 0.27083 | 0.142 | 0.109 | 1 | 6 | Cks2 | 0.033 | Young |
| 0.15628 | 0.16304 | 0.254 | 0.214 | 1 | 6 | Impdh2 | 0.04 | Young |
| 0.156619 | 0.803414 | 0.107 | 0.08 | 1 | 6 | Ccdc170 | 0.027 | Young |
| 0.15671 | 0.201676 | 0.58 | 0.529 | 1 | 6 | Ostf1 | 0.051 | Young |
| 0.157614 | 0.136845 | 0.497 | 0.449 | 1 | 6 | Batf3 | 0.048 | Young |
| 0.157878 | 0.002872 | 0.162 | 0.129 | 1 | 6 | Nup62 | 0.033 | Young |
| 0.15798 | 0.176901 | 0.211 | 0.171 | 1 | 6 | Ypel3 | 0.04 | Young |
| 0.15806 | 1.320733 | 0.054 | 0.034 | 1 | 6 | Haus6 | 0.02 | Young |
| 0.158078 | 0.567018 | 0.075 | 0.051 | 1 | 6 | Nemp1 | 0.024 | Young |
| 0.158432 | 0.298077 | 0.293 | 0.257 | 1 | 6 | Gspt1 | 0.036 | Young |
| 0.158447 | 0.273867 | 0.689 | 0.654 | 1 | 6 | Zfand5 | 0.035 | Young |
| 0.158674 | 0.449517 | 0.274 | 0.237 | 1 | 6 | D16Ertd47 | 0.037 | Young |
| 0.158979 | 0.259601 | 0.403 | 0.374 | 1 | 6 | Prkx | 0.029 | Young |
| 0.159425 | 1.037041 | 0.05 | 0.031 | 1 | 6 | 4930503L1 | 0.019 | Young |
| 0.159949 | 0.637342 | 0.179 | 0.149 | 1 | 6 | Rnf115 | 0.03 | Young |
| 0.160231 | 0.52747 | 0.04 | 0.023 | 1 | 6 | Tmem107 | 0.017 | Young |
| 0.160435 | 0.459438 | 0.072 | 0.049 | 1 | 6 | Paxip1 | 0.023 | Young |
| 0.160441 | 0.123564 | 0.791 | 0.743 | 1 | 6 | Gpr132 | 0.048 | Young |
| 0.160543 | 0.847867 | 0.09 | 0.066 | 1 | 6 | Tmem63b | 0.024 | Young |
| 0.160567 | 0.152914 | 0.697 | 0.66 | 1 | 6 | Nrros | 0.037 | Young |
| 0.160592 | 0.240634 | 0.085 | 0.06 | 1 | 6 | Rasgrp2 | 0.025 | Young |
| 0.160675 | 0.122955 | 0.587 | 0.569 | 1 | 6 | Calm3 | 0.018 | Young |
| 0.160822 | 0.932717 | 0.064 | 0.043 | 1 | 6 | Trim65 | 0.021 | Young |
| 0.160877 | 1.001681 | 0.05 | 0.031 | 1 | 6 | Letmd1 | 0.019 | Young |
| 0.161561 | 0.409451 | 0.043 | 0.026 | 1 | 6 | Tapt1 | 0.017 | Young |
| 0.161944 | 0.161339 | 0.366 | 0.32 | 1 | 6 | Commd8 | 0.046 | Young |
| 0.162021 | 0.391435 | 0.154 | 0.123 | 1 | 6 | Smim27 | 0.031 | Young |
| 0.162258 | 0.336207 | 0.144 | 0.114 | 1 | 6 | Trappc6a | 0.03 | Young |
| 0.162495 | 0.994149 | 0.028 | 0.014 | 1 | 6 | Ttc27 | 0.014 | Young |
| 0.162607 | 0.088965 | 0.826 | 0.806 | 1 | 6 | Ucp2 | 0.02 | Young |
| 0.162759 | 1.035317 | 0.05 | 0.031 | 1 | 6 | Flnb | 0.019 | Young |
| 0.162808 | 0.452221 | 0.069 | 0.046 | 1 | 6 | Gnl2 | 0.023 | Young |
| 0.163063 | 1.121887 | 0.028 | 0.014 | 1 | 6 | Fam151b | 0.014 | Young |
| 0.163227 | 0.302678 | 0.373 | 0.337 | 1 | 6 | Snap23 | 0.036 | Young |
| 0.163373 | 0.329442 | 0.263 | 0.226 | 1 | 6 | Samd9l | 0.037 | Young |
| 0.163556 | 0.131911 | 0.885 | 0.811 | 1 | 6 | Dazap2 | 0.074 | Young |
| 0.163617 | 0.476085 | 0.256 | 0.22 | 1 | 6 | Sertad2 | 0.036 | Young |
| 0.163662 | 0.84061 | 0.057 | 0.037 | 1 | 6 | Klhl21 | 0.02 | Young |
| 0.163676 | 0.515105 | 0.156 | 0.126 | 1 | 6 | Atp7a | 0.03 | Young |
| 0.163686 | 0.33954 | 0.318 | 0.291 | 1 | 6 | Xrn2 | 0.027 | Young |
| 0.163717 | 0.154854 | 0.393 | 0.357 | 1 | 6 | Uba1 | 0.036 | Young |
| 0.163799 | 0.252977 | 0.308 | 0.266 | 1 | 6 | Zmat2 | 0.042 | Young |
| 0.163912 | 0.143561 | 0.42 | 0.369 | 1 | 6 | Rnf149 | 0.051 | Young |

| | | | | | | | | |
|---|---|---|---|---|---|---|---|---|
| 0.164046 | 0.207674 | 0.378 | 0.34 | 1 | 6 | Mier1 | 0.038 | Young |
| 0.164318 | 0.627971 | 0.107 | 0.08 | 1 | 6 | Il11ra1 | 0.027 | Young |
| 0.164589 | 0.460796 | 0.256 | 0.223 | 1 | 6 | St3gal1 | 0.033 | Young |
| 0.164702 | 0.218294 | 0.403 | 0.363 | 1 | 6 | Snu13 | 0.04 | Young |
| 0.164729 | 0.320686 | 0.135 | 0.106 | 1 | 6 | 2310009B1 | 0.029 | Young |
| 0.165105 | 0.189514 | 0.246 | 0.203 | 1 | 6 | Ssfa2 | 0.043 | Young |
| 0.165242 | 0.216689 | 0.176 | 0.143 | 1 | 6 | Mtmr6 | 0.033 | Young |
| 0.165389 | 0.195498 | 0.365 | 0.326 | 1 | 6 | Sbds | 0.039 | Young |
| 0.165635 | 0.117195 | 0.07 | 0.097 | 1 | 6 | Fam117a | -0.027 | Young |
| 0.165866 | 1.925751 | 0.02 | 0.009 | 1 | 6 | Gm29340 | 0.011 | Young |
| 0.165932 | 1.444096 | 0.032 | 0.017 | 1 | 6 | Ocstamp | 0.015 | Young |
| 0.16594 | 0.289724 | 0.124 | 0.094 | 1 | 6 | Ndrg3 | 0.03 | Young |
| 0.166086 | 0.269015 | 0.189 | 0.154 | 1 | 6 | Stard4 | 0.035 | Young |
| 0.166158 | 0.894109 | 0.054 | 0.034 | 1 | 6 | Mitf | 0.02 | Young |
| 0.166423 | 0.752485 | 0.064 | 0.043 | 1 | 6 | Pgm2 | 0.021 | Young |
| 0.166799 | 0.144792 | 0.157 | 0.123 | 1 | 6 | Sos2 | 0.034 | Young |
| 0.166915 | 0.060234 | 0.5 | 0.449 | 1 | 6 | Arhgap45 | 0.051 | Young |
| 0.167278 | 0.299608 | 0.082 | 0.057 | 1 | 6 | Anxa3 | 0.025 | Young |
| 0.167477 | 0.039668 | 0.219 | 0.183 | 1 | 6 | Mien1 | 0.036 | Young |
| 0.167715 | 0.695398 | 0.084 | 0.06 | 1 | 6 | Cx3cr1 | 0.024 | Young |
| 0.167879 | 0.03467 | 0.824 | 0.766 | 1 | 6 | Fosl2 | 0.058 | Young |
| 0.168362 | 0.326637 | 0.242 | 0.206 | 1 | 6 | Dennd4b | 0.036 | Young |
| 0.168617 | 0.48879 | 0.067 | 0.046 | 1 | 6 | Tnip1 | 0.021 | Young |
| 0.168865 | 0.203611 | 0.107 | 0.08 | 1 | 6 | Med29 | 0.027 | Young |
| 0.169034 | 0.423612 | 0.212 | 0.18 | 1 | 6 | Zfp385a | 0.032 | Young |
| 0.169099 | 0.609665 | 0.074 | 0.051 | 1 | 6 | Tprkb | 0.023 | Young |
| 0.170074 | 1.269593 | 0.02 | 0.009 | 1 | 6 | Gm48066 | 0.011 | Young |
| 0.170202 | 0.018752 | 0.079 | 0.106 | 1 | 6 | Ints1 | -0.027 | Young |
| 0.17065 | 0.017993 | 0.405 | 0.366 | 1 | 6 | Arhgap17 | 0.039 | Young |
| 0.170691 | 0.917114 | 0.057 | 0.037 | 1 | 6 | Adam22 | 0.02 | Young |
| 0.170718 | 0.88614 | 0.064 | 0.043 | 1 | 6 | Rab9 | 0.021 | Young |
| 0.170828 | 0.126028 | 0.135 | 0.103 | 1 | 6 | Lcorl | 0.032 | Young |
| 0.170984 | 0.489973 | 0.172 | 0.143 | 1 | 6 | Utp14a | 0.029 | Young |
| 0.171139 | 0.777063 | 0.02 | 0.009 | 1 | 6 | Nsun5 | 0.011 | Young |
| 0.171163 | 0.327391 | 0.313 | 0.28 | 1 | 6 | Rps27rt | 0.033 | Young |
| 0.171386 | 0.318231 | 0.085 | 0.06 | 1 | 6 | E4f1 | 0.025 | Young |
| 0.171494 | 0.736587 | 0.194 | 0.163 | 1 | 6 | Cebpd | 0.031 | Young |
| 0.171495 | 0.937855 | 0.02 | 0.009 | 1 | 6 | Shpk | 0.011 | Young |
| 0.171532 | 0.485688 | 0.047 | 0.029 | 1 | 6 | Kdm1b | 0.018 | Young |
| 0.171688 | 0.88014 | 0.032 | 0.017 | 1 | 6 | Zmym1 | 0.015 | Young |
| 0.17221 | 0.453124 | 0.11 | 0.083 | 1 | 6 | 9930021J0 | 0.027 | Young |
| 0.172641 | 0.159867 | 0.202 | 0.166 | 1 | 6 | Cetn3 | 0.036 | Young |
| 0.17298 | 0.066009 | 0.413 | 0.371 | 1 | 6 | Ube3a | 0.042 | Young |
| 0.173546 | 0.278949 | 0.284 | 0.249 | 1 | 6 | Gas7 | 0.035 | Young |
| 0.173642 | 0.636156 | 0.02 | 0.009 | 1 | 6 | Icam2 | 0.011 | Young |

| | | | | | | | | |
|---|---|---|---|---|---|---|---|---|
| 0.173961 | 0.012111 | 0.05 | 0.031 | 1 | 6 | Adgrg6 | 0.019 | Young |
| 0.174453 | 0.351344 | 0.169 | 0.137 | 1 | 6 | Rybp | 0.032 | Young |
| 0.174471 | 0.035782 | 0.672 | 0.64 | 1 | 6 | Mtdh | 0.032 | Young |
| 0.174498 | 0.111284 | 0.166 | 0.131 | 1 | 6 | Mthfsl | 0.035 | Young |
| 0.174595 | 0.408627 | 0.09 | 0.066 | 1 | 6 | Rnf170 | 0.024 | Young |
| 0.174637 | 0.046226 | 0.656 | 0.583 | 1 | 6 | Gsn | 0.073 | Young |
| 0.174723 | 0.186552 | 0.02 | 0.009 | 1 | 6 | Srfbp1 | 0.011 | Young |
| 0.174795 | 0.379865 | 0.107 | 0.08 | 1 | 6 | Cul4a | 0.027 | Young |
| 0.174996 | 0.149593 | 0.448 | 0.409 | 1 | 6 | Anp32a | 0.039 | Young |
| 0.175044 | 0.580723 | 0.032 | 0.017 | 1 | 6 | Nat6 | 0.015 | Young |
| 0.175326 | 0.885825 | 0.032 | 0.017 | 1 | 6 | Olr1 | 0.015 | Young |
| 0.175497 | 0.404054 | 0.132 | 0.103 | 1 | 6 | Dock1 | 0.029 | Young |
| 0.175995 | 0.726006 | 0.13 | 0.103 | 1 | 6 | Gan | 0.027 | Young |
| 0.176964 | 0.027585 | 0.321 | 0.28 | 1 | 6 | Dap | 0.041 | Young |
| 0.177215 | 0.116121 | 0.351 | 0.311 | 1 | 6 | Susd6 | 0.04 | Young |
| 0.177266 | 0.132533 | 0.02 | 0.009 | 1 | 6 | Pwwp2b | 0.011 | Young |
| 0.177454 | 0.054599 | 0.764 | 0.743 | 1 | 6 | Sod1 | 0.021 | Young |
| 0.177659 | 0.745573 | 0.151 | 0.123 | 1 | 6 | Cebpa | 0.028 | Young |
| 0.178141 | 0.330083 | 0.368 | 0.337 | 1 | 6 | Ccnt1 | 0.031 | Young |
| 0.178414 | 0.703901 | 0.09 | 0.066 | 1 | 6 | Mxi1 | 0.024 | Young |
| 0.178442 | 0.069555 | 0.167 | 0.131 | 1 | 6 | Pom121 | 0.036 | Young |
| 0.178445 | 0.091842 | 0.565 | 0.534 | 1 | 6 | Snx20 | 0.031 | Young |
| 0.178692 | 0.093021 | 0.383 | 0.349 | 1 | 6 | Lpp | 0.034 | Young |
| 0.178697 | 0.671164 | 0.067 | 0.046 | 1 | 6 | Lsm10 | 0.021 | Young |
| 0.178947 | 0.118375 | 0.13 | 0.1 | 1 | 6 | Ero1l | 0.03 | Young |
| 0.179148 | 0.580894 | 0.104 | 0.077 | 1 | 6 | Adamts10 | 0.027 | Young |
| 0.179923 | 0.236279 | 0.211 | 0.171 | 1 | 6 | Rnasel | 0.04 | Young |
| 0.179947 | 1.011122 | 0.035 | 0.02 | 1 | 6 | Stard10 | 0.015 | Young |
| 0.18026 | 0.497081 | 0.149 | 0.12 | 1 | 6 | 0610030E2 | 0.029 | Young |
| 0.180273 | 0.2116 | 0.281 | 0.246 | 1 | 6 | Fam126a | 0.035 | Young |
| 0.180345 | 0.688185 | 0.092 | 0.069 | 1 | 6 | Mrps5 | 0.023 | Young |
| 0.1804 | 0.362661 | 0.221 | 0.186 | 1 | 6 | Errfi1 | 0.035 | Young |
| 0.180584 | 0.262224 | 0.324 | 0.289 | 1 | 6 | Me2 | 0.035 | Young |
| 0.180687 | 0.125974 | 0.921 | 0.883 | 1 | 6 | Nr4a1 | 0.038 | Young |
| 0.180832 | 0.230391 | 0.365 | 0.334 | 1 | 6 | Map1lc3b | 0.031 | Young |
| 0.180936 | 0.12221 | 0.161 | 0.126 | 1 | 6 | Plxna4 | 0.035 | Young |
| 0.181199 | 0.214041 | 0.604 | 0.566 | 1 | 6 | Cbl | 0.038 | Young |
| 0.181311 | 1.029121 | 0.035 | 0.02 | 1 | 6 | Gemin2 | 0.015 | Young |
| 0.181433 | 0.476759 | 0.067 | 0.046 | 1 | 6 | Mylip | 0.021 | Young |
| 0.181435 | 0.281173 | 0.087 | 0.063 | 1 | 6 | Sart3 | 0.024 | Young |
| 0.181504 | 0.037916 | 0.386 | 0.343 | 1 | 6 | Sfxn3 | 0.043 | Young |
| 0.181511 | 0.213231 | 0.293 | 0.257 | 1 | 6 | Ppp1r12c | 0.036 | Young |
| 0.181564 | 0.351674 | 0.084 | 0.06 | 1 | 6 | Ctnnd2 | 0.024 | Young |
| 0.182118 | 0.600154 | 0.038 | 0.023 | 1 | 6 | Smim24 | 0.015 | Young |
| 0.182134 | 1.080811 | 0.035 | 0.02 | 1 | 6 | Suv39h1 | 0.015 | Young |

| | | | | | | | | |
|---|---|---|---|---|---|---|---|---|
| 0.182404 | 0.126654 | 0.336 | 0.303 | 1 | 6 | Rin3 | 0.033 | Young |
| 0.182449 | 0.048312 | 0.336 | 0.291 | 1 | 6 | Baz1b | 0.045 | Young |
| 0.18258 | 0.034991 | 0.137 | 0.109 | 1 | 6 | Tax1bp3 | 0.028 | Young |
| 0.18292 | 0.679476 | 0.109 | 0.083 | 1 | 6 | Zfp120 | 0.026 | Young |
| 0.183526 | 0.167425 | 0.094 | 0.069 | 1 | 6 | Syap1 | 0.025 | Young |
| 0.183672 | 0.473535 | 0.224 | 0.197 | 1 | 6 | Myo1c | 0.027 | Young |
| 0.183771 | 0.229234 | 0.303 | 0.263 | 1 | 6 | Rela | 0.04 | Young |
| 0.183843 | 0.0916 | 0.253 | 0.217 | 1 | 6 | Ilk | 0.036 | Young |
| 0.1841 | 0.239273 | 0.222 | 0.189 | 1 | 6 | Fuca2 | 0.033 | Young |
| 0.184277 | 0.15893 | 0.296 | 0.254 | 1 | 6 | Smc1a | 0.042 | Young |
| 0.184381 | 0.419317 | 0.092 | 0.069 | 1 | 6 | Ciz1 | 0.023 | Young |
| 0.184729 | 0.189883 | 0.256 | 0.22 | 1 | 6 | Chchd1 | 0.036 | Young |
| 0.185813 | 0.30989 | 0.187 | 0.157 | 1 | 6 | Ccndbp1 | 0.03 | Young |
| 0.185855 | 0.168297 | 0.234 | 0.194 | 1 | 6 | Gadd45g | 0.04 | Young |
| 0.186167 | 1.404388 | 0.023 | 0.011 | 1 | 6 | Nxn | 0.012 | Young |
| 0.186319 | 0.55477 | 0.159 | 0.129 | 1 | 6 | Tspan3 | 0.03 | Young |
| 0.186455 | 0.199722 | 0.08 | 0.057 | 1 | 6 | Nmd3 | 0.023 | Young |
| 0.18653 | 0.005846 | 0.281 | 0.24 | 1 | 6 | Cpne3 | 0.041 | Young |
| 0.186565 | 0.396476 | 0.167 | 0.137 | 1 | 6 | Epb41l4aos | 0.03 | Young |
| 0.186973 | 0.236742 | 0.064 | 0.043 | 1 | 6 | Zscan29 | 0.021 | Young |
| 0.187063 | 0.092782 | 0.468 | 0.431 | 1 | 6 | Nabp1 | 0.037 | Young |
| 0.187162 | 0.826577 | 0.075 | 0.054 | 1 | 6 | Aqp9 | 0.021 | Young |
| 0.18726 | 0.048293 | 0.221 | 0.189 | 1 | 6 | Ascc3 | 0.032 | Young |
| 0.187331 | 0.412655 | 0.194 | 0.163 | 1 | 6 | Idh1 | 0.031 | Young |
| 0.187556 | 1.612723 | 0.023 | 0.011 | 1 | 6 | Taf1c | 0.012 | Young |
| 0.187624 | 0.176204 | 0.426 | 0.386 | 1 | 6 | Fam117b | 0.04 | Young |
| 0.187848 | 0.110491 | 0.868 | 0.82 | 1 | 6 | Atp5e | 0.048 | Young |
| 0.188087 | 0.457989 | 0.289 | 0.26 | 1 | 6 | Tnks2 | 0.029 | Young |
| 0.188134 | 0.069489 | 0.849 | 0.823 | 1 | 6 | Npm1 | 0.026 | Young |
| 0.188385 | 0.296282 | 0.142 | 0.114 | 1 | 6 | Lemd3 | 0.028 | Young |
| 0.188425 | 0.417281 | 0.172 | 0.14 | 1 | 6 | Cmip | 0.032 | Young |
| 0.188531 | 0.540813 | 0.035 | 0.02 | 1 | 6 | Ptges2 | 0.015 | Young |
| 0.189351 | 0.062214 | 0.222 | 0.186 | 1 | 6 | Gramd1a | 0.036 | Young |
| 0.189463 | 0.053744 | 0.139 | 0.177 | 1 | 6 | Hypk | -0.038 | Young |
| 0.189542 | 0.05045 | 0.661 | 0.589 | 1 | 6 | Il6ra | 0.072 | Young |
| 0.189983 | 0.550282 | 0.124 | 0.097 | 1 | 6 | Lrrc42 | 0.027 | Young |
| 0.190356 | 1.013484 | 0.023 | 0.011 | 1 | 6 | Tubd1 | 0.012 | Young |
| 0.190369 | 0.293684 | 0.186 | 0.154 | 1 | 6 | Nasp | 0.032 | Young |
| 0.190625 | 1.098163 | 0.038 | 0.023 | 1 | 6 | Cnbd2 | 0.015 | Young |
| 0.190692 | 0.34789 | 0.112 | 0.086 | 1 | 6 | Kdm3b | 0.026 | Young |
| 0.190766 | 0.271438 | 0.161 | 0.131 | 1 | 6 | Fam129b | 0.03 | Young |
| 0.190831 | 0.926796 | 0.042 | 0.026 | 1 | 6 | Lpcat2 | 0.016 | Young |
| 0.190895 | 0.611542 | 0.038 | 0.023 | 1 | 6 | Trim37 | 0.015 | Young |
| 0.191418 | 0.237985 | 0.256 | 0.22 | 1 | 6 | Secisbp2l | 0.036 | Young |
| 0.191596 | 1.576503 | 0.015 | 0.006 | 1 | 6 | Spata1 | 0.009 | Young |

| | | | | | | | | |
|---|---|---|---|---|---|---|---|---|
| 0.191631 | 0.181333 | 0.758 | 0.7 | 1 | 6 | Ly86 | 0.058 | Young |
| 0.192026 | 0.430261 | 0.102 | 0.077 | 1 | 6 | Bckdha | 0.025 | Young |
| 0.192444 | 0.102255 | 0.43 | 0.383 | 1 | 6 | Txndc17 | 0.047 | Young |
| 0.192556 | 0.04312 | 0.124 | 0.097 | 1 | 6 | Ipmk | 0.027 | Young |
| 0.192806 | 0.027988 | 0.366 | 0.337 | 1 | 6 | Agpat4 | 0.029 | Young |
| 0.192831 | 0.761495 | 0.023 | 0.011 | 1 | 6 | Lrp5 | 0.012 | Young |
| 0.192951 | 1.288952 | 0.015 | 0.006 | 1 | 6 | Nfe2 | 0.009 | Young |
| 0.192951 | 1.276249 | 0.015 | 0.006 | 1 | 6 | Slc25a35 | 0.009 | Young |
| 0.193076 | 0.267161 | 0.279 | 0.246 | 1 | 6 | Abcg1 | 0.033 | Young |
| 0.193383 | 0.132089 | 0.104 | 0.077 | 1 | 6 | Stab1 | 0.027 | Young |
| 0.193405 | 1.173093 | 0.015 | 0.006 | 1 | 6 | Gm1673 | 0.009 | Young |
| 0.193414 | 0.693537 | 0.129 | 0.103 | 1 | 6 | Ccdc9 | 0.026 | Young |
| 0.193859 | 1.514564 | 0.015 | 0.006 | 1 | 6 | Tctn3 | 0.009 | Young |
| 0.193859 | 1.383614 | 0.015 | 0.006 | 1 | 6 | Nr1h3 | 0.009 | Young |
| 0.193984 | 0.0467 | 0.567 | 0.509 | 1 | 6 | Skap2 | 0.058 | Young |
| 0.194163 | 0.79161 | 0.038 | 0.023 | 1 | 6 | Zfp101 | 0.015 | Young |
| 0.194303 | 0.511651 | 0.169 | 0.14 | 1 | 6 | Hmox1 | 0.029 | Young |
| 0.194314 | 1.284934 | 0.015 | 0.006 | 1 | 6 | Epha4 | 0.009 | Young |
| 0.194674 | 0.542847 | 0.092 | 0.069 | 1 | 6 | Rpf1 | 0.023 | Young |
| 0.19477 | 1.347726 | 0.015 | 0.006 | 1 | 6 | Zmynd15 | 0.009 | Young |
| 0.194794 | 0.245666 | 0.074 | 0.051 | 1 | 6 | Slc35f5 | 0.023 | Young |
| 0.194855 | 0.905185 | 0.048 | 0.031 | 1 | 6 | Uqcc1 | 0.017 | Young |
| 0.194873 | 0.840448 | 0.042 | 0.026 | 1 | 6 | Rbm12b1 | 0.016 | Young |
| 0.195226 | 1.129562 | 0.015 | 0.006 | 1 | 6 | Plekhg1 | 0.009 | Young |
| 0.195226 | 1.113671 | 0.015 | 0.006 | 1 | 6 | Tfb1m | 0.009 | Young |
| 0.195267 | 0.388028 | 0.284 | 0.257 | 1 | 6 | Nuak2 | 0.027 | Young |
| 0.195399 | 0.219245 | 0.042 | 0.026 | 1 | 6 | Mapk7 | 0.016 | Young |
| 0.195684 | 1.011384 | 0.015 | 0.006 | 1 | 6 | Ppp2r5b | 0.009 | Young |
| 0.196379 | 0.035982 | 0.533 | 0.491 | 1 | 6 | Snrpf | 0.042 | Young |
| 0.196454 | 0.570451 | 0.042 | 0.026 | 1 | 6 | Dgkg | 0.016 | Young |
| 0.196493 | 0.295346 | 0.189 | 0.157 | 1 | 6 | Nub1 | 0.032 | Young |
| 0.196546 | 0.071007 | 0.554 | 0.509 | 1 | 6 | Dstn | 0.045 | Young |
| 0.196601 | 0.787111 | 0.015 | 0.006 | 1 | 6 | Mrm3 | 0.009 | Young |
| 0.196769 | 0.462277 | 0.023 | 0.011 | 1 | 6 | Osbpl1a | 0.012 | Young |
| 0.197009 | 0.176306 | 0.784 | 0.731 | 1 | 6 | Wnk1 | 0.053 | Young |
| 0.19706 | 0.878961 | 0.015 | 0.006 | 1 | 6 | Pex11a | 0.009 | Young |
| 0.197262 | 0.43457 | 0.187 | 0.16 | 1 | 6 | Clptm1 | 0.027 | Young |
| 0.197521 | 0.596151 | 0.015 | 0.006 | 1 | 6 | Gdpd5 | 0.009 | Young |
| 0.197778 | 0.135082 | 0.199 | 0.16 | 1 | 6 | Sgk1 | 0.039 | Young |
| 0.197834 | 0.085652 | 0.839 | 0.817 | 1 | 6 | Zfp36l1 | 0.022 | Young |
| 0.197982 | 0.517446 | 0.015 | 0.006 | 1 | 6 | 0610040J0 | 0.009 | Young |
| 0.198618 | 0.151351 | 0.336 | 0.3 | 1 | 6 | Wdr89 | 0.036 | Young |
| 0.199059 | 0.909441 | 0.059 | 0.04 | 1 | 6 | Hsf2 | 0.019 | Young |
| 0.199371 | 0.32382 | 0.015 | 0.006 | 1 | 6 | Popdc2 | 0.009 | Young |
| 0.19944 | 0.168683 | 0.104 | 0.077 | 1 | 6 | Rps6ka5 | 0.027 | Young |

| | | | | | | | | |
|---|---|---|---|---|---|---|---|---|
| 0.199594 | 0.052153 | 0.134 | 0.103 | 1 | 6 | Capn7 | 0.031 | Young |
| 0.199609 | 0.57979 | 0.082 | 0.06 | 1 | 6 | Trp53inp2 | 0.022 | Young |
| 0.199862 | 0.024763 | 0.995 | 0.994 | 1 | 6 | Rpl37a | 0.001 | Young |
| 0.19991 | 1.45643 | 0.027 | 0.014 | 1 | 6 | 9930104L0 | 0.013 | Young |
| 0.199957 | 0.534356 | 0.059 | 0.04 | 1 | 6 | Scai | 0.019 | Young |
| 0.20023 | 0.176747 | 0.298 | 0.263 | 1 | 6 | Stub1 | 0.035 | Young |
| 0.200275 | 0.320413 | 0.288 | 0.257 | 1 | 6 | Lmo2 | 0.031 | Young |
| 0.200401 | 0.09881 | 0.067 | 0.046 | 1 | 6 | Tnfaip8l1 | 0.021 | Young |
| 0.200682 | 0.032211 | 0.296 | 0.254 | 1 | 6 | Tes | 0.042 | Young |
| 0.200866 | 0.160083 | 0.127 | 0.097 | 1 | 6 | Vkorc1 | 0.03 | Young |
| 0.201369 | 0.542272 | 0.104 | 0.08 | 1 | 6 | Nsun6 | 0.024 | Young |
| 0.201606 | 0.262867 | 0.197 | 0.166 | 1 | 6 | Dennd1c | 0.031 | Young |
| 0.201767 | 0.517711 | 0.147 | 0.12 | 1 | 6 | Cops2 | 0.027 | Young |
| 0.201988 | 0.843277 | 0.059 | 0.04 | 1 | 6 | Ndufaf3 | 0.019 | Young |
| 0.202071 | 0.087945 | 0.652 | 0.6 | 1 | 6 | S100a11 | 0.052 | Young |
| 0.202091 | 0.538218 | 0.082 | 0.06 | 1 | 6 | Ctnnbl1 | 0.022 | Young |
| 0.202134 | 0.346145 | 0.298 | 0.263 | 1 | 6 | Ddrgk1 | 0.035 | Young |
| 0.202263 | 0.502154 | 0.114 | 0.089 | 1 | 6 | Casp1 | 0.025 | Young |
| 0.202299 | 0.741244 | 0.045 | 0.029 | 1 | 6 | Utp14b | 0.016 | Young |
| 0.202308 | 0.699865 | 0.095 | 0.071 | 1 | 6 | Zadh2 | 0.024 | Young |
| 0.202542 | 0.393293 | 0.182 | 0.154 | 1 | 6 | Gclc | 0.028 | Young |
| 0.202559 | 0.726656 | 0.045 | 0.029 | 1 | 6 | Ampd3 | 0.016 | Young |
| 0.202577 | 0.133484 | 0.214 | 0.18 | 1 | 6 | Ints6 | 0.034 | Young |
| 0.20263 | 1.219616 | 0.027 | 0.014 | 1 | 6 | Batf2 | 0.013 | Young |
| 0.202707 | 0.577337 | 0.062 | 0.043 | 1 | 6 | Coq8b | 0.019 | Young |
| 0.202873 | 0.429655 | 0.042 | 0.026 | 1 | 6 | Zfp667 | 0.016 | Young |
| 0.202917 | 0.063241 | 0.064 | 0.043 | 1 | 6 | Tirap | 0.021 | Young |
| 0.203329 | 0.093117 | 0.726 | 0.657 | 1 | 6 | Cox6c | 0.069 | Young |
| 0.203882 | 0.065559 | 0.555 | 0.503 | 1 | 6 | Dusp5 | 0.052 | Young |
| 0.203958 | 0.431954 | 0.042 | 0.026 | 1 | 6 | Rab2b | 0.016 | Young |
| 0.204024 | 0.10821 | 0.192 | 0.157 | 1 | 6 | AC163018. | 0.035 | Young |
| 0.204381 | 0.948753 | 0.045 | 0.029 | 1 | 6 | Carmil1 | 0.016 | Young |
| 0.204687 | 0.888016 | 0.027 | 0.014 | 1 | 6 | Ripor1 | 0.013 | Young |
| 0.204687 | 0.587499 | 0.027 | 0.014 | 1 | 6 | D030056L2 | 0.013 | Young |
| 0.205002 | 0.053095 | 0.209 | 0.174 | 1 | 6 | Naa15 | 0.035 | Young |
| 0.205084 | 0.547017 | 0.094 | 0.071 | 1 | 6 | Hmgn2 | 0.023 | Young |
| 0.20529 | 0.714895 | 0.079 | 0.057 | 1 | 6 | Arhgef7 | 0.022 | Young |
| 0.20538 | 0.107121 | 0.95 | 0.914 | 1 | 6 | Zfp36 | 0.036 | Young |
| 0.205885 | 0.173186 | 0.079 | 0.057 | 1 | 6 | Phb | 0.022 | Young |
| 0.206102 | 0.370852 | 0.132 | 0.106 | 1 | 6 | Nipsnap3b | 0.026 | Young |
| 0.206302 | 0.101346 | 0.671 | 0.637 | 1 | 6 | Diaph1 | 0.034 | Young |
| 0.206679 | 0.925266 | 0.069 | 0.049 | 1 | 6 | Ndufaf2 | 0.02 | Young |
| 0.206844 | 0.160999 | 0.01 | 0.02 | 1 | 6 | Gm45871 | -0.01 | Young |
| 0.20689 | 0.18409 | 0.194 | 0.16 | 1 | 6 | Aldh1a2 | 0.034 | Young |
| 0.207089 | 0.60422 | 0.206 | 0.177 | 1 | 6 | C1qb | 0.029 | Young |

| | | | | | | | | |
|---|---|---|---|---|---|---|---|---|
| 0.207113 | 0.818635 | 0.084 | 0.063 | 1 | 6 | Lmbr1l | 0.021 | Young |
| 0.207305 | 0.197965 | 0.222 | 0.2 | 1 | 6 | Crtc3 | 0.022 | Young |
| 0.207355 | 0.569769 | 0.149 | 0.123 | 1 | 6 | Gm12840 | 0.026 | Young |
| 0.207652 | 0.233667 | 0.089 | 0.066 | 1 | 6 | Tor3a | 0.023 | Young |
| 0.207711 | 0.139425 | 0.431 | 0.394 | 1 | 6 | Drap1 | 0.037 | Young |
| 0.207723 | 0.120876 | 0.01 | 0.02 | 1 | 6 | Zfp637 | -0.01 | Young |
| 0.207997 | 0.232041 | 0.289 | 0.254 | 1 | 6 | Pacsin2 | 0.035 | Young |
| 0.208195 | 0.005186 | 0.07 | 0.094 | 1 | 6 | Rad54l2 | -0.024 | Young |
| 0.208232 | 0.202667 | 0.45 | 0.411 | 1 | 6 | Rilpl2 | 0.039 | Young |
| 0.208427 | 0.145126 | 0.453 | 0.414 | 1 | 6 | Sdf4 | 0.039 | Young |
| 0.208448 | 0.294149 | 0.104 | 0.08 | 1 | 6 | Cd63 | 0.024 | Young |
| 0.208635 | 0.244433 | 0.279 | 0.246 | 1 | 6 | Hsbp1 | 0.033 | Young |
| 0.209091 | 0.472496 | 0.052 | 0.034 | 1 | 6 | Map3k20 | 0.018 | Young |
| 0.209871 | 0.145027 | 0.4 | 0.363 | 1 | 6 | Sqstm1 | 0.037 | Young |
| 0.210024 | 0.674168 | 0.184 | 0.157 | 1 | 6 | Hsph1 | 0.027 | Young |
| 0.21003 | 0.146399 | 0.512 | 0.483 | 1 | 6 | Ndufa13 | 0.029 | Young |
| 0.210168 | 0.415388 | 0.099 | 0.074 | 1 | 6 | Usp39 | 0.025 | Young |
| 0.210248 | 0.507404 | 0.027 | 0.014 | 1 | 6 | Cdc42ep2 | 0.013 | Young |
| 0.211099 | 0.262766 | 0.294 | 0.266 | 1 | 6 | Phb2 | 0.028 | Young |
| 0.211311 | 0.240196 | 0.232 | 0.203 | 1 | 6 | Uxs1 | 0.029 | Young |
| 0.211576 | 0.036446 | 0.756 | 0.697 | 1 | 6 | Sh3bgrl | 0.059 | Young |
| 0.211672 | 1.284206 | 0.01 | 0.003 | 1 | 6 | Tfpi | 0.007 | Young |
| 0.211672 | 1.265678 | 0.01 | 0.003 | 1 | 6 | Paqr3 | 0.007 | Young |
| 0.211672 | 0.900012 | 0.01 | 0.003 | 1 | 6 | Taco1 | 0.007 | Young |
| 0.211713 | 0.11676 | 0.232 | 0.194 | 1 | 6 | Qpct | 0.038 | Young |
| 0.21205 | 0.117225 | 0.132 | 0.106 | 1 | 6 | Mgat4b | 0.026 | Young |
| 0.21216 | 0.537934 | 0.14 | 0.117 | 1 | 6 | Bag3 | 0.023 | Young |
| 0.212279 | 1.541573 | 0.01 | 0.003 | 1 | 6 | Kif18a | 0.007 | Young |
| 0.212279 | 1.500673 | 0.01 | 0.003 | 1 | 6 | Tmem17 | 0.007 | Young |
| 0.212279 | 1.20407 | 0.01 | 0.003 | 1 | 6 | Mir9-3hg | 0.007 | Young |
| 0.212279 | 1.082688 | 0.01 | 0.003 | 1 | 6 | Klhl42 | 0.007 | Young |
| 0.212279 | 1.081239 | 0.01 | 0.003 | 1 | 6 | Tgm1 | 0.007 | Young |
| 0.212279 | 0.964567 | 0.01 | 0.003 | 1 | 6 | AC125351. | 0.007 | Young |
| 0.212887 | 1.410737 | 0.01 | 0.003 | 1 | 6 | Sirt4 | 0.007 | Young |
| 0.212887 | 1.296768 | 0.01 | 0.003 | 1 | 6 | Xpo5 | 0.007 | Young |
| 0.212887 | 1.268316 | 0.01 | 0.003 | 1 | 6 | Gm26526 | 0.007 | Young |
| 0.212887 | 1.196634 | 0.01 | 0.003 | 1 | 6 | 1810009A1 | 0.007 | Young |
| 0.212887 | 1.079678 | 0.01 | 0.003 | 1 | 6 | Taco1os | 0.007 | Young |
| 0.212887 | 1.074994 | 0.01 | 0.003 | 1 | 6 | C230037L1 | 0.007 | Young |
| 0.212887 | 1.042809 | 0.01 | 0.003 | 1 | 6 | Mtus2 | 0.007 | Young |
| 0.212887 | 0.758085 | 0.01 | 0.003 | 1 | 6 | Nudt8.1 | 0.007 | Young |
| 0.213059 | 1.380246 | 0.03 | 0.017 | 1 | 6 | Slc2a8 | 0.013 | Young |
| 0.213132 | 0.408418 | 0.069 | 0.049 | 1 | 6 | Rbbp5 | 0.02 | Young |
| 0.213298 | 0.507215 | 0.144 | 0.117 | 1 | 6 | Gm11808 | 0.027 | Young |
| 0.213304 | 0.366479 | 0.231 | 0.2 | 1 | 6 | Atxn7 | 0.031 | Young |

| | | | | | | | | |
|---|---|---|---|---|---|---|---|---|
| 0.213338 | 0.491621 | 0.13 | 0.106 | 1 | 6 | Mospd3 | 0.024 | Young |
| 0.213348 | 0.283411 | 0.13 | 0.103 | 1 | 6 | Pfdn1 | 0.027 | Young |
| 0.213739 | 0.163092 | 0.609 | 0.571 | 1 | 6 | Tgfbi | 0.038 | Young |
| 0.213744 | 0.114969 | 0.831 | 0.769 | 1 | 6 | Gnb2 | 0.062 | Young |
| 0.213929 | 0.312654 | 0.079 | 0.057 | 1 | 6 | Fcor | 0.022 | Young |
| 0.214057 | 1.011705 | 0.03 | 0.017 | 1 | 6 | BC147527 | 0.013 | Young |
| 0.214107 | 0.819619 | 0.01 | 0.003 | 1 | 6 | Gm1965 | 0.007 | Young |
| 0.214107 | 0.801021 | 0.01 | 0.003 | 1 | 6 | Oxld1 | 0.007 | Young |
| 0.214107 | 0.776465 | 0.01 | 0.003 | 1 | 6 | Apbb1 | 0.007 | Young |
| 0.214107 | 0.767031 | 0.01 | 0.003 | 1 | 6 | Pemt | 0.007 | Young |
| 0.214107 | 0.762111 | 0.01 | 0.003 | 1 | 6 | F5 | 0.007 | Young |
| 0.214295 | 0.448038 | 0.11 | 0.086 | 1 | 6 | Acox1 | 0.024 | Young |
| 0.214864 | 0.385317 | 0.137 | 0.111 | 1 | 6 | Cdip1 | 0.026 | Young |
| 0.215332 | 0.515556 | 0.01 | 0.003 | 1 | 6 | Snx24 | 0.007 | Young |
| 0.215332 | 0.027896 | 0.01 | 0.003 | 1 | 6 | Frmd5 | 0.007 | Young |
| 0.215631 | 0.325878 | 0.319 | 0.291 | 1 | 6 | Ube2q1 | 0.028 | Young |
| 0.215727 | 0.919537 | 0.03 | 0.017 | 1 | 6 | Kif9 | 0.013 | Young |
| 0.2159 | 0.187792 | 0.415 | 0.377 | 1 | 6 | Tmod3 | 0.038 | Young |
| 0.216584 | 0.41232 | 0.075 | 0.054 | 1 | 6 | Ptchd1 | 0.021 | Young |
| 0.21686 | 0.100766 | 0.579 | 0.54 | 1 | 6 | Ppp2r5c | 0.039 | Young |
| 0.217105 | 0.469954 | 0.059 | 0.04 | 1 | 6 | Stam | 0.019 | Young |
| 0.21741 | 0.190764 | 0.043 | 0.063 | 1 | 6 | Tpgs1 | -0.02 | Young |
| 0.217468 | 0.213706 | 0.418 | 0.391 | 1 | 6 | Map2k3 | 0.027 | Young |
| 0.217611 | 0.318129 | 0.089 | 0.066 | 1 | 6 | Cnep1r1 | 0.023 | Young |
| 0.217691 | 0.065945 | 0.256 | 0.22 | 1 | 6 | Cers5 | 0.036 | Young |
| 0.217743 | 0.78874 | 0.03 | 0.017 | 1 | 6 | Ift122 | 0.013 | Young |
| 0.217755 | 0.117752 | 0.507 | 0.477 | 1 | 6 | Malt1 | 0.03 | Young |
| 0.217857 | 0.095043 | 0.082 | 0.06 | 1 | 6 | Brms1 | 0.022 | Young |
| 0.217923 | 0.25643 | 0.219 | 0.189 | 1 | 6 | Kmt2d | 0.03 | Young |
| 0.218191 | 0.069365 | 0.253 | 0.214 | 1 | 6 | Washc4 | 0.039 | Young |
| 0.218262 | 0.419564 | 0.082 | 0.06 | 1 | 6 | Abhd13 | 0.022 | Young |
| 0.218335 | 0.270347 | 0.216 | 0.183 | 1 | 6 | Wdfy2 | 0.033 | Young |
| 0.218686 | 0.111234 | 0.065 | 0.046 | 1 | 6 | Zfp865 | 0.019 | Young |
| 0.218835 | 0.232758 | 0.174 | 0.146 | 1 | 6 | Mrps25 | 0.028 | Young |
| 0.218906 | 0.24569 | 0.092 | 0.069 | 1 | 6 | Cfl2 | 0.023 | Young |
| 0.21918 | 0.166956 | 0.36 | 0.317 | 1 | 6 | Fli1 | 0.043 | Young |
| 0.219295 | 0.608628 | 0.062 | 0.043 | 1 | 6 | Gm15417 | 0.019 | Young |
| 0.220021 | 0.189061 | 0.129 | 0.1 | 1 | 6 | Fbxw11 | 0.029 | Young |
| 0.220476 | 0.315649 | 0.326 | 0.303 | 1 | 6 | Cd68 | 0.023 | Young |
| 0.220834 | 0.154762 | 0.559 | 0.52 | 1 | 6 | Map4k4 | 0.039 | Young |
| 0.221111 | 0.225663 | 0.095 | 0.071 | 1 | 6 | Apex1 | 0.024 | Young |
| 0.221134 | 0.638095 | 0.03 | 0.017 | 1 | 6 | Gsdme | 0.013 | Young |
| 0.22131 | 0.3261 | 0.059 | 0.04 | 1 | 6 | Lin37 | 0.019 | Young |
| 0.221424 | 0.275657 | 0.065 | 0.046 | 1 | 6 | Zfp729a | 0.019 | Young |
| 0.221591 | 0.070636 | 0.808 | 0.783 | 1 | 6 | Ccnl1 | 0.025 | Young |

| | | | | | | | | |
|---|---|---|---|---|---|---|---|---|
| 0.221804 | 0.370823 | 0.171 | 0.143 | 1 | 6 | Nab2 | 0.028 | Young |
| 0.22185 | 0.425078 | 0.237 | 0.211 | 1 | 6 | Aldh2 | 0.026 | Young |
| 0.222278 | 0.503644 | 0.172 | 0.146 | 1 | 6 | Sema4c | 0.026 | Young |
| 0.22277 | 0.305251 | 0.244 | 0.211 | 1 | 6 | Gxylt1 | 0.033 | Young |
| 0.222845 | 0.114259 | 0.478 | 0.434 | 1 | 6 | Cox5b | 0.044 | Young |
| 0.223668 | 0.024639 | 0.079 | 0.057 | 1 | 6 | Pacs1 | 0.022 | Young |
| 0.224269 | 1.199454 | 0.018 | 0.009 | 1 | 6 | BC051226 | 0.009 | Young |
| 0.224269 | 1.113481 | 0.018 | 0.009 | 1 | 6 | Gm32856 | 0.009 | Young |
| 0.224366 | 0.347921 | 0.261 | 0.231 | 1 | 6 | H2-Oa | 0.03 | Young |
| 0.224496 | 0.063838 | 0.935 | 0.849 | 1 | 6 | Ccr2 | 0.086 | Young |
| 0.224532 | 1.035645 | 0.037 | 0.023 | 1 | 6 | Ipo4 | 0.014 | Young |
| 0.224707 | 0.376222 | 0.169 | 0.143 | 1 | 6 | G6pdx | 0.026 | Young |
| 0.224963 | 0.207838 | 0.11 | 0.086 | 1 | 6 | Traf6 | 0.024 | Young |
| 0.225062 | 0.02297 | 0.656 | 0.614 | 1 | 6 | Rbms1 | 0.042 | Young |
| 0.225167 | 0.526901 | 0.018 | 0.009 | 1 | 6 | Gm14326 | 0.009 | Young |
| 0.225307 | 0.718653 | 0.033 | 0.02 | 1 | 6 | Sh3rf1 | 0.013 | Young |
| 0.225505 | 0.162247 | 0.329 | 0.297 | 1 | 6 | Rwdd1 | 0.032 | Young |
| 0.225752 | 0.391935 | 0.112 | 0.089 | 1 | 6 | Kbtbd2 | 0.023 | Young |
| 0.225992 | 0.054607 | 0.122 | 0.094 | 1 | 6 | Dgcr2 | 0.028 | Young |
| 0.226068 | 0.828957 | 0.018 | 0.009 | 1 | 6 | Cacnb1 | 0.009 | Young |
| 0.226068 | 0.602665 | 0.018 | 0.009 | 1 | 6 | Gm16638 | 0.009 | Young |
| 0.226299 | 0.336886 | 0.132 | 0.106 | 1 | 6 | Rbbp8 | 0.026 | Young |
| 0.226388 | 0.448993 | 0.062 | 0.043 | 1 | 6 | Wdsub1 | 0.019 | Young |
| 0.22652 | 0.945273 | 0.018 | 0.009 | 1 | 6 | Adra2a | 0.009 | Young |
| 0.226705 | 1.021142 | 0.037 | 0.023 | 1 | 6 | Cnksr3 | 0.014 | Young |
| 0.227057 | 1.167424 | 0.06 | 0.043 | 1 | 6 | Gypc | 0.017 | Young |
| 0.227094 | 0.437155 | 0.253 | 0.229 | 1 | 6 | Flii | 0.024 | Young |
| 0.227118 | 0.343015 | 0.008 | 0.017 | 1 | 6 | Fadd | -0.009 | Young |
| 0.227792 | 0.337108 | 0.065 | 0.046 | 1 | 6 | Slc25a44 | 0.019 | Young |
| 0.227928 | 0.880051 | 0.033 | 0.02 | 1 | 6 | Gna11 | 0.013 | Young |
| 0.227942 | 0.235338 | 0.226 | 0.194 | 1 | 6 | Atp6v1h | 0.032 | Young |
| 0.228332 | 0.593782 | 0.018 | 0.009 | 1 | 6 | Akap1 | 0.009 | Young |
| 0.228332 | 0.485764 | 0.018 | 0.009 | 1 | 6 | Tubg1 | 0.009 | Young |
| 0.228917 | 0.19702 | 0.033 | 0.02 | 1 | 6 | Plcb4 | 0.013 | Young |
| 0.229076 | 0.229562 | 0.03 | 0.017 | 1 | 6 | Cetn4 | 0.013 | Young |
| 0.229115 | 0.205881 | 0.244 | 0.209 | 1 | 6 | Kmo | 0.035 | Young |
| 0.229584 | 0.290571 | 0.296 | 0.269 | 1 | 6 | Insig1 | 0.027 | Young |
| 0.229634 | 0.072976 | 0.485 | 0.434 | 1 | 6 | Spi1 | 0.051 | Young |
| 0.229777 | 0.208467 | 0.38 | 0.351 | 1 | 6 | Dync1i2 | 0.029 | Young |
| 0.229814 | 0.42146 | 0.119 | 0.094 | 1 | 6 | Csad | 0.025 | Young |
| 0.230271 | 0.087808 | 0.035 | 0.051 | 1 | 6 | Zbtb6 | -0.016 | Young |
| 0.23049 | 0.058897 | 0.264 | 0.229 | 1 | 6 | Phf21a | 0.035 | Young |
| 0.230582 | 0.269987 | 0.289 | 0.257 | 1 | 6 | Osbpl9 | 0.032 | Young |
| 0.230786 | 0.234259 | 0.129 | 0.103 | 1 | 6 | Nceh1 | 0.026 | Young |
| 0.23107 | 0.135589 | 0.018 | 0.009 | 1 | 6 | Dynlt1a | 0.009 | Young |

| | | | | | | | | |
|---|---|---|---|---|---|---|---|---|
| 0.231181 | 0.127068 | 0.326 | 0.294 | 1 | 6 | Ik | 0.032 | Young |
| 0.231411 | 1.182591 | 0.037 | 0.023 | 1 | 6 | Gprc5c | 0.014 | Young |
| 0.231533 | 0.110128 | 0.03 | 0.017 | 1 | 6 | Dnajb5 | 0.013 | Young |
| 0.23155 | 0.186179 | 0.117 | 0.091 | 1 | 6 | Pi4k2b | 0.026 | Young |
| 0.231557 | 0.675006 | 0.114 | 0.091 | 1 | 6 | Psd4 | 0.023 | Young |
| 0.23171 | 0.142746 | 0.03 | 0.046 | 1 | 6 | Ak8 | -0.016 | Young |
| 0.232582 | 0.399563 | 0.164 | 0.137 | 1 | 6 | Cep120 | 0.027 | Young |
| 0.232695 | 0.06084 | 0.395 | 0.346 | 1 | 6 | Mapk1ip1l | 0.049 | Young |
| 0.232773 | 0.355295 | 0.174 | 0.149 | 1 | 6 | Ramp3 | 0.025 | Young |
| 0.232887 | 0.241414 | 0.231 | 0.203 | 1 | 6 | Cfap43 | 0.028 | Young |
| 0.233085 | 0.570572 | 0.099 | 0.077 | 1 | 6 | Tdrd7 | 0.022 | Young |
| 0.233544 | 0.117038 | 0.187 | 0.157 | 1 | 6 | Sptan1 | 0.03 | Young |
| 0.23361 | 0.22641 | 0.403 | 0.374 | 1 | 6 | Mkrn1 | 0.029 | Young |
| 0.233727 | 0.224654 | 0.458 | 0.42 | 1 | 6 | D8Ertd738 | 0.038 | Young |
| 0.23432 | 0.290037 | 0.084 | 0.063 | 1 | 6 | Dnajc9 | 0.021 | Young |
| 0.234485 | 0.243836 | 0.177 | 0.149 | 1 | 6 | Mybbp1a | 0.028 | Young |
| 0.234659 | 0.227009 | 0.179 | 0.149 | 1 | 6 | Esyt2 | 0.03 | Young |
| 0.234677 | 0.188875 | 0.273 | 0.243 | 1 | 6 | Cpt1a | 0.03 | Young |
| 0.234825 | 0.168632 | 0.189 | 0.157 | 1 | 6 | Arl1 | 0.032 | Young |
| 0.234869 | 0.108911 | 0.249 | 0.217 | 1 | 6 | Chtop | 0.032 | Young |
| 0.234908 | 0.464617 | 0.08 | 0.06 | 1 | 6 | Fkbp1b | 0.02 | Young |
| 0.235146 | 0.033199 | 0.908 | 0.9 | 1 | 6 | Ppia | 0.008 | Young |
| 0.23515 | 0.245498 | 0.174 | 0.143 | 1 | 6 | Ptdss1 | 0.031 | Young |
| 0.235233 | 0.951932 | 0.04 | 0.026 | 1 | 6 | Plekhg5 | 0.014 | Young |
| 0.235537 | 0.017958 | 0.114 | 0.089 | 1 | 6 | Lrrc8a | 0.025 | Young |
| 0.235538 | 0.992608 | 0.04 | 0.026 | 1 | 6 | Rprd1a | 0.014 | Young |
| 0.235603 | 0.207489 | 0.361 | 0.337 | 1 | 6 | Gpbp1 | 0.024 | Young |
| 0.235696 | 0.082116 | 0.727 | 0.677 | 1 | 6 | Irf2bp2 | 0.05 | Young |
| 0.235791 | 0.049209 | 0.059 | 0.04 | 1 | 6 | Dvl3 | 0.019 | Young |
| 0.235922 | 0.319096 | 0.033 | 0.02 | 1 | 6 | Tlr1 | 0.013 | Young |
| 0.236152 | 0.897646 | 0.047 | 0.031 | 1 | 6 | Rubcn | 0.016 | Young |
| 0.236884 | 0.520356 | 0.043 | 0.029 | 1 | 6 | Rfc4 | 0.014 | Young |
| 0.236913 | 0.026204 | 0.279 | 0.243 | 1 | 6 | Tmx3 | 0.036 | Young |
| 0.236929 | 1.115563 | 0.05 | 0.034 | 1 | 6 | Leo1 | 0.016 | Young |
| 0.236935 | 0.328935 | 0.033 | 0.02 | 1 | 6 | Cad | 0.013 | Young |
| 0.236935 | 0.038241 | 0.023 | 0.037 | 1 | 6 | Zfp952 | -0.014 | Young |
| 0.23694 | 0.225113 | 0.242 | 0.211 | 1 | 6 | Idh3g | 0.031 | Young |
| 0.236995 | 0.488968 | 0.094 | 0.071 | 1 | 6 | Ak6 | 0.023 | Young |
| 0.237012 | 0.43583 | 0.119 | 0.094 | 1 | 6 | Ints9 | 0.025 | Young |
| 0.237793 | 0.647915 | 0.037 | 0.023 | 1 | 6 | Dhx58 | 0.014 | Young |
| 0.238037 | 0.403539 | 0.216 | 0.189 | 1 | 6 | 5031425E2 | 0.027 | Young |
| 0.23804 | 0.088595 | 0.587 | 0.543 | 1 | 6 | Taok3 | 0.044 | Young |
| 0.238085 | 0.241149 | 0.304 | 0.277 | 1 | 6 | Stk40 | 0.027 | Young |
| 0.23895 | 0.938013 | 0.043 | 0.029 | 1 | 6 | 6330418K0 | 0.014 | Young |
| 0.239411 | 0.418083 | 0.084 | 0.063 | 1 | 6 | Ccdc102a | 0.021 | Young |

| | | | | | | | | |
|---|---|---|---|---|---|---|---|---|
| 0.239731 | 0.362757 | 0.037 | 0.023 | 1 | 6 | Mon1a | 0.014 | Young |
| 0.240119 | 0.740712 | 0.089 | 0.069 | 1 | 6 | Nit1 | 0.02 | Young |
| 0.240434 | 0.601097 | 0.043 | 0.029 | 1 | 6 | Slc52a3 | 0.014 | Young |
| 0.240555 | 0.04641 | 0.763 | 0.749 | 1 | 6 | Fnbp1 | 0.014 | Young |
| 0.240741 | 0.314843 | 0.13 | 0.106 | 1 | 6 | Mepce | 0.024 | Young |
| 0.240785 | 0.248063 | 0.247 | 0.214 | 1 | 6 | AI314180 | 0.033 | Young |
| 0.24103 | 0.646512 | 0.037 | 0.023 | 1 | 6 | Sec22a | 0.014 | Young |
| 0.241045 | 0.505749 | 0.191 | 0.163 | 1 | 6 | Relb | 0.028 | Young |
| 0.241204 | 0.688387 | 0.057 | 0.04 | 1 | 6 | Clpp | 0.017 | Young |
| 0.241433 | 1.37647 | 0.022 | 0.011 | 1 | 6 | Hectd2 | 0.011 | Young |
| 0.241671 | 0.294307 | 0.249 | 0.22 | 1 | 6 | Ndfip1 | 0.029 | Young |
| 0.241863 | 1.173176 | 0.022 | 0.011 | 1 | 6 | Gm7694 | 0.011 | Young |
| 0.242385 | 0.19504 | 0.395 | 0.363 | 1 | 6 | Zmiz1 | 0.032 | Young |
| 0.242435 | 0.123445 | 0.241 | 0.206 | 1 | 6 | Smarce1 | 0.035 | Young |
| 0.242662 | 0.322756 | 0.244 | 0.214 | 1 | 6 | Heca | 0.03 | Young |
| 0.243157 | 0.424946 | 0.142 | 0.117 | 1 | 6 | Thap12 | 0.025 | Young |
| 0.243193 | 0.007276 | 0.082 | 0.106 | 1 | 6 | Ap1g2 | -0.024 | Young |
| 0.24342 | 0.015412 | 0.789 | 0.731 | 1 | 6 | Ier5 | 0.058 | Young |
| 0.2436 | 0.128368 | 0.269 | 0.231 | 1 | 6 | Aup1 | 0.038 | Young |
| 0.243746 | 0.365343 | 0.142 | 0.117 | 1 | 6 | Pwwp2a | 0.025 | Young |
| 0.243892 | 0.510801 | 0.04 | 0.026 | 1 | 6 | Rab3ip | 0.014 | Young |
| 0.243917 | 0.183594 | 0.231 | 0.206 | 1 | 6 | Kdm5c | 0.025 | Young |
| 0.244453 | 1.118441 | 0.022 | 0.011 | 1 | 6 | Gm14286 | 0.011 | Young |
| 0.244852 | 0.181594 | 0.264 | 0.231 | 1 | 6 | Vav1 | 0.033 | Young |
| 0.245457 | 0.155012 | 0.647 | 0.614 | 1 | 6 | Smdt1 | 0.033 | Young |
| 0.245756 | 1.111374 | 0.022 | 0.011 | 1 | 6 | Plagl1 | 0.011 | Young |
| 0.245756 | 0.879362 | 0.022 | 0.011 | 1 | 6 | Dusp28 | 0.011 | Young |
| 0.245828 | 1.16062 | 0.043 | 0.029 | 1 | 6 | Gm15232 | 0.014 | Young |
| 0.24633 | 0.125745 | 0.164 | 0.134 | 1 | 6 | Tgfbr2 | 0.03 | Young |
| 0.246541 | 0.55146 | 0.102 | 0.08 | 1 | 6 | Kctd17 | 0.022 | Young |
| 0.246627 | 0.744743 | 0.022 | 0.011 | 1 | 6 | Ubxn11 | 0.011 | Young |
| 0.246727 | 0.144141 | 0.525 | 0.491 | 1 | 6 | Marveld1 | 0.034 | Young |
| 0.246821 | 0.412171 | 0.132 | 0.109 | 1 | 6 | Dck | 0.023 | Young |
| 0.247038 | 0.41373 | 0.043 | 0.029 | 1 | 6 | Plekhn1 | 0.014 | Young |
| 0.247692 | 0.140164 | 0.164 | 0.137 | 1 | 6 | Tlnrd1 | 0.027 | Young |
| 0.247913 | 0.090465 | 0.602 | 0.569 | 1 | 6 | Mat2a | 0.033 | Young |
| 0.248132 | 0.060991 | 0.428 | 0.389 | 1 | 6 | Pdlim5 | 0.039 | Young |
| 0.248376 | 1.016069 | 0.022 | 0.011 | 1 | 6 | Cnnm2 | 0.011 | Young |
| 0.248376 | 0.583988 | 0.022 | 0.011 | 1 | 6 | Tbc1d8b | 0.011 | Young |
| 0.248814 | 0.768927 | 0.022 | 0.011 | 1 | 6 | Smg8 | 0.011 | Young |
| 0.249014 | 0.092135 | 0.413 | 0.38 | 1 | 6 | Adgre1 | 0.033 | Young |
| 0.249053 | 0.6305 | 0.06 | 0.043 | 1 | 6 | Rnf8 | 0.017 | Young |
| 0.249114 | 0.27223 | 0.028 | 0.043 | 1 | 6 | Plcxd2 | -0.015 | Young |
| 0.249522 | 0.322976 | 0.316 | 0.291 | 1 | 6 | Nfe2l1 | 0.025 | Young |
| 0.249677 | 0.229058 | 0.07 | 0.051 | 1 | 6 | Rbmx | 0.019 | Young |

| | | | | | | | | |
|---|---|---|---|---|---|---|---|---|
| 0.249925 | 0.174504 | 0.475 | 0.449 | 1 | 6 | Ndufc2 | 0.026 | Young |
| 0.250133 | 0.549028 | 0.022 | 0.011 | 1 | 6 | Tspyl3 | 0.011 | Young |
| 0.250313 | 0.082571 | 0.125 | 0.1 | 1 | 6 | Ezh1 | 0.025 | Young |
| 0.250313 | 0.055805 | 0.125 | 0.1 | 1 | 6 | H2-Ke6 | 0.025 | Young |
| 0.250514 | 0.664258 | 0.054 | 0.037 | 1 | 6 | 1700019L1 | 0.017 | Young |
| 0.250574 | 0.635055 | 0.022 | 0.011 | 1 | 6 | Elmod3 | 0.011 | Young |
| 0.250935 | 0.317103 | 0.234 | 0.206 | 1 | 6 | Fndc3a | 0.028 | Young |
| 0.250957 | 0.18842 | 0.007 | 0.014 | 1 | 6 | Smo | -0.007 | Young |
| 0.251015 | 0.380695 | 0.022 | 0.011 | 1 | 6 | Ccdc73 | 0.011 | Young |
| 0.25137 | 0.311239 | 0.084 | 0.063 | 1 | 6 | Nudt16 | 0.021 | Young |
| 0.251579 | 0.303136 | 0.077 | 0.057 | 1 | 6 | Dnlz | 0.02 | Young |
| 0.251646 | 0.407136 | 0.214 | 0.189 | 1 | 6 | Imp3 | 0.025 | Young |
| 0.251899 | 0.675841 | 0.022 | 0.011 | 1 | 6 | Cx3cl1 | 0.011 | Young |
| 0.252073 | 0.228825 | 0.289 | 0.26 | 1 | 6 | Ryr1 | 0.029 | Young |
| 0.252503 | 0.339525 | 0.152 | 0.126 | 1 | 6 | Fam98c | 0.026 | Young |
| 0.252923 | 0.491893 | 0.105 | 0.083 | 1 | 6 | Strip2 | 0.022 | Young |
| 0.253093 | 0.483801 | 0.057 | 0.04 | 1 | 6 | Il1a | 0.017 | Young |
| 0.254077 | 0.33594 | 0.104 | 0.083 | 1 | 6 | Tstd3 | 0.021 | Young |
| 0.254392 | 0.097687 | 0.122 | 0.097 | 1 | 6 | Sc5d | 0.025 | Young |
| 0.254397 | 0.03008 | 0.278 | 0.246 | 1 | 6 | Bin1 | 0.032 | Young |
| 0.254481 | 0.199288 | 0.015 | 0.026 | 1 | 6 | Smyd2 | -0.011 | Young |
| 0.254643 | 0.290817 | 0.164 | 0.137 | 1 | 6 | Nmt1 | 0.027 | Young |
| 0.254816 | 0.257447 | 0.157 | 0.131 | 1 | 6 | Fxr2 | 0.026 | Young |
| 0.254982 | 0.164661 | 0.142 | 0.114 | 1 | 6 | Oser1 | 0.028 | Young |
| 0.255024 | 0.21949 | 0.047 | 0.031 | 1 | 6 | Sfmbt1 | 0.016 | Young |
| 0.255226 | 0.11114 | 0.351 | 0.32 | 1 | 6 | Itgax | 0.031 | Young |
| 0.255228 | 0.230617 | 0.117 | 0.094 | 1 | 6 | Stk38l | 0.023 | Young |
| 0.255355 | 0.218808 | 0.293 | 0.269 | 1 | 6 | Ndufv2 | 0.024 | Young |
| 0.255524 | 0.026672 | 0.443 | 0.406 | 1 | 6 | Camk1d | 0.037 | Young |
| 0.255938 | 0.172355 | 0.538 | 0.506 | 1 | 6 | Eef1d | 0.032 | Young |
| 0.256033 | 0.295618 | 0.05 | 0.034 | 1 | 6 | Atg4c | 0.016 | Young |
| 0.256396 | 0.332437 | 0.204 | 0.18 | 1 | 6 | Dpp8 | 0.024 | Young |
| 0.256569 | 1.214085 | 0.025 | 0.014 | 1 | 6 | Ift80 | 0.011 | Young |
| 0.256572 | 0.446576 | 0.212 | 0.189 | 1 | 6 | Synrg | 0.023 | Young |
| 0.256984 | 1.2178 | 0.025 | 0.014 | 1 | 6 | Zbtb45 | 0.011 | Young |
| 0.25706 | 0.215465 | 0.365 | 0.329 | 1 | 6 | Atp13a3 | 0.036 | Young |
| 0.257065 | 0.135481 | 0.094 | 0.071 | 1 | 6 | Scmh1 | 0.023 | Young |
| 0.258 | 0.143159 | 0.316 | 0.28 | 1 | 6 | Crk | 0.036 | Young |
| 0.258113 | 0.430925 | 0.199 | 0.171 | 1 | 6 | Tbl1x | 0.028 | Young |
| 0.258318 | 0.023957 | 0.505 | 0.474 | 1 | 6 | Tardbp | 0.031 | Young |
| 0.258717 | 0.135564 | 0.431 | 0.4 | 1 | 6 | Azin1 | 0.031 | Young |
| 0.259794 | 0.110328 | 0.06 | 0.043 | 1 | 6 | Ctbs | 0.017 | Young |
| 0.260204 | 0.297677 | 0.271 | 0.246 | 1 | 6 | Pigx | 0.025 | Young |
| 0.260314 | 1.622812 | 0.025 | 0.014 | 1 | 6 | Prdm15 | 0.011 | Young |
| 0.260314 | 0.893862 | 0.025 | 0.014 | 1 | 6 | Det1 | 0.011 | Young |

| | | | | | | | | |
|---|---|---|---|---|---|---|---|---|
| 0.260732 | 0.837181 | 0.025 | 0.014 | 1 | 6 | Mpi | 0.011 | Young |
| 0.261136 | 0.237486 | 0.268 | 0.24 | 1 | 6 | Tmbim1 | 0.028 | Young |
| 0.261151 | 0.830276 | 0.025 | 0.014 | 1 | 6 | Pigf | 0.011 | Young |
| 0.261425 | 1.445341 | 0.013 | 0.006 | 1 | 6 | Gja1 | 0.007 | Young |
| 0.261895 | 0.357847 | 0.288 | 0.266 | 1 | 6 | Phlda1 | 0.022 | Young |
| 0.262016 | 1.00795 | 0.013 | 0.006 | 1 | 6 | Lilra5 | 0.007 | Young |
| 0.262027 | 0.448435 | 0.172 | 0.149 | 1 | 6 | Wdr3 | 0.023 | Young |
| 0.26232 | 0.211111 | 0.226 | 0.2 | 1 | 6 | Actr1a | 0.026 | Young |
| 0.26241 | 0.760269 | 0.025 | 0.014 | 1 | 6 | Mocs1 | 0.011 | Young |
| 0.262549 | 0.017397 | 0.187 | 0.157 | 1 | 6 | Frrs1 | 0.03 | Young |
| 0.262573 | 0.124037 | 0.478 | 0.457 | 1 | 6 | Ndufb11 | 0.021 | Young |
| 0.262743 | 0.484999 | 0.102 | 0.08 | 1 | 6 | Strn | 0.022 | Young |
| 0.262832 | 0.189726 | 0.196 | 0.166 | 1 | 6 | Nfkb2 | 0.03 | Young |
| 0.262876 | 0.648405 | 0.075 | 0.057 | 1 | 6 | Hspbap1 | 0.018 | Young |
| 0.26304 | 0.398757 | 0.202 | 0.18 | 1 | 6 | Skiv2l | 0.022 | Young |
| 0.263083 | 0.032349 | 0.139 | 0.114 | 1 | 6 | Ccdc6 | 0.025 | Young |
| 0.263083 | 0.278 | 0.139 | 0.114 | 1 | 6 | Frmd8 | 0.025 | Young |
| 0.263199 | 1.613372 | 0.013 | 0.006 | 1 | 6 | Dpys | 0.007 | Young |
| 0.263199 | 1.146987 | 0.013 | 0.006 | 1 | 6 | Gm26756 | 0.007 | Young |
| 0.263199 | 1.077409 | 0.013 | 0.006 | 1 | 6 | Tmem201 | 0.007 | Young |
| 0.263283 | 0.038647 | 0.258 | 0.226 | 1 | 6 | Mllt10 | 0.032 | Young |
| 0.263304 | 0.13968 | 0.236 | 0.206 | 1 | 6 | Zfp800 | 0.03 | Young |
| 0.26354 | 0.148 | 0.366 | 0.337 | 1 | 6 | Ppig | 0.029 | Young |
| 0.263892 | 0.272875 | 0.074 | 0.054 | 1 | 6 | Urod | 0.02 | Young |
| 0.264309 | 0.048403 | 0.9 | 0.846 | 1 | 6 | Sh3bgrl3 | 0.054 | Young |
| 0.264386 | 1.143888 | 0.013 | 0.006 | 1 | 6 | Cracr2b | 0.007 | Young |
| 0.264386 | 1.141522 | 0.013 | 0.006 | 1 | 6 | Prmt6 | 0.007 | Young |
| 0.264386 | 0.826817 | 0.013 | 0.006 | 1 | 6 | Nhej1 | 0.007 | Young |
| 0.264518 | 0.30736 | 0.025 | 0.014 | 1 | 6 | Rbm4 | 0.011 | Young |
| 0.264817 | 0.201652 | 0.054 | 0.037 | 1 | 6 | Runx2 | 0.017 | Young |
| 0.264837 | 0.309178 | 0.157 | 0.131 | 1 | 6 | Mrps9 | 0.026 | Young |
| 0.264981 | 1.440339 | 0.013 | 0.006 | 1 | 6 | 9430015G1 | 0.007 | Young |
| 0.264981 | 1.345246 | 0.013 | 0.006 | 1 | 6 | Ldlrad3 | 0.007 | Young |
| 0.264981 | 0.968289 | 0.013 | 0.006 | 1 | 6 | 1700056N1 | 0.007 | Young |
| 0.264987 | 0.944058 | 0.065 | 0.049 | 1 | 6 | Miip | 0.016 | Young |
| 0.265144 | 0.252302 | 0.095 | 0.074 | 1 | 6 | Hist3h2a | 0.021 | Young |
| 0.265258 | 0.043585 | 0.092 | 0.071 | 1 | 6 | Amz2 | 0.021 | Young |
| 0.265577 | 0.594619 | 0.013 | 0.006 | 1 | 6 | Piwil2 | 0.007 | Young |
| 0.265716 | 0.539012 | 0.082 | 0.063 | 1 | 6 | Scaf1 | 0.019 | Young |
| 0.265788 | 0.517213 | 0.025 | 0.014 | 1 | 6 | Gtf2h2 | 0.011 | Young |
| 0.266114 | 0.230056 | 0.166 | 0.14 | 1 | 6 | Tmem57 | 0.026 | Young |
| 0.266174 | 0.746144 | 0.013 | 0.006 | 1 | 6 | Ccdc138 | 0.007 | Young |
| 0.266174 | 0.71417 | 0.013 | 0.006 | 1 | 6 | Fahd2a | 0.007 | Young |
| 0.266174 | 0.639544 | 0.013 | 0.006 | 1 | 6 | Pdlim7 | 0.007 | Young |
| 0.266281 | 0.240253 | 0.204 | 0.174 | 1 | 6 | Unc119b | 0.03 | Young |

| | | | | | | | | |
|---|---|---|---|---|---|---|---|---|
| 0.266335 | 0.187201 | 0.436 | 0.406 | 1 | 6 | Gpcpd1 | 0.03 | Young |
| 0.267006 | 0.199185 | 0.199 | 0.171 | 1 | 6 | Crybg3 | 0.028 | Young |
| 0.267633 | 0.131295 | 0.159 | 0.194 | 1 | 6 | Cnp | -0.035 | Young |
| 0.26769 | 0.555765 | 0.075 | 0.057 | 1 | 6 | Prkaa1 | 0.018 | Young |
| 0.267966 | 0.257722 | 0.304 | 0.277 | 1 | 6 | Map3k8 | 0.027 | Young |
| 0.26797 | 0.737133 | 0.013 | 0.006 | 1 | 6 | Zfp446 | 0.007 | Young |
| 0.26797 | 0.324244 | 0.013 | 0.006 | 1 | 6 | Ift43 | 0.007 | Young |
| 0.268773 | 0.555209 | 0.122 | 0.1 | 1 | 6 | Zdhhc5 | 0.022 | Young |
| 0.268805 | 0.316625 | 0.072 | 0.054 | 1 | 6 | Stx17 | 0.018 | Young |
| 0.268917 | 0.017979 | 0.119 | 0.146 | 1 | 6 | Sucla2 | -0.027 | Young |
| 0.268941 | 0.132121 | 0.06 | 0.043 | 1 | 6 | Scamp1 | 0.017 | Young |
| 0.26903 | 0.902462 | 0.028 | 0.017 | 1 | 6 | Tmod1 | 0.011 | Young |
| 0.269166 | 0.060167 | 0.617 | 0.563 | 1 | 6 | Ndel1 | 0.054 | Young |
| 0.269197 | 0.288959 | 0.025 | 0.014 | 1 | 6 | Cul7 | 0.011 | Young |
| 0.269461 | 0.44004 | 0.125 | 0.103 | 1 | 6 | Plpp2 | 0.022 | Young |
| 0.26979 | 0.464598 | 0.289 | 0.266 | 1 | 6 | Golph3 | 0.023 | Young |
| 0.270053 | 0.169093 | 0.025 | 0.014 | 1 | 6 | Diras2 | 0.011 | Young |
| 0.270067 | 0.266766 | 0.286 | 0.257 | 1 | 6 | Entpd1 | 0.029 | Young |
| 0.270089 | 0.057259 | 0.132 | 0.106 | 1 | 6 | Rbm12 | 0.026 | Young |
| 0.270097 | 1.794118 | 0.035 | 0.023 | 1 | 6 | Mad1l1 | 0.012 | Young |
| 0.270286 | 0.375072 | 0.182 | 0.157 | 1 | 6 | Stim2 | 0.025 | Young |
| 0.270406 | 0.193421 | 0.181 | 0.151 | 1 | 6 | Impact | 0.03 | Young |
| 0.270482 | 0.123798 | 0.025 | 0.014 | 1 | 6 | Megf9 | 0.011 | Young |
| 0.270688 | 0.118649 | 0.269 | 0.237 | 1 | 6 | Npm3 | 0.032 | Young |
| 0.270983 | 0.402536 | 0.154 | 0.131 | 1 | 6 | Dnajc14 | 0.023 | Young |
| 0.271355 | 0.038072 | 0.806 | 0.76 | 1 | 6 | S100a10 | 0.046 | Young |
| 0.272612 | 0.302151 | 0.11 | 0.089 | 1 | 6 | Ak3 | 0.021 | Young |
| 0.273582 | 0.066757 | 0.122 | 0.151 | 1 | 6 | Atraid | -0.029 | Young |
| 0.273775 | 0.406275 | 0.157 | 0.134 | 1 | 6 | Bag6 | 0.023 | Young |
| 0.274184 | 0.422125 | 0.109 | 0.089 | 1 | 6 | Mafk | 0.02 | Young |
| 0.274424 | 1.001603 | 0.032 | 0.02 | 1 | 6 | Kat8 | 0.012 | Young |
| 0.274874 | 0.339572 | 0.097 | 0.077 | 1 | 6 | Tmem43 | 0.02 | Young |
| 0.274936 | 0.314263 | 0.177 | 0.154 | 1 | 6 | Lsm7 | 0.023 | Young |
| 0.275133 | 0.049774 | 0.358 | 0.32 | 1 | 6 | Arih2 | 0.038 | Young |
| 0.275251 | 0.219447 | 0.452 | 0.42 | 1 | 6 | Elob | 0.032 | Young |
| 0.27557 | 0.752137 | 0.032 | 0.02 | 1 | 6 | Mvb12b | 0.012 | Young |
| 0.27562 | 0.213797 | 0.182 | 0.154 | 1 | 6 | Afdn | 0.028 | Young |
| 0.275665 | 0.016455 | 0.135 | 0.111 | 1 | 6 | Virma | 0.024 | Young |
| 0.276209 | 0.348566 | 0.137 | 0.114 | 1 | 6 | Babam1 | 0.023 | Young |
| 0.276247 | 0.41788 | 0.179 | 0.154 | 1 | 6 | Diaph2 | 0.025 | Young |
| 0.276693 | 0.638545 | 0.028 | 0.017 | 1 | 6 | Cox11 | 0.011 | Young |
| 0.276925 | 0.230601 | 0.11 | 0.089 | 1 | 6 | Usp32 | 0.021 | Young |
| 0.276958 | 0.290421 | 0.236 | 0.211 | 1 | 6 | Cndp2 | 0.025 | Young |
| 0.277033 | 0.264342 | 0.117 | 0.094 | 1 | 6 | Foxk2 | 0.023 | Young |
| 0.277262 | 0.137587 | 0.179 | 0.151 | 1 | 6 | Ireb2 | 0.028 | Young |

| | | | | | | | | |
|---|---|---|---|---|---|---|---|---|
| 0.277309 | 0.854409 | 0.055 | 0.04 | 1 | 6 | Pccb | 0.015 | Young |
| 0.277429 | 0.099218 | 0.154 | 0.129 | 1 | 6 | Fam133b | 0.025 | Young |
| 0.277611 | 0.250974 | 0.079 | 0.06 | 1 | 6 | Pogk | 0.019 | Young |
| 0.277856 | 0.301302 | 0.11 | 0.089 | 1 | 6 | Zfyve16 | 0.021 | Young |
| 0.278428 | 0.446068 | 0.129 | 0.106 | 1 | 6 | Fibp | 0.023 | Young |
| 0.278436 | 0.151576 | 0.612 | 0.574 | 1 | 6 | Cct5 | 0.038 | Young |
| 0.278911 | 0.155484 | 0.271 | 0.237 | 1 | 6 | Aimp1 | 0.034 | Young |
| 0.279414 | 1.173105 | 0.032 | 0.02 | 1 | 6 | Acvr2b | 0.012 | Young |
| 0.279512 | 1.027373 | 0.035 | 0.023 | 1 | 6 | Cd209b | 0.012 | Young |
| 0.279691 | 0.189769 | 0.254 | 0.229 | 1 | 6 | Arl2bp | 0.025 | Young |
| 0.279864 | 0.993291 | 0.048 | 0.034 | 1 | 6 | Zmym3 | 0.014 | Young |
| 0.280187 | 0.753538 | 0.032 | 0.02 | 1 | 6 | Fam206a | 0.012 | Young |
| 0.280572 | 0.39094 | 0.013 | 0.023 | 1 | 6 | Mdm1 | -0.01 | Young |
| 0.280803 | 0.568955 | 0.097 | 0.077 | 1 | 6 | Abca1 | 0.02 | Young |
| 0.281011 | 0.113897 | 0.666 | 0.617 | 1 | 6 | Tspo | 0.049 | Young |
| 0.281039 | 0.675911 | 0.065 | 0.049 | 1 | 6 | Mcee | 0.016 | Young |
| 0.281442 | 0.871688 | 0.038 | 0.026 | 1 | 6 | Slc38a7 | 0.012 | Young |
| 0.281476 | 0.061773 | 0.831 | 0.829 | 1 | 6 | Ybx1 | 0.002 | Young |
| 0.282084 | 0.850132 | 0.035 | 0.023 | 1 | 6 | Srrd | 0.012 | Young |
| 0.282115 | 0.56047 | 0.065 | 0.049 | 1 | 6 | Nfxl1 | 0.016 | Young |
| 0.282126 | 1.05293 | 0.042 | 0.029 | 1 | 6 | Abhd5 | 0.013 | Young |
| 0.282453 | 1.112274 | 0.035 | 0.023 | 1 | 6 | Gm44751 | 0.012 | Young |
| 0.283024 | 0.038536 | 0.368 | 0.323 | 1 | 6 | Nectin1 | 0.045 | Young |
| 0.283602 | 0.299958 | 0.206 | 0.18 | 1 | 6 | Alkbh1 | 0.026 | Young |
| 0.283675 | 0.252573 | 0.028 | 0.017 | 1 | 6 | Nlrp10 | 0.011 | Young |
| 0.283684 | 0.327903 | 0.032 | 0.02 | 1 | 6 | Ercc1 | 0.012 | Young |
| 0.283762 | 0.927168 | 0.048 | 0.034 | 1 | 6 | Myo7a | 0.014 | Young |
| 0.283776 | 0.260703 | 0.314 | 0.283 | 1 | 6 | Csnk1d | 0.031 | Young |
| 0.28417 | 0.717716 | 0.069 | 0.051 | 1 | 6 | Agfg2 | 0.018 | Young |
| 0.284519 | 0.286555 | 0.013 | 0.023 | 1 | 6 | Hemk1 | -0.01 | Young |
| 0.284646 | 0.067231 | 0.5 | 0.471 | 1 | 6 | Nop53 | 0.029 | Young |
| 0.284909 | 0.482083 | 0.082 | 0.063 | 1 | 6 | Fech | 0.019 | Young |
| 0.284967 | 0.955937 | 0.038 | 0.026 | 1 | 6 | Dcun1d2 | 0.012 | Young |
| 0.285247 | 0.484466 | 0.032 | 0.02 | 1 | 6 | Sox12 | 0.012 | Young |
| 0.285615 | 0.062471 | 0.391 | 0.351 | 1 | 6 | Nufip2 | 0.04 | Young |
| 0.285791 | 0.814958 | 0.052 | 0.037 | 1 | 6 | Riok2 | 0.015 | Young |
| 0.286385 | 0.64423 | 0.08 | 0.063 | 1 | 6 | Smox | 0.017 | Young |
| 0.286445 | 0.568536 | 0.151 | 0.126 | 1 | 6 | Lpar1 | 0.025 | Young |
| 0.286644 | 0.22228 | 0.162 | 0.137 | 1 | 6 | Snrk | 0.025 | Young |
| 0.28667 | 0.744493 | 0.059 | 0.043 | 1 | 6 | Hps1 | 0.016 | Young |
| 0.287616 | 0.343148 | 0.069 | 0.051 | 1 | 6 | Grasp | 0.018 | Young |
| 0.28764 | 0.142308 | 0.403 | 0.363 | 1 | 6 | Ndufb3 | 0.04 | Young |
| 0.287651 | 0.105291 | 0.176 | 0.149 | 1 | 6 | Erap1 | 0.027 | Young |
| 0.288031 | 0.685277 | 0.045 | 0.031 | 1 | 6 | Mthfr | 0.014 | Young |
| 0.28831 | 0.112298 | 0.503 | 0.474 | 1 | 6 | Mef2a | 0.029 | Young |

| | | | | | | | | |
|---|---|---|---|---|---|---|---|---|
| 0.28836 | 0.836048 | 0.045 | 0.031 | 1 | 6 | Slc27a4 | 0.014 | Young |
| 0.288436 | 0.31847 | 0.122 | 0.1 | 1 | 6 | Kpna3 | 0.022 | Young |
| 0.288565 | 0.09776 | 0.609 | 0.586 | 1 | 6 | Atp6v0b | 0.023 | Young |
| 0.288617 | 0.168411 | 0.37 | 0.337 | 1 | 6 | Naa50 | 0.033 | Young |
| 0.288982 | 0.299413 | 0.283 | 0.254 | 1 | 6 | Dcun1d5 | 0.029 | Young |
| 0.289452 | 0.12058 | 0.323 | 0.294 | 1 | 6 | Sh3kbp1 | 0.029 | Young |
| 0.289819 | 1.166666 | 0.017 | 0.009 | 1 | 6 | Csrp2 | 0.008 | Young |
| 0.289851 | 0.412414 | 0.152 | 0.129 | 1 | 6 | Btaf1 | 0.023 | Young |
| 0.289983 | 0.18065 | 0.119 | 0.097 | 1 | 6 | Swt1 | 0.022 | Young |
| 0.290009 | 0.121357 | 0.555 | 0.534 | 1 | 6 | Hmgb1 | 0.021 | Young |
| 0.290258 | 0.164683 | 0.189 | 0.163 | 1 | 6 | Gtf2b | 0.026 | Young |
| 0.290284 | 0.824134 | 0.042 | 0.029 | 1 | 6 | Mmgt2 | 0.013 | Young |
| 0.290359 | 0.130406 | 0.403 | 0.377 | 1 | 6 | Etfb | 0.026 | Young |
| 0.290553 | 0.509315 | 0.069 | 0.051 | 1 | 6 | Zfp597 | 0.018 | Young |
| 0.290707 | 0.522478 | 0.059 | 0.043 | 1 | 6 | Clec4n | 0.016 | Young |
| 0.290783 | 0.151173 | 0.095 | 0.074 | 1 | 6 | Ice1 | 0.021 | Young |
| 0.29087 | 0.717958 | 0.048 | 0.034 | 1 | 6 | Nipal3 | 0.014 | Young |
| 0.291613 | 0.302953 | 0.052 | 0.037 | 1 | 6 | Akr1c13 | 0.015 | Young |
| 0.291745 | 0.676136 | 0.038 | 0.026 | 1 | 6 | St7 | 0.012 | Young |
| 0.292314 | 0.499272 | 0.055 | 0.04 | 1 | 6 | Plxna4os1 | 0.015 | Young |
| 0.292325 | 0.173135 | 0.361 | 0.34 | 1 | 6 | Fam168b | 0.021 | Young |
| 0.292575 | 0.733257 | 0.13 | 0.111 | 1 | 6 | Tmem243 | 0.019 | Young |
| 0.293269 | 0.210906 | 0.202 | 0.174 | 1 | 6 | Celf4 | 0.028 | Young |
| 0.293301 | 0.591176 | 0.08 | 0.063 | 1 | 6 | Rnf167 | 0.017 | Young |
| 0.293802 | 0.155383 | 0.393 | 0.374 | 1 | 6 | Ppp1cb | 0.019 | Young |
| 0.293908 | 0.352772 | 0.038 | 0.026 | 1 | 6 | Engase | 0.012 | Young |
| 0.29468 | 0.428101 | 0.065 | 0.049 | 1 | 6 | Hsdl1 | 0.016 | Young |
| 0.295116 | 0.762309 | 0.042 | 0.029 | 1 | 6 | Arhgap22 | 0.013 | Young |
| 0.295219 | 0.770607 | 0.115 | 0.097 | 1 | 6 | Srd5a3 | 0.018 | Young |
| 0.295286 | 0.082934 | 0.666 | 0.614 | 1 | 6 | Sgms1 | 0.052 | Young |
| 0.295331 | 0.560096 | 0.052 | 0.037 | 1 | 6 | Adsl | 0.015 | Young |
| 0.295355 | 0.143263 | 0.217 | 0.189 | 1 | 6 | Brd1 | 0.028 | Young |
| 0.295415 | 1.141886 | 0.017 | 0.009 | 1 | 6 | Kcnc3 | 0.008 | Young |
| 0.295515 | 0.012649 | 0.062 | 0.046 | 1 | 6 | Aatf | 0.016 | Young |
| 0.295553 | 0.134693 | 0.032 | 0.02 | 1 | 6 | Bahcc1 | 0.012 | Young |
| 0.295706 | 0.477522 | 0.144 | 0.123 | 1 | 6 | Sae1 | 0.021 | Young |
| 0.29572 | 0.339671 | 0.331 | 0.311 | 1 | 6 | Gnai3 | 0.02 | Young |
| 0.295951 | 0.355807 | 0.333 | 0.311 | 1 | 6 | Pdcd5 | 0.022 | Young |
| 0.29607 | 0.099931 | 0.122 | 0.149 | 1 | 6 | Mdp1 | -0.027 | Young |
| 0.296375 | 0.250692 | 0.349 | 0.317 | 1 | 6 | Utrn | 0.032 | Young |
| 0.296456 | 0.268125 | 0.09 | 0.071 | 1 | 6 | Umad1 | 0.019 | Young |
| 0.296542 | 0.610464 | 0.017 | 0.009 | 1 | 6 | Rbks | 0.008 | Young |
| 0.297054 | 0.210216 | 0.144 | 0.12 | 1 | 6 | Spn | 0.024 | Young |
| 0.297202 | 0.35659 | 0.052 | 0.037 | 1 | 6 | Rcan3 | 0.015 | Young |
| 0.297371 | 0.265352 | 0.202 | 0.177 | 1 | 6 | Stk11 | 0.025 | Young |

| | | | | | | | | |
|---|---|---|---|---|---|---|---|---|
| 0.297473 | 0.327667 | 0.035 | 0.023 | 1 | 6 | Slc35e3 | 0.012 | Young |
| 0.297673 | 0.655243 | 0.017 | 0.009 | 1 | 6 | Prag1 | 0.008 | Young |
| 0.297919 | 0.494703 | 0.117 | 0.097 | 1 | 6 | Gtf2e2 | 0.02 | Young |
| 0.297978 | 0.245913 | 0.244 | 0.22 | 1 | 6 | F13a1 | 0.024 | Young |
| 0.298194 | 0.074114 | 0.769 | 0.726 | 1 | 6 | Sec61b | 0.043 | Young |
| 0.29824 | 0.777123 | 0.017 | 0.009 | 1 | 6 | Crocc | 0.008 | Young |
| 0.298807 | 0.743443 | 0.017 | 0.009 | 1 | 6 | Tmem38a | 0.008 | Young |
| 0.2993 | 0.469827 | 0.042 | 0.029 | 1 | 6 | Urgcp | 0.013 | Young |
| 0.299556 | 0.554083 | 0.048 | 0.034 | 1 | 6 | Iffo1 | 0.014 | Young |
| 0.299567 | 0.156228 | 0.468 | 0.44 | 1 | 6 | Arih1 | 0.028 | Young |
| 0.299939 | 0.039014 | 0.662 | 0.6 | 1 | 6 | Itgam | 0.062 | Young |
| 0.299943 | 0.536135 | 0.017 | 0.009 | 1 | 6 | Hnmt | 0.008 | Young |
| 0.299949 | 0.242658 | 0.44 | 0.414 | 1 | 6 | Pepd | 0.026 | Young |
| 0.300028 | 0.049639 | 0.182 | 0.154 | 1 | 6 | Gripap1 | 0.028 | Young |
| 0.300328 | 0.127535 | 0.109 | 0.089 | 1 | 6 | Stoml2 | 0.02 | Young |
| 0.300366 | 0.363546 | 0.197 | 0.174 | 1 | 6 | Pak1 | 0.023 | Young |
| 0.300541 | 0.566928 | 0.102 | 0.083 | 1 | 6 | Cask | 0.019 | Young |
| 0.300627 | 0.266694 | 0.09 | 0.071 | 1 | 6 | Golt1b | 0.019 | Young |
| 0.300671 | 0.16558 | 0.059 | 0.043 | 1 | 6 | Hic2 | 0.016 | Young |
| 0.300781 | 0.486094 | 0.055 | 0.04 | 1 | 6 | Aamdc | 0.015 | Young |
| 0.301668 | 0.238015 | 0.391 | 0.366 | 1 | 6 | Gns | 0.025 | Young |
| 0.301731 | 0.445633 | 0.087 | 0.069 | 1 | 6 | Lcmt1 | 0.018 | Young |
| 0.301834 | 0.119269 | 0.025 | 0.037 | 1 | 6 | Cbx8 | -0.012 | Young |
| 0.301853 | 0.023474 | 0.565 | 0.523 | 1 | 6 | Mcub | 0.042 | Young |
| 0.301985 | 0.022142 | 0.569 | 0.54 | 1 | 6 | Uvrag | 0.029 | Young |
| 0.302085 | 0.376975 | 0.109 | 0.089 | 1 | 6 | Ufd1 | 0.02 | Young |
| 0.302355 | 0.430165 | 0.129 | 0.109 | 1 | 6 | Polr2i | 0.02 | Young |
| 0.302612 | 0.014375 | 0.067 | 0.086 | 1 | 6 | Rab10os | -0.019 | Young |
| 0.303311 | 0.154223 | 0.237 | 0.209 | 1 | 6 | Rb1 | 0.028 | Young |
| 0.303961 | 0.162873 | 0.214 | 0.189 | 1 | 6 | Vti1b | 0.025 | Young |
| 0.304142 | 0.097759 | 0.038 | 0.026 | 1 | 6 | Pld2 | 0.012 | Young |
| 0.304784 | 0.405385 | 0.048 | 0.034 | 1 | 6 | Nbn | 0.014 | Young |
| 0.304808 | 0.226036 | 0.281 | 0.257 | 1 | 6 | Maff | 0.024 | Young |
| 0.3053 | 0.074131 | 0.244 | 0.211 | 1 | 6 | Pik3r5 | 0.033 | Young |
| 0.305451 | 0.023923 | 0.184 | 0.214 | 1 | 6 | Pik3ca | -0.03 | Young |
| 0.305668 | 0.013737 | 0.017 | 0.009 | 1 | 6 | Zmat1 | 0.008 | Young |
| 0.305924 | 0.16196 | 0.515 | 0.503 | 1 | 6 | Klf3 | 0.012 | Young |
| 0.306592 | 0.026106 | 0.413 | 0.377 | 1 | 6 | Top2b | 0.036 | Young |
| 0.306847 | 0.1488 | 0.226 | 0.194 | 1 | 6 | Zfp703 | 0.032 | Young |
| 0.306921 | 0.192445 | 0.137 | 0.114 | 1 | 6 | Arfrp1 | 0.023 | Young |
| 0.307359 | 0.428499 | 0.037 | 0.051 | 1 | 6 | Klri2 | -0.014 | Young |
| 0.307509 | 0.266015 | 0.172 | 0.149 | 1 | 6 | Prdm2 | 0.023 | Young |
| 0.307584 | 0.226061 | 0.151 | 0.126 | 1 | 6 | Fam114a2 | 0.025 | Young |
| 0.307627 | 0.590829 | 0.169 | 0.149 | 1 | 6 | Apobec1 | 0.02 | Young |
| 0.308691 | 0.30118 | 0.346 | 0.323 | 1 | 6 | Ptpro | 0.023 | Young |

| | | | | | | | | |
|---|---|---|---|---|---|---|---|---|
| 0.308894 | 0.00701 | 0.047 | 0.063 | 1 | 6 | Mbtps2 | -0.016 | Young |
| 0.308976 | 0.082487 | 0.515 | 0.483 | 1 | 6 | Snx3 | 0.032 | Young |
| 0.308993 | 0.076912 | 0.219 | 0.189 | 1 | 6 | Nap1l4 | 0.03 | Young |
| 0.309408 | 0.206993 | 0.048 | 0.034 | 1 | 6 | Isca1 | 0.014 | Young |
| 0.309701 | 0.235747 | 0.062 | 0.046 | 1 | 6 | Fnip2 | 0.016 | Young |
| 0.310038 | 1.212651 | 0.02 | 0.011 | 1 | 6 | Zfp995 | 0.009 | Young |
| 0.310162 | 0.14463 | 0.605 | 0.557 | 1 | 6 | Scand1 | 0.048 | Young |
| 0.310534 | 0.018172 | 0.739 | 0.689 | 1 | 6 | Cfp | 0.05 | Young |
| 0.310975 | 0.303918 | 0.125 | 0.103 | 1 | 6 | Spink2 | 0.022 | Young |
| 0.311523 | 0.049277 | 0.851 | 0.743 | 1 | 6 | Syngr2 | 0.108 | Young |
| 0.311729 | 0.142104 | 0.042 | 0.029 | 1 | 6 | Ncapg2 | 0.013 | Young |
| 0.31286 | 0.249534 | 0.087 | 0.069 | 1 | 6 | Gm9493 | 0.018 | Young |
| 0.312994 | 0.069881 | 0.018 | 0.029 | 1 | 6 | Gm12089 | -0.011 | Young |
| 0.31323 | 0.081574 | 0.324 | 0.297 | 1 | 6 | Cisd2 | 0.027 | Young |
| 0.31361 | 0.25294 | 0.222 | 0.197 | 1 | 6 | Tia1 | 0.025 | Young |
| 0.313728 | 1.213317 | 0.02 | 0.011 | 1 | 6 | 5031425F1 | 0.009 | Young |
| 0.313743 | 0.220751 | 0.09 | 0.071 | 1 | 6 | Zfp322a | 0.019 | Young |
| 0.314874 | 0.38766 | 0.161 | 0.137 | 1 | 6 | Efcab14 | 0.024 | Young |
| 0.31498 | 0.165835 | 0.134 | 0.111 | 1 | 6 | Xpa | 0.023 | Young |
| 0.315093 | 0.376768 | 0.11 | 0.091 | 1 | 6 | AU040320 | 0.019 | Young |
| 0.315768 | 0.07953 | 0.674 | 0.62 | 1 | 6 | Sirpa | 0.054 | Young |
| 0.315849 | 1.148474 | 0.02 | 0.011 | 1 | 6 | Kif3a | 0.009 | Young |
| 0.316308 | 0.229076 | 0.142 | 0.12 | 1 | 6 | Copg1 | 0.022 | Young |
| 0.316381 | 1.113306 | 0.02 | 0.011 | 1 | 6 | Gramd1c | 0.009 | Young |
| 0.316523 | 0.107732 | 0.363 | 0.34 | 1 | 6 | Nek7 | 0.023 | Young |
| 0.316675 | 0.077701 | 0.052 | 0.069 | 1 | 6 | Bet1l | -0.017 | Young |
| 0.316757 | 0.597383 | 0.012 | 0.02 | 1 | 6 | Irf7 | -0.008 | Young |
| 0.31687 | 0.470368 | 0.067 | 0.051 | 1 | 6 | Ppih | 0.016 | Young |
| 0.317056 | 0.105885 | 0.59 | 0.554 | 1 | 6 | Psmb1 | 0.036 | Young |
| 0.317148 | 0.03675 | 0.042 | 0.057 | 1 | 6 | Spred2 | -0.015 | Young |
| 0.31754 | 0.498048 | 0.07 | 0.054 | 1 | 6 | Txlng | 0.016 | Young |
| 0.3181 | 0.865381 | 0.07 | 0.054 | 1 | 6 | Trim3 | 0.016 | Young |
| 0.318108 | 0.20487 | 0.018 | 0.029 | 1 | 6 | Ankrd13d | -0.011 | Young |
| 0.318207 | 0.287577 | 0.436 | 0.411 | 1 | 6 | Sdc4 | 0.025 | Young |
| 0.318477 | 0.127261 | 0.355 | 0.326 | 1 | 6 | Rassf2 | 0.029 | Young |
| 0.318514 | 0.752833 | 0.02 | 0.011 | 1 | 6 | Szt2 | 0.009 | Young |
| 0.318739 | 0.795714 | 0.127 | 0.109 | 1 | 6 | Mink1 | 0.018 | Young |
| 0.318789 | 0.272983 | 0.132 | 0.111 | 1 | 6 | Phf1 | 0.021 | Young |
| 0.319584 | 0.034952 | 0.02 | 0.011 | 1 | 6 | Tnni3 | 0.009 | Young |
| 0.320052 | 0.496887 | 0.074 | 0.057 | 1 | 6 | Bmf | 0.017 | Young |
| 0.320147 | 0.41638 | 0.077 | 0.06 | 1 | 6 | Chmp6 | 0.017 | Young |
| 0.320656 | 1.099274 | 0.02 | 0.011 | 1 | 6 | Frat1 | 0.009 | Young |
| 0.321167 | 0.133369 | 0.301 | 0.277 | 1 | 6 | Swap70 | 0.024 | Young |
| 0.321798 | 0.104091 | 0.46 | 0.42 | 1 | 6 | Ythdc1 | 0.04 | Young |
| 0.322269 | 0.04747 | 0.02 | 0.011 | 1 | 6 | Abcb6 | 0.009 | Young |

| | | | | | | | | |
|---|---|---|---|---|---|---|---|---|
| 0.322372 | 0.082941 | 0.808 | 0.766 | 1 | 6 | Rbm3 | 0.042 | Young |
| 0.322697 | 0.128204 | 0.124 | 0.103 | 1 | 6 | Fam96b | 0.021 | Young |
| 0.323173 | 0.097012 | 0.105 | 0.086 | 1 | 6 | Wwc2 | 0.019 | Young |
| 0.32329 | 1.344724 | 0.023 | 0.014 | 1 | 6 | Slc30a6 | 0.009 | Young |
| 0.323721 | 0.186801 | 0.095 | 0.077 | 1 | 6 | Slc2a6 | 0.018 | Young |
| 0.323828 | 0.209212 | 0.115 | 0.094 | 1 | 6 | Immp1l | 0.021 | Young |
| 0.324835 | 0.140599 | 0.234 | 0.209 | 1 | 6 | Madd | 0.025 | Young |
| 0.3249 | 0.097131 | 0.293 | 0.271 | 1 | 6 | Il6st | 0.022 | Young |
| 0.325154 | 0.428847 | 0.07 | 0.054 | 1 | 6 | Wdr46 | 0.016 | Young |
| 0.325268 | 0.138356 | 0.276 | 0.249 | 1 | 6 | Med28 | 0.027 | Young |
| 0.325678 | 0.07587 | 0.202 | 0.177 | 1 | 6 | Bsdc1 | 0.025 | Young |
| 0.325725 | 0.009536 | 0.077 | 0.097 | 1 | 6 | Med17 | -0.02 | Young |
| 0.32614 | 0.095205 | 0.074 | 0.057 | 1 | 6 | Gtpbp6 | 0.017 | Young |
| 0.326244 | 0.067556 | 0.293 | 0.266 | 1 | 6 | Ccdc88b | 0.027 | Young |
| 0.326919 | 0.265552 | 0.105 | 0.086 | 1 | 6 | Map3k5 | 0.019 | Young |
| 0.326945 | 0.118146 | 0.761 | 0.714 | 1 | 6 | Unc93b1 | 0.047 | Young |
| 0.327906 | 0.948838 | 0.043 | 0.031 | 1 | 6 | Btf3l4 | 0.012 | Young |
| 0.328097 | 0.368149 | 0.089 | 0.071 | 1 | 6 | Ifitm5 | 0.018 | Young |
| 0.328723 | 0.036781 | 0.164 | 0.14 | 1 | 6 | Foxj2 | 0.024 | Young |
| 0.329159 | 0.391877 | 0.06 | 0.046 | 1 | 6 | Kifc3 | 0.014 | Young |
| 0.329244 | 0.323215 | 0.194 | 0.171 | 1 | 6 | Lacc1 | 0.023 | Young |
| 0.329422 | 0.263773 | 0.385 | 0.374 | 1 | 6 | Dapp1 | 0.011 | Young |
| 0.329582 | 0.04517 | 0.426 | 0.403 | 1 | 6 | Klf13 | 0.023 | Young |
| 0.329665 | 0.214958 | 0.162 | 0.14 | 1 | 6 | Fam76a | 0.022 | Young |
| 0.329846 | 0.655134 | 0.079 | 0.063 | 1 | 6 | Cep128 | 0.016 | Young |
| 0.330215 | 0.460288 | 0.122 | 0.106 | 1 | 6 | Tshz1 | 0.016 | Young |
| 0.33027 | 0.146146 | 0.119 | 0.097 | 1 | 6 | Mpp1 | 0.022 | Young |
| 0.33035 | 0.300116 | 0.077 | 0.06 | 1 | 6 | Prkra | 0.017 | Young |
| 0.330575 | 0.148571 | 0.284 | 0.254 | 1 | 6 | Sod2 | 0.03 | Young |
| 0.331047 | 0.080095 | 0.326 | 0.3 | 1 | 6 | Pptc7 | 0.026 | Young |
| 0.331202 | 0.074352 | 0.766 | 0.771 | 1 | 6 | Dock10 | -0.005 | Young |
| 0.331257 | 0.573152 | 0.057 | 0.043 | 1 | 6 | Saysd1 | 0.014 | Young |
| 0.331733 | 0.325009 | 0.07 | 0.054 | 1 | 6 | Nfia | 0.016 | Young |
| 0.331794 | 0.772027 | 0.023 | 0.014 | 1 | 6 | Klrg2 | 0.009 | Young |
| 0.33186 | 0.119834 | 0.064 | 0.049 | 1 | 6 | Rfx5 | 0.015 | Young |
| 0.331887 | 0.148607 | 0.41 | 0.394 | 1 | 6 | Bri3bp | 0.016 | Young |
| 0.332299 | 0.519812 | 0.023 | 0.014 | 1 | 6 | Cdan1 | 0.009 | Young |
| 0.332998 | 0.19198 | 0.102 | 0.083 | 1 | 6 | Ubxn7 | 0.019 | Young |
| 0.333043 | 0.487702 | 0.147 | 0.129 | 1 | 6 | Oaz2 | 0.018 | Young |
| 0.333108 | 0.162288 | 0.309 | 0.286 | 1 | 6 | Ddx21 | 0.023 | Young |
| 0.333146 | 0.01703 | 0.222 | 0.191 | 1 | 6 | Stxbp6 | 0.031 | Young |
| 0.333173 | 0.776214 | 0.057 | 0.043 | 1 | 6 | Sirt3 | 0.014 | Young |
| 0.333192 | 0.065742 | 0.035 | 0.049 | 1 | 6 | Helz2 | -0.014 | Young |
| 0.33329 | 0.274178 | 0.074 | 0.057 | 1 | 6 | Comtd1 | 0.017 | Young |
| 0.333797 | 0.010191 | 0.271 | 0.234 | 1 | 6 | Zfp644 | 0.037 | Young |

| | | | | | | | | |
|---|---|---|---|---|---|---|---|---|
| 0.333928 | 0.230275 | 0.214 | 0.191 | 1 | 6 | Cbx1 | 0.023 | Young |
| 0.334284 | 0.201882 | 0.064 | 0.049 | 1 | 6 | Xlr4b | 0.015 | Young |
| 0.334323 | 0.225078 | 0.023 | 0.014 | 1 | 6 | Gstt3 | 0.009 | Young |
| 0.334588 | 0.422255 | 0.064 | 0.049 | 1 | 6 | Pag1 | 0.015 | Young |
| 0.33462 | 0.104782 | 0.467 | 0.434 | 1 | 6 | Glg1 | 0.033 | Young |
| 0.334851 | 0.74577 | 0.145 | 0.129 | 1 | 6 | Tmem131 | 0.016 | Young |
| 0.335171 | 0.29479 | 0.082 | 0.066 | 1 | 6 | Zfp260 | 0.016 | Young |
| 0.335338 | 0.578065 | 0.023 | 0.014 | 1 | 6 | Tubb2b | 0.009 | Young |
| 0.335371 | 0.283039 | 0.154 | 0.131 | 1 | 6 | Ms4a4c | 0.023 | Young |
| 0.335579 | 0.584281 | 0.057 | 0.043 | 1 | 6 | Dus1l | 0.014 | Young |
| 0.336354 | 0.659457 | 0.023 | 0.014 | 1 | 6 | Selp | 0.009 | Young |
| 0.336566 | 0.216682 | 0.115 | 0.097 | 1 | 6 | Ninj1 | 0.018 | Young |
| 0.33661 | 0.014428 | 0.04 | 0.054 | 1 | 6 | Fbxl15 | -0.014 | Young |
| 0.336627 | 0.502605 | 0.177 | 0.154 | 1 | 6 | Irf9 | 0.023 | Young |
| 0.336854 | 0.086477 | 0.478 | 0.446 | 1 | 6 | Ppib | 0.032 | Young |
| 0.336867 | 0.049687 | 0.109 | 0.089 | 1 | 6 | Fam126b | 0.02 | Young |
| 0.337401 | 0.929266 | 0.037 | 0.026 | 1 | 6 | Phf7 | 0.011 | Young |
| 0.337463 | 0.214648 | 0.202 | 0.18 | 1 | 6 | Fnip1 | 0.022 | Young |
| 0.337637 | 0.4998 | 0.097 | 0.08 | 1 | 6 | Rab28 | 0.017 | Young |
| 0.33771 | 0.098325 | 0.156 | 0.134 | 1 | 6 | Tmem135 | 0.022 | Young |
| 0.338064 | 0.133766 | 0.393 | 0.374 | 1 | 6 | Huwe1 | 0.019 | Young |
| 0.338466 | 0.138159 | 0.05 | 0.066 | 1 | 6 | Ebag9 | -0.016 | Young |
| 0.339607 | 0.370887 | 0.06 | 0.046 | 1 | 6 | Atg2b | 0.014 | Young |
| 0.339765 | 0.468343 | 0.06 | 0.046 | 1 | 6 | Cog1 | 0.014 | Young |
| 0.339772 | 0.199731 | 0.196 | 0.169 | 1 | 6 | CAAA0111 | 0.027 | Young |
| 0.339881 | 0.054973 | 0.122 | 0.1 | 1 | 6 | Tada1 | 0.022 | Young |
| 0.340181 | 0.007686 | 0.639 | 0.58 | 1 | 6 | Tm6sf1 | 0.059 | Young |
| 0.340394 | 0.450945 | 0.052 | 0.069 | 1 | 6 | Ndufaf7 | -0.017 | Young |
| 0.340993 | 0.410674 | 0.251 | 0.231 | 1 | 6 | Cd300lg | 0.02 | Young |
| 0.341259 | 0.255418 | 0.209 | 0.186 | 1 | 6 | Cyp4f18 | 0.023 | Young |
| 0.341283 | 0.92475 | 0.033 | 0.023 | 1 | 6 | Gm9733 | 0.01 | Young |
| 0.341805 | 0.734003 | 0.03 | 0.02 | 1 | 6 | Rbm48 | 0.01 | Young |
| 0.341975 | 0.255662 | 0.06 | 0.046 | 1 | 6 | Clp1 | 0.014 | Young |
| 0.342351 | 0.663708 | 0.027 | 0.017 | 1 | 6 | Snx7 | 0.01 | Young |
| 0.343157 | 0.020648 | 0.52 | 0.494 | 1 | 6 | Eif4ebp2 | 0.026 | Young |
| 0.34331 | 0.630114 | 0.027 | 0.017 | 1 | 6 | Procr | 0.01 | Young |
| 0.34331 | 0.515606 | 0.027 | 0.017 | 1 | 6 | Opn3 | 0.01 | Young |
| 0.343919 | 0.369072 | 0.095 | 0.077 | 1 | 6 | Mmd | 0.018 | Young |
| 0.344271 | 0.746672 | 0.027 | 0.017 | 1 | 6 | Map4k3 | 0.01 | Young |
| 0.34428 | 0.79615 | 0.099 | 0.083 | 1 | 6 | Washc1 | 0.016 | Young |
| 0.344836 | 0.275274 | 0.181 | 0.16 | 1 | 6 | Pip5k1a | 0.021 | Young |
| 0.344902 | 0.66566 | 0.05 | 0.037 | 1 | 6 | Ttf1 | 0.013 | Young |
| 0.345702 | 0.187031 | 0.095 | 0.077 | 1 | 6 | Fzr1 | 0.018 | Young |
| 0.345806 | 0.226055 | 0.057 | 0.043 | 1 | 6 | Fastk | 0.014 | Young |
| 0.345868 | 1.006328 | 0.03 | 0.02 | 1 | 6 | Gcat | 0.01 | Young |

| | | | | | | | | |
|---|---|---|---|---|---|---|---|---|
| 0.345988 | 0.298015 | 0.054 | 0.04 | 1 | 6 | Rbm45 | 0.014 | Young |
| 0.346009 | 0.32303 | 0.13 | 0.111 | 1 | 6 | Ulk2 | 0.019 | Young |
| 0.34668 | 0.34354 | 0.027 | 0.017 | 1 | 6 | Mapk1ip1 | 0.01 | Young |
| 0.347027 | 0.344331 | 0.129 | 0.109 | 1 | 6 | Dok1 | 0.02 | Young |
| 0.347243 | 0.22005 | 0.109 | 0.089 | 1 | 6 | Agpat1 | 0.02 | Young |
| 0.347647 | 0.020439 | 0.027 | 0.017 | 1 | 6 | Zfp874b | 0.01 | Young |
| 0.347649 | 0.157711 | 0.279 | 0.26 | 1 | 6 | Add1 | 0.019 | Young |
| 0.348075 | 0.214066 | 0.122 | 0.103 | 1 | 6 | Gab2 | 0.019 | Young |
| 0.34822 | 0.668132 | 0.043 | 0.031 | 1 | 6 | Tbc1d32 | 0.012 | Young |
| 0.348396 | 0.264378 | 0.05 | 0.037 | 1 | 6 | Pdcd2 | 0.013 | Young |
| 0.348448 | 0.325359 | 0.11 | 0.091 | 1 | 6 | Scyl2 | 0.019 | Young |
| 0.34856 | 0.272691 | 0.033 | 0.023 | 1 | 6 | Dis3l | 0.01 | Young |
| 0.348593 | 0.742573 | 0.03 | 0.02 | 1 | 6 | Mad2l1bp | 0.01 | Young |
| 0.348593 | 0.523315 | 0.03 | 0.02 | 1 | 6 | Tab1 | 0.01 | Young |
| 0.348953 | 0.222445 | 0.085 | 0.069 | 1 | 6 | Tnfaip8l2 | 0.016 | Young |
| 0.349727 | 0.252672 | 0.043 | 0.031 | 1 | 6 | Pola2 | 0.012 | Young |
| 0.349961 | 0.452408 | 0.03 | 0.02 | 1 | 6 | Thap4 | 0.01 | Young |
| 0.350104 | 0.265645 | 0.104 | 0.086 | 1 | 6 | Nolc1 | 0.018 | Young |
| 0.350719 | 0.557206 | 0.033 | 0.023 | 1 | 6 | Lrrc28 | 0.01 | Young |
| 0.350892 | 0.113026 | 0.328 | 0.3 | 1 | 6 | Vps35 | 0.028 | Young |
| 0.351152 | 0.135514 | 0.033 | 0.023 | 1 | 6 | Nsl1 | 0.01 | Young |
| 0.351163 | 0.000778 | 0.01 | 0.017 | 1 | 6 | Sap130 | -0.007 | Young |
| 0.351231 | 0.32654 | 0.06 | 0.046 | 1 | 6 | Fzd7 | 0.014 | Young |
| 0.35157 | 0.274769 | 0.162 | 0.14 | 1 | 6 | Commd4 | 0.022 | Young |
| 0.351652 | 0.500098 | 0.037 | 0.026 | 1 | 6 | Ctps | 0.011 | Young |
| 0.351758 | 0.057614 | 0.425 | 0.391 | 1 | 6 | Degs1 | 0.034 | Young |
| 0.352248 | 0.718622 | 0.03 | 0.02 | 1 | 6 | A530017D[illegible] | 0.01 | Young |
| 0.35292 | 0.20344 | 0.038 | 0.051 | 1 | 6 | Ly96 | -0.013 | Young |
| 0.353134 | 0.022472 | 0.01 | 0.017 | 1 | 6 | Eif3j2 | -0.007 | Young |
| 0.353265 | 0.164494 | 0.493 | 0.477 | 1 | 6 | Eid1 | 0.016 | Young |
| 0.353288 | 0.042556 | 0.179 | 0.157 | 1 | 6 | Hnrnpll | 0.022 | Young |
| 0.353692 | 0.223559 | 0.187 | 0.166 | 1 | 6 | Wwp2 | 0.021 | Young |
| 0.353706 | 0.146668 | 0.304 | 0.283 | 1 | 6 | Pin4 | 0.021 | Young |
| 0.35372 | 0.506489 | 0.04 | 0.029 | 1 | 6 | Rps6kb2 | 0.011 | Young |
| 0.353897 | 1.367373 | 0.012 | 0.006 | 1 | 6 | Retn | 0.006 | Young |
| 0.353897 | 0.838426 | 0.012 | 0.006 | 1 | 6 | Gm11131 | 0.006 | Young |
| 0.354012 | 0.176509 | 0.202 | 0.177 | 1 | 6 | Mob3b | 0.025 | Young |
| 0.354395 | 0.536496 | 0.057 | 0.043 | 1 | 6 | Gstm1 | 0.014 | Young |
| 0.354545 | 0.212943 | 0.03 | 0.02 | 1 | 6 | Cdc7 | 0.01 | Young |
| 0.354657 | 0.477097 | 0.043 | 0.031 | 1 | 6 | Usp18 | 0.012 | Young |
| 0.354657 | 1.186734 | 0.012 | 0.006 | 1 | 6 | Ogg1 | 0.006 | Young |
| 0.354991 | 0.248572 | 0.159 | 0.14 | 1 | 6 | 0610012G[illegible] | 0.019 | Young |
| 0.355225 | 0.000933 | 0.293 | 0.257 | 1 | 6 | Cdc5l | 0.036 | Young |
| 0.355419 | 1.255796 | 0.012 | 0.006 | 1 | 6 | C1qtnf12 | 0.006 | Young |
| 0.356181 | 1.424766 | 0.012 | 0.006 | 1 | 6 | Uprt | 0.006 | Young |

| | | | | | | | | |
|---|---|---|---|---|---|---|---|---|
| 0.356181 | 1.202129 | 0.012 | 0.006 | 1 | 6 | Nup43 | 0.006 | Young |
| 0.356181 | 1.199924 | 0.012 | 0.006 | 1 | 6 | 9130221H1 | 0.006 | Young |
| 0.356181 | 0.84021 | 0.012 | 0.006 | 1 | 6 | Kbtbd8 | 0.006 | Young |
| 0.356225 | 0.432398 | 0.097 | 0.08 | 1 | 6 | Zbtb21 | 0.017 | Young |
| 0.356573 | 0.086469 | 0.329 | 0.309 | 1 | 6 | Edf1 | 0.02 | Young |
| 0.35678 | 0.210273 | 0.129 | 0.109 | 1 | 6 | Mrpl43 | 0.02 | Young |
| 0.356888 | 0.291867 | 0.258 | 0.237 | 1 | 6 | Vwa5a | 0.021 | Young |
| 0.356944 | 0.927928 | 0.012 | 0.006 | 1 | 6 | Cald1 | 0.006 | Young |
| 0.357191 | 0.116199 | 0.416 | 0.38 | 1 | 6 | Med13l | 0.036 | Young |
| 0.357295 | 0.481174 | 0.04 | 0.029 | 1 | 6 | Prkci | 0.011 | Young |
| 0.357313 | 0.08992 | 0.03 | 0.02 | 1 | 6 | Tmem241 | 0.01 | Young |
| 0.357597 | 0.472062 | 0.054 | 0.04 | 1 | 6 | 6720427I0 | 0.014 | Young |
| 0.357709 | 0.770821 | 0.012 | 0.006 | 1 | 6 | Znhit3 | 0.006 | Young |
| 0.358279 | 0.213773 | 0.125 | 0.106 | 1 | 6 | Brd9 | 0.019 | Young |
| 0.358291 | 0.068018 | 0.037 | 0.026 | 1 | 6 | Fas | 0.011 | Young |
| 0.358474 | 0.987702 | 0.012 | 0.006 | 1 | 6 | Kdelc2 | 0.006 | Young |
| 0.358474 | 0.942792 | 0.012 | 0.006 | 1 | 6 | Tagln | 0.006 | Young |
| 0.358474 | 0.897072 | 0.012 | 0.006 | 1 | 6 | Ttll1 | 0.006 | Young |
| 0.358474 | 0.821146 | 0.012 | 0.006 | 1 | 6 | Nme4 | 0.006 | Young |
| 0.358474 | 0.731782 | 0.012 | 0.006 | 1 | 6 | A430106G1 | 0.006 | Young |
| 0.358474 | 0.730031 | 0.012 | 0.006 | 1 | 6 | Mapre3 | 0.006 | Young |
| 0.358474 | 0.71762 | 0.012 | 0.006 | 1 | 6 | E430018J2 | 0.006 | Young |
| 0.358474 | 0.600792 | 0.012 | 0.006 | 1 | 6 | Usp11 | 0.006 | Young |
| 0.358861 | 0.065779 | 0.043 | 0.031 | 1 | 6 | Rasgrf2 | 0.012 | Young |
| 0.35899 | 0.705301 | 0.062 | 0.049 | 1 | 6 | Strada | 0.013 | Young |
| 0.35924 | 0.643685 | 0.012 | 0.006 | 1 | 6 | Gm11696 | 0.006 | Young |
| 0.35924 | 0.567848 | 0.012 | 0.006 | 1 | 6 | Pnma1 | 0.006 | Young |
| 0.35924 | 0.5355 | 0.012 | 0.006 | 1 | 6 | Adamtsl4 | 0.006 | Young |
| 0.35924 | 0.34521 | 0.012 | 0.006 | 1 | 6 | Zfp652os | 0.006 | Young |
| 0.359428 | 0.304281 | 0.232 | 0.211 | 1 | 6 | Rsf1 | 0.021 | Young |
| 0.359585 | 0.097638 | 0.274 | 0.254 | 1 | 6 | AC160336. | 0.02 | Young |
| 0.359927 | 0.165814 | 0.162 | 0.14 | 1 | 6 | Klhl18 | 0.022 | Young |
| 0.360008 | 0.813171 | 0.012 | 0.006 | 1 | 6 | AC113595. | 0.006 | Young |
| 0.360008 | 0.591324 | 0.012 | 0.006 | 1 | 6 | Tmem41a | 0.006 | Young |
| 0.360048 | 0.440295 | 0.109 | 0.091 | 1 | 6 | Zeb2os | 0.018 | Young |
| 0.36007 | 0.368987 | 0.157 | 0.14 | 1 | 6 | Slc25a39 | 0.017 | Young |
| 0.36016 | 0.216252 | 0.236 | 0.211 | 1 | 6 | Pros1 | 0.025 | Young |
| 0.36027 | 0.029862 | 0.483 | 0.426 | 1 | 6 | Cysltr1 | 0.057 | Young |
| 0.360438 | 0.016096 | 0.114 | 0.094 | 1 | 6 | Ing3 | 0.02 | Young |
| 0.360471 | 0.015087 | 0.125 | 0.149 | 1 | 6 | Trmt1 | -0.024 | Young |
| 0.360654 | 0.654859 | 0.079 | 0.063 | 1 | 6 | Daxx | 0.016 | Young |
| 0.360776 | 0.420734 | 0.012 | 0.006 | 1 | 6 | Mocs3 | 0.006 | Young |
| 0.361293 | 0.360431 | 0.201 | 0.183 | 1 | 6 | Tfg | 0.018 | Young |
| 0.361546 | 0.206999 | 0.012 | 0.006 | 1 | 6 | Dact3 | 0.006 | Young |
| 0.361576 | 0.008333 | 0.152 | 0.174 | 1 | 6 | Luc7l | -0.022 | Young |

| | | | | | | | | |
|---|---|---|---|---|---|---|---|---|
| 0.361937 | 0.134377 | 0.172 | 0.149 | 1 | 6 | Erg28 | 0.023 | Young |
| 0.361938 | 0.371191 | 0.206 | 0.183 | 1 | 6 | Mtmr3 | 0.023 | Young |
| 0.362317 | 0.096634 | 0.012 | 0.006 | 1 | 6 | Pkd1l2 | 0.006 | Young |
| 0.362697 | 0.089826 | 0.542 | 0.511 | 1 | 6 | Sumo2 | 0.031 | Young |
| 0.36272 | 0.080001 | 0.386 | 0.363 | 1 | 6 | Morf4l2 | 0.023 | Young |
| 0.363088 | 0.338787 | 0.012 | 0.006 | 1 | 6 | Neo1 | 0.006 | Young |
| 0.363119 | 0.421273 | 0.162 | 0.14 | 1 | 6 | Prkcb | 0.022 | Young |
| 0.363441 | 0.138717 | 0.263 | 0.234 | 1 | 6 | Eif3g | 0.029 | Young |
| 0.3635 | 0.287783 | 0.038 | 0.051 | 1 | 6 | Slc12a4 | -0.013 | Young |
| 0.363617 | 0.009996 | 0.222 | 0.194 | 1 | 6 | Tob1 | 0.028 | Young |
| 0.364142 | 0.785444 | 0.062 | 0.049 | 1 | 6 | Phf11a | 0.013 | Young |
| 0.364157 | 0.123046 | 0.333 | 0.306 | 1 | 6 | Arfgef1 | 0.027 | Young |
| 0.364233 | 0.451708 | 0.109 | 0.091 | 1 | 6 | Scn2b | 0.018 | Young |
| 0.364499 | 0.099403 | 0.119 | 0.1 | 1 | 6 | Polr2j | 0.019 | Young |
| 0.364667 | 0.322193 | 0.199 | 0.18 | 1 | 6 | Cenpx | 0.019 | Young |
| 0.364736 | 0.315906 | 0.033 | 0.023 | 1 | 6 | Zxdb | 0.01 | Young |
| 0.364988 | 0.117523 | 0.351 | 0.314 | 1 | 6 | Rell1 | 0.037 | Young |
| 0.365323 | 0.092502 | 0.328 | 0.297 | 1 | 6 | Arhgap26 | 0.031 | Young |
| 0.365469 | 0.499894 | 0.156 | 0.137 | 1 | 6 | Polr1c | 0.019 | Young |
| 0.365615 | 0.363664 | 0.043 | 0.031 | 1 | 6 | Rpa2 | 0.012 | Young |
| 0.36581 | 0.270341 | 0.043 | 0.031 | 1 | 6 | Gna15 | 0.012 | Young |
| 0.365983 | 0.256143 | 0.145 | 0.126 | 1 | 6 | Dirc2 | 0.019 | Young |
| 0.366562 | 0.405837 | 0.079 | 0.063 | 1 | 6 | Cxxc5 | 0.016 | Young |
| 0.366579 | 0.185109 | 0.176 | 0.154 | 1 | 6 | Dram2 | 0.022 | Young |
| 0.366603 | 0.028412 | 0.426 | 0.4 | 1 | 6 | Gatad2b | 0.026 | Young |
| 0.36717 | 0.241063 | 0.221 | 0.197 | 1 | 6 | Trpv2 | 0.024 | Young |
| 0.367248 | 0.100989 | 0.39 | 0.351 | 1 | 6 | Uqcc2 | 0.039 | Young |
| 0.367529 | 0.29755 | 0.127 | 0.109 | 1 | 6 | Rnd3 | 0.018 | Young |
| 0.367645 | 0.004362 | 0.05 | 0.037 | 1 | 6 | Cep68 | 0.013 | Young |
| 0.367942 | 0.295716 | 0.099 | 0.083 | 1 | 6 | Ginm1 | 0.016 | Young |
| 0.367944 | 0.163117 | 0.294 | 0.269 | 1 | 6 | Lmo1 | 0.025 | Young |
| 0.368194 | 0.045051 | 0.085 | 0.069 | 1 | 6 | Rab3gap2 | 0.016 | Young |
| 0.368696 | 0.239973 | 0.087 | 0.071 | 1 | 6 | Snx13 | 0.016 | Young |
| 0.368822 | 0.045047 | 0.037 | 0.026 | 1 | 6 | Kdelc1 | 0.011 | Young |
| 0.368986 | 0.628916 | 0.119 | 0.103 | 1 | 6 | Desi2 | 0.016 | Young |
| 0.369384 | 0.264285 | 0.144 | 0.123 | 1 | 6 | Rpl10-ps3 | 0.021 | Young |
| 0.369569 | 0.334898 | 0.211 | 0.191 | 1 | 6 | Vezf1 | 0.02 | Young |
| 0.369687 | 0.248351 | 0.122 | 0.103 | 1 | 6 | Glod4 | 0.019 | Young |
| 0.370182 | 0.037209 | 0.54 | 0.517 | 1 | 6 | Eif4b | 0.023 | Young |
| 0.370441 | 0.106454 | 0.388 | 0.363 | 1 | 6 | BC028528 | 0.025 | Young |
| 0.370829 | 0.029683 | 0.059 | 0.074 | 1 | 6 | Tmed4 | -0.015 | Young |
| 0.371143 | 0.086225 | 0.135 | 0.117 | 1 | 6 | Cab39l | 0.018 | Young |
| 0.371473 | 0.37025 | 0.157 | 0.14 | 1 | 6 | Cnpy3 | 0.017 | Young |
| 0.371855 | 0.19595 | 0.14 | 0.12 | 1 | 6 | Aip | 0.02 | Young |
| 0.372059 | 0.164513 | 0.179 | 0.157 | 1 | 6 | Lsm8 | 0.022 | Young |

| | | | | | | | | |
|---|---|---|---|---|---|---|---|---|
| 0.372625 | 0.257167 | 0.043 | 0.057 | 1 | 6 | Recql | -0.014 | Young |
| 0.372627 | 0.134843 | 0.251 | 0.223 | 1 | 6 | Bcl7c | 0.028 | Young |
| 0.372841 | 0.070806 | 0.043 | 0.031 | 1 | 6 | Cavin1 | 0.012 | Young |
| 0.373048 | 0.046033 | 0.881 | 0.834 | 1 | 6 | Atp5l | 0.047 | Young |
| 0.374089 | 0.232066 | 0.038 | 0.051 | 1 | 6 | Decr2 | -0.013 | Young |
| 0.37421 | 0.257867 | 0.027 | 0.037 | 1 | 6 | Plpp6 | -0.01 | Young |
| 0.374756 | 0.080459 | 0.186 | 0.163 | 1 | 6 | Micu1 | 0.023 | Young |
| 0.375931 | 0.001398 | 0.085 | 0.069 | 1 | 6 | Fam193b | 0.016 | Young |
| 0.376068 | 0.487818 | 0.062 | 0.049 | 1 | 6 | Gon7 | 0.013 | Young |
| 0.376105 | 0.256951 | 0.104 | 0.086 | 1 | 6 | Tmem87b | 0.018 | Young |
| 0.376392 | 0.099404 | 0.179 | 0.157 | 1 | 6 | Nfic | 0.022 | Young |
| 0.376433 | 0.373153 | 0.114 | 0.097 | 1 | 6 | Adrb2 | 0.017 | Young |
| 0.376503 | 0.290827 | 0.09 | 0.074 | 1 | 6 | Acads | 0.016 | Young |
| 0.377042 | 0.237611 | 0.097 | 0.08 | 1 | 6 | Thoc3 | 0.017 | Young |
| 0.377046 | 0.424933 | 0.087 | 0.071 | 1 | 6 | Wrn | 0.016 | Young |
| 0.377154 | 0.061945 | 0.711 | 0.651 | 1 | 6 | Zyx | 0.06 | Young |
| 0.377305 | 0.009541 | 0.313 | 0.28 | 1 | 6 | Ext1 | 0.033 | Young |
| 0.377315 | 0.043671 | 0.861 | 0.76 | 1 | 6 | Cotl1 | 0.101 | Young |
| 0.377403 | 0.18694 | 0.176 | 0.154 | 1 | 6 | Got2 | 0.022 | Young |
| 0.377967 | 0.199303 | 0.079 | 0.063 | 1 | 6 | Creb5 | 0.016 | Young |
| 0.378903 | 0.015992 | 0.548 | 0.517 | 1 | 6 | Lyn | 0.031 | Young |
| 0.379048 | 0.475127 | 0.065 | 0.051 | 1 | 6 | Dnajc30 | 0.014 | Young |
| 0.380271 | 0.178115 | 0.144 | 0.126 | 1 | 6 | Pou2f1 | 0.018 | Young |
| 0.380422 | 1.732339 | 0.015 | 0.009 | 1 | 6 | Gne | 0.006 | Young |
| 0.380634 | 0.190358 | 0.323 | 0.303 | 1 | 6 | Ap1b1 | 0.02 | Young |
| 0.380982 | 0.311576 | 0.087 | 0.071 | 1 | 6 | Slc14a1 | 0.016 | Young |
| 0.381112 | 1.258056 | 0.015 | 0.009 | 1 | 6 | Trmo | 0.006 | Young |
| 0.38129 | 0.051814 | 0.176 | 0.151 | 1 | 6 | Srgap2 | 0.025 | Young |
| 0.381326 | 0.073783 | 0.082 | 0.066 | 1 | 6 | Ccdc90b | 0.016 | Young |
| 0.381519 | 0.136651 | 0.221 | 0.194 | 1 | 6 | Rbm42 | 0.027 | Young |
| 0.381793 | 0.230047 | 0.117 | 0.143 | 1 | 6 | Ccdc115 | -0.026 | Young |
| 0.382395 | 0.058221 | 0.145 | 0.123 | 1 | 6 | Rnf2 | 0.022 | Young |
| 0.382913 | 0.197654 | 0.192 | 0.169 | 1 | 6 | Ahcyl1 | 0.023 | Young |
| 0.382925 | 0.034625 | 0.597 | 0.566 | 1 | 6 | Tab2 | 0.031 | Young |
| 0.383186 | 1.395199 | 0.015 | 0.009 | 1 | 6 | 1700056E2 | 0.006 | Young |
| 0.383879 | 1.193106 | 0.015 | 0.009 | 1 | 6 | Chac2 | 0.006 | Young |
| 0.383879 | 1.036772 | 0.015 | 0.009 | 1 | 6 | Tmem202 | 0.006 | Young |
| 0.383916 | 0.28598 | 0.065 | 0.051 | 1 | 6 | Gaa | 0.014 | Young |
| 0.384074 | 0.216846 | 0.112 | 0.094 | 1 | 6 | Xpnpep1 | 0.018 | Young |
| 0.384188 | 0.207211 | 0.278 | 0.257 | 1 | 6 | Plxnb2 | 0.021 | Young |
| 0.384206 | 0.752336 | 0.055 | 0.043 | 1 | 6 | Ikbip | 0.012 | Young |
| 0.384234 | 0.269708 | 0.069 | 0.054 | 1 | 6 | Hs2st1 | 0.015 | Young |
| 0.384256 | 0.475489 | 0.08 | 0.066 | 1 | 6 | Fkrp | 0.014 | Young |
| 0.385267 | 1.053698 | 0.015 | 0.009 | 1 | 6 | Armc9 | 0.006 | Young |
| 0.385775 | 0.156434 | 0.328 | 0.306 | 1 | 6 | Jup | 0.022 | Young |

| | | | | | | | | |
|---|---|---|---|---|---|---|---|---|
| 0.385788 | 0.021946 | 0.12 | 0.1 | 1 | 6 | Foxo3 | 0.02 | Young |
| 0.385962 | 1.016616 | 0.015 | 0.009 | 1 | 6 | Enc1 | 0.006 | Young |
| 0.385962 | 0.79895 | 0.015 | 0.009 | 1 | 6 | Cryab | 0.006 | Young |
| 0.386126 | 0.200443 | 0.015 | 0.023 | 1 | 6 | Gm10138 | -0.008 | Young |
| 0.386658 | 0.795024 | 0.015 | 0.009 | 1 | 6 | P2ry12 | 0.006 | Young |
| 0.386714 | 0.151864 | 0.125 | 0.106 | 1 | 6 | Utp11 | 0.019 | Young |
| 0.387355 | 0.857165 | 0.015 | 0.009 | 1 | 6 | Mtrr | 0.006 | Young |
| 0.387355 | 0.830772 | 0.015 | 0.009 | 1 | 6 | Gm26881 | 0.006 | Young |
| 0.387387 | 0.172753 | 0.097 | 0.08 | 1 | 6 | Nup50 | 0.017 | Young |
| 0.387758 | 0.019708 | 0.032 | 0.043 | 1 | 6 | Qsox2 | -0.011 | Young |
| 0.388052 | 0.668325 | 0.015 | 0.009 | 1 | 6 | Gm31597 | 0.006 | Young |
| 0.388209 | 0.028273 | 0.436 | 0.409 | 1 | 6 | Uqcrb | 0.027 | Young |
| 0.38875 | 0.755782 | 0.015 | 0.009 | 1 | 6 | Tspyl4 | 0.006 | Young |
| 0.388973 | 0.196725 | 0.373 | 0.351 | 1 | 6 | Fam174a | 0.022 | Young |
| 0.389449 | 0.673702 | 0.015 | 0.009 | 1 | 6 | Snap47 | 0.006 | Young |
| 0.389449 | 0.520157 | 0.015 | 0.009 | 1 | 6 | Ctsf | 0.006 | Young |
| 0.389572 | 0.91034 | 0.048 | 0.037 | 1 | 6 | Map4k5 | 0.011 | Young |
| 0.390309 | 0.46901 | 0.077 | 0.063 | 1 | 6 | Ptpn9 | 0.014 | Young |
| 0.390849 | 0.340169 | 0.015 | 0.009 | 1 | 6 | Tesk2 | 0.006 | Young |
| 0.391423 | 0.234763 | 0.015 | 0.023 | 1 | 6 | Mms22l | -0.008 | Young |
| 0.391495 | 0.224859 | 0.171 | 0.149 | 1 | 6 | Ralgapa1 | 0.022 | Young |
| 0.391677 | 0.042494 | 0.61 | 0.574 | 1 | 6 | Tiam1 | 0.036 | Young |
| 0.392184 | 0.183345 | 0.239 | 0.22 | 1 | 6 | Ets2 | 0.019 | Young |
| 0.392196 | 0.082482 | 0.008 | 0.014 | 1 | 6 | Tarm1 | -0.006 | Young |
| 0.392253 | 0.107648 | 0.015 | 0.009 | 1 | 6 | Zgrf1 | 0.006 | Young |
| 0.39236 | 0.480156 | 0.134 | 0.117 | 1 | 6 | Irak2 | 0.017 | Young |
| 0.392376 | 0.117038 | 0.089 | 0.109 | 1 | 6 | Kat6b | -0.02 | Young |
| 0.392606 | 0.123409 | 0.015 | 0.023 | 1 | 6 | Map7 | -0.008 | Young |
| 0.392856 | 0.267605 | 0.229 | 0.209 | 1 | 6 | Mcur1 | 0.02 | Young |
| 0.39294 | 0.33682 | 0.065 | 0.051 | 1 | 6 | Frmd6 | 0.014 | Young |
| 0.393587 | 0.347806 | 0.212 | 0.194 | 1 | 6 | Adprh | 0.018 | Young |
| 0.393618 | 0.24355 | 0.231 | 0.209 | 1 | 6 | Psmg4 | 0.022 | Young |
| 0.393713 | 0.510445 | 0.052 | 0.04 | 1 | 6 | Lrrc61 | 0.012 | Young |
| 0.393735 | 0.089259 | 0.008 | 0.014 | 1 | 6 | Fam161a | -0.006 | Young |
| 0.394363 | 0.570164 | 0.015 | 0.009 | 1 | 6 | Ntng2 | 0.006 | Young |
| 0.394363 | 0.139643 | 0.015 | 0.009 | 1 | 6 | 1700037C1 | 0.006 | Young |
| 0.39496 | 0.202086 | 0.261 | 0.243 | 1 | 6 | Glrx3 | 0.018 | Young |
| 0.395169 | 0.967486 | 0.038 | 0.029 | 1 | 6 | Zfp704 | 0.009 | Young |
| 0.395635 | 0.073459 | 0.55 | 0.523 | 1 | 6 | Jmjd1c | 0.027 | Young |
| 0.397237 | 0.408851 | 0.052 | 0.04 | 1 | 6 | Tmem199 | 0.012 | Young |
| 0.397506 | 0.897502 | 0.045 | 0.034 | 1 | 6 | Gss | 0.011 | Young |
| 0.397599 | 0.003159 | 0.008 | 0.014 | 1 | 6 | Adcy4 | -0.006 | Young |
| 0.397705 | 0.147293 | 0.119 | 0.1 | 1 | 6 | Upf3a | 0.019 | Young |
| 0.39785 | 1.199977 | 0.018 | 0.011 | 1 | 6 | Tox3 | 0.007 | Young |
| 0.397861 | 0.533736 | 0.042 | 0.031 | 1 | 6 | Nprl3 | 0.011 | Young |

| | | | | | | | | |
|---|---|---|---|---|---|---|---|---|
| 0.398004 | 0.572603 | 0.084 | 0.069 | 1 | 6 | Gopc | 0.015 | Young |
| 0.39809 | 0.104425 | 0.157 | 0.183 | 1 | 6 | Slc35b1 | -0.026 | Young |
| 0.398529 | 0.077495 | 0.681 | 0.654 | 1 | 6 | Traf1 | 0.027 | Young |
| 0.398885 | 0.082192 | 0.276 | 0.249 | 1 | 6 | Rab1b | 0.027 | Young |
| 0.399049 | 0.374682 | 0.182 | 0.163 | 1 | 6 | Eapp | 0.019 | Young |
| 0.399121 | 1.073798 | 0.018 | 0.011 | 1 | 6 | Gm42715 | 0.007 | Young |
| 0.399184 | 0.792968 | 0.064 | 0.051 | 1 | 6 | Ccdc85b | 0.013 | Young |
| 0.399238 | 0.037223 | 0.09 | 0.109 | 1 | 6 | Sh3bp2 | -0.019 | Young |
| 0.399521 | 0.241252 | 0.099 | 0.083 | 1 | 6 | Phf20 | 0.016 | Young |
| 0.400207 | 0.604823 | 0.089 | 0.074 | 1 | 6 | Mettl7a1 | 0.015 | Young |
| 0.400792 | 0.389597 | 0.059 | 0.046 | 1 | 6 | Myg1 | 0.013 | Young |
| 0.400829 | 0.059847 | 0.157 | 0.137 | 1 | 6 | Zfp622 | 0.02 | Young |
| 0.40104 | 0.267143 | 0.055 | 0.043 | 1 | 6 | Cep164 | 0.012 | Young |
| 0.401529 | 0.227926 | 0.052 | 0.04 | 1 | 6 | Tmem70 | 0.012 | Young |
| 0.401529 | 0.118481 | 0.052 | 0.04 | 1 | 6 | Parp12 | 0.012 | Young |
| 0.401532 | 0.789375 | 0.035 | 0.026 | 1 | 6 | Plekha8 | 0.009 | Young |
| 0.4016 | 0.917994 | 0.042 | 0.031 | 1 | 6 | Zfp738 | 0.011 | Young |
| 0.401927 | 0.116241 | 0.125 | 0.106 | 1 | 6 | Hras | 0.019 | Young |
| 0.402096 | 0.256187 | 0.156 | 0.137 | 1 | 6 | Phf10 | 0.019 | Young |
| 0.402201 | 0.050637 | 0.109 | 0.091 | 1 | 6 | Phrf1 | 0.018 | Young |
| 0.402308 | 0.769734 | 0.018 | 0.011 | 1 | 6 | Dcaf4 | 0.007 | Young |
| 0.402433 | 0.95837 | 0.042 | 0.031 | 1 | 6 | Ap3m2 | 0.011 | Young |
| 0.402556 | 0.26038 | 0.059 | 0.046 | 1 | 6 | Eid3 | 0.013 | Young |
| 0.402854 | 0.326213 | 0.105 | 0.089 | 1 | 6 | Nudt4 | 0.016 | Young |
| 0.402875 | 0.707941 | 0.079 | 0.066 | 1 | 6 | Smpd1 | 0.013 | Young |
| 0.403585 | 0.377249 | 0.055 | 0.043 | 1 | 6 | Kif1bp | 0.012 | Young |
| 0.403587 | 0.751003 | 0.018 | 0.011 | 1 | 6 | Sumf2 | 0.007 | Young |
| 0.403587 | 0.575798 | 0.018 | 0.011 | 1 | 6 | Tchp | 0.007 | Young |
| 0.403794 | 0.027613 | 0.304 | 0.329 | 1 | 6 | Mlec | -0.025 | Young |
| 0.403831 | 0.918108 | 0.038 | 0.029 | 1 | 6 | Wdr47 | 0.009 | Young |
| 0.404036 | 0.289318 | 0.174 | 0.154 | 1 | 6 | Hbp1 | 0.02 | Young |
| 0.40434 | 0.564036 | 0.104 | 0.089 | 1 | 6 | Herc6 | 0.015 | Young |
| 0.405134 | 0.025469 | 0.776 | 0.743 | 1 | 6 | Emp3 | 0.033 | Young |
| 0.40527 | 0.254775 | 0.124 | 0.106 | 1 | 6 | Dmtf1 | 0.018 | Young |
| 0.40551 | 0.619071 | 0.018 | 0.011 | 1 | 6 | Cep131 | 0.007 | Young |
| 0.405606 | 0.922703 | 0.064 | 0.051 | 1 | 6 | Ctps2 | 0.013 | Young |
| 0.405693 | 0.162581 | 0.237 | 0.22 | 1 | 6 | Mthfd2 | 0.017 | Young |
| 0.406601 | 0.045986 | 0.176 | 0.151 | 1 | 6 | Mboat7 | 0.025 | Young |
| 0.406734 | 0.330672 | 0.045 | 0.034 | 1 | 6 | Ints14 | 0.011 | Young |
| 0.407213 | 0.08306 | 0.177 | 0.154 | 1 | 6 | Spata2 | 0.023 | Young |
| 0.407849 | 0.109412 | 0.119 | 0.103 | 1 | 6 | Dhx30 | 0.016 | Young |
| 0.408249 | 0.148863 | 0.398 | 0.374 | 1 | 6 | Kpna4 | 0.024 | Young |
| 0.408727 | 0.472756 | 0.018 | 0.011 | 1 | 6 | Gm3448 | 0.007 | Young |
| 0.40915 | 0.270217 | 0.107 | 0.091 | 1 | 6 | Naip5 | 0.016 | Young |
| 0.409284 | 0.252743 | 0.224 | 0.209 | 1 | 6 | Rnf145 | 0.015 | Young |

| | | | | | | | | |
|---|---|---|---|---|---|---|---|---|
| 0.409418 | 0.157062 | 0.219 | 0.2 | 1 | 6 | Arid1b | 0.019 | Young |
| 0.409562 | 0.780404 | 0.042 | 0.031 | 1 | 6 | Stk32c | 0.011 | Young |
| 0.40977 | 0.881042 | 0.035 | 0.026 | 1 | 6 | 2700038G2 | 0.009 | Young |
| 0.410348 | 0.259705 | 0.092 | 0.077 | 1 | 6 | Clcn7 | 0.015 | Young |
| 0.410399 | 0.345897 | 0.156 | 0.137 | 1 | 6 | Yrdc | 0.019 | Young |
| 0.410809 | 0.020048 | 0.084 | 0.069 | 1 | 6 | Irak3 | 0.015 | Young |
| 0.411123 | 0.956258 | 0.025 | 0.017 | 1 | 6 | Zfp143 | 0.008 | Young |
| 0.411246 | 0.131596 | 0.386 | 0.36 | 1 | 6 | Ube2n | 0.026 | Young |
| 0.41125 | 0.576521 | 0.042 | 0.031 | 1 | 6 | Snip1 | 0.011 | Young |
| 0.411312 | 0.564386 | 0.018 | 0.011 | 1 | 6 | Nos1ap | 0.007 | Young |
| 0.411594 | 0.022351 | 0.308 | 0.28 | 1 | 6 | Mfsd1 | 0.028 | Young |
| 0.411713 | 0.313906 | 0.052 | 0.04 | 1 | 6 | Mettl6 | 0.012 | Young |
| 0.412094 | 0.065513 | 0.052 | 0.04 | 1 | 6 | Taf1b | 0.012 | Young |
| 0.412164 | 0.063622 | 0.059 | 0.046 | 1 | 6 | Tars2 | 0.013 | Young |
| 0.412289 | 0.269991 | 0.127 | 0.111 | 1 | 6 | Washc3 | 0.016 | Young |
| 0.4123 | 0.594731 | 0.097 | 0.083 | 1 | 6 | Golga5 | 0.014 | Young |
| 0.412518 | 0.237628 | 0.323 | 0.309 | 1 | 6 | Tank | 0.014 | Young |
| 0.412727 | 0.21429 | 0.217 | 0.2 | 1 | 6 | Smim10l1 | 0.017 | Young |
| 0.412751 | 0.359296 | 0.055 | 0.043 | 1 | 6 | Jmy | 0.012 | Young |
| 0.412893 | 0.036628 | 0.351 | 0.343 | 1 | 6 | Hnrnpul1 | 0.008 | Young |
| 0.412943 | 0.565174 | 0.042 | 0.031 | 1 | 6 | Slc46a3 | 0.011 | Young |
| 0.413235 | 0.302998 | 0.052 | 0.04 | 1 | 6 | Agap1 | 0.012 | Young |
| 0.413257 | 0.185235 | 0.018 | 0.011 | 1 | 6 | Helq | 0.007 | Young |
| 0.414119 | 0.49849 | 0.11 | 0.094 | 1 | 6 | Gatd1 | 0.016 | Young |
| 0.414397 | 0.846465 | 0.022 | 0.014 | 1 | 6 | Slc22a5 | 0.008 | Young |
| 0.414423 | 0.844962 | 0.025 | 0.017 | 1 | 6 | Cby1 | 0.008 | Young |
| 0.414728 | 0.191159 | 0.084 | 0.069 | 1 | 6 | Ano6 | 0.015 | Young |
| 0.414853 | 0.736023 | 0.035 | 0.026 | 1 | 6 | Map3k6 | 0.009 | Young |
| 0.414973 | 0.235633 | 0.186 | 0.169 | 1 | 6 | Ssr2 | 0.017 | Young |
| 0.414975 | 0.755645 | 0.025 | 0.017 | 1 | 6 | Lrrc32 | 0.008 | Young |
| 0.415075 | 0.170771 | 0.254 | 0.229 | 1 | 6 | Mob4 | 0.025 | Young |
| 0.415201 | 0.875815 | 0.028 | 0.02 | 1 | 6 | Cnpy4 | 0.008 | Young |
| 0.415538 | 0.737078 | 0.032 | 0.023 | 1 | 6 | Rab26os | 0.009 | Young |
| 0.415588 | 0.854583 | 0.022 | 0.014 | 1 | 6 | Thg1l | 0.008 | Young |
| 0.4156 | 0.832965 | 0.075 | 0.063 | 1 | 6 | Dhdds | 0.012 | Young |
| 0.415633 | 0.560935 | 0.085 | 0.071 | 1 | 6 | Slc16a7 | 0.014 | Young |
| 0.415944 | 0.220041 | 0.145 | 0.126 | 1 | 6 | Anpep | 0.019 | Young |
| 0.416088 | 0.576695 | 0.045 | 0.034 | 1 | 6 | Ralgds | 0.011 | Young |
| 0.416711 | 0.910286 | 0.035 | 0.026 | 1 | 6 | Ptgr1 | 0.009 | Young |
| 0.417191 | 0.421038 | 0.042 | 0.031 | 1 | 6 | Ppat | 0.011 | Young |
| 0.417377 | 0.562435 | 0.022 | 0.014 | 1 | 6 | Zfp426 | 0.008 | Young |
| 0.417641 | 0.480236 | 0.035 | 0.026 | 1 | 6 | Ncapd2 | 0.009 | Young |
| 0.417739 | 0.086998 | 0.025 | 0.017 | 1 | 6 | Wrb | 0.008 | Young |
| 0.418435 | 0.038247 | 0.164 | 0.143 | 1 | 6 | Zdhhc23 | 0.021 | Young |
| 0.418518 | 0.525523 | 0.129 | 0.114 | 1 | 6 | Nacc1 | 0.015 | Young |

| | | | | | | | | |
|---|---|---|---|---|---|---|---|---|
| 0.418607 | 0.110026 | 0.159 | 0.14 | 1 | 6 | Ccpg1 | 0.019 | Young |
| 0.418885 | 0.113839 | 0.211 | 0.191 | 1 | 6 | Kif21b | 0.02 | Young |
| 0.419325 | 0.224364 | 0.042 | 0.031 | 1 | 6 | Acss2 | 0.011 | Young |
| 0.419337 | 0.158373 | 0.047 | 0.06 | 1 | 6 | Psmf1 | -0.013 | Young |
| 0.419457 | 1.063779 | 0.032 | 0.023 | 1 | 6 | Mrps2 | 0.009 | Young |
| 0.41993 | 0.081451 | 0.284 | 0.263 | 1 | 6 | Gapvd1 | 0.021 | Young |
| 0.419985 | 0.245251 | 0.115 | 0.1 | 1 | 6 | Sh2d1b1 | 0.015 | Young |
| 0.420218 | 0.125579 | 0.316 | 0.291 | 1 | 6 | Msl2 | 0.025 | Young |
| 0.420745 | 0.116519 | 0.085 | 0.103 | 1 | 6 | Farsa | -0.018 | Young |
| 0.420879 | 0.018479 | 0.04 | 0.051 | 1 | 6 | Narfl | -0.011 | Young |
| 0.421071 | 0.234698 | 0.025 | 0.017 | 1 | 6 | Ppp1r14a | 0.008 | Young |
| 0.42149 | 0.502305 | 0.038 | 0.029 | 1 | 6 | Bex6 | 0.009 | Young |
| 0.421502 | 0.657801 | 0.09 | 0.077 | 1 | 6 | Ddt | 0.013 | Young |
| 0.421568 | 0.232742 | 0.022 | 0.014 | 1 | 6 | Elp4 | 0.008 | Young |
| 0.421676 | 0.107115 | 0.04 | 0.051 | 1 | 6 | Adap2 | -0.011 | Young |
| 0.421953 | 0.418594 | 0.1 | 0.086 | 1 | 6 | Pgrmc2 | 0.014 | Young |
| 0.421988 | 0.178765 | 0.361 | 0.34 | 1 | 6 | Mef2d | 0.021 | Young |
| 0.422185 | 0.798673 | 0.025 | 0.017 | 1 | 6 | Slc25a15 | 0.008 | Young |
| 0.422215 | 0.169538 | 0.202 | 0.183 | 1 | 6 | Amz1 | 0.019 | Young |
| 0.422753 | 0.455478 | 0.042 | 0.031 | 1 | 6 | Scn4b | 0.011 | Young |
| 0.422782 | 0.374205 | 0.092 | 0.077 | 1 | 6 | Rps6ka2 | 0.015 | Young |
| 0.422905 | 0.541457 | 0.032 | 0.023 | 1 | 6 | 1190007I0 | 0.009 | Young |
| 0.42322 | 0.35475 | 0.092 | 0.077 | 1 | 6 | Borcs7 | 0.015 | Young |
| 0.423736 | 0.015356 | 0.508 | 0.469 | 1 | 6 | Aes | 0.039 | Young |
| 0.423859 | 0.38983 | 0.025 | 0.017 | 1 | 6 | Ptar1 | 0.008 | Young |
| 0.424047 | 0.09198 | 0.236 | 0.22 | 1 | 6 | Rbm17 | 0.016 | Young |
| 0.424147 | 0.105457 | 0.074 | 0.06 | 1 | 6 | Nagpa | 0.014 | Young |
| 0.424281 | 0.108064 | 0.212 | 0.194 | 1 | 6 | 1810022K0 | 0.018 | Young |
| 0.424627 | 0.171183 | 0.038 | 0.029 | 1 | 6 | Snx16 | 0.009 | Young |
| 0.42466 | 0.375028 | 0.035 | 0.026 | 1 | 6 | Rhbdd2 | 0.009 | Young |
| 0.424769 | 0.23493 | 0.089 | 0.074 | 1 | 6 | Gucd1 | 0.015 | Young |
| 0.424863 | 0.115006 | 0.154 | 0.134 | 1 | 6 | Pnrc2 | 0.02 | Young |
| 0.424882 | 0.222372 | 0.025 | 0.034 | 1 | 6 | Ccdc71 | -0.009 | Young |
| 0.424977 | 0.430441 | 0.025 | 0.017 | 1 | 6 | Gm10036 | 0.008 | Young |
| 0.425389 | 0.143435 | 0.084 | 0.069 | 1 | 6 | Mrpl16 | 0.015 | Young |
| 0.425524 | 0.064742 | 0.211 | 0.189 | 1 | 6 | Vps13b | 0.022 | Young |
| 0.425601 | 0.440187 | 0.035 | 0.026 | 1 | 6 | B230118H0 | 0.009 | Young |
| 0.42562 | 0.240601 | 0.028 | 0.02 | 1 | 6 | Map3k12 | 0.008 | Young |
| 0.42572 | 0.016972 | 0.135 | 0.117 | 1 | 6 | Senp7 | 0.018 | Young |
| 0.425748 | 0.106884 | 0.125 | 0.109 | 1 | 6 | 1110038F1 | 0.016 | Young |
| 0.425766 | 0.471059 | 0.042 | 0.031 | 1 | 6 | Rpp30 | 0.011 | Young |
| 0.425827 | 0.185835 | 0.259 | 0.24 | 1 | 6 | Ccni | 0.019 | Young |
| 0.426023 | 0.065693 | 0.139 | 0.16 | 1 | 6 | Mcfd2 | -0.021 | Young |
| 0.426544 | 0.278323 | 0.035 | 0.026 | 1 | 6 | Btla | 0.009 | Young |
| 0.426658 | 0.456156 | 0.025 | 0.017 | 1 | 6 | Tmem35b | 0.008 | Young |

| | | | | | | | | |
|---|---|---|---|---|---|---|---|---|
| 0.426909 | 0.360453 | 0.212 | 0.194 | 1 | 6 | Ubr2 | 0.018 | Young |
| 0.427125 | 0.124607 | 0.403 | 0.374 | 1 | 6 | Phip | 0.029 | Young |
| 0.427196 | 0.608297 | 0.028 | 0.02 | 1 | 6 | Wdr4 | 0.008 | Young |
| 0.427488 | 0.142162 | 0.274 | 0.251 | 1 | 6 | Emd | 0.023 | Young |
| 0.427493 | 0.284297 | 0.042 | 0.031 | 1 | 6 | Slc7a11 | 0.011 | Young |
| 0.427657 | 0.024849 | 0.154 | 0.137 | 1 | 6 | Arhgap1 | 0.017 | Young |
| 0.427903 | 0.007294 | 0.657 | 0.64 | 1 | 6 | Rpl22l1 | 0.017 | Young |
| 0.427959 | 0.451625 | 0.035 | 0.026 | 1 | 6 | Zbtb42 | 0.009 | Young |
| 0.428067 | 0.138338 | 0.065 | 0.08 | 1 | 6 | Tbrg4 | -0.015 | Young |
| 0.428479 | 0.012132 | 0.045 | 0.034 | 1 | 6 | Ddx60 | 0.011 | Young |
| 0.428745 | 0.11504 | 0.365 | 0.343 | 1 | 6 | Arpp19 | 0.022 | Young |
| 0.428827 | 0.038157 | 0.575 | 0.554 | 1 | 6 | Calm2 | 0.021 | Young |
| 0.429156 | 0.332762 | 0.09 | 0.077 | 1 | 6 | Rrm1 | 0.013 | Young |
| 0.429351 | 0.28529 | 0.032 | 0.023 | 1 | 6 | Dimt1 | 0.009 | Young |
| 0.430026 | 0.080761 | 0.274 | 0.311 | 1 | 6 | Nars | -0.037 | Young |
| 0.430091 | 0.17531 | 0.042 | 0.031 | 1 | 6 | Mettl21a | 0.011 | Young |
| 0.430228 | 0.266228 | 0.189 | 0.174 | 1 | 6 | Mrpl48 | 0.015 | Young |
| 0.430308 | 0.040228 | 0.105 | 0.089 | 1 | 6 | Cdk8 | 0.016 | Young |
| 0.430586 | 0.145445 | 0.393 | 0.377 | 1 | 6 | Stk10 | 0.016 | Young |
| 0.430799 | 0.040182 | 0.035 | 0.026 | 1 | 6 | Gpatch4 | 0.009 | Young |
| 0.430863 | 1.438002 | 0.047 | 0.037 | 1 | 6 | Gpr85 | 0.01 | Young |
| 0.431387 | 0.565206 | 0.164 | 0.149 | 1 | 6 | Galnt10 | 0.015 | Young |
| 0.431411 | 0.040772 | 0.592 | 0.549 | 1 | 6 | Ppp1ca | 0.043 | Young |
| 0.431737 | 0.344771 | 0.125 | 0.109 | 1 | 6 | Dnttip2 | 0.016 | Young |
| 0.431847 | 0.001923 | 0.038 | 0.029 | 1 | 6 | Lin9 | 0.009 | Young |
| 0.431951 | 0.375691 | 0.13 | 0.114 | 1 | 6 | Zfp280d | 0.016 | Young |
| 0.432595 | 0.491014 | 0.075 | 0.063 | 1 | 6 | Panx1 | 0.012 | Young |
| 0.432883 | 0.633881 | 0.099 | 0.086 | 1 | 6 | Parp2 | 0.013 | Young |
| 0.433004 | 0.078149 | 0.013 | 0.02 | 1 | 6 | Btbd19 | -0.007 | Young |
| 0.433201 | 0.305649 | 0.06 | 0.049 | 1 | 6 | Ankrd49 | 0.011 | Young |
| 0.433672 | 0.204308 | 0.013 | 0.02 | 1 | 6 | Fads6 | -0.007 | Young |
| 0.433971 | 0.13586 | 0.406 | 0.383 | 1 | 6 | Pycard | 0.023 | Young |
| 0.434291 | 0.632891 | 0.054 | 0.043 | 1 | 6 | Man2b2 | 0.011 | Young |
| 0.434291 | 0.06399 | 0.045 | 0.057 | 1 | 6 | Gfm2 | -0.012 | Young |
| 0.434828 | 0.032821 | 0.147 | 0.129 | 1 | 6 | Htatsf1 | 0.018 | Young |
| 0.435086 | 0.283893 | 0.064 | 0.051 | 1 | 6 | Fgd3 | 0.013 | Young |
| 0.435105 | 0.112977 | 0.119 | 0.103 | 1 | 6 | Scrib | 0.016 | Young |
| 0.435658 | 0.275034 | 0.028 | 0.02 | 1 | 6 | Rhobtb1 | 0.008 | Young |
| 0.435678 | 0.000892 | 0.013 | 0.02 | 1 | 6 | 1810044D( | -0.007 | Young |
| 0.435849 | 0.067191 | 0.237 | 0.266 | 1 | 6 | Mib1 | -0.029 | Young |
| 0.435924 | 0.252026 | 0.102 | 0.089 | 1 | 6 | Aspscr1 | 0.013 | Young |
| 0.436103 | 0.232263 | 0.137 | 0.12 | 1 | 6 | Igf1r | 0.017 | Young |
| 0.43619 | 0.003177 | 0.028 | 0.02 | 1 | 6 | Natd1 | 0.008 | Young |
| 0.436214 | 0.622614 | 0.054 | 0.043 | 1 | 6 | Zfp943 | 0.011 | Young |
| 0.436859 | 0.047695 | 0.032 | 0.023 | 1 | 6 | Grwd1 | 0.009 | Young |

| | | | | | | | | |
|---|---|---|---|---|---|---|---|---|
| 0.437098 | 0.290574 | 0.264 | 0.249 | 1 | 6 | Smap1 | 0.015 | Young |
| 0.43756 | 0.248796 | 0.067 | 0.054 | 1 | 6 | Fhod1 | 0.013 | Young |
| 0.437813 | 0.215619 | 0.092 | 0.077 | 1 | 6 | Dhx8 | 0.015 | Young |
| 0.437978 | 0.104908 | 0.089 | 0.074 | 1 | 6 | Cbr2 | 0.015 | Young |
| 0.438093 | 0.35695 | 0.117 | 0.103 | 1 | 6 | Pdk3 | 0.014 | Young |
| 0.43837 | 0.037881 | 0.032 | 0.023 | 1 | 6 | Eps8l1 | 0.009 | Young |
| 0.439706 | 0.041049 | 0.013 | 0.02 | 1 | 6 | Slc25a14 | -0.007 | Young |
| 0.439857 | 0.08206 | 0.823 | 0.791 | 1 | 6 | Ndufa6 | 0.032 | Young |
| 0.439959 | 0.199771 | 0.085 | 0.071 | 1 | 6 | Tlr11 | 0.014 | Young |
| 0.441054 | 0.236974 | 0.013 | 0.02 | 1 | 6 | Nup35 | -0.007 | Young |
| 0.441054 | 0.205611 | 0.013 | 0.02 | 1 | 6 | Slc25a26 | -0.007 | Young |
| 0.441054 | 0.125492 | 0.013 | 0.02 | 1 | 6 | Nudt7 | -0.007 | Young |
| 0.441095 | 0.389588 | 0.092 | 0.077 | 1 | 6 | Mri1 | 0.015 | Young |
| 0.441127 | 0.096045 | 0.035 | 0.046 | 1 | 6 | Klf8 | -0.011 | Young |
| 0.441477 | 0.079232 | 0.677 | 0.651 | 1 | 6 | Hexa | 0.026 | Young |
| 0.44158 | 0.049877 | 0.261 | 0.243 | 1 | 6 | Nrbp1 | 0.018 | Young |
| 0.4417 | 0.051944 | 0.227 | 0.206 | 1 | 6 | Spata6 | 0.021 | Young |
| 0.44173 | 0.019904 | 0.47 | 0.437 | 1 | 6 | Prex1 | 0.033 | Young |
| 0.441888 | 0.096661 | 0.202 | 0.183 | 1 | 6 | Prr14 | 0.019 | Young |
| 0.441912 | 0.129488 | 0.11 | 0.094 | 1 | 6 | Snap29 | 0.016 | Young |
| 0.442325 | 0.263021 | 0.11 | 0.094 | 1 | 6 | Cyp27a1 | 0.016 | Young |
| 0.442791 | 0.308453 | 0.391 | 0.391 | 1 | 6 | Mrps21 | 0 | Young |
| 0.443 | 0.158235 | 0.007 | 0.011 | 1 | 6 | Fxyd1 | -0.004 | Young |
| 0.44322 | 0.42615 | 0.206 | 0.189 | 1 | 6 | Akap9 | 0.017 | Young |
| 0.443646 | 0.385976 | 0.099 | 0.086 | 1 | 6 | Lsm2 | 0.013 | Young |
| 0.443842 | 0.091497 | 0.119 | 0.103 | 1 | 6 | Gar1 | 0.016 | Young |
| 0.444036 | 0.516534 | 0.075 | 0.063 | 1 | 6 | Fbxo22 | 0.012 | Young |
| 0.444082 | 1.143147 | 0.062 | 0.051 | 1 | 6 | Fam8a1 | 0.011 | Young |
| 0.444517 | 0.259923 | 0.06 | 0.049 | 1 | 6 | Smarcd1 | 0.011 | Young |
| 0.444562 | 0.13592 | 0.055 | 0.069 | 1 | 6 | Cpped1 | -0.014 | Young |
| 0.444885 | 0.549329 | 0.06 | 0.049 | 1 | 6 | Tgfbr1 | 0.011 | Young |
| 0.444971 | 0.2176 | 0.197 | 0.183 | 1 | 6 | Uba2 | 0.014 | Young |
| 0.445025 | 0.432981 | 0.075 | 0.063 | 1 | 6 | Snx14 | 0.012 | Young |
| 0.445484 | 0.234306 | 0.035 | 0.046 | 1 | 6 | Pi4kb | -0.011 | Young |
| 0.445772 | 0.576526 | 0.007 | 0.011 | 1 | 6 | Vangl2 | -0.004 | Young |
| 0.445772 | 0.225705 | 0.007 | 0.011 | 1 | 6 | Zfp768 | -0.004 | Young |
| 0.445788 | 0.370805 | 0.013 | 0.02 | 1 | 6 | Simc1 | -0.007 | Young |
| 0.446021 | 0.326054 | 0.187 | 0.166 | 1 | 6 | Prr5l | 0.021 | Young |
| 0.446087 | 0.059757 | 0.069 | 0.083 | 1 | 6 | Mcm4 | -0.014 | Young |
| 0.446591 | 0.048274 | 0.45 | 0.426 | 1 | 6 | Il10ra | 0.024 | Young |
| 0.44662 | 0.129335 | 0.109 | 0.094 | 1 | 6 | Tk2 | 0.015 | Young |
| 0.446698 | 0.157532 | 0.007 | 0.011 | 1 | 6 | Sfn | -0.004 | Young |
| 0.446813 | 0.2316 | 0.291 | 0.28 | 1 | 6 | Kpnb1 | 0.011 | Young |
| 0.446977 | 0.342216 | 0.169 | 0.151 | 1 | 6 | Spg7 | 0.018 | Young |
| 0.447552 | 0.030007 | 0.117 | 0.1 | 1 | 6 | Pnpla7 | 0.017 | Young |

| | | | | | | | | |
|---|---|---|---|---|---|---|---|---|
| 0.447616 | 0.057279 | 0.256 | 0.237 | 1 | 6 | Bod1l | 0.019 | Young |
| 0.447625 | 1.036911 | 0.007 | 0.011 | 1 | 6 | Gm33104 | -0.004 | Young |
| 0.447635 | 0.056996 | 0.104 | 0.089 | 1 | 6 | Pold3 | 0.015 | Young |
| 0.448029 | 0.29405 | 0.094 | 0.08 | 1 | 6 | Thap2 | 0.014 | Young |
| 0.448313 | 0.211718 | 0.405 | 0.38 | 1 | 6 | Rnase6 | 0.025 | Young |
| 0.448554 | 0.867467 | 0.007 | 0.011 | 1 | 6 | Jade3 | -0.004 | Young |
| 0.448554 | 0.378917 | 0.007 | 0.011 | 1 | 6 | 3010003L2 | -0.004 | Young |
| 0.448849 | 0.208361 | 0.105 | 0.091 | 1 | 6 | Foxk1 | 0.014 | Young |
| 0.449268 | 0.071961 | 0.079 | 0.094 | 1 | 6 | Necap1 | -0.015 | Young |
| 0.449333 | 0.042867 | 0.11 | 0.129 | 1 | 6 | Maml1 | -0.019 | Young |
| 0.449369 | 0.058028 | 0.249 | 0.226 | 1 | 6 | Suz12 | 0.023 | Young |
| 0.449877 | 0.185888 | 0.373 | 0.369 | 1 | 6 | Prdx5 | 0.004 | Young |
| 0.45127 | 0.1968 | 0.132 | 0.114 | 1 | 6 | Pctp | 0.018 | Young |
| 0.452587 | 0.159939 | 0.268 | 0.249 | 1 | 6 | Cript | 0.019 | Young |
| 0.452639 | 0.068228 | 0.057 | 0.046 | 1 | 6 | Etv5 | 0.011 | Young |
| 0.45277 | 0.084208 | 0.247 | 0.226 | 1 | 6 | Usp19 | 0.021 | Young |
| 0.453773 | 0.501103 | 0.084 | 0.071 | 1 | 6 | Triobp | 0.013 | Young |
| 0.453979 | 0.400059 | 0.075 | 0.063 | 1 | 6 | Pyurf | 0.012 | Young |
| 0.454745 | 0.00855 | 0.266 | 0.24 | 1 | 6 | Csgalnact2 | 0.026 | Young |
| 0.455199 | 0.324848 | 0.05 | 0.04 | 1 | 6 | Acvr2a | 0.01 | Young |
| 0.455731 | 0.03721 | 0.187 | 0.166 | 1 | 6 | Arhgap39 | 0.021 | Young |
| 0.456488 | 0.23253 | 0.151 | 0.134 | 1 | 6 | Rlf | 0.017 | Young |
| 0.45688 | 0.637625 | 0.09 | 0.077 | 1 | 6 | Stk19 | 0.013 | Young |
| 0.457005 | 0.158868 | 0.443 | 0.429 | 1 | 6 | Cerk | 0.014 | Young |
| 0.457154 | 0.151448 | 0.283 | 0.266 | 1 | 6 | 2410015M | 0.017 | Young |
| 0.457306 | 0.272737 | 0.097 | 0.083 | 1 | 6 | Clec2i | 0.014 | Young |
| 0.45753 | 0.447403 | 0.109 | 0.094 | 1 | 6 | Lpcat3 | 0.015 | Young |
| 0.458124 | 0.100176 | 0.129 | 0.149 | 1 | 6 | Cops3 | -0.02 | Young |
| 0.45847 | 0.313656 | 0.189 | 0.171 | 1 | 6 | Itpk1 | 0.018 | Young |
| 0.458569 | 0.14397 | 0.085 | 0.071 | 1 | 6 | Ubr1 | 0.014 | Young |
| 0.459519 | 0.170589 | 0.085 | 0.071 | 1 | 6 | Wdr77 | 0.014 | Young |
| 0.459739 | 0.123359 | 0.06 | 0.049 | 1 | 6 | Slc11a1 | 0.011 | Young |
| 0.460119 | 0.025446 | 0.064 | 0.077 | 1 | 6 | Saal1 | -0.013 | Young |
| 0.460131 | 0.065549 | 0.764 | 0.726 | 1 | 6 | Gabarap | 0.038 | Young |
| 0.460147 | 0.379857 | 0.099 | 0.086 | 1 | 6 | Arl10 | 0.013 | Young |
| 0.460249 | 0.128343 | 0.172 | 0.157 | 1 | 6 | Hnrnph3 | 0.015 | Young |
| 0.460466 | 0.084349 | 0.023 | 0.031 | 1 | 6 | Tbl2 | -0.008 | Young |
| 0.460557 | 0.093407 | 0.181 | 0.166 | 1 | 6 | Mtmr14 | 0.015 | Young |
| 0.460792 | 0.095869 | 0.199 | 0.183 | 1 | 6 | Abca3 | 0.016 | Young |
| 0.460817 | 0.448485 | 0.047 | 0.037 | 1 | 6 | Rcn3 | 0.01 | Young |
| 0.460858 | 0.194656 | 0.054 | 0.043 | 1 | 6 | Mllt3 | 0.011 | Young |
| 0.461063 | 0.211871 | 0.495 | 0.474 | 1 | 6 | Ptpn1 | 0.021 | Young |
| 0.461086 | 0.147366 | 0.278 | 0.263 | 1 | 6 | Phactr2 | 0.015 | Young |
| 0.461588 | 0.131661 | 0.167 | 0.151 | 1 | 6 | Mir142hg | 0.016 | Young |
| 0.461789 | 0.727598 | 0.043 | 0.034 | 1 | 6 | Spg20 | 0.009 | Young |

| | | | | | | | | |
|---|---|---|---|---|---|---|---|---|
| 0.461906 | 0.052748 | 0.269 | 0.249 | 1 | 6 | Isy1 | 0.02 | Young |
| 0.462112 | 0.060164 | 0.124 | 0.143 | 1 | 6 | Mrpl15 | -0.019 | Young |
| 0.463011 | 0.217541 | 0.142 | 0.126 | 1 | 6 | Mid1 | 0.016 | Young |
| 0.463467 | 0.348629 | 0.109 | 0.094 | 1 | 6 | Snapin | 0.015 | Young |
| 0.46355 | 0.387297 | 0.107 | 0.094 | 1 | 6 | Nip7 | 0.013 | Young |
| 0.46389 | 0.064309 | 0.082 | 0.069 | 1 | 6 | Tex10 | 0.013 | Young |
| 0.464003 | 0.206857 | 0.132 | 0.117 | 1 | 6 | Brox | 0.015 | Young |
| 0.464183 | 0.198261 | 0.226 | 0.203 | 1 | 6 | Zfp445 | 0.023 | Young |
| 0.466022 | 0.046695 | 0.614 | 0.6 | 1 | 6 | Atpif1 | 0.014 | Young |
| 0.466252 | 0.024678 | 0.197 | 0.18 | 1 | 6 | Fbxl20 | 0.017 | Young |
| 0.466254 | 0.180413 | 0.119 | 0.103 | 1 | 6 | Rtca | 0.016 | Young |
| 0.466413 | 0.266211 | 0.075 | 0.063 | 1 | 6 | C5ar1 | 0.012 | Young |
| 0.466494 | 0.083138 | 0.306 | 0.331 | 1 | 6 | Cycs | -0.025 | Young |
| 0.466502 | 0.071253 | 0.064 | 0.051 | 1 | 6 | Kif13a | 0.013 | Young |
| 0.466773 | 0.471127 | 0.047 | 0.037 | 1 | 6 | Fbxw8 | 0.01 | Young |
| 0.468089 | 0.359624 | 0.149 | 0.134 | 1 | 6 | Ahsa1 | 0.015 | Young |
| 0.468418 | 0.34672 | 0.043 | 0.034 | 1 | 6 | Lnx2 | 0.009 | Young |
| 0.468556 | 0.114936 | 0.279 | 0.26 | 1 | 6 | Pqlc1 | 0.019 | Young |
| 0.468644 | 0.214797 | 0.057 | 0.046 | 1 | 6 | Paqr7 | 0.011 | Young |
| 0.469541 | 0.123379 | 0.187 | 0.169 | 1 | 6 | Hltf | 0.018 | Young |
| 0.471003 | 0.08484 | 0.134 | 0.154 | 1 | 6 | Sra1 | -0.02 | Young |
| 0.471083 | 0.317927 | 0.152 | 0.14 | 1 | 6 | Pxk | 0.012 | Young |
| 0.471562 | 0.479275 | 0.079 | 0.066 | 1 | 6 | Nup214 | 0.013 | Young |
| 0.471645 | 0.06028 | 0.085 | 0.071 | 1 | 6 | Mcoln1 | 0.014 | Young |
| 0.472529 | 0.285199 | 0.075 | 0.063 | 1 | 6 | Chchd4 | 0.012 | Young |
| 0.472582 | 0.329565 | 0.072 | 0.06 | 1 | 6 | Cwc25 | 0.012 | Young |
| 0.472667 | 0.153117 | 0.043 | 0.054 | 1 | 6 | Zfp715 | -0.011 | Young |
| 0.473088 | 0.003354 | 0.207 | 0.191 | 1 | 6 | Smim11 | 0.016 | Young |
| 0.473658 | 0.105508 | 0.321 | 0.297 | 1 | 6 | Trpc4ap | 0.024 | Young |
| 0.474376 | 0.10712 | 0.296 | 0.277 | 1 | 6 | Igbp1 | 0.019 | Young |
| 0.474778 | 0.282482 | 0.104 | 0.091 | 1 | 6 | Snapc5 | 0.013 | Young |
| 0.475438 | 0.223617 | 0.187 | 0.171 | 1 | 6 | Tmem128 | 0.016 | Young |
| 0.475566 | 0.402901 | 0.069 | 0.057 | 1 | 6 | Clcf1 | 0.012 | Young |
| 0.475812 | 0.095445 | 0.102 | 0.089 | 1 | 6 | Timm44 | 0.013 | Young |
| 0.476188 | 0.019572 | 0.028 | 0.037 | 1 | 6 | AU022252 | -0.009 | Young |
| 0.476271 | 0.01848 | 0.057 | 0.046 | 1 | 6 | I830077J02 | 0.011 | Young |
| 0.476539 | 0.105203 | 0.097 | 0.083 | 1 | 6 | Znfx1 | 0.014 | Young |
| 0.476924 | 0.400697 | 0.065 | 0.054 | 1 | 6 | Ptpn23 | 0.011 | Young |
| 0.477144 | 0.125676 | 0.306 | 0.291 | 1 | 6 | Ctsd | 0.015 | Young |
| 0.477338 | 0.501724 | 0.043 | 0.034 | 1 | 6 | Igip | 0.009 | Young |
| 0.477338 | 0.405726 | 0.043 | 0.034 | 1 | 6 | Carf | 0.009 | Young |
| 0.477773 | 0.513947 | 0.04 | 0.031 | 1 | 6 | 1810032O( | 0.009 | Young |
| 0.477867 | 1.779027 | 0.01 | 0.006 | 1 | 6 | Zfp438 | 0.004 | Young |
| 0.478055 | 0.203942 | 0.164 | 0.149 | 1 | 6 | Hcfc1 | 0.015 | Young |
| 0.478116 | 0.068105 | 0.054 | 0.043 | 1 | 6 | Pkdcc | 0.011 | Young |

| | | | | | | | | |
|---|---|---|---|---|---|---|---|---|
| 0.478149 | 0.125675 | 0.279 | 0.263 | 1 | 6 | Anapc13 | 0.016 | Young |
| 0.478235 | 0.261999 | 0.043 | 0.034 | 1 | 6 | Rbmx2 | 0.009 | Young |
| 0.47839 | 0.404533 | 0.119 | 0.106 | 1 | 6 | 1110008P1 | 0.013 | Young |
| 0.478482 | 0.295005 | 0.102 | 0.089 | 1 | 6 | Slc25a46 | 0.013 | Young |
| 0.478684 | 0.330959 | 0.161 | 0.146 | 1 | 6 | Lysmd3 | 0.015 | Young |
| 0.478735 | 0.070357 | 0.301 | 0.28 | 1 | 6 | Dna2 | 0.021 | Young |
| 0.47883 | 1.18721 | 0.01 | 0.006 | 1 | 6 | Gm26522 | 0.004 | Young |
| 0.47883 | 1.185553 | 0.01 | 0.006 | 1 | 6 | Cd34 | 0.004 | Young |
| 0.47883 | 1.093074 | 0.01 | 0.006 | 1 | 6 | Chtf18 | 0.004 | Young |
| 0.47883 | 0.97563 | 0.01 | 0.006 | 1 | 6 | Gm527 | 0.004 | Young |
| 0.479043 | 0.217884 | 0.094 | 0.08 | 1 | 6 | Ccdc34 | 0.014 | Young |
| 0.479177 | 0.884301 | 0.055 | 0.046 | 1 | 6 | Bcas3 | 0.009 | Young |
| 0.479463 | 0.056105 | 0.212 | 0.237 | 1 | 6 | Npepps | -0.025 | Young |
| 0.479794 | 1.234606 | 0.01 | 0.006 | 1 | 6 | Map7d3 | 0.004 | Young |
| 0.479794 | 1.001068 | 0.01 | 0.006 | 1 | 6 | Rec8 | 0.004 | Young |
| 0.479794 | 0.943747 | 0.01 | 0.006 | 1 | 6 | Wdpcp | 0.004 | Young |
| 0.480445 | 0.040776 | 0.346 | 0.326 | 1 | 6 | Aff4 | 0.02 | Young |
| 0.480759 | 1.703797 | 0.01 | 0.006 | 1 | 6 | Acta2 | 0.004 | Young |
| 0.480759 | 0.885678 | 0.01 | 0.006 | 1 | 6 | Wee1 | 0.004 | Young |
| 0.480802 | 0.212891 | 0.162 | 0.149 | 1 | 6 | Gpalpp1 | 0.013 | Young |
| 0.480976 | 0.029639 | 0.828 | 0.774 | 1 | 6 | Ubl3 | 0.054 | Young |
| 0.481725 | 0.625427 | 0.01 | 0.006 | 1 | 6 | Gba2 | 0.004 | Young |
| 0.481725 | 0.105412 | 0.028 | 0.037 | 1 | 6 | Slc12a2 | -0.009 | Young |
| 0.482691 | 0.675443 | 0.01 | 0.006 | 1 | 6 | Gm29367 | 0.004 | Young |
| 0.482691 | 0.635325 | 0.01 | 0.006 | 1 | 6 | Ttc8 | 0.004 | Young |
| 0.482691 | 0.593027 | 0.01 | 0.006 | 1 | 6 | Tceanc | 0.004 | Young |
| 0.482842 | 0.060349 | 0.105 | 0.123 | 1 | 6 | 5-Mar | -0.018 | Young |
| 0.482882 | 0.163672 | 0.074 | 0.063 | 1 | 6 | Acot8 | 0.011 | Young |
| 0.48293 | 0.078996 | 0.05 | 0.04 | 1 | 6 | Dhx57 | 0.01 | Young |
| 0.483161 | 0.105312 | 0.087 | 0.074 | 1 | 6 | Cttnbp2nl | 0.013 | Young |
| 0.483472 | 0.33224 | 0.105 | 0.091 | 1 | 6 | Mzt1 | 0.014 | Young |
| 0.483647 | 0.141837 | 0.177 | 0.163 | 1 | 6 | Dguok | 0.014 | Young |
| 0.483659 | 1.122554 | 0.01 | 0.006 | 1 | 6 | Ciart | 0.004 | Young |
| 0.483659 | 0.853976 | 0.01 | 0.006 | 1 | 6 | Mrm2 | 0.004 | Young |
| 0.483659 | 0.81949 | 0.01 | 0.006 | 1 | 6 | Morn2 | 0.004 | Young |
| 0.483659 | 0.799802 | 0.01 | 0.006 | 1 | 6 | 4930524J0 | 0.004 | Young |
| 0.483659 | 0.747042 | 0.01 | 0.006 | 1 | 6 | Zfand4 | 0.004 | Young |
| 0.483659 | 0.455544 | 0.01 | 0.006 | 1 | 6 | Galnt12 | 0.004 | Young |
| 0.483659 | 0.453512 | 0.01 | 0.006 | 1 | 6 | Gm11423 | 0.004 | Young |
| 0.483659 | 0.447363 | 0.01 | 0.006 | 1 | 6 | Htra3 | 0.004 | Young |
| 0.483659 | 0.27732 | 0.01 | 0.006 | 1 | 6 | Zfp605 | 0.004 | Young |
| 0.484629 | 0.710447 | 0.01 | 0.006 | 1 | 6 | Jmjd7 | 0.004 | Young |
| 0.484629 | 0.316763 | 0.01 | 0.006 | 1 | 6 | Pomc | 0.004 | Young |
| 0.484771 | 0.3524 | 0.087 | 0.074 | 1 | 6 | Zfp512 | 0.013 | Young |
| 0.484784 | 0.28903 | 0.226 | 0.211 | 1 | 6 | Pcmt1 | 0.015 | Young |

| | | | | | | | | |
|---|---|---|---|---|---|---|---|---|
| 0.485271 | 0.002754 | 0.5 | 0.477 | 1 | 6 | Add3 | 0.023 | Young |
| 0.485599 | 0.327158 | 0.01 | 0.006 | 1 | 6 | Palb2 | 0.004 | Young |
| 0.485599 | 0.243001 | 0.01 | 0.006 | 1 | 6 | Wdr62 | 0.004 | Young |
| 0.485599 | 0.186495 | 0.01 | 0.006 | 1 | 6 | Oas1c | 0.004 | Young |
| 0.48573 | 0.267009 | 0.095 | 0.083 | 1 | 6 | Isg20l2 | 0.012 | Young |
| 0.485741 | 0.439032 | 0.04 | 0.031 | 1 | 6 | Tenm4 | 0.009 | Young |
| 0.486206 | 0.034122 | 0.047 | 0.037 | 1 | 6 | Ccnyl1 | 0.01 | Young |
| 0.486309 | 0.247461 | 0.214 | 0.197 | 1 | 6 | Nop58 | 0.017 | Young |
| 0.48635 | 0.390321 | 0.043 | 0.034 | 1 | 6 | Frat2 | 0.009 | Young |
| 0.486442 | 0.231021 | 0.314 | 0.3 | 1 | 6 | Zfp263 | 0.014 | Young |
| 0.486545 | 0.828448 | 0.033 | 0.026 | 1 | 6 | Exd2 | 0.007 | Young |
| 0.48657 | 0.281946 | 0.01 | 0.006 | 1 | 6 | Gins3 | 0.004 | Young |
| 0.48657 | 0.102247 | 0.01 | 0.006 | 1 | 6 | Rgs12 | 0.004 | Young |
| 0.487016 | 0.016904 | 0.232 | 0.214 | 1 | 6 | Lpxn | 0.018 | Young |
| 0.487174 | 0.172665 | 0.192 | 0.177 | 1 | 6 | Aurkaip1 | 0.015 | Young |
| 0.487542 | 0.132244 | 0.01 | 0.006 | 1 | 6 | Nat8f1 | 0.004 | Young |
| 0.487542 | 0.080209 | 0.01 | 0.006 | 1 | 6 | Dmxl2 | 0.004 | Young |
| 0.487758 | 0.227012 | 0.309 | 0.297 | 1 | 6 | Scarb2 | 0.012 | Young |
| 0.48912 | 0.262965 | 0.214 | 0.2 | 1 | 6 | Eif4e | 0.014 | Young |
| 0.489516 | 0.430249 | 0.04 | 0.031 | 1 | 6 | Fancg | 0.009 | Young |
| 0.489577 | 0.172523 | 0.072 | 0.086 | 1 | 6 | Uvssa | -0.014 | Young |
| 0.489701 | 0.066946 | 0.177 | 0.16 | 1 | 6 | Atp6v0a1 | 0.017 | Young |
| 0.489753 | 0.102986 | 0.253 | 0.231 | 1 | 6 | Net1 | 0.022 | Young |
| 0.489944 | 0.053237 | 0.137 | 0.123 | 1 | 6 | Osgin2 | 0.014 | Young |
| 0.490113 | 0.075395 | 0.273 | 0.251 | 1 | 6 | Rnf166 | 0.022 | Young |
| 0.490151 | 0.022979 | 0.137 | 0.12 | 1 | 6 | Gcc1 | 0.017 | Young |
| 0.490165 | 0.433258 | 0.033 | 0.026 | 1 | 6 | Ttc38 | 0.007 | Young |
| 0.490206 | 0.188137 | 0.177 | 0.163 | 1 | 6 | Zfp131 | 0.014 | Young |
| 0.490461 | 0.585399 | 0.059 | 0.049 | 1 | 6 | Pparg | 0.01 | Young |
| 0.490519 | 0.415753 | 0.062 | 0.051 | 1 | 6 | Selenon | 0.011 | Young |
| 0.490683 | 0.641017 | 0.033 | 0.026 | 1 | 6 | Ednrb | 0.007 | Young |
| 0.491201 | 0.615523 | 0.033 | 0.026 | 1 | 6 | Zfp53 | 0.007 | Young |
| 0.491243 | 0.375892 | 0.087 | 0.074 | 1 | 6 | Ckap5 | 0.013 | Young |
| 0.491572 | 0.46434 | 0.037 | 0.029 | 1 | 6 | Mllt11 | 0.008 | Young |
| 0.492084 | 0.326079 | 0.065 | 0.054 | 1 | 6 | Hectd3 | 0.011 | Young |
| 0.492325 | 0.357792 | 0.12 | 0.109 | 1 | 6 | C1galt1 | 0.011 | Young |
| 0.492486 | 0.552251 | 0.012 | 0.017 | 1 | 6 | Gm38394 | -0.005 | Young |
| 0.492613 | 0.12939 | 0.4 | 0.383 | 1 | 6 | Ctdnep1 | 0.017 | Young |
| 0.492631 | 0.073863 | 0.503 | 0.491 | 1 | 6 | Selenok | 0.012 | Young |
| 0.492994 | 0.102045 | 0.151 | 0.134 | 1 | 6 | Dda1 | 0.017 | Young |
| 0.492995 | 0.041946 | 0.585 | 0.563 | 1 | 6 | Uqcrq | 0.022 | Young |
| 0.493072 | 0.18127 | 0.134 | 0.12 | 1 | 6 | Ddx23 | 0.014 | Young |
| 0.493082 | 0.219574 | 0.226 | 0.211 | 1 | 6 | Hmgcr | 0.015 | Young |
| 0.493082 | 0.288414 | 0.05 | 0.04 | 1 | 6 | Nup155 | 0.01 | Young |
| 0.493252 | 0.608843 | 0.07 | 0.06 | 1 | 6 | Styx | 0.01 | Young |

| | | | | | | | | |
|---|---|---|---|---|---|---|---|---|
| 0.493428 | 0.518952 | 0.182 | 0.171 | 1 | 6 | Mdm2 | 0.011 | Young |
| 0.493844 | 0.099258 | 0.077 | 0.091 | 1 | 6 | Lmf1 | -0.014 | Young |
| 0.494026 | 0.316217 | 0.012 | 0.017 | 1 | 6 | Tbc1d7 | -0.005 | Young |
| 0.494324 | 0.283162 | 0.065 | 0.054 | 1 | 6 | Lrrcc1 | 0.011 | Young |
| 0.494539 | 0.290922 | 0.043 | 0.034 | 1 | 6 | Tnfsf14 | 0.009 | Young |
| 0.49484 | 0.504091 | 0.033 | 0.026 | 1 | 6 | Gtf2h3 | 0.007 | Young |
| 0.495207 | 0.313341 | 0.04 | 0.031 | 1 | 6 | Sec16b | 0.009 | Young |
| 0.495569 | 0.163065 | 0.012 | 0.017 | 1 | 6 | Zfp287 | -0.005 | Young |
| 0.495629 | 0.091201 | 0.222 | 0.209 | 1 | 6 | Tmem50b | 0.013 | Young |
| 0.495806 | 0.205469 | 0.187 | 0.171 | 1 | 6 | Slc25a36 | 0.016 | Young |
| 0.496159 | 0.142225 | 0.04 | 0.031 | 1 | 6 | Ppp1r16a | 0.009 | Young |
| 0.496231 | 1.522343 | 0.013 | 0.009 | 1 | 6 | Nadsyn1 | 0.004 | Young |
| 0.497051 | 0.284211 | 0.089 | 0.077 | 1 | 6 | Tvp23b | 0.012 | Young |
| 0.497262 | 0.035165 | 0.44 | 0.417 | 1 | 6 | Ndufc1 | 0.023 | Young |
| 0.497696 | 0.221541 | 0.102 | 0.089 | 1 | 6 | Pmpca | 0.013 | Young |
| 0.497891 | 0.160606 | 0.303 | 0.283 | 1 | 6 | Rheb | 0.02 | Young |
| 0.497962 | 0.127283 | 0.047 | 0.057 | 1 | 6 | Mbd4 | -0.01 | Young |
| 0.498202 | 0.135176 | 0.043 | 0.034 | 1 | 6 | Pygl | 0.009 | Young |
| 0.498401 | 0.489554 | 0.142 | 0.129 | 1 | 6 | Snrnp48 | 0.013 | Young |
| 0.498455 | 0.005127 | 0.993 | 0.957 | 1 | 6 | Laptm5 | 0.036 | Young |
| 0.498746 | 0.510149 | 0.069 | 0.057 | 1 | 6 | Ptgr2 | 0.012 | Young |
| 0.498751 | 0.965428 | 0.013 | 0.009 | 1 | 6 | Meis3 | 0.004 | Young |
| 0.499102 | 0.016706 | 0.077 | 0.066 | 1 | 6 | Slc39a6 | 0.011 | Young |
| 0.499371 | 0.727661 | 0.03 | 0.023 | 1 | 6 | Echdc1 | 0.007 | Young |
| 0.499374 | 0.162703 | 0.592 | 0.591 | 1 | 6 | Sh3glb1 | 0.001 | Young |
| 0.499463 | 0.129615 | 0.072 | 0.06 | 1 | 6 | Gramd1b | 0.012 | Young |
| 0.499523 | 0.171524 | 0.037 | 0.029 | 1 | 6 | Zmynd19 | 0.008 | Young |
| 0.499544 | 0.146264 | 0.209 | 0.191 | 1 | 6 | Safb | 0.018 | Young |
| 0.499592 | 1.05938 | 0.013 | 0.009 | 1 | 6 | Slx4 | 0.004 | Young |
| 0.499705 | 0.083095 | 0.4 | 0.38 | 1 | 6 | Runx3 | 0.02 | Young |
| 0.499887 | 0.007418 | 0.127 | 0.111 | 1 | 6 | Hist1h2bc | 0.016 | Young |
| 0.49992 | 0.930936 | 0.027 | 0.02 | 1 | 6 | Dcun1d4 | 0.007 | Young |
| 0.49992 | 0.860801 | 0.027 | 0.02 | 1 | 6 | Tor4a | 0.007 | Young |
| 0.49992 | 0.856588 | 0.027 | 0.02 | 1 | 6 | E130311K1 | 0.007 | Young |
| 0.499923 | 0.779243 | 0.03 | 0.023 | 1 | 6 | Tex9 | 0.007 | Young |
| 0.499923 | 0.349116 | 0.03 | 0.023 | 1 | 6 | Rwdd2a | 0.007 | Young |
| 0.499989 | 0.157414 | 0.308 | 0.291 | 1 | 6 | Cd244 | 0.017 | Young |
| 0.50052 | 0.354003 | 0.055 | 0.046 | 1 | 6 | C2cd2l | 0.009 | Young |
| 0.500586 | 0.419696 | 0.033 | 0.026 | 1 | 6 | Ppp1r3e | 0.007 | Young |
| 0.50072 | 0.001498 | 0.495 | 0.477 | 1 | 6 | Pbrm1 | 0.018 | Young |
| 0.501029 | 0.29857 | 0.124 | 0.109 | 1 | 6 | F630028O1 | 0.015 | Young |
| 0.501111 | 0.163809 | 0.189 | 0.177 | 1 | 6 | Lrrc8d | 0.012 | Young |
| 0.501192 | 0.00452 | 0.127 | 0.146 | 1 | 6 | Commd6 | -0.019 | Young |
| 0.501277 | 0.589889 | 0.013 | 0.009 | 1 | 6 | Ugt1a7c | 0.004 | Young |
| 0.501577 | 0.205675 | 0.259 | 0.246 | 1 | 6 | Aff1 | 0.013 | Young |

| | | | | | | | | |
|---|---|---|---|---|---|---|---|---|
| 0.501588 | 0.221655 | 0.139 | 0.126 | 1 | 6 | Epc2 | 0.013 | Young |
| 0.502113 | 0.112781 | 0.42 | 0.4 | 1 | 6 | Ehbp1l1 | 0.02 | Young |
| 0.502117 | 0.024273 | 0.14 | 0.123 | 1 | 6 | Gm20696 | 0.017 | Young |
| 0.502121 | 0.695388 | 0.013 | 0.009 | 1 | 6 | Ifi214 | 0.004 | Young |
| 0.502134 | 0.45328 | 0.03 | 0.023 | 1 | 6 | Hmga1 | 0.007 | Young |
| 0.502138 | 0.401528 | 0.124 | 0.111 | 1 | 6 | Pdcd2l | 0.013 | Young |
| 0.502174 | 0.071666 | 0.408 | 0.403 | 1 | 6 | Eps15 | 0.005 | Young |
| 0.502687 | 0.638522 | 0.03 | 0.023 | 1 | 6 | A430046D: | 0.007 | Young |
| 0.502687 | 0.605675 | 0.03 | 0.023 | 1 | 6 | Tlcd2 | 0.007 | Young |
| 0.50292 | 0.006041 | 0.201 | 0.183 | 1 | 6 | Rmnd5a | 0.018 | Young |
| 0.502999 | 0.152131 | 0.197 | 0.183 | 1 | 6 | Lamtor5 | 0.014 | Young |
| 0.503484 | 0.271458 | 0.334 | 0.329 | 1 | 6 | Mvp | 0.005 | Young |
| 0.50381 | 0.87193 | 0.013 | 0.009 | 1 | 6 | 4833418N( | 0.004 | Young |
| 0.504246 | 0.119814 | 0.137 | 0.123 | 1 | 6 | Dscr3 | 0.014 | Young |
| 0.504656 | 0.499419 | 0.013 | 0.009 | 1 | 6 | Hibch | 0.004 | Young |
| 0.504656 | 0.488667 | 0.013 | 0.009 | 1 | 6 | Cmpk2 | 0.004 | Young |
| 0.504656 | 0.345142 | 0.013 | 0.009 | 1 | 6 | Zfp459 | 0.004 | Young |
| 0.505233 | 0.121699 | 0.161 | 0.146 | 1 | 6 | Gbp3 | 0.015 | Young |
| 0.50579 | 0.027269 | 0.047 | 0.057 | 1 | 6 | Plcl1 | -0.01 | Young |
| 0.505933 | 0.50999 | 0.074 | 0.063 | 1 | 6 | Znrf1 | 0.011 | Young |
| 0.506123 | 0.185118 | 0.08 | 0.069 | 1 | 6 | Chst12 | 0.011 | Young |
| 0.50635 | 0.216534 | 0.013 | 0.009 | 1 | 6 | Gstm5 | 0.004 | Young |
| 0.506631 | 0.019237 | 0.033 | 0.026 | 1 | 6 | Msr1 | 0.007 | Young |
| 0.506795 | 0.235505 | 0.172 | 0.157 | 1 | 6 | Nsfl1c | 0.015 | Young |
| 0.507003 | 0.466654 | 0.07 | 0.06 | 1 | 6 | Atmin | 0.01 | Young |
| 0.50704 | 0.11703 | 0.294 | 0.274 | 1 | 6 | Sft2d2 | 0.02 | Young |
| 0.507066 | 0.006686 | 0.059 | 0.049 | 1 | 6 | Cnot11 | 0.01 | Young |
| 0.507212 | 0.06671 | 0.42 | 0.4 | 1 | 6 | Eif3m | 0.02 | Young |
| 0.507382 | 0.923949 | 0.017 | 0.011 | 1 | 6 | Tmem38b | 0.006 | Young |
| 0.507539 | 0.200346 | 0.239 | 0.226 | 1 | 6 | Smim7 | 0.013 | Young |
| 0.507626 | 0.087434 | 0.172 | 0.154 | 1 | 6 | Tomm34 | 0.018 | Young |
| 0.507684 | 0.522219 | 0.03 | 0.023 | 1 | 6 | Slc12a7 | 0.007 | Young |
| 0.507697 | 0.749473 | 0.02 | 0.014 | 1 | 6 | Slc17a9 | 0.006 | Young |
| 0.507853 | 0.168968 | 0.179 | 0.166 | 1 | 6 | Atp11b | 0.013 | Young |
| 0.507933 | 0.790942 | 0.023 | 0.017 | 1 | 6 | Ston2 | 0.006 | Young |
| 0.508566 | 0.636948 | 0.023 | 0.017 | 1 | 6 | Ankrd37 | 0.006 | Young |
| 0.508648 | 0.024968 | 0.421 | 0.4 | 1 | 6 | Cdc37 | 0.021 | Young |
| 0.508685 | 0.025014 | 0.435 | 0.414 | 1 | 6 | Rab5c | 0.021 | Young |
| 0.50889 | 0.876584 | 0.017 | 0.011 | 1 | 6 | Gm15446 | 0.006 | Young |
| 0.509068 | 1.280012 | 0.02 | 0.014 | 1 | 6 | Macrod1 | 0.006 | Young |
| 0.509068 | 0.606236 | 0.02 | 0.014 | 1 | 6 | Ccdc77 | 0.006 | Young |
| 0.5092 | 0.053464 | 0.134 | 0.12 | 1 | 6 | Pitpnb | 0.014 | Young |
| 0.509424 | 0.267525 | 0.117 | 0.103 | 1 | 6 | Erh | 0.014 | Young |
| 0.509456 | 0.470766 | 0.132 | 0.12 | 1 | 6 | Racgap1 | 0.012 | Young |
| 0.509623 | 0.112996 | 0.099 | 0.086 | 1 | 6 | Sqle | 0.013 | Young |

| | | | | | | | | |
|---|---|---|---|---|---|---|---|---|
| 0.509913 | 0.519424 | 0.03 | 0.023 | 1 | 6 | Tacc3 | 0.007 | Young |
| 0.510441 | 0.775296 | 0.02 | 0.014 | 1 | 6 | Ss18l1 | 0.006 | Young |
| 0.510441 | 0.572448 | 0.02 | 0.014 | 1 | 6 | Chaf1a | 0.006 | Young |
| 0.510528 | 0.142927 | 0.027 | 0.02 | 1 | 6 | Gstp3 | 0.007 | Young |
| 0.510594 | 0.123378 | 0.033 | 0.026 | 1 | 6 | 3110062M | 0.007 | Young |
| 0.511123 | 0.05047 | 0.033 | 0.026 | 1 | 6 | Zfp317 | 0.007 | Young |
| 0.511157 | 0.675929 | 0.017 | 0.011 | 1 | 6 | Kazald1 | 0.006 | Young |
| 0.511157 | 0.579787 | 0.017 | 0.011 | 1 | 6 | Rdh13 | 0.006 | Young |
| 0.511157 | 0.572617 | 0.017 | 0.011 | 1 | 6 | Fastkd1 | 0.006 | Young |
| 0.511369 | 0.034326 | 0.192 | 0.174 | 1 | 6 | Dnajc21 | 0.018 | Young |
| 0.511555 | 0.274203 | 0.206 | 0.194 | 1 | 6 | Ubxn6 | 0.012 | Young |
| 0.511588 | 0.195856 | 0.03 | 0.023 | 1 | 6 | Zfand1 | 0.007 | Young |
| 0.511816 | 0.565259 | 0.02 | 0.014 | 1 | 6 | Gpsm1 | 0.006 | Young |
| 0.511816 | 0.112446 | 0.02 | 0.014 | 1 | 6 | Tmem129 | 0.006 | Young |
| 0.512409 | 0.315565 | 0.055 | 0.046 | 1 | 6 | Sntb2 | 0.009 | Young |
| 0.512459 | 0.283576 | 0.12 | 0.109 | 1 | 6 | Brf1 | 0.011 | Young |
| 0.512671 | 0.596187 | 0.017 | 0.011 | 1 | 6 | Zfp579 | 0.006 | Young |
| 0.512902 | 0.026058 | 0.022 | 0.029 | 1 | 6 | Asb13 | -0.007 | Young |
| 0.513372 | 0.023136 | 0.089 | 0.103 | 1 | 6 | Dohh | -0.014 | Young |
| 0.513429 | 0.680671 | 0.017 | 0.011 | 1 | 6 | Wdr73 | 0.006 | Young |
| 0.513718 | 0.010831 | 0.214 | 0.231 | 1 | 6 | Mnt | -0.017 | Young |
| 0.513776 | 0.00428 | 0.033 | 0.026 | 1 | 6 | Hbegf | 0.007 | Young |
| 0.513803 | 0.223604 | 0.314 | 0.306 | 1 | 6 | Sdhd | 0.008 | Young |
| 0.513882 | 0.625613 | 0.02 | 0.014 | 1 | 6 | 4930430F0 | 0.006 | Young |
| 0.514094 | 0.003413 | 0.156 | 0.14 | 1 | 6 | Anapc1 | 0.016 | Young |
| 0.514187 | 0.809278 | 0.017 | 0.011 | 1 | 6 | Slc26a11 | 0.006 | Young |
| 0.514187 | 0.196916 | 0.017 | 0.011 | 1 | 6 | Spout1 | 0.006 | Young |
| 0.514504 | 0.203103 | 0.062 | 0.074 | 1 | 6 | Rhbdd1 | -0.012 | Young |
| 0.514613 | 0.103507 | 0.037 | 0.029 | 1 | 6 | Ccdc91 | 0.008 | Young |
| 0.514799 | 0.103394 | 0.017 | 0.023 | 1 | 6 | Fahd1 | -0.006 | Young |
| 0.514868 | 0.288971 | 0.166 | 0.151 | 1 | 6 | Zfp451 | 0.015 | Young |
| 0.515402 | 0.026126 | 0.04 | 0.031 | 1 | 6 | Spred3 | 0.009 | Young |
| 0.515776 | 0.190236 | 0.14 | 0.126 | 1 | 6 | Mrpl32 | 0.014 | Young |
| 0.515844 | 0.19823 | 0.129 | 0.149 | 1 | 6 | Mrps28 | -0.02 | Young |
| 0.515876 | 0.570735 | 0.027 | 0.02 | 1 | 6 | Zc3h6 | 0.007 | Young |
| 0.51588 | 0.188767 | 0.059 | 0.049 | 1 | 6 | Efl1 | 0.01 | Young |
| 0.515943 | 0.711253 | 0.052 | 0.043 | 1 | 6 | Ccdc122 | 0.009 | Young |
| 0.516141 | 0.247046 | 0.017 | 0.023 | 1 | 6 | L3mbtl2 | -0.006 | Young |
| 0.516143 | 0.343277 | 0.069 | 0.057 | 1 | 6 | Llgl1 | 0.012 | Young |
| 0.516264 | 0.181499 | 0.037 | 0.046 | 1 | 6 | Il1rap | -0.009 | Young |
| 0.516644 | 0.743489 | 0.045 | 0.037 | 1 | 6 | Rfxap | 0.008 | Young |
| 0.516644 | 0.392694 | 0.02 | 0.014 | 1 | 6 | Mturn | 0.006 | Young |
| 0.516977 | 0.161126 | 0.318 | 0.303 | 1 | 6 | Tex261 | 0.015 | Young |
| 0.517069 | 0.259775 | 0.027 | 0.02 | 1 | 6 | Polr1e | 0.007 | Young |
| 0.517449 | 0.160652 | 0.344 | 0.32 | 1 | 6 | Emilin2 | 0.024 | Young |

| | | | | | | | | |
|---|---|---|---|---|---|---|---|---|
| 0.517471 | 0.516204 | 0.023 | 0.017 | 1 | 6 | Ccp110 | 0.006 | Young |
| 0.517471 | 0.481651 | 0.023 | 0.017 | 1 | 6 | Wdr92 | 0.006 | Young |
| 0.517471 | 0.152267 | 0.023 | 0.017 | 1 | 6 | Iba57 | 0.006 | Young |
| 0.517486 | 0.11208 | 0.017 | 0.023 | 1 | 6 | Praf2 | -0.006 | Young |
| 0.517821 | 0.125026 | 0.032 | 0.04 | 1 | 6 | Ifi211 | -0.008 | Young |
| 0.517968 | 0.163098 | 0.117 | 0.103 | 1 | 6 | Setdb1 | 0.014 | Young |
| 0.517988 | 0.141621 | 0.017 | 0.011 | 1 | 6 | Gm13012 | 0.006 | Young |
| 0.518262 | 0.206093 | 0.022 | 0.029 | 1 | 6 | Ass1 | -0.007 | Young |
| 0.518484 | 0.331841 | 0.172 | 0.16 | 1 | 6 | Chtf8 | 0.012 | Young |
| 0.5187 | 0.133732 | 0.047 | 0.057 | 1 | 6 | Exosc1 | -0.01 | Young |
| 0.51875 | 0.326358 | 0.023 | 0.017 | 1 | 6 | Rangrf | 0.006 | Young |
| 0.518863 | 0.25559 | 0.159 | 0.146 | 1 | 6 | Ei24 | 0.013 | Young |
| 0.518975 | 0.141334 | 0.209 | 0.197 | 1 | 6 | Klhl5 | 0.012 | Young |
| 0.519288 | 0.00293 | 0.341 | 0.314 | 1 | 6 | Fam111a | 0.027 | Young |
| 0.519413 | 0.101333 | 0.02 | 0.014 | 1 | 6 | Ttl | 0.006 | Young |
| 0.519443 | 0.092123 | 0.03 | 0.023 | 1 | 6 | Acad11 | 0.007 | Young |
| 0.519513 | 0.489692 | 0.017 | 0.011 | 1 | 6 | Col1a2 | 0.006 | Young |
| 0.519566 | 0.30444 | 0.065 | 0.077 | 1 | 6 | Cib2 | -0.012 | Young |
| 0.519946 | 0.376735 | 0.112 | 0.1 | 1 | 6 | Imp4 | 0.012 | Young |
| 0.520107 | 0.128409 | 0.02 | 0.014 | 1 | 6 | Tmem126b | 0.006 | Young |
| 0.520181 | 0.822009 | 0.017 | 0.023 | 1 | 6 | B3gnt8 | -0.006 | Young |
| 0.520184 | 0.392109 | 0.201 | 0.186 | 1 | 6 | Stx12 | 0.015 | Young |
| 0.520276 | 0.26025 | 0.017 | 0.011 | 1 | 6 | Zdhhc1 | 0.006 | Young |
| 0.520654 | 0.254473 | 0.027 | 0.02 | 1 | 6 | Fam26f | 0.007 | Young |
| 0.521108 | 0.33359 | 0.169 | 0.157 | 1 | 6 | Rab24 | 0.012 | Young |
| 0.521188 | 0.138888 | 0.11 | 0.097 | 1 | 6 | Mnat1 | 0.013 | Young |
| 0.521446 | 0.252122 | 0.052 | 0.063 | 1 | 6 | Tbc1d31 | -0.011 | Young |
| 0.521492 | 0.234488 | 0.085 | 0.074 | 1 | 6 | Icmt | 0.011 | Young |
| 0.521495 | 0.223916 | 0.02 | 0.014 | 1 | 6 | Pdia5 | 0.006 | Young |
| 0.522075 | 0.060037 | 0.279 | 0.266 | 1 | 6 | Casc3 | 0.013 | Young |
| 0.522802 | 0.069484 | 0.027 | 0.034 | 1 | 6 | Entpd5 | -0.007 | Young |
| 0.523016 | 0.238268 | 0.212 | 0.197 | 1 | 6 | Prmt1 | 0.015 | Young |
| 0.523471 | 0.073857 | 0.179 | 0.166 | 1 | 6 | Kctd10 | 0.013 | Young |
| 0.523652 | 0.106032 | 0.027 | 0.02 | 1 | 6 | Ccl25 | 0.007 | Young |
| 0.524164 | 0.350545 | 0.047 | 0.057 | 1 | 6 | Pex14 | -0.01 | Young |
| 0.524637 | 0.167308 | 0.065 | 0.077 | 1 | 6 | Acot11 | -0.012 | Young |
| 0.525022 | 0.068697 | 0.089 | 0.103 | 1 | 6 | Cyhr1 | -0.014 | Young |
| 0.525532 | 0.671316 | 0.027 | 0.034 | 1 | 6 | Gm37305 | -0.007 | Young |
| 0.525696 | 0.05527 | 0.055 | 0.046 | 1 | 6 | Surf1 | 0.009 | Young |
| 0.525869 | 0.135657 | 0.032 | 0.04 | 1 | 6 | Vps33b | -0.008 | Young |
| 0.525908 | 0.181829 | 0.074 | 0.063 | 1 | 6 | Cox19 | 0.011 | Young |
| 0.525939 | 0.115032 | 0.105 | 0.091 | 1 | 6 | Nomo1 | 0.014 | Young |
| 0.526158 | 0.248513 | 0.189 | 0.174 | 1 | 6 | Suds3 | 0.015 | Young |
| 0.526358 | 0.086458 | 0.156 | 0.137 | 1 | 6 | Arl13b | 0.019 | Young |
| 0.52684 | 0.125618 | 0.055 | 0.066 | 1 | 6 | Mlh1 | -0.011 | Young |

| | | | | | | | | |
|---|---|---|---|---|---|---|---|---|
| 0.52695 | 0.670954 | 0.064 | 0.054 | 1 | 6 | Tpra1 | 0.01 | Young |
| 0.52732 | 0.046805 | 0.204 | 0.189 | 1 | 6 | Dcaf8 | 0.015 | Young |
| 0.527679 | 0.38882 | 0.097 | 0.086 | 1 | 6 | Eif2b4 | 0.011 | Young |
| 0.528202 | 0.258275 | 0.241 | 0.229 | 1 | 6 | Nfya | 0.012 | Young |
| 0.528342 | 0.337447 | 0.137 | 0.126 | 1 | 6 | Ms4a8a | 0.011 | Young |
| 0.528342 | 0.210622 | 0.329 | 0.311 | 1 | 6 | Nup210 | 0.018 | Young |
| 0.528434 | 0.122322 | 0.059 | 0.049 | 1 | 6 | Fam13b | 0.01 | Young |
| 0.528532 | 0.233404 | 0.115 | 0.103 | 1 | 6 | Pmvk | 0.012 | Young |
| 0.528564 | 0.06914 | 0.237 | 0.217 | 1 | 6 | Socs6 | 0.02 | Young |
| 0.528988 | 0.005134 | 0.124 | 0.14 | 1 | 6 | Ythdf2 | -0.016 | Young |
| 0.529132 | 0.026615 | 0.284 | 0.266 | 1 | 6 | Ddx50 | 0.018 | Young |
| 0.529298 | 0.067971 | 0.401 | 0.383 | 1 | 6 | Pdlim1 | 0.018 | Young |
| 0.52955 | 0.312315 | 0.074 | 0.063 | 1 | 6 | Stk35 | 0.011 | Young |
| 0.529584 | 0.191385 | 0.207 | 0.194 | 1 | 6 | Fam234a | 0.013 | Young |
| 0.530216 | 0.216956 | 0.229 | 0.217 | 1 | 6 | Acat1 | 0.012 | Young |
| 0.530301 | 0.552848 | 0.067 | 0.057 | 1 | 6 | Ap4m1 | 0.01 | Young |
| 0.530425 | 0.43823 | 0.032 | 0.04 | 1 | 6 | Nmral1 | -0.008 | Young |
| 0.530433 | 0.268976 | 0.176 | 0.16 | 1 | 6 | Rnft1 | 0.016 | Young |
| 0.530934 | 0.411422 | 0.117 | 0.106 | 1 | 6 | Hccs | 0.011 | Young |
| 0.53191 | 0.051078 | 0.06 | 0.051 | 1 | 6 | Ints2 | 0.009 | Young |
| 0.532304 | 0.255346 | 0.207 | 0.191 | 1 | 6 | Map3k2 | 0.016 | Young |
| 0.53274 | 0.301174 | 0.045 | 0.037 | 1 | 6 | Bco2 | 0.008 | Young |
| 0.532826 | 0.281733 | 0.055 | 0.046 | 1 | 6 | Smim13 | 0.009 | Young |
| 0.53318 | 0.152345 | 0.239 | 0.223 | 1 | 6 | Fh1 | 0.016 | Young |
| 0.533668 | 0.348983 | 0.045 | 0.037 | 1 | 6 | Dnajc4 | 0.008 | Young |
| 0.533868 | 0.020567 | 0.283 | 0.263 | 1 | 6 | Polb | 0.02 | Young |
| 0.533982 | 0.251671 | 0.082 | 0.071 | 1 | 6 | Ivd | 0.011 | Young |
| 0.53407 | 0.088846 | 0.234 | 0.26 | 1 | 6 | Pstpip1 | -0.026 | Young |
| 0.534107 | 0.152378 | 0.177 | 0.163 | 1 | 6 | Ubac2 | 0.014 | Young |
| 0.534767 | 0.474773 | 0.087 | 0.077 | 1 | 6 | Phf6 | 0.01 | Young |
| 0.53493 | 0.232505 | 0.11 | 0.129 | 1 | 6 | N4bp2 | -0.019 | Young |
| 0.53506 | 0.55946 | 0.045 | 0.037 | 1 | 6 | Btg3 | 0.008 | Young |
| 0.535064 | 0.001433 | 0.224 | 0.243 | 1 | 6 | Atp6v0a2 | -0.019 | Young |
| 0.535162 | 0.030016 | 0.505 | 0.489 | 1 | 6 | Rabac1 | 0.016 | Young |
| 0.535173 | 0.01965 | 0.446 | 0.417 | 1 | 6 | Ids | 0.029 | Young |
| 0.535525 | 0.60025 | 0.045 | 0.037 | 1 | 6 | Ccdc28a | 0.008 | Young |
| 0.536254 | 0.153601 | 0.159 | 0.146 | 1 | 6 | Ube3b | 0.013 | Young |
| 0.536881 | 0.190895 | 0.099 | 0.114 | 1 | 6 | Fbxo28 | -0.015 | Young |
| 0.537946 | 0.418284 | 0.201 | 0.191 | 1 | 6 | Id3 | 0.01 | Young |
| 0.538674 | 0.131514 | 0.261 | 0.243 | 1 | 6 | Trafd1 | 0.018 | Young |
| 0.538688 | 0.093124 | 0.197 | 0.186 | 1 | 6 | Taf6l | 0.011 | Young |
| 0.538735 | 0.654353 | 0.027 | 0.034 | 1 | 6 | Mpst | -0.007 | Young |
| 0.539295 | 0.041795 | 0.395 | 0.386 | 1 | 6 | Gm15987 | 0.009 | Young |
| 0.539476 | 0.028794 | 0.478 | 0.449 | 1 | 6 | Wasf2 | 0.029 | Young |
| 0.53965 | 0.107369 | 0.119 | 0.134 | 1 | 6 | Tm9sf4 | -0.015 | Young |

| | | | | | | | | |
|---|---|---|---|---|---|---|---|---|
| 0.540378 | 0.786767 | 0.06 | 0.051 | 1 | 6 | Ino80c | 0.009 | Young |
| 0.540521 | 0.21751 | 0.217 | 0.203 | 1 | 6 | Cd302 | 0.014 | Young |
| 0.540914 | 0.131794 | 0.092 | 0.08 | 1 | 6 | Ranbp3 | 0.012 | Young |
| 0.540986 | 0.108506 | 0.052 | 0.043 | 1 | 6 | Smarcal1 | 0.009 | Young |
| 0.541368 | 0.016607 | 0.05 | 0.06 | 1 | 6 | Pgghg | -0.01 | Young |
| 0.541486 | 0.042211 | 0.206 | 0.186 | 1 | 6 | Max | 0.02 | Young |
| 0.541986 | 0.331292 | 0.094 | 0.083 | 1 | 6 | Fut11 | 0.011 | Young |
| 0.542258 | 0.390027 | 0.064 | 0.054 | 1 | 6 | Smim5 | 0.01 | Young |
| 0.54245 | 0.173789 | 0.224 | 0.211 | 1 | 6 | Usp3 | 0.013 | Young |
| 0.542777 | 0.165344 | 0.311 | 0.291 | 1 | 6 | Cd200r1 | 0.02 | Young |
| 0.54324 | 0.53405 | 0.072 | 0.063 | 1 | 6 | Tbkbp1 | 0.009 | Young |
| 0.543333 | 0.178109 | 0.127 | 0.114 | 1 | 6 | Yeats2 | 0.013 | Young |
| 0.543781 | 0.234338 | 0.171 | 0.157 | 1 | 6 | Ints6l | 0.014 | Young |
| 0.543909 | 0.086648 | 0.129 | 0.117 | 1 | 6 | Kcnk6 | 0.012 | Young |
| 0.544488 | 0.061795 | 0.052 | 0.043 | 1 | 6 | Vwf | 0.009 | Young |
| 0.544815 | 0.573647 | 0.154 | 0.149 | 1 | 6 | Sart1 | 0.005 | Young |
| 0.54505 | 0.013735 | 0.38 | 0.363 | 1 | 6 | Lmna | 0.017 | Young |
| 0.545056 | 0.218554 | 0.333 | 0.32 | 1 | 6 | Zfp710 | 0.013 | Young |
| 0.545114 | 0.186434 | 0.079 | 0.069 | 1 | 6 | Kansl2 | 0.01 | Young |
| 0.545372 | 0.508257 | 0.084 | 0.074 | 1 | 6 | Sdhaf2 | 0.01 | Young |
| 0.545686 | 0.261813 | 0.082 | 0.071 | 1 | 6 | Fbxl17 | 0.011 | Young |
| 0.545778 | 0.132829 | 0.338 | 0.317 | 1 | 6 | Acss1 | 0.021 | Young |
| 0.546674 | 0.910341 | 0.038 | 0.031 | 1 | 6 | Ttll4 | 0.007 | Young |
| 0.546879 | 0.018444 | 0.99 | 0.983 | 1 | 6 | Rpl41 | 0.007 | Young |
| 0.547429 | 0.367035 | 0.048 | 0.04 | 1 | 6 | Tceal8 | 0.008 | Young |
| 0.547716 | 0.111709 | 0.196 | 0.183 | 1 | 6 | Pcgf5 | 0.013 | Young |
| 0.547866 | 0.176096 | 0.167 | 0.151 | 1 | 6 | Prpf8 | 0.016 | Young |
| 0.548115 | 0.079698 | 0.681 | 0.654 | 1 | 6 | Pltp | 0.027 | Young |
| 0.548476 | 0.149308 | 0.069 | 0.08 | 1 | 6 | Rrp15 | -0.011 | Young |
| 0.548692 | 0.023319 | 0.079 | 0.091 | 1 | 6 | Nol12 | -0.012 | Young |
| 0.549488 | 0.004294 | 0.151 | 0.134 | 1 | 6 | Ncoa6 | 0.017 | Young |
| 0.549711 | 0.070391 | 0.064 | 0.074 | 1 | 6 | Nob1 | -0.01 | Young |
| 0.550404 | 0.082009 | 0.174 | 0.194 | 1 | 6 | Pigs | -0.02 | Young |
| 0.55041 | 0.149418 | 0.401 | 0.386 | 1 | 6 | Wdfy4 | 0.015 | Young |
| 0.55079 | 0.191272 | 0.226 | 0.214 | 1 | 6 | Pip5k1c | 0.012 | Young |
| 0.550873 | 0.104371 | 0.12 | 0.109 | 1 | 6 | Ccdc174 | 0.011 | Young |
| 0.551156 | 0.000119 | 0.567 | 0.566 | 1 | 6 | Minos1 | 0.001 | Young |
| 0.551225 | 0.699427 | 0.069 | 0.06 | 1 | 6 | Gtf2e1 | 0.009 | Young |
| 0.551354 | 0.261517 | 0.189 | 0.177 | 1 | 6 | Sdhb | 0.012 | Young |
| 0.551442 | 0.142151 | 0.045 | 0.037 | 1 | 6 | Cd1d1 | 0.008 | Young |
| 0.551704 | 0.436484 | 0.042 | 0.034 | 1 | 6 | Snta1 | 0.008 | Young |
| 0.551712 | 0.213439 | 0.07 | 0.06 | 1 | 6 | Fam3a | 0.01 | Young |
| 0.552484 | 0.081212 | 0.211 | 0.197 | 1 | 6 | Setd7 | 0.014 | Young |
| 0.552564 | 0.029083 | 0.279 | 0.306 | 1 | 6 | Myo1f | -0.027 | Young |
| 0.552573 | 0.03183 | 0.145 | 0.131 | 1 | 6 | Rbmxl1 | 0.014 | Young |

| | | | | | | | | |
|---|---|---|---|---|---|---|---|---|
| 0.552712 | 0.003712 | 0.788 | 0.751 | 1 | 6 | Msn | 0.037 | Young |
| 0.552921 | 0.153466 | 0.082 | 0.094 | 1 | 6 | Zdhhc7 | -0.012 | Young |
| 0.553382 | 0.016262 | 0.151 | 0.134 | 1 | 6 | Cbx4 | 0.017 | Young |
| 0.553581 | 0.413514 | 0.164 | 0.154 | 1 | 6 | Stx11 | 0.01 | Young |
| 0.554268 | 0.064378 | 0.092 | 0.08 | 1 | 6 | Ctr9 | 0.012 | Young |
| 0.555146 | 0.195248 | 0.497 | 0.494 | 1 | 6 | Spop | 0.003 | Young |
| 0.555567 | 0.212738 | 0.293 | 0.28 | 1 | 6 | B4galt6 | 0.013 | Young |
| 0.556174 | 0.050044 | 0.472 | 0.451 | 1 | 6 | Tcf25 | 0.021 | Young |
| 0.556593 | 0.289023 | 0.278 | 0.274 | 1 | 6 | Rnf19b | 0.004 | Young |
| 0.55675 | 0.122671 | 0.01 | 0.014 | 1 | 6 | Serpinb2 | -0.004 | Young |
| 0.556798 | 0.146183 | 0.085 | 0.074 | 1 | 6 | Fam160b2 | 0.011 | Young |
| 0.556836 | 0.033006 | 0.331 | 0.317 | 1 | 6 | Zmiz2 | 0.014 | Young |
| 0.556847 | 0.020883 | 0.171 | 0.16 | 1 | 6 | Xrn1 | 0.011 | Young |
| 0.556907 | 0.410371 | 0.156 | 0.146 | 1 | 6 | Impad1 | 0.01 | Young |
| 0.557229 | 0.184084 | 0.1 | 0.089 | 1 | 6 | Slf2 | 0.011 | Young |
| 0.557595 | 0.373991 | 0.042 | 0.034 | 1 | 6 | Fbxo25 | 0.008 | Young |
| 0.557662 | 0.069738 | 0.184 | 0.169 | 1 | 6 | Idnk | 0.015 | Young |
| 0.557863 | 0.128088 | 0.336 | 0.323 | 1 | 6 | Ndufv3 | 0.013 | Young |
| 0.558145 | 0.143013 | 0.057 | 0.049 | 1 | 6 | Rpp14 | 0.008 | Young |
| 0.558417 | 0.035877 | 0.125 | 0.14 | 1 | 6 | Clip1 | -0.015 | Young |
| 0.55858 | 0.153761 | 0.518 | 0.52 | 1 | 6 | Eif5 | -0.002 | Young |
| 0.558648 | 0.508018 | 0.087 | 0.077 | 1 | 6 | Hikeshi | 0.01 | Young |
| 0.558995 | 0.383126 | 0.099 | 0.089 | 1 | 6 | Gid4 | 0.01 | Young |
| 0.55901 | 0.106779 | 0.261 | 0.249 | 1 | 6 | Abcc1 | 0.012 | Young |
| 0.559513 | 0.912087 | 0.035 | 0.029 | 1 | 6 | Smyd3 | 0.006 | Young |
| 0.559513 | 0.667801 | 0.035 | 0.029 | 1 | 6 | Mapkapk5 | 0.006 | Young |
| 0.559584 | 0.166502 | 0.161 | 0.149 | 1 | 6 | Zc3h18 | 0.012 | Young |
| 0.559648 | 0.180234 | 0.082 | 0.071 | 1 | 6 | Hadh | 0.011 | Young |
| 0.559901 | 0.287937 | 0.207 | 0.197 | 1 | 6 | Bach2 | 0.01 | Young |
| 0.560001 | 0.483129 | 0.06 | 0.071 | 1 | 6 | Haghl | -0.011 | Young |
| 0.560058 | 0.179328 | 0.042 | 0.034 | 1 | 6 | Slc35d2 | 0.008 | Young |
| 0.560314 | 0.025569 | 0.01 | 0.014 | 1 | 6 | Atg9b | -0.004 | Young |
| 0.560628 | 0.116694 | 0.125 | 0.114 | 1 | 6 | Hdac3 | 0.011 | Young |
| 0.560662 | 0.062845 | 0.095 | 0.109 | 1 | 6 | Ppp1r7 | -0.014 | Young |
| 0.56068 | 0.200446 | 0.072 | 0.063 | 1 | 6 | Dlat | 0.009 | Young |
| 0.561233 | 0.278849 | 0.102 | 0.091 | 1 | 6 | Ltbr | 0.011 | Young |
| 0.561519 | 0.299023 | 0.04 | 0.049 | 1 | 6 | Lyrm9 | -0.009 | Young |
| 0.561624 | 0.013385 | 0.316 | 0.303 | 1 | 6 | Foxn3 | 0.013 | Young |
| 0.562816 | 0.156538 | 0.045 | 0.037 | 1 | 6 | Aktip | 0.008 | Young |
| 0.563425 | 0.097371 | 0.254 | 0.24 | 1 | 6 | Etfa | 0.014 | Young |
| 0.563996 | 0.101079 | 0.142 | 0.129 | 1 | 6 | Sf3a2 | 0.013 | Young |
| 0.564543 | 0.07698 | 0.408 | 0.4 | 1 | 6 | Cd48 | 0.008 | Young |
| 0.565048 | 0.196573 | 0.209 | 0.197 | 1 | 6 | Pik3c2a | 0.012 | Young |
| 0.565173 | 0.152515 | 0.082 | 0.071 | 1 | 6 | Aco1 | 0.011 | Young |
| 0.565206 | 0.04254 | 0.196 | 0.217 | 1 | 6 | Gnpda1 | -0.021 | Young |

| | | | | | | | | |
|---|---|---|---|---|---|---|---|---|
| 0.565612 | 0.069155 | 0.236 | 0.223 | 1 | 6 | Krcc1 | 0.013 | Young |
| 0.565837 | 0.096605 | 0.217 | 0.203 | 1 | 6 | Ccny | 0.014 | Young |
| 0.565845 | 0.094011 | 0.261 | 0.28 | 1 | 6 | Ugcg | -0.019 | Young |
| 0.566044 | 0.162044 | 0.219 | 0.206 | 1 | 6 | Crebrf | 0.013 | Young |
| 0.566188 | 0.078322 | 0.192 | 0.177 | 1 | 6 | Vat1 | 0.015 | Young |
| 0.566279 | 0.351862 | 0.065 | 0.077 | 1 | 6 | Rcbtb1 | -0.012 | Young |
| 0.566488 | 0.214311 | 0.042 | 0.034 | 1 | 6 | Fbxl18 | 0.008 | Young |
| 0.566547 | 0.171105 | 0.219 | 0.209 | 1 | 6 | Mrpl14 | 0.01 | Young |
| 0.566592 | 0.013989 | 0.054 | 0.063 | 1 | 6 | Kifap3 | -0.009 | Young |
| 0.567089 | 0.305026 | 0.144 | 0.131 | 1 | 6 | Akap11 | 0.013 | Young |
| 0.567164 | 0.179143 | 0.038 | 0.031 | 1 | 6 | Primpol | 0.007 | Young |
| 0.56718 | 0.248917 | 0.02 | 0.026 | 1 | 6 | Ipp | -0.006 | Young |
| 0.567498 | 0.098375 | 0.239 | 0.223 | 1 | 6 | Acadm | 0.016 | Young |
| 0.567578 | 0.226671 | 0.048 | 0.04 | 1 | 6 | Alg6 | 0.008 | Young |
| 0.56762 | 0.170606 | 0.135 | 0.123 | 1 | 6 | Pold4 | 0.012 | Young |
| 0.568606 | 0.056408 | 0.263 | 0.243 | 1 | 6 | Plin3 | 0.02 | Young |
| 0.569153 | 0.183727 | 0.363 | 0.343 | 1 | 6 | Sin3b | 0.02 | Young |
| 0.569271 | 1.560975 | 0.01 | 0.014 | 1 | 6 | Ikbke | -0.004 | Young |
| 0.569594 | 0.053037 | 0.095 | 0.109 | 1 | 6 | Nde1 | -0.014 | Young |
| 0.56961 | 0.393721 | 0.05 | 0.06 | 1 | 6 | Sp4 | -0.01 | Young |
| 0.569769 | 0.111304 | 0.069 | 0.08 | 1 | 6 | Mgst3 | -0.011 | Young |
| 0.569993 | 0.107982 | 0.288 | 0.269 | 1 | 6 | Timp2 | 0.019 | Young |
| 0.570144 | 0.108687 | 0.137 | 0.126 | 1 | 6 | Ago4 | 0.011 | Young |
| 0.570227 | 0.02398 | 0.259 | 0.283 | 1 | 6 | Jtb | -0.024 | Young |
| 0.570495 | 0.117592 | 0.227 | 0.211 | 1 | 6 | Retreg2 | 0.016 | Young |
| 0.570597 | 0.28497 | 0.145 | 0.163 | 1 | 6 | Tmc6 | -0.018 | Young |
| 0.571169 | 0.021161 | 0.085 | 0.074 | 1 | 6 | Gprc5a | 0.011 | Young |
| 0.571301 | 0.106687 | 0.092 | 0.106 | 1 | 6 | Parp11 | -0.014 | Young |
| 0.571315 | 0.282225 | 0.13 | 0.12 | 1 | 6 | Acbd6 | 0.01 | Young |
| 0.571518 | 0.03107 | 0.142 | 0.129 | 1 | 6 | Rfc1 | 0.013 | Young |
| 0.571663 | 0.136221 | 0.142 | 0.131 | 1 | 6 | Mrps6 | 0.011 | Young |
| 0.571689 | 0.054212 | 0.328 | 0.311 | 1 | 6 | Mindy2 | 0.017 | Young |
| 0.571927 | 0.2774 | 0.035 | 0.029 | 1 | 6 | Nudt22 | 0.006 | Young |
| 0.571956 | 0.027351 | 0.042 | 0.034 | 1 | 6 | Habp4 | 0.008 | Young |
| 0.572179 | 0.329638 | 0.232 | 0.226 | 1 | 6 | Stat5a | 0.006 | Young |
| 0.573012 | 0.598739 | 0.035 | 0.029 | 1 | 6 | Abcd3 | 0.006 | Young |
| 0.573329 | 0.15049 | 0.064 | 0.074 | 1 | 6 | Commd10 | -0.01 | Young |
| 0.573392 | 0.14722 | 0.181 | 0.169 | 1 | 6 | Timm8b | 0.012 | Young |
| 0.573555 | 0.218607 | 0.03 | 0.037 | 1 | 6 | Dusp8 | -0.007 | Young |
| 0.574099 | 0.239696 | 0.035 | 0.029 | 1 | 6 | Bpgm | 0.006 | Young |
| 0.574111 | 0.072615 | 0.348 | 0.334 | 1 | 6 | Wapl | 0.014 | Young |
| 0.574546 | 0.142353 | 0.167 | 0.186 | 1 | 6 | Idh3b | -0.019 | Young |
| 0.574637 | 0.003284 | 0.025 | 0.031 | 1 | 6 | Pdik1l | -0.006 | Young |
| 0.574742 | 0.33314 | 0.057 | 0.049 | 1 | 6 | Trip10 | 0.008 | Young |
| 0.574908 | 0.388347 | 0.129 | 0.117 | 1 | 6 | Cant1 | 0.012 | Young |

| | | | | | | | | |
|---|---|---|---|---|---|---|---|---|
| 0.574942 | 0.227847 | 0.038 | 0.031 | 1 | 6 | Numb | 0.007 | Young |
| 0.575449 | 0.245558 | 0.042 | 0.034 | 1 | 6 | Srbd1 | 0.008 | Young |
| 0.57552 | 0.09586 | 0.329 | 0.314 | 1 | 6 | Etv6 | 0.015 | Young |
| 0.575885 | 0.010441 | 0.952 | 0.891 | 1 | 6 | Coro1a | 0.061 | Young |
| 0.576796 | 0.406608 | 0.069 | 0.06 | 1 | 6 | Plscr3 | 0.009 | Young |
| 0.577887 | 0.121893 | 0.077 | 0.089 | 1 | 6 | Cyp2s1 | -0.012 | Young |
| 0.578318 | 0.352152 | 0.117 | 0.109 | 1 | 6 | Axin1 | 0.008 | Young |
| 0.578409 | 0.270483 | 0.064 | 0.054 | 1 | 6 | Pus10 | 0.01 | Young |
| 0.578455 | 1.06205 | 0.032 | 0.026 | 1 | 6 | Utp4 | 0.006 | Young |
| 0.579738 | 0.463493 | 0.11 | 0.1 | 1 | 6 | Nudcd2 | 0.01 | Young |
| 0.580174 | 0.864178 | 0.062 | 0.054 | 1 | 6 | Hmbs | 0.008 | Young |
| 0.580482 | 0.014629 | 0.654 | 0.626 | 1 | 6 | Vamp8 | 0.028 | Young |
| 0.580794 | 0.169048 | 0.264 | 0.254 | 1 | 6 | Slc29a3 | 0.01 | Young |
| 0.5812 | 0.206842 | 0.038 | 0.031 | 1 | 6 | Tmem184c | 0.007 | Young |
| 0.581576 | 0.032265 | 0.331 | 0.314 | 1 | 6 | Arrb1 | 0.017 | Young |
| 0.581898 | 0.303409 | 0.069 | 0.06 | 1 | 6 | Creb3 | 0.009 | Young |
| 0.582116 | 0.135631 | 0.202 | 0.189 | 1 | 6 | Gpatch8 | 0.013 | Young |
| 0.582344 | 0.066225 | 0.411 | 0.394 | 1 | 6 | Abi1 | 0.017 | Young |
| 0.583301 | 0.186805 | 0.099 | 0.089 | 1 | 6 | Hectd4 | 0.01 | Young |
| 0.583329 | 0.258793 | 0.067 | 0.077 | 1 | 6 | Cln8 | -0.01 | Young |
| 0.58338 | 0.285803 | 0.144 | 0.131 | 1 | 6 | Tpcn1 | 0.013 | Young |
| 0.583669 | 0.397761 | 0.05 | 0.043 | 1 | 6 | Capn5 | 0.007 | Young |
| 0.583853 | 0.129957 | 0.119 | 0.134 | 1 | 6 | Chrac1 | -0.015 | Young |
| 0.583922 | 0.62768 | 0.03 | 0.037 | 1 | 6 | Pus7 | -0.007 | Young |
| 0.584119 | 0.223105 | 0.119 | 0.109 | 1 | 6 | Zcchc9 | 0.01 | Young |
| 0.58447 | 0.094184 | 0.03 | 0.037 | 1 | 6 | Kctd11 | -0.007 | Young |
| 0.584736 | 0.193642 | 0.176 | 0.163 | 1 | 6 | Antxr2 | 0.013 | Young |
| 0.584746 | 0.234018 | 0.025 | 0.031 | 1 | 6 | Etv1 | -0.006 | Young |
| 0.58481 | 0.188755 | 0.269 | 0.257 | 1 | 6 | Hk2 | 0.012 | Young |
| 0.584821 | 0.111376 | 0.072 | 0.083 | 1 | 6 | Fam219b | -0.011 | Young |
| 0.584943 | 0.260869 | 0.02 | 0.026 | 1 | 6 | Dmap1 | -0.006 | Young |
| 0.585606 | 0.208209 | 0.02 | 0.026 | 1 | 6 | Prkcg | -0.006 | Young |
| 0.585927 | 0.513127 | 0.032 | 0.026 | 1 | 6 | Gm35154 | 0.006 | Young |
| 0.586644 | 0.095332 | 0.054 | 0.063 | 1 | 6 | Nxt1 | -0.009 | Young |
| 0.587372 | 0.058648 | 0.418 | 0.406 | 1 | 6 | P2rx4 | 0.012 | Young |
| 0.587417 | 0.044539 | 0.161 | 0.146 | 1 | 6 | Lilra6 | 0.015 | Young |
| 0.587748 | 0.312607 | 0.059 | 0.069 | 1 | 6 | Wdr12 | -0.01 | Young |
| 0.587797 | 0.313303 | 0.125 | 0.114 | 1 | 6 | Btbd9 | 0.011 | Young |
| 0.587989 | 0.283994 | 0.075 | 0.066 | 1 | 6 | C3ar1 | 0.009 | Young |
| 0.588102 | 0.507264 | 0.085 | 0.077 | 1 | 6 | Bola1 | 0.008 | Young |
| 0.588235 | 0.539101 | 0.032 | 0.026 | 1 | 6 | Cdpf1 | 0.006 | Young |
| 0.588532 | 0.223455 | 0.057 | 0.049 | 1 | 6 | Slc25a22 | 0.008 | Young |
| 0.588663 | 0.247151 | 0.112 | 0.103 | 1 | 6 | Lpin2 | 0.009 | Young |
| 0.588926 | 0.206026 | 0.02 | 0.026 | 1 | 6 | Ap4e1 | -0.006 | Young |
| 0.588976 | 0.281021 | 0.135 | 0.126 | 1 | 6 | Man1c1 | 0.009 | Young |

| | | | | | | | | |
|---|---|---|---|---|---|---|---|---|
| 0.588985 | 0.159215 | 0.507 | 0.497 | 1 | 6 | Pfkp | 0.01 | Young |
| 0.5891 | 0.074006 | 0.251 | 0.237 | 1 | 6 | Ino80d | 0.014 | Young |
| 0.589438 | 0.028278 | 0.038 | 0.046 | 1 | 6 | 1300002E1 | -0.008 | Young |
| 0.590256 | 0.247551 | 0.02 | 0.026 | 1 | 6 | Pcgf6 | -0.006 | Young |
| 0.590258 | 0.026279 | 0.114 | 0.103 | 1 | 6 | Cpeb2 | 0.011 | Young |
| 0.590263 | 0.052531 | 0.075 | 0.066 | 1 | 6 | Gas6 | 0.009 | Young |
| 0.590513 | 0.382647 | 0.035 | 0.029 | 1 | 6 | Mphosph9 | 0.006 | Young |
| 0.590828 | 0.155938 | 0.269 | 0.254 | 1 | 6 | Map4k1 | 0.015 | Young |
| 0.591687 | 0.168954 | 0.229 | 0.254 | 1 | 6 | Mrpl57 | -0.025 | Young |
| 0.592531 | 0.373935 | 0.025 | 0.031 | 1 | 6 | Taf5 | -0.006 | Young |
| 0.592757 | 0.039182 | 0.326 | 0.351 | 1 | 6 | Ttc14 | -0.025 | Young |
| 0.59305 | 0.184407 | 0.072 | 0.063 | 1 | 6 | Atp1a3 | 0.009 | Young |
| 0.593153 | 0.341122 | 0.032 | 0.026 | 1 | 6 | Map3k15 | 0.006 | Young |
| 0.593387 | 0.231494 | 0.157 | 0.146 | 1 | 6 | Rcor1 | 0.011 | Young |
| 0.593751 | 0.271931 | 0.069 | 0.06 | 1 | 6 | Anks3 | 0.009 | Young |
| 0.593829 | 0.088426 | 0.201 | 0.217 | 1 | 6 | Ubr5 | -0.016 | Young |
| 0.594283 | 0.599013 | 0.028 | 0.023 | 1 | 6 | Btbd3 | 0.005 | Young |
| 0.594335 | 0.661791 | 0.025 | 0.031 | 1 | 6 | Qpctl | -0.006 | Young |
| 0.594603 | 0.022521 | 0.032 | 0.026 | 1 | 6 | Surf2 | 0.006 | Young |
| 0.594789 | 0.214118 | 0.072 | 0.083 | 1 | 6 | Glb1 | -0.011 | Young |
| 0.594927 | 0.003567 | 0.035 | 0.029 | 1 | 6 | Prrg2 | 0.006 | Young |
| 0.595202 | 0.016794 | 0.079 | 0.069 | 1 | 6 | Katnbl1 | 0.01 | Young |
| 0.595458 | 0.106654 | 0.184 | 0.203 | 1 | 6 | Gga1 | -0.019 | Young |
| 0.595518 | 0.27269 | 0.179 | 0.169 | 1 | 6 | Rnf38 | 0.01 | Young |
| 0.595765 | 0.261874 | 0.032 | 0.026 | 1 | 6 | Hcar2 | 0.006 | Young |
| 0.595997 | 0.247951 | 0.132 | 0.12 | 1 | 6 | Carhsp1 | 0.012 | Young |
| 0.596205 | 0.103578 | 0.09 | 0.08 | 1 | 6 | Ppp4r1 | 0.01 | Young |
| 0.596346 | 0.455873 | 0.032 | 0.026 | 1 | 6 | Kctd13 | 0.006 | Young |
| 0.596548 | 0.168097 | 0.072 | 0.063 | 1 | 6 | Cacfd1 | 0.009 | Young |
| 0.596899 | 0.086925 | 0.505 | 0.497 | 1 | 6 | Kif5b | 0.008 | Young |
| 0.597593 | 0.25114 | 0.219 | 0.211 | 1 | 6 | Ctdsp1 | 0.008 | Young |
| 0.597679 | 0.403378 | 0.08 | 0.071 | 1 | 6 | Ctc1 | 0.009 | Young |
| 0.597967 | 0.776755 | 0.028 | 0.023 | 1 | 6 | 5730480H( | 0.005 | Young |
| 0.598719 | 0.111014 | 0.229 | 0.223 | 1 | 6 | Larp4 | 0.006 | Young |
| 0.598962 | 0.795366 | 0.025 | 0.02 | 1 | 6 | Pigo | 0.005 | Young |
| 0.599721 | 0.000124 | 0.13 | 0.146 | 1 | 6 | Ikzf5 | -0.016 | Young |
| 0.59976 | 0.095345 | 0.344 | 0.326 | 1 | 6 | Gusb | 0.018 | Young |
| 0.599783 | 0.050082 | 0.42 | 0.4 | 1 | 6 | Atp5o.1 | 0.02 | Young |
| 0.59981 | 0.119297 | 0.229 | 0.214 | 1 | 6 | Uqcc3 | 0.015 | Young |
| 0.600089 | 0.029838 | 0.408 | 0.426 | 1 | 6 | Hivep2 | -0.018 | Young |
| 0.600394 | 0.231036 | 0.115 | 0.106 | 1 | 6 | Crnkl1 | 0.009 | Young |
| 0.600667 | 0.116353 | 0.316 | 0.314 | 1 | 6 | AI504432 | 0.002 | Young |
| 0.600759 | 0.006554 | 0.403 | 0.434 | 1 | 6 | Rab1a | -0.031 | Young |
| 0.601307 | 0.025083 | 0.075 | 0.066 | 1 | 6 | Nbdy | 0.009 | Young |
| 0.601317 | 0.310818 | 0.038 | 0.046 | 1 | 6 | AI597479 | -0.008 | Young |

| | | | | | | | | |
|---|---|---|---|---|---|---|---|---|
| 0.601495 | 0.117941 | 0.515 | 0.509 | 1 | 6 | Tpr | 0.006 | Young |
| 0.601676 | 0.014435 | 0.326 | 0.346 | 1 | 6 | Trappc4 | -0.02 | Young |
| 0.602362 | 0.04122 | 0.145 | 0.134 | 1 | 6 | Msl3 | 0.011 | Young |
| 0.602589 | 0.346428 | 0.05 | 0.043 | 1 | 6 | Asna1 | 0.007 | Young |
| 0.60269 | 0.149037 | 0.351 | 0.34 | 1 | 6 | Fgr | 0.011 | Young |
| 0.602822 | 0.000463 | 0.122 | 0.109 | 1 | 6 | Map3k4 | 0.013 | Young |
| 0.602899 | 0.254184 | 0.314 | 0.309 | 1 | 6 | Tprgl | 0.005 | Young |
| 0.603205 | 0.3453 | 0.059 | 0.069 | 1 | 6 | Smim8 | -0.01 | Young |
| 0.603205 | 0.12816 | 0.065 | 0.057 | 1 | 6 | Zfp219 | 0.008 | Young |
| 0.603662 | 0.174657 | 0.132 | 0.12 | 1 | 6 | Sigirr | 0.012 | Young |
| 0.604477 | 0.122474 | 0.156 | 0.146 | 1 | 6 | Helz | 0.01 | Young |
| 0.604774 | 0.125131 | 0.067 | 0.077 | 1 | 6 | Dis3l2 | -0.01 | Young |
| 0.605577 | 0.087313 | 0.281 | 0.266 | 1 | 6 | Was | 0.015 | Young |
| 0.605714 | 0.083952 | 0.062 | 0.071 | 1 | 6 | Nup160 | -0.009 | Young |
| 0.605973 | 0.17548 | 0.077 | 0.069 | 1 | 6 | Mtf1 | 0.008 | Young |
| 0.606227 | 0.112219 | 0.221 | 0.211 | 1 | 6 | Tcf12 | 0.01 | Young |
| 0.606601 | 0.014987 | 0.251 | 0.237 | 1 | 6 | Tial1 | 0.014 | Young |
| 0.606605 | 0.096627 | 0.373 | 0.357 | 1 | 6 | Cwc15 | 0.016 | Young |
| 0.607224 | 0.05068 | 0.355 | 0.343 | 1 | 6 | Scaf11 | 0.012 | Young |
| 0.607295 | 0.009498 | 0.507 | 0.471 | 1 | 6 | Itgb7 | 0.036 | Young |
| 0.607503 | 0.406159 | 0.025 | 0.02 | 1 | 6 | Klhl36 | 0.005 | Young |
| 0.609236 | 0.089825 | 0.527 | 0.506 | 1 | 6 | Adgre5 | 0.021 | Young |
| 0.609481 | 0.138347 | 0.025 | 0.02 | 1 | 6 | Reps2 | 0.005 | Young |
| 0.6097 | 0.018445 | 0.028 | 0.023 | 1 | 6 | Pomgnt1 | 0.005 | Young |
| 0.610093 | 0.090499 | 0.269 | 0.263 | 1 | 6 | Map4 | 0.006 | Young |
| 0.610551 | 0.185181 | 0.033 | 0.04 | 1 | 6 | Pcca | -0.007 | Young |
| 0.611562 | 0.733972 | 0.028 | 0.023 | 1 | 6 | Arrdc2 | 0.005 | Young |
| 0.611735 | 0.071862 | 0.069 | 0.06 | 1 | 6 | Bmi1 | 0.009 | Young |
| 0.612785 | 0.406411 | 0.025 | 0.02 | 1 | 6 | Dhx37 | 0.005 | Young |
| 0.612785 | 0.209759 | 0.025 | 0.02 | 1 | 6 | Atad5 | 0.005 | Young |
| 0.612805 | 0.175758 | 0.028 | 0.023 | 1 | 6 | Otud1 | 0.005 | Young |
| 0.612905 | 0.053308 | 0.589 | 0.574 | 1 | 6 | Mrpl52 | 0.015 | Young |
| 0.613559 | 0.622917 | 0.047 | 0.04 | 1 | 6 | Asgr2 | 0.007 | Young |
| 0.613939 | 0.094485 | 0.254 | 0.243 | 1 | 6 | Sumo3 | 0.011 | Young |
| 0.613977 | 0.162454 | 0.062 | 0.054 | 1 | 6 | F2rl2 | 0.008 | Young |
| 0.614299 | 0.041693 | 0.033 | 0.04 | 1 | 6 | Rrp8 | -0.007 | Young |
| 0.614304 | 0.077388 | 0.176 | 0.163 | 1 | 6 | Msmo1 | 0.013 | Young |
| 0.614387 | 0.100432 | 0.244 | 0.231 | 1 | 6 | Zcrb1 | 0.013 | Young |
| 0.614728 | 0.143473 | 0.05 | 0.043 | 1 | 6 | Tbc1d22b | 0.007 | Young |
| 0.615273 | 0.04107 | 0.199 | 0.186 | 1 | 6 | Usp47 | 0.013 | Young |
| 0.615449 | 0.082087 | 0.452 | 0.437 | 1 | 6 | Rassf5 | 0.015 | Young |
| 0.615498 | 0.562889 | 0.047 | 0.04 | 1 | 6 | Dxo | 0.007 | Young |
| 0.616017 | 0.2416 | 0.067 | 0.077 | 1 | 6 | Sil1 | -0.01 | Young |
| 0.616135 | 0.127916 | 0.05 | 0.043 | 1 | 6 | Traf2 | 0.007 | Young |
| 0.616198 | 0.012094 | 0.115 | 0.129 | 1 | 6 | Bin2 | -0.014 | Young |

| | | | | | | | | |
|---|---|---|---|---|---|---|---|---|
| 0.616253 | 0.37304 | 0.177 | 0.171 | 1 | 6 | Ddx42 | 0.006 | Young |
| 0.616316 | 0.851324 | 0.043 | 0.037 | 1 | 6 | Tmem159 | 0.006 | Young |
| 0.61676 | 0.405967 | 0.025 | 0.02 | 1 | 6 | Nek4 | 0.005 | Young |
| 0.616975 | 0.020838 | 0.069 | 0.06 | 1 | 6 | Usb1 | 0.009 | Young |
| 0.617299 | 0.4613 | 0.022 | 0.017 | 1 | 6 | Chst7 | 0.005 | Young |
| 0.617489 | 0.45041 | 0.055 | 0.049 | 1 | 6 | Rabep2 | 0.006 | Young |
| 0.617756 | 1.235417 | 0.018 | 0.014 | 1 | 6 | Ppp1r18os | 0.004 | Young |
| 0.617783 | 0.246328 | 0.199 | 0.189 | 1 | 6 | Slc38a9 | 0.01 | Young |
| 0.617836 | 0.000327 | 0.047 | 0.054 | 1 | 6 | Naa16 | -0.007 | Young |
| 0.617924 | 0.22441 | 0.11 | 0.103 | 1 | 6 | Apmap | 0.007 | Young |
| 0.619417 | 0.264371 | 0.025 | 0.02 | 1 | 6 | Shq1 | 0.005 | Young |
| 0.620026 | 0.040781 | 0.279 | 0.257 | 1 | 6 | Oat | 0.022 | Young |
| 0.620154 | 0.859565 | 0.022 | 0.017 | 1 | 6 | Rhebl1 | 0.005 | Young |
| 0.621086 | 0.304951 | 0.074 | 0.066 | 1 | 6 | Rab20 | 0.008 | Young |
| 0.621507 | 0.350578 | 0.366 | 0.354 | 1 | 6 | Ccl4 | 0.012 | Young |
| 0.6216 | 0.142947 | 0.587 | 0.594 | 1 | 6 | Ccr5 | -0.007 | Young |
| 0.622001 | 0.275738 | 0.119 | 0.109 | 1 | 6 | Pdcd11 | 0.01 | Young |
| 0.622005 | 0.182252 | 0.06 | 0.069 | 1 | 6 | Rnaset2a | -0.009 | Young |
| 0.6223 | 0.388456 | 0.022 | 0.017 | 1 | 6 | Lhpp | 0.005 | Young |
| 0.62242 | 1.047659 | 0.018 | 0.014 | 1 | 6 | Pigb | 0.004 | Young |
| 0.623422 | 0.026538 | 0.154 | 0.169 | 1 | 6 | Zdhhc3 | -0.015 | Young |
| 0.623595 | 0.231445 | 0.075 | 0.086 | 1 | 6 | Ccdc93 | -0.011 | Young |
| 0.623731 | 0.032272 | 0.022 | 0.017 | 1 | 6 | Gab3 | 0.005 | Young |
| 0.623754 | 0.152177 | 0.095 | 0.086 | 1 | 6 | 9430038I0 | 0.009 | Young |
| 0.623972 | 0.029785 | 0.411 | 0.383 | 1 | 6 | Stag2 | 0.028 | Young |
| 0.624035 | 0.113665 | 0.115 | 0.106 | 1 | 6 | Nudt16l1 | 0.009 | Young |
| 0.624263 | 0.262388 | 0.069 | 0.06 | 1 | 6 | Mob2 | 0.009 | Young |
| 0.624478 | 0.137803 | 0.296 | 0.28 | 1 | 6 | Lpcat1 | 0.016 | Young |
| 0.625065 | 0.076686 | 0.065 | 0.057 | 1 | 6 | Tmem55a | 0.008 | Young |
| 0.625538 | 1.552148 | 0.018 | 0.014 | 1 | 6 | Zfp788 | 0.004 | Young |
| 0.625557 | 0.299554 | 0.055 | 0.049 | 1 | 6 | Rbl1 | 0.006 | Young |
| 0.625579 | 0.058533 | 0.192 | 0.18 | 1 | 6 | Cers6 | 0.012 | Young |
| 0.626318 | 0.616491 | 0.018 | 0.014 | 1 | 6 | Rdh10 | 0.004 | Young |
| 0.626821 | 0.037895 | 0.508 | 0.483 | 1 | 6 | Bola2 | 0.025 | Young |
| 0.627317 | 0.323377 | 0.022 | 0.017 | 1 | 6 | Ift27 | 0.005 | Young |
| 0.627317 | 0.31017 | 0.022 | 0.017 | 1 | 6 | Dusp18 | 0.005 | Young |
| 0.627623 | 0.421363 | 0.04 | 0.034 | 1 | 6 | 1110032A0 | 0.006 | Young |
| 0.627742 | 0.067479 | 0.074 | 0.066 | 1 | 6 | Sacm1l | 0.008 | Young |
| 0.62788 | 0.555617 | 0.018 | 0.014 | 1 | 6 | Cdc45 | 0.004 | Young |
| 0.627885 | 0.040407 | 0.263 | 0.283 | 1 | 6 | Ndufs7 | -0.02 | Young |
| 0.628431 | 0.442295 | 0.043 | 0.037 | 1 | 6 | Xlr | 0.006 | Young |
| 0.628859 | 0.083235 | 0.199 | 0.186 | 1 | 6 | Mogs | 0.013 | Young |
| 0.628898 | 0.454546 | 0.149 | 0.14 | 1 | 6 | Lman2l | 0.009 | Young |
| 0.628913 | 0.096771 | 0.147 | 0.137 | 1 | 6 | Vrk3 | 0.01 | Young |
| 0.628917 | 0.072182 | 0.313 | 0.303 | 1 | 6 | Phf3 | 0.01 | Young |

| | | | | | | | | |
|---|---|---|---|---|---|---|---|---|
| 0.628918 | 0.080964 | 0.186 | 0.171 | 1 | 6 | Luzp1 | 0.015 | Young |
| 0.628978 | 0.050847 | 0.249 | 0.237 | 1 | 6 | Gosr2 | 0.012 | Young |
| 0.629444 | 0.450377 | 0.018 | 0.014 | 1 | 6 | G430095P1 | 0.004 | Young |
| 0.629511 | 0.195776 | 0.028 | 0.034 | 1 | 6 | Rgmb | -0.006 | Young |
| 0.629545 | 0.168988 | 0.249 | 0.271 | 1 | 6 | 2610507B1 | -0.022 | Young |
| 0.629575 | 0.011209 | 0.717 | 0.729 | 1 | 6 | Ddx6 | -0.012 | Young |
| 0.629934 | 0.20699 | 0.08 | 0.091 | 1 | 6 | Nsmce2 | -0.011 | Young |
| 0.630035 | 0.222188 | 0.077 | 0.069 | 1 | 6 | Rab11fip5 | 0.008 | Young |
| 0.630101 | 0.069823 | 0.381 | 0.369 | 1 | 6 | Lasp1 | 0.012 | Young |
| 0.630192 | 0.054552 | 0.022 | 0.017 | 1 | 6 | Gm42031 | 0.005 | Young |
| 0.630192 | 0.012096 | 0.022 | 0.017 | 1 | 6 | Mrm1 | 0.005 | Young |
| 0.630256 | 0.938855 | 0.015 | 0.011 | 1 | 6 | Gm12689 | 0.004 | Young |
| 0.630539 | 0.120717 | 0.259 | 0.286 | 1 | 6 | Ube2e1 | -0.027 | Young |
| 0.630548 | 0.007166 | 0.552 | 0.551 | 1 | 6 | Jak1 | 0.001 | Young |
| 0.63067 | 0.388248 | 0.082 | 0.074 | 1 | 6 | Alg5 | 0.008 | Young |
| 0.63101 | 0.913831 | 0.018 | 0.014 | 1 | 6 | Gpr4 | 0.004 | Young |
| 0.631016 | 0.046749 | 0.247 | 0.234 | 1 | 6 | Eri1 | 0.013 | Young |
| 0.631072 | 0.035638 | 0.179 | 0.194 | 1 | 6 | Atp8b2 | -0.015 | Young |
| 0.631116 | 0.05079 | 0.328 | 0.314 | 1 | 6 | Pet100 | 0.014 | Young |
| 0.631237 | 0.226382 | 0.199 | 0.189 | 1 | 6 | Ing2 | 0.01 | Young |
| 0.631538 | 0.277424 | 0.206 | 0.2 | 1 | 6 | Magoh | 0.006 | Young |
| 0.631793 | 0.522321 | 0.018 | 0.014 | 1 | 6 | Kbtbd7 | 0.004 | Young |
| 0.631934 | 0.071669 | 0.443 | 0.423 | 1 | 6 | Ssr1 | 0.02 | Young |
| 0.632369 | 0.253361 | 0.04 | 0.034 | 1 | 6 | Sord | 0.006 | Young |
| 0.632445 | 0.241965 | 0.234 | 0.254 | 1 | 6 | Smc4 | -0.02 | Young |
| 0.632673 | 0.658138 | 0.037 | 0.031 | 1 | 6 | Ift140 | 0.006 | Young |
| 0.632992 | 0.306523 | 0.059 | 0.051 | 1 | 6 | Elac2 | 0.008 | Young |
| 0.633045 | 0.069299 | 0.286 | 0.303 | 1 | 6 | Thoc2 | -0.017 | Young |
| 0.633048 | 0.180412 | 0.14 | 0.131 | 1 | 6 | Snrnp27 | 0.009 | Young |
| 0.633294 | 0.199908 | 0.242 | 0.237 | 1 | 6 | Mrpl30 | 0.005 | Young |
| 0.633361 | 0.361259 | 0.018 | 0.014 | 1 | 6 | Uhrf1 | 0.004 | Young |
| 0.633361 | 0.311159 | 0.018 | 0.014 | 1 | 6 | Tctn1 | 0.004 | Young |
| 0.633794 | 0.247325 | 0.022 | 0.017 | 1 | 6 | Dnajc17 | 0.005 | Young |
| 0.634146 | 0.460965 | 0.018 | 0.014 | 1 | 6 | F10 | 0.004 | Young |
| 0.634971 | 0.103774 | 0.047 | 0.054 | 1 | 6 | Sass6 | -0.007 | Young |
| 0.63536 | 0.066636 | 0.416 | 0.406 | 1 | 6 | Chmp2a | 0.01 | Young |
| 0.635418 | 0.035118 | 0.055 | 0.063 | 1 | 6 | Eng | -0.008 | Young |
| 0.635431 | 0.166541 | 0.156 | 0.143 | 1 | 6 | Dph5 | 0.013 | Young |
| 0.635716 | 0.389338 | 0.018 | 0.014 | 1 | 6 | Nt5c3b | 0.004 | Young |
| 0.635944 | 0.366105 | 0.097 | 0.089 | 1 | 6 | Ints3 | 0.008 | Young |
| 0.636009 | 0.096095 | 0.047 | 0.04 | 1 | 6 | Snx21 | 0.007 | Young |
| 0.636301 | 0.50503 | 0.023 | 0.029 | 1 | 6 | Gzf1 | -0.006 | Young |
| 0.636319 | 0.479156 | 0.135 | 0.129 | 1 | 6 | Klf16 | 0.006 | Young |
| 0.637209 | 1.110567 | 0.015 | 0.011 | 1 | 6 | Tha1 | 0.004 | Young |
| 0.637209 | 0.599676 | 0.015 | 0.011 | 1 | 6 | Cbr1 | 0.004 | Young |

| | | | | | | | | |
|---|---|---|---|---|---|---|---|---|
| 0.637246 | 0.263204 | 0.052 | 0.06 | 1 | 6 | Fbxo30 | -0.008 | Young |
| 0.637527 | 0.292714 | 0.112 | 0.103 | 1 | 6 | Flot2 | 0.009 | Young |
| 0.637591 | 0.205222 | 0.023 | 0.029 | 1 | 6 | Pdp1 | -0.006 | Young |
| 0.637628 | 0.13962 | 0.151 | 0.14 | 1 | 6 | Slc31a2 | 0.011 | Young |
| 0.638017 | 0.076142 | 0.07 | 0.063 | 1 | 6 | Lonp2 | 0.007 | Young |
| 0.638075 | 0.972159 | 0.018 | 0.014 | 1 | 6 | Tanc1 | 0.004 | Young |
| 0.638093 | 0.062637 | 0.169 | 0.186 | 1 | 6 | Mrpl36 | -0.017 | Young |
| 0.638123 | 0.100736 | 0.069 | 0.077 | 1 | 6 | Ddit4 | -0.008 | Young |
| 0.638128 | 0.422632 | 0.018 | 0.023 | 1 | 6 | Tanc2 | -0.005 | Young |
| 0.638718 | 1.440223 | 0.012 | 0.009 | 1 | 6 | Bfsp1 | 0.003 | Young |
| 0.638881 | 0.14752 | 0.023 | 0.029 | 1 | 6 | Maged1 | -0.006 | Young |
| 0.638952 | 0.676967 | 0.015 | 0.011 | 1 | 6 | Pex11g | 0.004 | Young |
| 0.63965 | 0.165399 | 0.018 | 0.014 | 1 | 6 | Fam84b | 0.004 | Young |
| 0.639907 | 0.030659 | 0.085 | 0.077 | 1 | 6 | Elof1 | 0.008 | Young |
| 0.640007 | 0.205134 | 0.089 | 0.08 | 1 | 6 | Hck | 0.009 | Young |
| 0.640104 | 0.468084 | 0.008 | 0.011 | 1 | 6 | A930033H1 | -0.003 | Young |
| 0.640107 | 0.269752 | 0.097 | 0.089 | 1 | 6 | Timm10 | 0.008 | Young |
| 0.640705 | 1.009847 | 0.012 | 0.009 | 1 | 6 | Scarf1 | 0.003 | Young |
| 0.641077 | 0.136011 | 0.161 | 0.151 | 1 | 6 | Cmc1 | 0.01 | Young |
| 0.641078 | 0.515966 | 0.084 | 0.077 | 1 | 6 | Map3k3 | 0.007 | Young |
| 0.64157 | 0.553658 | 0.015 | 0.011 | 1 | 6 | Tmem143 | 0.004 | Young |
| 0.641699 | 0.888704 | 0.012 | 0.009 | 1 | 6 | Zfp948 | 0.003 | Young |
| 0.641907 | 0.539761 | 0.04 | 0.034 | 1 | 6 | Polr3f | 0.006 | Young |
| 0.642015 | 0.199113 | 0.018 | 0.014 | 1 | 6 | Msh3 | 0.004 | Young |
| 0.642199 | 0.164216 | 0.008 | 0.011 | 1 | 6 | Cd320 | -0.003 | Young |
| 0.642199 | 0.134815 | 0.008 | 0.011 | 1 | 6 | Siah1b | -0.003 | Young |
| 0.642199 | 0.110952 | 0.008 | 0.011 | 1 | 6 | Hist1h1d | -0.003 | Young |
| 0.642468 | 0.333688 | 0.094 | 0.086 | 1 | 6 | Casp2 | 0.008 | Young |
| 0.642694 | 1.015015 | 0.012 | 0.009 | 1 | 6 | Eef2k | 0.003 | Young |
| 0.642694 | 0.867385 | 0.012 | 0.009 | 1 | 6 | Trip13 | 0.003 | Young |
| 0.642708 | 0.382193 | 0.064 | 0.057 | 1 | 6 | Inafm1 | 0.007 | Young |
| 0.642956 | 0.120279 | 0.157 | 0.174 | 1 | 6 | Vcpip1 | -0.017 | Young |
| 0.643158 | 0.00711 | 0.187 | 0.174 | 1 | 6 | Cdk13 | 0.013 | Young |
| 0.643319 | 0.425594 | 0.015 | 0.011 | 1 | 6 | Lrsam1 | 0.004 | Young |
| 0.643319 | 0.331703 | 0.015 | 0.011 | 1 | 6 | Gm16124 | 0.004 | Young |
| 0.643319 | 0.318489 | 0.015 | 0.011 | 1 | 6 | Cpt1c | 0.004 | Young |
| 0.643407 | 0.0284 | 0.023 | 0.029 | 1 | 6 | C330018D2 | -0.006 | Young |
| 0.64369 | 0.808468 | 0.012 | 0.009 | 1 | 6 | Ank2 | 0.003 | Young |
| 0.643732 | 0.745885 | 0.037 | 0.031 | 1 | 6 | Lair1 | 0.006 | Young |
| 0.643925 | 0.382342 | 0.018 | 0.023 | 1 | 6 | Mettl8 | -0.005 | Young |
| 0.643925 | 0.153214 | 0.018 | 0.023 | 1 | 6 | Mertk | -0.005 | Young |
| 0.644193 | 0.499381 | 0.015 | 0.011 | 1 | 6 | 2510046G1 | 0.004 | Young |
| 0.644193 | 0.064416 | 0.015 | 0.011 | 1 | 6 | Rwdd2b | 0.004 | Young |
| 0.644297 | 0.26265 | 0.008 | 0.011 | 1 | 6 | Ptrh1 | -0.003 | Young |
| 0.644297 | 0.257105 | 0.008 | 0.011 | 1 | 6 | Pdk2 | -0.003 | Young |

| | | | | | | | | |
|---|---|---|---|---|---|---|---|---|
| 0.644424 | 0.53205 | 0.084 | 0.077 | 1 | 6 | Spns1 | 0.007 | Young |
| 0.644686 | 0.858434 | 0.012 | 0.009 | 1 | 6 | Tiam2 | 0.003 | Young |
| 0.644686 | 0.673921 | 0.012 | 0.009 | 1 | 6 | Lsmem1 | 0.003 | Young |
| 0.644938 | 0.013969 | 0.355 | 0.383 | 1 | 6 | Psenen | -0.028 | Young |
| 0.6451 | 0.052946 | 0.04 | 0.034 | 1 | 6 | Dgcr14 | 0.006 | Young |
| 0.645105 | 0.190751 | 0.013 | 0.017 | 1 | 6 | Sh2b2 | -0.004 | Young |
| 0.645127 | 0.001777 | 0.112 | 0.123 | 1 | 6 | Snx27 | -0.011 | Young |
| 0.645347 | 0.122475 | 0.008 | 0.011 | 1 | 6 | 6430531B1 | -0.003 | Young |
| 0.645683 | 0.796207 | 0.012 | 0.009 | 1 | 6 | Ophn1 | 0.003 | Young |
| 0.645791 | 0.187529 | 0.059 | 0.051 | 1 | 6 | Cpsf4 | 0.008 | Young |
| 0.645945 | 0.361433 | 0.015 | 0.011 | 1 | 6 | Aqp3 | 0.004 | Young |
| 0.646045 | 0.031232 | 0.241 | 0.263 | 1 | 6 | Pdcl3 | -0.022 | Young |
| 0.646105 | 0.363438 | 0.018 | 0.023 | 1 | 6 | Brat1 | -0.005 | Young |
| 0.646221 | 0.301339 | 0.142 | 0.134 | 1 | 6 | Utp3 | 0.008 | Young |
| 0.646266 | 0.181763 | 0.089 | 0.1 | 1 | 6 | Extl3 | -0.011 | Young |
| 0.646281 | 0.258428 | 0.112 | 0.126 | 1 | 6 | Gon4l | -0.014 | Young |
| 0.64665 | 0.055711 | 0.706 | 0.683 | 1 | 6 | Eif4a1 | 0.023 | Young |
| 0.646656 | 0.001143 | 0.115 | 0.126 | 1 | 6 | Ube2g1 | -0.011 | Young |
| 0.646767 | 0.083494 | 0.589 | 0.56 | 1 | 6 | Tra2a | 0.029 | Young |
| 0.64677 | 0.133775 | 0.122 | 0.134 | 1 | 6 | Zufsp | -0.012 | Young |
| 0.646794 | 0.463159 | 0.013 | 0.017 | 1 | 6 | H2-Eb2 | -0.004 | Young |
| 0.646794 | 0.34349 | 0.013 | 0.017 | 1 | 6 | Gm8369 | -0.004 | Young |
| 0.646821 | 0.23818 | 0.015 | 0.011 | 1 | 6 | Lrp4 | 0.004 | Young |
| 0.646832 | 0.242228 | 0.018 | 0.023 | 1 | 6 | Etfbkmt | -0.005 | Young |
| 0.646834 | 0.399286 | 0.055 | 0.049 | 1 | 6 | Pmf1 | 0.006 | Young |
| 0.647121 | 0.137492 | 0.059 | 0.051 | 1 | 6 | Btbd10 | 0.008 | Young |
| 0.647188 | 0.066526 | 0.331 | 0.32 | 1 | 6 | Rala | 0.011 | Young |
| 0.647246 | 0.213499 | 0.094 | 0.086 | 1 | 6 | Myef2 | 0.008 | Young |
| 0.647254 | 0.148155 | 0.316 | 0.311 | 1 | 6 | Nsf | 0.005 | Young |
| 0.647279 | 0.239565 | 0.137 | 0.129 | 1 | 6 | Eif4ebp1 | 0.008 | Young |
| 0.647281 | 0.126114 | 0.127 | 0.12 | 1 | 6 | Lyst | 0.007 | Young |
| 0.64761 | 0.216375 | 0.042 | 0.049 | 1 | 6 | Snhg20 | -0.007 | Young |
| 0.647639 | 0.151832 | 0.013 | 0.017 | 1 | 6 | Phpt1 | -0.004 | Young |
| 0.647678 | 0.692825 | 0.012 | 0.009 | 1 | 6 | Nthl1 | 0.003 | Young |
| 0.647678 | 0.549907 | 0.012 | 0.009 | 1 | 6 | Hacl1 | 0.003 | Young |
| 0.647678 | 0.421725 | 0.012 | 0.009 | 1 | 6 | Efemp2 | 0.003 | Young |
| 0.647878 | 0.176197 | 0.135 | 0.126 | 1 | 6 | Polr3c | 0.009 | Young |
| 0.648026 | 0.099351 | 0.084 | 0.094 | 1 | 6 | Rnf14 | -0.01 | Young |
| 0.64812 | 0.001631 | 0.156 | 0.169 | 1 | 6 | Herpud2 | -0.013 | Young |
| 0.648501 | 0.311638 | 0.008 | 0.011 | 1 | 6 | Gm26857 | -0.003 | Young |
| 0.648677 | 0.091993 | 0.012 | 0.009 | 1 | 6 | Cep152 | 0.003 | Young |
| 0.648677 | 0.083849 | 0.012 | 0.009 | 1 | 6 | Thada | 0.003 | Young |
| 0.648887 | 0.263286 | 0.055 | 0.063 | 1 | 6 | 1110059E2 | -0.008 | Young |
| 0.649015 | 0.121072 | 0.018 | 0.023 | 1 | 6 | Pcgf1 | -0.005 | Young |
| 0.649554 | 0.749524 | 0.008 | 0.011 | 1 | 6 | Kif15 | -0.003 | Young |

| | | | | | | | | |
|---|---|---|---|---|---|---|---|---|
| 0.649676 | 0.745696 | 0.012 | 0.009 | 1 | 6 | Nat2 | 0.003 | Young |
| 0.650134 | 0.041779 | 0.127 | 0.14 | 1 | 6 | Mapkap1 | -0.013 | Young |
| 0.650292 | 0.03715 | 0.527 | 0.509 | 1 | 6 | Atp5d | 0.018 | Young |
| 0.650331 | 0.079317 | 0.015 | 0.011 | 1 | 6 | Atp6v0c | 0.004 | Young |
| 0.650473 | 0.308306 | 0.018 | 0.023 | 1 | 6 | Pde6d | -0.005 | Young |
| 0.650473 | 0.170341 | 0.018 | 0.023 | 1 | 6 | Bphl | -0.005 | Young |
| 0.650597 | 0.194061 | 0.443 | 0.44 | 1 | 6 | Fcrls | 0.003 | Young |
| 0.650674 | 0.061165 | 0.059 | 0.051 | 1 | 6 | Phlpp2 | 0.008 | Young |
| 0.650676 | 0.280988 | 0.012 | 0.009 | 1 | 6 | Gm26698 | 0.003 | Young |
| 0.650828 | 0.191392 | 0.187 | 0.18 | 1 | 6 | Ap1s3 | 0.007 | Young |
| 0.651025 | 0.033771 | 0.013 | 0.017 | 1 | 6 | Zfp553 | -0.004 | Young |
| 0.651129 | 0.037148 | 0.251 | 0.243 | 1 | 6 | Tox4 | 0.008 | Young |
| 0.651366 | 0.661201 | 0.033 | 0.029 | 1 | 6 | Med19 | 0.004 | Young |
| 0.651677 | 0.750531 | 0.012 | 0.009 | 1 | 6 | Trim68 | 0.003 | Young |
| 0.651677 | 0.127742 | 0.012 | 0.009 | 1 | 6 | Tmem51os | 0.003 | Young |
| 0.651873 | 0.179801 | 0.013 | 0.017 | 1 | 6 | 2310040G2 | -0.004 | Young |
| 0.651945 | 0.039958 | 0.055 | 0.063 | 1 | 6 | Rrp12 | -0.008 | Young |
| 0.652078 | 0.668048 | 0.037 | 0.031 | 1 | 6 | 3110001I2 | 0.006 | Young |
| 0.65233 | 0.021204 | 0.206 | 0.191 | 1 | 6 | Plekhg3 | 0.015 | Young |
| 0.652535 | 1.010124 | 0.033 | 0.029 | 1 | 6 | Mcm2 | 0.004 | Young |
| 0.652662 | 0.588653 | 0.018 | 0.023 | 1 | 6 | Gm4070 | -0.005 | Young |
| 0.652678 | 0.24098 | 0.012 | 0.009 | 1 | 6 | Slc5a6 | 0.003 | Young |
| 0.652678 | 0.19151 | 0.012 | 0.009 | 1 | 6 | Tnnt3 | 0.003 | Young |
| 0.652678 | 0.062367 | 0.012 | 0.009 | 1 | 6 | Tmem62 | 0.003 | Young |
| 0.652678 | 0.01368 | 0.012 | 0.009 | 1 | 6 | Cd46 | 0.003 | Young |
| 0.652721 | 0.171629 | 0.013 | 0.017 | 1 | 6 | Rasal2 | -0.004 | Young |
| 0.653645 | 0.112716 | 0.04 | 0.034 | 1 | 6 | Yaf2 | 0.006 | Young |
| 0.653877 | 0.055635 | 0.144 | 0.134 | 1 | 6 | Mkl1 | 0.01 | Young |
| 0.654241 | 0.023094 | 0.391 | 0.377 | 1 | 6 | Uqcrfs1 | 0.014 | Young |
| 0.654897 | 0.020795 | 0.314 | 0.306 | 1 | 6 | Usp34 | 0.008 | Young |
| 0.655023 | 0.287739 | 0.152 | 0.146 | 1 | 6 | Cnot8 | 0.006 | Young |
| 0.655067 | 0.273776 | 0.074 | 0.066 | 1 | 6 | Kctd5 | 0.008 | Young |
| 0.655199 | 0.267184 | 0.134 | 0.126 | 1 | 6 | Adh5 | 0.008 | Young |
| 0.655659 | 0.133645 | 0.13 | 0.12 | 1 | 6 | Gpr137b | 0.01 | Young |
| 0.655767 | 0.134668 | 0.037 | 0.043 | 1 | 6 | Zdhhc12 | -0.006 | Young |
| 0.656086 | 0.172646 | 0.074 | 0.083 | 1 | 6 | Tesk1 | -0.009 | Young |
| 0.656313 | 0.00946 | 0.229 | 0.246 | 1 | 6 | Snf8 | -0.017 | Young |
| 0.656445 | 0.009055 | 0.318 | 0.303 | 1 | 6 | Arrdc1 | 0.015 | Young |
| 0.656579 | 0.096547 | 0.227 | 0.217 | 1 | 6 | Vamp3 | 0.01 | Young |
| 0.656676 | 0.213279 | 0.378 | 0.383 | 1 | 6 | Laptm4a | -0.005 | Young |
| 0.656833 | 0.097305 | 0.192 | 0.209 | 1 | 6 | Akirin2 | -0.017 | Young |
| 0.656862 | 0.105733 | 0.151 | 0.14 | 1 | 6 | Zhx1 | 0.011 | Young |
| 0.656862 | 0.128755 | 0.04 | 0.034 | 1 | 6 | Ppargc1b | 0.006 | Young |
| 0.657803 | 0.675767 | 0.033 | 0.029 | 1 | 6 | Sema7a | 0.004 | Young |
| 0.657813 | 0.198524 | 0.055 | 0.049 | 1 | 6 | Phtf2 | 0.006 | Young |

| | | | | | | | | |
|---|---|---|---|---|---|---|---|---|
| 0.657912 | 0.283918 | 0.114 | 0.106 | 1 | 6 | Snx15 | 0.008 | Young |
| 0.658341 | 0.114494 | 0.147 | 0.16 | 1 | 6 | Usp6nl | -0.013 | Young |
| 0.658763 | 0.220838 | 0.236 | 0.226 | 1 | 6 | Cldnd1 | 0.01 | Young |
| 0.659034 | 0.021556 | 0.179 | 0.197 | 1 | 6 | Snx6 | -0.018 | Young |
| 0.659779 | 0.080626 | 0.112 | 0.103 | 1 | 6 | Rbpms | 0.009 | Young |
| 0.660006 | 0.028374 | 0.037 | 0.043 | 1 | 6 | Ntpcr | -0.006 | Young |
| 0.660024 | 0.229504 | 0.151 | 0.166 | 1 | 6 | Dlg1 | -0.015 | Young |
| 0.6602 | 0.512003 | 0.048 | 0.043 | 1 | 6 | Nod2 | 0.005 | Young |
| 0.66216 | 0.012956 | 0.064 | 0.071 | 1 | 6 | Tmub2 | -0.007 | Young |
| 0.662626 | 0.354242 | 0.191 | 0.186 | 1 | 6 | Ptpn7 | 0.005 | Young |
| 0.662722 | 0.184439 | 0.059 | 0.051 | 1 | 6 | Ggt5 | 0.008 | Young |
| 0.66299 | 0.022027 | 0.164 | 0.154 | 1 | 6 | Mrps36 | 0.01 | Young |
| 0.663133 | 0.00727 | 0.137 | 0.149 | 1 | 6 | Evi2 | -0.012 | Young |
| 0.663275 | 0.497761 | 0.037 | 0.031 | 1 | 6 | Vps72 | 0.006 | Young |
| 0.663642 | 0.07053 | 0.067 | 0.06 | 1 | 6 | Man2c1 | 0.007 | Young |
| 0.663642 | 0.297121 | 0.344 | 0.351 | 1 | 6 | Tuba1c | -0.007 | Young |
| 0.664019 | 0.0195 | 0.064 | 0.071 | 1 | 6 | Tmem126a | -0.007 | Young |
| 0.664551 | 0.080569 | 0.452 | 0.434 | 1 | 6 | Ppp3r1 | 0.018 | Young |
| 0.664892 | 0.010354 | 0.289 | 0.283 | 1 | 6 | Coa5 | 0.006 | Young |
| 0.665388 | 0.247078 | 0.254 | 0.251 | 1 | 6 | Ube2z | 0.003 | Young |
| 0.665785 | 0.113895 | 0.401 | 0.391 | 1 | 6 | Jak2 | 0.01 | Young |
| 0.665822 | 0.061106 | 0.241 | 0.229 | 1 | 6 | Kxd1 | 0.012 | Young |
| 0.665989 | 0.250777 | 0.134 | 0.126 | 1 | 6 | Ten1 | 0.008 | Young |
| 0.665993 | 0.115934 | 0.283 | 0.266 | 1 | 6 | R3hdm2 | 0.017 | Young |
| 0.666649 | 0.166547 | 0.037 | 0.031 | 1 | 6 | Depdc7 | 0.006 | Young |
| 0.666964 | 0.343449 | 0.075 | 0.069 | 1 | 6 | Rad17 | 0.006 | Young |
| 0.66721 | 0.375981 | 0.033 | 0.029 | 1 | 6 | Phtf1 | 0.004 | Young |
| 0.667315 | 0.016347 | 0.403 | 0.38 | 1 | 6 | Srp14 | 0.023 | Young |
| 0.667706 | 0.018918 | 0.112 | 0.103 | 1 | 6 | Wdr13 | 0.009 | Young |
| 0.668517 | 0.232117 | 0.037 | 0.043 | 1 | 6 | Tmem39b | -0.006 | Young |
| 0.668582 | 0.153807 | 0.079 | 0.071 | 1 | 6 | Casp9 | 0.008 | Young |
| 0.668635 | 0.005442 | 0.05 | 0.057 | 1 | 6 | Zcchc8 | -0.007 | Young |
| 0.668721 | 0.119867 | 0.072 | 0.08 | 1 | 6 | Arhgap11a | -0.008 | Young |
| 0.66883 | 0.012713 | 0.279 | 0.271 | 1 | 6 | Epsti1 | 0.008 | Young |
| 0.669135 | 0.196182 | 0.184 | 0.174 | 1 | 6 | Appbp2 | 0.01 | Young |
| 0.66932 | 0.030772 | 0.085 | 0.077 | 1 | 6 | Wdr41 | 0.008 | Young |
| 0.669803 | 0.159204 | 0.274 | 0.266 | 1 | 6 | Piezo1 | 0.008 | Young |
| 0.671342 | 0.439663 | 0.033 | 0.029 | 1 | 6 | Tsr2 | 0.004 | Young |
| 0.67192 | 0.895723 | 0.03 | 0.026 | 1 | 6 | Ccar2 | 0.004 | Young |
| 0.672091 | 0.258715 | 0.079 | 0.071 | 1 | 6 | Ptrhd1 | 0.008 | Young |
| 0.672209 | 0.232065 | 0.052 | 0.046 | 1 | 6 | Acad9 | 0.006 | Young |
| 0.672287 | 0.071005 | 0.037 | 0.031 | 1 | 6 | Dynlt1f | 0.006 | Young |
| 0.672287 | 0.103507 | 0.037 | 0.031 | 1 | 6 | Syce2 | 0.006 | Young |
| 0.672438 | 0.246923 | 0.107 | 0.1 | 1 | 6 | Gpd1l | 0.007 | Young |
| 0.673116 | 0.358883 | 0.033 | 0.029 | 1 | 6 | Nkiras1 | 0.004 | Young |

| | | | | | | | | |
|---|---|---|---|---|---|---|---|---|
| 0.673438 | 0.056954 | 0.187 | 0.177 | 1 | 6 | Gigyf1 | 0.01 | Young |
| 0.67344 | 0.392038 | 0.048 | 0.043 | 1 | 6 | Nlrp1b | 0.005 | Young |
| 0.673654 | 0.205843 | 0.079 | 0.071 | 1 | 6 | Lactb2 | 0.008 | Young |
| 0.673875 | 0.046775 | 0.11 | 0.12 | 1 | 6 | Dnajc10 | -0.01 | Young |
| 0.673907 | 0.052662 | 0.247 | 0.237 | 1 | 6 | Hps3 | 0.01 | Young |
| 0.674007 | 0.050208 | 0.204 | 0.197 | 1 | 6 | Sipa1l3 | 0.007 | Young |
| 0.674232 | 0.067407 | 0.221 | 0.209 | 1 | 6 | Acvrl1 | 0.012 | Young |
| 0.674243 | 0.07695 | 0.152 | 0.143 | 1 | 6 | Cenpa | 0.009 | Young |
| 0.6743 | 0.322064 | 0.033 | 0.029 | 1 | 6 | Trim59 | 0.004 | Young |
| 0.674508 | 0.028796 | 0.206 | 0.197 | 1 | 6 | Xpo1 | 0.009 | Young |
| 0.675484 | 0.223498 | 0.033 | 0.029 | 1 | 6 | Poc5 | 0.004 | Young |
| 0.67599 | 0.051099 | 0.048 | 0.054 | 1 | 6 | Glce | -0.006 | Young |
| 0.676008 | 0.274324 | 0.184 | 0.177 | 1 | 6 | Yipf1 | 0.007 | Young |
| 0.67607 | 0.355713 | 0.087 | 0.08 | 1 | 6 | Klf11 | 0.007 | Young |
| 0.676256 | 0.087256 | 0.064 | 0.071 | 1 | 6 | Syt11 | -0.007 | Young |
| 0.676754 | 0.223917 | 0.09 | 0.083 | 1 | 6 | Spast | 0.007 | Young |
| 0.676899 | 0.461935 | 0.045 | 0.04 | 1 | 6 | Rpain | 0.005 | Young |
| 0.676966 | 0.166754 | 0.174 | 0.166 | 1 | 6 | Zfp326 | 0.008 | Young |
| 0.677133 | 0.112206 | 0.032 | 0.037 | 1 | 6 | Zcchc10 | -0.005 | Young |
| 0.677939 | 0.076569 | 0.045 | 0.051 | 1 | 6 | Agpat2 | -0.006 | Young |
| 0.677966 | 0.192121 | 0.112 | 0.123 | 1 | 6 | Usf3 | -0.011 | Young |
| 0.678094 | 0.411499 | 0.08 | 0.074 | 1 | 6 | Zfp362 | 0.006 | Young |
| 0.678283 | 0.103977 | 0.032 | 0.037 | 1 | 6 | Kif19a | -0.005 | Young |
| 0.678584 | 0.021772 | 0.145 | 0.137 | 1 | 6 | Kbtbd11 | 0.008 | Young |
| 0.678589 | 0.057398 | 0.09 | 0.083 | 1 | 6 | Pdcd7 | 0.007 | Young |
| 0.67874 | 0.03533 | 0.079 | 0.071 | 1 | 6 | Pex19 | 0.008 | Young |
| 0.678864 | 0.320083 | 0.048 | 0.043 | 1 | 6 | Cep44 | 0.005 | Young |
| 0.679617 | 0.047524 | 0.115 | 0.126 | 1 | 6 | Srpk1 | -0.011 | Young |
| 0.679658 | 0.220408 | 0.067 | 0.06 | 1 | 6 | Cgnl1 | 0.007 | Young |
| 0.679723 | 0.141291 | 0.06 | 0.054 | 1 | 6 | Dcaf1 | 0.006 | Young |
| 0.680214 | 0.008246 | 0.097 | 0.089 | 1 | 6 | Peak1 | 0.008 | Young |
| 0.680475 | 0.224838 | 0.147 | 0.14 | 1 | 6 | Plekhg2 | 0.007 | Young |
| 0.680824 | 0.196362 | 0.033 | 0.029 | 1 | 6 | S1pr2 | 0.004 | Young |
| 0.681049 | 0.137459 | 0.274 | 0.266 | 1 | 6 | Cyp51 | 0.008 | Young |
| 0.681279 | 0.56674 | 0.03 | 0.026 | 1 | 6 | Map3k10 | 0.004 | Young |
| 0.68177 | 0.106742 | 0.089 | 0.097 | 1 | 6 | Smim19 | -0.008 | Young |
| 0.682264 | 0.192607 | 0.1 | 0.111 | 1 | 6 | Eepd1 | -0.011 | Young |
| 0.682811 | 0.139692 | 0.338 | 0.329 | 1 | 6 | Cfh | 0.009 | Young |
| 0.683677 | 0.002145 | 0.054 | 0.06 | 1 | 6 | Megf8 | -0.006 | Young |
| 0.684145 | 0.122693 | 0.064 | 0.057 | 1 | 6 | Spg11 | 0.007 | Young |
| 0.684304 | 0.062772 | 0.584 | 0.603 | 1 | 6 | Tmed5 | -0.019 | Young |
| 0.684667 | 0.073096 | 0.657 | 0.634 | 1 | 6 | Trib1 | 0.023 | Young |
| 0.6848 | 0.33154 | 0.048 | 0.043 | 1 | 6 | C4b | 0.005 | Young |
| 0.685036 | 0.010693 | 0.027 | 0.031 | 1 | 6 | Coil | -0.004 | Young |
| 0.685257 | 0.019032 | 0.326 | 0.323 | 1 | 6 | Cmtm7 | 0.003 | Young |

| | | | | | | | | |
|---|---|---|---|---|---|---|---|---|
| 0.685307 | 0.070945 | 0.415 | 0.411 | 1 | 6 | Cox14 | 0.004 | Young |
| 0.685663 | 0.307785 | 0.03 | 0.026 | 1 | 6 | Rabif | 0.004 | Young |
| 0.685711 | 0.290488 | 0.177 | 0.171 | 1 | 6 | Zdhhc9 | 0.006 | Young |
| 0.685801 | 0.024147 | 0.256 | 0.274 | 1 | 6 | Vdac1 | -0.018 | Young |
| 0.687084 | 0.214887 | 0.192 | 0.186 | 1 | 6 | Rap2c | 0.006 | Young |
| 0.687393 | 0.396081 | 0.105 | 0.1 | 1 | 6 | Zpr1 | 0.005 | Young |
| 0.688271 | 0.05659 | 0.048 | 0.043 | 1 | 6 | Kti12 | 0.005 | Young |
| 0.688531 | 0.014411 | 0.256 | 0.271 | 1 | 6 | Lilrb4a | -0.015 | Young |
| 0.68864 | 0.048433 | 0.214 | 0.229 | 1 | 6 | Gatm | -0.015 | Young |
| 0.68879 | 0.503747 | 0.069 | 0.063 | 1 | 6 | Arsk | 0.006 | Young |
| 0.688801 | 0.116562 | 0.027 | 0.031 | 1 | 6 | Tmem104 | -0.004 | Young |
| 0.688845 | 0.049985 | 0.226 | 0.214 | 1 | 6 | Tcf20 | 0.012 | Young |
| 0.689056 | 0.016726 | 0.222 | 0.217 | 1 | 6 | Ufsp2 | 0.005 | Young |
| 0.690036 | 1.128109 | 0.027 | 0.023 | 1 | 6 | Gpatch1 | 0.004 | Young |
| 0.690096 | 0.226014 | 0.1 | 0.111 | 1 | 6 | Pex2 | -0.011 | Young |
| 0.690212 | 0.121969 | 0.217 | 0.211 | 1 | 6 | Nsmaf | 0.006 | Young |
| 0.690293 | 0.034008 | 0.247 | 0.24 | 1 | 6 | Gm8186 | 0.007 | Young |
| 0.690459 | 0.180403 | 0.132 | 0.126 | 1 | 6 | Cetn2 | 0.006 | Young |
| 0.690533 | 0.040689 | 0.39 | 0.417 | 1 | 6 | Nono | -0.027 | Young |
| 0.690837 | 1.31898 | 0.022 | 0.026 | 1 | 6 | Cxcl9 | -0.004 | Young |
| 0.691019 | 0.110198 | 0.328 | 0.32 | 1 | 6 | Nab1 | 0.008 | Young |
| 0.691074 | 0.034372 | 0.371 | 0.354 | 1 | 6 | Prelid1 | 0.017 | Young |
| 0.691659 | 0.081645 | 0.324 | 0.311 | 1 | 6 | Myof | 0.013 | Young |
| 0.691664 | 0.016161 | 0.045 | 0.051 | 1 | 6 | Polr3d | -0.006 | Young |
| 0.692567 | 0.098018 | 0.064 | 0.071 | 1 | 6 | Bicra | -0.007 | Young |
| 0.692823 | 0.073885 | 0.316 | 0.311 | 1 | 6 | Pten | 0.005 | Young |
| 0.693131 | 0.332286 | 0.137 | 0.149 | 1 | 6 | Atg5 | -0.012 | Young |
| 0.693203 | 0.272611 | 0.03 | 0.026 | 1 | 6 | 1110004E0 | 0.004 | Young |
| 0.693305 | 0.001131 | 0.139 | 0.149 | 1 | 6 | Pdgfb | -0.01 | Young |
| 0.693573 | 0.099424 | 0.181 | 0.174 | 1 | 6 | Nin | 0.007 | Young |
| 0.693895 | 0.660195 | 0.085 | 0.08 | 1 | 6 | Slc39a11 | 0.005 | Young |
| 0.693998 | 0.15488 | 0.072 | 0.08 | 1 | 6 | Bpnt1 | -0.008 | Young |
| 0.694087 | 0.407166 | 0.147 | 0.143 | 1 | 6 | Tmem165 | 0.004 | Young |
| 0.694463 | 0.430983 | 0.03 | 0.026 | 1 | 6 | Utp23 | 0.004 | Young |
| 0.694524 | 0.647715 | 0.042 | 0.037 | 1 | 6 | Zbtb33 | 0.005 | Young |
| 0.69453 | 0.076215 | 0.04 | 0.046 | 1 | 6 | Tbc1d10a | -0.006 | Young |
| 0.694741 | 0.151123 | 0.431 | 0.429 | 1 | 6 | Uggt1 | 0.002 | Young |
| 0.695111 | 0.563136 | 0.045 | 0.051 | 1 | 6 | Ccr7 | -0.006 | Young |
| 0.695234 | 0.144306 | 0.048 | 0.043 | 1 | 6 | Fam217b | 0.005 | Young |
| 0.695243 | 0.183206 | 0.202 | 0.194 | 1 | 6 | Ankle2 | 0.008 | Young |
| 0.695369 | 0.760924 | 0.027 | 0.023 | 1 | 6 | Negr1 | 0.004 | Young |
| 0.695369 | 0.540789 | 0.027 | 0.023 | 1 | 6 | C130050O: | 0.004 | Young |
| 0.695505 | 0.024522 | 0.145 | 0.16 | 1 | 6 | Zbtb44 | -0.015 | Young |
| 0.695551 | 0.30409 | 0.104 | 0.097 | 1 | 6 | Fam241a | 0.007 | Young |
| 0.695593 | 0.892192 | 0.042 | 0.037 | 1 | 6 | Ppp1r12b | 0.005 | Young |

| | | | | | | | | |
|---|---|---|---|---|---|---|---|---|
| 0.696159 | 0.213095 | 0.172 | 0.189 | 1 | 6 | Exoc4 | -0.017 | Young |
| 0.696581 | 0.125502 | 0.207 | 0.197 | 1 | 6 | Nhsl2 | 0.01 | Young |
| 0.697799 | 0.17867 | 0.186 | 0.177 | 1 | 6 | Dnajc13 | 0.009 | Young |
| 0.697805 | 0.067358 | 0.266 | 0.26 | 1 | 6 | Cyb5r4 | 0.006 | Young |
| 0.697939 | 0.380024 | 0.045 | 0.04 | 1 | 6 | Ncapd3 | 0.005 | Young |
| 0.698079 | 0.094921 | 0.169 | 0.157 | 1 | 6 | Slc30a7 | 0.012 | Young |
| 0.698475 | 0.11142 | 0.048 | 0.043 | 1 | 6 | Med6 | 0.005 | Young |
| 0.698605 | 0.307975 | 0.075 | 0.069 | 1 | 6 | Gdpd3 | 0.006 | Young |
| 0.698895 | 0.11576 | 0.333 | 0.329 | 1 | 6 | Rbm8a | 0.004 | Young |
| 0.69901 | 0.003567 | 0.092 | 0.1 | 1 | 6 | Srpk2 | -0.008 | Young |
| 0.699014 | 0.268542 | 0.161 | 0.151 | 1 | 6 | Fundc2 | 0.01 | Young |
| 0.699127 | 0.120354 | 0.177 | 0.171 | 1 | 6 | Plcb2 | 0.006 | Young |
| 0.699158 | 0.587044 | 0.05 | 0.057 | 1 | 6 | Atat1 | -0.007 | Young |
| 0.699196 | 0.286405 | 0.135 | 0.129 | 1 | 6 | Bmyc | 0.006 | Young |
| 0.699378 | 0.669138 | 0.027 | 0.023 | 1 | 6 | Thap1 | 0.004 | Young |
| 0.699489 | 0.075276 | 0.216 | 0.209 | 1 | 6 | Kdm1a | 0.007 | Young |
| 0.699749 | 0.560331 | 0.04 | 0.046 | 1 | 6 | Ints10 | -0.006 | Young |
| 0.700018 | 0.074522 | 0.057 | 0.063 | 1 | 6 | Trove2 | -0.006 | Young |
| 0.700511 | 0.265362 | 0.06 | 0.054 | 1 | 6 | Mis12 | 0.006 | Young |
| 0.701213 | 0.09738 | 0.054 | 0.06 | 1 | 6 | Ptp4a3 | -0.006 | Young |
| 0.701215 | 0.116792 | 0.079 | 0.071 | 1 | 6 | Ermp1 | 0.008 | Young |
| 0.701288 | 0.033707 | 0.022 | 0.026 | 1 | 6 | Snapc4 | -0.004 | Young |
| 0.701466 | 0.059077 | 0.102 | 0.111 | 1 | 6 | Hspbp1 | -0.009 | Young |
| 0.701506 | 0.2892 | 0.132 | 0.126 | 1 | 6 | Apopt1 | 0.006 | Young |
| 0.702356 | 0.200317 | 0.087 | 0.08 | 1 | 6 | Zfp407 | 0.007 | Young |
| 0.702671 | 0.277114 | 0.03 | 0.026 | 1 | 6 | Ptgs2os2 | 0.004 | Young |
| 0.703097 | 0.131161 | 0.067 | 0.06 | 1 | 6 | Ell | 0.007 | Young |
| 0.703311 | 0.000569 | 0.206 | 0.2 | 1 | 6 | Ssbp4 | 0.006 | Young |
| 0.704066 | 0.114541 | 0.027 | 0.023 | 1 | 6 | Mrpl49 | 0.004 | Young |
| 0.704291 | 0.207813 | 0.12 | 0.131 | 1 | 6 | Traf5 | -0.011 | Young |
| 0.704409 | 0.261389 | 0.159 | 0.151 | 1 | 6 | Rabgef1 | 0.008 | Young |
| 0.704459 | 0.072555 | 0.04 | 0.046 | 1 | 6 | Rbm19 | -0.006 | Young |
| 0.704736 | 0.138419 | 0.318 | 0.323 | 1 | 6 | Trmt112 | -0.005 | Young |
| 0.704786 | 0.252584 | 0.06 | 0.054 | 1 | 6 | Pdp2 | 0.006 | Young |
| 0.705394 | 0.25344 | 0.154 | 0.149 | 1 | 6 | Acot7 | 0.005 | Young |
| 0.705508 | 0.094048 | 0.04 | 0.046 | 1 | 6 | Stambpl1 | -0.006 | Young |
| 0.705838 | 0.012367 | 0.03 | 0.026 | 1 | 6 | Gm20342 | 0.004 | Young |
| 0.705889 | 0.009811 | 0.107 | 0.117 | 1 | 6 | Psen2 | -0.01 | Young |
| 0.706079 | 0.299781 | 0.027 | 0.023 | 1 | 6 | Omp | 0.004 | Young |
| 0.706391 | 0.004388 | 0.237 | 0.229 | 1 | 6 | 2-Mar | 0.008 | Young |
| 0.706472 | 0.088825 | 0.03 | 0.026 | 1 | 6 | Osbpl7 | 0.004 | Young |
| 0.706721 | 0.24451 | 0.107 | 0.1 | 1 | 6 | Klhdc4 | 0.007 | Young |
| 0.707274 | 0.102587 | 0.304 | 0.323 | 1 | 6 | Map3k1 | -0.019 | Young |
| 0.707387 | 0.020588 | 0.209 | 0.22 | 1 | 6 | Polr2g | -0.011 | Young |
| 0.707389 | 0.578417 | 0.042 | 0.037 | 1 | 6 | Trp53i13 | 0.005 | Young |

| | | | | | | | | |
|---|---|---|---|---|---|---|---|---|
| 0.707422 | 0.295966 | 0.027 | 0.023 | 1 | 6 | Rbm43 | 0.004 | Young |
| 0.707422 | 0.039015 | 0.027 | 0.023 | 1 | 6 | Tef | 0.004 | Young |
| 0.707437 | 0.035155 | 0.142 | 0.134 | 1 | 6 | Mrpl21 | 0.008 | Young |
| 0.707993 | 0.253316 | 0.167 | 0.163 | 1 | 6 | Sec13 | 0.004 | Young |
| 0.708288 | 0.16208 | 0.045 | 0.04 | 1 | 6 | Cebpe | 0.005 | Young |
| 0.708989 | 0.162518 | 0.022 | 0.026 | 1 | 6 | Pex11b | -0.004 | Young |
| 0.709438 | 0.612669 | 0.027 | 0.023 | 1 | 6 | Pop1 | 0.004 | Young |
| 0.709579 | 0.179738 | 0.147 | 0.14 | 1 | 6 | Klhl9 | 0.007 | Young |
| 0.709622 | 0.192763 | 0.122 | 0.114 | 1 | 6 | Usf1 | 0.008 | Young |
| 0.709966 | 0.169589 | 0.059 | 0.066 | 1 | 6 | Mvd | -0.007 | Young |
| 0.7104 | 0.064916 | 0.062 | 0.069 | 1 | 6 | Rbm38 | -0.007 | Young |
| 0.710829 | 0.01209 | 0.161 | 0.151 | 1 | 6 | Yipf5 | 0.01 | Young |
| 0.710836 | 0.037255 | 0.162 | 0.174 | 1 | 6 | Cuedc2 | -0.012 | Young |
| 0.711578 | 0.279555 | 0.072 | 0.08 | 1 | 6 | Rnf34 | -0.008 | Young |
| 0.711661 | 0.131593 | 0.487 | 0.491 | 1 | 6 | Nol7 | -0.004 | Young |
| 0.712381 | 0.405739 | 0.077 | 0.071 | 1 | 6 | Hsdl2 | 0.006 | Young |
| 0.712899 | 0.238845 | 0.157 | 0.151 | 1 | 6 | Gm2000 | 0.006 | Young |
| 0.713035 | 0.467085 | 0.07 | 0.066 | 1 | 6 | Cops7a | 0.004 | Young |
| 0.713245 | 0.391931 | 0.065 | 0.06 | 1 | 6 | Renbp | 0.005 | Young |
| 0.713415 | 0.196952 | 0.062 | 0.069 | 1 | 6 | Zfp266 | -0.007 | Young |
| 0.713477 | 0.121142 | 0.027 | 0.023 | 1 | 6 | Arhgef40 | 0.004 | Young |
| 0.71373 | 0.72833 | 0.023 | 0.02 | 1 | 6 | Snapc2 | 0.003 | Young |
| 0.713772 | 0.27432 | 0.147 | 0.14 | 1 | 6 | Tmem131l | 0.007 | Young |
| 0.714151 | 0.169156 | 0.027 | 0.023 | 1 | 6 | Cep76 | 0.004 | Young |
| 0.7142 | 0.104348 | 0.099 | 0.091 | 1 | 6 | Zfp277 | 0.008 | Young |
| 0.71433 | 0.021541 | 0.199 | 0.191 | 1 | 6 | Szrd1 | 0.008 | Young |
| 0.71445 | 0.767632 | 0.023 | 0.02 | 1 | 6 | Mthfd2l | 0.003 | Young |
| 0.714822 | 0.179758 | 0.089 | 0.097 | 1 | 6 | Nosip | -0.008 | Young |
| 0.715019 | 0.122782 | 0.109 | 0.103 | 1 | 6 | Taf2 | 0.006 | Young |
| 0.715327 | 0.00633 | 0.289 | 0.3 | 1 | 6 | Epb41l2 | -0.011 | Young |
| 0.715499 | 0.129704 | 0.027 | 0.023 | 1 | 6 | Phka1 | 0.004 | Young |
| 0.715811 | 0.275489 | 0.069 | 0.063 | 1 | 6 | Bysl | 0.006 | Young |
| 0.715811 | 0.222153 | 0.069 | 0.063 | 1 | 6 | Dusp10 | 0.006 | Young |
| 0.715912 | 0.076321 | 0.054 | 0.06 | 1 | 6 | Polr2d | -0.006 | Young |
| 0.716018 | 0.005952 | 0.035 | 0.04 | 1 | 6 | Dedd | -0.005 | Young |
| 0.71625 | 0.35121 | 0.057 | 0.051 | 1 | 6 | Zdhhc8 | 0.006 | Young |
| 0.716394 | 0.162986 | 0.08 | 0.074 | 1 | 6 | Usp36 | 0.006 | Young |
| 0.716494 | 0.141245 | 0.157 | 0.149 | 1 | 6 | Ap3m1 | 0.008 | Young |
| 0.716517 | 0.072348 | 0.254 | 0.246 | 1 | 6 | Smndc1 | 0.008 | Young |
| 0.71733 | 0.372322 | 0.023 | 0.02 | 1 | 6 | Umps | 0.003 | Young |
| 0.717367 | 0.042755 | 0.244 | 0.24 | 1 | 6 | Sec63 | 0.004 | Young |
| 0.718045 | 0.003977 | 0.234 | 0.22 | 1 | 6 | Tmem55b | 0.014 | Young |
| 0.718213 | 0.24765 | 0.117 | 0.129 | 1 | 6 | Ust | -0.012 | Young |
| 0.71963 | 0.013586 | 0.247 | 0.237 | 1 | 6 | Rps6ka4 | 0.01 | Young |
| 0.719662 | 0.039207 | 0.495 | 0.489 | 1 | 6 | Anp32e | 0.006 | Young |

| | | | | | | | | |
|---|---|---|---|---|---|---|---|---|
| 0.720392 | 0.353087 | 0.085 | 0.08 | 1 | 6 | Klra2 | 0.005 | Young |
| 0.720427 | 0.21913 | 0.095 | 0.089 | 1 | 6 | Pygo2 | 0.006 | Young |
| 0.720542 | 0.028715 | 0.353 | 0.34 | 1 | 6 | Atp2a2 | 0.013 | Young |
| 0.720846 | 0.253347 | 0.171 | 0.166 | 1 | 6 | Eps15l1 | 0.005 | Young |
| 0.721439 | 0.249282 | 0.054 | 0.049 | 1 | 6 | Klhl15 | 0.005 | Young |
| 0.721657 | 0.569678 | 0.023 | 0.02 | 1 | 6 | Scn1b | 0.003 | Young |
| 0.721735 | 0.257561 | 0.085 | 0.08 | 1 | 6 | Dcaf5 | 0.005 | Young |
| 0.722264 | 0.067683 | 0.06 | 0.066 | 1 | 6 | Rffl | -0.006 | Young |
| 0.722986 | 0.253519 | 0.017 | 0.02 | 1 | 6 | Zfp846 | -0.003 | Young |
| 0.722986 | 0.079056 | 0.017 | 0.02 | 1 | 6 | Per3 | -0.003 | Young |
| 0.723023 | 0.240943 | 0.082 | 0.077 | 1 | 6 | Mrpl58 | 0.005 | Young |
| 0.723173 | 0.093317 | 0.129 | 0.123 | 1 | 6 | Fam192a | 0.006 | Young |
| 0.723709 | 0.217313 | 0.075 | 0.083 | 1 | 6 | Vps33a | -0.008 | Young |
| 0.723824 | 0.046024 | 0.023 | 0.02 | 1 | 6 | Zfp709 | 0.003 | Young |
| 0.724172 | 0.203108 | 0.231 | 0.226 | 1 | 6 | Foxn2 | 0.005 | Young |
| 0.724285 | 0.039704 | 0.027 | 0.023 | 1 | 6 | Pgpep1 | 0.004 | Young |
| 0.724435 | 0.050909 | 0.132 | 0.123 | 1 | 6 | Atxn7l3 | 0.009 | Young |
| 0.724482 | 0.696475 | 0.059 | 0.054 | 1 | 6 | 0610010K1 | 0.005 | Young |
| 0.724547 | 0.267731 | 0.023 | 0.02 | 1 | 6 | Vwa8 | 0.003 | Young |
| 0.724822 | 0.048981 | 0.396 | 0.377 | 1 | 6 | Atxn2l | 0.019 | Young |
| 0.725576 | 0.119055 | 0.12 | 0.114 | 1 | 6 | Magohb | 0.006 | Young |
| 0.725723 | 0.384521 | 0.074 | 0.069 | 1 | 6 | Agl | 0.005 | Young |
| 0.726193 | 0.289032 | 0.017 | 0.02 | 1 | 6 | Gins2 | -0.003 | Young |
| 0.726241 | 0.127143 | 0.054 | 0.049 | 1 | 6 | Aifm1 | 0.005 | Young |
| 0.726352 | 0.293734 | 0.171 | 0.166 | 1 | 6 | Gtf2a2 | 0.005 | Young |
| 0.72713 | 0.330873 | 0.094 | 0.089 | 1 | 6 | Rnf103 | 0.005 | Young |
| 0.727192 | 0.149205 | 0.065 | 0.06 | 1 | 6 | Ppp5c | 0.005 | Young |
| 0.727785 | 0.326276 | 0.074 | 0.069 | 1 | 6 | Clpx | 0.005 | Young |
| 0.727867 | 0.004936 | 0.052 | 0.057 | 1 | 6 | Tgif2 | -0.005 | Young |
| 0.727877 | 0.18727 | 0.112 | 0.106 | 1 | 6 | Pnpo | 0.006 | Young |
| 0.728099 | 0.336007 | 0.077 | 0.071 | 1 | 6 | Abcf3 | 0.006 | Young |
| 0.728165 | 0.633859 | 0.054 | 0.06 | 1 | 6 | Zbtb2 | -0.006 | Young |
| 0.72857 | 0.448314 | 0.087 | 0.083 | 1 | 6 | Il18 | 0.004 | Young |
| 0.729324 | 0.287186 | 0.054 | 0.06 | 1 | 6 | Riox2 | -0.006 | Young |
| 0.729376 | 0.082695 | 0.157 | 0.169 | 1 | 6 | Arhgap25 | -0.012 | Young |
| 0.730108 | 0.021017 | 0.559 | 0.586 | 1 | 6 | Abracl | -0.027 | Young |
| 0.730294 | 0.066939 | 0.296 | 0.291 | 1 | 6 | Trim8 | 0.005 | Young |
| 0.730431 | 0.330354 | 0.179 | 0.174 | 1 | 6 | Mysm1 | 0.005 | Young |
| 0.730679 | 0.409459 | 0.035 | 0.04 | 1 | 6 | Lrch4 | -0.005 | Young |
| 0.73072 | 0.187346 | 0.045 | 0.04 | 1 | 6 | Maml3 | 0.005 | Young |
| 0.731231 | 0.002241 | 0.057 | 0.063 | 1 | 6 | Zfp110 | -0.006 | Young |
| 0.731245 | 0.359093 | 0.035 | 0.04 | 1 | 6 | Pyroxd1 | -0.005 | Young |
| 0.731459 | 0.221344 | 0.308 | 0.311 | 1 | 6 | Ndufa11 | -0.003 | Young |
| 0.731633 | 0.051275 | 0.164 | 0.174 | 1 | 6 | Nedd9 | -0.01 | Young |
| 0.731867 | 0.134234 | 0.124 | 0.117 | 1 | 6 | Apba1 | 0.007 | Young |

| | | | | | | | | |
|---|---|---|---|---|---|---|---|---|
| 0.732411 | 0.494473 | 0.105 | 0.1 | 1 | 6 | Gtf2f1 | 0.005 | Young |
| 0.732501 | 0.222606 | 0.054 | 0.049 | 1 | 6 | Slc35e2 | 0.005 | Young |
| 0.732514 | 0.103569 | 0.023 | 0.02 | 1 | 6 | Gfi1b | 0.003 | Young |
| 0.733549 | 0.001448 | 0.565 | 0.56 | 1 | 6 | Syk | 0.005 | Young |
| 0.733847 | 0.037286 | 0.03 | 0.034 | 1 | 6 | Tspan9 | -0.004 | Young |
| 0.734178 | 0.056316 | 0.179 | 0.189 | 1 | 6 | Ric8a | -0.01 | Young |
| 0.734368 | 0.251874 | 0.1 | 0.094 | 1 | 6 | Ddx47 | 0.006 | Young |
| 0.734651 | 0.086807 | 0.239 | 0.24 | 1 | 6 | Cnppd1 | -0.001 | Young |
| 0.734692 | 0.487412 | 0.023 | 0.02 | 1 | 6 | Map2k6 | 0.003 | Young |
| 0.735127 | 0.076801 | 0.186 | 0.2 | 1 | 6 | Ankfy1 | -0.014 | Young |
| 0.73521 | 0.423355 | 0.038 | 0.034 | 1 | 6 | Rbsn | 0.004 | Young |
| 0.735519 | 0.229808 | 0.174 | 0.169 | 1 | 6 | Kif2a | 0.005 | Young |
| 0.735777 | 0.391502 | 0.035 | 0.04 | 1 | 6 | Nsdhl | -0.005 | Young |
| 0.735824 | 1.190573 | 0.02 | 0.017 | 1 | 6 | Fancc | 0.003 | Young |
| 0.735824 | 0.954457 | 0.02 | 0.017 | 1 | 6 | Rsph3b | 0.003 | Young |
| 0.735959 | 0.294657 | 0.122 | 0.117 | 1 | 6 | Ncor2 | 0.005 | Young |
| 0.736061 | 0.086938 | 0.268 | 0.257 | 1 | 6 | Comt | 0.011 | Young |
| 0.736587 | 0.048469 | 0.012 | 0.014 | 1 | 6 | Fam20a | -0.002 | Young |
| 0.736637 | 0.096768 | 0.042 | 0.037 | 1 | 6 | Osm | 0.005 | Young |
| 0.736901 | 0.023986 | 0.03 | 0.034 | 1 | 6 | Ppil1 | -0.004 | Young |
| 0.737075 | 0.020182 | 0.289 | 0.283 | 1 | 6 | Dpm3 | 0.006 | Young |
| 0.737493 | 0.247711 | 0.105 | 0.1 | 1 | 6 | Tmem192 | 0.005 | Young |
| 0.737544 | 0.166892 | 0.012 | 0.014 | 1 | 6 | 8030462N1 | -0.002 | Young |
| 0.738123 | 0.175661 | 0.03 | 0.034 | 1 | 6 | Zcchc4 | -0.004 | Young |
| 0.738327 | 0.041993 | 0.023 | 0.02 | 1 | 6 | 2900089D1 | 0.003 | Young |
| 0.738467 | 0.100337 | 0.084 | 0.077 | 1 | 6 | Trappc11 | 0.007 | Young |
| 0.738477 | 0.189171 | 0.186 | 0.18 | 1 | 6 | Tcerg1 | 0.006 | Young |
| 0.738502 | 0.544134 | 0.012 | 0.014 | 1 | 6 | Hs1bp3 | -0.002 | Young |
| 0.738502 | 0.328875 | 0.012 | 0.014 | 1 | 6 | Ttll5 | -0.002 | Young |
| 0.738502 | 0.218309 | 0.012 | 0.014 | 1 | 6 | Ybey | -0.002 | Young |
| 0.738502 | 0.184232 | 0.012 | 0.014 | 1 | 6 | Adnp | -0.002 | Young |
| 0.738502 | 0.146026 | 0.012 | 0.014 | 1 | 6 | Tex14 | -0.002 | Young |
| 0.739054 | 0.126945 | 0.023 | 0.02 | 1 | 6 | 1110008L1 | 0.003 | Young |
| 0.739336 | 0.863766 | 0.043 | 0.04 | 1 | 6 | Rnf185 | 0.003 | Young |
| 0.739365 | 0.040306 | 0.042 | 0.037 | 1 | 6 | Oxsm | 0.005 | Young |
| 0.73946 | 0.369634 | 0.012 | 0.014 | 1 | 6 | Zfp51 | -0.002 | Young |
| 0.739591 | 0.000649 | 0.227 | 0.243 | 1 | 6 | Nt5c | -0.016 | Young |
| 0.739962 | 0.134308 | 0.274 | 0.266 | 1 | 6 | Hlcs | 0.008 | Young |
| 0.740279 | 0.121917 | 0.209 | 0.223 | 1 | 6 | Zc3h13 | -0.014 | Young |
| 0.740418 | 0.212379 | 0.012 | 0.014 | 1 | 6 | Sphk1 | -0.002 | Young |
| 0.740494 | 0.31599 | 0.067 | 0.074 | 1 | 6 | Slc25a19 | -0.007 | Young |
| 0.740987 | 0.037557 | 0.09 | 0.097 | 1 | 6 | Rabggtb | -0.007 | Young |
| 0.741246 | 0.044076 | 0.306 | 0.3 | 1 | 6 | Ndufb6 | 0.006 | Young |
| 0.741345 | 0.191882 | 0.124 | 0.117 | 1 | 6 | Armcx3 | 0.007 | Young |
| 0.741623 | 0.035838 | 0.273 | 0.269 | 1 | 6 | N4bp2l2 | 0.004 | Young |

| | | | | | | | | |
|---|---|---|---|---|---|---|---|---|
| 0.741833 | 0.230626 | 0.05 | 0.046 | 1 | 6 | Tmem2 | 0.004 | Young |
| 0.74189 | 0.120482 | 0.057 | 0.063 | 1 | 6 | 6030458C1 | -0.006 | Young |
| 0.742102 | 0.101608 | 0.02 | 0.017 | 1 | 6 | Cdyl2 | 0.003 | Young |
| 0.742408 | 0.037908 | 0.03 | 0.034 | 1 | 6 | Iqcb1 | -0.004 | Young |
| 0.742451 | 0.193597 | 0.172 | 0.166 | 1 | 6 | Zbtb8os | 0.006 | Young |
| 0.742656 | 0.901721 | 0.032 | 0.029 | 1 | 6 | Sco2 | 0.003 | Young |
| 0.742757 | 0.134846 | 0.152 | 0.143 | 1 | 6 | Vrk2 | 0.009 | Young |
| 0.742914 | 0.038955 | 0.151 | 0.143 | 1 | 6 | Mau2 | 0.008 | Young |
| 0.743129 | 0.175779 | 0.196 | 0.214 | 1 | 6 | Denr | -0.018 | Young |
| 0.743674 | 0.655832 | 0.02 | 0.017 | 1 | 6 | Fam234b | 0.003 | Young |
| 0.743876 | 0.162066 | 0.092 | 0.1 | 1 | 6 | Hdgfl2 | -0.008 | Young |
| 0.744125 | 0.175037 | 0.149 | 0.143 | 1 | 6 | Taok2 | 0.006 | Young |
| 0.744461 | 0.490699 | 0.02 | 0.017 | 1 | 6 | Trdmt1 | 0.003 | Young |
| 0.744601 | 0.189842 | 0.114 | 0.123 | 1 | 6 | Dnpep | -0.009 | Young |
| 0.74519 | 0.216448 | 0.246 | 0.243 | 1 | 6 | Pnpla8 | 0.003 | Young |
| 0.745948 | 0.113666 | 0.145 | 0.14 | 1 | 6 | Bcat2 | 0.005 | Young |
| 0.746077 | 0.241542 | 0.05 | 0.046 | 1 | 6 | Mavs | 0.004 | Young |
| 0.746364 | 0.074198 | 0.184 | 0.197 | 1 | 6 | Otud5 | -0.013 | Young |
| 0.746545 | 0.100837 | 0.167 | 0.16 | 1 | 6 | Mfsd5 | 0.007 | Young |
| 0.74656 | 0.147632 | 0.102 | 0.111 | 1 | 6 | Shkbp1 | -0.009 | Young |
| 0.746742 | 0.101758 | 0.094 | 0.1 | 1 | 6 | Abi2 | -0.006 | Young |
| 0.747044 | 0.028339 | 0.191 | 0.183 | 1 | 6 | Mbd3 | 0.008 | Young |
| 0.747139 | 0.611269 | 0.012 | 0.014 | 1 | 6 | 4632404H1 | -0.002 | Young |
| 0.74717 | 0.02198 | 0.145 | 0.14 | 1 | 6 | Axl | 0.005 | Young |
| 0.747357 | 0.192801 | 0.065 | 0.071 | 1 | 6 | Brpf3 | -0.006 | Young |
| 0.748543 | 0.397743 | 0.03 | 0.034 | 1 | 6 | Bud13 | -0.004 | Young |
| 0.749117 | 0.085294 | 0.564 | 0.554 | 1 | 6 | Anp32b | 0.01 | Young |
| 0.749185 | 0.561306 | 0.02 | 0.017 | 1 | 6 | Twnk | 0.003 | Young |
| 0.749434 | 0.215552 | 0.047 | 0.043 | 1 | 6 | Chd1l | 0.004 | Young |
| 0.749579 | 0.205094 | 0.05 | 0.046 | 1 | 6 | Haus2 | 0.004 | Young |
| 0.749748 | 0.044566 | 0.333 | 0.349 | 1 | 6 | Plcg2 | -0.016 | Young |
| 0.74978 | 0.35295 | 0.156 | 0.151 | 1 | 6 | Anxa4 | 0.005 | Young |
| 0.750792 | 0.396704 | 0.059 | 0.054 | 1 | 6 | Oma1 | 0.005 | Young |
| 0.750796 | 0.176249 | 0.137 | 0.131 | 1 | 6 | Gemin7 | 0.006 | Young |
| 0.751016 | 0.054277 | 0.097 | 0.091 | 1 | 6 | Edc4 | 0.006 | Young |
| 0.75103 | 0.280315 | 0.172 | 0.169 | 1 | 6 | Pilrb2 | 0.003 | Young |
| 0.751039 | 0.259166 | 0.162 | 0.157 | 1 | 6 | Naa10 | 0.005 | Young |
| 0.751551 | 0.218442 | 0.02 | 0.017 | 1 | 6 | Rsl1 | 0.003 | Young |
| 0.751573 | 0.099473 | 0.129 | 0.123 | 1 | 6 | 3830406C1 | 0.006 | Young |
| 0.751635 | 0.092268 | 0.062 | 0.057 | 1 | 6 | Amd1 | 0.005 | Young |
| 0.751795 | 0.079178 | 0.207 | 0.2 | 1 | 6 | Cdk12 | 0.007 | Young |
| 0.752229 | 0.261147 | 0.144 | 0.14 | 1 | 6 | Eftud2 | 0.004 | Young |
| 0.752302 | 0.074074 | 0.082 | 0.089 | 1 | 6 | Mycbp | -0.007 | Young |
| 0.752481 | 0.140796 | 0.127 | 0.123 | 1 | 6 | Trak2 | 0.004 | Young |
| 0.75265 | 0.442059 | 0.059 | 0.054 | 1 | 6 | Chst15 | 0.005 | Young |

| | | | | | | | | |
|---|---|---|---|---|---|---|---|---|
| 0.752741 | 0.012928 | 0.425 | 0.42 | 1 | 6 | Eif3i | 0.005 | Young |
| 0.752994 | 0.277202 | 0.062 | 0.057 | 1 | 6 | Tmem167b | 0.005 | Young |
| 0.753475 | 0.202964 | 0.253 | 0.251 | 1 | 6 | Stip1 | 0.002 | Young |
| 0.753611 | 0.125218 | 0.174 | 0.186 | 1 | 6 | Txndc9 | -0.012 | Young |
| 0.753929 | 0.416139 | 0.035 | 0.031 | 1 | 6 | Yae1d1 | 0.004 | Young |
| 0.754114 | 0.084734 | 0.154 | 0.166 | 1 | 6 | Cyb5r3 | -0.012 | Young |
| 0.754675 | 0.161868 | 0.095 | 0.089 | 1 | 6 | Trmt6 | 0.006 | Young |
| 0.754868 | 0.19562 | 0.166 | 0.16 | 1 | 6 | Aida | 0.006 | Young |
| 0.755711 | 0.003984 | 0.038 | 0.034 | 1 | 6 | Abt1 | 0.004 | Young |
| 0.755865 | 0.514439 | 0.1 | 0.097 | 1 | 6 | Polr2h | 0.003 | Young |
| 0.756594 | 0.021247 | 0.433 | 0.446 | 1 | 6 | Atp6v1b2 | -0.013 | Young |
| 0.756642 | 0.038305 | 0.112 | 0.106 | 1 | 6 | Ube3c | 0.006 | Young |
| 0.75695 | 0.063201 | 0.025 | 0.029 | 1 | 6 | Wdr83 | -0.004 | Young |
| 0.757074 | 0.22947 | 0.062 | 0.057 | 1 | 6 | Tpm3-rs7 | 0.005 | Young |
| 0.757301 | 0.417877 | 0.059 | 0.054 | 1 | 6 | Dubr | 0.005 | Young |
| 0.757331 | 0.196228 | 0.112 | 0.106 | 1 | 6 | Zcchc2 | 0.006 | Young |
| 0.757336 | 0.012785 | 0.804 | 0.774 | 1 | 6 | Eef1g | 0.03 | Young |
| 0.757344 | 0.023238 | 0.211 | 0.223 | 1 | 6 | Coro1b | -0.012 | Young |
| 0.757428 | 0.007194 | 0.038 | 0.034 | 1 | 6 | Rmdn3 | 0.004 | Young |
| 0.757623 | 0.147369 | 0.025 | 0.029 | 1 | 6 | Slc22a4 | -0.004 | Young |
| 0.757872 | 0.027606 | 0.02 | 0.017 | 1 | 6 | Pdlim3 | 0.003 | Young |
| 0.758858 | 0.218911 | 0.07 | 0.077 | 1 | 6 | Pigyl | -0.007 | Young |
| 0.75889 | 0.322397 | 0.062 | 0.057 | 1 | 6 | Cyp4v3 | 0.005 | Young |
| 0.75939 | 0.441686 | 0.171 | 0.166 | 1 | 6 | Pilrb1 | 0.005 | Young |
| 0.759442 | 0.026033 | 0.08 | 0.074 | 1 | 6 | Phactr4 | 0.006 | Young |
| 0.759723 | 0.086043 | 0.052 | 0.057 | 1 | 6 | Men1 | -0.005 | Young |
| 0.759905 | 0.379493 | 0.035 | 0.031 | 1 | 6 | Supv3l1 | 0.004 | Young |
| 0.759908 | 0.270624 | 0.137 | 0.134 | 1 | 6 | Zfp608 | 0.003 | Young |
| 0.760051 | 0.031885 | 0.104 | 0.097 | 1 | 6 | 1600010M | 0.007 | Young |
| 0.760064 | 0.002026 | 0.298 | 0.309 | 1 | 6 | Vapb | -0.011 | Young |
| 0.760275 | 0.112583 | 0.191 | 0.189 | 1 | 6 | Slc7a1 | 0.002 | Young |
| 0.760319 | 0.436634 | 0.025 | 0.029 | 1 | 6 | Tatdn1 | -0.004 | Young |
| 0.760986 | 0.562698 | 0.017 | 0.014 | 1 | 6 | Harbi1 | 0.003 | Young |
| 0.761018 | 0.126699 | 0.237 | 0.237 | 1 | 6 | Mydgf | 0 | Young |
| 0.761381 | 0.076699 | 0.151 | 0.16 | 1 | 6 | Cops5 | -0.009 | Young |
| 0.761871 | 0.375537 | 0.047 | 0.043 | 1 | 6 | Zc3h10 | 0.004 | Young |
| 0.76306 | 0.109116 | 0.202 | 0.197 | 1 | 6 | Ift20 | 0.005 | Young |
| 0.763189 | 0.632164 | 0.092 | 0.103 | 1 | 6 | Vamp7 | -0.011 | Young |
| 0.763467 | 0.106226 | 0.14 | 0.134 | 1 | 6 | Rasa4 | 0.006 | Young |
| 0.763826 | 0.179078 | 0.059 | 0.054 | 1 | 6 | Thra | 0.005 | Young |
| 0.763875 | 0.145196 | 0.232 | 0.231 | 1 | 6 | Sms | 0.001 | Young |
| 0.76403 | 0.53478 | 0.052 | 0.049 | 1 | 6 | Chkb | 0.003 | Young |
| 0.764165 | 0.004705 | 0.247 | 0.257 | 1 | 6 | Usp8 | -0.01 | Young |
| 0.764454 | 0.611641 | 0.017 | 0.014 | 1 | 6 | Ube2e2 | 0.003 | Young |
| 0.764454 | 0.583697 | 0.017 | 0.014 | 1 | 6 | Cenpw | 0.003 | Young |

| | | | | | | | | |
|---|---|---|---|---|---|---|---|---|
| 0.7648 | 0.096057 | 0.062 | 0.057 | 1 | 6 | Folr2 | 0.005 | Young |
| 0.764859 | 0.170442 | 0.124 | 0.134 | 1 | 6 | Eif2ak1 | -0.01 | Young |
| 0.765294 | 0.073792 | 0.035 | 0.031 | 1 | 6 | 2810454H( | 0.004 | Young |
| 0.765954 | 0.206072 | 0.052 | 0.057 | 1 | 6 | Etfrf1 | -0.005 | Young |
| 0.766045 | 0.396079 | 0.219 | 0.217 | 1 | 6 | C1qa | 0.002 | Young |
| 0.766535 | 0.010027 | 0.284 | 0.28 | 1 | 6 | Guk1 | 0.004 | Young |
| 0.767073 | 0.168303 | 0.025 | 0.029 | 1 | 6 | Gemin5 | -0.004 | Young |
| 0.767723 | 0.246818 | 0.172 | 0.169 | 1 | 6 | Gpat4 | 0.003 | Young |
| 0.768677 | 0.175745 | 0.142 | 0.137 | 1 | 6 | Trnt1 | 0.005 | Young |
| 0.768767 | 0.000311 | 0.289 | 0.277 | 1 | 6 | Tnfrsf13b | 0.012 | Young |
| 0.769083 | 0.06442 | 0.47 | 0.466 | 1 | 6 | Ehd4 | 0.004 | Young |
| 0.770189 | 0.127302 | 0.047 | 0.051 | 1 | 6 | Tspyl2 | -0.004 | Young |
| 0.770455 | 0.043392 | 0.02 | 0.023 | 1 | 6 | Zfp84 | -0.003 | Young |
| 0.770533 | 0.32943 | 0.017 | 0.014 | 1 | 6 | Pex26 | 0.003 | Young |
| 0.770533 | 0.074835 | 0.017 | 0.014 | 1 | 6 | Usp30 | 0.003 | Young |
| 0.770701 | 0.285439 | 0.095 | 0.103 | 1 | 6 | Akr7a5 | -0.008 | Young |
| 0.771123 | 0.068684 | 0.348 | 0.34 | 1 | 6 | Trim30a | 0.008 | Young |
| 0.771403 | 0.307174 | 0.017 | 0.014 | 1 | 6 | Maoa | 0.003 | Young |
| 0.771637 | 0.033037 | 0.062 | 0.057 | 1 | 6 | Arhgap23 | 0.005 | Young |
| 0.77224 | 0.175108 | 0.059 | 0.054 | 1 | 6 | Cyb561d2 | 0.005 | Young |
| 0.772742 | 0.516961 | 0.032 | 0.029 | 1 | 6 | 8-Mar | 0.003 | Young |
| 0.773143 | 0.606089 | 0.017 | 0.014 | 1 | 6 | Tspoap1 | 0.003 | Young |
| 0.773677 | 0.335015 | 0.043 | 0.04 | 1 | 6 | Oraov1 | 0.003 | Young |
| 0.773836 | 0.007139 | 0.625 | 0.626 | 1 | 6 | Taldo1 | -0.001 | Young |
| 0.774555 | 0.092069 | 0.294 | 0.286 | 1 | 6 | Dhrs3 | 0.008 | Young |
| 0.774647 | 0.24581 | 0.122 | 0.117 | 1 | 6 | Mier3 | 0.005 | Young |
| 0.775094 | 0.014271 | 0.124 | 0.131 | 1 | 6 | Letm1 | -0.007 | Young |
| 0.775123 | 0.138963 | 0.145 | 0.14 | 1 | 6 | Dgkz | 0.005 | Young |
| 0.775266 | 0.151143 | 0.032 | 0.029 | 1 | 6 | Invs | 0.003 | Young |
| 0.776236 | 0.224108 | 0.117 | 0.111 | 1 | 6 | Guca1a | 0.006 | Young |
| 0.776339 | 0.734384 | 0.028 | 0.026 | 1 | 6 | Sh2d3c | 0.002 | Young |
| 0.77713 | 0.23321 | 0.07 | 0.066 | 1 | 6 | Tmem120a | 0.004 | Young |
| 0.777383 | 0.227446 | 0.368 | 0.371 | 1 | 6 | Irf1 | -0.003 | Young |
| 0.777462 | 0.856891 | 0.043 | 0.04 | 1 | 6 | Cxcl1 | 0.003 | Young |
| 0.777462 | 0.171835 | 0.043 | 0.04 | 1 | 6 | Orc6 | 0.003 | Young |
| 0.778186 | 0.089326 | 0.161 | 0.154 | 1 | 6 | Ranbp9 | 0.007 | Young |
| 0.778275 | 0.248099 | 0.125 | 0.12 | 1 | 6 | Ube2q2 | 0.005 | Young |
| 0.778337 | 0.268857 | 0.028 | 0.026 | 1 | 6 | Fpgt | 0.002 | Young |
| 0.77837 | 0.000402 | 0.017 | 0.014 | 1 | 6 | Ccdc130 | 0.003 | Young |
| 0.778451 | 0.131569 | 0.187 | 0.18 | 1 | 6 | Gsdmd | 0.007 | Young |
| 0.778773 | 0.006551 | 0.02 | 0.023 | 1 | 6 | Slc37a1 | -0.003 | Young |
| 0.77905 | 0.03891 | 0.142 | 0.134 | 1 | 6 | Tbc1d22a | 0.008 | Young |
| 0.779055 | 0.419034 | 0.032 | 0.029 | 1 | 6 | Fam131a | 0.003 | Young |
| 0.779055 | 0.337112 | 0.032 | 0.029 | 1 | 6 | Tom1 | 0.003 | Young |
| 0.779243 | 0.081538 | 0.017 | 0.014 | 1 | 6 | Ccdc171 | 0.003 | Young |

| | | | | | | | | |
|---|---|---|---|---|---|---|---|---|
| 0.779243 | 0.017276 | 0.017 | 0.014 | 1 | 6 | Wfs1 | 0.003 | Young |
| 0.779665 | 0.186028 | 0.264 | 0.266 | 1 | 6 | Nhp2 | -0.002 | Young |
| 0.780288 | 0.031548 | 0.02 | 0.023 | 1 | 6 | Evi5l | -0.003 | Young |
| 0.78128 | 0.033915 | 0.36 | 0.351 | 1 | 6 | Clec4a1 | 0.009 | Young |
| 0.781293 | 0.042913 | 0.157 | 0.166 | 1 | 6 | Prelid3b | -0.009 | Young |
| 0.781354 | 0.129746 | 0.055 | 0.051 | 1 | 6 | Cyp2ab1 | 0.004 | Young |
| 0.781585 | 0.185019 | 0.032 | 0.029 | 1 | 6 | Tgds | 0.003 | Young |
| 0.781793 | 0.436958 | 0.043 | 0.04 | 1 | 6 | Golph3l | 0.003 | Young |
| 0.781837 | 0.086199 | 0.288 | 0.286 | 1 | 6 | Dock11 | 0.002 | Young |
| 0.781861 | 0.070054 | 0.017 | 0.014 | 1 | 6 | Mdga1 | 0.003 | Young |
| 0.782054 | 0.061439 | 0.092 | 0.097 | 1 | 6 | Vps11 | -0.005 | Young |
| 0.782217 | 0.289762 | 0.032 | 0.029 | 1 | 6 | Pradc1 | 0.003 | Young |
| 0.782734 | 0.044391 | 0.085 | 0.091 | 1 | 6 | Itpripl1 | -0.006 | Young |
| 0.783148 | 0.151756 | 0.043 | 0.04 | 1 | 6 | Olfr1167 | 0.003 | Young |
| 0.783208 | 0.02055 | 0.196 | 0.206 | 1 | 6 | Abcd1 | -0.01 | Young |
| 0.783321 | 0.049848 | 0.02 | 0.023 | 1 | 6 | Zdhhc14 | -0.003 | Young |
| 0.784007 | 0.003055 | 0.311 | 0.32 | 1 | 6 | Dynlrb1 | -0.009 | Young |
| 0.784748 | 0.129865 | 0.147 | 0.14 | 1 | 6 | Gpr146 | 0.007 | Young |
| 0.784838 | 0.046387 | 0.02 | 0.023 | 1 | 6 | Rnf25 | -0.003 | Young |
| 0.785098 | 0.022131 | 0.415 | 0.434 | 1 | 6 | Larp1 | -0.019 | Young |
| 0.7851 | 0.130355 | 0.197 | 0.191 | 1 | 6 | Irf2bpl | 0.006 | Young |
| 0.785263 | 0.037821 | 0.453 | 0.434 | 1 | 6 | Lilr4b | 0.019 | Young |
| 0.785355 | 0.189176 | 0.047 | 0.043 | 1 | 6 | Bud23 | 0.004 | Young |
| 0.785385 | 0.07572 | 0.45 | 0.443 | 1 | 6 | Rsrc2 | 0.007 | Young |
| 0.786132 | 0.050474 | 0.043 | 0.04 | 1 | 6 | Xpc | 0.003 | Young |
| 0.786177 | 0.064941 | 0.07 | 0.066 | 1 | 6 | Trappc13 | 0.004 | Young |
| 0.786357 | 0.335005 | 0.02 | 0.023 | 1 | 6 | Prmt9 | -0.003 | Young |
| 0.786387 | 0.910337 | 0.013 | 0.011 | 1 | 6 | Dnmt3b | 0.002 | Young |
| 0.786387 | 0.776727 | 0.013 | 0.011 | 1 | 6 | Repin1 | 0.002 | Young |
| 0.786542 | 0.004964 | 0.187 | 0.18 | 1 | 6 | Rnf216 | 0.007 | Young |
| 0.787217 | 0.275993 | 0.043 | 0.04 | 1 | 6 | Tmem242 | 0.003 | Young |
| 0.787227 | 0.038953 | 0.355 | 0.351 | 1 | 6 | Atp5g1 | 0.004 | Young |
| 0.787363 | 0.731885 | 0.013 | 0.011 | 1 | 6 | Samd10 | 0.002 | Young |
| 0.787636 | 0.111367 | 0.167 | 0.163 | 1 | 6 | Egln2 | 0.004 | Young |
| 0.787681 | 0.569598 | 0.028 | 0.026 | 1 | 6 | Zfp619 | 0.002 | Young |
| 0.78875 | 0.048353 | 0.137 | 0.131 | 1 | 6 | Mical1 | 0.006 | Young |
| 0.789396 | 0.390304 | 0.02 | 0.023 | 1 | 6 | Pqlc2 | -0.003 | Young |
| 0.789572 | 0.000301 | 0.938 | 0.903 | 1 | 6 | Arpc2 | 0.035 | Young |
| 0.790295 | 0.924663 | 0.013 | 0.011 | 1 | 6 | Ddx20 | 0.002 | Young |
| 0.790295 | 0.542622 | 0.013 | 0.011 | 1 | 6 | Sema6d | 0.002 | Young |
| 0.790384 | 0.172997 | 0.117 | 0.111 | 1 | 6 | Pmpcb | 0.006 | Young |
| 0.790557 | 0.689499 | 0.06 | 0.057 | 1 | 6 | Scnm1 | 0.003 | Young |
| 0.790683 | 0.017842 | 0.386 | 0.377 | 1 | 6 | Psmd8 | 0.009 | Young |
| 0.790718 | 0.083472 | 0.08 | 0.086 | 1 | 6 | Lrch1 | -0.006 | Young |
| 0.790751 | 0.079422 | 0.166 | 0.163 | 1 | 6 | Mrpl4 | 0.003 | Young |

| | | | | | | | | |
|---|---|---|---|---|---|---|---|---|
| 0.791011 | 0.46601 | 0.072 | 0.069 | 1 | 6 | Vps18 | 0.003 | Young |
| 0.79102 | 0.10567 | 0.393 | 0.391 | 1 | 6 | Arcn1 | 0.002 | Young |
| 0.791272 | 0.788384 | 0.013 | 0.011 | 1 | 6 | Ifit3 | 0.002 | Young |
| 0.791408 | 0.167578 | 0.069 | 0.066 | 1 | 6 | Il16 | 0.003 | Young |
| 0.791439 | 0.23721 | 0.124 | 0.131 | 1 | 6 | Kars | -0.007 | Young |
| 0.79201 | 0.07512 | 0.288 | 0.306 | 1 | 6 | Plekhj1 | -0.018 | Young |
| 0.792108 | 0.180559 | 0.043 | 0.04 | 1 | 6 | Cep135 | 0.003 | Young |
| 0.792364 | 0.508515 | 0.028 | 0.026 | 1 | 6 | Ears2 | 0.002 | Young |
| 0.793229 | 0.638757 | 0.013 | 0.011 | 1 | 6 | Rtp4 | 0.002 | Young |
| 0.793229 | 0.228617 | 0.013 | 0.011 | 1 | 6 | Wdr59 | 0.002 | Young |
| 0.793474 | 0.056115 | 0.334 | 0.331 | 1 | 6 | 1110008F1 | 0.003 | Young |
| 0.793704 | 0.76443 | 0.028 | 0.026 | 1 | 6 | Fam20c | 0.002 | Young |
| 0.793956 | 0.039862 | 0.042 | 0.046 | 1 | 6 | Leng1 | -0.004 | Young |
| 0.794208 | 0.267162 | 0.013 | 0.011 | 1 | 6 | Slc25a10 | 0.002 | Young |
| 0.794494 | 0.335536 | 0.042 | 0.046 | 1 | 6 | Cactin | -0.004 | Young |
| 0.794535 | 0.599829 | 0.04 | 0.037 | 1 | 6 | Ddx19b | 0.003 | Young |
| 0.794535 | 0.380407 | 0.04 | 0.037 | 1 | 6 | Prcc | 0.003 | Young |
| 0.79515 | 0.005402 | 0.156 | 0.163 | 1 | 6 | Psen1 | -0.007 | Young |
| 0.795187 | 0.27158 | 0.013 | 0.011 | 1 | 6 | Cox7a1 | 0.002 | Young |
| 0.795619 | 0.172941 | 0.221 | 0.22 | 1 | 6 | Os9 | 0.001 | Young |
| 0.795821 | 0.231774 | 0.134 | 0.143 | 1 | 6 | Zfp593 | -0.009 | Young |
| 0.796268 | 0.112746 | 0.104 | 0.111 | 1 | 6 | Zcchc17 | -0.007 | Young |
| 0.796485 | 0.072672 | 0.528 | 0.534 | 1 | 6 | Arhgap15 | -0.006 | Young |
| 0.796677 | 0.003916 | 0.351 | 0.363 | 1 | 6 | Gramd4 | -0.012 | Young |
| 0.797055 | 0.232839 | 0.028 | 0.026 | 1 | 6 | Pmm1 | 0.002 | Young |
| 0.797146 | 0.215063 | 0.013 | 0.011 | 1 | 6 | Dnah2 | 0.002 | Young |
| 0.797146 | 0.084602 | 0.013 | 0.011 | 1 | 6 | Zc4h2 | 0.002 | Young |
| 0.797146 | 0.082288 | 0.013 | 0.011 | 1 | 6 | Retsat | 0.002 | Young |
| 0.797146 | 0.053101 | 0.013 | 0.011 | 1 | 6 | Gm40645 | 0.002 | Young |
| 0.797189 | 0.446308 | 0.099 | 0.094 | 1 | 6 | Cog3 | 0.005 | Young |
| 0.797294 | 0.070993 | 0.11 | 0.106 | 1 | 6 | Gse1 | 0.004 | Young |
| 0.798913 | 0.311059 | 0.043 | 0.04 | 1 | 6 | Spsb3 | 0.003 | Young |
| 0.799068 | 0.242837 | 0.028 | 0.026 | 1 | 6 | Mvk | 0.002 | Young |
| 0.799106 | 0.107687 | 0.013 | 0.011 | 1 | 6 | Arsi | 0.002 | Young |
| 0.799106 | 0.020643 | 0.013 | 0.011 | 1 | 6 | 9330020H( | 0.002 | Young |
| 0.799339 | 0.383344 | 0.042 | 0.046 | 1 | 6 | Appl2 | -0.004 | Young |
| 0.799526 | 0.10113 | 0.331 | 0.331 | 1 | 6 | Supt16 | 0 | Young |
| 0.800106 | 0.149347 | 0.127 | 0.134 | 1 | 6 | Ldah | -0.007 | Young |
| 0.800143 | 0.193172 | 0.189 | 0.2 | 1 | 6 | Cept1 | -0.011 | Young |
| 0.800326 | 0.012782 | 0.978 | 0.971 | 1 | 6 | Rps26 | 0.007 | Young |
| 0.800539 | 0.082284 | 0.059 | 0.063 | 1 | 6 | Foxred1 | -0.004 | Young |
| 0.800819 | 0.055343 | 0.289 | 0.286 | 1 | 6 | Gm10073 | 0.003 | Young |
| 0.801015 | 0.158612 | 0.109 | 0.117 | 1 | 6 | Kmt2b | -0.008 | Young |
| 0.801141 | 0.107024 | 0.134 | 0.129 | 1 | 6 | Sav1 | 0.005 | Young |
| 0.801296 | 0.321195 | 0.052 | 0.049 | 1 | 6 | Sirpb1c | 0.003 | Young |

| | | | | | | | | |
|---|---|---|---|---|---|---|---|---|
| 0.801361 | 0.118153 | 0.186 | 0.18 | 1 | 6 | Clasp2 | 0.006 | Young |
| 0.801591 | 0.506117 | 0.08 | 0.077 | 1 | 6 | Mrps7 | 0.003 | Young |
| 0.801835 | 0.071468 | 0.283 | 0.28 | 1 | 6 | Paf1 | 0.003 | Young |
| 0.802184 | 0.119757 | 0.055 | 0.051 | 1 | 6 | Nubp2 | 0.004 | Young |
| 0.802425 | 0.327847 | 0.028 | 0.026 | 1 | 6 | Thbs1 | 0.002 | Young |
| 0.803022 | 0.276692 | 0.04 | 0.037 | 1 | 6 | Mrps10 | 0.003 | Young |
| 0.803063 | 0.038764 | 0.413 | 0.414 | 1 | 6 | Tmem258 | -0.001 | Young |
| 0.803652 | 0.174455 | 0.268 | 0.269 | 1 | 6 | Ssu72 | -0.001 | Young |
| 0.803872 | 0.137217 | 0.266 | 0.269 | 1 | 6 | Pxn | -0.003 | Young |
| 0.804027 | 0.121816 | 0.045 | 0.049 | 1 | 6 | Vezt | -0.004 | Young |
| 0.804129 | 0.173521 | 0.055 | 0.051 | 1 | 6 | Dph6 | 0.004 | Young |
| 0.804293 | 0.182883 | 0.276 | 0.277 | 1 | 6 | Pum1 | -0.001 | Young |
| 0.804607 | 0.075084 | 0.236 | 0.243 | 1 | 6 | Zc3h15 | -0.007 | Young |
| 0.804654 | 0.331534 | 0.015 | 0.017 | 1 | 6 | Platr17 | -0.002 | Young |
| 0.805674 | 0.024399 | 0.097 | 0.091 | 1 | 6 | Lnpk | 0.006 | Young |
| 0.806148 | 0.462345 | 0.109 | 0.106 | 1 | 6 | Traf7 | 0.003 | Young |
| 0.806397 | 0.190773 | 0.064 | 0.06 | 1 | 6 | Mrpl27 | 0.004 | Young |
| 0.806623 | 0.171708 | 0.176 | 0.171 | 1 | 6 | Bbx | 0.005 | Young |
| 0.807052 | 0.125195 | 0.084 | 0.08 | 1 | 6 | Thumpd1 | 0.004 | Young |
| 0.807991 | 0.094052 | 0.137 | 0.146 | 1 | 6 | Ammecr1l | -0.009 | Young |
| 0.808875 | 0.46985 | 0.069 | 0.066 | 1 | 6 | Rgs3 | 0.003 | Young |
| 0.809126 | 0.44352 | 0.064 | 0.06 | 1 | 6 | Timm8a1 | 0.004 | Young |
| 0.809289 | 0.105852 | 0.328 | 0.346 | 1 | 6 | Tnpo3 | -0.018 | Young |
| 0.80965 | 0.230228 | 0.084 | 0.08 | 1 | 6 | Tmub1 | 0.004 | Young |
| 0.809938 | 0.131112 | 0.015 | 0.017 | 1 | 6 | C5ar2 | -0.002 | Young |
| 0.810128 | 0.21523 | 0.08 | 0.077 | 1 | 6 | Zmym4 | 0.003 | Young |
| 0.810547 | 0.003024 | 0.473 | 0.497 | 1 | 6 | Sppl2a | -0.024 | Young |
| 0.810719 | 0.482086 | 0.097 | 0.094 | 1 | 6 | Wipi2 | 0.003 | Young |
| 0.81082 | 0.117381 | 0.015 | 0.017 | 1 | 6 | Apoo | -0.002 | Young |
| 0.811504 | 0.014199 | 0.184 | 0.18 | 1 | 6 | Mon2 | 0.004 | Young |
| 0.811701 | 0.09143 | 0.015 | 0.017 | 1 | 6 | Ccdc180 | -0.002 | Young |
| 0.811794 | 0.434948 | 0.186 | 0.189 | 1 | 6 | Pdhb | -0.003 | Young |
| 0.812398 | 0.468992 | 0.047 | 0.051 | 1 | 6 | Ofd1 | -0.004 | Young |
| 0.812414 | 0.011136 | 0.227 | 0.22 | 1 | 6 | Clcn4 | 0.007 | Young |
| 0.812521 | 0.288984 | 0.028 | 0.031 | 1 | 6 | Eif2b3 | -0.003 | Young |
| 0.81254 | 0.021201 | 0.53 | 0.523 | 1 | 6 | 2900097C1 | 0.007 | Young |
| 0.812583 | 0.586065 | 0.015 | 0.017 | 1 | 6 | Dgat2 | -0.002 | Young |
| 0.812583 | 0.418415 | 0.015 | 0.017 | 1 | 6 | AC130815. | -0.002 | Young |
| 0.812651 | 0.208644 | 0.12 | 0.114 | 1 | 6 | Gyg | 0.006 | Young |
| 0.812833 | 0.032489 | 0.306 | 0.303 | 1 | 6 | Rex1bd | 0.003 | Young |
| 0.813003 | 0.394636 | 0.025 | 0.023 | 1 | 6 | Zfp729b | 0.002 | Young |
| 0.813237 | 0.46109 | 0.04 | 0.037 | 1 | 6 | Cog4 | 0.003 | Young |
| 0.813354 | 0.077152 | 0.356 | 0.349 | 1 | 6 | Eno1 | 0.007 | Young |
| 0.813384 | 0.024959 | 0.055 | 0.051 | 1 | 6 | Rbms2 | 0.004 | Young |
| 0.813418 | 0.09025 | 0.067 | 0.063 | 1 | 6 | Ctns | 0.004 | Young |

| | | | | | | | | |
|---|---|---|---|---|---|---|---|---|
| 0.814087 | 0.332794 | 0.048 | 0.046 | 1 | 6 | Pik3r2 | 0.002 | Young |
| 0.815151 | 0.606971 | 0.025 | 0.023 | 1 | 6 | Rpap1 | 0.002 | Young |
| 0.815512 | 0.116879 | 0.261 | 0.283 | 1 | 6 | Psmb7 | -0.022 | Young |
| 0.815867 | 0.462021 | 0.025 | 0.023 | 1 | 6 | Tsen54 | 0.002 | Young |
| 0.815981 | 0.058336 | 0.135 | 0.131 | 1 | 6 | Sft2d1 | 0.004 | Young |
| 0.81612 | 0.072189 | 0.142 | 0.149 | 1 | 6 | Mapk8 | -0.007 | Young |
| 0.816337 | 0.118232 | 0.194 | 0.189 | 1 | 6 | Pira2 | 0.005 | Young |
| 0.81635 | 0.10191 | 0.102 | 0.109 | 1 | 6 | Ctnnbip1 | -0.007 | Young |
| 0.816458 | 0.001964 | 0.478 | 0.477 | 1 | 6 | Orai1 | 0.001 | Young |
| 0.816579 | 0.764423 | 0.01 | 0.009 | 1 | 6 | 1810034E1 | 0.001 | Young |
| 0.816579 | 0.372363 | 0.01 | 0.009 | 1 | 6 | Nudt6 | 0.001 | Young |
| 0.817137 | 0.047558 | 0.515 | 0.511 | 1 | 6 | Atp6v1f | 0.004 | Young |
| 0.817146 | 0.161942 | 0.037 | 0.04 | 1 | 6 | Cry1 | -0.003 | Young |
| 0.817327 | 0.156523 | 0.219 | 0.22 | 1 | 6 | Tns3 | -0.001 | Young |
| 0.817843 | 0.13035 | 0.07 | 0.066 | 1 | 6 | Malsu1 | 0.004 | Young |
| 0.817966 | 0.121175 | 0.164 | 0.163 | 1 | 6 | Atp11c | 0.001 | Young |
| 0.817969 | 0.203603 | 0.124 | 0.12 | 1 | 6 | Ube4a | 0.004 | Young |
| 0.818016 | 0.393239 | 0.025 | 0.023 | 1 | 6 | Abcg2 | 0.002 | Young |
| 0.818022 | 0.245404 | 0.109 | 0.106 | 1 | 6 | Gpaa1 | 0.003 | Young |
| 0.818853 | 0.809549 | 0.01 | 0.009 | 1 | 6 | Chp2 | 0.001 | Young |
| 0.818853 | 0.447806 | 0.01 | 0.009 | 1 | 6 | Fntb | 0.001 | Young |
| 0.819109 | 0.026047 | 0.202 | 0.197 | 1 | 6 | Sik1 | 0.005 | Young |
| 0.819421 | 0.067629 | 0.192 | 0.189 | 1 | 6 | Slc25a11 | 0.003 | Young |
| 0.819991 | 0.654658 | 0.01 | 0.009 | 1 | 6 | Sparcl1 | 0.001 | Young |
| 0.819991 | 0.367302 | 0.01 | 0.009 | 1 | 6 | 4833445I0 | 0.001 | Young |
| 0.819991 | 0.288804 | 0.01 | 0.009 | 1 | 6 | Csrnp2 | 0.001 | Young |
| 0.819991 | 0.116562 | 0.01 | 0.009 | 1 | 6 | 6720489N1 | 0.001 | Young |
| 0.820122 | 0.148926 | 0.351 | 0.354 | 1 | 6 | Tbcb | -0.003 | Young |
| 0.820345 | 0.065138 | 0.072 | 0.069 | 1 | 6 | Lgals8 | 0.003 | Young |
| 0.820532 | 0.766126 | 0.015 | 0.017 | 1 | 6 | Traf3ip1 | -0.002 | Young |
| 0.820699 | 0.067499 | 0.184 | 0.183 | 1 | 6 | Emc3 | 0.001 | Young |
| 0.820767 | 0.039453 | 0.622 | 0.629 | 1 | 6 | Wipf1 | -0.007 | Young |
| 0.820883 | 0.367651 | 0.025 | 0.023 | 1 | 6 | Nlrc4 | 0.002 | Young |
| 0.820883 | 0.102883 | 0.025 | 0.023 | 1 | 6 | Otof | 0.002 | Young |
| 0.821129 | 0.344877 | 0.01 | 0.009 | 1 | 6 | Zfp850 | 0.001 | Young |
| 0.821129 | 0.196358 | 0.01 | 0.009 | 1 | 6 | C030006K1 | 0.001 | Young |
| 0.821417 | 0.578582 | 0.015 | 0.017 | 1 | 6 | Ghdc | -0.002 | Young |
| 0.821501 | 0.101438 | 0.216 | 0.211 | 1 | 6 | Pgd | 0.005 | Young |
| 0.821774 | 0.155647 | 0.127 | 0.123 | 1 | 6 | Ctdspl2 | 0.004 | Young |
| 0.822267 | 0.187691 | 0.01 | 0.009 | 1 | 6 | Gm15853 | 0.001 | Young |
| 0.822267 | 0.119735 | 0.01 | 0.009 | 1 | 6 | Enkur | 0.001 | Young |
| 0.822695 | 0.049902 | 0.06 | 0.057 | 1 | 6 | Ggta1 | 0.003 | Young |
| 0.823373 | 0.166151 | 0.095 | 0.091 | 1 | 6 | Casd1 | 0.004 | Young |
| 0.823388 | 0.209993 | 0.045 | 0.049 | 1 | 6 | Exosc4 | -0.004 | Young |
| 0.823406 | 0.181534 | 0.01 | 0.009 | 1 | 6 | Tacc2 | 0.001 | Young |

| | | | | | | | | |
|---|---|---|---|---|---|---|---|---|
| 0.823419 | 0.422512 | 0.048 | 0.046 | 1 | 6 | Ric8b | 0.002 | Young |
| 0.82349 | 0.013955 | 0.475 | 0.48 | 1 | 6 | Efr3a | -0.005 | Young |
| 0.824393 | 0.000889 | 0.062 | 0.066 | 1 | 6 | Trappc12 | -0.004 | Young |
| 0.824471 | 0.178157 | 0.025 | 0.023 | 1 | 6 | Gstz1 | 0.002 | Young |
| 0.824471 | 0.173416 | 0.023 | 0.026 | 1 | 6 | Ckap2l | -0.003 | Young |
| 0.824628 | 0.169605 | 0.064 | 0.06 | 1 | 6 | Nup205 | 0.004 | Young |
| 0.824761 | 0.218277 | 0.042 | 0.046 | 1 | 6 | Trpm4 | -0.004 | Young |
| 0.824826 | 0.070762 | 0.067 | 0.071 | 1 | 6 | Tsta3 | -0.004 | Young |
| 0.824977 | 0.298024 | 0.048 | 0.046 | 1 | 6 | Anapc7 | 0.002 | Young |
| 0.825423 | 0.248575 | 0.166 | 0.174 | 1 | 6 | Bbip1 | -0.008 | Young |
| 0.825684 | 0.420251 | 0.01 | 0.009 | 1 | 6 | Fastkd3 | 0.001 | Young |
| 0.825684 | 0.220197 | 0.01 | 0.009 | 1 | 6 | Slfn9 | 0.001 | Young |
| 0.825684 | 0.127591 | 0.01 | 0.009 | 1 | 6 | Zfp658 | 0.001 | Young |
| 0.825705 | 0.161838 | 0.182 | 0.194 | 1 | 6 | Fbxo33 | -0.012 | Young |
| 0.825908 | 0.174029 | 0.025 | 0.023 | 1 | 6 | Btbd17 | 0.002 | Young |
| 0.825908 | 0.115432 | 0.025 | 0.023 | 1 | 6 | Lss | 0.002 | Young |
| 0.826008 | 0.042127 | 0.125 | 0.131 | 1 | 6 | Gcn1l1 | -0.006 | Young |
| 0.826121 | 0.094273 | 0.1 | 0.097 | 1 | 6 | Dctn5 | 0.003 | Young |
| 0.826824 | 0.419677 | 0.01 | 0.009 | 1 | 6 | Rsad1 | 0.001 | Young |
| 0.827372 | 0.065602 | 0.064 | 0.06 | 1 | 6 | Pde4dip | 0.004 | Young |
| 0.827811 | 0.264991 | 0.05 | 0.054 | 1 | 6 | 5430427O1 | -0.004 | Young |
| 0.829206 | 0.266928 | 0.087 | 0.083 | 1 | 6 | Pogz | 0.004 | Young |
| 0.829931 | 0.012386 | 0.452 | 0.451 | 1 | 6 | Rab7 | 0.001 | Young |
| 0.830292 | 0.042624 | 0.062 | 0.066 | 1 | 6 | R3hcc1l | -0.004 | Young |
| 0.830742 | 0.10245 | 0.045 | 0.049 | 1 | 6 | Rcc1 | -0.004 | Young |
| 0.83092 | 0.000102 | 0.465 | 0.466 | 1 | 6 | Tmem160 | -0.001 | Young |
| 0.831007 | 0.150343 | 0.105 | 0.111 | 1 | 6 | Pafah1b3 | -0.006 | Young |
| 0.831636 | 0.148491 | 0.032 | 0.034 | 1 | 6 | Micu3 | -0.002 | Young |
| 0.831658 | 0.07774 | 0.023 | 0.026 | 1 | 6 | Kiz | -0.003 | Young |
| 0.831794 | 0.143802 | 0.045 | 0.049 | 1 | 6 | Arfgef2 | -0.004 | Young |
| 0.831987 | 0.266107 | 0.14 | 0.137 | 1 | 6 | Nrp2 | 0.003 | Young |
| 0.832105 | 0.049288 | 0.219 | 0.229 | 1 | 6 | Chd1 | -0.01 | Young |
| 0.832258 | 0.018786 | 0.032 | 0.034 | 1 | 6 | Paox | -0.002 | Young |
| 0.832287 | 0.188782 | 0.125 | 0.131 | 1 | 6 | Rprd2 | -0.006 | Young |
| 0.832683 | 0.126591 | 0.164 | 0.16 | 1 | 6 | Ngdn | 0.004 | Young |
| 0.832772 | 0.075547 | 0.1 | 0.097 | 1 | 6 | Get4 | 0.003 | Young |
| 0.83294 | 0.041037 | 0.246 | 0.257 | 1 | 6 | Eif2a | -0.011 | Young |
| 0.833097 | 0.013637 | 0.023 | 0.026 | 1 | 6 | Gm45606 | -0.003 | Young |
| 0.83339 | 0.303217 | 0.079 | 0.083 | 1 | 6 | Zfp598 | -0.004 | Young |
| 0.833521 | 0.072413 | 0.084 | 0.08 | 1 | 6 | Pign | 0.004 | Young |
| 0.833817 | 0.158825 | 0.025 | 0.023 | 1 | 6 | Cc2d1a | 0.002 | Young |
| 0.834157 | 0.149625 | 0.187 | 0.2 | 1 | 6 | Psmd6 | -0.013 | Young |
| 0.834326 | 0.192872 | 0.139 | 0.137 | 1 | 6 | Slc25a45 | 0.002 | Young |
| 0.834531 | 0.699326 | 0.045 | 0.043 | 1 | 6 | Rtl8b | 0.002 | Young |
| 0.835094 | 0.029128 | 0.099 | 0.094 | 1 | 6 | Car5b | 0.005 | Young |

| | | | | | | | | |
|---|---|---|---|---|---|---|---|---|
| 0.835198 | 0.193123 | 0.114 | 0.111 | 1 | 6 | Gle1 | 0.003 | Young |
| 0.83658 | 0.102394 | 0.01 | 0.011 | 1 | 6 | Gm5431 | -0.001 | Young |
| 0.837418 | 0.042825 | 0.087 | 0.091 | 1 | 6 | Slc4a1ap | -0.004 | Young |
| 0.837864 | 0.910836 | 0.033 | 0.031 | 1 | 6 | Rhbdf2 | 0.002 | Young |
| 0.838138 | 0.571205 | 0.023 | 0.026 | 1 | 6 | Acp6 | -0.003 | Young |
| 0.83875 | 0.052051 | 0.01 | 0.011 | 1 | 6 | Mettl25 | -0.001 | Young |
| 0.839177 | 0.117485 | 0.196 | 0.191 | 1 | 6 | Tmf1 | 0.005 | Young |
| 0.839429 | 0.220476 | 0.045 | 0.049 | 1 | 6 | Coq10a | -0.004 | Young |
| 0.839825 | 0.04514 | 0.222 | 0.234 | 1 | 6 | Ergic3 | -0.012 | Young |
| 0.840922 | 0.215065 | 0.01 | 0.011 | 1 | 6 | Rgs5 | -0.001 | Young |
| 0.841133 | 0.75265 | 0.022 | 0.02 | 1 | 6 | Tfap4 | 0.002 | Young |
| 0.841705 | 0.428861 | 0.057 | 0.054 | 1 | 6 | Fbxo45 | 0.003 | Young |
| 0.842008 | 0.518236 | 0.01 | 0.011 | 1 | 6 | Lrfn1 | -0.001 | Young |
| 0.842119 | 0.277768 | 0.062 | 0.066 | 1 | 6 | Crtap | -0.004 | Young |
| 0.842188 | 0.094928 | 0.065 | 0.063 | 1 | 6 | Mlx | 0.002 | Young |
| 0.842575 | 0.211562 | 0.075 | 0.08 | 1 | 6 | Mrs2 | -0.005 | Young |
| 0.842642 | 0.268234 | 0.134 | 0.131 | 1 | 6 | Brwd3 | 0.003 | Young |
| 0.842677 | 0.597955 | 0.022 | 0.02 | 1 | 6 | Cbr4 | 0.002 | Young |
| 0.842677 | 0.457436 | 0.022 | 0.02 | 1 | 6 | Hspa2 | 0.002 | Young |
| 0.843095 | 0.255737 | 0.01 | 0.011 | 1 | 6 | Mgmt | -0.001 | Young |
| 0.843095 | 0.217246 | 0.01 | 0.011 | 1 | 6 | 1700001L0 | -0.001 | Young |
| 0.843495 | 0.588052 | 0.082 | 0.08 | 1 | 6 | Stn1 | 0.002 | Young |
| 0.844182 | 0.441025 | 0.01 | 0.011 | 1 | 6 | Casz1 | -0.001 | Young |
| 0.844182 | 0.338948 | 0.01 | 0.011 | 1 | 6 | Tldc1 | -0.001 | Young |
| 0.844182 | 0.26187 | 0.01 | 0.011 | 1 | 6 | Ngfr | -0.001 | Young |
| 0.844182 | 0.135471 | 0.01 | 0.011 | 1 | 6 | Glmn | -0.001 | Young |
| 0.844182 | 0.133009 | 0.01 | 0.011 | 1 | 6 | Acy1 | -0.001 | Young |
| 0.844182 | 0.056471 | 0.01 | 0.011 | 1 | 6 | Ddx11 | -0.001 | Young |
| 0.844194 | 0.114284 | 0.054 | 0.057 | 1 | 6 | Tfb2m | -0.003 | Young |
| 0.844603 | 0.128071 | 0.057 | 0.054 | 1 | 6 | Naf1 | 0.003 | Young |
| 0.844746 | 0.026623 | 0.192 | 0.197 | 1 | 6 | Ap1s1 | -0.005 | Young |
| 0.845269 | 0.268187 | 0.01 | 0.011 | 1 | 6 | Bcl2l14 | -0.001 | Young |
| 0.845269 | 0.027257 | 0.01 | 0.011 | 1 | 6 | Gng10 | -0.001 | Young |
| 0.845434 | 0.159379 | 0.162 | 0.16 | 1 | 6 | Ube2a | 0.002 | Young |
| 0.845976 | 0.435842 | 0.033 | 0.031 | 1 | 6 | Ankrd46 | 0.002 | Young |
| 0.845998 | 0.01035 | 0.207 | 0.2 | 1 | 6 | Cstf3 | 0.007 | Young |
| 0.846182 | 0.031438 | 0.112 | 0.109 | 1 | 6 | Pkd1 | 0.003 | Young |
| 0.846356 | 0.720375 | 0.01 | 0.011 | 1 | 6 | Mterf4 | -0.001 | Young |
| 0.846356 | 0.539933 | 0.01 | 0.011 | 1 | 6 | Gm6793 | -0.001 | Young |
| 0.846531 | 0.272468 | 0.142 | 0.143 | 1 | 6 | Tbc1d15 | -0.001 | Young |
| 0.846542 | 0.334747 | 0.022 | 0.02 | 1 | 6 | Gm16973 | 0.002 | Young |
| 0.8466 | 0.459329 | 0.033 | 0.031 | 1 | 6 | D17H6S53I | 0.002 | Young |
| 0.846918 | 0.139479 | 0.114 | 0.111 | 1 | 6 | Fem1b | 0.003 | Young |
| 0.847117 | 0.19921 | 0.06 | 0.057 | 1 | 6 | Nln | 0.003 | Young |
| 0.847718 | 0.165214 | 0.07 | 0.074 | 1 | 6 | Pygb | -0.004 | Young |

| | | | | | | | | |
|---|---|---|---|---|---|---|---|---|
| 0.847733 | 0.064857 | 0.057 | 0.06 | 1 | 6 | Slc13a3 | -0.003 | Young |
| 0.847951 | 0.005563 | 0.087 | 0.091 | 1 | 6 | Tmem250- | -0.004 | Young |
| 0.847987 | 0.282365 | 0.057 | 0.054 | 1 | 6 | Zdhhc13 | 0.003 | Young |
| 0.847987 | 0.000997 | 0.057 | 0.054 | 1 | 6 | Etfdh | 0.003 | Young |
| 0.848148 | 0.157238 | 0.07 | 0.074 | 1 | 6 | Mrps35 | -0.004 | Young |
| 0.848191 | 0.062414 | 0.147 | 0.154 | 1 | 6 | Nae1 | -0.007 | Young |
| 0.848299 | 0.462628 | 0.157 | 0.157 | 1 | 6 | Sap18 | 0 | Young |
| 0.848335 | 0.038919 | 0.154 | 0.16 | 1 | 6 | Dctn2 | -0.006 | Young |
| 0.848768 | 0.244079 | 0.054 | 0.051 | 1 | 6 | Hps4 | 0.003 | Young |
| 0.848862 | 0.411465 | 0.022 | 0.02 | 1 | 6 | 1110012L1 | 0.002 | Young |
| 0.848862 | 0.293343 | 0.022 | 0.02 | 1 | 6 | Zbtb5 | 0.002 | Young |
| 0.849141 | 0.116653 | 0.12 | 0.126 | 1 | 6 | Ern1 | -0.006 | Young |
| 0.849441 | 0.005615 | 0.048 | 0.051 | 1 | 6 | Mark4 | -0.003 | Young |
| 0.84963 | 0.269567 | 0.045 | 0.043 | 1 | 6 | Xaf1 | 0.002 | Young |
| 0.849842 | 0.093837 | 0.069 | 0.066 | 1 | 6 | Mfsd11 | 0.003 | Young |
| 0.849896 | 0.086884 | 0.107 | 0.103 | 1 | 6 | Med27 | 0.004 | Young |
| 0.85035 | 0.212107 | 0.033 | 0.031 | 1 | 6 | Dgkq | 0.002 | Young |
| 0.850709 | 0.864153 | 0.01 | 0.011 | 1 | 6 | Fitm2 | -0.001 | Young |
| 0.851958 | 0.19838 | 0.022 | 0.02 | 1 | 6 | 5430416N( | 0.002 | Young |
| 0.852002 | 0.731937 | 0.05 | 0.049 | 1 | 6 | Tpk1 | 0.001 | Young |
| 0.852312 | 0.229245 | 0.147 | 0.146 | 1 | 6 | 4833439L1 | 0.001 | Young |
| 0.852656 | 0.044777 | 0.149 | 0.154 | 1 | 6 | Prpf18 | -0.005 | Young |
| 0.852732 | 0.27237 | 0.022 | 0.02 | 1 | 6 | Amer1 | 0.002 | Young |
| 0.852873 | 0.49048 | 0.045 | 0.043 | 1 | 6 | Mfsd7a | 0.002 | Young |
| 0.853207 | 0.102154 | 0.281 | 0.297 | 1 | 6 | Rnf4 | -0.016 | Young |
| 0.853574 | 0.027371 | 0.341 | 0.334 | 1 | 6 | Blnk | 0.007 | Young |
| 0.853921 | 0.13738 | 0.194 | 0.191 | 1 | 6 | Ubqln1 | 0.003 | Young |
| 0.854065 | 0.017481 | 0.082 | 0.08 | 1 | 6 | Sik2 | 0.002 | Young |
| 0.854356 | 0.020145 | 0.659 | 0.623 | 1 | 6 | Cox6b1 | 0.036 | Young |
| 0.855982 | 0.4041 | 0.032 | 0.034 | 1 | 6 | Ccdc191 | -0.002 | Young |
| 0.856528 | 0.133089 | 0.149 | 0.154 | 1 | 6 | Prkag2 | -0.005 | Young |
| 0.857029 | 0.129829 | 0.1 | 0.106 | 1 | 6 | Stau1 | -0.006 | Young |
| 0.857389 | 0.22145 | 0.095 | 0.1 | 1 | 6 | Cds2 | -0.005 | Young |
| 0.858156 | 0.009515 | 0.022 | 0.02 | 1 | 6 | Oas2 | 0.002 | Young |
| 0.858357 | 0.016216 | 0.084 | 0.08 | 1 | 6 | Rgl1 | 0.004 | Young |
| 0.858553 | 0.849459 | 0.042 | 0.04 | 1 | 6 | Kcnd1 | 0.002 | Young |
| 0.858553 | 0.276382 | 0.04 | 0.043 | 1 | 6 | Prickle3 | -0.003 | Young |
| 0.858854 | 0.035242 | 0.998 | 0.963 | 1 | 6 | Actg1 | 0.035 | Young |
| 0.858886 | 0.106764 | 0.018 | 0.02 | 1 | 6 | Gm43660 | -0.002 | Young |
| 0.858932 | 0.187127 | 0.022 | 0.02 | 1 | 6 | Tyw5 | 0.002 | Young |
| 0.859114 | 0.325586 | 0.033 | 0.031 | 1 | 6 | Cox18 | 0.002 | Young |
| 0.859158 | 0.016871 | 0.525 | 0.529 | 1 | 6 | Polr1d | -0.004 | Young |
| 0.860337 | 0.220548 | 0.057 | 0.054 | 1 | 6 | Csf2rb2 | 0.003 | Young |
| 0.860882 | 0.216302 | 0.077 | 0.074 | 1 | 6 | Sp3os | 0.003 | Young |
| 0.860919 | 0.111399 | 0.075 | 0.08 | 1 | 6 | BC003965 | -0.005 | Young |

| | | | | | | | | |
|---|---|---|---|---|---|---|---|---|
| 0.861781 | 0.779012 | 0.054 | 0.057 | 1 | 6 | As3mt | -0.003 | Young |
| 0.861794 | 0.035089 | 0.08 | 0.077 | 1 | 6 | Bicral | 0.003 | Young |
| 0.862967 | 0.067725 | 0.057 | 0.06 | 1 | 6 | Hscb | -0.003 | Young |
| 0.863035 | 0.227379 | 0.065 | 0.063 | 1 | 6 | Adgra2 | 0.002 | Young |
| 0.86332 | 0.077877 | 0.127 | 0.131 | 1 | 6 | Chm | -0.004 | Young |
| 0.863365 | 0.037099 | 0.329 | 0.346 | 1 | 6 | Kdelr1 | -0.017 | Young |
| 0.863614 | 0.022028 | 0.251 | 0.254 | 1 | 6 | Chmp3 | -0.003 | Young |
| 0.864241 | 0.000587 | 0.936 | 0.923 | 1 | 6 | Oaz1 | 0.013 | Young |
| 0.864363 | 0.063436 | 0.022 | 0.02 | 1 | 6 | Hebp2 | 0.002 | Young |
| 0.864682 | 0.078237 | 0.214 | 0.209 | 1 | 6 | Ppp6c | 0.005 | Young |
| 0.864727 | 0.313139 | 0.04 | 0.043 | 1 | 6 | Ttll3 | -0.003 | Young |
| 0.864756 | 0.264506 | 0.032 | 0.034 | 1 | 6 | Fam78a | -0.002 | Young |
| 0.86515 | 0.036431 | 0.314 | 0.32 | 1 | 6 | Camkk2 | -0.006 | Young |
| 0.865271 | 0.394114 | 0.159 | 0.157 | 1 | 6 | Spopl | 0.002 | Young |
| 0.865627 | 0.082411 | 0.089 | 0.086 | 1 | 6 | Diablo | 0.003 | Young |
| 0.865763 | 0.065488 | 0.097 | 0.094 | 1 | 6 | Larp7 | 0.003 | Young |
| 0.865875 | 0.159727 | 0.164 | 0.163 | 1 | 6 | Ntan1 | 0.001 | Young |
| 0.866557 | 0.881403 | 0.03 | 0.029 | 1 | 6 | Srxn1 | 0.001 | Young |
| 0.866942 | 0.002036 | 0.669 | 0.677 | 1 | 6 | Ppp3ca | -0.008 | Young |
| 0.867059 | 0.163571 | 0.018 | 0.02 | 1 | 6 | Zfp688 | -0.002 | Young |
| 0.867266 | 0.086793 | 0.405 | 0.411 | 1 | 6 | Llph | -0.006 | Young |
| 0.867878 | 0.479613 | 0.018 | 0.02 | 1 | 6 | Ankzf1 | -0.002 | Young |
| 0.869255 | 0.047118 | 0.291 | 0.294 | 1 | 6 | Rb1cc1 | -0.003 | Young |
| 0.869915 | 0.085979 | 0.027 | 0.029 | 1 | 6 | Cenpv | -0.002 | Young |
| 0.869926 | 0.040356 | 0.1 | 0.097 | 1 | 6 | Pdxdc1 | 0.003 | Young |
| 0.869973 | 0.212242 | 0.048 | 0.051 | 1 | 6 | Gcsh | -0.003 | Young |
| 0.869977 | 0.032327 | 0.246 | 0.249 | 1 | 6 | Anapc5 | -0.003 | Young |
| 0.870348 | 0.534116 | 0.042 | 0.04 | 1 | 6 | Aff3 | 0.002 | Young |
| 0.870536 | 0.464785 | 0.122 | 0.12 | 1 | 6 | Mxd1 | 0.002 | Young |
| 0.870543 | 0.024701 | 0.217 | 0.226 | 1 | 6 | Eif6 | -0.009 | Young |
| 0.87094 | 0.127513 | 0.159 | 0.169 | 1 | 6 | Cc2d1b | -0.01 | Young |
| 0.871157 | 0.053253 | 0.535 | 0.537 | 1 | 6 | Cdkn1b | -0.002 | Young |
| 0.871165 | 0.010938 | 0.55 | 0.543 | 1 | 6 | Ptpre | 0.007 | Young |
| 0.871307 | 0.92079 | 0.018 | 0.017 | 1 | 6 | AC163032. | 0.001 | Young |
| 0.872149 | 0.700992 | 0.018 | 0.017 | 1 | 6 | Gm38843 | 0.001 | Young |
| 0.872231 | 0.347752 | 0.065 | 0.069 | 1 | 6 | Ctu2 | -0.004 | Young |
| 0.872248 | 0.238295 | 0.077 | 0.074 | 1 | 6 | Taf12 | 0.003 | Young |
| 0.872317 | 0.697708 | 0.042 | 0.04 | 1 | 6 | Slc36a3os | 0.002 | Young |
| 0.872477 | 0.303416 | 0.145 | 0.151 | 1 | 6 | Mlxip | -0.006 | Young |
| 0.873608 | 0.277354 | 0.018 | 0.02 | 1 | 6 | 9530059O1 | -0.002 | Young |
| 0.873647 | 0.144193 | 0.159 | 0.157 | 1 | 6 | Pdha1 | 0.002 | Young |
| 0.874723 | 0.012303 | 0.105 | 0.109 | 1 | 6 | Gm37494 | -0.004 | Young |
| 0.874805 | 0.435525 | 0.032 | 0.034 | 1 | 6 | Ankrd52 | -0.002 | Young |
| 0.875247 | 0.713954 | 0.018 | 0.02 | 1 | 6 | Siglecf | -0.002 | Young |
| 0.875266 | 0.052044 | 0.094 | 0.091 | 1 | 6 | Pgm2l1 | 0.003 | Young |

| | | | | | | | | |
|---|---|---|---|---|---|---|---|---|
| 0.875443 | 0.024279 | 0.107 | 0.103 | 1 | 6 | Uchl5 | 0.004 | Young |
| 0.875621 | 0.173686 | 0.077 | 0.074 | 1 | 6 | Smim12 | 0.003 | Young |
| 0.875807 | 0.638974 | 0.105 | 0.106 | 1 | 6 | Slc25a17 | -0.001 | Young |
| 0.875892 | 0.07493 | 0.234 | 0.229 | 1 | 6 | Ddx54 | 0.005 | Young |
| 0.876172 | 0.044761 | 0.09 | 0.094 | 1 | 6 | Arfip1 | -0.004 | Young |
| 0.876253 | 0.092781 | 0.132 | 0.131 | 1 | 6 | Cul5 | 0.001 | Young |
| 0.876601 | 0.190568 | 0.139 | 0.137 | 1 | 6 | Ncbp3 | 0.002 | Young |
| 0.876649 | 0.329469 | 0.07 | 0.074 | 1 | 6 | Pigq | -0.004 | Young |
| 0.876886 | 0.903496 | 0.018 | 0.02 | 1 | 6 | Pdlim4 | -0.002 | Young |
| 0.8772 | 0.014296 | 0.052 | 0.054 | 1 | 6 | Ppm1d | -0.002 | Young |
| 0.877543 | 0.060585 | 0.043 | 0.046 | 1 | 6 | Acad8 | -0.003 | Young |
| 0.877615 | 0.445813 | 0.067 | 0.066 | 1 | 6 | Aga | 0.001 | Young |
| 0.877728 | 0.193367 | 0.062 | 0.06 | 1 | 6 | Dhrs4 | 0.002 | Young |
| 0.877778 | 0.228055 | 0.03 | 0.029 | 1 | 6 | B430306N( | 0.001 | Young |
| 0.87802 | 0.112808 | 0.149 | 0.149 | 1 | 6 | Hspb11 | 0 | Young |
| 0.878048 | 0.459002 | 0.018 | 0.017 | 1 | 6 | Nars2 | 0.001 | Young |
| 0.878455 | 0.527084 | 0.054 | 0.057 | 1 | 6 | Mcm9 | -0.003 | Young |
| 0.878892 | 0.40588 | 0.018 | 0.017 | 1 | 6 | Ssh3 | 0.001 | Young |
| 0.879004 | 0.179574 | 0.164 | 0.171 | 1 | 6 | Mrps18c | -0.007 | Young |
| 0.87941 | 0.192741 | 0.074 | 0.071 | 1 | 6 | Adgrg5 | 0.003 | Young |
| 0.879899 | 0.263293 | 0.149 | 0.149 | 1 | 6 | Cass4 | 0 | Young |
| 0.880166 | 0.596377 | 0.018 | 0.02 | 1 | 6 | Hdac8 | -0.002 | Young |
| 0.880179 | 0.038812 | 0.085 | 0.089 | 1 | 6 | Iars | -0.004 | Young |
| 0.880198 | 0.260081 | 0.1 | 0.097 | 1 | 6 | Dpep2 | 0.003 | Young |
| 0.880523 | 0.172186 | 0.035 | 0.037 | 1 | 6 | Gpr18 | -0.002 | Young |
| 0.881162 | 0.102788 | 0.293 | 0.303 | 1 | 6 | Fam96a | -0.01 | Young |
| 0.881308 | 0.017191 | 0.508 | 0.5 | 1 | 6 | Plekho1 | 0.008 | Young |
| 0.881422 | 0.371484 | 0.018 | 0.017 | 1 | 6 | Arl15 | 0.001 | Young |
| 0.881845 | 0.021171 | 0.102 | 0.106 | 1 | 6 | Hif1an | -0.004 | Young |
| 0.881924 | 0.041567 | 0.234 | 0.234 | 1 | 6 | Rab18 | 0 | Young |
| 0.882335 | 0.35724 | 0.05 | 0.049 | 1 | 6 | Msantd2 | 0.001 | Young |
| 0.882339 | 0.019339 | 0.5 | 0.497 | 1 | 6 | Sf3b2 | 0.003 | Young |
| 0.882559 | 0.100724 | 0.135 | 0.134 | 1 | 6 | Map2k4 | 0.001 | Young |
| 0.882736 | 0.351484 | 0.04 | 0.043 | 1 | 6 | Nsun4 | -0.003 | Young |
| 0.88276 | 0.157289 | 0.11 | 0.109 | 1 | 6 | Prpf6 | 0.001 | Young |
| 0.88311 | 0.247496 | 0.018 | 0.017 | 1 | 6 | Urb2 | 0.001 | Young |
| 0.88311 | 0.135685 | 0.018 | 0.017 | 1 | 6 | Fut7 | 0.001 | Young |
| 0.883523 | 0.070523 | 0.065 | 0.063 | 1 | 6 | Ltbp3 | 0.002 | Young |
| 0.883641 | 0.440181 | 0.087 | 0.091 | 1 | 6 | Nelfa | -0.004 | Young |
| 0.883729 | 0.316848 | 0.03 | 0.029 | 1 | 6 | B3glct | 0.001 | Young |
| 0.883884 | 0.042551 | 0.161 | 0.157 | 1 | 6 | Ppm1m | 0.004 | Young |
| 0.8839 | 0.115354 | 0.186 | 0.183 | 1 | 6 | Smg7 | 0.003 | Young |
| 0.883954 | 0.208159 | 0.018 | 0.017 | 1 | 6 | Klri1 | 0.001 | Young |
| 0.883959 | 0.195885 | 0.052 | 0.054 | 1 | 6 | Pigc | -0.002 | Young |
| 0.884065 | 0.05433 | 0.181 | 0.189 | 1 | 6 | Kctd20 | -0.008 | Young |

| | | | | | | | | |
|---|---|---|---|---|---|---|---|---|
| 0.884699 | 0.167945 | 0.099 | 0.097 | 1 | 6 | Gm38399 | 0.002 | Young |
| 0.884991 | 0.22841 | 0.042 | 0.04 | 1 | 6 | Psmg1 | 0.002 | Young |
| 0.885514 | 0.231842 | 0.082 | 0.08 | 1 | 6 | Bcl2l2 | 0.002 | Young |
| 0.885928 | 0.017753 | 0.137 | 0.137 | 1 | 6 | Psmc2 | 0 | Young |
| 0.885951 | 0.01163 | 0.176 | 0.174 | 1 | 6 | Mrpl12 | 0.002 | Young |
| 0.886391 | 0.000922 | 0.212 | 0.217 | 1 | 6 | Plxnc1 | -0.005 | Young |
| 0.886478 | 0.390825 | 0.132 | 0.131 | 1 | 6 | Tdg | 0.001 | Young |
| 0.886834 | 0.011696 | 0.06 | 0.063 | 1 | 6 | Wdtc1 | -0.003 | Young |
| 0.887332 | 0.093192 | 0.018 | 0.017 | 1 | 6 | Ift74 | 0.001 | Young |
| 0.887553 | 1.171734 | 0.018 | 0.02 | 1 | 6 | Rad1 | -0.002 | Young |
| 0.8877 | 0.184145 | 0.03 | 0.029 | 1 | 6 | Gprasp1 | 0.001 | Young |
| 0.887812 | 0.080096 | 0.042 | 0.04 | 1 | 6 | Pigv | 0.002 | Young |
| 0.887923 | 0.349665 | 0.079 | 0.083 | 1 | 6 | Yif1a | -0.004 | Young |
| 0.888011 | 0.211037 | 0.05 | 0.049 | 1 | 6 | Slc22a15 | 0.001 | Young |
| 0.888176 | 0.229349 | 0.018 | 0.017 | 1 | 6 | Slc9a3r2 | 0.001 | Young |
| 0.888441 | 0.032736 | 0.134 | 0.137 | 1 | 6 | Mettl9 | -0.003 | Young |
| 0.888485 | 0.126315 | 0.122 | 0.12 | 1 | 6 | Ythdc2 | 0.002 | Young |
| 0.888716 | 0.228884 | 0.217 | 0.226 | 1 | 6 | Rbm6 | -0.009 | Young |
| 0.88875 | 0.112787 | 0.117 | 0.114 | 1 | 6 | Ugdh | 0.003 | Young |
| 0.888941 | 0.056241 | 0.042 | 0.04 | 1 | 6 | Timm21 | 0.002 | Young |
| 0.889013 | 0.170905 | 0.038 | 0.04 | 1 | 6 | Slc35b4 | -0.002 | Young |
| 0.889203 | 0.222317 | 0.082 | 0.08 | 1 | 6 | Timm50 | 0.002 | Young |
| 0.889232 | 0.043742 | 0.247 | 0.257 | 1 | 6 | Tmem106a | -0.01 | Young |
| 0.889591 | 0.092398 | 0.187 | 0.194 | 1 | 6 | Hadha | -0.007 | Young |
| 0.889686 | 0.383497 | 0.03 | 0.029 | 1 | 6 | Kcnk13 | 0.001 | Young |
| 0.889723 | 0.138697 | 0.054 | 0.051 | 1 | 6 | Rpl36a-ps1 | 0.003 | Young |
| 0.890123 | 0.305503 | 0.196 | 0.2 | 1 | 6 | Stat5b | -0.004 | Young |
| 0.890277 | 0.171917 | 0.132 | 0.131 | 1 | 6 | Eed | 0.001 | Young |
| 0.890471 | 0.103745 | 0.207 | 0.206 | 1 | 6 | Fam172a | 0.001 | Young |
| 0.890765 | 0.135579 | 0.035 | 0.037 | 1 | 6 | Gatc | -0.002 | Young |
| 0.890984 | 0.210118 | 0.224 | 0.22 | 1 | 6 | Glrx5 | 0.004 | Young |
| 0.89118 | 0.375712 | 0.119 | 0.117 | 1 | 6 | Adam9 | 0.002 | Young |
| 0.892401 | 0.398783 | 0.05 | 0.049 | 1 | 6 | Bnip3 | 0.001 | Young |
| 0.892631 | 0.278749 | 0.166 | 0.166 | 1 | 6 | Gatad2a | 0 | Young |
| 0.893381 | 0.182031 | 0.06 | 0.063 | 1 | 6 | Cpsf1 | -0.003 | Young |
| 0.893674 | 0.085961 | 0.157 | 0.154 | 1 | 6 | Sfxn2 | 0.003 | Young |
| 0.893736 | 0.114896 | 0.054 | 0.051 | 1 | 6 | Siah1a | 0.003 | Young |
| 0.89443 | 0.48085 | 0.127 | 0.129 | 1 | 6 | Pcbd2 | -0.002 | Young |
| 0.895089 | 0.027847 | 0.375 | 0.38 | 1 | 6 | Zc3h7a | -0.005 | Young |
| 0.895759 | 0.442541 | 0.082 | 0.086 | 1 | 6 | Lrrc47 | -0.004 | Young |
| 0.89622 | 0.101533 | 0.241 | 0.234 | 1 | 6 | Cd180 | 0.007 | Young |
| 0.896222 | 0.218984 | 0.059 | 0.057 | 1 | 6 | Pole3 | 0.002 | Young |
| 0.896313 | 0.060338 | 0.03 | 0.029 | 1 | 6 | Serpinb9 | 0.001 | Young |
| 0.896489 | 0.36679 | 0.119 | 0.123 | 1 | 6 | Zswim8 | -0.004 | Young |
| 0.896939 | 0.180687 | 0.074 | 0.077 | 1 | 6 | Orai2 | -0.003 | Young |

| | | | | | | | | |
|---|---|---|---|---|---|---|---|---|
| 0.897945 | 0.148054 | 0.187 | 0.189 | 1 | 6 | Blmh | -0.002 | Young |
| 0.898955 | 0.020352 | 0.271 | 0.277 | 1 | 6 | Atg3 | -0.006 | Young |
| 0.899462 | 0.024222 | 0.082 | 0.08 | 1 | 6 | Eif1ad | 0.002 | Young |
| 0.901202 | 0.278527 | 0.074 | 0.077 | 1 | 6 | Mre11a | -0.003 | Young |
| 0.901871 | 0.0496 | 0.286 | 0.294 | 1 | 6 | Eif3d | -0.008 | Young |
| 0.902468 | 0.137403 | 0.055 | 0.057 | 1 | 6 | N6amt1 | -0.002 | Young |
| 0.902484 | 0.540276 | 0.048 | 0.051 | 1 | 6 | Med31 | -0.003 | Young |
| 0.904243 | 0.281462 | 0.043 | 0.046 | 1 | 6 | Siae | -0.003 | Young |
| 0.905089 | 0.126294 | 0.169 | 0.169 | 1 | 6 | Tbc1d10b | 0 | Young |
| 0.90519 | 0.085612 | 0.105 | 0.109 | 1 | 6 | Prmt3 | -0.004 | Young |
| 0.905194 | 0.527635 | 0.027 | 0.026 | 1 | 6 | Vps51 | 0.001 | Young |
| 0.905321 | 0.248492 | 0.07 | 0.069 | 1 | 6 | Ndst2 | 0.001 | Young |
| 0.905463 | 0.591904 | 0.038 | 0.037 | 1 | 6 | Esr1 | 0.001 | Young |
| 0.905722 | 0.003876 | 0.43 | 0.426 | 1 | 6 | Rp2 | 0.004 | Young |
| 0.905794 | 0.092637 | 0.06 | 0.063 | 1 | 6 | Pitrm1 | -0.003 | Young |
| 0.9058 | 0.46064 | 0.047 | 0.046 | 1 | 6 | Hist1h4i | 0.001 | Young |
| 0.906341 | 0.057668 | 0.145 | 0.149 | 1 | 6 | Esf1 | -0.004 | Young |
| 0.906622 | 0.04594 | 0.493 | 0.506 | 1 | 6 | Irf2 | -0.013 | Young |
| 0.907234 | 0.254059 | 0.013 | 0.014 | 1 | 6 | Cbr3 | -0.001 | Young |
| 0.907283 | 1.093879 | 0.015 | 0.014 | 1 | 6 | Dync2h1 | 0.001 | Young |
| 0.907725 | 0.339457 | 0.112 | 0.111 | 1 | 6 | Mbd1 | 0.001 | Young |
| 0.90777 | 0.257368 | 0.067 | 0.066 | 1 | 6 | Fam110a | 0.001 | Young |
| 0.908054 | 0.059516 | 0.052 | 0.054 | 1 | 6 | Dzip3 | -0.002 | Young |
| 0.908222 | 0.179637 | 0.067 | 0.066 | 1 | 6 | E2f2 | 0.001 | Young |
| 0.908323 | 0.08668 | 0.194 | 0.203 | 1 | 6 | Slc26a2 | -0.009 | Young |
| 0.908422 | 0.092268 | 0.102 | 0.1 | 1 | 6 | Naip2 | 0.002 | Young |
| 0.908493 | 0.025573 | 0.03 | 0.031 | 1 | 6 | 1700021F0 | -0.001 | Young |
| 0.908623 | 0.087745 | 0.268 | 0.266 | 1 | 6 | Ifngr2 | 0.002 | Young |
| 0.908703 | 0.548047 | 0.027 | 0.026 | 1 | 6 | Tubgcp6 | 0.001 | Young |
| 0.908725 | 0.168667 | 0.038 | 0.04 | 1 | 6 | Mllt1 | -0.002 | Young |
| 0.908863 | 0.158366 | 0.117 | 0.114 | 1 | 6 | Aph1b | 0.003 | Young |
| 0.909147 | 1.016376 | 0.015 | 0.014 | 1 | 6 | Clec5a | 0.001 | Young |
| 0.909405 | 0.299438 | 0.027 | 0.026 | 1 | 6 | Ints5.1 | 0.001 | Young |
| 0.909946 | 0.17364 | 0.097 | 0.1 | 1 | 6 | Eif2ak2 | -0.003 | Young |
| 0.91008 | 0.782449 | 0.015 | 0.014 | 1 | 6 | Osgepl1 | 0.001 | Young |
| 0.910135 | 0.021549 | 0.013 | 0.014 | 1 | 6 | Snx33 | -0.001 | Young |
| 0.910453 | 0.147215 | 0.03 | 0.031 | 1 | 6 | Mad2l2 | -0.001 | Young |
| 0.910691 | 0.188456 | 0.167 | 0.174 | 1 | 6 | Gtf3c6 | -0.007 | Young |
| 0.911224 | 0.053745 | 0.079 | 0.077 | 1 | 6 | Odf2 | 0.002 | Young |
| 0.911282 | 0.043973 | 0.07 | 0.069 | 1 | 6 | Sssca1 | 0.001 | Young |
| 0.911832 | 0.334428 | 0.065 | 0.069 | 1 | 6 | Nelfb | -0.004 | Young |
| 0.911946 | 0.832789 | 0.015 | 0.014 | 1 | 6 | Gstm4 | 0.001 | Young |
| 0.911946 | 0.558469 | 0.015 | 0.014 | 1 | 6 | Tmem115 | 0.001 | Young |
| 0.912059 | 0.0912 | 0.132 | 0.137 | 1 | 6 | Wars | -0.005 | Young |
| 0.912071 | 0.086807 | 0.487 | 0.483 | 1 | 6 | Eif3c | 0.004 | Young |

| | | | | | | | | |
|---|---|---|---|---|---|---|---|---|
| 0.912171 | 0.4609 | 0.035 | 0.034 | 1 | 6 | Kcnq1 | 0.001 | Young |
| 0.912785 | 0.585989 | 0.035 | 0.034 | 1 | 6 | D930016D( | 0.001 | Young |
| 0.912858 | 0.113713 | 0.318 | 0.323 | 1 | 6 | Tmed3 | -0.005 | Young |
| 0.912868 | 0.03401 | 0.478 | 0.494 | 1 | 6 | Limd1 | -0.016 | Young |
| 0.912916 | 0.219099 | 0.027 | 0.026 | 1 | 6 | Fam114a1 | 0.001 | Young |
| 0.913066 | 0.214876 | 0.03 | 0.031 | 1 | 6 | AW554918 | -0.001 | Young |
| 0.913284 | 0.209927 | 0.059 | 0.057 | 1 | 6 | Arntl | 0.002 | Young |
| 0.913315 | 0.007849 | 0.533 | 0.537 | 1 | 6 | Cebpb | -0.004 | Young |
| 0.913689 | 0.016129 | 0.181 | 0.186 | 1 | 6 | Rras | -0.005 | Young |
| 0.913812 | 0.135826 | 0.015 | 0.014 | 1 | 6 | Phldb3 | 0.001 | Young |
| 0.914392 | 0.320553 | 0.022 | 0.023 | 1 | 6 | Def8 | -0.001 | Young |
| 0.914676 | 0.480498 | 0.055 | 0.054 | 1 | 6 | Tomm40l | 0.001 | Young |
| 0.914974 | 0.90106 | 0.013 | 0.014 | 1 | 6 | Mks1 | -0.001 | Young |
| 0.914974 | 0.30116 | 0.013 | 0.014 | 1 | 6 | Fcrl1 | -0.001 | Young |
| 0.915118 | 0.108784 | 0.192 | 0.2 | 1 | 6 | Gatad1 | -0.008 | Young |
| 0.915249 | 0.03836 | 0.156 | 0.157 | 1 | 6 | Polr1a | -0.001 | Young |
| 0.915428 | 0.0804 | 0.094 | 0.091 | 1 | 6 | Fmc1 | 0.003 | Young |
| 0.91548 | 0.033274 | 0.07 | 0.069 | 1 | 6 | Cdc14b | 0.001 | Young |
| 0.915484 | 0.329532 | 0.085 | 0.089 | 1 | 6 | Pot1b | -0.004 | Young |
| 0.915496 | 0.03522 | 0.102 | 0.1 | 1 | 6 | Tbc1d13 | 0.002 | Young |
| 0.915678 | 0.448148 | 0.015 | 0.014 | 1 | 6 | Zfp113 | 0.001 | Young |
| 0.915873 | 0.094841 | 0.115 | 0.117 | 1 | 6 | Slc39a14 | -0.002 | Young |
| 0.915922 | 0.567674 | 0.022 | 0.023 | 1 | 6 | Gtf3c4 | -0.001 | Young |
| 0.915942 | 0.442479 | 0.013 | 0.014 | 1 | 6 | Arhgap19 | -0.001 | Young |
| 0.915942 | 0.381779 | 0.013 | 0.014 | 1 | 6 | Arhgap27o | -0.001 | Young |
| 0.916109 | 0.123185 | 0.11 | 0.109 | 1 | 6 | Ech1 | 0.001 | Young |
| 0.916335 | 0.258344 | 0.03 | 0.031 | 1 | 6 | Fbxl19 | -0.001 | Young |
| 0.916836 | 0.288242 | 0.105 | 0.109 | 1 | 6 | Nfx1 | -0.004 | Young |
| 0.916888 | 0.052527 | 0.214 | 0.211 | 1 | 6 | Ankrd13c | 0.003 | Young |
| 0.916998 | 0.1329 | 0.075 | 0.074 | 1 | 6 | Dhx35 | 0.001 | Young |
| 0.917131 | 0.692854 | 0.027 | 0.026 | 1 | 6 | Gm28707 | 0.001 | Young |
| 0.917131 | 0.190248 | 0.027 | 0.026 | 1 | 6 | Zfp777 | 0.001 | Young |
| 0.917545 | 0.460891 | 0.015 | 0.014 | 1 | 6 | Fgfbp3 | 0.001 | Young |
| 0.917884 | 0.028733 | 0.087 | 0.086 | 1 | 6 | Gm17018 | 0.001 | Young |
| 0.918259 | 0.040774 | 0.411 | 0.4 | 1 | 6 | Ybx3 | 0.011 | Young |
| 0.918479 | 0.703138 | 0.015 | 0.014 | 1 | 6 | Tm9sf1 | 0.001 | Young |
| 0.918847 | 1.056974 | 0.013 | 0.014 | 1 | 6 | Slc19a1 | -0.001 | Young |
| 0.918983 | 0.344525 | 0.022 | 0.023 | 1 | 6 | Gm38251 | -0.001 | Young |
| 0.919024 | 0.011036 | 0.253 | 0.243 | 1 | 6 | Il6 | 0.01 | Young |
| 0.919212 | 0.007701 | 0.064 | 0.066 | 1 | 6 | Micall1 | -0.002 | Young |
| 0.919314 | 0.263084 | 0.167 | 0.166 | 1 | 6 | Slc9a3r1 | 0.001 | Young |
| 0.919413 | 0.203087 | 0.134 | 0.137 | 1 | 6 | 4833420G1 | -0.003 | Young |
| 0.919413 | 0.392267 | 0.015 | 0.014 | 1 | 6 | Cfd | 0.001 | Young |
| 0.919413 | 0.362057 | 0.015 | 0.014 | 1 | 6 | Zfp775 | 0.001 | Young |
| 0.919413 | 0.24951 | 0.015 | 0.014 | 1 | 6 | Fam120c | 0.001 | Young |

| | | | | | | | | |
|---|---|---|---|---|---|---|---|---|
| 0.919459 | 0.052102 | 0.129 | 0.126 | 1 | 6 | Rnf213 | 0.003 | Young |
| 0.919816 | 0.499217 | 0.013 | 0.014 | 1 | 6 | Zfp52 | -0.001 | Young |
| 0.919816 | 0.214671 | 0.013 | 0.014 | 1 | 6 | Gbp2b | -0.001 | Young |
| 0.920155 | 0.320165 | 0.035 | 0.034 | 1 | 6 | Slc28a2 | 0.001 | Young |
| 0.920347 | 1.06497 | 0.015 | 0.014 | 1 | 6 | Mmaa | 0.001 | Young |
| 0.920645 | 0.229042 | 0.027 | 0.026 | 1 | 6 | Bnip1 | 0.001 | Young |
| 0.921464 | 0.102839 | 0.435 | 0.44 | 1 | 6 | Epn1 | -0.005 | Young |
| 0.921753 | 0.960936 | 0.013 | 0.014 | 1 | 6 | Gins1 | -0.001 | Young |
| 0.921819 | 0.082421 | 0.356 | 0.371 | 1 | 6 | Snrpb2 | -0.015 | Young |
| 0.921822 | 0.28441 | 0.092 | 0.094 | 1 | 6 | BC005561 | -0.002 | Young |
| 0.921846 | 0.030622 | 0.047 | 0.046 | 1 | 6 | Coq7 | 0.001 | Young |
| 0.921963 | 0.03578 | 0.072 | 0.074 | 1 | 6 | Aasdhppt | -0.002 | Young |
| 0.922215 | 0.31186 | 0.015 | 0.014 | 1 | 6 | Nlrp1a | 0.001 | Young |
| 0.922556 | 0.253213 | 0.075 | 0.074 | 1 | 6 | 9330160F1 | 0.001 | Young |
| 0.922614 | 0.069017 | 0.251 | 0.249 | 1 | 6 | Col4a3bp | 0.002 | Young |
| 0.922642 | 0.091446 | 0.378 | 0.386 | 1 | 6 | Sgpp1 | -0.008 | Young |
| 0.922754 | 0.091876 | 0.089 | 0.091 | 1 | 6 | Nr2f6 | -0.002 | Young |
| 0.923099 | 0.224941 | 0.229 | 0.234 | 1 | 6 | Tmem173 | -0.005 | Young |
| 0.923265 | 0.042208 | 0.075 | 0.077 | 1 | 6 | Rhoq | -0.002 | Young |
| 0.923397 | 0.045383 | 0.062 | 0.06 | 1 | 6 | Sqor | 0.002 | Young |
| 0.923458 | 0.146093 | 0.027 | 0.026 | 1 | 6 | Zfp958 | 0.001 | Young |
| 0.923482 | 0.056383 | 0.236 | 0.237 | 1 | 6 | Hipk3 | -0.001 | Young |
| 0.923576 | 0.439286 | 0.022 | 0.023 | 1 | 6 | Cry2 | -0.001 | Young |
| 0.92369 | 0.000779 | 0.075 | 0.077 | 1 | 6 | Rnf19a | -0.002 | Young |
| 0.923911 | 0.043566 | 0.189 | 0.194 | 1 | 6 | Snrnp200 | -0.005 | Young |
| 0.924523 | 0.231797 | 0.047 | 0.046 | 1 | 6 | Nedd1 | 0.001 | Young |
| 0.924838 | 0.316969 | 0.03 | 0.031 | 1 | 6 | Ankrd50 | -0.001 | Young |
| 0.925018 | 0.105018 | 0.015 | 0.014 | 1 | 6 | Acsf2 | 0.001 | Young |
| 0.926136 | 0.000367 | 0.324 | 0.323 | 1 | 6 | Aph1a | 0.001 | Young |
| 0.92626 | 0.053387 | 0.194 | 0.197 | 1 | 6 | Twistnb | -0.003 | Young |
| 0.926465 | 0.293531 | 0.055 | 0.057 | 1 | 6 | Ppp2r1b | -0.002 | Young |
| 0.926503 | 0.22582 | 0.286 | 0.303 | 1 | 6 | Myd88 | -0.017 | Young |
| 0.926628 | 0.109692 | 0.234 | 0.24 | 1 | 6 | Hsd17b4 | -0.006 | Young |
| 0.926887 | 0.100587 | 0.015 | 0.014 | 1 | 6 | Dennd2c | 0.001 | Young |
| 0.92697 | 0.182507 | 0.212 | 0.211 | 1 | 6 | Nenf | 0.001 | Young |
| 0.926975 | 0.416164 | 0.027 | 0.026 | 1 | 6 | Nubpl | 0.001 | Young |
| 0.92699 | 0.089848 | 0.124 | 0.123 | 1 | 6 | Uap1l1 | 0.001 | Young |
| 0.927071 | 0.33545 | 0.092 | 0.091 | 1 | 6 | Yif1b | 0.001 | Young |
| 0.927092 | 0.321381 | 0.087 | 0.086 | 1 | 6 | Zfand2b | 0.001 | Young |
| 0.927202 | 0.033813 | 0.047 | 0.046 | 1 | 6 | Hacd2 | 0.001 | Young |
| 0.927373 | 0.322958 | 0.077 | 0.08 | 1 | 6 | Mrpl40 | -0.003 | Young |
| 0.927534 | 0.803171 | 0.023 | 0.023 | 1 | 6 | Endov | 0 | Young |
| 0.928034 | 0.220573 | 0.109 | 0.109 | 1 | 6 | Sec16a | 0 | Young |
| 0.928171 | 0.428806 | 0.022 | 0.023 | 1 | 6 | Spsb2 | -0.001 | Young |
| 0.92819 | 0.08036 | 0.206 | 0.211 | 1 | 6 | Dstyk | -0.005 | Young |

| | | | | | | | | |
|---|---|---|---|---|---|---|---|---|
| 0.928495 | 0.136514 | 0.038 | 0.04 | 1 | 6 | Hpse | -0.002 | Young |
| 0.928633 | 0.357365 | 0.077 | 0.08 | 1 | 6 | Ccdc117 | -0.003 | Young |
| 0.928763 | 0.143515 | 0.035 | 0.034 | 1 | 6 | Ogfod2 | 0.001 | Young |
| 0.929455 | 0.172207 | 0.115 | 0.114 | 1 | 6 | Ddx10 | 0.001 | Young |
| 0.931109 | 0.15571 | 0.241 | 0.246 | 1 | 6 | Hadhb | -0.005 | Young |
| 0.931258 | 0.166674 | 0.072 | 0.071 | 1 | 6 | Pla2g16 | 0.001 | Young |
| 0.932162 | 0.094351 | 0.187 | 0.194 | 1 | 6 | Rnps1 | -0.007 | Young |
| 0.932481 | 0.206361 | 0.047 | 0.049 | 1 | 6 | Tmcc2 | -0.002 | Young |
| 0.93273 | 0.033627 | 0.117 | 0.12 | 1 | 6 | Sbf1 | -0.003 | Young |
| 0.932778 | 0.708249 | 0.023 | 0.023 | 1 | 6 | Cep19 | 0 | Young |
| 0.933027 | 0.220052 | 0.043 | 0.043 | 1 | 6 | Zfp180 | 0 | Young |
| 0.933061 | 0.188809 | 0.14 | 0.143 | 1 | 6 | Stard7 | -0.003 | Young |
| 0.933351 | 0.326673 | 0.03 | 0.031 | 1 | 6 | Xpnpep3 | -0.001 | Young |
| 0.933527 | 0.366689 | 0.144 | 0.146 | 1 | 6 | Lsm3 | -0.002 | Young |
| 0.933822 | 0.074726 | 0.104 | 0.103 | 1 | 6 | Zeb1 | 0.001 | Young |
| 0.933825 | 0.310267 | 0.055 | 0.057 | 1 | 6 | Pla2g15 | -0.002 | Young |
| 0.934277 | 0.579838 | 0.023 | 0.023 | 1 | 6 | Oxnad1 | 0 | Young |
| 0.934661 | 0.548018 | 0.03 | 0.031 | 1 | 6 | 5830432E0 | -0.001 | Young |
| 0.935953 | 0.020447 | 0.055 | 0.054 | 1 | 6 | Ccdc43 | 0.001 | Young |
| 0.936314 | 0.063993 | 0.047 | 0.046 | 1 | 6 | Pstk | 0.001 | Young |
| 0.936434 | 0.083032 | 0.092 | 0.091 | 1 | 6 | Oxa1l | 0.001 | Young |
| 0.936645 | 0.17685 | 0.159 | 0.16 | 1 | 6 | Tbc1d20 | -0.001 | Young |
| 0.937164 | 0.319313 | 0.177 | 0.183 | 1 | 6 | Bcap29 | -0.006 | Young |
| 0.93734 | 0.103317 | 0.089 | 0.091 | 1 | 6 | Ppp2cb | -0.002 | Young |
| 0.93786 | 0.110225 | 0.05 | 0.051 | 1 | 6 | Cinp | -0.001 | Young |
| 0.937935 | 0.026394 | 0.099 | 0.097 | 1 | 6 | Abcg3 | 0.002 | Young |
| 0.938024 | 0.2686 | 0.023 | 0.023 | 1 | 6 | Zc3h8 | 0 | Young |
| 0.939814 | 0.266243 | 0.105 | 0.109 | 1 | 6 | Fnta | -0.004 | Young |
| 0.939843 | 0.343751 | 0.06 | 0.06 | 1 | 6 | Coq2 | 0 | Young |
| 0.939992 | 0.457877 | 0.052 | 0.051 | 1 | 6 | Kat5 | 0.001 | Young |
| 0.940667 | 0.118267 | 0.102 | 0.106 | 1 | 6 | Nqo2 | -0.004 | Young |
| 0.941486 | 0.323429 | 0.072 | 0.074 | 1 | 6 | Slc35e4 | -0.002 | Young |
| 0.941494 | 0.159211 | 0.042 | 0.043 | 1 | 6 | Dglucy | -0.001 | Young |
| 0.942056 | 0.179095 | 0.042 | 0.043 | 1 | 6 | Gngt2 | -0.001 | Young |
| 0.942107 | 0.247215 | 0.094 | 0.094 | 1 | 6 | Prrc1 | 0 | Young |
| 0.942458 | 0.347318 | 0.043 | 0.043 | 1 | 6 | Manbal | 0 | Young |
| 0.943494 | 0.034384 | 0.194 | 0.197 | 1 | 6 | Cdk2ap1 | -0.003 | Young |
| 0.944321 | 0.048988 | 0.157 | 0.163 | 1 | 6 | Gna12 | -0.006 | Young |
| 0.944333 | 0.059525 | 0.087 | 0.086 | 1 | 6 | Ipo11 | 0.001 | Young |
| 0.945665 | 0.197528 | 0.102 | 0.106 | 1 | 6 | Faap20 | -0.004 | Young |
| 0.945964 | 0.081386 | 0.099 | 0.1 | 1 | 6 | Rexo4 | -0.001 | Young |
| 0.94655 | 0.060374 | 0.042 | 0.043 | 1 | 6 | Diexf | -0.001 | Young |
| 0.946899 | 0.337562 | 0.043 | 0.043 | 1 | 6 | Fam35a | 0 | Young |
| 0.946962 | 0.085209 | 0.202 | 0.203 | 1 | 6 | Tm2d2 | -0.001 | Young |
| 0.947365 | 0.082183 | 0.239 | 0.24 | 1 | 6 | Atg2a | -0.001 | Young |

| | | | | | | | | |
|---|---|---|---|---|---|---|---|---|
| 0.948009 | 0.160973 | 0.043 | 0.043 | 1 | 6 | Eri3 | 0 | Young |
| 0.948169 | 0.220221 | 0.097 | 0.1 | 1 | 6 | Dctpp1 | -0.003 | Young |
| 0.948683 | 0.368862 | 0.075 | 0.074 | 1 | 6 | Fads1 | 0.001 | Young |
| 0.949276 | 0.0407 | 0.023 | 0.023 | 1 | 6 | Rab39 | 0 | Young |
| 0.949451 | 0.687117 | 0.09 | 0.091 | 1 | 6 | 2410002F2 | -0.001 | Young |
| 0.950334 | 0.153427 | 0.224 | 0.229 | 1 | 6 | Glmp | -0.005 | Young |
| 0.950401 | 0.226115 | 0.264 | 0.271 | 1 | 6 | Kat2b | -0.007 | Young |
| 0.950777 | 0.046833 | 0.023 | 0.023 | 1 | 6 | Mansc1 | 0 | Young |
| 0.951005 | 0.065176 | 0.064 | 0.063 | 1 | 6 | Dedd2 | 0.001 | Young |
| 0.95129 | 0.015084 | 0.418 | 0.414 | 1 | 6 | Wac | 0.004 | Young |
| 0.951368 | 0.303059 | 0.072 | 0.071 | 1 | 6 | Trim11 | 0.001 | Young |
| 0.951382 | 0.094323 | 0.348 | 0.357 | 1 | 6 | Eif3j1 | -0.009 | Young |
| 0.951628 | 0.120842 | 0.14 | 0.143 | 1 | 6 | Dnase1l1 | -0.003 | Young |
| 0.952103 | 0.041641 | 0.142 | 0.143 | 1 | 6 | Dapk3 | -0.001 | Young |
| 0.952826 | 0.119864 | 0.122 | 0.123 | 1 | 6 | Cox6b2 | -0.001 | Young |
| 0.952927 | 0.053748 | 0.254 | 0.257 | 1 | 6 | Cdkn2d | -0.003 | Young |
| 0.953253 | 0.143705 | 0.052 | 0.051 | 1 | 6 | Bbc3 | 0.001 | Young |
| 0.95331 | 0.116989 | 0.05 | 0.051 | 1 | 6 | Oaf | -0.001 | Young |
| 0.954011 | 0.065449 | 0.452 | 0.466 | 1 | 6 | Nedd8 | -0.014 | Young |
| 0.954116 | 0.055516 | 0.109 | 0.111 | 1 | 6 | Rpa3 | -0.002 | Young |
| 0.95417 | 0.07909 | 0.266 | 0.263 | 1 | 6 | Rnf111 | 0.003 | Young |
| 0.954755 | 0.396039 | 0.032 | 0.031 | 1 | 6 | Foxo4 | 0.001 | Young |
| 0.954779 | 0.03802 | 0.07 | 0.071 | 1 | 6 | Nostrin | -0.001 | Young |
| 0.954953 | 0.550297 | 0.055 | 0.057 | 1 | 6 | Cdk5 | -0.002 | Young |
| 0.955248 | 0.072193 | 0.035 | 0.034 | 1 | 6 | Lrig2 | 0.001 | Young |
| 0.956058 | 0.102953 | 0.025 | 0.026 | 1 | 6 | Alyref2 | -0.001 | Young |
| 0.956108 | 0.078264 | 0.064 | 0.063 | 1 | 6 | Timm9 | 0.001 | Young |
| 0.956235 | 0.18984 | 0.079 | 0.08 | 1 | 6 | Chid1 | -0.001 | Young |
| 0.956463 | 0.119185 | 0.207 | 0.209 | 1 | 6 | Tmem208 | -0.002 | Young |
| 0.95739 | 0.011556 | 0.017 | 0.017 | 1 | 6 | B230217C1 | 0 | Young |
| 0.957718 | 0.241944 | 0.129 | 0.129 | 1 | 6 | Vps13d | 0 | Young |
| 0.958268 | 0.038356 | 0.017 | 0.017 | 1 | 6 | Chd3os | 0 | Young |
| 0.958269 | 0.007102 | 0.055 | 0.054 | 1 | 6 | Vps8 | 0.001 | Young |
| 0.958657 | 0.36294 | 0.122 | 0.126 | 1 | 6 | Fam98b | -0.004 | Young |
| 0.958979 | 0.189966 | 0.197 | 0.203 | 1 | 6 | Ppm1a | -0.006 | Young |
| 0.959123 | 0.129442 | 0.219 | 0.22 | 1 | 6 | Uba6 | -0.001 | Young |
| 0.959313 | 1.102421 | 0.012 | 0.011 | 1 | 6 | Zfp809 | 0.001 | Young |
| 0.959349 | 0.010889 | 0.291 | 0.3 | 1 | 6 | Slc35c2 | -0.009 | Young |
| 0.960115 | 0.196786 | 0.097 | 0.097 | 1 | 6 | Faf1 | 0 | Young |
| 0.960369 | 1.00891 | 0.012 | 0.011 | 1 | 6 | Cdca7 | 0.001 | Young |
| 0.960369 | 0.649724 | 0.012 | 0.011 | 1 | 6 | Dip2c | 0.001 | Young |
| 0.961719 | 0.030159 | 0.462 | 0.48 | 1 | 6 | Rab11fip1 | -0.018 | Young |
| 0.96178 | 0.176368 | 0.017 | 0.017 | 1 | 6 | Pigw | 0 | Young |
| 0.961816 | 0.117704 | 0.025 | 0.026 | 1 | 6 | Ppip5k1 | -0.001 | Young |
| 0.962006 | 0.10339 | 0.201 | 0.2 | 1 | 6 | Ube2m | 0.001 | Young |

| | | | | | | | | |
|---|---|---|---|---|---|---|---|---|
| 0.962089 | 0.649765 | 0.048 | 0.049 | 1 | 6 | Ring1 | -0.001 | Young |
| 0.962141 | 0.018962 | 0.064 | 0.063 | 1 | 6 | Kitl | 0.001 | Young |
| 0.962289 | 0.249 | 0.042 | 0.043 | 1 | 6 | Slc35b3 | -0.001 | Young |
| 0.962298 | 0.026125 | 0.166 | 0.166 | 1 | 6 | Cyc1 | 0 | Young |
| 0.96329 | 0.163298 | 0.144 | 0.146 | 1 | 6 | Pdzd8 | -0.002 | Young |
| 0.96362 | 0.288137 | 0.05 | 0.051 | 1 | 6 | Supt3 | -0.001 | Young |
| 0.96431 | 0.188582 | 0.097 | 0.097 | 1 | 6 | Slc29a1 | 0 | Young |
| 0.964593 | 0.785717 | 0.012 | 0.011 | 1 | 6 | 6530409C1 | 0.001 | Young |
| 0.964593 | 0.763056 | 0.012 | 0.011 | 1 | 6 | Ubox5 | 0.001 | Young |
| 0.964596 | 0.567968 | 0.033 | 0.034 | 1 | 6 | Tjap1 | -0.001 | Young |
| 0.96519 | 0.129317 | 0.107 | 0.109 | 1 | 6 | Stx8 | -0.002 | Young |
| 0.965292 | 0.126468 | 0.017 | 0.017 | 1 | 6 | Vmac | 0 | Young |
| 0.965339 | 0.288986 | 0.092 | 0.094 | 1 | 6 | Pdcl | -0.002 | Young |
| 0.965387 | 0.524791 | 0.04 | 0.04 | 1 | 6 | Dcxr | 0 | Young |
| 0.965649 | 0.533739 | 0.012 | 0.011 | 1 | 6 | Gm13205 | 0.001 | Young |
| 0.966027 | 0.206484 | 0.097 | 0.097 | 1 | 6 | Fbxo38 | 0 | Young |
| 0.966242 | 0.085045 | 0.07 | 0.071 | 1 | 6 | Hinfp | -0.001 | Young |
| 0.966706 | 1.054321 | 0.012 | 0.011 | 1 | 6 | Gpt | 0.001 | Young |
| 0.966706 | 0.304364 | 0.012 | 0.011 | 1 | 6 | Selenbp1 | 0.001 | Young |
| 0.9668 | 0.145225 | 0.145 | 0.151 | 1 | 6 | Por | -0.006 | Young |
| 0.967048 | 0.28581 | 0.017 | 0.017 | 1 | 6 | Tmem191c | 0 | Young |
| 0.967497 | 0.296254 | 0.082 | 0.083 | 1 | 6 | Knop1 | -0.001 | Young |
| 0.967762 | 0.142265 | 0.012 | 0.011 | 1 | 6 | Hhat | 0.001 | Young |
| 0.967879 | 0.121723 | 0.06 | 0.06 | 1 | 6 | Entpd7 | 0 | Young |
| 0.967926 | 0.464202 | 0.017 | 0.017 | 1 | 6 | Nlrx1 | 0 | Young |
| 0.967926 | 0.095704 | 0.017 | 0.017 | 1 | 6 | Zfp212 | 0 | Young |
| 0.968017 | 0.163985 | 0.1 | 0.1 | 1 | 6 | Cyth2 | 0 | Young |
| 0.968154 | 0.145521 | 0.132 | 0.131 | 1 | 6 | Smarcad1 | 0.001 | Young |
| 0.968805 | 0.462591 | 0.017 | 0.017 | 1 | 6 | Cdca7l | 0 | Young |
| 0.968805 | 0.022438 | 0.017 | 0.017 | 1 | 6 | Pdzd4 | 0 | Young |
| 0.968819 | 0.201734 | 0.012 | 0.011 | 1 | 6 | E130102H2 | 0.001 | Young |
| 0.969016 | 0.23819 | 0.025 | 0.026 | 1 | 6 | Fig4 | -0.001 | Young |
| 0.969197 | 0.013208 | 0.107 | 0.109 | 1 | 6 | Foxj3 | -0.002 | Young |
| 0.969629 | 0.108419 | 0.079 | 0.08 | 1 | 6 | Pop7 | -0.001 | Young |
| 0.969683 | 0.515906 | 0.017 | 0.017 | 1 | 6 | Nxpe4 | 0 | Young |
| 0.969765 | 0.225341 | 0.142 | 0.146 | 1 | 6 | Spr | -0.004 | Young |
| 0.969875 | 0.77613 | 0.012 | 0.011 | 1 | 6 | Rpgr | 0.001 | Young |
| 0.97007 | 0.120429 | 0.127 | 0.129 | 1 | 6 | Hsd17b10 | -0.002 | Young |
| 0.970457 | 0.502803 | 0.025 | 0.026 | 1 | 6 | Wrap53 | -0.001 | Young |
| 0.970561 | 0.382883 | 0.017 | 0.017 | 1 | 6 | Nif3l1 | 0 | Young |
| 0.970718 | 0.447925 | 0.065 | 0.066 | 1 | 6 | Coa6 | -0.001 | Young |
| 0.970932 | 0.031813 | 0.012 | 0.011 | 1 | 6 | Ica1l | 0.001 | Young |
| 0.971034 | 0.057627 | 0.249 | 0.254 | 1 | 6 | Kat6a | -0.005 | Young |
| 0.971335 | 0.037361 | 0.498 | 0.514 | 1 | 6 | Dynll1 | -0.016 | Young |
| 0.971384 | 0.261442 | 0.212 | 0.214 | 1 | 6 | Napa | -0.002 | Young |

| | | | | | | | | |
|---|---|---|---|---|---|---|---|---|
| 0.971873 | 0.134396 | 0.05 | 0.051 | 1 | 6 | Inpp5a | -0.001 | Young |
| 0.971891 | 0.193482 | 0.149 | 0.151 | 1 | 6 | Cib1 | -0.002 | Young |
| 0.971988 | 0.030527 | 0.012 | 0.011 | 1 | 6 | C1s1 | 0.001 | Young |
| 0.972869 | 0.002327 | 0.779 | 0.749 | 1 | 6 | Anxa5 | 0.03 | Young |
| 0.972978 | 0.322755 | 0.042 | 0.043 | 1 | 6 | Galc | -0.001 | Young |
| 0.973045 | 0.226654 | 0.012 | 0.011 | 1 | 6 | Ly75 | 0.001 | Young |
| 0.973065 | 0.165786 | 0.077 | 0.077 | 1 | 6 | Mroh1 | 0 | Young |
| 0.973111 | 0.244884 | 0.06 | 0.06 | 1 | 6 | Neurl4 | 0 | Young |
| 0.973171 | 0.19664 | 0.04 | 0.04 | 1 | 6 | Cops7b | 0 | Young |
| 0.97352 | 0.141139 | 0.105 | 0.106 | 1 | 6 | Rif1 | -0.001 | Young |
| 0.973849 | 0.011047 | 0.097 | 0.097 | 1 | 6 | Fbxl6 | 0 | Young |
| 0.97442 | 0.077445 | 0.227 | 0.231 | 1 | 6 | Timm10b | -0.004 | Young |
| 0.974653 | 0.287928 | 0.199 | 0.197 | 1 | 6 | Mgat1 | 0.002 | Young |
| 0.975505 | 0.192876 | 0.07 | 0.071 | 1 | 6 | Ganc | -0.001 | Young |
| 0.975774 | 0.521018 | 0.048 | 0.049 | 1 | 6 | Galk1 | -0.001 | Young |
| 0.976005 | 0.331079 | 0.059 | 0.06 | 1 | 6 | Trappc2 | -0.001 | Young |
| 0.976055 | 0.183345 | 0.04 | 0.04 | 1 | 6 | Mmp19 | 0 | Young |
| 0.976171 | 0.477309 | 0.02 | 0.02 | 1 | 6 | Zfp1 | 0 | Young |
| 0.976869 | 0.041124 | 0.293 | 0.297 | 1 | 6 | Kcmf1 | -0.004 | Young |
| 0.977248 | 0.145557 | 0.453 | 0.471 | 1 | 6 | Ubald2 | -0.018 | Young |
| 0.977564 | 0.185384 | 0.105 | 0.106 | 1 | 6 | Ltn1 | -0.001 | Young |
| 0.977593 | 0.062891 | 0.174 | 0.174 | 1 | 6 | Ube2v2 | 0 | Young |
| 0.977836 | 0.01538 | 0.94 | 0.889 | 1 | 6 | Ctss | 0.051 | Young |
| 0.978289 | 0.000758 | 0.104 | 0.103 | 1 | 6 | Synj2 | 0.001 | Young |
| 0.978468 | 0.054204 | 0.254 | 0.254 | 1 | 6 | Esd | 0 | Young |
| 0.978569 | 0.108361 | 0.119 | 0.12 | 1 | 6 | Paip1 | -0.001 | Young |
| 0.978591 | 0.086925 | 0.074 | 0.074 | 1 | 6 | Cars2 | 0 | Young |
| 0.978983 | 0.098282 | 0.329 | 0.343 | 1 | 6 | Stx4a | -0.014 | Young |
| 0.979096 | 0.465682 | 0.05 | 0.051 | 1 | 6 | Dgka | -0.001 | Young |
| 0.979101 | 0.790959 | 0.025 | 0.026 | 1 | 6 | Slx1b | -0.001 | Young |
| 0.980093 | 0.031677 | 0.04 | 0.04 | 1 | 6 | Zfp335 | 0 | Young |
| 0.980228 | 0.213521 | 0.057 | 0.057 | 1 | 6 | Kif13b | 0 | Young |
| 0.980512 | 0.392406 | 0.048 | 0.049 | 1 | 6 | Dcun1d3 | -0.001 | Young |
| 0.980545 | 0.010989 | 0.278 | 0.283 | 1 | 6 | Rab8a | -0.005 | Young |
| 0.980692 | 0.100243 | 0.045 | 0.046 | 1 | 6 | Med9 | -0.001 | Young |
| 0.981204 | 0.207007 | 0.057 | 0.057 | 1 | 6 | Lmf2 | 0 | Young |
| 0.981285 | 0.113371 | 0.028 | 0.029 | 1 | 6 | 5730455P1 | -0.001 | Young |
| 0.981692 | 0.711011 | 0.037 | 0.037 | 1 | 6 | Klhl12 | 0 | Young |
| 0.982033 | 0.18492 | 0.232 | 0.237 | 1 | 6 | Rnf7 | -0.005 | Young |
| 0.982331 | 0.008324 | 0.403 | 0.406 | 1 | 6 | Lbr | -0.003 | Young |
| 0.982384 | 0.131045 | 0.176 | 0.174 | 1 | 6 | Chordc1 | 0.002 | Young |
| 0.982431 | 0.039026 | 0.164 | 0.169 | 1 | 6 | Qars | -0.005 | Young |
| 0.982632 | 0.574204 | 0.02 | 0.02 | 1 | 6 | Gdf15 | 0 | Young |
| 0.982915 | 0.334511 | 0.074 | 0.074 | 1 | 6 | Epm2aip1 | 0 | Young |
| 0.983093 | 0.100233 | 0.224 | 0.226 | 1 | 6 | Plp2 | -0.002 | Young |

| | | | | | | | | |
|---|---|---|---|---|---|---|---|---|
| 0.983602 | 0.06173 | 0.266 | 0.266 | 1 | 6 | Ptprs | 0 | Young |
| 0.984374 | 0.107572 | 0.258 | 0.266 | 1 | 6 | Cnih1 | -0.008 | Young |
| 0.984632 | 0.032409 | 0.473 | 0.48 | 1 | 6 | S100a13 | -0.007 | Young |
| 0.985411 | 0.03047 | 0.363 | 0.36 | 1 | 6 | Hdlbp | 0.003 | Young |
| 0.985459 | 0.450732 | 0.02 | 0.02 | 1 | 6 | Kremen1 | 0 | Young |
| 0.986085 | 0.285156 | 0.057 | 0.057 | 1 | 6 | Cryl1 | 0 | Young |
| 0.986335 | 0.024355 | 0.192 | 0.194 | 1 | 6 | Spg21 | -0.002 | Young |
| 0.986652 | 0.268847 | 0.062 | 0.063 | 1 | 6 | Trim30b | -0.001 | Young |
| 0.986671 | 0.340584 | 0.02 | 0.02 | 1 | 6 | Nme7 | 0 | Young |
| 0.986729 | 0.154264 | 0.028 | 0.029 | 1 | 6 | 2010320M | -0.001 | Young |
| 0.986811 | 0.168442 | 0.204 | 0.209 | 1 | 6 | Gmip | -0.005 | Young |
| 0.987521 | 0.543021 | 0.059 | 0.06 | 1 | 6 | Rufy1 | -0.001 | Young |
| 0.987558 | 0.08206 | 0.274 | 0.28 | 1 | 6 | Abhd12 | -0.006 | Young |
| 0.987927 | 0.124492 | 0.331 | 0.357 | 1 | 6 | Wdr83os | -0.026 | Young |
| 0.988771 | 0.245037 | 0.028 | 0.029 | 1 | 6 | Lin52 | -0.001 | Young |
| 0.988898 | 0.048465 | 0.309 | 0.317 | 1 | 6 | Cdk9 | -0.008 | Young |
| 0.989342 | 0.165587 | 0.094 | 0.094 | 1 | 6 | Polr2b | 0 | Young |
| 0.990046 | 0.040037 | 0.38 | 0.397 | 1 | 6 | Tbl1xr1 | -0.017 | Young |
| 0.990199 | 0.006704 | 0.087 | 0.089 | 1 | 6 | Ethe1 | -0.002 | Young |
| 0.990242 | 0.249161 | 0.06 | 0.06 | 1 | 6 | 0610037L1 | 0 | Young |
| 0.99039 | 0.280072 | 0.065 | 0.066 | 1 | 6 | Agap3 | -0.001 | Young |
| 0.990421 | 0.055962 | 0.219 | 0.226 | 1 | 6 | Cops4 | -0.007 | Young |
| 0.990481 | 0.620318 | 0.045 | 0.046 | 1 | 6 | Ampd2 | -0.001 | Young |
| 0.990953 | 0.117213 | 0.256 | 0.263 | 1 | 6 | Rad21 | -0.007 | Young |
| 0.991475 | 0.050324 | 0.144 | 0.143 | 1 | 6 | Cnot2 | 0.001 | Young |
| 0.991901 | 0.111818 | 0.11 | 0.111 | 1 | 6 | Secisbp2 | -0.001 | Young |
| 0.992488 | 0.156572 | 0.194 | 0.197 | 1 | 6 | Ganab | -0.003 | Young |
| 0.992667 | 0.136594 | 0.079 | 0.08 | 1 | 6 | Sap30l | -0.001 | Young |
| 0.992678 | 0.051348 | 0.065 | 0.066 | 1 | 6 | Tbcd | -0.001 | Young |
| 0.992859 | 0.011782 | 0.172 | 0.174 | 1 | 6 | Nfatc2 | -0.002 | Young |
| 0.993679 | 0.038595 | 0.048 | 0.049 | 1 | 6 | Pan2 | -0.001 | Young |
| 0.993941 | 0.073119 | 0.02 | 0.02 | 1 | 6 | Alkbh2 | 0 | Young |
| 0.994297 | 0.174387 | 0.037 | 0.037 | 1 | 6 | Pank4 | 0 | Young |
| 0.994897 | 0.253755 | 0.037 | 0.037 | 1 | 6 | Xrcc6 | 0 | Young |
| 0.995292 | 0.127472 | 0.164 | 0.166 | 1 | 6 | Pi4ka | -0.002 | Young |
| 0.995478 | 0.0339 | 0.06 | 0.06 | 1 | 6 | Klhdc10 | 0 | Young |
| 0.995891 | 0.207834 | 0.074 | 0.074 | 1 | 6 | Dym | 0 | Young |
| 0.996059 | 0.727859 | 0.042 | 0.043 | 1 | 6 | Fbxo42 | -0.001 | Young |
| 0.996313 | 0.295081 | 0.048 | 0.049 | 1 | 6 | Gm43848 | -0.001 | Young |
| 0.996338 | 0.010276 | 0.057 | 0.057 | 1 | 6 | Mfn1 | 0 | Young |
| 0.99649 | 0.216144 | 0.104 | 0.106 | 1 | 6 | Acot9 | -0.002 | Young |
| 0.996543 | 0.130764 | 0.206 | 0.211 | 1 | 6 | Ythdf1 | -0.005 | Young |
| 0.996899 | 0.000646 | 0.122 | 0.123 | 1 | 6 | Rfc2 | -0.001 | Young |
| 0.997008 | 0.411168 | 0.045 | 0.046 | 1 | 6 | Rpl9-ps6 | -0.001 | Young |
| 0.997299 | 0.269477 | 0.037 | 0.037 | 1 | 6 | Golm1 | 0 | Young |

| | | | | | | | | |
|---|---|---|---|---|---|---|---|---|
| 0.997879 | 0.458595 | 0.094 | 0.097 | 1 | 6 | Mrpl35 | -0.003 | Young |
| 0.998311 | 0.010219 | 0.161 | 0.16 | 1 | 6 | Trim35 | 0.001 | Young |
| 0.998664 | 0.040795 | 0.097 | 0.097 | 1 | 6 | Pdlim2 | 0 | Young |
| 0.998881 | 0.072752 | 0.069 | 0.069 | 1 | 6 | Wdr11 | 0 | Young |
| 0.998952 | 0.269005 | 0.079 | 0.08 | 1 | 6 | Stard3 | -0.001 | Young |
| 0.999116 | 0.199238 | 0.176 | 0.18 | 1 | 6 | Trpm7 | -0.004 | Young |
| 0.999728 | 0.276317 | 0.045 | 0.046 | 1 | 6 | Shb | -0.001 | Young |
| 0.999806 | 0.005563 | 0.094 | 0.094 | 1 | 6 | Zfp984 | 0 | Young |
| 1.81E-17 | 4.643376 | 0.251 | 0.027 | 5.63E-13 | 7 | Cxcl13 | 0.224 | Old |
| 2.90E-15 | 4.610491 | 0.192 | 0.009 | 9.02E-11 | 7 | Cd55 | 0.183 | Old |
| 1.63E-10 | 2.241837 | 0.259 | 0.081 | 5.06E-06 | 7 | C3 | 0.178 | Old |
| 1.91E-10 | 0.366225 | 0.982 | 0.97 | 5.93E-06 | 7 | Gpx1 | 0.012 | Old |
| 7.61E-10 | 4.657582 | 0.108 | 0 | 2.36E-05 | 7 | Colq | 0.108 | Old |
| 2.38E-08 | 1.82722 | 0.395 | 0.217 | 0.00074 | 7 | Ccl8 | 0.178 | Old |
| 5.81E-08 | 2.305973 | 0.359 | 0.205 | 0.001803 | 7 | Mmp9 | 0.154 | Old |
| 9.24E-08 | 0.453655 | 0.931 | 0.864 | 0.002869 | 7 | Ccl9 | 0.067 | Old |
| 1.62E-07 | 0.209803 | 1 | 1 | 0.005026 | 7 | mt-Co1 | 0 | Old |
| 4.40E-07 | 0.273563 | 0.987 | 0.97 | 0.013678 | 7 | mt-Cytb | 0.017 | Old |
| 6.03E-07 | 1.446267 | 0.362 | 0.208 | 0.018721 | 7 | Fabp5 | 0.154 | Old |
| 2.62E-06 | 1.828587 | 0.126 | 0.03 | 0.081253 | 7 | Actn1 | 0.096 | Old |
| 4.38E-06 | 6.292065 | 0.062 | 0 | 0.136029 | 7 | Ctsk | 0.062 | Old |
| 4.41E-06 | 0.619474 | 0.631 | 0.47 | 0.136987 | 7 | Cdk2ap2 | 0.161 | Old |
| 4.98E-06 | 0.573602 | 0.762 | 0.63 | 0.154693 | 7 | Lmna | 0.132 | Old |
| 5.29E-06 | 0.382786 | 0.864 | 0.789 | 0.164316 | 7 | Anxa2 | 0.075 | Old |
| 6.31E-06 | 3.491268 | 0.126 | 0.033 | 0.195822 | 7 | Acp5 | 0.093 | Old |
| 8.34E-06 | 0.412604 | 0.895 | 0.834 | 0.258968 | 7 | Atp5a1 | 0.061 | Old |
| 9.79E-06 | 0.253792 | 0.954 | 0.94 | 0.303924 | 7 | mt-Nd3 | 0.014 | Old |
| 1.13E-05 | 2.996772 | 0.056 | 0 | 0.350786 | 7 | Vldlr | 0.056 | Old |
| 1.36E-05 | 0.49894 | 0.81 | 0.693 | 0.421696 | 7 | Samhd1 | 0.117 | Old |
| 1.45E-05 | 0.183052 | 0.997 | 0.994 | 0.451087 | 7 | mt-Co3 | 0.003 | Old |
| 1.47E-05 | 2.348386 | 0.11 | 0.027 | 0.457091 | 7 | Nr1h3 | 0.083 | Old |
| 2.22E-05 | 1.206534 | 0.197 | 0.084 | 0.689423 | 7 | Usp39 | 0.113 | Old |
| 2.22E-05 | 2.01428 | 0.087 | 0.015 | 0.690195 | 7 | Igkc | 0.072 | Old |
| 2.36E-05 | 1.078889 | 0.369 | 0.223 | 0.733705 | 7 | Hist1h1c | 0.146 | Old |
| 2.68E-05 | 0.718989 | 0.626 | 0.497 | 0.832018 | 7 | Anxa1 | 0.129 | Old |
| 2.75E-05 | 1.765455 | 0.108 | 0.027 | 0.853073 | 7 | Gdf3 | 0.081 | Old |
| 2.75E-05 | 1.521205 | 0.238 | 0.12 | 0.855364 | 7 | Slfn5 | 0.118 | Old |
| 3.09E-05 | 2.812826 | 0.072 | 0.009 | 0.959555 | 7 | Mcoln2 | 0.063 | Old |
| 3.14E-05 | 0.53586 | 0.787 | 0.699 | 0.973729 | 7 | Lpl | 0.088 | Old |
| 3.45E-05 | 0.272172 | 0.954 | 0.925 | 1 | 7 | Vim | 0.029 | Old |
| 3.69E-05 | 0.385806 | 0.823 | 0.747 | 1 | 7 | Ndufa4 | 0.076 | Old |
| 4.08E-05 | 0.078449 | 1 | 1 | 1 | 7 | Gm42418 | 0 | Old |
| 4.08E-05 | 0.728788 | 0.608 | 0.467 | 1 | 7 | Atp6v0d1 | 0.141 | Old |
| 5.24E-05 | 2.154516 | 0.092 | 0.021 | 1 | 7 | Gpr68 | 0.071 | Old |
| 5.46E-05 | 0.369252 | 0.805 | 0.732 | 1 | 7 | Gda | 0.073 | Old |

| | | | | | | | | |
|---|---|---|---|---|---|---|---|---|
| 6.77E-05 | 0.735346 | 0.431 | 0.292 | 1 | 7 | Ppp4r2 | 0.139 | Old |
| 7.14E-05 | 1.752594 | 0.062 | 0.006 | 1 | 7 | B4galt7 | 0.056 | Old |
| 7.48E-05 | 4.271138 | 0.046 | 0 | 1 | 7 | Gpnmb | 0.046 | Old |
| 7.66E-05 | 0.761976 | 0.259 | 0.139 | 1 | 7 | Abcb7 | 0.12 | Old |
| 8.25E-05 | 1.096248 | 0.269 | 0.148 | 1 | 7 | Tsfm | 0.121 | Old |
| 8.49E-05 | 0.317062 | 0.962 | 0.961 | 1 | 7 | H2-K1 | 0.001 | Old |
| 8.67E-05 | 0.963963 | 0.4 | 0.259 | 1 | 7 | Lemd2 | 0.141 | Old |
| 8.68E-05 | 0.708746 | 0.303 | 0.169 | 1 | 7 | Lgals3bp | 0.134 | Old |
| 8.82E-05 | 1.33381 | 0.182 | 0.081 | 1 | 7 | Dusp10 | 0.101 | Old |
| 9.53E-05 | 1.006651 | 0.272 | 0.154 | 1 | 7 | Ugdh | 0.118 | Old |
| 9.98E-05 | 0.318821 | 0.544 | 0.38 | 1 | 7 | Zranb2 | 0.164 | Old |
| 0.000106 | 2.849378 | 0.059 | 0.006 | 1 | 7 | Dmpk | 0.053 | Old |
| 0.000107 | 1.879257 | 0.131 | 0.048 | 1 | 7 | Igf2r | 0.083 | Old |
| 0.000107 | 0.323191 | 0.851 | 0.783 | 1 | 7 | Ctsz | 0.068 | Old |
| 0.00011 | 0.590491 | 0.59 | 0.452 | 1 | 7 | Sdhd | 0.138 | Old |
| 0.00012 | 0.382997 | 0.815 | 0.741 | 1 | 7 | Prkcd | 0.074 | Old |
| 0.000168 | 0.163283 | 0.992 | 0.991 | 1 | 7 | mt-Atp6 | 0.001 | Old |
| 0.000169 | 0.47815 | 0.436 | 0.289 | 1 | 7 | Mvp | 0.147 | Old |
| 0.000171 | 0.809849 | 0.3 | 0.178 | 1 | 7 | Plod3 | 0.122 | Old |
| 0.000174 | 0.865867 | 0.428 | 0.292 | 1 | 7 | Ckap4 | 0.136 | Old |
| 0.000178 | 1.621617 | 0.169 | 0.078 | 1 | 7 | Dtx2 | 0.091 | Old |
| 0.000187 | 3.037745 | 0.049 | 0.003 | 1 | 7 | Spint2 | 0.046 | Old |
| 0.000193 | 3.752378 | 0.041 | 0 | 1 | 7 | Dclk1 | 0.041 | Old |
| 0.000215 | 1.708137 | 0.085 | 0.021 | 1 | 7 | Tgds | 0.064 | Old |
| 0.000238 | 1.468059 | 0.09 | 0.024 | 1 | 7 | Cdc42ep2 | 0.066 | Old |
| 0.000247 | 0.989152 | 0.195 | 0.096 | 1 | 7 | Pofut2 | 0.099 | Old |
| 0.000262 | 0.663011 | 0.369 | 0.247 | 1 | 7 | Agpat5 | 0.122 | Old |
| 0.000276 | 2.217628 | 0.077 | 0.018 | 1 | 7 | Myo10 | 0.059 | Old |
| 0.000293 | 0.532234 | 0.7 | 0.581 | 1 | 7 | H2-T23 | 0.119 | Old |
| 0.000296 | 0.863427 | 0.228 | 0.123 | 1 | 7 | Dph5 | 0.105 | Old |
| 0.0003 | 0.849517 | 0.231 | 0.123 | 1 | 7 | Myo1e | 0.108 | Old |
| 0.000307 | 0.210907 | 0.992 | 0.997 | 1 | 7 | Actg1 | -0.005 | Old |
| 0.000309 | 1.388533 | 0.077 | 0.018 | 1 | 7 | Adcy3 | 0.059 | Old |
| 0.000314 | 1.172287 | 0.174 | 0.084 | 1 | 7 | Caap1 | 0.09 | Old |
| 0.000325 | 0.601568 | 0.426 | 0.295 | 1 | 7 | Eif2a | 0.131 | Old |
| 0.000339 | 0.143158 | 1 | 0.997 | 1 | 7 | mt-Co2 | 0.003 | Old |
| 0.000349 | 0.448556 | 0.603 | 0.479 | 1 | 7 | Stxbp3 | 0.124 | Old |
| 0.00035 | 0.404549 | 0.592 | 0.455 | 1 | 7 | Atp6v1e1 | 0.137 | Old |
| 0.000375 | 1.796493 | 0.118 | 0.045 | 1 | 7 | Cd274 | 0.073 | Old |
| 0.000389 | 1.447805 | 0.11 | 0.039 | 1 | 7 | Snupn | 0.071 | Old |
| 0.000396 | 0.462111 | 0.651 | 0.53 | 1 | 7 | Apobec3 | 0.121 | Old |
| 0.000402 | 2.146061 | 0.064 | 0.012 | 1 | 7 | Mycl | 0.052 | Old |
| 0.000409 | 0.185422 | 0.995 | 0.991 | 1 | 7 | Psap | 0.004 | Old |
| 0.000412 | 0.636202 | 0.259 | 0.148 | 1 | 7 | Fkbp5 | 0.111 | Old |
| 0.000423 | 0.957311 | 0.146 | 0.063 | 1 | 7 | Dhx35 | 0.083 | Old |

| | | | | | | | | |
|---|---|---|---|---|---|---|---|---|
| 0.000427 | 0.450784 | 0.69 | 0.572 | 1 | 7 | Supt16 | 0.118 | Old |
| 0.000431 | 0.663578 | 0.292 | 0.178 | 1 | 7 | Rab20 | 0.114 | Old |
| 0.000438 | 1.072984 | 0.133 | 0.054 | 1 | 7 | Acad9 | 0.079 | Old |
| 0.000443 | 0.665484 | 0.333 | 0.22 | 1 | 7 | Tcf20 | 0.113 | Old |
| 0.00048 | 0.58266 | 0.521 | 0.383 | 1 | 7 | Sar1a | 0.138 | Old |
| 0.000493 | 0.871406 | 0.279 | 0.169 | 1 | 7 | Hccs | 0.11 | Old |
| 0.000544 | 0.367436 | 0.579 | 0.455 | 1 | 7 | Csk | 0.124 | Old |
| 0.000567 | 1.257208 | 0.115 | 0.045 | 1 | 7 | Fam78b | 0.07 | Old |
| 0.00058 | 0.339661 | 0.597 | 0.464 | 1 | 7 | Cnot1 | 0.133 | Old |
| 0.000591 | 0.253434 | 0.903 | 0.904 | 1 | 7 | Eef1b2 | -0.001 | Old |
| 0.000594 | 0.530586 | 0.272 | 0.166 | 1 | 7 | Arid2 | 0.106 | Old |
| 0.000647 | 0.567835 | 0.495 | 0.367 | 1 | 7 | Rnf19b | 0.128 | Old |
| 0.000661 | 0.615305 | 0.49 | 0.38 | 1 | 7 | Anxa4 | 0.11 | Old |
| 0.000686 | 0.965573 | 0.233 | 0.136 | 1 | 7 | Fnta | 0.097 | Old |
| 0.000704 | 0.37272 | 0.592 | 0.476 | 1 | 7 | Cnot6 | 0.116 | Old |
| 0.000721 | 1.71443 | 0.049 | 0.006 | 1 | 7 | Tubd1 | 0.043 | Old |
| 0.000735 | 0.310799 | 0.9 | 0.852 | 1 | 7 | Efhd2 | 0.048 | Old |
| 0.00074 | 0.737945 | 0.159 | 0.075 | 1 | 7 | Kpna2 | 0.084 | Old |
| 0.00079 | 0.334065 | 0.828 | 0.762 | 1 | 7 | Atp5f1 | 0.066 | Old |
| 0.000791 | 0.521298 | 0.477 | 0.349 | 1 | 7 | 2610001J0 | 0.128 | Old |
| 0.000823 | 0.412947 | 0.703 | 0.581 | 1 | 7 | Rab1a | 0.122 | Old |
| 0.00083 | 1.094843 | 0.172 | 0.087 | 1 | 7 | Ccnc | 0.085 | Old |
| 0.000837 | 0.182962 | 0.992 | 0.982 | 1 | 7 | mt-Nd4 | 0.01 | Old |
| 0.000843 | 0.59016 | 0.297 | 0.187 | 1 | 7 | Tmem128 | 0.11 | Old |
| 0.00088 | 0.487536 | 0.236 | 0.136 | 1 | 7 | Abhd17c | 0.1 | Old |
| 0.000895 | 0.474863 | 0.464 | 0.337 | 1 | 7 | R3hdm2 | 0.127 | Old |
| 0.000896 | 0.710752 | 0.179 | 0.09 | 1 | 7 | Mrpl37 | 0.089 | Old |
| 0.000899 | 1.157737 | 0.154 | 0.075 | 1 | 7 | Cnpy4 | 0.079 | Old |
| 0.000949 | 1.384322 | 0.059 | 0.012 | 1 | 7 | Src | 0.047 | Old |
| 0.000965 | 0.497143 | 0.567 | 0.452 | 1 | 7 | Eno1 | 0.115 | Old |
| 0.000986 | 1.258863 | 0.138 | 0.063 | 1 | 7 | Gnpnat1 | 0.075 | Old |
| 0.001011 | 0.222145 | 0.905 | 0.858 | 1 | 7 | Actr2 | 0.047 | Old |
| 0.001014 | 0.589393 | 0.374 | 0.262 | 1 | 7 | Snrpa1 | 0.112 | Old |
| 0.001015 | 2.06768 | 0.074 | 0.021 | 1 | 7 | C6 | 0.053 | Old |
| 0.001029 | 0.412683 | 0.685 | 0.59 | 1 | 7 | Psmd2 | 0.095 | Old |
| 0.001103 | 0.953724 | 0.144 | 0.066 | 1 | 7 | Pam | 0.078 | Old |
| 0.001107 | 0.418091 | 0.241 | 0.139 | 1 | 7 | Abhd16a | 0.102 | Old |
| 0.001134 | 2.66273 | 0.082 | 0.027 | 1 | 7 | Atp6v0d2 | 0.055 | Old |
| 0.001176 | 0.217116 | 0.908 | 0.883 | 1 | 7 | Hnrnpk | 0.025 | Old |
| 0.001197 | 0.513552 | 0.272 | 0.169 | 1 | 7 | Sestd1 | 0.103 | Old |
| 0.001231 | 0.620302 | 0.395 | 0.274 | 1 | 7 | B4galt5 | 0.121 | Old |
| 0.00126 | 0.770694 | 0.231 | 0.136 | 1 | 7 | Taf2 | 0.095 | Old |
| 0.001275 | 1.51995 | 0.118 | 0.051 | 1 | 7 | Gm41307 | 0.067 | Old |
| 0.001283 | 2.543094 | 0.031 | 0 | 1 | 7 | Htra1 | 0.031 | Old |
| 0.001283 | 2.400362 | 0.031 | 0 | 1 | 7 | Wdr25 | 0.031 | Old |

| | | | | | | | | |
|---|---|---|---|---|---|---|---|---|
| 0.001287 | 0.624327 | 0.492 | 0.386 | 1 | 7 | Tiam1 | 0.106 | Old |
| 0.001329 | 0.488009 | 0.305 | 0.19 | 1 | 7 | Acaa1a | 0.115 | Old |
| 0.00135 | 2.293101 | 0.051 | 0.009 | 1 | 7 | Ly6a | 0.042 | Old |
| 0.001356 | 0.384535 | 0.205 | 0.114 | 1 | 7 | Nup160 | 0.091 | Old |
| 0.001383 | 0.488678 | 0.331 | 0.22 | 1 | 7 | Abhd17b | 0.111 | Old |
| 0.001404 | 2.136952 | 0.056 | 0.012 | 1 | 7 | Sh3bgrl2 | 0.044 | Old |
| 0.001436 | 1.707504 | 0.056 | 0.012 | 1 | 7 | Gprc5b | 0.044 | Old |
| 0.001476 | 0.346011 | 0.603 | 0.494 | 1 | 7 | Tm9sf2 | 0.109 | Old |
| 0.001493 | 0.529333 | 0.379 | 0.271 | 1 | 7 | Hars | 0.108 | Old |
| 0.001562 | 0.480035 | 0.454 | 0.325 | 1 | 7 | Abce1 | 0.129 | Old |
| 0.001638 | 0.633566 | 0.251 | 0.157 | 1 | 7 | Dlg1 | 0.094 | Old |
| 0.001642 | 0.280055 | 0.818 | 0.783 | 1 | 7 | Vamp8 | 0.035 | Old |
| 0.001682 | 0.842237 | 0.179 | 0.096 | 1 | 7 | Snx25 | 0.083 | Old |
| 0.001686 | 0.584635 | 0.308 | 0.199 | 1 | 7 | Ccdc22 | 0.109 | Old |
| 0.001693 | 0.640385 | 0.131 | 0.06 | 1 | 7 | Fiz1 | 0.071 | Old |
| 0.001735 | 0.470239 | 0.254 | 0.16 | 1 | 7 | Rpe | 0.094 | Old |
| 0.001744 | 0.219401 | 0.992 | 0.973 | 1 | 7 | Crip1 | 0.019 | Old |
| 0.001747 | 0.786783 | 0.177 | 0.096 | 1 | 7 | Mief1 | 0.081 | Old |
| 0.001765 | 0.245592 | 0.71 | 0.593 | 1 | 7 | Plekho2 | 0.117 | Old |
| 0.001789 | 0.666624 | 0.062 | 0.015 | 1 | 7 | Dennd2a | 0.047 | Old |
| 0.001807 | 0.62206 | 0.331 | 0.223 | 1 | 7 | Tyk2 | 0.108 | Old |
| 0.00181 | 1.536365 | 0.195 | 0.117 | 1 | 7 | Cavin1 | 0.078 | Old |
| 0.001825 | 0.202683 | 0.362 | 0.235 | 1 | 7 | C1d | 0.127 | Old |
| 0.001851 | 0.505663 | 0.249 | 0.154 | 1 | 7 | Gsap | 0.095 | Old |
| 0.001852 | 0.327411 | 0.374 | 0.25 | 1 | 7 | Uchl5 | 0.124 | Old |
| 0.001882 | 0.485219 | 0.697 | 0.59 | 1 | 7 | Tuba1c | 0.107 | Old |
| 0.001898 | 0.180188 | 0.987 | 0.994 | 1 | 7 | mt-Nd1 | -0.007 | Old |
| 0.001911 | 0.553839 | 0.338 | 0.235 | 1 | 7 | Drg1 | 0.103 | Old |
| 0.001933 | 0.185074 | 0.977 | 0.97 | 1 | 7 | mt-Nd2 | 0.007 | Old |
| 0.001948 | 0.427967 | 0.687 | 0.62 | 1 | 7 | Cstb | 0.067 | Old |
| 0.001953 | 0.400259 | 0.651 | 0.584 | 1 | 7 | Ccr5 | 0.067 | Old |
| 0.001954 | 0.355103 | 0.59 | 0.458 | 1 | 7 | Cab39 | 0.132 | Old |
| 0.00199 | 0.297949 | 0.767 | 0.672 | 1 | 7 | M6pr | 0.095 | Old |
| 0.002016 | 1.048994 | 0.049 | 0.009 | 1 | 7 | F7 | 0.04 | Old |
| 0.002019 | 2.995786 | 0.036 | 0.003 | 1 | 7 | Ttn | 0.033 | Old |
| 0.002035 | 0.448259 | 0.179 | 0.096 | 1 | 7 | Nubp2 | 0.083 | Old |
| 0.002052 | 0.740053 | 0.118 | 0.051 | 1 | 7 | Fbxl15 | 0.067 | Old |
| 0.002056 | 2.580311 | 0.054 | 0.012 | 1 | 7 | Sema6a | 0.042 | Old |
| 0.002059 | 2.245253 | 0.036 | 0.003 | 1 | 7 | Sh2d4a | 0.033 | Old |
| 0.002066 | 2.298648 | 0.028 | 0 | 1 | 7 | Serpinb6b | 0.028 | Old |
| 0.002075 | 0.57573 | 0.392 | 0.28 | 1 | 7 | Plin3 | 0.112 | Old |
| 0.002086 | 0.201473 | 0.344 | 0.238 | 1 | 7 | Il1rn | 0.106 | Old |
| 0.002105 | 0.483572 | 0.603 | 0.497 | 1 | 7 | Trim25 | 0.106 | Old |
| 0.002169 | 0.192425 | 0.967 | 0.955 | 1 | 7 | Pfn1 | 0.012 | Old |
| 0.002243 | 0.090065 | 0.995 | 0.997 | 1 | 7 | Ptma | -0.002 | Old |

| | | | | | | | | |
|---|---|---|---|---|---|---|---|---|
| 0.002368 | 0.931747 | 0.21 | 0.123 | 1 | 7 | Slc25a17 | 0.087 | Old |
| 0.002379 | 0.300181 | 0.744 | 0.669 | 1 | 7 | Far1 | 0.075 | Old |
| 0.002388 | 0.936307 | 0.079 | 0.027 | 1 | 7 | Nudt22 | 0.052 | Old |
| 0.002419 | 0.385613 | 0.751 | 0.693 | 1 | 7 | Ifi27l2a | 0.058 | Old |
| 0.002462 | 0.771143 | 0.233 | 0.142 | 1 | 7 | Wsb2 | 0.091 | Old |
| 0.00248 | 0.627907 | 0.185 | 0.102 | 1 | 7 | Tnip1 | 0.083 | Old |
| 0.002481 | 0.447338 | 0.413 | 0.292 | 1 | 7 | Tbc1d10b | 0.121 | Old |
| 0.002493 | 0.439644 | 0.595 | 0.506 | 1 | 7 | Stub1 | 0.089 | Old |
| 0.002504 | 0.394112 | 0.385 | 0.271 | 1 | 7 | Trafd1 | 0.114 | Old |
| 0.002576 | 0.821224 | 0.2 | 0.12 | 1 | 7 | Ndufaf4 | 0.08 | Old |
| 0.002597 | 0.705464 | 0.151 | 0.078 | 1 | 7 | Tmem2 | 0.073 | Old |
| 0.002606 | 0.362624 | 0.592 | 0.479 | 1 | 7 | Cct3 | 0.113 | Old |
| 0.002628 | 0.991946 | 0.069 | 0.021 | 1 | 7 | Uaca | 0.048 | Old |
| 0.002629 | 0.405844 | 0.728 | 0.642 | 1 | 7 | Plin2 | 0.086 | Old |
| 0.00263 | 0.34536 | 0.662 | 0.545 | 1 | 7 | Zfp207 | 0.117 | Old |
| 0.002708 | 0.318543 | 0.777 | 0.723 | 1 | 7 | Cops9 | 0.054 | Old |
| 0.002722 | 0.28147 | 0.831 | 0.783 | 1 | 7 | 2010107E0 | 0.048 | Old |
| 0.002722 | 0.178179 | 0.967 | 0.928 | 1 | 7 | Ahnak | 0.039 | Old |
| 0.00273 | 1.7271 | 0.041 | 0.006 | 1 | 7 | Yars2 | 0.035 | Old |
| 0.00279 | 0.62654 | 0.246 | 0.157 | 1 | 7 | Cul2 | 0.089 | Old |
| 0.002815 | 0.382889 | 0.582 | 0.476 | 1 | 7 | Mdh2 | 0.106 | Old |
| 0.002883 | 0.224907 | 0.951 | 0.934 | 1 | 7 | H2-D1 | 0.017 | Old |
| 0.002991 | 0.667574 | 0.351 | 0.25 | 1 | 7 | Rcn2 | 0.101 | Old |
| 0.003001 | 0.508725 | 0.192 | 0.108 | 1 | 7 | Ranbp10 | 0.084 | Old |
| 0.003008 | 0.289535 | 0.821 | 0.801 | 1 | 7 | Cfp | 0.02 | Old |
| 0.003093 | 0.596328 | 0.262 | 0.169 | 1 | 7 | Dpf2 | 0.093 | Old |
| 0.003131 | 0.317488 | 0.828 | 0.801 | 1 | 7 | Cox6a1 | 0.027 | Old |
| 0.003145 | 0.579704 | 0.305 | 0.205 | 1 | 7 | Ulbp1 | 0.1 | Old |
| 0.003165 | 0.615378 | 0.264 | 0.169 | 1 | 7 | Ubap2 | 0.095 | Old |
| 0.003176 | 0.442651 | 0.487 | 0.377 | 1 | 7 | Bhlhe40 | 0.11 | Old |
| 0.00319 | 0.4296 | 0.687 | 0.596 | 1 | 7 | Selplg | 0.091 | Old |
| 0.003203 | 0.78616 | 0.1 | 0.042 | 1 | 7 | Slc37a4 | 0.058 | Old |
| 0.003205 | 0.99524 | 0.082 | 0.03 | 1 | 7 | Abcb10 | 0.052 | Old |
| 0.003205 | 0.438825 | 0.254 | 0.16 | 1 | 7 | Zpr1 | 0.094 | Old |
| 0.00325 | 0.40751 | 0.331 | 0.223 | 1 | 7 | Derl2 | 0.108 | Old |
| 0.003254 | 0.512509 | 0.313 | 0.217 | 1 | 7 | Babam1 | 0.096 | Old |
| 0.003272 | 0.72158 | 0.272 | 0.184 | 1 | 7 | Evi2 | 0.088 | Old |
| 0.003298 | 0.879631 | 0.154 | 0.081 | 1 | 7 | Cfap97 | 0.073 | Old |
| 0.003313 | 0.923133 | 0.213 | 0.133 | 1 | 7 | Dnajc11 | 0.08 | Old |
| 0.003314 | 0.390248 | 0.582 | 0.47 | 1 | 7 | Spcs2 | 0.112 | Old |
| 0.003335 | 1.717806 | 0.026 | 0 | 1 | 7 | Fkbpl | 0.026 | Old |
| 0.003399 | 0.340793 | 0.449 | 0.331 | 1 | 7 | 6-Mar | 0.118 | Old |
| 0.003419 | 0.492626 | 0.718 | 0.639 | 1 | 7 | Klf4 | 0.079 | Old |
| 0.003435 | 1.625389 | 0.062 | 0.018 | 1 | 7 | Klhl7 | 0.044 | Old |
| 0.00345 | 0.576029 | 0.19 | 0.108 | 1 | 7 | Ifi47 | 0.082 | Old |

| | | | | | | | | |
|---|---|---|---|---|---|---|---|---|
| 0.003528 | 0.744021 | 0.287 | 0.199 | 1 | 7 | Pira2 | 0.088 | Old |
| 0.003531 | 0.217364 | 0.287 | 0.184 | 1 | 7 | Pfdn4 | 0.103 | Old |
| 0.003543 | 1.572169 | 0.067 | 0.021 | 1 | 7 | Cipc | 0.046 | Old |
| 0.003643 | 0.28619 | 0.726 | 0.645 | 1 | 7 | Prrc2c | 0.081 | Old |
| 0.003655 | 0.650155 | 0.341 | 0.241 | 1 | 7 | 2310035C2 | 0.1 | Old |
| 0.003668 | 0.677795 | 0.344 | 0.247 | 1 | 7 | Tomm34 | 0.097 | Old |
| 0.003725 | 1.506014 | 0.085 | 0.033 | 1 | 7 | Adprm | 0.052 | Old |
| 0.003974 | 0.716146 | 0.29 | 0.199 | 1 | 7 | Sec13 | 0.091 | Old |
| 0.003999 | 0.299122 | 0.405 | 0.289 | 1 | 7 | Abcc1 | 0.116 | Old |
| 0.004016 | 2.082476 | 0.038 | 0.006 | 1 | 7 | Med18 | 0.032 | Old |
| 0.004031 | 1.341171 | 0.067 | 0.021 | 1 | 7 | Ccdc18 | 0.046 | Old |
| 0.004033 | 0.485824 | 0.426 | 0.322 | 1 | 7 | Acsl5 | 0.104 | Old |
| 0.004069 | 1.847531 | 0.038 | 0.006 | 1 | 7 | St18 | 0.032 | Old |
| 0.004096 | 0.439485 | 0.251 | 0.163 | 1 | 7 | Klhl28 | 0.088 | Old |
| 0.004097 | 0.58742 | 0.395 | 0.292 | 1 | 7 | Slc25a39 | 0.103 | Old |
| 0.004134 | 0.248575 | 0.241 | 0.145 | 1 | 7 | Bop1 | 0.096 | Old |
| 0.004139 | 2.241022 | 0.038 | 0.006 | 1 | 7 | AW146154 | 0.032 | Old |
| 0.004175 | 2.018331 | 0.038 | 0.006 | 1 | 7 | Epop | 0.032 | Old |
| 0.004193 | 1.816407 | 0.038 | 0.006 | 1 | 7 | Zfp456 | 0.032 | Old |
| 0.004218 | 0.697286 | 0.346 | 0.25 | 1 | 7 | Vti1b | 0.096 | Old |
| 0.004392 | 0.297412 | 0.767 | 0.675 | 1 | 7 | Ctsd | 0.092 | Old |
| 0.004402 | 0.576254 | 0.426 | 0.334 | 1 | 7 | Cyld | 0.092 | Old |
| 0.004434 | 0.740991 | 0.228 | 0.145 | 1 | 7 | Minpp1 | 0.083 | Old |
| 0.004497 | 0.321723 | 0.277 | 0.187 | 1 | 7 | Pqlc3 | 0.09 | Old |
| 0.004598 | 0.451618 | 0.508 | 0.416 | 1 | 7 | Ldlr | 0.092 | Old |
| 0.004658 | 0.69119 | 0.174 | 0.099 | 1 | 7 | Zfp608 | 0.075 | Old |
| 0.00474 | 0.322704 | 0.303 | 0.202 | 1 | 7 | Arl6ip4 | 0.101 | Old |
| 0.004744 | 0.581454 | 0.141 | 0.075 | 1 | 7 | Mrps18b | 0.066 | Old |
| 0.004748 | 0.86558 | 0.1 | 0.045 | 1 | 7 | Tmem184c | 0.055 | Old |
| 0.004755 | 1.038885 | 0.069 | 0.024 | 1 | 7 | Aspa | 0.045 | Old |
| 0.004758 | 0.848486 | 0.074 | 0.027 | 1 | 7 | Rusc2 | 0.047 | Old |
| 0.004787 | 2.762672 | 0.044 | 0.009 | 1 | 7 | Plac8 | 0.035 | Old |
| 0.004817 | 0.68753 | 0.41 | 0.322 | 1 | 7 | Adam8 | 0.088 | Old |
| 0.004855 | 0.224498 | 0.979 | 0.961 | 1 | 7 | B2m | 0.018 | Old |
| 0.004877 | 0.76824 | 0.195 | 0.117 | 1 | 7 | Gpr137b | 0.078 | Old |
| 0.004921 | 1.321246 | 0.044 | 0.009 | 1 | 7 | Zbtb26 | 0.035 | Old |
| 0.004923 | 1.02018 | 0.105 | 0.048 | 1 | 7 | Sigirr | 0.057 | Old |
| 0.00496 | 0.750348 | 0.138 | 0.072 | 1 | 7 | Hpse | 0.066 | Old |
| 0.004985 | 0.7687 | 0.256 | 0.169 | 1 | 7 | 2310022A1 | 0.087 | Old |
| 0.004987 | 0.873545 | 0.238 | 0.157 | 1 | 7 | Iars2 | 0.081 | Old |
| 0.005008 | 0.289286 | 0.508 | 0.404 | 1 | 7 | Myo9b | 0.104 | Old |
| 0.005058 | 0.437481 | 0.146 | 0.078 | 1 | 7 | Dpep2 | 0.068 | Old |
| 0.005099 | 1.797649 | 0.054 | 0.015 | 1 | 7 | Cyp2j6 | 0.039 | Old |
| 0.005134 | 0.516797 | 0.397 | 0.301 | 1 | 7 | Spc24 | 0.096 | Old |
| 0.005175 | 0.376236 | 0.618 | 0.536 | 1 | 7 | Cct2 | 0.082 | Old |

| | | | | | | | | |
|---|---|---|---|---|---|---|---|---|
| 0.005181 | 1.795152 | 0.049 | 0.012 | 1 | 7 | Gm17745 | 0.037 | Old |
| 0.00519 | 0.66964 | 0.249 | 0.163 | 1 | 7 | Snhg6 | 0.086 | Old |
| 0.005216 | 1.049006 | 0.069 | 0.024 | 1 | 7 | Tarsl2 | 0.045 | Old |
| 0.00523 | 0.155314 | 0.962 | 0.97 | 1 | 7 | Hsp90b1 | -0.008 | Old |
| 0.005296 | 0.145421 | 0.141 | 0.075 | 1 | 7 | Mlh1 | 0.066 | Old |
| 0.005331 | 1.314722 | 0.054 | 0.015 | 1 | 7 | Atp23 | 0.039 | Old |
| 0.005395 | 1.753449 | 0.023 | 0 | 1 | 7 | Adgra3 | 0.023 | Old |
| 0.005395 | 1.67695 | 0.023 | 0 | 1 | 7 | Cpeb1 | 0.023 | Old |
| 0.005555 | 0.7763 | 0.172 | 0.099 | 1 | 7 | Ppa2 | 0.073 | Old |
| 0.005557 | 1.050548 | 0.213 | 0.136 | 1 | 7 | Naip2 | 0.077 | Old |
| 0.005565 | 0.430515 | 0.533 | 0.434 | 1 | 7 | Nadk | 0.099 | Old |
| 0.005613 | 0.564626 | 0.287 | 0.193 | 1 | 7 | Cops6 | 0.094 | Old |
| 0.005624 | 0.451478 | 0.505 | 0.401 | 1 | 7 | Rwdd1 | 0.104 | Old |
| 0.005627 | 0.482674 | 0.528 | 0.437 | 1 | 7 | Mapre2 | 0.091 | Old |
| 0.005657 | 0.726762 | 0.113 | 0.054 | 1 | 7 | Smyd5 | 0.059 | Old |
| 0.00566 | 0.801289 | 0.105 | 0.048 | 1 | 7 | Urgcp | 0.057 | Old |
| 0.005673 | 0.696827 | 0.287 | 0.202 | 1 | 7 | Bub1 | 0.085 | Old |
| 0.005676 | 0.773845 | 0.087 | 0.036 | 1 | 7 | Tfec | 0.051 | Old |
| 0.005721 | 0.342278 | 0.774 | 0.705 | 1 | 7 | Colgalt1 | 0.069 | Old |
| 0.005743 | 1.293206 | 0.136 | 0.075 | 1 | 7 | Ppargc1b | 0.061 | Old |
| 0.00575 | 1.973905 | 0.09 | 0.039 | 1 | 7 | Rbm45 | 0.051 | Old |
| 0.005825 | 0.884147 | 0.054 | 0.015 | 1 | 7 | Akr1c12 | 0.039 | Old |
| 0.005825 | 0.548004 | 0.054 | 0.015 | 1 | 7 | Zmat1 | 0.039 | Old |
| 0.005847 | 0.288191 | 0.395 | 0.286 | 1 | 7 | Il2rg | 0.109 | Old |
| 0.005902 | 0.694144 | 0.174 | 0.102 | 1 | 7 | Rnf103 | 0.072 | Old |
| 0.005944 | 0.492095 | 0.385 | 0.286 | 1 | 7 | Mfhas1 | 0.099 | Old |
| 0.006037 | 0.059852 | 0.241 | 0.151 | 1 | 7 | Nsrp1 | 0.09 | Old |
| 0.00607 | 0.612891 | 0.197 | 0.123 | 1 | 7 | Grk5 | 0.074 | Old |
| 0.006102 | 0.312477 | 0.323 | 0.226 | 1 | 7 | Ddx18 | 0.097 | Old |
| 0.00618 | 0.265236 | 0.574 | 0.458 | 1 | 7 | Tmem123 | 0.116 | Old |
| 0.006246 | 0.866067 | 0.121 | 0.06 | 1 | 7 | Primpol | 0.061 | Old |
| 0.006246 | 0.559304 | 0.21 | 0.13 | 1 | 7 | Ttc13 | 0.08 | Old |
| 0.006259 | 1.037554 | 0.103 | 0.048 | 1 | 7 | Nup35 | 0.055 | Old |
| 0.006341 | 0.561224 | 0.344 | 0.256 | 1 | 7 | Sec24b | 0.088 | Old |
| 0.006365 | 0.279514 | 0.333 | 0.241 | 1 | 7 | Ndufs1 | 0.092 | Old |
| 0.006457 | 0.311452 | 0.377 | 0.274 | 1 | 7 | Tpm1 | 0.103 | Old |
| 0.006472 | 0.746958 | 0.203 | 0.127 | 1 | 7 | Ago1 | 0.076 | Old |
| 0.006486 | 0.314074 | 0.623 | 0.521 | 1 | 7 | Ddost | 0.102 | Old |
| 0.006509 | 1.128408 | 0.159 | 0.093 | 1 | 7 | Rnf121 | 0.066 | Old |
| 0.006614 | 0.376589 | 0.392 | 0.301 | 1 | 7 | Adssl1 | 0.091 | Old |
| 0.006625 | 0.674776 | 0.169 | 0.099 | 1 | 7 | Pex16 | 0.07 | Old |
| 0.006643 | 0.388059 | 0.456 | 0.352 | 1 | 7 | Slc25a24 | 0.104 | Old |
| 0.00666 | 0.487467 | 0.133 | 0.069 | 1 | 7 | Zc3hc1 | 0.064 | Old |
| 0.006686 | 0.128867 | 0.928 | 0.925 | 1 | 7 | Tuba1b | 0.003 | Old |
| 0.006693 | 0.264032 | 0.685 | 0.572 | 1 | 7 | Vdac2 | 0.113 | Old |

| | | | | | | | | |
|---|---|---|---|---|---|---|---|---|
| 0.006698 | 0.727792 | 0.159 | 0.09 | 1 | 7 | Spred2 | 0.069 | Old |
| 0.006712 | 0.513236 | 0.308 | 0.214 | 1 | 7 | Ubqln2 | 0.094 | Old |
| 0.006745 | 0.482029 | 0.538 | 0.452 | 1 | 7 | Arhgap9 | 0.086 | Old |
| 0.006831 | 0.342538 | 0.036 | 0.006 | 1 | 7 | Mief2 | 0.03 | Old |
| 0.006845 | 0.376714 | 0.641 | 0.551 | 1 | 7 | Usmg5 | 0.09 | Old |
| 0.007025 | 0.718896 | 0.2 | 0.123 | 1 | 7 | Brwd3 | 0.077 | Old |
| 0.007071 | 0.354101 | 0.579 | 0.452 | 1 | 7 | Efr3a | 0.127 | Old |
| 0.007087 | 0.185454 | 0.841 | 0.795 | 1 | 7 | mt-Nd4l | 0.046 | Old |
| 0.007089 | 0.96491 | 0.11 | 0.054 | 1 | 7 | 3110040N1 | 0.056 | Old |
| 0.007199 | 0.330447 | 0.577 | 0.464 | 1 | 7 | Mtch1 | 0.113 | Old |
| 0.007245 | 0.910013 | 0.095 | 0.042 | 1 | 7 | Gpx3 | 0.053 | Old |
| 0.007259 | 0.693871 | 0.187 | 0.114 | 1 | 7 | Arsa | 0.073 | Old |
| 0.007334 | 0.620303 | 0.267 | 0.181 | 1 | 7 | Vps37a | 0.086 | Old |
| 0.007351 | 0.551063 | 0.323 | 0.229 | 1 | 7 | Ube2v2 | 0.094 | Old |
| 0.007355 | 0.364777 | 0.526 | 0.434 | 1 | 7 | Ubtf | 0.092 | Old |
| 0.007369 | 0.401339 | 0.741 | 0.669 | 1 | 7 | Klf2 | 0.072 | Old |
| 0.007415 | 0.281496 | 0.659 | 0.551 | 1 | 7 | Hif1a | 0.108 | Old |
| 0.007422 | 1.018191 | 0.187 | 0.114 | 1 | 7 | Urm1 | 0.073 | Old |
| 0.00753 | 0.229733 | 0.938 | 0.886 | 1 | 7 | Anxa5 | 0.052 | Old |
| 0.007548 | 1.698249 | 0.051 | 0.015 | 1 | 7 | Tasp1 | 0.036 | Old |
| 0.007609 | 0.184365 | 0.851 | 0.81 | 1 | 7 | Ywhah | 0.041 | Old |
| 0.00764 | 0.890379 | 0.223 | 0.145 | 1 | 7 | Stx11 | 0.078 | Old |
| 0.007654 | 0.36863 | 0.238 | 0.154 | 1 | 7 | Mfsd12 | 0.084 | Old |
| 0.007677 | 0.720348 | 0.167 | 0.099 | 1 | 7 | Znrf1 | 0.068 | Old |
| 0.00769 | 0.272479 | 0.262 | 0.172 | 1 | 7 | Tbc1d2b | 0.09 | Old |
| 0.007697 | 0.624833 | 0.31 | 0.22 | 1 | 7 | Cdc26 | 0.09 | Old |
| 0.007703 | 0.644915 | 0.282 | 0.193 | 1 | 7 | Rps27rt | 0.089 | Old |
| 0.007709 | 0.226131 | 0.6 | 0.494 | 1 | 7 | Lcp2 | 0.106 | Old |
| 0.00771 | 0.628945 | 0.159 | 0.09 | 1 | 7 | Rhot2 | 0.069 | Old |
| 0.007742 | 0.380945 | 0.503 | 0.395 | 1 | 7 | Trim30a | 0.108 | Old |
| 0.007812 | 0.651919 | 0.097 | 0.045 | 1 | 7 | Spg11 | 0.052 | Old |
| 0.007887 | 0.289274 | 0.351 | 0.253 | 1 | 7 | Frrs1 | 0.098 | Old |
| 0.007984 | 0.34259 | 0.418 | 0.319 | 1 | 7 | AC160336. | 0.099 | Old |
| 0.008023 | 0.880636 | 0.113 | 0.057 | 1 | 7 | Tpcn2 | 0.056 | Old |
| 0.008042 | 0.300235 | 0.597 | 0.512 | 1 | 7 | Slc39a1 | 0.085 | Old |
| 0.00816 | 0.881113 | 0.056 | 0.018 | 1 | 7 | Rnf123 | 0.038 | Old |
| 0.008167 | 0.189475 | 0.969 | 0.934 | 1 | 7 | Myl6 | 0.035 | Old |
| 0.008167 | 2.383894 | 0.028 | 0.003 | 1 | 7 | Bmpr1a | 0.025 | Old |
| 0.008176 | 0.365922 | 0.251 | 0.166 | 1 | 7 | Dcps | 0.085 | Old |
| 0.008207 | 0.501697 | 0.223 | 0.148 | 1 | 7 | Fam43a | 0.075 | Old |
| 0.008213 | 0.976357 | 0.136 | 0.075 | 1 | 7 | Ttl | 0.061 | Old |
| 0.008236 | 0.979895 | 0.1 | 0.048 | 1 | 7 | Noc3l | 0.052 | Old |
| 0.008266 | 0.028907 | 0.351 | 0.244 | 1 | 7 | Ss18 | 0.107 | Old |
| 0.008281 | 0.745213 | 0.162 | 0.09 | 1 | 7 | Heatr6 | 0.072 | Old |
| 0.008325 | 0.671091 | 0.195 | 0.12 | 1 | 7 | Ahsa2 | 0.075 | Old |

| | | | | | | | | |
|---|---|---|---|---|---|---|---|---|
| 0.008393 | 0.97314 | 0.128 | 0.069 | 1 | 7 | Sertad3 | 0.059 | Old |
| 0.008396 | 0.880163 | 0.318 | 0.238 | 1 | 7 | Timm50 | 0.08 | Old |
| 0.008424 | 0.963149 | 0.062 | 0.021 | 1 | 7 | Bahcc1 | 0.041 | Old |
| 0.008444 | 1.827925 | 0.028 | 0.003 | 1 | 7 | D130040H: | 0.025 | Old |
| 0.008541 | 0.372147 | 0.159 | 0.093 | 1 | 7 | Rrp12 | 0.066 | Old |
| 0.008572 | 0.814398 | 0.051 | 0.015 | 1 | 7 | Tefm | 0.036 | Old |
| 0.008686 | 0.325678 | 0.446 | 0.346 | 1 | 7 | Tsc22d1 | 0.1 | Old |
| 0.008729 | 0.265684 | 0.718 | 0.608 | 1 | 7 | Slk | 0.11 | Old |
| 0.008747 | 0.217647 | 0.413 | 0.313 | 1 | 7 | Cpne3 | 0.1 | Old |
| 0.008757 | 0.518442 | 0.213 | 0.136 | 1 | 7 | P2ry10b | 0.077 | Old |
| 0.008758 | 2.402322 | 0.021 | 0 | 1 | 7 | Klra3 | 0.021 | Old |
| 0.008758 | 2.343068 | 0.021 | 0 | 1 | 7 | Gm15675 | 0.021 | Old |
| 0.008758 | 1.724075 | 0.021 | 0 | 1 | 7 | 2200002D( | 0.021 | Old |
| 0.008758 | 1.686788 | 0.021 | 0 | 1 | 7 | Anxa9 | 0.021 | Old |
| 0.008794 | 0.496232 | 0.313 | 0.22 | 1 | 7 | Sfxn1 | 0.093 | Old |
| 0.008797 | 0.263368 | 0.39 | 0.286 | 1 | 7 | Crybg1 | 0.104 | Old |
| 0.008801 | 0.541993 | 0.197 | 0.123 | 1 | 7 | Selenon | 0.074 | Old |
| 0.008876 | 0.283441 | 0.633 | 0.539 | 1 | 7 | Prdx2 | 0.094 | Old |
| 0.008978 | 0.579196 | 0.541 | 0.494 | 1 | 7 | Cd63 | 0.047 | Old |
| 0.009018 | 1.024853 | 0.095 | 0.045 | 1 | 7 | Stk39 | 0.05 | Old |
| 0.009095 | 0.290003 | 0.505 | 0.389 | 1 | 7 | Ccna2 | 0.116 | Old |
| 0.009102 | 0.142829 | 0.882 | 0.877 | 1 | 7 | Slc25a3 | 0.005 | Old |
| 0.009109 | 0.25067 | 0.136 | 0.072 | 1 | 7 | Fam53a | 0.064 | Old |
| 0.009194 | 0.234697 | 0.836 | 0.801 | 1 | 7 | Atp5g3 | 0.035 | Old |
| 0.00936 | 0.348612 | 0.423 | 0.328 | 1 | 7 | Tmem168 | 0.095 | Old |
| 0.009476 | 0.667523 | 0.323 | 0.238 | 1 | 7 | Parl | 0.085 | Old |
| 0.009524 | 0.566038 | 0.246 | 0.163 | 1 | 7 | Pabpc4 | 0.083 | Old |
| 0.00955 | 0.571115 | 0.231 | 0.154 | 1 | 7 | Thoc3 | 0.077 | Old |
| 0.009552 | 0.412097 | 0.503 | 0.395 | 1 | 7 | Psmb10 | 0.108 | Old |
| 0.009555 | 1.364264 | 0.069 | 0.027 | 1 | 7 | Gstt2 | 0.042 | Old |
| 0.009557 | 0.551096 | 0.354 | 0.262 | 1 | 7 | Dr1 | 0.092 | Old |
| 0.009699 | 0.622426 | 0.203 | 0.127 | 1 | 7 | Acot9 | 0.076 | Old |
| 0.0097 | 0.538358 | 0.374 | 0.28 | 1 | 7 | Nudcd2 | 0.094 | Old |
| 0.009759 | 0.146466 | 0.9 | 0.834 | 1 | 7 | Arpc5 | 0.066 | Old |
| 0.009764 | 1.333906 | 0.082 | 0.036 | 1 | 7 | Klhl12 | 0.046 | Old |
| 0.009774 | 0.693185 | 0.303 | 0.211 | 1 | 7 | Nup93 | 0.092 | Old |
| 0.009797 | 1.603949 | 0.077 | 0.033 | 1 | 7 | Rad9a | 0.044 | Old |
| 0.009821 | 0.143591 | 0.169 | 0.096 | 1 | 7 | Trmt2b | 0.073 | Old |
| 0.01002 | 0.507408 | 0.436 | 0.352 | 1 | 7 | Mrpl12 | 0.084 | Old |
| 0.010093 | 0.498137 | 0.354 | 0.268 | 1 | 7 | Chmp2b | 0.086 | Old |
| 0.010106 | 1.353621 | 0.059 | 0.021 | 1 | 7 | Mcrip2 | 0.038 | Old |
| 0.010124 | 0.393547 | 0.544 | 0.458 | 1 | 7 | Parvb | 0.086 | Old |
| 0.010176 | 0.302363 | 0.446 | 0.355 | 1 | 7 | Fam104a | 0.091 | Old |
| 0.010219 | 0.80974 | 0.087 | 0.039 | 1 | 7 | Riox1 | 0.048 | Old |
| 0.010245 | 0.419962 | 0.513 | 0.446 | 1 | 7 | Ear2 | 0.067 | Old |

| | | | | | | | | |
|---|---|---|---|---|---|---|---|---|
| 0.010283 | 0.569735 | 0.285 | 0.202 | 1 | 7 | Phf23 | 0.083 | Old |
| 0.010305 | 0.870221 | 0.115 | 0.06 | 1 | 7 | Acot11 | 0.055 | Old |
| 0.010376 | 0.21747 | 0.374 | 0.271 | 1 | 7 | Rab18 | 0.103 | Old |
| 0.0104 | 0.576289 | 0.228 | 0.151 | 1 | 7 | Mrpl16 | 0.077 | Old |
| 0.010407 | 0.342187 | 0.533 | 0.428 | 1 | 7 | Rab31 | 0.105 | Old |
| 0.010502 | 0.389058 | 0.574 | 0.503 | 1 | 7 | Dnmt3a | 0.071 | Old |
| 0.010525 | 0.590578 | 0.185 | 0.114 | 1 | 7 | Orc2 | 0.071 | Old |
| 0.010575 | 0.269547 | 0.764 | 0.666 | 1 | 7 | Lgals3 | 0.098 | Old |
| 0.010655 | 0.274828 | 0.374 | 0.271 | 1 | 7 | Hcfc1 | 0.103 | Old |
| 0.01077 | 0.676294 | 0.185 | 0.114 | 1 | 7 | Amn1 | 0.071 | Old |
| 0.01078 | 0.229236 | 0.231 | 0.148 | 1 | 7 | Gpd2 | 0.083 | Old |
| 0.010817 | 0.542701 | 0.2 | 0.127 | 1 | 7 | Gsto1 | 0.073 | Old |
| 0.010822 | 0.260494 | 0.531 | 0.449 | 1 | 7 | Aco2 | 0.082 | Old |
| 0.010847 | 0.518771 | 0.308 | 0.226 | 1 | 7 | Itpr1 | 0.082 | Old |
| 0.010894 | 0.735791 | 0.082 | 0.036 | 1 | 7 | Zfp160 | 0.046 | Old |
| 0.010959 | 0.576103 | 0.321 | 0.244 | 1 | 7 | Capn1 | 0.077 | Old |
| 0.011013 | 0.38119 | 0.405 | 0.313 | 1 | 7 | B4galnt1 | 0.092 | Old |
| 0.011088 | 0.809408 | 0.254 | 0.175 | 1 | 7 | Kctd10 | 0.079 | Old |
| 0.011109 | 0.46871 | 0.333 | 0.244 | 1 | 7 | Tmem147 | 0.089 | Old |
| 0.011116 | 0.688185 | 0.362 | 0.28 | 1 | 7 | Hopx | 0.082 | Old |
| 0.011233 | 0.445596 | 0.305 | 0.214 | 1 | 7 | Mob1b | 0.091 | Old |
| 0.011274 | 2.1191 | 0.054 | 0.018 | 1 | 7 | Chst15 | 0.036 | Old |
| 0.011303 | 0.320816 | 0.492 | 0.373 | 1 | 7 | Pole4 | 0.119 | Old |
| 0.011305 | 0.309309 | 0.156 | 0.09 | 1 | 7 | Elp3 | 0.066 | Old |
| 0.011312 | 0.497892 | 0.405 | 0.31 | 1 | 7 | Lrrc8c | 0.095 | Old |
| 0.011328 | 0.529228 | 0.377 | 0.289 | 1 | 7 | Gga1 | 0.088 | Old |
| 0.011332 | 0.373819 | 0.531 | 0.434 | 1 | 7 | Sf3b3 | 0.097 | Old |
| 0.011351 | 0.348307 | 0.144 | 0.078 | 1 | 7 | Cenpp | 0.066 | Old |
| 0.011428 | 1.544098 | 0.038 | 0.009 | 1 | 7 | Slc6a8 | 0.029 | Old |
| 0.01143 | 2.165862 | 0.044 | 0.012 | 1 | 7 | Coq4 | 0.032 | Old |
| 0.011435 | 0.344675 | 0.608 | 0.536 | 1 | 7 | Etv3 | 0.072 | Old |
| 0.011456 | 0.283418 | 0.515 | 0.428 | 1 | 7 | Anxa7 | 0.087 | Old |
| 0.011609 | 1.171226 | 0.154 | 0.093 | 1 | 7 | Pitpnm1 | 0.061 | Old |
| 0.011634 | 0.339691 | 0.254 | 0.169 | 1 | 7 | Cdk17 | 0.085 | Old |
| 0.011657 | 0.349062 | 0.392 | 0.292 | 1 | 7 | Egln1 | 0.1 | Old |
| 0.011737 | 0.770765 | 0.167 | 0.102 | 1 | 7 | Kifc1 | 0.065 | Old |
| 0.011754 | 0.475452 | 0.292 | 0.211 | 1 | 7 | Tug1 | 0.081 | Old |
| 0.011768 | 1.433343 | 0.049 | 0.015 | 1 | 7 | Galm | 0.034 | Old |
| 0.011845 | 0.256869 | 0.344 | 0.253 | 1 | 7 | Tnfaip2 | 0.091 | Old |
| 0.011905 | 0.384025 | 0.379 | 0.286 | 1 | 7 | Vps13b | 0.093 | Old |
| 0.011969 | 0.565794 | 0.274 | 0.196 | 1 | 7 | Aip | 0.078 | Old |
| 0.011991 | 0.517519 | 0.3 | 0.223 | 1 | 7 | Gigyf2 | 0.077 | Old |
| 0.012145 | 0.928734 | 0.077 | 0.033 | 1 | 7 | Surf6 | 0.044 | Old |
| 0.012198 | 0.2215 | 0.823 | 0.738 | 1 | 7 | Rpn1 | 0.085 | Old |
| 0.012239 | 0.195615 | 0.392 | 0.301 | 1 | 7 | Mtch2 | 0.091 | Old |

| | | | | | | | | |
|---|---|---|---|---|---|---|---|---|
| 0.012391 | 0.374747 | 0.69 | 0.633 | 1 | 7 | Hspd1 | 0.057 | Old |
| 0.012477 | 0.559595 | 0.377 | 0.289 | 1 | 7 | Grk3 | 0.088 | Old |
| 0.012534 | 1.10923 | 0.092 | 0.045 | 1 | 7 | Pdpr | 0.047 | Old |
| 0.012574 | 0.55502 | 0.269 | 0.187 | 1 | 7 | Rars | 0.082 | Old |
| 0.012635 | 0.44338 | 0.418 | 0.322 | 1 | 7 | Ttyh3 | 0.096 | Old |
| 0.012763 | 0.542485 | 0.133 | 0.075 | 1 | 7 | Rpp14 | 0.058 | Old |
| 0.012771 | 0.370325 | 0.372 | 0.277 | 1 | 7 | Sec22b | 0.095 | Old |
| 0.012875 | 0.339482 | 0.808 | 0.744 | 1 | 7 | Jpt1 | 0.064 | Old |
| 0.012884 | 0.565661 | 0.251 | 0.175 | 1 | 7 | 4833439L1 | 0.076 | Old |
| 0.012958 | 0.52831 | 0.282 | 0.202 | 1 | 7 | Rnf220 | 0.08 | Old |
| 0.012984 | 0.414515 | 0.487 | 0.38 | 1 | 7 | Bax | 0.107 | Old |
| 0.013002 | 0.438993 | 0.233 | 0.157 | 1 | 7 | Camk2g | 0.076 | Old |
| 0.013009 | 0.330057 | 0.338 | 0.256 | 1 | 7 | Noct | 0.082 | Old |
| 0.013034 | 0.178147 | 0.418 | 0.313 | 1 | 7 | Ugp2 | 0.105 | Old |
| 0.013096 | 1.02781 | 0.077 | 0.033 | 1 | 7 | Slc25a40 | 0.044 | Old |
| 0.013175 | 0.75887 | 0.29 | 0.214 | 1 | 7 | Prdx3 | 0.076 | Old |
| 0.013202 | 0.250318 | 0.185 | 0.111 | 1 | 7 | Wasl | 0.074 | Old |
| 0.013343 | 0.129771 | 0.603 | 0.521 | 1 | 7 | Ogdh | 0.082 | Old |
| 0.013406 | 0.607547 | 0.203 | 0.133 | 1 | 7 | Cdc45 | 0.07 | Old |
| 0.013573 | 0.497812 | 0.336 | 0.256 | 1 | 7 | Uap1l1 | 0.08 | Old |
| 0.01362 | 0.319905 | 0.118 | 0.063 | 1 | 7 | Rrp1b | 0.055 | Old |
| 0.01366 | 0.774192 | 0.136 | 0.078 | 1 | 7 | St8sia6 | 0.058 | Old |
| 0.013747 | 0.331823 | 0.203 | 0.13 | 1 | 7 | Tmem120a | 0.073 | Old |
| 0.013785 | 0.428722 | 0.485 | 0.395 | 1 | 7 | Tap1 | 0.09 | Old |
| 0.013834 | 0.847208 | 0.162 | 0.099 | 1 | 7 | Lztfl1 | 0.063 | Old |
| 0.013845 | 0.794219 | 0.092 | 0.045 | 1 | 7 | Slx4 | 0.047 | Old |
| 0.013898 | 0.398079 | 0.462 | 0.361 | 1 | 7 | Malt1 | 0.101 | Old |
| 0.013911 | 1.776522 | 0.108 | 0.057 | 1 | 7 | Dcn | 0.051 | Old |
| 0.013958 | 0.264262 | 0.441 | 0.331 | 1 | 7 | Tmx1 | 0.11 | Old |
| 0.013997 | 0.313847 | 0.154 | 0.09 | 1 | 7 | Hmgb3 | 0.064 | Old |
| 0.014052 | 0.699772 | 0.195 | 0.127 | 1 | 7 | Mmd | 0.068 | Old |
| 0.014052 | 0.50625 | 0.362 | 0.28 | 1 | 7 | Gfpt1 | 0.082 | Old |
| 0.014204 | 0.521978 | 0.323 | 0.244 | 1 | 7 | Snrnp25 | 0.079 | Old |
| 0.014245 | 0.333092 | 0.618 | 0.536 | 1 | 7 | Lims1 | 0.082 | Old |
| 0.014246 | 0.281114 | 0.567 | 0.473 | 1 | 7 | Plekhb2 | 0.094 | Old |
| 0.014248 | 0.100684 | 0.949 | 0.937 | 1 | 7 | Pabpc1 | 0.012 | Old |
| 0.014256 | 0.682459 | 0.387 | 0.295 | 1 | 7 | Ctsl | 0.092 | Old |
| 0.014266 | 0.269489 | 0.336 | 0.247 | 1 | 7 | Bcat2 | 0.089 | Old |
| 0.014272 | 1.529361 | 0.062 | 0.024 | 1 | 7 | BC002059 | 0.038 | Old |
| 0.014277 | 3.120221 | 0.018 | 0 | 1 | 7 | Ccl11 | 0.018 | Old |
| 0.014277 | 2.091744 | 0.018 | 0 | 1 | 7 | Eln | 0.018 | Old |
| 0.014277 | 1.371794 | 0.018 | 0 | 1 | 7 | Ltb | 0.018 | Old |
| 0.014277 | 1.307196 | 0.018 | 0 | 1 | 7 | Dusp9 | 0.018 | Old |
| 0.014277 | 1.307033 | 0.018 | 0 | 1 | 7 | Hsd17b14 | 0.018 | Old |
| 0.014304 | 0.454339 | 0.215 | 0.145 | 1 | 7 | Rpf1 | 0.07 | Old |

| | | | | | | | | |
|---|---|---|---|---|---|---|---|---|
| 0.014343 | 0.287102 | 0.428 | 0.331 | 1 | 7 | Uso1 | 0.097 | Old |
| 0.014409 | 0.350229 | 0.305 | 0.217 | 1 | 7 | Dld | 0.088 | Old |
| 0.014414 | 0.284234 | 0.592 | 0.521 | 1 | 7 | Rnf149 | 0.071 | Old |
| 0.014644 | 0.267756 | 0.59 | 0.491 | 1 | 7 | Aldoa | 0.099 | Old |
| 0.014771 | 1.311004 | 0.087 | 0.042 | 1 | 7 | Irgm2 | 0.045 | Old |
| 0.014772 | 0.17642 | 0.972 | 0.946 | 1 | 7 | H2afz | 0.026 | Old |
| 0.014821 | 0.570984 | 0.192 | 0.123 | 1 | 7 | Gpsm2 | 0.069 | Old |
| 0.014848 | 0.686813 | 0.313 | 0.235 | 1 | 7 | Naa20 | 0.078 | Old |
| 0.014876 | 0.803176 | 0.113 | 0.06 | 1 | 7 | Dpp7 | 0.053 | Old |
| 0.014884 | 0.091489 | 0.692 | 0.587 | 1 | 7 | Actn4 | 0.105 | Old |
| 0.014968 | 0.745476 | 0.079 | 0.036 | 1 | 7 | Fdxacb1 | 0.043 | Old |
| 0.014974 | 1.970392 | 0.051 | 0.018 | 1 | 7 | Chchd10 | 0.033 | Old |
| 0.01503 | 0.215597 | 0.513 | 0.404 | 1 | 7 | Pnn | 0.109 | Old |
| 0.015044 | 0.389502 | 0.167 | 0.102 | 1 | 7 | Rbmx | 0.065 | Old |
| 0.015054 | 0.305761 | 0.721 | 0.633 | 1 | 7 | Atp6v1g1 | 0.088 | Old |
| 0.015134 | 0.386275 | 0.39 | 0.31 | 1 | 7 | Cops4 | 0.08 | Old |
| 0.015155 | 0.660101 | 0.146 | 0.087 | 1 | 7 | E130309D( | 0.059 | Old |
| 0.015169 | 0.38796 | 0.492 | 0.413 | 1 | 7 | Kras | 0.079 | Old |
| 0.015175 | 0.257763 | 0.231 | 0.157 | 1 | 7 | Terf1 | 0.074 | Old |
| 0.015238 | 0.210887 | 0.531 | 0.419 | 1 | 7 | Gna13 | 0.112 | Old |
| 0.015322 | 0.254569 | 0.708 | 0.636 | 1 | 7 | Csf2ra | 0.072 | Old |
| 0.01536 | 0.621591 | 0.113 | 0.06 | 1 | 7 | Mettl3 | 0.053 | Old |
| 0.015454 | 0.40518 | 0.323 | 0.238 | 1 | 7 | Adrm1 | 0.085 | Old |
| 0.015461 | 0.813397 | 0.2 | 0.136 | 1 | 7 | Scfd1 | 0.064 | Old |
| 0.015517 | 0.278975 | 0.841 | 0.804 | 1 | 7 | Eif3a | 0.037 | Old |
| 0.01553 | 0.29104 | 0.479 | 0.38 | 1 | 7 | Ubn1 | 0.099 | Old |
| 0.015639 | 0.498263 | 0.244 | 0.169 | 1 | 7 | Ppp1r35 | 0.075 | Old |
| 0.015702 | 0.29628 | 0.644 | 0.572 | 1 | 7 | Skap2 | 0.072 | Old |
| 0.015758 | 0.275908 | 0.533 | 0.437 | 1 | 7 | Mrpl28 | 0.096 | Old |
| 0.015762 | 0.422931 | 0.479 | 0.392 | 1 | 7 | Etfa | 0.087 | Old |
| 0.015771 | 2.044232 | 0.051 | 0.018 | 1 | 7 | Dmwd | 0.033 | Old |
| 0.015841 | 1.782233 | 0.031 | 0.006 | 1 | 7 | Adal | 0.025 | Old |
| 0.015855 | 1.397849 | 0.087 | 0.042 | 1 | 7 | Edrf1 | 0.045 | Old |
| 0.01586 | 0.275521 | 0.379 | 0.283 | 1 | 7 | Snrnp200 | 0.096 | Old |
| 0.015909 | 0.200802 | 0.469 | 0.364 | 1 | 7 | Uqcrc2 | 0.105 | Old |
| 0.015972 | 0.15119 | 0.259 | 0.178 | 1 | 7 | Stk3 | 0.081 | Old |
| 0.016015 | 1.548584 | 0.056 | 0.021 | 1 | 7 | Akr1b8 | 0.035 | Old |
| 0.016102 | 0.338721 | 0.418 | 0.325 | 1 | 7 | Hat1 | 0.093 | Old |
| 0.016207 | 0.630187 | 0.128 | 0.072 | 1 | 7 | Fdxr | 0.056 | Old |
| 0.01623 | 0.336913 | 0.503 | 0.407 | 1 | 7 | Map2k2 | 0.096 | Old |
| 0.016272 | 0.244819 | 0.715 | 0.648 | 1 | 7 | G3bp1 | 0.067 | Old |
| 0.016283 | 0.433044 | 0.428 | 0.337 | 1 | 7 | Akirin2 | 0.091 | Old |
| 0.016341 | 0.822145 | 0.056 | 0.021 | 1 | 7 | Mettl17 | 0.035 | Old |
| 0.016437 | 1.307883 | 0.031 | 0.006 | 1 | 7 | Arl4d | 0.025 | Old |
| 0.016589 | 0.565065 | 0.205 | 0.136 | 1 | 7 | Tdp2 | 0.069 | Old |

| | | | | | | | | |
|---|---|---|---|---|---|---|---|---|
| 0.016613 | 0.734563 | 0.138 | 0.081 | 1 | 7 | Dnph1 | 0.057 | Old |
| 0.016647 | 0.430273 | 0.164 | 0.102 | 1 | 7 | Chmp6 | 0.062 | Old |
| 0.016673 | 0.902282 | 0.056 | 0.021 | 1 | 7 | Cbwd1 | 0.035 | Old |
| 0.016772 | 1.003326 | 0.09 | 0.045 | 1 | 7 | Dus3l | 0.045 | Old |
| 0.016776 | 1.14577 | 0.031 | 0.006 | 1 | 7 | Zfp12 | 0.025 | Old |
| 0.016785 | 0.315055 | 0.482 | 0.392 | 1 | 7 | Dnase2a | 0.09 | Old |
| 0.016876 | 0.457541 | 0.31 | 0.235 | 1 | 7 | Zdhhc9 | 0.075 | Old |
| 0.016881 | 1.08062 | 0.162 | 0.102 | 1 | 7 | Dgat1 | 0.06 | Old |
| 0.016943 | 0.34549 | 0.331 | 0.247 | 1 | 7 | Cdk6 | 0.084 | Old |
| 0.01706 | 1.164583 | 0.056 | 0.021 | 1 | 7 | Bloc1s3 | 0.035 | Old |
| 0.017089 | 1.685199 | 0.036 | 0.009 | 1 | 7 | Gm16104 | 0.027 | Old |
| 0.017122 | 0.700239 | 0.031 | 0.006 | 1 | 7 | Gm10269 | 0.025 | Old |
| 0.017168 | 0.472483 | 0.241 | 0.169 | 1 | 7 | Ppp4r1 | 0.072 | Old |
| 0.01721 | 1.010426 | 0.095 | 0.048 | 1 | 7 | Inpp5a | 0.047 | Old |
| 0.017422 | 0.431186 | 0.454 | 0.364 | 1 | 7 | Ptpn12 | 0.09 | Old |
| 0.017482 | 0.320092 | 0.636 | 0.551 | 1 | 7 | Abi1 | 0.085 | Old |
| 0.017489 | 0.276373 | 0.341 | 0.253 | 1 | 7 | Kpna3 | 0.088 | Old |
| 0.017583 | 0.298444 | 0.303 | 0.217 | 1 | 7 | Tm2d1 | 0.086 | Old |
| 0.017691 | 1.09722 | 0.051 | 0.018 | 1 | 7 | Amer1 | 0.033 | Old |
| 0.017714 | 1.046796 | 0.169 | 0.108 | 1 | 7 | Lrrc8b | 0.061 | Old |
| 0.01773 | 0.764281 | 0.159 | 0.099 | 1 | 7 | Coq2 | 0.06 | Old |
| 0.017838 | 0.232266 | 0.231 | 0.16 | 1 | 7 | Rabgef1 | 0.071 | Old |
| 0.017864 | 1.395393 | 0.036 | 0.009 | 1 | 7 | Wdr59 | 0.027 | Old |
| 0.01793 | 0.228715 | 0.654 | 0.575 | 1 | 7 | Ptges3 | 0.079 | Old |
| 0.017937 | 0.632138 | 0.231 | 0.16 | 1 | 7 | Zfp592 | 0.071 | Old |
| 0.017948 | 0.874401 | 0.097 | 0.051 | 1 | 7 | Scfd2 | 0.046 | Old |
| 0.017961 | 0.475912 | 0.531 | 0.461 | 1 | 7 | Atp6v1a | 0.07 | Old |
| 0.018018 | 0.310985 | 0.705 | 0.599 | 1 | 7 | Psmd13 | 0.106 | Old |
| 0.018037 | 1.424209 | 0.046 | 0.015 | 1 | 7 | Txnl4b | 0.031 | Old |
| 0.018059 | 0.475546 | 0.441 | 0.352 | 1 | 7 | Pdxk | 0.089 | Old |
| 0.01807 | 0.220211 | 0.287 | 0.202 | 1 | 7 | Tsnax | 0.085 | Old |
| 0.018234 | 1.09857 | 0.069 | 0.03 | 1 | 7 | Tmem138 | 0.039 | Old |
| 0.018347 | 0.491229 | 0.141 | 0.084 | 1 | 7 | Akr1e1 | 0.057 | Old |
| 0.018431 | 0.768203 | 0.236 | 0.163 | 1 | 7 | Bin3 | 0.073 | Old |
| 0.018439 | 0.506894 | 0.418 | 0.34 | 1 | 7 | Fgd2 | 0.078 | Old |
| 0.0185 | 0.54033 | 0.146 | 0.087 | 1 | 7 | Pym1 | 0.059 | Old |
| 0.018503 | 0.40217 | 0.679 | 0.59 | 1 | 7 | Eif4b | 0.089 | Old |
| 0.018659 | 0.978821 | 0.121 | 0.069 | 1 | 7 | Atg4c | 0.052 | Old |
| 0.01868 | 0.287389 | 0.454 | 0.358 | 1 | 7 | Slc4a7 | 0.096 | Old |
| 0.018752 | 0.328358 | 0.108 | 0.057 | 1 | 7 | Zfp738 | 0.051 | Old |
| 0.018845 | 0.200612 | 0.854 | 0.801 | 1 | 7 | Cox5a | 0.053 | Old |
| 0.018871 | 0.22311 | 0.313 | 0.229 | 1 | 7 | Galnt7 | 0.084 | Old |
| 0.018876 | 0.965764 | 0.174 | 0.114 | 1 | 7 | Rab10os | 0.06 | Old |
| 0.018913 | 0.904172 | 0.097 | 0.051 | 1 | 7 | Rft1 | 0.046 | Old |
| 0.018963 | 0.236361 | 0.592 | 0.506 | 1 | 7 | Ppig | 0.086 | Old |

| | | | | | | | | |
|---|---|---|---|---|---|---|---|---|
| 0.018979 | 0.5661 | 0.182 | 0.117 | 1 | 7 | Rgs3 | 0.065 | Old |
| 0.01899 | 0.571877 | 0.259 | 0.184 | 1 | 7 | Flvcr1 | 0.075 | Old |
| 0.019059 | 0.207267 | 0.715 | 0.633 | 1 | 7 | Ccdc88a | 0.082 | Old |
| 0.01911 | 0.367354 | 0.623 | 0.542 | 1 | 7 | Degs1 | 0.081 | Old |
| 0.019222 | 0.348612 | 0.395 | 0.316 | 1 | 7 | Fgr | 0.079 | Old |
| 0.019232 | 0.243346 | 0.303 | 0.22 | 1 | 7 | Arhgap19 | 0.083 | Old |
| 0.019336 | 0.967432 | 0.113 | 0.063 | 1 | 7 | Gnptab | 0.05 | Old |
| 0.019342 | 0.404429 | 0.285 | 0.208 | 1 | 7 | Dap3 | 0.077 | Old |
| 0.019392 | 0.367536 | 0.746 | 0.675 | 1 | 7 | Tapbp | 0.071 | Old |
| 0.019468 | 0.220931 | 0.713 | 0.645 | 1 | 7 | Cmpk1 | 0.068 | Old |
| 0.019524 | 0.464573 | 0.308 | 0.232 | 1 | 7 | Gramd4 | 0.076 | Old |
| 0.019663 | 0.307721 | 0.533 | 0.443 | 1 | 7 | Cers2 | 0.09 | Old |
| 0.019692 | 0.29511 | 0.292 | 0.211 | 1 | 7 | Rfc5 | 0.081 | Old |
| 0.019732 | 0.395427 | 0.21 | 0.142 | 1 | 7 | Terf2 | 0.068 | Old |
| 0.019769 | 0.484487 | 0.262 | 0.187 | 1 | 7 | Igsf8 | 0.075 | Old |
| 0.019776 | 0.234321 | 0.721 | 0.63 | 1 | 7 | Atp6v0e | 0.091 | Old |
| 0.019779 | 0.154281 | 0.833 | 0.777 | 1 | 7 | Gnb1 | 0.056 | Old |
| 0.019866 | 0.60539 | 0.144 | 0.087 | 1 | 7 | Gcc1 | 0.057 | Old |
| 0.019985 | 0.665946 | 0.128 | 0.075 | 1 | 7 | Parn | 0.053 | Old |
| 0.020062 | 0.409258 | 0.195 | 0.133 | 1 | 7 | Psmg2 | 0.062 | Old |
| 0.020104 | 0.43118 | 0.174 | 0.111 | 1 | 7 | Zfp260 | 0.063 | Old |
| 0.020179 | 0.179206 | 0.144 | 0.084 | 1 | 7 | Sil1 | 0.06 | Old |
| 0.020203 | 0.21072 | 0.603 | 0.521 | 1 | 7 | Itgb2 | 0.082 | Old |
| 0.020207 | 0.47642 | 0.269 | 0.19 | 1 | 7 | Nudt16l1 | 0.079 | Old |
| 0.020365 | 0.376932 | 0.474 | 0.395 | 1 | 7 | Stk4 | 0.079 | Old |
| 0.020369 | 0.658883 | 0.149 | 0.093 | 1 | 7 | Gucd1 | 0.056 | Old |
| 0.02044 | 0.910683 | 0.141 | 0.087 | 1 | 7 | Fblim1 | 0.054 | Old |
| 0.020605 | 0.452198 | 0.264 | 0.187 | 1 | 7 | Zc3h14 | 0.077 | Old |
| 0.02064 | 0.385271 | 0.341 | 0.25 | 1 | 7 | Tmem51 | 0.091 | Old |
| 0.020641 | 0.666247 | 0.241 | 0.172 | 1 | 7 | Setdb1 | 0.069 | Old |
| 0.020701 | 0.083602 | 0.187 | 0.123 | 1 | 7 | Exoc1 | 0.064 | Old |
| 0.020703 | 0.402224 | 0.167 | 0.102 | 1 | 7 | Med11 | 0.065 | Old |
| 0.020746 | 0.410286 | 0.19 | 0.123 | 1 | 7 | Hcfc2 | 0.067 | Old |
| 0.020801 | 0.536472 | 0.213 | 0.142 | 1 | 7 | Nomo1 | 0.071 | Old |
| 0.021055 | 0.470199 | 0.149 | 0.09 | 1 | 7 | Entpd7 | 0.059 | Old |
| 0.021109 | 0.405496 | 0.441 | 0.355 | 1 | 7 | Knstrn | 0.086 | Old |
| 0.021174 | 0.530816 | 0.136 | 0.081 | 1 | 7 | Scly | 0.055 | Old |
| 0.021344 | 0.151175 | 0.413 | 0.313 | 1 | 7 | Emd | 0.1 | Old |
| 0.021391 | 0.196092 | 0.295 | 0.211 | 1 | 7 | Ankrd40 | 0.084 | Old |
| 0.021447 | 0.224652 | 0.333 | 0.247 | 1 | 7 | C330027C( | 0.086 | Old |
| 0.021449 | 0.098126 | 1 | 1 | 1 | 7 | Actb | 0 | Old |
| 0.021461 | 0.240006 | 0.854 | 0.786 | 1 | 7 | Flna | 0.068 | Old |
| 0.021467 | 0.390055 | 0.628 | 0.551 | 1 | 7 | Timm13 | 0.077 | Old |
| 0.0215 | 0.626769 | 0.064 | 0.027 | 1 | 7 | Mccc1 | 0.037 | Old |
| 0.021561 | 0.555685 | 0.215 | 0.151 | 1 | 7 | Slc27a1 | 0.064 | Old |

| | | | | | | | | |
|---|---|---|---|---|---|---|---|---|
| 0.021571 | 0.51809 | 0.497 | 0.422 | 1 | 7 | Copa | 0.075 | Old |
| 0.021581 | 0.838683 | 0.146 | 0.09 | 1 | 7 | Lsm14b | 0.056 | Old |
| 0.021591 | 1.32204 | 0.023 | 0.003 | 1 | 7 | 9330175E1 | 0.02 | Old |
| 0.021614 | 0.332353 | 0.138 | 0.084 | 1 | 7 | Troap | 0.054 | Old |
| 0.021663 | 0.865886 | 0.085 | 0.042 | 1 | 7 | Abhd11 | 0.043 | Old |
| 0.021679 | 0.15024 | 0.485 | 0.392 | 1 | 7 | Gm21188 | 0.093 | Old |
| 0.021692 | 1.995858 | 0.023 | 0.003 | 1 | 7 | Zfp51 | 0.02 | Old |
| 0.021692 | 1.334029 | 0.023 | 0.003 | 1 | 7 | Cttn | 0.02 | Old |
| 0.021742 | 0.333567 | 0.418 | 0.328 | 1 | 7 | Apobec1 | 0.09 | Old |
| 0.021835 | 0.377174 | 0.146 | 0.09 | 1 | 7 | Pip4k2c | 0.056 | Old |
| 0.021841 | 0.232137 | 0.715 | 0.623 | 1 | 7 | Tmem30a | 0.092 | Old |
| 0.021841 | 0.201747 | 0.118 | 0.066 | 1 | 7 | Mettl6 | 0.052 | Old |
| 0.021973 | 0.735457 | 0.3 | 0.226 | 1 | 7 | Cops3 | 0.074 | Old |
| 0.021998 | 1.079753 | 0.023 | 0.003 | 1 | 7 | Rpp40 | 0.02 | Old |
| 0.022213 | 0.239882 | 0.203 | 0.136 | 1 | 7 | Zcchc8 | 0.067 | Old |
| 0.022219 | 0.368391 | 0.297 | 0.22 | 1 | 7 | Idh3b | 0.077 | Old |
| 0.022224 | 0.142536 | 0.946 | 0.922 | 1 | 7 | Mcl1 | 0.024 | Old |
| 0.022245 | 0.176303 | 0.828 | 0.768 | 1 | 7 | Akr1a1 | 0.06 | Old |
| 0.022248 | 0.637382 | 0.221 | 0.154 | 1 | 7 | Tspan5 | 0.067 | Old |
| 0.022389 | 0.4705 | 0.128 | 0.075 | 1 | 7 | Adprhl2 | 0.053 | Old |
| 0.022426 | 0.186206 | 0.297 | 0.217 | 1 | 7 | Slc16a10 | 0.08 | Old |
| 0.022433 | 0.671121 | 0.108 | 0.06 | 1 | 7 | AI987944 | 0.048 | Old |
| 0.022487 | 0.464849 | 0.133 | 0.078 | 1 | 7 | Rbm10 | 0.055 | Old |
| 0.022556 | 0.714112 | 0.059 | 0.024 | 1 | 7 | Nmnat3 | 0.035 | Old |
| 0.022568 | 0.682122 | 0.126 | 0.072 | 1 | 7 | Foxred1 | 0.054 | Old |
| 0.02262 | 0.360287 | 0.254 | 0.181 | 1 | 7 | Mvb12a | 0.073 | Old |
| 0.022634 | 0.336074 | 0.474 | 0.383 | 1 | 7 | Eif1b | 0.091 | Old |
| 0.022643 | 0.165451 | 0.09 | 0.045 | 1 | 7 | Lrrc61 | 0.045 | Old |
| 0.022688 | 0.167828 | 0.092 | 0.048 | 1 | 7 | Otud7b | 0.044 | Old |
| 0.022761 | 0.517848 | 0.238 | 0.169 | 1 | 7 | Mrpl2 | 0.069 | Old |
| 0.0228 | 0.229093 | 0.651 | 0.575 | 1 | 7 | Ptpn1 | 0.076 | Old |
| 0.022822 | 0.00827 | 0.097 | 0.051 | 1 | 7 | Rps6kb2 | 0.046 | Old |
| 0.02286 | 0.752868 | 0.059 | 0.024 | 1 | 7 | Irf7 | 0.035 | Old |
| 0.022901 | 0.751081 | 0.141 | 0.087 | 1 | 7 | Prmt7 | 0.054 | Old |
| 0.022906 | 0.697314 | 0.156 | 0.096 | 1 | 7 | Cog8 | 0.06 | Old |
| 0.022941 | 0.092605 | 0.385 | 0.292 | 1 | 7 | Ncstn | 0.093 | Old |
| 0.023018 | 0.147231 | 0.659 | 0.539 | 1 | 7 | Baz1b | 0.12 | Old |
| 0.023054 | 0.190837 | 0.315 | 0.229 | 1 | 7 | Pqbp1 | 0.086 | Old |
| 0.023166 | 0.290844 | 0.528 | 0.434 | 1 | 7 | Sbno2 | 0.094 | Old |
| 0.023177 | 0.019504 | 0.61 | 0.506 | 1 | 7 | Gpr132 | 0.104 | Old |
| 0.023191 | 0.9468 | 0.085 | 0.042 | 1 | 7 | Zfp513 | 0.043 | Old |
| 0.023247 | 0.39011 | 0.31 | 0.235 | 1 | 7 | Vps13c | 0.075 | Old |
| 0.023272 | 0.476297 | 0.174 | 0.111 | 1 | 7 | Zdhhc6 | 0.063 | Old |
| 0.023363 | 0.301067 | 0.472 | 0.38 | 1 | 7 | Rad23a | 0.092 | Old |
| 0.023368 | 0.391844 | 0.351 | 0.268 | 1 | 7 | Pik3r5 | 0.083 | Old |

| | | | | | | | | |
|---|---|---|---|---|---|---|---|---|
| 0.023371 | 0.428328 | 0.179 | 0.117 | 1 | 7 | Tinf2 | 0.062 | Old |
| 0.023394 | 0.440922 | 0.356 | 0.271 | 1 | 7 | Safb | 0.085 | Old |
| 0.023406 | 2.466284 | 0.015 | 0 | 1 | 7 | Cerkl | 0.015 | Old |
| 0.023406 | 2.362864 | 0.015 | 0 | 1 | 7 | Cdh5 | 0.015 | Old |
| 0.023406 | 1.849653 | 0.015 | 0 | 1 | 7 | Gm43796 | 0.015 | Old |
| 0.023406 | 1.827067 | 0.015 | 0 | 1 | 7 | 9430002A1 | 0.015 | Old |
| 0.023406 | 1.826836 | 0.015 | 0 | 1 | 7 | Neil2 | 0.015 | Old |
| 0.023406 | 1.693149 | 0.015 | 0 | 1 | 7 | Gm10384 | 0.015 | Old |
| 0.023406 | 1.646101 | 0.015 | 0 | 1 | 7 | Fam189a2 | 0.015 | Old |
| 0.023406 | 1.540526 | 0.015 | 0 | 1 | 7 | AL591952.: | 0.015 | Old |
| 0.023406 | 1.38439 | 0.015 | 0 | 1 | 7 | Gpr82 | 0.015 | Old |
| 0.023406 | 1.256856 | 0.015 | 0 | 1 | 7 | Ceacam16 | 0.015 | Old |
| 0.023406 | 1.202621 | 0.015 | 0 | 1 | 7 | Gm49207 | 0.015 | Old |
| 0.023406 | 1.164988 | 0.015 | 0 | 1 | 7 | Gnal | 0.015 | Old |
| 0.023415 | 0.434887 | 0.182 | 0.12 | 1 | 7 | Mkl2 | 0.062 | Old |
| 0.023638 | 0.382485 | 0.562 | 0.479 | 1 | 7 | Eif3i | 0.083 | Old |
| 0.023698 | 0.48586 | 0.177 | 0.117 | 1 | 7 | Ndrg3 | 0.06 | Old |
| 0.023698 | 0.379032 | 0.777 | 0.744 | 1 | 7 | Txn1 | 0.033 | Old |
| 0.023706 | 0.459381 | 0.385 | 0.307 | 1 | 7 | Trp53 | 0.078 | Old |
| 0.023852 | 0.179854 | 0.849 | 0.864 | 1 | 7 | Arhgdib | -0.015 | Old |
| 0.02388 | 0.428307 | 0.305 | 0.229 | 1 | 7 | Cep192 | 0.076 | Old |
| 0.023931 | 0.318085 | 0.759 | 0.69 | 1 | 7 | Nme1 | 0.069 | Old |
| 0.024069 | 0.237306 | 0.351 | 0.268 | 1 | 7 | Scamp3 | 0.083 | Old |
| 0.024139 | 0.773519 | 0.118 | 0.069 | 1 | 7 | Uqcc1 | 0.049 | Old |
| 0.024229 | 0.550505 | 0.108 | 0.06 | 1 | 7 | Tmem87a | 0.048 | Old |
| 0.024271 | 0.184557 | 0.662 | 0.602 | 1 | 7 | Naa50 | 0.06 | Old |
| 0.024297 | 0.410078 | 0.149 | 0.093 | 1 | 7 | Tomm6 | 0.056 | Old |
| 0.024348 | 0.202277 | 0.359 | 0.271 | 1 | 7 | Zfp516 | 0.088 | Old |
| 0.024358 | 0.535141 | 0.192 | 0.127 | 1 | 7 | Arl13b | 0.065 | Old |
| 0.024474 | 0.249741 | 0.249 | 0.169 | 1 | 7 | Exoc2 | 0.08 | Old |
| 0.024481 | 0.439516 | 0.464 | 0.38 | 1 | 7 | Arhgap11a | 0.084 | Old |
| 0.024613 | 1.099221 | 0.067 | 0.03 | 1 | 7 | L3mbtl2 | 0.037 | Old |
| 0.024646 | 0.319737 | 0.321 | 0.235 | 1 | 7 | Bfar | 0.086 | Old |
| 0.024715 | 0.427851 | 0.213 | 0.148 | 1 | 7 | Hdac4 | 0.065 | Old |
| 0.024821 | 0.291873 | 0.121 | 0.069 | 1 | 7 | Tor2a | 0.052 | Old |
| 0.024842 | 0.541405 | 0.087 | 0.045 | 1 | 7 | Btrc | 0.042 | Old |
| 0.024901 | 0.178714 | 0.356 | 0.274 | 1 | 7 | Ap3d1 | 0.082 | Old |
| 0.024905 | 0.297273 | 0.821 | 0.771 | 1 | 7 | Srsf2 | 0.05 | Old |
| 0.024907 | 0.348815 | 0.315 | 0.238 | 1 | 7 | Dbf4 | 0.077 | Old |
| 0.024934 | 0.284668 | 0.772 | 0.717 | 1 | 7 | Arpc4 | 0.055 | Old |
| 0.025007 | 0.979488 | 0.159 | 0.105 | 1 | 7 | Ptgir | 0.054 | Old |
| 0.025059 | 1.518763 | 0.062 | 0.027 | 1 | 7 | Fbxl12 | 0.035 | Old |
| 0.02506 | 0.672085 | 0.136 | 0.081 | 1 | 7 | Rfxap | 0.055 | Old |
| 0.025118 | 0.316636 | 0.244 | 0.175 | 1 | 7 | Mcm10 | 0.069 | Old |
| 0.025207 | 0.412565 | 0.203 | 0.136 | 1 | 7 | Cdk5rap2 | 0.067 | Old |

| | | | | | | | | |
|---|---|---|---|---|---|---|---|---|
| 0.025238 | 0.085029 | 0.379 | 0.286 | 1 | 7 | Ccdc82 | 0.093 | Old |
| 0.02526 | 0.120199 | 0.531 | 0.437 | 1 | 7 | Nxf1 | 0.094 | Old |
| 0.02535 | 0.845234 | 0.067 | 0.03 | 1 | 7 | Oas3 | 0.037 | Old |
| 0.025397 | 0.717475 | 0.251 | 0.181 | 1 | 7 | Ccnk | 0.07 | Old |
| 0.025399 | 0.358223 | 0.274 | 0.199 | 1 | 7 | Irf2bpl | 0.075 | Old |
| 0.025456 | 0.811569 | 0.136 | 0.084 | 1 | 7 | Aga | 0.052 | Old |
| 0.025475 | 1.244849 | 0.067 | 0.03 | 1 | 7 | D2hgdh | 0.037 | Old |
| 0.025489 | 0.196073 | 0.136 | 0.081 | 1 | 7 | Mrpl22 | 0.055 | Old |
| 0.025519 | 0.110511 | 0.428 | 0.34 | 1 | 7 | Cyb5b | 0.088 | Old |
| 0.02556 | 0.740504 | 0.123 | 0.072 | 1 | 7 | Ipo13 | 0.051 | Old |
| 0.025563 | 1.140284 | 0.103 | 0.057 | 1 | 7 | Fam58b | 0.046 | Old |
| 0.0256 | 0.863003 | 0.067 | 0.03 | 1 | 7 | Sirt3 | 0.037 | Old |
| 0.025682 | 0.31468 | 0.631 | 0.545 | 1 | 7 | Birc3 | 0.086 | Old |
| 0.025782 | 0.318838 | 0.295 | 0.22 | 1 | 7 | Cldnd1 | 0.075 | Old |
| 0.025805 | 0.231207 | 0.182 | 0.117 | 1 | 7 | Fbxo34 | 0.065 | Old |
| 0.025952 | 0.735072 | 0.044 | 0.015 | 1 | 7 | Dntt | 0.029 | Old |
| 0.02597 | 0.223769 | 0.777 | 0.681 | 1 | 7 | Rab14 | 0.096 | Old |
| 0.026034 | 0.024993 | 0.438 | 0.331 | 1 | 7 | Zfp148 | 0.107 | Old |
| 0.026238 | 0.407328 | 0.344 | 0.271 | 1 | 7 | Psmd7 | 0.073 | Old |
| 0.026252 | 0.236398 | 0.89 | 0.904 | 1 | 7 | Sdc3 | -0.014 | Old |
| 0.026275 | 0.71889 | 0.274 | 0.196 | 1 | 7 | Cnot2 | 0.078 | Old |
| 0.026438 | 1.264043 | 0.044 | 0.015 | 1 | 7 | Ahnak2 | 0.029 | Old |
| 0.026446 | 0.401298 | 0.177 | 0.12 | 1 | 7 | Hist1h4i | 0.057 | Old |
| 0.026584 | 0.717748 | 0.318 | 0.247 | 1 | 7 | Tollip | 0.071 | Old |
| 0.026689 | 0.331387 | 0.613 | 0.521 | 1 | 7 | Snrpd3 | 0.092 | Old |
| 0.0267 | 0.582025 | 0.028 | 0.006 | 1 | 7 | Mrm3 | 0.022 | Old |
| 0.026714 | 1.551521 | 0.033 | 0.009 | 1 | 7 | Gm10521 | 0.024 | Old |
| 0.026736 | 0.350084 | 0.564 | 0.5 | 1 | 7 | Myof | 0.064 | Old |
| 0.026751 | 1.039509 | 0.097 | 0.054 | 1 | 7 | Lncpint | 0.043 | Old |
| 0.026808 | 0.153518 | 0.923 | 0.898 | 1 | 7 | Actr3 | 0.025 | Old |
| 0.026967 | 1.247769 | 0.082 | 0.042 | 1 | 7 | Aig1 | 0.04 | Old |
| 0.027029 | 0.199454 | 0.39 | 0.295 | 1 | 7 | Adprh | 0.095 | Old |
| 0.027071 | 0.546276 | 0.249 | 0.181 | 1 | 7 | Plod1 | 0.068 | Old |
| 0.027098 | 1.122568 | 0.044 | 0.015 | 1 | 7 | Ap5b1 | 0.029 | Old |
| 0.027116 | 0.245419 | 0.387 | 0.298 | 1 | 7 | Btbd1 | 0.089 | Old |
| 0.02712 | 0.250961 | 0.336 | 0.25 | 1 | 7 | Cers6 | 0.086 | Old |
| 0.027224 | 0.263616 | 0.5 | 0.413 | 1 | 7 | Nin | 0.087 | Old |
| 0.027262 | 0.362528 | 0.505 | 0.404 | 1 | 7 | Nfe2l1 | 0.101 | Old |
| 0.027278 | 0.502991 | 0.213 | 0.151 | 1 | 7 | Ift52 | 0.062 | Old |
| 0.027297 | 0.985399 | 0.033 | 0.009 | 1 | 7 | 9530036M | 0.024 | Old |
| 0.027369 | 0.669552 | 0.215 | 0.157 | 1 | 7 | Akr1c13 | 0.058 | Old |
| 0.027376 | 0.3743 | 0.251 | 0.181 | 1 | 7 | Elp2 | 0.07 | Old |
| 0.027475 | 1.344491 | 0.038 | 0.012 | 1 | 7 | Il4i1 | 0.026 | Old |
| 0.027502 | 0.597245 | 0.215 | 0.154 | 1 | 7 | Atp2a3 | 0.061 | Old |
| 0.027533 | 0.41736 | 0.197 | 0.133 | 1 | 7 | Ero1lb | 0.064 | Old |

| | | | | | | | | |
|---|---|---|---|---|---|---|---|---|
| 0.027534 | 0.309192 | 0.282 | 0.205 | 1 | 7 | Tbc1d20 | 0.077 | Old |
| 0.027542 | 0.367312 | 0.605 | 0.497 | 1 | 7 | Vdac3 | 0.108 | Old |
| 0.027542 | 0.794227 | 0.187 | 0.13 | 1 | 7 | Gsr | 0.057 | Old |
| 0.027655 | 0.693148 | 0.182 | 0.12 | 1 | 7 | Nt5c3 | 0.062 | Old |
| 0.027657 | 0.326264 | 0.5 | 0.407 | 1 | 7 | Ist1 | 0.093 | Old |
| 0.027723 | 0.323195 | 0.11 | 0.063 | 1 | 7 | Rars2 | 0.047 | Old |
| 0.02807 | 0.825705 | 0.208 | 0.145 | 1 | 7 | Rassf8 | 0.063 | Old |
| 0.028209 | 0.449828 | 0.187 | 0.123 | 1 | 7 | Mrpl19 | 0.064 | Old |
| 0.028306 | 0.485344 | 0.441 | 0.364 | 1 | 7 | Sms | 0.077 | Old |
| 0.028321 | 0.754928 | 0.074 | 0.036 | 1 | 7 | Gpc1 | 0.038 | Old |
| 0.028333 | 0.663877 | 0.195 | 0.133 | 1 | 7 | Fuom | 0.062 | Old |
| 0.028401 | 0.31722 | 0.336 | 0.253 | 1 | 7 | Srsf4 | 0.083 | Old |
| 0.028418 | 0.198601 | 0.231 | 0.16 | 1 | 7 | Vta1 | 0.071 | Old |
| 0.028461 | 0.228493 | 0.685 | 0.593 | 1 | 7 | Srsf6 | 0.092 | Old |
| 0.028492 | 0.89149 | 0.09 | 0.048 | 1 | 7 | Carf | 0.042 | Old |
| 0.028529 | 0.269777 | 0.513 | 0.422 | 1 | 7 | Adss | 0.091 | Old |
| 0.028645 | 0.642999 | 0.074 | 0.036 | 1 | 7 | Rsad2 | 0.038 | Old |
| 0.028649 | 0.155119 | 0.356 | 0.286 | 1 | 7 | Mrps17 | 0.07 | Old |
| 0.028667 | 0.483919 | 0.277 | 0.208 | 1 | 7 | Metrn | 0.069 | Old |
| 0.028684 | 0.309516 | 0.19 | 0.123 | 1 | 7 | 2610002M | 0.067 | Old |
| 0.028769 | 0.744566 | 0.038 | 0.012 | 1 | 7 | Tmem177 | 0.026 | Old |
| 0.028774 | 0.208997 | 0.869 | 0.828 | 1 | 7 | Emp3 | 0.041 | Old |
| 0.028882 | 0.434937 | 0.474 | 0.392 | 1 | 7 | Sltm | 0.082 | Old |
| 0.029013 | 0.615606 | 0.118 | 0.069 | 1 | 7 | Mecr | 0.049 | Old |
| 0.029065 | 0.395005 | 0.118 | 0.069 | 1 | 7 | Rabggta | 0.049 | Old |
| 0.029075 | 0.276463 | 0.397 | 0.31 | 1 | 7 | Pgrmc1 | 0.087 | Old |
| 0.02913 | 0.278107 | 0.544 | 0.461 | 1 | 7 | Birc5 | 0.083 | Old |
| 0.029142 | 0.429488 | 0.169 | 0.111 | 1 | 7 | Tusc2 | 0.058 | Old |
| 0.029169 | 0.287674 | 0.459 | 0.373 | 1 | 7 | Washc2 | 0.086 | Old |
| 0.029279 | 0.236367 | 0.159 | 0.102 | 1 | 7 | Gfm1 | 0.057 | Old |
| 0.029339 | 0.1347 | 0.497 | 0.407 | 1 | 7 | Arl8b | 0.09 | Old |
| 0.02951 | 0.321517 | 0.403 | 0.319 | 1 | 7 | Ncf4 | 0.084 | Old |
| 0.029513 | 0.29681 | 0.315 | 0.238 | 1 | 7 | Ckap2 | 0.077 | Old |
| 0.029518 | 0.321223 | 0.192 | 0.13 | 1 | 7 | Gpn1 | 0.062 | Old |
| 0.029533 | 0.447138 | 0.118 | 0.069 | 1 | 7 | Kat14 | 0.049 | Old |
| 0.029748 | 0.40341 | 0.144 | 0.09 | 1 | 7 | Stard5 | 0.054 | Old |
| 0.02982 | 0.804686 | 0.133 | 0.084 | 1 | 7 | Ifih1 | 0.049 | Old |
| 0.029965 | 0.030398 | 0.095 | 0.051 | 1 | 7 | Ankle1 | 0.044 | Old |
| 0.030059 | 0.72235 | 0.272 | 0.211 | 1 | 7 | Net1 | 0.061 | Old |
| 0.030069 | 0.179228 | 0.908 | 0.889 | 1 | 7 | Chchd2 | 0.019 | Old |
| 0.030075 | 0.338399 | 0.208 | 0.142 | 1 | 7 | Fbxo22 | 0.066 | Old |
| 0.030151 | 0.257157 | 0.708 | 0.642 | 1 | 7 | Dynll1 | 0.066 | Old |
| 0.030181 | 0.974762 | 0.115 | 0.069 | 1 | 7 | Tmem167b | 0.046 | Old |
| 0.030216 | 1.316142 | 0.085 | 0.045 | 1 | 7 | Tmem38b | 0.04 | Old |
| 0.030251 | 0.412563 | 0.174 | 0.114 | 1 | 7 | Ssh1 | 0.06 | Old |

| | | | | | | | | |
|---|---|---|---|---|---|---|---|---|
| 0.030315 | 0.471389 | 0.618 | 0.581 | 1 | 7 | 9-Sep | 0.037 | Old |
| 0.03033 | 0.954841 | 0.056 | 0.024 | 1 | 7 | Acad11 | 0.032 | Old |
| 0.030402 | 0.420204 | 0.497 | 0.422 | 1 | 7 | Mia3 | 0.075 | Old |
| 0.030415 | 0.699094 | 0.09 | 0.048 | 1 | 7 | 3110062M | 0.042 | Old |
| 0.030458 | 0.320934 | 0.644 | 0.554 | 1 | 7 | Srsf1 | 0.09 | Old |
| 0.030787 | 0.943424 | 0.09 | 0.048 | 1 | 7 | Rmdn3 | 0.042 | Old |
| 0.030879 | 0.276994 | 0.338 | 0.265 | 1 | 7 | Pgp | 0.073 | Old |
| 0.030923 | 0.518916 | 0.351 | 0.283 | 1 | 7 | Sarnp | 0.068 | Old |
| 0.030941 | 0.213616 | 0.39 | 0.316 | 1 | 7 | Usp16 | 0.074 | Old |
| 0.031043 | 0.112063 | 0.238 | 0.166 | 1 | 7 | Srp68 | 0.072 | Old |
| 0.031146 | 0.859691 | 0.105 | 0.06 | 1 | 7 | Fsd1l | 0.045 | Old |
| 0.031191 | 0.146428 | 0.367 | 0.28 | 1 | 7 | Atic | 0.087 | Old |
| 0.031254 | 0.162368 | 0.262 | 0.187 | 1 | 7 | Mre11a | 0.075 | Old |
| 0.031454 | 0.649005 | 0.128 | 0.078 | 1 | 7 | Mier2 | 0.05 | Old |
| 0.031548 | 0.412172 | 0.351 | 0.274 | 1 | 7 | Rbck1 | 0.077 | Old |
| 0.031597 | 0.993135 | 0.077 | 0.039 | 1 | 7 | Stk32c | 0.038 | Old |
| 0.031614 | 0.453667 | 0.128 | 0.078 | 1 | 7 | Bst1 | 0.05 | Old |
| 0.031627 | 0.429668 | 0.267 | 0.19 | 1 | 7 | Arfgap2 | 0.077 | Old |
| 0.031727 | 0.35414 | 0.533 | 0.452 | 1 | 7 | P2rx4 | 0.081 | Old |
| 0.03173 | 1.016943 | 0.138 | 0.087 | 1 | 7 | Tomm40l | 0.051 | Old |
| 0.031839 | 0.973783 | 0.097 | 0.054 | 1 | 7 | Pgap1 | 0.043 | Old |
| 0.031848 | 1.218988 | 0.051 | 0.021 | 1 | 7 | Tsen54 | 0.03 | Old |
| 0.031919 | 0.87406 | 0.092 | 0.051 | 1 | 7 | Paxx | 0.041 | Old |
| 0.032122 | 0.154428 | 0.908 | 0.886 | 1 | 7 | Ctss | 0.022 | Old |
| 0.032124 | 0.147485 | 0.462 | 0.386 | 1 | 7 | Psmd6 | 0.076 | Old |
| 0.032277 | 0.466391 | 0.082 | 0.042 | 1 | 7 | Sufu | 0.04 | Old |
| 0.032288 | 0.224371 | 0.231 | 0.163 | 1 | 7 | Itpa | 0.068 | Old |
| 0.032352 | 0.219276 | 0.659 | 0.539 | 1 | 7 | Tsn | 0.12 | Old |
| 0.032398 | 0.268746 | 0.531 | 0.437 | 1 | 7 | Smarcc1 | 0.094 | Old |
| 0.032434 | 0.393662 | 0.433 | 0.364 | 1 | 7 | H2-Q7 | 0.069 | Old |
| 0.03244 | 0.246456 | 0.364 | 0.283 | 1 | 7 | Fchsd2 | 0.081 | Old |
| 0.032442 | 0.428359 | 0.454 | 0.37 | 1 | 7 | Hadhb | 0.084 | Old |
| 0.032479 | 0.510525 | 0.146 | 0.093 | 1 | 7 | Akr7a5 | 0.053 | Old |
| 0.032549 | 1.051511 | 0.051 | 0.021 | 1 | 7 | Rarb | 0.03 | Old |
| 0.032615 | 0.328214 | 0.328 | 0.262 | 1 | 7 | Lrrc59 | 0.066 | Old |
| 0.032686 | 0.060609 | 0.367 | 0.289 | 1 | 7 | Odc1 | 0.078 | Old |
| 0.032736 | 0.486988 | 0.164 | 0.108 | 1 | 7 | Fam20c | 0.056 | Old |
| 0.032796 | 0.347669 | 0.595 | 0.53 | 1 | 7 | Prpf40a | 0.065 | Old |
| 0.03281 | 0.519079 | 0.205 | 0.145 | 1 | 7 | Gm42047 | 0.06 | Old |
| 0.032831 | 0.345259 | 0.377 | 0.292 | 1 | 7 | Bcl7c | 0.085 | Old |
| 0.032833 | 0.250869 | 0.744 | 0.702 | 1 | 7 | Atp5k | 0.042 | Old |
| 0.03288 | 0.903351 | 0.092 | 0.051 | 1 | 7 | Tmem129 | 0.041 | Old |
| 0.032895 | 0.571612 | 0.182 | 0.123 | 1 | 7 | Xab2 | 0.059 | Old |
| 0.032896 | 0.091526 | 0.367 | 0.289 | 1 | 7 | Hsd17b4 | 0.078 | Old |
| 0.033071 | 0.26067 | 0.195 | 0.136 | 1 | 7 | Ino80c | 0.059 | Old |

| | | | | | | | | |
|---|---|---|---|---|---|---|---|---|
| 0.033114 | 0.307199 | 0.256 | 0.19 | 1 | 7 | Ncapd3 | 0.066 | Old |
| 0.033135 | 0.25066 | 0.728 | 0.654 | 1 | 7 | Eif2s2 | 0.074 | Old |
| 0.03314 | 0.996065 | 0.092 | 0.051 | 1 | 7 | Alpk1 | 0.041 | Old |
| 0.033166 | 1.539265 | 0.095 | 0.054 | 1 | 7 | Ifi211 | 0.041 | Old |
| 0.033177 | 1.158619 | 0.085 | 0.045 | 1 | 7 | Agk | 0.04 | Old |
| 0.033183 | 0.455641 | 0.136 | 0.084 | 1 | 7 | Scap | 0.052 | Old |
| 0.033191 | 0.405003 | 0.126 | 0.075 | 1 | 7 | Alg14 | 0.051 | Old |
| 0.033239 | 0.542428 | 0.09 | 0.048 | 1 | 7 | Abcb8 | 0.042 | Old |
| 0.033385 | 0.585266 | 0.082 | 0.042 | 1 | 7 | Kctd3 | 0.04 | Old |
| 0.033431 | 0.279925 | 0.185 | 0.123 | 1 | 7 | Rnf126 | 0.062 | Old |
| 0.033538 | 0.403901 | 0.226 | 0.16 | 1 | 7 | Acsl1 | 0.066 | Old |
| 0.033566 | 0.144162 | 0.844 | 0.807 | 1 | 7 | Arl6ip1 | 0.037 | Old |
| 0.033717 | 0.945107 | 0.051 | 0.021 | 1 | 7 | Tnfsf13os | 0.03 | Old |
| 0.033783 | 0.481857 | 0.336 | 0.259 | 1 | 7 | Smpdl3a | 0.077 | Old |
| 0.033883 | 0.656804 | 0.121 | 0.072 | 1 | 7 | Engase | 0.049 | Old |
| 0.033947 | 0.547506 | 0.156 | 0.102 | 1 | 7 | Sephs2 | 0.054 | Old |
| 0.034083 | 0.921916 | 0.051 | 0.021 | 1 | 7 | 2010320M | 0.03 | Old |
| 0.034315 | 0.283826 | 0.374 | 0.298 | 1 | 7 | Slc25a37 | 0.076 | Old |
| 0.034339 | 0.227419 | 0.582 | 0.518 | 1 | 7 | Tpi1 | 0.064 | Old |
| 0.034361 | 0.195935 | 0.197 | 0.133 | 1 | 7 | Ankrd10 | 0.064 | Old |
| 0.034382 | 0.473087 | 0.351 | 0.271 | 1 | 7 | Ppm1g | 0.08 | Old |
| 0.034389 | 0.095497 | 0.09 | 0.048 | 1 | 7 | Mpi | 0.042 | Old |
| 0.034625 | 0.364608 | 0.118 | 0.072 | 1 | 7 | Hps1 | 0.046 | Old |
| 0.034678 | 0.217501 | 0.595 | 0.506 | 1 | 7 | Vdac1 | 0.089 | Old |
| 0.034709 | 3.391974 | 0.021 | 0.003 | 1 | 7 | Chil3 | 0.018 | Old |
| 0.034795 | 0.371809 | 0.146 | 0.093 | 1 | 7 | Zbed4 | 0.053 | Old |
| 0.034814 | 0.62618 | 0.392 | 0.322 | 1 | 7 | Zmiz2 | 0.07 | Old |
| 0.034842 | 0.310411 | 0.482 | 0.389 | 1 | 7 | Tacc3 | 0.093 | Old |
| 0.034869 | 1.354874 | 0.021 | 0.003 | 1 | 7 | Card10 | 0.018 | Old |
| 0.034918 | 0.167808 | 0.238 | 0.166 | 1 | 7 | Usp5 | 0.072 | Old |
| 0.034938 | 0.375673 | 0.164 | 0.108 | 1 | 7 | Sepsecs | 0.056 | Old |
| 0.034954 | 0.270181 | 0.29 | 0.214 | 1 | 7 | Crtc3 | 0.076 | Old |
| 0.034969 | 0.696372 | 0.113 | 0.066 | 1 | 7 | Cpt2 | 0.047 | Old |
| 0.03503 | 1.168904 | 0.021 | 0.003 | 1 | 7 | Lbhd1 | 0.018 | Old |
| 0.03508 | 0.456764 | 0.187 | 0.127 | 1 | 7 | Kmt2b | 0.06 | Old |
| 0.035127 | 0.268461 | 0.364 | 0.286 | 1 | 7 | Marveld1 | 0.078 | Old |
| 0.035191 | 1.314396 | 0.021 | 0.003 | 1 | 7 | Lrp4 | 0.018 | Old |
| 0.035191 | 1.310165 | 0.021 | 0.003 | 1 | 7 | AC130815. | 0.018 | Old |
| 0.035191 | 0.994823 | 0.021 | 0.003 | 1 | 7 | Micall2 | 0.018 | Old |
| 0.035353 | 1.100138 | 0.021 | 0.003 | 1 | 7 | Zfp28 | 0.018 | Old |
| 0.035494 | 0.503142 | 0.182 | 0.123 | 1 | 7 | Apobr | 0.059 | Old |
| 0.035516 | 0.765706 | 0.021 | 0.003 | 1 | 7 | Mageh1 | 0.018 | Old |
| 0.035516 | 0.544334 | 0.021 | 0.003 | 1 | 7 | Cdk18 | 0.018 | Old |
| 0.035608 | 0.52486 | 0.121 | 0.072 | 1 | 7 | Clk2 | 0.049 | Old |
| 0.035609 | 0.9006 | 0.495 | 0.416 | 1 | 7 | Nrp2 | 0.079 | Old |

| | | | | | | | | |
|---|---|---|---|---|---|---|---|---|
| 0.035634 | 0.654387 | 0.1 | 0.057 | 1 | 7 | Tmem242 | 0.043 | Old |
| 0.035663 | 0.292455 | 0.246 | 0.178 | 1 | 7 | Vps16 | 0.068 | Old |
| 0.035802 | 0.987584 | 0.115 | 0.069 | 1 | 7 | Gnl2 | 0.046 | Old |
| 0.035823 | 0.304985 | 0.446 | 0.364 | 1 | 7 | Copb2 | 0.082 | Old |
| 0.036081 | 0.583945 | 0.141 | 0.09 | 1 | 7 | Pgm1 | 0.051 | Old |
| 0.036087 | 0.548603 | 0.085 | 0.045 | 1 | 7 | Ercc1 | 0.04 | Old |
| 0.036169 | 0.575618 | 0.151 | 0.099 | 1 | 7 | Parp12 | 0.052 | Old |
| 0.036234 | 0.454442 | 0.349 | 0.277 | 1 | 7 | Nfil3 | 0.072 | Old |
| 0.036261 | 0.100668 | 0.385 | 0.307 | 1 | 7 | Napa | 0.078 | Old |
| 0.036263 | 0.312073 | 0.2 | 0.136 | 1 | 7 | Zfyve16 | 0.064 | Old |
| 0.036273 | 0.089582 | 0.559 | 0.479 | 1 | 7 | Tmem167 | 0.08 | Old |
| 0.036415 | 0.72111 | 0.241 | 0.178 | 1 | 7 | Sash3 | 0.063 | Old |
| 0.036511 | 0.240693 | 0.418 | 0.343 | 1 | 7 | Sdhb | 0.075 | Old |
| 0.036523 | 0.244106 | 0.231 | 0.163 | 1 | 7 | Ino80e | 0.068 | Old |
| 0.036635 | 0.289428 | 0.372 | 0.292 | 1 | 7 | Exoc4 | 0.08 | Old |
| 0.036677 | 0.724474 | 0.072 | 0.036 | 1 | 7 | Gnb4 | 0.036 | Old |
| 0.03676 | 0.427773 | 0.397 | 0.316 | 1 | 7 | Cop1 | 0.081 | Old |
| 0.036811 | 0.19446 | 0.503 | 0.404 | 1 | 7 | Nudt21 | 0.099 | Old |
| 0.037045 | 1.359515 | 0.041 | 0.015 | 1 | 7 | Tmem267 | 0.026 | Old |
| 0.037183 | 0.282343 | 0.69 | 0.645 | 1 | 7 | Atp5b | 0.045 | Old |
| 0.037668 | 0.175088 | 0.487 | 0.392 | 1 | 7 | Srebf2 | 0.095 | Old |
| 0.037714 | 0.339692 | 0.354 | 0.277 | 1 | 7 | Slc25a11 | 0.077 | Old |
| 0.037759 | 0.415913 | 0.141 | 0.09 | 1 | 7 | Fam117a | 0.051 | Old |
| 0.037833 | 0.333121 | 0.192 | 0.133 | 1 | 7 | Mrpl38 | 0.059 | Old |
| 0.037842 | 0.281897 | 0.103 | 0.06 | 1 | 7 | Dock5 | 0.043 | Old |
| 0.037911 | 0.770493 | 0.103 | 0.06 | 1 | 7 | Itpkc | 0.043 | Old |
| 0.038022 | 0.468362 | 0.485 | 0.428 | 1 | 7 | Spred1 | 0.057 | Old |
| 0.038046 | 0.140417 | 0.374 | 0.283 | 1 | 7 | Slc1a5 | 0.091 | Old |
| 0.038079 | 0.786538 | 0.087 | 0.048 | 1 | 7 | Rad1 | 0.039 | Old |
| 0.038113 | 0.373851 | 0.379 | 0.304 | 1 | 7 | Ccnb2 | 0.075 | Old |
| 0.038148 | 0.327142 | 0.436 | 0.352 | 1 | 7 | Kat2b | 0.084 | Old |
| 0.038176 | 0.127131 | 0.385 | 0.304 | 1 | 7 | Rbm47 | 0.081 | Old |
| 0.038198 | 0.425822 | 0.287 | 0.223 | 1 | 7 | Dguok | 0.064 | Old |
| 0.038239 | 0.12344 | 0.046 | 0.018 | 1 | 7 | Nck2 | 0.028 | Old |
| 0.038343 | 0.186339 | 0.731 | 0.669 | 1 | 7 | Rpn2 | 0.062 | Old |
| 0.0384 | 0.30988 | 0.192 | 0.133 | 1 | 7 | Blm | 0.059 | Old |
| 0.038404 | 1.187995 | 0.041 | 0.015 | 1 | 7 | Tspoap1 | 0.026 | Old |
| 0.038407 | 0.338345 | 0.562 | 0.494 | 1 | 7 | Ubald2 | 0.068 | Old |
| 0.038429 | 0.654042 | 0.085 | 0.045 | 1 | 7 | Zfp426 | 0.04 | Old |
| 0.038482 | 0.620272 | 0.151 | 0.099 | 1 | 7 | Lsm1 | 0.052 | Old |
| 0.038653 | 0.157311 | 0.141 | 0.09 | 1 | 7 | Srbd1 | 0.051 | Old |
| 0.038678 | 3.325653 | 0.013 | 0 | 1 | 7 | Ly6c2 | 0.013 | Old |
| 0.038678 | 2.98846 | 0.013 | 0 | 1 | 7 | Gm7967 | 0.013 | Old |
| 0.038678 | 2.273402 | 0.013 | 0 | 1 | 7 | Bcl6b | 0.013 | Old |
| 0.038678 | 2.263434 | 0.013 | 0 | 1 | 7 | Dmkn | 0.013 | Old |

| | | | | | | | | |
|---|---|---|---|---|---|---|---|---|
| 0.038678 | 1.900434 | 0.013 | 0 | 1 | 7 | Car2 | 0.013 | Old |
| 0.038678 | 1.824666 | 0.013 | 0 | 1 | 7 | Fam71e1 | 0.013 | Old |
| 0.038678 | 1.808662 | 0.013 | 0 | 1 | 7 | Cystm1 | 0.013 | Old |
| 0.038678 | 1.775485 | 0.013 | 0 | 1 | 7 | Hcn2 | 0.013 | Old |
| 0.038678 | 1.687657 | 0.013 | 0 | 1 | 7 | Six5 | 0.013 | Old |
| 0.038678 | 1.647468 | 0.013 | 0 | 1 | 7 | Pdcd1lg2 | 0.013 | Old |
| 0.038678 | 1.541288 | 0.013 | 0 | 1 | 7 | Rxrg | 0.013 | Old |
| 0.038678 | 1.37888 | 0.013 | 0 | 1 | 7 | Farp2 | 0.013 | Old |
| 0.038678 | 1.371205 | 0.013 | 0 | 1 | 7 | Hist1h4k | 0.013 | Old |
| 0.038678 | 1.348911 | 0.013 | 0 | 1 | 7 | Mipol1 | 0.013 | Old |
| 0.038678 | 1.301408 | 0.013 | 0 | 1 | 7 | Efna2 | 0.013 | Old |
| 0.038678 | 1.247126 | 0.013 | 0 | 1 | 7 | Ip6k3 | 0.013 | Old |
| 0.038678 | 1.188761 | 0.013 | 0 | 1 | 7 | Srcin1 | 0.013 | Old |
| 0.038678 | 1.129584 | 0.013 | 0 | 1 | 7 | Vegfc | 0.013 | Old |
| 0.038678 | 1.052609 | 0.013 | 0 | 1 | 7 | Il4 | 0.013 | Old |
| 0.038678 | 0.955014 | 0.013 | 0 | 1 | 7 | Exoc3l4 | 0.013 | Old |
| 0.038678 | 0.84768 | 0.013 | 0 | 1 | 7 | St5 | 0.013 | Old |
| 0.038678 | 0.817408 | 0.013 | 0 | 1 | 7 | Ccpg1os | 0.013 | Old |
| 0.038695 | 0.349926 | 0.121 | 0.072 | 1 | 7 | Ints2 | 0.049 | Old |
| 0.038705 | 0.569542 | 0.3 | 0.238 | 1 | 7 | Vcl | 0.062 | Old |
| 0.03873 | 0.351063 | 0.313 | 0.238 | 1 | 7 | Fuca2 | 0.075 | Old |
| 0.038766 | 0.230095 | 0.203 | 0.142 | 1 | 7 | Tmem209 | 0.061 | Old |
| 0.039091 | 0.53155 | 0.128 | 0.078 | 1 | 7 | Oasl2 | 0.05 | Old |
| 0.039094 | 0.510693 | 0.313 | 0.247 | 1 | 7 | Gpalpp1 | 0.066 | Old |
| 0.039153 | 0.108682 | 0.905 | 0.861 | 1 | 7 | Rhoa | 0.044 | Old |
| 0.039317 | 1.417008 | 0.103 | 0.06 | 1 | 7 | Trp53bp1 | 0.043 | Old |
| 0.039439 | 0.153344 | 0.782 | 0.705 | 1 | 7 | Grb2 | 0.077 | Old |
| 0.039449 | 0.6681 | 0.133 | 0.084 | 1 | 7 | Uckl1 | 0.049 | Old |
| 0.039465 | 0.366905 | 0.1 | 0.057 | 1 | 7 | Phtf1 | 0.043 | Old |
| 0.039549 | 0.342678 | 0.197 | 0.139 | 1 | 7 | Tubgcp2 | 0.058 | Old |
| 0.039635 | 0.207807 | 0.785 | 0.738 | 1 | 7 | Mt1 | 0.047 | Old |
| 0.039666 | 0.454939 | 0.182 | 0.123 | 1 | 7 | Plpp2 | 0.059 | Old |
| 0.039679 | 0.306583 | 0.31 | 0.241 | 1 | 7 | Asxl2 | 0.069 | Old |
| 0.039686 | 0.264075 | 0.213 | 0.154 | 1 | 7 | Clcn7 | 0.059 | Old |
| 0.039749 | 0.239453 | 0.141 | 0.09 | 1 | 7 | Ankrd16 | 0.051 | Old |
| 0.039839 | 1.271006 | 0.036 | 0.012 | 1 | 7 | Sh3d19 | 0.024 | Old |
| 0.039873 | 0.141657 | 0.536 | 0.443 | 1 | 7 | Mndal | 0.093 | Old |
| 0.039875 | 0.377577 | 0.192 | 0.133 | 1 | 7 | Vopp1 | 0.059 | Old |
| 0.040102 | 1.870391 | 0.031 | 0.009 | 1 | 7 | Pcdh7 | 0.022 | Old |
| 0.040118 | 1.180627 | 0.026 | 0.006 | 1 | 7 | Slc22a4 | 0.02 | Old |
| 0.040203 | 0.527088 | 0.19 | 0.133 | 1 | 7 | Pkp4 | 0.057 | Old |
| 0.040347 | 0.954705 | 0.169 | 0.12 | 1 | 7 | Hilpda | 0.049 | Old |
| 0.040403 | 0.66698 | 0.041 | 0.015 | 1 | 7 | Ptrh1 | 0.026 | Old |
| 0.040426 | 1.375976 | 0.074 | 0.039 | 1 | 7 | Dnaaf3 | 0.035 | Old |
| 0.040433 | 1.454507 | 0.026 | 0.006 | 1 | 7 | Hspa12a | 0.02 | Old |

| | | | | | | | | |
|---|---|---|---|---|---|---|---|---|
| 0.040518 | 0.147389 | 0.608 | 0.503 | 1 | 7 | Ube3a | 0.105 | Old |
| 0.040525 | 1.763928 | 0.031 | 0.009 | 1 | 7 | Gm31243 | 0.022 | Old |
| 0.040592 | 1.257062 | 0.026 | 0.006 | 1 | 7 | Jcad | 0.02 | Old |
| 0.040593 | 0.281399 | 0.81 | 0.765 | 1 | 7 | Tkt | 0.045 | Old |
| 0.040619 | 0.25357 | 0.287 | 0.217 | 1 | 7 | Tnfrsf21 | 0.07 | Old |
| 0.040694 | 0.145498 | 0.759 | 0.69 | 1 | 7 | Eif5a | 0.069 | Old |
| 0.040767 | 0.467945 | 0.408 | 0.334 | 1 | 7 | Hmox2 | 0.074 | Old |
| 0.040886 | 0.501727 | 0.041 | 0.015 | 1 | 7 | Tnni3 | 0.026 | Old |
| 0.040959 | 0.193448 | 0.292 | 0.226 | 1 | 7 | Hypk | 0.066 | Old |
| 0.041134 | 0.208945 | 0.508 | 0.425 | 1 | 7 | Mrpl23 | 0.083 | Old |
| 0.041232 | 0.31258 | 0.256 | 0.19 | 1 | 7 | Gabpa | 0.066 | Old |
| 0.041279 | 0.742866 | 0.113 | 0.069 | 1 | 7 | Naa25 | 0.044 | Old |
| 0.041305 | 0.23305 | 0.826 | 0.759 | 1 | 7 | Sub1 | 0.067 | Old |
| 0.04131 | 0.063308 | 0.177 | 0.117 | 1 | 7 | Tbcel | 0.06 | Old |
| 0.041329 | 0.993436 | 0.054 | 0.024 | 1 | 7 | Camk2a | 0.03 | Old |
| 0.041346 | 0.828234 | 0.079 | 0.042 | 1 | 7 | Ndufaf5 | 0.037 | Old |
| 0.041391 | 0.755604 | 0.026 | 0.006 | 1 | 7 | Spata7 | 0.02 | Old |
| 0.041409 | 1.486039 | 0.09 | 0.051 | 1 | 7 | Ccdc163 | 0.039 | Old |
| 0.041433 | 0.901809 | 0.103 | 0.06 | 1 | 7 | Wrap73 | 0.043 | Old |
| 0.041674 | 2.065395 | 0.031 | 0.009 | 1 | 7 | Il18bp | 0.022 | Old |
| 0.041688 | 0.366275 | 0.079 | 0.042 | 1 | 7 | Cep89 | 0.037 | Old |
| 0.041695 | 0.613624 | 0.203 | 0.145 | 1 | 7 | Ly9 | 0.058 | Old |
| 0.041716 | 0.297992 | 0.495 | 0.407 | 1 | 7 | Immt | 0.088 | Old |
| 0.04182 | 1.240953 | 0.031 | 0.009 | 1 | 7 | Pcbp4 | 0.022 | Old |
| 0.041832 | 0.223005 | 0.269 | 0.202 | 1 | 7 | Cdk13 | 0.067 | Old |
| 0.041856 | 1.40749 | 0.054 | 0.024 | 1 | 7 | Cd4 | 0.03 | Old |
| 0.041957 | 0.341315 | 0.11 | 0.066 | 1 | 7 | Ring1 | 0.044 | Old |
| 0.042059 | 0.30843 | 0.577 | 0.542 | 1 | 7 | Shtn1 | 0.035 | Old |
| 0.042065 | 0.558215 | 0.036 | 0.012 | 1 | 7 | Ick | 0.024 | Old |
| 0.042118 | 0.335923 | 0.079 | 0.042 | 1 | 7 | Dhrs13 | 0.037 | Old |
| 0.042406 | 0.649363 | 0.031 | 0.009 | 1 | 7 | Pigl | 0.022 | Old |
| 0.042604 | 0.596584 | 0.036 | 0.012 | 1 | 7 | Tarbp1 | 0.024 | Old |
| 0.042674 | 0.238448 | 0.664 | 0.578 | 1 | 7 | Alyref | 0.086 | Old |
| 0.042714 | 1.720079 | 0.049 | 0.021 | 1 | 7 | Fzd1 | 0.028 | Old |
| 0.042733 | 0.583114 | 0.067 | 0.033 | 1 | 7 | Siah1b | 0.034 | Old |
| 0.04284 | 0.599442 | 0.315 | 0.25 | 1 | 7 | Rsl24d1 | 0.065 | Old |
| 0.042845 | 0.514905 | 0.072 | 0.036 | 1 | 7 | Ears2 | 0.036 | Old |
| 0.042897 | 0.715733 | 0.2 | 0.145 | 1 | 7 | Sumf1 | 0.055 | Old |
| 0.042926 | 0.617049 | 0.208 | 0.151 | 1 | 7 | Zfp984 | 0.057 | Old |
| 0.042941 | 0.743348 | 0.169 | 0.114 | 1 | 7 | Dnpep | 0.055 | Old |
| 0.043183 | 0.799343 | 0.087 | 0.048 | 1 | 7 | Zscan21 | 0.039 | Old |
| 0.043225 | 0.387879 | 0.228 | 0.166 | 1 | 7 | Zfp330 | 0.062 | Old |
| 0.043237 | 0.297945 | 0.474 | 0.398 | 1 | 7 | Cycs | 0.076 | Old |
| 0.043242 | 0.289798 | 0.133 | 0.084 | 1 | 7 | Stk16 | 0.049 | Old |
| 0.043244 | 0.284986 | 0.433 | 0.352 | 1 | 7 | Nhp2 | 0.081 | Old |

| | | | | | | | | |
|---|---|---|---|---|---|---|---|---|
| 0.043294 | 0.290358 | 0.244 | 0.175 | 1 | 7 | Dgcr8 | 0.069 | Old |
| 0.043308 | 0.353661 | 0.315 | 0.238 | 1 | 7 | Cept1 | 0.077 | Old |
| 0.043428 | 0.362981 | 0.177 | 0.12 | 1 | 7 | Ddx49 | 0.057 | Old |
| 0.043597 | 0.099054 | 0.092 | 0.051 | 1 | 7 | 2010315B0 | 0.041 | Old |
| 0.043634 | 0.826743 | 0.074 | 0.039 | 1 | 7 | Dnal4 | 0.035 | Old |
| 0.043669 | 0.238852 | 0.164 | 0.108 | 1 | 7 | Rtca | 0.056 | Old |
| 0.043894 | 0.264334 | 0.349 | 0.274 | 1 | 7 | Pabpn1 | 0.075 | Old |
| 0.043939 | 0.564153 | 0.277 | 0.214 | 1 | 7 | Mt2 | 0.063 | Old |
| 0.044063 | 0.38827 | 0.131 | 0.084 | 1 | 7 | Bcl2l13 | 0.047 | Old |
| 0.044079 | 0.266288 | 0.374 | 0.289 | 1 | 7 | Suds3 | 0.085 | Old |
| 0.044134 | 0.060073 | 0.133 | 0.084 | 1 | 7 | Tdrkh | 0.049 | Old |
| 0.044149 | 0.064802 | 0.421 | 0.337 | 1 | 7 | Kcnn4 | 0.084 | Old |
| 0.044233 | 0.206024 | 0.454 | 0.373 | 1 | 7 | Psmc5 | 0.081 | Old |
| 0.044256 | 0.38508 | 0.459 | 0.38 | 1 | 7 | Phf20l1 | 0.079 | Old |
| 0.044336 | 0.399514 | 0.118 | 0.072 | 1 | 7 | Sugp1 | 0.046 | Old |
| 0.044348 | 0.061078 | 0.226 | 0.163 | 1 | 7 | Mtr | 0.063 | Old |
| 0.044574 | 0.286423 | 0.638 | 0.566 | 1 | 7 | Morf4l2 | 0.072 | Old |
| 0.044672 | 0.342315 | 0.379 | 0.301 | 1 | 7 | Casp8ap2 | 0.078 | Old |
| 0.044726 | 0.341881 | 0.174 | 0.12 | 1 | 7 | Rnf157 | 0.054 | Old |
| 0.04474 | 0.137033 | 0.141 | 0.09 | 1 | 7 | Rnf8 | 0.051 | Old |
| 0.044826 | 0.380486 | 0.297 | 0.226 | 1 | 7 | Mrpl32 | 0.071 | Old |
| 0.044866 | 0.439122 | 0.408 | 0.334 | 1 | 7 | Snap23 | 0.074 | Old |
| 0.044901 | 0.662659 | 0.151 | 0.102 | 1 | 7 | Tirap | 0.049 | Old |
| 0.045105 | 0.565509 | 0.154 | 0.102 | 1 | 7 | Dnaja3 | 0.052 | Old |
| 0.04519 | 0.577187 | 0.067 | 0.033 | 1 | 7 | C130050O1 | 0.034 | Old |
| 0.045274 | 1.188653 | 0.049 | 0.021 | 1 | 7 | Dcp1b | 0.028 | Old |
| 0.045375 | 0.373205 | 0.297 | 0.232 | 1 | 7 | Sars | 0.065 | Old |
| 0.045389 | 0.39589 | 0.349 | 0.271 | 1 | 7 | Rgs1 | 0.078 | Old |
| 0.045551 | 0.54574 | 0.144 | 0.096 | 1 | 7 | Rnf41 | 0.048 | Old |
| 0.045612 | 0.670907 | 0.179 | 0.127 | 1 | 7 | Cdc25b | 0.052 | Old |
| 0.045649 | 0.706899 | 0.182 | 0.13 | 1 | 7 | Ptpn9 | 0.052 | Old |
| 0.045719 | 0.480525 | 0.39 | 0.328 | 1 | 7 | Tomm70a | 0.062 | Old |
| 0.045938 | 0.177255 | 0.497 | 0.401 | 1 | 7 | Eif1ax | 0.096 | Old |
| 0.046017 | 1.1697 | 0.062 | 0.03 | 1 | 7 | Gfi1b | 0.032 | Old |
| 0.046086 | 0.349452 | 0.115 | 0.069 | 1 | 7 | Eif2b3 | 0.046 | Old |
| 0.04623 | 0.839396 | 0.062 | 0.03 | 1 | 7 | Adnp | 0.032 | Old |
| 0.046296 | 0.557877 | 0.315 | 0.25 | 1 | 7 | Ms4a8a | 0.065 | Old |
| 0.046314 | 0.33036 | 0.523 | 0.434 | 1 | 7 | Ppp1r9b | 0.089 | Old |
| 0.046335 | 0.251882 | 0.305 | 0.232 | 1 | 7 | Cant1 | 0.073 | Old |
| 0.046406 | 0.243494 | 0.677 | 0.633 | 1 | 7 | Hnrnpc | 0.044 | Old |
| 0.046536 | 0.192172 | 0.582 | 0.497 | 1 | 7 | Elavl1 | 0.085 | Old |
| 0.046606 | 0.543192 | 0.233 | 0.175 | 1 | 7 | Gapt | 0.058 | Old |
| 0.046654 | 0.386913 | 0.162 | 0.108 | 1 | 7 | Gosr1 | 0.054 | Old |
| 0.046751 | 0.102034 | 0.846 | 0.798 | 1 | 7 | Hnrnpab | 0.048 | Old |
| 0.046851 | 0.296194 | 0.338 | 0.262 | 1 | 7 | Tcf3 | 0.076 | Old |

| | | | | | | | | |
|---|---|---|---|---|---|---|---|---|
| 0.046859 | 0.195811 | 0.851 | 0.807 | 1 | 7 | Pcbp1 | 0.044 | Old |
| 0.046862 | 0.439845 | 0.377 | 0.316 | 1 | 7 | Eif2s1 | 0.061 | Old |
| 0.046952 | 0.093907 | 0.59 | 0.5 | 1 | 7 | Rp2 | 0.09 | Old |
| 0.046994 | 0.127541 | 0.99 | 0.985 | 1 | 7 | Rps26 | 0.005 | Old |
| 0.047089 | 0.749085 | 0.164 | 0.114 | 1 | 7 | Prkaa1 | 0.05 | Old |
| 0.047151 | 0.841727 | 0.074 | 0.039 | 1 | 7 | Ube2o | 0.035 | Old |
| 0.047431 | 0.250355 | 0.526 | 0.443 | 1 | 7 | Mapk14 | 0.083 | Old |
| 0.047449 | 0.132181 | 0.067 | 0.033 | 1 | 7 | Shq1 | 0.034 | Old |
| 0.047601 | 0.418077 | 0.118 | 0.072 | 1 | 7 | Fbxo42 | 0.046 | Old |
| 0.047983 | 0.047257 | 0.167 | 0.111 | 1 | 7 | Smg6 | 0.056 | Old |
| 0.04802 | 0.210482 | 0.836 | 0.795 | 1 | 7 | Btg1 | 0.041 | Old |
| 0.048022 | 0.239335 | 0.556 | 0.473 | 1 | 7 | Rcc2 | 0.083 | Old |
| 0.048036 | 0.829758 | 0.123 | 0.078 | 1 | 7 | Gm12840 | 0.045 | Old |
| 0.048042 | 0.621563 | 0.074 | 0.039 | 1 | 7 | Zfp143 | 0.035 | Old |
| 0.048167 | 0.390225 | 0.192 | 0.136 | 1 | 7 | Ubqln4 | 0.056 | Old |
| 0.048349 | 0.700479 | 0.231 | 0.175 | 1 | 7 | Anapc4 | 0.056 | Old |
| 0.048614 | 0.365852 | 0.438 | 0.361 | 1 | 7 | Papd4 | 0.077 | Old |
| 0.048663 | 0.140851 | 0.628 | 0.542 | 1 | 7 | Map4k4 | 0.086 | Old |
| 0.048721 | 0.35269 | 0.41 | 0.334 | 1 | 7 | Rgl1 | 0.076 | Old |
| 0.048745 | 0.17526 | 0.121 | 0.075 | 1 | 7 | Gabpb1 | 0.046 | Old |
| 0.048769 | 0.2696 | 0.138 | 0.09 | 1 | 7 | Zfp101 | 0.048 | Old |
| 0.049054 | 0.277624 | 0.387 | 0.307 | 1 | 7 | Guk1 | 0.08 | Old |
| 0.049093 | 0.183493 | 0.472 | 0.404 | 1 | 7 | Psmb5 | 0.068 | Old |
| 0.049409 | 0.496344 | 0.128 | 0.081 | 1 | 7 | Tmem237 | 0.047 | Old |
| 0.049441 | 0.368114 | 0.364 | 0.298 | 1 | 7 | Lgals9 | 0.066 | Old |
| 0.049581 | 1.450123 | 0.044 | 0.018 | 1 | 7 | Eral1 | 0.026 | Old |
| 0.049591 | 0.302427 | 0.405 | 0.328 | 1 | 7 | Acadl | 0.077 | Old |
| 0.049598 | 0.068244 | 0.156 | 0.105 | 1 | 7 | Dok3 | 0.051 | Old |
| 0.049703 | 0.293603 | 0.621 | 0.566 | 1 | 7 | Pycard | 0.055 | Old |
| 0.049706 | 0.783783 | 0.159 | 0.111 | 1 | 7 | Cherp | 0.048 | Old |
| 0.049781 | 0.440566 | 0.251 | 0.19 | 1 | 7 | D5Ertd579 | 0.061 | Old |
| 0.049785 | 0.322208 | 0.272 | 0.205 | 1 | 7 | Fam133b | 0.067 | Old |
| 0.049844 | 0.351786 | 0.279 | 0.211 | 1 | 7 | Snrnp48 | 0.068 | Old |
| 0.04989 | 0.332277 | 0.641 | 0.602 | 1 | 7 | Ssr1 | 0.039 | Old |
| 0.05004 | 0.220249 | 0.2 | 0.142 | 1 | 7 | Esrra | 0.058 | Old |
| 0.050225 | 0.319927 | 0.659 | 0.587 | 1 | 7 | Ube2s | 0.072 | Old |
| 0.050342 | 0.289704 | 0.305 | 0.235 | 1 | 7 | Acaa2 | 0.07 | Old |
| 0.050387 | 0.284033 | 0.367 | 0.295 | 1 | 7 | Ube2g2 | 0.072 | Old |
| 0.050389 | 0.275052 | 0.251 | 0.187 | 1 | 7 | Mrpl35 | 0.064 | Old |
| 0.050408 | 0.279429 | 0.674 | 0.575 | 1 | 7 | Selenof | 0.099 | Old |
| 0.050477 | 0.247603 | 0.528 | 0.461 | 1 | 7 | 1110008F1 | 0.067 | Old |
| 0.05063 | 0.550051 | 0.123 | 0.078 | 1 | 7 | Zfand2b | 0.045 | Old |
| 0.050637 | 0.201181 | 0.708 | 0.687 | 1 | 7 | Atp6v1b2 | 0.021 | Old |
| 0.050923 | 0.131254 | 0.664 | 0.602 | 1 | 7 | Gusb | 0.062 | Old |
| 0.050924 | 0.306137 | 0.285 | 0.217 | 1 | 7 | Znrd1 | 0.068 | Old |

| | | | | | | | | |
|---|---|---|---|---|---|---|---|---|
| 0.051042 | 0.131998 | 0.646 | 0.581 | 1 | 7 | Shisa5 | 0.065 | Old |
| 0.051106 | 0.32452 | 0.244 | 0.184 | 1 | 7 | Ube4a | 0.06 | Old |
| 0.051133 | 0.387519 | 0.136 | 0.087 | 1 | 7 | Thoc1 | 0.049 | Old |
| 0.051158 | 0.73188 | 0.1 | 0.06 | 1 | 7 | Naf1 | 0.04 | Old |
| 0.051169 | 0.60494 | 0.156 | 0.105 | 1 | 7 | Riok2 | 0.051 | Old |
| 0.051187 | 0.520608 | 0.162 | 0.108 | 1 | 7 | Cdc23 | 0.054 | Old |
| 0.051211 | 0.232904 | 0.815 | 0.804 | 1 | 7 | Slc25a5 | 0.011 | Old |
| 0.051323 | 0.135721 | 0.541 | 0.455 | 1 | 7 | Tmem258 | 0.086 | Old |
| 0.051367 | 0.190234 | 0.259 | 0.19 | 1 | 7 | Manba | 0.069 | Old |
| 0.0514 | 0.191486 | 0.618 | 0.527 | 1 | 7 | Sgpl1 | 0.091 | Old |
| 0.051452 | 0.52336 | 0.146 | 0.096 | 1 | 7 | Rrp8 | 0.05 | Old |
| 0.051468 | 0.247481 | 0.659 | 0.587 | 1 | 7 | Capza1 | 0.072 | Old |
| 0.051514 | 0.413097 | 0.415 | 0.346 | 1 | 7 | Vars | 0.069 | Old |
| 0.051624 | 0.437215 | 0.182 | 0.127 | 1 | 7 | Sfxn2 | 0.055 | Old |
| 0.051683 | 0.446508 | 0.085 | 0.048 | 1 | 7 | Dolk | 0.037 | Old |
| 0.051881 | 0.699558 | 0.085 | 0.048 | 1 | 7 | Rnf113a2 | 0.037 | Old |
| 0.051887 | 0.352603 | 0.331 | 0.271 | 1 | 7 | Chchd3 | 0.06 | Old |
| 0.051907 | 0.263964 | 0.744 | 0.708 | 1 | 7 | S100a4 | 0.036 | Old |
| 0.052115 | 0.399196 | 0.105 | 0.063 | 1 | 7 | Tma16 | 0.042 | Old |
| 0.052368 | 0.46267 | 0.328 | 0.256 | 1 | 7 | Tars | 0.072 | Old |
| 0.052472 | 0.253118 | 0.236 | 0.175 | 1 | 7 | Sucla2 | 0.061 | Old |
| 0.052546 | 0.28334 | 0.674 | 0.633 | 1 | 7 | Hspe1 | 0.041 | Old |
| 0.05277 | 0.498976 | 0.223 | 0.169 | 1 | 7 | Lpar1 | 0.054 | Old |
| 0.052814 | 0.340211 | 0.354 | 0.286 | 1 | 7 | Cmas | 0.068 | Old |
| 0.053034 | 0.166694 | 0.249 | 0.181 | 1 | 7 | Dlgap5 | 0.068 | Old |
| 0.053055 | 0.641189 | 0.192 | 0.136 | 1 | 7 | Mtx1 | 0.056 | Old |
| 0.053158 | 0.285315 | 0.444 | 0.355 | 1 | 7 | Lamtor3 | 0.089 | Old |
| 0.053296 | 1.49692 | 0.051 | 0.024 | 1 | 7 | Fntb | 0.027 | Old |
| 0.05344 | 0.122147 | 0.385 | 0.31 | 1 | 7 | Ube2j2 | 0.075 | Old |
| 0.053477 | 0.26228 | 0.677 | 0.605 | 1 | 7 | Nktr | 0.072 | Old |
| 0.053501 | 0.43722 | 0.062 | 0.03 | 1 | 7 | Hmga1 | 0.032 | Old |
| 0.053517 | 0.37364 | 0.185 | 0.133 | 1 | 7 | Adar | 0.052 | Old |
| 0.053603 | 0.195733 | 0.277 | 0.208 | 1 | 7 | Nrf1 | 0.069 | Old |
| 0.053676 | 0.197282 | 0.187 | 0.133 | 1 | 7 | Spns1 | 0.054 | Old |
| 0.053693 | 0.319909 | 0.151 | 0.102 | 1 | 7 | Borcs5 | 0.049 | Old |
| 0.053761 | 0.164368 | 0.308 | 0.232 | 1 | 7 | Raf1 | 0.076 | Old |
| 0.053829 | 0.511078 | 0.195 | 0.139 | 1 | 7 | Psmd5 | 0.056 | Old |
| 0.053877 | 0.216756 | 0.851 | 0.849 | 1 | 7 | Cebpb | 0.002 | Old |
| 0.054063 | 0.295478 | 0.333 | 0.265 | 1 | 7 | Parp14 | 0.068 | Old |
| 0.054156 | 1.933915 | 0.038 | 0.015 | 1 | 7 | Ccdc73 | 0.023 | Old |
| 0.054184 | 0.218048 | 0.162 | 0.108 | 1 | 7 | Sec16a | 0.054 | Old |
| 0.054218 | 0.805835 | 0.126 | 0.081 | 1 | 7 | Bnip3 | 0.045 | Old |
| 0.054313 | 2.024684 | 0.038 | 0.015 | 1 | 7 | Zfp951 | 0.023 | Old |
| 0.054386 | 0.530609 | 0.126 | 0.081 | 1 | 7 | Cmss1 | 0.045 | Old |
| 0.054411 | 0.473461 | 0.044 | 0.018 | 1 | 7 | Kdelc1 | 0.026 | Old |

| | | | | | | | | |
|---|---|---|---|---|---|---|---|---|
| 0.054481 | 0.036158 | 0.444 | 0.343 | 1 | 7 | Ccnt1 | 0.101 | Old |
| 0.054495 | 0.518865 | 0.092 | 0.054 | 1 | 7 | Vps8 | 0.038 | Old |
| 0.054562 | 0.179031 | 0.71 | 0.654 | 1 | 7 | Atp1b3 | 0.056 | Old |
| 0.054732 | 0.546732 | 0.326 | 0.259 | 1 | 7 | Plgrkt | 0.067 | Old |
| 0.054814 | 0.297578 | 0.295 | 0.232 | 1 | 7 | Nudt5 | 0.063 | Old |
| 0.054901 | 0.588959 | 0.303 | 0.244 | 1 | 7 | Rnft1 | 0.059 | Old |
| 0.054908 | 1.078663 | 0.087 | 0.051 | 1 | 7 | Cd200r4 | 0.036 | Old |
| 0.055027 | 0.549632 | 0.085 | 0.048 | 1 | 7 | Pus7 | 0.037 | Old |
| 0.055063 | 0.349445 | 0.249 | 0.187 | 1 | 7 | Pmpca | 0.062 | Old |
| 0.055063 | 0.358717 | 0.272 | 0.205 | 1 | 7 | Dph3 | 0.067 | Old |
| 0.055219 | 0.781311 | 0.11 | 0.069 | 1 | 7 | Acap3 | 0.041 | Old |
| 0.055307 | 0.424971 | 0.2 | 0.145 | 1 | 7 | Eif2ak2 | 0.055 | Old |
| 0.055334 | 0.18723 | 0.749 | 0.693 | 1 | 7 | Zfand5 | 0.056 | Old |
| 0.055438 | 1.447496 | 0.018 | 0.003 | 1 | 7 | 4930518I1! | 0.015 | Old |
| 0.055445 | 0.503261 | 0.2 | 0.145 | 1 | 7 | Ecd | 0.055 | Old |
| 0.055527 | 0.555878 | 0.254 | 0.196 | 1 | 7 | Nsmaf | 0.058 | Old |
| 0.055602 | 0.306928 | 0.274 | 0.211 | 1 | 7 | Snx12 | 0.063 | Old |
| 0.055664 | 0.461222 | 0.205 | 0.151 | 1 | 7 | Nr2f6 | 0.054 | Old |
| 0.055682 | 0.317122 | 0.418 | 0.358 | 1 | 7 | Kif15 | 0.06 | Old |
| 0.05569 | 2.173755 | 0.018 | 0.003 | 1 | 7 | Tnxb | 0.015 | Old |
| 0.055696 | 0.307674 | 0.336 | 0.259 | 1 | 7 | Pi4k2a | 0.077 | Old |
| 0.055705 | 0.140607 | 0.626 | 0.566 | 1 | 7 | Pirb | 0.06 | Old |
| 0.055717 | 0.366607 | 0.449 | 0.392 | 1 | 7 | Stt3b | 0.057 | Old |
| 0.0558 | 0.6082 | 0.21 | 0.157 | 1 | 7 | Mid1 | 0.053 | Old |
| 0.05588 | 0.204162 | 0.467 | 0.377 | 1 | 7 | Mrpl18 | 0.09 | Old |
| 0.055944 | 1.771092 | 0.018 | 0.003 | 1 | 7 | AC129328. | 0.015 | Old |
| 0.055944 | 1.51198 | 0.018 | 0.003 | 1 | 7 | Igf2bp2 | 0.015 | Old |
| 0.055944 | 1.107064 | 0.018 | 0.003 | 1 | 7 | Gm16701.: | 0.015 | Old |
| 0.055969 | 0.759214 | 0.085 | 0.048 | 1 | 7 | Ttc7b | 0.037 | Old |
| 0.056026 | 0.515889 | 0.056 | 0.027 | 1 | 7 | Tmem35b | 0.029 | Old |
| 0.056072 | 1.082445 | 0.038 | 0.015 | 1 | 7 | Piwil2 | 0.023 | Old |
| 0.056075 | 0.936493 | 0.085 | 0.048 | 1 | 7 | Dmac2 | 0.037 | Old |
| 0.056185 | 0.192164 | 0.762 | 0.69 | 1 | 7 | Snrpb | 0.072 | Old |
| 0.056273 | 0.121689 | 0.01 | 0.03 | 1 | 7 | Intu | -0.02 | Old |
| 0.056453 | 2.168294 | 0.018 | 0.003 | 1 | 7 | Zfp532 | 0.015 | Old |
| 0.056667 | 0.192088 | 0.346 | 0.274 | 1 | 7 | Med1 | 0.072 | Old |
| 0.056825 | 0.481692 | 0.115 | 0.072 | 1 | 7 | Nat10 | 0.043 | Old |
| 0.056838 | 0.594684 | 0.1 | 0.06 | 1 | 7 | C2cd2 | 0.04 | Old |
| 0.056887 | 0.243503 | 0.1 | 0.06 | 1 | 7 | Slc2a1 | 0.04 | Old |
| 0.056901 | 0.499941 | 0.241 | 0.181 | 1 | 7 | Ten1 | 0.06 | Old |
| 0.056985 | 0.526916 | 0.1 | 0.06 | 1 | 7 | Zdhhc4 | 0.04 | Old |
| 0.057032 | 0.599267 | 0.108 | 0.066 | 1 | 7 | Gcat | 0.042 | Old |
| 0.057032 | 0.517363 | 0.085 | 0.048 | 1 | 7 | Polr3d | 0.037 | Old |
| 0.057073 | 0.36766 | 0.177 | 0.123 | 1 | 7 | Eif2b4 | 0.054 | Old |
| 0.057193 | 0.273473 | 0.131 | 0.084 | 1 | 7 | Appl2 | 0.047 | Old |

| | | | | | | | | |
|---|---|---|---|---|---|---|---|---|
| 0.057223 | 0.782184 | 0.018 | 0.003 | 1 | 7 | 1700061G1 | 0.015 | Old |
| 0.057223 | 0.709218 | 0.018 | 0.003 | 1 | 7 | Car6 | 0.015 | Old |
| 0.057223 | 0.565015 | 0.018 | 0.003 | 1 | 7 | Napb | 0.015 | Old |
| 0.057223 | 0.427292 | 0.018 | 0.003 | 1 | 7 | CAAA0114 | 0.015 | Old |
| 0.057313 | 0.482285 | 0.523 | 0.47 | 1 | 7 | Fgl2 | 0.053 | Old |
| 0.05732 | 0.377634 | 0.136 | 0.09 | 1 | 7 | Zfp410 | 0.046 | Old |
| 0.057396 | 0.373548 | 0.208 | 0.151 | 1 | 7 | Traf3 | 0.057 | Old |
| 0.057409 | 0.850794 | 0.123 | 0.078 | 1 | 7 | Abcc4 | 0.045 | Old |
| 0.057423 | 0.61212 | 0.179 | 0.127 | 1 | 7 | Pcyox1l | 0.052 | Old |
| 0.057452 | 0.457236 | 0.385 | 0.313 | 1 | 7 | Usp14 | 0.072 | Old |
| 0.057697 | 0.478905 | 0.487 | 0.434 | 1 | 7 | Tcirg1 | 0.053 | Old |
| 0.057796 | 0.149942 | 0.349 | 0.274 | 1 | 7 | G2e3 | 0.075 | Old |
| 0.057947 | 0.426052 | 0.351 | 0.292 | 1 | 7 | Idi1 | 0.059 | Old |
| 0.058117 | 0.387013 | 0.577 | 0.512 | 1 | 7 | Prkar2a | 0.065 | Old |
| 0.058291 | 0.575025 | 0.056 | 0.027 | 1 | 7 | Rgs12 | 0.029 | Old |
| 0.058391 | 0.580487 | 0.154 | 0.105 | 1 | 7 | C1galt1c1 | 0.049 | Old |
| 0.058493 | 0.107128 | 0.579 | 0.518 | 1 | 7 | Pik3cd | 0.061 | Old |
| 0.058513 | 1.117114 | 0.051 | 0.024 | 1 | 7 | Dgkq | 0.027 | Old |
| 0.058527 | 0.3543 | 0.277 | 0.22 | 1 | 7 | Gtf3c6 | 0.057 | Old |
| 0.058554 | 0.319818 | 0.185 | 0.133 | 1 | 7 | Coq5 | 0.052 | Old |
| 0.058688 | 0.020514 | 0.423 | 0.343 | 1 | 7 | Klhl9 | 0.08 | Old |
| 0.058768 | 0.581721 | 0.085 | 0.048 | 1 | 7 | Papolg | 0.037 | Old |
| 0.058857 | 0.254535 | 0.146 | 0.099 | 1 | 7 | Ttf1 | 0.047 | Old |
| 0.059155 | 0.436325 | 0.141 | 0.093 | 1 | 7 | Rhbdf2 | 0.048 | Old |
| 0.05945 | 0.587478 | 0.203 | 0.148 | 1 | 7 | Nol12 | 0.055 | Old |
| 0.059454 | 0.476347 | 0.421 | 0.355 | 1 | 7 | Snw1 | 0.066 | Old |
| 0.059561 | 0.665675 | 0.113 | 0.072 | 1 | 7 | Gpr137 | 0.041 | Old |
| 0.059615 | 0.738697 | 0.115 | 0.075 | 1 | 7 | Bnip1 | 0.04 | Old |
| 0.059651 | 0.035395 | 0.274 | 0.205 | 1 | 7 | Atxn3 | 0.069 | Old |
| 0.059656 | 0.320793 | 0.397 | 0.328 | 1 | 7 | Mrpl30 | 0.069 | Old |
| 0.059668 | 0.32755 | 0.218 | 0.163 | 1 | 7 | Bloc1s2 | 0.055 | Old |
| 0.05972 | 0.108453 | 0.364 | 0.289 | 1 | 7 | Atf1 | 0.075 | Old |
| 0.059808 | 0.751331 | 0.051 | 0.024 | 1 | 7 | Cdc42bpg | 0.027 | Old |
| 0.059903 | 0.389244 | 0.038 | 0.015 | 1 | 7 | Runx2 | 0.023 | Old |
| 0.059926 | 0.056405 | 0.315 | 0.241 | 1 | 7 | Snrnp40 | 0.074 | Old |
| 0.059936 | 0.852777 | 0.095 | 0.057 | 1 | 7 | Parp10 | 0.038 | Old |
| 0.060184 | 0.489864 | 0.169 | 0.117 | 1 | 7 | Sntb2 | 0.052 | Old |
| 0.060375 | 0.1664 | 0.756 | 0.672 | 1 | 7 | Mrpl33 | 0.084 | Old |
| 0.060606 | 0.322685 | 0.305 | 0.238 | 1 | 7 | Mrpl15 | 0.067 | Old |
| 0.06063 | 0.1758 | 0.454 | 0.386 | 1 | 7 | Sec31a | 0.068 | Old |
| 0.060826 | 0.419578 | 0.341 | 0.274 | 1 | 7 | Senp2 | 0.067 | Old |
| 0.060873 | 1.203752 | 0.059 | 0.03 | 1 | 7 | Nkrf | 0.029 | Old |
| 0.06092 | 0.67927 | 0.128 | 0.084 | 1 | 7 | Med9 | 0.044 | Old |
| 0.061055 | 0.445674 | 0.403 | 0.346 | 1 | 7 | C1qbp | 0.057 | Old |
| 0.061057 | 0.448189 | 0.351 | 0.283 | 1 | 7 | Ube2a | 0.068 | Old |

| | | | | | | | | |
|---|---|---|---|---|---|---|---|---|
| 0.06114 | 0.4264 | 0.185 | 0.133 | 1 | 7 | Vamp7 | 0.052 | Old |
| 0.061165 | 2.414966 | 0.023 | 0.006 | 1 | 7 | Zbtb48 | 0.017 | Old |
| 0.061201 | 0.277044 | 0.351 | 0.28 | 1 | 7 | Prpf8 | 0.071 | Old |
| 0.061207 | 0.367945 | 0.095 | 0.057 | 1 | 7 | Pgpep1 | 0.038 | Old |
| 0.061326 | 0.503637 | 0.321 | 0.256 | 1 | 7 | Cdc34 | 0.065 | Old |
| 0.061399 | 1.589146 | 0.023 | 0.006 | 1 | 7 | A630089N( | 0.017 | Old |
| 0.061423 | 0.853782 | 0.087 | 0.051 | 1 | 7 | Rnf219 | 0.036 | Old |
| 0.061546 | 0.155033 | 0.215 | 0.16 | 1 | 7 | Mmgt1 | 0.055 | Old |
| 0.061772 | 0.313416 | 0.19 | 0.139 | 1 | 7 | Pdk3 | 0.051 | Old |
| 0.061805 | 2.297995 | 0.028 | 0.009 | 1 | 7 | Dyrk3 | 0.019 | Old |
| 0.061872 | 0.538126 | 0.087 | 0.051 | 1 | 7 | Gm13373 | 0.036 | Old |
| 0.06191 | 0.432004 | 0.187 | 0.133 | 1 | 7 | Copg2 | 0.054 | Old |
| 0.062064 | 0.49775 | 0.123 | 0.081 | 1 | 7 | 0610009B2 | 0.042 | Old |
| 0.06208 | 0.279807 | 0.356 | 0.283 | 1 | 7 | Prkx | 0.073 | Old |
| 0.06212 | 0.694446 | 0.033 | 0.012 | 1 | 7 | Mks1 | 0.021 | Old |
| 0.062222 | 0.341696 | 0.423 | 0.349 | 1 | 7 | Fip1l1 | 0.074 | Old |
| 0.062232 | 0.143715 | 0.308 | 0.238 | 1 | 7 | Igsf6 | 0.07 | Old |
| 0.062268 | 0.380108 | 0.118 | 0.075 | 1 | 7 | Ctu2 | 0.043 | Old |
| 0.062274 | 0.288966 | 0.41 | 0.331 | 1 | 7 | Pld3 | 0.079 | Old |
| 0.062394 | 0.274278 | 0.6 | 0.533 | 1 | 7 | Racgap1 | 0.067 | Old |
| 0.062434 | 1.390244 | 0.028 | 0.009 | 1 | 7 | Bdh2 | 0.019 | Old |
| 0.062503 | 0.437329 | 0.318 | 0.253 | 1 | 7 | Churc1 | 0.065 | Old |
| 0.062527 | 1.210695 | 0.059 | 0.03 | 1 | 7 | Mus81 | 0.029 | Old |
| 0.062586 | 0.345569 | 0.146 | 0.099 | 1 | 7 | Sc5d | 0.047 | Old |
| 0.062785 | 0.637941 | 0.113 | 0.072 | 1 | 7 | Msi2 | 0.041 | Old |
| 0.062817 | 0.355444 | 0.023 | 0.006 | 1 | 7 | Insl6 | 0.017 | Old |
| 0.062916 | 0.391805 | 0.597 | 0.533 | 1 | 7 | Rpl13a | 0.064 | Old |
| 0.063201 | 0.162081 | 0.636 | 0.56 | 1 | 7 | Scp2 | 0.076 | Old |
| 0.063256 | 0.139157 | 0.518 | 0.434 | 1 | 7 | Cnih1 | 0.084 | Old |
| 0.063352 | 0.530117 | 0.195 | 0.142 | 1 | 7 | Mrpl40 | 0.053 | Old |
| 0.063357 | 0.215026 | 0.238 | 0.175 | 1 | 7 | Fundc1 | 0.063 | Old |
| 0.0634 | 0.604575 | 0.318 | 0.25 | 1 | 7 | Samd1 | 0.068 | Old |
| 0.063473 | 0.268346 | 0.126 | 0.081 | 1 | 7 | Map2k5 | 0.045 | Old |
| 0.063506 | 0.398019 | 0.367 | 0.301 | 1 | 7 | Spg21 | 0.066 | Old |
| 0.063521 | 0.853829 | 0.079 | 0.045 | 1 | 7 | Dusp18 | 0.034 | Old |
| 0.063536 | 0.942711 | 0.023 | 0.006 | 1 | 7 | Syngr1 | 0.017 | Old |
| 0.063536 | 0.891405 | 0.023 | 0.006 | 1 | 7 | Irgc1 | 0.017 | Old |
| 0.063814 | 0.604314 | 0.128 | 0.084 | 1 | 7 | Cog6 | 0.044 | Old |
| 0.063966 | 0.300341 | 0.221 | 0.166 | 1 | 7 | Gtf2f1 | 0.055 | Old |
| 0.064195 | 0.881173 | 0.128 | 0.084 | 1 | 7 | Gmppa | 0.044 | Old |
| 0.064356 | 1.55601 | 0.028 | 0.009 | 1 | 7 | Kcnd1 | 0.019 | Old |
| 0.064356 | 1.010362 | 0.028 | 0.009 | 1 | 7 | Ift88 | 0.019 | Old |
| 0.064356 | 0.891064 | 0.028 | 0.009 | 1 | 7 | Tm7sf2 | 0.019 | Old |
| 0.064421 | 0.321639 | 0.144 | 0.099 | 1 | 7 | G6pc3 | 0.045 | Old |
| 0.064489 | 0.600005 | 0.2 | 0.148 | 1 | 7 | Casp1 | 0.052 | Old |

| | | | | | | | | |
|---|---|---|---|---|---|---|---|---|
| 0.064502 | 0.554999 | 0.059 | 0.03 | 1 | 7 | Zfp518b | 0.029 | Old |
| 0.064572 | 0.828789 | 0.028 | 0.009 | 1 | 7 | Taco1 | 0.019 | Old |
| 0.064591 | 0.300881 | 0.318 | 0.25 | 1 | 7 | Nemf | 0.068 | Old |
| 0.064604 | 0.121204 | 0.459 | 0.392 | 1 | 7 | Eif3d | 0.067 | Old |
| 0.064615 | 0.693081 | 0.079 | 0.045 | 1 | 7 | Zbtb33 | 0.034 | Old |
| 0.064639 | 1.991504 | 0.01 | 0 | 1 | 7 | Hoxb6 | 0.01 | Old |
| 0.064639 | 1.862585 | 0.01 | 0 | 1 | 7 | Wdr31 | 0.01 | Old |
| 0.064639 | 1.827133 | 0.01 | 0 | 1 | 7 | Cdh6 | 0.01 | Old |
| 0.064639 | 1.700678 | 0.01 | 0 | 1 | 7 | Nbl1 | 0.01 | Old |
| 0.064639 | 1.699481 | 0.01 | 0 | 1 | 7 | Akr1c19 | 0.01 | Old |
| 0.064639 | 1.675604 | 0.01 | 0 | 1 | 7 | Rnf128 | 0.01 | Old |
| 0.064639 | 1.548365 | 0.01 | 0 | 1 | 7 | Kcnc3 | 0.01 | Old |
| 0.064639 | 1.50833 | 0.01 | 0 | 1 | 7 | Dpyd | 0.01 | Old |
| 0.064639 | 1.478733 | 0.01 | 0 | 1 | 7 | Guca1b | 0.01 | Old |
| 0.064639 | 1.447717 | 0.01 | 0 | 1 | 7 | Ace | 0.01 | Old |
| 0.064639 | 1.394564 | 0.01 | 0 | 1 | 7 | 4833413G1 | 0.01 | Old |
| 0.064639 | 1.343087 | 0.01 | 0 | 1 | 7 | Il12a | 0.01 | Old |
| 0.064639 | 1.337477 | 0.01 | 0 | 1 | 7 | Zfp869 | 0.01 | Old |
| 0.064639 | 1.282507 | 0.01 | 0 | 1 | 7 | Peg12 | 0.01 | Old |
| 0.064639 | 1.218974 | 0.01 | 0 | 1 | 7 | Nhlrc1 | 0.01 | Old |
| 0.064639 | 1.206403 | 0.01 | 0 | 1 | 7 | Pnp2 | 0.01 | Old |
| 0.064639 | 1.18598 | 0.01 | 0 | 1 | 7 | 1700029I1 | 0.01 | Old |
| 0.064639 | 1.160611 | 0.01 | 0 | 1 | 7 | Morc4 | 0.01 | Old |
| 0.064639 | 1.156822 | 0.01 | 0 | 1 | 7 | Cyb5d1 | 0.01 | Old |
| 0.064639 | 1.081875 | 0.01 | 0 | 1 | 7 | Camkk1 | 0.01 | Old |
| 0.064639 | 1.072594 | 0.01 | 0 | 1 | 7 | Nat14 | 0.01 | Old |
| 0.064639 | 1.061219 | 0.01 | 0 | 1 | 7 | Ltf | 0.01 | Old |
| 0.064639 | 1.051969 | 0.01 | 0 | 1 | 7 | Art2b | 0.01 | Old |
| 0.064639 | 1.040018 | 0.01 | 0 | 1 | 7 | Mcam | 0.01 | Old |
| 0.064639 | 1.032483 | 0.01 | 0 | 1 | 7 | Zfp54 | 0.01 | Old |
| 0.064639 | 0.983182 | 0.01 | 0 | 1 | 7 | Hoxa1 | 0.01 | Old |
| 0.064639 | 0.955985 | 0.01 | 0 | 1 | 7 | Plekhf1 | 0.01 | Old |
| 0.064639 | 0.950776 | 0.01 | 0 | 1 | 7 | Unc5b | 0.01 | Old |
| 0.064639 | 0.862359 | 0.01 | 0 | 1 | 7 | Tmem79 | 0.01 | Old |
| 0.064639 | 0.856788 | 0.01 | 0 | 1 | 7 | Gm26733 | 0.01 | Old |
| 0.064639 | 0.837241 | 0.01 | 0 | 1 | 7 | Gdap1l1 | 0.01 | Old |
| 0.064639 | 0.836932 | 0.01 | 0 | 1 | 7 | Pemt | 0.01 | Old |
| 0.064639 | 0.828176 | 0.01 | 0 | 1 | 7 | Vps25 | 0.01 | Old |
| 0.064639 | 0.819301 | 0.01 | 0 | 1 | 7 | Oit3 | 0.01 | Old |
| 0.064639 | 0.797481 | 0.01 | 0 | 1 | 7 | Gm8797 | 0.01 | Old |
| 0.064639 | 0.776713 | 0.01 | 0 | 1 | 7 | Plppr2 | 0.01 | Old |
| 0.064639 | 0.759333 | 0.01 | 0 | 1 | 7 | Gm26912 | 0.01 | Old |
| 0.064639 | 0.724355 | 0.01 | 0 | 1 | 7 | Cmbl | 0.01 | Old |
| 0.064639 | 0.704329 | 0.01 | 0 | 1 | 7 | Xylb | 0.01 | Old |
| 0.064639 | 0.658642 | 0.01 | 0 | 1 | 7 | Krt18 | 0.01 | Old |

| | | | | | | | | |
|---|---|---|---|---|---|---|---|---|
| 0.064639 | 0.63233 | 0.01 | 0 | 1 | 7 | Nek3 | 0.01 | Old |
| 0.064722 | 0.409651 | 0.421 | 0.355 | 1 | 7 | Rab11fip1 | 0.066 | Old |
| 0.06474 | 0.112384 | 0.549 | 0.473 | 1 | 7 | Ppp2r1a | 0.076 | Old |
| 0.064771 | 0.154412 | 0.128 | 0.084 | 1 | 7 | Slc25a16 | 0.044 | Old |
| 0.064932 | 0.807027 | 0.059 | 0.03 | 1 | 7 | 9930104L0 | 0.029 | Old |
| 0.064999 | 0.194566 | 0.5 | 0.422 | 1 | 7 | Lypla2 | 0.078 | Old |
| 0.065007 | 0.461715 | 0.028 | 0.009 | 1 | 7 | Jmjd7 | 0.019 | Old |
| 0.065102 | 0.157682 | 0.477 | 0.392 | 1 | 7 | Dot1l | 0.085 | Old |
| 0.065225 | 0.332912 | 0.028 | 0.009 | 1 | 7 | Ropn1l | 0.019 | Old |
| 0.065234 | 0.550795 | 0.308 | 0.253 | 1 | 7 | Atg5 | 0.055 | Old |
| 0.065351 | 0.354621 | 0.123 | 0.081 | 1 | 7 | Depdc5 | 0.042 | Old |
| 0.065365 | 0.792738 | 0.059 | 0.03 | 1 | 7 | Gm26870 | 0.029 | Old |
| 0.065387 | 0.338315 | 0.051 | 0.024 | 1 | 7 | Macrod1 | 0.027 | Old |
| 0.065547 | 0.462749 | 0.179 | 0.13 | 1 | 7 | Dlat | 0.049 | Old |
| 0.065649 | 0.66281 | 0.067 | 0.036 | 1 | 7 | Tapt1 | 0.031 | Old |
| 0.065671 | 0.201204 | 0.164 | 0.114 | 1 | 7 | Tmem97 | 0.05 | Old |
| 0.065728 | 0.306426 | 0.297 | 0.232 | 1 | 7 | Slc11a1 | 0.065 | Old |
| 0.066127 | 0.178957 | 0.449 | 0.377 | 1 | 7 | Chtop | 0.072 | Old |
| 0.066171 | 0.56709 | 0.095 | 0.057 | 1 | 7 | Panx1 | 0.038 | Old |
| 0.066194 | 0.825008 | 0.067 | 0.036 | 1 | 7 | Sh2b2 | 0.031 | Old |
| 0.06622 | 0.280239 | 0.29 | 0.232 | 1 | 7 | H2-T22 | 0.058 | Old |
| 0.066252 | 0.574964 | 0.144 | 0.099 | 1 | 7 | Las1l | 0.045 | Old |
| 0.066483 | 0.68294 | 0.046 | 0.021 | 1 | 7 | Ankzf1 | 0.025 | Old |
| 0.066502 | 0.58243 | 0.126 | 0.081 | 1 | 7 | Kif19a | 0.045 | Old |
| 0.066536 | 0.755321 | 0.11 | 0.072 | 1 | 7 | Lmf1 | 0.038 | Old |
| 0.066583 | 0.502675 | 0.174 | 0.123 | 1 | 7 | Bad | 0.051 | Old |
| 0.066716 | 0.29073 | 0.179 | 0.13 | 1 | 7 | Tbck | 0.049 | Old |
| 0.066782 | 0.206329 | 0.346 | 0.274 | 1 | 7 | Cd37 | 0.072 | Old |
| 0.066845 | 0.252607 | 0.395 | 0.322 | 1 | 7 | Mff | 0.073 | Old |
| 0.066981 | 0.909414 | 0.097 | 0.06 | 1 | 7 | Fanca | 0.037 | Old |
| 0.066987 | 1.164945 | 0.046 | 0.021 | 1 | 7 | Fam109b | 0.025 | Old |
| 0.067382 | 0.095897 | 0.733 | 0.66 | 1 | 7 | Plxnd1 | 0.073 | Old |
| 0.067566 | 0.484438 | 0.069 | 0.039 | 1 | 7 | Prdm10 | 0.03 | Old |
| 0.067585 | 0.056932 | 0.415 | 0.343 | 1 | 7 | Wdr89 | 0.072 | Old |
| 0.06774 | 0.082487 | 0.013 | 0.033 | 1 | 7 | Mccc2 | -0.02 | Old |
| 0.067746 | 0.463864 | 0.282 | 0.223 | 1 | 7 | Sdccag3 | 0.059 | Old |
| 0.067834 | 1.052997 | 0.046 | 0.021 | 1 | 7 | Bcs1l | 0.025 | Old |
| 0.067888 | 0.399619 | 0.172 | 0.123 | 1 | 7 | Slc39a6 | 0.049 | Old |
| 0.067923 | 0.170598 | 0.515 | 0.437 | 1 | 7 | 1110004F1 | 0.078 | Old |
| 0.068083 | 0.270319 | 0.41 | 0.334 | 1 | 7 | Tmbim1 | 0.076 | Old |
| 0.068093 | 0.156426 | 0.631 | 0.557 | 1 | 7 | Metap2 | 0.074 | Old |
| 0.068219 | 0.123422 | 0.174 | 0.123 | 1 | 7 | Tmem60 | 0.051 | Old |
| 0.068441 | 0.192765 | 0.156 | 0.108 | 1 | 7 | Rnf170 | 0.048 | Old |
| 0.068458 | 0.438915 | 0.059 | 0.03 | 1 | 7 | Pigh | 0.029 | Old |
| 0.068549 | 0.738613 | 0.067 | 0.036 | 1 | 7 | Kat8 | 0.031 | Old |

| | | | | | | | | |
|---|---|---|---|---|---|---|---|---|
| 0.068943 | 0.586344 | 0.192 | 0.142 | 1 | 7 | Sdc1 | 0.05 | Old |
| 0.069047 | 0.005801 | 0.259 | 0.19 | 1 | 7 | Vps11 | 0.069 | Old |
| 0.069061 | 0.67197 | 0.059 | 0.03 | 1 | 7 | Sac3d1 | 0.029 | Old |
| 0.069067 | 0.17458 | 0.295 | 0.226 | 1 | 7 | Gripap1 | 0.069 | Old |
| 0.06909 | 0.572839 | 0.185 | 0.136 | 1 | 7 | Vps52 | 0.049 | Old |
| 0.069255 | 0.968554 | 0.067 | 0.036 | 1 | 7 | Mlh3 | 0.031 | Old |
| 0.069352 | 0.506316 | 0.218 | 0.163 | 1 | 7 | Sigmar1 | 0.055 | Old |
| 0.06938 | 0.737591 | 0.046 | 0.021 | 1 | 7 | Gfod1 | 0.025 | Old |
| 0.069487 | 0.187112 | 0.318 | 0.253 | 1 | 7 | Rpl7l1 | 0.065 | Old |
| 0.069552 | 0.789138 | 0.092 | 0.057 | 1 | 7 | Hebp1 | 0.035 | Old |
| 0.069622 | 0.411644 | 0.438 | 0.386 | 1 | 7 | Cd24a | 0.052 | Old |
| 0.069708 | 0.144716 | 0.718 | 0.639 | 1 | 7 | Xbp1 | 0.079 | Old |
| 0.069728 | 0.562149 | 0.046 | 0.021 | 1 | 7 | Slc2a9 | 0.025 | Old |
| 0.069985 | 0.336615 | 0.356 | 0.289 | 1 | 7 | Atp6v0a1 | 0.067 | Old |
| 0.070026 | 0.31775 | 0.354 | 0.283 | 1 | 7 | Dars | 0.071 | Old |
| 0.070056 | 0.174008 | 0.323 | 0.253 | 1 | 7 | Yipf5 | 0.07 | Old |
| 0.07011 | 0.170578 | 0.067 | 0.036 | 1 | 7 | Rad51b | 0.031 | Old |
| 0.070164 | 0.263773 | 0.236 | 0.178 | 1 | 7 | Mocs2 | 0.058 | Old |
| 0.070253 | 0.094818 | 0.4 | 0.322 | 1 | 7 | Nans | 0.078 | Old |
| 0.070638 | 0.466728 | 0.215 | 0.16 | 1 | 7 | Utp6 | 0.055 | Old |
| 0.070753 | 0.531934 | 0.133 | 0.09 | 1 | 7 | Nat9 | 0.043 | Old |
| 0.070829 | 0.449761 | 0.067 | 0.036 | 1 | 7 | Sgcb | 0.031 | Old |
| 0.070847 | 0.298542 | 0.326 | 0.265 | 1 | 7 | Atp8b4 | 0.061 | Old |
| 0.07089 | 0.397884 | 0.082 | 0.048 | 1 | 7 | Gtf2e1 | 0.034 | Old |
| 0.070959 | 0.554532 | 0.09 | 0.054 | 1 | 7 | Mtg1 | 0.036 | Old |
| 0.070998 | 0.772184 | 0.213 | 0.16 | 1 | 7 | Abl2 | 0.053 | Old |
| 0.071024 | 0.2118 | 0.195 | 0.142 | 1 | 7 | Noc4l | 0.053 | Old |
| 0.071237 | 0.094777 | 0.359 | 0.289 | 1 | 7 | Spag7 | 0.07 | Old |
| 0.071255 | 0.024753 | 0.572 | 0.494 | 1 | 7 | Plec | 0.078 | Old |
| 0.071298 | 0.215537 | 0.774 | 0.738 | 1 | 7 | Naaa | 0.036 | Old |
| 0.071354 | 0.126973 | 0.462 | 0.373 | 1 | 7 | Coa5 | 0.089 | Old |
| 0.071361 | 0.17439 | 0.244 | 0.187 | 1 | 7 | Tnpo2 | 0.057 | Old |
| 0.071377 | 1.08029 | 0.041 | 0.018 | 1 | 7 | Creb3l1 | 0.023 | Old |
| 0.071743 | 0.565916 | 0.128 | 0.087 | 1 | 7 | Pde4dip | 0.041 | Old |
| 0.072135 | 1.228753 | 0.041 | 0.018 | 1 | 7 | Col4a1 | 0.023 | Old |
| 0.072138 | 0.095889 | 0.218 | 0.166 | 1 | 7 | Coq7 | 0.052 | Old |
| 0.072234 | 0.180397 | 0.151 | 0.105 | 1 | 7 | Trim65 | 0.046 | Old |
| 0.072506 | 0.296587 | 0.408 | 0.343 | 1 | 7 | Kif11 | 0.065 | Old |
| 0.072517 | 1.793307 | 0.041 | 0.018 | 1 | 7 | Adgrg1 | 0.023 | Old |
| 0.072658 | 0.608362 | 0.1 | 0.063 | 1 | 7 | Ogfod1 | 0.037 | Old |
| 0.0728 | 0.164027 | 0.8 | 0.78 | 1 | 7 | Hdgf | 0.02 | Old |
| 0.072814 | 0.173052 | 0.544 | 0.467 | 1 | 7 | Mfsd14b | 0.077 | Old |
| 0.072879 | 0.343601 | 0.231 | 0.175 | 1 | 7 | Slc12a7 | 0.056 | Old |
| 0.072901 | 0.716251 | 0.041 | 0.018 | 1 | 7 | Pigo | 0.023 | Old |
| 0.072907 | 0.243814 | 0.279 | 0.217 | 1 | 7 | Socs4 | 0.062 | Old |

| | | | | | | | | |
|---|---|---|---|---|---|---|---|---|
| 0.072936 | 0.305383 | 0.254 | 0.196 | 1 | 7 | Ppdpf | 0.058 | Old |
| 0.072947 | 0.186502 | 0.297 | 0.232 | 1 | 7 | Rest | 0.065 | Old |
| 0.073479 | 1.576064 | 0.041 | 0.018 | 1 | 7 | Arhgap27 | 0.023 | Old |
| 0.073479 | 1.120064 | 0.041 | 0.018 | 1 | 7 | Lhfpl2 | 0.023 | Old |
| 0.073623 | 0.225672 | 0.279 | 0.217 | 1 | 7 | Brd9 | 0.062 | Old |
| 0.073877 | 0.321258 | 0.254 | 0.196 | 1 | 7 | Gsdmd | 0.058 | Old |
| 0.073924 | 0.205301 | 0.223 | 0.169 | 1 | 7 | Mdn1 | 0.054 | Old |
| 0.073965 | 0.253248 | 0.279 | 0.217 | 1 | 7 | Cpeb4 | 0.062 | Old |
| 0.074268 | 0.233129 | 0.451 | 0.398 | 1 | 7 | Ap1s1 | 0.053 | Old |
| 0.074322 | 0.402839 | 0.185 | 0.136 | 1 | 7 | Gtf2e2 | 0.049 | Old |
| 0.07448 | 0.089224 | 0.3 | 0.235 | 1 | 7 | Por | 0.065 | Old |
| 0.074513 | 0.184527 | 0.282 | 0.22 | 1 | 7 | Anapc16 | 0.062 | Old |
| 0.074543 | 0.148233 | 0.126 | 0.081 | 1 | 7 | Tsen15 | 0.045 | Old |
| 0.07455 | 0.18022 | 0.951 | 0.919 | 1 | 7 | BC005537 | 0.032 | Old |
| 0.074604 | 0.66846 | 0.105 | 0.066 | 1 | 7 | Slc39a13 | 0.039 | Old |
| 0.074726 | 0.34151 | 0.213 | 0.157 | 1 | 7 | Jmjd6 | 0.056 | Old |
| 0.074732 | 0.73023 | 0.054 | 0.027 | 1 | 7 | Ercc8 | 0.027 | Old |
| 0.074784 | 0.290158 | 0.146 | 0.102 | 1 | 7 | Rai1 | 0.044 | Old |
| 0.074841 | 0.999911 | 0.179 | 0.136 | 1 | 7 | Abhd2 | 0.043 | Old |
| 0.074967 | 0.036945 | 0.067 | 0.036 | 1 | 7 | Acy1 | 0.031 | Old |
| 0.075013 | 0.507562 | 0.223 | 0.172 | 1 | 7 | Uba5 | 0.051 | Old |
| 0.07513 | 0.408305 | 0.238 | 0.187 | 1 | 7 | Mapk8 | 0.051 | Old |
| 0.075181 | 0.470121 | 0.446 | 0.383 | 1 | 7 | Chmp5 | 0.063 | Old |
| 0.075184 | 0.981494 | 0.062 | 0.033 | 1 | 7 | Wdr62 | 0.029 | Old |
| 0.075253 | 0.279072 | 0.221 | 0.16 | 1 | 7 | Cdip1 | 0.061 | Old |
| 0.075401 | 0.659852 | 0.136 | 0.093 | 1 | 7 | 1600012H( | 0.043 | Old |
| 0.075679 | 0.235229 | 0.146 | 0.099 | 1 | 7 | Tbrg4 | 0.047 | Old |
| 0.075758 | 0.165275 | 0.523 | 0.452 | 1 | 7 | Baz1a | 0.071 | Old |
| 0.075816 | 0.795623 | 0.062 | 0.033 | 1 | 7 | Tube1 | 0.029 | Old |
| 0.075907 | 0.440812 | 0.315 | 0.262 | 1 | 7 | Tns3 | 0.053 | Old |
| 0.076044 | 0.412029 | 0.256 | 0.199 | 1 | 7 | Wdr61 | 0.057 | Old |
| 0.076125 | 0.633831 | 0.11 | 0.072 | 1 | 7 | Bag5 | 0.038 | Old |
| 0.076522 | 0.428369 | 0.341 | 0.274 | 1 | 7 | Spn | 0.067 | Old |
| 0.076597 | 0.361752 | 0.405 | 0.337 | 1 | 7 | Atp6v1c1 | 0.068 | Old |
| 0.076753 | 0.177671 | 0.231 | 0.172 | 1 | 7 | Grsf1 | 0.059 | Old |
| 0.076782 | 0.125689 | 0.787 | 0.741 | 1 | 7 | Psmb4 | 0.046 | Old |
| 0.076851 | 0.267044 | 0.326 | 0.265 | 1 | 7 | Ptpn22 | 0.061 | Old |
| 0.076876 | 0.163754 | 0.197 | 0.142 | 1 | 7 | Rgs14 | 0.055 | Old |
| 0.076943 | 0.158513 | 0.264 | 0.202 | 1 | 7 | Lats2 | 0.062 | Old |
| 0.07705 | 0.015403 | 0.144 | 0.099 | 1 | 7 | Med27 | 0.045 | Old |
| 0.077072 | 0.908614 | 0.103 | 0.066 | 1 | 7 | Cd82 | 0.037 | Old |
| 0.077535 | 0.396609 | 0.546 | 0.494 | 1 | 7 | Psma2 | 0.052 | Old |
| 0.077587 | 0.309161 | 0.318 | 0.253 | 1 | 7 | Vps41 | 0.065 | Old |
| 0.077903 | 0.386982 | 0.115 | 0.075 | 1 | 7 | Gpatch4 | 0.04 | Old |
| 0.077914 | 0.691331 | 0.162 | 0.117 | 1 | 7 | Tgfbrap1 | 0.045 | Old |

| | | | | | | | | |
|---|---|---|---|---|---|---|---|---|
| 0.077973 | 0.392167 | 0.169 | 0.12 | 1 | 7 | Ccpg1 | 0.049 | Old |
| 0.077985 | 0.152962 | 0.362 | 0.292 | 1 | 7 | Kdm5a | 0.07 | Old |
| 0.078009 | 0.178451 | 0.5 | 0.431 | 1 | 7 | Ybx3 | 0.069 | Old |
| 0.078148 | 0.758398 | 0.11 | 0.072 | 1 | 7 | Narfl | 0.038 | Old |
| 0.078157 | 0.443735 | 0.174 | 0.123 | 1 | 7 | Cdk10 | 0.051 | Old |
| 0.078217 | 0.427147 | 0.118 | 0.078 | 1 | 7 | Tbl3 | 0.04 | Old |
| 0.078511 | 0.362014 | 0.246 | 0.196 | 1 | 7 | Ano6 | 0.05 | Old |
| 0.078546 | 0.493374 | 0.333 | 0.274 | 1 | 7 | Phf5a | 0.059 | Old |
| 0.078567 | 0.26936 | 0.254 | 0.202 | 1 | 7 | Gpr35 | 0.052 | Old |
| 0.078713 | 0.338809 | 0.29 | 0.229 | 1 | 7 | Cd226 | 0.061 | Old |
| 0.078998 | 0.272867 | 0.231 | 0.175 | 1 | 7 | Plcl2 | 0.056 | Old |
| 0.079195 | 1.46659 | 0.036 | 0.015 | 1 | 7 | Pxylp1 | 0.021 | Old |
| 0.079234 | 0.261975 | 0.282 | 0.223 | 1 | 7 | Golgb1 | 0.059 | Old |
| 0.079496 | 0.450625 | 0.403 | 0.355 | 1 | 7 | Samsn1 | 0.048 | Old |
| 0.079802 | 0.817269 | 0.156 | 0.111 | 1 | 7 | B3glct | 0.045 | Old |
| 0.07986 | 1.323854 | 0.036 | 0.015 | 1 | 7 | Zbtb25 | 0.021 | Old |
| 0.079959 | 0.944102 | 0.049 | 0.024 | 1 | 7 | 5730480H( | 0.025 | Old |
| 0.080147 | 0.049733 | 0.19 | 0.142 | 1 | 7 | Dera | 0.048 | Old |
| 0.080168 | 0.146083 | 0.264 | 0.205 | 1 | 7 | Dennd4c | 0.059 | Old |
| 0.080175 | 0.20568 | 0.587 | 0.527 | 1 | 7 | Ywhag | 0.06 | Old |
| 0.0802 | 0.336797 | 0.077 | 0.045 | 1 | 7 | Utp4 | 0.032 | Old |
| 0.08029 | 0.327411 | 0.167 | 0.117 | 1 | 7 | Fam160b2 | 0.05 | Old |
| 0.080435 | 0.319311 | 0.585 | 0.554 | 1 | 7 | Tnip3 | 0.031 | Old |
| 0.080457 | 0.15155 | 0.513 | 0.434 | 1 | 7 | Arih1 | 0.079 | Old |
| 0.080483 | 0.194762 | 0.433 | 0.361 | 1 | 7 | Srp19 | 0.072 | Old |
| 0.080485 | 0.195018 | 0.238 | 0.184 | 1 | 7 | Gm15987 | 0.054 | Old |
| 0.080604 | 0.026604 | 0.126 | 0.084 | 1 | 7 | Rbm15b | 0.042 | Old |
| 0.080685 | 0.421197 | 0.336 | 0.277 | 1 | 7 | Cd200r1 | 0.059 | Old |
| 0.080696 | 0.142253 | 0.559 | 0.488 | 1 | 7 | Copz1 | 0.071 | Old |
| 0.080735 | 0.422499 | 0.226 | 0.178 | 1 | 7 | Ak6 | 0.048 | Old |
| 0.080791 | 0.073828 | 0.349 | 0.28 | 1 | 7 | Txndc9 | 0.069 | Old |
| 0.080852 | 0.458025 | 0.203 | 0.157 | 1 | 7 | Nupr1 | 0.046 | Old |
| 0.08101 | 0.566543 | 0.115 | 0.075 | 1 | 7 | Akap7 | 0.04 | Old |
| 0.081028 | 0.156414 | 0.282 | 0.217 | 1 | 7 | Fam192a | 0.065 | Old |
| 0.081091 | 0.737721 | 0.049 | 0.024 | 1 | 7 | Ndufaf6 | 0.025 | Old |
| 0.08127 | 0.161804 | 0.918 | 0.883 | 1 | 7 | Lamp1 | 0.035 | Old |
| 0.081311 | 0.238715 | 0.249 | 0.19 | 1 | 7 | Epb41l1 | 0.059 | Old |
| 0.08142 | 0.970454 | 0.113 | 0.075 | 1 | 7 | Mfsd7a | 0.038 | Old |
| 0.081837 | 0.185747 | 0.077 | 0.045 | 1 | 7 | Tti2 | 0.032 | Old |
| 0.08188 | 0.842448 | 0.036 | 0.015 | 1 | 7 | Tmem186 | 0.021 | Old |
| 0.081881 | 0.457343 | 0.167 | 0.12 | 1 | 7 | Sh3tc1 | 0.047 | Old |
| 0.082044 | 0.922844 | 0.049 | 0.024 | 1 | 7 | Letm2 | 0.025 | Old |
| 0.082055 | 0.517964 | 0.1 | 0.063 | 1 | 7 | Kif16b | 0.037 | Old |
| 0.082531 | 0.338611 | 0.154 | 0.108 | 1 | 7 | Phb | 0.046 | Old |
| 0.082661 | 1.089508 | 0.087 | 0.054 | 1 | 7 | Hectd3 | 0.033 | Old |

| | | | | | | | | |
|---|---|---|---|---|---|---|---|---|
| 0.083092 | 0.304877 | 0.397 | 0.325 | 1 | 7 | Slc35b2 | 0.072 | Old |
| 0.083227 | 0.198639 | 0.377 | 0.304 | 1 | 7 | Cd180 | 0.073 | Old |
| 0.083567 | 0.520979 | 0.069 | 0.039 | 1 | 7 | Chchd6 | 0.03 | Old |
| 0.083589 | 1.062444 | 0.049 | 0.024 | 1 | 7 | 2610021A0 | 0.025 | Old |
| 0.083722 | 0.54726 | 0.085 | 0.051 | 1 | 7 | Tamm41 | 0.034 | Old |
| 0.083759 | 0.163224 | 0.546 | 0.455 | 1 | 7 | Atxn7l3b | 0.091 | Old |
| 0.083795 | 0.24415 | 0.533 | 0.461 | 1 | 7 | Dbnl | 0.072 | Old |
| 0.08382 | 0.090266 | 0.528 | 0.437 | 1 | 7 | Dhx9 | 0.091 | Old |
| 0.083942 | 0.727884 | 0.036 | 0.015 | 1 | 7 | Slc37a1 | 0.021 | Old |
| 0.083979 | 0.811158 | 0.049 | 0.024 | 1 | 7 | Det1 | 0.025 | Old |
| 0.084023 | 0.179838 | 0.533 | 0.458 | 1 | 7 | Sdhc | 0.075 | Old |
| 0.084056 | 0.33812 | 0.144 | 0.099 | 1 | 7 | Zwilch | 0.045 | Old |
| 0.084059 | 0.36608 | 0.177 | 0.13 | 1 | 7 | Ptgr1 | 0.047 | Old |
| 0.084088 | 0.18247 | 0.29 | 0.226 | 1 | 7 | Casc3 | 0.064 | Old |
| 0.084138 | 0.254874 | 0.218 | 0.166 | 1 | 7 | Nkap | 0.052 | Old |
| 0.084182 | 0.772507 | 0.179 | 0.136 | 1 | 7 | Gm36161 | 0.043 | Old |
| 0.084333 | 0.346247 | 0.092 | 0.057 | 1 | 7 | Mrps22 | 0.035 | Old |
| 0.084338 | 0.373002 | 0.444 | 0.395 | 1 | 7 | Emilin2 | 0.049 | Old |
| 0.084402 | 0.205448 | 0.208 | 0.16 | 1 | 7 | Nifk | 0.048 | Old |
| 0.084406 | 0.674535 | 0.036 | 0.015 | 1 | 7 | Aldoc | 0.021 | Old |
| 0.084457 | 0.454219 | 0.151 | 0.105 | 1 | 7 | Mrps11 | 0.046 | Old |
| 0.08458 | 0.385641 | 0.354 | 0.292 | 1 | 7 | Cdk2ap1 | 0.062 | Old |
| 0.084582 | 0.168165 | 0.097 | 0.06 | 1 | 7 | Wdr83 | 0.037 | Old |
| 0.084617 | 0.2929 | 0.062 | 0.033 | 1 | 7 | Tgm2 | 0.029 | Old |
| 0.084754 | 0.284245 | 0.179 | 0.13 | 1 | 7 | Smg9 | 0.049 | Old |
| 0.084799 | 0.09008 | 0.595 | 0.512 | 1 | 7 | Slc43a2 | 0.083 | Old |
| 0.084851 | 0.433263 | 0.251 | 0.199 | 1 | 7 | Hk1 | 0.052 | Old |
| 0.084871 | 0.42643 | 0.036 | 0.015 | 1 | 7 | Ppm1l | 0.021 | Old |
| 0.085116 | 0.431753 | 0.221 | 0.172 | 1 | 7 | Rit1 | 0.049 | Old |
| 0.085137 | 0.082365 | 0.121 | 0.081 | 1 | 7 | 2310022B0 | 0.04 | Old |
| 0.085301 | 0.338319 | 0.141 | 0.099 | 1 | 7 | Rttn | 0.042 | Old |
| 0.085342 | 0.357221 | 0.208 | 0.154 | 1 | 7 | Prpf4 | 0.054 | Old |
| 0.085367 | 0.232086 | 0.287 | 0.223 | 1 | 7 | Uimc1 | 0.064 | Old |
| 0.085552 | 0.624983 | 0.079 | 0.048 | 1 | 7 | Fam160a2 | 0.031 | Old |
| 0.085649 | 0.379608 | 0.344 | 0.277 | 1 | 7 | Elf4 | 0.067 | Old |
| 0.085755 | 1.11413 | 0.121 | 0.081 | 1 | 7 | Dopey2 | 0.04 | Old |
| 0.0859 | 0.752218 | 0.11 | 0.072 | 1 | 7 | Ndufaf7 | 0.038 | Old |
| 0.085906 | 0.60396 | 0.056 | 0.03 | 1 | 7 | Qrsl1 | 0.026 | Old |
| 0.085989 | 0.083798 | 0.356 | 0.283 | 1 | 7 | Hnrnpll | 0.073 | Old |
| 0.086015 | 0.497472 | 0.162 | 0.117 | 1 | 7 | Dram1 | 0.045 | Old |
| 0.086045 | 0.05122 | 0.036 | 0.015 | 1 | 7 | Slc30a6 | 0.021 | Old |
| 0.086053 | 0.169668 | 0.318 | 0.262 | 1 | 7 | Wdr82 | 0.056 | Old |
| 0.086122 | 0.406626 | 0.213 | 0.163 | 1 | 7 | Ell2 | 0.05 | Old |
| 0.086252 | 0.315684 | 0.262 | 0.205 | 1 | 7 | Thoc6 | 0.057 | Old |
| 0.086307 | 0.119821 | 0.108 | 0.069 | 1 | 7 | Vamp5 | 0.039 | Old |

| | | | | | | | | |
|---|---|---|---|---|---|---|---|---|
| 0.086423 | 0.341924 | 0.585 | 0.515 | 1 | 7 | Stat3 | 0.07 | Old |
| 0.086502 | 0.296362 | 0.121 | 0.081 | 1 | 7 | Slc25a44 | 0.04 | Old |
| 0.086538 | 0.377489 | 0.156 | 0.111 | 1 | 7 | Ptrhd1 | 0.045 | Old |
| 0.086599 | 0.164502 | 0.197 | 0.145 | 1 | 7 | Mrps30 | 0.052 | Old |
| 0.08663 | 0.551721 | 0.259 | 0.202 | 1 | 7 | Slc25a28 | 0.057 | Old |
| 0.08663 | 0.494299 | 0.251 | 0.199 | 1 | 7 | Iqgap3 | 0.052 | Old |
| 0.086691 | 0.094686 | 0.226 | 0.172 | 1 | 7 | Pdcd11 | 0.054 | Old |
| 0.086783 | 0.131768 | 0.418 | 0.352 | 1 | 7 | Sun2 | 0.066 | Old |
| 0.087076 | 0.177756 | 0.569 | 0.491 | 1 | 7 | Blnk | 0.078 | Old |
| 0.087087 | 0.185405 | 0.338 | 0.283 | 1 | 7 | Hdac2 | 0.055 | Old |
| 0.087142 | 0.34756 | 0.205 | 0.157 | 1 | 7 | Hspb11 | 0.048 | Old |
| 0.087176 | 0.1489 | 0.069 | 0.039 | 1 | 7 | Ftsj1 | 0.03 | Old |
| 0.08728 | 0.127623 | 0.705 | 0.627 | 1 | 7 | Gm26917 | 0.078 | Old |
| 0.08776 | 0.391496 | 0.156 | 0.111 | 1 | 7 | Bahd1 | 0.045 | Old |
| 0.087942 | 1.132533 | 0.056 | 0.03 | 1 | 7 | Lrrc49 | 0.026 | Old |
| 0.087975 | 0.27787 | 0.195 | 0.145 | 1 | 7 | Srm | 0.05 | Old |
| 0.088247 | 1.162436 | 0.072 | 0.042 | 1 | 7 | Bcas3 | 0.03 | Old |
| 0.088386 | 0.906416 | 0.064 | 0.036 | 1 | 7 | Zfp943 | 0.028 | Old |
| 0.088392 | 0.236044 | 0.121 | 0.081 | 1 | 7 | Isca1 | 0.04 | Old |
| 0.088521 | 0.169427 | 0.656 | 0.572 | 1 | 7 | Fam120a | 0.084 | Old |
| 0.088576 | 0.04517 | 0.382 | 0.313 | 1 | 7 | Cers5 | 0.069 | Old |
| 0.088661 | 0.881784 | 0.187 | 0.145 | 1 | 7 | Ube2l6 | 0.042 | Old |
| 0.08894 | 0.634892 | 0.308 | 0.256 | 1 | 7 | Arnt | 0.052 | Old |
| 0.088972 | 0.909259 | 0.115 | 0.078 | 1 | 7 | Cops7b | 0.037 | Old |
| 0.089011 | 0.182764 | 0.567 | 0.5 | 1 | 7 | Pigt | 0.067 | Old |
| 0.089035 | 0.184457 | 0.408 | 0.334 | 1 | 7 | Mrpl14 | 0.074 | Old |
| 0.089123 | 0.374689 | 0.177 | 0.13 | 1 | 7 | Pctp | 0.047 | Old |
| 0.089125 | 0.190415 | 0.728 | 0.696 | 1 | 7 | Ndufa6 | 0.032 | Old |
| 0.089181 | 0.314896 | 0.326 | 0.265 | 1 | 7 | Herc2 | 0.061 | Old |
| 0.089193 | 0.17799 | 0.487 | 0.425 | 1 | 7 | Sec61a1 | 0.062 | Old |
| 0.089287 | 0.061791 | 0.618 | 0.53 | 1 | 7 | Cast | 0.088 | Old |
| 0.089297 | 0.203619 | 0.267 | 0.208 | 1 | 7 | Aaed1 | 0.059 | Old |
| 0.089395 | 0.748149 | 0.131 | 0.09 | 1 | 7 | Mob2 | 0.041 | Old |
| 0.089724 | 0.902125 | 0.031 | 0.012 | 1 | 7 | Syp | 0.019 | Old |
| 0.08973 | 0.290034 | 0.441 | 0.364 | 1 | 7 | Srp72 | 0.077 | Old |
| 0.089826 | 0.911719 | 0.056 | 0.03 | 1 | 7 | Cenpv | 0.026 | Old |
| 0.089893 | 0.247064 | 0.538 | 0.476 | 1 | 7 | Eif3l | 0.062 | Old |
| 0.089905 | 1.161505 | 0.079 | 0.048 | 1 | 7 | Ppp1r12b | 0.031 | Old |
| 0.090021 | 0.036547 | 0.231 | 0.172 | 1 | 7 | Larp1b | 0.059 | Old |
| 0.090073 | 1.360574 | 0.015 | 0.003 | 1 | 7 | H2-M2 | 0.012 | Old |
| 0.090212 | 0.474465 | 0.144 | 0.102 | 1 | 7 | Neurl4 | 0.042 | Old |
| 0.090285 | 0.145743 | 0.238 | 0.184 | 1 | 7 | Grpel1 | 0.054 | Old |
| 0.090335 | 0.666274 | 0.156 | 0.114 | 1 | 7 | Mkl1 | 0.042 | Old |
| 0.090408 | 0.427344 | 0.11 | 0.072 | 1 | 7 | St7 | 0.038 | Old |
| 0.090435 | 0.280495 | 0.359 | 0.301 | 1 | 7 | Dctn3 | 0.058 | Old |

| | | | | | | | | |
|---|---|---|---|---|---|---|---|---|
| 0.090474 | 1.133721 | 0.015 | 0.003 | 1 | 7 | Spib | 0.012 | Old |
| 0.090586 | 0.636589 | 0.087 | 0.054 | 1 | 7 | Coq6 | 0.033 | Old |
| 0.090758 | 0.388931 | 0.133 | 0.093 | 1 | 7 | Fbxl6 | 0.04 | Old |
| 0.090787 | 1.0823 | 0.044 | 0.021 | 1 | 7 | Ccdc171 | 0.023 | Old |
| 0.090889 | 0.14226 | 0.303 | 0.241 | 1 | 7 | Kif1c | 0.062 | Old |
| 0.090979 | 0.572028 | 0.192 | 0.145 | 1 | 7 | Usp50 | 0.047 | Old |
| 0.091047 | 0.675799 | 0.095 | 0.06 | 1 | 7 | Nol10 | 0.035 | Old |
| 0.091376 | 0.229894 | 0.359 | 0.295 | 1 | 7 | Sbf2 | 0.064 | Old |
| 0.091685 | 1.767251 | 0.015 | 0.003 | 1 | 7 | Atg9b | 0.012 | Old |
| 0.091685 | 0.701976 | 0.015 | 0.003 | 1 | 7 | Stxbp4 | 0.012 | Old |
| 0.091767 | 0.270264 | 0.256 | 0.196 | 1 | 7 | Agap1 | 0.06 | Old |
| 0.091839 | 0.332272 | 0.415 | 0.352 | 1 | 7 | Ppp4r3a | 0.063 | Old |
| 0.092017 | 0.244338 | 0.228 | 0.175 | 1 | 7 | Mbd1 | 0.053 | Old |
| 0.092092 | 0.958435 | 0.015 | 0.003 | 1 | 7 | C330011M | 0.012 | Old |
| 0.092092 | 0.892545 | 0.015 | 0.003 | 1 | 7 | Tbc1d10c | 0.012 | Old |
| 0.092092 | 0.843499 | 0.015 | 0.003 | 1 | 7 | 9130230L2 | 0.012 | Old |
| 0.092115 | 0.949483 | 0.044 | 0.021 | 1 | 7 | 2410131K1 | 0.023 | Old |
| 0.092124 | 0.335641 | 0.064 | 0.036 | 1 | 7 | Mvk | 0.028 | Old |
| 0.092314 | 0.317297 | 0.118 | 0.078 | 1 | 7 | Commd9 | 0.04 | Old |
| 0.092338 | 1.313675 | 0.044 | 0.021 | 1 | 7 | AW209491 | 0.023 | Old |
| 0.092414 | 0.229818 | 0.636 | 0.581 | 1 | 7 | Pgam1 | 0.055 | Old |
| 0.092434 | 1.317111 | 0.026 | 0.009 | 1 | 7 | Sbsn | 0.017 | Old |
| 0.0925 | 0.171441 | 0.015 | 0.003 | 1 | 7 | Pars2 | 0.012 | Old |
| 0.0925 | 0.076433 | 0.015 | 0.003 | 1 | 7 | A730063M | 0.012 | Old |
| 0.09257 | 0.245892 | 0.459 | 0.404 | 1 | 7 | Cs | 0.055 | Old |
| 0.092735 | 1.559047 | 0.026 | 0.009 | 1 | 7 | Gpr162 | 0.017 | Old |
| 0.092892 | 0.26466 | 0.223 | 0.172 | 1 | 7 | Melk | 0.051 | Old |
| 0.092909 | 0.462852 | 0.056 | 0.03 | 1 | 7 | Coq3 | 0.026 | Old |
| 0.092983 | 0.288455 | 0.177 | 0.13 | 1 | 7 | Senp5 | 0.047 | Old |
| 0.093051 | 0.054954 | 0.103 | 0.066 | 1 | 7 | Acad8 | 0.037 | Old |
| 0.093077 | 0.108749 | 0.356 | 0.28 | 1 | 7 | Dna2 | 0.076 | Old |
| 0.093575 | 0.263463 | 0.305 | 0.241 | 1 | 7 | Egln2 | 0.064 | Old |
| 0.093937 | 0.187686 | 0.146 | 0.102 | 1 | 7 | Bora | 0.044 | Old |
| 0.093949 | 1.695535 | 0.026 | 0.009 | 1 | 7 | Naprt | 0.017 | Old |
| 0.093981 | 0.261365 | 0.208 | 0.157 | 1 | 7 | Psmg3 | 0.051 | Old |
| 0.094054 | 0.033339 | 0.031 | 0.012 | 1 | 7 | Car13 | 0.019 | Old |
| 0.094092 | 0.00365 | 0.423 | 0.328 | 1 | 7 | Vbp1 | 0.095 | Old |
| 0.094108 | 0.250521 | 0.423 | 0.358 | 1 | 7 | Mrpl57 | 0.065 | Old |
| 0.094173 | 0.467369 | 0.274 | 0.22 | 1 | 7 | Cstf3 | 0.054 | Old |
| 0.094255 | 0.981042 | 0.026 | 0.009 | 1 | 7 | Laptm4b | 0.017 | Old |
| 0.094293 | 0.028193 | 0.208 | 0.157 | 1 | 7 | Lrrcc1 | 0.051 | Old |
| 0.094303 | 0.265863 | 0.146 | 0.102 | 1 | 7 | Rngtt | 0.044 | Old |
| 0.094368 | 0.377241 | 0.472 | 0.398 | 1 | 7 | Gmfb | 0.074 | Old |
| 0.09447 | 0.230677 | 0.364 | 0.301 | 1 | 7 | Snx4 | 0.063 | Old |
| 0.0945 | 0.227994 | 0.21 | 0.16 | 1 | 7 | Nvl | 0.05 | Old |

| | | | | | | | | |
|---|---|---|---|---|---|---|---|---|
| 0.09459 | 0.127723 | 0.351 | 0.289 | 1 | 7 | Pdhb | 0.062 | Old |
| 0.09497 | 1.612461 | 0.051 | 0.027 | 1 | 7 | Rsph3b | 0.024 | Old |
| 0.095037 | 0.128946 | 0.349 | 0.286 | 1 | 7 | Chtf8 | 0.063 | Old |
| 0.095157 | 0.334409 | 0.254 | 0.196 | 1 | 7 | Suclg2 | 0.058 | Old |
| 0.095211 | 0.442902 | 0.072 | 0.042 | 1 | 7 | Gng11 | 0.03 | Old |
| 0.095347 | 0.135085 | 0.433 | 0.361 | 1 | 7 | Cse1l | 0.072 | Old |
| 0.095484 | 1.033945 | 0.026 | 0.009 | 1 | 7 | Btbd11 | 0.017 | Old |
| 0.095767 | 0.199947 | 0.677 | 0.62 | 1 | 7 | Gls | 0.057 | Old |
| 0.095778 | 0.35082 | 0.462 | 0.419 | 1 | 7 | Tubb4b | 0.043 | Old |
| 0.095794 | 1.030852 | 0.026 | 0.009 | 1 | 7 | Timp3 | 0.017 | Old |
| 0.095803 | 0.209104 | 0.551 | 0.473 | 1 | 7 | Rsrc2 | 0.078 | Old |
| 0.095876 | 1.071852 | 0.021 | 0.006 | 1 | 7 | Rnd2 | 0.015 | Old |
| 0.095985 | 0.054074 | 0.156 | 0.111 | 1 | 7 | Jagn1 | 0.045 | Old |
| 0.096027 | 0.19063 | 0.69 | 0.614 | 1 | 7 | Rad21 | 0.076 | Old |
| 0.096028 | 0.18213 | 0.644 | 0.593 | 1 | 7 | Oxct1 | 0.051 | Old |
| 0.096074 | 0.462008 | 0.215 | 0.166 | 1 | 7 | H2-Q6 | 0.049 | Old |
| 0.096104 | 1.254143 | 0.026 | 0.009 | 1 | 7 | Ttyh2 | 0.017 | Old |
| 0.096104 | 0.918332 | 0.026 | 0.009 | 1 | 7 | 4933417E1 | 0.017 | Old |
| 0.096374 | 0.379682 | 0.292 | 0.235 | 1 | 7 | Tmem131l | 0.057 | Old |
| 0.096415 | 0.351072 | 0.113 | 0.075 | 1 | 7 | 1190007I0 | 0.038 | Old |
| 0.096447 | 0.238473 | 0.431 | 0.355 | 1 | 7 | Clptm1l | 0.076 | Old |
| 0.096468 | 0.424671 | 0.303 | 0.253 | 1 | 7 | Dnttip2 | 0.05 | Old |
| 0.096649 | 0.728601 | 0.051 | 0.027 | 1 | 7 | Kctd11 | 0.024 | Old |
| 0.096915 | 0.237811 | 0.359 | 0.292 | 1 | 7 | Arl5a | 0.067 | Old |
| 0.096927 | 0.202924 | 0.064 | 0.036 | 1 | 7 | Pcgf6 | 0.028 | Old |
| 0.096939 | 1.148294 | 0.021 | 0.006 | 1 | 7 | Zfp963 | 0.015 | Old |
| 0.096939 | 1.005474 | 0.021 | 0.006 | 1 | 7 | Nutf2 | 0.015 | Old |
| 0.096961 | 0.085902 | 0.462 | 0.383 | 1 | 7 | Rab5a | 0.079 | Old |
| 0.097083 | 0.149637 | 0.267 | 0.205 | 1 | 7 | Suclg1 | 0.062 | Old |
| 0.097305 | 0.354702 | 0.064 | 0.036 | 1 | 7 | Prrg1 | 0.028 | Old |
| 0.097352 | 0.207789 | 0.172 | 0.123 | 1 | 7 | Klhdc2 | 0.049 | Old |
| 0.097353 | 0.690677 | 0.026 | 0.009 | 1 | 7 | Zfp235 | 0.017 | Old |
| 0.097353 | 0.566209 | 0.026 | 0.009 | 1 | 7 | Top1mt | 0.017 | Old |
| 0.097494 | 0.479407 | 0.064 | 0.036 | 1 | 7 | Fbxo21 | 0.028 | Old |
| 0.097503 | 0.290858 | 0.408 | 0.325 | 1 | 7 | Lrp6 | 0.083 | Old |
| 0.097564 | 0.281572 | 0.131 | 0.09 | 1 | 7 | Il27ra | 0.041 | Old |
| 0.097647 | 0.183602 | 0.303 | 0.244 | 1 | 7 | Plk4 | 0.059 | Old |
| 0.097652 | 1.66635 | 0.021 | 0.006 | 1 | 7 | Hspg2 | 0.015 | Old |
| 0.097652 | 1.094318 | 0.021 | 0.006 | 1 | 7 | Pcp4l1 | 0.015 | Old |
| 0.097668 | 0.248751 | 0.451 | 0.373 | 1 | 7 | Vapb | 0.078 | Old |
| 0.097713 | 0.326847 | 0.567 | 0.488 | 1 | 7 | Chmp2a | 0.079 | Old |
| 0.097726 | 0.485317 | 0.154 | 0.111 | 1 | 7 | Kif13b | 0.043 | Old |
| 0.097764 | 0.093744 | 0.251 | 0.196 | 1 | 7 | Atf2 | 0.055 | Old |
| 0.097951 | 0.241296 | 0.162 | 0.114 | 1 | 7 | Tmem238 | 0.048 | Old |
| 0.097952 | 0.161328 | 0.177 | 0.13 | 1 | 7 | Rad18 | 0.047 | Old |

| | | | | | | | | |
|---|---|---|---|---|---|---|---|---|
| 0.098011 | 1.115207 | 0.021 | 0.006 | 1 | 7 | Atp1b1 | 0.015 | Old |
| 0.098011 | 0.860756 | 0.021 | 0.006 | 1 | 7 | Rhou | 0.015 | Old |
| 0.098227 | 0.335975 | 0.131 | 0.09 | 1 | 7 | Bysl | 0.041 | Old |
| 0.098254 | 0.366105 | 0.064 | 0.036 | 1 | 7 | Ndufaf1 | 0.028 | Old |
| 0.098298 | 0.401997 | 0.026 | 0.009 | 1 | 7 | Slc17a9 | 0.017 | Old |
| 0.098371 | 1.191707 | 0.021 | 0.006 | 1 | 7 | AC139347. | 0.015 | Old |
| 0.098371 | 1.014117 | 0.021 | 0.006 | 1 | 7 | Gm29562 | 0.015 | Old |
| 0.098409 | 0.21426 | 0.151 | 0.108 | 1 | 7 | Urod | 0.043 | Old |
| 0.098442 | 0.174927 | 0.364 | 0.304 | 1 | 7 | Cntrl | 0.06 | Old |
| 0.098458 | 0.406948 | 0.087 | 0.054 | 1 | 7 | Srpyd7 | 0.033 | Old |
| 0.09852 | 0.158689 | 0.326 | 0.262 | 1 | 7 | Tiprl | 0.064 | Old |
| 0.098572 | 0.267368 | 0.454 | 0.392 | 1 | 7 | Txnrd1 | 0.062 | Old |
| 0.098691 | 0.546309 | 0.2 | 0.157 | 1 | 7 | Chd6 | 0.043 | Old |
| 0.098731 | 0.374168 | 0.021 | 0.006 | 1 | 7 | Gm36266 | 0.015 | Old |
| 0.098842 | 0.136186 | 0.823 | 0.798 | 1 | 7 | Ywhab | 0.025 | Old |
| 0.098868 | 0.305116 | 0.431 | 0.37 | 1 | 7 | Ubash3b | 0.061 | Old |
| 0.099156 | 0.188575 | 0.151 | 0.108 | 1 | 7 | Pxmp4 | 0.043 | Old |
| 0.099201 | 0.244287 | 0.2 | 0.151 | 1 | 7 | Polr3gl | 0.049 | Old |
| 0.099292 | 0.224837 | 0.203 | 0.154 | 1 | 7 | Pdcl | 0.049 | Old |
| 0.099311 | 0.197182 | 0.451 | 0.377 | 1 | 7 | Polr2k | 0.074 | Old |
| 0.099419 | 0.742958 | 0.082 | 0.051 | 1 | 7 | Vdr | 0.031 | Old |
| 0.099455 | 0.601155 | 0.021 | 0.006 | 1 | 7 | Ank2 | 0.015 | Old |
| 0.099471 | 0.785669 | 0.067 | 0.039 | 1 | 7 | Mrps27 | 0.028 | Old |
| 0.099726 | 0.130509 | 0.685 | 0.596 | 1 | 7 | Syk | 0.089 | Old |
| 0.099819 | 0.477392 | 0.021 | 0.006 | 1 | 7 | Ttc30b | 0.015 | Old |
| 0.099885 | 0.356992 | 0.115 | 0.078 | 1 | 7 | Med24 | 0.037 | Old |
| 0.099916 | 0.556455 | 0.1 | 0.066 | 1 | 7 | P2ry1 | 0.034 | Old |
| 0.099956 | 0.176599 | 0.556 | 0.494 | 1 | 7 | Usp1 | 0.062 | Old |
| 0.100054 | 0.422974 | 0.369 | 0.301 | 1 | 7 | Mfsd5 | 0.068 | Old |
| 0.100134 | 0.212532 | 0.359 | 0.289 | 1 | 7 | Tmem9b | 0.07 | Old |
| 0.100397 | 0.108199 | 0.928 | 0.931 | 1 | 7 | Iqgap1 | -0.003 | Old |
| 0.100409 | 0.112596 | 0.849 | 0.804 | 1 | 7 | Hnrnpa0 | 0.045 | Old |
| 0.100462 | 0.486851 | 0.059 | 0.033 | 1 | 7 | Kif3a | 0.026 | Old |
| 0.100502 | 0.183304 | 0.192 | 0.145 | 1 | 7 | Ttpal | 0.047 | Old |
| 0.100561 | 0.369274 | 0.295 | 0.244 | 1 | 7 | Fam49a | 0.051 | Old |
| 0.100958 | 0.172406 | 0.882 | 0.861 | 1 | 7 | Ywhaz | 0.021 | Old |
| 0.101022 | 0.603039 | 0.236 | 0.187 | 1 | 7 | Pex7 | 0.049 | Old |
| 0.101023 | 0.169764 | 0.531 | 0.461 | 1 | 7 | Jarid2 | 0.07 | Old |
| 0.101102 | 0.277912 | 0.438 | 0.377 | 1 | 7 | Ube2z | 0.061 | Old |
| 0.101325 | 0.200003 | 0.354 | 0.286 | 1 | 7 | Vps13a | 0.068 | Old |
| 0.101387 | 0.768454 | 0.051 | 0.027 | 1 | 7 | 1600014C1 | 0.024 | Old |
| 0.101507 | 1.180629 | 0.038 | 0.018 | 1 | 7 | Gm5150 | 0.02 | Old |
| 0.101606 | 0.551405 | 0.051 | 0.027 | 1 | 7 | Taf9b | 0.024 | Old |
| 0.10165 | 0.488927 | 0.105 | 0.069 | 1 | 7 | 1600002K0 | 0.036 | Old |
| 0.10195 | 0.28933 | 0.228 | 0.178 | 1 | 7 | Stxbp5 | 0.05 | Old |

| | | | | | | | | |
|---|---|---|---|---|---|---|---|---|
| 0.102179 | 0.249174 | 0.349 | 0.286 | 1 | 7 | Ogfr | 0.063 | Old |
| 0.10226 | 0.09398 | 0.246 | 0.19 | 1 | 7 | Srpk2 | 0.056 | Old |
| 0.102518 | 0.090672 | 0.174 | 0.127 | 1 | 7 | Pnpla2 | 0.047 | Old |
| 0.103038 | 0.307548 | 0.318 | 0.256 | 1 | 7 | Phc3 | 0.062 | Old |
| 0.103043 | 0.193257 | 0.164 | 0.12 | 1 | 7 | Haus3 | 0.044 | Old |
| 0.103073 | 0.123691 | 0.413 | 0.331 | 1 | 7 | Lsm14a | 0.082 | Old |
| 0.103133 | 0.283682 | 0.172 | 0.127 | 1 | 7 | Proser1 | 0.045 | Old |
| 0.103136 | 0.261716 | 0.523 | 0.449 | 1 | 7 | Nasp | 0.074 | Old |
| 0.103262 | 0.187686 | 0.405 | 0.331 | 1 | 7 | Pcmtd1 | 0.074 | Old |
| 0.103314 | 0.192196 | 0.549 | 0.455 | 1 | 7 | Slc20a1 | 0.094 | Old |
| 0.103793 | 0.494579 | 0.262 | 0.208 | 1 | 7 | Foxm1 | 0.054 | Old |
| 0.104028 | 1.174468 | 0.103 | 0.069 | 1 | 7 | Wdr7 | 0.034 | Old |
| 0.104058 | 0.266377 | 0.067 | 0.039 | 1 | 7 | Nbas | 0.028 | Old |
| 0.104175 | 0.229367 | 0.418 | 0.349 | 1 | 7 | Abcf1 | 0.069 | Old |
| 0.104339 | 0.146255 | 0.236 | 0.187 | 1 | 7 | 0610010F0 | 0.049 | Old |
| 0.104431 | 0.136938 | 0.287 | 0.229 | 1 | 7 | Htra2 | 0.058 | Old |
| 0.104447 | 0.49888 | 0.095 | 0.06 | 1 | 7 | Alkbh7 | 0.035 | Old |
| 0.104467 | 0.139904 | 0.241 | 0.19 | 1 | 7 | Dolpp1 | 0.051 | Old |
| 0.104476 | 0.61085 | 0.208 | 0.16 | 1 | 7 | Ndnf | 0.048 | Old |
| 0.104512 | 0.345897 | 0.195 | 0.145 | 1 | 7 | Brf1 | 0.05 | Old |
| 0.104521 | 0.577573 | 0.172 | 0.13 | 1 | 7 | Prmt3 | 0.042 | Old |
| 0.104576 | 0.217927 | 0.538 | 0.464 | 1 | 7 | Ehbp1l1 | 0.074 | Old |
| 0.104604 | 0.171984 | 0.326 | 0.259 | 1 | 7 | Map1s | 0.067 | Old |
| 0.10467 | 0.295408 | 0.613 | 0.551 | 1 | 7 | Id2 | 0.062 | Old |
| 0.104886 | 0.521992 | 0.118 | 0.081 | 1 | 7 | Cog4 | 0.037 | Old |
| 0.10501 | 0.074497 | 0.128 | 0.087 | 1 | 7 | Psmg1 | 0.041 | Old |
| 0.105173 | 0.105136 | 0.051 | 0.027 | 1 | 7 | Rdh13 | 0.024 | Old |
| 0.105297 | 0.417031 | 0.187 | 0.142 | 1 | 7 | Med16 | 0.045 | Old |
| 0.105412 | 0.15058 | 0.149 | 0.108 | 1 | 7 | Pphln1 | 0.041 | Old |
| 0.105416 | 0.634387 | 0.038 | 0.018 | 1 | 7 | Unc5a | 0.02 | Old |
| 0.105609 | 0.526522 | 0.423 | 0.37 | 1 | 7 | Foxn2 | 0.053 | Old |
| 0.105681 | 0.580247 | 0.038 | 0.018 | 1 | 7 | Hyal2 | 0.02 | Old |
| 0.105907 | 0.18726 | 0.136 | 0.096 | 1 | 7 | Rrp9 | 0.04 | Old |
| 0.106348 | 0.293722 | 0.1 | 0.066 | 1 | 7 | Cyp4v3 | 0.034 | Old |
| 0.106398 | 0.126715 | 0.456 | 0.373 | 1 | 7 | Psmb9 | 0.083 | Old |
| 0.106434 | 1.014325 | 0.046 | 0.024 | 1 | 7 | Mettl25 | 0.022 | Old |
| 0.106521 | 0.014768 | 0.087 | 0.054 | 1 | 7 | Fars2 | 0.033 | Old |
| 0.106756 | 0.098736 | 0.095 | 0.06 | 1 | 7 | Dcun1d2 | 0.035 | Old |
| 0.106764 | 1.023438 | 0.051 | 0.027 | 1 | 7 | Gys1 | 0.024 | Old |
| 0.10693 | 0.443058 | 0.115 | 0.078 | 1 | 7 | Scnm1 | 0.037 | Old |
| 0.107172 | 0.424 | 0.213 | 0.163 | 1 | 7 | E2f4 | 0.05 | Old |
| 0.107624 | 0.384468 | 0.262 | 0.208 | 1 | 7 | Hnrnph3 | 0.054 | Old |
| 0.107652 | 0.318326 | 0.123 | 0.084 | 1 | 7 | Ubtd1 | 0.039 | Old |
| 0.107835 | 0.234781 | 0.354 | 0.289 | 1 | 7 | Baiap2 | 0.065 | Old |
| 0.107884 | 0.865318 | 0.046 | 0.024 | 1 | 7 | Tnfrsf25 | 0.022 | Old |

| | | | | | | | | |
|---|---|---|---|---|---|---|---|---|
| 0.107935 | 0.269795 | 0.485 | 0.428 | 1 | 7 | Fis1 | 0.057 | Old |
| 0.108025 | 0.235272 | 0.277 | 0.22 | 1 | 7 | Trim27 | 0.057 | Old |
| 0.108082 | 0.255305 | 0.538 | 0.47 | 1 | 7 | Tomm5 | 0.068 | Old |
| 0.108112 | 0.156202 | 0.164 | 0.123 | 1 | 7 | Pggt1b | 0.041 | Old |
| 0.108145 | 0.25815 | 0.59 | 0.518 | 1 | 7 | Azin1 | 0.072 | Old |
| 0.108295 | 0.270243 | 0.441 | 0.38 | 1 | 7 | Cmtm3 | 0.061 | Old |
| 0.108316 | 0.170434 | 0.605 | 0.536 | 1 | 7 | Ndufa3 | 0.069 | Old |
| 0.108504 | 0.140959 | 0.377 | 0.313 | 1 | 7 | Rere | 0.064 | Old |
| 0.108648 | 0.870726 | 0.095 | 0.063 | 1 | 7 | Ercc5 | 0.032 | Old |
| 0.108705 | 0.187405 | 0.6 | 0.539 | 1 | 7 | Tmem14c | 0.061 | Old |
| 0.108713 | 0.27708 | 0.279 | 0.223 | 1 | 7 | Erg28 | 0.056 | Old |
| 0.108849 | 0.109577 | 0.254 | 0.199 | 1 | 7 | Napg | 0.055 | Old |
| 0.108914 | 0.31047 | 0.5 | 0.431 | 1 | 7 | Ttc7 | 0.069 | Old |
| 0.108985 | 0.199484 | 0.197 | 0.148 | 1 | 7 | Usp38 | 0.049 | Old |
| 0.109011 | 0.231224 | 0.5 | 0.428 | 1 | 7 | Khsrp | 0.072 | Old |
| 0.109017 | 0.55655 | 0.059 | 0.033 | 1 | 7 | Cnnm2 | 0.026 | Old |
| 0.109027 | 0.269419 | 0.572 | 0.506 | 1 | 7 | Ikzf1 | 0.066 | Old |
| 0.109222 | 0.095952 | 0.395 | 0.334 | 1 | 7 | Arf2 | 0.061 | Old |
| 0.109259 | 0.137219 | 0.464 | 0.383 | 1 | 7 | Usp4 | 0.081 | Old |
| 0.109388 | 0.105826 | 0.59 | 0.53 | 1 | 7 | Ncor1 | 0.06 | Old |
| 0.10953 | 0.183896 | 0.467 | 0.389 | 1 | 7 | Dapk1 | 0.078 | Old |
| 0.109553 | 0.041158 | 0.238 | 0.181 | 1 | 7 | Apex1 | 0.057 | Old |
| 0.109596 | 1.493182 | 0.046 | 0.024 | 1 | 7 | Zfp874b | 0.022 | Old |
| 0.109627 | 0.032526 | 0.105 | 0.069 | 1 | 7 | Dvl3 | 0.036 | Old |
| 0.109767 | 0.138092 | 0.269 | 0.214 | 1 | 7 | Selenoi | 0.055 | Old |
| 0.11018 | 0.088182 | 0.162 | 0.117 | 1 | 7 | Slc39a14 | 0.045 | Old |
| 0.110231 | 0.578016 | 0.067 | 0.039 | 1 | 7 | Fzd7 | 0.028 | Old |
| 0.110318 | 0.094562 | 0.61 | 0.533 | 1 | 7 | Lmnb1 | 0.077 | Old |
| 0.110497 | 0.093797 | 0.877 | 0.867 | 1 | 7 | Canx | 0.01 | Old |
| 0.11074 | 0.407039 | 0.3 | 0.232 | 1 | 7 | Oxr1 | 0.068 | Old |
| 0.110905 | 0.907958 | 0.115 | 0.081 | 1 | 7 | A4galt | 0.034 | Old |
| 0.111044 | 0.286199 | 0.338 | 0.28 | 1 | 7 | Iscu | 0.058 | Old |
| 0.111048 | 0.119707 | 0.308 | 0.244 | 1 | 7 | Txndc5 | 0.064 | Old |
| 0.111107 | 0.128496 | 0.879 | 0.889 | 1 | 7 | Btf3 | -0.01 | Old |
| 0.111119 | 0.139465 | 0.074 | 0.045 | 1 | 7 | Zbtb42 | 0.029 | Old |
| 0.111166 | 0.346588 | 0.128 | 0.09 | 1 | 7 | Ptcd3 | 0.038 | Old |
| 0.111252 | 0.375687 | 0.146 | 0.105 | 1 | 7 | Rpusd4 | 0.041 | Old |
| 0.111256 | 0.532644 | 0.11 | 0.075 | 1 | 7 | Trip10 | 0.035 | Old |
| 0.111301 | 0.391554 | 0.349 | 0.289 | 1 | 7 | 9030624J0 | 0.06 | Old |
| 0.111802 | 0.072609 | 0.672 | 0.575 | 1 | 7 | Atp6v1f | 0.097 | Old |
| 0.111859 | 0.100854 | 0.672 | 0.599 | 1 | 7 | St13 | 0.073 | Old |
| 0.112152 | 0.054474 | 0.369 | 0.31 | 1 | 7 | Prelid3b | 0.059 | Old |
| 0.112407 | 0.174774 | 0.167 | 0.12 | 1 | 7 | Sp3os | 0.047 | Old |
| 0.112563 | 0.32068 | 0.185 | 0.139 | 1 | 7 | Dnajc25 | 0.046 | Old |
| 0.11258 | 0.518417 | 0.046 | 0.024 | 1 | 7 | Brf2 | 0.022 | Old |

| | | | | | | | | |
|---|---|---|---|---|---|---|---|---|
| 0.112714 | 0.518588 | 0.274 | 0.22 | 1 | 7 | Idh3a | 0.054 | Old |
| 0.112811 | 0.095665 | 0.182 | 0.139 | 1 | 7 | Zfp593 | 0.043 | Old |
| 0.112975 | 0.125607 | 0.677 | 0.599 | 1 | 7 | Larp4b | 0.078 | Old |
| 0.113519 | 0.18838 | 0.223 | 0.169 | 1 | 7 | Ppih | 0.054 | Old |
| 0.113589 | 0.870378 | 0.046 | 0.024 | 1 | 7 | P2ry14 | 0.022 | Old |
| 0.113744 | 0.235434 | 0.292 | 0.232 | 1 | 7 | Esf1 | 0.06 | Old |
| 0.113765 | 1.123005 | 0.033 | 0.015 | 1 | 7 | Szt2 | 0.018 | Old |
| 0.113772 | 0.32643 | 0.146 | 0.105 | 1 | 7 | Nav2 | 0.041 | Old |
| 0.113835 | 0.252925 | 0.682 | 0.672 | 1 | 7 | Fn1 | 0.01 | Old |
| 0.113999 | 0.194258 | 0.418 | 0.352 | 1 | 7 | H2afx | 0.066 | Old |
| 0.114096 | 0.145378 | 0.046 | 0.024 | 1 | 7 | Usp18 | 0.022 | Old |
| 0.114109 | 1.141178 | 0.095 | 0.063 | 1 | 7 | Sgsm2 | 0.032 | Old |
| 0.114285 | 0.746804 | 0.113 | 0.078 | 1 | 7 | 2810006K2 | 0.035 | Old |
| 0.114372 | 1.023849 | 0.033 | 0.015 | 1 | 7 | Zrsr1 | 0.018 | Old |
| 0.114605 | 1.265114 | 0.046 | 0.024 | 1 | 7 | Mmp12 | 0.022 | Old |
| 0.114962 | 1.052001 | 0.062 | 0.036 | 1 | 7 | Adgrl2 | 0.026 | Old |
| 0.115127 | 0.175012 | 0.821 | 0.78 | 1 | 7 | Ptafr | 0.041 | Old |
| 0.115145 | 0.163311 | 0.503 | 0.434 | 1 | 7 | Psmd4 | 0.069 | Old |
| 0.115287 | 1.116804 | 0.033 | 0.015 | 1 | 7 | Srd5a1 | 0.018 | Old |
| 0.115359 | 0.322861 | 0.395 | 0.334 | 1 | 7 | Asf1b | 0.061 | Old |
| 0.115412 | 0.157283 | 0.287 | 0.229 | 1 | 7 | Ndufv1 | 0.058 | Old |
| 0.115442 | 0.727815 | 0.069 | 0.042 | 1 | 7 | Clmp | 0.027 | Old |
| 0.115508 | 0.310232 | 0.454 | 0.395 | 1 | 7 | Trappc4 | 0.059 | Old |
| 0.115535 | 0.262261 | 0.454 | 0.389 | 1 | 7 | Psmc1 | 0.065 | Old |
| 0.115726 | 0.460722 | 0.103 | 0.069 | 1 | 7 | Fbxo6 | 0.034 | Old |
| 0.1159 | 1.152892 | 0.033 | 0.015 | 1 | 7 | Pgap3 | 0.018 | Old |
| 0.115904 | 0.35396 | 0.085 | 0.054 | 1 | 7 | Mfsd8 | 0.031 | Old |
| 0.11591 | 0.459916 | 0.174 | 0.133 | 1 | 7 | Malsu1 | 0.041 | Old |
| 0.116257 | 0.370497 | 0.279 | 0.226 | 1 | 7 | Vcam1 | 0.053 | Old |
| 0.116457 | 0.327078 | 0.485 | 0.422 | 1 | 7 | Psmb7 | 0.063 | Old |
| 0.116516 | 0.372545 | 0.033 | 0.015 | 1 | 7 | Tcim | 0.018 | Old |
| 0.116658 | 0.873468 | 0.046 | 0.024 | 1 | 7 | Osgepl1 | 0.022 | Old |
| 0.116722 | 0.524181 | 0.623 | 0.627 | 1 | 7 | Ly6e | -0.004 | Old |
| 0.117035 | 0.20132 | 0.413 | 0.352 | 1 | 7 | Arf6 | 0.061 | Old |
| 0.117087 | 0.369439 | 0.103 | 0.069 | 1 | 7 | Mlkl | 0.034 | Old |
| 0.117135 | 0.793225 | 0.033 | 0.015 | 1 | 7 | Abcb4 | 0.018 | Old |
| 0.117457 | 0.077614 | 0.413 | 0.355 | 1 | 7 | Ndufv2 | 0.058 | Old |
| 0.117522 | 1.441824 | 0.054 | 0.03 | 1 | 7 | Sparc | 0.024 | Old |
| 0.117648 | 0.270967 | 0.128 | 0.09 | 1 | 7 | Usb1 | 0.038 | Old |
| 0.117911 | 0.269049 | 0.092 | 0.06 | 1 | 7 | Cenpo | 0.032 | Old |
| 0.118124 | 0.529984 | 0.328 | 0.283 | 1 | 7 | Nop56 | 0.045 | Old |
| 0.11823 | 0.290296 | 0.451 | 0.392 | 1 | 7 | Smarcd2 | 0.059 | Old |
| 0.118503 | 1.052764 | 0.062 | 0.036 | 1 | 7 | Nop2 | 0.026 | Old |
| 0.118683 | 0.217839 | 0.31 | 0.259 | 1 | 7 | Vrk2 | 0.051 | Old |
| 0.118692 | 0.681729 | 0.033 | 0.015 | 1 | 7 | Gm10053 | 0.018 | Old |

| | | | | | | | | |
|---|---|---|---|---|---|---|---|---|
| 0.118923 | 0.223616 | 0.385 | 0.319 | 1 | 7 | Tmem219 | 0.066 | Old |
| 0.119206 | 0.344859 | 0.392 | 0.34 | 1 | 7 | Ube2q1 | 0.052 | Old |
| 0.119228 | 0.457251 | 0.231 | 0.184 | 1 | 7 | Lypla1 | 0.047 | Old |
| 0.11932 | 0.528316 | 0.033 | 0.015 | 1 | 7 | Zcchc3 | 0.018 | Old |
| 0.119445 | 0.24808 | 0.059 | 0.09 | 1 | 7 | Areg | -0.031 | Old |
| 0.119626 | 0.282635 | 0.069 | 0.042 | 1 | 7 | Zfp952 | 0.027 | Old |
| 0.119635 | 0.743349 | 0.033 | 0.015 | 1 | 7 | Nudt15 | 0.018 | Old |
| 0.11993 | 1.306474 | 0.054 | 0.03 | 1 | 7 | 1810043G( | 0.024 | Old |
| 0.120196 | 0.462287 | 0.203 | 0.157 | 1 | 7 | Eif2ak1 | 0.046 | Old |
| 0.120288 | 0.497248 | 0.092 | 0.06 | 1 | 7 | Homer3 | 0.032 | Old |
| 0.120329 | 0.433314 | 0.256 | 0.205 | 1 | 7 | Nup54 | 0.051 | Old |
| 0.120365 | 0.263506 | 0.254 | 0.205 | 1 | 7 | Tufm | 0.049 | Old |
| 0.120491 | 0.300981 | 0.305 | 0.247 | 1 | 7 | 4933434E2 | 0.058 | Old |
| 0.120572 | 0.08764 | 0.39 | 0.322 | 1 | 7 | Fam173a | 0.068 | Old |
| 0.12077 | 0.168376 | 0.223 | 0.172 | 1 | 7 | Ergic1 | 0.051 | Old |
| 0.120815 | 0.173387 | 0.333 | 0.271 | 1 | 7 | BC005624 | 0.062 | Old |
| 0.120824 | 0.502197 | 0.2 | 0.154 | 1 | 7 | Fopnl | 0.046 | Old |
| 0.120884 | 0.244763 | 0.587 | 0.518 | 1 | 7 | Nras | 0.069 | Old |
| 0.1209 | 0.523533 | 0.033 | 0.015 | 1 | 7 | BC049352 | 0.018 | Old |
| 0.121082 | 0.37146 | 0.182 | 0.139 | 1 | 7 | Heatr1 | 0.043 | Old |
| 0.121118 | 0.928031 | 0.046 | 0.024 | 1 | 7 | Zfp825 | 0.022 | Old |
| 0.12119 | 0.504269 | 0.077 | 0.048 | 1 | 7 | Ttc27 | 0.029 | Old |
| 0.12119 | 0.245965 | 0.077 | 0.048 | 1 | 7 | Fam83d | 0.029 | Old |
| 0.121393 | 0.151508 | 0.344 | 0.277 | 1 | 7 | Plaa | 0.067 | Old |
| 0.121513 | 0.575922 | 0.136 | 0.099 | 1 | 7 | Traf2 | 0.037 | Old |
| 0.121515 | 0.36505 | 0.192 | 0.148 | 1 | 7 | Zmym2 | 0.044 | Old |
| 0.121561 | 0.290451 | 0.279 | 0.232 | 1 | 7 | Seh1l | 0.047 | Old |
| 0.121566 | 0.004431 | 0.213 | 0.163 | 1 | 7 | Cdc20 | 0.05 | Old |
| 0.121617 | 0.469818 | 0.1 | 0.066 | 1 | 7 | Mrps2 | 0.034 | Old |
| 0.12169 | 0.307553 | 0.431 | 0.367 | 1 | 7 | Strap | 0.064 | Old |
| 0.121707 | 0.320614 | 0.313 | 0.253 | 1 | 7 | Pik3ca | 0.06 | Old |
| 0.121873 | 1.701763 | 0.041 | 0.021 | 1 | 7 | Gbp2b | 0.02 | Old |
| 0.121926 | 0.258124 | 0.185 | 0.139 | 1 | 7 | Wwc2 | 0.046 | Old |
| 0.12218 | 0.409569 | 0.208 | 0.166 | 1 | 7 | Fubp3 | 0.042 | Old |
| 0.122497 | 0.895678 | 0.033 | 0.015 | 1 | 7 | Fahd2a | 0.018 | Old |
| 0.122497 | 0.363405 | 0.033 | 0.015 | 1 | 7 | Mfsd3 | 0.018 | Old |
| 0.122524 | 0.508002 | 0.174 | 0.136 | 1 | 7 | Ints3 | 0.038 | Old |
| 0.122582 | 0.135791 | 0.19 | 0.148 | 1 | 7 | Ralgapa1 | 0.042 | Old |
| 0.122756 | 0.221741 | 0.256 | 0.202 | 1 | 7 | Polr2b | 0.054 | Old |
| 0.123018 | 0.25792 | 0.118 | 0.081 | 1 | 7 | Nbdy | 0.037 | Old |
| 0.123185 | 0.291094 | 0.118 | 0.081 | 1 | 7 | Dcp1a | 0.037 | Old |
| 0.123515 | 0.70449 | 0.238 | 0.199 | 1 | 7 | Gtf3a | 0.039 | Old |
| 0.123517 | 0.058857 | 0.405 | 0.34 | 1 | 7 | Tcof1 | 0.065 | Old |
| 0.123565 | 0.153548 | 0.497 | 0.428 | 1 | 7 | Ppp4c | 0.069 | Old |
| 0.123636 | 0.288341 | 0.087 | 0.057 | 1 | 7 | Neurl2 | 0.03 | Old |

| | | | | | | | | |
|---|---|---|---|---|---|---|---|---|
| 0.124059 | 0.191468 | 0.077 | 0.048 | 1 | 7 | Sesn2 | 0.029 | Old |
| 0.124147 | 0.44738 | 0.069 | 0.042 | 1 | 7 | Ranbp6 | 0.027 | Old |
| 0.124403 | 0.437497 | 0.121 | 0.084 | 1 | 7 | Grcc10 | 0.037 | Old |
| 0.124403 | 0.253685 | 0.121 | 0.084 | 1 | 7 | Ccdc69 | 0.037 | Old |
| 0.124604 | 0.13198 | 0.81 | 0.774 | 1 | 7 | Snrnp70 | 0.036 | Old |
| 0.125016 | 0.063621 | 0.169 | 0.127 | 1 | 7 | Pus1 | 0.042 | Old |
| 0.125299 | 0.199133 | 0.251 | 0.199 | 1 | 7 | Pspc1 | 0.052 | Old |
| 0.125438 | 0.68221 | 0.072 | 0.045 | 1 | 7 | Hdac10 | 0.027 | Old |
| 0.125606 | 0.791695 | 0.062 | 0.036 | 1 | 7 | Atat1 | 0.026 | Old |
| 0.125612 | 0.226276 | 0.559 | 0.497 | 1 | 7 | Cct7 | 0.062 | Old |
| 0.125771 | 0.204961 | 0.546 | 0.476 | 1 | 7 | Pkig | 0.07 | Old |
| 0.126145 | 0.344852 | 0.228 | 0.184 | 1 | 7 | Ccnb1 | 0.044 | Old |
| 0.126226 | 0.5667 | 0.105 | 0.072 | 1 | 7 | Lrr1 | 0.033 | Old |
| 0.126231 | 0.37672 | 0.236 | 0.19 | 1 | 7 | Yrdc | 0.046 | Old |
| 0.126236 | 0.244716 | 0.121 | 0.084 | 1 | 7 | Atg9a | 0.037 | Old |
| 0.126366 | 0.044036 | 0.341 | 0.274 | 1 | 7 | Dnajb9 | 0.067 | Old |
| 0.126494 | 0.390544 | 0.156 | 0.117 | 1 | 7 | Cp | 0.039 | Old |
| 0.126546 | 0.26335 | 0.323 | 0.256 | 1 | 7 | Kpna1 | 0.067 | Old |
| 0.126642 | 0.341333 | 0.269 | 0.214 | 1 | 7 | Nubp1 | 0.055 | Old |
| 0.126706 | 0.35619 | 0.154 | 0.114 | 1 | 7 | Ubac1 | 0.04 | Old |
| 0.126891 | 0.19187 | 0.562 | 0.482 | 1 | 7 | Pmaip1 | 0.08 | Old |
| 0.127061 | 0.057156 | 0.233 | 0.187 | 1 | 7 | Katnbl1 | 0.046 | Old |
| 0.127214 | 0.311747 | 0.569 | 0.527 | 1 | 7 | Tubb2a | 0.042 | Old |
| 0.127313 | 0.164249 | 0.382 | 0.313 | 1 | 7 | Gbp7 | 0.069 | Old |
| 0.127456 | 0.062305 | 0.003 | 0.012 | 1 | 7 | Gm20522 | -0.009 | Old |
| 0.127464 | 0.115737 | 0.231 | 0.181 | 1 | 7 | Cdc42ep3 | 0.05 | Old |
| 0.127478 | 0.240475 | 0.459 | 0.386 | 1 | 7 | Rab6a | 0.073 | Old |
| 0.127567 | 1.134446 | 0.028 | 0.012 | 1 | 7 | Tulp3 | 0.016 | Old |
| 0.127718 | 0.468558 | 0.205 | 0.157 | 1 | 7 | Prep | 0.048 | Old |
| 0.127828 | 0.246996 | 0.131 | 0.093 | 1 | 7 | Rrn3 | 0.038 | Old |
| 0.127951 | 0.217941 | 0.144 | 0.105 | 1 | 7 | Fbxo30 | 0.039 | Old |
| 0.128011 | 0.231984 | 0.303 | 0.253 | 1 | 7 | Asxl1 | 0.05 | Old |
| 0.128118 | 0.057276 | 0.544 | 0.476 | 1 | 7 | Fam96a | 0.068 | Old |
| 0.128181 | 0.316719 | 0.123 | 0.087 | 1 | 7 | Nol6 | 0.036 | Old |
| 0.128293 | 1.772745 | 0.028 | 0.012 | 1 | 7 | Cxcl9 | 0.016 | Old |
| 0.128352 | 0.13267 | 0.305 | 0.25 | 1 | 7 | Eapp | 0.055 | Old |
| 0.128491 | 0.240061 | 0.179 | 0.136 | 1 | 7 | Gpd1l | 0.043 | Old |
| 0.128509 | 0.344905 | 0.213 | 0.166 | 1 | 7 | Mrps7 | 0.047 | Old |
| 0.128591 | 0.273701 | 0.172 | 0.13 | 1 | 7 | Dst | 0.042 | Old |
| 0.128657 | 0.731793 | 0.028 | 0.012 | 1 | 7 | Sirt4 | 0.016 | Old |
| 0.128857 | 0.460799 | 0.251 | 0.205 | 1 | 7 | Jkamp | 0.046 | Old |
| 0.128922 | 0.408104 | 0.277 | 0.232 | 1 | 7 | Abca3 | 0.045 | Old |
| 0.129137 | 0.203364 | 0.305 | 0.244 | 1 | 7 | Htatsf1 | 0.061 | Old |
| 0.129436 | 0.050916 | 0.054 | 0.03 | 1 | 7 | Gpr157 | 0.024 | Old |
| 0.129557 | 0.09286 | 0.39 | 0.331 | 1 | 7 | Skp1a | 0.059 | Old |

| | | | | | | | | |
|---|---|---|---|---|---|---|---|---|
| 0.129561 | 0.37996 | 0.418 | 0.358 | 1 | 7 | Nab1 | 0.06 | Old |
| 0.129601 | 0.259933 | 0.256 | 0.205 | 1 | 7 | Fbxw11 | 0.051 | Old |
| 0.129614 | 0.278155 | 0.318 | 0.265 | 1 | 7 | Ssfa2 | 0.053 | Old |
| 0.130281 | 0.662761 | 0.167 | 0.13 | 1 | 7 | Skp2 | 0.037 | Old |
| 0.130323 | 0.219151 | 0.041 | 0.021 | 1 | 7 | Trip6 | 0.02 | Old |
| 0.130351 | 0.131358 | 0.167 | 0.12 | 1 | 7 | Rnpc3 | 0.047 | Old |
| 0.130386 | 0.294317 | 0.182 | 0.139 | 1 | 7 | Ints11 | 0.043 | Old |
| 0.13053 | 0.294419 | 0.236 | 0.184 | 1 | 7 | Pigm | 0.052 | Old |
| 0.130672 | 0.185271 | 0.585 | 0.518 | 1 | 7 | Srsf7 | 0.067 | Old |
| 0.130731 | 0.719378 | 0.138 | 0.102 | 1 | 7 | Pogk | 0.036 | Old |
| 0.130749 | 0.572639 | 0.108 | 0.075 | 1 | 7 | Tecpr1 | 0.033 | Old |
| 0.130832 | 0.196305 | 0.415 | 0.352 | 1 | 7 | Idh1 | 0.063 | Old |
| 0.130872 | 0.267206 | 0.262 | 0.214 | 1 | 7 | Haus4 | 0.048 | Old |
| 0.13093 | 0.459371 | 0.108 | 0.075 | 1 | 7 | Nuak1 | 0.033 | Old |
| 0.131027 | 0.286056 | 0.285 | 0.232 | 1 | 7 | Emc2 | 0.053 | Old |
| 0.1311 | 0.522557 | 0.079 | 0.051 | 1 | 7 | Alg3 | 0.028 | Old |
| 0.131123 | 0.948475 | 0.09 | 0.06 | 1 | 7 | Kcnq1 | 0.03 | Old |
| 0.131228 | 0.823149 | 0.028 | 0.012 | 1 | 7 | Lrfn1 | 0.016 | Old |
| 0.131394 | 0.173332 | 0.831 | 0.825 | 1 | 7 | Ran | 0.006 | Old |
| 0.131497 | 0.548029 | 0.167 | 0.127 | 1 | 7 | Mpg | 0.04 | Old |
| 0.131599 | 0.69385 | 0.028 | 0.012 | 1 | 7 | Slc22a5 | 0.016 | Old |
| 0.131599 | 0.382924 | 0.197 | 0.154 | 1 | 7 | Irf3 | 0.043 | Old |
| 0.131668 | 0.124948 | 0.246 | 0.196 | 1 | 7 | Hibadh | 0.05 | Old |
| 0.131668 | 0.187003 | 0.141 | 0.102 | 1 | 7 | Ammecr1 | 0.039 | Old |
| 0.131727 | 0.105908 | 0.892 | 0.895 | 1 | 7 | Ptp4a2 | -0.003 | Old |
| 0.131816 | 0.15551 | 0.259 | 0.214 | 1 | 7 | Trappc8 | 0.045 | Old |
| 0.131825 | 0.517327 | 0.041 | 0.021 | 1 | 7 | Trmt61a | 0.02 | Old |
| 0.131854 | 0.186033 | 0.505 | 0.443 | 1 | 7 | Ak2 | 0.062 | Old |
| 0.13204 | 0.158047 | 0.295 | 0.241 | 1 | 7 | Cfap43 | 0.054 | Old |
| 0.132388 | 0.418386 | 0.246 | 0.199 | 1 | 7 | CAAA0111 | 0.047 | Old |
| 0.132406 | 0.1716 | 0.2 | 0.154 | 1 | 7 | Nip7 | 0.046 | Old |
| 0.13243 | 0.259622 | 0.274 | 0.223 | 1 | 7 | Ncor2 | 0.051 | Old |
| 0.132669 | 0.038603 | 0.29 | 0.232 | 1 | 7 | Kif20b | 0.058 | Old |
| 0.132732 | 0.637288 | 0.041 | 0.021 | 1 | 7 | AC163032. | 0.02 | Old |
| 0.132814 | 0.201814 | 0.369 | 0.313 | 1 | 7 | Ppm1a | 0.056 | Old |
| 0.132884 | 0.100984 | 0.656 | 0.593 | 1 | 7 | Prpf38b | 0.063 | Old |
| 0.132925 | 0.02749 | 0.321 | 0.256 | 1 | 7 | Kif2a | 0.065 | Old |
| 0.132936 | 0.143109 | 0.349 | 0.292 | 1 | 7 | Grk6 | 0.057 | Old |
| 0.133021 | 0.238675 | 0.433 | 0.361 | 1 | 7 | Mob4 | 0.072 | Old |
| 0.133165 | 0.282238 | 0.2 | 0.16 | 1 | 7 | Dkc1 | 0.04 | Old |
| 0.133184 | 0.156243 | 0.426 | 0.355 | 1 | 7 | Pfdn2 | 0.071 | Old |
| 0.133328 | 0.214312 | 0.264 | 0.211 | 1 | 7 | Eif4ebp1 | 0.053 | Old |
| 0.13334 | 0.655083 | 0.041 | 0.021 | 1 | 7 | Gabbr1 | 0.02 | Old |
| 0.133444 | 0.430697 | 0.079 | 0.051 | 1 | 7 | Btg3 | 0.028 | Old |
| 0.133464 | 0.151568 | 0.028 | 0.012 | 1 | 7 | Tmem229a | 0.016 | Old |

| | | | | | | | | |
|---|---|---|---|---|---|---|---|---|
| 0.133643 | 0.726006 | 0.113 | 0.081 | 1 | 7 | Anpep | 0.032 | Old |
| 0.133644 | 0.468023 | 0.041 | 0.021 | 1 | 7 | A130010J1 | 0.02 | Old |
| 0.133653 | 0.670976 | 0.064 | 0.039 | 1 | 7 | Hacd1 | 0.025 | Old |
| 0.133745 | 0.209525 | 0.241 | 0.187 | 1 | 7 | Nmt2 | 0.054 | Old |
| 0.133749 | 0.036603 | 0.7 | 0.63 | 1 | 7 | Ube2d2a | 0.07 | Old |
| 0.133812 | 0.441676 | 0.251 | 0.211 | 1 | 7 | Rnd3 | 0.04 | Old |
| 0.13384 | 0.293018 | 0.028 | 0.012 | 1 | 7 | Bbs9 | 0.016 | Old |
| 0.133927 | 0.402775 | 0.09 | 0.06 | 1 | 7 | Zmym3 | 0.03 | Old |
| 0.133957 | 0.557509 | 0.228 | 0.184 | 1 | 7 | Otud6b | 0.044 | Old |
| 0.13402 | 0.25478 | 0.062 | 0.036 | 1 | 7 | Alyref2 | 0.026 | Old |
| 0.134129 | 0.153722 | 0.574 | 0.497 | 1 | 7 | Wdr83os | 0.077 | Old |
| 0.134216 | 0.36668 | 0.028 | 0.012 | 1 | 7 | Coq8a | 0.016 | Old |
| 0.134336 | 0.194424 | 0.167 | 0.127 | 1 | 7 | Gla | 0.04 | Old |
| 0.134381 | 0.143049 | 0.81 | 0.759 | 1 | 7 | Eif4a1 | 0.051 | Old |
| 0.134408 | 0.283916 | 0.446 | 0.386 | 1 | 7 | Cdca8 | 0.06 | Old |
| 0.134556 | 0.188754 | 0.803 | 0.756 | 1 | 7 | Atp6v0b | 0.047 | Old |
| 0.134561 | 0.680904 | 0.041 | 0.021 | 1 | 7 | Flrt2 | 0.02 | Old |
| 0.134777 | 0.159437 | 0.292 | 0.235 | 1 | 7 | Ncbp1 | 0.057 | Old |
| 0.134856 | 0.242196 | 0.497 | 0.452 | 1 | 7 | Ndufb6 | 0.045 | Old |
| 0.134867 | 0.152919 | 0.715 | 0.627 | 1 | 7 | Ddx39b | 0.088 | Old |
| 0.134868 | 0.444779 | 0.041 | 0.021 | 1 | 7 | Tiam2 | 0.02 | Old |
| 0.135128 | 0.0718 | 0.326 | 0.262 | 1 | 7 | Cltb | 0.064 | Old |
| 0.135451 | 0.188262 | 0.726 | 0.693 | 1 | 7 | Mapre1 | 0.033 | Old |
| 0.135453 | 0.193173 | 0.664 | 0.648 | 1 | 7 | Ppp1ca | 0.016 | Old |
| 0.135482 | 0.002643 | 0.108 | 0.072 | 1 | 7 | Yae1d1 | 0.036 | Old |
| 0.135483 | 0.463452 | 0.041 | 0.021 | 1 | 7 | Lhpp | 0.02 | Old |
| 0.135492 | 0.328397 | 0.336 | 0.277 | 1 | 7 | Gabpb2 | 0.059 | Old |
| 0.135502 | 0.843258 | 0.103 | 0.072 | 1 | 7 | Hes6 | 0.031 | Old |
| 0.135518 | 0.137514 | 0.567 | 0.497 | 1 | 7 | Arcn1 | 0.07 | Old |
| 0.135547 | 0.189926 | 0.123 | 0.087 | 1 | 7 | Med7 | 0.036 | Old |
| 0.135566 | 0.616146 | 0.056 | 0.033 | 1 | 7 | Zfp87 | 0.023 | Old |
| 0.13565 | 0.419269 | 0.072 | 0.045 | 1 | 7 | Mettl21a | 0.027 | Old |
| 0.13565 | 0.16611 | 0.349 | 0.289 | 1 | 7 | Hells | 0.06 | Old |
| 0.135683 | 0.824321 | 0.049 | 0.027 | 1 | 7 | Endov | 0.022 | Old |
| 0.135729 | 0.239934 | 0.028 | 0.012 | 1 | 7 | Tcaf1 | 0.016 | Old |
| 0.135792 | 0.195406 | 0.041 | 0.021 | 1 | 7 | Prnp | 0.02 | Old |
| 0.135975 | 0.210817 | 0.241 | 0.193 | 1 | 7 | Rab29 | 0.048 | Old |
| 0.136126 | 0.275693 | 0.341 | 0.28 | 1 | 7 | Chd1 | 0.061 | Old |
| 0.136537 | 0.186887 | 0.31 | 0.253 | 1 | 7 | Gpbp1l1 | 0.057 | Old |
| 0.136761 | 0.264903 | 0.097 | 0.066 | 1 | 7 | Gmppb | 0.031 | Old |
| 0.136953 | 0.172047 | 0.351 | 0.298 | 1 | 7 | Usp47 | 0.053 | Old |
| 0.13732 | 0.589417 | 0.082 | 0.054 | 1 | 7 | Mrpl47 | 0.028 | Old |
| 0.137322 | 1.877689 | 0.023 | 0.009 | 1 | 7 | Ifit3 | 0.014 | Old |
| 0.137476 | 0.220321 | 0.467 | 0.398 | 1 | 7 | Ppp6r3 | 0.069 | Old |
| 0.137696 | 0.471198 | 0.318 | 0.271 | 1 | 7 | Naga | 0.047 | Old |

| | | | | | | | | |
|---|---|---|---|---|---|---|---|---|
| 0.137768 | 0.237934 | 0.192 | 0.148 | 1 | 7 | Ska1 | 0.044 | Old |
| 0.13791 | 0.536941 | 0.128 | 0.093 | 1 | 7 | Smyd2 | 0.035 | Old |
| 0.138001 | 0.179538 | 0.472 | 0.404 | 1 | 7 | Mkrn1 | 0.068 | Old |
| 0.138024 | 0.142863 | 0.028 | 0.012 | 1 | 7 | Aars2 | 0.016 | Old |
| 0.13834 | 0.043219 | 0.454 | 0.401 | 1 | 7 | Stip1 | 0.053 | Old |
| 0.13834 | 0.131807 | 0.687 | 0.596 | 1 | 7 | Dok2 | 0.091 | Old |
| 0.139005 | 0.499575 | 0.128 | 0.096 | 1 | 7 | Zdhhc8 | 0.032 | Old |
| 0.139367 | 0.405947 | 0.19 | 0.148 | 1 | 7 | Ints9 | 0.042 | Old |
| 0.13944 | 1.237365 | 0.036 | 0.018 | 1 | 7 | Gm10184 | 0.018 | Old |
| 0.139442 | 0.131571 | 0.221 | 0.172 | 1 | 7 | 1810013L2 | 0.049 | Old |
| 0.139729 | 0.178766 | 0.192 | 0.148 | 1 | 7 | Sssca1 | 0.044 | Old |
| 0.139802 | 0.029426 | 0.118 | 0.081 | 1 | 7 | Zc3h7b | 0.037 | Old |
| 0.139833 | 0.096885 | 0.295 | 0.232 | 1 | 7 | Ap1g1 | 0.063 | Old |
| 0.139893 | 0.380201 | 0.331 | 0.277 | 1 | 7 | Cdkn2d | 0.054 | Old |
| 0.139942 | 0.463975 | 0.049 | 0.027 | 1 | 7 | Gdpd1 | 0.022 | Old |
| 0.140033 | 0.164794 | 0.364 | 0.307 | 1 | 7 | Tank | 0.057 | Old |
| 0.140341 | 0.945061 | 0.023 | 0.009 | 1 | 7 | 6430531B1 | 0.014 | Old |
| 0.140446 | 0.110591 | 0.877 | 0.828 | 1 | 7 | Irf2bp2 | 0.049 | Old |
| 0.14064 | 0.437217 | 0.679 | 0.657 | 1 | 7 | Gsn | 0.022 | Old |
| 0.140777 | 1.07107 | 0.023 | 0.009 | 1 | 7 | Ehd2 | 0.014 | Old |
| 0.140825 | 0.375783 | 0.162 | 0.12 | 1 | 7 | Trap1 | 0.042 | Old |
| 0.140908 | 0.33145 | 0.3 | 0.259 | 1 | 7 | Pgap2 | 0.041 | Old |
| 0.14091 | 0.038891 | 0.469 | 0.392 | 1 | 7 | Irak1 | 0.077 | Old |
| 0.141 | 0.001466 | 0.297 | 0.241 | 1 | 7 | Skiv2l | 0.056 | Old |
| 0.141098 | 0.086564 | 0.485 | 0.416 | 1 | 7 | Cdk9 | 0.069 | Old |
| 0.141134 | 0.923679 | 0.036 | 0.018 | 1 | 7 | Mppe1 | 0.018 | Old |
| 0.141265 | 0.323994 | 0.374 | 0.319 | 1 | 7 | Rnh1 | 0.055 | Old |
| 0.141629 | 0.653132 | 0.056 | 0.033 | 1 | 7 | Pif1 | 0.023 | Old |
| 0.141636 | 0.118965 | 0.579 | 0.503 | 1 | 7 | Srsf9 | 0.076 | Old |
| 0.141739 | 0.112156 | 0.321 | 0.268 | 1 | 7 | Rabl6 | 0.053 | Old |
| 0.141807 | 0.458093 | 0.154 | 0.117 | 1 | 7 | Plekhg5 | 0.037 | Old |
| 0.142221 | 0.375038 | 0.162 | 0.123 | 1 | 7 | Comtd1 | 0.039 | Old |
| 0.142404 | 0.453764 | 0.169 | 0.13 | 1 | 7 | Atp8b2 | 0.039 | Old |
| 0.142704 | 0.9537 | 0.056 | 0.033 | 1 | 7 | Zkscan14 | 0.023 | Old |
| 0.14297 | 0.383878 | 0.023 | 0.009 | 1 | 7 | Kcng2 | 0.014 | Old |
| 0.143118 | 0.713217 | 0.3 | 0.262 | 1 | 7 | Il6 | 0.038 | Old |
| 0.143167 | 0.614802 | 0.241 | 0.199 | 1 | 7 | Tbc1d1 | 0.042 | Old |
| 0.143226 | 0.813104 | 0.082 | 0.054 | 1 | 7 | Poc5 | 0.028 | Old |
| 0.143412 | 0.722931 | 0.023 | 0.009 | 1 | 7 | Rnd1 | 0.014 | Old |
| 0.143417 | 0.263581 | 0.197 | 0.154 | 1 | 7 | Skiv2l2 | 0.043 | Old |
| 0.143461 | 0.457159 | 0.21 | 0.163 | 1 | 7 | Ppp1r8 | 0.047 | Old |
| 0.143567 | 0.16535 | 0.449 | 0.389 | 1 | 7 | Tspan3 | 0.06 | Old |
| 0.143766 | 0.284439 | 0.444 | 0.392 | 1 | 7 | Eri1 | 0.052 | Old |
| 0.143805 | 0.191807 | 0.195 | 0.151 | 1 | 7 | Rap1gap2 | 0.044 | Old |
| 0.144205 | 0.367873 | 0.544 | 0.485 | 1 | 7 | Ghitm | 0.059 | Old |

| | | | | | | | | |
|---|---|---|---|---|---|---|---|---|
| 0.14429 | 0.209021 | 0.338 | 0.295 | 1 | 7 | Zfp292 | 0.043 | Old |
| 0.144299 | 0.213804 | 0.267 | 0.214 | 1 | 7 | Copg1 | 0.053 | Old |
| 0.144345 | 0.219827 | 0.187 | 0.145 | 1 | 7 | Nup107 | 0.042 | Old |
| 0.144476 | 0.543912 | 0.221 | 0.178 | 1 | 7 | Mrpl41 | 0.043 | Old |
| 0.144507 | 0.296928 | 0.151 | 0.111 | 1 | 7 | Cgnl1 | 0.04 | Old |
| 0.144563 | 0.170251 | 0.421 | 0.361 | 1 | 7 | Gnl3 | 0.06 | Old |
| 0.144743 | 1.238219 | 0.023 | 0.009 | 1 | 7 | Prps1l3 | 0.014 | Old |
| 0.14475 | 0.204677 | 0.313 | 0.259 | 1 | 7 | Plekha1 | 0.054 | Old |
| 0.144892 | 0.163275 | 0.049 | 0.027 | 1 | 7 | Gm17092 | 0.022 | Old |
| 0.145071 | 0.364615 | 0.067 | 0.042 | 1 | 7 | Zfp866 | 0.025 | Old |
| 0.145263 | 0.963431 | 0.036 | 0.018 | 1 | 7 | Rwdd2b | 0.018 | Old |
| 0.145314 | 0.255103 | 0.231 | 0.184 | 1 | 7 | Tmem87b | 0.047 | Old |
| 0.14545 | 0.325337 | 0.144 | 0.105 | 1 | 7 | Ruvbl2 | 0.039 | Old |
| 0.145572 | 0.905717 | 0.044 | 0.024 | 1 | 7 | BC017158 | 0.02 | Old |
| 0.145636 | 0.366128 | 0.023 | 0.009 | 1 | 7 | Pstpip2 | 0.014 | Old |
| 0.145969 | 0.398396 | 0.351 | 0.301 | 1 | 7 | Cyc1 | 0.05 | Old |
| 0.146047 | 0.215971 | 0.379 | 0.322 | 1 | 7 | Stk11 | 0.057 | Old |
| 0.146079 | 0.522203 | 0.131 | 0.096 | 1 | 7 | Drg2 | 0.035 | Old |
| 0.146084 | 0.170053 | 0.023 | 0.009 | 1 | 7 | Mettl4 | 0.014 | Old |
| 0.146199 | 1.266532 | 0.044 | 0.024 | 1 | 7 | Ppcs | 0.02 | Old |
| 0.146207 | 0.036343 | 0.485 | 0.407 | 1 | 7 | Nr4a2 | 0.078 | Old |
| 0.146427 | 0.677238 | 0.085 | 0.057 | 1 | 7 | Unkl | 0.028 | Old |
| 0.146663 | 0.182857 | 0.338 | 0.271 | 1 | 7 | Tmem165 | 0.067 | Old |
| 0.146698 | 0.318174 | 0.421 | 0.367 | 1 | 7 | Tsen34 | 0.054 | Old |
| 0.146797 | 0.08784 | 0.292 | 0.238 | 1 | 7 | Fzr1 | 0.054 | Old |
| 0.146937 | 1.101761 | 0.013 | 0.003 | 1 | 7 | Hsd11b1 | 0.01 | Old |
| 0.146937 | 1.041647 | 0.013 | 0.003 | 1 | 7 | Shank1 | 0.01 | Old |
| 0.146937 | 0.871014 | 0.013 | 0.003 | 1 | 7 | Ankrd55 | 0.01 | Old |
| 0.146956 | 1.197808 | 0.018 | 0.006 | 1 | 7 | Gm16740 | 0.012 | Old |
| 0.147281 | 0.255463 | 0.197 | 0.154 | 1 | 7 | Abcg3 | 0.043 | Old |
| 0.147296 | 0.177982 | 0.341 | 0.277 | 1 | 7 | Ilkap | 0.064 | Old |
| 0.147527 | 1.069598 | 0.092 | 0.063 | 1 | 7 | AC163354. | 0.029 | Old |
| 0.147556 | 0.306419 | 0.321 | 0.271 | 1 | 7 | Ddx39 | 0.05 | Old |
| 0.147572 | 1.842437 | 0.013 | 0.003 | 1 | 7 | Gm11696 | 0.01 | Old |
| 0.147572 | 1.562 | 0.013 | 0.003 | 1 | 7 | Amotl1 | 0.01 | Old |
| 0.147572 | 1.416381 | 0.013 | 0.003 | 1 | 7 | Galnt15 | 0.01 | Old |
| 0.147572 | 1.223626 | 0.013 | 0.003 | 1 | 7 | Sntg1 | 0.01 | Old |
| 0.147572 | 1.174024 | 0.013 | 0.003 | 1 | 7 | Morn1 | 0.01 | Old |
| 0.147572 | 1.066891 | 0.013 | 0.003 | 1 | 7 | Zfp689 | 0.01 | Old |
| 0.147572 | 0.941944 | 0.013 | 0.003 | 1 | 7 | Ubxn11 | 0.01 | Old |
| 0.147572 | 0.914082 | 0.013 | 0.003 | 1 | 7 | Ccdc80 | 0.01 | Old |
| 0.147877 | 0.107648 | 0.513 | 0.467 | 1 | 7 | Hook3 | 0.046 | Old |
| 0.147887 | 0.063045 | 0.023 | 0.009 | 1 | 7 | Zfp382 | 0.014 | Old |
| 0.147926 | 0.251146 | 0.172 | 0.133 | 1 | 7 | Mtap | 0.039 | Old |
| 0.14821 | 1.218218 | 0.013 | 0.003 | 1 | 7 | Adm | 0.01 | Old |

| | | | | | | | | |
|---|---|---|---|---|---|---|---|---|
| 0.14821 | 1.02036 | 0.013 | 0.003 | 1 | 7 | Synpo2 | 0.01 | Old |
| 0.14821 | 0.979901 | 0.013 | 0.003 | 1 | 7 | Aqp7 | 0.01 | Old |
| 0.14821 | 0.928073 | 0.013 | 0.003 | 1 | 7 | Gpr21 | 0.01 | Old |
| 0.14821 | 0.925528 | 0.013 | 0.003 | 1 | 7 | Galnt2 | 0.01 | Old |
| 0.14821 | 0.862346 | 0.013 | 0.003 | 1 | 7 | AI463229 | 0.01 | Old |
| 0.148232 | 0.182812 | 0.249 | 0.202 | 1 | 7 | Rbm15 | 0.047 | Old |
| 0.148282 | 0.196976 | 0.259 | 0.217 | 1 | 7 | Ncoa1 | 0.042 | Old |
| 0.148362 | 0.056049 | 0.521 | 0.458 | 1 | 7 | Psma1 | 0.063 | Old |
| 0.148365 | 0.495383 | 0.097 | 0.066 | 1 | 7 | Cdan1 | 0.031 | Old |
| 0.148571 | 0.413756 | 0.133 | 0.096 | 1 | 7 | Traf6 | 0.037 | Old |
| 0.14875 | 0.550522 | 0.144 | 0.108 | 1 | 7 | Gpatch2 | 0.036 | Old |
| 0.148755 | 0.457041 | 0.182 | 0.139 | 1 | 7 | Mphosph6 | 0.043 | Old |
| 0.14885 | 1.269552 | 0.013 | 0.003 | 1 | 7 | Bicdl1 | 0.01 | Old |
| 0.14885 | 1.172558 | 0.013 | 0.003 | 1 | 7 | Gm49201 | 0.01 | Old |
| 0.14885 | 0.677442 | 0.013 | 0.003 | 1 | 7 | Cpe | 0.01 | Old |
| 0.14885 | 0.603858 | 0.013 | 0.003 | 1 | 7 | Airn | 0.01 | Old |
| 0.14885 | 0.599077 | 0.013 | 0.003 | 1 | 7 | Gm614 | 0.01 | Old |
| 0.148888 | 0.443514 | 0.172 | 0.133 | 1 | 7 | Pold2 | 0.039 | Old |
| 0.148968 | 0.084168 | 0.81 | 0.804 | 1 | 7 | Glud1 | 0.006 | Old |
| 0.149005 | 0.038166 | 0.331 | 0.265 | 1 | 7 | Myo1c | 0.066 | Old |
| 0.149045 | 1.304943 | 0.018 | 0.006 | 1 | 7 | Gm29994 | 0.012 | Old |
| 0.149197 | 0.798815 | 0.113 | 0.081 | 1 | 7 | Glt8d1 | 0.032 | Old |
| 0.14926 | 0.256079 | 0.482 | 0.446 | 1 | 7 | Esyt1 | 0.036 | Old |
| 0.149492 | 1.222503 | 0.013 | 0.003 | 1 | 7 | Ccdc141 | 0.01 | Old |
| 0.149492 | 0.99988 | 0.013 | 0.003 | 1 | 7 | 6330562C2 | 0.01 | Old |
| 0.149492 | 0.82163 | 0.013 | 0.003 | 1 | 7 | Kirrel | 0.01 | Old |
| 0.149492 | 0.643163 | 0.013 | 0.003 | 1 | 7 | C1s1 | 0.01 | Old |
| 0.149492 | 0.621027 | 0.013 | 0.003 | 1 | 7 | Gm20939 | 0.01 | Old |
| 0.149492 | 0.515099 | 0.013 | 0.003 | 1 | 7 | Ksr2 | 0.01 | Old |
| 0.149512 | 0.138364 | 0.595 | 0.515 | 1 | 7 | Ppp1cb | 0.08 | Old |
| 0.14957 | 1.240765 | 0.018 | 0.006 | 1 | 7 | Gm29336 | 0.012 | Old |
| 0.14957 | 1.001393 | 0.018 | 0.006 | 1 | 7 | Trim6 | 0.012 | Old |
| 0.149611 | 1.298146 | 0.051 | 0.03 | 1 | 7 | Cbr3 | 0.021 | Old |
| 0.149673 | 0.022969 | 0.072 | 0.045 | 1 | 7 | Zfp429 | 0.027 | Old |
| 0.149746 | 0.44124 | 0.156 | 0.117 | 1 | 7 | Stard3 | 0.039 | Old |
| 0.149759 | 0.170279 | 0.259 | 0.208 | 1 | 7 | Unc50 | 0.051 | Old |
| 0.149785 | 0.059438 | 0.851 | 0.84 | 1 | 7 | Celf2 | 0.011 | Old |
| 0.149978 | 0.096388 | 0.362 | 0.301 | 1 | 7 | Dnajc9 | 0.061 | Old |
| 0.150136 | 0.64604 | 0.013 | 0.003 | 1 | 7 | Dlg3 | 0.01 | Old |
| 0.150136 | 0.367146 | 0.013 | 0.003 | 1 | 7 | Zfp85 | 0.01 | Old |
| 0.150136 | 0.366566 | 0.013 | 0.003 | 1 | 7 | Brdt | 0.01 | Old |
| 0.150136 | 0.137771 | 0.013 | 0.003 | 1 | 7 | Bace1 | 0.01 | Old |
| 0.150136 | 0.050531 | 0.013 | 0.003 | 1 | 7 | Fabp7 | 0.01 | Old |
| 0.150142 | 0.279534 | 0.269 | 0.22 | 1 | 7 | Nck1 | 0.049 | Old |
| 0.150189 | 0.506291 | 0.123 | 0.09 | 1 | 7 | Caml | 0.033 | Old |

| | | | | | | | | |
|---|---|---|---|---|---|---|---|---|
| 0.150196 | 0.838973 | 0.036 | 0.018 | 1 | 7 | Tmc8 | 0.018 | Old |
| 0.150196 | 0.098051 | 0.036 | 0.018 | 1 | 7 | Xpo5 | 0.018 | Old |
| 0.150341 | 0.130406 | 0.359 | 0.304 | 1 | 7 | Ociad1 | 0.055 | Old |
| 0.150429 | 0.330891 | 0.328 | 0.28 | 1 | 7 | Zfp263 | 0.048 | Old |
| 0.150547 | 0.441532 | 0.092 | 0.063 | 1 | 7 | 0610009O2 | 0.029 | Old |
| 0.150604 | 0.171591 | 0.162 | 0.12 | 1 | 7 | 4930503L1 | 0.042 | Old |
| 0.150626 | 0.7805 | 0.018 | 0.006 | 1 | 7 | Gm48420 | 0.012 | Old |
| 0.150626 | 0.626508 | 0.018 | 0.006 | 1 | 7 | Csprs | 0.012 | Old |
| 0.150669 | 0.318696 | 0.185 | 0.142 | 1 | 7 | Parp9 | 0.043 | Old |
| 0.15067 | 0.230311 | 0.3 | 0.25 | 1 | 7 | Pik3r6 | 0.05 | Old |
| 0.150696 | 0.170458 | 0.336 | 0.28 | 1 | 7 | Efcab14 | 0.056 | Old |
| 0.151286 | 1.18959 | 0.044 | 0.024 | 1 | 7 | Mill2 | 0.02 | Old |
| 0.151541 | 0.373646 | 0.097 | 0.066 | 1 | 7 | Kcne3 | 0.031 | Old |
| 0.151607 | 0.930656 | 0.095 | 0.066 | 1 | 7 | Pi16 | 0.029 | Old |
| 0.151629 | 0.395396 | 0.036 | 0.018 | 1 | 7 | Zfp867 | 0.018 | Old |
| 0.151687 | 0.79631 | 0.018 | 0.006 | 1 | 7 | Zfp772 | 0.012 | Old |
| 0.15183 | 0.272247 | 0.138 | 0.102 | 1 | 7 | Syne3 | 0.036 | Old |
| 0.151989 | 0.24569 | 0.036 | 0.018 | 1 | 7 | Cwf19l1 | 0.018 | Old |
| 0.152094 | 0.23053 | 0.308 | 0.259 | 1 | 7 | Kcnab2 | 0.049 | Old |
| 0.152119 | 0.331981 | 0.21 | 0.169 | 1 | 7 | Kat7 | 0.041 | Old |
| 0.152348 | 0.378264 | 0.113 | 0.081 | 1 | 7 | Mir142hg | 0.032 | Old |
| 0.152382 | 0.117643 | 0.249 | 0.202 | 1 | 7 | Wdr33 | 0.047 | Old |
| 0.152409 | 0.019563 | 0.185 | 0.139 | 1 | 7 | Alg10b | 0.046 | Old |
| 0.152452 | 0.00035 | 0.223 | 0.175 | 1 | 7 | Nup155 | 0.048 | Old |
| 0.152704 | 0.419388 | 0.141 | 0.105 | 1 | 7 | C2cd2l | 0.036 | Old |
| 0.152956 | 0.778214 | 0.067 | 0.042 | 1 | 7 | Zfp146 | 0.025 | Old |
| 0.153145 | 0.348383 | 0.144 | 0.108 | 1 | 7 | 4930453N2 | 0.036 | Old |
| 0.153289 | 0.526858 | 0.018 | 0.006 | 1 | 7 | Tgfbr3 | 0.012 | Old |
| 0.153393 | 0.219488 | 0.092 | 0.063 | 1 | 7 | Tmem126b | 0.029 | Old |
| 0.153495 | 0.32604 | 0.2 | 0.157 | 1 | 7 | Mrpl58 | 0.043 | Old |
| 0.153515 | 0.409055 | 0.059 | 0.036 | 1 | 7 | Dhrs9 | 0.023 | Old |
| 0.153754 | 0.147653 | 0.415 | 0.377 | 1 | 7 | Nus1 | 0.038 | Old |
| 0.153792 | 0.704929 | 0.059 | 0.036 | 1 | 7 | Tab3 | 0.023 | Old |
| 0.153826 | 0.526048 | 0.018 | 0.006 | 1 | 7 | Ppic | 0.012 | Old |
| 0.153826 | 0.290215 | 0.018 | 0.006 | 1 | 7 | Zfp189 | 0.012 | Old |
| 0.153934 | 0.211231 | 0.497 | 0.434 | 1 | 7 | Dnajc8 | 0.063 | Old |
| 0.15407 | 0.576793 | 0.059 | 0.036 | 1 | 7 | Sumf2 | 0.023 | Old |
| 0.154072 | 0.128109 | 0.221 | 0.178 | 1 | 7 | Fcna | 0.043 | Old |
| 0.154074 | 0.167407 | 0.449 | 0.413 | 1 | 7 | Atg3 | 0.036 | Old |
| 0.15432 | 0.201864 | 0.669 | 0.627 | 1 | 7 | Hnrnpa1 | 0.042 | Old |
| 0.154338 | 0.012898 | 0.292 | 0.235 | 1 | 7 | Cryl1 | 0.057 | Old |
| 0.154364 | 0.474363 | 0.018 | 0.006 | 1 | 7 | Als2cl | 0.012 | Old |
| 0.154627 | 0.1235 | 0.133 | 0.096 | 1 | 7 | Nadk2 | 0.037 | Old |
| 0.154811 | 0.412587 | 0.305 | 0.253 | 1 | 7 | Rasa3 | 0.052 | Old |
| 0.154871 | 0.097935 | 0.156 | 0.199 | 1 | 7 | Xpa | -0.043 | Old |

| | | | | | | | | |
|---|---|---|---|---|---|---|---|---|
| 0.154943 | 0.341712 | 0.249 | 0.205 | 1 | 7 | Cnep1r1 | 0.044 | Old |
| 0.155069 | 0.207665 | 0.51 | 0.449 | 1 | 7 | Psmc6 | 0.061 | Old |
| 0.155221 | 0.152032 | 0.808 | 0.75 | 1 | 7 | Ifitm3 | 0.058 | Old |
| 0.155244 | 0.143817 | 0.331 | 0.271 | 1 | 7 | Ccdc115 | 0.06 | Old |
| 0.155339 | 0.370082 | 0.379 | 0.325 | 1 | 7 | Elf2 | 0.054 | Old |
| 0.155551 | 0.844998 | 0.051 | 0.03 | 1 | 7 | Atp10a | 0.021 | Old |
| 0.15574 | 0.189386 | 0.536 | 0.47 | 1 | 7 | Cul3 | 0.066 | Old |
| 0.155928 | 0.856367 | 0.087 | 0.06 | 1 | 7 | Gtf3c5 | 0.027 | Old |
| 0.155985 | 0.58684 | 0.105 | 0.075 | 1 | 7 | Gm43848 | 0.03 | Old |
| 0.156017 | 0.189896 | 0.679 | 0.611 | 1 | 7 | Anp32a | 0.068 | Old |
| 0.156154 | 1.062919 | 0.051 | 0.03 | 1 | 7 | Lamc1 | 0.021 | Old |
| 0.156177 | 0.581268 | 0.044 | 0.024 | 1 | 7 | Itfg2 | 0.02 | Old |
| 0.156382 | 0.143544 | 0.136 | 0.099 | 1 | 7 | Arhgef39 | 0.037 | Old |
| 0.156425 | 0.297417 | 0.197 | 0.154 | 1 | 7 | Hist1h2bc | 0.043 | Old |
| 0.15644 | 0.267742 | 0.359 | 0.31 | 1 | 7 | Msl3 | 0.049 | Old |
| 0.156702 | 0.637646 | 0.077 | 0.051 | 1 | 7 | Cacna1f | 0.026 | Old |
| 0.15676 | 0.546334 | 0.051 | 0.03 | 1 | 7 | Cdc42bpa | 0.021 | Old |
| 0.156838 | 0.271013 | 0.162 | 0.127 | 1 | 7 | Exosc7 | 0.035 | Old |
| 0.157045 | 0.346361 | 0.262 | 0.223 | 1 | 7 | Rnf213 | 0.039 | Old |
| 0.157377 | 0.169627 | 0.359 | 0.301 | 1 | 7 | Taf6l | 0.058 | Old |
| 0.157475 | 0.201586 | 0.218 | 0.172 | 1 | 7 | Phrf1 | 0.046 | Old |
| 0.157892 | 0.521584 | 0.115 | 0.084 | 1 | 7 | Pole | 0.031 | Old |
| 0.158022 | 0.092224 | 0.513 | 0.461 | 1 | 7 | Hectd1 | 0.052 | Old |
| 0.158126 | 0.184972 | 0.362 | 0.307 | 1 | 7 | Fen1 | 0.055 | Old |
| 0.1582 | 0.145634 | 0.282 | 0.232 | 1 | 7 | Smg7 | 0.05 | Old |
| 0.158265 | 0.1668 | 0.126 | 0.09 | 1 | 7 | Hic2 | 0.036 | Old |
| 0.158295 | 0.141899 | 0.462 | 0.398 | 1 | 7 | Retreg2 | 0.064 | Old |
| 0.15848 | 0.351359 | 0.067 | 0.042 | 1 | 7 | Srrd | 0.025 | Old |
| 0.158494 | 1.960626 | 0.031 | 0.015 | 1 | 7 | Amotl2 | 0.016 | Old |
| 0.1585 | 0.327579 | 0.044 | 0.024 | 1 | 7 | Exo5 | 0.02 | Old |
| 0.158523 | 0.165923 | 0.923 | 0.898 | 1 | 7 | Pcbp2 | 0.025 | Old |
| 0.158703 | 0.515785 | 0.074 | 0.048 | 1 | 7 | Frat2 | 0.026 | Old |
| 0.158893 | 0.798869 | 0.051 | 0.03 | 1 | 7 | Uros | 0.021 | Old |
| 0.158897 | 0.142225 | 0.546 | 0.491 | 1 | 7 | Rbbp4 | 0.055 | Old |
| 0.159023 | 0.238309 | 0.323 | 0.265 | 1 | 7 | Gmip | 0.058 | Old |
| 0.159056 | 0.797374 | 0.095 | 0.066 | 1 | 7 | Trim23 | 0.029 | Old |
| 0.159241 | 0.300253 | 0.297 | 0.25 | 1 | 7 | Nae1 | 0.047 | Old |
| 0.159619 | 0.03345 | 0.492 | 0.416 | 1 | 7 | Ube2k | 0.076 | Old |
| 0.159929 | 0.361364 | 0.082 | 0.054 | 1 | 7 | Rad54b | 0.028 | Old |
| 0.160098 | 0.397332 | 0.138 | 0.102 | 1 | 7 | Cd40 | 0.036 | Old |
| 0.160132 | 0.364141 | 0.19 | 0.148 | 1 | 7 | Med4 | 0.042 | Old |
| 0.160363 | 0.517091 | 0.174 | 0.136 | 1 | 7 | Acads | 0.038 | Old |
| 0.160517 | 0.426308 | 0.146 | 0.111 | 1 | 7 | Recql | 0.035 | Old |
| 0.160565 | 0.374646 | 0.059 | 0.036 | 1 | 7 | Ankrd37 | 0.023 | Old |
| 0.160566 | 0.26325 | 0.367 | 0.313 | 1 | 7 | Cln8 | 0.054 | Old |

| | | | | | | | | |
|---|---|---|---|---|---|---|---|---|
| 0.160609 | 0.11243 | 0.687 | 0.605 | 1 | 7 | Psme1 | 0.082 | Old |
| 0.160632 | 0.89084 | 0.105 | 0.075 | 1 | 7 | Man2b2 | 0.03 | Old |
| 0.160738 | 1.314866 | 0.051 | 0.03 | 1 | 7 | Cryz | 0.021 | Old |
| 0.160849 | 0.355289 | 0.044 | 0.024 | 1 | 7 | Bex6 | 0.02 | Old |
| 0.16119 | 0.311943 | 0.156 | 0.117 | 1 | 7 | Cpsf1 | 0.039 | Old |
| 0.161218 | 0.265583 | 0.205 | 0.163 | 1 | 7 | Ppfibp1 | 0.042 | Old |
| 0.161302 | 0.194043 | 0.559 | 0.503 | 1 | 7 | Ago2 | 0.056 | Old |
| 0.161313 | 0.161906 | 0.541 | 0.479 | 1 | 7 | Emg1 | 0.062 | Old |
| 0.161441 | 0.37667 | 0.344 | 0.289 | 1 | 7 | Fam32a | 0.055 | Old |
| 0.161482 | 0.620165 | 0.113 | 0.081 | 1 | 7 | Hectd4 | 0.032 | Old |
| 0.161646 | 0.137076 | 0.515 | 0.455 | 1 | 7 | Itch | 0.06 | Old |
| 0.161738 | 0.810724 | 0.031 | 0.015 | 1 | 7 | Itgb3 | 0.016 | Old |
| 0.161755 | 0.24666 | 0.662 | 0.633 | 1 | 7 | Atp6ap2 | 0.029 | Old |
| 0.162135 | 0.211073 | 0.323 | 0.28 | 1 | 7 | Plpbp | 0.043 | Old |
| 0.162204 | 0.298152 | 0.1 | 0.069 | 1 | 7 | Aaas | 0.031 | Old |
| 0.162283 | 0.1033 | 0.431 | 0.358 | 1 | 7 | Noc2l | 0.073 | Old |
| 0.162422 | 0.282361 | 0.249 | 0.205 | 1 | 7 | Asnsd1 | 0.044 | Old |
| 0.162671 | 0.211398 | 0.118 | 0.084 | 1 | 7 | Nos1ap | 0.034 | Old |
| 0.162967 | 0.984926 | 0.031 | 0.015 | 1 | 7 | Krtcap3 | 0.016 | Old |
| 0.163049 | 0.10424 | 0.59 | 0.509 | 1 | 7 | Rangap1 | 0.081 | Old |
| 0.16309 | 0.827878 | 0.133 | 0.102 | 1 | 7 | N6amt1 | 0.031 | Old |
| 0.163379 | 0.751791 | 0.031 | 0.015 | 1 | 7 | Mocos | 0.016 | Old |
| 0.163446 | 0.272281 | 0.29 | 0.235 | 1 | 7 | Rnaseh2b | 0.055 | Old |
| 0.163606 | 0.058733 | 0.333 | 0.274 | 1 | 7 | Rasa1 | 0.059 | Old |
| 0.163761 | 0.051346 | 0.408 | 0.34 | 1 | 7 | Mcts1 | 0.068 | Old |
| 0.163791 | 1.751542 | 0.031 | 0.015 | 1 | 7 | Dpt | 0.016 | Old |
| 0.163962 | 0.13587 | 0.182 | 0.142 | 1 | 7 | Fam53b | 0.04 | Old |
| 0.164391 | 0.410695 | 0.185 | 0.148 | 1 | 7 | Dok1 | 0.037 | Old |
| 0.164698 | 0.160483 | 0.11 | 0.078 | 1 | 7 | Akr1b10 | 0.032 | Old |
| 0.165038 | 0.441256 | 0.113 | 0.081 | 1 | 7 | Ncdn | 0.032 | Old |
| 0.165102 | 0.05944 | 0.344 | 0.295 | 1 | 7 | 1500011K1 | 0.049 | Old |
| 0.165129 | 0.838076 | 0.146 | 0.111 | 1 | 7 | Ubl7 | 0.035 | Old |
| 0.1652 | 0.693972 | 0.11 | 0.081 | 1 | 7 | Mpv17 | 0.029 | Old |
| 0.165389 | 0.110262 | 0.164 | 0.123 | 1 | 7 | Rnf181 | 0.041 | Old |
| 0.165447 | 1.463496 | 0.031 | 0.015 | 1 | 7 | Tanc1 | 0.016 | Old |
| 0.165472 | 0.319131 | 0.19 | 0.151 | 1 | 7 | Prmt5 | 0.039 | Old |
| 0.165574 | 0.036899 | 0.367 | 0.307 | 1 | 7 | Wdfy3 | 0.06 | Old |
| 0.165658 | 0.212934 | 0.131 | 0.096 | 1 | 7 | Dbr1 | 0.035 | Old |
| 0.165807 | 0.10138 | 0.533 | 0.497 | 1 | 7 | Uqcrfs1 | 0.036 | Old |
| 0.165838 | 0.154933 | 0.328 | 0.277 | 1 | 7 | Pank2 | 0.051 | Old |
| 0.165863 | 0.805196 | 0.031 | 0.015 | 1 | 7 | Slc26a6 | 0.016 | Old |
| 0.166334 | 0.714007 | 0.21 | 0.169 | 1 | 7 | Cdca5 | 0.041 | Old |
| 0.166698 | 0.958696 | 0.031 | 0.015 | 1 | 7 | Abi3 | 0.016 | Old |
| 0.166746 | 0.161526 | 0.326 | 0.283 | 1 | 7 | Snd1 | 0.043 | Old |
| 0.167082 | 0.116396 | 0.103 | 0.072 | 1 | 7 | Decr2 | 0.031 | Old |

| | | | | | | | | |
|---|---|---|---|---|---|---|---|---|
| 0.167116 | 0.699046 | 0.031 | 0.015 | 1 | 7 | Gm15564 | 0.016 | Old |
| 0.167222 | 0.113718 | 0.708 | 0.639 | 1 | 7 | Fosl2 | 0.069 | Old |
| 0.167258 | 0.185864 | 0.379 | 0.316 | 1 | 7 | 4932438A1 | 0.063 | Old |
| 0.167452 | 0.941463 | 0.087 | 0.06 | 1 | 7 | Dhx34 | 0.027 | Old |
| 0.167471 | 0.589736 | 0.085 | 0.057 | 1 | 7 | Ddr1 | 0.028 | Old |
| 0.167536 | 0.520489 | 0.031 | 0.015 | 1 | 7 | Tut1 | 0.016 | Old |
| 0.167536 | 0.491975 | 0.031 | 0.015 | 1 | 7 | AU041133 | 0.016 | Old |
| 0.167647 | 0.080523 | 0.887 | 0.87 | 1 | 7 | Gm10076 | 0.017 | Old |
| 0.167707 | 0.093305 | 0.656 | 0.596 | 1 | 7 | Rrm1 | 0.06 | Old |
| 0.167752 | 0.235269 | 0.103 | 0.072 | 1 | 7 | Asb7 | 0.031 | Old |
| 0.167783 | 0.147564 | 0.359 | 0.307 | 1 | 7 | Senp6 | 0.052 | Old |
| 0.167956 | 0.303215 | 0.031 | 0.015 | 1 | 7 | Fastkd3 | 0.016 | Old |
| 0.167999 | 0.786976 | 0.167 | 0.133 | 1 | 7 | Cln5 | 0.034 | Old |
| 0.168003 | 0.674356 | 0.156 | 0.123 | 1 | 7 | Serpinb1a | 0.033 | Old |
| 0.168038 | 0.356533 | 0.251 | 0.208 | 1 | 7 | Wdr43 | 0.043 | Old |
| 0.168154 | 0.28436 | 0.059 | 0.036 | 1 | 7 | 1110012L1 | 0.023 | Old |
| 0.168241 | 0.501361 | 0.092 | 0.063 | 1 | 7 | Zfp938 | 0.029 | Old |
| 0.168376 | 0.494322 | 0.031 | 0.015 | 1 | 7 | Nedd4 | 0.016 | Old |
| 0.168401 | 0.108845 | 0.285 | 0.232 | 1 | 7 | Rprd2 | 0.053 | Old |
| 0.168787 | 0.186752 | 0.246 | 0.196 | 1 | 7 | Rfc4 | 0.05 | Old |
| 0.168824 | 0.064991 | 0.274 | 0.226 | 1 | 7 | Themis2 | 0.048 | Old |
| 0.168945 | 1.040808 | 0.038 | 0.021 | 1 | 7 | H2-T24 | 0.017 | Old |
| 0.168947 | 0.166534 | 0.195 | 0.154 | 1 | 7 | Sec24a | 0.041 | Old |
| 0.169775 | 0.566418 | 0.103 | 0.072 | 1 | 7 | Lzic | 0.031 | Old |
| 0.169777 | 0.433607 | 0.128 | 0.096 | 1 | 7 | Fam219a | 0.032 | Old |
| 0.169786 | 0.528133 | 0.144 | 0.108 | 1 | 7 | Slc35c1 | 0.036 | Old |
| 0.169842 | 0.268353 | 0.077 | 0.051 | 1 | 7 | Bcdin3d | 0.026 | Old |
| 0.169854 | 0.391272 | 0.244 | 0.199 | 1 | 7 | Atf6b | 0.045 | Old |
| 0.169912 | 0.085591 | 0.479 | 0.41 | 1 | 7 | Tial1 | 0.069 | Old |
| 0.17011 | 0.511547 | 0.074 | 0.048 | 1 | 7 | Odf2l | 0.026 | Old |
| 0.170153 | 0.264735 | 0.231 | 0.19 | 1 | 7 | Oard1 | 0.041 | Old |
| 0.170271 | 0.545085 | 0.062 | 0.039 | 1 | 7 | Trdmt1 | 0.023 | Old |
| 0.170272 | 0.139404 | 0.241 | 0.196 | 1 | 7 | Mrpl4 | 0.045 | Old |
| 0.170492 | 0.017007 | 0.031 | 0.015 | 1 | 7 | Dennd2d | 0.016 | Old |
| 0.170671 | 0.149551 | 0.613 | 0.542 | 1 | 7 | St8sia4 | 0.071 | Old |
| 0.170806 | 0.391702 | 0.121 | 0.09 | 1 | 7 | Xrcc4 | 0.031 | Old |
| 0.171091 | 0.75567 | 0.231 | 0.19 | 1 | 7 | Prps1 | 0.041 | Old |
| 0.171581 | 0.458997 | 0.105 | 0.075 | 1 | 7 | Mpp5 | 0.03 | Old |
| 0.171646 | 0.288768 | 0.2 | 0.16 | 1 | 7 | Nagk | 0.04 | Old |
| 0.171771 | 0.679265 | 0.031 | 0.015 | 1 | 7 | Dph7 | 0.016 | Old |
| 0.171771 | 0.277367 | 0.031 | 0.015 | 1 | 7 | Arhgef5 | 0.016 | Old |
| 0.171856 | 0.096114 | 0.572 | 0.485 | 1 | 7 | Lsm6 | 0.087 | Old |
| 0.171908 | 0.079704 | 0.926 | 0.907 | 1 | 7 | Tagln2 | 0.019 | Old |
| 0.171912 | 0.26766 | 0.567 | 0.524 | 1 | 7 | Git2 | 0.043 | Old |
| 0.172088 | 0.326508 | 0.415 | 0.355 | 1 | 7 | Kif1b | 0.06 | Old |

| | | | | | | | | |
|---|---|---|---|---|---|---|---|---|
| 0.172157 | 0.077196 | 0.444 | 0.386 | 1 | 7 | Med13l | 0.058 | Old |
| 0.172454 | 0.246515 | 0.528 | 0.464 | 1 | 7 | 2810474O1 | 0.064 | Old |
| 0.172695 | 0.198756 | 0.069 | 0.045 | 1 | 7 | Fmn1 | 0.024 | Old |
| 0.172884 | 0.143567 | 0.513 | 0.458 | 1 | 7 | Ppp1r15b | 0.055 | Old |
| 0.172951 | 0.576731 | 0.118 | 0.087 | 1 | 7 | Ppil3 | 0.031 | Old |
| 0.173205 | 0.164065 | 0.136 | 0.099 | 1 | 7 | Tbcd | 0.037 | Old |
| 0.173337 | 0.149011 | 0.277 | 0.232 | 1 | 7 | Aggf1 | 0.045 | Old |
| 0.173365 | 0.76098 | 0.054 | 0.033 | 1 | 7 | Fadd | 0.021 | Old |
| 0.173396 | 0.000986 | 0.244 | 0.199 | 1 | 7 | Slc16a3 | 0.045 | Old |
| 0.173485 | 0.733271 | 0.079 | 0.054 | 1 | 7 | Otof | 0.025 | Old |
| 0.17353 | 0.641707 | 0.069 | 0.045 | 1 | 7 | Cenpj | 0.024 | Old |
| 0.173683 | 0.352307 | 0.054 | 0.033 | 1 | 7 | Micu3 | 0.021 | Old |
| 0.173839 | 1.027409 | 0.038 | 0.021 | 1 | 7 | Gstm4 | 0.017 | Old |
| 0.173963 | 0.187926 | 0.905 | 0.861 | 1 | 7 | Npm1 | 0.044 | Old |
| 0.174047 | 0.555675 | 0.077 | 0.051 | 1 | 7 | Aldh18a1 | 0.026 | Old |
| 0.17418 | 0.332384 | 0.377 | 0.334 | 1 | 7 | Glrx | 0.043 | Old |
| 0.174181 | 0.155472 | 0.336 | 0.28 | 1 | 7 | Spg7 | 0.056 | Old |
| 0.174564 | 0.203958 | 0.462 | 0.425 | 1 | 7 | Bak1 | 0.037 | Old |
| 0.174798 | 0.26557 | 0.431 | 0.389 | 1 | 7 | Notch1 | 0.042 | Old |
| 0.175093 | 0.008747 | 0.146 | 0.108 | 1 | 7 | Hacd3 | 0.038 | Old |
| 0.175223 | 0.334227 | 0.136 | 0.099 | 1 | 7 | Slc25a32 | 0.037 | Old |
| 0.175439 | 0.459583 | 0.251 | 0.217 | 1 | 7 | Cd300lg | 0.034 | Old |
| 0.175754 | 0.117576 | 0.133 | 0.099 | 1 | 7 | Cactin | 0.034 | Old |
| 0.175772 | 0.339257 | 0.179 | 0.139 | 1 | 7 | Crls1 | 0.04 | Old |
| 0.175872 | 0.086953 | 0.472 | 0.404 | 1 | 7 | Uqcc2 | 0.068 | Old |
| 0.175883 | 0.074963 | 0.313 | 0.259 | 1 | 7 | Gba | 0.054 | Old |
| 0.175885 | 0.8548 | 0.105 | 0.075 | 1 | 7 | Cacfd1 | 0.03 | Old |
| 0.175886 | 0.092136 | 0.226 | 0.178 | 1 | 7 | Zbtb18 | 0.048 | Old |
| 0.176101 | 0.205749 | 0.574 | 0.518 | 1 | 7 | Tomm22 | 0.056 | Old |
| 0.176135 | 0.123619 | 0.151 | 0.114 | 1 | 7 | Mapk8ip3 | 0.037 | Old |
| 0.176177 | 0.172313 | 0.51 | 0.443 | 1 | 7 | Hpcal1 | 0.067 | Old |
| 0.176603 | 0.734292 | 0.118 | 0.087 | 1 | 7 | Dnajc30 | 0.031 | Old |
| 0.176618 | 0.513068 | 0.069 | 0.045 | 1 | 7 | Pomgnt1 | 0.024 | Old |
| 0.176667 | 0.286324 | 0.482 | 0.425 | 1 | 7 | Gmnn | 0.057 | Old |
| 0.176801 | 0.125477 | 0.295 | 0.25 | 1 | 7 | Arhgap26 | 0.045 | Old |
| 0.176904 | 0.708749 | 0.038 | 0.021 | 1 | 7 | Gm16973 | 0.017 | Old |
| 0.176918 | 0.203992 | 0.444 | 0.383 | 1 | 7 | Ap3s1 | 0.061 | Old |
| 0.177117 | 0.012988 | 0.103 | 0.072 | 1 | 7 | Snx16 | 0.031 | Old |
| 0.17726 | 1.202517 | 0.072 | 0.048 | 1 | 7 | Lrp5 | 0.024 | Old |
| 0.177303 | 0.05781 | 0.251 | 0.205 | 1 | 7 | Prpf31 | 0.046 | Old |
| 0.17731 | 0.291276 | 0.374 | 0.319 | 1 | 7 | Kdelr2 | 0.055 | Old |
| 0.177378 | 0.20739 | 0.5 | 0.452 | 1 | 7 | Aebp2 | 0.048 | Old |
| 0.177498 | 0.036773 | 0.123 | 0.09 | 1 | 7 | Acaca | 0.033 | Old |
| 0.177524 | 0.087743 | 0.054 | 0.033 | 1 | 7 | Map3k15 | 0.021 | Old |
| 0.177585 | 0.229493 | 0.272 | 0.22 | 1 | 7 | Bcap29 | 0.052 | Old |

| | | | | | | | | |
|---|---|---|---|---|---|---|---|---|
| 0.177643 | 0.253375 | 0.323 | 0.265 | 1 | 7 | Slc35c2 | 0.058 | Old |
| 0.177686 | 0.758338 | 0.046 | 0.027 | 1 | 7 | Isoc2b | 0.019 | Old |
| 0.177688 | 0.756017 | 0.062 | 0.039 | 1 | 7 | Sirpb1b | 0.023 | Old |
| 0.177815 | 0.969071 | 0.072 | 0.048 | 1 | 7 | Mpp7 | 0.024 | Old |
| 0.177887 | 0.302745 | 0.572 | 0.518 | 1 | 7 | Cks1b | 0.054 | Old |
| 0.177995 | 0.092642 | 0.292 | 0.244 | 1 | 7 | Mrps36 | 0.048 | Old |
| 0.178042 | 1.338434 | 0.026 | 0.012 | 1 | 7 | Kyat1 | 0.014 | Old |
| 0.178077 | 0.125017 | 0.864 | 0.816 | 1 | 7 | Anp32b | 0.048 | Old |
| 0.178081 | 0.55891 | 0.2 | 0.16 | 1 | 7 | Agap3 | 0.04 | Old |
| 0.17828 | 0.282617 | 0.385 | 0.334 | 1 | 7 | Incenp | 0.051 | Old |
| 0.178495 | 0.541577 | 0.054 | 0.033 | 1 | 7 | Alkbh2 | 0.021 | Old |
| 0.178495 | 0.443662 | 0.054 | 0.033 | 1 | 7 | Mettl8 | 0.021 | Old |
| 0.178839 | 0.471845 | 0.038 | 0.021 | 1 | 7 | Nupl2 | 0.017 | Old |
| 0.178866 | 0.022193 | 0.241 | 0.19 | 1 | 7 | Glb1 | 0.051 | Old |
| 0.178979 | 0.236725 | 0.251 | 0.202 | 1 | 7 | Dnajc14 | 0.049 | Old |
| 0.179019 | 0.304257 | 0.203 | 0.163 | 1 | 7 | Nol8 | 0.04 | Old |
| 0.179176 | 0.215058 | 0.415 | 0.355 | 1 | 7 | Srrt | 0.06 | Old |
| 0.179238 | 0.692112 | 0.103 | 0.075 | 1 | 7 | Polr3f | 0.028 | Old |
| 0.179378 | 0.497989 | 0.249 | 0.211 | 1 | 7 | Gcn1l1 | 0.038 | Old |
| 0.179469 | 0.433203 | 0.054 | 0.033 | 1 | 7 | Trub1 | 0.021 | Old |
| 0.179762 | 0.434687 | 0.136 | 0.102 | 1 | 7 | Fhl3 | 0.034 | Old |
| 0.180063 | 0.421767 | 0.079 | 0.054 | 1 | 7 | Vcpkmt | 0.025 | Old |
| 0.180151 | 0.6523 | 0.046 | 0.027 | 1 | 7 | Tnfrsf13c | 0.019 | Old |
| 0.180662 | 0.200418 | 0.308 | 0.259 | 1 | 7 | Ebna1bp2 | 0.049 | Old |
| 0.180722 | 0.49298 | 0.062 | 0.039 | 1 | 7 | Smco4 | 0.023 | Old |
| 0.180927 | 0.214974 | 0.526 | 0.47 | 1 | 7 | Uba2 | 0.056 | Old |
| 0.180973 | 0.084867 | 0.226 | 0.181 | 1 | 7 | Kdm6a | 0.045 | Old |
| 0.181014 | 0.657966 | 0.082 | 0.057 | 1 | 7 | Hars2 | 0.025 | Old |
| 0.181094 | 0.865665 | 0.138 | 0.108 | 1 | 7 | Mafk | 0.03 | Old |
| 0.181191 | 0.238355 | 0.405 | 0.352 | 1 | 7 | Polr2f | 0.053 | Old |
| 0.181575 | 0.208137 | 0.203 | 0.16 | 1 | 7 | Lpcat3 | 0.043 | Old |
| 0.181637 | 0.125128 | 0.097 | 0.069 | 1 | 7 | Tbccd1 | 0.028 | Old |
| 0.181871 | 0.685197 | 0.092 | 0.066 | 1 | 7 | 1700021F0 | 0.026 | Old |
| 0.181923 | 0.182994 | 0.203 | 0.16 | 1 | 7 | Exosc10 | 0.043 | Old |
| 0.18195 | 0.239889 | 0.174 | 0.136 | 1 | 7 | Scaf1 | 0.038 | Old |
| 0.182251 | 0.048375 | 0.149 | 0.111 | 1 | 7 | Rev1 | 0.038 | Old |
| 0.182284 | 0.334201 | 0.046 | 0.027 | 1 | 7 | Recql4 | 0.019 | Old |
| 0.182729 | 0.366632 | 0.497 | 0.476 | 1 | 7 | Ube2c | 0.021 | Old |
| 0.182745 | 0.376876 | 0.079 | 0.054 | 1 | 7 | B230307C2 | 0.025 | Old |
| 0.183062 | 0.021259 | 0.223 | 0.178 | 1 | 7 | Prkab2 | 0.045 | Old |
| 0.183118 | 0.207328 | 0.344 | 0.304 | 1 | 7 | Ufm1 | 0.04 | Old |
| 0.183152 | 0.439299 | 0.038 | 0.021 | 1 | 7 | Ttll1 | 0.017 | Old |
| 0.18326 | 0.047947 | 0.349 | 0.298 | 1 | 7 | Ing1 | 0.051 | Old |
| 0.183295 | 0.126832 | 0.426 | 0.37 | 1 | 7 | Mib1 | 0.056 | Old |
| 0.183358 | 0.598994 | 0.046 | 0.027 | 1 | 7 | Adck5 | 0.019 | Old |

| | | | | | | | | |
|---|---|---|---|---|---|---|---|---|
| 0.183556 | 0.82839 | 0.079 | 0.054 | 1 | 7 | Suv39h2 | 0.025 | Old |
| 0.18366 | 0.350493 | 0.292 | 0.25 | 1 | 7 | Ntan1 | 0.042 | Old |
| 0.183669 | 0.31133 | 0.585 | 0.545 | 1 | 7 | Add3 | 0.04 | Old |
| 0.183723 | 0.452352 | 0.072 | 0.048 | 1 | 7 | Tenm4 | 0.024 | Old |
| 0.183727 | 0.802057 | 0.064 | 0.042 | 1 | 7 | Ube2e2 | 0.022 | Old |
| 0.183802 | 0.134117 | 0.815 | 0.819 | 1 | 7 | Set | -0.004 | Old |
| 0.184 | 0.628542 | 0.09 | 0.063 | 1 | 7 | Zcchc10 | 0.027 | Old |
| 0.184336 | 0.361181 | 0.482 | 0.44 | 1 | 7 | Snx10 | 0.042 | Old |
| 0.184412 | 0.26896 | 0.274 | 0.232 | 1 | 7 | Micu1 | 0.042 | Old |
| 0.184657 | 0.184745 | 0.649 | 0.578 | 1 | 7 | Atp5c1 | 0.071 | Old |
| 0.184797 | 0.453692 | 0.046 | 0.027 | 1 | 7 | Tarm1 | 0.019 | Old |
| 0.18527 | 0.626526 | 0.087 | 0.06 | 1 | 7 | Cox18 | 0.027 | Old |
| 0.185701 | 0.257583 | 0.364 | 0.319 | 1 | 7 | Map3k11 | 0.045 | Old |
| 0.185724 | 0.544879 | 0.072 | 0.048 | 1 | 7 | Lyrm4 | 0.024 | Old |
| 0.186082 | 0.435052 | 0.251 | 0.211 | 1 | 7 | Mpdu1 | 0.04 | Old |
| 0.186392 | 0.547284 | 0.026 | 0.012 | 1 | 7 | Tyw3 | 0.014 | Old |
| 0.186481 | 0.244136 | 0.379 | 0.325 | 1 | 7 | Samm50 | 0.054 | Old |
| 0.186534 | 0.264939 | 0.262 | 0.22 | 1 | 7 | 1700123O2 | 0.042 | Old |
| 0.18657 | 0.256381 | 0.21 | 0.172 | 1 | 7 | Rinl | 0.038 | Old |
| 0.18668 | 0.109903 | 0.351 | 0.295 | 1 | 7 | Dctn4 | 0.056 | Old |
| 0.186696 | 0.031837 | 0.113 | 0.084 | 1 | 7 | Asb3 | 0.029 | Old |
| 0.186838 | 0.093454 | 0.087 | 0.06 | 1 | 7 | Grwd1 | 0.027 | Old |
| 0.186899 | 0.125567 | 0.444 | 0.392 | 1 | 7 | Cpsf2 | 0.052 | Old |
| 0.186972 | 0.14502 | 0.033 | 0.054 | 1 | 7 | Xlr | -0.021 | Old |
| 0.187405 | 0.415855 | 0.054 | 0.033 | 1 | 7 | Zfp74 | 0.021 | Old |
| 0.187699 | 0.528672 | 0.046 | 0.027 | 1 | 7 | Thrb | 0.019 | Old |
| 0.187862 | 0.134139 | 0.149 | 0.114 | 1 | 7 | Nudt14 | 0.035 | Old |
| 0.187921 | 0.278124 | 0.313 | 0.256 | 1 | 7 | Mrpl17 | 0.057 | Old |
| 0.188224 | 0.058965 | 0.331 | 0.271 | 1 | 7 | Arl6ip5 | 0.06 | Old |
| 0.188756 | 0.593914 | 0.072 | 0.048 | 1 | 7 | Zfp617 | 0.024 | Old |
| 0.188901 | 0.312272 | 0.072 | 0.048 | 1 | 7 | Rhbdd2 | 0.024 | Old |
| 0.188903 | 0.699132 | 0.026 | 0.012 | 1 | 7 | Senp8 | 0.014 | Old |
| 0.18897 | 0.255355 | 0.187 | 0.148 | 1 | 7 | Sipa1l3 | 0.039 | Old |
| 0.189014 | 0.18615 | 0.236 | 0.19 | 1 | 7 | P4ha1 | 0.046 | Old |
| 0.189149 | 0.831469 | 0.09 | 0.063 | 1 | 7 | Zfyve1 | 0.027 | Old |
| 0.189597 | 0.423801 | 0.21 | 0.178 | 1 | 7 | Jdp2 | 0.032 | Old |
| 0.189615 | 0.13271 | 0.108 | 0.078 | 1 | 7 | Ptcd2 | 0.03 | Old |
| 0.189703 | 0.781692 | 0.115 | 0.087 | 1 | 7 | Rfx1 | 0.028 | Old |
| 0.189772 | 0.191526 | 0.2 | 0.16 | 1 | 7 | Dhx16 | 0.04 | Old |
| 0.190367 | 0.188432 | 0.572 | 0.509 | 1 | 7 | Pgls | 0.063 | Old |
| 0.190369 | 0.129485 | 0.062 | 0.039 | 1 | 7 | Ccdc91 | 0.023 | Old |
| 0.190421 | 0.415957 | 0.026 | 0.012 | 1 | 7 | D11Wsu47 | 0.014 | Old |
| 0.190591 | 0.136108 | 0.446 | 0.389 | 1 | 7 | Ddx21 | 0.057 | Old |
| 0.1906 | 0.494656 | 0.121 | 0.09 | 1 | 7 | Gemin6 | 0.031 | Old |
| 0.19084 | 0.054577 | 0.454 | 0.383 | 1 | 7 | Dock7 | 0.071 | Old |

| | | | | | | | | |
|---|---|---|---|---|---|---|---|---|
| 0.191438 | 0.519794 | 0.026 | 0.012 | 1 | 7 | Msrb2 | 0.014 | Old |
| 0.191438 | 0.096147 | 0.026 | 0.012 | 1 | 7 | Pigz | 0.014 | Old |
| 0.191468 | 0.164008 | 0.277 | 0.229 | 1 | 7 | Ctbp2 | 0.048 | Old |
| 0.191528 | 0.348062 | 0.079 | 0.054 | 1 | 7 | Snx19 | 0.025 | Old |
| 0.19208 | 0.100398 | 0.497 | 0.428 | 1 | 7 | Tmod3 | 0.069 | Old |
| 0.192104 | 0.492778 | 0.133 | 0.102 | 1 | 7 | Uck2 | 0.031 | Old |
| 0.192459 | 0.521905 | 0.026 | 0.012 | 1 | 7 | Zfp820 | 0.014 | Old |
| 0.192649 | 0.308068 | 0.272 | 0.226 | 1 | 7 | Pdlim4 | 0.046 | Old |
| 0.192793 | 0.209423 | 0.39 | 0.328 | 1 | 7 | Naa15 | 0.062 | Old |
| 0.192962 | 0.122105 | 0.49 | 0.434 | 1 | 7 | Eif2s3x | 0.056 | Old |
| 0.192967 | 0.169695 | 0.408 | 0.34 | 1 | 7 | Plp2 | 0.068 | Old |
| 0.193091 | 0.337606 | 0.064 | 0.042 | 1 | 7 | Tmem241 | 0.022 | Old |
| 0.193243 | 0.786021 | 0.067 | 0.045 | 1 | 7 | Stk11ip | 0.022 | Old |
| 0.193484 | 0.26891 | 0.026 | 0.012 | 1 | 7 | Tmem41a | 0.014 | Old |
| 0.193704 | 0.119325 | 0.631 | 0.596 | 1 | 7 | Pebp1 | 0.035 | Old |
| 0.193994 | 0.151989 | 0.474 | 0.416 | 1 | 7 | Rab8a | 0.058 | Old |
| 0.194056 | 0.200453 | 0.549 | 0.494 | 1 | 7 | Ezr | 0.055 | Old |
| 0.194512 | 0.520712 | 0.074 | 0.051 | 1 | 7 | Aqp9 | 0.023 | Old |
| 0.194669 | 0.516134 | 0.31 | 0.262 | 1 | 7 | Wbp1l | 0.048 | Old |
| 0.194709 | 0.124856 | 0.315 | 0.259 | 1 | 7 | Ncapg | 0.056 | Old |
| 0.194786 | 0.265489 | 0.231 | 0.19 | 1 | 7 | Usf1 | 0.041 | Old |
| 0.195113 | 0.248614 | 0.456 | 0.395 | 1 | 7 | Ube2i | 0.061 | Old |
| 0.195142 | 0.053266 | 0.092 | 0.066 | 1 | 7 | Gm20186 | 0.026 | Old |
| 0.195171 | 0.199001 | 0.062 | 0.039 | 1 | 7 | Rdh10 | 0.023 | Old |
| 0.195376 | 0.271425 | 0.072 | 0.048 | 1 | 7 | Gm7694 | 0.024 | Old |
| 0.195431 | 0.178335 | 0.431 | 0.367 | 1 | 7 | Riok3 | 0.064 | Old |
| 0.195572 | 0.068236 | 0.795 | 0.753 | 1 | 7 | Reep5 | 0.042 | Old |
| 0.195899 | 0.223507 | 0.69 | 0.657 | 1 | 7 | Eps8 | 0.033 | Old |
| 0.195967 | 1.48641 | 0.033 | 0.018 | 1 | 7 | Nthl1 | 0.015 | Old |
| 0.195974 | 0.296352 | 0.141 | 0.108 | 1 | 7 | Fbxw8 | 0.033 | Old |
| 0.196045 | 0.127625 | 0.405 | 0.361 | 1 | 7 | Hspa14 | 0.044 | Old |
| 0.196197 | 0.130685 | 0.49 | 0.44 | 1 | 7 | Lfng | 0.05 | Old |
| 0.196263 | 0.567954 | 0.074 | 0.051 | 1 | 7 | Adora2a | 0.023 | Old |
| 0.196344 | 0.440318 | 0.159 | 0.12 | 1 | 7 | Tstd3 | 0.039 | Old |
| 0.196414 | 1.084873 | 0.033 | 0.018 | 1 | 7 | Zfp788 | 0.015 | Old |
| 0.196558 | 0.040391 | 0.421 | 0.352 | 1 | 7 | Klc1 | 0.069 | Old |
| 0.196768 | 0.362011 | 0.249 | 0.211 | 1 | 7 | Taz | 0.038 | Old |
| 0.197022 | 0.163265 | 0.272 | 0.226 | 1 | 7 | Sf3a2 | 0.046 | Old |
| 0.197066 | 0.1158 | 0.549 | 0.494 | 1 | 7 | Rad23b | 0.055 | Old |
| 0.197291 | 0.509022 | 0.308 | 0.271 | 1 | 7 | Ltb4r1 | 0.037 | Old |
| 0.197301 | 0.279247 | 0.213 | 0.172 | 1 | 7 | Gon4l | 0.041 | Old |
| 0.197315 | 0.263857 | 0.377 | 0.328 | 1 | 7 | Gtpbp4 | 0.049 | Old |
| 0.197458 | 0.002686 | 0.079 | 0.054 | 1 | 7 | Coa4 | 0.025 | Old |
| 0.197463 | 0.15432 | 0.692 | 0.648 | 1 | 7 | Cct5 | 0.044 | Old |
| 0.197593 | 0.050801 | 0.418 | 0.352 | 1 | 7 | Erbin | 0.066 | Old |

| | | | | | | | | |
|---|---|---|---|---|---|---|---|---|
| 0.197706 | 0.088909 | 0.315 | 0.265 | 1 | 7 | Ccdc71l | 0.05 | Old |
| 0.197939 | 0.13944 | 0.579 | 0.515 | 1 | 7 | Zmiz1 | 0.064 | Old |
| 0.198209 | 0.737254 | 0.033 | 0.018 | 1 | 7 | Ift43 | 0.015 | Old |
| 0.198255 | 0.33795 | 0.285 | 0.241 | 1 | 7 | Kidins220 | 0.044 | Old |
| 0.198355 | 0.28327 | 0.308 | 0.262 | 1 | 7 | Sh2b3 | 0.046 | Old |
| 0.198431 | 0.095273 | 0.872 | 0.867 | 1 | 7 | Ccl6 | 0.005 | Old |
| 0.198436 | 0.273974 | 0.554 | 0.527 | 1 | 7 | Ppp1r14b | 0.027 | Old |
| 0.198443 | 0.081228 | 0.249 | 0.202 | 1 | 7 | Sirt7 | 0.047 | Old |
| 0.198483 | 0.155078 | 0.836 | 0.834 | 1 | 7 | Pitpna | 0.002 | Old |
| 0.198647 | 0.471009 | 0.121 | 0.09 | 1 | 7 | Prkci | 0.031 | Old |
| 0.198724 | 0.188442 | 0.641 | 0.608 | 1 | 7 | Top2a | 0.033 | Old |
| 0.19888 | 0.082596 | 0.356 | 0.307 | 1 | 7 | Polr2g | 0.049 | Old |
| 0.198947 | 0.182861 | 0.277 | 0.232 | 1 | 7 | Stard4 | 0.045 | Old |
| 0.199022 | 0.153459 | 0.169 | 0.133 | 1 | 7 | Ccne2 | 0.036 | Old |
| 0.199114 | 0.428845 | 0.121 | 0.09 | 1 | 7 | Agl | 0.031 | Old |
| 0.199579 | 0.526296 | 0.064 | 0.042 | 1 | 7 | Magi3 | 0.022 | Old |
| 0.199691 | 0.948196 | 0.049 | 0.03 | 1 | 7 | Fastkd1 | 0.019 | Old |
| 0.199773 | 0.092306 | 0.056 | 0.036 | 1 | 7 | Mtbp | 0.02 | Old |
| 0.19986 | 0.099778 | 0.908 | 0.901 | 1 | 7 | Capza2 | 0.007 | Old |
| 0.199927 | 0.470399 | 0.333 | 0.295 | 1 | 7 | Cyp51 | 0.038 | Old |
| 0.199932 | 0.069909 | 0.19 | 0.151 | 1 | 7 | Ccdc9 | 0.039 | Old |
| 0.200069 | 0.23772 | 0.421 | 0.383 | 1 | 7 | Ninj1 | 0.038 | Old |
| 0.200227 | 0.225033 | 0.182 | 0.145 | 1 | 7 | Rpia | 0.037 | Old |
| 0.20073 | 0.151186 | 0.515 | 0.443 | 1 | 7 | Mdfic | 0.072 | Old |
| 0.200733 | 0.183975 | 0.333 | 0.277 | 1 | 7 | Strn4 | 0.056 | Old |
| 0.200793 | 0.13997 | 0.072 | 0.048 | 1 | 7 | Tti1 | 0.024 | Old |
| 0.200924 | 0.633988 | 0.033 | 0.018 | 1 | 7 | Lins1 | 0.015 | Old |
| 0.200924 | 0.543713 | 0.033 | 0.018 | 1 | 7 | Rangrf | 0.015 | Old |
| 0.20099 | 0.211287 | 0.544 | 0.482 | 1 | 7 | Sh3kbp1 | 0.062 | Old |
| 0.201077 | 0.326743 | 0.274 | 0.235 | 1 | 7 | Fyn | 0.039 | Old |
| 0.201105 | 0.764354 | 0.113 | 0.084 | 1 | 7 | Tpm3-rs7 | 0.029 | Old |
| 0.20113 | 0.426086 | 0.105 | 0.075 | 1 | 7 | Dsn1 | 0.03 | Old |
| 0.20125 | 0.534697 | 0.192 | 0.154 | 1 | 7 | Wdr3 | 0.038 | Old |
| 0.201584 | 0.162883 | 0.549 | 0.506 | 1 | 7 | Eif3h | 0.043 | Old |
| 0.201607 | 0.21687 | 0.228 | 0.19 | 1 | 7 | Ddx56 | 0.038 | Old |
| 0.20162 | 0.973345 | 0.087 | 0.063 | 1 | 7 | F10 | 0.024 | Old |
| 0.201693 | 0.327293 | 0.151 | 0.117 | 1 | 7 | Hif1an | 0.034 | Old |
| 0.201832 | 0.028489 | 0.097 | 0.069 | 1 | 7 | Eef2k | 0.028 | Old |
| 0.201902 | 0.054831 | 0.564 | 0.509 | 1 | 7 | Surf4 | 0.055 | Old |
| 0.20193 | 0.015821 | 0.123 | 0.09 | 1 | 7 | Uck1 | 0.033 | Old |
| 0.202253 | 0.969061 | 0.067 | 0.045 | 1 | 7 | Nlrx1 | 0.022 | Old |
| 0.202486 | 0.04545 | 0.426 | 0.364 | 1 | 7 | Kmt5a | 0.062 | Old |
| 0.202565 | 0.349983 | 0.113 | 0.084 | 1 | 7 | Sec24d | 0.029 | Old |
| 0.20262 | 0.319355 | 0.097 | 0.069 | 1 | 7 | Pskh1 | 0.028 | Old |
| 0.202749 | 0.693811 | 0.033 | 0.018 | 1 | 7 | Mocs3 | 0.015 | Old |

| | | | | | | | | |
|---|---|---|---|---|---|---|---|---|
| 0.202934 | 0.222008 | 0.141 | 0.108 | 1 | 7 | A530064D( | 0.033 | Old |
| 0.202941 | 0.062595 | 0.205 | 0.163 | 1 | 7 | Ubr7 | 0.042 | Old |
| 0.203049 | 0.205269 | 0.559 | 0.5 | 1 | 7 | Etfb | 0.059 | Old |
| 0.203126 | 0.151739 | 0.287 | 0.238 | 1 | 7 | Lrpap1 | 0.049 | Old |
| 0.203207 | 0.973985 | 0.033 | 0.018 | 1 | 7 | Itga2 | 0.015 | Old |
| 0.203219 | 0.623988 | 0.103 | 0.075 | 1 | 7 | Abt1 | 0.028 | Old |
| 0.203535 | 1.235446 | 0.021 | 0.009 | 1 | 7 | Ddr2 | 0.012 | Old |
| 0.203542 | 0.081641 | 0.072 | 0.048 | 1 | 7 | Rtn4ip1 | 0.024 | Old |
| 0.203586 | 0.09303 | 0.697 | 0.605 | 1 | 7 | Vmp1 | 0.092 | Old |
| 0.203715 | 1.173233 | 0.041 | 0.024 | 1 | 7 | Tctn3 | 0.017 | Old |
| 0.204125 | 1.534108 | 0.033 | 0.018 | 1 | 7 | Fbxo46 | 0.015 | Old |
| 0.204231 | 0.165529 | 0.397 | 0.334 | 1 | 7 | Phf14 | 0.063 | Old |
| 0.204259 | 0.309842 | 0.351 | 0.304 | 1 | 7 | Vat1 | 0.047 | Old |
| 0.204415 | 0.29264 | 0.282 | 0.241 | 1 | 7 | C1galt1 | 0.041 | Old |
| 0.204569 | 0.360954 | 0.146 | 0.114 | 1 | 7 | Pdzd11 | 0.032 | Old |
| 0.204585 | 0.204554 | 0.033 | 0.018 | 1 | 7 | Ankrd24 | 0.015 | Old |
| 0.204684 | 0.010248 | 0.082 | 0.057 | 1 | 7 | Tmem18 | 0.025 | Old |
| 0.204737 | 0.253511 | 0.092 | 0.066 | 1 | 7 | Rcc1l | 0.026 | Old |
| 0.205137 | 0.360336 | 0.182 | 0.145 | 1 | 7 | BC003965 | 0.037 | Old |
| 0.205341 | 1.540702 | 0.021 | 0.009 | 1 | 7 | Ifit1 | 0.012 | Old |
| 0.205341 | 1.485892 | 0.021 | 0.009 | 1 | 7 | Tmem216 | 0.012 | Old |
| 0.205356 | 0.594827 | 0.077 | 0.054 | 1 | 7 | Chpf | 0.023 | Old |
| 0.205393 | 0.421808 | 0.128 | 0.096 | 1 | 7 | Wdyhv1 | 0.032 | Old |
| 0.205509 | 0.421194 | 0.033 | 0.018 | 1 | 7 | Mmachc | 0.015 | Old |
| 0.205544 | 0.208689 | 0.395 | 0.34 | 1 | 7 | Prdx4 | 0.055 | Old |
| 0.205662 | 0.421408 | 0.262 | 0.226 | 1 | 7 | Mrps6 | 0.036 | Old |
| 0.205758 | 0.093533 | 0.444 | 0.377 | 1 | 7 | Vamp3 | 0.067 | Old |
| 0.205773 | 0.564801 | 0.041 | 0.024 | 1 | 7 | Rbfox2 | 0.017 | Old |
| 0.205865 | 0.569927 | 0.138 | 0.108 | 1 | 7 | Tsta3 | 0.03 | Old |
| 0.205884 | 0.169619 | 0.623 | 0.566 | 1 | 7 | Sgms1 | 0.057 | Old |
| 0.20594 | 0.522999 | 0.179 | 0.142 | 1 | 7 | Xrcc1 | 0.037 | Old |
| 0.206003 | 0.142218 | 0.397 | 0.352 | 1 | 7 | Ddx46 | 0.045 | Old |
| 0.206032 | 0.363396 | 0.246 | 0.208 | 1 | 7 | Cnp | 0.038 | Old |
| 0.206036 | 0.988947 | 0.049 | 0.03 | 1 | 7 | Rida | 0.019 | Old |
| 0.206289 | 0.017901 | 0.292 | 0.241 | 1 | 7 | C2cd5 | 0.051 | Old |
| 0.20635 | 0.480262 | 0.056 | 0.036 | 1 | 7 | Klhl22 | 0.02 | Old |
| 0.206403 | 0.022056 | 0.074 | 0.051 | 1 | 7 | 2700097O( | 0.023 | Old |
| 0.206477 | 0.263059 | 0.377 | 0.325 | 1 | 7 | Krit1 | 0.052 | Old |
| 0.20654 | 0.644182 | 0.144 | 0.111 | 1 | 7 | Med25 | 0.033 | Old |
| 0.206645 | 0.139728 | 0.608 | 0.545 | 1 | 7 | Rps6ka3 | 0.063 | Old |
| 0.207158 | 1.816281 | 0.021 | 0.009 | 1 | 7 | Zfp605 | 0.012 | Old |
| 0.207254 | 0.102698 | 0.8 | 0.768 | 1 | 7 | Aplp2 | 0.032 | Old |
| 0.207479 | 0.065697 | 0.138 | 0.105 | 1 | 7 | Ap4m1 | 0.033 | Old |
| 0.207536 | 0.2105 | 0.728 | 0.672 | 1 | 7 | Ppp2r5c | 0.056 | Old |
| 0.207766 | 1.177339 | 0.021 | 0.009 | 1 | 7 | Glis2 | 0.012 | Old |

| | | | | | | | | |
|---|---|---|---|---|---|---|---|---|
| 0.207769 | 0.064947 | 0.551 | 0.488 | 1 | 7 | Bub3 | 0.063 | Old |
| 0.208123 | 1.020842 | 0.11 | 0.084 | 1 | 7 | Ngrn | 0.026 | Old |
| 0.208296 | 0.110532 | 0.033 | 0.018 | 1 | 7 | Ccdc137 | 0.015 | Old |
| 0.20835 | 0.102298 | 0.346 | 0.295 | 1 | 7 | Alkbh5 | 0.051 | Old |
| 0.208376 | 1.28675 | 0.021 | 0.009 | 1 | 7 | Isg20 | 0.012 | Old |
| 0.208691 | 0.482722 | 0.049 | 0.03 | 1 | 7 | Ogg1 | 0.019 | Old |
| 0.208748 | 0.43751 | 0.141 | 0.111 | 1 | 7 | Tusc1 | 0.03 | Old |
| 0.20901 | 0.391542 | 0.238 | 0.199 | 1 | 7 | Idnk | 0.039 | Old |
| 0.209443 | 0.085088 | 0.569 | 0.512 | 1 | 7 | Epb41l2 | 0.057 | Old |
| 0.209454 | 0.786702 | 0.049 | 0.03 | 1 | 7 | Ppp1r9a | 0.019 | Old |
| 0.209496 | 0.086271 | 0.254 | 0.205 | 1 | 7 | Polr3c | 0.049 | Old |
| 0.209599 | 0.908811 | 0.021 | 0.009 | 1 | 7 | Zswim3 | 0.012 | Old |
| 0.209599 | 0.665443 | 0.021 | 0.009 | 1 | 7 | Aldh1l1 | 0.012 | Old |
| 0.209759 | 0.31772 | 0.174 | 0.142 | 1 | 7 | Gtf3c1 | 0.032 | Old |
| 0.209895 | 0.160707 | 0.308 | 0.268 | 1 | 7 | Lin54 | 0.04 | Old |
| 0.209915 | 0.281685 | 0.313 | 0.268 | 1 | 7 | Adam9 | 0.045 | Old |
| 0.21003 | 0.257557 | 0.123 | 0.09 | 1 | 7 | Mtmr9 | 0.033 | Old |
| 0.210142 | 0.247804 | 0.077 | 0.054 | 1 | 7 | Fam122a | 0.023 | Old |
| 0.210212 | 0.762011 | 0.021 | 0.009 | 1 | 7 | Nadsyn1 | 0.012 | Old |
| 0.210212 | 0.726385 | 0.021 | 0.009 | 1 | 7 | Gm11131 | 0.012 | Old |
| 0.210291 | 0.136943 | 0.618 | 0.548 | 1 | 7 | 2900097C1 | 0.07 | Old |
| 0.210362 | 0.041341 | 0.697 | 0.627 | 1 | 7 | Rtn3 | 0.07 | Old |
| 0.210709 | 0.062492 | 0.567 | 0.497 | 1 | 7 | Mia2 | 0.07 | Old |
| 0.210774 | 0.218601 | 0.041 | 0.024 | 1 | 7 | Zfp579 | 0.017 | Old |
| 0.210774 | 0.169492 | 0.041 | 0.024 | 1 | 7 | Pus3 | 0.017 | Old |
| 0.210827 | 1.015844 | 0.021 | 0.009 | 1 | 7 | Rab34 | 0.012 | Old |
| 0.210827 | 0.630743 | 0.021 | 0.009 | 1 | 7 | Xrcc3 | 0.012 | Old |
| 0.210897 | 0.235576 | 0.367 | 0.316 | 1 | 7 | Wdfy2 | 0.051 | Old |
| 0.210992 | 0.100514 | 0.372 | 0.322 | 1 | 7 | Dpy30 | 0.05 | Old |
| 0.211045 | 0.114669 | 0.613 | 0.56 | 1 | 7 | Cdv3 | 0.053 | Old |
| 0.211051 | 0.10366 | 0.262 | 0.217 | 1 | 7 | Ipo9 | 0.045 | Old |
| 0.21116 | 0.415888 | 0.123 | 0.093 | 1 | 7 | Champ1 | 0.03 | Old |
| 0.211472 | 0.322746 | 0.262 | 0.214 | 1 | 7 | Opa1 | 0.048 | Old |
| 0.211964 | 0.324927 | 0.244 | 0.208 | 1 | 7 | Nub1 | 0.036 | Old |
| 0.212061 | 1.439911 | 0.021 | 0.009 | 1 | 7 | 9130011E1 | 0.012 | Old |
| 0.212061 | 0.797502 | 0.021 | 0.009 | 1 | 7 | D130017N( | 0.012 | Old |
| 0.212061 | 0.629441 | 0.021 | 0.009 | 1 | 7 | Tmem14a | 0.012 | Old |
| 0.212061 | 0.371263 | 0.021 | 0.009 | 1 | 7 | Armcx1 | 0.012 | Old |
| 0.21214 | 0.399445 | 0.049 | 0.03 | 1 | 7 | Elp4 | 0.019 | Old |
| 0.21222 | 0.732859 | 0.067 | 0.045 | 1 | 7 | Aven | 0.022 | Old |
| 0.212679 | 0.396555 | 0.021 | 0.009 | 1 | 7 | A530032D1 | 0.012 | Old |
| 0.212832 | 0.718754 | 0.087 | 0.063 | 1 | 7 | Gemin2 | 0.024 | Old |
| 0.212906 | 0.241866 | 0.382 | 0.343 | 1 | 7 | Pilra | 0.039 | Old |
| 0.21292 | 0.295145 | 0.131 | 0.099 | 1 | 7 | Kdm5d | 0.032 | Old |
| 0.212933 | 0.127863 | 0.128 | 0.096 | 1 | 7 | Spice1 | 0.032 | Old |

| | | | | | | | | |
|---|---|---|---|---|---|---|---|---|
| 0.213008 | 0.115461 | 0.315 | 0.268 | 1 | 7 | Chmp1a | 0.047 | Old |
| 0.213288 | 0.535782 | 0.279 | 0.244 | 1 | 7 | Bmt2 | 0.035 | Old |
| 0.213299 | 1.433997 | 0.021 | 0.009 | 1 | 7 | Ispd | 0.012 | Old |
| 0.213299 | 0.783841 | 0.021 | 0.009 | 1 | 7 | Gm26756 | 0.012 | Old |
| 0.213417 | 0.063104 | 0.415 | 0.352 | 1 | 7 | Rab35 | 0.063 | Old |
| 0.213479 | 0.206567 | 0.077 | 0.054 | 1 | 7 | Tuft1 | 0.023 | Old |
| 0.213565 | 0.105804 | 0.321 | 0.268 | 1 | 7 | Knl1 | 0.053 | Old |
| 0.213801 | 0.565364 | 0.056 | 0.036 | 1 | 7 | Ppp1r3e | 0.02 | Old |
| 0.213872 | 0.483393 | 0.174 | 0.142 | 1 | 7 | Pop7 | 0.032 | Old |
| 0.213921 | 0.709993 | 0.021 | 0.009 | 1 | 7 | B3galt4 | 0.012 | Old |
| 0.21397 | 0.64615 | 0.172 | 0.142 | 1 | 7 | Nt5c2 | 0.03 | Old |
| 0.21409 | 0.213309 | 0.282 | 0.232 | 1 | 7 | Kif22 | 0.05 | Old |
| 0.214459 | 0.451983 | 0.167 | 0.136 | 1 | 7 | Faap20 | 0.031 | Old |
| 0.214543 | 0.269383 | 0.021 | 0.009 | 1 | 7 | Pafah2 | 0.012 | Old |
| 0.214548 | 0.437908 | 0.162 | 0.13 | 1 | 7 | Usf3 | 0.032 | Old |
| 0.214752 | 1.037618 | 0.154 | 0.127 | 1 | 7 | Aimp2 | 0.027 | Old |
| 0.214804 | 0.063559 | 0.382 | 0.319 | 1 | 7 | Lman1 | 0.063 | Old |
| 0.215027 | 0.244906 | 0.185 | 0.151 | 1 | 7 | Swt1 | 0.034 | Old |
| 0.215149 | 0.072848 | 0.456 | 0.401 | 1 | 7 | Atf6 | 0.055 | Old |
| 0.215202 | 0.095534 | 0.318 | 0.271 | 1 | 7 | Luc7l | 0.047 | Old |
| 0.215428 | 0.534169 | 0.146 | 0.114 | 1 | 7 | Galns | 0.032 | Old |
| 0.215781 | 0.209788 | 0.418 | 0.37 | 1 | 7 | N4bp1 | 0.048 | Old |
| 0.215884 | 0.300296 | 0.169 | 0.136 | 1 | 7 | E2f3 | 0.033 | Old |
| 0.215944 | 0.330629 | 0.208 | 0.169 | 1 | 7 | Lilra6 | 0.039 | Old |
| 0.21629 | 0.00168 | 0.041 | 0.024 | 1 | 7 | Pxmp2 | 0.017 | Old |
| 0.216419 | 0.309563 | 0.021 | 0.009 | 1 | 7 | Lysmd4 | 0.012 | Old |
| 0.21653 | 0.080021 | 0.597 | 0.545 | 1 | 7 | Pdcd5 | 0.052 | Old |
| 0.216835 | 1.128434 | 0.067 | 0.045 | 1 | 7 | Tbc1d32 | 0.022 | Old |
| 0.216963 | 0.183723 | 0.131 | 0.099 | 1 | 7 | Yif1a | 0.032 | Old |
| 0.217057 | 0.154183 | 0.479 | 0.428 | 1 | 7 | Trmt112 | 0.051 | Old |
| 0.217234 | 0.068751 | 0.074 | 0.051 | 1 | 7 | Zfyve21 | 0.023 | Old |
| 0.217275 | 0.355967 | 0.156 | 0.123 | 1 | 7 | Lrch1 | 0.033 | Old |
| 0.217564 | 0.103055 | 0.231 | 0.187 | 1 | 7 | Hgsnat | 0.044 | Old |
| 0.217599 | 0.070008 | 0.338 | 0.286 | 1 | 7 | Cyb5r3 | 0.052 | Old |
| 0.217605 | 0.131767 | 0.515 | 0.452 | 1 | 7 | Stx4a | 0.063 | Old |
| 0.218402 | 0.629697 | 0.077 | 0.054 | 1 | 7 | Fbxl4 | 0.023 | Old |
| 0.218598 | 0.6038 | 0.059 | 0.039 | 1 | 7 | Cul9 | 0.02 | Old |
| 0.219073 | 0.226609 | 0.244 | 0.202 | 1 | 7 | Hyou1 | 0.042 | Old |
| 0.219748 | 0.467109 | 0.231 | 0.199 | 1 | 7 | Ncln | 0.032 | Old |
| 0.219965 | 0.310006 | 0.128 | 0.099 | 1 | 7 | Cep70 | 0.029 | Old |
| 0.220084 | 0.514392 | 0.177 | 0.145 | 1 | 7 | Tmem68 | 0.032 | Old |
| 0.22017 | 0.279933 | 0.041 | 0.024 | 1 | 7 | Polrmt | 0.017 | Old |
| 0.220192 | 0.213724 | 0.464 | 0.407 | 1 | 7 | Phb2 | 0.057 | Old |
| 0.220388 | 0.150797 | 0.8 | 0.747 | 1 | 7 | Csnk1a1 | 0.053 | Old |
| 0.220528 | 0.318289 | 0.159 | 0.127 | 1 | 7 | Trmt2a | 0.032 | Old |

| | | | | | | | | |
|---|---|---|---|---|---|---|---|---|
| 0.22082 | 0.296712 | 0.305 | 0.265 | 1 | 7 | Fam76a | 0.04 | Old |
| 0.220854 | 0.52979 | 0.141 | 0.111 | 1 | 7 | Msantd4 | 0.03 | Old |
| 0.220916 | 0.044645 | 0.472 | 0.407 | 1 | 7 | Tmed3 | 0.065 | Old |
| 0.220993 | 0.318541 | 0.205 | 0.166 | 1 | 7 | Pigq | 0.039 | Old |
| 0.221421 | 0.059029 | 0.538 | 0.47 | 1 | 7 | Atxn2l | 0.068 | Old |
| 0.221743 | 0.258266 | 0.313 | 0.265 | 1 | 7 | Rbm42 | 0.048 | Old |
| 0.221872 | 0.045452 | 0.233 | 0.19 | 1 | 7 | Klhdc4 | 0.043 | Old |
| 0.221938 | 0.16853 | 0.049 | 0.03 | 1 | 7 | Cradd | 0.019 | Old |
| 0.221961 | 0.206461 | 0.162 | 0.127 | 1 | 7 | Matk | 0.035 | Old |
| 0.222155 | 0.534685 | 0.228 | 0.196 | 1 | 7 | Ccdc88c | 0.032 | Old |
| 0.222436 | 0.0842 | 0.228 | 0.187 | 1 | 7 | Enox2 | 0.041 | Old |
| 0.22302 | 0.276398 | 0.482 | 0.452 | 1 | 7 | Cenpa | 0.03 | Old |
| 0.223123 | 0.208438 | 0.656 | 0.587 | 1 | 7 | Taok3 | 0.069 | Old |
| 0.223138 | 0.068294 | 0.456 | 0.389 | 1 | 7 | Dnajc2 | 0.067 | Old |
| 0.223193 | 0.023811 | 0.023 | 0.039 | 1 | 7 | Zfp553 | -0.016 | Old |
| 0.223398 | 0.123321 | 0.341 | 0.298 | 1 | 7 | Lpcat1 | 0.043 | Old |
| 0.223914 | 0.25005 | 0.103 | 0.075 | 1 | 7 | Zfp729a | 0.028 | Old |
| 0.224017 | 0.113486 | 0.282 | 0.241 | 1 | 7 | Rnf44 | 0.041 | Old |
| 0.224164 | 0.514884 | 0.164 | 0.133 | 1 | 7 | Dnlz | 0.031 | Old |
| 0.224469 | 0.107894 | 0.19 | 0.151 | 1 | 7 | Nuf2 | 0.039 | Old |
| 0.224573 | 0.319317 | 0.067 | 0.045 | 1 | 7 | Ankrd39 | 0.022 | Old |
| 0.224973 | 0.574004 | 0.154 | 0.127 | 1 | 7 | Gen1 | 0.027 | Old |
| 0.225021 | 0.260687 | 0.397 | 0.349 | 1 | 7 | Cdk1 | 0.048 | Old |
| 0.225154 | 0.00864 | 0.231 | 0.187 | 1 | 7 | 3110043O2 | 0.044 | Old |
| 0.225293 | 0.298483 | 0.367 | 0.322 | 1 | 7 | Poldip3 | 0.045 | Old |
| 0.225379 | 0.949988 | 0.044 | 0.027 | 1 | 7 | Snap47 | 0.017 | Old |
| 0.225592 | 0.324805 | 0.126 | 0.096 | 1 | 7 | Tars2 | 0.03 | Old |
| 0.22569 | 0.536723 | 0.051 | 0.033 | 1 | 7 | Inafm2 | 0.018 | Old |
| 0.225817 | 0.163355 | 0.715 | 0.687 | 1 | 7 | Ldha | 0.028 | Old |
| 0.225825 | 0.124846 | 0.362 | 0.319 | 1 | 7 | Wtap | 0.043 | Old |
| 0.2261 | 0.32892 | 0.287 | 0.241 | 1 | 7 | Commd4 | 0.046 | Old |
| 0.226536 | 0.218849 | 0.541 | 0.479 | 1 | 7 | Hpf1 | 0.062 | Old |
| 0.226653 | 0.272911 | 0.354 | 0.31 | 1 | 7 | Npepps | 0.044 | Old |
| 0.22669 | 0.887417 | 0.028 | 0.015 | 1 | 7 | Fam234b | 0.013 | Old |
| 0.226719 | 0.16573 | 0.105 | 0.078 | 1 | 7 | Dis3 | 0.027 | Old |
| 0.226777 | 0.115795 | 0.695 | 0.66 | 1 | 7 | Rock1 | 0.035 | Old |
| 0.227237 | 1.59013 | 0.015 | 0.006 | 1 | 7 | Zfp775 | 0.009 | Old |
| 0.227337 | 0.075118 | 0.213 | 0.178 | 1 | 7 | Cnot7 | 0.035 | Old |
| 0.227463 | 0.237089 | 0.29 | 0.25 | 1 | 7 | Tfrc | 0.04 | Old |
| 0.227468 | 0.424823 | 0.254 | 0.211 | 1 | 7 | Agtpbp1 | 0.043 | Old |
| 0.227508 | 0.951511 | 0.044 | 0.027 | 1 | 7 | Spryd4 | 0.017 | Old |
| 0.227635 | 0.427775 | 0.146 | 0.114 | 1 | 7 | Cebpe | 0.032 | Old |
| 0.227699 | 0.282808 | 0.233 | 0.193 | 1 | 7 | Tbc1d13 | 0.04 | Old |
| 0.228103 | 0.178982 | 0.7 | 0.66 | 1 | 7 | 7-Sep | 0.04 | Old |
| 0.228141 | 0.185376 | 0.074 | 0.051 | 1 | 7 | Fpgs | 0.023 | Old |

| | | | | | | | | |
|---|---|---|---|---|---|---|---|---|
| 0.228191 | 1.116992 | 0.082 | 0.06 | 1 | 7 | Cryzl2 | 0.022 | Old |
| 0.228313 | 0.793001 | 0.028 | 0.015 | 1 | 7 | Hectd2 | 0.013 | Old |
| 0.228435 | 0.626754 | 0.051 | 0.033 | 1 | 7 | Mtrr | 0.018 | Old |
| 0.228495 | 0.049076 | 0.451 | 0.407 | 1 | 7 | Mrpl20 | 0.044 | Old |
| 0.228797 | 0.289763 | 0.272 | 0.229 | 1 | 7 | Sptlc1 | 0.043 | Old |
| 0.22901 | 0.08656 | 0.541 | 0.467 | 1 | 7 | Clec2d | 0.074 | Old |
| 0.229181 | 0.095548 | 0.6 | 0.551 | 1 | 7 | Psma3 | 0.049 | Old |
| 0.229189 | 0.132422 | 0.105 | 0.078 | 1 | 7 | Pdk1 | 0.027 | Old |
| 0.229399 | 0.70481 | 0.028 | 0.015 | 1 | 7 | Procr | 0.013 | Old |
| 0.229416 | 0.319314 | 0.369 | 0.331 | 1 | 7 | Cyth1 | 0.038 | Old |
| 0.229428 | 0.495099 | 0.231 | 0.199 | 1 | 7 | Atg16l1 | 0.032 | Old |
| 0.229464 | 0.018029 | 0.428 | 0.358 | 1 | 7 | Dpy19l1 | 0.07 | Old |
| 0.229525 | 1.088804 | 0.015 | 0.006 | 1 | 7 | Bmp1 | 0.009 | Old |
| 0.229679 | 0.325988 | 0.033 | 0.051 | 1 | 7 | Rpl9-ps6 | -0.018 | Old |
| 0.229942 | 0.29067 | 0.356 | 0.316 | 1 | 7 | Rras2 | 0.04 | Old |
| 0.229967 | 0.225237 | 0.385 | 0.343 | 1 | 7 | Sod2 | 0.042 | Old |
| 0.230047 | 1.189325 | 0.082 | 0.06 | 1 | 7 | Elovl6 | 0.022 | Old |
| 0.230235 | 1.3982 | 0.036 | 0.021 | 1 | 7 | Cfap45 | 0.015 | Old |
| 0.230291 | 1.160126 | 0.015 | 0.006 | 1 | 7 | Ceacam19 | 0.009 | Old |
| 0.230291 | 1.10917 | 0.015 | 0.006 | 1 | 7 | Lrig3 | 0.009 | Old |
| 0.230291 | 0.939322 | 0.015 | 0.006 | 1 | 7 | B3gnt7 | 0.009 | Old |
| 0.230291 | 0.745792 | 0.015 | 0.006 | 1 | 7 | Cdon | 0.009 | Old |
| 0.230345 | 0.208709 | 0.09 | 0.066 | 1 | 7 | Rbm38 | 0.024 | Old |
| 0.230416 | 0.41419 | 0.172 | 0.142 | 1 | 7 | Acadsb | 0.03 | Old |
| 0.230453 | 0.003547 | 0.203 | 0.238 | 1 | 7 | Milr1 | -0.035 | Old |
| 0.230477 | 0.309674 | 0.097 | 0.072 | 1 | 7 | Tfap4 | 0.025 | Old |
| 0.230484 | 0.183623 | 0.087 | 0.063 | 1 | 7 | E130307A1 | 0.024 | Old |
| 0.23079 | 0.363811 | 0.172 | 0.139 | 1 | 7 | Atg7 | 0.033 | Old |
| 0.231035 | 0.626571 | 0.028 | 0.015 | 1 | 7 | Ccdc94 | 0.013 | Old |
| 0.23106 | 0.931116 | 0.015 | 0.006 | 1 | 7 | Rasip1 | 0.009 | Old |
| 0.23106 | 0.87325 | 0.015 | 0.006 | 1 | 7 | Trim13 | 0.009 | Old |
| 0.23106 | 0.833509 | 0.015 | 0.006 | 1 | 7 | Gca | 0.009 | Old |
| 0.231177 | 0.059499 | 0.146 | 0.114 | 1 | 7 | Irgm1 | 0.032 | Old |
| 0.231376 | 1.118384 | 0.044 | 0.027 | 1 | 7 | Nim1k | 0.017 | Old |
| 0.231393 | 0.541675 | 0.249 | 0.214 | 1 | 7 | Supt6 | 0.035 | Old |
| 0.231541 | 0.432352 | 0.205 | 0.169 | 1 | 7 | Ruvbl1 | 0.036 | Old |
| 0.231582 | 0.843289 | 0.028 | 0.015 | 1 | 7 | E030030I0 | 0.013 | Old |
| 0.231582 | 0.801681 | 0.028 | 0.015 | 1 | 7 | Cdk5rap1 | 0.013 | Old |
| 0.231829 | 1.501067 | 0.015 | 0.006 | 1 | 7 | Gm34590 | 0.009 | Old |
| 0.231829 | 1.221275 | 0.015 | 0.006 | 1 | 7 | Frg2f1 | 0.009 | Old |
| 0.231829 | 0.997278 | 0.015 | 0.006 | 1 | 7 | Trpt1 | 0.009 | Old |
| 0.231829 | 0.944728 | 0.015 | 0.006 | 1 | 7 | Lgals4 | 0.009 | Old |
| 0.231829 | 0.602532 | 0.015 | 0.006 | 1 | 7 | Cd5l | 0.009 | Old |
| 0.231848 | 0.35898 | 0.177 | 0.145 | 1 | 7 | Trappc9 | 0.032 | Old |
| 0.232131 | 1.176798 | 0.028 | 0.015 | 1 | 7 | Sell | 0.013 | Old |

| | | | | | | | | |
|---|---|---|---|---|---|---|---|---|
| 0.232131 | 0.711517 | 0.028 | 0.015 | 1 | 7 | Pla2g6 | 0.013 | Old |
| 0.232287 | 0.09236 | 0.372 | 0.325 | 1 | 7 | Rbm7 | 0.047 | Old |
| 0.232601 | 0.792509 | 0.015 | 0.006 | 1 | 7 | Rims3 | 0.009 | Old |
| 0.232601 | 0.715533 | 0.015 | 0.006 | 1 | 7 | Mettl22 | 0.009 | Old |
| 0.232601 | 0.31193 | 0.015 | 0.006 | 1 | 7 | Mcts2 | 0.009 | Old |
| 0.232906 | 0.599672 | 0.108 | 0.081 | 1 | 7 | Tmem214 | 0.027 | Old |
| 0.233102 | 0.185524 | 0.144 | 0.114 | 1 | 7 | Pdhx | 0.03 | Old |
| 0.23323 | 1.486791 | 0.028 | 0.015 | 1 | 7 | Gm10031 | 0.013 | Old |
| 0.23323 | 0.597465 | 0.028 | 0.015 | 1 | 7 | 1700052K1 | 0.013 | Old |
| 0.233375 | 0.665911 | 0.015 | 0.006 | 1 | 7 | Sult1e1 | 0.009 | Old |
| 0.233375 | 0.515068 | 0.015 | 0.006 | 1 | 7 | Ankrd9 | 0.009 | Old |
| 0.233375 | 0.435096 | 0.015 | 0.006 | 1 | 7 | Klrb1f | 0.009 | Old |
| 0.233375 | 0.314324 | 0.015 | 0.006 | 1 | 7 | 4930404N1 | 0.009 | Old |
| 0.233477 | 0.605325 | 0.092 | 0.066 | 1 | 7 | Traip | 0.026 | Old |
| 0.233597 | 1.113839 | 0.036 | 0.021 | 1 | 7 | Izumo4 | 0.015 | Old |
| 0.233719 | 0.550428 | 0.11 | 0.084 | 1 | 7 | Zscan29 | 0.026 | Old |
| 0.233807 | 0.231173 | 0.103 | 0.078 | 1 | 7 | AI837181 | 0.025 | Old |
| 0.233921 | 0.11544 | 0.421 | 0.373 | 1 | 7 | Samd4b | 0.048 | Old |
| 0.234045 | 0.237268 | 0.267 | 0.226 | 1 | 7 | Naa10 | 0.041 | Old |
| 0.23415 | 1.169352 | 0.015 | 0.006 | 1 | 7 | Smpd5 | 0.009 | Old |
| 0.23415 | 0.711015 | 0.015 | 0.006 | 1 | 7 | Rhox8 | 0.009 | Old |
| 0.23415 | 0.460992 | 0.015 | 0.006 | 1 | 7 | Gne | 0.009 | Old |
| 0.23415 | 0.185562 | 0.015 | 0.006 | 1 | 7 | Eml5 | 0.009 | Old |
| 0.234183 | 0.559867 | 0.141 | 0.111 | 1 | 7 | Tjp2 | 0.03 | Old |
| 0.234202 | 0.025804 | 0.328 | 0.271 | 1 | 7 | Tspyl1 | 0.057 | Old |
| 0.234415 | 0.221358 | 0.413 | 0.37 | 1 | 7 | Ipo5 | 0.043 | Old |
| 0.234417 | 0.68349 | 0.044 | 0.027 | 1 | 7 | Gm36445 | 0.017 | Old |
| 0.234699 | 0.122864 | 0.328 | 0.283 | 1 | 7 | Slc9a3r1 | 0.045 | Old |
| 0.234771 | 0.422409 | 0.213 | 0.178 | 1 | 7 | Aldh3a2 | 0.035 | Old |
| 0.234886 | 0.661848 | 0.028 | 0.015 | 1 | 7 | 9030025P2 | 0.013 | Old |
| 0.234928 | 0.498798 | 0.015 | 0.006 | 1 | 7 | 2510017J1 | 0.009 | Old |
| 0.235194 | 0.080592 | 0.559 | 0.503 | 1 | 7 | Psma6 | 0.056 | Old |
| 0.235235 | 0.296269 | 0.233 | 0.199 | 1 | 7 | Arhgap39 | 0.034 | Old |
| 0.235503 | 0.214449 | 0.241 | 0.199 | 1 | 7 | Ing4 | 0.042 | Old |
| 0.235551 | 0.140083 | 0.095 | 0.069 | 1 | 7 | Ints5.1 | 0.026 | Old |
| 0.235591 | 0.314726 | 0.228 | 0.196 | 1 | 7 | Gas2l1 | 0.032 | Old |
| 0.235698 | 0.316168 | 0.069 | 0.048 | 1 | 7 | Wdr4 | 0.021 | Old |
| 0.235707 | 0.244982 | 0.015 | 0.006 | 1 | 7 | Cryab | 0.009 | Old |
| 0.23571 | 0.203682 | 0.197 | 0.163 | 1 | 7 | Cdk16 | 0.034 | Old |
| 0.235775 | 0.057256 | 0.246 | 0.199 | 1 | 7 | Paxip1 | 0.047 | Old |
| 0.23579 | 0.672691 | 0.072 | 0.051 | 1 | 7 | Mrrf | 0.021 | Old |
| 0.236382 | 0.10474 | 0.379 | 0.322 | 1 | 7 | Osbpl8 | 0.057 | Old |
| 0.236424 | 0.164039 | 0.174 | 0.142 | 1 | 7 | Heatr3 | 0.032 | Old |
| 0.236463 | 0.890066 | 0.062 | 0.042 | 1 | 7 | Copz2 | 0.02 | Old |
| 0.236488 | 0.183755 | 0.015 | 0.006 | 1 | 7 | Ggnbp1 | 0.009 | Old |

| | | | | | | | | |
|---|---|---|---|---|---|---|---|---|
| 0.23669 | 0.233188 | 0.272 | 0.226 | 1 | 7 | Psmc4 | 0.046 | Old |
| 0.236735 | 0.32318 | 0.095 | 0.069 | 1 | 7 | Mto1 | 0.026 | Old |
| 0.236819 | 0.162649 | 0.318 | 0.274 | 1 | 7 | Sap18 | 0.044 | Old |
| 0.23683 | 0.273456 | 0.4 | 0.361 | 1 | 7 | Qpct | 0.039 | Old |
| 0.237112 | 0.395028 | 0.141 | 0.111 | 1 | 7 | Prr12 | 0.03 | Old |
| 0.237236 | 0.251145 | 0.433 | 0.389 | 1 | 7 | Me2 | 0.044 | Old |
| 0.23731 | 0.085682 | 0.159 | 0.127 | 1 | 7 | Rbm4b | 0.032 | Old |
| 0.237505 | 0.277323 | 0.238 | 0.199 | 1 | 7 | Inpp5f | 0.039 | Old |
| 0.237625 | 0.128211 | 0.095 | 0.069 | 1 | 7 | Pwp2 | 0.026 | Old |
| 0.237626 | 0.000314 | 0.103 | 0.078 | 1 | 7 | Pnpt1 | 0.025 | Old |
| 0.237641 | 0.454589 | 0.059 | 0.039 | 1 | 7 | Pmm1 | 0.02 | Old |
| 0.237731 | 0.516701 | 0.185 | 0.157 | 1 | 7 | Dbt | 0.028 | Old |
| 0.238007 | 0.005945 | 0.223 | 0.181 | 1 | 7 | Sacm1l | 0.042 | Old |
| 0.238028 | 0.212684 | 0.154 | 0.123 | 1 | 7 | Fam213b | 0.031 | Old |
| 0.238267 | 0.219054 | 0.287 | 0.25 | 1 | 7 | Fto | 0.037 | Old |
| 0.238276 | 0.077534 | 0.654 | 0.602 | 1 | 7 | Tram1 | 0.052 | Old |
| 0.238299 | 0.038437 | 0.444 | 0.488 | 1 | 7 | Man1a | -0.044 | Old |
| 0.238542 | 0.180823 | 0.354 | 0.304 | 1 | 7 | Tap2 | 0.05 | Old |
| 0.238781 | 0.174387 | 0.226 | 0.187 | 1 | 7 | Eef1e1 | 0.039 | Old |
| 0.238785 | 0.738393 | 0.09 | 0.066 | 1 | 7 | Trmt10a | 0.024 | Old |
| 0.238879 | 0.137173 | 0.231 | 0.19 | 1 | 7 | Mrpl9 | 0.041 | Old |
| 0.239513 | 0.626971 | 0.069 | 0.048 | 1 | 7 | Rfx2 | 0.021 | Old |
| 0.239776 | 0.440432 | 0.062 | 0.042 | 1 | 7 | BC052040 | 0.02 | Old |
| 0.239899 | 0.189537 | 0.077 | 0.054 | 1 | 7 | Nit2 | 0.023 | Old |
| 0.240036 | 0.831947 | 0.11 | 0.084 | 1 | 7 | Snip1 | 0.026 | Old |
| 0.240132 | 0.21322 | 0.108 | 0.081 | 1 | 7 | Gbe1 | 0.027 | Old |
| 0.240254 | 0.012415 | 0.587 | 0.521 | 1 | 7 | Cggbp1 | 0.066 | Old |
| 0.240336 | 0.112755 | 0.292 | 0.253 | 1 | 7 | Prpf18 | 0.039 | Old |
| 0.24034 | 0.400477 | 0.115 | 0.09 | 1 | 7 | Bcar3 | 0.025 | Old |
| 0.240547 | 0.395654 | 0.203 | 0.169 | 1 | 7 | Zfp322a | 0.034 | Old |
| 0.240893 | 2.48769 | 0.01 | 0.003 | 1 | 7 | Pla1a | 0.007 | Old |
| 0.240893 | 1.520615 | 0.01 | 0.003 | 1 | 7 | Gpr27 | 0.007 | Old |
| 0.240893 | 1.307398 | 0.01 | 0.003 | 1 | 7 | Pde7b | 0.007 | Old |
| 0.240893 | 1.28281 | 0.01 | 0.003 | 1 | 7 | Gm28874 | 0.007 | Old |
| 0.240893 | 1.211359 | 0.01 | 0.003 | 1 | 7 | Mybpc3 | 0.007 | Old |
| 0.240893 | 1.200813 | 0.01 | 0.003 | 1 | 7 | Ablim3 | 0.007 | Old |
| 0.240893 | 1.016927 | 0.01 | 0.003 | 1 | 7 | 4-Sep | 0.007 | Old |
| 0.240893 | 0.948374 | 0.01 | 0.003 | 1 | 7 | Gm9796 | 0.007 | Old |
| 0.240893 | 0.910165 | 0.01 | 0.003 | 1 | 7 | Dnase1l2 | 0.007 | Old |
| 0.241094 | 0.03468 | 0.205 | 0.169 | 1 | 7 | Esco2 | 0.036 | Old |
| 0.241474 | 0.566029 | 0.044 | 0.027 | 1 | 7 | Asb4 | 0.017 | Old |
| 0.241496 | 0.151922 | 0.197 | 0.166 | 1 | 7 | Pex6 | 0.031 | Old |
| 0.241503 | 0.243519 | 0.451 | 0.416 | 1 | 7 | Lrmp | 0.035 | Old |
| 0.241895 | 2.043575 | 0.01 | 0.003 | 1 | 7 | Prss23 | 0.007 | Old |
| 0.241895 | 1.659096 | 0.01 | 0.003 | 1 | 7 | Cd200r3 | 0.007 | Old |

| | | | | | | | | |
|---|---|---|---|---|---|---|---|---|
| 0.241895 | 1.293726 | 0.01 | 0.003 | 1 | 7 | Adgrf5 | 0.007 | Old |
| 0.241895 | 1.149604 | 0.01 | 0.003 | 1 | 7 | Hist1h2bh | 0.007 | Old |
| 0.241895 | 0.974639 | 0.01 | 0.003 | 1 | 7 | 1700030J2 | 0.007 | Old |
| 0.241895 | 0.862688 | 0.01 | 0.003 | 1 | 7 | Nupr1l | 0.007 | Old |
| 0.241895 | 0.809459 | 0.01 | 0.003 | 1 | 7 | Fcrla | 0.007 | Old |
| 0.241895 | 0.798581 | 0.01 | 0.003 | 1 | 7 | Vwa1 | 0.007 | Old |
| 0.241895 | 0.791486 | 0.01 | 0.003 | 1 | 7 | Hist1h4j | 0.007 | Old |
| 0.241895 | 0.660908 | 0.01 | 0.003 | 1 | 7 | Hist2h2bb | 0.007 | Old |
| 0.241971 | 0.420898 | 0.179 | 0.148 | 1 | 7 | Trpv2 | 0.031 | Old |
| 0.242169 | 0.216453 | 0.449 | 0.413 | 1 | 7 | Map4 | 0.036 | Old |
| 0.242457 | 0.080483 | 0.597 | 0.518 | 1 | 7 | Rnf13 | 0.079 | Old |
| 0.24273 | 0.109446 | 0.403 | 0.355 | 1 | 7 | Idh2 | 0.048 | Old |
| 0.242899 | 1.23345 | 0.01 | 0.003 | 1 | 7 | Gm15326 | 0.007 | Old |
| 0.242899 | 1.159686 | 0.01 | 0.003 | 1 | 7 | Zfp418 | 0.007 | Old |
| 0.242899 | 1.159019 | 0.01 | 0.003 | 1 | 7 | Pmel | 0.007 | Old |
| 0.242899 | 1.079735 | 0.01 | 0.003 | 1 | 7 | Tst | 0.007 | Old |
| 0.242899 | 1.024666 | 0.01 | 0.003 | 1 | 7 | Rasl2-9 | 0.007 | Old |
| 0.242899 | 0.732934 | 0.01 | 0.003 | 1 | 7 | Zfp69 | 0.007 | Old |
| 0.242899 | 0.724517 | 0.01 | 0.003 | 1 | 7 | Ank1 | 0.007 | Old |
| 0.242899 | 0.660145 | 0.01 | 0.003 | 1 | 7 | G730003C1 | 0.007 | Old |
| 0.242899 | 0.589146 | 0.01 | 0.003 | 1 | 7 | Pde8b | 0.007 | Old |
| 0.242899 | 0.586388 | 0.01 | 0.003 | 1 | 7 | Col4a5 | 0.007 | Old |
| 0.242899 | 0.559517 | 0.01 | 0.003 | 1 | 7 | Gm15228 | 0.007 | Old |
| 0.242899 | 0.504983 | 0.01 | 0.003 | 1 | 7 | Sat2 | 0.007 | Old |
| 0.242899 | 0.481662 | 0.01 | 0.003 | 1 | 7 | Cldn12 | 0.007 | Old |
| 0.242899 | 0.441002 | 0.01 | 0.003 | 1 | 7 | Zfp759 | 0.007 | Old |
| 0.243264 | 0.167164 | 0.328 | 0.28 | 1 | 7 | Rab3d | 0.048 | Old |
| 0.243419 | 0.132662 | 0.577 | 0.533 | 1 | 7 | Mdh1 | 0.044 | Old |
| 0.243673 | 0.102054 | 0.492 | 0.446 | 1 | 7 | Sp100 | 0.046 | Old |
| 0.243709 | 0.244203 | 0.228 | 0.193 | 1 | 7 | Slc39a9 | 0.035 | Old |
| 0.24371 | 0.373367 | 0.044 | 0.027 | 1 | 7 | Med26 | 0.017 | Old |
| 0.24371 | 0.340872 | 0.044 | 0.027 | 1 | 7 | Gale | 0.017 | Old |
| 0.243851 | 0.5029 | 0.246 | 0.22 | 1 | 7 | A430005L1 | 0.026 | Old |
| 0.24387 | 0.693114 | 0.062 | 0.042 | 1 | 7 | Abcb6 | 0.02 | Old |
| 0.243906 | 0.72004 | 0.01 | 0.003 | 1 | 7 | Lrp11 | 0.007 | Old |
| 0.243906 | 0.588561 | 0.01 | 0.003 | 1 | 7 | Gnmt | 0.007 | Old |
| 0.243906 | 0.524373 | 0.01 | 0.003 | 1 | 7 | Klre1 | 0.007 | Old |
| 0.243906 | 0.502806 | 0.01 | 0.003 | 1 | 7 | Gm17690 | 0.007 | Old |
| 0.243906 | 0.501099 | 0.01 | 0.003 | 1 | 7 | Ly6c1 | 0.007 | Old |
| 0.243906 | 0.384194 | 0.01 | 0.003 | 1 | 7 | Mfap3l | 0.007 | Old |
| 0.244163 | 0.152718 | 0.051 | 0.033 | 1 | 7 | Lmln | 0.018 | Old |
| 0.244184 | 0.309593 | 0.131 | 0.102 | 1 | 7 | Cenpn | 0.029 | Old |
| 0.244359 | 0.315199 | 0.233 | 0.199 | 1 | 7 | Itfg1 | 0.034 | Old |
| 0.244389 | 0.350355 | 0.036 | 0.021 | 1 | 7 | Rab23 | 0.015 | Old |
| 0.244916 | 0.447866 | 0.01 | 0.003 | 1 | 7 | Serpina3g | 0.007 | Old |

| | | | | | | | | |
|---|---|---|---|---|---|---|---|---|
| 0.244916 | 0.40008 | 0.01 | 0.003 | 1 | 7 | Col15a1 | 0.007 | Old |
| 0.244916 | 0.387615 | 0.01 | 0.003 | 1 | 7 | Gm11110 | 0.007 | Old |
| 0.244916 | 0.385209 | 0.01 | 0.003 | 1 | 7 | 5730422E0 | 0.007 | Old |
| 0.244916 | 0.035744 | 0.01 | 0.003 | 1 | 7 | Cd300e | 0.007 | Old |
| 0.24504 | 0.344383 | 0.079 | 0.057 | 1 | 7 | Tmem185a | 0.022 | Old |
| 0.245187 | 0.170163 | 0.121 | 0.093 | 1 | 7 | Pdgfa | 0.028 | Old |
| 0.245387 | 0.350028 | 0.036 | 0.021 | 1 | 7 | Abhd14b | 0.015 | Old |
| 0.245748 | 0.286802 | 0.262 | 0.229 | 1 | 7 | Acot7 | 0.033 | Old |
| 0.245848 | 0.127875 | 0.069 | 0.048 | 1 | 7 | Aifm2 | 0.021 | Old |
| 0.245887 | 0.688925 | 0.036 | 0.021 | 1 | 7 | Tnfsf14 | 0.015 | Old |
| 0.245887 | 0.349749 | 0.036 | 0.021 | 1 | 7 | Mrm1 | 0.015 | Old |
| 0.24619 | 0.980555 | 0.09 | 0.066 | 1 | 7 | Ap5z1 | 0.024 | Old |
| 0.246289 | 0.280289 | 0.205 | 0.172 | 1 | 7 | Plekha3 | 0.033 | Old |
| 0.246421 | 0.445709 | 0.072 | 0.051 | 1 | 7 | Vwa8 | 0.021 | Old |
| 0.246552 | 0.038865 | 0.131 | 0.099 | 1 | 7 | Cdkn2aip | 0.032 | Old |
| 0.246728 | 0.242643 | 0.195 | 0.16 | 1 | 7 | Taf5l | 0.035 | Old |
| 0.246905 | 0.483821 | 0.082 | 0.06 | 1 | 7 | Ccdc77 | 0.022 | Old |
| 0.246986 | 0.370238 | 0.138 | 0.108 | 1 | 7 | Nob1 | 0.03 | Old |
| 0.247091 | 0.115661 | 0.728 | 0.663 | 1 | 7 | Tspo | 0.065 | Old |
| 0.247217 | 0.513485 | 0.121 | 0.093 | 1 | 7 | Rnf215 | 0.028 | Old |
| 0.247453 | 0.073714 | 0.226 | 0.184 | 1 | 7 | Fam234a | 0.042 | Old |
| 0.247724 | 0.142113 | 0.356 | 0.31 | 1 | 7 | Hdac1 | 0.046 | Old |
| 0.247801 | 0.22202 | 0.108 | 0.081 | 1 | 7 | Polq | 0.027 | Old |
| 0.248103 | 0.063256 | 0.646 | 0.584 | 1 | 7 | Vsir | 0.062 | Old |
| 0.248199 | 0.28008 | 0.331 | 0.298 | 1 | 7 | Nfkbie | 0.033 | Old |
| 0.248225 | 0.038315 | 0.044 | 0.027 | 1 | 7 | Cox7a1 | 0.017 | Old |
| 0.248234 | 0.142244 | 0.631 | 0.587 | 1 | 7 | Nol7 | 0.044 | Old |
| 0.248358 | 0.258625 | 0.079 | 0.057 | 1 | 7 | Nptxr | 0.022 | Old |
| 0.248823 | 0.483891 | 0.262 | 0.223 | 1 | 7 | AW112010 | 0.039 | Old |
| 0.249152 | 0.059338 | 0.051 | 0.033 | 1 | 7 | Slc9a4 | 0.018 | Old |
| 0.249315 | 0.231532 | 0.659 | 0.62 | 1 | 7 | Lbr | 0.039 | Old |
| 0.249733 | 0.190904 | 0.71 | 0.678 | 1 | 7 | Olfm1 | 0.032 | Old |
| 0.249862 | 0.167715 | 0.608 | 0.557 | 1 | 7 | Cnn2 | 0.051 | Old |
| 0.25018 | 0.675447 | 0.082 | 0.06 | 1 | 7 | Trmt10c | 0.022 | Old |
| 0.250634 | 0.330135 | 0.185 | 0.151 | 1 | 7 | Calhm2 | 0.034 | Old |
| 0.250764 | 0.221609 | 0.364 | 0.313 | 1 | 7 | Dennd6a | 0.051 | Old |
| 0.251216 | 0.1184 | 0.069 | 0.048 | 1 | 7 | Hook2 | 0.021 | Old |
| 0.25125 | 0.169534 | 0.408 | 0.364 | 1 | 7 | Smarca4 | 0.044 | Old |
| 0.251339 | 0.225577 | 0.295 | 0.256 | 1 | 7 | Slc35a3 | 0.039 | Old |
| 0.251429 | 0.145801 | 0.241 | 0.205 | 1 | 7 | Mrpl10 | 0.036 | Old |
| 0.25166 | 0.115266 | 0.362 | 0.31 | 1 | 7 | Ripk1 | 0.052 | Old |
| 0.251699 | 0.054266 | 0.387 | 0.346 | 1 | 7 | Ythdf1 | 0.041 | Old |
| 0.251743 | 0.201627 | 0.038 | 0.057 | 1 | 7 | Asgr2 | -0.019 | Old |
| 0.251749 | 0.97092 | 0.085 | 0.063 | 1 | 7 | Gbp5 | 0.022 | Old |
| 0.251886 | 0.129253 | 0.844 | 0.816 | 1 | 7 | Hmgb1 | 0.028 | Old |

| | | | | | | | | |
|---|---|---|---|---|---|---|---|---|
| 0.251896 | 0.171488 | 0.413 | 0.361 | 1 | 7 | Sel1l | 0.052 | Old |
| 0.251961 | 0.161856 | 0.462 | 0.422 | 1 | 7 | Socs6 | 0.04 | Old |
| 0.252278 | 0.103465 | 0.162 | 0.127 | 1 | 7 | Eif2b2 | 0.035 | Old |
| 0.252335 | 0.277083 | 0.197 | 0.166 | 1 | 7 | Tex10 | 0.031 | Old |
| 0.252546 | 0.03742 | 0.269 | 0.223 | 1 | 7 | Ncbp3 | 0.046 | Old |
| 0.252572 | 0.020868 | 0.115 | 0.087 | 1 | 7 | Tbce | 0.028 | Old |
| 0.252606 | 0.183787 | 0.5 | 0.461 | 1 | 7 | Tbcb | 0.039 | Old |
| 0.252796 | 0.080502 | 0.872 | 0.88 | 1 | 7 | Atp5l | -0.008 | Old |
| 0.252876 | 0.030173 | 0.554 | 0.491 | 1 | 7 | Spcs1 | 0.063 | Old |
| 0.252943 | 0.257516 | 0.336 | 0.298 | 1 | 7 | Pim3 | 0.038 | Old |
| 0.253265 | 0.033067 | 0.308 | 0.271 | 1 | 7 | Gnl1 | 0.037 | Old |
| 0.253869 | 0.389399 | 0.241 | 0.205 | 1 | 7 | Gid8 | 0.036 | Old |
| 0.25415 | 0.556125 | 0.082 | 0.06 | 1 | 7 | Hdac6 | 0.022 | Old |
| 0.254454 | 0.379517 | 0.21 | 0.175 | 1 | 7 | Zcchc9 | 0.035 | Old |
| 0.254515 | 0.24543 | 0.097 | 0.072 | 1 | 7 | Rpgrip1 | 0.025 | Old |
| 0.254657 | 0.139292 | 0.849 | 0.828 | 1 | 7 | Gabarap | 0.021 | Old |
| 0.255099 | 0.123666 | 0.905 | 0.892 | 1 | 7 | Hmgb2 | 0.013 | Old |
| 0.255241 | 0.086246 | 0.149 | 0.117 | 1 | 7 | Zbtb21 | 0.032 | Old |
| 0.255315 | 0.43258 | 0.121 | 0.093 | 1 | 7 | Nup37 | 0.028 | Old |
| 0.255647 | 0.393454 | 0.313 | 0.274 | 1 | 7 | Mrps16 | 0.039 | Old |
| 0.255693 | 0.109171 | 0.244 | 0.205 | 1 | 7 | Ppfia1 | 0.039 | Old |
| 0.255768 | 0.360539 | 0.079 | 0.057 | 1 | 7 | Btf3l4 | 0.022 | Old |
| 0.255952 | 0.101503 | 0.169 | 0.136 | 1 | 7 | Pigu | 0.033 | Old |
| 0.256027 | 0.052815 | 0.333 | 0.28 | 1 | 7 | Evi5 | 0.053 | Old |
| 0.256053 | 0.385142 | 0.151 | 0.123 | 1 | 7 | Cenpl | 0.028 | Old |
| 0.25628 | 0.105237 | 0.428 | 0.377 | 1 | 7 | Mbtd1 | 0.051 | Old |
| 0.256331 | 0.413061 | 0.272 | 0.235 | 1 | 7 | Clec4e | 0.037 | Old |
| 0.256481 | 0.212342 | 0.503 | 0.455 | 1 | 7 | Ptbp1 | 0.048 | Old |
| 0.256649 | 0.060968 | 0.172 | 0.139 | 1 | 7 | Nrbf2 | 0.033 | Old |
| 0.256806 | 0.397619 | 0.013 | 0.024 | 1 | 7 | Sall2 | -0.011 | Old |
| 0.25702 | 0.018571 | 0.423 | 0.37 | 1 | 7 | Apaf1 | 0.053 | Old |
| 0.257023 | 0.12842 | 0.105 | 0.078 | 1 | 7 | Lyrm9 | 0.027 | Old |
| 0.257065 | 0.341127 | 0.054 | 0.036 | 1 | 7 | Egfr | 0.018 | Old |
| 0.25721 | 0.596555 | 0.154 | 0.127 | 1 | 7 | Il15 | 0.027 | Old |
| 0.257221 | 0.330173 | 0.062 | 0.042 | 1 | 7 | Invs | 0.02 | Old |
| 0.257426 | 0.496376 | 0.318 | 0.277 | 1 | 7 | Stat1 | 0.041 | Old |
| 0.25767 | 0.072127 | 0.982 | 0.988 | 1 | 7 | Eef1a1 | -0.006 | Old |
| 0.257737 | 0.148537 | 0.277 | 0.238 | 1 | 7 | Sart1 | 0.039 | Old |
| 0.257754 | 0.021275 | 0.192 | 0.157 | 1 | 7 | Mdp1 | 0.035 | Old |
| 0.258142 | 0.584146 | 0.121 | 0.096 | 1 | 7 | Nudt1 | 0.025 | Old |
| 0.258148 | 0.078923 | 0.603 | 0.542 | 1 | 7 | Acer3 | 0.061 | Old |
| 0.258285 | 0.287061 | 0.103 | 0.078 | 1 | 7 | Dmap1 | 0.025 | Old |
| 0.258456 | 0.083855 | 0.51 | 0.455 | 1 | 7 | Nufip2 | 0.055 | Old |
| 0.258569 | 0.211108 | 0.105 | 0.081 | 1 | 7 | Gtpbp6 | 0.024 | Old |
| 0.259033 | 0.609333 | 0.046 | 0.03 | 1 | 7 | Polg2 | 0.016 | Old |

| | | | | | | | | |
|---|---|---|---|---|---|---|---|---|
| 0.259452 | 0.051535 | 0.154 | 0.123 | 1 | 7 | Isoc1 | 0.031 | Old |
| 0.259536 | 0.004371 | 0.172 | 0.136 | 1 | 7 | Prrc1 | 0.036 | Old |
| 0.259565 | 0.210914 | 0.185 | 0.151 | 1 | 7 | Srebf1 | 0.034 | Old |
| 0.259572 | 0.049928 | 0.554 | 0.506 | 1 | 7 | Usp25 | 0.048 | Old |
| 0.260125 | 0.447098 | 0.149 | 0.12 | 1 | 7 | Daxx | 0.029 | Old |
| 0.260387 | 0.096658 | 0.544 | 0.494 | 1 | 7 | Tnpo1 | 0.05 | Old |
| 0.260739 | 1.591942 | 0.023 | 0.012 | 1 | 7 | Tns2 | 0.011 | Old |
| 0.260983 | 0.231245 | 0.179 | 0.151 | 1 | 7 | Rpf2 | 0.028 | Old |
| 0.26152 | 0.088432 | 0.062 | 0.042 | 1 | 7 | Polr1b | 0.02 | Old |
| 0.261762 | 0.089709 | 0.356 | 0.31 | 1 | 7 | Fkbp4 | 0.046 | Old |
| 0.261841 | 0.336259 | 0.164 | 0.133 | 1 | 7 | Creb3 | 0.031 | Old |
| 0.261861 | 0.546262 | 0.172 | 0.142 | 1 | 7 | Smim12 | 0.03 | Old |
| 0.261973 | 0.180977 | 0.551 | 0.506 | 1 | 7 | Pcna | 0.045 | Old |
| 0.261997 | 0.111933 | 0.364 | 0.325 | 1 | 7 | Cdca3 | 0.039 | Old |
| 0.262183 | 0.21293 | 0.344 | 0.295 | 1 | 7 | Mybbp1a | 0.049 | Old |
| 0.262407 | 0.307073 | 0.115 | 0.09 | 1 | 7 | Stambp | 0.025 | Old |
| 0.262483 | 0.023117 | 0.89 | 0.867 | 1 | 7 | Ywhae | 0.023 | Old |
| 0.262637 | 0.07905 | 0.331 | 0.286 | 1 | 7 | Bcl3 | 0.045 | Old |
| 0.262953 | 0.039665 | 0.431 | 0.38 | 1 | 7 | Tmem164 | 0.051 | Old |
| 0.263384 | 1.356836 | 0.023 | 0.012 | 1 | 7 | Oasl1 | 0.011 | Old |
| 0.264048 | 0.232461 | 0.013 | 0.024 | 1 | 7 | Ehbp1 | -0.011 | Old |
| 0.264307 | 0.101655 | 0.251 | 0.211 | 1 | 7 | Rnf114 | 0.04 | Old |
| 0.26436 | 0.130476 | 0.203 | 0.166 | 1 | 7 | Scaf8 | 0.037 | Old |
| 0.26452 | 0.330619 | 0.428 | 0.395 | 1 | 7 | Fam168a | 0.033 | Old |
| 0.264572 | 0.280876 | 0.105 | 0.081 | 1 | 7 | Slc35e3 | 0.024 | Old |
| 0.264622 | 0.037054 | 0.421 | 0.361 | 1 | 7 | Ccni | 0.06 | Old |
| 0.264713 | 0.815009 | 0.023 | 0.012 | 1 | 7 | Zhx2 | 0.011 | Old |
| 0.264713 | 0.683919 | 0.023 | 0.012 | 1 | 7 | A330069E1 | 0.011 | Old |
| 0.26498 | 0.131929 | 0.746 | 0.72 | 1 | 7 | Pomp | 0.026 | Old |
| 0.265075 | 0.196801 | 0.09 | 0.066 | 1 | 7 | Coq10a | 0.024 | Old |
| 0.265093 | 0.640129 | 0.064 | 0.045 | 1 | 7 | Usp30 | 0.019 | Old |
| 0.265105 | 0.162378 | 0.323 | 0.274 | 1 | 7 | Trio | 0.049 | Old |
| 0.26538 | 0.065787 | 0.013 | 0.024 | 1 | 7 | Klhl26 | -0.011 | Old |
| 0.265481 | 0.210973 | 0.064 | 0.045 | 1 | 7 | Galnt9 | 0.019 | Old |
| 0.26576 | 1.1017 | 0.031 | 0.018 | 1 | 7 | Ehf | 0.013 | Old |
| 0.266047 | 0.5765 | 0.023 | 0.012 | 1 | 7 | Lcmt2 | 0.011 | Old |
| 0.266061 | 0.160102 | 0.09 | 0.066 | 1 | 7 | Pot1b | 0.024 | Old |
| 0.266072 | 0.130675 | 0.146 | 0.117 | 1 | 7 | Ccdc43 | 0.029 | Old |
| 0.266298 | 0.364763 | 0.236 | 0.202 | 1 | 7 | Commd2 | 0.034 | Old |
| 0.266536 | 0.00437 | 0.156 | 0.187 | 1 | 7 | Rsbn1 | -0.031 | Old |
| 0.266591 | 0.084004 | 0.849 | 0.828 | 1 | 7 | Erp29 | 0.021 | Old |
| 0.266716 | 0.976938 | 0.023 | 0.012 | 1 | 7 | Ctif | 0.011 | Old |
| 0.266716 | 0.368012 | 0.023 | 0.012 | 1 | 7 | Mansc1 | 0.011 | Old |
| 0.266916 | 0.313581 | 0.477 | 0.44 | 1 | 7 | Sulf2 | 0.037 | Old |
| 0.267126 | 0.375491 | 0.113 | 0.087 | 1 | 7 | 2310001H1 | 0.026 | Old |

| | | | | | | | | |
|---|---|---|---|---|---|---|---|---|
| 0.267157 | 0.328378 | 0.3 | 0.265 | 1 | 7 | Rreb1 | 0.035 | Old |
| 0.267261 | 0.135088 | 0.382 | 0.34 | 1 | 7 | Erp44 | 0.042 | Old |
| 0.267378 | 0.097532 | 0.721 | 0.675 | 1 | 7 | Ppp2ca | 0.046 | Old |
| 0.267386 | 0.30356 | 0.023 | 0.012 | 1 | 7 | Sdr42e1 | 0.011 | Old |
| 0.267571 | 0.089103 | 0.072 | 0.051 | 1 | 7 | Heatr5b | 0.021 | Old |
| 0.267736 | 0.043312 | 0.367 | 0.319 | 1 | 7 | Imp3 | 0.048 | Old |
| 0.267977 | 0.063556 | 0.597 | 0.533 | 1 | 7 | Rbm8a | 0.064 | Old |
| 0.268057 | 0.376992 | 0.136 | 0.108 | 1 | 7 | Ddt | 0.028 | Old |
| 0.268143 | 0.585071 | 0.133 | 0.108 | 1 | 7 | Xist | 0.025 | Old |
| 0.268162 | 0.320937 | 0.372 | 0.334 | 1 | 7 | Pin4 | 0.038 | Old |
| 0.268315 | 0.035298 | 0.354 | 0.298 | 1 | 7 | Gmps | 0.056 | Old |
| 0.268372 | 0.150941 | 0.103 | 0.078 | 1 | 7 | Ralgapa2 | 0.025 | Old |
| 0.26867 | 0.212945 | 0.413 | 0.352 | 1 | 7 | Mrps15 | 0.061 | Old |
| 0.268682 | 0.539783 | 0.103 | 0.078 | 1 | 7 | Slc38a7 | 0.025 | Old |
| 0.268729 | 0.550351 | 0.023 | 0.012 | 1 | 7 | AI480526 | 0.011 | Old |
| 0.269102 | 0.580073 | 0.046 | 0.03 | 1 | 7 | Cables1 | 0.016 | Old |
| 0.269195 | 0.206809 | 0.221 | 0.19 | 1 | 7 | Dctn5 | 0.031 | Old |
| 0.269196 | 0.162491 | 0.556 | 0.503 | 1 | 7 | Vapa | 0.053 | Old |
| 0.269413 | 0.339209 | 0.274 | 0.235 | 1 | 7 | Chaf1a | 0.039 | Old |
| 0.269551 | 0.136389 | 0.318 | 0.283 | 1 | 7 | Slc39a7 | 0.035 | Old |
| 0.269734 | 0.113925 | 0.054 | 0.036 | 1 | 7 | Ccdc28a | 0.018 | Old |
| 0.269749 | 0.094014 | 0.192 | 0.16 | 1 | 7 | Ppt2 | 0.032 | Old |
| 0.270031 | 0.497515 | 0.046 | 0.03 | 1 | 7 | Eml2 | 0.016 | Old |
| 0.270413 | 0.941947 | 0.038 | 0.024 | 1 | 7 | E130311K1 | 0.014 | Old |
| 0.270475 | 0.935709 | 0.087 | 0.066 | 1 | 7 | Pgghg | 0.021 | Old |
| 0.270617 | 0.074494 | 0.146 | 0.114 | 1 | 7 | Zfp407 | 0.032 | Old |
| 0.270794 | 0.548857 | 0.138 | 0.111 | 1 | 7 | Apip | 0.027 | Old |
| 0.270817 | 0.30217 | 0.082 | 0.06 | 1 | 7 | Pparg | 0.022 | Old |
| 0.270904 | 0.116143 | 0.072 | 0.051 | 1 | 7 | Lin52 | 0.021 | Old |
| 0.270919 | 0.888992 | 0.031 | 0.018 | 1 | 7 | A430106G1 | 0.013 | Old |
| 0.270925 | 0.679018 | 0.038 | 0.024 | 1 | 7 | Vars2 | 0.014 | Old |
| 0.271145 | 0.121994 | 0.323 | 0.28 | 1 | 7 | Otud4 | 0.043 | Old |
| 0.271753 | 0.351344 | 0.262 | 0.223 | 1 | 7 | Hivep1 | 0.039 | Old |
| 0.271852 | 0.050228 | 0.141 | 0.111 | 1 | 7 | Slc9a1 | 0.03 | Old |
| 0.272074 | 0.586171 | 0.031 | 0.018 | 1 | 7 | 1700056E2 | 0.013 | Old |
| 0.272074 | 0.545568 | 0.031 | 0.018 | 1 | 7 | 9130221H1 | 0.013 | Old |
| 0.272079 | 0.496677 | 0.156 | 0.127 | 1 | 7 | Mtmr1 | 0.029 | Old |
| 0.272197 | 0.254482 | 0.19 | 0.157 | 1 | 7 | Lgals8 | 0.033 | Old |
| 0.272408 | 0.065237 | 0.221 | 0.184 | 1 | 7 | Cep55 | 0.037 | Old |
| 0.272653 | 0.637268 | 0.031 | 0.018 | 1 | 7 | Bend3 | 0.013 | Old |
| 0.272664 | 0.249978 | 0.274 | 0.235 | 1 | 7 | Rgl2 | 0.039 | Old |
| 0.272829 | 0.210867 | 0.046 | 0.03 | 1 | 7 | Klc4 | 0.016 | Old |
| 0.273048 | 0.063801 | 0.679 | 0.611 | 1 | 7 | Raph1 | 0.068 | Old |
| 0.273088 | 0.119587 | 0.351 | 0.295 | 1 | 7 | Frg1 | 0.056 | Old |
| 0.273091 | 0.014598 | 0.118 | 0.09 | 1 | 7 | Paip2b | 0.028 | Old |

| | | | | | | | | |
|---|---|---|---|---|---|---|---|---|
| 0.273257 | 0.227909 | 0.082 | 0.06 | 1 | 7 | Hus1 | 0.022 | Old |
| 0.273334 | 0.202638 | 0.272 | 0.232 | 1 | 7 | Arhgap1 | 0.04 | Old |
| 0.273372 | 0.086221 | 0.364 | 0.307 | 1 | 7 | Zfp644 | 0.057 | Old |
| 0.273451 | 0.272598 | 0.318 | 0.286 | 1 | 7 | Cit | 0.032 | Old |
| 0.273507 | 0.237889 | 0.482 | 0.425 | 1 | 7 | Psma5 | 0.057 | Old |
| 0.273687 | 0.175939 | 0.231 | 0.193 | 1 | 7 | Ddx19a | 0.038 | Old |
| 0.273967 | 0.166279 | 0.426 | 0.37 | 1 | 7 | Sp3 | 0.056 | Old |
| 0.274063 | 0.143394 | 0.136 | 0.108 | 1 | 7 | Lztr1 | 0.028 | Old |
| 0.274396 | 0.591649 | 0.031 | 0.018 | 1 | 7 | Atp6v0c | 0.013 | Old |
| 0.274828 | 0.132751 | 0.362 | 0.319 | 1 | 7 | Armc8 | 0.043 | Old |
| 0.274978 | 1.374497 | 0.031 | 0.018 | 1 | 7 | Pdia5 | 0.013 | Old |
| 0.274978 | 0.551398 | 0.031 | 0.018 | 1 | 7 | Stbd1 | 0.013 | Old |
| 0.275156 | 0.155648 | 0.341 | 0.292 | 1 | 7 | Nsun2 | 0.049 | Old |
| 0.275344 | 0.125844 | 0.9 | 0.889 | 1 | 7 | Npc2 | 0.011 | Old |
| 0.275392 | 0.448022 | 0.072 | 0.051 | 1 | 7 | Oip5 | 0.021 | Old |
| 0.275576 | 0.17139 | 0.21 | 0.175 | 1 | 7 | Braf | 0.035 | Old |
| 0.275638 | 0.035088 | 0.192 | 0.157 | 1 | 7 | Trim12a | 0.035 | Old |
| 0.275942 | 0.267357 | 0.421 | 0.364 | 1 | 7 | Ipo7 | 0.057 | Old |
| 0.275986 | 0.707147 | 0.136 | 0.111 | 1 | 7 | Scyl1 | 0.025 | Old |
| 0.276281 | 0.312906 | 0.644 | 0.596 | 1 | 7 | Cd84 | 0.048 | Old |
| 0.276433 | 0.011029 | 0.474 | 0.416 | 1 | 7 | Pde4b | 0.058 | Old |
| 0.27665 | 0.051893 | 0.174 | 0.139 | 1 | 7 | Adpgk | 0.035 | Old |
| 0.276875 | 0.053506 | 0.428 | 0.361 | 1 | 7 | Gas2l3 | 0.067 | Old |
| 0.276932 | 0.405574 | 0.085 | 0.063 | 1 | 7 | Alkbh8 | 0.022 | Old |
| 0.276988 | 0.152416 | 0.615 | 0.605 | 1 | 7 | Hspa9 | 0.01 | Old |
| 0.277285 | 0.214819 | 0.064 | 0.045 | 1 | 7 | 3110002H1 | 0.019 | Old |
| 0.277737 | 0.315823 | 0.192 | 0.163 | 1 | 7 | Ipo8 | 0.029 | Old |
| 0.27779 | 0.199808 | 0.267 | 0.229 | 1 | 7 | Nfyb | 0.038 | Old |
| 0.277832 | 0.476415 | 0.149 | 0.12 | 1 | 7 | Cfap36 | 0.029 | Old |
| 0.277903 | 0.486777 | 0.031 | 0.018 | 1 | 7 | Neto2 | 0.013 | Old |
| 0.277903 | 0.450695 | 0.031 | 0.018 | 1 | 7 | Ldlrad3 | 0.013 | Old |
| 0.27806 | 0.117369 | 0.126 | 0.099 | 1 | 7 | Zfp991 | 0.027 | Old |
| 0.278095 | 0.007763 | 0.582 | 0.527 | 1 | 7 | Dusp3 | 0.055 | Old |
| 0.278208 | 0.069267 | 0.259 | 0.22 | 1 | 7 | Tbc1d31 | 0.039 | Old |
| 0.278622 | 0.197089 | 0.318 | 0.277 | 1 | 7 | Snrpa | 0.041 | Old |
| 0.278654 | 0.037494 | 0.131 | 0.102 | 1 | 7 | Trim11 | 0.029 | Old |
| 0.27965 | 0.18123 | 0.615 | 0.554 | 1 | 7 | Tubb6 | 0.061 | Old |
| 0.279698 | 0.175943 | 0.19 | 0.16 | 1 | 7 | Slc25a30 | 0.03 | Old |
| 0.2799 | 0.100187 | 0.705 | 0.669 | 1 | 7 | Prkar1a | 0.036 | Old |
| 0.279904 | 0.286822 | 0.292 | 0.253 | 1 | 7 | Rara | 0.039 | Old |
| 0.279968 | 0.033689 | 0.6 | 0.533 | 1 | 7 | Apbb1ip | 0.067 | Old |
| 0.279999 | 0.533919 | 0.105 | 0.081 | 1 | 7 | Flnb | 0.024 | Old |
| 0.28008 | 0.249683 | 0.367 | 0.322 | 1 | 7 | Syf2 | 0.045 | Old |
| 0.280258 | 0.58529 | 0.031 | 0.018 | 1 | 7 | 1810055G( | 0.013 | Old |
| 0.280845 | 0.131439 | 0.177 | 0.145 | 1 | 7 | Slc30a9 | 0.032 | Old |

| | | | | | | | | |
|---|---|---|---|---|---|---|---|---|
| 0.280849 | 0.393027 | 0.031 | 0.018 | 1 | 7 | Zfp606 | 0.013 | Old |
| 0.281347 | 0.205148 | 0.144 | 0.114 | 1 | 7 | Rrp15 | 0.03 | Old |
| 0.281613 | 0.401405 | 0.323 | 0.286 | 1 | 7 | Evl | 0.037 | Old |
| 0.281787 | 0.223318 | 0.172 | 0.139 | 1 | 7 | Zbtb7b | 0.033 | Old |
| 0.28183 | 0.322133 | 0.205 | 0.175 | 1 | 7 | Sympk | 0.03 | Old |
| 0.282032 | 0.234994 | 0.344 | 0.304 | 1 | 7 | Sdf2l1 | 0.04 | Old |
| 0.282034 | 0.192776 | 0.031 | 0.018 | 1 | 7 | Fbxo32 | 0.013 | Old |
| 0.28215 | 0.21529 | 0.308 | 0.271 | 1 | 7 | Stom | 0.037 | Old |
| 0.282165 | 0.074212 | 0.377 | 0.331 | 1 | 7 | Mgea5 | 0.046 | Old |
| 0.282504 | 0.218457 | 0.174 | 0.145 | 1 | 7 | Mtmr2 | 0.029 | Old |
| 0.282577 | 0.535095 | 0.067 | 0.048 | 1 | 7 | Sco1 | 0.019 | Old |
| 0.283221 | 1.276397 | 0.031 | 0.018 | 1 | 7 | Tmtc2 | 0.013 | Old |
| 0.28341 | 0.209388 | 0.038 | 0.024 | 1 | 7 | Hyal1 | 0.014 | Old |
| 0.283565 | 0.080421 | 0.479 | 0.431 | 1 | 7 | Tmem189 | 0.048 | Old |
| 0.283651 | 0.023031 | 0.315 | 0.262 | 1 | 7 | Bicd2 | 0.053 | Old |
| 0.283656 | 0.51383 | 0.141 | 0.114 | 1 | 7 | Gpatch11 | 0.027 | Old |
| 0.283938 | 0.284198 | 0.038 | 0.024 | 1 | 7 | Gpr19 | 0.014 | Old |
| 0.284115 | 0.770508 | 0.138 | 0.111 | 1 | 7 | Gbp2 | 0.027 | Old |
| 0.284148 | 0.31141 | 0.236 | 0.205 | 1 | 7 | Sgo2a | 0.031 | Old |
| 0.284281 | 0.641845 | 0.085 | 0.063 | 1 | 7 | Tnfaip8l1 | 0.022 | Old |
| 0.284383 | 0.096549 | 0.2 | 0.166 | 1 | 7 | Mrpl45 | 0.034 | Old |
| 0.284997 | 0.118009 | 0.038 | 0.024 | 1 | 7 | Syn1 | 0.014 | Old |
| 0.285099 | 0.240762 | 0.197 | 0.169 | 1 | 7 | Fbxw7 | 0.028 | Old |
| 0.285128 | 0.065254 | 0.579 | 0.53 | 1 | 7 | Tax1bp1 | 0.049 | Old |
| 0.285221 | 0.053975 | 0.123 | 0.096 | 1 | 7 | Ccdc167 | 0.027 | Old |
| 0.285326 | 0.579562 | 0.082 | 0.06 | 1 | 7 | Ogfod2 | 0.022 | Old |
| 0.285359 | 0.147246 | 0.246 | 0.208 | 1 | 7 | Rad51 | 0.038 | Old |
| 0.285607 | 0.08582 | 0.031 | 0.018 | 1 | 7 | C130013H( | 0.013 | Old |
| 0.285615 | 0.10676 | 0.126 | 0.099 | 1 | 7 | Mbtps2 | 0.027 | Old |
| 0.286202 | 0.260939 | 0.126 | 0.099 | 1 | 7 | Ddx55 | 0.027 | Old |
| 0.286238 | 0.076232 | 0.246 | 0.208 | 1 | 7 | Hirip3 | 0.038 | Old |
| 0.286493 | 0.366518 | 0.295 | 0.256 | 1 | 7 | Erh | 0.039 | Old |
| 0.286788 | 0.419527 | 0.079 | 0.06 | 1 | 7 | Ddx51 | 0.019 | Old |
| 0.28709 | 0.394446 | 0.056 | 0.039 | 1 | 7 | Aunip | 0.017 | Old |
| 0.287307 | 0.135388 | 0.274 | 0.235 | 1 | 7 | Arpc5l | 0.039 | Old |
| 0.287329 | 0.0844 | 0.81 | 0.789 | 1 | 7 | Prcp | 0.021 | Old |
| 0.28752 | 0.143594 | 0.603 | 0.539 | 1 | 7 | Cox17 | 0.064 | Old |
| 0.287622 | 0.03938 | 0.577 | 0.53 | 1 | 7 | Serinc1 | 0.047 | Old |
| 0.288192 | 0.123835 | 0.554 | 0.482 | 1 | 7 | Snrpd2 | 0.072 | Old |
| 0.288836 | 0.26395 | 0.056 | 0.039 | 1 | 7 | Polm | 0.017 | Old |
| 0.28918 | 0.059525 | 0.39 | 0.343 | 1 | 7 | Tspan32 | 0.047 | Old |
| 0.28926 | 0.275507 | 0.038 | 0.024 | 1 | 7 | Spred3 | 0.014 | Old |
| 0.289293 | 0.293844 | 0.108 | 0.084 | 1 | 7 | Lancl2 | 0.024 | Old |
| 0.289888 | 0.922734 | 0.059 | 0.042 | 1 | 7 | Atf5 | 0.017 | Old |
| 0.289893 | 0.101397 | 0.862 | 0.858 | 1 | 7 | Atp5e | 0.004 | Old |

| | | | | | | | | |
|---|---|---|---|---|---|---|---|---|
| 0.289988 | 0.165086 | 0.213 | 0.178 | 1 | 7 | Ezh1 | 0.035 | Old |
| 0.290006 | 0.110277 | 0.408 | 0.358 | 1 | 7 | Rnps1 | 0.05 | Old |
| 0.290008 | 0.058831 | 0.464 | 0.413 | 1 | 7 | Prkcsh | 0.051 | Old |
| 0.290329 | 0.089395 | 0.141 | 0.111 | 1 | 7 | Patz1 | 0.03 | Old |
| 0.290339 | 0.199643 | 0.318 | 0.283 | 1 | 7 | Rap2c | 0.035 | Old |
| 0.290589 | 0.142967 | 0.056 | 0.039 | 1 | 7 | Cby1 | 0.017 | Old |
| 0.291155 | 0.241406 | 0.105 | 0.081 | 1 | 7 | Piga | 0.024 | Old |
| 0.291202 | 0.533784 | 0.108 | 0.084 | 1 | 7 | Arhgef18 | 0.024 | Old |
| 0.291222 | 0.171257 | 0.39 | 0.337 | 1 | 7 | Ilf3 | 0.053 | Old |
| 0.291965 | 0.255556 | 0.223 | 0.19 | 1 | 7 | Tmem5 | 0.033 | Old |
| 0.292076 | 0.108203 | 0.2 | 0.169 | 1 | 7 | Bin2 | 0.031 | Old |
| 0.292272 | 0.411474 | 0.077 | 0.057 | 1 | 7 | Fancm | 0.02 | Old |
| 0.292291 | 0.760806 | 0.049 | 0.033 | 1 | 7 | Msl3l2 | 0.016 | Old |
| 0.292349 | 0.418264 | 0.056 | 0.039 | 1 | 7 | Fpr2 | 0.017 | Old |
| 0.292376 | 0.132415 | 0.095 | 0.072 | 1 | 7 | Nt5dc1 | 0.023 | Old |
| 0.2929 | 0.000646 | 0.241 | 0.205 | 1 | 7 | Timm44 | 0.036 | Old |
| 0.293063 | 0.236412 | 0.154 | 0.127 | 1 | 7 | Mxd3 | 0.027 | Old |
| 0.293273 | 0.321481 | 0.203 | 0.169 | 1 | 7 | Vav2 | 0.034 | Old |
| 0.293318 | 0.663347 | 0.059 | 0.042 | 1 | 7 | Zfp53 | 0.017 | Old |
| 0.293462 | 0.054466 | 0.262 | 0.223 | 1 | 7 | Znfx1 | 0.039 | Old |
| 0.294103 | 0.102453 | 0.19 | 0.16 | 1 | 7 | Map1lc3a | 0.03 | Old |
| 0.294197 | 0.164028 | 0.262 | 0.22 | 1 | 7 | 2210016F1 | 0.042 | Old |
| 0.294311 | 0.142763 | 0.71 | 0.651 | 1 | 7 | Il6ra | 0.059 | Old |
| 0.294508 | 0.203426 | 0.197 | 0.169 | 1 | 7 | S100pbp | 0.028 | Old |
| 0.294651 | 0.245612 | 0.3 | 0.265 | 1 | 7 | Meaf6 | 0.035 | Old |
| 0.29467 | 0.883047 | 0.049 | 0.033 | 1 | 7 | Papss2 | 0.016 | Old |
| 0.29469 | 0.09227 | 0.195 | 0.16 | 1 | 7 | Fbxo28 | 0.035 | Old |
| 0.295094 | 0.061022 | 0.231 | 0.196 | 1 | 7 | Lrrc8d | 0.035 | Old |
| 0.295576 | 0.50713 | 0.138 | 0.114 | 1 | 7 | Emc4 | 0.024 | Old |
| 0.295783 | 0.005802 | 0.374 | 0.325 | 1 | 7 | Plekha2 | 0.049 | Old |
| 0.295968 | 0.029091 | 0.285 | 0.238 | 1 | 7 | Pigk | 0.047 | Old |
| 0.296252 | 0.185889 | 0.228 | 0.193 | 1 | 7 | Hps5 | 0.035 | Old |
| 0.296264 | 0.473048 | 0.192 | 0.16 | 1 | 7 | Nsl1 | 0.032 | Old |
| 0.296303 | 0.246525 | 0.126 | 0.099 | 1 | 7 | Kat2a | 0.027 | Old |
| 0.29665 | 0.133734 | 0.226 | 0.193 | 1 | 7 | Kctd2 | 0.033 | Old |
| 0.296681 | 0.126527 | 0.603 | 0.572 | 1 | 7 | Srrm1 | 0.031 | Old |
| 0.297122 | 0.109608 | 0.062 | 0.081 | 1 | 7 | Wdr45 | -0.019 | Old |
| 0.297205 | 0.205823 | 0.126 | 0.099 | 1 | 7 | Srgap3 | 0.027 | Old |
| 0.297242 | 0.256835 | 0.103 | 0.078 | 1 | 7 | Hbegf | 0.025 | Old |
| 0.297461 | 0.490083 | 0.067 | 0.048 | 1 | 7 | Vps33b | 0.019 | Old |
| 0.297942 | 0.116625 | 0.541 | 0.5 | 1 | 7 | Rpl36al | 0.041 | Old |
| 0.298213 | 0.142182 | 0.677 | 0.639 | 1 | 7 | P4hb | 0.038 | Old |
| 0.298518 | 0.083306 | 0.097 | 0.123 | 1 | 7 | Alg13 | -0.026 | Old |
| 0.298642 | 0.015708 | 0.085 | 0.063 | 1 | 7 | Osbpl1a | 0.022 | Old |
| 0.298767 | 0.584621 | 0.172 | 0.142 | 1 | 7 | Insig2 | 0.03 | Old |

| | | | | | | | | |
|---|---|---|---|---|---|---|---|---|
| 0.298825 | 0.147336 | 0.164 | 0.133 | 1 | 7 | 1110008P1 | 0.031 | Old |
| 0.298889 | 0.238935 | 0.095 | 0.072 | 1 | 7 | Ccdc122 | 0.023 | Old |
| 0.299197 | 0.117351 | 0.118 | 0.093 | 1 | 7 | Tmem104 | 0.025 | Old |
| 0.299381 | 0.185925 | 0.41 | 0.367 | 1 | 7 | Ppp6r1 | 0.043 | Old |
| 0.299461 | 0.046384 | 0.056 | 0.039 | 1 | 7 | Hmgn3 | 0.017 | Old |
| 0.299724 | 0.07127 | 0.433 | 0.398 | 1 | 7 | Rnasel | 0.035 | Old |
| 0.300237 | 0.09593 | 0.244 | 0.208 | 1 | 7 | Sqor | 0.036 | Old |
| 0.300277 | 0.136751 | 0.492 | 0.443 | 1 | 7 | Gm16286 | 0.049 | Old |
| 0.300396 | 0.190567 | 0.467 | 0.416 | 1 | 7 | Rp9 | 0.051 | Old |
| 0.300759 | 0.196131 | 0.397 | 0.358 | 1 | 7 | Aurkaip1 | 0.039 | Old |
| 0.300815 | 0.19159 | 0.167 | 0.136 | 1 | 7 | Diablo | 0.031 | Old |
| 0.301159 | 0.187652 | 0.067 | 0.048 | 1 | 7 | Rabif | 0.019 | Old |
| 0.301221 | 0.076243 | 0.374 | 0.331 | 1 | 7 | Ubr2 | 0.043 | Old |
| 0.301274 | 0.005875 | 0.285 | 0.241 | 1 | 7 | Zfc3h1 | 0.044 | Old |
| 0.301319 | 0.470582 | 0.11 | 0.087 | 1 | 7 | Pigc | 0.023 | Old |
| 0.301418 | 0.009299 | 0.285 | 0.244 | 1 | 7 | Eed | 0.041 | Old |
| 0.301569 | 0.006772 | 0.423 | 0.364 | 1 | 7 | Arrdc1 | 0.059 | Old |
| 0.301666 | 0.130095 | 0.213 | 0.178 | 1 | 7 | Ddx47 | 0.035 | Old |
| 0.302165 | 0.162881 | 0.077 | 0.057 | 1 | 7 | Slc22a15 | 0.02 | Old |
| 0.302355 | 0.098642 | 0.095 | 0.072 | 1 | 7 | Etfrf1 | 0.023 | Old |
| 0.302673 | 0.198156 | 0.195 | 0.163 | 1 | 7 | Cask | 0.032 | Old |
| 0.303034 | 0.222047 | 0.369 | 0.322 | 1 | 7 | Nucb1 | 0.047 | Old |
| 0.303063 | 0.41794 | 0.354 | 0.319 | 1 | 7 | Zcchc24 | 0.035 | Old |
| 0.303571 | 0.033524 | 0.969 | 0.988 | 1 | 7 | Cfl1 | -0.019 | Old |
| 0.303573 | 0.233928 | 0.118 | 0.093 | 1 | 7 | Polr3b | 0.025 | Old |
| 0.304124 | 0.111953 | 0.308 | 0.268 | 1 | 7 | Polr2c | 0.04 | Old |
| 0.304144 | 1.224275 | 0.018 | 0.009 | 1 | 7 | Fgfrl1 | 0.009 | Old |
| 0.304327 | 0.089021 | 0.731 | 0.699 | 1 | 7 | Cnbp | 0.032 | Old |
| 0.304496 | 0.036338 | 0.608 | 0.548 | 1 | 7 | Ier3ip1 | 0.06 | Old |
| 0.304707 | 1.034839 | 0.033 | 0.021 | 1 | 7 | Kazald1 | 0.012 | Old |
| 0.304977 | 0.758313 | 0.018 | 0.009 | 1 | 7 | Cyp39a1 | 0.009 | Old |
| 0.305092 | 0.178909 | 0.151 | 0.127 | 1 | 7 | Dpm2 | 0.024 | Old |
| 0.305169 | 0.043167 | 0.974 | 0.967 | 1 | 7 | H2-Ab1 | 0.007 | Old |
| 0.305321 | 0.142686 | 0.759 | 0.684 | 1 | 7 | Rap1a | 0.075 | Old |
| 0.305368 | 0.088458 | 0.069 | 0.051 | 1 | 7 | Clec4d | 0.018 | Old |
| 0.305461 | 0.118345 | 0.338 | 0.295 | 1 | 7 | Srp54a | 0.043 | Old |
| 0.305468 | 0.250135 | 0.167 | 0.139 | 1 | 7 | Pfkfb4 | 0.028 | Old |
| 0.305481 | 0.096722 | 0.469 | 0.422 | 1 | 7 | Rnf4 | 0.047 | Old |
| 0.305494 | 0.055247 | 0.308 | 0.268 | 1 | 7 | Pola1 | 0.04 | Old |
| 0.305812 | 1.196548 | 0.018 | 0.009 | 1 | 7 | Uprt | 0.009 | Old |
| 0.305812 | 0.539828 | 0.018 | 0.009 | 1 | 7 | Serping1 | 0.009 | Old |
| 0.30595 | 0.147899 | 0.608 | 0.56 | 1 | 7 | Mapk1 | 0.048 | Old |
| 0.306012 | 0.076977 | 0.9 | 0.922 | 1 | 7 | Eif4g2 | -0.022 | Old |
| 0.306243 | 0.07301 | 0.156 | 0.13 | 1 | 7 | Cnnm3 | 0.026 | Old |
| 0.306458 | 0.205185 | 0.29 | 0.256 | 1 | 7 | Rad50 | 0.034 | Old |

| | | | | | | | | |
|---|---|---|---|---|---|---|---|---|
| 0.306648 | 0.45994 | 0.018 | 0.009 | 1 | 7 | Mmab | 0.009 | Old |
| 0.306836 | 0.210835 | 0.279 | 0.25 | 1 | 7 | Tlnrd1 | 0.029 | Old |
| 0.306865 | 0.497376 | 0.059 | 0.042 | 1 | 7 | Scimp | 0.017 | Old |
| 0.307221 | 0.092588 | 0.103 | 0.081 | 1 | 7 | Rgcc | 0.022 | Old |
| 0.30732 | 0.10509 | 0.262 | 0.226 | 1 | 7 | Cdk12 | 0.036 | Old |
| 0.307481 | 0.165184 | 0.628 | 0.593 | 1 | 7 | Ywhaq | 0.035 | Old |
| 0.307486 | 0.86232 | 0.018 | 0.009 | 1 | 7 | Hist1h3a | 0.009 | Old |
| 0.307486 | 0.411515 | 0.018 | 0.009 | 1 | 7 | Pcgf2 | 0.009 | Old |
| 0.307532 | 0.081434 | 0.172 | 0.145 | 1 | 7 | Exoc6b | 0.027 | Old |
| 0.307654 | 1.191131 | 0.041 | 0.027 | 1 | 7 | Bcorl1 | 0.014 | Old |
| 0.307708 | 0.062748 | 0.251 | 0.22 | 1 | 7 | Aim2 | 0.031 | Old |
| 0.307831 | 0.060662 | 0.233 | 0.202 | 1 | 7 | Imp4 | 0.031 | Old |
| 0.307944 | 0.142284 | 0.128 | 0.102 | 1 | 7 | Slc35a2 | 0.026 | Old |
| 0.308004 | 0.143552 | 0.567 | 0.497 | 1 | 7 | Hspa4 | 0.07 | Old |
| 0.30819 | 0.832738 | 0.041 | 0.027 | 1 | 7 | Nod1 | 0.014 | Old |
| 0.308197 | 0.176666 | 0.059 | 0.042 | 1 | 7 | Tmed1 | 0.017 | Old |
| 0.308281 | 1.052042 | 0.033 | 0.021 | 1 | 7 | Ttc30a1 | 0.012 | Old |
| 0.308325 | 0.510155 | 0.018 | 0.009 | 1 | 7 | Erbb3 | 0.009 | Old |
| 0.308325 | 0.387937 | 0.018 | 0.009 | 1 | 7 | Zscan22 | 0.009 | Old |
| 0.308375 | 0.557382 | 0.09 | 0.069 | 1 | 7 | Dgcr14 | 0.021 | Old |
| 0.308695 | 0.035647 | 0.277 | 0.235 | 1 | 7 | Polr2l | 0.042 | Old |
| 0.309002 | 0.196524 | 0.779 | 0.783 | 1 | 7 | H2afy | -0.004 | Old |
| 0.309165 | 0.581458 | 0.018 | 0.009 | 1 | 7 | Mpzl3 | 0.009 | Old |
| 0.309165 | 0.197083 | 0.018 | 0.009 | 1 | 7 | 1110002L0 | 0.009 | Old |
| 0.30931 | 0.513558 | 0.005 | 0.012 | 1 | 7 | E430024P1 | -0.007 | Old |
| 0.309508 | 0.284178 | 0.192 | 0.163 | 1 | 7 | Slc13a3 | 0.029 | Old |
| 0.309932 | 0.283556 | 0.249 | 0.217 | 1 | 7 | Pbx2 | 0.032 | Old |
| 0.310075 | 0.248248 | 0.262 | 0.235 | 1 | 7 | Nfkb2 | 0.027 | Old |
| 0.310149 | 0.07821 | 0.838 | 0.777 | 1 | 7 | S100a6 | 0.061 | Old |
| 0.310429 | 0.358752 | 0.018 | 0.009 | 1 | 7 | Gm10642 | 0.009 | Old |
| 0.31077 | 0.403276 | 0.079 | 0.06 | 1 | 7 | Clpb | 0.019 | Old |
| 0.310907 | 1.094573 | 0.09 | 0.069 | 1 | 7 | Dffa | 0.021 | Old |
| 0.310965 | 0.171875 | 0.105 | 0.081 | 1 | 7 | Bpnt1 | 0.024 | Old |
| 0.310965 | 0.090061 | 0.105 | 0.081 | 1 | 7 | Il7r | 0.024 | Old |
| 0.31097 | 0.259509 | 0.121 | 0.096 | 1 | 7 | Coq8b | 0.025 | Old |
| 0.311084 | 0.04735 | 0.11 | 0.087 | 1 | 7 | Ttc37 | 0.023 | Old |
| 0.311322 | 0.66959 | 0.059 | 0.042 | 1 | 7 | Tor4a | 0.017 | Old |
| 0.311482 | 1.260004 | 0.005 | 0.012 | 1 | 7 | Kif5a | -0.007 | Old |
| 0.311612 | 0.121354 | 0.646 | 0.623 | 1 | 7 | Cct8 | 0.023 | Old |
| 0.311696 | 0.299088 | 0.018 | 0.009 | 1 | 7 | Satb1 | 0.009 | Old |
| 0.311696 | 0.282157 | 0.018 | 0.009 | 1 | 7 | Spry2 | 0.009 | Old |
| 0.31174 | 0.468079 | 0.133 | 0.108 | 1 | 7 | Mastl | 0.025 | Old |
| 0.311821 | 0.036349 | 0.19 | 0.157 | 1 | 7 | Naa30 | 0.033 | Old |
| 0.311853 | 0.189867 | 0.2 | 0.169 | 1 | 7 | Ankib1 | 0.031 | Old |
| 0.311883 | 0.8666 | 0.033 | 0.021 | 1 | 7 | Pecam1 | 0.012 | Old |

| | | | | | | | | |
|---|---|---|---|---|---|---|---|---|
| 0.311949 | 0.195724 | 0.228 | 0.196 | 1 | 7 | Rab3gap2 | 0.032 | Old |
| 0.312031 | 0.13095 | 0.367 | 0.322 | 1 | 7 | H2-M3 | 0.045 | Old |
| 0.312118 | 0.008799 | 0.597 | 0.548 | 1 | 7 | Trip12 | 0.049 | Old |
| 0.312265 | 0.255097 | 0.177 | 0.151 | 1 | 7 | Bid | 0.026 | Old |
| 0.312543 | 0.209426 | 0.018 | 0.009 | 1 | 7 | Gm45894 | 0.009 | Old |
| 0.312543 | 0.061884 | 0.018 | 0.009 | 1 | 7 | Abca5 | 0.009 | Old |
| 0.312572 | 0.865651 | 0.005 | 0.012 | 1 | 7 | Bicc1 | -0.007 | Old |
| 0.312572 | 0.14027 | 0.005 | 0.012 | 1 | 7 | Gm26590 | -0.007 | Old |
| 0.312572 | 0.066085 | 0.005 | 0.012 | 1 | 7 | Rom1 | -0.007 | Old |
| 0.312823 | 0.006676 | 0.915 | 0.901 | 1 | 7 | Dazap2 | 0.014 | Old |
| 0.312837 | 0.152901 | 0.177 | 0.148 | 1 | 7 | Dhps | 0.029 | Old |
| 0.312913 | 0.076674 | 0.095 | 0.072 | 1 | 7 | Mthfs | 0.023 | Old |
| 0.313269 | 0.097302 | 0.272 | 0.235 | 1 | 7 | Phf20 | 0.037 | Old |
| 0.313587 | 0.382066 | 0.041 | 0.027 | 1 | 7 | Pex12 | 0.014 | Old |
| 0.313667 | 0.157476 | 0.618 | 0.563 | 1 | 7 | Ndufa13 | 0.055 | Old |
| 0.313735 | 0.006657 | 0.118 | 0.093 | 1 | 7 | Crbn | 0.025 | Old |
| 0.313798 | 0.194976 | 0.256 | 0.226 | 1 | 7 | Ube2d1 | 0.03 | Old |
| 0.314211 | 0.900437 | 0.026 | 0.015 | 1 | 7 | 4930555A( | 0.011 | Old |
| 0.314674 | 0.363125 | 0.041 | 0.027 | 1 | 7 | Fut4 | 0.014 | Old |
| 0.314905 | 0.759131 | 0.033 | 0.021 | 1 | 7 | Pter | 0.012 | Old |
| 0.314907 | 0.383629 | 0.118 | 0.096 | 1 | 7 | Gramd1b | 0.022 | Old |
| 0.314977 | 0.168441 | 0.356 | 0.313 | 1 | 7 | Plaur | 0.043 | Old |
| 0.315218 | 0.035232 | 0.041 | 0.027 | 1 | 7 | Eef1akmt4 | 0.014 | Old |
| 0.316258 | 0.070726 | 0.067 | 0.048 | 1 | 7 | Cbr4 | 0.019 | Old |
| 0.31627 | 0.114117 | 0.582 | 0.536 | 1 | 7 | S100a11 | 0.046 | Old |
| 0.316306 | 0.697909 | 0.026 | 0.015 | 1 | 7 | Otud3 | 0.011 | Old |
| 0.316776 | 0.180534 | 0.479 | 0.434 | 1 | 7 | Cebpg | 0.045 | Old |
| 0.317006 | 0.839536 | 0.026 | 0.015 | 1 | 7 | Slc25a42 | 0.011 | Old |
| 0.317022 | 0.499146 | 0.077 | 0.057 | 1 | 7 | Oas1a | 0.02 | Old |
| 0.317059 | 0.888757 | 0.051 | 0.036 | 1 | 7 | Mylk | 0.015 | Old |
| 0.317109 | 0.059786 | 0.933 | 0.961 | 1 | 7 | Arpc2 | -0.028 | Old |
| 0.317572 | 0.148487 | 0.795 | 0.795 | 1 | 7 | Csde1 | 0 | Old |
| 0.317708 | 0.823078 | 0.026 | 0.015 | 1 | 7 | Gdpgp1 | 0.011 | Old |
| 0.318059 | 0.650578 | 0.026 | 0.015 | 1 | 7 | Nme2 | 0.011 | Old |
| 0.318585 | 0.325052 | 0.09 | 0.069 | 1 | 7 | Chst7 | 0.021 | Old |
| 0.318719 | 0.0424 | 0.159 | 0.13 | 1 | 7 | Zfp24 | 0.029 | Old |
| 0.319006 | 0.545221 | 0.051 | 0.036 | 1 | 7 | Zfp953 | 0.015 | Old |
| 0.319114 | 0.697235 | 0.026 | 0.015 | 1 | 7 | Rpgr | 0.011 | Old |
| 0.319169 | 0.861871 | 0.033 | 0.021 | 1 | 7 | Cpne8 | 0.012 | Old |
| 0.319262 | 0.088775 | 0.187 | 0.154 | 1 | 7 | Rbm34 | 0.033 | Old |
| 0.319451 | 0.103363 | 0.374 | 0.334 | 1 | 7 | Cuedc2 | 0.04 | Old |
| 0.319511 | 0.522925 | 0.103 | 0.081 | 1 | 7 | D430042O( | 0.022 | Old |
| 0.319781 | 0.451268 | 0.033 | 0.021 | 1 | 7 | Tbc1d19 | 0.012 | Old |
| 0.320023 | 0.092534 | 0.179 | 0.148 | 1 | 7 | Specc1 | 0.031 | Old |
| 0.320025 | 0.399415 | 0.156 | 0.13 | 1 | 7 | Armc5 | 0.026 | Old |

| | | | | | | | | |
|---|---|---|---|---|---|---|---|---|
| 0.320171 | 0.143099 | 0.254 | 0.22 | 1 | 7 | Bola3 | 0.034 | Old |
| 0.320487 | 0.782767 | 0.113 | 0.09 | 1 | 7 | Rrs1 | 0.023 | Old |
| 0.32056 | 0.937613 | 0.072 | 0.054 | 1 | 7 | Wdr92 | 0.018 | Old |
| 0.320751 | 0.129395 | 0.231 | 0.196 | 1 | 7 | Mut | 0.035 | Old |
| 0.320819 | 0.355015 | 0.336 | 0.304 | 1 | 7 | Ifitm6 | 0.032 | Old |
| 0.321055 | 0.601473 | 0.11 | 0.087 | 1 | 7 | Sorl1 | 0.023 | Old |
| 0.32123 | 0.543818 | 0.026 | 0.015 | 1 | 7 | Leng9 | 0.011 | Old |
| 0.321406 | 0.670402 | 0.126 | 0.105 | 1 | 7 | Slc35f5 | 0.021 | Old |
| 0.321461 | 0.151482 | 0.592 | 0.536 | 1 | 7 | Cnot6l | 0.056 | Old |
| 0.322431 | 0.320413 | 0.051 | 0.036 | 1 | 7 | Unc13d | 0.015 | Old |
| 0.322556 | 0.144693 | 0.233 | 0.196 | 1 | 7 | Atad1 | 0.037 | Old |
| 0.322597 | 0.225502 | 0.177 | 0.145 | 1 | 7 | Cdc7 | 0.032 | Old |
| 0.322666 | 0.04694 | 0.513 | 0.443 | 1 | 7 | Fcho2 | 0.07 | Old |
| 0.323258 | 0.061819 | 0.59 | 0.548 | 1 | 7 | Prr13 | 0.042 | Old |
| 0.323562 | 0.301093 | 0.213 | 0.184 | 1 | 7 | Slc36a4 | 0.029 | Old |
| 0.324052 | 0.065863 | 0.685 | 0.651 | 1 | 7 | Uqcr10 | 0.034 | Old |
| 0.324077 | 0.337936 | 0.092 | 0.072 | 1 | 7 | Lrch4 | 0.02 | Old |
| 0.324154 | 0.157022 | 0.582 | 0.554 | 1 | 7 | Eif4ebp2 | 0.028 | Old |
| 0.324186 | 0.377847 | 0.185 | 0.154 | 1 | 7 | Isg20l2 | 0.031 | Old |
| 0.324216 | 0.221475 | 0.321 | 0.286 | 1 | 7 | Ndfip1 | 0.035 | Old |
| 0.324579 | 0.208187 | 0.095 | 0.075 | 1 | 7 | Pigv | 0.02 | Old |
| 0.324654 | 0.007758 | 0.233 | 0.193 | 1 | 7 | Taf13 | 0.04 | Old |
| 0.324705 | 0.059606 | 0.187 | 0.157 | 1 | 7 | Phkg2 | 0.03 | Old |
| 0.324806 | 0.061613 | 0.21 | 0.181 | 1 | 7 | Plekhf2 | 0.029 | Old |
| 0.325058 | 0.170548 | 0.121 | 0.096 | 1 | 7 | Usp31 | 0.025 | Old |
| 0.325073 | 0.406736 | 0.167 | 0.193 | 1 | 7 | Vps53 | -0.026 | Old |
| 0.325231 | 0.166565 | 0.641 | 0.587 | 1 | 7 | Mcub | 0.054 | Old |
| 0.325596 | 0.896746 | 0.044 | 0.03 | 1 | 7 | Aldh1b1 | 0.014 | Old |
| 0.325715 | 0.179277 | 0.264 | 0.226 | 1 | 7 | Ubp1 | 0.038 | Old |
| 0.326037 | 0.077198 | 0.344 | 0.298 | 1 | 7 | Isy1 | 0.046 | Old |
| 0.326203 | 0.396731 | 0.026 | 0.015 | 1 | 7 | Fut10 | 0.011 | Old |
| 0.326203 | 0.25186 | 0.026 | 0.015 | 1 | 7 | Nhej1 | 0.011 | Old |
| 0.326283 | 0.467531 | 0.092 | 0.072 | 1 | 7 | Slc2a8 | 0.02 | Old |
| 0.326429 | 0.082676 | 0.592 | 0.539 | 1 | 7 | Lbh | 0.053 | Old |
| 0.326817 | 0.437654 | 0.287 | 0.259 | 1 | 7 | Ube2j1 | 0.028 | Old |
| 0.326904 | 0.135991 | 0.238 | 0.211 | 1 | 7 | Cmip | 0.027 | Old |
| 0.327168 | 0.110396 | 0.905 | 0.855 | 1 | 7 | Arf5 | 0.05 | Old |
| 0.327355 | 0.13058 | 0.541 | 0.503 | 1 | 7 | U2surp | 0.038 | Old |
| 0.327365 | 0.35389 | 0.051 | 0.036 | 1 | 7 | Cutc | 0.015 | Old |
| 0.327634 | 1.358054 | 0.026 | 0.015 | 1 | 7 | Myo1b | 0.011 | Old |
| 0.328039 | 0.047581 | 0.349 | 0.304 | 1 | 7 | Tulp4 | 0.045 | Old |
| 0.328203 | 0.072127 | 0.444 | 0.47 | 1 | 7 | Pik3r1 | -0.026 | Old |
| 0.328285 | 0.284551 | 0.097 | 0.075 | 1 | 7 | Zfp942 | 0.022 | Old |
| 0.328358 | 0.720912 | 0.051 | 0.036 | 1 | 7 | Kbtbd3 | 0.015 | Old |
| 0.329142 | 0.761905 | 0.074 | 0.057 | 1 | 7 | Gm20342 | 0.017 | Old |

| | | | | | | | | |
|---|---|---|---|---|---|---|---|---|
| 0.329233 | 0.645532 | 0.062 | 0.045 | 1 | 7 | Xlr4b | 0.017 | Old |
| 0.3293 | 0.057976 | 0.215 | 0.184 | 1 | 7 | Nop16 | 0.031 | Old |
| 0.329382 | 0.773777 | 0.231 | 0.208 | 1 | 7 | Pla2g2d | 0.023 | Old |
| 0.329534 | 0.010387 | 0.756 | 0.696 | 1 | 7 | Pbrm1 | 0.06 | Old |
| 0.329624 | 0.209605 | 0.187 | 0.16 | 1 | 7 | Tmem11 | 0.027 | Old |
| 0.329708 | 0.087078 | 0.072 | 0.054 | 1 | 7 | Jpx | 0.018 | Old |
| 0.329761 | 0.623714 | 0.123 | 0.099 | 1 | 7 | Tcp11l2 | 0.024 | Old |
| 0.329973 | 0.222381 | 0.295 | 0.25 | 1 | 7 | Ptdss1 | 0.045 | Old |
| 0.329974 | 0.173979 | 0.508 | 0.476 | 1 | 7 | Vma21 | 0.032 | Old |
| 0.330002 | 0.089069 | 0.246 | 0.214 | 1 | 7 | Rnaseh2a | 0.032 | Old |
| 0.3303 | 0.053597 | 0.551 | 0.506 | 1 | 7 | Znrf2 | 0.045 | Old |
| 0.330383 | 0.150214 | 0.418 | 0.373 | 1 | 7 | Aprt | 0.045 | Old |
| 0.330848 | 0.338136 | 0.051 | 0.036 | 1 | 7 | Etfbkmt | 0.015 | Old |
| 0.331198 | 0.207201 | 0.164 | 0.136 | 1 | 7 | Rnf14 | 0.028 | Old |
| 0.331439 | 0.12135 | 0.6 | 0.557 | 1 | 7 | Mfsd1 | 0.043 | Old |
| 0.331553 | 0.311841 | 0.244 | 0.211 | 1 | 7 | Ddx52 | 0.033 | Old |
| 0.331892 | 0.026731 | 0.649 | 0.611 | 1 | 7 | Reep3 | 0.038 | Old |
| 0.33207 | 0.660356 | 0.044 | 0.03 | 1 | 7 | Wdr53 | 0.014 | Old |
| 0.332312 | 0.109207 | 0.551 | 0.494 | 1 | 7 | Hp1bp3 | 0.057 | Old |
| 0.332425 | 0.162582 | 0.221 | 0.19 | 1 | 7 | Akap8 | 0.031 | Old |
| 0.332715 | 0.080878 | 0.154 | 0.127 | 1 | 7 | Pex5 | 0.027 | Old |
| 0.332722 | 0.333352 | 0.208 | 0.178 | 1 | 7 | Nop9 | 0.03 | Old |
| 0.33287 | 0.450972 | 0.062 | 0.045 | 1 | 7 | Dusp19 | 0.017 | Old |
| 0.332874 | 0.107524 | 0.179 | 0.151 | 1 | 7 | Nom1 | 0.028 | Old |
| 0.333033 | 0.25138 | 0.149 | 0.175 | 1 | 7 | Tmem88 | -0.026 | Old |
| 0.333189 | 0.111715 | 0.362 | 0.316 | 1 | 7 | Birc2 | 0.046 | Old |
| 0.333281 | 0.291798 | 0.462 | 0.425 | 1 | 7 | Pdcd10 | 0.037 | Old |
| 0.333286 | 0.141687 | 0.313 | 0.277 | 1 | 7 | Cops2 | 0.036 | Old |
| 0.333291 | 0.101288 | 0.208 | 0.175 | 1 | 7 | Hspa13 | 0.033 | Old |
| 0.333335 | 0.155981 | 0.518 | 0.488 | 1 | 7 | Notch2 | 0.03 | Old |
| 0.333537 | 0.111311 | 0.749 | 0.696 | 1 | 7 | Pdcd6ip | 0.053 | Old |
| 0.333896 | 0.302474 | 0.503 | 0.482 | 1 | 7 | Ahcyl2 | 0.021 | Old |
| 0.333934 | 0.123609 | 0.569 | 0.524 | 1 | 7 | Lsm5 | 0.045 | Old |
| 0.333967 | 0.090685 | 0.136 | 0.111 | 1 | 7 | Adora2b | 0.025 | Old |
| 0.333986 | 0.301921 | 0.205 | 0.175 | 1 | 7 | Fbxo18 | 0.03 | Old |
| 0.334131 | 0.447091 | 0.254 | 0.22 | 1 | 7 | Hmg20b | 0.034 | Old |
| 0.334422 | 0.328041 | 0.079 | 0.06 | 1 | 7 | Nsun3 | 0.019 | Old |
| 0.334792 | 0.707659 | 0.044 | 0.03 | 1 | 7 | Slc7a5 | 0.014 | Old |
| 0.334924 | 0.2512 | 0.182 | 0.154 | 1 | 7 | Vps54 | 0.028 | Old |
| 0.334931 | 0.033836 | 0.162 | 0.136 | 1 | 7 | Mfn2 | 0.026 | Old |
| 0.335084 | 0.261855 | 0.295 | 0.265 | 1 | 7 | Zranb1 | 0.03 | Old |
| 0.335151 | 0.135182 | 0.1 | 0.078 | 1 | 7 | Arl11 | 0.022 | Old |
| 0.33534 | 0.065962 | 0.654 | 0.608 | 1 | 7 | Ap2m1 | 0.046 | Old |
| 0.335884 | 0.776273 | 0.044 | 0.03 | 1 | 7 | Plcxd2 | 0.014 | Old |
| 0.335918 | 0.163476 | 0.464 | 0.425 | 1 | 7 | Ivns1abp | 0.039 | Old |

| | | | | | | | | |
|---|---|---|---|---|---|---|---|---|
| 0.336145 | 0.06019 | 0.295 | 0.259 | 1 | 7 | Atp11b | 0.036 | Old |
| 0.336699 | 0.196978 | 0.515 | 0.473 | 1 | 7 | Snx9 | 0.042 | Old |
| 0.336837 | 0.192768 | 0.149 | 0.123 | 1 | 7 | Ddx58 | 0.026 | Old |
| 0.337374 | 0.582942 | 0.085 | 0.066 | 1 | 7 | Rad51d | 0.019 | Old |
| 0.337414 | 0.399173 | 0.082 | 0.063 | 1 | 7 | Acot13 | 0.019 | Old |
| 0.338038 | 0.080817 | 0.869 | 0.843 | 1 | 7 | Mob1a | 0.026 | Old |
| 0.338187 | 0.064998 | 0.11 | 0.133 | 1 | 7 | Mlycd | -0.023 | Old |
| 0.338213 | 0.390707 | 0.082 | 0.063 | 1 | 7 | Qpctl | 0.019 | Old |
| 0.338496 | 0.263106 | 0.149 | 0.123 | 1 | 7 | Brms1 | 0.026 | Old |
| 0.33852 | 0.402006 | 0.095 | 0.075 | 1 | 7 | Wiz | 0.02 | Old |
| 0.338682 | 0.128265 | 0.069 | 0.051 | 1 | 7 | Capn5 | 0.018 | Old |
| 0.338909 | 0.292969 | 0.251 | 0.217 | 1 | 7 | Nab2 | 0.034 | Old |
| 0.338921 | 0.154982 | 0.123 | 0.099 | 1 | 7 | Hmces | 0.024 | Old |
| 0.339769 | 0.146799 | 0.297 | 0.265 | 1 | 7 | Pnrc2 | 0.032 | Old |
| 0.339915 | 0.054626 | 0.051 | 0.036 | 1 | 7 | Gins3 | 0.015 | Old |
| 0.340162 | 0.249598 | 0.108 | 0.087 | 1 | 7 | Hps4 | 0.021 | Old |
| 0.340264 | 0.127978 | 0.403 | 0.361 | 1 | 7 | Gtf2a1 | 0.042 | Old |
| 0.340512 | 0.136892 | 0.079 | 0.06 | 1 | 7 | Gin1 | 0.019 | Old |
| 0.340614 | 0.090269 | 0.082 | 0.063 | 1 | 7 | Fam136a | 0.019 | Old |
| 0.341202 | 0.140407 | 0.115 | 0.093 | 1 | 7 | Tmem107 | 0.022 | Old |
| 0.341722 | 0.572056 | 0.2 | 0.175 | 1 | 7 | Ppfibp2 | 0.025 | Old |
| 0.342451 | 1.077665 | 0.036 | 0.024 | 1 | 7 | Prmt6 | 0.012 | Old |
| 0.342542 | 0.065543 | 0.062 | 0.045 | 1 | 7 | Prkca | 0.017 | Old |
| 0.343341 | 0.157804 | 0.097 | 0.075 | 1 | 7 | Nlrc5 | 0.022 | Old |
| 0.343624 | 0.13611 | 0.338 | 0.298 | 1 | 7 | Znhit1 | 0.04 | Old |
| 0.343813 | 0.133632 | 0.431 | 0.401 | 1 | 7 | Ccnd3 | 0.03 | Old |
| 0.344907 | 0.185868 | 0.254 | 0.226 | 1 | 7 | Pofut1 | 0.028 | Old |
| 0.345217 | 0.315267 | 0.133 | 0.111 | 1 | 7 | Cpped1 | 0.022 | Old |
| 0.345432 | 0.049123 | 0.315 | 0.28 | 1 | 7 | Tm2d2 | 0.035 | Old |
| 0.345683 | 0.018398 | 0.454 | 0.407 | 1 | 7 | Ppp1r21 | 0.047 | Old |
| 0.345806 | 0.077251 | 0.244 | 0.211 | 1 | 7 | Tmem222 | 0.033 | Old |
| 0.346261 | 0.342915 | 0.082 | 0.063 | 1 | 7 | Elac1 | 0.019 | Old |
| 0.34638 | 0.193722 | 0.044 | 0.03 | 1 | 7 | Rpusd1 | 0.014 | Old |
| 0.346664 | 0.360296 | 0.113 | 0.09 | 1 | 7 | Mfng | 0.023 | Old |
| 0.347151 | 0.33009 | 0.185 | 0.157 | 1 | 7 | Mink1 | 0.028 | Old |
| 0.347242 | 0.12242 | 0.167 | 0.142 | 1 | 7 | Slf1 | 0.025 | Old |
| 0.347497 | 0.332541 | 0.044 | 0.03 | 1 | 7 | Cd300lf | 0.014 | Old |
| 0.347499 | 0.310926 | 0.221 | 0.193 | 1 | 7 | Rhoq | 0.028 | Old |
| 0.348404 | 0.953157 | 0.013 | 0.006 | 1 | 7 | Tcea2 | 0.007 | Old |
| 0.348464 | 0.541047 | 0.085 | 0.066 | 1 | 7 | Pkp3 | 0.019 | Old |
| 0.348688 | 0.133508 | 0.103 | 0.081 | 1 | 7 | Cd79b | 0.022 | Old |
| 0.348833 | 0.409699 | 0.128 | 0.105 | 1 | 7 | Ksr1 | 0.023 | Old |
| 0.349365 | 0.026684 | 0.769 | 0.708 | 1 | 7 | Tpr | 0.061 | Old |
| 0.349638 | 0.177918 | 0.159 | 0.136 | 1 | 7 | Psd4 | 0.023 | Old |
| 0.349738 | 0.140675 | 0.044 | 0.03 | 1 | 7 | Them4 | 0.014 | Old |

| | | | | | | | | |
|---|---|---|---|---|---|---|---|---|
| 0.349809 | 0.018765 | 0.464 | 0.416 | 1 | 7 | Dusp11 | 0.048 | Old |
| 0.35003 | 0.025663 | 0.062 | 0.045 | 1 | 7 | Tmem115 | 0.017 | Old |
| 0.350458 | 0.885362 | 0.036 | 0.024 | 1 | 7 | Ftx | 0.012 | Old |
| 0.350566 | 0.051329 | 0.138 | 0.114 | 1 | 7 | Def6 | 0.024 | Old |
| 0.350577 | 1.382814 | 0.013 | 0.006 | 1 | 7 | Bvht | 0.007 | Old |
| 0.350577 | 0.767742 | 0.013 | 0.006 | 1 | 7 | Zfp808 | 0.007 | Old |
| 0.350969 | 0.109139 | 0.326 | 0.286 | 1 | 7 | Sys1 | 0.04 | Old |
| 0.35119 | 0.030955 | 0.221 | 0.19 | 1 | 7 | Ssbp1 | 0.031 | Old |
| 0.351199 | 0.243436 | 0.092 | 0.072 | 1 | 7 | Extl2 | 0.02 | Old |
| 0.351204 | 0.199125 | 0.485 | 0.449 | 1 | 7 | Bcas2 | 0.036 | Old |
| 0.35125 | 0.030441 | 0.536 | 0.497 | 1 | 7 | Larp1 | 0.039 | Old |
| 0.351332 | 0.134637 | 0.477 | 0.425 | 1 | 7 | Huwe1 | 0.052 | Old |
| 0.351571 | 0.207574 | 0.736 | 0.726 | 1 | 7 | Ifitm2 | 0.01 | Old |
| 0.351667 | 0.760798 | 0.013 | 0.006 | 1 | 7 | Gm38336 | 0.007 | Old |
| 0.351679 | 0.160776 | 0.569 | 0.521 | 1 | 7 | Cdc42se2 | 0.048 | Old |
| 0.352255 | 0.046581 | 0.146 | 0.12 | 1 | 7 | Abcd2 | 0.026 | Old |
| 0.352512 | 0.088053 | 0.721 | 0.705 | 1 | 7 | Fam111a | 0.016 | Old |
| 0.352759 | 0.969091 | 0.013 | 0.006 | 1 | 7 | Spa17 | 0.007 | Old |
| 0.352759 | 0.90646 | 0.013 | 0.006 | 1 | 7 | 9330020H( | 0.007 | Old |
| 0.352759 | 0.903943 | 0.013 | 0.006 | 1 | 7 | Kdm8 | 0.007 | Old |
| 0.352759 | 0.702908 | 0.013 | 0.006 | 1 | 7 | Dync2li1 | 0.007 | Old |
| 0.352759 | 0.66248 | 0.013 | 0.006 | 1 | 7 | Icos | 0.007 | Old |
| 0.352759 | 0.443469 | 0.013 | 0.006 | 1 | 7 | Ssc4d | 0.007 | Old |
| 0.353194 | 0.559959 | 0.087 | 0.069 | 1 | 7 | Sirpb1c | 0.018 | Old |
| 0.353298 | 0.035335 | 0.492 | 0.449 | 1 | 7 | Api5 | 0.043 | Old |
| 0.353569 | 0.688374 | 0.036 | 0.024 | 1 | 7 | Ifi213 | 0.012 | Old |
| 0.353569 | 0.358544 | 0.036 | 0.024 | 1 | 7 | Rbak | 0.012 | Old |
| 0.353853 | 0.662281 | 0.013 | 0.006 | 1 | 7 | Spns2 | 0.007 | Old |
| 0.353853 | 0.437863 | 0.013 | 0.006 | 1 | 7 | Peak1os | 0.007 | Old |
| 0.353853 | 0.420115 | 0.013 | 0.006 | 1 | 7 | Spns3 | 0.007 | Old |
| 0.353853 | 0.253832 | 0.013 | 0.006 | 1 | 7 | Gm30025 | 0.007 | Old |
| 0.354385 | 0.840149 | 0.008 | 0.015 | 1 | 7 | Dnal1 | -0.007 | Old |
| 0.354576 | 0.1395 | 0.226 | 0.196 | 1 | 7 | Dhfr | 0.03 | Old |
| 0.354949 | 0.390743 | 0.013 | 0.006 | 1 | 7 | Mtfp1 | 0.007 | Old |
| 0.35568 | 0.08454 | 0.123 | 0.099 | 1 | 7 | Rnf31 | 0.024 | Old |
| 0.35597 | 0.60283 | 0.067 | 0.051 | 1 | 7 | Zfyve19 | 0.016 | Old |
| 0.356047 | 0.472304 | 0.013 | 0.006 | 1 | 7 | 5033406O( | 0.007 | Old |
| 0.356047 | 0.463363 | 0.013 | 0.006 | 1 | 7 | Ahrr | 0.007 | Old |
| 0.356047 | 0.35343 | 0.013 | 0.006 | 1 | 7 | Gm13012 | 0.007 | Old |
| 0.356047 | 0.322015 | 0.013 | 0.006 | 1 | 7 | Dner | 0.007 | Old |
| 0.356047 | 0.28931 | 0.013 | 0.006 | 1 | 7 | Gm15834 | 0.007 | Old |
| 0.356047 | 0.166091 | 0.013 | 0.006 | 1 | 7 | Plagl1 | 0.007 | Old |
| 0.356047 | 0.121691 | 0.013 | 0.006 | 1 | 7 | Zfp316 | 0.007 | Old |
| 0.35607 | 0.489203 | 0.036 | 0.024 | 1 | 7 | Thg1l | 0.012 | Old |
| 0.356199 | 0.02746 | 0.303 | 0.262 | 1 | 7 | Il6st | 0.041 | Old |

| | | | | | | | | |
|---|---|---|---|---|---|---|---|---|
| 0.356201 | 0.268666 | 0.459 | 0.416 | 1 | 7 | Sae1 | 0.043 | Old |
| 0.356206 | 0.028444 | 0.341 | 0.301 | 1 | 7 | Brd3 | 0.04 | Old |
| 0.356537 | 0.558109 | 0.162 | 0.139 | 1 | 7 | Sos1 | 0.023 | Old |
| 0.356902 | 0.360005 | 0.218 | 0.19 | 1 | 7 | Aldh16a1 | 0.028 | Old |
| 0.356986 | 0.107225 | 0.074 | 0.057 | 1 | 7 | Nprl2 | 0.017 | Old |
| 0.357083 | 0.402105 | 0.044 | 0.03 | 1 | 7 | Rpgrip1l | 0.014 | Old |
| 0.357126 | 0.199962 | 0.582 | 0.548 | 1 | 7 | Rtraf | 0.034 | Old |
| 0.357148 | 0.509602 | 0.013 | 0.006 | 1 | 7 | Zfp11 | 0.007 | Old |
| 0.357148 | 0.376108 | 0.013 | 0.006 | 1 | 7 | Tspan7 | 0.007 | Old |
| 0.357325 | 0.749718 | 0.036 | 0.024 | 1 | 7 | Pms1 | 0.012 | Old |
| 0.357325 | 0.380195 | 0.036 | 0.024 | 1 | 7 | Numbl | 0.012 | Old |
| 0.357813 | 0.031255 | 0.203 | 0.172 | 1 | 7 | Ppil1 | 0.031 | Old |
| 0.357918 | 0.101012 | 0.521 | 0.473 | 1 | 7 | Cisd2 | 0.048 | Old |
| 0.35834 | 0.038684 | 0.297 | 0.259 | 1 | 7 | Tk1 | 0.038 | Old |
| 0.358379 | 0.408707 | 0.105 | 0.084 | 1 | 7 | Ddx20 | 0.021 | Old |
| 0.358406 | 0.090134 | 0.679 | 0.627 | 1 | 7 | Matr3 | 0.052 | Old |
| 0.358457 | 0.131609 | 0.331 | 0.292 | 1 | 7 | Cr1l | 0.039 | Old |
| 0.358523 | 0.078623 | 0.115 | 0.093 | 1 | 7 | Mtfr2 | 0.022 | Old |
| 0.358968 | 0.202486 | 0.085 | 0.066 | 1 | 7 | Ublcp1 | 0.019 | Old |
| 0.359031 | 0.417939 | 0.238 | 0.208 | 1 | 7 | Lrrk1 | 0.03 | Old |
| 0.35913 | 0.290838 | 0.149 | 0.123 | 1 | 7 | Hsf2 | 0.026 | Old |
| 0.359355 | 0.120041 | 0.013 | 0.006 | 1 | 7 | Gm21988 | 0.007 | Old |
| 0.359355 | 0.043137 | 0.013 | 0.006 | 1 | 7 | Timm23 | 0.007 | Old |
| 0.359465 | 0.240903 | 0.274 | 0.244 | 1 | 7 | Fbxo3 | 0.03 | Old |
| 0.359529 | 0.124278 | 0.428 | 0.41 | 1 | 7 | Uhmk1 | 0.018 | Old |
| 0.359822 | 0.113359 | 0.679 | 0.648 | 1 | 7 | Syncrip | 0.031 | Old |
| 0.359843 | 0.733639 | 0.036 | 0.024 | 1 | 7 | Gm20703 | 0.012 | Old |
| 0.360289 | 0.190061 | 0.282 | 0.247 | 1 | 7 | Dpagt1 | 0.035 | Old |
| 0.360474 | 0.81567 | 0.036 | 0.024 | 1 | 7 | Gorasp1 | 0.012 | Old |
| 0.360626 | 0.0899 | 0.156 | 0.133 | 1 | 7 | Phf19 | 0.023 | Old |
| 0.36065 | 0.307829 | 0.141 | 0.117 | 1 | 7 | Ginm1 | 0.024 | Old |
| 0.360716 | 0.113485 | 0.159 | 0.133 | 1 | 7 | Tmem19 | 0.026 | Old |
| 0.361078 | 0.028335 | 0.144 | 0.117 | 1 | 7 | Dhdds | 0.027 | Old |
| 0.361488 | 0.029441 | 0.421 | 0.364 | 1 | 7 | Park7 | 0.057 | Old |
| 0.362044 | 0.129817 | 0.431 | 0.398 | 1 | 7 | Cacybp | 0.033 | Old |
| 0.362052 | 0.115957 | 0.115 | 0.093 | 1 | 7 | Pcnx3 | 0.022 | Old |
| 0.362163 | 0.790211 | 0.028 | 0.018 | 1 | 7 | Trim16 | 0.01 | Old |
| 0.362167 | 0.235493 | 0.138 | 0.114 | 1 | 7 | Ccdc86 | 0.024 | Old |
| 0.362372 | 0.504572 | 0.036 | 0.024 | 1 | 7 | Mtif3 | 0.012 | Old |
| 0.362882 | 1.525504 | 0.028 | 0.018 | 1 | 7 | Tmem38a | 0.01 | Old |
| 0.362885 | 0.005817 | 0.931 | 0.94 | 1 | 7 | Itm2b | -0.009 | Old |
| 0.363078 | 0.106419 | 0.074 | 0.057 | 1 | 7 | Ipcef1 | 0.017 | Old |
| 0.363151 | 0.021208 | 0.405 | 0.355 | 1 | 7 | Cd302 | 0.05 | Old |
| 0.36386 | 0.044643 | 0.321 | 0.277 | 1 | 7 | Fam162a | 0.044 | Old |
| 0.363893 | 0.017662 | 0.977 | 0.976 | 1 | 7 | Rpl38 | 0.001 | Old |

| | | | | | | | | |
|---|---|---|---|---|---|---|---|---|
| 0.363975 | 0.21284 | 0.21 | 0.184 | 1 | 7 | Mak16 | 0.026 | Old |
| 0.364021 | 0.226195 | 0.151 | 0.127 | 1 | 7 | Eml3 | 0.024 | Old |
| 0.364276 | 0.313651 | 0.036 | 0.024 | 1 | 7 | Lysmd1 | 0.012 | Old |
| 0.364847 | 0.092327 | 0.246 | 0.217 | 1 | 7 | Gtf2h1 | 0.029 | Old |
| 0.365007 | 0.126189 | 0.295 | 0.262 | 1 | 7 | Yipf4 | 0.033 | Old |
| 0.365123 | 0.297784 | 0.149 | 0.123 | 1 | 7 | Mta3 | 0.026 | Old |
| 0.365141 | 0.008489 | 0.482 | 0.425 | 1 | 7 | Cxcr4 | 0.057 | Old |
| 0.365285 | 0.249991 | 0.074 | 0.093 | 1 | 7 | Gm44751 | -0.019 | Old |
| 0.365574 | 0.160517 | 0.546 | 0.515 | 1 | 7 | Mcm5 | 0.031 | Old |
| 0.365632 | 0.015151 | 0.233 | 0.199 | 1 | 7 | Mplkip | 0.034 | Old |
| 0.365678 | 1.434575 | 0.021 | 0.012 | 1 | 7 | Ikzf2 | 0.009 | Old |
| 0.365678 | 0.969093 | 0.021 | 0.012 | 1 | 7 | Gm20513 | 0.009 | Old |
| 0.365873 | 0.105898 | 0.128 | 0.105 | 1 | 7 | Ttc33 | 0.023 | Old |
| 0.365991 | 0.243863 | 0.459 | 0.428 | 1 | 7 | Vwa5a | 0.031 | Old |
| 0.366171 | 0.126808 | 0.113 | 0.09 | 1 | 7 | Adsl | 0.023 | Old |
| 0.366388 | 0.20287 | 0.108 | 0.087 | 1 | 7 | Cwc27 | 0.021 | Old |
| 0.366722 | 0.107303 | 0.618 | 0.563 | 1 | 7 | Xiap | 0.055 | Old |
| 0.366893 | 0.096735 | 0.092 | 0.114 | 1 | 7 | Zfp219 | -0.022 | Old |
| 0.366914 | 0.130256 | 0.054 | 0.039 | 1 | 7 | Acsf3 | 0.015 | Old |
| 0.367042 | 0.325026 | 0.064 | 0.048 | 1 | 7 | Alg2 | 0.016 | Old |
| 0.367314 | 0.114868 | 0.777 | 0.795 | 1 | 7 | Uqcr11 | -0.018 | Old |
| 0.367394 | 1.107537 | 0.021 | 0.012 | 1 | 7 | Mcf2l | 0.009 | Old |
| 0.367619 | 0.176007 | 0.2 | 0.172 | 1 | 7 | Eif2b5 | 0.028 | Old |
| 0.367737 | 0.027741 | 0.646 | 0.62 | 1 | 7 | Top1 | 0.026 | Old |
| 0.36821 | 0.016836 | 0.449 | 0.401 | 1 | 7 | Ddx24 | 0.048 | Old |
| 0.368252 | 0.273247 | 0.128 | 0.105 | 1 | 7 | Phtf2 | 0.023 | Old |
| 0.368326 | 0.089209 | 0.379 | 0.343 | 1 | 7 | Pikfyve | 0.036 | Old |
| 0.368429 | 0.29294 | 0.505 | 0.464 | 1 | 7 | Mif | 0.041 | Old |
| 0.3686 | 0.14707 | 0.772 | 0.753 | 1 | 7 | Rpl22l1 | 0.019 | Old |
| 0.368715 | 0.056214 | 0.138 | 0.114 | 1 | 7 | Vti1a | 0.024 | Old |
| 0.368812 | 0.256283 | 0.097 | 0.078 | 1 | 7 | Trove2 | 0.019 | Old |
| 0.36885 | 0.37206 | 0.067 | 0.051 | 1 | 7 | Gpatch1 | 0.016 | Old |
| 0.369297 | 0.030782 | 0.962 | 0.958 | 1 | 7 | Rack1 | 0.004 | Old |
| 0.369456 | 0.158912 | 0.487 | 0.446 | 1 | 7 | Calu | 0.041 | Old |
| 0.369486 | 0.258866 | 0.285 | 0.253 | 1 | 7 | Senp1 | 0.032 | Old |
| 0.369596 | 0.738169 | 0.046 | 0.033 | 1 | 7 | Slc52a3 | 0.013 | Old |
| 0.369616 | 0.061304 | 0.128 | 0.105 | 1 | 7 | Zbp1 | 0.023 | Old |
| 0.369937 | 0.067756 | 0.251 | 0.22 | 1 | 7 | Rapgef2 | 0.031 | Old |
| 0.369977 | 1.597122 | 0.021 | 0.012 | 1 | 7 | Klhl8 | 0.009 | Old |
| 0.369977 | 1.134612 | 0.021 | 0.012 | 1 | 7 | 2610318N( | 0.009 | Old |
| 0.369991 | 0.074064 | 0.156 | 0.13 | 1 | 7 | Pam16 | 0.026 | Old |
| 0.370116 | 0.541913 | 0.028 | 0.018 | 1 | 7 | Dnm1 | 0.01 | Old |
| 0.370148 | 0.075372 | 0.451 | 0.407 | 1 | 7 | Rbm5 | 0.044 | Old |
| 0.370499 | 0.555949 | 0.085 | 0.066 | 1 | 7 | Mrpl39 | 0.019 | Old |
| 0.370811 | 0.376286 | 0.208 | 0.187 | 1 | 7 | Gadd45gip | 0.021 | Old |

| | | | | | | | | |
|---|---|---|---|---|---|---|---|---|
| 0.370841 | 0.693917 | 0.021 | 0.012 | 1 | 7 | Gm4070 | 0.009 | Old |
| 0.371171 | 0.131101 | 0.146 | 0.12 | 1 | 7 | Thap11 | 0.026 | Old |
| 0.371382 | 0.528047 | 0.079 | 0.063 | 1 | 7 | Zbtb17 | 0.016 | Old |
| 0.371706 | 0.96262 | 0.021 | 0.012 | 1 | 7 | Rhoj | 0.009 | Old |
| 0.372572 | 0.863305 | 0.021 | 0.012 | 1 | 7 | Dlc1 | 0.009 | Old |
| 0.372582 | 0.018923 | 0.085 | 0.066 | 1 | 7 | Thop1 | 0.019 | Old |
| 0.372772 | 0.01189 | 0.254 | 0.22 | 1 | 7 | Rabggtb | 0.034 | Old |
| 0.372916 | 0.203734 | 0.538 | 0.503 | 1 | 7 | Fbl | 0.035 | Old |
| 0.373035 | 1.056954 | 0.028 | 0.018 | 1 | 7 | Nicn1 | 0.01 | Old |
| 0.373101 | 0.289078 | 0.405 | 0.38 | 1 | 7 | Nusap1 | 0.025 | Old |
| 0.373106 | 0.058366 | 0.397 | 0.355 | 1 | 7 | Rnf187 | 0.042 | Old |
| 0.373191 | 0.321928 | 0.167 | 0.142 | 1 | 7 | Eepd1 | 0.025 | Old |
| 0.374267 | 0.510979 | 0.295 | 0.265 | 1 | 7 | Eif1ad | 0.03 | Old |
| 0.374309 | 0.708487 | 0.021 | 0.012 | 1 | 7 | Amot | 0.009 | Old |
| 0.374314 | 0.060145 | 0.238 | 0.211 | 1 | 7 | Tmem141 | 0.027 | Old |
| 0.374557 | 0.289113 | 0.11 | 0.09 | 1 | 7 | Zdhhc13 | 0.02 | Old |
| 0.374601 | 0.031352 | 0.264 | 0.226 | 1 | 7 | Dhrs1 | 0.038 | Old |
| 0.37484 | 0.414421 | 0.123 | 0.102 | 1 | 7 | Slc30a1 | 0.021 | Old |
| 0.374972 | 0.205355 | 0.482 | 0.452 | 1 | 7 | Mrps33 | 0.03 | Old |
| 0.37538 | 0.013992 | 0.436 | 0.38 | 1 | 7 | Usp8 | 0.056 | Old |
| 0.375526 | 0.21524 | 0.105 | 0.084 | 1 | 7 | Tdp1 | 0.021 | Old |
| 0.375818 | 0.03446 | 0.328 | 0.292 | 1 | 7 | Gars | 0.036 | Old |
| 0.375864 | 0.112152 | 0.251 | 0.22 | 1 | 7 | Utp14a | 0.031 | Old |
| 0.375991 | 0.28945 | 0.182 | 0.157 | 1 | 7 | Mmp19 | 0.025 | Old |
| 0.37605 | 0.556178 | 0.021 | 0.012 | 1 | 7 | Cntd1 | 0.009 | Old |
| 0.37605 | 0.132684 | 0.021 | 0.012 | 1 | 7 | 2510046G1 | 0.009 | Old |
| 0.376293 | 0.143713 | 0.259 | 0.229 | 1 | 7 | Smarcad1 | 0.03 | Old |
| 0.376332 | 0.303178 | 0.077 | 0.06 | 1 | 7 | Nek1 | 0.017 | Old |
| 0.3767 | 0.2927 | 0.295 | 0.265 | 1 | 7 | Usp22 | 0.03 | Old |
| 0.37708 | 0.120321 | 0.81 | 0.816 | 1 | 7 | Rac1 | -0.006 | Old |
| 0.377116 | 0.210766 | 0.156 | 0.133 | 1 | 7 | Mapk9 | 0.023 | Old |
| 0.378158 | 0.302761 | 0.228 | 0.202 | 1 | 7 | Ryr1 | 0.026 | Old |
| 0.378207 | 0.1034 | 0.731 | 0.672 | 1 | 7 | Dock2 | 0.059 | Old |
| 0.378351 | 0.289308 | 0.087 | 0.069 | 1 | 7 | Stn1 | 0.018 | Old |
| 0.378531 | 0.10121 | 0.379 | 0.334 | 1 | 7 | Krcc1 | 0.045 | Old |
| 0.378671 | 0.213267 | 0.021 | 0.012 | 1 | 7 | Foxd2 | 0.009 | Old |
| 0.378671 | 0.009579 | 0.021 | 0.012 | 1 | 7 | 4931428F0 | 0.009 | Old |
| 0.378914 | 0.127641 | 0.028 | 0.018 | 1 | 7 | Cacna1a | 0.01 | Old |
| 0.378991 | 0.464235 | 0.11 | 0.09 | 1 | 7 | Ccne1 | 0.02 | Old |
| 0.37906 | 0.467809 | 0.3 | 0.271 | 1 | 7 | Rasgrp4 | 0.029 | Old |
| 0.379601 | 0.106742 | 0.179 | 0.154 | 1 | 7 | Fbxo45 | 0.025 | Old |
| 0.379892 | 0.22078 | 0.233 | 0.205 | 1 | 7 | Spen | 0.028 | Old |
| 0.379907 | 0.101984 | 0.118 | 0.096 | 1 | 7 | Manbal | 0.022 | Old |
| 0.380011 | 0.289194 | 0.087 | 0.069 | 1 | 7 | Rbm19 | 0.018 | Old |
| 0.380072 | 0.248813 | 0.272 | 0.241 | 1 | 7 | Bag6 | 0.031 | Old |

| | | | | | | | | |
|---|---|---|---|---|---|---|---|---|
| 0.38036 | 0.138958 | 0.046 | 0.033 | 1 | 7 | Zfp1 | 0.013 | Old |
| 0.380365 | 0.071311 | 0.777 | 0.771 | 1 | 7 | Sod1 | 0.006 | Old |
| 0.380425 | 0.589517 | 0.021 | 0.012 | 1 | 7 | Abtb2 | 0.009 | Old |
| 0.380509 | 0.265065 | 0.215 | 0.187 | 1 | 7 | Tbpl1 | 0.028 | Old |
| 0.380614 | 0.708928 | 0.074 | 0.057 | 1 | 7 | Zranb3 | 0.017 | Old |
| 0.380687 | 0.102196 | 0.541 | 0.506 | 1 | 7 | Irf5 | 0.035 | Old |
| 0.380752 | 0.028855 | 0.626 | 0.566 | 1 | 7 | Pds5a | 0.06 | Old |
| 0.381071 | 0.148372 | 0.518 | 0.479 | 1 | 7 | Mbnl2 | 0.039 | Old |
| 0.38109 | 0.022217 | 0.849 | 0.81 | 1 | 7 | Ssr3 | 0.039 | Old |
| 0.381158 | 0.081645 | 0.469 | 0.428 | 1 | 7 | Gnpda1 | 0.041 | Old |
| 0.381303 | 0.290188 | 0.021 | 0.012 | 1 | 7 | Polr3g | 0.009 | Old |
| 0.381355 | 0.512163 | 0.069 | 0.054 | 1 | 7 | Cdkal1 | 0.015 | Old |
| 0.381364 | 0.047625 | 0.987 | 0.991 | 1 | 7 | Rpsa | -0.004 | Old |
| 0.381504 | 1.060474 | 0.046 | 0.033 | 1 | 7 | Nmnat1 | 0.013 | Old |
| 0.381698 | 0.104547 | 0.713 | 0.672 | 1 | 7 | Rbm25 | 0.041 | Old |
| 0.381808 | 0.042889 | 0.221 | 0.19 | 1 | 7 | Lmbrd2 | 0.031 | Old |
| 0.38189 | 0.107103 | 0.733 | 0.729 | 1 | 7 | Arf1 | 0.004 | Old |
| 0.382038 | 0.11078 | 0.3 | 0.274 | 1 | 7 | Eif4e2 | 0.026 | Old |
| 0.382479 | 0.237838 | 0.169 | 0.142 | 1 | 7 | Tmem185b | 0.027 | Old |
| 0.382875 | 0.202169 | 0.126 | 0.105 | 1 | 7 | Etfdh | 0.021 | Old |
| 0.382927 | 0.141538 | 0.2 | 0.172 | 1 | 7 | Polr1a | 0.028 | Old |
| 0.383443 | 0.080732 | 0.067 | 0.051 | 1 | 7 | Zfp821 | 0.016 | Old |
| 0.383639 | 0.106312 | 0.164 | 0.139 | 1 | 7 | Pvr | 0.025 | Old |
| 0.384873 | 0.243607 | 0.067 | 0.051 | 1 | 7 | Foxp4 | 0.016 | Old |
| 0.384985 | 0.166491 | 0.492 | 0.44 | 1 | 7 | Lipa | 0.052 | Old |
| 0.3852 | 0.040846 | 0.262 | 0.232 | 1 | 7 | Pxk | 0.03 | Old |
| 0.385849 | 0.082835 | 0.197 | 0.169 | 1 | 7 | Pigp | 0.028 | Old |
| 0.385911 | 0.043625 | 0.479 | 0.431 | 1 | 7 | Ap2b1 | 0.048 | Old |
| 0.386181 | 0.326654 | 0.223 | 0.196 | 1 | 7 | Tubgcp3 | 0.027 | Old |
| 0.386207 | 0.237803 | 0.249 | 0.214 | 1 | 7 | Impa1 | 0.035 | Old |
| 0.386371 | 0.020383 | 0.056 | 0.042 | 1 | 7 | Tecpr2 | 0.014 | Old |
| 0.38675 | 0.354918 | 0.205 | 0.178 | 1 | 7 | Wdr48 | 0.027 | Old |
| 0.38723 | 0.117018 | 0.359 | 0.325 | 1 | 7 | Med15 | 0.034 | Old |
| 0.387366 | 0.058803 | 0.085 | 0.066 | 1 | 7 | AC127341. | 0.019 | Old |
| 0.387606 | 0.114405 | 0.641 | 0.62 | 1 | 7 | Metrnl | 0.021 | Old |
| 0.388224 | 0.285036 | 0.067 | 0.051 | 1 | 7 | Def8 | 0.016 | Old |
| 0.388284 | 0.012912 | 0.672 | 0.63 | 1 | 7 | Bsg | 0.042 | Old |
| 0.38835 | 0.222357 | 0.346 | 0.319 | 1 | 7 | Fndc3a | 0.027 | Old |
| 0.388476 | 0.084159 | 0.621 | 0.569 | 1 | 7 | Tep1 | 0.052 | Old |
| 0.388559 | 0.067354 | 0.169 | 0.145 | 1 | 7 | Scd2 | 0.024 | Old |
| 0.388705 | 0.221289 | 0.067 | 0.051 | 1 | 7 | 2700038G2 | 0.016 | Old |
| 0.388819 | 0.112417 | 0.295 | 0.262 | 1 | 7 | Dyrk1a | 0.033 | Old |
| 0.388956 | 0.038104 | 0.521 | 0.482 | 1 | 7 | Eif1a | 0.039 | Old |
| 0.389168 | 0.02195 | 0.151 | 0.127 | 1 | 7 | Fnip2 | 0.024 | Old |
| 0.390092 | 0.005015 | 0.128 | 0.105 | 1 | 7 | Trim5 | 0.023 | Old |

| | | | | | | | | |
|---|---|---|---|---|---|---|---|---|
| 0.390203 | 0.340051 | 0.144 | 0.123 | 1 | 7 | Atad3a | 0.021 | Old |
| 0.390402 | 0.97039 | 0.038 | 0.027 | 1 | 7 | Gbp4 | 0.011 | Old |
| 0.391278 | 0.048582 | 0.326 | 0.28 | 1 | 7 | Smc5 | 0.046 | Old |
| 0.391334 | 0.188814 | 0.379 | 0.343 | 1 | 7 | Ncoa3 | 0.036 | Old |
| 0.391587 | 0.062871 | 0.162 | 0.139 | 1 | 7 | Ttll12 | 0.023 | Old |
| 0.392105 | 0.179169 | 0.09 | 0.072 | 1 | 7 | Uba7 | 0.018 | Old |
| 0.392579 | 0.204329 | 0.487 | 0.458 | 1 | 7 | Arglu1 | 0.029 | Old |
| 0.392952 | 0.631486 | 0.038 | 0.027 | 1 | 7 | Ttc21b | 0.011 | Old |
| 0.392992 | 0.083981 | 0.854 | 0.837 | 1 | 7 | Pim1 | 0.017 | Old |
| 0.393747 | 0.152314 | 0.403 | 0.37 | 1 | 7 | Pla2g7 | 0.033 | Old |
| 0.393866 | 0.045625 | 0.774 | 0.744 | 1 | 7 | Anp32e | 0.03 | Old |
| 0.394299 | 0.164302 | 0.469 | 0.428 | 1 | 7 | Asap1 | 0.041 | Old |
| 0.39498 | 0.091474 | 0.067 | 0.051 | 1 | 7 | Pcca | 0.016 | Old |
| 0.395082 | 0.136944 | 0.013 | 0.021 | 1 | 7 | Tjp1 | -0.008 | Old |
| 0.39512 | 0.092287 | 0.485 | 0.434 | 1 | 7 | Psmb3 | 0.051 | Old |
| 0.395169 | 0.203267 | 0.113 | 0.133 | 1 | 7 | Pygl | -0.02 | Old |
| 0.395459 | 0.341124 | 0.146 | 0.123 | 1 | 7 | Atg4d | 0.023 | Old |
| 0.395838 | 0.001442 | 0.713 | 0.711 | 1 | 7 | Brk1 | 0.002 | Old |
| 0.396004 | 0.043224 | 0.087 | 0.069 | 1 | 7 | Tdrd3 | 0.018 | Old |
| 0.396602 | 0.330989 | 0.044 | 0.057 | 1 | 7 | Gm30978 | -0.013 | Old |
| 0.396866 | 0.177348 | 0.056 | 0.042 | 1 | 7 | Usp49 | 0.014 | Old |
| 0.397104 | 0.153053 | 0.259 | 0.229 | 1 | 7 | Hsph1 | 0.03 | Old |
| 0.39713 | 0.044869 | 0.064 | 0.081 | 1 | 7 | Snta1 | -0.017 | Old |
| 0.39722 | 0.535068 | 0.049 | 0.036 | 1 | 7 | 4930522L1 | 0.013 | Old |
| 0.397232 | 0.017159 | 0.518 | 0.476 | 1 | 7 | Uhrf2 | 0.042 | Old |
| 0.397437 | 0.39749 | 0.038 | 0.027 | 1 | 7 | Sec16b | 0.011 | Old |
| 0.397437 | 0.37502 | 0.038 | 0.027 | 1 | 7 | Tmem206 | 0.011 | Old |
| 0.397475 | 0.256958 | 0.1 | 0.081 | 1 | 7 | Dhcr7 | 0.019 | Old |
| 0.397639 | 0.778004 | 0.082 | 0.066 | 1 | 7 | Ifrd2 | 0.016 | Old |
| 0.397769 | 0.218228 | 0.264 | 0.232 | 1 | 7 | Dync1li2 | 0.032 | Old |
| 0.397784 | 1.1983 | 0.013 | 0.021 | 1 | 7 | Dock9 | -0.008 | Old |
| 0.397938 | 0.115066 | 0.39 | 0.355 | 1 | 7 | Rbm17 | 0.035 | Old |
| 0.398304 | 0.1868 | 0.231 | 0.202 | 1 | 7 | Ube3c | 0.029 | Old |
| 0.398455 | 0.166907 | 0.056 | 0.042 | 1 | 7 | Gorab | 0.014 | Old |
| 0.398569 | 0.077653 | 0.369 | 0.322 | 1 | 7 | Htt | 0.047 | Old |
| 0.398753 | 0.09671 | 0.203 | 0.175 | 1 | 7 | Xdh | 0.028 | Old |
| 0.398811 | 0.249597 | 0.09 | 0.072 | 1 | 7 | Mon1b | 0.018 | Old |
| 0.399067 | 0.040789 | 0.887 | 0.88 | 1 | 7 | Rap1b | 0.007 | Old |
| 0.399096 | 0.187644 | 0.136 | 0.114 | 1 | 7 | Mri1 | 0.022 | Old |
| 0.399127 | 0.105222 | 0.195 | 0.22 | 1 | 7 | Akna | -0.025 | Old |
| 0.399275 | 0.16511 | 0.197 | 0.172 | 1 | 7 | Cd2bp2 | 0.025 | Old |
| 0.399591 | 0.690472 | 0.013 | 0.021 | 1 | 7 | Gm13091 | -0.008 | Old |
| 0.400014 | 0.224241 | 0.197 | 0.172 | 1 | 7 | Eef1akmt1 | 0.025 | Old |
| 0.400019 | 0.082914 | 0.108 | 0.087 | 1 | 7 | Chek2 | 0.021 | Old |
| 0.400086 | 0.011812 | 0.367 | 0.328 | 1 | 7 | Zfand3 | 0.039 | Old |

| | | | | | | | | |
|---|---|---|---|---|---|---|---|---|
| 0.400096 | 0.088935 | 0.859 | 0.861 | 1 | 7 | Serbp1 | -0.002 | Old |
| 0.400199 | 0.09601 | 0.141 | 0.117 | 1 | 7 | Thumpd3 | 0.024 | Old |
| 0.400302 | 0.583493 | 0.154 | 0.133 | 1 | 7 | Apeh | 0.021 | Old |
| 0.400555 | 0.586268 | 0.092 | 0.075 | 1 | 7 | Utp23 | 0.017 | Old |
| 0.400817 | 0.205217 | 0.015 | 0.024 | 1 | 7 | Rad52 | -0.009 | Old |
| 0.40084 | 0.553302 | 0.133 | 0.114 | 1 | 7 | Rapgef1 | 0.019 | Old |
| 0.400873 | 0.057881 | 0.249 | 0.22 | 1 | 7 | Mms19 | 0.029 | Old |
| 0.401019 | 0.258838 | 0.272 | 0.241 | 1 | 7 | Atxn2 | 0.031 | Old |
| 0.401217 | 0.040128 | 0.651 | 0.596 | 1 | 7 | Selenot | 0.055 | Old |
| 0.401307 | 0.288536 | 0.038 | 0.027 | 1 | 7 | Zfp52 | 0.011 | Old |
| 0.40141 | 0.089723 | 0.449 | 0.41 | 1 | 7 | Polr2m | 0.039 | Old |
| 0.401645 | 0.069979 | 0.428 | 0.392 | 1 | 7 | AI314180 | 0.036 | Old |
| 0.401954 | 0.346715 | 0.038 | 0.027 | 1 | 7 | Phf11a | 0.011 | Old |
| 0.402034 | 0.054584 | 0.908 | 0.898 | 1 | 7 | Cyba | 0.01 | Old |
| 0.402047 | 0.365644 | 0.079 | 0.063 | 1 | 7 | Gpr180 | 0.016 | Old |
| 0.402419 | 0.044115 | 0.574 | 0.527 | 1 | 7 | Limd1 | 0.047 | Old |
| 0.40272 | 0.493976 | 0.136 | 0.117 | 1 | 7 | Fam98a | 0.019 | Old |
| 0.402789 | 0.175213 | 0.251 | 0.223 | 1 | 7 | Galnt10 | 0.028 | Old |
| 0.402928 | 0.790483 | 0.049 | 0.036 | 1 | 7 | Fam198b | 0.013 | Old |
| 0.403113 | 0.009715 | 0.103 | 0.12 | 1 | 7 | Prkrip1 | -0.017 | Old |
| 0.40313 | 0.112576 | 0.108 | 0.087 | 1 | 7 | Bag4 | 0.021 | Old |
| 0.40325 | 0.372988 | 0.038 | 0.027 | 1 | 7 | Rexo5 | 0.011 | Old |
| 0.403395 | 0.266868 | 0.079 | 0.063 | 1 | 7 | Acvr2a | 0.016 | Old |
| 0.403737 | 0.034566 | 0.244 | 0.214 | 1 | 7 | Zkscan3 | 0.03 | Old |
| 0.403926 | 0.205862 | 0.415 | 0.383 | 1 | 7 | Cks2 | 0.032 | Old |
| 0.404136 | 0.013057 | 0.451 | 0.41 | 1 | 7 | Ncapd2 | 0.041 | Old |
| 0.404241 | 0.060674 | 0.91 | 0.892 | 1 | 7 | Wnk1 | 0.018 | Old |
| 0.40426 | 0.161038 | 0.067 | 0.051 | 1 | 7 | Zcchc4 | 0.016 | Old |
| 0.40434 | 0.112283 | 0.254 | 0.223 | 1 | 7 | Nup205 | 0.031 | Old |
| 0.404549 | 0.355848 | 0.038 | 0.027 | 1 | 7 | Hgh1 | 0.011 | Old |
| 0.404679 | 0.041823 | 0.795 | 0.756 | 1 | 7 | Pdia6 | 0.039 | Old |
| 0.404859 | 0.02874 | 0.7 | 0.636 | 1 | 7 | Dnaja1 | 0.064 | Old |
| 0.404885 | 0.238134 | 0.351 | 0.319 | 1 | 7 | Atp6v1d | 0.032 | Old |
| 0.404925 | 0.241059 | 0.441 | 0.416 | 1 | 7 | Bud31 | 0.025 | Old |
| 0.4051 | 0.122061 | 0.279 | 0.247 | 1 | 7 | Atraid | 0.032 | Old |
| 0.405302 | 0.341468 | 0.108 | 0.09 | 1 | 7 | Med19 | 0.018 | Old |
| 0.405331 | 0.111256 | 0.379 | 0.352 | 1 | 7 | Magoh | 0.027 | Old |
| 0.40585 | 0.192812 | 0.038 | 0.027 | 1 | 7 | Cep131 | 0.011 | Old |
| 0.407598 | 0.139826 | 0.149 | 0.127 | 1 | 7 | Zfp524 | 0.022 | Old |
| 0.408004 | 0.074462 | 0.023 | 0.033 | 1 | 7 | Dcun1d4 | -0.01 | Old |
| 0.408132 | 0.467414 | 0.059 | 0.045 | 1 | 7 | Pomt1 | 0.014 | Old |
| 0.408192 | 0.211612 | 0.513 | 0.476 | 1 | 7 | Casp8 | 0.037 | Old |
| 0.408369 | 0.1131 | 0.315 | 0.28 | 1 | 7 | Ssr2 | 0.035 | Old |
| 0.408744 | 0.10463 | 0.682 | 0.645 | 1 | 7 | Capns1 | 0.037 | Old |
| 0.409072 | 0.204264 | 0.231 | 0.205 | 1 | 7 | Pak1ip1 | 0.026 | Old |

| | | | | | | | | |
|---|---|---|---|---|---|---|---|---|
| 0.409139 | 0.360583 | 0.133 | 0.111 | 1 | 7 | Arid5b | 0.022 | Old |
| 0.40918 | 0.088203 | 0.821 | 0.801 | 1 | 7 | Prdx1 | 0.02 | Old |
| 0.409571 | 0.195578 | 0.428 | 0.38 | 1 | 7 | Dcaf12 | 0.048 | Old |
| 0.409698 | 0.77453 | 0.031 | 0.021 | 1 | 7 | A930033H1 | 0.01 | Old |
| 0.409814 | 0.074983 | 0.497 | 0.467 | 1 | 7 | Adgre5 | 0.03 | Old |
| 0.409863 | 0.712335 | 0.069 | 0.054 | 1 | 7 | Zfp595 | 0.015 | Old |
| 0.409863 | 0.366416 | 0.069 | 0.054 | 1 | 7 | Gm9844 | 0.015 | Old |
| 0.410083 | 0.300298 | 0.041 | 0.054 | 1 | 7 | Plscr1 | -0.013 | Old |
| 0.410186 | 0.143386 | 0.1 | 0.12 | 1 | 7 | Abcb1b | -0.02 | Old |
| 0.410332 | 0.035963 | 0.236 | 0.208 | 1 | 7 | Ammecr1l | 0.028 | Old |
| 0.410359 | 0.163668 | 0.121 | 0.099 | 1 | 7 | Umps | 0.022 | Old |
| 0.410432 | 0.034988 | 0.031 | 0.021 | 1 | 7 | Gm31718 | 0.01 | Old |
| 0.410438 | 0.457974 | 0.213 | 0.19 | 1 | 7 | Steap3 | 0.023 | Old |
| 0.410792 | 0.162105 | 0.269 | 0.235 | 1 | 7 | Dnm1l | 0.034 | Old |
| 0.410939 | 0.597171 | 0.131 | 0.111 | 1 | 7 | Ltbp3 | 0.02 | Old |
| 0.41122 | 0.070689 | 0.167 | 0.142 | 1 | 7 | Oxa1l | 0.025 | Old |
| 0.411235 | 0.187198 | 0.097 | 0.078 | 1 | 7 | Zbtb45 | 0.019 | Old |
| 0.411813 | 0.279406 | 0.059 | 0.045 | 1 | 7 | Hexdc | 0.014 | Old |
| 0.412074 | 0.097394 | 0.241 | 0.211 | 1 | 7 | Paxbp1 | 0.03 | Old |
| 0.41258 | 0.373868 | 0.246 | 0.223 | 1 | 7 | Mrpl21 | 0.023 | Old |
| 0.412744 | 0.252946 | 0.049 | 0.036 | 1 | 7 | Glb1l | 0.013 | Old |
| 0.413021 | 0.34095 | 0.018 | 0.027 | 1 | 7 | Cavin2 | -0.009 | Old |
| 0.413046 | 0.039932 | 0.279 | 0.256 | 1 | 7 | Zc3h4 | 0.023 | Old |
| 0.413442 | 0.273039 | 0.167 | 0.145 | 1 | 7 | Tceanc2 | 0.022 | Old |
| 0.413908 | 0.387627 | 0.049 | 0.036 | 1 | 7 | Yipf2 | 0.013 | Old |
| 0.413908 | 0.305907 | 0.049 | 0.036 | 1 | 7 | Kremen1 | 0.013 | Old |
| 0.41412 | 0.231984 | 0.126 | 0.105 | 1 | 7 | Cstf1 | 0.021 | Old |
| 0.414124 | 0.102574 | 0.208 | 0.181 | 1 | 7 | Mpnd | 0.027 | Old |
| 0.415168 | 0.005393 | 0.067 | 0.051 | 1 | 7 | Wfs1 | 0.016 | Old |
| 0.415524 | 0.045009 | 0.203 | 0.226 | 1 | 7 | Ranbp9 | -0.023 | Old |
| 0.415841 | 0.160345 | 0.449 | 0.413 | 1 | 7 | Sri | 0.036 | Old |
| 0.415871 | 0.13325 | 0.079 | 0.063 | 1 | 7 | Snx11 | 0.016 | Old |
| 0.416646 | 0.201061 | 0.026 | 0.036 | 1 | 7 | Mblac1 | -0.01 | Old |
| 0.416797 | 0.125197 | 0.564 | 0.524 | 1 | 7 | Usp9x | 0.04 | Old |
| 0.416827 | 0.419629 | 0.049 | 0.036 | 1 | 7 | Ankrd28 | 0.013 | Old |
| 0.417343 | 0.191362 | 0.023 | 0.033 | 1 | 7 | Bfsp1 | -0.01 | Old |
| 0.417403 | 0.072256 | 0.331 | 0.301 | 1 | 7 | Syne2 | 0.03 | Old |
| 0.417404 | 0.270115 | 0.172 | 0.148 | 1 | 7 | Gcsh | 0.024 | Old |
| 0.41766 | 1.51E-05 | 0.503 | 0.509 | 1 | 7 | Ost4 | -0.006 | Old |
| 0.41879 | 0.075822 | 0.546 | 0.494 | 1 | 7 | Wac | 0.052 | Old |
| 0.418867 | 0.008938 | 0.059 | 0.072 | 1 | 7 | Mpst | -0.013 | Old |
| 0.419063 | 0.044226 | 0.031 | 0.042 | 1 | 7 | Zfhx2 | -0.011 | Old |
| 0.419097 | 0.465144 | 0.133 | 0.114 | 1 | 7 | 4930455G( | 0.019 | Old |
| 0.419193 | 0.288208 | 0.341 | 0.31 | 1 | 7 | Pank3 | 0.031 | Old |
| 0.419702 | 0.448707 | 0.041 | 0.03 | 1 | 7 | Iqcc | 0.011 | Old |

| | | | | | | | | |
|---|---|---|---|---|---|---|---|---|
| 0.419982 | 0.066176 | 0.172 | 0.148 | 1 | 7 | Hax1 | 0.024 | Old |
| 0.419982 | 0.016746 | 0.172 | 0.148 | 1 | 7 | Tmem205 | 0.024 | Old |
| 0.420346 | 0.051231 | 0.049 | 0.036 | 1 | 7 | L2hgdh | 0.013 | Old |
| 0.420347 | 0.081042 | 0.19 | 0.166 | 1 | 7 | Fam208b | 0.024 | Old |
| 0.420368 | 0.039813 | 0.397 | 0.361 | 1 | 7 | Tspan14 | 0.036 | Old |
| 0.420691 | 0.053433 | 0.162 | 0.181 | 1 | 7 | Hira | -0.019 | Old |
| 0.420826 | 0.052618 | 0.044 | 0.057 | 1 | 7 | Aasdh | -0.013 | Old |
| 0.420934 | 0.258556 | 0.049 | 0.036 | 1 | 7 | Gm4951 | 0.013 | Old |
| 0.421129 | 0.176005 | 0.226 | 0.199 | 1 | 7 | Ccndbp1 | 0.027 | Old |
| 0.421273 | 0.126305 | 0.141 | 0.12 | 1 | 7 | S1pr1 | 0.021 | Old |
| 0.421541 | 0.089045 | 0.308 | 0.277 | 1 | 7 | Rps6ka4 | 0.031 | Old |
| 0.421616 | 0.058035 | 0.197 | 0.169 | 1 | 7 | Ranbp3 | 0.028 | Old |
| 0.421711 | 0.23981 | 0.082 | 0.066 | 1 | 7 | Oaf | 0.016 | Old |
| 0.42173 | 0.121365 | 0.151 | 0.13 | 1 | 7 | Clpp | 0.021 | Old |
| 0.422278 | 0.13684 | 0.031 | 0.021 | 1 | 7 | Zdhhc24 | 0.01 | Old |
| 0.422316 | 0.943713 | 0.062 | 0.048 | 1 | 7 | Adat2 | 0.014 | Old |
| 0.422619 | 0.279533 | 0.082 | 0.066 | 1 | 7 | Cd244 | 0.016 | Old |
| 0.422819 | 0.003535 | 0.09 | 0.072 | 1 | 7 | Mtrf1l | 0.018 | Old |
| 0.422968 | 0.102277 | 0.479 | 0.446 | 1 | 7 | Mlf2 | 0.033 | Old |
| 0.423601 | 0.41169 | 0.131 | 0.111 | 1 | 7 | Plcb1 | 0.02 | Old |
| 0.423979 | 0.030395 | 0.815 | 0.795 | 1 | 7 | S100a10 | 0.02 | Old |
| 0.424784 | 0.024109 | 0.187 | 0.166 | 1 | 7 | Immp1l | 0.021 | Old |
| 0.424832 | 0.447439 | 0.041 | 0.03 | 1 | 7 | Cnr2 | 0.011 | Old |
| 0.425232 | 0.032937 | 0.749 | 0.735 | 1 | 7 | Mrfap1 | 0.014 | Old |
| 0.425271 | 0.310615 | 0.031 | 0.021 | 1 | 7 | Rwdd2a | 0.01 | Old |
| 0.425317 | 0.2438 | 0.079 | 0.063 | 1 | 7 | Dcun1d3 | 0.016 | Old |
| 0.425327 | 0.164673 | 0.526 | 0.503 | 1 | 7 | Ctdnep1 | 0.023 | Old |
| 0.425349 | 0.214694 | 0.082 | 0.066 | 1 | 7 | 1600010M | 0.016 | Old |
| 0.425445 | 0.358899 | 0.149 | 0.13 | 1 | 7 | Pja1 | 0.019 | Old |
| 0.425554 | 0.055066 | 0.192 | 0.169 | 1 | 7 | Fignl1 | 0.023 | Old |
| 0.425805 | 0.787331 | 0.082 | 0.066 | 1 | 7 | Itgb3bp | 0.016 | Old |
| 0.425916 | 0.002846 | 0.564 | 0.518 | 1 | 7 | Llph | 0.046 | Old |
| 0.426021 | 0.520136 | 0.031 | 0.021 | 1 | 7 | Tagap1 | 0.01 | Old |
| 0.426729 | 0.336156 | 0.059 | 0.045 | 1 | 7 | Cluap1 | 0.014 | Old |
| 0.427523 | 0.118088 | 0.031 | 0.021 | 1 | 7 | Adamtsl4 | 0.01 | Old |
| 0.427569 | 0.107374 | 0.49 | 0.44 | 1 | 7 | Cdc37 | 0.05 | Old |
| 0.427995 | 0.153622 | 0.146 | 0.123 | 1 | 7 | Fdft1 | 0.023 | Old |
| 0.428001 | 0.247884 | 0.203 | 0.178 | 1 | 7 | Ubl4a | 0.025 | Old |
| 0.428052 | 0.23972 | 0.367 | 0.337 | 1 | 7 | Svil | 0.03 | Old |
| 0.428301 | 0.026661 | 0.715 | 0.702 | 1 | 7 | Wdr26 | 0.013 | Old |
| 0.428357 | 0.09785 | 0.328 | 0.295 | 1 | 7 | Cpsf7 | 0.033 | Old |
| 0.428569 | 0.060077 | 0.262 | 0.235 | 1 | 7 | Ift20 | 0.027 | Old |
| 0.428683 | 1.747787 | 0.023 | 0.015 | 1 | 7 | Snx33 | 0.008 | Old |
| 0.428743 | 0.083622 | 0.138 | 0.117 | 1 | 7 | Mphosph9 | 0.021 | Old |
| 0.428886 | 0.055675 | 0.059 | 0.045 | 1 | 7 | Haao | 0.014 | Old |

| | | | | | | | | |
|---|---|---|---|---|---|---|---|---|
| 0.429082 | 0.102499 | 0.244 | 0.214 | 1 | 7 | Clns1a | 0.03 | Old |
| 0.429119 | 0.014861 | 0.39 | 0.355 | 1 | 7 | Psmb2 | 0.035 | Old |
| 0.42943 | 0.069102 | 0.282 | 0.25 | 1 | 7 | Cebpa | 0.032 | Old |
| 0.429577 | 0.165703 | 0.033 | 0.045 | 1 | 7 | Lss | -0.012 | Old |
| 0.429634 | 0.33645 | 0.218 | 0.19 | 1 | 7 | Slu7 | 0.028 | Old |
| 0.429702 | 0.025458 | 0.423 | 0.37 | 1 | 7 | Was | 0.053 | Old |
| 0.429884 | 0.073557 | 0.803 | 0.774 | 1 | 7 | Cd44 | 0.029 | Old |
| 0.43 | 0.37088 | 0.041 | 0.03 | 1 | 7 | Clcf1 | 0.011 | Old |
| 0.430666 | 0.347599 | 0.151 | 0.13 | 1 | 7 | B9d2 | 0.021 | Old |
| 0.431574 | 0.043684 | 0.187 | 0.16 | 1 | 7 | Usp45 | 0.027 | Old |
| 0.43159 | 0.441862 | 0.059 | 0.045 | 1 | 7 | 2610008E1 | 0.014 | Old |
| 0.431607 | 0.076707 | 0.092 | 0.075 | 1 | 7 | Dtx4 | 0.017 | Old |
| 0.431704 | 0.330192 | 0.151 | 0.13 | 1 | 7 | Mrs2 | 0.021 | Old |
| 0.431996 | 0.176844 | 0.387 | 0.355 | 1 | 7 | Rfc2 | 0.032 | Old |
| 0.432076 | 0.275849 | 0.328 | 0.298 | 1 | 7 | Ahcyl1 | 0.03 | Old |
| 0.432189 | 1.099912 | 0.023 | 0.015 | 1 | 7 | Cyfip2 | 0.008 | Old |
| 0.432257 | 0.098909 | 0.103 | 0.084 | 1 | 7 | Dscc1 | 0.019 | Old |
| 0.432455 | 0.086303 | 0.903 | 0.901 | 1 | 7 | Capzb | 0.002 | Old |
| 0.432757 | 0.362952 | 0.159 | 0.139 | 1 | 7 | Spdl1 | 0.02 | Old |
| 0.432831 | 0.014917 | 0.59 | 0.542 | 1 | 7 | Scaf11 | 0.048 | Old |
| 0.433135 | 0.236707 | 0.203 | 0.181 | 1 | 7 | Trmt6 | 0.022 | Old |
| 0.433214 | 0.060865 | 0.269 | 0.238 | 1 | 7 | Slc25a51 | 0.031 | Old |
| 0.433468 | 0.14347 | 0.292 | 0.259 | 1 | 7 | Trnau1ap | 0.033 | Old |
| 0.433916 | 0.16301 | 0.103 | 0.084 | 1 | 7 | Ripor1 | 0.019 | Old |
| 0.434159 | 0.016327 | 0.328 | 0.289 | 1 | 7 | Zwint | 0.039 | Old |
| 0.434529 | 1.010486 | 0.077 | 0.063 | 1 | 7 | Slc7a11 | 0.014 | Old |
| 0.434587 | 0.042981 | 0.664 | 0.636 | 1 | 7 | Smc4 | 0.028 | Old |
| 0.434829 | 0.766227 | 0.023 | 0.015 | 1 | 7 | Eya4 | 0.008 | Old |
| 0.434985 | 0.596662 | 0.082 | 0.066 | 1 | 7 | Isg15 | 0.016 | Old |
| 0.435063 | 0.084778 | 0.069 | 0.054 | 1 | 7 | Slc25a15 | 0.015 | Old |
| 0.435476 | 0.323373 | 0.108 | 0.09 | 1 | 7 | Tada3 | 0.018 | Old |
| 0.435667 | 0.168965 | 0.141 | 0.12 | 1 | 7 | Phf6 | 0.021 | Old |
| 0.435711 | 0.408361 | 0.023 | 0.015 | 1 | 7 | Kctd7 | 0.008 | Old |
| 0.435972 | 0.267527 | 0.133 | 0.111 | 1 | 7 | Dvl1 | 0.022 | Old |
| 0.436272 | 0.118955 | 0.29 | 0.262 | 1 | 7 | Chfr | 0.028 | Old |
| 0.436927 | 0.067561 | 0.426 | 0.392 | 1 | 7 | Zfp703 | 0.034 | Old |
| 0.437164 | 0.917225 | 0.041 | 0.03 | 1 | 7 | A230050P2 | 0.011 | Old |
| 0.437819 | 0.099298 | 0.041 | 0.03 | 1 | 7 | Arl16 | 0.011 | Old |
| 0.438117 | 0.081908 | 0.667 | 0.608 | 1 | 7 | Ap2s1 | 0.059 | Old |
| 0.438304 | 0.163224 | 0.326 | 0.295 | 1 | 7 | Stat5b | 0.031 | Old |
| 0.43881 | 0.13208 | 0.051 | 0.039 | 1 | 7 | AU022252 | 0.012 | Old |
| 0.439012 | 0.080286 | 0.087 | 0.105 | 1 | 7 | Slc14a1 | -0.018 | Old |
| 0.439292 | 0.266685 | 0.438 | 0.416 | 1 | 7 | Coro2a | 0.022 | Old |
| 0.439395 | 0.39655 | 0.051 | 0.039 | 1 | 7 | Esr1 | 0.012 | Old |
| 0.439439 | 0.026866 | 0.349 | 0.31 | 1 | 7 | Actl6a | 0.039 | Old |

| | | | | | | | | |
|---|---|---|---|---|---|---|---|---|
| 0.439635 | 0.093795 | 0.628 | 0.584 | 1 | 7 | Hnrnpul1 | 0.044 | Old |
| 0.439654 | 0.292722 | 0.031 | 0.021 | 1 | 7 | Gimap6 | 0.01 | Old |
| 0.439765 | 0.288209 | 0.182 | 0.16 | 1 | 7 | Tango2 | 0.022 | Old |
| 0.439982 | 1.229322 | 0.015 | 0.009 | 1 | 7 | Efr3b | 0.006 | Old |
| 0.439982 | 1.113392 | 0.015 | 0.009 | 1 | 7 | Mb21d2 | 0.006 | Old |
| 0.439982 | 0.791311 | 0.015 | 0.009 | 1 | 7 | Ppp1r3d | 0.006 | Old |
| 0.440138 | 0.142333 | 0.023 | 0.015 | 1 | 7 | Fam212a | 0.008 | Old |
| 0.440443 | 0.389605 | 0.041 | 0.03 | 1 | 7 | Pick1 | 0.011 | Old |
| 0.440565 | 0.087843 | 0.297 | 0.271 | 1 | 7 | St6galnac6 | 0.026 | Old |
| 0.440575 | 0.132984 | 0.203 | 0.178 | 1 | 7 | Hdac9 | 0.025 | Old |
| 0.440735 | 0.1754 | 0.303 | 0.28 | 1 | 7 | Cisd1 | 0.023 | Old |
| 0.441086 | 1.497624 | 0.015 | 0.009 | 1 | 7 | Tespa1 | 0.006 | Old |
| 0.441086 | 0.640206 | 0.015 | 0.009 | 1 | 7 | 1-Sep | 0.006 | Old |
| 0.441314 | 0.143456 | 0.226 | 0.199 | 1 | 7 | Fry | 0.027 | Old |
| 0.441915 | 0.294172 | 0.023 | 0.015 | 1 | 7 | Ablim1 | 0.008 | Old |
| 0.441915 | 0.077402 | 0.023 | 0.015 | 1 | 7 | Nectin2 | 0.008 | Old |
| 0.441915 | 0.024996 | 0.023 | 0.015 | 1 | 7 | Fitm2 | 0.008 | Old |
| 0.442242 | 0.116086 | 0.618 | 0.563 | 1 | 7 | Ndufc2 | 0.055 | Old |
| 0.442417 | 0.198554 | 0.041 | 0.03 | 1 | 7 | B930036N1 | 0.011 | Old |
| 0.442598 | 0.049307 | 0.331 | 0.295 | 1 | 7 | Acss1 | 0.036 | Old |
| 0.442927 | 0.218028 | 0.228 | 0.205 | 1 | 7 | Rad51ap1 | 0.023 | Old |
| 0.443298 | 1.183331 | 0.015 | 0.009 | 1 | 7 | Samd14 | 0.006 | Old |
| 0.443298 | 0.904275 | 0.015 | 0.009 | 1 | 7 | Wwtr1 | 0.006 | Old |
| 0.443298 | 0.869693 | 0.015 | 0.009 | 1 | 7 | Tlr11 | 0.006 | Old |
| 0.443298 | 0.644648 | 0.015 | 0.009 | 1 | 7 | Slc39a8 | 0.006 | Old |
| 0.443298 | 0.593819 | 0.015 | 0.009 | 1 | 7 | Msrb3 | 0.006 | Old |
| 0.443298 | 0.460284 | 0.015 | 0.009 | 1 | 7 | Gm31597 | 0.006 | Old |
| 0.443298 | 0.341433 | 0.015 | 0.009 | 1 | 7 | Tmsb15b1 | 0.006 | Old |
| 0.443298 | 0.251461 | 0.015 | 0.009 | 1 | 7 | Casp12 | 0.006 | Old |
| 0.443298 | 0.081948 | 0.015 | 0.009 | 1 | 7 | Gpr18 | 0.006 | Old |
| 0.443383 | 0.230207 | 0.254 | 0.226 | 1 | 7 | Clcn3 | 0.028 | Old |
| 0.443384 | 0.008178 | 0.605 | 0.569 | 1 | 7 | Ssrp1 | 0.036 | Old |
| 0.443471 | 0.05204 | 0.385 | 0.337 | 1 | 7 | Gatad1 | 0.048 | Old |
| 0.444397 | 0.133595 | 0.041 | 0.03 | 1 | 7 | Usp11 | 0.011 | Old |
| 0.444407 | 0.63251 | 0.015 | 0.009 | 1 | 7 | Phtf1os | 0.006 | Old |
| 0.444407 | 0.313304 | 0.015 | 0.009 | 1 | 7 | Gm38251 | 0.006 | Old |
| 0.444489 | 0.327704 | 0.192 | 0.172 | 1 | 7 | Crtc2 | 0.02 | Old |
| 0.444701 | 0.286798 | 0.346 | 0.319 | 1 | 7 | Rrm2 | 0.027 | Old |
| 0.444705 | 0.072237 | 0.641 | 0.596 | 1 | 7 | Dync1h1 | 0.045 | Old |
| 0.445061 | 0.162783 | 0.659 | 0.614 | 1 | 7 | Eif3c | 0.045 | Old |
| 0.445378 | 0.822031 | 0.033 | 0.024 | 1 | 7 | Gm42715 | 0.009 | Old |
| 0.445483 | 0.326361 | 0.023 | 0.015 | 1 | 7 | Adrb1 | 0.008 | Old |
| 0.445517 | 0.32554 | 0.015 | 0.009 | 1 | 7 | Gvin1 | 0.006 | Old |
| 0.445517 | 0.224029 | 0.015 | 0.009 | 1 | 7 | Osbpl5 | 0.006 | Old |
| 0.445854 | 0.139804 | 0.559 | 0.53 | 1 | 7 | Batf3 | 0.029 | Old |

| | | | | | | | | |
|---|---|---|---|---|---|---|---|---|
| 0.446041 | 0.07448 | 0.503 | 0.47 | 1 | 7 | Sin3b | 0.033 | Old |
| 0.446102 | 0.410931 | 0.169 | 0.151 | 1 | 7 | Slc28a2 | 0.018 | Old |
| 0.446381 | 0.100765 | 0.041 | 0.03 | 1 | 7 | Mamld1 | 0.011 | Old |
| 0.446854 | 0.224112 | 0.182 | 0.157 | 1 | 7 | Acsl3 | 0.025 | Old |
| 0.446863 | 0.022903 | 0.267 | 0.238 | 1 | 7 | Ttc39b | 0.029 | Old |
| 0.447179 | 0.103112 | 0.21 | 0.187 | 1 | 7 | Ctr9 | 0.023 | Old |
| 0.447541 | 0.001456 | 0.254 | 0.223 | 1 | 7 | Bsdc1 | 0.031 | Old |
| 0.447743 | 0.560495 | 0.015 | 0.009 | 1 | 7 | Zfp97 | 0.006 | Old |
| 0.447743 | 0.106885 | 0.015 | 0.009 | 1 | 7 | 1700056N1 | 0.006 | Old |
| 0.447795 | 0.020048 | 0.518 | 0.467 | 1 | 7 | Rer1 | 0.051 | Old |
| 0.447799 | 0.112272 | 0.562 | 0.509 | 1 | 7 | Atp2a2 | 0.053 | Old |
| 0.447833 | 0.449859 | 0.141 | 0.123 | 1 | 7 | Lrwd1 | 0.018 | Old |
| 0.447968 | 0.184512 | 0.272 | 0.25 | 1 | 7 | Carhsp1 | 0.022 | Old |
| 0.44803 | 0.034946 | 0.638 | 0.575 | 1 | 7 | Cox5b | 0.063 | Old |
| 0.44817 | 0.128536 | 0.023 | 0.015 | 1 | 7 | A530040E1 | 0.008 | Old |
| 0.448547 | 0.338112 | 0.215 | 0.193 | 1 | 7 | Nacc2 | 0.022 | Old |
| 0.448858 | 0.540491 | 0.015 | 0.009 | 1 | 7 | Ifi208 | 0.006 | Old |
| 0.448858 | 0.274271 | 0.015 | 0.009 | 1 | 7 | Gm6793 | 0.006 | Old |
| 0.449387 | 0.06592 | 0.264 | 0.241 | 1 | 7 | Mfsd11 | 0.023 | Old |
| 0.449412 | 0.120163 | 0.051 | 0.039 | 1 | 7 | Mmp14 | 0.012 | Old |
| 0.449493 | 0.061971 | 0.554 | 0.518 | 1 | 7 | Suz12 | 0.036 | Old |
| 0.44977 | 0.058813 | 0.892 | 0.883 | 1 | 7 | Sh3bgrl3 | 0.009 | Old |
| 0.44979 | 0.778097 | 0.033 | 0.024 | 1 | 7 | 9530059O1 | 0.009 | Old |
| 0.449879 | 0.001875 | 0.556 | 0.539 | 1 | 7 | Ubxn1 | 0.017 | Old |
| 0.449975 | 0.207015 | 0.015 | 0.009 | 1 | 7 | Gas2 | 0.006 | Old |
| 0.450031 | 0.108157 | 0.354 | 0.319 | 1 | 7 | Col4a3bp | 0.035 | Old |
| 0.450396 | 0.044047 | 0.146 | 0.127 | 1 | 7 | Decr1 | 0.019 | Old |
| 0.45073 | 0.492554 | 0.172 | 0.151 | 1 | 7 | Pygo2 | 0.021 | Old |
| 0.450944 | 0.050205 | 0.674 | 0.654 | 1 | 7 | Bola2 | 0.02 | Old |
| 0.451006 | 0.149064 | 0.569 | 0.524 | 1 | 7 | Nav1 | 0.045 | Old |
| 0.451588 | 0.144119 | 0.395 | 0.37 | 1 | 7 | Dap | 0.025 | Old |
| 0.451688 | 0.491638 | 0.074 | 0.06 | 1 | 7 | Rbm12b2 | 0.014 | Old |
| 0.451885 | 0.304878 | 0.349 | 0.328 | 1 | 7 | B4galt1 | 0.021 | Old |
| 0.451926 | 0.23662 | 0.208 | 0.184 | 1 | 7 | Smim27 | 0.024 | Old |
| 0.451984 | 0.38341 | 0.162 | 0.145 | 1 | 7 | D6Wsu163 | 0.017 | Old |
| 0.452213 | 0.063312 | 0.015 | 0.009 | 1 | 7 | Slc22a18 | 0.006 | Old |
| 0.452213 | 0.019699 | 0.015 | 0.009 | 1 | 7 | Gm42670 | 0.006 | Old |
| 0.452407 | 0.364421 | 0.21 | 0.19 | 1 | 7 | Aars | 0.02 | Old |
| 0.452884 | 0.275334 | 0.072 | 0.057 | 1 | 7 | Mbip | 0.015 | Old |
| 0.453043 | 0.129521 | 0.59 | 0.53 | 1 | 7 | Smc3 | 0.06 | Old |
| 0.453303 | 0.061306 | 0.151 | 0.13 | 1 | 7 | Rasa2 | 0.021 | Old |
| 0.453326 | 0.051034 | 0.879 | 0.873 | 1 | 7 | Hnrnpa3 | 0.006 | Old |
| 0.453665 | 0.383407 | 0.074 | 0.06 | 1 | 7 | Ttll4 | 0.014 | Old |
| 0.45416 | 0.166657 | 0.074 | 0.06 | 1 | 7 | Wdr55 | 0.014 | Old |
| 0.454228 | 0.625939 | 0.033 | 0.024 | 1 | 7 | Helz2 | 0.009 | Old |

| | | | | | | | | |
|---|---|---|---|---|---|---|---|---|
| 0.454243 | 0.196078 | 0.285 | 0.253 | 1 | 7 | Eif4enif1 | 0.032 | Old |
| 0.454369 | 0.193541 | 0.041 | 0.03 | 1 | 7 | Adra2a | 0.011 | Old |
| 0.454592 | 0.376123 | 0.136 | 0.117 | 1 | 7 | Pla2g16 | 0.019 | Old |
| 0.454625 | 0.042815 | 0.287 | 0.256 | 1 | 7 | Ngdn | 0.031 | Old |
| 0.45487 | 0.162075 | 0.062 | 0.048 | 1 | 7 | Pvt1 | 0.014 | Old |
| 0.45509 | 0.14114 | 0.279 | 0.25 | 1 | 7 | Dhx38 | 0.029 | Old |
| 0.455547 | 0.235263 | 0.105 | 0.087 | 1 | 7 | Tmem161b | 0.018 | Old |
| 0.455712 | 0.408864 | 0.033 | 0.024 | 1 | 7 | Acss2 | 0.009 | Old |
| 0.455796 | 0.351437 | 0.103 | 0.087 | 1 | 7 | Prr3 | 0.016 | Old |
| 0.455837 | 0.134342 | 0.456 | 0.434 | 1 | 7 | Nsf | 0.022 | Old |
| 0.45592 | 0.229265 | 0.251 | 0.232 | 1 | 7 | Atp6v0a2 | 0.019 | Old |
| 0.456567 | 0.064807 | 0.472 | 0.41 | 1 | 7 | Crlf3 | 0.062 | Old |
| 0.456816 | 0.132915 | 0.579 | 0.548 | 1 | 7 | Tcf25 | 0.031 | Old |
| 0.457172 | 0.009825 | 0.259 | 0.277 | 1 | 7 | Klhl6 | -0.018 | Old |
| 0.457244 | 0.05531 | 0.077 | 0.093 | 1 | 7 | P3h2 | -0.016 | Old |
| 0.457758 | 0.166315 | 0.051 | 0.039 | 1 | 7 | Flad1 | 0.012 | Old |
| 0.457936 | 0.002651 | 0.287 | 0.256 | 1 | 7 | Cops5 | 0.031 | Old |
| 0.457978 | 0.020576 | 0.146 | 0.127 | 1 | 7 | Zfyve27 | 0.019 | Old |
| 0.45821 | 0.197371 | 0.167 | 0.148 | 1 | 7 | Isca2 | 0.019 | Old |
| 0.459103 | 0.16644 | 0.274 | 0.241 | 1 | 7 | Hdac3 | 0.033 | Old |
| 0.459211 | 0.445058 | 0.118 | 0.102 | 1 | 7 | Pnkp | 0.016 | Old |
| 0.459763 | 0.059825 | 0.226 | 0.199 | 1 | 7 | Oaz2 | 0.027 | Old |
| 0.459781 | 0.083828 | 0.956 | 0.937 | 1 | 7 | Zeb2 | 0.019 | Old |
| 0.46005 | 0.340923 | 0.085 | 0.069 | 1 | 7 | Creb3l2 | 0.016 | Old |
| 0.460071 | 0.06509 | 0.218 | 0.187 | 1 | 7 | Stoml2 | 0.031 | Old |
| 0.460217 | 0.402171 | 0.313 | 0.289 | 1 | 7 | Fxyd2 | 0.024 | Old |
| 0.460749 | 0.254038 | 0.346 | 0.325 | 1 | 7 | Dnajc15 | 0.021 | Old |
| 0.46093 | 0.707526 | 0.033 | 0.024 | 1 | 7 | Aldh4a1 | 0.009 | Old |
| 0.46093 | 0.468984 | 0.033 | 0.024 | 1 | 7 | Hist1h4c | 0.009 | Old |
| 0.460989 | 0.038324 | 0.085 | 0.069 | 1 | 7 | Tmem175 | 0.016 | Old |
| 0.461024 | 0.581926 | 0.092 | 0.078 | 1 | 7 | Rem1 | 0.014 | Old |
| 0.461613 | 0.175662 | 0.4 | 0.373 | 1 | 7 | Dip2b | 0.027 | Old |
| 0.461678 | 0.792849 | 0.033 | 0.024 | 1 | 7 | Usp20 | 0.009 | Old |
| 0.462007 | 0.05149 | 0.072 | 0.057 | 1 | 7 | Clybl | 0.015 | Old |
| 0.462136 | 0.230531 | 0.638 | 0.63 | 1 | 7 | Sdha | 0.008 | Old |
| 0.462153 | 0.24035 | 0.195 | 0.172 | 1 | 7 | Stx5a | 0.023 | Old |
| 0.462427 | 0.185488 | 0.033 | 0.024 | 1 | 7 | Pde5a | 0.009 | Old |
| 0.46269 | 0.186926 | 0.138 | 0.12 | 1 | 7 | Cars | 0.018 | Old |
| 0.463121 | 0.062537 | 0.297 | 0.265 | 1 | 7 | Espl1 | 0.032 | Old |
| 0.463453 | 0.046007 | 0.19 | 0.166 | 1 | 7 | 9430038I0 | 0.024 | Old |
| 0.463574 | 0.149738 | 0.085 | 0.069 | 1 | 7 | Card9 | 0.016 | Old |
| 0.463879 | 0.057763 | 0.505 | 0.464 | 1 | 7 | Pdia4 | 0.041 | Old |
| 0.464137 | 0.038707 | 0.374 | 0.343 | 1 | 7 | Derl1 | 0.031 | Old |
| 0.464264 | 0.036406 | 0.231 | 0.199 | 1 | 7 | Pes1 | 0.032 | Old |
| 0.464677 | 0.335173 | 0.033 | 0.024 | 1 | 7 | Batf2 | 0.009 | Old |

| | | | | | | | | |
|---|---|---|---|---|---|---|---|---|
| 0.464832 | 0.011979 | 0.756 | 0.711 | 1 | 7 | Tardbp | 0.045 | Old |
| 0.465035 | 0.038789 | 0.487 | 0.443 | 1 | 7 | Dynlrb1 | 0.044 | Old |
| 0.465358 | 0.208349 | 0.177 | 0.154 | 1 | 7 | Arfip1 | 0.023 | Old |
| 0.465702 | 0.045166 | 0.513 | 0.467 | 1 | 7 | Nop53 | 0.046 | Old |
| 0.466058 | 0.028296 | 0.474 | 0.434 | 1 | 7 | Slc29a1 | 0.04 | Old |
| 0.466303 | 0.197883 | 0.154 | 0.133 | 1 | 7 | Gpatch2l | 0.021 | Old |
| 0.466896 | 0.498837 | 0.238 | 0.214 | 1 | 7 | Rnf168 | 0.024 | Old |
| 0.466913 | 0.524809 | 0.067 | 0.054 | 1 | 7 | Bcl7a | 0.013 | Old |
| 0.46702 | 0.290796 | 0.154 | 0.133 | 1 | 7 | Ap5m1 | 0.021 | Old |
| 0.467026 | 0.335793 | 0.087 | 0.072 | 1 | 7 | Cd151 | 0.015 | Old |
| 0.467462 | 0.04285 | 0.185 | 0.205 | 1 | 7 | Tbc1d9 | -0.02 | Old |
| 0.468026 | 0.199202 | 0.044 | 0.033 | 1 | 7 | Brat1 | 0.011 | Old |
| 0.468053 | 0.265093 | 0.177 | 0.16 | 1 | 7 | Ece1 | 0.017 | Old |
| 0.468173 | 0.117874 | 0.108 | 0.09 | 1 | 7 | Riox2 | 0.018 | Old |
| 0.468269 | 0.070073 | 0.636 | 0.627 | 1 | 7 | Tcp1 | 0.009 | Old |
| 0.468441 | 0.330916 | 0.033 | 0.024 | 1 | 7 | Zbtb32 | 0.009 | Old |
| 0.468682 | 0.592812 | 0.044 | 0.033 | 1 | 7 | Nradd | 0.011 | Old |
| 0.468845 | 0.107876 | 0.192 | 0.169 | 1 | 7 | Chchd5 | 0.023 | Old |
| 0.469358 | 0.03724 | 0.156 | 0.136 | 1 | 7 | Nxt2 | 0.02 | Old |
| 0.469695 | 0.113144 | 0.61 | 0.566 | 1 | 7 | Ubap2l | 0.044 | Old |
| 0.470275 | 0.254319 | 0.054 | 0.042 | 1 | 7 | Smyd4 | 0.012 | Old |
| 0.470483 | 0.170245 | 0.415 | 0.373 | 1 | 7 | Unc119 | 0.042 | Old |
| 0.470998 | 0.114318 | 0.238 | 0.205 | 1 | 7 | Ppip5k2 | 0.033 | Old |
| 0.471025 | 0.347705 | 0.192 | 0.172 | 1 | 7 | Rhoh | 0.02 | Old |
| 0.471058 | 0.177736 | 0.241 | 0.214 | 1 | 7 | Vps50 | 0.027 | Old |
| 0.471605 | 0.019706 | 0.423 | 0.389 | 1 | 7 | Usp15 | 0.034 | Old |
| 0.471826 | 0.078991 | 0.462 | 0.437 | 1 | 7 | Pdap1 | 0.025 | Old |
| 0.472241 | 0.10881 | 0.5 | 0.461 | 1 | 7 | Psmd11 | 0.039 | Old |
| 0.472392 | 0.084075 | 0.49 | 0.455 | 1 | 7 | Ndufb4 | 0.035 | Old |
| 0.472763 | 0.034872 | 0.531 | 0.485 | 1 | 7 | Tnpo3 | 0.046 | Old |
| 0.472763 | 0.097657 | 0.09 | 0.075 | 1 | 7 | Dcxr | 0.015 | Old |
| 0.472878 | 0.01177 | 0.259 | 0.229 | 1 | 7 | Rnf20 | 0.03 | Old |
| 0.472981 | 0.148 | 0.167 | 0.145 | 1 | 7 | Wdr77 | 0.022 | Old |
| 0.47319 | 0.07987 | 0.546 | 0.503 | 1 | 7 | Srsf11 | 0.043 | Old |
| 0.473542 | 0.153387 | 0.621 | 0.623 | 1 | 7 | Clec10a | -0.002 | Old |
| 0.474273 | 0.6997 | 0.121 | 0.105 | 1 | 7 | Pde8a | 0.016 | Old |
| 0.47433 | 0.407229 | 0.059 | 0.072 | 1 | 7 | Wrap53 | -0.013 | Old |
| 0.474417 | 0.003059 | 0.667 | 0.645 | 1 | 7 | Tra2a | 0.022 | Old |
| 0.47502 | 0.431921 | 0.054 | 0.042 | 1 | 7 | Smtn | 0.012 | Old |
| 0.475103 | 0.169024 | 0.103 | 0.087 | 1 | 7 | Eif2b1 | 0.016 | Old |
| 0.475206 | 0.34598 | 0.087 | 0.072 | 1 | 7 | Wdfy1 | 0.015 | Old |
| 0.475424 | 0.101026 | 0.613 | 0.557 | 1 | 7 | U2af2 | 0.056 | Old |
| 0.475526 | 0.16531 | 0.323 | 0.295 | 1 | 7 | Ext1 | 0.028 | Old |
| 0.475588 | 0.045342 | 0.156 | 0.139 | 1 | 7 | Hr | 0.017 | Old |
| 0.475917 | 0.264351 | 0.182 | 0.166 | 1 | 7 | Klrk1 | 0.016 | Old |

| | | | | | | | | |
|---|---|---|---|---|---|---|---|---|
| 0.475938 | 0.485847 | 0.044 | 0.033 | 1 | 7 | 1110019D1 | 0.011 | Old |
| 0.476101 | 0.161743 | 0.256 | 0.232 | 1 | 7 | Gemin7 | 0.024 | Old |
| 0.477042 | 0.139129 | 0.377 | 0.349 | 1 | 7 | Ep400 | 0.028 | Old |
| 0.477439 | 0.047562 | 0.059 | 0.072 | 1 | 7 | Rbm12b1 | -0.013 | Old |
| 0.478024 | 0.113463 | 0.238 | 0.214 | 1 | 7 | Mmadhc | 0.024 | Old |
| 0.478091 | 0.467207 | 0.067 | 0.054 | 1 | 7 | Kctd18 | 0.013 | Old |
| 0.478333 | 0.276272 | 0.351 | 0.325 | 1 | 7 | Adipor2 | 0.026 | Old |
| 0.478504 | 0.028714 | 0.326 | 0.292 | 1 | 7 | Hipk3 | 0.034 | Old |
| 0.478559 | 0.117797 | 0.397 | 0.361 | 1 | 7 | Ube2e1 | 0.036 | Old |
| 0.478617 | 0.049843 | 0.205 | 0.184 | 1 | 7 | Gyg | 0.021 | Old |
| 0.478624 | 0.116131 | 0.318 | 0.283 | 1 | 7 | Rpa1 | 0.035 | Old |
| 0.478653 | 0.133102 | 0.6 | 0.566 | 1 | 7 | Bag1 | 0.034 | Old |
| 0.479454 | 0.427705 | 0.103 | 0.087 | 1 | 7 | Fra10ac1 | 0.016 | Old |
| 0.479475 | 0.045923 | 0.428 | 0.386 | 1 | 7 | Cebpz | 0.042 | Old |
| 0.479601 | 0.014026 | 0.556 | 0.524 | 1 | 7 | Cd68 | 0.032 | Old |
| 0.479757 | 1.120377 | 0.036 | 0.027 | 1 | 7 | Plcg1 | 0.009 | Old |
| 0.479834 | 0.028203 | 0.033 | 0.024 | 1 | 7 | Cd101 | 0.009 | Old |
| 0.480512 | 0.123157 | 0.344 | 0.319 | 1 | 7 | Zc3h12a | 0.025 | Old |
| 0.480636 | 0.401836 | 0.126 | 0.108 | 1 | 7 | Atxn1l | 0.018 | Old |
| 0.481494 | 0.094548 | 0.705 | 0.654 | 1 | 7 | Ncf2 | 0.051 | Old |
| 0.481717 | 0.041357 | 0.321 | 0.289 | 1 | 7 | Slc26a2 | 0.032 | Old |
| 0.482219 | 0.555939 | 0.079 | 0.066 | 1 | 7 | Kdm1b | 0.013 | Old |
| 0.482456 | 0.104374 | 0.123 | 0.105 | 1 | 7 | Dcstamp | 0.018 | Old |
| 0.482517 | 0.290548 | 0.087 | 0.072 | 1 | 7 | Sec22a | 0.015 | Old |
| 0.483091 | 0.175649 | 0.187 | 0.166 | 1 | 7 | Cenps | 0.021 | Old |
| 0.483151 | 0.030842 | 0.256 | 0.229 | 1 | 7 | Arl2bp | 0.027 | Old |
| 0.483375 | 0.041204 | 0.849 | 0.813 | 1 | 7 | Hnrnpu | 0.036 | Old |
| 0.483819 | 0.472942 | 0.09 | 0.105 | 1 | 7 | Rtl8b | -0.015 | Old |
| 0.483826 | 1.154183 | 0.082 | 0.069 | 1 | 7 | Slc46a3 | 0.013 | Old |
| 0.484435 | 0.238373 | 0.182 | 0.163 | 1 | 7 | Mrpl27 | 0.019 | Old |
| 0.484519 | 0.138431 | 0.149 | 0.13 | 1 | 7 | Stam | 0.019 | Old |
| 0.484944 | 0.023509 | 0.395 | 0.364 | 1 | 7 | Tmx3 | 0.031 | Old |
| 0.485116 | 0.116127 | 0.215 | 0.187 | 1 | 7 | Pknox1 | 0.028 | Old |
| 0.485718 | 0.59817 | 0.026 | 0.018 | 1 | 7 | Bcr | 0.008 | Old |
| 0.48598 | 0.135898 | 0.5 | 0.458 | 1 | 7 | Ndufa5 | 0.042 | Old |
| 0.485999 | 0.030213 | 0.09 | 0.105 | 1 | 7 | Syne1 | -0.015 | Old |
| 0.48646 | 0.063807 | 0.441 | 0.413 | 1 | 7 | Hsd17b11 | 0.028 | Old |
| 0.486465 | 0.094861 | 0.062 | 0.075 | 1 | 7 | Gpr141 | -0.013 | Old |
| 0.486714 | 0.251572 | 0.169 | 0.148 | 1 | 7 | Rev3l | 0.021 | Old |
| 0.487102 | 0.992811 | 0.036 | 0.027 | 1 | 7 | Olfr1167 | 0.009 | Old |
| 0.487306 | 0.31701 | 0.123 | 0.105 | 1 | 7 | Arhgap23 | 0.018 | Old |
| 0.487993 | 0.120975 | 0.213 | 0.19 | 1 | 7 | Timm10 | 0.023 | Old |
| 0.488235 | 0.29281 | 0.131 | 0.114 | 1 | 7 | Iffo2 | 0.017 | Old |
| 0.488467 | 0.129874 | 0.205 | 0.184 | 1 | 7 | Pygb | 0.021 | Old |
| 0.488505 | 0.207178 | 0.379 | 0.352 | 1 | 7 | Mthfd2 | 0.027 | Old |

| | | | | | | | | |
|---|---|---|---|---|---|---|---|---|
| 0.488652 | 0.103046 | 0.082 | 0.096 | 1 | 7 | Gm1673 | -0.014 | Old |
| 0.488871 | 0.318315 | 0.067 | 0.054 | 1 | 7 | Tmem39b | 0.013 | Old |
| 0.489056 | 0.182676 | 0.241 | 0.22 | 1 | 7 | Pigs | 0.021 | Old |
| 0.489274 | 0.020883 | 0.662 | 0.602 | 1 | 7 | Phip | 0.06 | Old |
| 0.490075 | 0.226139 | 0.164 | 0.145 | 1 | 7 | Mea1 | 0.019 | Old |
| 0.490122 | 0.662338 | 0.026 | 0.018 | 1 | 7 | Slc18b1 | 0.008 | Old |
| 0.490458 | 0.157582 | 0.128 | 0.108 | 1 | 7 | Leo1 | 0.02 | Old |
| 0.490474 | 0.27896 | 0.103 | 0.12 | 1 | 7 | Fcgr4 | -0.017 | Old |
| 0.490658 | 0.25569 | 0.126 | 0.111 | 1 | 7 | Tprkb | 0.015 | Old |
| 0.490774 | 0.045659 | 0.651 | 0.614 | 1 | 7 | Zfp706 | 0.037 | Old |
| 0.491595 | 0.073743 | 0.756 | 0.726 | 1 | 7 | Dstn | 0.03 | Old |
| 0.49189 | 0.80644 | 0.026 | 0.018 | 1 | 7 | Gja1 | 0.008 | Old |
| 0.49189 | 0.046399 | 0.026 | 0.018 | 1 | 7 | Slc7a4 | 0.008 | Old |
| 0.492145 | 0.517548 | 0.079 | 0.066 | 1 | 7 | Dhx57 | 0.013 | Old |
| 0.492304 | 0.037235 | 0.479 | 0.44 | 1 | 7 | Ostc | 0.039 | Old |
| 0.492399 | 0.092072 | 0.731 | 0.705 | 1 | 7 | Myadm | 0.026 | Old |
| 0.49248 | 0.083773 | 0.056 | 0.045 | 1 | 7 | Fancc | 0.011 | Old |
| 0.492617 | 0.343379 | 0.082 | 0.069 | 1 | 7 | Dhx33 | 0.013 | Old |
| 0.492705 | 0.030411 | 0.487 | 0.452 | 1 | 7 | Tpp1 | 0.035 | Old |
| 0.492775 | 0.465895 | 0.026 | 0.018 | 1 | 7 | Nipsnap1 | 0.008 | Old |
| 0.492775 | 0.238975 | 0.026 | 0.018 | 1 | 7 | Gm14305 | 0.008 | Old |
| 0.492775 | 0.226156 | 0.026 | 0.018 | 1 | 7 | Wdr5b | 0.008 | Old |
| 0.492812 | 0.049286 | 0.195 | 0.175 | 1 | 7 | Slc45a4 | 0.02 | Old |
| 0.492912 | 0.07743 | 0.736 | 0.711 | 1 | 7 | Tm9sf3 | 0.025 | Old |
| 0.493219 | 0.135559 | 0.067 | 0.054 | 1 | 7 | Supt7l | 0.013 | Old |
| 0.493275 | 0.275221 | 0.377 | 0.349 | 1 | 7 | Mpc1 | 0.028 | Old |
| 0.493353 | 0.09282 | 0.451 | 0.416 | 1 | 7 | Smg1 | 0.035 | Old |
| 0.493758 | 0.035759 | 0.546 | 0.557 | 1 | 7 | Psmb8 | -0.011 | Old |
| 0.494108 | 0.094337 | 0.644 | 0.617 | 1 | 7 | Taldo1 | 0.027 | Old |
| 0.494387 | 0.282192 | 0.069 | 0.057 | 1 | 7 | Pyurf | 0.012 | Old |
| 0.494416 | 0.032981 | 0.292 | 0.265 | 1 | 7 | Slamf8 | 0.027 | Old |
| 0.49453 | 0.045765 | 0.064 | 0.051 | 1 | 7 | Plpp5 | 0.013 | Old |
| 0.494547 | 0.830947 | 0.026 | 0.018 | 1 | 7 | Gtf2h3 | 0.008 | Old |
| 0.494682 | 0.05024 | 0.044 | 0.033 | 1 | 7 | Serpine1 | 0.011 | Old |
| 0.49501 | 0.047172 | 0.508 | 0.479 | 1 | 7 | Dlst | 0.029 | Old |
| 0.495265 | 0.062672 | 0.615 | 0.569 | 1 | 7 | Ndufb3 | 0.046 | Old |
| 0.495295 | 0.131541 | 0.644 | 0.599 | 1 | 7 | Tmco1 | 0.045 | Old |
| 0.495439 | 0.18283 | 0.056 | 0.045 | 1 | 7 | Phykpl | 0.011 | Old |
| 0.495904 | 0.234515 | 0.179 | 0.157 | 1 | 7 | Wdr36 | 0.022 | Old |
| 0.496675 | 0.422705 | 0.115 | 0.099 | 1 | 7 | B3galnt2 | 0.016 | Old |
| 0.496942 | 0.107613 | 0.146 | 0.127 | 1 | 7 | Wdr18 | 0.019 | Old |
| 0.497281 | 0.238773 | 0.046 | 0.036 | 1 | 7 | Aste1 | 0.01 | Old |
| 0.497389 | 0.387774 | 0.2 | 0.181 | 1 | 7 | 5-Mar | 0.019 | Old |
| 0.497545 | 0.106871 | 0.621 | 0.587 | 1 | 7 | Lsm4 | 0.034 | Old |
| 0.497598 | 0.313744 | 0.069 | 0.057 | 1 | 7 | Rufy2 | 0.012 | Old |

| | | | | | | | | |
|---|---|---|---|---|---|---|---|---|
| 0.49764 | 0.077685 | 0.179 | 0.157 | 1 | 7 | Kri1 | 0.022 | Old |
| 0.497863 | 0.064767 | 0.656 | 0.614 | 1 | 7 | Creg1 | 0.042 | Old |
| 0.497884 | 0.150985 | 0.51 | 0.47 | 1 | 7 | Pa2g4 | 0.04 | Old |
| 0.498407 | 0.269063 | 0.056 | 0.045 | 1 | 7 | Pole2 | 0.011 | Old |
| 0.498516 | 0.074213 | 0.251 | 0.22 | 1 | 7 | Gpr108 | 0.031 | Old |
| 0.49889 | 0.398912 | 0.208 | 0.19 | 1 | 7 | Psmd9 | 0.018 | Old |
| 0.499058 | 0.054269 | 0.105 | 0.09 | 1 | 7 | Dxo | 0.015 | Old |
| 0.499339 | 0.096406 | 0.777 | 0.765 | 1 | 7 | Wdr1 | 0.012 | Old |
| 0.499419 | 0.200017 | 0.241 | 0.217 | 1 | 7 | Fem1b | 0.024 | Old |
| 0.499517 | 0.033549 | 0.156 | 0.136 | 1 | 7 | Tmem250- | 0.02 | Old |
| 0.49978 | 0.185005 | 0.067 | 0.054 | 1 | 7 | Tfip11 | 0.013 | Old |
| 0.499849 | 0.152327 | 0.836 | 0.825 | 1 | 7 | Ncl | 0.011 | Old |
| 0.499932 | 0.261857 | 0.351 | 0.331 | 1 | 7 | Lta4h | 0.02 | Old |
| 0.500194 | 0.017587 | 0.228 | 0.247 | 1 | 7 | Mms22l | -0.019 | Old |
| 0.500277 | 0.115878 | 0.387 | 0.407 | 1 | 7 | Cmtm7 | -0.02 | Old |
| 0.500905 | 0.137942 | 0.149 | 0.133 | 1 | 7 | Pigyl | 0.016 | Old |
| 0.501167 | 0.079255 | 0.126 | 0.108 | 1 | 7 | Foxo1 | 0.018 | Old |
| 0.50248 | 0.138879 | 0.269 | 0.247 | 1 | 7 | Zfp451 | 0.022 | Old |
| 0.502698 | 0.110577 | 0.823 | 0.792 | 1 | 7 | Gltp | 0.031 | Old |
| 0.503479 | 0.224179 | 0.438 | 0.422 | 1 | 7 | Bnip3l | 0.016 | Old |
| 0.503514 | 0.422169 | 0.069 | 0.057 | 1 | 7 | Rab13 | 0.012 | Old |
| 0.503873 | 0.104286 | 0.313 | 0.289 | 1 | 7 | Zdhhc3 | 0.024 | Old |
| 0.504261 | 0.149158 | 0.097 | 0.114 | 1 | 7 | Fancl | -0.017 | Old |
| 0.504678 | 0.130212 | 0.567 | 0.551 | 1 | 7 | Srp14 | 0.016 | Old |
| 0.504934 | 0.061812 | 0.21 | 0.187 | 1 | 7 | Mknk1 | 0.023 | Old |
| 0.504949 | 0.130636 | 0.169 | 0.151 | 1 | 7 | Saal1 | 0.018 | Old |
| 0.504969 | 0.46058 | 0.082 | 0.069 | 1 | 7 | 6030458C1 | 0.013 | Old |
| 0.505217 | 0.556372 | 0.082 | 0.069 | 1 | 7 | Cadm1 | 0.013 | Old |
| 0.50549 | 0.197445 | 0.295 | 0.274 | 1 | 7 | Yipf1 | 0.021 | Old |
| 0.505628 | 0.23599 | 0.182 | 0.199 | 1 | 7 | Kansl2 | -0.017 | Old |
| 0.505756 | 0.07886 | 0.382 | 0.346 | 1 | 7 | Mrps24 | 0.036 | Old |
| 0.505776 | 0.125576 | 0.405 | 0.367 | 1 | 7 | Ubxn4 | 0.038 | Old |
| 0.5062 | 0.046104 | 0.597 | 0.563 | 1 | 7 | Csrnp1 | 0.034 | Old |
| 0.506223 | 0.187428 | 0.079 | 0.066 | 1 | 7 | Nat6 | 0.013 | Old |
| 0.506236 | 0.197746 | 0.182 | 0.16 | 1 | 7 | Slc25a12 | 0.022 | Old |
| 0.506481 | 0.349488 | 0.036 | 0.027 | 1 | 7 | Praf2 | 0.009 | Old |
| 0.506481 | 0.082195 | 0.036 | 0.027 | 1 | 7 | Srr | 0.009 | Old |
| 0.506513 | 0.179681 | 0.046 | 0.036 | 1 | 7 | Hist1h2ab | 0.01 | Old |
| 0.506535 | 0.248697 | 0.154 | 0.136 | 1 | 7 | Ncoa5 | 0.018 | Old |
| 0.506626 | 0.00015 | 0.4 | 0.352 | 1 | 7 | Gorasp2 | 0.048 | Old |
| 0.50663 | 0.737442 | 0.133 | 0.117 | 1 | 7 | Triobp | 0.016 | Old |
| 0.50753 | 0.065953 | 0.2 | 0.181 | 1 | 7 | Cfap20 | 0.019 | Old |
| 0.507713 | 0.002371 | 0.041 | 0.051 | 1 | 7 | Tmem9 | -0.01 | Old |
| 0.508201 | 0.19506 | 0.318 | 0.292 | 1 | 7 | Yme1l1 | 0.026 | Old |
| 0.508456 | 0.307625 | 0.177 | 0.157 | 1 | 7 | Zswim8 | 0.02 | Old |

| | | | | | | | | |
|---|---|---|---|---|---|---|---|---|
| 0.508495 | 0.442878 | 0.085 | 0.072 | 1 | 7 | Hvcn1 | 0.013 | Old |
| 0.508943 | 0.015596 | 0.362 | 0.328 | 1 | 7 | Timm10b | 0.034 | Old |
| 0.509547 | 0.249559 | 0.182 | 0.163 | 1 | 7 | Ccdc174 | 0.019 | Old |
| 0.509912 | 0.738997 | 0.018 | 0.012 | 1 | 7 | Trem1 | 0.006 | Old |
| 0.509973 | 0.02541 | 0.497 | 0.449 | 1 | 7 | Ddi2 | 0.048 | Old |
| 0.51008 | 0.122945 | 0.277 | 0.298 | 1 | 7 | Hmgcs1 | -0.021 | Old |
| 0.510281 | 0.13506 | 0.172 | 0.154 | 1 | 7 | Crebzf | 0.018 | Old |
| 0.510523 | 0.046516 | 0.21 | 0.187 | 1 | 7 | Sart3 | 0.023 | Old |
| 0.510565 | 0.165096 | 0.095 | 0.081 | 1 | 7 | Rffl | 0.014 | Old |
| 0.510778 | 0.114758 | 0.551 | 0.521 | 1 | 7 | Rhog | 0.03 | Old |
| 0.510796 | 0.677989 | 0.059 | 0.048 | 1 | 7 | Trit1 | 0.011 | Old |
| 0.511265 | 0.061279 | 0.123 | 0.108 | 1 | 7 | Mybl2 | 0.015 | Old |
| 0.511289 | 0.054445 | 0.533 | 0.488 | 1 | 7 | Sypl | 0.045 | Old |
| 0.511299 | 0.16928 | 0.449 | 0.44 | 1 | 7 | Dctn1 | 0.009 | Old |
| 0.512549 | 0.275387 | 0.131 | 0.114 | 1 | 7 | Fam8a1 | 0.017 | Old |
| 0.513106 | 0.158217 | 0.4 | 0.373 | 1 | 7 | Tcf12 | 0.027 | Old |
| 0.513168 | 0.525461 | 0.018 | 0.012 | 1 | 7 | Lyplal1 | 0.006 | Old |
| 0.513255 | 0.355726 | 0.236 | 0.256 | 1 | 7 | Fam50a | -0.02 | Old |
| 0.51328 | 0.365591 | 0.146 | 0.13 | 1 | 7 | Smim26 | 0.016 | Old |
| 0.513737 | 0.491078 | 0.059 | 0.048 | 1 | 7 | Fam110b | 0.011 | Old |
| 0.513737 | 0.779563 | 0.059 | 0.048 | 1 | 7 | Rnaseh1 | 0.011 | Old |
| 0.513894 | 0.181169 | 0.246 | 0.22 | 1 | 7 | Pgrmc2 | 0.026 | Old |
| 0.514172 | 0.120694 | 0.256 | 0.232 | 1 | 7 | Ube3b | 0.024 | Old |
| 0.514293 | 0.157129 | 0.469 | 0.431 | 1 | 7 | Zmat2 | 0.038 | Old |
| 0.514425 | 0.147597 | 0.487 | 0.458 | 1 | 7 | Arhgef1 | 0.029 | Old |
| 0.514464 | 0.459527 | 0.082 | 0.069 | 1 | 7 | Amd1 | 0.013 | Old |
| 0.514684 | 0.265207 | 0.262 | 0.232 | 1 | 7 | Ykt6 | 0.03 | Old |
| 0.514857 | 0.099821 | 0.397 | 0.358 | 1 | 7 | Mrpl24 | 0.039 | Old |
| 0.515345 | 1.472633 | 0.018 | 0.012 | 1 | 7 | Kdr | 0.006 | Old |
| 0.515345 | 0.91714 | 0.018 | 0.012 | 1 | 7 | AC162302. | 0.006 | Old |
| 0.515345 | 0.266321 | 0.018 | 0.012 | 1 | 7 | Ocstamp | 0.006 | Old |
| 0.515382 | 0.031853 | 0.556 | 0.569 | 1 | 7 | Eif4g1 | -0.013 | Old |
| 0.516003 | 0.346491 | 0.069 | 0.057 | 1 | 7 | Olr1 | 0.012 | Old |
| 0.516119 | 0.028159 | 0.431 | 0.398 | 1 | 7 | Blmh | 0.033 | Old |
| 0.516291 | 0.007787 | 0.559 | 0.527 | 1 | 7 | Lasp1 | 0.032 | Old |
| 0.516977 | 0.022035 | 0.544 | 0.548 | 1 | 7 | Psmc3 | -0.004 | Old |
| 0.517044 | 0.075791 | 0.372 | 0.34 | 1 | 7 | 2510039O1 | 0.032 | Old |
| 0.517297 | 0.182499 | 0.103 | 0.087 | 1 | 7 | Ccs | 0.016 | Old |
| 0.517527 | 0.69142 | 0.018 | 0.012 | 1 | 7 | Cavin3 | 0.006 | Old |
| 0.517527 | 0.1331 | 0.018 | 0.012 | 1 | 7 | Mettl27 | 0.006 | Old |
| 0.517607 | 0.023676 | 0.418 | 0.395 | 1 | 7 | Rlim | 0.023 | Old |
| 0.517612 | 0.054313 | 0.21 | 0.232 | 1 | 7 | Tmem106b | -0.022 | Old |
| 0.517892 | 0.576438 | 0.049 | 0.039 | 1 | 7 | Nxpe4 | 0.01 | Old |
| 0.518134 | 0.103782 | 0.418 | 0.386 | 1 | 7 | Exoc5 | 0.032 | Old |
| 0.518619 | 0.683402 | 0.018 | 0.012 | 1 | 7 | Immp2l | 0.006 | Old |

| | | | | | | | | |
|---|---|---|---|---|---|---|---|---|
| 0.518619 | 0.347683 | 0.018 | 0.012 | 1 | 7 | 1700109H( | 0.006 | Old |
| 0.518667 | 0.070465 | 0.582 | 0.611 | 1 | 7 | Zfp91 | -0.029 | Old |
| 0.518742 | 0.160562 | 0.259 | 0.238 | 1 | 7 | Camta2 | 0.021 | Old |
| 0.518788 | 0.306727 | 0.113 | 0.099 | 1 | 7 | Ormdl3 | 0.014 | Old |
| 0.51971 | 0.125178 | 0.192 | 0.172 | 1 | 7 | Rwdd4a | 0.02 | Old |
| 0.519713 | 0.145231 | 0.018 | 0.012 | 1 | 7 | Prodh | 0.006 | Old |
| 0.520615 | 0.070419 | 0.272 | 0.247 | 1 | 7 | Gpaa1 | 0.025 | Old |
| 0.520808 | 0.45353 | 0.018 | 0.012 | 1 | 7 | Hist4h4 | 0.006 | Old |
| 0.520808 | 0.26287 | 0.018 | 0.012 | 1 | 7 | Cdkl2 | 0.006 | Old |
| 0.520808 | 0.056966 | 0.018 | 0.012 | 1 | 7 | Dpf3 | 0.006 | Old |
| 0.520938 | 0.122139 | 0.831 | 0.804 | 1 | 7 | Cd47 | 0.027 | Old |
| 0.521045 | 0.103347 | 0.569 | 0.515 | 1 | 7 | Ranbp2 | 0.054 | Old |
| 0.521312 | 0.11309 | 0.141 | 0.123 | 1 | 7 | Snx14 | 0.018 | Old |
| 0.522253 | 0.085993 | 0.264 | 0.238 | 1 | 7 | Bri3bp | 0.026 | Old |
| 0.522392 | 0.024568 | 0.151 | 0.133 | 1 | 7 | Ppp2cb | 0.018 | Old |
| 0.522797 | 0.021011 | 0.39 | 0.364 | 1 | 7 | Cyb5r4 | 0.026 | Old |
| 0.523804 | 0.40929 | 0.059 | 0.048 | 1 | 7 | Gstk1 | 0.011 | Old |
| 0.5241 | 0.229081 | 0.018 | 0.012 | 1 | 7 | Fads3 | 0.006 | Old |
| 0.5241 | 0.039446 | 0.018 | 0.012 | 1 | 7 | Poli | 0.006 | Old |
| 0.524424 | 0.111433 | 0.115 | 0.099 | 1 | 7 | Wrnip1 | 0.016 | Old |
| 0.524518 | 0.025141 | 0.085 | 0.096 | 1 | 7 | Chkb | -0.011 | Old |
| 0.524574 | 1.091412 | 0.028 | 0.021 | 1 | 7 | Rnf217 | 0.007 | Old |
| 0.524848 | 0.29584 | 0.305 | 0.283 | 1 | 7 | Max | 0.022 | Old |
| 0.525029 | 0.033622 | 0.803 | 0.774 | 1 | 7 | Dbi | 0.029 | Old |
| 0.525063 | 0.330059 | 0.203 | 0.184 | 1 | 7 | Plk1 | 0.019 | Old |
| 0.526161 | 0.214462 | 0.187 | 0.172 | 1 | 7 | Bcl6 | 0.015 | Old |
| 0.526301 | 0.058108 | 0.018 | 0.012 | 1 | 7 | Cd34 | 0.006 | Old |
| 0.526612 | 0.090864 | 0.151 | 0.133 | 1 | 7 | Nagpa | 0.018 | Old |
| 0.526706 | 0.002325 | 0.346 | 0.31 | 1 | 7 | Clptm1 | 0.036 | Old |
| 0.526725 | 0.472778 | 0.072 | 0.06 | 1 | 7 | Ppif | 0.012 | Old |
| 0.52678 | 0.300253 | 0.131 | 0.114 | 1 | 7 | Kntc1 | 0.017 | Old |
| 0.527115 | 0.119363 | 0.241 | 0.217 | 1 | 7 | Ap1ar | 0.024 | Old |
| 0.527477 | 1.16916 | 0.01 | 0.006 | 1 | 7 | Gas1 | 0.004 | Old |
| 0.527477 | 0.868252 | 0.01 | 0.006 | 1 | 7 | Zfp976 | 0.004 | Old |
| 0.527477 | 0.738195 | 0.01 | 0.006 | 1 | 7 | 4930558J1 | 0.004 | Old |
| 0.527477 | 0.615321 | 0.01 | 0.006 | 1 | 7 | Tmem120b | 0.004 | Old |
| 0.527609 | 0.270146 | 0.179 | 0.16 | 1 | 7 | Haus6 | 0.019 | Old |
| 0.528391 | 0.149524 | 0.474 | 0.431 | 1 | 7 | Adcy7 | 0.043 | Old |
| 0.528451 | 0.001567 | 0.379 | 0.358 | 1 | 7 | Pan3 | 0.021 | Old |
| 0.528506 | 0.096154 | 0.018 | 0.012 | 1 | 7 | Pced1b | 0.006 | Old |
| 0.528877 | 0.029629 | 0.303 | 0.274 | 1 | 7 | Nde1 | 0.029 | Old |
| 0.528961 | 0.114647 | 0.146 | 0.13 | 1 | 7 | Arl4a | 0.016 | Old |
| 0.529191 | 0.105507 | 0.513 | 0.485 | 1 | 7 | Dnm2 | 0.028 | Old |
| 0.529196 | 0.083008 | 0.238 | 0.217 | 1 | 7 | Nol11 | 0.021 | Old |
| 0.529278 | 0.263379 | 0.118 | 0.102 | 1 | 7 | Slc7a6os | 0.016 | Old |

| | | | | | | | | |
|---|---|---|---|---|---|---|---|---|
| 0.52947 | 0.012404 | 0.113 | 0.096 | 1 | 7 | Cpsf4 | 0.017 | Old |
| 0.529748 | 0.025887 | 0.133 | 0.117 | 1 | 7 | Ocel1 | 0.016 | Old |
| 0.530457 | 1.729369 | 0.01 | 0.006 | 1 | 7 | Cldn5 | 0.004 | Old |
| 0.530457 | 1.461502 | 0.01 | 0.006 | 1 | 7 | Sapcd1 | 0.004 | Old |
| 0.530457 | 1.355136 | 0.01 | 0.006 | 1 | 7 | Ccdc17 | 0.004 | Old |
| 0.530457 | 1.155733 | 0.01 | 0.006 | 1 | 7 | Tsku | 0.004 | Old |
| 0.530457 | 1.100541 | 0.01 | 0.006 | 1 | 7 | Tlr9 | 0.004 | Old |
| 0.530457 | 0.979211 | 0.01 | 0.006 | 1 | 7 | Fbxl22 | 0.004 | Old |
| 0.530457 | 0.859634 | 0.01 | 0.006 | 1 | 7 | Adamts5 | 0.004 | Old |
| 0.530457 | 0.522959 | 0.01 | 0.006 | 1 | 7 | Tox2 | 0.004 | Old |
| 0.530457 | 0.501568 | 0.01 | 0.006 | 1 | 7 | Gm10734 | 0.004 | Old |
| 0.530457 | 0.335191 | 0.01 | 0.006 | 1 | 7 | Padi3 | 0.004 | Old |
| 0.530468 | 0.155758 | 0.156 | 0.175 | 1 | 7 | Upf2 | -0.019 | Old |
| 0.530523 | 0.424172 | 0.146 | 0.13 | 1 | 7 | Map3k3 | 0.016 | Old |
| 0.530866 | 0.093859 | 0.069 | 0.057 | 1 | 7 | Fkbp9 | 0.012 | Old |
| 0.530976 | 0.535785 | 0.049 | 0.039 | 1 | 7 | Calcoco1 | 0.01 | Old |
| 0.531217 | 0.069016 | 0.228 | 0.247 | 1 | 7 | Baz2a | -0.019 | Old |
| 0.531352 | 0.263696 | 0.128 | 0.111 | 1 | 7 | Ctps | 0.017 | Old |
| 0.531368 | 0.073039 | 0.074 | 0.087 | 1 | 7 | Abhd5 | -0.013 | Old |
| 0.531635 | 0.417367 | 0.049 | 0.039 | 1 | 7 | Rnf135 | 0.01 | Old |
| 0.53164 | 0.242632 | 0.187 | 0.166 | 1 | 7 | Ms4a6d | 0.021 | Old |
| 0.531951 | 0.847918 | 0.01 | 0.006 | 1 | 7 | 4930539J0 | 0.004 | Old |
| 0.531951 | 0.639023 | 0.01 | 0.006 | 1 | 7 | 2810405F1 | 0.004 | Old |
| 0.531951 | 0.540085 | 0.01 | 0.006 | 1 | 7 | Npr1 | 0.004 | Old |
| 0.531951 | 0.430829 | 0.01 | 0.006 | 1 | 7 | Zfp296 | 0.004 | Old |
| 0.531951 | 0.355419 | 0.01 | 0.006 | 1 | 7 | Gm29394 | 0.004 | Old |
| 0.532113 | 0.113578 | 0.305 | 0.277 | 1 | 7 | Tfdp1 | 0.028 | Old |
| 0.532154 | 0.522021 | 0.072 | 0.06 | 1 | 7 | Ppp1r16a | 0.012 | Old |
| 0.532262 | 0.327254 | 0.121 | 0.105 | 1 | 7 | Thap3 | 0.016 | Old |
| 0.532508 | 0.177202 | 0.136 | 0.117 | 1 | 7 | Amz2 | 0.019 | Old |
| 0.532702 | 0.555932 | 0.1 | 0.087 | 1 | 7 | Strbp | 0.013 | Old |
| 0.533191 | 0.005372 | 0.897 | 0.861 | 1 | 7 | Rbm3 | 0.036 | Old |
| 0.533446 | 1.30002 | 0.01 | 0.006 | 1 | 7 | Hoxb7 | 0.004 | Old |
| 0.533446 | 1.033353 | 0.01 | 0.006 | 1 | 7 | Gm47815 | 0.004 | Old |
| 0.533446 | 0.97451 | 0.01 | 0.006 | 1 | 7 | Gm15013 | 0.004 | Old |
| 0.533446 | 0.611973 | 0.01 | 0.006 | 1 | 7 | Zfp59 | 0.004 | Old |
| 0.533446 | 0.402268 | 0.01 | 0.006 | 1 | 7 | Gm42893 | 0.004 | Old |
| 0.533446 | 0.372413 | 0.01 | 0.006 | 1 | 7 | Cx3cl1 | 0.004 | Old |
| 0.533446 | 0.352453 | 0.01 | 0.006 | 1 | 7 | Gm13822 | 0.004 | Old |
| 0.533446 | 0.321507 | 0.01 | 0.006 | 1 | 7 | Tmem158 | 0.004 | Old |
| 0.533446 | 0.293225 | 0.01 | 0.006 | 1 | 7 | Gm15964 | 0.004 | Old |
| 0.533446 | 0.281719 | 0.01 | 0.006 | 1 | 7 | Gm12166 | 0.004 | Old |
| 0.533446 | 0.264801 | 0.01 | 0.006 | 1 | 7 | Gm15050 | 0.004 | Old |
| 0.533446 | 0.264107 | 0.01 | 0.006 | 1 | 7 | 4930590J0 | 0.004 | Old |
| 0.533446 | 0.260816 | 0.01 | 0.006 | 1 | 7 | Hist1h2bm | 0.004 | Old |

| | | | | | | | | |
|---|---|---|---|---|---|---|---|---|
| 0.533446 | 0.071353 | 0.01 | 0.006 | 1 | 7 | Zfp975 | 0.004 | Old |
| 0.534393 | 0.097127 | 0.038 | 0.048 | 1 | 7 | Poll | -0.01 | Old |
| 0.534698 | 0.171442 | 0.179 | 0.163 | 1 | 7 | Sun1 | 0.016 | Old |
| 0.534784 | 0.11642 | 0.392 | 0.364 | 1 | 7 | Ilf2 | 0.028 | Old |
| 0.534933 | 0.621548 | 0.049 | 0.039 | 1 | 7 | Zfp688 | 0.01 | Old |
| 0.534944 | 1.122641 | 0.01 | 0.006 | 1 | 7 | Srl | 0.004 | Old |
| 0.534944 | 0.475311 | 0.01 | 0.006 | 1 | 7 | Slpi | 0.004 | Old |
| 0.534944 | 0.248191 | 0.01 | 0.006 | 1 | 7 | 9130604C2 | 0.004 | Old |
| 0.534944 | 0.139357 | 0.01 | 0.006 | 1 | 7 | Adh1 | 0.004 | Old |
| 0.534944 | 0.085503 | 0.01 | 0.006 | 1 | 7 | Rnls | 0.004 | Old |
| 0.534944 | 0.029954 | 0.01 | 0.006 | 1 | 7 | 4930477G( | 0.004 | Old |
| 0.535129 | 0.175392 | 0.095 | 0.081 | 1 | 7 | Paqr7 | 0.014 | Old |
| 0.535274 | 0.13379 | 0.492 | 0.455 | 1 | 7 | Itsn2 | 0.037 | Old |
| 0.535712 | 0.171512 | 0.338 | 0.316 | 1 | 7 | Tmbim4 | 0.022 | Old |
| 0.535736 | 0.076824 | 0.092 | 0.078 | 1 | 7 | Zdhhc12 | 0.014 | Old |
| 0.535773 | 0.394492 | 0.2 | 0.187 | 1 | 7 | Map2k4 | 0.013 | Old |
| 0.536033 | 0.297696 | 0.197 | 0.181 | 1 | 7 | Arhgap4 | 0.016 | Old |
| 0.536202 | 0.138462 | 0.313 | 0.274 | 1 | 7 | Ptov1 | 0.039 | Old |
| 0.536351 | 0.053384 | 0.11 | 0.096 | 1 | 7 | Vipas39 | 0.014 | Old |
| 0.536376 | 0.535733 | 0.085 | 0.072 | 1 | 7 | Hist1h3c | 0.013 | Old |
| 0.536443 | 1.290905 | 0.01 | 0.006 | 1 | 7 | Fam212b | 0.004 | Old |
| 0.536443 | 0.471627 | 0.01 | 0.006 | 1 | 7 | Asb2 | 0.004 | Old |
| 0.536443 | 0.338194 | 0.01 | 0.006 | 1 | 7 | Rhof | 0.004 | Old |
| 0.536443 | 0.31614 | 0.01 | 0.006 | 1 | 7 | Adam4 | 0.004 | Old |
| 0.536443 | 0.250152 | 0.01 | 0.006 | 1 | 7 | Htr1b | 0.004 | Old |
| 0.536443 | 0.223412 | 0.01 | 0.006 | 1 | 7 | Gm38832 | 0.004 | Old |
| 0.536443 | 0.20894 | 0.01 | 0.006 | 1 | 7 | Hist2h3c2 | 0.004 | Old |
| 0.536443 | 0.172074 | 0.01 | 0.006 | 1 | 7 | Dusp4 | 0.004 | Old |
| 0.536443 | 0.115455 | 0.01 | 0.006 | 1 | 7 | Prss30 | 0.004 | Old |
| 0.536443 | 0.020109 | 0.01 | 0.006 | 1 | 7 | Tnfrsf14 | 0.004 | Old |
| 0.536625 | 0.230911 | 0.223 | 0.211 | 1 | 7 | Tyms | 0.012 | Old |
| 0.536918 | 0.400988 | 0.049 | 0.039 | 1 | 7 | Dhx37 | 0.01 | Old |
| 0.537152 | 0.531207 | 0.146 | 0.133 | 1 | 7 | Zeb2os | 0.013 | Old |
| 0.537614 | 0.258956 | 0.028 | 0.021 | 1 | 7 | Eif3j2 | 0.007 | Old |
| 0.537861 | 0.121875 | 0.436 | 0.416 | 1 | 7 | Gfra2 | 0.02 | Old |
| 0.537892 | 0.109358 | 0.472 | 0.443 | 1 | 7 | Sumo1 | 0.029 | Old |
| 0.537945 | 0.005307 | 0.01 | 0.006 | 1 | 7 | 2700016F2 | 0.004 | Old |
| 0.538052 | 0.067523 | 0.869 | 0.849 | 1 | 7 | Cotl1 | 0.02 | Old |
| 0.539204 | 0.001335 | 0.285 | 0.259 | 1 | 7 | H2-Q4 | 0.026 | Old |
| 0.539449 | 0.026396 | 0.01 | 0.006 | 1 | 7 | Dqx1 | 0.004 | Old |
| 0.539692 | 0.158591 | 0.128 | 0.111 | 1 | 7 | Gm5547 | 0.017 | Old |
| 0.539708 | 0.280369 | 0.162 | 0.142 | 1 | 7 | Cfl2 | 0.02 | Old |
| 0.539989 | 0.309933 | 0.474 | 0.443 | 1 | 7 | Elmo2 | 0.031 | Old |
| 0.540513 | 0.028398 | 0.141 | 0.157 | 1 | 7 | Guca1a | -0.016 | Old |
| 0.540604 | 0.070352 | 0.762 | 0.741 | 1 | 7 | Tomm7 | 0.021 | Old |

| | | | | | | | | |
|---|---|---|---|---|---|---|---|---|
| 0.540877 | 0.008578 | 0.259 | 0.232 | 1 | 7 | Polr3k | 0.027 | Old |
| 0.540939 | 0.360289 | 0.044 | 0.054 | 1 | 7 | Arhgef10 | -0.01 | Old |
| 0.542089 | 0.015598 | 0.79 | 0.792 | 1 | 7 | Hnrnpm | -0.002 | Old |
| 0.542853 | 0.043023 | 0.29 | 0.262 | 1 | 7 | Katna1 | 0.028 | Old |
| 0.542912 | 0.277516 | 0.182 | 0.166 | 1 | 7 | Sdhaf2 | 0.016 | Old |
| 0.543413 | 0.163117 | 0.121 | 0.105 | 1 | 7 | Nmi | 0.016 | Old |
| 0.543754 | 0.239476 | 0.028 | 0.021 | 1 | 7 | Enpep | 0.007 | Old |
| 0.543822 | 0.129553 | 0.156 | 0.142 | 1 | 7 | Paox | 0.014 | Old |
| 0.544018 | 0.1313 | 0.036 | 0.045 | 1 | 7 | Fam72a | -0.009 | Old |
| 0.544195 | 0.305784 | 0.072 | 0.06 | 1 | 7 | Btbd17 | 0.012 | Old |
| 0.544609 | 0.179723 | 0.249 | 0.226 | 1 | 7 | Zcchc17 | 0.023 | Old |
| 0.544634 | 1.011602 | 0.028 | 0.021 | 1 | 7 | Pxdc1 | 0.007 | Old |
| 0.545101 | 0.640592 | 0.113 | 0.099 | 1 | 7 | Wdr75 | 0.014 | Old |
| 0.545362 | 0.139751 | 0.308 | 0.283 | 1 | 7 | Dync1li1 | 0.025 | Old |
| 0.545459 | 0.119007 | 0.269 | 0.247 | 1 | 7 | Gle1 | 0.022 | Old |
| 0.545515 | 0.058875 | 0.59 | 0.563 | 1 | 7 | Siva1 | 0.027 | Old |
| 0.545559 | 0.381852 | 0.049 | 0.039 | 1 | 7 | Zgpat | 0.01 | Old |
| 0.545559 | 0.083483 | 0.049 | 0.039 | 1 | 7 | Slc25a26 | 0.01 | Old |
| 0.545764 | 0.038679 | 0.218 | 0.199 | 1 | 7 | Txndc11 | 0.019 | Old |
| 0.545892 | 0.079899 | 0.156 | 0.139 | 1 | 7 | Akt3 | 0.017 | Old |
| 0.545923 | 0.151498 | 0.062 | 0.051 | 1 | 7 | Spic | 0.011 | Old |
| 0.546474 | 0.253696 | 0.038 | 0.03 | 1 | 7 | Hspa2 | 0.008 | Old |
| 0.547726 | 0.170345 | 0.074 | 0.063 | 1 | 7 | Wwox | 0.011 | Old |
| 0.547929 | 0.198287 | 0.121 | 0.108 | 1 | 7 | 1110059E2 | 0.013 | Old |
| 0.547988 | 0.280656 | 0.028 | 0.036 | 1 | 7 | Kiz | -0.008 | Old |
| 0.548161 | 0.133674 | 0.028 | 0.021 | 1 | 7 | Lrmda | 0.007 | Old |
| 0.548562 | 0.060706 | 0.549 | 0.524 | 1 | 7 | Psmd8 | 0.025 | Old |
| 0.548597 | 0.446448 | 0.1 | 0.087 | 1 | 7 | Cdc42ep4 | 0.013 | Old |
| 0.548732 | 0.463777 | 0.038 | 0.03 | 1 | 7 | Hist2h2aa1 | 0.008 | Old |
| 0.548732 | 0.100146 | 0.038 | 0.03 | 1 | 7 | Zfp111 | 0.008 | Old |
| 0.548832 | 0.05127 | 0.282 | 0.256 | 1 | 7 | Inppl1 | 0.026 | Old |
| 0.548958 | 0.293283 | 0.382 | 0.358 | 1 | 7 | Ptprs | 0.024 | Old |
| 0.54928 | 0.149858 | 0.154 | 0.136 | 1 | 7 | Kbtbd11 | 0.018 | Old |
| 0.549418 | 0.106162 | 0.323 | 0.301 | 1 | 7 | Coa3 | 0.022 | Old |
| 0.549455 | 0.371899 | 0.149 | 0.133 | 1 | 7 | Ppp2r3d | 0.016 | Old |
| 0.549528 | 0.131229 | 0.023 | 0.03 | 1 | 7 | Col1a2 | -0.007 | Old |
| 0.549929 | 0.273623 | 0.028 | 0.021 | 1 | 7 | 4732440D( | 0.007 | Old |
| 0.55024 | 0.280037 | 0.031 | 0.039 | 1 | 7 | Hpn | -0.008 | Old |
| 0.550469 | 0.202364 | 0.762 | 0.765 | 1 | 7 | Tmem176a | -0.003 | Old |
| 0.550722 | 0.175089 | 0.144 | 0.13 | 1 | 7 | Ulk1 | 0.014 | Old |
| 0.55081 | 0.068474 | 0.221 | 0.199 | 1 | 7 | Cenpq | 0.022 | Old |
| 0.550819 | 0.007077 | 0.072 | 0.06 | 1 | 7 | Pla2g12a | 0.012 | Old |
| 0.550913 | 0.123094 | 0.438 | 0.407 | 1 | 7 | Thoc7 | 0.031 | Old |
| 0.551212 | 0.634167 | 0.008 | 0.012 | 1 | 7 | Saa3 | -0.004 | Old |
| 0.55152 | 0.03836 | 0.097 | 0.084 | 1 | 7 | Idua | 0.013 | Old |

| | | | | | | | | |
|---|---|---|---|---|---|---|---|---|
| 0.551681 | 0.115639 | 0.164 | 0.181 | 1 | 7 | Slc35a5 | -0.017 | Old |
| 0.551762 | 0.234522 | 0.121 | 0.139 | 1 | 7 | Gdf15 | -0.018 | Old |
| 0.551889 | 0.129314 | 0.387 | 0.358 | 1 | 7 | Med28 | 0.029 | Old |
| 0.552107 | 0.025266 | 0.141 | 0.123 | 1 | 7 | Pik3cb | 0.018 | Old |
| 0.552622 | 0.160358 | 0.008 | 0.012 | 1 | 7 | Acsbg1 | -0.004 | Old |
| 0.553068 | 0.345252 | 0.062 | 0.051 | 1 | 7 | BC147527 | 0.011 | Old |
| 0.55367 | 0.060685 | 0.151 | 0.133 | 1 | 7 | Atn1 | 0.018 | Old |
| 0.55368 | 0.078807 | 0.944 | 0.931 | 1 | 7 | Klf6 | 0.013 | Old |
| 0.554151 | 0.049318 | 0.544 | 0.497 | 1 | 7 | Tmem160 | 0.047 | Old |
| 0.554223 | 0.374183 | 0.21 | 0.193 | 1 | 7 | Tbrg1 | 0.017 | Old |
| 0.554421 | 0.011796 | 0.503 | 0.524 | 1 | 7 | Asah1 | -0.021 | Old |
| 0.554551 | 0.097105 | 0.682 | 0.636 | 1 | 7 | Ubl5 | 0.046 | Old |
| 0.554775 | 0.515105 | 0.051 | 0.042 | 1 | 7 | Zbtb24 | 0.009 | Old |
| 0.555363 | 0.178585 | 0.233 | 0.214 | 1 | 7 | Plekhm3 | 0.019 | Old |
| 0.555429 | 0.335405 | 0.051 | 0.042 | 1 | 7 | Pced1a | 0.009 | Old |
| 0.555448 | 1.576189 | 0.008 | 0.012 | 1 | 7 | Zfp341 | -0.004 | Old |
| 0.555448 | 0.384866 | 0.008 | 0.012 | 1 | 7 | Cd27 | -0.004 | Old |
| 0.556046 | 0.162688 | 0.041 | 0.033 | 1 | 7 | Serpinb9 | 0.008 | Old |
| 0.556333 | 0.067943 | 0.595 | 0.569 | 1 | 7 | Bclaf1 | 0.026 | Old |
| 0.556365 | 0.2739 | 0.087 | 0.075 | 1 | 7 | Zbtb2 | 0.012 | Old |
| 0.556645 | 0.26155 | 0.028 | 0.036 | 1 | 7 | Mtmr11 | -0.008 | Old |
| 0.556806 | 0.005953 | 0.092 | 0.078 | 1 | 7 | Gstp1 | 0.014 | Old |
| 0.556864 | 0.367846 | 0.008 | 0.012 | 1 | 7 | Fam129c | -0.004 | Old |
| 0.556864 | 0.236854 | 0.008 | 0.012 | 1 | 7 | Gm40773 | -0.004 | Old |
| 0.557054 | 0.066641 | 0.423 | 0.386 | 1 | 7 | Ube2e3 | 0.037 | Old |
| 0.557101 | 0.045126 | 0.469 | 0.443 | 1 | 7 | Comt | 0.026 | Old |
| 0.557815 | 0.087136 | 0.069 | 0.057 | 1 | 7 | Mul1 | 0.012 | Old |
| 0.557857 | 0.513617 | 0.062 | 0.051 | 1 | 7 | Ift74 | 0.011 | Old |
| 0.557876 | 0.000642 | 0.123 | 0.105 | 1 | 7 | Styx | 0.018 | Old |
| 0.557943 | 0.019016 | 0.297 | 0.271 | 1 | 7 | Slc35a4 | 0.026 | Old |
| 0.558002 | 0.240999 | 0.308 | 0.289 | 1 | 7 | Nfu1 | 0.019 | Old |
| 0.558175 | 0.184218 | 0.023 | 0.03 | 1 | 7 | Cbarp | -0.007 | Old |
| 0.558281 | 0.480418 | 0.008 | 0.012 | 1 | 7 | Thrsp | -0.004 | Old |
| 0.558281 | 0.35259 | 0.008 | 0.012 | 1 | 7 | Cdkl3 | -0.004 | Old |
| 0.558281 | 0.316456 | 0.008 | 0.012 | 1 | 7 | Gm527 | -0.004 | Old |
| 0.558721 | 0.357239 | 0.097 | 0.084 | 1 | 7 | Ampd3 | 0.013 | Old |
| 0.55884 | 0.127348 | 0.151 | 0.136 | 1 | 7 | Cdc27 | 0.015 | Old |
| 0.558852 | 0.002749 | 0.433 | 0.389 | 1 | 7 | Mydgf | 0.044 | Old |
| 0.559225 | 0.17222 | 0.133 | 0.117 | 1 | 7 | Lxn | 0.016 | Old |
| 0.5595 | 0.330606 | 0.323 | 0.298 | 1 | 7 | Otub1 | 0.025 | Old |
| 0.559644 | 0.124885 | 0.208 | 0.19 | 1 | 7 | Dtnbp1 | 0.018 | Old |
| 0.5597 | 0.424849 | 0.008 | 0.012 | 1 | 7 | Gm16124 | -0.004 | Old |
| 0.5597 | 0.186388 | 0.008 | 0.012 | 1 | 7 | E430024I0! | -0.004 | Old |
| 0.559864 | 0.309108 | 0.177 | 0.16 | 1 | 7 | Atmin | 0.017 | Old |
| 0.560259 | 0.695087 | 0.062 | 0.051 | 1 | 7 | Kbtbd4 | 0.011 | Old |

| | | | | | | | | |
|---|---|---|---|---|---|---|---|---|
| 0.560259 | 0.248135 | 0.062 | 0.051 | 1 | 7 | Ccdc61 | 0.011 | Old |
| 0.560259 | 0.035372 | 0.062 | 0.051 | 1 | 7 | Sh3bp5l | 0.011 | Old |
| 0.560673 | 0.750065 | 0.051 | 0.042 | 1 | 7 | C1qtnf12 | 0.009 | Old |
| 0.561121 | 0.518794 | 0.008 | 0.012 | 1 | 7 | Des | -0.004 | Old |
| 0.561121 | 0.43894 | 0.008 | 0.012 | 1 | 7 | Tmem51os | -0.004 | Old |
| 0.561384 | 0.924337 | 0.1 | 0.087 | 1 | 7 | Usp21 | 0.013 | Old |
| 0.561499 | 0.022548 | 0.162 | 0.142 | 1 | 7 | Abl1 | 0.02 | Old |
| 0.561615 | 0.10013 | 0.038 | 0.03 | 1 | 7 | Btd | 0.008 | Old |
| 0.5619 | 0.308799 | 0.074 | 0.063 | 1 | 7 | Fmnl3 | 0.011 | Old |
| 0.562019 | 0.08755 | 0.733 | 0.732 | 1 | 7 | Ctsa | 0.001 | Old |
| 0.562148 | 0.058125 | 0.544 | 0.491 | 1 | 7 | Furin | 0.053 | Old |
| 0.562484 | 0.057837 | 0.064 | 0.054 | 1 | 7 | Katnal1 | 0.01 | Old |
| 0.562582 | 0.043353 | 0.269 | 0.247 | 1 | 7 | Cfdp1 | 0.022 | Old |
| 0.562646 | 0.340562 | 0.051 | 0.042 | 1 | 7 | Zfand1 | 0.009 | Old |
| 0.562947 | 0.075595 | 0.01 | 0.015 | 1 | 7 | Ces1d | -0.005 | Old |
| 0.56339 | 0.117835 | 0.179 | 0.16 | 1 | 7 | Memo1 | 0.019 | Old |
| 0.563608 | 0.025812 | 0.19 | 0.169 | 1 | 7 | Ppp2r5d | 0.021 | Old |
| 0.564446 | 0.245887 | 0.251 | 0.235 | 1 | 7 | Abcg1 | 0.016 | Old |
| 0.564875 | 0.116004 | 0.628 | 0.578 | 1 | 7 | Twf2 | 0.05 | Old |
| 0.565078 | 0.178409 | 0.062 | 0.051 | 1 | 7 | B4galt4 | 0.011 | Old |
| 0.565298 | 0.06244 | 0.205 | 0.187 | 1 | 7 | Slc25a45 | 0.018 | Old |
| 0.565562 | 0.69164 | 0.013 | 0.018 | 1 | 7 | Olfml3 | -0.005 | Old |
| 0.565563 | 0.152923 | 0.136 | 0.154 | 1 | 7 | Sema4d | -0.018 | Old |
| 0.565625 | 0.020067 | 0.172 | 0.151 | 1 | 7 | Prpsap2 | 0.021 | Old |
| 0.565646 | 0.069411 | 0.964 | 0.961 | 1 | 7 | H3f3b | 0.003 | Old |
| 0.565932 | 0.094418 | 0.11 | 0.096 | 1 | 7 | Prcc | 0.014 | Old |
| 0.566156 | 0.240282 | 0.1 | 0.087 | 1 | 7 | Cblb | 0.013 | Old |
| 0.566196 | 0.686465 | 0.031 | 0.039 | 1 | 7 | Fggy | -0.008 | Old |
| 0.566302 | 0.277216 | 0.074 | 0.087 | 1 | 7 | Ift27 | -0.013 | Old |
| 0.566427 | 0.015041 | 0.659 | 0.669 | 1 | 7 | Diaph1 | -0.01 | Old |
| 0.566536 | 0.15146 | 0.108 | 0.093 | 1 | 7 | Slc25a19 | 0.015 | Old |
| 0.566541 | 0.029392 | 0.521 | 0.524 | 1 | 7 | Drap1 | -0.003 | Old |
| 0.566879 | 0.317015 | 0.182 | 0.163 | 1 | 7 | Mrps5 | 0.019 | Old |
| 0.567317 | 0.007166 | 0.469 | 0.434 | 1 | 7 | Csf2rb | 0.035 | Old |
| 0.567536 | 0.189999 | 0.018 | 0.024 | 1 | 7 | Gm9530 | -0.006 | Old |
| 0.567892 | 0.24729 | 0.128 | 0.114 | 1 | 7 | Grap2 | 0.014 | Old |
| 0.567988 | 0.268461 | 0.01 | 0.015 | 1 | 7 | Ccl5 | -0.005 | Old |
| 0.567995 | 0.014965 | 0.241 | 0.226 | 1 | 7 | Pou2f1 | 0.015 | Old |
| 0.568148 | 0.009927 | 0.533 | 0.497 | 1 | 7 | Zbtb20 | 0.036 | Old |
| 0.568319 | 0.277831 | 0.128 | 0.114 | 1 | 7 | Mcm9 | 0.014 | Old |
| 0.568525 | 0.194694 | 0.146 | 0.133 | 1 | 7 | Gpr146 | 0.013 | Old |
| 0.568781 | 0.048721 | 0.944 | 0.952 | 1 | 7 | Rpl8 | -0.008 | Old |
| 0.568791 | 0.038808 | 0.372 | 0.337 | 1 | 7 | Gatad2a | 0.035 | Old |
| 0.569189 | 0.018738 | 0.272 | 0.253 | 1 | 7 | Emc8 | 0.019 | Old |
| 0.569294 | 0.384674 | 0.133 | 0.117 | 1 | 7 | Mrpl50 | 0.016 | Old |

| | | | | | | | | |
|---|---|---|---|---|---|---|---|---|
| 0.569501 | 0.027402 | 0.018 | 0.024 | 1 | 7 | Gm4610 | -0.006 | Old |
| 0.569501 | 0.023856 | 0.018 | 0.024 | 1 | 7 | E230032D2 | -0.006 | Old |
| 0.569572 | 0.27385 | 0.064 | 0.054 | 1 | 7 | Gm26782 | 0.01 | Old |
| 0.569576 | 0.100739 | 0.485 | 0.461 | 1 | 7 | Ddb1 | 0.024 | Old |
| 0.569852 | 0.036216 | 0.097 | 0.084 | 1 | 7 | 5730455P1 | 0.013 | Old |
| 0.57021 | 0.583769 | 0.123 | 0.111 | 1 | 7 | Npl | 0.012 | Old |
| 0.570622 | 0.109501 | 0.441 | 0.419 | 1 | 7 | Wls | 0.022 | Old |
| 0.571232 | 0.031438 | 0.608 | 0.566 | 1 | 7 | Ndufa1 | 0.042 | Old |
| 0.571266 | 0.141247 | 0.023 | 0.03 | 1 | 7 | Card6 | -0.007 | Old |
| 0.571465 | 0.068885 | 0.428 | 0.401 | 1 | 7 | Zcrb1 | 0.027 | Old |
| 0.572235 | 0.42857 | 0.041 | 0.033 | 1 | 7 | Fbxl8 | 0.008 | Old |
| 0.572456 | 0.2807 | 0.018 | 0.024 | 1 | 7 | Ugt1a7c | -0.006 | Old |
| 0.572468 | 0.597865 | 0.079 | 0.069 | 1 | 7 | Rpap3 | 0.01 | Old |
| 0.57305 | 0.221518 | 0.01 | 0.015 | 1 | 7 | Met | -0.005 | Old |
| 0.573102 | 1.624177 | 0.021 | 0.015 | 1 | 7 | Thnsl1 | 0.006 | Old |
| 0.573102 | 0.353572 | 0.015 | 0.021 | 1 | 7 | Sfxn5 | -0.006 | Old |
| 0.573343 | 0.254102 | 0.292 | 0.283 | 1 | 7 | Zfp800 | 0.009 | Old |
| 0.573423 | 0.372404 | 0.113 | 0.099 | 1 | 7 | Sirt1 | 0.014 | Old |
| 0.57358 | 0.015133 | 0.013 | 0.018 | 1 | 7 | Zfp58 | -0.005 | Old |
| 0.574163 | 0.016229 | 0.444 | 0.428 | 1 | 7 | Psmd14 | 0.016 | Old |
| 0.574336 | 0.242517 | 0.446 | 0.428 | 1 | 7 | Oat | 0.018 | Old |
| 0.574485 | 0.007713 | 0.192 | 0.172 | 1 | 7 | Ints1 | 0.02 | Old |
| 0.574543 | 0.055429 | 0.364 | 0.337 | 1 | 7 | Pih1d1 | 0.027 | Old |
| 0.57476 | 0.160769 | 0.123 | 0.108 | 1 | 7 | Tstd2 | 0.015 | Old |
| 0.574796 | 0.030898 | 0.677 | 0.654 | 1 | 7 | Ewsr1 | 0.023 | Old |
| 0.575026 | 0.055535 | 0.805 | 0.768 | 1 | 7 | 1810037I1 | 0.037 | Old |
| 0.575277 | 0.475066 | 0.1 | 0.087 | 1 | 7 | Tbc1d8 | 0.013 | Old |
| 0.5753 | 0.17594 | 0.41 | 0.377 | 1 | 7 | Kat6a | 0.033 | Old |
| 0.575343 | 0.061728 | 0.131 | 0.114 | 1 | 7 | Usp10 | 0.017 | Old |
| 0.575836 | 0.033366 | 0.449 | 0.416 | 1 | 7 | Zdhhc20 | 0.033 | Old |
| 0.576036 | 0.080746 | 0.177 | 0.16 | 1 | 7 | Ube2w | 0.017 | Old |
| 0.576076 | 0.478223 | 0.056 | 0.066 | 1 | 7 | Mllt3 | -0.01 | Old |
| 0.576272 | 0.167606 | 0.318 | 0.298 | 1 | 7 | Mien1 | 0.02 | Old |
| 0.57681 | 0.030398 | 0.756 | 0.729 | 1 | 7 | Atp1a1 | 0.027 | Old |
| 0.576965 | 0.219444 | 0.09 | 0.078 | 1 | 7 | Setdb2 | 0.012 | Old |
| 0.577342 | 1.195154 | 0.021 | 0.015 | 1 | 7 | Clec9a | 0.006 | Old |
| 0.578223 | 0.037394 | 0.115 | 0.102 | 1 | 7 | Lonp2 | 0.013 | Old |
| 0.578269 | 0.471522 | 0.236 | 0.223 | 1 | 7 | Gdap10 | 0.013 | Old |
| 0.578481 | 0.026091 | 0.254 | 0.235 | 1 | 7 | Relt | 0.019 | Old |
| 0.579588 | 0.156939 | 0.077 | 0.066 | 1 | 7 | Pycrl | 0.011 | Old |
| 0.579726 | 0.203269 | 0.118 | 0.105 | 1 | 7 | Gon7 | 0.013 | Old |
| 0.579906 | 0.044235 | 0.826 | 0.813 | 1 | 7 | Sfpq | 0.013 | Old |
| 0.580436 | 0.479589 | 0.021 | 0.027 | 1 | 7 | Ptgs2os2 | -0.006 | Old |
| 0.580506 | 0.25228 | 0.069 | 0.081 | 1 | 7 | Slc25a22 | -0.012 | Old |
| 0.580682 | 0.042232 | 0.077 | 0.066 | 1 | 7 | Phf11d | 0.011 | Old |

| | | | | | | | | |
|---|---|---|---|---|---|---|---|---|
| 0.581077 | 0.0369 | 0.726 | 0.705 | 1 | 7 | Ostf1 | 0.021 | Old |
| 0.58116 | 0.21709 | 0.041 | 0.033 | 1 | 7 | Rasal3 | 0.008 | Old |
| 0.581416 | 0.02653 | 0.077 | 0.087 | 1 | 7 | Tedc1 | -0.01 | Old |
| 0.581505 | 0.097199 | 0.367 | 0.349 | 1 | 7 | Rnf111 | 0.018 | Old |
| 0.581772 | 0.056512 | 0.328 | 0.298 | 1 | 7 | Coro1b | 0.03 | Old |
| 0.581772 | 0.146726 | 0.331 | 0.295 | 1 | 7 | Pkmyt1 | 0.036 | Old |
| 0.582376 | 0.522194 | 0.156 | 0.142 | 1 | 7 | Banp | 0.014 | Old |
| 0.582513 | 0.497031 | 0.031 | 0.024 | 1 | 7 | Ggcx | 0.007 | Old |
| 0.582663 | 0.759215 | 0.021 | 0.015 | 1 | 7 | Rab38 | 0.006 | Old |
| 0.582966 | 0.15465 | 0.131 | 0.117 | 1 | 7 | Alg6 | 0.014 | Old |
| 0.5834 | 0.034844 | 0.338 | 0.313 | 1 | 7 | Rtf2 | 0.025 | Old |
| 0.58373 | 0.139755 | 0.021 | 0.015 | 1 | 7 | Cxcl3 | 0.006 | Old |
| 0.584083 | 0.289079 | 0.09 | 0.078 | 1 | 7 | Nfrkb | 0.012 | Old |
| 0.58425 | 0.100923 | 0.249 | 0.232 | 1 | 7 | Ccnh | 0.017 | Old |
| 0.584395 | 0.449805 | 0.138 | 0.127 | 1 | 7 | Parp11 | 0.011 | Old |
| 0.584703 | 0.091259 | 0.136 | 0.12 | 1 | 7 | Abcf3 | 0.016 | Old |
| 0.584798 | 0.494209 | 0.021 | 0.015 | 1 | 7 | Shpk | 0.006 | Old |
| 0.584895 | 0.085863 | 0.556 | 0.512 | 1 | 7 | Mpp6 | 0.044 | Old |
| 0.584954 | 0.026056 | 0.438 | 0.41 | 1 | 7 | Mrps14 | 0.028 | Old |
| 0.585544 | 0.106211 | 0.444 | 0.407 | 1 | 7 | Agps | 0.037 | Old |
| 0.585593 | 0.065111 | 0.423 | 0.389 | 1 | 7 | Sec14l1 | 0.034 | Old |
| 0.585851 | 0.208675 | 0.267 | 0.244 | 1 | 7 | Agpat4 | 0.023 | Old |
| 0.586416 | 0.360377 | 0.318 | 0.301 | 1 | 7 | BC031181 | 0.017 | Old |
| 0.58664 | 0.09208 | 0.638 | 0.602 | 1 | 7 | Atp5o.1 | 0.036 | Old |
| 0.58684 | 0.070006 | 0.031 | 0.024 | 1 | 7 | Rccd1 | 0.007 | Old |
| 0.58689 | 0.023845 | 0.228 | 0.211 | 1 | 7 | 2310009B1 | 0.017 | Old |
| 0.586936 | 0.418512 | 0.021 | 0.015 | 1 | 7 | Treml2 | 0.006 | Old |
| 0.586936 | 0.38873 | 0.021 | 0.015 | 1 | 7 | Mfsd6l | 0.006 | Old |
| 0.586936 | 0.147409 | 0.021 | 0.015 | 1 | 7 | 4930481A1 | 0.006 | Old |
| 0.586936 | 0.114936 | 0.021 | 0.015 | 1 | 7 | Dnmt3b | 0.006 | Old |
| 0.586936 | 0.110911 | 0.021 | 0.015 | 1 | 7 | Cd109 | 0.006 | Old |
| 0.587008 | 0.028755 | 0.387 | 0.355 | 1 | 7 | Golph3 | 0.032 | Old |
| 0.587504 | 0.13741 | 0.256 | 0.238 | 1 | 7 | 1810030O0 | 0.018 | Old |
| 0.587524 | 0.009094 | 0.279 | 0.262 | 1 | 7 | Wdr37 | 0.017 | Old |
| 0.587657 | 0.005083 | 0.09 | 0.078 | 1 | 7 | Mllt1 | 0.012 | Old |
| 0.588007 | 0.220569 | 0.021 | 0.015 | 1 | 7 | Kbtbd8 | 0.006 | Old |
| 0.588007 | 0.093709 | 0.021 | 0.015 | 1 | 7 | Gm39469 | 0.006 | Old |
| 0.588007 | 0.040599 | 0.021 | 0.015 | 1 | 7 | 2810021J2 | 0.006 | Old |
| 0.588213 | 0.034866 | 0.297 | 0.268 | 1 | 7 | Map2k7 | 0.029 | Old |
| 0.588978 | 0.117954 | 0.282 | 0.301 | 1 | 7 | Ppm1b | -0.019 | Old |
| 0.589075 | 0.232198 | 0.618 | 0.617 | 1 | 7 | Sec11c | 0.001 | Old |
| 0.589078 | 0.213066 | 0.021 | 0.015 | 1 | 7 | P3h1 | 0.006 | Old |
| 0.589192 | 0.227787 | 0.09 | 0.078 | 1 | 7 | Oas2 | 0.012 | Old |
| 0.589297 | 0.082541 | 0.167 | 0.151 | 1 | 7 | Hmg20a | 0.016 | Old |
| 0.589755 | 0.337324 | 0.054 | 0.045 | 1 | 7 | Rab3ip | 0.009 | Old |

| | | | | | | | | |
|---|---|---|---|---|---|---|---|---|
| 0.589897 | 0.058772 | 0.367 | 0.325 | 1 | 7 | Nudt4 | 0.042 | Old |
| 0.589958 | 0.219777 | 0.144 | 0.13 | 1 | 7 | Flcn | 0.014 | Old |
| 0.591041 | 0.088208 | 0.205 | 0.187 | 1 | 7 | Glipr1 | 0.018 | Old |
| 0.591044 | 0.174767 | 0.174 | 0.157 | 1 | 7 | Fbxl20 | 0.017 | Old |
| 0.591224 | 0.002153 | 0.021 | 0.015 | 1 | 7 | Bhlhb9 | 0.006 | Old |
| 0.591308 | 0.104056 | 0.138 | 0.154 | 1 | 7 | S1pr2 | -0.016 | Old |
| 0.592089 | 0.245848 | 0.372 | 0.358 | 1 | 7 | Ndufb2 | 0.014 | Old |
| 0.592916 | 0.054842 | 0.359 | 0.328 | 1 | 7 | Akirin1 | 0.031 | Old |
| 0.59333 | 0.235962 | 0.077 | 0.066 | 1 | 7 | Acvr1b | 0.011 | Old |
| 0.59347 | 0.281479 | 0.236 | 0.22 | 1 | 7 | Upb1 | 0.016 | Old |
| 0.593546 | 0.06249 | 0.156 | 0.145 | 1 | 7 | Cog5 | 0.011 | Old |
| 0.593582 | 0.105315 | 0.064 | 0.075 | 1 | 7 | 1110032A( | -0.011 | Old |
| 0.593753 | 0.006354 | 0.123 | 0.108 | 1 | 7 | Siae | 0.015 | Old |
| 0.593977 | 0.412767 | 0.054 | 0.063 | 1 | 7 | Xrcc2 | -0.009 | Old |
| 0.59421 | 0.076071 | 0.659 | 0.611 | 1 | 7 | Wipf1 | 0.048 | Old |
| 0.594305 | 0.163667 | 0.213 | 0.193 | 1 | 7 | Golt1b | 0.02 | Old |
| 0.594413 | 0.322672 | 0.103 | 0.09 | 1 | 7 | Dnajc24 | 0.013 | Old |
| 0.594786 | 0.198836 | 0.279 | 0.262 | 1 | 7 | Ltn1 | 0.017 | Old |
| 0.594989 | 0.314146 | 0.077 | 0.066 | 1 | 7 | Habp4 | 0.011 | Old |
| 0.595107 | 0.134322 | 0.079 | 0.069 | 1 | 7 | Gatc | 0.01 | Old |
| 0.595398 | 0.056157 | 0.392 | 0.373 | 1 | 7 | 1810058I2 | 0.019 | Old |
| 0.595723 | 0.025617 | 0.349 | 0.325 | 1 | 7 | Rbm22 | 0.024 | Old |
| 0.596162 | 0.484075 | 0.182 | 0.166 | 1 | 7 | Tax1bp3 | 0.016 | Old |
| 0.596311 | 0.58082 | 0.056 | 0.048 | 1 | 7 | Gm49368 | 0.008 | Old |
| 0.596461 | 0.043004 | 0.754 | 0.705 | 1 | 7 | Pafah1b1 | 0.049 | Old |
| 0.596844 | 0.087043 | 0.236 | 0.214 | 1 | 7 | Zfp131 | 0.022 | Old |
| 0.597285 | 0.292911 | 0.031 | 0.024 | 1 | 7 | Sag | 0.007 | Old |
| 0.597556 | 0.019775 | 0.562 | 0.533 | 1 | 7 | Anapc11 | 0.029 | Old |
| 0.598222 | 0.106924 | 0.597 | 0.566 | 1 | 7 | Jmjd1c | 0.031 | Old |
| 0.598433 | 0.000693 | 0.382 | 0.343 | 1 | 7 | Fam126a | 0.039 | Old |
| 0.598467 | 0.125435 | 0.592 | 0.569 | 1 | 7 | Snrpg | 0.023 | Old |
| 0.598624 | 0.049867 | 0.438 | 0.416 | 1 | 7 | Sowahc | 0.022 | Old |
| 0.598698 | 0.060698 | 0.618 | 0.584 | 1 | 7 | Arf4 | 0.034 | Old |
| 0.598919 | 0.102207 | 0.079 | 0.069 | 1 | 7 | Zcchc14 | 0.01 | Old |
| 0.599776 | 0.213515 | 0.105 | 0.093 | 1 | 7 | Gpr89 | 0.012 | Old |
| 0.59991 | 0.020029 | 0.031 | 0.024 | 1 | 7 | Afg1l | 0.007 | Old |
| 0.599976 | 0.001612 | 0.062 | 0.051 | 1 | 7 | Prickle3 | 0.011 | Old |
| 0.600277 | 0.141091 | 0.164 | 0.148 | 1 | 7 | Sp2 | 0.016 | Old |
| 0.601073 | 0.427196 | 0.056 | 0.066 | 1 | 7 | Ramp3 | -0.01 | Old |
| 0.601422 | 0.043335 | 0.646 | 0.633 | 1 | 7 | Ranbp1 | 0.013 | Old |
| 0.601616 | 0.077873 | 0.333 | 0.316 | 1 | 7 | Plcg2 | 0.017 | Old |
| 0.601877 | 0.027108 | 0.197 | 0.178 | 1 | 7 | Itgb7 | 0.019 | Old |
| 0.601987 | 0.049985 | 0.064 | 0.054 | 1 | 7 | Rfesd | 0.01 | Old |
| 0.602464 | 0.131415 | 0.187 | 0.169 | 1 | 7 | Stau1 | 0.018 | Old |
| 0.602592 | 0.018472 | 0.267 | 0.241 | 1 | 7 | Mecp2 | 0.026 | Old |

| | | | | | | | | |
|---|---|---|---|---|---|---|---|---|
| 0.602743 | 0.213929 | 0.056 | 0.048 | 1 | 7 | Zfp729b | 0.008 | Old |
| 0.603071 | 0.033231 | 0.246 | 0.226 | 1 | 7 | Hist1h1b | 0.02 | Old |
| 0.603102 | 0.626171 | 0.067 | 0.057 | 1 | 7 | Retsat | 0.01 | Old |
| 0.603187 | 0.367665 | 0.164 | 0.151 | 1 | 7 | Tmem55a | 0.013 | Old |
| 0.603548 | 0.25069 | 0.285 | 0.268 | 1 | 7 | Dnajc1 | 0.017 | Old |
| 0.60364 | 0.104641 | 0.141 | 0.127 | 1 | 7 | Aen | 0.014 | Old |
| 0.603647 | 0.025381 | 0.233 | 0.217 | 1 | 7 | Chchd7 | 0.016 | Old |
| 0.604555 | 0.376357 | 0.131 | 0.117 | 1 | 7 | Bbc3 | 0.014 | Old |
| 0.605037 | 0.037974 | 0.851 | 0.846 | 1 | 7 | Arhgdia | 0.005 | Old |
| 0.605089 | 0.093355 | 0.19 | 0.172 | 1 | 7 | Ilvbl | 0.018 | Old |
| 0.605207 | 0.537345 | 0.072 | 0.063 | 1 | 7 | Dgkg | 0.009 | Old |
| 0.605813 | 0.123422 | 0.408 | 0.392 | 1 | 7 | Nop58 | 0.016 | Old |
| 0.605877 | 0.044288 | 0.79 | 0.75 | 1 | 7 | Ap2a2 | 0.04 | Old |
| 0.606054 | 0.031513 | 0.031 | 0.024 | 1 | 7 | Ppp2r5b | 0.007 | Old |
| 0.606084 | 0.660854 | 0.067 | 0.057 | 1 | 7 | Tjap1 | 0.01 | Old |
| 0.606622 | 0.010436 | 0.113 | 0.099 | 1 | 7 | Lix1l | 0.014 | Old |
| 0.607043 | 0.094405 | 0.874 | 0.843 | 1 | 7 | Sdcbp | 0.031 | Old |
| 0.607158 | 0.042014 | 0.454 | 0.413 | 1 | 7 | Glyr1 | 0.041 | Old |
| 0.607434 | 0.095348 | 0.351 | 0.328 | 1 | 7 | Psme4 | 0.023 | Old |
| 0.607683 | 0.053421 | 0.408 | 0.373 | 1 | 7 | Cenpx | 0.035 | Old |
| 0.6077 | 0.218012 | 0.056 | 0.066 | 1 | 7 | Cry2 | -0.01 | Old |
| 0.607739 | 0.087368 | 0.941 | 0.931 | 1 | 7 | Tubb5 | 0.01 | Old |
| 0.607795 | 0.147269 | 0.108 | 0.096 | 1 | 7 | Sugp2 | 0.012 | Old |
| 0.607975 | 0.276289 | 0.082 | 0.072 | 1 | 7 | Golga1 | 0.01 | Old |
| 0.608035 | 0.110472 | 0.077 | 0.087 | 1 | 7 | Socs5 | -0.01 | Old |
| 0.608209 | 0.128579 | 0.133 | 0.12 | 1 | 7 | Mettl1 | 0.013 | Old |
| 0.608311 | 0.1378 | 0.249 | 0.229 | 1 | 7 | Ncaph | 0.02 | Old |
| 0.608857 | 0.131618 | 0.41 | 0.401 | 1 | 7 | R3hdm1 | 0.009 | Old |
| 0.609206 | 0.658295 | 0.056 | 0.048 | 1 | 7 | Ddx28 | 0.008 | Old |
| 0.609324 | 0.133863 | 0.172 | 0.154 | 1 | 7 | Tmx2 | 0.018 | Old |
| 0.609777 | 0.134808 | 0.269 | 0.25 | 1 | 7 | Get4 | 0.019 | Old |
| 0.60984 | 0.15526 | 0.041 | 0.033 | 1 | 7 | Phlda3 | 0.008 | Old |
| 0.611166 | 0.065322 | 0.687 | 0.657 | 1 | 7 | Syngr2 | 0.03 | Old |
| 0.611179 | 0.216995 | 0.123 | 0.136 | 1 | 7 | Frmd6 | -0.013 | Old |
| 0.61139 | 0.261079 | 0.09 | 0.078 | 1 | 7 | Mex3d | 0.012 | Old |
| 0.611851 | 0.633429 | 0.033 | 0.027 | 1 | 7 | Armcx5 | 0.006 | Old |
| 0.613319 | 0.105032 | 0.205 | 0.187 | 1 | 7 | Fut11 | 0.018 | Old |
| 0.613423 | 0.109041 | 0.121 | 0.108 | 1 | 7 | Mitd1 | 0.013 | Old |
| 0.614123 | 0.02224 | 0.538 | 0.5 | 1 | 7 | Trim28 | 0.038 | Old |
| 0.614222 | 0.457049 | 0.126 | 0.117 | 1 | 7 | Slc37a2 | 0.009 | Old |
| 0.615043 | 0.043114 | 0.377 | 0.355 | 1 | 7 | Tbl1x | 0.022 | Old |
| 0.615206 | 0.135706 | 0.221 | 0.238 | 1 | 7 | Mctp1 | -0.017 | Old |
| 0.615383 | 0.090313 | 0.079 | 0.069 | 1 | 7 | Bcl9 | 0.01 | Old |
| 0.615397 | 0.128459 | 0.397 | 0.373 | 1 | 7 | Smim7 | 0.024 | Old |
| 0.615666 | 0.029111 | 0.333 | 0.307 | 1 | 7 | Ssbp3 | 0.026 | Old |

| | | | | | | | | |
|---|---|---|---|---|---|---|---|---|
| 0.616087 | 0.165439 | 0.169 | 0.154 | 1 | 7 | Bard1 | 0.015 | Old |
| 0.616559 | 0.068124 | 0.467 | 0.434 | 1 | 7 | Maz | 0.033 | Old |
| 0.616568 | 0.255897 | 0.149 | 0.136 | 1 | 7 | Lpin1 | 0.013 | Old |
| 0.616924 | 0.036457 | 0.495 | 0.455 | 1 | 7 | Csnk2b | 0.04 | Old |
| 0.617066 | 0.001318 | 0.467 | 0.443 | 1 | 7 | Rab21 | 0.024 | Old |
| 0.618103 | 0.075168 | 0.267 | 0.277 | 1 | 7 | Chn2 | -0.01 | Old |
| 0.618302 | 0.082934 | 0.097 | 0.108 | 1 | 7 | Kctd17 | -0.011 | Old |
| 0.618625 | 0.285299 | 0.133 | 0.12 | 1 | 7 | Ino80 | 0.013 | Old |
| 0.619223 | 0.294882 | 0.405 | 0.383 | 1 | 7 | Smarcc2 | 0.022 | Old |
| 0.619635 | 0.111641 | 0.167 | 0.184 | 1 | 7 | Lipo3 | -0.017 | Old |
| 0.619726 | 0.225914 | 0.044 | 0.036 | 1 | 7 | Irgq | 0.008 | Old |
| 0.619785 | 0.026325 | 0.287 | 0.265 | 1 | 7 | Slc38a10 | 0.022 | Old |
| 0.620148 | 1.924987 | 0.013 | 0.009 | 1 | 7 | Pcdhga12 | 0.004 | Old |
| 0.620263 | 0.317966 | 0.056 | 0.048 | 1 | 7 | Tbc1d7 | 0.008 | Old |
| 0.620561 | 0.398549 | 0.095 | 0.084 | 1 | 7 | Bud23 | 0.011 | Old |
| 0.620916 | 0.796682 | 0.056 | 0.048 | 1 | 7 | 4933427D1 | 0.008 | Old |
| 0.62132 | 0.002214 | 0.759 | 0.843 | 1 | 7 | Ccr2 | -0.084 | Old |
| 0.621542 | 0.728992 | 0.013 | 0.009 | 1 | 7 | Bbs5 | 0.004 | Old |
| 0.621542 | 0.325928 | 0.013 | 0.009 | 1 | 7 | Xkr5 | 0.004 | Old |
| 0.62158 | 0.117462 | 0.344 | 0.316 | 1 | 7 | Mnt | 0.028 | Old |
| 0.621929 | 0.011705 | 0.713 | 0.708 | 1 | 7 | Chd4 | 0.005 | Old |
| 0.622089 | 0.713127 | 0.095 | 0.084 | 1 | 7 | Taf1b | 0.011 | Old |
| 0.62228 | 0.095913 | 0.213 | 0.196 | 1 | 7 | Pop5 | 0.017 | Old |
| 0.622524 | 0.008544 | 0.636 | 0.596 | 1 | 7 | Smc2 | 0.04 | Old |
| 0.622937 | 1.211172 | 0.013 | 0.009 | 1 | 7 | Gpihbp1 | 0.004 | Old |
| 0.622937 | 1.012483 | 0.013 | 0.009 | 1 | 7 | Enkur | 0.004 | Old |
| 0.62341 | 0.099223 | 0.077 | 0.087 | 1 | 7 | Gmeb2 | -0.01 | Old |
| 0.624334 | 0.653153 | 0.013 | 0.009 | 1 | 7 | Gm9856 | 0.004 | Old |
| 0.624334 | 0.459794 | 0.013 | 0.009 | 1 | 7 | Gm14295 | 0.004 | Old |
| 0.624783 | 0.783839 | 0.023 | 0.018 | 1 | 7 | Mpv17l | 0.005 | Old |
| 0.625056 | 0.125713 | 0.372 | 0.337 | 1 | 7 | Lrrfip1 | 0.035 | Old |
| 0.625332 | 0.430796 | 0.097 | 0.087 | 1 | 7 | Fyco1 | 0.01 | Old |
| 0.625732 | 1.263818 | 0.013 | 0.009 | 1 | 7 | Ildr1 | 0.004 | Old |
| 0.625732 | 0.493787 | 0.013 | 0.009 | 1 | 7 | Hus1b | 0.004 | Old |
| 0.62585 | 0.004941 | 0.221 | 0.205 | 1 | 7 | Dusp16 | 0.016 | Old |
| 0.626264 | 0.180411 | 0.033 | 0.027 | 1 | 7 | Helq | 0.006 | Old |
| 0.626378 | 0.034073 | 0.259 | 0.244 | 1 | 7 | Gng12 | 0.015 | Old |
| 0.627089 | 0.037924 | 0.118 | 0.105 | 1 | 7 | Plrg1 | 0.013 | Old |
| 0.627116 | 0.505012 | 0.033 | 0.027 | 1 | 7 | Nrg2 | 0.006 | Old |
| 0.627116 | 0.239867 | 0.033 | 0.027 | 1 | 7 | Oxld1 | 0.006 | Old |
| 0.627132 | 0.428869 | 0.013 | 0.009 | 1 | 7 | Vsig10l | 0.004 | Old |
| 0.627132 | 0.408934 | 0.013 | 0.009 | 1 | 7 | Creg2 | 0.004 | Old |
| 0.627132 | 0.379311 | 0.013 | 0.009 | 1 | 7 | Rpp38 | 0.004 | Old |
| 0.627132 | 0.334261 | 0.013 | 0.009 | 1 | 7 | Tchh | 0.004 | Old |
| 0.627775 | 0.023021 | 0.556 | 0.584 | 1 | 7 | Adam10 | -0.028 | Old |

| | | | | | | | | |
|---|---|---|---|---|---|---|---|---|
| 0.627864 | 0.503346 | 0.023 | 0.018 | 1 | 7 | Hacl1 | 0.005 | Old |
| 0.62853 | 0.178658 | 0.208 | 0.193 | 1 | 7 | Erap1 | 0.015 | Old |
| 0.628533 | 0.260543 | 0.013 | 0.009 | 1 | 7 | C77080 | 0.004 | Old |
| 0.628533 | 0.236581 | 0.013 | 0.009 | 1 | 7 | Cysltr2 | 0.004 | Old |
| 0.628593 | 0.027742 | 0.495 | 0.452 | 1 | 7 | Add1 | 0.043 | Old |
| 0.628892 | 0.788668 | 0.023 | 0.018 | 1 | 7 | Eml6 | 0.005 | Old |
| 0.629935 | 0.645152 | 0.013 | 0.009 | 1 | 7 | Glcci1 | 0.004 | Old |
| 0.629935 | 0.281492 | 0.013 | 0.009 | 1 | 7 | Tmem171 | 0.004 | Old |
| 0.629935 | 0.176154 | 0.013 | 0.009 | 1 | 7 | Klhl11 | 0.004 | Old |
| 0.630078 | 0.699322 | 0.072 | 0.063 | 1 | 7 | Exo1 | 0.009 | Old |
| 0.630282 | 0.357318 | 0.177 | 0.163 | 1 | 7 | Timm9 | 0.014 | Old |
| 0.630322 | 0.15473 | 0.11 | 0.099 | 1 | 7 | Camsap1 | 0.011 | Old |
| 0.630556 | 0.026177 | 0.264 | 0.241 | 1 | 7 | Sik1 | 0.023 | Old |
| 0.630951 | 0.533646 | 0.09 | 0.081 | 1 | 7 | Rnf144b | 0.009 | Old |
| 0.631099 | 0.041468 | 0.659 | 0.639 | 1 | 7 | 11-Sep | 0.02 | Old |
| 0.631184 | 0.093844 | 0.205 | 0.187 | 1 | 7 | Ext2 | 0.018 | Old |
| 0.631339 | 0.216961 | 0.013 | 0.009 | 1 | 7 | Gm10320 | 0.004 | Old |
| 0.631339 | 0.210365 | 0.013 | 0.009 | 1 | 7 | Smoc2 | 0.004 | Old |
| 0.631339 | 0.208901 | 0.013 | 0.009 | 1 | 7 | 1700003F1 | 0.004 | Old |
| 0.631339 | 0.157969 | 0.013 | 0.009 | 1 | 7 | Necab3 | 0.004 | Old |
| 0.631339 | 0.123716 | 0.013 | 0.009 | 1 | 7 | Stard10 | 0.004 | Old |
| 0.631339 | 0.117874 | 0.013 | 0.009 | 1 | 7 | Gm15853 | 0.004 | Old |
| 0.631646 | 0.090136 | 0.136 | 0.123 | 1 | 7 | Gnl3l | 0.013 | Old |
| 0.631735 | 0.337146 | 0.049 | 0.057 | 1 | 7 | Asb6 | -0.008 | Old |
| 0.632042 | 0.389063 | 0.182 | 0.169 | 1 | 7 | Gskip | 0.013 | Old |
| 0.632223 | 0.709832 | 0.046 | 0.039 | 1 | 7 | 3-Sep | 0.007 | Old |
| 0.632297 | 0.176926 | 0.295 | 0.28 | 1 | 7 | Srf | 0.015 | Old |
| 0.632515 | 0.050646 | 0.577 | 0.551 | 1 | 7 | Eif5 | 0.026 | Old |
| 0.632744 | 0.15948 | 0.013 | 0.009 | 1 | 7 | Gm17655 | 0.004 | Old |
| 0.632744 | 0.078227 | 0.013 | 0.009 | 1 | 7 | Fam13a | 0.004 | Old |
| 0.632827 | 0.060129 | 0.274 | 0.253 | 1 | 7 | Rcc1 | 0.021 | Old |
| 0.633013 | 0.551681 | 0.023 | 0.018 | 1 | 7 | Gm47675 | 0.005 | Old |
| 0.633013 | 0.507287 | 0.023 | 0.018 | 1 | 7 | Hes2 | 0.005 | Old |
| 0.633208 | 0.206437 | 0.305 | 0.286 | 1 | 7 | Agtrap | 0.019 | Old |
| 0.633223 | 0.280813 | 0.256 | 0.25 | 1 | 7 | Timm17a | 0.006 | Old |
| 0.633486 | 0.158457 | 0.6 | 0.59 | 1 | 7 | Slbp | 0.01 | Old |
| 0.634115 | 0.299266 | 0.162 | 0.148 | 1 | 7 | Pttg1 | 0.014 | Old |
| 0.634665 | 0.085578 | 0.677 | 0.666 | 1 | 7 | Tomm20 | 0.011 | Old |
| 0.63472 | 0.026537 | 0.423 | 0.383 | 1 | 7 | mt-Atp8 | 0.04 | Old |
| 0.635002 | 0.243411 | 0.036 | 0.042 | 1 | 7 | Hotairm1 | -0.006 | Old |
| 0.635078 | 0.368765 | 0.023 | 0.018 | 1 | 7 | Gemin8 | 0.005 | Old |
| 0.635667 | 0.030296 | 0.033 | 0.027 | 1 | 7 | Als2 | 0.006 | Old |
| 0.635678 | 0.115283 | 0.313 | 0.292 | 1 | 7 | Mrps12 | 0.021 | Old |
| 0.636111 | 0.577835 | 0.023 | 0.018 | 1 | 7 | Rgl3 | 0.005 | Old |
| 0.636111 | 0.246293 | 0.023 | 0.018 | 1 | 7 | Gm48529 | 0.005 | Old |

| | | | | | | | | |
|---|---|---|---|---|---|---|---|---|
| 0.636155 | 0.054643 | 0.044 | 0.051 | 1 | 7 | Hist1h1a | -0.007 | Old |
| 0.636166 | 0.242012 | 0.123 | 0.111 | 1 | 7 | Ttc19 | 0.012 | Old |
| 0.636299 | 0.045755 | 0.515 | 0.503 | 1 | 7 | Prelid1 | 0.012 | Old |
| 0.637096 | 0.864364 | 0.072 | 0.063 | 1 | 7 | AI597479 | 0.009 | Old |
| 0.637145 | 0.11301 | 0.023 | 0.018 | 1 | 7 | Samd10 | 0.005 | Old |
| 0.637728 | 0.212943 | 0.4 | 0.373 | 1 | 7 | Tspan31 | 0.027 | Old |
| 0.638027 | 0.553595 | 0.049 | 0.042 | 1 | 7 | Sgsm3 | 0.007 | Old |
| 0.638234 | 0.028466 | 0.259 | 0.241 | 1 | 7 | Trim47 | 0.018 | Old |
| 0.639216 | 0.414938 | 0.023 | 0.018 | 1 | 7 | B130006D( | 0.005 | Old |
| 0.639253 | 0.041196 | 0.885 | 0.867 | 1 | 7 | Nucks1 | 0.018 | Old |
| 0.639301 | 0.821114 | 0.3 | 0.28 | 1 | 7 | Fabp4 | 0.02 | Old |
| 0.639391 | 0.03414 | 0.418 | 0.398 | 1 | 7 | Zc3h15 | 0.02 | Old |
| 0.639658 | 0.309459 | 0.156 | 0.145 | 1 | 7 | Ttc32 | 0.011 | Old |
| 0.639906 | 0.138213 | 0.567 | 0.545 | 1 | 7 | Nipa2 | 0.022 | Old |
| 0.640223 | 0.439201 | 0.046 | 0.039 | 1 | 7 | Repin1 | 0.007 | Old |
| 0.640252 | 0.260073 | 0.023 | 0.018 | 1 | 7 | Jmjd4 | 0.005 | Old |
| 0.640918 | 0.149591 | 0.438 | 0.416 | 1 | 7 | Serpinb8 | 0.022 | Old |
| 0.641289 | 0.062937 | 0.023 | 0.018 | 1 | 7 | BC051226 | 0.005 | Old |
| 0.641308 | 0.101526 | 0.205 | 0.19 | 1 | 7 | Cd209g | 0.015 | Old |
| 0.641626 | 0.039105 | 0.723 | 0.708 | 1 | 7 | Gm2a | 0.015 | Old |
| 0.641682 | 0.161255 | 0.046 | 0.039 | 1 | 7 | Cep95 | 0.007 | Old |
| 0.642784 | 0.1459 | 0.133 | 0.12 | 1 | 7 | Ciz1 | 0.013 | Old |
| 0.64309 | 0.21126 | 0.187 | 0.172 | 1 | 7 | Gps2 | 0.015 | Old |
| 0.64331 | 0.360093 | 0.074 | 0.066 | 1 | 7 | Psph | 0.008 | Old |
| 0.643365 | 0.044053 | 0.023 | 0.018 | 1 | 7 | Gm4349 | 0.005 | Old |
| 0.643545 | 0.058246 | 0.105 | 0.093 | 1 | 7 | Haus7 | 0.012 | Old |
| 0.643699 | 0.023222 | 0.505 | 0.488 | 1 | 7 | Dmac1 | 0.017 | Old |
| 0.644215 | 0.028015 | 0.349 | 0.331 | 1 | 7 | Dcaf8 | 0.018 | Old |
| 0.644604 | 0.062529 | 0.046 | 0.039 | 1 | 7 | R3hcc1 | 0.007 | Old |
| 0.64495 | 0.158305 | 0.41 | 0.38 | 1 | 7 | Ddx50 | 0.03 | Old |
| 0.645288 | 0.063727 | 0.556 | 0.56 | 1 | 7 | Alcam | -0.004 | Old |
| 0.645288 | 0.141091 | 0.056 | 0.048 | 1 | 7 | Faap24 | 0.008 | Old |
| 0.645678 | 0.190528 | 0.41 | 0.398 | 1 | 7 | Fgd4 | 0.012 | Old |
| 0.646153 | 0.039372 | 0.741 | 0.693 | 1 | 7 | Pip4k2a | 0.048 | Old |
| 0.646484 | 0.262106 | 0.023 | 0.018 | 1 | 7 | Lamb2 | 0.005 | Old |
| 0.646761 | 0.705297 | 0.036 | 0.03 | 1 | 7 | Gm47754 | 0.006 | Old |
| 0.646933 | 0.073537 | 0.046 | 0.054 | 1 | 7 | Rab2b | -0.008 | Old |
| 0.647013 | 0.131551 | 0.413 | 0.401 | 1 | 7 | Pum1 | 0.012 | Old |
| 0.647526 | 0.197342 | 0.023 | 0.018 | 1 | 7 | Slc19a2 | 0.005 | Old |
| 0.648762 | 0.142942 | 0.405 | 0.389 | 1 | 7 | Mknk2 | 0.016 | Old |
| 0.649241 | 0.126904 | 0.379 | 0.352 | 1 | 7 | Csgalnact2 | 0.027 | Old |
| 0.649373 | 0.099124 | 0.246 | 0.229 | 1 | 7 | Reep4 | 0.017 | Old |
| 0.650198 | 0.746023 | 0.085 | 0.075 | 1 | 7 | Dexi | 0.01 | Old |
| 0.65266 | 0.088704 | 0.254 | 0.232 | 1 | 7 | Ftsj3 | 0.022 | Old |
| 0.653226 | 0.109164 | 0.646 | 0.605 | 1 | 7 | Mapkapk2 | 0.041 | Old |

| | | | | | | | | |
|---|---|---|---|---|---|---|---|---|
| 0.653272 | 0.041948 | 0.213 | 0.196 | 1 | 7 | Fibp | 0.017 | Old |
| 0.653662 | 0.100161 | 0.505 | 0.524 | 1 | 7 | Amfr | -0.019 | Old |
| 0.654843 | 0.205461 | 0.646 | 0.614 | 1 | 7 | Ndufc1 | 0.032 | Old |
| 0.65489 | 0.067567 | 0.677 | 0.639 | 1 | 7 | Sfr1 | 0.038 | Old |
| 0.655127 | 0.103523 | 0.279 | 0.262 | 1 | 7 | Lage3 | 0.017 | Old |
| 0.655854 | 0.133451 | 0.408 | 0.377 | 1 | 7 | Bscl2 | 0.031 | Old |
| 0.656348 | 0.075016 | 0.046 | 0.039 | 1 | 7 | Ccdc191 | 0.007 | Old |
| 0.656516 | 0.071029 | 0.036 | 0.042 | 1 | 7 | Selenoo | -0.006 | Old |
| 0.657072 | 0.050117 | 0.462 | 0.482 | 1 | 7 | Pros1 | -0.02 | Old |
| 0.6572 | 0.19668 | 0.121 | 0.108 | 1 | 7 | Exosc4 | 0.013 | Old |
| 0.657776 | 0.212128 | 0.221 | 0.208 | 1 | 7 | Tm9sf4 | 0.013 | Old |
| 0.6589 | 0.011096 | 0.292 | 0.268 | 1 | 7 | Echs1 | 0.024 | Old |
| 0.659233 | 1.092551 | 0.036 | 0.03 | 1 | 7 | Tmem53 | 0.006 | Old |
| 0.659233 | 0.289571 | 0.036 | 0.03 | 1 | 7 | Zfat | 0.006 | Old |
| 0.659613 | 0.171437 | 0.036 | 0.042 | 1 | 7 | Evi5l | -0.006 | Old |
| 0.660082 | 0.265969 | 0.123 | 0.111 | 1 | 7 | Pkd2 | 0.012 | Old |
| 0.660142 | 0.209846 | 0.344 | 0.316 | 1 | 7 | Nudcd3 | 0.028 | Old |
| 0.660626 | 0.640833 | 0.1 | 0.09 | 1 | 7 | Borcs8 | 0.01 | Old |
| 0.660723 | 0.251011 | 0.072 | 0.063 | 1 | 7 | Csf3r | 0.009 | Old |
| 0.661009 | 0.127483 | 0.123 | 0.111 | 1 | 7 | Cep135 | 0.012 | Old |
| 0.66119 | 0.072385 | 0.287 | 0.274 | 1 | 7 | Mrpl3 | 0.013 | Old |
| 0.661337 | 0.008813 | 0.185 | 0.169 | 1 | 7 | Pdxdc1 | 0.016 | Old |
| 0.661969 | 0.346839 | 0.228 | 0.211 | 1 | 7 | Rhno1 | 0.017 | Old |
| 0.662575 | 0.18115 | 0.036 | 0.03 | 1 | 7 | Trmt5 | 0.006 | Old |
| 0.66296 | 0.421062 | 0.049 | 0.042 | 1 | 7 | Strip2 | 0.007 | Old |
| 0.66296 | 0.417401 | 0.049 | 0.042 | 1 | 7 | Akap1 | 0.007 | Old |
| 0.663595 | 0.341982 | 0.115 | 0.105 | 1 | 7 | Gm45902 | 0.01 | Old |
| 0.663732 | 0.019856 | 0.046 | 0.039 | 1 | 7 | Cnst | 0.007 | Old |
| 0.664203 | 0.280027 | 0.085 | 0.075 | 1 | 7 | 2810402E2 | 0.01 | Old |
| 0.664385 | 0.025057 | 0.472 | 0.482 | 1 | 7 | Rassf3 | -0.01 | Old |
| 0.664427 | 0.011303 | 0.297 | 0.271 | 1 | 7 | Kitl | 0.026 | Old |
| 0.664616 | 0.339858 | 0.105 | 0.096 | 1 | 7 | Bmp2 | 0.009 | Old |
| 0.664856 | 0.023204 | 0.526 | 0.503 | 1 | 7 | Cnih4 | 0.023 | Old |
| 0.664949 | 0.203096 | 0.246 | 0.232 | 1 | 7 | Tusc3 | 0.014 | Old |
| 0.665076 | 0.02977 | 0.29 | 0.268 | 1 | 7 | Dennd4b | 0.022 | Old |
| 0.665488 | 0.227118 | 0.167 | 0.154 | 1 | 7 | Hmbs | 0.013 | Old |
| 0.665579 | 0.30934 | 0.179 | 0.169 | 1 | 7 | Batf | 0.01 | Old |
| 0.665753 | 0.016749 | 0.277 | 0.259 | 1 | 7 | Ppp1r11 | 0.018 | Old |
| 0.665883 | 0.638784 | 0.038 | 0.045 | 1 | 7 | Cdk20 | -0.007 | Old |
| 0.667281 | 0.476663 | 0.072 | 0.063 | 1 | 7 | Mcm8 | 0.009 | Old |
| 0.6676 | 0.40639 | 0.036 | 0.03 | 1 | 7 | Pik3ap1 | 0.006 | Old |
| 0.667678 | 0.034468 | 0.167 | 0.178 | 1 | 7 | Thyn1 | -0.011 | Old |
| 0.667802 | 0.269986 | 0.064 | 0.057 | 1 | 7 | B230118H( | 0.007 | Old |
| 0.667878 | 0.174862 | 0.072 | 0.063 | 1 | 7 | Lmtk2 | 0.009 | Old |
| 0.668113 | 0.337399 | 0.277 | 0.268 | 1 | 7 | Rchy1 | 0.009 | Old |

| | | | | | | | | |
|---|---|---|---|---|---|---|---|---|
| 0.668621 | 0.031734 | 0.759 | 0.756 | 1 | 7 | Tmed10 | 0.003 | Old |
| 0.669279 | 0.688066 | 0.036 | 0.03 | 1 | 7 | Spata24 | 0.006 | Old |
| 0.669554 | 0.093236 | 0.677 | 0.675 | 1 | 7 | Rab8b | 0.002 | Old |
| 0.669573 | 0.042258 | 0.454 | 0.434 | 1 | 7 | Nfix | 0.02 | Old |
| 0.670118 | 0.247694 | 0.036 | 0.03 | 1 | 7 | Mapk1ip1 | 0.006 | Old |
| 0.670193 | 0.218672 | 0.608 | 0.581 | 1 | 7 | Psma4 | 0.027 | Old |
| 0.670462 | 0.011332 | 0.215 | 0.196 | 1 | 7 | Spast | 0.019 | Old |
| 0.670959 | 0.143132 | 0.036 | 0.03 | 1 | 7 | Txnrd3 | 0.006 | Old |
| 0.671108 | 0.012995 | 0.351 | 0.325 | 1 | 7 | Pdcl3 | 0.026 | Old |
| 0.671457 | 0.598504 | 0.182 | 0.199 | 1 | 7 | Cxcl10 | -0.017 | Old |
| 0.671468 | 0.303504 | 0.072 | 0.063 | 1 | 7 | Gm14548 | 0.009 | Old |
| 0.671577 | 0.610702 | 0.064 | 0.057 | 1 | 7 | Fam114a1 | 0.007 | Old |
| 0.67187 | 0.023713 | 0.277 | 0.256 | 1 | 7 | Med30 | 0.021 | Old |
| 0.67213 | 0.808327 | 0.097 | 0.09 | 1 | 7 | Gnptg | 0.007 | Old |
| 0.672338 | 0.178873 | 0.087 | 0.078 | 1 | 7 | Pms2 | 0.009 | Old |
| 0.673072 | 0.164237 | 0.085 | 0.093 | 1 | 7 | Zfp654 | -0.008 | Old |
| 0.673107 | 0.863474 | 0.026 | 0.021 | 1 | 7 | Ubtd2 | 0.005 | Old |
| 0.673207 | 0.315791 | 0.059 | 0.051 | 1 | 7 | Ap4e1 | 0.008 | Old |
| 0.673305 | 0.108819 | 0.197 | 0.184 | 1 | 7 | Arhgap25 | 0.013 | Old |
| 0.673482 | 0.520354 | 0.036 | 0.03 | 1 | 7 | Enc1 | 0.006 | Old |
| 0.673538 | 0.051842 | 0.215 | 0.202 | 1 | 7 | Zfp871 | 0.013 | Old |
| 0.673707 | 0.008923 | 0.267 | 0.247 | 1 | 7 | Odf2 | 0.02 | Old |
| 0.673782 | 0.059599 | 0.21 | 0.226 | 1 | 7 | Gcc2 | -0.016 | Old |
| 0.674486 | 0.060158 | 0.049 | 0.042 | 1 | 7 | Gstz1 | 0.007 | Old |
| 0.676098 | 0.001 | 0.585 | 0.596 | 1 | 7 | Emb | -0.011 | Old |
| 0.676837 | 0.161681 | 0.169 | 0.157 | 1 | 7 | Golga2 | 0.012 | Old |
| 0.676869 | 0.494483 | 0.133 | 0.123 | 1 | 7 | Smarcal1 | 0.01 | Old |
| 0.677314 | 0.154832 | 0.123 | 0.111 | 1 | 7 | Dclre1b | 0.012 | Old |
| 0.677379 | 0.266361 | 0.049 | 0.042 | 1 | 7 | Sars2 | 0.007 | Old |
| 0.677871 | 0.183344 | 0.128 | 0.117 | 1 | 7 | Utp14b | 0.011 | Old |
| 0.678089 | 0.166742 | 0.026 | 0.021 | 1 | 7 | Meis1 | 0.005 | Old |
| 0.679076 | 0.647306 | 0.077 | 0.069 | 1 | 7 | Bclaf3 | 0.008 | Old |
| 0.679436 | 0.365561 | 0.167 | 0.157 | 1 | 7 | AY036118 | 0.01 | Old |
| 0.680159 | 0.715812 | 0.051 | 0.045 | 1 | 7 | Ppm1f | 0.006 | Old |
| 0.6802 | 0.033226 | 0.164 | 0.175 | 1 | 7 | Kin | -0.011 | Old |
| 0.680347 | 0.10868 | 0.3 | 0.28 | 1 | 7 | Cln6 | 0.02 | Old |
| 0.680605 | 0.008982 | 0.472 | 0.449 | 1 | 7 | Kpna4 | 0.023 | Old |
| 0.680698 | 0.146032 | 0.038 | 0.033 | 1 | 7 | Gm45606 | 0.005 | Old |
| 0.680878 | 0.105989 | 0.103 | 0.093 | 1 | 7 | Oxsm | 0.01 | Old |
| 0.681085 | 0.421179 | 0.026 | 0.021 | 1 | 7 | Chst11 | 0.005 | Old |
| 0.681897 | 0.276876 | 0.074 | 0.066 | 1 | 7 | AC163018. | 0.008 | Old |
| 0.682 | 0.552753 | 0.131 | 0.123 | 1 | 7 | Fam173b | 0.008 | Old |
| 0.683089 | 0.103151 | 0.569 | 0.518 | 1 | 7 | Fermt3 | 0.051 | Old |
| 0.683956 | 0.060859 | 0.587 | 0.554 | 1 | 7 | Atp5d | 0.033 | Old |
| 0.683992 | 0.138824 | 0.492 | 0.464 | 1 | 7 | Vasp | 0.028 | Old |

| | | | | | | | | |
|---|---|---|---|---|---|---|---|---|
| 0.684086 | 0.586363 | 0.026 | 0.021 | 1 | 7 | Gm26601 | 0.005 | Old |
| 0.684184 | 0.453768 | 0.1 | 0.09 | 1 | 7 | Elac2 | 0.01 | Old |
| 0.684425 | 0.095201 | 0.444 | 0.419 | 1 | 7 | Ndufs7 | 0.025 | Old |
| 0.684458 | 0.047323 | 0.031 | 0.036 | 1 | 7 | Apold1 | -0.005 | Old |
| 0.684626 | 0.06539 | 0.367 | 0.34 | 1 | 7 | Man1c1 | 0.027 | Old |
| 0.685451 | 0.63008 | 0.236 | 0.232 | 1 | 7 | Ash2l | 0.004 | Old |
| 0.685564 | 0.455136 | 0.038 | 0.045 | 1 | 7 | Gm46430 | -0.007 | Old |
| 0.685569 | 0.034018 | 0.538 | 0.509 | 1 | 7 | Ppp1cc | 0.029 | Old |
| 0.686052 | 0.218308 | 0.141 | 0.154 | 1 | 7 | Setd1a | -0.013 | Old |
| 0.686843 | 0.01827 | 0.992 | 0.997 | 1 | 7 | Rpl41 | -0.005 | Old |
| 0.687422 | 0.141412 | 0.267 | 0.25 | 1 | 7 | Cd300lb | 0.017 | Old |
| 0.687962 | 0.183306 | 0.146 | 0.133 | 1 | 7 | Yipf6 | 0.013 | Old |
| 0.688597 | 0.695049 | 0.026 | 0.021 | 1 | 7 | Harbi1 | 0.005 | Old |
| 0.689099 | 0.305872 | 0.026 | 0.021 | 1 | 7 | Timp1 | 0.005 | Old |
| 0.690103 | 0.053411 | 0.026 | 0.021 | 1 | 7 | A430046D1 | 0.005 | Old |
| 0.690163 | 0.215658 | 0.079 | 0.087 | 1 | 7 | Slc41a1 | -0.008 | Old |
| 0.690397 | 0.082415 | 0.031 | 0.036 | 1 | 7 | Rbm41 | -0.005 | Old |
| 0.690767 | 0.011712 | 0.051 | 0.045 | 1 | 7 | Nxn | 0.006 | Old |
| 0.690851 | 0.06455 | 0.21 | 0.199 | 1 | 7 | Prkaca | 0.011 | Old |
| 0.691316 | 0.384747 | 0.067 | 0.06 | 1 | 7 | Nme4 | 0.007 | Old |
| 0.691494 | 0.658622 | 0.054 | 0.048 | 1 | 7 | Apoo | 0.006 | Old |
| 0.691767 | 0.023743 | 0.726 | 0.72 | 1 | 7 | Tm6sf1 | 0.006 | Old |
| 0.692114 | 0.146133 | 0.026 | 0.021 | 1 | 7 | Alg12 | 0.005 | Old |
| 0.692286 | 0.016614 | 0.633 | 0.59 | 1 | 7 | Myl12a | 0.043 | Old |
| 0.692626 | 1.1548 | 0.015 | 0.012 | 1 | 7 | Epas1 | 0.003 | Old |
| 0.692938 | 0.030731 | 0.5 | 0.473 | 1 | 7 | Uba1 | 0.027 | Old |
| 0.69409 | 0.037535 | 0.303 | 0.283 | 1 | 7 | 2810403A0 | 0.02 | Old |
| 0.694126 | 0.309582 | 0.026 | 0.021 | 1 | 7 | Gm26669 | 0.005 | Old |
| 0.694568 | 0.083111 | 0.356 | 0.334 | 1 | 7 | Gm8186 | 0.022 | Old |
| 0.694796 | 0.140128 | 0.254 | 0.238 | 1 | 7 | Cdca4 | 0.016 | Old |
| 0.694798 | 0.191483 | 0.185 | 0.169 | 1 | 7 | Myo18a | 0.016 | Old |
| 0.695133 | 0.04164 | 0.026 | 0.021 | 1 | 7 | H2-Q10 | 0.005 | Old |
| 0.69527 | 0.026929 | 0.485 | 0.458 | 1 | 7 | Prex1 | 0.027 | Old |
| 0.695502 | 0.193501 | 0.031 | 0.036 | 1 | 7 | Rhobtb2 | -0.005 | Old |
| 0.695552 | 0.022424 | 0.385 | 0.361 | 1 | 7 | Ckap5 | 0.024 | Old |
| 0.695633 | 0.088513 | 0.462 | 0.44 | 1 | 7 | Wbp11 | 0.022 | Old |
| 0.696546 | 0.459829 | 0.015 | 0.012 | 1 | 7 | Rpusd3 | 0.003 | Old |
| 0.69674 | 0.064547 | 0.074 | 0.066 | 1 | 7 | Ano10 | 0.008 | Old |
| 0.696838 | 0.183699 | 0.146 | 0.136 | 1 | 7 | Ndst2 | 0.01 | Old |
| 0.697275 | 0.01966 | 0.146 | 0.136 | 1 | 7 | Cog7 | 0.01 | Old |
| 0.697801 | 0.03461 | 0.09 | 0.081 | 1 | 7 | Zfp994 | 0.009 | Old |
| 0.697854 | 0.641161 | 0.015 | 0.012 | 1 | 7 | Rad9b | 0.003 | Old |
| 0.699804 | 0.562408 | 0.054 | 0.048 | 1 | 7 | Zbtb40 | 0.006 | Old |
| 0.700283 | 0.11253 | 0.059 | 0.066 | 1 | 7 | Nt5c3b | -0.007 | Old |
| 0.700372 | 0.015489 | 0.4 | 0.373 | 1 | 7 | Ddrgk1 | 0.027 | Old |

| | | | | | | | | |
|---|---|---|---|---|---|---|---|---|
| 0.700473 | 0.289355 | 0.015 | 0.012 | 1 | 7 | Zfp677 | 0.003 | Old |
| 0.700473 | 0.185443 | 0.015 | 0.012 | 1 | 7 | Lrrc1 | 0.003 | Old |
| 0.700545 | 0.093022 | 0.608 | 0.569 | 1 | 7 | Smc6 | 0.039 | Old |
| 0.700722 | 0.555389 | 0.051 | 0.045 | 1 | 7 | Adck1 | 0.006 | Old |
| 0.700967 | 0.115383 | 0.182 | 0.172 | 1 | 7 | Helb | 0.01 | Old |
| 0.701141 | 0.221208 | 0.038 | 0.033 | 1 | 7 | Gm5086 | 0.005 | Old |
| 0.70115 | 0.051274 | 0.487 | 0.464 | 1 | 7 | Hmox1 | 0.023 | Old |
| 0.701476 | 0.093488 | 0.031 | 0.036 | 1 | 7 | Omp | -0.005 | Old |
| 0.701543 | 0.04522 | 0.108 | 0.117 | 1 | 7 | Tnfrsf12a | -0.009 | Old |
| 0.701674 | 0.171238 | 0.144 | 0.133 | 1 | 7 | 4931406P1 | 0.011 | Old |
| 0.701784 | 0.524341 | 0.015 | 0.012 | 1 | 7 | Plxna2 | 0.003 | Old |
| 0.701784 | 0.303831 | 0.015 | 0.012 | 1 | 7 | Plcb4 | 0.003 | Old |
| 0.701784 | 0.128067 | 0.015 | 0.012 | 1 | 7 | Tle6 | 0.003 | Old |
| 0.702106 | 0.017785 | 0.164 | 0.154 | 1 | 7 | Fam160b1 | 0.01 | Old |
| 0.702435 | 0.035566 | 0.3 | 0.277 | 1 | 7 | Kdm4a | 0.023 | Old |
| 0.702524 | 0.238145 | 0.103 | 0.111 | 1 | 7 | Chpf2 | -0.008 | Old |
| 0.702558 | 0.145941 | 0.144 | 0.133 | 1 | 7 | Zfp992 | 0.011 | Old |
| 0.702774 | 0.652314 | 0.041 | 0.036 | 1 | 7 | C1rl | 0.005 | Old |
| 0.702855 | 0.053952 | 0.203 | 0.211 | 1 | 7 | Dhx30 | -0.008 | Old |
| 0.703096 | 0.498562 | 0.015 | 0.012 | 1 | 7 | Ubox5 | 0.003 | Old |
| 0.703282 | 0.43045 | 0.151 | 0.145 | 1 | 7 | Gt(ROSA)2( | 0.006 | Old |
| 0.703576 | 0.040394 | 0.051 | 0.045 | 1 | 7 | Ydjc | 0.006 | Old |
| 0.704409 | 0.535076 | 0.015 | 0.012 | 1 | 7 | Hist1h4a | 0.003 | Old |
| 0.704409 | 0.264248 | 0.015 | 0.012 | 1 | 7 | Etl4 | 0.003 | Old |
| 0.704409 | 0.147333 | 0.015 | 0.012 | 1 | 7 | Frs3 | 0.003 | Old |
| 0.704696 | 0.031407 | 0.559 | 0.548 | 1 | 7 | 9530068E0 | 0.011 | Old |
| 0.705722 | 0.212448 | 0.015 | 0.012 | 1 | 7 | Ltbp1 | 0.003 | Old |
| 0.70602 | 0.040121 | 0.695 | 0.648 | 1 | 7 | Lnpep | 0.047 | Old |
| 0.707037 | 0.024819 | 0.015 | 0.012 | 1 | 7 | BC030499 | 0.003 | Old |
| 0.707037 | 0.012886 | 0.015 | 0.012 | 1 | 7 | Matn2 | 0.003 | Old |
| 0.70726 | 0.394414 | 0.059 | 0.066 | 1 | 7 | Mbnl3 | -0.007 | Old |
| 0.707593 | 0.612933 | 0.028 | 0.024 | 1 | 7 | Rnf144a | 0.004 | Old |
| 0.70764 | 0.107669 | 0.09 | 0.081 | 1 | 7 | Dars2 | 0.009 | Old |
| 0.707895 | 0.001023 | 0.059 | 0.066 | 1 | 7 | H2-Ob | -0.007 | Old |
| 0.708256 | 0.006121 | 0.123 | 0.133 | 1 | 7 | Ddx60 | -0.01 | Old |
| 0.708352 | 0.117016 | 0.015 | 0.012 | 1 | 7 | AI427809 | 0.003 | Old |
| 0.708352 | 0.039341 | 0.015 | 0.012 | 1 | 7 | Gm21887 | 0.003 | Old |
| 0.708414 | 0.094889 | 0.331 | 0.325 | 1 | 7 | Idh3g | 0.006 | Old |
| 0.708519 | 0.039524 | 0.095 | 0.102 | 1 | 7 | Dpp4 | -0.007 | Old |
| 0.708684 | 0.20046 | 0.249 | 0.235 | 1 | 7 | Armc1 | 0.014 | Old |
| 0.709225 | 0.04943 | 0.1 | 0.108 | 1 | 7 | Mbd5 | -0.008 | Old |
| 0.710312 | 0.066931 | 0.149 | 0.139 | 1 | 7 | Mthfd1l | 0.01 | Old |
| 0.711032 | 0.177455 | 0.038 | 0.033 | 1 | 7 | Cptp | 0.005 | Old |
| 0.711557 | 0.22254 | 0.113 | 0.102 | 1 | 7 | Git1 | 0.011 | Old |
| 0.711743 | 0.257496 | 0.169 | 0.157 | 1 | 7 | AU019823 | 0.012 | Old |

| | | | | | | | | |
|---|---|---|---|---|---|---|---|---|
| 0.711777 | 0.297454 | 0.118 | 0.108 | 1 | 7 | Abhd13 | 0.01 | Old |
| 0.71178 | 0.065128 | 0.603 | 0.569 | 1 | 7 | Sf3b5 | 0.034 | Old |
| 0.711804 | 0.120015 | 0.246 | 0.262 | 1 | 7 | Rbbp8 | -0.016 | Old |
| 0.711857 | 0.06192 | 0.062 | 0.054 | 1 | 7 | Lrrc28 | 0.008 | Old |
| 0.712492 | 0.001094 | 0.774 | 0.765 | 1 | 7 | Snrpe | 0.009 | Old |
| 0.713538 | 0.384763 | 0.026 | 0.03 | 1 | 7 | 2810455O( | -0.004 | Old |
| 0.71354 | 0.075365 | 0.092 | 0.084 | 1 | 7 | Ankmy2 | 0.008 | Old |
| 0.713733 | 0.007255 | 0.536 | 0.491 | 1 | 7 | B230219D2 | 0.045 | Old |
| 0.713735 | 0.568269 | 0.054 | 0.048 | 1 | 7 | Tepsin | 0.006 | Old |
| 0.714041 | 0.23304 | 0.054 | 0.06 | 1 | 7 | Iigp1 | -0.006 | Old |
| 0.714053 | 0.021201 | 0.864 | 0.895 | 1 | 7 | Tpm3 | -0.031 | Old |
| 0.714226 | 0.042963 | 0.09 | 0.081 | 1 | 7 | Cdk5 | 0.009 | Old |
| 0.714317 | 0.096029 | 0.746 | 0.705 | 1 | 7 | Itgam | 0.041 | Old |
| 0.714474 | 0.003304 | 0.579 | 0.545 | 1 | 7 | Rdx | 0.034 | Old |
| 0.715605 | 0.026381 | 0.162 | 0.151 | 1 | 7 | Maml1 | 0.011 | Old |
| 0.71568 | 0.066987 | 0.841 | 0.825 | 1 | 7 | Pltp | 0.016 | Old |
| 0.716379 | 0.706789 | 0.056 | 0.051 | 1 | 7 | Mtx3 | 0.005 | Old |
| 0.717097 | 0.182471 | 0.079 | 0.072 | 1 | 7 | Trp53inp2 | 0.007 | Old |
| 0.717161 | 0.116175 | 0.303 | 0.286 | 1 | 7 | Tmem248 | 0.017 | Old |
| 0.717261 | 0.02908 | 0.221 | 0.205 | 1 | 7 | Spink2 | 0.016 | Old |
| 0.717292 | 0.016461 | 0.303 | 0.28 | 1 | 7 | Tor1b | 0.023 | Old |
| 0.717584 | 0.180833 | 0.105 | 0.096 | 1 | 7 | Ndrg1 | 0.009 | Old |
| 0.717689 | 0.430781 | 0.067 | 0.06 | 1 | 7 | AL590144.: | 0.007 | Old |
| 0.718183 | 0.575958 | 0.028 | 0.024 | 1 | 7 | Optn | 0.004 | Old |
| 0.718188 | 0.255542 | 0.251 | 0.241 | 1 | 7 | D10Jhu81e | 0.01 | Old |
| 0.718268 | 0.402163 | 0.313 | 0.301 | 1 | 7 | Cyb5a | 0.012 | Old |
| 0.718421 | 0.067665 | 0.579 | 0.572 | 1 | 7 | Mrps21 | 0.007 | Old |
| 0.718627 | 0.524306 | 0.051 | 0.045 | 1 | 7 | Fcho1 | 0.006 | Old |
| 0.719111 | 0.020731 | 0.972 | 0.967 | 1 | 7 | Jund | 0.005 | Old |
| 0.719365 | 0.089133 | 0.477 | 0.482 | 1 | 7 | Pias1 | -0.005 | Old |
| 0.720087 | 0.072939 | 0.497 | 0.473 | 1 | 7 | Spcs3 | 0.024 | Old |
| 0.720108 | 0.0487 | 0.128 | 0.117 | 1 | 7 | Gga3 | 0.011 | Old |
| 0.721723 | 0.049451 | 0.19 | 0.175 | 1 | 7 | Washc5 | 0.015 | Old |
| 0.721862 | 0.07556 | 0.141 | 0.133 | 1 | 7 | Anapc10 | 0.008 | Old |
| 0.721918 | 0.04151 | 0.656 | 0.614 | 1 | 7 | Nono | 0.042 | Old |
| 0.723015 | 1.490465 | 0.028 | 0.024 | 1 | 7 | Mfsd4b4 | 0.004 | Old |
| 0.723502 | 0.06999 | 0.256 | 0.235 | 1 | 7 | Rnf216 | 0.021 | Old |
| 0.723856 | 0.045877 | 0.267 | 0.25 | 1 | 7 | Sertad2 | 0.017 | Old |
| 0.724329 | 0.358405 | 0.041 | 0.036 | 1 | 7 | Mapkbp1 | 0.005 | Old |
| 0.724869 | 0.235812 | 0.023 | 0.027 | 1 | 7 | 1700006J1 | -0.004 | Old |
| 0.725365 | 0.037859 | 0.931 | 0.922 | 1 | 7 | Hnrnpa2b1 | 0.009 | Old |
| 0.725502 | 0.241228 | 0.028 | 0.033 | 1 | 7 | Hspb6 | -0.005 | Old |
| 0.726186 | 0.144415 | 0.377 | 0.358 | 1 | 7 | Sidt2 | 0.019 | Old |
| 0.726529 | 0.014744 | 0.174 | 0.16 | 1 | 7 | Il17ra | 0.014 | Old |
| 0.726731 | 0.529587 | 0.026 | 0.03 | 1 | 7 | Fam20a | -0.004 | Old |

| | | | | | | | | |
|---|---|---|---|---|---|---|---|---|
| 0.726889 | 0.395441 | 0.028 | 0.024 | 1 | 7 | Pkd1l2 | 0.004 | Old |
| 0.727884 | 0.185934 | 0.167 | 0.157 | 1 | 7 | Mllt6 | 0.01 | Old |
| 0.72821 | 0.313511 | 0.028 | 0.033 | 1 | 7 | Klf8 | -0.005 | Old |
| 0.728345 | 0.227287 | 0.041 | 0.036 | 1 | 7 | Zfp874a | 0.005 | Old |
| 0.730374 | 0.018906 | 0.403 | 0.38 | 1 | 7 | Mef2d | 0.023 | Old |
| 0.730579 | 0.445202 | 0.054 | 0.048 | 1 | 7 | Nudt7 | 0.006 | Old |
| 0.731071 | 0.466393 | 0.062 | 0.069 | 1 | 7 | Dnajc4 | -0.007 | Old |
| 0.73137 | 0.11918 | 0.292 | 0.271 | 1 | 7 | Prkag1 | 0.021 | Old |
| 0.731458 | 0.086383 | 0.413 | 0.392 | 1 | 7 | Cenpb | 0.021 | Old |
| 0.731666 | 0.007862 | 0.179 | 0.169 | 1 | 7 | Tarbp2 | 0.01 | Old |
| 0.73174 | 0.217185 | 0.028 | 0.024 | 1 | 7 | 1700019L1 | 0.004 | Old |
| 0.732369 | 0.311397 | 0.041 | 0.036 | 1 | 7 | Gm26881 | 0.005 | Old |
| 0.732922 | 0.116442 | 0.4 | 0.377 | 1 | 7 | Dennd1b | 0.023 | Old |
| 0.733445 | 0.084141 | 0.079 | 0.072 | 1 | 7 | Hscb | 0.007 | Old |
| 0.734282 | 0.122148 | 0.615 | 0.608 | 1 | 7 | Eef1d | 0.007 | Old |
| 0.734958 | 0.005729 | 0.777 | 0.768 | 1 | 7 | Cyth4 | 0.009 | Old |
| 0.735791 | 0.079352 | 0.079 | 0.072 | 1 | 7 | Asap2 | 0.007 | Old |
| 0.735917 | 0.058574 | 0.762 | 0.759 | 1 | 7 | Slc6a6 | 0.003 | Old |
| 0.73614 | 0.098594 | 0.228 | 0.211 | 1 | 7 | Upf3a | 0.017 | Old |
| 0.736205 | 0.189238 | 0.026 | 0.03 | 1 | 7 | Mterf1a | -0.004 | Old |
| 0.736614 | 0.061051 | 0.774 | 0.774 | 1 | 7 | H2-DMb1 | 0 | Old |
| 0.736762 | 0.074061 | 0.328 | 0.316 | 1 | 7 | Rab11fip5 | 0.012 | Old |
| 0.737594 | 0.173608 | 0.095 | 0.102 | 1 | 7 | Apool | -0.007 | Old |
| 0.739022 | 0.392004 | 0.223 | 0.211 | 1 | 7 | Cnot10 | 0.012 | Old |
| 0.739116 | 0.052282 | 0.377 | 0.358 | 1 | 7 | Rassf5 | 0.019 | Old |
| 0.739378 | 0.168027 | 0.387 | 0.383 | 1 | 7 | Il1rl1 | 0.004 | Old |
| 0.739652 | 0.003878 | 0.282 | 0.265 | 1 | 7 | Hlcs | 0.017 | Old |
| 0.740006 | 0.677321 | 0.026 | 0.03 | 1 | 7 | Serpinb2 | -0.004 | Old |
| 0.740095 | 0.399607 | 0.097 | 0.09 | 1 | 7 | Elmod2 | 0.007 | Old |
| 0.740329 | 0.30542 | 0.126 | 0.117 | 1 | 7 | Ddx41 | 0.009 | Old |
| 0.740405 | 0.045472 | 0.131 | 0.139 | 1 | 7 | Slc10a7 | -0.008 | Old |
| 0.740693 | 0.259446 | 0.141 | 0.154 | 1 | 7 | Cyth3 | -0.013 | Old |
| 0.741263 | 0.03205 | 0.926 | 0.919 | 1 | 7 | Rpl24 | 0.007 | Old |
| 0.741784 | 0.053364 | 0.536 | 0.56 | 1 | 7 | Capn2 | -0.024 | Old |
| 0.742112 | 0.099853 | 0.067 | 0.072 | 1 | 7 | Synpo | -0.005 | Old |
| 0.742462 | 0.401676 | 0.121 | 0.111 | 1 | 7 | Snhg12 | 0.01 | Old |
| 0.744171 | 0.189685 | 0.136 | 0.148 | 1 | 7 | Cxcl12 | -0.012 | Old |
| 0.744365 | 0.021033 | 0.762 | 0.774 | 1 | 7 | H2-DMa | -0.012 | Old |
| 0.744484 | 0.547628 | 0.074 | 0.069 | 1 | 7 | Csf2rb2 | 0.005 | Old |
| 0.744484 | 0.353333 | 0.041 | 0.036 | 1 | 7 | Ift172 | 0.005 | Old |
| 0.744695 | 0.209635 | 0.469 | 0.428 | 1 | 7 | Taf15 | 0.041 | Old |
| 0.744781 | 0.319414 | 0.174 | 0.163 | 1 | 7 | Tmem154 | 0.011 | Old |
| 0.744972 | 0.02093 | 0.118 | 0.108 | 1 | 7 | Tfdp2 | 0.01 | Old |
| 0.745119 | 0.231728 | 0.336 | 0.334 | 1 | 7 | AI467606 | 0.002 | Old |
| 0.745196 | 0.400797 | 0.1 | 0.093 | 1 | 7 | Wdr90 | 0.007 | Old |

| | | | | | | | | |
|---|---|---|---|---|---|---|---|---|
| 0.745606 | 0.019898 | 0.505 | 0.527 | 1 | 7 | Ap1b1 | -0.022 | Old |
| 0.745941 | 0.132176 | 0.203 | 0.19 | 1 | 7 | Phf12 | 0.013 | Old |
| 0.746077 | 0.679797 | 0.077 | 0.072 | 1 | 7 | Exoc8 | 0.005 | Old |
| 0.746673 | 0.005854 | 0.356 | 0.328 | 1 | 7 | Stx12 | 0.028 | Old |
| 0.746843 | 0.211009 | 0.049 | 0.054 | 1 | 7 | Dalrd3 | -0.005 | Old |
| 0.746939 | 0.006824 | 0.523 | 0.491 | 1 | 7 | Pdlim1 | 0.032 | Old |
| 0.747097 | 0.289511 | 0.108 | 0.117 | 1 | 7 | Rfng | -0.009 | Old |
| 0.747261 | 0.020974 | 0.174 | 0.166 | 1 | 7 | Ccdc186 | 0.008 | Old |
| 0.74727 | 0.136438 | 0.021 | 0.024 | 1 | 7 | A430078G2 | -0.003 | Old |
| 0.748631 | 0.017725 | 0.082 | 0.075 | 1 | 7 | Efl1 | 0.007 | Old |
| 0.748836 | 0.005967 | 0.51 | 0.476 | 1 | 7 | Tnfrsf1b | 0.034 | Old |
| 0.749165 | 0.107486 | 0.333 | 0.316 | 1 | 7 | Myd88 | 0.017 | Old |
| 0.749424 | 0.034826 | 0.374 | 0.352 | 1 | 7 | Piezo1 | 0.022 | Old |
| 0.749545 | 0.043777 | 0.187 | 0.178 | 1 | 7 | Slc25a10 | 0.009 | Old |
| 0.749609 | 1.879653 | 0.031 | 0.027 | 1 | 7 | Mgp | 0.004 | Old |
| 0.749826 | 0.005188 | 0.5 | 0.47 | 1 | 7 | Top2b | 0.03 | Old |
| 0.750531 | 0.029975 | 0.769 | 0.753 | 1 | 7 | Sh3glb1 | 0.016 | Old |
| 0.750736 | 0.408668 | 0.072 | 0.066 | 1 | 7 | Tmed8 | 0.006 | Old |
| 0.752552 | 1.074984 | 0.018 | 0.015 | 1 | 7 | Slc12a5 | 0.003 | Old |
| 0.752552 | 0.674169 | 0.018 | 0.015 | 1 | 7 | Padi4 | 0.003 | Old |
| 0.752552 | 0.636235 | 0.018 | 0.015 | 1 | 7 | Napepld | 0.003 | Old |
| 0.75286 | 0.087882 | 0.813 | 0.768 | 1 | 7 | Cox6b1 | 0.045 | Old |
| 0.753329 | 0.054282 | 0.095 | 0.102 | 1 | 7 | Gamt | -0.007 | Old |
| 0.753697 | 0.777165 | 0.059 | 0.054 | 1 | 7 | Rab27a | 0.005 | Old |
| 0.75378 | 0.031606 | 0.018 | 0.015 | 1 | 7 | Meis3 | 0.003 | Old |
| 0.753908 | 0.067574 | 0.251 | 0.238 | 1 | 7 | Coro7 | 0.013 | Old |
| 0.754206 | 0.138784 | 0.264 | 0.253 | 1 | 7 | Etv6 | 0.011 | Old |
| 0.754216 | 0.13687 | 0.438 | 0.428 | 1 | 7 | Ppp4r3b | 0.01 | Old |
| 0.754378 | 0.104991 | 0.336 | 0.319 | 1 | 7 | Ocrl | 0.017 | Old |
| 0.754538 | 0.071687 | 0.679 | 0.66 | 1 | 7 | Rassf4 | 0.019 | Old |
| 0.754736 | 0.091044 | 0.264 | 0.253 | 1 | 7 | Rspry1 | 0.011 | Old |
| 0.75474 | 0.085098 | 0.382 | 0.355 | 1 | 7 | Nfatc3 | 0.027 | Old |
| 0.755008 | 0.255261 | 0.018 | 0.015 | 1 | 7 | Lix1 | 0.003 | Old |
| 0.755207 | 0.196439 | 0.297 | 0.283 | 1 | 7 | Dram2 | 0.014 | Old |
| 0.755571 | 0.066828 | 0.695 | 0.669 | 1 | 7 | Nsa2 | 0.026 | Old |
| 0.755676 | 0.067483 | 0.782 | 0.816 | 1 | 7 | Tmem176b | -0.034 | Old |
| 0.756237 | 0.645914 | 0.018 | 0.015 | 1 | 7 | Cd59a | 0.003 | Old |
| 0.757074 | 0.092181 | 0.562 | 0.539 | 1 | 7 | Aamp | 0.023 | Old |
| 0.757396 | 0.104456 | 0.382 | 0.364 | 1 | 7 | Zfp638 | 0.018 | Old |
| 0.75793 | 0.322396 | 0.021 | 0.024 | 1 | 7 | Mgll | -0.003 | Old |
| 0.75793 | 1.04598 | 0.021 | 0.024 | 1 | 7 | Phgdh | -0.003 | Old |
| 0.757935 | 0.093157 | 0.459 | 0.44 | 1 | 7 | Ap1s2 | 0.019 | Old |
| 0.758049 | 0.125685 | 0.526 | 0.497 | 1 | 7 | Map7d1 | 0.029 | Old |
| 0.758724 | 0.168419 | 0.19 | 0.184 | 1 | 7 | Igsf9 | 0.006 | Old |
| 0.758801 | 0.09126 | 0.133 | 0.127 | 1 | 7 | Zfp646 | 0.006 | Old |

| | | | | | | | | |
|---|---|---|---|---|---|---|---|---|
| 0.759767 | 0.306607 | 0.018 | 0.021 | 1 | 7 | Cav1 | -0.003 | Old |
| 0.759853 | 0.024489 | 0.133 | 0.142 | 1 | 7 | Chchd4 | -0.009 | Old |
| 0.760005 | 0.11659 | 0.044 | 0.039 | 1 | 7 | Gm29340 | 0.005 | Old |
| 0.760068 | 0.247827 | 0.021 | 0.024 | 1 | 7 | F5 | -0.003 | Old |
| 0.760452 | 0.428863 | 0.315 | 0.304 | 1 | 7 | Twistnb | 0.011 | Old |
| 0.762392 | 0.377539 | 0.018 | 0.015 | 1 | 7 | Mast4 | 0.003 | Old |
| 0.762602 | 0.100992 | 0.056 | 0.051 | 1 | 7 | Smyd3 | 0.005 | Old |
| 0.7627 | 0.47726 | 0.031 | 0.027 | 1 | 7 | Zmym6 | 0.004 | Old |
| 0.762977 | 0.090049 | 0.251 | 0.244 | 1 | 7 | Gm6377 | 0.007 | Old |
| 0.763294 | 0.659414 | 0.074 | 0.069 | 1 | 7 | Tcf7l2 | 0.005 | Old |
| 0.763638 | 0.251337 | 0.031 | 0.027 | 1 | 7 | Adck2 | 0.004 | Old |
| 0.764858 | 0.676266 | 0.018 | 0.015 | 1 | 7 | Gm14296 | 0.003 | Old |
| 0.765481 | 0.171918 | 0.018 | 0.021 | 1 | 7 | Lepr | -0.003 | Old |
| 0.765514 | 0.073428 | 0.031 | 0.027 | 1 | 7 | Fndc7 | 0.004 | Old |
| 0.765592 | 0.321063 | 0.072 | 0.066 | 1 | 7 | Rad51c | 0.006 | Old |
| 0.765932 | 0.471086 | 0.059 | 0.054 | 1 | 7 | Mgmt | 0.005 | Old |
| 0.766068 | 0.076966 | 0.141 | 0.151 | 1 | 7 | Gkap1 | -0.01 | Old |
| 0.767326 | 0.018067 | 0.018 | 0.015 | 1 | 7 | Gm28557 | 0.003 | Old |
| 0.767605 | 0.061227 | 0.438 | 0.44 | 1 | 7 | Creb1 | -0.002 | Old |
| 0.768177 | 0.180635 | 0.136 | 0.145 | 1 | 7 | Tfam | -0.009 | Old |
| 0.768561 | 0.111294 | 0.018 | 0.015 | 1 | 7 | 2610301B2 | 0.003 | Old |
| 0.768561 | 0.103289 | 0.018 | 0.015 | 1 | 7 | Atxn7l2 | 0.003 | Old |
| 0.768616 | 0.005656 | 0.318 | 0.301 | 1 | 7 | Aldh3b1 | 0.017 | Old |
| 0.768621 | 0.184624 | 0.221 | 0.211 | 1 | 7 | Txndc15 | 0.01 | Old |
| 0.769151 | 0.092931 | 0.146 | 0.139 | 1 | 7 | Dcaf10 | 0.007 | Old |
| 0.769272 | 0.549034 | 0.031 | 0.027 | 1 | 7 | Ssh3 | 0.004 | Old |
| 0.769324 | 0.16968 | 0.372 | 0.37 | 1 | 7 | Glipr2 | 0.002 | Old |
| 0.77006 | 0.080789 | 0.018 | 0.021 | 1 | 7 | Gprin3 | -0.003 | Old |
| 0.770339 | 0.048134 | 0.795 | 0.744 | 1 | 7 | Clic1 | 0.051 | Old |
| 0.770427 | 0.076592 | 0.762 | 0.777 | 1 | 7 | Gnas | -0.015 | Old |
| 0.771026 | 0.040045 | 0.041 | 0.045 | 1 | 7 | Ulk3 | -0.004 | Old |
| 0.771207 | 0.203206 | 0.018 | 0.021 | 1 | 7 | Nmb | -0.003 | Old |
| 0.771519 | 0.152672 | 0.044 | 0.048 | 1 | 7 | Cep112 | -0.004 | Old |
| 0.771861 | 0.224953 | 0.044 | 0.039 | 1 | 7 | Coa7 | 0.005 | Old |
| 0.772123 | 0.068874 | 0.603 | 0.575 | 1 | 7 | Rab7 | 0.028 | Old |
| 0.772353 | 0.276176 | 0.018 | 0.021 | 1 | 7 | Kifc5b | -0.003 | Old |
| 0.772555 | 0.073479 | 0.105 | 0.111 | 1 | 7 | Cep164 | -0.006 | Old |
| 0.772823 | 0.159058 | 0.462 | 0.425 | 1 | 7 | Pxn | 0.037 | Old |
| 0.773034 | 0.257394 | 0.031 | 0.027 | 1 | 7 | Bphl | 0.004 | Old |
| 0.773528 | 0.452057 | 0.138 | 0.133 | 1 | 7 | Cstf2 | 0.005 | Old |
| 0.774126 | 0.116706 | 0.015 | 0.018 | 1 | 7 | Hp | -0.003 | Old |
| 0.774768 | 0.015205 | 0.756 | 0.744 | 1 | 7 | Lgals1 | 0.012 | Old |
| 0.775645 | 0.114323 | 0.454 | 0.428 | 1 | 7 | Nek7 | 0.026 | Old |
| 0.776603 | 0.115009 | 0.015 | 0.018 | 1 | 7 | Abcc10 | -0.003 | Old |
| 0.776603 | 0.104157 | 0.015 | 0.018 | 1 | 7 | Zfp790 | -0.003 | Old |

| | | | | | | | | |
|---|---|---|---|---|---|---|---|---|
| 0.776801 | 0.084181 | 0.379 | 0.383 | 1 | 7 | Puf60 | -0.004 | Old |
| 0.776805 | 0.06244 | 0.318 | 0.301 | 1 | 7 | Mbd3 | 0.017 | Old |
| 0.776929 | 0.035562 | 0.169 | 0.157 | 1 | 7 | Tor3a | 0.012 | Old |
| 0.776945 | 0.228224 | 0.018 | 0.021 | 1 | 7 | Gm38394 | -0.003 | Old |
| 0.77764 | 0.18687 | 0.479 | 0.461 | 1 | 7 | Rassf2 | 0.018 | Old |
| 0.778508 | 0.063387 | 0.49 | 0.467 | 1 | 7 | Mafg | 0.023 | Old |
| 0.779169 | 0.172117 | 0.249 | 0.238 | 1 | 7 | Hcst | 0.011 | Old |
| 0.779363 | 0.050251 | 0.097 | 0.09 | 1 | 7 | Daam1 | 0.007 | Old |
| 0.780405 | 0.230667 | 0.069 | 0.075 | 1 | 7 | Zfp568 | -0.006 | Old |
| 0.781233 | 0.261425 | 0.159 | 0.154 | 1 | 7 | D10Wsu10 | 0.005 | Old |
| 0.782245 | 0.450014 | 0.074 | 0.069 | 1 | 7 | Tbl2 | 0.005 | Old |
| 0.782405 | 0.036755 | 0.131 | 0.127 | 1 | 7 | Slc9a8 | 0.004 | Old |
| 0.782447 | 0.149985 | 0.267 | 0.262 | 1 | 7 | Upf1 | 0.005 | Old |
| 0.782806 | 0.316433 | 0.015 | 0.018 | 1 | 7 | Gm45051 | -0.003 | Old |
| 0.782806 | 0.274362 | 0.015 | 0.018 | 1 | 7 | Lum | -0.003 | Old |
| 0.782806 | 0.07161 | 0.015 | 0.018 | 1 | 7 | Adarb1 | -0.003 | Old |
| 0.783705 | 0.024783 | 0.449 | 0.416 | 1 | 7 | Mrpl51 | 0.033 | Old |
| 0.783822 | 0.149099 | 0.362 | 0.355 | 1 | 7 | Sdf2 | 0.007 | Old |
| 0.784349 | 0.212196 | 0.11 | 0.102 | 1 | 7 | Prkab1 | 0.008 | Old |
| 0.784938 | 0.04234 | 0.344 | 0.325 | 1 | 7 | Exosc8 | 0.019 | Old |
| 0.785217 | 0.062193 | 0.164 | 0.169 | 1 | 7 | Pfkl | -0.005 | Old |
| 0.785291 | 0.495764 | 0.015 | 0.018 | 1 | 7 | Tspyl4 | -0.003 | Old |
| 0.785291 | 0.235747 | 0.015 | 0.018 | 1 | 7 | Sirt6 | -0.003 | Old |
| 0.785291 | 0.213633 | 0.015 | 0.018 | 1 | 7 | Gm15706 | -0.003 | Old |
| 0.785524 | 0.118975 | 0.138 | 0.133 | 1 | 7 | Dusp23 | 0.005 | Old |
| 0.785634 | 0.046467 | 0.156 | 0.148 | 1 | 7 | Timm22 | 0.008 | Old |
| 0.7864 | 0.036809 | 0.769 | 0.729 | 1 | 7 | Sf3b1 | 0.04 | Old |
| 0.786549 | 0.051071 | 0.272 | 0.28 | 1 | 7 | Cd9 | -0.008 | Old |
| 0.787778 | 0.101485 | 0.015 | 0.018 | 1 | 7 | Slc9a7 | -0.003 | Old |
| 0.787825 | 0.137839 | 0.059 | 0.054 | 1 | 7 | Hnmt | 0.005 | Old |
| 0.787846 | 0.266825 | 0.238 | 0.232 | 1 | 7 | BC003331 | 0.006 | Old |
| 0.788 | 0.0035 | 0.149 | 0.142 | 1 | 7 | Vps33a | 0.007 | Old |
| 0.788734 | 0.126526 | 0.267 | 0.253 | 1 | 7 | Dda1 | 0.014 | Old |
| 0.788982 | 0.025817 | 0.044 | 0.048 | 1 | 7 | Fam210b | -0.004 | Old |
| 0.789022 | 0.452463 | 0.015 | 0.018 | 1 | 7 | Btnl2 | -0.003 | Old |
| 0.789329 | 0.271508 | 0.103 | 0.096 | 1 | 7 | Dis3l2 | 0.007 | Old |
| 0.790085 | 0.192176 | 0.362 | 0.349 | 1 | 7 | Apc | 0.013 | Old |
| 0.790875 | 0.011276 | 0.623 | 0.59 | 1 | 7 | Arhgef6 | 0.033 | Old |
| 0.792345 | 0.458627 | 0.226 | 0.217 | 1 | 7 | Pole3 | 0.009 | Old |
| 0.792814 | 0.021831 | 0.274 | 0.265 | 1 | 7 | Ccng1 | 0.009 | Old |
| 0.792943 | 0.093728 | 0.041 | 0.045 | 1 | 7 | Gtpbp3 | -0.004 | Old |
| 0.793481 | 0.14352 | 0.11 | 0.105 | 1 | 7 | Pde4a | 0.005 | Old |
| 0.794005 | 0.273806 | 0.015 | 0.018 | 1 | 7 | Fam120c | -0.003 | Old |
| 0.794071 | 0.197527 | 0.174 | 0.166 | 1 | 7 | Slc35e1 | 0.008 | Old |
| 0.794307 | 0.526835 | 0.069 | 0.066 | 1 | 7 | Zfp142 | 0.003 | Old |

| | | | | | | | | |
|---|---|---|---|---|---|---|---|---|
| 0.794703 | 0.004247 | 0.059 | 0.054 | 1 | 7 | Chtf18 | 0.005 | Old |
| 0.79487 | 0.275624 | 0.013 | 0.015 | 1 | 7 | Lrrc56 | -0.002 | Old |
| 0.795017 | 0.03423 | 0.192 | 0.184 | 1 | 7 | Mum1 | 0.008 | Old |
| 0.795528 | 0.026374 | 0.144 | 0.136 | 1 | 7 | Camsap2 | 0.008 | Old |
| 0.795893 | 0.191502 | 0.113 | 0.105 | 1 | 7 | Wdtc1 | 0.008 | Old |
| 0.795986 | 0.405935 | 0.144 | 0.136 | 1 | 7 | 2310033P0 | 0.008 | Old |
| 0.796766 | 0.26865 | 0.046 | 0.042 | 1 | 7 | Phf7 | 0.004 | Old |
| 0.797124 | 1.57172 | 0.021 | 0.018 | 1 | 7 | Fignl2 | 0.003 | Old |
| 0.797983 | 0.256807 | 0.2 | 0.196 | 1 | 7 | Sh3gl1 | 0.004 | Old |
| 0.798228 | 0.079194 | 0.087 | 0.093 | 1 | 7 | Prpf3 | -0.006 | Old |
| 0.798888 | 0.288838 | 0.033 | 0.03 | 1 | 7 | Tkfc | 0.003 | Old |
| 0.798966 | 0.227935 | 0.013 | 0.015 | 1 | 7 | Kif9 | -0.002 | Old |
| 0.799291 | 0.00434 | 0.659 | 0.611 | 1 | 7 | Nfam1 | 0.048 | Old |
| 0.79944 | 0.573791 | 0.021 | 0.018 | 1 | 7 | Bend6 | 0.003 | Old |
| 0.800043 | 0.164397 | 0.144 | 0.151 | 1 | 7 | Arhgef7 | -0.007 | Old |
| 0.800332 | 0.032535 | 0.013 | 0.015 | 1 | 7 | Rabl2 | -0.002 | Old |
| 0.800598 | 0.648218 | 0.021 | 0.018 | 1 | 7 | Zfp473 | 0.003 | Old |
| 0.800643 | 0.623614 | 0.046 | 0.042 | 1 | 7 | Arrdc2 | 0.004 | Old |
| 0.801377 | 0.103055 | 0.31 | 0.298 | 1 | 7 | Bbip1 | 0.012 | Old |
| 0.801699 | 0.273554 | 0.013 | 0.015 | 1 | 7 | Sphk1 | -0.002 | Old |
| 0.801699 | 0.164297 | 0.013 | 0.015 | 1 | 7 | Fam120aos | -0.002 | Old |
| 0.801699 | 0.099408 | 0.013 | 0.015 | 1 | 7 | Kif5c | -0.002 | Old |
| 0.802196 | 1.103055 | 0.046 | 0.042 | 1 | 7 | Hyls1 | 0.004 | Old |
| 0.802565 | 0.094004 | 0.328 | 0.313 | 1 | 7 | Ganab | 0.015 | Old |
| 0.802916 | 0.695378 | 0.021 | 0.018 | 1 | 7 | Ift122 | 0.003 | Old |
| 0.803686 | 0.065797 | 0.477 | 0.455 | 1 | 7 | Dynll2 | 0.022 | Old |
| 0.804075 | 0.460255 | 0.021 | 0.018 | 1 | 7 | Prelid3a | 0.003 | Old |
| 0.804435 | 0.334318 | 0.013 | 0.015 | 1 | 7 | Tspan2 | -0.002 | Old |
| 0.804435 | 0.308697 | 0.013 | 0.015 | 1 | 7 | Phactr1 | -0.002 | Old |
| 0.804435 | 0.245743 | 0.013 | 0.015 | 1 | 7 | Hist1h3b | -0.002 | Old |
| 0.807122 | 0.037791 | 0.059 | 0.054 | 1 | 7 | Ints4 | 0.005 | Old |
| 0.807598 | 0.027101 | 0.29 | 0.28 | 1 | 7 | Samd9l | 0.01 | Old |
| 0.80779 | 0.025653 | 0.126 | 0.133 | 1 | 7 | Bicral | -0.007 | Old |
| 0.808098 | 0.19553 | 0.049 | 0.045 | 1 | 7 | Socs1 | 0.004 | Old |
| 0.808544 | 0.502401 | 0.013 | 0.015 | 1 | 7 | Sparcl1 | -0.002 | Old |
| 0.808871 | 0.240362 | 0.19 | 0.184 | 1 | 7 | Ikzf5 | 0.006 | Old |
| 0.808894 | 0.023591 | 0.974 | 0.967 | 1 | 7 | Rpl7 | 0.007 | Old |
| 0.809135 | 0.144998 | 0.038 | 0.042 | 1 | 7 | Cdkn2a | -0.004 | Old |
| 0.809253 | 0.007181 | 0.077 | 0.072 | 1 | 7 | Mad2l2 | 0.005 | Old |
| 0.809354 | 0.273544 | 0.205 | 0.196 | 1 | 7 | Mtmr12 | 0.009 | Old |
| 0.809879 | 0.711894 | 0.021 | 0.018 | 1 | 7 | Hist1h2be | 0.003 | Old |
| 0.809879 | 0.403581 | 0.021 | 0.018 | 1 | 7 | Fam185a | 0.003 | Old |
| 0.810728 | 0.246839 | 0.033 | 0.03 | 1 | 7 | Tigar | 0.003 | Old |
| 0.810807 | 0.527805 | 0.087 | 0.081 | 1 | 7 | Tmem86a | 0.006 | Old |
| 0.811122 | 0.070414 | 0.049 | 0.045 | 1 | 7 | Hes1 | 0.004 | Old |

| | | | | | | | | |
|---|---|---|---|---|---|---|---|---|
| 0.811152 | 0.002607 | 0.244 | 0.232 | 1 | 7 | Wdr5 | 0.012 | Old |
| 0.811452 | 0.073205 | 0.372 | 0.358 | 1 | 7 | Pmf1 | 0.014 | Old |
| 0.812657 | 1.592959 | 0.013 | 0.015 | 1 | 7 | Gstp2 | -0.002 | Old |
| 0.813126 | 0.178128 | 0.141 | 0.136 | 1 | 7 | Nudt16 | 0.005 | Old |
| 0.813367 | 1.025477 | 0.021 | 0.018 | 1 | 7 | Ddit4 | 0.003 | Old |
| 0.813367 | 0.411593 | 0.021 | 0.018 | 1 | 7 | Zfp446 | 0.003 | Old |
| 0.813466 | 0.20749 | 0.033 | 0.03 | 1 | 7 | Hibch | 0.003 | Old |
| 0.814316 | 0.089446 | 0.538 | 0.497 | 1 | 7 | Ube2r2 | 0.041 | Old |
| 0.814393 | 0.042071 | 0.249 | 0.241 | 1 | 7 | Actr10 | 0.008 | Old |
| 0.814631 | 0.068375 | 0.074 | 0.069 | 1 | 7 | Pls3 | 0.005 | Old |
| 0.814976 | 0.346695 | 0.059 | 0.063 | 1 | 7 | Pdp1 | -0.004 | Old |
| 0.81522 | 0.640065 | 0.064 | 0.06 | 1 | 7 | Eif2ak4 | 0.004 | Old |
| 0.815656 | 0.035066 | 0.651 | 0.648 | 1 | 7 | Usf2 | 0.003 | Old |
| 0.815694 | 0.033387 | 0.021 | 0.018 | 1 | 7 | Gnb1l | 0.003 | Old |
| 0.815785 | 0.098139 | 0.526 | 0.521 | 1 | 7 | Ndufab1 | 0.005 | Old |
| 0.816627 | 0.075032 | 0.162 | 0.166 | 1 | 7 | Tnfaip8l2 | -0.004 | Old |
| 0.816856 | 0.034677 | 0.131 | 0.127 | 1 | 7 | Cwc22 | 0.004 | Old |
| 0.816925 | 0.031945 | 0.09 | 0.084 | 1 | 7 | Pef1 | 0.006 | Old |
| 0.818054 | 0.016314 | 0.01 | 0.012 | 1 | 7 | Usp27x | -0.002 | Old |
| 0.818201 | 0.145512 | 0.203 | 0.196 | 1 | 7 | Nedd9 | 0.007 | Old |
| 0.818937 | 0.159149 | 0.038 | 0.042 | 1 | 7 | Elmod3 | -0.004 | Old |
| 0.819497 | 0.180709 | 0.036 | 0.039 | 1 | 7 | Suox | -0.003 | Old |
| 0.819589 | 0.249899 | 0.01 | 0.012 | 1 | 7 | Slc46a1 | -0.002 | Old |
| 0.819589 | 0.182192 | 0.01 | 0.012 | 1 | 7 | Dact3 | -0.002 | Old |
| 0.819858 | 0.320527 | 0.079 | 0.075 | 1 | 7 | Zxdc | 0.004 | Old |
| 0.819933 | 0.019876 | 0.831 | 0.852 | 1 | 7 | Alox5ap | -0.021 | Old |
| 0.820233 | 0.107625 | 0.197 | 0.19 | 1 | 7 | Taf11 | 0.007 | Old |
| 0.820274 | 0.036243 | 0.105 | 0.099 | 1 | 7 | Arhgef11 | 0.006 | Old |
| 0.821124 | 0.923785 | 0.01 | 0.012 | 1 | 7 | Arhgap29 | -0.002 | Old |
| 0.821124 | 0.157679 | 0.01 | 0.012 | 1 | 7 | Dbn1 | -0.002 | Old |
| 0.821124 | 0.066708 | 0.01 | 0.012 | 1 | 7 | Zdhhc2 | -0.002 | Old |
| 0.821124 | 0.002781 | 0.01 | 0.012 | 1 | 7 | B230303O1 | -0.002 | Old |
| 0.821654 | 0.116384 | 0.19 | 0.193 | 1 | 7 | Ak3 | -0.003 | Old |
| 0.822157 | 0.161773 | 0.195 | 0.19 | 1 | 7 | Ap4s1 | 0.005 | Old |
| 0.822326 | 0.117682 | 0.172 | 0.175 | 1 | 7 | Lyrm2 | -0.003 | Old |
| 0.822586 | 0.045428 | 0.079 | 0.084 | 1 | 7 | Ttc39c | -0.005 | Old |
| 0.822626 | 0.036274 | 0.103 | 0.108 | 1 | 7 | Pask | -0.005 | Old |
| 0.822657 | 0.014273 | 0.969 | 0.979 | 1 | 7 | H2-Aa | -0.01 | Old |
| 0.82266 | 0.609393 | 0.01 | 0.012 | 1 | 7 | Jag1 | -0.002 | Old |
| 0.82266 | 0.036427 | 0.01 | 0.012 | 1 | 7 | Gm12689 | -0.002 | Old |
| 0.823113 | 0.052018 | 0.223 | 0.226 | 1 | 7 | Micu2 | -0.003 | Old |
| 0.823354 | 0.052266 | 0.356 | 0.349 | 1 | 7 | Man2a2 | 0.007 | Old |
| 0.824197 | 0.067368 | 0.01 | 0.012 | 1 | 7 | Ccdc173 | -0.002 | Old |
| 0.824554 | 0.279232 | 0.246 | 0.235 | 1 | 7 | Naa35 | 0.011 | Old |
| 0.824852 | 0.052065 | 0.415 | 0.407 | 1 | 7 | Cdk4 | 0.008 | Old |

| | | | | | | | | |
|---|---|---|---|---|---|---|---|---|
| 0.824883 | 0.396451 | 0.182 | 0.178 | 1 | 7 | Erlec1 | 0.004 | Old |
| 0.825735 | 0.359971 | 0.01 | 0.012 | 1 | 7 | Fam131a | -0.002 | Old |
| 0.82586 | 0.198594 | 0.113 | 0.105 | 1 | 7 | Lrrc20 | 0.008 | Old |
| 0.826277 | 0.090536 | 0.036 | 0.039 | 1 | 7 | Slc25a14 | -0.003 | Old |
| 0.826346 | 0.046451 | 0.131 | 0.123 | 1 | 7 | Dubr | 0.008 | Old |
| 0.826863 | 0.45577 | 0.036 | 0.033 | 1 | 7 | 1500015A( | 0.003 | Old |
| 0.827086 | 0.099244 | 0.056 | 0.06 | 1 | 7 | Gm42031 | -0.004 | Old |
| 0.827273 | 0.315347 | 0.01 | 0.012 | 1 | 7 | Klk8 | -0.002 | Old |
| 0.828423 | 0.017798 | 0.662 | 0.633 | 1 | 7 | Eif4h | 0.029 | Old |
| 0.828778 | 0.140819 | 0.077 | 0.072 | 1 | 7 | 1110034G2 | 0.005 | Old |
| 0.828812 | 0.540402 | 0.01 | 0.012 | 1 | 7 | Nid1 | -0.002 | Old |
| 0.828812 | 0.335511 | 0.01 | 0.012 | 1 | 7 | 4930513N1 | -0.002 | Old |
| 0.82994 | 0.361347 | 0.077 | 0.081 | 1 | 7 | Ptgfrn | -0.004 | Old |
| 0.830351 | 0.54866 | 0.01 | 0.012 | 1 | 7 | Gm5431 | -0.002 | Old |
| 0.830387 | 0.375761 | 0.036 | 0.033 | 1 | 7 | Wdr73 | 0.003 | Old |
| 0.83108 | 0.138441 | 0.244 | 0.235 | 1 | 7 | Cul5 | 0.009 | Old |
| 0.832151 | 0.546638 | 0.036 | 0.033 | 1 | 7 | Zfp251 | 0.003 | Old |
| 0.832282 | 0.089928 | 0.179 | 0.172 | 1 | 7 | Smpd1 | 0.007 | Old |
| 0.832566 | 0.461602 | 0.067 | 0.063 | 1 | 7 | Gprc5a | 0.004 | Old |
| 0.832575 | 0.023252 | 0.292 | 0.277 | 1 | 7 | Btbd7 | 0.015 | Old |
| 0.832797 | 0.080613 | 0.682 | 0.657 | 1 | 7 | Nptn | 0.025 | Old |
| 0.833006 | 0.120954 | 0.426 | 0.428 | 1 | 7 | Prc1 | -0.002 | Old |
| 0.833238 | 0.192853 | 0.228 | 0.223 | 1 | 7 | Ddx1 | 0.005 | Old |
| 0.83399 | 0.067291 | 0.592 | 0.575 | 1 | 7 | Ndufb8 | 0.017 | Old |
| 0.834124 | 0.014448 | 0.187 | 0.19 | 1 | 7 | Rab43 | -0.003 | Old |
| 0.834235 | 0.248625 | 0.082 | 0.078 | 1 | 7 | Dtd2 | 0.004 | Old |
| 0.834235 | 0.090349 | 0.082 | 0.078 | 1 | 7 | Clp1 | 0.004 | Old |
| 0.83468 | 0.410377 | 0.313 | 0.31 | 1 | 7 | Palld | 0.003 | Old |
| 0.835107 | 0.03049 | 0.631 | 0.63 | 1 | 7 | Fkbp1a | 0.001 | Old |
| 0.835149 | 0.522053 | 0.064 | 0.069 | 1 | 7 | Cd209b | -0.005 | Old |
| 0.83568 | 0.558649 | 0.036 | 0.033 | 1 | 7 | Focad | 0.003 | Old |
| 0.836071 | 0.010475 | 0.603 | 0.587 | 1 | 7 | Snu13 | 0.016 | Old |
| 0.836476 | 0.599296 | 0.023 | 0.021 | 1 | 7 | Eps8l1 | 0.002 | Old |
| 0.836868 | 0.101569 | 0.09 | 0.084 | 1 | 7 | Gtf3c4 | 0.006 | Old |
| 0.837031 | 0.062369 | 0.177 | 0.172 | 1 | 7 | Dcun1d1 | 0.005 | Old |
| 0.837312 | 0.009925 | 0.972 | 0.97 | 1 | 7 | Ftl1 | 0.002 | Old |
| 0.837551 | 0.271905 | 0.115 | 0.12 | 1 | 7 | Abr | -0.005 | Old |
| 0.837624 | 0.002233 | 0.579 | 0.563 | 1 | 7 | Ube2l3 | 0.016 | Old |
| 0.837692 | 0.030077 | 0.144 | 0.139 | 1 | 7 | Bmi1 | 0.005 | Old |
| 0.837913 | 0.012985 | 0.531 | 0.509 | 1 | 7 | Cd48 | 0.022 | Old |
| 0.838407 | 0.018179 | 0.395 | 0.38 | 1 | 7 | Rnaseh2c | 0.015 | Old |
| 0.838807 | 0.025803 | 0.218 | 0.211 | 1 | 7 | Pum3 | 0.007 | Old |
| 0.839073 | 0.199712 | 0.351 | 0.34 | 1 | 7 | Golga4 | 0.011 | Old |
| 0.840156 | 0.028412 | 0.818 | 0.807 | 1 | 7 | Sh3bgrl | 0.011 | Old |
| 0.840331 | 0.265071 | 0.115 | 0.111 | 1 | 7 | Nr2c2ap | 0.004 | Old |

| | | | | | | | | |
|---|---|---|---|---|---|---|---|---|
| 0.840606 | 0.069826 | 0.731 | 0.723 | 1 | 7 | Ppp3ca | 0.008 | Old |
| 0.840675 | 0.001721 | 0.241 | 0.232 | 1 | 7 | Rnf10 | 0.009 | Old |
| 0.840785 | 0.067234 | 0.136 | 0.139 | 1 | 7 | Poldip2 | -0.003 | Old |
| 0.841661 | 0.042202 | 0.169 | 0.16 | 1 | 7 | 1700037H( | 0.009 | Old |
| 0.841789 | 0.076034 | 0.133 | 0.13 | 1 | 7 | Tbc1d17 | 0.003 | Old |
| 0.841948 | 0.038319 | 0.2 | 0.193 | 1 | 7 | Mnat1 | 0.007 | Old |
| 0.841964 | 0.844051 | 0.023 | 0.021 | 1 | 7 | Cald1 | 0.002 | Old |
| 0.842398 | 0.263963 | 0.172 | 0.166 | 1 | 7 | Smim13 | 0.006 | Old |
| 0.842633 | 0.131543 | 0.354 | 0.349 | 1 | 7 | Vav1 | 0.005 | Old |
| 0.842748 | 0.094042 | 0.036 | 0.033 | 1 | 7 | Arl15 | 0.003 | Old |
| 0.84314 | 0.164673 | 0.082 | 0.078 | 1 | 7 | Vps72 | 0.004 | Old |
| 0.843217 | 0.039882 | 0.054 | 0.057 | 1 | 7 | Ccdc102a | -0.003 | Old |
| 0.843471 | 0.212543 | 0.172 | 0.166 | 1 | 7 | Fbxo4 | 0.006 | Old |
| 0.843553 | 0.048567 | 0.121 | 0.114 | 1 | 7 | Ubxn7 | 0.007 | Old |
| 0.84357 | 0.512586 | 0.056 | 0.054 | 1 | 7 | Eva1b | 0.002 | Old |
| 0.843626 | 0.204975 | 0.056 | 0.06 | 1 | 7 | Sv2b | -0.004 | Old |
| 0.843957 | 0.193687 | 0.062 | 0.057 | 1 | 7 | Polr3h | 0.005 | Old |
| 0.844137 | 0.93015 | 0.038 | 0.036 | 1 | 7 | Zfp317 | 0.002 | Old |
| 0.844999 | 0.056945 | 0.067 | 0.063 | 1 | 7 | Tpk1 | 0.004 | Old |
| 0.845286 | 0.149233 | 0.177 | 0.172 | 1 | 7 | Fut8 | 0.005 | Old |
| 0.847232 | 0.062104 | 0.074 | 0.078 | 1 | 7 | Bet1l | -0.004 | Old |
| 0.847315 | 0.140703 | 0.177 | 0.178 | 1 | 7 | Ap1m1 | -0.001 | Old |
| 0.847497 | 0.139823 | 0.608 | 0.572 | 1 | 7 | Smc1a | 0.036 | Old |
| 0.847548 | 0.968502 | 0.038 | 0.036 | 1 | 7 | Eri2 | 0.002 | Old |
| 0.847563 | 0.033804 | 0.051 | 0.048 | 1 | 7 | Zfp809 | 0.003 | Old |
| 0.84763 | 0.095008 | 0.133 | 0.139 | 1 | 7 | Lrrc8a | -0.006 | Old |
| 0.848176 | 0.263578 | 0.079 | 0.075 | 1 | 7 | Phlpp2 | 0.004 | Old |
| 0.848378 | 0.028135 | 0.323 | 0.313 | 1 | 7 | Slc15a3 | 0.01 | Old |
| 0.848604 | 0.041054 | 0.423 | 0.407 | 1 | 7 | Ndufa8 | 0.016 | Old |
| 0.849108 | 0.133318 | 0.162 | 0.157 | 1 | 7 | Trappc3 | 0.005 | Old |
| 0.849146 | 0.0229 | 0.049 | 0.045 | 1 | 7 | Setd6 | 0.004 | Old |
| 0.849434 | 0.234611 | 0.203 | 0.208 | 1 | 7 | Ddx23 | -0.005 | Old |
| 0.849873 | 0.156412 | 0.067 | 0.069 | 1 | 7 | Tonsl | -0.002 | Old |
| 0.849965 | 0.13603 | 0.244 | 0.235 | 1 | 7 | Tmem183a | 0.009 | Old |
| 0.849983 | 0.030154 | 0.397 | 0.395 | 1 | 7 | Csrp1 | 0.002 | Old |
| 0.850078 | 0.025974 | 0.069 | 0.072 | 1 | 7 | Ankrd26 | -0.003 | Old |
| 0.850369 | 0.043422 | 0.687 | 0.708 | 1 | 7 | Lsp1 | -0.021 | Old |
| 0.85076 | 0.240192 | 0.023 | 0.021 | 1 | 7 | Dnah2 | 0.002 | Old |
| 0.850777 | 0.052944 | 0.292 | 0.274 | 1 | 7 | Ing2 | 0.018 | Old |
| 0.851015 | 0.084292 | 0.318 | 0.313 | 1 | 7 | 4930523C( | 0.005 | Old |
| 0.851575 | 0.053601 | 0.192 | 0.184 | 1 | 7 | Lrif1 | 0.008 | Old |
| 0.852403 | 0.125188 | 0.292 | 0.292 | 1 | 7 | Chd3 | 0 | Old |
| 0.852962 | 0.287685 | 0.023 | 0.021 | 1 | 7 | Ifi206 | 0.002 | Old |
| 0.853374 | 0.186196 | 0.036 | 0.033 | 1 | 7 | Efemp2 | 0.003 | Old |
| 0.853414 | 0.066622 | 0.092 | 0.096 | 1 | 7 | Heg1 | -0.004 | Old |

| | | | | | | | | |
|---|---|---|---|---|---|---|---|---|
| 0.853849 | 0.235612 | 0.179 | 0.175 | 1 | 7 | Rdm1 | 0.004 | Old |
| 0.854063 | 0.14322 | 0.023 | 0.021 | 1 | 7 | Hist1h3f | 0.002 | Old |
| 0.854277 | 0.311366 | 0.051 | 0.054 | 1 | 7 | 5830432E0 | -0.003 | Old |
| 0.854475 | 0.01698 | 0.159 | 0.163 | 1 | 7 | Trim26 | -0.004 | Old |
| 0.855295 | 0.038491 | 0.474 | 0.452 | 1 | 7 | Ndufs2 | 0.022 | Old |
| 0.855722 | 0.405036 | 0.031 | 0.033 | 1 | 7 | Fuz | -0.002 | Old |
| 0.856267 | 0.086295 | 0.023 | 0.021 | 1 | 7 | Gm16867 | 0.002 | Old |
| 0.856472 | 0.226471 | 0.387 | 0.386 | 1 | 7 | Ccr1 | 0.001 | Old |
| 0.856833 | 0.047322 | 0.082 | 0.078 | 1 | 7 | Gm6563 | 0.004 | Old |
| 0.85715 | 0.183692 | 0.831 | 0.783 | 1 | 7 | Ptms | 0.048 | Old |
| 0.857199 | 0.324731 | 0.177 | 0.175 | 1 | 7 | Phka2 | 0.002 | Old |
| 0.857758 | 0.073272 | 0.533 | 0.509 | 1 | 7 | Gns | 0.024 | Old |
| 0.858051 | 0.091213 | 0.515 | 0.518 | 1 | 7 | Banf1 | -0.003 | Old |
| 0.858522 | 0.805676 | 0.01 | 0.009 | 1 | 7 | Vsig8 | 0.001 | Old |
| 0.859093 | 0.008824 | 0.272 | 0.259 | 1 | 7 | Zfp652 | 0.013 | Old |
| 0.859404 | 0.125788 | 0.21 | 0.214 | 1 | 7 | Cyp20a1 | -0.004 | Old |
| 0.859574 | 0.166271 | 0.023 | 0.021 | 1 | 7 | Gas8 | 0.002 | Old |
| 0.860017 | 0.120493 | 0.187 | 0.178 | 1 | 7 | Papss1 | 0.009 | Old |
| 0.86006 | 0.044593 | 0.354 | 0.337 | 1 | 7 | Jup | 0.017 | Old |
| 0.860179 | 0.606496 | 0.01 | 0.009 | 1 | 7 | Gm4924 | 0.001 | Old |
| 0.860365 | 0.025962 | 0.038 | 0.036 | 1 | 7 | Dgke | 0.002 | Old |
| 0.860828 | 0.199667 | 0.069 | 0.066 | 1 | 7 | Ppp2r3a | 0.003 | Old |
| 0.861416 | 0.000283 | 0.067 | 0.063 | 1 | 7 | Cdc25c | 0.004 | Old |
| 0.861757 | 0.281485 | 0.146 | 0.142 | 1 | 7 | Yif1b | 0.004 | Old |
| 0.861836 | 0.72102 | 0.01 | 0.009 | 1 | 7 | Ccno | 0.001 | Old |
| 0.861836 | 0.387685 | 0.01 | 0.009 | 1 | 7 | Cacnb4 | 0.001 | Old |
| 0.861836 | 0.291959 | 0.01 | 0.009 | 1 | 7 | Lgi4 | 0.001 | Old |
| 0.86244 | 0.084293 | 0.328 | 0.328 | 1 | 7 | Tgs1 | 0 | Old |
| 0.862933 | 0.245274 | 0.038 | 0.036 | 1 | 7 | Cd320 | 0.002 | Old |
| 0.863495 | 0.890442 | 0.01 | 0.009 | 1 | 7 | Zfp652os | 0.001 | Old |
| 0.863495 | 0.318584 | 0.01 | 0.009 | 1 | 7 | Llgl2 | 0.001 | Old |
| 0.864043 | 0.659583 | 0.031 | 0.033 | 1 | 7 | Efnb2 | -0.002 | Old |
| 0.864478 | 0.028673 | 0.544 | 0.53 | 1 | 7 | Brd4 | 0.014 | Old |
| 0.865153 | 1.068818 | 0.01 | 0.009 | 1 | 7 | Colca2 | 0.001 | Old |
| 0.865153 | 0.497542 | 0.01 | 0.009 | 1 | 7 | Gm15952 | 0.001 | Old |
| 0.865153 | 0.496125 | 0.01 | 0.009 | 1 | 7 | Hist1h2af | 0.001 | Old |
| 0.865153 | 0.404521 | 0.01 | 0.009 | 1 | 7 | Kit | 0.001 | Old |
| 0.865153 | 0.400228 | 0.01 | 0.009 | 1 | 7 | Map1b | 0.001 | Old |
| 0.865153 | 0.333166 | 0.01 | 0.009 | 1 | 7 | Gm29650 | 0.001 | Old |
| 0.865153 | 0.283576 | 0.01 | 0.009 | 1 | 7 | Gm6133 | 0.001 | Old |
| 0.865153 | 0.211202 | 0.01 | 0.009 | 1 | 7 | Gm7879 | 0.001 | Old |
| 0.86554 | 0.13258 | 0.238 | 0.235 | 1 | 7 | Cd209f | 0.003 | Old |
| 0.865792 | 0.056477 | 0.159 | 0.154 | 1 | 7 | Prkag2 | 0.005 | Old |
| 0.866096 | 0.154981 | 0.174 | 0.172 | 1 | 7 | Ate1 | 0.002 | Old |
| 0.866123 | 0.115734 | 0.128 | 0.123 | 1 | 7 | Rps6kc1 | 0.005 | Old |

| | | | | | | | | |
|---|---|---|---|---|---|---|---|---|
| 0.866196 | 0.342752 | 0.023 | 0.021 | 1 | 7 | Gm44777 | 0.002 | Old |
| 0.866379 | 0.355034 | 0.11 | 0.114 | 1 | 7 | Mtmr4 | -0.004 | Old |
| 0.866813 | 0.629924 | 0.01 | 0.009 | 1 | 7 | Lefty1 | 0.001 | Old |
| 0.866813 | 0.238244 | 0.01 | 0.009 | 1 | 7 | Ak1 | 0.001 | Old |
| 0.866813 | 0.223651 | 0.01 | 0.009 | 1 | 7 | Tex14 | 0.001 | Old |
| 0.866813 | 0.155877 | 0.01 | 0.009 | 1 | 7 | Dpys | 0.001 | Old |
| 0.866813 | 0.140802 | 0.01 | 0.009 | 1 | 7 | Hist1h2ak | 0.001 | Old |
| 0.866813 | 0.11735 | 0.01 | 0.009 | 1 | 7 | Xcr1 | 0.001 | Old |
| 0.866813 | 0.102694 | 0.01 | 0.009 | 1 | 7 | Frat1 | 0.001 | Old |
| 0.866813 | 0.035807 | 0.01 | 0.009 | 1 | 7 | Tlr12 | 0.001 | Old |
| 0.867063 | 0.01437 | 0.959 | 0.937 | 1 | 7 | Hsp90ab1 | 0.022 | Old |
| 0.867745 | 0.923512 | 0.031 | 0.033 | 1 | 7 | Dcaf6 | -0.002 | Old |
| 0.867802 | 0.090868 | 0.651 | 0.605 | 1 | 7 | Edem1 | 0.046 | Old |
| 0.868073 | 0.459455 | 0.038 | 0.036 | 1 | 7 | Nars2 | 0.002 | Old |
| 0.868073 | 0.237909 | 0.038 | 0.036 | 1 | 7 | Taf4b | 0.002 | Old |
| 0.868473 | 0.190386 | 0.01 | 0.009 | 1 | 7 | Tstd1 | 0.001 | Old |
| 0.868473 | 0.032446 | 0.01 | 0.009 | 1 | 7 | Paqr8 | 0.001 | Old |
| 0.869067 | 0.031964 | 0.3 | 0.286 | 1 | 7 | Azi2 | 0.014 | Old |
| 0.869511 | 0.058612 | 0.023 | 0.021 | 1 | 7 | Snapc4 | 0.002 | Old |
| 0.869766 | 0.085711 | 0.462 | 0.455 | 1 | 7 | Aftph | 0.007 | Old |
| 0.870022 | 0.232021 | 0.064 | 0.06 | 1 | 7 | Trim30c | 0.004 | Old |
| 0.870119 | 0.091825 | 0.579 | 0.548 | 1 | 7 | Rac2 | 0.031 | Old |
| 0.870133 | 0.647518 | 0.01 | 0.009 | 1 | 7 | Gm45238 | 0.001 | Old |
| 0.870133 | 0.004479 | 0.01 | 0.009 | 1 | 7 | Zfp27 | 0.001 | Old |
| 0.870483 | 0.02288 | 0.585 | 0.554 | 1 | 7 | Scamp2 | 0.031 | Old |
| 0.87064 | 0.280482 | 0.054 | 0.051 | 1 | 7 | Rnf152 | 0.003 | Old |
| 0.870645 | 0.193081 | 0.038 | 0.036 | 1 | 7 | Miga1 | 0.002 | Old |
| 0.871007 | 0.155024 | 0.351 | 0.34 | 1 | 7 | Aurkb | 0.011 | Old |
| 0.871794 | 0.048931 | 0.01 | 0.009 | 1 | 7 | Bbs12 | 0.001 | Old |
| 0.872463 | 0.220259 | 0.105 | 0.099 | 1 | 7 | Zfp771 | 0.006 | Old |
| 0.872602 | 0.001314 | 0.218 | 0.22 | 1 | 7 | Pbxip1 | -0.002 | Old |
| 0.872975 | 0.114384 | 0.367 | 0.358 | 1 | 7 | Stk24 | 0.009 | Old |
| 0.873116 | 0.011146 | 0.574 | 0.557 | 1 | 7 | Ube2n | 0.017 | Old |
| 0.873305 | 0.489864 | 0.031 | 0.033 | 1 | 7 | Lrp8 | -0.002 | Old |
| 0.873315 | 0.372465 | 0.041 | 0.039 | 1 | 7 | Tfcp2 | 0.002 | Old |
| 0.873939 | 0.039693 | 0.19 | 0.181 | 1 | 7 | Lrrc47 | 0.009 | Old |
| 0.873976 | 0.094309 | 0.151 | 0.154 | 1 | 7 | Orc4 | -0.003 | Old |
| 0.874149 | 0.020787 | 0.421 | 0.425 | 1 | 7 | Arl8a | -0.004 | Old |
| 0.874284 | 0.279374 | 0.054 | 0.051 | 1 | 7 | Gde1 | 0.003 | Old |
| 0.874565 | 0.074303 | 0.295 | 0.301 | 1 | 7 | Mettl9 | -0.006 | Old |
| 0.876471 | 0.015355 | 0.054 | 0.051 | 1 | 7 | Mex3b | 0.003 | Old |
| 0.876658 | 0.112623 | 0.051 | 0.048 | 1 | 7 | Zxdb | 0.003 | Old |
| 0.877916 | 0.006417 | 0.592 | 0.587 | 1 | 7 | Tmed7 | 0.005 | Old |
| 0.878006 | 0.065399 | 0.603 | 0.581 | 1 | 7 | Coro1c | 0.022 | Old |
| 0.87822 | 0.084383 | 0.151 | 0.145 | 1 | 7 | Atp11a | 0.006 | Old |

| | | | | | | | | |
|---|---|---|---|---|---|---|---|---|
| 0.878416 | 0.000221 | 0.738 | 0.696 | 1 | 7 | Serp1 | 0.042 | Old |
| 0.879273 | 0.009505 | 0.028 | 0.03 | 1 | 7 | Rnf122 | -0.002 | Old |
| 0.879402 | 0.031223 | 0.674 | 0.633 | 1 | 7 | Psmb1 | 0.041 | Old |
| 0.880523 | 0.230275 | 0.085 | 0.087 | 1 | 7 | Rfx5 | -0.002 | Old |
| 0.881243 | 0.04214 | 0.138 | 0.136 | 1 | 7 | Ccp110 | 0.002 | Old |
| 0.881409 | 0.045513 | 0.146 | 0.148 | 1 | 7 | Pdlim2 | -0.002 | Old |
| 0.882439 | 0.126402 | 0.233 | 0.235 | 1 | 7 | Mapkap1 | -0.002 | Old |
| 0.884113 | 0.274628 | 0.067 | 0.069 | 1 | 7 | Dnajc18 | -0.002 | Old |
| 0.884454 | 0.046557 | 0.123 | 0.117 | 1 | 7 | Aatf | 0.006 | Old |
| 0.885633 | 0.260121 | 0.026 | 0.024 | 1 | 7 | Maged2 | 0.002 | Old |
| 0.886525 | 0.311321 | 0.087 | 0.09 | 1 | 7 | Tmem63b | -0.003 | Old |
| 0.887039 | 0.103158 | 0.028 | 0.03 | 1 | 7 | Zfp931 | -0.002 | Old |
| 0.887102 | 0.00778 | 0.19 | 0.187 | 1 | 7 | Btaf1 | 0.003 | Old |
| 0.887972 | 0.055929 | 0.608 | 0.587 | 1 | 7 | Selenow | 0.021 | Old |
| 0.888318 | 0.094702 | 0.056 | 0.054 | 1 | 7 | Atpaf1 | 0.002 | Old |
| 0.888688 | 0.0191 | 0.554 | 0.542 | 1 | 7 | Necap2 | 0.012 | Old |
| 0.888774 | 0.483137 | 0.026 | 0.024 | 1 | 7 | Slfn1 | 0.002 | Old |
| 0.889096 | 0.070589 | 0.123 | 0.127 | 1 | 7 | Fam207a | -0.004 | Old |
| 0.889106 | 0.177763 | 0.09 | 0.087 | 1 | 7 | Tspan33 | 0.003 | Old |
| 0.890545 | 0.01586 | 0.085 | 0.081 | 1 | 7 | Ano8 | 0.004 | Old |
| 0.892563 | 0.071763 | 0.138 | 0.142 | 1 | 7 | 2210016L2 | -0.004 | Old |
| 0.892595 | 0.154178 | 0.169 | 0.163 | 1 | 7 | Cass4 | 0.006 | Old |
| 0.892615 | 0.114778 | 0.092 | 0.09 | 1 | 7 | E2f5 | 0.002 | Old |
| 0.892965 | 0.168185 | 0.026 | 0.024 | 1 | 7 | Fcor | 0.002 | Old |
| 0.893053 | 0.010429 | 0.964 | 0.973 | 1 | 7 | Junb | -0.009 | Old |
| 0.89319 | 0.0319 | 0.146 | 0.142 | 1 | 7 | Cyth2 | 0.004 | Old |
| 0.893845 | 0.212668 | 0.067 | 0.069 | 1 | 7 | Pyroxd1 | -0.002 | Old |
| 0.894029 | 0.07117 | 0.056 | 0.054 | 1 | 7 | 2-Sep | 0.002 | Old |
| 0.894743 | 0.257303 | 0.056 | 0.054 | 1 | 7 | Vegfa | 0.002 | Old |
| 0.895113 | 0.085203 | 0.074 | 0.072 | 1 | 7 | Fam149b | 0.002 | Old |
| 0.895245 | 0.010965 | 0.849 | 0.828 | 1 | 7 | Nap1l1 | 0.021 | Old |
| 0.895457 | 0.269768 | 0.056 | 0.054 | 1 | 7 | Stambpl1 | 0.002 | Old |
| 0.895476 | 0.129419 | 0.233 | 0.241 | 1 | 7 | Pstpip1 | -0.008 | Old |
| 0.896077 | 0.140248 | 0.418 | 0.404 | 1 | 7 | Lsm3 | 0.014 | Old |
| 0.896488 | 0.064739 | 0.431 | 0.413 | 1 | 7 | Mapk6 | 0.018 | Old |
| 0.896631 | 0.009347 | 0.579 | 0.566 | 1 | 7 | Ppp2r5a | 0.013 | Old |
| 0.897087 | 0.124571 | 0.179 | 0.187 | 1 | 7 | Tmco3 | -0.008 | Old |
| 0.897159 | 0.076463 | 0.026 | 0.024 | 1 | 7 | Itpr3 | 0.002 | Old |
| 0.897359 | 0.218447 | 0.192 | 0.187 | 1 | 7 | Samd8 | 0.005 | Old |
| 0.897488 | 0.186828 | 0.151 | 0.154 | 1 | 7 | Ppm1d | -0.003 | Old |
| 0.899257 | 0.018093 | 0.19 | 0.184 | 1 | 7 | Atg12 | 0.006 | Old |
| 0.900307 | 0.112291 | 0.026 | 0.024 | 1 | 7 | Gm21781 | 0.002 | Old |
| 0.900892 | 0.638776 | 0.069 | 0.072 | 1 | 7 | Snhg9 | -0.003 | Old |
| 0.90094 | 0.127071 | 0.123 | 0.127 | 1 | 7 | Txndc12 | -0.004 | Old |
| 0.901035 | 0.022932 | 0.649 | 0.642 | 1 | 7 | Lamtor4 | 0.007 | Old |

| | | | | | | | | |
|---|---|---|---|---|---|---|---|---|
| 0.901146 | 0.127502 | 0.103 | 0.099 | 1 | 7 | Nol4l | 0.004 | Old |
| 0.901354 | 0.006639 | 0.631 | 0.605 | 1 | 7 | Ndufb10 | 0.026 | Old |
| 0.901407 | 0.109627 | 0.303 | 0.304 | 1 | 7 | Uqcc3 | -0.001 | Old |
| 0.901722 | 0.040554 | 0.151 | 0.148 | 1 | 7 | Btbd9 | 0.003 | Old |
| 0.90188 | 0.031476 | 0.467 | 0.458 | 1 | 7 | Crk | 0.009 | Old |
| 0.901952 | 0.012781 | 0.849 | 0.849 | 1 | 7 | Hspa5 | 0 | Old |
| 0.901999 | 0.251128 | 0.218 | 0.226 | 1 | 7 | Naa40 | -0.008 | Old |
| 0.902714 | 0.178382 | 0.121 | 0.123 | 1 | 7 | Usp40 | -0.002 | Old |
| 0.903089 | 0.030846 | 0.718 | 0.681 | 1 | 7 | Hcls1 | 0.037 | Old |
| 0.903184 | 0.256864 | 0.313 | 0.322 | 1 | 7 | Elovl5 | -0.009 | Old |
| 0.903268 | 0.082538 | 0.474 | 0.449 | 1 | 7 | Fkbp8 | 0.025 | Old |
| 0.90332 | 0.101393 | 0.056 | 0.054 | 1 | 7 | Whamm | 0.002 | Old |
| 0.903856 | 0.037854 | 0.103 | 0.099 | 1 | 7 | Gpkow | 0.004 | Old |
| 0.904783 | 0.059964 | 0.913 | 0.913 | 1 | 7 | Grn | 0 | Old |
| 0.905735 | 0.060342 | 0.523 | 0.5 | 1 | 7 | Zbtb7a | 0.023 | Old |
| 0.906307 | 0.009666 | 0.841 | 0.828 | 1 | 7 | Rbms1 | 0.013 | Old |
| 0.906912 | 0.142803 | 0.092 | 0.096 | 1 | 7 | Fbxo7 | -0.004 | Old |
| 0.907417 | 0.083232 | 0.041 | 0.039 | 1 | 7 | Hebp2 | 0.002 | Old |
| 0.907923 | 0.953031 | 0.067 | 0.066 | 1 | 7 | Zfp444 | 0.001 | Old |
| 0.908251 | 0.011824 | 0.041 | 0.039 | 1 | 7 | C730034F0 | 0.002 | Old |
| 0.908605 | 0.175298 | 0.287 | 0.283 | 1 | 7 | Tfg | 0.004 | Old |
| 0.908701 | 0.096415 | 0.513 | 0.509 | 1 | 7 | Mier1 | 0.004 | Old |
| 0.909023 | 0.32638 | 0.026 | 0.027 | 1 | 7 | Mical3 | -0.001 | Old |
| 0.909023 | 0.028813 | 0.026 | 0.027 | 1 | 7 | 2700081O1 | -0.001 | Old |
| 0.909563 | 0.035975 | 0.541 | 0.53 | 1 | 7 | Ikbkb | 0.011 | Old |
| 0.909822 | 0.038967 | 0.749 | 0.726 | 1 | 7 | Tpm4 | 0.023 | Old |
| 0.911403 | 0.03774 | 0.067 | 0.069 | 1 | 7 | Ssx2ip | -0.002 | Old |
| 0.911613 | 0.034469 | 0.09 | 0.093 | 1 | 7 | Capn15 | -0.003 | Old |
| 0.912754 | 0.54051 | 0.028 | 0.027 | 1 | 7 | Pdxp | 0.001 | Old |
| 0.912853 | 0.170099 | 0.059 | 0.057 | 1 | 7 | 1500011B0 | 0.002 | Old |
| 0.913145 | 0.010554 | 0.7 | 0.675 | 1 | 7 | Bri3 | 0.025 | Old |
| 0.914272 | 0.111734 | 0.446 | 0.44 | 1 | 7 | Krtcap2 | 0.006 | Old |
| 0.914691 | 0.764406 | 0.013 | 0.012 | 1 | 7 | Fmo2 | 0.001 | Old |
| 0.915865 | 0.104832 | 0.51 | 0.497 | 1 | 7 | Lat2 | 0.013 | Old |
| 0.916169 | 0.251098 | 0.013 | 0.012 | 1 | 7 | Klc2 | 0.001 | Old |
| 0.916192 | 0.291828 | 0.026 | 0.027 | 1 | 7 | Sorbs3 | -0.001 | Old |
| 0.91675 | 0.100329 | 0.028 | 0.027 | 1 | 7 | Cyp2ab1 | 0.001 | Old |
| 0.916824 | 0.072423 | 0.218 | 0.217 | 1 | 7 | Gab2 | 0.001 | Old |
| 0.917272 | 0.002112 | 0.418 | 0.419 | 1 | 7 | Cdipt | -0.001 | Old |
| 0.917592 | 0.419312 | 0.044 | 0.042 | 1 | 7 | Mtfmt | 0.002 | Old |
| 0.91764 | 0.31451 | 0.274 | 0.274 | 1 | 7 | Cope | 0 | Old |
| 0.917647 | 0.503028 | 0.013 | 0.012 | 1 | 7 | A530010L1 | 0.001 | Old |
| 0.917731 | 0.016463 | 0.992 | 0.988 | 1 | 7 | Rps8 | 0.004 | Old |
| 0.918401 | 0.461836 | 0.044 | 0.042 | 1 | 7 | 4930430F0 | 0.002 | Old |
| 0.918928 | 0.017793 | 0.251 | 0.244 | 1 | 7 | Nipsnap3b | 0.007 | Old |

| | | | | | | | | |
|---|---|---|---|---|---|---|---|---|
| 0.919126 | 1.249454 | 0.013 | 0.012 | 1 | 7 | Mpzl1 | 0.001 | Old |
| 0.919126 | 0.563013 | 0.013 | 0.012 | 1 | 7 | Gm12258 | 0.001 | Old |
| 0.919788 | 0.067114 | 0.787 | 0.798 | 1 | 7 | Eef2 | -0.011 | Old |
| 0.919873 | 0.018806 | 0.082 | 0.084 | 1 | 7 | Alkbh6 | -0.002 | Old |
| 0.919908 | 0.136533 | 0.336 | 0.334 | 1 | 7 | Vezf1 | 0.002 | Old |
| 0.920569 | 0.07724 | 0.613 | 0.602 | 1 | 7 | Chmp4b | 0.011 | Old |
| 0.920605 | 0.726182 | 0.013 | 0.012 | 1 | 7 | Fam161a | 0.001 | Old |
| 0.920749 | 0.186888 | 0.028 | 0.027 | 1 | 7 | Gm6169 | 0.001 | Old |
| 0.922084 | 2.168655 | 0.013 | 0.012 | 1 | 7 | Fam171a1 | 0.001 | Old |
| 0.922084 | 0.346527 | 0.013 | 0.012 | 1 | 7 | S100a16 | 0.001 | Old |
| 0.922084 | 0.14157 | 0.013 | 0.012 | 1 | 7 | Rab26os | 0.001 | Old |
| 0.922084 | 0.021473 | 0.013 | 0.012 | 1 | 7 | Wdr78 | 0.001 | Old |
| 0.9221 | 0.635654 | 0.074 | 0.072 | 1 | 7 | Zfp746 | 0.002 | Old |
| 0.922284 | 0.046563 | 0.192 | 0.193 | 1 | 7 | Aph1c | -0.001 | Old |
| 0.922358 | 0.013672 | 0.023 | 0.024 | 1 | 7 | Morn2 | -0.001 | Old |
| 0.922446 | 0.560631 | 0.041 | 0.039 | 1 | 7 | Ipo4 | 0.002 | Old |
| 0.923564 | 0.23662 | 0.013 | 0.012 | 1 | 7 | Tmem17 | 0.001 | Old |
| 0.923564 | 0.122378 | 0.013 | 0.012 | 1 | 7 | Gm5914 | 0.001 | Old |
| 0.923564 | 0.118753 | 0.013 | 0.012 | 1 | 7 | Klhl23 | 0.001 | Old |
| 0.923564 | 0.080339 | 0.013 | 0.012 | 1 | 7 | Thsd4 | 0.001 | Old |
| 0.923568 | 0.03854 | 0.197 | 0.205 | 1 | 7 | Ppm1m | -0.008 | Old |
| 0.924669 | 0.46742 | 0.062 | 0.06 | 1 | 7 | Klhl15 | 0.002 | Old |
| 0.925686 | 0.088559 | 0.044 | 0.042 | 1 | 7 | Ccdc130 | 0.002 | Old |
| 0.92575 | 0.319955 | 0.028 | 0.027 | 1 | 7 | Zbtb39 | 0.001 | Old |
| 0.926338 | 0.357592 | 0.131 | 0.13 | 1 | 7 | BC029722 | 0.001 | Old |
| 0.926523 | 0.370273 | 0.013 | 0.012 | 1 | 7 | Oplah | 0.001 | Old |
| 0.926523 | 0.326394 | 0.013 | 0.012 | 1 | 7 | Pard3b | 0.001 | Old |
| 0.926523 | 0.081182 | 0.013 | 0.012 | 1 | 7 | A930007I1 | 0.001 | Old |
| 0.926766 | 0.029961 | 0.208 | 0.211 | 1 | 7 | Tob1 | -0.003 | Old |
| 0.927821 | 0.021803 | 0.949 | 0.955 | 1 | 7 | Rpl37 | -0.006 | Old |
| 0.927844 | 0.005102 | 0.492 | 0.476 | 1 | 7 | Gnai3 | 0.016 | Old |
| 0.928004 | 0.01102 | 0.013 | 0.012 | 1 | 7 | Abhd15 | 0.001 | Old |
| 0.928628 | 0.0889 | 0.041 | 0.042 | 1 | 7 | Zfp319 | -0.001 | Old |
| 0.928751 | 0.225811 | 0.028 | 0.027 | 1 | 7 | Pou6f1 | 0.001 | Old |
| 0.928751 | 0.127254 | 0.028 | 0.027 | 1 | 7 | Neil1 | 0.001 | Old |
| 0.929522 | 0.504309 | 0.026 | 0.027 | 1 | 7 | Fn3krp | -0.001 | Old |
| 0.929553 | 0.065694 | 0.338 | 0.331 | 1 | 7 | Xpr1 | 0.007 | Old |
| 0.929567 | 0.132038 | 0.077 | 0.078 | 1 | 7 | Serinc5 | -0.001 | Old |
| 0.930099 | 0.115285 | 0.138 | 0.136 | 1 | 7 | Tk2 | 0.002 | Old |
| 0.930287 | 0.089519 | 0.169 | 0.169 | 1 | 7 | Herc6 | 0 | Old |
| 0.93101 | 0.033496 | 0.723 | 0.672 | 1 | 7 | Clk1 | 0.051 | Old |
| 0.931016 | 0.032392 | 0.746 | 0.72 | 1 | 7 | Rpl29 | 0.026 | Old |
| 0.931366 | 0.126541 | 0.097 | 0.096 | 1 | 7 | Il21r | 0.001 | Old |
| 0.93182 | 0.243488 | 0.138 | 0.139 | 1 | 7 | Pkib | -0.001 | Old |
| 0.931922 | 0.078966 | 0.144 | 0.139 | 1 | 7 | Ascc1 | 0.005 | Old |

| | | | | | | | | |
|---|---|---|---|---|---|---|---|---|
| 0.932997 | 0.030629 | 0.721 | 0.702 | 1 | 7 | Tra2b | 0.019 | Old |
| 0.933023 | 0.163586 | 0.269 | 0.271 | 1 | 7 | Bmpr2 | -0.002 | Old |
| 0.934158 | 0.21377 | 0.046 | 0.045 | 1 | 7 | Zkscan6 | 0.001 | Old |
| 0.93461 | 0.19072 | 0.126 | 0.123 | 1 | 7 | Chid1 | 0.003 | Old |
| 0.934618 | 0.206879 | 0.064 | 0.063 | 1 | 7 | Itga1 | 0.001 | Old |
| 0.934967 | 0.214526 | 0.11 | 0.108 | 1 | 7 | Cnn3 | 0.002 | Old |
| 0.935182 | 0.269935 | 0.044 | 0.045 | 1 | 7 | Plekhg1 | -0.001 | Old |
| 0.935214 | 0.171499 | 0.041 | 0.042 | 1 | 7 | Hemk1 | -0.001 | Old |
| 0.935603 | 0.200048 | 0.064 | 0.066 | 1 | 7 | Zkscan17 | -0.002 | Old |
| 0.935731 | 0.352408 | 0.046 | 0.045 | 1 | 7 | Coro2b | 0.001 | Old |
| 0.936076 | 0.012652 | 0.956 | 0.949 | 1 | 7 | Calm1 | 0.007 | Old |
| 0.936228 | 0.012067 | 0.485 | 0.482 | 1 | 7 | Cuta | 0.003 | Old |
| 0.936267 | 0.241345 | 0.031 | 0.03 | 1 | 7 | Rtn4rl1 | 0.001 | Old |
| 0.936442 | 0.307677 | 0.023 | 0.024 | 1 | 7 | 6330403L0 | -0.001 | Old |
| 0.936676 | 0.112166 | 0.103 | 0.105 | 1 | 7 | Fam222b | -0.002 | Old |
| 0.936688 | 0.017866 | 0.059 | 0.057 | 1 | 7 | Ppp1r16b | 0.002 | Old |
| 0.937263 | 0.037834 | 0.341 | 0.325 | 1 | 7 | Ppp2r2d | 0.016 | Old |
| 0.937526 | 0.179675 | 0.023 | 0.024 | 1 | 7 | Tcaim | -0.001 | Old |
| 0.93761 | 0.080507 | 0.144 | 0.145 | 1 | 7 | Narf | -0.001 | Old |
| 0.938136 | 0.010986 | 0.187 | 0.184 | 1 | 7 | Senp7 | 0.003 | Old |
| 0.938161 | 0.023614 | 0.077 | 0.078 | 1 | 7 | Ahi1 | -0.001 | Old |
| 0.93821 | 0.097524 | 0.203 | 0.199 | 1 | 7 | Man1b1 | 0.004 | Old |
| 0.938273 | 0.089526 | 0.382 | 0.373 | 1 | 7 | Cd86 | 0.009 | Old |
| 0.938572 | 0.524717 | 0.038 | 0.039 | 1 | 7 | Tgm1 | -0.001 | Old |
| 0.93861 | 0.31213 | 0.023 | 0.024 | 1 | 7 | Nek8 | -0.001 | Old |
| 0.938764 | 0.083601 | 0.172 | 0.172 | 1 | 7 | Neurl1b | 0 | Old |
| 0.938765 | 0.562639 | 0.028 | 0.027 | 1 | 7 | Gadd45a | 0.001 | Old |
| 0.939049 | 0.017823 | 0.092 | 0.093 | 1 | 7 | L1cam | -0.001 | Old |
| 0.939225 | 0.055216 | 0.531 | 0.512 | 1 | 7 | Rab11a | 0.019 | Old |
| 0.939767 | 0.075397 | 0.028 | 0.027 | 1 | 7 | Arpin | 0.001 | Old |
| 0.94087 | 0.123398 | 0.405 | 0.404 | 1 | 7 | H1f0 | 0.001 | Old |
| 0.941117 | 0.066544 | 0.264 | 0.259 | 1 | 7 | Nfatc1 | 0.005 | Old |
| 0.941481 | 0.134841 | 0.213 | 0.211 | 1 | 7 | Mcrs1 | 0.002 | Old |
| 0.94307 | 0.133973 | 0.174 | 0.172 | 1 | 7 | Nfkbib | 0.002 | Old |
| 0.943143 | 0.09309 | 0.154 | 0.157 | 1 | 7 | Celf4 | -0.003 | Old |
| 0.94392 | 0.435287 | 0.031 | 0.03 | 1 | 7 | Sgce | 0.001 | Old |
| 0.945713 | 2.44E-05 | 0.577 | 0.563 | 1 | 7 | Dnajc3 | 0.014 | Old |
| 0.946112 | 0.018077 | 0.844 | 0.87 | 1 | 7 | Cox7c | -0.026 | Old |
| 0.946275 | 0.675491 | 0.021 | 0.021 | 1 | 7 | Kif3c | 0 | Old |
| 0.947576 | 0.054403 | 0.203 | 0.199 | 1 | 7 | Syvn1 | 0.004 | Old |
| 0.94898 | 0.056204 | 0.467 | 0.464 | 1 | 7 | 9930111J2 | 0.003 | Old |
| 0.949109 | 0.0108 | 0.344 | 0.328 | 1 | 7 | Mb21d1 | 0.016 | Old |
| 0.94945 | 0.336751 | 0.174 | 0.178 | 1 | 7 | Mertk | -0.004 | Old |
| 0.949859 | 0.006187 | 0.5 | 0.491 | 1 | 7 | Eif3b | 0.009 | Old |
| 0.949997 | 0.016009 | 0.246 | 0.247 | 1 | 7 | Trim44 | -0.001 | Old |

| | | | | | | | | |
|---|---|---|---|---|---|---|---|---|
| 0.950241 | 0.055168 | 0.079 | 0.078 | 1 | 7 | Gm15417 | 0.001 | Old |
| 0.95025 | 0.042603 | 0.126 | 0.123 | 1 | 7 | Bicra | 0.003 | Old |
| 0.95148 | 0.076003 | 0.367 | 0.352 | 1 | 7 | Mta2 | 0.015 | Old |
| 0.951579 | 0.263826 | 0.031 | 0.03 | 1 | 7 | Zfp619 | 0.001 | Old |
| 0.951835 | 0.032514 | 0.474 | 0.461 | 1 | 7 | Fam102b | 0.013 | Old |
| 0.952045 | 0.364447 | 0.021 | 0.021 | 1 | 7 | Amigo3 | 0 | Old |
| 0.952645 | 0.382675 | 0.069 | 0.069 | 1 | 7 | Thap1 | 0 | Old |
| 0.954192 | 0.1352 | 0.215 | 0.217 | 1 | 7 | Dcaf13 | -0.002 | Old |
| 0.954601 | 0.01069 | 0.956 | 0.955 | 1 | 7 | Ppia | 0.001 | Old |
| 0.955236 | 0.050711 | 0.449 | 0.434 | 1 | 7 | Slc29a3 | 0.015 | Old |
| 0.955442 | 0.085684 | 0.136 | 0.133 | 1 | 7 | Hk3 | 0.003 | Old |
| 0.955583 | 0.426841 | 0.062 | 0.06 | 1 | 7 | Zfp932 | 0.002 | Old |
| 0.955903 | 0.344721 | 0.054 | 0.054 | 1 | 7 | Rfxank | 0 | Old |
| 0.956029 | 0.05091 | 0.167 | 0.163 | 1 | 7 | Plscr3 | 0.004 | Old |
| 0.956645 | 0.861618 | 0.041 | 0.042 | 1 | 7 | Gm10282 | -0.001 | Old |
| 0.95746 | 0.002549 | 0.346 | 0.337 | 1 | 7 | C330007P( | 0.009 | Old |
| 0.95747 | 0.355378 | 0.041 | 0.042 | 1 | 7 | Ice2 | -0.001 | Old |
| 0.958383 | 0.04989 | 0.536 | 0.524 | 1 | 7 | Usp7 | 0.012 | Old |
| 0.959119 | 0.002856 | 0.185 | 0.181 | 1 | 7 | Farsb | 0.004 | Old |
| 0.959963 | 0.058025 | 0.372 | 0.358 | 1 | 7 | Klhl24 | 0.014 | Old |
| 0.960193 | 0.079992 | 0.29 | 0.295 | 1 | 7 | 2-Mar | -0.005 | Old |
| 0.960201 | 0.130839 | 0.031 | 0.03 | 1 | 7 | 1110051M | 0.001 | Old |
| 0.960296 | 1.787492 | 0.015 | 0.015 | 1 | 7 | Alkbh4 | 0 | Old |
| 0.960731 | 0.379105 | 0.038 | 0.039 | 1 | 7 | Ddx59 | -0.001 | Old |
| 0.960939 | 0.310604 | 0.046 | 0.045 | 1 | 7 | Atp2b4 | 0.001 | Old |
| 0.961621 | 0.098388 | 0.251 | 0.253 | 1 | 7 | Trim56 | -0.002 | Old |
| 0.96191 | 0.104382 | 0.056 | 0.057 | 1 | 7 | Bckdhb | -0.001 | Old |
| 0.962147 | 0.076576 | 0.2 | 0.199 | 1 | 7 | Vac14 | 0.001 | Old |
| 0.962293 | 0.148162 | 0.156 | 0.157 | 1 | 7 | Cenpi | -0.001 | Old |
| 0.962436 | 0.097904 | 0.021 | 0.021 | 1 | 7 | Rsl1 | 0 | Old |
| 0.962649 | 0.050121 | 0.379 | 0.364 | 1 | 7 | Eif4a3 | 0.015 | Old |
| 0.962685 | 0.152924 | 0.382 | 0.377 | 1 | 7 | Cript | 0.005 | Old |
| 0.962741 | 0.007208 | 0.338 | 0.328 | 1 | 7 | Eftud2 | 0.01 | Old |
| 0.96275 | 0.03495 | 0.077 | 0.078 | 1 | 7 | Dynlt1a | -0.001 | Old |
| 0.963149 | 0.02883 | 0.287 | 0.286 | 1 | 7 | Sav1 | 0.001 | Old |
| 0.963916 | 0.06897 | 0.226 | 0.226 | 1 | 7 | Dpp9 | 0 | Old |
| 0.964093 | 0.214461 | 0.046 | 0.045 | 1 | 7 | Mnd1 | 0.001 | Old |
| 0.964207 | 0.064711 | 0.305 | 0.292 | 1 | 7 | Zrsr2 | 0.013 | Old |
| 0.964446 | 0.033663 | 0.385 | 0.37 | 1 | 7 | St3gal5 | 0.015 | Old |
| 0.965403 | 0.283755 | 0.231 | 0.232 | 1 | 7 | C4b | -0.001 | Old |
| 0.965862 | 0.185788 | 0.428 | 0.419 | 1 | 7 | Osgep | 0.009 | Old |
| 0.965902 | 0.010468 | 0.482 | 0.47 | 1 | 7 | Cd164 | 0.012 | Old |
| 0.965972 | 0.048018 | 0.415 | 0.413 | 1 | 7 | Rgs18 | 0.002 | Old |
| 0.966233 | 0.089432 | 0.128 | 0.127 | 1 | 7 | Kdsr | 0.001 | Old |
| 0.967022 | 0.438283 | 0.015 | 0.015 | 1 | 7 | Fgf11 | 0 | Old |

| | | | | | | | | |
|---|---|---|---|---|---|---|---|---|
| 0.96705 | 0.001809 | 0.626 | 0.599 | 1 | 7 | Tmed5 | 0.027 | Old |
| 0.967604 | 0.016913 | 0.197 | 0.193 | 1 | 7 | Tbk1 | 0.004 | Old |
| 0.967609 | 0.05553 | 0.208 | 0.202 | 1 | 7 | Fbxo11 | 0.006 | Old |
| 0.969007 | 0.13262 | 0.551 | 0.551 | 1 | 7 | Herpud1 | 0 | Old |
| 0.969317 | 0.480639 | 0.074 | 0.075 | 1 | 7 | Mapk7 | -0.001 | Old |
| 0.969506 | 0.517888 | 0.085 | 0.084 | 1 | 7 | Pik3r2 | 0.001 | Old |
| 0.969712 | 0.189943 | 0.015 | 0.015 | 1 | 7 | Adgra2 | 0 | Old |
| 0.969746 | 0.113824 | 0.223 | 0.22 | 1 | 7 | Il13ra1 | 0.003 | Old |
| 0.970378 | 0.054781 | 0.397 | 0.392 | 1 | 7 | Chmp3 | 0.005 | Old |
| 0.97043 | 0.093064 | 0.238 | 0.232 | 1 | 7 | Mif4gd | 0.006 | Old |
| 0.971063 | 0.057361 | 0.185 | 0.184 | 1 | 7 | Tspan4 | 0.001 | Old |
| 0.972306 | 0.104809 | 0.6 | 0.617 | 1 | 7 | Rab7b | -0.017 | Old |
| 0.972677 | 0.056074 | 0.038 | 0.039 | 1 | 7 | Mrvi1 | -0.001 | Old |
| 0.972745 | 0.078587 | 0.11 | 0.111 | 1 | 7 | Arl10 | -0.001 | Old |
| 0.973749 | 0.144525 | 0.015 | 0.015 | 1 | 7 | Enah | 0 | Old |
| 0.973749 | 0.105151 | 0.015 | 0.015 | 1 | 7 | Il15ra | 0 | Old |
| 0.973989 | 1.332632 | 0.021 | 0.021 | 1 | 7 | Mapre3 | 0 | Old |
| 0.974481 | 0.067224 | 0.213 | 0.214 | 1 | 7 | Plekhm2 | -0.001 | Old |
| 0.974547 | 0.148714 | 0.303 | 0.304 | 1 | 7 | Dyrk2 | -0.001 | Old |
| 0.975056 | 0.255685 | 0.049 | 0.048 | 1 | 7 | Zfp709 | 0.001 | Old |
| 0.975224 | 0.013262 | 0.036 | 0.036 | 1 | 7 | Wdr24 | 0 | Old |
| 0.975238 | 0.189057 | 0.038 | 0.039 | 1 | 7 | Pard6a | -0.001 | Old |
| 0.975858 | 0.266476 | 0.067 | 0.066 | 1 | 7 | Nr2c1 | 0.001 | Old |
| 0.975989 | 0.005466 | 0.197 | 0.199 | 1 | 7 | Orc6 | -0.002 | Old |
| 0.976413 | 0.208372 | 0.11 | 0.111 | 1 | 7 | Tns4 | -0.001 | Old |
| 0.977246 | 0.076585 | 0.733 | 0.759 | 1 | 7 | Egr1 | -0.026 | Old |
| 0.977378 | 0.013252 | 0.192 | 0.193 | 1 | 7 | Ing5 | -0.001 | Old |
| 0.977903 | 0.000571 | 0.8 | 0.777 | 1 | 7 | Cox7a2l | 0.023 | Old |
| 0.977968 | 0.042323 | 0.549 | 0.548 | 1 | 7 | Tmem59 | 0.001 | Old |
| 0.978814 | 0.041037 | 0.631 | 0.593 | 1 | 7 | Ctdsp2 | 0.038 | Old |
| 0.978991 | 0.2595 | 0.056 | 0.057 | 1 | 7 | Ecsit | -0.001 | Old |
| 0.979012 | 0.026612 | 0.392 | 0.383 | 1 | 7 | Rrp1 | 0.009 | Old |
| 0.979199 | 0.099603 | 0.205 | 0.208 | 1 | 7 | Wdr44 | -0.003 | Old |
| 0.979984 | 0.061424 | 0.177 | 0.175 | 1 | 7 | Ccdc127 | 0.002 | Old |
| 0.98005 | 0.490435 | 0.267 | 0.271 | 1 | 7 | Irf4 | -0.004 | Old |
| 0.980284 | 0.082225 | 0.244 | 0.244 | 1 | 7 | St6galnac4 | 0 | Old |
| 0.980285 | 0.126646 | 0.336 | 0.334 | 1 | 7 | Ubn2 | 0.002 | Old |
| 0.980782 | 0.105551 | 0.267 | 0.271 | 1 | 7 | Borcs6 | -0.004 | Old |
| 0.981275 | 0.008781 | 0.177 | 0.175 | 1 | 7 | Rarg | 0.002 | Old |
| 0.981828 | 0.445697 | 0.069 | 0.069 | 1 | 7 | Ppp1r13b | 0 | Old |
| 0.982447 | 0.266498 | 0.077 | 0.078 | 1 | 7 | Mcph1 | -0.001 | Old |
| 0.983208 | 0.021309 | 0.431 | 0.425 | 1 | 7 | Smim11 | 0.006 | Old |
| 0.983593 | 0.114193 | 0.054 | 0.054 | 1 | 7 | Fbxl19 | 0 | Old |
| 0.98444 | 0.119472 | 0.221 | 0.22 | 1 | 7 | Tpst2 | 0.001 | Old |
| 0.984498 | 0.359494 | 0.018 | 0.018 | 1 | 7 | Xlr3b | 0 | Old |

| | | | | | | | | |
|---|---|---|---|---|---|---|---|---|
| 0.985058 | 0.017551 | 0.787 | 0.756 | 1 | 7 | Purb | 0.031 | Old |
| 0.985269 | 0.011818 | 0.841 | 0.883 | 1 | 7 | Naca | -0.042 | Old |
| 0.985363 | 0.132871 | 0.31 | 0.31 | 1 | 7 | Vps28 | 0 | Old |
| 0.987948 | 0.025635 | 0.338 | 0.334 | 1 | 7 | Fkbp3 | 0.004 | Old |
| 0.988276 | 0.162227 | 0.131 | 0.13 | 1 | 7 | Ascc2 | 0.001 | Old |
| 0.989244 | 0.058368 | 0.226 | 0.223 | 1 | 7 | Mob3c | 0.003 | Old |
| 0.990435 | 0.076031 | 0.282 | 0.277 | 1 | 7 | Supt4a | 0.005 | Old |
| 0.990684 | 0.021537 | 0.174 | 0.172 | 1 | 7 | Cbx4 | 0.002 | Old |
| 0.990758 | 0.01084 | 0.072 | 0.072 | 1 | 7 | H6pd | 0 | Old |
| 0.991743 | 0.091802 | 0.364 | 0.373 | 1 | 7 | Slc48a1 | -0.009 | Old |
| 0.992334 | 0.127859 | 0.362 | 0.352 | 1 | 7 | Commd7 | 0.01 | Old |
| 0.992484 | 0.074972 | 0.074 | 0.075 | 1 | 7 | Nlrc4 | -0.001 | Old |
| 0.992844 | 0.116312 | 0.513 | 0.5 | 1 | 7 | Fli1 | 0.013 | Old |
| 0.993179 | 0.213345 | 0.018 | 0.018 | 1 | 7 | Rassf7 | 0 | Old |
| 0.993179 | 0.180142 | 0.018 | 0.018 | 1 | 7 | Gm48066 | 0 | Old |
| 0.993306 | 0.02933 | 0.174 | 0.175 | 1 | 7 | Zfp518a | -0.001 | Old |
| 0.993863 | 0.039844 | 0.559 | 0.545 | 1 | 7 | Nipbl | 0.014 | Old |
| 0.9941 | 0.066193 | 0.382 | 0.373 | 1 | 7 | Sec63 | 0.009 | Old |
| 0.994419 | 0.983391 | 0.018 | 0.018 | 1 | 7 | Gm13431 | 0 | Old |
| 0.994419 | 0.13184 | 0.018 | 0.018 | 1 | 7 | Cdiptos | 0 | Old |
| 0.994582 | 0.370661 | 0.072 | 0.072 | 1 | 7 | Tysnd1 | 0 | Old |
| 0.995143 | 0.029132 | 0.928 | 0.937 | 1 | 7 | Rpl6 | -0.009 | Old |
| 0.99566 | 0.03239 | 0.154 | 0.151 | 1 | 7 | Nfyc | 0.003 | Old |
| 0.995778 | 0.120944 | 0.049 | 0.048 | 1 | 7 | Ubxn8 | 0.001 | Old |
| 0.996375 | 0.216568 | 0.179 | 0.181 | 1 | 7 | Pla2g4a | -0.002 | Old |
| 0.996899 | 1.082723 | 0.018 | 0.018 | 1 | 7 | Mmp13 | 0 | Old |
| 0.997132 | 0.06235 | 0.072 | 0.072 | 1 | 7 | Ankrd50 | 0 | Old |
| 0.997249 | 0.045942 | 0.223 | 0.22 | 1 | 7 | 1700017B0 | 0.003 | Old |
| 0.997621 | 0.187105 | 0.572 | 0.578 | 1 | 7 | Cflar | -0.006 | Old |
| 0.998003 | 0.443991 | 0.164 | 0.163 | 1 | 7 | Carnmt1 | 0.001 | Old |
| 0.99814 | 0.601692 | 0.018 | 0.018 | 1 | 7 | Gm26510 | 0 | Old |
| 0.998219 | 0.446816 | 0.056 | 0.057 | 1 | 7 | Gdpd3 | -0.001 | Old |
| 0.998346 | 0.214432 | 0.067 | 0.066 | 1 | 7 | Arl14ep | 0.001 | Old |
| 0.999634 | 0.083276 | 0.269 | 0.268 | 1 | 7 | Pla2g15 | 0.001 | Old |
| 2.29E-18 | 1.305077 | 0.608 | 0.313 | 7.12E-14 | 7 | Hspa1a | 0.295 | Young |
| 1.27E-16 | 1.330612 | 0.575 | 0.29 | 3.93E-12 | 7 | Hspa1b | 0.285 | Young |
| 6.32E-11 | 0.825298 | 0.849 | 0.705 | 1.96E-06 | 7 | Apoe | 0.144 | Young |
| 1.11E-10 | 1.399166 | 0.419 | 0.203 | 3.43E-06 | 7 | Cx3cr1 | 0.216 | Young |
| 1.34E-10 | 1.049165 | 0.434 | 0.218 | 4.17E-06 | 7 | Cd33 | 0.216 | Young |
| 2.05E-10 | 1.228903 | 0.389 | 0.185 | 6.37E-06 | 7 | Hpgd | 0.204 | Young |
| 1.22E-09 | 1.701744 | 0.346 | 0.164 | 3.80E-05 | 7 | Slco2b1 | 0.182 | Young |
| 4.00E-09 | 2.72314 | 0.184 | 0.049 | 0.000124 | 7 | Itga6 | 0.135 | Young |
| 1.18E-08 | 0.678065 | 0.822 | 0.741 | 0.000366 | 7 | Dab2 | 0.081 | Young |
| 5.97E-08 | 1.122062 | 0.63 | 0.485 | 0.001855 | 7 | Gatm | 0.145 | Young |
| 2.56E-07 | 0.340496 | 0.973 | 0.959 | 0.007935 | 7 | Cst3 | 0.014 | Young |

| | | | | | | | | |
|---|---|---|---|---|---|---|---|---|
| 3.00E-07 | 0.754506 | 0.524 | 0.354 | 0.009319 | 7 | Stab1 | 0.17 | Young |
| 4.42E-07 | 0.961257 | 0.593 | 0.474 | 0.01371 | 7 | Igfbp4 | 0.119 | Young |
| 4.67E-07 | 0.761656 | 0.681 | 0.61 | 0.0145 | 7 | Mef2c | 0.071 | Young |
| 5.22E-07 | 0.498565 | 0.889 | 0.777 | 0.016214 | 7 | Nfkbiz | 0.112 | Young |
| 6.52E-07 | 2.361994 | 0.099 | 0.015 | 0.020242 | 7 | Col14a1 | 0.084 | Young |
| 6.77E-07 | 1.616115 | 0.235 | 0.103 | 0.021034 | 7 | Plxdc2 | 0.132 | Young |
| 8.85E-07 | 0.595969 | 0.795 | 0.649 | 0.027486 | 7 | Pf4 | 0.146 | Young |
| 9.41E-07 | 0.281622 | 0.976 | 0.964 | 0.029211 | 7 | Rps14 | 0.012 | Young |
| 9.92E-07 | 0.928859 | 0.431 | 0.272 | 0.0308 | 7 | Pou2f2 | 0.159 | Young |
| 1.14E-06 | 1.29856 | 0.422 | 0.272 | 0.035381 | 7 | Ccl4 | 0.15 | Young |
| 1.30E-06 | 1.635252 | 0.196 | 0.077 | 0.040522 | 7 | Serpinf1 | 0.119 | Young |
| 2.97E-06 | 0.573607 | 0.78 | 0.708 | 0.092107 | 7 | Csf1r | 0.072 | Young |
| 3.15E-06 | 1.013806 | 0.398 | 0.264 | 0.097884 | 7 | Ccl24 | 0.134 | Young |
| 3.50E-06 | 0.515898 | 0.786 | 0.71 | 0.108587 | 7 | Zfp36l1 | 0.076 | Young |
| 9.58E-06 | 0.166467 | 0.991 | 0.995 | 0.297391 | 7 | Rps29 | -0.004 | Young |
| 1.02E-05 | 0.37065 | 0.792 | 0.664 | 0.31582 | 7 | C1qb | 0.128 | Young |
| 1.88E-05 | 0.240024 | 0.925 | 0.915 | 0.585223 | 7 | Fcer1g | 0.01 | Young |
| 2.06E-05 | 0.225865 | 0.964 | 0.951 | 0.639863 | 7 | Rps4x | 0.013 | Young |
| 2.09E-05 | 0.569887 | 0.717 | 0.608 | 0.649811 | 7 | Maf | 0.109 | Young |
| 2.89E-05 | 0.451077 | 0.69 | 0.592 | 0.897403 | 7 | Trf | 0.098 | Young |
| 3.18E-05 | 1.78446 | 0.111 | 0.033 | 0.987362 | 7 | Pmepa1 | 0.078 | Young |
| 3.47E-05 | 0.467282 | 0.87 | 0.815 | 1 | 7 | Cd83 | 0.055 | Young |
| 3.81E-05 | 0.214225 | 0.976 | 0.985 | 1 | 7 | Rps24 | -0.009 | Young |
| 3.95E-05 | 1.337121 | 0.133 | 0.046 | 1 | 7 | Rasgrp3 | 0.087 | Young |
| 4.42E-05 | 0.337898 | 0.822 | 0.697 | 1 | 7 | C1qa | 0.125 | Young |
| 4.64E-05 | 0.728287 | 0.482 | 0.351 | 1 | 7 | Abca1 | 0.131 | Young |
| 7.60E-05 | 1.136616 | 0.274 | 0.162 | 1 | 7 | Susd3 | 0.112 | Young |
| 8.38E-05 | 1.146419 | 0.187 | 0.09 | 1 | 7 | Tmem8 | 0.097 | Young |
| 8.41E-05 | 0.499098 | 0.389 | 0.254 | 1 | 7 | Nlrp3 | 0.135 | Young |
| 0.000117 | 1.504888 | 0.229 | 0.126 | 1 | 7 | Gpr34 | 0.103 | Young |
| 0.000129 | 0.395135 | 0.822 | 0.795 | 1 | 7 | Jun | 0.027 | Young |
| 0.000155 | 0.502586 | 0.581 | 0.467 | 1 | 7 | Fcgrt | 0.114 | Young |
| 0.000172 | 0.597463 | 0.578 | 0.477 | 1 | 7 | Zfhx3 | 0.101 | Young |
| 0.000213 | 0.531889 | 0.705 | 0.631 | 1 | 7 | Cxcl2 | 0.074 | Young |
| 0.000217 | 0.106179 | 0.991 | 0.992 | 1 | 7 | Rplp1 | -0.001 | Young |
| 0.000243 | 0.650276 | 0.392 | 0.277 | 1 | 7 | Ms4a7 | 0.115 | Young |
| 0.000258 | 0.36353 | 0.735 | 0.672 | 1 | 7 | Nfkbid | 0.063 | Young |
| 0.000267 | 0.78494 | 0.401 | 0.274 | 1 | 7 | Cd163 | 0.127 | Young |
| 0.000306 | 0.451174 | 0.726 | 0.656 | 1 | 7 | Ptpn18 | 0.07 | Young |
| 0.000396 | 0.958767 | 0.328 | 0.215 | 1 | 7 | Tnf | 0.113 | Young |
| 0.000415 | 0.281198 | 0.958 | 0.944 | 1 | 7 | Marcks | 0.014 | Young |
| 0.00043 | 1.98505 | 0.066 | 0.015 | 1 | 7 | Flt3 | 0.051 | Young |
| 0.000478 | 1.715308 | 0.211 | 0.121 | 1 | 7 | Ccl3 | 0.09 | Young |
| 0.000488 | 0.17881 | 0.967 | 0.962 | 1 | 7 | Rpl27a | 0.005 | Young |
| 0.000567 | 0.204516 | 0.94 | 0.936 | 1 | 7 | Rps10 | 0.004 | Young |

| | | | | | | | | |
|---|---|---|---|---|---|---|---|---|
| 0.000585 | 1.276544 | 0.114 | 0.046 | 1 | 7 | Lpcat2 | 0.068 | Young |
| 0.000653 | 0.190804 | 0.961 | 0.941 | 1 | 7 | Rps13 | 0.02 | Young |
| 0.000666 | 0.725197 | 0.313 | 0.208 | 1 | 7 | Dusp6 | 0.105 | Young |
| 0.00089 | 1.126718 | 0.193 | 0.108 | 1 | 7 | Sh3bp5 | 0.085 | Young |
| 0.000908 | 0.954835 | 0.545 | 0.485 | 1 | 7 | Hexb | 0.06 | Young |
| 0.00091 | 2.177835 | 0.066 | 0.018 | 1 | 7 | Nid2 | 0.048 | Young |
| 0.000992 | 0.734494 | 0.346 | 0.249 | 1 | 7 | Nr4a3 | 0.097 | Young |
| 0.001006 | 0.661709 | 0.545 | 0.441 | 1 | 7 | Ier3 | 0.104 | Young |
| 0.001048 | 0.764423 | 0.349 | 0.251 | 1 | 7 | Rnase4 | 0.098 | Young |
| 0.001058 | 0.493955 | 0.648 | 0.551 | 1 | 7 | Marcksl1 | 0.097 | Young |
| 0.001087 | 0.460057 | 0.771 | 0.692 | 1 | 7 | Txnip | 0.079 | Young |
| 0.001138 | 0.436766 | 0.651 | 0.569 | 1 | 7 | Ptprj | 0.082 | Young |
| 0.001142 | 0.517524 | 0.551 | 0.495 | 1 | 7 | Ighm | 0.056 | Young |
| 0.001185 | 0.896131 | 0.283 | 0.185 | 1 | 7 | Msr1 | 0.098 | Young |
| 0.001203 | 0.23476 | 0.925 | 0.892 | 1 | 7 | Rpl26 | 0.033 | Young |
| 0.001223 | 0.114059 | 1 | 1 | 1 | 7 | Tmsb4x | 0 | Young |
| 0.001247 | 0.423317 | 0.572 | 0.485 | 1 | 7 | Lamtor2 | 0.087 | Young |
| 0.001252 | 0.792795 | 0.334 | 0.241 | 1 | 7 | Ltc4s | 0.093 | Young |
| 0.001369 | 1.277967 | 0.19 | 0.11 | 1 | 7 | St3gal6 | 0.08 | Young |
| 0.001402 | 0.566995 | 0.536 | 0.436 | 1 | 7 | Gas6 | 0.1 | Young |
| 0.001426 | 0.684029 | 0.389 | 0.3 | 1 | 7 | Smim14 | 0.089 | Young |
| 0.001429 | 1.069589 | 0.31 | 0.221 | 1 | 7 | Sesn1 | 0.089 | Young |
| 0.001443 | 0.454716 | 0.666 | 0.603 | 1 | 7 | Rel | 0.063 | Young |
| 0.001455 | 0.331903 | 0.925 | 0.921 | 1 | 7 | Ctsc | 0.004 | Young |
| 0.001651 | 0.745458 | 0.108 | 0.046 | 1 | 7 | Slc40a1 | 0.062 | Young |
| 0.001745 | 0.438465 | 0.729 | 0.669 | 1 | 7 | Ehd4 | 0.06 | Young |
| 0.001851 | 0.274376 | 0.837 | 0.797 | 1 | 7 | Gdi2 | 0.04 | Young |
| 0.00193 | 0.273446 | 0.955 | 0.897 | 1 | 7 | Mrc1 | 0.058 | Young |
| 0.002057 | 1.076781 | 0.184 | 0.105 | 1 | 7 | Smpdl3b | 0.079 | Young |
| 0.002109 | 0.236832 | 0.88 | 0.849 | 1 | 7 | Rps3 | 0.031 | Young |
| 0.002381 | 0.198568 | 0.985 | 0.99 | 1 | 7 | Fth1 | -0.005 | Young |
| 0.002413 | 0.789884 | 0.322 | 0.231 | 1 | 7 | Slc9a9 | 0.091 | Young |
| 0.002556 | 1.098922 | 0.114 | 0.054 | 1 | 7 | Tmem37 | 0.06 | Young |
| 0.002721 | 0.220401 | 0.913 | 0.872 | 1 | 7 | Rps6 | 0.041 | Young |
| 0.002743 | 1.741291 | 0.102 | 0.046 | 1 | 7 | Ttc28 | 0.056 | Young |
| 0.002875 | 0.950047 | 0.205 | 0.126 | 1 | 7 | Retreg1 | 0.079 | Young |
| 0.002894 | 0.660712 | 0.485 | 0.403 | 1 | 7 | Pmp22 | 0.082 | Young |
| 0.002986 | 0.154308 | 0.976 | 0.951 | 1 | 7 | Rps16 | 0.025 | Young |
| 0.003236 | 1.06703 | 0.12 | 0.059 | 1 | 7 | Dusp8 | 0.061 | Young |
| 0.003284 | 0.330215 | 0.837 | 0.779 | 1 | 7 | Nr4a1 | 0.058 | Young |
| 0.003299 | 0.226305 | 0.877 | 0.844 | 1 | 7 | Rpl11 | 0.033 | Young |
| 0.003309 | 0.191316 | 0.91 | 0.91 | 1 | 7 | Rpl34 | 0 | Young |
| 0.003313 | 0.751107 | 0.096 | 0.041 | 1 | 7 | Diexf | 0.055 | Young |
| 0.003402 | 1.709198 | 0.054 | 0.015 | 1 | 7 | Cxxc5 | 0.039 | Young |
| 0.003424 | 1.052192 | 0.286 | 0.205 | 1 | 7 | Pitpnc1 | 0.081 | Young |

| | | | | | | | | |
|---|---|---|---|---|---|---|---|---|
| 0.003602 | 0.754585 | 0.392 | 0.31 | 1 | 7 | Tnfsf9 | 0.082 | Young |
| 0.003667 | 0.438934 | 0.265 | 0.172 | 1 | 7 | Slamf7 | 0.093 | Young |
| 0.003675 | 1.58583 | 0.075 | 0.028 | 1 | 7 | Smad7 | 0.047 | Young |
| 0.003788 | 1.058844 | 0.34 | 0.251 | 1 | 7 | Cd209a | 0.089 | Young |
| 0.003813 | 0.990083 | 0.12 | 0.059 | 1 | 7 | Pag1 | 0.061 | Young |
| 0.003913 | 0.359943 | 0.783 | 0.741 | 1 | 7 | F13a1 | 0.042 | Young |
| 0.003914 | 1.217337 | 0.111 | 0.054 | 1 | 7 | Mamdc2 | 0.057 | Young |
| 0.003923 | 0.84726 | 0.217 | 0.138 | 1 | 7 | Tanc2 | 0.079 | Young |
| 0.003949 | 0.847434 | 0.063 | 0.021 | 1 | 7 | AA467197 | 0.042 | Young |
| 0.003996 | 2.210544 | 0.021 | 0 | 1 | 7 | Kcnip3 | 0.021 | Young |
| 0.003996 | 1.712033 | 0.021 | 0 | 1 | 7 | Lzts3 | 0.021 | Young |
| 0.004012 | 0.354289 | 0.681 | 0.649 | 1 | 7 | Adgre1 | 0.032 | Young |
| 0.004043 | 0.236661 | 0.937 | 0.908 | 1 | 7 | Serinc3 | 0.029 | Young |
| 0.004077 | 2.296616 | 0.066 | 0.023 | 1 | 7 | Cfd | 0.043 | Young |
| 0.0041 | 0.193013 | 0.961 | 0.933 | 1 | 7 | Dusp1 | 0.028 | Young |
| 0.0041 | 0.168182 | 0.982 | 0.977 | 1 | 7 | Fau | 0.005 | Young |
| 0.004111 | 0.129151 | 0.967 | 0.956 | 1 | 7 | Rps12 | 0.011 | Young |
| 0.004121 | 0.563242 | 0.256 | 0.164 | 1 | 7 | Creb5 | 0.092 | Young |
| 0.004409 | 0.508585 | 0.524 | 0.482 | 1 | 7 | Prdx5 | 0.042 | Young |
| 0.004413 | 0.298108 | 0.831 | 0.769 | 1 | 7 | Ly86 | 0.062 | Young |
| 0.004611 | 1.835868 | 0.048 | 0.013 | 1 | 7 | Ass1 | 0.035 | Young |
| 0.004652 | 0.514898 | 0.455 | 0.385 | 1 | 7 | Tpp2 | 0.07 | Young |
| 0.004775 | 0.558737 | 0.389 | 0.305 | 1 | 7 | Sirt2 | 0.084 | Young |
| 0.004775 | 0.279231 | 0.807 | 0.767 | 1 | 7 | Selenop | 0.04 | Young |
| 0.004806 | 0.396339 | 0.687 | 0.628 | 1 | 7 | Atf3 | 0.059 | Young |
| 0.004867 | 2.718774 | 0.027 | 0.003 | 1 | 7 | Tesmin | 0.024 | Young |
| 0.00487 | 0.282352 | 0.783 | 0.756 | 1 | 7 | Qk | 0.027 | Young |
| 0.004894 | 2.412734 | 0.027 | 0.003 | 1 | 7 | Zfp229 | 0.024 | Young |
| 0.005025 | 0.48279 | 0.521 | 0.433 | 1 | 7 | C3ar1 | 0.088 | Young |
| 0.005058 | 1.207621 | 0.027 | 0.003 | 1 | 7 | Tbc1d2 | 0.024 | Young |
| 0.005175 | 0.24696 | 0.687 | 0.605 | 1 | 7 | Cd81 | 0.082 | Young |
| 0.005533 | 0.158474 | 0.97 | 0.962 | 1 | 7 | Rpl13 | 0.008 | Young |
| 0.005918 | 0.46893 | 0.175 | 0.105 | 1 | 7 | Smagp | 0.07 | Young |
| 0.006415 | 1.12364 | 0.196 | 0.126 | 1 | 7 | Cd72 | 0.07 | Young |
| 0.006454 | 1.538657 | 0.114 | 0.059 | 1 | 7 | Arid3b | 0.055 | Young |
| 0.006652 | 0.778804 | 0.253 | 0.179 | 1 | 7 | Trem2 | 0.074 | Young |
| 0.006693 | 0.278027 | 0.828 | 0.79 | 1 | 7 | Atp5h | 0.038 | Young |
| 0.006798 | 0.233037 | 0.916 | 0.89 | 1 | 7 | Cltc | 0.026 | Young |
| 0.007025 | 0.332761 | 0.696 | 0.662 | 1 | 7 | Plek | 0.034 | Young |
| 0.007103 | 2.272557 | 0.054 | 0.018 | 1 | 7 | BC024978 | 0.036 | Young |
| 0.007276 | 0.497428 | 0.38 | 0.297 | 1 | 7 | Rnf150 | 0.083 | Young |
| 0.007508 | 0.210625 | 0.515 | 0.41 | 1 | 7 | Cbr2 | 0.105 | Young |
| 0.007746 | 1.450611 | 0.018 | 0 | 1 | 7 | Gm47389 | 0.018 | Young |
| 0.007841 | 1.131294 | 0.172 | 0.108 | 1 | 7 | Bmyc | 0.064 | Young |
| 0.007855 | 2.595318 | 0.057 | 0.021 | 1 | 7 | Gbp9 | 0.036 | Young |

| | | | | | | | | |
|---|---|---|---|---|---|---|---|---|
| 0.008247 | 0.374847 | 0.708 | 0.649 | 1 | 7 | Cd14 | 0.059 | Young |
| 0.008726 | 0.534274 | 0.416 | 0.333 | 1 | 7 | Basp1 | 0.083 | Young |
| 0.008751 | 2.039605 | 0.03 | 0.005 | 1 | 7 | B630019K0 | 0.025 | Young |
| 0.008871 | 0.38278 | 0.62 | 0.574 | 1 | 7 | Pea15a | 0.046 | Young |
| 0.008876 | 2.005075 | 0.03 | 0.005 | 1 | 7 | 2900026A0 | 0.025 | Young |
| 0.008945 | 1.574361 | 0.069 | 0.028 | 1 | 7 | Phc1 | 0.041 | Young |
| 0.009064 | 0.402327 | 0.804 | 0.767 | 1 | 7 | Lgmn | 0.037 | Young |
| 0.009123 | 0.24855 | 0.467 | 0.382 | 1 | 7 | Stap1 | 0.085 | Young |
| 0.009131 | 0.393194 | 0.581 | 0.526 | 1 | 7 | Ctnna1 | 0.055 | Young |
| 0.009178 | 0.925018 | 0.175 | 0.11 | 1 | 7 | F630028O1 | 0.065 | Young |
| 0.009196 | 0.191331 | 0.759 | 0.677 | 1 | 7 | C1qc | 0.082 | Young |
| 0.009226 | 2.114513 | 0.024 | 0.003 | 1 | 7 | Nccrp1 | 0.021 | Young |
| 0.009226 | 1.88795 | 0.024 | 0.003 | 1 | 7 | Cd300ld2 | 0.021 | Young |
| 0.009272 | 0.386094 | 0.581 | 0.533 | 1 | 7 | Rgs10 | 0.048 | Young |
| 0.009276 | 1.870185 | 0.024 | 0.003 | 1 | 7 | Zfp438 | 0.021 | Young |
| 0.009377 | 1.914891 | 0.024 | 0.003 | 1 | 7 | Ppp1r3f | 0.021 | Young |
| 0.00956 | 0.687346 | 0.196 | 0.128 | 1 | 7 | Ntpcr | 0.068 | Young |
| 0.009583 | 1.171506 | 0.024 | 0.003 | 1 | 7 | D830025C0 | 0.021 | Young |
| 0.009733 | 1.183341 | 0.133 | 0.077 | 1 | 7 | Mcemp1 | 0.056 | Young |
| 0.009832 | 0.594578 | 0.247 | 0.172 | 1 | 7 | Clcn5 | 0.075 | Young |
| 0.009913 | 0.246451 | 0.732 | 0.697 | 1 | 7 | Pfdn5 | 0.035 | Young |
| 0.009976 | 0.378978 | 0.756 | 0.736 | 1 | 7 | Ier5 | 0.02 | Young |
| 0.01 | 0.136043 | 0.961 | 0.941 | 1 | 7 | Rpl39 | 0.02 | Young |
| 0.01006 | 0.192983 | 0.91 | 0.9 | 1 | 7 | Rpl21 | 0.01 | Young |
| 0.010139 | 0.829685 | 0.084 | 0.038 | 1 | 7 | Zfp629 | 0.046 | Young |
| 0.010159 | 1.504459 | 0.075 | 0.033 | 1 | 7 | Maml3 | 0.042 | Young |
| 0.010345 | 1.29851 | 0.133 | 0.077 | 1 | 7 | Gpr155 | 0.056 | Young |
| 0.010693 | 0.680044 | 0.449 | 0.397 | 1 | 7 | Ssh2 | 0.052 | Young |
| 0.010702 | 0.465443 | 0.431 | 0.359 | 1 | 7 | Lacc1 | 0.072 | Young |
| 0.010866 | 0.711175 | 0.437 | 0.385 | 1 | 7 | Dgkd | 0.052 | Young |
| 0.011178 | 0.455519 | 0.373 | 0.292 | 1 | 7 | Cd93 | 0.081 | Young |
| 0.011226 | 0.387319 | 0.711 | 0.672 | 1 | 7 | Snx2 | 0.039 | Young |
| 0.0115 | 0.235607 | 0.813 | 0.756 | 1 | 7 | Ctsh | 0.057 | Young |
| 0.011584 | 0.470109 | 0.28 | 0.203 | 1 | 7 | Bank1 | 0.077 | Young |
| 0.011717 | 0.586814 | 0.313 | 0.233 | 1 | 7 | Peak1 | 0.08 | Young |
| 0.012032 | 0.163736 | 0.943 | 0.91 | 1 | 7 | Rps25 | 0.033 | Young |
| 0.012796 | 0.970696 | 0.154 | 0.095 | 1 | 7 | Slc15a2 | 0.059 | Young |
| 0.012837 | 3.184855 | 0.033 | 0.008 | 1 | 7 | Cxcl14 | 0.025 | Young |
| 0.013273 | 1.133616 | 0.051 | 0.018 | 1 | 7 | Il1a | 0.033 | Young |
| 0.013332 | 1.86415 | 0.033 | 0.008 | 1 | 7 | Tmem119 | 0.025 | Young |
| 0.013439 | 0.784763 | 0.145 | 0.087 | 1 | 7 | Xylt2 | 0.058 | Young |
| 0.013643 | 0.194641 | 0.877 | 0.849 | 1 | 7 | Rpl35 | 0.028 | Young |
| 0.013672 | 1.616487 | 0.033 | 0.008 | 1 | 7 | Lrrc75a | 0.025 | Young |
| 0.0137 | 0.858238 | 0.078 | 0.036 | 1 | 7 | Bmf | 0.042 | Young |
| 0.013731 | 0.558461 | 0.488 | 0.423 | 1 | 7 | Phlda1 | 0.065 | Young |

| | | | | | | | | |
|---|---|---|---|---|---|---|---|---|
| 0.013735 | 1.04833 | 0.078 | 0.036 | 1 | 7 | Gm20707 | 0.042 | Young |
| 0.014198 | 0.523375 | 0.497 | 0.456 | 1 | 7 | Snx6 | 0.041 | Young |
| 0.014415 | 0.921618 | 0.196 | 0.136 | 1 | 7 | AU020206 | 0.06 | Young |
| 0.014729 | 0.767944 | 0.31 | 0.246 | 1 | 7 | Mrpl48 | 0.064 | Young |
| 0.014842 | 0.35513 | 0.557 | 0.515 | 1 | 7 | Ndufa7 | 0.042 | Young |
| 0.014848 | 0.043944 | 0.051 | 0.103 | 1 | 7 | Smad3 | -0.052 | Young |
| 0.01492 | 0.466271 | 0.334 | 0.274 | 1 | 7 | Dnajb1 | 0.06 | Young |
| 0.015091 | 0.463083 | 0.235 | 0.169 | 1 | 7 | Tagap | 0.066 | Young |
| 0.01514 | 2.604983 | 0.015 | 0 | 1 | 7 | Csrnp2 | 0.015 | Young |
| 0.01514 | 2.022914 | 0.015 | 0 | 1 | 7 | Hs3st1 | 0.015 | Young |
| 0.01514 | 1.528047 | 0.015 | 0 | 1 | 7 | Arhgap24 | 0.015 | Young |
| 0.01514 | 1.456658 | 0.015 | 0 | 1 | 7 | Bik | 0.015 | Young |
| 0.01514 | 1.317887 | 0.015 | 0 | 1 | 7 | Dnhd1 | 0.015 | Young |
| 0.015199 | 0.405635 | 0.301 | 0.226 | 1 | 7 | Ulk2 | 0.075 | Young |
| 0.015997 | 0.12191 | 0.967 | 0.936 | 1 | 7 | Eif1 | 0.031 | Young |
| 0.016298 | 2.339503 | 0.027 | 0.005 | 1 | 7 | Cyr61 | 0.022 | Young |
| 0.016373 | 1.936456 | 0.027 | 0.005 | 1 | 7 | Acta2 | 0.022 | Young |
| 0.016381 | 0.802287 | 0.232 | 0.164 | 1 | 7 | Pdcd2l | 0.068 | Young |
| 0.016474 | 0.278537 | 0.666 | 0.638 | 1 | 7 | Slc38a2 | 0.028 | Young |
| 0.016692 | 0.718327 | 0.25 | 0.185 | 1 | 7 | Anks1 | 0.065 | Young |
| 0.016726 | 1.188947 | 0.072 | 0.033 | 1 | 7 | Flywch1 | 0.039 | Young |
| 0.017012 | 0.684556 | 0.373 | 0.305 | 1 | 7 | Lyz1 | 0.068 | Young |
| 0.017346 | 0.209135 | 0.907 | 0.854 | 1 | 7 | Tgfbi | 0.053 | Young |
| 0.01753 | 1.093676 | 0.093 | 0.049 | 1 | 7 | Ang | 0.044 | Young |
| 0.017713 | 2.43804 | 0.021 | 0.003 | 1 | 7 | Ctla2b | 0.018 | Young |
| 0.017713 | 1.474365 | 0.021 | 0.003 | 1 | 7 | 3222401L1 | 0.018 | Young |
| 0.017842 | 0.248786 | 0.319 | 0.241 | 1 | 7 | Ebi3 | 0.078 | Young |
| 0.017903 | 1.894899 | 0.021 | 0.003 | 1 | 7 | 2900060B1 | 0.018 | Young |
| 0.017905 | 1.289304 | 0.099 | 0.054 | 1 | 7 | Wtip | 0.045 | Young |
| 0.018077 | 0.32922 | 0.723 | 0.703 | 1 | 7 | Gmfg | 0.02 | Young |
| 0.018104 | 0.628745 | 0.325 | 0.254 | 1 | 7 | Tnfsf12 | 0.071 | Young |
| 0.018191 | 1.182494 | 0.021 | 0.003 | 1 | 7 | Dynlt1c | 0.018 | Young |
| 0.018269 | 1.283988 | 0.057 | 0.023 | 1 | 7 | Ccl22 | 0.034 | Young |
| 0.018806 | 0.843931 | 0.175 | 0.115 | 1 | 7 | Tsc1 | 0.06 | Young |
| 0.019174 | 0.855272 | 0.214 | 0.149 | 1 | 7 | Cmah | 0.065 | Young |
| 0.019269 | 0.186569 | 0.988 | 0.962 | 1 | 7 | Cd74 | 0.026 | Young |
| 0.019289 | 0.627304 | 0.47 | 0.405 | 1 | 7 | Cited2 | 0.065 | Young |
| 0.019616 | 0.587505 | 0.256 | 0.19 | 1 | 7 | Crkl | 0.066 | Young |
| 0.019671 | 0.496166 | 0.331 | 0.264 | 1 | 7 | Tes | 0.067 | Young |
| 0.019806 | 0.118127 | 0.108 | 0.059 | 1 | 7 | Nfkbil1 | 0.049 | Young |
| 0.019893 | 0.649239 | 0.253 | 0.19 | 1 | 7 | Ypel3 | 0.063 | Young |
| 0.020069 | 0.083357 | 0.991 | 0.997 | 1 | 7 | Rps9 | -0.006 | Young |
| 0.020132 | 0.611333 | 0.193 | 0.133 | 1 | 7 | Ddit3 | 0.06 | Young |
| 0.020172 | 0.709669 | 0.283 | 0.215 | 1 | 7 | Rftn1 | 0.068 | Young |
| 0.020684 | 0.568767 | 0.386 | 0.31 | 1 | 7 | Igf1 | 0.076 | Young |

| | | | | | | | | |
|---|---|---|---|---|---|---|---|---|
| 0.020868 | 0.592985 | 0.169 | 0.11 | 1 | 7 | Hist1h1e | 0.059 | Young |
| 0.020902 | 0.134709 | 0.943 | 0.933 | 1 | 7 | Rpl17 | 0.01 | Young |
| 0.021309 | 0.449346 | 0.41 | 0.331 | 1 | 7 | Axl | 0.079 | Young |
| 0.021574 | 1.154284 | 0.048 | 0.018 | 1 | 7 | Rubcnl | 0.03 | Young |
| 0.021667 | 0.16987 | 0.024 | 0.059 | 1 | 7 | C130026I2 | -0.035 | Young |
| 0.021757 | 0.791497 | 0.235 | 0.174 | 1 | 7 | Osm | 0.061 | Young |
| 0.021949 | 1.228771 | 0.084 | 0.044 | 1 | 7 | Sp4 | 0.04 | Young |
| 0.022165 | 0.240964 | 0.205 | 0.144 | 1 | 7 | Timm17b | 0.061 | Young |
| 0.022724 | 2.083686 | 0.03 | 0.008 | 1 | 7 | Neurl1a | 0.022 | Young |
| 0.02289 | 0.138552 | 0.94 | 0.928 | 1 | 7 | Rpl19 | 0.012 | Young |
| 0.022917 | 2.122968 | 0.039 | 0.013 | 1 | 7 | Hspb1 | 0.026 | Young |
| 0.023097 | 2.30759 | 0.03 | 0.008 | 1 | 7 | Spata13 | 0.022 | Young |
| 0.023108 | 0.16682 | 0.904 | 0.915 | 1 | 7 | Atp2b1 | -0.011 | Young |
| 0.023351 | 0.415488 | 0.66 | 0.618 | 1 | 7 | Ogt | 0.042 | Young |
| 0.02337 | 0.100778 | 0.901 | 0.879 | 1 | 7 | Rps2 | 0.022 | Young |
| 0.023393 | 1.369212 | 0.051 | 0.021 | 1 | 7 | Negr1 | 0.03 | Young |
| 0.02345 | 0.263044 | 0.184 | 0.121 | 1 | 7 | Adgre4 | 0.063 | Young |
| 0.023486 | 0.41006 | 0.503 | 0.433 | 1 | 7 | Fxr1 | 0.07 | Young |
| 0.023666 | 1.903514 | 0.03 | 0.008 | 1 | 7 | Platr17 | 0.022 | Young |
| 0.023741 | 1.256781 | 0.051 | 0.021 | 1 | 7 | Rmdn1 | 0.03 | Young |
| 0.023972 | 1.221298 | 0.039 | 0.013 | 1 | 7 | Rnf180 | 0.026 | Young |
| 0.023972 | 1.218727 | 0.039 | 0.013 | 1 | 7 | Nnt | 0.026 | Young |
| 0.024024 | 1.116382 | 0.196 | 0.138 | 1 | 7 | Fgfr1op | 0.058 | Young |
| 0.024053 | 1.401777 | 0.03 | 0.008 | 1 | 7 | Gm8369 | 0.022 | Young |
| 0.024096 | 0.104632 | 0.988 | 0.979 | 1 | 7 | Rpl35a | 0.009 | Young |
| 0.024163 | 1.451345 | 0.051 | 0.021 | 1 | 7 | Dtx3 | 0.03 | Young |
| 0.024352 | 1.252883 | 0.096 | 0.054 | 1 | 7 | Drosha | 0.042 | Young |
| 0.024444 | 0.578734 | 0.199 | 0.138 | 1 | 7 | Il16 | 0.061 | Young |
| 0.024702 | 0.381486 | 0.578 | 0.538 | 1 | 7 | Ncf1 | 0.04 | Young |
| 0.025171 | 0.66492 | 0.253 | 0.19 | 1 | 7 | Hpgds | 0.063 | Young |
| 0.025788 | 0.206648 | 0.687 | 0.664 | 1 | 7 | Fcgr2b | 0.023 | Young |
| 0.025909 | 1.556712 | 0.072 | 0.036 | 1 | 7 | Cnrip1 | 0.036 | Young |
| 0.025912 | 0.634728 | 0.166 | 0.11 | 1 | 7 | Hhex | 0.056 | Young |
| 0.026163 | 0.15812 | 0.925 | 0.931 | 1 | 7 | Kctd12 | -0.006 | Young |
| 0.026397 | 0.699205 | 0.283 | 0.221 | 1 | 7 | Snx30 | 0.062 | Young |
| 0.026413 | 0.477611 | 0.672 | 0.654 | 1 | 7 | Il1b | 0.018 | Young |
| 0.026459 | 0.473192 | 0.346 | 0.282 | 1 | 7 | Ssbp4 | 0.064 | Young |
| 0.026554 | 0.456429 | 0.503 | 0.451 | 1 | 7 | Il10ra | 0.052 | Young |
| 0.027012 | 1.192819 | 0.114 | 0.069 | 1 | 7 | Dock4 | 0.045 | Young |
| 0.027266 | 1.829184 | 0.042 | 0.015 | 1 | 7 | Map3k12 | 0.027 | Young |
| 0.027526 | 0.297546 | 0.651 | 0.6 | 1 | 7 | Lrp1 | 0.051 | Young |
| 0.02769 | 0.192497 | 0.533 | 0.438 | 1 | 7 | C5ar1 | 0.095 | Young |
| 0.027734 | 0.662496 | 0.352 | 0.295 | 1 | 7 | Slc7a8 | 0.057 | Young |
| 0.027761 | 0.373961 | 0.617 | 0.595 | 1 | 7 | Gadd45b | 0.022 | Young |
| 0.02817 | 0.962238 | 0.208 | 0.151 | 1 | 7 | Tgfbr1 | 0.057 | Young |

| | | | | | | | | |
|---|---|---|---|---|---|---|---|---|
| 0.028322 | 0.42835 | 0.578 | 0.531 | 1 | 7 | Rgs2 | 0.047 | Young |
| 0.028356 | 1.624153 | 0.108 | 0.064 | 1 | 7 | Dhx32 | 0.044 | Young |
| 0.028597 | 0.069819 | 0.997 | 0.997 | 1 | 7 | Tpt1 | 0 | Young |
| 0.028657 | 1.468512 | 0.042 | 0.015 | 1 | 7 | Znrf3 | 0.027 | Young |
| 0.028733 | 0.860528 | 0.16 | 0.108 | 1 | 7 | Dynlt1f | 0.052 | Young |
| 0.028896 | 0.454033 | 0.123 | 0.074 | 1 | 7 | Gprc5c | 0.049 | Young |
| 0.028918 | 1.052406 | 0.196 | 0.144 | 1 | 7 | Trim24 | 0.052 | Young |
| 0.029021 | 0.144607 | 0.886 | 0.862 | 1 | 7 | Nfkbia | 0.024 | Young |
| 0.029137 | 1.83395 | 0.024 | 0.005 | 1 | 7 | Lsmem1 | 0.019 | Young |
| 0.029268 | 2.292121 | 0.024 | 0.005 | 1 | 7 | Mettl18 | 0.019 | Young |
| 0.029399 | 1.105519 | 0.042 | 0.015 | 1 | 7 | C030034L1 | 0.027 | Young |
| 0.029399 | 1.025836 | 0.042 | 0.015 | 1 | 7 | Ccl17 | 0.027 | Young |
| 0.029626 | 0.693734 | 0.157 | 0.103 | 1 | 7 | Nt5dc2 | 0.054 | Young |
| 0.029706 | 0.651233 | 0.181 | 0.123 | 1 | 7 | Pcmtd2 | 0.058 | Young |
| 0.029947 | 1.599928 | 0.012 | 0 | 1 | 7 | Gjc1 | 0.012 | Young |
| 0.029947 | 1.295394 | 0.012 | 0 | 1 | 7 | Fah | 0.012 | Young |
| 0.029947 | 1.247478 | 0.012 | 0 | 1 | 7 | Afmid | 0.012 | Young |
| 0.029947 | 1.233333 | 0.012 | 0 | 1 | 7 | 3-Mar | 0.012 | Young |
| 0.029947 | 0.922241 | 0.012 | 0 | 1 | 7 | Rdh5 | 0.012 | Young |
| 0.029947 | 0.893305 | 0.012 | 0 | 1 | 7 | Glis3 | 0.012 | Young |
| 0.029947 | 0.850614 | 0.012 | 0 | 1 | 7 | Ccdc149 | 0.012 | Young |
| 0.029947 | 0.795817 | 0.012 | 0 | 1 | 7 | Gm11273 | 0.012 | Young |
| 0.029947 | 0.773891 | 0.012 | 0 | 1 | 7 | Spats2l | 0.012 | Young |
| 0.029947 | 0.651319 | 0.012 | 0 | 1 | 7 | Plin4 | 0.012 | Young |
| 0.030605 | 0.971419 | 0.024 | 0.005 | 1 | 7 | Scml2 | 0.019 | Young |
| 0.030745 | 0.325761 | 0.476 | 0.4 | 1 | 7 | Cfh | 0.076 | Young |
| 0.030878 | 1.021119 | 0.024 | 0.005 | 1 | 7 | H2-Oa | 0.019 | Young |
| 0.030975 | 0.673993 | 0.111 | 0.067 | 1 | 7 | Uvssa | 0.044 | Young |
| 0.030981 | 0.56385 | 0.193 | 0.138 | 1 | 7 | Dnajb4 | 0.055 | Young |
| 0.031053 | 1.926971 | 0.063 | 0.031 | 1 | 7 | Pald1 | 0.032 | Young |
| 0.031178 | 2.105644 | 0.045 | 0.018 | 1 | 7 | Tmem201 | 0.027 | Young |
| 0.03143 | 0.070572 | 0.994 | 0.99 | 1 | 7 | Rplp2 | 0.004 | Young |
| 0.031517 | 0.129444 | 0.81 | 0.803 | 1 | 7 | Gng5 | 0.007 | Young |
| 0.031584 | 0.756099 | 0.175 | 0.123 | 1 | 7 | Atg4b | 0.052 | Young |
| 0.031623 | 0.156553 | 0.205 | 0.287 | 1 | 7 | Itpk1 | -0.082 | Young |
| 0.031949 | 0.417056 | 0.455 | 0.403 | 1 | 7 | Ppp1r12c | 0.052 | Young |
| 0.032112 | 0.15085 | 0.925 | 0.933 | 1 | 7 | Rps11 | -0.008 | Young |
| 0.032233 | 1.043682 | 0.081 | 0.044 | 1 | 7 | Gm20604 | 0.037 | Young |
| 0.03245 | 0.436518 | 0.41 | 0.344 | 1 | 7 | Tgif1 | 0.066 | Young |
| 0.032855 | 1.142783 | 0.033 | 0.01 | 1 | 7 | Cbx2 | 0.023 | Young |
| 0.033503 | 0.368834 | 0.533 | 0.497 | 1 | 7 | Tet3 | 0.036 | Young |
| 0.033798 | 0.83719 | 0.334 | 0.282 | 1 | 7 | Blvra | 0.052 | Young |
| 0.034027 | 0.504433 | 0.419 | 0.372 | 1 | 7 | Lmo2 | 0.047 | Young |
| 0.034088 | 1.853503 | 0.018 | 0.003 | 1 | 7 | Hes7 | 0.015 | Young |
| 0.034266 | 1.212849 | 0.018 | 0.003 | 1 | 7 | 1700001L0 | 0.015 | Young |

| | | | | | | | | |
|---|---|---|---|---|---|---|---|---|
| 0.034276 | 0.834739 | 0.226 | 0.172 | 1 | 7 | Fam98c | 0.054 | Young |
| 0.034446 | 1.294012 | 0.018 | 0.003 | 1 | 7 | Prdm16 | 0.015 | Young |
| 0.034446 | 1.186408 | 0.018 | 0.003 | 1 | 7 | Aplnr | 0.015 | Young |
| 0.034446 | 1.016831 | 0.018 | 0.003 | 1 | 7 | Naip1 | 0.015 | Young |
| 0.034626 | 0.976337 | 0.018 | 0.003 | 1 | 7 | Porcn | 0.015 | Young |
| 0.034635 | 0.524499 | 0.271 | 0.21 | 1 | 7 | Stim1 | 0.061 | Young |
| 0.034807 | 0.83708 | 0.018 | 0.003 | 1 | 7 | Clic5 | 0.015 | Young |
| 0.034879 | 0.191063 | 0.958 | 0.962 | 1 | 7 | Zfp36 | -0.004 | Young |
| 0.034933 | 0.559976 | 0.431 | 0.377 | 1 | 7 | Ptpn11 | 0.054 | Young |
| 0.034936 | 0.839705 | 0.045 | 0.018 | 1 | 7 | Zfp628 | 0.027 | Young |
| 0.03505 | 0.560273 | 0.235 | 0.177 | 1 | 7 | E2f2 | 0.058 | Young |
| 0.035344 | 0.562733 | 0.22 | 0.162 | 1 | 7 | Acox3 | 0.058 | Young |
| 0.03543 | 1.692606 | 0.048 | 0.021 | 1 | 7 | Zfp609 | 0.027 | Young |
| 0.036392 | 1.027296 | 0.151 | 0.1 | 1 | 7 | Abcd4 | 0.051 | Young |
| 0.036431 | 0.861151 | 0.063 | 0.031 | 1 | 7 | Btbd3 | 0.032 | Young |
| 0.036437 | 0.194245 | 0.03 | 0.064 | 1 | 7 | Urb1 | -0.034 | Young |
| 0.036823 | 1.113061 | 0.178 | 0.126 | 1 | 7 | Ssbp2 | 0.052 | Young |
| 0.036833 | 0.47321 | 0.37 | 0.31 | 1 | 7 | Fam208a | 0.06 | Young |
| 0.037103 | 0.842909 | 0.211 | 0.156 | 1 | 7 | Etv1 | 0.055 | Young |
| 0.037732 | 1.212894 | 0.093 | 0.054 | 1 | 7 | Mns1 | 0.039 | Young |
| 0.038595 | 1.09779 | 0.069 | 0.036 | 1 | 7 | Klrd1 | 0.033 | Young |
| 0.038813 | 1.209007 | 0.127 | 0.082 | 1 | 7 | Slc39a3 | 0.045 | Young |
| 0.038994 | 0.473649 | 0.395 | 0.331 | 1 | 7 | Fnip1 | 0.064 | Young |
| 0.038997 | 0.145759 | 0.018 | 0.046 | 1 | 7 | Cmpk2 | -0.028 | Young |
| 0.039163 | 0.62456 | 0.226 | 0.169 | 1 | 7 | Itgb5 | 0.057 | Young |
| 0.039512 | 1.01939 | 0.099 | 0.059 | 1 | 7 | Pkdcc | 0.04 | Young |
| 0.039785 | 1.509766 | 0.072 | 0.038 | 1 | 7 | Mir155hg | 0.034 | Young |
| 0.039827 | 0.865655 | 0.12 | 0.077 | 1 | 7 | Hepacam2 | 0.043 | Young |
| 0.039867 | 0.531296 | 0.102 | 0.062 | 1 | 7 | Rab6b | 0.04 | Young |
| 0.039899 | 0.130669 | 0.154 | 0.218 | 1 | 7 | Crtap | -0.064 | Young |
| 0.040054 | 0.925878 | 0.096 | 0.056 | 1 | 7 | Rrad | 0.04 | Young |
| 0.040197 | 1.790263 | 0.027 | 0.008 | 1 | 7 | Cyb5rl | 0.019 | Young |
| 0.040684 | 0.596741 | 0.211 | 0.154 | 1 | 7 | Naxe | 0.057 | Young |
| 0.04083 | 1.593348 | 0.027 | 0.008 | 1 | 7 | Gm26825 | 0.019 | Young |
| 0.040953 | 0.778474 | 0.036 | 0.013 | 1 | 7 | Park2 | 0.023 | Young |
| 0.041068 | 1.493945 | 0.075 | 0.041 | 1 | 7 | Fmo5 | 0.034 | Young |
| 0.041102 | 0.329723 | 0.524 | 0.474 | 1 | 7 | Tsc22d3 | 0.05 | Young |
| 0.041369 | 1.07803 | 0.051 | 0.023 | 1 | 7 | Msh5 | 0.028 | Young |
| 0.041397 | 0.376036 | 0.443 | 0.385 | 1 | 7 | Kcmf1 | 0.058 | Young |
| 0.041432 | 0.083363 | 0.229 | 0.167 | 1 | 7 | Cd209d | 0.062 | Young |
| 0.041474 | 0.028425 | 0.036 | 0.072 | 1 | 7 | Mettl26 | -0.036 | Young |
| 0.04179 | 0.911021 | 0.111 | 0.069 | 1 | 7 | Gm20696 | 0.042 | Young |
| 0.041914 | 0.129459 | 0.922 | 0.913 | 1 | 7 | Rpl9 | 0.009 | Young |
| 0.041958 | 0.956592 | 0.027 | 0.008 | 1 | 7 | Gm29367 | 0.019 | Young |
| 0.042078 | 0.130364 | 0.867 | 0.874 | 1 | 7 | Tyrobp | -0.007 | Young |

| | | | | | | | | |
|---|---|---|---|---|---|---|---|---|
| 0.042602 | 0.457696 | 0.184 | 0.128 | 1 | 7 | Irak2 | 0.056 | Young |
| 0.043018 | 0.635401 | 0.211 | 0.159 | 1 | 7 | Arid3a | 0.052 | Young |
| 0.043076 | 0.099278 | 0.253 | 0.326 | 1 | 7 | Cc2d1b | -0.073 | Young |
| 0.043193 | 0.560152 | 0.392 | 0.328 | 1 | 7 | Fam129a | 0.064 | Young |
| 0.043698 | 0.15411 | 0.196 | 0.267 | 1 | 7 | Dhrs7b | -0.071 | Young |
| 0.04384 | 0.813674 | 0.114 | 0.072 | 1 | 7 | Ebag9 | 0.042 | Young |
| 0.043878 | 0.365219 | 0.395 | 0.333 | 1 | 7 | Tns1 | 0.062 | Young |
| 0.043992 | 0.121509 | 0.961 | 0.959 | 1 | 7 | Rps23 | 0.002 | Young |
| 0.044434 | 0.371344 | 0.428 | 0.379 | 1 | 7 | Pacsin2 | 0.049 | Young |
| 0.04467 | 0.21238 | 0.596 | 0.546 | 1 | 7 | Ms4a6c | 0.05 | Young |
| 0.044724 | 0.290319 | 0.614 | 0.559 | 1 | 7 | Ptpre | 0.055 | Young |
| 0.04479 | 0.695359 | 0.434 | 0.395 | 1 | 7 | Skil | 0.039 | Young |
| 0.044921 | 0.432073 | 0.684 | 0.664 | 1 | 7 | Cd36 | 0.02 | Young |
| 0.045161 | 1.887527 | 0.054 | 0.026 | 1 | 7 | Ch25h | 0.028 | Young |
| 0.045282 | 0.749844 | 0.139 | 0.092 | 1 | 7 | P2ry12 | 0.047 | Young |
| 0.046211 | 0.631906 | 0.151 | 0.1 | 1 | 7 | Cish | 0.051 | Young |
| 0.046387 | 0.339596 | 0.358 | 0.3 | 1 | 7 | Mxd4 | 0.058 | Young |
| 0.046678 | 1.274914 | 0.057 | 0.028 | 1 | 7 | Zfp523 | 0.029 | Young |
| 0.046692 | 1.327649 | 0.039 | 0.015 | 1 | 7 | Sap130 | 0.024 | Young |
| 0.047029 | 0.216446 | 0.536 | 0.485 | 1 | 7 | Hipk1 | 0.051 | Young |
| 0.047311 | 0.563653 | 0.214 | 0.159 | 1 | 7 | Tbc1d22a | 0.055 | Young |
| 0.047791 | 0.28329 | 0.235 | 0.174 | 1 | 7 | Plxnc1 | 0.061 | Young |
| 0.047859 | 0.374217 | 0.59 | 0.567 | 1 | 7 | Ptpra | 0.023 | Young |
| 0.047919 | 0.324359 | 0.566 | 0.521 | 1 | 7 | Mycbp2 | 0.045 | Young |
| 0.048276 | 0.225157 | 0.765 | 0.7 | 1 | 7 | Khdrbs1 | 0.065 | Young |
| 0.048437 | 1.258524 | 0.102 | 0.064 | 1 | 7 | Car3 | 0.038 | Young |
| 0.048573 | 0.350878 | 0.337 | 0.277 | 1 | 7 | Cat | 0.06 | Young |
| 0.049261 | 0.609085 | 0.265 | 0.21 | 1 | 7 | AI504432 | 0.055 | Young |
| 0.04928 | 0.730777 | 0.078 | 0.044 | 1 | 7 | Zc3h10 | 0.034 | Young |
| 0.049611 | 0.379901 | 0.096 | 0.056 | 1 | 7 | Rgp1 | 0.04 | Young |
| 0.050506 | 0.369398 | 0.681 | 0.656 | 1 | 7 | Fnbp1 | 0.025 | Young |
| 0.050665 | 0.592405 | 0.373 | 0.328 | 1 | 7 | Mcm2 | 0.045 | Young |
| 0.050738 | 0.147824 | 0.148 | 0.213 | 1 | 7 | Rbm18 | -0.065 | Young |
| 0.051501 | 0.211921 | 0.813 | 0.8 | 1 | 7 | Rsrp1 | 0.013 | Young |
| 0.052134 | 0.171882 | 0.608 | 0.572 | 1 | 7 | Fcgr3 | 0.036 | Young |
| 0.052445 | 0.44794 | 0.184 | 0.133 | 1 | 7 | Klhl18 | 0.051 | Young |
| 0.052792 | 1.935425 | 0.021 | 0.005 | 1 | 7 | Dnajc28 | 0.016 | Young |
| 0.053064 | 1.644927 | 0.03 | 0.01 | 1 | 7 | Colec12 | 0.02 | Young |
| 0.053064 | 1.278686 | 0.03 | 0.01 | 1 | 7 | Tnks1bp1 | 0.02 | Young |
| 0.05325 | 1.502519 | 0.021 | 0.005 | 1 | 7 | Gm26542 | 0.016 | Young |
| 0.053413 | 0.807829 | 0.208 | 0.156 | 1 | 7 | Ralgps2 | 0.052 | Young |
| 0.05348 | 1.422525 | 0.021 | 0.005 | 1 | 7 | Kif24 | 0.016 | Young |
| 0.054037 | 0.616024 | 0.163 | 0.113 | 1 | 7 | Syt11 | 0.05 | Young |
| 0.054117 | 0.569592 | 0.018 | 0.044 | 1 | 7 | Nipa1 | -0.026 | Young |
| 0.054161 | 0.722153 | 0.081 | 0.046 | 1 | 7 | Trp53inp1 | 0.035 | Young |

| | | | | | | | | |
|---|---|---|---|---|---|---|---|---|
| 0.054175 | 1.241817 | 0.021 | 0.005 | 1 | 7 | Gm10501 | 0.016 | Young |
| 0.054408 | 0.534815 | 0.021 | 0.005 | 1 | 7 | Gm12159 | 0.016 | Young |
| 0.054807 | 0.975719 | 0.108 | 0.069 | 1 | 7 | Nxpe5 | 0.039 | Young |
| 0.054877 | 1.012364 | 0.021 | 0.005 | 1 | 7 | Gm20655 | 0.016 | Young |
| 0.054945 | 0.292828 | 0.304 | 0.256 | 1 | 7 | Rcor1 | 0.048 | Young |
| 0.055113 | 0.73603 | 0.021 | 0.005 | 1 | 7 | Sorbs1 | 0.016 | Young |
| 0.055303 | 1.859704 | 0.045 | 0.021 | 1 | 7 | Arsi | 0.024 | Young |
| 0.055331 | 0.261019 | 0.575 | 0.546 | 1 | 7 | Runx1 | 0.029 | Young |
| 0.05568 | 0.485634 | 0.437 | 0.39 | 1 | 7 | Bcl2a1b | 0.047 | Young |
| 0.055962 | 0.161554 | 0.114 | 0.167 | 1 | 7 | E2f7 | -0.053 | Young |
| 0.055982 | 0.425646 | 0.328 | 0.272 | 1 | 7 | Sash1 | 0.056 | Young |
| 0.056623 | 0.240675 | 0.042 | 0.077 | 1 | 7 | Pan2 | -0.035 | Young |
| 0.057161 | 0.711908 | 0.063 | 0.033 | 1 | 7 | Klrb1b | 0.03 | Young |
| 0.057296 | 0.521566 | 0.355 | 0.297 | 1 | 7 | P2ry6 | 0.058 | Young |
| 0.05751 | 0.02891 | 0.136 | 0.197 | 1 | 7 | Arhgap21 | -0.061 | Young |
| 0.058516 | 0.109442 | 0.705 | 0.636 | 1 | 7 | Mgl2 | 0.069 | Young |
| 0.058754 | 0.376587 | 0.627 | 0.597 | 1 | 7 | Myl12b | 0.03 | Young |
| 0.059147 | 0.319219 | 0.497 | 0.446 | 1 | 7 | Arrb2 | 0.051 | Young |
| 0.059689 | 1.405535 | 0.127 | 0.087 | 1 | 7 | Mgme1 | 0.04 | Young |
| 0.060035 | 0.565659 | 0.337 | 0.282 | 1 | 7 | Stx16 | 0.055 | Young |
| 0.060442 | 0.348172 | 0.298 | 0.236 | 1 | 7 | Map3k2 | 0.062 | Young |
| 0.06059 | 1.110662 | 0.045 | 0.021 | 1 | 7 | Pde1b | 0.024 | Young |
| 0.060707 | 0.122852 | 0.94 | 0.931 | 1 | 7 | Rps5 | 0.009 | Young |
| 0.061155 | 0.105872 | 0.967 | 0.959 | 1 | 7 | Rpl23 | 0.008 | Young |
| 0.061253 | 2.137481 | 0.033 | 0.013 | 1 | 7 | Klhl13 | 0.02 | Young |
| 0.061397 | 0.960965 | 0.193 | 0.149 | 1 | 7 | Zfp704 | 0.044 | Young |
| 0.061661 | 0.691004 | 0.377 | 0.349 | 1 | 7 | Ecm1 | 0.028 | Young |
| 0.062383 | 0.939341 | 0.072 | 0.041 | 1 | 7 | Tslp | 0.031 | Young |
| 0.062694 | 0.463735 | 0.16 | 0.113 | 1 | 7 | Ap3s2 | 0.047 | Young |
| 0.062714 | 0.276518 | 0.346 | 0.295 | 1 | 7 | Cnot4 | 0.051 | Young |
| 0.062775 | 0.129911 | 0.913 | 0.897 | 1 | 7 | Serf2 | 0.016 | Young |
| 0.063066 | 0.181473 | 0.991 | 0.969 | 1 | 7 | Malat1 | 0.022 | Young |
| 0.063545 | 0.070188 | 0.563 | 0.513 | 1 | 7 | Arl4c | 0.05 | Young |
| 0.064025 | 0.288405 | 0.033 | 0.013 | 1 | 7 | Fam149a | 0.02 | Young |
| 0.06406 | 0.464419 | 0.045 | 0.021 | 1 | 7 | Inpp4b | 0.024 | Young |
| 0.064583 | 1.066558 | 0.075 | 0.044 | 1 | 7 | Rbm48 | 0.031 | Young |
| 0.064826 | 1.017454 | 0.048 | 0.023 | 1 | 7 | Sgms2 | 0.025 | Young |
| 0.064839 | 0.631277 | 0.12 | 0.079 | 1 | 7 | Gtdc1 | 0.041 | Young |
| 0.064888 | 0.084611 | 0.117 | 0.172 | 1 | 7 | Chm | -0.055 | Young |
| 0.065132 | 1.592581 | 0.015 | 0.003 | 1 | 7 | Tagln | 0.012 | Young |
| 0.065132 | 1.33674 | 0.015 | 0.003 | 1 | 7 | Cd46 | 0.012 | Young |
| 0.065132 | 1.324712 | 0.015 | 0.003 | 1 | 7 | Armcx2 | 0.012 | Young |
| 0.06526 | 0.491424 | 0.364 | 0.315 | 1 | 7 | Gsk3a | 0.049 | Young |
| 0.065464 | 1.45491 | 0.015 | 0.003 | 1 | 7 | Gm3336 | 0.012 | Young |
| 0.065464 | 0.996995 | 0.015 | 0.003 | 1 | 7 | Gm34006 | 0.012 | Young |

| | | | | | | | | |
|---|---|---|---|---|---|---|---|---|
| 0.065476 | 0.978567 | 0.081 | 0.049 | 1 | 7 | Slc10a3 | 0.032 | Young |
| 0.065609 | 0.12826 | 0.87 | 0.849 | 1 | 7 | Rpl12 | 0.021 | Young |
| 0.065619 | 0.148537 | 0.13 | 0.185 | 1 | 7 | Vps45 | -0.055 | Young |
| 0.065798 | 1.0234 | 0.015 | 0.003 | 1 | 7 | Nt5e | 0.012 | Young |
| 0.065893 | 1.525163 | 0.051 | 0.026 | 1 | 7 | Gm38843 | 0.025 | Young |
| 0.065949 | 0.141938 | 0.711 | 0.654 | 1 | 7 | Ckb | 0.057 | Young |
| 0.066025 | 0.056375 | 0.063 | 0.103 | 1 | 7 | Rbm43 | -0.04 | Young |
| 0.066073 | 0.880648 | 0.175 | 0.131 | 1 | 7 | Wdr12 | 0.044 | Young |
| 0.066131 | 0.506511 | 0.229 | 0.182 | 1 | 7 | Etv5 | 0.047 | Young |
| 0.066133 | 1.136902 | 0.015 | 0.003 | 1 | 7 | Gm42722 | 0.012 | Young |
| 0.066288 | 0.987212 | 0.072 | 0.041 | 1 | 7 | Zdhhc23 | 0.031 | Young |
| 0.06634 | 0.546113 | 0.271 | 0.226 | 1 | 7 | Nfxl1 | 0.045 | Young |
| 0.066394 | 0.400967 | 0.28 | 0.226 | 1 | 7 | Pcif1 | 0.054 | Young |
| 0.066409 | 0.034311 | 0.187 | 0.256 | 1 | 7 | Polg | -0.069 | Young |
| 0.066469 | 0.458372 | 0.015 | 0.003 | 1 | 7 | Tspan9 | 0.012 | Young |
| 0.066734 | 0.125904 | 0.922 | 0.913 | 1 | 7 | Rpl28 | 0.009 | Young |
| 0.066912 | 0.220645 | 0.648 | 0.621 | 1 | 7 | Kdm6b | 0.027 | Young |
| 0.067696 | 0.233147 | 0.78 | 0.767 | 1 | 7 | Cd52 | 0.013 | Young |
| 0.067989 | 0.049418 | 0.045 | 0.079 | 1 | 7 | Ptpn4 | -0.034 | Young |
| 0.067992 | 0.079756 | 0.136 | 0.187 | 1 | 7 | Ubap1 | -0.051 | Young |
| 0.068511 | 0.024442 | 0.226 | 0.29 | 1 | 7 | Dnajc10 | -0.064 | Young |
| 0.068652 | 0.026411 | 0.09 | 0.136 | 1 | 7 | Asf1a | -0.046 | Young |
| 0.068675 | 0.097281 | 0.241 | 0.313 | 1 | 7 | Stard7 | -0.072 | Young |
| 0.069329 | 0.09115 | 0.928 | 0.928 | 1 | 7 | Rps27 | 0 | Young |
| 0.069792 | 0.41298 | 0.401 | 0.356 | 1 | 7 | Arsb | 0.045 | Young |
| 0.069902 | 0.204415 | 0.021 | 0.046 | 1 | 7 | Magee1 | -0.025 | Young |
| 0.070056 | 3.010862 | 0.024 | 0.008 | 1 | 7 | 4732471J0 | 0.016 | Young |
| 0.070099 | 0.477608 | 0.482 | 0.436 | 1 | 7 | Neurl3 | 0.046 | Young |
| 0.070139 | 0.005133 | 0.057 | 0.095 | 1 | 7 | Lclat1 | -0.038 | Young |
| 0.070167 | 1.071158 | 0.051 | 0.026 | 1 | 7 | Svip | 0.025 | Young |
| 0.0703 | 0.25905 | 0.44 | 0.39 | 1 | 7 | Adipor1 | 0.05 | Young |
| 0.070356 | 0.206116 | 0.53 | 0.479 | 1 | 7 | Mcm7 | 0.051 | Young |
| 0.07044 | 0.356513 | 0.503 | 0.477 | 1 | 7 | Nsd3 | 0.026 | Young |
| 0.070778 | 0.340393 | 0.307 | 0.251 | 1 | 7 | Cdc37l1 | 0.056 | Young |
| 0.070836 | 0.246197 | 0.599 | 0.569 | 1 | 7 | Man2b1 | 0.03 | Young |
| 0.071106 | 2.010651 | 0.024 | 0.008 | 1 | 7 | Tal1 | 0.016 | Young |
| 0.07112 | 0.276969 | 0.431 | 0.379 | 1 | 7 | Rfc1 | 0.052 | Young |
| 0.071318 | 0.384954 | 0.313 | 0.264 | 1 | 7 | Dctpp1 | 0.049 | Young |
| 0.07172 | 1.343607 | 0.036 | 0.015 | 1 | 7 | Slc25a23 | 0.021 | Young |
| 0.071822 | 0.555611 | 0.054 | 0.028 | 1 | 7 | Ntmt1 | 0.026 | Young |
| 0.071902 | 2.000824 | 0.024 | 0.008 | 1 | 7 | Myo19 | 0.016 | Young |
| 0.071902 | 1.347934 | 0.024 | 0.008 | 1 | 7 | Nrep | 0.016 | Young |
| 0.072095 | 0.221538 | 0.373 | 0.326 | 1 | 7 | Crlf2 | 0.047 | Young |
| 0.072169 | 0.927204 | 0.024 | 0.008 | 1 | 7 | Igfbp6 | 0.016 | Young |
| 0.07282 | 0.425229 | 0.205 | 0.154 | 1 | 7 | Fbrsl1 | 0.051 | Young |

| | | | | | | | | |
|---|---|---|---|---|---|---|---|---|
| 0.073064 | 0.674279 | 0.136 | 0.095 | 1 | 7 | Calcrl | 0.041 | Young |
| 0.073257 | 1.155928 | 0.051 | 0.026 | 1 | 7 | Pqlc2 | 0.025 | Young |
| 0.073458 | 0.260854 | 0.202 | 0.154 | 1 | 7 | Nemp1 | 0.048 | Young |
| 0.073475 | 0.578906 | 0.184 | 0.138 | 1 | 7 | Eci2 | 0.046 | Young |
| 0.073516 | 0.972199 | 0.024 | 0.008 | 1 | 7 | Wdpcp | 0.016 | Young |
| 0.073702 | 0.04404 | 0.096 | 0.144 | 1 | 7 | Xylt1 | -0.048 | Young |
| 0.073915 | 0.211917 | 0.726 | 0.664 | 1 | 7 | Fcrls | 0.062 | Young |
| 0.074029 | 0.763532 | 0.087 | 0.054 | 1 | 7 | Emilin1 | 0.033 | Young |
| 0.074112 | 1.047377 | 0.12 | 0.082 | 1 | 7 | Phf1 | 0.038 | Young |
| 0.074695 | 1.025729 | 0.036 | 0.015 | 1 | 7 | Arvcf | 0.021 | Young |
| 0.074784 | 1.078227 | 0.09 | 0.056 | 1 | 7 | Ints8 | 0.034 | Young |
| 0.074852 | 0.938628 | 0.057 | 0.031 | 1 | 7 | 2900076A( | 0.026 | Young |
| 0.075051 | 0.170913 | 0.102 | 0.149 | 1 | 7 | Oma1 | -0.047 | Young |
| 0.075078 | 0.434356 | 0.452 | 0.405 | 1 | 7 | Klf10 | 0.047 | Young |
| 0.075128 | 0.824028 | 0.036 | 0.015 | 1 | 7 | Mdga1 | 0.021 | Young |
| 0.075176 | 0.294562 | 0.09 | 0.056 | 1 | 7 | Men1 | 0.034 | Young |
| 0.075753 | 0.131124 | 0.807 | 0.792 | 1 | 7 | Atpif1 | 0.015 | Young |
| 0.076128 | 0.011637 | 0.268 | 0.331 | 1 | 7 | Mgrn1 | -0.063 | Young |
| 0.076274 | 0.037973 | 0.169 | 0.228 | 1 | 7 | Nbr1 | -0.059 | Young |
| 0.07628 | 0.049581 | 0.184 | 0.241 | 1 | 7 | Lcmt1 | -0.057 | Young |
| 0.076457 | 0.639697 | 0.163 | 0.118 | 1 | 7 | Ophn1 | 0.045 | Young |
| 0.077439 | 0.318034 | 0.633 | 0.628 | 1 | 7 | Zcchc6 | 0.005 | Young |
| 0.077508 | 0.295955 | 0.298 | 0.246 | 1 | 7 | Ccl12 | 0.052 | Young |
| 0.077662 | 0.091017 | 0.232 | 0.303 | 1 | 7 | Rfk | -0.071 | Young |
| 0.077672 | 0.199586 | 0.395 | 0.344 | 1 | 7 | Blvrb | 0.051 | Young |
| 0.077688 | 0.782412 | 0.217 | 0.174 | 1 | 7 | Acadvl | 0.043 | Young |
| 0.077883 | 0.730294 | 0.226 | 0.182 | 1 | 7 | Rxra | 0.044 | Young |
| 0.078065 | 0.610616 | 0.202 | 0.159 | 1 | 7 | Krr1 | 0.043 | Young |
| 0.078074 | 0.178151 | 0.834 | 0.821 | 1 | 7 | Clta | 0.013 | Young |
| 0.078101 | 0.504708 | 0.09 | 0.056 | 1 | 7 | Dennd2c | 0.034 | Young |
| 0.078198 | 0.199859 | 0.593 | 0.554 | 1 | 7 | Tcf4 | 0.039 | Young |
| 0.07857 | 0.507534 | 0.139 | 0.097 | 1 | 7 | Manea | 0.042 | Young |
| 0.078818 | 0.375846 | 0.361 | 0.305 | 1 | 7 | Chd2 | 0.056 | Young |
| 0.079027 | 0.187886 | 0.575 | 0.554 | 1 | 7 | Eps15 | 0.021 | Young |
| 0.079108 | 0.931009 | 0.057 | 0.031 | 1 | 7 | Dsel | 0.026 | Young |
| 0.079558 | 0.747674 | 0.139 | 0.097 | 1 | 7 | Zfp282 | 0.042 | Young |
| 0.079685 | 0.065671 | 0.364 | 0.459 | 1 | 7 | Traf1 | -0.095 | Young |
| 0.080115 | 0.08059 | 0.584 | 0.518 | 1 | 7 | Clec4b1 | 0.066 | Young |
| 0.080762 | 0.392948 | 0.37 | 0.328 | 1 | 7 | Ramp1 | 0.042 | Young |
| 0.081067 | 1.057911 | 0.16 | 0.118 | 1 | 7 | Tle1 | 0.042 | Young |
| 0.081284 | 0.52221 | 0.488 | 0.454 | 1 | 7 | Cdkn1b | 0.034 | Young |
| 0.081698 | 0.527002 | 0.193 | 0.144 | 1 | 7 | C5ar2 | 0.049 | Young |
| 0.081897 | 0.269311 | 0.681 | 0.662 | 1 | 7 | Tmed2 | 0.019 | Young |
| 0.082814 | 0.112918 | 0.024 | 0.049 | 1 | 7 | Dnajc12 | -0.025 | Young |
| 0.082924 | 0.595872 | 0.139 | 0.097 | 1 | 7 | Farsa | 0.042 | Young |

| | | | | | | | | |
|---|---|---|---|---|---|---|---|---|
| 0.083048 | 0.890306 | 0.145 | 0.105 | 1 | 7 | Rnf40 | 0.04 | Young |
| 0.083092 | 1.142702 | 0.066 | 0.038 | 1 | 7 | Nanos1 | 0.028 | Young |
| 0.08338 | 0.137981 | 0.81 | 0.756 | 1 | 7 | Btg2 | 0.054 | Young |
| 0.083463 | 0.046703 | 0.277 | 0.351 | 1 | 7 | Slc25a36 | -0.074 | Young |
| 0.083669 | 0.061531 | 0.331 | 0.418 | 1 | 7 | Ankfy1 | -0.087 | Young |
| 0.084289 | 0.512429 | 0.057 | 0.031 | 1 | 7 | Ttll3 | 0.026 | Young |
| 0.084301 | 0.132725 | 0.241 | 0.303 | 1 | 7 | Psat1 | -0.062 | Young |
| 0.085363 | 0.153344 | 0.506 | 0.451 | 1 | 7 | Tlr2 | 0.055 | Young |
| 0.085952 | 0.105313 | 0.925 | 0.903 | 1 | 7 | Rps17 | 0.022 | Young |
| 0.086369 | 0.512137 | 0.259 | 0.208 | 1 | 7 | Tcn2 | 0.051 | Young |
| 0.086784 | 0.228238 | 0.675 | 0.656 | 1 | 7 | Rab5c | 0.019 | Young |
| 0.086854 | 2.674182 | 0.027 | 0.01 | 1 | 7 | Fam187b | 0.017 | Young |
| 0.087141 | 1.645756 | 0.027 | 0.01 | 1 | 7 | Retn | 0.017 | Young |
| 0.087833 | 0.0205 | 0.081 | 0.123 | 1 | 7 | BC030867 | -0.042 | Young |
| 0.087882 | 0.163026 | 0.193 | 0.144 | 1 | 7 | Cd38 | 0.049 | Young |
| 0.087951 | 0.759142 | 0.145 | 0.105 | 1 | 7 | Golim4 | 0.04 | Young |
| 0.088023 | 0.932292 | 0.069 | 0.041 | 1 | 7 | B230354K1 | 0.028 | Young |
| 0.088302 | 1.123303 | 0.12 | 0.085 | 1 | 7 | Timeless | 0.035 | Young |
| 0.088877 | 0.795589 | 0.069 | 0.041 | 1 | 7 | Arap3 | 0.028 | Young |
| 0.089144 | 1.161193 | 0.099 | 0.067 | 1 | 7 | Ddah2 | 0.032 | Young |
| 0.089407 | 0.429081 | 0.226 | 0.179 | 1 | 7 | Kbtbd2 | 0.047 | Young |
| 0.08951 | 0.087069 | 0.262 | 0.333 | 1 | 7 | B3gat3 | -0.071 | Young |
| 0.089566 | 1.114174 | 0.069 | 0.041 | 1 | 7 | Emc9 | 0.028 | Young |
| 0.089756 | 1.08219 | 0.027 | 0.01 | 1 | 7 | Dclk2 | 0.017 | Young |
| 0.09022 | 0.122965 | 0.907 | 0.877 | 1 | 7 | Rps7 | 0.03 | Young |
| 0.090287 | 0.223775 | 0.539 | 0.495 | 1 | 7 | Paics | 0.044 | Young |
| 0.090507 | 0.901452 | 0.066 | 0.038 | 1 | 7 | Otud1 | 0.028 | Young |
| 0.090507 | 0.334539 | 0.066 | 0.038 | 1 | 7 | Ms4a4c | 0.028 | Young |
| 0.090903 | 0.272479 | 0.542 | 0.492 | 1 | 7 | Taok1 | 0.05 | Young |
| 0.091287 | 0.053083 | 0.28 | 0.351 | 1 | 7 | Zcchc7 | -0.071 | Young |
| 0.091305 | 1.574115 | 0.069 | 0.041 | 1 | 7 | Tppp | 0.028 | Young |
| 0.091506 | 0.188446 | 0.596 | 0.556 | 1 | 7 | Luc7l3 | 0.04 | Young |
| 0.091833 | 0.610531 | 0.027 | 0.01 | 1 | 7 | Arhgap32 | 0.017 | Young |
| 0.091947 | 1.03837 | 0.169 | 0.128 | 1 | 7 | Mylip | 0.041 | Young |
| 0.092257 | 0.456815 | 0.464 | 0.441 | 1 | 7 | Nrp1 | 0.023 | Young |
| 0.092765 | 0.667732 | 0.367 | 0.331 | 1 | 7 | Camk1 | 0.036 | Young |
| 0.093599 | 0.597133 | 0.268 | 0.223 | 1 | 7 | Nuak2 | 0.045 | Young |
| 0.093817 | 1.141638 | 0.084 | 0.054 | 1 | 7 | Serf1 | 0.03 | Young |
| 0.093943 | 0.493035 | 0.253 | 0.208 | 1 | 7 | Klf9 | 0.045 | Young |
| 0.094049 | 0.08797 | 0.825 | 0.823 | 1 | 7 | Ube2d3 | 0.002 | Young |
| 0.09405 | 1.365757 | 0.042 | 0.021 | 1 | 7 | Chst1 | 0.021 | Young |
| 0.094679 | 0.762828 | 0.108 | 0.074 | 1 | 7 | Cd99l2 | 0.034 | Young |
| 0.094782 | 0.533158 | 0.316 | 0.272 | 1 | 7 | 1-Mar | 0.044 | Young |
| 0.094839 | 0.027757 | 0.295 | 0.367 | 1 | 7 | Nampt | -0.072 | Young |
| 0.094885 | 0.487973 | 0.012 | 0.031 | 1 | 7 | Dhrs11 | -0.019 | Young |

| | | | | | | | | |
|---|---|---|---|---|---|---|---|---|
| 0.095096 | 0.929351 | 0.081 | 0.051 | 1 | 7 | Rnf185 | 0.03 | Young |
| 0.095787 | 1.690607 | 0.018 | 0.005 | 1 | 7 | Cdo1 | 0.013 | Young |
| 0.096181 | 2.399108 | 0.018 | 0.005 | 1 | 7 | Gm47469 | 0.013 | Young |
| 0.096462 | 0.70867 | 0.093 | 0.062 | 1 | 7 | Ggta1 | 0.031 | Young |
| 0.096577 | 0.798923 | 0.018 | 0.005 | 1 | 7 | Nphp4 | 0.013 | Young |
| 0.097264 | 0.524231 | 0.485 | 0.438 | 1 | 7 | Kif23 | 0.047 | Young |
| 0.097372 | 1.688648 | 0.018 | 0.005 | 1 | 7 | Gm29291 | 0.013 | Young |
| 0.097372 | 1.092341 | 0.018 | 0.005 | 1 | 7 | Rapsn | 0.013 | Young |
| 0.097604 | 0.825926 | 0.048 | 0.026 | 1 | 7 | Casz1 | 0.022 | Young |
| 0.097771 | 1.333199 | 0.018 | 0.005 | 1 | 7 | Grk4 | 0.013 | Young |
| 0.098543 | 0.081842 | 0.988 | 0.982 | 1 | 7 | Rps3a1 | 0.006 | Young |
| 0.098592 | 0.506725 | 0.262 | 0.215 | 1 | 7 | Hbp1 | 0.047 | Young |
| 0.09913 | 0.010098 | 0.377 | 0.441 | 1 | 7 | Zmynd8 | -0.064 | Young |
| 0.099382 | 0.859896 | 0.018 | 0.005 | 1 | 7 | Lrrc32 | 0.013 | Young |
| 0.099382 | 0.699007 | 0.018 | 0.005 | 1 | 7 | Hist1h3i | 0.013 | Young |
| 0.099382 | 0.621039 | 0.018 | 0.005 | 1 | 7 | Serpinh1 | 0.013 | Young |
| 0.099456 | 0.63991 | 0.157 | 0.118 | 1 | 7 | Bap1 | 0.039 | Young |
| 0.099788 | 0.413293 | 0.295 | 0.241 | 1 | 7 | Psme2b | 0.054 | Young |
| 0.099788 | 0.121358 | 0.018 | 0.005 | 1 | 7 | Gm15247 | 0.013 | Young |
| 0.099794 | 0.178407 | 0.747 | 0.749 | 1 | 7 | Eif3f | -0.002 | Young |
| 0.099917 | 0.52819 | 0.108 | 0.074 | 1 | 7 | Ripor2 | 0.034 | Young |
| 0.100184 | 0.476992 | 0.006 | 0.021 | 1 | 7 | Lgi3 | -0.015 | Young |
| 0.100529 | 0.506053 | 0.413 | 0.364 | 1 | 7 | R3hdm4 | 0.049 | Young |
| 0.100767 | 1.307517 | 0.03 | 0.013 | 1 | 7 | Hdac8 | 0.017 | Young |
| 0.10083 | 1.76861 | 0.051 | 0.028 | 1 | 7 | Large1 | 0.023 | Young |
| 0.100872 | 0.882139 | 0.081 | 0.051 | 1 | 7 | Bcam | 0.03 | Young |
| 0.101443 | 0.645242 | 0.172 | 0.133 | 1 | 7 | Chp2 | 0.039 | Young |
| 0.101498 | 0.601708 | 0.208 | 0.162 | 1 | 7 | Stx6 | 0.046 | Young |
| 0.101531 | 0.100025 | 0.666 | 0.618 | 1 | 7 | Slamf9 | 0.048 | Young |
| 0.102032 | 0.316969 | 0.608 | 0.579 | 1 | 7 | Wwp1 | 0.029 | Young |
| 0.10228 | 1.628434 | 0.03 | 0.013 | 1 | 7 | Fbxo10 | 0.017 | Young |
| 0.102319 | 1.04269 | 0.193 | 0.154 | 1 | 7 | Ing3 | 0.039 | Young |
| 0.102485 | 0.130199 | 0.684 | 0.664 | 1 | 7 | Stk17b | 0.02 | Young |
| 0.102836 | 0.802647 | 0.084 | 0.054 | 1 | 7 | Siglec1 | 0.03 | Young |
| 0.102875 | 1.108293 | 0.048 | 0.026 | 1 | 7 | Tesk2 | 0.022 | Young |
| 0.10289 | 1.413794 | 0.03 | 0.013 | 1 | 7 | Rapgef5 | 0.017 | Young |
| 0.102906 | 0.001949 | 0.247 | 0.31 | 1 | 7 | 2310011J0 | -0.063 | Young |
| 0.103475 | 0.073414 | 0.949 | 0.954 | 1 | 7 | Rps19 | -0.005 | Young |
| 0.103621 | 0.112922 | 0.892 | 0.882 | 1 | 7 | Rpl36 | 0.01 | Young |
| 0.103676 | 0.430964 | 0.467 | 0.433 | 1 | 7 | Per1 | 0.034 | Young |
| 0.103824 | 0.001327 | 0.148 | 0.197 | 1 | 7 | Snapc1 | -0.049 | Young |
| 0.103862 | 0.369836 | 0.377 | 0.331 | 1 | 7 | Lair1 | 0.046 | Young |
| 0.103918 | 1.960182 | 0.066 | 0.041 | 1 | 7 | Cers4 | 0.025 | Young |
| 0.104208 | 0.217174 | 0.515 | 0.485 | 1 | 7 | Ppib | 0.03 | Young |
| 0.104421 | 0.810865 | 0.087 | 0.056 | 1 | 7 | Atg10 | 0.031 | Young |

| | | | | | | | | |
|---|---|---|---|---|---|---|---|---|
| 0.104737 | 1.022746 | 0.03 | 0.013 | 1 | 7 | Gm340 | 0.017 | Young |
| 0.104799 | 0.089185 | 0.054 | 0.087 | 1 | 7 | Znhit3 | -0.033 | Young |
| 0.104968 | 0.578932 | 0.172 | 0.133 | 1 | 7 | Qser1 | 0.039 | Young |
| 0.105629 | 0.093545 | 0.072 | 0.108 | 1 | 7 | Tpgs2 | -0.036 | Young |
| 0.105742 | 0.22076 | 0.729 | 0.695 | 1 | 7 | Akap13 | 0.034 | Young |
| 0.105866 | 0.214379 | 0.777 | 0.764 | 1 | 7 | Atox1 | 0.013 | Young |
| 0.105978 | 0.041933 | 0.316 | 0.374 | 1 | 7 | Nabp1 | -0.058 | Young |
| 0.106074 | 0.384278 | 0.509 | 0.477 | 1 | 7 | Elf1 | 0.032 | Young |
| 0.106125 | 0.148533 | 0.792 | 0.79 | 1 | 7 | Hexa | 0.002 | Young |
| 0.106442 | 0.419216 | 0.241 | 0.197 | 1 | 7 | Polr2j | 0.044 | Young |
| 0.106509 | 0.39392 | 0.051 | 0.028 | 1 | 7 | Akr1c18 | 0.023 | Young |
| 0.106966 | 0.044272 | 0.41 | 0.482 | 1 | 7 | Pfkp | -0.072 | Young |
| 0.107035 | 0.581049 | 0.172 | 0.131 | 1 | 7 | Tmem243 | 0.041 | Young |
| 0.107237 | 1.065454 | 0.111 | 0.077 | 1 | 7 | Mdm1 | 0.034 | Young |
| 0.107675 | 0.986207 | 0.051 | 0.028 | 1 | 7 | Pdlim7 | 0.023 | Young |
| 0.108 | 0.063848 | 0.467 | 0.413 | 1 | 7 | Cxcl16 | 0.054 | Young |
| 0.109281 | 1.075195 | 0.057 | 0.033 | 1 | 7 | Map3k10 | 0.024 | Young |
| 0.10953 | 0.315743 | 0.506 | 0.462 | 1 | 7 | Slc38a1 | 0.044 | Young |
| 0.109685 | 0.37765 | 0.274 | 0.231 | 1 | 7 | Atf7 | 0.043 | Young |
| 0.110437 | 0.794958 | 0.054 | 0.031 | 1 | 7 | Natd1 | 0.023 | Young |
| 0.110675 | 0.78667 | 0.355 | 0.326 | 1 | 7 | Cpd | 0.029 | Young |
| 0.111047 | 0.201136 | 0.286 | 0.231 | 1 | 7 | Aldh1a2 | 0.055 | Young |
| 0.111278 | 0.55414 | 0.404 | 0.369 | 1 | 7 | Sugt1 | 0.035 | Young |
| 0.111426 | 0.557566 | 0.102 | 0.069 | 1 | 7 | Tnni2 | 0.033 | Young |
| 0.111829 | 0.177752 | 0.13 | 0.09 | 1 | 7 | Mical1 | 0.04 | Young |
| 0.11216 | 0.03119 | 0.208 | 0.272 | 1 | 7 | Dapk3 | -0.064 | Young |
| 0.112717 | 0.945161 | 0.151 | 0.113 | 1 | 7 | Agpat3 | 0.038 | Young |
| 0.113022 | 0.251354 | 0.557 | 0.521 | 1 | 7 | Pacs2 | 0.036 | Young |
| 0.113466 | 0.312025 | 0.247 | 0.2 | 1 | 7 | Dtl | 0.047 | Young |
| 0.113515 | 0.45274 | 0.346 | 0.305 | 1 | 7 | Tnrc6c | 0.041 | Young |
| 0.113522 | 0.381523 | 0.331 | 0.282 | 1 | 7 | Ms4a6b | 0.049 | Young |
| 0.11367 | 1.444827 | 0.033 | 0.015 | 1 | 7 | Arhgef40 | 0.018 | Young |
| 0.114344 | 0.271978 | 0.355 | 0.308 | 1 | 7 | Entpd1 | 0.047 | Young |
| 0.114415 | 0.641689 | 0.172 | 0.136 | 1 | 7 | Ndufaf2 | 0.036 | Young |
| 0.114819 | 0.18371 | 0.702 | 0.7 | 1 | 7 | Snx5 | 0.002 | Young |
| 0.114847 | 1.146923 | 0.06 | 0.036 | 1 | 7 | Zfp777 | 0.024 | Young |
| 0.114921 | 1.174265 | 0.033 | 0.015 | 1 | 7 | 2610020C0 | 0.018 | Young |
| 0.115182 | 0.125996 | 0.759 | 0.736 | 1 | 7 | Snx3 | 0.023 | Young |
| 0.115749 | 0.458979 | 0.166 | 0.128 | 1 | 7 | Ung | 0.038 | Young |
| 0.115774 | 0.17142 | 0.229 | 0.292 | 1 | 7 | Galk2 | -0.063 | Young |
| 0.115888 | 0.431807 | 0.377 | 0.328 | 1 | 7 | Tgfbr2 | 0.049 | Young |
| 0.115913 | 0.269825 | 0.37 | 0.315 | 1 | 7 | Ptger4 | 0.055 | Young |
| 0.116182 | 1.192198 | 0.033 | 0.015 | 1 | 7 | Pdk2 | 0.018 | Young |
| 0.116398 | 0.592696 | 0.172 | 0.133 | 1 | 7 | Thra | 0.039 | Young |
| 0.11677 | 0.516038 | 0.268 | 0.228 | 1 | 7 | Adrb2 | 0.04 | Young |

| | | | | | | | | |
|---|---|---|---|---|---|---|---|---|
| 0.116918 | 0.841217 | 0.111 | 0.079 | 1 | 7 | 2510009E0 | 0.032 | Young |
| 0.116948 | 0.264097 | 0.578 | 0.533 | 1 | 7 | Dync1i2 | 0.045 | Young |
| 0.117109 | 0.008762 | 0.22 | 0.274 | 1 | 7 | Sfswap | -0.054 | Young |
| 0.117135 | 0.940689 | 0.033 | 0.015 | 1 | 7 | Mturn | 0.018 | Young |
| 0.117143 | 0.745791 | 0.217 | 0.182 | 1 | 7 | Mgat4a | 0.035 | Young |
| 0.117454 | 1.082139 | 0.033 | 0.015 | 1 | 7 | Cd300ld3 | 0.018 | Young |
| 0.117879 | 0.682359 | 0.13 | 0.095 | 1 | 7 | Maged1 | 0.035 | Young |
| 0.118226 | 0.103282 | 0.913 | 0.895 | 1 | 7 | Ddx5 | 0.018 | Young |
| 0.118449 | 0.372043 | 0.214 | 0.169 | 1 | 7 | Parp8 | 0.045 | Young |
| 0.118507 | 0.107614 | 0.181 | 0.236 | 1 | 7 | Ddx10 | -0.055 | Young |
| 0.118769 | 0.443447 | 0.19 | 0.149 | 1 | 7 | Ermp1 | 0.041 | Young |
| 0.118871 | 1.19276 | 0.066 | 0.041 | 1 | 7 | Smo | 0.025 | Young |
| 0.119623 | 0.491074 | 0.325 | 0.285 | 1 | 7 | Ppp1r10 | 0.04 | Young |
| 0.119854 | 0.289331 | 0.515 | 0.492 | 1 | 7 | Cmtm6 | 0.023 | Young |
| 0.120251 | 1.983115 | 0.081 | 0.054 | 1 | 7 | Spp1 | 0.027 | Young |
| 0.120522 | 0.358999 | 0.166 | 0.126 | 1 | 7 | Fam193b | 0.04 | Young |
| 0.120607 | 0.450758 | 0.133 | 0.095 | 1 | 7 | Ripk2 | 0.038 | Young |
| 0.120663 | 0.234619 | 0.744 | 0.728 | 1 | 7 | Son | 0.016 | Young |
| 0.120728 | 0.240878 | 0.645 | 0.613 | 1 | 7 | Hnrnpd | 0.032 | Young |
| 0.121157 | 0.115858 | 0.123 | 0.169 | 1 | 7 | Dhx29 | -0.046 | Young |
| 0.121726 | 1.544471 | 0.021 | 0.008 | 1 | 7 | Tmc4 | 0.013 | Young |
| 0.121823 | 0.863878 | 0.172 | 0.136 | 1 | 7 | Arsk | 0.036 | Young |
| 0.122204 | 0.664349 | 0.235 | 0.192 | 1 | 7 | Arrdc3 | 0.043 | Young |
| 0.122378 | 0.147088 | 0.349 | 0.297 | 1 | 7 | Zfp395 | 0.052 | Young |
| 0.122475 | 0.328859 | 0.114 | 0.159 | 1 | 7 | Cdyl | -0.045 | Young |
| 0.122589 | 0.091876 | 0.69 | 0.644 | 1 | 7 | Tpd52 | 0.046 | Young |
| 0.122792 | 0.040836 | 0.22 | 0.277 | 1 | 7 | Plxna1 | -0.057 | Young |
| 0.122843 | 0.718558 | 0.256 | 0.215 | 1 | 7 | Fam204a | 0.041 | Young |
| 0.122853 | 0.53432 | 0.377 | 0.341 | 1 | 7 | Rab3il1 | 0.036 | Young |
| 0.122977 | 0.28349 | 0.105 | 0.149 | 1 | 7 | Alad | -0.044 | Young |
| 0.12312 | 0.055531 | 0.916 | 0.913 | 1 | 7 | Cox4i1 | 0.003 | Young |
| 0.123288 | 0.783641 | 0.117 | 0.085 | 1 | 7 | Pdik1l | 0.032 | Young |
| 0.123541 | 0.048344 | 0.078 | 0.115 | 1 | 7 | Ints13 | -0.037 | Young |
| 0.123755 | 0.163586 | 0.569 | 0.549 | 1 | 7 | Sdc4 | 0.02 | Young |
| 0.123786 | 0.110159 | 0.015 | 0.033 | 1 | 7 | Zscan12 | -0.018 | Young |
| 0.123787 | 0.06722 | 0.735 | 0.697 | 1 | 7 | Pld4 | 0.038 | Young |
| 0.123871 | 1.081698 | 0.021 | 0.008 | 1 | 7 | Gm33973 | 0.013 | Young |
| 0.124125 | 0.829985 | 0.142 | 0.105 | 1 | 7 | Gphn | 0.037 | Young |
| 0.124737 | 1.481435 | 0.021 | 0.008 | 1 | 7 | Mss51 | 0.013 | Young |
| 0.124904 | 0.079372 | 0.036 | 0.062 | 1 | 7 | Tyw5 | -0.026 | Young |
| 0.124985 | 1.691413 | 0.012 | 0.003 | 1 | 7 | Gm14391 | 0.009 | Young |
| 0.124985 | 1.376677 | 0.012 | 0.003 | 1 | 7 | B4galt2 | 0.009 | Young |
| 0.125172 | 1.394384 | 0.021 | 0.008 | 1 | 7 | Zfp763 | 0.013 | Young |
| 0.125337 | 0.29588 | 0.491 | 0.464 | 1 | 7 | Fuca1 | 0.027 | Young |
| 0.125489 | 1.071213 | 0.063 | 0.038 | 1 | 7 | Slc26a11 | 0.025 | Young |

| | | | | | | | | |
|---|---|---|---|---|---|---|---|---|
| 0.1256 | 4.037914 | 0.012 | 0.003 | 1 | 7 | Mucl1 | 0.009 | Young |
| 0.1256 | 1.155807 | 0.012 | 0.003 | 1 | 7 | Sema7a | 0.009 | Young |
| 0.1256 | 0.846409 | 0.012 | 0.003 | 1 | 7 | 1700102H2 | 0.009 | Young |
| 0.125608 | 1.194149 | 0.021 | 0.008 | 1 | 7 | Zfp503 | 0.013 | Young |
| 0.125608 | 1.110081 | 0.021 | 0.008 | 1 | 7 | Ms4a4a | 0.013 | Young |
| 0.125608 | 0.792085 | 0.021 | 0.008 | 1 | 7 | Ccdc106 | 0.013 | Young |
| 0.125754 | 0.183728 | 0.536 | 0.505 | 1 | 7 | Numa1 | 0.031 | Young |
| 0.125901 | 0.090416 | 0.084 | 0.121 | 1 | 7 | Pafah1b3 | -0.037 | Young |
| 0.126216 | 1.474138 | 0.012 | 0.003 | 1 | 7 | 6230400D1 | 0.009 | Young |
| 0.126216 | 1.294142 | 0.012 | 0.003 | 1 | 7 | Gm13479 | 0.009 | Young |
| 0.126216 | 1.198428 | 0.012 | 0.003 | 1 | 7 | Ggact | 0.009 | Young |
| 0.126216 | 1.075452 | 0.012 | 0.003 | 1 | 7 | Med12l | 0.009 | Young |
| 0.126216 | 0.872462 | 0.012 | 0.003 | 1 | 7 | Gsg1 | 0.009 | Young |
| 0.126216 | 0.768172 | 0.012 | 0.003 | 1 | 7 | Spaca1 | 0.009 | Young |
| 0.126484 | 0.783035 | 0.021 | 0.008 | 1 | 7 | Gm7160 | 0.013 | Young |
| 0.126835 | 0.710709 | 0.012 | 0.003 | 1 | 7 | Gm8013 | 0.009 | Young |
| 0.126835 | 0.628378 | 0.012 | 0.003 | 1 | 7 | Rsph9 | 0.009 | Young |
| 0.126835 | 0.598222 | 0.012 | 0.003 | 1 | 7 | H2-Eb2 | 0.009 | Young |
| 0.126924 | 1.248085 | 0.021 | 0.008 | 1 | 7 | Plpp1 | 0.013 | Young |
| 0.127456 | 0.504943 | 0.012 | 0.003 | 1 | 7 | Wdr19 | 0.009 | Young |
| 0.127456 | 0.129735 | 0.012 | 0.003 | 1 | 7 | Tceal1 | 0.009 | Young |
| 0.128426 | 0.169906 | 0.166 | 0.126 | 1 | 7 | Ago4 | 0.04 | Young |
| 0.12849 | 0.26923 | 0.557 | 0.526 | 1 | 7 | Etf1 | 0.031 | Young |
| 0.128737 | 0.938505 | 0.154 | 0.121 | 1 | 7 | Cebpzos | 0.033 | Young |
| 0.12875 | 0.162492 | 0.455 | 0.397 | 1 | 7 | Pura | 0.058 | Young |
| 0.129098 | 0.553542 | 0.19 | 0.151 | 1 | 7 | Prr14 | 0.039 | Young |
| 0.129765 | 0.315495 | 0.154 | 0.115 | 1 | 7 | Lpar6 | 0.039 | Young |
| 0.129879 | 0.068075 | 0.386 | 0.459 | 1 | 7 | Stard3nl | -0.073 | Young |
| 0.129934 | 0.396098 | 0.247 | 0.21 | 1 | 7 | Ncoa2 | 0.037 | Young |
| 0.129972 | 0.612256 | 0.175 | 0.138 | 1 | 7 | Erf | 0.037 | Young |
| 0.130076 | 0.578802 | 0.13 | 0.095 | 1 | 7 | Mfn1 | 0.035 | Young |
| 0.13055 | 0.709678 | 0.175 | 0.138 | 1 | 7 | Cdca7 | 0.037 | Young |
| 0.130565 | 0.800833 | 0.084 | 0.056 | 1 | 7 | Coq9 | 0.028 | Young |
| 0.130795 | 0.788836 | 0.169 | 0.133 | 1 | 7 | Hivep3 | 0.036 | Young |
| 0.131124 | 1.075476 | 0.039 | 0.021 | 1 | 7 | Gpt | 0.018 | Young |
| 0.131142 | 0.453767 | 0.325 | 0.287 | 1 | 7 | Pcf11 | 0.038 | Young |
| 0.131811 | 0.25982 | 0.084 | 0.056 | 1 | 7 | Aacs | 0.028 | Young |
| 0.132023 | 0.143291 | 0.744 | 0.751 | 1 | 7 | Picalm | -0.007 | Young |
| 0.132742 | 0.697057 | 0.075 | 0.049 | 1 | 7 | Lmbr1l | 0.026 | Young |
| 0.133537 | 0.112424 | 0.06 | 0.092 | 1 | 7 | Rrnad1 | -0.032 | Young |
| 0.133623 | 0.534873 | 0.235 | 0.195 | 1 | 7 | Elof1 | 0.04 | Young |
| 0.133833 | 0.21426 | 0.069 | 0.044 | 1 | 7 | Rac3 | 0.025 | Young |
| 0.134232 | 0.240193 | 0.47 | 0.441 | 1 | 7 | Ski | 0.029 | Young |
| 0.134664 | 0.165789 | 0.096 | 0.136 | 1 | 7 | Gm46224 | -0.04 | Young |
| 0.135092 | 0.3316 | 0.398 | 0.359 | 1 | 7 | Cyp4f18 | 0.039 | Young |

| | | | | | | | | |
|---|---|---|---|---|---|---|---|---|
| 0.135439 | 0.007066 | 0.22 | 0.272 | 1 | 7 | Tmem245 | -0.052 | Young |
| 0.135461 | 0.122988 | 0.623 | 0.572 | 1 | 7 | Aif1 | 0.051 | Young |
| 0.13582 | 0.251617 | 0.051 | 0.079 | 1 | 7 | Ccl25 | -0.028 | Young |
| 0.136113 | 0.854773 | 0.075 | 0.049 | 1 | 7 | Pcyt1b | 0.026 | Young |
| 0.13636 | 0.392437 | 0.232 | 0.192 | 1 | 7 | Rhoc | 0.04 | Young |
| 0.136549 | 0.152804 | 0.09 | 0.062 | 1 | 7 | Tom1l2 | 0.028 | Young |
| 0.136871 | 0.47733 | 0.039 | 0.021 | 1 | 7 | Mcc | 0.018 | Young |
| 0.136905 | 0.250118 | 0.069 | 0.044 | 1 | 7 | Gm49359 | 0.025 | Young |
| 0.137382 | 0.662618 | 0.069 | 0.044 | 1 | 7 | Cox6b2 | 0.025 | Young |
| 0.137402 | 0.071825 | 0.922 | 0.913 | 1 | 7 | Gnai2 | 0.009 | Young |
| 0.138379 | 0.066429 | 0.732 | 0.721 | 1 | 7 | Minos1 | 0.011 | Young |
| 0.138387 | 0.346663 | 0.063 | 0.038 | 1 | 7 | Wdr6 | 0.025 | Young |
| 0.138645 | 0.510663 | 0.099 | 0.069 | 1 | 7 | Abtb1 | 0.03 | Young |
| 0.138705 | 0.054796 | 0.834 | 0.823 | 1 | 7 | Rpl10 | 0.011 | Young |
| 0.138746 | 0.252367 | 0.274 | 0.226 | 1 | 7 | Tbc1d5 | 0.048 | Young |
| 0.138756 | 0.035242 | 0.069 | 0.103 | 1 | 7 | C87436 | -0.034 | Young |
| 0.138846 | 0.617493 | 0.096 | 0.067 | 1 | 7 | 2310061I0 | 0.029 | Young |
| 0.138978 | 0.059774 | 0.25 | 0.31 | 1 | 7 | Hmbox1 | -0.06 | Young |
| 0.139077 | 0.887013 | 0.169 | 0.133 | 1 | 7 | Tlr7 | 0.036 | Young |
| 0.139157 | 0.554969 | 0.039 | 0.021 | 1 | 7 | Slc25a53 | 0.018 | Young |
| 0.139467 | 0.779695 | 0.09 | 0.062 | 1 | 7 | Lin37 | 0.028 | Young |
| 0.139638 | 0.16732 | 0.105 | 0.074 | 1 | 7 | Morc2a | 0.031 | Young |
| 0.140025 | 1.352183 | 0.045 | 0.026 | 1 | 7 | Stat4 | 0.019 | Young |
| 0.141027 | 0.175214 | 0.274 | 0.226 | 1 | 7 | Ndfip2 | 0.048 | Young |
| 0.141198 | 0.144637 | 0.654 | 0.641 | 1 | 7 | Romo1 | 0.013 | Young |
| 0.141356 | 0.006806 | 0.247 | 0.303 | 1 | 7 | Aqr | -0.056 | Young |
| 0.141989 | 0.447506 | 0.105 | 0.074 | 1 | 7 | Aldh7a1 | 0.031 | Young |
| 0.142343 | 0.449692 | 0.229 | 0.19 | 1 | 7 | Adap2 | 0.039 | Young |
| 0.142448 | 0.250099 | 0.16 | 0.126 | 1 | 7 | Klf16 | 0.034 | Young |
| 0.142833 | 0.59814 | 0.13 | 0.097 | 1 | 7 | Gnpda2 | 0.033 | Young |
| 0.1431 | 0.101783 | 0.837 | 0.787 | 1 | 7 | Cd53 | 0.05 | Young |
| 0.143118 | 0.061489 | 0.361 | 0.331 | 1 | 7 | Heca | 0.03 | Young |
| 0.143405 | 0.089438 | 0.777 | 0.782 | 1 | 7 | Cox8a | -0.005 | Young |
| 0.143633 | 1.720275 | 0.024 | 0.01 | 1 | 7 | Htra3 | 0.014 | Young |
| 0.144703 | 0.213395 | 0.633 | 0.618 | 1 | 7 | Eid1 | 0.015 | Young |
| 0.144747 | 0.08635 | 0.937 | 0.897 | 1 | 7 | Rpl22 | 0.04 | Young |
| 0.145111 | 0.322265 | 0.38 | 0.344 | 1 | 7 | Anxa3 | 0.036 | Young |
| 0.145143 | 0.138803 | 0.873 | 0.872 | 1 | 7 | Rtn4 | 0.001 | Young |
| 0.145646 | 0.474745 | 0.211 | 0.172 | 1 | 7 | Tada1 | 0.039 | Young |
| 0.145656 | 0.059096 | 0.34 | 0.413 | 1 | 7 | Acsl4 | -0.073 | Young |
| 0.14589 | 0.002764 | 0.099 | 0.138 | 1 | 7 | Cds1 | -0.039 | Young |
| 0.145917 | 0.102044 | 0.87 | 0.854 | 1 | 7 | Atp5g2 | 0.016 | Young |
| 0.146084 | 0.758066 | 0.009 | 0.023 | 1 | 7 | 2210408I2 | -0.014 | Young |
| 0.146949 | 0.050822 | 0.596 | 0.582 | 1 | 7 | Snrpb2 | 0.014 | Young |
| 0.147099 | 0.032861 | 0.169 | 0.215 | 1 | 7 | Dhdh | -0.046 | Young |

| | | | | | | | | |
|---|---|---|---|---|---|---|---|---|
| 0.147209 | 1.413474 | 0.024 | 0.01 | 1 | 7 | Cuedc1 | 0.014 | Young |
| 0.147334 | 0.265847 | 0.193 | 0.154 | 1 | 7 | Trim30d | 0.039 | Young |
| 0.147429 | 0.121837 | 0.563 | 0.536 | 1 | 7 | Nop10 | 0.027 | Young |
| 0.147566 | 0.236121 | 0.62 | 0.6 | 1 | 7 | Raly | 0.02 | Young |
| 0.14771 | 0.904764 | 0.105 | 0.077 | 1 | 7 | Rprd1a | 0.028 | Young |
| 0.148046 | 0.099511 | 0.111 | 0.151 | 1 | 7 | Smpd4 | -0.04 | Young |
| 0.148091 | 0.880299 | 0.045 | 0.026 | 1 | 7 | Mical2 | 0.019 | Young |
| 0.148408 | 0.912776 | 0.045 | 0.026 | 1 | 7 | Dtd1 | 0.019 | Young |
| 0.148539 | 0.016825 | 0.081 | 0.115 | 1 | 7 | Il10 | -0.034 | Young |
| 0.149478 | 0.557304 | 0.024 | 0.01 | 1 | 7 | Hist1h2bb | 0.014 | Young |
| 0.149727 | 0.836862 | 0.048 | 0.028 | 1 | 7 | Ankrd13d | 0.02 | Young |
| 0.149867 | 0.090357 | 0.988 | 0.985 | 1 | 7 | Rps21 | 0.003 | Young |
| 0.150136 | 0.175196 | 0.003 | 0.013 | 1 | 7 | C920021L1 | -0.01 | Young |
| 0.150393 | 0.342909 | 0.024 | 0.01 | 1 | 7 | Tmem240 | 0.014 | Young |
| 0.150644 | 0.194447 | 0.819 | 0.797 | 1 | 7 | Srrm2 | 0.022 | Young |
| 0.150881 | 0.41325 | 0.123 | 0.09 | 1 | 7 | Sfi1 | 0.033 | Young |
| 0.151214 | 0.563997 | 0.229 | 0.187 | 1 | 7 | Hdgfl2 | 0.042 | Young |
| 0.151313 | 0.104862 | 0.223 | 0.277 | 1 | 7 | Slc8b1 | -0.054 | Young |
| 0.151727 | 0.125383 | 0.759 | 0.736 | 1 | 7 | Ubb | 0.023 | Young |
| 0.152332 | 0.709619 | 0.166 | 0.131 | 1 | 7 | Pml | 0.035 | Young |
| 0.152377 | 0.201529 | 0.449 | 0.431 | 1 | 7 | Camk1d | 0.018 | Young |
| 0.153053 | 0.29648 | 0.386 | 0.349 | 1 | 7 | Srpr | 0.037 | Young |
| 0.153382 | 0.174049 | 0.123 | 0.09 | 1 | 7 | Cyp27a1 | 0.033 | Young |
| 0.153405 | 0.378643 | 0.139 | 0.105 | 1 | 7 | Scmh1 | 0.034 | Young |
| 0.154124 | 0.164583 | 0.587 | 0.574 | 1 | 7 | Nedd8 | 0.013 | Young |
| 0.154367 | 0.132731 | 0.102 | 0.138 | 1 | 7 | Nln | -0.036 | Young |
| 0.155763 | 0.329461 | 0.554 | 0.564 | 1 | 7 | Nr3c1 | -0.01 | Young |
| 0.156087 | 0.81525 | 0.117 | 0.087 | 1 | 7 | Rtl8a | 0.03 | Young |
| 0.156307 | 0.093503 | 0.229 | 0.282 | 1 | 7 | Med8 | -0.053 | Young |
| 0.156439 | 1.128846 | 0.051 | 0.031 | 1 | 7 | Hist2h2ac | 0.02 | Young |
| 0.156587 | 0.962159 | 0.06 | 0.038 | 1 | 7 | Ubiad1 | 0.022 | Young |
| 0.156611 | 0.205214 | 0.16 | 0.208 | 1 | 7 | Ubac2 | -0.048 | Young |
| 0.157051 | 0.887136 | 0.123 | 0.092 | 1 | 7 | Smim3 | 0.031 | Young |
| 0.157303 | 0.713181 | 0.235 | 0.2 | 1 | 7 | Snx13 | 0.035 | Young |
| 0.157932 | 0.17554 | 0.364 | 0.323 | 1 | 7 | Golga7 | 0.041 | Young |
| 0.158293 | 0.60281 | 0.051 | 0.031 | 1 | 7 | Zmynd15 | 0.02 | Young |
| 0.158446 | 0.633969 | 0.054 | 0.033 | 1 | 7 | Shb | 0.021 | Young |
| 0.158594 | 0.245946 | 0.726 | 0.728 | 1 | 7 | Cdkn1a | -0.002 | Young |
| 0.158833 | 0.971011 | 0.096 | 0.069 | 1 | 7 | Msh3 | 0.027 | Young |
| 0.158833 | 0.911724 | 0.096 | 0.069 | 1 | 7 | Il1rap | 0.027 | Young |
| 0.158959 | 0.41349 | 0.253 | 0.213 | 1 | 7 | Arhgef3 | 0.04 | Young |
| 0.159401 | 0.178568 | 0.62 | 0.613 | 1 | 7 | Mef2a | 0.007 | Young |
| 0.160116 | 0.896062 | 0.172 | 0.138 | 1 | 7 | Pmvk | 0.034 | Young |
| 0.160144 | 0.225291 | 0.44 | 0.41 | 1 | 7 | Foxn3 | 0.03 | Young |
| 0.16082 | 0.08371 | 0.178 | 0.223 | 1 | 7 | N4bp2l1 | -0.045 | Young |

| | | | | | | | | |
|---|---|---|---|---|---|---|---|---|
| 0.161058 | 1.368815 | 0.027 | 0.013 | 1 | 7 | Ctla2a | 0.014 | Young |
| 0.161271 | 0.591251 | 0.295 | 0.259 | 1 | 7 | Man1a2 | 0.036 | Young |
| 0.161378 | 0.561132 | 0.175 | 0.141 | 1 | 7 | Mcee | 0.034 | Young |
| 0.161587 | 0.873597 | 0.232 | 0.2 | 1 | 7 | Itgb1bp1 | 0.032 | Young |
| 0.161677 | 0.040121 | 0.226 | 0.269 | 1 | 7 | Irs2 | -0.043 | Young |
| 0.161835 | 0.630952 | 0.367 | 0.341 | 1 | 7 | Tsc22d2 | 0.026 | Young |
| 0.161866 | 0.867074 | 0.099 | 0.072 | 1 | 7 | Katnb1 | 0.027 | Young |
| 0.162012 | 0.609201 | 0.048 | 0.074 | 1 | 7 | Wrb | -0.026 | Young |
| 0.162105 | 0.167925 | 0.081 | 0.113 | 1 | 7 | Eef2kmt | -0.032 | Young |
| 0.162638 | 0.73216 | 0.093 | 0.067 | 1 | 7 | Rgmb | 0.026 | Young |
| 0.162737 | 0.729629 | 0.108 | 0.079 | 1 | 7 | Chd3os | 0.029 | Young |
| 0.16274 | 0.265753 | 0.084 | 0.121 | 1 | 7 | Tigd2 | -0.037 | Young |
| 0.162742 | 0.557668 | 0.337 | 0.313 | 1 | 7 | Stxbp6 | 0.024 | Young |
| 0.164095 | 0.677934 | 0.145 | 0.113 | 1 | 7 | Kctd12b | 0.032 | Young |
| 0.164242 | 0.015835 | 0.117 | 0.156 | 1 | 7 | Orc3 | -0.039 | Young |
| 0.164248 | 0.564541 | 0.181 | 0.146 | 1 | 7 | Deptor | 0.035 | Young |
| 0.164311 | 0.074877 | 0.904 | 0.885 | 1 | 7 | Rpl36a | 0.019 | Young |
| 0.164672 | 1.026477 | 0.027 | 0.013 | 1 | 7 | Thumpd2 | 0.014 | Young |
| 0.164743 | 0.09895 | 0.265 | 0.323 | 1 | 7 | Usp48 | -0.058 | Young |
| 0.165175 | 0.070057 | 0.916 | 0.905 | 1 | 7 | Ybx1 | 0.011 | Young |
| 0.165396 | 0.716136 | 0.096 | 0.069 | 1 | 7 | Nudcd1 | 0.027 | Young |
| 0.165428 | 0.649466 | 0.217 | 0.179 | 1 | 7 | Lmo1 | 0.038 | Young |
| 0.16562 | 0.239801 | 0.392 | 0.341 | 1 | 7 | Fam117b | 0.051 | Young |
| 0.165833 | 0.613213 | 0.102 | 0.074 | 1 | 7 | Cracr2b | 0.028 | Young |
| 0.166042 | 1.138226 | 0.027 | 0.013 | 1 | 7 | Agbl3 | 0.014 | Young |
| 0.166083 | 0.302424 | 0.352 | 0.315 | 1 | 7 | Polb | 0.037 | Young |
| 0.166501 | 0.627187 | 0.027 | 0.013 | 1 | 7 | Ms4a14 | 0.014 | Young |
| 0.166969 | 1.083556 | 0.06 | 0.038 | 1 | 7 | Gm26518 | 0.022 | Young |
| 0.167179 | 0.641222 | 0.166 | 0.131 | 1 | 7 | Gm37494 | 0.035 | Young |
| 0.16741 | 0.13279 | 0.151 | 0.195 | 1 | 7 | Pi4k2b | -0.044 | Young |
| 0.167605 | 0.105235 | 0.828 | 0.818 | 1 | 7 | Rrbp1 | 0.01 | Young |
| 0.167668 | 1.261128 | 0.054 | 0.033 | 1 | 7 | Ahdc1 | 0.021 | Young |
| 0.168068 | 0.722765 | 0.105 | 0.077 | 1 | 7 | Selenbp1 | 0.028 | Young |
| 0.168237 | 1.660076 | 0.063 | 0.041 | 1 | 7 | Tnfsf13b | 0.022 | Young |
| 0.168323 | 0.354112 | 0.241 | 0.2 | 1 | 7 | Cntln | 0.041 | Young |
| 0.16847 | 0.874357 | 0.084 | 0.059 | 1 | 7 | Thap4 | 0.025 | Young |
| 0.168507 | 0.153399 | 0.202 | 0.246 | 1 | 7 | Prpf38a | -0.044 | Young |
| 0.16879 | 0.078781 | 0.051 | 0.031 | 1 | 7 | Fam92a | 0.02 | Young |
| 0.169743 | 0.532013 | 0.31 | 0.279 | 1 | 7 | Hltf | 0.031 | Young |
| 0.169908 | 0.115466 | 0.298 | 0.349 | 1 | 7 | Yipf3 | -0.051 | Young |
| 0.169909 | 0.138737 | 0.735 | 0.713 | 1 | 7 | Gapdh | 0.022 | Young |
| 0.170366 | 0.710855 | 0.172 | 0.138 | 1 | 7 | Serhl | 0.034 | Young |
| 0.170673 | 0.923874 | 0.027 | 0.013 | 1 | 7 | Palm | 0.014 | Young |
| 0.170697 | 0.391092 | 0.127 | 0.095 | 1 | 7 | Casp2 | 0.032 | Young |
| 0.171122 | 0.350984 | 0.494 | 0.472 | 1 | 7 | Irf1 | 0.022 | Young |

| | | | | | | | | |
|---|---|---|---|---|---|---|---|---|
| 0.171194 | 0.366399 | 0.193 | 0.154 | 1 | 7 | Mosmo | 0.039 | Young |
| 0.171246 | 0.068124 | 0.861 | 0.872 | 1 | 7 | Ubc | -0.011 | Young |
| 0.171444 | 0.287546 | 0.036 | 0.059 | 1 | 7 | Nsun4 | -0.023 | Young |
| 0.171591 | 0.473252 | 0.22 | 0.187 | 1 | 7 | Mxd1 | 0.033 | Young |
| 0.171747 | 0.169155 | 0.789 | 0.762 | 1 | 7 | Fxyd5 | 0.027 | Young |
| 0.171862 | 0.573551 | 0.256 | 0.218 | 1 | 7 | Ip6k1 | 0.038 | Young |
| 0.171906 | 0.813404 | 0.123 | 0.095 | 1 | 7 | Rab40c | 0.028 | Young |
| 0.171962 | 1.722508 | 0.015 | 0.005 | 1 | 7 | Ypel1 | 0.01 | Young |
| 0.171986 | 1.244039 | 0.084 | 0.059 | 1 | 7 | Gna11 | 0.025 | Young |
| 0.172218 | 0.494707 | 0.268 | 0.236 | 1 | 7 | Pde7a | 0.032 | Young |
| 0.172394 | 0.167565 | 0.325 | 0.377 | 1 | 7 | Ndc1 | -0.052 | Young |
| 0.172606 | 0.303352 | 0.166 | 0.131 | 1 | 7 | Rabgap1l | 0.035 | Young |
| 0.172625 | 1.611331 | 0.015 | 0.005 | 1 | 7 | Ccdc125 | 0.01 | Young |
| 0.172984 | 1.442976 | 0.03 | 0.015 | 1 | 7 | Gm15446 | 0.015 | Young |
| 0.173427 | 1.358225 | 0.03 | 0.015 | 1 | 7 | 1700030KC | 0.015 | Young |
| 0.173957 | 1.895906 | 0.015 | 0.005 | 1 | 7 | Me1 | 0.01 | Young |
| 0.174369 | 0.678786 | 0.13 | 0.1 | 1 | 7 | Polk | 0.03 | Young |
| 0.174486 | 0.215898 | 0.133 | 0.174 | 1 | 7 | Map3k20 | -0.041 | Young |
| 0.174626 | 1.279004 | 0.015 | 0.005 | 1 | 7 | Gm6710 | 0.01 | Young |
| 0.174713 | 0.522502 | 0.211 | 0.177 | 1 | 7 | Snapin | 0.034 | Young |
| 0.175297 | 1.111389 | 0.015 | 0.005 | 1 | 7 | Wnt4 | 0.01 | Young |
| 0.175297 | 0.668355 | 0.015 | 0.005 | 1 | 7 | Gng10 | 0.01 | Young |
| 0.175297 | 0.355852 | 0.015 | 0.005 | 1 | 7 | Gm26699 | 0.01 | Young |
| 0.175418 | 0.441774 | 0.102 | 0.074 | 1 | 7 | Mrpl49 | 0.028 | Young |
| 0.175813 | 0.985497 | 0.084 | 0.059 | 1 | 7 | Cog1 | 0.025 | Young |
| 0.17597 | 2.023818 | 0.015 | 0.005 | 1 | 7 | Slc7a2 | 0.01 | Young |
| 0.17597 | 0.862466 | 0.015 | 0.005 | 1 | 7 | Clec4g | 0.01 | Young |
| 0.176205 | 0.304542 | 0.434 | 0.392 | 1 | 7 | Slc25a4 | 0.042 | Young |
| 0.176387 | 0.226943 | 0.491 | 0.467 | 1 | 7 | Tfe3 | 0.024 | Young |
| 0.176421 | 0.375401 | 0.262 | 0.226 | 1 | 7 | Anapc15 | 0.036 | Young |
| 0.176606 | 0.030203 | 0.069 | 0.1 | 1 | 7 | Pde6d | -0.031 | Young |
| 0.176714 | 0.014855 | 0.051 | 0.077 | 1 | 7 | Ptges2 | -0.026 | Young |
| 0.177276 | 0.163256 | 0.25 | 0.208 | 1 | 7 | Cd300c2 | 0.042 | Young |
| 0.177324 | 0.299987 | 0.419 | 0.395 | 1 | 7 | Arid4a | 0.024 | Young |
| 0.177942 | 0.402078 | 0.334 | 0.3 | 1 | 7 | Clcn4 | 0.034 | Young |
| 0.178193 | 0.383103 | 0.25 | 0.21 | 1 | 7 | Zfp385a | 0.04 | Young |
| 0.178327 | 0.433895 | 0.386 | 0.356 | 1 | 7 | Ehmt2 | 0.03 | Young |
| 0.178382 | 0.097115 | 0.873 | 0.874 | 1 | 7 | Zfp36l2 | -0.001 | Young |
| 0.17852 | 0.096702 | 0.316 | 0.269 | 1 | 7 | Lsm7 | 0.047 | Young |
| 0.178583 | 0.071736 | 0.277 | 0.338 | 1 | 7 | Hexim1 | -0.061 | Young |
| 0.178657 | 0.462131 | 0.066 | 0.044 | 1 | 7 | Pianp | 0.022 | Young |
| 0.178874 | 0.189611 | 0.259 | 0.223 | 1 | 7 | Creld2 | 0.036 | Young |
| 0.179065 | 0.725756 | 0.133 | 0.103 | 1 | 7 | Stk35 | 0.03 | Young |
| 0.179265 | 0.791943 | 0.03 | 0.015 | 1 | 7 | Tlr5 | 0.015 | Young |
| 0.179265 | 0.665589 | 0.03 | 0.015 | 1 | 7 | Vcan | 0.015 | Young |

| | | | | | | | | |
|---|---|---|---|---|---|---|---|---|
| 0.180003 | 0.678965 | 0.133 | 0.103 | 1 | 7 | Ganc | 0.03 | Young |
| 0.180135 | 0.881331 | 0.117 | 0.087 | 1 | 7 | Brpf3 | 0.03 | Young |
| 0.180346 | 0.194314 | 0.44 | 0.392 | 1 | 7 | Frmd4b | 0.048 | Young |
| 0.180633 | 0.829141 | 0.03 | 0.015 | 1 | 7 | Slc9a5 | 0.015 | Young |
| 0.180747 | 0.136615 | 0.702 | 0.7 | 1 | 7 | Myh9 | 0.002 | Young |
| 0.180977 | 0.387119 | 0.06 | 0.038 | 1 | 7 | Tmco6 | 0.022 | Young |
| 0.181549 | 0.694608 | 0.03 | 0.015 | 1 | 7 | Zfp287 | 0.015 | Young |
| 0.181797 | 0.218523 | 0.364 | 0.326 | 1 | 7 | Ndufa12 | 0.038 | Young |
| 0.181822 | 0.342829 | 0.325 | 0.287 | 1 | 7 | Gosr2 | 0.038 | Young |
| 0.181959 | 0.055217 | 0.973 | 0.979 | 1 | 7 | Rps20 | -0.006 | Young |
| 0.182936 | 0.458538 | 0.154 | 0.121 | 1 | 7 | Kifap3 | 0.033 | Young |
| 0.183176 | 0.133373 | 0.208 | 0.251 | 1 | 7 | Oxsr1 | -0.043 | Young |
| 0.183206 | 1.317914 | 0.033 | 0.018 | 1 | 7 | Gm10762 | 0.015 | Young |
| 0.183467 | 0.122086 | 0.274 | 0.328 | 1 | 7 | Vps26a | -0.054 | Young |
| 0.183712 | 0.174075 | 0.06 | 0.087 | 1 | 7 | Ccdc28b | -0.027 | Young |
| 0.18379 | 0.081636 | 0.045 | 0.069 | 1 | 7 | Ift140 | -0.024 | Young |
| 0.183806 | 0.313901 | 0.075 | 0.051 | 1 | 7 | Mybl1 | 0.024 | Young |
| 0.183855 | 0.667893 | 0.03 | 0.015 | 1 | 7 | Fam84b | 0.015 | Young |
| 0.184064 | 0.65246 | 0.265 | 0.226 | 1 | 7 | Brpf1 | 0.039 | Young |
| 0.18429 | 0.279216 | 0.331 | 0.292 | 1 | 7 | Fam105a | 0.039 | Young |
| 0.185137 | 0.230094 | 0.66 | 0.646 | 1 | 7 | Cbl | 0.014 | Young |
| 0.185621 | 0.011708 | 0.163 | 0.208 | 1 | 7 | Sh3glb2 | -0.045 | Young |
| 0.18672 | 1.353661 | 0.033 | 0.018 | 1 | 7 | Ildr2 | 0.015 | Young |
| 0.18672 | 1.298987 | 0.033 | 0.018 | 1 | 7 | F830016B0 | 0.015 | Young |
| 0.186917 | 0.868329 | 0.123 | 0.097 | 1 | 7 | Extl3 | 0.026 | Young |
| 0.187069 | 0.261704 | 0.545 | 0.528 | 1 | 7 | Rab10 | 0.017 | Young |
| 0.187208 | 0.686791 | 0.169 | 0.136 | 1 | 7 | Shprh | 0.033 | Young |
| 0.187983 | 0.710534 | 0.066 | 0.044 | 1 | 7 | Zfp719 | 0.022 | Young |
| 0.188025 | 0.788308 | 0.12 | 0.092 | 1 | 7 | Pias3 | 0.028 | Young |
| 0.188077 | 0.038313 | 0.033 | 0.054 | 1 | 7 | Rundc1 | -0.021 | Young |
| 0.188714 | 0.626208 | 0.127 | 0.097 | 1 | 7 | Snx24 | 0.03 | Young |
| 0.189004 | 0.480478 | 0.114 | 0.087 | 1 | 7 | Trp53bp2 | 0.027 | Young |
| 0.189232 | 0.341414 | 0.416 | 0.39 | 1 | 7 | Arhgap45 | 0.026 | Young |
| 0.189408 | 0.248069 | 0.012 | 0.026 | 1 | 7 | Armc9 | -0.014 | Young |
| 0.189511 | 0.07205 | 0.747 | 0.705 | 1 | 7 | Mpeg1 | 0.042 | Young |
| 0.189577 | 0.162113 | 0.286 | 0.244 | 1 | 7 | Id3 | 0.042 | Young |
| 0.190054 | 0.810969 | 0.199 | 0.172 | 1 | 7 | Bcl2l1 | 0.027 | Young |
| 0.190232 | 0.079477 | 0.187 | 0.231 | 1 | 7 | Rnmt | -0.044 | Young |
| 0.190277 | 0.284194 | 0.045 | 0.069 | 1 | 7 | Tbc1d12 | -0.024 | Young |
| 0.190347 | 0.067009 | 0.145 | 0.187 | 1 | 7 | Kctd5 | -0.042 | Young |
| 0.190446 | 0.3331 | 0.033 | 0.054 | 1 | 7 | Mon1a | -0.021 | Young |
| 0.190873 | 0.069316 | 0.12 | 0.159 | 1 | 7 | Txnl4a | -0.039 | Young |
| 0.190899 | 0.255782 | 0.244 | 0.208 | 1 | 7 | Arhgap5 | 0.036 | Young |
| 0.191103 | 0.091063 | 0.822 | 0.821 | 1 | 7 | Rps18 | 0.001 | Young |
| 0.191769 | 0.124824 | 0.136 | 0.172 | 1 | 7 | Mpv17l2 | -0.036 | Young |

| | | | | | | | | |
|---|---|---|---|---|---|---|---|---|
| 0.192157 | 0.42904 | 0.319 | 0.285 | 1 | 7 | Chordc1 | 0.034 | Young |
| 0.1925 | 1.015858 | 0.151 | 0.121 | 1 | 7 | Fhod1 | 0.03 | Young |
| 0.192667 | 0.200116 | 0.096 | 0.128 | 1 | 7 | Tpra1 | -0.032 | Young |
| 0.192691 | 0.187317 | 0.28 | 0.244 | 1 | 7 | Klhl5 | 0.036 | Young |
| 0.192971 | 0.040907 | 0.012 | 0.026 | 1 | 7 | Tlr8 | -0.014 | Young |
| 0.193409 | 0.348166 | 0.084 | 0.059 | 1 | 7 | Bcl11a | 0.025 | Young |
| 0.193426 | 0.176876 | 0.723 | 0.721 | 1 | 7 | Psma7 | 0.002 | Young |
| 0.193536 | 0.041525 | 0.127 | 0.164 | 1 | 7 | Miip | -0.037 | Young |
| 0.193693 | 0.085701 | 0.919 | 0.887 | 1 | 7 | Gnb2 | 0.032 | Young |
| 0.193945 | 0.062404 | 0.244 | 0.297 | 1 | 7 | Dnajc21 | -0.053 | Young |
| 0.194451 | 0.294792 | 0.389 | 0.349 | 1 | 7 | Npm3 | 0.04 | Young |
| 0.1955 | 0.180481 | 0.428 | 0.379 | 1 | 7 | Arl5b | 0.049 | Young |
| 0.195662 | 1.239559 | 0.036 | 0.021 | 1 | 7 | Zbtb10 | 0.015 | Young |
| 0.195933 | 0.261943 | 0.533 | 0.51 | 1 | 7 | Epn1 | 0.023 | Young |
| 0.195953 | 0.167343 | 0.108 | 0.144 | 1 | 7 | Cep44 | -0.036 | Young |
| 0.19623 | 0.266523 | 0.551 | 0.536 | 1 | 7 | Dennd4a | 0.015 | Young |
| 0.196583 | 0.524679 | 0.012 | 0.026 | 1 | 7 | AC163040. | -0.014 | Young |
| 0.196641 | 0.69835 | 0.033 | 0.018 | 1 | 7 | Tnfrsf17 | 0.015 | Young |
| 0.197229 | 1.89779 | 0.042 | 0.026 | 1 | 7 | Nle1 | 0.016 | Young |
| 0.197679 | 0.031371 | 0.151 | 0.187 | 1 | 7 | Gins4 | -0.036 | Young |
| 0.198001 | 0.277601 | 0.5 | 0.485 | 1 | 7 | Hacd4 | 0.015 | Young |
| 0.198101 | 0.067007 | 0.123 | 0.162 | 1 | 7 | Dohh | -0.039 | Young |
| 0.198516 | 0.017291 | 0.166 | 0.21 | 1 | 7 | Mtmr14 | -0.044 | Young |
| 0.198553 | 0.312834 | 0.467 | 0.451 | 1 | 7 | Nfic | 0.016 | Young |
| 0.199118 | 0.744493 | 0.105 | 0.079 | 1 | 7 | Hint3 | 0.026 | Young |
| 0.199346 | 0.18052 | 0.696 | 0.682 | 1 | 7 | Snrpf | 0.014 | Young |
| 0.199361 | 0.437128 | 0.349 | 0.328 | 1 | 7 | Sla | 0.021 | Young |
| 0.199594 | 0.122817 | 0.789 | 0.772 | 1 | 7 | Lamp2 | 0.017 | Young |
| 0.199606 | 0.73426 | 0.036 | 0.021 | 1 | 7 | Gm4673 | 0.015 | Young |
| 0.200012 | 0.194504 | 0.672 | 0.664 | 1 | 7 | Jak1 | 0.008 | Young |
| 0.201191 | 0.538239 | 0.295 | 0.267 | 1 | 7 | Nfatc2 | 0.028 | Young |
| 0.201224 | 0.446237 | 0.169 | 0.136 | 1 | 7 | Pot1a | 0.033 | Young |
| 0.201768 | 0.179526 | 0.283 | 0.338 | 1 | 7 | Slc30a5 | -0.055 | Young |
| 0.201965 | 0.350956 | 0.322 | 0.282 | 1 | 7 | D1Ertd622 | 0.04 | Young |
| 0.203553 | 0.012131 | 0.425 | 0.49 | 1 | 7 | Ppp1r12a | -0.065 | Young |
| 0.203607 | 0.824621 | 0.036 | 0.021 | 1 | 7 | Zfp213 | 0.015 | Young |
| 0.203748 | 0.039651 | 0.205 | 0.254 | 1 | 7 | Faf1 | -0.049 | Young |
| 0.203801 | 0.360136 | 0.175 | 0.141 | 1 | 7 | Zbtb4 | 0.034 | Young |
| 0.204056 | 0.605036 | 0.036 | 0.021 | 1 | 7 | Gpsm1 | 0.015 | Young |
| 0.204772 | 0.141086 | 0.093 | 0.126 | 1 | 7 | Cables2 | -0.033 | Young |
| 0.204931 | 0.401208 | 0.28 | 0.244 | 1 | 7 | Cmklr1 | 0.036 | Young |
| 0.205515 | 0.181329 | 0.711 | 0.695 | 1 | 7 | Smarca5 | 0.016 | Young |
| 0.206404 | 0.068519 | 0.178 | 0.223 | 1 | 7 | Rab5b | -0.045 | Young |
| 0.206877 | 1.386352 | 0.018 | 0.008 | 1 | 7 | Cacnb1 | 0.01 | Young |
| 0.207051 | 0.327821 | 0.151 | 0.118 | 1 | 7 | Tspan13 | 0.033 | Young |

| | | | | | | | | |
|---|---|---|---|---|---|---|---|---|
| 0.207164 | 1.496098 | 0.048 | 0.031 | 1 | 7 | Mblac2 | 0.017 | Young |
| 0.207176 | 0.581692 | 0.386 | 0.364 | 1 | 7 | Mpp1 | 0.022 | Young |
| 0.207203 | 0.066805 | 0.81 | 0.803 | 1 | 7 | Msn | 0.007 | Young |
| 0.207256 | 0.262225 | 0.494 | 0.49 | 1 | 7 | Acat1 | 0.004 | Young |
| 0.207448 | 0.072499 | 0.22 | 0.272 | 1 | 7 | Mis18bp1 | -0.052 | Young |
| 0.207846 | 0.412039 | 0.042 | 0.026 | 1 | 7 | Mrm2 | 0.016 | Young |
| 0.207934 | 0.356906 | 0.054 | 0.079 | 1 | 7 | Ercc2 | -0.025 | Young |
| 0.208125 | 0.161788 | 0.145 | 0.182 | 1 | 7 | Zfp120 | -0.037 | Young |
| 0.20822 | 1.483047 | 0.018 | 0.008 | 1 | 7 | Mypop | 0.01 | Young |
| 0.20822 | 0.865995 | 0.018 | 0.008 | 1 | 7 | Rbp1 | 0.01 | Young |
| 0.208798 | 0.270552 | 0.096 | 0.126 | 1 | 7 | Yod1 | -0.03 | Young |
| 0.208881 | 0.165543 | 0.569 | 0.533 | 1 | 7 | Trib1 | 0.036 | Young |
| 0.20915 | 0.398584 | 0.075 | 0.054 | 1 | 7 | Mettl7a1 | 0.021 | Young |
| 0.209326 | 0.430181 | 0.364 | 0.338 | 1 | 7 | Secisbp2l | 0.026 | Young |
| 0.209423 | 0.086891 | 0.271 | 0.321 | 1 | 7 | Fbxl5 | -0.05 | Young |
| 0.20949 | 0.260625 | 0.036 | 0.021 | 1 | 7 | Hist1h2ac | 0.015 | Young |
| 0.209518 | 0.616861 | 0.16 | 0.128 | 1 | 7 | Bcl2 | 0.032 | Young |
| 0.209618 | 0.264705 | 0.404 | 0.364 | 1 | 7 | Gpr65 | 0.04 | Young |
| 0.209618 | 0.086141 | 0.057 | 0.082 | 1 | 7 | 1110004E0 | -0.025 | Young |
| 0.209927 | 0.033312 | 0.157 | 0.197 | 1 | 7 | Pex2 | -0.04 | Young |
| 0.210223 | 0.62356 | 0.039 | 0.023 | 1 | 7 | Fancb | 0.016 | Young |
| 0.21023 | 0.477508 | 0.114 | 0.087 | 1 | 7 | Pcnx | 0.027 | Young |
| 0.210247 | 1.107884 | 0.018 | 0.008 | 1 | 7 | C920006O1 | 0.01 | Young |
| 0.210305 | 0.005724 | 0.25 | 0.295 | 1 | 7 | Dctn2 | -0.045 | Young |
| 0.210378 | 0.071142 | 0.175 | 0.218 | 1 | 7 | Lmnb2 | -0.043 | Young |
| 0.210661 | 0.881579 | 0.039 | 0.023 | 1 | 7 | Gcnt2 | 0.016 | Young |
| 0.210774 | 1.279592 | 0.042 | 0.026 | 1 | 7 | Creld1 | 0.016 | Young |
| 0.210926 | 0.963749 | 0.018 | 0.008 | 1 | 7 | Fam222a | 0.01 | Young |
| 0.210926 | 0.694458 | 0.018 | 0.008 | 1 | 7 | Gm5127 | 0.01 | Young |
| 0.211373 | 0.088939 | 0.063 | 0.09 | 1 | 7 | Gga2 | -0.027 | Young |
| 0.211616 | 0.034 | 0.024 | 0.041 | 1 | 7 | Eno3 | -0.017 | Young |
| 0.211726 | 0.123762 | 0.148 | 0.19 | 1 | 7 | Rce1 | -0.042 | Young |
| 0.212289 | 0.845267 | 0.018 | 0.008 | 1 | 7 | Prdm9 | 0.01 | Young |
| 0.212543 | 0.258289 | 0.051 | 0.074 | 1 | 7 | Zfp335 | -0.023 | Young |
| 0.212546 | 0.170624 | 0.542 | 0.492 | 1 | 7 | Tmem109 | 0.05 | Young |
| 0.212571 | 0.482217 | 0.145 | 0.115 | 1 | 7 | Itgal | 0.03 | Young |
| 0.212832 | 0.111209 | 0.795 | 0.79 | 1 | 7 | Rpl31 | 0.005 | Young |
| 0.212861 | 0.598463 | 0.039 | 0.023 | 1 | 7 | Taf1c | 0.016 | Young |
| 0.212947 | 1.577534 | 0.054 | 0.036 | 1 | 7 | Tchp | 0.018 | Young |
| 0.212973 | 0.218191 | 0.018 | 0.008 | 1 | 7 | Crybb1 | 0.01 | Young |
| 0.213745 | 0.800562 | 0.039 | 0.023 | 1 | 7 | 1700096K1 | 0.016 | Young |
| 0.213745 | 0.600537 | 0.039 | 0.023 | 1 | 7 | Macrod2 | 0.016 | Young |
| 0.21382 | 0.200163 | 0.316 | 0.277 | 1 | 7 | Ncoa7 | 0.039 | Young |
| 0.213958 | 0.009302 | 0.38 | 0.444 | 1 | 7 | Tnks2 | -0.064 | Young |
| 0.214156 | 0.97271 | 0.042 | 0.026 | 1 | 7 | Cbr1 | 0.016 | Young |

| | | | | | | | | |
|---|---|---|---|---|---|---|---|---|
| 0.214263 | 0.603407 | 0.178 | 0.149 | 1 | 7 | Msh6 | 0.029 | Young |
| 0.214756 | 0.028651 | 0.187 | 0.149 | 1 | 7 | Itga9 | 0.038 | Young |
| 0.214963 | 1.265586 | 0.111 | 0.087 | 1 | 7 | 1110065P2 | 0.024 | Young |
| 0.21499 | 0.249305 | 0.259 | 0.223 | 1 | 7 | Ndc80 | 0.036 | Young |
| 0.214995 | 0.320303 | 0.111 | 0.085 | 1 | 7 | Timm21 | 0.026 | Young |
| 0.215229 | 0.042759 | 0.301 | 0.362 | 1 | 7 | Smndc1 | -0.061 | Young |
| 0.215289 | 1.077611 | 0.069 | 0.049 | 1 | 7 | Sox12 | 0.02 | Young |
| 0.215801 | 0.119402 | 0.316 | 0.379 | 1 | 7 | Rfwd3 | -0.063 | Young |
| 0.216184 | 0.122047 | 0.28 | 0.246 | 1 | 7 | Il4ra | 0.034 | Young |
| 0.21629 | 0.812473 | 0.042 | 0.026 | 1 | 7 | Rab39 | 0.016 | Young |
| 0.217512 | 0.546887 | 0.358 | 0.341 | 1 | 7 | Rps6ka1 | 0.017 | Young |
| 0.217653 | 0.25098 | 0.563 | 0.546 | 1 | 7 | Dennd5a | 0.017 | Young |
| 0.217819 | 0.227179 | 0.581 | 0.549 | 1 | 7 | Atad2 | 0.032 | Young |
| 0.218201 | 0.021623 | 0.392 | 0.446 | 1 | 7 | Uhrf1 | -0.054 | Young |
| 0.218501 | 0.537791 | 0.395 | 0.382 | 1 | 7 | Setx | 0.013 | Young |
| 0.21895 | 0.099587 | 0.178 | 0.223 | 1 | 7 | Glo1 | -0.045 | Young |
| 0.219058 | 0.060698 | 0.184 | 0.228 | 1 | 7 | Tmem184b | -0.044 | Young |
| 0.219105 | 0.069149 | 0.636 | 0.626 | 1 | 7 | Ifngr1 | 0.01 | Young |
| 0.219645 | 0.937097 | 0.078 | 0.056 | 1 | 7 | Tnfrsf18 | 0.022 | Young |
| 0.219718 | 1.173622 | 0.018 | 0.033 | 1 | 7 | Gpr84 | -0.015 | Young |
| 0.220223 | 0.062343 | 0.97 | 0.951 | 1 | 7 | Rpl23a | 0.019 | Young |
| 0.22029 | 0.969692 | 0.072 | 0.051 | 1 | 7 | Agpat2 | 0.021 | Young |
| 0.220364 | 0.124399 | 0.075 | 0.103 | 1 | 7 | Znhit6 | -0.028 | Young |
| 0.220472 | 0.158362 | 0.093 | 0.123 | 1 | 7 | Rpain | -0.03 | Young |
| 0.22061 | 0.348451 | 0.259 | 0.226 | 1 | 7 | Jak2 | 0.033 | Young |
| 0.220724 | 0.627546 | 0.111 | 0.085 | 1 | 7 | Alg9 | 0.026 | Young |
| 0.221143 | 0.465164 | 0.081 | 0.059 | 1 | 7 | Scd1 | 0.022 | Young |
| 0.221163 | 0.438675 | 0.045 | 0.028 | 1 | 7 | Glmn | 0.017 | Young |
| 0.221181 | 0.149064 | 0.38 | 0.351 | 1 | 7 | Bin1 | 0.029 | Young |
| 0.221321 | 0.272173 | 0.545 | 0.521 | 1 | 7 | Myo5a | 0.024 | Young |
| 0.22145 | 0.867694 | 0.081 | 0.059 | 1 | 7 | Mettl13 | 0.022 | Young |
| 0.221552 | 1.404709 | 0.054 | 0.036 | 1 | 7 | Sh3bp4 | 0.018 | Young |
| 0.22191 | 0.492519 | 0.042 | 0.026 | 1 | 7 | 4933433G1 | 0.016 | Young |
| 0.22195 | 0.373618 | 0.401 | 0.374 | 1 | 7 | Trpm7 | 0.027 | Young |
| 0.222384 | 0.371781 | 0.491 | 0.487 | 1 | 7 | Dnajb14 | 0.004 | Young |
| 0.222458 | 0.306656 | 0.205 | 0.174 | 1 | 7 | Lrch3 | 0.031 | Young |
| 0.222942 | 0.164791 | 0.563 | 0.549 | 1 | 7 | Psme2 | 0.014 | Young |
| 0.223052 | 0.028923 | 0.253 | 0.303 | 1 | 7 | Arid1b | -0.05 | Young |
| 0.223155 | 0.4914 | 0.105 | 0.079 | 1 | 7 | Nek6 | 0.026 | Young |
| 0.2234 | 0.699747 | 0.057 | 0.082 | 1 | 7 | Shmt1 | -0.025 | Young |
| 0.223814 | 0.696412 | 0.048 | 0.031 | 1 | 7 | Tlcd2 | 0.017 | Young |
| 0.224221 | 0.876741 | 0.057 | 0.038 | 1 | 7 | Ahcy | 0.019 | Young |
| 0.225011 | 0.747269 | 0.142 | 0.118 | 1 | 7 | Sgf29 | 0.024 | Young |
| 0.225128 | 0.431428 | 0.295 | 0.259 | 1 | 7 | Clasp2 | 0.036 | Young |
| 0.225368 | 0.263917 | 0.042 | 0.064 | 1 | 7 | Pop1 | -0.022 | Young |

| | | | | | | | | |
|---|---|---|---|---|---|---|---|---|
| 0.225791 | 0.078306 | 0.771 | 0.726 | 1 | 7 | Mafb | 0.045 | Young |
| 0.225804 | 0.138601 | 0.127 | 0.097 | 1 | 7 | R3hcc1l | 0.03 | Young |
| 0.225949 | 0.42513 | 0.214 | 0.182 | 1 | 7 | Clpx | 0.032 | Young |
| 0.226085 | 0.779248 | 0.051 | 0.033 | 1 | 7 | Zbed3 | 0.018 | Young |
| 0.226446 | 0.302443 | 0.078 | 0.105 | 1 | 7 | Pramef8 | -0.027 | Young |
| 0.226489 | 0.124319 | 0.831 | 0.838 | 1 | 7 | Fam49b | -0.007 | Young |
| 0.226598 | 0.66319 | 0.13 | 0.103 | 1 | 7 | 2610524H( | 0.027 | Young |
| 0.227539 | 0.431286 | 0.105 | 0.079 | 1 | 7 | Cib2 | 0.026 | Young |
| 0.227829 | 0.361145 | 0.163 | 0.133 | 1 | 7 | Sfmbt1 | 0.03 | Young |
| 0.228189 | 0.324313 | 0.133 | 0.169 | 1 | 7 | Yaf2 | -0.036 | Young |
| 0.228258 | 0.206095 | 0.398 | 0.369 | 1 | 7 | Safb2 | 0.029 | Young |
| 0.229509 | 0.112699 | 0.295 | 0.359 | 1 | 7 | Avpi1 | -0.064 | Young |
| 0.229679 | 0.071342 | 0.051 | 0.033 | 1 | 7 | Ptcd1 | 0.018 | Young |
| 0.229762 | 0.521801 | 0.099 | 0.074 | 1 | 7 | Plxna4os1 | 0.025 | Young |
| 0.229959 | 0.292922 | 0.473 | 0.456 | 1 | 7 | Ggh | 0.017 | Young |
| 0.230006 | 0.260332 | 0.358 | 0.338 | 1 | 7 | Nsmce1 | 0.02 | Young |
| 0.230215 | 0.030493 | 0.066 | 0.092 | 1 | 7 | Trmt13 | -0.026 | Young |
| 0.230483 | 0.421635 | 0.051 | 0.033 | 1 | 7 | Ccdc14 | 0.018 | Young |
| 0.230827 | 0.494924 | 0.25 | 0.223 | 1 | 7 | Alox5 | 0.027 | Young |
| 0.231235 | 0.275592 | 0.048 | 0.031 | 1 | 7 | Tcea3 | 0.017 | Young |
| 0.231301 | 0.089595 | 0.175 | 0.221 | 1 | 7 | Fam96b | -0.046 | Young |
| 0.23135 | 0.006365 | 0.292 | 0.338 | 1 | 7 | Otulin | -0.046 | Young |
| 0.231389 | 0.816061 | 0.069 | 0.049 | 1 | 7 | I830077J02 | 0.02 | Young |
| 0.231405 | 0.212116 | 0.175 | 0.144 | 1 | 7 | Tlr4 | 0.031 | Young |
| 0.232121 | 0.166493 | 0.102 | 0.133 | 1 | 7 | Rnf19a | -0.031 | Young |
| 0.232217 | 0.778251 | 0.06 | 0.041 | 1 | 7 | Mov10 | 0.019 | Young |
| 0.232307 | 0.042975 | 0.319 | 0.367 | 1 | 7 | Bub1b | -0.048 | Young |
| 0.232394 | 0.196147 | 0.491 | 0.451 | 1 | 7 | Ctcf | 0.04 | Young |
| 0.23251 | 1.22813 | 0.021 | 0.01 | 1 | 7 | Ttc12 | 0.011 | Young |
| 0.232754 | 0.797338 | 0.054 | 0.036 | 1 | 7 | Gm37233 | 0.018 | Young |
| 0.23289 | 0.639783 | 0.057 | 0.038 | 1 | 7 | Nectin4 | 0.019 | Young |
| 0.233343 | 0.083376 | 0.325 | 0.374 | 1 | 7 | Eif6 | -0.049 | Young |
| 0.233781 | 0.201156 | 0.015 | 0.028 | 1 | 7 | Cpt1c | -0.013 | Young |
| 0.233833 | 1.491851 | 0.021 | 0.01 | 1 | 7 | Il27 | 0.011 | Young |
| 0.234534 | 0.527518 | 0.051 | 0.033 | 1 | 7 | Tgif2 | 0.018 | Young |
| 0.235025 | 0.239658 | 0.114 | 0.087 | 1 | 7 | Slc35b3 | 0.027 | Young |
| 0.235351 | 0.373439 | 0.313 | 0.295 | 1 | 7 | Antxr2 | 0.018 | Young |
| 0.235707 | 0.249335 | 0.006 | 0.015 | 1 | 7 | Dpep3 | -0.009 | Young |
| 0.235827 | 0.466528 | 0.021 | 0.01 | 1 | 7 | Tsacc | 0.011 | Young |
| 0.235914 | 0.638281 | 0.063 | 0.044 | 1 | 7 | D930030I0 | 0.019 | Young |
| 0.236461 | 0.03979 | 0.289 | 0.341 | 1 | 7 | Ythdf2 | -0.052 | Young |
| 0.237002 | 0.002607 | 0.262 | 0.3 | 1 | 7 | Tor1a | -0.038 | Young |
| 0.237063 | 0.304453 | 0.503 | 0.479 | 1 | 7 | Crebbp | 0.024 | Young |
| 0.237164 | 0.484165 | 0.021 | 0.01 | 1 | 7 | Trmu | 0.011 | Young |
| 0.23718 | 0.078865 | 0.557 | 0.533 | 1 | 7 | Evi2a | 0.024 | Young |

| | | | | | | | | |
|---|---|---|---|---|---|---|---|---|
| 0.237912 | 0.682742 | 0.151 | 0.123 | 1 | 7 | Pex13 | 0.028 | Young |
| 0.238025 | 0.106912 | 0.084 | 0.113 | 1 | 7 | Txnrd2 | -0.029 | Young |
| 0.238117 | 0.290372 | 0.497 | 0.487 | 1 | 7 | Ppp1r15a | 0.01 | Young |
| 0.238181 | 0.804703 | 0.117 | 0.092 | 1 | 7 | Slc33a1 | 0.025 | Young |
| 0.238208 | 0.917982 | 0.06 | 0.041 | 1 | 7 | Zfp65 | 0.019 | Young |
| 0.238226 | 0.177842 | 0.13 | 0.103 | 1 | 7 | Jaml | 0.027 | Young |
| 0.238453 | 0.006752 | 0.37 | 0.431 | 1 | 7 | Lima1 | -0.061 | Young |
| 0.238505 | 0.947012 | 0.021 | 0.01 | 1 | 7 | Hist3h2ba | 0.011 | Young |
| 0.238505 | 0.499341 | 0.021 | 0.01 | 1 | 7 | Flt1 | 0.011 | Young |
| 0.238691 | 0.588432 | 0.072 | 0.051 | 1 | 7 | Abcc3 | 0.021 | Young |
| 0.238832 | 0.091931 | 0.196 | 0.238 | 1 | 7 | Rfc3 | -0.042 | Young |
| 0.239123 | 0.332132 | 0.111 | 0.085 | 1 | 7 | Eef1akmt2 | 0.026 | Young |
| 0.239447 | 0.474241 | 0.108 | 0.141 | 1 | 7 | Fech | -0.033 | Young |
| 0.239558 | 0.055102 | 0.313 | 0.374 | 1 | 7 | Rtcb | -0.061 | Young |
| 0.239654 | 0.134163 | 0.127 | 0.1 | 1 | 7 | Cnot11 | 0.027 | Young |
| 0.239754 | 0.007257 | 0.271 | 0.323 | 1 | 7 | Dhx36 | -0.052 | Young |
| 0.24001 | 0.165285 | 0.069 | 0.095 | 1 | 7 | Armt1 | -0.026 | Young |
| 0.240181 | 0.066391 | 0.202 | 0.246 | 1 | 7 | Kif3b | -0.044 | Young |
| 0.240192 | 0.578706 | 0.142 | 0.113 | 1 | 7 | Tpgs1 | 0.029 | Young |
| 0.240365 | 0.137193 | 0.437 | 0.413 | 1 | 7 | Lst1 | 0.024 | Young |
| 0.240528 | 0.410581 | 0.021 | 0.01 | 1 | 7 | Tmem221 | 0.011 | Young |
| 0.241543 | 0.866775 | 0.09 | 0.069 | 1 | 7 | Cog2 | 0.021 | Young |
| 0.241728 | 0.061098 | 0.527 | 0.462 | 1 | 7 | Cysltr1 | 0.065 | Young |
| 0.242156 | 0.292437 | 0.015 | 0.028 | 1 | 7 | Lzts2 | -0.013 | Young |
| 0.242158 | 0.199703 | 0.639 | 0.628 | 1 | 7 | Clec4a2 | 0.011 | Young |
| 0.242419 | 0.073173 | 0.133 | 0.169 | 1 | 7 | Asb8 | -0.036 | Young |
| 0.242883 | 0.324087 | 0.196 | 0.167 | 1 | 7 | Pola2 | 0.029 | Young |
| 0.243523 | 0.047497 | 0.262 | 0.313 | 1 | 7 | Mark3 | -0.051 | Young |
| 0.243589 | 0.331882 | 0.199 | 0.167 | 1 | 7 | Rufy1 | 0.032 | Young |
| 0.244407 | 0.615323 | 0.105 | 0.082 | 1 | 7 | Snx7 | 0.023 | Young |
| 0.244661 | 0.194063 | 0.416 | 0.39 | 1 | 7 | Tecr | 0.026 | Young |
| 0.244916 | 0.069706 | 0.003 | 0.01 | 1 | 7 | E130208F1 | -0.007 | Young |
| 0.245198 | 0.122305 | 0.133 | 0.169 | 1 | 7 | Esco1 | -0.036 | Young |
| 0.245211 | 1.036621 | 0.181 | 0.156 | 1 | 7 | Brca1 | 0.025 | Young |
| 0.245744 | 0.424562 | 0.205 | 0.179 | 1 | 7 | Cpne2 | 0.026 | Young |
| 0.245881 | 0.166327 | 0.551 | 0.533 | 1 | 7 | Mcm6 | 0.018 | Young |
| 0.245992 | 0.196405 | 0.476 | 0.456 | 1 | 7 | Cd300ld | 0.02 | Young |
| 0.24604 | 0.042052 | 0.166 | 0.2 | 1 | 7 | Naxd | -0.034 | Young |
| 0.246076 | 0.522667 | 0.202 | 0.174 | 1 | 7 | Rabgap1 | 0.028 | Young |
| 0.246277 | 0.024608 | 0.437 | 0.503 | 1 | 7 | Uggt1 | -0.066 | Young |
| 0.246955 | 0.321854 | 0.09 | 0.121 | 1 | 7 | Polr2d | -0.031 | Young |
| 0.247138 | 0.059779 | 0.383 | 0.426 | 1 | 7 | Cenpf | -0.043 | Young |
| 0.247187 | 0.130298 | 0.319 | 0.287 | 1 | 7 | Bcl2a1d | 0.032 | Young |
| 0.24722 | 0.004836 | 0.208 | 0.254 | 1 | 7 | Dag1 | -0.046 | Young |
| 0.247687 | 0.224089 | 0.253 | 0.223 | 1 | 7 | Pwwp2a | 0.03 | Young |

| | | | | | | | | |
|---|---|---|---|---|---|---|---|---|
| 0.247789 | 1.140692 | 0.024 | 0.013 | 1 | 7 | Zfp707 | 0.011 | Young |
| 0.247864 | 0.361094 | 0.22 | 0.19 | 1 | 7 | Runx3 | 0.03 | Young |
| 0.247867 | 0.003233 | 0.208 | 0.249 | 1 | 7 | Lrpprc | -0.041 | Young |
| 0.248004 | 0.126927 | 0.777 | 0.705 | 1 | 7 | Fosb | 0.072 | Young |
| 0.248087 | 0.331964 | 0.087 | 0.115 | 1 | 7 | Slc35a1 | -0.028 | Young |
| 0.248125 | 0.478205 | 0.226 | 0.197 | 1 | 7 | G6pdx | 0.029 | Young |
| 0.248795 | 0.985349 | 0.084 | 0.064 | 1 | 7 | Rabepk | 0.02 | Young |
| 0.248903 | 0.559727 | 0.021 | 0.036 | 1 | 7 | Rab3a | -0.015 | Young |
| 0.248952 | 0.255924 | 0.06 | 0.041 | 1 | 7 | Armc7 | 0.019 | Young |
| 0.249259 | 0.43255 | 0.096 | 0.072 | 1 | 7 | Ryk | 0.024 | Young |
| 0.249418 | 0.027532 | 0.048 | 0.069 | 1 | 7 | Rai14 | -0.021 | Young |
| 0.249596 | 0.55546 | 0.235 | 0.21 | 1 | 7 | Hmgcl | 0.025 | Young |
| 0.249698 | 0.713923 | 0.25 | 0.228 | 1 | 7 | Fam98b | 0.022 | Young |
| 0.249719 | 0.719287 | 0.214 | 0.19 | 1 | 7 | Msh2 | 0.024 | Young |
| 0.249806 | 0.057116 | 0.087 | 0.064 | 1 | 7 | Ddx31 | 0.023 | Young |
| 0.250178 | 0.248818 | 0.136 | 0.174 | 1 | 7 | Slc25a13 | -0.038 | Young |
| 0.250239 | 0.740275 | 0.286 | 0.262 | 1 | 7 | Zmym5 | 0.024 | Young |
| 0.25041 | 0.206171 | 0.033 | 0.051 | 1 | 7 | Ccdc15 | -0.018 | Young |
| 0.250956 | 0.300968 | 0.461 | 0.459 | 1 | 7 | Dnajc19 | 0.002 | Young |
| 0.251537 | 0.349692 | 0.437 | 0.415 | 1 | 7 | Cul1 | 0.022 | Young |
| 0.251545 | 1.069461 | 0.081 | 0.062 | 1 | 7 | Zhx3 | 0.019 | Young |
| 0.25177 | 0.194041 | 0.756 | 0.782 | 1 | 7 | Hint1 | -0.026 | Young |
| 0.251838 | 0.187899 | 0.714 | 0.708 | 1 | 7 | Foxp1 | 0.006 | Young |
| 0.252046 | 0.402024 | 0.268 | 0.244 | 1 | 7 | Mospd2 | 0.024 | Young |
| 0.25243 | 0.066331 | 0.919 | 0.938 | 1 | 7 | Rpl18a | -0.019 | Young |
| 0.252444 | 0.563389 | 0.331 | 0.31 | 1 | 7 | Kansl1 | 0.021 | Young |
| 0.252512 | 0.245324 | 0.211 | 0.179 | 1 | 7 | Sec23ip | 0.032 | Young |
| 0.252562 | 0.529049 | 0.214 | 0.179 | 1 | 7 | Smad1 | 0.035 | Young |
| 0.252673 | 0.116542 | 0.289 | 0.249 | 1 | 7 | Phactr2 | 0.04 | Young |
| 0.253121 | 0.123224 | 0.069 | 0.095 | 1 | 7 | Ap1s3 | -0.026 | Young |
| 0.25326 | 0.41361 | 0.238 | 0.208 | 1 | 7 | Desi2 | 0.03 | Young |
| 0.253987 | 0.93324 | 0.021 | 0.036 | 1 | 7 | Tada2a | -0.015 | Young |
| 0.254439 | 0.321753 | 0.53 | 0.518 | 1 | 7 | 2610507B1 | 0.012 | Young |
| 0.25463 | 0.950662 | 0.087 | 0.067 | 1 | 7 | Ndufaf3 | 0.02 | Young |
| 0.254969 | 0.147982 | 0.102 | 0.131 | 1 | 7 | Mrpl46 | -0.029 | Young |
| 0.254974 | 0.441279 | 0.205 | 0.172 | 1 | 7 | Vkorc1l1 | 0.033 | Young |
| 0.255063 | 0.180023 | 0.771 | 0.792 | 1 | 7 | Tln1 | -0.021 | Young |
| 0.255237 | 0.524505 | 0.169 | 0.141 | 1 | 7 | Gopc | 0.028 | Young |
| 0.255352 | 0.136541 | 0.274 | 0.238 | 1 | 7 | Sgta | 0.036 | Young |
| 0.255564 | 0.291338 | 0.377 | 0.359 | 1 | 7 | Etnk1 | 0.018 | Young |
| 0.255746 | 0.664031 | 0.148 | 0.123 | 1 | 7 | Svbp | 0.025 | Young |
| 0.255912 | 0.53809 | 0.307 | 0.282 | 1 | 7 | Ccny | 0.025 | Young |
| 0.255935 | 0.089254 | 0.081 | 0.108 | 1 | 7 | Nup188 | -0.027 | Young |
| 0.256154 | 0.868349 | 0.024 | 0.013 | 1 | 7 | Tshz2 | 0.011 | Young |
| 0.256271 | 0.097342 | 0.208 | 0.251 | 1 | 7 | Timm8a1 | -0.043 | Young |

| | | | | | | | | |
|---|---|---|---|---|---|---|---|---|
| 0.256538 | 0.41212 | 0.346 | 0.328 | 1 | 7 | Pik3cg | 0.018 | Young |
| 0.256796 | 0.089932 | 0.117 | 0.146 | 1 | 7 | Lias | -0.029 | Young |
| 0.256806 | 0.517506 | 0.024 | 0.013 | 1 | 7 | Usp35 | 0.011 | Young |
| 0.256809 | 0.243512 | 0.617 | 0.613 | 1 | 7 | Tsc22d4 | 0.004 | Young |
| 0.257041 | 0.468494 | 0.163 | 0.136 | 1 | 7 | Rtn1 | 0.027 | Young |
| 0.257429 | 0.189798 | 0.506 | 0.487 | 1 | 7 | Clint1 | 0.019 | Young |
| 0.257473 | 0.238354 | 0.057 | 0.079 | 1 | 7 | Znhit2 | -0.022 | Young |
| 0.257585 | 0.009372 | 0.114 | 0.146 | 1 | 7 | Zfp397 | -0.032 | Young |
| 0.257836 | 0.092136 | 0.843 | 0.831 | 1 | 7 | Rpl7a | 0.012 | Young |
| 0.258034 | 1.631187 | 0.027 | 0.015 | 1 | 7 | Zfp799 | 0.012 | Young |
| 0.258628 | 0.142837 | 0.054 | 0.077 | 1 | 7 | Numb | -0.023 | Young |
| 0.259842 | 0.775581 | 0.102 | 0.079 | 1 | 7 | Adamts10 | 0.023 | Young |
| 0.260142 | 0.51479 | 0.099 | 0.077 | 1 | 7 | Fam118a | 0.022 | Young |
| 0.260153 | 0.016621 | 0.154 | 0.19 | 1 | 7 | Agfg2 | -0.036 | Young |
| 0.260908 | 0.102762 | 0.193 | 0.236 | 1 | 7 | Rras | -0.043 | Young |
| 0.261179 | 0.214115 | 0.099 | 0.074 | 1 | 7 | Golm1 | 0.025 | Young |
| 0.262016 | 0.050594 | 0.44 | 0.485 | 1 | 7 | Gm26532 | -0.045 | Young |
| 0.262328 | 2.094054 | 0.027 | 0.015 | 1 | 7 | Lpcat4 | 0.012 | Young |
| 0.262359 | 0.646133 | 0.199 | 0.172 | 1 | 7 | Qrich1 | 0.027 | Young |
| 0.262956 | 0.622001 | 0.166 | 0.141 | 1 | 7 | Kdm3a | 0.025 | Young |
| 0.263141 | 0.416285 | 0.253 | 0.223 | 1 | 7 | Slc35f6 | 0.03 | Young |
| 0.263541 | 0.011511 | 0.105 | 0.136 | 1 | 7 | Cbx7 | -0.031 | Young |
| 0.263563 | 1.178251 | 0.027 | 0.015 | 1 | 7 | Zfp850 | 0.012 | Young |
| 0.263849 | 0.086706 | 0.163 | 0.2 | 1 | 7 | Wdr91 | -0.037 | Young |
| 0.264024 | 0.408322 | 0.084 | 0.113 | 1 | 7 | Crcp | -0.029 | Young |
| 0.264164 | 0.094159 | 0.063 | 0.087 | 1 | 7 | Alkbh3 | -0.024 | Young |
| 0.264183 | 1.418468 | 0.027 | 0.015 | 1 | 7 | Asah2 | 0.012 | Young |
| 0.264278 | 0.240741 | 0.593 | 0.595 | 1 | 7 | Galnt1 | -0.002 | Young |
| 0.264342 | 0.111427 | 0.843 | 0.859 | 1 | 7 | Tmem50a | -0.016 | Young |
| 0.264543 | 0.085988 | 0.919 | 0.923 | 1 | 7 | Rpl32 | -0.004 | Young |
| 0.264937 | 0.278676 | 0.449 | 0.421 | 1 | 7 | Stag1 | 0.028 | Young |
| 0.265086 | 0.337186 | 0.352 | 0.333 | 1 | 7 | Fbxl3 | 0.019 | Young |
| 0.265425 | 1.725303 | 0.027 | 0.015 | 1 | 7 | Stag3 | 0.012 | Young |
| 0.265457 | 0.089978 | 0.401 | 0.454 | 1 | 7 | Txnl1 | -0.053 | Young |
| 0.265497 | 0.242506 | 0.163 | 0.133 | 1 | 7 | Plk3 | 0.03 | Young |
| 0.265821 | 0.313474 | 0.217 | 0.259 | 1 | 7 | Vps4a | -0.042 | Young |
| 0.265854 | 0.212087 | 0.277 | 0.236 | 1 | 7 | Lcorl | 0.041 | Young |
| 0.266007 | 0.131225 | 0.75 | 0.71 | 1 | 7 | Mat2a | 0.04 | Young |
| 0.266645 | 0.096353 | 0.045 | 0.064 | 1 | 7 | Sergef | -0.019 | Young |
| 0.26668 | 0.072432 | 0.172 | 0.205 | 1 | 7 | Ero1l | -0.033 | Young |
| 0.26669 | 0.159543 | 0.364 | 0.331 | 1 | 7 | Ebp | 0.033 | Young |
| 0.266735 | 0.324371 | 0.383 | 0.372 | 1 | 7 | Tmf1 | 0.011 | Young |
| 0.266952 | 0.077108 | 0.184 | 0.223 | 1 | 7 | Amdhd2 | -0.039 | Young |
| 0.267099 | 0.053476 | 0.127 | 0.156 | 1 | 7 | Bcl2l12 | -0.029 | Young |
| 0.267642 | 0.021018 | 0.202 | 0.246 | 1 | 7 | Reps1 | -0.044 | Young |

| | | | | | | | | |
|---|---|---|---|---|---|---|---|---|
| 0.267845 | 0.133492 | 0.298 | 0.356 | 1 | 7 | Tle4 | -0.058 | Young |
| 0.26792 | 0.979161 | 0.027 | 0.015 | 1 | 7 | Atp6v1g2 | 0.012 | Young |
| 0.267989 | 0.030366 | 0.217 | 0.256 | 1 | 7 | Wipf2 | -0.039 | Young |
| 0.268016 | 0.589206 | 0.307 | 0.285 | 1 | 7 | Prrc2b | 0.022 | Young |
| 0.268148 | 0.721577 | 0.172 | 0.146 | 1 | 7 | Fem1a | 0.026 | Young |
| 0.26816 | 0.043083 | 0.283 | 0.251 | 1 | 7 | BC028528 | 0.032 | Young |
| 0.268167 | 0.130732 | 0.28 | 0.336 | 1 | 7 | Zfr | -0.056 | Young |
| 0.268235 | 0.550345 | 0.105 | 0.082 | 1 | 7 | Fanci | 0.023 | Young |
| 0.268625 | 0.035664 | 0.69 | 0.649 | 1 | 7 | Wfdc17 | 0.041 | Young |
| 0.26917 | 0.424539 | 0.304 | 0.277 | 1 | 7 | Cep83 | 0.027 | Young |
| 0.269391 | 0.009143 | 0.148 | 0.182 | 1 | 7 | Brap | -0.034 | Young |
| 0.26955 | 0.278883 | 0.181 | 0.149 | 1 | 7 | Prkar2b | 0.032 | Young |
| 0.269628 | 0.604665 | 0.235 | 0.205 | 1 | 7 | Paip1 | 0.03 | Young |
| 0.269696 | 0.145509 | 0.488 | 0.469 | 1 | 7 | Ndufs5 | 0.019 | Young |
| 0.26993 | 0.283949 | 0.358 | 0.333 | 1 | 7 | Mon2 | 0.025 | Young |
| 0.270208 | 0.163384 | 0.127 | 0.159 | 1 | 7 | Cops7a | -0.032 | Young |
| 0.270305 | 0.066271 | 0.081 | 0.108 | 1 | 7 | Nedd4l | -0.027 | Young |
| 0.270542 | 0.117571 | 0.256 | 0.305 | 1 | 7 | Rnf5 | -0.049 | Young |
| 0.270569 | 0.060903 | 0.795 | 0.749 | 1 | 7 | Ier2 | 0.046 | Young |
| 0.270665 | 0.02746 | 0.298 | 0.272 | 1 | 7 | Glrx2 | 0.026 | Young |
| 0.271044 | 0.890422 | 0.081 | 0.062 | 1 | 7 | Polr3e | 0.019 | Young |
| 0.271063 | 0.454511 | 0.027 | 0.015 | 1 | 7 | Tsix | 0.012 | Young |
| 0.271063 | 0.303022 | 0.027 | 0.015 | 1 | 7 | Dpy19l3 | 0.012 | Young |
| 0.271148 | 0.797676 | 0.181 | 0.156 | 1 | 7 | Plau | 0.025 | Young |
| 0.271469 | 0.702764 | 0.096 | 0.074 | 1 | 7 | Chd1l | 0.022 | Young |
| 0.271966 | 0.21715 | 0.714 | 0.721 | 1 | 7 | Ddx6 | -0.007 | Young |
| 0.272108 | 0.010405 | 0.012 | 0.023 | 1 | 7 | 4833445I0 | -0.011 | Young |
| 0.272327 | 0.633419 | 0.027 | 0.015 | 1 | 7 | Zfp60 | 0.012 | Young |
| 0.272664 | 0.582686 | 0.16 | 0.133 | 1 | 7 | Hinfp | 0.027 | Young |
| 0.272787 | 0.156303 | 0.012 | 0.023 | 1 | 7 | Swsap1 | -0.011 | Young |
| 0.272937 | 0.440516 | 0.105 | 0.136 | 1 | 7 | Cdkn3 | -0.031 | Young |
| 0.273033 | 0.001503 | 0.157 | 0.192 | 1 | 7 | Zfp512 | -0.035 | Young |
| 0.273122 | 0.983171 | 0.069 | 0.051 | 1 | 7 | Iqsec2 | 0.018 | Young |
| 0.273744 | 0.035439 | 0.175 | 0.213 | 1 | 7 | Xpo4 | -0.038 | Young |
| 0.274191 | 0.05036 | 0.458 | 0.423 | 1 | 7 | Tceal9 | 0.035 | Young |
| 0.274193 | 0.195236 | 0.196 | 0.244 | 1 | 7 | Nelfcd | -0.048 | Young |
| 0.274284 | 0.669483 | 0.087 | 0.067 | 1 | 7 | Dnase1l3 | 0.02 | Young |
| 0.274329 | 1.10565 | 0.108 | 0.087 | 1 | 7 | Maoa | 0.021 | Young |
| 0.27438 | 0.170647 | 0.072 | 0.097 | 1 | 7 | Hist1h1d | -0.025 | Young |
| 0.274652 | 0.300815 | 0.211 | 0.182 | 1 | 7 | Slc35b1 | 0.029 | Young |
| 0.275001 | 0.059706 | 0.19 | 0.228 | 1 | 7 | Phf11b | -0.038 | Young |
| 0.275031 | 0.167431 | 0.53 | 0.508 | 1 | 7 | Mapk1ip1l | 0.022 | Young |
| 0.275031 | 0.045667 | 0.602 | 0.572 | 1 | 7 | Cdc42se1 | 0.03 | Young |
| 0.275467 | 0.136425 | 0.669 | 0.715 | 1 | 7 | Ifrd1 | -0.046 | Young |
| 0.275504 | 0.380486 | 0.027 | 0.015 | 1 | 7 | Lamb3 | 0.012 | Young |

| | | | | | | | | |
|---|---|---|---|---|---|---|---|---|
| 0.275504 | 0.346041 | 0.027 | 0.015 | 1 | 7 | Siglecf | 0.012 | Young |
| 0.275923 | 0.046496 | 0.217 | 0.185 | 1 | 7 | Tdrd7 | 0.032 | Young |
| 0.27609 | 0.881037 | 0.084 | 0.064 | 1 | 7 | Gfm2 | 0.02 | Young |
| 0.276126 | 1.710609 | 0.036 | 0.023 | 1 | 7 | Zfp580 | 0.013 | Young |
| 0.276899 | 0.702596 | 0.072 | 0.054 | 1 | 7 | Snx21 | 0.018 | Young |
| 0.277082 | 0.726151 | 0.117 | 0.095 | 1 | 7 | Mkrn2 | 0.022 | Young |
| 0.277123 | 0.473071 | 0.06 | 0.082 | 1 | 7 | Gm16536 | -0.022 | Young |
| 0.277563 | 0.199026 | 0.494 | 0.479 | 1 | 7 | Ssr4 | 0.015 | Young |
| 0.278265 | 0.468436 | 0.367 | 0.351 | 1 | 7 | Fndc3b | 0.016 | Young |
| 0.278803 | 0.41047 | 0.241 | 0.215 | 1 | 7 | Rab12 | 0.026 | Young |
| 0.278902 | 0.545554 | 0.123 | 0.1 | 1 | 7 | Kat6b | 0.023 | Young |
| 0.279126 | 0.720741 | 0.151 | 0.126 | 1 | 7 | Gmeb1 | 0.025 | Young |
| 0.279489 | 0.227515 | 0.554 | 0.528 | 1 | 7 | Sec62 | 0.026 | Young |
| 0.279665 | 0.907673 | 0.078 | 0.059 | 1 | 7 | Ttc38 | 0.019 | Young |
| 0.279871 | 0.13636 | 0.148 | 0.185 | 1 | 7 | Tbc1d9b | -0.037 | Young |
| 0.279915 | 0.681115 | 0.081 | 0.062 | 1 | 7 | Slc44a2 | 0.019 | Young |
| 0.280073 | 0.109029 | 0.919 | 0.887 | 1 | 7 | Fos | 0.032 | Young |
| 0.280411 | 0.346403 | 0.105 | 0.082 | 1 | 7 | Bach2 | 0.023 | Young |
| 0.280606 | 0.060748 | 0.151 | 0.182 | 1 | 7 | Snx15 | -0.031 | Young |
| 0.280634 | 0.566606 | 0.093 | 0.072 | 1 | 7 | Erich1 | 0.021 | Young |
| 0.2807 | 0.213291 | 0.274 | 0.244 | 1 | 7 | Hspa4l | 0.03 | Young |
| 0.280759 | 0.517692 | 0.515 | 0.523 | 1 | 7 | Hmgn1 | -0.008 | Young |
| 0.280816 | 0.523686 | 0.09 | 0.069 | 1 | 7 | Lilra5 | 0.021 | Young |
| 0.280867 | 0.89623 | 0.03 | 0.046 | 1 | 7 | Fam206a | -0.016 | Young |
| 0.281607 | 0.121155 | 0.343 | 0.315 | 1 | 7 | Phc2 | 0.028 | Young |
| 0.281843 | 1.127774 | 0.03 | 0.018 | 1 | 7 | 2610044O1 | 0.012 | Young |
| 0.281897 | 0.079947 | 0.154 | 0.187 | 1 | 7 | Commd10 | -0.033 | Young |
| 0.281912 | 0.006442 | 0.19 | 0.226 | 1 | 7 | Srek1ip1 | -0.036 | Young |
| 0.281964 | 0.12851 | 0.741 | 0.764 | 1 | 7 | Cyfip1 | -0.023 | Young |
| 0.282008 | 0.706814 | 0.199 | 0.172 | 1 | 7 | Klf11 | 0.027 | Young |
| 0.282219 | 0.768236 | 0.051 | 0.072 | 1 | 7 | Lrrc45 | -0.021 | Young |
| 0.282483 | 0.170958 | 0.593 | 0.585 | 1 | 7 | Arhgap18 | 0.008 | Young |
| 0.282534 | 0.659428 | 0.211 | 0.19 | 1 | 7 | Trip13 | 0.021 | Young |
| 0.282719 | 0.739234 | 0.196 | 0.174 | 1 | 7 | Abcc5 | 0.022 | Young |
| 0.283635 | 0.470552 | 0.175 | 0.149 | 1 | 7 | Sp140 | 0.026 | Young |
| 0.283681 | 0.383317 | 0.328 | 0.297 | 1 | 7 | Tnfrsf13b | 0.031 | Young |
| 0.283751 | 0.604169 | 0.072 | 0.054 | 1 | 7 | Ap3m2 | 0.018 | Young |
| 0.284342 | 1.189139 | 0.036 | 0.023 | 1 | 7 | Klri2 | 0.013 | Young |
| 0.284373 | 0.98118 | 0.06 | 0.044 | 1 | 7 | Crocc | 0.016 | Young |
| 0.28513 | 0.214989 | 0.169 | 0.203 | 1 | 7 | Trmt1l | -0.034 | Young |
| 0.285626 | 0.459851 | 0.084 | 0.064 | 1 | 7 | BC055324 | 0.02 | Young |
| 0.285695 | 0.579408 | 0.081 | 0.062 | 1 | 7 | Leng1 | 0.019 | Young |
| 0.285895 | 0.91017 | 0.033 | 0.021 | 1 | 7 | Galnt12 | 0.012 | Young |
| 0.285947 | 0.031799 | 0.19 | 0.231 | 1 | 7 | Trak2 | -0.041 | Young |
| 0.286005 | 0.940978 | 0.036 | 0.023 | 1 | 7 | Zfp949 | 0.013 | Young |

| | | | | | | | | |
|---|---|---|---|---|---|---|---|---|
| 0.286131 | 0.46348 | 0.03 | 0.018 | 1 | 7 | Trim14 | 0.012 | Young |
| 0.286471 | 0.527943 | 0.178 | 0.154 | 1 | 7 | Tpst1 | 0.024 | Young |
| 0.286548 | 0.575036 | 0.09 | 0.118 | 1 | 7 | Hsdl2 | -0.028 | Young |
| 0.286713 | 0.18466 | 0.163 | 0.197 | 1 | 7 | Orai3 | -0.034 | Young |
| 0.286757 | 0.075676 | 0.181 | 0.221 | 1 | 7 | Kif2c | -0.04 | Young |
| 0.287035 | 0.762624 | 0.078 | 0.059 | 1 | 7 | Zbtb46 | 0.019 | Young |
| 0.287243 | 0.558393 | 0.105 | 0.136 | 1 | 7 | Gtf3c3 | -0.031 | Young |
| 0.287582 | 0.129886 | 0.642 | 0.646 | 1 | 7 | Glg1 | -0.004 | Young |
| 0.287804 | 0.265729 | 0.151 | 0.123 | 1 | 7 | Zfp281 | 0.028 | Young |
| 0.287994 | 0.321358 | 0.133 | 0.108 | 1 | 7 | Trim36 | 0.025 | Young |
| 0.288104 | 0.109867 | 0.166 | 0.203 | 1 | 7 | Ubxn2a | -0.037 | Young |
| 0.28818 | 0.343771 | 0.256 | 0.236 | 1 | 7 | Bet1 | 0.02 | Young |
| 0.288234 | 0.880284 | 0.033 | 0.021 | 1 | 7 | Lig4 | 0.012 | Young |
| 0.288377 | 0.157158 | 0.075 | 0.097 | 1 | 7 | Pstk | -0.022 | Young |
| 0.28934 | 0.666228 | 0.075 | 0.056 | 1 | 7 | Gngt2 | 0.019 | Young |
| 0.289608 | 0.267061 | 0.575 | 0.569 | 1 | 7 | Kpnb1 | 0.006 | Young |
| 0.289842 | 0.608286 | 0.03 | 0.018 | 1 | 7 | Gm19951 | 0.012 | Young |
| 0.290056 | 0.018593 | 0.407 | 0.456 | 1 | 7 | Copb1 | -0.049 | Young |
| 0.29006 | 0.12967 | 0.045 | 0.064 | 1 | 7 | Plekha8 | -0.019 | Young |
| 0.290352 | 0.986166 | 0.066 | 0.049 | 1 | 7 | Taf8 | 0.017 | Young |
| 0.291568 | 0.409179 | 0.247 | 0.223 | 1 | 7 | Ankle2 | 0.024 | Young |
| 0.29162 | 0.141646 | 0.295 | 0.346 | 1 | 7 | Mdm2 | -0.051 | Young |
| 0.29171 | 0.763493 | 0.03 | 0.018 | 1 | 7 | Grasp | 0.012 | Young |
| 0.29171 | 0.389945 | 0.03 | 0.018 | 1 | 7 | Grhpr | 0.012 | Young |
| 0.291804 | 0.012792 | 0.157 | 0.19 | 1 | 7 | Lrrc41 | -0.033 | Young |
| 0.291913 | 0.332769 | 0.377 | 0.354 | 1 | 7 | Mat2b | 0.023 | Young |
| 0.291937 | 0.326039 | 0.377 | 0.351 | 1 | 7 | Aida | 0.026 | Young |
| 0.292159 | 0.795459 | 0.036 | 0.023 | 1 | 7 | Trim7 | 0.013 | Young |
| 0.292293 | 0.796456 | 0.081 | 0.062 | 1 | 7 | Arfgap3 | 0.019 | Young |
| 0.29274 | 0.536668 | 0.205 | 0.177 | 1 | 7 | 4833420G1 | 0.028 | Young |
| 0.292764 | 0.043856 | 0.693 | 0.713 | 1 | 7 | Ccdc50 | -0.02 | Young |
| 0.293051 | 0.242321 | 0.434 | 0.415 | 1 | 7 | Pkn1 | 0.019 | Young |
| 0.293284 | 0.375434 | 0.328 | 0.305 | 1 | 7 | Araf | 0.023 | Young |
| 0.293542 | 0.377669 | 0.033 | 0.021 | 1 | 7 | Hdhd5 | 0.012 | Young |
| 0.293743 | 0.005716 | 0.262 | 0.295 | 1 | 7 | Rbmxl1 | -0.033 | Young |
| 0.293853 | 0.867991 | 0.036 | 0.023 | 1 | 7 | Wdr34 | 0.013 | Young |
| 0.294041 | 0.041593 | 0.295 | 0.341 | 1 | 7 | Pptc7 | -0.046 | Young |
| 0.294956 | 0.244359 | 0.464 | 0.444 | 1 | 7 | Ccdc12 | 0.02 | Young |
| 0.294967 | 1.053397 | 0.045 | 0.031 | 1 | 7 | Chst14 | 0.014 | Young |
| 0.294989 | 0.195966 | 0.093 | 0.072 | 1 | 7 | Pank1 | 0.021 | Young |
| 0.295258 | 0.350883 | 0.081 | 0.062 | 1 | 7 | Zbtb16 | 0.019 | Young |
| 0.29529 | 0.163604 | 0.316 | 0.282 | 1 | 7 | Uap1 | 0.034 | Young |
| 0.29533 | 0.717989 | 0.178 | 0.154 | 1 | 7 | Snx29 | 0.024 | Young |
| 0.295436 | 0.126493 | 0.536 | 0.523 | 1 | 7 | Hmgn2 | 0.013 | Young |
| 0.295496 | 0.223997 | 0.166 | 0.141 | 1 | 7 | Brcc3 | 0.025 | Young |

| | | | | | | | | |
|---|---|---|---|---|---|---|---|---|
| 0.295626 | 0.463996 | 0.31 | 0.297 | 1 | 7 | Mtmr6 | 0.013 | Young |
| 0.295867 | 0.470649 | 0.075 | 0.056 | 1 | 7 | Hmgxb3 | 0.019 | Young |
| 0.295891 | 0.56499 | 0.039 | 0.056 | 1 | 7 | Aptx | -0.017 | Young |
| 0.296108 | 0.208853 | 0.229 | 0.274 | 1 | 7 | Mta1 | -0.045 | Young |
| 0.296112 | 0.196656 | 0.148 | 0.121 | 1 | 7 | 1190002N1 | 0.027 | Young |
| 0.296336 | 0.270127 | 0.039 | 0.056 | 1 | 7 | Ip6k2 | -0.017 | Young |
| 0.296342 | 0.851605 | 0.054 | 0.038 | 1 | 7 | Palb2 | 0.016 | Young |
| 0.296476 | 0.61112 | 0.045 | 0.031 | 1 | 7 | Dbp | 0.014 | Young |
| 0.296823 | 0.929257 | 0.039 | 0.026 | 1 | 7 | Uhrf1bp1 | 0.013 | Young |
| 0.297925 | 0.315021 | 0.133 | 0.108 | 1 | 7 | Ccdc117 | 0.025 | Young |
| 0.298107 | 0.896228 | 0.16 | 0.136 | 1 | 7 | Gmcl1 | 0.024 | Young |
| 0.298119 | 0.261745 | 0.178 | 0.149 | 1 | 7 | Hist1h2ae | 0.029 | Young |
| 0.298295 | 0.209062 | 0.151 | 0.123 | 1 | 7 | Mfap1b | 0.028 | Young |
| 0.298318 | 0.690011 | 0.105 | 0.085 | 1 | 7 | Zfp398 | 0.02 | Young |
| 0.298548 | 0.15923 | 0.63 | 0.633 | 1 | 7 | Dock10 | -0.003 | Young |
| 0.298663 | 0.192004 | 0.524 | 0.51 | 1 | 7 | Mindy2 | 0.014 | Young |
| 0.298734 | 0.206128 | 0.322 | 0.3 | 1 | 7 | Gtf2a2 | 0.022 | Young |
| 0.298894 | 0.26028 | 0.208 | 0.251 | 1 | 7 | Cops8 | -0.043 | Young |
| 0.298918 | 0.019528 | 0.316 | 0.359 | 1 | 7 | Rsbn1l | -0.043 | Young |
| 0.29897 | 0.4398 | 0.313 | 0.292 | 1 | 7 | Sft2d2 | 0.021 | Young |
| 0.298973 | 0.216841 | 0.036 | 0.023 | 1 | 7 | Arl6 | 0.013 | Young |
| 0.299037 | 0.865072 | 0.099 | 0.079 | 1 | 7 | C1qtnf1 | 0.02 | Young |
| 0.29951 | 0.063497 | 0.217 | 0.256 | 1 | 7 | Tssc4 | -0.039 | Young |
| 0.29975 | 0.128153 | 0.22 | 0.19 | 1 | 7 | Scarb1 | 0.03 | Young |
| 0.299897 | 0.317963 | 0.334 | 0.303 | 1 | 7 | Uba52 | 0.031 | Young |
| 0.29995 | 0.090621 | 0.033 | 0.049 | 1 | 7 | Kdm5b | -0.016 | Young |
| 0.300105 | 1.045541 | 0.039 | 0.026 | 1 | 7 | Nod2 | 0.013 | Young |
| 0.300118 | 0.156228 | 0.033 | 0.021 | 1 | 7 | Sdk1 | 0.012 | Young |
| 0.300128 | 0.619136 | 0.142 | 0.118 | 1 | 7 | Ttk | 0.024 | Young |
| 0.300526 | 0.829961 | 0.045 | 0.031 | 1 | 7 | Zswim7 | 0.014 | Young |
| 0.300605 | 0.042442 | 0.181 | 0.218 | 1 | 7 | Slc41a3 | -0.037 | Young |
| 0.300693 | 0.393479 | 0.036 | 0.023 | 1 | 7 | Cep97 | 0.013 | Young |
| 0.300871 | 0.385477 | 0.108 | 0.087 | 1 | 7 | Crip2 | 0.021 | Young |
| 0.300941 | 0.650333 | 0.054 | 0.038 | 1 | 7 | Zfp944 | 0.016 | Young |
| 0.301266 | 0.742186 | 0.042 | 0.028 | 1 | 7 | Plekhh3 | 0.014 | Young |
| 0.301359 | 0.390527 | 0.256 | 0.228 | 1 | 7 | Dcaf15 | 0.028 | Young |
| 0.301594 | 0.02348 | 0.419 | 0.482 | 1 | 7 | Aph1a | -0.063 | Young |
| 0.30179 | 0.217241 | 0.458 | 0.441 | 1 | 7 | Myo1f | 0.017 | Young |
| 0.301924 | 0.119558 | 0.295 | 0.336 | 1 | 7 | Agfg1 | -0.041 | Young |
| 0.302388 | 0.081829 | 0.503 | 0.551 | 1 | 7 | Zfp106 | -0.048 | Young |
| 0.302549 | 0.151628 | 0.172 | 0.21 | 1 | 7 | Mitf | -0.038 | Young |
| 0.302609 | 0.521958 | 0.057 | 0.041 | 1 | 7 | Iqce | 0.016 | Young |
| 0.302609 | 0.290837 | 0.057 | 0.041 | 1 | 7 | Stau2 | 0.016 | Young |
| 0.302715 | 0.022767 | 0.928 | 0.923 | 1 | 7 | Hnrnpf | 0.005 | Young |
| 0.30297 | 0.519324 | 0.346 | 0.315 | 1 | 7 | Ap3b1 | 0.031 | Young |

| | | | | | | | | |
|---|---|---|---|---|---|---|---|---|
| 0.302981 | 0.486703 | 0.072 | 0.054 | 1 | 7 | Mib2 | 0.018 | Young |
| 0.303201 | 0.34776 | 0.151 | 0.126 | 1 | 7 | Fasn | 0.025 | Young |
| 0.303602 | 0.08506 | 0.178 | 0.213 | 1 | 7 | Tubgcp4 | -0.035 | Young |
| 0.303633 | 0.296237 | 0.5 | 0.467 | 1 | 7 | Sgk3 | 0.033 | Young |
| 0.303702 | 0.006129 | 0.211 | 0.246 | 1 | 7 | Pafah1b2 | -0.035 | Young |
| 0.303923 | 1.736811 | 0.012 | 0.005 | 1 | 7 | 1700010I1 | 0.007 | Young |
| 0.303923 | 1.185634 | 0.012 | 0.005 | 1 | 7 | Tnnt3 | 0.007 | Young |
| 0.303923 | 1.006249 | 0.012 | 0.005 | 1 | 7 | Il17rc | 0.007 | Young |
| 0.304307 | 0.543196 | 0.157 | 0.133 | 1 | 7 | Sdccag8 | 0.024 | Young |
| 0.304373 | 0.261332 | 0.584 | 0.587 | 1 | 7 | Slc3a2 | -0.003 | Young |
| 0.304417 | 0.026791 | 0.039 | 0.056 | 1 | 7 | Sh3rf1 | -0.017 | Young |
| 0.304713 | 0.719512 | 0.187 | 0.162 | 1 | 7 | Pilrb1 | 0.025 | Young |
| 0.304804 | 0.04967 | 0.232 | 0.267 | 1 | 7 | Mrps9 | -0.035 | Young |
| 0.30526 | 0.004883 | 0.515 | 0.549 | 1 | 7 | Dck | -0.034 | Young |
| 0.30527 | 0.710046 | 0.069 | 0.051 | 1 | 7 | Parp6 | 0.018 | Young |
| 0.30529 | 0.213028 | 0.178 | 0.21 | 1 | 7 | Zufsp | -0.032 | Young |
| 0.305415 | 0.175044 | 0.322 | 0.364 | 1 | 7 | Elk4 | -0.042 | Young |
| 0.305562 | 0.240471 | 0.048 | 0.033 | 1 | 7 | Zfp946 | 0.015 | Young |
| 0.305785 | 0.077327 | 0.084 | 0.108 | 1 | 7 | Zswim4 | -0.024 | Young |
| 0.305853 | 0.047537 | 0.301 | 0.351 | 1 | 7 | Oser1 | -0.05 | Young |
| 0.30607 | 1.502563 | 0.012 | 0.005 | 1 | 7 | Aatk | 0.007 | Young |
| 0.30607 | 1.318464 | 0.012 | 0.005 | 1 | 7 | Gm27003 | 0.007 | Young |
| 0.30607 | 1.295329 | 0.012 | 0.005 | 1 | 7 | Ccdc36 | 0.007 | Young |
| 0.30607 | 1.013445 | 0.012 | 0.005 | 1 | 7 | Gm3604 | 0.007 | Young |
| 0.30607 | 0.650863 | 0.012 | 0.005 | 1 | 7 | Zfp30 | 0.007 | Young |
| 0.306114 | 0.098932 | 0.187 | 0.223 | 1 | 7 | Lats1 | -0.036 | Young |
| 0.306176 | 0.487322 | 0.111 | 0.09 | 1 | 7 | Lrrc57 | 0.021 | Young |
| 0.306382 | 0.685292 | 0.157 | 0.133 | 1 | 7 | Pogz | 0.024 | Young |
| 0.306439 | 0.152388 | 0.142 | 0.172 | 1 | 7 | Lactb2 | -0.03 | Young |
| 0.306579 | 0.524049 | 0.19 | 0.167 | 1 | 7 | Vps9d1 | 0.023 | Young |
| 0.306717 | 0.194022 | 0.72 | 0.715 | 1 | 7 | Tnfaip3 | 0.005 | Young |
| 0.307118 | 0.63123 | 0.042 | 0.028 | 1 | 7 | Zfp40 | 0.014 | Young |
| 0.307148 | 1.084821 | 0.012 | 0.005 | 1 | 7 | Gm7324 | 0.007 | Young |
| 0.307148 | 0.642082 | 0.012 | 0.005 | 1 | 7 | Noxo1 | 0.007 | Young |
| 0.307148 | 0.555887 | 0.012 | 0.005 | 1 | 7 | Chrna5 | 0.007 | Young |
| 0.307249 | 0.425694 | 0.265 | 0.238 | 1 | 7 | Clip1 | 0.027 | Young |
| 0.307305 | 0.041272 | 0.259 | 0.303 | 1 | 7 | Tmem43 | -0.044 | Young |
| 0.307708 | 0.131248 | 0.096 | 0.123 | 1 | 7 | Fbxo8 | -0.027 | Young |
| 0.307777 | 0.303839 | 0.404 | 0.39 | 1 | 7 | Mbd2 | 0.014 | Young |
| 0.307847 | 0.476571 | 0.265 | 0.244 | 1 | 7 | Atxn7 | 0.021 | Young |
| 0.307858 | 0.295962 | 0.515 | 0.518 | 1 | 7 | Fgfr1 | -0.003 | Young |
| 0.307883 | 0.1674 | 0.208 | 0.246 | 1 | 7 | Ptbp2 | -0.038 | Young |
| 0.307965 | 0.235722 | 0.157 | 0.128 | 1 | 7 | Gan | 0.029 | Young |
| 0.308227 | 1.172177 | 0.012 | 0.005 | 1 | 7 | Atp1a2 | 0.007 | Young |
| 0.308227 | 0.846339 | 0.012 | 0.005 | 1 | 7 | Gm49087 | 0.007 | Young |

| | | | | | | | | |
|---|---|---|---|---|---|---|---|---|
| 0.308227 | 0.836417 | 0.012 | 0.005 | 1 | 7 | Ppbp | 0.007 | Young |
| 0.308227 | 0.8226 | 0.012 | 0.005 | 1 | 7 | Zfp169 | 0.007 | Young |
| 0.308227 | 0.795621 | 0.012 | 0.005 | 1 | 7 | Rbm44 | 0.007 | Young |
| 0.308227 | 0.773397 | 0.012 | 0.005 | 1 | 7 | Gm13205 | 0.007 | Young |
| 0.308227 | 0.693451 | 0.012 | 0.005 | 1 | 7 | Gm26947 | 0.007 | Young |
| 0.308227 | 0.582688 | 0.012 | 0.005 | 1 | 7 | Rdh12 | 0.007 | Young |
| 0.308227 | 0.524015 | 0.012 | 0.005 | 1 | 7 | 3010003L2 | 0.007 | Young |
| 0.308227 | 0.482345 | 0.012 | 0.005 | 1 | 7 | Gm40304 | 0.007 | Young |
| 0.308514 | 0.042573 | 0.316 | 0.369 | 1 | 7 | Rif1 | -0.053 | Young |
| 0.308806 | 0.139388 | 0.036 | 0.023 | 1 | 7 | Susd1 | 0.013 | Young |
| 0.308855 | 0.330831 | 0.178 | 0.151 | 1 | 7 | Nosip | 0.027 | Young |
| 0.308991 | 0.304084 | 0.069 | 0.051 | 1 | 7 | Nsdhl | 0.018 | Young |
| 0.309053 | 0.265494 | 0.048 | 0.033 | 1 | 7 | 2310040G2 | 0.015 | Young |
| 0.30931 | 0.421963 | 0.012 | 0.005 | 1 | 7 | Gm47230 | 0.007 | Young |
| 0.30931 | 0.418311 | 0.012 | 0.005 | 1 | 7 | Cfap126 | 0.007 | Young |
| 0.30931 | 0.284828 | 0.012 | 0.005 | 1 | 7 | Paqr3 | 0.007 | Young |
| 0.309495 | 0.88522 | 0.087 | 0.069 | 1 | 7 | Kank2 | 0.018 | Young |
| 0.309921 | 0.396666 | 0.259 | 0.231 | 1 | 7 | Ehmt1 | 0.028 | Young |
| 0.309964 | 0.7032 | 0.163 | 0.138 | 1 | 7 | Actr1b | 0.025 | Young |
| 0.310123 | 0.517848 | 0.244 | 0.223 | 1 | 7 | Fads1 | 0.021 | Young |
| 0.310344 | 0.422329 | 0.039 | 0.056 | 1 | 7 | Rpap2 | -0.017 | Young |
| 0.310394 | 0.574061 | 0.012 | 0.005 | 1 | 7 | Ogn | 0.007 | Young |
| 0.310394 | 0.513662 | 0.012 | 0.005 | 1 | 7 | Il2ra | 0.007 | Young |
| 0.310394 | 0.446185 | 0.012 | 0.005 | 1 | 7 | Adamts6 | 0.007 | Young |
| 0.310881 | 0.517909 | 0.042 | 0.028 | 1 | 7 | Ddias | 0.014 | Young |
| 0.310959 | 0.085688 | 0.09 | 0.115 | 1 | 7 | Top3b | -0.025 | Young |
| 0.311176 | 0.483434 | 0.051 | 0.036 | 1 | 7 | 5730409E0 | 0.015 | Young |
| 0.311482 | 0.11361 | 0.012 | 0.005 | 1 | 7 | Il1r2 | 0.007 | Young |
| 0.312045 | 0.333569 | 0.175 | 0.215 | 1 | 7 | Mrps26 | -0.04 | Young |
| 0.312297 | 0.422082 | 0.154 | 0.187 | 1 | 7 | 2610037D0 | -0.033 | Young |
| 0.312419 | 0.638559 | 0.108 | 0.087 | 1 | 7 | Zdhhc14 | 0.021 | Young |
| 0.312503 | 0.453631 | 0.042 | 0.028 | 1 | 7 | Cmtr2 | 0.014 | Young |
| 0.312904 | 0.310182 | 0.066 | 0.049 | 1 | 7 | Wdr47 | 0.017 | Young |
| 0.312977 | 0.282555 | 0.25 | 0.226 | 1 | 7 | Rasgef1b | 0.024 | Young |
| 0.313455 | 1.072359 | 0.099 | 0.079 | 1 | 7 | Accs | 0.02 | Young |
| 0.313561 | 0.90576 | 0.133 | 0.11 | 1 | 7 | Rcor3 | 0.023 | Young |
| 0.313593 | 0.038325 | 0.214 | 0.249 | 1 | 7 | Clasp1 | -0.035 | Young |
| 0.313841 | 0.054624 | 0.925 | 0.921 | 1 | 7 | Rplp0 | 0.004 | Young |
| 0.314073 | 0.401246 | 0.054 | 0.038 | 1 | 7 | Sh2d3c | 0.016 | Young |
| 0.31413 | 0.002701 | 0.042 | 0.028 | 1 | 7 | Zfp958 | 0.014 | Young |
| 0.314218 | 0.096478 | 0.232 | 0.277 | 1 | 7 | Slfn9 | -0.045 | Young |
| 0.314234 | 0.151928 | 0.304 | 0.349 | 1 | 7 | Utp11 | -0.045 | Young |
| 0.314234 | 0.191815 | 0.229 | 0.205 | 1 | 7 | Pde3b | 0.024 | Young |
| 0.314241 | 0.981941 | 0.009 | 0.018 | 1 | 7 | Cped1 | -0.009 | Young |
| 0.314298 | 0.332444 | 0.096 | 0.123 | 1 | 7 | Slc39a11 | -0.027 | Young |

| | | | | | | | | |
|---|---|---|---|---|---|---|---|---|
| 0.314478 | 0.32604 | 0.352 | 0.336 | 1 | 7 | Rb1 | 0.016 | Young |
| 0.314662 | 0.163954 | 0.111 | 0.141 | 1 | 7 | Fam110a | -0.03 | Young |
| 0.314767 | 0.068943 | 0.13 | 0.159 | 1 | 7 | Cpox | -0.029 | Young |
| 0.314824 | 0.034499 | 0.611 | 0.582 | 1 | 7 | Aes | 0.029 | Young |
| 0.314836 | 0.087571 | 0.078 | 0.103 | 1 | 7 | Slc27a4 | -0.025 | Young |
| 0.314914 | 0.401603 | 0.205 | 0.185 | 1 | 7 | Prkce | 0.02 | Young |
| 0.315218 | 0.160447 | 0.027 | 0.041 | 1 | 7 | Fbxw9 | -0.014 | Young |
| 0.315367 | 0.519956 | 0.13 | 0.108 | 1 | 7 | Rcbtb2 | 0.022 | Young |
| 0.315371 | 0.225024 | 0.47 | 0.456 | 1 | 7 | Sqstm1 | 0.014 | Young |
| 0.315872 | 0.411375 | 0.361 | 0.346 | 1 | 7 | Tprgl | 0.015 | Young |
| 0.315891 | 0.375958 | 0.205 | 0.177 | 1 | 7 | Asl | 0.028 | Young |
| 0.315919 | 0.087799 | 0.383 | 0.451 | 1 | 7 | Rhob | -0.068 | Young |
| 0.315945 | 0.364125 | 0.009 | 0.018 | 1 | 7 | 1110035H1 | -0.009 | Young |
| 0.316622 | 0.099781 | 0.352 | 0.397 | 1 | 7 | Prdx6 | -0.045 | Young |
| 0.316684 | 0.053816 | 0.048 | 0.067 | 1 | 7 | Mtrf1 | -0.019 | Young |
| 0.317057 | 0.334142 | 0.051 | 0.036 | 1 | 7 | Lnx2 | 0.015 | Young |
| 0.317087 | 0.174457 | 0.084 | 0.064 | 1 | 7 | Spata5 | 0.02 | Young |
| 0.317147 | 0.314408 | 0.063 | 0.046 | 1 | 7 | Dnmbp | 0.017 | Young |
| 0.317274 | 0.066316 | 0.518 | 0.585 | 1 | 7 | Gnaq | -0.067 | Young |
| 0.31728 | 0.206527 | 0.458 | 0.444 | 1 | 7 | Ggnbp2 | 0.014 | Young |
| 0.317399 | 0.006827 | 0.265 | 0.3 | 1 | 7 | Mtmr3 | -0.035 | Young |
| 0.317709 | 0.310367 | 0.111 | 0.09 | 1 | 7 | Lsm11 | 0.021 | Young |
| 0.317933 | 0.206582 | 0.539 | 0.526 | 1 | 7 | Ndufv3 | 0.013 | Young |
| 0.318044 | 0.275162 | 0.051 | 0.036 | 1 | 7 | Hoxb4 | 0.015 | Young |
| 0.318104 | 0.566026 | 0.105 | 0.085 | 1 | 7 | Osbpl2 | 0.02 | Young |
| 0.318132 | 0.029032 | 0.084 | 0.108 | 1 | 7 | Impa2 | -0.024 | Young |
| 0.318661 | 0.050222 | 0.048 | 0.033 | 1 | 7 | Myb | 0.015 | Young |
| 0.318807 | 0.030135 | 0.364 | 0.418 | 1 | 7 | Lactb | -0.054 | Young |
| 0.318911 | 0.283081 | 0.163 | 0.136 | 1 | 7 | Khnyn | 0.027 | Young |
| 0.319143 | 0.385951 | 0.057 | 0.041 | 1 | 7 | Twnk | 0.016 | Young |
| 0.319154 | 0.186161 | 0.539 | 0.531 | 1 | 7 | Cox14 | 0.008 | Young |
| 0.320696 | 0.572544 | 0.027 | 0.041 | 1 | 7 | Nsun5 | -0.014 | Young |
| 0.320802 | 0.139826 | 0.069 | 0.09 | 1 | 7 | Traf3ip3 | -0.021 | Young |
| 0.32126 | 0.443423 | 0.142 | 0.118 | 1 | 7 | Zdhhc18 | 0.024 | Young |
| 0.321391 | 0.456492 | 0.111 | 0.09 | 1 | 7 | Pnpo | 0.021 | Young |
| 0.321827 | 0.710886 | 0.117 | 0.097 | 1 | 7 | Toe1 | 0.02 | Young |
| 0.321912 | 0.430379 | 0.139 | 0.172 | 1 | 7 | Gpn3 | -0.033 | Young |
| 0.321954 | 0.233891 | 0.051 | 0.069 | 1 | 7 | Snapc2 | -0.018 | Young |
| 0.32267 | 0.093359 | 0.599 | 0.61 | 1 | 7 | Tmem256 | -0.011 | Young |
| 0.32267 | 0.247234 | 0.117 | 0.146 | 1 | 7 | Fam60a | -0.029 | Young |
| 0.322737 | 0.407622 | 0.111 | 0.09 | 1 | 7 | Dip2a | 0.021 | Young |
| 0.322781 | 0.323321 | 0.334 | 0.308 | 1 | 7 | Uba6 | 0.026 | Young |
| 0.323557 | 0.05079 | 0.12 | 0.149 | 1 | 7 | Fam220a | -0.029 | Young |
| 0.323776 | 0.051377 | 0.075 | 0.056 | 1 | 7 | Sco2 | 0.019 | Young |
| 0.324323 | 0.193747 | 0.657 | 0.644 | 1 | 7 | Pepd | 0.013 | Young |

| | | | | | | | | |
|---|---|---|---|---|---|---|---|---|
| 0.324487 | 0.054951 | 0.042 | 0.059 | 1 | 7 | Nme6 | -0.017 | Young |
| 0.324777 | 0.790185 | 0.015 | 0.026 | 1 | 7 | Gm26982 | -0.011 | Young |
| 0.325103 | 0.192085 | 0.337 | 0.385 | 1 | 7 | Zzef1 | -0.048 | Young |
| 0.325152 | 0.269945 | 0.265 | 0.323 | 1 | 7 | Rbm28 | -0.058 | Young |
| 0.325294 | 0.223248 | 0.313 | 0.367 | 1 | 7 | Mrps18c | -0.054 | Young |
| 0.325677 | 0.26875 | 0.139 | 0.169 | 1 | 7 | Slc9a6 | -0.03 | Young |
| 0.325749 | 0.021183 | 0.705 | 0.736 | 1 | 7 | Tmpo | -0.031 | Young |
| 0.326204 | 0.246762 | 0.057 | 0.041 | 1 | 7 | Pde4d | 0.016 | Young |
| 0.326475 | 0.288283 | 0.265 | 0.233 | 1 | 7 | Smarca2 | 0.032 | Young |
| 0.326514 | 0.285287 | 0.479 | 0.472 | 1 | 7 | Map1lc3b | 0.007 | Young |
| 0.326517 | 0.310644 | 0.214 | 0.19 | 1 | 7 | Dgcr2 | 0.024 | Young |
| 0.327529 | 0.932805 | 0.102 | 0.085 | 1 | 7 | Gna15 | 0.017 | Young |
| 0.327627 | 0.161883 | 0.509 | 0.487 | 1 | 7 | Ezh2 | 0.022 | Young |
| 0.32783 | 0.190804 | 0.542 | 0.523 | 1 | 7 | Tcea1 | 0.019 | Young |
| 0.328005 | 0.201335 | 0.271 | 0.246 | 1 | 7 | Ago3 | 0.025 | Young |
| 0.32827 | 0.093617 | 0.057 | 0.077 | 1 | 7 | Zfp934 | -0.02 | Young |
| 0.328728 | 0.0613 | 0.148 | 0.179 | 1 | 7 | Ap1g2 | -0.031 | Young |
| 0.329003 | 0.29132 | 0.181 | 0.151 | 1 | 7 | Casd1 | 0.03 | Young |
| 0.329028 | 0.353543 | 0.027 | 0.041 | 1 | 7 | Map2k6 | -0.014 | Young |
| 0.329475 | 0.145912 | 0.786 | 0.751 | 1 | 7 | Itgb1 | 0.035 | Young |
| 0.32981 | 0.551407 | 0.151 | 0.128 | 1 | 7 | Tbc1d4 | 0.023 | Young |
| 0.330145 | 0.114413 | 0.069 | 0.09 | 1 | 7 | Cbfa2t2 | -0.021 | Young |
| 0.330239 | 0.587523 | 0.105 | 0.085 | 1 | 7 | Fbxw5 | 0.02 | Young |
| 0.33029 | 0.067783 | 0.244 | 0.285 | 1 | 7 | Mrps23 | -0.041 | Young |
| 0.330395 | 0.440363 | 0.13 | 0.108 | 1 | 7 | Npat | 0.022 | Young |
| 0.330478 | 0.334968 | 0.214 | 0.192 | 1 | 7 | Yeats2 | 0.022 | Young |
| 0.330513 | 0.003683 | 0.142 | 0.172 | 1 | 7 | Stat2 | -0.03 | Young |
| 0.331355 | 0.098632 | 0.172 | 0.146 | 1 | 7 | Nmral1 | 0.026 | Young |
| 0.331362 | 0.08638 | 0.497 | 0.469 | 1 | 7 | Ankrd44 | 0.028 | Young |
| 0.331791 | 0.175451 | 0.31 | 0.369 | 1 | 7 | Zc3h13 | -0.059 | Young |
| 0.332671 | 0.336803 | 0.015 | 0.026 | 1 | 7 | Apbb2 | -0.011 | Young |
| 0.332683 | 0.084037 | 0.608 | 0.592 | 1 | 7 | Ralbp1 | 0.016 | Young |
| 0.333028 | 0.147693 | 0.831 | 0.805 | 1 | 7 | Pid1 | 0.026 | Young |
| 0.333688 | 0.774857 | 0.181 | 0.159 | 1 | 7 | Cpeb2 | 0.022 | Young |
| 0.333875 | 0.36372 | 0.232 | 0.272 | 1 | 7 | Ncbp2 | -0.04 | Young |
| 0.334121 | 0.003574 | 0.226 | 0.262 | 1 | 7 | Tmem65 | -0.036 | Young |
| 0.334251 | 0.221173 | 0.482 | 0.482 | 1 | 7 | Zfand6 | 0 | Young |
| 0.334751 | 0.269933 | 0.048 | 0.033 | 1 | 7 | Trim32 | 0.015 | Young |
| 0.334802 | 0.228965 | 0.148 | 0.179 | 1 | 7 | Rexo4 | -0.031 | Young |
| 0.335131 | 0.615593 | 0.193 | 0.228 | 1 | 7 | Chrac1 | -0.035 | Young |
| 0.335729 | 0.42447 | 0.307 | 0.287 | 1 | 7 | Usp37 | 0.02 | Young |
| 0.336228 | 0.356468 | 0.398 | 0.385 | 1 | 7 | 2410015M | 0.013 | Young |
| 0.33647 | 0.131907 | 0.148 | 0.179 | 1 | 7 | Pelp1 | -0.031 | Young |
| 0.3366 | 0.417659 | 0.021 | 0.033 | 1 | 7 | 5430416N( | -0.012 | Young |
| 0.336641 | 0.331633 | 0.166 | 0.141 | 1 | 7 | Rab28 | 0.025 | Young |

| | | | | | | | | |
|---|---|---|---|---|---|---|---|---|
| 0.337234 | 0.487258 | 0.021 | 0.033 | 1 | 7 | Gm8113 | -0.012 | Young |
| 0.337249 | 0.052339 | 0.488 | 0.556 | 1 | 7 | Gspt1 | -0.068 | Young |
| 0.337324 | 0.558539 | 0.313 | 0.297 | 1 | 7 | Rufy3 | 0.016 | Young |
| 0.337439 | 0.016514 | 0.111 | 0.09 | 1 | 7 | Slc35b4 | 0.021 | Young |
| 0.337523 | 0.217332 | 0.172 | 0.208 | 1 | 7 | Ppa1 | -0.036 | Young |
| 0.337852 | 0.015214 | 0.31 | 0.356 | 1 | 7 | Ibtk | -0.046 | Young |
| 0.337989 | 0.276196 | 0.599 | 0.597 | 1 | 7 | Nfe2l2 | 0.002 | Young |
| 0.338371 | 0.651065 | 0.226 | 0.208 | 1 | 7 | Slc23a2 | 0.018 | Young |
| 0.33894 | 0.075877 | 0.145 | 0.174 | 1 | 7 | Slc35e4 | -0.029 | Young |
| 0.339259 | 0.131997 | 0.081 | 0.103 | 1 | 7 | Dhx58 | -0.022 | Young |
| 0.339994 | 0.950871 | 0.084 | 0.067 | 1 | 7 | Fktn | 0.017 | Young |
| 0.339997 | 0.048689 | 0.136 | 0.164 | 1 | 7 | Desi1 | -0.028 | Young |
| 0.340648 | 0.300826 | 0.145 | 0.123 | 1 | 7 | Nelfb | 0.022 | Young |
| 0.341463 | 0.087248 | 0.151 | 0.182 | 1 | 7 | Psen2 | -0.031 | Young |
| 0.341968 | 0.215697 | 0.301 | 0.279 | 1 | 7 | Ankrd13c | 0.022 | Young |
| 0.342207 | 1.506267 | 0.015 | 0.008 | 1 | 7 | Pnpla1 | 0.007 | Young |
| 0.342207 | 1.345671 | 0.015 | 0.008 | 1 | 7 | Ighd | 0.007 | Young |
| 0.342207 | 0.922491 | 0.015 | 0.008 | 1 | 7 | Poln | 0.007 | Young |
| 0.342328 | 0.010403 | 0.295 | 0.264 | 1 | 7 | Arid4b | 0.031 | Young |
| 0.342558 | 0.016223 | 0.06 | 0.079 | 1 | 7 | Mthfd2l | -0.019 | Young |
| 0.342815 | 0.056808 | 0.973 | 0.964 | 1 | 7 | Hspa8 | 0.009 | Young |
| 0.342921 | 0.033231 | 0.334 | 0.379 | 1 | 7 | Cdk14 | -0.045 | Young |
| 0.34303 | 0.452098 | 0.105 | 0.085 | 1 | 7 | Dglucy | 0.02 | Young |
| 0.343211 | 1.732602 | 0.015 | 0.008 | 1 | 7 | Slc4a11 | 0.007 | Young |
| 0.343211 | 1.127166 | 0.015 | 0.008 | 1 | 7 | Kcnb1 | 0.007 | Young |
| 0.343211 | 1.092118 | 0.015 | 0.008 | 1 | 7 | Ldlrad1 | 0.007 | Young |
| 0.343211 | 0.925749 | 0.015 | 0.008 | 1 | 7 | Spata2l | 0.007 | Young |
| 0.343315 | 0.17653 | 0.425 | 0.4 | 1 | 7 | Folr2 | 0.025 | Young |
| 0.343641 | 0.402603 | 0.099 | 0.079 | 1 | 7 | Vegfb | 0.02 | Young |
| 0.343738 | 0.20489 | 0.352 | 0.333 | 1 | 7 | Gpat4 | 0.019 | Young |
| 0.343745 | 0.114391 | 0.292 | 0.254 | 1 | 7 | H2-DMb2 | 0.038 | Young |
| 0.343789 | 0.367316 | 0.06 | 0.079 | 1 | 7 | Epg5 | -0.019 | Young |
| 0.343982 | 0.831562 | 0.084 | 0.067 | 1 | 7 | Per3 | 0.017 | Young |
| 0.344218 | 1.025787 | 0.015 | 0.008 | 1 | 7 | Eid2 | 0.007 | Young |
| 0.344218 | 0.878254 | 0.015 | 0.008 | 1 | 7 | Snn | 0.007 | Young |
| 0.344218 | 0.65628 | 0.015 | 0.008 | 1 | 7 | Gm43569 | 0.007 | Young |
| 0.344835 | 0.268121 | 0.458 | 0.446 | 1 | 7 | Psmd1 | 0.012 | Young |
| 0.345226 | 1.28362 | 0.015 | 0.008 | 1 | 7 | Wdr27 | 0.007 | Young |
| 0.345226 | 1.012584 | 0.015 | 0.008 | 1 | 7 | Gm26522 | 0.007 | Young |
| 0.345226 | 0.774761 | 0.015 | 0.008 | 1 | 7 | Acot1 | 0.007 | Young |
| 0.345226 | 0.564796 | 0.015 | 0.008 | 1 | 7 | Etnk2 | 0.007 | Young |
| 0.345474 | 0.188232 | 0.434 | 0.408 | 1 | 7 | Dcun1d5 | 0.026 | Young |
| 0.345474 | 0.495845 | 0.111 | 0.092 | 1 | 7 | Pgm2 | 0.019 | Young |
| 0.345476 | 0.262077 | 0.087 | 0.11 | 1 | 7 | Mad2l1bp | -0.023 | Young |
| 0.345856 | 0.236852 | 0.063 | 0.082 | 1 | 7 | Kctd13 | -0.019 | Young |

| | | | | | | | | |
|---|---|---|---|---|---|---|---|---|
| 0.345931 | 0.238462 | 0.169 | 0.144 | 1 | 7 | Wdr20 | 0.025 | Young |
| 0.345967 | 0.084219 | 0.09 | 0.113 | 1 | 7 | Zfp830 | -0.023 | Young |
| 0.346128 | 0.185996 | 0.416 | 0.403 | 1 | 7 | Elmo1 | 0.013 | Young |
| 0.346181 | 0.055026 | 0.087 | 0.11 | 1 | 7 | Hacd2 | -0.023 | Young |
| 0.346196 | 0.109198 | 0.241 | 0.215 | 1 | 7 | Maff | 0.026 | Young |
| 0.346236 | 1.204667 | 0.015 | 0.008 | 1 | 7 | Gm14286 | 0.007 | Young |
| 0.346236 | 0.48931 | 0.015 | 0.008 | 1 | 7 | Fam167b | 0.007 | Young |
| 0.346429 | 0.771812 | 0.09 | 0.072 | 1 | 7 | Btbd2 | 0.018 | Young |
| 0.346496 | 0.026069 | 0.123 | 0.151 | 1 | 7 | Triap1 | -0.028 | Young |
| 0.346951 | 0.047313 | 0.154 | 0.185 | 1 | 7 | Exosc9 | -0.031 | Young |
| 0.346955 | 0.009238 | 0.096 | 0.121 | 1 | 7 | Dus1l | -0.025 | Young |
| 0.347107 | 0.096225 | 0.259 | 0.3 | 1 | 7 | Snx27 | -0.041 | Young |
| 0.347204 | 0.897479 | 0.072 | 0.056 | 1 | 7 | Galt | 0.016 | Young |
| 0.347204 | 0.71334 | 0.072 | 0.056 | 1 | 7 | Arfip2 | 0.016 | Young |
| 0.347248 | 0.349225 | 0.015 | 0.008 | 1 | 7 | Gm5141 | 0.007 | Young |
| 0.347248 | 0.274437 | 0.015 | 0.008 | 1 | 7 | Fchsd1 | 0.007 | Young |
| 0.347467 | 0.255641 | 0.361 | 0.41 | 1 | 7 | Cenpe | -0.049 | Young |
| 0.347874 | 0.092467 | 0.172 | 0.208 | 1 | 7 | Xpnpep1 | -0.036 | Young |
| 0.348262 | 1.046491 | 0.015 | 0.008 | 1 | 7 | Vangl2 | 0.007 | Young |
| 0.348292 | 0.043372 | 0.063 | 0.082 | 1 | 7 | Zfp84 | -0.019 | Young |
| 0.348493 | 0.00614 | 0.208 | 0.182 | 1 | 7 | Tmem50b | 0.026 | Young |
| 0.348592 | 0.354844 | 0.226 | 0.205 | 1 | 7 | Nploc4 | 0.021 | Young |
| 0.348622 | 0.268036 | 0.036 | 0.051 | 1 | 7 | Abraxas1 | -0.015 | Young |
| 0.348876 | 0.260531 | 0.066 | 0.087 | 1 | 7 | 2300009A0 | -0.021 | Young |
| 0.349085 | 0.118399 | 0.247 | 0.29 | 1 | 7 | Clspn | -0.043 | Young |
| 0.349251 | 0.037874 | 0.099 | 0.123 | 1 | 7 | Tm7sf3 | -0.024 | Young |
| 0.349278 | 0.32597 | 0.015 | 0.008 | 1 | 7 | Traf4 | 0.007 | Young |
| 0.349278 | 0.119795 | 0.015 | 0.008 | 1 | 7 | Zfp112 | 0.007 | Young |
| 0.349457 | 0.226032 | 0.392 | 0.377 | 1 | 7 | Mcm4 | 0.015 | Young |
| 0.349817 | 0.326917 | 0.142 | 0.121 | 1 | 7 | 0610037L1 | 0.021 | Young |
| 0.349861 | 0.146946 | 0.319 | 0.292 | 1 | 7 | Map2k1 | 0.027 | Young |
| 0.349965 | 0.043205 | 0.756 | 0.713 | 1 | 7 | Timp2 | 0.043 | Young |
| 0.349994 | 0.000915 | 0.13 | 0.156 | 1 | 7 | Gm13212 | -0.026 | Young |
| 0.350295 | 0.258075 | 0.015 | 0.008 | 1 | 7 | AC113595. | 0.007 | Young |
| 0.350463 | 0.041349 | 0.358 | 0.4 | 1 | 7 | Asph | -0.042 | Young |
| 0.350512 | 0.030439 | 0.617 | 0.672 | 1 | 7 | Mki67 | -0.055 | Young |
| 0.350751 | 0.266177 | 0.235 | 0.213 | 1 | 7 | Ednrb | 0.022 | Young |
| 0.351083 | 0.405517 | 0.127 | 0.105 | 1 | 7 | Sema6d | 0.022 | Young |
| 0.3512 | 0.533361 | 0.154 | 0.133 | 1 | 7 | Cwc25 | 0.021 | Young |
| 0.351243 | 0.454363 | 0.334 | 0.323 | 1 | 7 | Sf3a1 | 0.011 | Young |
| 0.351269 | 0.000863 | 0.19 | 0.223 | 1 | 7 | Nedd1 | -0.033 | Young |
| 0.352117 | 0.520395 | 0.169 | 0.149 | 1 | 7 | Trim37 | 0.02 | Young |
| 0.352269 | 0.291137 | 0.09 | 0.113 | 1 | 7 | Slco3a1 | -0.023 | Young |
| 0.352472 | 0.875481 | 0.066 | 0.051 | 1 | 7 | Rtel1 | 0.015 | Young |
| 0.352487 | 0.056087 | 0.184 | 0.218 | 1 | 7 | Zbtb1 | -0.034 | Young |

| | | | | | | | | |
|---|---|---|---|---|---|---|---|---|
| 0.352491 | 0.726817 | 0.142 | 0.123 | 1 | 7 | Borcs7 | 0.019 | Young |
| 0.352948 | 0.333158 | 0.123 | 0.151 | 1 | 7 | Vps26b | -0.028 | Young |
| 0.353025 | 0.010704 | 0.13 | 0.156 | 1 | 7 | Rasgrp1 | -0.026 | Young |
| 0.35375 | 0.048196 | 0.054 | 0.072 | 1 | 7 | Clasrp | -0.018 | Young |
| 0.354293 | 0.639281 | 0.111 | 0.092 | 1 | 7 | Cul4b | 0.019 | Young |
| 0.354432 | 0.056752 | 0.298 | 0.274 | 1 | 7 | Rab32 | 0.024 | Young |
| 0.3548 | 0.06536 | 0.259 | 0.303 | 1 | 7 | Rnf145 | -0.044 | Young |
| 0.354908 | 0.002295 | 0.446 | 0.495 | 1 | 7 | Bst2 | -0.049 | Young |
| 0.354944 | 0.288049 | 0.081 | 0.064 | 1 | 7 | Gtpbp10 | 0.017 | Young |
| 0.355015 | 0.10376 | 0.117 | 0.095 | 1 | 7 | Cdc14b | 0.022 | Young |
| 0.355116 | 0.36878 | 0.301 | 0.274 | 1 | 7 | Atxn10 | 0.027 | Young |
| 0.355379 | 0.076947 | 0.03 | 0.044 | 1 | 7 | Apba3 | -0.014 | Young |
| 0.355627 | 0.441274 | 0.148 | 0.126 | 1 | 7 | Ifi35 | 0.022 | Young |
| 0.355957 | 1.325761 | 0.069 | 0.054 | 1 | 7 | Haspin | 0.015 | Young |
| 0.356047 | 0.047772 | 0.006 | 0.013 | 1 | 7 | Zfp882 | -0.007 | Young |
| 0.356132 | 0.19422 | 0.105 | 0.085 | 1 | 7 | Haus5 | 0.02 | Young |
| 0.356757 | 0.317954 | 0.105 | 0.131 | 1 | 7 | Gk | -0.026 | Young |
| 0.356831 | 0.357849 | 0.093 | 0.074 | 1 | 7 | Fas | 0.019 | Young |
| 0.357148 | 0.060403 | 0.006 | 0.013 | 1 | 7 | Elavl4 | -0.007 | Young |
| 0.357459 | 0.00432 | 0.108 | 0.133 | 1 | 7 | Epm2aip1 | -0.025 | Young |
| 0.357699 | 0.366734 | 0.148 | 0.177 | 1 | 7 | Rae1 | -0.029 | Young |
| 0.357966 | 0.320503 | 0.491 | 0.485 | 1 | 7 | Scpep1 | 0.006 | Young |
| 0.35825 | 0.11294 | 0.006 | 0.013 | 1 | 7 | Ss18l1 | -0.007 | Young |
| 0.35846 | 0.392334 | 0.13 | 0.108 | 1 | 7 | Prr5l | 0.022 | Young |
| 0.358791 | 0.264246 | 0.03 | 0.044 | 1 | 7 | Ggct | -0.014 | Young |
| 0.358823 | 0.213898 | 0.069 | 0.09 | 1 | 7 | Gss | -0.021 | Young |
| 0.359038 | 0.366196 | 0.114 | 0.095 | 1 | 7 | Slx4ip | 0.019 | Young |
| 0.359355 | 0.06829 | 0.006 | 0.013 | 1 | 7 | Mindy4 | -0.007 | Young |
| 0.359564 | 0.009164 | 0.114 | 0.136 | 1 | 7 | Arhgap6 | -0.022 | Young |
| 0.359895 | 0.203511 | 0.455 | 0.451 | 1 | 7 | Stard8 | 0.004 | Young |
| 0.359952 | 0.584676 | 0.081 | 0.064 | 1 | 7 | Wdcp | 0.017 | Young |
| 0.360417 | 0.06848 | 0.069 | 0.09 | 1 | 7 | Thap7 | -0.021 | Young |
| 0.360494 | 0.065979 | 0.045 | 0.062 | 1 | 7 | Zc3h12c | -0.017 | Young |
| 0.360552 | 0.18359 | 0.307 | 0.282 | 1 | 7 | Stxbp2 | 0.025 | Young |
| 0.360627 | 0.1639 | 0.452 | 0.421 | 1 | 7 | Ifi207 | 0.031 | Young |
| 0.360663 | 0.158921 | 0.142 | 0.169 | 1 | 7 | Dcaf11 | -0.027 | Young |
| 0.361738 | 0.183505 | 0.024 | 0.036 | 1 | 7 | Dctd | -0.012 | Young |
| 0.361837 | 0.486968 | 0.069 | 0.054 | 1 | 7 | Nes | 0.015 | Young |
| 0.361856 | 0.036209 | 0.304 | 0.277 | 1 | 7 | Fnbp1l | 0.027 | Young |
| 0.361906 | 1.236897 | 0.018 | 0.01 | 1 | 7 | Klhdc1 | 0.008 | Young |
| 0.36288 | 0.352693 | 0.184 | 0.162 | 1 | 7 | Josd1 | 0.022 | Young |
| 0.363284 | 0.001105 | 0.145 | 0.172 | 1 | 7 | Dhx8 | -0.027 | Young |
| 0.363327 | 0.164429 | 0.202 | 0.177 | 1 | 7 | Mfap3 | 0.025 | Young |
| 0.363361 | 0.057446 | 0.759 | 0.759 | 1 | 7 | H2afv | 0 | Young |
| 0.363516 | 0.783945 | 0.075 | 0.059 | 1 | 7 | Nudt18 | 0.016 | Young |

| | | | | | | | | |
|---|---|---|---|---|---|---|---|---|
| 0.363743 | 0.124607 | 0.247 | 0.223 | 1 | 7 | Pigx | 0.024 | Young |
| 0.364276 | 0.785122 | 0.024 | 0.036 | 1 | 7 | Nme7 | -0.012 | Young |
| 0.364451 | 0.473553 | 0.136 | 0.113 | 1 | 7 | Ctns | 0.023 | Young |
| 0.364706 | 0.828605 | 0.018 | 0.01 | 1 | 7 | Trim46 | 0.008 | Young |
| 0.364943 | 0.448285 | 0.127 | 0.105 | 1 | 7 | Cyp4f13 | 0.022 | Young |
| 0.365732 | 0.056861 | 0.759 | 0.738 | 1 | 7 | Sat1 | 0.021 | Young |
| 0.365783 | 0.454764 | 0.214 | 0.19 | 1 | 7 | Cir1 | 0.024 | Young |
| 0.365959 | 0.383307 | 0.093 | 0.115 | 1 | 7 | Irf2bp1 | -0.022 | Young |
| 0.366023 | 0.033905 | 0.148 | 0.123 | 1 | 7 | Shmt2 | 0.025 | Young |
| 0.36638 | 0.140047 | 0.202 | 0.236 | 1 | 7 | Wdhd1 | -0.034 | Young |
| 0.366581 | 0.899475 | 0.018 | 0.01 | 1 | 7 | Ppp1r18os | 0.008 | Young |
| 0.36659 | 0.019983 | 0.488 | 0.536 | 1 | 7 | Cd2ap | -0.048 | Young |
| 0.366985 | 0.111855 | 0.139 | 0.167 | 1 | 7 | Irak3 | -0.028 | Young |
| 0.36735 | 0.006827 | 0.265 | 0.244 | 1 | 7 | Med14 | 0.021 | Young |
| 0.36752 | 1.021679 | 0.018 | 0.01 | 1 | 7 | Zc4h2 | 0.008 | Young |
| 0.367887 | 0.222826 | 0.413 | 0.395 | 1 | 7 | Fes | 0.018 | Young |
| 0.368213 | 0.581359 | 0.093 | 0.118 | 1 | 7 | Zfp780b | -0.025 | Young |
| 0.368461 | 1.274766 | 0.018 | 0.01 | 1 | 7 | Phf11c | 0.008 | Young |
| 0.368461 | 1.150264 | 0.018 | 0.01 | 1 | 7 | E430018J2 | 0.008 | Young |
| 0.368676 | 0.130405 | 0.268 | 0.249 | 1 | 7 | Aagab | 0.019 | Young |
| 0.368702 | 0.012739 | 0.09 | 0.113 | 1 | 7 | Kat5 | -0.023 | Young |
| 0.369299 | 0.857777 | 0.093 | 0.077 | 1 | 7 | Kantr | 0.016 | Young |
| 0.369331 | 0.310601 | 0.355 | 0.338 | 1 | 7 | Mgat1 | 0.017 | Young |
| 0.369348 | 0.144065 | 0.494 | 0.482 | 1 | 7 | Kmt2e | 0.012 | Young |
| 0.369384 | 0.247348 | 0.024 | 0.036 | 1 | 7 | Pusl1 | -0.012 | Young |
| 0.369404 | 0.955312 | 0.018 | 0.01 | 1 | 7 | Tnfrsf26 | 0.008 | Young |
| 0.369418 | 0.270616 | 0.048 | 0.064 | 1 | 7 | Cela1 | -0.016 | Young |
| 0.369446 | 0.061803 | 0.111 | 0.133 | 1 | 7 | Ptp4a3 | -0.022 | Young |
| 0.370025 | 0.320722 | 0.024 | 0.036 | 1 | 7 | Ptges3l | -0.012 | Young |
| 0.370135 | 0.141025 | 0.229 | 0.264 | 1 | 7 | Ube4b | -0.035 | Young |
| 0.370291 | 1.057694 | 0.054 | 0.041 | 1 | 7 | Dnajb2 | 0.013 | Young |
| 0.370535 | 0.018016 | 0.087 | 0.069 | 1 | 7 | Orc5 | 0.018 | Young |
| 0.370582 | 0.499584 | 0.066 | 0.051 | 1 | 7 | Fzd5 | 0.015 | Young |
| 0.370701 | 0.158399 | 0.596 | 0.597 | 1 | 7 | Tmcc1 | -0.001 | Young |
| 0.371152 | 0.606963 | 0.045 | 0.062 | 1 | 7 | Zfp760 | -0.017 | Young |
| 0.371293 | 0.536954 | 0.018 | 0.01 | 1 | 7 | Trim62 | 0.008 | Young |
| 0.371758 | 0.105342 | 0.127 | 0.151 | 1 | 7 | Mrps34 | -0.024 | Young |
| 0.371807 | 0.003955 | 0.163 | 0.195 | 1 | 7 | Pmpcb | -0.032 | Young |
| 0.372038 | 0.117183 | 0.449 | 0.51 | 1 | 7 | Prpf4b | -0.061 | Young |
| 0.372145 | 0.144618 | 0.678 | 0.679 | 1 | 7 | Gpi1 | -0.001 | Young |
| 0.37224 | 0.345487 | 0.018 | 0.01 | 1 | 7 | Col3a1 | 0.008 | Young |
| 0.372337 | 0.124196 | 0.057 | 0.074 | 1 | 7 | Cdk7 | -0.017 | Young |
| 0.372793 | 0.349466 | 0.148 | 0.126 | 1 | 7 | Fam199x | 0.022 | Young |
| 0.372854 | 0.071494 | 0.988 | 0.977 | 1 | 7 | Laptm5 | 0.011 | Young |
| 0.373051 | 0.216648 | 0.337 | 0.313 | 1 | 7 | Lmbrd1 | 0.024 | Young |

| | | | | | | | | |
|---|---|---|---|---|---|---|---|---|
| 0.373432 | 0.096818 | 0.09 | 0.113 | 1 | 7 | Dgcr6 | -0.023 | Young |
| 0.373676 | 0.130681 | 0.226 | 0.267 | 1 | 7 | Irf9 | -0.041 | Young |
| 0.374107 | 0.170719 | 0.142 | 0.167 | 1 | 7 | Nufip1 | -0.025 | Young |
| 0.374117 | 0.321398 | 0.075 | 0.059 | 1 | 7 | Tmem263 | 0.016 | Young |
| 0.374579 | 0.47092 | 0.078 | 0.062 | 1 | 7 | Fance | 0.016 | Young |
| 0.374743 | 1.601879 | 0.021 | 0.013 | 1 | 7 | Gm30329 | 0.008 | Young |
| 0.374772 | 0.018137 | 0.148 | 0.174 | 1 | 7 | Btbd10 | -0.026 | Young |
| 0.374885 | 0.064966 | 0.913 | 0.905 | 1 | 7 | Rps15a | 0.008 | Young |
| 0.37509 | 0.362305 | 0.018 | 0.01 | 1 | 7 | Gm43936 | 0.008 | Young |
| 0.37509 | 0.211953 | 0.018 | 0.01 | 1 | 7 | Catip | 0.008 | Young |
| 0.375118 | 0.11774 | 0.434 | 0.505 | 1 | 7 | Rilpl2 | -0.071 | Young |
| 0.375155 | 0.332081 | 0.202 | 0.185 | 1 | 7 | Cpq | 0.017 | Young |
| 0.375217 | 0.126689 | 0.12 | 0.149 | 1 | 7 | Dnttip1 | -0.029 | Young |
| 0.375636 | 0.04299 | 0.97 | 0.962 | 1 | 7 | Rps15 | 0.008 | Young |
| 0.375672 | 0.194442 | 0.072 | 0.056 | 1 | 7 | Ppox | 0.016 | Young |
| 0.375784 | 0.009761 | 0.313 | 0.364 | 1 | 7 | Cdk19 | -0.051 | Young |
| 0.375852 | 0.319894 | 0.133 | 0.113 | 1 | 7 | Med29 | 0.02 | Young |
| 0.376816 | 0.858006 | 0.057 | 0.044 | 1 | 7 | Slc35g1 | 0.013 | Young |
| 0.376868 | 0.519797 | 0.229 | 0.21 | 1 | 7 | Gabarapl1 | 0.019 | Young |
| 0.377359 | 1.521561 | 0.021 | 0.013 | 1 | 7 | Tln2 | 0.008 | Young |
| 0.37751 | 0.141923 | 0.648 | 0.613 | 1 | 7 | Glul | 0.035 | Young |
| 0.377519 | 0.680052 | 0.205 | 0.19 | 1 | 7 | Trak1 | 0.015 | Young |
| 0.377563 | 0.196763 | 0.142 | 0.169 | 1 | 7 | Ly96 | -0.027 | Young |
| 0.377809 | 0.339427 | 0.343 | 0.328 | 1 | 7 | Sfxn3 | 0.015 | Young |
| 0.377954 | 0.031783 | 0.018 | 0.01 | 1 | 7 | Gm17435 | 0.008 | Young |
| 0.378128 | 0.193055 | 0.608 | 0.613 | 1 | 7 | Elob | -0.005 | Young |
| 0.378233 | 1.174755 | 0.021 | 0.013 | 1 | 7 | Tmem202 | 0.008 | Young |
| 0.378843 | 0.083421 | 0.611 | 0.587 | 1 | 7 | Gng2 | 0.024 | Young |
| 0.378914 | 0.072386 | 0.018 | 0.028 | 1 | 7 | Snx32 | -0.01 | Young |
| 0.378938 | 0.080155 | 0.262 | 0.295 | 1 | 7 | Hmmr | -0.033 | Young |
| 0.379042 | 1.178092 | 0.048 | 0.036 | 1 | 7 | 1700066M | 0.012 | Young |
| 0.379109 | 1.393821 | 0.021 | 0.013 | 1 | 7 | Zfp623 | 0.008 | Young |
| 0.379355 | 0.017664 | 0.684 | 0.728 | 1 | 7 | Mtpn | -0.044 | Young |
| 0.379492 | 0.077922 | 0.283 | 0.323 | 1 | 7 | Mtm1 | -0.04 | Young |
| 0.379899 | 1.035844 | 0.057 | 0.044 | 1 | 7 | Pex1 | 0.013 | Young |
| 0.379942 | 0.613235 | 0.142 | 0.121 | 1 | 7 | Cdc14a | 0.021 | Young |
| 0.380096 | 0.235594 | 0.461 | 0.449 | 1 | 7 | Acap2 | 0.012 | Young |
| 0.380272 | 1.171157 | 0.054 | 0.041 | 1 | 7 | Parvg | 0.013 | Young |
| 0.380626 | 0.7854 | 0.117 | 0.144 | 1 | 7 | Aifm1 | -0.027 | Young |
| 0.381 | 0.017383 | 0.235 | 0.272 | 1 | 7 | Ifi27 | -0.037 | Young |
| 0.381439 | 0.69577 | 0.048 | 0.064 | 1 | 7 | Mars2 | -0.016 | Young |
| 0.381504 | 0.652366 | 0.033 | 0.046 | 1 | 7 | Miga2 | -0.013 | Young |
| 0.381622 | 0.163772 | 0.06 | 0.077 | 1 | 7 | Oxnad1 | -0.017 | Young |
| 0.381743 | 0.711021 | 0.021 | 0.013 | 1 | 7 | Gm2415 | 0.008 | Young |
| 0.38175 | 0.353339 | 0.316 | 0.303 | 1 | 7 | Ifnar2 | 0.013 | Young |

| | | | | | | | | |
|---|---|---|---|---|---|---|---|---|
| 0.382171 | 0.188612 | 0.142 | 0.169 | 1 | 7 | Ppp5c | -0.027 | Young |
| 0.382587 | 1.914155 | 0.024 | 0.015 | 1 | 7 | 5830416I1! | 0.009 | Young |
| 0.38264 | 0.092708 | 0.172 | 0.2 | 1 | 7 | Slc25a46 | -0.028 | Young |
| 0.382672 | 0.117803 | 0.678 | 0.715 | 1 | 7 | Srp9 | -0.037 | Young |
| 0.382755 | 0.291308 | 0.425 | 0.405 | 1 | 7 | Bmp2k | 0.02 | Young |
| 0.38291 | 0.233453 | 0.431 | 0.421 | 1 | 7 | Orai1 | 0.01 | Young |
| 0.383176 | 0.421829 | 0.06 | 0.046 | 1 | 7 | Smim5 | 0.014 | Young |
| 0.383506 | 0.715974 | 0.021 | 0.013 | 1 | 7 | Tbc1d30 | 0.008 | Young |
| 0.383573 | 0.038065 | 0.946 | 0.949 | 1 | 7 | H2-Eb1 | -0.003 | Young |
| 0.384792 | 0.384612 | 0.133 | 0.11 | 1 | 7 | Hace1 | 0.023 | Young |
| 0.385057 | 0.927681 | 0.054 | 0.041 | 1 | 7 | Nkiras1 | 0.013 | Young |
| 0.385826 | 0.070295 | 0.428 | 0.474 | 1 | 7 | Icam1 | -0.046 | Young |
| 0.38588 | 0.069935 | 0.849 | 0.833 | 1 | 7 | Unc93b1 | 0.016 | Young |
| 0.386159 | 1.576859 | 0.021 | 0.013 | 1 | 7 | Aqp1 | 0.008 | Young |
| 0.386159 | 0.86812 | 0.021 | 0.013 | 1 | 7 | Ift81 | 0.008 | Young |
| 0.386204 | 0.109389 | 0.398 | 0.446 | 1 | 7 | Casp4 | -0.048 | Young |
| 0.38631 | 1.136994 | 0.027 | 0.018 | 1 | 7 | Stk26 | 0.009 | Young |
| 0.386403 | 0.01925 | 0.386 | 0.431 | 1 | 7 | Plekhj1 | -0.045 | Young |
| 0.38695 | 1.269603 | 0.036 | 0.026 | 1 | 7 | Afap1l1 | 0.01 | Young |
| 0.38697 | 0.78855 | 0.093 | 0.077 | 1 | 7 | Rhbdd1 | 0.016 | Young |
| 0.386998 | 0.379847 | 0.352 | 0.336 | 1 | 7 | Mark2 | 0.016 | Young |
| 0.387489 | 0.92974 | 0.012 | 0.021 | 1 | 7 | Acap1 | -0.009 | Young |
| 0.387489 | 0.735771 | 0.012 | 0.021 | 1 | 7 | Tceanc | -0.009 | Young |
| 0.387651 | 0.102397 | 0.28 | 0.321 | 1 | 7 | Rnpepl1 | -0.041 | Young |
| 0.387933 | 0.488923 | 0.021 | 0.013 | 1 | 7 | Slfn3 | 0.008 | Young |
| 0.387989 | 0.528699 | 0.093 | 0.077 | 1 | 7 | Cep76 | 0.016 | Young |
| 0.388066 | 0.345518 | 0.208 | 0.187 | 1 | 7 | Prdm2 | 0.021 | Young |
| 0.388134 | 0.151253 | 0.238 | 0.29 | 1 | 7 | Mrpl43 | -0.052 | Young |
| 0.388379 | 0.078143 | 0.289 | 0.262 | 1 | 7 | Synrg | 0.027 | Young |
| 0.3888 | 1.036979 | 0.087 | 0.072 | 1 | 7 | Pbx1 | 0.015 | Young |
| 0.389084 | 0.492241 | 0.111 | 0.092 | 1 | 7 | Scaper | 0.019 | Young |
| 0.389225 | 0.992722 | 0.048 | 0.036 | 1 | 7 | Zfp276 | 0.012 | Young |
| 0.38924 | 0.713849 | 0.057 | 0.044 | 1 | 7 | Vill | 0.013 | Young |
| 0.389982 | 1.139866 | 0.024 | 0.015 | 1 | 7 | Tex9 | 0.009 | Young |
| 0.389982 | 0.821076 | 0.024 | 0.015 | 1 | 7 | Gm4631 | 0.009 | Young |
| 0.390091 | 0.430915 | 0.018 | 0.028 | 1 | 7 | Ovca2 | -0.01 | Young |
| 0.390147 | 0.164495 | 0.051 | 0.067 | 1 | 7 | Inpp5e | -0.016 | Young |
| 0.390183 | 0.438598 | 0.238 | 0.218 | 1 | 7 | Slc44a1 | 0.02 | Young |
| 0.390283 | 0.094205 | 0.545 | 0.538 | 1 | 7 | Hprt | 0.007 | Young |
| 0.390429 | 0.022883 | 0.744 | 0.751 | 1 | 7 | Grk2 | -0.007 | Young |
| 0.390506 | 0.309404 | 0.087 | 0.11 | 1 | 7 | Tbc1d24 | -0.023 | Young |
| 0.390662 | 0.85027 | 0.045 | 0.033 | 1 | 7 | Abhd14a | 0.012 | Young |
| 0.390734 | 0.163905 | 0.048 | 0.064 | 1 | 7 | Arel1 | -0.016 | Young |
| 0.391498 | 0.995296 | 0.021 | 0.013 | 1 | 7 | Efcab2 | 0.008 | Young |
| 0.391498 | 0.034826 | 0.021 | 0.013 | 1 | 7 | Gstt1 | 0.008 | Young |

| | | | | | | | | |
|---|---|---|---|---|---|---|---|---|
| 0.391625 | 0.331498 | 0.184 | 0.164 | 1 | 7 | Pink1 | 0.02 | Young |
| 0.391971 | 0.177476 | 0.081 | 0.103 | 1 | 7 | Cenpt | -0.022 | Young |
| 0.39231 | 0.475084 | 0.051 | 0.038 | 1 | 7 | Adgrl1 | 0.013 | Young |
| 0.392359 | 0.166855 | 0.262 | 0.303 | 1 | 7 | Fbxw2 | -0.041 | Young |
| 0.392419 | 0.05589 | 0.458 | 0.433 | 1 | 7 | Dpm3 | 0.025 | Young |
| 0.392467 | 1.149476 | 0.024 | 0.015 | 1 | 7 | Peg10 | 0.009 | Young |
| 0.392607 | 0.582883 | 0.099 | 0.082 | 1 | 7 | Dnajc16 | 0.017 | Young |
| 0.392817 | 0.63931 | 0.274 | 0.262 | 1 | 7 | Gch1 | 0.012 | Young |
| 0.393028 | 0.851407 | 0.087 | 0.072 | 1 | 7 | Orai2 | 0.015 | Young |
| 0.393288 | 0.496021 | 0.021 | 0.013 | 1 | 7 | Nr6a1 | 0.008 | Young |
| 0.393297 | 0.588221 | 0.024 | 0.015 | 1 | 7 | Gm28707 | 0.009 | Young |
| 0.393297 | 0.531973 | 0.024 | 0.015 | 1 | 7 | Gm15859 | 0.009 | Young |
| 0.393334 | 0.089073 | 0.623 | 0.597 | 1 | 7 | Rpl27 | 0.026 | Young |
| 0.39337 | 0.490084 | 0.271 | 0.256 | 1 | 7 | Ssna1 | 0.015 | Young |
| 0.393461 | 0.209549 | 0.22 | 0.197 | 1 | 7 | Prpf19 | 0.023 | Young |
| 0.39391 | 0.17721 | 0.253 | 0.228 | 1 | 7 | Snrk | 0.025 | Young |
| 0.393983 | 0.077609 | 0.193 | 0.223 | 1 | 7 | Cwf19l2 | -0.03 | Young |
| 0.394128 | 0.851584 | 0.024 | 0.015 | 1 | 7 | 4933404O1 | 0.009 | Young |
| 0.394215 | 0.355171 | 0.205 | 0.182 | 1 | 7 | Polr2h | 0.023 | Young |
| 0.394242 | 0.075038 | 0.066 | 0.085 | 1 | 7 | Gpn2 | -0.019 | Young |
| 0.394609 | 0.259739 | 0.108 | 0.09 | 1 | 7 | Thbd | 0.018 | Young |
| 0.394628 | 0.185634 | 0.066 | 0.051 | 1 | 7 | Cep41 | 0.015 | Young |
| 0.394633 | 0.463192 | 0.325 | 0.315 | 1 | 7 | Luzp1 | 0.01 | Young |
| 0.394681 | 0.669136 | 0.117 | 0.1 | 1 | 7 | Fbxl17 | 0.017 | Young |
| 0.394706 | 0.21629 | 0.223 | 0.254 | 1 | 7 | Fam69a | -0.031 | Young |
| 0.394772 | 0.083969 | 0.075 | 0.095 | 1 | 7 | Aar2 | -0.02 | Young |
| 0.394813 | 0.295944 | 0.033 | 0.046 | 1 | 7 | Dppa3 | -0.013 | Young |
| 0.394834 | 1.646041 | 0.027 | 0.018 | 1 | 7 | Prkcg | 0.009 | Young |
| 0.394918 | 1.371347 | 0.036 | 0.026 | 1 | 7 | Trim30b | 0.01 | Young |
| 0.394961 | 0.158199 | 0.024 | 0.015 | 1 | 7 | Znrd1as | 0.009 | Young |
| 0.395079 | 0.255219 | 0.123 | 0.103 | 1 | 7 | Nrde2 | 0.02 | Young |
| 0.395102 | 1.233641 | 0.03 | 0.021 | 1 | 7 | Slc22a21 | 0.009 | Young |
| 0.395107 | 0.120282 | 0.066 | 0.085 | 1 | 7 | Zfp715 | -0.019 | Young |
| 0.395173 | 0.441859 | 0.355 | 0.336 | 1 | 7 | Rbm26 | 0.019 | Young |
| 0.395256 | 0.245253 | 0.142 | 0.169 | 1 | 7 | Usp6nl | -0.027 | Young |
| 0.395281 | 0.909687 | 0.033 | 0.023 | 1 | 7 | Hps6 | 0.01 | Young |
| 0.395406 | 0.750824 | 0.045 | 0.033 | 1 | 7 | 1810011H1 | 0.012 | Young |
| 0.395614 | 0.90472 | 0.027 | 0.018 | 1 | 7 | Zfp692 | 0.009 | Young |
| 0.395674 | 0.199061 | 0.358 | 0.351 | 1 | 7 | Smap1 | 0.007 | Young |
| 0.395727 | 0.278675 | 0.578 | 0.574 | 1 | 7 | Mcm3 | 0.004 | Young |
| 0.395764 | 0.116016 | 0.108 | 0.09 | 1 | 7 | Slc43a3 | 0.018 | Young |
| 0.395783 | 0.105207 | 0.88 | 0.872 | 1 | 7 | Retnla | 0.008 | Young |
| 0.395886 | 0.160592 | 0.075 | 0.059 | 1 | 7 | Stk38l | 0.016 | Young |
| 0.395981 | 0.5463 | 0.021 | 0.013 | 1 | 7 | Gm16287 | 0.008 | Young |
| 0.395981 | 0.033947 | 0.021 | 0.013 | 1 | 7 | Hoxb3 | 0.008 | Young |

| | | | | | | | | |
|---|---|---|---|---|---|---|---|---|
| 0.396045 | 0.215131 | 0.108 | 0.133 | 1 | 7 | Rad17 | -0.025 | Young |
| 0.396061 | 0.042441 | 0.373 | 0.354 | 1 | 7 | Ddx42 | 0.019 | Young |
| 0.39652 | 0.362958 | 0.102 | 0.128 | 1 | 7 | 2310039H( | -0.026 | Young |
| 0.396628 | 1.27499 | 0.024 | 0.015 | 1 | 7 | Nlrp10 | 0.009 | Young |
| 0.396628 | 0.514371 | 0.024 | 0.015 | 1 | 7 | Nynrin | 0.009 | Young |
| 0.396628 | 0.352324 | 0.024 | 0.015 | 1 | 7 | Il18r1 | 0.009 | Young |
| 0.396992 | 0.603684 | 0.298 | 0.282 | 1 | 7 | Lockd | 0.016 | Young |
| 0.397024 | 0.110975 | 0.262 | 0.305 | 1 | 7 | Utrn | -0.043 | Young |
| 0.397068 | 0.872282 | 0.042 | 0.031 | 1 | 7 | Ccdc51 | 0.011 | Young |
| 0.397358 | 0.203886 | 0.404 | 0.451 | 1 | 7 | Cpt1a | -0.047 | Young |
| 0.397378 | 0.173853 | 0.151 | 0.128 | 1 | 7 | Rrp7a | 0.023 | Young |
| 0.397984 | 0.770928 | 0.06 | 0.046 | 1 | 7 | Siglech | 0.014 | Young |
| 0.39817 | 0.271003 | 0.13 | 0.156 | 1 | 7 | Sik3 | -0.026 | Young |
| 0.398293 | 0.463469 | 0.455 | 0.474 | 1 | 7 | Prkacb | -0.019 | Young |
| 0.398301 | 0.023839 | 0.024 | 0.015 | 1 | 7 | Zfp39 | 0.009 | Young |
| 0.398384 | 0.212561 | 0.051 | 0.067 | 1 | 7 | Eogt | -0.016 | Young |
| 0.39845 | 0.39203 | 0.051 | 0.038 | 1 | 7 | Pidd1 | 0.013 | Young |
| 0.398876 | 0.018414 | 0.099 | 0.121 | 1 | 7 | Arfgap1 | -0.022 | Young |
| 0.398877 | 0.074108 | 0.075 | 0.095 | 1 | 7 | Tbp | -0.02 | Young |
| 0.399096 | 0.813501 | 0.114 | 0.097 | 1 | 7 | Thoc5 | 0.017 | Young |
| 0.399138 | 0.606389 | 0.024 | 0.015 | 1 | 7 | Fndc4 | 0.009 | Young |
| 0.399316 | 0.09974 | 0.226 | 0.205 | 1 | 7 | Acbd6 | 0.021 | Young |
| 0.399369 | 0.134039 | 0.027 | 0.038 | 1 | 7 | H1fx | -0.011 | Young |
| 0.399611 | 0.902408 | 0.036 | 0.026 | 1 | 7 | Trmo | 0.01 | Young |
| 0.399996 | 0.219876 | 0.175 | 0.154 | 1 | 7 | Cxcl1 | 0.021 | Young |
| 0.400017 | 0.234216 | 0.542 | 0.541 | 1 | 7 | Cndp2 | 0.001 | Young |
| 0.400076 | 0.362076 | 0.286 | 0.267 | 1 | 7 | Syce2 | 0.019 | Young |
| 0.40016 | 0.06438 | 0.193 | 0.223 | 1 | 7 | Rnf11 | -0.03 | Young |
| 0.400184 | 0.981961 | 0.045 | 0.033 | 1 | 7 | Tbc1d8b | 0.012 | Young |
| 0.4002 | 0.931858 | 0.033 | 0.023 | 1 | 7 | Asrgl1 | 0.01 | Young |
| 0.400313 | 0.227599 | 0.069 | 0.054 | 1 | 7 | Mtus1 | 0.015 | Young |
| 0.400317 | 0.70233 | 0.027 | 0.018 | 1 | 7 | Ugt1a6a | 0.009 | Young |
| 0.400358 | 0.138428 | 0.343 | 0.4 | 1 | 7 | Tbc1d14 | -0.057 | Young |
| 0.400817 | 0.347838 | 0.024 | 0.015 | 1 | 7 | Mtcp1 | 0.009 | Young |
| 0.401246 | 0.351488 | 0.208 | 0.187 | 1 | 7 | Kansl3 | 0.021 | Young |
| 0.401261 | 0.292491 | 0.051 | 0.038 | 1 | 7 | Tlr3 | 0.013 | Young |
| 0.401307 | 0.950579 | 0.039 | 0.028 | 1 | 7 | Trmt12 | 0.011 | Young |
| 0.401384 | 0.683892 | 0.045 | 0.033 | 1 | 7 | Zfand4 | 0.012 | Young |
| 0.401574 | 0.182694 | 0.367 | 0.349 | 1 | 7 | Elmsan1 | 0.018 | Young |
| 0.401626 | 0.189412 | 0.443 | 0.426 | 1 | 7 | Iqgap2 | 0.017 | Young |
| 0.401658 | 0.326528 | 0.024 | 0.015 | 1 | 7 | Usp2 | 0.009 | Young |
| 0.401862 | 0.351666 | 0.199 | 0.177 | 1 | 7 | Aak1 | 0.022 | Young |
| 0.401895 | 0.402604 | 0.048 | 0.036 | 1 | 7 | Diras2 | 0.012 | Young |
| 0.402292 | 0.259695 | 0.145 | 0.174 | 1 | 7 | Mrpl55 | -0.029 | Young |
| 0.402493 | 0.416783 | 0.358 | 0.349 | 1 | 7 | Pdpk1 | 0.009 | Young |

| | | | | | | | | |
|---|---|---|---|---|---|---|---|---|
| 0.402602 | 0.598653 | 0.039 | 0.028 | 1 | 7 | Slc25a33 | 0.011 | Young |
| 0.402984 | 0.960726 | 0.036 | 0.026 | 1 | 7 | Pigb | 0.01 | Young |
| 0.40328 | 0.469975 | 0.042 | 0.031 | 1 | 7 | Epor | 0.011 | Young |
| 0.403289 | 0.541866 | 0.087 | 0.072 | 1 | 7 | Zfp945 | 0.015 | Young |
| 0.403343 | 0.389356 | 0.024 | 0.015 | 1 | 7 | Gm16638 | 0.009 | Young |
| 0.403378 | 0.973173 | 0.084 | 0.069 | 1 | 7 | Xpc | 0.015 | Young |
| 0.403721 | 0.286034 | 0.37 | 0.351 | 1 | 7 | Arl1 | 0.019 | Young |
| 0.404002 | 0.708892 | 0.03 | 0.021 | 1 | 7 | Slc5a6 | 0.009 | Young |
| 0.404549 | 0.095197 | 0.039 | 0.028 | 1 | 7 | Ccdc126 | 0.011 | Young |
| 0.404633 | 0.177321 | 0.307 | 0.344 | 1 | 7 | Erlin1 | -0.037 | Young |
| 0.40475 | 1.140046 | 0.03 | 0.021 | 1 | 7 | Pfkm | 0.009 | Young |
| 0.405053 | 0.499826 | 0.027 | 0.018 | 1 | 7 | Mmaa | 0.009 | Young |
| 0.405053 | 0.430499 | 0.027 | 0.018 | 1 | 7 | Tmem67 | 0.009 | Young |
| 0.405082 | 0.117094 | 0.087 | 0.108 | 1 | 7 | Ddx19b | -0.021 | Young |
| 0.405215 | 0.377082 | 0.051 | 0.038 | 1 | 7 | Fahd1 | 0.013 | Young |
| 0.405756 | 0.060087 | 0.214 | 0.246 | 1 | 7 | Ddhd2 | -0.032 | Young |
| 0.40585 | 0.713668 | 0.039 | 0.028 | 1 | 7 | Ttc8 | 0.011 | Young |
| 0.40585 | 0.459208 | 0.039 | 0.028 | 1 | 7 | Trmt44 | 0.011 | Young |
| 0.405913 | 0.047076 | 0.108 | 0.131 | 1 | 7 | Atrn | -0.023 | Young |
| 0.406013 | 0.244526 | 0.145 | 0.123 | 1 | 7 | Armc10 | 0.022 | Young |
| 0.406017 | 0.568042 | 0.033 | 0.046 | 1 | 7 | Tbkbp1 | -0.013 | Young |
| 0.406206 | 0.62005 | 0.045 | 0.033 | 1 | 7 | Smpd2 | 0.012 | Young |
| 0.406771 | 0.110503 | 0.181 | 0.159 | 1 | 7 | Zfp560 | 0.022 | Young |
| 0.407253 | 0.398068 | 0.25 | 0.287 | 1 | 7 | Lin7c | -0.037 | Young |
| 0.407434 | 0.140481 | 0.027 | 0.018 | 1 | 7 | Gm45871 | 0.009 | Young |
| 0.407611 | 0.575922 | 0.087 | 0.072 | 1 | 7 | Micall1 | 0.015 | Young |
| 0.407786 | 0.205771 | 0.117 | 0.141 | 1 | 7 | Crot | -0.024 | Young |
| 0.408028 | 0.676795 | 0.13 | 0.11 | 1 | 7 | Gtpbp1 | 0.02 | Young |
| 0.408229 | 0.282448 | 0.027 | 0.018 | 1 | 7 | Ccr7 | 0.009 | Young |
| 0.408291 | 0.908488 | 0.042 | 0.031 | 1 | 7 | Arhgap33 | 0.011 | Young |
| 0.408459 | 0.321161 | 0.027 | 0.038 | 1 | 7 | Upp2 | -0.011 | Young |
| 0.408623 | 0.367541 | 0.051 | 0.038 | 1 | 7 | Cul7 | 0.013 | Young |
| 0.40863 | 0.461369 | 0.045 | 0.033 | 1 | 7 | Nipal3 | 0.012 | Young |
| 0.408637 | 0.428761 | 0.166 | 0.146 | 1 | 7 | Scamp1 | 0.02 | Young |
| 0.408778 | 0.289089 | 0.419 | 0.405 | 1 | 7 | Nudc | 0.014 | Young |
| 0.40892 | 0.636666 | 0.042 | 0.031 | 1 | 7 | Fzd4 | 0.011 | Young |
| 0.409224 | 0.302766 | 0.048 | 0.064 | 1 | 7 | Dus2 | -0.016 | Young |
| 0.409292 | 0.030376 | 0.175 | 0.197 | 1 | 7 | Smim19 | -0.022 | Young |
| 0.409433 | 1.066364 | 0.033 | 0.023 | 1 | 7 | Dtwd2 | 0.01 | Young |
| 0.409841 | 0.058524 | 0.633 | 0.628 | 1 | 7 | Ndufb9 | 0.005 | Young |
| 0.41048 | 0.180518 | 0.223 | 0.256 | 1 | 7 | Preb | -0.033 | Young |
| 0.410557 | 0.502016 | 0.13 | 0.11 | 1 | 7 | Arntl | 0.02 | Young |
| 0.410658 | 0.39414 | 0.093 | 0.115 | 1 | 7 | Zdhhc16 | -0.022 | Young |
| 0.410756 | 0.305561 | 0.03 | 0.021 | 1 | 7 | Fastkd5 | 0.009 | Young |
| 0.410922 | 0.4555 | 0.241 | 0.226 | 1 | 7 | Fam103a1 | 0.015 | Young |

| | | | | | | | | |
|---|---|---|---|---|---|---|---|---|
| 0.411326 | 0.275865 | 0.295 | 0.277 | 1 | 7 | Marf1 | 0.018 | Young |
| 0.411582 | 0.081599 | 0.036 | 0.049 | 1 | 7 | Zc3h6 | -0.013 | Young |
| 0.411708 | 0.135068 | 0.148 | 0.126 | 1 | 7 | Mgst3 | 0.022 | Young |
| 0.411902 | 0.076475 | 0.277 | 0.254 | 1 | 7 | Zhx1 | 0.023 | Young |
| 0.412037 | 0.42593 | 0.479 | 0.485 | 1 | 7 | Ankrd17 | -0.006 | Young |
| 0.41208 | 0.063869 | 0.277 | 0.31 | 1 | 7 | Spc25 | -0.033 | Young |
| 0.412299 | 0.419606 | 0.033 | 0.023 | 1 | 7 | Tmem218 | 0.01 | Young |
| 0.412384 | 0.078171 | 0.798 | 0.797 | 1 | 7 | Ndufa2 | 0.001 | Young |
| 0.412392 | 0.482326 | 0.039 | 0.028 | 1 | 7 | Gtf2ird2 | 0.011 | Young |
| 0.412587 | 0.478483 | 0.078 | 0.064 | 1 | 7 | Zcwpw1 | 0.014 | Young |
| 0.412608 | 0.033153 | 0.316 | 0.354 | 1 | 7 | Gps1 | -0.038 | Young |
| 0.412796 | 0.05572 | 0.142 | 0.169 | 1 | 7 | Ppp2r3c | -0.027 | Young |
| 0.412901 | 0.323287 | 0.063 | 0.079 | 1 | 7 | Atg4a | -0.016 | Young |
| 0.413639 | 0.141751 | 0.22 | 0.197 | 1 | 7 | Dpp3 | 0.023 | Young |
| 0.413817 | 0.063191 | 0.373 | 0.418 | 1 | 7 | Rc3h1 | -0.045 | Young |
| 0.413985 | 0.133616 | 0.572 | 0.549 | 1 | 7 | Snx20 | 0.023 | Young |
| 0.414116 | 0.155385 | 0.413 | 0.392 | 1 | 7 | Thoc2 | 0.021 | Young |
| 0.414189 | 0.121793 | 0.06 | 0.046 | 1 | 7 | Abhd18 | 0.014 | Young |
| 0.414571 | 0.244245 | 0.229 | 0.21 | 1 | 7 | Ostm1 | 0.019 | Young |
| 0.41508 | 0.031079 | 0.081 | 0.1 | 1 | 7 | Ccdc112 | -0.019 | Young |
| 0.41512 | 0.008446 | 0.443 | 0.485 | 1 | 7 | Nars | -0.042 | Young |
| 0.415137 | 0.027563 | 0.205 | 0.231 | 1 | 7 | Fam129b | -0.026 | Young |
| 0.415872 | 0.150474 | 0.111 | 0.136 | 1 | 7 | Xrcc6 | -0.025 | Young |
| 0.415995 | 0.164554 | 0.048 | 0.036 | 1 | 7 | Hrh1 | 0.012 | Young |
| 0.416127 | 0.097321 | 0.867 | 0.838 | 1 | 7 | Morf4l1 | 0.029 | Young |
| 0.416348 | 0.106752 | 0.039 | 0.028 | 1 | 7 | Lrrk2 | 0.011 | Young |
| 0.416532 | 0.350939 | 0.169 | 0.151 | 1 | 7 | Ccdc90b | 0.018 | Young |
| 0.416877 | 0.602654 | 0.151 | 0.131 | 1 | 7 | Rpl27-ps3 | 0.02 | Young |
| 0.416943 | 0.266568 | 0.325 | 0.313 | 1 | 7 | Xpo6 | 0.012 | Young |
| 0.417291 | 0.063283 | 0.187 | 0.215 | 1 | 7 | Sap30 | -0.028 | Young |
| 0.417387 | 0.161531 | 0.717 | 0.751 | 1 | 7 | Ubl3 | -0.034 | Young |
| 0.417741 | 0.041505 | 0.858 | 0.831 | 1 | 7 | Ptbp3 | 0.027 | Young |
| 0.41776 | 0.869871 | 0.123 | 0.108 | 1 | 7 | Twsg1 | 0.015 | Young |
| 0.417795 | 0.388327 | 0.045 | 0.033 | 1 | 7 | Trmt11 | 0.012 | Young |
| 0.418338 | 0.274845 | 0.03 | 0.021 | 1 | 7 | Zfp141 | 0.009 | Young |
| 0.418481 | 0.215166 | 0.289 | 0.272 | 1 | 7 | Nhsl2 | 0.017 | Young |
| 0.418539 | 0.303299 | 0.084 | 0.105 | 1 | 7 | St3gal2 | -0.021 | Young |
| 0.418814 | 0.245689 | 0.142 | 0.123 | 1 | 7 | Ints10 | 0.019 | Young |
| 0.418914 | 0.020235 | 0.105 | 0.126 | 1 | 7 | Casp6 | -0.021 | Young |
| 0.419062 | 0.508405 | 0.081 | 0.067 | 1 | 7 | Cd1d1 | 0.014 | Young |
| 0.419342 | 0.209309 | 0.238 | 0.218 | 1 | 7 | Sult1a1 | 0.02 | Young |
| 0.419517 | 0.783658 | 0.078 | 0.064 | 1 | 7 | Exd2 | 0.014 | Young |
| 0.419857 | 0.353062 | 0.211 | 0.192 | 1 | 7 | Vps37b | 0.019 | Young |
| 0.419899 | 0.223572 | 0.322 | 0.305 | 1 | 7 | Ndufs4 | 0.017 | Young |
| 0.419913 | 0.026852 | 0.262 | 0.295 | 1 | 7 | Npepl1 | -0.033 | Young |

| | | | | | | | | |
|---|---|---|---|---|---|---|---|---|
| 0.420105 | 0.012077 | 0.051 | 0.038 | 1 | 7 | Ddx11 | 0.013 | Young |
| 0.42013 | 0.417829 | 0.154 | 0.136 | 1 | 7 | Pck2 | 0.018 | Young |
| 0.420152 | 0.119697 | 0.184 | 0.215 | 1 | 7 | Adap1 | -0.031 | Young |
| 0.420242 | 0.169454 | 0.033 | 0.023 | 1 | 7 | Nat8l | 0.01 | Young |
| 0.421264 | 0.266657 | 0.051 | 0.038 | 1 | 7 | Echdc1 | 0.013 | Young |
| 0.421484 | 0.576769 | 0.123 | 0.108 | 1 | 7 | Tapbpl | 0.015 | Young |
| 0.421545 | 0.11657 | 0.256 | 0.292 | 1 | 7 | Ube2f | -0.036 | Young |
| 0.421729 | 0.261535 | 0.259 | 0.241 | 1 | 7 | Stk40 | 0.018 | Young |
| 0.421891 | 0.222551 | 0.295 | 0.272 | 1 | 7 | Commd3 | 0.023 | Young |
| 0.422521 | 0.058612 | 0.133 | 0.156 | 1 | 7 | Slc16a6 | -0.023 | Young |
| 0.422525 | 0.068939 | 0.268 | 0.305 | 1 | 7 | Abhd17a | -0.037 | Young |
| 0.422733 | 0.022615 | 0.943 | 0.951 | 1 | 7 | Rpl30 | -0.008 | Young |
| 0.423084 | 0.304615 | 0.271 | 0.251 | 1 | 7 | Ormdl2 | 0.02 | Young |
| 0.423753 | 0.246928 | 0.048 | 0.036 | 1 | 7 | Rasal2 | 0.012 | Young |
| 0.424333 | 0.464464 | 0.202 | 0.185 | 1 | 7 | Cdc25a | 0.017 | Young |
| 0.424419 | 0.049307 | 0.214 | 0.244 | 1 | 7 | Apmap | -0.03 | Young |
| 0.424457 | 0.038531 | 0.229 | 0.267 | 1 | 7 | Suco | -0.038 | Young |
| 0.425048 | 0.205567 | 0.37 | 0.351 | 1 | 7 | Ubr5 | 0.019 | Young |
| 0.425082 | 0.385679 | 0.145 | 0.126 | 1 | 7 | Prdm4 | 0.019 | Young |
| 0.425182 | 0.142193 | 0.5 | 0.49 | 1 | 7 | Hsbp1 | 0.01 | Young |
| 0.425333 | 0.38131 | 0.051 | 0.038 | 1 | 7 | 6330418K0 | 0.013 | Young |
| 0.425746 | 0.050619 | 0.211 | 0.249 | 1 | 7 | Mras | -0.038 | Young |
| 0.426708 | 0.170948 | 0.47 | 0.456 | 1 | 7 | Dse | 0.014 | Young |
| 0.427193 | 0.572827 | 0.09 | 0.074 | 1 | 7 | Metap1d | 0.016 | Young |
| 0.427471 | 0.086904 | 0.09 | 0.11 | 1 | 7 | Pnpla7 | -0.02 | Young |
| 0.427727 | 0.351561 | 0.175 | 0.203 | 1 | 7 | Frmd8 | -0.028 | Young |
| 0.428572 | 0.293886 | 0.063 | 0.079 | 1 | 7 | Parp3 | -0.016 | Young |
| 0.428698 | 0.318218 | 0.431 | 0.436 | 1 | 7 | Zc3h7a | -0.005 | Young |
| 0.429249 | 0.193754 | 0.62 | 0.633 | 1 | 7 | Trps1 | -0.013 | Young |
| 0.429352 | 0.309531 | 0.042 | 0.031 | 1 | 7 | Gstp3 | 0.011 | Young |
| 0.429528 | 0.03594 | 0.307 | 0.344 | 1 | 7 | Sf3a3 | -0.037 | Young |
| 0.4301 | 0.736316 | 0.12 | 0.105 | 1 | 7 | Smarcd1 | 0.015 | Young |
| 0.430819 | 0.637999 | 0.133 | 0.115 | 1 | 7 | Ska3 | 0.018 | Young |
| 0.430909 | 0.01575 | 0.304 | 0.346 | 1 | 7 | Ide | -0.042 | Young |
| 0.430977 | 0.032908 | 0.244 | 0.277 | 1 | 7 | Lmf2 | -0.033 | Young |
| 0.43108 | 0.607309 | 0.105 | 0.09 | 1 | 7 | Aamdc | 0.015 | Young |
| 0.43144 | 0.036947 | 0.117 | 0.138 | 1 | 7 | Fam241a | -0.021 | Young |
| 0.432239 | 0.172568 | 0.169 | 0.2 | 1 | 7 | 1110038F1 | -0.031 | Young |
| 0.432403 | 0.503129 | 0.081 | 0.067 | 1 | 7 | Cenpu | 0.014 | Young |
| 0.432405 | 0.038855 | 0.123 | 0.105 | 1 | 7 | Rab11fip2 | 0.018 | Young |
| 0.432837 | 0.137205 | 0.654 | 0.644 | 1 | 7 | Prrc2a | 0.01 | Young |
| 0.433219 | 0.567083 | 0.087 | 0.072 | 1 | 7 | Zfp626 | 0.015 | Young |
| 0.433273 | 0.405078 | 0.139 | 0.164 | 1 | 7 | Slc2a6 | -0.025 | Young |
| 0.433434 | 0.303018 | 0.13 | 0.156 | 1 | 7 | Zscan26 | -0.026 | Young |
| 0.43352 | 0.050274 | 0.44 | 0.485 | 1 | 7 | Strn3 | -0.045 | Young |

| | | | | | | | | |
|---|---|---|---|---|---|---|---|---|
| 0.433631 | 0.482309 | 0.19 | 0.172 | 1 | 7 | Slc25a1 | 0.018 | Young |
| 0.433715 | 0.128287 | 0.063 | 0.079 | 1 | 7 | Depdc7 | -0.016 | Young |
| 0.434277 | 0.188958 | 0.084 | 0.069 | 1 | 7 | Pdgfc | 0.015 | Young |
| 0.434278 | 0.147863 | 0.09 | 0.11 | 1 | 7 | D030056L2 | -0.02 | Young |
| 0.434306 | 0.651525 | 0.151 | 0.133 | 1 | 7 | C2cd3 | 0.018 | Young |
| 0.434452 | 0.023522 | 0.072 | 0.059 | 1 | 7 | Rmi2 | 0.013 | Young |
| 0.43451 | 0.069708 | 0.849 | 0.826 | 1 | 7 | Rps27l | 0.023 | Young |
| 0.434572 | 0.183585 | 0.117 | 0.1 | 1 | 7 | Clec5a | 0.017 | Young |
| 0.434619 | 0.044359 | 0.139 | 0.164 | 1 | 7 | Yars | -0.025 | Young |
| 0.434767 | 0.772995 | 0.093 | 0.115 | 1 | 7 | Prrg2 | -0.022 | Young |
| 0.43509 | 0.029412 | 0.467 | 0.526 | 1 | 7 | Cox20 | -0.059 | Young |
| 0.435308 | 0.054814 | 0.313 | 0.341 | 1 | 7 | Ptpn2 | -0.028 | Young |
| 0.435741 | 0.310589 | 0.172 | 0.203 | 1 | 7 | Nkiras2 | -0.031 | Young |
| 0.43606 | 0.478842 | 0.241 | 0.221 | 1 | 7 | Itgax | 0.02 | Young |
| 0.436233 | 0.024612 | 0.289 | 0.321 | 1 | 7 | Plekhg3 | -0.032 | Young |
| 0.43635 | 0.240427 | 0.205 | 0.187 | 1 | 7 | Sft2d1 | 0.018 | Young |
| 0.436364 | 0.46846 | 0.136 | 0.118 | 1 | 7 | H2-Ke6 | 0.018 | Young |
| 0.436388 | 0.078029 | 0.145 | 0.126 | 1 | 7 | Xxylt1 | 0.019 | Young |
| 0.436473 | 0.088329 | 0.181 | 0.162 | 1 | 7 | Ppme1 | 0.019 | Young |
| 0.436823 | 0.009734 | 0.069 | 0.087 | 1 | 7 | Kti12 | -0.018 | Young |
| 0.437042 | 0.440408 | 0.256 | 0.241 | 1 | 7 | Apopt1 | 0.015 | Young |
| 0.437212 | 0.188124 | 0.123 | 0.149 | 1 | 7 | Sephs1 | -0.026 | Young |
| 0.437479 | 0.308835 | 0.015 | 0.023 | 1 | 7 | Gm12355 | -0.008 | Young |
| 0.437776 | 0.093631 | 0.283 | 0.318 | 1 | 7 | Fmr1 | -0.035 | Young |
| 0.438198 | 0.218086 | 0.036 | 0.049 | 1 | 7 | Apex2 | -0.013 | Young |
| 0.438452 | 0.070695 | 0.925 | 0.913 | 1 | 7 | Rpl18 | 0.012 | Young |
| 0.438652 | 0.002722 | 0.316 | 0.295 | 1 | 7 | Sec23b | 0.021 | Young |
| 0.438761 | 0.558946 | 0.172 | 0.154 | 1 | 7 | AU040320 | 0.018 | Young |
| 0.438794 | 0.067779 | 0.325 | 0.297 | 1 | 7 | Cib1 | 0.028 | Young |
| 0.438858 | 0.056132 | 0.136 | 0.156 | 1 | 7 | Usp32 | -0.02 | Young |
| 0.438934 | 0.073173 | 0.301 | 0.274 | 1 | 7 | Jade1 | 0.027 | Young |
| 0.439686 | 0.400096 | 0.163 | 0.144 | 1 | 7 | Ctbs | 0.019 | Young |
| 0.439817 | 0.440694 | 0.093 | 0.113 | 1 | 7 | Smim8 | -0.02 | Young |
| 0.439854 | 0.151138 | 0.211 | 0.19 | 1 | 7 | Slc31a2 | 0.021 | Young |
| 0.439891 | 0.078568 | 0.169 | 0.197 | 1 | 7 | Shc1 | -0.028 | Young |
| 0.439953 | 0.280353 | 0.178 | 0.162 | 1 | 7 | Enoph1 | 0.016 | Young |
| 0.439956 | 0.003693 | 0.178 | 0.203 | 1 | 7 | Fam126b | -0.025 | Young |
| 0.440158 | 0.13822 | 0.136 | 0.159 | 1 | 7 | Atr | -0.023 | Young |
| 0.441157 | 0.032634 | 0.208 | 0.236 | 1 | 7 | Rbl2 | -0.028 | Young |
| 0.441193 | 0.269167 | 0.202 | 0.182 | 1 | 7 | Rnf146 | 0.02 | Young |
| 0.441283 | 0.153313 | 0.304 | 0.346 | 1 | 7 | Mdm4 | -0.042 | Young |
| 0.441627 | 0.486208 | 0.069 | 0.056 | 1 | 7 | Pinx1 | 0.013 | Young |
| 0.441646 | 0.120317 | 0.413 | 0.377 | 1 | 7 | Ndufa10 | 0.036 | Young |
| 0.441659 | 0.08334 | 0.226 | 0.256 | 1 | 7 | Gar1 | -0.03 | Young |
| 0.442255 | 0.348937 | 0.181 | 0.213 | 1 | 7 | Lman2l | -0.032 | Young |

| | | | | | | | | |
|---|---|---|---|---|---|---|---|---|
| 0.442506 | 0.383545 | 0.045 | 0.059 | 1 | 7 | Map4k5 | -0.014 | Young |
| 0.442806 | 0.479319 | 0.015 | 0.023 | 1 | 7 | Lgalsl | -0.008 | Young |
| 0.443022 | 0.095397 | 0.53 | 0.523 | 1 | 7 | Rnf141 | 0.007 | Young |
| 0.443133 | 0.090035 | 0.413 | 0.39 | 1 | 7 | Vps36 | 0.023 | Young |
| 0.443156 | 0.342145 | 0.081 | 0.067 | 1 | 7 | Ogfod3 | 0.014 | Young |
| 0.443218 | 0.24206 | 0.108 | 0.092 | 1 | 7 | Slc41a2 | 0.016 | Young |
| 0.443719 | 0.278984 | 0.136 | 0.118 | 1 | 7 | Anapc7 | 0.018 | Young |
| 0.443943 | 0.683661 | 0.102 | 0.087 | 1 | 7 | Pou5f2 | 0.015 | Young |
| 0.444397 | 0.076459 | 0.03 | 0.041 | 1 | 7 | Ctu1 | -0.011 | Young |
| 0.44468 | 0.090118 | 0.431 | 0.479 | 1 | 7 | Bptf | -0.048 | Young |
| 0.445464 | 0.311806 | 0.479 | 0.49 | 1 | 7 | Fam168b | -0.011 | Young |
| 0.44581 | 0.594827 | 0.358 | 0.346 | 1 | 7 | Rbm6 | 0.012 | Young |
| 0.446378 | 0.170042 | 0.015 | 0.023 | 1 | 7 | Slc37a3 | -0.008 | Young |
| 0.446536 | 0.011727 | 0.117 | 0.138 | 1 | 7 | Ubr1 | -0.021 | Young |
| 0.446626 | 0.032835 | 0.166 | 0.19 | 1 | 7 | Dennd1c | -0.024 | Young |
| 0.446629 | 0.42256 | 0.009 | 0.015 | 1 | 7 | Tmem170 | -0.006 | Young |
| 0.447056 | 0.056752 | 0.142 | 0.123 | 1 | 7 | Inafm1 | 0.019 | Young |
| 0.447144 | 0.08963 | 0.123 | 0.144 | 1 | 7 | Nop14 | -0.021 | Young |
| 0.447249 | 0.077548 | 0.623 | 0.628 | 1 | 7 | Sf1 | -0.005 | Young |
| 0.447267 | 0.208019 | 0.163 | 0.192 | 1 | 7 | Ppm1h | -0.029 | Young |
| 0.447749 | 0.373039 | 0.075 | 0.062 | 1 | 7 | Hcar2 | 0.013 | Young |
| 0.448047 | 0.427777 | 0.187 | 0.174 | 1 | 7 | Ints7 | 0.013 | Young |
| 0.448093 | 0.609588 | 0.123 | 0.108 | 1 | 7 | Etaa1 | 0.015 | Young |
| 0.44817 | 0.441602 | 0.015 | 0.023 | 1 | 7 | 4931414P1 | -0.008 | Young |
| 0.44817 | 0.163952 | 0.015 | 0.023 | 1 | 7 | 1700037C1 | -0.008 | Young |
| 0.44837 | 0.330211 | 0.03 | 0.041 | 1 | 7 | Sapcd2 | -0.011 | Young |
| 0.448371 | 0.054131 | 0.265 | 0.3 | 1 | 7 | Synj2bp | -0.035 | Young |
| 0.448594 | 0.14831 | 0.28 | 0.256 | 1 | 7 | Rnf38 | 0.024 | Young |
| 0.448753 | 0.377008 | 0.075 | 0.092 | 1 | 7 | Fam13b | -0.017 | Young |
| 0.448858 | 0.265935 | 0.009 | 0.015 | 1 | 7 | Zfp105 | -0.006 | Young |
| 0.448984 | 0.050874 | 0.455 | 0.436 | 1 | 7 | Rassf1 | 0.019 | Young |
| 0.448996 | 0.633184 | 0.154 | 0.138 | 1 | 7 | Klhl20 | 0.016 | Young |
| 0.449028 | 0.338513 | 0.211 | 0.192 | 1 | 7 | Pcyox1 | 0.019 | Young |
| 0.449034 | 0.162377 | 0.533 | 0.546 | 1 | 7 | Klf13 | -0.013 | Young |
| 0.449067 | 0.389065 | 0.015 | 0.023 | 1 | 7 | Homez | -0.008 | Young |
| 0.449224 | 0.831936 | 0.075 | 0.062 | 1 | 7 | Acot2 | 0.013 | Young |
| 0.449525 | 0.340045 | 0.084 | 0.069 | 1 | 7 | Eci1 | 0.015 | Young |
| 0.449863 | 0.178483 | 0.13 | 0.154 | 1 | 7 | Ppat | -0.024 | Young |
| 0.449966 | 0.270161 | 0.015 | 0.023 | 1 | 7 | Gstm5 | -0.008 | Young |
| 0.450308 | 0.146819 | 0.169 | 0.197 | 1 | 7 | Nectin1 | -0.028 | Young |
| 0.450983 | 0.227689 | 0.617 | 0.623 | 1 | 7 | Selenok | -0.006 | Young |
| 0.451375 | 0.384749 | 0.217 | 0.246 | 1 | 7 | Ipmk | -0.029 | Young |
| 0.451446 | 0.033427 | 0.277 | 0.303 | 1 | 7 | Nup153 | -0.026 | Young |
| 0.451488 | 0.005593 | 0.123 | 0.144 | 1 | 7 | Depdc1a | -0.021 | Young |
| 0.451536 | 0.127102 | 0.289 | 0.267 | 1 | 7 | Gabarapl2 | 0.022 | Young |

| | | | | | | | | |
|---|---|---|---|---|---|---|---|---|
| 0.451679 | 0.202698 | 0.133 | 0.113 | 1 | 7 | Lrp12 | 0.02 | Young |
| 0.451835 | 0.43882 | 0.069 | 0.056 | 1 | 7 | Ifitm5 | 0.013 | Young |
| 0.452014 | 0.6732 | 0.063 | 0.051 | 1 | 7 | Tpbgl | 0.012 | Young |
| 0.452213 | 0.973405 | 0.009 | 0.015 | 1 | 7 | Tnfsf10 | -0.006 | Young |
| 0.452213 | 0.4551 | 0.009 | 0.015 | 1 | 7 | Tha1 | -0.006 | Young |
| 0.452213 | 0.316081 | 0.009 | 0.015 | 1 | 7 | Zfp607b | -0.006 | Young |
| 0.452296 | 0.174983 | 0.44 | 0.441 | 1 | 7 | Aldh9a1 | -0.001 | Young |
| 0.452358 | 0.643827 | 0.136 | 0.121 | 1 | 7 | Zfp236 | 0.015 | Young |
| 0.45247 | 0.132575 | 0.136 | 0.162 | 1 | 7 | Ect2 | -0.026 | Young |
| 0.452564 | 0.463371 | 0.208 | 0.192 | 1 | 7 | P2rx7 | 0.016 | Young |
| 0.452598 | 0.050486 | 0.169 | 0.195 | 1 | 7 | Mthfd1 | -0.026 | Young |
| 0.452701 | 0.089508 | 0.127 | 0.149 | 1 | 7 | Tcp11l1 | -0.022 | Young |
| 0.452978 | 0.708576 | 0.039 | 0.051 | 1 | 7 | Rhebl1 | -0.012 | Young |
| 0.453124 | 0.034026 | 0.503 | 0.562 | 1 | 7 | Gsk3b | -0.059 | Young |
| 0.453149 | 0.346739 | 0.199 | 0.182 | 1 | 7 | Zfp664 | 0.017 | Young |
| 0.453334 | 1.200594 | 0.009 | 0.015 | 1 | 7 | Plppr4 | -0.006 | Young |
| 0.453334 | 0.861727 | 0.009 | 0.015 | 1 | 7 | Mafa | -0.006 | Young |
| 0.453557 | 0.089989 | 0.178 | 0.205 | 1 | 7 | Kdm5c | -0.027 | Young |
| 0.454256 | 0.339662 | 0.211 | 0.192 | 1 | 7 | Lyst | 0.019 | Young |
| 0.454277 | 0.029546 | 0.533 | 0.577 | 1 | 7 | Sumo3 | -0.044 | Young |
| 0.454293 | 0.283612 | 0.238 | 0.274 | 1 | 7 | Exosc3 | -0.036 | Young |
| 0.454459 | 0.008474 | 0.274 | 0.308 | 1 | 7 | Psmc2 | -0.034 | Young |
| 0.455025 | 0.106777 | 0.102 | 0.123 | 1 | 7 | Tsga10 | -0.021 | Young |
| 0.455042 | 0.569579 | 0.199 | 0.182 | 1 | 7 | Smim20 | 0.017 | Young |
| 0.455098 | 0.239034 | 0.395 | 0.379 | 1 | 7 | Map3k1 | 0.016 | Young |
| 0.455207 | 0.05547 | 0.943 | 0.933 | 1 | 7 | Ctsb | 0.01 | Young |
| 0.455239 | 0.426091 | 0.151 | 0.177 | 1 | 7 | Coa6 | -0.026 | Young |
| 0.455365 | 0.049392 | 0.039 | 0.051 | 1 | 7 | Dgka | -0.012 | Young |
| 0.455493 | 0.242851 | 0.47 | 0.454 | 1 | 7 | Psenen | 0.016 | Young |
| 0.456258 | 0.17196 | 0.123 | 0.108 | 1 | 7 | Rasgrp2 | 0.015 | Young |
| 0.456415 | 0.265671 | 0.295 | 0.272 | 1 | 7 | Arid5a | 0.023 | Young |
| 0.456558 | 0.141007 | 0.142 | 0.121 | 1 | 7 | Slc7a7 | 0.021 | Young |
| 0.45664 | 0.019789 | 0.06 | 0.074 | 1 | 7 | Ifitm1 | -0.014 | Young |
| 0.456767 | 0.241147 | 0.142 | 0.123 | 1 | 7 | Nbn | 0.019 | Young |
| 0.457146 | 0.020369 | 0.142 | 0.164 | 1 | 7 | Tex264 | -0.022 | Young |
| 0.457324 | 0.189258 | 0.325 | 0.308 | 1 | 7 | Ccdc47 | 0.017 | Young |
| 0.457327 | 0.546243 | 0.229 | 0.215 | 1 | 7 | Zfp280d | 0.014 | Young |
| 0.457713 | 1.04207 | 0.081 | 0.069 | 1 | 7 | D230025D1 | 0.012 | Young |
| 0.457725 | 0.02093 | 0.244 | 0.277 | 1 | 7 | Ppp2r5e | -0.033 | Young |
| 0.457758 | 0.355229 | 0.039 | 0.051 | 1 | 7 | Ccdc181 | -0.012 | Young |
| 0.457759 | 0.165163 | 0.358 | 0.405 | 1 | 7 | Arhgap31 | -0.047 | Young |
| 0.45796 | 0.098371 | 0.458 | 0.513 | 1 | 7 | Emc7 | -0.055 | Young |
| 0.458152 | 0.026433 | 0.428 | 0.472 | 1 | 7 | Dnajc7 | -0.044 | Young |
| 0.458182 | 0.025662 | 0.259 | 0.287 | 1 | 7 | Sp110 | -0.028 | Young |
| 0.45903 | 0.067496 | 0.485 | 0.533 | 1 | 7 | Akt1 | -0.048 | Young |

| | | | | | | | | |
|---|---|---|---|---|---|---|---|---|
| 0.45928 | 0.152254 | 0.295 | 0.336 | 1 | 7 | Sike1 | -0.041 | Young |
| 0.459352 | 0.262417 | 0.108 | 0.092 | 1 | 7 | Scai | 0.016 | Young |
| 0.459434 | 0.105588 | 0.175 | 0.2 | 1 | 7 | Poc1b | -0.025 | Young |
| 0.459476 | 0.298831 | 0.199 | 0.182 | 1 | 7 | Nmd3 | 0.017 | Young |
| 0.459494 | 0.188033 | 0.298 | 0.272 | 1 | 7 | Arl6ip6 | 0.026 | Young |
| 0.459767 | 0.241062 | 0.268 | 0.303 | 1 | 7 | Ireb2 | -0.035 | Young |
| 0.459793 | 0.193412 | 0.148 | 0.128 | 1 | 7 | Bckdha | 0.02 | Young |
| 0.460661 | 0.064856 | 0.292 | 0.326 | 1 | 7 | Trip11 | -0.034 | Young |
| 0.460901 | 0.220781 | 0.488 | 0.495 | 1 | 7 | Tmed9 | -0.007 | Young |
| 0.46093 | 0.094641 | 0.024 | 0.033 | 1 | 7 | Gm15448 | -0.009 | Young |
| 0.461121 | 0.047745 | 0.145 | 0.169 | 1 | 7 | Golga3 | -0.024 | Young |
| 0.461272 | 0.527607 | 0.078 | 0.097 | 1 | 7 | Zfp26 | -0.019 | Young |
| 0.461323 | 0.073202 | 0.28 | 0.31 | 1 | 7 | Mrpl34 | -0.03 | Young |
| 0.461361 | 0.036609 | 0.039 | 0.051 | 1 | 7 | Vsig10 | -0.012 | Young |
| 0.461458 | 0.065545 | 0.292 | 0.326 | 1 | 7 | Rheb | -0.034 | Young |
| 0.461601 | 0.138941 | 0.346 | 0.387 | 1 | 7 | Dazap1 | -0.041 | Young |
| 0.461903 | 0.618125 | 0.148 | 0.131 | 1 | 7 | Taf12 | 0.017 | Young |
| 0.461963 | 0.196986 | 0.039 | 0.051 | 1 | 7 | D3Ertd751 | -0.012 | Young |
| 0.462095 | 0.246304 | 0.41 | 0.397 | 1 | 7 | Rap1gds1 | 0.013 | Young |
| 0.462187 | 0.197826 | 0.181 | 0.159 | 1 | 7 | Slc4a1ap | 0.022 | Young |
| 0.462243 | 0.114144 | 0.648 | 0.641 | 1 | 7 | Lyn | 0.007 | Young |
| 0.462462 | 0.092621 | 0.241 | 0.269 | 1 | 7 | Snrnp27 | -0.028 | Young |
| 0.462566 | 0.409802 | 0.039 | 0.051 | 1 | 7 | Fads6 | -0.012 | Young |
| 0.462675 | 0.279604 | 0.13 | 0.11 | 1 | 7 | Icmt | 0.02 | Young |
| 0.462938 | 0.171291 | 0.256 | 0.295 | 1 | 7 | Smad4 | -0.039 | Young |
| 0.463016 | 0.392286 | 0.199 | 0.187 | 1 | 7 | Gadd45g | 0.012 | Young |
| 0.463191 | 0.05883 | 0.587 | 0.636 | 1 | 7 | Ehd1 | -0.049 | Young |
| 0.463478 | 0.440718 | 0.178 | 0.203 | 1 | 7 | Cep128 | -0.025 | Young |
| 0.463863 | 0.146129 | 0.34 | 0.372 | 1 | 7 | Emc6 | -0.032 | Young |
| 0.464917 | 0.021947 | 0.093 | 0.077 | 1 | 7 | Mzt2 | 0.016 | Young |
| 0.465071 | 0.088996 | 0.072 | 0.059 | 1 | 7 | Vwf | 0.013 | Young |
| 0.465081 | 0.428381 | 0.178 | 0.208 | 1 | 7 | Nek2 | -0.03 | Young |
| 0.46529 | 0.261618 | 0.066 | 0.054 | 1 | 7 | Usp42 | 0.012 | Young |
| 0.465463 | 0.70792 | 0.108 | 0.095 | 1 | 7 | Zfp90 | 0.013 | Young |
| 0.465514 | 0.077643 | 0.325 | 0.308 | 1 | 7 | Bdp1 | 0.017 | Young |
| 0.46556 | 0.024044 | 0.404 | 0.446 | 1 | 7 | Serpinb6a | -0.042 | Young |
| 0.465958 | 0.099951 | 0.078 | 0.064 | 1 | 7 | Mark4 | 0.014 | Young |
| 0.466056 | 0.018066 | 0.41 | 0.444 | 1 | 7 | Eprs | -0.034 | Young |
| 0.466486 | 0.104607 | 0.09 | 0.108 | 1 | 7 | Ptk2 | -0.018 | Young |
| 0.467024 | 1.45588 | 0.048 | 0.038 | 1 | 7 | Rbbp9 | 0.01 | Young |
| 0.467401 | 0.113645 | 0.039 | 0.051 | 1 | 7 | Inpp5b | -0.012 | Young |
| 0.468173 | 0.072678 | 0.09 | 0.108 | 1 | 7 | Mrpl1 | -0.018 | Young |
| 0.468195 | 0.398222 | 0.072 | 0.09 | 1 | 7 | Nt5dc3 | -0.018 | Young |
| 0.468356 | 0.03692 | 0.524 | 0.505 | 1 | 7 | Dnmt1 | 0.019 | Young |
| 0.46839 | 0.159924 | 0.425 | 0.413 | 1 | 7 | Akt2 | 0.012 | Young |

| | | | | | | | | |
|---|---|---|---|---|---|---|---|---|
| 0.468425 | 0.348329 | 0.202 | 0.185 | 1 | 7 | Tmem135 | 0.017 | Young |
| 0.468462 | 0.161532 | 0.102 | 0.121 | 1 | 7 | Fdps | -0.019 | Young |
| 0.468597 | 0.198327 | 0.334 | 0.387 | 1 | 7 | Nup62 | -0.053 | Young |
| 0.468637 | 0.076289 | 0.554 | 0.549 | 1 | 7 | Lman2 | 0.005 | Young |
| 0.468759 | 0.230484 | 0.319 | 0.308 | 1 | 7 | Trappc10 | 0.011 | Young |
| 0.468961 | 0.025093 | 0.066 | 0.082 | 1 | 7 | Tspyl2 | -0.016 | Young |
| 0.469033 | 0.350946 | 0.066 | 0.054 | 1 | 7 | Clstn1 | 0.012 | Young |
| 0.46926 | 0.004735 | 0.202 | 0.226 | 1 | 7 | Mettl23 | -0.024 | Young |
| 0.469913 | 0.67153 | 0.16 | 0.146 | 1 | 7 | Paqr4 | 0.014 | Young |
| 0.469963 | 0.003603 | 0.16 | 0.185 | 1 | 7 | Washc1 | -0.025 | Young |
| 0.470022 | 0.041362 | 0.545 | 0.603 | 1 | 7 | Lamtor1 | -0.058 | Young |
| 0.470054 | 0.204526 | 0.111 | 0.095 | 1 | 7 | Sema4b | 0.016 | Young |
| 0.470664 | 0.001038 | 0.06 | 0.074 | 1 | 7 | Zkscan1 | -0.014 | Young |
| 0.470975 | 0.465762 | 0.259 | 0.244 | 1 | 7 | Homer1 | 0.015 | Young |
| 0.471025 | 0.036646 | 0.367 | 0.403 | 1 | 7 | Glrx5 | -0.036 | Young |
| 0.471121 | 0.054668 | 0.13 | 0.151 | 1 | 7 | Gm17018 | -0.021 | Young |
| 0.471399 | 0.138066 | 0.377 | 0.354 | 1 | 7 | Ccnt2 | 0.023 | Young |
| 0.472535 | 0.301725 | 0.277 | 0.264 | 1 | 7 | Ints6l | 0.013 | Young |
| 0.47254 | 0.47015 | 0.253 | 0.236 | 1 | 7 | Gpr160 | 0.017 | Young |
| 0.472836 | 0.52481 | 0.096 | 0.082 | 1 | 7 | Thbs1 | 0.014 | Young |
| 0.473662 | 0.088044 | 0.099 | 0.118 | 1 | 7 | Donson | -0.019 | Young |
| 0.473739 | 0.588401 | 0.024 | 0.033 | 1 | 7 | Sirt5 | -0.009 | Young |
| 0.474007 | 0.138358 | 0.449 | 0.438 | 1 | 7 | Arrb1 | 0.011 | Young |
| 0.474036 | 0.056558 | 0.075 | 0.092 | 1 | 7 | Letmd1 | -0.017 | Young |
| 0.474077 | 0.213567 | 0.13 | 0.113 | 1 | 7 | Tspan17 | 0.017 | Young |
| 0.474295 | 0.604247 | 0.108 | 0.095 | 1 | 7 | Fam78a | 0.013 | Young |
| 0.474715 | 0.113894 | 0.075 | 0.062 | 1 | 7 | Zfp386 | 0.013 | Young |
| 0.474852 | 0.234471 | 0.235 | 0.221 | 1 | 7 | Lpin2 | 0.014 | Young |
| 0.475556 | 0.156039 | 0.355 | 0.4 | 1 | 7 | Appl1 | -0.045 | Young |
| 0.475852 | 0.074771 | 0.271 | 0.3 | 1 | 7 | Map3k7 | -0.029 | Young |
| 0.475854 | 0.464337 | 0.301 | 0.287 | 1 | 7 | Igbp1 | 0.014 | Young |
| 0.475931 | 0.805603 | 0.039 | 0.051 | 1 | 7 | Depdc1b | -0.012 | Young |
| 0.476176 | 0.198784 | 0.123 | 0.144 | 1 | 7 | Agpat1 | -0.021 | Young |
| 0.476475 | 0.279395 | 0.259 | 0.241 | 1 | 7 | Trim12c | 0.018 | Young |
| 0.47685 | 0.079792 | 0.705 | 0.69 | 1 | 7 | Uqcrq | 0.015 | Young |
| 0.476853 | 0.754325 | 0.114 | 0.1 | 1 | 7 | Tmem29 | 0.014 | Young |
| 0.476943 | 0.235441 | 0.172 | 0.154 | 1 | 7 | Icosl | 0.018 | Young |
| 0.477262 | 0.052519 | 0.904 | 0.867 | 1 | 7 | Cybb | 0.037 | Young |
| 0.477544 | 0.109685 | 0.024 | 0.033 | 1 | 7 | Fam57a | -0.009 | Young |
| 0.477613 | 0.178822 | 0.093 | 0.11 | 1 | 7 | Hdac7 | -0.017 | Young |
| 0.477729 | 0.047656 | 0.223 | 0.244 | 1 | 7 | Abi2 | -0.021 | Young |
| 0.478061 | 0.087258 | 0.13 | 0.151 | 1 | 7 | Hmgn5 | -0.021 | Young |
| 0.478407 | 0.287788 | 0.154 | 0.136 | 1 | 7 | Sclt1 | 0.018 | Young |
| 0.47846 | 0.127272 | 0.069 | 0.056 | 1 | 7 | Fam46c | 0.013 | Young |
| 0.478946 | 0.144406 | 0.446 | 0.433 | 1 | 7 | Prkd3 | 0.013 | Young |

| | | | | | | | | |
|---|---|---|---|---|---|---|---|---|
| 0.478972 | 0.519127 | 0.087 | 0.074 | 1 | 7 | Ticam1 | 0.013 | Young |
| 0.478995 | 0.835621 | 0.057 | 0.046 | 1 | 7 | Zfp35 | 0.011 | Young |
| 0.479023 | 0.45778 | 0.199 | 0.187 | 1 | 7 | Slc30a7 | 0.012 | Young |
| 0.479215 | 0.408827 | 0.256 | 0.238 | 1 | 7 | Cdkn2c | 0.018 | Young |
| 0.479269 | 0.077625 | 0.976 | 0.964 | 1 | 7 | Lyz2 | 0.012 | Young |
| 0.479316 | 0.461299 | 0.211 | 0.192 | 1 | 7 | Cd80 | 0.019 | Young |
| 0.479713 | 0.327984 | 0.127 | 0.113 | 1 | 7 | Carm1 | 0.014 | Young |
| 0.479813 | 0.032376 | 0.316 | 0.362 | 1 | 7 | Lpp | -0.046 | Young |
| 0.480306 | 0.057686 | 0.217 | 0.195 | 1 | 7 | Foxj2 | 0.022 | Young |
| 0.480421 | 0.071978 | 0.081 | 0.097 | 1 | 7 | Jade2 | -0.016 | Young |
| 0.480456 | 0.048566 | 0.069 | 0.085 | 1 | 7 | Trub2 | -0.016 | Young |
| 0.480599 | 0.168272 | 0.024 | 0.033 | 1 | 7 | Loxl3 | -0.009 | Young |
| 0.480814 | 0.299962 | 0.238 | 0.277 | 1 | 7 | Pi4ka | -0.039 | Young |
| 0.480871 | 0.881468 | 0.048 | 0.038 | 1 | 7 | 2700062CC | 0.01 | Young |
| 0.481052 | 0.348007 | 0.193 | 0.174 | 1 | 7 | Ltbr | 0.019 | Young |
| 0.481446 | 0.320349 | 0.117 | 0.103 | 1 | 7 | Phf13 | 0.014 | Young |
| 0.481768 | 0.068226 | 0.274 | 0.305 | 1 | 7 | Ddx54 | -0.031 | Young |
| 0.48182 | 0.000352 | 0.452 | 0.49 | 1 | 7 | Ttc14 | -0.038 | Young |
| 0.482118 | 0.255631 | 0.093 | 0.079 | 1 | 7 | Tubgcp6 | 0.014 | Young |
| 0.482138 | 1.12786 | 0.084 | 0.072 | 1 | 7 | Rnaset2b | 0.012 | Young |
| 0.482314 | 0.699757 | 0.154 | 0.141 | 1 | 7 | Ticam2 | 0.013 | Young |
| 0.482489 | 0.385131 | 0.319 | 0.315 | 1 | 7 | Mid1ip1 | 0.004 | Young |
| 0.483078 | 0.571109 | 0.057 | 0.046 | 1 | 7 | Armc6 | 0.011 | Young |
| 0.483202 | 0.475689 | 0.09 | 0.077 | 1 | 7 | Jmy | 0.013 | Young |
| 0.483321 | 0.554213 | 0.078 | 0.067 | 1 | 7 | Gprasp1 | 0.011 | Young |
| 0.4834 | 0.371134 | 0.346 | 0.338 | 1 | 7 | Cdk2 | 0.008 | Young |
| 0.483578 | 0.160771 | 0.19 | 0.215 | 1 | 7 | Usp12 | -0.025 | Young |
| 0.483661 | 0.379119 | 0.286 | 0.269 | 1 | 7 | Diaph2 | 0.017 | Young |
| 0.48378 | 0.23567 | 0.395 | 0.39 | 1 | 7 | Stk10 | 0.005 | Young |
| 0.483893 | 0.183562 | 0.078 | 0.064 | 1 | 7 | Oraov1 | 0.014 | Young |
| 0.48392 | 0.735834 | 0.114 | 0.1 | 1 | 7 | Pibf1 | 0.014 | Young |
| 0.483944 | 0.037952 | 0.681 | 0.672 | 1 | 7 | Eif3k | 0.009 | Young |
| 0.483987 | 0.705373 | 0.054 | 0.044 | 1 | 7 | Sec61a2 | 0.01 | Young |
| 0.484196 | 0.135076 | 0.172 | 0.197 | 1 | 7 | Faf2 | -0.025 | Young |
| 0.484258 | 0.288101 | 0.361 | 0.349 | 1 | 7 | Nrip1 | 0.012 | Young |
| 0.484421 | 0.115553 | 0.25 | 0.277 | 1 | 7 | Glod4 | -0.027 | Young |
| 0.485021 | 0.603186 | 0.193 | 0.179 | 1 | 7 | Spata2 | 0.014 | Young |
| 0.48507 | 0.220477 | 0.202 | 0.182 | 1 | 7 | Spag5 | 0.02 | Young |
| 0.485419 | 0.53275 | 0.057 | 0.046 | 1 | 7 | Nubpl | 0.011 | Young |
| 0.485591 | 0.555638 | 0.114 | 0.1 | 1 | 7 | Wdr81 | 0.014 | Young |
| 0.485826 | 0.127597 | 0.235 | 0.272 | 1 | 7 | Pdha1 | -0.037 | Young |
| 0.486402 | 1.150025 | 0.051 | 0.041 | 1 | 7 | Ttbk2 | 0.01 | Young |
| 0.486514 | 0.084357 | 0.783 | 0.795 | 1 | 7 | Srgn | -0.012 | Young |
| 0.486893 | 0.623212 | 0.25 | 0.244 | 1 | 7 | Cebpd | 0.006 | Young |
| 0.48707 | 0.611836 | 0.166 | 0.151 | 1 | 7 | Aspscr1 | 0.015 | Young |

| | | | | | | | | |
|---|---|---|---|---|---|---|---|---|
| 0.487409 | 0.199911 | 0.34 | 0.323 | 1 | 7 | Snx8 | 0.017 | Young |
| 0.487528 | 0.462443 | 0.361 | 0.367 | 1 | 7 | Osbpl9 | -0.006 | Young |
| 0.4876 | 0.780028 | 0.054 | 0.044 | 1 | 7 | Ccdc138 | 0.01 | Young |
| 0.487753 | 0.130749 | 0.139 | 0.159 | 1 | 7 | Foxred2 | -0.02 | Young |
| 0.488107 | 0.325055 | 0.093 | 0.079 | 1 | 7 | Slc30a4 | 0.014 | Young |
| 0.488343 | 0.124318 | 0.482 | 0.469 | 1 | 7 | Msl1 | 0.013 | Young |
| 0.488378 | 0.024172 | 0.943 | 0.926 | 1 | 7 | Rpl3 | 0.017 | Young |
| 0.488378 | 0.182542 | 0.178 | 0.208 | 1 | 7 | Faim | -0.03 | Young |
| 0.488983 | 0.00647 | 0.955 | 0.954 | 1 | 7 | Oaz1 | 0.001 | Young |
| 0.489026 | 0.202272 | 0.169 | 0.192 | 1 | 7 | Upf3b | -0.023 | Young |
| 0.489028 | 0.506578 | 0.181 | 0.167 | 1 | 7 | Pop4 | 0.014 | Young |
| 0.489157 | 1.035316 | 0.048 | 0.038 | 1 | 7 | Spef1 | 0.01 | Young |
| 0.489274 | 0.193988 | 0.211 | 0.236 | 1 | 7 | Trabd | -0.025 | Young |
| 0.489658 | 0.141285 | 0.108 | 0.128 | 1 | 7 | Kif18b | -0.02 | Young |
| 0.489797 | 0.881681 | 0.048 | 0.038 | 1 | 7 | C030034I2 | 0.01 | Young |
| 0.490513 | 0.115008 | 0.268 | 0.254 | 1 | 7 | Ccz1 | 0.014 | Young |
| 0.490632 | 0.555629 | 0.033 | 0.044 | 1 | 7 | Spaca6 | -0.011 | Young |
| 0.490795 | 0.462819 | 0.12 | 0.105 | 1 | 7 | Ofd1 | 0.015 | Young |
| 0.490966 | 0.177434 | 0.527 | 0.533 | 1 | 7 | Arpp19 | -0.006 | Young |
| 0.491014 | 0.090071 | 0.889 | 0.877 | 1 | 7 | Arpc1b | 0.012 | Young |
| 0.491214 | 0.233299 | 0.093 | 0.11 | 1 | 7 | Slc20a2 | -0.017 | Young |
| 0.491227 | 0.270246 | 0.054 | 0.044 | 1 | 7 | Zfp668 | 0.01 | Young |
| 0.491834 | 0.285886 | 0.244 | 0.231 | 1 | 7 | Vkorc1 | 0.013 | Young |
| 0.492029 | 0.604784 | 0.12 | 0.105 | 1 | 7 | Fam122b | 0.015 | Young |
| 0.492161 | 0.211465 | 0.081 | 0.097 | 1 | 7 | Pdcd2 | -0.016 | Young |
| 0.492284 | 0.49926 | 0.235 | 0.223 | 1 | 7 | Dclre1c | 0.012 | Young |
| 0.492401 | 0.003929 | 0.127 | 0.146 | 1 | 7 | Ell | -0.019 | Young |
| 0.492439 | 0.034043 | 0.042 | 0.054 | 1 | 7 | Disc1 | -0.012 | Young |
| 0.492757 | 0.44099 | 0.289 | 0.277 | 1 | 7 | Ldb1 | 0.012 | Young |
| 0.492902 | 0.128591 | 0.361 | 0.408 | 1 | 7 | Dnajc13 | -0.047 | Young |
| 0.493046 | 0.083162 | 0.042 | 0.054 | 1 | 7 | Qtrt2 | -0.012 | Young |
| 0.493164 | 0.540539 | 0.142 | 0.128 | 1 | 7 | Ephx1 | 0.014 | Young |
| 0.49385 | 0.30194 | 0.105 | 0.126 | 1 | 7 | Utp15 | -0.021 | Young |
| 0.494403 | 0.037428 | 0.593 | 0.608 | 1 | 7 | Ankrd11 | -0.015 | Young |
| 0.49507 | 0.375226 | 0.133 | 0.118 | 1 | 7 | Klra2 | 0.015 | Young |
| 0.495073 | 0.035909 | 0.274 | 0.254 | 1 | 7 | Fam114a2 | 0.02 | Young |
| 0.495476 | 0.753515 | 0.054 | 0.044 | 1 | 7 | Zfp930 | 0.01 | Young |
| 0.4966 | 0.487853 | 0.142 | 0.128 | 1 | 7 | Zbtb22 | 0.014 | Young |
| 0.496774 | 0.548399 | 0.187 | 0.174 | 1 | 7 | Lcor | 0.013 | Young |
| 0.496777 | 0.085379 | 0.163 | 0.187 | 1 | 7 | Scaf4 | -0.024 | Young |
| 0.496869 | 0.495671 | 0.154 | 0.138 | 1 | 7 | Wipi2 | 0.016 | Young |
| 0.497052 | 0.026554 | 0.145 | 0.167 | 1 | 7 | Cbll1 | -0.022 | Young |
| 0.497204 | 0.330589 | 0.175 | 0.2 | 1 | 7 | Kmt5c | -0.025 | Young |
| 0.497289 | 0.090307 | 0.108 | 0.128 | 1 | 7 | Kdm4c | -0.02 | Young |
| 0.497588 | 0.029202 | 0.054 | 0.067 | 1 | 7 | Cep72 | -0.013 | Young |

| | | | | | | | | |
|---|---|---|---|---|---|---|---|---|
| 0.498103 | 0.031814 | 0.018 | 0.026 | 1 | 7 | 8030462N1 | -0.008 | Young |
| 0.498525 | 0.579021 | 0.087 | 0.074 | 1 | 7 | Bcl9l | 0.013 | Young |
| 0.498707 | 0.127548 | 0.102 | 0.087 | 1 | 7 | Sik2 | 0.015 | Young |
| 0.499123 | 0.141573 | 0.133 | 0.151 | 1 | 7 | Rbbp5 | -0.018 | Young |
| 0.499256 | 0.228928 | 0.217 | 0.251 | 1 | 7 | Pimreg | -0.034 | Young |
| 0.499544 | 0.002405 | 0.196 | 0.226 | 1 | 7 | Uhrf1bp1l | -0.03 | Young |
| 0.499886 | 0.576703 | 0.018 | 0.026 | 1 | 7 | Inpp1 | -0.008 | Young |
| 0.500305 | 0.824203 | 0.145 | 0.133 | 1 | 7 | Gnpat | 0.012 | Young |
| 0.500357 | 0.582512 | 0.054 | 0.044 | 1 | 7 | Plscr4 | 0.01 | Young |
| 0.500627 | 0.126828 | 0.12 | 0.138 | 1 | 7 | Maip1 | -0.018 | Young |
| 0.500689 | 0.317514 | 0.28 | 0.264 | 1 | 7 | Tec | 0.016 | Young |
| 0.500755 | 1.257442 | 0.048 | 0.038 | 1 | 7 | Gm10138 | 0.01 | Young |
| 0.500755 | 0.745552 | 0.048 | 0.038 | 1 | 7 | Tatdn3 | 0.01 | Young |
| 0.500759 | 0.166722 | 0.304 | 0.292 | 1 | 7 | Fbxo33 | 0.012 | Young |
| 0.500778 | 0.04369 | 0.018 | 0.026 | 1 | 7 | Xlr4a | -0.008 | Young |
| 0.500945 | 0.001385 | 0.136 | 0.121 | 1 | 7 | Qsox1 | 0.015 | Young |
| 0.501525 | 0.614418 | 0.265 | 0.256 | 1 | 7 | Rab22a | 0.009 | Young |
| 0.501807 | 0.236695 | 0.051 | 0.064 | 1 | 7 | Gpt2 | -0.013 | Young |
| 0.502068 | 0.535369 | 0.151 | 0.179 | 1 | 7 | Zadh2 | -0.028 | Young |
| 0.502472 | 0.517169 | 0.229 | 0.215 | 1 | 7 | Lyl1 | 0.014 | Young |
| 0.502688 | 0.14853 | 0.202 | 0.231 | 1 | 7 | Pin1 | -0.029 | Young |
| 0.502875 | 0.286317 | 0.199 | 0.179 | 1 | 7 | Mboat7 | 0.02 | Young |
| 0.502926 | 0.075209 | 0.497 | 0.492 | 1 | 7 | Lrrc58 | 0.005 | Young |
| 0.503519 | 0.634298 | 0.265 | 0.259 | 1 | 7 | Nr2c2 | 0.006 | Young |
| 0.503682 | 0.289118 | 0.313 | 0.295 | 1 | 7 | Ergic2 | 0.018 | Young |
| 0.503994 | 0.441377 | 0.078 | 0.067 | 1 | 7 | Trp53rka | 0.011 | Young |
| 0.504184 | 0.012425 | 0.599 | 0.615 | 1 | 7 | Caprin1 | -0.016 | Young |
| 0.504291 | 0.369552 | 0.298 | 0.285 | 1 | 7 | Vamp2 | 0.013 | Young |
| 0.504449 | 0.328226 | 0.09 | 0.077 | 1 | 7 | Dcakd | 0.013 | Young |
| 0.504485 | 0.057506 | 0.94 | 0.936 | 1 | 7 | Lcp1 | 0.004 | Young |
| 0.504658 | 0.280986 | 0.295 | 0.336 | 1 | 7 | Ckap2l | -0.041 | Young |
| 0.504749 | 0.323288 | 0.096 | 0.082 | 1 | 7 | Tmtc3 | 0.014 | Young |
| 0.504883 | 0.268321 | 0.25 | 0.279 | 1 | 7 | Mepce | -0.029 | Young |
| 0.504901 | 0.266139 | 0.093 | 0.079 | 1 | 7 | Ppard | 0.014 | Young |
| 0.505005 | 0.208425 | 0.425 | 0.421 | 1 | 7 | Eny2 | 0.004 | Young |
| 0.505098 | 0.228315 | 0.208 | 0.192 | 1 | 7 | Letm1 | 0.016 | Young |
| 0.505283 | 0.083733 | 0.054 | 0.067 | 1 | 7 | Fam213a | -0.013 | Young |
| 0.505568 | 0.276441 | 0.057 | 0.046 | 1 | 7 | Thada | 0.011 | Young |
| 0.505633 | 0.340554 | 0.298 | 0.292 | 1 | 7 | Iws1 | 0.006 | Young |
| 0.505694 | 0.520274 | 0.12 | 0.141 | 1 | 7 | Poglut1 | -0.021 | Young |
| 0.50607 | 0.317019 | 0.256 | 0.238 | 1 | 7 | Hist1h2ap | 0.018 | Young |
| 0.506107 | 0.064389 | 0.06 | 0.049 | 1 | 7 | Fbf1 | 0.011 | Young |
| 0.506285 | 0.648283 | 0.045 | 0.036 | 1 | 7 | Mypopos | 0.009 | Young |
| 0.506558 | 0.001394 | 0.184 | 0.21 | 1 | 7 | Sh3bp2 | -0.026 | Young |
| 0.506609 | 0.557905 | 0.048 | 0.038 | 1 | 7 | Ipp | 0.01 | Young |

| | | | | | | | | |
|---|---|---|---|---|---|---|---|---|
| 0.506774 | 0.262602 | 0.247 | 0.274 | 1 | 7 | Kcnq1ot1 | -0.027 | Young |
| 0.50705 | 0.218566 | 0.322 | 0.308 | 1 | 7 | Lilrb4a | 0.014 | Young |
| 0.50747 | 0.046167 | 0.331 | 0.372 | 1 | 7 | Rapgef6 | -0.041 | Young |
| 0.507634 | 0.07189 | 0.271 | 0.3 | 1 | 7 | Commd1 | -0.029 | Young |
| 0.508161 | 0.681328 | 0.042 | 0.033 | 1 | 7 | Gm10036 | 0.009 | Young |
| 0.508251 | 0.085696 | 0.139 | 0.123 | 1 | 7 | Nsun6 | 0.016 | Young |
| 0.508505 | 0.424663 | 0.193 | 0.177 | 1 | 7 | Gatd1 | 0.016 | Young |
| 0.508568 | 0.37891 | 0.048 | 0.038 | 1 | 7 | 2810454H( | 0.01 | Young |
| 0.508921 | 0.206081 | 0.123 | 0.144 | 1 | 7 | Ptger2 | -0.021 | Young |
| 0.509086 | 0.443944 | 0.111 | 0.097 | 1 | 7 | Csad | 0.014 | Young |
| 0.509203 | 0.33089 | 0.178 | 0.203 | 1 | 7 | Zfp62 | -0.025 | Young |
| 0.509308 | 0.349824 | 0.096 | 0.115 | 1 | 7 | Chek1 | -0.019 | Young |
| 0.509429 | 0.234347 | 0.172 | 0.192 | 1 | 7 | Aurka | -0.02 | Young |
| 0.50955 | 0.337468 | 0.178 | 0.164 | 1 | 7 | Hck | 0.014 | Young |
| 0.509622 | 0.128962 | 0.232 | 0.264 | 1 | 7 | Zmpste24 | -0.032 | Young |
| 0.509758 | 0.119564 | 0.346 | 0.392 | 1 | 7 | Rbm27 | -0.046 | Young |
| 0.510496 | 0.151685 | 0.398 | 0.364 | 1 | 7 | Ciita | 0.034 | Young |
| 0.510896 | 0.204629 | 0.072 | 0.087 | 1 | 7 | Ccdc93 | -0.015 | Young |
| 0.510982 | 0.285486 | 0.081 | 0.097 | 1 | 7 | Dpysl3 | -0.016 | Young |
| 0.511485 | 0.034823 | 0.063 | 0.077 | 1 | 7 | Tmod1 | -0.014 | Young |
| 0.51205 | 0.424885 | 0.054 | 0.044 | 1 | 7 | 6720489N1 | 0.01 | Young |
| 0.51218 | 0.008327 | 0.343 | 0.377 | 1 | 7 | Cep350 | -0.034 | Young |
| 0.512495 | 0.124526 | 0.325 | 0.305 | 1 | 7 | Dpp8 | 0.02 | Young |
| 0.512679 | 0.140636 | 0.34 | 0.321 | 1 | 7 | Tia1 | 0.019 | Young |
| 0.513041 | 0.697984 | 0.045 | 0.036 | 1 | 7 | Bbs7 | 0.009 | Young |
| 0.513124 | 0.68353 | 0.033 | 0.044 | 1 | 7 | Hist2h2be | -0.011 | Young |
| 0.513164 | 0.815356 | 0.039 | 0.031 | 1 | 7 | Phpt1 | 0.008 | Young |
| 0.513233 | 0.373728 | 0.301 | 0.287 | 1 | 7 | Ankrd12 | 0.014 | Young |
| 0.51332 | 0.566604 | 0.208 | 0.195 | 1 | 7 | Uri1 | 0.013 | Young |
| 0.513387 | 0.083259 | 0.045 | 0.056 | 1 | 7 | Gm26853 | -0.011 | Young |
| 0.513839 | 0.169554 | 0.133 | 0.154 | 1 | 7 | Prpsap1 | -0.021 | Young |
| 0.513855 | 0.170167 | 0.307 | 0.287 | 1 | 7 | Flot2 | 0.02 | Young |
| 0.514502 | 0.236112 | 0.12 | 0.141 | 1 | 7 | Cep290 | -0.021 | Young |
| 0.514584 | 0.623771 | 0.16 | 0.149 | 1 | 7 | Rcan3 | 0.011 | Young |
| 0.514622 | 0.870035 | 0.039 | 0.031 | 1 | 7 | Klrb1a | 0.008 | Young |
| 0.514647 | 0.131131 | 0.142 | 0.162 | 1 | 7 | Hikeshi | -0.02 | Young |
| 0.514711 | 0.151604 | 0.051 | 0.041 | 1 | 7 | Ccdc66 | 0.01 | Young |
| 0.514872 | 0.259601 | 0.066 | 0.079 | 1 | 7 | 9430015G1 | -0.013 | Young |
| 0.51496 | 0.277329 | 0.608 | 0.644 | 1 | 7 | Tob2 | -0.036 | Young |
| 0.515342 | 0.651742 | 0.139 | 0.123 | 1 | 7 | Hsdl1 | 0.016 | Young |
| 0.51545 | 0.58055 | 0.084 | 0.072 | 1 | 7 | Amd2 | 0.012 | Young |
| 0.515741 | 0.062802 | 0.482 | 0.528 | 1 | 7 | Pnisr | -0.046 | Young |
| 0.515762 | 0.272403 | 0.319 | 0.313 | 1 | 7 | Nrd1 | 0.006 | Young |
| 0.515877 | 0.085941 | 0.13 | 0.151 | 1 | 7 | Afg3l2 | -0.021 | Young |
| 0.516308 | 0.032705 | 0.428 | 0.477 | 1 | 7 | U2af1 | -0.049 | Young |

| | | | | | | | | |
|---|---|---|---|---|---|---|---|---|
| 0.516888 | 0.791421 | 0.217 | 0.208 | 1 | 7 | Lyve1 | 0.009 | Young |
| 0.517217 | 0.21179 | 0.09 | 0.108 | 1 | 7 | Cgrrf1 | -0.018 | Young |
| 0.518171 | 0.077959 | 0.13 | 0.151 | 1 | 7 | Spg20 | -0.021 | Young |
| 0.518368 | 0.739815 | 0.117 | 0.105 | 1 | 7 | Asna1 | 0.012 | Young |
| 0.518458 | 0.109568 | 0.256 | 0.29 | 1 | 7 | Crybg3 | -0.034 | Young |
| 0.518546 | 0.328809 | 0.145 | 0.128 | 1 | 7 | Cdc42bpb | 0.017 | Young |
| 0.518663 | 0.548005 | 0.211 | 0.2 | 1 | 7 | Kif4 | 0.011 | Young |
| 0.519027 | 0.518512 | 0.114 | 0.136 | 1 | 7 | Klhl2 | -0.022 | Young |
| 0.519168 | 0.359105 | 0.322 | 0.313 | 1 | 7 | Chd9 | 0.009 | Young |
| 0.519198 | 0.187878 | 0.196 | 0.179 | 1 | 7 | Taf3 | 0.017 | Young |
| 0.519265 | 0.040073 | 0.422 | 0.454 | 1 | 7 | Arhgef2 | -0.032 | Young |
| 0.519329 | 0.156291 | 0.702 | 0.692 | 1 | 7 | Tab2 | 0.01 | Young |
| 0.519367 | 0.234021 | 0.027 | 0.036 | 1 | 7 | Pus7l | -0.009 | Young |
| 0.519397 | 1.019834 | 0.042 | 0.033 | 1 | 7 | Adgrg5 | 0.009 | Young |
| 0.519833 | 0.338369 | 0.142 | 0.162 | 1 | 7 | Exosc2 | -0.02 | Young |
| 0.520098 | 0.341644 | 0.099 | 0.118 | 1 | 7 | Eng | -0.019 | Young |
| 0.52051 | 0.217321 | 0.066 | 0.079 | 1 | 7 | Ccar2 | -0.013 | Young |
| 0.52125 | 0.174203 | 0.154 | 0.177 | 1 | 7 | AI413582 | -0.023 | Young |
| 0.52153 | 0.100815 | 0.127 | 0.146 | 1 | 7 | Galk1 | -0.019 | Young |
| 0.521584 | 0.129915 | 0.25 | 0.231 | 1 | 7 | Neil3 | 0.019 | Young |
| 0.52168 | 0.849983 | 0.051 | 0.064 | 1 | 7 | Pkn3 | -0.013 | Young |
| 0.52176 | 0.246998 | 0.304 | 0.295 | 1 | 7 | Itsn1 | 0.009 | Young |
| 0.521762 | 0.837185 | 0.105 | 0.092 | 1 | 7 | Ctdp1 | 0.013 | Young |
| 0.521857 | 0.280278 | 0.563 | 0.549 | 1 | 7 | Vps35 | 0.014 | Young |
| 0.521904 | 0.254395 | 0.012 | 0.018 | 1 | 7 | Gpr137c | -0.006 | Young |
| 0.522191 | 0.404085 | 0.072 | 0.087 | 1 | 7 | Naip6 | -0.015 | Young |
| 0.522263 | 0.036355 | 0.355 | 0.397 | 1 | 7 | Bod1l | -0.042 | Young |
| 0.522392 | 0.106749 | 0.133 | 0.151 | 1 | 7 | Ceacam1 | -0.018 | Young |
| 0.522432 | 0.169658 | 0.063 | 0.077 | 1 | 7 | Zfp212 | -0.014 | Young |
| 0.522486 | 0.147682 | 0.045 | 0.056 | 1 | 7 | Slc35d2 | -0.011 | Young |
| 0.52291 | 0.167619 | 0.31 | 0.29 | 1 | 7 | Atxn7l1 | 0.02 | Young |
| 0.522922 | 0.307453 | 0.41 | 0.413 | 1 | 7 | Ptpn7 | -0.003 | Young |
| 0.52293 | 0.044981 | 0.208 | 0.233 | 1 | 7 | Arhgap10 | -0.025 | Young |
| 0.523216 | 0.387115 | 0.117 | 0.136 | 1 | 7 | Fez2 | -0.019 | Young |
| 0.523297 | 0.056281 | 0.229 | 0.256 | 1 | 7 | Msmo1 | -0.027 | Young |
| 0.523565 | 0.127956 | 0.105 | 0.092 | 1 | 7 | Lrba | 0.013 | Young |
| 0.52374 | 0.197145 | 0.09 | 0.077 | 1 | 7 | Ust | 0.013 | Young |
| 0.524048 | 0.833132 | 0.096 | 0.085 | 1 | 7 | Osbpl3 | 0.011 | Young |
| 0.524097 | 0.14306 | 0.428 | 0.431 | 1 | 7 | Il10rb | -0.003 | Young |
| 0.524264 | 0.004801 | 0.901 | 0.887 | 1 | 7 | Rpl10a | 0.014 | Young |
| 0.524544 | 0.371461 | 0.123 | 0.144 | 1 | 7 | Coasy | -0.021 | Young |
| 0.524625 | 0.24186 | 0.22 | 0.197 | 1 | 7 | Adam19 | 0.023 | Young |
| 0.524677 | 0.804308 | 0.075 | 0.064 | 1 | 7 | Ikbkg | 0.011 | Young |
| 0.524722 | 0.800677 | 0.036 | 0.028 | 1 | 7 | Gm20300 | 0.008 | Young |
| 0.524722 | 0.166628 | 0.027 | 0.036 | 1 | 7 | Fam221a | -0.009 | Young |

| | | | | | | | | |
|---|---|---|---|---|---|---|---|---|
| 0.524785 | 0.137364 | 0.057 | 0.069 | 1 | 7 | Urb2 | -0.012 | Young |
| 0.525059 | 0.0202 | 0.232 | 0.251 | 1 | 7 | Ythdc2 | -0.019 | Young |
| 0.525073 | 0.195196 | 0.102 | 0.087 | 1 | 7 | Snrnp35 | 0.015 | Young |
| 0.525138 | 0.346447 | 0.054 | 0.044 | 1 | 7 | Slc9a3r2 | 0.01 | Young |
| 0.5252 | 0.582088 | 0.117 | 0.105 | 1 | 7 | Fancd2 | 0.012 | Young |
| 0.5252 | 0.213116 | 0.012 | 0.018 | 1 | 7 | Zbtb7c | -0.006 | Young |
| 0.525387 | 0.346378 | 0.181 | 0.208 | 1 | 7 | Uspl1 | -0.027 | Young |
| 0.525536 | 0.343857 | 0.084 | 0.1 | 1 | 7 | Akip1 | -0.016 | Young |
| 0.525589 | 0.095953 | 0.093 | 0.108 | 1 | 7 | Myc | -0.015 | Young |
| 0.52578 | 0.053346 | 0.136 | 0.154 | 1 | 7 | Tm2d3 | -0.018 | Young |
| 0.526248 | 0.535511 | 0.199 | 0.19 | 1 | 7 | Polr1c | 0.009 | Young |
| 0.526301 | 1.259462 | 0.012 | 0.018 | 1 | 7 | Zfp455 | -0.006 | Young |
| 0.526483 | 0.741823 | 0.042 | 0.033 | 1 | 7 | Gm26778 | 0.009 | Young |
| 0.526545 | 0.398851 | 0.208 | 0.197 | 1 | 7 | Ldah | 0.011 | Young |
| 0.526991 | 0.213278 | 0.145 | 0.131 | 1 | 7 | Fam214b | 0.014 | Young |
| 0.527241 | 0.211586 | 0.145 | 0.164 | 1 | 7 | Prkra | -0.019 | Young |
| 0.527403 | 0.333067 | 0.012 | 0.018 | 1 | 7 | Prkar1b | -0.006 | Young |
| 0.52763 | 0.069213 | 0.202 | 0.228 | 1 | 7 | Atad5 | -0.026 | Young |
| 0.527688 | 0.29846 | 0.063 | 0.077 | 1 | 7 | Ccdc58 | -0.014 | Young |
| 0.527731 | 0.039631 | 0.066 | 0.079 | 1 | 7 | Ptdss2 | -0.013 | Young |
| 0.527843 | 0.083072 | 0.34 | 0.379 | 1 | 7 | Nenf | -0.039 | Young |
| 0.528004 | 0.086504 | 0.105 | 0.121 | 1 | 7 | Siglecg | -0.016 | Young |
| 0.528108 | 0.047503 | 0.175 | 0.197 | 1 | 7 | Abcf2 | -0.022 | Young |
| 0.528266 | 0.064899 | 0.346 | 0.387 | 1 | 7 | Cd300a | -0.041 | Young |
| 0.52837 | 0.205906 | 0.404 | 0.382 | 1 | 7 | Psme3 | 0.022 | Young |
| 0.528598 | 0.135427 | 0.045 | 0.056 | 1 | 7 | Rabep2 | -0.011 | Young |
| 0.528598 | 0.077617 | 0.045 | 0.056 | 1 | 7 | Hspbap1 | -0.011 | Young |
| 0.528674 | 0.036163 | 0.678 | 0.674 | 1 | 7 | Tbca | 0.004 | Young |
| 0.528994 | 0.372887 | 0.268 | 0.262 | 1 | 7 | Kmt2d | 0.006 | Young |
| 0.529611 | 0.316598 | 0.012 | 0.018 | 1 | 7 | Zfp449 | -0.006 | Young |
| 0.529788 | 0.051155 | 0.343 | 0.385 | 1 | 7 | Zmynd11 | -0.042 | Young |
| 0.530056 | 0.188659 | 0.039 | 0.031 | 1 | 7 | Cd69 | 0.008 | Young |
| 0.530131 | 1.063135 | 0.111 | 0.1 | 1 | 7 | Gypc | 0.011 | Young |
| 0.530133 | 0.023031 | 0.217 | 0.241 | 1 | 7 | Nolc1 | -0.024 | Young |
| 0.530569 | 0.07011 | 0.102 | 0.118 | 1 | 7 | Rrm2b | -0.016 | Young |
| 0.530648 | 0.142723 | 0.663 | 0.703 | 1 | 7 | Pnrc1 | -0.04 | Young |
| 0.530927 | 0.100725 | 0.259 | 0.287 | 1 | 7 | Polr2i | -0.028 | Young |
| 0.531473 | 0.276664 | 0.042 | 0.033 | 1 | 7 | Dtnb | 0.009 | Young |
| 0.531517 | 0.339915 | 0.045 | 0.036 | 1 | 7 | 3110001I2 | 0.009 | Young |
| 0.531567 | 0.431057 | 0.066 | 0.056 | 1 | 7 | Tbcc | 0.01 | Young |
| 0.531842 | 0.105364 | 0.178 | 0.159 | 1 | 7 | Stx3 | 0.019 | Young |
| 0.531904 | 0.307257 | 0.178 | 0.2 | 1 | 7 | Uchl3 | -0.022 | Young |
| 0.53228 | 0.502092 | 0.039 | 0.031 | 1 | 7 | Gk5 | 0.008 | Young |
| 0.532383 | 0.013448 | 0.048 | 0.038 | 1 | 7 | Zfp971 | 0.01 | Young |
| 0.532986 | 0.107486 | 0.084 | 0.1 | 1 | 7 | Gstcd | -0.016 | Young |

| | | | | | | | | |
|---|---|---|---|---|---|---|---|---|
| 0.533033 | 0.540623 | 0.226 | 0.21 | 1 | 7 | Riok1 | 0.016 | Young |
| 0.533243 | 0.486932 | 0.148 | 0.133 | 1 | 7 | Slc12a9 | 0.015 | Young |
| 0.533326 | 0.905831 | 0.033 | 0.026 | 1 | 7 | Kbtbd7 | 0.007 | Young |
| 0.533354 | 0.164763 | 0.271 | 0.308 | 1 | 7 | Vcpip1 | -0.037 | Young |
| 0.533652 | 0.015618 | 0.364 | 0.403 | 1 | 7 | Sppl3 | -0.039 | Young |
| 0.53368 | 0.184504 | 0.093 | 0.11 | 1 | 7 | Arl2 | -0.017 | Young |
| 0.533894 | 0.386837 | 0.181 | 0.167 | 1 | 7 | Tmem260 | 0.014 | Young |
| 0.533968 | 0.778187 | 0.036 | 0.028 | 1 | 7 | Dync2h1 | 0.008 | Young |
| 0.534038 | 0.125799 | 0.093 | 0.079 | 1 | 7 | Usp46 | 0.014 | Young |
| 0.534292 | 0.044475 | 0.313 | 0.292 | 1 | 7 | Pcbd2 | 0.021 | Young |
| 0.534612 | 0.430211 | 0.072 | 0.087 | 1 | 7 | Platr25 | -0.015 | Young |
| 0.534742 | 1.005875 | 0.036 | 0.028 | 1 | 7 | Sccpdh | 0.008 | Young |
| 0.534821 | 0.016193 | 0.166 | 0.187 | 1 | 7 | Ptgr2 | -0.021 | Young |
| 0.534917 | 0.548652 | 0.175 | 0.162 | 1 | 7 | Stat5a | 0.013 | Young |
| 0.534944 | 0.493938 | 0.006 | 0.01 | 1 | 7 | Rap1gap | -0.004 | Young |
| 0.535163 | 0.384066 | 0.102 | 0.09 | 1 | 7 | Dvl2 | 0.012 | Young |
| 0.535353 | 0.268801 | 0.117 | 0.136 | 1 | 7 | Cep68 | -0.019 | Young |
| 0.535516 | 0.558147 | 0.036 | 0.028 | 1 | 7 | Vamp1 | 0.008 | Young |
| 0.535792 | 0.410804 | 0.205 | 0.192 | 1 | 7 | Bcl2a1a | 0.013 | Young |
| 0.536061 | 0.294277 | 0.22 | 0.208 | 1 | 7 | Gbp3 | 0.012 | Young |
| 0.536263 | 0.794929 | 0.117 | 0.108 | 1 | 7 | Edem3 | 0.009 | Young |
| 0.536443 | 0.036487 | 0.006 | 0.01 | 1 | 7 | Rbpms2 | -0.004 | Young |
| 0.536552 | 0.038142 | 0.151 | 0.172 | 1 | 7 | Mphosph1 | -0.021 | Young |
| 0.536799 | 0.131938 | 0.123 | 0.141 | 1 | 7 | Pnkd | -0.018 | Young |
| 0.537208 | 0.227911 | 0.34 | 0.336 | 1 | 7 | Chd8 | 0.004 | Young |
| 0.537488 | 0.69069 | 0.03 | 0.038 | 1 | 7 | Elp6 | -0.008 | Young |
| 0.537567 | 0.115136 | 0.392 | 0.444 | 1 | 7 | Scarb2 | -0.052 | Young |
| 0.53758 | 0.340353 | 0.039 | 0.049 | 1 | 7 | C330018D2 | -0.01 | Young |
| 0.537894 | 0.71706 | 0.13 | 0.118 | 1 | 7 | Akap8l | 0.012 | Young |
| 0.537945 | 0.28099 | 0.006 | 0.01 | 1 | 7 | Zfp661 | -0.004 | Young |
| 0.537945 | 0.06491 | 0.006 | 0.01 | 1 | 7 | Gm17767 | -0.004 | Young |
| 0.538186 | 0.698861 | 0.033 | 0.026 | 1 | 7 | Mcat | 0.007 | Young |
| 0.538256 | 0.976483 | 0.027 | 0.021 | 1 | 7 | Bbs2 | 0.006 | Young |
| 0.538345 | 0.126227 | 0.069 | 0.082 | 1 | 7 | Zc3h8 | -0.013 | Young |
| 0.538375 | 0.117593 | 0.244 | 0.272 | 1 | 7 | Psen1 | -0.028 | Young |
| 0.538721 | 0.110068 | 0.151 | 0.174 | 1 | 7 | Trappc11 | -0.023 | Young |
| 0.538752 | 0.065054 | 0.286 | 0.313 | 1 | 7 | Nbeal1 | -0.027 | Young |
| 0.539237 | 0.444085 | 0.349 | 0.346 | 1 | 7 | Dusp22 | 0.003 | Young |
| 0.539449 | 1.566186 | 0.006 | 0.01 | 1 | 7 | Gm34961 | -0.004 | Young |
| 0.539449 | 1.013967 | 0.006 | 0.01 | 1 | 7 | Parp16 | -0.004 | Young |
| 0.539449 | 0.724593 | 0.006 | 0.01 | 1 | 7 | 2010110K1 | -0.004 | Young |
| 0.539449 | 0.659612 | 0.006 | 0.01 | 1 | 7 | Dagla | -0.004 | Young |
| 0.539449 | 0.040321 | 0.006 | 0.01 | 1 | 7 | Bgn | -0.004 | Young |
| 0.539449 | 0.03921 | 0.006 | 0.01 | 1 | 7 | Epha4 | -0.004 | Young |
| 0.539667 | 0.154524 | 0.584 | 0.608 | 1 | 7 | Wasf2 | -0.024 | Young |

| | | | | | | | | |
|---|---|---|---|---|---|---|---|---|
| 0.539835 | 0.117388 | 0.045 | 0.036 | 1 | 7 | Gm26890 | 0.009 | Young |
| 0.540058 | 2.121523 | 0.027 | 0.021 | 1 | 7 | Mr1 | 0.006 | Young |
| 0.540371 | 0.067148 | 0.873 | 0.841 | 1 | 7 | mt-Nd5 | 0.032 | Young |
| 0.540385 | 0.118962 | 0.202 | 0.228 | 1 | 7 | Gpr107 | -0.026 | Young |
| 0.540673 | 0.295947 | 0.392 | 0.385 | 1 | 7 | Brd8 | 0.007 | Young |
| 0.54106 | 0.031699 | 0.358 | 0.387 | 1 | 7 | Acp1 | -0.029 | Young |
| 0.541188 | 0.230281 | 0.22 | 0.251 | 1 | 7 | Irak4 | -0.031 | Young |
| 0.541447 | 0.198727 | 0.271 | 0.259 | 1 | 7 | Slmap | 0.012 | Young |
| 0.541734 | 0.34198 | 0.036 | 0.028 | 1 | 7 | Zfp472 | 0.008 | Young |
| 0.542514 | 0.063992 | 0.036 | 0.028 | 1 | 7 | Gm14326 | 0.008 | Young |
| 0.542585 | 0.168713 | 0.172 | 0.156 | 1 | 7 | Fbrs | 0.016 | Young |
| 0.543402 | 0.314752 | 0.123 | 0.11 | 1 | 7 | Cox10 | 0.013 | Young |
| 0.543403 | 0.103907 | 0.22 | 0.246 | 1 | 7 | Mzt1 | -0.026 | Young |
| 0.543414 | 0.023358 | 0.084 | 0.097 | 1 | 7 | Haghl | -0.013 | Young |
| 0.54344 | 0.558581 | 0.172 | 0.159 | 1 | 7 | Cln3 | 0.013 | Young |
| 0.543471 | 0.456694 | 0.039 | 0.031 | 1 | 7 | Zdhhc1 | 0.008 | Young |
| 0.543854 | 0.078384 | 0.404 | 0.449 | 1 | 7 | Pon2 | -0.045 | Young |
| 0.543915 | 0.121253 | 0.244 | 0.272 | 1 | 7 | Clcc1 | -0.028 | Young |
| 0.544011 | 0.04199 | 0.13 | 0.115 | 1 | 7 | Megf8 | 0.015 | Young |
| 0.544188 | 1.052973 | 0.012 | 0.008 | 1 | 7 | Azin2 | 0.004 | Young |
| 0.54444 | 0.049366 | 0.386 | 0.421 | 1 | 7 | Arfgef1 | -0.035 | Young |
| 0.54449 | 0.152449 | 0.265 | 0.295 | 1 | 7 | Mgat4b | -0.03 | Young |
| 0.544851 | 0.169587 | 0.226 | 0.246 | 1 | 7 | Mindy1 | -0.02 | Young |
| 0.545109 | 0.735946 | 0.117 | 0.133 | 1 | 7 | Ift46 | -0.016 | Young |
| 0.545238 | 0.399419 | 0.349 | 0.346 | 1 | 7 | Usp19 | 0.003 | Young |
| 0.545381 | 0.226145 | 0.057 | 0.069 | 1 | 7 | Tsen2 | -0.012 | Young |
| 0.545589 | 0.941276 | 0.012 | 0.008 | 1 | 7 | Inhba | 0.004 | Young |
| 0.545589 | 0.654748 | 0.012 | 0.008 | 1 | 7 | Shisa8 | 0.004 | Young |
| 0.545635 | 0.122249 | 0.078 | 0.067 | 1 | 7 | Tacc2 | 0.011 | Young |
| 0.545723 | 0.963391 | 0.039 | 0.031 | 1 | 7 | Gm48604 | 0.008 | Young |
| 0.545732 | 0.190831 | 0.193 | 0.221 | 1 | 7 | Ado | -0.028 | Young |
| 0.545747 | 0.095156 | 0.521 | 0.554 | 1 | 7 | Cbfb | -0.033 | Young |
| 0.545837 | 0.175419 | 0.139 | 0.123 | 1 | 7 | Ccser2 | 0.016 | Young |
| 0.545892 | 0.01631 | 0.139 | 0.156 | 1 | 7 | Plcb3 | -0.017 | Young |
| 0.546017 | 0.510637 | 0.142 | 0.128 | 1 | 7 | Hsf1 | 0.014 | Young |
| 0.546336 | 0.521633 | 0.033 | 0.026 | 1 | 7 | Zfyve9 | 0.007 | Young |
| 0.546341 | 0.143969 | 0.524 | 0.518 | 1 | 7 | Eloc | 0.006 | Young |
| 0.546396 | 0.802535 | 0.021 | 0.028 | 1 | 7 | Wars2 | -0.007 | Young |
| 0.546507 | 0.29936 | 0.166 | 0.182 | 1 | 7 | Tmem251 | -0.016 | Young |
| 0.546992 | 1.08929 | 0.012 | 0.008 | 1 | 7 | Ddx43 | 0.004 | Young |
| 0.546992 | 0.978618 | 0.012 | 0.008 | 1 | 7 | Xk | 0.004 | Young |
| 0.546992 | 0.471655 | 0.012 | 0.008 | 1 | 7 | Pou5f1 | 0.004 | Young |
| 0.547155 | 1.177467 | 0.033 | 0.026 | 1 | 7 | Map2k3os | 0.007 | Young |
| 0.547294 | 0.979788 | 0.027 | 0.021 | 1 | 7 | Steap4 | 0.006 | Young |
| 0.547709 | 0.115994 | 0.593 | 0.592 | 1 | 7 | Clec12a | 0.001 | Young |

| | | | | | | | | |
|---|---|---|---|---|---|---|---|---|
| 0.548161 | 0.409417 | 0.021 | 0.028 | 1 | 7 | Nat2 | -0.007 | Young |
| 0.548267 | 0.192411 | 0.075 | 0.064 | 1 | 7 | Aff3 | 0.011 | Young |
| 0.548397 | 0.826996 | 0.012 | 0.008 | 1 | 7 | Zfp119b | 0.004 | Young |
| 0.548397 | 0.748421 | 0.012 | 0.008 | 1 | 7 | Tpm2 | 0.004 | Young |
| 0.548397 | 0.745391 | 0.012 | 0.008 | 1 | 7 | Gm26698 | 0.004 | Young |
| 0.548397 | 0.682292 | 0.012 | 0.008 | 1 | 7 | Gm10827 | 0.004 | Young |
| 0.548397 | 0.657604 | 0.012 | 0.008 | 1 | 7 | Clmn | 0.004 | Young |
| 0.548399 | 0.39887 | 0.286 | 0.282 | 1 | 7 | Trappc1 | 0.004 | Young |
| 0.548409 | 0.112179 | 0.163 | 0.179 | 1 | 7 | Clec2i | -0.016 | Young |
| 0.548587 | 0.177209 | 0.617 | 0.615 | 1 | 7 | Ctnnb1 | 0.002 | Young |
| 0.548793 | 0.578439 | 0.033 | 0.026 | 1 | 7 | Pcbp3 | 0.007 | Young |
| 0.548793 | 0.1772 | 0.033 | 0.026 | 1 | 7 | 1700008J0 | 0.007 | Young |
| 0.548951 | 0.231437 | 0.099 | 0.115 | 1 | 7 | Ivd | -0.016 | Young |
| 0.548962 | 0.212572 | 0.473 | 0.469 | 1 | 7 | Edf1 | 0.004 | Young |
| 0.549433 | 0.029287 | 0.181 | 0.203 | 1 | 7 | Mettl16 | -0.022 | Young |
| 0.549487 | 0.138075 | 0.042 | 0.033 | 1 | 7 | Tspyl3 | 0.009 | Young |
| 0.549504 | 0.540206 | 0.175 | 0.162 | 1 | 7 | Prim1 | 0.013 | Young |
| 0.54969 | 1.429484 | 0.018 | 0.013 | 1 | 7 | Qprt | 0.005 | Young |
| 0.549804 | 1.717072 | 0.012 | 0.008 | 1 | 7 | Spsb1 | 0.004 | Young |
| 0.549804 | 0.930205 | 0.012 | 0.008 | 1 | 7 | 4930532G1 | 0.004 | Young |
| 0.549804 | 0.733186 | 0.012 | 0.008 | 1 | 7 | Pdzd4 | 0.004 | Young |
| 0.549804 | 0.512201 | 0.012 | 0.008 | 1 | 7 | M1ap | 0.004 | Young |
| 0.54981 | 0.093513 | 0.316 | 0.346 | 1 | 7 | Nfat5 | -0.03 | Young |
| 0.550019 | 0.625377 | 0.027 | 0.021 | 1 | 7 | Pdk4 | 0.006 | Young |
| 0.550426 | 0.309406 | 0.389 | 0.372 | 1 | 7 | Zfp710 | 0.017 | Young |
| 0.550765 | 0.006112 | 0.196 | 0.177 | 1 | 7 | Zmym4 | 0.019 | Young |
| 0.551008 | 0.089773 | 0.337 | 0.377 | 1 | 7 | Slc12a6 | -0.04 | Young |
| 0.551144 | 0.315228 | 0.13 | 0.115 | 1 | 7 | Csrp2 | 0.015 | Young |
| 0.551153 | 0.065899 | 0.247 | 0.231 | 1 | 7 | Bcor | 0.016 | Young |
| 0.551212 | 0.787466 | 0.012 | 0.008 | 1 | 7 | Tubb3 | 0.004 | Young |
| 0.551212 | 0.67386 | 0.012 | 0.008 | 1 | 7 | Pard6g | 0.004 | Young |
| 0.551212 | 0.524847 | 0.012 | 0.008 | 1 | 7 | 1810034E1 | 0.004 | Young |
| 0.551212 | 0.496299 | 0.012 | 0.008 | 1 | 7 | 1700113A1 | 0.004 | Young |
| 0.551237 | 0.309037 | 0.163 | 0.151 | 1 | 7 | Gm11808 | 0.012 | Young |
| 0.551252 | 0.539754 | 0.03 | 0.023 | 1 | 7 | Fam216a | 0.007 | Young |
| 0.551252 | 0.290638 | 0.03 | 0.023 | 1 | 7 | Pim2 | 0.007 | Young |
| 0.551252 | 0.043499 | 0.03 | 0.023 | 1 | 7 | Gm42595 | 0.007 | Young |
| 0.551281 | 0.24398 | 0.066 | 0.079 | 1 | 7 | Sbk1 | -0.013 | Young |
| 0.55132 | 0.254575 | 0.139 | 0.162 | 1 | 7 | Arrdc4 | -0.023 | Young |
| 0.55134 | 0.138773 | 0.148 | 0.133 | 1 | 7 | Avl9 | 0.015 | Young |
| 0.55156 | 0.239723 | 0.422 | 0.421 | 1 | 7 | Tet2 | 0.001 | Young |
| 0.551835 | 0.425475 | 0.114 | 0.103 | 1 | 7 | Amz1 | 0.011 | Young |
| 0.551886 | 0.37602 | 0.09 | 0.079 | 1 | 7 | Terf2ip | 0.011 | Young |
| 0.55196 | 0.31567 | 0.151 | 0.138 | 1 | 7 | Foxk1 | 0.013 | Young |
| 0.552082 | 0.064401 | 0.602 | 0.59 | 1 | 7 | Zyx | 0.012 | Young |

| | | | | | | | | |
|---|---|---|---|---|---|---|---|---|
| 0.552115 | 0.245129 | 0.03 | 0.023 | 1 | 7 | Pcnx4 | 0.007 | Young |
| 0.552154 | 0.145869 | 0.048 | 0.059 | 1 | 7 | Scarf1 | -0.011 | Young |
| 0.552365 | 0.081211 | 0.386 | 0.421 | 1 | 7 | Kmt2a | -0.035 | Young |
| 0.552413 | 0.325817 | 0.163 | 0.149 | 1 | 7 | Wee1 | 0.014 | Young |
| 0.552568 | 0.342619 | 0.238 | 0.226 | 1 | 7 | Inpp5k | 0.012 | Young |
| 0.552622 | 1.123233 | 0.012 | 0.008 | 1 | 7 | Stac2 | 0.004 | Young |
| 0.552622 | 1.095859 | 0.012 | 0.008 | 1 | 7 | Camk2n1 | 0.004 | Young |
| 0.552622 | 0.924634 | 0.012 | 0.008 | 1 | 7 | Plxna3 | 0.004 | Young |
| 0.552622 | 0.615666 | 0.012 | 0.008 | 1 | 7 | Cyp2r1 | 0.004 | Young |
| 0.552622 | 0.430926 | 0.012 | 0.008 | 1 | 7 | Gm26759 | 0.004 | Young |
| 0.552622 | 0.385572 | 0.012 | 0.008 | 1 | 7 | Oas1g | 0.004 | Young |
| 0.552622 | 0.294686 | 0.012 | 0.008 | 1 | 7 | Slc24a5 | 0.004 | Young |
| 0.552622 | 0.031363 | 0.012 | 0.008 | 1 | 7 | Ly75 | 0.004 | Young |
| 0.552736 | 0.634011 | 0.066 | 0.056 | 1 | 7 | Crebl2 | 0.01 | Young |
| 0.552899 | 0.896135 | 0.024 | 0.018 | 1 | 7 | E130102H2 | 0.006 | Young |
| 0.5529 | 0.359103 | 0.033 | 0.026 | 1 | 7 | Slamf6 | 0.007 | Young |
| 0.552931 | 1.281933 | 0.015 | 0.01 | 1 | 7 | Osgin1 | 0.005 | Young |
| 0.552931 | 1.262401 | 0.015 | 0.01 | 1 | 7 | Ifi214 | 0.005 | Young |
| 0.552931 | 1.043805 | 0.015 | 0.01 | 1 | 7 | Ankrd33b | 0.005 | Young |
| 0.553073 | 1.262377 | 0.018 | 0.013 | 1 | 7 | Map3k9 | 0.005 | Young |
| 0.553263 | 0.118029 | 0.039 | 0.031 | 1 | 7 | Lmbr1 | 0.008 | Young |
| 0.553664 | 1.017066 | 0.027 | 0.021 | 1 | 7 | Hist1h3d | 0.006 | Young |
| 0.553832 | 0.242963 | 0.373 | 0.377 | 1 | 7 | Tnrc6b | -0.004 | Young |
| 0.553843 | 0.474789 | 0.03 | 0.023 | 1 | 7 | Tpmt | 0.007 | Young |
| 0.553844 | 0.54359 | 0.069 | 0.059 | 1 | 7 | Myh10 | 0.01 | Young |
| 0.554016 | 0.425161 | 0.178 | 0.164 | 1 | 7 | Alg5 | 0.014 | Young |
| 0.554034 | 0.425206 | 0.012 | 0.008 | 1 | 7 | 9130019O2 | 0.004 | Young |
| 0.554034 | 0.330019 | 0.012 | 0.008 | 1 | 7 | Hspa1l | 0.004 | Young |
| 0.554034 | 0.268791 | 0.012 | 0.008 | 1 | 7 | Il20rb | 0.004 | Young |
| 0.554034 | 0.16949 | 0.012 | 0.008 | 1 | 7 | Mustn1 | 0.004 | Young |
| 0.554034 | 0.126728 | 0.012 | 0.008 | 1 | 7 | Gm5113 | 0.004 | Young |
| 0.554111 | 0.429577 | 0.196 | 0.185 | 1 | 7 | Prune2 | 0.011 | Young |
| 0.554123 | 0.221129 | 0.331 | 0.369 | 1 | 7 | Acp2 | -0.038 | Young |
| 0.554284 | 0.035318 | 0.325 | 0.315 | 1 | 7 | Trappc6b | 0.01 | Young |
| 0.554476 | 0.078701 | 0.422 | 0.415 | 1 | 7 | Ifi203 | 0.007 | Young |
| 0.554497 | 0.132882 | 0.512 | 0.495 | 1 | 7 | Leprot | 0.017 | Young |
| 0.554547 | 0.86045 | 0.033 | 0.026 | 1 | 7 | Cpeb3 | 0.007 | Young |
| 0.554829 | 0.303086 | 0.078 | 0.092 | 1 | 7 | Spsb3 | -0.014 | Young |
| 0.555129 | 0.30791 | 0.127 | 0.146 | 1 | 7 | Fdx1 | -0.019 | Young |
| 0.555311 | 0.094981 | 0.259 | 0.277 | 1 | 7 | Gigyf1 | -0.018 | Young |
| 0.555427 | 1.299203 | 0.015 | 0.01 | 1 | 7 | Jag2 | 0.005 | Young |
| 0.555427 | 1.233457 | 0.015 | 0.01 | 1 | 7 | Fgfbp3 | 0.005 | Young |
| 0.555438 | 0.313918 | 0.069 | 0.082 | 1 | 7 | Gtpbp8 | -0.013 | Young |
| 0.555448 | 0.604633 | 0.012 | 0.008 | 1 | 7 | Psd | 0.004 | Young |
| 0.555448 | 0.317988 | 0.012 | 0.008 | 1 | 7 | Lrfn4 | 0.004 | Young |

| | | | | | | | | |
|---|---|---|---|---|---|---|---|---|
| 0.555535 | 0.353709 | 0.187 | 0.172 | 1 | 7 | Pdcd7 | 0.015 | Young |
| 0.55565 | 0.078617 | 0.377 | 0.413 | 1 | 7 | Chic2 | -0.036 | Young |
| 0.555757 | 0.497319 | 0.154 | 0.146 | 1 | 7 | Ormdl1 | 0.008 | Young |
| 0.555958 | 0.232761 | 0.202 | 0.228 | 1 | 7 | Wdr41 | -0.026 | Young |
| 0.555972 | 0.009505 | 0.506 | 0.554 | 1 | 7 | Ndufb5 | -0.048 | Young |
| 0.55629 | 0.898856 | 0.021 | 0.015 | 1 | 7 | Gm20632 | 0.006 | Young |
| 0.556299 | 0.157743 | 0.202 | 0.187 | 1 | 7 | Siglece | 0.015 | Young |
| 0.55638 | 0.037081 | 0.217 | 0.236 | 1 | 7 | Papd5 | -0.019 | Young |
| 0.556405 | 0.335275 | 0.027 | 0.021 | 1 | 7 | Trim68 | 0.006 | Young |
| 0.556576 | 0.167047 | 0.512 | 0.562 | 1 | 7 | Hnrnph2 | -0.05 | Young |
| 0.556629 | 0.128878 | 0.087 | 0.103 | 1 | 7 | Zfp865 | -0.016 | Young |
| 0.556677 | 0.726116 | 0.015 | 0.01 | 1 | 7 | Cdh24 | 0.005 | Young |
| 0.556784 | 0.385943 | 0.024 | 0.018 | 1 | 7 | Tmem62 | 0.006 | Young |
| 0.556864 | 0.030795 | 0.012 | 0.008 | 1 | 7 | Hddc3 | 0.004 | Young |
| 0.556864 | 0.029605 | 0.012 | 0.008 | 1 | 7 | Tmem236 | 0.004 | Young |
| 0.556926 | 0.058595 | 0.19 | 0.21 | 1 | 7 | Renbp | -0.02 | Young |
| 0.557158 | 0.627453 | 0.166 | 0.154 | 1 | 7 | Fam53c | 0.012 | Young |
| 0.557334 | 0.736671 | 0.021 | 0.015 | 1 | 7 | Hrh2 | 0.006 | Young |
| 0.557334 | 0.584959 | 0.021 | 0.015 | 1 | 7 | Tssk6 | 0.006 | Young |
| 0.557378 | 0.250568 | 0.087 | 0.077 | 1 | 7 | Trpm4 | 0.01 | Young |
| 0.557758 | 0.635248 | 0.024 | 0.018 | 1 | 7 | Gm15232 | 0.006 | Young |
| 0.557829 | 0.02503 | 0.13 | 0.146 | 1 | 7 | Rptor | -0.016 | Young |
| 0.557876 | 0.28785 | 0.214 | 0.2 | 1 | 7 | Vps13d | 0.014 | Young |
| 0.557928 | 1.21992 | 0.015 | 0.01 | 1 | 7 | Ulk4 | 0.005 | Young |
| 0.557928 | 0.61577 | 0.015 | 0.01 | 1 | 7 | Gm38037 | 0.005 | Young |
| 0.557928 | 0.556772 | 0.015 | 0.01 | 1 | 7 | Gm9938 | 0.005 | Young |
| 0.558043 | 0.480688 | 0.142 | 0.131 | 1 | 7 | Dcaf1 | 0.011 | Young |
| 0.558291 | 0.447269 | 0.078 | 0.092 | 1 | 7 | Atg2b | -0.014 | Young |
| 0.558443 | 0.615867 | 0.063 | 0.054 | 1 | 7 | Tfpi | 0.009 | Young |
| 0.558567 | 0.006495 | 0.654 | 0.656 | 1 | 7 | Adam15 | -0.002 | Young |
| 0.558583 | 0.24827 | 0.259 | 0.244 | 1 | 7 | Elp5 | 0.015 | Young |
| 0.558943 | 0.041432 | 0.148 | 0.167 | 1 | 7 | Setd7 | -0.019 | Young |
| 0.558975 | 0.216301 | 0.524 | 0.546 | 1 | 7 | Nsd2 | -0.022 | Young |
| 0.559043 | 0.695353 | 0.03 | 0.023 | 1 | 7 | Usp54 | 0.007 | Young |
| 0.559361 | 0.471109 | 0.172 | 0.159 | 1 | 7 | 6-Sep | 0.013 | Young |
| 0.560071 | 0.752413 | 0.027 | 0.021 | 1 | 7 | Clec1a | 0.006 | Young |
| 0.560206 | 0.086465 | 0.102 | 0.121 | 1 | 7 | Unc119b | -0.019 | Young |
| 0.560211 | 0.628172 | 0.28 | 0.267 | 1 | 7 | Ptgs2 | 0.013 | Young |
| 0.56032 | 0.047722 | 0.569 | 0.623 | 1 | 7 | Bach1 | -0.054 | Young |
| 0.560435 | 0.647245 | 0.015 | 0.01 | 1 | 7 | Zbtb49 | 0.005 | Young |
| 0.560435 | 0.459485 | 0.015 | 0.01 | 1 | 7 | Cygb | 0.005 | Young |
| 0.560682 | 0.480466 | 0.024 | 0.018 | 1 | 7 | Lekr1 | 0.006 | Young |
| 0.560842 | 0.67376 | 0.081 | 0.097 | 1 | 7 | Sipa1l2 | -0.016 | Young |
| 0.561013 | 0.150532 | 0.536 | 0.541 | 1 | 7 | 1700020I1 | -0.005 | Young |
| 0.56114 | 0.60995 | 0.06 | 0.051 | 1 | 7 | Smpd3 | 0.009 | Young |

| | | | | | | | | |
|---|---|---|---|---|---|---|---|---|
| 0.561141 | 0.554842 | 0.078 | 0.067 | 1 | 7 | Mlst8 | 0.011 | Young |
| 0.561518 | 0.267716 | 0.021 | 0.015 | 1 | 7 | Plpp3 | 0.006 | Young |
| 0.561651 | 0.438853 | 0.03 | 0.023 | 1 | 7 | Gm37305 | 0.007 | Young |
| 0.56169 | 0.545058 | 0.015 | 0.01 | 1 | 7 | Gm26826 | 0.005 | Young |
| 0.56169 | 0.238431 | 0.015 | 0.01 | 1 | 7 | A530013C2 | 0.005 | Young |
| 0.561824 | 0.251828 | 0.087 | 0.103 | 1 | 7 | Rnf214 | -0.016 | Young |
| 0.561972 | 0.286896 | 0.479 | 0.479 | 1 | 7 | Tnrc18 | 0 | Young |
| 0.561988 | 0.08629 | 0.033 | 0.026 | 1 | 7 | Zfp667 | 0.007 | Young |
| 0.562142 | 0.539511 | 0.018 | 0.013 | 1 | 7 | Gm48653 | 0.005 | Young |
| 0.562142 | 0.174029 | 0.018 | 0.013 | 1 | 7 | Gm12589 | 0.005 | Young |
| 0.562156 | 0.269074 | 0.223 | 0.256 | 1 | 7 | Slc15a4 | -0.033 | Young |
| 0.562855 | 0.631136 | 0.09 | 0.079 | 1 | 7 | Trim3 | 0.011 | Young |
| 0.562929 | 0.059784 | 0.373 | 0.364 | 1 | 7 | Mgat2 | 0.009 | Young |
| 0.562985 | 0.06112 | 0.19 | 0.215 | 1 | 7 | Cep250 | -0.025 | Young |
| 0.562993 | 0.209665 | 0.154 | 0.138 | 1 | 7 | Cdc16 | 0.016 | Young |
| 0.563281 | 0.726027 | 0.018 | 0.013 | 1 | 7 | Tjp3 | 0.005 | Young |
| 0.56336 | 0.559764 | 0.154 | 0.144 | 1 | 7 | 4930402H2 | 0.01 | Young |
| 0.563387 | 0.425309 | 0.151 | 0.141 | 1 | 7 | Cmtm4 | 0.01 | Young |
| 0.563387 | 0.413524 | 0.136 | 0.154 | 1 | 7 | Sipa1l1 | -0.018 | Young |
| 0.563639 | 0.098348 | 0.524 | 0.518 | 1 | 7 | Fgfr1op2 | 0.006 | Young |
| 0.563674 | 0.493968 | 0.111 | 0.1 | 1 | 7 | Nf2 | 0.011 | Young |
| 0.563765 | 0.255937 | 0.479 | 0.49 | 1 | 7 | Rex1bd | -0.011 | Young |
| 0.563766 | 0.255114 | 0.151 | 0.136 | 1 | 7 | Ppm1k | 0.015 | Young |
| 0.563853 | 0.154553 | 0.114 | 0.131 | 1 | 7 | Fgd6 | -0.017 | Young |
| 0.563903 | 0.599542 | 0.03 | 0.038 | 1 | 7 | Ndst1 | -0.008 | Young |
| 0.564164 | 0.326807 | 0.151 | 0.136 | 1 | 7 | Tmc6 | 0.015 | Young |
| 0.564214 | 0.14424 | 0.34 | 0.323 | 1 | 7 | Lamtor5 | 0.017 | Young |
| 0.564249 | 0.317804 | 0.133 | 0.121 | 1 | 7 | Klhl21 | 0.012 | Young |
| 0.564348 | 0.663021 | 0.066 | 0.056 | 1 | 7 | Dcbld2 | 0.01 | Young |
| 0.564366 | 0.498752 | 0.105 | 0.095 | 1 | 7 | Med22 | 0.01 | Young |
| 0.564421 | 0.509504 | 0.018 | 0.013 | 1 | 7 | Arhgef4 | 0.005 | Young |
| 0.564421 | 0.197568 | 0.018 | 0.013 | 1 | 7 | Fam109a | 0.005 | Young |
| 0.564571 | 0.309379 | 0.136 | 0.154 | 1 | 7 | Umad1 | -0.018 | Young |
| 0.564594 | 0.56329 | 0.024 | 0.018 | 1 | 7 | 4632404H1 | 0.006 | Young |
| 0.565436 | 0.082539 | 0.449 | 0.454 | 1 | 7 | Impdh2 | -0.005 | Young |
| 0.565465 | 0.136312 | 0.015 | 0.01 | 1 | 7 | Tle2 | 0.005 | Young |
| 0.565468 | 0.150284 | 0.19 | 0.177 | 1 | 7 | Axin1 | 0.013 | Young |
| 0.565562 | 0.254409 | 0.018 | 0.013 | 1 | 7 | Clec2g | 0.005 | Young |
| 0.565574 | 0.222129 | 0.024 | 0.018 | 1 | 7 | Dph2 | 0.006 | Young |
| 0.565704 | 0.221535 | 0.111 | 0.097 | 1 | 7 | Slc27a3 | 0.014 | Young |
| 0.565704 | 0.217769 | 0.111 | 0.097 | 1 | 7 | Pccb | 0.014 | Young |
| 0.566026 | 0.308045 | 0.205 | 0.192 | 1 | 7 | Atp8a1 | 0.013 | Young |
| 0.566155 | 0.40546 | 0.151 | 0.136 | 1 | 7 | Rdh11 | 0.015 | Young |
| 0.566397 | 0.309519 | 0.205 | 0.195 | 1 | 7 | Mdc1 | 0.01 | Young |
| 0.566554 | 0.585937 | 0.024 | 0.018 | 1 | 7 | Ciart | 0.006 | Young |

| | | | | | | | | |
|---|---|---|---|---|---|---|---|---|
| 0.566698 | 0.180091 | 0.395 | 0.428 | 1 | 7 | Swap70 | -0.033 | Young |
| 0.566704 | 0.260672 | 0.018 | 0.013 | 1 | 7 | Bbs4 | 0.005 | Young |
| 0.566791 | 0.069507 | 0.599 | 0.592 | 1 | 7 | H2afj | 0.007 | Young |
| 0.566991 | 0.081241 | 0.346 | 0.341 | 1 | 7 | Eif3g | 0.005 | Young |
| 0.567462 | 0.228068 | 0.416 | 0.418 | 1 | 7 | Pkn2 | -0.002 | Young |
| 0.567536 | 0.247221 | 0.024 | 0.018 | 1 | 7 | G430095P1 | 0.006 | Young |
| 0.567637 | 0.055711 | 0.175 | 0.195 | 1 | 7 | Prelid2 | -0.02 | Young |
| 0.567733 | 0.052747 | 0.214 | 0.238 | 1 | 7 | Fnbp4 | -0.024 | Young |
| 0.567822 | 0.268936 | 0.021 | 0.015 | 1 | 7 | Spsb4 | 0.006 | Young |
| 0.567822 | 0.10607 | 0.021 | 0.015 | 1 | 7 | Polh | 0.006 | Young |
| 0.567847 | 0.154334 | 0.018 | 0.013 | 1 | 7 | Eaf2 | 0.005 | Young |
| 0.567847 | 0.080194 | 0.018 | 0.013 | 1 | 7 | Nlrp1a | 0.005 | Young |
| 0.567854 | 0.175263 | 0.066 | 0.056 | 1 | 7 | Cry1 | 0.01 | Young |
| 0.567988 | 0.056412 | 0.015 | 0.01 | 1 | 7 | Pcx | 0.005 | Young |
| 0.567988 | 0.031457 | 0.015 | 0.01 | 1 | 7 | Gm4117 | 0.005 | Young |
| 0.568114 | 0.504497 | 0.072 | 0.062 | 1 | 7 | Slc35e2 | 0.01 | Young |
| 0.568115 | 0.086571 | 0.072 | 0.085 | 1 | 7 | Ccdc85b | -0.013 | Young |
| 0.568143 | 0.219705 | 0.307 | 0.3 | 1 | 7 | Ccnd2 | 0.007 | Young |
| 0.568387 | 0.074045 | 0.253 | 0.285 | 1 | 7 | Tifa | -0.032 | Young |
| 0.568772 | 0.328906 | 0.274 | 0.264 | 1 | 7 | Phf21a | 0.01 | Young |
| 0.568876 | 0.313032 | 0.021 | 0.015 | 1 | 7 | Cldn34c1 | 0.006 | Young |
| 0.568876 | 0.263014 | 0.021 | 0.015 | 1 | 7 | Gm11772 | 0.006 | Young |
| 0.568992 | 0.023328 | 0.018 | 0.013 | 1 | 7 | Gm12089 | 0.005 | Young |
| 0.569335 | 0.00336 | 0.774 | 0.779 | 1 | 7 | Tma7 | -0.005 | Young |
| 0.569366 | 0.414441 | 0.241 | 0.233 | 1 | 7 | Tbc1d15 | 0.008 | Young |
| 0.569946 | 0.446973 | 0.151 | 0.133 | 1 | 7 | Dnajb12 | 0.018 | Young |
| 0.570137 | 0.140757 | 0.018 | 0.013 | 1 | 7 | Slc8a1 | 0.005 | Young |
| 0.570263 | 0.283316 | 0.325 | 0.326 | 1 | 7 | Mga | -0.001 | Young |
| 0.570316 | 0.22504 | 0.271 | 0.262 | 1 | 7 | Ftl1-ps1 | 0.009 | Young |
| 0.570987 | 0.142185 | 0.021 | 0.015 | 1 | 7 | Vmac | 0.006 | Young |
| 0.570987 | 0.11379 | 0.021 | 0.015 | 1 | 7 | BC048403 | 0.006 | Young |
| 0.571136 | 0.125152 | 0.027 | 0.021 | 1 | 7 | P2ry13 | 0.006 | Young |
| 0.571136 | 0.044957 | 0.027 | 0.021 | 1 | 7 | Gm3448 | 0.006 | Young |
| 0.57147 | 0.378388 | 0.024 | 0.018 | 1 | 7 | Calml4 | 0.006 | Young |
| 0.571572 | 0.656362 | 0.063 | 0.054 | 1 | 7 | Hsd17b7 | 0.009 | Young |
| 0.571955 | 0.100272 | 0.331 | 0.323 | 1 | 7 | Maea | 0.008 | Young |
| 0.572306 | 0.091559 | 0.157 | 0.174 | 1 | 7 | Stk19 | -0.017 | Young |
| 0.572431 | 0.080651 | 0.018 | 0.013 | 1 | 7 | Nfib | 0.005 | Young |
| 0.572483 | 0.028831 | 0.801 | 0.797 | 1 | 7 | Arpc3 | 0.004 | Young |
| 0.572529 | 0.246416 | 0.361 | 0.356 | 1 | 7 | Tsg101 | 0.005 | Young |
| 0.572743 | 0.020431 | 0.443 | 0.495 | 1 | 7 | Ppp1r2 | -0.052 | Young |
| 0.573051 | 0.050813 | 0.265 | 0.287 | 1 | 7 | Atad2b | -0.022 | Young |
| 0.573102 | 0.493149 | 0.021 | 0.015 | 1 | 7 | Scn2b | 0.006 | Young |
| 0.573135 | 0.050118 | 0.44 | 0.487 | 1 | 7 | Nsd1 | -0.047 | Young |
| 0.573536 | 0.091043 | 0.069 | 0.079 | 1 | 7 | Dimt1 | -0.01 | Young |

| | | | | | | | | |
|---|---|---|---|---|---|---|---|---|
| 0.573627 | 0.131632 | 0.542 | 0.546 | 1 | 7 | Fkbp2 | -0.004 | Young |
| 0.573691 | 0.199803 | 0.087 | 0.077 | 1 | 7 | Zbtb14 | 0.01 | Young |
| 0.573717 | 0.351866 | 0.087 | 0.103 | 1 | 7 | Frs2 | -0.016 | Young |
| 0.57418 | 0.251195 | 0.136 | 0.123 | 1 | 7 | Rnf26 | 0.013 | Young |
| 0.574236 | 0.109638 | 0.262 | 0.287 | 1 | 7 | Fh1 | -0.025 | Young |
| 0.57436 | 0.190676 | 0.072 | 0.085 | 1 | 7 | Nol9 | -0.013 | Young |
| 0.574644 | 0.282076 | 0.114 | 0.133 | 1 | 7 | Eri3 | -0.019 | Young |
| 0.57473 | 0.153655 | 0.018 | 0.013 | 1 | 7 | Nat8f1 | 0.005 | Young |
| 0.574775 | 0.195696 | 0.051 | 0.062 | 1 | 7 | Phldb1 | -0.011 | Young |
| 0.574795 | 0.167971 | 0.087 | 0.1 | 1 | 7 | Ikbke | -0.013 | Young |
| 0.575369 | 0.164226 | 0.386 | 0.372 | 1 | 7 | Tpx2 | 0.014 | Young |
| 0.575391 | 0.023751 | 0.407 | 0.433 | 1 | 7 | Got1 | -0.026 | Young |
| 0.575434 | 0.339229 | 0.322 | 0.313 | 1 | 7 | Tmem208 | 0.009 | Young |
| 0.575579 | 0.374916 | 0.148 | 0.136 | 1 | 7 | Rpp30 | 0.012 | Young |
| 0.575917 | 0.009693 | 0.25 | 0.233 | 1 | 7 | Crebrf | 0.017 | Young |
| 0.57601 | 0.218163 | 0.355 | 0.405 | 1 | 7 | Rin2 | -0.05 | Young |
| 0.576039 | 0.011064 | 0.548 | 0.528 | 1 | 7 | Hnrnpdl | 0.02 | Young |
| 0.576067 | 0.494913 | 0.169 | 0.159 | 1 | 7 | Gid4 | 0.01 | Young |
| 0.57628 | 0.124143 | 0.021 | 0.015 | 1 | 7 | Zfyve28 | 0.006 | Young |
| 0.576318 | 0.04632 | 0.386 | 0.408 | 1 | 7 | Twf1 | -0.022 | Young |
| 0.576438 | 0.040975 | 0.485 | 0.521 | 1 | 7 | Vps4b | -0.036 | Young |
| 0.577148 | 0.626814 | 0.12 | 0.11 | 1 | 7 | Rcbtb1 | 0.01 | Young |
| 0.577211 | 0.300164 | 0.051 | 0.062 | 1 | 7 | Fpgt | -0.011 | Young |
| 0.577532 | 0.200032 | 0.093 | 0.082 | 1 | 7 | Ccdc32 | 0.011 | Young |
| 0.57764 | 0.522316 | 0.102 | 0.118 | 1 | 7 | Rbpms | -0.016 | Young |
| 0.577846 | 0.249277 | 0.066 | 0.056 | 1 | 7 | Mllt11 | 0.01 | Young |
| 0.577894 | 0.165241 | 0.262 | 0.256 | 1 | 7 | Ylpm1 | 0.006 | Young |
| 0.578084 | 0.013761 | 0.208 | 0.231 | 1 | 7 | Nabp2 | -0.023 | Young |
| 0.578191 | 0.706349 | 0.063 | 0.054 | 1 | 7 | Spout1 | 0.009 | Young |
| 0.578391 | 0.006959 | 0.349 | 0.387 | 1 | 7 | Ttc3 | -0.038 | Young |
| 0.578455 | 0.474062 | 0.069 | 0.059 | 1 | 7 | Socs2 | 0.01 | Young |
| 0.578794 | 0.546498 | 0.114 | 0.103 | 1 | 7 | Ptpn23 | 0.011 | Young |
| 0.578841 | 0.271775 | 0.166 | 0.151 | 1 | 7 | Soga1 | 0.015 | Young |
| 0.579066 | 0.495054 | 0.084 | 0.074 | 1 | 7 | Clec4n | 0.01 | Young |
| 0.579299 | 0.484934 | 0.208 | 0.195 | 1 | 7 | Vps18 | 0.013 | Young |
| 0.580279 | 0.123821 | 0.244 | 0.272 | 1 | 7 | Pbx3 | -0.028 | Young |
| 0.580322 | 0.037731 | 0.524 | 0.51 | 1 | 7 | Arap1 | 0.014 | Young |
| 0.580671 | 0.466884 | 0.208 | 0.197 | 1 | 7 | Skint3 | 0.011 | Young |
| 0.580705 | 0.070439 | 0.361 | 0.392 | 1 | 7 | Lrrc25 | -0.031 | Young |
| 0.580723 | 0.064977 | 0.949 | 0.928 | 1 | 7 | H3f3a | 0.021 | Young |
| 0.580946 | 0.42602 | 0.31 | 0.305 | 1 | 7 | Pom121 | 0.005 | Young |
| 0.58123 | 0.174244 | 0.497 | 0.485 | 1 | 7 | Cbx1 | 0.012 | Young |
| 0.581426 | 0.344735 | 0.205 | 0.195 | 1 | 7 | Eipr1 | 0.01 | Young |
| 0.581494 | 0.571404 | 0.081 | 0.095 | 1 | 7 | Rtl5 | -0.014 | Young |
| 0.581593 | 0.072053 | 0.271 | 0.3 | 1 | 7 | Clk3 | -0.029 | Young |

| | | | | | | | | |
|---|---|---|---|---|---|---|---|---|
| 0.58168 | 0.569145 | 0.06 | 0.072 | 1 | 7 | Rmnd1 | -0.012 | Young |
| 0.581903 | 0.187525 | 0.256 | 0.246 | 1 | 7 | Rnf166 | 0.01 | Young |
| 0.582471 | 0.133133 | 0.458 | 0.497 | 1 | 7 | Acly | -0.039 | Young |
| 0.582563 | 0.034294 | 0.684 | 0.682 | 1 | 7 | Abracl | 0.002 | Young |
| 0.582663 | 0.778285 | 0.096 | 0.113 | 1 | 7 | Ampd2 | -0.017 | Young |
| 0.582682 | 0.336812 | 0.211 | 0.2 | 1 | 7 | Fcgr1 | 0.011 | Young |
| 0.582743 | 0.334813 | 0.401 | 0.405 | 1 | 7 | Rtf1 | -0.004 | Young |
| 0.582752 | 0.361307 | 0.133 | 0.118 | 1 | 7 | Usp33 | 0.015 | Young |
| 0.582793 | 0.240824 | 0.22 | 0.244 | 1 | 7 | Gm28875 | -0.024 | Young |
| 0.583097 | 0.09107 | 0.289 | 0.305 | 1 | 7 | Herc4 | -0.016 | Young |
| 0.58358 | 0.101309 | 0.127 | 0.144 | 1 | 7 | Cog3 | -0.017 | Young |
| 0.58389 | 0.096138 | 0.383 | 0.379 | 1 | 7 | Rpa2 | 0.004 | Young |
| 0.584141 | 0.050399 | 0.286 | 0.31 | 1 | 7 | Psmd12 | -0.024 | Young |
| 0.584154 | 0.035382 | 0.145 | 0.162 | 1 | 7 | Golga5 | -0.017 | Young |
| 0.584366 | 0.237703 | 0.145 | 0.133 | 1 | 7 | 10-Sep | 0.012 | Young |
| 0.584505 | 0.142201 | 0.343 | 0.387 | 1 | 7 | Cnppd1 | -0.044 | Young |
| 0.584662 | 0.080169 | 0.292 | 0.331 | 1 | 7 | Sf3b4 | -0.039 | Young |
| 0.584798 | 0.229091 | 0.015 | 0.021 | 1 | 7 | Tnfrsf22 | -0.006 | Young |
| 0.584845 | 0.145285 | 0.41 | 0.459 | 1 | 7 | Midn | -0.049 | Young |
| 0.585358 | 0.499977 | 0.151 | 0.138 | 1 | 7 | Plekhm1 | 0.013 | Young |
| 0.585726 | 0.090552 | 0.063 | 0.074 | 1 | 7 | Qtrt1 | -0.011 | Young |
| 0.586301 | 0.046068 | 0.232 | 0.215 | 1 | 7 | Ncoa6 | 0.017 | Young |
| 0.586322 | 0.055464 | 0.575 | 0.597 | 1 | 7 | Rbx1 | -0.022 | Young |
| 0.586404 | 0.672095 | 0.057 | 0.049 | 1 | 7 | Atrip | 0.008 | Young |
| 0.586567 | 0.231821 | 0.262 | 0.29 | 1 | 7 | Alkbh1 | -0.028 | Young |
| 0.58676 | 0.127505 | 0.639 | 0.649 | 1 | 7 | Plekho1 | -0.01 | Young |
| 0.586903 | 0.031807 | 0.383 | 0.421 | 1 | 7 | Peli1 | -0.038 | Young |
| 0.587398 | 0.061036 | 0.735 | 0.731 | 1 | 7 | Calm3 | 0.004 | Young |
| 0.587475 | 0.0124 | 0.072 | 0.085 | 1 | 7 | Anks3 | -0.013 | Young |
| 0.587689 | 0.257902 | 0.47 | 0.482 | 1 | 7 | Nup98 | -0.012 | Young |
| 0.587869 | 0.953979 | 0.099 | 0.09 | 1 | 7 | Cox19 | 0.009 | Young |
| 0.588007 | 0.174342 | 0.015 | 0.021 | 1 | 7 | Arg1 | -0.006 | Young |
| 0.588773 | 0.396357 | 0.054 | 0.046 | 1 | 7 | Ccnyl1 | 0.008 | Young |
| 0.588892 | 0.024421 | 0.358 | 0.395 | 1 | 7 | Csnk1g2 | -0.037 | Young |
| 0.589296 | 0.307586 | 0.193 | 0.179 | 1 | 7 | Tlk2 | 0.014 | Young |
| 0.589687 | 0.195684 | 0.145 | 0.167 | 1 | 7 | Rhot1 | -0.022 | Young |
| 0.589819 | 0.823689 | 0.072 | 0.064 | 1 | 7 | Xaf1 | 0.008 | Young |
| 0.589856 | 0.101637 | 0.096 | 0.085 | 1 | 7 | Zfp329 | 0.011 | Young |
| 0.58994 | 0.043776 | 0.042 | 0.051 | 1 | 7 | Dnajc27 | -0.009 | Young |
| 0.590092 | 0.240274 | 0.06 | 0.051 | 1 | 7 | Adam33 | 0.009 | Young |
| 0.590715 | 0.102204 | 0.06 | 0.051 | 1 | 7 | Zfp157 | 0.009 | Young |
| 0.590717 | 0.138667 | 0.193 | 0.215 | 1 | 7 | Zfp622 | -0.022 | Young |
| 0.590846 | 0.113394 | 0.494 | 0.49 | 1 | 7 | Arf3 | 0.004 | Young |
| 0.5909 | 0.109159 | 0.033 | 0.041 | 1 | 7 | Agrn | -0.008 | Young |
| 0.590989 | 0.259269 | 0.25 | 0.233 | 1 | 7 | Pbdc1 | 0.017 | Young |

| | | | | | | | | |
|---|---|---|---|---|---|---|---|---|
| 0.591292 | 0.316545 | 0.136 | 0.126 | 1 | 7 | Khk | 0.01 | Young |
| 0.591333 | 0.096836 | 0.063 | 0.074 | 1 | 7 | Gab3 | -0.011 | Young |
| 0.591956 | 0.072048 | 0.142 | 0.159 | 1 | 7 | Endod1 | -0.017 | Young |
| 0.592147 | 0.583648 | 0.057 | 0.049 | 1 | 7 | Nemp2 | 0.008 | Young |
| 0.592298 | 0.319027 | 0.015 | 0.021 | 1 | 7 | Zfp566 | -0.006 | Young |
| 0.592409 | 0.665881 | 0.16 | 0.149 | 1 | 7 | Spidr | 0.011 | Young |
| 0.592493 | 0.873085 | 0.093 | 0.085 | 1 | 7 | Dmxl2 | 0.008 | Young |
| 0.592536 | 0.600451 | 0.19 | 0.177 | 1 | 7 | Mob3b | 0.013 | Young |
| 0.592614 | 0.200905 | 0.105 | 0.095 | 1 | 7 | Tmem199 | 0.01 | Young |
| 0.592727 | 0.061068 | 0.675 | 0.703 | 1 | 7 | Sf3b2 | -0.028 | Young |
| 0.593042 | 0.121567 | 0.148 | 0.136 | 1 | 7 | Gcnt1 | 0.012 | Young |
| 0.593068 | 0.422541 | 0.133 | 0.118 | 1 | 7 | Josd2 | 0.015 | Young |
| 0.593178 | 0.164924 | 0.142 | 0.162 | 1 | 7 | Atm | -0.02 | Young |
| 0.593729 | 0.136787 | 0.485 | 0.5 | 1 | 7 | Ptp4a1 | -0.015 | Young |
| 0.593924 | 0.045786 | 0.235 | 0.256 | 1 | 7 | Dido1 | -0.021 | Young |
| 0.593977 | 0.151747 | 0.063 | 0.054 | 1 | 7 | Sort1 | 0.009 | Young |
| 0.594708 | 0.219704 | 0.057 | 0.049 | 1 | 7 | Ctnnd2 | 0.008 | Young |
| 0.595374 | 0.022589 | 0.389 | 0.426 | 1 | 7 | Ptgs1 | -0.037 | Young |
| 0.595538 | 0.048817 | 0.024 | 0.031 | 1 | 7 | Ccdc180 | -0.007 | Young |
| 0.59556 | 0.494048 | 0.163 | 0.151 | 1 | 7 | Stil | 0.012 | Young |
| 0.595963 | 0.022126 | 0.145 | 0.162 | 1 | 7 | Zmat5 | -0.017 | Young |
| 0.596161 | 0.180393 | 0.145 | 0.162 | 1 | 7 | Akap10 | -0.017 | Young |
| 0.596411 | 0.668122 | 0.024 | 0.031 | 1 | 7 | Ttll5 | -0.007 | Young |
| 0.596563 | 0.107188 | 0.151 | 0.141 | 1 | 7 | Syap1 | 0.01 | Young |
| 0.596599 | 0.1662 | 0.328 | 0.313 | 1 | 7 | Npc1 | 0.015 | Young |
| 0.596603 | 0.032703 | 0.015 | 0.021 | 1 | 7 | Zfp61 | -0.006 | Young |
| 0.59689 | 0.176647 | 0.44 | 0.438 | 1 | 7 | Esd | 0.002 | Young |
| 0.59698 | 0.102801 | 0.045 | 0.054 | 1 | 7 | B430306N( | -0.009 | Young |
| 0.597028 | 0.058535 | 0.41 | 0.449 | 1 | 7 | Vrk1 | -0.039 | Young |
| 0.59704 | 0.276861 | 0.256 | 0.238 | 1 | 7 | Cnpy3 | 0.018 | Young |
| 0.597194 | 0.048758 | 0.512 | 0.559 | 1 | 7 | Ep300 | -0.047 | Young |
| 0.597283 | 0.037473 | 0.774 | 0.79 | 1 | 7 | App | -0.016 | Young |
| 0.597424 | 0.125556 | 0.133 | 0.121 | 1 | 7 | AI606181 | 0.012 | Young |
| 0.597482 | 0.010919 | 0.506 | 0.541 | 1 | 7 | Psmb6 | -0.035 | Young |
| 0.597529 | 0.266228 | 0.352 | 0.392 | 1 | 7 | Emc3 | -0.04 | Young |
| 0.597682 | 0.186197 | 0.015 | 0.021 | 1 | 7 | Nphp1 | -0.006 | Young |
| 0.597738 | 0.033302 | 0.054 | 0.064 | 1 | 7 | Supv3l1 | -0.01 | Young |
| 0.597862 | 0.292266 | 0.232 | 0.221 | 1 | 7 | Fcf1 | 0.011 | Young |
| 0.598042 | 0.281932 | 0.042 | 0.051 | 1 | 7 | Nek4 | -0.009 | Young |
| 0.598132 | 0.46553 | 0.099 | 0.09 | 1 | 7 | Mvd | 0.009 | Young |
| 0.598444 | 0.37081 | 0.033 | 0.041 | 1 | 7 | Mx1 | -0.008 | Young |
| 0.598642 | 0.087401 | 0.286 | 0.267 | 1 | 7 | Ccdc59 | 0.019 | Young |
| 0.598761 | 0.337596 | 0.015 | 0.021 | 1 | 7 | Prr15 | -0.006 | Young |
| 0.599047 | 0.311127 | 0.105 | 0.121 | 1 | 7 | Cars2 | -0.016 | Young |
| 0.599736 | 0.508136 | 0.102 | 0.092 | 1 | 7 | Gaa | 0.01 | Young |

| | | | | | | | | |
|---|---|---|---|---|---|---|---|---|
| 0.599958 | 0.862421 | 0.033 | 0.041 | 1 | 7 | Rnase2a | -0.008 | Young |
| 0.600502 | 0.331575 | 0.096 | 0.11 | 1 | 7 | Dclre1a | -0.014 | Young |
| 0.600656 | 0.168406 | 0.491 | 0.495 | 1 | 7 | Rabac1 | -0.004 | Young |
| 0.600749 | 0.3854 | 0.286 | 0.313 | 1 | 7 | Angel2 | -0.027 | Young |
| 0.601082 | 0.116898 | 0.542 | 0.554 | 1 | 7 | Tmem106a | -0.012 | Young |
| 0.601524 | 0.052418 | 0.524 | 0.582 | 1 | 7 | Tgoln1 | -0.058 | Young |
| 0.601536 | 0.395271 | 0.09 | 0.079 | 1 | 7 | Scyl3 | 0.011 | Young |
| 0.60157 | 0.367546 | 0.099 | 0.09 | 1 | 7 | Rmnd5b | 0.009 | Young |
| 0.601628 | 0.285401 | 0.211 | 0.2 | 1 | 7 | Slc5a3 | 0.011 | Young |
| 0.601946 | 0.126072 | 0.111 | 0.1 | 1 | 7 | Kdelc2 | 0.011 | Young |
| 0.602029 | 0.00166 | 0.578 | 0.633 | 1 | 7 | Capg | -0.055 | Young |
| 0.602078 | 0.313516 | 0.247 | 0.236 | 1 | 7 | Itga5 | 0.011 | Young |
| 0.602352 | 0.003676 | 0.241 | 0.226 | 1 | 7 | Epc1 | 0.015 | Young |
| 0.602582 | 0.363059 | 0.196 | 0.215 | 1 | 7 | Rbm12 | -0.019 | Young |
| 0.602814 | 0.090302 | 0.313 | 0.336 | 1 | 7 | Anln | -0.023 | Young |
| 0.603066 | 0.297641 | 0.28 | 0.269 | 1 | 7 | Rnase6 | 0.011 | Young |
| 0.603102 | 0.070179 | 0.057 | 0.067 | 1 | 7 | Elk1 | -0.01 | Young |
| 0.603353 | 0.273834 | 0.123 | 0.138 | 1 | 7 | Tyw1 | -0.015 | Young |
| 0.603413 | 0.041911 | 0.614 | 0.667 | 1 | 7 | Smchd1 | -0.053 | Young |
| 0.603477 | 0.207596 | 0.404 | 0.4 | 1 | 7 | Lsm12 | 0.004 | Young |
| 0.6037 | 0.103026 | 0.139 | 0.154 | 1 | 7 | Pde12 | -0.015 | Young |
| 0.603752 | 0.0643 | 0.623 | 0.636 | 1 | 7 | Uqcrb | -0.013 | Young |
| 0.603812 | 0.528447 | 0.051 | 0.044 | 1 | 7 | Samd4 | 0.007 | Young |
| 0.604122 | 0.673167 | 0.217 | 0.215 | 1 | 7 | Cttnbp2nl | 0.002 | Young |
| 0.604245 | 0.189691 | 0.045 | 0.054 | 1 | 7 | Fam26f | -0.009 | Young |
| 0.60442 | 0.175813 | 0.054 | 0.064 | 1 | 7 | Zfp839 | -0.01 | Young |
| 0.604754 | 0.318481 | 0.133 | 0.123 | 1 | 7 | Prr14l | 0.01 | Young |
| 0.60489 | 0.097602 | 0.063 | 0.074 | 1 | 7 | Epn2 | -0.011 | Young |
| 0.604927 | 0.596956 | 0.12 | 0.113 | 1 | 7 | Cdadc1 | 0.007 | Young |
| 0.604973 | 0.123389 | 0.163 | 0.182 | 1 | 7 | Ube2t | -0.019 | Young |
| 0.605173 | 0.797658 | 0.051 | 0.044 | 1 | 7 | Adamtsl5 | 0.007 | Young |
| 0.605174 | 0.712879 | 0.024 | 0.031 | 1 | 7 | Tmem81 | -0.007 | Young |
| 0.605286 | 0.055846 | 0.259 | 0.285 | 1 | 7 | Clock | -0.026 | Young |
| 0.605425 | 0.172178 | 0.157 | 0.174 | 1 | 7 | Atrnl1 | -0.017 | Young |
| 0.605457 | 0.165003 | 0.063 | 0.074 | 1 | 7 | Lig3 | -0.011 | Young |
| 0.60553 | 0.139863 | 0.47 | 0.469 | 1 | 7 | Herc1 | 0.001 | Young |
| 0.605717 | 0.242585 | 0.226 | 0.221 | 1 | 7 | Itpripl1 | 0.005 | Young |
| 0.605749 | 0.152492 | 0.111 | 0.126 | 1 | 7 | Wdr46 | -0.015 | Young |
| 0.606104 | 0.086316 | 0.066 | 0.077 | 1 | 7 | Il1rl2 | -0.011 | Young |
| 0.606514 | 0.161145 | 0.446 | 0.436 | 1 | 7 | Rbbp6 | 0.01 | Young |
| 0.606619 | 0.081648 | 0.081 | 0.072 | 1 | 7 | Mthfr | 0.009 | Young |
| 0.606828 | 0.004176 | 0.34 | 0.367 | 1 | 7 | Hadha | -0.027 | Young |
| 0.606981 | 0.282525 | 0.075 | 0.087 | 1 | 7 | Fam210a | -0.012 | Young |
| 0.607028 | 0.477742 | 0.199 | 0.187 | 1 | 7 | Fbxw4 | 0.012 | Young |
| 0.607138 | 0.042372 | 0.304 | 0.333 | 1 | 7 | Pqlc1 | -0.029 | Young |

| | | | | | | | | |
|---|---|---|---|---|---|---|---|---|
| 0.607484 | 0.739078 | 0.069 | 0.062 | 1 | 7 | Ccdc71 | 0.007 | Young |
| 0.607834 | 0.349064 | 0.256 | 0.292 | 1 | 7 | Zzz3 | -0.036 | Young |
| 0.607916 | 0.731939 | 0.075 | 0.067 | 1 | 7 | Mfge8 | 0.008 | Young |
| 0.608069 | 0.052192 | 0.467 | 0.505 | 1 | 7 | Camkk2 | -0.038 | Young |
| 0.608082 | 0.399337 | 0.102 | 0.118 | 1 | 7 | Ppan | -0.016 | Young |
| 0.608475 | 0.221601 | 0.087 | 0.1 | 1 | 7 | Selenom | -0.013 | Young |
| 0.60848 | 0.458873 | 0.075 | 0.067 | 1 | 7 | Msto1 | 0.008 | Young |
| 0.608673 | 0.255455 | 0.53 | 0.546 | 1 | 7 | Ndel1 | -0.016 | Young |
| 0.608863 | 0.234074 | 0.25 | 0.274 | 1 | 7 | Gm47283 | -0.024 | Young |
| 0.608882 | 0.004722 | 0.057 | 0.049 | 1 | 7 | 9330160F1 | 0.008 | Young |
| 0.609078 | 0.450366 | 0.033 | 0.041 | 1 | 7 | Dtwd1 | -0.008 | Young |
| 0.609098 | 0.466409 | 0.277 | 0.277 | 1 | 7 | Trappc5 | 0 | Young |
| 0.60916 | 0.030026 | 0.849 | 0.867 | 1 | 7 | Ucp2 | -0.018 | Young |
| 0.609543 | 0.279422 | 0.578 | 0.61 | 1 | 7 | Ptpa | -0.032 | Young |
| 0.609731 | 0.179418 | 0.223 | 0.244 | 1 | 7 | Hint2 | -0.021 | Young |
| 0.609773 | 0.307123 | 0.286 | 0.315 | 1 | 7 | BC004004 | -0.029 | Young |
| 0.609832 | 0.02759 | 0.434 | 0.459 | 1 | 7 | Msl2 | -0.025 | Young |
| 0.609886 | 0.245872 | 0.377 | 0.372 | 1 | 7 | Qdpr | 0.005 | Young |
| 0.610402 | 0.104908 | 0.304 | 0.338 | 1 | 7 | Dnajb11 | -0.034 | Young |
| 0.610751 | 0.235322 | 0.25 | 0.238 | 1 | 7 | Rybp | 0.012 | Young |
| 0.610923 | 0.442987 | 0.196 | 0.187 | 1 | 7 | Bbx | 0.009 | Young |
| 0.611135 | 0.035391 | 0.066 | 0.077 | 1 | 7 | Tsr2 | -0.011 | Young |
| 0.611414 | 0.052059 | 0.178 | 0.197 | 1 | 7 | Kif20a | -0.019 | Young |
| 0.611439 | 0.2527 | 0.518 | 0.528 | 1 | 7 | Tbxas1 | -0.01 | Young |
| 0.611584 | 0.235893 | 0.349 | 0.331 | 1 | 7 | Wbp2 | 0.018 | Young |
| 0.611748 | 0.480188 | 0.054 | 0.064 | 1 | 7 | Zfp182 | -0.01 | Young |
| 0.611918 | 0.389947 | 0.265 | 0.259 | 1 | 7 | 1810022K0 | 0.006 | Young |
| 0.611961 | 0.068601 | 0.084 | 0.074 | 1 | 7 | Zfp369 | 0.01 | Young |
| 0.612104 | 0.508012 | 0.072 | 0.064 | 1 | 7 | Mmgt2 | 0.008 | Young |
| 0.612267 | 0.255098 | 0.295 | 0.287 | 1 | 7 | Phf10 | 0.008 | Young |
| 0.612271 | 0.069253 | 0.578 | 0.567 | 1 | 7 | D8Ertd738 | 0.011 | Young |
| 0.612432 | 0.174408 | 0.078 | 0.09 | 1 | 7 | Uevld | -0.012 | Young |
| 0.612489 | 0.034608 | 0.093 | 0.082 | 1 | 7 | Hagh | 0.011 | Young |
| 0.612893 | 0.207886 | 0.277 | 0.303 | 1 | 7 | Zdhhc21 | -0.026 | Young |
| 0.613038 | 0.049691 | 0.096 | 0.108 | 1 | 7 | Srcap | -0.012 | Young |
| 0.613074 | 0.024851 | 0.111 | 0.123 | 1 | 7 | Aasdhppt | -0.012 | Young |
| 0.614214 | 0.25999 | 0.045 | 0.054 | 1 | 7 | Abca2 | -0.009 | Young |
| 0.614235 | 0.023841 | 0.226 | 0.244 | 1 | 7 | Siah2 | -0.018 | Young |
| 0.614598 | 0.087868 | 0.581 | 0.59 | 1 | 7 | Clec4a1 | -0.009 | Young |
| 0.614786 | 0.116136 | 0.172 | 0.156 | 1 | 7 | Gtf2f2 | 0.016 | Young |
| 0.614811 | 0.304338 | 0.211 | 0.236 | 1 | 7 | Mex3c | -0.025 | Young |
| 0.61526 | 0.105177 | 0.06 | 0.051 | 1 | 7 | Gm27010 | 0.009 | Young |
| 0.615529 | 0.147655 | 0.166 | 0.154 | 1 | 7 | AW549877 | 0.012 | Young |
| 0.615668 | 0.330071 | 0.105 | 0.095 | 1 | 7 | Xndc1 | 0.01 | Young |
| 0.616009 | 0.689602 | 0.048 | 0.041 | 1 | 7 | Gatb | 0.007 | Young |

| | | | | | | | | |
|---|---|---|---|---|---|---|---|---|
| 0.616271 | 0.014268 | 0.627 | 0.674 | 1 | 7 | Bnip2 | -0.047 | Young |
| 0.616565 | 0.158894 | 0.352 | 0.351 | 1 | 7 | Sbds | 0.001 | Young |
| 0.616606 | 0.209965 | 0.181 | 0.205 | 1 | 7 | Fbxl14 | -0.024 | Young |
| 0.616993 | 0.322242 | 0.081 | 0.072 | 1 | 7 | Ankrd52 | 0.009 | Young |
| 0.617128 | 0.527133 | 0.202 | 0.192 | 1 | 7 | Rnf2 | 0.01 | Young |
| 0.617169 | 0.246814 | 0.268 | 0.29 | 1 | 7 | Exosc5 | -0.022 | Young |
| 0.617553 | 0.02031 | 0.045 | 0.054 | 1 | 7 | Fastkd2 | -0.009 | Young |
| 0.617969 | 0.167808 | 0.139 | 0.154 | 1 | 7 | Dpm1 | -0.015 | Young |
| 0.617984 | 0.000327 | 0.373 | 0.392 | 1 | 7 | Prmt1 | -0.019 | Young |
| 0.618591 | 0.192727 | 0.627 | 0.644 | 1 | 7 | Nfkb1 | -0.017 | Young |
| 0.618778 | 0.367234 | 0.102 | 0.092 | 1 | 7 | Tsr1 | 0.01 | Young |
| 0.619008 | 0.158731 | 0.482 | 0.469 | 1 | 7 | Susd6 | 0.013 | Young |
| 0.620254 | 0.706765 | 0.048 | 0.041 | 1 | 7 | Erc1 | 0.007 | Young |
| 0.621368 | 0.331622 | 0.253 | 0.279 | 1 | 7 | Aspm | -0.026 | Young |
| 0.621576 | 0.580707 | 0.087 | 0.079 | 1 | 7 | Acd | 0.008 | Young |
| 0.62161 | 0.421125 | 0.238 | 0.236 | 1 | 7 | Trmt1 | 0.002 | Young |
| 0.62192 | 0.01772 | 0.642 | 0.679 | 1 | 7 | Kif5b | -0.037 | Young |
| 0.621944 | 0.280837 | 0.298 | 0.287 | 1 | 7 | Atp2c1 | 0.011 | Young |
| 0.62237 | 0.372662 | 0.313 | 0.308 | 1 | 7 | Pnpla8 | 0.005 | Young |
| 0.622381 | 0.257885 | 0.048 | 0.041 | 1 | 7 | Crtc1 | 0.007 | Young |
| 0.622581 | 0.223979 | 0.169 | 0.156 | 1 | 7 | Mxi1 | 0.013 | Young |
| 0.622889 | 0.278802 | 0.078 | 0.09 | 1 | 7 | Recql5 | -0.012 | Young |
| 0.622939 | 0.523001 | 0.139 | 0.131 | 1 | 7 | Dhrs4 | 0.008 | Young |
| 0.622992 | 0.27929 | 0.051 | 0.044 | 1 | 7 | Slc25a38 | 0.007 | Young |
| 0.623002 | 0.011858 | 0.53 | 0.515 | 1 | 7 | Anapc5 | 0.015 | Young |
| 0.623369 | 0.108804 | 0.205 | 0.192 | 1 | 7 | Dtx3l | 0.013 | Young |
| 0.623911 | 0.466241 | 0.196 | 0.215 | 1 | 7 | Anapc2 | -0.019 | Young |
| 0.62403 | 0.299735 | 0.117 | 0.105 | 1 | 7 | Pld1 | 0.012 | Young |
| 0.624163 | 0.182431 | 0.449 | 0.454 | 1 | 7 | Uvrag | -0.005 | Young |
| 0.624468 | 0.353647 | 0.142 | 0.131 | 1 | 7 | Ern1 | 0.011 | Young |
| 0.624512 | 0.161297 | 0.048 | 0.041 | 1 | 7 | Icam2 | 0.007 | Young |
| 0.624661 | 0.152748 | 0.416 | 0.415 | 1 | 7 | Pdlim5 | 0.001 | Young |
| 0.624817 | 0.039157 | 0.072 | 0.082 | 1 | 7 | Bambi | -0.01 | Young |
| 0.625234 | 0.316466 | 0.425 | 0.433 | 1 | 7 | Tmem234 | -0.008 | Young |
| 0.625309 | 0.034769 | 0.515 | 0.51 | 1 | 7 | Emc10 | 0.005 | Young |
| 0.625505 | 0.59273 | 0.075 | 0.067 | 1 | 7 | Mxra7 | 0.008 | Young |
| 0.625893 | 0.197271 | 0.139 | 0.128 | 1 | 7 | Dym | 0.011 | Young |
| 0.62619 | 0.027954 | 0.205 | 0.187 | 1 | 7 | Tuba4a | 0.018 | Young |
| 0.626302 | 0.459718 | 0.187 | 0.182 | 1 | 7 | Trappc13 | 0.005 | Young |
| 0.626359 | 0.17995 | 0.28 | 0.303 | 1 | 7 | Utp3 | -0.023 | Young |
| 0.626401 | 0.026299 | 0.316 | 0.297 | 1 | 7 | Ctdspl2 | 0.019 | Young |
| 0.626544 | 0.191609 | 0.283 | 0.318 | 1 | 7 | Mrpl36 | -0.035 | Young |
| 0.626682 | 0.161315 | 0.084 | 0.095 | 1 | 7 | AC165953. | -0.011 | Young |
| 0.626945 | 0.021504 | 0.452 | 0.495 | 1 | 7 | Mlec | -0.043 | Young |
| 0.62724 | 0.187085 | 0.509 | 0.495 | 1 | 7 | Mcrip1 | 0.014 | Young |

| | | | | | | | | |
|---|---|---|---|---|---|---|---|---|
| 0.627592 | 0.022739 | 0.66 | 0.692 | 1 | 7 | Fubp1 | -0.032 | Young |
| 0.62762 | 0.082415 | 0.078 | 0.09 | 1 | 7 | Stx18 | -0.012 | Young |
| 0.627739 | 0.749214 | 0.066 | 0.059 | 1 | 7 | Mboat1 | 0.007 | Young |
| 0.627763 | 0.078489 | 0.355 | 0.354 | 1 | 7 | Eif4g3 | 0.001 | Young |
| 0.627885 | 0.216344 | 0.145 | 0.159 | 1 | 7 | Zcchc2 | -0.014 | Young |
| 0.628054 | 0.281894 | 0.488 | 0.513 | 1 | 7 | Zc3hav1 | -0.025 | Young |
| 0.628069 | 0.500731 | 0.048 | 0.041 | 1 | 7 | Ralgps1 | 0.007 | Young |
| 0.628533 | 0.202335 | 0.009 | 0.013 | 1 | 7 | Cks1brt | -0.004 | Young |
| 0.628554 | 0.270602 | 0.084 | 0.097 | 1 | 7 | Limk1 | -0.013 | Young |
| 0.628663 | 0.117837 | 0.036 | 0.044 | 1 | 7 | Fam177a | -0.008 | Young |
| 0.628733 | 0.058147 | 0.145 | 0.131 | 1 | 7 | Fbxo9 | 0.014 | Young |
| 0.628823 | 0.149146 | 0.027 | 0.033 | 1 | 7 | Cmtr1 | -0.006 | Young |
| 0.629139 | 0.03511 | 0.681 | 0.662 | 1 | 7 | Bzw1 | 0.019 | Young |
| 0.629182 | 0.287005 | 0.105 | 0.121 | 1 | 7 | Abcd3 | -0.016 | Young |
| 0.629187 | 0.247489 | 0.205 | 0.223 | 1 | 7 | Sra1 | -0.018 | Young |
| 0.629203 | 0.230464 | 0.187 | 0.208 | 1 | 7 | Brix1 | -0.021 | Young |
| 0.629663 | 0.018859 | 0.114 | 0.126 | 1 | 7 | Srxn1 | -0.012 | Young |
| 0.62971 | 0.196759 | 0.37 | 0.369 | 1 | 7 | Rragc | 0.001 | Young |
| 0.629722 | 0.281617 | 0.349 | 0.346 | 1 | 7 | Kxd1 | 0.003 | Young |
| 0.629912 | 0.684388 | 0.069 | 0.062 | 1 | 7 | Zfp687 | 0.007 | Young |
| 0.630009 | 0.401477 | 0.193 | 0.187 | 1 | 7 | Rnf139 | 0.006 | Young |
| 0.630122 | 0.02569 | 0.172 | 0.187 | 1 | 7 | Scyl2 | -0.015 | Young |
| 0.630365 | 0.849464 | 0.084 | 0.077 | 1 | 7 | Pex14 | 0.007 | Young |
| 0.630922 | 0.33057 | 0.127 | 0.118 | 1 | 7 | Nckipsd | 0.009 | Young |
| 0.630944 | 0.026913 | 0.181 | 0.169 | 1 | 7 | Aoah | 0.012 | Young |
| 0.631102 | 0.764211 | 0.069 | 0.062 | 1 | 7 | Surf2 | 0.007 | Young |
| 0.631194 | 0.104333 | 0.241 | 0.259 | 1 | 7 | Arhgef12 | -0.018 | Young |
| 0.631294 | 0.125938 | 0.151 | 0.138 | 1 | 7 | Rpl10-ps3 | 0.013 | Young |
| 0.631339 | 0.13832 | 0.009 | 0.013 | 1 | 7 | Egfl8 | -0.004 | Young |
| 0.631371 | 0.320546 | 0.187 | 0.177 | 1 | 7 | Knop1 | 0.01 | Young |
| 0.631464 | 0.03715 | 0.175 | 0.195 | 1 | 7 | Ercc6l | -0.02 | Young |
| 0.631547 | 0.025974 | 0.901 | 0.879 | 1 | 7 | Ptprc | 0.022 | Young |
| 0.632355 | 0.013512 | 0.515 | 0.559 | 1 | 7 | Dhx15 | -0.044 | Young |
| 0.632512 | 0.33304 | 0.298 | 0.292 | 1 | 7 | Pdcd4 | 0.006 | Young |
| 0.632594 | 0.034014 | 0.151 | 0.164 | 1 | 7 | Ctps2 | -0.013 | Young |
| 0.632633 | 0.42346 | 0.133 | 0.123 | 1 | 7 | Cbx6 | 0.01 | Young |
| 0.632723 | 0.020703 | 0.048 | 0.056 | 1 | 7 | 1810032O( | -0.008 | Young |
| 0.632727 | 0.169537 | 0.506 | 0.505 | 1 | 7 | Laptm4a | 0.001 | Young |
| 0.632744 | 0.274203 | 0.009 | 0.013 | 1 | 7 | Gm12184 | -0.004 | Young |
| 0.633073 | 0.550911 | 0.133 | 0.123 | 1 | 7 | Nr1d2 | 0.01 | Young |
| 0.633097 | 0.084083 | 0.027 | 0.033 | 1 | 7 | 1110008L1 | -0.006 | Young |
| 0.633416 | 0.058351 | 0.099 | 0.09 | 1 | 7 | Sema4c | 0.009 | Young |
| 0.633524 | 0.133807 | 0.268 | 0.256 | 1 | 7 | Atg101 | 0.012 | Young |
| 0.63415 | 2.048602 | 0.009 | 0.013 | 1 | 7 | Sfn | -0.004 | Young |
| 0.63415 | 0.629952 | 0.009 | 0.013 | 1 | 7 | Afap1 | -0.004 | Young |

| | | | | | | | | |
|---|---|---|---|---|---|---|---|---|
| 0.63415 | 0.068412 | 0.009 | 0.013 | 1 | 7 | 9130023H2 | -0.004 | Young |
| 0.63415 | 0.058667 | 0.009 | 0.013 | 1 | 7 | Gm10643 | -0.004 | Young |
| 0.634317 | 0.382773 | 0.16 | 0.151 | 1 | 7 | Ska2 | 0.009 | Young |
| 0.63447 | 0.013195 | 0.253 | 0.274 | 1 | 7 | 4931406CC | -0.021 | Young |
| 0.635256 | 0.058298 | 0.639 | 0.651 | 1 | 7 | Slfn2 | -0.012 | Young |
| 0.635339 | 0.214564 | 0.208 | 0.231 | 1 | 7 | Mospd3 | -0.023 | Young |
| 0.635501 | 0.06127 | 0.401 | 0.436 | 1 | 7 | Ik | -0.035 | Young |
| 0.635557 | 0.368625 | 0.009 | 0.013 | 1 | 7 | Kbtbd6 | -0.004 | Young |
| 0.63571 | 0.10638 | 0.66 | 0.654 | 1 | 7 | Ccl2 | 0.006 | Young |
| 0.6367 | 0.114726 | 0.657 | 0.667 | 1 | 7 | Fam46a | -0.01 | Young |
| 0.636941 | 0.117954 | 0.708 | 0.695 | 1 | 7 | Rnf130 | 0.013 | Young |
| 0.636966 | 0.537917 | 0.009 | 0.013 | 1 | 7 | Ptprm | -0.004 | Young |
| 0.637112 | 0.234204 | 0.139 | 0.126 | 1 | 7 | Ciao1 | 0.013 | Young |
| 0.63737 | 0.233844 | 0.422 | 0.428 | 1 | 7 | Trpc4ap | -0.006 | Young |
| 0.637992 | 0.178335 | 0.072 | 0.082 | 1 | 7 | Isyna1 | -0.01 | Young |
| 0.638372 | 0.144575 | 0.452 | 0.451 | 1 | 7 | Taf10 | 0.001 | Young |
| 0.638376 | 0.716839 | 0.009 | 0.013 | 1 | 7 | Col27a1 | -0.004 | Young |
| 0.639151 | 0.002519 | 0.467 | 0.454 | 1 | 7 | Dnajc5 | 0.013 | Young |
| 0.639432 | 0.029269 | 0.349 | 0.382 | 1 | 7 | Ufsp2 | -0.033 | Young |
| 0.639576 | 0.006616 | 0.289 | 0.279 | 1 | 7 | Keap1 | 0.01 | Young |
| 0.639608 | 0.116979 | 0.268 | 0.259 | 1 | 7 | Cenpw | 0.009 | Young |
| 0.639645 | 0.470165 | 0.214 | 0.208 | 1 | 7 | Tmem229b | 0.006 | Young |
| 0.639906 | 0.169454 | 0.127 | 0.141 | 1 | 7 | Cep162 | -0.014 | Young |
| 0.640194 | 0.547656 | 0.063 | 0.056 | 1 | 7 | Gem | 0.007 | Young |
| 0.640657 | 0.368153 | 0.06 | 0.069 | 1 | 7 | Tbc1d22b | -0.009 | Young |
| 0.640754 | 0.212748 | 0.205 | 0.195 | 1 | 7 | Parp2 | 0.01 | Young |
| 0.64076 | 0.206565 | 0.133 | 0.121 | 1 | 7 | Kifc3 | 0.012 | Young |
| 0.641121 | 0.271202 | 0.193 | 0.182 | 1 | 7 | Retreg3 | 0.011 | Young |
| 0.642097 | 0.182794 | 0.205 | 0.226 | 1 | 7 | Bccip | -0.021 | Young |
| 0.642239 | 0.011252 | 0.16 | 0.177 | 1 | 7 | Smn1 | -0.017 | Young |
| 0.642347 | 0.503609 | 0.093 | 0.085 | 1 | 7 | Gzf1 | 0.008 | Young |
| 0.642502 | 0.339599 | 0.117 | 0.131 | 1 | 7 | Nfs1 | -0.014 | Young |
| 0.642544 | 0.053329 | 0.027 | 0.033 | 1 | 7 | Dyrk1b | -0.006 | Young |
| 0.642567 | 0.352858 | 0.099 | 0.113 | 1 | 7 | Qsox2 | -0.014 | Young |
| 0.642606 | 0.03879 | 0.235 | 0.256 | 1 | 7 | Nelfe | -0.021 | Young |
| 0.642608 | 0.048592 | 0.41 | 0.449 | 1 | 7 | Myo1g | -0.039 | Young |
| 0.643253 | 0.143074 | 0.154 | 0.144 | 1 | 7 | Vgll4 | 0.01 | Young |
| 0.643263 | 0.138531 | 0.217 | 0.241 | 1 | 7 | Ube2q2 | -0.024 | Young |
| 0.643308 | 0.027265 | 0.5 | 0.536 | 1 | 7 | Itpkb | -0.036 | Young |
| 0.643365 | 0.057536 | 0.018 | 0.023 | 1 | 7 | 1110046J0 | -0.005 | Young |
| 0.64369 | 0.057643 | 0.587 | 0.6 | 1 | 7 | B4galt6 | -0.013 | Young |
| 0.643942 | 0.076173 | 0.271 | 0.264 | 1 | 7 | Sptlc2 | 0.007 | Young |
| 0.643943 | 0.272154 | 0.404 | 0.405 | 1 | 7 | Mis18a | -0.001 | Young |
| 0.644061 | 0.189866 | 0.178 | 0.167 | 1 | 7 | Cd2 | 0.011 | Young |
| 0.644199 | 0.942458 | 0.045 | 0.038 | 1 | 7 | Timd4 | 0.007 | Young |

| | | | | | | | | |
|---|---|---|---|---|---|---|---|---|
| 0.644267 | 0.555115 | 0.027 | 0.033 | 1 | 7 | Car9 | -0.006 | Young |
| 0.64431 | 0.338979 | 0.5 | 0.531 | 1 | 7 | Acin1 | -0.031 | Young |
| 0.64438 | 0.185431 | 0.133 | 0.146 | 1 | 7 | Nlk | -0.013 | Young |
| 0.644467 | 0.01048 | 0.452 | 0.474 | 1 | 7 | Fam174a | -0.022 | Young |
| 0.644553 | 0.105322 | 0.157 | 0.144 | 1 | 7 | Slc25a20 | 0.013 | Young |
| 0.644751 | 0.035172 | 0.642 | 0.636 | 1 | 7 | Itm2c | 0.006 | Young |
| 0.645364 | 0.229379 | 0.139 | 0.128 | 1 | 7 | Rnf167 | 0.011 | Young |
| 0.64544 | 0.767437 | 0.099 | 0.092 | 1 | 7 | Mgst1 | 0.007 | Young |
| 0.645444 | 0.295744 | 0.018 | 0.023 | 1 | 7 | Dnaaf5 | -0.005 | Young |
| 0.645487 | 0.010971 | 0.211 | 0.231 | 1 | 7 | Trim59 | -0.02 | Young |
| 0.646429 | 0.947501 | 0.045 | 0.038 | 1 | 7 | Scrn3 | 0.007 | Young |
| 0.646761 | 0.351503 | 0.03 | 0.036 | 1 | 7 | Ifit2 | -0.006 | Young |
| 0.646801 | 0.285149 | 0.154 | 0.172 | 1 | 7 | Cinp | -0.018 | Young |
| 0.647075 | 0.078736 | 0.16 | 0.144 | 1 | 7 | Rhobtb1 | 0.016 | Young |
| 0.647167 | 0.136394 | 0.494 | 0.485 | 1 | 7 | Abhd12 | 0.009 | Young |
| 0.647187 | 0.213006 | 0.232 | 0.254 | 1 | 7 | Herpud2 | -0.022 | Young |
| 0.647221 | 0.087162 | 0.139 | 0.149 | 1 | 7 | Trappc12 | -0.01 | Young |
| 0.647526 | 0.48092 | 0.018 | 0.023 | 1 | 7 | Cdkl5 | -0.005 | Young |
| 0.647526 | 0.138046 | 0.018 | 0.023 | 1 | 7 | Card11 | -0.005 | Young |
| 0.647526 | 0.056354 | 0.018 | 0.023 | 1 | 7 | Gm17586 | -0.005 | Young |
| 0.647608 | 0.174375 | 0.217 | 0.21 | 1 | 7 | Pold1 | 0.007 | Young |
| 0.647791 | 0.034872 | 0.407 | 0.428 | 1 | 7 | Phf3 | -0.021 | Young |
| 0.647863 | 0.118204 | 0.06 | 0.069 | 1 | 7 | Cox15 | -0.009 | Young |
| 0.647975 | 0.047728 | 0.702 | 0.71 | 1 | 7 | Celf1 | -0.008 | Young |
| 0.64803 | 0.260769 | 0.377 | 0.377 | 1 | 7 | Med13 | 0 | Young |
| 0.648275 | 0.355962 | 0.072 | 0.064 | 1 | 7 | Mroh1 | 0.008 | Young |
| 0.648554 | 0.367668 | 0.151 | 0.141 | 1 | 7 | Ctso | 0.01 | Young |
| 0.648567 | 0.127558 | 0.018 | 0.023 | 1 | 7 | Ifi205 | -0.005 | Young |
| 0.648652 | 0.040764 | 0.419 | 0.415 | 1 | 7 | Aldh2 | 0.004 | Young |
| 0.648947 | 0.181828 | 0.114 | 0.128 | 1 | 7 | Cep85 | -0.014 | Young |
| 0.649372 | 0.237851 | 0.19 | 0.179 | 1 | 7 | Tmem131 | 0.011 | Young |
| 0.64961 | 0.232764 | 0.018 | 0.023 | 1 | 7 | Acsf2 | -0.005 | Young |
| 0.649885 | 0.031627 | 0.12 | 0.133 | 1 | 7 | Senp3 | -0.013 | Young |
| 0.649951 | 0.111339 | 0.401 | 0.397 | 1 | 7 | Brd1 | 0.004 | Young |
| 0.650073 | 1.33798 | 0.039 | 0.033 | 1 | 7 | Zkscan8 | 0.006 | Young |
| 0.650087 | 0.324561 | 0.277 | 0.264 | 1 | 7 | Dhrs7 | 0.013 | Young |
| 0.650119 | 0.12778 | 0.117 | 0.131 | 1 | 7 | Med12 | -0.014 | Young |
| 0.650316 | 0.311838 | 0.027 | 0.033 | 1 | 7 | Zfp974 | -0.006 | Young |
| 0.650527 | 0.098004 | 0.244 | 0.259 | 1 | 7 | Fxr2 | -0.015 | Young |
| 0.65087 | 0.019261 | 0.331 | 0.351 | 1 | 7 | Rap2b | -0.02 | Young |
| 0.651181 | 0.09483 | 0.596 | 0.608 | 1 | 7 | Sptssa | -0.012 | Young |
| 0.651334 | 0.197081 | 0.256 | 0.285 | 1 | 7 | Nup50 | -0.029 | Young |
| 0.651434 | 0.024673 | 0.238 | 0.256 | 1 | 7 | Ppie | -0.018 | Young |
| 0.651757 | 0.060171 | 0.241 | 0.256 | 1 | 7 | Rbm14 | -0.015 | Young |
| 0.651821 | 0.193104 | 0.072 | 0.064 | 1 | 7 | Ttc17 | 0.008 | Young |

| | | | | | | | | |
|---|---|---|---|---|---|---|---|---|
| 0.651934 | 0.097834 | 0.039 | 0.046 | 1 | 7 | Pdss1 | -0.007 | Young |
| 0.652635 | 0.293034 | 0.136 | 0.128 | 1 | 7 | Ctnnbl1 | 0.008 | Young |
| 0.652736 | 0.042105 | 0.172 | 0.159 | 1 | 7 | Cdk8 | 0.013 | Young |
| 0.652742 | 0.13624 | 0.018 | 0.023 | 1 | 7 | 9130008F2 | -0.005 | Young |
| 0.652915 | 0.310256 | 0.027 | 0.033 | 1 | 7 | Cbx8 | -0.006 | Young |
| 0.653005 | 0.220408 | 0.232 | 0.226 | 1 | 7 | Cab39l | 0.006 | Young |
| 0.65327 | 0.884413 | 0.039 | 0.033 | 1 | 7 | Mettl2 | 0.006 | Young |
| 0.653523 | 0.137201 | 0.229 | 0.254 | 1 | 7 | Zbtb44 | -0.025 | Young |
| 0.653656 | 0.640479 | 0.111 | 0.126 | 1 | 7 | Ttc4 | -0.015 | Young |
| 0.653895 | 0.187658 | 0.142 | 0.156 | 1 | 7 | Trip4 | -0.014 | Young |
| 0.65392 | 0.44888 | 0.136 | 0.149 | 1 | 7 | Pitrm1 | -0.013 | Young |
| 0.65414 | 0.058975 | 0.039 | 0.046 | 1 | 7 | Cep19 | -0.007 | Young |
| 0.654183 | 0.02467 | 0.334 | 0.364 | 1 | 7 | Anapc13 | -0.03 | Young |
| 0.654542 | 0.225531 | 0.084 | 0.095 | 1 | 7 | Nudt2 | -0.011 | Young |
| 0.654833 | 1.198508 | 0.018 | 0.023 | 1 | 7 | Fancf | -0.005 | Young |
| 0.654833 | 0.561269 | 0.018 | 0.023 | 1 | 7 | Cc2d2a | -0.005 | Young |
| 0.65492 | 0.201618 | 0.136 | 0.146 | 1 | 7 | Lipe | -0.01 | Young |
| 0.655449 | 0.089591 | 0.16 | 0.172 | 1 | 7 | Gbf1 | -0.012 | Young |
| 0.655549 | 0.20002 | 0.346 | 0.341 | 1 | 7 | Ctnnd1 | 0.005 | Young |
| 0.655584 | 0.073431 | 0.084 | 0.095 | 1 | 7 | Ankrd46 | -0.011 | Young |
| 0.655646 | 0.569667 | 0.202 | 0.2 | 1 | 7 | Cep295 | 0.002 | Young |
| 0.655698 | 0.036616 | 0.283 | 0.3 | 1 | 7 | Kdm1a | -0.017 | Young |
| 0.655716 | 0.264299 | 0.099 | 0.09 | 1 | 7 | Gab1 | 0.009 | Young |
| 0.655767 | 0.627529 | 0.208 | 0.205 | 1 | 7 | Tshz1 | 0.003 | Young |
| 0.655781 | 0.234947 | 0.172 | 0.187 | 1 | 7 | Pmm2 | -0.015 | Young |
| 0.655945 | 0.175528 | 0.244 | 0.269 | 1 | 7 | Fyttd1 | -0.025 | Young |
| 0.656055 | 0.030653 | 0.247 | 0.277 | 1 | 7 | Ktn1 | -0.03 | Young |
| 0.656086 | 0.296846 | 0.277 | 0.31 | 1 | 7 | Lrp10 | -0.033 | Young |
| 0.656141 | 0.28236 | 0.367 | 0.362 | 1 | 7 | Taf1 | 0.005 | Young |
| 0.65642 | 0.03985 | 0.5 | 0.51 | 1 | 7 | Abca9 | -0.01 | Young |
| 0.656493 | 0.060417 | 0.593 | 0.564 | 1 | 7 | Ccnd1 | 0.029 | Young |
| 0.656592 | 0.357458 | 0.157 | 0.149 | 1 | 7 | Stx8 | 0.008 | Young |
| 0.657015 | 0.253029 | 0.142 | 0.154 | 1 | 7 | Nsmf | -0.012 | Young |
| 0.657275 | 0.696465 | 0.039 | 0.033 | 1 | 7 | Arhgap35 | 0.006 | Young |
| 0.657289 | 0.093348 | 0.042 | 0.036 | 1 | 7 | Nxpe3 | 0.006 | Young |
| 0.657545 | 0.341649 | 0.172 | 0.164 | 1 | 7 | 9130401M | 0.008 | Young |
| 0.657553 | 0.009427 | 0.404 | 0.438 | 1 | 7 | Lsm2 | -0.034 | Young |
| 0.658033 | 0.657746 | 0.172 | 0.167 | 1 | 7 | Atp7a | 0.005 | Young |
| 0.658135 | 0.358374 | 0.09 | 0.103 | 1 | 7 | Ccdc97 | -0.013 | Young |
| 0.658174 | 0.154018 | 0.136 | 0.149 | 1 | 7 | Cdc6 | -0.013 | Young |
| 0.65819 | 0.101475 | 0.084 | 0.095 | 1 | 7 | Mars | -0.011 | Young |
| 0.658308 | 0.052343 | 0.623 | 0.618 | 1 | 7 | Mtss1 | 0.005 | Young |
| 0.658515 | 0.148895 | 0.208 | 0.228 | 1 | 7 | Washc3 | -0.02 | Young |
| 0.658897 | 0.122002 | 0.172 | 0.19 | 1 | 7 | Edc4 | -0.018 | Young |
| 0.659738 | 0.084958 | 0.554 | 0.592 | 1 | 7 | Sf3b6 | -0.038 | Young |

| | | | | | | | | |
|---|---|---|---|---|---|---|---|---|
| 0.659986 | 0.172058 | 0.211 | 0.208 | 1 | 7 | Mob3a | 0.003 | Young |
| 0.660072 | 0.178354 | 0.051 | 0.059 | 1 | 7 | Iqcb1 | -0.008 | Young |
| 0.660224 | 0.36005 | 0.069 | 0.079 | 1 | 7 | Dph6 | -0.01 | Young |
| 0.660441 | 0.216376 | 0.172 | 0.19 | 1 | 7 | Ebpl | -0.018 | Young |
| 0.660581 | 0.183257 | 0.398 | 0.444 | 1 | 7 | Mcmbp | -0.046 | Young |
| 0.660694 | 0.707404 | 0.066 | 0.077 | 1 | 7 | Ric8b | -0.011 | Young |
| 0.660843 | 0.058711 | 0.178 | 0.192 | 1 | 7 | Ppid | -0.014 | Young |
| 0.661035 | 0.389106 | 0.108 | 0.1 | 1 | 7 | Sgsh | 0.008 | Young |
| 0.661159 | 0.103882 | 0.069 | 0.062 | 1 | 7 | Gemin5 | 0.007 | Young |
| 0.661315 | 0.087228 | 0.292 | 0.315 | 1 | 7 | Rbm33 | -0.023 | Young |
| 0.661526 | 0.225535 | 0.096 | 0.11 | 1 | 7 | Pigf | -0.014 | Young |
| 0.661913 | 0.227663 | 0.072 | 0.064 | 1 | 7 | Tmco4 | 0.008 | Young |
| 0.662632 | 0.095401 | 0.5 | 0.495 | 1 | 7 | Arhgap15 | 0.005 | Young |
| 0.663104 | 0.167524 | 0.072 | 0.064 | 1 | 7 | Zfp948 | 0.008 | Young |
| 0.6633 | 0.112511 | 0.265 | 0.254 | 1 | 7 | Akap9 | 0.011 | Young |
| 0.663572 | 0.522593 | 0.229 | 0.228 | 1 | 7 | Epc2 | 0.001 | Young |
| 0.663747 | 0.209169 | 0.343 | 0.367 | 1 | 7 | Vav3 | -0.024 | Young |
| 0.66402 | 0.023248 | 0.316 | 0.344 | 1 | 7 | Tox4 | -0.028 | Young |
| 0.664139 | 0.195453 | 0.105 | 0.118 | 1 | 7 | Mtif2 | -0.013 | Young |
| 0.664263 | 0.306756 | 0.214 | 0.208 | 1 | 7 | Fam193a | 0.006 | Young |
| 0.66469 | 0.012375 | 0.801 | 0.818 | 1 | 7 | Dek | -0.017 | Young |
| 0.664725 | 0.453356 | 0.172 | 0.164 | 1 | 7 | Dpy19l4 | 0.008 | Young |
| 0.66515 | 0.250918 | 0.214 | 0.238 | 1 | 7 | Slc36a1 | -0.024 | Young |
| 0.665299 | 0.089334 | 0.256 | 0.249 | 1 | 7 | Rsrc1 | 0.007 | Young |
| 0.665316 | 0.489326 | 0.039 | 0.033 | 1 | 7 | Tmem140 | 0.006 | Young |
| 0.665358 | 0.022899 | 0.524 | 0.562 | 1 | 7 | Ndufs6 | -0.038 | Young |
| 0.665485 | 0.298939 | 0.169 | 0.159 | 1 | 7 | Tcf19 | 0.01 | Young |
| 0.66585 | 0.123004 | 0.596 | 0.613 | 1 | 7 | Cct4 | -0.017 | Young |
| 0.666038 | 0.251983 | 0.117 | 0.108 | 1 | 7 | Cep57l1 | 0.009 | Young |
| 0.666071 | 0.030109 | 0.47 | 0.505 | 1 | 7 | Atp5g1 | -0.035 | Young |
| 0.666684 | 0.460696 | 0.072 | 0.064 | 1 | 7 | Pi4kb | 0.008 | Young |
| 0.666693 | 0.227994 | 0.37 | 0.377 | 1 | 7 | Cand1 | -0.007 | Young |
| 0.666744 | 0.208189 | 0.133 | 0.121 | 1 | 7 | Ints12 | 0.012 | Young |
| 0.667063 | 0.074252 | 0.533 | 0.541 | 1 | 7 | Lifr | -0.008 | Young |
| 0.667125 | 0.090991 | 0.16 | 0.174 | 1 | 7 | Tmem63a | -0.014 | Young |
| 0.668132 | 0.275867 | 0.114 | 0.108 | 1 | 7 | Uty | 0.006 | Young |
| 0.668412 | 0.23633 | 0.343 | 0.338 | 1 | 7 | Elk3 | 0.005 | Young |
| 0.668458 | 0.114267 | 0.06 | 0.069 | 1 | 7 | Igip | -0.009 | Young |
| 0.669307 | 0.095829 | 0.307 | 0.287 | 1 | 7 | Dhx40 | 0.02 | Young |
| 0.669347 | 0.354403 | 0.069 | 0.079 | 1 | 7 | Cd3eap | -0.01 | Young |
| 0.66935 | 0.473066 | 0.039 | 0.033 | 1 | 7 | Zfp768 | 0.006 | Young |
| 0.669547 | 0.028862 | 0.383 | 0.4 | 1 | 7 | Rsl1d1 | -0.017 | Young |
| 0.669706 | 0.238604 | 0.151 | 0.138 | 1 | 7 | Hist1h4d | 0.013 | Young |
| 0.669952 | 0.223777 | 0.199 | 0.19 | 1 | 7 | Tmem230 | 0.009 | Young |
| 0.669993 | 0.182437 | 0.389 | 0.374 | 1 | 7 | Wdr45b | 0.015 | Young |

| | | | | | | | | |
|---|---|---|---|---|---|---|---|---|
| 0.670154 | 0.087828 | 0.042 | 0.049 | 1 | 7 | Chac2 | -0.007 | Young |
| 0.670158 | 0.44899 | 0.039 | 0.033 | 1 | 7 | Tfb1m | 0.006 | Young |
| 0.670158 | 0.20299 | 0.039 | 0.033 | 1 | 7 | Ankrd13b | 0.006 | Young |
| 0.670328 | 0.164155 | 0.169 | 0.185 | 1 | 7 | Stam2 | -0.016 | Young |
| 0.671232 | 0.570092 | 0.06 | 0.054 | 1 | 7 | A530017D: | 0.006 | Young |
| 0.67206 | 0.139202 | 0.042 | 0.036 | 1 | 7 | Aplf | 0.006 | Young |
| 0.672202 | 0.18679 | 0.253 | 0.249 | 1 | 7 | Osbpl11 | 0.004 | Young |
| 0.672233 | 0.05103 | 0.175 | 0.19 | 1 | 7 | Sgo1 | -0.015 | Young |
| 0.672435 | 0.019913 | 0.217 | 0.203 | 1 | 7 | Prdm1 | 0.014 | Young |
| 0.672508 | 0.082302 | 0.352 | 0.338 | 1 | 7 | Nfya | 0.014 | Young |
| 0.672697 | 0.173204 | 0.12 | 0.11 | 1 | 7 | Ggt5 | 0.01 | Young |
| 0.67273 | 0.225545 | 0.06 | 0.069 | 1 | 7 | 2010111I0: | -0.009 | Young |
| 0.673187 | 0.215342 | 0.244 | 0.264 | 1 | 7 | Ahr | -0.02 | Young |
| 0.673379 | 0.099345 | 0.039 | 0.046 | 1 | 7 | Acat2 | -0.007 | Young |
| 0.67339 | 0.224657 | 0.084 | 0.095 | 1 | 7 | Zgrf1 | -0.011 | Young |
| 0.673562 | 0.144296 | 0.467 | 0.469 | 1 | 7 | Ets2 | -0.002 | Young |
| 0.674226 | 0.010252 | 0.075 | 0.067 | 1 | 7 | Tmem101 | 0.008 | Young |
| 0.674458 | 0.089447 | 0.274 | 0.303 | 1 | 7 | Rab24 | -0.029 | Young |
| 0.67481 | 0.042139 | 0.265 | 0.254 | 1 | 7 | Slc38a9 | 0.011 | Young |
| 0.674873 | 0.210423 | 0.199 | 0.19 | 1 | 7 | Hs2st1 | 0.009 | Young |
| 0.674908 | 0.326721 | 0.169 | 0.164 | 1 | 7 | Eaf1 | 0.005 | Young |
| 0.674988 | 0.23717 | 0.127 | 0.118 | 1 | 7 | Sh3pxd2a | 0.009 | Young |
| 0.675268 | 0.271327 | 0.187 | 0.21 | 1 | 7 | Tmx4 | -0.023 | Young |
| 0.675302 | 0.0502 | 0.849 | 0.877 | 1 | 7 | Neat1 | -0.028 | Young |
| 0.67532 | 0.639767 | 0.114 | 0.108 | 1 | 7 | Zfp362 | 0.006 | Young |
| 0.675469 | 0.0878 | 0.202 | 0.221 | 1 | 7 | Zbtb8os | -0.019 | Young |
| 0.676186 | 0.570832 | 0.063 | 0.056 | 1 | 7 | Rhbdd3 | 0.007 | Young |
| 0.676223 | 0.264985 | 0.163 | 0.179 | 1 | 7 | A630001G: | -0.016 | Young |
| 0.676506 | 0.095743 | 0.051 | 0.059 | 1 | 7 | B3gnt8 | -0.008 | Young |
| 0.676637 | 0.773818 | 0.039 | 0.033 | 1 | 7 | Mfsd4a | 0.006 | Young |
| 0.676637 | 0.222421 | 0.039 | 0.033 | 1 | 7 | Fam151b | 0.006 | Young |
| 0.676852 | 0.293463 | 0.03 | 0.036 | 1 | 7 | Traf3ip1 | -0.006 | Young |
| 0.676852 | 0.147416 | 0.03 | 0.036 | 1 | 7 | Dnajc17 | -0.006 | Young |
| 0.676889 | 0.013096 | 0.175 | 0.19 | 1 | 7 | Nqo2 | -0.015 | Young |
| 0.677545 | 0.037155 | 0.172 | 0.187 | 1 | 7 | Ddhd1 | -0.015 | Young |
| 0.678067 | 0.118371 | 0.063 | 0.072 | 1 | 7 | B4gat1 | -0.009 | Young |
| 0.678264 | 0.063664 | 0.663 | 0.685 | 1 | 7 | Eif3e | -0.022 | Young |
| 0.67848 | 0.392612 | 0.352 | 0.364 | 1 | 7 | Snx17 | -0.012 | Young |
| 0.678631 | 0.070116 | 0.087 | 0.097 | 1 | 7 | Ino80b | -0.01 | Young |
| 0.679214 | 0.206645 | 0.485 | 0.49 | 1 | 7 | Stk38 | -0.005 | Young |
| 0.679381 | 0.04196 | 0.063 | 0.056 | 1 | 7 | Cmc4 | 0.007 | Young |
| 0.679574 | 0.225761 | 0.205 | 0.226 | 1 | 7 | Mysm1 | -0.021 | Young |
| 0.679788 | 0.223021 | 0.108 | 0.121 | 1 | 7 | Exoc7 | -0.013 | Young |
| 0.679793 | 0.032605 | 0.184 | 0.197 | 1 | 7 | Ralgds | -0.013 | Young |
| 0.680383 | 0.095493 | 0.352 | 0.338 | 1 | 7 | Stat6 | 0.014 | Young |

| | | | | | | | | |
|---|---|---|---|---|---|---|---|---|
| 0.680698 | 0.308509 | 0.039 | 0.033 | 1 | 7 | Hist1h3e | 0.006 | Young |
| 0.680754 | 0.010258 | 0.539 | 0.559 | 1 | 7 | H13 | -0.02 | Young |
| 0.680804 | 0.222362 | 0.163 | 0.182 | 1 | 7 | Pias2 | -0.019 | Young |
| 0.681103 | 0.441475 | 0.178 | 0.197 | 1 | 7 | Slc31a1 | -0.019 | Young |
| 0.681356 | 0.097463 | 0.392 | 0.418 | 1 | 7 | Ifngr2 | -0.026 | Young |
| 0.681729 | 0.169773 | 0.461 | 0.477 | 1 | 7 | Cdk11b | -0.016 | Young |
| 0.681904 | 0.157716 | 0.123 | 0.115 | 1 | 7 | Gdap2 | 0.008 | Young |
| 0.682503 | 0.204249 | 0.084 | 0.077 | 1 | 7 | Psmf1 | 0.007 | Young |
| 0.682745 | 0.366414 | 0.154 | 0.172 | 1 | 7 | Hlx | -0.018 | Young |
| 0.682863 | 0.46985 | 0.22 | 0.215 | 1 | 7 | Spr | 0.005 | Young |
| 0.68304 | 0.065833 | 0.256 | 0.241 | 1 | 7 | Gramd1a | 0.015 | Young |
| 0.68312 | 0.422003 | 0.166 | 0.159 | 1 | 7 | Tmem71 | 0.007 | Young |
| 0.683383 | 0.008858 | 0.096 | 0.087 | 1 | 7 | Exoc6 | 0.009 | Young |
| 0.683465 | 0.017725 | 0.13 | 0.141 | 1 | 7 | Htatip2 | -0.011 | Young |
| 0.683572 | 0.402825 | 0.081 | 0.074 | 1 | 7 | Abcg2 | 0.007 | Young |
| 0.68361 | 0.095905 | 0.491 | 0.503 | 1 | 7 | Yy1 | -0.012 | Young |
| 0.683762 | 0.204072 | 0.548 | 0.572 | 1 | 7 | Wsb1 | -0.024 | Young |
| 0.684428 | 0.020662 | 0.033 | 0.028 | 1 | 7 | Klhl25 | 0.005 | Young |
| 0.684465 | 0.121019 | 0.28 | 0.267 | 1 | 7 | Madd | 0.013 | Young |
| 0.684491 | 0.075868 | 0.286 | 0.313 | 1 | 7 | Esyt2 | -0.027 | Young |
| 0.684655 | 0.102139 | 0.521 | 0.518 | 1 | 7 | Gapvd1 | 0.003 | Young |
| 0.684768 | 0.117108 | 0.117 | 0.128 | 1 | 7 | Zfp266 | -0.011 | Young |
| 0.68479 | 0.016228 | 0.88 | 0.895 | 1 | 7 | Uqcrh | -0.015 | Young |
| 0.684852 | 0.085236 | 0.383 | 0.382 | 1 | 7 | Btk | 0.001 | Young |
| 0.68502 | 0.104404 | 0.53 | 0.577 | 1 | 7 | Atp13a3 | -0.047 | Young |
| 0.685192 | 0.232922 | 0.313 | 0.349 | 1 | 7 | Mrpl13 | -0.036 | Young |
| 0.685921 | 0.231201 | 0.202 | 0.218 | 1 | 7 | Adh5 | -0.016 | Young |
| 0.686028 | 0.135211 | 0.199 | 0.213 | 1 | 7 | Slc39a10 | -0.014 | Young |
| 0.686172 | 0.328257 | 0.148 | 0.162 | 1 | 7 | Eya3 | -0.014 | Young |
| 0.686402 | 0.211887 | 0.343 | 0.338 | 1 | 7 | Snf8 | 0.005 | Young |
| 0.686514 | 0.313509 | 0.117 | 0.11 | 1 | 7 | Pgam5 | 0.007 | Young |
| 0.687197 | 0.042383 | 0.711 | 0.738 | 1 | 7 | Gpx4 | -0.027 | Young |
| 0.687216 | 0.11637 | 0.208 | 0.231 | 1 | 7 | Tnks | -0.023 | Young |
| 0.687248 | 0.061059 | 0.123 | 0.133 | 1 | 7 | Plekhg2 | -0.01 | Young |
| 0.687399 | 0.001437 | 0.229 | 0.246 | 1 | 7 | Mtf2 | -0.017 | Young |
| 0.687644 | 0.38194 | 0.187 | 0.185 | 1 | 7 | Smim4 | 0.002 | Young |
| 0.687765 | 0.063857 | 0.518 | 0.526 | 1 | 7 | Tex261 | -0.008 | Young |
| 0.687849 | 0.241465 | 0.036 | 0.031 | 1 | 7 | 2900089D1 | 0.005 | Young |
| 0.687881 | 0.283785 | 0.184 | 0.205 | 1 | 7 | Ethe1 | -0.021 | Young |
| 0.688068 | 0.389812 | 0.142 | 0.133 | 1 | 7 | Papd7 | 0.009 | Young |
| 0.688186 | 0.026961 | 0.572 | 0.608 | 1 | 7 | Akr1b3 | -0.036 | Young |
| 0.688338 | 1.104997 | 0.03 | 0.026 | 1 | 7 | 5430403G1 | 0.004 | Young |
| 0.688466 | 0.241529 | 0.12 | 0.113 | 1 | 7 | Tubg1 | 0.007 | Young |
| 0.688498 | 0.571438 | 0.196 | 0.19 | 1 | 7 | Pex19 | 0.006 | Young |
| 0.688579 | 0.029459 | 0.274 | 0.295 | 1 | 7 | Rpa3 | -0.021 | Young |

| | | | | | | | | |
|---|---|---|---|---|---|---|---|---|
| 0.688586 | 0.213061 | 0.057 | 0.051 | 1 | 7 | Slc36a3os | 0.006 | Young |
| 0.689099 | 0.418849 | 0.021 | 0.026 | 1 | 7 | Dus4l | -0.005 | Young |
| 0.689241 | 0.081953 | 0.292 | 0.279 | 1 | 7 | Ap3m1 | 0.013 | Young |
| 0.689365 | 0.111612 | 0.042 | 0.049 | 1 | 7 | Bivm | -0.007 | Young |
| 0.68947 | 0.000115 | 0.572 | 0.587 | 1 | 7 | Cbx5 | -0.015 | Young |
| 0.689551 | 0.721973 | 0.057 | 0.067 | 1 | 7 | Vps37c | -0.01 | Young |
| 0.6896 | 0.09346 | 0.066 | 0.074 | 1 | 7 | Hist1h4h | -0.008 | Young |
| 0.689602 | 0.276594 | 0.093 | 0.105 | 1 | 7 | Haus2 | -0.012 | Young |
| 0.689729 | 0.26627 | 0.042 | 0.049 | 1 | 7 | Zc2hc1a | -0.007 | Young |
| 0.689864 | 0.086877 | 0.09 | 0.1 | 1 | 7 | Uxt | -0.01 | Young |
| 0.68992 | 0.073803 | 0.214 | 0.228 | 1 | 7 | E2f8 | -0.014 | Young |
| 0.690327 | 0.557366 | 0.22 | 0.218 | 1 | 7 | Ipo11 | 0.002 | Young |
| 0.690401 | 0.043043 | 0.34 | 0.372 | 1 | 7 | Ndufaf8 | -0.032 | Young |
| 0.690458 | 0.009255 | 0.042 | 0.049 | 1 | 7 | Acrbp | -0.007 | Young |
| 0.690706 | 0.05214 | 0.214 | 0.203 | 1 | 7 | Magohb | 0.011 | Young |
| 0.690875 | 0.095134 | 0.554 | 0.577 | 1 | 7 | Ptpn6 | -0.023 | Young |
| 0.690914 | 0.017068 | 0.205 | 0.223 | 1 | 7 | Tomm40 | -0.018 | Young |
| 0.69104 | 0.283475 | 0.346 | 0.341 | 1 | 7 | Parp1 | 0.005 | Young |
| 0.691188 | 0.31551 | 0.042 | 0.049 | 1 | 7 | Cep78 | -0.007 | Young |
| 0.691238 | 0.009897 | 0.271 | 0.29 | 1 | 7 | Plbd2 | -0.019 | Young |
| 0.691564 | 0.127144 | 0.148 | 0.159 | 1 | 7 | Rps19bp1 | -0.011 | Young |
| 0.692009 | 0.196877 | 0.199 | 0.215 | 1 | 7 | Mtx2 | -0.016 | Young |
| 0.692086 | 0.312699 | 0.09 | 0.082 | 1 | 7 | Zdhhc17 | 0.008 | Young |
| 0.692124 | 0.074419 | 0.217 | 0.236 | 1 | 7 | Ccm2 | -0.019 | Young |
| 0.692452 | 0.151932 | 0.494 | 0.508 | 1 | 7 | Topbp1 | -0.014 | Young |
| 0.692914 | 0.168092 | 0.253 | 0.249 | 1 | 7 | Galc | 0.004 | Young |
| 0.693119 | 0.284033 | 0.021 | 0.026 | 1 | 7 | Gstt3 | -0.005 | Young |
| 0.693799 | 0.180202 | 0.036 | 0.031 | 1 | 7 | Enkd1 | 0.005 | Young |
| 0.693909 | 0.946447 | 0.03 | 0.026 | 1 | 7 | Acad12 | 0.004 | Young |
| 0.693934 | 0.143273 | 0.166 | 0.182 | 1 | 7 | Dhcr24 | -0.016 | Young |
| 0.693963 | 0.130509 | 0.066 | 0.059 | 1 | 7 | Usp28 | 0.007 | Young |
| 0.694099 | 0.028488 | 0.985 | 0.977 | 1 | 7 | Rps28 | 0.008 | Young |
| 0.694126 | 0.086756 | 0.542 | 0.569 | 1 | 7 | Swi5 | -0.027 | Young |
| 0.694147 | 0.181119 | 0.105 | 0.115 | 1 | 7 | Mad1l1 | -0.01 | Young |
| 0.694171 | 1.121784 | 0.033 | 0.028 | 1 | 7 | B3gnt3 | 0.005 | Young |
| 0.694209 | 0.698692 | 0.166 | 0.162 | 1 | 7 | Pilrb2 | 0.004 | Young |
| 0.694372 | 0.337169 | 0.139 | 0.131 | 1 | 7 | Zfp110 | 0.008 | Young |
| 0.694586 | 0.124336 | 0.241 | 0.256 | 1 | 7 | Bms1 | -0.015 | Young |
| 0.695036 | 0.023873 | 0.461 | 0.485 | 1 | 7 | Rnf7 | -0.024 | Young |
| 0.695196 | 0.035134 | 0.181 | 0.174 | 1 | 7 | Ppfia4 | 0.007 | Young |
| 0.695269 | 0.092369 | 0.651 | 0.646 | 1 | 7 | Tnfaip8 | 0.005 | Young |
| 0.695448 | 0.086214 | 0.47 | 0.469 | 1 | 7 | Eif3m | 0.001 | Young |
| 0.695566 | 0.074613 | 0.337 | 0.349 | 1 | 7 | Csnk2a2 | -0.012 | Young |
| 0.696003 | 0.73388 | 0.057 | 0.051 | 1 | 7 | Mtg2 | 0.006 | Young |
| 0.696477 | 0.408218 | 0.235 | 0.233 | 1 | 7 | Camk2d | 0.002 | Young |

| | | | | | | | | |
|---|---|---|---|---|---|---|---|---|
| 0.696701 | 0.921405 | 0.03 | 0.026 | 1 | 7 | Caprin2 | 0.004 | Young |
| 0.696803 | 0.191202 | 0.12 | 0.131 | 1 | 7 | Fam120b | -0.011 | Young |
| 0.696899 | 0.032189 | 0.304 | 0.336 | 1 | 7 | Nmt1 | -0.032 | Young |
| 0.696921 | 0.019369 | 0.53 | 0.531 | 1 | 7 | Hdlbp | -0.001 | Young |
| 0.697086 | 0.639988 | 0.072 | 0.067 | 1 | 7 | Rint1 | 0.005 | Young |
| 0.697148 | 0.212027 | 0.021 | 0.026 | 1 | 7 | Prdm11 | -0.005 | Young |
| 0.697152 | 0.029637 | 0.253 | 0.274 | 1 | 7 | Otud5 | -0.021 | Young |
| 0.697262 | 0.187061 | 0.181 | 0.172 | 1 | 7 | Foxo3 | 0.009 | Young |
| 0.697329 | 0.003714 | 0.214 | 0.231 | 1 | 7 | Tpd52l2 | -0.017 | Young |
| 0.697588 | 0.148526 | 0.542 | 0.569 | 1 | 7 | Inpp5d | -0.027 | Young |
| 0.697713 | 0.28167 | 0.25 | 0.267 | 1 | 7 | Cspp1 | -0.017 | Young |
| 0.697855 | 0.176353 | 0.033 | 0.038 | 1 | 7 | Tcta | -0.005 | Young |
| 0.697872 | 0.129533 | 0.395 | 0.423 | 1 | 7 | Bcl2l11 | -0.028 | Young |
| 0.697958 | 0.060898 | 0.292 | 0.318 | 1 | 7 | Dynlt3 | -0.026 | Young |
| 0.698063 | 0.184941 | 0.136 | 0.128 | 1 | 7 | Nit1 | 0.008 | Young |
| 0.698488 | 0.195253 | 0.202 | 0.223 | 1 | 7 | Zbtb11 | -0.021 | Young |
| 0.698565 | 0.730693 | 0.03 | 0.026 | 1 | 7 | Per2 | 0.004 | Young |
| 0.698608 | 0.230086 | 0.398 | 0.395 | 1 | 7 | Hjurp | 0.003 | Young |
| 0.699037 | 0.063461 | 0.217 | 0.233 | 1 | 7 | Map4k1 | -0.016 | Young |
| 0.699166 | 0.355322 | 0.021 | 0.026 | 1 | 7 | Tldc1 | -0.005 | Young |
| 0.699333 | 0.245211 | 0.337 | 0.333 | 1 | 7 | Lpgat1 | 0.004 | Young |
| 0.699759 | 0.418034 | 0.157 | 0.149 | 1 | 7 | Hmgxb4 | 0.008 | Young |
| 0.700246 | 0.180162 | 0.373 | 0.362 | 1 | 7 | Nup210 | 0.011 | Young |
| 0.700771 | 0.165823 | 0.127 | 0.136 | 1 | 7 | Mtpap | -0.009 | Young |
| 0.701081 | 0.053314 | 0.268 | 0.262 | 1 | 7 | Aff1 | 0.006 | Young |
| 0.701133 | 0.109079 | 0.259 | 0.282 | 1 | 7 | Ctc1 | -0.023 | Young |
| 0.701186 | 0.015327 | 0.021 | 0.026 | 1 | 7 | Ddb2 | -0.005 | Young |
| 0.701289 | 0.19454 | 0.033 | 0.028 | 1 | 7 | Limk2 | 0.005 | Young |
| 0.702028 | 0.406113 | 0.111 | 0.103 | 1 | 7 | Pfkfb3 | 0.008 | Young |
| 0.702149 | 0.119593 | 0.045 | 0.051 | 1 | 7 | Mios | -0.006 | Young |
| 0.702196 | 0.127566 | 0.021 | 0.026 | 1 | 7 | Gba2 | -0.005 | Young |
| 0.702272 | 0.194552 | 0.063 | 0.072 | 1 | 7 | Entpd6 | -0.009 | Young |
| 0.702292 | 0.042795 | 0.623 | 0.618 | 1 | 7 | Spop | 0.005 | Young |
| 0.702611 | 0.198906 | 0.28 | 0.279 | 1 | 7 | Dennd1a | 0.001 | Young |
| 0.702638 | 0.199706 | 0.337 | 0.341 | 1 | 7 | Irf8 | -0.004 | Young |
| 0.702751 | 0.519397 | 0.06 | 0.054 | 1 | 7 | 2700049A( | 0.006 | Young |
| 0.703015 | 0.164663 | 0.334 | 0.338 | 1 | 7 | Ankhd1 | -0.004 | Young |
| 0.70302 | 0.022412 | 0.455 | 0.487 | 1 | 7 | Cytip | -0.032 | Young |
| 0.703056 | 0.060601 | 0.208 | 0.197 | 1 | 7 | Wnt11 | 0.011 | Young |
| 0.703112 | 0.215639 | 0.175 | 0.19 | 1 | 7 | Foxk2 | -0.015 | Young |
| 0.703351 | 0.086446 | 0.169 | 0.179 | 1 | 7 | Pisd | -0.01 | Young |
| 0.703363 | 0.604236 | 0.108 | 0.103 | 1 | 7 | Lrig2 | 0.005 | Young |
| 0.703413 | 0.209797 | 0.06 | 0.054 | 1 | 7 | Alg11 | 0.006 | Young |
| 0.703593 | 0.141039 | 0.355 | 0.349 | 1 | 7 | Flii | 0.006 | Young |
| 0.703598 | 0.021653 | 0.069 | 0.077 | 1 | 7 | Unc45a | -0.008 | Young |

| | | | | | | | | |
|---|---|---|---|---|---|---|---|---|
| 0.703941 | 0.209959 | 0.377 | 0.377 | 1 | 7 | N4bp2l2 | 0 | Young |
| 0.703943 | 0.007562 | 0.545 | 0.592 | 1 | 7 | Cox7b | -0.047 | Young |
| 0.704575 | 0.093831 | 0.434 | 0.459 | 1 | 7 | Selenoh | -0.025 | Young |
| 0.704922 | 0.414502 | 0.09 | 0.1 | 1 | 7 | Gmpr2 | -0.01 | Young |
| 0.70497 | 0.026746 | 0.184 | 0.195 | 1 | 7 | Acvrl1 | -0.011 | Young |
| 0.705029 | 0.166422 | 0.096 | 0.108 | 1 | 7 | Smcr8 | -0.012 | Young |
| 0.705444 | 0.142238 | 0.09 | 0.1 | 1 | 7 | Naa16 | -0.01 | Young |
| 0.70566 | 0.052105 | 0.413 | 0.454 | 1 | 7 | Tcerg1 | -0.041 | Young |
| 0.706186 | 0.07509 | 0.307 | 0.308 | 1 | 7 | Hfe | -0.001 | Young |
| 0.706629 | 0.212387 | 0.084 | 0.077 | 1 | 7 | Zfp280c | 0.007 | Young |
| 0.706902 | 0.164506 | 0.259 | 0.282 | 1 | 7 | Slc7a1 | -0.023 | Young |
| 0.706943 | 0.0267 | 0.801 | 0.818 | 1 | 7 | Cox6c | -0.017 | Young |
| 0.707037 | 0.062683 | 0.012 | 0.015 | 1 | 7 | Ica1l | -0.003 | Young |
| 0.707037 | 0.021978 | 0.012 | 0.015 | 1 | 7 | Gm32856 | -0.003 | Young |
| 0.707206 | 0.017115 | 0.283 | 0.297 | 1 | 7 | Trim41 | -0.014 | Young |
| 0.707258 | 0.521537 | 0.021 | 0.026 | 1 | 7 | Adora3 | -0.005 | Young |
| 0.707258 | 0.09175 | 0.139 | 0.149 | 1 | 7 | Pprc1 | -0.01 | Young |
| 0.707314 | 0.224954 | 0.199 | 0.218 | 1 | 7 | Auh | -0.019 | Young |
| 0.707601 | 0.015831 | 0.711 | 0.692 | 1 | 7 | Cox7a2 | 0.019 | Young |
| 0.707692 | 0.138919 | 0.41 | 0.433 | 1 | 7 | Ids | -0.023 | Young |
| 0.707714 | 0.212885 | 0.262 | 0.256 | 1 | 7 | Fam118b | 0.006 | Young |
| 0.707787 | 0.147439 | 0.214 | 0.228 | 1 | 7 | Cox16 | -0.014 | Young |
| 0.707865 | 1.411207 | 0.051 | 0.046 | 1 | 7 | Naif1 | 0.005 | Young |
| 0.707976 | 0.189452 | 0.166 | 0.182 | 1 | 7 | Atxn7l3 | -0.016 | Young |
| 0.708011 | 0.256953 | 0.127 | 0.136 | 1 | 7 | Tmem161a | -0.009 | Young |
| 0.708108 | 0.025405 | 0.741 | 0.777 | 1 | 7 | Tacc1 | -0.036 | Young |
| 0.708285 | 0.136496 | 0.533 | 0.597 | 1 | 7 | Cpsf6 | -0.064 | Young |
| 0.708847 | 0.247931 | 0.03 | 0.026 | 1 | 7 | Gm44067 | 0.004 | Young |
| 0.709067 | 0.349529 | 0.117 | 0.131 | 1 | 7 | Mcm3ap | -0.014 | Young |
| 0.709228 | 0.448841 | 0.054 | 0.062 | 1 | 7 | Tmem268 | -0.008 | Young |
| 0.709286 | 0.608176 | 0.021 | 0.026 | 1 | 7 | Kcnj2 | -0.005 | Young |
| 0.70938 | 0.085328 | 0.033 | 0.038 | 1 | 7 | Cyp2s1 | -0.005 | Young |
| 0.709483 | 0.131298 | 0.069 | 0.077 | 1 | 7 | Rcl1 | -0.008 | Young |
| 0.709669 | 0.863953 | 0.012 | 0.015 | 1 | 7 | Nrg1 | -0.003 | Young |
| 0.709669 | 0.144549 | 0.012 | 0.015 | 1 | 7 | Slc25a43 | -0.003 | Young |
| 0.70974 | 0.106943 | 0.142 | 0.154 | 1 | 7 | Pdrg1 | -0.012 | Young |
| 0.709755 | 0.096541 | 0.145 | 0.156 | 1 | 7 | Mier3 | -0.011 | Young |
| 0.709764 | 0.71329 | 0.072 | 0.067 | 1 | 7 | Grpel2 | 0.005 | Young |
| 0.710151 | 0.035491 | 0.087 | 0.097 | 1 | 7 | Blcap | -0.01 | Young |
| 0.710229 | 0.074912 | 0.343 | 0.374 | 1 | 7 | Atf7ip | -0.031 | Young |
| 0.710423 | 0.342891 | 0.313 | 0.303 | 1 | 7 | Mrps25 | 0.01 | Young |
| 0.710539 | 0.258582 | 0.148 | 0.164 | 1 | 7 | Wdr11 | -0.016 | Young |
| 0.710542 | 0.193074 | 0.054 | 0.062 | 1 | 7 | Ghdc | -0.008 | Young |
| 0.710611 | 0.284153 | 0.154 | 0.146 | 1 | 7 | Adgrg6 | 0.008 | Young |
| 0.710729 | 0.574189 | 0.051 | 0.046 | 1 | 7 | Blzf1 | 0.005 | Young |

| | | | | | | | | |
|---|---|---|---|---|---|---|---|---|
| 0.710986 | 0.833772 | 0.012 | 0.015 | 1 | 7 | Gm9733 | -0.003 | Young |
| 0.710986 | 0.744637 | 0.012 | 0.015 | 1 | 7 | Gnb5 | -0.003 | Young |
| 0.710986 | 0.743526 | 0.012 | 0.015 | 1 | 7 | Nlrc3 | -0.003 | Young |
| 0.710986 | 0.711208 | 0.012 | 0.015 | 1 | 7 | Slc22a17 | -0.003 | Young |
| 0.710986 | 0.206532 | 0.012 | 0.015 | 1 | 7 | Rgs5 | -0.003 | Young |
| 0.710986 | 0.099257 | 0.012 | 0.015 | 1 | 7 | Pglyrp1 | -0.003 | Young |
| 0.71149 | 0.766736 | 0.027 | 0.023 | 1 | 7 | Spon1 | 0.004 | Young |
| 0.711996 | 0.521351 | 0.274 | 0.287 | 1 | 7 | Dusp5 | -0.013 | Young |
| 0.712025 | 0.198761 | 0.187 | 0.203 | 1 | 7 | Crem | -0.016 | Young |
| 0.712044 | 0.126999 | 0.06 | 0.054 | 1 | 7 | Fbxo31 | 0.006 | Young |
| 0.712084 | 0.135358 | 0.075 | 0.069 | 1 | 7 | Coprs | 0.006 | Young |
| 0.712287 | 0.298597 | 0.057 | 0.051 | 1 | 7 | Tram2 | 0.006 | Young |
| 0.712304 | 0.486405 | 0.012 | 0.015 | 1 | 7 | P3h3 | -0.003 | Young |
| 0.712304 | 0.253701 | 0.012 | 0.015 | 1 | 7 | D130020LC | -0.003 | Young |
| 0.71232 | 0.356706 | 0.13 | 0.121 | 1 | 7 | Tmem70 | 0.009 | Young |
| 0.712339 | 0.264183 | 0.211 | 0.226 | 1 | 7 | Tmem33 | -0.015 | Young |
| 0.712402 | 0.097183 | 0.19 | 0.205 | 1 | 7 | Topors | -0.015 | Young |
| 0.712471 | 0.042554 | 0.575 | 0.626 | 1 | 7 | Sppl2a | -0.051 | Young |
| 0.712599 | 0.581044 | 0.03 | 0.026 | 1 | 7 | Enpp1 | 0.004 | Young |
| 0.712599 | 0.103387 | 0.03 | 0.026 | 1 | 7 | Zfp512b | 0.004 | Young |
| 0.713314 | 0.185155 | 0.319 | 0.326 | 1 | 7 | Rap2a | -0.007 | Young |
| 0.713467 | 0.612192 | 0.027 | 0.023 | 1 | 7 | Jmjd8 | 0.004 | Young |
| 0.71356 | 0.132897 | 0.163 | 0.177 | 1 | 7 | Lin9 | -0.014 | Young |
| 0.714617 | 0.227527 | 0.316 | 0.346 | 1 | 7 | Acbd5 | -0.03 | Young |
| 0.714942 | 0.423252 | 0.012 | 0.015 | 1 | 7 | Zfp428 | -0.003 | Young |
| 0.714942 | 0.232247 | 0.012 | 0.015 | 1 | 7 | Zfp420 | -0.003 | Young |
| 0.715307 | 0.315288 | 0.066 | 0.074 | 1 | 7 | Eefsec | -0.008 | Young |
| 0.715445 | 0.757841 | 0.027 | 0.023 | 1 | 7 | Dnajb5 | 0.004 | Young |
| 0.715602 | 0.224465 | 0.431 | 0.485 | 1 | 7 | Cdc5l | -0.054 | Young |
| 0.715794 | 0.212519 | 0.06 | 0.067 | 1 | 7 | Ypel2 | -0.007 | Young |
| 0.715806 | 0.014813 | 0.554 | 0.592 | 1 | 7 | Pclaf | -0.038 | Young |
| 0.715853 | 0.264839 | 0.479 | 0.487 | 1 | 7 | Ensa | -0.008 | Young |
| 0.715948 | 0.053739 | 0.831 | 0.836 | 1 | 7 | Hsp90aa1 | -0.005 | Young |
| 0.716555 | 0.045669 | 0.078 | 0.072 | 1 | 7 | Tada2b | 0.006 | Young |
| 0.716812 | 0.172185 | 0.066 | 0.074 | 1 | 7 | Txlng | -0.008 | Young |
| 0.716962 | 0.292413 | 0.114 | 0.126 | 1 | 7 | Chpt1 | -0.012 | Young |
| 0.717025 | 0.244479 | 0.151 | 0.164 | 1 | 7 | Mfap1a | -0.013 | Young |
| 0.717448 | 0.316772 | 0.184 | 0.2 | 1 | 7 | Gm2000 | -0.016 | Young |
| 0.717492 | 0.227031 | 0.437 | 0.446 | 1 | 7 | Tiparp | -0.009 | Young |
| 0.71804 | 0.271507 | 0.06 | 0.054 | 1 | 7 | Megf9 | 0.006 | Young |
| 0.719216 | 0.005516 | 0.157 | 0.169 | 1 | 7 | Ap2a1 | -0.012 | Young |
| 0.719279 | 0.164768 | 0.319 | 0.346 | 1 | 7 | Mbp | -0.027 | Young |
| 0.719461 | 0.512338 | 0.28 | 0.279 | 1 | 7 | Arih2 | 0.001 | Young |
| 0.719639 | 0.088345 | 0.117 | 0.128 | 1 | 7 | BC005561 | -0.011 | Young |
| 0.719809 | 0.004931 | 0.756 | 0.754 | 1 | 7 | Ccnl1 | 0.002 | Young |

| | | | | | | | | |
|---|---|---|---|---|---|---|---|---|
| 0.720124 | 0.399355 | 0.03 | 0.026 | 1 | 7 | Enpp4 | 0.004 | Young |
| 0.720313 | 0.120182 | 0.208 | 0.2 | 1 | 7 | Mrpl11 | 0.008 | Young |
| 0.720327 | 0.301176 | 0.099 | 0.108 | 1 | 7 | Pus10 | -0.009 | Young |
| 0.720576 | 0.260459 | 0.133 | 0.126 | 1 | 7 | Alms1 | 0.007 | Young |
| 0.720898 | 0.111729 | 0.509 | 0.497 | 1 | 7 | Ccdc34 | 0.012 | Young |
| 0.721022 | 0.104521 | 0.274 | 0.272 | 1 | 7 | Pip5k1a | 0.002 | Young |
| 0.721261 | 0.443832 | 0.096 | 0.108 | 1 | 7 | Iffo1 | -0.012 | Young |
| 0.721379 | 0.437377 | 0.235 | 0.231 | 1 | 7 | Sass6 | 0.004 | Young |
| 0.721848 | 0.182159 | 0.057 | 0.051 | 1 | 7 | Tmtc4 | 0.006 | Young |
| 0.721955 | 0.276076 | 0.244 | 0.262 | 1 | 7 | Ndufs8 | -0.018 | Young |
| 0.722355 | 0.536642 | 0.148 | 0.141 | 1 | 7 | Cdca2 | 0.007 | Young |
| 0.722492 | 0.734774 | 0.069 | 0.077 | 1 | 7 | Zfand2a | -0.008 | Young |
| 0.722556 | 0.13357 | 0.127 | 0.118 | 1 | 7 | Kansl1l | 0.009 | Young |
| 0.723182 | 0.001888 | 0.247 | 0.262 | 1 | 7 | Smad2 | -0.015 | Young |
| 0.723669 | 0.274149 | 0.045 | 0.051 | 1 | 7 | Plekhn1 | -0.006 | Young |
| 0.723693 | 0.531905 | 0.256 | 0.259 | 1 | 7 | Dscr3 | -0.003 | Young |
| 0.723833 | 0.332917 | 0.256 | 0.282 | 1 | 7 | Ccng2 | -0.026 | Young |
| 0.723931 | 0.221809 | 0.211 | 0.205 | 1 | 7 | Chsy1 | 0.006 | Young |
| 0.72399 | 1.566146 | 0.024 | 0.021 | 1 | 7 | Rilp | 0.003 | Young |
| 0.724262 | 0.059267 | 0.226 | 0.215 | 1 | 7 | Med10 | 0.011 | Young |
| 0.724372 | 0.392362 | 0.027 | 0.023 | 1 | 7 | Olfr920 | 0.004 | Young |
| 0.724451 | 0.1368 | 0.136 | 0.131 | 1 | 7 | Faap100 | 0.005 | Young |
| 0.724579 | 0.696384 | 0.084 | 0.079 | 1 | 7 | Rdh14 | 0.005 | Young |
| 0.724727 | 0.090763 | 0.482 | 0.513 | 1 | 7 | Snrpc | -0.031 | Young |
| 0.725278 | 0.231776 | 0.319 | 0.318 | 1 | 7 | Ergic3 | 0.001 | Young |
| 0.725366 | 0.549783 | 0.027 | 0.023 | 1 | 7 | Ccdc152 | 0.004 | Young |
| 0.725472 | 0.184117 | 0.358 | 0.367 | 1 | 7 | Fundc2 | -0.009 | Young |
| 0.725716 | 0.605247 | 0.093 | 0.087 | 1 | 7 | Mtfr1 | 0.006 | Young |
| 0.725883 | 0.200398 | 0.175 | 0.169 | 1 | 7 | Akap11 | 0.006 | Young |
| 0.726059 | 0.146366 | 0.265 | 0.285 | 1 | 7 | Txlna | -0.02 | Young |
| 0.726199 | 0.137024 | 0.5 | 0.505 | 1 | 7 | Mkln1 | -0.005 | Young |
| 0.727103 | 0.251281 | 0.289 | 0.285 | 1 | 7 | Pitpnb | 0.004 | Young |
| 0.72715 | 1.746768 | 0.024 | 0.021 | 1 | 7 | Carmil1 | 0.003 | Young |
| 0.727495 | 0.038829 | 0.41 | 0.433 | 1 | 7 | Trim8 | -0.023 | Young |
| 0.728014 | 0.136749 | 0.072 | 0.067 | 1 | 7 | Kptn | 0.005 | Young |
| 0.728016 | 0.221849 | 0.063 | 0.072 | 1 | 7 | Mospd1 | -0.009 | Young |
| 0.728032 | 0.135862 | 0.184 | 0.195 | 1 | 7 | Ppwd1 | -0.011 | Young |
| 0.728205 | 0.715862 | 0.024 | 0.021 | 1 | 7 | Gm14005 | 0.003 | Young |
| 0.72904 | 0.178046 | 0.111 | 0.123 | 1 | 7 | Zfp511 | -0.012 | Young |
| 0.729459 | 0.084576 | 0.377 | 0.413 | 1 | 7 | Bckdk | -0.036 | Young |
| 0.729497 | 0.651508 | 0.048 | 0.044 | 1 | 7 | Osbpl7 | 0.004 | Young |
| 0.729732 | 0.433958 | 0.196 | 0.19 | 1 | 7 | Ric8a | 0.006 | Young |
| 0.729937 | 0.110524 | 0.223 | 0.244 | 1 | 7 | Mfsd10 | -0.021 | Young |
| 0.730266 | 0.072428 | 0.133 | 0.126 | 1 | 7 | Tctex1d2 | 0.007 | Young |
| 0.730765 | 0.357711 | 0.057 | 0.051 | 1 | 7 | 1300002E1 | 0.006 | Young |

| | | | | | | | | |
|---|---|---|---|---|---|---|---|---|
| 0.730959 | 0.778523 | 0.033 | 0.038 | 1 | 7 | Jade3 | -0.005 | Young |
| 0.731371 | 1.016697 | 0.024 | 0.021 | 1 | 7 | 2610035D1 | 0.003 | Young |
| 0.731371 | 1.010613 | 0.024 | 0.021 | 1 | 7 | Prr5 | 0.003 | Young |
| 0.731373 | 0.309248 | 0.114 | 0.108 | 1 | 7 | Zbtb37 | 0.006 | Young |
| 0.731575 | 0.178413 | 0.232 | 0.223 | 1 | 7 | Bex3 | 0.009 | Young |
| 0.732278 | 0.338959 | 0.06 | 0.067 | 1 | 7 | Cyb561d1 | -0.007 | Young |
| 0.73234 | 0.353434 | 0.045 | 0.051 | 1 | 7 | Arhgap22 | -0.006 | Young |
| 0.732428 | 0.85253 | 0.024 | 0.021 | 1 | 7 | Adam22 | 0.003 | Young |
| 0.732428 | 0.626887 | 0.024 | 0.021 | 1 | 7 | Pex11g | 0.003 | Young |
| 0.733008 | 0.323664 | 0.187 | 0.2 | 1 | 7 | Mrto4 | -0.013 | Young |
| 0.733009 | 0.121237 | 0.16 | 0.154 | 1 | 7 | Rfx7 | 0.006 | Young |
| 0.733358 | 0.190611 | 0.03 | 0.026 | 1 | 7 | Tmem42 | 0.004 | Young |
| 0.733363 | 0.041764 | 0.892 | 0.908 | 1 | 7 | Rpl14 | -0.016 | Young |
| 0.733778 | 0.027625 | 0.099 | 0.108 | 1 | 7 | Cchcr1 | -0.009 | Young |
| 0.733788 | 0.065256 | 0.045 | 0.051 | 1 | 7 | Zfp770 | -0.006 | Young |
| 0.734105 | 0.311264 | 0.048 | 0.054 | 1 | 7 | Trim21 | -0.006 | Young |
| 0.73414 | 0.156827 | 0.16 | 0.172 | 1 | 7 | Lemd3 | -0.012 | Young |
| 0.734221 | 0.593754 | 0.078 | 0.072 | 1 | 7 | Actr5 | 0.006 | Young |
| 0.734542 | 0.876291 | 0.024 | 0.021 | 1 | 7 | Fam189b | 0.003 | Young |
| 0.734542 | 0.051855 | 0.024 | 0.021 | 1 | 7 | Akap17b | 0.003 | Young |
| 0.734656 | 0.090553 | 0.307 | 0.305 | 1 | 7 | Saraf | 0.002 | Young |
| 0.735333 | 0.064658 | 0.027 | 0.023 | 1 | 7 | N4bp3 | 0.004 | Young |
| 0.735511 | 0.255203 | 0.175 | 0.169 | 1 | 7 | Vrk3 | 0.006 | Young |
| 0.735971 | 0.083501 | 0.072 | 0.067 | 1 | 7 | Cisd3 | 0.005 | Young |
| 0.7364 | 0.048545 | 0.036 | 0.041 | 1 | 7 | Zbtb8a | -0.005 | Young |
| 0.736492 | 0.279883 | 0.361 | 0.372 | 1 | 7 | Trappc2l | -0.011 | Young |
| 0.736521 | 0.14717 | 0.265 | 0.29 | 1 | 7 | Sde2 | -0.025 | Young |
| 0.736923 | 0.172663 | 0.262 | 0.259 | 1 | 7 | Gclc | 0.003 | Young |
| 0.737181 | 0.261891 | 0.139 | 0.154 | 1 | 7 | Aph1b | -0.015 | Young |
| 0.737312 | 0.255687 | 0.114 | 0.108 | 1 | 7 | Ppp2r1b | 0.006 | Young |
| 0.737441 | 0.039148 | 0.645 | 0.687 | 1 | 7 | Clic4 | -0.042 | Young |
| 0.737565 | 0.288735 | 0.166 | 0.177 | 1 | 7 | 3830406C1 | -0.011 | Young |
| 0.737643 | 0.370926 | 0.033 | 0.038 | 1 | 7 | Il18rap | -0.005 | Young |
| 0.737714 | 0.024249 | 0.127 | 0.118 | 1 | 7 | Stx17 | 0.009 | Young |
| 0.737717 | 0.481372 | 0.024 | 0.021 | 1 | 7 | Nudt6 | 0.003 | Young |
| 0.737845 | 0.033525 | 0.34 | 0.359 | 1 | 7 | Mogs | -0.019 | Young |
| 0.73794 | 0.015245 | 0.539 | 0.569 | 1 | 7 | Cep170 | -0.03 | Young |
| 0.738015 | 0.265942 | 0.036 | 0.041 | 1 | 7 | Btbd6 | -0.005 | Young |
| 0.738139 | 0.107292 | 0.072 | 0.079 | 1 | 7 | Usp53 | -0.007 | Young |
| 0.73828 | 0.121925 | 0.304 | 0.328 | 1 | 7 | Mad2l1 | -0.024 | Young |
| 0.738289 | 0.233493 | 0.196 | 0.21 | 1 | 7 | Supt20 | -0.014 | Young |
| 0.738771 | 0.164562 | 0.437 | 0.482 | 1 | 7 | Pten | -0.045 | Young |
| 0.738865 | 0.531209 | 0.051 | 0.046 | 1 | 7 | Zfp346 | 0.005 | Young |
| 0.738933 | 0.035002 | 0.226 | 0.215 | 1 | 7 | Smad5 | 0.011 | Young |
| 0.739074 | 0.069646 | 0.289 | 0.29 | 1 | 7 | Tmcc3 | -0.001 | Young |

| | | | | | | | | |
|---|---|---|---|---|---|---|---|---|
| 0.739498 | 0.113515 | 0.404 | 0.397 | 1 | 7 | Aimp1 | 0.007 | Young |
| 0.739619 | 0.586554 | 0.09 | 0.1 | 1 | 7 | Mterf3 | -0.01 | Young |
| 0.739631 | 0.779069 | 0.036 | 0.041 | 1 | 7 | Eid2b | -0.005 | Young |
| 0.739773 | 0.303129 | 0.325 | 0.333 | 1 | 7 | Msrb1 | -0.008 | Young |
| 0.739942 | 0.137584 | 0.13 | 0.121 | 1 | 7 | Bloc1s6 | 0.009 | Young |
| 0.740043 | 0.179329 | 0.154 | 0.146 | 1 | 7 | Ralgapb | 0.008 | Young |
| 0.740349 | 0.663201 | 0.148 | 0.144 | 1 | 7 | Nfatc2ip | 0.004 | Young |
| 0.740373 | 0.044279 | 0.681 | 0.71 | 1 | 7 | Smdt1 | -0.029 | Young |
| 0.740734 | 0.578625 | 0.078 | 0.072 | 1 | 7 | Dip2c | 0.006 | Young |
| 0.740774 | 0.188006 | 0.449 | 0.479 | 1 | 7 | Tppp3 | -0.03 | Young |
| 0.740897 | 0.719407 | 0.024 | 0.021 | 1 | 7 | Zfp764 | 0.003 | Young |
| 0.740897 | 0.610987 | 0.024 | 0.021 | 1 | 7 | Fscn1 | 0.003 | Young |
| 0.740915 | 0.004615 | 0.157 | 0.167 | 1 | 7 | Pcgf3 | -0.01 | Young |
| 0.740918 | 0.517337 | 0.093 | 0.087 | 1 | 7 | Pacs1 | 0.006 | Young |
| 0.741063 | 0.275352 | 0.187 | 0.205 | 1 | 7 | Spopl | -0.018 | Young |
| 0.741215 | 0.259791 | 0.437 | 0.467 | 1 | 7 | Man2a1 | -0.03 | Young |
| 0.741318 | 0.733401 | 0.136 | 0.133 | 1 | 7 | Ctsf | 0.003 | Young |
| 0.741322 | 0.300211 | 0.443 | 0.469 | 1 | 7 | Nudt3 | -0.026 | Young |
| 0.741574 | 0.600124 | 0.021 | 0.018 | 1 | 7 | Cobll1 | 0.003 | Young |
| 0.741636 | 0.350674 | 0.096 | 0.09 | 1 | 7 | Kctd9 | 0.006 | Young |
| 0.741639 | 0.215732 | 0.389 | 0.385 | 1 | 7 | Mfsd14a | 0.004 | Young |
| 0.741902 | 0.15868 | 0.142 | 0.154 | 1 | 7 | Llgl1 | -0.012 | Young |
| 0.742259 | 0.000382 | 0.515 | 0.569 | 1 | 7 | Synj1 | -0.054 | Young |
| 0.742344 | 0.641974 | 0.045 | 0.041 | 1 | 7 | Ccdc170 | 0.004 | Young |
| 0.742426 | 0.000219 | 0.503 | 0.531 | 1 | 7 | Gatad2b | -0.028 | Young |
| 0.74262 | 0.222699 | 0.172 | 0.185 | 1 | 7 | Nacc1 | -0.013 | Young |
| 0.742707 | 0.138876 | 0.163 | 0.174 | 1 | 7 | Pik3c3 | -0.011 | Young |
| 0.74302 | 0.198574 | 0.024 | 0.021 | 1 | 7 | Tmem69 | 0.003 | Young |
| 0.743531 | 0.309958 | 0.277 | 0.274 | 1 | 7 | Pcyt1a | 0.003 | Young |
| 0.743826 | 0.815104 | 0.042 | 0.038 | 1 | 7 | Gm20045 | 0.004 | Young |
| 0.74399 | 0.583724 | 0.199 | 0.2 | 1 | 7 | Hspbp1 | -0.001 | Young |
| 0.744061 | 0.091007 | 0.44 | 0.467 | 1 | 7 | Cbfa2t3 | -0.027 | Young |
| 0.744082 | 0.232175 | 0.024 | 0.021 | 1 | 7 | Rtkn2 | 0.003 | Young |
| 0.744116 | 0.434886 | 0.057 | 0.064 | 1 | 7 | Zfp467 | -0.007 | Young |
| 0.744118 | 0.094366 | 0.151 | 0.162 | 1 | 7 | 6430548M | -0.011 | Young |
| 0.744132 | 0.359795 | 0.193 | 0.19 | 1 | 7 | Iars | 0.003 | Young |
| 0.744405 | 0.396886 | 0.024 | 0.028 | 1 | 7 | Slc7a13 | -0.004 | Young |
| 0.744492 | 0.505254 | 0.238 | 0.231 | 1 | 7 | Myef2 | 0.007 | Young |
| 0.744622 | 0.080639 | 0.464 | 0.487 | 1 | 7 | Pnp | -0.023 | Young |
| 0.745144 | 0.571514 | 0.024 | 0.021 | 1 | 7 | 4930512J1 | 0.003 | Young |
| 0.745444 | 0.040695 | 0.506 | 0.528 | 1 | 7 | Dad1 | -0.022 | Young |
| 0.746077 | 0.196365 | 0.078 | 0.072 | 1 | 7 | Zeb1 | 0.006 | Young |
| 0.746207 | 0.161034 | 0.024 | 0.021 | 1 | 7 | Sox4 | 0.003 | Young |
| 0.74636 | 0.241173 | 0.024 | 0.028 | 1 | 7 | 3110009E1 | -0.004 | Young |
| 0.746406 | 0.226093 | 0.425 | 0.433 | 1 | 7 | Rsu1 | -0.008 | Young |

| | | | | | | | | |
|---|---|---|---|---|---|---|---|---|
| 0.746482 | 0.496884 | 0.123 | 0.136 | 1 | 7 | Stk25 | -0.013 | Young |
| 0.746609 | 0.172624 | 0.241 | 0.256 | 1 | 7 | Dnase1l1 | -0.015 | Young |
| 0.746664 | 0.405983 | 0.226 | 0.246 | 1 | 7 | Gtf2i | -0.02 | Young |
| 0.746867 | 0.196336 | 0.045 | 0.051 | 1 | 7 | Stradb | -0.006 | Young |
| 0.746898 | 0.06028 | 0.437 | 0.441 | 1 | 7 | Smim15 | -0.004 | Young |
| 0.747244 | 1.056991 | 0.021 | 0.018 | 1 | 7 | Pyroxd2 | 0.003 | Young |
| 0.747269 | 0.307373 | 0.349 | 0.362 | 1 | 7 | Pcm1 | -0.013 | Young |
| 0.747337 | 0.359969 | 0.024 | 0.028 | 1 | 7 | 1810009A1 | -0.004 | Young |
| 0.747553 | 0.128215 | 0.048 | 0.054 | 1 | 7 | Clcn6 | -0.006 | Young |
| 0.748201 | 0.279844 | 0.479 | 0.487 | 1 | 7 | Srsf10 | -0.008 | Young |
| 0.748263 | 0.003646 | 0.054 | 0.049 | 1 | 7 | Amigo1 | 0.005 | Young |
| 0.748361 | 0.269203 | 0.352 | 0.372 | 1 | 7 | Aup1 | -0.02 | Young |
| 0.74838 | 0.668655 | 0.021 | 0.018 | 1 | 7 | Trp53rkb | 0.003 | Young |
| 0.74838 | 0.456097 | 0.021 | 0.018 | 1 | 7 | Gtf2ird1 | 0.003 | Young |
| 0.748445 | 0.199852 | 0.259 | 0.256 | 1 | 7 | Atl3 | 0.003 | Young |
| 0.748631 | 0.655161 | 0.081 | 0.077 | 1 | 7 | Ercc4 | 0.004 | Young |
| 0.748836 | 0.036968 | 0.111 | 0.123 | 1 | 7 | Eif2ak3 | -0.012 | Young |
| 0.749162 | 0.472523 | 0.169 | 0.185 | 1 | 7 | Rps6ka5 | -0.016 | Young |
| 0.749349 | 0.087 | 0.036 | 0.041 | 1 | 7 | Spats2 | -0.005 | Young |
| 0.749516 | 0.877926 | 0.021 | 0.018 | 1 | 7 | Ficd | 0.003 | Young |
| 0.749684 | 0.501045 | 0.048 | 0.054 | 1 | 7 | Phka1 | -0.006 | Young |
| 0.749739 | 0.186823 | 0.364 | 0.364 | 1 | 7 | Cux1 | 0 | Young |
| 0.749922 | 0.326551 | 0.461 | 0.472 | 1 | 7 | Pdcd6 | -0.011 | Young |
| 0.750103 | 0.117632 | 0.166 | 0.179 | 1 | 7 | Nelfa | -0.013 | Young |
| 0.75016 | 0.35961 | 0.036 | 0.041 | 1 | 7 | Prr7 | -0.005 | Young |
| 0.750203 | 0.476387 | 0.108 | 0.121 | 1 | 7 | Exosc1 | -0.013 | Young |
| 0.75055 | 0.432779 | 0.069 | 0.077 | 1 | 7 | Nlrp1b | -0.008 | Young |
| 0.750992 | 0.320427 | 0.295 | 0.297 | 1 | 7 | Kctd20 | -0.002 | Young |
| 0.751059 | 0.121241 | 0.09 | 0.097 | 1 | 7 | Lancl1 | -0.007 | Young |
| 0.751122 | 0.041827 | 0.238 | 0.256 | 1 | 7 | Rasa4 | -0.018 | Young |
| 0.751253 | 0.5512 | 0.024 | 0.028 | 1 | 7 | Col4a2 | -0.004 | Young |
| 0.751312 | 0.392927 | 0.057 | 0.064 | 1 | 7 | Ap5s1 | -0.007 | Young |
| 0.751418 | 0.09774 | 0.307 | 0.328 | 1 | 7 | Fam91a1 | -0.021 | Young |
| 0.751744 | 0.226485 | 0.265 | 0.295 | 1 | 7 | Nup85 | -0.03 | Young |
| 0.751765 | 0.011607 | 0.193 | 0.205 | 1 | 7 | Ubald1 | -0.012 | Young |
| 0.752133 | 0.068404 | 0.437 | 0.441 | 1 | 7 | Paip2 | -0.004 | Young |
| 0.752739 | 0.230732 | 0.123 | 0.118 | 1 | 7 | St7l | 0.005 | Young |
| 0.752876 | 0.034358 | 0.84 | 0.864 | 1 | 7 | Rpl15 | -0.024 | Young |
| 0.752929 | 0.263901 | 0.021 | 0.018 | 1 | 7 | Myom1 | 0.003 | Young |
| 0.753206 | 0.07287 | 0.325 | 0.341 | 1 | 7 | Pcgf5 | -0.016 | Young |
| 0.753264 | 0.124147 | 0.545 | 0.541 | 1 | 7 | Lilr4b | 0.004 | Young |
| 0.753289 | 0.253272 | 0.232 | 0.233 | 1 | 7 | Opa3 | -0.001 | Young |
| 0.753422 | 0.173307 | 0.373 | 0.408 | 1 | 7 | Ppp6c | -0.035 | Young |
| 0.753463 | 0.1588 | 0.358 | 0.395 | 1 | 7 | Lsm8 | -0.037 | Young |
| 0.753586 | 0.172865 | 0.229 | 0.244 | 1 | 7 | Trnt1 | -0.015 | Young |

| | | | | | | | | |
|---|---|---|---|---|---|---|---|---|
| 0.754002 | 0.325099 | 0.223 | 0.244 | 1 | 7 | Atp11c | -0.021 | Young |
| 0.754132 | 0.008092 | 0.13 | 0.138 | 1 | 7 | Strip1 | -0.008 | Young |
| 0.754555 | 0.42498 | 0.084 | 0.092 | 1 | 7 | P2ry10 | -0.008 | Young |
| 0.755023 | 0.00847 | 0.518 | 0.546 | 1 | 7 | Kdm7a | -0.028 | Young |
| 0.755206 | 0.488315 | 0.021 | 0.018 | 1 | 7 | 1110002J0 | 0.003 | Young |
| 0.755375 | 0.082554 | 0.054 | 0.049 | 1 | 7 | Cnksr3 | 0.005 | Young |
| 0.755457 | 0.161495 | 0.151 | 0.144 | 1 | 7 | Med20 | 0.007 | Young |
| 0.755573 | 0.180312 | 0.202 | 0.213 | 1 | 7 | Cenph | -0.011 | Young |
| 0.755622 | 0.186116 | 0.241 | 0.244 | 1 | 7 | Slc50a1 | -0.003 | Young |
| 0.755678 | 0.217586 | 0.072 | 0.067 | 1 | 7 | Prmt2 | 0.005 | Young |
| 0.755796 | 0.221315 | 0.187 | 0.179 | 1 | 7 | 9930021J0 | 0.008 | Young |
| 0.755839 | 0.158106 | 0.25 | 0.244 | 1 | 7 | Pbk | 0.006 | Young |
| 0.75586 | 0.14434 | 0.346 | 0.351 | 1 | 7 | Ufc1 | -0.005 | Young |
| 0.756153 | 0.23027 | 0.163 | 0.156 | 1 | 7 | Fbxo5 | 0.007 | Young |
| 0.756156 | 0.190141 | 0.355 | 0.356 | 1 | 7 | Lrrfip2 | -0.001 | Young |
| 0.756346 | 0.298476 | 0.021 | 0.018 | 1 | 7 | Tnnt1 | 0.003 | Young |
| 0.756346 | 0.264807 | 0.021 | 0.018 | 1 | 7 | Hhat | 0.003 | Young |
| 0.75639 | 0.089383 | 0.825 | 0.818 | 1 | 7 | Srsf5 | 0.007 | Young |
| 0.756594 | 0.198478 | 0.557 | 0.569 | 1 | 7 | Eif3j1 | -0.012 | Young |
| 0.756663 | 0.561986 | 0.036 | 0.041 | 1 | 7 | Ppp3cc | -0.005 | Young |
| 0.756862 | 0.291963 | 0.235 | 0.259 | 1 | 7 | Epb41 | -0.024 | Young |
| 0.756984 | 0.01542 | 0.509 | 0.544 | 1 | 7 | Stag2 | -0.035 | Young |
| 0.75699 | 0.481822 | 0.163 | 0.162 | 1 | 7 | 0610010K1 | 0.001 | Young |
| 0.757031 | 0.882695 | 0.045 | 0.041 | 1 | 7 | E330009J0 | 0.004 | Young |
| 0.757113 | 0.035474 | 0.268 | 0.287 | 1 | 7 | Dmxl1 | -0.019 | Young |
| 0.757262 | 0.004127 | 0.211 | 0.226 | 1 | 7 | Pycr2 | -0.015 | Young |
| 0.757377 | 0.187921 | 0.166 | 0.177 | 1 | 7 | Edem2 | -0.011 | Young |
| 0.757486 | 1.120942 | 0.021 | 0.018 | 1 | 7 | Cd209e | 0.003 | Young |
| 0.758018 | 0.009469 | 0.256 | 0.274 | 1 | 7 | Gart | -0.018 | Young |
| 0.75828 | 0.328135 | 0.163 | 0.159 | 1 | 7 | Gins2 | 0.004 | Young |
| 0.75916 | 0.04055 | 0.404 | 0.405 | 1 | 7 | Gpsm3 | -0.001 | Young |
| 0.759319 | 0.215381 | 0.292 | 0.292 | 1 | 7 | Gpatch8 | 0 | Young |
| 0.759358 | 0.302227 | 0.045 | 0.041 | 1 | 7 | Tmem86b | 0.004 | Young |
| 0.759767 | 0.133464 | 0.021 | 0.018 | 1 | 7 | Arsg | 0.003 | Young |
| 0.759801 | 1.303981 | 0.042 | 0.038 | 1 | 7 | Aldh6a1 | 0.004 | Young |
| 0.759921 | 0.562513 | 0.036 | 0.041 | 1 | 7 | Gm35154 | -0.005 | Young |
| 0.760151 | 0.053779 | 0.566 | 0.562 | 1 | 7 | Hnrnpul2 | 0.004 | Young |
| 0.760278 | 0.32264 | 0.244 | 0.244 | 1 | 7 | Cirbp | 0 | Young |
| 0.760625 | 0.341196 | 0.256 | 0.279 | 1 | 7 | Ube2h | -0.023 | Young |
| 0.760909 | 0.220987 | 0.021 | 0.018 | 1 | 7 | Prepl | 0.003 | Young |
| 0.760917 | 0.133355 | 0.063 | 0.069 | 1 | 7 | Fam35a | -0.006 | Young |
| 0.760926 | 0.37405 | 0.123 | 0.118 | 1 | 7 | Camta1 | 0.005 | Young |
| 0.761461 | 0.282281 | 0.313 | 0.313 | 1 | 7 | Gak | 0 | Young |
| 0.761835 | 0.001314 | 0.145 | 0.136 | 1 | 7 | Atp1a3 | 0.009 | Young |
| 0.762572 | 0.169607 | 0.154 | 0.164 | 1 | 7 | Rogdi | -0.01 | Young |

| | | | | | | | | |
|---|---|---|---|---|---|---|---|---|
| 0.762739 | 0.235293 | 0.581 | 0.592 | 1 | 7 | Eif4a2 | -0.011 | Young |
| 0.762748 | 0.321615 | 0.271 | 0.274 | 1 | 7 | Snx1 | -0.003 | Young |
| 0.762882 | 0.204972 | 0.286 | 0.315 | 1 | 7 | Ndufs3 | -0.029 | Young |
| 0.763008 | 1.15923 | 0.018 | 0.015 | 1 | 7 | Wdr60 | 0.003 | Young |
| 0.763109 | 0.207825 | 0.202 | 0.221 | 1 | 7 | Utp20 | -0.019 | Young |
| 0.763124 | 0.177242 | 0.142 | 0.136 | 1 | 7 | Zfp598 | 0.006 | Young |
| 0.763194 | 0.776271 | 0.021 | 0.018 | 1 | 7 | 9830107B1 | 0.003 | Young |
| 0.763292 | 0.352138 | 0.244 | 0.246 | 1 | 7 | Srpk1 | -0.002 | Young |
| 0.763624 | 0.242062 | 0.015 | 0.018 | 1 | 7 | B230217C1 | -0.003 | Young |
| 0.763624 | 0.061867 | 0.015 | 0.018 | 1 | 7 | B3gntl1 | -0.003 | Young |
| 0.763677 | 0.076184 | 0.06 | 0.067 | 1 | 7 | Pwwp2b | -0.007 | Young |
| 0.763855 | 0.431316 | 0.12 | 0.128 | 1 | 7 | Pak4 | -0.008 | Young |
| 0.764126 | 0.026405 | 0.699 | 0.69 | 1 | 7 | 5031439G( | 0.009 | Young |
| 0.764176 | 0.212616 | 0.178 | 0.19 | 1 | 7 | Dirc2 | -0.012 | Young |
| 0.764207 | 0.063603 | 0.148 | 0.138 | 1 | 7 | Kif14 | 0.01 | Young |
| 0.764337 | 0.072227 | 0.021 | 0.018 | 1 | 7 | Dnaaf2 | 0.003 | Young |
| 0.764351 | 0.581994 | 0.075 | 0.082 | 1 | 7 | Bloc1s1 | -0.007 | Young |
| 0.765217 | 0.750498 | 0.093 | 0.09 | 1 | 7 | Strada | 0.003 | Young |
| 0.765422 | 0.10334 | 0.25 | 0.249 | 1 | 7 | Sec24c | 0.001 | Young |
| 0.765621 | 0.171765 | 0.108 | 0.103 | 1 | 7 | Mbd4 | 0.005 | Young |
| 0.765662 | 0.077619 | 0.247 | 0.236 | 1 | 7 | Sptan1 | 0.011 | Young |
| 0.765714 | 0.025356 | 0.346 | 0.364 | 1 | 7 | Acadm | -0.018 | Young |
| 0.766103 | 0.072868 | 0.199 | 0.208 | 1 | 7 | Phactr4 | -0.009 | Young |
| 0.76645 | 0.571725 | 0.036 | 0.041 | 1 | 7 | Lrsam1 | -0.005 | Young |
| 0.7665 | 0.590621 | 0.078 | 0.074 | 1 | 7 | Rcan1 | 0.004 | Young |
| 0.766618 | 0.14852 | 0.346 | 0.367 | 1 | 7 | Ppp2r2a | -0.021 | Young |
| 0.766641 | 0.086159 | 0.485 | 0.503 | 1 | 7 | Ralb | -0.018 | Young |
| 0.766709 | 0.807665 | 0.018 | 0.015 | 1 | 7 | Zc3hav1l | 0.003 | Young |
| 0.766709 | 0.716735 | 0.018 | 0.015 | 1 | 7 | Phldb3 | 0.003 | Young |
| 0.766904 | 0.21353 | 0.06 | 0.067 | 1 | 7 | F2rl2 | -0.007 | Young |
| 0.767191 | 0.145741 | 0.286 | 0.287 | 1 | 7 | Hk2 | -0.001 | Young |
| 0.767385 | 0.147776 | 0.383 | 0.39 | 1 | 7 | Ythdf3 | -0.007 | Young |
| 0.767559 | 0.273385 | 0.383 | 0.408 | 1 | 7 | Gxylt1 | -0.025 | Young |
| 0.767565 | 0.200805 | 0.286 | 0.29 | 1 | 7 | Slf2 | -0.004 | Young |
| 0.767769 | 0.039248 | 0.021 | 0.018 | 1 | 7 | Zfp324 | 0.003 | Young |
| 0.767786 | 0.398658 | 0.184 | 0.2 | 1 | 7 | Rexo1 | -0.016 | Young |
| 0.76783 | 0.197788 | 0.139 | 0.149 | 1 | 7 | Ttf2 | -0.01 | Young |
| 0.767937 | 0.184131 | 0.139 | 0.131 | 1 | 7 | Chaf1b | 0.008 | Young |
| 0.768091 | 0.382522 | 0.274 | 0.269 | 1 | 7 | Gm49336 | 0.005 | Young |
| 0.768133 | 0.091126 | 0.184 | 0.197 | 1 | 7 | Brca2 | -0.013 | Young |
| 0.768507 | 0.409587 | 0.063 | 0.069 | 1 | 7 | Ccsap | -0.006 | Young |
| 0.76853 | 0.359275 | 0.286 | 0.285 | 1 | 7 | Nceh1 | 0.001 | Young |
| 0.768688 | 0.447159 | 0.045 | 0.041 | 1 | 7 | Acp6 | 0.004 | Young |
| 0.768688 | 0.210033 | 0.214 | 0.231 | 1 | 7 | Tmem106c | -0.017 | Young |
| 0.768694 | 0.051025 | 0.039 | 0.044 | 1 | 7 | Smug1 | -0.005 | Young |

| | | | | | | | | |
|---|---|---|---|---|---|---|---|---|
| 0.7687 | 0.298416 | 0.301 | 0.338 | 1 | 7 | Glmp | -0.037 | Young |
| 0.768735 | 0.511922 | 0.145 | 0.159 | 1 | 7 | Zfp639 | -0.014 | Young |
| 0.768973 | 0.674497 | 0.114 | 0.11 | 1 | 7 | Zfp68 | 0.004 | Young |
| 0.769068 | 0.443996 | 0.063 | 0.059 | 1 | 7 | Mag | 0.004 | Young |
| 0.769179 | 1.186876 | 0.018 | 0.015 | 1 | 7 | Tmem156 | 0.003 | Young |
| 0.769215 | 0.059826 | 0.289 | 0.3 | 1 | 7 | Zfx | -0.011 | Young |
| 0.769561 | 0.102057 | 0.566 | 0.577 | 1 | 7 | Txndc17 | -0.011 | Young |
| 0.769797 | 0.153589 | 0.015 | 0.018 | 1 | 7 | Disp1 | -0.003 | Young |
| 0.770097 | 0.017924 | 0.127 | 0.136 | 1 | 7 | Naglu | -0.009 | Young |
| 0.770144 | 0.060653 | 0.163 | 0.177 | 1 | 7 | Wdr13 | -0.014 | Young |
| 0.770164 | 0.085478 | 0.09 | 0.097 | 1 | 7 | Zfp64 | -0.007 | Young |
| 0.770227 | 0.60269 | 0.093 | 0.103 | 1 | 7 | Acyp1 | -0.01 | Young |
| 0.770247 | 0.163816 | 0.078 | 0.085 | 1 | 7 | Kmo | -0.007 | Young |
| 0.770446 | 0.534087 | 0.142 | 0.138 | 1 | 7 | Ube2g1 | 0.004 | Young |
| 0.770688 | 0.580187 | 0.093 | 0.09 | 1 | 7 | Casp9 | 0.003 | Young |
| 0.770924 | 0.23232 | 0.328 | 0.359 | 1 | 7 | Acbd3 | -0.031 | Young |
| 0.771033 | 0.315623 | 0.015 | 0.018 | 1 | 7 | Ankrd66 | -0.003 | Young |
| 0.771258 | 0.788568 | 0.072 | 0.079 | 1 | 7 | Atg13 | -0.007 | Young |
| 0.77164 | 0.009811 | 0.645 | 0.654 | 1 | 7 | Spi1 | -0.009 | Young |
| 0.771651 | 0.709394 | 0.018 | 0.015 | 1 | 7 | Prpf40b | 0.003 | Young |
| 0.771651 | 0.328895 | 0.018 | 0.015 | 1 | 7 | Srfbp1 | 0.003 | Young |
| 0.771768 | 0.762573 | 0.045 | 0.051 | 1 | 7 | Proser3 | -0.006 | Young |
| 0.77227 | 0.394921 | 0.015 | 0.018 | 1 | 7 | Gatsl3 | -0.003 | Young |
| 0.772326 | 0.367526 | 0.09 | 0.097 | 1 | 7 | Sema6b | -0.007 | Young |
| 0.772485 | 0.017886 | 0.214 | 0.228 | 1 | 7 | Ddx27 | -0.014 | Young |
| 0.772888 | 0.608137 | 0.018 | 0.015 | 1 | 7 | Xkr8 | 0.003 | Young |
| 0.772888 | 0.516779 | 0.018 | 0.015 | 1 | 7 | Havcr2 | 0.003 | Young |
| 0.772888 | 0.495174 | 0.018 | 0.015 | 1 | 7 | Dnah8 | 0.003 | Young |
| 0.773018 | 0.182939 | 0.199 | 0.192 | 1 | 7 | Mesd | 0.007 | Young |
| 0.773507 | 0.98444 | 0.015 | 0.018 | 1 | 7 | Capn10 | -0.003 | Young |
| 0.773507 | 0.240772 | 0.015 | 0.018 | 1 | 7 | Zfp442 | -0.003 | Young |
| 0.773515 | 0.436182 | 0.105 | 0.115 | 1 | 7 | Ric1 | -0.01 | Young |
| 0.773566 | 0.188344 | 0.208 | 0.223 | 1 | 7 | Prim2 | -0.015 | Young |
| 0.773578 | 0.066163 | 0.386 | 0.397 | 1 | 7 | Coq10b | -0.011 | Young |
| 0.773685 | 0.133619 | 0.274 | 0.295 | 1 | 7 | Chka | -0.021 | Young |
| 0.773707 | 0.347546 | 0.084 | 0.079 | 1 | 7 | Telo2 | 0.005 | Young |
| 0.773802 | 0.258529 | 0.175 | 0.169 | 1 | 7 | Ubfd1 | 0.006 | Young |
| 0.774079 | 0.100535 | 0.416 | 0.421 | 1 | 7 | Tuba1a | -0.005 | Young |
| 0.774371 | 0.070207 | 0.464 | 0.479 | 1 | 7 | Bcap31 | -0.015 | Young |
| 0.774713 | 0.082214 | 0.166 | 0.164 | 1 | 7 | Tex30 | 0.002 | Young |
| 0.774745 | 0.591885 | 0.015 | 0.018 | 1 | 7 | Dennd5b | -0.003 | Young |
| 0.774745 | 0.495378 | 0.015 | 0.018 | 1 | 7 | Pdpn | -0.003 | Young |
| 0.77486 | 0.307947 | 0.386 | 0.421 | 1 | 7 | Setd3 | -0.035 | Young |
| 0.774922 | 0.022545 | 0.066 | 0.072 | 1 | 7 | Wipi1 | -0.006 | Young |
| 0.775364 | 0.467307 | 0.018 | 0.015 | 1 | 7 | Mrnip | 0.003 | Young |

| | | | | | | | | |
|---|---|---|---|---|---|---|---|---|
| 0.775364 | 0.246398 | 0.018 | 0.015 | 1 | 7 | Clec3b | 0.003 | Young |
| 0.775473 | 0.267579 | 0.075 | 0.082 | 1 | 7 | Zbtb43 | -0.007 | Young |
| 0.775625 | 0.0249 | 0.217 | 0.218 | 1 | 7 | Scoc | -0.001 | Young |
| 0.775667 | 0.190158 | 0.253 | 0.272 | 1 | 7 | Rictor | -0.019 | Young |
| 0.775875 | 0.090619 | 0.151 | 0.144 | 1 | 7 | Pgs1 | 0.007 | Young |
| 0.77617 | 0.278971 | 0.277 | 0.282 | 1 | 7 | Psmd3 | -0.005 | Young |
| 0.776473 | 0.030136 | 1 | 0.997 | 1 | 7 | Rpl37a | 0.003 | Young |
| 0.776603 | 0.327057 | 0.018 | 0.015 | 1 | 7 | Gm43672 | 0.003 | Young |
| 0.776603 | 0.157273 | 0.018 | 0.015 | 1 | 7 | Zfp933 | 0.003 | Young |
| 0.776603 | 0.031965 | 0.018 | 0.015 | 1 | 7 | Cep170b | 0.003 | Young |
| 0.776752 | 0.13536 | 0.139 | 0.133 | 1 | 7 | Ice1 | 0.006 | Young |
| 0.777117 | 0.245971 | 0.127 | 0.121 | 1 | 7 | AI839979 | 0.006 | Young |
| 0.777252 | 0.444213 | 0.178 | 0.177 | 1 | 7 | Txndc16 | 0.001 | Young |
| 0.777343 | 0.005082 | 0.509 | 0.533 | 1 | 7 | Kdm2a | -0.024 | Young |
| 0.777412 | 0.537655 | 0.039 | 0.044 | 1 | 7 | Pdss2 | -0.005 | Young |
| 0.778031 | 0.055727 | 0.127 | 0.136 | 1 | 7 | Trpv4 | -0.009 | Young |
| 0.77809 | 0.353603 | 0.13 | 0.126 | 1 | 7 | Tmem126a | 0.004 | Young |
| 0.778099 | 0.359767 | 0.401 | 0.426 | 1 | 7 | Rb1cc1 | -0.025 | Young |
| 0.778123 | 0.281442 | 0.268 | 0.295 | 1 | 7 | Cdc73 | -0.027 | Young |
| 0.778168 | 0.064827 | 0.608 | 0.636 | 1 | 7 | Mrpl52 | -0.028 | Young |
| 0.778411 | 0.070731 | 0.226 | 0.223 | 1 | 7 | Mau2 | 0.003 | Young |
| 0.778598 | 0.196541 | 0.148 | 0.156 | 1 | 7 | Mtf1 | -0.008 | Young |
| 0.778662 | 0.488402 | 0.069 | 0.064 | 1 | 7 | 2510002D2 | 0.005 | Young |
| 0.778833 | 0.112428 | 0.045 | 0.041 | 1 | 7 | Ippk | 0.004 | Young |
| 0.778835 | 0.150869 | 0.19 | 0.185 | 1 | 7 | Gm34084 | 0.005 | Young |
| 0.779083 | 0.520538 | 0.018 | 0.015 | 1 | 7 | Zfp873 | 0.003 | Young |
| 0.779083 | 0.146962 | 0.018 | 0.015 | 1 | 7 | Wdr74 | 0.003 | Young |
| 0.779506 | 0.929718 | 0.057 | 0.054 | 1 | 7 | Eif5a2 | 0.003 | Young |
| 0.779642 | 0.290165 | 0.268 | 0.274 | 1 | 7 | Cenpc1 | -0.006 | Young |
| 0.779915 | 0.627345 | 0.042 | 0.038 | 1 | 7 | Tedc2 | 0.004 | Young |
| 0.780002 | 0.216317 | 0.127 | 0.121 | 1 | 7 | Sap30bp | 0.006 | Young |
| 0.780112 | 0.013838 | 0.142 | 0.151 | 1 | 7 | Nudt19 | -0.009 | Young |
| 0.780507 | 0.150448 | 0.28 | 0.279 | 1 | 7 | Ncoa4 | 0.001 | Young |
| 0.780722 | 1.130047 | 0.042 | 0.038 | 1 | 7 | F8a | 0.004 | Young |
| 0.780949 | 0.245075 | 0.193 | 0.203 | 1 | 7 | Cenpm | -0.01 | Young |
| 0.781045 | 0.321078 | 0.274 | 0.269 | 1 | 7 | Mis12 | 0.005 | Young |
| 0.781097 | 0.142321 | 0.229 | 0.226 | 1 | 7 | Mtfr1l | 0.003 | Young |
| 0.781194 | 0.057988 | 0.461 | 0.451 | 1 | 7 | Pcnp | 0.01 | Young |
| 0.78159 | 0.309086 | 0.364 | 0.374 | 1 | 7 | Usp34 | -0.01 | Young |
| 0.781822 | 0.250143 | 0.163 | 0.169 | 1 | 7 | Mtmr10 | -0.006 | Young |
| 0.781872 | 0.550555 | 0.163 | 0.177 | 1 | 7 | Itpr2 | -0.014 | Young |
| 0.781878 | 0.062745 | 0.169 | 0.179 | 1 | 7 | Taf4 | -0.01 | Young |
| 0.781894 | 0.11825 | 0.09 | 0.085 | 1 | 7 | Trim34a | 0.005 | Young |
| 0.781988 | 0.581722 | 0.108 | 0.115 | 1 | 7 | Mgat5 | -0.007 | Young |
| 0.782183 | 0.204391 | 0.051 | 0.056 | 1 | 7 | Ercc6 | -0.005 | Young |

| | | | | | | | | |
|---|---|---|---|---|---|---|---|---|
| 0.782185 | 0.7728 | 0.015 | 0.018 | 1 | 7 | Gm28151 | -0.003 | Young |
| 0.782277 | 0.209644 | 0.117 | 0.113 | 1 | 7 | Abhd4 | 0.004 | Young |
| 0.782745 | 0.294313 | 0.045 | 0.041 | 1 | 7 | Ppp1r3b | 0.004 | Young |
| 0.782751 | 0.59176 | 0.139 | 0.136 | 1 | 7 | Golph3l | 0.003 | Young |
| 0.782945 | 0.118044 | 0.53 | 0.541 | 1 | 7 | Papola | -0.011 | Young |
| 0.783063 | 0.366027 | 0.31 | 0.321 | 1 | 7 | Rgs19 | -0.011 | Young |
| 0.78316 | 0.058342 | 0.569 | 0.592 | 1 | 7 | Clec4a3 | -0.023 | Young |
| 0.783205 | 0.225811 | 0.277 | 0.272 | 1 | 7 | Hmgcr | 0.005 | Young |
| 0.783483 | 0.236448 | 0.217 | 0.215 | 1 | 7 | Nxt1 | 0.002 | Young |
| 0.783505 | 0.465183 | 0.093 | 0.103 | 1 | 7 | Prorsd1 | -0.01 | Young |
| 0.783754 | 0.033475 | 0.063 | 0.069 | 1 | 7 | Spryd3 | -0.006 | Young |
| 0.78403 | 0.138879 | 0.093 | 0.1 | 1 | 7 | Pdp2 | -0.007 | Young |
| 0.784045 | 0.432501 | 0.187 | 0.182 | 1 | 7 | Nipsnap2 | 0.005 | Young |
| 0.784048 | 0.085669 | 0.018 | 0.015 | 1 | 7 | Plat | 0.003 | Young |
| 0.784357 | 0.182108 | 0.449 | 0.474 | 1 | 7 | Eea1 | -0.025 | Young |
| 0.784359 | 0.142719 | 0.617 | 0.638 | 1 | 7 | Snrpd1 | -0.021 | Young |
| 0.784671 | 0.298327 | 0.114 | 0.126 | 1 | 7 | Gipc1 | -0.012 | Young |
| 0.784764 | 0.284033 | 0.042 | 0.038 | 1 | 7 | F3 | 0.004 | Young |
| 0.784882 | 0.256249 | 0.211 | 0.233 | 1 | 7 | Hadh | -0.022 | Young |
| 0.784918 | 0.081739 | 0.322 | 0.321 | 1 | 7 | Hps3 | 0.001 | Young |
| 0.785096 | 0.035911 | 0.045 | 0.041 | 1 | 7 | Rbm4 | 0.004 | Young |
| 0.785111 | 0.093844 | 0.569 | 0.569 | 1 | 7 | Stx7 | 0 | Young |
| 0.785177 | 0.127233 | 0.048 | 0.044 | 1 | 7 | Tango6 | 0.004 | Young |
| 0.78521 | 0.017592 | 0.274 | 0.287 | 1 | 7 | Osbp | -0.013 | Young |
| 0.785263 | 0.032799 | 0.654 | 0.667 | 1 | 7 | Pum2 | -0.013 | Young |
| 0.785291 | 0.200699 | 0.018 | 0.015 | 1 | 7 | Gm17259 | 0.003 | Young |
| 0.785312 | 0.339689 | 0.202 | 0.223 | 1 | 7 | Cstf2t | -0.021 | Young |
| 0.785336 | 0.714705 | 0.015 | 0.013 | 1 | 7 | AC125149. | 0.002 | Young |
| 0.785336 | 0.641008 | 0.015 | 0.013 | 1 | 7 | 4933406I1 | 0.002 | Young |
| 0.785495 | 0.067985 | 0.636 | 0.641 | 1 | 7 | Arhgap30 | -0.005 | Young |
| 0.786005 | 0.122995 | 0.163 | 0.174 | 1 | 7 | Snap29 | -0.011 | Young |
| 0.786158 | 0.254663 | 0.13 | 0.126 | 1 | 7 | Cyb5r1 | 0.004 | Young |
| 0.786203 | 0.632648 | 0.084 | 0.092 | 1 | 7 | Zbtb6 | -0.008 | Young |
| 0.786228 | 0.161872 | 0.431 | 0.451 | 1 | 7 | Lig1 | -0.02 | Young |
| 0.786534 | 0.497531 | 0.018 | 0.015 | 1 | 7 | Gm11423 | 0.003 | Young |
| 0.786643 | 0.631629 | 0.039 | 0.036 | 1 | 7 | Krt10 | 0.003 | Young |
| 0.786696 | 0.883923 | 0.015 | 0.013 | 1 | 7 | Wdr35 | 0.002 | Young |
| 0.786696 | 0.684086 | 0.015 | 0.013 | 1 | 7 | A430033K0 | 0.002 | Young |
| 0.786918 | 0.091659 | 0.102 | 0.11 | 1 | 7 | Taf6 | -0.008 | Young |
| 0.787192 | 0.169488 | 0.042 | 0.038 | 1 | 7 | Cetn4 | 0.004 | Young |
| 0.787208 | 0.063715 | 0.16 | 0.151 | 1 | 7 | Psmd10 | 0.009 | Young |
| 0.787208 | 0.776743 | 0.087 | 0.097 | 1 | 7 | Vps39 | -0.01 | Young |
| 0.787579 | 0.307151 | 0.063 | 0.069 | 1 | 7 | Actr6 | -0.006 | Young |
| 0.787723 | 0.44455 | 0.178 | 0.192 | 1 | 7 | Lap3 | -0.014 | Young |
| 0.787752 | 0.190594 | 0.039 | 0.044 | 1 | 7 | Zfp46 | -0.005 | Young |

| | | | | | | | | |
|---|---|---|---|---|---|---|---|---|
| 0.787804 | 0.175054 | 0.051 | 0.056 | 1 | 7 | Spaca9 | -0.005 | Young |
| 0.787815 | 0.196703 | 0.133 | 0.141 | 1 | 7 | Usp36 | -0.008 | Young |
| 0.787876 | 0.049767 | 0.298 | 0.321 | 1 | 7 | Lars2 | -0.023 | Young |
| 0.788004 | 0.002829 | 0.419 | 0.415 | 1 | 7 | Ppp3cb | 0.004 | Young |
| 0.788137 | 0.198876 | 0.027 | 0.031 | 1 | 7 | Cox11 | -0.004 | Young |
| 0.788157 | 0.079789 | 0.142 | 0.151 | 1 | 7 | Sec23a | -0.009 | Young |
| 0.788208 | 0.014474 | 0.434 | 0.451 | 1 | 7 | Ilk | -0.017 | Young |
| 0.788549 | 0.472741 | 0.039 | 0.044 | 1 | 7 | Zc3h12d | -0.005 | Young |
| 0.788665 | 0.478638 | 0.142 | 0.138 | 1 | 7 | Kpna6 | 0.004 | Young |
| 0.788703 | 0.249191 | 0.075 | 0.082 | 1 | 7 | Ticrr | -0.007 | Young |
| 0.788763 | 0.132032 | 0.139 | 0.133 | 1 | 7 | Klhdc10 | 0.006 | Young |
| 0.788951 | 0.307958 | 0.066 | 0.062 | 1 | 7 | Zmat3 | 0.004 | Young |
| 0.788997 | 0.507492 | 0.09 | 0.087 | 1 | 7 | Mrpl44 | 0.003 | Young |
| 0.789346 | 0.026823 | 0.039 | 0.044 | 1 | 7 | Pcgf1 | -0.005 | Young |
| 0.789418 | 0.04772 | 0.015 | 0.013 | 1 | 7 | Magi2 | 0.002 | Young |
| 0.789457 | 0.197708 | 0.139 | 0.133 | 1 | 7 | Rad54l2 | 0.006 | Young |
| 0.78959 | 0.32334 | 0.202 | 0.2 | 1 | 7 | Epb41l4aos | 0.002 | Young |
| 0.789816 | 0.013805 | 0.304 | 0.29 | 1 | 7 | Rnf6 | 0.014 | Young |
| 0.790124 | 0.209869 | 0.102 | 0.097 | 1 | 7 | Zfp180 | 0.005 | Young |
| 0.790465 | 0.341206 | 0.292 | 0.321 | 1 | 7 | Mcfd2 | -0.029 | Young |
| 0.790468 | 0.415368 | 0.099 | 0.095 | 1 | 7 | Ccdc134 | 0.004 | Young |
| 0.790527 | 0.218125 | 0.331 | 0.331 | 1 | 7 | Tmem259 | 0 | Young |
| 0.790639 | 0.106397 | 0.361 | 0.359 | 1 | 7 | Rala | 0.002 | Young |
| 0.79078 | 0.671827 | 0.015 | 0.013 | 1 | 7 | Ppp1r13l | 0.002 | Young |
| 0.79078 | 0.188962 | 0.015 | 0.013 | 1 | 7 | Arhgap27o | 0.002 | Young |
| 0.79079 | 0.388459 | 0.105 | 0.1 | 1 | 7 | Hddc2 | 0.005 | Young |
| 0.790941 | 0.408072 | 0.039 | 0.044 | 1 | 7 | Zfp748 | -0.005 | Young |
| 0.791872 | 0.004667 | 0.087 | 0.082 | 1 | 7 | Rbms2 | 0.005 | Young |
| 0.791926 | 0.721136 | 0.027 | 0.031 | 1 | 7 | Zfp113 | -0.004 | Young |
| 0.792143 | 0.723014 | 0.015 | 0.013 | 1 | 7 | Apbb1 | 0.002 | Young |
| 0.792143 | 0.530161 | 0.015 | 0.013 | 1 | 7 | Adat1 | 0.002 | Young |
| 0.792143 | 0.422712 | 0.015 | 0.013 | 1 | 7 | C230037L1 | 0.002 | Young |
| 0.792162 | 0.333692 | 0.265 | 0.292 | 1 | 7 | S100a1 | -0.027 | Young |
| 0.792814 | 0.328492 | 0.084 | 0.079 | 1 | 7 | D730003I1 | 0.005 | Young |
| 0.792837 | 0.105713 | 0.611 | 0.662 | 1 | 7 | Trir | -0.051 | Young |
| 0.792838 | 0.159419 | 0.398 | 0.4 | 1 | 7 | Nsmce4a | -0.002 | Young |
| 0.79297 | 0.207187 | 0.096 | 0.105 | 1 | 7 | Ankrd54 | -0.009 | Young |
| 0.793113 | 0.081979 | 0.31 | 0.328 | 1 | 7 | Sertad1 | -0.018 | Young |
| 0.793152 | 0.207032 | 0.346 | 0.354 | 1 | 7 | Rbl1 | -0.008 | Young |
| 0.793155 | 0.041219 | 0.292 | 0.287 | 1 | 7 | Diaph3 | 0.005 | Young |
| 0.793395 | 0.29571 | 0.084 | 0.079 | 1 | 7 | Ift22 | 0.005 | Young |
| 0.793506 | 0.565385 | 0.015 | 0.013 | 1 | 7 | Id1 | 0.002 | Young |
| 0.793506 | 0.149812 | 0.015 | 0.013 | 1 | 7 | Scml4 | 0.002 | Young |
| 0.793732 | 0.017491 | 0.87 | 0.828 | 1 | 7 | Coro1a | 0.042 | Young |
| 0.79377 | 0.079462 | 0.431 | 0.438 | 1 | 7 | Hivep2 | -0.007 | Young |

| | | | | | | | | |
|---|---|---|---|---|---|---|---|---|
| 0.793958 | 0.179204 | 0.169 | 0.179 | 1 | 7 | Cenpk | -0.01 | Young |
| 0.794141 | 0.385356 | 0.051 | 0.056 | 1 | 7 | Smim24 | -0.005 | Young |
| 0.794347 | 0.845755 | 0.03 | 0.033 | 1 | 7 | Trmt10b | -0.003 | Young |
| 0.79487 | 0.598281 | 0.015 | 0.013 | 1 | 7 | Them6 | 0.002 | Young |
| 0.794898 | 0.254628 | 0.081 | 0.077 | 1 | 7 | Mefv | 0.004 | Young |
| 0.794926 | 0.048659 | 0.349 | 0.379 | 1 | 7 | Tipin | -0.03 | Young |
| 0.795393 | 0.165422 | 0.358 | 0.385 | 1 | 7 | Emp1 | -0.027 | Young |
| 0.795799 | 0.244157 | 0.355 | 0.359 | 1 | 7 | Eloa | -0.004 | Young |
| 0.796113 | 0.065802 | 0.042 | 0.038 | 1 | 7 | C1ra | 0.004 | Young |
| 0.796235 | 0.691694 | 0.015 | 0.013 | 1 | 7 | Bhlhe41 | 0.002 | Young |
| 0.796235 | 0.515982 | 0.015 | 0.013 | 1 | 7 | Cdk5r1 | 0.002 | Young |
| 0.796277 | 0.126677 | 0.127 | 0.138 | 1 | 7 | Fancg | -0.011 | Young |
| 0.796459 | 0.060788 | 0.38 | 0.379 | 1 | 7 | Dock11 | 0.001 | Young |
| 0.79649 | 0.203352 | 0.367 | 0.369 | 1 | 7 | Mllt10 | -0.002 | Young |
| 0.796517 | 0.025746 | 0.654 | 0.667 | 1 | 7 | Gas7 | -0.013 | Young |
| 0.796794 | 0.012819 | 0.292 | 0.315 | 1 | 7 | Rin3 | -0.023 | Young |
| 0.79712 | 0.051049 | 0.244 | 0.256 | 1 | 7 | Klhdc3 | -0.012 | Young |
| 0.797229 | 0.015076 | 0.669 | 0.669 | 1 | 7 | Pgk1 | 0 | Young |
| 0.797351 | 0.047375 | 0.163 | 0.159 | 1 | 7 | Ldlrap1 | 0.004 | Young |
| 0.797458 | 0.319753 | 0.054 | 0.059 | 1 | 7 | Zfp358 | -0.005 | Young |
| 0.797534 | 0.130208 | 0.202 | 0.197 | 1 | 7 | Phf8 | 0.005 | Young |
| 0.797548 | 0.147309 | 0.238 | 0.254 | 1 | 7 | Mphosph8 | -0.016 | Young |
| 0.7976 | 0.134128 | 0.015 | 0.013 | 1 | 7 | Trim45 | 0.002 | Young |
| 0.7976 | 0.045929 | 0.015 | 0.013 | 1 | 7 | Gm16845 | 0.002 | Young |
| 0.797717 | 0.182585 | 0.205 | 0.218 | 1 | 7 | Cluh | -0.013 | Young |
| 0.797717 | 0.178031 | 0.205 | 0.218 | 1 | 7 | Ctnnbip1 | -0.013 | Young |
| 0.797764 | 0.261014 | 0.277 | 0.305 | 1 | 7 | Dapp1 | -0.028 | Young |
| 0.797917 | 0.160296 | 0.136 | 0.131 | 1 | 7 | Ppp1r37 | 0.005 | Young |
| 0.798303 | 0.278651 | 0.12 | 0.133 | 1 | 7 | Sphk2 | -0.013 | Young |
| 0.798324 | 0.2046 | 0.178 | 0.19 | 1 | 7 | Snapc5 | -0.012 | Young |
| 0.798462 | 0.225486 | 0.247 | 0.246 | 1 | 7 | Atp13a1 | 0.001 | Young |
| 0.798555 | 0.044739 | 0.407 | 0.436 | 1 | 7 | Becn1 | -0.029 | Young |
| 0.798791 | 0.013886 | 0.361 | 0.385 | 1 | 7 | Mrpl42 | -0.024 | Young |
| 0.798966 | 0.148426 | 0.015 | 0.013 | 1 | 7 | AC139671. | 0.002 | Young |
| 0.799106 | 0.330403 | 0.223 | 0.244 | 1 | 7 | Slc4a2 | -0.021 | Young |
| 0.799153 | 0.501788 | 0.211 | 0.21 | 1 | 7 | Sar1b | 0.001 | Young |
| 0.799221 | 0.07908 | 0.512 | 0.549 | 1 | 7 | Sema4a | -0.037 | Young |
| 0.799867 | 0.55781 | 0.093 | 0.09 | 1 | 7 | Wdsub1 | 0.003 | Young |
| 0.800082 | 0.238915 | 0.127 | 0.136 | 1 | 7 | Mrps10 | -0.009 | Young |
| 0.800436 | 0.248916 | 0.199 | 0.21 | 1 | 7 | Filip1l | -0.011 | Young |
| 0.800528 | 0.196115 | 0.039 | 0.044 | 1 | 7 | Mapkapk5 | -0.005 | Young |
| 0.800583 | 0.227715 | 0.09 | 0.097 | 1 | 7 | Aktip | -0.007 | Young |
| 0.800691 | 0.274238 | 0.262 | 0.254 | 1 | 7 | Dusp2 | 0.008 | Young |
| 0.800707 | 0.153871 | 0.03 | 0.033 | 1 | 7 | Fmnl2 | -0.003 | Young |
| 0.801183 | 0.163512 | 0.455 | 0.467 | 1 | 7 | Xpo1 | -0.012 | Young |

| | | | | | | | | |
|---|---|---|---|---|---|---|---|---|
| 0.801314 | 0.180676 | 0.355 | 0.367 | 1 | 7 | Ino80d | -0.012 | Young |
| 0.801385 | 0.298717 | 0.223 | 0.218 | 1 | 7 | Ccdc25 | 0.005 | Young |
| 0.801653 | 0.689924 | 0.054 | 0.051 | 1 | 7 | Slc7a6 | 0.003 | Young |
| 0.802655 | 0.714813 | 0.06 | 0.067 | 1 | 7 | Mipep | -0.007 | Young |
| 0.802955 | 0.168114 | 0.286 | 0.308 | 1 | 7 | Pcnt | -0.022 | Young |
| 0.80317 | 1.021277 | 0.036 | 0.033 | 1 | 7 | Ighmbp2 | 0.003 | Young |
| 0.803185 | 0.277881 | 0.235 | 0.254 | 1 | 7 | Ppil2 | -0.019 | Young |
| 0.803431 | 0.126394 | 0.042 | 0.038 | 1 | 7 | Zfp868 | 0.004 | Young |
| 0.803548 | 0.187643 | 0.163 | 0.172 | 1 | 7 | Myo7a | -0.009 | Young |
| 0.803755 | 0.381478 | 0.223 | 0.221 | 1 | 7 | Insr | 0.002 | Young |
| 0.803755 | 0.269525 | 0.223 | 0.221 | 1 | 7 | Med21 | 0.002 | Young |
| 0.803783 | 0.16776 | 0.169 | 0.179 | 1 | 7 | Fmc1 | -0.01 | Young |
| 0.803863 | 0.00191 | 0.678 | 0.705 | 1 | 7 | Ssb | -0.027 | Young |
| 0.803883 | 0.200487 | 0.084 | 0.09 | 1 | 7 | Zfp954 | -0.006 | Young |
| 0.804297 | 0.122531 | 0.087 | 0.095 | 1 | 7 | Prkch | -0.008 | Young |
| 0.804473 | 0.275234 | 0.087 | 0.085 | 1 | 7 | Fgd3 | 0.002 | Young |
| 0.804759 | 0.124246 | 0.271 | 0.285 | 1 | 7 | Uxs1 | -0.014 | Young |
| 0.804935 | 1.002028 | 0.063 | 0.059 | 1 | 7 | Jak3 | 0.004 | Young |
| 0.804935 | 0.450797 | 0.063 | 0.059 | 1 | 7 | Zfp41 | 0.004 | Young |
| 0.805048 | 0.023036 | 0.054 | 0.059 | 1 | 7 | Sec22c | -0.005 | Young |
| 0.805461 | 0.327545 | 0.111 | 0.121 | 1 | 7 | Sdhaf1 | -0.01 | Young |
| 0.805545 | 0.295563 | 0.244 | 0.267 | 1 | 7 | Xpo7 | -0.023 | Young |
| 0.805609 | 0.456604 | 0.187 | 0.187 | 1 | 7 | Tmem192 | 0 | Young |
| 0.805739 | 0.031696 | 0.054 | 0.059 | 1 | 7 | 5031425F1 | -0.005 | Young |
| 0.805765 | 0.226145 | 0.181 | 0.195 | 1 | 7 | Lysmd3 | -0.014 | Young |
| 0.805885 | 0.359113 | 0.123 | 0.133 | 1 | 7 | Fam3a | -0.01 | Young |
| 0.80608 | 0.116526 | 0.042 | 0.046 | 1 | 7 | Prdm15 | -0.004 | Young |
| 0.806114 | 0.352903 | 0.337 | 0.349 | 1 | 7 | Gtf3c2 | -0.012 | Young |
| 0.80643 | 0.454816 | 0.054 | 0.059 | 1 | 7 | Pank4 | -0.005 | Young |
| 0.80679 | 0.372327 | 0.084 | 0.079 | 1 | 7 | Cep104 | 0.005 | Young |
| 0.80682 | 0.407016 | 0.039 | 0.036 | 1 | 7 | Ptpn21 | 0.003 | Young |
| 0.806857 | 0.313845 | 0.042 | 0.046 | 1 | 7 | Klhl36 | -0.004 | Young |
| 0.807147 | 0.364143 | 0.123 | 0.118 | 1 | 7 | Brwd1 | 0.005 | Young |
| 0.807206 | 0.353962 | 0.111 | 0.118 | 1 | 7 | Ppp6r2 | -0.007 | Young |
| 0.807287 | 0.062695 | 0.392 | 0.436 | 1 | 7 | Tnrc6a | -0.044 | Young |
| 0.807301 | 0.385576 | 0.169 | 0.185 | 1 | 7 | Dopey1 | -0.016 | Young |
| 0.807415 | 0.031374 | 0.395 | 0.428 | 1 | 7 | Sh3bp1 | -0.033 | Young |
| 0.807459 | 0.009248 | 0.873 | 0.874 | 1 | 7 | Sirpa | -0.001 | Young |
| 0.807593 | 0.116146 | 0.352 | 0.377 | 1 | 7 | Ccnl2 | -0.025 | Young |
| 0.807604 | 0.124159 | 0.105 | 0.1 | 1 | 7 | Med31 | 0.005 | Young |
| 0.807663 | 0.293466 | 0.039 | 0.036 | 1 | 7 | 1810024B0 | 0.003 | Young |
| 0.807813 | 0.155299 | 0.054 | 0.059 | 1 | 7 | Zfp959 | -0.005 | Young |
| 0.807847 | 0.088206 | 0.084 | 0.09 | 1 | 7 | Tef | -0.006 | Young |
| 0.808405 | 0.293928 | 0.175 | 0.19 | 1 | 7 | Zfp384 | -0.015 | Young |
| 0.808429 | 0.003059 | 0.898 | 0.936 | 1 | 7 | Mbnl1 | -0.038 | Young |

| | | | | | | | | |
|---|---|---|---|---|---|---|---|---|
| 0.808462 | 0.051107 | 0.614 | 0.654 | 1 | 7 | Atp6ap1 | -0.04 | Young |
| 0.808606 | 0.266313 | 0.057 | 0.062 | 1 | 7 | Parpbp | -0.005 | Young |
| 0.808743 | 0.253809 | 0.096 | 0.103 | 1 | 7 | Taf5 | -0.007 | Young |
| 0.80886 | 1.955915 | 0.012 | 0.01 | 1 | 7 | Smim1 | 0.002 | Young |
| 0.808875 | 0.064363 | 0.066 | 0.062 | 1 | 7 | Tatdn1 | 0.004 | Young |
| 0.80897 | 0.05816 | 0.271 | 0.285 | 1 | 7 | Stim2 | -0.014 | Young |
| 0.809108 | 0.237448 | 0.238 | 0.238 | 1 | 7 | Kcnk6 | 0 | Young |
| 0.809297 | 0.137723 | 0.187 | 0.195 | 1 | 7 | Parg | -0.008 | Young |
| 0.809428 | 0.278712 | 0.205 | 0.221 | 1 | 7 | Pgm2l1 | -0.016 | Young |
| 0.809888 | 0.729793 | 0.054 | 0.059 | 1 | 7 | Pomt2 | -0.005 | Young |
| 0.809944 | 0.468272 | 0.117 | 0.115 | 1 | 7 | Mlx | 0.002 | Young |
| 0.809969 | 0.037974 | 0.042 | 0.046 | 1 | 7 | Coil | -0.004 | Young |
| 0.8101 | 0.096301 | 0.301 | 0.313 | 1 | 7 | Paf1 | -0.012 | Young |
| 0.810391 | 1.373391 | 0.012 | 0.01 | 1 | 7 | Tnfrsf23 | 0.002 | Young |
| 0.810508 | 0.10504 | 0.467 | 0.505 | 1 | 7 | Ncaph2 | -0.038 | Young |
| 0.810603 | 0.114386 | 0.069 | 0.074 | 1 | 7 | Slc19a1 | -0.005 | Young |
| 0.810891 | 0.119908 | 0.12 | 0.128 | 1 | 7 | Atxn1 | -0.008 | Young |
| 0.811258 | 0.126906 | 0.373 | 0.4 | 1 | 7 | Setd2 | -0.027 | Young |
| 0.811375 | 0.179478 | 0.127 | 0.123 | 1 | 7 | Pcsk7 | 0.004 | Young |
| 0.811523 | 0.091475 | 0.316 | 0.333 | 1 | 7 | Casp3 | -0.017 | Young |
| 0.811723 | 0.034861 | 0.44 | 0.459 | 1 | 7 | Ankrd13a | -0.019 | Young |
| 0.811898 | 0.077601 | 0.458 | 0.464 | 1 | 7 | Ddx3y | -0.006 | Young |
| 0.811922 | 1.267285 | 0.012 | 0.01 | 1 | 7 | C030006K1 | 0.002 | Young |
| 0.811922 | 0.948665 | 0.012 | 0.01 | 1 | 7 | Tlcd1 | 0.002 | Young |
| 0.811922 | 0.433485 | 0.012 | 0.01 | 1 | 7 | Bcl2a1c | 0.002 | Young |
| 0.81195 | 0.265178 | 0.352 | 0.387 | 1 | 7 | Anxa6 | -0.035 | Young |
| 0.812215 | 0.34268 | 0.108 | 0.105 | 1 | 7 | Glce | 0.003 | Young |
| 0.81291 | 0.0195 | 0.28 | 0.295 | 1 | 7 | Tnfrsf11a | -0.015 | Young |
| 0.813155 | 0.785599 | 0.075 | 0.072 | 1 | 7 | Mettl5 | 0.003 | Young |
| 0.813408 | 0.138235 | 0.127 | 0.133 | 1 | 7 | Pno1 | -0.006 | Young |
| 0.813422 | 0.026079 | 0.259 | 0.272 | 1 | 7 | Lnpk | -0.013 | Young |
| 0.813454 | 0.792931 | 0.012 | 0.01 | 1 | 7 | Plxdc1 | 0.002 | Young |
| 0.813454 | 0.778128 | 0.012 | 0.01 | 1 | 7 | Zfp956 | 0.002 | Young |
| 0.813454 | 0.715547 | 0.012 | 0.01 | 1 | 7 | Gm14403 | 0.002 | Young |
| 0.813454 | 0.56389 | 0.012 | 0.01 | 1 | 7 | Anks6 | 0.002 | Young |
| 0.813454 | 0.326235 | 0.012 | 0.01 | 1 | 7 | Kif21a | 0.002 | Young |
| 0.813588 | 0.318244 | 0.114 | 0.123 | 1 | 7 | Med23 | -0.009 | Young |
| 0.813998 | 0.18579 | 0.286 | 0.287 | 1 | 7 | Wbp4 | -0.001 | Young |
| 0.814042 | 0.05928 | 0.105 | 0.11 | 1 | 7 | Dcaf5 | -0.005 | Young |
| 0.814098 | 0.227633 | 0.133 | 0.138 | 1 | 7 | Ikbkap | -0.005 | Young |
| 0.814589 | 0.008061 | 0.994 | 0.985 | 1 | 7 | Rps27a | 0.009 | Young |
| 0.814852 | 0.323224 | 0.175 | 0.187 | 1 | 7 | Inf2 | -0.012 | Young |
| 0.814971 | 0.152285 | 0.084 | 0.079 | 1 | 7 | Zer1 | 0.005 | Young |
| 0.814987 | 0.952754 | 0.012 | 0.01 | 1 | 7 | Ptpn3 | 0.002 | Young |
| 0.814987 | 0.828188 | 0.012 | 0.01 | 1 | 7 | Gm16062 | 0.002 | Young |

| | | | | | | | | |
|---|---|---|---|---|---|---|---|---|
| 0.814999 | 0.289818 | 0.075 | 0.072 | 1 | 7 | Zbtb34 | 0.003 | Young |
| 0.815293 | 0.876338 | 0.033 | 0.031 | 1 | 7 | Tmem143 | 0.002 | Young |
| 0.815361 | 0.874658 | 0.09 | 0.097 | 1 | 7 | Galnt4 | -0.007 | Young |
| 0.815421 | 0.216578 | 0.042 | 0.046 | 1 | 7 | Lonrf3 | -0.004 | Young |
| 0.815556 | 0.204559 | 0.078 | 0.085 | 1 | 7 | Xpnpep3 | -0.007 | Young |
| 0.815687 | 0.511931 | 0.096 | 0.103 | 1 | 7 | Trappc2 | -0.007 | Young |
| 0.815694 | 0.068686 | 0.018 | 0.021 | 1 | 7 | Cdyl2 | -0.003 | Young |
| 0.815782 | 0.544104 | 0.084 | 0.082 | 1 | 7 | Hs1bp3 | 0.002 | Young |
| 0.815783 | 0.350651 | 0.154 | 0.164 | 1 | 7 | Gse1 | -0.01 | Young |
| 0.816233 | 0.163857 | 0.34 | 0.351 | 1 | 7 | Mrpl54 | -0.011 | Young |
| 0.816317 | 0.069075 | 0.063 | 0.059 | 1 | 7 | Foxo4 | 0.004 | Young |
| 0.816477 | 0.048474 | 0.325 | 0.341 | 1 | 7 | Lyar | -0.016 | Young |
| 0.81652 | 0.181679 | 0.012 | 0.01 | 1 | 7 | Pcna-ps2 | 0.002 | Young |
| 0.81652 | 0.160964 | 0.012 | 0.01 | 1 | 7 | Pitpnm2 | 0.002 | Young |
| 0.816645 | 0.28052 | 0.157 | 0.154 | 1 | 7 | Crim1 | 0.003 | Young |
| 0.816757 | 0.029289 | 0.096 | 0.103 | 1 | 7 | Gm9493 | -0.007 | Young |
| 0.816858 | 0.004356 | 0.018 | 0.021 | 1 | 7 | Prag1 | -0.003 | Young |
| 0.816867 | 0.069802 | 0.105 | 0.113 | 1 | 7 | Kif18a | -0.008 | Young |
| 0.817131 | 0.310634 | 0.241 | 0.241 | 1 | 7 | Dgkz | 0 | Young |
| 0.817541 | 0.190816 | 0.114 | 0.123 | 1 | 7 | Zfpl1 | -0.009 | Young |
| 0.817804 | 0.164788 | 0.039 | 0.036 | 1 | 7 | D3Ertd254 | 0.003 | Young |
| 0.817866 | 0.427968 | 0.102 | 0.1 | 1 | 7 | Bcl2l2 | 0.002 | Young |
| 0.817966 | 0.190347 | 0.157 | 0.162 | 1 | 7 | Ciapin1 | -0.005 | Young |
| 0.818054 | 0.163955 | 0.012 | 0.01 | 1 | 7 | Lag3 | 0.002 | Young |
| 0.818054 | 0.129818 | 0.012 | 0.01 | 1 | 7 | Aoc2 | 0.002 | Young |
| 0.818071 | 0.412851 | 0.286 | 0.292 | 1 | 7 | Rps12-ps3 | -0.006 | Young |
| 0.818127 | 0.180271 | 0.13 | 0.136 | 1 | 7 | Arl3 | -0.006 | Young |
| 0.818434 | 0.146265 | 0.524 | 0.556 | 1 | 7 | Irf2 | -0.032 | Young |
| 0.818465 | 0.006677 | 0.268 | 0.287 | 1 | 7 | Dicer1 | -0.019 | Young |
| 0.818526 | 0.232305 | 0.298 | 0.318 | 1 | 7 | Chp1 | -0.02 | Young |
| 0.818895 | 0.532413 | 0.054 | 0.059 | 1 | 7 | Stoml1 | -0.005 | Young |
| 0.818901 | 0.2563 | 0.087 | 0.092 | 1 | 7 | Mir17hg | -0.005 | Young |
| 0.819187 | 0.318598 | 0.018 | 0.021 | 1 | 7 | Tbc1d25 | -0.003 | Young |
| 0.819187 | 0.257691 | 0.018 | 0.021 | 1 | 7 | Zkscan5 | -0.003 | Young |
| 0.819273 | 0.070355 | 0.723 | 0.754 | 1 | 7 | Atf4 | -0.031 | Young |
| 0.819303 | 0.291494 | 0.108 | 0.118 | 1 | 7 | Lsg1 | -0.01 | Young |
| 0.819373 | 0.036748 | 0.533 | 0.533 | 1 | 7 | Cwc15 | 0 | Young |
| 0.819589 | 0.301011 | 0.012 | 0.01 | 1 | 7 | Kcnj10 | 0.002 | Young |
| 0.819728 | 0.138756 | 0.464 | 0.477 | 1 | 7 | Commd8 | -0.013 | Young |
| 0.819767 | 0.401975 | 0.084 | 0.09 | 1 | 7 | Zfp597 | -0.006 | Young |
| 0.819809 | 0.168015 | 0.467 | 0.503 | 1 | 7 | Pttg1ip | -0.036 | Young |
| 0.819903 | 0.316477 | 0.298 | 0.328 | 1 | 7 | Pfdn1 | -0.03 | Young |
| 0.820127 | 0.206005 | 0.187 | 0.192 | 1 | 7 | Zyg11b | -0.005 | Young |
| 0.820198 | 0.031881 | 0.392 | 0.395 | 1 | 7 | Glrx3 | -0.003 | Young |
| 0.820353 | 0.632631 | 0.018 | 0.021 | 1 | 7 | Klhl33 | -0.003 | Young |

| | | | | | | | | |
|---|---|---|---|---|---|---|---|---|
| 0.820381 | 0.008739 | 0.298 | 0.313 | 1 | 7 | Rsf1 | -0.015 | Young |
| 0.820418 | 0.339288 | 0.337 | 0.336 | 1 | 7 | Cdt1 | 0.001 | Young |
| 0.82094 | 0.105508 | 0.407 | 0.41 | 1 | 7 | Clk4 | -0.003 | Young |
| 0.821021 | 0.426691 | 0.069 | 0.067 | 1 | 7 | Bud13 | 0.002 | Young |
| 0.821124 | 0.07339 | 0.012 | 0.01 | 1 | 7 | Nkapl | 0.002 | Young |
| 0.82159 | 0.388131 | 0.06 | 0.056 | 1 | 7 | Sppl2b | 0.004 | Young |
| 0.82188 | 0.122313 | 0.244 | 0.262 | 1 | 7 | Kmt5b | -0.018 | Young |
| 0.822038 | 0.02027 | 0.039 | 0.036 | 1 | 7 | Fam102a | 0.003 | Young |
| 0.822078 | 0.310858 | 0.087 | 0.095 | 1 | 7 | Acvr1 | -0.008 | Young |
| 0.82266 | 0.051777 | 0.012 | 0.01 | 1 | 7 | Gdpd5 | 0.002 | Young |
| 0.82266 | 0.013826 | 0.012 | 0.01 | 1 | 7 | 0610040B1 | 0.002 | Young |
| 0.823024 | 0.090501 | 0.223 | 0.238 | 1 | 7 | Jpt2 | -0.015 | Young |
| 0.823088 | 0.196837 | 0.25 | 0.264 | 1 | 7 | Thap12 | -0.014 | Young |
| 0.823099 | 0.645561 | 0.096 | 0.095 | 1 | 7 | Rps6ka2 | 0.001 | Young |
| 0.823502 | 0.040029 | 0.16 | 0.169 | 1 | 7 | Fbxo38 | -0.009 | Young |
| 0.823851 | 0.764231 | 0.018 | 0.021 | 1 | 7 | Pex11a | -0.003 | Young |
| 0.823851 | 0.214227 | 0.018 | 0.021 | 1 | 7 | Oas1c | -0.003 | Young |
| 0.823851 | 0.043299 | 0.018 | 0.021 | 1 | 7 | Zfp870 | -0.003 | Young |
| 0.82394 | 0.198197 | 0.123 | 0.131 | 1 | 7 | Slc11a2 | -0.008 | Young |
| 0.823985 | 0.026645 | 0.461 | 0.477 | 1 | 7 | Tbl1xr1 | -0.016 | Young |
| 0.824197 | 0.081425 | 0.012 | 0.01 | 1 | 7 | Sox13 | 0.002 | Young |
| 0.824878 | 0.145825 | 0.069 | 0.074 | 1 | 7 | Med6 | -0.005 | Young |
| 0.825018 | 0.257466 | 0.018 | 0.021 | 1 | 7 | Camk2n2 | -0.003 | Young |
| 0.825018 | 0.172223 | 0.018 | 0.021 | 1 | 7 | Rab11fip3 | -0.003 | Young |
| 0.825753 | 0.133606 | 0.316 | 0.341 | 1 | 7 | Naa38 | -0.025 | Young |
| 0.82587 | 0.045322 | 0.154 | 0.162 | 1 | 7 | Prr11 | -0.008 | Young |
| 0.825983 | 0.45295 | 0.033 | 0.036 | 1 | 7 | Rflnb | -0.003 | Young |
| 0.826009 | 0.2611 | 0.229 | 0.226 | 1 | 7 | Ints6 | 0.003 | Young |
| 0.826259 | 0.331277 | 0.093 | 0.09 | 1 | 7 | Abhd10 | 0.003 | Young |
| 0.826281 | 0.200282 | 0.241 | 0.262 | 1 | 7 | Smarcb1 | -0.021 | Young |
| 0.826283 | 0.141061 | 0.045 | 0.049 | 1 | 7 | Pomk | -0.004 | Young |
| 0.826441 | 0.101041 | 0.105 | 0.11 | 1 | 7 | Fkbp1b | -0.005 | Young |
| 0.826701 | 0.31571 | 0.172 | 0.169 | 1 | 7 | Naa60 | 0.003 | Young |
| 0.826918 | 0.266799 | 0.057 | 0.062 | 1 | 7 | Igtp | -0.005 | Young |
| 0.82707 | 0.139897 | 0.12 | 0.115 | 1 | 7 | As3mt | 0.005 | Young |
| 0.827086 | 0.078297 | 0.06 | 0.056 | 1 | 7 | Eid3 | 0.004 | Young |
| 0.827157 | 0.110445 | 0.099 | 0.105 | 1 | 7 | Eif2d | -0.006 | Young |
| 0.827319 | 0.077686 | 0.087 | 0.092 | 1 | 7 | Casp7 | -0.005 | Young |
| 0.827544 | 0.457419 | 0.181 | 0.2 | 1 | 7 | Cul4a | -0.019 | Young |
| 0.828229 | 0.811364 | 0.054 | 0.051 | 1 | 7 | Map4k3 | 0.003 | Young |
| 0.828286 | 0.150259 | 0.461 | 0.472 | 1 | 7 | Use1 | -0.011 | Young |
| 0.828355 | 0.145186 | 0.386 | 0.385 | 1 | 7 | Kdm2b | 0.001 | Young |
| 0.828648 | 0.649531 | 0.051 | 0.049 | 1 | 7 | Trpm2 | 0.002 | Young |
| 0.829051 | 0.352933 | 0.102 | 0.1 | 1 | 7 | Tmem39a | 0.002 | Young |
| 0.829154 | 0.234796 | 0.292 | 0.295 | 1 | 7 | Ahctf1 | -0.003 | Young |

| | | | | | | | | |
|---|---|---|---|---|---|---|---|---|
| 0.829228 | 0.317262 | 0.256 | 0.272 | 1 | 7 | Prpf39 | -0.016 | Young |
| 0.829256 | 0.213222 | 0.172 | 0.169 | 1 | 7 | Foxj3 | 0.003 | Young |
| 0.829379 | 0.069591 | 0.563 | 0.592 | 1 | 7 | Itpripl2 | -0.029 | Young |
| 0.829781 | 0.476268 | 0.13 | 0.128 | 1 | 7 | Pon3 | 0.002 | Young |
| 0.830092 | 0.218209 | 0.205 | 0.2 | 1 | 7 | Ubr3 | 0.005 | Young |
| 0.83054 | 0.088027 | 0.148 | 0.141 | 1 | 7 | Tmem181a | 0.007 | Young |
| 0.831004 | 0.06006 | 0.256 | 0.251 | 1 | 7 | Scrib | 0.005 | Young |
| 0.831531 | 0.235241 | 0.196 | 0.21 | 1 | 7 | Mast3 | -0.014 | Young |
| 0.831666 | 0.054685 | 0.19 | 0.187 | 1 | 7 | Nr1h2 | 0.003 | Young |
| 0.831928 | 0.401492 | 0.057 | 0.054 | 1 | 7 | Polr3a | 0.003 | Young |
| 0.832059 | 0.474786 | 0.151 | 0.149 | 1 | 7 | Pold4 | 0.002 | Young |
| 0.832556 | 0.248733 | 0.208 | 0.218 | 1 | 7 | Map3k5 | -0.01 | Young |
| 0.832698 | 0.142516 | 0.03 | 0.033 | 1 | 7 | Tbc1d16 | -0.003 | Young |
| 0.833033 | 0.512843 | 0.036 | 0.033 | 1 | 7 | Lipt2 | 0.003 | Young |
| 0.83309 | 0.175928 | 0.051 | 0.049 | 1 | 7 | Nfe2 | 0.002 | Young |
| 0.833301 | 0.159844 | 0.133 | 0.141 | 1 | 7 | Plxna4 | -0.008 | Young |
| 0.833328 | 0.109067 | 0.172 | 0.185 | 1 | 7 | Secisbp2 | -0.013 | Young |
| 0.833545 | 0.21011 | 0.235 | 0.238 | 1 | 7 | Kars | -0.003 | Young |
| 0.83394 | 0.306409 | 0.208 | 0.221 | 1 | 7 | Prps2 | -0.013 | Young |
| 0.834406 | 0.213909 | 0.057 | 0.062 | 1 | 7 | Timmdc1 | -0.005 | Young |
| 0.834535 | 1.006922 | 0.03 | 0.033 | 1 | 7 | Efnb1 | -0.003 | Young |
| 0.834535 | 0.804 | 0.033 | 0.031 | 1 | 7 | Cd276 | 0.002 | Young |
| 0.83463 | 0.068891 | 0.476 | 0.479 | 1 | 7 | Sbno1 | -0.003 | Young |
| 0.834876 | 0.187288 | 0.111 | 0.108 | 1 | 7 | Hist3h2a | 0.003 | Young |
| 0.835127 | 0.088094 | 0.166 | 0.177 | 1 | 7 | Kif13a | -0.011 | Young |
| 0.835373 | 0.028305 | 0.235 | 0.244 | 1 | 7 | Zfp326 | -0.009 | Young |
| 0.835541 | 0.303055 | 0.13 | 0.128 | 1 | 7 | Ints14 | 0.002 | Young |
| 0.835647 | 0.167511 | 0.398 | 0.423 | 1 | 7 | Supt5 | -0.025 | Young |
| 0.836157 | 0.286351 | 0.057 | 0.054 | 1 | 7 | Dusp12 | 0.003 | Young |
| 0.836783 | 0.029176 | 0.431 | 0.464 | 1 | 7 | Taf1d | -0.033 | Young |
| 0.836818 | 0.132395 | 0.383 | 0.397 | 1 | 7 | Emc1 | -0.014 | Young |
| 0.836825 | 0.097633 | 0.34 | 0.344 | 1 | 7 | 1700025G( | -0.004 | Young |
| 0.836881 | 0.302577 | 0.214 | 0.226 | 1 | 7 | Afdn | -0.012 | Young |
| 0.837948 | 0.087777 | 0.536 | 0.587 | 1 | 7 | Brd2 | -0.051 | Young |
| 0.837964 | 0.349103 | 0.081 | 0.087 | 1 | 7 | Cad | -0.006 | Young |
| 0.838073 | 0.181734 | 0.229 | 0.241 | 1 | 7 | Gtf2b | -0.012 | Young |
| 0.838083 | 0.041942 | 0.419 | 0.438 | 1 | 7 | Pet100 | -0.019 | Young |
| 0.838116 | 0.053849 | 0.346 | 0.364 | 1 | 7 | Atp6v1h | -0.018 | Young |
| 0.838211 | 0.481016 | 0.033 | 0.031 | 1 | 7 | Ada | 0.002 | Young |
| 0.838513 | 0.018101 | 0.786 | 0.79 | 1 | 7 | Tmsb10 | -0.004 | Young |
| 0.838711 | 0.728209 | 0.151 | 0.162 | 1 | 7 | B4galt3 | -0.011 | Young |
| 0.839071 | 0.370091 | 0.172 | 0.169 | 1 | 7 | Ufd1 | 0.003 | Young |
| 0.839109 | 0.000961 | 0.181 | 0.174 | 1 | 7 | Uba3 | 0.007 | Young |
| 0.839119 | 0.049846 | 0.485 | 0.497 | 1 | 7 | Rbbp7 | -0.012 | Young |
| 0.839154 | 0.502764 | 0.084 | 0.09 | 1 | 7 | Ppcdc | -0.006 | Young |

| | | | | | | | | |
|---|---|---|---|---|---|---|---|---|
| 0.839182 | 0.066997 | 0.25 | 0.259 | 1 | 7 | Spata6 | -0.009 | Young |
| 0.839269 | 0.238245 | 0.145 | 0.154 | 1 | 7 | Cnnm4 | -0.009 | Young |
| 0.839314 | 0.002822 | 0.078 | 0.074 | 1 | 7 | Cc2d1a | 0.004 | Young |
| 0.839365 | 0.447522 | 0.12 | 0.118 | 1 | 7 | Ercc3 | 0.002 | Young |
| 0.83972 | 0.127447 | 0.145 | 0.154 | 1 | 7 | Apba1 | -0.009 | Young |
| 0.839866 | 0.35794 | 0.123 | 0.133 | 1 | 7 | Dctn6 | -0.01 | Young |
| 0.839983 | 0.171567 | 0.072 | 0.069 | 1 | 7 | Angptl4 | 0.003 | Young |
| 0.840219 | 0.303898 | 0.157 | 0.164 | 1 | 7 | Ss18l2 | -0.007 | Young |
| 0.840371 | 0.194525 | 0.096 | 0.103 | 1 | 7 | 2410004B1 | -0.007 | Young |
| 0.840725 | 0.098801 | 0.268 | 0.269 | 1 | 7 | Rraga | -0.001 | Young |
| 0.840793 | 0.029163 | 0.455 | 0.482 | 1 | 7 | Pgd | -0.027 | Young |
| 0.84101 | 0.578445 | 0.166 | 0.169 | 1 | 7 | Prpf6 | -0.003 | Young |
| 0.841288 | 0.22066 | 0.199 | 0.21 | 1 | 7 | Ufl1 | -0.011 | Young |
| 0.841364 | 0.121586 | 0.099 | 0.095 | 1 | 7 | Gm38399 | 0.004 | Young |
| 0.841494 | 0.054017 | 0.31 | 0.321 | 1 | 7 | Os9 | -0.011 | Young |
| 0.841619 | 0.260922 | 0.048 | 0.051 | 1 | 7 | Cdpf1 | -0.003 | Young |
| 0.841864 | 0.215449 | 0.033 | 0.036 | 1 | 7 | Fut7 | -0.003 | Young |
| 0.842117 | 0.239519 | 0.217 | 0.215 | 1 | 7 | Reps2 | 0.002 | Young |
| 0.842249 | 0.733445 | 0.081 | 0.079 | 1 | 7 | Dis3l | 0.002 | Young |
| 0.842681 | 0.129221 | 0.41 | 0.436 | 1 | 7 | Mpc2 | -0.026 | Young |
| 0.842729 | 0.193484 | 0.13 | 0.136 | 1 | 7 | Rnf138 | -0.006 | Young |
| 0.84289 | 0.144202 | 0.241 | 0.236 | 1 | 7 | Mindy3 | 0.005 | Young |
| 0.843127 | 0.09118 | 0.114 | 0.113 | 1 | 7 | Kif21b | 0.001 | Young |
| 0.843632 | 0.09188 | 0.033 | 0.036 | 1 | 7 | Herc3 | -0.003 | Young |
| 0.843882 | 0.17189 | 0.208 | 0.218 | 1 | 7 | Tdg | -0.01 | Young |
| 0.844128 | 0.363462 | 0.117 | 0.123 | 1 | 7 | Padi2 | -0.006 | Young |
| 0.844148 | 0.050868 | 0.377 | 0.397 | 1 | 7 | Pcmt1 | -0.02 | Young |
| 0.844331 | 0.107926 | 0.114 | 0.11 | 1 | 7 | Rcn3 | 0.004 | Young |
| 0.84468 | 0.181278 | 0.096 | 0.103 | 1 | 7 | Atpaf2 | -0.007 | Young |
| 0.844876 | 0.440506 | 0.084 | 0.09 | 1 | 7 | Man2c1 | -0.006 | Young |
| 0.844997 | 0.069096 | 0.193 | 0.203 | 1 | 7 | Hsd17b10 | -0.01 | Young |
| 0.845139 | 0.106112 | 0.22 | 0.226 | 1 | 7 | Atg2a | -0.006 | Young |
| 0.845391 | 0.07358 | 0.596 | 0.618 | 1 | 7 | Hnrnpl | -0.022 | Young |
| 0.845402 | 0.524031 | 0.033 | 0.036 | 1 | 7 | Dcaf4 | -0.003 | Young |
| 0.845835 | 0.253026 | 0.316 | 0.321 | 1 | 7 | Actr1a | -0.005 | Young |
| 0.84612 | 0.287074 | 0.157 | 0.154 | 1 | 7 | Fam219b | 0.003 | Young |
| 0.846174 | 0.174189 | 0.139 | 0.136 | 1 | 7 | Vezt | 0.003 | Young |
| 0.846253 | 0.053227 | 0.208 | 0.203 | 1 | 7 | Nsmce2 | 0.005 | Young |
| 0.846845 | 0.143966 | 0.178 | 0.185 | 1 | 7 | Kdm3b | -0.007 | Young |
| 0.846928 | 0.042005 | 0.087 | 0.085 | 1 | 7 | Jrkl | 0.002 | Young |
| 0.846931 | 0.207503 | 0.274 | 0.272 | 1 | 7 | Ccdc88b | 0.002 | Young |
| 0.846985 | 0.455511 | 0.154 | 0.151 | 1 | 7 | Necap1 | 0.003 | Young |
| 0.847059 | 0.254575 | 0.114 | 0.121 | 1 | 7 | Togaram1 | -0.007 | Young |
| 0.84712 | 0.460008 | 0.169 | 0.169 | 1 | 7 | Traf7 | 0 | Young |
| 0.847172 | 0.099929 | 0.033 | 0.036 | 1 | 7 | Map3k6 | -0.003 | Young |

| | | | | | | | | |
|---|---|---|---|---|---|---|---|---|
| 0.847291 | 0.273976 | 0.108 | 0.103 | 1 | 7 | Ercc6l2 | 0.005 | Young |
| 0.8474 | 0.358531 | 0.19 | 0.203 | 1 | 7 | Zdhhc7 | -0.013 | Young |
| 0.847674 | 0.112354 | 0.518 | 0.546 | 1 | 7 | Rab2a | -0.028 | Young |
| 0.847729 | 0.101049 | 0.31 | 0.338 | 1 | 7 | Rmnd5a | -0.028 | Young |
| 0.847877 | 0.324679 | 0.06 | 0.064 | 1 | 7 | Cyb561d2 | -0.004 | Young |
| 0.848 | 0.866816 | 0.048 | 0.046 | 1 | 7 | Galnt6 | 0.002 | Young |
| 0.848128 | 0.09718 | 0.328 | 0.359 | 1 | 7 | Vps29 | -0.031 | Young |
| 0.848175 | 0.066183 | 0.681 | 0.705 | 1 | 7 | Sumo2 | -0.024 | Young |
| 0.848253 | 0.275619 | 0.169 | 0.179 | 1 | 7 | Pias4 | -0.01 | Young |
| 0.848296 | 0.034118 | 0.669 | 0.679 | 1 | 7 | Macf1 | -0.01 | Young |
| 0.84834 | 0.492569 | 0.033 | 0.031 | 1 | 7 | Arap2 | 0.002 | Young |
| 0.848382 | 0.557839 | 0.045 | 0.049 | 1 | 7 | Sft2d3 | -0.004 | Young |
| 0.848484 | 0.108069 | 0.422 | 0.436 | 1 | 7 | Daglb | -0.014 | Young |
| 0.848824 | 0.132439 | 0.199 | 0.197 | 1 | 7 | Lsr | 0.002 | Young |
| 0.848841 | 0.24252 | 0.232 | 0.246 | 1 | 7 | Helz | -0.014 | Young |
| 0.848943 | 0.027981 | 0.033 | 0.036 | 1 | 7 | Ptch1 | -0.003 | Young |
| 0.849378 | 0.536274 | 0.102 | 0.1 | 1 | 7 | Fam214a | 0.002 | Young |
| 0.849413 | 0.136791 | 0.139 | 0.146 | 1 | 7 | Smurf1 | -0.007 | Young |
| 0.849659 | 0.093803 | 0.021 | 0.023 | 1 | 7 | Gm15445 | -0.002 | Young |
| 0.849766 | 0.063403 | 0.328 | 0.338 | 1 | 7 | Nrbp1 | -0.01 | Young |
| 0.849885 | 0.498466 | 0.06 | 0.064 | 1 | 7 | E130308A1 | -0.004 | Young |
| 0.849962 | 0.573913 | 0.096 | 0.095 | 1 | 7 | Tmem110 | 0.001 | Young |
| 0.850494 | 0.157392 | 0.247 | 0.256 | 1 | 7 | Usp24 | -0.009 | Young |
| 0.850584 | 0.111535 | 0.307 | 0.328 | 1 | 7 | AB124611 | -0.021 | Young |
| 0.850676 | 0.315963 | 0.157 | 0.156 | 1 | 7 | 1110059G1 | 0.001 | Young |
| 0.850705 | 0.09847 | 0.575 | 0.592 | 1 | 7 | Dnajb6 | -0.017 | Young |
| 0.850823 | 0.134212 | 0.449 | 0.454 | 1 | 7 | Socs3 | -0.005 | Young |
| 0.850964 | 0.003054 | 0.84 | 0.869 | 1 | 7 | Calm2 | -0.029 | Young |
| 0.851601 | 0.109285 | 0.033 | 0.036 | 1 | 7 | Sdhaf3 | -0.003 | Young |
| 0.851845 | 0.017897 | 0.696 | 0.715 | 1 | 7 | Atrx | -0.019 | Young |
| 0.851965 | 0.04876 | 0.136 | 0.141 | 1 | 7 | Dusp7 | -0.005 | Young |
| 0.851975 | 0.20967 | 0.395 | 0.408 | 1 | 7 | Larp4 | -0.013 | Young |
| 0.852209 | 0.093347 | 0.244 | 0.244 | 1 | 7 | Stard9 | 0 | Young |
| 0.852403 | 0.039528 | 0.286 | 0.297 | 1 | 7 | Ccdc124 | -0.011 | Young |
| 0.852587 | 1.011368 | 0.048 | 0.046 | 1 | 7 | Slc2a3 | 0.002 | Young |
| 0.852633 | 0.339096 | 0.16 | 0.159 | 1 | 7 | Map3k14 | 0.001 | Young |
| 0.852822 | 0.083757 | 0.175 | 0.169 | 1 | 7 | Zw10 | 0.006 | Young |
| 0.852899 | 0.078644 | 0.084 | 0.09 | 1 | 7 | Gm45716 | -0.006 | Young |
| 0.853093 | 0.222031 | 0.069 | 0.067 | 1 | 7 | Agmo | 0.002 | Young |
| 0.853253 | 0.115611 | 0.087 | 0.092 | 1 | 7 | Gtf2h4 | -0.005 | Young |
| 0.853374 | 0.280782 | 0.033 | 0.036 | 1 | 7 | Zfp995 | -0.003 | Young |
| 0.853392 | 0.430046 | 0.178 | 0.192 | 1 | 7 | Nudt9 | -0.014 | Young |
| 0.853614 | 0.154001 | 0.145 | 0.151 | 1 | 7 | Frmd4a | -0.006 | Young |
| 0.853876 | 0.177038 | 0.033 | 0.031 | 1 | 7 | Kdm4b | 0.002 | Young |
| 0.853928 | 0.113052 | 0.093 | 0.1 | 1 | 7 | Il18 | -0.007 | Young |

| | | | | | | | | |
|---|---|---|---|---|---|---|---|---|
| 0.854031 | 0.454619 | 0.099 | 0.097 | 1 | 7 | Pfas | 0.002 | Young |
| 0.854046 | 0.106704 | 0.084 | 0.09 | 1 | 7 | 2410089E0 | -0.006 | Young |
| 0.854222 | 0.247885 | 0.485 | 0.495 | 1 | 7 | Zcchc11 | -0.01 | Young |
| 0.854447 | 0.109075 | 0.536 | 0.579 | 1 | 7 | Dock8 | -0.043 | Young |
| 0.854721 | 0.237154 | 0.226 | 0.241 | 1 | 7 | 5031425E2 | -0.015 | Young |
| 0.854816 | 0.114561 | 0.37 | 0.372 | 1 | 7 | Dcaf7 | -0.002 | Young |
| 0.855013 | 0.338027 | 0.184 | 0.182 | 1 | 7 | Sord | 0.002 | Young |
| 0.856894 | 0.19246 | 0.337 | 0.349 | 1 | 7 | Nap1l4 | -0.012 | Young |
| 0.856922 | 0.995972 | 0.033 | 0.036 | 1 | 7 | A830010M | -0.003 | Young |
| 0.857013 | 0.651217 | 0.03 | 0.028 | 1 | 7 | B3gnt5 | 0.002 | Young |
| 0.857027 | 0.143784 | 0.072 | 0.077 | 1 | 7 | Ift80 | -0.005 | Young |
| 0.857037 | 0.057593 | 0.373 | 0.382 | 1 | 7 | Leprotl1 | -0.009 | Young |
| 0.857312 | 0.58029 | 0.148 | 0.156 | 1 | 7 | Ptrh2 | -0.008 | Young |
| 0.857603 | 0.22227 | 0.069 | 0.067 | 1 | 7 | Phlpp1 | 0.002 | Young |
| 0.857613 | 0.665602 | 0.051 | 0.049 | 1 | 7 | Gramd3 | 0.002 | Young |
| 0.857667 | 0.096342 | 0.202 | 0.2 | 1 | 7 | Tmem170b | 0.002 | Young |
| 0.857755 | 0.126486 | 0.346 | 0.351 | 1 | 7 | Kmt2c | -0.005 | Young |
| 0.857809 | 0.174329 | 0.033 | 0.036 | 1 | 7 | Polr1e | -0.003 | Young |
| 0.858092 | 0.01719 | 0.175 | 0.182 | 1 | 7 | Med17 | -0.007 | Young |
| 0.858517 | 0.276411 | 0.226 | 0.233 | 1 | 7 | Tmem57 | -0.007 | Young |
| 0.858616 | 0.12908 | 0.145 | 0.151 | 1 | 7 | Rpp25l | -0.006 | Young |
| 0.858735 | 0.534225 | 0.123 | 0.131 | 1 | 7 | Naip5 | -0.008 | Young |
| 0.858925 | 0.276413 | 0.214 | 0.215 | 1 | 7 | Spin1 | -0.001 | Young |
| 0.859076 | 0.297093 | 0.283 | 0.285 | 1 | 7 | Slirp | -0.002 | Young |
| 0.859145 | 0.317676 | 0.256 | 0.259 | 1 | 7 | Chml | -0.003 | Young |
| 0.859585 | 0.745253 | 0.033 | 0.036 | 1 | 7 | Btbd19 | -0.003 | Young |
| 0.859959 | 0.246019 | 0.301 | 0.308 | 1 | 7 | Pja2 | -0.007 | Young |
| 0.86007 | 0.000591 | 0.151 | 0.149 | 1 | 7 | Inip | 0.002 | Young |
| 0.860233 | 0.089122 | 0.105 | 0.113 | 1 | 7 | Fastk | -0.008 | Young |
| 0.860363 | 0.541103 | 0.084 | 0.09 | 1 | 7 | Adi1 | -0.006 | Young |
| 0.860412 | 0.026148 | 0.813 | 0.849 | 1 | 7 | Stmn1 | -0.036 | Young |
| 0.860602 | 0.052172 | 0.702 | 0.733 | 1 | 7 | Rock2 | -0.031 | Young |
| 0.860677 | 0.117794 | 0.021 | 0.023 | 1 | 7 | Spry1 | -0.002 | Young |
| 0.860758 | 0.423021 | 0.066 | 0.064 | 1 | 7 | Gtse1 | 0.002 | Young |
| 0.860779 | 0.164078 | 0.09 | 0.087 | 1 | 7 | Stxbp1 | 0.003 | Young |
| 0.860924 | 0.26281 | 0.307 | 0.31 | 1 | 7 | Hs6st1 | -0.003 | Young |
| 0.861029 | 0.124778 | 0.142 | 0.151 | 1 | 7 | Phyh | -0.009 | Young |
| 0.861267 | 0.044194 | 0.033 | 0.031 | 1 | 7 | Tm9sf1 | 0.002 | Young |
| 0.861473 | 0.089132 | 0.114 | 0.113 | 1 | 7 | Tmem203 | 0.001 | Young |
| 0.861625 | 0.005285 | 0.633 | 0.659 | 1 | 7 | Dpysl2 | -0.026 | Young |
| 0.861849 | 0.352623 | 0.187 | 0.195 | 1 | 7 | Mrps28 | -0.008 | Young |
| 0.862074 | 0.955544 | 0.066 | 0.064 | 1 | 7 | Bod1 | 0.002 | Young |
| 0.86217 | 0.316903 | 0.157 | 0.154 | 1 | 7 | Rnf169 | 0.003 | Young |
| 0.862185 | 0.263449 | 0.226 | 0.238 | 1 | 7 | Elovl1 | -0.012 | Young |
| 0.862192 | 0.306804 | 0.099 | 0.097 | 1 | 7 | Scn1b | 0.002 | Young |

| | | | | | | | | |
|---|---|---|---|---|---|---|---|---|
| 0.86231 | 0.455256 | 0.105 | 0.113 | 1 | 7 | Cdr2 | -0.008 | Young |
| 0.862809 | 0.898636 | 0.03 | 0.028 | 1 | 7 | Lif | 0.002 | Young |
| 0.862829 | 0.041343 | 0.111 | 0.108 | 1 | 7 | Rubcn | 0.003 | Young |
| 0.862836 | 0.04775 | 0.807 | 0.821 | 1 | 7 | Eef1g | -0.014 | Young |
| 0.862884 | 0.135055 | 0.259 | 0.259 | 1 | 7 | Eif4e3 | 0 | Young |
| 0.86289 | 0.073284 | 0.557 | 0.597 | 1 | 7 | Arid1a | -0.04 | Young |
| 0.862939 | 0.155676 | 0.178 | 0.187 | 1 | 7 | Wdr76 | -0.009 | Young |
| 0.862976 | 0.073255 | 0.467 | 0.492 | 1 | 7 | Ndufa11 | -0.025 | Young |
| 0.863579 | 0.30606 | 0.051 | 0.049 | 1 | 7 | Pnpla6 | 0.002 | Young |
| 0.863579 | 0.169114 | 0.051 | 0.049 | 1 | 7 | Plpp6 | 0.002 | Young |
| 0.863776 | 0.319069 | 0.03 | 0.028 | 1 | 7 | Lppos | 0.002 | Young |
| 0.863843 | 0.156915 | 0.31 | 0.326 | 1 | 7 | Cklf | -0.016 | Young |
| 0.863923 | 0.024526 | 0.66 | 0.656 | 1 | 7 | Ndufb11 | 0.004 | Young |
| 0.863925 | 0.214201 | 0.307 | 0.318 | 1 | 7 | Ogfrl1 | -0.011 | Young |
| 0.864043 | 0.615644 | 0.033 | 0.031 | 1 | 7 | Zfp747 | 0.002 | Young |
| 0.864202 | 0.08841 | 0.307 | 0.331 | 1 | 7 | Gdi1 | -0.024 | Young |
| 0.864331 | 0.24768 | 0.386 | 0.387 | 1 | 7 | Hip1 | -0.001 | Young |
| 0.864554 | 0.090315 | 0.166 | 0.172 | 1 | 7 | Rab9 | -0.006 | Young |
| 0.864579 | 0.06354 | 0.355 | 0.379 | 1 | 7 | Ube2m | -0.024 | Young |
| 0.864622 | 0.103106 | 0.304 | 0.305 | 1 | 7 | Napsa | -0.001 | Young |
| 0.864699 | 0.218429 | 0.048 | 0.051 | 1 | 7 | Gtf2h2 | -0.003 | Young |
| 0.864778 | 0.118501 | 0.22 | 0.233 | 1 | 7 | Iah1 | -0.013 | Young |
| 0.864844 | 0.403104 | 0.048 | 0.046 | 1 | 7 | Fig4 | 0.002 | Young |
| 0.865352 | 0.518842 | 0.111 | 0.113 | 1 | 7 | Fkrp | -0.002 | Young |
| 0.865366 | 0.30842 | 0.066 | 0.064 | 1 | 7 | Bloc1s5 | 0.002 | Young |
| 0.865587 | 0.002817 | 0.256 | 0.267 | 1 | 7 | Cacul1 | -0.011 | Young |
| 0.865675 | 0.018503 | 0.145 | 0.144 | 1 | 7 | Lonp1 | 0.001 | Young |
| 0.86627 | 0.140194 | 0.111 | 0.118 | 1 | 7 | Arfgef2 | -0.007 | Young |
| 0.866338 | 0.033773 | 0.292 | 0.3 | 1 | 7 | Ubqln1 | -0.008 | Young |
| 0.866374 | 0.237807 | 0.277 | 0.292 | 1 | 7 | Spty2d1 | -0.015 | Young |
| 0.866406 | 0.157578 | 0.136 | 0.133 | 1 | 7 | Bend4 | 0.003 | Young |
| 0.866637 | 0.003482 | 0.199 | 0.195 | 1 | 7 | Armcx3 | 0.004 | Young |
| 0.866639 | 0.179657 | 0.066 | 0.069 | 1 | 7 | Mrps31 | -0.003 | Young |
| 0.866678 | 0.267501 | 0.03 | 0.028 | 1 | 7 | Zfp658 | 0.002 | Young |
| 0.866813 | 0.40843 | 0.009 | 0.01 | 1 | 7 | Zfp3 | -0.001 | Young |
| 0.866813 | 0.076542 | 0.009 | 0.01 | 1 | 7 | Klhdc8b | -0.001 | Young |
| 0.866899 | 0.306662 | 0.072 | 0.077 | 1 | 7 | 6720427I0 | -0.005 | Young |
| 0.866931 | 0.010387 | 0.819 | 0.81 | 1 | 7 | Rpl4 | 0.009 | Young |
| 0.866962 | 0.086138 | 0.066 | 0.069 | 1 | 7 | Tab1 | -0.003 | Young |
| 0.866986 | 0.166972 | 0.346 | 0.362 | 1 | 7 | Tnfrsf1a | -0.016 | Young |
| 0.867238 | 0.014314 | 0.22 | 0.228 | 1 | 7 | Tpcn1 | -0.008 | Young |
| 0.867301 | 0.157737 | 0.021 | 0.023 | 1 | 7 | Zc3h11a | -0.002 | Young |
| 0.867583 | 0.251945 | 0.033 | 0.036 | 1 | 7 | Sap18b | -0.003 | Young |
| 0.868473 | 0.682624 | 0.009 | 0.01 | 1 | 7 | Gm49339 | -0.001 | Young |
| 0.868473 | 0.290441 | 0.009 | 0.01 | 1 | 7 | Gm14325 | -0.001 | Young |

| | | | | | | | | |
|---|---|---|---|---|---|---|---|---|
| 0.868476 | 0.547609 | 0.081 | 0.079 | 1 | 7 | Mrgbp | 0.002 | Young |
| 0.868616 | 0.04467 | 0.22 | 0.233 | 1 | 7 | Relb | -0.013 | Young |
| 0.868663 | 0.130374 | 0.377 | 0.379 | 1 | 7 | Wdfy4 | -0.002 | Young |
| 0.868677 | 0.073422 | 0.06 | 0.064 | 1 | 7 | Nt5m | -0.004 | Young |
| 0.869636 | 0.206155 | 0.223 | 0.223 | 1 | 7 | Hras | 0 | Young |
| 0.869699 | 0.549062 | 0.078 | 0.077 | 1 | 7 | Phospho2 | 0.001 | Young |
| 0.870133 | 1.003148 | 0.009 | 0.01 | 1 | 7 | Dzip1 | -0.001 | Young |
| 0.870133 | 0.083292 | 0.009 | 0.01 | 1 | 7 | Gm27029 | -0.001 | Young |
| 0.870195 | 0.110257 | 0.31 | 0.305 | 1 | 7 | Ppt1 | 0.005 | Young |
| 0.870216 | 0.435958 | 0.048 | 0.046 | 1 | 7 | Noa1 | 0.002 | Young |
| 0.870295 | 0.296724 | 0.244 | 0.249 | 1 | 7 | Rpp21 | -0.005 | Young |
| 0.870435 | 0.325797 | 0.066 | 0.072 | 1 | 7 | Fbxl18 | -0.006 | Young |
| 0.870541 | 0.102996 | 0.386 | 0.415 | 1 | 7 | Ssu72 | -0.029 | Young |
| 0.870697 | 0.238511 | 0.325 | 0.326 | 1 | 7 | Rela | -0.001 | Young |
| 0.870849 | 0.386215 | 0.111 | 0.118 | 1 | 7 | Siah1a | -0.007 | Young |
| 0.871156 | 0.166792 | 0.214 | 0.226 | 1 | 7 | BC035044 | -0.012 | Young |
| 0.871211 | 0.189833 | 0.301 | 0.31 | 1 | 7 | Ubr4 | -0.009 | Young |
| 0.871322 | 0.006229 | 0.467 | 0.469 | 1 | 7 | AC149090. | -0.002 | Young |
| 0.871412 | 0.081702 | 0.401 | 0.418 | 1 | 7 | Gna12 | -0.017 | Young |
| 0.871436 | 0.19138 | 0.355 | 0.354 | 1 | 7 | Gpcpd1 | 0.001 | Young |
| 0.871653 | 0.120896 | 0.169 | 0.177 | 1 | 7 | Cetn2 | -0.008 | Young |
| 0.871722 | 0.134655 | 0.295 | 0.295 | 1 | 7 | Krt80 | 0 | Young |
| 0.871794 | 0.168158 | 0.009 | 0.01 | 1 | 7 | Tmlhe | -0.001 | Young |
| 0.871794 | 0.099587 | 0.009 | 0.01 | 1 | 7 | Adap2os | -0.001 | Young |
| 0.871955 | 0.368527 | 0.111 | 0.115 | 1 | 7 | Tex2 | -0.004 | Young |
| 0.872134 | 0.166549 | 0.184 | 0.182 | 1 | 7 | Cpsf3 | 0.002 | Young |
| 0.872205 | 0.150651 | 0.09 | 0.087 | 1 | 7 | Zbtb41 | 0.003 | Young |
| 0.872487 | 0.259397 | 0.03 | 0.028 | 1 | 7 | Gm30074 | 0.002 | Young |
| 0.872821 | 0.107535 | 0.286 | 0.308 | 1 | 7 | Fam76b | -0.022 | Young |
| 0.873004 | 0.270167 | 0.181 | 0.182 | 1 | 7 | Bzw2 | -0.001 | Young |
| 0.873063 | 0.102482 | 0.133 | 0.136 | 1 | 7 | Vhl | -0.003 | Young |
| 0.873289 | 0.321183 | 0.048 | 0.046 | 1 | 7 | Efcab11 | 0.002 | Young |
| 0.873456 | 0.722108 | 0.009 | 0.01 | 1 | 7 | Gm9774 | -0.001 | Young |
| 0.873456 | 0.59117 | 0.009 | 0.01 | 1 | 7 | 2010015M | -0.001 | Young |
| 0.873456 | 0.373462 | 0.009 | 0.01 | 1 | 7 | Taco1os | -0.001 | Young |
| 0.873456 | 0.362428 | 0.009 | 0.01 | 1 | 7 | Ffar4 | -0.001 | Young |
| 0.873456 | 0.227794 | 0.009 | 0.01 | 1 | 7 | Rsph3a | -0.001 | Young |
| 0.873456 | 0.10016 | 0.009 | 0.01 | 1 | 7 | Mst1r | -0.001 | Young |
| 0.874285 | 0.137613 | 0.226 | 0.226 | 1 | 7 | Tmem64 | 0 | Young |
| 0.874414 | 0.52982 | 0.048 | 0.051 | 1 | 7 | 2900093K2 | -0.003 | Young |
| 0.874425 | 0.255047 | 0.03 | 0.028 | 1 | 7 | Zbtb11os1 | 0.002 | Young |
| 0.874611 | 0.059307 | 0.114 | 0.11 | 1 | 7 | Kif1bp | 0.004 | Young |
| 0.874624 | 0.119843 | 0.283 | 0.297 | 1 | 7 | Cnpy2 | -0.014 | Young |
| 0.874779 | 0.019351 | 0.087 | 0.09 | 1 | 7 | Alg1 | -0.003 | Young |
| 0.875118 | 0.917899 | 0.009 | 0.01 | 1 | 7 | Gm49085 | -0.001 | Young |

| | | | | | | | | |
|---|---|---|---|---|---|---|---|---|
| 0.875118 | 0.111194 | 0.009 | 0.01 | 1 | 7 | Gpd1 | -0.001 | Young |
| 0.875133 | 0.404085 | 0.169 | 0.169 | 1 | 7 | Strn | 0 | Young |
| 0.875217 | 0.30491 | 0.262 | 0.269 | 1 | 7 | Lars | -0.007 | Young |
| 0.875295 | 0.362884 | 0.087 | 0.085 | 1 | 7 | AW554918 | 0.002 | Young |
| 0.875386 | 0.172871 | 0.154 | 0.162 | 1 | 7 | Mast2 | -0.008 | Young |
| 0.875532 | 0.154129 | 0.476 | 0.513 | 1 | 7 | Dnaja2 | -0.037 | Young |
| 0.875537 | 0.904504 | 0.045 | 0.044 | 1 | 7 | Dhodh | 0.001 | Young |
| 0.875657 | 0.192416 | 0.271 | 0.277 | 1 | 7 | Fryl | -0.006 | Young |
| 0.875662 | 0.220977 | 0.172 | 0.179 | 1 | 7 | Shcbp1 | -0.007 | Young |
| 0.875778 | 0.310571 | 0.193 | 0.2 | 1 | 7 | Map3k4 | -0.007 | Young |
| 0.875794 | 0.047945 | 0.036 | 0.038 | 1 | 7 | Taf1a | -0.002 | Young |
| 0.875828 | 0.584368 | 0.096 | 0.1 | 1 | 7 | Rnf34 | -0.004 | Young |
| 0.875979 | 0.449083 | 0.045 | 0.049 | 1 | 7 | Mkks | -0.004 | Young |
| 0.876147 | 0.409052 | 0.021 | 0.023 | 1 | 7 | Klhl42 | -0.002 | Young |
| 0.876343 | 0.026192 | 0.102 | 0.108 | 1 | 7 | Smox | -0.006 | Young |
| 0.876364 | 1.050314 | 0.03 | 0.028 | 1 | 7 | Rbks | 0.002 | Young |
| 0.876364 | 0.291524 | 0.03 | 0.028 | 1 | 7 | Dffb | 0.002 | Young |
| 0.876412 | 0.054601 | 0.268 | 0.282 | 1 | 7 | Atp13a2 | -0.014 | Young |
| 0.876491 | 0.429354 | 0.093 | 0.092 | 1 | 7 | Atg14 | 0.001 | Young |
| 0.876646 | 0.073453 | 0.172 | 0.167 | 1 | 7 | Ankra2 | 0.005 | Young |
| 0.876728 | 0.011255 | 0.13 | 0.126 | 1 | 7 | Ikbip | 0.004 | Young |
| 0.876781 | 0.965813 | 0.009 | 0.01 | 1 | 7 | Tnfaip6 | -0.001 | Young |
| 0.876939 | 0.277724 | 0.154 | 0.159 | 1 | 7 | Gclm | -0.005 | Young |
| 0.876951 | 0.116989 | 0.407 | 0.413 | 1 | 7 | Morc3 | -0.006 | Young |
| 0.877032 | 0.538047 | 0.051 | 0.049 | 1 | 7 | Unk | 0.002 | Young |
| 0.877043 | 0.012353 | 0.136 | 0.131 | 1 | 7 | 2410002F2 | 0.005 | Young |
| 0.87709 | 0.015463 | 0.563 | 0.569 | 1 | 7 | Fyb | -0.006 | Young |
| 0.877095 | 0.245414 | 0.142 | 0.138 | 1 | 7 | Mavs | 0.004 | Young |
| 0.877103 | 0.094537 | 0.467 | 0.487 | 1 | 7 | Sptbn1 | -0.02 | Young |
| 0.87712 | 0.772386 | 0.045 | 0.044 | 1 | 7 | Zfp653 | 0.001 | Young |
| 0.877451 | 0.686843 | 0.078 | 0.082 | 1 | 7 | 3110082I1 | -0.004 | Young |
| 0.87766 | 0.102257 | 0.5 | 0.526 | 1 | 7 | Nisch | -0.026 | Young |
| 0.877912 | 0.524234 | 0.045 | 0.044 | 1 | 7 | Nepro | 0.001 | Young |
| 0.878303 | 0.225904 | 0.03 | 0.028 | 1 | 7 | Peli2 | 0.002 | Young |
| 0.878361 | 0.755176 | 0.021 | 0.023 | 1 | 7 | Cnbd2 | -0.002 | Young |
| 0.878393 | 0.262961 | 0.027 | 0.026 | 1 | 7 | F11r | 0.001 | Young |
| 0.878398 | 0.219698 | 0.485 | 0.51 | 1 | 7 | Cerk | -0.025 | Young |
| 0.878422 | 0.318588 | 0.265 | 0.274 | 1 | 7 | Vamp4 | -0.009 | Young |
| 0.878444 | 0.592973 | 0.009 | 0.01 | 1 | 7 | Zfp202 | -0.001 | Young |
| 0.878492 | 0.277842 | 0.19 | 0.2 | 1 | 7 | Nup133 | -0.01 | Young |
| 0.878623 | 0.152488 | 0.328 | 0.336 | 1 | 7 | Hcfc1r1 | -0.008 | Young |
| 0.879406 | 0.019636 | 0.852 | 0.841 | 1 | 7 | Tmbim6 | 0.011 | Young |
| 0.879533 | 0.020001 | 0.196 | 0.208 | 1 | 7 | Aco1 | -0.012 | Young |
| 0.879556 | 0.033935 | 0.16 | 0.167 | 1 | 7 | Espn | -0.007 | Young |
| 0.87971 | 0.054858 | 0.355 | 0.351 | 1 | 7 | Ugcg | 0.004 | Young |

| | | | | | | | | |
|---|---|---|---|---|---|---|---|---|
| 0.879807 | 0.106374 | 0.765 | 0.787 | 1 | 7 | Ddx3x | -0.022 | Young |
| 0.880129 | 0.511435 | 0.099 | 0.1 | 1 | 7 | Map4k2 | -0.001 | Young |
| 0.880287 | 0.066952 | 0.623 | 0.672 | 1 | 7 | Dhrs3 | -0.049 | Young |
| 0.880287 | 0.589502 | 0.045 | 0.044 | 1 | 7 | Eme1 | 0.001 | Young |
| 0.880311 | 0.207008 | 0.211 | 0.213 | 1 | 7 | Mbd6 | -0.002 | Young |
| 0.880328 | 0.131116 | 0.367 | 0.397 | 1 | 7 | Washc4 | -0.03 | Young |
| 0.880376 | 0.061615 | 0.286 | 0.287 | 1 | 7 | Pdzd8 | -0.001 | Young |
| 0.880393 | 0.238078 | 0.13 | 0.136 | 1 | 7 | Ankrd27 | -0.006 | Young |
| 0.880418 | 0.646463 | 0.093 | 0.092 | 1 | 7 | Zmym1 | 0.001 | Young |
| 0.880421 | 0.22151 | 0.334 | 0.341 | 1 | 7 | Fam172a | -0.007 | Young |
| 0.880448 | 0.234702 | 0.581 | 0.6 | 1 | 7 | Tor1aip1 | -0.019 | Young |
| 0.880912 | 0.052049 | 0.819 | 0.849 | 1 | 7 | Rbm39 | -0.03 | Young |
| 0.881384 | 0.148366 | 0.229 | 0.228 | 1 | 7 | Arl5c | 0.001 | Young |
| 0.88152 | 0.231143 | 0.069 | 0.067 | 1 | 7 | Cep152 | 0.002 | Young |
| 0.881661 | 0.180371 | 0.458 | 0.479 | 1 | 7 | Hnrnpr | -0.021 | Young |
| 0.88231 | 0.175796 | 0.051 | 0.054 | 1 | 7 | Dedd | -0.003 | Young |
| 0.882519 | 0.365352 | 0.048 | 0.046 | 1 | 7 | Flrt3 | 0.002 | Young |
| 0.882838 | 0.029211 | 0.672 | 0.692 | 1 | 7 | Atp5j | -0.02 | Young |
| 0.883301 | 0.114707 | 0.244 | 0.246 | 1 | 7 | Cmc1 | -0.002 | Young |
| 0.883405 | 0.182545 | 0.084 | 0.09 | 1 | 7 | Nbeal2 | -0.006 | Young |
| 0.883479 | 0.151324 | 0.178 | 0.177 | 1 | 7 | Cds2 | 0.001 | Young |
| 0.883487 | 0.862844 | 0.027 | 0.026 | 1 | 7 | Setd4 | 0.001 | Young |
| 0.88371 | 0.122013 | 0.19 | 0.19 | 1 | 7 | Dlgap4 | 0 | Young |
| 0.883744 | 0.179694 | 0.078 | 0.082 | 1 | 7 | Ubxn2b | -0.004 | Young |
| 0.883838 | 0.186147 | 0.295 | 0.313 | 1 | 7 | Mprip | -0.018 | Young |
| 0.883954 | 0.076763 | 0.117 | 0.123 | 1 | 7 | Mfsd6 | -0.006 | Young |
| 0.883966 | 0.145977 | 0.551 | 0.585 | 1 | 7 | Manf | -0.034 | Young |
| 0.884102 | 0.188218 | 0.319 | 0.328 | 1 | 7 | Smu1 | -0.009 | Young |
| 0.88413 | 0.195731 | 0.289 | 0.292 | 1 | 7 | St6gal1 | -0.003 | Young |
| 0.884199 | 0.10406 | 0.262 | 0.262 | 1 | 7 | 8-Sep | 0 | Young |
| 0.884392 | 0.519692 | 0.093 | 0.1 | 1 | 7 | Snapc3 | -0.007 | Young |
| 0.884444 | 0.147832 | 0.175 | 0.174 | 1 | 7 | Parp4 | 0.001 | Young |
| 0.884522 | 0.070438 | 0.051 | 0.049 | 1 | 7 | Rfx3 | 0.002 | Young |
| 0.884643 | 0.039997 | 0.476 | 0.492 | 1 | 7 | Egr2 | -0.016 | Young |
| 0.885097 | 0.008034 | 0.03 | 0.028 | 1 | 7 | Trim2 | 0.002 | Young |
| 0.885229 | 0.148107 | 0.196 | 0.195 | 1 | 7 | Sdad1 | 0.001 | Young |
| 0.88525 | 0.232197 | 0.214 | 0.228 | 1 | 7 | Qars | -0.014 | Young |
| 0.886117 | 0.199546 | 0.235 | 0.236 | 1 | 7 | Tmem173 | -0.001 | Young |
| 0.886131 | 0.102668 | 0.5 | 0.528 | 1 | 7 | Ctbp1 | -0.028 | Young |
| 0.886229 | 0.283317 | 0.096 | 0.092 | 1 | 7 | Tradd | 0.004 | Young |
| 0.887124 | 0.66324 | 0.12 | 0.128 | 1 | 7 | Tom1 | -0.008 | Young |
| 0.887134 | 0.094304 | 0.292 | 0.315 | 1 | 7 | Pold3 | -0.023 | Young |
| 0.887145 | 0.328939 | 0.048 | 0.051 | 1 | 7 | Lrrc14 | -0.003 | Young |
| 0.887349 | 0.202576 | 0.307 | 0.336 | 1 | 7 | Exoc3 | -0.029 | Young |
| 0.887629 | 0.065638 | 0.437 | 0.449 | 1 | 7 | Higd1a | -0.012 | Young |

| | | | | | | | | |
|---|---|---|---|---|---|---|---|---|
| 0.888251 | 0.044237 | 0.127 | 0.133 | 1 | 7 | Cxxc1 | -0.006 | Young |
| 0.888455 | 0.248542 | 0.117 | 0.123 | 1 | 7 | Mthfsl | -0.006 | Young |
| 0.888544 | 0.264823 | 0.241 | 0.249 | 1 | 7 | Plagl2 | -0.008 | Young |
| 0.888634 | 0.125204 | 0.093 | 0.097 | 1 | 7 | Pld2 | -0.004 | Young |
| 0.888841 | 0.064944 | 0.09 | 0.095 | 1 | 7 | Gatsl2 | -0.005 | Young |
| 0.889508 | 0.328535 | 0.346 | 0.356 | 1 | 7 | Rc3h2 | -0.01 | Young |
| 0.88998 | 0.25128 | 0.154 | 0.154 | 1 | 7 | Mycbp | 0 | Young |
| 0.890003 | 0.435807 | 0.169 | 0.179 | 1 | 7 | Suv39h1 | -0.01 | Young |
| 0.890097 | 0.29876 | 0.187 | 0.187 | 1 | 7 | Zbtb38 | 0 | Young |
| 0.890158 | 0.105644 | 0.148 | 0.144 | 1 | 7 | Taf7 | 0.004 | Young |
| 0.890233 | 0.056328 | 0.328 | 0.338 | 1 | 7 | Specc1l | -0.01 | Young |
| 0.89024 | 0.23158 | 0.304 | 0.31 | 1 | 7 | Srek1 | -0.006 | Young |
| 0.890409 | 0.237628 | 0.036 | 0.038 | 1 | 7 | Zfp408 | -0.002 | Young |
| 0.890594 | 0.442049 | 0.045 | 0.044 | 1 | 7 | Ift57 | 0.001 | Young |
| 0.890704 | 0.151701 | 0.446 | 0.49 | 1 | 7 | Adam17 | -0.044 | Young |
| 0.890852 | 0.162691 | 0.331 | 0.341 | 1 | 7 | Szrd1 | -0.01 | Young |
| 0.890926 | 0.079299 | 0.03 | 0.028 | 1 | 7 | Bco2 | 0.002 | Young |
| 0.89095 | 0.188774 | 0.265 | 0.285 | 1 | 7 | Trim35 | -0.02 | Young |
| 0.890955 | 0.420907 | 0.214 | 0.231 | 1 | 7 | Cramp1l | -0.017 | Young |
| 0.891628 | 0.011231 | 0.102 | 0.105 | 1 | 7 | Zfp414 | -0.003 | Young |
| 0.892129 | 0.219621 | 0.133 | 0.131 | 1 | 7 | N4bp2 | 0.002 | Young |
| 0.892181 | 0.667764 | 0.045 | 0.044 | 1 | 7 | Zfp637 | 0.001 | Young |
| 0.892294 | 0.124424 | 0.467 | 0.485 | 1 | 7 | 2-Mar | -0.018 | Young |
| 0.892438 | 0.091571 | 0.063 | 0.067 | 1 | 7 | Zbtb5 | -0.004 | Young |
| 0.892521 | 0.36552 | 0.139 | 0.146 | 1 | 7 | Fam45a | -0.007 | Young |
| 0.892536 | 0.055873 | 0.048 | 0.046 | 1 | 7 | Acpp | 0.002 | Young |
| 0.892566 | 0.110205 | 0.295 | 0.3 | 1 | 7 | Ola1 | -0.005 | Young |
| 0.892649 | 0.200518 | 0.298 | 0.318 | 1 | 7 | Crat | -0.02 | Young |
| 0.892846 | 0.216956 | 0.325 | 0.333 | 1 | 7 | Cnot3 | -0.008 | Young |
| 0.893628 | 0.531842 | 0.096 | 0.095 | 1 | 7 | Tfb2m | 0.001 | Young |
| 0.893629 | 0.215889 | 0.38 | 0.385 | 1 | 7 | Ncapg2 | -0.005 | Young |
| 0.893762 | 0.320039 | 0.066 | 0.064 | 1 | 7 | Tfpt | 0.002 | Young |
| 0.893769 | 0.672505 | 0.045 | 0.044 | 1 | 7 | 1600020E0 | 0.001 | Young |
| 0.894431 | 0.053702 | 0.096 | 0.1 | 1 | 7 | Snhg20 | -0.004 | Young |
| 0.894563 | 0.168946 | 0.045 | 0.044 | 1 | 7 | Zfp961 | 0.001 | Young |
| 0.894569 | 0.355502 | 0.187 | 0.19 | 1 | 7 | Patl1 | -0.003 | Young |
| 0.894595 | 0.093771 | 0.762 | 0.79 | 1 | 7 | Ddx17 | -0.028 | Young |
| 0.894759 | 0.24247 | 0.295 | 0.315 | 1 | 7 | Grina | -0.02 | Young |
| 0.895062 | 0.099074 | 0.024 | 0.026 | 1 | 7 | Zswim1 | -0.002 | Young |
| 0.895576 | 0.523155 | 0.036 | 0.038 | 1 | 7 | Opn3 | -0.002 | Young |
| 0.895735 | 0.123423 | 0.211 | 0.221 | 1 | 7 | Phf2 | -0.01 | Young |
| 0.895749 | 0.01867 | 0.078 | 0.082 | 1 | 7 | Slc35d1 | -0.004 | Young |
| 0.89586 | 0.302306 | 0.205 | 0.2 | 1 | 7 | Lrrc42 | 0.005 | Young |
| 0.895897 | 0.242084 | 0.256 | 0.264 | 1 | 7 | Rabep1 | -0.008 | Young |
| 0.896003 | 0.120293 | 0.093 | 0.095 | 1 | 7 | Ppip5k1 | -0.002 | Young |

| | | | | | | | | |
|---|---|---|---|---|---|---|---|---|
| 0.896404 | 0.080945 | 0.093 | 0.097 | 1 | 7 | E4f1 | -0.004 | Young |
| 0.896425 | 0.930164 | 0.024 | 0.023 | 1 | 7 | Ccl27a | 0.001 | Young |
| 0.896646 | 0.471734 | 0.108 | 0.108 | 1 | 7 | Timm29 | 0 | Young |
| 0.896691 | 0.160849 | 0.13 | 0.136 | 1 | 7 | Shkbp1 | -0.006 | Young |
| 0.896753 | 0.269795 | 0.027 | 0.026 | 1 | 7 | Pex26 | 0.001 | Young |
| 0.896753 | 0.077108 | 0.027 | 0.026 | 1 | 7 | Exog | 0.001 | Young |
| 0.896774 | 0.005422 | 0.747 | 0.751 | 1 | 7 | Vcp | -0.004 | Young |
| 0.896901 | 0.399213 | 0.048 | 0.051 | 1 | 7 | Abhd8 | -0.003 | Young |
| 0.896995 | 0.070482 | 0.298 | 0.315 | 1 | 7 | St3gal1 | -0.017 | Young |
| 0.897278 | 0.257095 | 0.09 | 0.095 | 1 | 7 | Tmcc2 | -0.005 | Young |
| 0.897473 | 0.62215 | 0.211 | 0.21 | 1 | 7 | Mlxip | 0.001 | Young |
| 0.897774 | 0.226944 | 0.027 | 0.026 | 1 | 7 | Lyrm1 | 0.001 | Young |
| 0.897836 | 0.315719 | 0.133 | 0.138 | 1 | 7 | Cdca7l | -0.005 | Young |
| 0.897936 | 0.241601 | 0.048 | 0.046 | 1 | 7 | Dedd2 | 0.002 | Young |
| 0.89817 | 0.199352 | 0.25 | 0.249 | 1 | 7 | Xpot | 0.001 | Young |
| 0.898208 | 0.136024 | 0.024 | 0.026 | 1 | 7 | Gmpr | -0.002 | Young |
| 0.898246 | 0.492077 | 0.06 | 0.059 | 1 | 7 | Tsr3 | 0.001 | Young |
| 0.898494 | 0.617521 | 0.072 | 0.077 | 1 | 7 | Tmub1 | -0.005 | Young |
| 0.899065 | 0.16132 | 0.744 | 0.785 | 1 | 7 | Hnrnph1 | -0.041 | Young |
| 0.89913 | 0.138785 | 0.217 | 0.218 | 1 | 7 | Ttc9c | -0.001 | Young |
| 0.899492 | 0.213698 | 0.247 | 0.254 | 1 | 7 | Chuk | -0.007 | Young |
| 0.899561 | 0.206576 | 0.25 | 0.267 | 1 | 7 | Pip4k2b | -0.017 | Young |
| 0.900063 | 0.071671 | 0.099 | 0.103 | 1 | 7 | Saysd1 | -0.004 | Young |
| 0.900111 | 0.198145 | 0.211 | 0.213 | 1 | 7 | Ptk2b | -0.002 | Young |
| 0.900134 | 0.093841 | 0.389 | 0.415 | 1 | 7 | Cmc2 | -0.026 | Young |
| 0.900227 | 0.06847 | 0.401 | 0.408 | 1 | 7 | 7-Mar | -0.007 | Young |
| 0.900292 | 0.322002 | 0.096 | 0.095 | 1 | 7 | Nup43 | 0.001 | Young |
| 0.900556 | 0.07089 | 0.163 | 0.167 | 1 | 7 | Plk2 | -0.004 | Young |
| 0.900594 | 0.770035 | 0.051 | 0.054 | 1 | 7 | Zfp846 | -0.003 | Young |
| 0.900659 | 0.166974 | 0.256 | 0.259 | 1 | 7 | Fam20b | -0.003 | Young |
| 0.900871 | 0.254288 | 0.127 | 0.131 | 1 | 7 | Mettl14 | -0.004 | Young |
| 0.900919 | 0.464867 | 0.045 | 0.044 | 1 | 7 | Nif3l1 | 0.001 | Young |
| 0.901004 | 0.113576 | 0.569 | 0.603 | 1 | 7 | Magt1 | -0.034 | Young |
| 0.901058 | 0.534694 | 0.06 | 0.064 | 1 | 7 | Zfp81 | -0.004 | Young |
| 0.901541 | 0.160418 | 0.349 | 0.367 | 1 | 7 | Cdc123 | -0.018 | Young |
| 0.901797 | 0.350661 | 0.048 | 0.046 | 1 | 7 | Prkd2 | 0.002 | Young |
| 0.901927 | 0.579835 | 0.081 | 0.082 | 1 | 7 | Gmds | -0.001 | Young |
| 0.902207 | 0.059885 | 0.19 | 0.2 | 1 | 7 | Capn7 | -0.01 | Young |
| 0.90221 | 0.193708 | 0.389 | 0.415 | 1 | 7 | BC017643 | -0.026 | Young |
| 0.902414 | 0.183967 | 0.039 | 0.041 | 1 | 7 | 4930524J0 | -0.002 | Young |
| 0.902451 | 0.213284 | 0.247 | 0.249 | 1 | 7 | Pfdn6 | -0.002 | Young |
| 0.902743 | 0.404748 | 0.202 | 0.213 | 1 | 7 | Errfi1 | -0.011 | Young |
| 0.903438 | 0.207382 | 0.123 | 0.123 | 1 | 7 | Mrps35 | 0 | Young |
| 0.903549 | 0.000226 | 0.584 | 0.626 | 1 | 7 | Rab11b | -0.042 | Young |
| 0.904025 | 0.035386 | 0.316 | 0.331 | 1 | 7 | Pip5k1c | -0.015 | Young |

| | | | | | | | | |
|---|---|---|---|---|---|---|---|---|
| 0.9041 | 0.219568 | 0.045 | 0.044 | 1 | 7 | Mvb12b | 0.001 | Young |
| 0.904258 | 0.067994 | 0.051 | 0.054 | 1 | 7 | Maml2 | -0.003 | Young |
| 0.904363 | 0.199728 | 0.127 | 0.126 | 1 | 7 | Ripk3 | 0.001 | Young |
| 0.904506 | 1.379659 | 0.024 | 0.026 | 1 | 7 | Zfp607a | -0.002 | Young |
| 0.904506 | 0.432005 | 0.024 | 0.026 | 1 | 7 | Slc16a13 | -0.002 | Young |
| 0.904606 | 0.104823 | 0.343 | 0.351 | 1 | 7 | Fkbp15 | -0.008 | Young |
| 0.904627 | 0.576078 | 0.099 | 0.1 | 1 | 7 | Tlr13 | -0.001 | Young |
| 0.904751 | 0.081624 | 0.054 | 0.056 | 1 | 7 | Deaf1 | -0.002 | Young |
| 0.904912 | 0.528394 | 0.13 | 0.133 | 1 | 7 | Flot1 | -0.003 | Young |
| 0.904931 | 0.431167 | 0.027 | 0.026 | 1 | 7 | Abca6 | 0.001 | Young |
| 0.904931 | 0.055973 | 0.027 | 0.026 | 1 | 7 | Tmem132a | 0.001 | Young |
| 0.904943 | 0.19083 | 0.208 | 0.208 | 1 | 7 | Ifi209 | 0 | Young |
| 0.905449 | 0.12893 | 0.22 | 0.218 | 1 | 7 | Ccdc6 | 0.002 | Young |
| 0.905556 | 0.431185 | 0.024 | 0.026 | 1 | 7 | Rcn1 | -0.002 | Young |
| 0.905556 | 0.107693 | 0.024 | 0.026 | 1 | 7 | Tmem220 | -0.002 | Young |
| 0.905643 | 0.584418 | 0.075 | 0.074 | 1 | 7 | Entpd5 | 0.001 | Young |
| 0.905672 | 0.331246 | 0.081 | 0.079 | 1 | 7 | Rnaset2a | 0.002 | Young |
| 0.905894 | 0.081843 | 0.346 | 0.367 | 1 | 7 | Rab1b | -0.021 | Young |
| 0.905934 | 0.117819 | 0.277 | 0.292 | 1 | 7 | Insig1 | -0.015 | Young |
| 0.906087 | 0.478439 | 0.111 | 0.118 | 1 | 7 | Mcoln1 | -0.007 | Young |
| 0.906121 | 0.0574 | 0.792 | 0.8 | 1 | 7 | Calr | -0.008 | Young |
| 0.906157 | 0.085437 | 0.157 | 0.164 | 1 | 7 | Tfeb | -0.007 | Young |
| 0.90658 | 0.186727 | 0.178 | 0.174 | 1 | 7 | 8-Mar | 0.004 | Young |
| 0.906751 | 0.125119 | 0.452 | 0.469 | 1 | 7 | Xrn2 | -0.017 | Young |
| 0.906788 | 0.548166 | 0.036 | 0.038 | 1 | 7 | Rhbdf1 | -0.002 | Young |
| 0.907149 | 0.602399 | 0.06 | 0.064 | 1 | 7 | Mocs1 | -0.004 | Young |
| 0.90722 | 0.052304 | 0.605 | 0.633 | 1 | 7 | Scand1 | -0.028 | Young |
| 0.907274 | 0.014511 | 0.587 | 0.605 | 1 | 7 | Spag9 | -0.018 | Young |
| 0.907451 | 0.090698 | 0.416 | 0.436 | 1 | 7 | Sec11a | -0.02 | Young |
| 0.90757 | 0.003106 | 0.672 | 0.672 | 1 | 7 | Plbd1 | 0 | Young |
| 0.907606 | 0.101083 | 0.464 | 0.487 | 1 | 7 | Rell1 | -0.023 | Young |
| 0.907652 | 0.012404 | 0.099 | 0.103 | 1 | 7 | Chmp7 | -0.004 | Young |
| 0.907892 | 0.099904 | 0.196 | 0.192 | 1 | 7 | Cav2 | 0.004 | Young |
| 0.907925 | 0.643356 | 0.06 | 0.059 | 1 | 7 | Asb1 | 0.001 | Young |
| 0.907944 | 0.103266 | 0.346 | 0.356 | 1 | 7 | Ctdsp1 | -0.01 | Young |
| 0.908 | 0.124215 | 0.416 | 0.438 | 1 | 7 | Tle3 | -0.022 | Young |
| 0.908077 | 0.581736 | 0.045 | 0.044 | 1 | 7 | Ier5l | 0.001 | Young |
| 0.908272 | 0.098819 | 0.395 | 0.405 | 1 | 7 | Stt3a | -0.01 | Young |
| 0.90901 | 0.309767 | 0.063 | 0.067 | 1 | 7 | Spsb2 | -0.004 | Young |
| 0.909137 | 0.145035 | 0.446 | 0.459 | 1 | 7 | Polr2a | -0.013 | Young |
| 0.909326 | 0.156672 | 0.331 | 0.344 | 1 | 7 | Rps6kb1 | -0.013 | Young |
| 0.909379 | 0.321755 | 0.187 | 0.192 | 1 | 7 | Socs7 | -0.005 | Young |
| 0.90938 | 0.811943 | 0.024 | 0.023 | 1 | 7 | Armc3 | 0.001 | Young |
| 0.909475 | 0.058728 | 0.202 | 0.208 | 1 | 7 | Il11ra1 | -0.006 | Young |
| 0.909537 | 0.066143 | 0.232 | 0.236 | 1 | 7 | Chd7 | -0.004 | Young |

| | | | | | | | | |
|---|---|---|---|---|---|---|---|---|
| 0.909828 | 0.125894 | 0.105 | 0.103 | 1 | 7 | Sharpin | 0.002 | Young |
| 0.909867 | 0.080448 | 0.681 | 0.7 | 1 | 7 | Ppp1r18 | -0.019 | Young |
| 0.90995 | 0.153852 | 0.298 | 0.308 | 1 | 7 | Larp7 | -0.01 | Young |
| 0.910083 | 0.022092 | 0.081 | 0.085 | 1 | 7 | 5430427O1 | -0.004 | Young |
| 0.910131 | 0.065912 | 0.13 | 0.128 | 1 | 7 | St3gal3 | 0.002 | Young |
| 0.910185 | 0.090816 | 0.259 | 0.269 | 1 | 7 | Arhgef10l | -0.01 | Young |
| 0.910251 | 0.047974 | 0.361 | 0.385 | 1 | 7 | Dcp2 | -0.024 | Young |
| 0.91046 | 0.988023 | 0.024 | 0.023 | 1 | 7 | Lca5 | 0.001 | Young |
| 0.91046 | 0.497236 | 0.024 | 0.023 | 1 | 7 | Nmrk1 | 0.001 | Young |
| 0.910609 | 0.114457 | 0.139 | 0.136 | 1 | 7 | Zfp787 | 0.003 | Young |
| 0.910785 | 0.138577 | 0.175 | 0.174 | 1 | 7 | Ccnf | 0.001 | Young |
| 0.91081 | 0.060766 | 0.024 | 0.026 | 1 | 7 | Purg | -0.002 | Young |
| 0.910991 | 0.261905 | 0.139 | 0.138 | 1 | 7 | Tmed4 | 0.001 | Young |
| 0.91101 | 0.196377 | 0.331 | 0.344 | 1 | 7 | Zfp367 | -0.013 | Young |
| 0.911039 | 0.450401 | 0.226 | 0.231 | 1 | 7 | Rab3gap1 | -0.005 | Young |
| 0.911756 | 0.235298 | 0.223 | 0.221 | 1 | 7 | Nfx1 | 0.002 | Young |
| 0.911811 | 0.114494 | 0.283 | 0.287 | 1 | 7 | Ahsa1 | -0.004 | Young |
| 0.911869 | 0.146697 | 0.304 | 0.323 | 1 | 7 | Sh3pxd2b | -0.019 | Young |
| 0.911929 | 0.0171 | 0.271 | 0.269 | 1 | 7 | Tbc1d23 | 0.002 | Young |
| 0.9123 | 0.042178 | 0.295 | 0.305 | 1 | 7 | Brip1os | -0.01 | Young |
| 0.912482 | 0.122393 | 0.301 | 0.31 | 1 | 7 | Zc3h18 | -0.009 | Young |
| 0.912582 | 0.075912 | 0.37 | 0.369 | 1 | 7 | Hsd17b12 | 0.001 | Young |
| 0.912734 | 0.204154 | 0.277 | 0.29 | 1 | 7 | Rnf115 | -0.013 | Young |
| 0.912749 | 0.641718 | 0.114 | 0.115 | 1 | 7 | Slc38a6 | -0.001 | Young |
| 0.913044 | 0.266589 | 0.148 | 0.149 | 1 | 7 | Srprb | -0.001 | Young |
| 0.913357 | 0.233997 | 0.066 | 0.069 | 1 | 7 | Pex11b | -0.003 | Young |
| 0.913492 | 0.057113 | 0.217 | 0.221 | 1 | 7 | Adk | -0.004 | Young |
| 0.913973 | 0.142129 | 0.343 | 0.344 | 1 | 7 | Csnk1e | -0.001 | Young |
| 0.914056 | 0.180849 | 0.241 | 0.254 | 1 | 7 | Eif2s3y | -0.013 | Young |
| 0.914092 | 0.037048 | 0.039 | 0.041 | 1 | 7 | Cyb5d2 | -0.002 | Young |
| 0.914157 | 0.125924 | 0.169 | 0.177 | 1 | 7 | Zfp318 | -0.008 | Young |
| 0.914191 | 0.264048 | 0.096 | 0.1 | 1 | 7 | Hdhd2 | -0.004 | Young |
| 0.914227 | 0.263984 | 0.364 | 0.397 | 1 | 7 | Csnk2a1 | -0.033 | Young |
| 0.91428 | 0.278718 | 0.205 | 0.203 | 1 | 7 | Ttc5 | 0.002 | Young |
| 0.914453 | 0.049048 | 0.518 | 0.531 | 1 | 7 | Nckap1l | -0.013 | Young |
| 0.914593 | 0.421617 | 0.163 | 0.159 | 1 | 7 | Nf1 | 0.004 | Young |
| 0.914775 | 0.188012 | 0.054 | 0.056 | 1 | 7 | Fbxw17 | -0.002 | Young |
| 0.914869 | 0.098558 | 0.536 | 0.559 | 1 | 7 | Itga4 | -0.023 | Young |
| 0.915134 | 0.618111 | 0.139 | 0.141 | 1 | 7 | Pex3 | -0.002 | Young |
| 0.915212 | 0.630978 | 0.084 | 0.087 | 1 | 7 | Prune1 | -0.003 | Young |
| 0.915356 | 0.200441 | 0.117 | 0.121 | 1 | 7 | Dzip3 | -0.004 | Young |
| 0.915669 | 0.238856 | 0.292 | 0.295 | 1 | 7 | Heatr5a | -0.003 | Young |
| 0.915762 | 0.445818 | 0.039 | 0.041 | 1 | 7 | Zfp275 | -0.002 | Young |
| 0.915781 | 0.186479 | 0.163 | 0.162 | 1 | 7 | Zfp655 | 0.001 | Young |
| 0.916208 | 0.316939 | 0.054 | 0.056 | 1 | 7 | Ermard | -0.002 | Young |

| | | | | | | | | |
|---|---|---|---|---|---|---|---|---|
| 0.916288 | 0.632453 | 0.145 | 0.146 | 1 | 7 | Acox1 | -0.001 | Young |
| 0.916924 | 0.20453 | 0.054 | 0.056 | 1 | 7 | Prmt9 | -0.002 | Young |
| 0.916948 | 0.634263 | 0.024 | 0.023 | 1 | 7 | Cd28 | 0.001 | Young |
| 0.917087 | 0.174961 | 0.389 | 0.39 | 1 | 7 | Baz2b | -0.001 | Young |
| 0.917408 | 0.276383 | 0.196 | 0.197 | 1 | 7 | Utp18 | -0.001 | Young |
| 0.917505 | 0.117703 | 0.069 | 0.072 | 1 | 7 | Poc1a | -0.003 | Young |
| 0.917591 | 0.027885 | 0.229 | 0.233 | 1 | 7 | Babam2 | -0.004 | Young |
| 0.917592 | 0.190213 | 0.042 | 0.044 | 1 | 7 | Orc1 | -0.002 | Young |
| 0.917992 | 0.233298 | 0.208 | 0.218 | 1 | 7 | Zfp950 | -0.01 | Young |
| 0.918135 | 0.18305 | 0.181 | 0.185 | 1 | 7 | Cyp4f16 | -0.004 | Young |
| 0.918171 | 0.728056 | 0.024 | 0.026 | 1 | 7 | Enho | -0.002 | Young |
| 0.918591 | 0.671777 | 0.057 | 0.056 | 1 | 7 | Gcdh | 0.001 | Young |
| 0.918659 | 0.019544 | 0.717 | 0.718 | 1 | 7 | Nrros | -0.001 | Young |
| 0.91898 | 0.133356 | 0.449 | 0.459 | 1 | 7 | Ccl7 | -0.01 | Young |
| 0.919221 | 0.436684 | 0.066 | 0.069 | 1 | 7 | Ak8 | -0.003 | Young |
| 0.919566 | 0.342549 | 0.202 | 0.215 | 1 | 7 | Bag3 | -0.013 | Young |
| 0.919768 | 0.264858 | 0.223 | 0.221 | 1 | 7 | Itgav | 0.002 | Young |
| 0.919808 | 0.090197 | 0.419 | 0.418 | 1 | 7 | Srgap2 | 0.001 | Young |
| 0.920121 | 0.165944 | 0.127 | 0.126 | 1 | 7 | Pradc1 | 0.001 | Young |
| 0.920144 | 0.454112 | 0.232 | 0.251 | 1 | 7 | Pak1 | -0.019 | Young |
| 0.920256 | 0.055251 | 0.657 | 0.7 | 1 | 7 | Sec61g | -0.043 | Young |
| 0.920257 | 0.280576 | 0.154 | 0.154 | 1 | 7 | Ccdc107 | 0 | Young |
| 0.920334 | 0.095518 | 0.346 | 0.354 | 1 | 7 | Sipa1 | -0.008 | Young |
| 0.920345 | 0.27283 | 0.108 | 0.108 | 1 | 7 | Adcy9 | 0 | Young |
| 0.920681 | 0.001054 | 0.099 | 0.103 | 1 | 7 | Tmem198b | -0.004 | Young |
| 0.920696 | 0.075136 | 0.226 | 0.231 | 1 | 7 | Tlk1 | -0.005 | Young |
| 0.920719 | 0.420239 | 0.22 | 0.231 | 1 | 7 | Hbs1l | -0.011 | Young |
| 0.920847 | 0.057342 | 0.572 | 0.597 | 1 | 7 | Txn2 | -0.025 | Young |
| 0.920868 | 0.272927 | 0.127 | 0.131 | 1 | 7 | Tesk1 | -0.004 | Young |
| 0.921354 | 0.300046 | 0.142 | 0.141 | 1 | 7 | Zfp277 | 0.001 | Young |
| 0.921421 | 0.05695 | 0.307 | 0.31 | 1 | 7 | Hipk2 | -0.003 | Young |
| 0.921719 | 0.169884 | 0.178 | 0.187 | 1 | 7 | Metap1 | -0.009 | Young |
| 0.922233 | 0.036545 | 0.771 | 0.769 | 1 | 7 | Luc7l2 | 0.002 | Young |
| 0.922274 | 0.260729 | 0.108 | 0.113 | 1 | 7 | Gm48065 | -0.005 | Young |
| 0.922481 | 0.312271 | 0.066 | 0.069 | 1 | 7 | Psmc3ip | -0.003 | Young |
| 0.923427 | 0.280761 | 0.151 | 0.156 | 1 | 7 | Msra | -0.005 | Young |
| 0.923564 | 0.014527 | 0.012 | 0.013 | 1 | 7 | Igfbp7 | -0.001 | Young |
| 0.923724 | 0.153164 | 0.283 | 0.297 | 1 | 7 | Setd5 | -0.014 | Young |
| 0.924032 | 0.204932 | 0.349 | 0.362 | 1 | 7 | Higd2a | -0.013 | Young |
| 0.924071 | 0.919824 | 0.036 | 0.038 | 1 | 7 | Zfp280b | -0.002 | Young |
| 0.924493 | 0.111078 | 0.105 | 0.105 | 1 | 7 | Zmynd19 | 0 | Young |
| 0.924523 | 0.333029 | 0.024 | 0.023 | 1 | 7 | Rasgrf2 | 0.001 | Young |
| 0.924692 | 0.175199 | 0.458 | 0.495 | 1 | 7 | Epsti1 | -0.037 | Young |
| 0.924776 | 0.411021 | 0.123 | 0.133 | 1 | 7 | Pik3r4 | -0.01 | Young |
| 0.925193 | 0.054683 | 0.099 | 0.097 | 1 | 7 | Fbxo25 | 0.002 | Young |

| | | | | | | | | |
|---|---|---|---|---|---|---|---|---|
| 0.92539 | 0.68309 | 0.102 | 0.105 | 1 | 7 | Edc3 | -0.003 | Young |
| 0.925547 | 0.043369 | 0.322 | 0.326 | 1 | 7 | Ascc3 | -0.004 | Young |
| 0.92579 | 0.339328 | 0.039 | 0.041 | 1 | 7 | 2810001G2 | -0.002 | Young |
| 0.926043 | 0.270548 | 0.093 | 0.092 | 1 | 7 | Slc12a4 | 0.001 | Young |
| 0.926175 | 0.037493 | 0.169 | 0.172 | 1 | 7 | Cnot9 | -0.003 | Young |
| 0.926274 | 0.060281 | 0.253 | 0.254 | 1 | 7 | 2810004N2 | -0.001 | Young |
| 0.926503 | 0.265466 | 0.319 | 0.331 | 1 | 7 | Nup88 | -0.012 | Young |
| 0.926579 | 0.334855 | 0.247 | 0.251 | 1 | 7 | Rprd1b | -0.004 | Young |
| 0.926601 | 0.049834 | 0.328 | 0.344 | 1 | 7 | Haus8 | -0.016 | Young |
| 0.926689 | 0.230993 | 0.024 | 0.023 | 1 | 7 | Mfsd13a | 0.001 | Young |
| 0.926689 | 0.170444 | 0.024 | 0.023 | 1 | 7 | Tctn1 | 0.001 | Young |
| 0.926767 | 0.117768 | 0.081 | 0.079 | 1 | 7 | Msantd2 | 0.002 | Young |
| 0.926934 | 0.028547 | 0.229 | 0.223 | 1 | 7 | Virma | 0.006 | Young |
| 0.927001 | 0.168632 | 0.102 | 0.105 | 1 | 7 | Tlr1 | -0.003 | Young |
| 0.927016 | 0.162432 | 0.096 | 0.095 | 1 | 7 | Cacna1d | 0.001 | Young |
| 0.927092 | 0.747372 | 0.087 | 0.09 | 1 | 7 | Brms1l | -0.003 | Young |
| 0.927106 | 0.35075 | 0.069 | 0.072 | 1 | 7 | Fer | -0.003 | Young |
| 0.927355 | 0.267522 | 0.145 | 0.144 | 1 | 7 | Rmi1 | 0.001 | Young |
| 0.927415 | 0.397259 | 0.181 | 0.185 | 1 | 7 | Nsmce3 | -0.004 | Young |
| 0.927762 | 0.473939 | 0.145 | 0.151 | 1 | 7 | Spire1 | -0.006 | Young |
| 0.928004 | 0.238191 | 0.012 | 0.013 | 1 | 7 | Satb2 | -0.001 | Young |
| 0.928004 | 0.182433 | 0.012 | 0.013 | 1 | 7 | Tnfsfm13 | -0.001 | Young |
| 0.928054 | 0.224662 | 0.142 | 0.146 | 1 | 7 | Nhlrc2 | -0.004 | Young |
| 0.92807 | 0.251461 | 0.19 | 0.195 | 1 | 7 | Crnkl1 | -0.005 | Young |
| 0.928086 | 0.063739 | 0.425 | 0.421 | 1 | 7 | Ccrl2 | 0.004 | Young |
| 0.928141 | 0.404653 | 0.151 | 0.154 | 1 | 7 | Rbmx2 | -0.003 | Young |
| 0.928298 | 0.566276 | 0.039 | 0.041 | 1 | 7 | Nphp3 | -0.002 | Young |
| 0.928298 | 0.361184 | 0.039 | 0.041 | 1 | 7 | Ptar1 | -0.002 | Young |
| 0.928668 | 0.642934 | 0.084 | 0.087 | 1 | 7 | Car5b | -0.003 | Young |
| 0.928811 | 0.021188 | 0.283 | 0.292 | 1 | 7 | Eml4 | -0.009 | Young |
| 0.929027 | 0.486572 | 0.069 | 0.072 | 1 | 7 | Tnk2 | -0.003 | Young |
| 0.929173 | 0.470989 | 0.114 | 0.115 | 1 | 7 | Cep63 | -0.001 | Young |
| 0.929318 | 0.183425 | 0.187 | 0.192 | 1 | 7 | Hdac5 | -0.005 | Young |
| 0.929376 | 0.231185 | 0.307 | 0.318 | 1 | 7 | Dock1 | -0.011 | Young |
| 0.929484 | 0.305301 | 0.012 | 0.013 | 1 | 7 | Rtl6 | -0.001 | Young |
| 0.929484 | 0.28801 | 0.012 | 0.013 | 1 | 7 | Rsad1 | -0.001 | Young |
| 0.929484 | 0.122367 | 0.012 | 0.013 | 1 | 7 | Ngfr | -0.001 | Young |
| 0.929484 | 0.081732 | 0.012 | 0.013 | 1 | 7 | Lgr4 | -0.001 | Young |
| 0.929544 | 0.163734 | 0.114 | 0.118 | 1 | 7 | Guf1 | -0.004 | Young |
| 0.929656 | 0.048481 | 0.377 | 0.392 | 1 | 7 | E2f1 | -0.015 | Young |
| 0.929708 | 0.157841 | 0.319 | 0.344 | 1 | 7 | Gtpbp2 | -0.025 | Young |
| 0.929739 | 0.254203 | 0.187 | 0.187 | 1 | 7 | Trappc6a | 0 | Young |
| 0.929971 | 0.604291 | 0.039 | 0.041 | 1 | 7 | Ap4b1 | -0.002 | Young |
| 0.930218 | 0.037727 | 0.657 | 0.674 | 1 | 7 | Pkm | -0.017 | Young |
| 0.930398 | 0.093701 | 0.22 | 0.233 | 1 | 7 | Tmem41b | -0.013 | Young |

| | | | | | | | | |
|---|---|---|---|---|---|---|---|---|
| 0.930618 | 0.259766 | 0.133 | 0.133 | 1 | 7 | Stx2 | 0 | Young |
| 0.930645 | 0.260439 | 0.057 | 0.056 | 1 | 7 | Rora | 0.001 | Young |
| 0.930666 | 0.106019 | 0.431 | 0.438 | 1 | 7 | Gpbp1 | -0.007 | Young |
| 0.930753 | 1.142478 | 0.027 | 0.028 | 1 | 7 | Dusp28 | -0.001 | Young |
| 0.930965 | 0.493714 | 0.012 | 0.013 | 1 | 7 | Akap12 | -0.001 | Young |
| 0.930965 | 0.321072 | 0.012 | 0.013 | 1 | 7 | Zfp72 | -0.001 | Young |
| 0.930965 | 0.052077 | 0.012 | 0.013 | 1 | 7 | Gm26624 | -0.001 | Young |
| 0.93101 | 0.370471 | 0.334 | 0.349 | 1 | 7 | Setd1b | -0.015 | Young |
| 0.931291 | 0.027029 | 0.849 | 0.872 | 1 | 7 | Fus | -0.023 | Young |
| 0.931486 | 0.034441 | 0.331 | 0.331 | 1 | 7 | Cetn3 | 0 | Young |
| 0.931812 | 0.160865 | 0.268 | 0.277 | 1 | 7 | Wwp2 | -0.009 | Young |
| 0.931958 | 0.232032 | 0.108 | 0.108 | 1 | 7 | Plcb2 | 0 | Young |
| 0.931972 | 0.099141 | 0.313 | 0.313 | 1 | 7 | Shoc2 | 0 | Young |
| 0.931988 | 0.411229 | 0.054 | 0.056 | 1 | 7 | Endog | -0.002 | Young |
| 0.932446 | 1.057914 | 0.012 | 0.013 | 1 | 7 | Il7 | -0.001 | Young |
| 0.932446 | 0.549834 | 0.012 | 0.013 | 1 | 7 | Layn | -0.001 | Young |
| 0.932446 | 0.535827 | 0.012 | 0.013 | 1 | 7 | Pecr | -0.001 | Young |
| 0.932446 | 0.281392 | 0.012 | 0.013 | 1 | 7 | Sardh | -0.001 | Young |
| 0.932446 | 0.09559 | 0.012 | 0.013 | 1 | 7 | Gm15545 | -0.001 | Young |
| 0.933304 | 0.323728 | 0.274 | 0.285 | 1 | 7 | Xrn1 | -0.011 | Young |
| 0.933551 | 0.120051 | 0.166 | 0.169 | 1 | 7 | Cep120 | -0.003 | Young |
| 0.933604 | 0.097217 | 0.627 | 0.644 | 1 | 7 | S100a13 | -0.017 | Young |
| 0.933634 | 0.992867 | 0.051 | 0.054 | 1 | 7 | Iba57 | -0.003 | Young |
| 0.933711 | 0.232647 | 0.093 | 0.097 | 1 | 7 | Pithd1 | -0.004 | Young |
| 0.933927 | 0.215237 | 0.012 | 0.013 | 1 | 7 | Gpr150 | -0.001 | Young |
| 0.933927 | 0.197837 | 0.012 | 0.013 | 1 | 7 | Txk | -0.001 | Young |
| 0.934031 | 0.177659 | 0.316 | 0.338 | 1 | 7 | Zfp445 | -0.022 | Young |
| 0.934262 | 0.05356 | 0.205 | 0.205 | 1 | 7 | Rxrb | 0 | Young |
| 0.934274 | 0.133124 | 0.024 | 0.023 | 1 | 7 | Plekha5 | 0.001 | Young |
| 0.934308 | 0.026894 | 0.41 | 0.428 | 1 | 7 | D17Wsu92 | -0.018 | Young |
| 0.934315 | 0.936804 | 0.039 | 0.038 | 1 | 7 | Zfp758 | 0.001 | Young |
| 0.934506 | 0.080659 | 0.211 | 0.213 | 1 | 7 | Nfia | -0.002 | Young |
| 0.934647 | 0.212581 | 0.09 | 0.092 | 1 | 7 | Trim39 | -0.002 | Young |
| 0.93467 | 0.202623 | 0.214 | 0.213 | 1 | 7 | Smg5 | 0.001 | Young |
| 0.934738 | 0.427904 | 0.127 | 0.128 | 1 | 7 | Tvp23b | -0.001 | Young |
| 0.9353 | 0.264438 | 0.166 | 0.167 | 1 | 7 | Abraxas2 | -0.001 | Young |
| 0.935358 | 0.236293 | 0.024 | 0.023 | 1 | 7 | Zswim9 | 0.001 | Young |
| 0.935409 | 1.270686 | 0.012 | 0.013 | 1 | 7 | Ndor1 | -0.001 | Young |
| 0.935409 | 0.086791 | 0.012 | 0.013 | 1 | 7 | AC146911. | -0.001 | Young |
| 0.935768 | 0.084136 | 0.482 | 0.485 | 1 | 7 | Lmo4 | -0.003 | Young |
| 0.935941 | 0.278939 | 0.199 | 0.205 | 1 | 7 | Trrap | -0.006 | Young |
| 0.936078 | 0.090022 | 0.693 | 0.741 | 1 | 7 | Pak2 | -0.048 | Young |
| 0.936252 | 0.14159 | 0.377 | 0.39 | 1 | 7 | Smarce1 | -0.013 | Young |
| 0.936299 | 0.094133 | 0.211 | 0.213 | 1 | 7 | Cep57 | -0.002 | Young |
| 0.936666 | 0.653149 | 0.039 | 0.041 | 1 | 7 | Ccdc84 | -0.002 | Young |

| | | | | | | | | |
|---|---|---|---|---|---|---|---|---|
| 0.936688 | 0.100189 | 0.057 | 0.059 | 1 | 7 | Slc12a2 | -0.002 | Young |
| 0.936705 | 0.476108 | 0.181 | 0.185 | 1 | 7 | Cdk5rap3 | -0.004 | Young |
| 0.936762 | 0.572687 | 0.027 | 0.028 | 1 | 7 | Amacr | -0.001 | Young |
| 0.936891 | 0.667309 | 0.012 | 0.013 | 1 | 7 | Garnl3 | -0.001 | Young |
| 0.936958 | 0.113261 | 0.117 | 0.115 | 1 | 7 | Slc16a1 | 0.002 | Young |
| 0.937217 | 0.049168 | 0.825 | 0.841 | 1 | 7 | Srsf3 | -0.016 | Young |
| 0.937335 | 0.604008 | 0.078 | 0.082 | 1 | 7 | Mthfsd | -0.004 | Young |
| 0.93761 | 0.232667 | 0.145 | 0.144 | 1 | 7 | Tceal8 | 0.001 | Young |
| 0.938034 | 0.235725 | 0.139 | 0.144 | 1 | 7 | Grap | -0.005 | Young |
| 0.938399 | 0.533921 | 0.069 | 0.069 | 1 | 7 | Rusc1 | 0 | Young |
| 0.93876 | 0.009777 | 0.895 | 0.918 | 1 | 7 | Sem1 | -0.023 | Young |
| 0.938765 | 0.054347 | 0.027 | 0.028 | 1 | 7 | Gm43660 | -0.001 | Young |
| 0.939534 | 0.369376 | 0.211 | 0.21 | 1 | 7 | Chmp1b | 0.001 | Young |
| 0.939855 | 1.287916 | 0.012 | 0.013 | 1 | 7 | Lbx2 | -0.001 | Young |
| 0.939855 | 0.959168 | 0.012 | 0.013 | 1 | 7 | Slc10a6 | -0.001 | Young |
| 0.940116 | 0.113231 | 0.401 | 0.403 | 1 | 7 | Uqcrc1 | -0.002 | Young |
| 0.940275 | 0.436354 | 0.042 | 0.044 | 1 | 7 | 4632415L0 | -0.002 | Young |
| 0.940458 | 0.018832 | 0.06 | 0.062 | 1 | 7 | Mterf4 | -0.002 | Young |
| 0.940655 | 0.157311 | 0.154 | 0.156 | 1 | 7 | Tmub2 | -0.002 | Young |
| 0.940686 | 0.078649 | 0.605 | 0.631 | 1 | 7 | Litaf | -0.026 | Young |
| 0.940887 | 0.163626 | 0.331 | 0.344 | 1 | 7 | Smim10l1 | -0.013 | Young |
| 0.940982 | 0.18859 | 0.277 | 0.29 | 1 | 7 | Dstyk | -0.013 | Young |
| 0.940993 | 0.062209 | 0.169 | 0.167 | 1 | 7 | Gpr183 | 0.002 | Young |
| 0.941042 | 0.24673 | 0.187 | 0.187 | 1 | 7 | D16Ertd47 | 0 | Young |
| 0.941337 | 0.653207 | 0.012 | 0.013 | 1 | 7 | Pex10 | -0.001 | Young |
| 0.941412 | 0.117681 | 0.325 | 0.344 | 1 | 7 | Arpc1a | -0.019 | Young |
| 0.94156 | 0.124336 | 0.41 | 0.426 | 1 | 7 | Rexo2 | -0.016 | Young |
| 0.941741 | 0.04523 | 0.386 | 0.387 | 1 | 7 | Card19 | -0.001 | Young |
| 0.941978 | 0.361194 | 0.039 | 0.038 | 1 | 7 | Clec16a | 0.001 | Young |
| 0.942028 | 0.098276 | 0.045 | 0.046 | 1 | 7 | B3galt6 | -0.001 | Young |
| 0.942057 | 0.144806 | 0.28 | 0.287 | 1 | 7 | Abca7 | -0.007 | Young |
| 0.942275 | 0.048643 | 0.651 | 0.682 | 1 | 7 | Atp5j2 | -0.031 | Young |
| 0.94246 | 0.097163 | 0.078 | 0.077 | 1 | 7 | Pgm3 | 0.001 | Young |
| 0.942593 | 0.143366 | 0.217 | 0.221 | 1 | 7 | Ppil4 | -0.004 | Young |
| 0.942648 | 0.328814 | 0.13 | 0.131 | 1 | 7 | Traf5 | -0.001 | Young |
| 0.942718 | 0.085126 | 0.184 | 0.19 | 1 | 7 | Rlf | -0.006 | Young |
| 0.942738 | 0.829031 | 0.139 | 0.144 | 1 | 7 | Sh2d1b1 | -0.005 | Young |
| 0.942763 | 0.374486 | 0.054 | 0.056 | 1 | 7 | Slx1b | -0.002 | Young |
| 0.94294 | 0.177159 | 0.178 | 0.179 | 1 | 7 | Mrps18a | -0.001 | Young |
| 0.942973 | 0.269275 | 0.145 | 0.146 | 1 | 7 | Scamp5 | -0.001 | Young |
| 0.942982 | 0.058493 | 0.307 | 0.31 | 1 | 7 | Ifi204 | -0.003 | Young |
| 0.943045 | 0.390467 | 0.416 | 0.449 | 1 | 7 | Leng8 | -0.033 | Young |
| 0.94322 | 0.527305 | 0.081 | 0.082 | 1 | 7 | Dlg4 | -0.001 | Young |
| 0.943365 | 0.531608 | 0.039 | 0.041 | 1 | 7 | Vps51 | -0.002 | Young |
| 0.943407 | 0.244949 | 0.148 | 0.154 | 1 | 7 | Impdh1 | -0.006 | Young |

| | | | | | | | | |
|---|---|---|---|---|---|---|---|---|
| 0.943448 | 0.177321 | 0.316 | 0.328 | 1 | 7 | Map3k8 | -0.012 | Young |
| 0.943528 | 0.033557 | 0.271 | 0.277 | 1 | 7 | Rbfa | -0.006 | Young |
| 0.943651 | 0.290493 | 0.226 | 0.241 | 1 | 7 | Galnt11 | -0.015 | Young |
| 0.943707 | 0.177817 | 0.247 | 0.249 | 1 | 7 | Wars | -0.002 | Young |
| 0.943776 | 0.372813 | 0.027 | 0.028 | 1 | 7 | Plcl1 | -0.001 | Young |
| 0.943869 | 0.160128 | 0.289 | 0.3 | 1 | 7 | Cdkn2aipnl | -0.011 | Young |
| 0.943888 | 0.035486 | 0.548 | 0.569 | 1 | 7 | Plxnb2 | -0.021 | Young |
| 0.944026 | 0.063584 | 0.452 | 0.477 | 1 | 7 | Map2k3 | -0.025 | Young |
| 0.944177 | 0.231761 | 0.226 | 0.228 | 1 | 7 | Pdgfb | -0.002 | Young |
| 0.94425 | 0.044883 | 0.476 | 0.482 | 1 | 7 | Arhgap17 | -0.006 | Young |
| 0.944268 | 0.131058 | 0.193 | 0.2 | 1 | 7 | Osgin2 | -0.007 | Young |
| 0.944295 | 0.162464 | 0.172 | 0.179 | 1 | 7 | Ltv1 | -0.007 | Young |
| 0.944413 | 0.01163 | 0.057 | 0.059 | 1 | 7 | Acvr2b | -0.002 | Young |
| 0.944451 | 0.058631 | 0.461 | 0.477 | 1 | 7 | Ptpro | -0.016 | Young |
| 0.94467 | 0.046605 | 0.256 | 0.259 | 1 | 7 | Anxa11 | -0.003 | Young |
| 0.9449 | 0.113866 | 0.175 | 0.179 | 1 | 7 | Ngly1 | -0.004 | Young |
| 0.9451 | 0.020547 | 0.042 | 0.041 | 1 | 7 | Map7d3 | 0.001 | Young |
| 0.945122 | 0.097525 | 0.021 | 0.021 | 1 | 7 | Abhd6 | 0 | Young |
| 0.945318 | 0.487897 | 0.142 | 0.144 | 1 | 7 | Cabin1 | -0.002 | Young |
| 0.945481 | 0.081692 | 0.108 | 0.11 | 1 | 7 | Sqle | -0.002 | Young |
| 0.945697 | 0.172477 | 0.069 | 0.072 | 1 | 7 | Tmem94 | -0.003 | Young |
| 0.94613 | 0.044742 | 0.211 | 0.21 | 1 | 7 | Brip1 | 0.001 | Young |
| 0.946732 | 0.061401 | 0.193 | 0.2 | 1 | 7 | Tatdn2 | -0.007 | Young |
| 0.946968 | 0.296836 | 0.307 | 0.315 | 1 | 7 | Neu1 | -0.008 | Young |
| 0.947011 | 0.533076 | 0.244 | 0.254 | 1 | 7 | Brox | -0.01 | Young |
| 0.947293 | 0.48752 | 0.099 | 0.1 | 1 | 7 | Pcid2 | -0.001 | Young |
| 0.947455 | 0.09651 | 0.072 | 0.072 | 1 | 7 | Cep85l | 0 | Young |
| 0.947621 | 0.136486 | 0.145 | 0.146 | 1 | 7 | Gins1 | -0.001 | Young |
| 0.947786 | 0.773758 | 0.027 | 0.028 | 1 | 7 | Rita1 | -0.001 | Young |
| 0.947786 | 0.402313 | 0.027 | 0.028 | 1 | 7 | C1qtnf6 | -0.001 | Young |
| 0.947941 | 0.337515 | 0.223 | 0.228 | 1 | 7 | Ppp1r7 | -0.005 | Young |
| 0.948055 | 0.211874 | 0.277 | 0.292 | 1 | 7 | Abcd1 | -0.015 | Young |
| 0.948092 | 0.089934 | 0.322 | 0.336 | 1 | 7 | Pds5b | -0.014 | Young |
| 0.948464 | 0.164796 | 0.253 | 0.259 | 1 | 7 | Usp3 | -0.006 | Young |
| 0.948876 | 0.16911 | 0.322 | 0.333 | 1 | 7 | Tifab | -0.011 | Young |
| 0.948984 | 0.124288 | 0.386 | 0.4 | 1 | 7 | Cic | -0.014 | Young |
| 0.949586 | 0.275646 | 0.139 | 0.141 | 1 | 7 | Cryzl1 | -0.002 | Young |
| 0.949626 | 0.052365 | 0.446 | 0.451 | 1 | 7 | Fmnl1 | -0.005 | Young |
| 0.949737 | 0.438183 | 0.021 | 0.021 | 1 | 7 | Ccdc166 | 0 | Young |
| 0.949737 | 0.292111 | 0.021 | 0.021 | 1 | 7 | Mttp | 0 | Young |
| 0.950005 | 0.041584 | 0.142 | 0.144 | 1 | 7 | Kcnk13 | -0.002 | Young |
| 0.95013 | 0.383946 | 0.102 | 0.105 | 1 | 7 | 4930579G2 | -0.003 | Young |
| 0.950419 | 0.169076 | 0.205 | 0.205 | 1 | 7 | Trim33 | 0 | Young |
| 0.950702 | 0.677732 | 0.069 | 0.069 | 1 | 7 | D930016D( | 0 | Young |
| 0.95073 | 0.015697 | 0.208 | 0.213 | 1 | 7 | Nucb2 | -0.005 | Young |

| | | | | | | | | |
|---|---|---|---|---|---|---|---|---|
| 0.950891 | 0.776067 | 0.021 | 0.021 | 1 | 7 | Gm4707 | 0 | Young |
| 0.951136 | 0.198187 | 0.304 | 0.318 | 1 | 7 | Lpxn | -0.014 | Young |
| 0.951198 | 0.009075 | 0.437 | 0.438 | 1 | 7 | Chchd1 | -0.001 | Young |
| 0.951283 | 0.452256 | 0.111 | 0.113 | 1 | 7 | Rbsn | -0.002 | Young |
| 0.95177 | 0.041861 | 0.16 | 0.162 | 1 | 7 | Ttc1 | -0.002 | Young |
| 0.951796 | 0.621044 | 0.036 | 0.036 | 1 | 7 | Ybey | 0 | Young |
| 0.951834 | 0.342183 | 0.202 | 0.208 | 1 | 7 | Tnip2 | -0.006 | Young |
| 0.951896 | 0.119292 | 0.334 | 0.349 | 1 | 7 | Myo9a | -0.015 | Young |
| 0.951897 | 0.680892 | 0.075 | 0.077 | 1 | 7 | Kctd6 | -0.002 | Young |
| 0.952227 | 0.154333 | 0.319 | 0.333 | 1 | 7 | 0610012G( | -0.014 | Young |
| 0.95238 | 0.153307 | 0.232 | 0.233 | 1 | 7 | Ctdspl | -0.001 | Young |
| 0.952521 | 0.643836 | 0.042 | 0.041 | 1 | 7 | D17H6S53I | 0.001 | Young |
| 0.952537 | 0.258437 | 0.03 | 0.031 | 1 | 7 | Simc1 | -0.001 | Young |
| 0.953056 | 0.12486 | 0.045 | 0.046 | 1 | 7 | Ets1 | -0.001 | Young |
| 0.953299 | 1.075247 | 0.051 | 0.051 | 1 | 7 | Psrc1 | 0 | Young |
| 0.953324 | 0.197615 | 0.117 | 0.118 | 1 | 7 | Synj2 | -0.001 | Young |
| 0.953371 | 0.211519 | 0.105 | 0.108 | 1 | 7 | Slc17a5 | -0.003 | Young |
| 0.953653 | 0.105957 | 0.205 | 0.205 | 1 | 7 | Ech1 | 0 | Young |
| 0.953775 | 0.229242 | 0.12 | 0.123 | 1 | 7 | Lrrc40 | -0.003 | Young |
| 0.953816 | 0.088113 | 0.072 | 0.072 | 1 | 7 | Rpl36a-ps1 | 0 | Young |
| 0.953898 | 0.313825 | 0.262 | 0.274 | 1 | 7 | Mbtps1 | -0.012 | Young |
| 0.954249 | 0.121589 | 0.087 | 0.09 | 1 | 7 | Mynn | -0.003 | Young |
| 0.954353 | 0.315799 | 0.021 | 0.021 | 1 | 7 | Atp5s | 0 | Young |
| 0.954402 | 0.042057 | 0.473 | 0.485 | 1 | 7 | Kdelr1 | -0.012 | Young |
| 0.954597 | 0.242293 | 0.09 | 0.092 | 1 | 7 | Adnp2 | -0.002 | Young |
| 0.954792 | 1.350951 | 0.051 | 0.051 | 1 | 7 | Nr1d1 | 0 | Young |
| 0.95517 | 0.057983 | 0.355 | 0.374 | 1 | 7 | Aff4 | -0.019 | Young |
| 0.955583 | 0.151286 | 0.06 | 0.062 | 1 | 7 | 2310057M | -0.002 | Young |
| 0.955614 | 0.040744 | 0.039 | 0.038 | 1 | 7 | Ptchd1 | 0.001 | Young |
| 0.955904 | 0.200554 | 0.62 | 0.669 | 1 | 7 | Mapk3 | -0.049 | Young |
| 0.956271 | 0.303762 | 0.06 | 0.062 | 1 | 7 | Trp53i13 | -0.002 | Young |
| 0.956662 | 0.309514 | 0.021 | 0.021 | 1 | 7 | 1810044D( | 0 | Young |
| 0.956662 | 0.088792 | 0.021 | 0.021 | 1 | 7 | Tmem191c | 0 | Young |
| 0.957055 | 0.109731 | 0.31 | 0.321 | 1 | 7 | Psd3 | -0.011 | Young |
| 0.957093 | 0.26052 | 0.289 | 0.305 | 1 | 7 | Ypel5 | -0.016 | Young |
| 0.957098 | 0.216116 | 0.105 | 0.108 | 1 | 7 | Nudt13 | -0.003 | Young |
| 0.95721 | 0.139726 | 0.196 | 0.197 | 1 | 7 | Taok2 | -0.001 | Young |
| 0.957222 | 0.107075 | 0.187 | 0.192 | 1 | 7 | Zfp740 | -0.005 | Young |
| 0.957228 | 0.170275 | 0.467 | 0.495 | 1 | 7 | Yeats4 | -0.028 | Young |
| 0.957596 | 0.277887 | 0.446 | 0.477 | 1 | 7 | Ccar1 | -0.031 | Young |
| 0.957598 | 0.218845 | 0.286 | 0.3 | 1 | 7 | Ei24 | -0.014 | Young |
| 0.958132 | 0.642004 | 0.042 | 0.044 | 1 | 7 | Tubb2b | -0.002 | Young |
| 0.958671 | 0.314552 | 0.307 | 0.326 | 1 | 7 | Slain2 | -0.019 | Young |
| 0.958671 | 0.006771 | 0.322 | 0.328 | 1 | 7 | Limd2 | -0.006 | Young |
| 0.958878 | 0.008903 | 0.943 | 0.936 | 1 | 7 | Cdc42 | 0.007 | Young |

| | | | | | | | | |
|---|---|---|---|---|---|---|---|---|
| 0.959025 | 0.298941 | 0.039 | 0.038 | 1 | 7 | Pigg | 0.001 | Young |
| 0.959328 | 0.047489 | 0.283 | 0.287 | 1 | 7 | Maf1 | -0.004 | Young |
| 0.959698 | 0.188728 | 0.515 | 0.538 | 1 | 7 | Thrap3 | -0.023 | Young |
| 0.959944 | 0.104871 | 0.16 | 0.164 | 1 | 7 | Chst12 | -0.004 | Young |
| 0.960019 | 0.022268 | 0.178 | 0.179 | 1 | 7 | Phax | -0.001 | Young |
| 0.960323 | 0.066336 | 0.142 | 0.144 | 1 | 7 | Kank3 | -0.002 | Young |
| 0.960462 | 1.221971 | 0.033 | 0.033 | 1 | 7 | Hgf | 0 | Young |
| 0.960742 | 0.245722 | 0.108 | 0.11 | 1 | 7 | Lsm10 | -0.002 | Young |
| 0.961079 | 0.389627 | 0.136 | 0.138 | 1 | 7 | Phkb | -0.002 | Young |
| 0.961156 | 0.267745 | 0.301 | 0.318 | 1 | 7 | Csnk1g3 | -0.017 | Young |
| 0.961199 | 0.099085 | 0.075 | 0.077 | 1 | 7 | Ncs1 | -0.002 | Young |
| 0.961284 | 0.158862 | 0.057 | 0.059 | 1 | 7 | Lonrf1 | -0.002 | Young |
| 0.961516 | 0.109618 | 0.19 | 0.197 | 1 | 7 | Alg8 | -0.007 | Young |
| 0.961532 | 0.349866 | 0.157 | 0.159 | 1 | 7 | Ankrd49 | -0.002 | Young |
| 0.96198 | 0.127726 | 0.509 | 0.546 | 1 | 7 | Gtf2h5 | -0.037 | Young |
| 0.962033 | 0.157136 | 0.253 | 0.259 | 1 | 7 | Cnot8 | -0.006 | Young |
| 0.962063 | 0.233384 | 0.241 | 0.251 | 1 | 7 | Tor1aip2 | -0.01 | Young |
| 0.962148 | 0.184453 | 0.617 | 0.631 | 1 | 7 | Polr1d | -0.014 | Young |
| 0.962178 | 0.257289 | 0.148 | 0.154 | 1 | 7 | Zfp422 | -0.006 | Young |
| 0.962609 | 0.483197 | 0.099 | 0.103 | 1 | 7 | Dgkh | -0.004 | Young |
| 0.962653 | 0.277542 | 0.163 | 0.167 | 1 | 7 | Cyhr1 | -0.004 | Young |
| 0.962952 | 0.114297 | 0.494 | 0.51 | 1 | 7 | Bcl10 | -0.016 | Young |
| 0.963076 | 0.383367 | 0.03 | 0.031 | 1 | 7 | Rnf24 | -0.001 | Young |
| 0.963157 | 0.299725 | 0.09 | 0.092 | 1 | 7 | Sesn3 | -0.002 | Young |
| 0.963285 | 0.321965 | 0.154 | 0.156 | 1 | 7 | Gstm1 | -0.002 | Young |
| 0.963453 | 0.153804 | 0.259 | 0.272 | 1 | 7 | Timm8b | -0.013 | Young |
| 0.963566 | 0.004566 | 0.428 | 0.441 | 1 | 7 | Eif4e | -0.013 | Young |
| 0.963589 | 0.2111 | 0.084 | 0.085 | 1 | 7 | Prkdc | -0.001 | Young |
| 0.963597 | 0.511463 | 0.102 | 0.105 | 1 | 7 | Tsc2 | -0.003 | Young |
| 0.96365 | 0.347117 | 0.114 | 0.115 | 1 | 7 | Ggps1 | -0.001 | Young |
| 0.963664 | 0.397879 | 0.069 | 0.069 | 1 | 7 | Zfp574 | 0 | Young |
| 0.963838 | 0.746053 | 0.027 | 0.028 | 1 | 7 | Gpx7 | -0.001 | Young |
| 0.964097 | 0.389896 | 0.117 | 0.115 | 1 | 7 | Ambra1 | 0.002 | Young |
| 0.964169 | 0.852856 | 0.036 | 0.036 | 1 | 7 | A630072M | 0 | Young |
| 0.964169 | 0.822645 | 0.036 | 0.036 | 1 | 7 | Pcolce | 0 | Young |
| 0.964312 | 0.162383 | 0.187 | 0.192 | 1 | 7 | Erlin2 | -0.005 | Young |
| 0.964532 | 0.169768 | 0.413 | 0.428 | 1 | 7 | Soat1 | -0.015 | Young |
| 0.964882 | 0.006305 | 0.045 | 0.046 | 1 | 7 | Zfp810 | -0.001 | Young |
| 0.964993 | 0.488512 | 0.03 | 0.031 | 1 | 7 | Zbtb12 | -0.001 | Young |
| 0.965013 | 0.240827 | 0.346 | 0.346 | 1 | 7 | Klf7 | 0 | Young |
| 0.965144 | 0.014492 | 0.401 | 0.41 | 1 | 7 | Csnk1d | -0.009 | Young |
| 0.965749 | 0.183612 | 0.22 | 0.233 | 1 | 7 | Commd6 | -0.013 | Young |
| 0.965852 | 0.009363 | 0.56 | 0.567 | 1 | 7 | Klf3 | -0.007 | Young |
| 0.96656 | 0.340026 | 0.211 | 0.223 | 1 | 7 | Tnfaip1 | -0.012 | Young |
| 0.967196 | 0.252157 | 0.319 | 0.328 | 1 | 7 | Emsy | -0.009 | Young |

| | | | | | | | | |
|---|---|---|---|---|---|---|---|---|
| 0.967487 | 0.588883 | 0.051 | 0.051 | 1 | 7 | Rnf25 | 0 | Young |
| 0.968293 | 0.028346 | 0.063 | 0.064 | 1 | 7 | Flt3l | -0.001 | Young |
| 0.968533 | 0.166396 | 0.295 | 0.308 | 1 | 7 | Nt5c | -0.013 | Young |
| 0.968544 | 0.10386 | 0.47 | 0.485 | 1 | 7 | Ash1l | -0.015 | Young |
| 0.968642 | 0.142587 | 0.108 | 0.11 | 1 | 7 | Gfer | -0.002 | Young |
| 0.968659 | 0.015661 | 0.06 | 0.062 | 1 | 7 | Smg8 | -0.002 | Young |
| 0.968708 | 0.146687 | 0.172 | 0.172 | 1 | 7 | Sdhaf4 | 0 | Young |
| 0.968866 | 0.297942 | 0.09 | 0.092 | 1 | 7 | Pcyt2 | -0.002 | Young |
| 0.968949 | 0.021858 | 0.666 | 0.7 | 1 | 7 | Tgfb1 | -0.034 | Young |
| 0.969263 | 0.058577 | 0.039 | 0.038 | 1 | 7 | Gcfc2 | 0.001 | Young |
| 0.969403 | 0.541167 | 0.133 | 0.136 | 1 | 7 | Dcaf17 | -0.003 | Young |
| 0.969592 | 0.354252 | 0.202 | 0.2 | 1 | 7 | Sos2 | 0.002 | Young |
| 0.969728 | 0.572782 | 0.051 | 0.051 | 1 | 7 | Nprl3 | 0 | Young |
| 0.969732 | 0.294557 | 0.148 | 0.149 | 1 | 7 | Pkd1 | -0.001 | Young |
| 0.970117 | 0.174597 | 0.039 | 0.038 | 1 | 7 | Gsdme | 0.001 | Young |
| 0.970471 | 0.0416 | 0.211 | 0.208 | 1 | 7 | Eps15l1 | 0.003 | Young |
| 0.970546 | 0.430135 | 0.205 | 0.215 | 1 | 7 | Atl2 | -0.01 | Young |
| 0.970554 | 0.362669 | 0.117 | 0.121 | 1 | 7 | Haus1 | -0.004 | Young |
| 0.970635 | 0.430675 | 0.123 | 0.126 | 1 | 7 | Rrp36 | -0.003 | Young |
| 0.970896 | 0.00975 | 0.723 | 0.754 | 1 | 7 | Pdia3 | -0.031 | Young |
| 0.971146 | 0.597775 | 0.163 | 0.169 | 1 | 7 | Arhgap12 | -0.006 | Young |
| 0.971168 | 0.152986 | 0.301 | 0.313 | 1 | 7 | Pik3c2a | -0.012 | Young |
| 0.971772 | 0.221357 | 0.154 | 0.159 | 1 | 7 | Tubgcp5 | -0.005 | Young |
| 0.972342 | 0.122531 | 0.509 | 0.538 | 1 | 7 | Birc6 | -0.029 | Young |
| 0.972403 | 0.452272 | 0.015 | 0.015 | 1 | 7 | AW011738 | 0 | Young |
| 0.972403 | 0.076706 | 0.015 | 0.015 | 1 | 7 | Ms4a4b | 0 | Young |
| 0.972662 | 0.346876 | 0.03 | 0.031 | 1 | 7 | Spindoc | -0.001 | Young |
| 0.97271 | 0.106858 | 0.331 | 0.344 | 1 | 7 | Il3ra | -0.013 | Young |
| 0.973559 | 0.156301 | 0.045 | 0.046 | 1 | 7 | Slc52a2 | -0.001 | Young |
| 0.97362 | 0.241354 | 0.322 | 0.331 | 1 | 7 | Anapc1 | -0.009 | Young |
| 0.973749 | 0.053423 | 0.015 | 0.015 | 1 | 7 | Map7 | 0 | Young |
| 0.973776 | 0.020556 | 0.738 | 0.741 | 1 | 7 | Sec61b | -0.003 | Young |
| 0.973875 | 0.235663 | 0.066 | 0.067 | 1 | 7 | Fxn | -0.001 | Young |
| 0.973897 | 0.3265 | 0.036 | 0.036 | 1 | 7 | L3mbtl3 | 0 | Young |
| 0.974153 | 0.014663 | 0.678 | 0.682 | 1 | 7 | Ndufb7 | -0.004 | Young |
| 0.974634 | 0.11193 | 0.392 | 0.403 | 1 | 7 | Polr2e | -0.011 | Young |
| 0.974768 | 0.05047 | 0.142 | 0.141 | 1 | 7 | Sbf1 | 0.001 | Young |
| 0.974936 | 0.005745 | 0.723 | 0.759 | 1 | 7 | Mtdh | -0.036 | Young |
| 0.975095 | 0.103098 | 0.015 | 0.015 | 1 | 7 | Gm10605 | 0 | Young |
| 0.975095 | 0.036072 | 0.015 | 0.015 | 1 | 7 | Fndc10 | 0 | Young |
| 0.975103 | 0.029164 | 0.127 | 0.128 | 1 | 7 | Sacs | -0.001 | Young |
| 0.975488 | 0.285253 | 0.28 | 0.292 | 1 | 7 | Fam107b | -0.012 | Young |
| 0.975539 | 0.628643 | 0.03 | 0.031 | 1 | 7 | Rmdn2 | -0.001 | Young |
| 0.975718 | 0.142287 | 0.217 | 0.221 | 1 | 7 | Zdhhc5 | -0.004 | Young |
| 0.975719 | 0.668731 | 0.09 | 0.092 | 1 | 7 | Surf1 | -0.002 | Young |

| | | | | | | | | |
|---|---|---|---|---|---|---|---|---|
| 0.975743 | 0.111154 | 0.105 | 0.108 | 1 | 7 | Mcu | -0.003 | Young |
| 0.975747 | 0.19609 | 0.313 | 0.331 | 1 | 7 | Prkcb | -0.018 | Young |
| 0.975794 | 0.064204 | 0.44 | 0.464 | 1 | 7 | Fem1c | -0.024 | Young |
| 0.975799 | 0.366722 | 0.096 | 0.097 | 1 | 7 | Tbc1d10a | -0.001 | Young |
| 0.975926 | 1.008041 | 0.045 | 0.046 | 1 | 7 | Bpgm | -0.001 | Young |
| 0.97603 | 0.067627 | 0.524 | 0.538 | 1 | 7 | Ube2b | -0.014 | Young |
| 0.976134 | 0.354873 | 0.184 | 0.192 | 1 | 7 | Bloc1s4 | -0.008 | Young |
| 0.976169 | 0.008943 | 0.521 | 0.538 | 1 | 7 | Ppp3r1 | -0.017 | Young |
| 0.976223 | 0.070008 | 0.229 | 0.233 | 1 | 7 | Ubxn6 | -0.004 | Young |
| 0.976312 | 0.763275 | 0.072 | 0.074 | 1 | 7 | Supt3 | -0.002 | Young |
| 0.976374 | 0.339565 | 0.157 | 0.159 | 1 | 7 | BC037034 | -0.002 | Young |
| 0.976441 | 0.241881 | 0.015 | 0.015 | 1 | 7 | Sirpb1a | 0 | Young |
| 0.976498 | 0.851826 | 0.03 | 0.031 | 1 | 7 | Tmem144 | -0.001 | Young |
| 0.976968 | 0.154138 | 0.244 | 0.249 | 1 | 7 | Atp9b | -0.005 | Young |
| 0.977457 | 0.748583 | 0.03 | 0.031 | 1 | 7 | Clip2 | -0.001 | Young |
| 0.977787 | 0.384061 | 0.015 | 0.015 | 1 | 7 | Mettl15 | 0 | Young |
| 0.977787 | 0.252867 | 0.015 | 0.015 | 1 | 7 | Mutyh | 0 | Young |
| 0.97786 | 0.014323 | 0.931 | 0.933 | 1 | 7 | Rpl5 | -0.002 | Young |
| 0.97789 | 0.132508 | 0.395 | 0.423 | 1 | 7 | Denr | -0.028 | Young |
| 0.977984 | 0.287846 | 0.087 | 0.09 | 1 | 7 | Tmem159 | -0.003 | Young |
| 0.978416 | 0.228631 | 0.03 | 0.031 | 1 | 7 | Zfp7 | -0.001 | Young |
| 0.978575 | 0.241167 | 0.09 | 0.092 | 1 | 7 | Tmem80 | -0.002 | Young |
| 0.978651 | 0.124716 | 0.072 | 0.072 | 1 | 7 | Rpap1 | 0 | Young |
| 0.979189 | 0.170092 | 0.295 | 0.303 | 1 | 7 | Impad1 | -0.008 | Young |
| 0.979199 | 0.254214 | 0.205 | 0.208 | 1 | 7 | Tmem127 | -0.003 | Young |
| 0.979293 | 0.037119 | 0.166 | 0.167 | 1 | 7 | Sap30l | -0.001 | Young |
| 0.979315 | 0.204623 | 0.229 | 0.233 | 1 | 7 | Sgk1 | -0.004 | Young |
| 0.979407 | 0.058757 | 0.349 | 0.354 | 1 | 7 | St3gal4 | -0.005 | Young |
| 0.979409 | 0.118083 | 0.325 | 0.341 | 1 | 7 | Jtb | -0.016 | Young |
| 0.979474 | 0.33273 | 0.105 | 0.108 | 1 | 7 | Gm5617 | -0.003 | Young |
| 0.979523 | 0.272022 | 0.157 | 0.159 | 1 | 7 | Slc16a7 | -0.002 | Young |
| 0.979538 | 0.781799 | 0.018 | 0.018 | 1 | 7 | Irak1bp1 | 0 | Young |
| 0.979644 | 0.072409 | 0.307 | 0.315 | 1 | 7 | Nsfl1c | -0.008 | Young |
| 0.979872 | 0.751584 | 0.045 | 0.046 | 1 | 7 | Top3a | -0.001 | Young |
| 0.980144 | 0.106325 | 0.12 | 0.123 | 1 | 7 | Wdr70 | -0.003 | Young |
| 0.980158 | 0.038963 | 0.235 | 0.244 | 1 | 7 | Rcsd1 | -0.009 | Young |
| 0.980685 | 0.012869 | 0.033 | 0.033 | 1 | 7 | Tmem150b | 0 | Young |
| 0.980778 | 0.456039 | 0.018 | 0.018 | 1 | 7 | Ccdc114 | 0 | Young |
| 0.980778 | 0.357932 | 0.018 | 0.018 | 1 | 7 | Zfp458 | 0 | Young |
| 0.980805 | 0.251023 | 0.096 | 0.097 | 1 | 7 | Gpr85 | -0.001 | Young |
| 0.981042 | 0.123173 | 0.151 | 0.154 | 1 | 7 | Bola1 | -0.003 | Young |
| 0.981148 | 0.256263 | 0.066 | 0.067 | 1 | 7 | Wbp1 | -0.001 | Young |
| 0.981825 | 0.407757 | 0.015 | 0.015 | 1 | 7 | Wfdc18 | 0 | Young |
| 0.981825 | 0.246639 | 0.015 | 0.015 | 1 | 7 | 1700086O( | 0 | Young |
| 0.981825 | 0.055109 | 0.015 | 0.015 | 1 | 7 | Fbxo48 | 0 | Young |

| | | | | | | | | |
|---|---|---|---|---|---|---|---|---|
| 0.982018 | 0.440751 | 0.018 | 0.018 | 1 | 7 | Gm4285 | 0 | Young |
| 0.982018 | 0.407609 | 0.018 | 0.018 | 1 | 7 | Cidec | 0 | Young |
| 0.982018 | 0.143354 | 0.018 | 0.018 | 1 | 7 | 3300005D( | 0 | Young |
| 0.982331 | 0.028936 | 0.238 | 0.246 | 1 | 7 | Srd5a3 | -0.008 | Young |
| 0.983073 | 0.197099 | 0.298 | 0.318 | 1 | 7 | Appbp2 | -0.02 | Young |
| 0.983125 | 0.07647 | 0.069 | 0.069 | 1 | 7 | Sh2b1 | 0 | Young |
| 0.983171 | 1.345636 | 0.015 | 0.015 | 1 | 7 | Hexim2 | 0 | Young |
| 0.983171 | 0.525794 | 0.015 | 0.015 | 1 | 7 | Arv1 | 0 | Young |
| 0.983401 | 0.150191 | 0.62 | 0.621 | 1 | 7 | Eif5b | -0.001 | Young |
| 0.983695 | 0.31142 | 0.238 | 0.249 | 1 | 7 | Rad54l | -0.011 | Young |
| 0.983698 | 0.161659 | 0.166 | 0.167 | 1 | 7 | Thap2 | -0.001 | Young |
| 0.983738 | 0.844241 | 0.105 | 0.108 | 1 | 7 | Nostrin | -0.003 | Young |
| 0.984174 | 0.125419 | 0.274 | 0.29 | 1 | 7 | Afg3l1 | -0.016 | Young |
| 0.984287 | 0.381158 | 0.09 | 0.092 | 1 | 7 | Rab33b | -0.002 | Young |
| 0.984472 | 0.119631 | 0.334 | 0.328 | 1 | 7 | Gm26740 | 0.006 | Young |
| 0.984498 | 1.106462 | 0.018 | 0.018 | 1 | 7 | Fsbp | 0 | Young |
| 0.984705 | 0.024798 | 0.883 | 0.882 | 1 | 7 | Rbpj | 0.001 | Young |
| 0.984707 | 0.412864 | 0.241 | 0.251 | 1 | 7 | Zfp217 | -0.01 | Young |
| 0.984977 | 0.325025 | 0.108 | 0.11 | 1 | 7 | Akt1s1 | -0.002 | Young |
| 0.985044 | 0.184644 | 0.343 | 0.351 | 1 | 7 | Inpp4a | -0.008 | Young |
| 0.985165 | 0.173428 | 0.102 | 0.105 | 1 | 7 | Cntrob | -0.003 | Young |
| 0.985194 | 0.227564 | 0.38 | 0.395 | 1 | 7 | Sp1 | -0.015 | Young |
| 0.985253 | 0.104061 | 0.554 | 0.579 | 1 | 7 | Wapl | -0.025 | Young |
| 0.985357 | 0.110452 | 0.542 | 0.572 | 1 | 7 | Selenos | -0.03 | Young |
| 0.985579 | 0.226697 | 0.175 | 0.177 | 1 | 7 | Dmtf1 | -0.002 | Young |
| 0.985721 | 0.096481 | 0.069 | 0.069 | 1 | 7 | Mtor | 0 | Young |
| 0.985778 | 0.458212 | 0.066 | 0.067 | 1 | 7 | Sprtn | -0.001 | Young |
| 0.985803 | 0.165305 | 0.163 | 0.164 | 1 | 7 | Slfn8 | -0.001 | Young |
| 0.985863 | 1.024168 | 0.015 | 0.015 | 1 | 7 | Fhit | 0 | Young |
| 0.986015 | 0.232855 | 0.184 | 0.182 | 1 | 7 | Alas1 | 0.002 | Young |
| 0.986025 | 0.044297 | 0.163 | 0.164 | 1 | 7 | Pign | -0.001 | Young |
| 0.986198 | 0.200673 | 0.383 | 0.397 | 1 | 7 | Dut | -0.014 | Young |
| 0.986265 | 0.301063 | 0.133 | 0.136 | 1 | 7 | Zfp672 | -0.003 | Young |
| 0.986307 | 0.053726 | 0.437 | 0.441 | 1 | 7 | Ank | -0.004 | Young |
| 0.98659 | 0.103822 | 0.434 | 0.436 | 1 | 7 | Rnpep | -0.002 | Young |
| 0.986871 | 0.21345 | 0.277 | 0.287 | 1 | 7 | Got2 | -0.01 | Young |
| 0.986957 | 0.212004 | 0.163 | 0.167 | 1 | 7 | Wrn | -0.004 | Young |
| 0.986978 | 0.097902 | 0.018 | 0.018 | 1 | 7 | Ptpdc1 | 0 | Young |
| 0.987101 | 0.094238 | 0.066 | 0.067 | 1 | 7 | Atg16l2 | -0.001 | Young |
| 0.987239 | 0.181528 | 0.054 | 0.054 | 1 | 7 | Zc3h3 | 0 | Young |
| 0.987487 | 0.021122 | 0.334 | 0.346 | 1 | 7 | Mcur1 | -0.012 | Young |
| 0.987762 | 0.536015 | 0.232 | 0.244 | 1 | 7 | Fam3c | -0.012 | Young |
| 0.987891 | 0.619507 | 0.196 | 0.2 | 1 | 7 | mt-Nd6 | -0.004 | Young |
| 0.988482 | 0.055036 | 0.38 | 0.403 | 1 | 7 | Gm10073 | -0.023 | Young |
| 0.988556 | 0.874724 | 0.015 | 0.015 | 1 | 7 | Egfl7 | 0 | Young |

| | | | | | | | | |
|---|---|---|---|---|---|---|---|---|
| 0.988571 | 0.298255 | 0.271 | 0.279 | 1 | 7 | Nek9 | -0.008 | Young |
| 0.988773 | 0.420123 | 0.373 | 0.4 | 1 | 7 | Snx18 | -0.027 | Young |
| 0.9888 | 0.356823 | 0.217 | 0.226 | 1 | 7 | 0610030E2 | -0.009 | Young |
| 0.988908 | 0.419778 | 0.16 | 0.167 | 1 | 7 | Slc25a25 | -0.007 | Young |
| 0.988967 | 0.477026 | 0.03 | 0.031 | 1 | 7 | Asb13 | -0.001 | Young |
| 0.989026 | 0.139944 | 0.117 | 0.118 | 1 | 7 | Iqsec1 | -0.001 | Young |
| 0.989458 | 0.038962 | 0.018 | 0.018 | 1 | 7 | Fkbp7 | 0 | Young |
| 0.989578 | 0.049756 | 0.295 | 0.3 | 1 | 7 | Ndufa9 | -0.005 | Young |
| 0.989823 | 0.047976 | 0.036 | 0.036 | 1 | 7 | Cdkn1c | 0 | Young |
| 0.989877 | 0.041768 | 0.717 | 0.703 | 1 | 7 | Cbx3 | 0.014 | Young |
| 0.989947 | 0.118165 | 0.178 | 0.179 | 1 | 7 | Csnk1g1 | -0.001 | Young |
| 0.990089 | 0.051364 | 0.461 | 0.477 | 1 | 7 | Dtymk | -0.016 | Young |
| 0.990168 | 0.108652 | 0.298 | 0.308 | 1 | 7 | Bcl7b | -0.01 | Young |
| 0.990408 | 0.31424 | 0.066 | 0.067 | 1 | 7 | Tlr6 | -0.001 | Young |
| 0.990458 | 0.29674 | 0.114 | 0.115 | 1 | 7 | 4921524J1 | -0.001 | Young |
| 0.990802 | 0.6861 | 0.033 | 0.033 | 1 | 7 | Gpatch3 | 0 | Young |
| 0.991113 | 0.206866 | 0.389 | 0.405 | 1 | 7 | Nrm | -0.016 | Young |
| 0.991149 | 0.037492 | 0.696 | 0.677 | 1 | 7 | G3bp2 | 0.019 | Young |
| 0.992121 | 0.346146 | 0.087 | 0.087 | 1 | 7 | Acot8 | 0 | Young |
| 0.992246 | 0.216057 | 0.123 | 0.123 | 1 | 7 | Actr8 | 0 | Young |
| 0.992547 | 0.282408 | 0.075 | 0.077 | 1 | 7 | Fam217b | -0.002 | Young |
| 0.992601 | 0.332933 | 0.16 | 0.159 | 1 | 7 | Mapkapk3 | 0.001 | Young |
| 0.992919 | 0.445428 | 0.175 | 0.179 | 1 | 7 | Nhlrc3 | -0.004 | Young |
| 0.993031 | 0.400503 | 0.123 | 0.126 | 1 | 7 | Myg1 | -0.003 | Young |
| 0.993282 | 0.045724 | 0.205 | 0.2 | 1 | 7 | Sin3a | 0.005 | Young |
| 0.993383 | 0.39802 | 0.09 | 0.09 | 1 | 7 | E2f6 | 0 | Young |
| 0.993561 | 0.342107 | 0.033 | 0.033 | 1 | 7 | Pigw | 0 | Young |
| 0.993561 | 0.313821 | 0.033 | 0.033 | 1 | 7 | Acyp2 | 0 | Young |
| 0.9936 | 0.081297 | 0.56 | 0.579 | 1 | 7 | Sgpp1 | -0.019 | Young |
| 0.993739 | 0.328764 | 0.208 | 0.205 | 1 | 7 | Egr3 | 0.003 | Young |
| 0.994602 | 0.178216 | 0.063 | 0.064 | 1 | 7 | Tprn | -0.001 | Young |
| 0.995184 | 0.119455 | 0.187 | 0.19 | 1 | 7 | Smurf2 | -0.003 | Young |
| 0.995372 | 0.047603 | 0.551 | 0.569 | 1 | 7 | Ythdc1 | -0.018 | Young |
| 0.99545 | 0.180562 | 0.172 | 0.174 | 1 | 7 | Thumpd1 | -0.002 | Young |
| 0.996092 | 0.207497 | 0.437 | 0.464 | 1 | 7 | Smap2 | -0.027 | Young |
| 0.99648 | 0.138247 | 0.22 | 0.228 | 1 | 7 | Igf1r | -0.008 | Young |
| 0.996494 | 0.036284 | 0.407 | 0.428 | 1 | 7 | Ifnar1 | -0.021 | Young |
| 0.996532 | 0.01747 | 0.428 | 0.444 | 1 | 7 | Psip1 | -0.016 | Young |
| 0.996743 | 0.128854 | 0.084 | 0.085 | 1 | 7 | B3galnt1 | -0.001 | Young |
| 0.996766 | 0.203511 | 0.325 | 0.341 | 1 | 7 | Cdc40 | -0.016 | Young |
| 0.997146 | 0.007625 | 0.283 | 0.295 | 1 | 7 | Impact | -0.012 | Young |
| 0.997647 | 0.379924 | 0.175 | 0.182 | 1 | 7 | Arfrp1 | -0.007 | Young |
| 0.998548 | 0.667733 | 0.13 | 0.136 | 1 | 7 | Zfyve26 | -0.006 | Young |
| 0.998747 | 0.286351 | 0.286 | 0.287 | 1 | 7 | Pts | -0.001 | Young |
| 0.999084 | 0.725215 | 0.078 | 0.079 | 1 | 7 | Gpank1 | -0.001 | Young |

| | | | | | | | | |
|---|---|---|---|---|---|---|---|---|
| 0.999112 | 0.198451 | 0.084 | 0.085 | 1 | 7 | Rtp4 | -0.001 | Young |
| 0.999118 | 0.32332 | 0.289 | 0.308 | 1 | 7 | Psmg4 | -0.019 | Young |
| 0.999267 | 0.023168 | 0.627 | 0.628 | 1 | 7 | Cap1 | -0.001 | Young |
| 0.999616 | 0.490523 | 0.048 | 0.049 | 1 | 7 | Rabl3 | -0.001 | Young |
| 0.999696 | 0.036254 | 0.503 | 0.521 | 1 | 7 | Sdf4 | -0.018 | Young |
| 7.15E-46 | 5.186957 | 0.592 | 0.065 | 2.22E-41 | 8 | Cxcl13 | 0.527 | Old |
| 2.15E-32 | 4.519714 | 0.391 | 0.021 | 6.66E-28 | 8 | Cd55 | 0.37 | Old |
| 2.73E-25 | 3.469252 | 0.43 | 0.078 | 8.48E-21 | 8 | C3 | 0.352 | Old |
| 5.47E-22 | 1.766206 | 0.788 | 0.46 | 1.70E-17 | 8 | Ccl8 | 0.328 | Old |
| 1.31E-20 | 0.573056 | 0.994 | 0.997 | 4.06E-16 | 8 | B2m | -0.003 | Old |
| 1.25E-17 | 0.907575 | 0.939 | 0.865 | 3.87E-13 | 8 | Gda | 0.074 | Old |
| 1.07E-15 | 1.318754 | 0.721 | 0.442 | 3.33E-11 | 8 | Mmp9 | 0.279 | Old |
| 4.72E-14 | 0.913171 | 0.883 | 0.777 | 1.47E-09 | 8 | Ly6e | 0.106 | Old |
| 9.62E-14 | 0.691113 | 1 | 0.932 | 2.99E-09 | 8 | Sdc3 | 0.068 | Old |
| 2.01E-13 | 1.100071 | 0.804 | 0.652 | 6.23E-09 | 8 | Cstb | 0.152 | Old |
| 7.50E-12 | 0.502869 | 1 | 0.995 | 2.33E-07 | 8 | H2-K1 | 0.005 | Old |
| 3.24E-11 | 0.306771 | 1 | 1 | 1.01E-06 | 8 | Psap | 0 | Old |
| 8.29E-11 | 4.666669 | 0.106 | 0 | 2.58E-06 | 8 | Ttn | 0.106 | Old |
| 8.86E-11 | 0.472215 | 1 | 1 | 2.75E-06 | 8 | Junb | 0 | Old |
| 1.13E-10 | 3.919805 | 0.123 | 0.005 | 3.50E-06 | 8 | Colq | 0.118 | Old |
| 1.34E-10 | 0.368777 | 1 | 1 | 4.15E-06 | 8 | mt-Co3 | 0 | Old |
| 2.20E-10 | 0.644976 | 0.983 | 0.979 | 6.83E-06 | 8 | mt-Nd3 | 0.004 | Old |
| 9.07E-10 | 0.592956 | 0.994 | 0.987 | 2.82E-05 | 8 | Hsp90b1 | 0.007 | Old |
| 1.16E-08 | 0.91974 | 0.754 | 0.603 | 0.000359 | 8 | Apobec3 | 0.151 | Old |
| 2.03E-08 | 0.535146 | 0.978 | 0.943 | 0.000629 | 8 | Samhd1 | 0.035 | Old |
| 2.66E-08 | 0.540145 | 0.972 | 0.943 | 0.000827 | 8 | BC005537 | 0.029 | Old |
| 4.36E-08 | 0.583654 | 0.978 | 0.953 | 0.001354 | 8 | Irf2bp2 | 0.025 | Old |
| 6.05E-08 | 1.638805 | 0.441 | 0.252 | 0.001878 | 8 | Fam43a | 0.189 | Old |
| 7.16E-08 | 0.816141 | 0.927 | 0.823 | 0.002224 | 8 | Lpl | 0.104 | Old |
| 1.04E-07 | 0.346709 | 1 | 1 | 0.003242 | 8 | mt-Co2 | 0 | Old |
| 1.19E-07 | 1.185335 | 0.497 | 0.309 | 0.003682 | 8 | Hist1h1c | 0.188 | Old |
| 1.23E-07 | 0.225998 | 1 | 1 | 0.00381 | 8 | mt-Co1 | 0 | Old |
| 1.36E-07 | 0.337908 | 0.966 | 0.969 | 0.004215 | 8 | Ctss | -0.003 | Old |
| 1.96E-07 | 3.482693 | 0.095 | 0.008 | 0.006081 | 8 | Dmpk | 0.087 | Old |
| 2.39E-07 | 0.187376 | 1 | 1 | 0.007418 | 8 | Actb | 0 | Old |
| 2.83E-07 | 0.381388 | 0.989 | 0.987 | 0.008794 | 8 | Ccl6 | 0.002 | Old |
| 3.08E-07 | 0.350702 | 0.989 | 0.99 | 0.009565 | 8 | Ccl9 | -0.001 | Old |
| 7.46E-07 | 0.367321 | 0.989 | 0.977 | 0.023177 | 8 | Pfn1 | 0.012 | Old |
| 8.73E-07 | 0.329946 | 1 | 1 | 0.027103 | 8 | mt-Cytb | 0 | Old |
| 9.05E-07 | 0.437136 | 0.944 | 0.906 | 0.028088 | 8 | Clec10a | 0.038 | Old |
| 9.07E-07 | 1.057906 | 0.592 | 0.413 | 0.028154 | 8 | Tspan3 | 0.179 | Old |
| 9.94E-07 | 0.374971 | 0.989 | 0.997 | 0.030869 | 8 | Ptma | -0.008 | Old |
| 1.78E-06 | 0.54724 | 0.888 | 0.823 | 0.055182 | 8 | Tuba1b | 0.065 | Old |
| 1.88E-06 | 0.725549 | 0.821 | 0.706 | 0.058489 | 8 | Arpc4 | 0.115 | Old |
| 2.35E-06 | 0.417505 | 0.983 | 0.971 | 0.073069 | 8 | Arf5 | 0.012 | Old |

| | | | | | | | | |
|---|---|---|---|---|---|---|---|---|
| 2.96E-06 | 3.553455 | 0.056 | 0 | 0.09183 | 8 | Vldlr | 0.056 | Old |
| 3.10E-06 | 1.740971 | 0.162 | 0.047 | 0.096258 | 8 | C6 | 0.115 | Old |
| 5.70E-06 | 0.636981 | 0.86 | 0.803 | 0.176993 | 8 | Xbp1 | 0.057 | Old |
| 8.25E-06 | 0.471314 | 0.888 | 0.883 | 0.256316 | 8 | Prkcd | 0.005 | Old |
| 8.28E-06 | 2.829562 | 0.106 | 0.021 | 0.257216 | 8 | Gm26870 | 0.085 | Old |
| 8.70E-06 | 0.423806 | 1 | 1 | 0.270028 | 8 | Jund | 0 | Old |
| 9.30E-06 | 0.300925 | 0.989 | 0.984 | 0.288751 | 8 | H3f3b | 0.005 | Old |
| 1.17E-05 | 0.648765 | 0.81 | 0.764 | 0.36311 | 8 | Cd63 | 0.046 | Old |
| 1.61E-05 | 0.489224 | 0.894 | 0.805 | 0.499597 | 8 | Fn1 | 0.089 | Old |
| 1.66E-05 | 0.197384 | 1 | 1 | 0.516173 | 8 | Actg1 | 0 | Old |
| 1.76E-05 | 1.45241 | 0.38 | 0.221 | 0.545905 | 8 | Bcat2 | 0.159 | Old |
| 2.27E-05 | 2.500065 | 0.084 | 0.013 | 0.705293 | 8 | Adgrg1 | 0.071 | Old |
| 2.58E-05 | 0.23765 | 1 | 1 | 0.800932 | 8 | mt-Atp6 | 0 | Old |
| 2.60E-05 | 0.599791 | 0.732 | 0.579 | 0.808609 | 8 | Sulf2 | 0.153 | Old |
| 2.85E-05 | 0.599206 | 0.877 | 0.816 | 0.88466 | 8 | Pdia6 | 0.061 | Old |
| 2.90E-05 | 0.667198 | 0.698 | 0.561 | 0.902019 | 8 | Nrp2 | 0.137 | Old |
| 2.99E-05 | 0.503308 | 0.872 | 0.868 | 0.929881 | 8 | Nrros | 0.004 | Old |
| 3.01E-05 | 0.46024 | 0.961 | 0.953 | 0.935948 | 8 | Cebpb | 0.008 | Old |
| 3.29E-05 | 0.551869 | 0.989 | 0.987 | 1 | 8 | Lgmn | 0.002 | Old |
| 3.45E-05 | 0.935811 | 0.43 | 0.268 | 1 | 8 | B4galnt1 | 0.162 | Old |
| 3.57E-05 | 0.447419 | 0.916 | 0.909 | 1 | 8 | Ier5 | 0.007 | Old |
| 4.05E-05 | 0.369723 | 0.927 | 0.935 | 1 | 8 | Anxa2 | -0.008 | Old |
| 5.03E-05 | 1.575406 | 0.207 | 0.088 | 1 | 8 | St8sia6 | 0.119 | Old |
| 5.76E-05 | 0.426047 | 0.972 | 0.979 | 1 | 8 | Grn | -0.007 | Old |
| 6.35E-05 | 0.51111 | 0.866 | 0.821 | 1 | 8 | Ssr3 | 0.045 | Old |
| 6.67E-05 | 2.737826 | 0.05 | 0.003 | 1 | 8 | Src | 0.047 | Old |
| 7.86E-05 | 0.886895 | 0.698 | 0.59 | 1 | 8 | Ddx39b | 0.108 | Old |
| 7.89E-05 | 0.613409 | 0.648 | 0.519 | 1 | 8 | Dnmt3a | 0.129 | Old |
| 8.64E-05 | 0.310914 | 0.983 | 0.992 | 1 | 8 | Ctsb | -0.009 | Old |
| 9.64E-05 | 3.020046 | 0.039 | 0 | 1 | 8 | Sh2d4a | 0.039 | Old |
| 9.90E-05 | 0.460264 | 0.866 | 0.868 | 1 | 8 | Ctsd | -0.002 | Old |
| 0.000102 | 1.416776 | 0.184 | 0.075 | 1 | 8 | Il27ra | 0.109 | Old |
| 0.000106 | 0.356072 | 0.816 | 0.829 | 1 | 8 | Gnb1 | -0.013 | Old |
| 0.00012 | 0.890559 | 0.402 | 0.249 | 1 | 8 | Rab20 | 0.153 | Old |
| 0.000123 | 0.402932 | 0.966 | 0.961 | 1 | 8 | Anxa5 | 0.005 | Old |
| 0.000127 | 0.81312 | 0.553 | 0.426 | 1 | 8 | Eif1b | 0.127 | Old |
| 0.000129 | 2.643389 | 0.061 | 0.008 | 1 | 8 | Kif2c | 0.053 | Old |
| 0.000137 | 0.658215 | 0.609 | 0.53 | 1 | 8 | Anxa1 | 0.079 | Old |
| 0.000147 | 1.479128 | 0.156 | 0.06 | 1 | 8 | Aar2 | 0.096 | Old |
| 0.000151 | 0.502184 | 0.804 | 0.735 | 1 | 8 | Pdcd6ip | 0.069 | Old |
| 0.000152 | 1.025335 | 0.425 | 0.281 | 1 | 8 | Stat1 | 0.144 | Old |
| 0.000161 | 0.840628 | 0.67 | 0.556 | 1 | 8 | Actn4 | 0.114 | Old |
| 0.000168 | 0.32458 | 0.944 | 0.891 | 1 | 8 | Klf2 | 0.053 | Old |
| 0.000168 | 1.177415 | 0.313 | 0.174 | 1 | 8 | Akr1c13 | 0.139 | Old |
| 0.000169 | 1.254494 | 0.201 | 0.091 | 1 | 8 | Dram1 | 0.11 | Old |

| | | | | | | | | |
|---|---|---|---|---|---|---|---|---|
| 0.000193 | 0.847219 | 0.207 | 0.091 | 1 | 8 | F10 | 0.116 | Old |
| 0.000211 | 0.286675 | 0.978 | 0.995 | 1 | 8 | Klf6 | -0.017 | Old |
| 0.000221 | 0.836101 | 0.525 | 0.364 | 1 | 8 | Vcam1 | 0.161 | Old |
| 0.000229 | 0.273 | 0.989 | 0.997 | 1 | 8 | Fos | -0.008 | Old |
| 0.000234 | 0.350409 | 0.894 | 0.883 | 1 | 8 | Mob1a | 0.011 | Old |
| 0.000248 | 0.522294 | 0.816 | 0.795 | 1 | 8 | Plin2 | 0.021 | Old |
| 0.000248 | 1.133343 | 0.324 | 0.192 | 1 | 8 | Ralgds | 0.132 | Old |
| 0.000252 | 0.299714 | 0.939 | 0.966 | 1 | 8 | Efhd2 | -0.027 | Old |
| 0.000306 | 0.261402 | 1 | 0.997 | 1 | 8 | C1qa | 0.003 | Old |
| 0.000316 | 0.944612 | 0.514 | 0.358 | 1 | 8 | Tfrc | 0.156 | Old |
| 0.00032 | 0.246259 | 1 | 1 | 1 | 8 | mt-Nd1 | 0 | Old |
| 0.000451 | 1.021122 | 0.408 | 0.281 | 1 | 8 | Tnfaip2 | 0.127 | Old |
| 0.000472 | 0.530265 | 0.698 | 0.584 | 1 | 8 | Cd209f | 0.114 | Old |
| 0.000507 | 0.312551 | 0.939 | 0.93 | 1 | 8 | Hspa5 | 0.009 | Old |
| 0.000563 | 1.061586 | 0.302 | 0.177 | 1 | 8 | Galnt7 | 0.125 | Old |
| 0.00058 | 1.830456 | 0.151 | 0.062 | 1 | 8 | Igf2r | 0.089 | Old |
| 0.000685 | 0.435689 | 0.95 | 0.899 | 1 | 8 | Chchd2 | 0.051 | Old |
| 0.000708 | 0.6917 | 0.553 | 0.436 | 1 | 8 | Armc8 | 0.117 | Old |
| 0.000709 | 0.461621 | 0.888 | 0.896 | 1 | 8 | Fam49b | -0.008 | Old |
| 0.00072 | 0.219561 | 0.944 | 0.987 | 1 | 8 | Lamp1 | -0.043 | Old |
| 0.00074 | 0.364395 | 0.877 | 0.896 | 1 | 8 | Tmem176a | -0.019 | Old |
| 0.000766 | 1.105806 | 0.134 | 0.052 | 1 | 8 | Rrp12 | 0.082 | Old |
| 0.000832 | 0.235035 | 0.955 | 0.969 | 1 | 8 | Actr3 | -0.014 | Old |
| 0.000953 | 2.362971 | 0.045 | 0.005 | 1 | 8 | Klra3 | 0.04 | Old |
| 0.000991 | 0.369197 | 0.894 | 0.842 | 1 | 8 | Cd44 | 0.052 | Old |
| 0.001004 | 2.227708 | 0.028 | 0 | 1 | 8 | Cdh6 | 0.028 | Old |
| 0.001009 | 0.999225 | 0.369 | 0.252 | 1 | 8 | Ckap4 | 0.117 | Old |
| 0.001083 | 2.119853 | 0.067 | 0.016 | 1 | 8 | Cpne8 | 0.051 | Old |
| 0.001086 | 0.91429 | 0.525 | 0.408 | 1 | 8 | Becn1 | 0.117 | Old |
| 0.001087 | 3.223113 | 0.05 | 0.008 | 1 | 8 | Xist | 0.042 | Old |
| 0.001092 | 0.386054 | 0.866 | 0.842 | 1 | 8 | Pcbp1 | 0.024 | Old |
| 0.001114 | 0.363642 | 0.754 | 0.712 | 1 | 8 | Hif1a | 0.042 | Old |
| 0.001143 | 0.25312 | 0.972 | 0.956 | 1 | 8 | Mafb | 0.016 | Old |
| 0.001148 | 1.893675 | 0.05 | 0.008 | 1 | 8 | Slfn1 | 0.042 | Old |
| 0.001153 | 0.493144 | 0.715 | 0.678 | 1 | 8 | Cmpk1 | 0.037 | Old |
| 0.001175 | 2.477592 | 0.05 | 0.008 | 1 | 8 | Gm15675 | 0.042 | Old |
| 0.001178 | 2.486401 | 0.067 | 0.016 | 1 | 8 | Sh3bgrl2 | 0.051 | Old |
| 0.001179 | 0.396894 | 0.788 | 0.751 | 1 | 8 | Ehd1 | 0.037 | Old |
| 0.001307 | 0.510494 | 0.676 | 0.618 | 1 | 8 | Cdc42se2 | 0.058 | Old |
| 0.00131 | 0.310795 | 0.905 | 0.904 | 1 | 8 | Gltp | 0.001 | Old |
| 0.001319 | 0.669762 | 0.575 | 0.465 | 1 | 8 | Spred1 | 0.11 | Old |
| 0.001355 | 0.464649 | 0.782 | 0.79 | 1 | 8 | Atp6v1b2 | -0.008 | Old |
| 0.001405 | 0.49567 | 0.821 | 0.782 | 1 | 8 | Gm26917 | 0.039 | Old |
| 0.001573 | 0.33636 | 0.955 | 0.964 | 1 | 8 | Arpc5 | -0.009 | Old |
| 0.001647 | 0.152422 | 0.983 | 1 | 1 | 8 | Gpx1 | -0.017 | Old |

| | | | | | | | | |
|---|---|---|---|---|---|---|---|---|
| 0.001681 | 1.03204 | 0.207 | 0.112 | 1 | 8 | Il21r | 0.095 | Old |
| 0.001775 | 0.393659 | 0.765 | 0.738 | 1 | 8 | Srsf2 | 0.027 | Old |
| 0.00182 | 5.879776 | 0.084 | 0.026 | 1 | 8 | Igkc | 0.058 | Old |
| 0.001865 | 0.808372 | 0.447 | 0.356 | 1 | 8 | Coro2a | 0.091 | Old |
| 0.001869 | 0.631551 | 0.497 | 0.382 | 1 | 8 | Ms4a8a | 0.115 | Old |
| 0.002009 | 2.880541 | 0.034 | 0.003 | 1 | 8 | Ctsk | 0.031 | Old |
| 0.002147 | 0.932651 | 0.101 | 0.036 | 1 | 8 | Sema6a | 0.065 | Old |
| 0.002185 | 0.489297 | 0.777 | 0.704 | 1 | 8 | Nme1 | 0.073 | Old |
| 0.002198 | 0.309185 | 0.821 | 0.849 | 1 | 8 | Plxnd1 | -0.028 | Old |
| 0.002228 | 0.527791 | 0.771 | 0.717 | 1 | 8 | Sgms1 | 0.054 | Old |
| 0.002245 | 0.661393 | 0.598 | 0.509 | 1 | 8 | Tuba1c | 0.089 | Old |
| 0.002304 | 0.903091 | 0.469 | 0.374 | 1 | 8 | Rgs1 | 0.095 | Old |
| 0.002419 | 0.192776 | 0.994 | 0.982 | 1 | 8 | Myl6 | 0.012 | Old |
| 0.002427 | 0.512062 | 0.709 | 0.657 | 1 | 8 | Cdk2ap2 | 0.052 | Old |
| 0.002482 | 0.544 | 0.642 | 0.553 | 1 | 8 | Hspa9 | 0.089 | Old |
| 0.002546 | 0.480032 | 0.659 | 0.561 | 1 | 8 | Ppp3r1 | 0.098 | Old |
| 0.002589 | 1.176242 | 0.212 | 0.119 | 1 | 8 | Ly9 | 0.093 | Old |
| 0.002783 | 0.714733 | 0.43 | 0.314 | 1 | 8 | Etfa | 0.116 | Old |
| 0.002791 | 2.252131 | 0.039 | 0.005 | 1 | 8 | C920006O: | 0.034 | Old |
| 0.002801 | 1.226524 | 0.156 | 0.075 | 1 | 8 | Hvcn1 | 0.081 | Old |
| 0.002809 | 0.528158 | 0.57 | 0.46 | 1 | 8 | Emilin2 | 0.11 | Old |
| 0.003025 | 2.173765 | 0.061 | 0.016 | 1 | 8 | Dmwd | 0.045 | Old |
| 0.003292 | 2.402319 | 0.022 | 0 | 1 | 8 | Rxrg | 0.022 | Old |
| 0.003434 | 1.740679 | 0.05 | 0.01 | 1 | 8 | Zfp867 | 0.04 | Old |
| 0.00352 | 0.793838 | 0.341 | 0.231 | 1 | 8 | Rasa4 | 0.11 | Old |
| 0.00387 | 0.423201 | 0.793 | 0.761 | 1 | 8 | Ccr5 | 0.032 | Old |
| 0.003885 | 1.247018 | 0.19 | 0.104 | 1 | 8 | Gpx3 | 0.086 | Old |
| 0.003998 | 0.355362 | 0.765 | 0.699 | 1 | 8 | Prkar1a | 0.066 | Old |
| 0.004004 | 0.931069 | 0.229 | 0.132 | 1 | 8 | Ifi47 | 0.097 | Old |
| 0.004043 | 0.67438 | 0.425 | 0.322 | 1 | 8 | Cers6 | 0.103 | Old |
| 0.004138 | 0.433668 | 0.793 | 0.779 | 1 | 8 | Rab8b | 0.014 | Old |
| 0.004214 | 1.087699 | 0.168 | 0.086 | 1 | 8 | Rnf40 | 0.082 | Old |
| 0.004219 | 2.182667 | 0.078 | 0.026 | 1 | 8 | Ippk | 0.052 | Old |
| 0.004302 | 0.34245 | 0.855 | 0.826 | 1 | 8 | Wdr26 | 0.029 | Old |
| 0.004608 | 0.227613 | 1 | 1 | 1 | 8 | Dusp1 | 0 | Old |
| 0.004741 | 0.360026 | 0.816 | 0.771 | 1 | 8 | Atp5g3 | 0.045 | Old |
| 0.004767 | 0.628411 | 0.598 | 0.512 | 1 | 8 | Tmem123 | 0.086 | Old |
| 0.004925 | 1.998134 | 0.073 | 0.023 | 1 | 8 | Rbm12b2 | 0.05 | Old |
| 0.00504 | 0.650827 | 0.598 | 0.556 | 1 | 8 | Ccr1 | 0.042 | Old |
| 0.005047 | 0.805755 | 0.156 | 0.078 | 1 | 8 | Nrde2 | 0.078 | Old |
| 0.005126 | 0.314006 | 0.911 | 0.961 | 1 | 8 | Sirpa | -0.05 | Old |
| 0.005296 | 0.899915 | 0.665 | 0.753 | 1 | 8 | Ighm | -0.088 | Old |
| 0.005322 | 0.657974 | 0.475 | 0.369 | 1 | 8 | Kcnn4 | 0.106 | Old |
| 0.00534 | 1.543714 | 0.112 | 0.049 | 1 | 8 | Vegfa | 0.063 | Old |
| 0.005486 | 0.762653 | 0.508 | 0.439 | 1 | 8 | Vat1 | 0.069 | Old |

| | | | | | | | | |
|---|---|---|---|---|---|---|---|---|
| 0.005559 | 0.165438 | 0.983 | 0.992 | 1 | 8 | Cfl1 | -0.009 | Old |
| 0.005834 | 0.688424 | 0.296 | 0.19 | 1 | 8 | Hr | 0.106 | Old |
| 0.005859 | 0.236521 | 0.955 | 0.961 | 1 | 8 | Timp2 | -0.006 | Old |
| 0.005864 | 1.161714 | 0.151 | 0.078 | 1 | 8 | Abcd4 | 0.073 | Old |
| 0.005901 | 0.40126 | 0.737 | 0.647 | 1 | 8 | Pmp22 | 0.09 | Old |
| 0.005961 | 0.45409 | 0.715 | 0.683 | 1 | 8 | Hdgf | 0.032 | Old |
| 0.006356 | 0.245449 | 0.922 | 0.951 | 1 | 8 | Actr2 | -0.029 | Old |
| 0.006425 | 1.574087 | 0.128 | 0.062 | 1 | 8 | Tmem110 | 0.066 | Old |
| 0.006519 | 0.938301 | 0.101 | 0.042 | 1 | 8 | Fpgt | 0.059 | Old |
| 0.006568 | 1.304716 | 0.028 | 0.003 | 1 | 8 | Clcn2 | 0.025 | Old |
| 0.006711 | 0.245658 | 0.966 | 0.964 | 1 | 8 | Pim1 | 0.002 | Old |
| 0.007021 | 0.272458 | 0.95 | 0.977 | 1 | 8 | Btg1 | -0.027 | Old |
| 0.007038 | 0.601313 | 0.475 | 0.361 | 1 | 8 | Srpr | 0.114 | Old |
| 0.007129 | 0.377356 | 0.86 | 0.839 | 1 | 8 | Arl6ip1 | 0.021 | Old |
| 0.007185 | 0.217399 | 0.927 | 0.935 | 1 | 8 | Sdcbp | -0.008 | Old |
| 0.007281 | 0.253186 | 0.821 | 0.914 | 1 | 8 | Ctsz | -0.093 | Old |
| 0.007362 | 0.550101 | 0.709 | 0.694 | 1 | 8 | Lmna | 0.015 | Old |
| 0.007544 | 0.7358 | 0.43 | 0.332 | 1 | 8 | Gorasp2 | 0.098 | Old |
| 0.007659 | 0.179273 | 0.95 | 0.969 | 1 | 8 | Capza2 | -0.019 | Old |
| 0.007765 | 0.870089 | 0.19 | 0.112 | 1 | 8 | Umad1 | 0.078 | Old |
| 0.007872 | 2.666211 | 0.034 | 0.005 | 1 | 8 | Rhou | 0.029 | Old |
| 0.007999 | 1.99176 | 0.034 | 0.005 | 1 | 8 | Cpeb1 | 0.029 | Old |
| 0.008064 | 2.130384 | 0.034 | 0.005 | 1 | 8 | 1700109H( | 0.029 | Old |
| 0.008064 | 1.780295 | 0.034 | 0.005 | 1 | 8 | Gm16094 | 0.029 | Old |
| 0.008079 | 1.758417 | 0.089 | 0.036 | 1 | 8 | Pgm3 | 0.053 | Old |
| 0.008326 | 1.051222 | 0.034 | 0.005 | 1 | 8 | 1700030KC | 0.029 | Old |
| 0.008387 | 1.188849 | 0.128 | 0.062 | 1 | 8 | Polr3f | 0.066 | Old |
| 0.008399 | 0.311315 | 0.855 | 0.852 | 1 | 8 | Dstn | 0.003 | Old |
| 0.008552 | 1.015164 | 0.335 | 0.242 | 1 | 8 | B4galt5 | 0.093 | Old |
| 0.008601 | 1.148258 | 0.134 | 0.068 | 1 | 8 | Dnaaf3 | 0.066 | Old |
| 0.00869 | 1.494228 | 0.045 | 0.01 | 1 | 8 | Parp16 | 0.035 | Old |
| 0.008782 | 1.979115 | 0.039 | 0.008 | 1 | 8 | Vsig8 | 0.031 | Old |
| 0.009095 | 1.204309 | 0.045 | 0.01 | 1 | 8 | Csprs | 0.035 | Old |
| 0.009112 | 0.41826 | 0.844 | 0.839 | 1 | 8 | Atp1a1 | 0.005 | Old |
| 0.009114 | 1.082368 | 0.324 | 0.229 | 1 | 8 | Odc1 | 0.095 | Old |
| 0.009358 | 0.459034 | 0.788 | 0.764 | 1 | 8 | Rpn1 | 0.024 | Old |
| 0.009413 | 0.627351 | 0.475 | 0.377 | 1 | 8 | Nsf | 0.098 | Old |
| 0.009471 | 0.217589 | 0.944 | 0.966 | 1 | 8 | Hnrnpa2b1 | -0.022 | Old |
| 0.009631 | 1.30057 | 0.156 | 0.086 | 1 | 8 | Mylk | 0.07 | Old |
| 0.009639 | 0.594423 | 0.447 | 0.353 | 1 | 8 | Gfpt1 | 0.094 | Old |
| 0.009657 | 0.556998 | 0.587 | 0.545 | 1 | 8 | Slc39a1 | 0.042 | Old |
| 0.009791 | 0.140239 | 0.994 | 0.997 | 1 | 8 | Ahnak | -0.003 | Old |
| 0.009867 | 0.363458 | 0.922 | 0.938 | 1 | 8 | Ap2a2 | -0.016 | Old |
| 0.009895 | 1.027635 | 0.201 | 0.119 | 1 | 8 | Thbs1 | 0.082 | Old |
| 0.009899 | 0.516505 | 0.687 | 0.665 | 1 | 8 | Plekhb2 | 0.022 | Old |

| | | | | | | | | |
|---|---|---|---|---|---|---|---|---|
| 0.009947 | 0.197633 | 0.961 | 0.956 | 1 | 8 | Ptp4a2 | 0.005 | Old |
| 0.009954 | 0.390181 | 0.754 | 0.722 | 1 | 8 | Mcub | 0.032 | Old |
| 0.010229 | 0.616733 | 0.385 | 0.294 | 1 | 8 | Rnh1 | 0.091 | Old |
| 0.010422 | 1.003534 | 0.123 | 0.06 | 1 | 8 | Hebp1 | 0.063 | Old |
| 0.010496 | 1.468615 | 0.084 | 0.034 | 1 | 8 | 10-Sep | 0.05 | Old |
| 0.010653 | 1.527737 | 0.067 | 0.023 | 1 | 8 | F7 | 0.044 | Old |
| 0.010757 | 1.2742 | 0.067 | 0.023 | 1 | 8 | Trim2 | 0.044 | Old |
| 0.011007 | 2.195413 | 0.017 | 0 | 1 | 8 | Tpx2 | 0.017 | Old |
| 0.011007 | 1.853194 | 0.017 | 0 | 1 | 8 | Rbm8a2 | 0.017 | Old |
| 0.011007 | 1.827504 | 0.017 | 0 | 1 | 8 | Cenph | 0.017 | Old |
| 0.011007 | 1.770933 | 0.017 | 0 | 1 | 8 | A930006I0 | 0.017 | Old |
| 0.011007 | 1.513558 | 0.017 | 0 | 1 | 8 | Ptpn3 | 0.017 | Old |
| 0.011007 | 1.499021 | 0.017 | 0 | 1 | 8 | Gm26582 | 0.017 | Old |
| 0.011007 | 1.466122 | 0.017 | 0 | 1 | 8 | Adm | 0.017 | Old |
| 0.011007 | 1.438956 | 0.017 | 0 | 1 | 8 | Brip1 | 0.017 | Old |
| 0.011007 | 1.253393 | 0.017 | 0 | 1 | 8 | Gm29650 | 0.017 | Old |
| 0.011007 | 1.213079 | 0.017 | 0 | 1 | 8 | Gm26867 | 0.017 | Old |
| 0.011059 | 1.400522 | 0.117 | 0.057 | 1 | 8 | Vdr | 0.06 | Old |
| 0.011436 | 0.719834 | 0.101 | 0.044 | 1 | 8 | Aig1 | 0.057 | Old |
| 0.011516 | 1.037516 | 0.307 | 0.218 | 1 | 8 | Utp11 | 0.089 | Old |
| 0.011799 | 0.693255 | 0.251 | 0.166 | 1 | 8 | Mmp19 | 0.085 | Old |
| 0.011869 | 0.588588 | 0.682 | 0.587 | 1 | 8 | Ctsl | 0.095 | Old |
| 0.012056 | 0.262006 | 0.916 | 0.94 | 1 | 8 | Pcbp2 | -0.024 | Old |
| 0.0123 | 1.452831 | 0.061 | 0.021 | 1 | 8 | Purg | 0.04 | Old |
| 0.012377 | 0.227736 | 0.899 | 0.919 | 1 | 8 | H2-DMb1 | -0.02 | Old |
| 0.012384 | 0.423953 | 0.698 | 0.665 | 1 | 8 | Gsn | 0.033 | Old |
| 0.012596 | 0.607216 | 0.24 | 0.151 | 1 | 8 | Klhdc4 | 0.089 | Old |
| 0.012644 | 1.627263 | 0.061 | 0.021 | 1 | 8 | Pop1 | 0.04 | Old |
| 0.012792 | 1.018298 | 0.151 | 0.083 | 1 | 8 | Ezh2 | 0.068 | Old |
| 0.012882 | 0.801694 | 0.352 | 0.257 | 1 | 8 | Prmt1 | 0.095 | Old |
| 0.013369 | 0.283805 | 0.872 | 0.881 | 1 | 8 | Atp5a1 | -0.009 | Old |
| 0.013439 | 0.912617 | 0.145 | 0.081 | 1 | 8 | Bcl2l13 | 0.064 | Old |
| 0.013521 | 1.381201 | 0.123 | 0.062 | 1 | 8 | Snhg9 | 0.061 | Old |
| 0.013594 | 0.477512 | 0.525 | 0.455 | 1 | 8 | Anxa4 | 0.07 | Old |
| 0.013856 | 1.988415 | 0.056 | 0.018 | 1 | 8 | Zfp764 | 0.038 | Old |
| 0.01387 | 0.876263 | 0.313 | 0.229 | 1 | 8 | Renbp | 0.084 | Old |
| 0.014545 | 1.16562 | 0.14 | 0.075 | 1 | 8 | Usp53 | 0.065 | Old |
| 0.014852 | 0.320613 | 0.687 | 0.626 | 1 | 8 | Eif5b | 0.061 | Old |
| 0.015136 | 0.999962 | 0.056 | 0.018 | 1 | 8 | Ifi213 | 0.038 | Old |
| 0.015363 | 1.125255 | 0.212 | 0.138 | 1 | 8 | Dpep2 | 0.074 | Old |
| 0.015383 | 0.254376 | 0.911 | 0.912 | 1 | 8 | Ptbp3 | -0.001 | Old |
| 0.01549 | 0.553341 | 0.536 | 0.447 | 1 | 8 | Epb41l2 | 0.089 | Old |
| 0.015634 | 1.049105 | 0.279 | 0.2 | 1 | 8 | Fabp5 | 0.079 | Old |
| 0.015759 | 1.325604 | 0.073 | 0.029 | 1 | 8 | Cd4 | 0.044 | Old |
| 0.015981 | 0.108133 | 0.955 | 0.99 | 1 | 8 | Cdc42 | -0.035 | Old |

| | | | | | | | | |
|---|---|---|---|---|---|---|---|---|
| 0.016101 | 0.51167 | 0.056 | 0.018 | 1 | 8 | Polr3h | 0.038 | Old |
| 0.016229 | 1.056606 | 0.112 | 0.055 | 1 | 8 | Kti12 | 0.057 | Old |
| 0.016819 | 0.24001 | 0.955 | 0.945 | 1 | 8 | Rap1b | 0.01 | Old |
| 0.016887 | 1.354056 | 0.084 | 0.036 | 1 | 8 | Asb1 | 0.048 | Old |
| 0.017004 | 0.884877 | 0.358 | 0.278 | 1 | 8 | Ccdc115 | 0.08 | Old |
| 0.017093 | 1.327019 | 0.095 | 0.044 | 1 | 8 | Zw10 | 0.051 | Old |
| 0.017493 | 0.922085 | 0.196 | 0.122 | 1 | 8 | Traf3 | 0.074 | Old |
| 0.017546 | 0.819858 | 0.302 | 0.223 | 1 | 8 | Rnd3 | 0.079 | Old |
| 0.01763 | 0.59309 | 0.497 | 0.403 | 1 | 8 | Ppp1r21 | 0.094 | Old |
| 0.017708 | 0.941344 | 0.173 | 0.104 | 1 | 8 | Ccdc127 | 0.069 | Old |
| 0.017721 | 0.949423 | 0.302 | 0.221 | 1 | 8 | 2310022A1 | 0.081 | Old |
| 0.017777 | 1.61587 | 0.05 | 0.016 | 1 | 8 | C130026I2 | 0.034 | Old |
| 0.017822 | 0.920719 | 0.151 | 0.086 | 1 | 8 | Dkc1 | 0.065 | Old |
| 0.017873 | 1.100987 | 0.05 | 0.016 | 1 | 8 | Hgh1 | 0.034 | Old |
| 0.017984 | 1.190225 | 0.084 | 0.036 | 1 | 8 | Thrb | 0.048 | Old |
| 0.018176 | 0.569093 | 0.598 | 0.582 | 1 | 8 | Pla2g7 | 0.016 | Old |
| 0.018182 | 0.453948 | 0.693 | 0.655 | 1 | 8 | Rnf149 | 0.038 | Old |
| 0.018262 | 0.989602 | 0.05 | 0.016 | 1 | 8 | Bend3 | 0.034 | Old |
| 0.019136 | 1.758519 | 0.045 | 0.013 | 1 | 8 | Rsad1 | 0.032 | Old |
| 0.019255 | 0.230449 | 0.872 | 0.87 | 1 | 8 | Calr | 0.002 | Old |
| 0.019286 | 0.358871 | 0.648 | 0.569 | 1 | 8 | Tnfrsf1b | 0.079 | Old |
| 0.019579 | 1.465781 | 0.045 | 0.013 | 1 | 8 | Slc25a43 | 0.032 | Old |
| 0.019579 | 1.426675 | 0.045 | 0.013 | 1 | 8 | Tnni3 | 0.032 | Old |
| 0.019885 | 1.614172 | 0.022 | 0.003 | 1 | 8 | Spint2 | 0.019 | Old |
| 0.019885 | 1.247272 | 0.022 | 0.003 | 1 | 8 | Gm10320 | 0.019 | Old |
| 0.01989 | 0.702176 | 0.374 | 0.309 | 1 | 8 | Tnfrsf21 | 0.065 | Old |
| 0.019988 | 0.465306 | 0.687 | 0.636 | 1 | 8 | Rhob | 0.051 | Old |
| 0.020108 | 0.246979 | 0.927 | 0.925 | 1 | 8 | Hnrnpf | 0.002 | Old |
| 0.020373 | 0.121804 | 0.994 | 0.997 | 1 | 8 | Itm2b | -0.003 | Old |
| 0.020375 | 1.170869 | 0.045 | 0.013 | 1 | 8 | Irgc1 | 0.032 | Old |
| 0.020427 | 0.212147 | 0.944 | 0.961 | 1 | 8 | Ptafr | -0.017 | Old |
| 0.020436 | 1.161523 | 0.022 | 0.003 | 1 | 8 | Stab2 | 0.019 | Old |
| 0.02096 | 1.151315 | 0.145 | 0.086 | 1 | 8 | Myo1e | 0.059 | Old |
| 0.0211 | 1.706624 | 0.061 | 0.023 | 1 | 8 | Zfp653 | 0.038 | Old |
| 0.021189 | 0.342784 | 0.575 | 0.481 | 1 | 8 | Supt16 | 0.094 | Old |
| 0.021745 | 0.807205 | 0.128 | 0.07 | 1 | 8 | Ppan | 0.058 | Old |
| 0.021965 | 0.281431 | 0.754 | 0.779 | 1 | 8 | 11-Sep | -0.025 | Old |
| 0.021966 | 1.19638 | 0.117 | 0.062 | 1 | 8 | Asap2 | 0.055 | Old |
| 0.021979 | 0.500121 | 0.642 | 0.595 | 1 | 8 | Sowahc | 0.047 | Old |
| 0.022194 | 1.030143 | 0.101 | 0.049 | 1 | 8 | Oas1a | 0.052 | Old |
| 0.022357 | 0.565135 | 0.525 | 0.462 | 1 | 8 | Cd209g | 0.063 | Old |
| 0.022379 | 0.788946 | 0.436 | 0.377 | 1 | 8 | Srsf7 | 0.059 | Old |
| 0.022449 | 1.100105 | 0.117 | 0.062 | 1 | 8 | Pam | 0.055 | Old |
| 0.022525 | 0.947569 | 0.168 | 0.101 | 1 | 8 | Chchd4 | 0.067 | Old |
| 0.022536 | 2.521546 | 0.028 | 0.005 | 1 | 8 | Aldh1l1 | 0.023 | Old |

| | | | | | | | | |
|---|---|---|---|---|---|---|---|---|
| 0.022881 | 1.959359 | 0.028 | 0.005 | 1 | 8 | Zfp532 | 0.023 | Old |
| 0.023172 | 1.516368 | 0.112 | 0.06 | 1 | 8 | Snhg20 | 0.052 | Old |
| 0.023214 | 0.404716 | 0.508 | 0.426 | 1 | 8 | Smc6 | 0.082 | Old |
| 0.02323 | 2.025841 | 0.028 | 0.005 | 1 | 8 | Gm10800 | 0.023 | Old |
| 0.02323 | 1.448256 | 0.028 | 0.005 | 1 | 8 | Tsacc | 0.023 | Old |
| 0.023249 | 0.325066 | 0.737 | 0.649 | 1 | 8 | Bhlhe40 | 0.088 | Old |
| 0.023406 | 1.844576 | 0.028 | 0.005 | 1 | 8 | Abraxas1 | 0.023 | Old |
| 0.023406 | 1.591895 | 0.028 | 0.005 | 1 | 8 | Hus1b | 0.023 | Old |
| 0.023406 | 1.488767 | 0.028 | 0.005 | 1 | 8 | Mmab | 0.023 | Old |
| 0.023584 | 1.401552 | 0.028 | 0.005 | 1 | 8 | Car13 | 0.023 | Old |
| 0.023584 | 1.336307 | 0.028 | 0.005 | 1 | 8 | Gm26533 | 0.023 | Old |
| 0.023604 | 1.405367 | 0.034 | 0.008 | 1 | 8 | Col16a1 | 0.026 | Old |
| 0.023762 | 1.500595 | 0.028 | 0.005 | 1 | 8 | Inhba | 0.023 | Old |
| 0.023762 | 1.111789 | 0.028 | 0.005 | 1 | 8 | Rpgrip1l | 0.023 | Old |
| 0.02391 | 0.960916 | 0.101 | 0.049 | 1 | 8 | Abhd18 | 0.052 | Old |
| 0.023954 | 0.943056 | 0.117 | 0.062 | 1 | 8 | 1110059E2 | 0.055 | Old |
| 0.024123 | 0.867327 | 0.028 | 0.005 | 1 | 8 | Arg1 | 0.023 | Old |
| 0.024225 | 0.491933 | 0.285 | 0.205 | 1 | 8 | Hnrnpll | 0.08 | Old |
| 0.024235 | 1.066154 | 0.179 | 0.109 | 1 | 8 | Mphosph6 | 0.07 | Old |
| 0.024329 | 0.842644 | 0.24 | 0.166 | 1 | 8 | A4galt | 0.074 | Old |
| 0.024386 | 1.004474 | 0.296 | 0.226 | 1 | 8 | Gpr146 | 0.07 | Old |
| 0.024797 | 0.566739 | 0.274 | 0.192 | 1 | 8 | Vps50 | 0.082 | Old |
| 0.024803 | 0.815286 | 0.302 | 0.229 | 1 | 8 | Rnft1 | 0.073 | Old |
| 0.025117 | 0.188134 | 0.905 | 0.94 | 1 | 8 | Cd53 | -0.035 | Old |
| 0.02559 | 0.268916 | 0.922 | 0.932 | 1 | 8 | Canx | -0.01 | Old |
| 0.025922 | 1.379752 | 0.128 | 0.073 | 1 | 8 | Coq7 | 0.055 | Old |
| 0.025967 | 0.133756 | 1 | 1 | 1 | 8 | mt-Nd4 | 0 | Old |
| 0.025992 | 0.338131 | 0.709 | 0.665 | 1 | 8 | Metrnl | 0.044 | Old |
| 0.02601 | 0.431285 | 0.296 | 0.203 | 1 | 8 | Rnf213 | 0.093 | Old |
| 0.026011 | 0.46804 | 0.687 | 0.675 | 1 | 8 | Atp6ap2 | 0.012 | Old |
| 0.026046 | 0.220895 | 0.804 | 0.826 | 1 | 8 | Itgb1 | -0.022 | Old |
| 0.026054 | 0.249224 | 0.838 | 0.842 | 1 | 8 | Ywhab | -0.004 | Old |
| 0.026209 | 0.141621 | 0.978 | 0.979 | 1 | 8 | C1qc | -0.001 | Old |
| 0.026282 | 0.600978 | 0.318 | 0.231 | 1 | 8 | Kif1c | 0.087 | Old |
| 0.026315 | 0.494023 | 0.458 | 0.384 | 1 | 8 | Nfil3 | 0.074 | Old |
| 0.026422 | 0.838951 | 0.341 | 0.26 | 1 | 8 | Rhoq | 0.081 | Old |
| 0.026854 | 0.46712 | 0.587 | 0.504 | 1 | 8 | Lypla2 | 0.083 | Old |
| 0.027096 | 0.325823 | 0.302 | 0.21 | 1 | 8 | 4930455G( | 0.092 | Old |
| 0.027448 | 1.887435 | 0.056 | 0.021 | 1 | 8 | Dhrs9 | 0.035 | Old |
| 0.027823 | 0.875212 | 0.184 | 0.119 | 1 | 8 | Nup93 | 0.065 | Old |
| 0.028062 | 0.257589 | 0.816 | 0.792 | 1 | 8 | Dok2 | 0.024 | Old |
| 0.028184 | 0.926583 | 0.179 | 0.109 | 1 | 8 | Nuak1 | 0.07 | Old |
| 0.028803 | 0.411068 | 0.374 | 0.283 | 1 | 8 | Il2rg | 0.091 | Old |
| 0.028979 | 0.461619 | 0.564 | 0.517 | 1 | 8 | Idh1 | 0.047 | Old |
| 0.029261 | 1.439878 | 0.162 | 0.104 | 1 | 8 | Ltbp3 | 0.058 | Old |

| | | | | | | | | |
|---|---|---|---|---|---|---|---|---|
| 0.029698 | 0.846711 | 0.056 | 0.021 | 1 | 8 | Klhl8 | 0.035 | Old |
| 0.029744 | 2.260742 | 0.05 | 0.018 | 1 | 8 | 2610021A( | 0.032 | Old |
| 0.029887 | 0.737728 | 0.391 | 0.319 | 1 | 8 | Sp3 | 0.072 | Old |
| 0.029926 | 0.245821 | 0.827 | 0.839 | 1 | 8 | Ddx17 | -0.012 | Old |
| 0.030313 | 1.638222 | 0.078 | 0.036 | 1 | 8 | Uck2 | 0.042 | Old |
| 0.030517 | 0.352958 | 0.721 | 0.706 | 1 | 8 | Ptpn1 | 0.015 | Old |
| 0.030764 | 0.395691 | 0.626 | 0.59 | 1 | 8 | Atp6v1e1 | 0.036 | Old |
| 0.030824 | 0.401772 | 0.665 | 0.626 | 1 | 8 | Tpp1 | 0.039 | Old |
| 0.03098 | 0.586501 | 0.151 | 0.088 | 1 | 8 | Rrp36 | 0.063 | Old |
| 0.031124 | 0.473586 | 0.609 | 0.538 | 1 | 8 | Bag1 | 0.071 | Old |
| 0.031251 | 0.821056 | 0.19 | 0.125 | 1 | 8 | Rbm34 | 0.065 | Old |
| 0.03149 | 0.309308 | 0.877 | 0.852 | 1 | 8 | Glul | 0.025 | Old |
| 0.031549 | 0.340667 | 0.726 | 0.66 | 1 | 8 | Hmox1 | 0.066 | Old |
| 0.032039 | 1.283064 | 0.134 | 0.078 | 1 | 8 | Cdca4 | 0.056 | Old |
| 0.032186 | 0.553136 | 0.318 | 0.242 | 1 | 8 | Plxna1 | 0.076 | Old |
| 0.032206 | 0.838388 | 0.352 | 0.291 | 1 | 8 | Cp | 0.061 | Old |
| 0.032208 | 0.793676 | 0.235 | 0.164 | 1 | 8 | Rars | 0.071 | Old |
| 0.032237 | 0.478073 | 0.698 | 0.655 | 1 | 8 | Pgk1 | 0.043 | Old |
| 0.032669 | 0.469077 | 0.251 | 0.171 | 1 | 8 | Eps15l1 | 0.08 | Old |
| 0.032734 | 0.793449 | 0.246 | 0.174 | 1 | 8 | Ccdc174 | 0.072 | Old |
| 0.032853 | 0.25238 | 0.955 | 0.943 | 1 | 8 | Ier2 | 0.012 | Old |
| 0.033148 | 1.351085 | 0.112 | 0.062 | 1 | 8 | Tmtc3 | 0.05 | Old |
| 0.033271 | 0.967804 | 0.067 | 0.029 | 1 | 8 | Flrt2 | 0.038 | Old |
| 0.033637 | 0.783111 | 0.514 | 0.475 | 1 | 8 | Pla2g2d | 0.039 | Old |
| 0.03369 | 0.94419 | 0.106 | 0.057 | 1 | 8 | Ggta1 | 0.049 | Old |
| 0.034033 | 0.232355 | 0.944 | 0.956 | 1 | 8 | Rhoa | -0.012 | Old |
| 0.034235 | 1.060191 | 0.078 | 0.036 | 1 | 8 | Mcoln2 | 0.042 | Old |
| 0.034414 | 0.737163 | 0.408 | 0.33 | 1 | 8 | Htt | 0.078 | Old |
| 0.034766 | 0.565635 | 0.503 | 0.455 | 1 | 8 | Pld3 | 0.048 | Old |
| 0.034792 | 1.106434 | 0.095 | 0.049 | 1 | 8 | Drg2 | 0.046 | Old |
| 0.034942 | 1.330651 | 0.184 | 0.127 | 1 | 8 | Klra2 | 0.057 | Old |
| 0.035174 | 0.221488 | 0.804 | 0.857 | 1 | 8 | Atp5f1 | -0.053 | Old |
| 0.035519 | 0.882388 | 0.268 | 0.195 | 1 | 8 | Brpf1 | 0.073 | Old |
| 0.036332 | 1.613538 | 0.073 | 0.034 | 1 | 8 | 2010111I0: | 0.039 | Old |
| 0.03661 | 0.410153 | 0.754 | 0.751 | 1 | 8 | Ppp2ca | 0.003 | Old |
| 0.036645 | 1.068226 | 0.112 | 0.062 | 1 | 8 | Epg5 | 0.05 | Old |
| 0.036683 | 0.864192 | 0.078 | 0.036 | 1 | 8 | Gm9844 | 0.042 | Old |
| 0.037041 | 0.675761 | 0.201 | 0.138 | 1 | 8 | Nolc1 | 0.063 | Old |
| 0.037165 | 0.012101 | 0.43 | 0.574 | 1 | 8 | Arl4c | -0.144 | Old |
| 0.037431 | 0.856217 | 0.151 | 0.094 | 1 | 8 | Eif1ad | 0.057 | Old |
| 0.037437 | 1.259696 | 0.073 | 0.034 | 1 | 8 | Rfc5 | 0.039 | Old |
| 0.037697 | 0.725917 | 0.313 | 0.242 | 1 | 8 | Epb41l1 | 0.071 | Old |
| 0.037844 | 0.211017 | 0.983 | 0.99 | 1 | 8 | Ctsc | -0.007 | Old |
| 0.037889 | 0.687779 | 0.33 | 0.252 | 1 | 8 | Yipf4 | 0.078 | Old |
| 0.03815 | 1.006291 | 0.045 | 0.016 | 1 | 8 | Oas3 | 0.029 | Old |

| | | | | | | | | |
|---|---|---|---|---|---|---|---|---|
| 0.038154 | 2.435065 | 0.011 | 0 | 1 | 8 | Mnd1 | 0.011 | Old |
| 0.038154 | 2.213947 | 0.011 | 0 | 1 | 8 | Rasl10a | 0.011 | Old |
| 0.038154 | 2.186578 | 0.011 | 0 | 1 | 8 | Syt7 | 0.011 | Old |
| 0.038154 | 2.119084 | 0.011 | 0 | 1 | 8 | Gm14302 | 0.011 | Old |
| 0.038154 | 2.099086 | 0.011 | 0 | 1 | 8 | Adamts14 | 0.011 | Old |
| 0.038154 | 2.0839 | 0.011 | 0 | 1 | 8 | Gm47439 | 0.011 | Old |
| 0.038154 | 2.032678 | 0.011 | 0 | 1 | 8 | Dpf1 | 0.011 | Old |
| 0.038154 | 1.953235 | 0.011 | 0 | 1 | 8 | Iglc2 | 0.011 | Old |
| 0.038154 | 1.862763 | 0.011 | 0 | 1 | 8 | Gm40787 | 0.011 | Old |
| 0.038154 | 1.844353 | 0.011 | 0 | 1 | 8 | Bdh2 | 0.011 | Old |
| 0.038154 | 1.833835 | 0.011 | 0 | 1 | 8 | Sfmbt2 | 0.011 | Old |
| 0.038154 | 1.797736 | 0.011 | 0 | 1 | 8 | Colec11 | 0.011 | Old |
| 0.038154 | 1.747555 | 0.011 | 0 | 1 | 8 | Gm47917 | 0.011 | Old |
| 0.038154 | 1.609197 | 0.011 | 0 | 1 | 8 | Cnga1 | 0.011 | Old |
| 0.038154 | 1.605435 | 0.011 | 0 | 1 | 8 | Cd79a | 0.011 | Old |
| 0.038154 | 1.596822 | 0.011 | 0 | 1 | 8 | 3100002H( | 0.011 | Old |
| 0.038154 | 1.587059 | 0.011 | 0 | 1 | 8 | Poln | 0.011 | Old |
| 0.038154 | 1.547063 | 0.011 | 0 | 1 | 8 | Gm47922 | 0.011 | Old |
| 0.038154 | 1.445981 | 0.011 | 0 | 1 | 8 | Hsd17b14 | 0.011 | Old |
| 0.038154 | 1.387966 | 0.011 | 0 | 1 | 8 | Prg4 | 0.011 | Old |
| 0.038154 | 1.378341 | 0.011 | 0 | 1 | 8 | Chic1 | 0.011 | Old |
| 0.038154 | 1.377742 | 0.011 | 0 | 1 | 8 | AK157302 | 0.011 | Old |
| 0.038154 | 1.356615 | 0.011 | 0 | 1 | 8 | Rbakdn | 0.011 | Old |
| 0.038154 | 1.301316 | 0.011 | 0 | 1 | 8 | Hif3a | 0.011 | Old |
| 0.038154 | 1.285292 | 0.011 | 0 | 1 | 8 | Vnn3 | 0.011 | Old |
| 0.038154 | 1.269246 | 0.011 | 0 | 1 | 8 | Gm30122 | 0.011 | Old |
| 0.038154 | 1.252418 | 0.011 | 0 | 1 | 8 | 4933424M | 0.011 | Old |
| 0.038154 | 1.248853 | 0.011 | 0 | 1 | 8 | Rwdd3 | 0.011 | Old |
| 0.038154 | 1.246733 | 0.011 | 0 | 1 | 8 | Gm35339 | 0.011 | Old |
| 0.038154 | 1.245758 | 0.011 | 0 | 1 | 8 | Dpyd | 0.011 | Old |
| 0.038154 | 1.242459 | 0.011 | 0 | 1 | 8 | Megf11 | 0.011 | Old |
| 0.038154 | 1.227808 | 0.011 | 0 | 1 | 8 | Lamb1 | 0.011 | Old |
| 0.038154 | 1.225851 | 0.011 | 0 | 1 | 8 | Prkg1 | 0.011 | Old |
| 0.038154 | 1.224573 | 0.011 | 0 | 1 | 8 | 1110020A2 | 0.011 | Old |
| 0.038154 | 1.19528 | 0.011 | 0 | 1 | 8 | Nrarp | 0.011 | Old |
| 0.038154 | 1.158755 | 0.011 | 0 | 1 | 8 | Gm15972 | 0.011 | Old |
| 0.038154 | 1.158731 | 0.011 | 0 | 1 | 8 | Gm10734 | 0.011 | Old |
| 0.038154 | 1.149342 | 0.011 | 0 | 1 | 8 | AC168977. | 0.011 | Old |
| 0.038154 | 1.144577 | 0.011 | 0 | 1 | 8 | Ptges | 0.011 | Old |
| 0.038154 | 1.143843 | 0.011 | 0 | 1 | 8 | Gm44777 | 0.011 | Old |
| 0.038154 | 1.138241 | 0.011 | 0 | 1 | 8 | Plin1 | 0.011 | Old |
| 0.038154 | 1.111249 | 0.011 | 0 | 1 | 8 | P2rx5 | 0.011 | Old |
| 0.038154 | 1.061151 | 0.011 | 0 | 1 | 8 | Gm29994 | 0.011 | Old |
| 0.038154 | 1.032501 | 0.011 | 0 | 1 | 8 | Fxyd4 | 0.011 | Old |
| 0.038154 | 1.030127 | 0.011 | 0 | 1 | 8 | Gm49249 | 0.011 | Old |

| | | | | | | | | |
|---|---|---|---|---|---|---|---|---|
| 0.038154 | 1.02844 | 0.011 | 0 | 1 | 8 | 4833427F1 | 0.011 | Old |
| 0.038154 | 1.028365 | 0.011 | 0 | 1 | 8 | Lox | 0.011 | Old |
| 0.038154 | 1.020259 | 0.011 | 0 | 1 | 8 | Pabpn1l | 0.011 | Old |
| 0.038154 | 1.015558 | 0.011 | 0 | 1 | 8 | Gm28535 | 0.011 | Old |
| 0.038154 | 1.007106 | 0.011 | 0 | 1 | 8 | AC150035. | 0.011 | Old |
| 0.038154 | 0.999456 | 0.011 | 0 | 1 | 8 | Ak4 | 0.011 | Old |
| 0.038154 | 0.983757 | 0.011 | 0 | 1 | 8 | Myl2 | 0.011 | Old |
| 0.038154 | 0.98292 | 0.011 | 0 | 1 | 8 | Gpr55 | 0.011 | Old |
| 0.038154 | 0.974017 | 0.011 | 0 | 1 | 8 | Gm14168 | 0.011 | Old |
| 0.038154 | 0.966093 | 0.011 | 0 | 1 | 8 | Exo1 | 0.011 | Old |
| 0.038154 | 0.921694 | 0.011 | 0 | 1 | 8 | Apol7d | 0.011 | Old |
| 0.038154 | 0.914513 | 0.011 | 0 | 1 | 8 | Anxa9 | 0.011 | Old |
| 0.038154 | 0.900871 | 0.011 | 0 | 1 | 8 | Mipol1 | 0.011 | Old |
| 0.038154 | 0.880385 | 0.011 | 0 | 1 | 8 | Gm10244 | 0.011 | Old |
| 0.038154 | 0.860347 | 0.011 | 0 | 1 | 8 | Slc25a35 | 0.011 | Old |
| 0.038154 | 0.814406 | 0.011 | 0 | 1 | 8 | Tnik | 0.011 | Old |
| 0.038154 | 0.810066 | 0.011 | 0 | 1 | 8 | Podn | 0.011 | Old |
| 0.038154 | 0.808142 | 0.011 | 0 | 1 | 8 | A430105J0 | 0.011 | Old |
| 0.038154 | 0.807106 | 0.011 | 0 | 1 | 8 | Ckm | 0.011 | Old |
| 0.038154 | 0.79795 | 0.011 | 0 | 1 | 8 | Dnmt3aos | 0.011 | Old |
| 0.038154 | 0.763323 | 0.011 | 0 | 1 | 8 | Pcdhgb5 | 0.011 | Old |
| 0.038362 | 0.650383 | 0.184 | 0.119 | 1 | 8 | Dubr | 0.065 | Old |
| 0.038535 | 1.18239 | 0.045 | 0.016 | 1 | 8 | Galm | 0.029 | Old |
| 0.038941 | 0.303052 | 0.721 | 0.691 | 1 | 8 | Kif5b | 0.03 | Old |
| 0.039025 | 1.236769 | 0.117 | 0.068 | 1 | 8 | Gng11 | 0.049 | Old |
| 0.039532 | 0.827398 | 0.117 | 0.065 | 1 | 8 | Traf2 | 0.052 | Old |
| 0.039575 | 0.748036 | 0.274 | 0.197 | 1 | 8 | Slc12a7 | 0.077 | Old |
| 0.039881 | 0.541955 | 0.408 | 0.338 | 1 | 8 | Dock7 | 0.07 | Old |
| 0.040208 | 0.49763 | 0.145 | 0.086 | 1 | 8 | Inip | 0.059 | Old |
| 0.040288 | 0.25212 | 0.939 | 0.971 | 1 | 8 | Lcp1 | -0.032 | Old |
| 0.040294 | 0.986173 | 0.084 | 0.042 | 1 | 8 | Tlr3 | 0.042 | Old |
| 0.04053 | 0.158749 | 0.955 | 0.956 | 1 | 8 | Cltc | -0.001 | Old |
| 0.041321 | 0.971753 | 0.196 | 0.135 | 1 | 8 | Mis12 | 0.061 | Old |
| 0.042275 | 0.620277 | 0.497 | 0.41 | 1 | 8 | 6-Mar | 0.087 | Old |
| 0.042296 | 0.597859 | 0.38 | 0.306 | 1 | 8 | Cyp51 | 0.074 | Old |
| 0.042596 | 0.000338 | 0.235 | 0.322 | 1 | 8 | Angel2 | -0.087 | Old |
| 0.042667 | 1.140867 | 0.184 | 0.122 | 1 | 8 | Ero1l | 0.062 | Old |
| 0.043073 | 0.783363 | 0.268 | 0.205 | 1 | 8 | Snrpa | 0.063 | Old |
| 0.043393 | 0.732308 | 0.156 | 0.099 | 1 | 8 | Prdm4 | 0.057 | Old |
| 0.043646 | 0.724136 | 0.391 | 0.338 | 1 | 8 | Ak2 | 0.053 | Old |
| 0.043806 | 0.662592 | 0.581 | 0.553 | 1 | 8 | Emc10 | 0.028 | Old |
| 0.043921 | 0.237283 | 0.849 | 0.836 | 1 | 8 | Vamp8 | 0.013 | Old |
| 0.044113 | 0.783676 | 0.151 | 0.094 | 1 | 8 | Yars | 0.057 | Old |
| 0.044478 | 0.473567 | 0.458 | 0.403 | 1 | 8 | Ugcg | 0.055 | Old |
| 0.044666 | 0.226483 | 0.855 | 0.86 | 1 | 8 | Wdr1 | -0.005 | Old |

| | | | | | | | | |
|---|---|---|---|---|---|---|---|---|
| 0.044768 | 0.703954 | 0.385 | 0.322 | 1 | 8 | Yipf3 | 0.063 | Old |
| 0.04513 | 1.61168 | 0.056 | 0.023 | 1 | 8 | Gm31718 | 0.033 | Old |
| 0.045142 | 0.156948 | 0.844 | 0.847 | 1 | 8 | Dock2 | -0.003 | Old |
| 0.045275 | 0.841407 | 0.151 | 0.096 | 1 | 8 | Ppargc1b | 0.055 | Old |
| 0.045487 | 0.480414 | 0.084 | 0.042 | 1 | 8 | Blzf1 | 0.042 | Old |
| 0.045626 | 1.49115 | 0.039 | 0.013 | 1 | 8 | Cacna1a | 0.026 | Old |
| 0.045704 | 1.069601 | 0.056 | 0.023 | 1 | 8 | Gm16104 | 0.033 | Old |
| 0.04619 | 0.249553 | 0.704 | 0.688 | 1 | 8 | 2900097C1 | 0.016 | Old |
| 0.046503 | 1.070173 | 0.095 | 0.052 | 1 | 8 | Cox10 | 0.043 | Old |
| 0.046821 | 0.396599 | 0.62 | 0.574 | 1 | 8 | Psmd13 | 0.046 | Old |
| 0.047069 | 1.320164 | 0.056 | 0.023 | 1 | 8 | E2f6 | 0.033 | Old |
| 0.047267 | 0.885653 | 0.056 | 0.023 | 1 | 8 | Tsen54 | 0.033 | Old |
| 0.047306 | 0.577756 | 0.441 | 0.361 | 1 | 8 | Eif3d | 0.08 | Old |
| 0.047384 | 0.223631 | 0.933 | 0.956 | 1 | 8 | Ywhaz | -0.023 | Old |
| 0.048079 | 1.012815 | 0.112 | 0.065 | 1 | 8 | Wrnip1 | 0.047 | Old |
| 0.048164 | 0.419117 | 0.665 | 0.636 | 1 | 8 | Ezr | 0.029 | Old |
| 0.048654 | 0.662381 | 0.291 | 0.221 | 1 | 8 | Katna1 | 0.07 | Old |
| 0.048698 | 0.303294 | 0.777 | 0.766 | 1 | 8 | H13 | 0.011 | Old |
| 0.049074 | 0.524423 | 0.542 | 0.509 | 1 | 8 | Gnpda1 | 0.033 | Old |
| 0.049103 | 0.862674 | 0.156 | 0.099 | 1 | 8 | Mri1 | 0.057 | Old |
| 0.049263 | 0.601323 | 0.458 | 0.397 | 1 | 8 | Evl | 0.061 | Old |
| 0.049364 | 1.762248 | 0.101 | 0.057 | 1 | 8 | Dnm1 | 0.044 | Old |
| 0.049804 | 0.243772 | 0.899 | 0.948 | 1 | 8 | Rtn4 | -0.049 | Old |
| 0.050129 | 0.316671 | 0.575 | 0.517 | 1 | 8 | Adam8 | 0.058 | Old |
| 0.050194 | 1.214 | 0.145 | 0.094 | 1 | 8 | Arhgap21 | 0.051 | Old |
| 0.050824 | 1.322617 | 0.101 | 0.057 | 1 | 8 | Parp10 | 0.044 | Old |
| 0.051083 | 0.281835 | 0.698 | 0.688 | 1 | 8 | Ldha | 0.01 | Old |
| 0.051305 | 1.287206 | 0.034 | 0.01 | 1 | 8 | 4931414P1 | 0.024 | Old |
| 0.051496 | 0.545954 | 0.659 | 0.631 | 1 | 8 | Selenos | 0.028 | Old |
| 0.051568 | 0.447683 | 0.553 | 0.486 | 1 | 8 | Mdfic | 0.067 | Old |
| 0.052181 | 1.086296 | 0.034 | 0.01 | 1 | 8 | Cd5l | 0.024 | Old |
| 0.052406 | 1.107608 | 0.117 | 0.07 | 1 | 8 | Nol6 | 0.047 | Old |
| 0.052476 | 1.283504 | 0.034 | 0.01 | 1 | 8 | Pcdh7 | 0.024 | Old |
| 0.052799 | 0.665734 | 0.078 | 0.039 | 1 | 8 | Phf7 | 0.039 | Old |
| 0.053112 | 0.499381 | 0.503 | 0.434 | 1 | 8 | Pdcd10 | 0.069 | Old |
| 0.053119 | 0.404455 | 0.62 | 0.613 | 1 | 8 | Pycard | 0.007 | Old |
| 0.05328 | 0.603785 | 0.374 | 0.294 | 1 | 8 | Cept1 | 0.08 | Old |
| 0.053438 | 0.385004 | 0.737 | 0.717 | 1 | 8 | Snrpb | 0.02 | Old |
| 0.05367 | 1.065603 | 0.034 | 0.01 | 1 | 8 | Ccdc17 | 0.024 | Old |
| 0.053998 | 2.095246 | 0.073 | 0.036 | 1 | 8 | Irf7 | 0.037 | Old |
| 0.054582 | 0.552304 | 0.179 | 0.117 | 1 | 8 | Tbck | 0.062 | Old |
| 0.05471 | 1.606909 | 0.05 | 0.021 | 1 | 8 | Exo5 | 0.029 | Old |
| 0.055136 | 0.390619 | 0.525 | 0.481 | 1 | 8 | Lfng | 0.044 | Old |
| 0.055396 | 0.810039 | 0.196 | 0.135 | 1 | 8 | Pla2g4a | 0.061 | Old |
| 0.055502 | 1.120112 | 0.034 | 0.01 | 1 | 8 | E030030I0 | 0.024 | Old |

| | | | | | | | | |
|---|---|---|---|---|---|---|---|---|
| 0.056753 | 0.273727 | 0.034 | 0.01 | 1 | 8 | Fsd1l | 0.024 | Old |
| 0.056865 | 1.431056 | 0.073 | 0.036 | 1 | 8 | Unk | 0.037 | Old |
| 0.057218 | 0.265023 | 0.844 | 0.823 | 1 | 8 | Sfpq | 0.021 | Old |
| 0.057241 | 0.870083 | 0.212 | 0.151 | 1 | 8 | Ddx58 | 0.061 | Old |
| 0.057324 | 0.846614 | 0.318 | 0.252 | 1 | 8 | Dync1li1 | 0.066 | Old |
| 0.057388 | 1.01779 | 0.151 | 0.099 | 1 | 8 | Neurl2 | 0.052 | Old |
| 0.057618 | 0.737761 | 0.313 | 0.249 | 1 | 8 | Tmem33 | 0.064 | Old |
| 0.057707 | 0.555991 | 0.486 | 0.439 | 1 | 8 | Apobec1 | 0.047 | Old |
| 0.057973 | 0.698328 | 0.251 | 0.184 | 1 | 8 | Dnm1l | 0.067 | Old |
| 0.058047 | 1.386753 | 0.073 | 0.036 | 1 | 8 | Zbtb24 | 0.037 | Old |
| 0.058102 | 1.076348 | 0.05 | 0.021 | 1 | 8 | Arhgap27 | 0.029 | Old |
| 0.058254 | 0.57899 | 0.24 | 0.174 | 1 | 8 | Slc7a1 | 0.066 | Old |
| 0.058533 | 1.926333 | 0.061 | 0.029 | 1 | 8 | Mmp12 | 0.032 | Old |
| 0.058851 | 0.701294 | 0.05 | 0.021 | 1 | 8 | Wars2 | 0.029 | Old |
| 0.058962 | 0.504527 | 0.469 | 0.423 | 1 | 8 | Yeats4 | 0.046 | Old |
| 0.058984 | 0.820296 | 0.179 | 0.125 | 1 | 8 | Rae1 | 0.054 | Old |
| 0.059238 | 1.850031 | 0.028 | 0.008 | 1 | 8 | Fam129c | 0.02 | Old |
| 0.059249 | 1.102359 | 0.073 | 0.036 | 1 | 8 | Timmdc1 | 0.037 | Old |
| 0.059345 | 0.245818 | 0.832 | 0.86 | 1 | 8 | Itgam | -0.028 | Old |
| 0.059599 | 0.089372 | 0.14 | 0.221 | 1 | 8 | Txndc16 | -0.081 | Old |
| 0.059604 | 1.557487 | 0.028 | 0.008 | 1 | 8 | Tmem267 | 0.02 | Old |
| 0.059604 | 1.34789 | 0.028 | 0.008 | 1 | 8 | Htra3 | 0.02 | Old |
| 0.059617 | 0.721517 | 0.201 | 0.14 | 1 | 8 | Nlk | 0.061 | Old |
| 0.059645 | 0.866017 | 0.061 | 0.029 | 1 | 8 | Ggct | 0.032 | Old |
| 0.059728 | 0.110188 | 0.966 | 0.984 | 1 | 8 | Arpc2 | -0.018 | Old |
| 0.059757 | 0.372361 | 0.57 | 0.496 | 1 | 8 | Rab11fip1 | 0.074 | Old |
| 0.060024 | 0.311337 | 0.592 | 0.551 | 1 | 8 | Tppp3 | 0.041 | Old |
| 0.060062 | 1.060579 | 0.073 | 0.036 | 1 | 8 | Trim30c | 0.037 | Old |
| 0.060118 | 0.55215 | 0.447 | 0.382 | 1 | 8 | Ythdf3 | 0.065 | Old |
| 0.060472 | 0.948641 | 0.073 | 0.036 | 1 | 8 | Tgds | 0.037 | Old |
| 0.060576 | 0.611808 | 0.151 | 0.096 | 1 | 8 | Arhgef11 | 0.055 | Old |
| 0.060742 | 0.866358 | 0.229 | 0.171 | 1 | 8 | Wdr91 | 0.058 | Old |
| 0.060888 | 0.252117 | 0.05 | 0.021 | 1 | 8 | Fggy | 0.029 | Old |
| 0.061003 | 1.170625 | 0.061 | 0.029 | 1 | 8 | Chac2 | 0.032 | Old |
| 0.061052 | 0.891226 | 0.101 | 0.057 | 1 | 8 | Pigz | 0.044 | Old |
| 0.061069 | 0.152625 | 0.911 | 0.925 | 1 | 8 | Cfp | -0.014 | Old |
| 0.061086 | 0.975622 | 0.028 | 0.008 | 1 | 8 | Fsd2 | 0.02 | Old |
| 0.061146 | 0.985682 | 0.05 | 0.021 | 1 | 8 | Chst14 | 0.029 | Old |
| 0.061224 | 0.905299 | 0.101 | 0.057 | 1 | 8 | Mtfr1 | 0.044 | Old |
| 0.061528 | 0.568175 | 0.235 | 0.169 | 1 | 8 | Parp9 | 0.066 | Old |
| 0.061659 | 0.092786 | 0.006 | 0.031 | 1 | 8 | A130077B1 | -0.025 | Old |
| 0.061803 | 2.179641 | 0.017 | 0.003 | 1 | 8 | Ifi44 | 0.014 | Old |
| 0.061803 | 1.721385 | 0.017 | 0.003 | 1 | 8 | Ccdc80 | 0.014 | Old |
| 0.061803 | 1.445149 | 0.017 | 0.003 | 1 | 8 | Defb25 | 0.014 | Old |
| 0.061803 | 1.376409 | 0.017 | 0.003 | 1 | 8 | Gprin3 | 0.014 | Old |

| | | | | | | | | |
|---|---|---|---|---|---|---|---|---|
| 0.061838 | 1.093904 | 0.028 | 0.008 | 1 | 8 | Ppp1r14a | 0.02 | Old |
| 0.061928 | 0.498784 | 0.184 | 0.122 | 1 | 8 | Cog3 | 0.062 | Old |
| 0.062038 | 0.174419 | 0.927 | 0.951 | 1 | 8 | Gnb2 | -0.024 | Old |
| 0.062066 | 0.982153 | 0.123 | 0.075 | 1 | 8 | Pramef8 | 0.048 | Old |
| 0.062338 | 2.418642 | 0.017 | 0.003 | 1 | 8 | Rnf128 | 0.014 | Old |
| 0.062338 | 1.364504 | 0.017 | 0.003 | 1 | 8 | Mss51 | 0.014 | Old |
| 0.062338 | 1.297534 | 0.017 | 0.003 | 1 | 8 | Gm6793 | 0.014 | Old |
| 0.062338 | 1.234277 | 0.017 | 0.003 | 1 | 8 | Gm30108 | 0.014 | Old |
| 0.062338 | 1.193934 | 0.017 | 0.003 | 1 | 8 | Gvin1 | 0.014 | Old |
| 0.062338 | 1.013755 | 0.017 | 0.003 | 1 | 8 | Gm5127 | 0.014 | Old |
| 0.06243 | 0.658787 | 0.453 | 0.39 | 1 | 8 | Api5 | 0.063 | Old |
| 0.062547 | 0.420468 | 0.084 | 0.044 | 1 | 8 | Atat1 | 0.04 | Old |
| 0.062715 | 0.781823 | 0.179 | 0.122 | 1 | 8 | Ttc33 | 0.057 | Old |
| 0.062765 | 0.403833 | 0.581 | 0.54 | 1 | 8 | Ikzf1 | 0.041 | Old |
| 0.062859 | 0.338032 | 0.592 | 0.558 | 1 | 8 | Stag2 | 0.034 | Old |
| 0.062876 | 1.129041 | 0.017 | 0.003 | 1 | 8 | Tex14 | 0.014 | Old |
| 0.062982 | 0.649521 | 0.028 | 0.008 | 1 | 8 | Osbpl5 | 0.02 | Old |
| 0.063185 | 0.348242 | 0.709 | 0.686 | 1 | 8 | Ran | 0.023 | Old |
| 0.063271 | 5.499683 | 0.022 | 0.005 | 1 | 8 | Jchain | 0.017 | Old |
| 0.063271 | 1.664177 | 0.022 | 0.005 | 1 | 8 | Suv39h2 | 0.017 | Old |
| 0.063418 | 1.092933 | 0.017 | 0.003 | 1 | 8 | Gm9856 | 0.014 | Old |
| 0.063418 | 0.961202 | 0.017 | 0.003 | 1 | 8 | Gm10605 | 0.014 | Old |
| 0.063418 | 0.884429 | 0.017 | 0.003 | 1 | 8 | 4932438H2 | 0.014 | Old |
| 0.063418 | 0.741339 | 0.017 | 0.003 | 1 | 8 | Creb3l4 | 0.014 | Old |
| 0.063504 | 0.238494 | 0.793 | 0.821 | 1 | 8 | Tm6sf1 | -0.028 | Old |
| 0.063716 | 1.173271 | 0.022 | 0.005 | 1 | 8 | Spib | 0.017 | Old |
| 0.063744 | 0.046697 | 0.061 | 0.114 | 1 | 8 | Rdh14 | -0.053 | Old |
| 0.063818 | 0.715436 | 0.223 | 0.161 | 1 | 8 | Anpep | 0.062 | Old |
| 0.064059 | 0.833083 | 0.162 | 0.112 | 1 | 8 | Psmg3 | 0.05 | Old |
| 0.064164 | 1.684971 | 0.022 | 0.005 | 1 | 8 | Zbtb49 | 0.017 | Old |
| 0.064164 | 1.312045 | 0.022 | 0.005 | 1 | 8 | Tsku | 0.017 | Old |
| 0.06419 | 0.884468 | 0.128 | 0.081 | 1 | 8 | Zxdc | 0.047 | Old |
| 0.064372 | 0.449506 | 0.274 | 0.208 | 1 | 8 | Smarcad1 | 0.066 | Old |
| 0.064615 | 1.359779 | 0.078 | 0.042 | 1 | 8 | Tmem38b | 0.036 | Old |
| 0.064615 | 1.419918 | 0.022 | 0.005 | 1 | 8 | H3f3aos | 0.017 | Old |
| 0.064615 | 1.281951 | 0.022 | 0.005 | 1 | 8 | Il7 | 0.017 | Old |
| 0.065039 | 0.43181 | 0.592 | 0.545 | 1 | 8 | Rhog | 0.047 | Old |
| 0.065068 | 1.21399 | 0.022 | 0.005 | 1 | 8 | Ereg | 0.017 | Old |
| 0.065068 | 1.082618 | 0.022 | 0.005 | 1 | 8 | Ifi205 | 0.017 | Old |
| 0.06508 | 0.916229 | 0.106 | 0.062 | 1 | 8 | Chpf | 0.044 | Old |
| 0.065141 | 0.336679 | 0.447 | 0.379 | 1 | 8 | Brd7 | 0.068 | Old |
| 0.065308 | 0.402496 | 0.536 | 0.475 | 1 | 8 | Sypl | 0.061 | Old |
| 0.065524 | 1.689712 | 0.022 | 0.005 | 1 | 8 | Pdlim3 | 0.017 | Old |
| 0.065596 | 1.054504 | 0.274 | 0.213 | 1 | 8 | Sf3a1 | 0.061 | Old |
| 0.065816 | 0.861879 | 0.184 | 0.13 | 1 | 8 | Ddx56 | 0.054 | Old |

| | | | | | | | | |
|---|---|---|---|---|---|---|---|---|
| 0.065856 | 0.716406 | 0.425 | 0.384 | 1 | 8 | Nus1 | 0.041 | Old |
| 0.065983 | 1.460879 | 0.022 | 0.005 | 1 | 8 | Fam216a | 0.017 | Old |
| 0.065983 | 1.024564 | 0.022 | 0.005 | 1 | 8 | AC174678. | 0.017 | Old |
| 0.065999 | 0.543923 | 0.112 | 0.065 | 1 | 8 | Snupn | 0.047 | Old |
| 0.066 | 1.310267 | 0.045 | 0.018 | 1 | 8 | Tyms | 0.027 | Old |
| 0.066021 | 0.457837 | 0.397 | 0.34 | 1 | 8 | Mgat4b | 0.057 | Old |
| 0.066411 | 0.203175 | 0.832 | 0.852 | 1 | 8 | Slc25a5 | -0.02 | Old |
| 0.066444 | 0.826581 | 0.022 | 0.005 | 1 | 8 | Rnd1 | 0.017 | Old |
| 0.066444 | 0.815639 | 0.022 | 0.005 | 1 | 8 | Gpr137c | 0.017 | Old |
| 0.066694 | 0.569737 | 0.408 | 0.343 | 1 | 8 | Ube2j2 | 0.065 | Old |
| 0.066765 | 0.497656 | 0.397 | 0.338 | 1 | 8 | Dnajb9 | 0.059 | Old |
| 0.066979 | 0.431361 | 0.687 | 0.655 | 1 | 8 | Ap1b1 | 0.032 | Old |
| 0.067541 | 0.553326 | 0.257 | 0.192 | 1 | 8 | Gdap10 | 0.065 | Old |
| 0.067591 | 0.485316 | 0.453 | 0.39 | 1 | 8 | Rnf4 | 0.063 | Old |
| 0.06776 | 0.521566 | 0.453 | 0.403 | 1 | 8 | Tspan31 | 0.05 | Old |
| 0.067944 | 1.544876 | 0.056 | 0.026 | 1 | 8 | Lins1 | 0.03 | Old |
| 0.067947 | 0.550422 | 0.302 | 0.229 | 1 | 8 | Msl3 | 0.073 | Old |
| 0.069292 | 1.031255 | 0.045 | 0.018 | 1 | 8 | Chd3os | 0.027 | Old |
| 0.06938 | 1.183313 | 0.151 | 0.104 | 1 | 8 | Ocel1 | 0.047 | Old |
| 0.069428 | 0.219747 | 0.777 | 0.758 | 1 | 8 | Atp1b3 | 0.019 | Old |
| 0.069649 | 0.368668 | 0.397 | 0.317 | 1 | 8 | Arnt | 0.08 | Old |
| 0.069674 | 0.542679 | 0.587 | 0.514 | 1 | 8 | Serpinb8 | 0.073 | Old |
| 0.069808 | 0.32066 | 0.972 | 0.961 | 1 | 8 | Dazap2 | 0.011 | Old |
| 0.070213 | 0.801485 | 0.045 | 0.018 | 1 | 8 | Snx33 | 0.027 | Old |
| 0.070375 | 0.069574 | 0.112 | 0.177 | 1 | 8 | Msi2 | -0.065 | Old |
| 0.070771 | 1.120409 | 0.145 | 0.096 | 1 | 8 | Rpa1 | 0.049 | Old |
| 0.070801 | 0.207032 | 0.983 | 0.969 | 1 | 8 | Neat1 | 0.014 | Old |
| 0.070842 | 0.618733 | 0.145 | 0.094 | 1 | 8 | Rfng | 0.051 | Old |
| 0.071089 | 1.047633 | 0.145 | 0.096 | 1 | 8 | Sdhaf1 | 0.049 | Old |
| 0.07185 | 0.629495 | 0.106 | 0.062 | 1 | 8 | Eif2b2 | 0.044 | Old |
| 0.071957 | 1.540096 | 0.056 | 0.026 | 1 | 8 | Ccdc180 | 0.03 | Old |
| 0.07242 | 0.729051 | 0.067 | 0.034 | 1 | 8 | Gin1 | 0.033 | Old |
| 0.072916 | 1.542393 | 0.067 | 0.034 | 1 | 8 | Akap1 | 0.033 | Old |
| 0.073951 | 0.713879 | 0.302 | 0.239 | 1 | 8 | Slc23a2 | 0.063 | Old |
| 0.074227 | 0.687071 | 0.318 | 0.257 | 1 | 8 | Rcor1 | 0.061 | Old |
| 0.074429 | 0.431651 | 0.307 | 0.244 | 1 | 8 | Sec23b | 0.063 | Old |
| 0.074569 | 0.437121 | 0.464 | 0.408 | 1 | 8 | Cd200r1 | 0.056 | Old |
| 0.074969 | 0.341103 | 0.67 | 0.644 | 1 | 8 | Cap1 | 0.026 | Old |
| 0.075302 | 0.70597 | 0.391 | 0.33 | 1 | 8 | Blmh | 0.061 | Old |
| 0.075425 | 0.854214 | 0.229 | 0.174 | 1 | 8 | Sfxn2 | 0.055 | Old |
| 0.07582 | 0.44115 | 0.587 | 0.517 | 1 | 8 | Psmd8 | 0.07 | Old |
| 0.075985 | 0.250688 | 0.838 | 0.826 | 1 | 8 | Colgalt1 | 0.012 | Old |
| 0.076146 | 0.445736 | 0.43 | 0.374 | 1 | 8 | Glyr1 | 0.056 | Old |
| 0.076652 | 0.615152 | 0.19 | 0.132 | 1 | 8 | Ccdc90b | 0.058 | Old |
| 0.076902 | 0.178008 | 0.626 | 0.561 | 1 | 8 | Man1a | 0.065 | Old |

| | | | | | | | | |
|---|---|---|---|---|---|---|---|---|
| 0.076944 | 0.542176 | 0.106 | 0.062 | 1 | 8 | Gnptab | 0.044 | Old |
| 0.077015 | 0.225299 | 0.832 | 0.868 | 1 | 8 | App | -0.036 | Old |
| 0.077212 | 0.849428 | 0.302 | 0.244 | 1 | 8 | Cdc26 | 0.058 | Old |
| 0.077246 | 0.842671 | 0.067 | 0.034 | 1 | 8 | Vill | 0.033 | Old |
| 0.077249 | 0.236922 | 0.827 | 0.813 | 1 | 8 | Eif5a | 0.014 | Old |
| 0.077507 | 0.631863 | 0.067 | 0.034 | 1 | 8 | Zfp456 | 0.033 | Old |
| 0.077622 | 0.705353 | 0.084 | 0.047 | 1 | 8 | Wwox | 0.037 | Old |
| 0.078527 | 0.051335 | 0.296 | 0.395 | 1 | 8 | Tnrc6c | -0.099 | Old |
| 0.078537 | 1.103966 | 0.095 | 0.055 | 1 | 8 | Gm14548 | 0.04 | Old |
| 0.078568 | 0.913318 | 0.128 | 0.083 | 1 | 8 | Slc25a13 | 0.045 | Old |
| 0.078782 | 0.288517 | 0.615 | 0.558 | 1 | 8 | Anxa7 | 0.057 | Old |
| 0.078801 | 1.465422 | 0.039 | 0.016 | 1 | 8 | Nptxr | 0.023 | Old |
| 0.079129 | 0.541016 | 0.263 | 0.2 | 1 | 8 | Vps13d | 0.063 | Old |
| 0.080251 | 0.057312 | 0.179 | 0.252 | 1 | 8 | Atp11c | -0.073 | Old |
| 0.080344 | 0.439839 | 0.246 | 0.179 | 1 | 8 | Fam214b | 0.067 | Old |
| 0.080401 | 0.361407 | 0.732 | 0.722 | 1 | 8 | Clec4a1 | 0.01 | Old |
| 0.081008 | 1.201437 | 0.039 | 0.016 | 1 | 8 | Nemp2 | 0.023 | Old |
| 0.082111 | 0.947713 | 0.128 | 0.083 | 1 | 8 | Tmem161b | 0.045 | Old |
| 0.082229 | 0.135073 | 0.989 | 0.987 | 1 | 8 | H2-D1 | 0.002 | Old |
| 0.082307 | 0.481822 | 0.134 | 0.086 | 1 | 8 | AI606181 | 0.048 | Old |
| 0.082354 | 0.404607 | 0.559 | 0.53 | 1 | 8 | Hpcal1 | 0.029 | Old |
| 0.082589 | 0.468204 | 0.464 | 0.397 | 1 | 8 | Rras2 | 0.067 | Old |
| 0.083204 | 0.090329 | 0.279 | 0.364 | 1 | 8 | Il13ra1 | -0.085 | Old |
| 0.083291 | 0.545916 | 0.358 | 0.296 | 1 | 8 | Ubr5 | 0.062 | Old |
| 0.083368 | 0.222916 | 0.816 | 0.857 | 1 | 8 | Serbp1 | -0.041 | Old |
| 0.083579 | 0.359225 | 0.715 | 0.725 | 1 | 8 | Dusp3 | -0.01 | Old |
| 0.083609 | 1.347548 | 0.073 | 0.039 | 1 | 8 | Urb1 | 0.034 | Old |
| 0.083739 | 0.347885 | 0.441 | 0.361 | 1 | 8 | Glmp | 0.08 | Old |
| 0.083872 | 0.422892 | 0.609 | 0.545 | 1 | 8 | Manf | 0.064 | Old |
| 0.084165 | 1.296839 | 0.061 | 0.031 | 1 | 8 | Dexi | 0.03 | Old |
| 0.084331 | 0.937176 | 0.268 | 0.213 | 1 | 8 | Sgk1 | 0.055 | Old |
| 0.084411 | 0.515999 | 0.039 | 0.016 | 1 | 8 | Hist2h3c2 | 0.023 | Old |
| 0.08494 | 0.249639 | 0.687 | 0.662 | 1 | 8 | Ifi207 | 0.025 | Old |
| 0.08509 | 0.403286 | 0.626 | 0.605 | 1 | 8 | AC149090. | 0.021 | Old |
| 0.085274 | 0.037955 | 0.168 | 0.239 | 1 | 8 | Tdrd7 | -0.071 | Old |
| 0.085499 | 0.489519 | 0.095 | 0.055 | 1 | 8 | F2rl2 | 0.04 | Old |
| 0.085503 | 1.091417 | 0.184 | 0.135 | 1 | 8 | Ccdc25 | 0.049 | Old |
| 0.085568 | 0.707114 | 0.134 | 0.088 | 1 | 8 | Nt5dc3 | 0.046 | Old |
| 0.085629 | 1.511833 | 0.061 | 0.031 | 1 | 8 | Twnk | 0.03 | Old |
| 0.085811 | 0.508436 | 0.302 | 0.231 | 1 | 8 | Sdccag3 | 0.071 | Old |
| 0.086107 | 0.432264 | 0.525 | 0.483 | 1 | 8 | Slc29a3 | 0.042 | Old |
| 0.086221 | 1.314326 | 0.061 | 0.031 | 1 | 8 | Mettl17 | 0.03 | Old |
| 0.086362 | 0.765747 | 0.246 | 0.195 | 1 | 8 | Cavin1 | 0.051 | Old |
| 0.08643 | 0.218396 | 0.866 | 0.862 | 1 | 8 | Sh3glb1 | 0.004 | Old |
| 0.086586 | 0.376033 | 0.14 | 0.091 | 1 | 8 | Zdhhc6 | 0.049 | Old |

| | | | | | | | | |
|---|---|---|---|---|---|---|---|---|
| 0.087017 | 0.784692 | 0.24 | 0.192 | 1 | 8 | H2-Q4 | 0.048 | Old |
| 0.087081 | 0.346993 | 0.732 | 0.706 | 1 | 8 | Clec4a3 | 0.026 | Old |
| 0.087737 | 0.580197 | 0.291 | 0.229 | 1 | 8 | 8-Mar | 0.062 | Old |
| 0.087841 | 1.061484 | 0.128 | 0.086 | 1 | 8 | Sqle | 0.042 | Old |
| 0.088101 | 0.377032 | 0.609 | 0.535 | 1 | 8 | Adam10 | 0.074 | Old |
| 0.088184 | 1.089704 | 0.151 | 0.106 | 1 | 8 | Ormdl1 | 0.045 | Old |
| 0.08829 | 0.021825 | 0.363 | 0.452 | 1 | 8 | Ypel5 | -0.089 | Old |
| 0.088631 | 0.203265 | 0.754 | 0.753 | 1 | 8 | Cops9 | 0.001 | Old |
| 0.089014 | 0.686415 | 0.073 | 0.039 | 1 | 8 | Rcc1 | 0.034 | Old |
| 0.089017 | 0.148552 | 0.899 | 0.958 | 1 | 8 | Alox5ap | -0.059 | Old |
| 0.08923 | 1.105764 | 0.061 | 0.031 | 1 | 8 | Mphosph9 | 0.03 | Old |
| 0.089531 | 0.200589 | 0.715 | 0.688 | 1 | 8 | Lyn | 0.027 | Old |
| 0.089942 | 0.138076 | 0.978 | 0.992 | 1 | 8 | Zeb2 | -0.014 | Old |
| 0.090407 | 0.049175 | 0.061 | 0.109 | 1 | 8 | Tmem126a | -0.048 | Old |
| 0.090489 | 0.449834 | 0.531 | 0.499 | 1 | 8 | Socs6 | 0.032 | Old |
| 0.090633 | 0.930267 | 0.358 | 0.301 | 1 | 8 | Mindy1 | 0.057 | Old |
| 0.092235 | 1.019918 | 0.101 | 0.062 | 1 | 8 | Mthfr | 0.039 | Old |
| 0.093004 | 0.645202 | 0.257 | 0.2 | 1 | 8 | Creld2 | 0.057 | Old |
| 0.093314 | 0.184278 | 0.687 | 0.603 | 1 | 8 | Ccl7 | 0.084 | Old |
| 0.093583 | 1.087738 | 0.061 | 0.031 | 1 | 8 | Pold2 | 0.03 | Old |
| 0.093639 | 0.219756 | 0.939 | 0.932 | 1 | 8 | Egr1 | 0.007 | Old |
| 0.093855 | 0.484135 | 0.14 | 0.094 | 1 | 8 | Glt8d1 | 0.046 | Old |
| 0.094031 | 0.137252 | 0.011 | 0.036 | 1 | 8 | Gstm4 | -0.025 | Old |
| 0.094094 | 0.338307 | 0.112 | 0.07 | 1 | 8 | Isg15 | 0.042 | Old |
| 0.094473 | 1.742029 | 0.067 | 0.036 | 1 | 8 | Nod1 | 0.031 | Old |
| 0.095008 | 0.42797 | 0.587 | 0.556 | 1 | 8 | Sdf4 | 0.031 | Old |
| 0.095282 | 0.605129 | 0.05 | 0.023 | 1 | 8 | Acp6 | 0.027 | Old |
| 0.095514 | 0.982177 | 0.078 | 0.044 | 1 | 8 | Rrnad1 | 0.034 | Old |
| 0.096282 | 0.512276 | 0.251 | 0.192 | 1 | 8 | Mmadhc | 0.059 | Old |
| 0.096425 | 0.347145 | 0.397 | 0.338 | 1 | 8 | Ppm1a | 0.059 | Old |
| 0.096784 | 0.677632 | 0.335 | 0.286 | 1 | 8 | Nsfl1c | 0.049 | Old |
| 0.096855 | 1.095086 | 0.14 | 0.096 | 1 | 8 | Vps39 | 0.044 | Old |
| 0.097076 | 0.586637 | 0.302 | 0.242 | 1 | 8 | Plekhm2 | 0.06 | Old |
| 0.097272 | 0.355936 | 0.302 | 0.239 | 1 | 8 | Mitf | 0.063 | Old |
| 0.097501 | 1.635063 | 0.034 | 0.013 | 1 | 8 | Klhl25 | 0.021 | Old |
| 0.098208 | 0.446789 | 0.274 | 0.213 | 1 | 8 | Nkiras2 | 0.061 | Old |
| 0.098212 | 0.314643 | 0.05 | 0.023 | 1 | 8 | Meis3 | 0.027 | Old |
| 0.098355 | 0.960185 | 0.134 | 0.091 | 1 | 8 | Fdx1 | 0.043 | Old |
| 0.099063 | 0.004445 | 0.011 | 0.036 | 1 | 8 | Dennd2d | -0.025 | Old |
| 0.099163 | 0.756626 | 0.229 | 0.177 | 1 | 8 | Pum3 | 0.052 | Old |
| 0.099389 | 1.197197 | 0.034 | 0.013 | 1 | 8 | Ddx59 | 0.021 | Old |
| 0.099704 | 0.484044 | 0.089 | 0.052 | 1 | 8 | Sprtn | 0.037 | Old |
| 0.099866 | 1.749399 | 0.034 | 0.013 | 1 | 8 | Hist1h4h | 0.021 | Old |
| 0.099866 | 1.413396 | 0.034 | 0.013 | 1 | 8 | Polrmt | 0.021 | Old |
| 0.100095 | 0.710772 | 0.061 | 0.031 | 1 | 8 | Fam198b | 0.03 | Old |

| | | | | | | | | |
|---|---|---|---|---|---|---|---|---|
| 0.100344 | 1.162276 | 0.034 | 0.013 | 1 | 8 | A830010M | 0.021 | Old |
| 0.100344 | 0.886157 | 0.034 | 0.013 | 1 | 8 | Gm5150 | 0.021 | Old |
| 0.100402 | 0.727234 | 0.184 | 0.135 | 1 | 8 | Pik3c3 | 0.049 | Old |
| 0.100413 | 0.438619 | 0.598 | 0.556 | 1 | 8 | Rgl1 | 0.042 | Old |
| 0.100472 | 0.361906 | 0.81 | 0.818 | 1 | 8 | Klf4 | -0.008 | Old |
| 0.100654 | 2.889502 | 0.045 | 0.021 | 1 | 8 | Serpine1 | 0.024 | Old |
| 0.100949 | 0.611494 | 0.425 | 0.4 | 1 | 8 | Man2a1 | 0.025 | Old |
| 0.100991 | 0.817843 | 0.196 | 0.145 | 1 | 8 | Pfdn4 | 0.051 | Old |
| 0.10125 | 0.33608 | 0.285 | 0.223 | 1 | 8 | Pxk | 0.062 | Old |
| 0.101307 | 1.350568 | 0.034 | 0.013 | 1 | 8 | Prr15 | 0.021 | Old |
| 0.101307 | 1.275542 | 0.034 | 0.013 | 1 | 8 | Klhl11 | 0.021 | Old |
| 0.101307 | 0.881036 | 0.034 | 0.013 | 1 | 8 | Eya4 | 0.021 | Old |
| 0.10132 | 0.465188 | 0.179 | 0.125 | 1 | 8 | Tmem185b | 0.054 | Old |
| 0.101767 | 0.360581 | 0.715 | 0.722 | 1 | 8 | Far1 | -0.007 | Old |
| 0.101883 | 0.396897 | 0.682 | 0.644 | 1 | 8 | G3bp1 | 0.038 | Old |
| 0.102513 | 0.202321 | 0.274 | 0.366 | 1 | 8 | N4bp2l1 | -0.092 | Old |
| 0.102547 | 0.595676 | 0.19 | 0.135 | 1 | 8 | Rnpc3 | 0.055 | Old |
| 0.102937 | 0.369928 | 0.168 | 0.114 | 1 | 8 | Nelfb | 0.054 | Old |
| 0.103254 | 0.664546 | 0.034 | 0.013 | 1 | 8 | Ddx11 | 0.021 | Old |
| 0.103254 | 0.557852 | 0.034 | 0.013 | 1 | 8 | Zfp408 | 0.021 | Old |
| 0.103696 | 0.937529 | 0.112 | 0.073 | 1 | 8 | Rpp14 | 0.039 | Old |
| 0.10387 | 0.285483 | 0.659 | 0.631 | 1 | 8 | Ier3ip1 | 0.028 | Old |
| 0.104114 | 0.230318 | 0.665 | 0.644 | 1 | 8 | Tmem106a | 0.021 | Old |
| 0.104172 | 0.126099 | 0.86 | 0.888 | 1 | 8 | Ccnl1 | -0.028 | Old |
| 0.104225 | 0.153477 | 0.162 | 0.223 | 1 | 8 | Pgm2l1 | -0.061 | Old |
| 0.105163 | 0.419172 | 0.581 | 0.556 | 1 | 8 | Smap2 | 0.025 | Old |
| 0.105883 | 0.660162 | 0.19 | 0.14 | 1 | 8 | Cops3 | 0.05 | Old |
| 0.105995 | 0.850835 | 0.128 | 0.088 | 1 | 8 | Parp3 | 0.04 | Old |
| 0.106263 | 0.739488 | 0.19 | 0.14 | 1 | 8 | Nelfcd | 0.05 | Old |
| 0.106511 | 0.445256 | 0.419 | 0.369 | 1 | 8 | Il1rn | 0.05 | Old |
| 0.106636 | 0.517103 | 0.436 | 0.374 | 1 | 8 | Skp1a | 0.062 | Old |
| 0.106675 | 0.378201 | 0.123 | 0.078 | 1 | 8 | Riox2 | 0.045 | Old |
| 0.10675 | 0.513052 | 0.441 | 0.384 | 1 | 8 | Larp4 | 0.057 | Old |
| 0.106886 | 0.468528 | 0.173 | 0.122 | 1 | 8 | Rnf138 | 0.051 | Old |
| 0.107044 | 0.473319 | 0.313 | 0.26 | 1 | 8 | Lgals3bp | 0.053 | Old |
| 0.107066 | 0.008111 | 0.531 | 0.605 | 1 | 8 | Dad1 | -0.074 | Old |
| 0.107104 | 0.328953 | 0.385 | 0.319 | 1 | 8 | Ap3b1 | 0.066 | Old |
| 0.107667 | 0.5746 | 0.324 | 0.27 | 1 | 8 | Pgap2 | 0.054 | Old |
| 0.107688 | 0.654379 | 0.078 | 0.044 | 1 | 8 | Pus3 | 0.034 | Old |
| 0.107851 | 0.462104 | 0.486 | 0.413 | 1 | 8 | Mapk6 | 0.073 | Old |
| 0.108677 | 0.416939 | 0.475 | 0.429 | 1 | 8 | Arf2 | 0.046 | Old |
| 0.108992 | 0.450172 | 0.056 | 0.029 | 1 | 8 | Ndufaf5 | 0.027 | Old |
| 0.109048 | 0.052238 | 0.022 | 0.052 | 1 | 8 | BC051226 | -0.03 | Old |
| 0.109274 | 0.96769 | 0.084 | 0.049 | 1 | 8 | 1700096K1 | 0.035 | Old |
| 0.109669 | 0.157038 | 0.799 | 0.834 | 1 | 8 | Tacc1 | -0.035 | Old |

| | | | | | | | | |
|---|---|---|---|---|---|---|---|---|
| 0.109763 | 0.430971 | 0.324 | 0.275 | 1 | 8 | Hmgcs1 | 0.049 | Old |
| 0.10977 | 1.043401 | 0.045 | 0.021 | 1 | 8 | Ugt1a7c | 0.024 | Old |
| 0.11021 | 0.347279 | 0.665 | 0.652 | 1 | 8 | Selenot | 0.013 | Old |
| 0.110248 | 0.419195 | 0.162 | 0.112 | 1 | 8 | Otud6b | 0.05 | Old |
| 0.110631 | 1.260776 | 0.045 | 0.021 | 1 | 8 | Dnmt3b | 0.024 | Old |
| 0.110873 | 0.714759 | 0.056 | 0.029 | 1 | 8 | C1qtnf12 | 0.027 | Old |
| 0.111439 | 0.20154 | 0.855 | 0.842 | 1 | 8 | Lgals3 | 0.013 | Old |
| 0.111477 | 0.098555 | 0.441 | 0.551 | 1 | 8 | Mndal | -0.11 | Old |
| 0.112129 | 0.725867 | 0.184 | 0.138 | 1 | 8 | Hilpda | 0.046 | Old |
| 0.112402 | 0.592963 | 0.218 | 0.164 | 1 | 8 | Acbd6 | 0.054 | Old |
| 0.112581 | 0.336881 | 0.587 | 0.556 | 1 | 8 | Itch | 0.031 | Old |
| 0.112638 | 0.296432 | 0.575 | 0.569 | 1 | 8 | Ppp1r14b | 0.006 | Old |
| 0.112804 | 0.713392 | 0.045 | 0.021 | 1 | 8 | Aldh18a1 | 0.024 | Old |
| 0.113136 | 0.406455 | 0.151 | 0.104 | 1 | 8 | Brpf3 | 0.047 | Old |
| 0.113313 | 0.184394 | 0.749 | 0.774 | 1 | 8 | Dhrs3 | -0.025 | Old |
| 0.113551 | 0.945723 | 0.056 | 0.029 | 1 | 8 | Mtfmt | 0.027 | Old |
| 0.113778 | 0.756615 | 0.128 | 0.088 | 1 | 8 | Eml3 | 0.04 | Old |
| 0.114003 | 0.507676 | 0.475 | 0.416 | 1 | 8 | Slc20a1 | 0.059 | Old |
| 0.114683 | 0.869076 | 0.156 | 0.109 | 1 | 8 | Sfi1 | 0.047 | Old |
| 0.114844 | 0.378633 | 0.128 | 0.083 | 1 | 8 | Fbxo45 | 0.045 | Old |
| 0.115173 | 0.813813 | 0.073 | 0.042 | 1 | 8 | Rhbdf1 | 0.031 | Old |
| 0.115261 | 0.214732 | 0.615 | 0.569 | 1 | 8 | Arf3 | 0.046 | Old |
| 0.11573 | 0.597194 | 0.067 | 0.036 | 1 | 8 | Sco1 | 0.031 | Old |
| 0.116033 | 0.418926 | 0.419 | 0.364 | 1 | 8 | Rc3h2 | 0.055 | Old |
| 0.11633 | 0.017063 | 0.743 | 0.836 | 1 | 8 | H2afy | -0.093 | Old |
| 0.116431 | 0.461858 | 0.425 | 0.374 | 1 | 8 | Noc2l | 0.051 | Old |
| 0.116673 | 0.427727 | 0.056 | 0.029 | 1 | 8 | Tmem101 | 0.027 | Old |
| 0.116919 | 1.379517 | 0.028 | 0.01 | 1 | 8 | Fcho1 | 0.018 | Old |
| 0.116964 | 0.264574 | 0.771 | 0.79 | 1 | 8 | Psmb4 | -0.019 | Old |
| 0.118158 | 0.457699 | 0.045 | 0.021 | 1 | 8 | Setd4 | 0.024 | Old |
| 0.118259 | 0.919668 | 0.056 | 0.029 | 1 | 8 | A230050P2 | 0.027 | Old |
| 0.118386 | 0.600387 | 0.302 | 0.249 | 1 | 8 | Galk2 | 0.053 | Old |
| 0.118462 | 1.554231 | 0.084 | 0.052 | 1 | 8 | Fahd1 | 0.032 | Old |
| 0.118592 | 0.464345 | 0.413 | 0.356 | 1 | 8 | Slc26a2 | 0.057 | Old |
| 0.118641 | 0.045398 | 0.028 | 0.06 | 1 | 8 | Ankrd39 | -0.032 | Old |
| 0.119001 | 0.641481 | 0.291 | 0.242 | 1 | 8 | Aco1 | 0.049 | Old |
| 0.119036 | 1.304094 | 0.061 | 0.034 | 1 | 8 | Ppp1r3e | 0.027 | Old |
| 0.119244 | 0.820873 | 0.073 | 0.042 | 1 | 8 | Dhx58 | 0.031 | Old |
| 0.119326 | 1.96644 | 0.028 | 0.01 | 1 | 8 | Ttc30a1 | 0.018 | Old |
| 0.119419 | 0.464954 | 0.268 | 0.208 | 1 | 8 | Slc27a1 | 0.06 | Old |
| 0.119487 | 0.398304 | 0.168 | 0.117 | 1 | 8 | Zfp608 | 0.051 | Old |
| 0.119768 | 0.30401 | 0.62 | 0.587 | 1 | 8 | Magt1 | 0.033 | Old |
| 0.120124 | 0.294941 | 0.754 | 0.797 | 1 | 8 | Ppp1r18 | -0.043 | Old |
| 0.120691 | 0.670481 | 0.207 | 0.156 | 1 | 8 | Crtap | 0.051 | Old |
| 0.121075 | 0.598351 | 0.056 | 0.029 | 1 | 8 | Cd300lf | 0.027 | Old |

| | | | | | | | | |
|---|---|---|---|---|---|---|---|---|
| 0.121075 | 0.290177 | 0.056 | 0.029 | 1 | 8 | Pdss2 | 0.027 | Old |
| 0.121508 | 0.49601 | 0.743 | 0.74 | 1 | 8 | Tmem30a | 0.003 | Old |
| 0.121689 | 1.112615 | 0.061 | 0.034 | 1 | 8 | Smg8 | 0.027 | Old |
| 0.121752 | 0.72611 | 0.341 | 0.278 | 1 | 8 | Chmp2b | 0.063 | Old |
| 0.121772 | 0.667233 | 0.028 | 0.01 | 1 | 8 | A530032D1 | 0.018 | Old |
| 0.122345 | 0.555216 | 0.168 | 0.122 | 1 | 8 | Rgs3 | 0.046 | Old |
| 0.122625 | 0.800864 | 0.196 | 0.145 | 1 | 8 | Snrnp40 | 0.051 | Old |
| 0.122686 | 0.411378 | 0.084 | 0.049 | 1 | 8 | Rnaseh1 | 0.035 | Old |
| 0.122693 | 0.429983 | 0.156 | 0.109 | 1 | 8 | Cd151 | 0.047 | Old |
| 0.122797 | 0.588737 | 0.19 | 0.14 | 1 | 8 | Dhx16 | 0.05 | Old |
| 0.122841 | 0.782583 | 0.061 | 0.034 | 1 | 8 | Fpr2 | 0.027 | Old |
| 0.122937 | 1.009392 | 0.123 | 0.083 | 1 | 8 | Tfb2m | 0.04 | Old |
| 0.12301 | 1.132153 | 0.028 | 0.01 | 1 | 8 | Amacr | 0.018 | Old |
| 0.12301 | 1.024608 | 0.028 | 0.01 | 1 | 8 | Il4i1 | 0.018 | Old |
| 0.123226 | 1.506426 | 0.061 | 0.034 | 1 | 8 | Gga2 | 0.027 | Old |
| 0.123427 | 0.58988 | 0.073 | 0.042 | 1 | 8 | Oaf | 0.031 | Old |
| 0.123613 | 1.109636 | 0.061 | 0.034 | 1 | 8 | Mms22l | 0.027 | Old |
| 0.123632 | 1.211322 | 0.028 | 0.01 | 1 | 8 | Zfp119a | 0.018 | Old |
| 0.123632 | 1.101205 | 0.028 | 0.01 | 1 | 8 | Tmem51os | 0.018 | Old |
| 0.123647 | 0.474697 | 0.486 | 0.449 | 1 | 8 | Srsf1 | 0.037 | Old |
| 0.124 | 0.481929 | 0.419 | 0.384 | 1 | 8 | Sec24b | 0.035 | Old |
| 0.124753 | 0.141066 | 0.894 | 0.93 | 1 | 8 | Slc25a3 | -0.036 | Old |
| 0.124894 | 1.499379 | 0.05 | 0.026 | 1 | 8 | Gadd45a | 0.024 | Old |
| 0.124912 | 0.387729 | 0.721 | 0.745 | 1 | 8 | Celf1 | -0.024 | Old |
| 0.125109 | 0.23302 | 0.011 | 0.034 | 1 | 8 | Map3k12 | -0.023 | Old |
| 0.125125 | 0.997047 | 0.251 | 0.205 | 1 | 8 | Med8 | 0.046 | Old |
| 0.125183 | 0.590203 | 0.263 | 0.21 | 1 | 8 | Rpe | 0.053 | Old |
| 0.125246 | 0.485509 | 0.168 | 0.119 | 1 | 8 | Fam20c | 0.049 | Old |
| 0.125514 | 0.692664 | 0.028 | 0.01 | 1 | 8 | Cavin3 | 0.018 | Old |
| 0.125848 | 0.31632 | 0.81 | 0.803 | 1 | 8 | Ccl2 | 0.007 | Old |
| 0.125878 | 0.380726 | 0.559 | 0.53 | 1 | 8 | Gm21188 | 0.029 | Old |
| 0.12682 | 1.09284 | 0.173 | 0.132 | 1 | 8 | Traf6 | 0.041 | Old |
| 0.127279 | 0.165993 | 0.676 | 0.655 | 1 | 8 | Bola2 | 0.021 | Old |
| 0.128351 | 0.228369 | 0.709 | 0.714 | 1 | 8 | Scamp2 | -0.005 | Old |
| 0.128585 | 0.354989 | 0.201 | 0.148 | 1 | 8 | Cebpe | 0.053 | Old |
| 0.128697 | 0.354126 | 0.659 | 0.662 | 1 | 8 | P2rx4 | -0.003 | Old |
| 0.129108 | 0.871448 | 0.14 | 0.099 | 1 | 8 | Mtmr2 | 0.041 | Old |
| 0.129172 | 0.507208 | 0.458 | 0.403 | 1 | 8 | Tmx3 | 0.055 | Old |
| 0.12924 | 0.548929 | 0.24 | 0.184 | 1 | 8 | Rab29 | 0.056 | Old |
| 0.129243 | 0.322724 | 0.531 | 0.504 | 1 | 8 | Samsn1 | 0.027 | Old |
| 0.129812 | 0.353746 | 0.626 | 0.603 | 1 | 8 | P4hb | 0.023 | Old |
| 0.129824 | 0.042999 | 0.039 | 0.073 | 1 | 8 | Zc3h6 | -0.034 | Old |
| 0.130331 | 0.942952 | 0.061 | 0.034 | 1 | 8 | Ccnyl1 | 0.027 | Old |
| 0.131041 | 0.862166 | 0.14 | 0.096 | 1 | 8 | Chid1 | 0.044 | Old |
| 0.131051 | 0.459326 | 0.106 | 0.068 | 1 | 8 | Tpcn2 | 0.038 | Old |

| | | | | | | | | |
|---|---|---|---|---|---|---|---|---|
| 0.131076 | 1.828122 | 0.039 | 0.018 | 1 | 8 | Tigar | 0.021 | Old |
| 0.131379 | 0.453307 | 0.408 | 0.345 | 1 | 8 | Vps13c | 0.063 | Old |
| 0.131405 | 0.407769 | 0.335 | 0.275 | 1 | 8 | Araf | 0.06 | Old |
| 0.131439 | 0.463439 | 0.302 | 0.244 | 1 | 8 | Pira2 | 0.058 | Old |
| 0.131545 | 0.683081 | 0.061 | 0.034 | 1 | 8 | Ercc8 | 0.027 | Old |
| 0.131602 | 1.308328 | 0.039 | 0.018 | 1 | 8 | Fancm | 0.021 | Old |
| 0.131711 | 0.46838 | 0.475 | 0.442 | 1 | 8 | Cd37 | 0.033 | Old |
| 0.132033 | 1.196814 | 0.05 | 0.026 | 1 | 8 | Ehf | 0.024 | Old |
| 0.132106 | 0.39087 | 0.609 | 0.605 | 1 | 8 | Rp2 | 0.004 | Old |
| 0.132131 | 1.15804 | 0.039 | 0.018 | 1 | 8 | Gm20300 | 0.021 | Old |
| 0.132394 | 0.626547 | 0.263 | 0.213 | 1 | 8 | Ddx10 | 0.05 | Old |
| 0.13266 | 1.060872 | 0.039 | 0.018 | 1 | 8 | Tnnt1 | 0.021 | Old |
| 0.133043 | 0.272637 | 0.743 | 0.764 | 1 | 8 | Arf1 | -0.021 | Old |
| 0.133405 | 0.551316 | 0.14 | 0.096 | 1 | 8 | Mterf3 | 0.044 | Old |
| 0.133866 | 1.128464 | 0.05 | 0.026 | 1 | 8 | B3gnt5 | 0.024 | Old |
| 0.134322 | 0.170787 | 0.872 | 0.891 | 1 | 8 | Atp6v0b | -0.019 | Old |
| 0.134473 | 0.104227 | 0.966 | 0.979 | 1 | 8 | Ccr2 | -0.013 | Old |
| 0.134809 | 0.616298 | 0.251 | 0.2 | 1 | 8 | Slc16a10 | 0.051 | Old |
| 0.135376 | 0.252898 | 0.626 | 0.59 | 1 | 8 | Fermt3 | 0.036 | Old |
| 0.135398 | 0.70818 | 0.223 | 0.174 | 1 | 8 | Dgcr2 | 0.049 | Old |
| 0.135874 | 1.121721 | 0.039 | 0.018 | 1 | 8 | Zdhhc24 | 0.021 | Old |
| 0.135941 | 0.88845 | 0.134 | 0.096 | 1 | 8 | Tenm4 | 0.038 | Old |
| 0.136208 | 0.522513 | 0.43 | 0.371 | 1 | 8 | Tbc1d14 | 0.059 | Old |
| 0.136346 | 0.428851 | 0.162 | 0.114 | 1 | 8 | Gpn1 | 0.048 | Old |
| 0.13668 | 0.517417 | 0.168 | 0.122 | 1 | 8 | Heatr1 | 0.046 | Old |
| 0.137004 | 0.552942 | 0.173 | 0.127 | 1 | 8 | Iars | 0.046 | Old |
| 0.137122 | 0.746384 | 0.05 | 0.026 | 1 | 8 | Ints5.1 | 0.024 | Old |
| 0.137503 | 1.078592 | 0.039 | 0.018 | 1 | 8 | Asf1b | 0.021 | Old |
| 0.137707 | 0.502439 | 0.369 | 0.322 | 1 | 8 | Ube2m | 0.047 | Old |
| 0.138049 | 0.780573 | 0.039 | 0.018 | 1 | 8 | Coq3 | 0.021 | Old |
| 0.138063 | 0.879714 | 0.05 | 0.026 | 1 | 8 | Nup107 | 0.024 | Old |
| 0.138121 | 0.282547 | 0.715 | 0.706 | 1 | 8 | Rab7b | 0.009 | Old |
| 0.138429 | 0.441556 | 0.553 | 0.538 | 1 | 8 | Vwa5a | 0.015 | Old |
| 0.138527 | 0.274971 | 0.698 | 0.699 | 1 | 8 | Trim25 | -0.001 | Old |
| 0.13857 | 0.981146 | 0.117 | 0.081 | 1 | 8 | Flt3l | 0.036 | Old |
| 0.139147 | 0.752859 | 0.039 | 0.018 | 1 | 8 | Top3a | 0.021 | Old |
| 0.139673 | 0.952139 | 0.179 | 0.138 | 1 | 8 | Mrpl10 | 0.041 | Old |
| 0.139739 | 0.483992 | 0.078 | 0.047 | 1 | 8 | Ipo4 | 0.031 | Old |
| 0.139792 | 0.059733 | 0.006 | 0.023 | 1 | 8 | A630089N( | -0.017 | Old |
| 0.140251 | 0.998106 | 0.039 | 0.018 | 1 | 8 | Zscan12 | 0.021 | Old |
| 0.140251 | 0.77374 | 0.039 | 0.018 | 1 | 8 | Atp1a2 | 0.021 | Old |
| 0.141298 | 0.858602 | 0.156 | 0.112 | 1 | 8 | Gtf2f1 | 0.044 | Old |
| 0.141396 | 0.795258 | 0.05 | 0.026 | 1 | 8 | Nmb | 0.024 | Old |
| 0.141397 | 0.301678 | 0.559 | 0.532 | 1 | 8 | Tmco1 | 0.027 | Old |
| 0.141535 | 0.309272 | 0.408 | 0.364 | 1 | 8 | 2610001J0 | 0.044 | Old |

| | | | | | | | | |
|---|---|---|---|---|---|---|---|---|
| 0.141878 | 0.599556 | 0.05 | 0.026 | 1 | 8 | Btd | 0.024 | Old |
| 0.142176 | 1.228724 | 0.022 | 0.008 | 1 | 8 | 3010003L2 | 0.014 | Old |
| 0.142362 | 0.104457 | 0.939 | 0.961 | 1 | 8 | Rbpj | -0.022 | Old |
| 0.142481 | 0.637014 | 0.039 | 0.018 | 1 | 8 | Colca2 | 0.021 | Old |
| 0.14253 | 0.289496 | 0.715 | 0.699 | 1 | 8 | Ube2d2a | 0.016 | Old |
| 0.142535 | 0.52807 | 0.201 | 0.153 | 1 | 8 | Tdg | 0.048 | Old |
| 0.14269 | 0.404327 | 0.335 | 0.275 | 1 | 8 | Smu1 | 0.06 | Old |
| 0.142803 | 0.359383 | 0.307 | 0.252 | 1 | 8 | Fbxw2 | 0.055 | Old |
| 0.142933 | 0.280351 | 0.598 | 0.548 | 1 | 8 | Galnt1 | 0.05 | Old |
| 0.142965 | 2.239823 | 0.022 | 0.008 | 1 | 8 | Fignl1 | 0.014 | Old |
| 0.143221 | 0.682426 | 0.251 | 0.205 | 1 | 8 | Csnk1g1 | 0.046 | Old |
| 0.143414 | 0.247835 | 0.034 | 0.065 | 1 | 8 | Mical2 | -0.031 | Old |
| 0.143517 | 0.316778 | 0.682 | 0.649 | 1 | 8 | Tomm20 | 0.033 | Old |
| 0.143756 | 1.083768 | 0.022 | 0.008 | 1 | 8 | Kif3c | 0.014 | Old |
| 0.143756 | 1.03575 | 0.022 | 0.008 | 1 | 8 | Hes2 | 0.014 | Old |
| 0.144249 | 0.796212 | 0.067 | 0.039 | 1 | 8 | Cstf1 | 0.028 | Old |
| 0.144249 | 0.718492 | 0.067 | 0.039 | 1 | 8 | Tbccd1 | 0.028 | Old |
| 0.14438 | 0.183563 | 0.894 | 0.886 | 1 | 8 | Sh3bgrl | 0.008 | Old |
| 0.144551 | 1.749273 | 0.022 | 0.008 | 1 | 8 | Sgo2a | 0.014 | Old |
| 0.144551 | 1.397577 | 0.022 | 0.008 | 1 | 8 | Gm48420 | 0.014 | Old |
| 0.144551 | 1.093303 | 0.022 | 0.008 | 1 | 8 | Harbi1 | 0.014 | Old |
| 0.14471 | 0.512178 | 0.212 | 0.169 | 1 | 8 | Ndufaf4 | 0.043 | Old |
| 0.144903 | 1.158989 | 0.095 | 0.062 | 1 | 8 | Cep70 | 0.033 | Old |
| 0.144965 | 0.79446 | 0.128 | 0.091 | 1 | 8 | Etfdh | 0.037 | Old |
| 0.145321 | 0.498628 | 0.274 | 0.218 | 1 | 8 | Jmjd6 | 0.056 | Old |
| 0.14535 | 1.315173 | 0.022 | 0.008 | 1 | 8 | Rtkn | 0.014 | Old |
| 0.14535 | 1.285871 | 0.022 | 0.008 | 1 | 8 | Rbpms2 | 0.014 | Old |
| 0.145357 | 0.17303 | 0.285 | 0.358 | 1 | 8 | Gtf2a1 | -0.073 | Old |
| 0.145362 | 0.79948 | 0.207 | 0.161 | 1 | 8 | Tmem68 | 0.046 | Old |
| 0.146152 | 1.004037 | 0.022 | 0.008 | 1 | 8 | St5 | 0.014 | Old |
| 0.146152 | 0.963358 | 0.022 | 0.008 | 1 | 8 | Smim1 | 0.014 | Old |
| 0.146643 | 0.39323 | 0.592 | 0.577 | 1 | 8 | Tm9sf2 | 0.015 | Old |
| 0.146946 | 0.333588 | 0.508 | 0.462 | 1 | 8 | Anxa3 | 0.046 | Old |
| 0.146957 | 1.0317 | 0.022 | 0.008 | 1 | 8 | Porcn | 0.014 | Old |
| 0.146957 | 0.915374 | 0.022 | 0.008 | 1 | 8 | Rexo5 | 0.014 | Old |
| 0.146989 | 0.708845 | 0.212 | 0.158 | 1 | 8 | Golga3 | 0.054 | Old |
| 0.147083 | 0.374647 | 0.682 | 0.675 | 1 | 8 | Rtn3 | 0.007 | Old |
| 0.147765 | 1.288035 | 0.022 | 0.008 | 1 | 8 | Xlr3b | 0.014 | Old |
| 0.147765 | 0.932826 | 0.022 | 0.008 | 1 | 8 | Gm38832 | 0.014 | Old |
| 0.147836 | 0.061424 | 0.33 | 0.397 | 1 | 8 | Ivns1abp | -0.067 | Old |
| 0.148577 | 0.805917 | 0.022 | 0.008 | 1 | 8 | Gemin8 | 0.014 | Old |
| 0.148577 | 0.760752 | 0.022 | 0.008 | 1 | 8 | 6430531B1 | 0.014 | Old |
| 0.148645 | 0.606067 | 0.212 | 0.161 | 1 | 8 | Ipo11 | 0.051 | Old |
| 0.149149 | 0.297189 | 0.62 | 0.6 | 1 | 8 | Srsf6 | 0.02 | Old |
| 0.149393 | 0.710427 | 0.022 | 0.008 | 1 | 8 | Msh5 | 0.014 | Old |

| | | | | | | | | |
|---|---|---|---|---|---|---|---|---|
| 0.149393 | 0.664728 | 0.022 | 0.008 | 1 | 8 | Fignl2 | 0.014 | Old |
| 0.149455 | 0.56969 | 0.184 | 0.14 | 1 | 8 | Golt1b | 0.044 | Old |
| 0.149912 | 0.402332 | 0.307 | 0.247 | 1 | 8 | P4ha1 | 0.06 | Old |
| 0.15014 | 1.235812 | 0.056 | 0.031 | 1 | 8 | Glis2 | 0.025 | Old |
| 0.150211 | 0.776149 | 0.022 | 0.008 | 1 | 8 | Cttn | 0.014 | Old |
| 0.150211 | 0.566625 | 0.022 | 0.008 | 1 | 8 | Tha1 | 0.014 | Old |
| 0.150211 | 0.541467 | 0.022 | 0.008 | 1 | 8 | Pdpn | 0.014 | Old |
| 0.150211 | 0.345731 | 0.022 | 0.008 | 1 | 8 | Tldc1 | 0.014 | Old |
| 0.150692 | 0.600994 | 0.24 | 0.195 | 1 | 8 | Fem1b | 0.045 | Old |
| 0.151063 | 0.799391 | 0.218 | 0.171 | 1 | 8 | Cdk17 | 0.047 | Old |
| 0.151409 | 0.331186 | 0.492 | 0.444 | 1 | 8 | Ap3s1 | 0.048 | Old |
| 0.151704 | 0.62287 | 0.274 | 0.231 | 1 | 8 | Smarcb1 | 0.043 | Old |
| 0.151859 | 0.28247 | 0.022 | 0.008 | 1 | 8 | Hist1h2be | 0.014 | Old |
| 0.151877 | 0.601319 | 0.24 | 0.19 | 1 | 8 | Ctso | 0.05 | Old |
| 0.152089 | 0.141696 | 0.933 | 0.938 | 1 | 8 | Tagln2 | -0.005 | Old |
| 0.153147 | 0.26664 | 0.084 | 0.052 | 1 | 8 | Dhx34 | 0.032 | Old |
| 0.153232 | 0.690007 | 0.223 | 0.179 | 1 | 8 | Naxd | 0.044 | Old |
| 0.153752 | 0.150704 | 0.771 | 0.795 | 1 | 8 | Reep5 | -0.024 | Old |
| 0.15393 | 0.278224 | 0.592 | 0.538 | 1 | 8 | Nadk | 0.054 | Old |
| 0.154098 | 0.879139 | 0.184 | 0.145 | 1 | 8 | mt-Nd6 | 0.039 | Old |
| 0.154242 | 0.183462 | 0.693 | 0.678 | 1 | 8 | Larp4b | 0.015 | Old |
| 0.154406 | 0.636404 | 0.056 | 0.031 | 1 | 8 | Fam26f | 0.025 | Old |
| 0.154924 | 0.835373 | 0.196 | 0.153 | 1 | 8 | Ddx39 | 0.043 | Old |
| 0.154979 | 0.364618 | 0.575 | 0.569 | 1 | 8 | Eif4g1 | 0.006 | Old |
| 0.155182 | 0.82591 | 0.162 | 0.119 | 1 | 8 | Atp8b2 | 0.043 | Old |
| 0.155339 | 0.591436 | 0.223 | 0.177 | 1 | 8 | Mtmr10 | 0.046 | Old |
| 0.155498 | 1.177562 | 0.073 | 0.044 | 1 | 8 | Abcg2 | 0.029 | Old |
| 0.155765 | 0.312383 | 0.196 | 0.143 | 1 | 8 | Ccnk | 0.053 | Old |
| 0.155952 | 0.549975 | 0.101 | 0.065 | 1 | 8 | Dcp1a | 0.036 | Old |
| 0.156529 | 0.411377 | 0.458 | 0.416 | 1 | 8 | Rcc2 | 0.042 | Old |
| 0.156591 | 0.425278 | 0.067 | 0.039 | 1 | 8 | Mtx3 | 0.028 | Old |
| 0.156633 | 0.69482 | 0.106 | 0.073 | 1 | 8 | Spg11 | 0.033 | Old |
| 0.156869 | 0.44972 | 0.19 | 0.14 | 1 | 8 | Lnpk | 0.05 | Old |
| 0.157646 | 0.316179 | 0.52 | 0.47 | 1 | 8 | Psma6 | 0.05 | Old |
| 0.158066 | 0.17539 | 0.922 | 0.951 | 1 | 8 | Tmem50a | -0.029 | Old |
| 0.158147 | 0.310186 | 0.737 | 0.787 | 1 | 8 | Jak1 | -0.05 | Old |
| 0.158421 | 1.065854 | 0.061 | 0.036 | 1 | 8 | Hmga1 | 0.025 | Old |
| 0.158763 | 0.780586 | 0.056 | 0.031 | 1 | 8 | 1810024B0 | 0.025 | Old |
| 0.158922 | 0.230754 | 0.872 | 0.904 | 1 | 8 | Ly86 | -0.032 | Old |
| 0.159732 | 0.812885 | 0.095 | 0.062 | 1 | 8 | Mtor | 0.033 | Old |
| 0.160777 | 0.834957 | 0.201 | 0.156 | 1 | 8 | Stat2 | 0.045 | Old |
| 0.160897 | 0.241231 | 0.598 | 0.577 | 1 | 8 | Copz1 | 0.021 | Old |
| 0.161065 | 0.139874 | 0.156 | 0.117 | 1 | 8 | Gdf15 | 0.039 | Old |
| 0.16122 | 0.665348 | 0.145 | 0.106 | 1 | 8 | Mid1 | 0.039 | Old |
| 0.161356 | 0.192365 | 0.682 | 0.688 | 1 | 8 | Clec4a2 | -0.006 | Old |

| | | | | | | | | |
|---|---|---|---|---|---|---|---|---|
| 0.161463 | 0.76968 | 0.223 | 0.182 | 1 | 8 | Usf1 | 0.041 | Old |
| 0.161725 | 0.094966 | 0.084 | 0.052 | 1 | 8 | Sacs | 0.032 | Old |
| 0.161727 | 0.975602 | 0.145 | 0.109 | 1 | 8 | Ccdc43 | 0.036 | Old |
| 0.161731 | 0.471693 | 0.363 | 0.309 | 1 | 8 | Commd7 | 0.054 | Old |
| 0.16272 | 0.504059 | 0.212 | 0.166 | 1 | 8 | Smurf1 | 0.046 | Old |
| 0.162786 | 0.081449 | 0.246 | 0.312 | 1 | 8 | Arrdc3 | -0.066 | Old |
| 0.16309 | 0.512047 | 0.168 | 0.127 | 1 | 8 | Arl4a | 0.041 | Old |
| 0.163431 | 0.572416 | 0.201 | 0.161 | 1 | 8 | Sc5d | 0.04 | Old |
| 0.163759 | 0.079324 | 0.034 | 0.062 | 1 | 8 | Mbip | -0.028 | Old |
| 0.163799 | 0.764176 | 0.095 | 0.062 | 1 | 8 | Polr3d | 0.033 | Old |
| 0.163805 | 0.877753 | 0.184 | 0.143 | 1 | 8 | Eci2 | 0.041 | Old |
| 0.164128 | 0.461273 | 0.503 | 0.462 | 1 | 8 | Smpdl3a | 0.041 | Old |
| 0.16419 | 0.687098 | 0.246 | 0.2 | 1 | 8 | Hk1 | 0.046 | Old |
| 0.164305 | 1.161749 | 0.045 | 0.023 | 1 | 8 | Lhpp | 0.022 | Old |
| 0.164364 | 0.570056 | 0.374 | 0.338 | 1 | 8 | Eloa | 0.036 | Old |
| 0.164398 | 0.565701 | 0.218 | 0.174 | 1 | 8 | Samd1 | 0.044 | Old |
| 0.164665 | 0.774675 | 0.101 | 0.068 | 1 | 8 | Pitrm1 | 0.033 | Old |
| 0.164716 | 0.223603 | 0.022 | 0.047 | 1 | 8 | Ccdc181 | -0.025 | Old |
| 0.164842 | 0.485894 | 0.073 | 0.044 | 1 | 8 | Ercc4 | 0.029 | Old |
| 0.164929 | 0.383232 | 0.19 | 0.148 | 1 | 8 | Capn7 | 0.042 | Old |
| 0.164997 | 0.430618 | 0.19 | 0.143 | 1 | 8 | Ralgapa1 | 0.047 | Old |
| 0.165103 | 0.034268 | 0.296 | 0.358 | 1 | 8 | Mff | -0.062 | Old |
| 0.165158 | 0.744672 | 0.223 | 0.179 | 1 | 8 | Pdcd11 | 0.044 | Old |
| 0.165445 | 1.270416 | 0.045 | 0.023 | 1 | 8 | Mrm1 | 0.022 | Old |
| 0.165592 | 0.353461 | 0.268 | 0.216 | 1 | 8 | Glb1 | 0.052 | Old |
| 0.166149 | 0.901851 | 0.089 | 0.06 | 1 | 8 | Yae1d1 | 0.029 | Old |
| 0.166511 | 0.395943 | 0.285 | 0.229 | 1 | 8 | Pqlc3 | 0.056 | Old |
| 0.166698 | 0.475157 | 0.162 | 0.119 | 1 | 8 | 2610002M | 0.043 | Old |
| 0.167169 | 0.267367 | 0.682 | 0.657 | 1 | 8 | Arf4 | 0.025 | Old |
| 0.167183 | 0.47327 | 0.179 | 0.135 | 1 | 8 | Spns1 | 0.044 | Old |
| 0.167297 | 0.344385 | 0.52 | 0.475 | 1 | 8 | Cul3 | 0.045 | Old |
| 0.167357 | 0.301031 | 0.38 | 0.327 | 1 | 8 | Cebpa | 0.053 | Old |
| 0.167433 | 0.552603 | 0.123 | 0.086 | 1 | 8 | Gfm1 | 0.037 | Old |
| 0.167462 | 0.465898 | 0.073 | 0.044 | 1 | 8 | Rhbdd3 | 0.029 | Old |
| 0.167604 | 0.50301 | 0.525 | 0.522 | 1 | 8 | Sumo3 | 0.003 | Old |
| 0.167653 | 1.151468 | 0.078 | 0.049 | 1 | 8 | Alyref2 | 0.029 | Old |
| 0.167707 | 0.151816 | 0.112 | 0.161 | 1 | 8 | Lrp12 | -0.049 | Old |
| 0.167813 | 1.120965 | 0.061 | 0.036 | 1 | 8 | Ydjc | 0.025 | Old |
| 0.168121 | 0.75879 | 0.223 | 0.179 | 1 | 8 | Dcaf15 | 0.044 | Old |
| 0.168177 | 1.09359 | 0.034 | 0.016 | 1 | 8 | Trmt44 | 0.018 | Old |
| 0.168774 | 1.080714 | 0.061 | 0.036 | 1 | 8 | Steap4 | 0.025 | Old |
| 0.168917 | 0.878274 | 0.078 | 0.049 | 1 | 8 | Zfp839 | 0.029 | Old |
| 0.169683 | 0.396116 | 0.223 | 0.177 | 1 | 8 | 6430548M | 0.046 | Old |
| 0.169898 | 1.207663 | 0.034 | 0.016 | 1 | 8 | Piwil2 | 0.018 | Old |
| 0.169982 | 1.284756 | 0.061 | 0.036 | 1 | 8 | C1ra | 0.025 | Old |

| | | | | | | | | |
|---|---|---|---|---|---|---|---|---|
| 0.170244 | 0.82835 | 0.034 | 0.016 | 1 | 8 | Enc1 | 0.018 | Old |
| 0.170244 | 0.623519 | 0.034 | 0.016 | 1 | 8 | Rnf217 | 0.018 | Old |
| 0.170338 | 2.007481 | 0.017 | 0.005 | 1 | 8 | Magi3 | 0.012 | Old |
| 0.170338 | 1.879504 | 0.017 | 0.005 | 1 | 8 | Dact1 | 0.012 | Old |
| 0.170755 | 0.29981 | 0.559 | 0.491 | 1 | 8 | Cd164 | 0.068 | Old |
| 0.170802 | 0.329173 | 0.458 | 0.41 | 1 | 8 | Ndufab1 | 0.048 | Old |
| 0.170817 | 1.125393 | 0.218 | 0.179 | 1 | 8 | Lmbrd2 | 0.039 | Old |
| 0.170937 | 0.837168 | 0.034 | 0.016 | 1 | 8 | Gm11423 | 0.018 | Old |
| 0.170943 | 1.17854 | 0.084 | 0.055 | 1 | 8 | Pla2g12a | 0.029 | Old |
| 0.171011 | 0.440096 | 0.788 | 0.805 | 1 | 8 | Mtss1 | -0.017 | Old |
| 0.171197 | 0.899045 | 0.061 | 0.036 | 1 | 8 | Ddx20 | 0.025 | Old |
| 0.171235 | 1.120496 | 0.045 | 0.023 | 1 | 8 | Gstm5 | 0.022 | Old |
| 0.171406 | 1.659371 | 0.017 | 0.005 | 1 | 8 | 4930579G2 | 0.012 | Old |
| 0.171415 | 0.191972 | 0.782 | 0.743 | 1 | 8 | Sdc4 | 0.039 | Old |
| 0.171574 | 0.135004 | 0.352 | 0.286 | 1 | 8 | Pts | 0.066 | Old |
| 0.171632 | 0.633324 | 0.034 | 0.016 | 1 | 8 | E230032D2 | 0.018 | Old |
| 0.171836 | 0.864103 | 0.151 | 0.117 | 1 | 8 | Dusp23 | 0.034 | Old |
| 0.172468 | 0.148633 | 0.313 | 0.384 | 1 | 8 | Guk1 | -0.071 | Old |
| 0.172479 | 1.971631 | 0.017 | 0.005 | 1 | 8 | Scn3b | 0.012 | Old |
| 0.172479 | 1.90668 | 0.017 | 0.005 | 1 | 8 | Camk2b | 0.012 | Old |
| 0.172479 | 1.832521 | 0.017 | 0.005 | 1 | 8 | Hspg2 | 0.012 | Old |
| 0.172479 | 1.599836 | 0.017 | 0.005 | 1 | 8 | Bcas1 | 0.012 | Old |
| 0.172479 | 1.491971 | 0.017 | 0.005 | 1 | 8 | Ppm1l | 0.012 | Old |
| 0.172479 | 1.325795 | 0.017 | 0.005 | 1 | 8 | Armcx1 | 0.012 | Old |
| 0.172479 | 1.227661 | 0.017 | 0.005 | 1 | 8 | Znrd1as | 0.012 | Old |
| 0.172479 | 0.997068 | 0.017 | 0.005 | 1 | 8 | Zfp937 | 0.012 | Old |
| 0.172479 | 0.94555 | 0.017 | 0.005 | 1 | 8 | Mmp23 | 0.012 | Old |
| 0.173021 | 0.464125 | 0.034 | 0.062 | 1 | 8 | Mmp14 | -0.028 | Old |
| 0.17314 | 0.313151 | 0.709 | 0.748 | 1 | 8 | G3bp2 | -0.039 | Old |
| 0.173557 | 1.426358 | 0.017 | 0.005 | 1 | 8 | Gm13091 | 0.012 | Old |
| 0.173557 | 1.264303 | 0.017 | 0.005 | 1 | 8 | Shmt1 | 0.012 | Old |
| 0.173557 | 0.862846 | 0.017 | 0.005 | 1 | 8 | S100a16 | 0.012 | Old |
| 0.173557 | 0.8195 | 0.017 | 0.005 | 1 | 8 | AC130815. | 0.012 | Old |
| 0.173896 | 0.56011 | 0.168 | 0.127 | 1 | 8 | Sp2 | 0.041 | Old |
| 0.174315 | 0.346795 | 0.603 | 0.569 | 1 | 8 | Camk1d | 0.034 | Old |
| 0.174451 | 0.447824 | 0.145 | 0.104 | 1 | 8 | Fbxo30 | 0.041 | Old |
| 0.174641 | 1.346223 | 0.017 | 0.005 | 1 | 8 | Fzd8 | 0.012 | Old |
| 0.174641 | 0.972397 | 0.017 | 0.005 | 1 | 8 | Fbxl22 | 0.012 | Old |
| 0.174641 | 0.967648 | 0.017 | 0.005 | 1 | 8 | AL591952.: | 0.012 | Old |
| 0.174641 | 0.799703 | 0.017 | 0.005 | 1 | 8 | Gm9465 | 0.012 | Old |
| 0.174641 | 0.794026 | 0.017 | 0.005 | 1 | 8 | Enkd1 | 0.012 | Old |
| 0.174641 | 0.727258 | 0.017 | 0.005 | 1 | 8 | Ccdc166 | 0.012 | Old |
| 0.174641 | 0.645985 | 0.017 | 0.005 | 1 | 8 | Vps25 | 0.012 | Old |
| 0.174781 | 0.526941 | 0.045 | 0.023 | 1 | 8 | Gm13431 | 0.022 | Old |
| 0.175055 | 0.354597 | 0.073 | 0.044 | 1 | 8 | Tmem39b | 0.029 | Old |

| | | | | | | | | |
|---|---|---|---|---|---|---|---|---|
| 0.175377 | 0.540305 | 0.045 | 0.023 | 1 | 8 | Zfp607a | 0.022 | Old |
| 0.175541 | 0.342051 | 0.123 | 0.086 | 1 | 8 | Ints2 | 0.037 | Old |
| 0.175549 | 0.517811 | 0.313 | 0.27 | 1 | 8 | Vbp1 | 0.043 | Old |
| 0.175583 | 0.369969 | 0.587 | 0.564 | 1 | 8 | Gm26532 | 0.023 | Old |
| 0.175729 | 0.708604 | 0.017 | 0.005 | 1 | 8 | Nphp1 | 0.012 | Old |
| 0.175729 | 0.600079 | 0.017 | 0.005 | 1 | 8 | Gm19412 | 0.012 | Old |
| 0.175729 | 0.545818 | 0.017 | 0.005 | 1 | 8 | Zfp72 | 0.012 | Old |
| 0.175847 | 0.153671 | 0.034 | 0.016 | 1 | 8 | 9030025P2 | 0.018 | Old |
| 0.175909 | 1.078909 | 0.084 | 0.055 | 1 | 8 | Fam122b | 0.029 | Old |
| 0.175984 | 1.626764 | 0.05 | 0.029 | 1 | 8 | Nr1h3 | 0.021 | Old |
| 0.176119 | 0.817268 | 0.061 | 0.036 | 1 | 8 | Mast4 | 0.025 | Old |
| 0.176617 | 0.800381 | 0.061 | 0.036 | 1 | 8 | 1810032O( | 0.025 | Old |
| 0.176687 | 0.49049 | 0.207 | 0.161 | 1 | 8 | Gm36161 | 0.046 | Old |
| 0.176747 | 1.0734 | 0.084 | 0.055 | 1 | 8 | Tbl3 | 0.029 | Old |
| 0.176949 | 0.022448 | 0.335 | 0.416 | 1 | 8 | Fabp4 | -0.081 | Old |
| 0.177386 | 0.673491 | 0.162 | 0.122 | 1 | 8 | Rrm2b | 0.04 | Old |
| 0.177399 | 0.529192 | 0.318 | 0.273 | 1 | 8 | Ube2d1 | 0.045 | Old |
| 0.17744 | 0.023292 | 0.469 | 0.538 | 1 | 8 | Hnrnpd | -0.069 | Old |
| 0.178052 | 0.149641 | 0.05 | 0.083 | 1 | 8 | E4f1 | -0.033 | Old |
| 0.17854 | 0.435148 | 0.291 | 0.247 | 1 | 8 | Nsun2 | 0.044 | Old |
| 0.178608 | 0.243586 | 0.14 | 0.099 | 1 | 8 | Atad3a | 0.041 | Old |
| 0.178924 | 0.385668 | 0.492 | 0.457 | 1 | 8 | Acsl4 | 0.035 | Old |
| 0.17934 | 0.182386 | 0.726 | 0.732 | 1 | 8 | Cnbp | -0.006 | Old |
| 0.179357 | 0.601281 | 0.196 | 0.151 | 1 | 8 | Trappc13 | 0.045 | Old |
| 0.179392 | 0.068059 | 0.972 | 1 | 1 | 8 | Mcl1 | -0.028 | Old |
| 0.179519 | 0.519432 | 0.168 | 0.125 | 1 | 8 | Adora2b | 0.043 | Old |
| 0.179571 | 0.281277 | 0.318 | 0.26 | 1 | 8 | Keap1 | 0.058 | Old |
| 0.179627 | 0.563231 | 0.061 | 0.036 | 1 | 8 | Zfp874b | 0.025 | Old |
| 0.180037 | 0.660753 | 0.212 | 0.169 | 1 | 8 | Sepsecs | 0.043 | Old |
| 0.180042 | 0.419283 | 0.318 | 0.268 | 1 | 8 | Nceh1 | 0.05 | Old |
| 0.180043 | 0.827422 | 0.324 | 0.291 | 1 | 8 | Mogs | 0.033 | Old |
| 0.180482 | 0.716953 | 0.251 | 0.21 | 1 | 8 | Atic | 0.041 | Old |
| 0.180744 | 0.36043 | 0.43 | 0.392 | 1 | 8 | Tmem168 | 0.038 | Old |
| 0.181478 | 0.570914 | 0.257 | 0.213 | 1 | 8 | Tmem184b | 0.044 | Old |
| 0.182248 | 0.28828 | 0.352 | 0.299 | 1 | 8 | 9030624J0 | 0.053 | Old |
| 0.182507 | 0.713259 | 0.095 | 0.065 | 1 | 8 | E2f3 | 0.03 | Old |
| 0.183023 | 0.722328 | 0.128 | 0.094 | 1 | 8 | Ptgr1 | 0.034 | Old |
| 0.183244 | 1.172068 | 0.156 | 0.119 | 1 | 8 | Gmppa | 0.037 | Old |
| 0.183244 | 0.114316 | 0.089 | 0.132 | 1 | 8 | Fam207a | -0.043 | Old |
| 0.183665 | 0.174555 | 0.201 | 0.268 | 1 | 8 | Aaed1 | -0.067 | Old |
| 0.184113 | 1.292749 | 0.067 | 0.042 | 1 | 8 | Actr6 | 0.025 | Old |
| 0.184251 | 0.464868 | 0.24 | 0.195 | 1 | 8 | Itgb1bp1 | 0.045 | Old |
| 0.18437 | 0.338435 | 0.006 | 0.021 | 1 | 8 | Plxnb3 | -0.015 | Old |
| 0.184626 | 0.814558 | 0.145 | 0.109 | 1 | 8 | Akr7a5 | 0.036 | Old |
| 0.185016 | 0.707218 | 0.184 | 0.145 | 1 | 8 | Nupr1 | 0.039 | Old |

| | | | | | | | | |
|---|---|---|---|---|---|---|---|---|
| 0.185022 | 0.831409 | 0.073 | 0.047 | 1 | 8 | Agk | 0.026 | Old |
| 0.185761 | 0.655628 | 0.061 | 0.036 | 1 | 8 | Nkiras1 | 0.025 | Old |
| 0.186199 | 0.22161 | 0.374 | 0.314 | 1 | 8 | Ap1s1 | 0.06 | Old |
| 0.186756 | 0.496273 | 0.296 | 0.26 | 1 | 8 | Clec4e | 0.036 | Old |
| 0.187026 | 0.837022 | 0.184 | 0.143 | 1 | 8 | Pmpcb | 0.041 | Old |
| 0.187313 | 1.307414 | 0.05 | 0.029 | 1 | 8 | Tanc1 | 0.021 | Old |
| 0.188267 | 0.762477 | 0.106 | 0.073 | 1 | 8 | Ampd2 | 0.033 | Old |
| 0.188795 | 0.327496 | 0.084 | 0.055 | 1 | 8 | Arl14ep | 0.029 | Old |
| 0.189057 | 1.774827 | 0.05 | 0.029 | 1 | 8 | Ddx51 | 0.021 | Old |
| 0.189254 | 0.288888 | 0.385 | 0.338 | 1 | 8 | Phf14 | 0.047 | Old |
| 0.189555 | 0.486236 | 0.486 | 0.462 | 1 | 8 | Map3k1 | 0.024 | Old |
| 0.191027 | 0.909161 | 0.067 | 0.042 | 1 | 8 | Pold1 | 0.025 | Old |
| 0.191055 | 0.220928 | 0.62 | 0.618 | 1 | 8 | Vdac2 | 0.002 | Old |
| 0.191615 | 0.019944 | 0.106 | 0.148 | 1 | 8 | Arfip1 | -0.042 | Old |
| 0.191758 | 0.421018 | 0.48 | 0.447 | 1 | 8 | Irf8 | 0.033 | Old |
| 0.191992 | 0.691118 | 0.05 | 0.029 | 1 | 8 | Zfp87 | 0.021 | Old |
| 0.192134 | 3.14337 | 0.011 | 0.003 | 1 | 8 | Gfod2 | 0.008 | Old |
| 0.192134 | 2.101803 | 0.011 | 0.003 | 1 | 8 | Medag | 0.008 | Old |
| 0.192134 | 1.904571 | 0.011 | 0.003 | 1 | 8 | Dennd6b | 0.008 | Old |
| 0.192134 | 1.796145 | 0.011 | 0.003 | 1 | 8 | Zfp37 | 0.008 | Old |
| 0.192134 | 1.766323 | 0.011 | 0.003 | 1 | 8 | 1700003F1 | 0.008 | Old |
| 0.192134 | 1.67711 | 0.011 | 0.003 | 1 | 8 | Hsf2bp | 0.008 | Old |
| 0.192134 | 1.554816 | 0.011 | 0.003 | 1 | 8 | Efcab5 | 0.008 | Old |
| 0.192134 | 1.523837 | 0.011 | 0.003 | 1 | 8 | B3gnt6 | 0.008 | Old |
| 0.192134 | 1.419762 | 0.011 | 0.003 | 1 | 8 | Txlnb | 0.008 | Old |
| 0.192134 | 1.403899 | 0.011 | 0.003 | 1 | 8 | 4930480K2 | 0.008 | Old |
| 0.192134 | 1.372522 | 0.011 | 0.003 | 1 | 8 | Exoc3l4 | 0.008 | Old |
| 0.192134 | 1.294415 | 0.011 | 0.003 | 1 | 8 | 1-Sep | 0.008 | Old |
| 0.192134 | 1.280972 | 0.011 | 0.003 | 1 | 8 | Gm48529 | 0.008 | Old |
| 0.192134 | 1.164654 | 0.011 | 0.003 | 1 | 8 | Gm26610 | 0.008 | Old |
| 0.192134 | 1.109964 | 0.011 | 0.003 | 1 | 8 | Pgf | 0.008 | Old |
| 0.192134 | 1.099975 | 0.011 | 0.003 | 1 | 8 | Bmp1 | 0.008 | Old |
| 0.192134 | 1.040671 | 0.011 | 0.003 | 1 | 8 | AU019990 | 0.008 | Old |
| 0.192134 | 1.031683 | 0.011 | 0.003 | 1 | 8 | 9930022D1 | 0.008 | Old |
| 0.192134 | 1.008347 | 0.011 | 0.003 | 1 | 8 | Gnai1 | 0.008 | Old |
| 0.192134 | 0.990824 | 0.011 | 0.003 | 1 | 8 | Ablim1 | 0.008 | Old |
| 0.192134 | 0.94372 | 0.011 | 0.003 | 1 | 8 | Atp10a | 0.008 | Old |
| 0.192134 | 0.925329 | 0.011 | 0.003 | 1 | 8 | Rab26 | 0.008 | Old |
| 0.192134 | 0.859145 | 0.011 | 0.003 | 1 | 8 | Cacna1b | 0.008 | Old |
| 0.192134 | 0.832116 | 0.011 | 0.003 | 1 | 8 | Dner | 0.008 | Old |
| 0.192134 | 0.783236 | 0.011 | 0.003 | 1 | 8 | Insl6 | 0.008 | Old |
| 0.192134 | 0.758188 | 0.011 | 0.003 | 1 | 8 | Fanca | 0.008 | Old |
| 0.192134 | 0.754843 | 0.011 | 0.003 | 1 | 8 | F730311O2 | 0.008 | Old |
| 0.192134 | 0.753861 | 0.011 | 0.003 | 1 | 8 | Nutf2 | 0.008 | Old |
| 0.192134 | 0.682877 | 0.011 | 0.003 | 1 | 8 | Gm29666 | 0.008 | Old |

| | | | | | | | | |
|---|---|---|---|---|---|---|---|---|
| 0.192134 | 0.681085 | 0.011 | 0.003 | 1 | 8 | Gm26514 | 0.008 | Old |
| 0.192977 | 0.39201 | 0.106 | 0.073 | 1 | 8 | Nt5dc1 | 0.033 | Old |
| 0.193126 | 0.293963 | 0.397 | 0.358 | 1 | 8 | Gga1 | 0.039 | Old |
| 0.19364 | 1.712759 | 0.011 | 0.003 | 1 | 8 | Ahnak2 | 0.008 | Old |
| 0.19364 | 1.63887 | 0.011 | 0.003 | 1 | 8 | Dusp4 | 0.008 | Old |
| 0.19364 | 1.590239 | 0.011 | 0.003 | 1 | 8 | Gm14325 | 0.008 | Old |
| 0.19364 | 1.520732 | 0.011 | 0.003 | 1 | 8 | Arl14epl | 0.008 | Old |
| 0.19364 | 1.47392 | 0.011 | 0.003 | 1 | 8 | Gm15965 | 0.008 | Old |
| 0.19364 | 1.408773 | 0.011 | 0.003 | 1 | 8 | Ccl11 | 0.008 | Old |
| 0.19364 | 1.32045 | 0.011 | 0.003 | 1 | 8 | Cdk18 | 0.008 | Old |
| 0.19364 | 1.259753 | 0.011 | 0.003 | 1 | 8 | Pdgfra | 0.008 | Old |
| 0.19364 | 1.228378 | 0.011 | 0.003 | 1 | 8 | Ankrd55 | 0.008 | Old |
| 0.19364 | 1.216761 | 0.011 | 0.003 | 1 | 8 | 9330175E1 | 0.008 | Old |
| 0.19364 | 1.03354 | 0.011 | 0.003 | 1 | 8 | Gm10260 | 0.008 | Old |
| 0.19364 | 0.922705 | 0.011 | 0.003 | 1 | 8 | Srgap1 | 0.008 | Old |
| 0.19364 | 0.909082 | 0.011 | 0.003 | 1 | 8 | Morn1 | 0.008 | Old |
| 0.19364 | 0.892555 | 0.011 | 0.003 | 1 | 8 | Stk26 | 0.008 | Old |
| 0.19364 | 0.888339 | 0.011 | 0.003 | 1 | 8 | Htra1 | 0.008 | Old |
| 0.19364 | 0.864026 | 0.011 | 0.003 | 1 | 8 | 9130604C2 | 0.008 | Old |
| 0.19364 | 0.834358 | 0.011 | 0.003 | 1 | 8 | Cyp2r1 | 0.008 | Old |
| 0.19364 | 0.827099 | 0.011 | 0.003 | 1 | 8 | Ptger1 | 0.008 | Old |
| 0.19364 | 0.825386 | 0.011 | 0.003 | 1 | 8 | Fbxo44 | 0.008 | Old |
| 0.19364 | 0.804193 | 0.011 | 0.003 | 1 | 8 | Hspb6 | 0.008 | Old |
| 0.19364 | 0.799159 | 0.011 | 0.003 | 1 | 8 | Afp | 0.008 | Old |
| 0.19364 | 0.788332 | 0.011 | 0.003 | 1 | 8 | Cngb1 | 0.008 | Old |
| 0.19364 | 0.78321 | 0.011 | 0.003 | 1 | 8 | Gm26526 | 0.008 | Old |
| 0.19364 | 0.765291 | 0.011 | 0.003 | 1 | 8 | Hic1 | 0.008 | Old |
| 0.19364 | 0.73584 | 0.011 | 0.003 | 1 | 8 | H2-Eb2 | 0.008 | Old |
| 0.19364 | 0.706751 | 0.011 | 0.003 | 1 | 8 | Zfp661 | 0.008 | Old |
| 0.19364 | 0.652782 | 0.011 | 0.003 | 1 | 8 | Gm42726 | 0.008 | Old |
| 0.19364 | 0.649167 | 0.011 | 0.003 | 1 | 8 | Prss23 | 0.008 | Old |
| 0.19364 | 0.57383 | 0.011 | 0.003 | 1 | 8 | 1810062O1 | 0.008 | Old |
| 0.193913 | 0.269537 | 0.704 | 0.704 | 1 | 8 | Clec2d | 0 | Old |
| 0.194209 | 0.524776 | 0.123 | 0.088 | 1 | 8 | Prkrip1 | 0.035 | Old |
| 0.194262 | 0.48972 | 0.335 | 0.288 | 1 | 8 | Nck1 | 0.047 | Old |
| 0.194312 | 0.358692 | 0.14 | 0.101 | 1 | 8 | Ubr7 | 0.039 | Old |
| 0.194578 | 0.70613 | 0.274 | 0.236 | 1 | 8 | Echs1 | 0.038 | Old |
| 0.194714 | 0.348411 | 0.061 | 0.036 | 1 | 8 | Ifrd2 | 0.025 | Old |
| 0.195055 | 0.376539 | 0.196 | 0.153 | 1 | 8 | Arhgap5 | 0.043 | Old |
| 0.195155 | 0.729196 | 0.011 | 0.003 | 1 | 8 | D930028M | 0.008 | Old |
| 0.195155 | 0.703571 | 0.011 | 0.003 | 1 | 8 | Gm14320 | 0.008 | Old |
| 0.195155 | 0.698482 | 0.011 | 0.003 | 1 | 8 | Gm16701.1 | 0.008 | Old |
| 0.195155 | 0.66304 | 0.011 | 0.003 | 1 | 8 | Hoxa4 | 0.008 | Old |
| 0.195155 | 0.649417 | 0.011 | 0.003 | 1 | 8 | Pcdhga7 | 0.008 | Old |
| 0.195155 | 0.632855 | 0.011 | 0.003 | 1 | 8 | Tmc3 | 0.008 | Old |

| | | | | | | | | |
|---|---|---|---|---|---|---|---|---|
| 0.195155 | 0.614747 | 0.011 | 0.003 | 1 | 8 | Zbed5 | 0.008 | Old |
| 0.195155 | 0.530134 | 0.011 | 0.003 | 1 | 8 | Met | 0.008 | Old |
| 0.195155 | 0.523733 | 0.011 | 0.003 | 1 | 8 | Lrrc63 | 0.008 | Old |
| 0.195155 | 0.512878 | 0.011 | 0.003 | 1 | 8 | En2 | 0.008 | Old |
| 0.195155 | 0.403089 | 0.011 | 0.003 | 1 | 8 | Rasgef1a | 0.008 | Old |
| 0.195155 | 0.40052 | 0.011 | 0.003 | 1 | 8 | Htr7 | 0.008 | Old |
| 0.195155 | 0.384494 | 0.011 | 0.003 | 1 | 8 | Kif5a | 0.008 | Old |
| 0.195155 | 0.38292 | 0.011 | 0.003 | 1 | 8 | 4930432K2 | 0.008 | Old |
| 0.195155 | 0.322821 | 0.011 | 0.003 | 1 | 8 | Phf11c | 0.008 | Old |
| 0.195155 | 0.285077 | 0.011 | 0.003 | 1 | 8 | Des | 0.008 | Old |
| 0.195155 | 0.277209 | 0.011 | 0.003 | 1 | 8 | Ank1 | 0.008 | Old |
| 0.195155 | 0.220645 | 0.011 | 0.003 | 1 | 8 | Pole | 0.008 | Old |
| 0.195155 | 0.195243 | 0.011 | 0.003 | 1 | 8 | Ms4a4b | 0.008 | Old |
| 0.195155 | 0.156162 | 0.011 | 0.003 | 1 | 8 | A330069E1 | 0.008 | Old |
| 0.195959 | 0.100314 | 0.14 | 0.19 | 1 | 8 | Taf5l | -0.05 | Old |
| 0.19626 | 0.17838 | 0.782 | 0.831 | 1 | 8 | Cox5a | -0.049 | Old |
| 0.196402 | 0.222559 | 0.615 | 0.597 | 1 | 8 | Snu13 | 0.018 | Old |
| 0.196498 | 0.816527 | 0.279 | 0.236 | 1 | 8 | Dnajc21 | 0.043 | Old |
| 0.196757 | 0.510255 | 0.263 | 0.218 | 1 | 8 | Asxl1 | 0.045 | Old |
| 0.196786 | 0.184482 | 0.86 | 0.862 | 1 | 8 | Rbm3 | -0.002 | Old |
| 0.196788 | 0.375745 | 0.514 | 0.506 | 1 | 8 | Larp1 | 0.008 | Old |
| 0.196792 | 0.520432 | 0.436 | 0.397 | 1 | 8 | Smarcd2 | 0.039 | Old |
| 0.197123 | 0.003568 | 0.352 | 0.413 | 1 | 8 | Eea1 | -0.061 | Old |
| 0.197961 | 0.917749 | 0.05 | 0.029 | 1 | 8 | Gemin2 | 0.021 | Old |
| 0.197963 | 0.453749 | 0.179 | 0.14 | 1 | 8 | Fbxo38 | 0.039 | Old |
| 0.198015 | 1.144343 | 0.078 | 0.052 | 1 | 8 | Dusp18 | 0.026 | Old |
| 0.198565 | 0.531171 | 0.05 | 0.029 | 1 | 8 | Gm46430 | 0.021 | Old |
| 0.198758 | 0.311607 | 0.056 | 0.088 | 1 | 8 | Yaf2 | -0.032 | Old |
| 0.198947 | 0.548957 | 0.38 | 0.335 | 1 | 8 | Anapc5 | 0.045 | Old |
| 0.199171 | 1.173933 | 0.05 | 0.029 | 1 | 8 | Polr1e | 0.021 | Old |
| 0.19945 | 0.859931 | 0.101 | 0.07 | 1 | 8 | Gtpbp1 | 0.031 | Old |
| 0.199778 | 1.375687 | 0.05 | 0.029 | 1 | 8 | Iqcc | 0.021 | Old |
| 0.199881 | 0.408896 | 0.581 | 0.574 | 1 | 8 | Llph | 0.007 | Old |
| 0.199885 | 0.876406 | 0.073 | 0.047 | 1 | 8 | Mrps2 | 0.026 | Old |
| 0.200063 | 0.646768 | 0.279 | 0.236 | 1 | 8 | Smad2 | 0.043 | Old |
| 0.200084 | 0.528468 | 0.212 | 0.169 | 1 | 8 | Aggf1 | 0.043 | Old |
| 0.200993 | 1.074323 | 0.039 | 0.021 | 1 | 8 | Zfp619 | 0.018 | Old |
| 0.200999 | 0.077714 | 0.117 | 0.081 | 1 | 8 | Mag | 0.036 | Old |
| 0.201425 | 0.554667 | 0.173 | 0.135 | 1 | 8 | C1galt1c1 | 0.038 | Old |
| 0.201769 | 0.548129 | 0.296 | 0.255 | 1 | 8 | Arhgef10l | 0.041 | Old |
| 0.202134 | 0.391302 | 0.458 | 0.418 | 1 | 8 | Wdr45b | 0.04 | Old |
| 0.202144 | 0.296814 | 0.693 | 0.678 | 1 | 8 | Id2 | 0.015 | Old |
| 0.20283 | 0.366241 | 0.145 | 0.106 | 1 | 8 | Flnb | 0.039 | Old |
| 0.202909 | 0.384853 | 0.061 | 0.096 | 1 | 8 | Zkscan1 | -0.035 | Old |
| 0.203172 | 1.47154 | 0.028 | 0.013 | 1 | 8 | Gm35363 | 0.015 | Old |

| | | | | | | | | |
|---|---|---|---|---|---|---|---|---|
| 0.203508 | 0.2247 | 0.447 | 0.421 | 1 | 8 | Cox20 | 0.026 | Old |
| 0.204102 | 0.362729 | 0.564 | 0.53 | 1 | 8 | 1110008F1 | 0.034 | Old |
| 0.204707 | 0.541317 | 0.207 | 0.166 | 1 | 8 | Cpsf3 | 0.041 | Old |
| 0.20516 | 0.308318 | 0.615 | 0.582 | 1 | 8 | Wsb1 | 0.033 | Old |
| 0.20529 | 0.156742 | 0.849 | 0.906 | 1 | 8 | Lamp2 | -0.057 | Old |
| 0.205306 | 0.537577 | 0.101 | 0.07 | 1 | 8 | Wdr45 | 0.031 | Old |
| 0.20538 | 0.208785 | 0.201 | 0.156 | 1 | 8 | Zfp560 | 0.045 | Old |
| 0.205531 | 0.444122 | 0.654 | 0.647 | 1 | 8 | Arap1 | 0.007 | Old |
| 0.20577 | 1.039825 | 0.028 | 0.013 | 1 | 8 | Tgfbr3 | 0.015 | Old |
| 0.20577 | 0.784087 | 0.028 | 0.013 | 1 | 8 | Cxcr3 | 0.015 | Old |
| 0.205948 | 1.172665 | 0.039 | 0.021 | 1 | 8 | Ckap2l | 0.018 | Old |
| 0.206002 | 0.520299 | 0.145 | 0.106 | 1 | 8 | Zfyve26 | 0.039 | Old |
| 0.206642 | 1.428327 | 0.028 | 0.013 | 1 | 8 | Adgrl2 | 0.015 | Old |
| 0.207383 | 0.669237 | 0.073 | 0.047 | 1 | 8 | Enpep | 0.026 | Old |
| 0.207516 | 1.455935 | 0.028 | 0.013 | 1 | 8 | Lgi4 | 0.015 | Old |
| 0.207731 | 0.367214 | 0.112 | 0.158 | 1 | 8 | Klhl21 | -0.046 | Old |
| 0.207817 | 0.530808 | 0.201 | 0.156 | 1 | 8 | Slc30a1 | 0.045 | Old |
| 0.208418 | 0.822315 | 0.011 | 0.029 | 1 | 8 | Pigw | -0.018 | Old |
| 0.208958 | 0.183635 | 0.816 | 0.868 | 1 | 8 | Eif3a | -0.052 | Old |
| 0.209191 | 0.13599 | 0.011 | 0.029 | 1 | 8 | Vsig10 | -0.018 | Old |
| 0.209221 | 0.423777 | 0.592 | 0.577 | 1 | 8 | Rab11a | 0.015 | Old |
| 0.209273 | 0.905244 | 0.028 | 0.013 | 1 | 8 | Tceal1 | 0.015 | Old |
| 0.209416 | 0.167338 | 0.888 | 0.943 | 1 | 8 | Hnrnpk | -0.055 | Old |
| 0.209508 | 0.013834 | 0.302 | 0.377 | 1 | 8 | Dmxl1 | -0.075 | Old |
| 0.209542 | 0.898173 | 0.039 | 0.021 | 1 | 8 | Lysmd1 | 0.018 | Old |
| 0.209567 | 0.616771 | 0.229 | 0.184 | 1 | 8 | Hadh | 0.045 | Old |
| 0.209993 | 0.655904 | 0.078 | 0.052 | 1 | 8 | Aven | 0.026 | Old |
| 0.210155 | 0.667172 | 0.028 | 0.013 | 1 | 8 | Pole2 | 0.015 | Old |
| 0.210266 | 1.066829 | 0.039 | 0.021 | 1 | 8 | Akr1c12 | 0.018 | Old |
| 0.210438 | 0.855688 | 0.073 | 0.047 | 1 | 8 | Ccdc163 | 0.026 | Old |
| 0.210764 | 0.002462 | 0.341 | 0.408 | 1 | 8 | Dclre1c | -0.067 | Old |
| 0.210848 | 0.472275 | 0.291 | 0.244 | 1 | 8 | Hist1h2bc | 0.047 | Old |
| 0.210992 | 0.729458 | 0.039 | 0.021 | 1 | 8 | Kcnj2 | 0.018 | Old |
| 0.210992 | 0.493171 | 0.039 | 0.021 | 1 | 8 | Creb3l1 | 0.018 | Old |
| 0.21104 | 0.783717 | 0.028 | 0.013 | 1 | 8 | Shank1 | 0.015 | Old |
| 0.211134 | 0.108187 | 0.33 | 0.268 | 1 | 8 | Fbxo33 | 0.062 | Old |
| 0.21119 | 0.515153 | 0.274 | 0.231 | 1 | 8 | Fbxw11 | 0.043 | Old |
| 0.211316 | 0.790974 | 0.19 | 0.153 | 1 | 8 | Psmd9 | 0.037 | Old |
| 0.211323 | 0.554667 | 0.257 | 0.213 | 1 | 8 | Bmp2 | 0.044 | Old |
| 0.21152 | 1.207959 | 0.112 | 0.083 | 1 | 8 | Gnptg | 0.029 | Old |
| 0.211569 | 0.228417 | 0.05 | 0.029 | 1 | 8 | Fancc | 0.021 | Old |
| 0.211803 | 0.21615 | 0.274 | 0.223 | 1 | 8 | Ola1 | 0.051 | Old |
| 0.211859 | 0.638813 | 0.168 | 0.127 | 1 | 8 | Slc25a12 | 0.041 | Old |
| 0.21202 | 0.083717 | 0.905 | 0.899 | 1 | 8 | Tnfaip3 | 0.006 | Old |
| 0.212238 | 0.53306 | 0.307 | 0.273 | 1 | 8 | Xpo6 | 0.034 | Old |

| | | | | | | | | |
|---|---|---|---|---|---|---|---|---|
| 0.212335 | 0.140649 | 0.777 | 0.8 | 1 | 8 | Tm9sf3 | -0.023 | Old |
| 0.213036 | 0.531406 | 0.095 | 0.065 | 1 | 8 | Zfp568 | 0.03 | Old |
| 0.213644 | 0.273332 | 0.676 | 0.699 | 1 | 8 | M6pr | -0.023 | Old |
| 0.213711 | 1.051041 | 0.028 | 0.013 | 1 | 8 | Tk1 | 0.015 | Old |
| 0.214607 | 0.910924 | 0.028 | 0.013 | 1 | 8 | Adat1 | 0.015 | Old |
| 0.214607 | 0.543913 | 0.028 | 0.013 | 1 | 8 | 2510046G1 | 0.015 | Old |
| 0.214607 | 0.210122 | 0.028 | 0.013 | 1 | 8 | Pla2g6 | 0.015 | Old |
| 0.214992 | 0.031023 | 0.43 | 0.509 | 1 | 8 | Trim30a | -0.079 | Old |
| 0.215081 | 0.794663 | 0.218 | 0.182 | 1 | 8 | Tmem65 | 0.036 | Old |
| 0.215085 | 0.519091 | 0.117 | 0.086 | 1 | 8 | Zbtb43 | 0.031 | Old |
| 0.215151 | 0.392309 | 0.089 | 0.06 | 1 | 8 | Fbxl4 | 0.029 | Old |
| 0.215375 | 0.069775 | 0.017 | 0.036 | 1 | 8 | Cep131 | -0.019 | Old |
| 0.215427 | 0.710655 | 0.078 | 0.052 | 1 | 8 | Rpl36a-ps1 | 0.026 | Old |
| 0.215491 | 0.41073 | 0.508 | 0.439 | 1 | 8 | Fcna | 0.069 | Old |
| 0.215604 | 0.274725 | 0.251 | 0.205 | 1 | 8 | Chsy1 | 0.046 | Old |
| 0.215805 | 0.327187 | 0.218 | 0.169 | 1 | 8 | Tmem205 | 0.049 | Old |
| 0.215883 | 0.803192 | 0.128 | 0.096 | 1 | 8 | Gpr68 | 0.032 | Old |
| 0.216058 | 0.386869 | 0.106 | 0.145 | 1 | 8 | Cd244 | -0.039 | Old |
| 0.216124 | 0.358431 | 0.039 | 0.021 | 1 | 8 | Tmem150b | 0.018 | Old |
| 0.216314 | 0.278299 | 0.391 | 0.335 | 1 | 8 | Eif4e | 0.056 | Old |
| 0.216725 | 0.45304 | 0.369 | 0.338 | 1 | 8 | Arhgef3 | 0.031 | Old |
| 0.21718 | 1.040128 | 0.084 | 0.057 | 1 | 8 | Gprc5b | 0.027 | Old |
| 0.217186 | 0.205238 | 0.62 | 0.566 | 1 | 8 | Rbx1 | 0.054 | Old |
| 0.217223 | 0.530326 | 0.318 | 0.275 | 1 | 8 | Rest | 0.043 | Old |
| 0.217495 | 0.463441 | 0.436 | 0.41 | 1 | 8 | Hip1 | 0.026 | Old |
| 0.217617 | 1.172644 | 0.056 | 0.034 | 1 | 8 | Ip6k2 | 0.022 | Old |
| 0.217765 | 0.420357 | 0.291 | 0.244 | 1 | 8 | Cse1l | 0.047 | Old |
| 0.218351 | 0.554818 | 0.039 | 0.021 | 1 | 8 | Wdr60 | 0.018 | Old |
| 0.218351 | 0.542954 | 0.039 | 0.021 | 1 | 8 | Timeless | 0.018 | Old |
| 0.218455 | 0.158053 | 0.832 | 0.868 | 1 | 8 | Pld4 | -0.036 | Old |
| 0.219127 | 0.039819 | 0.028 | 0.013 | 1 | 8 | Dnajc28 | 0.015 | Old |
| 0.2192 | 0.069753 | 0.402 | 0.34 | 1 | 8 | Ccl12 | 0.062 | Old |
| 0.219238 | 0.452504 | 0.128 | 0.094 | 1 | 8 | Daam1 | 0.034 | Old |
| 0.219436 | 1.048031 | 0.056 | 0.034 | 1 | 8 | Vwa8 | 0.022 | Old |
| 0.220054 | 0.251036 | 0.715 | 0.683 | 1 | 8 | Ms4a6c | 0.032 | Old |
| 0.22029 | 0.363119 | 0.335 | 0.294 | 1 | 8 | Acadl | 0.041 | Old |
| 0.220417 | 0.681276 | 0.229 | 0.195 | 1 | 8 | Paip1 | 0.034 | Old |
| 0.22044 | 0.235659 | 0.888 | 0.899 | 1 | 8 | Sat1 | -0.011 | Old |
| 0.220798 | 0.512133 | 0.123 | 0.088 | 1 | 8 | Nudt14 | 0.035 | Old |
| 0.221266 | 0.439019 | 0.056 | 0.034 | 1 | 8 | Srrd | 0.022 | Old |
| 0.221497 | 0.155999 | 0.816 | 0.818 | 1 | 8 | Srsf3 | -0.002 | Old |
| 0.221552 | 0.044872 | 0.626 | 0.727 | 1 | 8 | Scp2 | -0.101 | Old |
| 0.221564 | 0.217449 | 0.503 | 0.444 | 1 | 8 | Ptpn12 | 0.059 | Old |
| 0.221593 | 0.878924 | 0.061 | 0.039 | 1 | 8 | Fam20a | 0.022 | Old |
| 0.22172 | 0.144231 | 0.285 | 0.226 | 1 | 8 | Gps1 | 0.059 | Old |

| | | | | | | | | |
|---|---|---|---|---|---|---|---|---|
| 0.221927 | 0.333666 | 0.391 | 0.343 | 1 | 8 | Ptprs | 0.048 | Old |
| 0.221965 | 0.266843 | 0.592 | 0.605 | 1 | 8 | Eif5 | -0.013 | Old |
| 0.221989 | 0.491908 | 0.251 | 0.21 | 1 | 8 | Manba | 0.041 | Old |
| 0.222007 | 0.384621 | 0.447 | 0.41 | 1 | 8 | Rer1 | 0.037 | Old |
| 0.222713 | 0.275527 | 0.151 | 0.112 | 1 | 8 | Pop7 | 0.039 | Old |
| 0.222821 | 0.44976 | 0.123 | 0.088 | 1 | 8 | Pgm2 | 0.035 | Old |
| 0.22331 | 0.555359 | 0.318 | 0.281 | 1 | 8 | Idi1 | 0.037 | Old |
| 0.223364 | 0.398427 | 0.285 | 0.236 | 1 | 8 | Flot2 | 0.049 | Old |
| 0.223849 | 0.500897 | 0.112 | 0.081 | 1 | 8 | Ankrd49 | 0.031 | Old |
| 0.224325 | 0.823567 | 0.117 | 0.083 | 1 | 8 | Uckl1 | 0.034 | Old |
| 0.224882 | 0.428535 | 0.603 | 0.569 | 1 | 8 | Cdv3 | 0.034 | Old |
| 0.224939 | 0.442586 | 0.089 | 0.127 | 1 | 8 | Nit1 | -0.038 | Old |
| 0.225191 | 0.485281 | 0.341 | 0.291 | 1 | 8 | Yipf5 | 0.05 | Old |
| 0.225203 | 0.643248 | 0.151 | 0.114 | 1 | 8 | Malsu1 | 0.037 | Old |
| 0.226199 | 0.534562 | 0.056 | 0.034 | 1 | 8 | Alg3 | 0.022 | Old |
| 0.226309 | 0.403994 | 0.391 | 0.351 | 1 | 8 | Psmd6 | 0.04 | Old |
| 0.226697 | 0.213148 | 0.547 | 0.514 | 1 | 8 | Cab39 | 0.033 | Old |
| 0.227044 | 0.20044 | 0.061 | 0.094 | 1 | 8 | Pef1 | -0.033 | Old |
| 0.227053 | 0.589745 | 0.106 | 0.078 | 1 | 8 | Btbd17 | 0.028 | Old |
| 0.227114 | 0.838486 | 0.045 | 0.026 | 1 | 8 | Rhobtb2 | 0.019 | Old |
| 0.227623 | 1.02985 | 0.101 | 0.073 | 1 | 8 | Etfrf1 | 0.028 | Old |
| 0.227693 | 0.464008 | 0.101 | 0.07 | 1 | 8 | Cacfd1 | 0.031 | Old |
| 0.227915 | 0.473591 | 0.196 | 0.158 | 1 | 8 | Alas1 | 0.038 | Old |
| 0.228003 | 0.397322 | 0.274 | 0.234 | 1 | 8 | Aim2 | 0.04 | Old |
| 0.228212 | 0.439156 | 0.179 | 0.138 | 1 | 8 | Specc1 | 0.041 | Old |
| 0.229031 | 0.942467 | 0.095 | 0.068 | 1 | 8 | Rars2 | 0.027 | Old |
| 0.229193 | 0.65482 | 0.061 | 0.039 | 1 | 8 | Dennd2a | 0.022 | Old |
| 0.22944 | 0.152816 | 0.821 | 0.87 | 1 | 8 | 2010107E0 | -0.049 | Old |
| 0.22952 | 0.494382 | 0.101 | 0.07 | 1 | 8 | Smad3 | 0.031 | Old |
| 0.230745 | 0.821384 | 0.14 | 0.109 | 1 | 8 | Gmcl1 | 0.031 | Old |
| 0.230851 | 0.464042 | 0.296 | 0.255 | 1 | 8 | Txndc5 | 0.041 | Old |
| 0.231212 | 0.595306 | 0.279 | 0.239 | 1 | 8 | Ncoa6 | 0.04 | Old |
| 0.231234 | 0.377456 | 0.318 | 0.275 | 1 | 8 | Ddx42 | 0.043 | Old |
| 0.231276 | 0.459215 | 0.279 | 0.231 | 1 | 8 | Rhoh | 0.048 | Old |
| 0.231286 | 0.450111 | 0.318 | 0.281 | 1 | 8 | Exoc3 | 0.037 | Old |
| 0.231983 | 0.326143 | 0.581 | 0.551 | 1 | 8 | Nras | 0.03 | Old |
| 0.232021 | 0.550612 | 0.24 | 0.205 | 1 | 8 | Slc44a1 | 0.035 | Old |
| 0.232116 | 0.259328 | 0.564 | 0.545 | 1 | 8 | Ubxn1 | 0.019 | Old |
| 0.232339 | 0.100654 | 0.894 | 0.943 | 1 | 8 | Eif4g2 | -0.049 | Old |
| 0.23234 | 0.129094 | 0.391 | 0.468 | 1 | 8 | Ppp1cc | -0.077 | Old |
| 0.232422 | 0.592441 | 0.341 | 0.309 | 1 | 8 | Tmem208 | 0.032 | Old |
| 0.232757 | 0.223859 | 0.609 | 0.603 | 1 | 8 | Ube2n | 0.006 | Old |
| 0.232784 | 0.438819 | 0.547 | 0.519 | 1 | 8 | Pias1 | 0.028 | Old |
| 0.232971 | 0.641862 | 0.279 | 0.242 | 1 | 8 | Ubr4 | 0.037 | Old |
| 0.233078 | 0.109957 | 0.173 | 0.13 | 1 | 8 | Cdk8 | 0.043 | Old |

| | | | | | | | | |
|---|---|---|---|---|---|---|---|---|
| 0.233294 | 0.365665 | 0.173 | 0.135 | 1 | 8 | Ankrd27 | 0.038 | Old |
| 0.233352 | 0.775675 | 0.134 | 0.101 | 1 | 8 | Lsm1 | 0.033 | Old |
| 0.233429 | 0.144187 | 0.274 | 0.226 | 1 | 8 | Trim44 | 0.048 | Old |
| 0.233434 | 0.120294 | 0.05 | 0.078 | 1 | 8 | Flywch1 | -0.028 | Old |
| 0.23382 | 0.006895 | 0.369 | 0.439 | 1 | 8 | Mgea5 | -0.07 | Old |
| 0.233896 | 0.088965 | 0.117 | 0.164 | 1 | 8 | Upf2 | -0.047 | Old |
| 0.234169 | 0.11192 | 0.905 | 0.93 | 1 | 8 | Gabarap | -0.025 | Old |
| 0.234205 | 1.1171 | 0.045 | 0.026 | 1 | 8 | Camk2a | 0.019 | Old |
| 0.234505 | 0.278555 | 0.559 | 0.522 | 1 | 8 | Tmem189 | 0.037 | Old |
| 0.234618 | 0.566979 | 0.385 | 0.356 | 1 | 8 | Hexim1 | 0.029 | Old |
| 0.23502 | 0.248337 | 0.056 | 0.034 | 1 | 8 | Asb4 | 0.022 | Old |
| 0.23561 | 0.665486 | 0.134 | 0.104 | 1 | 8 | Akr1e1 | 0.03 | Old |
| 0.235637 | 0.608818 | 0.162 | 0.125 | 1 | 8 | Hbegf | 0.037 | Old |
| 0.235659 | 0.245608 | 0.056 | 0.034 | 1 | 8 | Hoxb4 | 0.022 | Old |
| 0.236329 | 0.024807 | 0.084 | 0.119 | 1 | 8 | Idua | -0.035 | Old |
| 0.236565 | 0.657814 | 0.179 | 0.145 | 1 | 8 | Dgat1 | 0.034 | Old |
| 0.23686 | 0.290461 | 0.196 | 0.153 | 1 | 8 | Usp45 | 0.043 | Old |
| 0.236941 | 0.109049 | 0.056 | 0.034 | 1 | 8 | Emilin1 | 0.022 | Old |
| 0.237085 | 0.939745 | 0.045 | 0.026 | 1 | 8 | Thap1 | 0.019 | Old |
| 0.23819 | 0.49366 | 0.14 | 0.106 | 1 | 8 | Acpp | 0.034 | Old |
| 0.238535 | 0.97593 | 0.045 | 0.026 | 1 | 8 | Mrm2 | 0.019 | Old |
| 0.238622 | 0.646473 | 0.084 | 0.057 | 1 | 8 | Fam151b | 0.027 | Old |
| 0.238631 | 0.233592 | 0.33 | 0.281 | 1 | 8 | Peak1 | 0.049 | Old |
| 0.238751 | 0.047356 | 0.38 | 0.449 | 1 | 8 | Marveld1 | -0.069 | Old |
| 0.238854 | 0.011146 | 0.028 | 0.049 | 1 | 8 | Tmed1 | -0.021 | Old |
| 0.238913 | 0.493191 | 0.229 | 0.187 | 1 | 8 | Gemin7 | 0.042 | Old |
| 0.239166 | 0.282956 | 0.609 | 0.592 | 1 | 8 | Sh3kbp1 | 0.017 | Old |
| 0.239262 | 0.439348 | 0.045 | 0.026 | 1 | 8 | Cby1 | 0.019 | Old |
| 0.239404 | 0.267095 | 0.212 | 0.164 | 1 | 8 | Ranbp10 | 0.048 | Old |
| 0.239954 | 0.269831 | 0.799 | 0.836 | 1 | 8 | Mat2a | -0.037 | Old |
| 0.239991 | 0.84598 | 0.045 | 0.026 | 1 | 8 | Rad18 | 0.019 | Old |
| 0.240374 | 0.257327 | 0.307 | 0.252 | 1 | 8 | Nfkb2 | 0.055 | Old |
| 0.240434 | 0.948977 | 0.095 | 0.068 | 1 | 8 | Alg13 | 0.027 | Old |
| 0.240815 | 0.110718 | 0.162 | 0.127 | 1 | 8 | B9d2 | 0.035 | Old |
| 0.24116 | 0.527823 | 0.067 | 0.044 | 1 | 8 | Uck1 | 0.023 | Old |
| 0.241187 | 0.514001 | 0.089 | 0.062 | 1 | 8 | Znhit2 | 0.027 | Old |
| 0.241357 | 0.204542 | 0.067 | 0.099 | 1 | 8 | Kif3a | -0.032 | Old |
| 0.241441 | 0.327912 | 0.413 | 0.384 | 1 | 8 | Slc25a37 | 0.029 | Old |
| 0.241452 | 1.11193 | 0.045 | 0.026 | 1 | 8 | Lmbr1 | 0.019 | Old |
| 0.241861 | 0.030028 | 0.285 | 0.351 | 1 | 8 | Acbd3 | -0.066 | Old |
| 0.241887 | 0.964524 | 0.095 | 0.068 | 1 | 8 | Tbp | 0.027 | Old |
| 0.242235 | 0.08793 | 0.123 | 0.169 | 1 | 8 | Tnfaip1 | -0.046 | Old |
| 0.242236 | 0.316412 | 0.223 | 0.182 | 1 | 8 | Esrra | 0.041 | Old |
| 0.242327 | 0.779674 | 0.067 | 0.044 | 1 | 8 | Rnf8 | 0.023 | Old |
| 0.242372 | 0.118425 | 0.056 | 0.086 | 1 | 8 | Adprm | -0.03 | Old |

| | | | | | | | | |
|---|---|---|---|---|---|---|---|---|
| 0.242474 | 0.452685 | 0.061 | 0.039 | 1 | 8 | Fam213a | 0.022 | Old |
| 0.242844 | 1.003235 | 0.101 | 0.073 | 1 | 8 | Efl1 | 0.028 | Old |
| 0.243528 | 0.478649 | 0.346 | 0.309 | 1 | 8 | Il3ra | 0.037 | Old |
| 0.243553 | 0.079769 | 0.061 | 0.091 | 1 | 8 | Serinc5 | -0.03 | Old |
| 0.243831 | 0.483408 | 0.006 | 0.018 | 1 | 8 | Oas1c | -0.012 | Old |
| 0.243831 | 0.40303 | 0.006 | 0.018 | 1 | 8 | A430078G2 | -0.012 | Old |
| 0.243956 | 1.376531 | 0.101 | 0.075 | 1 | 8 | Dcn | 0.026 | Old |
| 0.24434 | 0.513541 | 0.084 | 0.057 | 1 | 8 | H2afx | 0.027 | Old |
| 0.244864 | 0.500959 | 0.084 | 0.057 | 1 | 8 | Galns | 0.027 | Old |
| 0.245185 | 0.837632 | 0.151 | 0.119 | 1 | 8 | Abcf3 | 0.032 | Old |
| 0.245435 | 0.405675 | 0.341 | 0.296 | 1 | 8 | Hadha | 0.045 | Old |
| 0.245658 | 0.302045 | 0.514 | 0.475 | 1 | 8 | 7-Mar | 0.039 | Old |
| 0.246037 | 0.188029 | 0.52 | 0.475 | 1 | 8 | Ubald2 | 0.045 | Old |
| 0.246115 | 0.322629 | 0.302 | 0.255 | 1 | 8 | Arhgap25 | 0.047 | Old |
| 0.246313 | 0.008256 | 0.665 | 0.732 | 1 | 8 | Rab1a | -0.067 | Old |
| 0.246533 | 0.656435 | 0.134 | 0.104 | 1 | 8 | Osbpl2 | 0.03 | Old |
| 0.246994 | 0.077873 | 0.196 | 0.236 | 1 | 8 | BC003331 | -0.04 | Old |
| 0.248254 | 0.394605 | 0.056 | 0.086 | 1 | 8 | Apool | -0.03 | Old |
| 0.24838 | 0.38784 | 0.659 | 0.655 | 1 | 8 | Herpud1 | 0.004 | Old |
| 0.248393 | 0.262526 | 0.715 | 0.73 | 1 | 8 | Rpn2 | -0.015 | Old |
| 0.248519 | 0.249441 | 0.24 | 0.192 | 1 | 8 | Ppm1b | 0.048 | Old |
| 0.248652 | 1.116638 | 0.201 | 0.177 | 1 | 8 | Osm | 0.024 | Old |
| 0.248665 | 0.370972 | 0.285 | 0.249 | 1 | 8 | Tor1b | 0.036 | Old |
| 0.249664 | 0.572148 | 0.106 | 0.078 | 1 | 8 | 1110065P2 | 0.028 | Old |
| 0.250465 | 0.291792 | 0.218 | 0.278 | 1 | 8 | Il11ra1 | -0.06 | Old |
| 0.250487 | 1.350707 | 0.05 | 0.031 | 1 | 8 | Nol10 | 0.019 | Old |
| 0.250553 | 1.308491 | 0.034 | 0.018 | 1 | 8 | Tspyl3 | 0.016 | Old |
| 0.250553 | 1.094532 | 0.034 | 0.018 | 1 | 8 | Gm15853 | 0.016 | Old |
| 0.250799 | 1.243302 | 0.095 | 0.07 | 1 | 8 | Slc25a22 | 0.025 | Old |
| 0.250844 | 0.455788 | 0.33 | 0.301 | 1 | 8 | Atp8b4 | 0.029 | Old |
| 0.251009 | 1.006223 | 0.073 | 0.049 | 1 | 8 | Bcar3 | 0.024 | Old |
| 0.251204 | 0.314109 | 0.067 | 0.044 | 1 | 8 | Ears2 | 0.023 | Old |
| 0.251242 | 0.131565 | 0.123 | 0.169 | 1 | 8 | Slc7a7 | -0.046 | Old |
| 0.251269 | 0.357323 | 0.637 | 0.649 | 1 | 8 | Serpinb6a | -0.012 | Old |
| 0.251434 | 1.018448 | 0.034 | 0.018 | 1 | 8 | Ctif | 0.016 | Old |
| 0.251528 | 0.169594 | 0.86 | 0.883 | 1 | 8 | Ndufa2 | -0.023 | Old |
| 0.252231 | 0.652333 | 0.095 | 0.068 | 1 | 8 | Brms1l | 0.027 | Old |
| 0.253028 | 0.261556 | 0.38 | 0.335 | 1 | 8 | Rb1 | 0.045 | Old |
| 0.253204 | 0.851977 | 0.034 | 0.018 | 1 | 8 | Spef1 | 0.016 | Old |
| 0.253443 | 0.523874 | 0.235 | 0.19 | 1 | 8 | Agtpbp1 | 0.045 | Old |
| 0.253584 | 0.427356 | 0.251 | 0.213 | 1 | 8 | Unc50 | 0.038 | Old |
| 0.253861 | 0.264692 | 0.676 | 0.706 | 1 | 8 | Bnip2 | -0.03 | Old |
| 0.253948 | 0.711538 | 0.089 | 0.065 | 1 | 8 | Btrc | 0.024 | Old |
| 0.253976 | 0.514514 | 0.117 | 0.086 | 1 | 8 | Morc2a | 0.031 | Old |
| 0.254069 | 0.113434 | 0.223 | 0.182 | 1 | 8 | Zpr1 | 0.041 | Old |

| | | | | | | | | |
|---|---|---|---|---|---|---|---|---|
| 0.254604 | 0.143839 | 0.022 | 0.042 | 1 | 8 | Zfp275 | -0.02 | Old |
| 0.254886 | 0.485878 | 0.486 | 0.491 | 1 | 8 | D17Wsu92 | -0.005 | Old |
| 0.254983 | 1.034634 | 0.034 | 0.018 | 1 | 8 | Ifi206 | 0.016 | Old |
| 0.254983 | 0.980282 | 0.034 | 0.018 | 1 | 8 | Gdf3 | 0.016 | Old |
| 0.25512 | 0.250982 | 0.168 | 0.216 | 1 | 8 | Lipe | -0.048 | Old |
| 0.255136 | 0.329619 | 0.128 | 0.096 | 1 | 8 | Cherp | 0.032 | Old |
| 0.25532 | 0.052312 | 0.475 | 0.548 | 1 | 8 | Uggt1 | -0.073 | Old |
| 0.255719 | 0.214804 | 0.486 | 0.429 | 1 | 8 | Cnot1 | 0.057 | Old |
| 0.255876 | 0.71704 | 0.034 | 0.018 | 1 | 8 | Aldh1b1 | 0.016 | Old |
| 0.255886 | 0.576538 | 0.285 | 0.244 | 1 | 8 | 1810030O( | 0.041 | Old |
| 0.256052 | 0.167296 | 0.022 | 0.042 | 1 | 8 | Dtd1 | -0.02 | Old |
| 0.25631 | 0.452559 | 0.101 | 0.073 | 1 | 8 | Ddx19b | 0.028 | Old |
| 0.256543 | 0.665155 | 0.089 | 0.062 | 1 | 8 | Nudt1 | 0.027 | Old |
| 0.25677 | 1.108001 | 0.034 | 0.018 | 1 | 8 | Btbd6 | 0.016 | Old |
| 0.25677 | 0.868619 | 0.034 | 0.018 | 1 | 8 | Zfp41 | 0.016 | Old |
| 0.25683 | 0.265315 | 0.598 | 0.592 | 1 | 8 | Cerk | 0.006 | Old |
| 0.256898 | 0.552287 | 0.128 | 0.099 | 1 | 8 | Noc4l | 0.029 | Old |
| 0.257246 | 0.705308 | 0.067 | 0.044 | 1 | 8 | Actn1 | 0.023 | Old |
| 0.257532 | 0.837692 | 0.05 | 0.031 | 1 | 8 | Ly6a | 0.019 | Old |
| 0.257557 | 0.256702 | 0.173 | 0.226 | 1 | 8 | Mcrs1 | -0.053 | Old |
| 0.257656 | 0.095792 | 1 | 0.995 | 1 | 8 | Laptm5 | 0.005 | Old |
| 0.257806 | 0.368837 | 0.134 | 0.104 | 1 | 8 | Isoc1 | 0.03 | Old |
| 0.257809 | 1.635277 | 0.022 | 0.01 | 1 | 8 | Dennd5b | 0.012 | Old |
| 0.257809 | 1.347746 | 0.022 | 0.01 | 1 | 8 | Gm14005 | 0.012 | Old |
| 0.257966 | 0.690089 | 0.201 | 0.169 | 1 | 8 | Taf13 | 0.032 | Old |
| 0.258102 | 0.275528 | 0.626 | 0.597 | 1 | 8 | Trir | 0.029 | Old |
| 0.258378 | 0.293753 | 0.028 | 0.049 | 1 | 8 | Nradd | -0.021 | Old |
| 0.258566 | 0.806085 | 0.034 | 0.018 | 1 | 8 | Gm15446 | 0.016 | Old |
| 0.258951 | 1.347799 | 0.022 | 0.01 | 1 | 8 | Zfp975 | 0.012 | Old |
| 0.259467 | 0.97928 | 0.034 | 0.018 | 1 | 8 | Trmt12 | 0.016 | Old |
| 0.259672 | 1.002019 | 0.05 | 0.031 | 1 | 8 | Sgsm3 | 0.019 | Old |
| 0.259696 | 0.093565 | 0.022 | 0.042 | 1 | 8 | Ccdc84 | -0.02 | Old |
| 0.259886 | 0.254955 | 0.453 | 0.405 | 1 | 8 | Alyref | 0.048 | Old |
| 0.259924 | 0.359706 | 0.257 | 0.213 | 1 | 8 | Tusc3 | 0.044 | Old |
| 0.260371 | 0.674906 | 0.034 | 0.018 | 1 | 8 | Nectin2 | 0.016 | Old |
| 0.260388 | 0.858315 | 0.05 | 0.031 | 1 | 8 | 2700062CC | 0.019 | Old |
| 0.260526 | 0.236147 | 0.307 | 0.257 | 1 | 8 | AW112010 | 0.05 | Old |
| 0.261068 | 0.634601 | 0.145 | 0.114 | 1 | 8 | Trp53inp2 | 0.031 | Old |
| 0.261164 | 0.043602 | 0.022 | 0.042 | 1 | 8 | Zfp579 | -0.02 | Old |
| 0.261245 | 1.109474 | 0.022 | 0.01 | 1 | 8 | Gpc1 | 0.012 | Old |
| 0.261245 | 0.934345 | 0.022 | 0.01 | 1 | 8 | Erbb3 | 0.012 | Old |
| 0.261245 | 0.853148 | 0.022 | 0.01 | 1 | 8 | Slc4a11 | 0.012 | Old |
| 0.261586 | 0.465228 | 0.196 | 0.158 | 1 | 8 | Ipo8 | 0.038 | Old |
| 0.262074 | 1.209562 | 0.056 | 0.036 | 1 | 8 | Scarf1 | 0.02 | Old |
| 0.262184 | 0.500215 | 0.034 | 0.018 | 1 | 8 | Zfp948 | 0.016 | Old |

| | | | | | | | | |
|---|---|---|---|---|---|---|---|---|
| 0.262446 | 0.094851 | 0.257 | 0.325 | 1 | 8 | Adam9 | -0.068 | Old |
| 0.262564 | 0.05192 | 0.184 | 0.234 | 1 | 8 | Pofut1 | -0.05 | Old |
| 0.26258 | 0.035059 | 0.346 | 0.403 | 1 | 8 | Hsd17b12 | -0.057 | Old |
| 0.262797 | 0.45089 | 0.067 | 0.099 | 1 | 8 | Spaca6 | -0.032 | Old |
| 0.263023 | 0.367922 | 0.419 | 0.379 | 1 | 8 | Ppp6c | 0.04 | Old |
| 0.263553 | 0.773255 | 0.022 | 0.01 | 1 | 8 | Cdkl3 | 0.012 | Old |
| 0.263553 | 0.761408 | 0.022 | 0.01 | 1 | 8 | Zfp61 | 0.012 | Old |
| 0.263553 | 0.530975 | 0.022 | 0.01 | 1 | 8 | Clec2g | 0.012 | Old |
| 0.263628 | 0.631831 | 0.05 | 0.031 | 1 | 8 | Msl3l2 | 0.019 | Old |
| 0.263628 | 0.472741 | 0.24 | 0.2 | 1 | 8 | Impa1 | 0.04 | Old |
| 0.263716 | 0.654499 | 0.223 | 0.184 | 1 | 8 | Ap1ar | 0.039 | Old |
| 0.263922 | 0.110137 | 0.67 | 0.634 | 1 | 8 | Rab5c | 0.036 | Old |
| 0.264394 | 0.214097 | 0.218 | 0.182 | 1 | 8 | Cds2 | 0.036 | Old |
| 0.264713 | 1.144784 | 0.022 | 0.01 | 1 | 8 | Dpf3 | 0.012 | Old |
| 0.265299 | 0.279619 | 0.346 | 0.304 | 1 | 8 | Chfr | 0.042 | Old |
| 0.26538 | 0.11665 | 0.017 | 0.034 | 1 | 8 | Dusp19 | -0.017 | Old |
| 0.265837 | 0.525686 | 0.034 | 0.018 | 1 | 8 | Ggcx | 0.016 | Old |
| 0.265875 | 0.924772 | 0.022 | 0.01 | 1 | 8 | Gm15013 | 0.012 | Old |
| 0.26612 | 0.109738 | 0.402 | 0.356 | 1 | 8 | Tap2 | 0.046 | Old |
| 0.266184 | 0.279722 | 0.453 | 0.416 | 1 | 8 | Ccnd2 | 0.037 | Old |
| 0.26621 | 0.373066 | 0.017 | 0.034 | 1 | 8 | Lzts2 | -0.017 | Old |
| 0.266897 | 0.048542 | 0.469 | 0.545 | 1 | 8 | Gnai3 | -0.076 | Old |
| 0.267278 | 0.487764 | 0.011 | 0.026 | 1 | 8 | Tm9sf1 | -0.015 | Old |
| 0.267676 | 0.107315 | 0.034 | 0.018 | 1 | 8 | Gne | 0.016 | Old |
| 0.267732 | 0.375241 | 0.229 | 0.195 | 1 | 8 | Cog5 | 0.034 | Old |
| 0.267904 | 0.205392 | 0.542 | 0.488 | 1 | 8 | Naa50 | 0.054 | Old |
| 0.268211 | 0.603845 | 0.022 | 0.01 | 1 | 8 | Gpnmb | 0.012 | Old |
| 0.268211 | 0.599159 | 0.022 | 0.01 | 1 | 8 | Ropn1l | 0.012 | Old |
| 0.268245 | 0.178145 | 0.922 | 0.878 | 1 | 8 | Marcksl1 | 0.044 | Old |
| 0.268599 | 0.365241 | 0.034 | 0.018 | 1 | 8 | Zfp119b | 0.016 | Old |
| 0.269185 | 0.455847 | 0.179 | 0.143 | 1 | 8 | Arhgap4 | 0.036 | Old |
| 0.269385 | 0.187983 | 0.022 | 0.01 | 1 | 8 | Gm49339 | 0.012 | Old |
| 0.269567 | 0.131611 | 0.469 | 0.545 | 1 | 8 | Ccrl2 | -0.076 | Old |
| 0.269664 | 0.026416 | 0.508 | 0.571 | 1 | 8 | Tns1 | -0.063 | Old |
| 0.269664 | 1.003465 | 0.084 | 0.06 | 1 | 8 | Gm7694 | 0.024 | Old |
| 0.269752 | 0.32703 | 0.676 | 0.688 | 1 | 8 | 7-Sep | -0.012 | Old |
| 0.270039 | 0.281941 | 0.257 | 0.21 | 1 | 8 | Cops5 | 0.047 | Old |
| 0.270308 | 0.001012 | 0.279 | 0.345 | 1 | 8 | Taf1d | -0.066 | Old |
| 0.27034 | 0.600531 | 0.112 | 0.083 | 1 | 8 | 4930453N2 | 0.029 | Old |
| 0.270357 | 0.340981 | 0.168 | 0.13 | 1 | 8 | Cyth2 | 0.038 | Old |
| 0.270557 | 0.533611 | 0.05 | 0.031 | 1 | 8 | Trmt10b | 0.019 | Old |
| 0.270726 | 0.280964 | 0.296 | 0.252 | 1 | 8 | Itpk1 | 0.044 | Old |
| 0.270789 | 0.40015 | 0.251 | 0.208 | 1 | 8 | Trappc8 | 0.043 | Old |
| 0.271256 | 0.354559 | 0.615 | 0.616 | 1 | 8 | Gusb | -0.001 | Old |
| 0.271498 | 0.229374 | 0.508 | 0.491 | 1 | 8 | Rbbp6 | 0.017 | Old |

| | | | | | | | | |
|---|---|---|---|---|---|---|---|---|
| 0.27169 | 0.341398 | 0.369 | 0.335 | 1 | 8 | Ube2f | 0.034 | Old |
| 0.272207 | 0.544047 | 0.268 | 0.236 | 1 | 8 | Msmo1 | 0.032 | Old |
| 0.272212 | 0.635878 | 0.218 | 0.184 | 1 | 8 | Sh3gl1 | 0.034 | Old |
| 0.27225 | 0.117648 | 0.095 | 0.068 | 1 | 8 | Ddx41 | 0.027 | Old |
| 0.272324 | 0.332199 | 0.101 | 0.073 | 1 | 8 | Mrpl22 | 0.028 | Old |
| 0.272578 | 0.277818 | 0.609 | 0.621 | 1 | 8 | Drap1 | -0.012 | Old |
| 0.272881 | 0.422012 | 0.151 | 0.119 | 1 | 8 | Prr14l | 0.032 | Old |
| 0.273015 | 0.209813 | 0.235 | 0.299 | 1 | 8 | Ccdc88b | -0.064 | Old |
| 0.273169 | 0.632658 | 0.123 | 0.094 | 1 | 8 | Fam222b | 0.029 | Old |
| 0.273512 | 1.325549 | 0.05 | 0.031 | 1 | 8 | A430106G1 | 0.019 | Old |
| 0.273512 | 0.174894 | 0.05 | 0.031 | 1 | 8 | Acy1 | 0.019 | Old |
| 0.27363 | 0.291001 | 0.726 | 0.753 | 1 | 8 | Gls | -0.027 | Old |
| 0.273805 | 0.360855 | 0.179 | 0.143 | 1 | 8 | Zfpl1 | 0.036 | Old |
| 0.274141 | 0.038591 | 0.43 | 0.499 | 1 | 8 | Rab21 | -0.069 | Old |
| 0.274591 | 0.461745 | 0.168 | 0.135 | 1 | 8 | Uty | 0.033 | Old |
| 0.27489 | 0.012956 | 0.134 | 0.169 | 1 | 8 | Traf5 | -0.035 | Old |
| 0.274988 | 0.215642 | 0.508 | 0.494 | 1 | 8 | Snrpd3 | 0.014 | Old |
| 0.27504 | 0.331671 | 0.151 | 0.117 | 1 | 8 | Sipa1l1 | 0.034 | Old |
| 0.275054 | 0.458182 | 0.218 | 0.182 | 1 | 8 | Thoc6 | 0.036 | Old |
| 0.275882 | 0.418371 | 0.391 | 0.361 | 1 | 8 | Arf6 | 0.03 | Old |
| 0.276113 | 0.041097 | 0.575 | 0.652 | 1 | 8 | Atxn7l3b | -0.077 | Old |
| 0.276646 | 0.736494 | 0.151 | 0.119 | 1 | 8 | Dnajc11 | 0.032 | Old |
| 0.276676 | 0.320025 | 0.598 | 0.564 | 1 | 8 | Nipa2 | 0.034 | Old |
| 0.276816 | 0.427753 | 0.196 | 0.161 | 1 | 8 | Cd2bp2 | 0.035 | Old |
| 0.277066 | 0.614419 | 0.156 | 0.125 | 1 | 8 | Secisbp2 | 0.031 | Old |
| 0.277926 | 0.062309 | 0.335 | 0.392 | 1 | 8 | Cat | -0.057 | Old |
| 0.278311 | 0.445658 | 0.196 | 0.161 | 1 | 8 | Atp1a3 | 0.035 | Old |
| 0.278622 | 0.497506 | 0.229 | 0.19 | 1 | 8 | Utp18 | 0.039 | Old |
| 0.278677 | 0.201791 | 0.14 | 0.177 | 1 | 8 | Zfp281 | -0.037 | Old |
| 0.278803 | 0.392236 | 0.385 | 0.356 | 1 | 8 | Nfat5 | 0.029 | Old |
| 0.278896 | 0.472436 | 0.19 | 0.156 | 1 | 8 | Shc1 | 0.034 | Old |
| 0.278918 | 0.989748 | 0.095 | 0.07 | 1 | 8 | AI597479 | 0.025 | Old |
| 0.278918 | 0.836766 | 0.095 | 0.07 | 1 | 8 | AC165953. | 0.025 | Old |
| 0.279159 | 0.116561 | 0.095 | 0.068 | 1 | 8 | Il1rap | 0.027 | Old |
| 0.279305 | 0.418345 | 0.235 | 0.195 | 1 | 8 | Hyou1 | 0.04 | Old |
| 0.27933 | 0.150646 | 0.145 | 0.19 | 1 | 8 | Immp1l | -0.045 | Old |
| 0.279752 | 0.248317 | 0.52 | 0.496 | 1 | 8 | Hectd1 | 0.024 | Old |
| 0.279763 | 0.257576 | 0.631 | 0.631 | 1 | 8 | Tram1 | 0 | Old |
| 0.280058 | 0.181652 | 0.804 | 0.826 | 1 | 8 | Pdia3 | -0.022 | Old |
| 0.280118 | 0.186166 | 0.598 | 0.569 | 1 | 8 | Necap2 | 0.029 | Old |
| 0.28013 | 0.611587 | 0.352 | 0.325 | 1 | 8 | Cand1 | 0.027 | Old |
| 0.280395 | 0.252447 | 0.425 | 0.361 | 1 | 8 | Ccni | 0.064 | Old |
| 0.280579 | 1.108017 | 0.039 | 0.023 | 1 | 8 | Hmgn3 | 0.016 | Old |
| 0.281014 | 0.265559 | 0.531 | 0.525 | 1 | 8 | Plec | 0.006 | Old |
| 0.281039 | 0.339695 | 0.291 | 0.242 | 1 | 8 | Gabpb2 | 0.049 | Old |

| | | | | | | | | |
|---|---|---|---|---|---|---|---|---|
| 0.281109 | 0.422505 | 0.145 | 0.114 | 1 | 8 | Trp53bp2 | 0.031 | Old |
| 0.281626 | 0.349232 | 0.101 | 0.073 | 1 | 8 | Slc25a32 | 0.028 | Old |
| 0.282005 | 0.48293 | 0.268 | 0.231 | 1 | 8 | Zbtb11 | 0.037 | Old |
| 0.282391 | 0.18734 | 0.894 | 0.868 | 1 | 8 | Rbms1 | 0.026 | Old |
| 0.282458 | 0.714293 | 0.101 | 0.075 | 1 | 8 | Elac2 | 0.026 | Old |
| 0.283588 | 0.885901 | 0.056 | 0.036 | 1 | 8 | Pde1b | 0.02 | Old |
| 0.283646 | 0.241212 | 0.162 | 0.208 | 1 | 8 | Hbs1l | -0.046 | Old |
| 0.283852 | 0.56059 | 0.151 | 0.119 | 1 | 8 | Evi2 | 0.032 | Old |
| 0.283945 | 0.439884 | 0.156 | 0.125 | 1 | 8 | Atg9a | 0.031 | Old |
| 0.284554 | 0.155068 | 0.095 | 0.068 | 1 | 8 | Il7r | 0.027 | Old |
| 0.284621 | 0.256178 | 0.134 | 0.169 | 1 | 8 | Trim5 | -0.035 | Old |
| 0.284948 | 0.516501 | 0.274 | 0.239 | 1 | 8 | Nudt16l1 | 0.035 | Old |
| 0.285006 | 0.346474 | 0.257 | 0.221 | 1 | 8 | Sae1 | 0.036 | Old |
| 0.28507 | 0.09312 | 0.173 | 0.221 | 1 | 8 | Spr | -0.048 | Old |
| 0.285588 | 0.360439 | 0.128 | 0.099 | 1 | 8 | Mex3d | 0.029 | Old |
| 0.285671 | 0.454392 | 0.397 | 0.387 | 1 | 8 | Appl1 | 0.01 | Old |
| 0.285716 | 0.400664 | 0.385 | 0.356 | 1 | 8 | Eny2 | 0.029 | Old |
| 0.286001 | 0.106086 | 0.229 | 0.275 | 1 | 8 | Mob1b | -0.046 | Old |
| 0.286209 | 0.486921 | 0.609 | 0.579 | 1 | 8 | Fam120a | 0.03 | Old |
| 0.286325 | 0.06539 | 0.19 | 0.148 | 1 | 8 | Enoph1 | 0.042 | Old |
| 0.286587 | 0.745322 | 0.067 | 0.047 | 1 | 8 | Klhl22 | 0.02 | Old |
| 0.286634 | 0.144686 | 0.201 | 0.242 | 1 | 8 | Atf6b | -0.041 | Old |
| 0.28665 | 0.175914 | 0.682 | 0.662 | 1 | 8 | Nucks1 | 0.02 | Old |
| 0.287054 | 0.352444 | 0.302 | 0.255 | 1 | 8 | Appbp2 | 0.047 | Old |
| 0.287061 | 0.974899 | 0.24 | 0.213 | 1 | 8 | Dck | 0.027 | Old |
| 0.287088 | 0.302092 | 0.285 | 0.239 | 1 | 8 | Twistnb | 0.046 | Old |
| 0.28752 | 0.761849 | 0.039 | 0.023 | 1 | 8 | Cacna1f | 0.016 | Old |
| 0.287719 | 0.402136 | 0.151 | 0.119 | 1 | 8 | Timm8a1 | 0.032 | Old |
| 0.288032 | 0.600381 | 0.335 | 0.322 | 1 | 8 | Tomm70a | 0.013 | Old |
| 0.288704 | 0.156862 | 0.229 | 0.187 | 1 | 8 | Tbc1d2b | 0.042 | Old |
| 0.288801 | 0.380523 | 0.078 | 0.055 | 1 | 8 | Pkp3 | 0.023 | Old |
| 0.288842 | 0.014437 | 0.073 | 0.049 | 1 | 8 | Rbm38 | 0.024 | Old |
| 0.289063 | 0.397835 | 0.117 | 0.088 | 1 | 8 | Aga | 0.029 | Old |
| 0.289329 | 0.874498 | 0.056 | 0.036 | 1 | 8 | Telo2 | 0.02 | Old |
| 0.289481 | 0.398031 | 0.296 | 0.255 | 1 | 8 | Ammecr1l | 0.041 | Old |
| 0.289902 | 0.140014 | 0.106 | 0.145 | 1 | 8 | Pdk3 | -0.039 | Old |
| 0.290329 | 0.705177 | 0.162 | 0.132 | 1 | 8 | Sh2d1b1 | 0.03 | Old |
| 0.290656 | 0.371756 | 0.184 | 0.151 | 1 | 8 | Lap3 | 0.033 | Old |
| 0.290986 | 0.679922 | 0.151 | 0.122 | 1 | 8 | Pcsk7 | 0.029 | Old |
| 0.291003 | 0.018479 | 0.117 | 0.086 | 1 | 8 | Coro2b | 0.031 | Old |
| 0.291246 | 0.005327 | 0.251 | 0.291 | 1 | 8 | Kif3b | -0.04 | Old |
| 0.291641 | 0.122082 | 0.927 | 0.982 | 1 | 8 | Ptprc | -0.055 | Old |
| 0.291917 | 0.883128 | 0.039 | 0.023 | 1 | 8 | Spout1 | 0.016 | Old |
| 0.291917 | 0.80924 | 0.039 | 0.023 | 1 | 8 | Arsg | 0.016 | Old |
| 0.292619 | 0.661542 | 0.207 | 0.179 | 1 | 8 | Arhgap10 | 0.028 | Old |

| | | | | | | | | |
|---|---|---|---|---|---|---|---|---|
| 0.29269 | 0.811282 | 0.073 | 0.052 | 1 | 8 | Alkbh8 | 0.021 | Old |
| 0.292836 | 0.070608 | 0.279 | 0.33 | 1 | 8 | Stk11 | -0.051 | Old |
| 0.292957 | 0.27432 | 0.056 | 0.036 | 1 | 8 | Bcdin3d | 0.02 | Old |
| 0.293001 | 0.537081 | 0.123 | 0.094 | 1 | 8 | Wdyhv1 | 0.029 | Old |
| 0.293516 | 0.203062 | 0.648 | 0.6 | 1 | 8 | Rilpl2 | 0.048 | Old |
| 0.294024 | 0.752398 | 0.101 | 0.075 | 1 | 8 | Bcl2l12 | 0.026 | Old |
| 0.294194 | 0.052526 | 0.034 | 0.055 | 1 | 8 | Trit1 | -0.021 | Old |
| 0.294494 | 0.366959 | 0.089 | 0.065 | 1 | 8 | Cdc23 | 0.024 | Old |
| 0.294557 | 0.807237 | 0.101 | 0.075 | 1 | 8 | Ipcef1 | 0.026 | Old |
| 0.295141 | 0.020047 | 0.564 | 0.657 | 1 | 8 | Ap2s1 | -0.093 | Old |
| 0.295409 | 0.519842 | 0.145 | 0.114 | 1 | 8 | Hinfp | 0.031 | Old |
| 0.295467 | 0.804687 | 0.039 | 0.023 | 1 | 8 | Dppa3 | 0.016 | Old |
| 0.295539 | 0.006594 | 0.743 | 0.844 | 1 | 8 | Ndufa4 | -0.101 | Old |
| 0.295924 | 0.655537 | 0.061 | 0.042 | 1 | 8 | Nudt18 | 0.019 | Old |
| 0.295994 | 0.001327 | 0.162 | 0.2 | 1 | 8 | Slc35a5 | -0.038 | Old |
| 0.295994 | 0.106977 | 0.559 | 0.522 | 1 | 8 | Ddost | 0.037 | Old |
| 0.296134 | 0.296631 | 0.112 | 0.083 | 1 | 8 | Urgcp | 0.029 | Old |
| 0.296271 | 0.723394 | 0.061 | 0.042 | 1 | 8 | Ccar2 | 0.019 | Old |
| 0.296351 | 0.307188 | 0.179 | 0.145 | 1 | 8 | Faap20 | 0.034 | Old |
| 0.297099 | 0.800112 | 0.106 | 0.081 | 1 | 8 | Rbsn | 0.025 | Old |
| 0.297424 | 0.557116 | 0.402 | 0.39 | 1 | 8 | H2-M3 | 0.012 | Old |
| 0.29795 | 0.025239 | 0.402 | 0.47 | 1 | 8 | Numa1 | -0.068 | Old |
| 0.298148 | 0.658725 | 0.039 | 0.023 | 1 | 8 | Zfp953 | 0.016 | Old |
| 0.298347 | 0.691822 | 0.218 | 0.19 | 1 | 8 | Myo7a | 0.028 | Old |
| 0.298468 | 0.090537 | 0.352 | 0.416 | 1 | 8 | Sod2 | -0.064 | Old |
| 0.298936 | 0.046988 | 0.156 | 0.197 | 1 | 8 | Tmc6 | -0.041 | Old |
| 0.298973 | 0.286944 | 0.201 | 0.166 | 1 | 8 | Rem1 | 0.035 | Old |
| 0.299944 | 1.141592 | 0.039 | 0.023 | 1 | 8 | Zfp873 | 0.016 | Old |
| 0.300087 | 0.16405 | 0.397 | 0.348 | 1 | 8 | Eif1ax | 0.049 | Old |
| 0.300324 | 0.106541 | 0.503 | 0.574 | 1 | 8 | Prkacb | -0.071 | Old |
| 0.300465 | 0.391043 | 0.101 | 0.075 | 1 | 8 | Lzic | 0.026 | Old |
| 0.300629 | 0.444614 | 0.475 | 0.465 | 1 | 8 | Aco2 | 0.01 | Old |
| 0.300999 | 0.286825 | 0.603 | 0.618 | 1 | 8 | Psmd2 | -0.015 | Old |
| 0.301468 | 0.2222 | 0.838 | 0.842 | 1 | 8 | Vcp | -0.004 | Old |
| 0.301696 | 0.359119 | 0.57 | 0.566 | 1 | 8 | Apbb1ip | 0.004 | Old |
| 0.301735 | 0.017624 | 0.263 | 0.304 | 1 | 8 | Xpo1 | -0.041 | Old |
| 0.302141 | 0.42027 | 0.391 | 0.353 | 1 | 8 | Med28 | 0.038 | Old |
| 0.302272 | 0.300903 | 0.324 | 0.281 | 1 | 8 | Hmbox1 | 0.043 | Old |
| 0.302347 | 0.702357 | 0.106 | 0.081 | 1 | 8 | Abt1 | 0.025 | Old |
| 0.302596 | 0.579419 | 0.464 | 0.468 | 1 | 8 | Dusp5 | -0.004 | Old |
| 0.302807 | 0.701485 | 0.173 | 0.143 | 1 | 8 | Hdac4 | 0.03 | Old |
| 0.303169 | 0.317671 | 0.196 | 0.161 | 1 | 8 | Smg9 | 0.035 | Old |
| 0.303365 | 0.135211 | 0.173 | 0.218 | 1 | 8 | Rnf2 | -0.045 | Old |
| 0.303409 | 0.007885 | 0.296 | 0.348 | 1 | 8 | Ahr | -0.052 | Old |
| 0.303559 | 0.345001 | 0.039 | 0.023 | 1 | 8 | Gorasp1 | 0.016 | Old |

| | | | | | | | | |
|---|---|---|---|---|---|---|---|---|
| 0.303676 | 0.009186 | 0.207 | 0.242 | 1 | 8 | Ahctf1 | -0.035 | Old |
| 0.303704 | 0.945585 | 0.067 | 0.047 | 1 | 8 | Tchp | 0.02 | Old |
| 0.303909 | 0.077586 | 0.274 | 0.327 | 1 | 8 | Kdm1a | -0.053 | Old |
| 0.304056 | 0.401463 | 0.402 | 0.377 | 1 | 8 | Id3 | 0.025 | Old |
| 0.304193 | 0.348808 | 0.525 | 0.504 | 1 | 8 | Nxf1 | 0.021 | Old |
| 0.304773 | 0.681689 | 0.179 | 0.148 | 1 | 8 | Golga2 | 0.031 | Old |
| 0.304937 | 0.291819 | 0.084 | 0.06 | 1 | 8 | Stk32c | 0.024 | Old |
| 0.305103 | 0.428149 | 0.061 | 0.088 | 1 | 8 | Snx19 | -0.027 | Old |
| 0.305816 | 0.438594 | 0.235 | 0.197 | 1 | 8 | Vps37a | 0.038 | Old |
| 0.305902 | 0.226759 | 0.592 | 0.595 | 1 | 8 | Lcp2 | -0.003 | Old |
| 0.306093 | 0.642687 | 0.061 | 0.042 | 1 | 8 | Mocs1 | 0.019 | Old |
| 0.306486 | 0.552075 | 0.229 | 0.195 | 1 | 8 | Lrrc8a | 0.034 | Old |
| 0.306702 | 0.214647 | 0.207 | 0.171 | 1 | 8 | Zfp280d | 0.036 | Old |
| 0.307202 | 0.316991 | 0.039 | 0.023 | 1 | 8 | Arhgef5 | 0.016 | Old |
| 0.307202 | 0.310627 | 0.039 | 0.023 | 1 | 8 | 2700081O1 | 0.016 | Old |
| 0.307668 | 0.563651 | 0.179 | 0.148 | 1 | 8 | Nfyb | 0.031 | Old |
| 0.307709 | 0.358103 | 0.503 | 0.457 | 1 | 8 | AC160336. | 0.046 | Old |
| 0.307776 | 0.454581 | 0.112 | 0.086 | 1 | 8 | Naf1 | 0.026 | Old |
| 0.308225 | 0.373862 | 0.061 | 0.042 | 1 | 8 | Rab39 | 0.019 | Old |
| 0.308444 | 0.459158 | 0.458 | 0.439 | 1 | 8 | Fam126a | 0.019 | Old |
| 0.308734 | 0.330071 | 0.531 | 0.499 | 1 | 8 | Notch1 | 0.032 | Old |
| 0.308764 | 0.240285 | 0.788 | 0.792 | 1 | 8 | Pafah1b1 | -0.004 | Old |
| 0.308831 | 0.457552 | 0.279 | 0.242 | 1 | 8 | Rasa3 | 0.037 | Old |
| 0.308906 | 0.445939 | 0.508 | 0.496 | 1 | 8 | Mia3 | 0.012 | Old |
| 0.308994 | 0.191759 | 0.123 | 0.094 | 1 | 8 | Acot13 | 0.029 | Old |
| 0.30904 | 0.648944 | 0.128 | 0.101 | 1 | 8 | Ints13 | 0.027 | Old |
| 0.309074 | 0.792119 | 0.128 | 0.104 | 1 | 8 | Dusp10 | 0.024 | Old |
| 0.309169 | 0.219119 | 0.575 | 0.584 | 1 | 8 | Cct4 | -0.009 | Old |
| 0.310249 | 0.263672 | 0.765 | 0.761 | 1 | 8 | Tapbp | 0.004 | Old |
| 0.310462 | 0.054433 | 0.095 | 0.125 | 1 | 8 | Dnajc24 | -0.03 | Old |
| 0.310582 | 0.626363 | 0.123 | 0.099 | 1 | 8 | Rfxap | 0.024 | Old |
| 0.310733 | 0.542695 | 0.14 | 0.109 | 1 | 8 | Fdft1 | 0.031 | Old |
| 0.311158 | 1.468423 | 0.028 | 0.016 | 1 | 8 | A430033K0 | 0.012 | Old |
| 0.311498 | 0.057893 | 0.335 | 0.387 | 1 | 8 | Acss1 | -0.052 | Old |
| 0.311781 | 0.291978 | 0.575 | 0.571 | 1 | 8 | Aftph | 0.004 | Old |
| 0.312512 | 0.53358 | 0.067 | 0.047 | 1 | 8 | Sema6b | 0.02 | Old |
| 0.312785 | 0.462022 | 0.207 | 0.171 | 1 | 8 | Strn | 0.036 | Old |
| 0.313006 | 0.701727 | 0.112 | 0.086 | 1 | 8 | Habp4 | 0.026 | Old |
| 0.313097 | 0.374701 | 0.14 | 0.109 | 1 | 8 | Med23 | 0.031 | Old |
| 0.313633 | 0.433721 | 0.369 | 0.338 | 1 | 8 | Csnk1g2 | 0.031 | Old |
| 0.313685 | 0.123054 | 0.804 | 0.818 | 1 | 8 | Hnrnpa0 | -0.014 | Old |
| 0.313911 | 0.096495 | 0.022 | 0.039 | 1 | 8 | Col27a1 | -0.017 | Old |
| 0.313918 | 0.036036 | 0.508 | 0.595 | 1 | 8 | Fbl | -0.087 | Old |
| 0.314363 | 0.27905 | 0.218 | 0.182 | 1 | 8 | Rbbp8 | 0.036 | Old |
| 0.314495 | 0.167546 | 0.235 | 0.283 | 1 | 8 | Scamp3 | -0.048 | Old |

| | | | | | | | | |
|---|---|---|---|---|---|---|---|---|
| 0.314821 | 0.676155 | 0.095 | 0.07 | 1 | 8 | Dxo | 0.025 | Old |
| 0.314855 | 0.436748 | 0.156 | 0.125 | 1 | 8 | Cpsf1 | 0.031 | Old |
| 0.315343 | 0.343623 | 0.397 | 0.358 | 1 | 8 | Ggh | 0.039 | Old |
| 0.315419 | 0.194958 | 0.089 | 0.065 | 1 | 8 | Ttc39c | 0.024 | Old |
| 0.315581 | 0.30339 | 0.402 | 0.384 | 1 | 8 | Copb1 | 0.018 | Old |
| 0.315621 | 1.183446 | 0.045 | 0.029 | 1 | 8 | Prrg1 | 0.016 | Old |
| 0.315809 | 0.939688 | 0.073 | 0.052 | 1 | 8 | Mcm2 | 0.021 | Old |
| 0.315897 | 0.576132 | 0.128 | 0.101 | 1 | 8 | Riok2 | 0.027 | Old |
| 0.3159 | 0.099125 | 0.128 | 0.164 | 1 | 8 | Atp2a3 | -0.036 | Old |
| 0.315908 | 0.443635 | 0.251 | 0.218 | 1 | 8 | Nfia | 0.033 | Old |
| 0.316853 | 0.306103 | 0.061 | 0.042 | 1 | 8 | Gm1673 | 0.019 | Old |
| 0.316975 | 0.598852 | 0.184 | 0.158 | 1 | 8 | Plcb1 | 0.026 | Old |
| 0.317088 | 0.27862 | 0.575 | 0.582 | 1 | 8 | Dse | -0.007 | Old |
| 0.317616 | 0.080561 | 0.106 | 0.14 | 1 | 8 | Kdm5d | -0.034 | Old |
| 0.31787 | 0.059308 | 0.145 | 0.184 | 1 | 8 | Ttll12 | -0.039 | Old |
| 0.318097 | 1.011126 | 0.095 | 0.073 | 1 | 8 | Ano10 | 0.022 | Old |
| 0.318344 | 0.351493 | 0.145 | 0.114 | 1 | 8 | Brix1 | 0.031 | Old |
| 0.318512 | 0.252713 | 0.24 | 0.203 | 1 | 8 | Cops6 | 0.037 | Old |
| 0.318707 | 0.436745 | 0.067 | 0.047 | 1 | 8 | Ift74 | 0.02 | Old |
| 0.319072 | 0.254223 | 0.173 | 0.216 | 1 | 8 | Gid8 | -0.043 | Old |
| 0.319745 | 0.1219 | 0.112 | 0.083 | 1 | 8 | Aifm2 | 0.029 | Old |
| 0.320021 | 0.291138 | 0.095 | 0.07 | 1 | 8 | Cep85l | 0.025 | Old |
| 0.320099 | 0.698295 | 0.028 | 0.016 | 1 | 8 | Zfp189 | 0.012 | Old |
| 0.320099 | 0.696125 | 0.028 | 0.016 | 1 | 8 | Frs3 | 0.012 | Old |
| 0.320521 | 1.003507 | 0.084 | 0.062 | 1 | 8 | Tmem268 | 0.022 | Old |
| 0.32079 | 0.48375 | 0.067 | 0.047 | 1 | 8 | Ripk3 | 0.02 | Old |
| 0.321079 | 0.054622 | 0.279 | 0.236 | 1 | 8 | Tob1 | 0.043 | Old |
| 0.321228 | 1.360173 | 0.028 | 0.016 | 1 | 8 | Sgms2 | 0.012 | Old |
| 0.321228 | 1.035992 | 0.028 | 0.016 | 1 | 8 | Zbtb39 | 0.012 | Old |
| 0.321491 | 0.429004 | 0.229 | 0.197 | 1 | 8 | Bccip | 0.032 | Old |
| 0.32168 | 0.707602 | 0.168 | 0.138 | 1 | 8 | Acot7 | 0.03 | Old |
| 0.32236 | 0.77435 | 0.028 | 0.016 | 1 | 8 | Apoo | 0.012 | Old |
| 0.32236 | 0.590895 | 0.028 | 0.016 | 1 | 8 | Gm4707 | 0.012 | Old |
| 0.32239 | 0.665952 | 0.145 | 0.119 | 1 | 8 | Olr1 | 0.026 | Old |
| 0.322522 | 0.076437 | 0.022 | 0.039 | 1 | 8 | Letm2 | -0.017 | Old |
| 0.32255 | 0.186721 | 0.458 | 0.429 | 1 | 8 | Mrpl20 | 0.029 | Old |
| 0.322677 | 0.531925 | 0.235 | 0.208 | 1 | 8 | Plagl2 | 0.027 | Old |
| 0.322727 | 0.297208 | 0.162 | 0.127 | 1 | 8 | Zdhhc18 | 0.035 | Old |
| 0.323218 | 0.724648 | 0.19 | 0.164 | 1 | 8 | Fut11 | 0.026 | Old |
| 0.323392 | 0.411275 | 0.045 | 0.029 | 1 | 8 | Gm16124 | 0.016 | Old |
| 0.323479 | 0.102101 | 0.983 | 0.987 | 1 | 8 | Retnla | -0.004 | Old |
| 0.323523 | 0.005764 | 0.263 | 0.314 | 1 | 8 | Cpsf7 | -0.051 | Old |
| 0.323557 | 0.883324 | 0.006 | 0.016 | 1 | 8 | Gm9733 | -0.01 | Old |
| 0.324003 | 0.213979 | 0.782 | 0.79 | 1 | 8 | Atf4 | -0.008 | Old |
| 0.324106 | 0.229442 | 0.089 | 0.119 | 1 | 8 | Cpsf4 | -0.03 | Old |

| | | | | | | | | |
|---|---|---|---|---|---|---|---|---|
| 0.324157 | 0.062926 | 0.067 | 0.094 | 1 | 8 | Bcl9l | -0.027 | Old |
| 0.324275 | 0.454514 | 0.19 | 0.161 | 1 | 8 | Ccdc6 | 0.029 | Old |
| 0.324483 | 0.59384 | 0.19 | 0.161 | 1 | 8 | Pex7 | 0.029 | Old |
| 0.324632 | 1.231237 | 0.028 | 0.016 | 1 | 8 | Birc5 | 0.012 | Old |
| 0.324632 | 0.570582 | 0.028 | 0.016 | 1 | 8 | Syp | 0.012 | Old |
| 0.325771 | 0.592095 | 0.028 | 0.016 | 1 | 8 | Adal | 0.012 | Old |
| 0.325771 | 0.379046 | 0.028 | 0.016 | 1 | 8 | Fam167b | 0.012 | Old |
| 0.325842 | 0.371236 | 0.341 | 0.306 | 1 | 8 | Ehmt2 | 0.035 | Old |
| 0.325943 | 0.340856 | 0.179 | 0.145 | 1 | 8 | Akap8l | 0.034 | Old |
| 0.326186 | 0.029144 | 0.609 | 0.725 | 1 | 8 | Asah1 | -0.116 | Old |
| 0.326914 | 1.06498 | 0.028 | 0.016 | 1 | 8 | Zfp449 | 0.012 | Old |
| 0.32706 | 0.498221 | 0.201 | 0.169 | 1 | 8 | Hccs | 0.032 | Old |
| 0.327256 | 0.566766 | 0.402 | 0.371 | 1 | 8 | Klc1 | 0.031 | Old |
| 0.327387 | 0.777606 | 0.078 | 0.057 | 1 | 8 | Ppp1r16a | 0.021 | Old |
| 0.327496 | 0.320631 | 0.397 | 0.366 | 1 | 8 | Ipo7 | 0.031 | Old |
| 0.32777 | 0.463224 | 0.246 | 0.208 | 1 | 8 | Dirc2 | 0.038 | Old |
| 0.328059 | 0.56993 | 0.028 | 0.016 | 1 | 8 | Cavin2 | 0.012 | Old |
| 0.328059 | 0.269236 | 0.028 | 0.016 | 1 | 8 | Zfp229 | 0.012 | Old |
| 0.328059 | 0.040571 | 0.028 | 0.016 | 1 | 8 | Prkar1b | 0.012 | Old |
| 0.328514 | 0.762288 | 0.145 | 0.119 | 1 | 8 | Paox | 0.026 | Old |
| 0.328813 | 0.326075 | 0.095 | 0.07 | 1 | 8 | F3 | 0.025 | Old |
| 0.328907 | 0.481795 | 0.19 | 0.156 | 1 | 8 | Prkra | 0.034 | Old |
| 0.329009 | 0.172216 | 0.592 | 0.579 | 1 | 8 | Rell1 | 0.013 | Old |
| 0.329391 | 0.252766 | 0.682 | 0.691 | 1 | 8 | Bzw1 | -0.009 | Old |
| 0.329521 | 0.436521 | 0.045 | 0.029 | 1 | 8 | Sh3d19 | 0.016 | Old |
| 0.329698 | 1.55249 | 0.017 | 0.008 | 1 | 8 | 1110046J0 | 0.009 | Old |
| 0.329698 | 1.346336 | 0.017 | 0.008 | 1 | 8 | Hrh2 | 0.009 | Old |
| 0.329698 | 1.289368 | 0.017 | 0.008 | 1 | 8 | Sptbn4 | 0.009 | Old |
| 0.329732 | 0.334369 | 0.201 | 0.169 | 1 | 8 | Slc4a2 | 0.032 | Old |
| 0.330238 | 0.532856 | 0.145 | 0.114 | 1 | 8 | Lrba | 0.031 | Old |
| 0.330465 | 0.140686 | 0.888 | 0.899 | 1 | 8 | Flna | -0.011 | Old |
| 0.330542 | 0.182984 | 0.084 | 0.114 | 1 | 8 | Rcor3 | -0.03 | Old |
| 0.33125 | 2.224443 | 0.017 | 0.008 | 1 | 8 | Rnls | 0.009 | Old |
| 0.33125 | 1.494041 | 0.017 | 0.008 | 1 | 8 | Gm12258 | 0.009 | Old |
| 0.33125 | 0.792203 | 0.017 | 0.008 | 1 | 8 | Rasl2-9 | 0.009 | Old |
| 0.33125 | 0.655084 | 0.017 | 0.008 | 1 | 8 | Timm23 | 0.009 | Old |
| 0.331757 | 0.204197 | 0.592 | 0.592 | 1 | 8 | Notch2 | 0 | Old |
| 0.331772 | 0.170439 | 0.374 | 0.426 | 1 | 8 | Arfgef1 | -0.052 | Old |
| 0.332175 | 0.396993 | 0.246 | 0.205 | 1 | 8 | Gatd1 | 0.041 | Old |
| 0.332582 | 0.432714 | 0.492 | 0.488 | 1 | 8 | Psmd4 | 0.004 | Old |
| 0.332664 | 0.329241 | 0.028 | 0.016 | 1 | 8 | Pstpip2 | 0.012 | Old |
| 0.33275 | 0.291059 | 0.201 | 0.166 | 1 | 8 | Adar | 0.035 | Old |
| 0.3328 | 0.182906 | 0.721 | 0.725 | 1 | 8 | Hnrnpm | -0.004 | Old |
| 0.332807 | 1.493321 | 0.017 | 0.008 | 1 | 8 | Mettl5os | 0.009 | Old |
| 0.332807 | 1.472365 | 0.017 | 0.008 | 1 | 8 | 4930477G0 | 0.009 | Old |

| | | | | | | | | |
|---|---|---|---|---|---|---|---|---|
| 0.332807 | 1.304087 | 0.017 | 0.008 | 1 | 8 | Art2b | 0.009 | Old |
| 0.332807 | 1.258958 | 0.017 | 0.008 | 1 | 8 | Ltb | 0.009 | Old |
| 0.332807 | 0.827679 | 0.017 | 0.008 | 1 | 8 | Gm4610 | 0.009 | Old |
| 0.332807 | 0.716929 | 0.017 | 0.008 | 1 | 8 | Lrig3 | 0.009 | Old |
| 0.332807 | 0.708529 | 0.017 | 0.008 | 1 | 8 | Dock6 | 0.009 | Old |
| 0.333057 | 0.460429 | 0.045 | 0.029 | 1 | 8 | Cyp2j6 | 0.016 | Old |
| 0.333136 | 0.431009 | 0.19 | 0.158 | 1 | 8 | Topors | 0.032 | Old |
| 0.33378 | 0.451189 | 0.05 | 0.034 | 1 | 8 | Hus1 | 0.016 | Old |
| 0.333822 | 0.085528 | 0.011 | 0.023 | 1 | 8 | Dgke | -0.012 | Old |
| 0.334256 | 0.046774 | 0.302 | 0.351 | 1 | 8 | Mrpl30 | -0.049 | Old |
| 0.334369 | 1.185994 | 0.017 | 0.008 | 1 | 8 | Car6 | 0.009 | Old |
| 0.334369 | 0.831439 | 0.017 | 0.008 | 1 | 8 | Lca5 | 0.009 | Old |
| 0.334369 | 0.775255 | 0.017 | 0.008 | 1 | 8 | Prdm16 | 0.009 | Old |
| 0.334369 | 0.751493 | 0.017 | 0.008 | 1 | 8 | Ccdc63 | 0.009 | Old |
| 0.334369 | 0.595112 | 0.017 | 0.008 | 1 | 8 | Ildr1 | 0.009 | Old |
| 0.334369 | 0.56271 | 0.017 | 0.008 | 1 | 8 | Ggn | 0.009 | Old |
| 0.334408 | 0.869365 | 0.106 | 0.083 | 1 | 8 | Pgpep1 | 0.023 | Old |
| 0.334492 | 0.134464 | 0.754 | 0.764 | 1 | 8 | Gpx4 | -0.01 | Old |
| 0.335282 | 0.27675 | 0.073 | 0.052 | 1 | 8 | 3110040N1 | 0.021 | Old |
| 0.335551 | 0.158396 | 0.156 | 0.125 | 1 | 8 | Phkg2 | 0.031 | Old |
| 0.335764 | 0.009587 | 0.631 | 0.701 | 1 | 8 | Dock10 | -0.07 | Old |
| 0.335936 | 0.64732 | 0.017 | 0.008 | 1 | 8 | Msantd1 | 0.009 | Old |
| 0.335936 | 0.635837 | 0.017 | 0.008 | 1 | 8 | Slc8a1 | 0.009 | Old |
| 0.335936 | 0.634633 | 0.017 | 0.008 | 1 | 8 | Top1mt | 0.009 | Old |
| 0.335936 | 0.573968 | 0.017 | 0.008 | 1 | 8 | Pde7b | 0.009 | Old |
| 0.335936 | 0.485306 | 0.017 | 0.008 | 1 | 8 | Plag1 | 0.009 | Old |
| 0.336336 | 0.430566 | 0.067 | 0.047 | 1 | 8 | Rmnd1 | 0.02 | Old |
| 0.336525 | 0.507738 | 0.374 | 0.338 | 1 | 8 | Psma5 | 0.036 | Old |
| 0.337311 | 0.490592 | 0.011 | 0.023 | 1 | 8 | Prkcg | -0.012 | Old |
| 0.337311 | 0.124106 | 0.028 | 0.016 | 1 | 8 | Ficd | 0.012 | Old |
| 0.337339 | 0.168831 | 0.536 | 0.522 | 1 | 8 | Copa | 0.014 | Old |
| 0.337339 | 0.423884 | 0.123 | 0.096 | 1 | 8 | Tex2 | 0.027 | Old |
| 0.33734 | 0.39624 | 0.073 | 0.052 | 1 | 8 | Txnl4b | 0.021 | Old |
| 0.337466 | 0.687417 | 0.134 | 0.106 | 1 | 8 | Txnrd2 | 0.028 | Old |
| 0.337507 | 1.14389 | 0.017 | 0.008 | 1 | 8 | Arv1 | 0.009 | Old |
| 0.337507 | 0.535725 | 0.017 | 0.008 | 1 | 8 | Gm11713 | 0.009 | Old |
| 0.337781 | 0.20623 | 0.117 | 0.091 | 1 | 8 | Srgap3 | 0.026 | Old |
| 0.338133 | 0.330663 | 0.039 | 0.06 | 1 | 8 | Mrpl39 | -0.021 | Old |
| 0.338311 | 1.108462 | 0.112 | 0.088 | 1 | 8 | Zbtb46 | 0.024 | Old |
| 0.338429 | 0.026118 | 0.067 | 0.094 | 1 | 8 | Tsta3 | -0.027 | Old |
| 0.338554 | 0.422508 | 0.263 | 0.234 | 1 | 8 | Dcstamp | 0.029 | Old |
| 0.339083 | 0.357146 | 0.017 | 0.008 | 1 | 8 | Rpusd3 | 0.009 | Old |
| 0.339083 | 0.312938 | 0.017 | 0.008 | 1 | 8 | Gm39469 | 0.009 | Old |
| 0.339196 | 0.166019 | 0.615 | 0.584 | 1 | 8 | Ube3a | 0.031 | Old |
| 0.33955 | 0.189364 | 0.24 | 0.205 | 1 | 8 | Vps4a | 0.035 | Old |

| | | | | | | | | |
|---|---|---|---|---|---|---|---|---|
| 0.339754 | 1.203249 | 0.078 | 0.06 | 1 | 8 | Skp2 | 0.018 | Old |
| 0.339958 | 0.129543 | 0.145 | 0.114 | 1 | 8 | B3glct | 0.031 | Old |
| 0.340664 | 0.160204 | 0.017 | 0.008 | 1 | 8 | AU022793 | 0.009 | Old |
| 0.341102 | 0.122598 | 0.045 | 0.029 | 1 | 8 | Chek2 | 0.016 | Old |
| 0.34116 | 0.705445 | 0.078 | 0.057 | 1 | 8 | Yipf2 | 0.021 | Old |
| 0.341214 | 0.043511 | 0.112 | 0.145 | 1 | 8 | BC005561 | -0.033 | Old |
| 0.341521 | 1.242713 | 0.056 | 0.039 | 1 | 8 | Klhl12 | 0.017 | Old |
| 0.341578 | 0.245881 | 0.173 | 0.138 | 1 | 8 | Selenon | 0.035 | Old |
| 0.341599 | 0.501994 | 0.179 | 0.148 | 1 | 8 | Med12 | 0.031 | Old |
| 0.34197 | 0.145955 | 0.095 | 0.127 | 1 | 8 | Cav2 | -0.032 | Old |
| 0.34225 | 0.14402 | 0.017 | 0.008 | 1 | 8 | Trim45 | 0.009 | Old |
| 0.342416 | 0.427024 | 0.145 | 0.114 | 1 | 8 | Bloc1s4 | 0.031 | Old |
| 0.342648 | 0.548762 | 0.207 | 0.179 | 1 | 8 | Slc25a1 | 0.028 | Old |
| 0.342935 | 0.343431 | 0.391 | 0.358 | 1 | 8 | Glrx5 | 0.033 | Old |
| 0.34308 | 0.461781 | 0.101 | 0.078 | 1 | 8 | Stk16 | 0.023 | Old |
| 0.343248 | 0.195031 | 0.123 | 0.158 | 1 | 8 | Pcnt | -0.035 | Old |
| 0.343684 | 0.239823 | 0.212 | 0.177 | 1 | 8 | Ext2 | 0.035 | Old |
| 0.343897 | 1.206973 | 0.056 | 0.039 | 1 | 8 | Ranbp6 | 0.017 | Old |
| 0.343897 | 0.701777 | 0.056 | 0.039 | 1 | 8 | Tnfsf13os | 0.017 | Old |
| 0.344729 | 0.29541 | 0.162 | 0.13 | 1 | 8 | Ptgr2 | 0.032 | Old |
| 0.345497 | 0.405917 | 0.05 | 0.034 | 1 | 8 | Socs1 | 0.016 | Old |
| 0.346088 | 0.454247 | 0.385 | 0.384 | 1 | 8 | Retreg2 | 0.001 | Old |
| 0.346164 | 0.004029 | 0.179 | 0.218 | 1 | 8 | Ltn1 | -0.039 | Old |
| 0.346217 | 0.262405 | 0.492 | 0.473 | 1 | 8 | Cpt1a | 0.019 | Old |
| 0.346344 | 0.268292 | 0.05 | 0.034 | 1 | 8 | Zfp637 | 0.016 | Old |
| 0.346479 | 0.36741 | 0.38 | 0.358 | 1 | 8 | Pdxk | 0.022 | Old |
| 0.346668 | 0.566194 | 0.235 | 0.205 | 1 | 8 | Dap3 | 0.03 | Old |
| 0.346883 | 0.641897 | 0.196 | 0.169 | 1 | 8 | Lrpprc | 0.027 | Old |
| 0.347054 | 0.096078 | 0.151 | 0.187 | 1 | 8 | Scd2 | -0.036 | Old |
| 0.347795 | 0.562229 | 0.285 | 0.26 | 1 | 8 | Yipf1 | 0.025 | Old |
| 0.348157 | 0.440173 | 0.207 | 0.174 | 1 | 8 | Ttpal | 0.033 | Old |
| 0.348329 | 0.172438 | 0.05 | 0.073 | 1 | 8 | I830077J02 | -0.023 | Old |
| 0.348566 | 0.036482 | 0.073 | 0.099 | 1 | 8 | Cmtm4 | -0.026 | Old |
| 0.348679 | 0.833728 | 0.056 | 0.039 | 1 | 8 | Zcchc4 | 0.017 | Old |
| 0.348747 | 0.116016 | 0.156 | 0.2 | 1 | 8 | Zfp330 | -0.044 | Old |
| 0.348777 | 0.217491 | 0.693 | 0.701 | 1 | 8 | Atp6ap1 | -0.008 | Old |
| 0.349132 | 0.197278 | 0.14 | 0.174 | 1 | 8 | Irak2 | -0.034 | Old |
| 0.349132 | 0.021102 | 0.615 | 0.699 | 1 | 8 | Myl12b | -0.084 | Old |
| 0.349142 | 0.10922 | 0.38 | 0.452 | 1 | 8 | Cbfb | -0.072 | Old |
| 0.349172 | 0.424611 | 0.469 | 0.444 | 1 | 8 | Rnpep | 0.025 | Old |
| 0.349637 | 0.149931 | 0.201 | 0.164 | 1 | 8 | Cul2 | 0.037 | Old |
| 0.349856 | 0.186572 | 0.073 | 0.052 | 1 | 8 | Eogt | 0.021 | Old |
| 0.350008 | 0.634915 | 0.095 | 0.073 | 1 | 8 | Plrg1 | 0.022 | Old |
| 0.350153 | 0.578339 | 0.156 | 0.13 | 1 | 8 | Pygo2 | 0.026 | Old |
| 0.350239 | 0.528859 | 0.123 | 0.099 | 1 | 8 | Iqsec1 | 0.024 | Old |

| | | | | | | | | |
|---|---|---|---|---|---|---|---|---|
| 0.350378 | 0.560396 | 0.229 | 0.2 | 1 | 8 | Cln6 | 0.029 | Old |
| 0.350384 | 0.179046 | 0.374 | 0.34 | 1 | 8 | Elk4 | 0.034 | Old |
| 0.35046 | 0.333777 | 0.151 | 0.119 | 1 | 8 | Bst1 | 0.032 | Old |
| 0.350561 | 0.024161 | 0.095 | 0.122 | 1 | 8 | Zfp639 | -0.027 | Old |
| 0.351199 | 0.48709 | 0.173 | 0.145 | 1 | 8 | Elp2 | 0.028 | Old |
| 0.351293 | 0.913342 | 0.034 | 0.021 | 1 | 8 | Jag1 | 0.013 | Old |
| 0.351417 | 0.460293 | 0.559 | 0.548 | 1 | 8 | 2610507B1 | 0.011 | Old |
| 0.351489 | 1.09632 | 0.056 | 0.039 | 1 | 8 | Map3k15 | 0.017 | Old |
| 0.351563 | 0.112296 | 0.246 | 0.301 | 1 | 8 | Cnot3 | -0.055 | Old |
| 0.351643 | 0.098592 | 0.184 | 0.229 | 1 | 8 | Washc3 | -0.045 | Old |
| 0.351868 | 0.317198 | 0.553 | 0.551 | 1 | 8 | Ubash3b | 0.002 | Old |
| 0.352044 | 0.411752 | 0.156 | 0.127 | 1 | 8 | Cdk2 | 0.029 | Old |
| 0.352228 | 0.553671 | 0.084 | 0.062 | 1 | 8 | Prune1 | 0.022 | Old |
| 0.352664 | 0.243077 | 0.268 | 0.314 | 1 | 8 | Vcl | -0.046 | Old |
| 0.352882 | 0.872438 | 0.084 | 0.062 | 1 | 8 | Spice1 | 0.022 | Old |
| 0.353307 | 0.53526 | 0.168 | 0.14 | 1 | 8 | Adpgk | 0.028 | Old |
| 0.353454 | 0.563854 | 0.201 | 0.177 | 1 | 8 | Uspl1 | 0.024 | Old |
| 0.353645 | 0.696583 | 0.089 | 0.068 | 1 | 8 | Polr3e | 0.021 | Old |
| 0.354226 | 0.419678 | 0.223 | 0.187 | 1 | 8 | Nudcd2 | 0.036 | Old |
| 0.354496 | 1.360219 | 0.034 | 0.021 | 1 | 8 | Gm42595 | 0.013 | Old |
| 0.354802 | 0.189499 | 0.073 | 0.052 | 1 | 8 | Dedd | 0.021 | Old |
| 0.355064 | 0.240249 | 0.145 | 0.179 | 1 | 8 | Yeats2 | -0.034 | Old |
| 0.355567 | 0.66747 | 0.034 | 0.021 | 1 | 8 | Tctn3 | 0.013 | Old |
| 0.355844 | 0.312516 | 0.268 | 0.236 | 1 | 8 | Abcc5 | 0.032 | Old |
| 0.355933 | 0.611082 | 0.056 | 0.039 | 1 | 8 | Evi5l | 0.017 | Old |
| 0.356476 | 0.0083 | 0.33 | 0.384 | 1 | 8 | Klf9 | -0.054 | Old |
| 0.35661 | 0.295645 | 0.05 | 0.034 | 1 | 8 | Dars2 | 0.016 | Old |
| 0.357293 | 0.34928 | 0.525 | 0.517 | 1 | 8 | U2af2 | 0.008 | Old |
| 0.357456 | 0.33394 | 0.089 | 0.068 | 1 | 8 | Atpaf2 | 0.021 | Old |
| 0.357662 | 0.091555 | 0.33 | 0.387 | 1 | 8 | Reps2 | -0.057 | Old |
| 0.357811 | 0.196299 | 0.497 | 0.465 | 1 | 8 | Dynlrb1 | 0.032 | Old |
| 0.358086 | 0.702745 | 0.034 | 0.052 | 1 | 8 | Tmed8 | -0.018 | Old |
| 0.358337 | 0.92045 | 0.067 | 0.049 | 1 | 8 | Wdr83 | 0.018 | Old |
| 0.358658 | 0.164257 | 0.363 | 0.332 | 1 | 8 | Zfc3h1 | 0.031 | Old |
| 0.358794 | 1.525724 | 0.034 | 0.021 | 1 | 8 | Hspb1 | 0.013 | Old |
| 0.359397 | 0.05704 | 0.156 | 0.19 | 1 | 8 | Cnot7 | -0.034 | Old |
| 0.359874 | 0.739767 | 0.034 | 0.021 | 1 | 8 | Papss2 | 0.013 | Old |
| 0.359896 | 0.150707 | 0.14 | 0.174 | 1 | 8 | Mrps26 | -0.034 | Old |
| 0.360365 | 0.048788 | 0.397 | 0.447 | 1 | 8 | Abcc1 | -0.05 | Old |
| 0.360432 | 0.390416 | 0.363 | 0.34 | 1 | 8 | Ubxn6 | 0.023 | Old |
| 0.360913 | 0.625451 | 0.184 | 0.156 | 1 | 8 | Gpd2 | 0.028 | Old |
| 0.361789 | 0.315598 | 0.38 | 0.351 | 1 | 8 | Abce1 | 0.029 | Old |
| 0.36204 | 0.381288 | 0.034 | 0.021 | 1 | 8 | Eps8l1 | 0.013 | Old |
| 0.362159 | 0.098432 | 0.078 | 0.057 | 1 | 8 | Pwp2 | 0.021 | Old |
| 0.362187 | 0.407065 | 0.581 | 0.577 | 1 | 8 | Morf4l2 | 0.004 | Old |

| | | | | | | | | |
|---|---|---|---|---|---|---|---|---|
| 0.362386 | 0.170648 | 0.123 | 0.158 | 1 | 8 | Pdrg1 | -0.035 | Old |
| 0.362453 | 0.021276 | 0.313 | 0.351 | 1 | 8 | Pole4 | -0.038 | Old |
| 0.362619 | 0.113945 | 0.14 | 0.174 | 1 | 8 | Mapk8 | -0.034 | Old |
| 0.362675 | 0.032315 | 0.123 | 0.153 | 1 | 8 | Mfap1a | -0.03 | Old |
| 0.362706 | 0.373768 | 0.229 | 0.195 | 1 | 8 | Cdc34 | 0.034 | Old |
| 0.362777 | 0.034068 | 0.173 | 0.21 | 1 | 8 | Prrc1 | -0.037 | Old |
| 0.36285 | 0.156396 | 0.078 | 0.057 | 1 | 8 | Abhd11 | 0.021 | Old |
| 0.36322 | 0.554225 | 0.089 | 0.068 | 1 | 8 | Mgmt | 0.021 | Old |
| 0.363376 | 0.711616 | 0.145 | 0.122 | 1 | 8 | Wdr77 | 0.023 | Old |
| 0.363835 | 1.215898 | 0.078 | 0.06 | 1 | 8 | Mill2 | 0.018 | Old |
| 0.363939 | 0.244658 | 0.246 | 0.216 | 1 | 8 | Naa35 | 0.03 | Old |
| 0.364014 | 0.232925 | 0.436 | 0.421 | 1 | 8 | Arhgef2 | 0.015 | Old |
| 0.364213 | 0.61489 | 0.034 | 0.021 | 1 | 8 | Fam177a | 0.013 | Old |
| 0.364213 | 0.523318 | 0.034 | 0.021 | 1 | 8 | Wdr34 | 0.013 | Old |
| 0.364824 | 0.562276 | 0.173 | 0.145 | 1 | 8 | Rpf1 | 0.028 | Old |
| 0.364986 | 0.261637 | 0.603 | 0.592 | 1 | 8 | Hspd1 | 0.011 | Old |
| 0.365016 | 0.335165 | 0.514 | 0.488 | 1 | 8 | Sdhd | 0.026 | Old |
| 0.365079 | 0.290473 | 0.162 | 0.135 | 1 | 8 | Atp11a | 0.027 | Old |
| 0.36531 | 0.251286 | 0.028 | 0.044 | 1 | 8 | Park2 | -0.016 | Old |
| 0.365367 | 0.748738 | 0.067 | 0.049 | 1 | 8 | Nudcd1 | 0.018 | Old |
| 0.365736 | 0.316652 | 0.335 | 0.304 | 1 | 8 | Slc25a36 | 0.031 | Old |
| 0.36575 | 0.43524 | 0.056 | 0.039 | 1 | 8 | Cald1 | 0.017 | Old |
| 0.366187 | 0.055353 | 0.028 | 0.044 | 1 | 8 | Anln | -0.016 | Old |
| 0.366203 | 0.233216 | 0.698 | 0.725 | 1 | 8 | Atp5k | -0.027 | Old |
| 0.366351 | 0.46902 | 0.246 | 0.216 | 1 | 8 | Syne2 | 0.03 | Old |
| 0.366444 | 0.402058 | 0.38 | 0.358 | 1 | 8 | Khsrp | 0.022 | Old |
| 0.366642 | 0.548321 | 0.106 | 0.083 | 1 | 8 | Ctu2 | 0.023 | Old |
| 0.367214 | 0.222423 | 0.318 | 0.275 | 1 | 8 | Mrpl34 | 0.043 | Old |
| 0.367359 | 0.154153 | 0.05 | 0.07 | 1 | 8 | Zfp367 | -0.02 | Old |
| 0.367459 | 0.396531 | 0.084 | 0.062 | 1 | 8 | Ogfod2 | 0.022 | Old |
| 0.367462 | 0.14753 | 0.397 | 0.348 | 1 | 8 | Ppp3cb | 0.049 | Old |
| 0.367465 | 0.495095 | 0.151 | 0.125 | 1 | 8 | Pbdc1 | 0.026 | Old |
| 0.367607 | 0.337415 | 0.196 | 0.166 | 1 | 8 | Yipf6 | 0.03 | Old |
| 0.367946 | 0.195722 | 0.05 | 0.034 | 1 | 8 | D17H6S53I | 0.016 | Old |
| 0.368003 | 0.44204 | 0.508 | 0.499 | 1 | 8 | Mef2d | 0.009 | Old |
| 0.368086 | 0.219365 | 0.804 | 0.826 | 1 | 8 | Prdx1 | -0.022 | Old |
| 0.368231 | 0.388377 | 0.056 | 0.039 | 1 | 8 | Chst11 | 0.017 | Old |
| 0.36851 | 0.089028 | 0.112 | 0.086 | 1 | 8 | Cd200r4 | 0.026 | Old |
| 0.368657 | 0.103954 | 0.413 | 0.46 | 1 | 8 | Ube2q1 | -0.047 | Old |
| 0.369284 | 0.181967 | 0.894 | 0.906 | 1 | 8 | Naaa | -0.012 | Old |
| 0.369665 | 0.772963 | 0.134 | 0.112 | 1 | 8 | Rufy1 | 0.022 | Old |
| 0.369683 | 0.554425 | 0.034 | 0.021 | 1 | 8 | Rtn4ip1 | 0.013 | Old |
| 0.369686 | 0.327318 | 0.145 | 0.114 | 1 | 8 | Nup205 | 0.031 | Old |
| 0.370207 | 0.550934 | 0.106 | 0.083 | 1 | 8 | Tbrg4 | 0.023 | Old |
| 0.370784 | 0.8249 | 0.034 | 0.021 | 1 | 8 | Eml2 | 0.013 | Old |

| | | | | | | | | |
|---|---|---|---|---|---|---|---|---|
| 0.370784 | 0.521213 | 0.034 | 0.021 | 1 | 8 | A930033H1 | 0.013 | Old |
| 0.370865 | 0.407585 | 0.173 | 0.148 | 1 | 8 | Kansl2 | 0.025 | Old |
| 0.371479 | 0.341639 | 0.151 | 0.125 | 1 | 8 | Glipr1 | 0.026 | Old |
| 0.371805 | 0.069778 | 0.179 | 0.145 | 1 | 8 | Ate1 | 0.034 | Old |
| 0.372105 | 0.344078 | 0.553 | 0.53 | 1 | 8 | Ttc7 | 0.023 | Old |
| 0.37227 | 0.007574 | 0.117 | 0.091 | 1 | 8 | 2410002F2 | 0.026 | Old |
| 0.372301 | 1.134789 | 0.039 | 0.026 | 1 | 8 | Gm20604 | 0.013 | Old |
| 0.372328 | 0.359377 | 0.302 | 0.275 | 1 | 8 | Tomm34 | 0.027 | Old |
| 0.37249 | 0.101201 | 0.067 | 0.091 | 1 | 8 | Kdm1b | -0.024 | Old |
| 0.372505 | 0.249974 | 0.307 | 0.278 | 1 | 8 | Anks1 | 0.029 | Old |
| 0.37299 | 0.466859 | 0.034 | 0.021 | 1 | 8 | Pcbp4 | 0.013 | Old |
| 0.374059 | 0.230995 | 0.056 | 0.039 | 1 | 8 | Tbc1d16 | 0.017 | Old |
| 0.374096 | 0.367568 | 0.034 | 0.021 | 1 | 8 | 1700056E2 | 0.013 | Old |
| 0.374103 | 0.258826 | 0.575 | 0.569 | 1 | 8 | Atp6v0d1 | 0.006 | Old |
| 0.374141 | 0.599677 | 0.028 | 0.044 | 1 | 8 | Twsg1 | -0.016 | Old |
| 0.374237 | 0.090269 | 0.218 | 0.265 | 1 | 8 | Bet1 | -0.047 | Old |
| 0.374406 | 0.203466 | 0.531 | 0.506 | 1 | 8 | Cyp4f18 | 0.025 | Old |
| 0.374429 | 0.622645 | 0.073 | 0.055 | 1 | 8 | Plpp6 | 0.018 | Old |
| 0.374649 | 0.386447 | 0.084 | 0.112 | 1 | 8 | Stmn1 | -0.028 | Old |
| 0.374787 | 0.206576 | 0.33 | 0.294 | 1 | 8 | BC031181 | 0.036 | Old |
| 0.375153 | 1.211337 | 0.073 | 0.055 | 1 | 8 | Lin52 | 0.018 | Old |
| 0.375245 | 0.239679 | 0.48 | 0.457 | 1 | 8 | Psmd11 | 0.023 | Old |
| 0.375488 | 0.426227 | 0.117 | 0.094 | 1 | 8 | Dip2a | 0.023 | Old |
| 0.375744 | 0.116681 | 0.397 | 0.452 | 1 | 8 | Thrap3 | -0.055 | Old |
| 0.37576 | 0.568422 | 0.112 | 0.088 | 1 | 8 | Pde4dip | 0.024 | Old |
| 0.37576 | 0.153682 | 0.385 | 0.434 | 1 | 8 | Ms4a6b | -0.049 | Old |
| 0.375883 | 0.592806 | 0.067 | 0.049 | 1 | 8 | Pomgnt1 | 0.018 | Old |
| 0.376405 | 0.055676 | 0.279 | 0.327 | 1 | 8 | Gatad2a | -0.048 | Old |
| 0.376762 | 0.037209 | 0.117 | 0.148 | 1 | 8 | Prpf6 | -0.031 | Old |
| 0.377231 | 0.297869 | 0.704 | 0.727 | 1 | 8 | Snx2 | -0.023 | Old |
| 0.37763 | 0.412078 | 0.184 | 0.156 | 1 | 8 | Tufm | 0.028 | Old |
| 0.377765 | 0.711295 | 0.061 | 0.044 | 1 | 8 | Wfdc18 | 0.017 | Old |
| 0.378112 | 0.511743 | 0.156 | 0.132 | 1 | 8 | Pik3r4 | 0.024 | Old |
| 0.37823 | 0.189397 | 0.76 | 0.792 | 1 | 8 | Rnf130 | -0.032 | Old |
| 0.3785 | 0.415033 | 0.134 | 0.109 | 1 | 8 | Agap1 | 0.025 | Old |
| 0.378634 | 0.489376 | 0.101 | 0.078 | 1 | 8 | Rdm1 | 0.023 | Old |
| 0.378644 | 0.628275 | 0.145 | 0.122 | 1 | 8 | Nufip1 | 0.023 | Old |
| 0.378712 | 0.355545 | 0.101 | 0.132 | 1 | 8 | Atg4c | -0.031 | Old |
| 0.379108 | 0.032344 | 0.162 | 0.13 | 1 | 8 | Gtf2f2 | 0.032 | Old |
| 0.379252 | 0.450036 | 0.263 | 0.239 | 1 | 8 | Kcnab2 | 0.024 | Old |
| 0.379857 | 0.660041 | 0.14 | 0.117 | 1 | 8 | Cfl2 | 0.023 | Old |
| 0.379877 | 0.370677 | 0.101 | 0.078 | 1 | 8 | Pcnx3 | 0.023 | Old |
| 0.380586 | 0.393412 | 0.235 | 0.208 | 1 | 8 | Dag1 | 0.027 | Old |
| 0.38059 | 0.39119 | 0.246 | 0.218 | 1 | 8 | Hsph1 | 0.028 | Old |
| 0.380681 | 0.115336 | 0.279 | 0.335 | 1 | 8 | Pa2g4 | -0.056 | Old |

| | | | | | | | | |
|---|---|---|---|---|---|---|---|---|
| 0.380789 | 0.397947 | 0.43 | 0.413 | 1 | 8 | Uhmk1 | 0.017 | Old |
| 0.381199 | 0.106821 | 0.123 | 0.153 | 1 | 8 | Wrn | -0.03 | Old |
| 0.381283 | 0.59349 | 0.151 | 0.127 | 1 | 8 | Fubp3 | 0.024 | Old |
| 0.381792 | 0.44178 | 0.525 | 0.519 | 1 | 8 | Fxyd2 | 0.006 | Old |
| 0.381945 | 0.473735 | 0.095 | 0.073 | 1 | 8 | Uxt | 0.022 | Old |
| 0.382077 | 0.490448 | 0.168 | 0.143 | 1 | 8 | Scrib | 0.025 | Old |
| 0.382263 | 0.031457 | 0.184 | 0.221 | 1 | 8 | Ctdspl2 | -0.037 | Old |
| 0.382452 | 0.329236 | 0.33 | 0.304 | 1 | 8 | Fbxl5 | 0.026 | Old |
| 0.38257 | 0.493241 | 0.19 | 0.166 | 1 | 8 | Eif4enif1 | 0.024 | Old |
| 0.382705 | 0.372991 | 0.123 | 0.099 | 1 | 8 | Agmo | 0.024 | Old |
| 0.382881 | 0.073826 | 0.827 | 0.87 | 1 | 8 | Arpc3 | -0.043 | Old |
| 0.382893 | 0.1168 | 0.335 | 0.294 | 1 | 8 | Tmem9b | 0.041 | Old |
| 0.383107 | 0.192125 | 0.564 | 0.545 | 1 | 8 | Dnajb14 | 0.019 | Old |
| 0.383333 | 0.115189 | 0.056 | 0.039 | 1 | 8 | 2900093K2 | 0.017 | Old |
| 0.383469 | 0.128139 | 0.073 | 0.096 | 1 | 8 | Sass6 | -0.023 | Old |
| 0.383507 | 0.616876 | 0.067 | 0.049 | 1 | 8 | Slc16a1 | 0.018 | Old |
| 0.383601 | 0.273925 | 0.279 | 0.247 | 1 | 8 | Cmas | 0.032 | Old |
| 0.383907 | 0.060552 | 0.123 | 0.151 | 1 | 8 | Ginm1 | -0.028 | Old |
| 0.384119 | 0.14554 | 0.257 | 0.309 | 1 | 8 | Elf4 | -0.052 | Old |
| 0.384179 | 0.322828 | 0.061 | 0.044 | 1 | 8 | Ttc7b | 0.017 | Old |
| 0.384696 | 0.627563 | 0.078 | 0.06 | 1 | 8 | Phospho2 | 0.018 | Old |
| 0.384712 | 0.160016 | 0.112 | 0.143 | 1 | 8 | Nol12 | -0.031 | Old |
| 0.384841 | 0.227103 | 0.559 | 0.558 | 1 | 8 | Pgam1 | 0.001 | Old |
| 0.385024 | 0.44453 | 0.05 | 0.07 | 1 | 8 | Rft1 | -0.02 | Old |
| 0.385228 | 0.154256 | 0.866 | 0.899 | 1 | 8 | Srsf5 | -0.033 | Old |
| 0.385996 | 0.100624 | 0.022 | 0.036 | 1 | 8 | Tmem263 | -0.014 | Old |
| 0.386858 | 0.73651 | 0.212 | 0.19 | 1 | 8 | Prkag2 | 0.022 | Old |
| 0.387012 | 0.436261 | 0.207 | 0.179 | 1 | 8 | Slc12a9 | 0.028 | Old |
| 0.387103 | 0.299843 | 0.447 | 0.416 | 1 | 8 | Gspt1 | 0.031 | Old |
| 0.387115 | 0.068668 | 0.346 | 0.387 | 1 | 8 | Polr2f | -0.041 | Old |
| 0.387259 | 0.359223 | 0.33 | 0.304 | 1 | 8 | Rara | 0.026 | Old |
| 0.387522 | 0.316554 | 0.246 | 0.21 | 1 | 8 | Oxr1 | 0.036 | Old |
| 0.387556 | 0.774974 | 0.039 | 0.026 | 1 | 8 | Trmt61a | 0.013 | Old |
| 0.388144 | 0.105178 | 0.207 | 0.252 | 1 | 8 | Arl2bp | -0.045 | Old |
| 0.388313 | 0.399208 | 0.279 | 0.252 | 1 | 8 | Herc2 | 0.027 | Old |
| 0.388509 | 0.368123 | 0.302 | 0.265 | 1 | 8 | Plod3 | 0.037 | Old |
| 0.389133 | 0.349048 | 0.453 | 0.421 | 1 | 8 | Pkig | 0.032 | Old |
| 0.389581 | 0.220722 | 0.33 | 0.403 | 1 | 8 | Tsen34 | -0.073 | Old |
| 0.390012 | 0.06977 | 0.24 | 0.275 | 1 | 8 | Rbm28 | -0.035 | Old |
| 0.390058 | 0.704558 | 0.173 | 0.153 | 1 | 8 | Tada1 | 0.02 | Old |
| 0.39037 | 0.568026 | 0.184 | 0.164 | 1 | 8 | Mrps18a | 0.02 | Old |
| 0.390397 | 0.005926 | 0.184 | 0.221 | 1 | 8 | Ethe1 | -0.037 | Old |
| 0.390653 | 0.54196 | 0.039 | 0.026 | 1 | 8 | Dhrs11 | 0.013 | Old |
| 0.390683 | 0.249692 | 0.24 | 0.281 | 1 | 8 | AI504432 | -0.041 | Old |
| 0.390778 | 0.157583 | 0.447 | 0.403 | 1 | 8 | Srgap2 | 0.044 | Old |

| | | | | | | | | |
|---|---|---|---|---|---|---|---|---|
| 0.391349 | 0.078347 | 0.374 | 0.426 | 1 | 8 | Snrpc | -0.052 | Old |
| 0.391473 | 0.300646 | 0.061 | 0.044 | 1 | 8 | Fam210b | 0.017 | Old |
| 0.391666 | 0.78034 | 0.089 | 0.07 | 1 | 8 | Ercc6l2 | 0.019 | Old |
| 0.391688 | 0.551491 | 0.039 | 0.026 | 1 | 8 | Tada2a | 0.013 | Old |
| 0.391774 | 0.027967 | 0.48 | 0.551 | 1 | 8 | Dnm2 | -0.071 | Old |
| 0.392326 | 0.035769 | 0.358 | 0.314 | 1 | 8 | Otulin | 0.044 | Old |
| 0.392479 | 0.049315 | 0.408 | 0.449 | 1 | 8 | Ncoa3 | -0.041 | Old |
| 0.392659 | 0.323004 | 0.134 | 0.109 | 1 | 8 | Tmem2 | 0.025 | Old |
| 0.392732 | 0.032665 | 0.587 | 0.655 | 1 | 8 | Myl12a | -0.068 | Old |
| 0.392791 | 0.653904 | 0.101 | 0.081 | 1 | 8 | Wdfy1 | 0.02 | Old |
| 0.39299 | 0.537815 | 0.291 | 0.265 | 1 | 8 | Alkbh1 | 0.026 | Old |
| 0.393647 | 0.151339 | 0.134 | 0.169 | 1 | 8 | Nkap | -0.035 | Old |
| 0.393764 | 0.46613 | 0.039 | 0.026 | 1 | 8 | Adnp | 0.013 | Old |
| 0.394106 | 0.196599 | 0.291 | 0.255 | 1 | 8 | Ibtk | 0.036 | Old |
| 0.394392 | 0.128606 | 0.453 | 0.475 | 1 | 8 | Ccar1 | -0.022 | Old |
| 0.394481 | 0.073922 | 0.302 | 0.353 | 1 | 8 | Cdc40 | -0.051 | Old |
| 0.394527 | 0.645616 | 0.358 | 0.33 | 1 | 8 | Trim12c | 0.028 | Old |
| 0.394667 | 0.00304 | 0.514 | 0.603 | 1 | 8 | Fuca1 | -0.089 | Old |
| 0.394762 | 0.214452 | 0.078 | 0.101 | 1 | 8 | Mynn | -0.023 | Old |
| 0.395239 | 0.413268 | 0.117 | 0.094 | 1 | 8 | Prkaa1 | 0.023 | Old |
| 0.395358 | 0.30357 | 0.363 | 0.335 | 1 | 8 | Dusp6 | 0.028 | Old |
| 0.395804 | 1.511327 | 0.022 | 0.013 | 1 | 8 | Tspan7 | 0.009 | Old |
| 0.395813 | 0.043846 | 0.179 | 0.208 | 1 | 8 | Tmem131 | -0.029 | Old |
| 0.396292 | 0.44914 | 0.38 | 0.348 | 1 | 8 | Cln8 | 0.032 | Old |
| 0.396296 | 0.018493 | 0.564 | 0.623 | 1 | 8 | Ewsr1 | -0.059 | Old |
| 0.396806 | 0.089518 | 0.184 | 0.216 | 1 | 8 | Rnf169 | -0.032 | Old |
| 0.39689 | 0.772731 | 0.039 | 0.026 | 1 | 8 | Phpt1 | 0.013 | Old |
| 0.397134 | 0.086704 | 0.073 | 0.096 | 1 | 8 | Gramd3 | -0.023 | Old |
| 0.397135 | 0.29634 | 0.196 | 0.171 | 1 | 8 | Cdc42ep3 | 0.025 | Old |
| 0.397616 | 0.254957 | 0.408 | 0.392 | 1 | 8 | Tmem55b | 0.016 | Old |
| 0.397876 | 0.314246 | 0.408 | 0.382 | 1 | 8 | Ssrp1 | 0.026 | Old |
| 0.397913 | 0.357379 | 0.341 | 0.312 | 1 | 8 | Bcl7b | 0.029 | Old |
| 0.397935 | 0.509648 | 0.039 | 0.026 | 1 | 8 | Aste1 | 0.013 | Old |
| 0.398071 | 0.023665 | 0.687 | 0.699 | 1 | 8 | Top1 | -0.012 | Old |
| 0.398252 | 0.556953 | 0.067 | 0.049 | 1 | 8 | Mettl3 | 0.018 | Old |
| 0.398323 | 0.318029 | 0.117 | 0.096 | 1 | 8 | Gpn3 | 0.021 | Old |
| 0.398475 | 0.118585 | 0.324 | 0.387 | 1 | 8 | Sec63 | -0.063 | Old |
| 0.398656 | 1.750155 | 0.022 | 0.013 | 1 | 8 | Hoxb3 | 0.009 | Old |
| 0.398703 | 0.412814 | 0.559 | 0.538 | 1 | 8 | Kdelr1 | 0.021 | Old |
| 0.399284 | 0.429287 | 0.251 | 0.231 | 1 | 8 | Atp11b | 0.02 | Old |
| 0.399535 | 0.700614 | 0.073 | 0.055 | 1 | 8 | Rcl1 | 0.018 | Old |
| 0.399815 | 0.294932 | 0.503 | 0.494 | 1 | 8 | Myo9b | 0.009 | Old |
| 0.400086 | 1.567931 | 0.022 | 0.013 | 1 | 8 | Aatk | 0.009 | Old |
| 0.400087 | 0.325417 | 0.184 | 0.156 | 1 | 8 | Hcfc2 | 0.028 | Old |
| 0.401122 | 0.286553 | 0.43 | 0.426 | 1 | 8 | Snx18 | 0.004 | Old |

| | | | | | | | | |
|---|---|---|---|---|---|---|---|---|
| 0.401151 | 0.574339 | 0.084 | 0.065 | 1 | 8 | Fam58b | 0.019 | Old |
| 0.40152 | 1.112281 | 0.022 | 0.013 | 1 | 8 | Gdpgp1 | 0.009 | Old |
| 0.40152 | 1.053783 | 0.022 | 0.013 | 1 | 8 | Rab26os | 0.009 | Old |
| 0.40152 | 0.703143 | 0.022 | 0.013 | 1 | 8 | Ovca2 | 0.009 | Old |
| 0.401794 | 0.016582 | 0.458 | 0.501 | 1 | 8 | Brd1 | -0.043 | Old |
| 0.401842 | 0.193744 | 0.151 | 0.122 | 1 | 8 | Rab9 | 0.029 | Old |
| 0.402134 | 0.302777 | 0.039 | 0.026 | 1 | 8 | Tprn | 0.013 | Old |
| 0.402345 | 0.164129 | 0.274 | 0.236 | 1 | 8 | Nup62 | 0.038 | Old |
| 0.402557 | 0.415154 | 0.48 | 0.455 | 1 | 8 | Nfix | 0.025 | Old |
| 0.402625 | 0.422861 | 0.106 | 0.083 | 1 | 8 | Mettl14 | 0.023 | Old |
| 0.402725 | 0.054208 | 0.48 | 0.512 | 1 | 8 | Ptgs1 | -0.032 | Old |
| 0.402957 | 1.004293 | 0.022 | 0.013 | 1 | 8 | Gm17586 | 0.009 | Old |
| 0.402957 | 0.831361 | 0.022 | 0.013 | 1 | 8 | C1qtnf6 | 0.009 | Old |
| 0.402957 | 0.830808 | 0.022 | 0.013 | 1 | 8 | Cyp2d22 | 0.009 | Old |
| 0.403056 | 0.079331 | 0.151 | 0.184 | 1 | 8 | Vta1 | -0.033 | Old |
| 0.403494 | 0.080572 | 0.145 | 0.179 | 1 | 8 | Rbm4b | -0.034 | Old |
| 0.403504 | 0.350769 | 0.151 | 0.125 | 1 | 8 | Hmbs | 0.026 | Old |
| 0.403592 | 0.346144 | 0.358 | 0.335 | 1 | 8 | Cop1 | 0.023 | Old |
| 0.403847 | 0.47483 | 0.112 | 0.091 | 1 | 8 | Nr2c2ap | 0.021 | Old |
| 0.404098 | 0.110466 | 0.38 | 0.436 | 1 | 8 | Plekhj1 | -0.056 | Old |
| 0.404132 | 0.289479 | 0.413 | 0.39 | 1 | 8 | Themis2 | 0.023 | Old |
| 0.404181 | 0.160043 | 0.52 | 0.488 | 1 | 8 | Hsd17b11 | 0.032 | Old |
| 0.404227 | 0.361257 | 0.123 | 0.101 | 1 | 8 | Ccdc107 | 0.022 | Old |
| 0.404333 | 0.025681 | 0.553 | 0.639 | 1 | 8 | Ptpra | -0.086 | Old |
| 0.404397 | 0.766955 | 0.022 | 0.013 | 1 | 8 | Zfp605 | 0.009 | Old |
| 0.404454 | 0.579232 | 0.184 | 0.161 | 1 | 8 | Ldlrap1 | 0.023 | Old |
| 0.404564 | 0.344069 | 0.151 | 0.125 | 1 | 8 | Nol8 | 0.026 | Old |
| 0.404982 | 0.249374 | 0.598 | 0.6 | 1 | 8 | Dapk1 | -0.002 | Old |
| 0.405134 | 0.07223 | 0.81 | 0.821 | 1 | 8 | Luc7l2 | -0.011 | Old |
| 0.405307 | 0.091474 | 0.017 | 0.029 | 1 | 8 | Lrp4 | -0.012 | Old |
| 0.405363 | 0.487577 | 0.179 | 0.151 | 1 | 8 | Vac14 | 0.028 | Old |
| 0.40584 | 1.05153 | 0.022 | 0.013 | 1 | 8 | Nhej1 | 0.009 | Old |
| 0.40584 | 0.975085 | 0.022 | 0.013 | 1 | 8 | Gm38394 | 0.009 | Old |
| 0.40584 | 0.738548 | 0.022 | 0.013 | 1 | 8 | Tle6 | 0.009 | Old |
| 0.406358 | 0.055775 | 0.039 | 0.026 | 1 | 8 | Mpv17l | 0.013 | Old |
| 0.406604 | 0.193398 | 0.235 | 0.283 | 1 | 8 | Igsf9 | -0.048 | Old |
| 0.407186 | 0.30266 | 0.24 | 0.21 | 1 | 8 | Snx12 | 0.03 | Old |
| 0.407276 | 0.239403 | 0.626 | 0.644 | 1 | 8 | Dnajb6 | -0.018 | Old |
| 0.407286 | 0.38268 | 0.022 | 0.013 | 1 | 8 | Gm13479 | 0.009 | Old |
| 0.407499 | 0.204415 | 0.218 | 0.184 | 1 | 8 | Polr3c | 0.034 | Old |
| 0.407726 | 0.56906 | 0.212 | 0.187 | 1 | 8 | Nup54 | 0.025 | Old |
| 0.407776 | 0.166047 | 0.235 | 0.265 | 1 | 8 | Ldah | -0.03 | Old |
| 0.407835 | 0.685493 | 0.156 | 0.135 | 1 | 8 | Eipr1 | 0.021 | Old |
| 0.408036 | 1.095979 | 0.045 | 0.031 | 1 | 8 | Ybey | 0.014 | Old |
| 0.40818 | 0.34868 | 0.559 | 0.54 | 1 | 8 | Arcn1 | 0.019 | Old |

| | | | | | | | | |
|---|---|---|---|---|---|---|---|---|
| 0.408736 | 0.999744 | 0.022 | 0.013 | 1 | 8 | Gimap1 | 0.009 | Old |
| 0.408736 | 0.482259 | 0.022 | 0.013 | 1 | 8 | Rab11fip3 | 0.009 | Old |
| 0.408889 | 0.233893 | 0.134 | 0.166 | 1 | 8 | Ssbp2 | -0.032 | Old |
| 0.408927 | 0.045864 | 0.106 | 0.083 | 1 | 8 | Rbmx | 0.023 | Old |
| 0.408961 | 0.455115 | 0.223 | 0.197 | 1 | 8 | Babam1 | 0.026 | Old |
| 0.409204 | 0.605205 | 0.078 | 0.06 | 1 | 8 | Zfp182 | 0.018 | Old |
| 0.409714 | 0.023431 | 0.994 | 0.997 | 1 | 8 | Lyz2 | -0.003 | Old |
| 0.410188 | 1.130695 | 0.022 | 0.013 | 1 | 8 | Eri2 | 0.009 | Old |
| 0.410188 | 0.83951 | 0.022 | 0.013 | 1 | 8 | Gm48653 | 0.009 | Old |
| 0.410188 | 0.457371 | 0.022 | 0.013 | 1 | 8 | Tceanc | 0.009 | Old |
| 0.410188 | 0.21492 | 0.022 | 0.013 | 1 | 8 | Bbs9 | 0.009 | Old |
| 0.410283 | 0.263089 | 0.223 | 0.195 | 1 | 8 | Syvn1 | 0.028 | Old |
| 0.410722 | 0.108906 | 0.235 | 0.27 | 1 | 8 | Txlna | -0.035 | Old |
| 0.410808 | 0.509263 | 0.229 | 0.21 | 1 | 8 | Tmem128 | 0.019 | Old |
| 0.411102 | 0.033188 | 0.034 | 0.049 | 1 | 8 | Copz2 | -0.015 | Old |
| 0.411114 | 0.285215 | 0.503 | 0.473 | 1 | 8 | Prkcsh | 0.03 | Old |
| 0.411571 | 0.210585 | 0.235 | 0.203 | 1 | 8 | Slc41a3 | 0.032 | Old |
| 0.411642 | 0.445345 | 0.223 | 0.2 | 1 | 8 | Plxnc1 | 0.023 | Old |
| 0.411643 | 0.987024 | 0.022 | 0.013 | 1 | 8 | Lig4 | 0.009 | Old |
| 0.411643 | 0.385853 | 0.022 | 0.013 | 1 | 8 | Gm15050 | 0.009 | Old |
| 0.411643 | 0.377246 | 0.022 | 0.013 | 1 | 8 | Sgtb | 0.009 | Old |
| 0.411729 | 0.45063 | 0.084 | 0.065 | 1 | 8 | Tmem175 | 0.019 | Old |
| 0.412237 | 0.260568 | 0.598 | 0.605 | 1 | 8 | Cnot6l | -0.007 | Old |
| 0.412459 | 0.294385 | 0.073 | 0.055 | 1 | 8 | Pycrl | 0.018 | Old |
| 0.41267 | 0.477575 | 0.207 | 0.182 | 1 | 8 | Zswim8 | 0.025 | Old |
| 0.41267 | 0.052277 | 0.168 | 0.2 | 1 | 8 | Med30 | -0.032 | Old |
| 0.412673 | 0.549404 | 0.089 | 0.07 | 1 | 8 | Smyd2 | 0.019 | Old |
| 0.41285 | 0.387512 | 0.05 | 0.07 | 1 | 8 | Rab3ip | -0.02 | Old |
| 0.413101 | 0.740855 | 0.022 | 0.013 | 1 | 8 | Lbx2 | 0.009 | Old |
| 0.413554 | 0.221188 | 0.408 | 0.374 | 1 | 8 | Zzef1 | 0.034 | Old |
| 0.41433 | 0.256516 | 0.581 | 0.595 | 1 | 8 | Ehbp1l1 | -0.014 | Old |
| 0.414363 | 0.491965 | 0.078 | 0.06 | 1 | 8 | Haus6 | 0.018 | Old |
| 0.414363 | 0.305994 | 0.078 | 0.06 | 1 | 8 | Lrrc57 | 0.018 | Old |
| 0.414563 | 0.651545 | 0.022 | 0.013 | 1 | 8 | Tpmt | 0.009 | Old |
| 0.414563 | 0.427646 | 0.022 | 0.013 | 1 | 8 | Gm11131 | 0.009 | Old |
| 0.414834 | 0.0719 | 0.525 | 0.577 | 1 | 8 | Sgpp1 | -0.052 | Old |
| 0.414943 | 0.937842 | 0.045 | 0.031 | 1 | 8 | Adcy3 | 0.014 | Old |
| 0.415194 | 0.390078 | 0.218 | 0.192 | 1 | 8 | Stat5a | 0.026 | Old |
| 0.415537 | 0.436324 | 0.14 | 0.117 | 1 | 8 | Pias2 | 0.023 | Old |
| 0.415817 | 0.39856 | 0.43 | 0.423 | 1 | 8 | R3hdm4 | 0.007 | Old |
| 0.416027 | 0.54262 | 0.022 | 0.013 | 1 | 8 | Zfp775 | 0.009 | Old |
| 0.41629 | 0.242571 | 0.743 | 0.805 | 1 | 8 | Pepd | -0.062 | Old |
| 0.416414 | 0.611062 | 0.397 | 0.395 | 1 | 8 | Immt | 0.002 | Old |
| 0.416695 | 0.178923 | 0.687 | 0.678 | 1 | 8 | Rock1 | 0.009 | Old |
| 0.416833 | 0.381985 | 0.346 | 0.335 | 1 | 8 | Dynlt3 | 0.011 | Old |

| | | | | | | | | |
|---|---|---|---|---|---|---|---|---|
| 0.416929 | 0.413198 | 0.045 | 0.031 | 1 | 8 | Rangrf | 0.014 | Old |
| 0.416929 | 0.271616 | 0.045 | 0.031 | 1 | 8 | Zkscan6 | 0.014 | Old |
| 0.416983 | 1.031916 | 0.05 | 0.036 | 1 | 8 | Mtrr | 0.014 | Old |
| 0.417112 | 0.396079 | 0.123 | 0.101 | 1 | 8 | Zcchc8 | 0.022 | Old |
| 0.417145 | 0.548627 | 0.101 | 0.081 | 1 | 8 | Clpp | 0.02 | Old |
| 0.417328 | 0.263227 | 0.531 | 0.553 | 1 | 8 | Atp6v1a | -0.022 | Old |
| 0.417444 | 0.239278 | 0.587 | 0.584 | 1 | 8 | Cndp2 | 0.003 | Old |
| 0.417495 | 0.06222 | 0.022 | 0.013 | 1 | 8 | Mcf2l | 0.009 | Old |
| 0.417548 | 0.101992 | 0.352 | 0.4 | 1 | 8 | Cbx1 | -0.048 | Old |
| 0.418201 | 0.541811 | 0.089 | 0.07 | 1 | 8 | Bloc1s5 | 0.019 | Old |
| 0.418457 | 0.355893 | 0.475 | 0.444 | 1 | 8 | Vamp3 | 0.031 | Old |
| 0.418626 | 0.268756 | 0.073 | 0.055 | 1 | 8 | Gm49368 | 0.018 | Old |
| 0.418808 | 0.074133 | 0.771 | 0.805 | 1 | 8 | Vmp1 | -0.034 | Old |
| 0.419419 | 0.395288 | 0.458 | 0.455 | 1 | 8 | Pum1 | 0.003 | Old |
| 0.41983 | 0.12364 | 0.33 | 0.377 | 1 | 8 | Agps | -0.047 | Old |
| 0.419929 | 0.221418 | 0.726 | 0.784 | 1 | 8 | Syk | -0.058 | Old |
| 0.420074 | 0.359759 | 0.324 | 0.299 | 1 | 8 | Rps12-ps3 | 0.025 | Old |
| 0.420344 | 0.111752 | 0.318 | 0.364 | 1 | 8 | Snf8 | -0.046 | Old |
| 0.420713 | 0.516706 | 0.05 | 0.036 | 1 | 8 | Cmpk2 | 0.014 | Old |
| 0.420917 | 0.185382 | 0.034 | 0.049 | 1 | 8 | Hemk1 | -0.015 | Old |
| 0.421439 | 0.305011 | 0.531 | 0.54 | 1 | 8 | Mcrip1 | -0.009 | Old |
| 0.421444 | 0.233154 | 0.257 | 0.221 | 1 | 8 | Hmg20b | 0.036 | Old |
| 0.42148 | 0.830511 | 0.106 | 0.088 | 1 | 8 | Adamtsl5 | 0.018 | Old |
| 0.421683 | 0.470021 | 0.268 | 0.242 | 1 | 8 | Dars | 0.026 | Old |
| 0.421748 | 0.452777 | 0.268 | 0.247 | 1 | 8 | Tars | 0.021 | Old |
| 0.421755 | 0.373797 | 0.223 | 0.2 | 1 | 8 | Hdac2 | 0.023 | Old |
| 0.421943 | 0.693952 | 0.123 | 0.104 | 1 | 8 | Zfp954 | 0.019 | Old |
| 0.421977 | 0.626504 | 0.235 | 0.21 | 1 | 8 | Fam204a | 0.025 | Old |
| 0.422183 | 0.459169 | 0.011 | 0.021 | 1 | 8 | Elp6 | -0.01 | Old |
| 0.422372 | 0.016761 | 0.872 | 0.961 | 1 | 8 | Ucp2 | -0.089 | Old |
| 0.422498 | 0.115068 | 0.101 | 0.125 | 1 | 8 | Zfp282 | -0.024 | Old |
| 0.422531 | 0.170004 | 0.553 | 0.574 | 1 | 8 | Epsti1 | -0.021 | Old |
| 0.422965 | 0.51709 | 0.184 | 0.164 | 1 | 8 | Pspc1 | 0.02 | Old |
| 0.423148 | 0.160203 | 0.156 | 0.187 | 1 | 8 | Soga1 | -0.031 | Old |
| 0.423179 | 0.39066 | 0.223 | 0.195 | 1 | 8 | Cep120 | 0.028 | Old |
| 0.423588 | 0.246724 | 0.011 | 0.021 | 1 | 8 | Ccdc138 | -0.01 | Old |
| 0.423703 | 0.301897 | 0.324 | 0.296 | 1 | 8 | Ulk2 | 0.028 | Old |
| 0.424251 | 0.597416 | 0.145 | 0.125 | 1 | 8 | Htatip2 | 0.02 | Old |
| 0.42446 | 0.305593 | 0.201 | 0.171 | 1 | 8 | Tomm40 | 0.03 | Old |
| 0.424507 | 0.303675 | 0.631 | 0.629 | 1 | 8 | Degs1 | 0.002 | Old |
| 0.424548 | 0.050169 | 0.285 | 0.335 | 1 | 8 | Gabarapl2 | -0.05 | Old |
| 0.424996 | 0.751779 | 0.011 | 0.021 | 1 | 8 | Rab34 | -0.01 | Old |
| 0.425015 | 0.14355 | 0.173 | 0.208 | 1 | 8 | Klf11 | -0.035 | Old |
| 0.425522 | 0.18761 | 0.14 | 0.114 | 1 | 8 | Fiz1 | 0.026 | Old |
| 0.426014 | 0.245635 | 0.542 | 0.535 | 1 | 8 | Lat2 | 0.007 | Old |

| | | | | | | | | |
|---|---|---|---|---|---|---|---|---|
| 0.426087 | 0.096746 | 0.084 | 0.065 | 1 | 8 | Cog2 | 0.019 | Old |
| 0.426093 | 0.13657 | 0.285 | 0.255 | 1 | 8 | Rap2b | 0.03 | Old |
| 0.426267 | 0.261198 | 0.453 | 0.434 | 1 | 8 | Tank | 0.019 | Old |
| 0.426652 | 0.312764 | 0.218 | 0.192 | 1 | 8 | Dtymk | 0.026 | Old |
| 0.426679 | 0.4733 | 0.128 | 0.106 | 1 | 8 | Ctnnbl1 | 0.022 | Old |
| 0.426866 | 0.119966 | 0.318 | 0.369 | 1 | 8 | Mcts1 | -0.051 | Old |
| 0.428049 | 2.450786 | 0.011 | 0.005 | 1 | 8 | Slpi | 0.006 | Old |
| 0.428049 | 1.965127 | 0.011 | 0.005 | 1 | 8 | Mcm10 | 0.006 | Old |
| 0.428049 | 1.619809 | 0.011 | 0.005 | 1 | 8 | Gm6710 | 0.006 | Old |
| 0.428049 | 1.535093 | 0.011 | 0.005 | 1 | 8 | Gm43796 | 0.006 | Old |
| 0.428049 | 1.494416 | 0.011 | 0.005 | 1 | 8 | Rnf144a | 0.006 | Old |
| 0.428049 | 1.479102 | 0.011 | 0.005 | 1 | 8 | Nav2 | 0.006 | Old |
| 0.428049 | 1.430927 | 0.011 | 0.005 | 1 | 8 | Cdon | 0.006 | Old |
| 0.428049 | 1.277803 | 0.011 | 0.005 | 1 | 8 | Rsph9 | 0.006 | Old |
| 0.428049 | 1.232627 | 0.011 | 0.005 | 1 | 8 | Zfp459 | 0.006 | Old |
| 0.428049 | 1.196421 | 0.011 | 0.005 | 1 | 8 | Gm12184 | 0.006 | Old |
| 0.428049 | 0.963822 | 0.011 | 0.005 | 1 | 8 | Gins3 | 0.006 | Old |
| 0.428049 | 0.919041 | 0.011 | 0.005 | 1 | 8 | Gm45733 | 0.006 | Old |
| 0.428049 | 0.806754 | 0.011 | 0.005 | 1 | 8 | Gm6169 | 0.006 | Old |
| 0.428049 | 0.758279 | 0.011 | 0.005 | 1 | 8 | Itga2 | 0.006 | Old |
| 0.428049 | 0.66202 | 0.011 | 0.005 | 1 | 8 | Aqp7 | 0.006 | Old |
| 0.428049 | 0.644477 | 0.011 | 0.005 | 1 | 8 | Pcx | 0.006 | Old |
| 0.428049 | 0.634186 | 0.011 | 0.005 | 1 | 8 | Zfp296 | 0.006 | Old |
| 0.428612 | 0.12021 | 0.246 | 0.21 | 1 | 8 | Actl6a | 0.036 | Old |
| 0.429326 | 0.007295 | 0.575 | 0.657 | 1 | 8 | Tmem59 | -0.082 | Old |
| 0.429463 | 0.391537 | 0.207 | 0.179 | 1 | 8 | Fam126b | 0.028 | Old |
| 0.429758 | 0.001597 | 0.318 | 0.353 | 1 | 8 | Pi4k2a | -0.035 | Old |
| 0.429918 | 0.022996 | 0.57 | 0.629 | 1 | 8 | Mindy2 | -0.059 | Old |
| 0.429974 | 0.007435 | 0.045 | 0.031 | 1 | 8 | Rnf24 | 0.014 | Old |
| 0.430121 | 0.481572 | 0.05 | 0.036 | 1 | 8 | Tmem229a | 0.014 | Old |
| 0.430279 | 1.50837 | 0.011 | 0.005 | 1 | 8 | Cdc45 | 0.006 | Old |
| 0.430279 | 1.381493 | 0.011 | 0.005 | 1 | 8 | Saa3 | 0.006 | Old |
| 0.430279 | 1.118511 | 0.011 | 0.005 | 1 | 8 | E430018J2 | 0.006 | Old |
| 0.430279 | 0.898278 | 0.011 | 0.005 | 1 | 8 | Tgfa | 0.006 | Old |
| 0.430279 | 0.854313 | 0.011 | 0.005 | 1 | 8 | Tmem158 | 0.006 | Old |
| 0.430279 | 0.724345 | 0.011 | 0.005 | 1 | 8 | Gm6277 | 0.006 | Old |
| 0.430279 | 0.680422 | 0.011 | 0.005 | 1 | 8 | D930020B1 | 0.006 | Old |
| 0.430279 | 0.676663 | 0.011 | 0.005 | 1 | 8 | Appbp2os | 0.006 | Old |
| 0.430279 | 0.670226 | 0.011 | 0.005 | 1 | 8 | Pkia | 0.006 | Old |
| 0.430279 | 0.655935 | 0.011 | 0.005 | 1 | 8 | Srpx | 0.006 | Old |
| 0.430279 | 0.65329 | 0.011 | 0.005 | 1 | 8 | Nup210l | 0.006 | Old |
| 0.430279 | 0.621644 | 0.011 | 0.005 | 1 | 8 | Zfp248 | 0.006 | Old |
| 0.430279 | 0.583154 | 0.011 | 0.005 | 1 | 8 | Zfp286 | 0.006 | Old |
| 0.430279 | 0.516712 | 0.011 | 0.005 | 1 | 8 | Prtn3 | 0.006 | Old |
| 0.430279 | 0.49996 | 0.011 | 0.005 | 1 | 8 | 1010001B2 | 0.006 | Old |

| | | | | | | | | |
|---|---|---|---|---|---|---|---|---|
| 0.430279 | 0.442218 | 0.011 | 0.005 | 1 | 8 | 4930518I1! | 0.006 | Old |
| 0.430417 | 0.095955 | 0.816 | 0.906 | 1 | 8 | Cd47 | -0.09 | Old |
| 0.430656 | 0.196516 | 0.011 | 0.021 | 1 | 8 | Cd69 | -0.01 | Old |
| 0.430656 | 0.079395 | 0.011 | 0.021 | 1 | 8 | Ifit3 | -0.01 | Old |
| 0.430916 | 0.161792 | 0.039 | 0.055 | 1 | 8 | Scrn3 | -0.016 | Old |
| 0.430919 | 0.612122 | 0.056 | 0.042 | 1 | 8 | Gatc | 0.014 | Old |
| 0.430996 | 0.293724 | 0.24 | 0.21 | 1 | 8 | Nae1 | 0.03 | Old |
| 0.43112 | 0.696979 | 0.006 | 0.013 | 1 | 8 | Mettl4 | -0.007 | Old |
| 0.43112 | 0.69075 | 0.006 | 0.013 | 1 | 8 | C230037L1 | -0.007 | Old |
| 0.43112 | 0.214239 | 0.006 | 0.013 | 1 | 8 | Sdk1 | -0.007 | Old |
| 0.431121 | 0.130138 | 0.073 | 0.055 | 1 | 8 | Diexf | 0.018 | Old |
| 0.431835 | 0.313612 | 0.184 | 0.156 | 1 | 8 | Dcun1d1 | 0.028 | Old |
| 0.432002 | 0.194101 | 0.045 | 0.031 | 1 | 8 | Zfp788 | 0.014 | Old |
| 0.432078 | 0.5232 | 0.011 | 0.021 | 1 | 8 | Tmem144 | -0.01 | Old |
| 0.432178 | 0.329107 | 0.274 | 0.229 | 1 | 8 | Ggt5 | 0.045 | Old |
| 0.432192 | 0.178656 | 0.782 | 0.823 | 1 | 8 | Bri3 | -0.041 | Old |
| 0.432217 | 0.316508 | 0.335 | 0.314 | 1 | 8 | Emc6 | 0.021 | Old |
| 0.432361 | 0.332859 | 0.112 | 0.091 | 1 | 8 | Ankrd54 | 0.021 | Old |
| 0.432515 | 1.727033 | 0.011 | 0.005 | 1 | 8 | Gm26947 | 0.006 | Old |
| 0.432515 | 1.492619 | 0.011 | 0.005 | 1 | 8 | Zfp94 | 0.006 | Old |
| 0.432515 | 0.992958 | 0.011 | 0.005 | 1 | 8 | Exd1 | 0.006 | Old |
| 0.432515 | 0.696239 | 0.011 | 0.005 | 1 | 8 | Gm9725 | 0.006 | Old |
| 0.432515 | 0.677405 | 0.011 | 0.005 | 1 | 8 | Mlxipl | 0.006 | Old |
| 0.432515 | 0.653566 | 0.011 | 0.005 | 1 | 8 | Entpd2 | 0.006 | Old |
| 0.432515 | 0.522477 | 0.011 | 0.005 | 1 | 8 | Pcdhgc4 | 0.006 | Old |
| 0.432515 | 0.511215 | 0.011 | 0.005 | 1 | 8 | Gm4924 | 0.006 | Old |
| 0.432515 | 0.50267 | 0.011 | 0.005 | 1 | 8 | Camk2n2 | 0.006 | Old |
| 0.432515 | 0.450057 | 0.011 | 0.005 | 1 | 8 | Pdxp | 0.006 | Old |
| 0.432515 | 0.406039 | 0.011 | 0.005 | 1 | 8 | Gm14698 | 0.006 | Old |
| 0.432709 | 0.650968 | 0.056 | 0.042 | 1 | 8 | Pou6f1 | 0.014 | Old |
| 0.433056 | 0.171329 | 0.603 | 0.597 | 1 | 8 | Cd300ld | 0.006 | Old |
| 0.43322 | 0.397854 | 0.397 | 0.371 | 1 | 8 | Krit1 | 0.026 | Old |
| 0.433232 | 0.606553 | 0.095 | 0.075 | 1 | 8 | Scly | 0.02 | Old |
| 0.433493 | 0.161141 | 0.419 | 0.377 | 1 | 8 | Vav3 | 0.042 | Old |
| 0.433503 | 0.812929 | 0.011 | 0.021 | 1 | 8 | Znrf3 | -0.01 | Old |
| 0.433504 | 0.598049 | 0.268 | 0.257 | 1 | 8 | St6galnac4 | 0.011 | Old |
| 0.433533 | 0.024083 | 0.358 | 0.387 | 1 | 8 | Usp16 | -0.029 | Old |
| 0.434275 | 0.340497 | 0.257 | 0.229 | 1 | 8 | Gpr107 | 0.028 | Old |
| 0.434511 | 0.091727 | 0.067 | 0.049 | 1 | 8 | B230354K1 | 0.018 | Old |
| 0.434516 | 0.531679 | 0.447 | 0.442 | 1 | 8 | Stub1 | 0.005 | Old |
| 0.434554 | 0.100243 | 0.436 | 0.486 | 1 | 8 | Adipor1 | -0.05 | Old |
| 0.434758 | 0.518722 | 0.011 | 0.005 | 1 | 8 | Gm15326 | 0.006 | Old |
| 0.434758 | 0.453353 | 0.011 | 0.005 | 1 | 8 | Cfap126 | 0.006 | Old |
| 0.434758 | 0.441181 | 0.011 | 0.005 | 1 | 8 | 2610318N( | 0.006 | Old |
| 0.434758 | 0.415834 | 0.011 | 0.005 | 1 | 8 | Recql4 | 0.006 | Old |

| | | | | | | | | |
|---|---|---|---|---|---|---|---|---|
| 0.434758 | 0.405836 | 0.011 | 0.005 | 1 | 8 | BC055324 | 0.006 | Old |
| 0.434758 | 0.387928 | 0.011 | 0.005 | 1 | 8 | Gm12166 | 0.006 | Old |
| 0.434758 | 0.383232 | 0.011 | 0.005 | 1 | 8 | Hap1 | 0.006 | Old |
| 0.434758 | 0.351548 | 0.011 | 0.005 | 1 | 8 | 1810021B2 | 0.006 | Old |
| 0.434758 | 0.299322 | 0.011 | 0.005 | 1 | 8 | Dync2li1 | 0.006 | Old |
| 0.434758 | 0.292678 | 0.011 | 0.005 | 1 | 8 | Gm10269 | 0.006 | Old |
| 0.434758 | 0.076066 | 0.011 | 0.005 | 1 | 8 | Zfp69 | 0.006 | Old |
| 0.435151 | 0.143513 | 0.732 | 0.756 | 1 | 8 | Hnrnpab | -0.024 | Old |
| 0.435622 | 0.365622 | 0.48 | 0.449 | 1 | 8 | Sdf2 | 0.031 | Old |
| 0.435708 | 0.038577 | 0.453 | 0.483 | 1 | 8 | Erbin | -0.03 | Old |
| 0.436316 | 0.143443 | 0.073 | 0.094 | 1 | 8 | Slc25a16 | -0.021 | Old |
| 0.43636 | 0.001511 | 0.687 | 0.748 | 1 | 8 | Nfam1 | -0.061 | Old |
| 0.436698 | 0.012099 | 0.14 | 0.166 | 1 | 8 | Dopey2 | -0.026 | Old |
| 0.436925 | 0.425534 | 0.196 | 0.171 | 1 | 8 | Stk19 | 0.025 | Old |
| 0.437008 | 0.316154 | 0.011 | 0.005 | 1 | 8 | Cep170b | 0.006 | Old |
| 0.437008 | 0.253827 | 0.011 | 0.005 | 1 | 8 | Gm15441 | 0.006 | Old |
| 0.437008 | 0.252831 | 0.011 | 0.005 | 1 | 8 | Gm26935 | 0.006 | Old |
| 0.437008 | 0.186204 | 0.011 | 0.005 | 1 | 8 | Kif15 | 0.006 | Old |
| 0.437008 | 0.169215 | 0.011 | 0.005 | 1 | 8 | Eme2 | 0.006 | Old |
| 0.437008 | 0.167412 | 0.011 | 0.005 | 1 | 8 | Cfap99 | 0.006 | Old |
| 0.437008 | 0.164944 | 0.011 | 0.005 | 1 | 8 | Gm34590 | 0.006 | Old |
| 0.437008 | 0.132599 | 0.011 | 0.005 | 1 | 8 | Olfr1033 | 0.006 | Old |
| 0.437008 | 0.10319 | 0.011 | 0.005 | 1 | 8 | Serping1 | 0.006 | Old |
| 0.437008 | 0.077984 | 0.011 | 0.005 | 1 | 8 | AC163040. | 0.006 | Old |
| 0.437008 | 0.074968 | 0.011 | 0.005 | 1 | 8 | Gm29107 | 0.006 | Old |
| 0.437447 | 0.356592 | 0.162 | 0.138 | 1 | 8 | Sf3a2 | 0.024 | Old |
| 0.437495 | 0.429021 | 0.123 | 0.101 | 1 | 8 | Wdr70 | 0.022 | Old |
| 0.437579 | 0.013355 | 0.235 | 0.275 | 1 | 8 | Rspry1 | -0.04 | Old |
| 0.437791 | 0.171737 | 0.777 | 0.818 | 1 | 8 | Tspo | -0.041 | Old |
| 0.438273 | 0.211149 | 0.117 | 0.096 | 1 | 8 | Rrp15 | 0.021 | Old |
| 0.438423 | 0.06483 | 0.078 | 0.099 | 1 | 8 | Usp21 | -0.021 | Old |
| 0.438424 | 0.204024 | 0.715 | 0.706 | 1 | 8 | Mapre1 | 0.009 | Old |
| 0.43871 | 0.268263 | 0.112 | 0.091 | 1 | 8 | Cnot9 | 0.021 | Old |
| 0.439256 | 0.565041 | 0.084 | 0.065 | 1 | 8 | Panx1 | 0.019 | Old |
| 0.439265 | 0.189622 | 0.676 | 0.717 | 1 | 8 | Slk | -0.041 | Old |
| 0.439624 | 0.278675 | 0.162 | 0.135 | 1 | 8 | Nagpa | 0.027 | Old |
| 0.440083 | 0.917076 | 0.084 | 0.068 | 1 | 8 | Etfbkmt | 0.016 | Old |
| 0.440346 | 0.124017 | 0.151 | 0.179 | 1 | 8 | Fam118a | -0.028 | Old |
| 0.440376 | 0.222165 | 0.14 | 0.169 | 1 | 8 | Trove2 | -0.029 | Old |
| 0.44061 | 0.372296 | 0.05 | 0.036 | 1 | 8 | Casp7 | 0.014 | Old |
| 0.44061 | 0.325321 | 0.05 | 0.036 | 1 | 8 | Tmem186 | 0.014 | Old |
| 0.440817 | 0.52041 | 0.056 | 0.042 | 1 | 8 | Rbm41 | 0.014 | Old |
| 0.440908 | 1.306611 | 0.028 | 0.018 | 1 | 8 | Kyat1 | 0.01 | Old |
| 0.44112 | 0.096665 | 0.927 | 0.969 | 1 | 8 | Zfp36l2 | -0.042 | Old |
| 0.441264 | 0.308749 | 0.028 | 0.042 | 1 | 8 | Ccdc73 | -0.014 | Old |

| | | | | | | | | |
|---|---|---|---|---|---|---|---|---|
| 0.441669 | 0.095211 | 0.112 | 0.088 | 1 | 8 | Narfl | 0.024 | Old |
| 0.441675 | 0.16044 | 0.251 | 0.288 | 1 | 8 | Prpf39 | -0.037 | Old |
| 0.441786 | 0.442417 | 0.128 | 0.106 | 1 | 8 | Mir17hg | 0.022 | Old |
| 0.442092 | 0.296153 | 0.575 | 0.571 | 1 | 8 | Eif1a | 0.004 | Old |
| 0.442392 | 0.245692 | 0.402 | 0.358 | 1 | 8 | Pfdn2 | 0.044 | Old |
| 0.442451 | 0.074432 | 0.944 | 0.979 | 1 | 8 | Unc93b1 | -0.035 | Old |
| 0.442533 | 0.289504 | 0.05 | 0.036 | 1 | 8 | Mars | 0.014 | Old |
| 0.442557 | 0.047968 | 0.324 | 0.387 | 1 | 8 | Fnip1 | -0.063 | Old |
| 0.443096 | 0.092263 | 0.039 | 0.055 | 1 | 8 | Dsel | -0.016 | Old |
| 0.443685 | 0.54113 | 0.235 | 0.216 | 1 | 8 | Tmem183a | 0.019 | Old |
| 0.443743 | 0.142393 | 0.682 | 0.681 | 1 | 8 | Batf3 | 0.001 | Old |
| 0.444587 | 0.984174 | 0.061 | 0.047 | 1 | 8 | Kbtbd3 | 0.014 | Old |
| 0.444587 | 0.588436 | 0.061 | 0.047 | 1 | 8 | Tor4a | 0.014 | Old |
| 0.444725 | 0.05692 | 0.369 | 0.418 | 1 | 8 | Qpct | -0.049 | Old |
| 0.444868 | 1.027797 | 0.028 | 0.018 | 1 | 8 | Pigb | 0.01 | Old |
| 0.445241 | 0.515881 | 0.073 | 0.057 | 1 | 8 | Kiz | 0.016 | Old |
| 0.445776 | 0.513622 | 0.162 | 0.138 | 1 | 8 | 2310001H1 | 0.024 | Old |
| 0.446033 | 0.831936 | 0.073 | 0.057 | 1 | 8 | Fer | 0.016 | Old |
| 0.446193 | 0.993813 | 0.028 | 0.018 | 1 | 8 | 4930481A1 | 0.01 | Old |
| 0.446271 | 0.53198 | 0.056 | 0.042 | 1 | 8 | Ttf2 | 0.014 | Old |
| 0.446317 | 0.669801 | 0.061 | 0.047 | 1 | 8 | Fam114a1 | 0.014 | Old |
| 0.446333 | 0.150289 | 0.704 | 0.709 | 1 | 8 | Pkm | -0.005 | Old |
| 0.446948 | 0.360624 | 0.291 | 0.268 | 1 | 8 | Slc8b1 | 0.023 | Old |
| 0.44752 | 0.515548 | 0.028 | 0.018 | 1 | 8 | Oasl1 | 0.01 | Old |
| 0.447688 | 0.425237 | 0.263 | 0.239 | 1 | 8 | Eif4a3 | 0.024 | Old |
| 0.44797 | 0.483218 | 0.302 | 0.265 | 1 | 8 | Dennd4c | 0.037 | Old |
| 0.448079 | 0.030443 | 0.061 | 0.081 | 1 | 8 | Pdp1 | -0.02 | Old |
| 0.448859 | 0.052717 | 0.235 | 0.273 | 1 | 8 | Tm2d1 | -0.038 | Old |
| 0.44897 | 0.503494 | 0.229 | 0.205 | 1 | 8 | Sdc1 | 0.024 | Old |
| 0.449085 | 0.269952 | 0.173 | 0.145 | 1 | 8 | Uba3 | 0.028 | Old |
| 0.449337 | 0.740851 | 0.028 | 0.042 | 1 | 8 | Tdp1 | -0.014 | Old |
| 0.449435 | 0.340592 | 0.196 | 0.169 | 1 | 8 | Tsfm | 0.027 | Old |
| 0.449442 | 0.057193 | 0.101 | 0.125 | 1 | 8 | Ell | -0.024 | Old |
| 0.449451 | 0.82826 | 0.134 | 0.114 | 1 | 8 | Trim11 | 0.02 | Old |
| 0.449483 | 0.444473 | 0.246 | 0.226 | 1 | 8 | Amdhd2 | 0.02 | Old |
| 0.449708 | 0.018192 | 0.263 | 0.299 | 1 | 8 | Ahcyl1 | -0.036 | Old |
| 0.449955 | 0.236069 | 0.609 | 0.616 | 1 | 8 | Taok1 | -0.007 | Old |
| 0.450453 | 0.349383 | 0.223 | 0.203 | 1 | 8 | Ubqln2 | 0.02 | Old |
| 0.450793 | 0.462056 | 0.14 | 0.117 | 1 | 8 | Fem1a | 0.023 | Old |
| 0.450845 | 0.56126 | 0.056 | 0.042 | 1 | 8 | Nsl1 | 0.014 | Old |
| 0.451514 | 0.649951 | 0.028 | 0.018 | 1 | 8 | Ccdc106 | 0.01 | Old |
| 0.451849 | 0.134979 | 0.352 | 0.403 | 1 | 8 | Mob4 | -0.051 | Old |
| 0.452016 | 0.234891 | 0.358 | 0.319 | 1 | 8 | Sesn1 | 0.039 | Old |
| 0.452096 | 0.066025 | 0.503 | 0.561 | 1 | 8 | Pfkp | -0.058 | Old |
| 0.45285 | 0.659395 | 0.028 | 0.018 | 1 | 8 | 9830107B1 | 0.01 | Old |

| | | | | | | | | |
|---|---|---|---|---|---|---|---|---|
| 0.45285 | 0.592985 | 0.028 | 0.018 | 1 | 8 | 5830416I1! | 0.01 | Old |
| 0.453155 | 0.571851 | 0.212 | 0.19 | 1 | 8 | Ptbp2 | 0.022 | Old |
| 0.453224 | 0.210798 | 0.464 | 0.447 | 1 | 8 | Cnot6 | 0.017 | Old |
| 0.453276 | 0.201267 | 0.061 | 0.047 | 1 | 8 | Nfatc2ip | 0.014 | Old |
| 0.453996 | 0.328533 | 0.05 | 0.068 | 1 | 8 | Zfp729b | -0.018 | Old |
| 0.453996 | 0.166802 | 0.05 | 0.068 | 1 | 8 | Zfp444 | -0.018 | Old |
| 0.454024 | 0.303754 | 0.279 | 0.249 | 1 | 8 | Thap12 | 0.03 | Old |
| 0.45415 | 0.230412 | 0.212 | 0.244 | 1 | 8 | Armcx3 | -0.032 | Old |
| 0.454207 | 0.168763 | 0.045 | 0.062 | 1 | 8 | Gpatch1 | -0.017 | Old |
| 0.454424 | 0.332994 | 0.028 | 0.042 | 1 | 8 | 2900089D1 | -0.014 | Old |
| 0.454443 | 1.01669 | 0.078 | 0.062 | 1 | 8 | Armt1 | 0.016 | Old |
| 0.454538 | 0.306039 | 0.402 | 0.384 | 1 | 8 | Rad23a | 0.018 | Old |
| 0.454588 | 0.47195 | 0.061 | 0.047 | 1 | 8 | Asb6 | 0.014 | Old |
| 0.454712 | 0.263733 | 0.285 | 0.255 | 1 | 8 | Zkscan3 | 0.03 | Old |
| 0.454991 | 0.015948 | 0.19 | 0.213 | 1 | 8 | Casd1 | -0.023 | Old |
| 0.455026 | 0.632294 | 0.061 | 0.047 | 1 | 8 | Map3k10 | 0.014 | Old |
| 0.455321 | 0.236586 | 0.302 | 0.268 | 1 | 8 | Gpalpp1 | 0.034 | Old |
| 0.455528 | 0.532133 | 0.028 | 0.018 | 1 | 8 | 6330562C2 | 0.01 | Old |
| 0.455821 | 0.016804 | 0.318 | 0.356 | 1 | 8 | Cycs | -0.038 | Old |
| 0.455902 | 0.308072 | 0.061 | 0.047 | 1 | 8 | Eva1b | 0.014 | Old |
| 0.455907 | 0.570802 | 0.14 | 0.119 | 1 | 8 | Neurl4 | 0.021 | Old |
| 0.45606 | 0.059492 | 0.385 | 0.431 | 1 | 8 | N4bp1 | -0.046 | Old |
| 0.456088 | 0.254024 | 0.732 | 0.758 | 1 | 8 | Lbh | -0.026 | Old |
| 0.456417 | 0.153533 | 0.793 | 0.868 | 1 | 8 | Csde1 | -0.075 | Old |
| 0.457287 | 0.358758 | 0.413 | 0.413 | 1 | 8 | Atp6v1h | 0 | Old |
| 0.45736 | 0.051479 | 0.073 | 0.094 | 1 | 8 | Zfp84 | -0.021 | Old |
| 0.457588 | 0.402452 | 0.296 | 0.273 | 1 | 8 | Tnfrsf11a | 0.023 | Old |
| 0.457588 | 0.149921 | 0.302 | 0.27 | 1 | 8 | Mtch2 | 0.032 | Old |
| 0.457655 | 0.232472 | 0.14 | 0.119 | 1 | 8 | Ptger2 | 0.021 | Old |
| 0.457819 | 0.69035 | 0.106 | 0.088 | 1 | 8 | Dcun1d3 | 0.018 | Old |
| 0.458215 | 0.811939 | 0.028 | 0.018 | 1 | 8 | Ldlrad4 | 0.01 | Old |
| 0.458215 | 0.411206 | 0.028 | 0.018 | 1 | 8 | Prdm11 | 0.01 | Old |
| 0.458229 | 0.072479 | 0.464 | 0.423 | 1 | 8 | Rexo2 | 0.041 | Old |
| 0.458306 | 0.059754 | 0.056 | 0.073 | 1 | 8 | Haus4 | -0.017 | Old |
| 0.458508 | 0.039517 | 0.151 | 0.125 | 1 | 8 | Zfp672 | 0.026 | Old |
| 0.45856 | 0.717393 | 0.084 | 0.068 | 1 | 8 | Clpb | 0.016 | Old |
| 0.458737 | 0.083074 | 0.073 | 0.094 | 1 | 8 | Tinf2 | -0.021 | Old |
| 0.459236 | 0.223501 | 0.162 | 0.14 | 1 | 8 | Snrnp25 | 0.022 | Old |
| 0.459256 | 0.231224 | 0.279 | 0.317 | 1 | 8 | Fam172a | -0.038 | Old |
| 0.459562 | 0.344144 | 0.028 | 0.018 | 1 | 8 | Gpsm1 | 0.01 | Old |
| 0.459941 | 0.387405 | 0.156 | 0.135 | 1 | 8 | Abhd2 | 0.021 | Old |
| 0.460475 | 0.399454 | 0.235 | 0.21 | 1 | 8 | Pdlim4 | 0.025 | Old |
| 0.460521 | 0.107206 | 0.134 | 0.161 | 1 | 8 | Foxk1 | -0.027 | Old |
| 0.460589 | 0.134764 | 0.218 | 0.255 | 1 | 8 | Copg1 | -0.037 | Old |
| 0.460605 | 0.024712 | 0.263 | 0.294 | 1 | 8 | Nsmce4a | -0.031 | Old |

| | | | | | | | | |
|---|---|---|---|---|---|---|---|---|
| 0.460706 | 0.032376 | 0.145 | 0.169 | 1 | 8 | Zfp120 | -0.024 | Old |
| 0.461277 | 0.274469 | 0.425 | 0.429 | 1 | 8 | Dnajc13 | -0.004 | Old |
| 0.461616 | 0.009466 | 0.19 | 0.223 | 1 | 8 | Rabgap1 | -0.033 | Old |
| 0.46193 | 0.289536 | 0.056 | 0.042 | 1 | 8 | Zfp146 | 0.014 | Old |
| 0.461933 | 0.198978 | 0.022 | 0.034 | 1 | 8 | Nup37 | -0.012 | Old |
| 0.46213 | 0.379666 | 0.486 | 0.465 | 1 | 8 | Man1c1 | 0.021 | Old |
| 0.462263 | 0.238271 | 0.028 | 0.018 | 1 | 8 | Gm10762 | 0.01 | Old |
| 0.462307 | 0.593539 | 0.084 | 0.068 | 1 | 8 | Mrpl1 | 0.016 | Old |
| 0.462396 | 0.165834 | 0.101 | 0.122 | 1 | 8 | Ammecr1 | -0.021 | Old |
| 0.462501 | 0.561636 | 0.134 | 0.114 | 1 | 8 | Cnn3 | 0.02 | Old |
| 0.462628 | 0.242543 | 0.039 | 0.055 | 1 | 8 | Rasal3 | -0.016 | Old |
| 0.463281 | 0.536875 | 0.156 | 0.135 | 1 | 8 | Dcaf10 | 0.021 | Old |
| 0.463527 | 0.03756 | 0.039 | 0.055 | 1 | 8 | Wdr47 | -0.016 | Old |
| 0.463616 | 0.156222 | 0.028 | 0.018 | 1 | 8 | B3galt6 | 0.01 | Old |
| 0.464034 | 0.076952 | 0.436 | 0.478 | 1 | 8 | Ist1 | -0.042 | Old |
| 0.464381 | 0.11052 | 0.497 | 0.561 | 1 | 8 | Csk | -0.064 | Old |
| 0.464527 | 0.333323 | 0.173 | 0.151 | 1 | 8 | Dpm2 | 0.022 | Old |
| 0.464542 | 0.090242 | 0.536 | 0.582 | 1 | 8 | Nipbl | -0.046 | Old |
| 0.464579 | 0.091293 | 0.173 | 0.145 | 1 | 8 | Cnnm3 | 0.028 | Old |
| 0.464871 | 0.290425 | 0.631 | 0.665 | 1 | 8 | Ssb | -0.034 | Old |
| 0.464924 | 0.188818 | 0.81 | 0.865 | 1 | 8 | Ifrd1 | -0.055 | Old |
| 0.464951 | 0.222218 | 0.48 | 0.481 | 1 | 8 | Stk4 | -0.001 | Old |
| 0.465255 | 0.557111 | 0.218 | 0.2 | 1 | 8 | Usp37 | 0.018 | Old |
| 0.465915 | 0.135067 | 0.134 | 0.112 | 1 | 8 | Vamp7 | 0.022 | Old |
| 0.466096 | 0.085203 | 0.201 | 0.239 | 1 | 8 | Srek1ip1 | -0.038 | Old |
| 0.466416 | 0.182637 | 0.486 | 0.478 | 1 | 8 | Ranbp1 | 0.008 | Old |
| 0.466593 | 0.103975 | 0.056 | 0.042 | 1 | 8 | Cep76 | 0.014 | Old |
| 0.466757 | 0.405531 | 0.207 | 0.244 | 1 | 8 | Eml4 | -0.037 | Old |
| 0.467038 | 0.416462 | 0.279 | 0.262 | 1 | 8 | Grk6 | 0.017 | Old |
| 0.467389 | 0.112319 | 0.235 | 0.27 | 1 | 8 | Tor1aip2 | -0.035 | Old |
| 0.467562 | 0.563006 | 0.168 | 0.145 | 1 | 8 | Naa30 | 0.023 | Old |
| 0.46769 | 0.043441 | 0.028 | 0.018 | 1 | 8 | Ppcs | 0.01 | Old |
| 0.467752 | 0.319977 | 0.682 | 0.699 | 1 | 8 | Sgpl1 | -0.017 | Old |
| 0.468037 | 0.166644 | 0.039 | 0.055 | 1 | 8 | Gpt2 | -0.016 | Old |
| 0.468917 | 0.021301 | 0.162 | 0.187 | 1 | 8 | Rps6kc1 | -0.025 | Old |
| 0.469021 | 0.243823 | 0.419 | 0.418 | 1 | 8 | Mcur1 | 0.001 | Old |
| 0.469153 | 0.068957 | 0.352 | 0.395 | 1 | 8 | Psmc5 | -0.043 | Old |
| 0.46926 | 0.224159 | 0.112 | 0.091 | 1 | 8 | Pphln1 | 0.021 | Old |
| 0.469517 | 0.250374 | 0.497 | 0.486 | 1 | 8 | Pgd | 0.011 | Old |
| 0.469874 | 0.24589 | 0.313 | 0.273 | 1 | 8 | Sav1 | 0.04 | Old |
| 0.470226 | 0.577201 | 0.067 | 0.052 | 1 | 8 | Rbm12b1 | 0.015 | Old |
| 0.470534 | 0.021692 | 0.022 | 0.034 | 1 | 8 | Herc3 | -0.012 | Old |
| 0.470999 | 0.451564 | 0.352 | 0.345 | 1 | 8 | Sdhb | 0.007 | Old |
| 0.471194 | 0.090621 | 0.263 | 0.234 | 1 | 8 | Cd300lb | 0.029 | Old |
| 0.471249 | 0.146893 | 0.112 | 0.091 | 1 | 8 | Prmt5 | 0.021 | Old |

| | | | | | | | | |
|---|---|---|---|---|---|---|---|---|
| 0.471338 | 0.490003 | 0.223 | 0.203 | 1 | 8 | Hibadh | 0.02 | Old |
| 0.471393 | 0.812046 | 0.184 | 0.166 | 1 | 8 | Fam199x | 0.018 | Old |
| 0.471782 | 0.502226 | 0.184 | 0.161 | 1 | 8 | Srebf1 | 0.023 | Old |
| 0.471906 | 0.187091 | 0.536 | 0.553 | 1 | 8 | Cers2 | -0.017 | Old |
| 0.47199 | 0.437513 | 0.095 | 0.078 | 1 | 8 | Snx21 | 0.017 | Old |
| 0.47211 | 0.169723 | 0.704 | 0.743 | 1 | 8 | Mtpn | -0.039 | Old |
| 0.472119 | 0.798983 | 0.034 | 0.023 | 1 | 8 | Uaca | 0.011 | Old |
| 0.472483 | 0.028818 | 0.078 | 0.099 | 1 | 8 | Rnf126 | -0.021 | Old |
| 0.472536 | 0.055194 | 0.363 | 0.413 | 1 | 8 | Pbxip1 | -0.05 | Old |
| 0.47264 | 0.123727 | 0.151 | 0.179 | 1 | 8 | Atad2 | -0.028 | Old |
| 0.472889 | 0.42097 | 0.084 | 0.068 | 1 | 8 | Tmem87a | 0.016 | Old |
| 0.472893 | 0.090864 | 0.508 | 0.595 | 1 | 8 | Fgfr1op2 | -0.087 | Old |
| 0.473236 | 0.41108 | 0.196 | 0.174 | 1 | 8 | Cracr2b | 0.022 | Old |
| 0.473638 | 0.324618 | 0.067 | 0.052 | 1 | 8 | Ptgfrn | 0.015 | Old |
| 0.473734 | 0.220814 | 0.067 | 0.088 | 1 | 8 | H6pd | -0.021 | Old |
| 0.47375 | 0.31872 | 0.156 | 0.138 | 1 | 8 | Rpl10-ps3 | 0.018 | Old |
| 0.473808 | 0.370496 | 0.385 | 0.369 | 1 | 8 | Adssl1 | 0.016 | Old |
| 0.474382 | 0.438962 | 0.559 | 0.574 | 1 | 8 | 9530068E0 | -0.015 | Old |
| 0.474438 | 0.075119 | 0.196 | 0.223 | 1 | 8 | Snap29 | -0.027 | Old |
| 0.474527 | 0.460988 | 0.123 | 0.104 | 1 | 8 | Haghl | 0.019 | Old |
| 0.474747 | 0.250508 | 0.14 | 0.119 | 1 | 8 | Tstd3 | 0.021 | Old |
| 0.474791 | 0.00992 | 0.173 | 0.2 | 1 | 8 | Larp7 | -0.027 | Old |
| 0.474822 | 0.265182 | 0.346 | 0.332 | 1 | 8 | Capn1 | 0.014 | Old |
| 0.474884 | 0.50192 | 0.218 | 0.197 | 1 | 8 | Ugdh | 0.021 | Old |
| 0.475093 | 0.005143 | 0.547 | 0.571 | 1 | 8 | Arih1 | -0.024 | Old |
| 0.475158 | 0.523746 | 0.022 | 0.034 | 1 | 8 | Armc6 | -0.012 | Old |
| 0.475224 | 0.370995 | 0.285 | 0.265 | 1 | 8 | Hivep1 | 0.02 | Old |
| 0.475316 | 0.023566 | 0.374 | 0.403 | 1 | 8 | Cnih1 | -0.029 | Old |
| 0.475418 | 0.299842 | 0.559 | 0.54 | 1 | 8 | Leprot | 0.019 | Old |
| 0.47548 | 0.202186 | 0.408 | 0.39 | 1 | 8 | U2surp | 0.018 | Old |
| 0.475809 | 1.234991 | 0.034 | 0.023 | 1 | 8 | Intu | 0.011 | Old |
| 0.475811 | 0.231047 | 0.391 | 0.374 | 1 | 8 | Aup1 | 0.017 | Old |
| 0.475918 | 0.39581 | 0.145 | 0.122 | 1 | 8 | Zfp180 | 0.023 | Old |
| 0.47606 | 0.254795 | 0.24 | 0.216 | 1 | 8 | Cnp | 0.024 | Old |
| 0.47606 | 0.233248 | 0.24 | 0.216 | 1 | 8 | Rassf8 | 0.024 | Old |
| 0.47624 | 0.055793 | 0.073 | 0.091 | 1 | 8 | Erich1 | -0.018 | Old |
| 0.476281 | 0.45437 | 0.095 | 0.078 | 1 | 8 | Tmem242 | 0.017 | Old |
| 0.476317 | 0.490711 | 0.022 | 0.034 | 1 | 8 | Arap3 | -0.012 | Old |
| 0.476362 | 0.266111 | 0.397 | 0.384 | 1 | 8 | Phf20l1 | 0.013 | Old |
| 0.4765 | 0.320965 | 0.514 | 0.532 | 1 | 8 | Ranbp2 | -0.018 | Old |
| 0.47695 | 0.066374 | 0.045 | 0.06 | 1 | 8 | Zfand1 | -0.015 | Old |
| 0.477042 | 0.962678 | 0.034 | 0.023 | 1 | 8 | Zfp933 | 0.011 | Old |
| 0.477063 | 0.413923 | 0.067 | 0.052 | 1 | 8 | Taf6 | 0.015 | Old |
| 0.477209 | 0.280351 | 0.061 | 0.047 | 1 | 8 | Zfp868 | 0.014 | Old |
| 0.477347 | 0.448493 | 0.156 | 0.135 | 1 | 8 | Il10 | 0.021 | Old |

| | | | | | | | | |
|---|---|---|---|---|---|---|---|---|
| 0.477569 | 0.723643 | 0.218 | 0.197 | 1 | 8 | Wdr82 | 0.021 | Old |
| 0.47774 | 0.698464 | 0.101 | 0.083 | 1 | 8 | Rhbdd1 | 0.018 | Old |
| 0.477759 | 0.169041 | 0.626 | 0.652 | 1 | 8 | Aldoa | -0.026 | Old |
| 0.478109 | 0.125901 | 0.061 | 0.047 | 1 | 8 | Zfp932 | 0.014 | Old |
| 0.478277 | 0.388957 | 0.034 | 0.023 | 1 | 8 | Wdr73 | 0.011 | Old |
| 0.478361 | 0.490969 | 0.073 | 0.057 | 1 | 8 | Angptl4 | 0.016 | Old |
| 0.478887 | 0.213453 | 0.52 | 0.514 | 1 | 8 | Dmac1 | 0.006 | Old |
| 0.478888 | 0.452367 | 0.307 | 0.299 | 1 | 8 | Mospd2 | 0.008 | Old |
| 0.478937 | 0.030157 | 0.346 | 0.384 | 1 | 8 | Clk4 | -0.038 | Old |
| 0.478995 | 0.640109 | 0.084 | 0.068 | 1 | 8 | Pvt1 | 0.016 | Old |
| 0.479514 | 0.861066 | 0.034 | 0.023 | 1 | 8 | Cfap45 | 0.011 | Old |
| 0.480122 | 0.236114 | 0.14 | 0.119 | 1 | 8 | Gtf2e2 | 0.021 | Old |
| 0.48013 | 0.166452 | 0.419 | 0.478 | 1 | 8 | Rnf19b | -0.059 | Old |
| 0.480492 | 0.255783 | 0.369 | 0.369 | 1 | 8 | Gm6377 | 0 | Old |
| 0.480752 | 0.298261 | 0.034 | 0.023 | 1 | 8 | Wdr76 | 0.011 | Old |
| 0.480765 | 0.135521 | 0.156 | 0.187 | 1 | 8 | Pnpla7 | -0.031 | Old |
| 0.481086 | 0.440098 | 0.156 | 0.138 | 1 | 8 | Tor3a | 0.018 | Old |
| 0.481375 | 0.265124 | 0.33 | 0.306 | 1 | 8 | Slc15a4 | 0.024 | Old |
| 0.481564 | 0.116093 | 0.402 | 0.439 | 1 | 8 | Ube2s | -0.037 | Old |
| 0.481934 | 0.300317 | 0.324 | 0.314 | 1 | 8 | Stx12 | 0.01 | Old |
| 0.482413 | 0.074268 | 0.034 | 0.047 | 1 | 8 | Prmt6 | -0.013 | Old |
| 0.482499 | 0.335708 | 0.123 | 0.101 | 1 | 8 | Desi1 | 0.022 | Old |
| 0.482577 | 0.130855 | 0.782 | 0.756 | 1 | 8 | Pea15a | 0.026 | Old |
| 0.483037 | 0.678646 | 0.156 | 0.135 | 1 | 8 | Vps33a | 0.021 | Old |
| 0.483235 | 0.451428 | 0.034 | 0.023 | 1 | 8 | E130308A1 | 0.011 | Old |
| 0.483235 | 0.415792 | 0.034 | 0.023 | 1 | 8 | Taf4b | 0.011 | Old |
| 0.483403 | 0.291175 | 0.034 | 0.047 | 1 | 8 | Atad5 | -0.013 | Old |
| 0.483461 | 0.147764 | 0.078 | 0.099 | 1 | 8 | Fam35a | -0.021 | Old |
| 0.483525 | 0.001749 | 0.061 | 0.047 | 1 | 8 | Nsdhl | 0.014 | Old |
| 0.483923 | 0.290181 | 0.251 | 0.223 | 1 | 8 | Hcfc1 | 0.028 | Old |
| 0.484143 | 0.30602 | 0.346 | 0.33 | 1 | 8 | Ube2h | 0.016 | Old |
| 0.484431 | 0.087248 | 0.061 | 0.047 | 1 | 8 | Mrpl44 | 0.014 | Old |
| 0.48497 | 0.518183 | 0.184 | 0.171 | 1 | 8 | Ppp2r3d | 0.013 | Old |
| 0.485051 | 0.630086 | 0.078 | 0.062 | 1 | 8 | Isca1 | 0.016 | Old |
| 0.485096 | 0.243676 | 0.492 | 0.481 | 1 | 8 | Ago2 | 0.011 | Old |
| 0.485126 | 0.278737 | 0.324 | 0.312 | 1 | 8 | Ccny | 0.012 | Old |
| 0.485651 | 0.050215 | 0.095 | 0.117 | 1 | 8 | Togaram1 | -0.022 | Old |
| 0.486007 | 0.147732 | 0.173 | 0.205 | 1 | 8 | Polr1a | -0.032 | Old |
| 0.486246 | 0.257881 | 0.061 | 0.047 | 1 | 8 | 9530059O1 | 0.014 | Old |
| 0.48665 | 0.223025 | 0.056 | 0.073 | 1 | 8 | Fbxo31 | -0.017 | Old |
| 0.486913 | 0.225453 | 0.358 | 0.332 | 1 | 8 | Ndufa10 | 0.026 | Old |
| 0.487297 | 0.987129 | 0.039 | 0.029 | 1 | 8 | Ccdc77 | 0.01 | Old |
| 0.487676 | 0.023324 | 0.073 | 0.091 | 1 | 8 | Fhod1 | -0.018 | Old |
| 0.487782 | 0.133765 | 0.682 | 0.714 | 1 | 8 | Bsg | -0.032 | Old |
| 0.488003 | 0.555318 | 0.134 | 0.117 | 1 | 8 | Zfp24 | 0.017 | Old |

| | | | | | | | | |
|---|---|---|---|---|---|---|---|---|
| 0.488196 | 0.600756 | 0.363 | 0.358 | 1 | 8 | Pank3 | 0.005 | Old |
| 0.48822 | 0.32644 | 0.034 | 0.023 | 1 | 8 | Exog | 0.011 | Old |
| 0.488252 | 0.354801 | 0.078 | 0.062 | 1 | 8 | Zfyve21 | 0.016 | Old |
| 0.488441 | 0.854427 | 0.039 | 0.029 | 1 | 8 | Lrmda | 0.01 | Old |
| 0.488441 | 0.573055 | 0.039 | 0.029 | 1 | 8 | Cryz | 0.01 | Old |
| 0.488548 | 0.484147 | 0.14 | 0.119 | 1 | 8 | Cdadc1 | 0.021 | Old |
| 0.489137 | 0.38876 | 0.073 | 0.057 | 1 | 8 | Ercc2 | 0.016 | Old |
| 0.489264 | 0.189155 | 0.184 | 0.161 | 1 | 8 | Batf | 0.023 | Old |
| 0.490011 | 0.37805 | 0.095 | 0.078 | 1 | 8 | Jak3 | 0.017 | Old |
| 0.490028 | 0.226891 | 0.067 | 0.052 | 1 | 8 | Ppp1r12b | 0.015 | Old |
| 0.490308 | 0.479517 | 0.184 | 0.166 | 1 | 8 | Crebzf | 0.018 | Old |
| 0.490569 | 0.597465 | 0.089 | 0.073 | 1 | 8 | Sirpb1c | 0.016 | Old |
| 0.49066 | 0.450167 | 0.078 | 0.062 | 1 | 8 | Slc2a1 | 0.016 | Old |
| 0.490699 | 0.132851 | 0.81 | 0.829 | 1 | 8 | Itm2c | -0.019 | Old |
| 0.490936 | 1.034075 | 0.089 | 0.075 | 1 | 8 | 1500011B0 | 0.014 | Old |
| 0.490978 | 0.087603 | 0.302 | 0.351 | 1 | 8 | Zmiz2 | -0.049 | Old |
| 0.490992 | 0.303481 | 0.296 | 0.275 | 1 | 8 | Wtap | 0.021 | Old |
| 0.491008 | 0.715637 | 0.106 | 0.088 | 1 | 8 | Cstf2 | 0.018 | Old |
| 0.491201 | 0.160678 | 0.715 | 0.751 | 1 | 8 | Tra2b | -0.036 | Old |
| 0.49133 | 0.043893 | 0.346 | 0.39 | 1 | 8 | Zfp148 | -0.044 | Old |
| 0.491542 | 0.260254 | 0.385 | 0.364 | 1 | 8 | Taf15 | 0.021 | Old |
| 0.492072 | 0.345164 | 0.089 | 0.073 | 1 | 8 | Exoc8 | 0.016 | Old |
| 0.49235 | 0.224692 | 0.587 | 0.592 | 1 | 8 | Sdha | -0.005 | Old |
| 0.49265 | 0.147262 | 0.358 | 0.319 | 1 | 8 | Fam32a | 0.039 | Old |
| 0.493171 | 0.382097 | 0.346 | 0.327 | 1 | 8 | Ubqln1 | 0.019 | Old |
| 0.49373 | 0.023383 | 0.263 | 0.286 | 1 | 8 | Spata6 | -0.023 | Old |
| 0.49424 | 0.248469 | 0.553 | 0.566 | 1 | 8 | Trip12 | -0.013 | Old |
| 0.494599 | 0.537835 | 0.285 | 0.268 | 1 | 8 | Esyt2 | 0.017 | Old |
| 0.495029 | 0.194737 | 0.547 | 0.556 | 1 | 8 | Tiparp | -0.009 | Old |
| 0.495267 | 0.214121 | 0.067 | 0.052 | 1 | 8 | Slc35d1 | 0.015 | Old |
| 0.495339 | 0.132693 | 0.101 | 0.122 | 1 | 8 | Adamts10 | -0.021 | Old |
| 0.495409 | 0.022429 | 0.067 | 0.086 | 1 | 8 | Tlr13 | -0.019 | Old |
| 0.495726 | 0.294115 | 0.48 | 0.468 | 1 | 8 | Atp6v0a1 | 0.012 | Old |
| 0.496014 | 0.16622 | 0.464 | 0.449 | 1 | 8 | Smg1 | 0.015 | Old |
| 0.496017 | 0.438772 | 0.14 | 0.122 | 1 | 8 | Aspscr1 | 0.018 | Old |
| 0.49603 | 0.115376 | 0.313 | 0.281 | 1 | 8 | Derl2 | 0.032 | Old |
| 0.49648 | 0.709647 | 0.173 | 0.156 | 1 | 8 | Ddx19a | 0.017 | Old |
| 0.496486 | 0.878969 | 0.039 | 0.029 | 1 | 8 | Gfi1b | 0.01 | Old |
| 0.496486 | 0.834717 | 0.039 | 0.029 | 1 | 8 | Slc37a1 | 0.01 | Old |
| 0.497323 | 0.406313 | 0.095 | 0.078 | 1 | 8 | Gabpb1 | 0.017 | Old |
| 0.497607 | 0.174975 | 0.128 | 0.153 | 1 | 8 | Anapc15 | -0.025 | Old |
| 0.497705 | 0.020986 | 0.156 | 0.182 | 1 | 8 | Vopp1 | -0.026 | Old |
| 0.498266 | 0.039349 | 0.235 | 0.268 | 1 | 8 | Mecp2 | -0.033 | Old |
| 0.498522 | 0.070531 | 0.816 | 0.862 | 1 | 8 | Hnrnpu | -0.046 | Old |
| 0.498593 | 0.01591 | 0.268 | 0.236 | 1 | 8 | Mrpl17 | 0.032 | Old |

| | | | | | | | | |
|---|---|---|---|---|---|---|---|---|
| 0.498734 | 0.211309 | 0.078 | 0.062 | 1 | 8 | Carf | 0.016 | Old |
| 0.498797 | 0.367521 | 0.039 | 0.029 | 1 | 8 | Rnf123 | 0.01 | Old |
| 0.498826 | 0.428279 | 0.156 | 0.138 | 1 | 8 | Cul4a | 0.018 | Old |
| 0.499001 | 0.445401 | 0.145 | 0.125 | 1 | 8 | Atr | 0.02 | Old |
| 0.499068 | 0.483776 | 0.291 | 0.273 | 1 | 8 | Ftl1-ps1 | 0.018 | Old |
| 0.499141 | 0.390336 | 0.207 | 0.187 | 1 | 8 | Stard7 | 0.02 | Old |
| 0.499146 | 0.311413 | 0.128 | 0.109 | 1 | 8 | Tmem167k | 0.019 | Old |
| 0.499933 | 0.026934 | 0.179 | 0.208 | 1 | 8 | Pcyox1 | -0.029 | Old |
| 0.500025 | 0.388249 | 0.106 | 0.088 | 1 | 8 | Ganc | 0.018 | Old |
| 0.500185 | 0.258165 | 0.078 | 0.099 | 1 | 8 | Sec22a | -0.021 | Old |
| 0.500447 | 0.871486 | 0.061 | 0.049 | 1 | 8 | Utp23 | 0.012 | Old |
| 0.501072 | 0.144218 | 0.101 | 0.125 | 1 | 8 | Iffo1 | -0.024 | Old |
| 0.501109 | 0.068715 | 0.017 | 0.026 | 1 | 8 | Kcng2 | -0.009 | Old |
| 0.501393 | 0.176456 | 0.039 | 0.052 | 1 | 8 | Zkscan8 | -0.013 | Old |
| 0.502677 | 0.290858 | 0.162 | 0.14 | 1 | 8 | Ggps1 | 0.022 | Old |
| 0.502931 | 1.03525 | 0.05 | 0.039 | 1 | 8 | Atp6v0d2 | 0.011 | Old |
| 0.502937 | 0.451594 | 0.106 | 0.091 | 1 | 8 | Iffo2 | 0.015 | Old |
| 0.503264 | 0.207577 | 0.352 | 0.332 | 1 | 8 | Slc11a1 | 0.02 | Old |
| 0.503553 | 0.314844 | 0.33 | 0.312 | 1 | 8 | Rbm17 | 0.018 | Old |
| 0.503575 | 0.075897 | 0.698 | 0.782 | 1 | 8 | Atp6v1g1 | -0.084 | Old |
| 0.503831 | 0.067221 | 0.017 | 0.026 | 1 | 8 | Grhpr | -0.009 | Old |
| 0.504131 | 0.016386 | 0.402 | 0.449 | 1 | 8 | Hfe | -0.047 | Old |
| 0.504917 | 0.484949 | 0.106 | 0.088 | 1 | 8 | AI837181 | 0.018 | Old |
| 0.505024 | 0.425109 | 0.263 | 0.244 | 1 | 8 | Prelid3b | 0.019 | Old |
| 0.50523 | 0.075343 | 0.212 | 0.239 | 1 | 8 | Fyn | -0.027 | Old |
| 0.505396 | 0.206458 | 0.33 | 0.377 | 1 | 8 | Psme4 | -0.047 | Old |
| 0.505968 | 0.583315 | 0.045 | 0.034 | 1 | 8 | Gm42715 | 0.011 | Old |
| 0.5065 | 0.309089 | 0.318 | 0.304 | 1 | 8 | Agfg1 | 0.014 | Old |
| 0.506707 | 0.167662 | 0.078 | 0.096 | 1 | 8 | Phf13 | -0.018 | Old |
| 0.507051 | 1.045136 | 0.045 | 0.034 | 1 | 8 | Hirip3 | 0.011 | Old |
| 0.507051 | 0.464931 | 0.045 | 0.034 | 1 | 8 | Arhgap35 | 0.011 | Old |
| 0.507142 | 1.048467 | 0.056 | 0.044 | 1 | 8 | Smpd3 | 0.012 | Old |
| 0.507604 | 0.094427 | 0.374 | 0.41 | 1 | 8 | Prkcb | -0.036 | Old |
| 0.507928 | 0.547877 | 0.017 | 0.026 | 1 | 8 | Pkn3 | -0.009 | Old |
| 0.508135 | 0.672774 | 0.045 | 0.034 | 1 | 8 | Elp4 | 0.011 | Old |
| 0.508313 | 0.114 | 0.676 | 0.722 | 1 | 8 | Abca1 | -0.046 | Old |
| 0.508682 | 0.210223 | 0.497 | 0.483 | 1 | 8 | Gm16286 | 0.014 | Old |
| 0.508985 | 0.141119 | 0.151 | 0.13 | 1 | 8 | Slc17a5 | 0.021 | Old |
| 0.509221 | 0.141598 | 0.084 | 0.104 | 1 | 8 | Mre11a | -0.02 | Old |
| 0.509722 | 0.104499 | 0.631 | 0.626 | 1 | 8 | Ldlr | 0.005 | Old |
| 0.509961 | 0.226178 | 0.24 | 0.218 | 1 | 8 | Lsm7 | 0.022 | Old |
| 0.510041 | 0.958132 | 0.05 | 0.039 | 1 | 8 | Cnnm2 | 0.011 | Old |
| 0.510058 | 0.578331 | 0.14 | 0.122 | 1 | 8 | Ppa1 | 0.018 | Old |
| 0.51018 | 0.09358 | 0.123 | 0.143 | 1 | 8 | Gm11808 | -0.02 | Old |
| 0.510258 | 0.437881 | 0.117 | 0.099 | 1 | 8 | Cdkn2aip | 0.018 | Old |

| | | | | | | | | |
|---|---|---|---|---|---|---|---|---|
| 0.510346 | 0.045041 | 0.296 | 0.335 | 1 | 8 | Myo1c | -0.039 | Old |
| 0.510441 | 0.467679 | 0.039 | 0.029 | 1 | 8 | Trim32 | 0.01 | Old |
| 0.510854 | 0.36449 | 0.419 | 0.4 | 1 | 8 | Emc1 | 0.019 | Old |
| 0.510981 | 0.575272 | 0.302 | 0.294 | 1 | 8 | Ccdc71l | 0.008 | Old |
| 0.51107 | 0.315316 | 0.089 | 0.073 | 1 | 8 | Mfsd8 | 0.016 | Old |
| 0.511288 | 0.662566 | 0.14 | 0.122 | 1 | 8 | Ppid | 0.018 | Old |
| 0.511395 | 1.045694 | 0.045 | 0.034 | 1 | 8 | Supv3l1 | 0.011 | Old |
| 0.511488 | 0.076821 | 0.626 | 0.686 | 1 | 8 | Tardbp | -0.06 | Old |
| 0.511838 | 0.065328 | 0.089 | 0.073 | 1 | 8 | Nudt13 | 0.016 | Old |
| 0.511845 | 0.149005 | 0.771 | 0.779 | 1 | 8 | Grk2 | -0.008 | Old |
| 0.511968 | 0.268457 | 0.246 | 0.286 | 1 | 8 | Anapc1 | -0.04 | Old |
| 0.512026 | 0.406789 | 0.207 | 0.184 | 1 | 8 | Rgs14 | 0.023 | Old |
| 0.512036 | 0.115509 | 0.402 | 0.452 | 1 | 8 | Ctbp1 | -0.05 | Old |
| 0.512673 | 0.136413 | 0.218 | 0.247 | 1 | 8 | Ccdc88c | -0.029 | Old |
| 0.512712 | 0.015357 | 0.207 | 0.242 | 1 | 8 | Pfdn1 | -0.035 | Old |
| 0.51286 | 0.331944 | 0.397 | 0.405 | 1 | 8 | Mfhas1 | -0.008 | Old |
| 0.512874 | 0.319701 | 0.469 | 0.449 | 1 | 8 | Nabp1 | 0.02 | Old |
| 0.512906 | 0.050624 | 0.419 | 0.374 | 1 | 8 | Ncoa4 | 0.045 | Old |
| 0.513139 | 0.111905 | 0.609 | 0.608 | 1 | 8 | Itpkb | 0.001 | Old |
| 0.513202 | 0.197276 | 0.073 | 0.091 | 1 | 8 | Mcm9 | -0.018 | Old |
| 0.513494 | 0.022368 | 0.318 | 0.343 | 1 | 8 | Elf2 | -0.025 | Old |
| 0.513691 | 0.073094 | 0.285 | 0.319 | 1 | 8 | Ptpn7 | -0.034 | Old |
| 0.513742 | 0.309185 | 0.257 | 0.236 | 1 | 8 | Cd2 | 0.021 | Old |
| 0.513892 | 0.852909 | 0.056 | 0.044 | 1 | 8 | Setdb2 | 0.012 | Old |
| 0.514292 | 0.047768 | 0.726 | 0.774 | 1 | 8 | Gas7 | -0.048 | Old |
| 0.514403 | 0.10667 | 0.285 | 0.325 | 1 | 8 | Zfp263 | -0.04 | Old |
| 0.51451 | 0.074147 | 0.095 | 0.114 | 1 | 8 | Mxra7 | -0.019 | Old |
| 0.514775 | 0.357728 | 0.106 | 0.088 | 1 | 8 | Abhd5 | 0.018 | Old |
| 0.514795 | 0.043663 | 0.017 | 0.026 | 1 | 8 | Cox11 | -0.009 | Old |
| 0.514862 | 0.390055 | 0.302 | 0.278 | 1 | 8 | Cdk19 | 0.024 | Old |
| 0.514915 | 0.153023 | 0.089 | 0.073 | 1 | 8 | Cnpy4 | 0.016 | Old |
| 0.514928 | 0.058094 | 0.279 | 0.306 | 1 | 8 | Eif4e2 | -0.027 | Old |
| 0.515025 | 0.021937 | 0.436 | 0.47 | 1 | 8 | Itsn1 | -0.034 | Old |
| 0.515137 | 0.163097 | 0.039 | 0.029 | 1 | 8 | Zfp958 | 0.01 | Old |
| 0.515442 | 0.053865 | 0.592 | 0.66 | 1 | 8 | Srp14 | -0.068 | Old |
| 0.515758 | 0.519592 | 0.045 | 0.034 | 1 | 8 | Hpn | 0.011 | Old |
| 0.516049 | 0.453966 | 0.263 | 0.244 | 1 | 8 | Rab3gap1 | 0.019 | Old |
| 0.516628 | 0.166874 | 0.525 | 0.532 | 1 | 8 | Srrm1 | -0.007 | Old |
| 0.516683 | 1.444059 | 0.017 | 0.01 | 1 | 8 | Smtn | 0.007 | Old |
| 0.516714 | 0.09224 | 0.251 | 0.281 | 1 | 8 | Gramd1a | -0.03 | Old |
| 0.516809 | 0.090187 | 0.246 | 0.216 | 1 | 8 | Mrps23 | 0.03 | Old |
| 0.517207 | 0.64379 | 0.123 | 0.106 | 1 | 8 | Ubap2 | 0.017 | Old |
| 0.517222 | 0.577252 | 0.084 | 0.07 | 1 | 8 | Micall1 | 0.014 | Old |
| 0.517316 | 0.110215 | 0.061 | 0.078 | 1 | 8 | Slf1 | -0.017 | Old |
| 0.51744 | 0.635625 | 0.218 | 0.205 | 1 | 8 | Atxn10 | 0.013 | Old |

| | | | | | | | | |
|---|---|---|---|---|---|---|---|---|
| 0.517483 | 0.372523 | 0.291 | 0.268 | 1 | 8 | Ccndbp1 | 0.023 | Old |
| 0.517504 | 0.069038 | 0.168 | 0.192 | 1 | 8 | Fam96b | -0.024 | Old |
| 0.517762 | 0.197484 | 0.385 | 0.361 | 1 | 8 | Ufm1 | 0.024 | Old |
| 0.51785 | 0.792543 | 0.061 | 0.049 | 1 | 8 | Cdk7 | 0.012 | Old |
| 0.517866 | 0.566056 | 0.162 | 0.145 | 1 | 8 | Parp11 | 0.017 | Old |
| 0.517937 | 0.13385 | 0.391 | 0.361 | 1 | 8 | Col4a3bp | 0.03 | Old |
| 0.517956 | 0.054744 | 0.922 | 0.971 | 1 | 8 | Capzb | -0.049 | Old |
| 0.517969 | 0.249983 | 0.246 | 0.291 | 1 | 8 | Sptlc1 | -0.045 | Old |
| 0.518094 | 0.577456 | 0.089 | 0.075 | 1 | 8 | Ddx55 | 0.014 | Old |
| 0.518231 | 0.655769 | 0.05 | 0.039 | 1 | 8 | Rilp | 0.011 | Old |
| 0.518237 | 0.227165 | 0.251 | 0.229 | 1 | 8 | Gtf3c6 | 0.022 | Old |
| 0.518395 | 0.052141 | 0.112 | 0.135 | 1 | 8 | Ttc32 | -0.023 | Old |
| 0.518557 | 1.170636 | 0.017 | 0.01 | 1 | 8 | Fkbp14 | 0.007 | Old |
| 0.518557 | 0.916573 | 0.017 | 0.01 | 1 | 8 | Gm10767 | 0.007 | Old |
| 0.518557 | 0.782172 | 0.017 | 0.01 | 1 | 8 | Procr | 0.007 | Old |
| 0.518557 | 0.693956 | 0.017 | 0.01 | 1 | 8 | Zfp956 | 0.007 | Old |
| 0.518557 | 0.644997 | 0.017 | 0.01 | 1 | 8 | Siah1b | 0.007 | Old |
| 0.518819 | 0.03524 | 0.151 | 0.171 | 1 | 8 | A430005L1 | -0.02 | Old |
| 0.518833 | 0.295629 | 0.263 | 0.239 | 1 | 8 | Hspa14 | 0.024 | Old |
| 0.519223 | 0.157711 | 0.184 | 0.216 | 1 | 8 | Rbm18 | -0.032 | Old |
| 0.519284 | 0.15254 | 0.687 | 0.719 | 1 | 8 | Prr13 | -0.032 | Old |
| 0.519463 | 0.788698 | 0.067 | 0.055 | 1 | 8 | Heatr5b | 0.012 | Old |
| 0.519658 | 0.116273 | 0.084 | 0.104 | 1 | 8 | Anks3 | -0.02 | Old |
| 0.519856 | 0.000372 | 0.039 | 0.029 | 1 | 8 | Enpp4 | 0.01 | Old |
| 0.519905 | 0.128022 | 0.112 | 0.132 | 1 | 8 | Sec24d | -0.02 | Old |
| 0.520434 | 1.536743 | 0.017 | 0.01 | 1 | 8 | Gbp8 | 0.007 | Old |
| 0.520434 | 0.945749 | 0.017 | 0.01 | 1 | 8 | Ciart | 0.007 | Old |
| 0.520434 | 0.870822 | 0.017 | 0.01 | 1 | 8 | Afap1 | 0.007 | Old |
| 0.520558 | 0.016564 | 0.525 | 0.566 | 1 | 8 | H2afv | -0.041 | Old |
| 0.520597 | 0.084418 | 0.369 | 0.343 | 1 | 8 | Il6st | 0.026 | Old |
| 0.520753 | 0.275168 | 0.218 | 0.2 | 1 | 8 | Ube3b | 0.018 | Old |
| 0.52086 | 0.111669 | 0.235 | 0.27 | 1 | 8 | Acvrl1 | -0.035 | Old |
| 0.521583 | 0.385454 | 0.162 | 0.143 | 1 | 8 | Ksr1 | 0.019 | Old |
| 0.521641 | 0.130094 | 0.134 | 0.117 | 1 | 8 | Acot8 | 0.017 | Old |
| 0.521662 | 0.621662 | 0.056 | 0.044 | 1 | 8 | Tmem107 | 0.012 | Old |
| 0.521816 | 0.959732 | 0.14 | 0.125 | 1 | 8 | Mroh1 | 0.015 | Old |
| 0.52217 | 0.608217 | 0.112 | 0.096 | 1 | 8 | Gtf3c3 | 0.016 | Old |
| 0.522314 | 1.336537 | 0.017 | 0.01 | 1 | 8 | Mmachc | 0.007 | Old |
| 0.522314 | 1.042079 | 0.017 | 0.01 | 1 | 8 | Cdh5 | 0.007 | Old |
| 0.522314 | 0.708023 | 0.017 | 0.01 | 1 | 8 | Ptprm | 0.007 | Old |
| 0.522314 | 0.467624 | 0.017 | 0.01 | 1 | 8 | Scnn1a | 0.007 | Old |
| 0.522314 | 0.461803 | 0.017 | 0.01 | 1 | 8 | Brf2 | 0.007 | Old |
| 0.52234 | 0.349972 | 0.045 | 0.034 | 1 | 8 | Nif3l1 | 0.011 | Old |
| 0.52234 | 0.14655 | 0.045 | 0.034 | 1 | 8 | 2410131K1 | 0.011 | Old |
| 0.522371 | 0.032631 | 0.486 | 0.535 | 1 | 8 | Tmod3 | -0.049 | Old |

| | | | | | | | | |
|---|---|---|---|---|---|---|---|---|
| 0.522481 | 0.591264 | 0.19 | 0.174 | 1 | 8 | Plekhf2 | 0.016 | Old |
| 0.522589 | 0.195652 | 0.028 | 0.039 | 1 | 8 | Cep78 | -0.011 | Old |
| 0.522675 | 0.098175 | 0.363 | 0.408 | 1 | 8 | Senp6 | -0.045 | Old |
| 0.523122 | 0.016963 | 0.687 | 0.777 | 1 | 8 | Litaf | -0.09 | Old |
| 0.523421 | 0.136765 | 0.391 | 0.356 | 1 | 8 | Stag1 | 0.035 | Old |
| 0.523434 | 0.118265 | 0.587 | 0.657 | 1 | 8 | Sf1 | -0.07 | Old |
| 0.523588 | 0.588863 | 0.145 | 0.127 | 1 | 8 | Nsmf | 0.018 | Old |
| 0.523663 | 0.10487 | 0.497 | 0.558 | 1 | 8 | Hnrnph2 | -0.061 | Old |
| 0.523709 | 0.336616 | 0.184 | 0.164 | 1 | 8 | Nol11 | 0.02 | Old |
| 0.523717 | 0.072135 | 0.028 | 0.039 | 1 | 8 | Cbarp | -0.011 | Old |
| 0.523765 | 0.326182 | 0.307 | 0.286 | 1 | 8 | Senp2 | 0.021 | Old |
| 0.52379 | 0.15452 | 0.408 | 0.384 | 1 | 8 | Sash1 | 0.024 | Old |
| 0.523863 | 0.792091 | 0.084 | 0.104 | 1 | 8 | Abcb1b | -0.02 | Old |
| 0.524199 | 1.688855 | 0.017 | 0.01 | 1 | 8 | B3galt2 | 0.007 | Old |
| 0.524199 | 1.6268 | 0.017 | 0.01 | 1 | 8 | Mrm3 | 0.007 | Old |
| 0.524199 | 0.448658 | 0.017 | 0.01 | 1 | 8 | Lgals4 | 0.007 | Old |
| 0.524199 | 0.366952 | 0.017 | 0.01 | 1 | 8 | Tubb3 | 0.007 | Old |
| 0.524713 | 0.698399 | 0.112 | 0.099 | 1 | 8 | AL590144.: | 0.013 | Old |
| 0.524923 | 0.532304 | 0.235 | 0.226 | 1 | 8 | Flvcr1 | 0.009 | Old |
| 0.52576 | 0.052906 | 0.089 | 0.073 | 1 | 8 | Hnmt | 0.016 | Old |
| 0.525869 | 0.752365 | 0.117 | 0.104 | 1 | 8 | Gon7 | 0.013 | Old |
| 0.525975 | 0.05922 | 0.028 | 0.039 | 1 | 8 | Zfp709 | -0.011 | Old |
| 0.525985 | 0.832675 | 0.073 | 0.06 | 1 | 8 | Ikzf2 | 0.013 | Old |
| 0.526086 | 1.563176 | 0.017 | 0.01 | 1 | 8 | Ccno | 0.007 | Old |
| 0.526086 | 1.099421 | 0.017 | 0.01 | 1 | 8 | Zfp951 | 0.007 | Old |
| 0.526086 | 0.918617 | 0.017 | 0.01 | 1 | 8 | Zscan20 | 0.007 | Old |
| 0.526356 | 0.117993 | 0.318 | 0.286 | 1 | 8 | Noct | 0.032 | Old |
| 0.526457 | 0.006447 | 0.24 | 0.275 | 1 | 8 | Slmap | -0.035 | Old |
| 0.526551 | 0.121449 | 0.341 | 0.306 | 1 | 8 | Tmem51 | 0.035 | Old |
| 0.526571 | 0.401634 | 0.184 | 0.166 | 1 | 8 | Asb8 | 0.018 | Old |
| 0.526655 | 0.24346 | 0.145 | 0.127 | 1 | 8 | Tusc1 | 0.018 | Old |
| 0.526724 | 0.095935 | 0.106 | 0.13 | 1 | 8 | Gtpbp6 | -0.024 | Old |
| 0.526742 | 0.226156 | 0.24 | 0.278 | 1 | 8 | Emc2 | -0.038 | Old |
| 0.526977 | 0.069629 | 0.039 | 0.029 | 1 | 8 | A630072M | 0.01 | Old |
| 0.527074 | 0.195918 | 0.257 | 0.299 | 1 | 8 | Srf | -0.042 | Old |
| 0.52708 | 0.190196 | 0.358 | 0.408 | 1 | 8 | Rbm26 | -0.05 | Old |
| 0.527121 | 0.009645 | 0.201 | 0.226 | 1 | 8 | Clcn7 | -0.025 | Old |
| 0.527368 | 0.167041 | 0.101 | 0.125 | 1 | 8 | Stam | -0.024 | Old |
| 0.527418 | 0.141631 | 0.19 | 0.221 | 1 | 8 | Spin1 | -0.031 | Old |
| 0.527427 | 0.350144 | 0.14 | 0.122 | 1 | 8 | AY036118 | 0.018 | Old |
| 0.527579 | 0.130824 | 0.397 | 0.46 | 1 | 8 | Scarb2 | -0.063 | Old |
| 0.527977 | 0.604657 | 0.017 | 0.01 | 1 | 8 | Fam13a | 0.007 | Old |
| 0.527977 | 0.488729 | 0.017 | 0.01 | 1 | 8 | Tcf19 | 0.007 | Old |
| 0.527977 | 0.35439 | 0.017 | 0.01 | 1 | 8 | Ccdc173 | 0.007 | Old |
| 0.527977 | 0.350599 | 0.017 | 0.01 | 1 | 8 | Ifi208 | 0.007 | Old |

| | | | | | | | | |
|---|---|---|---|---|---|---|---|---|
| 0.527977 | 0.072405 | 0.017 | 0.01 | 1 | 8 | Taco1os | 0.007 | Old |
| 0.527977 | 0.048848 | 0.017 | 0.01 | 1 | 8 | Wdr27 | 0.007 | Old |
| 0.528374 | 0.063772 | 0.078 | 0.062 | 1 | 8 | Fancl | 0.016 | Old |
| 0.528468 | 0.588282 | 0.179 | 0.213 | 1 | 8 | Htra2 | -0.034 | Old |
| 0.528789 | 0.334734 | 0.207 | 0.244 | 1 | 8 | Preb | -0.037 | Old |
| 0.528964 | 0.659141 | 0.045 | 0.034 | 1 | 8 | Dpt | 0.011 | Old |
| 0.5291 | 0.583362 | 0.101 | 0.086 | 1 | 8 | Gfer | 0.015 | Old |
| 0.529145 | 0.10459 | 0.246 | 0.281 | 1 | 8 | Nfu1 | -0.035 | Old |
| 0.529485 | 0.033386 | 0.123 | 0.143 | 1 | 8 | Ivd | -0.02 | Old |
| 0.529702 | 0.25295 | 0.48 | 0.494 | 1 | 8 | Derl1 | -0.014 | Old |
| 0.529793 | 0.050053 | 0.296 | 0.332 | 1 | 8 | Suz12 | -0.036 | Old |
| 0.529872 | 0.513612 | 0.017 | 0.01 | 1 | 8 | Pard3b | 0.007 | Old |
| 0.529872 | 0.303387 | 0.017 | 0.01 | 1 | 8 | Il17rd | 0.007 | Old |
| 0.529872 | 0.269496 | 0.017 | 0.01 | 1 | 8 | Ankrd24 | 0.007 | Old |
| 0.529872 | 0.111523 | 0.017 | 0.01 | 1 | 8 | Kcnc3 | 0.007 | Old |
| 0.530034 | 0.105314 | 0.659 | 0.67 | 1 | 8 | Pbrm1 | -0.011 | Old |
| 0.530072 | 0.619785 | 0.045 | 0.034 | 1 | 8 | Nme6 | 0.011 | Old |
| 0.530269 | 0.137055 | 0.145 | 0.169 | 1 | 8 | Hhex | -0.024 | Old |
| 0.530809 | 0.245243 | 0.291 | 0.273 | 1 | 8 | Ide | 0.018 | Old |
| 0.531234 | 0.262979 | 0.134 | 0.158 | 1 | 8 | E2f1 | -0.024 | Old |
| 0.531326 | 0.082805 | 0.179 | 0.208 | 1 | 8 | Sumf1 | -0.029 | Old |
| 0.531433 | 0.363329 | 0.246 | 0.229 | 1 | 8 | Ndufs1 | 0.017 | Old |
| 0.531438 | 0.450056 | 0.19 | 0.177 | 1 | 8 | Afdn | 0.013 | Old |
| 0.531747 | 0.334225 | 0.385 | 0.371 | 1 | 8 | Fam49a | 0.014 | Old |
| 0.531769 | 0.183188 | 0.017 | 0.01 | 1 | 8 | Klri1 | 0.007 | Old |
| 0.531769 | 0.089717 | 0.017 | 0.01 | 1 | 8 | Gm13205 | 0.007 | Old |
| 0.531769 | 0.088692 | 0.017 | 0.01 | 1 | 8 | Il18bp | 0.007 | Old |
| 0.531934 | 0.428773 | 0.184 | 0.161 | 1 | 8 | Clns1a | 0.023 | Old |
| 0.532557 | 0.032426 | 0.184 | 0.216 | 1 | 8 | Dhdh | -0.032 | Old |
| 0.532726 | 0.299934 | 0.05 | 0.039 | 1 | 8 | Nat2 | 0.011 | Old |
| 0.533113 | 0.087229 | 0.145 | 0.169 | 1 | 8 | Crnkl1 | -0.024 | Old |
| 0.533204 | 0.074548 | 0.123 | 0.104 | 1 | 8 | Bzw2 | 0.019 | Old |
| 0.53324 | 0.191356 | 0.19 | 0.21 | 1 | 8 | Map1s | -0.02 | Old |
| 0.533287 | 0.034596 | 0.413 | 0.439 | 1 | 8 | Soat1 | -0.026 | Old |
| 0.533671 | 0.153565 | 0.017 | 0.01 | 1 | 8 | Tmem150a | 0.007 | Old |
| 0.533671 | 0.09748 | 0.017 | 0.01 | 1 | 8 | Zfp493 | 0.007 | Old |
| 0.533673 | 0.293071 | 0.369 | 0.356 | 1 | 8 | Msrb1 | 0.013 | Old |
| 0.533807 | 0.325362 | 0.056 | 0.07 | 1 | 8 | Rcc1l | -0.014 | Old |
| 0.534352 | 0.279931 | 0.318 | 0.301 | 1 | 8 | Tspyl1 | 0.017 | Old |
| 0.534418 | 0.635293 | 0.056 | 0.044 | 1 | 8 | Tmem241 | 0.012 | Old |
| 0.534589 | 0.713446 | 0.207 | 0.197 | 1 | 8 | Fam133b | 0.01 | Old |
| 0.535067 | 0.283141 | 0.296 | 0.281 | 1 | 8 | Nek9 | 0.015 | Old |
| 0.535413 | 0.021212 | 0.207 | 0.231 | 1 | 8 | Retreg3 | -0.024 | Old |
| 0.535415 | 0.016354 | 0.24 | 0.265 | 1 | 8 | Fry | -0.025 | Old |
| 0.535453 | 0.32767 | 0.285 | 0.27 | 1 | 8 | Mga | 0.015 | Old |

| | | | | | | | | |
|---|---|---|---|---|---|---|---|---|
| 0.535616 | 0.083981 | 0.223 | 0.257 | 1 | 8 | Lilra6 | -0.034 | Old |
| 0.535814 | 0.026822 | 0.14 | 0.161 | 1 | 8 | Mkl1 | -0.021 | Old |
| 0.536201 | 0.443844 | 0.028 | 0.039 | 1 | 8 | Cep57l1 | -0.011 | Old |
| 0.536492 | 0.211897 | 0.246 | 0.223 | 1 | 8 | Grk5 | 0.023 | Old |
| 0.536612 | 0.170386 | 0.547 | 0.54 | 1 | 8 | Ube2r2 | 0.007 | Old |
| 0.536954 | 0.235212 | 0.223 | 0.197 | 1 | 8 | 3830406C1 | 0.026 | Old |
| 0.537429 | 0.405115 | 0.162 | 0.143 | 1 | 8 | Cacna1d | 0.019 | Old |
| 0.537565 | 0.045667 | 0.961 | 0.979 | 1 | 8 | Nr4a1 | -0.018 | Old |
| 0.537635 | 0.009742 | 0.011 | 0.018 | 1 | 8 | Slamf6 | -0.007 | Old |
| 0.537771 | 0.21979 | 0.145 | 0.127 | 1 | 8 | Nfrkb | 0.018 | Old |
| 0.5382 | 0.095753 | 0.536 | 0.569 | 1 | 8 | Lbr | -0.033 | Old |
| 0.538206 | 0.209949 | 0.559 | 0.574 | 1 | 8 | Gsk3b | -0.015 | Old |
| 0.538765 | 0.294654 | 0.335 | 0.322 | 1 | 8 | Lpar1 | 0.013 | Old |
| 0.538844 | 0.208056 | 0.24 | 0.221 | 1 | 8 | Ap1g1 | 0.019 | Old |
| 0.53888 | 0.750817 | 0.112 | 0.099 | 1 | 8 | Fhl3 | 0.013 | Old |
| 0.53899 | 0.14769 | 0.24 | 0.265 | 1 | 8 | Tmem230 | -0.025 | Old |
| 0.539365 | 0.491389 | 0.056 | 0.044 | 1 | 8 | Orai2 | 0.012 | Old |
| 0.53943 | 0.133907 | 0.693 | 0.686 | 1 | 8 | Atp5b | 0.007 | Old |
| 0.53965 | 0.294486 | 0.19 | 0.171 | 1 | 8 | Mafk | 0.019 | Old |
| 0.539802 | 0.347558 | 0.385 | 0.361 | 1 | 8 | Mrpl28 | 0.024 | Old |
| 0.539806 | 0.151288 | 0.101 | 0.083 | 1 | 8 | Eef1akmt2 | 0.018 | Old |
| 0.539806 | 0.069567 | 0.078 | 0.094 | 1 | 8 | Ptrh2 | -0.016 | Old |
| 0.540049 | 0.560287 | 0.05 | 0.039 | 1 | 8 | Zfp518b | 0.011 | Old |
| 0.540064 | 0.019449 | 0.324 | 0.351 | 1 | 8 | Acaa2 | -0.027 | Old |
| 0.540211 | 0.382012 | 0.207 | 0.249 | 1 | 8 | Txndc15 | -0.042 | Old |
| 0.540628 | 0.019673 | 0.654 | 0.725 | 1 | 8 | Nfe2l2 | -0.071 | Old |
| 0.540649 | 0.546322 | 0.14 | 0.122 | 1 | 8 | Ppp1r37 | 0.018 | Old |
| 0.541477 | 0.141266 | 0.151 | 0.177 | 1 | 8 | Fam160b2 | -0.026 | Old |
| 0.541555 | 0.307643 | 0.391 | 0.444 | 1 | 8 | Acp2 | -0.053 | Old |
| 0.541807 | 0.025632 | 0.302 | 0.335 | 1 | 8 | Dpp8 | -0.033 | Old |
| 0.542088 | 0.58842 | 0.089 | 0.075 | 1 | 8 | Taf1b | 0.014 | Old |
| 0.542569 | 0.476662 | 0.196 | 0.177 | 1 | 8 | Atg16l1 | 0.019 | Old |
| 0.542612 | 0.044266 | 0.469 | 0.525 | 1 | 8 | Tcea1 | -0.056 | Old |
| 0.543057 | 0.323855 | 0.302 | 0.281 | 1 | 8 | Mras | 0.021 | Old |
| 0.543129 | 0.02858 | 0.536 | 0.605 | 1 | 8 | Hnrnpul2 | -0.069 | Old |
| 0.543203 | 0.720666 | 0.05 | 0.039 | 1 | 8 | Txnrd3 | 0.011 | Old |
| 0.54321 | 0.257841 | 0.413 | 0.436 | 1 | 8 | Fndc3a | -0.023 | Old |
| 0.54334 | 0.241421 | 0.056 | 0.044 | 1 | 8 | Card6 | 0.012 | Old |
| 0.543429 | 0.01803 | 0.246 | 0.281 | 1 | 8 | Adprh | -0.035 | Old |
| 0.543666 | 0.215115 | 0.581 | 0.577 | 1 | 8 | Metap2 | 0.004 | Old |
| 0.544407 | 0.312617 | 0.011 | 0.018 | 1 | 8 | Samd10 | -0.007 | Old |
| 0.544407 | 0.034521 | 0.011 | 0.018 | 1 | 8 | Kdelc1 | -0.007 | Old |
| 0.544439 | 0.364323 | 0.089 | 0.075 | 1 | 8 | Primpol | 0.014 | Old |
| 0.545253 | 0.117222 | 0.397 | 0.444 | 1 | 8 | Wls | -0.047 | Old |
| 0.545842 | 0.472069 | 0.358 | 0.348 | 1 | 8 | Tug1 | 0.01 | Old |

| | | | | | | | | |
|---|---|---|---|---|---|---|---|---|
| 0.54587 | 0.108871 | 0.324 | 0.361 | 1 | 8 | Irf2bpl | -0.037 | Old |
| 0.546378 | 0.084682 | 0.307 | 0.348 | 1 | 8 | Alox5 | -0.041 | Old |
| 0.546405 | 0.053524 | 0.19 | 0.221 | 1 | 8 | Itpripl1 | -0.031 | Old |
| 0.546469 | 0.018273 | 0.385 | 0.358 | 1 | 8 | Rmnd5a | 0.027 | Old |
| 0.546515 | 0.540256 | 0.123 | 0.106 | 1 | 8 | Gtf2e1 | 0.017 | Old |
| 0.546693 | 0.411279 | 0.078 | 0.065 | 1 | 8 | Mtmr9 | 0.013 | Old |
| 0.546707 | 0.648775 | 0.073 | 0.06 | 1 | 8 | Haus3 | 0.013 | Old |
| 0.546765 | 0.035075 | 0.128 | 0.145 | 1 | 8 | Strip1 | -0.017 | Old |
| 0.546882 | 0.306447 | 0.307 | 0.294 | 1 | 8 | Pdpk1 | 0.013 | Old |
| 0.547207 | 0.141071 | 0.168 | 0.197 | 1 | 8 | Ufl1 | -0.029 | Old |
| 0.547512 | 0.283752 | 0.207 | 0.231 | 1 | 8 | Hars | -0.024 | Old |
| 0.547532 | 0.608771 | 0.117 | 0.143 | 1 | 8 | AI839979 | -0.026 | Old |
| 0.547579 | 0.447616 | 0.073 | 0.06 | 1 | 8 | Ercc6 | 0.013 | Old |
| 0.548293 | 0.746671 | 0.184 | 0.166 | 1 | 8 | Gbp2 | 0.018 | Old |
| 0.548407 | 0.006235 | 0.251 | 0.294 | 1 | 8 | Polr2g | -0.043 | Old |
| 0.548604 | 0.672258 | 0.134 | 0.117 | 1 | 8 | Fkbp5 | 0.017 | Old |
| 0.54868 | 0.327642 | 0.508 | 0.491 | 1 | 8 | Sp100 | 0.017 | Old |
| 0.548918 | 0.133722 | 0.374 | 0.351 | 1 | 8 | Stt3a | 0.023 | Old |
| 0.549308 | 0.599798 | 0.073 | 0.091 | 1 | 8 | Gk | -0.018 | Old |
| 0.549426 | 0.328007 | 0.14 | 0.164 | 1 | 8 | Rif1 | -0.024 | Old |
| 0.549513 | 0.739815 | 0.011 | 0.018 | 1 | 8 | Mansc1 | -0.007 | Old |
| 0.549513 | 0.57623 | 0.011 | 0.018 | 1 | 8 | Zfp341 | -0.007 | Old |
| 0.549523 | 0.116337 | 0.251 | 0.283 | 1 | 8 | Plaa | -0.032 | Old |
| 0.549631 | 0.002907 | 0.844 | 0.909 | 1 | 8 | Rac1 | -0.065 | Old |
| 0.549888 | 0.165329 | 0.358 | 0.332 | 1 | 8 | Rbm7 | 0.026 | Old |
| 0.55005 | 0.050592 | 0.291 | 0.319 | 1 | 8 | Crtc3 | -0.028 | Old |
| 0.550084 | 0.236482 | 0.073 | 0.088 | 1 | 8 | Cgrrf1 | -0.015 | Old |
| 0.550168 | 0.304624 | 0.117 | 0.101 | 1 | 8 | Pdgfa | 0.016 | Old |
| 0.550674 | 0.506803 | 0.128 | 0.114 | 1 | 8 | B3galnt2 | 0.014 | Old |
| 0.55124 | 0.455488 | 0.101 | 0.086 | 1 | 8 | Eri3 | 0.015 | Old |
| 0.551371 | 0.106392 | 0.547 | 0.532 | 1 | 8 | Papola | 0.015 | Old |
| 0.5515 | 0.117924 | 0.441 | 0.501 | 1 | 8 | Hopx | -0.06 | Old |
| 0.551674 | 0.560999 | 0.151 | 0.135 | 1 | 8 | Tm7sf3 | 0.016 | Old |
| 0.551834 | 0.191361 | 0.212 | 0.19 | 1 | 8 | Cluh | 0.022 | Old |
| 0.55195 | 0.431631 | 0.073 | 0.06 | 1 | 8 | Trmt10c | 0.013 | Old |
| 0.552166 | 0.514761 | 0.078 | 0.065 | 1 | 8 | Mtrf1l | 0.013 | Old |
| 0.552512 | 0.682098 | 0.106 | 0.13 | 1 | 8 | Ryk | -0.024 | Old |
| 0.552536 | 0.341466 | 0.346 | 0.335 | 1 | 8 | Ube2g2 | 0.011 | Old |
| 0.552917 | 0.104919 | 0.33 | 0.366 | 1 | 8 | Mcmbp | -0.036 | Old |
| 0.553023 | 0.231276 | 0.207 | 0.234 | 1 | 8 | Slc45a4 | -0.027 | Old |
| 0.553234 | 0.386706 | 0.173 | 0.156 | 1 | 8 | Med17 | 0.017 | Old |
| 0.553319 | 0.242127 | 0.201 | 0.179 | 1 | 8 | Atad1 | 0.022 | Old |
| 0.553419 | 0.043185 | 0.251 | 0.273 | 1 | 8 | Socs4 | -0.022 | Old |
| 0.553642 | 0.285412 | 0.067 | 0.055 | 1 | 8 | Rbm45 | 0.012 | Old |
| 0.554297 | 0.055251 | 0.034 | 0.044 | 1 | 8 | Gm37305 | -0.01 | Old |

| | | | | | | | | |
|---|---|---|---|---|---|---|---|---|
| 0.554451 | 0.073762 | 0.106 | 0.088 | 1 | 8 | Ankrd50 | 0.018 | Old |
| 0.55512 | 0.24835 | 0.073 | 0.088 | 1 | 8 | Noc3l | -0.015 | Old |
| 0.55605 | 0.222719 | 0.279 | 0.26 | 1 | 8 | Wars | 0.019 | Old |
| 0.556301 | 0.363382 | 0.223 | 0.203 | 1 | 8 | Wsb2 | 0.02 | Old |
| 0.556321 | 0.355387 | 0.106 | 0.091 | 1 | 8 | Cfap97 | 0.015 | Old |
| 0.556329 | 0.120159 | 0.207 | 0.234 | 1 | 8 | Kctd10 | -0.027 | Old |
| 0.556619 | 0.555149 | 0.084 | 0.07 | 1 | 8 | Snta1 | 0.014 | Old |
| 0.55675 | 0.17761 | 0.346 | 0.319 | 1 | 8 | Abcg1 | 0.027 | Old |
| 0.556911 | 0.470144 | 0.391 | 0.384 | 1 | 8 | Tial1 | 0.007 | Old |
| 0.556963 | 0.036412 | 0.14 | 0.161 | 1 | 8 | Avl9 | -0.021 | Old |
| 0.557062 | 0.871611 | 0.089 | 0.075 | 1 | 8 | Tmem18 | 0.014 | Old |
| 0.5574 | 0.661149 | 0.061 | 0.049 | 1 | 8 | Def8 | 0.012 | Old |
| 0.557541 | 1.342688 | 0.022 | 0.016 | 1 | 8 | Dagla | 0.006 | Old |
| 0.55781 | 0.026635 | 0.095 | 0.112 | 1 | 8 | Lrig2 | -0.017 | Old |
| 0.557858 | 0.103865 | 0.207 | 0.234 | 1 | 8 | Zfp740 | -0.027 | Old |
| 0.55797 | 0.417862 | 0.101 | 0.086 | 1 | 8 | Slc9a4 | 0.015 | Old |
| 0.558452 | 0.438585 | 0.145 | 0.127 | 1 | 8 | Mcm6 | 0.018 | Old |
| 0.558561 | 0.247076 | 0.358 | 0.343 | 1 | 8 | Ppp4r3a | 0.015 | Old |
| 0.559964 | 0.560741 | 0.156 | 0.143 | 1 | 8 | Tmco3 | 0.013 | Old |
| 0.56007 | 0.110393 | 0.095 | 0.081 | 1 | 8 | Spsb3 | 0.014 | Old |
| 0.56024 | 0.481972 | 0.089 | 0.075 | 1 | 8 | Fas | 0.014 | Old |
| 0.560392 | 0.201378 | 0.056 | 0.044 | 1 | 8 | Nprl3 | 0.012 | Old |
| 0.56055 | 0.02364 | 0.888 | 0.917 | 1 | 8 | Aplp2 | -0.029 | Old |
| 0.560637 | 0.381393 | 0.078 | 0.065 | 1 | 8 | Aifm1 | 0.013 | Old |
| 0.560718 | 0.095312 | 0.626 | 0.704 | 1 | 8 | Gns | -0.078 | Old |
| 0.560804 | 0.125656 | 0.464 | 0.436 | 1 | 8 | Mvp | 0.028 | Old |
| 0.561014 | 0.058474 | 0.38 | 0.434 | 1 | 8 | Srsf10 | -0.054 | Old |
| 0.561301 | 0.243052 | 0.251 | 0.234 | 1 | 8 | Pdhb | 0.017 | Old |
| 0.561424 | 0.224707 | 0.196 | 0.174 | 1 | 8 | Lgals8 | 0.022 | Old |
| 0.561633 | 0.113392 | 0.475 | 0.551 | 1 | 8 | Fam111a | -0.076 | Old |
| 0.562339 | 0.389521 | 0.078 | 0.065 | 1 | 8 | Pdk1 | 0.013 | Old |
| 0.562389 | 0.511915 | 0.095 | 0.081 | 1 | 8 | Stx18 | 0.014 | Old |
| 0.562446 | 0.639545 | 0.022 | 0.016 | 1 | 8 | Zgrf1 | 0.006 | Old |
| 0.562467 | 0.004004 | 0.145 | 0.164 | 1 | 8 | Cabin1 | -0.019 | Old |
| 0.562571 | 0.085357 | 0.972 | 0.971 | 1 | 8 | Atp2b1 | 0.001 | Old |
| 0.562589 | 0.206455 | 0.212 | 0.239 | 1 | 8 | Kdm6a | -0.027 | Old |
| 0.562952 | 0.207588 | 0.212 | 0.242 | 1 | 8 | Pwwp2a | -0.03 | Old |
| 0.564064 | 0.130452 | 0.134 | 0.156 | 1 | 8 | Sgsh | -0.022 | Old |
| 0.564337 | 0.169761 | 0.458 | 0.517 | 1 | 8 | Map2k3 | -0.059 | Old |
| 0.564398 | 0.266363 | 0.112 | 0.096 | 1 | 8 | Coq10a | 0.016 | Old |
| 0.564635 | 0.168896 | 0.514 | 0.494 | 1 | 8 | Wapl | 0.02 | Old |
| 0.56466 | 0.020559 | 0.05 | 0.062 | 1 | 8 | Ahdc1 | -0.012 | Old |
| 0.564947 | 0.266773 | 0.24 | 0.278 | 1 | 8 | S1pr1 | -0.038 | Old |
| 0.564969 | 0.395303 | 0.291 | 0.273 | 1 | 8 | Mfsd11 | 0.018 | Old |
| 0.565111 | 0.262261 | 0.106 | 0.091 | 1 | 8 | Ptk2 | 0.015 | Old |

| | | | | | | | | |
|---|---|---|---|---|---|---|---|---|
| 0.565405 | 0.030867 | 0.296 | 0.325 | 1 | 8 | Hipk3 | -0.029 | Old |
| 0.565537 | 0.219988 | 0.497 | 0.488 | 1 | 8 | Gmfb | 0.009 | Old |
| 0.565727 | 1.14068 | 0.022 | 0.016 | 1 | 8 | Vsig4 | 0.006 | Old |
| 0.565727 | 0.712938 | 0.022 | 0.016 | 1 | 8 | Ccdc51 | 0.006 | Old |
| 0.56587 | 0.256389 | 0.257 | 0.234 | 1 | 8 | Tbc1d13 | 0.023 | Old |
| 0.566061 | 0.100339 | 0.151 | 0.171 | 1 | 8 | Stx3 | -0.02 | Old |
| 0.566312 | 0.366 | 0.268 | 0.249 | 1 | 8 | Pih1d1 | 0.019 | Old |
| 0.566323 | 0.637235 | 0.179 | 0.161 | 1 | 8 | Mms19 | 0.018 | Old |
| 0.566403 | 0.564734 | 0.24 | 0.226 | 1 | 8 | Arid2 | 0.014 | Old |
| 0.566427 | 0.439832 | 0.201 | 0.184 | 1 | 8 | Matk | 0.017 | Old |
| 0.566551 | 0.310798 | 0.112 | 0.096 | 1 | 8 | 5031425F1 | 0.016 | Old |
| 0.566664 | 0.50705 | 0.14 | 0.125 | 1 | 8 | Synj2 | 0.015 | Old |
| 0.567041 | 0.256689 | 0.067 | 0.083 | 1 | 8 | Sco2 | -0.016 | Old |
| 0.567087 | 0.039447 | 0.704 | 0.797 | 1 | 8 | Snx3 | -0.093 | Old |
| 0.56732 | 0.197812 | 0.084 | 0.101 | 1 | 8 | Cnot11 | -0.017 | Old |
| 0.567372 | 1.242742 | 0.022 | 0.016 | 1 | 8 | Spon1 | 0.006 | Old |
| 0.567372 | 1.224247 | 0.022 | 0.016 | 1 | 8 | Cptp | 0.006 | Old |
| 0.567372 | 0.700358 | 0.022 | 0.016 | 1 | 8 | Pim2 | 0.006 | Old |
| 0.567372 | 0.455711 | 0.022 | 0.016 | 1 | 8 | Tmlhe | 0.006 | Old |
| 0.567372 | 0.348344 | 0.022 | 0.016 | 1 | 8 | Ctla2a | 0.006 | Old |
| 0.56741 | 0.038912 | 0.251 | 0.27 | 1 | 8 | Tsnax | -0.019 | Old |
| 0.567596 | 0.228342 | 0.408 | 0.395 | 1 | 8 | Clptm1l | 0.013 | Old |
| 0.567657 | 0.214675 | 0.447 | 0.439 | 1 | 8 | Rbm8a | 0.008 | Old |
| 0.567717 | 0.161059 | 0.201 | 0.234 | 1 | 8 | Taok2 | -0.033 | Old |
| 0.567957 | 0.060243 | 0.106 | 0.125 | 1 | 8 | Cep63 | -0.019 | Old |
| 0.568114 | 0.006569 | 0.296 | 0.322 | 1 | 8 | Aff1 | -0.026 | Old |
| 0.568217 | 0.01114 | 0.246 | 0.281 | 1 | 8 | Vrk2 | -0.035 | Old |
| 0.56831 | 0.11173 | 0.436 | 0.403 | 1 | 8 | Syf2 | 0.033 | Old |
| 0.568376 | 0.564576 | 0.251 | 0.236 | 1 | 8 | Clcn3 | 0.015 | Old |
| 0.568404 | 0.235034 | 0.419 | 0.4 | 1 | 8 | Plin3 | 0.019 | Old |
| 0.568493 | 0.205824 | 0.034 | 0.044 | 1 | 8 | Gm47754 | -0.01 | Old |
| 0.568747 | 0.031283 | 0.039 | 0.049 | 1 | 8 | Ccp110 | -0.01 | Old |
| 0.568949 | 0.300359 | 0.352 | 0.332 | 1 | 8 | Rsl1d1 | 0.02 | Old |
| 0.569018 | 0.600032 | 0.022 | 0.016 | 1 | 8 | 1110002L0 | 0.006 | Old |
| 0.569018 | 0.36678 | 0.022 | 0.016 | 1 | 8 | Xrcc2 | 0.006 | Old |
| 0.569021 | 0.382557 | 0.151 | 0.135 | 1 | 8 | Fbxo28 | 0.016 | Old |
| 0.569047 | 0.73486 | 0.106 | 0.094 | 1 | 8 | Wdr18 | 0.012 | Old |
| 0.569603 | 0.27349 | 0.073 | 0.06 | 1 | 8 | 0610009B2 | 0.013 | Old |
| 0.569801 | 0.282883 | 0.559 | 0.603 | 1 | 8 | Surf4 | -0.044 | Old |
| 0.569864 | 0.120895 | 0.067 | 0.081 | 1 | 8 | E130307A1 | -0.014 | Old |
| 0.569875 | 0.335266 | 0.184 | 0.166 | 1 | 8 | Irf3 | 0.018 | Old |
| 0.569969 | 0.187179 | 0.285 | 0.26 | 1 | 8 | Usp22 | 0.025 | Old |
| 0.570321 | 0.339586 | 0.162 | 0.14 | 1 | 8 | Snhg12 | 0.022 | Old |
| 0.570336 | 0.372689 | 0.251 | 0.234 | 1 | 8 | Mthfd2 | 0.017 | Old |
| 0.570418 | 0.048426 | 0.045 | 0.057 | 1 | 8 | Nup35 | -0.012 | Old |

| | | | | | | | | |
|---|---|---|---|---|---|---|---|---|
| 0.570667 | 0.276129 | 0.022 | 0.016 | 1 | 8 | Zfp458 | 0.006 | Old |
| 0.57081 | 0.075998 | 0.592 | 0.629 | 1 | 8 | Inpp5d | -0.037 | Old |
| 0.570885 | 0.176423 | 0.078 | 0.065 | 1 | 8 | Siglec1 | 0.013 | Old |
| 0.571216 | 0.10416 | 0.413 | 0.392 | 1 | 8 | Zmynd8 | 0.021 | Old |
| 0.571768 | 0.392802 | 0.145 | 0.13 | 1 | 8 | Mrpl16 | 0.015 | Old |
| 0.571873 | 0.002975 | 0.207 | 0.179 | 1 | 8 | Nop56 | 0.028 | Old |
| 0.571885 | 0.194771 | 0.168 | 0.148 | 1 | 8 | Klhl20 | 0.02 | Old |
| 0.571896 | 0.230376 | 0.196 | 0.177 | 1 | 8 | Prpf38a | 0.019 | Old |
| 0.572055 | 0.219114 | 0.162 | 0.145 | 1 | 8 | Trappc12 | 0.017 | Old |
| 0.572318 | 0.969631 | 0.022 | 0.016 | 1 | 8 | Proser3 | 0.006 | Old |
| 0.572318 | 0.890712 | 0.022 | 0.016 | 1 | 8 | Cxcl3 | 0.006 | Old |
| 0.572318 | 0.538787 | 0.022 | 0.016 | 1 | 8 | Map7 | 0.006 | Old |
| 0.572318 | 0.448898 | 0.022 | 0.016 | 1 | 8 | Tbc1d30 | 0.006 | Old |
| 0.572318 | 0.385813 | 0.022 | 0.016 | 1 | 8 | Adarb1 | 0.006 | Old |
| 0.572318 | 0.366658 | 0.022 | 0.016 | 1 | 8 | Mtbp | 0.006 | Old |
| 0.572318 | 0.340921 | 0.022 | 0.016 | 1 | 8 | Gm12589 | 0.006 | Old |
| 0.572318 | 0.227929 | 0.022 | 0.016 | 1 | 8 | Prpf40b | 0.006 | Old |
| 0.572318 | 0.149884 | 0.022 | 0.016 | 1 | 8 | Klk8 | 0.006 | Old |
| 0.572493 | 0.208784 | 0.095 | 0.081 | 1 | 8 | Aff3 | 0.014 | Old |
| 0.572709 | 0.466882 | 0.179 | 0.161 | 1 | 8 | Nudt5 | 0.018 | Old |
| 0.573249 | 0.675329 | 0.117 | 0.104 | 1 | 8 | Mecr | 0.013 | Old |
| 0.573867 | 0.413153 | 0.201 | 0.184 | 1 | 8 | Stk3 | 0.017 | Old |
| 0.573971 | 0.334623 | 0.022 | 0.016 | 1 | 8 | Pou5f1 | 0.006 | Old |
| 0.574475 | 0.142556 | 0.335 | 0.374 | 1 | 8 | Tap1 | -0.039 | Old |
| 0.574536 | 0.362203 | 0.497 | 0.491 | 1 | 8 | Hmgb2 | 0.006 | Old |
| 0.574894 | 0.231111 | 0.603 | 0.623 | 1 | 8 | Ubap2l | -0.02 | Old |
| 0.574963 | 0.108777 | 0.006 | 0.01 | 1 | 8 | Chek1 | -0.004 | Old |
| 0.575056 | 0.162809 | 0.358 | 0.338 | 1 | 8 | Mdm2 | 0.02 | Old |
| 0.575206 | 0.17429 | 0.089 | 0.106 | 1 | 8 | Patz1 | -0.017 | Old |
| 0.575441 | 0.132384 | 0.62 | 0.618 | 1 | 8 | Ifitm6 | 0.002 | Old |
| 0.575627 | 0.193276 | 0.022 | 0.016 | 1 | 8 | Cox6a2 | 0.006 | Old |
| 0.575627 | 0.179373 | 0.022 | 0.016 | 1 | 8 | Tmem201 | 0.006 | Old |
| 0.575627 | 0.169335 | 0.022 | 0.016 | 1 | 8 | Catip | 0.006 | Old |
| 0.575627 | 0.158578 | 0.022 | 0.016 | 1 | 8 | Tm7sf2 | 0.006 | Old |
| 0.575711 | 0.172441 | 0.564 | 0.577 | 1 | 8 | Plxnb2 | -0.013 | Old |
| 0.576034 | 0.002745 | 0.341 | 0.358 | 1 | 8 | Bub3 | -0.017 | Old |
| 0.57629 | 0.295227 | 0.022 | 0.031 | 1 | 8 | Xlr4a | -0.009 | Old |
| 0.576713 | 0.223297 | 0.05 | 0.062 | 1 | 8 | Rasgrf2 | -0.012 | Old |
| 0.577297 | 1.128463 | 0.006 | 0.01 | 1 | 8 | Gm12195 | -0.004 | Old |
| 0.577297 | 0.895857 | 0.006 | 0.01 | 1 | 8 | Cenpm | -0.004 | Old |
| 0.577297 | 0.403004 | 0.006 | 0.01 | 1 | 8 | Ndor1 | -0.004 | Old |
| 0.577297 | 0.383364 | 0.006 | 0.01 | 1 | 8 | Kctd14 | -0.004 | Old |
| 0.577297 | 0.340993 | 0.006 | 0.01 | 1 | 8 | Fancd2 | -0.004 | Old |
| 0.577297 | 0.317093 | 0.006 | 0.01 | 1 | 8 | B3gnt7 | -0.004 | Old |
| 0.577297 | 0.20059 | 0.006 | 0.01 | 1 | 8 | L3hypdh | -0.004 | Old |

| | | | | | | | | |
|---|---|---|---|---|---|---|---|---|
| 0.577297 | 0.081815 | 0.006 | 0.01 | 1 | 8 | Edn1 | -0.004 | Old |
| 0.577297 | 0.080823 | 0.006 | 0.01 | 1 | 8 | Polq | -0.004 | Old |
| 0.577297 | 0.045074 | 0.006 | 0.01 | 1 | 8 | AC151836. | -0.004 | Old |
| 0.577297 | 0.031093 | 0.006 | 0.01 | 1 | 8 | Ttc12 | -0.004 | Old |
| 0.577425 | 0.113807 | 0.173 | 0.195 | 1 | 8 | Nmt2 | -0.022 | Old |
| 0.577634 | 0.172804 | 0.073 | 0.06 | 1 | 8 | Tbc1d32 | 0.013 | Old |
| 0.577832 | 0.172716 | 0.162 | 0.145 | 1 | 8 | Faf1 | 0.017 | Old |
| 0.578146 | 0.648455 | 0.145 | 0.13 | 1 | 8 | Timm50 | 0.015 | Old |
| 0.578156 | 0.05548 | 0.313 | 0.335 | 1 | 8 | Gpatch8 | -0.022 | Old |
| 0.578428 | 0.129019 | 0.631 | 0.699 | 1 | 8 | Ninj1 | -0.068 | Old |
| 0.57853 | 0.155241 | 0.073 | 0.06 | 1 | 8 | Ddx28 | 0.013 | Old |
| 0.578701 | 0.51095 | 0.028 | 0.021 | 1 | 8 | Cdiptos | 0.007 | Old |
| 0.578945 | 0.10014 | 0.022 | 0.016 | 1 | 8 | C730034F0 | 0.006 | Old |
| 0.579095 | 0.065439 | 0.318 | 0.345 | 1 | 8 | Rabep1 | -0.027 | Old |
| 0.579512 | 0.202703 | 0.05 | 0.062 | 1 | 8 | Gm45902 | -0.012 | Old |
| 0.5803 | 0.07104 | 0.089 | 0.075 | 1 | 8 | Jrkl | 0.014 | Old |
| 0.580607 | 0.111096 | 0.022 | 0.016 | 1 | 8 | Lrrc32 | 0.006 | Old |
| 0.581131 | 0.162198 | 0.045 | 0.057 | 1 | 8 | Mccc2 | -0.012 | Old |
| 0.581415 | 0.426542 | 0.318 | 0.304 | 1 | 8 | Safb | 0.014 | Old |
| 0.581451 | 0.313349 | 0.101 | 0.086 | 1 | 8 | Alg14 | 0.015 | Old |
| 0.581569 | 0.498791 | 0.196 | 0.182 | 1 | 8 | Ddx1 | 0.014 | Old |
| 0.581685 | 0.24365 | 0.385 | 0.366 | 1 | 8 | Dhx9 | 0.019 | Old |
| 0.581943 | 0.195141 | 0.24 | 0.27 | 1 | 8 | Rsl24d1 | -0.03 | Old |
| 0.582119 | 0.013645 | 0.073 | 0.06 | 1 | 8 | Mul1 | 0.013 | Old |
| 0.582218 | 0.03391 | 0.061 | 0.075 | 1 | 8 | Ubxn8 | -0.014 | Old |
| 0.582272 | 0.143226 | 0.022 | 0.016 | 1 | 8 | Zbtb7c | 0.006 | Old |
| 0.582859 | 0.187982 | 0.112 | 0.099 | 1 | 8 | Nf2 | 0.013 | Old |
| 0.583129 | 0.058686 | 0.939 | 0.943 | 1 | 8 | Tpm3 | -0.004 | Old |
| 0.583197 | 0.383524 | 0.128 | 0.117 | 1 | 8 | Atn1 | 0.011 | Old |
| 0.583409 | 0.135395 | 0.341 | 0.374 | 1 | 8 | Pptc7 | -0.033 | Old |
| 0.58361 | 0.235619 | 0.369 | 0.366 | 1 | 8 | Igsf6 | 0.003 | Old |
| 0.583945 | 0.221075 | 0.43 | 0.426 | 1 | 8 | Rbm47 | 0.004 | Old |
| 0.584154 | 0.063569 | 0.413 | 0.4 | 1 | 8 | Rela | 0.013 | Old |
| 0.584191 | 0.264739 | 0.05 | 0.062 | 1 | 8 | Lrrc28 | -0.012 | Old |
| 0.58443 | 0.294176 | 0.385 | 0.426 | 1 | 8 | Rad21 | -0.041 | Old |
| 0.584558 | 0.924775 | 0.028 | 0.021 | 1 | 8 | Grk4 | 0.007 | Old |
| 0.584558 | 0.630607 | 0.028 | 0.021 | 1 | 8 | Gcat | 0.007 | Old |
| 0.584672 | 0.007534 | 0.587 | 0.631 | 1 | 8 | Znrf2 | -0.044 | Old |
| 0.585037 | 0.034053 | 0.207 | 0.229 | 1 | 8 | Utp14a | -0.022 | Old |
| 0.585114 | 0.185677 | 0.196 | 0.223 | 1 | 8 | Afg3l1 | -0.027 | Old |
| 0.585378 | 0.134044 | 0.078 | 0.094 | 1 | 8 | Dvl2 | -0.016 | Old |
| 0.585607 | 0.02694 | 0.022 | 0.016 | 1 | 8 | Cbwd1 | 0.006 | Old |
| 0.585722 | 0.063045 | 0.341 | 0.377 | 1 | 8 | Gxylt1 | -0.036 | Old |
| 0.585781 | 0.177356 | 0.162 | 0.145 | 1 | 8 | Ahsa2 | 0.017 | Old |
| 0.586026 | 1.472141 | 0.028 | 0.021 | 1 | 8 | Dffb | 0.007 | Old |

| | | | | | | | | |
|---|---|---|---|---|---|---|---|---|
| 0.586196 | 0.330046 | 0.073 | 0.088 | 1 | 8 | Gprasp1 | -0.015 | Old |
| 0.586678 | 0.231733 | 0.603 | 0.621 | 1 | 8 | Chmp4b | -0.018 | Old |
| 0.586822 | 0.535233 | 0.251 | 0.239 | 1 | 8 | Mepce | 0.012 | Old |
| 0.586877 | 0.115994 | 0.196 | 0.179 | 1 | 8 | Fibp | 0.017 | Old |
| 0.587309 | 0.082179 | 0.453 | 0.488 | 1 | 8 | Cmtm3 | -0.035 | Old |
| 0.587455 | 0.141788 | 0.128 | 0.151 | 1 | 8 | Spg20 | -0.023 | Old |
| 0.587496 | 0.885095 | 0.028 | 0.021 | 1 | 8 | Gm21781 | 0.007 | Old |
| 0.587892 | 0.242209 | 0.128 | 0.145 | 1 | 8 | Gclm | -0.017 | Old |
| 0.587919 | 0.304651 | 0.101 | 0.119 | 1 | 8 | Msantd4 | -0.018 | Old |
| 0.587935 | 0.035521 | 0.296 | 0.325 | 1 | 8 | Anxa11 | -0.029 | Old |
| 0.587966 | 0.009846 | 0.43 | 0.486 | 1 | 8 | Vdac1 | -0.056 | Old |
| 0.588029 | 0.204185 | 0.447 | 0.431 | 1 | 8 | Dlst | 0.016 | Old |
| 0.588376 | 0.24605 | 0.497 | 0.496 | 1 | 8 | Ap2b1 | 0.001 | Old |
| 0.588568 | 0.023379 | 0.754 | 0.761 | 1 | 8 | Selenok | -0.007 | Old |
| 0.588753 | 0.174997 | 0.285 | 0.265 | 1 | 8 | Lrrc59 | 0.02 | Old |
| 0.588968 | 0.761267 | 0.028 | 0.021 | 1 | 8 | Timp1 | 0.007 | Old |
| 0.589359 | 0.075274 | 0.19 | 0.216 | 1 | 8 | Trappc1 | -0.026 | Old |
| 0.589686 | 0.122659 | 0.095 | 0.109 | 1 | 8 | Irf2bp1 | -0.014 | Old |
| 0.589897 | 0.005282 | 0.24 | 0.26 | 1 | 8 | Zfp451 | -0.02 | Old |
| 0.589993 | 0.34677 | 0.564 | 0.577 | 1 | 8 | Pros1 | -0.013 | Old |
| 0.590182 | 0.240606 | 0.609 | 0.621 | 1 | 8 | Cct8 | -0.012 | Old |
| 0.590265 | 0.321799 | 0.268 | 0.252 | 1 | 8 | Etv5 | 0.016 | Old |
| 0.590418 | 0.225202 | 0.358 | 0.348 | 1 | 8 | Stxbp2 | 0.01 | Old |
| 0.590515 | 0.373329 | 0.067 | 0.081 | 1 | 8 | Wrap73 | -0.014 | Old |
| 0.590809 | 0.767746 | 0.123 | 0.112 | 1 | 8 | Ppp2r3c | 0.011 | Old |
| 0.590866 | 0.154998 | 0.503 | 0.501 | 1 | 8 | Esyt1 | 0.002 | Old |
| 0.590986 | 0.223945 | 0.073 | 0.088 | 1 | 8 | Map3k20 | -0.015 | Old |
| 0.591607 | 0.112292 | 0.302 | 0.327 | 1 | 8 | Tle4 | -0.025 | Old |
| 0.591625 | 0.414183 | 0.128 | 0.114 | 1 | 8 | St7l | 0.014 | Old |
| 0.591801 | 0.008158 | 0.637 | 0.691 | 1 | 8 | Anp32a | -0.054 | Old |
| 0.592929 | 0.159667 | 0.24 | 0.268 | 1 | 8 | Ms4a6d | -0.028 | Old |
| 0.592973 | 0.191186 | 0.151 | 0.135 | 1 | 8 | Timm22 | 0.016 | Old |
| 0.593147 | 0.46248 | 0.218 | 0.203 | 1 | 8 | Ddx60 | 0.015 | Old |
| 0.593199 | 0.15107 | 0.084 | 0.07 | 1 | 8 | Ints7 | 0.014 | Old |
| 0.5932 | 0.318331 | 0.235 | 0.216 | 1 | 8 | Zc3h4 | 0.019 | Old |
| 0.593393 | 0.055587 | 0.028 | 0.021 | 1 | 8 | Paqr8 | 0.007 | Old |
| 0.59356 | 0.068877 | 0.709 | 0.735 | 1 | 8 | Ear2 | -0.026 | Old |
| 0.593725 | 0.078171 | 0.246 | 0.226 | 1 | 8 | Zbtb18 | 0.02 | Old |
| 0.593875 | 0.248765 | 0.039 | 0.049 | 1 | 8 | Phka1 | -0.01 | Old |
| 0.593898 | 0.795724 | 0.056 | 0.047 | 1 | 8 | Gstt2 | 0.009 | Old |
| 0.593923 | 0.240297 | 0.078 | 0.094 | 1 | 8 | Tmem29 | -0.016 | Old |
| 0.594164 | 0.380953 | 0.385 | 0.382 | 1 | 8 | Psmc1 | 0.003 | Old |
| 0.595148 | 0.135905 | 0.296 | 0.325 | 1 | 8 | Abcd1 | -0.029 | Old |
| 0.595264 | 0.508705 | 0.179 | 0.166 | 1 | 8 | Usp5 | 0.013 | Old |
| 0.595312 | 0.284302 | 0.43 | 0.434 | 1 | 8 | Pdia4 | -0.004 | Old |

| | | | | | | | | |
|---|---|---|---|---|---|---|---|---|
| 0.59557 | 0.018946 | 0.274 | 0.291 | 1 | 8 | Gripap1 | -0.017 | Old |
| 0.595676 | 0.166233 | 0.106 | 0.094 | 1 | 8 | Sh2b1 | 0.012 | Old |
| 0.59575 | 0.228671 | 0.313 | 0.345 | 1 | 8 | Nampt | -0.032 | Old |
| 0.59586 | 0.179535 | 0.503 | 0.504 | 1 | 8 | Arl8a | -0.001 | Old |
| 0.596317 | 0.299867 | 0.184 | 0.169 | 1 | 8 | Sfxn1 | 0.015 | Old |
| 0.596352 | 0.175101 | 0.028 | 0.021 | 1 | 8 | Zfp799 | 0.007 | Old |
| 0.596374 | 0.055068 | 0.089 | 0.106 | 1 | 8 | Aktip | -0.017 | Old |
| 0.59653 | 0.080524 | 0.095 | 0.081 | 1 | 8 | Tex30 | 0.014 | Old |
| 0.596579 | 0.061465 | 0.084 | 0.07 | 1 | 8 | Dusp12 | 0.014 | Old |
| 0.596838 | 0.076544 | 0.603 | 0.662 | 1 | 8 | Spop | -0.059 | Old |
| 0.597351 | 0.492252 | 0.067 | 0.057 | 1 | 8 | Rabif | 0.01 | Old |
| 0.597435 | 0.223625 | 0.151 | 0.135 | 1 | 8 | Ak6 | 0.016 | Old |
| 0.597549 | 1.375336 | 0.034 | 0.026 | 1 | 8 | Fitm2 | 0.008 | Old |
| 0.597631 | 0.038161 | 0.341 | 0.356 | 1 | 8 | Insig1 | -0.015 | Old |
| 0.597656 | 0.685047 | 0.073 | 0.062 | 1 | 8 | Saal1 | 0.011 | Old |
| 0.597748 | 0.539974 | 0.134 | 0.161 | 1 | 8 | Nup214 | -0.027 | Old |
| 0.597834 | 0.38918 | 0.028 | 0.021 | 1 | 8 | Rccd1 | 0.007 | Old |
| 0.597834 | 0.297821 | 0.028 | 0.021 | 1 | 8 | Tmem81 | 0.007 | Old |
| 0.597834 | 0.182297 | 0.028 | 0.021 | 1 | 8 | Tgm1 | 0.007 | Old |
| 0.597845 | 1.004287 | 0.039 | 0.031 | 1 | 8 | Zfp280b | 0.008 | Old |
| 0.5982 | 0.032172 | 0.525 | 0.571 | 1 | 8 | Smchd1 | -0.046 | Old |
| 0.598205 | 0.219704 | 0.156 | 0.177 | 1 | 8 | Pgrmc2 | -0.021 | Old |
| 0.598226 | 0.186439 | 0.089 | 0.075 | 1 | 8 | Tmem70 | 0.014 | Old |
| 0.598228 | 0.173645 | 0.313 | 0.288 | 1 | 8 | Gba | 0.025 | Old |
| 0.5984 | 0.233139 | 0.324 | 0.371 | 1 | 8 | Ccnt2 | -0.047 | Old |
| 0.598587 | 0.261378 | 0.179 | 0.161 | 1 | 8 | Ecd | 0.018 | Old |
| 0.598636 | 0.186423 | 0.089 | 0.075 | 1 | 8 | Zfp280c | 0.014 | Old |
| 0.598672 | 0.481367 | 0.162 | 0.148 | 1 | 8 | 1600012H( | 0.014 | Old |
| 0.598844 | 0.103228 | 0.24 | 0.275 | 1 | 8 | Lin7c | -0.035 | Old |
| 0.599131 | 0.017238 | 0.352 | 0.382 | 1 | 8 | Palld | -0.03 | Old |
| 0.599219 | 0.239848 | 0.38 | 0.364 | 1 | 8 | Mfsd5 | 0.016 | Old |
| 0.599238 | 0.34993 | 0.469 | 0.478 | 1 | 8 | Hnrnpdl | -0.009 | Old |
| 0.599317 | 0.804818 | 0.028 | 0.021 | 1 | 8 | Nup43 | 0.007 | Old |
| 0.599317 | 0.446196 | 0.028 | 0.021 | 1 | 8 | Sphk1 | 0.007 | Old |
| 0.599708 | 0.051851 | 0.235 | 0.262 | 1 | 8 | Commd6 | -0.027 | Old |
| 0.599813 | 0.18714 | 0.525 | 0.592 | 1 | 8 | Pik3cd | -0.067 | Old |
| 0.600212 | 0.017689 | 0.726 | 0.758 | 1 | 8 | Ptpre | -0.032 | Old |
| 0.600232 | 0.814908 | 0.034 | 0.026 | 1 | 8 | Akr1b8 | 0.008 | Old |
| 0.600411 | 0.06893 | 0.101 | 0.117 | 1 | 8 | Golph3l | -0.016 | Old |
| 0.600428 | 0.278296 | 0.62 | 0.681 | 1 | 8 | Txn2 | -0.061 | Old |
| 0.600476 | 0.214189 | 0.464 | 0.47 | 1 | 8 | Stx4a | -0.006 | Old |
| 0.600641 | 0.031457 | 0.117 | 0.135 | 1 | 8 | Rtl5 | -0.018 | Old |
| 0.600804 | 0.222779 | 0.223 | 0.208 | 1 | 8 | Ctbp2 | 0.015 | Old |
| 0.601123 | 0.002712 | 0.229 | 0.249 | 1 | 8 | Trnau1ap | -0.02 | Old |
| 0.601301 | 0.356718 | 0.218 | 0.203 | 1 | 8 | Adh5 | 0.015 | Old |

| | | | | | | | | |
|---|---|---|---|---|---|---|---|---|
| 0.601616 | 0.35071 | 0.078 | 0.068 | 1 | 8 | Cd79b | 0.01 | Old |
| 0.601757 | 0.355743 | 0.168 | 0.153 | 1 | 8 | Bahd1 | 0.015 | Old |
| 0.601788 | 0.306158 | 0.112 | 0.099 | 1 | 8 | Slc35c1 | 0.013 | Old |
| 0.602004 | 0.237327 | 0.615 | 0.631 | 1 | 8 | Tmed7 | -0.016 | Old |
| 0.602289 | 0.250985 | 0.028 | 0.021 | 1 | 8 | Car9 | 0.007 | Old |
| 0.602362 | 0.071752 | 0.039 | 0.049 | 1 | 8 | Lonrf1 | -0.01 | Old |
| 0.602787 | 0.99461 | 0.039 | 0.031 | 1 | 8 | Adck2 | 0.008 | Old |
| 0.603158 | 0.105953 | 0.089 | 0.075 | 1 | 8 | D230025D: | 0.014 | Old |
| 0.603218 | 0.085595 | 0.128 | 0.148 | 1 | 8 | Slc4a1ap | -0.02 | Old |
| 0.603255 | 0.705424 | 0.112 | 0.099 | 1 | 8 | Fasn | 0.013 | Old |
| 0.603385 | 0.068876 | 0.358 | 0.392 | 1 | 8 | Usp19 | -0.034 | Old |
| 0.603446 | 0.018613 | 0.145 | 0.166 | 1 | 8 | Ddx27 | -0.021 | Old |
| 0.603527 | 0.050726 | 0.134 | 0.151 | 1 | 8 | Mtmr1 | -0.017 | Old |
| 0.603778 | 0.445547 | 0.028 | 0.021 | 1 | 8 | Pdk2 | 0.007 | Old |
| 0.603789 | 0.01486 | 0.061 | 0.075 | 1 | 8 | Lrp5 | -0.014 | Old |
| 0.603865 | 0.036694 | 0.525 | 0.582 | 1 | 8 | Lrrc58 | -0.057 | Old |
| 0.603985 | 0.367246 | 0.145 | 0.13 | 1 | 8 | Coq2 | 0.015 | Old |
| 0.604186 | 0.154424 | 0.33 | 0.314 | 1 | 8 | Etv6 | 0.016 | Old |
| 0.604265 | 0.392903 | 0.034 | 0.026 | 1 | 8 | Gja1 | 0.008 | Old |
| 0.604297 | 0.189512 | 0.101 | 0.088 | 1 | 8 | Prcc | 0.013 | Old |
| 0.604714 | 0.139182 | 0.095 | 0.112 | 1 | 8 | Mkrn2 | -0.017 | Old |
| 0.604719 | 0.065024 | 0.084 | 0.099 | 1 | 8 | Ndufaf3 | -0.015 | Old |
| 0.604933 | 0.204292 | 0.732 | 0.784 | 1 | 8 | Trps1 | -0.052 | Old |
| 0.605133 | 0.201778 | 0.128 | 0.148 | 1 | 8 | Cdc42bpb | -0.02 | Old |
| 0.605232 | 0.264409 | 0.285 | 0.265 | 1 | 8 | Rufy3 | 0.02 | Old |
| 0.605284 | 0.139379 | 0.117 | 0.135 | 1 | 8 | 0610037L1 | -0.018 | Old |
| 0.60543 | 0.608486 | 0.106 | 0.094 | 1 | 8 | Txnl4a | 0.012 | Old |
| 0.605613 | 0.754073 | 0.034 | 0.026 | 1 | 8 | Pgap3 | 0.008 | Old |
| 0.605847 | 0.030701 | 0.57 | 0.6 | 1 | 8 | Pebp1 | -0.03 | Old |
| 0.605917 | 0.042586 | 0.084 | 0.07 | 1 | 8 | Aptx | 0.014 | Old |
| 0.605986 | 0.060316 | 0.905 | 0.945 | 1 | 8 | Pitpna | -0.04 | Old |
| 0.606013 | 0.12231 | 0.19 | 0.174 | 1 | 8 | Ftsj3 | 0.016 | Old |
| 0.606108 | 0.14879 | 0.106 | 0.119 | 1 | 8 | Ifih1 | -0.013 | Old |
| 0.606375 | 0.180264 | 0.017 | 0.023 | 1 | 8 | Tnfrsf26 | -0.006 | Old |
| 0.607067 | 0.046816 | 0.156 | 0.177 | 1 | 8 | Irak3 | -0.021 | Old |
| 0.607432 | 0.045611 | 0.14 | 0.156 | 1 | 8 | Creb3 | -0.016 | Old |
| 0.607569 | 0.280695 | 0.302 | 0.291 | 1 | 8 | Lars2 | 0.011 | Old |
| 0.607592 | 0.106094 | 0.374 | 0.361 | 1 | 8 | Gnl3 | 0.013 | Old |
| 0.607644 | 0.354695 | 0.207 | 0.192 | 1 | 8 | Tmem43 | 0.015 | Old |
| 0.607928 | 0.416534 | 0.017 | 0.023 | 1 | 8 | Nek8 | -0.006 | Old |
| 0.608253 | 0.289999 | 0.028 | 0.021 | 1 | 8 | Tmem143 | 0.007 | Old |
| 0.60831 | 0.027227 | 0.156 | 0.138 | 1 | 8 | Smg6 | 0.018 | Old |
| 0.608968 | 0.305881 | 0.067 | 0.081 | 1 | 8 | Nfkbil1 | -0.014 | Old |
| 0.609087 | 0.704815 | 0.095 | 0.083 | 1 | 8 | Phtf2 | 0.012 | Old |
| 0.609168 | 0.087628 | 0.335 | 0.379 | 1 | 8 | Nhsl2 | -0.044 | Old |

| | | | | | | | | |
|---|---|---|---|---|---|---|---|---|
| 0.609662 | 0.379858 | 0.034 | 0.026 | 1 | 8 | Ift57 | 0.008 | Old |
| 0.609795 | 0.075446 | 0.324 | 0.361 | 1 | 8 | Agpat4 | -0.037 | Old |
| 0.610093 | 0.128563 | 0.654 | 0.688 | 1 | 8 | Zfp706 | -0.034 | Old |
| 0.610322 | 0.222757 | 0.43 | 0.442 | 1 | 8 | Kcmf1 | -0.012 | Old |
| 0.610333 | 0.303397 | 0.101 | 0.117 | 1 | 8 | Cd99l2 | -0.016 | Old |
| 0.610433 | 0.154187 | 0.112 | 0.096 | 1 | 8 | Ppwd1 | 0.016 | Old |
| 0.611015 | 0.588238 | 0.034 | 0.026 | 1 | 8 | Tns2 | 0.008 | Old |
| 0.61103 | 0.153065 | 0.207 | 0.231 | 1 | 8 | Nab2 | -0.024 | Old |
| 0.612351 | 0.218055 | 0.084 | 0.073 | 1 | 8 | Usp31 | 0.011 | Old |
| 0.612498 | 0.247839 | 0.436 | 0.429 | 1 | 8 | Ppp4c | 0.007 | Old |
| 0.612724 | 0.410862 | 0.028 | 0.036 | 1 | 8 | Btbd19 | -0.008 | Old |
| 0.612761 | 0.181191 | 0.073 | 0.088 | 1 | 8 | Zfp626 | -0.015 | Old |
| 0.612844 | 0.005758 | 0.978 | 0.995 | 1 | 8 | Kctd12 | -0.017 | Old |
| 0.612878 | 0.40463 | 0.218 | 0.205 | 1 | 8 | Setd7 | 0.013 | Old |
| 0.612909 | 0.445218 | 0.603 | 0.621 | 1 | 8 | Rassf2 | -0.018 | Old |
| 0.613059 | 0.012918 | 0.179 | 0.197 | 1 | 8 | Lrch3 | -0.018 | Old |
| 0.613071 | 0.551254 | 0.095 | 0.083 | 1 | 8 | Dus1l | 0.012 | Old |
| 0.61312 | 0.799533 | 0.123 | 0.112 | 1 | 8 | Med4 | 0.011 | Old |
| 0.613384 | 0.227928 | 0.212 | 0.187 | 1 | 8 | Slu7 | 0.025 | Old |
| 0.613405 | 0.153833 | 0.324 | 0.366 | 1 | 8 | Luzp1 | -0.042 | Old |
| 0.613463 | 0.254986 | 0.151 | 0.138 | 1 | 8 | Rab28 | 0.013 | Old |
| 0.614295 | 0.385332 | 0.045 | 0.036 | 1 | 8 | Lrrc45 | 0.009 | Old |
| 0.614394 | 0.356034 | 0.112 | 0.132 | 1 | 8 | Daxx | -0.02 | Old |
| 0.614913 | 0.03759 | 0.341 | 0.319 | 1 | 8 | Snx4 | 0.022 | Old |
| 0.615221 | 0.220373 | 0.453 | 0.475 | 1 | 8 | Trpc4ap | -0.022 | Old |
| 0.615284 | 0.026391 | 0.268 | 0.296 | 1 | 8 | Tbc1d23 | -0.028 | Old |
| 0.615378 | 0.083637 | 0.966 | 0.987 | 1 | 8 | Calm1 | -0.021 | Old |
| 0.615459 | 0.425428 | 0.045 | 0.036 | 1 | 8 | Nprl2 | 0.009 | Old |
| 0.616197 | 0.028632 | 0.106 | 0.122 | 1 | 8 | Cars | -0.016 | Old |
| 0.616439 | 0.477587 | 0.034 | 0.026 | 1 | 8 | Crocc | 0.008 | Old |
| 0.616439 | 0.444961 | 0.034 | 0.026 | 1 | 8 | Gm16638 | 0.008 | Old |
| 0.616498 | 0.458067 | 0.212 | 0.2 | 1 | 8 | Naa20 | 0.012 | Old |
| 0.616502 | 0.084225 | 0.447 | 0.418 | 1 | 8 | Riok3 | 0.029 | Old |
| 0.616638 | 0.683313 | 0.128 | 0.117 | 1 | 8 | Ikbkap | 0.011 | Old |
| 0.616729 | 0.185042 | 0.128 | 0.145 | 1 | 8 | Scyl2 | -0.017 | Old |
| 0.617278 | 0.970122 | 0.05 | 0.042 | 1 | 8 | Sccpdh | 0.008 | Old |
| 0.61772 | 0.667773 | 0.039 | 0.031 | 1 | 8 | Gm26756 | 0.008 | Old |
| 0.617788 | 0.976089 | 0.045 | 0.036 | 1 | 8 | Mapre3 | 0.009 | Old |
| 0.617798 | 0.080717 | 0.034 | 0.026 | 1 | 8 | Gorab | 0.008 | Old |
| 0.617866 | 0.185834 | 0.078 | 0.091 | 1 | 8 | Pik3r2 | -0.013 | Old |
| 0.618258 | 0.477927 | 0.101 | 0.088 | 1 | 8 | Psat1 | 0.013 | Old |
| 0.618305 | 0.223216 | 0.335 | 0.384 | 1 | 8 | Qdpr | -0.049 | Old |
| 0.618372 | 0.526354 | 0.045 | 0.036 | 1 | 8 | Esr1 | 0.009 | Old |
| 0.618469 | 0.040855 | 0.257 | 0.281 | 1 | 8 | Pbx3 | -0.024 | Old |
| 0.618627 | 0.61191 | 0.056 | 0.047 | 1 | 8 | Mllt11 | 0.009 | Old |

| | | | | | | | | |
|---|---|---|---|---|---|---|---|---|
| 0.618711 | 0.308518 | 0.24 | 0.223 | 1 | 8 | Ehmt1 | 0.017 | Old |
| 0.618955 | 0.841582 | 0.045 | 0.036 | 1 | 8 | Nes | 0.009 | Old |
| 0.618972 | 0.910405 | 0.039 | 0.031 | 1 | 8 | Abhd14a | 0.008 | Old |
| 0.619028 | 0.035277 | 0.056 | 0.068 | 1 | 8 | Trip10 | -0.012 | Old |
| 0.619315 | 0.12591 | 0.207 | 0.19 | 1 | 8 | Tbc1d22a | 0.017 | Old |
| 0.619318 | 0.245761 | 0.201 | 0.187 | 1 | 8 | Tmem154 | 0.014 | Old |
| 0.620181 | 0.471802 | 0.291 | 0.291 | 1 | 8 | Cyb5r3 | 0 | Old |
| 0.620412 | 0.246785 | 0.017 | 0.023 | 1 | 8 | Chchd10 | -0.006 | Old |
| 0.620412 | 0.11901 | 0.017 | 0.023 | 1 | 8 | Tmem41a | -0.006 | Old |
| 0.62086 | 0.014976 | 0.268 | 0.286 | 1 | 8 | Pi4ka | -0.018 | Old |
| 0.620959 | 0.311792 | 0.212 | 0.192 | 1 | 8 | Pop5 | 0.02 | Old |
| 0.621479 | 0.032076 | 0.028 | 0.036 | 1 | 8 | Mettl2 | -0.008 | Old |
| 0.621591 | 0.018173 | 0.073 | 0.086 | 1 | 8 | Coprs | -0.013 | Old |
| 0.62174 | 0.570108 | 0.061 | 0.075 | 1 | 8 | Slc35e2 | -0.014 | Old |
| 0.621802 | 0.179448 | 0.056 | 0.068 | 1 | 8 | Pdik1l | -0.012 | Old |
| 0.621883 | 0.165723 | 0.034 | 0.026 | 1 | 8 | Ube2t | 0.008 | Old |
| 0.621883 | 0.12132 | 0.034 | 0.026 | 1 | 8 | Gas8 | 0.008 | Old |
| 0.621949 | 0.333967 | 0.156 | 0.179 | 1 | 8 | Mthfd1 | -0.023 | Old |
| 0.622277 | 0.142901 | 0.38 | 0.416 | 1 | 8 | Hmox2 | -0.036 | Old |
| 0.622457 | 0.185559 | 0.207 | 0.184 | 1 | 8 | Tssc4 | 0.023 | Old |
| 0.622734 | 0.572066 | 0.039 | 0.031 | 1 | 8 | Rdh13 | 0.008 | Old |
| 0.622743 | 0.007785 | 0.179 | 0.197 | 1 | 8 | Tmem131l | -0.018 | Old |
| 0.622818 | 0.245881 | 0.291 | 0.273 | 1 | 8 | Insr | 0.018 | Old |
| 0.622831 | 0.176984 | 0.089 | 0.078 | 1 | 8 | 2410004B1 | 0.011 | Old |
| 0.623034 | 0.697513 | 0.073 | 0.062 | 1 | 8 | Tmem126b | 0.011 | Old |
| 0.623498 | 0.287007 | 0.575 | 0.618 | 1 | 8 | Aamp | -0.043 | Old |
| 0.623725 | 0.214759 | 0.101 | 0.088 | 1 | 8 | Acaca | 0.013 | Old |
| 0.62399 | 0.43991 | 0.039 | 0.031 | 1 | 8 | Usp18 | 0.008 | Old |
| 0.624386 | 0.795783 | 0.078 | 0.068 | 1 | 8 | Scfd2 | 0.01 | Old |
| 0.624627 | 0.206724 | 0.24 | 0.221 | 1 | 8 | Dld | 0.019 | Old |
| 0.624913 | 0.271275 | 0.313 | 0.335 | 1 | 8 | Smarcc2 | -0.022 | Old |
| 0.624973 | 0.528459 | 0.05 | 0.042 | 1 | 8 | Clec4d | 0.008 | Old |
| 0.625021 | 0.128049 | 0.123 | 0.14 | 1 | 8 | Slc35e4 | -0.017 | Old |
| 0.625122 | 0.543673 | 0.017 | 0.023 | 1 | 8 | Dlg3 | -0.006 | Old |
| 0.625122 | 0.033144 | 0.017 | 0.023 | 1 | 8 | Zfp60 | -0.006 | Old |
| 0.625291 | 0.176647 | 0.19 | 0.174 | 1 | 8 | Ppp2cb | 0.016 | Old |
| 0.625788 | 0.344577 | 0.184 | 0.169 | 1 | 8 | Trpv4 | 0.015 | Old |
| 0.625856 | 0.11112 | 0.335 | 0.325 | 1 | 8 | Dr1 | 0.01 | Old |
| 0.625938 | 0.038865 | 0.346 | 0.374 | 1 | 8 | Dram2 | -0.028 | Old |
| 0.62619 | 0.782657 | 0.173 | 0.161 | 1 | 8 | Necap1 | 0.012 | Old |
| 0.626467 | 0.322891 | 0.101 | 0.088 | 1 | 8 | Armc7 | 0.013 | Old |
| 0.626505 | 0.13926 | 0.028 | 0.036 | 1 | 8 | Cetn4 | -0.008 | Old |
| 0.626668 | 0.231612 | 0.162 | 0.19 | 1 | 8 | Tnpo2 | -0.028 | Old |
| 0.626696 | 0.192441 | 0.017 | 0.023 | 1 | 8 | Sh3rf1 | -0.006 | Old |
| 0.626892 | 0.117083 | 0.112 | 0.096 | 1 | 8 | Elmod2 | 0.016 | Old |

| | | | | | | | | |
|---|---|---|---|---|---|---|---|---|
| 0.626901 | 0.271582 | 0.162 | 0.145 | 1 | 8 | Adap1 | 0.017 | Old |
| 0.627037 | 0.405741 | 0.078 | 0.068 | 1 | 8 | Hars2 | 0.01 | Old |
| 0.627162 | 0.657834 | 0.201 | 0.197 | 1 | 8 | Mkl2 | 0.004 | Old |
| 0.627167 | 0.098353 | 0.24 | 0.273 | 1 | 8 | Gng12 | -0.033 | Old |
| 0.627262 | 0.306429 | 0.207 | 0.19 | 1 | 8 | Csrp1 | 0.017 | Old |
| 0.627431 | 0.201042 | 0.603 | 0.621 | 1 | 8 | Ube2l3 | -0.018 | Old |
| 0.627439 | 0.183975 | 0.564 | 0.574 | 1 | 8 | Susd6 | -0.01 | Old |
| 0.628029 | 0.060814 | 0.045 | 0.055 | 1 | 8 | Wdcp | -0.01 | Old |
| 0.628143 | 0.054014 | 0.447 | 0.465 | 1 | 8 | Ifngr2 | -0.018 | Old |
| 0.628558 | 0.066112 | 0.682 | 0.756 | 1 | 8 | Ppp2r5c | -0.074 | Old |
| 0.628873 | 0.102152 | 0.48 | 0.525 | 1 | 8 | Usp25 | -0.045 | Old |
| 0.628972 | 0.676458 | 0.106 | 0.094 | 1 | 8 | Tapbpl | 0.012 | Old |
| 0.629025 | 0.223074 | 0.039 | 0.031 | 1 | 8 | Rwdd2a | 0.008 | Old |
| 0.629172 | 0.29572 | 0.156 | 0.143 | 1 | 8 | Fam120b | 0.013 | Old |
| 0.629496 | 0.547294 | 0.045 | 0.036 | 1 | 8 | Cenpl | 0.009 | Old |
| 0.629547 | 0.216591 | 0.503 | 0.504 | 1 | 8 | Klhl9 | -0.001 | Old |
| 0.629647 | 0.126036 | 0.413 | 0.395 | 1 | 8 | Dap | 0.018 | Old |
| 0.629659 | 0.008891 | 0.263 | 0.239 | 1 | 8 | Fam118b | 0.024 | Old |
| 0.629848 | 0.384807 | 0.017 | 0.023 | 1 | 8 | AI480526 | -0.006 | Old |
| 0.629848 | 0.381594 | 0.017 | 0.023 | 1 | 8 | Nkapl | -0.006 | Old |
| 0.629897 | 0.046598 | 0.385 | 0.405 | 1 | 8 | Cnppd1 | -0.02 | Old |
| 0.630325 | 0.032514 | 0.184 | 0.166 | 1 | 8 | Gm34084 | 0.018 | Old |
| 0.630618 | 0.661792 | 0.128 | 0.119 | 1 | 8 | Hira | 0.009 | Old |
| 0.630672 | 0.238342 | 0.045 | 0.036 | 1 | 8 | Rbak | 0.009 | Old |
| 0.630676 | 0.131626 | 0.173 | 0.192 | 1 | 8 | Mcm3 | -0.019 | Old |
| 0.630727 | 0.399121 | 0.061 | 0.052 | 1 | 8 | Trim21 | 0.009 | Old |
| 0.630799 | 0.033863 | 0.408 | 0.431 | 1 | 8 | Napa | -0.023 | Old |
| 0.63101 | 0.292625 | 0.246 | 0.229 | 1 | 8 | Rit1 | 0.017 | Old |
| 0.631029 | 0.198873 | 0.525 | 0.53 | 1 | 8 | Ralb | -0.005 | Old |
| 0.63112 | 0.147621 | 0.045 | 0.055 | 1 | 8 | Upp2 | -0.01 | Old |
| 0.63129 | 0.498731 | 0.073 | 0.062 | 1 | 8 | Rhbdd2 | 0.011 | Old |
| 0.631461 | 0.070205 | 0.034 | 0.026 | 1 | 8 | 1810044D( | 0.008 | Old |
| 0.631526 | 0.587064 | 0.179 | 0.169 | 1 | 8 | Ipmk | 0.01 | Old |
| 0.631708 | 0.213721 | 0.408 | 0.392 | 1 | 8 | Camk1 | 0.016 | Old |
| 0.631848 | 0.619382 | 0.045 | 0.036 | 1 | 8 | Brat1 | 0.009 | Old |
| 0.63191 | 0.316773 | 0.078 | 0.068 | 1 | 8 | Dclre1b | 0.01 | Old |
| 0.631944 | 0.362234 | 0.235 | 0.221 | 1 | 8 | Ly96 | 0.014 | Old |
| 0.632144 | 0.164401 | 0.067 | 0.078 | 1 | 8 | Gm16536 | -0.011 | Old |
| 0.632156 | 0.041415 | 0.235 | 0.262 | 1 | 8 | Ncoa1 | -0.027 | Old |
| 0.632207 | 0.40615 | 0.056 | 0.047 | 1 | 8 | Ttc4 | 0.009 | Old |
| 0.632207 | 0.325788 | 0.056 | 0.047 | 1 | 8 | Cry1 | 0.009 | Old |
| 0.632217 | 0.014713 | 0.291 | 0.314 | 1 | 8 | Ccng1 | -0.023 | Old |
| 0.63223 | 0.699732 | 0.123 | 0.112 | 1 | 8 | Itpa | 0.011 | Old |
| 0.632294 | 0.088059 | 0.324 | 0.358 | 1 | 8 | Stip1 | -0.034 | Old |
| 0.632661 | 0.132685 | 0.33 | 0.312 | 1 | 8 | Aida | 0.018 | Old |

| | | | | | | | | |
|---|---|---|---|---|---|---|---|---|
| 0.632767 | 0.083384 | 0.14 | 0.161 | 1 | 8 | Scn1b | -0.021 | Old |
| 0.632825 | 0.090706 | 0.145 | 0.161 | 1 | 8 | Ccnh | -0.016 | Old |
| 0.632898 | 0.53796 | 0.168 | 0.156 | 1 | 8 | Nipsnap2 | 0.012 | Old |
| 0.63313 | 0.099191 | 0.073 | 0.062 | 1 | 8 | Leng1 | 0.011 | Old |
| 0.633326 | 0.2143 | 0.084 | 0.099 | 1 | 8 | Pex16 | -0.015 | Old |
| 0.633478 | 0.002506 | 0.296 | 0.325 | 1 | 8 | Tmx1 | -0.029 | Old |
| 0.633535 | 0.046559 | 0.302 | 0.332 | 1 | 8 | Adam19 | -0.03 | Old |
| 0.63363 | 0.257799 | 0.341 | 0.327 | 1 | 8 | Nudt21 | 0.014 | Old |
| 0.633727 | 0.346537 | 0.061 | 0.052 | 1 | 8 | Wrb | 0.009 | Old |
| 0.633742 | 0.122118 | 0.291 | 0.312 | 1 | 8 | Lamtor5 | -0.021 | Old |
| 0.633796 | 0.101271 | 0.24 | 0.273 | 1 | 8 | Rnf38 | -0.033 | Old |
| 0.634052 | 0.525984 | 0.073 | 0.062 | 1 | 8 | P2ry1 | 0.011 | Old |
| 0.634057 | 0.282033 | 0.346 | 0.39 | 1 | 8 | Atf7ip | -0.044 | Old |
| 0.634117 | 0.38097 | 0.212 | 0.203 | 1 | 8 | Znfx1 | 0.009 | Old |
| 0.634124 | 0.483646 | 0.246 | 0.242 | 1 | 8 | Ndnf | 0.004 | Old |
| 0.63439 | 0.723243 | 0.162 | 0.153 | 1 | 8 | 0610010F0 | 0.009 | Old |
| 0.634412 | 0.048546 | 0.799 | 0.868 | 1 | 8 | Olfm1 | -0.069 | Old |
| 0.634475 | 0.124441 | 0.475 | 0.522 | 1 | 8 | Tmem167 | -0.047 | Old |
| 0.634729 | 0.525514 | 0.061 | 0.052 | 1 | 8 | Gm20707 | 0.009 | Old |
| 0.634837 | 0.043867 | 1 | 0.995 | 1 | 8 | Rps28 | 0.005 | Old |
| 0.634961 | 0.282591 | 0.268 | 0.255 | 1 | 8 | Impad1 | 0.013 | Old |
| 0.635359 | 0.519935 | 0.056 | 0.047 | 1 | 8 | Cdan1 | 0.009 | Old |
| 0.635359 | 0.185222 | 0.056 | 0.047 | 1 | 8 | Svip | 0.009 | Old |
| 0.635372 | 0.033764 | 0.179 | 0.197 | 1 | 8 | Ppp4r1 | -0.018 | Old |
| 0.635467 | 0.098761 | 0.19 | 0.208 | 1 | 8 | Fbrs | -0.018 | Old |
| 0.635924 | 0.026576 | 0.179 | 0.161 | 1 | 8 | Abcd2 | 0.018 | Old |
| 0.636256 | 0.058256 | 0.067 | 0.081 | 1 | 8 | Rrs1 | -0.014 | Old |
| 0.636291 | 0.362666 | 0.151 | 0.135 | 1 | 8 | Sh3tc1 | 0.016 | Old |
| 0.636683 | 0.231496 | 0.179 | 0.203 | 1 | 8 | Arl6ip6 | -0.024 | Old |
| 0.636733 | 0.275265 | 0.061 | 0.052 | 1 | 8 | Pi16 | 0.009 | Old |
| 0.636756 | 0.036592 | 0.251 | 0.229 | 1 | 8 | Tbc1d1 | 0.022 | Old |
| 0.636769 | 0.142717 | 0.251 | 0.234 | 1 | 8 | Kcnq1ot1 | 0.017 | Old |
| 0.637164 | 0.166043 | 0.279 | 0.257 | 1 | 8 | Asxl2 | 0.022 | Old |
| 0.637583 | 0.002636 | 0.117 | 0.132 | 1 | 8 | Tlr1 | -0.015 | Old |
| 0.637879 | 0.317456 | 0.039 | 0.031 | 1 | 8 | Itfg2 | 0.008 | Old |
| 0.638108 | 0.635453 | 0.067 | 0.057 | 1 | 8 | Gpr180 | 0.01 | Old |
| 0.638129 | 0.148967 | 0.173 | 0.19 | 1 | 8 | Rnf14 | -0.017 | Old |
| 0.638244 | 0.030564 | 0.302 | 0.317 | 1 | 8 | Emc3 | -0.015 | Old |
| 0.638357 | 0.173391 | 0.045 | 0.055 | 1 | 8 | Mon1a | -0.01 | Old |
| 0.638517 | 0.309862 | 0.056 | 0.047 | 1 | 8 | Mllt1 | 0.009 | Old |
| 0.638518 | 0.096071 | 0.285 | 0.317 | 1 | 8 | Trio | -0.032 | Old |
| 0.638673 | 0.01432 | 0.061 | 0.073 | 1 | 8 | Nlrx1 | -0.012 | Old |
| 0.638876 | 0.492447 | 0.101 | 0.119 | 1 | 8 | Ubtd1 | -0.018 | Old |
| 0.638929 | 0.227844 | 0.045 | 0.036 | 1 | 8 | Ap4b1 | 0.009 | Old |
| 0.639096 | 0.036002 | 0.151 | 0.166 | 1 | 8 | Acox1 | -0.015 | Old |

| | | | | | | | | |
|---|---|---|---|---|---|---|---|---|
| 0.639164 | 0.215061 | 0.24 | 0.221 | 1 | 8 | Ykt6 | 0.019 | Old |
| 0.639473 | 0.293831 | 0.078 | 0.068 | 1 | 8 | Rmdn3 | 0.01 | Old |
| 0.639613 | 0.127223 | 0.425 | 0.46 | 1 | 8 | Csnk1e | -0.035 | Old |
| 0.639842 | 0.327371 | 0.464 | 0.506 | 1 | 8 | Tmem14c | -0.042 | Old |
| 0.640058 | 0.358226 | 0.151 | 0.138 | 1 | 8 | Nvl | 0.013 | Old |
| 0.640249 | 0.412518 | 0.061 | 0.052 | 1 | 8 | Zfp629 | 0.009 | Old |
| 0.640347 | 0.13636 | 0.246 | 0.229 | 1 | 8 | Mrps9 | 0.017 | Old |
| 0.640569 | 0.175451 | 0.257 | 0.278 | 1 | 8 | Skiv2l | -0.021 | Old |
| 0.640732 | 0.284976 | 0.156 | 0.143 | 1 | 8 | Slc35a2 | 0.013 | Old |
| 0.640875 | 0.09911 | 0.117 | 0.104 | 1 | 8 | Urod | 0.013 | Old |
| 0.641119 | 0.425267 | 0.173 | 0.164 | 1 | 8 | Ptgir | 0.009 | Old |
| 0.641324 | 0.216864 | 0.453 | 0.457 | 1 | 8 | Ostc | -0.004 | Old |
| 0.641418 | 0.092325 | 0.598 | 0.634 | 1 | 8 | Bclaf1 | -0.036 | Old |
| 0.641435 | 0.04187 | 0.229 | 0.255 | 1 | 8 | Cdk4 | -0.026 | Old |
| 0.641444 | 0.122902 | 0.101 | 0.114 | 1 | 8 | Tmem37 | -0.013 | Old |
| 0.641612 | 0.037989 | 0.117 | 0.104 | 1 | 8 | Nos1ap | 0.013 | Old |
| 0.641689 | 0.408643 | 0.039 | 0.031 | 1 | 8 | Pced1b | 0.008 | Old |
| 0.641731 | 0.582473 | 0.134 | 0.122 | 1 | 8 | Gm17018 | 0.012 | Old |
| 0.642089 | 0.308899 | 0.246 | 0.231 | 1 | 8 | Xpot | 0.015 | Old |
| 0.642299 | 0.277385 | 0.335 | 0.327 | 1 | 8 | U2af1 | 0.008 | Old |
| 0.643067 | 0.561958 | 0.302 | 0.296 | 1 | 8 | C1d | 0.006 | Old |
| 0.643135 | 0.150333 | 0.128 | 0.117 | 1 | 8 | Crls1 | 0.011 | Old |
| 0.643449 | 0.281384 | 0.229 | 0.21 | 1 | 8 | Glod4 | 0.019 | Old |
| 0.643772 | 0.556277 | 0.061 | 0.052 | 1 | 8 | Sbk1 | 0.009 | Old |
| 0.643936 | 0.196925 | 0.19 | 0.174 | 1 | 8 | Itga9 | 0.016 | Old |
| 0.644261 | 0.257415 | 0.045 | 0.036 | 1 | 8 | Spindoc | 0.009 | Old |
| 0.644538 | 0.454404 | 0.145 | 0.132 | 1 | 8 | Entpd7 | 0.013 | Old |
| 0.644801 | 0.235155 | 0.564 | 0.587 | 1 | 8 | Pigt | -0.023 | Old |
| 0.644855 | 0.268665 | 0.034 | 0.042 | 1 | 8 | Plcg1 | -0.008 | Old |
| 0.644855 | 0.05615 | 0.045 | 0.036 | 1 | 8 | Miga2 | 0.009 | Old |
| 0.645062 | 0.049865 | 0.112 | 0.096 | 1 | 8 | Cdk5 | 0.016 | Old |
| 0.645172 | 0.101675 | 0.821 | 0.86 | 1 | 8 | Tubb5 | -0.039 | Old |
| 0.645402 | 0.070281 | 0.084 | 0.096 | 1 | 8 | Rdh11 | -0.012 | Old |
| 0.645495 | 0.046591 | 0.335 | 0.361 | 1 | 8 | Btk | -0.026 | Old |
| 0.645624 | 0.39914 | 0.374 | 0.408 | 1 | 8 | Smim7 | -0.034 | Old |
| 0.645687 | 0.099812 | 0.307 | 0.335 | 1 | 8 | Iscu | -0.028 | Old |
| 0.646175 | 0.060084 | 0.123 | 0.138 | 1 | 8 | Aars | -0.015 | Old |
| 0.646221 | 0.163406 | 0.134 | 0.122 | 1 | 8 | Psmd5 | 0.012 | Old |
| 0.646567 | 0.1162 | 0.134 | 0.122 | 1 | 8 | Pex3 | 0.012 | Old |
| 0.646588 | 0.52275 | 0.073 | 0.086 | 1 | 8 | Alg2 | -0.013 | Old |
| 0.646716 | 0.078689 | 0.05 | 0.06 | 1 | 8 | Ier5l | -0.01 | Old |
| 0.646903 | 0.035206 | 0.24 | 0.257 | 1 | 8 | Rnmt | -0.017 | Old |
| 0.646977 | 0.249532 | 0.078 | 0.091 | 1 | 8 | Spred2 | -0.013 | Old |
| 0.647027 | 0.263633 | 0.128 | 0.143 | 1 | 8 | Rnf168 | -0.015 | Old |
| 0.647231 | 0.288816 | 0.034 | 0.042 | 1 | 8 | Rbm4 | -0.008 | Old |

| | | | | | | | | |
|---|---|---|---|---|---|---|---|---|
| 0.647265 | 0.018885 | 0.106 | 0.119 | 1 | 8 | Scai | -0.013 | Old |
| 0.647366 | 0.2669 | 0.207 | 0.197 | 1 | 8 | Lrrc47 | 0.01 | Old |
| 0.647399 | 0.112759 | 0.246 | 0.27 | 1 | 8 | Rbm22 | -0.024 | Old |
| 0.647499 | 0.078803 | 0.318 | 0.351 | 1 | 8 | Gdi1 | -0.033 | Old |
| 0.647761 | 0.111978 | 0.257 | 0.234 | 1 | 8 | Napg | 0.023 | Old |
| 0.648173 | 0.060825 | 0.24 | 0.273 | 1 | 8 | Ift20 | -0.033 | Old |
| 0.648421 | 0.030017 | 0.045 | 0.036 | 1 | 8 | Chchd6 | 0.009 | Old |
| 0.648489 | 0.023118 | 0.559 | 0.592 | 1 | 8 | Rab2a | -0.033 | Old |
| 0.648638 | 0.034192 | 0.201 | 0.184 | 1 | 8 | Taf11 | 0.017 | Old |
| 0.648719 | 0.285099 | 0.067 | 0.057 | 1 | 8 | Zbtb17 | 0.01 | Old |
| 0.649421 | 0.319631 | 0.05 | 0.042 | 1 | 8 | Zfp319 | 0.008 | Old |
| 0.649427 | 0.175921 | 0.341 | 0.317 | 1 | 8 | Plekha2 | 0.024 | Old |
| 0.649476 | 0.39073 | 0.268 | 0.262 | 1 | 8 | Mrpl43 | 0.006 | Old |
| 0.650173 | 0.209535 | 0.084 | 0.096 | 1 | 8 | Snx24 | -0.012 | Old |
| 0.650516 | 0.277306 | 0.469 | 0.465 | 1 | 8 | Gna12 | 0.004 | Old |
| 0.650567 | 0.111067 | 0.212 | 0.234 | 1 | 8 | Tmem147 | -0.022 | Old |
| 0.650803 | 0.452469 | 0.034 | 0.042 | 1 | 8 | Pced1a | -0.008 | Old |
| 0.650806 | 0.100319 | 0.441 | 0.481 | 1 | 8 | Top2b | -0.04 | Old |
| 0.651348 | 0.212659 | 0.363 | 0.392 | 1 | 8 | Aprt | -0.029 | Old |
| 0.651539 | 0.00019 | 0.151 | 0.166 | 1 | 8 | Pfkfb4 | -0.015 | Old |
| 0.651951 | 0.099967 | 0.279 | 0.299 | 1 | 8 | Dcp2 | -0.02 | Old |
| 0.652008 | 0.168098 | 0.14 | 0.122 | 1 | 8 | Nr1d2 | 0.018 | Old |
| 0.652226 | 0.049055 | 0.475 | 0.545 | 1 | 8 | Gfra2 | -0.07 | Old |
| 0.65226 | 0.199765 | 0.285 | 0.268 | 1 | 8 | Uap1l1 | 0.017 | Old |
| 0.652314 | 0.202082 | 0.43 | 0.436 | 1 | 8 | Mapk1ip1l | -0.006 | Old |
| 0.652483 | 0.039143 | 0.631 | 0.673 | 1 | 8 | Tor1aip1 | -0.042 | Old |
| 0.652593 | 0.447417 | 0.229 | 0.21 | 1 | 8 | Eed | 0.019 | Old |
| 0.652626 | 0.088852 | 0.123 | 0.109 | 1 | 8 | Zfp414 | 0.014 | Old |
| 0.652827 | 0.079351 | 0.196 | 0.218 | 1 | 8 | Cramp1l | -0.022 | Old |
| 0.652838 | 0.264593 | 0.179 | 0.166 | 1 | 8 | Mcoln1 | 0.013 | Old |
| 0.652868 | 0.03752 | 0.061 | 0.052 | 1 | 8 | Wdr55 | 0.009 | Old |
| 0.652935 | 0.293064 | 0.045 | 0.055 | 1 | 8 | Zfp212 | -0.01 | Old |
| 0.653083 | 0.210668 | 0.184 | 0.208 | 1 | 8 | Clasp1 | -0.024 | Old |
| 0.653189 | 0.057823 | 0.045 | 0.036 | 1 | 8 | Det1 | 0.009 | Old |
| 0.653194 | 0.298169 | 0.257 | 0.249 | 1 | 8 | C330007P0 | 0.008 | Old |
| 0.653545 | 0.001101 | 0.95 | 0.982 | 1 | 8 | Mbnl1 | -0.032 | Old |
| 0.653845 | 0.066496 | 0.698 | 0.732 | 1 | 8 | Tpd52 | -0.034 | Old |
| 0.654342 | 0.475506 | 0.095 | 0.083 | 1 | 8 | E2f5 | 0.012 | Old |
| 0.654342 | 0.502949 | 0.156 | 0.145 | 1 | 8 | Fam220a | 0.011 | Old |
| 0.654971 | 0.097365 | 0.302 | 0.322 | 1 | 8 | Hpf1 | -0.02 | Old |
| 0.655032 | 0.09644 | 0.05 | 0.042 | 1 | 8 | Snapc2 | 0.008 | Old |
| 0.655042 | 0.011368 | 0.659 | 0.735 | 1 | 8 | Dynll1 | -0.076 | Old |
| 0.655827 | 0.186646 | 0.391 | 0.444 | 1 | 8 | Cmklr1 | -0.053 | Old |
| 0.656043 | 0.128456 | 0.218 | 0.244 | 1 | 8 | Pisd | -0.026 | Old |
| 0.656087 | 0.187779 | 0.168 | 0.153 | 1 | 8 | Slfn8 | 0.015 | Old |

| | | | | | | | | |
|---|---|---|---|---|---|---|---|---|
| 0.656182 | 0.37173 | 0.084 | 0.073 | 1 | 8 | Pex14 | 0.011 | Old |
| 0.656637 | 0.034142 | 0.067 | 0.078 | 1 | 8 | Zbtb41 | -0.011 | Old |
| 0.657037 | 0.238556 | 0.777 | 0.842 | 1 | 8 | Ifi27l2a | -0.065 | Old |
| 0.657282 | 0.322128 | 0.05 | 0.042 | 1 | 8 | Mettl13 | 0.008 | Old |
| 0.657326 | 0.344772 | 0.318 | 0.306 | 1 | 8 | Nfatc3 | 0.012 | Old |
| 0.658556 | 0.202081 | 0.101 | 0.088 | 1 | 8 | Exoc7 | 0.013 | Old |
| 0.658556 | 0.112438 | 0.101 | 0.088 | 1 | 8 | Dym | 0.013 | Old |
| 0.658773 | 0.00234 | 0.564 | 0.623 | 1 | 8 | Map4k4 | -0.059 | Old |
| 0.658804 | 0.392755 | 0.173 | 0.161 | 1 | 8 | Wdr5 | 0.012 | Old |
| 0.658804 | 0.405253 | 0.151 | 0.171 | 1 | 8 | Pdcl | -0.02 | Old |
| 0.659219 | 0.487713 | 0.078 | 0.068 | 1 | 8 | Mad2l1bp | 0.01 | Old |
| 0.659248 | 0.11697 | 0.765 | 0.829 | 1 | 8 | Ctsa | -0.064 | Old |
| 0.659364 | 0.050191 | 0.346 | 0.371 | 1 | 8 | Trim41 | -0.025 | Old |
| 0.659399 | 0.417708 | 0.128 | 0.117 | 1 | 8 | Pgm1 | 0.011 | Old |
| 0.659535 | 0.10692 | 0.05 | 0.042 | 1 | 8 | Lmln | 0.008 | Old |
| 0.660108 | 0.346519 | 0.101 | 0.091 | 1 | 8 | Gramd1b | 0.01 | Old |
| 0.660367 | 0.015218 | 0.034 | 0.042 | 1 | 8 | 2310040G2 | -0.008 | Old |
| 0.660481 | 0.272503 | 0.151 | 0.138 | 1 | 8 | Prpf19 | 0.013 | Old |
| 0.660935 | 0.342797 | 0.168 | 0.158 | 1 | 8 | Gsap | 0.01 | Old |
| 0.66112 | 0.373813 | 0.302 | 0.283 | 1 | 8 | Arhgap1 | 0.019 | Old |
| 0.661329 | 0.357 | 0.095 | 0.083 | 1 | 8 | Pou5f2 | 0.012 | Old |
| 0.661366 | 0.040028 | 0.168 | 0.187 | 1 | 8 | Polr1c | -0.019 | Old |
| 0.661491 | 0.014375 | 0.682 | 0.738 | 1 | 8 | Sf3b2 | -0.056 | Old |
| 0.661578 | 0.057405 | 0.994 | 1 | 1 | 8 | Crip1 | -0.006 | Old |
| 0.661678 | 0.048899 | 0.525 | 0.577 | 1 | 8 | Ptpn6 | -0.052 | Old |
| 0.661893 | 0.042799 | 0.285 | 0.309 | 1 | 8 | Blvra | -0.024 | Old |
| 0.661975 | 0.168479 | 0.52 | 0.543 | 1 | 8 | Dhx15 | -0.023 | Old |
| 0.662436 | 0.45437 | 0.184 | 0.174 | 1 | 8 | Gyg | 0.01 | Old |
| 0.662565 | 0.006031 | 0.078 | 0.091 | 1 | 8 | Foxo4 | -0.013 | Old |
| 0.662643 | 0.067319 | 0.497 | 0.535 | 1 | 8 | Ndufs6 | -0.038 | Old |
| 0.662827 | 0.003033 | 0.397 | 0.434 | 1 | 8 | Ddx24 | -0.037 | Old |
| 0.662842 | 0.281639 | 0.536 | 0.53 | 1 | 8 | Casp8 | 0.006 | Old |
| 0.66285 | 0.11529 | 0.212 | 0.195 | 1 | 8 | Atxn7l3 | 0.017 | Old |
| 0.66292 | 0.308218 | 0.05 | 0.042 | 1 | 8 | Gm26881 | 0.008 | Old |
| 0.662936 | 0.353045 | 0.201 | 0.195 | 1 | 8 | Katnbl1 | 0.006 | Old |
| 0.662995 | 0.111866 | 0.754 | 0.797 | 1 | 8 | Ywhah | -0.043 | Old |
| 0.663054 | 0.245159 | 0.291 | 0.283 | 1 | 8 | Slc25a51 | 0.008 | Old |
| 0.663582 | 0.265349 | 0.341 | 0.366 | 1 | 8 | Chchd1 | -0.025 | Old |
| 0.663715 | 0.029237 | 0.05 | 0.06 | 1 | 8 | 1500015A0 | -0.01 | Old |
| 0.664136 | 0.105839 | 0.022 | 0.029 | 1 | 8 | Gm31597 | -0.007 | Old |
| 0.66427 | 0.201936 | 0.145 | 0.13 | 1 | 8 | Mtx2 | 0.015 | Old |
| 0.664432 | 0.01222 | 0.168 | 0.151 | 1 | 8 | Gtf3a | 0.017 | Old |
| 0.664487 | 0.307764 | 0.257 | 0.244 | 1 | 8 | Frrs1 | 0.013 | Old |
| 0.664879 | 0.282297 | 0.458 | 0.439 | 1 | 8 | Lmo4 | 0.019 | Old |
| 0.665077 | 0.255551 | 0.061 | 0.052 | 1 | 8 | Timm21 | 0.009 | Old |

| | | | | | | | | |
|---|---|---|---|---|---|---|---|---|
| 0.665181 | 0.272199 | 0.291 | 0.319 | 1 | 8 | Cib1 | -0.028 | Old |
| 0.665449 | 0.174454 | 0.212 | 0.236 | 1 | 8 | Ckap5 | -0.024 | Old |
| 0.665513 | 0.031156 | 0.112 | 0.099 | 1 | 8 | Manbal | 0.013 | Old |
| 0.665583 | 0.120225 | 0.022 | 0.029 | 1 | 8 | Mmaa | -0.007 | Old |
| 0.666404 | 0.184113 | 0.33 | 0.366 | 1 | 8 | Dctn4 | -0.036 | Old |
| 0.666598 | 0.433469 | 0.168 | 0.158 | 1 | 8 | Fech | 0.01 | Old |
| 0.666731 | 0.405402 | 0.156 | 0.143 | 1 | 8 | Acsl3 | 0.013 | Old |
| 0.666863 | 0.349766 | 0.162 | 0.148 | 1 | 8 | Eif2b4 | 0.014 | Old |
| 0.666986 | 0.02521 | 0.413 | 0.429 | 1 | 8 | Cdc5l | -0.016 | Old |
| 0.667239 | 0.105827 | 0.212 | 0.192 | 1 | 8 | Nploc4 | 0.02 | Old |
| 0.667458 | 0.175856 | 0.302 | 0.288 | 1 | 8 | Mgat2 | 0.014 | Old |
| 0.667734 | 0.089213 | 0.369 | 0.397 | 1 | 8 | Trafd1 | -0.028 | Old |
| 0.667788 | 0.190199 | 0.084 | 0.096 | 1 | 8 | Sugp2 | -0.012 | Old |
| 0.667899 | 0.191451 | 0.307 | 0.338 | 1 | 8 | Casc3 | -0.031 | Old |
| 0.667934 | 0.351667 | 0.436 | 0.442 | 1 | 8 | Vma21 | -0.006 | Old |
| 0.668005 | 0.061711 | 0.073 | 0.083 | 1 | 8 | F8a | -0.01 | Old |
| 0.668258 | 0.063198 | 0.603 | 0.616 | 1 | 8 | Rnasel | -0.013 | Old |
| 0.668292 | 0.217505 | 0.324 | 0.366 | 1 | 8 | Puf60 | -0.042 | Old |
| 0.668426 | 0.103443 | 0.419 | 0.457 | 1 | 8 | Tet2 | -0.038 | Old |
| 0.668703 | 0.607112 | 0.106 | 0.096 | 1 | 8 | Rbbp5 | 0.01 | Old |
| 0.668761 | 0.015233 | 0.324 | 0.364 | 1 | 8 | Lrrfip2 | -0.04 | Old |
| 0.668764 | 0.240746 | 0.559 | 0.527 | 1 | 8 | Got1 | 0.032 | Old |
| 0.668869 | 0.361946 | 0.101 | 0.091 | 1 | 8 | Cd274 | 0.01 | Old |
| 0.66946 | 0.101059 | 0.291 | 0.309 | 1 | 8 | Synj2bp | -0.018 | Old |
| 0.669727 | 0.067701 | 0.112 | 0.099 | 1 | 8 | Utp14b | 0.013 | Old |
| 0.669757 | 0.262133 | 0.05 | 0.06 | 1 | 8 | Creld1 | -0.01 | Old |
| 0.669947 | 0.192117 | 0.324 | 0.364 | 1 | 8 | Csnk2a2 | -0.04 | Old |
| 0.670001 | 0.263753 | 0.095 | 0.083 | 1 | 8 | Hps1 | 0.012 | Old |
| 0.670318 | 0.21238 | 0.642 | 0.714 | 1 | 8 | Cd84 | -0.072 | Old |
| 0.67039 | 0.417185 | 0.246 | 0.231 | 1 | 8 | Srd5a3 | 0.015 | Old |
| 0.670456 | 0.026493 | 0.994 | 0.995 | 1 | 8 | Zfp36 | -0.001 | Old |
| 0.670816 | 0.304374 | 0.045 | 0.055 | 1 | 8 | Acvr2b | -0.01 | Old |
| 0.671002 | 0.154566 | 0.078 | 0.068 | 1 | 8 | Mrps18b | 0.01 | Old |
| 0.671042 | 0.115485 | 0.089 | 0.104 | 1 | 8 | Dmxl2 | -0.015 | Old |
| 0.671058 | 0.002037 | 0.609 | 0.66 | 1 | 8 | Furin | -0.051 | Old |
| 0.67107 | 0.353991 | 0.084 | 0.073 | 1 | 8 | Clk2 | 0.011 | Old |
| 0.671561 | 0.30302 | 0.201 | 0.184 | 1 | 8 | Slc39a9 | 0.017 | Old |
| 0.671621 | 0.022138 | 0.503 | 0.517 | 1 | 8 | Ubtf | -0.014 | Old |
| 0.671657 | 0.008182 | 0.078 | 0.091 | 1 | 8 | Trmt13 | -0.013 | Old |
| 0.671775 | 0.112707 | 0.05 | 0.06 | 1 | 8 | Clmp | -0.01 | Old |
| 0.672091 | 0.298752 | 0.112 | 0.101 | 1 | 8 | Tmem198b | 0.011 | Old |
| 0.672241 | 0.103866 | 0.128 | 0.117 | 1 | 8 | Heatr3 | 0.011 | Old |
| 0.672581 | 0.025507 | 0.067 | 0.078 | 1 | 8 | Cog1 | -0.011 | Old |
| 0.672929 | 0.123816 | 0.318 | 0.345 | 1 | 8 | Paf1 | -0.027 | Old |
| 0.673186 | 0.115423 | 0.76 | 0.797 | 1 | 8 | Plekho2 | -0.037 | Old |

| | | | | | | | | |
|---|---|---|---|---|---|---|---|---|
| 0.673289 | 0.244199 | 0.117 | 0.135 | 1 | 8 | Nip7 | -0.018 | Old |
| 0.674247 | 0.994595 | 0.073 | 0.065 | 1 | 8 | Ifi211 | 0.008 | Old |
| 0.67531 | 0.306834 | 0.095 | 0.086 | 1 | 8 | Csf3r | 0.009 | Old |
| 0.675421 | 0.294483 | 0.229 | 0.257 | 1 | 8 | Foxo3 | -0.028 | Old |
| 0.676114 | 0.522433 | 0.073 | 0.065 | 1 | 8 | Alad | 0.008 | Old |
| 0.676301 | 0.24942 | 0.106 | 0.119 | 1 | 8 | Sbf1 | -0.013 | Old |
| 0.676393 | 0.22764 | 0.235 | 0.268 | 1 | 8 | Orai3 | -0.033 | Old |
| 0.676419 | 0.006721 | 0.229 | 0.205 | 1 | 8 | Ncln | 0.024 | Old |
| 0.676785 | 0.10666 | 0.575 | 0.571 | 1 | 8 | Ctdnep1 | 0.004 | Old |
| 0.676866 | 0.026935 | 0.291 | 0.314 | 1 | 8 | 2810403A( | -0.023 | Old |
| 0.676873 | 0.584921 | 0.101 | 0.091 | 1 | 8 | Slc35e3 | 0.01 | Old |
| 0.677202 | 0.375995 | 0.022 | 0.029 | 1 | 8 | Acsf2 | -0.007 | Old |
| 0.677202 | 0.361215 | 0.022 | 0.029 | 1 | 8 | Slc46a1 | -0.007 | Old |
| 0.677291 | 0.203675 | 0.257 | 0.236 | 1 | 8 | Fam192a | 0.021 | Old |
| 0.677324 | 0.185342 | 0.313 | 0.332 | 1 | 8 | Abca7 | -0.019 | Old |
| 0.677363 | 0.174045 | 0.318 | 0.301 | 1 | 8 | Ndufa12 | 0.017 | Old |
| 0.678239 | 0.242813 | 0.123 | 0.138 | 1 | 8 | Pctp | -0.015 | Old |
| 0.67826 | 0.072751 | 0.358 | 0.387 | 1 | 8 | Gtpbp4 | -0.029 | Old |
| 0.678501 | 0.05954 | 0.313 | 0.33 | 1 | 8 | Setd3 | -0.017 | Old |
| 0.67866 | 0.973363 | 0.022 | 0.029 | 1 | 8 | Gpsm2 | -0.007 | Old |
| 0.678718 | 0.137786 | 0.056 | 0.065 | 1 | 8 | Rbm10 | -0.009 | Old |
| 0.679139 | 0.306796 | 0.492 | 0.506 | 1 | 8 | Rad23b | -0.014 | Old |
| 0.679211 | 0.192772 | 0.117 | 0.106 | 1 | 8 | Map3k3 | 0.011 | Old |
| 0.679258 | 0.169204 | 0.497 | 0.488 | 1 | 8 | Smim15 | 0.009 | Old |
| 0.679716 | 8.22E-05 | 0.335 | 0.353 | 1 | 8 | Zfp652 | -0.018 | Old |
| 0.679748 | 0.288415 | 0.207 | 0.195 | 1 | 8 | Prkaca | 0.012 | Old |
| 0.679874 | 0.901859 | 0.05 | 0.06 | 1 | 8 | Ogfod3 | -0.01 | Old |
| 0.679884 | 0.216244 | 0.134 | 0.153 | 1 | 8 | Impdh1 | -0.019 | Old |
| 0.680531 | 0.218952 | 0.106 | 0.122 | 1 | 8 | Tbce | -0.016 | Old |
| 0.680819 | 0.275045 | 0.196 | 0.184 | 1 | 8 | Nfx1 | 0.012 | Old |
| 0.681136 | 0.145249 | 0.089 | 0.101 | 1 | 8 | Edem3 | -0.012 | Old |
| 0.681298 | 1.164892 | 0.078 | 0.07 | 1 | 8 | Ahi1 | 0.008 | Old |
| 0.681349 | 0.092014 | 0.011 | 0.016 | 1 | 8 | AW011738 | -0.005 | Old |
| 0.681434 | 0.424761 | 0.123 | 0.112 | 1 | 8 | Gcsh | 0.011 | Old |
| 0.68147 | 0.234717 | 0.173 | 0.192 | 1 | 8 | Flcn | -0.019 | Old |
| 0.6816 | 0.141351 | 0.14 | 0.156 | 1 | 8 | Pex19 | -0.016 | Old |
| 0.682014 | 0.264771 | 0.196 | 0.184 | 1 | 8 | Armc1 | 0.012 | Old |
| 0.682063 | 0.111722 | 0.218 | 0.239 | 1 | 8 | Smim27 | -0.021 | Old |
| 0.682313 | 0.184009 | 0.207 | 0.221 | 1 | 8 | Zbtb38 | -0.014 | Old |
| 0.682422 | 0.24569 | 0.246 | 0.236 | 1 | 8 | Tmem170b | 0.01 | Old |
| 0.682695 | 0.286815 | 0.251 | 0.234 | 1 | 8 | Spen | 0.017 | Old |
| 0.682835 | 0.235979 | 0.341 | 0.338 | 1 | 8 | Myd88 | 0.003 | Old |
| 0.683062 | 0.093718 | 0.251 | 0.278 | 1 | 8 | Sec22b | -0.027 | Old |
| 0.683101 | 0.346892 | 0.391 | 0.364 | 1 | 8 | Zfp516 | 0.027 | Old |
| 0.683338 | 0.310951 | 0.011 | 0.016 | 1 | 8 | Zfp790 | -0.005 | Old |

| | | | | | | | | |
|---|---|---|---|---|---|---|---|---|
| 0.683338 | 0.200321 | 0.011 | 0.016 | 1 | 8 | Rasip1 | -0.005 | Old |
| 0.68335 | 0.05232 | 0.682 | 0.691 | 1 | 8 | Romo1 | -0.009 | Old |
| 0.68351 | 0.023883 | 0.089 | 0.101 | 1 | 8 | Cdc27 | -0.012 | Old |
| 0.683831 | 0.087291 | 0.397 | 0.387 | 1 | 8 | Erp44 | 0.01 | Old |
| 0.684084 | 0.155995 | 0.469 | 0.455 | 1 | 8 | Sltm | 0.014 | Old |
| 0.684088 | 1.52787 | 0.011 | 0.008 | 1 | 8 | Hyls1 | 0.003 | Old |
| 0.684088 | 1.490734 | 0.011 | 0.008 | 1 | 8 | Srcin1 | 0.003 | Old |
| 0.684088 | 1.41443 | 0.011 | 0.008 | 1 | 8 | Gimap9 | 0.003 | Old |
| 0.684088 | 0.768089 | 0.011 | 0.008 | 1 | 8 | Hist2h2aa1 | 0.003 | Old |
| 0.684374 | 0.074799 | 0.106 | 0.119 | 1 | 8 | Qsox1 | -0.013 | Old |
| 0.684504 | 0.345738 | 0.022 | 0.029 | 1 | 8 | Pdk4 | -0.007 | Old |
| 0.684512 | 0.209873 | 0.156 | 0.143 | 1 | 8 | Nln | 0.013 | Old |
| 0.684954 | 0.742758 | 0.061 | 0.055 | 1 | 8 | Cdkal1 | 0.006 | Old |
| 0.685091 | 0.35003 | 0.313 | 0.304 | 1 | 8 | Atp6v0a2 | 0.009 | Old |
| 0.685138 | 0.158037 | 0.106 | 0.096 | 1 | 8 | Smim3 | 0.01 | Old |
| 0.685481 | 0.898472 | 0.073 | 0.065 | 1 | 8 | Otud1 | 0.008 | Old |
| 0.685537 | 0.135892 | 0.056 | 0.065 | 1 | 8 | Pigf | -0.009 | Old |
| 0.68554 | 0.060506 | 0.385 | 0.421 | 1 | 8 | Gbp7 | -0.036 | Old |
| 0.6859 | 0.50904 | 0.162 | 0.179 | 1 | 8 | Plekhm1 | -0.017 | Old |
| 0.685916 | 0.069885 | 0.067 | 0.057 | 1 | 8 | Bclaf3 | 0.01 | Old |
| 0.686339 | 0.417657 | 0.117 | 0.106 | 1 | 8 | Fbxo7 | 0.011 | Old |
| 0.686601 | 1.299161 | 0.011 | 0.008 | 1 | 8 | Bloc1s6os | 0.003 | Old |
| 0.686601 | 1.235947 | 0.011 | 0.008 | 1 | 8 | G0s2 | 0.003 | Old |
| 0.686601 | 0.822594 | 0.011 | 0.008 | 1 | 8 | Bmpr1a | 0.003 | Old |
| 0.686601 | 0.746948 | 0.011 | 0.008 | 1 | 8 | Sugct | 0.003 | Old |
| 0.686601 | 0.699707 | 0.011 | 0.008 | 1 | 8 | Zscan18 | 0.003 | Old |
| 0.686601 | 0.669637 | 0.011 | 0.008 | 1 | 8 | 2810414N( | 0.003 | Old |
| 0.686601 | 0.59837 | 0.011 | 0.008 | 1 | 8 | Lrrc51 | 0.003 | Old |
| 0.686601 | 0.499139 | 0.011 | 0.008 | 1 | 8 | Gm15964 | 0.003 | Old |
| 0.686601 | 0.497664 | 0.011 | 0.008 | 1 | 8 | Phtf1os | 0.003 | Old |
| 0.686621 | 0.003657 | 0.279 | 0.268 | 1 | 8 | Trappc6b | 0.011 | Old |
| 0.68667 | 0.046199 | 0.33 | 0.356 | 1 | 8 | Wbp1l | -0.026 | Old |
| 0.68721 | 0.044433 | 0.24 | 0.221 | 1 | 8 | Supt20 | 0.019 | Old |
| 0.687322 | 0.108625 | 0.011 | 0.016 | 1 | 8 | Timp4 | -0.005 | Old |
| 0.687433 | 0.019204 | 0.022 | 0.029 | 1 | 8 | Mtmr11 | -0.007 | Old |
| 0.687689 | 0.46332 | 0.117 | 0.109 | 1 | 8 | Zfand2b | 0.008 | Old |
| 0.687878 | 0.448614 | 0.24 | 0.236 | 1 | 8 | Ythdf2 | 0.004 | Old |
| 0.688202 | 0.563394 | 0.123 | 0.114 | 1 | 8 | Ambra1 | 0.009 | Old |
| 0.688352 | 0.287081 | 0.156 | 0.174 | 1 | 8 | Brd9 | -0.018 | Old |
| 0.688599 | 0.374298 | 0.123 | 0.138 | 1 | 8 | Bnip3 | -0.015 | Old |
| 0.688775 | 0.358987 | 0.307 | 0.304 | 1 | 8 | Tmem165 | 0.003 | Old |
| 0.688821 | 0.033356 | 0.559 | 0.582 | 1 | 8 | Spcs2 | -0.023 | Old |
| 0.689118 | 1.11961 | 0.011 | 0.008 | 1 | 8 | Lrrc29 | 0.003 | Old |
| 0.689118 | 0.70998 | 0.011 | 0.008 | 1 | 8 | Nuf2 | 0.003 | Old |
| 0.689118 | 0.65236 | 0.011 | 0.008 | 1 | 8 | Tmc4 | 0.003 | Old |

| | | | | | | | | |
|---|---|---|---|---|---|---|---|---|
| 0.689118 | 0.581365 | 0.011 | 0.008 | 1 | 8 | Gm12355 | 0.003 | Old |
| 0.689118 | 0.569696 | 0.011 | 0.008 | 1 | 8 | Trem1 | 0.003 | Old |
| 0.689118 | 0.535916 | 0.011 | 0.008 | 1 | 8 | E430024P1 | 0.003 | Old |
| 0.689118 | 0.496667 | 0.011 | 0.008 | 1 | 8 | Gm9484 | 0.003 | Old |
| 0.689118 | 0.474564 | 0.011 | 0.008 | 1 | 8 | Tnfrsf10b | 0.003 | Old |
| 0.689118 | 0.449565 | 0.011 | 0.008 | 1 | 8 | Ebf2 | 0.003 | Old |
| 0.689118 | 0.383456 | 0.011 | 0.008 | 1 | 8 | 2010109A1 | 0.003 | Old |
| 0.689118 | 0.353163 | 0.011 | 0.008 | 1 | 8 | Rcn1 | 0.003 | Old |
| 0.689118 | 0.342381 | 0.011 | 0.008 | 1 | 8 | Fgf10 | 0.003 | Old |
| 0.689118 | 0.335853 | 0.011 | 0.008 | 1 | 8 | Mctp2 | 0.003 | Old |
| 0.689118 | 0.288438 | 0.011 | 0.008 | 1 | 8 | 6230400D1 | 0.003 | Old |
| 0.689118 | 0.266744 | 0.011 | 0.008 | 1 | 8 | Syt3 | 0.003 | Old |
| 0.689118 | 0.123681 | 0.011 | 0.008 | 1 | 8 | 5830454E0 | 0.003 | Old |
| 0.689318 | 0.186686 | 0.011 | 0.016 | 1 | 8 | Amot | -0.005 | Old |
| 0.689318 | 0.175659 | 0.011 | 0.016 | 1 | 8 | 9530082P2 | -0.005 | Old |
| 0.689475 | 0.11736 | 0.302 | 0.33 | 1 | 8 | R3hdm1 | -0.028 | Old |
| 0.689787 | 0.290854 | 0.218 | 0.203 | 1 | 8 | Ddhd1 | 0.015 | Old |
| 0.690208 | 0.141353 | 0.587 | 0.631 | 1 | 8 | Dync1i2 | -0.044 | Old |
| 0.690232 | 0.11491 | 0.687 | 0.727 | 1 | 8 | Add3 | -0.04 | Old |
| 0.690267 | 0.020848 | 0.106 | 0.119 | 1 | 8 | Pibf1 | -0.013 | Old |
| 0.690334 | 0.656586 | 0.246 | 0.236 | 1 | 8 | Dapk3 | 0.01 | Old |
| 0.690366 | 0.171705 | 0.687 | 0.701 | 1 | 8 | Dync1h1 | -0.014 | Old |
| 0.690998 | 0.093891 | 0.212 | 0.2 | 1 | 8 | Suco | 0.012 | Old |
| 0.691315 | 0.536889 | 0.011 | 0.016 | 1 | 8 | Phactr1 | -0.005 | Old |
| 0.691385 | 0.343667 | 0.128 | 0.119 | 1 | 8 | Pxmp4 | 0.009 | Old |
| 0.691638 | 1.214881 | 0.011 | 0.008 | 1 | 8 | Bhlhb9 | 0.003 | Old |
| 0.691638 | 0.77527 | 0.011 | 0.008 | 1 | 8 | Zfp273 | 0.003 | Old |
| 0.691638 | 0.772348 | 0.011 | 0.008 | 1 | 8 | Gpd1 | 0.003 | Old |
| 0.691638 | 0.665859 | 0.011 | 0.008 | 1 | 8 | Ptpn13 | 0.003 | Old |
| 0.691638 | 0.620554 | 0.011 | 0.008 | 1 | 8 | Gm20234 | 0.003 | Old |
| 0.691638 | 0.351454 | 0.011 | 0.008 | 1 | 8 | Cdca3 | 0.003 | Old |
| 0.691638 | 0.232609 | 0.011 | 0.008 | 1 | 8 | Creb3l3 | 0.003 | Old |
| 0.691638 | 0.221489 | 0.011 | 0.008 | 1 | 8 | D730045B0 | 0.003 | Old |
| 0.691638 | 0.216149 | 0.011 | 0.008 | 1 | 8 | Gm36839 | 0.003 | Old |
| 0.691638 | 0.195374 | 0.011 | 0.008 | 1 | 8 | Cenpu | 0.003 | Old |
| 0.691638 | 0.184989 | 0.011 | 0.008 | 1 | 8 | B230369F2 | 0.003 | Old |
| 0.691748 | 0.433828 | 0.302 | 0.288 | 1 | 8 | Dennd1a | 0.014 | Old |
| 0.692064 | 0.218853 | 0.184 | 0.171 | 1 | 8 | Zfyve16 | 0.013 | Old |
| 0.692975 | 0.377304 | 0.151 | 0.143 | 1 | 8 | Fblim1 | 0.008 | Old |
| 0.693008 | 0.101988 | 0.212 | 0.229 | 1 | 8 | Ncbp3 | -0.017 | Old |
| 0.69307 | 0.128144 | 0.402 | 0.392 | 1 | 8 | Cyb5r4 | 0.01 | Old |
| 0.693113 | 0.084341 | 0.05 | 0.06 | 1 | 8 | Gtpbp10 | -0.01 | Old |
| 0.693416 | 0.255121 | 0.441 | 0.436 | 1 | 8 | Uhrf2 | 0.005 | Old |
| 0.693753 | 0.21879 | 0.291 | 0.322 | 1 | 8 | Rps6ka1 | -0.031 | Old |
| 0.693849 | 0.029165 | 0.067 | 0.057 | 1 | 8 | Ice2 | 0.01 | Old |

| | | | | | | | | |
|---|---|---|---|---|---|---|---|---|
| 0.693945 | 0.387777 | 0.223 | 0.213 | 1 | 8 | Mdp1 | 0.01 | Old |
| 0.694071 | 0.289362 | 0.067 | 0.078 | 1 | 8 | Smim24 | -0.011 | Old |
| 0.694162 | 0.289152 | 0.011 | 0.008 | 1 | 8 | Gpihbp1 | 0.003 | Old |
| 0.694162 | 0.13529 | 0.011 | 0.008 | 1 | 8 | Agbl5 | 0.003 | Old |
| 0.694162 | 0.088623 | 0.011 | 0.008 | 1 | 8 | Hoxd8 | 0.003 | Old |
| 0.694162 | 0.054696 | 0.011 | 0.008 | 1 | 8 | Fam185a | 0.003 | Old |
| 0.694162 | 0.035804 | 0.011 | 0.008 | 1 | 8 | Lockd | 0.003 | Old |
| 0.694477 | 0.233336 | 0.441 | 0.452 | 1 | 8 | Sel1l | -0.011 | Old |
| 0.694712 | 0.002167 | 0.374 | 0.392 | 1 | 8 | Pnp | -0.018 | Old |
| 0.694811 | 0.454371 | 0.117 | 0.104 | 1 | 8 | Sssca1 | 0.013 | Old |
| 0.695056 | 0.259296 | 0.274 | 0.262 | 1 | 8 | Suds3 | 0.012 | Old |
| 0.695138 | 0.015534 | 0.542 | 0.574 | 1 | 8 | Atp5o.1 | -0.032 | Old |
| 0.695171 | 0.216533 | 0.095 | 0.086 | 1 | 8 | Ogfod1 | 0.009 | Old |
| 0.695345 | 0.020524 | 0.268 | 0.291 | 1 | 8 | Strn4 | -0.023 | Old |
| 0.695539 | 0.473208 | 0.14 | 0.13 | 1 | 8 | Coq8b | 0.01 | Old |
| 0.695832 | 0.107643 | 0.458 | 0.491 | 1 | 8 | Cbx3 | -0.033 | Old |
| 0.69584 | 0.215615 | 0.073 | 0.065 | 1 | 8 | 2810402E2 | 0.008 | Old |
| 0.695877 | 0.072239 | 0.162 | 0.174 | 1 | 8 | Fbxw7 | -0.012 | Old |
| 0.695882 | 0.077681 | 0.246 | 0.268 | 1 | 8 | Gpr108 | -0.022 | Old |
| 0.695989 | 0.017003 | 0.039 | 0.047 | 1 | 8 | Cluap1 | -0.008 | Old |
| 0.696041 | 0.19564 | 0.128 | 0.14 | 1 | 8 | Fam45a | -0.012 | Old |
| 0.696303 | 0.181622 | 0.056 | 0.065 | 1 | 8 | Sgf29 | -0.009 | Old |
| 0.696517 | 0.277179 | 0.101 | 0.114 | 1 | 8 | Lmf2 | -0.013 | Old |
| 0.696688 | 0.09328 | 0.011 | 0.008 | 1 | 8 | Gm47469 | 0.003 | Old |
| 0.696762 | 0.463531 | 0.117 | 0.132 | 1 | 8 | Zfp524 | -0.015 | Old |
| 0.697087 | 0.309022 | 0.346 | 0.343 | 1 | 8 | Pik3r5 | 0.003 | Old |
| 0.697338 | 0.043068 | 0.844 | 0.901 | 1 | 8 | Ywhae | -0.057 | Old |
| 0.697595 | 0.354308 | 0.212 | 0.205 | 1 | 8 | Fam193a | 0.007 | Old |
| 0.697805 | 0.21297 | 0.24 | 0.231 | 1 | 8 | Dhx38 | 0.009 | Old |
| 0.697864 | 0.14696 | 0.302 | 0.327 | 1 | 8 | Vps13b | -0.025 | Old |
| 0.699145 | 0.412013 | 0.324 | 0.325 | 1 | 8 | Frg1 | -0.001 | Old |
| 0.699219 | 0.089037 | 0.011 | 0.008 | 1 | 8 | Scn4b | 0.003 | Old |
| 0.699503 | 0.071311 | 0.246 | 0.27 | 1 | 8 | Meaf6 | -0.024 | Old |
| 0.699621 | 0.338865 | 0.073 | 0.065 | 1 | 8 | Rusc2 | 0.008 | Old |
| 0.699641 | 0.313005 | 0.078 | 0.091 | 1 | 8 | Kat14 | -0.013 | Old |
| 0.700155 | 0.452333 | 0.196 | 0.182 | 1 | 8 | Arfrp1 | 0.014 | Old |
| 0.700159 | 0.178317 | 0.246 | 0.239 | 1 | 8 | Hmgcr | 0.007 | Old |
| 0.700281 | 0.964975 | 0.061 | 0.055 | 1 | 8 | Tlr5 | 0.006 | Old |
| 0.700392 | 0.080495 | 0.291 | 0.314 | 1 | 8 | Vcpip1 | -0.023 | Old |
| 0.700498 | 0.286056 | 0.196 | 0.187 | 1 | 8 | Micu2 | 0.009 | Old |
| 0.700632 | 0.089561 | 0.307 | 0.338 | 1 | 8 | Crat | -0.031 | Old |
| 0.701098 | 0.268495 | 0.251 | 0.268 | 1 | 8 | Fam50a | -0.017 | Old |
| 0.701275 | 0.173455 | 0.637 | 0.668 | 1 | 8 | C3ar1 | -0.031 | Old |
| 0.701393 | 0.26972 | 0.229 | 0.213 | 1 | 8 | Mtmr12 | 0.016 | Old |
| 0.701551 | 0.320223 | 0.151 | 0.14 | 1 | 8 | Abcf2 | 0.011 | Old |

| | | | | | | | | |
|---|---|---|---|---|---|---|---|---|
| 0.701996 | 0.031572 | 0.184 | 0.197 | 1 | 8 | Man1b1 | -0.013 | Old |
| 0.702214 | 0.315205 | 0.19 | 0.179 | 1 | 8 | Bin3 | 0.011 | Old |
| 0.702587 | 0.188035 | 0.235 | 0.223 | 1 | 8 | Enox2 | 0.012 | Old |
| 0.702588 | 0.396873 | 0.391 | 0.413 | 1 | 8 | Tcirg1 | -0.022 | Old |
| 0.702847 | 0.210304 | 0.19 | 0.2 | 1 | 8 | Narf | -0.01 | Old |
| 0.702913 | 0.261848 | 0.106 | 0.096 | 1 | 8 | Pigv | 0.01 | Old |
| 0.7031 | 0.121941 | 0.24 | 0.229 | 1 | 8 | Supt6 | 0.011 | Old |
| 0.703192 | 0.237039 | 0.374 | 0.413 | 1 | 8 | H1f0 | -0.039 | Old |
| 0.703414 | 0.049072 | 0.089 | 0.101 | 1 | 8 | Ercc5 | -0.012 | Old |
| 0.703454 | 0.122688 | 0.508 | 0.545 | 1 | 8 | Arrb2 | -0.037 | Old |
| 0.703459 | 0.185714 | 0.145 | 0.135 | 1 | 8 | Lsm2 | 0.01 | Old |
| 0.703657 | 0.139834 | 0.374 | 0.413 | 1 | 8 | Apaf1 | -0.039 | Old |
| 0.703978 | 0.083079 | 0.14 | 0.156 | 1 | 8 | Elof1 | -0.016 | Old |
| 0.704098 | 0.304311 | 0.218 | 0.208 | 1 | 8 | Ppfibp2 | 0.01 | Old |
| 0.704148 | 0.0567 | 0.335 | 0.361 | 1 | 8 | Idh2 | -0.026 | Old |
| 0.704185 | 0.099099 | 0.223 | 0.21 | 1 | 8 | 1700123O2 | 0.013 | Old |
| 0.70458 | 0.030144 | 0.346 | 0.319 | 1 | 8 | Azi2 | 0.027 | Old |
| 0.704682 | 0.088271 | 0.503 | 0.512 | 1 | 8 | Ppp1r15b | -0.009 | Old |
| 0.704895 | 0.075989 | 0.318 | 0.299 | 1 | 8 | Uxs1 | 0.019 | Old |
| 0.705054 | 0.163396 | 0.469 | 0.452 | 1 | 8 | Plaur | 0.017 | Old |
| 0.705061 | 0.07639 | 0.369 | 0.41 | 1 | 8 | Lamtor3 | -0.041 | Old |
| 0.705425 | 0.155819 | 0.123 | 0.135 | 1 | 8 | Vezt | -0.012 | Old |
| 0.706038 | 0.060615 | 0.084 | 0.094 | 1 | 8 | Ppp1r8 | -0.01 | Old |
| 0.706256 | 0.184612 | 0.073 | 0.065 | 1 | 8 | Zfp101 | 0.008 | Old |
| 0.706256 | 0.726197 | 0.073 | 0.065 | 1 | 8 | Iigp1 | 0.008 | Old |
| 0.706446 | 0.646948 | 0.061 | 0.055 | 1 | 8 | Igtp | 0.006 | Old |
| 0.706492 | 0.119715 | 0.162 | 0.177 | 1 | 8 | Arl3 | -0.015 | Old |
| 0.706588 | 0.138865 | 0.615 | 0.616 | 1 | 8 | Ppp2r5a | -0.001 | Old |
| 0.706714 | 0.194207 | 0.235 | 0.262 | 1 | 8 | Ahsa1 | -0.027 | Old |
| 0.706817 | 0.090886 | 0.212 | 0.231 | 1 | 8 | Ppp2r5e | -0.019 | Old |
| 0.707125 | 0.237463 | 0.073 | 0.083 | 1 | 8 | Gab3 | -0.01 | Old |
| 0.70735 | 0.255632 | 0.296 | 0.291 | 1 | 8 | Mrps17 | 0.005 | Old |
| 0.707386 | 0.269955 | 0.145 | 0.138 | 1 | 8 | Poglut1 | 0.007 | Old |
| 0.707715 | 0.475796 | 0.078 | 0.07 | 1 | 8 | Bud23 | 0.008 | Old |
| 0.708025 | 0.392343 | 0.168 | 0.161 | 1 | 8 | Faf2 | 0.007 | Old |
| 0.708083 | 0.015463 | 0.101 | 0.088 | 1 | 8 | Gmpr2 | 0.013 | Old |
| 0.708198 | 0.078313 | 0.508 | 0.532 | 1 | 8 | Vps4b | -0.024 | Old |
| 0.70833 | 0.024431 | 0.078 | 0.086 | 1 | 8 | Acyp1 | -0.008 | Old |
| 0.708494 | 0.216829 | 0.084 | 0.096 | 1 | 8 | Slc35b4 | -0.012 | Old |
| 0.708683 | 0.25073 | 0.24 | 0.27 | 1 | 8 | Zfp950 | -0.03 | Old |
| 0.70869 | 0.120079 | 0.039 | 0.047 | 1 | 8 | Clip2 | -0.008 | Old |
| 0.708942 | 0.002184 | 0.123 | 0.135 | 1 | 8 | Mycbp | -0.012 | Old |
| 0.709106 | 0.379635 | 0.056 | 0.065 | 1 | 8 | BC002059 | -0.009 | Old |
| 0.709563 | 0.246385 | 0.156 | 0.145 | 1 | 8 | Tubgcp3 | 0.011 | Old |
| 0.70973 | 0.217446 | 0.263 | 0.255 | 1 | 8 | Galnt10 | 0.008 | Old |

| | | | | | | | | |
|---|---|---|---|---|---|---|---|---|
| 0.710269 | 0.163582 | 0.246 | 0.262 | 1 | 8 | Gigyf2 | -0.016 | Old |
| 0.710398 | 0.946621 | 0.017 | 0.013 | 1 | 8 | Bcl7a | 0.004 | Old |
| 0.710398 | 0.85011 | 0.017 | 0.013 | 1 | 8 | Klhdc1 | 0.004 | Old |
| 0.710471 | 0.243812 | 0.313 | 0.34 | 1 | 8 | Limd2 | -0.027 | Old |
| 0.710567 | 0.653174 | 0.061 | 0.055 | 1 | 8 | Repin1 | 0.006 | Old |
| 0.711065 | 0.478731 | 0.184 | 0.177 | 1 | 8 | Psip1 | 0.007 | Old |
| 0.711083 | 0.199226 | 0.067 | 0.06 | 1 | 8 | Dgkq | 0.007 | Old |
| 0.711261 | 0.186293 | 0.067 | 0.078 | 1 | 8 | Rad51d | -0.011 | Old |
| 0.711399 | 0.403906 | 0.095 | 0.109 | 1 | 8 | Ppa2 | -0.014 | Old |
| 0.711812 | 0.14039 | 0.229 | 0.252 | 1 | 8 | Prpf18 | -0.023 | Old |
| 0.712192 | 0.234526 | 0.358 | 0.348 | 1 | 8 | Morc3 | 0.01 | Old |
| 0.712341 | 1.649856 | 0.028 | 0.023 | 1 | 8 | Trim68 | 0.005 | Old |
| 0.712417 | 0.992025 | 0.017 | 0.013 | 1 | 8 | Fzd1 | 0.004 | Old |
| 0.712534 | 0.053047 | 0.553 | 0.608 | 1 | 8 | Zfp207 | -0.055 | Old |
| 0.712748 | 0.150927 | 0.587 | 0.592 | 1 | 8 | Cd24a | -0.005 | Old |
| 0.713215 | 0.249947 | 0.168 | 0.158 | 1 | 8 | Timm44 | 0.01 | Old |
| 0.713294 | 0.062807 | 0.145 | 0.161 | 1 | 8 | Slc39a14 | -0.016 | Old |
| 0.713867 | 0.06527 | 0.073 | 0.065 | 1 | 8 | Eid3 | 0.008 | Old |
| 0.714205 | 0.432042 | 0.235 | 0.234 | 1 | 8 | Mif4gd | 0.001 | Old |
| 0.714334 | 0.171356 | 0.587 | 0.603 | 1 | 8 | Ahcyl2 | -0.016 | Old |
| 0.714415 | 0.169801 | 0.134 | 0.148 | 1 | 8 | Pign | -0.014 | Old |
| 0.714439 | 1.572843 | 0.017 | 0.013 | 1 | 8 | Pkmyt1 | 0.004 | Old |
| 0.714833 | 0.264956 | 0.33 | 0.309 | 1 | 8 | Npc1 | 0.021 | Old |
| 0.715146 | 0.390001 | 0.503 | 0.522 | 1 | 8 | Birc6 | -0.019 | Old |
| 0.715221 | 0.395515 | 0.251 | 0.252 | 1 | 8 | Arhgap12 | -0.001 | Old |
| 0.715235 | 0.23198 | 0.089 | 0.081 | 1 | 8 | Orc3 | 0.008 | Old |
| 0.71539 | 0.176414 | 0.028 | 0.034 | 1 | 8 | Usp35 | -0.006 | Old |
| 0.715475 | 0.322357 | 0.268 | 0.265 | 1 | 8 | Sgta | 0.003 | Old |
| 0.715577 | 0.242283 | 0.156 | 0.145 | 1 | 8 | Gpatch2l | 0.011 | Old |
| 0.715928 | 0.254181 | 0.179 | 0.169 | 1 | 8 | Tmem251 | 0.01 | Old |
| 0.716042 | 0.167851 | 0.324 | 0.322 | 1 | 8 | Sqor | 0.002 | Old |
| 0.716314 | 0.034618 | 0.156 | 0.169 | 1 | 8 | Slc25a30 | -0.013 | Old |
| 0.716468 | 0.356362 | 0.363 | 0.371 | 1 | 8 | Trim28 | -0.008 | Old |
| 0.716738 | 0.098302 | 0.24 | 0.265 | 1 | 8 | Rnaseh2c | -0.025 | Old |
| 0.716907 | 0.691718 | 0.123 | 0.117 | 1 | 8 | Nrm | 0.006 | Old |
| 0.717051 | 0.214631 | 0.117 | 0.109 | 1 | 8 | Zfp329 | 0.008 | Old |
| 0.71783 | 0.116457 | 0.067 | 0.075 | 1 | 8 | Entpd6 | -0.008 | Old |
| 0.71783 | 0.188372 | 0.089 | 0.081 | 1 | 8 | Calcrl | 0.008 | Old |
| 0.717883 | 0.069402 | 0.369 | 0.392 | 1 | 8 | Gramd4 | -0.023 | Old |
| 0.717949 | 0.309169 | 0.056 | 0.049 | 1 | 8 | Dcp1b | 0.007 | Old |
| 0.717978 | 0.132038 | 0.039 | 0.047 | 1 | 8 | Pcgf6 | -0.008 | Old |
| 0.718488 | 0.618894 | 0.017 | 0.013 | 1 | 8 | Zfp503 | 0.004 | Old |
| 0.718512 | 0.313071 | 0.251 | 0.275 | 1 | 8 | Rangap1 | -0.024 | Old |
| 0.718514 | 0.092885 | 0.123 | 0.112 | 1 | 8 | Mief1 | 0.011 | Old |
| 0.7187 | 0.201858 | 0.48 | 0.53 | 1 | 8 | Hspa4 | -0.05 | Old |

| | | | | | | | | |
|---|---|---|---|---|---|---|---|---|
| 0.71935 | 0.081568 | 0.084 | 0.094 | 1 | 8 | Unkl | -0.01 | Old |
| 0.719423 | 1.332919 | 0.022 | 0.018 | 1 | 8 | Zfp949 | 0.004 | Old |
| 0.719423 | 1.271449 | 0.022 | 0.018 | 1 | 8 | Rhebl1 | 0.004 | Old |
| 0.719557 | 0.493007 | 0.05 | 0.044 | 1 | 8 | Prmt9 | 0.006 | Old |
| 0.719627 | 0.077539 | 0.285 | 0.317 | 1 | 8 | Glrx | -0.032 | Old |
| 0.719887 | 0.150048 | 0.101 | 0.091 | 1 | 8 | Tradd | 0.01 | Old |
| 0.71994 | 0.229387 | 0.413 | 0.423 | 1 | 8 | Usp34 | -0.01 | Old |
| 0.720236 | 0.024372 | 0.413 | 0.426 | 1 | 8 | Ndufa5 | -0.013 | Old |
| 0.720515 | 0.74345 | 0.017 | 0.013 | 1 | 8 | L2hgdh | 0.004 | Old |
| 0.720515 | 0.47498 | 0.017 | 0.013 | 1 | 8 | Gm47232 | 0.004 | Old |
| 0.720515 | 0.311998 | 0.017 | 0.013 | 1 | 8 | 1110008L1 | 0.004 | Old |
| 0.720515 | 0.261497 | 0.017 | 0.013 | 1 | 8 | Jade3 | 0.004 | Old |
| 0.7207 | 0.606238 | 0.05 | 0.044 | 1 | 8 | Rttn | 0.006 | Old |
| 0.720863 | 0.295805 | 0.128 | 0.117 | 1 | 8 | Ublcp1 | 0.011 | Old |
| 0.721204 | 0.636179 | 0.056 | 0.049 | 1 | 8 | Fance | 0.007 | Old |
| 0.721435 | 0.169904 | 0.123 | 0.114 | 1 | 8 | Tirap | 0.009 | Old |
| 0.721521 | 0.301731 | 0.078 | 0.07 | 1 | 8 | Dph6 | 0.008 | Old |
| 0.721528 | 0.357197 | 0.101 | 0.091 | 1 | 8 | Map2k5 | 0.01 | Old |
| 0.721547 | 0.020818 | 0.112 | 0.101 | 1 | 8 | Bet1l | 0.011 | Old |
| 0.721605 | 0.10694 | 0.279 | 0.306 | 1 | 8 | Psmd3 | -0.027 | Old |
| 0.721672 | 0.074313 | 0.207 | 0.195 | 1 | 8 | Slc30a9 | 0.012 | Old |
| 0.722042 | 0.175075 | 0.302 | 0.288 | 1 | 8 | Fnbp1l | 0.014 | Old |
| 0.722269 | 0.239648 | 0.207 | 0.195 | 1 | 8 | Slc16a7 | 0.012 | Old |
| 0.722545 | 0.712328 | 0.017 | 0.013 | 1 | 8 | Fam189b | 0.004 | Old |
| 0.722545 | 0.552175 | 0.017 | 0.013 | 1 | 8 | 4930524J0 | 0.004 | Old |
| 0.722545 | 0.522986 | 0.017 | 0.013 | 1 | 8 | Flt1 | 0.004 | Old |
| 0.722545 | 0.334018 | 0.017 | 0.013 | 1 | 8 | Gm4285 | 0.004 | Old |
| 0.722545 | 0.282548 | 0.017 | 0.013 | 1 | 8 | Epas1 | 0.004 | Old |
| 0.722545 | 0.259897 | 0.017 | 0.013 | 1 | 8 | Xk | 0.004 | Old |
| 0.722545 | 0.117657 | 0.017 | 0.013 | 1 | 8 | Mgll | 0.004 | Old |
| 0.722635 | 0.27503 | 0.374 | 0.408 | 1 | 8 | Cript | -0.034 | Old |
| 0.722638 | 0.737546 | 0.039 | 0.047 | 1 | 8 | Grpel2 | -0.008 | Old |
| 0.722638 | 0.23032 | 0.039 | 0.047 | 1 | 8 | Gm26778 | -0.008 | Old |
| 0.722716 | 0.375551 | 0.089 | 0.101 | 1 | 8 | Igip | -0.012 | Old |
| 0.72325 | 0.434916 | 0.196 | 0.19 | 1 | 8 | Slc25a28 | 0.006 | Old |
| 0.723302 | 0.293417 | 0.14 | 0.132 | 1 | 8 | Mpv17 | 0.008 | Old |
| 0.723363 | 0.33976 | 0.089 | 0.099 | 1 | 8 | Zfp738 | -0.01 | Old |
| 0.723481 | 0.091672 | 0.207 | 0.221 | 1 | 8 | Ndufa9 | -0.014 | Old |
| 0.724132 | 0.124007 | 0.173 | 0.19 | 1 | 8 | Nsrp1 | -0.017 | Old |
| 0.724134 | 0.816729 | 0.05 | 0.044 | 1 | 8 | Lclat1 | 0.006 | Old |
| 0.724576 | 0.481504 | 0.017 | 0.013 | 1 | 8 | Zfp105 | 0.004 | Old |
| 0.724576 | 0.370536 | 0.017 | 0.013 | 1 | 8 | Rnd2 | 0.004 | Old |
| 0.724576 | 0.245247 | 0.017 | 0.013 | 1 | 8 | Gm43672 | 0.004 | Old |
| 0.724576 | 0.21426 | 0.017 | 0.013 | 1 | 8 | Oxld1 | 0.004 | Old |
| 0.724689 | 0.141717 | 0.117 | 0.106 | 1 | 8 | Egfr | 0.011 | Old |

| | | | | | | | | |
|---|---|---|---|---|---|---|---|---|
| 0.724932 | 0.350299 | 0.134 | 0.125 | 1 | 8 | Vps52 | 0.009 | Old |
| 0.724939 | 0.817499 | 0.028 | 0.034 | 1 | 8 | Usp20 | -0.006 | Old |
| 0.725006 | 0.037052 | 0.045 | 0.052 | 1 | 8 | Cnst | -0.007 | Old |
| 0.725056 | 0.623678 | 0.061 | 0.055 | 1 | 8 | Alg11 | 0.006 | Old |
| 0.725297 | 0.336985 | 0.24 | 0.226 | 1 | 8 | Erg28 | 0.014 | Old |
| 0.725336 | 0.019797 | 0.061 | 0.07 | 1 | 8 | Trpm2 | -0.009 | Old |
| 0.725363 | 0.337475 | 0.19 | 0.182 | 1 | 8 | Atmin | 0.008 | Old |
| 0.725546 | 0.044567 | 0.263 | 0.283 | 1 | 8 | Pcm1 | -0.02 | Old |
| 0.725563 | 0.446918 | 0.179 | 0.171 | 1 | 8 | Ift52 | 0.008 | Old |
| 0.725618 | 0.391176 | 0.419 | 0.426 | 1 | 8 | Eif2s3x | -0.007 | Old |
| 0.725618 | 0.264121 | 0.43 | 0.421 | 1 | 8 | Pcnp | 0.009 | Old |
| 0.725942 | 0.28001 | 0.145 | 0.138 | 1 | 8 | Rcan3 | 0.007 | Old |
| 0.725985 | 0.074809 | 0.531 | 0.595 | 1 | 8 | Arhgap17 | -0.064 | Old |
| 0.726207 | 0.666906 | 0.028 | 0.023 | 1 | 8 | Gm47675 | 0.005 | Old |
| 0.726307 | 0.260984 | 0.028 | 0.034 | 1 | 8 | Gm14326 | -0.006 | Old |
| 0.726369 | 1.641631 | 0.022 | 0.018 | 1 | 8 | A730063M | 0.004 | Old |
| 0.726369 | 0.529884 | 0.022 | 0.018 | 1 | 8 | Myo10 | 0.004 | Old |
| 0.726384 | 1.043251 | 0.045 | 0.039 | 1 | 8 | Usp11 | 0.006 | Old |
| 0.726396 | 0.159336 | 0.263 | 0.247 | 1 | 8 | Stard9 | 0.016 | Old |
| 0.726408 | 0.051364 | 0.117 | 0.127 | 1 | 8 | Cpped1 | -0.01 | Old |
| 0.72661 | 0.323007 | 0.017 | 0.013 | 1 | 8 | Hk1os | 0.004 | Old |
| 0.72661 | 0.1505 | 0.017 | 0.013 | 1 | 8 | Rom1 | 0.004 | Old |
| 0.72661 | 0.031187 | 0.017 | 0.013 | 1 | 8 | Homez | 0.004 | Old |
| 0.72661 | 0.028247 | 0.017 | 0.013 | 1 | 8 | Lpcat4 | 0.004 | Old |
| 0.726625 | 0.253426 | 0.089 | 0.099 | 1 | 8 | Prrg2 | -0.01 | Old |
| 0.726672 | 0.066005 | 0.14 | 0.151 | 1 | 8 | Josd1 | -0.011 | Old |
| 0.726814 | 0.36359 | 0.363 | 0.377 | 1 | 8 | Dennd6a | -0.014 | Old |
| 0.726985 | 0.150409 | 0.184 | 0.2 | 1 | 8 | Pilrb1 | -0.016 | Old |
| 0.727006 | 0.099107 | 0.134 | 0.145 | 1 | 8 | D6Wsu163 | -0.011 | Old |
| 0.727026 | 0.08394 | 0.101 | 0.112 | 1 | 8 | Klhdc10 | -0.011 | Old |
| 0.727219 | 0.138619 | 0.045 | 0.052 | 1 | 8 | Urb2 | -0.007 | Old |
| 0.727373 | 0.476653 | 0.034 | 0.029 | 1 | 8 | Osbpl3 | 0.005 | Old |
| 0.727676 | 0.399548 | 0.028 | 0.034 | 1 | 8 | Ppif | -0.006 | Old |
| 0.727676 | 0.353949 | 0.028 | 0.034 | 1 | 8 | Slc25a14 | -0.006 | Old |
| 0.728072 | 0.311466 | 0.106 | 0.096 | 1 | 8 | Zfp362 | 0.01 | Old |
| 0.72831 | 0.375166 | 0.268 | 0.257 | 1 | 8 | 1700017B0 | 0.011 | Old |
| 0.728555 | 0.145834 | 0.218 | 0.203 | 1 | 8 | Atl3 | 0.015 | Old |
| 0.728572 | 0.083454 | 0.101 | 0.112 | 1 | 8 | Naa25 | -0.011 | Old |
| 0.728645 | 0.705075 | 0.017 | 0.013 | 1 | 8 | Asb2 | 0.004 | Old |
| 0.728645 | 0.152561 | 0.017 | 0.013 | 1 | 8 | Pomc | 0.004 | Old |
| 0.728645 | 0.089015 | 0.017 | 0.013 | 1 | 8 | Arl4d | 0.004 | Old |
| 0.728645 | 0.050294 | 0.017 | 0.013 | 1 | 8 | Fads3 | 0.004 | Old |
| 0.728752 | 0.025137 | 0.106 | 0.117 | 1 | 8 | Socs5 | -0.011 | Old |
| 0.728781 | 0.767248 | 0.034 | 0.029 | 1 | 8 | Dpy19l3 | 0.005 | Old |
| 0.728895 | 0.33435 | 0.123 | 0.114 | 1 | 8 | Rpusd4 | 0.009 | Old |

| | | | | | | | | |
|---|---|---|---|---|---|---|---|---|
| 0.72894 | 0.468745 | 0.039 | 0.034 | 1 | 8 | Ldlrad3 | 0.005 | Old |
| 0.729013 | 0.213487 | 0.179 | 0.197 | 1 | 8 | Ube4a | -0.018 | Old |
| 0.729045 | 0.568411 | 0.028 | 0.034 | 1 | 8 | Mars2 | -0.006 | Old |
| 0.729173 | 0.338957 | 0.073 | 0.065 | 1 | 8 | Gpn2 | 0.008 | Old |
| 0.729605 | 0.543203 | 0.117 | 0.132 | 1 | 8 | Pip4k2c | -0.015 | Old |
| 0.729667 | 0.139966 | 0.162 | 0.179 | 1 | 8 | Hspa13 | -0.017 | Old |
| 0.729851 | 0.75436 | 0.022 | 0.018 | 1 | 8 | Mis18bp1 | 0.004 | Old |
| 0.729965 | 0.368006 | 0.134 | 0.125 | 1 | 8 | P2ry10b | 0.009 | Old |
| 0.730005 | 0.656589 | 0.084 | 0.096 | 1 | 8 | Tmem161a | -0.012 | Old |
| 0.730563 | 0.303072 | 0.095 | 0.106 | 1 | 8 | Pacs1 | -0.011 | Old |
| 0.730682 | 0.486337 | 0.017 | 0.013 | 1 | 8 | Srd5a1 | 0.004 | Old |
| 0.730682 | 0.283413 | 0.017 | 0.013 | 1 | 8 | Zfp784 | 0.004 | Old |
| 0.730682 | 0.272951 | 0.017 | 0.013 | 1 | 8 | Cpt1c | 0.004 | Old |
| 0.730682 | 0.203774 | 0.017 | 0.013 | 1 | 8 | Ubox5 | 0.004 | Old |
| 0.730682 | 0.04203 | 0.017 | 0.013 | 1 | 8 | Atg9b | 0.004 | Old |
| 0.730787 | 0.302178 | 0.223 | 0.244 | 1 | 8 | Arrdc4 | -0.021 | Old |
| 0.731069 | 0.244684 | 0.084 | 0.094 | 1 | 8 | Fam136a | -0.01 | Old |
| 0.731253 | 0.301825 | 0.045 | 0.039 | 1 | 8 | Bex6 | 0.006 | Old |
| 0.731281 | 0.322243 | 0.324 | 0.325 | 1 | 8 | Usp14 | -0.001 | Old |
| 0.731449 | 0.048258 | 0.754 | 0.779 | 1 | 8 | Folr2 | -0.025 | Old |
| 0.731466 | 0.013994 | 0.134 | 0.148 | 1 | 8 | Wdr20 | -0.014 | Old |
| 0.731669 | 0.962084 | 0.095 | 0.091 | 1 | 8 | Ramp3 | 0.004 | Old |
| 0.732055 | 0.340019 | 0.061 | 0.07 | 1 | 8 | Pcyox1l | -0.009 | Old |
| 0.732164 | 0.140274 | 0.151 | 0.138 | 1 | 8 | Prpsap2 | 0.013 | Old |
| 0.732328 | 0.984973 | 0.095 | 0.088 | 1 | 8 | Rusc1 | 0.007 | Old |
| 0.732561 | 0.155108 | 0.609 | 0.626 | 1 | 8 | Jarid2 | -0.017 | Old |
| 0.732604 | 0.209262 | 0.246 | 0.275 | 1 | 8 | Pbx2 | -0.029 | Old |
| 0.732721 | 0.003207 | 0.017 | 0.013 | 1 | 8 | Aqp1 | 0.004 | Old |
| 0.733008 | 0.148116 | 0.425 | 0.41 | 1 | 8 | Pcmtd1 | 0.015 | Old |
| 0.733338 | 0.588204 | 0.022 | 0.018 | 1 | 8 | Slc6a8 | 0.004 | Old |
| 0.733338 | 0.184073 | 0.022 | 0.018 | 1 | 8 | Polg2 | 0.004 | Old |
| 0.733712 | 0.062398 | 0.603 | 0.642 | 1 | 8 | Zfp106 | -0.039 | Old |
| 0.734063 | 0.078049 | 0.14 | 0.153 | 1 | 8 | Mxd1 | -0.013 | Old |
| 0.734294 | 0.116726 | 0.123 | 0.112 | 1 | 8 | Dgcr8 | 0.011 | Old |
| 0.734461 | 0.013902 | 0.978 | 0.984 | 1 | 8 | Iqgap1 | -0.006 | Old |
| 0.734467 | 0.154503 | 0.05 | 0.044 | 1 | 8 | Polr1b | 0.006 | Old |
| 0.734532 | 0.274711 | 0.028 | 0.034 | 1 | 8 | Vcpkmt | -0.006 | Old |
| 0.734757 | 0.069157 | 0.408 | 0.436 | 1 | 8 | 1-Mar | -0.028 | Old |
| 0.734779 | 0.013095 | 0.078 | 0.086 | 1 | 8 | Scnm1 | -0.008 | Old |
| 0.734986 | 0.420432 | 0.257 | 0.252 | 1 | 8 | Galc | 0.005 | Old |
| 0.735083 | 0.590048 | 0.022 | 0.018 | 1 | 8 | Ndufaf6 | 0.004 | Old |
| 0.735083 | 0.575579 | 0.022 | 0.018 | 1 | 8 | Cdc7 | 0.004 | Old |
| 0.735083 | 0.453327 | 0.022 | 0.018 | 1 | 8 | Gm43660 | 0.004 | Old |
| 0.73537 | 0.298884 | 0.134 | 0.125 | 1 | 8 | Zdhhc14 | 0.009 | Old |
| 0.735405 | 0.107788 | 0.469 | 0.449 | 1 | 8 | Exoc5 | 0.02 | Old |

| | | | | | | | | |
|---|---|---|---|---|---|---|---|---|
| 0.735459 | 1.00062 | 0.039 | 0.034 | 1 | 8 | Cenpt | 0.005 | Old |
| 0.735459 | 0.307377 | 0.039 | 0.034 | 1 | 8 | Zfp930 | 0.005 | Old |
| 0.735502 | 1.203504 | 0.028 | 0.023 | 1 | 8 | Etl4 | 0.005 | Old |
| 0.735502 | 0.360411 | 0.028 | 0.023 | 1 | 8 | Dand5 | 0.005 | Old |
| 0.736099 | 0.484322 | 0.179 | 0.174 | 1 | 8 | 9130401M | 0.005 | Old |
| 0.736154 | 0.371982 | 0.201 | 0.226 | 1 | 8 | Parp4 | -0.025 | Old |
| 0.736191 | 0.109162 | 0.497 | 0.535 | 1 | 8 | Map7d1 | -0.038 | Old |
| 0.736317 | 0.024938 | 0.302 | 0.291 | 1 | 8 | Atg2a | 0.011 | Old |
| 0.736322 | 0.44723 | 0.128 | 0.145 | 1 | 8 | Tmem55a | -0.017 | Old |
| 0.736463 | 0.470075 | 0.056 | 0.049 | 1 | 8 | Tfcp2 | 0.007 | Old |
| 0.736765 | 0.008487 | 0.039 | 0.034 | 1 | 8 | Pigo | 0.005 | Old |
| 0.73683 | 0.765627 | 0.022 | 0.018 | 1 | 8 | Morn2 | 0.004 | Old |
| 0.73683 | 0.414427 | 0.022 | 0.018 | 1 | 8 | Tubb2b | 0.004 | Old |
| 0.73683 | 0.217149 | 0.022 | 0.018 | 1 | 8 | Gm38251 | 0.004 | Old |
| 0.73683 | 0.17587 | 0.022 | 0.018 | 1 | 8 | Ift43 | 0.004 | Old |
| 0.737055 | 0.218883 | 0.028 | 0.023 | 1 | 8 | Ankrd33b | 0.005 | Old |
| 0.737083 | 0.098151 | 0.274 | 0.309 | 1 | 8 | Clcn5 | -0.035 | Old |
| 0.737247 | 0.294067 | 0.034 | 0.029 | 1 | 8 | A530017D2 | 0.005 | Old |
| 0.73728 | 0.453132 | 0.028 | 0.034 | 1 | 8 | Tmem218 | -0.006 | Old |
| 0.737473 | 0.152463 | 0.296 | 0.314 | 1 | 8 | Atp6v1d | -0.018 | Old |
| 0.737557 | 0.489753 | 0.056 | 0.049 | 1 | 8 | Fbxo21 | 0.007 | Old |
| 0.737557 | 0.443831 | 0.056 | 0.049 | 1 | 8 | Rfc3 | 0.007 | Old |
| 0.737761 | 0.361192 | 0.101 | 0.114 | 1 | 8 | Aamdc | -0.013 | Old |
| 0.73796 | 0.219561 | 0.056 | 0.065 | 1 | 8 | Fbxl18 | -0.009 | Old |
| 0.738177 | 0.334316 | 0.117 | 0.13 | 1 | 8 | Gcnt1 | -0.013 | Old |
| 0.738329 | 0.013165 | 0.743 | 0.813 | 1 | 8 | Dnaja1 | -0.07 | Old |
| 0.738577 | 0.572861 | 0.045 | 0.039 | 1 | 8 | B430306N( | 0.006 | Old |
| 0.738579 | 0.417 | 0.022 | 0.018 | 1 | 8 | Clec3b | 0.004 | Old |
| 0.7386 | 0.198302 | 0.061 | 0.055 | 1 | 8 | Flad1 | 0.006 | Old |
| 0.738619 | 0.318081 | 0.134 | 0.125 | 1 | 8 | Chpt1 | 0.009 | Old |
| 0.7387 | 0.021919 | 0.385 | 0.395 | 1 | 8 | Herc4 | -0.01 | Old |
| 0.738903 | 0.343107 | 0.101 | 0.094 | 1 | 8 | Pskh1 | 0.007 | Old |
| 0.739098 | 0.198956 | 0.145 | 0.138 | 1 | 8 | Taf4 | 0.007 | Old |
| 0.739139 | 0.010662 | 0.067 | 0.075 | 1 | 8 | Asna1 | -0.008 | Old |
| 0.739511 | 0.170161 | 0.324 | 0.358 | 1 | 8 | Vps36 | -0.034 | Old |
| 0.739759 | 0.174332 | 0.061 | 0.07 | 1 | 8 | Cdca8 | -0.009 | Old |
| 0.739886 | 0.229765 | 0.073 | 0.083 | 1 | 8 | Gm41307 | -0.01 | Old |
| 0.739937 | 0.070446 | 0.492 | 0.527 | 1 | 8 | Tmed9 | -0.035 | Old |
| 0.739964 | 0.213809 | 0.067 | 0.06 | 1 | 8 | Nek4 | 0.007 | Old |
| 0.740164 | 1.447882 | 0.028 | 0.023 | 1 | 8 | Slfn9 | 0.005 | Old |
| 0.740328 | 0.20635 | 0.022 | 0.018 | 1 | 8 | Zfyve28 | 0.004 | Old |
| 0.740328 | 0.13763 | 0.022 | 0.018 | 1 | 8 | Lyrm1 | 0.004 | Old |
| 0.740674 | 0.066522 | 0.207 | 0.218 | 1 | 8 | Fbxo4 | -0.011 | Old |
| 0.740857 | 0.081169 | 0.525 | 0.551 | 1 | 8 | C4b | -0.026 | Old |
| 0.740919 | 0.102843 | 0.369 | 0.397 | 1 | 8 | Vav1 | -0.028 | Old |

| | | | | | | | | |
|---|---|---|---|---|---|---|---|---|
| 0.741034 | 0.021833 | 0.849 | 0.922 | 1 | 8 | Tmem176b | -0.073 | Old |
| 0.741036 | 0.055691 | 0.168 | 0.156 | 1 | 8 | Pgs1 | 0.012 | Old |
| 0.741151 | 0.465473 | 0.19 | 0.184 | 1 | 8 | Cstf2t | 0.006 | Old |
| 0.741269 | 0.142147 | 0.257 | 0.273 | 1 | 8 | Ptdss1 | -0.016 | Old |
| 0.741384 | 0.76319 | 0.05 | 0.044 | 1 | 8 | Ap5s1 | 0.006 | Old |
| 0.741384 | 0.457722 | 0.05 | 0.044 | 1 | 8 | Tpbgl | 0.006 | Old |
| 0.741384 | 0.289882 | 0.05 | 0.044 | 1 | 8 | 2610044O1 | 0.006 | Old |
| 0.741492 | 0.422783 | 0.034 | 0.029 | 1 | 8 | Zfp580 | 0.005 | Old |
| 0.74172 | 0.31033 | 0.028 | 0.023 | 1 | 8 | Hgf | 0.005 | Old |
| 0.741938 | 0.256709 | 0.056 | 0.049 | 1 | 8 | Noa1 | 0.007 | Old |
| 0.74208 | 0.284686 | 0.022 | 0.018 | 1 | 8 | 4933417E1 | 0.004 | Old |
| 0.74208 | 0.086509 | 0.022 | 0.018 | 1 | 8 | Mblac1 | 0.004 | Old |
| 0.742166 | 0.105339 | 0.575 | 0.629 | 1 | 8 | Klf13 | -0.054 | Old |
| 0.742538 | 0.203377 | 0.05 | 0.044 | 1 | 8 | Slc44a2 | 0.006 | Old |
| 0.742742 | 0.353524 | 0.156 | 0.148 | 1 | 8 | Psmd10 | 0.008 | Old |
| 0.742841 | 0.567195 | 1 | 1 | 1 | 8 | Gm42418 | 0 | Old |
| 0.742908 | 0.388383 | 0.034 | 0.029 | 1 | 8 | Slc17a9 | 0.005 | Old |
| 0.742908 | 0.289599 | 0.034 | 0.029 | 1 | 8 | Dimt1 | 0.005 | Old |
| 0.743035 | 0.685019 | 0.056 | 0.049 | 1 | 8 | Nup188 | 0.007 | Old |
| 0.743277 | 1.017401 | 0.028 | 0.023 | 1 | 8 | Trub1 | 0.005 | Old |
| 0.743277 | 0.461936 | 0.028 | 0.023 | 1 | 8 | Spdl1 | 0.005 | Old |
| 0.743292 | 0.016129 | 0.263 | 0.281 | 1 | 8 | Chuk | -0.018 | Old |
| 0.743319 | 0.156034 | 0.24 | 0.255 | 1 | 8 | Nfatc1 | -0.015 | Old |
| 0.743694 | 0.716239 | 0.05 | 0.044 | 1 | 8 | Mettl8 | 0.006 | Old |
| 0.743798 | 0.07455 | 0.067 | 0.075 | 1 | 8 | Commd9 | -0.008 | Old |
| 0.743831 | 0.107363 | 0.061 | 0.055 | 1 | 8 | Echdc1 | 0.006 | Old |
| 0.743832 | 0.443008 | 0.022 | 0.018 | 1 | 8 | Agrn | 0.004 | Old |
| 0.743894 | 0.381363 | 0.307 | 0.34 | 1 | 8 | Tollip | -0.033 | Old |
| 0.744085 | 0.152174 | 0.156 | 0.169 | 1 | 8 | Trmt1l | -0.013 | Old |
| 0.744326 | 0.477442 | 0.034 | 0.029 | 1 | 8 | Ppp3cc | 0.005 | Old |
| 0.744326 | 0.070435 | 0.034 | 0.029 | 1 | 8 | Hectd2 | 0.005 | Old |
| 0.744545 | 0.121275 | 0.419 | 0.465 | 1 | 8 | Chmp3 | -0.046 | Old |
| 0.744618 | 0.79859 | 0.039 | 0.034 | 1 | 8 | Zfp628 | 0.005 | Old |
| 0.744632 | 0.221162 | 0.268 | 0.26 | 1 | 8 | Iah1 | 0.008 | Old |
| 0.744835 | 0.681483 | 0.028 | 0.023 | 1 | 8 | Snx32 | 0.005 | Old |
| 0.744835 | 0.264469 | 0.028 | 0.023 | 1 | 8 | Rtel1 | 0.005 | Old |
| 0.74485 | 0.25319 | 0.05 | 0.044 | 1 | 8 | Iba57 | 0.006 | Old |
| 0.744889 | 0.316936 | 0.251 | 0.247 | 1 | 8 | Polr2l | 0.004 | Old |
| 0.745025 | 0.35805 | 0.112 | 0.104 | 1 | 8 | Tdp2 | 0.008 | Old |
| 0.745544 | 0.157349 | 0.564 | 0.574 | 1 | 8 | Tubb6 | -0.01 | Old |
| 0.745586 | 0.155176 | 0.017 | 0.021 | 1 | 8 | Patj | -0.004 | Old |
| 0.745586 | 0.100868 | 0.022 | 0.018 | 1 | 8 | Ldlrad1 | 0.004 | Old |
| 0.745586 | 0.003355 | 0.022 | 0.018 | 1 | 8 | Nsun5 | 0.004 | Old |
| 0.745657 | 0.345482 | 0.101 | 0.112 | 1 | 8 | Triap1 | -0.011 | Old |
| 0.745689 | 0.022664 | 0.469 | 0.496 | 1 | 8 | Cct3 | -0.027 | Old |

| | | | | | | | | |
|---|---|---|---|---|---|---|---|---|
| 0.745725 | 0.721827 | 0.184 | 0.179 | 1 | 8 | Gon4l | 0.005 | Old |
| 0.745745 | 0.114516 | 0.737 | 0.784 | 1 | 8 | Lnpep | -0.047 | Old |
| 0.74581 | 0.350984 | 0.112 | 0.104 | 1 | 8 | Slc27a4 | 0.008 | Old |
| 0.745891 | 0.062492 | 0.413 | 0.431 | 1 | 8 | Rap1gds1 | -0.018 | Old |
| 0.745929 | 0.211895 | 0.039 | 0.034 | 1 | 8 | Tti2 | 0.005 | Old |
| 0.74625 | 0.407552 | 0.229 | 0.221 | 1 | 8 | Gm49336 | 0.008 | Old |
| 0.746307 | 0.123856 | 0.073 | 0.081 | 1 | 8 | Gm30978 | -0.008 | Old |
| 0.746395 | 0.056499 | 0.028 | 0.023 | 1 | 8 | Lrrc1 | 0.005 | Old |
| 0.746451 | 0.051409 | 0.536 | 0.558 | 1 | 8 | Spcs1 | -0.022 | Old |
| 0.746457 | 0.352307 | 0.095 | 0.106 | 1 | 8 | Jmy | -0.011 | Old |
| 0.746724 | 0.147601 | 0.531 | 0.504 | 1 | 8 | Mfsd14b | 0.027 | Old |
| 0.746944 | 0.005996 | 0.134 | 0.125 | 1 | 8 | Abhd17c | 0.009 | Old |
| 0.746992 | 0.70682 | 0.067 | 0.06 | 1 | 8 | Foxp4 | 0.007 | Old |
| 0.747073 | 0.084063 | 0.084 | 0.094 | 1 | 8 | Tspyl2 | -0.01 | Old |
| 0.747482 | 0.128053 | 0.151 | 0.161 | 1 | 8 | Rprd1b | -0.01 | Old |
| 0.74751 | 0.240177 | 0.151 | 0.143 | 1 | 8 | Scfd1 | 0.008 | Old |
| 0.747605 | 0.253474 | 0.128 | 0.122 | 1 | 8 | Cpox | 0.006 | Old |
| 0.747687 | 0.082914 | 0.564 | 0.61 | 1 | 8 | Bcl10 | -0.046 | Old |
| 0.747893 | 0.157625 | 0.341 | 0.332 | 1 | 8 | Usp47 | 0.009 | Old |
| 0.748024 | 0.225739 | 0.061 | 0.055 | 1 | 8 | Plscr4 | 0.006 | Old |
| 0.748024 | 0.206802 | 0.061 | 0.055 | 1 | 8 | Cutc | 0.006 | Old |
| 0.748024 | 0.127257 | 0.061 | 0.055 | 1 | 8 | Tram2 | 0.006 | Old |
| 0.748062 | 0.249292 | 0.212 | 0.205 | 1 | 8 | Tspan5 | 0.007 | Old |
| 0.748089 | 0.032253 | 0.207 | 0.195 | 1 | 8 | Dido1 | 0.012 | Old |
| 0.748308 | 1.234194 | 0.028 | 0.034 | 1 | 8 | Krt10 | -0.006 | Old |
| 0.748489 | 0.101274 | 0.609 | 0.675 | 1 | 8 | Capns1 | -0.066 | Old |
| 0.748565 | 0.108119 | 0.095 | 0.104 | 1 | 8 | Acad8 | -0.009 | Old |
| 0.748821 | 0.177327 | 0.48 | 0.47 | 1 | 8 | Trappc4 | 0.01 | Old |
| 0.749296 | 0.53179 | 0.117 | 0.112 | 1 | 8 | Cnnm4 | 0.005 | Old |
| 0.749369 | 0.348701 | 0.106 | 0.099 | 1 | 8 | Fbxo25 | 0.007 | Old |
| 0.749479 | 0.323119 | 0.05 | 0.044 | 1 | 8 | Smpd2 | 0.006 | Old |
| 0.749516 | 0.219728 | 0.028 | 0.023 | 1 | 8 | Tmem138 | 0.005 | Old |
| 0.749516 | 0.086437 | 0.028 | 0.023 | 1 | 8 | Naif1 | 0.005 | Old |
| 0.749532 | 0.164567 | 0.363 | 0.348 | 1 | 8 | Cd226 | 0.015 | Old |
| 0.749783 | 0.060331 | 0.156 | 0.143 | 1 | 8 | Thap11 | 0.013 | Old |
| 0.749825 | 0.038956 | 0.475 | 0.519 | 1 | 8 | Rab6a | -0.044 | Old |
| 0.750458 | 0.419653 | 0.14 | 0.132 | 1 | 8 | Ino80c | 0.008 | Old |
| 0.750638 | 0.228879 | 0.05 | 0.044 | 1 | 8 | Zmym6 | 0.006 | Old |
| 0.750855 | 0.337557 | 0.017 | 0.021 | 1 | 8 | Zfp12 | -0.004 | Old |
| 0.751307 | 0.286379 | 0.419 | 0.413 | 1 | 8 | Bud31 | 0.006 | Old |
| 0.751427 | 0.257292 | 0.034 | 0.029 | 1 | 8 | Slc25a40 | 0.005 | Old |
| 0.751495 | 0.158413 | 0.302 | 0.286 | 1 | 8 | Cops4 | 0.016 | Old |
| 0.751543 | 0.13778 | 0.078 | 0.088 | 1 | 8 | Nlrc5 | -0.01 | Old |
| 0.751811 | 0.365047 | 0.05 | 0.057 | 1 | 8 | Cpt2 | -0.007 | Old |
| 0.75204 | 0.064933 | 0.497 | 0.532 | 1 | 8 | Comt | -0.035 | Old |

| | | | | | | | | |
|---|---|---|---|---|---|---|---|---|
| 0.752065 | 0.073045 | 0.045 | 0.039 | 1 | 8 | Rrp1b | 0.006 | Old |
| 0.752497 | 0.310599 | 0.039 | 0.034 | 1 | 8 | Paqr4 | 0.005 | Old |
| 0.752525 | 0.362494 | 0.196 | 0.19 | 1 | 8 | Dpy19l4 | 0.006 | Old |
| 0.752662 | 0.114239 | 0.682 | 0.719 | 1 | 8 | Ifngr1 | -0.037 | Old |
| 0.752868 | 0.226419 | 0.196 | 0.187 | 1 | 8 | Sp140 | 0.009 | Old |
| 0.753295 | 0.169645 | 0.045 | 0.039 | 1 | 8 | Tysnd1 | 0.006 | Old |
| 0.753413 | 0.010517 | 0.408 | 0.403 | 1 | 8 | Fam208a | 0.005 | Old |
| 0.753817 | 0.101432 | 0.391 | 0.439 | 1 | 8 | Sar1a | -0.048 | Old |
| 0.753844 | 0.327272 | 0.084 | 0.094 | 1 | 8 | Mettl1 | -0.01 | Old |
| 0.754015 | 0.436375 | 0.112 | 0.109 | 1 | 8 | Poldip2 | 0.003 | Old |
| 0.754036 | 0.078906 | 0.14 | 0.148 | 1 | 8 | Manea | -0.008 | Old |
| 0.754193 | 0.086959 | 0.229 | 0.218 | 1 | 8 | Imp4 | 0.011 | Old |
| 0.754206 | 0.131209 | 0.028 | 0.023 | 1 | 8 | 4732440D( | 0.005 | Old |
| 0.754273 | 0.148189 | 0.034 | 0.029 | 1 | 8 | Ccdc171 | 0.005 | Old |
| 0.754403 | 0.437943 | 0.078 | 0.073 | 1 | 8 | Card9 | 0.005 | Old |
| 0.754477 | 0.425611 | 0.168 | 0.179 | 1 | 8 | Gm28875 | -0.011 | Old |
| 0.754526 | 0.251434 | 0.045 | 0.039 | 1 | 8 | Zfp512b | 0.006 | Old |
| 0.754593 | 0.174705 | 0.503 | 0.504 | 1 | 8 | Snx10 | -0.001 | Old |
| 0.754724 | 0.415274 | 0.151 | 0.143 | 1 | 8 | Alg5 | 0.008 | Old |
| 0.755278 | 0.733117 | 0.05 | 0.044 | 1 | 8 | Zfp141 | 0.006 | Old |
| 0.75538 | 0.21796 | 0.061 | 0.055 | 1 | 8 | 2610008E1 | 0.006 | Old |
| 0.755507 | 0.079732 | 0.128 | 0.135 | 1 | 8 | Gpaa1 | -0.007 | Old |
| 0.75554 | 0.073834 | 0.67 | 0.719 | 1 | 8 | Cct5 | -0.049 | Old |
| 0.755757 | 0.107314 | 0.045 | 0.039 | 1 | 8 | Omp | 0.006 | Old |
| 0.755994 | 0.303094 | 0.19 | 0.21 | 1 | 8 | Pou2f1 | -0.02 | Old |
| 0.756094 | 0.172446 | 0.184 | 0.2 | 1 | 8 | Phyh | -0.016 | Old |
| 0.756135 | 0.252916 | 0.017 | 0.021 | 1 | 8 | Mocs3 | -0.004 | Old |
| 0.756198 | 0.323627 | 0.045 | 0.052 | 1 | 8 | Rbm19 | -0.007 | Old |
| 0.756433 | 0.174023 | 0.128 | 0.119 | 1 | 8 | Zfp991 | 0.009 | Old |
| 0.75683 | 0.048024 | 0.101 | 0.109 | 1 | 8 | Rnf121 | -0.008 | Old |
| 0.756899 | 0.310352 | 0.084 | 0.078 | 1 | 8 | Hotairm1 | 0.006 | Old |
| 0.756936 | 0.213468 | 0.296 | 0.294 | 1 | 8 | Tcp11l2 | 0.002 | Old |
| 0.756989 | 0.021139 | 0.045 | 0.039 | 1 | 8 | Zmynd15 | 0.006 | Old |
| 0.757324 | 0.091547 | 0.788 | 0.836 | 1 | 8 | Cox6a1 | -0.048 | Old |
| 0.757457 | 0.0056 | 0.531 | 0.582 | 1 | 8 | Ndufb8 | -0.051 | Old |
| 0.757625 | 0.460422 | 0.117 | 0.109 | 1 | 8 | Fam78b | 0.008 | Old |
| 0.757753 | 0.108194 | 0.525 | 0.548 | 1 | 8 | Malt1 | -0.023 | Old |
| 0.757765 | 0.262878 | 0.039 | 0.034 | 1 | 8 | Itgb3bp | 0.005 | Old |
| 0.757833 | 0.173656 | 0.067 | 0.075 | 1 | 8 | Ppat | -0.008 | Old |
| 0.757898 | 0.279716 | 0.017 | 0.021 | 1 | 8 | Fut10 | -0.004 | Old |
| 0.757916 | 0.091267 | 0.43 | 0.462 | 1 | 8 | Emg1 | -0.032 | Old |
| 0.758372 | 0.101919 | 0.48 | 0.517 | 1 | 8 | Eif3l | -0.037 | Old |
| 0.758394 | 0.285653 | 0.196 | 0.208 | 1 | 8 | Zscan26 | -0.012 | Old |
| 0.758724 | 0.079101 | 0.43 | 0.418 | 1 | 8 | Txnrd1 | 0.012 | Old |
| 0.758849 | 0.284053 | 0.24 | 0.234 | 1 | 8 | Chchd3 | 0.006 | Old |

| | | | | | | | | |
|---|---|---|---|---|---|---|---|---|
| 0.759454 | 0.195474 | 0.045 | 0.039 | 1 | 8 | Spaca9 | 0.006 | Old |
| 0.759519 | 0.161219 | 0.218 | 0.239 | 1 | 8 | Smad5 | -0.021 | Old |
| 0.759531 | 0.139327 | 0.123 | 0.135 | 1 | 8 | Dhdds | -0.012 | Old |
| 0.759653 | 0.793088 | 0.067 | 0.062 | 1 | 8 | Slx1b | 0.005 | Old |
| 0.759662 | 0.190301 | 0.017 | 0.021 | 1 | 8 | Fscn1 | -0.004 | Old |
| 0.75977 | 0.089603 | 0.369 | 0.361 | 1 | 8 | Cenpb | 0.008 | Old |
| 0.760403 | 0.790644 | 0.039 | 0.034 | 1 | 8 | Mvk | 0.005 | Old |
| 0.760403 | 0.030988 | 0.039 | 0.034 | 1 | 8 | Fam102a | 0.005 | Old |
| 0.760641 | 0.594061 | 0.034 | 0.039 | 1 | 8 | D2hgdh | -0.005 | Old |
| 0.760662 | 0.456889 | 0.106 | 0.099 | 1 | 8 | Cwc27 | 0.007 | Old |
| 0.760925 | 0.269502 | 0.151 | 0.143 | 1 | 8 | Dst | 0.008 | Old |
| 0.761441 | 0.416302 | 0.162 | 0.156 | 1 | 8 | Bcam | 0.006 | Old |
| 0.761703 | 0.250317 | 0.061 | 0.055 | 1 | 8 | Them4 | 0.006 | Old |
| 0.761703 | 0.17931 | 0.061 | 0.055 | 1 | 8 | Usp42 | 0.006 | Old |
| 0.761715 | 0.312647 | 0.257 | 0.291 | 1 | 8 | Plpbp | -0.034 | Old |
| 0.761755 | 0.242312 | 0.056 | 0.049 | 1 | 8 | Tgm2 | 0.007 | Old |
| 0.761782 | 0.02059 | 0.922 | 0.951 | 1 | 8 | Cotl1 | -0.029 | Old |
| 0.762092 | 0.064959 | 0.453 | 0.481 | 1 | 8 | Mdh1 | -0.028 | Old |
| 0.762187 | 0.10746 | 0.369 | 0.405 | 1 | 8 | Lrp10 | -0.036 | Old |
| 0.762502 | 0.026671 | 0.196 | 0.184 | 1 | 8 | Vps9d1 | 0.012 | Old |
| 0.762948 | 0.108584 | 0.508 | 0.553 | 1 | 8 | Paip2 | -0.045 | Old |
| 0.763156 | 0.21669 | 0.112 | 0.122 | 1 | 8 | Ngrn | -0.01 | Old |
| 0.763631 | 0.159541 | 0.425 | 0.416 | 1 | 8 | Aebp2 | 0.009 | Old |
| 0.764511 | 0.135427 | 0.034 | 0.039 | 1 | 8 | Tmem38a | -0.005 | Old |
| 0.764644 | 0.19846 | 0.531 | 0.584 | 1 | 8 | Tsn | -0.053 | Old |
| 0.764751 | 0.032872 | 0.307 | 0.317 | 1 | 8 | Atxn7 | -0.01 | Old |
| 0.76486 | 0.45346 | 0.134 | 0.153 | 1 | 8 | Ing5 | -0.019 | Old |
| 0.764962 | 0.739941 | 0.017 | 0.021 | 1 | 8 | Ccnf | -0.004 | Old |
| 0.76542 | 0.190391 | 0.453 | 0.504 | 1 | 8 | Ifnar1 | -0.051 | Old |
| 0.765545 | 0.003235 | 0.872 | 0.935 | 1 | 8 | Arhgdia | -0.063 | Old |
| 0.765595 | 0.103857 | 0.33 | 0.345 | 1 | 8 | Ryr1 | -0.015 | Old |
| 0.766181 | 0.057069 | 0.056 | 0.049 | 1 | 8 | Adsl | 0.007 | Old |
| 0.76635 | 0.097968 | 0.324 | 0.332 | 1 | 8 | Spty2d1 | -0.008 | Old |
| 0.766404 | 0.354847 | 0.128 | 0.122 | 1 | 8 | Mlycd | 0.006 | Old |
| 0.766829 | 0.108214 | 0.497 | 0.514 | 1 | 8 | Tnpo1 | -0.017 | Old |
| 0.766916 | 0.008796 | 0.291 | 0.314 | 1 | 8 | Mob3c | -0.023 | Old |
| 0.767039 | 0.03874 | 0.374 | 0.392 | 1 | 8 | Phf21a | -0.018 | Old |
| 0.767095 | 0.277963 | 0.034 | 0.039 | 1 | 8 | Fadd | -0.005 | Old |
| 0.767138 | 0.093952 | 0.106 | 0.099 | 1 | 8 | Slc10a7 | 0.007 | Old |
| 0.767234 | 0.047695 | 0.145 | 0.156 | 1 | 8 | Ascc1 | -0.011 | Old |
| 0.767259 | 0.080132 | 0.162 | 0.179 | 1 | 8 | Trak2 | -0.017 | Old |
| 0.767383 | 0.140889 | 0.324 | 0.314 | 1 | 8 | Dennd4b | 0.01 | Old |
| 0.767656 | 0.229507 | 0.358 | 0.384 | 1 | 8 | Lpgat1 | -0.026 | Old |
| 0.767698 | 0.061544 | 0.408 | 0.403 | 1 | 8 | Hadhb | 0.005 | Old |
| 0.767732 | 0.05037 | 0.078 | 0.086 | 1 | 8 | Arfgap3 | -0.008 | Old |

| | | | | | | | | |
|---|---|---|---|---|---|---|---|---|
| 0.767807 | 0.271411 | 0.101 | 0.094 | 1 | 8 | Rabggta | 0.007 | Old |
| 0.767816 | 0.281677 | 0.268 | 0.26 | 1 | 8 | Ankib1 | 0.008 | Old |
| 0.767972 | 0.410459 | 0.324 | 0.325 | 1 | 8 | Trappc2l | -0.001 | Old |
| 0.76816 | 0.246752 | 0.514 | 0.548 | 1 | 8 | Ids | -0.034 | Old |
| 0.768387 | 0.163351 | 0.034 | 0.039 | 1 | 8 | Mdm1 | -0.005 | Old |
| 0.768479 | 0.438722 | 0.106 | 0.117 | 1 | 8 | Nedd1 | -0.011 | Old |
| 0.768637 | 0.178794 | 0.089 | 0.099 | 1 | 8 | Polr2d | -0.01 | Old |
| 0.768785 | 0.360446 | 0.162 | 0.156 | 1 | 8 | Aak1 | 0.006 | Old |
| 0.7691 | 0.03665 | 0.061 | 0.055 | 1 | 8 | Dhx32 | 0.006 | Old |
| 0.769532 | 0.322042 | 0.464 | 0.468 | 1 | 8 | Fgl2 | -0.004 | Old |
| 0.76955 | 0.144663 | 0.156 | 0.166 | 1 | 8 | Rnf139 | -0.01 | Old |
| 0.769555 | 0.132781 | 0.162 | 0.174 | 1 | 8 | Tmem71 | -0.012 | Old |
| 0.769657 | 0.035684 | 0.039 | 0.034 | 1 | 8 | Cd320 | 0.005 | Old |
| 0.769681 | 0.056798 | 0.034 | 0.039 | 1 | 8 | Prelid3a | -0.005 | Old |
| 0.769736 | 0.177479 | 0.117 | 0.127 | 1 | 8 | Bbc3 | -0.01 | Old |
| 0.77028 | 0.059438 | 1 | 0.987 | 1 | 8 | Marcks | 0.013 | Old |
| 0.770331 | 0.130457 | 0.358 | 0.338 | 1 | 8 | Mgrn1 | 0.02 | Old |
| 0.7707 | 0.317373 | 0.112 | 0.122 | 1 | 8 | Borcs7 | -0.01 | Old |
| 0.770864 | 0.334605 | 0.218 | 0.205 | 1 | 8 | Nabp2 | 0.013 | Old |
| 0.770877 | 0.263911 | 0.156 | 0.171 | 1 | 8 | Ube2g1 | -0.015 | Old |
| 0.771168 | 0.274967 | 0.229 | 0.221 | 1 | 8 | Gclc | 0.008 | Old |
| 0.771519 | 0.597921 | 0.112 | 0.106 | 1 | 8 | Gnpat | 0.006 | Old |
| 0.77152 | 0.034497 | 0.419 | 0.397 | 1 | 8 | Ndufa8 | 0.022 | Old |
| 0.771582 | 0.168852 | 0.615 | 0.688 | 1 | 8 | Glg1 | -0.073 | Old |
| 0.771781 | 0.245453 | 0.173 | 0.187 | 1 | 8 | Fam193b | -0.014 | Old |
| 0.771914 | 0.137867 | 0.156 | 0.148 | 1 | 8 | Cryzl1 | 0.008 | Old |
| 0.771917 | 0.262122 | 0.38 | 0.379 | 1 | 8 | Cep350 | 0.001 | Old |
| 0.771939 | 0.003254 | 0.682 | 0.797 | 1 | 8 | Lrp1 | -0.115 | Old |
| 0.771947 | 0.391152 | 0.045 | 0.052 | 1 | 8 | Ccl25 | -0.007 | Old |
| 0.771963 | 0.549669 | 0.089 | 0.099 | 1 | 8 | Ctdp1 | -0.01 | Old |
| 0.771963 | 0.284554 | 0.089 | 0.099 | 1 | 8 | Poc1b | -0.01 | Old |
| 0.772008 | 0.047519 | 0.14 | 0.13 | 1 | 8 | Ap4m1 | 0.01 | Old |
| 0.772464 | 0.65015 | 0.162 | 0.156 | 1 | 8 | Stk25 | 0.006 | Old |
| 0.77281 | 0.193676 | 0.542 | 0.574 | 1 | 8 | Lasp1 | -0.032 | Old |
| 0.772833 | 0.055024 | 0.056 | 0.049 | 1 | 8 | Ptges2 | 0.007 | Old |
| 0.773046 | 0.181332 | 0.078 | 0.086 | 1 | 8 | Hsf2 | -0.008 | Old |
| 0.773436 | 0.085423 | 0.145 | 0.161 | 1 | 8 | Mmgt1 | -0.016 | Old |
| 0.773689 | 0.277051 | 0.341 | 0.348 | 1 | 8 | N4bp2l2 | -0.007 | Old |
| 0.774411 | 0.18703 | 0.179 | 0.197 | 1 | 8 | Trpv2 | -0.018 | Old |
| 0.774712 | 0.751511 | 0.056 | 0.052 | 1 | 8 | Cdpf1 | 0.004 | Old |
| 0.774788 | 0.319929 | 0.335 | 0.34 | 1 | 8 | Slc39a7 | -0.005 | Old |
| 0.775216 | 0.144787 | 0.318 | 0.34 | 1 | 8 | Stt3b | -0.022 | Old |
| 0.775456 | 0.280833 | 0.162 | 0.153 | 1 | 8 | Senp5 | 0.009 | Old |
| 0.775469 | 0.245585 | 0.385 | 0.384 | 1 | 8 | Tspan32 | 0.001 | Old |
| 0.775491 | 0.127901 | 0.659 | 0.66 | 1 | 8 | Polr1d | -0.001 | Old |

| | | | | | | | | |
|---|---|---|---|---|---|---|---|---|
| 0.775852 | 0.003913 | 0.073 | 0.078 | 1 | 8 | Recql5 | -0.005 | Old |
| 0.775906 | 0.315016 | 0.117 | 0.13 | 1 | 8 | Dnajc16 | -0.013 | Old |
| 0.776135 | 0.338983 | 0.425 | 0.447 | 1 | 8 | Arrdc1 | -0.022 | Old |
| 0.776154 | 0.111738 | 0.156 | 0.166 | 1 | 8 | Mrpl2 | -0.01 | Old |
| 0.776468 | 0.472909 | 0.05 | 0.057 | 1 | 8 | Mterf4 | -0.007 | Old |
| 0.776468 | 0.333779 | 0.05 | 0.057 | 1 | 8 | Taf8 | -0.007 | Old |
| 0.776712 | 0.127391 | 0.469 | 0.478 | 1 | 8 | Fam168a | -0.009 | Old |
| 0.776738 | 0.419469 | 0.335 | 0.332 | 1 | 8 | Dnajb11 | 0.003 | Old |
| 0.77682 | 0.115924 | 0.151 | 0.161 | 1 | 8 | Eaf1 | -0.01 | Old |
| 0.77691 | 0.099119 | 0.313 | 0.325 | 1 | 8 | Rbbp4 | -0.012 | Old |
| 0.777414 | 0.292407 | 0.318 | 0.317 | 1 | 8 | Hs6st1 | 0.001 | Old |
| 0.777707 | 0.073512 | 0.134 | 0.145 | 1 | 8 | Thap2 | -0.011 | Old |
| 0.777739 | 0.142539 | 0.642 | 0.655 | 1 | 8 | Rgs2 | -0.013 | Old |
| 0.777802 | 0.416324 | 0.196 | 0.21 | 1 | 8 | Tbk1 | -0.014 | Old |
| 0.778005 | 0.158394 | 0.19 | 0.205 | 1 | 8 | Riok1 | -0.015 | Old |
| 0.778242 | 0.186388 | 0.38 | 0.418 | 1 | 8 | Ccnl2 | -0.038 | Old |
| 0.77837 | 0.045508 | 0.078 | 0.086 | 1 | 8 | Med31 | -0.008 | Old |
| 0.778623 | 0.041773 | 0.05 | 0.057 | 1 | 8 | Fbxl19 | -0.007 | Old |
| 0.778779 | 0.254296 | 0.128 | 0.14 | 1 | 8 | Krr1 | -0.012 | Old |
| 0.779042 | 0.019156 | 0.251 | 0.27 | 1 | 8 | Mrpl3 | -0.019 | Old |
| 0.779053 | 0.181455 | 0.24 | 0.231 | 1 | 8 | Tmem57 | 0.009 | Old |
| 0.779186 | 0.396645 | 0.201 | 0.205 | 1 | 8 | Gcc1 | -0.004 | Old |
| 0.779234 | 0.095261 | 0.525 | 0.577 | 1 | 8 | Orai1 | -0.052 | Old |
| 0.779275 | 0.307682 | 0.335 | 0.332 | 1 | 8 | Crybg1 | 0.003 | Old |
| 0.779681 | 0.279699 | 0.151 | 0.145 | 1 | 8 | Brf1 | 0.006 | Old |
| 0.779812 | 0.403381 | 0.101 | 0.109 | 1 | 8 | Ints11 | -0.008 | Old |
| 0.779997 | 0.1437 | 0.469 | 0.46 | 1 | 8 | Ybx3 | 0.009 | Old |
| 0.780315 | 0.18581 | 0.257 | 0.247 | 1 | 8 | Idh3b | 0.01 | Old |
| 0.78051 | 0.193351 | 0.358 | 0.358 | 1 | 8 | Ascc3 | 0 | Old |
| 0.780787 | 0.008093 | 0.263 | 0.257 | 1 | 8 | Mettl23 | 0.006 | Old |
| 0.781033 | 0.02454 | 0.112 | 0.119 | 1 | 8 | Nob1 | -0.007 | Old |
| 0.781167 | 0.025494 | 0.408 | 0.426 | 1 | 8 | Dnajc7 | -0.018 | Old |
| 0.781235 | 0.020985 | 0.464 | 0.496 | 1 | 8 | Cd302 | -0.032 | Old |
| 0.781245 | 0.212234 | 0.128 | 0.122 | 1 | 8 | Parp12 | 0.006 | Old |
| 0.781572 | 0.279469 | 0.162 | 0.174 | 1 | 8 | Med10 | -0.012 | Old |
| 0.781815 | 0.02181 | 0.553 | 0.571 | 1 | 8 | Ptp4a1 | -0.018 | Old |
| 0.781927 | 0.167414 | 0.38 | 0.366 | 1 | 8 | Dcaf7 | 0.014 | Old |
| 0.781979 | 0.165742 | 0.475 | 0.488 | 1 | 8 | Sin3b | -0.013 | Old |
| 0.782057 | 0.124728 | 0.112 | 0.122 | 1 | 8 | Mir142hg | -0.01 | Old |
| 0.782104 | 0.031352 | 0.14 | 0.13 | 1 | 8 | Kat7 | 0.01 | Old |
| 0.782271 | 0.044861 | 0.089 | 0.096 | 1 | 8 | Tmem199 | -0.007 | Old |
| 0.782436 | 0.057141 | 0.112 | 0.122 | 1 | 8 | Tor2a | -0.01 | Old |
| 0.782649 | 0.604163 | 0.034 | 0.039 | 1 | 8 | Lrrc49 | -0.005 | Old |
| 0.78283 | 0.299343 | 0.095 | 0.088 | 1 | 8 | Ubac1 | 0.007 | Old |
| 0.783607 | 0.022372 | 0.039 | 0.044 | 1 | 8 | BC024978 | -0.005 | Old |

| | | | | | | | | |
|---|---|---|---|---|---|---|---|---|
| 0.783677 | 0.898716 | 0.145 | 0.14 | 1 | 8 | Ttc13 | 0.005 | Old |
| 0.783843 | 0.026332 | 0.19 | 0.2 | 1 | 8 | Tmem63a | -0.01 | Old |
| 0.784019 | 0.486232 | 0.05 | 0.057 | 1 | 8 | Trmt11 | -0.007 | Old |
| 0.784029 | 0.292174 | 0.14 | 0.132 | 1 | 8 | Fbxo34 | 0.008 | Old |
| 0.78508 | 0.033358 | 0.274 | 0.291 | 1 | 8 | Ccdc124 | -0.017 | Old |
| 0.78555 | 0.064408 | 0.318 | 0.314 | 1 | 8 | Cers5 | 0.004 | Old |
| 0.785653 | 0.172736 | 0.385 | 0.382 | 1 | 8 | Psmd14 | 0.003 | Old |
| 0.785687 | 0.081685 | 0.76 | 0.803 | 1 | 8 | Adam15 | -0.043 | Old |
| 0.785869 | 0.342733 | 0.279 | 0.281 | 1 | 8 | Ube2v2 | -0.002 | Old |
| 0.785939 | 0.119584 | 0.156 | 0.148 | 1 | 8 | Hmg20a | 0.008 | Old |
| 0.786302 | 0.173527 | 0.201 | 0.195 | 1 | 8 | Map2k4 | 0.006 | Old |
| 0.786394 | 0.036672 | 0.425 | 0.397 | 1 | 8 | Mfsd14a | 0.028 | Old |
| 0.786845 | 0.17817 | 0.302 | 0.322 | 1 | 8 | Lpxn | -0.02 | Old |
| 0.787247 | 0.048097 | 0.553 | 0.558 | 1 | 8 | Midn | -0.005 | Old |
| 0.787382 | 0.118678 | 0.559 | 0.579 | 1 | 8 | Mapk1 | -0.02 | Old |
| 0.787552 | 0.446128 | 0.196 | 0.213 | 1 | 8 | Ash2l | -0.017 | Old |
| 0.787842 | 0.202101 | 0.084 | 0.091 | 1 | 8 | Tomm40l | -0.007 | Old |
| 0.788036 | 0.503289 | 0.067 | 0.075 | 1 | 8 | Hes1 | -0.008 | Old |
| 0.788348 | 0.208707 | 0.358 | 0.353 | 1 | 8 | Myo9a | 0.005 | Old |
| 0.789049 | 0.067013 | 0.078 | 0.086 | 1 | 8 | Ndc1 | -0.008 | Old |
| 0.789111 | 0.329842 | 0.246 | 0.239 | 1 | 8 | Fam3c | 0.007 | Old |
| 0.789555 | 0.065022 | 0.274 | 0.286 | 1 | 8 | Vamp4 | -0.012 | Old |
| 0.789586 | 0.3135 | 0.067 | 0.06 | 1 | 8 | Ap5z1 | 0.007 | Old |
| 0.789849 | 0.64284 | 0.123 | 0.119 | 1 | 8 | Smarcal1 | 0.004 | Old |
| 0.789913 | 0.356433 | 0.57 | 0.587 | 1 | 8 | Mfsd1 | -0.017 | Old |
| 0.790208 | 0.092167 | 0.749 | 0.769 | 1 | 8 | Tab2 | -0.02 | Old |
| 0.790329 | 0.350471 | 0.089 | 0.099 | 1 | 8 | Crebl2 | -0.01 | Old |
| 0.790625 | 0.230978 | 0.43 | 0.426 | 1 | 8 | Csnk1d | 0.004 | Old |
| 0.790807 | 0.140849 | 0.257 | 0.27 | 1 | 8 | Mrps12 | -0.013 | Old |
| 0.791147 | 0.026946 | 0.257 | 0.268 | 1 | 8 | Gtf3c2 | -0.011 | Old |
| 0.791159 | 0.41718 | 0.385 | 0.4 | 1 | 8 | Tmem109 | -0.015 | Old |
| 0.791465 | 0.187564 | 0.296 | 0.288 | 1 | 8 | Dnajc2 | 0.008 | Old |
| 0.791772 | 0.026989 | 0.916 | 0.943 | 1 | 8 | Wnk1 | -0.027 | Old |
| 0.791886 | 0.063661 | 0.168 | 0.156 | 1 | 8 | Npl | 0.012 | Old |
| 0.792107 | 0.108278 | 0.257 | 0.247 | 1 | 8 | Polr2c | 0.01 | Old |
| 0.792399 | 0.280922 | 0.425 | 0.475 | 1 | 8 | Baz1a | -0.05 | Old |
| 0.792502 | 0.264487 | 0.547 | 0.584 | 1 | 8 | Wac | -0.037 | Old |
| 0.79287 | 0.310294 | 0.112 | 0.106 | 1 | 8 | Psmg2 | 0.006 | Old |
| 0.79308 | 0.146323 | 0.33 | 0.345 | 1 | 8 | 2510039O1 | -0.015 | Old |
| 0.793376 | 0.23735 | 0.101 | 0.109 | 1 | 8 | Ddx49 | -0.008 | Old |
| 0.793635 | 0.015038 | 0.503 | 0.525 | 1 | 8 | Map2k2 | -0.022 | Old |
| 0.793826 | 0.131853 | 0.38 | 0.403 | 1 | 8 | Tspan14 | -0.023 | Old |
| 0.793848 | 0.276218 | 0.369 | 0.361 | 1 | 8 | Sppl3 | 0.008 | Old |
| 0.794041 | 0.172846 | 0.112 | 0.119 | 1 | 8 | Prkci | -0.007 | Old |
| 0.794815 | 0.562468 | 0.291 | 0.294 | 1 | 8 | Znhit1 | -0.003 | Old |

| | | | | | | | | |
|---|---|---|---|---|---|---|---|---|
| 0.795203 | 0.286678 | 0.352 | 0.361 | 1 | 8 | Wdfy2 | -0.009 | Old |
| 0.79543 | 0.032892 | 0.274 | 0.294 | 1 | 8 | Eif6 | -0.02 | Old |
| 0.795624 | 0.449297 | 0.151 | 0.145 | 1 | 8 | Phax | 0.006 | Old |
| 0.795717 | 0.214585 | 0.095 | 0.088 | 1 | 8 | C2cd2l | 0.007 | Old |
| 0.79582 | 0.049692 | 0.073 | 0.068 | 1 | 8 | Shb | 0.005 | Old |
| 0.796276 | 0.253496 | 0.14 | 0.132 | 1 | 8 | Scarb1 | 0.008 | Old |
| 0.796307 | 0.047872 | 0.229 | 0.218 | 1 | 8 | Tcf3 | 0.011 | Old |
| 0.796443 | 0.19166 | 0.039 | 0.044 | 1 | 8 | Gpr84 | -0.005 | Old |
| 0.796503 | 0.187386 | 0.291 | 0.286 | 1 | 8 | Ndufs4 | 0.005 | Old |
| 0.79653 | 0.332169 | 0.123 | 0.135 | 1 | 8 | Mfsd12 | -0.012 | Old |
| 0.796627 | 0.174934 | 0.056 | 0.062 | 1 | 8 | Dbf4 | -0.006 | Old |
| 0.796884 | 0.449959 | 0.106 | 0.099 | 1 | 8 | Aasdhppt | 0.007 | Old |
| 0.797187 | 0.418953 | 0.19 | 0.187 | 1 | 8 | Senp1 | 0.003 | Old |
| 0.797209 | 0.087743 | 0.369 | 0.403 | 1 | 8 | Csgalnact2 | -0.034 | Old |
| 0.797224 | 0.0349 | 0.24 | 0.252 | 1 | 8 | Ano6 | -0.012 | Old |
| 0.798013 | 0.238142 | 0.201 | 0.195 | 1 | 8 | Pds5b | 0.006 | Old |
| 0.798299 | 0.06079 | 0.246 | 0.234 | 1 | 8 | Slc16a3 | 0.012 | Old |
| 0.798401 | 0.109428 | 0.676 | 0.727 | 1 | 8 | Shisa5 | -0.051 | Old |
| 0.798663 | 0.143215 | 0.425 | 0.465 | 1 | 8 | Card19 | -0.04 | Old |
| 0.799193 | 0.221701 | 0.022 | 0.026 | 1 | 8 | Ctu1 | -0.004 | Old |
| 0.799284 | 0.165024 | 0.073 | 0.081 | 1 | 8 | Mrps35 | -0.008 | Old |
| 0.799344 | 0.460288 | 0.173 | 0.166 | 1 | 8 | Topbp1 | 0.007 | Old |
| 0.799507 | 0.201955 | 0.039 | 0.044 | 1 | 8 | Dcun1d4 | -0.005 | Old |
| 0.799556 | 0.037865 | 0.101 | 0.094 | 1 | 8 | Alkbh6 | 0.007 | Old |
| 0.799839 | 0.077534 | 0.151 | 0.143 | 1 | 8 | Rpp25l | 0.008 | Old |
| 0.799887 | 0.051049 | 0.128 | 0.138 | 1 | 8 | Cdk10 | -0.01 | Old |
| 0.79995 | 0.025582 | 0.793 | 0.823 | 1 | 8 | Csf2ra | -0.03 | Old |
| 0.799951 | 0.083113 | 0.369 | 0.353 | 1 | 8 | Ddx46 | 0.016 | Old |
| 0.800183 | 0.006703 | 0.112 | 0.119 | 1 | 8 | Carnmt1 | -0.007 | Old |
| 0.800277 | 0.190731 | 0.223 | 0.247 | 1 | 8 | Snrpa1 | -0.024 | Old |
| 0.800358 | 0.312904 | 0.274 | 0.265 | 1 | 8 | Tfg | 0.009 | Old |
| 0.80054 | 0.108655 | 0.201 | 0.21 | 1 | 8 | Mtr | -0.009 | Old |
| 0.800686 | 0.29524 | 0.531 | 0.574 | 1 | 8 | Gapvd1 | -0.043 | Old |
| 0.800689 | 0.267966 | 0.179 | 0.169 | 1 | 8 | H2-Q6 | 0.01 | Old |
| 0.800699 | 0.141186 | 0.235 | 0.229 | 1 | 8 | Uimc1 | 0.006 | Old |
| 0.800813 | 0.199064 | 0.346 | 0.343 | 1 | 8 | 2-Mar | 0.003 | Old |
| 0.800949 | 0.112877 | 0.128 | 0.135 | 1 | 8 | Hax1 | -0.007 | Old |
| 0.80096 | 0.13851 | 0.698 | 0.756 | 1 | 8 | Atp6v1f | -0.058 | Old |
| 0.801486 | 0.50056 | 0.128 | 0.125 | 1 | 8 | Polk | 0.003 | Old |
| 0.801745 | 0.060959 | 0.279 | 0.294 | 1 | 8 | Nop58 | -0.015 | Old |
| 0.802499 | 0.182744 | 0.229 | 0.221 | 1 | 8 | Utp3 | 0.008 | Old |
| 0.80322 | 0.524036 | 0.291 | 0.291 | 1 | 8 | Wdr37 | 0 | Old |
| 0.803275 | 0.622648 | 0.05 | 0.047 | 1 | 8 | 2-Sep | 0.003 | Old |
| 0.803348 | 0.129387 | 0.201 | 0.218 | 1 | 8 | Xpa | -0.017 | Old |
| 0.80376 | 0.138804 | 0.251 | 0.265 | 1 | 8 | Gsk3a | -0.014 | Old |

| | | | | | | | | |
|---|---|---|---|---|---|---|---|---|
| 0.803862 | 0.477426 | 0.089 | 0.083 | 1 | 8 | Cep85 | 0.006 | Old |
| 0.804031 | 0.109218 | 0.117 | 0.127 | 1 | 8 | Rnf19a | -0.01 | Old |
| 0.804208 | 0.166489 | 0.246 | 0.262 | 1 | 8 | Plgrkt | -0.016 | Old |
| 0.804246 | 0.258273 | 0.078 | 0.086 | 1 | 8 | 6030458C1 | -0.008 | Old |
| 0.804332 | 0.105151 | 0.128 | 0.119 | 1 | 8 | Tsga10 | 0.009 | Old |
| 0.80434 | 0.133043 | 0.095 | 0.088 | 1 | 8 | Xpc | 0.007 | Old |
| 0.804348 | 0.061887 | 0.726 | 0.753 | 1 | 8 | Syngr2 | -0.027 | Old |
| 0.804351 | 0.332149 | 0.084 | 0.091 | 1 | 8 | Ciao1 | -0.007 | Old |
| 0.804422 | 0.274278 | 0.067 | 0.073 | 1 | 8 | Eif2ak4 | -0.006 | Old |
| 0.80455 | 0.022701 | 0.318 | 0.301 | 1 | 8 | Smad4 | 0.017 | Old |
| 0.804765 | 0.290263 | 0.268 | 0.268 | 1 | 8 | Plekhg5 | 0 | Old |
| 0.804815 | 0.318883 | 0.335 | 0.345 | 1 | 8 | Pdap1 | -0.01 | Old |
| 0.805564 | 0.032112 | 0.022 | 0.026 | 1 | 8 | Htr1b | -0.004 | Old |
| 0.805675 | 0.17839 | 0.235 | 0.226 | 1 | 8 | Eif4ebp1 | 0.009 | Old |
| 0.805855 | 0.092522 | 0.061 | 0.068 | 1 | 8 | Cnksr3 | -0.007 | Old |
| 0.80654 | 0.372342 | 0.475 | 0.475 | 1 | 8 | Cebpd | 0 | Old |
| 0.806597 | 0.50178 | 0.056 | 0.052 | 1 | 8 | Msh6 | 0.004 | Old |
| 0.806597 | 0.071065 | 0.117 | 0.125 | 1 | 8 | Ifi35 | -0.008 | Old |
| 0.806955 | 0.161375 | 0.067 | 0.062 | 1 | 8 | Dnaja3 | 0.005 | Old |
| 0.807064 | 0.368104 | 0.419 | 0.481 | 1 | 8 | Zfp703 | -0.062 | Old |
| 0.807159 | 0.853831 | 0.022 | 0.026 | 1 | 8 | Klhl26 | -0.004 | Old |
| 0.807159 | 0.202043 | 0.022 | 0.026 | 1 | 8 | Slc37a3 | -0.004 | Old |
| 0.807346 | 0.218616 | 0.425 | 0.423 | 1 | 8 | Fam96a | 0.002 | Old |
| 0.807436 | 0.146612 | 0.38 | 0.403 | 1 | 8 | Uba2 | -0.023 | Old |
| 0.807918 | 0.207069 | 0.279 | 0.294 | 1 | 8 | Golgb1 | -0.015 | Old |
| 0.808072 | 0.370671 | 0.168 | 0.161 | 1 | 8 | Arntl | 0.007 | Old |
| 0.808104 | 0.449677 | 0.039 | 0.044 | 1 | 8 | Tubgcp6 | -0.005 | Old |
| 0.808176 | 0.189463 | 0.486 | 0.514 | 1 | 8 | Vapa | -0.028 | Old |
| 0.808755 | 0.262912 | 0.022 | 0.026 | 1 | 8 | 3-Mar | -0.004 | Old |
| 0.808783 | 0.05109 | 0.48 | 0.499 | 1 | 8 | Crk | -0.019 | Old |
| 0.808902 | 0.486892 | 0.128 | 0.14 | 1 | 8 | Dhx8 | -0.012 | Old |
| 0.808958 | 0.034559 | 0.128 | 0.135 | 1 | 8 | Pkd2 | -0.007 | Old |
| 0.809024 | 0.117403 | 0.212 | 0.223 | 1 | 8 | Casp8ap2 | -0.011 | Old |
| 0.809117 | 0.08395 | 0.061 | 0.057 | 1 | 8 | Gtpbp8 | 0.004 | Old |
| 0.809334 | 0.247063 | 0.039 | 0.044 | 1 | 8 | C330018D2 | -0.005 | Old |
| 0.80971 | 0.0775 | 0.134 | 0.127 | 1 | 8 | Arfgef2 | 0.007 | Old |
| 0.809798 | 0.31304 | 0.235 | 0.26 | 1 | 8 | Psmc4 | -0.025 | Old |
| 0.80984 | 0.252169 | 0.302 | 0.296 | 1 | 8 | Ilf3 | 0.006 | Old |
| 0.809941 | 0.150561 | 0.419 | 0.429 | 1 | 8 | Pon2 | -0.01 | Old |
| 0.81016 | 0.285148 | 0.056 | 0.062 | 1 | 8 | Dus3l | -0.006 | Old |
| 0.810219 | 0.261452 | 0.067 | 0.073 | 1 | 8 | Cep128 | -0.006 | Old |
| 0.810228 | 0.005777 | 0.201 | 0.208 | 1 | 8 | Cwf19l2 | -0.007 | Old |
| 0.810351 | 0.064344 | 0.022 | 0.026 | 1 | 8 | Rpusd1 | -0.004 | Old |
| 0.810351 | 0.064282 | 0.022 | 0.026 | 1 | 8 | Tedc2 | -0.004 | Old |
| 0.81039 | 0.053233 | 0.095 | 0.088 | 1 | 8 | Acot2 | 0.007 | Old |

| | | | | | | | | |
|---|---|---|---|---|---|---|---|---|
| 0.810533 | 0.246701 | 0.207 | 0.205 | 1 | 8 | Ptk2b | 0.002 | Old |
| 0.810787 | 0.437092 | 0.117 | 0.127 | 1 | 8 | Cenpc1 | -0.01 | Old |
| 0.811327 | 0.011108 | 1 | 1 | 1 | 8 | H2-Aa | 0 | Old |
| 0.811864 | 0.103626 | 0.279 | 0.304 | 1 | 8 | Sertad2 | -0.025 | Old |
| 0.811875 | 0.078514 | 0.128 | 0.135 | 1 | 8 | Lsm14b | -0.007 | Old |
| 0.812304 | 0.034078 | 0.86 | 0.914 | 1 | 8 | Btf3 | -0.054 | Old |
| 0.812422 | 0.239425 | 0.073 | 0.068 | 1 | 8 | C87436 | 0.005 | Old |
| 0.813088 | 0.075152 | 0.525 | 0.54 | 1 | 8 | Casp4 | -0.015 | Old |
| 0.813501 | 0.100553 | 0.615 | 0.636 | 1 | 8 | Matr3 | -0.021 | Old |
| 0.813562 | 0.252207 | 0.24 | 0.236 | 1 | 8 | Nup153 | 0.004 | Old |
| 0.813739 | 0.157334 | 0.045 | 0.049 | 1 | 8 | Aldh6a1 | -0.004 | Old |
| 0.8138 | 0.137303 | 0.564 | 0.639 | 1 | 8 | Eif4ebp2 | -0.075 | Old |
| 0.814004 | 0.118844 | 0.073 | 0.078 | 1 | 8 | Rrp9 | -0.005 | Old |
| 0.814181 | 0.220076 | 0.173 | 0.166 | 1 | 8 | Fbxl14 | 0.007 | Old |
| 0.814482 | 0.102425 | 0.615 | 0.657 | 1 | 8 | Eif4b | -0.042 | Old |
| 0.815145 | 0.029261 | 0.022 | 0.026 | 1 | 8 | D3Ertd751 | -0.004 | Old |
| 0.815403 | 0.151374 | 0.196 | 0.205 | 1 | 8 | Nf1 | -0.009 | Old |
| 0.81542 | 0.018278 | 0.089 | 0.083 | 1 | 8 | Cdc42ep2 | 0.006 | Old |
| 0.815546 | 0.233617 | 0.408 | 0.442 | 1 | 8 | Kat2b | -0.034 | Old |
| 0.815761 | 0.150591 | 0.497 | 0.496 | 1 | 8 | Zfand6 | 0.001 | Old |
| 0.815822 | 0.152838 | 0.419 | 0.447 | 1 | 8 | mt-Atp8 | -0.028 | Old |
| 0.815855 | 0.161144 | 0.33 | 0.325 | 1 | 8 | Tmf1 | 0.005 | Old |
| 0.816431 | 0.181737 | 0.128 | 0.119 | 1 | 8 | Slc41a2 | 0.009 | Old |
| 0.816491 | 0.263659 | 0.168 | 0.177 | 1 | 8 | Cln3 | -0.009 | Old |
| 0.816658 | 0.151136 | 0.112 | 0.104 | 1 | 8 | N6amt1 | 0.008 | Old |
| 0.81689 | 0.219415 | 0.33 | 0.325 | 1 | 8 | Plekhg3 | 0.005 | Old |
| 0.817235 | 0.525532 | 0.05 | 0.047 | 1 | 8 | Ndst1 | 0.003 | Old |
| 0.81755 | 0.085677 | 0.804 | 0.839 | 1 | 8 | Ppp3ca | -0.035 | Old |
| 0.817555 | 0.128593 | 0.57 | 0.621 | 1 | 8 | Itgb2 | -0.051 | Old |
| 0.817644 | 0.122031 | 0.179 | 0.19 | 1 | 8 | Polr3gl | -0.011 | Old |
| 0.817723 | 0.084337 | 0.771 | 0.847 | 1 | 8 | Cd81 | -0.076 | Old |
| 0.817906 | 0.132537 | 0.397 | 0.397 | 1 | 8 | Slfn5 | 0 | Old |
| 0.817912 | 0.54966 | 0.117 | 0.13 | 1 | 8 | Banp | -0.013 | Old |
| 0.817965 | 0.263104 | 0.067 | 0.073 | 1 | 8 | Dcakd | -0.006 | Old |
| 0.818003 | 0.225779 | 0.156 | 0.148 | 1 | 8 | Gaa | 0.008 | Old |
| 0.818302 | 0.29792 | 0.073 | 0.068 | 1 | 8 | Exosc7 | 0.005 | Old |
| 0.818345 | 0.637588 | 0.022 | 0.026 | 1 | 8 | Ccne1 | -0.004 | Old |
| 0.818401 | 0.686759 | 0.05 | 0.047 | 1 | 8 | B4galt7 | 0.003 | Old |
| 0.818696 | 0.254848 | 0.162 | 0.156 | 1 | 8 | Zfp384 | 0.006 | Old |
| 0.818787 | 0.111449 | 0.151 | 0.143 | 1 | 8 | Rad54l2 | 0.008 | Old |
| 0.81913 | 0.242956 | 0.475 | 0.509 | 1 | 8 | Pxn | -0.034 | Old |
| 0.819145 | 0.360999 | 0.101 | 0.096 | 1 | 8 | Nup160 | 0.005 | Old |
| 0.819324 | 0.457025 | 0.089 | 0.086 | 1 | 8 | Rev3l | 0.003 | Old |
| 0.819628 | 0.099276 | 0.145 | 0.135 | 1 | 8 | Utp20 | 0.01 | Old |
| 0.819653 | 0.0838 | 0.307 | 0.309 | 1 | 8 | St6galnac6 | -0.002 | Old |

| | | | | | | | | |
|---|---|---|---|---|---|---|---|---|
| 0.819704 | 0.09805 | 0.335 | 0.358 | 1 | 8 | Pcmt1 | -0.023 | Old |
| 0.819838 | 0.504078 | 0.179 | 0.174 | 1 | 8 | Papss1 | 0.005 | Old |
| 0.819889 | 0.570785 | 0.056 | 0.052 | 1 | 8 | Clybl | 0.004 | Old |
| 0.819889 | 0.373997 | 0.056 | 0.052 | 1 | 8 | Orc6 | 0.004 | Old |
| 0.819914 | 0.227911 | 0.156 | 0.169 | 1 | 8 | Spink2 | -0.013 | Old |
| 0.820208 | 0.092449 | 0.453 | 0.481 | 1 | 8 | Irak1 | -0.028 | Old |
| 0.820265 | 0.406145 | 0.073 | 0.068 | 1 | 8 | Kantr | 0.005 | Old |
| 0.820396 | 0.261845 | 0.374 | 0.403 | 1 | 8 | Nrbp1 | -0.029 | Old |
| 0.820398 | 0.220257 | 0.352 | 0.343 | 1 | 8 | 4931406C0 | 0.009 | Old |
| 0.820458 | 0.083867 | 0.207 | 0.218 | 1 | 8 | Pfkfb3 | -0.011 | Old |
| 0.820626 | 0.073624 | 0.19 | 0.197 | 1 | 8 | Reps1 | -0.007 | Old |
| 0.820707 | 0.141701 | 0.246 | 0.257 | 1 | 8 | Luc7l | -0.011 | Old |
| 0.820884 | 0.145614 | 0.453 | 0.457 | 1 | 8 | Tmed3 | -0.004 | Old |
| 0.820999 | 0.690356 | 0.056 | 0.052 | 1 | 8 | Dhx37 | 0.004 | Old |
| 0.821164 | 0.187545 | 0.24 | 0.234 | 1 | 8 | Uhrf1bp1l | 0.006 | Old |
| 0.821392 | 0.130448 | 0.168 | 0.177 | 1 | 8 | Aldh3a2 | -0.009 | Old |
| 0.821738 | 0.338861 | 0.117 | 0.112 | 1 | 8 | Tbpl1 | 0.005 | Old |
| 0.822109 | 0.300355 | 0.056 | 0.052 | 1 | 8 | Mgme1 | 0.004 | Old |
| 0.822228 | 0.135216 | 0.073 | 0.068 | 1 | 8 | Zc3hc1 | 0.005 | Old |
| 0.822474 | 0.119238 | 0.391 | 0.423 | 1 | 8 | Bak1 | -0.032 | Old |
| 0.822549 | 0.328243 | 0.112 | 0.119 | 1 | 8 | Dcaf1 | -0.007 | Old |
| 0.822631 | 0.052493 | 0.318 | 0.332 | 1 | 8 | Mnt | -0.014 | Old |
| 0.822923 | 0.28331 | 0.061 | 0.068 | 1 | 8 | Golm1 | -0.007 | Old |
| 0.823074 | 0.025036 | 0.184 | 0.19 | 1 | 8 | Cyp20a1 | -0.006 | Old |
| 0.82327 | 0.115524 | 0.285 | 0.299 | 1 | 8 | Gpat4 | -0.014 | Old |
| 0.823289 | 0.150626 | 0.486 | 0.527 | 1 | 8 | Rpl36al | -0.041 | Old |
| 0.823322 | 0.175094 | 0.112 | 0.119 | 1 | 8 | Rbm15 | -0.007 | Old |
| 0.82395 | 0.189309 | 0.67 | 0.719 | 1 | 8 | B4galt6 | -0.049 | Old |
| 0.824193 | 0.475954 | 0.073 | 0.068 | 1 | 8 | Mpi | 0.005 | Old |
| 0.824265 | 1.67E-05 | 0.642 | 0.688 | 1 | 8 | Slc3a2 | -0.046 | Old |
| 0.824331 | 0.543085 | 0.056 | 0.052 | 1 | 8 | Gdpd1 | 0.004 | Old |
| 0.824675 | 0.150498 | 0.603 | 0.655 | 1 | 8 | Sppl2a | -0.052 | Old |
| 0.825034 | 0.121813 | 0.24 | 0.231 | 1 | 8 | Ccdc22 | 0.009 | Old |
| 0.825036 | 0.290412 | 0.24 | 0.252 | 1 | 8 | Med1 | -0.012 | Old |
| 0.825619 | 0.087544 | 0.123 | 0.117 | 1 | 8 | Vps26b | 0.006 | Old |
| 0.825845 | 0.619449 | 0.056 | 0.062 | 1 | 8 | Dnajc12 | -0.006 | Old |
| 0.825877 | 0.19614 | 0.134 | 0.13 | 1 | 8 | Cwc22 | 0.004 | Old |
| 0.825892 | 0.430006 | 0.173 | 0.19 | 1 | 8 | Lman2l | -0.017 | Old |
| 0.826316 | 0.137394 | 0.341 | 0.366 | 1 | 8 | Psd3 | -0.025 | Old |
| 0.82633 | 0.10507 | 0.156 | 0.148 | 1 | 8 | Rap1gap2 | 0.008 | Old |
| 0.826554 | 0.393788 | 0.089 | 0.099 | 1 | 8 | Pianp | -0.01 | Old |
| 0.826971 | 0.20427 | 0.229 | 0.239 | 1 | 8 | 1810013L2 | -0.01 | Old |
| 0.827881 | 0.242364 | 0.028 | 0.031 | 1 | 8 | Pola2 | -0.003 | Old |
| 0.828073 | 0.055937 | 0.095 | 0.101 | 1 | 8 | Fam78a | -0.006 | Old |
| 0.828379 | 0.11624 | 0.101 | 0.096 | 1 | 8 | Capn5 | 0.005 | Old |

| | | | | | | | | |
|---|---|---|---|---|---|---|---|---|
| 0.82876 | 0.29286 | 0.268 | 0.286 | 1 | 8 | Stim2 | -0.018 | Old |
| 0.828817 | 0.268535 | 0.201 | 0.21 | 1 | 8 | Fmr1 | -0.009 | Old |
| 0.82905 | 0.094918 | 0.251 | 0.257 | 1 | 8 | Ppil2 | -0.006 | Old |
| 0.829342 | 0.243431 | 0.028 | 0.031 | 1 | 8 | Ankrd37 | -0.003 | Old |
| 0.829657 | 0.10083 | 0.14 | 0.132 | 1 | 8 | Slc25a25 | 0.008 | Old |
| 0.830056 | 0.120085 | 0.374 | 0.395 | 1 | 8 | Akirin2 | -0.021 | Old |
| 0.830826 | 0.317447 | 0.089 | 0.086 | 1 | 8 | Guf1 | 0.003 | Old |
| 0.830973 | 0.045582 | 0.542 | 0.584 | 1 | 8 | Cdc42se1 | -0.042 | Old |
| 0.830973 | 0.302576 | 0.145 | 0.156 | 1 | 8 | Rnf103 | -0.011 | Old |
| 0.831005 | 0.506893 | 0.056 | 0.052 | 1 | 8 | Mlst8 | 0.004 | Old |
| 0.831005 | 0.177963 | 0.056 | 0.052 | 1 | 8 | Lppos | 0.004 | Old |
| 0.831782 | 0.480875 | 0.374 | 0.395 | 1 | 8 | Sertad1 | -0.021 | Old |
| 0.831923 | 0.008536 | 0.782 | 0.865 | 1 | 8 | Eif4a1 | -0.083 | Old |
| 0.831932 | 0.247529 | 0.285 | 0.301 | 1 | 8 | Parl | -0.016 | Old |
| 0.831959 | 0.356559 | 0.179 | 0.192 | 1 | 8 | Pigm | -0.013 | Old |
| 0.832229 | 0.034699 | 0.173 | 0.177 | 1 | 8 | Ube2w | -0.004 | Old |
| 0.832623 | 0.038708 | 0.352 | 0.34 | 1 | 8 | Tecr | 0.012 | Old |
| 0.832698 | 0.175622 | 0.352 | 0.39 | 1 | 8 | Csnk2a1 | -0.038 | Old |
| 0.832823 | 0.162144 | 0.179 | 0.169 | 1 | 8 | Ppip5k2 | 0.01 | Old |
| 0.832912 | 0.059443 | 0.665 | 0.694 | 1 | 8 | Eif2s2 | -0.029 | Old |
| 0.832958 | 0.511933 | 0.14 | 0.138 | 1 | 8 | Exoc1 | 0.002 | Old |
| 0.833047 | 0.138885 | 0.073 | 0.068 | 1 | 8 | Nbn | 0.005 | Old |
| 0.833082 | 0.070467 | 0.575 | 0.592 | 1 | 8 | Ctnnb1 | -0.017 | Old |
| 0.833137 | 0.308043 | 0.117 | 0.125 | 1 | 8 | Cog6 | -0.008 | Old |
| 0.833399 | 0.075073 | 0.112 | 0.119 | 1 | 8 | Agl | -0.007 | Old |
| 0.833502 | 0.559925 | 0.156 | 0.153 | 1 | 8 | Zmym4 | 0.003 | Old |
| 0.833542 | 0.042038 | 0.603 | 0.558 | 1 | 8 | Cct2 | 0.045 | Old |
| 0.833728 | 0.051542 | 0.028 | 0.031 | 1 | 8 | 1600014C1 | -0.003 | Old |
| 0.834218 | 0.029537 | 0.179 | 0.184 | 1 | 8 | Mtf2 | -0.005 | Old |
| 0.835181 | 0.283969 | 0.168 | 0.182 | 1 | 8 | Phf1 | -0.014 | Old |
| 0.835196 | 0.139882 | 0.156 | 0.164 | 1 | 8 | Gpr137b | -0.008 | Old |
| 0.835243 | 0.66165 | 0.106 | 0.104 | 1 | 8 | Ppp1r13b | 0.002 | Old |
| 0.835456 | 0.619478 | 0.067 | 0.073 | 1 | 8 | Arel1 | -0.006 | Old |
| 0.836003 | 0.280127 | 0.073 | 0.068 | 1 | 8 | Usp28 | 0.005 | Old |
| 0.836059 | 0.188153 | 0.101 | 0.106 | 1 | 8 | Nubp2 | -0.005 | Old |
| 0.836794 | 0.427865 | 0.101 | 0.096 | 1 | 8 | Thoc1 | 0.005 | Old |
| 0.836805 | 0.279291 | 0.168 | 0.179 | 1 | 8 | Gse1 | -0.011 | Old |
| 0.836991 | 0.36893 | 0.095 | 0.091 | 1 | 8 | Cul9 | 0.004 | Old |
| 0.83711 | 0.838913 | 0.045 | 0.049 | 1 | 8 | Tfap4 | -0.004 | Old |
| 0.83711 | 0.435761 | 0.045 | 0.049 | 1 | 8 | Ncs1 | -0.004 | Old |
| 0.837188 | 0.044462 | 0.106 | 0.112 | 1 | 8 | Sik3 | -0.006 | Old |
| 0.837842 | 0.098194 | 0.151 | 0.143 | 1 | 8 | Eef2k | 0.008 | Old |
| 0.838119 | 0.002336 | 0.028 | 0.031 | 1 | 8 | Sap130 | -0.003 | Old |
| 0.838126 | 0.138218 | 0.078 | 0.073 | 1 | 8 | Mettl26 | 0.005 | Old |
| 0.838248 | 0.088123 | 0.05 | 0.055 | 1 | 8 | Cep104 | -0.005 | Old |

| | | | | | | | | |
|---|---|---|---|---|---|---|---|---|
| 0.838673 | 0.297099 | 0.402 | 0.439 | 1 | 8 | Golph3 | -0.037 | Old |
| 0.838723 | 0.07909 | 0.184 | 0.182 | 1 | 8 | Zfp318 | 0.002 | Old |
| 0.838961 | 0.103797 | 0.464 | 0.494 | 1 | 8 | Ppp2r1a | -0.03 | Old |
| 0.839126 | 0.083146 | 0.559 | 0.584 | 1 | 8 | Arhgap9 | -0.025 | Old |
| 0.839428 | 0.22144 | 0.447 | 0.483 | 1 | 8 | Rab11fip5 | -0.036 | Old |
| 0.839614 | 0.189751 | 0.296 | 0.291 | 1 | 8 | Kpna1 | 0.005 | Old |
| 0.839778 | 0.352625 | 0.162 | 0.153 | 1 | 8 | Slc25a17 | 0.009 | Old |
| 0.839793 | 0.084526 | 0.346 | 0.335 | 1 | 8 | Arpc1a | 0.011 | Old |
| 0.839887 | 0.120818 | 0.084 | 0.078 | 1 | 8 | Galt | 0.006 | Old |
| 0.839927 | 0.165142 | 0.061 | 0.057 | 1 | 8 | Zfp160 | 0.004 | Old |
| 0.839948 | 0.077563 | 0.073 | 0.068 | 1 | 8 | Atg13 | 0.005 | Old |
| 0.840322 | 0.052737 | 0.073 | 0.078 | 1 | 8 | Uqcc1 | -0.005 | Old |
| 0.840629 | 0.665408 | 0.045 | 0.049 | 1 | 8 | Eif2b3 | -0.004 | Old |
| 0.840994 | 0.176 | 0.056 | 0.06 | 1 | 8 | Etaa1 | -0.004 | Old |
| 0.841012 | 0.058309 | 0.291 | 0.312 | 1 | 8 | Nin | -0.021 | Old |
| 0.841101 | 0.040625 | 0.251 | 0.255 | 1 | 8 | Cirbp | -0.004 | Old |
| 0.841185 | 0.01314 | 0.128 | 0.135 | 1 | 8 | Arhgef7 | -0.007 | Old |
| 0.841477 | 0.126806 | 0.089 | 0.086 | 1 | 8 | Pot1b | 0.003 | Old |
| 0.841477 | 0.546546 | 0.151 | 0.145 | 1 | 8 | Zfp787 | 0.006 | Old |
| 0.841659 | 0.233991 | 0.223 | 0.234 | 1 | 8 | Lrrk1 | -0.011 | Old |
| 0.841682 | 0.007046 | 0.369 | 0.358 | 1 | 8 | Map2k7 | 0.011 | Old |
| 0.841709 | 0.275365 | 0.039 | 0.036 | 1 | 8 | Cul7 | 0.003 | Old |
| 0.841802 | 0.292743 | 0.05 | 0.047 | 1 | 8 | Fdxacb1 | 0.003 | Old |
| 0.842535 | 0.081436 | 0.436 | 0.447 | 1 | 8 | Ndufa11 | -0.011 | Old |
| 0.842688 | 0.201618 | 0.453 | 0.462 | 1 | 8 | Dip2b | -0.009 | Old |
| 0.842894 | 0.216508 | 0.078 | 0.073 | 1 | 8 | Lyrm9 | 0.005 | Old |
| 0.843035 | 0.036078 | 0.676 | 0.701 | 1 | 8 | Mapkapk2 | -0.025 | Old |
| 0.843044 | 0.320219 | 0.095 | 0.091 | 1 | 8 | Ncdn | 0.004 | Old |
| 0.843045 | 0.136072 | 0.615 | 0.618 | 1 | 8 | Prrc2a | -0.003 | Old |
| 0.843152 | 0.555518 | 0.145 | 0.145 | 1 | 8 | Terf2 | 0 | Old |
| 0.843153 | 0.260271 | 0.073 | 0.078 | 1 | 8 | Zbtb2 | -0.005 | Old |
| 0.843191 | 0.154131 | 0.665 | 0.717 | 1 | 8 | Sema4a | -0.052 | Old |
| 0.843367 | 0.209986 | 0.179 | 0.171 | 1 | 8 | Ttc5 | 0.008 | Old |
| 0.843506 | 0.213312 | 0.14 | 0.148 | 1 | 8 | Hsdl2 | -0.008 | Old |
| 0.843535 | 0.682613 | 0.089 | 0.099 | 1 | 8 | Tlr6 | -0.01 | Old |
| 0.843582 | 0.082525 | 0.106 | 0.112 | 1 | 8 | Kin | -0.006 | Old |
| 0.843869 | 0.303201 | 0.196 | 0.203 | 1 | 8 | Gtf2h1 | -0.007 | Old |
| 0.843982 | 0.353013 | 0.028 | 0.031 | 1 | 8 | Opn3 | -0.003 | Old |
| 0.844051 | 0.088797 | 0.162 | 0.171 | 1 | 8 | Fam129b | -0.009 | Old |
| 0.844575 | 0.439288 | 0.045 | 0.042 | 1 | 8 | Pomk | 0.003 | Old |
| 0.844887 | 0.023642 | 0.134 | 0.14 | 1 | 8 | Dnpep | -0.006 | Old |
| 0.845007 | 0.018718 | 0.86 | 0.927 | 1 | 8 | Tmbim6 | -0.067 | Old |
| 0.845191 | 0.166785 | 0.223 | 0.234 | 1 | 8 | Ten1 | -0.011 | Old |
| 0.845285 | 0.088733 | 0.112 | 0.106 | 1 | 8 | Arfgap1 | 0.006 | Old |
| 0.845325 | 0.510091 | 0.05 | 0.047 | 1 | 8 | Atf5 | 0.003 | Old |

| | | | | | | | | |
|---|---|---|---|---|---|---|---|---|
| 0.845325 | 0.174207 | 0.05 | 0.047 | 1 | 8 | Zfp846 | 0.003 | Old |
| 0.84545 | 0.004997 | 0.341 | 0.34 | 1 | 8 | Phf23 | 0.001 | Old |
| 0.845644 | 0.057728 | 0.268 | 0.262 | 1 | 8 | Sarnp | 0.006 | Old |
| 0.845934 | 0.10205 | 0.698 | 0.743 | 1 | 8 | Nptn | -0.045 | Old |
| 0.846003 | 0.049955 | 0.391 | 0.423 | 1 | 8 | Usp4 | -0.032 | Old |
| 0.846372 | 0.286004 | 0.419 | 0.423 | 1 | 8 | Mif | -0.004 | Old |
| 0.8465 | 0.365491 | 0.05 | 0.047 | 1 | 8 | Rpap1 | 0.003 | Old |
| 0.846813 | 0.439394 | 0.089 | 0.086 | 1 | 8 | Cep164 | 0.003 | Old |
| 0.846876 | 0.120548 | 0.263 | 0.26 | 1 | 8 | Tmem248 | 0.003 | Old |
| 0.846917 | 0.340569 | 0.028 | 0.031 | 1 | 8 | Il1r2 | -0.003 | Old |
| 0.847067 | 0.434083 | 0.168 | 0.164 | 1 | 8 | Sdhaf2 | 0.004 | Old |
| 0.847133 | 0.118525 | 0.436 | 0.452 | 1 | 8 | Mlf2 | -0.016 | Old |
| 0.847301 | 0.183766 | 0.112 | 0.106 | 1 | 8 | Mthfsl | 0.006 | Old |
| 0.847315 | 0.071067 | 0.084 | 0.091 | 1 | 8 | Pdcd2 | -0.007 | Old |
| 0.847369 | 0.445825 | 0.19 | 0.182 | 1 | 8 | Dhrs7b | 0.008 | Old |
| 0.847502 | 0.311044 | 0.458 | 0.499 | 1 | 8 | Tnks2 | -0.041 | Old |
| 0.847756 | 0.186942 | 0.307 | 0.314 | 1 | 8 | Ing2 | -0.007 | Old |
| 0.84788 | 0.314076 | 0.084 | 0.081 | 1 | 8 | Gm45716 | 0.003 | Old |
| 0.847896 | 0.011822 | 0.972 | 0.992 | 1 | 8 | Rack1 | -0.02 | Old |
| 0.848217 | 0.035983 | 0.497 | 0.496 | 1 | 8 | Cxcl16 | 0.001 | Old |
| 0.848472 | 0.27658 | 0.061 | 0.057 | 1 | 8 | Tmem237 | 0.004 | Old |
| 0.848733 | 0.220024 | 0.123 | 0.127 | 1 | 8 | Tusc2 | -0.004 | Old |
| 0.848821 | 0.04942 | 0.073 | 0.078 | 1 | 8 | Cdt1 | -0.005 | Old |
| 0.848851 | 0.064768 | 0.045 | 0.049 | 1 | 8 | Ccdc61 | -0.004 | Old |
| 0.848879 | 0.04676 | 0.492 | 0.519 | 1 | 8 | Fgd4 | -0.027 | Old |
| 0.848904 | 0.748707 | 0.034 | 0.031 | 1 | 8 | Enpp1 | 0.003 | Old |
| 0.848934 | 0.053272 | 0.145 | 0.151 | 1 | 8 | Adgrg6 | -0.006 | Old |
| 0.848938 | 0.157022 | 0.358 | 0.384 | 1 | 8 | Psmd1 | -0.026 | Old |
| 0.849298 | 0.230491 | 0.19 | 0.182 | 1 | 8 | Fnta | 0.008 | Old |
| 0.849475 | 0.217927 | 0.268 | 0.281 | 1 | 8 | Tmem141 | -0.013 | Old |
| 0.849739 | 0.061273 | 0.229 | 0.208 | 1 | 8 | Klrk1 | 0.021 | Old |
| 0.849991 | 0.286036 | 0.173 | 0.171 | 1 | 8 | Polr2j | 0.002 | Old |
| 0.85013 | 0.128733 | 0.637 | 0.694 | 1 | 8 | Tmed5 | -0.057 | Old |
| 0.850202 | 0.470223 | 0.061 | 0.068 | 1 | 8 | Myc | -0.007 | Old |
| 0.85068 | 0.071194 | 0.38 | 0.387 | 1 | 8 | Ddx21 | -0.007 | Old |
| 0.850792 | 0.178264 | 0.045 | 0.042 | 1 | 8 | H2-T24 | 0.003 | Old |
| 0.851203 | 0.712844 | 0.045 | 0.049 | 1 | 8 | Sgcb | -0.004 | Old |
| 0.851203 | 0.323879 | 0.05 | 0.047 | 1 | 8 | Gk5 | 0.003 | Old |
| 0.851943 | 0.090474 | 0.486 | 0.481 | 1 | 8 | Tpi1 | 0.005 | Old |
| 0.852046 | 0.118405 | 0.503 | 0.501 | 1 | 8 | Dbnl | 0.002 | Old |
| 0.852219 | 0.122973 | 0.341 | 0.358 | 1 | 8 | H2-Q7 | -0.017 | Old |
| 0.852449 | 0.083052 | 0.112 | 0.114 | 1 | 8 | Nup85 | -0.002 | Old |
| 0.852534 | 0.707782 | 0.173 | 0.177 | 1 | 8 | Fam53c | -0.004 | Old |
| 0.852595 | 0.041177 | 0.285 | 0.296 | 1 | 8 | Egln1 | -0.011 | Old |
| 0.852798 | 0.104263 | 0.073 | 0.068 | 1 | 8 | Med19 | 0.005 | Old |

| | | | | | | | | |
|---|---|---|---|---|---|---|---|---|
| 0.853003 | 0.281122 | 0.251 | 0.262 | 1 | 8 | Etv1 | -0.011 | Old |
| 0.853238 | 0.224926 | 0.101 | 0.106 | 1 | 8 | Trim3 | -0.005 | Old |
| 0.85355 | 0.282629 | 0.073 | 0.078 | 1 | 8 | Ralgapa2 | -0.005 | Old |
| 0.854009 | 0.324567 | 0.067 | 0.073 | 1 | 8 | Acap3 | -0.006 | Old |
| 0.854517 | 0.188646 | 0.123 | 0.13 | 1 | 8 | Rab33b | -0.007 | Old |
| 0.854527 | 0.121385 | 0.045 | 0.042 | 1 | 8 | Phf11a | 0.003 | Old |
| 0.854533 | 0.269368 | 0.128 | 0.138 | 1 | 8 | Cdyl | -0.01 | Old |
| 0.854798 | 0.021394 | 0.106 | 0.112 | 1 | 8 | Atrn | -0.006 | Old |
| 0.854882 | 0.166092 | 0.52 | 0.543 | 1 | 8 | Pdcd5 | -0.023 | Old |
| 0.85508 | 0.174451 | 0.24 | 0.236 | 1 | 8 | Acox3 | 0.004 | Old |
| 0.855545 | 0.085144 | 0.547 | 0.564 | 1 | 8 | Tnrc18 | -0.017 | Old |
| 0.8558 | 0.115529 | 0.525 | 0.54 | 1 | 8 | Prpf40a | -0.015 | Old |
| 0.855846 | 0.226987 | 0.279 | 0.299 | 1 | 8 | Plbd2 | -0.02 | Old |
| 0.855923 | 0.292319 | 0.235 | 0.231 | 1 | 8 | Dnajc14 | 0.004 | Old |
| 0.856126 | 0.238368 | 0.14 | 0.138 | 1 | 8 | AU019823 | 0.002 | Old |
| 0.856283 | 0.197552 | 0.486 | 0.483 | 1 | 8 | Huwe1 | 0.003 | Old |
| 0.85629 | 0.707756 | 0.067 | 0.065 | 1 | 8 | Acvr2a | 0.002 | Old |
| 0.856306 | 0.089914 | 0.771 | 0.823 | 1 | 8 | Jpt1 | -0.052 | Old |
| 0.856577 | 0.136539 | 0.145 | 0.14 | 1 | 8 | 1110059G1 | 0.005 | Old |
| 0.856866 | 0.159107 | 0.268 | 0.262 | 1 | 8 | Sec24c | 0.006 | Old |
| 0.857019 | 0.405413 | 0.045 | 0.042 | 1 | 8 | 2900076A0 | 0.003 | Old |
| 0.85703 | 0.175455 | 0.413 | 0.447 | 1 | 8 | Rlim | -0.034 | Old |
| 0.857047 | 0.189364 | 0.324 | 0.348 | 1 | 8 | Rbck1 | -0.024 | Old |
| 0.857052 | 0.356726 | 0.084 | 0.081 | 1 | 8 | Mtap | 0.003 | Old |
| 0.85709 | 0.560618 | 0.045 | 0.049 | 1 | 8 | Slc7a13 | -0.004 | Old |
| 0.857218 | 0.322109 | 0.419 | 0.439 | 1 | 8 | Ankrd13a | -0.02 | Old |
| 0.857556 | 0.715741 | 0.011 | 0.013 | 1 | 8 | Cep89 | -0.002 | Old |
| 0.857556 | 0.610666 | 0.011 | 0.013 | 1 | 8 | Klc2 | -0.002 | Old |
| 0.857556 | 0.559366 | 0.011 | 0.013 | 1 | 8 | Hes7 | -0.002 | Old |
| 0.857556 | 0.290608 | 0.011 | 0.013 | 1 | 8 | Dnph1 | -0.002 | Old |
| 0.857799 | 0.320482 | 0.112 | 0.106 | 1 | 8 | Slc25a44 | 0.006 | Old |
| 0.858188 | 0.247028 | 0.374 | 0.397 | 1 | 8 | Polr2k | -0.023 | Old |
| 0.858467 | 0.241022 | 0.335 | 0.369 | 1 | 8 | Stx16 | -0.034 | Old |
| 0.858592 | 0.0576 | 0.726 | 0.823 | 1 | 8 | Grb2 | -0.097 | Old |
| 0.858758 | 0.414805 | 0.101 | 0.096 | 1 | 8 | Cops7b | 0.005 | Old |
| 0.858915 | 0.565095 | 0.034 | 0.031 | 1 | 8 | Zfp952 | 0.003 | Old |
| 0.859079 | 0.15503 | 0.341 | 0.343 | 1 | 8 | Tulp4 | -0.002 | Old |
| 0.859112 | 0.076127 | 0.223 | 0.216 | 1 | 8 | Tlk1 | 0.007 | Old |
| 0.859178 | 0.3235 | 0.056 | 0.06 | 1 | 8 | Zfp157 | -0.004 | Old |
| 0.859261 | 0.137635 | 0.201 | 0.197 | 1 | 8 | Mut | 0.004 | Old |
| 0.859286 | 0.357602 | 0.257 | 0.262 | 1 | 8 | Hdac1 | -0.005 | Old |
| 0.859765 | 0.116134 | 0.542 | 0.548 | 1 | 8 | Tiam1 | -0.006 | Old |
| 0.859829 | 0.958253 | 0.011 | 0.013 | 1 | 8 | Pfkfb2 | -0.002 | Old |
| 0.859829 | 0.89772 | 0.011 | 0.013 | 1 | 8 | Ppef2 | -0.002 | Old |
| 0.859829 | 0.612523 | 0.011 | 0.013 | 1 | 8 | Zkscan7 | -0.002 | Old |

| | | | | | | | | |
|---|---|---|---|---|---|---|---|---|
| 0.859829 | 0.584297 | 0.011 | 0.013 | 1 | 8 | Tmem14a | -0.002 | Old |
| 0.859829 | 0.509395 | 0.011 | 0.013 | 1 | 8 | Mb21d2 | -0.002 | Old |
| 0.859829 | 0.302839 | 0.011 | 0.013 | 1 | 8 | Tmem240 | -0.002 | Old |
| 0.859829 | 0.044915 | 0.011 | 0.013 | 1 | 8 | Gm10053 | -0.002 | Old |
| 0.860233 | 0.032939 | 0.184 | 0.177 | 1 | 8 | Zmym2 | 0.007 | Old |
| 0.860416 | 0.072697 | 0.106 | 0.112 | 1 | 8 | Tmub1 | -0.006 | Old |
| 0.860495 | 0.234573 | 0.307 | 0.309 | 1 | 8 | Chp2 | -0.002 | Old |
| 0.860542 | 0.006265 | 0.201 | 0.213 | 1 | 8 | Mex3c | -0.012 | Old |
| 0.860585 | 0.14129 | 0.251 | 0.26 | 1 | 8 | Rhoc | -0.009 | Old |
| 0.860799 | 0.098801 | 0.615 | 0.621 | 1 | 8 | Lims1 | -0.006 | Old |
| 0.860985 | 0.080933 | 0.777 | 0.797 | 1 | 8 | Csrnp1 | -0.02 | Old |
| 0.861122 | 0.081037 | 0.408 | 0.436 | 1 | 8 | Baiap2 | -0.028 | Old |
| 0.861186 | 0.065549 | 0.078 | 0.083 | 1 | 8 | Abcc4 | -0.005 | Old |
| 0.861617 | 0.120114 | 0.24 | 0.231 | 1 | 8 | Gskip | 0.009 | Old |
| 0.861621 | 0.566277 | 0.028 | 0.031 | 1 | 8 | Btc | -0.003 | Old |
| 0.86166 | 0.677881 | 0.128 | 0.127 | 1 | 8 | Il15 | 0.001 | Old |
| 0.861707 | 0.07067 | 0.38 | 0.374 | 1 | 8 | Setd1b | 0.006 | Old |
| 0.861727 | 0.574272 | 0.123 | 0.132 | 1 | 8 | Triobp | -0.009 | Old |
| 0.861779 | 0.616766 | 0.034 | 0.031 | 1 | 8 | Atp5s | 0.003 | Old |
| 0.861879 | 0.191157 | 0.575 | 0.59 | 1 | 8 | Sf3b6 | -0.015 | Old |
| 0.862104 | 0.192828 | 0.011 | 0.013 | 1 | 8 | Gm45871 | -0.002 | Old |
| 0.862104 | 0.086224 | 0.011 | 0.013 | 1 | 8 | Smkr-ps | -0.002 | Old |
| 0.862395 | 0.104866 | 0.056 | 0.06 | 1 | 8 | Zfp386 | -0.004 | Old |
| 0.862523 | 0.310804 | 0.358 | 0.382 | 1 | 8 | Dcun1d5 | -0.024 | Old |
| 0.862627 | 0.207127 | 0.145 | 0.153 | 1 | 8 | Rcbtb2 | -0.008 | Old |
| 0.862827 | 0.196655 | 0.106 | 0.112 | 1 | 8 | Rps6ka2 | -0.006 | Old |
| 0.862943 | 0.455816 | 0.039 | 0.036 | 1 | 8 | Mad2l2 | 0.003 | Old |
| 0.862998 | 0.365696 | 0.095 | 0.091 | 1 | 8 | Depdc5 | 0.004 | Old |
| 0.863927 | 0.277524 | 0.117 | 0.112 | 1 | 8 | Zfp771 | 0.005 | Old |
| 0.864031 | 0.073607 | 0.324 | 0.327 | 1 | 8 | Plekha1 | -0.003 | Old |
| 0.86438 | 0.779866 | 0.011 | 0.013 | 1 | 8 | Abcc10 | -0.002 | Old |
| 0.864401 | 0.152878 | 0.246 | 0.255 | 1 | 8 | Slc5a3 | -0.009 | Old |
| 0.864516 | 0.030925 | 0.542 | 0.569 | 1 | 8 | Rtraf | -0.027 | Old |
| 0.864531 | 0.166281 | 0.106 | 0.101 | 1 | 8 | Abcd3 | 0.005 | Old |
| 0.864541 | 0.156091 | 0.061 | 0.057 | 1 | 8 | Epn2 | 0.004 | Old |
| 0.8647 | 0.069871 | 0.274 | 0.281 | 1 | 8 | Fip1l1 | -0.007 | Old |
| 0.865011 | 0.20073 | 0.212 | 0.205 | 1 | 8 | Lrrc41 | 0.007 | Old |
| 0.865245 | 0.398114 | 0.179 | 0.177 | 1 | 8 | Fgd6 | 0.002 | Old |
| 0.865276 | 0.286942 | 0.212 | 0.21 | 1 | 8 | Psen2 | 0.002 | Old |
| 0.865948 | 0.260433 | 0.458 | 0.481 | 1 | 8 | Supt5 | -0.023 | Old |
| 0.866375 | 0.096089 | 0.045 | 0.042 | 1 | 8 | Strbp | 0.003 | Old |
| 0.86645 | 0.180477 | 0.235 | 0.229 | 1 | 8 | Gars | 0.006 | Old |
| 0.866524 | 0.022374 | 0.05 | 0.047 | 1 | 8 | Klhl7 | 0.003 | Old |
| 0.866549 | 0.133708 | 0.358 | 0.361 | 1 | 8 | Dcaf8 | -0.003 | Old |
| 0.866657 | 0.787102 | 0.011 | 0.013 | 1 | 8 | Stox2 | -0.002 | Old |

| | | | | | | | | |
|---|---|---|---|---|---|---|---|---|
| 0.866657 | 0.604004 | 0.011 | 0.013 | 1 | 8 | Zfp963 | -0.002 | Old |
| 0.866657 | 0.450025 | 0.011 | 0.013 | 1 | 8 | Mbnl3 | -0.002 | Old |
| 0.867036 | 0.161433 | 0.207 | 0.205 | 1 | 8 | Trak1 | 0.002 | Old |
| 0.867189 | 0.266411 | 0.391 | 0.41 | 1 | 8 | St3gal4 | -0.019 | Old |
| 0.867514 | 0.114364 | 0.034 | 0.031 | 1 | 8 | Kbtbd4 | 0.003 | Old |
| 0.867768 | 0.036889 | 0.073 | 0.078 | 1 | 8 | Ak8 | -0.005 | Old |
| 0.867885 | 0.068313 | 0.369 | 0.364 | 1 | 8 | Map4 | 0.005 | Old |
| 0.868018 | 0.018771 | 0.101 | 0.106 | 1 | 8 | Jagn1 | -0.005 | Old |
| 0.868478 | 0.573877 | 0.05 | 0.055 | 1 | 8 | Tatdn3 | -0.005 | Old |
| 0.868479 | 0.265048 | 0.112 | 0.119 | 1 | 8 | Mrps30 | -0.007 | Old |
| 0.868483 | 0.342544 | 0.128 | 0.132 | 1 | 8 | Slc39a11 | -0.004 | Old |
| 0.868513 | 0.105045 | 0.19 | 0.2 | 1 | 8 | Ppil4 | -0.01 | Old |
| 0.868594 | 0.18648 | 0.246 | 0.236 | 1 | 8 | Nucb2 | 0.01 | Old |
| 0.868935 | 1.639432 | 0.011 | 0.013 | 1 | 8 | Rnase2a | -0.002 | Old |
| 0.868935 | 1.130596 | 0.011 | 0.013 | 1 | 8 | 1700099I0 | -0.002 | Old |
| 0.868949 | 0.601012 | 0.034 | 0.031 | 1 | 8 | Pfas | 0.003 | Old |
| 0.869014 | 0.043474 | 0.117 | 0.122 | 1 | 8 | Cln5 | -0.005 | Old |
| 0.869033 | 0.353755 | 0.229 | 0.229 | 1 | 8 | Pak1ip1 | 0 | Old |
| 0.869161 | 0.197796 | 0.218 | 0.229 | 1 | 8 | Sec24a | -0.011 | Old |
| 0.869357 | 0.021592 | 0.436 | 0.447 | 1 | 8 | Lsm12 | -0.011 | Old |
| 0.869377 | 0.184602 | 0.184 | 0.177 | 1 | 8 | Srp68 | 0.007 | Old |
| 0.869695 | 0.404945 | 0.067 | 0.073 | 1 | 8 | Rsad2 | -0.006 | Old |
| 0.870747 | 0.119038 | 0.045 | 0.042 | 1 | 8 | 5830432E0 | 0.003 | Old |
| 0.870932 | 0.093732 | 0.039 | 0.036 | 1 | 8 | F5 | 0.003 | Old |
| 0.870935 | 0.08534 | 0.207 | 0.203 | 1 | 8 | Rnf220 | 0.004 | Old |
| 0.870983 | 0.005311 | 0.207 | 0.21 | 1 | 8 | Stx11 | -0.003 | Old |
| 0.871085 | 0.048317 | 0.659 | 0.766 | 1 | 8 | Ptprj | -0.107 | Old |
| 0.871612 | 0.217204 | 0.268 | 0.283 | 1 | 8 | Lars | -0.015 | Old |
| 0.87183 | 1.440916 | 0.028 | 0.026 | 1 | 8 | Lrsam1 | 0.002 | Old |
| 0.872132 | 0.108381 | 0.101 | 0.106 | 1 | 8 | Epm2aip1 | -0.005 | Old |
| 0.872265 | 0.204638 | 0.039 | 0.036 | 1 | 8 | Slc5a6 | 0.003 | Old |
| 0.872331 | 0.103492 | 0.095 | 0.099 | 1 | 8 | Top3b | -0.004 | Old |
| 0.87248 | 0.080929 | 0.106 | 0.112 | 1 | 8 | Slc39a6 | -0.006 | Old |
| 0.872715 | 0.186967 | 0.279 | 0.275 | 1 | 8 | Chordc1 | 0.004 | Old |
| 0.872902 | 0.022718 | 0.324 | 0.33 | 1 | 8 | Dock1 | -0.006 | Old |
| 0.873272 | 0.196067 | 0.268 | 0.283 | 1 | 8 | Itgax | -0.015 | Old |
| 0.87418 | 0.247381 | 0.52 | 0.582 | 1 | 8 | Camkk2 | -0.062 | Old |
| 0.87421 | 0.307406 | 0.056 | 0.06 | 1 | 8 | Mettl5 | -0.004 | Old |
| 0.874602 | 0.201918 | 0.101 | 0.106 | 1 | 8 | Extl3 | -0.005 | Old |
| 0.875991 | 0.208294 | 0.128 | 0.122 | 1 | 8 | Tmem214 | 0.006 | Old |
| 0.876092 | 0.096483 | 0.385 | 0.384 | 1 | 8 | Thoc2 | 0.001 | Old |
| 0.876099 | 0.089793 | 0.061 | 0.065 | 1 | 8 | Invs | -0.004 | Old |
| 0.876129 | 0.111923 | 0.034 | 0.031 | 1 | 8 | Zfp523 | 0.003 | Old |
| 0.876131 | 0.106942 | 0.48 | 0.488 | 1 | 8 | Elf1 | -0.008 | Old |
| 0.876235 | 0.167702 | 0.246 | 0.239 | 1 | 8 | Foxj2 | 0.007 | Old |

| | | | | | | | | |
|---|---|---|---|---|---|---|---|---|
| 0.87625 | 0.605823 | 0.106 | 0.104 | 1 | 8 | Zfp746 | 0.002 | Old |
| 0.876276 | 0.150621 | 0.246 | 0.242 | 1 | 8 | Cul5 | 0.004 | Old |
| 0.876404 | 0.476815 | 0.039 | 0.042 | 1 | 8 | Ints4 | -0.003 | Old |
| 0.876495 | 0.165031 | 0.089 | 0.094 | 1 | 8 | Bpnt1 | -0.005 | Old |
| 0.876538 | 0.663168 | 0.028 | 0.026 | 1 | 8 | Lag3 | 0.002 | Old |
| 0.876574 | 0.60084 | 0.067 | 0.073 | 1 | 8 | Tmco6 | -0.006 | Old |
| 0.876714 | 0.126194 | 0.475 | 0.506 | 1 | 8 | Pten | -0.031 | Old |
| 0.876831 | 0.298619 | 0.263 | 0.26 | 1 | 8 | Cc2d1b | 0.003 | Old |
| 0.877065 | 0.096277 | 0.508 | 0.514 | 1 | 8 | Tomm5 | -0.006 | Old |
| 0.877521 | 0.530607 | 0.123 | 0.119 | 1 | 8 | Utp6 | 0.004 | Old |
| 0.877524 | 0.183375 | 0.117 | 0.125 | 1 | 8 | Naa40 | -0.008 | Old |
| 0.87756 | 0.127056 | 0.196 | 0.203 | 1 | 8 | AI413582 | -0.007 | Old |
| 0.877574 | 0.572177 | 0.101 | 0.109 | 1 | 8 | Zfp407 | -0.008 | Old |
| 0.878108 | 0.747158 | 0.028 | 0.026 | 1 | 8 | Dtnb | 0.002 | Old |
| 0.878108 | 0.722848 | 0.028 | 0.026 | 1 | 8 | Msrb2 | 0.002 | Old |
| 0.878709 | 0.161263 | 0.307 | 0.322 | 1 | 8 | Gosr2 | -0.015 | Old |
| 0.878917 | 0.054266 | 0.413 | 0.426 | 1 | 8 | Inpp4a | -0.013 | Old |
| 0.878936 | 0.182934 | 0.441 | 0.418 | 1 | 8 | Cs | 0.023 | Old |
| 0.879143 | 0.179912 | 0.084 | 0.081 | 1 | 8 | Chst7 | 0.003 | Old |
| 0.879198 | 0.026897 | 0.061 | 0.065 | 1 | 8 | Hmgxb3 | -0.004 | Old |
| 0.879558 | 0.103644 | 0.592 | 0.613 | 1 | 8 | Dpysl2 | -0.021 | Old |
| 0.879798 | 0.258596 | 0.436 | 0.46 | 1 | 8 | Acap2 | -0.024 | Old |
| 0.87995 | 0.071786 | 0.76 | 0.782 | 1 | 8 | H2-T23 | -0.022 | Old |
| 0.88027 | 0.65316 | 0.039 | 0.036 | 1 | 8 | Zfp931 | 0.003 | Old |
| 0.880442 | 1.006607 | 0.034 | 0.031 | 1 | 8 | Ankzf1 | 0.003 | Old |
| 0.880757 | 0.302551 | 0.045 | 0.042 | 1 | 8 | Tarm1 | 0.003 | Old |
| 0.88079 | 0.050429 | 0.508 | 0.514 | 1 | 8 | Acin1 | -0.006 | Old |
| 0.880862 | 0.356658 | 0.034 | 0.036 | 1 | 8 | Miga1 | -0.002 | Old |
| 0.881074 | 0.37771 | 0.268 | 0.273 | 1 | 8 | Acaa1a | -0.005 | Old |
| 0.881712 | 0.185177 | 0.112 | 0.106 | 1 | 8 | 2310022B0 | 0.006 | Old |
| 0.881735 | 0.134529 | 0.212 | 0.218 | 1 | 8 | Pde7a | -0.006 | Old |
| 0.881881 | 0.055677 | 0.034 | 0.031 | 1 | 8 | Hdac10 | 0.003 | Old |
| 0.882209 | 0.45809 | 0.101 | 0.099 | 1 | 8 | 2510002D2 | 0.002 | Old |
| 0.882213 | 0.283674 | 0.257 | 0.257 | 1 | 8 | Pitpnb | 0 | Old |
| 0.882572 | 0.412466 | 0.19 | 0.205 | 1 | 8 | Lypla1 | -0.015 | Old |
| 0.882586 | 1.253276 | 0.022 | 0.021 | 1 | 8 | Serpinb2 | 0.001 | Old |
| 0.882602 | 0.176008 | 0.089 | 0.094 | 1 | 8 | Crbn | -0.005 | Old |
| 0.88308 | 0.048229 | 0.05 | 0.047 | 1 | 8 | Kctd9 | 0.003 | Old |
| 0.883308 | 0.150893 | 0.441 | 0.444 | 1 | 8 | Chmp5 | -0.003 | Old |
| 0.883452 | 0.434665 | 0.084 | 0.088 | 1 | 8 | Snx25 | -0.004 | Old |
| 0.883861 | 0.144067 | 0.19 | 0.197 | 1 | 8 | Itgav | -0.007 | Old |
| 0.883934 | 0.496024 | 0.078 | 0.075 | 1 | 8 | Rrp8 | 0.003 | Old |
| 0.884122 | 0.266442 | 0.352 | 0.364 | 1 | 8 | Ptpn2 | -0.012 | Old |
| 0.884352 | 0.032897 | 0.944 | 0.964 | 1 | 8 | Pltp | -0.02 | Old |
| 0.884395 | 0.641011 | 0.028 | 0.026 | 1 | 8 | Ift80 | 0.002 | Old |

| | | | | | | | | |
|---|---|---|---|---|---|---|---|---|
| 0.884431 | 0.937395 | 0.084 | 0.083 | 1 | 8 | Pogk | 0.001 | Old |
| 0.884905 | 0.343616 | 0.196 | 0.197 | 1 | 8 | Smad1 | -0.001 | Old |
| 0.88494 | 0.481089 | 0.101 | 0.109 | 1 | 8 | Ascc2 | -0.008 | Old |
| 0.885639 | 0.165205 | 0.14 | 0.145 | 1 | 8 | Ppp1r7 | -0.005 | Old |
| 0.885967 | 0.184483 | 0.028 | 0.026 | 1 | 8 | Kpna2 | 0.002 | Old |
| 0.886009 | 0.221291 | 0.095 | 0.101 | 1 | 8 | Rtp4 | -0.006 | Old |
| 0.886041 | 0.462516 | 0.117 | 0.122 | 1 | 8 | Snx14 | -0.005 | Old |
| 0.886052 | 0.827145 | 0.056 | 0.06 | 1 | 8 | Prpf3 | -0.004 | Old |
| 0.886096 | 0.182011 | 0.089 | 0.094 | 1 | 8 | Dvl1 | -0.005 | Old |
| 0.886151 | 0.072116 | 0.112 | 0.117 | 1 | 8 | Rngtt | -0.005 | Old |
| 0.886198 | 0.051917 | 0.14 | 0.132 | 1 | 8 | Ino80 | 0.008 | Old |
| 0.886209 | 0.138087 | 0.475 | 0.496 | 1 | 8 | Sdhc | -0.021 | Old |
| 0.886239 | 0.24443 | 0.162 | 0.171 | 1 | 8 | Tmem106c | -0.009 | Old |
| 0.886341 | 0.31823 | 0.369 | 0.379 | 1 | 8 | Mrpl14 | -0.01 | Old |
| 0.886588 | 0.059325 | 0.525 | 0.545 | 1 | 8 | Mafg | -0.02 | Old |
| 0.887275 | 0.102783 | 0.458 | 0.462 | 1 | 8 | Adam17 | -0.004 | Old |
| 0.887499 | 0.414751 | 0.089 | 0.096 | 1 | 8 | Zfp511 | -0.007 | Old |
| 0.887541 | 0.254588 | 0.028 | 0.026 | 1 | 8 | Lgi3 | 0.002 | Old |
| 0.887541 | 0.235691 | 0.028 | 0.026 | 1 | 8 | Cntrob | 0.002 | Old |
| 0.887701 | 0.343785 | 0.034 | 0.036 | 1 | 8 | Gm19951 | -0.002 | Old |
| 0.887939 | 0.201003 | 0.084 | 0.088 | 1 | 8 | Cinp | -0.004 | Old |
| 0.888017 | 0.455867 | 0.145 | 0.145 | 1 | 8 | Rab6b | 0 | Old |
| 0.888024 | 0.007062 | 0.575 | 0.603 | 1 | 8 | Frmd4b | -0.028 | Old |
| 0.888029 | 0.307084 | 0.201 | 0.218 | 1 | 8 | Tgfbrap1 | -0.017 | Old |
| 0.888486 | 0.425626 | 0.24 | 0.239 | 1 | 8 | Sec13 | 0.001 | Old |
| 0.888534 | 0.189622 | 0.173 | 0.171 | 1 | 8 | Kif19a | 0.002 | Old |
| 0.889029 | 0.296139 | 0.179 | 0.179 | 1 | 8 | Kmt5c | 0 | Old |
| 0.889039 | 0.249491 | 0.268 | 0.286 | 1 | 8 | Ilf2 | -0.018 | Old |
| 0.889114 | 0.188958 | 0.028 | 0.026 | 1 | 8 | Ttll4 | 0.002 | Old |
| 0.889114 | 0.132729 | 0.028 | 0.026 | 1 | 8 | Trdmt1 | 0.002 | Old |
| 0.889191 | 0.627041 | 0.039 | 0.042 | 1 | 8 | Zfand2a | -0.003 | Old |
| 0.889327 | 0.136685 | 0.196 | 0.197 | 1 | 8 | Larp1b | -0.001 | Old |
| 0.889531 | 0.119644 | 0.045 | 0.042 | 1 | 8 | Jpx | 0.003 | Old |
| 0.889542 | 0.212681 | 0.061 | 0.065 | 1 | 8 | Lig3 | -0.004 | Old |
| 0.889571 | 0.072835 | 0.067 | 0.07 | 1 | 8 | Spsb2 | -0.003 | Old |
| 0.889602 | 0.829159 | 0.022 | 0.021 | 1 | 8 | Dnaaf5 | 0.001 | Old |
| 0.889602 | 0.404304 | 0.022 | 0.021 | 1 | 8 | Mttp | 0.001 | Old |
| 0.889662 | 0.314676 | 0.173 | 0.174 | 1 | 8 | Arid5b | -0.001 | Old |
| 0.88998 | 0.039386 | 0.698 | 0.792 | 1 | 8 | Set | -0.094 | Old |
| 0.890122 | 0.139474 | 0.173 | 0.184 | 1 | 8 | Bin2 | -0.011 | Old |
| 0.890439 | 0.324169 | 0.034 | 0.036 | 1 | 8 | Taf1c | -0.002 | Old |
| 0.890474 | 0.168772 | 0.257 | 0.26 | 1 | 8 | Fxr2 | -0.003 | Old |
| 0.89056 | 0.107792 | 0.173 | 0.179 | 1 | 8 | Pmpca | -0.006 | Old |
| 0.890814 | 0.384508 | 0.168 | 0.166 | 1 | 8 | Fnbp4 | 0.002 | Old |
| 0.890832 | 0.384505 | 0.179 | 0.192 | 1 | 8 | Wdr43 | -0.013 | Old |

| | | | | | | | | |
|---|---|---|---|---|---|---|---|---|
| 0.891646 | 0.024677 | 0.268 | 0.278 | 1 | 8 | Brox | -0.01 | Old |
| 0.892057 | 0.035548 | 0.503 | 0.504 | 1 | 8 | Kras | -0.001 | Old |
| 0.892194 | 0.302316 | 0.201 | 0.21 | 1 | 8 | Vps11 | -0.009 | Old |
| 0.892263 | 0.301152 | 0.028 | 0.026 | 1 | 8 | Fam222a | 0.002 | Old |
| 0.892351 | 0.106254 | 0.19 | 0.195 | 1 | 8 | Ppp1r35 | -0.005 | Old |
| 0.892538 | 0.193009 | 0.173 | 0.171 | 1 | 8 | Snapc1 | 0.002 | Old |
| 0.892917 | 0.009988 | 0.201 | 0.187 | 1 | 8 | Cnot10 | 0.014 | Old |
| 0.893113 | 0.257877 | 0.022 | 0.021 | 1 | 8 | Neil1 | 0.001 | Old |
| 0.893394 | 0.078631 | 0.346 | 0.327 | 1 | 8 | Phc3 | 0.019 | Old |
| 0.893751 | 0.397361 | 0.151 | 0.148 | 1 | 8 | Rasa2 | 0.003 | Old |
| 0.893838 | 0.333662 | 0.028 | 0.026 | 1 | 8 | Slc25a53 | 0.002 | Old |
| 0.893838 | 0.082798 | 0.028 | 0.026 | 1 | 8 | Mrps27 | 0.002 | Old |
| 0.893838 | 0.074771 | 0.028 | 0.026 | 1 | 8 | Nlrp1a | 0.002 | Old |
| 0.893838 | 0.063855 | 0.028 | 0.026 | 1 | 8 | Tmem67 | 0.002 | Old |
| 0.893853 | 0.560249 | 0.089 | 0.088 | 1 | 8 | Arl2 | 0.001 | Old |
| 0.893963 | 0.152682 | 0.089 | 0.094 | 1 | 8 | Otud7b | -0.005 | Old |
| 0.894125 | 0.549638 | 0.196 | 0.2 | 1 | 8 | Cops8 | -0.004 | Old |
| 0.894314 | 0.497024 | 0.039 | 0.042 | 1 | 8 | Tarsl2 | -0.003 | Old |
| 0.894705 | 0.171895 | 0.503 | 0.548 | 1 | 8 | Elavl1 | -0.045 | Old |
| 0.894724 | 0.148235 | 0.078 | 0.081 | 1 | 8 | Tpm3-rs7 | -0.003 | Old |
| 0.894749 | 0.07022 | 0.179 | 0.174 | 1 | 8 | Smc5 | 0.005 | Old |
| 0.89487 | 0.763191 | 0.022 | 0.021 | 1 | 8 | Nrep | 0.001 | Old |
| 0.895049 | 0.232589 | 0.425 | 0.444 | 1 | 8 | Rab3il1 | -0.019 | Old |
| 0.895125 | 0.232391 | 0.089 | 0.086 | 1 | 8 | Cbfa2t2 | 0.003 | Old |
| 0.89592 | 0.27401 | 0.034 | 0.036 | 1 | 8 | Traf3ip1 | -0.002 | Old |
| 0.896202 | 0.405934 | 0.156 | 0.164 | 1 | 8 | Cyp4f13 | -0.008 | Old |
| 0.89623 | 0.306656 | 0.19 | 0.197 | 1 | 8 | Rab3gap2 | -0.007 | Old |
| 0.896741 | 0.110279 | 0.358 | 0.358 | 1 | 8 | Gnl1 | 0 | Old |
| 0.897073 | 0.402229 | 0.274 | 0.296 | 1 | 8 | D1Ertd622 | -0.022 | Old |
| 0.897789 | 0.331548 | 0.419 | 0.481 | 1 | 8 | Nudt3 | -0.062 | Old |
| 0.898099 | 0.231101 | 0.067 | 0.065 | 1 | 8 | Tuft1 | 0.002 | Old |
| 0.898145 | 0.134415 | 0.179 | 0.177 | 1 | 8 | Pold3 | 0.002 | Old |
| 0.898384 | 0.192241 | 0.022 | 0.021 | 1 | 8 | Efcab2 | 0.001 | Old |
| 0.898384 | 0.150063 | 0.022 | 0.021 | 1 | 8 | Bhlhe41 | 0.001 | Old |
| 0.898384 | 0.05537 | 0.022 | 0.021 | 1 | 8 | Zfp692 | 0.001 | Old |
| 0.898483 | 0.025705 | 0.045 | 0.047 | 1 | 8 | Polm | -0.002 | Old |
| 0.898624 | 0.078097 | 0.263 | 0.281 | 1 | 8 | Mrpl42 | -0.018 | Old |
| 0.899098 | 0.26585 | 0.218 | 0.216 | 1 | 8 | Slc25a45 | 0.002 | Old |
| 0.899194 | 0.294419 | 0.078 | 0.075 | 1 | 8 | Zfp994 | 0.003 | Old |
| 0.899312 | 0.214733 | 0.128 | 0.135 | 1 | 8 | Zmat5 | -0.007 | Old |
| 0.899332 | 0.107077 | 0.475 | 0.501 | 1 | 8 | Ghitm | -0.026 | Old |
| 0.899424 | 0.161936 | 0.151 | 0.148 | 1 | 8 | Chmp6 | 0.003 | Old |
| 0.899506 | 0.111924 | 0.017 | 0.018 | 1 | 8 | Selp | -0.001 | Old |
| 0.899518 | 0.277514 | 0.162 | 0.158 | 1 | 8 | Kif13b | 0.004 | Old |
| 0.899579 | 0.082444 | 0.123 | 0.127 | 1 | 8 | Armc5 | -0.004 | Old |

| | | | | | | | | |
|---|---|---|---|---|---|---|---|---|
| 0.899611 | 0.036778 | 0.436 | 0.444 | 1 | 8 | Stat6 | -0.008 | Old |
| 0.899698 | 0.297146 | 0.313 | 0.327 | 1 | 8 | Egln2 | -0.014 | Old |
| 0.899833 | 0.199361 | 0.14 | 0.148 | 1 | 8 | Med16 | -0.008 | Old |
| 0.899956 | 0.048039 | 0.324 | 0.33 | 1 | 8 | Gpbp1l1 | -0.006 | Old |
| 0.900016 | 0.084285 | 0.592 | 0.618 | 1 | 8 | Zmiz1 | -0.026 | Old |
| 0.900114 | 0.126813 | 0.57 | 0.608 | 1 | 8 | Prdx2 | -0.038 | Old |
| 0.900134 | 0.071174 | 0.151 | 0.148 | 1 | 8 | Thbd | 0.003 | Old |
| 0.900141 | 0.642973 | 0.028 | 0.026 | 1 | 8 | Gins1 | 0.002 | Old |
| 0.900142 | 0.375237 | 0.022 | 0.021 | 1 | 8 | Pigl | 0.001 | Old |
| 0.900157 | 0.068456 | 0.693 | 0.732 | 1 | 8 | Pirb | -0.039 | Old |
| 0.900642 | 0.262075 | 0.475 | 0.522 | 1 | 8 | AI467606 | -0.047 | Old |
| 0.900792 | 0.089665 | 0.268 | 0.268 | 1 | 8 | Ube2a | 0 | Old |
| 0.900859 | 0.208157 | 0.251 | 0.255 | 1 | 8 | Psen1 | -0.004 | Old |
| 0.900954 | 0.130647 | 0.335 | 0.34 | 1 | 8 | Phc2 | -0.005 | Old |
| 0.901021 | 0.244008 | 0.106 | 0.104 | 1 | 8 | Hcar2 | 0.002 | Old |
| 0.901427 | 0.189106 | 0.648 | 0.745 | 1 | 8 | Skap2 | -0.097 | Old |
| 0.901429 | 0.124959 | 0.017 | 0.018 | 1 | 8 | Gm15952 | -0.001 | Old |
| 0.901833 | 0.268414 | 0.251 | 0.247 | 1 | 8 | Tiprl | 0.004 | Old |
| 0.9019 | 0.175141 | 0.022 | 0.021 | 1 | 8 | Numbl | 0.001 | Old |
| 0.9019 | 0.1198 | 0.022 | 0.021 | 1 | 8 | Bivm | 0.001 | Old |
| 0.902058 | 0.080197 | 0.034 | 0.031 | 1 | 8 | Sirt4 | 0.003 | Old |
| 0.902083 | 0.381658 | 0.14 | 0.138 | 1 | 8 | Bop1 | 0.002 | Old |
| 0.902157 | 0.102574 | 0.581 | 0.59 | 1 | 8 | Ptges3 | -0.009 | Old |
| 0.902189 | 0.292167 | 0.123 | 0.122 | 1 | 8 | Casp6 | 0.001 | Old |
| 0.902475 | 0.502584 | 0.179 | 0.195 | 1 | 8 | Ptpn9 | -0.016 | Old |
| 0.902622 | 0.529908 | 0.223 | 0.242 | 1 | 8 | Plk2 | -0.019 | Old |
| 0.902821 | 0.211075 | 0.547 | 0.569 | 1 | 8 | Akt1 | -0.022 | Old |
| 0.903155 | 0.022693 | 0.168 | 0.171 | 1 | 8 | Nacc1 | -0.003 | Old |
| 0.903221 | 0.106828 | 0.089 | 0.086 | 1 | 8 | Asb7 | 0.003 | Old |
| 0.903322 | 0.064956 | 0.045 | 0.047 | 1 | 8 | Cmc4 | -0.002 | Old |
| 0.903685 | 0.284097 | 0.073 | 0.07 | 1 | 8 | Dnajc9 | 0.003 | Old |
| 0.903897 | 0.435239 | 0.173 | 0.171 | 1 | 8 | Pex2 | 0.002 | Old |
| 0.903901 | 0.139876 | 0.24 | 0.249 | 1 | 8 | Gmps | -0.009 | Old |
| 0.904069 | 0.374629 | 0.14 | 0.14 | 1 | 8 | Gucd1 | 0 | Old |
| 0.904287 | 0.044734 | 0.123 | 0.119 | 1 | 8 | Slc30a4 | 0.004 | Old |
| 0.904288 | 0.013913 | 0.207 | 0.21 | 1 | 8 | Abl2 | -0.003 | Old |
| 0.904317 | 0.814762 | 0.017 | 0.016 | 1 | 8 | Chrna5 | 0.001 | Old |
| 0.905093 | 0.097003 | 0.061 | 0.065 | 1 | 8 | Rnaset2b | -0.004 | Old |
| 0.9051 | 0.158583 | 0.128 | 0.132 | 1 | 8 | Ttf1 | -0.004 | Old |
| 0.905345 | 0.027759 | 0.749 | 0.79 | 1 | 8 | Myadm | -0.041 | Old |
| 0.905414 | 0.239834 | 0.218 | 0.231 | 1 | 8 | Srsf4 | -0.013 | Old |
| 0.905454 | 0.204555 | 0.117 | 0.122 | 1 | 8 | Pon3 | -0.005 | Old |
| 0.905527 | 0.627104 | 0.134 | 0.143 | 1 | 8 | Slc9a6 | -0.009 | Old |
| 0.905702 | 0.016429 | 0.553 | 0.584 | 1 | 8 | Egr2 | -0.031 | Old |
| 0.905734 | 0.15423 | 0.112 | 0.106 | 1 | 8 | Trim34a | 0.006 | Old |

| | | | | | | | | |
|---|---|---|---|---|---|---|---|---|
| 0.905859 | 0.124944 | 0.469 | 0.501 | 1 | 8 | Azin1 | -0.032 | Old |
| 0.90625 | 0.197997 | 0.123 | 0.13 | 1 | 8 | Siah1a | -0.007 | Old |
| 0.906343 | 0.902611 | 0.017 | 0.016 | 1 | 8 | Cep112 | 0.001 | Old |
| 0.906343 | 0.351688 | 0.017 | 0.016 | 1 | 8 | Icam2 | 0.001 | Old |
| 0.906455 | 0.109445 | 0.212 | 0.205 | 1 | 8 | Washc1 | 0.007 | Old |
| 0.906455 | 0.077306 | 0.201 | 0.21 | 1 | 8 | Pttg1 | -0.009 | Old |
| 0.906522 | 0.109048 | 0.14 | 0.145 | 1 | 8 | Cep250 | -0.005 | Old |
| 0.906629 | 0.407035 | 0.123 | 0.13 | 1 | 8 | Mlx | -0.007 | Old |
| 0.906715 | 0.210234 | 0.212 | 0.221 | 1 | 8 | Fam160b1 | -0.009 | Old |
| 0.906975 | 0.078563 | 0.425 | 0.439 | 1 | 8 | Rragc | -0.014 | Old |
| 0.907019 | 0.082007 | 0.212 | 0.221 | 1 | 8 | Adrm1 | -0.009 | Old |
| 0.907142 | 0.02496 | 0.469 | 0.473 | 1 | 8 | Arglu1 | -0.004 | Old |
| 0.907178 | 0.260431 | 0.022 | 0.021 | 1 | 8 | Trim59 | 0.001 | Old |
| 0.9072 | 0.473662 | 0.017 | 0.018 | 1 | 8 | 2210408I2 | -0.001 | Old |
| 0.9072 | 0.023569 | 0.017 | 0.018 | 1 | 8 | Tkfc | -0.001 | Old |
| 0.90746 | 0.062087 | 0.475 | 0.473 | 1 | 8 | Dnase2a | 0.002 | Old |
| 0.907672 | 0.072054 | 0.162 | 0.166 | 1 | 8 | Oxa1l | -0.004 | Old |
| 0.908117 | 0.232239 | 0.352 | 0.353 | 1 | 8 | Ocrl | -0.001 | Old |
| 0.90814 | 0.071919 | 0.156 | 0.161 | 1 | 8 | Tceal8 | -0.005 | Old |
| 0.90837 | 0.814243 | 0.017 | 0.016 | 1 | 8 | Paqr9 | 0.001 | Old |
| 0.90873 | 0.390647 | 0.056 | 0.06 | 1 | 8 | Rgmb | -0.004 | Old |
| 0.908741 | 0.238989 | 0.084 | 0.081 | 1 | 8 | Phlpp2 | 0.003 | Old |
| 0.908943 | 0.477437 | 0.117 | 0.117 | 1 | 8 | Fam214a | 0 | Old |
| 0.909125 | 0.856815 | 0.017 | 0.018 | 1 | 8 | Dhrs13 | -0.001 | Old |
| 0.909125 | 0.287187 | 0.017 | 0.018 | 1 | 8 | Rdh5 | -0.001 | Old |
| 0.909233 | 0.059279 | 0.575 | 0.621 | 1 | 8 | Rnf141 | -0.046 | Old |
| 0.909246 | 0.165443 | 0.061 | 0.065 | 1 | 8 | Adnp2 | -0.004 | Old |
| 0.909499 | 0.103844 | 0.123 | 0.127 | 1 | 8 | Sap30 | -0.004 | Old |
| 0.909708 | 0.374513 | 0.039 | 0.042 | 1 | 8 | Izumo4 | -0.003 | Old |
| 0.909881 | 0.14109 | 0.196 | 0.19 | 1 | 8 | Gbf1 | 0.006 | Old |
| 0.910284 | 0.538368 | 0.106 | 0.114 | 1 | 8 | AC163354. | -0.008 | Old |
| 0.910398 | 0.942996 | 0.017 | 0.016 | 1 | 8 | Poli | 0.001 | Old |
| 0.910398 | 0.09238 | 0.017 | 0.016 | 1 | 8 | Ift140 | 0.001 | Old |
| 0.910691 | 0.765924 | 0.117 | 0.125 | 1 | 8 | Llgl1 | -0.008 | Old |
| 0.910699 | 0.092854 | 0.022 | 0.021 | 1 | 8 | Zfp324 | 0.001 | Old |
| 0.910834 | 0.215077 | 0.151 | 0.161 | 1 | 8 | Rpa3 | -0.01 | Old |
| 0.91105 | 0.153828 | 0.017 | 0.018 | 1 | 8 | Gm20513 | -0.001 | Old |
| 0.911405 | 0.216103 | 0.212 | 0.208 | 1 | 8 | Lyar | 0.004 | Old |
| 0.911553 | 0.089766 | 0.492 | 0.519 | 1 | 8 | Cdk9 | -0.027 | Old |
| 0.911765 | 0.019329 | 0.145 | 0.148 | 1 | 8 | Rasgrp1 | -0.003 | Old |
| 0.911923 | 0.192014 | 0.212 | 0.218 | 1 | 8 | Ccng2 | -0.006 | Old |
| 0.911999 | 0.146905 | 0.33 | 0.335 | 1 | 8 | Taf1 | -0.005 | Old |
| 0.912023 | 0.026115 | 0.777 | 0.849 | 1 | 8 | Hsp90aa1 | -0.072 | Old |
| 0.912165 | 0.062243 | 0.263 | 0.257 | 1 | 8 | Rnf11 | 0.006 | Old |
| 0.912426 | 0.482729 | 0.017 | 0.016 | 1 | 8 | Ifi214 | 0.001 | Old |

| | | | | | | | | |
|---|---|---|---|---|---|---|---|---|
| 0.912426 | 0.299389 | 0.017 | 0.016 | 1 | 8 | Bco2 | 0.001 | Old |
| 0.912426 | 0.172551 | 0.017 | 0.016 | 1 | 8 | Pcbp3 | 0.001 | Old |
| 0.912442 | 0.128801 | 0.196 | 0.195 | 1 | 8 | Yrdc | 0.001 | Old |
| 0.91246 | 0.342209 | 0.022 | 0.021 | 1 | 8 | Tiam2 | 0.001 | Old |
| 0.912479 | 0.043036 | 0.145 | 0.148 | 1 | 8 | 1110008P1 | -0.003 | Old |
| 0.912633 | 0.297337 | 0.318 | 0.319 | 1 | 8 | Rbm6 | -0.001 | Old |
| 0.912749 | 0.296432 | 0.14 | 0.148 | 1 | 8 | Mdn1 | -0.008 | Old |
| 0.912953 | 0.101279 | 0.207 | 0.21 | 1 | 8 | Vps54 | -0.003 | Old |
| 0.912976 | 0.73763 | 0.017 | 0.018 | 1 | 8 | Rad52 | -0.001 | Old |
| 0.912976 | 0.355365 | 0.017 | 0.018 | 1 | 8 | Fn3krp | -0.001 | Old |
| 0.913006 | 0.299715 | 0.335 | 0.356 | 1 | 8 | Npepps | -0.021 | Old |
| 0.91301 | 0.112254 | 0.045 | 0.047 | 1 | 8 | Hspa2 | -0.002 | Old |
| 0.91301 | 0.688665 | 0.168 | 0.182 | 1 | 8 | Arsk | -0.014 | Old |
| 0.913129 | 0.127923 | 0.693 | 0.743 | 1 | 8 | Tmed2 | -0.05 | Old |
| 0.913445 | 0.409226 | 0.101 | 0.106 | 1 | 8 | Zdhhc8 | -0.005 | Old |
| 0.913445 | 0.354404 | 0.101 | 0.106 | 1 | 8 | Slc7a11 | -0.005 | Old |
| 0.913598 | 0.286352 | 0.38 | 0.379 | 1 | 8 | Adss | 0.001 | Old |
| 0.913791 | 0.059591 | 0.218 | 0.213 | 1 | 8 | Zc3h18 | 0.005 | Old |
| 0.914014 | 0.377889 | 0.117 | 0.122 | 1 | 8 | Tsc2 | -0.005 | Old |
| 0.914052 | 0.460012 | 0.184 | 0.187 | 1 | 8 | Gcn1l1 | -0.003 | Old |
| 0.914058 | 0.449129 | 0.151 | 0.151 | 1 | 8 | Cog7 | 0 | Old |
| 0.914178 | 0.004037 | 0.877 | 0.896 | 1 | 8 | Fosl2 | -0.019 | Old |
| 0.914221 | 0.085996 | 0.045 | 0.047 | 1 | 8 | Smyd3 | -0.002 | Old |
| 0.914321 | 0.104936 | 0.615 | 0.662 | 1 | 8 | Serinc1 | -0.047 | Old |
| 0.914455 | 0.636975 | 0.017 | 0.016 | 1 | 8 | Zfp287 | 0.001 | Old |
| 0.914455 | 0.019041 | 0.017 | 0.016 | 1 | 8 | Gm26520 | 0.001 | Old |
| 0.914935 | 0.250271 | 0.335 | 0.348 | 1 | 8 | Tns3 | -0.013 | Old |
| 0.91496 | 0.126431 | 0.743 | 0.766 | 1 | 8 | Gpr132 | -0.023 | Old |
| 0.915525 | 0.148747 | 0.391 | 0.39 | 1 | 8 | Ndfip1 | 0.001 | Old |
| 0.915879 | 0.0679 | 0.642 | 0.717 | 1 | 8 | Eif3c | -0.075 | Old |
| 0.916082 | 0.183444 | 0.162 | 0.158 | 1 | 8 | Rce1 | 0.004 | Old |
| 0.916173 | 0.242336 | 0.201 | 0.203 | 1 | 8 | Mpnd | -0.002 | Old |
| 0.916309 | 0.044588 | 0.173 | 0.177 | 1 | 8 | Rfc2 | -0.004 | Old |
| 0.916364 | 0.1223 | 0.603 | 0.649 | 1 | 8 | Rab7 | -0.046 | Old |
| 0.916484 | 0.126246 | 0.017 | 0.016 | 1 | 8 | Zfp7 | 0.001 | Old |
| 0.916484 | 0.071027 | 0.017 | 0.016 | 1 | 8 | Dyrk3 | 0.001 | Old |
| 0.916542 | 0.305539 | 0.112 | 0.109 | 1 | 8 | Aen | 0.003 | Old |
| 0.91683 | 0.513049 | 0.017 | 0.018 | 1 | 8 | Map10 | -0.001 | Old |
| 0.916834 | 0.157961 | 0.128 | 0.135 | 1 | 8 | Dnajc25 | -0.007 | Old |
| 0.917313 | 0.344355 | 0.179 | 0.182 | 1 | 8 | Ppm1m | -0.003 | Old |
| 0.917362 | 0.006535 | 0.078 | 0.075 | 1 | 8 | Ric8b | 0.003 | Old |
| 0.917643 | 0.044228 | 0.687 | 0.722 | 1 | 8 | Capg | -0.035 | Old |
| 0.917654 | 0.090525 | 0.458 | 0.47 | 1 | 8 | Rnf7 | -0.012 | Old |
| 0.917815 | 0.049399 | 0.508 | 0.53 | 1 | 8 | Spcs3 | -0.022 | Old |
| 0.918053 | 0.271109 | 0.335 | 0.343 | 1 | 8 | Ctcf | -0.008 | Old |

| | | | | | | | | |
|---|---|---|---|---|---|---|---|---|
| 0.918275 | 0.345617 | 0.056 | 0.055 | 1 | 8 | Sh3bp5l | 0.001 | Old |
| 0.918514 | 0.374747 | 0.017 | 0.016 | 1 | 8 | Ocstamp | 0.001 | Old |
| 0.918514 | 0.136626 | 0.017 | 0.016 | 1 | 8 | Zfp446 | 0.001 | Old |
| 0.918538 | 0.223859 | 0.084 | 0.088 | 1 | 8 | Nol4l | -0.004 | Old |
| 0.918601 | 0.057886 | 0.592 | 0.673 | 1 | 8 | Macf1 | -0.081 | Old |
| 0.918704 | 0.445107 | 0.039 | 0.042 | 1 | 8 | Rpl9-ps6 | -0.003 | Old |
| 0.918757 | 0.136284 | 0.017 | 0.018 | 1 | 8 | E430024I08 | -0.001 | Old |
| 0.918976 | 0.375528 | 0.112 | 0.109 | 1 | 8 | Vps45 | 0.003 | Old |
| 0.919027 | 0.418518 | 0.106 | 0.106 | 1 | 8 | Hmces | 0 | Old |
| 0.919071 | 0.099503 | 0.045 | 0.047 | 1 | 8 | 9930104L0 | -0.002 | Old |
| 0.919282 | 0.165491 | 0.179 | 0.182 | 1 | 8 | Stxbp5 | -0.003 | Old |
| 0.919502 | 0.296316 | 0.19 | 0.192 | 1 | 8 | Hdac3 | -0.002 | Old |
| 0.919655 | 0.051779 | 0.564 | 0.592 | 1 | 8 | Fam174a | -0.028 | Old |
| 0.919665 | 0.443637 | 0.101 | 0.106 | 1 | 8 | Smcr8 | -0.005 | Old |
| 0.919703 | 0.128267 | 0.542 | 0.564 | 1 | 8 | Ppp1r9b | -0.022 | Old |
| 0.920429 | 0.073226 | 0.564 | 0.569 | 1 | 8 | Syncrip | -0.005 | Old |
| 0.920506 | 0.140861 | 0.682 | 0.719 | 1 | 8 | Khdrbs1 | -0.037 | Old |
| 0.920544 | 0.647113 | 0.017 | 0.016 | 1 | 8 | Kbtbd8 | 0.001 | Old |
| 0.920685 | 0.781042 | 0.017 | 0.018 | 1 | 8 | Mocos | -0.001 | Old |
| 0.920685 | 0.734481 | 0.017 | 0.018 | 1 | 8 | Zfp652os | -0.001 | Old |
| 0.920685 | 0.223537 | 0.017 | 0.018 | 1 | 8 | C1rl | -0.001 | Old |
| 0.920755 | 0.29713 | 0.112 | 0.112 | 1 | 8 | Rffl | 0 | Old |
| 0.920926 | 0.211614 | 0.201 | 0.2 | 1 | 8 | Zc3h14 | 0.001 | Old |
| 0.921247 | 0.084342 | 0.156 | 0.158 | 1 | 8 | Ints1 | -0.002 | Old |
| 0.921347 | 0.02855 | 0.464 | 0.481 | 1 | 8 | Ifi204 | -0.017 | Old |
| 0.921476 | 0.108288 | 0.626 | 0.652 | 1 | 8 | Prpf38b | -0.026 | Old |
| 0.921516 | 0.450157 | 0.089 | 0.088 | 1 | 8 | Dclre1a | 0.001 | Old |
| 0.921575 | 0.140682 | 0.536 | 0.53 | 1 | 8 | Sptssa | 0.006 | Old |
| 0.921714 | 0.060222 | 0.19 | 0.195 | 1 | 8 | Zbtb8os | -0.005 | Old |
| 0.921773 | 0.217271 | 0.151 | 0.145 | 1 | 8 | Rbpms | 0.006 | Old |
| 0.921781 | 0.183172 | 0.184 | 0.192 | 1 | 8 | Axin1 | -0.008 | Old |
| 0.92188 | 0.154715 | 0.078 | 0.081 | 1 | 8 | 3110082I1 | -0.003 | Old |
| 0.921888 | 0.040891 | 0.263 | 0.265 | 1 | 8 | Dgkz | -0.002 | Old |
| 0.922493 | 0.305559 | 0.123 | 0.127 | 1 | 8 | Csf2rb2 | -0.004 | Old |
| 0.922575 | 0.128402 | 0.017 | 0.016 | 1 | 8 | Tnfrsf22 | 0.001 | Old |
| 0.922575 | 0.002586 | 0.017 | 0.016 | 1 | 8 | Sdr42e1 | 0.001 | Old |
| 0.922675 | 0.163881 | 0.223 | 0.234 | 1 | 8 | Ccdc82 | -0.011 | Old |
| 0.922864 | 0.060143 | 0.413 | 0.423 | 1 | 8 | Ube2i | -0.01 | Old |
| 0.923075 | 0.321065 | 0.123 | 0.122 | 1 | 8 | Tmem97 | 0.001 | Old |
| 0.92385 | 1.171663 | 0.039 | 0.042 | 1 | 8 | 2610035D1 | -0.003 | Old |
| 0.923878 | 0.422459 | 0.061 | 0.06 | 1 | 8 | Fpgs | 0.001 | Old |
| 0.924005 | 0.191568 | 0.156 | 0.153 | 1 | 8 | Ankra2 | 0.003 | Old |
| 0.924088 | 1.302391 | 0.011 | 0.01 | 1 | 8 | Gspt2 | 0.001 | Old |
| 0.924088 | 1.272497 | 0.011 | 0.01 | 1 | 8 | Ptges3l | 0.001 | Old |
| 0.924088 | 0.786482 | 0.011 | 0.01 | 1 | 8 | Slc16a12 | 0.001 | Old |

| | | | | | | | | |
|---|---|---|---|---|---|---|---|---|
| 0.924088 | 0.703724 | 0.011 | 0.01 | 1 | 8 | Hs3st3b1 | 0.001 | Old |
| 0.924088 | 0.431451 | 0.011 | 0.01 | 1 | 8 | Gm20633 | 0.001 | Old |
| 0.924169 | 0.035108 | 0.235 | 0.229 | 1 | 8 | Ormdl2 | 0.006 | Old |
| 0.92421 | 0.220556 | 0.073 | 0.075 | 1 | 8 | Zfyve1 | -0.002 | Old |
| 0.924343 | 0.295297 | 0.073 | 0.073 | 1 | 8 | Msto1 | 0 | Old |
| 0.924456 | 0.027787 | 0.771 | 0.797 | 1 | 8 | Tmed10 | -0.026 | Old |
| 0.924652 | 0.283457 | 0.089 | 0.094 | 1 | 8 | Zdhhc13 | -0.005 | Old |
| 0.924879 | 0.07711 | 0.05 | 0.052 | 1 | 8 | Tma16 | -0.002 | Old |
| 0.92522 | 0.306384 | 0.335 | 0.345 | 1 | 8 | Btbd1 | -0.01 | Old |
| 0.92542 | 0.194669 | 0.101 | 0.099 | 1 | 8 | Akt1s1 | 0.002 | Old |
| 0.92553 | 0.128182 | 0.089 | 0.094 | 1 | 8 | Heg1 | -0.005 | Old |
| 0.925598 | 0.280341 | 0.48 | 0.535 | 1 | 8 | Pik3r1 | -0.055 | Old |
| 0.925759 | 0.422928 | 0.156 | 0.153 | 1 | 8 | Espl1 | 0.003 | Old |
| 0.926016 | 0.494285 | 0.061 | 0.06 | 1 | 8 | Slc22a15 | 0.001 | Old |
| 0.926103 | 0.089089 | 0.782 | 0.823 | 1 | 8 | Tkt | -0.041 | Old |
| 0.92657 | 0.857612 | 0.011 | 0.01 | 1 | 8 | Tcea2 | 0.001 | Old |
| 0.92657 | 0.795432 | 0.011 | 0.01 | 1 | 8 | Spry1 | 0.001 | Old |
| 0.92657 | 0.752309 | 0.011 | 0.01 | 1 | 8 | Dtl | 0.001 | Old |
| 0.92657 | 0.694037 | 0.011 | 0.01 | 1 | 8 | 3830403N1 | 0.001 | Old |
| 0.92657 | 0.451389 | 0.011 | 0.01 | 1 | 8 | Cryba4 | 0.001 | Old |
| 0.926761 | 0.332639 | 0.078 | 0.078 | 1 | 8 | Emc9 | 0 | Old |
| 0.92679 | 0.101413 | 0.413 | 0.408 | 1 | 8 | Cxcr4 | 0.005 | Old |
| 0.926937 | 0.166903 | 0.358 | 0.369 | 1 | 8 | B4galt1 | -0.011 | Old |
| 0.926942 | 0.025984 | 0.078 | 0.075 | 1 | 8 | Mlh1 | 0.003 | Old |
| 0.927587 | 0.438867 | 0.067 | 0.07 | 1 | 8 | Pcid2 | -0.003 | Old |
| 0.927779 | 0.166797 | 0.179 | 0.184 | 1 | 8 | Ubxn2a | -0.005 | Old |
| 0.927834 | 0.208588 | 0.251 | 0.257 | 1 | 8 | Dicer1 | -0.006 | Old |
| 0.928133 | 0.101269 | 0.799 | 0.831 | 1 | 8 | Csnk1a1 | -0.032 | Old |
| 0.928136 | 0.084466 | 0.48 | 0.499 | 1 | 8 | Lrmp | -0.019 | Old |
| 0.928215 | 0.070081 | 0.212 | 0.205 | 1 | 8 | Gapt | 0.007 | Old |
| 0.928253 | 0.303927 | 0.101 | 0.104 | 1 | 8 | Fuom | -0.003 | Old |
| 0.928281 | 0.241009 | 0.19 | 0.187 | 1 | 8 | 1110038F1 | 0.003 | Old |
| 0.928559 | 0.309498 | 0.123 | 0.125 | 1 | 8 | Farsa | -0.002 | Old |
| 0.928588 | 0.14391 | 0.251 | 0.26 | 1 | 8 | Slc36a1 | -0.009 | Old |
| 0.928824 | 0.225627 | 0.352 | 0.361 | 1 | 8 | Hmgn1 | -0.009 | Old |
| 0.928959 | 0.218885 | 0.24 | 0.242 | 1 | 8 | Dhrs1 | -0.002 | Old |
| 0.929047 | 0.047028 | 0.263 | 0.27 | 1 | 8 | Mtmr3 | -0.007 | Old |
| 0.929052 | 1.366138 | 0.011 | 0.01 | 1 | 8 | Prox2 | 0.001 | Old |
| 0.929052 | 1.159984 | 0.011 | 0.01 | 1 | 8 | Tcaf1 | 0.001 | Old |
| 0.929052 | 1.062393 | 0.011 | 0.01 | 1 | 8 | Cacnb4 | 0.001 | Old |
| 0.929052 | 0.620732 | 0.011 | 0.01 | 1 | 8 | Iqgap3 | 0.001 | Old |
| 0.929052 | 0.467986 | 0.011 | 0.01 | 1 | 8 | Gm26624 | 0.001 | Old |
| 0.929052 | 0.380337 | 0.011 | 0.01 | 1 | 8 | 9430002A1 | 0.001 | Old |
| 0.929052 | 0.309546 | 0.011 | 0.01 | 1 | 8 | Tnfrsf14 | 0.001 | Old |
| 0.929052 | 0.296787 | 0.011 | 0.01 | 1 | 8 | Ninl | 0.001 | Old |

| | | | | | | | | |
|---|---|---|---|---|---|---|---|---|
| 0.929052 | 0.155451 | 0.011 | 0.01 | 1 | 8 | Angptl6 | 0.001 | Old |
| 0.929224 | 0.343545 | 0.207 | 0.205 | 1 | 8 | Hdac9 | 0.002 | Old |
| 0.929243 | 0.113703 | 0.721 | 0.745 | 1 | 8 | Birc3 | -0.024 | Old |
| 0.929653 | 0.259983 | 0.363 | 0.405 | 1 | 8 | Neu1 | -0.042 | Old |
| 0.929934 | 0.137382 | 0.056 | 0.057 | 1 | 8 | Gm9530 | -0.001 | Old |
| 0.929994 | 0.200448 | 0.045 | 0.047 | 1 | 8 | Umps | -0.002 | Old |
| 0.930028 | 0.129802 | 0.441 | 0.429 | 1 | 8 | Cpne3 | 0.012 | Old |
| 0.930081 | 0.201014 | 0.257 | 0.262 | 1 | 8 | Epc1 | -0.005 | Old |
| 0.930381 | 0.068879 | 0.279 | 0.27 | 1 | 8 | Cfap43 | 0.009 | Old |
| 0.930394 | 0.284435 | 0.207 | 0.208 | 1 | 8 | BC037034 | -0.001 | Old |
| 0.930652 | 0.071846 | 0.408 | 0.423 | 1 | 8 | Vapb | -0.015 | Old |
| 0.930782 | 0.274086 | 0.184 | 0.179 | 1 | 8 | Auh | 0.005 | Old |
| 0.930857 | 0.20195 | 0.168 | 0.164 | 1 | 8 | Nup50 | 0.004 | Old |
| 0.93123 | 0.239231 | 0.218 | 0.223 | 1 | 8 | Wdr41 | -0.005 | Old |
| 0.931536 | 0.370285 | 0.011 | 0.01 | 1 | 8 | BC048403 | 0.001 | Old |
| 0.931536 | 0.208583 | 0.011 | 0.01 | 1 | 8 | Gm13822 | 0.001 | Old |
| 0.931536 | 0.144924 | 0.011 | 0.01 | 1 | 8 | Slc43a3 | 0.001 | Old |
| 0.931536 | 0.118293 | 0.011 | 0.01 | 1 | 8 | Amigo3 | 0.001 | Old |
| 0.931536 | 0.089148 | 0.011 | 0.01 | 1 | 8 | Bbs1 | 0.001 | Old |
| 0.931536 | 0.084731 | 0.011 | 0.01 | 1 | 8 | Gm9888 | 0.001 | Old |
| 0.931536 | 0.032101 | 0.011 | 0.01 | 1 | 8 | Thrsp | 0.001 | Old |
| 0.931789 | 0.091317 | 0.246 | 0.242 | 1 | 8 | Fbxo3 | 0.004 | Old |
| 0.9321 | 0.369857 | 0.095 | 0.099 | 1 | 8 | Utp15 | -0.004 | Old |
| 0.932443 | 0.286952 | 0.123 | 0.127 | 1 | 8 | Gdap2 | -0.004 | Old |
| 0.932887 | 0.203525 | 0.43 | 0.431 | 1 | 8 | Mib1 | -0.001 | Old |
| 0.932973 | 0.125541 | 0.285 | 0.288 | 1 | 8 | Ogfr | -0.003 | Old |
| 0.933114 | 0.00356 | 0.123 | 0.119 | 1 | 8 | Mrpl38 | 0.004 | Old |
| 0.933254 | 0.037746 | 0.218 | 0.226 | 1 | 8 | Hnrnph3 | -0.008 | Old |
| 0.933334 | 0.195411 | 0.162 | 0.161 | 1 | 8 | Vps53 | 0.001 | Old |
| 0.933504 | 0.12666 | 0.117 | 0.122 | 1 | 8 | Msra | -0.005 | Old |
| 0.93354 | 0.392071 | 0.173 | 0.179 | 1 | 8 | Tspan17 | -0.006 | Old |
| 0.933858 | 0.026514 | 0.117 | 0.119 | 1 | 8 | Tarbp2 | -0.002 | Old |
| 0.933932 | 0.139847 | 0.52 | 0.553 | 1 | 8 | Mdh2 | -0.033 | Old |
| 0.933968 | 0.123839 | 0.436 | 0.423 | 1 | 8 | Smim10l1 | 0.013 | Old |
| 0.934019 | 1.017419 | 0.011 | 0.01 | 1 | 8 | Gas1 | 0.001 | Old |
| 0.934019 | 0.273879 | 0.011 | 0.01 | 1 | 8 | Fgfrl1 | 0.001 | Old |
| 0.934019 | 0.132629 | 0.011 | 0.01 | 1 | 8 | Jam2 | 0.001 | Old |
| 0.934019 | 0.120492 | 0.011 | 0.01 | 1 | 8 | Sorbs1 | 0.001 | Old |
| 0.934019 | 0.068312 | 0.011 | 0.01 | 1 | 8 | Nrg1 | 0.001 | Old |
| 0.934019 | 0.023105 | 0.011 | 0.01 | 1 | 8 | Wnt4 | 0.001 | Old |
| 0.934019 | 0.022947 | 0.011 | 0.01 | 1 | 8 | Sfn | 0.001 | Old |
| 0.934019 | 0.011829 | 0.011 | 0.01 | 1 | 8 | Gnal | 0.001 | Old |
| 0.934102 | 0.568191 | 0.05 | 0.052 | 1 | 8 | Acad12 | -0.002 | Old |
| 0.934366 | 0.496815 | 0.307 | 0.322 | 1 | 8 | Gtpbp2 | -0.015 | Old |
| 0.934418 | 0.185251 | 0.302 | 0.314 | 1 | 8 | Net1 | -0.012 | Old |

| | | | | | | | | |
|---|---|---|---|---|---|---|---|---|
| 0.934448 | 0.363868 | 0.095 | 0.094 | 1 | 8 | Naip6 | 0.001 | Old |
| 0.934534 | 0.016142 | 0.564 | 0.574 | 1 | 8 | Csf2rb | -0.01 | Old |
| 0.935026 | 0.42268 | 0.056 | 0.055 | 1 | 8 | Zbtb5 | 0.001 | Old |
| 0.935317 | 0.17422 | 0.022 | 0.023 | 1 | 8 | Nfe2 | -0.001 | Old |
| 0.935317 | 0.007619 | 0.022 | 0.023 | 1 | 8 | Gstt3 | -0.001 | Old |
| 0.935644 | 0.53379 | 0.061 | 0.06 | 1 | 8 | Abhd10 | 0.001 | Old |
| 0.936518 | 0.210359 | 0.05 | 0.049 | 1 | 8 | Depdc7 | 0.001 | Old |
| 0.936558 | 0.341952 | 0.268 | 0.27 | 1 | 8 | Psme3 | -0.002 | Old |
| 0.936673 | 0.226921 | 0.374 | 0.4 | 1 | 8 | Chtop | -0.026 | Old |
| 0.937016 | 0.06452 | 0.022 | 0.023 | 1 | 8 | Atp1b1 | -0.001 | Old |
| 0.937123 | 0.04426 | 0.263 | 0.273 | 1 | 8 | Gbp3 | -0.01 | Old |
| 0.937227 | 0.009188 | 0.687 | 0.675 | 1 | 8 | Fubp1 | 0.012 | Old |
| 0.937327 | 0.235473 | 0.223 | 0.234 | 1 | 8 | Fundc2 | -0.011 | Old |
| 0.937328 | 0.47945 | 0.14 | 0.138 | 1 | 8 | Cfap36 | 0.002 | Old |
| 0.937464 | 0.236809 | 0.402 | 0.405 | 1 | 8 | Dynll2 | -0.003 | Old |
| 0.937501 | 0.322111 | 0.084 | 0.088 | 1 | 8 | Endod1 | -0.004 | Old |
| 0.937821 | 0.15034 | 0.173 | 0.169 | 1 | 8 | Mrps28 | 0.004 | Old |
| 0.937864 | 0.193128 | 0.296 | 0.304 | 1 | 8 | Sars | -0.008 | Old |
| 0.937958 | 0.425344 | 0.045 | 0.044 | 1 | 8 | Tjap1 | 0.001 | Old |
| 0.938264 | 0.053748 | 0.486 | 0.494 | 1 | 8 | Msl2 | -0.008 | Old |
| 0.938498 | 0.717683 | 0.045 | 0.047 | 1 | 8 | Rab38 | -0.002 | Old |
| 0.938715 | 0.277389 | 0.022 | 0.023 | 1 | 8 | 9130008F2 | -0.001 | Old |
| 0.938715 | 0.029259 | 0.022 | 0.023 | 1 | 8 | Zfp809 | -0.001 | Old |
| 0.938717 | 0.289244 | 0.05 | 0.052 | 1 | 8 | Top2a | -0.002 | Old |
| 0.938884 | 0.224631 | 0.291 | 0.294 | 1 | 8 | Ing4 | -0.003 | Old |
| 0.93918 | 0.034279 | 0.609 | 0.644 | 1 | 8 | Psma3 | -0.035 | Old |
| 0.939322 | 0.039395 | 0.095 | 0.096 | 1 | 8 | Wdr75 | -0.001 | Old |
| 0.939324 | 0.294144 | 0.123 | 0.13 | 1 | 8 | Gnpda2 | -0.007 | Old |
| 0.939642 | 0.035742 | 0.581 | 0.59 | 1 | 8 | Pgls | -0.009 | Old |
| 0.939935 | 0.154199 | 0.503 | 0.543 | 1 | 8 | Tmbim1 | -0.04 | Old |
| 0.940414 | 0.986078 | 0.022 | 0.023 | 1 | 8 | Tnk2 | -0.001 | Old |
| 0.940475 | 0.04703 | 0.207 | 0.205 | 1 | 8 | Fuca2 | 0.002 | Old |
| 0.940593 | 0.560443 | 0.039 | 0.042 | 1 | 8 | Coa7 | -0.003 | Old |
| 0.940647 | 0.121184 | 0.162 | 0.158 | 1 | 8 | Kars | 0.004 | Old |
| 0.94075 | 0.191276 | 0.352 | 0.364 | 1 | 8 | Xpr1 | -0.012 | Old |
| 0.940857 | 0.232675 | 0.156 | 0.153 | 1 | 8 | Wdr11 | 0.003 | Old |
| 0.940871 | 0.233313 | 0.123 | 0.127 | 1 | 8 | Cops7a | -0.004 | Old |
| 0.940906 | 0.060792 | 0.698 | 0.725 | 1 | 8 | Pum2 | -0.027 | Old |
| 0.941011 | 0.554143 | 0.073 | 0.073 | 1 | 8 | Zfp26 | 0 | Old |
| 0.941026 | 0.274479 | 0.05 | 0.052 | 1 | 8 | Cipc | -0.002 | Old |
| 0.941052 | 0.15601 | 0.385 | 0.408 | 1 | 8 | Fndc3b | -0.023 | Old |
| 0.941086 | 0.078012 | 0.251 | 0.257 | 1 | 8 | D5Ertd579 | -0.006 | Old |
| 0.94118 | 0.25192 | 0.207 | 0.21 | 1 | 8 | Fbrsl1 | -0.003 | Old |
| 0.941578 | 0.387336 | 0.128 | 0.13 | 1 | 8 | Ubr1 | -0.002 | Old |
| 0.941581 | 0.133439 | 0.101 | 0.099 | 1 | 8 | Slc38a7 | 0.002 | Old |

| | | | | | | | | |
|---|---|---|---|---|---|---|---|---|
| 0.941685 | 0.183215 | 0.123 | 0.122 | 1 | 8 | Stard5 | 0.001 | Old |
| 0.941826 | 0.307598 | 0.145 | 0.148 | 1 | 8 | Mettl16 | -0.003 | Old |
| 0.942113 | 0.145343 | 0.022 | 0.023 | 1 | 8 | Dhodh | -0.001 | Old |
| 0.942123 | 0.27338 | 0.117 | 0.117 | 1 | 8 | Lrch1 | 0 | Old |
| 0.942145 | 0.43619 | 0.045 | 0.047 | 1 | 8 | Epor | -0.002 | Old |
| 0.942205 | 0.191804 | 0.358 | 0.366 | 1 | 8 | Phb2 | -0.008 | Old |
| 0.942519 | 0.0141 | 0.246 | 0.249 | 1 | 8 | Sf3b4 | -0.003 | Old |
| 0.942545 | 0.051563 | 0.078 | 0.081 | 1 | 8 | Dhx57 | -0.003 | Old |
| 0.942902 | 0.019008 | 0.134 | 0.138 | 1 | 8 | Mrps5 | -0.004 | Old |
| 0.943312 | 0.159614 | 0.073 | 0.07 | 1 | 8 | Edrf1 | 0.003 | Old |
| 0.943374 | 0.624531 | 0.078 | 0.078 | 1 | 8 | Sppl2b | 0 | Old |
| 0.943395 | 0.397313 | 0.151 | 0.153 | 1 | 8 | Comtd1 | -0.002 | Old |
| 0.943419 | 0.152621 | 0.156 | 0.161 | 1 | 8 | Taf2 | -0.005 | Old |
| 0.944179 | 0.130362 | 0.128 | 0.132 | 1 | 8 | Rrp7a | -0.004 | Old |
| 0.944363 | 0.511981 | 0.123 | 0.125 | 1 | 8 | Mrpl45 | -0.002 | Old |
| 0.944489 | 0.106439 | 0.05 | 0.052 | 1 | 8 | Stradb | -0.002 | Old |
| 0.944494 | 0.14574 | 0.397 | 0.41 | 1 | 8 | Smarcc1 | -0.013 | Old |
| 0.944961 | 0.177951 | 0.061 | 0.062 | 1 | 8 | Pex11b | -0.001 | Old |
| 0.944961 | 0.046057 | 0.061 | 0.062 | 1 | 8 | Wipi1 | -0.001 | Old |
| 0.94538 | 0.464061 | 0.095 | 0.096 | 1 | 8 | Pelp1 | -0.001 | Old |
| 0.945637 | 0.107084 | 0.346 | 0.366 | 1 | 8 | Rnf187 | -0.02 | Old |
| 0.946182 | 0.248011 | 0.201 | 0.2 | 1 | 8 | Nagk | 0.001 | Old |
| 0.946675 | 0.075824 | 0.307 | 0.301 | 1 | 8 | Irf9 | 0.006 | Old |
| 0.947145 | 0.15989 | 0.413 | 0.431 | 1 | 8 | Chn2 | -0.018 | Old |
| 0.947213 | 0.718265 | 0.022 | 0.023 | 1 | 8 | Shq1 | -0.001 | Old |
| 0.947259 | 0.204325 | 0.173 | 0.177 | 1 | 8 | Gm13212 | -0.004 | Old |
| 0.947507 | 0.109357 | 0.458 | 0.478 | 1 | 8 | Sbno1 | -0.02 | Old |
| 0.947592 | 0.144204 | 0.296 | 0.309 | 1 | 8 | Uap1 | -0.013 | Old |
| 0.947665 | 0.091621 | 0.117 | 0.117 | 1 | 8 | Stx2 | 0 | Old |
| 0.947707 | 0.035869 | 0.564 | 0.579 | 1 | 8 | Myof | -0.015 | Old |
| 0.948179 | 0.096106 | 0.508 | 0.512 | 1 | 8 | Lima1 | -0.004 | Old |
| 0.948536 | 0.784705 | 0.039 | 0.039 | 1 | 8 | Nxpe4 | 0 | Old |
| 0.948751 | 0.184736 | 0.19 | 0.192 | 1 | 8 | Sestd1 | -0.002 | Old |
| 0.948911 | 0.045996 | 0.134 | 0.135 | 1 | 8 | Usp50 | -0.001 | Old |
| 0.948914 | 0.665255 | 0.022 | 0.023 | 1 | 8 | Tmem62 | -0.001 | Old |
| 0.948914 | 0.601911 | 0.022 | 0.023 | 1 | 8 | Arl6 | -0.001 | Old |
| 0.948914 | 0.183628 | 0.022 | 0.023 | 1 | 8 | Gm42047 | -0.001 | Old |
| 0.948914 | 0.075376 | 0.022 | 0.023 | 1 | 8 | Gm45051 | -0.001 | Old |
| 0.949076 | 0.013944 | 0.156 | 0.158 | 1 | 8 | Pmvk | -0.002 | Old |
| 0.94925 | 0.100614 | 0.441 | 0.462 | 1 | 8 | Zcchc24 | -0.021 | Old |
| 0.949259 | 0.014227 | 0.173 | 0.177 | 1 | 8 | Mplkip | -0.004 | Old |
| 0.949312 | 0.174624 | 0.162 | 0.158 | 1 | 8 | Slc38a6 | 0.004 | Old |
| 0.949862 | 0.189801 | 0.346 | 0.361 | 1 | 8 | Psmb7 | -0.015 | Old |
| 0.949872 | 0.167347 | 0.441 | 0.47 | 1 | 8 | Emc7 | -0.029 | Old |
| 0.950049 | 0.055867 | 0.821 | 0.816 | 1 | 8 | Ddx6 | 0.005 | Old |

| | | | | | | | | |
|---|---|---|---|---|---|---|---|---|
| 0.950614 | 0.85765 | 0.022 | 0.023 | 1 | 8 | Mageh1 | -0.001 | Old |
| 0.950638 | 0.484868 | 0.061 | 0.06 | 1 | 8 | Gemin6 | 0.001 | Old |
| 0.950949 | 0.138374 | 0.101 | 0.099 | 1 | 8 | Gzf1 | 0.002 | Old |
| 0.951021 | 0.178533 | 0.48 | 0.509 | 1 | 8 | Dusp11 | -0.029 | Old |
| 0.951111 | 0.574397 | 0.14 | 0.143 | 1 | 8 | Cfap20 | -0.003 | Old |
| 0.951116 | 0.022103 | 0.615 | 0.631 | 1 | 8 | Rab11b | -0.016 | Old |
| 0.951686 | 0.260793 | 0.134 | 0.14 | 1 | 8 | C2cd3 | -0.006 | Old |
| 0.952315 | 0.297759 | 0.179 | 0.179 | 1 | 8 | Slc31a1 | 0 | Old |
| 0.952315 | 0.538174 | 0.022 | 0.023 | 1 | 8 | Zfp566 | -0.001 | Old |
| 0.952315 | 0.420858 | 0.022 | 0.023 | 1 | 8 | Zfp251 | -0.001 | Old |
| 0.95253 | 0.059116 | 0.788 | 0.86 | 1 | 8 | Plbd1 | -0.072 | Old |
| 0.952635 | 0.028937 | 0.134 | 0.135 | 1 | 8 | Agpat1 | -0.001 | Old |
| 0.952973 | 0.065341 | 0.112 | 0.114 | 1 | 8 | Akap7 | -0.002 | Old |
| 0.95365 | 0.17279 | 0.095 | 0.094 | 1 | 8 | Rab11fip2 | 0.001 | Old |
| 0.95371 | 0.108061 | 0.352 | 0.348 | 1 | 8 | 2310035C2 | 0.004 | Old |
| 0.95375 | 0.093956 | 0.19 | 0.195 | 1 | 8 | Guca1a | -0.005 | Old |
| 0.953877 | 0.097471 | 0.313 | 0.327 | 1 | 8 | Lrrc8c | -0.014 | Old |
| 0.954016 | 0.148507 | 0.022 | 0.023 | 1 | 8 | Tagap1 | -0.001 | Old |
| 0.954045 | 0.238682 | 0.056 | 0.055 | 1 | 8 | Spats2 | 0.001 | Old |
| 0.954128 | 0.427083 | 0.05 | 0.049 | 1 | 8 | Pls3 | 0.001 | Old |
| 0.954246 | 0.055613 | 0.201 | 0.203 | 1 | 8 | Mrpl21 | -0.002 | Old |
| 0.954264 | 0.048809 | 0.14 | 0.138 | 1 | 8 | Ruvbl1 | 0.002 | Old |
| 0.955009 | 0.355355 | 0.19 | 0.184 | 1 | 8 | Ss18l2 | 0.006 | Old |
| 0.955507 | 0.052237 | 0.983 | 0.992 | 1 | 8 | Hsp90ab1 | -0.009 | Old |
| 0.955538 | 0.097004 | 0.402 | 0.413 | 1 | 8 | Calu | -0.011 | Old |
| 0.955741 | 0.266801 | 0.296 | 0.301 | 1 | 8 | Zdhhc21 | -0.005 | Old |
| 0.956129 | 0.379978 | 0.179 | 0.182 | 1 | 8 | Magohb | -0.003 | Old |
| 0.956441 | 0.186068 | 0.123 | 0.122 | 1 | 8 | Tnip1 | 0.001 | Old |
| 0.95663 | 0.484739 | 0.196 | 0.197 | 1 | 8 | Edc4 | -0.001 | Old |
| 0.956909 | 0.148289 | 0.693 | 0.766 | 1 | 8 | Rbm25 | -0.073 | Old |
| 0.957166 | 0.033938 | 0.101 | 0.104 | 1 | 8 | Cxxc1 | -0.003 | Old |
| 0.957215 | 0.145496 | 0.168 | 0.169 | 1 | 8 | Naip5 | -0.001 | Old |
| 0.957448 | 0.564309 | 0.207 | 0.218 | 1 | 8 | Srpk2 | -0.011 | Old |
| 0.957532 | 0.074102 | 0.145 | 0.143 | 1 | 8 | Ctbs | 0.002 | Old |
| 0.957736 | 0.123655 | 0.156 | 0.153 | 1 | 8 | Zfp984 | 0.003 | Old |
| 0.957794 | 0.294783 | 0.207 | 0.213 | 1 | 8 | Rpl7l1 | -0.006 | Old |
| 0.958198 | 0.094661 | 0.291 | 0.304 | 1 | 8 | Atxn2 | -0.013 | Old |
| 0.958401 | 0.262507 | 0.14 | 0.143 | 1 | 8 | Txndc12 | -0.003 | Old |
| 0.958425 | 0.222021 | 0.173 | 0.174 | 1 | 8 | Rhot1 | -0.001 | Old |
| 0.958516 | 0.290106 | 0.201 | 0.2 | 1 | 8 | Eif2ak2 | 0.001 | Old |
| 0.959095 | 0.437216 | 0.173 | 0.174 | 1 | 8 | Scaf8 | -0.001 | Old |
| 0.959121 | 0.897495 | 0.022 | 0.023 | 1 | 8 | Cep95 | -0.001 | Old |
| 0.959121 | 0.815052 | 0.022 | 0.023 | 1 | 8 | Slc25a23 | -0.001 | Old |
| 0.959121 | 0.339562 | 0.022 | 0.023 | 1 | 8 | Pusl1 | -0.001 | Old |
| 0.959167 | 0.165002 | 0.391 | 0.4 | 1 | 8 | Tcn2 | -0.009 | Old |

| | | | | | | | | |
|---|---|---|---|---|---|---|---|---|
| 0.959211 | 0.127507 | 0.156 | 0.164 | 1 | 8 | Haus8 | -0.008 | Old |
| 0.959386 | 0.2204 | 0.492 | 0.512 | 1 | 8 | Atp13a3 | -0.02 | Old |
| 0.959764 | 0.119912 | 0.235 | 0.242 | 1 | 8 | Tlnrd1 | -0.007 | Old |
| 0.959785 | 0.213127 | 0.207 | 0.216 | 1 | 8 | Dnajb4 | -0.009 | Old |
| 0.959954 | 0.170143 | 0.346 | 0.34 | 1 | 8 | Slc25a39 | 0.006 | Old |
| 0.960593 | 0.446459 | 0.196 | 0.2 | 1 | 8 | Sh3bp2 | -0.004 | Old |
| 0.960601 | 0.060217 | 0.117 | 0.119 | 1 | 8 | Snip1 | -0.002 | Old |
| 0.96077 | 0.001828 | 0.067 | 0.068 | 1 | 8 | Chkb | -0.001 | Old |
| 0.960823 | 0.461842 | 0.022 | 0.023 | 1 | 8 | Map2k6 | -0.001 | Old |
| 0.960839 | 0.447926 | 0.089 | 0.091 | 1 | 8 | 6720427I0 | -0.002 | Old |
| 0.960942 | 0.067573 | 0.101 | 0.104 | 1 | 8 | Zbp1 | -0.003 | Old |
| 0.961308 | 0.213771 | 0.603 | 0.631 | 1 | 8 | Pttg1ip | -0.028 | Old |
| 0.961743 | 0.03215 | 0.397 | 0.416 | 1 | 8 | Snx8 | -0.019 | Old |
| 0.961788 | 0.246472 | 0.067 | 0.068 | 1 | 8 | Terf2ip | -0.001 | Old |
| 0.961898 | 1.089406 | 0.056 | 0.057 | 1 | 8 | Mrrf | -0.001 | Old |
| 0.962075 | 0.062349 | 0.525 | 0.553 | 1 | 8 | Phlda1 | -0.028 | Old |
| 0.962081 | 0.206363 | 0.659 | 0.722 | 1 | 8 | Sbno2 | -0.063 | Old |
| 0.962332 | 0.045775 | 0.43 | 0.421 | 1 | 8 | Slc12a6 | 0.009 | Old |
| 0.962466 | 0.237979 | 0.145 | 0.151 | 1 | 8 | Isg20l2 | -0.006 | Old |
| 0.962482 | 0.042097 | 0.151 | 0.153 | 1 | 8 | Dlat | -0.002 | Old |
| 0.962803 | 0.077129 | 0.145 | 0.145 | 1 | 8 | Ap1g2 | 0 | Old |
| 0.962804 | 0.355968 | 0.212 | 0.218 | 1 | 8 | Ap4s1 | -0.006 | Old |
| 0.962898 | 0.215276 | 0.223 | 0.236 | 1 | 8 | Chmp1b | -0.013 | Old |
| 0.963386 | 0.514143 | 0.089 | 0.094 | 1 | 8 | Exosc2 | -0.005 | Old |
| 0.963402 | 0.009575 | 0.324 | 0.319 | 1 | 8 | Kdm4a | 0.005 | Old |
| 0.963719 | 0.331582 | 0.151 | 0.151 | 1 | 8 | Pdcd7 | 0 | Old |
| 0.964316 | 0.548258 | 0.084 | 0.086 | 1 | 8 | Parn | -0.002 | Old |
| 0.964379 | 0.017061 | 0.369 | 0.366 | 1 | 8 | Agtrap | 0.003 | Old |
| 0.964542 | 0.158078 | 0.475 | 0.473 | 1 | 8 | 1810058I2 | 0.002 | Old |
| 0.965141 | 0.116683 | 0.151 | 0.151 | 1 | 8 | Tvp23b | 0 | Old |
| 0.96529 | 0.384687 | 0.101 | 0.101 | 1 | 8 | Btbd10 | 0 | Old |
| 0.965596 | 0.281462 | 0.123 | 0.125 | 1 | 8 | Dnajb12 | -0.002 | Old |
| 0.965826 | 0.225503 | 0.112 | 0.114 | 1 | 8 | Ccdc93 | -0.002 | Old |
| 0.965837 | 0.273398 | 0.128 | 0.13 | 1 | 8 | Hacd3 | -0.002 | Old |
| 0.96618 | 0.157441 | 0.285 | 0.288 | 1 | 8 | Vps26a | -0.003 | Old |
| 0.966197 | 0.365513 | 0.078 | 0.078 | 1 | 8 | Ttl | 0 | Old |
| 0.966258 | 0.168003 | 0.385 | 0.392 | 1 | 8 | Pim3 | -0.007 | Old |
| 0.966881 | 0.015977 | 0.207 | 0.208 | 1 | 8 | Rbmxl1 | -0.001 | Old |
| 0.96733 | 0.171101 | 0.33 | 0.351 | 1 | 8 | Zc3h13 | -0.021 | Old |
| 0.967425 | 0.115811 | 0.374 | 0.377 | 1 | 8 | Nenf | -0.003 | Old |
| 0.967467 | 0.069492 | 0.151 | 0.148 | 1 | 8 | Ado | 0.003 | Old |
| 0.967546 | 0.339083 | 0.128 | 0.132 | 1 | 8 | Med20 | -0.004 | Old |
| 0.967666 | 0.167044 | 0.028 | 0.029 | 1 | 8 | Slc26a6 | -0.001 | Old |
| 0.968074 | 0.267654 | 0.168 | 0.169 | 1 | 8 | Mnat1 | -0.001 | Old |
| 0.968301 | 0.347548 | 0.24 | 0.247 | 1 | 8 | Cyc1 | -0.007 | Old |

| | | | | | | | | |
|---|---|---|---|---|---|---|---|---|
| 0.968497 | 0.147478 | 0.101 | 0.104 | 1 | 8 | Nt5c3 | -0.003 | Old |
| 0.968931 | 0.293366 | 0.218 | 0.223 | 1 | 8 | Zyg11b | -0.005 | Old |
| 0.969052 | 0.056347 | 0.078 | 0.078 | 1 | 8 | Coq9 | 0 | Old |
| 0.969378 | 0.312241 | 0.196 | 0.197 | 1 | 8 | 5-Mar | -0.001 | Old |
| 0.969387 | 0.339736 | 0.263 | 0.273 | 1 | 8 | Rab12 | -0.01 | Old |
| 0.969631 | 0.417114 | 0.128 | 0.13 | 1 | 8 | 4930402H2 | -0.002 | Old |
| 0.969808 | 0.161295 | 0.318 | 0.335 | 1 | 8 | Ociad1 | -0.017 | Old |
| 0.969907 | 0.14381 | 0.413 | 0.444 | 1 | 8 | Fgr | -0.031 | Old |
| 0.969983 | 0.212141 | 0.492 | 0.509 | 1 | 8 | Nsd3 | -0.017 | Old |
| 0.970197 | 0.120295 | 0.045 | 0.044 | 1 | 8 | Aspa | 0.001 | Old |
| 0.970744 | 0.001124 | 0.028 | 0.029 | 1 | 8 | Syn1 | -0.001 | Old |
| 0.970955 | 0.656354 | 0.039 | 0.039 | 1 | 8 | Efemp2 | 0 | Old |
| 0.970992 | 0.049496 | 0.659 | 0.701 | 1 | 8 | Raph1 | -0.042 | Old |
| 0.971071 | 0.201975 | 0.05 | 0.052 | 1 | 8 | Hsd17b7 | -0.002 | Old |
| 0.971131 | 0.425815 | 0.095 | 0.094 | 1 | 8 | Zfp597 | 0.001 | Old |
| 0.971469 | 0.185207 | 0.168 | 0.169 | 1 | 8 | Abraxas2 | -0.001 | Old |
| 0.97168 | 0.318136 | 0.14 | 0.145 | 1 | 8 | Vrk3 | -0.005 | Old |
| 0.971755 | 0.302879 | 0.38 | 0.41 | 1 | 8 | Sec31a | -0.03 | Old |
| 0.972221 | 0.04661 | 0.788 | 0.816 | 1 | 8 | Hnrnph1 | -0.028 | Old |
| 0.972285 | 0.139187 | 0.274 | 0.27 | 1 | 8 | Inppl1 | 0.004 | Old |
| 0.972334 | 0.148009 | 0.38 | 0.397 | 1 | 8 | Siva1 | -0.017 | Old |
| 0.972487 | 0.161849 | 0.106 | 0.104 | 1 | 8 | Ptpn23 | 0.002 | Old |
| 0.972604 | 0.143349 | 0.397 | 0.41 | 1 | 8 | Rab18 | -0.013 | Old |
| 0.97266 | 0.264923 | 0.453 | 0.481 | 1 | 8 | Rassf3 | -0.028 | Old |
| 0.972943 | 0.241074 | 0.168 | 0.171 | 1 | 8 | Mrps7 | -0.003 | Old |
| 0.973484 | 0.485651 | 0.196 | 0.205 | 1 | 8 | Cyp4f16 | -0.009 | Old |
| 0.973595 | 0.574321 | 0.039 | 0.039 | 1 | 8 | Tbc1d22b | 0 | Old |
| 0.973664 | 0.157824 | 0.207 | 0.205 | 1 | 8 | Dtx3l | 0.002 | Old |
| 0.973871 | 0.010802 | 0.927 | 0.987 | 1 | 8 | Vim | -0.06 | Old |
| 0.973958 | 0.385626 | 0.168 | 0.171 | 1 | 8 | Lactb2 | -0.003 | Old |
| 0.974091 | 0.145276 | 0.212 | 0.216 | 1 | 8 | Bms1 | -0.004 | Old |
| 0.974101 | 0.156365 | 0.285 | 0.296 | 1 | 8 | Clptm1 | -0.011 | Old |
| 0.974114 | 0.421365 | 0.05 | 0.049 | 1 | 8 | Lnx2 | 0.001 | Old |
| 0.974184 | 0.740736 | 0.089 | 0.091 | 1 | 8 | Fam210a | -0.002 | Old |
| 0.974395 | 0.118623 | 0.324 | 0.338 | 1 | 8 | Lgals9 | -0.014 | Old |
| 0.974526 | 0.282521 | 0.313 | 0.319 | 1 | 8 | Tsg101 | -0.006 | Old |
| 0.974627 | 0.381134 | 0.207 | 0.216 | 1 | 8 | Cdk6 | -0.009 | Old |
| 0.975073 | 0.595901 | 0.089 | 0.091 | 1 | 8 | Hdhd2 | -0.002 | Old |
| 0.975362 | 0.304005 | 0.028 | 0.029 | 1 | 8 | Slc10a6 | -0.001 | Old |
| 0.975362 | 0.294425 | 0.028 | 0.029 | 1 | 8 | Pus7l | -0.001 | Old |
| 0.97553 | 0.258464 | 0.229 | 0.236 | 1 | 8 | Nudt16 | -0.007 | Old |
| 0.975687 | 0.017906 | 0.313 | 0.309 | 1 | 8 | Ndufb2 | 0.004 | Old |
| 0.9761 | 0.173866 | 0.179 | 0.187 | 1 | 8 | Timm17a | -0.008 | Old |
| 0.97616 | 0.247889 | 0.145 | 0.145 | 1 | 8 | Dbt | 0 | Old |
| 0.976333 | 0.14009 | 0.553 | 0.553 | 1 | 8 | Uba1 | 0 | Old |

| | | | | | | | | |
|---|---|---|---|---|---|---|---|---|
| 0.976488 | 0.495131 | 0.156 | 0.164 | 1 | 8 | Taf7 | -0.008 | Old |
| 0.976507 | 0.509152 | 0.117 | 0.122 | 1 | 8 | Mavs | -0.005 | Old |
| 0.976562 | 0.285451 | 0.162 | 0.171 | 1 | 8 | Hist3h2a | -0.009 | Old |
| 0.976657 | 0.306317 | 0.229 | 0.234 | 1 | 8 | Camk2d | -0.005 | Old |
| 0.976815 | 0.232354 | 0.296 | 0.304 | 1 | 8 | Slamf8 | -0.008 | Old |
| 0.977016 | 0.228108 | 0.492 | 0.538 | 1 | 8 | Il10rb | -0.046 | Old |
| 0.977178 | 0.108717 | 0.291 | 0.296 | 1 | 8 | Ganab | -0.005 | Old |
| 0.977324 | 0.113398 | 0.112 | 0.112 | 1 | 8 | Tmem104 | 0 | Old |
| 0.977623 | 0.231113 | 0.257 | 0.262 | 1 | 8 | Atp9b | -0.005 | Old |
| 0.977656 | 0.011427 | 0.106 | 0.106 | 1 | 8 | Pnpo | 0 | Old |
| 0.977708 | 0.20814 | 0.268 | 0.275 | 1 | 8 | Nup88 | -0.007 | Old |
| 0.977962 | 0.145993 | 0.123 | 0.125 | 1 | 8 | Ikbip | -0.002 | Old |
| 0.97805 | 0.054246 | 0.313 | 0.322 | 1 | 8 | Ugp2 | -0.009 | Old |
| 0.978133 | 0.447692 | 0.218 | 0.226 | 1 | 8 | Rnf5 | -0.008 | Old |
| 0.978263 | 0.096113 | 0.302 | 0.306 | 1 | 8 | Gpr35 | -0.004 | Old |
| 0.978459 | 0.690549 | 0.056 | 0.057 | 1 | 8 | Msh3 | -0.001 | Old |
| 0.978505 | 0.302814 | 0.268 | 0.273 | 1 | 8 | Ap3m1 | -0.005 | Old |
| 0.978535 | 0.074206 | 0.173 | 0.174 | 1 | 8 | Scoc | -0.001 | Old |
| 0.978624 | 0.101232 | 0.173 | 0.177 | 1 | 8 | 9430038I0 | -0.004 | Old |
| 0.979009 | 0.063379 | 0.486 | 0.519 | 1 | 8 | Mpp1 | -0.033 | Old |
| 0.979466 | 0.18293 | 0.151 | 0.153 | 1 | 8 | Atg4b | -0.002 | Old |
| 0.979478 | 0.271705 | 0.134 | 0.135 | 1 | 8 | Ap3s2 | -0.001 | Old |
| 0.979996 | 0.15326 | 0.05 | 0.049 | 1 | 8 | Amd2 | 0.001 | Old |
| 0.980129 | 0.155997 | 0.045 | 0.044 | 1 | 8 | Zfp276 | 0.001 | Old |
| 0.980589 | 0.09199 | 0.458 | 0.481 | 1 | 8 | Snrpd2 | -0.023 | Old |
| 0.980627 | 0.106838 | 0.173 | 0.177 | 1 | 8 | Pnpla2 | -0.004 | Old |
| 0.980846 | 0.14665 | 0.235 | 0.231 | 1 | 8 | Gas2l1 | 0.004 | Old |
| 0.98152 | 0.714041 | 0.028 | 0.029 | 1 | 8 | Fcor | -0.001 | Old |
| 0.981579 | 0.138159 | 0.034 | 0.034 | 1 | 8 | Mlkl | 0 | Old |
| 0.981752 | 0.135689 | 0.201 | 0.205 | 1 | 8 | Glo1 | -0.004 | Old |
| 0.982011 | 0.057062 | 0.581 | 0.569 | 1 | 8 | Hnrnpl | 0.012 | Old |
| 0.982228 | 0.171359 | 0.441 | 0.444 | 1 | 8 | Peli1 | -0.003 | Old |
| 0.982612 | 0.024371 | 0.045 | 0.044 | 1 | 8 | Btf3l4 | 0.001 | Old |
| 0.982877 | 0.164458 | 0.056 | 0.057 | 1 | 8 | Nat6 | -0.001 | Old |
| 0.98288 | 0.033531 | 0.123 | 0.122 | 1 | 8 | Sil1 | 0.001 | Old |
| 0.982996 | 0.213152 | 0.034 | 0.034 | 1 | 8 | Cmtr1 | 0 | Old |
| 0.983026 | 0.033446 | 0.14 | 0.138 | 1 | 8 | Med27 | 0.002 | Old |
| 0.983033 | 0.390962 | 0.078 | 0.081 | 1 | 8 | Gmeb2 | -0.003 | Old |
| 0.983053 | 0.255381 | 0.061 | 0.062 | 1 | 8 | Cables1 | -0.001 | Old |
| 0.98306 | 0.258206 | 0.028 | 0.029 | 1 | 8 | Dus2 | -0.001 | Old |
| 0.983116 | 0.201256 | 0.43 | 0.436 | 1 | 8 | Man2a2 | -0.006 | Old |
| 0.983179 | 0.049229 | 0.056 | 0.055 | 1 | 8 | Ncapg2 | 0.001 | Old |
| 0.9832 | 0.172282 | 0.374 | 0.377 | 1 | 8 | Sirt2 | -0.003 | Old |
| 0.983469 | 0.162802 | 0.196 | 0.2 | 1 | 8 | Slc2a6 | -0.004 | Old |
| 0.983526 | 0.402616 | 0.084 | 0.086 | 1 | 8 | Zfp513 | -0.002 | Old |

| | | | | | | | | |
|---|---|---|---|---|---|---|---|---|
| 0.983854 | 0.453796 | 0.045 | 0.044 | 1 | 8 | Lrrc14 | 0.001 | Old |
| 0.984285 | 0.063892 | 0.078 | 0.078 | 1 | 8 | Foxred1 | 0 | Old |
| 0.98439 | 0.324728 | 0.106 | 0.109 | 1 | 8 | Actr8 | -0.003 | Old |
| 0.984451 | 0.071999 | 0.419 | 0.421 | 1 | 8 | Vdac3 | -0.002 | Old |
| 0.9846 | 0.892214 | 0.028 | 0.029 | 1 | 8 | 4933427D1 | -0.001 | Old |
| 0.984629 | 0.804857 | 0.123 | 0.127 | 1 | 8 | Usp39 | -0.004 | Old |
| 0.984686 | 0.066705 | 0.196 | 0.197 | 1 | 8 | Scamp1 | -0.001 | Old |
| 0.984796 | 0.114811 | 0.33 | 0.319 | 1 | 8 | Alkbh5 | 0.011 | Old |
| 0.984817 | 0.234679 | 0.559 | 0.592 | 1 | 8 | Edem1 | -0.033 | Old |
| 0.984894 | 0.64485 | 0.067 | 0.07 | 1 | 8 | Tfpi | -0.003 | Old |
| 0.985026 | 0.319384 | 0.212 | 0.223 | 1 | 8 | Hras | -0.011 | Old |
| 0.985096 | 0.448875 | 0.045 | 0.044 | 1 | 8 | Zfp810 | 0.001 | Old |
| 0.98524 | 0.171166 | 0.073 | 0.073 | 1 | 8 | 1700021F0 | 0 | Old |
| 0.985755 | 0.548754 | 0.089 | 0.091 | 1 | 8 | Ccdc32 | -0.002 | Old |
| 0.985843 | 0.216251 | 0.302 | 0.304 | 1 | 8 | Rnpepl1 | -0.002 | Old |
| 0.9859 | 0.061871 | 0.134 | 0.138 | 1 | 8 | Mapk9 | -0.004 | Old |
| 0.985983 | 0.103923 | 0.391 | 0.403 | 1 | 8 | Aldh1a2 | -0.012 | Old |
| 0.986118 | 0.359011 | 0.095 | 0.096 | 1 | 8 | Pwp1 | -0.001 | Old |
| 0.986338 | 0.102153 | 0.045 | 0.044 | 1 | 8 | Inpp4b | 0.001 | Old |
| 0.986591 | 0.137599 | 0.313 | 0.319 | 1 | 8 | Gch1 | -0.006 | Old |
| 0.986715 | 0.069221 | 0.229 | 0.226 | 1 | 8 | Mrpl13 | 0.003 | Old |
| 0.987059 | 0.191635 | 0.453 | 0.483 | 1 | 8 | Dgkd | -0.03 | Old |
| 0.987295 | 0.11265 | 0.056 | 0.057 | 1 | 8 | Ccdc58 | -0.001 | Old |
| 0.987568 | 0.099963 | 0.341 | 0.361 | 1 | 8 | Ube2z | -0.02 | Old |
| 0.988116 | 0.279229 | 0.039 | 0.039 | 1 | 8 | Zfp770 | 0 | Old |
| 0.988349 | 0.323619 | 0.246 | 0.252 | 1 | 8 | Pigk | -0.006 | Old |
| 0.988663 | 0.16705 | 0.034 | 0.034 | 1 | 8 | Rgl3 | 0 | Old |
| 0.988688 | 0.829552 | 0.078 | 0.081 | 1 | 8 | Fam217b | -0.003 | Old |
| 0.988991 | 0.242053 | 0.101 | 0.101 | 1 | 8 | Ccnc | 0 | Old |
| 0.989218 | 0.256132 | 0.508 | 0.532 | 1 | 8 | Pura | -0.024 | Old |
| 0.989298 | 0.144963 | 0.43 | 0.452 | 1 | 8 | Cct7 | -0.022 | Old |
| 0.989326 | 0.051898 | 0.151 | 0.151 | 1 | 8 | Ints9 | 0 | Old |
| 0.989378 | 0.234736 | 0.151 | 0.153 | 1 | 8 | Farsb | -0.002 | Old |
| 0.989436 | 0.011735 | 0.039 | 0.039 | 1 | 8 | Rai14 | 0 | Old |
| 0.989852 | 0.272292 | 0.229 | 0.236 | 1 | 8 | Slc35a3 | -0.007 | Old |
| 0.989915 | 0.489696 | 0.212 | 0.223 | 1 | 8 | Rabgef1 | -0.011 | Old |
| 0.990229 | 0.095369 | 0.587 | 0.652 | 1 | 8 | Mapk3 | -0.065 | Old |
| 0.990309 | 0.282011 | 0.218 | 0.229 | 1 | 8 | Galnt11 | -0.011 | Old |
| 0.990467 | 0.566667 | 0.061 | 0.062 | 1 | 8 | Klc4 | -0.001 | Old |
| 0.990604 | 0.139672 | 0.453 | 0.468 | 1 | 8 | Grina | -0.015 | Old |
| 0.990655 | 0.147982 | 0.145 | 0.148 | 1 | 8 | Hif1an | -0.003 | Old |
| 0.990759 | 0.530042 | 0.028 | 0.029 | 1 | 8 | AC163032. | -0.001 | Old |
| 0.991018 | 0.206895 | 0.179 | 0.177 | 1 | 8 | Tspan4 | 0.002 | Old |
| 0.991324 | 0.500085 | 0.095 | 0.096 | 1 | 8 | Rhno1 | -0.001 | Old |
| 0.991335 | 0.03634 | 0.212 | 0.213 | 1 | 8 | Kdm5c | -0.001 | Old |

| | | | | | | | | |
|---|---|---|---|---|---|---|---|---|
| 0.992419 | 0.056573 | 0.721 | 0.782 | 1 | 8 | Zyx | -0.061 | Old |
| 0.992738 | 0.111258 | 0.162 | 0.161 | 1 | 8 | Tesk1 | 0.001 | Old |
| 0.992739 | 0.089467 | 0.637 | 0.639 | 1 | 8 | Cflar | -0.002 | Old |
| 0.992802 | 0.184284 | 0.101 | 0.101 | 1 | 8 | Tom1 | 0 | Old |
| 0.993101 | 0.159451 | 0.33 | 0.327 | 1 | 8 | Ifnar2 | 0.003 | Old |
| 0.993176 | 0.067071 | 0.436 | 0.444 | 1 | 8 | Eif4g3 | -0.008 | Old |
| 0.993387 | 0.104792 | 0.291 | 0.294 | 1 | 8 | Sik1 | -0.003 | Old |
| 0.994096 | 0.130135 | 0.073 | 0.073 | 1 | 8 | Fars2 | 0 | Old |
| 0.994128 | 0.20653 | 0.24 | 0.247 | 1 | 8 | Rnf115 | -0.007 | Old |
| 0.994327 | 0.096566 | 0.352 | 0.364 | 1 | 8 | Zfp292 | -0.012 | Old |
| 0.994336 | 0.409995 | 0.201 | 0.205 | 1 | 8 | Cfdp1 | -0.004 | Old |
| 0.994393 | 0.095995 | 0.52 | 0.54 | 1 | 8 | Ank | -0.02 | Old |
| 0.994853 | 0.108129 | 0.508 | 0.54 | 1 | 8 | Ddi2 | -0.032 | Old |
| 0.994919 | 0.36214 | 0.101 | 0.101 | 1 | 8 | Tfdp1 | 0 | Old |
| 0.995 | 0.27078 | 0.324 | 0.343 | 1 | 8 | Dhrs7 | -0.019 | Old |
| 0.995243 | 0.095797 | 0.631 | 0.655 | 1 | 8 | Stxbp3 | -0.024 | Old |
| 0.995299 | 0.040153 | 0.704 | 0.732 | 1 | 8 | Minos1 | -0.028 | Old |
| 0.99538 | 0.884447 | 0.028 | 0.029 | 1 | 8 | Pkd1l2 | -0.001 | Old |
| 0.99561 | 0.295051 | 0.207 | 0.205 | 1 | 8 | Zfp622 | 0.002 | Old |
| 0.995749 | 0.252935 | 0.034 | 0.034 | 1 | 8 | Nle1 | 0 | Old |
| 0.995792 | 0.159241 | 0.156 | 0.156 | 1 | 8 | Ric1 | 0 | Old |
| 0.996006 | 0.111456 | 0.318 | 0.327 | 1 | 8 | Fchsd2 | -0.009 | Old |
| 0.996059 | 0.068778 | 0.346 | 0.335 | 1 | 8 | Fam91a1 | 0.011 | Old |
| 0.996454 | 0.041834 | 0.296 | 0.304 | 1 | 8 | Pcna | -0.008 | Old |
| 0.996613 | 0.107368 | 0.101 | 0.101 | 1 | 8 | Snx11 | 0 | Old |
| 0.997031 | 0.529631 | 0.134 | 0.138 | 1 | 8 | Nsmce3 | -0.004 | Old |
| 0.997166 | 0.796859 | 0.034 | 0.034 | 1 | 8 | Pdlim7 | 0 | Old |
| 0.997359 | 0.182032 | 0.039 | 0.039 | 1 | 8 | Thada | 0 | Old |
| 0.997359 | 0.12546 | 0.039 | 0.039 | 1 | 8 | Arrdc2 | 0 | Old |
| 0.99741 | 0.389233 | 0.184 | 0.192 | 1 | 8 | Cd209b | -0.008 | Old |
| 0.997706 | 0.080604 | 0.369 | 0.377 | 1 | 8 | Zc3h15 | -0.008 | Old |
| 0.998067 | 0.133105 | 0.123 | 0.125 | 1 | 8 | Lipo3 | -0.002 | Old |
| 0.998169 | 0.443449 | 0.084 | 0.086 | 1 | 8 | Fbxl15 | -0.002 | Old |
| 0.998393 | 0.085119 | 0.196 | 0.192 | 1 | 8 | Zadh2 | 0.004 | Old |
| 0.998442 | 0.067 | 0.749 | 0.808 | 1 | 8 | Tnfaip8 | -0.059 | Old |
| 0.998623 | 0.187821 | 0.162 | 0.164 | 1 | 8 | Ppfibp1 | -0.002 | Old |
| 0.998858 | 0.097489 | 0.263 | 0.265 | 1 | 8 | Fyttd1 | -0.002 | Old |
| 0.998922 | 0.211901 | 0.145 | 0.148 | 1 | 8 | Ciz1 | -0.003 | Old |
| 0.999051 | 0.151821 | 0.475 | 0.504 | 1 | 8 | Sgk3 | -0.029 | Old |
| 0.999076 | 0.072709 | 0.52 | 0.592 | 1 | 8 | Ptpa | -0.072 | Old |
| 0.999337 | 0.045528 | 0.179 | 0.179 | 1 | 8 | Stard4 | 0 | Old |
| 3.16E-20 | 0.617323 | 0.99 | 0.955 | 9.82E-16 | 8 | Rps14 | 0.035 | Young |
| 4.27E-20 | 0.550356 | 0.995 | 0.916 | 1.33E-15 | 8 | Rps23 | 0.079 | Young |
| 7.45E-20 | 0.652027 | 0.987 | 0.933 | 2.31E-15 | 8 | Rps10 | 0.054 | Young |
| 1.25E-19 | 0.548045 | 0.997 | 0.989 | 3.87E-15 | 8 | Rpl13 | 0.008 | Young |

| | | | | | | | | |
|---|---|---|---|---|---|---|---|---|
| 5.35E-19 | 0.707936 | 0.982 | 0.894 | 1.66E-14 | 8 | Rpl18a | 0.088 | Young |
| 6.01E-19 | 0.435691 | 0.997 | 0.994 | 1.87E-14 | 8 | Rpl35a | 0.003 | Young |
| 1.00E-18 | 0.77916 | 0.961 | 0.877 | 3.11E-14 | 8 | Rpl32 | 0.084 | Young |
| 2.65E-18 | 0.653122 | 0.99 | 0.894 | 8.22E-14 | 8 | Rps13 | 0.096 | Young |
| 2.67E-18 | 0.465757 | 1 | 1 | 8.29E-14 | 8 | Fau | 0 | Young |
| 1.98E-17 | 0.56193 | 0.992 | 0.95 | 6.15E-13 | 8 | Rps4x | 0.042 | Young |
| 4.47E-17 | 0.639286 | 0.987 | 0.899 | 1.39E-12 | 8 | Rpl26 | 0.088 | Young |
| 5.52E-17 | 0.786245 | 0.938 | 0.782 | 1.72E-12 | 8 | Rpl11 | 0.156 | Young |
| 7.45E-17 | 0.577804 | 0.997 | 0.983 | 2.31E-12 | 8 | Cst3 | 0.014 | Young |
| 7.90E-17 | 0.549152 | 0.99 | 0.978 | 2.45E-12 | 8 | Rps16 | 0.012 | Young |
| 8.73E-17 | 0.724485 | 0.735 | 0.341 | 2.71E-12 | 8 | Hspa1b | 0.394 | Young |
| 1.18E-16 | 0.612019 | 0.977 | 0.888 | 3.67E-12 | 8 | Rpl9 | 0.089 | Young |
| 1.22E-16 | 0.61105 | 0.982 | 0.927 | 3.80E-12 | 8 | Rps5 | 0.055 | Young |
| 2.82E-16 | 0.464105 | 1 | 0.983 | 8.74E-12 | 8 | Rps24 | 0.017 | Young |
| 4.43E-16 | 0.557378 | 0.987 | 0.939 | 1.38E-11 | 8 | Rpl30 | 0.048 | Young |
| 1.08E-15 | 0.645228 | 0.974 | 0.899 | 3.35E-11 | 8 | Rpl28 | 0.075 | Young |
| 1.58E-15 | 0.579856 | 0.969 | 0.95 | 4.91E-11 | 8 | Rpl34 | 0.019 | Young |
| 4.53E-15 | 0.498195 | 0.992 | 0.978 | 1.41E-10 | 8 | Rpl23 | 0.014 | Young |
| 4.74E-15 | 0.739119 | 0.997 | 0.966 | 1.47E-10 | 8 | Apoe | 0.031 | Young |
| 5.34E-15 | 0.406696 | 1 | 0.994 | 1.66E-10 | 8 | Rplp1 | 0.006 | Young |
| 2.27E-14 | 0.429777 | 0.997 | 0.978 | 7.04E-10 | 8 | Rpl27a | 0.019 | Young |
| 2.29E-14 | 0.508256 | 0.997 | 0.955 | 7.12E-10 | 8 | Rps19 | 0.042 | Young |
| 2.61E-14 | 0.447555 | 0.997 | 0.961 | 8.12E-10 | 8 | Rps12 | 0.036 | Young |
| 2.93E-14 | 0.526983 | 0.987 | 0.944 | 9.09E-10 | 8 | Rps27 | 0.043 | Young |
| 5.08E-14 | 0.497209 | 0.987 | 0.955 | 1.58E-09 | 8 | Rpl39 | 0.032 | Young |
| 8.90E-14 | 1.50247 | 0.584 | 0.24 | 2.76E-09 | 8 | Hpgd | 0.344 | Young |
| 3.78E-13 | 0.558027 | 0.969 | 0.855 | 1.17E-08 | 8 | Tyrobp | 0.114 | Young |
| 2.02E-12 | 0.602241 | 0.945 | 0.86 | 6.28E-08 | 8 | Rps3 | 0.085 | Young |
| 4.04E-12 | 0.587673 | 0.948 | 0.816 | 1.25E-07 | 8 | Rpl12 | 0.132 | Young |
| 5.24E-12 | 0.396082 | 1 | 0.972 | 1.63E-07 | 8 | Rps20 | 0.028 | Young |
| 5.70E-12 | 0.63082 | 0.938 | 0.771 | 1.77E-07 | 8 | Rpl35 | 0.167 | Young |
| 9.64E-12 | 0.555405 | 0.958 | 0.883 | 2.99E-07 | 8 | Rps7 | 0.075 | Young |
| 1.03E-11 | 0.65565 | 0.935 | 0.788 | 3.19E-07 | 8 | Rpl10 | 0.147 | Young |
| 1.73E-11 | 0.516784 | 0.971 | 0.922 | 5.39E-07 | 8 | Rpl18 | 0.049 | Young |
| 1.96E-11 | 0.435397 | 1 | 1 | 6.09E-07 | 8 | Cd74 | 0 | Young |
| 2.00E-11 | 0.658425 | 0.73 | 0.453 | 6.22E-07 | 8 | Hspa1a | 0.277 | Young |
| 3.13E-11 | 0.484948 | 0.987 | 0.939 | 9.71E-07 | 8 | Rpl19 | 0.048 | Young |
| 3.65E-11 | 0.495905 | 0.99 | 0.933 | 1.13E-06 | 8 | Rps11 | 0.057 | Young |
| 4.61E-11 | 0.426907 | 0.984 | 0.927 | 1.43E-06 | 8 | Rpl22 | 0.057 | Young |
| 6.81E-11 | 0.39407 | 0.987 | 0.966 | 2.12E-06 | 8 | Rpl17 | 0.021 | Young |
| 1.37E-10 | 0.530358 | 0.958 | 0.86 | 4.26E-06 | 8 | Rps6 | 0.098 | Young |
| 1.39E-10 | 0.448116 | 0.992 | 0.916 | 4.31E-06 | 8 | Rpl5 | 0.076 | Young |
| 1.85E-10 | 0.466229 | 0.971 | 0.888 | 5.75E-06 | 8 | Rpl3 | 0.083 | Young |
| 4.76E-10 | 0.315544 | 1 | 0.989 | 1.48E-05 | 8 | Rps8 | 0.011 | Young |
| 9.09E-10 | 0.301243 | 0.997 | 0.994 | 2.82E-05 | 8 | Rps21 | 0.003 | Young |

| | | | | | | | | |
|---|---|---|---|---|---|---|---|---|
| 1.24E-09 | 0.998995 | 0.647 | 0.363 | 3.85E-05 | 8 | Gatm | 0.284 | Young |
| 1.73E-09 | 0.229234 | 1 | 0.994 | 5.39E-05 | 8 | Rps29 | 0.006 | Young |
| 2.04E-09 | 0.476552 | 0.969 | 0.877 | 6.33E-05 | 8 | Rps15a | 0.092 | Young |
| 7.03E-09 | 0.474581 | 0.951 | 0.866 | 0.000218 | 8 | Rpl21 | 0.085 | Young |
| 7.66E-09 | 0.754166 | 0.647 | 0.391 | 0.000238 | 8 | Cd209a | 0.256 | Young |
| 7.81E-09 | 0.431004 | 0.966 | 0.883 | 0.000243 | 8 | Rpl24 | 0.083 | Young |
| 8.18E-09 | 0.403182 | 0.987 | 0.922 | 0.000254 | 8 | Rps25 | 0.065 | Young |
| 9.56E-09 | 0.346065 | 0.995 | 0.961 | 0.000297 | 8 | Rps15 | 0.034 | Young |
| 1.72E-08 | 0.212231 | 1 | 1 | 0.000534 | 8 | Tpt1 | 0 | Young |
| 2.09E-08 | 0.564887 | 0.888 | 0.737 | 0.00065 | 8 | Rpl31 | 0.151 | Young |
| 3.30E-08 | 0.559279 | 0.93 | 0.816 | 0.001024 | 8 | Wfdc17 | 0.114 | Young |
| 3.52E-08 | 0.553029 | 0.914 | 0.788 | 0.001093 | 8 | Rps18 | 0.126 | Young |
| 4.98E-08 | 0.538369 | 0.901 | 0.777 | 0.001546 | 8 | Rpl15 | 0.124 | Young |
| 6.00E-08 | 0.387734 | 0.966 | 0.927 | 0.001863 | 8 | Rplp0 | 0.039 | Young |
| 6.12E-08 | 0.534908 | 0.865 | 0.721 | 0.001902 | 8 | Cox8a | 0.144 | Young |
| 7.74E-08 | 0.347224 | 0.979 | 0.933 | 0.002403 | 8 | Rpl6 | 0.046 | Young |
| 1.18E-07 | 2.305362 | 0.239 | 0.056 | 0.003667 | 8 | Slc15a2 | 0.183 | Young |
| 1.27E-07 | 0.650535 | 0.751 | 0.508 | 0.00393 | 8 | Tsc22d3 | 0.243 | Young |
| 1.88E-07 | 0.559417 | 0.86 | 0.715 | 0.005838 | 8 | Ptpn18 | 0.145 | Young |
| 4.30E-07 | 0.343532 | 0.977 | 0.905 | 0.013354 | 8 | Serf2 | 0.072 | Young |
| 4.64E-07 | 0.298196 | 0.995 | 0.972 | 0.01442 | 8 | Rpl23a | 0.023 | Young |
| 5.56E-07 | 0.37853 | 0.961 | 0.894 | 0.017278 | 8 | Rpl36 | 0.067 | Young |
| 6.22E-07 | 0.485712 | 0.883 | 0.737 | 0.0193 | 8 | Rpl4 | 0.146 | Young |
| 9.31E-07 | 0.774942 | 0.603 | 0.391 | 0.028919 | 8 | Fam129a | 0.212 | Young |
| 1.10E-06 | 1.034049 | 0.538 | 0.352 | 0.034177 | 8 | Napsa | 0.186 | Young |
| 1.15E-06 | 1.336024 | 0.294 | 0.106 | 0.035648 | 8 | Sh3bp5 | 0.188 | Young |
| 1.22E-06 | 0.404356 | 0.966 | 0.872 | 0.037833 | 8 | Rpl10a | 0.094 | Young |
| 1.79E-06 | 0.387529 | 0.953 | 0.849 | 0.055591 | 8 | Rpl36a | 0.104 | Young |
| 2.02E-06 | 0.901664 | 0.629 | 0.453 | 0.062753 | 8 | Bcl2a1b | 0.176 | Young |
| 2.57E-06 | 0.39835 | 0.938 | 0.832 | 0.07991 | 8 | Rps2 | 0.106 | Young |
| 3.09E-06 | 1.00786 | 0.475 | 0.279 | 0.095969 | 8 | Bcl2a1d | 0.196 | Young |
| 3.10E-06 | 0.197822 | 1 | 0.994 | 0.096393 | 8 | Rps3a1 | 0.006 | Young |
| 3.22E-06 | 2.069171 | 0.218 | 0.061 | 0.099914 | 8 | Slc40a1 | 0.157 | Young |
| 3.33E-06 | 0.553759 | 0.86 | 0.743 | 0.103286 | 8 | Gadd45b | 0.117 | Young |
| 3.54E-06 | 0.657291 | 0.686 | 0.525 | 0.109827 | 8 | Ankrd44 | 0.161 | Young |
| 3.54E-06 | 0.296008 | 0.992 | 0.944 | 0.109968 | 8 | Fcer1g | 0.048 | Young |
| 3.76E-06 | 1.066523 | 0.387 | 0.19 | 0.116693 | 8 | Cmah | 0.197 | Young |
| 3.79E-06 | 1.062724 | 0.468 | 0.257 | 0.117658 | 8 | Ccl4 | 0.211 | Young |
| 3.84E-06 | 0.487498 | 0.847 | 0.698 | 0.119182 | 8 | Atpif1 | 0.149 | Young |
| 3.86E-06 | 0.553163 | 0.587 | 0.369 | 0.119835 | 8 | Blvrb | 0.218 | Young |
| 3.99E-06 | 1.372916 | 0.296 | 0.117 | 0.12404 | 8 | Plxdc2 | 0.179 | Young |
| 5.33E-06 | 0.143653 | 1 | 0.994 | 0.165491 | 8 | Rps9 | 0.006 | Young |
| 5.44E-06 | 0.298208 | 0.992 | 0.95 | 0.16878 | 8 | Rpl37 | 0.042 | Young |
| 5.67E-06 | 0.698212 | 0.642 | 0.464 | 0.176148 | 8 | Eif3h | 0.178 | Young |
| 5.72E-06 | 1.818421 | 0.262 | 0.101 | 0.177599 | 8 | Bmyc | 0.161 | Young |

| | | | | | | | | |
|---|---|---|---|---|---|---|---|---|
| 5.76E-06 | 0.525446 | 0.8 | 0.665 | 0.178748 | 8 | Eps15 | 0.135 | Young |
| 6.45E-06 | 1.847637 | 0.244 | 0.084 | 0.200379 | 8 | Ccl3 | 0.16 | Young |
| 6.85E-06 | 0.421931 | 0.901 | 0.804 | 0.212755 | 8 | Atp5h | 0.097 | Young |
| 7.99E-06 | 4.387944 | 0.104 | 0 | 0.248145 | 8 | Colec12 | 0.104 | Young |
| 8.06E-06 | 0.42396 | 0.925 | 0.838 | 0.250263 | 8 | Rpl7a | 0.087 | Young |
| 8.17E-06 | 0.741554 | 0.587 | 0.397 | 0.253646 | 8 | Nr4a3 | 0.19 | Young |
| 8.79E-06 | 0.461621 | 0.857 | 0.743 | 0.272816 | 8 | Ubb | 0.114 | Young |
| 9.86E-06 | 0.65631 | 0.655 | 0.48 | 0.306038 | 8 | Rpl13a | 0.175 | Young |
| 1.19E-05 | 0.24197 | 0.992 | 0.961 | 0.368772 | 8 | Hspa8 | 0.031 | Young |
| 1.26E-05 | 0.745697 | 0.603 | 0.441 | 0.391089 | 8 | Ost4 | 0.162 | Young |
| 1.50E-05 | 0.364581 | 0.984 | 0.955 | 0.464372 | 8 | Pf4 | 0.029 | Young |
| 1.57E-05 | 0.51946 | 0.8 | 0.654 | 0.486035 | 8 | Rpl29 | 0.146 | Young |
| 1.67E-05 | 1.604895 | 0.242 | 0.089 | 0.517204 | 8 | Klrb1b | 0.153 | Young |
| 1.75E-05 | 2.416133 | 0.187 | 0.056 | 0.543484 | 8 | Dpp4 | 0.131 | Young |
| 1.97E-05 | 0.347152 | 0.974 | 0.922 | 0.612058 | 8 | Sh3bgrl3 | 0.052 | Young |
| 2.27E-05 | 0.439461 | 0.6 | 0.413 | 0.70476 | 8 | Iqgap2 | 0.187 | Young |
| 2.49E-05 | 0.498082 | 0.823 | 0.687 | 0.772968 | 8 | Mef2c | 0.136 | Young |
| 2.60E-05 | 0.426761 | 0.704 | 0.486 | 0.808095 | 8 | Eef1d | 0.218 | Young |
| 2.73E-05 | 0.481233 | 0.906 | 0.877 | 0.847185 | 8 | Il1b | 0.029 | Young |
| 2.93E-05 | 0.528936 | 0.706 | 0.559 | 0.910861 | 8 | Dek | 0.147 | Young |
| 2.95E-05 | 1.098703 | 0.343 | 0.173 | 0.914567 | 8 | Bri3bp | 0.17 | Young |
| 3.67E-05 | 0.400059 | 0.917 | 0.816 | 1 | 8 | Cox7a2l | 0.101 | Young |
| 3.72E-05 | 0.596831 | 0.701 | 0.531 | 1 | 8 | Rabac1 | 0.17 | Young |
| 4.06E-05 | 0.19446 | 1 | 0.994 | 1 | 8 | Eef1a1 | 0.006 | Young |
| 4.35E-05 | 1.086409 | 0.291 | 0.134 | 1 | 8 | Crim1 | 0.157 | Young |
| 4.50E-05 | 0.531893 | 0.579 | 0.397 | 1 | 8 | Brd4 | 0.182 | Young |
| 4.68E-05 | 0.706884 | 0.54 | 0.346 | 1 | 8 | Myo1g | 0.194 | Young |
| 4.80E-05 | 0.316172 | 0.966 | 0.866 | 1 | 8 | Rps17 | 0.1 | Young |
| 4.83E-05 | 0.728964 | 0.319 | 0.145 | 1 | 8 | Cx3cr1 | 0.174 | Young |
| 5.20E-05 | 0.369102 | 0.87 | 0.726 | 1 | 8 | Akap13 | 0.144 | Young |
| 5.23E-05 | 0.800728 | 0.543 | 0.374 | 1 | 8 | Rnf150 | 0.169 | Young |
| 5.42E-05 | 0.645693 | 0.545 | 0.346 | 1 | 8 | Trmt112 | 0.199 | Young |
| 5.55E-05 | 1.040236 | 0.348 | 0.184 | 1 | 8 | Cxcl12 | 0.164 | Young |
| 5.70E-05 | 0.846284 | 0.366 | 0.196 | 1 | 8 | Mid1ip1 | 0.17 | Young |
| 5.76E-05 | 0.191278 | 1 | 1 | 1 | 8 | Fth1 | 0 | Young |
| 5.95E-05 | 0.308554 | 0.969 | 0.899 | 1 | 8 | Rpl14 | 0.07 | Young |
| 6.16E-05 | 0.406807 | 0.865 | 0.76 | 1 | 8 | Gng5 | 0.105 | Young |
| 6.29E-05 | 0.447985 | 0.779 | 0.626 | 1 | 8 | Tgfb1 | 0.153 | Young |
| 7.27E-05 | 0.432005 | 0.787 | 0.615 | 1 | 8 | Man2b1 | 0.172 | Young |
| 7.31E-05 | 0.61253 | 0.543 | 0.38 | 1 | 8 | Fkbp2 | 0.163 | Young |
| 7.68E-05 | 0.956324 | 0.291 | 0.134 | 1 | 8 | Trem2 | 0.157 | Young |
| 8.30E-05 | 0.528927 | 0.683 | 0.547 | 1 | 8 | Aes | 0.136 | Young |
| 8.70E-05 | 1.700634 | 0.145 | 0.034 | 1 | 8 | Pkp4 | 0.111 | Young |
| 8.89E-05 | 1.417168 | 0.203 | 0.073 | 1 | 8 | Mxi1 | 0.13 | Young |
| 9.32E-05 | 0.305008 | 0.938 | 0.849 | 1 | 8 | Csf1r | 0.089 | Young |

| | | | | | | | | |
|---|---|---|---|---|---|---|---|---|
| 9.49E-05 | 0.974236 | 0.444 | 0.285 | 1 | 8 | Slamf7 | 0.159 | Young |
| 9.68E-05 | 2.202481 | 0.132 | 0.028 | 1 | 8 | AU020206 | 0.104 | Young |
| 0.000104 | 0.359421 | 0.925 | 0.793 | 1 | 8 | Cd52 | 0.132 | Young |
| 0.000108 | 0.745067 | 0.291 | 0.134 | 1 | 8 | Hepacam2 | 0.157 | Young |
| 0.000119 | 0.304736 | 0.974 | 0.866 | 1 | 8 | Coro1a | 0.108 | Young |
| 0.00012 | 0.289509 | 0.969 | 0.899 | 1 | 8 | Arpc1b | 0.07 | Young |
| 0.000122 | 0.55188 | 0.566 | 0.402 | 1 | 8 | Arhgap45 | 0.164 | Young |
| 0.000151 | 0.725334 | 0.338 | 0.179 | 1 | 8 | Itgb7 | 0.159 | Young |
| 0.000156 | 0.584704 | 0.61 | 0.475 | 1 | 8 | Ppib | 0.135 | Young |
| 0.000165 | 0.673549 | 0.758 | 0.62 | 1 | 8 | Ccl24 | 0.138 | Young |
| 0.000166 | 0.620944 | 0.436 | 0.274 | 1 | 8 | Arsb | 0.162 | Young |
| 0.000177 | 1.42944 | 0.179 | 0.061 | 1 | 8 | C1qtnf1 | 0.118 | Young |
| 0.000177 | 0.744124 | 0.41 | 0.257 | 1 | 8 | Mctp1 | 0.153 | Young |
| 0.000178 | 1.967334 | 0.119 | 0.022 | 1 | 8 | Ttc28 | 0.097 | Young |
| 0.000184 | 0.942091 | 0.247 | 0.112 | 1 | 8 | Ntpcr | 0.135 | Young |
| 0.000186 | 0.840532 | 0.348 | 0.19 | 1 | 8 | Ranbp9 | 0.158 | Young |
| 0.000189 | 0.244628 | 0.997 | 0.955 | 1 | 8 | S100a6 | 0.042 | Young |
| 0.00019 | 0.414749 | 0.777 | 0.581 | 1 | 8 | Sfr1 | 0.196 | Young |
| 0.00019 | 0.764845 | 0.366 | 0.207 | 1 | 8 | Snrnp200 | 0.159 | Young |
| 0.000192 | 0.749814 | 0.369 | 0.207 | 1 | 8 | Trappc5 | 0.162 | Young |
| 0.000193 | 0.913544 | 0.33 | 0.184 | 1 | 8 | Casp3 | 0.146 | Young |
| 0.0002 | 0.568523 | 0.748 | 0.631 | 1 | 8 | Dennd4a | 0.117 | Young |
| 0.000204 | 0.701281 | 0.361 | 0.201 | 1 | 8 | Dctn2 | 0.16 | Young |
| 0.000205 | 0.341402 | 0.917 | 0.838 | 1 | 8 | Gm2a | 0.079 | Young |
| 0.000213 | 0.461928 | 0.491 | 0.296 | 1 | 8 | Zfp710 | 0.195 | Young |
| 0.000233 | 0.458281 | 0.649 | 0.492 | 1 | 8 | Gng2 | 0.157 | Young |
| 0.000237 | 0.502205 | 0.566 | 0.391 | 1 | 8 | Atf6 | 0.175 | Young |
| 0.000249 | 1.308412 | 0.221 | 0.095 | 1 | 8 | Vgll4 | 0.126 | Young |
| 0.000252 | 0.19843 | 1 | 0.994 | 1 | 8 | Ftl1 | 0.006 | Young |
| 0.00026 | 0.655498 | 0.442 | 0.279 | 1 | 8 | Ppp1r10 | 0.163 | Young |
| 0.000286 | 0.326432 | 0.953 | 0.905 | 1 | 8 | Rsrp1 | 0.048 | Young |
| 0.0003 | 0.254943 | 1 | 1 | 1 | 8 | Malat1 | 0 | Young |
| 0.000318 | 0.373648 | 0.826 | 0.721 | 1 | 8 | Pfdn5 | 0.105 | Young |
| 0.00032 | 0.671569 | 0.548 | 0.397 | 1 | 8 | Bptf | 0.151 | Young |
| 0.000332 | 0.629885 | 0.577 | 0.425 | 1 | 8 | Skil | 0.152 | Young |
| 0.000334 | 0.604352 | 0.512 | 0.363 | 1 | 8 | Arhgap31 | 0.149 | Young |
| 0.000341 | 0.454619 | 0.719 | 0.581 | 1 | 8 | Ndel1 | 0.138 | Young |
| 0.000354 | 0.849519 | 0.405 | 0.257 | 1 | 8 | Aimp1 | 0.148 | Young |
| 0.000358 | 0.669158 | 0.509 | 0.346 | 1 | 8 | Cd33 | 0.163 | Young |
| 0.000372 | 0.64477 | 0.462 | 0.296 | 1 | 8 | Bod1l | 0.166 | Young |
| 0.000373 | 0.426229 | 0.706 | 0.542 | 1 | 8 | Ralbp1 | 0.164 | Young |
| 0.000375 | 0.450723 | 0.623 | 0.475 | 1 | 8 | Tomm22 | 0.148 | Young |
| 0.000379 | 0.390119 | 0.771 | 0.598 | 1 | 8 | H2afj | 0.173 | Young |
| 0.000399 | 0.394665 | 0.839 | 0.693 | 1 | 8 | Ier3 | 0.146 | Young |
| 0.000426 | 0.493199 | 0.449 | 0.285 | 1 | 8 | Lacc1 | 0.164 | Young |

| | | | | | | | | |
|---|---|---|---|---|---|---|---|---|
| 0.000438 | 0.662501 | 0.47 | 0.318 | 1 | 8 | Rwdd1 | 0.152 | Young |
| 0.000443 | 0.909472 | 0.366 | 0.229 | 1 | 8 | Hnrnpr | 0.137 | Young |
| 0.000444 | 0.458578 | 0.626 | 0.453 | 1 | 8 | Bnip3l | 0.173 | Young |
| 0.000456 | 1.488382 | 0.13 | 0.034 | 1 | 8 | Bckdhb | 0.096 | Young |
| 0.000462 | 0.405649 | 0.779 | 0.603 | 1 | 8 | Stab1 | 0.176 | Young |
| 0.000486 | 0.495814 | 0.504 | 0.324 | 1 | 8 | Fmnl1 | 0.18 | Young |
| 0.000494 | 0.306411 | 0.475 | 0.307 | 1 | 8 | Nrip1 | 0.168 | Young |
| 0.000501 | 2.235974 | 0.099 | 0.017 | 1 | 8 | Cd209c | 0.082 | Young |
| 0.00051 | 0.594083 | 0.429 | 0.274 | 1 | 8 | Tnfrsf13b | 0.155 | Young |
| 0.000525 | 0.766109 | 0.364 | 0.223 | 1 | 8 | Nfxl1 | 0.141 | Young |
| 0.000553 | 0.491453 | 0.655 | 0.542 | 1 | 8 | Clec4b1 | 0.113 | Young |
| 0.000558 | 0.466553 | 0.678 | 0.52 | 1 | 8 | Nr3c1 | 0.158 | Young |
| 0.000578 | 0.323009 | 0.522 | 0.38 | 1 | 8 | Fgd2 | 0.142 | Young |
| 0.000579 | 2.887955 | 0.086 | 0.011 | 1 | 8 | Ms4a4c | 0.075 | Young |
| 0.000583 | 0.797379 | 0.4 | 0.257 | 1 | 8 | Ccnd3 | 0.143 | Young |
| 0.00059 | 0.32875 | 0.86 | 0.721 | 1 | 8 | Gmfg | 0.139 | Young |
| 0.000591 | 0.48056 | 0.455 | 0.296 | 1 | 8 | Usp8 | 0.159 | Young |
| 0.000613 | 0.449145 | 0.722 | 0.592 | 1 | 8 | Cd163 | 0.13 | Young |
| 0.000618 | 0.855932 | 0.317 | 0.184 | 1 | 8 | Tbc1d9 | 0.133 | Young |
| 0.000624 | 0.736676 | 0.332 | 0.19 | 1 | 8 | Btbd7 | 0.142 | Young |
| 0.000629 | 0.278873 | 0.94 | 0.804 | 1 | 8 | Cox7c | 0.136 | Young |
| 0.000647 | 1.355631 | 0.221 | 0.106 | 1 | 8 | P2ry12 | 0.115 | Young |
| 0.000649 | 0.302504 | 0.922 | 0.788 | 1 | 8 | Rbm39 | 0.134 | Young |
| 0.000663 | 2.145204 | 0.114 | 0.028 | 1 | 8 | Lilra5 | 0.086 | Young |
| 0.000672 | 1.318786 | 0.203 | 0.089 | 1 | 8 | Zfp131 | 0.114 | Young |
| 0.000681 | 0.561557 | 0.556 | 0.43 | 1 | 8 | Tnfsf9 | 0.126 | Young |
| 0.000686 | 0.337558 | 0.873 | 0.765 | 1 | 8 | Cox6c | 0.108 | Young |
| 0.000695 | 0.705136 | 0.377 | 0.235 | 1 | 8 | Tanc2 | 0.142 | Young |
| 0.000766 | 0.955722 | 0.247 | 0.123 | 1 | 8 | Plxna4 | 0.124 | Young |
| 0.000768 | 0.367006 | 0.769 | 0.631 | 1 | 8 | Chd4 | 0.138 | Young |
| 0.000784 | 0.499086 | 0.499 | 0.335 | 1 | 8 | Slco2b1 | 0.164 | Young |
| 0.000793 | 2.104786 | 0.112 | 0.028 | 1 | 8 | Cfd | 0.084 | Young |
| 0.000794 | 0.401283 | 0.852 | 0.749 | 1 | 8 | Rassf4 | 0.103 | Young |
| 0.000803 | 0.426345 | 0.592 | 0.458 | 1 | 8 | Mapk14 | 0.134 | Young |
| 0.000804 | 0.621345 | 0.361 | 0.218 | 1 | 8 | Pik3c2a | 0.143 | Young |
| 0.000823 | 0.413895 | 0.704 | 0.531 | 1 | 8 | Ddx3y | 0.173 | Young |
| 0.000852 | 0.191561 | 0.966 | 0.899 | 1 | 8 | Gnai2 | 0.067 | Young |
| 0.00087 | 0.385417 | 0.696 | 0.547 | 1 | 8 | Ndufb10 | 0.149 | Young |
| 0.000872 | 0.402568 | 0.499 | 0.346 | 1 | 8 | Mpc2 | 0.153 | Young |
| 0.000874 | 0.46261 | 0.566 | 0.402 | 1 | 8 | Hipk1 | 0.164 | Young |
| 0.000904 | 0.808915 | 0.382 | 0.251 | 1 | 8 | BC028528 | 0.131 | Young |
| 0.000916 | 0.696598 | 0.335 | 0.201 | 1 | 8 | Arl5c | 0.134 | Young |
| 0.000943 | 0.999653 | 0.21 | 0.095 | 1 | 8 | Serpinf1 | 0.115 | Young |
| 0.000955 | 0.426602 | 0.626 | 0.48 | 1 | 8 | Arhgap15 | 0.146 | Young |
| 0.000969 | 0.331221 | 0.875 | 0.76 | 1 | 8 | Eif3f | 0.115 | Young |

| | | | | | | | | |
|---|---|---|---|---|---|---|---|---|
| 0.001003 | 0.513226 | 0.59 | 0.458 | 1 | 8 | Ltc4s | 0.132 | Young |
| 0.001003 | 1.232669 | 0.19 | 0.084 | 1 | 8 | Exosc8 | 0.106 | Young |
| 0.001026 | 0.319258 | 0.847 | 0.698 | 1 | 8 | Snx5 | 0.149 | Young |
| 0.001044 | 0.309001 | 0.753 | 0.631 | 1 | 8 | Zbtb20 | 0.122 | Young |
| 0.001064 | 0.341142 | 0.808 | 0.687 | 1 | 8 | Smdt1 | 0.121 | Young |
| 0.001065 | 0.30003 | 0.917 | 0.799 | 1 | 8 | Fxyd5 | 0.118 | Young |
| 0.001083 | 0.683222 | 0.317 | 0.179 | 1 | 8 | Runx3 | 0.138 | Young |
| 0.001111 | 1.874584 | 0.117 | 0.034 | 1 | 8 | Ift46 | 0.083 | Young |
| 0.001133 | 0.752849 | 0.387 | 0.251 | 1 | 8 | Dapp1 | 0.136 | Young |
| 0.001173 | 0.420683 | 0.59 | 0.447 | 1 | 8 | Coq10b | 0.143 | Young |
| 0.001195 | 1.323692 | 0.081 | 0.011 | 1 | 8 | B4galt4 | 0.07 | Young |
| 0.00122 | 0.350559 | 0.792 | 0.676 | 1 | 8 | Kdm7a | 0.116 | Young |
| 0.001264 | 0.315531 | 0.938 | 0.883 | 1 | 8 | Npm1 | 0.055 | Young |
| 0.001267 | 1.12522 | 0.177 | 0.073 | 1 | 8 | Bcl11a | 0.104 | Young |
| 0.001278 | 0.520607 | 0.403 | 0.257 | 1 | 8 | Os9 | 0.146 | Young |
| 0.0013 | 2.847467 | 0.078 | 0.011 | 1 | 8 | Fmnl2 | 0.067 | Young |
| 0.001303 | 2.135029 | 0.106 | 0.028 | 1 | 8 | Myh10 | 0.078 | Young |
| 0.001323 | 0.523014 | 0.304 | 0.168 | 1 | 8 | Ogfrl1 | 0.136 | Young |
| 0.00133 | 0.237566 | 0.878 | 0.782 | 1 | 8 | Fcgr2b | 0.096 | Young |
| 0.001348 | 1.02928 | 0.239 | 0.123 | 1 | 8 | Mta1 | 0.116 | Young |
| 0.001367 | 1.811287 | 0.088 | 0.017 | 1 | 8 | Lrrk2 | 0.071 | Young |
| 0.001416 | 0.904226 | 0.223 | 0.112 | 1 | 8 | Kmo | 0.111 | Young |
| 0.001419 | 1.938448 | 0.138 | 0.05 | 1 | 8 | Flt3 | 0.088 | Young |
| 0.001443 | 1.190367 | 0.151 | 0.056 | 1 | 8 | Ccdc97 | 0.095 | Young |
| 0.001454 | 1.276947 | 0.231 | 0.123 | 1 | 8 | Rasgrp3 | 0.108 | Young |
| 0.001458 | 0.231345 | 0.974 | 0.899 | 1 | 8 | H2-DMa | 0.075 | Young |
| 0.00146 | 0.385318 | 0.597 | 0.436 | 1 | 8 | Nup98 | 0.161 | Young |
| 0.001479 | 0.299303 | 0.595 | 0.43 | 1 | 8 | Ash1l | 0.165 | Young |
| 0.001517 | 0.568019 | 0.447 | 0.33 | 1 | 8 | P2ry6 | 0.117 | Young |
| 0.001529 | 0.341169 | 0.73 | 0.564 | 1 | 8 | Ndufa13 | 0.166 | Young |
| 0.001627 | 0.939475 | 0.262 | 0.145 | 1 | 8 | Nbr1 | 0.117 | Young |
| 0.001663 | 0.268189 | 0.966 | 0.899 | 1 | 8 | Uqcrh | 0.067 | Young |
| 0.00168 | 0.222316 | 0.969 | 0.933 | 1 | 8 | Cox4i1 | 0.036 | Young |
| 0.0017 | 0.704465 | 0.262 | 0.145 | 1 | 8 | Chchd7 | 0.117 | Young |
| 0.00171 | 0.32772 | 0.86 | 0.788 | 1 | 8 | Mpeg1 | 0.072 | Young |
| 0.001725 | 0.158588 | 0.995 | 0.989 | 1 | 8 | Rpl7 | 0.006 | Young |
| 0.001733 | 0.756494 | 0.426 | 0.307 | 1 | 8 | Park7 | 0.119 | Young |
| 0.00174 | 1.167386 | 0.203 | 0.101 | 1 | 8 | P2ry10 | 0.102 | Young |
| 0.001771 | 0.816513 | 0.27 | 0.156 | 1 | 8 | St3gal6 | 0.114 | Young |
| 0.001773 | 0.507306 | 0.569 | 0.441 | 1 | 8 | Il1rl1 | 0.128 | Young |
| 0.001818 | 0.924841 | 0.14 | 0.05 | 1 | 8 | Siae | 0.09 | Young |
| 0.001853 | 0.370837 | 0.883 | 0.782 | 1 | 8 | Tmsb10 | 0.101 | Young |
| 0.00186 | 1.099301 | 0.236 | 0.128 | 1 | 8 | Susd3 | 0.108 | Young |
| 0.001861 | 0.509903 | 0.478 | 0.352 | 1 | 8 | Ltb4r1 | 0.126 | Young |
| 0.001874 | 0.381422 | 0.439 | 0.279 | 1 | 8 | Dhx40 | 0.16 | Young |

| | | | | | | | | |
|---|---|---|---|---|---|---|---|---|
| 0.001886 | 0.302486 | 0.787 | 0.648 | 1 | 8 | Arhgap30 | 0.139 | Young |
| 0.001895 | 0.237833 | 0.948 | 0.844 | 1 | 8 | Ctsh | 0.104 | Young |
| 0.001897 | 0.417014 | 0.574 | 0.453 | 1 | 8 | Psme2 | 0.121 | Young |
| 0.001946 | 0.529588 | 0.332 | 0.196 | 1 | 8 | Igf1r | 0.136 | Young |
| 0.001968 | 2.035403 | 0.075 | 0.011 | 1 | 8 | Prkd2 | 0.064 | Young |
| 0.001992 | 1.006565 | 0.255 | 0.145 | 1 | 8 | Mvb12a | 0.11 | Young |
| 0.002022 | 1.464793 | 0.114 | 0.034 | 1 | 8 | Dock4 | 0.08 | Young |
| 0.002031 | 0.654674 | 0.41 | 0.285 | 1 | 8 | Skint3 | 0.125 | Young |
| 0.002054 | 0.628231 | 0.273 | 0.151 | 1 | 8 | Bbx | 0.122 | Young |
| 0.002074 | 0.499432 | 0.47 | 0.335 | 1 | 8 | Taf10 | 0.135 | Young |
| 0.002098 | 0.745539 | 0.314 | 0.196 | 1 | 8 | Mob3a | 0.118 | Young |
| 0.002109 | 0.655325 | 0.268 | 0.151 | 1 | 8 | Vps37b | 0.117 | Young |
| 0.002154 | 0.505187 | 0.494 | 0.358 | 1 | 8 | Rbm5 | 0.136 | Young |
| 0.00219 | 0.540238 | 0.455 | 0.318 | 1 | 8 | Rassf5 | 0.137 | Young |
| 0.002246 | 0.29301 | 0.748 | 0.615 | 1 | 8 | Selenow | 0.133 | Young |
| 0.002266 | 1.299099 | 0.205 | 0.106 | 1 | 8 | Tfeb | 0.099 | Young |
| 0.002298 | 1.167227 | 0.216 | 0.112 | 1 | 8 | Spire1 | 0.104 | Young |
| 0.002306 | 0.951881 | 0.182 | 0.084 | 1 | 8 | Tmem203 | 0.098 | Young |
| 0.002306 | 1.401063 | 0.119 | 0.039 | 1 | 8 | Eepd1 | 0.08 | Young |
| 0.002313 | 0.424573 | 0.569 | 0.43 | 1 | 8 | Ssr4 | 0.139 | Young |
| 0.002337 | 0.239085 | 0.961 | 0.888 | 1 | 8 | Tgfbi | 0.073 | Young |
| 0.002398 | 0.9452 | 0.226 | 0.117 | 1 | 8 | Ero1lb | 0.109 | Young |
| 0.002402 | 0.415906 | 0.818 | 0.726 | 1 | 8 | Etv3 | 0.092 | Young |
| 0.002405 | 1.010607 | 0.221 | 0.117 | 1 | 8 | Slc14a1 | 0.104 | Young |
| 0.002449 | 1.50964 | 0.135 | 0.05 | 1 | 8 | Zdhhc23 | 0.085 | Young |
| 0.002461 | 0.38423 | 0.509 | 0.374 | 1 | 8 | Washc4 | 0.135 | Young |
| 0.002473 | 0.313551 | 0.691 | 0.587 | 1 | 8 | Atp5j | 0.104 | Young |
| 0.002478 | 0.678997 | 0.421 | 0.291 | 1 | 8 | Rab1b | 0.13 | Young |
| 0.002533 | 3.26671 | 0.049 | 0 | 1 | 8 | F11r | 0.049 | Young |
| 0.002536 | 0.268312 | 0.8 | 0.648 | 1 | 8 | Eif3e | 0.152 | Young |
| 0.002546 | 0.438918 | 0.683 | 0.581 | 1 | 8 | Coro1c | 0.102 | Young |
| 0.002556 | 0.25958 | 0.87 | 0.793 | 1 | 8 | Igfbp4 | 0.077 | Young |
| 0.00258 | 0.252013 | 0.673 | 0.514 | 1 | 8 | Rnase4 | 0.159 | Young |
| 0.002587 | 2.186112 | 0.091 | 0.022 | 1 | 8 | Zfp142 | 0.069 | Young |
| 0.0026 | 1.218035 | 0.119 | 0.039 | 1 | 8 | Gm4951 | 0.08 | Young |
| 0.002624 | 0.264158 | 0.878 | 0.76 | 1 | 8 | Pnrc1 | 0.118 | Young |
| 0.002624 | 0.653143 | 0.395 | 0.285 | 1 | 8 | 4930523C( | 0.11 | Young |
| 0.002652 | 0.250334 | 0.8 | 0.693 | 1 | 8 | Atp6v0e | 0.107 | Young |
| 0.002661 | 1.051352 | 0.166 | 0.073 | 1 | 8 | Appl2 | 0.093 | Young |
| 0.002674 | 0.240277 | 0.956 | 0.888 | 1 | 8 | H2afz | 0.068 | Young |
| 0.00273 | 1.367197 | 0.135 | 0.05 | 1 | 8 | Mgat5 | 0.085 | Young |
| 0.002732 | 0.323398 | 0.603 | 0.469 | 1 | 8 | Zfp91 | 0.134 | Young |
| 0.002766 | 0.552096 | 0.465 | 0.346 | 1 | 8 | Gm10073 | 0.119 | Young |
| 0.002793 | 0.348233 | 0.686 | 0.553 | 1 | 8 | Ikbkb | 0.133 | Young |
| 0.002794 | 0.421679 | 0.771 | 0.637 | 1 | 8 | Gapdh | 0.134 | Young |

| | | | | | | | | |
|---|---|---|---|---|---|---|---|---|
| 0.00281 | 0.303932 | 0.756 | 0.581 | 1 | 8 | St8sia4 | 0.175 | Young |
| 0.00282 | 0.837152 | 0.301 | 0.184 | 1 | 8 | Rcsd1 | 0.117 | Young |
| 0.002927 | 1.321881 | 0.119 | 0.039 | 1 | 8 | Mzt2 | 0.08 | Young |
| 0.002978 | 0.709984 | 0.431 | 0.307 | 1 | 8 | Zfp644 | 0.124 | Young |
| 0.002993 | 0.163907 | 0.439 | 0.291 | 1 | 8 | Bcl2l11 | 0.148 | Young |
| 0.002997 | 0.462839 | 0.501 | 0.374 | 1 | 8 | Tmcc3 | 0.127 | Young |
| 0.003015 | 0.303908 | 0.722 | 0.57 | 1 | 8 | Tsc22d4 | 0.152 | Young |
| 0.003049 | 0.704026 | 0.345 | 0.223 | 1 | 8 | Gpr34 | 0.122 | Young |
| 0.003065 | 0.892973 | 0.192 | 0.095 | 1 | 8 | Cpne2 | 0.097 | Young |
| 0.003087 | 0.303739 | 0.691 | 0.57 | 1 | 8 | Clec12a | 0.121 | Young |
| 0.003135 | 2.250391 | 0.06 | 0.006 | 1 | 8 | Gm26510 | 0.054 | Young |
| 0.003152 | 1.005933 | 0.218 | 0.117 | 1 | 8 | Creb5 | 0.101 | Young |
| 0.00318 | 0.542287 | 0.556 | 0.419 | 1 | 8 | Eif3i | 0.137 | Young |
| 0.003187 | 0.765401 | 0.179 | 0.084 | 1 | 8 | Uchl3 | 0.095 | Young |
| 0.003236 | 0.363908 | 0.566 | 0.413 | 1 | 8 | Hexb | 0.153 | Young |
| 0.003288 | 1.03902 | 0.192 | 0.095 | 1 | 8 | Phka2 | 0.097 | Young |
| 0.003305 | 0.40331 | 0.577 | 0.441 | 1 | 8 | Swi5 | 0.136 | Young |
| 0.003365 | 0.250824 | 0.829 | 0.726 | 1 | 8 | Ogt | 0.103 | Young |
| 0.003413 | 0.530722 | 0.468 | 0.341 | 1 | 8 | Ramp1 | 0.127 | Young |
| 0.003415 | 1.262273 | 0.205 | 0.106 | 1 | 8 | Siglech | 0.099 | Young |
| 0.003457 | 0.214746 | 0.958 | 0.877 | 1 | 8 | Ybx1 | 0.081 | Young |
| 0.003526 | 0.355197 | 0.779 | 0.642 | 1 | 8 | Tra2a | 0.137 | Young |
| 0.00357 | 0.576735 | 0.283 | 0.173 | 1 | 8 | Ankle2 | 0.11 | Young |
| 0.003571 | 0.742539 | 0.301 | 0.184 | 1 | 8 | Trp53 | 0.117 | Young |
| 0.003602 | 2.2067 | 0.078 | 0.017 | 1 | 8 | Ptpn4 | 0.061 | Young |
| 0.003609 | 0.3598 | 0.59 | 0.441 | 1 | 8 | Ccnd1 | 0.149 | Young |
| 0.003664 | 0.204733 | 0.99 | 0.927 | 1 | 8 | Rpl8 | 0.063 | Young |
| 0.003687 | 0.696413 | 0.353 | 0.235 | 1 | 8 | Cd9 | 0.118 | Young |
| 0.003696 | 0.243154 | 0.927 | 0.838 | 1 | 8 | Atp5e | 0.089 | Young |
| 0.003723 | 0.969591 | 0.145 | 0.061 | 1 | 8 | Cnrip1 | 0.084 | Young |
| 0.003726 | 0.750375 | 0.182 | 0.084 | 1 | 8 | Mefv | 0.098 | Young |
| 0.003742 | 0.353357 | 0.582 | 0.453 | 1 | 8 | Foxn3 | 0.129 | Young |
| 0.00375 | 0.296687 | 0.847 | 0.698 | 1 | 8 | Atox1 | 0.149 | Young |
| 0.003769 | 0.946603 | 0.174 | 0.084 | 1 | 8 | Siglecg | 0.09 | Young |
| 0.003856 | 0.265414 | 0.725 | 0.615 | 1 | 8 | Alcam | 0.11 | Young |
| 0.003862 | 0.404464 | 0.579 | 0.464 | 1 | 8 | Lamtor2 | 0.115 | Young |
| 0.003889 | 0.469951 | 0.457 | 0.324 | 1 | 8 | Kmt2a | 0.133 | Young |
| 0.003891 | 0.273343 | 0.605 | 0.453 | 1 | 8 | Ppig | 0.152 | Young |
| 0.003916 | 0.266881 | 0.361 | 0.229 | 1 | 8 | Dnajb1 | 0.132 | Young |
| 0.003932 | 0.441228 | 0.636 | 0.497 | 1 | 8 | Usmg5 | 0.139 | Young |
| 0.003934 | 0.71014 | 0.377 | 0.268 | 1 | 8 | Nt5c | 0.109 | Young |
| 0.003976 | 0.328481 | 0.694 | 0.553 | 1 | 8 | D8Ertd738 | 0.141 | Young |
| 0.003999 | 1.293706 | 0.166 | 0.078 | 1 | 8 | Mea1 | 0.088 | Young |
| 0.004002 | 0.554156 | 0.294 | 0.179 | 1 | 8 | Cntrl | 0.115 | Young |
| 0.004018 | 0.426617 | 0.395 | 0.268 | 1 | 8 | Pcf11 | 0.127 | Young |

| | | | | | | | | |
|---|---|---|---|---|---|---|---|---|
| 0.004025 | 0.345328 | 0.509 | 0.358 | 1 | 8 | Ccdc12 | 0.151 | Young |
| 0.004083 | 0.538911 | 0.506 | 0.38 | 1 | 8 | Dpm3 | 0.126 | Young |
| 0.004107 | 1.205612 | 0.138 | 0.056 | 1 | 8 | Nup133 | 0.082 | Young |
| 0.00415 | 0.440743 | 0.171 | 0.078 | 1 | 8 | Ubxn7 | 0.093 | Young |
| 0.004153 | 0.460479 | 0.426 | 0.302 | 1 | 8 | Slbp | 0.124 | Young |
| 0.004162 | 0.3266 | 0.727 | 0.631 | 1 | 8 | Tbca | 0.096 | Young |
| 0.004169 | 0.485497 | 0.496 | 0.391 | 1 | 8 | Edf1 | 0.105 | Young |
| 0.004181 | 0.571901 | 0.34 | 0.218 | 1 | 8 | Cenpx | 0.122 | Young |
| 0.004181 | 0.426987 | 0.387 | 0.263 | 1 | 8 | Tox4 | 0.124 | Young |
| 0.004242 | 0.592934 | 0.434 | 0.318 | 1 | 8 | Hcfc1r1 | 0.116 | Young |
| 0.004274 | 0.393307 | 0.647 | 0.525 | 1 | 8 | Elob | 0.122 | Young |
| 0.004294 | 0.330128 | 0.774 | 0.631 | 1 | 8 | Vsir | 0.143 | Young |
| 0.004413 | 0.496566 | 0.384 | 0.257 | 1 | 8 | Cux1 | 0.127 | Young |
| 0.004453 | 0.645924 | 0.192 | 0.095 | 1 | 8 | Mier3 | 0.097 | Young |
| 0.004463 | 1.048586 | 0.262 | 0.162 | 1 | 8 | Mrpl15 | 0.1 | Young |
| 0.004488 | 0.758759 | 0.319 | 0.212 | 1 | 8 | Pstpip1 | 0.107 | Young |
| 0.004541 | 0.795536 | 0.247 | 0.145 | 1 | 8 | Ang | 0.102 | Young |
| 0.004556 | 0.664887 | 0.364 | 0.246 | 1 | 8 | Gadd45g | 0.118 | Young |
| 0.004559 | 0.351108 | 0.566 | 0.419 | 1 | 8 | Mkln1 | 0.147 | Young |
| 0.004572 | 0.497601 | 0.319 | 0.212 | 1 | 8 | Gigyf1 | 0.107 | Young |
| 0.004577 | 0.568384 | 0.226 | 0.117 | 1 | 8 | Prkag1 | 0.109 | Young |
| 0.004589 | 0.836952 | 0.236 | 0.134 | 1 | 8 | Mfap3 | 0.102 | Young |
| 0.00461 | 0.335374 | 0.805 | 0.693 | 1 | 8 | Eif3k | 0.112 | Young |
| 0.004717 | 0.209711 | 0.984 | 0.95 | 1 | 8 | Nfkbiz | 0.034 | Young |
| 0.004787 | 0.682995 | 0.317 | 0.207 | 1 | 8 | Abca3 | 0.11 | Young |
| 0.00483 | 1.485542 | 0.119 | 0.045 | 1 | 8 | Ap3m2 | 0.074 | Young |
| 0.004831 | 0.477193 | 0.405 | 0.274 | 1 | 8 | Bfar | 0.131 | Young |
| 0.004848 | 0.384333 | 0.452 | 0.318 | 1 | 8 | Cyb5a | 0.134 | Young |
| 0.004907 | 0.5807 | 0.322 | 0.207 | 1 | 8 | Mrps18c | 0.115 | Young |
| 0.005003 | 0.276662 | 0.797 | 0.687 | 1 | 8 | Myh9 | 0.11 | Young |
| 0.005026 | 0.600821 | 0.358 | 0.24 | 1 | 8 | Arid4b | 0.118 | Young |
| 0.00504 | 1.423253 | 0.086 | 0.022 | 1 | 8 | P2ry13 | 0.064 | Young |
| 0.005052 | 1.844905 | 0.101 | 0.034 | 1 | 8 | Ercc1 | 0.067 | Young |
| 0.005079 | 0.544507 | 0.47 | 0.341 | 1 | 8 | Krtcap2 | 0.129 | Young |
| 0.005101 | 1.386486 | 0.086 | 0.022 | 1 | 8 | Lmbr1l | 0.064 | Young |
| 0.005116 | 0.39734 | 0.494 | 0.374 | 1 | 8 | Was | 0.12 | Young |
| 0.005181 | 0.803803 | 0.291 | 0.19 | 1 | 8 | Uba52 | 0.101 | Young |
| 0.005228 | 0.693238 | 0.182 | 0.089 | 1 | 8 | Itgal | 0.093 | Young |
| 0.005231 | 1.365185 | 0.112 | 0.039 | 1 | 8 | Bambi | 0.073 | Young |
| 0.005272 | 0.291867 | 0.956 | 0.899 | 1 | 8 | Srgn | 0.057 | Young |
| 0.005275 | 0.349419 | 0.662 | 0.525 | 1 | 8 | Adgre5 | 0.137 | Young |
| 0.005289 | 0.176658 | 0.969 | 0.939 | 1 | 8 | Eef1b2 | 0.03 | Young |
| 0.005311 | 0.719659 | 0.19 | 0.095 | 1 | 8 | Abl1 | 0.095 | Young |
| 0.005358 | 0.917755 | 0.182 | 0.089 | 1 | 8 | Kdsr | 0.093 | Young |
| 0.005429 | 0.466742 | 0.475 | 0.341 | 1 | 8 | Elmo1 | 0.134 | Young |

| | | | | | | | | |
|---|---|---|---|---|---|---|---|---|
| 0.005449 | 2.074416 | 0.065 | 0.011 | 1 | 8 | Tln2 | 0.054 | Young |
| 0.005449 | 2.020081 | 0.065 | 0.011 | 1 | 8 | Trmt10a | 0.054 | Young |
| 0.005459 | 0.598217 | 0.413 | 0.318 | 1 | 8 | Stk24 | 0.095 | Young |
| 0.005476 | 0.713945 | 0.205 | 0.106 | 1 | 8 | Foxj3 | 0.099 | Young |
| 0.005539 | 0.218316 | 0.519 | 0.374 | 1 | 8 | Bcap31 | 0.145 | Young |
| 0.005566 | 0.38603 | 0.81 | 0.754 | 1 | 8 | Rel | 0.056 | Young |
| 0.005587 | 0.401679 | 0.356 | 0.235 | 1 | 8 | Lair1 | 0.121 | Young |
| 0.005617 | 0.318308 | 0.668 | 0.542 | 1 | 8 | Cbfa2t3 | 0.126 | Young |
| 0.005625 | 0.58987 | 0.332 | 0.218 | 1 | 8 | Tbc1d5 | 0.114 | Young |
| 0.005698 | 0.556219 | 0.405 | 0.285 | 1 | 8 | Wdfy4 | 0.12 | Young |
| 0.005704 | 0.521041 | 0.34 | 0.218 | 1 | 8 | Eif3g | 0.122 | Young |
| 0.005716 | 1.226214 | 0.161 | 0.078 | 1 | 8 | Bcl2l2 | 0.083 | Young |
| 0.005757 | 0.286558 | 0.571 | 0.441 | 1 | 8 | Ski | 0.13 | Young |
| 0.005763 | 0.318565 | 0.777 | 0.631 | 1 | 8 | Spag9 | 0.146 | Young |
| 0.005768 | 0.350492 | 0.758 | 0.659 | 1 | 8 | Cytip | 0.099 | Young |
| 0.005803 | 0.866007 | 0.179 | 0.089 | 1 | 8 | Upf3b | 0.09 | Young |
| 0.005872 | 0.936543 | 0.197 | 0.106 | 1 | 8 | Bicral | 0.091 | Young |
| 0.005952 | 0.905656 | 0.242 | 0.145 | 1 | 8 | Ago4 | 0.097 | Young |
| 0.005955 | 0.525436 | 0.431 | 0.307 | 1 | 8 | Ypel3 | 0.124 | Young |
| 0.005973 | 0.217561 | 0.958 | 0.872 | 1 | 8 | Lsp1 | 0.086 | Young |
| 0.006047 | 0.523833 | 0.332 | 0.212 | 1 | 8 | Cttnbp2nl | 0.12 | Young |
| 0.006054 | 0.475301 | 0.361 | 0.24 | 1 | 8 | Prrc2b | 0.121 | Young |
| 0.006067 | 1.46891 | 0.158 | 0.078 | 1 | 8 | Scyl1 | 0.08 | Young |
| 0.006124 | 1.064272 | 0.174 | 0.089 | 1 | 8 | Gm46224 | 0.085 | Young |
| 0.006216 | 0.454109 | 0.444 | 0.324 | 1 | 8 | Dennd1b | 0.12 | Young |
| 0.00625 | 0.34226 | 0.681 | 0.559 | 1 | 8 | Rgs10 | 0.122 | Young |
| 0.0063 | 0.752207 | 0.294 | 0.184 | 1 | 8 | Mrps16 | 0.11 | Young |
| 0.006319 | 1.473812 | 0.109 | 0.039 | 1 | 8 | Dip2c | 0.07 | Young |
| 0.006338 | 1.269587 | 0.153 | 0.073 | 1 | 8 | Camta1 | 0.08 | Young |
| 0.006363 | 1.375532 | 0.109 | 0.039 | 1 | 8 | H2-Oa | 0.07 | Young |
| 0.006374 | 0.198903 | 0.839 | 0.743 | 1 | 8 | Zcchc6 | 0.096 | Young |
| 0.006377 | 0.141242 | 0.997 | 0.994 | 1 | 8 | Rpsa | 0.003 | Young |
| 0.006397 | 0.587393 | 0.21 | 0.112 | 1 | 8 | Lysmd3 | 0.098 | Young |
| 0.006409 | 1.086659 | 0.169 | 0.084 | 1 | 8 | Slc27a3 | 0.085 | Young |
| 0.006554 | 0.226415 | 0.969 | 0.916 | 1 | 8 | Dab2 | 0.053 | Young |
| 0.00669 | 0.337438 | 0.727 | 0.62 | 1 | 8 | Tep1 | 0.107 | Young |
| 0.006697 | 0.399033 | 0.494 | 0.385 | 1 | 8 | Atp5g1 | 0.109 | Young |
| 0.006812 | 0.561848 | 0.405 | 0.291 | 1 | 8 | Adap2 | 0.114 | Young |
| 0.00688 | 0.320349 | 0.647 | 0.52 | 1 | 8 | Dennd5a | 0.127 | Young |
| 0.006884 | 0.449803 | 0.39 | 0.279 | 1 | 8 | Phactr2 | 0.111 | Young |
| 0.00694 | 0.580593 | 0.395 | 0.285 | 1 | 8 | Pik3cg | 0.11 | Young |
| 0.006978 | 0.594673 | 0.249 | 0.145 | 1 | 8 | Tmem64 | 0.104 | Young |
| 0.007101 | 0.345272 | 0.639 | 0.531 | 1 | 8 | Ssh2 | 0.108 | Young |
| 0.007127 | 1.362924 | 0.091 | 0.028 | 1 | 8 | Rint1 | 0.063 | Young |
| 0.007167 | 1.284617 | 0.099 | 0.034 | 1 | 8 | St3gal2 | 0.065 | Young |

| | | | | | | | | |
|---|---|---|---|---|---|---|---|---|
| 0.007352 | 0.382805 | 0.213 | 0.117 | 1 | 8 | Rptor | 0.096 | Young |
| 0.007438 | 0.243345 | 0.655 | 0.497 | 1 | 8 | Snx20 | 0.158 | Young |
| 0.007479 | 0.137794 | 1 | 0.989 | 1 | 8 | Rps27a | 0.011 | Young |
| 0.007518 | 2.323052 | 0.039 | 0 | 1 | 8 | Ighd | 0.039 | Young |
| 0.007536 | 1.856835 | 0.088 | 0.028 | 1 | 8 | Ift22 | 0.06 | Young |
| 0.007692 | 0.740321 | 0.288 | 0.184 | 1 | 8 | Mprip | 0.104 | Young |
| 0.007735 | 0.99851 | 0.091 | 0.028 | 1 | 8 | Nme4 | 0.063 | Young |
| 0.007748 | 0.234056 | 0.966 | 0.95 | 1 | 8 | F13a1 | 0.016 | Young |
| 0.007807 | 0.297294 | 0.683 | 0.547 | 1 | 8 | Cd2ap | 0.136 | Young |
| 0.007808 | 0.20165 | 0.805 | 0.654 | 1 | 8 | Ctdsp2 | 0.151 | Young |
| 0.007817 | 1.367027 | 0.127 | 0.056 | 1 | 8 | Sipa1l2 | 0.071 | Young |
| 0.007907 | 0.628428 | 0.317 | 0.218 | 1 | 8 | Tmem127 | 0.099 | Young |
| 0.007935 | 0.6759 | 0.27 | 0.168 | 1 | 8 | Cnpy3 | 0.102 | Young |
| 0.007989 | 0.370511 | 0.556 | 0.43 | 1 | 8 | Pdlim1 | 0.126 | Young |
| 0.008004 | 0.341628 | 0.696 | 0.615 | 1 | 8 | Runx1 | 0.081 | Young |
| 0.008206 | 0.455313 | 0.319 | 0.212 | 1 | 8 | Snx13 | 0.107 | Young |
| 0.008235 | 0.511366 | 0.249 | 0.14 | 1 | 8 | Tmem8 | 0.109 | Young |
| 0.008275 | 0.333213 | 0.325 | 0.218 | 1 | 8 | Ebi3 | 0.107 | Young |
| 0.008323 | 0.266686 | 0.608 | 0.492 | 1 | 8 | Aldh2 | 0.116 | Young |
| 0.008331 | 0.359987 | 0.766 | 0.62 | 1 | 8 | Nsa2 | 0.146 | Young |
| 0.00836 | 0.808622 | 0.138 | 0.061 | 1 | 8 | Mtus1 | 0.077 | Young |
| 0.008376 | 0.232205 | 0.748 | 0.626 | 1 | 8 | Zfhx3 | 0.122 | Young |
| 0.008418 | 0.206494 | 0.974 | 0.933 | 1 | 8 | Jun | 0.041 | Young |
| 0.008446 | 0.18161 | 0.608 | 0.464 | 1 | 8 | Sf3b5 | 0.144 | Young |
| 0.008505 | 1.552462 | 0.096 | 0.034 | 1 | 8 | Diras2 | 0.062 | Young |
| 0.008538 | 0.420963 | 0.169 | 0.084 | 1 | 8 | Kifap3 | 0.085 | Young |
| 0.008577 | 2.698753 | 0.049 | 0.006 | 1 | 8 | Slc2a3 | 0.043 | Young |
| 0.008608 | 0.38938 | 0.208 | 0.112 | 1 | 8 | Atxn1 | 0.096 | Young |
| 0.008648 | 1.186038 | 0.104 | 0.039 | 1 | 8 | Cxxc5 | 0.065 | Young |
| 0.008655 | 1.390671 | 0.127 | 0.056 | 1 | 8 | AC163018. | 0.071 | Young |
| 0.008667 | 0.59215 | 0.301 | 0.201 | 1 | 8 | Mrpl51 | 0.1 | Young |
| 0.008719 | 0.27051 | 0.571 | 0.425 | 1 | 8 | Srsf11 | 0.146 | Young |
| 0.008719 | 0.828737 | 0.203 | 0.112 | 1 | 8 | Arl10 | 0.091 | Young |
| 0.00875 | 1.967356 | 0.088 | 0.028 | 1 | 8 | Ccl17 | 0.06 | Young |
| 0.008796 | 0.224244 | 0.865 | 0.777 | 1 | 8 | Stk17b | 0.088 | Young |
| 0.008894 | 0.417487 | 0.54 | 0.441 | 1 | 8 | Pou2f2 | 0.099 | Young |
| 0.008933 | 1.537122 | 0.112 | 0.045 | 1 | 8 | Arhgef10 | 0.067 | Young |
| 0.008957 | 0.355236 | 0.618 | 0.531 | 1 | 8 | Scaf11 | 0.087 | Young |
| 0.008961 | 0.27757 | 0.647 | 0.531 | 1 | 8 | Irf1 | 0.116 | Young |
| 0.009043 | 0.756657 | 0.21 | 0.123 | 1 | 8 | Commd1 | 0.087 | Young |
| 0.009055 | 0.710915 | 0.174 | 0.089 | 1 | 8 | Tnfrsf12a | 0.085 | Young |
| 0.00918 | 0.418867 | 0.574 | 0.453 | 1 | 8 | Cdkn1b | 0.121 | Young |
| 0.009203 | 1.271569 | 0.06 | 0.011 | 1 | 8 | Ctnnd2 | 0.049 | Young |
| 0.009256 | 0.193645 | 0.745 | 0.609 | 1 | 8 | Itga4 | 0.136 | Young |
| 0.009275 | 0.250077 | 0.818 | 0.693 | 1 | 8 | Fgfr1 | 0.125 | Young |

| | | | | | | | | |
|---|---|---|---|---|---|---|---|---|
| 0.009296 | 0.316281 | 0.426 | 0.296 | 1 | 8 | Apc | 0.13 | Young |
| 0.009368 | 0.246081 | 0.47 | 0.341 | 1 | 8 | Nucb1 | 0.129 | Young |
| 0.009417 | 0.282397 | 0.906 | 0.754 | 1 | 8 | Trf | 0.152 | Young |
| 0.009444 | 1.214011 | 0.049 | 0.006 | 1 | 8 | Tmem129 | 0.043 | Young |
| 0.009708 | 0.264605 | 0.826 | 0.721 | 1 | 8 | Tma7 | 0.105 | Young |
| 0.009847 | 0.616183 | 0.252 | 0.151 | 1 | 8 | Trim33 | 0.101 | Young |
| 0.00985 | 1.514035 | 0.06 | 0.011 | 1 | 8 | Dbp | 0.049 | Young |
| 0.009871 | 1.309307 | 0.14 | 0.067 | 1 | 8 | Zfyve27 | 0.073 | Young |
| 0.009881 | 2.806799 | 0.036 | 0 | 1 | 8 | Fam109b | 0.036 | Young |
| 0.009908 | 0.679353 | 0.171 | 0.089 | 1 | 8 | Hjurp | 0.082 | Young |
| 0.009933 | 1.364676 | 0.104 | 0.039 | 1 | 8 | Abcc3 | 0.065 | Young |
| 0.010183 | 0.189688 | 0.59 | 0.458 | 1 | 8 | Cited2 | 0.132 | Young |
| 0.010341 | 1.633626 | 0.117 | 0.05 | 1 | 8 | Slco3a1 | 0.067 | Young |
| 0.01037 | 0.25184 | 0.553 | 0.436 | 1 | 8 | Fam168b | 0.117 | Young |
| 0.010401 | 0.264387 | 0.655 | 0.547 | 1 | 8 | Ndufa7 | 0.108 | Young |
| 0.010445 | 0.317126 | 0.61 | 0.475 | 1 | 8 | Kmt2e | 0.135 | Young |
| 0.010467 | 0.214168 | 0.727 | 0.626 | 1 | 8 | Slc43a2 | 0.101 | Young |
| 0.010475 | 0.338794 | 0.917 | 0.76 | 1 | 8 | Zfand5 | 0.157 | Young |
| 0.010509 | 0.837193 | 0.184 | 0.101 | 1 | 8 | Tmem120a | 0.083 | Young |
| 0.010636 | 0.8803 | 0.221 | 0.128 | 1 | 8 | Nasp | 0.093 | Young |
| 0.010675 | 0.307181 | 0.748 | 0.587 | 1 | 8 | Tgoln1 | 0.161 | Young |
| 0.010689 | 0.618311 | 0.351 | 0.251 | 1 | 8 | Dna2 | 0.1 | Young |
| 0.010727 | 0.458513 | 0.47 | 0.38 | 1 | 8 | Ik | 0.09 | Young |
| 0.010765 | 0.320255 | 0.475 | 0.369 | 1 | 8 | Krt80 | 0.106 | Young |
| 0.010842 | 0.576089 | 0.514 | 0.425 | 1 | 8 | Rp9 | 0.089 | Young |
| 0.010962 | 0.214194 | 0.67 | 0.564 | 1 | 8 | Cmtm6 | 0.106 | Young |
| 0.01108 | 0.278412 | 0.849 | 0.76 | 1 | 8 | Wwp1 | 0.089 | Young |
| 0.011088 | 1.409186 | 0.094 | 0.034 | 1 | 8 | Scimp | 0.06 | Young |
| 0.011166 | 1.101498 | 0.078 | 0.022 | 1 | 8 | Cep44 | 0.056 | Young |
| 0.011327 | 0.478599 | 0.353 | 0.24 | 1 | 8 | Vezf1 | 0.113 | Young |
| 0.011377 | 1.45219 | 0.086 | 0.028 | 1 | 8 | Gcnt2 | 0.058 | Young |
| 0.0114 | 0.799786 | 0.223 | 0.128 | 1 | 8 | Grsf1 | 0.095 | Young |
| 0.011429 | 0.707155 | 0.283 | 0.19 | 1 | 8 | Cmip | 0.093 | Young |
| 0.011458 | 0.506158 | 0.205 | 0.117 | 1 | 8 | Itgb5 | 0.088 | Young |
| 0.011484 | 0.355014 | 0.603 | 0.503 | 1 | 8 | Use1 | 0.1 | Young |
| 0.011507 | 0.527799 | 0.527 | 0.425 | 1 | 8 | Ednrb | 0.102 | Young |
| 0.011566 | 0.36296 | 0.483 | 0.358 | 1 | 8 | Uqcc2 | 0.125 | Young |
| 0.011577 | 2.130676 | 0.057 | 0.011 | 1 | 8 | Arap2 | 0.046 | Young |
| 0.011788 | 2.27616 | 0.057 | 0.011 | 1 | 8 | Grasp | 0.046 | Young |
| 0.011809 | 0.207121 | 0.784 | 0.693 | 1 | 8 | Snrpe | 0.091 | Young |
| 0.011825 | 0.267803 | 0.849 | 0.726 | 1 | 8 | Psma7 | 0.123 | Young |
| 0.012038 | 0.698076 | 0.2 | 0.112 | 1 | 8 | Rad50 | 0.088 | Young |
| 0.012086 | 0.770467 | 0.236 | 0.145 | 1 | 8 | Gm47283 | 0.091 | Young |
| 0.012148 | 0.868183 | 0.169 | 0.089 | 1 | 8 | Tmem260 | 0.08 | Young |
| 0.012224 | 0.583825 | 0.338 | 0.235 | 1 | 8 | Ankhd1 | 0.103 | Young |

| | | | | | | | | |
|---|---|---|---|---|---|---|---|---|
| 0.012225 | 0.5463 | 0.377 | 0.274 | 1 | 8 | Ppt1 | 0.103 | Young |
| 0.012294 | 0.196956 | 0.821 | 0.715 | 1 | 8 | Hint1 | 0.106 | Young |
| 0.012295 | 0.938034 | 0.174 | 0.095 | 1 | 8 | Usp32 | 0.079 | Young |
| 0.012315 | 1.568106 | 0.075 | 0.022 | 1 | 8 | Katnal1 | 0.053 | Young |
| 0.01235 | 0.564224 | 0.369 | 0.263 | 1 | 8 | Upb1 | 0.106 | Young |
| 0.012424 | 0.510779 | 0.221 | 0.128 | 1 | 8 | Plcl2 | 0.093 | Young |
| 0.012442 | 0.291481 | 0.301 | 0.184 | 1 | 8 | Fh1 | 0.117 | Young |
| 0.012488 | 0.502452 | 0.299 | 0.196 | 1 | 8 | Phf5a | 0.103 | Young |
| 0.012621 | 0.794472 | 0.247 | 0.156 | 1 | 8 | Qars | 0.091 | Young |
| 0.012749 | 0.909955 | 0.221 | 0.134 | 1 | 8 | Frmd6 | 0.087 | Young |
| 0.012778 | 0.750785 | 0.094 | 0.034 | 1 | 8 | Mthfd2l | 0.06 | Young |
| 0.012779 | 0.223642 | 0.699 | 0.587 | 1 | 8 | Mrpl52 | 0.112 | Young |
| 0.01278 | 0.280122 | 0.797 | 0.693 | 1 | 8 | Fcgr3 | 0.104 | Young |
| 0.012803 | 0.448062 | 0.119 | 0.05 | 1 | 8 | Slx4ip | 0.069 | Young |
| 0.012899 | 0.253069 | 0.2 | 0.112 | 1 | 8 | Tmed4 | 0.088 | Young |
| 0.012902 | 0.722382 | 0.229 | 0.14 | 1 | 8 | Icosl | 0.089 | Young |
| 0.012998 | 2.793259 | 0.034 | 0 | 1 | 8 | Als2cl | 0.034 | Young |
| 0.012998 | 2.513902 | 0.034 | 0 | 1 | 8 | Fndc4 | 0.034 | Young |
| 0.012998 | 2.220959 | 0.034 | 0 | 1 | 8 | Ccdc137 | 0.034 | Young |
| 0.013008 | 1.22462 | 0.156 | 0.084 | 1 | 8 | Pafah1b3 | 0.072 | Young |
| 0.013009 | 0.510449 | 0.452 | 0.358 | 1 | 8 | Bax | 0.094 | Young |
| 0.013068 | 0.847519 | 0.216 | 0.128 | 1 | 8 | Thra | 0.088 | Young |
| 0.013083 | 0.28859 | 0.766 | 0.626 | 1 | 8 | Mef2a | 0.14 | Young |
| 0.013138 | 0.42542 | 0.109 | 0.045 | 1 | 8 | Timd4 | 0.064 | Young |
| 0.013152 | 0.631456 | 0.125 | 0.056 | 1 | 8 | Fkrp | 0.069 | Young |
| 0.013221 | 0.857105 | 0.109 | 0.045 | 1 | 8 | Med22 | 0.064 | Young |
| 0.013367 | 0.34468 | 0.613 | 0.503 | 1 | 8 | Mpp6 | 0.11 | Young |
| 0.013402 | 0.318024 | 0.699 | 0.587 | 1 | 8 | Myo5a | 0.112 | Young |
| 0.013431 | 1.087181 | 0.109 | 0.045 | 1 | 8 | Hscb | 0.064 | Young |
| 0.013441 | 0.267912 | 0.429 | 0.307 | 1 | 8 | Oat | 0.122 | Young |
| 0.013498 | 0.52946 | 0.312 | 0.212 | 1 | 8 | Fbxw4 | 0.1 | Young |
| 0.013519 | 0.786346 | 0.252 | 0.156 | 1 | 8 | Pdgfb | 0.096 | Young |
| 0.013574 | 0.265575 | 0.925 | 0.844 | 1 | 8 | S100a10 | 0.081 | Young |
| 0.013611 | 0.359521 | 0.457 | 0.335 | 1 | 8 | Egr3 | 0.122 | Young |
| 0.013712 | 0.195005 | 0.686 | 0.542 | 1 | 8 | Rps6ka3 | 0.144 | Young |
| 0.01374 | 0.517797 | 0.377 | 0.279 | 1 | 8 | Fam173a | 0.098 | Young |
| 0.013812 | 0.258956 | 0.899 | 0.849 | 1 | 8 | Eef2 | 0.05 | Young |
| 0.013826 | 0.376063 | 0.291 | 0.19 | 1 | 8 | Rftn1 | 0.101 | Young |
| 0.013859 | 0.626942 | 0.27 | 0.173 | 1 | 8 | Zbtb4 | 0.097 | Young |
| 0.014083 | 0.397376 | 0.652 | 0.553 | 1 | 8 | Hacd4 | 0.099 | Young |
| 0.014087 | 0.230773 | 0.543 | 0.425 | 1 | 8 | Sec14l1 | 0.118 | Young |
| 0.014102 | 0.420793 | 0.506 | 0.419 | 1 | 8 | Cd86 | 0.087 | Young |
| 0.014217 | 0.478325 | 0.551 | 0.475 | 1 | 8 | Cebpg | 0.076 | Young |
| 0.014223 | 0.326873 | 0.491 | 0.374 | 1 | 8 | Specc1l | 0.117 | Young |
| 0.014232 | 0.417421 | 0.512 | 0.408 | 1 | 8 | Bin1 | 0.104 | Young |

| | | | | | | | | |
|---|---|---|---|---|---|---|---|---|
| 0.014266 | 0.149281 | 0.421 | 0.302 | 1 | 8 | Irf4 | 0.119 | Young |
| 0.014289 | 0.716577 | 0.229 | 0.14 | 1 | 8 | Mphosph8 | 0.089 | Young |
| 0.014359 | 0.500407 | 0.294 | 0.201 | 1 | 8 | Klhl6 | 0.093 | Young |
| 0.014369 | 0.628887 | 0.278 | 0.184 | 1 | 8 | Parp8 | 0.094 | Young |
| 0.014455 | 0.753981 | 0.101 | 0.039 | 1 | 8 | Eci1 | 0.062 | Young |
| 0.014545 | 0.330788 | 0.278 | 0.184 | 1 | 8 | Kctd2 | 0.094 | Young |
| 0.014551 | 0.398326 | 0.231 | 0.145 | 1 | 8 | Tbc1d15 | 0.086 | Young |
| 0.01462 | 0.540679 | 0.268 | 0.173 | 1 | 8 | Akap9 | 0.095 | Young |
| 0.014687 | 0.539507 | 0.387 | 0.291 | 1 | 8 | Szrd1 | 0.096 | Young |
| 0.014714 | 0.769193 | 0.205 | 0.123 | 1 | 8 | Tubgcp5 | 0.082 | Young |
| 0.014751 | 0.739775 | 0.197 | 0.117 | 1 | 8 | Samd4 | 0.08 | Young |
| 0.014761 | 0.423596 | 0.229 | 0.134 | 1 | 8 | Ppfia1 | 0.095 | Young |
| 0.014784 | 1.210432 | 0.177 | 0.101 | 1 | 8 | Zfp277 | 0.076 | Young |
| 0.014788 | 2.444739 | 0.044 | 0.006 | 1 | 8 | Bpgm | 0.038 | Young |
| 0.014833 | 1.247662 | 0.106 | 0.045 | 1 | 8 | Vps72 | 0.061 | Young |
| 0.014946 | 0.813108 | 0.252 | 0.168 | 1 | 8 | Mrpl4 | 0.084 | Young |
| 0.015001 | 0.410862 | 0.382 | 0.274 | 1 | 8 | Ino80d | 0.108 | Young |
| 0.015161 | 1.779776 | 0.055 | 0.011 | 1 | 8 | Stat4 | 0.044 | Young |
| 0.015192 | 0.20461 | 0.164 | 0.084 | 1 | 8 | Mical1 | 0.08 | Young |
| 0.015205 | 0.295758 | 0.478 | 0.363 | 1 | 8 | Slc48a1 | 0.115 | Young |
| 0.015229 | 1.801604 | 0.055 | 0.011 | 1 | 8 | Serpinb9 | 0.044 | Young |
| 0.015397 | 0.789141 | 0.151 | 0.078 | 1 | 8 | Borcs5 | 0.073 | Young |
| 0.015406 | 0.833572 | 0.213 | 0.128 | 1 | 8 | Naa10 | 0.085 | Young |
| 0.015437 | 0.88777 | 0.13 | 0.061 | 1 | 8 | Icmt | 0.069 | Young |
| 0.015504 | 0.573664 | 0.275 | 0.179 | 1 | 8 | Bcl6 | 0.096 | Young |
| 0.015572 | 0.510304 | 0.322 | 0.223 | 1 | 8 | Cd300c2 | 0.099 | Young |
| 0.015584 | 0.093636 | 1 | 1 | 1 | 8 | Rplp2 | 0 | Young |
| 0.015604 | 1.182696 | 0.127 | 0.061 | 1 | 8 | Ppil3 | 0.066 | Young |
| 0.015692 | 0.218412 | 0.6 | 0.486 | 1 | 8 | Epn1 | 0.114 | Young |
| 0.015707 | 2.055953 | 0.044 | 0.006 | 1 | 8 | Mmp13 | 0.038 | Young |
| 0.01575 | 0.409461 | 0.597 | 0.508 | 1 | 8 | Nol7 | 0.089 | Young |
| 0.015776 | 0.356854 | 0.379 | 0.279 | 1 | 8 | Vps29 | 0.1 | Young |
| 0.015818 | 0.489949 | 0.2 | 0.117 | 1 | 8 | Rasgrp2 | 0.083 | Young |
| 0.015834 | 0.233105 | 0.696 | 0.547 | 1 | 8 | Jmjd1c | 0.149 | Young |
| 0.015871 | 0.224756 | 0.481 | 0.346 | 1 | 8 | Pikfyve | 0.135 | Young |
| 0.015957 | 0.625083 | 0.091 | 0.034 | 1 | 8 | Tsen15 | 0.057 | Young |
| 0.016007 | 0.16785 | 0.353 | 0.246 | 1 | 8 | Hbp1 | 0.107 | Young |
| 0.016213 | 0.451487 | 0.356 | 0.263 | 1 | 8 | Bmpr2 | 0.093 | Young |
| 0.016267 | 0.279248 | 0.805 | 0.682 | 1 | 8 | Mrfap1 | 0.123 | Young |
| 0.016333 | 0.690827 | 0.278 | 0.19 | 1 | 8 | Gstm1 | 0.088 | Young |
| 0.016401 | 0.559342 | 0.252 | 0.162 | 1 | 8 | Siglece | 0.09 | Young |
| 0.016451 | 0.257636 | 0.678 | 0.564 | 1 | 8 | Ndufb11 | 0.114 | Young |
| 0.016463 | 0.242786 | 0.545 | 0.419 | 1 | 8 | Raly | 0.126 | Young |
| 0.016578 | 0.35092 | 0.395 | 0.291 | 1 | 8 | Rapgef6 | 0.104 | Young |
| 0.016649 | 0.356892 | 0.216 | 0.128 | 1 | 8 | Ssbp1 | 0.088 | Young |

| | | | | | | | | |
|---|---|---|---|---|---|---|---|---|
| 0.016676 | 1.369754 | 0.081 | 0.028 | 1 | 8 | Micu3 | 0.053 | Young |
| 0.016736 | 0.765885 | 0.223 | 0.14 | 1 | 8 | Tmem243 | 0.083 | Young |
| 0.0168 | 1.127437 | 0.119 | 0.056 | 1 | 8 | Xaf1 | 0.063 | Young |
| 0.016906 | 0.31104 | 0.655 | 0.542 | 1 | 8 | Rac2 | 0.113 | Young |
| 0.016946 | 1.014617 | 0.073 | 0.022 | 1 | 8 | Stambpl1 | 0.051 | Young |
| 0.016997 | 0.243181 | 0.655 | 0.508 | 1 | 8 | Nckap1l | 0.147 | Young |
| 0.017017 | 0.940871 | 0.114 | 0.05 | 1 | 8 | Recql | 0.064 | Young |
| 0.017023 | 0.264006 | 0.751 | 0.67 | 1 | 8 | Hmgb1 | 0.081 | Young |
| 0.017034 | 1.23467 | 0.112 | 0.05 | 1 | 8 | 0610010K1 | 0.062 | Young |
| 0.017118 | 2.463472 | 0.031 | 0 | 1 | 8 | Nudt6 | 0.031 | Young |
| 0.017118 | 2.222516 | 0.031 | 0 | 1 | 8 | Zbtb10 | 0.031 | Young |
| 0.017118 | 2.222441 | 0.031 | 0 | 1 | 8 | Gm29340 | 0.031 | Young |
| 0.017299 | 0.183816 | 0.738 | 0.609 | 1 | 8 | Mycbp2 | 0.129 | Young |
| 0.017426 | 0.432396 | 0.265 | 0.168 | 1 | 8 | Desi2 | 0.097 | Young |
| 0.017449 | 0.371742 | 0.423 | 0.33 | 1 | 8 | Chd2 | 0.093 | Young |
| 0.017462 | 0.353915 | 0.382 | 0.274 | 1 | 8 | Mark2 | 0.108 | Young |
| 0.017567 | 1.61306 | 0.088 | 0.034 | 1 | 8 | Ston2 | 0.054 | Young |
| 0.017696 | 1.192926 | 0.158 | 0.089 | 1 | 8 | Fam3a | 0.069 | Young |
| 0.017767 | 0.187575 | 0.722 | 0.642 | 1 | 8 | Selplg | 0.08 | Young |
| 0.017863 | 1.437932 | 0.096 | 0.039 | 1 | 8 | Sirpb1b | 0.057 | Young |
| 0.017872 | 1.295035 | 0.112 | 0.05 | 1 | 8 | Klrd1 | 0.062 | Young |
| 0.017908 | 0.20082 | 0.738 | 0.682 | 1 | 8 | Eid1 | 0.056 | Young |
| 0.018034 | 0.305174 | 0.171 | 0.095 | 1 | 8 | Abhd4 | 0.076 | Young |
| 0.018099 | 0.24377 | 0.421 | 0.307 | 1 | 8 | Dock11 | 0.114 | Young |
| 0.018147 | 0.833081 | 0.164 | 0.089 | 1 | 8 | Tmub2 | 0.075 | Young |
| 0.018147 | 0.766598 | 0.26 | 0.173 | 1 | 8 | Sra1 | 0.087 | Young |
| 0.018251 | 0.765555 | 0.223 | 0.145 | 1 | 8 | Dcaf11 | 0.078 | Young |
| 0.018326 | 0.363296 | 0.364 | 0.257 | 1 | 8 | Zcchc7 | 0.107 | Young |
| 0.018444 | 0.218179 | 0.488 | 0.358 | 1 | 8 | Crlf2 | 0.13 | Young |
| 0.018448 | 0.250004 | 0.4 | 0.291 | 1 | 8 | Arl1 | 0.109 | Young |
| 0.018503 | 1.244241 | 0.14 | 0.073 | 1 | 8 | Gpkow | 0.067 | Young |
| 0.018545 | 0.503044 | 0.345 | 0.251 | 1 | 8 | Sdf2l1 | 0.094 | Young |
| 0.01908 | 0.157257 | 0.618 | 0.492 | 1 | 8 | Clint1 | 0.126 | Young |
| 0.019159 | 0.336764 | 0.392 | 0.274 | 1 | 8 | Basp1 | 0.118 | Young |
| 0.019231 | 2.722631 | 0.052 | 0.011 | 1 | 8 | Cyr61 | 0.041 | Young |
| 0.019336 | 0.266624 | 0.374 | 0.263 | 1 | 8 | Mrpl57 | 0.111 | Young |
| 0.019424 | 1.072722 | 0.112 | 0.05 | 1 | 8 | Dis3l2 | 0.062 | Young |
| 0.019445 | 0.275793 | 0.512 | 0.391 | 1 | 8 | Dnaja2 | 0.121 | Young |
| 0.019575 | 2.057876 | 0.052 | 0.011 | 1 | 8 | Col14a1 | 0.041 | Young |
| 0.019606 | 0.518688 | 0.288 | 0.201 | 1 | 8 | Smagp | 0.087 | Young |
| 0.019615 | 0.535669 | 0.335 | 0.24 | 1 | 8 | Rybp | 0.095 | Young |
| 0.01962 | 0.165693 | 0.304 | 0.196 | 1 | 8 | Slc25a11 | 0.108 | Young |
| 0.019625 | 0.773121 | 0.088 | 0.034 | 1 | 8 | Slc26a11 | 0.054 | Young |
| 0.019709 | 1.924719 | 0.07 | 0.022 | 1 | 8 | Iqce | 0.048 | Young |
| 0.019725 | 0.395615 | 0.59 | 0.48 | 1 | 8 | Dot1l | 0.11 | Young |

| | | | | | | | | |
|---|---|---|---|---|---|---|---|---|
| 0.019813 | 0.748122 | 0.262 | 0.179 | 1 | 8 | Cenpa | 0.083 | Young |
| 0.019814 | 0.520616 | 0.2 | 0.117 | 1 | 8 | A630001G: | 0.083 | Young |
| 0.019818 | 2.037876 | 0.042 | 0.006 | 1 | 8 | Barx2 | 0.036 | Young |
| 0.019923 | 0.913939 | 0.127 | 0.061 | 1 | 8 | Svbp | 0.066 | Young |
| 0.019964 | 0.364271 | 0.332 | 0.223 | 1 | 8 | Samm50 | 0.109 | Young |
| 0.019975 | 0.531561 | 0.335 | 0.24 | 1 | 8 | Nuak2 | 0.095 | Young |
| 0.019998 | 0.358602 | 0.561 | 0.436 | 1 | 8 | Luc7l3 | 0.125 | Young |
| 0.020091 | 1.632639 | 0.078 | 0.028 | 1 | 8 | Tnni2 | 0.05 | Young |
| 0.020097 | 0.53563 | 0.294 | 0.207 | 1 | 8 | Gtf2a2 | 0.087 | Young |
| 0.020105 | 0.604958 | 0.296 | 0.207 | 1 | 8 | Brd3 | 0.089 | Young |
| 0.020121 | 0.773618 | 0.088 | 0.034 | 1 | 8 | Slc10a3 | 0.054 | Young |
| 0.020232 | 0.777037 | 0.208 | 0.128 | 1 | 8 | Cnep1r1 | 0.08 | Young |
| 0.020269 | 0.571191 | 0.252 | 0.168 | 1 | 8 | Tpst2 | 0.084 | Young |
| 0.020408 | 0.519947 | 0.231 | 0.145 | 1 | 8 | Cdip1 | 0.086 | Young |
| 0.020421 | 2.25664 | 0.042 | 0.006 | 1 | 8 | Gm26522 | 0.036 | Young |
| 0.020463 | 0.305567 | 0.517 | 0.408 | 1 | 8 | Crlf3 | 0.109 | Young |
| 0.020615 | 1.217858 | 0.07 | 0.022 | 1 | 8 | Zbtb45 | 0.048 | Young |
| 0.020655 | 0.0942 | 0.483 | 0.341 | 1 | 8 | Cd93 | 0.142 | Young |
| 0.020661 | 0.240447 | 0.358 | 0.263 | 1 | 8 | Zfx | 0.095 | Young |
| 0.020706 | 0.996564 | 0.138 | 0.073 | 1 | 8 | Med7 | 0.065 | Young |
| 0.020709 | 0.966905 | 0.226 | 0.151 | 1 | 8 | Plk3 | 0.075 | Young |
| 0.020779 | 0.29537 | 0.888 | 0.816 | 1 | 8 | Cd14 | 0.072 | Young |
| 0.020932 | 0.100122 | 0.397 | 0.296 | 1 | 8 | Zfp395 | 0.101 | Young |
| 0.020935 | 1.306391 | 0.042 | 0.006 | 1 | 8 | Olfr920 | 0.036 | Young |
| 0.020994 | 0.453279 | 0.335 | 0.24 | 1 | 8 | Nemf | 0.095 | Young |
| 0.021019 | 0.972529 | 0.156 | 0.084 | 1 | 8 | Pot1a | 0.072 | Young |
| 0.021039 | 0.201545 | 0.912 | 0.81 | 1 | 8 | Tln1 | 0.102 | Young |
| 0.021056 | 0.470161 | 0.14 | 0.073 | 1 | 8 | Ppp6r2 | 0.067 | Young |
| 0.021139 | 0.307867 | 0.564 | 0.43 | 1 | 8 | Msl1 | 0.134 | Young |
| 0.021145 | 0.281369 | 0.688 | 0.559 | 1 | 8 | Rpl27 | 0.129 | Young |
| 0.021224 | 0.311993 | 0.439 | 0.346 | 1 | 8 | Ube2e1 | 0.093 | Young |
| 0.021279 | 0.433841 | 0.481 | 0.391 | 1 | 8 | Csnk2b | 0.09 | Young |
| 0.021281 | 1.224373 | 0.052 | 0.011 | 1 | 8 | Tsr3 | 0.041 | Young |
| 0.021359 | 0.329968 | 0.608 | 0.48 | 1 | 8 | Cox14 | 0.128 | Young |
| 0.021408 | 0.481736 | 0.348 | 0.257 | 1 | 8 | Hpgds | 0.091 | Young |
| 0.021476 | 1.347724 | 0.07 | 0.022 | 1 | 8 | Lrrc8b | 0.048 | Young |
| 0.021528 | 0.504103 | 0.371 | 0.279 | 1 | 8 | Arl6ip5 | 0.092 | Young |
| 0.02157 | 0.777171 | 0.171 | 0.101 | 1 | 8 | Ndrg3 | 0.07 | Young |
| 0.021584 | 0.258941 | 0.652 | 0.575 | 1 | 8 | Xiap | 0.077 | Young |
| 0.021635 | 1.321513 | 0.101 | 0.045 | 1 | 8 | Ap1s3 | 0.056 | Young |
| 0.021683 | 0.368416 | 0.304 | 0.201 | 1 | 8 | Cant1 | 0.103 | Young |
| 0.021718 | 0.518475 | 0.112 | 0.05 | 1 | 8 | Slc9a3r2 | 0.062 | Young |
| 0.021923 | 0.249465 | 0.857 | 0.782 | 1 | 8 | Purb | 0.075 | Young |
| 0.021988 | 0.160363 | 0.875 | 0.793 | 1 | 8 | Cyth4 | 0.082 | Young |
| 0.021991 | 0.170647 | 0.519 | 0.402 | 1 | 8 | Add1 | 0.117 | Young |

| | | | | | | | | |
|---|---|---|---|---|---|---|---|---|
| 0.021996 | 0.797897 | 0.174 | 0.101 | 1 | 8 | Casp1 | 0.073 | Young |
| 0.022093 | 0.592084 | 0.294 | 0.212 | 1 | 8 | Rras | 0.082 | Young |
| 0.022135 | 0.333221 | 0.538 | 0.436 | 1 | 8 | Tbl1xr1 | 0.102 | Young |
| 0.022187 | 0.430366 | 0.143 | 0.073 | 1 | 8 | Mgst3 | 0.07 | Young |
| 0.022329 | 0.57744 | 0.171 | 0.095 | 1 | 8 | Tslp | 0.076 | Young |
| 0.022388 | 0.189469 | 0.725 | 0.642 | 1 | 8 | Plekho1 | 0.083 | Young |
| 0.022576 | 2.326151 | 0.029 | 0 | 1 | 8 | Ubtd2 | 0.029 | Young |
| 0.022576 | 2.144478 | 0.029 | 0 | 1 | 8 | 1810034E1 | 0.029 | Young |
| 0.022576 | 2.037865 | 0.029 | 0 | 1 | 8 | 1810055G( | 0.029 | Young |
| 0.022576 | 1.995495 | 0.029 | 0 | 1 | 8 | Dpys | 0.029 | Young |
| 0.022784 | 0.91569 | 0.101 | 0.045 | 1 | 8 | Ccl22 | 0.056 | Young |
| 0.022856 | 1.205638 | 0.086 | 0.034 | 1 | 8 | Fam110a | 0.052 | Young |
| 0.022921 | 0.165736 | 0.561 | 0.447 | 1 | 8 | Arid1a | 0.114 | Young |
| 0.022923 | 1.243845 | 0.101 | 0.045 | 1 | 8 | Styx | 0.056 | Young |
| 0.022972 | 0.832936 | 0.132 | 0.067 | 1 | 8 | Cyp4v3 | 0.065 | Young |
| 0.023009 | 0.371156 | 0.223 | 0.14 | 1 | 8 | Foxred2 | 0.083 | Young |
| 0.023089 | 1.770457 | 0.06 | 0.017 | 1 | 8 | Mex3b | 0.043 | Young |
| 0.023194 | 0.488223 | 0.403 | 0.302 | 1 | 8 | Cr1l | 0.101 | Young |
| 0.023231 | 0.243164 | 0.571 | 0.453 | 1 | 8 | Lsm4 | 0.118 | Young |
| 0.023248 | 0.473671 | 0.252 | 0.168 | 1 | 8 | Tspan13 | 0.084 | Young |
| 0.023292 | 0.247066 | 0.73 | 0.603 | 1 | 8 | S100a13 | 0.127 | Young |
| 0.023556 | 0.710201 | 0.112 | 0.05 | 1 | 8 | Arhgef18 | 0.062 | Young |
| 0.023602 | 1.245398 | 0.114 | 0.056 | 1 | 8 | Gamt | 0.058 | Young |
| 0.023683 | 0.988402 | 0.078 | 0.028 | 1 | 8 | Zfp574 | 0.05 | Young |
| 0.023729 | 0.363572 | 0.288 | 0.19 | 1 | 8 | Klhdc3 | 0.098 | Young |
| 0.023788 | 0.2612 | 0.67 | 0.559 | 1 | 8 | Ppp1r15a | 0.111 | Young |
| 0.023822 | 0.456509 | 0.426 | 0.335 | 1 | 8 | Mpc1 | 0.091 | Young |
| 0.02392 | 0.866135 | 0.117 | 0.056 | 1 | 8 | Fbxw5 | 0.061 | Young |
| 0.02393 | 0.844041 | 0.078 | 0.028 | 1 | 8 | Sec61a2 | 0.05 | Young |
| 0.024116 | 0.958825 | 0.109 | 0.05 | 1 | 8 | Lxn | 0.059 | Young |
| 0.024135 | 0.366393 | 0.294 | 0.196 | 1 | 8 | Cope | 0.098 | Young |
| 0.024175 | 2.283843 | 0.049 | 0.011 | 1 | 8 | Adgrg5 | 0.038 | Young |
| 0.024189 | 0.046429 | 0.455 | 0.33 | 1 | 8 | Rasgrp4 | 0.125 | Young |
| 0.024204 | 0.7121 | 0.101 | 0.045 | 1 | 8 | Klrb1a | 0.056 | Young |
| 0.024238 | 1.58946 | 0.068 | 0.022 | 1 | 8 | Zfp959 | 0.046 | Young |
| 0.024477 | 0.815497 | 0.197 | 0.123 | 1 | 8 | Khnyn | 0.074 | Young |
| 0.024569 | 0.620726 | 0.268 | 0.184 | 1 | 8 | Fryl | 0.084 | Young |
| 0.02465 | 0.314075 | 0.455 | 0.363 | 1 | 8 | Cic | 0.092 | Young |
| 0.024652 | 0.620631 | 0.179 | 0.106 | 1 | 8 | Gar1 | 0.073 | Young |
| 0.024712 | 2.220198 | 0.049 | 0.011 | 1 | 8 | Gpr141 | 0.038 | Young |
| 0.024712 | 2.030621 | 0.049 | 0.011 | 1 | 8 | Adam22 | 0.038 | Young |
| 0.024884 | 1.169304 | 0.122 | 0.061 | 1 | 8 | Lncpint | 0.061 | Young |
| 0.024912 | 0.633334 | 0.117 | 0.056 | 1 | 8 | Msh2 | 0.061 | Young |
| 0.024913 | 0.274784 | 0.571 | 0.464 | 1 | 8 | Prex1 | 0.107 | Young |
| 0.025235 | 0.503499 | 0.377 | 0.291 | 1 | 8 | Gm15987 | 0.086 | Young |

| | | | | | | | | |
|---|---|---|---|---|---|---|---|---|
| 0.02527 | 0.277506 | 0.535 | 0.447 | 1 | 8 | Fam102b | 0.088 | Young |
| 0.025305 | 0.448325 | 0.291 | 0.196 | 1 | 8 | Mrps6 | 0.095 | Young |
| 0.025427 | 0.250912 | 0.571 | 0.486 | 1 | 8 | Efr3a | 0.085 | Young |
| 0.025451 | 0.267872 | 0.525 | 0.413 | 1 | 8 | Ubn2 | 0.112 | Young |
| 0.025526 | 0.264891 | 0.732 | 0.615 | 1 | 8 | Ppp1ca | 0.117 | Young |
| 0.025655 | 2.175436 | 0.039 | 0.006 | 1 | 8 | Cep19 | 0.033 | Young |
| 0.025841 | 1.508223 | 0.075 | 0.028 | 1 | 8 | Mvb12b | 0.047 | Young |
| 0.025861 | 0.138138 | 0.387 | 0.279 | 1 | 8 | Lsm14a | 0.108 | Young |
| 0.025901 | 0.38045 | 0.249 | 0.162 | 1 | 8 | Tnks | 0.087 | Young |
| 0.025906 | 0.534246 | 0.192 | 0.112 | 1 | 8 | Pkd1 | 0.08 | Young |
| 0.025912 | 1.91668 | 0.039 | 0.006 | 1 | 8 | Apbb1 | 0.033 | Young |
| 0.025937 | 0.240897 | 0.727 | 0.637 | 1 | 8 | Arhgap18 | 0.09 | Young |
| 0.025967 | 0.399762 | 0.286 | 0.201 | 1 | 8 | Rasgef1b | 0.085 | Young |
| 0.026043 | 0.261351 | 0.408 | 0.302 | 1 | 8 | Evi5 | 0.106 | Young |
| 0.026049 | 0.975148 | 0.114 | 0.056 | 1 | 8 | Zscan29 | 0.058 | Young |
| 0.026123 | 0.795245 | 0.223 | 0.151 | 1 | 8 | Mfsd6 | 0.072 | Young |
| 0.026308 | 0.223803 | 0.483 | 0.374 | 1 | 8 | Klhl24 | 0.109 | Young |
| 0.026357 | 0.497692 | 0.34 | 0.257 | 1 | 8 | Bsdc1 | 0.083 | Young |
| 0.026382 | 0.401404 | 0.299 | 0.207 | 1 | 8 | Naa38 | 0.092 | Young |
| 0.026411 | 0.26449 | 0.784 | 0.704 | 1 | 8 | Trib1 | 0.08 | Young |
| 0.026489 | 0.934374 | 0.122 | 0.061 | 1 | 8 | Dnase1l3 | 0.061 | Young |
| 0.026606 | 1.448271 | 0.083 | 0.034 | 1 | 8 | Cd1d1 | 0.049 | Young |
| 0.026606 | 1.311717 | 0.083 | 0.034 | 1 | 8 | Dmac2 | 0.049 | Young |
| 0.026608 | 0.605121 | 0.262 | 0.179 | 1 | 8 | Il16 | 0.083 | Young |
| 0.026689 | 1.073327 | 0.143 | 0.078 | 1 | 8 | Akr1b10 | 0.065 | Young |
| 0.026695 | 0.16815 | 0.655 | 0.503 | 1 | 8 | Lipa | 0.152 | Young |
| 0.026843 | 1.109927 | 0.075 | 0.028 | 1 | 8 | Ccdc28b | 0.047 | Young |
| 0.026915 | 0.185292 | 0.631 | 0.503 | 1 | 8 | Itsn2 | 0.128 | Young |
| 0.027004 | 0.278281 | 0.345 | 0.246 | 1 | 8 | Hipk2 | 0.099 | Young |
| 0.027144 | 0.558924 | 0.358 | 0.274 | 1 | 8 | Tifa | 0.084 | Young |
| 0.027181 | 0.76805 | 0.148 | 0.084 | 1 | 8 | Pam16 | 0.064 | Young |
| 0.027226 | 0.289637 | 0.384 | 0.296 | 1 | 8 | Dctn3 | 0.088 | Young |
| 0.027245 | 0.574292 | 0.226 | 0.151 | 1 | 8 | Tnfaip8l2 | 0.075 | Young |
| 0.027292 | 0.726189 | 0.171 | 0.101 | 1 | 8 | Xylt2 | 0.07 | Young |
| 0.027363 | 1.142565 | 0.039 | 0.006 | 1 | 8 | Casz1 | 0.033 | Young |
| 0.027446 | 0.368938 | 0.301 | 0.207 | 1 | 8 | Dnase1l1 | 0.094 | Young |
| 0.027456 | 0.674226 | 0.244 | 0.162 | 1 | 8 | Ccz1 | 0.082 | Young |
| 0.027465 | 0.37408 | 0.325 | 0.223 | 1 | 8 | Snx29 | 0.102 | Young |
| 0.027593 | 0.630127 | 0.242 | 0.162 | 1 | 8 | Opa1 | 0.08 | Young |
| 0.027704 | 0.355445 | 0.273 | 0.179 | 1 | 8 | Slc31a2 | 0.094 | Young |
| 0.027795 | 0.999958 | 0.049 | 0.011 | 1 | 8 | Psph | 0.038 | Young |
| 0.027795 | 0.914317 | 0.177 | 0.106 | 1 | 8 | Cep192 | 0.071 | Young |
| 0.027853 | 0.267725 | 0.47 | 0.374 | 1 | 8 | Ccnt1 | 0.096 | Young |
| 0.027872 | 1.119175 | 0.068 | 0.022 | 1 | 8 | Sufu | 0.046 | Young |
| 0.027878 | 0.221745 | 0.652 | 0.575 | 1 | 8 | Eif4a2 | 0.077 | Young |

| | | | | | | | | |
|---|---|---|---|---|---|---|---|---|
| 0.02793 | 0.198332 | 0.408 | 0.296 | 1 | 8 | Tmem259 | 0.112 | Young |
| 0.028178 | 1.051809 | 0.068 | 0.022 | 1 | 8 | Akip1 | 0.046 | Young |
| 0.028317 | 1.067612 | 0.114 | 0.056 | 1 | 8 | Ebag9 | 0.058 | Young |
| 0.028403 | 0.30825 | 0.392 | 0.291 | 1 | 8 | Otub1 | 0.101 | Young |
| 0.028614 | 2.044183 | 0.057 | 0.017 | 1 | 8 | Thsd4 | 0.04 | Young |
| 0.028614 | 1.97246 | 0.057 | 0.017 | 1 | 8 | C030034I2: | 0.04 | Young |
| 0.028629 | 0.589867 | 0.192 | 0.117 | 1 | 8 | Erf | 0.075 | Young |
| 0.028772 | 1.230006 | 0.083 | 0.034 | 1 | 8 | Agpat2 | 0.049 | Young |
| 0.02878 | 0.332353 | 0.553 | 0.436 | 1 | 8 | Ndufb5 | 0.117 | Young |
| 0.028821 | 0.989096 | 0.148 | 0.084 | 1 | 8 | Sik2 | 0.064 | Young |
| 0.028973 | 0.212047 | 0.301 | 0.207 | 1 | 8 | 4933434E2 | 0.094 | Young |
| 0.029346 | 0.643324 | 0.208 | 0.134 | 1 | 8 | Nrbf2 | 0.074 | Young |
| 0.02944 | 0.2875 | 0.14 | 0.073 | 1 | 8 | Mcemp1 | 0.067 | Young |
| 0.029456 | 0.919764 | 0.099 | 0.045 | 1 | 8 | Syne1 | 0.054 | Young |
| 0.029486 | 0.728047 | 0.2 | 0.128 | 1 | 8 | Tax1bp3 | 0.072 | Young |
| 0.029711 | 0.310293 | 0.509 | 0.413 | 1 | 8 | Fkbp8 | 0.096 | Young |
| 0.029717 | 0.456353 | 0.416 | 0.335 | 1 | 8 | Smap1 | 0.081 | Young |
| 0.029829 | 1.801457 | 0.026 | 0 | 1 | 8 | Bcl2a1c | 0.026 | Young |
| 0.029829 | 2.226689 | 0.026 | 0 | 1 | 8 | Frat1 | 0.026 | Young |
| 0.029829 | 2.21534 | 0.026 | 0 | 1 | 8 | G430095P1 | 0.026 | Young |
| 0.029829 | 2.168947 | 0.026 | 0 | 1 | 8 | B230217C1 | 0.026 | Young |
| 0.029829 | 2.031388 | 0.026 | 0 | 1 | 8 | Slc22a4 | 0.026 | Young |
| 0.029829 | 1.993836 | 0.026 | 0 | 1 | 8 | Ica1l | 0.026 | Young |
| 0.029829 | 1.944092 | 0.026 | 0 | 1 | 8 | Cygb | 0.026 | Young |
| 0.029829 | 1.93813 | 0.026 | 0 | 1 | 8 | 1110002J0 | 0.026 | Young |
| 0.029956 | 0.559629 | 0.2 | 0.128 | 1 | 8 | Klhl18 | 0.072 | Young |
| 0.029975 | 0.469652 | 0.247 | 0.168 | 1 | 8 | Cnot8 | 0.079 | Young |
| 0.030053 | 0.339413 | 0.286 | 0.201 | 1 | 8 | Pdha1 | 0.085 | Young |
| 0.030075 | 0.423594 | 0.299 | 0.201 | 1 | 8 | Rnf20 | 0.098 | Young |
| 0.030201 | 0.960547 | 0.145 | 0.084 | 1 | 8 | Oma1 | 0.061 | Young |
| 0.030412 | 0.443374 | 0.358 | 0.279 | 1 | 8 | Psmg4 | 0.079 | Young |
| 0.030789 | 0.537847 | 0.216 | 0.134 | 1 | 8 | Retreg1 | 0.082 | Young |
| 0.030881 | 0.2685 | 0.169 | 0.101 | 1 | 8 | Tuba4a | 0.068 | Young |
| 0.030885 | 0.346681 | 0.371 | 0.268 | 1 | 8 | Uqcrc1 | 0.103 | Young |
| 0.030893 | 0.188474 | 0.371 | 0.279 | 1 | 8 | Lilrb4a | 0.092 | Young |
| 0.031264 | 0.122311 | 0.995 | 0.983 | 1 | 8 | Rpl38 | 0.012 | Young |
| 0.03128 | 0.283315 | 0.387 | 0.285 | 1 | 8 | Smndc1 | 0.102 | Young |
| 0.03142 | 0.599507 | 0.208 | 0.134 | 1 | 8 | Lcorl | 0.074 | Young |
| 0.031675 | 0.315161 | 0.306 | 0.207 | 1 | 8 | Aph1c | 0.099 | Young |
| 0.031796 | 0.162882 | 0.795 | 0.704 | 1 | 8 | Cbl | 0.091 | Young |
| 0.031936 | 0.488789 | 0.413 | 0.318 | 1 | 8 | Foxn2 | 0.095 | Young |
| 0.03208 | 0.824959 | 0.174 | 0.106 | 1 | 8 | Pigq | 0.068 | Young |
| 0.032093 | 0.188422 | 0.706 | 0.581 | 1 | 8 | Spi1 | 0.125 | Young |
| 0.032198 | 0.225509 | 0.696 | 0.615 | 1 | 8 | Ap2m1 | 0.081 | Young |
| 0.032375 | 0.348267 | 0.626 | 0.547 | 1 | 8 | Cox7b | 0.079 | Young |

| | | | | | | | | |
|---|---|---|---|---|---|---|---|---|
| 0.032383 | 0.195909 | 0.569 | 0.436 | 1 | 8 | Ensa | 0.133 | Young |
| 0.032406 | 0.696055 | 0.184 | 0.112 | 1 | 8 | Sema4d | 0.072 | Young |
| 0.032436 | 2.00575 | 0.047 | 0.011 | 1 | 8 | Gm5431 | 0.036 | Young |
| 0.032571 | 1.137246 | 0.088 | 0.039 | 1 | 8 | Ptdss2 | 0.049 | Young |
| 0.032648 | 0.232821 | 0.836 | 0.737 | 1 | 8 | Tmcc1 | 0.099 | Young |
| 0.032719 | 0.19266 | 0.613 | 0.52 | 1 | 8 | 9930111J2 | 0.093 | Young |
| 0.032809 | 1.066803 | 0.119 | 0.061 | 1 | 8 | Gm9493 | 0.058 | Young |
| 0.032954 | 0.440353 | 0.273 | 0.184 | 1 | 8 | Zfp326 | 0.089 | Young |
| 0.033075 | 0.576829 | 0.057 | 0.017 | 1 | 8 | C030034L1 | 0.04 | Young |
| 0.033346 | 0.285305 | 0.353 | 0.268 | 1 | 8 | Ap3d1 | 0.085 | Young |
| 0.033431 | 1.86799 | 0.047 | 0.011 | 1 | 8 | Zfp946 | 0.036 | Young |
| 0.033452 | 1.327784 | 0.081 | 0.034 | 1 | 8 | Tmtc2 | 0.047 | Young |
| 0.033473 | 0.381927 | 0.403 | 0.302 | 1 | 8 | Fam104a | 0.101 | Young |
| 0.033524 | 0.233568 | 0.652 | 0.536 | 1 | 8 | Rab10 | 0.116 | Young |
| 0.033565 | 0.327689 | 0.488 | 0.385 | 1 | 8 | Gpr65 | 0.103 | Young |
| 0.03372 | 1.70673 | 0.047 | 0.011 | 1 | 8 | Arfip2 | 0.036 | Young |
| 0.033724 | 0.355137 | 0.449 | 0.369 | 1 | 8 | Slc9a9 | 0.08 | Young |
| 0.033952 | 0.6889 | 0.2 | 0.128 | 1 | 8 | Senp7 | 0.072 | Young |
| 0.034062 | 1.527635 | 0.065 | 0.022 | 1 | 8 | Zmym1 | 0.043 | Young |
| 0.034068 | 1.806217 | 0.036 | 0.006 | 1 | 8 | Tchh | 0.03 | Young |
| 0.034081 | 0.233215 | 0.725 | 0.648 | 1 | 8 | Calm3 | 0.077 | Young |
| 0.034236 | 0.258283 | 0.558 | 0.453 | 1 | 8 | Mkrn1 | 0.105 | Young |
| 0.034336 | 0.277677 | 0.506 | 0.402 | 1 | 8 | Prelid1 | 0.104 | Young |
| 0.034359 | 0.184746 | 0.852 | 0.749 | 1 | 8 | Son | 0.103 | Young |
| 0.034406 | 2.040127 | 0.036 | 0.006 | 1 | 8 | Trip13 | 0.03 | Young |
| 0.034446 | 0.360268 | 0.309 | 0.223 | 1 | 8 | Emsy | 0.086 | Young |
| 0.034475 | 1.034198 | 0.145 | 0.084 | 1 | 8 | Tmod1 | 0.061 | Young |
| 0.034576 | 1.793954 | 0.036 | 0.006 | 1 | 8 | Spata13 | 0.03 | Young |
| 0.034579 | 0.284925 | 0.397 | 0.296 | 1 | 8 | Kansl1 | 0.101 | Young |
| 0.03461 | 0.106227 | 1 | 0.994 | 1 | 8 | Rpl41 | 0.006 | Young |
| 0.034679 | 0.991659 | 0.065 | 0.022 | 1 | 8 | Gbp2b | 0.043 | Young |
| 0.034746 | 1.427405 | 0.073 | 0.028 | 1 | 8 | Fbxo32 | 0.045 | Young |
| 0.034746 | 1.321699 | 0.073 | 0.028 | 1 | 8 | Phldb1 | 0.045 | Young |
| 0.034864 | 0.935695 | 0.073 | 0.028 | 1 | 8 | Acrbp | 0.045 | Young |
| 0.034875 | 1.379815 | 0.081 | 0.034 | 1 | 8 | Zfp715 | 0.047 | Young |
| 0.034933 | 0.658431 | 0.195 | 0.123 | 1 | 8 | Gan | 0.072 | Young |
| 0.035006 | 0.634284 | 0.182 | 0.112 | 1 | 8 | Rsrc1 | 0.07 | Young |
| 0.035206 | 0.526571 | 0.299 | 0.218 | 1 | 8 | P2rx7 | 0.081 | Young |
| 0.03524 | 0.107373 | 0.574 | 0.458 | 1 | 8 | Mier1 | 0.116 | Young |
| 0.035308 | 0.12633 | 0.509 | 0.38 | 1 | 8 | Icam1 | 0.129 | Young |
| 0.035328 | 0.657125 | 0.301 | 0.223 | 1 | 8 | Irak4 | 0.078 | Young |
| 0.035355 | 1.014191 | 0.096 | 0.045 | 1 | 8 | Parp6 | 0.051 | Young |
| 0.035437 | 1.541718 | 0.036 | 0.006 | 1 | 8 | Sh3bp4 | 0.03 | Young |
| 0.035567 | 0.098954 | 0.332 | 0.223 | 1 | 8 | Wnt11 | 0.109 | Young |
| 0.0356 | 0.308474 | 0.525 | 0.436 | 1 | 8 | Acly | 0.089 | Young |

| | | | | | | | | |
|---|---|---|---|---|---|---|---|---|
| 0.035611 | 1.422884 | 0.036 | 0.006 | 1 | 8 | Ptgs2os2 | 0.03 | Young |
| 0.035707 | 0.479673 | 0.351 | 0.268 | 1 | 8 | Vrk1 | 0.083 | Young |
| 0.035795 | 1.413726 | 0.094 | 0.045 | 1 | 8 | Zbtb14 | 0.049 | Young |
| 0.035821 | 0.619165 | 0.099 | 0.045 | 1 | 8 | Gm20342 | 0.054 | Young |
| 0.035944 | 0.141121 | 0.571 | 0.458 | 1 | 8 | Tgfbr2 | 0.113 | Young |
| 0.036036 | 0.065728 | 0.299 | 0.207 | 1 | 8 | Slc38a10 | 0.092 | Young |
| 0.03612 | 1.130842 | 0.081 | 0.034 | 1 | 8 | Syce2 | 0.047 | Young |
| 0.0362 | 0.248908 | 0.649 | 0.525 | 1 | 8 | Nisch | 0.124 | Young |
| 0.036239 | 0.789805 | 0.145 | 0.084 | 1 | 8 | Sh3pxd2a | 0.061 | Young |
| 0.036266 | 0.315756 | 0.405 | 0.313 | 1 | 8 | Churc1 | 0.092 | Young |
| 0.036355 | 1.225269 | 0.088 | 0.039 | 1 | 8 | Rufy2 | 0.049 | Young |
| 0.036499 | 0.348996 | 0.569 | 0.492 | 1 | 8 | Lmo2 | 0.077 | Young |
| 0.036583 | 0.303439 | 0.543 | 0.464 | 1 | 8 | B230219D2 | 0.079 | Young |
| 0.036587 | 0.549632 | 0.19 | 0.117 | 1 | 8 | Nmd3 | 0.073 | Young |
| 0.036595 | 0.347854 | 0.478 | 0.402 | 1 | 8 | Ilk | 0.076 | Young |
| 0.036606 | 0.565456 | 0.169 | 0.101 | 1 | 8 | Ice1 | 0.068 | Young |
| 0.036629 | 0.330147 | 0.418 | 0.33 | 1 | 8 | Secisbp2l | 0.088 | Young |
| 0.0368 | 0.48803 | 0.213 | 0.14 | 1 | 8 | Anapc2 | 0.073 | Young |
| 0.036988 | 0.493427 | 0.273 | 0.19 | 1 | 8 | Snrk | 0.083 | Young |
| 0.036995 | 0.32957 | 0.21 | 0.134 | 1 | 8 | Cbx5 | 0.076 | Young |
| 0.037017 | 0.756553 | 0.125 | 0.067 | 1 | 8 | Cebpzos | 0.058 | Young |
| 0.037056 | 0.442885 | 0.281 | 0.207 | 1 | 8 | Ubr3 | 0.074 | Young |
| 0.037076 | 0.227819 | 0.294 | 0.196 | 1 | 8 | Mob3b | 0.098 | Young |
| 0.037115 | 0.411551 | 0.517 | 0.441 | 1 | 8 | Gpsm3 | 0.076 | Young |
| 0.037116 | 0.271566 | 0.691 | 0.609 | 1 | 8 | Psmb1 | 0.082 | Young |
| 0.037148 | 0.718229 | 0.166 | 0.101 | 1 | 8 | Jaml | 0.065 | Young |
| 0.037263 | 0.844649 | 0.177 | 0.112 | 1 | 8 | Trappc3 | 0.065 | Young |
| 0.037284 | 0.801594 | 0.081 | 0.034 | 1 | 8 | D930030I0 | 0.047 | Young |
| 0.03757 | 0.256614 | 0.631 | 0.536 | 1 | 8 | Il10ra | 0.095 | Young |
| 0.03772 | 0.552061 | 0.244 | 0.168 | 1 | 8 | Ngdn | 0.076 | Young |
| 0.038011 | 1.620009 | 0.055 | 0.017 | 1 | 8 | Pde4d | 0.038 | Young |
| 0.038096 | 0.265091 | 0.571 | 0.486 | 1 | 8 | Pnisr | 0.085 | Young |
| 0.038158 | 1.146377 | 0.055 | 0.017 | 1 | 8 | Cers4 | 0.038 | Young |
| 0.03817 | 0.681039 | 0.21 | 0.14 | 1 | 8 | Tagap | 0.07 | Young |
| 0.038237 | 0.850368 | 0.143 | 0.084 | 1 | 8 | Cds1 | 0.059 | Young |
| 0.038358 | 0.261668 | 0.345 | 0.251 | 1 | 8 | Pcyt1a | 0.094 | Young |
| 0.038591 | 0.464879 | 0.283 | 0.201 | 1 | 8 | Idnk | 0.082 | Young |
| 0.038709 | 0.641072 | 0.2 | 0.128 | 1 | 8 | Ttc9c | 0.072 | Young |
| 0.038753 | 1.493694 | 0.055 | 0.017 | 1 | 8 | Slc18b1 | 0.038 | Young |
| 0.038862 | 0.52417 | 0.234 | 0.156 | 1 | 8 | Cd40 | 0.078 | Young |
| 0.038901 | 0.278683 | 0.379 | 0.279 | 1 | 8 | Hk2 | 0.1 | Young |
| 0.039196 | 0.270977 | 0.322 | 0.218 | 1 | 8 | Ubr2 | 0.104 | Young |
| 0.039239 | 0.759183 | 0.19 | 0.123 | 1 | 8 | Snapc5 | 0.067 | Young |
| 0.039274 | 0.379207 | 0.273 | 0.19 | 1 | 8 | Mlxip | 0.083 | Young |
| 0.039395 | 0.100093 | 0.652 | 0.553 | 1 | 8 | Nrp1 | 0.099 | Young |

| | | | | | | | | |
|---|---|---|---|---|---|---|---|---|
| 0.039496 | 0.451678 | 0.164 | 0.095 | 1 | 8 | Zbtb37 | 0.069 | Young |
| 0.039503 | 2.724842 | 0.023 | 0 | 1 | 8 | Cxcl9 | 0.023 | Young |
| 0.039503 | 2.327737 | 0.023 | 0 | 1 | 8 | Zfand4 | 0.023 | Young |
| 0.039503 | 2.218632 | 0.023 | 0 | 1 | 8 | B3gnt2 | 0.023 | Young |
| 0.039503 | 2.086534 | 0.023 | 0 | 1 | 8 | 0610040J0 | 0.023 | Young |
| 0.039503 | 1.947657 | 0.023 | 0 | 1 | 8 | Hp | 0.023 | Young |
| 0.039503 | 1.917697 | 0.023 | 0 | 1 | 8 | Epha4 | 0.023 | Young |
| 0.039503 | 1.824013 | 0.023 | 0 | 1 | 8 | Kif20b | 0.023 | Young |
| 0.039503 | 1.818844 | 0.023 | 0 | 1 | 8 | Tmem8b | 0.023 | Young |
| 0.039503 | 1.803291 | 0.023 | 0 | 1 | 8 | Tcaim | 0.023 | Young |
| 0.039513 | 0.177233 | 0.548 | 0.436 | 1 | 8 | Stk38 | 0.112 | Young |
| 0.03974 | 0.489763 | 0.262 | 0.179 | 1 | 8 | Pcbd2 | 0.083 | Young |
| 0.039778 | 0.203048 | 0.621 | 0.536 | 1 | 8 | Ets2 | 0.085 | Young |
| 0.040062 | 0.304577 | 0.21 | 0.134 | 1 | 8 | Ngly1 | 0.076 | Young |
| 0.040276 | 0.670513 | 0.223 | 0.156 | 1 | 8 | Rogdi | 0.067 | Young |
| 0.040323 | 0.379691 | 0.345 | 0.257 | 1 | 8 | Rnase6 | 0.088 | Young |
| 0.040352 | 0.498207 | 0.257 | 0.179 | 1 | 8 | Chd6 | 0.078 | Young |
| 0.040505 | 0.035036 | 0.145 | 0.078 | 1 | 8 | Scap | 0.067 | Young |
| 0.040521 | 0.645944 | 0.234 | 0.162 | 1 | 8 | Nsd2 | 0.072 | Young |
| 0.040573 | 0.297213 | 0.73 | 0.631 | 1 | 8 | Socs3 | 0.099 | Young |
| 0.040672 | 0.542782 | 0.216 | 0.145 | 1 | 8 | Plscr3 | 0.071 | Young |
| 0.04074 | 0.875401 | 0.055 | 0.017 | 1 | 8 | Afap1l1 | 0.038 | Young |
| 0.040767 | 0.442812 | 0.369 | 0.291 | 1 | 8 | St6gal1 | 0.078 | Young |
| 0.040853 | 1.577281 | 0.062 | 0.022 | 1 | 8 | Rfesd | 0.04 | Young |
| 0.0411 | 0.367032 | 0.135 | 0.073 | 1 | 8 | Xylt1 | 0.062 | Young |
| 0.041169 | 0.14791 | 0.714 | 0.581 | 1 | 8 | Cox7a2 | 0.133 | Young |
| 0.041316 | 0.163229 | 0.745 | 0.626 | 1 | 8 | Lamtor4 | 0.119 | Young |
| 0.041345 | 0.561392 | 0.377 | 0.302 | 1 | 8 | Chd7 | 0.075 | Young |
| 0.041454 | 1.992396 | 0.044 | 0.011 | 1 | 8 | Asah2 | 0.033 | Young |
| 0.04164 | 0.287467 | 0.203 | 0.134 | 1 | 8 | Akap11 | 0.069 | Young |
| 0.041726 | 0.44534 | 0.309 | 0.229 | 1 | 8 | Upf1 | 0.08 | Young |
| 0.041757 | 0.300658 | 0.525 | 0.413 | 1 | 8 | Etfb | 0.112 | Young |
| 0.041772 | 0.564883 | 0.125 | 0.067 | 1 | 8 | Galnt4 | 0.058 | Young |
| 0.041868 | 0.276909 | 0.509 | 0.408 | 1 | 8 | Ppp1r12a | 0.101 | Young |
| 0.041971 | 0.179487 | 0.304 | 0.218 | 1 | 8 | Ube2q2 | 0.086 | Young |
| 0.042005 | 0.86499 | 0.055 | 0.017 | 1 | 8 | Mypopos | 0.038 | Young |
| 0.042069 | 0.185825 | 0.496 | 0.397 | 1 | 8 | Sft2d2 | 0.099 | Young |
| 0.042087 | 0.809436 | 0.117 | 0.061 | 1 | 8 | Usp10 | 0.056 | Young |
| 0.042179 | 1.523631 | 0.062 | 0.022 | 1 | 8 | Trim30b | 0.04 | Young |
| 0.042192 | 0.174537 | 0.709 | 0.598 | 1 | 8 | Sumo2 | 0.111 | Young |
| 0.042286 | 0.514797 | 0.174 | 0.106 | 1 | 8 | 2210016L2 | 0.068 | Young |
| 0.042348 | 1.644713 | 0.044 | 0.011 | 1 | 8 | Fastkd3 | 0.033 | Young |
| 0.042497 | 0.873537 | 0.158 | 0.095 | 1 | 8 | Pnkd | 0.063 | Young |
| 0.04271 | 1.630084 | 0.044 | 0.011 | 1 | 8 | Nid2 | 0.033 | Young |
| 0.042828 | 0.310728 | 0.205 | 0.134 | 1 | 8 | Slc35b1 | 0.071 | Young |

| | | | | | | | | |
|---|---|---|---|---|---|---|---|---|
| 0.04283 | 0.787005 | 0.109 | 0.056 | 1 | 8 | Xpnpep3 | 0.053 | Young |
| 0.042925 | 0.510572 | 0.332 | 0.257 | 1 | 8 | Rin3 | 0.075 | Young |
| 0.042943 | 0.112745 | 0.236 | 0.151 | 1 | 8 | Dlg1 | 0.085 | Young |
| 0.042987 | 0.273891 | 0.288 | 0.201 | 1 | 8 | Erlec1 | 0.087 | Young |
| 0.043213 | 0.501777 | 0.19 | 0.123 | 1 | 8 | Ankrd10 | 0.067 | Young |
| 0.043245 | 0.653283 | 0.177 | 0.112 | 1 | 8 | Sord | 0.065 | Young |
| 0.043279 | 0.297369 | 0.468 | 0.374 | 1 | 8 | Ppp6r3 | 0.094 | Young |
| 0.043501 | 1.170952 | 0.07 | 0.028 | 1 | 8 | Dnmbp | 0.042 | Young |
| 0.043671 | 1.39598 | 0.112 | 0.061 | 1 | 8 | Rps6ka5 | 0.051 | Young |
| 0.043792 | 0.333185 | 0.125 | 0.067 | 1 | 8 | Dcxr | 0.058 | Young |
| 0.043935 | 0.73091 | 0.086 | 0.039 | 1 | 8 | Pxdc1 | 0.047 | Young |
| 0.044024 | 0.303843 | 0.509 | 0.425 | 1 | 8 | Hprt | 0.084 | Young |
| 0.04411 | 0.468956 | 0.361 | 0.285 | 1 | 8 | Mbd2 | 0.076 | Young |
| 0.044127 | 0.797322 | 0.156 | 0.095 | 1 | 8 | Lonp2 | 0.061 | Young |
| 0.04417 | 2.25995 | 0.034 | 0.006 | 1 | 8 | Gm8369 | 0.028 | Young |
| 0.04421 | 0.23056 | 0.584 | 0.497 | 1 | 8 | Nop53 | 0.087 | Young |
| 0.044307 | 0.163958 | 0.447 | 0.358 | 1 | 8 | Uso1 | 0.089 | Young |
| 0.044426 | 0.462182 | 0.319 | 0.235 | 1 | 8 | Pigs | 0.084 | Young |
| 0.044471 | 0.983568 | 0.062 | 0.022 | 1 | 8 | Zfp617 | 0.04 | Young |
| 0.044638 | 0.714069 | 0.117 | 0.061 | 1 | 8 | Rpa2 | 0.056 | Young |
| 0.044648 | 0.566852 | 0.273 | 0.196 | 1 | 8 | Helz | 0.077 | Young |
| 0.044653 | 0.198514 | 0.522 | 0.419 | 1 | 8 | Fam117b | 0.103 | Young |
| 0.04475 | 0.720377 | 0.044 | 0.011 | 1 | 8 | Zbtb11os1 | 0.033 | Young |
| 0.044775 | 0.259153 | 0.392 | 0.302 | 1 | 8 | Itpr1 | 0.09 | Young |
| 0.044783 | 0.299596 | 0.358 | 0.268 | 1 | 8 | Pip5k1c | 0.09 | Young |
| 0.044792 | 0.98769 | 0.078 | 0.034 | 1 | 8 | Peli2 | 0.044 | Young |
| 0.044824 | 2.27253 | 0.034 | 0.006 | 1 | 8 | Ildr2 | 0.028 | Young |
| 0.044857 | 0.358512 | 0.223 | 0.151 | 1 | 8 | AW549877 | 0.072 | Young |
| 0.044864 | 0.089833 | 0.148 | 0.084 | 1 | 8 | Uvssa | 0.064 | Young |
| 0.04494 | 1.352696 | 0.044 | 0.011 | 1 | 8 | Ptcd1 | 0.033 | Young |
| 0.044963 | 0.297321 | 0.473 | 0.374 | 1 | 8 | Rab5a | 0.099 | Young |
| 0.045048 | 0.601648 | 0.319 | 0.24 | 1 | 8 | Mbd3 | 0.079 | Young |
| 0.045271 | 0.493763 | 0.314 | 0.235 | 1 | 8 | Abhd17a | 0.079 | Young |
| 0.045316 | 0.374021 | 0.335 | 0.251 | 1 | 8 | Strap | 0.084 | Young |
| 0.045421 | 0.329129 | 0.252 | 0.173 | 1 | 8 | Ppm1g | 0.079 | Young |
| 0.045441 | 0.574968 | 0.174 | 0.112 | 1 | 8 | Ufd1 | 0.062 | Young |
| 0.045526 | 0.162716 | 0.655 | 0.57 | 1 | 8 | Nr4a2 | 0.085 | Young |
| 0.045536 | 0.242094 | 0.345 | 0.257 | 1 | 8 | Rgs19 | 0.088 | Young |
| 0.045539 | 0.35033 | 0.223 | 0.151 | 1 | 8 | Smim4 | 0.072 | Young |
| 0.045657 | 0.006747 | 0.314 | 0.218 | 1 | 8 | Rab22a | 0.096 | Young |
| 0.045684 | 1.042811 | 0.122 | 0.067 | 1 | 8 | Chmp7 | 0.055 | Young |
| 0.04571 | 1.870445 | 0.034 | 0.006 | 1 | 8 | Cenpj | 0.028 | Young |
| 0.04571 | 1.760922 | 0.034 | 0.006 | 1 | 8 | Zswim1 | 0.028 | Young |
| 0.045803 | 0.111513 | 0.982 | 0.922 | 1 | 8 | H3f3a | 0.06 | Young |
| 0.045904 | 0.209437 | 0.761 | 0.715 | 1 | 8 | Clic4 | 0.046 | Young |

| | | | | | | | | |
|---|---|---|---|---|---|---|---|---|
| 0.046013 | 0.424777 | 0.223 | 0.151 | 1 | 8 | Hgsnat | 0.072 | Young |
| 0.046086 | 1.067017 | 0.127 | 0.073 | 1 | 8 | Cars2 | 0.054 | Young |
| 0.04635 | 0.539537 | 0.078 | 0.034 | 1 | 8 | Rasal2 | 0.044 | Young |
| 0.046407 | 0.61803 | 0.281 | 0.212 | 1 | 8 | Ndufs3 | 0.069 | Young |
| 0.046627 | 0.438705 | 0.262 | 0.179 | 1 | 8 | Lats2 | 0.083 | Young |
| 0.046639 | 0.273638 | 0.327 | 0.246 | 1 | 8 | Vps28 | 0.081 | Young |
| 0.046689 | 0.207657 | 0.732 | 0.631 | 1 | 8 | Ubl5 | 0.101 | Young |
| 0.046838 | 1.025511 | 0.034 | 0.006 | 1 | 8 | Fam161a | 0.028 | Young |
| 0.047067 | 0.537205 | 0.034 | 0.006 | 1 | 8 | Gnb1l | 0.028 | Young |
| 0.047067 | 0.4746 | 0.034 | 0.006 | 1 | 8 | Rtl6 | 0.028 | Young |
| 0.047204 | 0.872793 | 0.114 | 0.061 | 1 | 8 | Fmo5 | 0.053 | Young |
| 0.047205 | 0.290434 | 0.335 | 0.246 | 1 | 8 | Chmp1a | 0.089 | Young |
| 0.047428 | 0.383497 | 0.504 | 0.436 | 1 | 8 | Nek7 | 0.068 | Young |
| 0.047513 | 0.190926 | 0.626 | 0.52 | 1 | 8 | Ms4a7 | 0.106 | Young |
| 0.047582 | 0.804946 | 0.101 | 0.05 | 1 | 8 | Dnajb2 | 0.051 | Young |
| 0.047626 | 0.882715 | 0.114 | 0.061 | 1 | 8 | Mrps10 | 0.053 | Young |
| 0.047644 | 0.791118 | 0.099 | 0.05 | 1 | 8 | Mbd5 | 0.049 | Young |
| 0.047713 | 0.340598 | 0.348 | 0.268 | 1 | 8 | Smarca4 | 0.08 | Young |
| 0.047718 | 0.69046 | 0.094 | 0.045 | 1 | 8 | Scd1 | 0.049 | Young |
| 0.047789 | 0.901637 | 0.203 | 0.14 | 1 | 8 | Prdm1 | 0.063 | Young |
| 0.047914 | 1.827156 | 0.052 | 0.017 | 1 | 8 | E2f2 | 0.035 | Young |
| 0.047975 | 0.629363 | 0.205 | 0.134 | 1 | 8 | Ssh1 | 0.071 | Young |
| 0.048143 | 0.773317 | 0.184 | 0.117 | 1 | 8 | Gpr183 | 0.067 | Young |
| 0.048168 | 0.690204 | 0.203 | 0.134 | 1 | 8 | Nectin1 | 0.069 | Young |
| 0.048658 | 0.596413 | 0.294 | 0.229 | 1 | 8 | Dguok | 0.065 | Young |
| 0.048735 | 0.389483 | 0.2 | 0.134 | 1 | 8 | Pla2g16 | 0.066 | Young |
| 0.048762 | 0.299156 | 0.273 | 0.19 | 1 | 8 | Rfc1 | 0.083 | Young |
| 0.048882 | 0.399156 | 0.281 | 0.196 | 1 | 8 | Fbxo11 | 0.085 | Young |
| 0.0489 | 0.179729 | 0.527 | 0.441 | 1 | 8 | Sfxn3 | 0.086 | Young |
| 0.04893 | 0.206931 | 0.377 | 0.285 | 1 | 8 | Wdr89 | 0.092 | Young |
| 0.04895 | 0.175877 | 0.387 | 0.296 | 1 | 8 | Prdx6 | 0.091 | Young |
| 0.048956 | 1.235083 | 0.091 | 0.045 | 1 | 8 | Myg1 | 0.046 | Young |
| 0.049016 | 0.571847 | 0.192 | 0.123 | 1 | 8 | Cyhr1 | 0.069 | Young |
| 0.049131 | 0.593628 | 0.234 | 0.162 | 1 | 8 | Nfkbib | 0.072 | Young |
| 0.04926 | 0.082329 | 1 | 1 | 1 | 8 | H2-Ab1 | 0 | Young |
| 0.049394 | 1.45826 | 0.052 | 0.017 | 1 | 8 | Thop1 | 0.035 | Young |
| 0.049416 | 0.334876 | 0.301 | 0.218 | 1 | 8 | Hltf | 0.083 | Young |
| 0.049628 | 0.476677 | 0.184 | 0.117 | 1 | 8 | Psd4 | 0.067 | Young |
| 0.049638 | 0.738899 | 0.192 | 0.128 | 1 | 8 | Zfp704 | 0.064 | Young |
| 0.049708 | 0.752885 | 0.153 | 0.095 | 1 | 8 | Gm15417 | 0.058 | Young |
| 0.049752 | 0.211375 | 0.439 | 0.341 | 1 | 8 | Mon2 | 0.098 | Young |
| 0.049807 | 0.460673 | 0.148 | 0.089 | 1 | 8 | Smpdl3b | 0.059 | Young |
| 0.049817 | 1.400919 | 0.083 | 0.039 | 1 | 8 | Mcm3ap | 0.044 | Young |
| 0.049913 | 0.625219 | 0.07 | 0.028 | 1 | 8 | Cyp2ab1 | 0.042 | Young |
| 0.049913 | 0.145166 | 0.865 | 0.732 | 1 | 8 | Mtdh | 0.133 | Young |

| | | | | | | | | |
|---|---|---|---|---|---|---|---|---|
| 0.049977 | 0.242536 | 0.506 | 0.413 | 1 | 8 | Sult1a1 | 0.093 | Young |
| 0.050069 | 0.39143 | 0.335 | 0.263 | 1 | 8 | Heatr5a | 0.072 | Young |
| 0.050263 | 0.147941 | 0.413 | 0.313 | 1 | 8 | Pja2 | 0.1 | Young |
| 0.050471 | 0.246493 | 0.203 | 0.128 | 1 | 8 | Tex264 | 0.075 | Young |
| 0.050619 | 0.618909 | 0.101 | 0.05 | 1 | 8 | Kif23 | 0.051 | Young |
| 0.050678 | 0.813137 | 0.078 | 0.034 | 1 | 8 | Bloc1s3 | 0.044 | Young |
| 0.050773 | 0.495717 | 0.257 | 0.184 | 1 | 8 | Zwint | 0.073 | Young |
| 0.050778 | 0.197507 | 0.288 | 0.201 | 1 | 8 | Map3k14 | 0.087 | Young |
| 0.050797 | 0.755977 | 0.166 | 0.106 | 1 | 8 | Smox | 0.06 | Young |
| 0.050848 | 0.675555 | 0.158 | 0.095 | 1 | 8 | Tmem39a | 0.063 | Young |
| 0.050901 | 0.206239 | 0.764 | 0.67 | 1 | 8 | Uqcrq | 0.094 | Young |
| 0.050913 | 1.164037 | 0.052 | 0.017 | 1 | 8 | Tcea3 | 0.035 | Young |
| 0.051118 | 0.37794 | 0.345 | 0.274 | 1 | 8 | Psmb9 | 0.071 | Young |
| 0.051118 | 0.655547 | 0.213 | 0.145 | 1 | 8 | Zfp62 | 0.068 | Young |
| 0.051366 | 0.52656 | 0.19 | 0.123 | 1 | 8 | Nxpe5 | 0.067 | Young |
| 0.051367 | 0.626937 | 0.106 | 0.056 | 1 | 8 | Pop4 | 0.05 | Young |
| 0.051443 | 0.893343 | 0.156 | 0.095 | 1 | 8 | 6-Sep | 0.061 | Young |
| 0.051505 | 0.346531 | 0.213 | 0.145 | 1 | 8 | Aoah | 0.068 | Young |
| 0.051632 | 0.341308 | 0.252 | 0.179 | 1 | 8 | Rcn3 | 0.073 | Young |
| 0.05172 | 0.778716 | 0.203 | 0.14 | 1 | 8 | Acads | 0.063 | Young |
| 0.051797 | 0.668674 | 0.135 | 0.078 | 1 | 8 | Dnttip1 | 0.057 | Young |
| 0.051881 | 0.276565 | 0.382 | 0.296 | 1 | 8 | AI314180 | 0.086 | Young |
| 0.051996 | 0.7519 | 0.249 | 0.184 | 1 | 8 | Mapkap1 | 0.065 | Young |
| 0.052009 | 0.13922 | 0.61 | 0.508 | 1 | 8 | Rsrc2 | 0.102 | Young |
| 0.052187 | 0.311372 | 0.444 | 0.346 | 1 | 8 | Ankfy1 | 0.098 | Young |
| 0.052468 | 2.27481 | 0.021 | 0 | 1 | 8 | Fgf9 | 0.021 | Young |
| 0.052468 | 1.95088 | 0.021 | 0 | 1 | 8 | Nr6a1 | 0.021 | Young |
| 0.052468 | 1.927275 | 0.021 | 0 | 1 | 8 | Gm20632 | 0.021 | Young |
| 0.052468 | 1.920533 | 0.021 | 0 | 1 | 8 | Uhrf1bp1 | 0.021 | Young |
| 0.052468 | 1.89321 | 0.021 | 0 | 1 | 8 | Dusp28 | 0.021 | Young |
| 0.052468 | 1.847546 | 0.021 | 0 | 1 | 8 | K230015D( | 0.021 | Young |
| 0.052468 | 1.805081 | 0.021 | 0 | 1 | 8 | BC052040 | 0.021 | Young |
| 0.052468 | 1.798043 | 0.021 | 0 | 1 | 8 | Tspan9 | 0.021 | Young |
| 0.052468 | 1.734135 | 0.021 | 0 | 1 | 8 | Zfp3 | 0.021 | Young |
| 0.052528 | 0.457589 | 0.148 | 0.089 | 1 | 8 | Zfp90 | 0.059 | Young |
| 0.052685 | 0.571754 | 0.309 | 0.235 | 1 | 8 | Zfp800 | 0.074 | Young |
| 0.052735 | 2.138088 | 0.042 | 0.011 | 1 | 8 | Neurl1a | 0.031 | Young |
| 0.053389 | 0.329624 | 0.213 | 0.14 | 1 | 8 | Dok1 | 0.073 | Young |
| 0.053457 | 0.220571 | 0.392 | 0.291 | 1 | 8 | Dstyk | 0.101 | Young |
| 0.053784 | 0.545636 | 0.138 | 0.078 | 1 | 8 | A530064D( | 0.06 | Young |
| 0.053801 | 1.429648 | 0.068 | 0.028 | 1 | 8 | 2810006K2 | 0.04 | Young |
| 0.053836 | 0.780713 | 0.127 | 0.073 | 1 | 8 | Cox19 | 0.054 | Young |
| 0.053895 | 1.357341 | 0.075 | 0.034 | 1 | 8 | Ccdc91 | 0.041 | Young |
| 0.054142 | 0.238372 | 0.431 | 0.352 | 1 | 8 | Tsc22d2 | 0.079 | Young |
| 0.054306 | 0.204183 | 0.2 | 0.134 | 1 | 8 | Agpat3 | 0.066 | Young |

| | | | | | | | | |
|---|---|---|---|---|---|---|---|---|
| 0.054439 | 0.290929 | 0.27 | 0.196 | 1 | 8 | Prune2 | 0.074 | Young |
| 0.054459 | 0.343441 | 0.519 | 0.458 | 1 | 8 | Unc119 | 0.061 | Young |
| 0.054542 | 0.943637 | 0.119 | 0.067 | 1 | 8 | Paxip1 | 0.052 | Young |
| 0.054555 | 1.103236 | 0.083 | 0.039 | 1 | 8 | Med9 | 0.044 | Young |
| 0.054561 | 1.33086 | 0.06 | 0.022 | 1 | 8 | Dcun1d2 | 0.038 | Young |
| 0.054597 | 0.55713 | 0.143 | 0.084 | 1 | 8 | Ptrhd1 | 0.059 | Young |
| 0.054958 | 0.191775 | 0.935 | 0.832 | 1 | 8 | Gm10076 | 0.103 | Young |
| 0.054961 | 0.269719 | 0.442 | 0.363 | 1 | 8 | Prkx | 0.079 | Young |
| 0.055086 | 0.249355 | 0.527 | 0.453 | 1 | 8 | Ttc14 | 0.074 | Young |
| 0.055114 | 0.56187 | 0.203 | 0.134 | 1 | 8 | Pom121 | 0.069 | Young |
| 0.055259 | 0.540766 | 0.099 | 0.05 | 1 | 8 | Mfge8 | 0.049 | Young |
| 0.055261 | 0.2303 | 0.509 | 0.402 | 1 | 8 | Fxr1 | 0.107 | Young |
| 0.05532 | 0.924556 | 0.06 | 0.022 | 1 | 8 | Tbl2 | 0.038 | Young |
| 0.055476 | 0.117638 | 0.966 | 0.905 | 1 | 8 | Ddx5 | 0.061 | Young |
| 0.055753 | 0.318958 | 0.421 | 0.335 | 1 | 8 | Acbd5 | 0.086 | Young |
| 0.055758 | 0.235626 | 0.468 | 0.363 | 1 | 8 | Tnfrsf1a | 0.105 | Young |
| 0.055895 | 0.965712 | 0.06 | 0.022 | 1 | 8 | Sec16b | 0.038 | Young |
| 0.055903 | 0.48549 | 0.127 | 0.073 | 1 | 8 | Pex13 | 0.054 | Young |
| 0.055932 | 0.38776 | 0.481 | 0.397 | 1 | 8 | Pdcd4 | 0.084 | Young |
| 0.05606 | 0.143651 | 0.616 | 0.525 | 1 | 8 | Asap1 | 0.091 | Young |
| 0.056065 | 0.468449 | 0.304 | 0.223 | 1 | 8 | Clock | 0.081 | Young |
| 0.056167 | 1.489191 | 0.042 | 0.011 | 1 | 8 | Bahcc1 | 0.031 | Young |
| 0.056245 | 0.997294 | 0.104 | 0.056 | 1 | 8 | Mettl7a1 | 0.048 | Young |
| 0.056304 | 0.146803 | 0.34 | 0.251 | 1 | 8 | Rab32 | 0.089 | Young |
| 0.056717 | 0.242493 | 0.335 | 0.257 | 1 | 8 | Fam76a | 0.078 | Young |
| 0.05673 | 0.41738 | 0.364 | 0.274 | 1 | 8 | Hcst | 0.09 | Young |
| 0.056743 | 0.464864 | 0.223 | 0.156 | 1 | 8 | Rpp21 | 0.067 | Young |
| 0.05686 | 0.095734 | 0.886 | 0.799 | 1 | 8 | Fcgrt | 0.087 | Young |
| 0.056875 | 0.235144 | 0.538 | 0.43 | 1 | 8 | Ndufv3 | 0.108 | Young |
| 0.056971 | 0.122488 | 0.894 | 0.816 | 1 | 8 | Qk | 0.078 | Young |
| 0.057004 | 0.215569 | 0.369 | 0.296 | 1 | 8 | Tmbim4 | 0.073 | Young |
| 0.057073 | 0.325305 | 0.408 | 0.324 | 1 | 8 | Jtb | 0.084 | Young |
| 0.057337 | 0.60694 | 0.153 | 0.095 | 1 | 8 | Fopnl | 0.058 | Young |
| 0.057364 | 0.383141 | 0.356 | 0.279 | 1 | 8 | Dnajc15 | 0.077 | Young |
| 0.057512 | 0.746564 | 0.177 | 0.117 | 1 | 8 | Tango2 | 0.06 | Young |
| 0.057591 | 0.112106 | 0.613 | 0.508 | 1 | 8 | Timm13 | 0.105 | Young |
| 0.057643 | 0.479724 | 0.197 | 0.134 | 1 | 8 | Tubb4b | 0.063 | Young |
| 0.057716 | 0.232932 | 0.678 | 0.592 | 1 | 8 | Neurl3 | 0.086 | Young |
| 0.057731 | 0.168839 | 0.19 | 0.123 | 1 | 8 | Klf16 | 0.067 | Young |
| 0.057799 | 0.268253 | 0.332 | 0.257 | 1 | 8 | 4932438A1 | 0.075 | Young |
| 0.057944 | 0.19094 | 0.14 | 0.084 | 1 | 8 | Herc6 | 0.056 | Young |
| 0.057991 | 1.063697 | 0.125 | 0.073 | 1 | 8 | Ptp4a3 | 0.052 | Young |
| 0.057996 | 0.198358 | 0.46 | 0.369 | 1 | 8 | Ddrgk1 | 0.091 | Young |
| 0.058047 | 0.157526 | 0.06 | 0.022 | 1 | 8 | Zbtb40 | 0.038 | Young |
| 0.058047 | 0.666066 | 0.171 | 0.112 | 1 | 8 | Cyth3 | 0.059 | Young |

| | | | | | | | | |
|---|---|---|---|---|---|---|---|---|
| 0.058073 | 0.716732 | 0.042 | 0.011 | 1 | 8 | Bbs7 | 0.031 | Young |
| 0.058079 | 0.115886 | 0.964 | 0.927 | 1 | 8 | C1qb | 0.037 | Young |
| 0.05817 | 0.222037 | 0.678 | 0.575 | 1 | 8 | Txndc17 | 0.103 | Young |
| 0.058176 | 2.132816 | 0.031 | 0.006 | 1 | 8 | Snn | 0.025 | Young |
| 0.058176 | 1.919619 | 0.031 | 0.006 | 1 | 8 | Gm36445 | 0.025 | Young |
| 0.058197 | 0.450186 | 0.4 | 0.324 | 1 | 8 | Usp15 | 0.076 | Young |
| 0.058294 | 0.376625 | 0.268 | 0.196 | 1 | 8 | Opa3 | 0.072 | Young |
| 0.058315 | 1.41081 | 0.042 | 0.011 | 1 | 8 | 4833445I0 | 0.031 | Young |
| 0.058352 | 0.308689 | 0.481 | 0.38 | 1 | 8 | Myo1f | 0.101 | Young |
| 0.05846 | 2.100046 | 0.031 | 0.006 | 1 | 8 | Zfp51 | 0.025 | Young |
| 0.05846 | 1.696942 | 0.031 | 0.006 | 1 | 8 | Ptpdc1 | 0.025 | Young |
| 0.058702 | 0.178249 | 0.325 | 0.235 | 1 | 8 | Lpin1 | 0.09 | Young |
| 0.058764 | 0.736627 | 0.151 | 0.095 | 1 | 8 | Lrrcc1 | 0.056 | Young |
| 0.058801 | 1.219564 | 0.042 | 0.011 | 1 | 8 | Zbtb26 | 0.031 | Young |
| 0.058814 | 0.283489 | 0.208 | 0.14 | 1 | 8 | Mrpl41 | 0.068 | Young |
| 0.058816 | 0.210919 | 0.816 | 0.765 | 1 | 8 | Tob2 | 0.051 | Young |
| 0.058862 | 1.677243 | 0.057 | 0.022 | 1 | 8 | BC147527 | 0.035 | Young |
| 0.059006 | 0.254009 | 0.452 | 0.363 | 1 | 8 | Esd | 0.089 | Young |
| 0.059015 | 0.191995 | 0.655 | 0.542 | 1 | 8 | Ndufc1 | 0.113 | Young |
| 0.059031 | 1.74729 | 0.031 | 0.006 | 1 | 8 | Nynrin | 0.025 | Young |
| 0.059031 | 1.73541 | 0.031 | 0.006 | 1 | 8 | Nlrp10 | 0.025 | Young |
| 0.059258 | 0.527514 | 0.068 | 0.028 | 1 | 8 | Dennd2c | 0.04 | Young |
| 0.059319 | 1.923883 | 0.031 | 0.006 | 1 | 8 | Zfp707 | 0.025 | Young |
| 0.059319 | 1.857244 | 0.031 | 0.006 | 1 | 8 | Coq8a | 0.025 | Young |
| 0.059319 | 1.769466 | 0.031 | 0.006 | 1 | 8 | 1700037C1 | 0.025 | Young |
| 0.059421 | 1.510062 | 0.049 | 0.017 | 1 | 8 | Ppp2r5b | 0.032 | Young |
| 0.059648 | 0.587325 | 0.197 | 0.134 | 1 | 8 | 2310009B1 | 0.063 | Young |
| 0.059659 | 0.278879 | 0.364 | 0.285 | 1 | 8 | Cnpy2 | 0.079 | Young |
| 0.059744 | 0.323518 | 0.462 | 0.369 | 1 | 8 | Gpbp1 | 0.093 | Young |
| 0.059774 | 0.171038 | 0.774 | 0.654 | 1 | 8 | Sec61g | 0.12 | Young |
| 0.059783 | 0.159322 | 0.41 | 0.307 | 1 | 8 | Nab1 | 0.103 | Young |
| 0.059791 | 0.527833 | 0.221 | 0.151 | 1 | 8 | Rlf | 0.07 | Young |
| 0.059869 | 0.51525 | 0.226 | 0.162 | 1 | 8 | Ebpl | 0.064 | Young |
| 0.059897 | 1.593506 | 0.031 | 0.006 | 1 | 8 | Ttyh2 | 0.025 | Young |
| 0.06009 | 0.199507 | 0.322 | 0.235 | 1 | 8 | Gmip | 0.087 | Young |
| 0.060095 | 1.72601 | 0.049 | 0.017 | 1 | 8 | Slc25a15 | 0.032 | Young |
| 0.060179 | 0.477762 | 0.27 | 0.201 | 1 | 8 | Kmt2d | 0.069 | Young |
| 0.060187 | 1.703907 | 0.031 | 0.006 | 1 | 8 | Gm15232 | 0.025 | Young |
| 0.060547 | 1.64835 | 0.049 | 0.017 | 1 | 8 | Cd300ld3 | 0.032 | Young |
| 0.061039 | 0.439575 | 0.158 | 0.095 | 1 | 8 | Plp2 | 0.063 | Young |
| 0.061051 | 0.392106 | 0.335 | 0.251 | 1 | 8 | Thoc7 | 0.084 | Young |
| 0.061067 | 0.694998 | 0.119 | 0.067 | 1 | 8 | Pkdcc | 0.052 | Young |
| 0.061068 | 0.254831 | 0.41 | 0.33 | 1 | 8 | Sipa1 | 0.08 | Young |
| 0.061178 | 0.27729 | 0.322 | 0.235 | 1 | 8 | Kcnk6 | 0.087 | Young |
| 0.06128 | 1.616522 | 0.065 | 0.028 | 1 | 8 | Gm10138 | 0.037 | Young |

| | | | | | | | | |
|---|---|---|---|---|---|---|---|---|
| 0.061312 | 0.257614 | 0.345 | 0.263 | 1 | 8 | Sike1 | 0.082 | Young |
| 0.061317 | 0.435308 | 0.195 | 0.128 | 1 | 8 | Atg12 | 0.067 | Young |
| 0.061452 | 0.525682 | 0.345 | 0.268 | 1 | 8 | Chd8 | 0.077 | Young |
| 0.061624 | 0.422512 | 0.299 | 0.223 | 1 | 8 | Fam162a | 0.076 | Young |
| 0.061659 | 0.845249 | 0.031 | 0.006 | 1 | 8 | Ptrh1 | 0.025 | Young |
| 0.061816 | 0.387933 | 0.343 | 0.257 | 1 | 8 | 1700025G( | 0.086 | Young |
| 0.062004 | 0.270068 | 0.312 | 0.223 | 1 | 8 | Tes | 0.089 | Young |
| 0.062145 | 0.466773 | 0.226 | 0.156 | 1 | 8 | Mocs2 | 0.07 | Young |
| 0.062245 | 0.263531 | 0.4 | 0.318 | 1 | 8 | Anxa6 | 0.082 | Young |
| 0.062309 | 0.942376 | 0.104 | 0.056 | 1 | 8 | Tmem80 | 0.048 | Young |
| 0.062427 | 0.260094 | 0.481 | 0.385 | 1 | 8 | Dpy19l1 | 0.096 | Young |
| 0.062662 | 0.498105 | 0.213 | 0.151 | 1 | 8 | Fcf1 | 0.062 | Young |
| 0.062677 | 1.432167 | 0.065 | 0.028 | 1 | 8 | Pnpla6 | 0.037 | Young |
| 0.062833 | 0.610465 | 0.119 | 0.067 | 1 | 8 | Gkap1 | 0.052 | Young |
| 0.062996 | 0.144049 | 0.41 | 0.318 | 1 | 8 | Setd2 | 0.092 | Young |
| 0.063147 | 0.423447 | 0.34 | 0.274 | 1 | 8 | Slc9a3r1 | 0.066 | Young |
| 0.063323 | 0.542664 | 0.088 | 0.045 | 1 | 8 | Gpr85 | 0.043 | Young |
| 0.063394 | 0.511437 | 0.286 | 0.212 | 1 | 8 | Prpf8 | 0.074 | Young |
| 0.063441 | 0.197941 | 0.434 | 0.363 | 1 | 8 | Pkn2 | 0.071 | Young |
| 0.063445 | 0.15894 | 0.361 | 0.285 | 1 | 8 | Sh3pxd2b | 0.076 | Young |
| 0.06351 | 1.374408 | 0.057 | 0.022 | 1 | 8 | Ecsit | 0.035 | Young |
| 0.063533 | 0.328042 | 0.309 | 0.229 | 1 | 8 | Ints6 | 0.08 | Young |
| 0.063553 | 0.189548 | 0.361 | 0.274 | 1 | 8 | Snx30 | 0.087 | Young |
| 0.063557 | 0.690988 | 0.049 | 0.017 | 1 | 8 | Pex11a | 0.032 | Young |
| 0.063601 | 0.161064 | 0.925 | 0.866 | 1 | 8 | Ifitm3 | 0.059 | Young |
| 0.063714 | 0.473834 | 0.205 | 0.14 | 1 | 8 | Stim1 | 0.065 | Young |
| 0.063867 | 0.354822 | 0.335 | 0.251 | 1 | 8 | Vamp2 | 0.084 | Young |
| 0.064137 | 0.92031 | 0.081 | 0.039 | 1 | 8 | Trim39 | 0.042 | Young |
| 0.06414 | 0.225035 | 0.748 | 0.682 | 1 | 8 | Nfkb1 | 0.066 | Young |
| 0.064166 | 1.304139 | 0.057 | 0.022 | 1 | 8 | 1600020E( | 0.035 | Young |
| 0.064225 | 0.795599 | 0.122 | 0.073 | 1 | 8 | Mark4 | 0.049 | Young |
| 0.064256 | 0.45381 | 0.174 | 0.112 | 1 | 8 | Mcee | 0.062 | Young |
| 0.064269 | 1.209294 | 0.049 | 0.017 | 1 | 8 | Cox7a1 | 0.032 | Young |
| 0.064433 | 0.454708 | 0.247 | 0.179 | 1 | 8 | Pex6 | 0.068 | Young |
| 0.064449 | 1.2992 | 0.094 | 0.05 | 1 | 8 | Zfp938 | 0.044 | Young |
| 0.064461 | 0.188422 | 0.665 | 0.547 | 1 | 8 | Ndufb3 | 0.118 | Young |
| 0.064508 | 0.547654 | 0.049 | 0.017 | 1 | 8 | Adamtsl4 | 0.032 | Young |
| 0.064561 | 0.506687 | 0.132 | 0.078 | 1 | 8 | Ifitm1 | 0.054 | Young |
| 0.064588 | 0.138527 | 0.855 | 0.76 | 1 | 8 | Sec61b | 0.095 | Young |
| 0.064593 | 0.947616 | 0.096 | 0.05 | 1 | 8 | Ripor1 | 0.046 | Young |
| 0.064735 | 0.271163 | 0.364 | 0.285 | 1 | 8 | Mien1 | 0.079 | Young |
| 0.064748 | 0.930921 | 0.049 | 0.017 | 1 | 8 | Zfp53 | 0.032 | Young |
| 0.064793 | 0.218393 | 0.119 | 0.067 | 1 | 8 | Wtip | 0.052 | Young |
| 0.064832 | 0.295844 | 0.314 | 0.235 | 1 | 8 | Mrpl48 | 0.079 | Young |
| 0.064858 | 0.146796 | 0.81 | 0.743 | 1 | 8 | Ckb | 0.067 | Young |

| | | | | | | | | |
|---|---|---|---|---|---|---|---|---|
| 0.065057 | 0.012853 | 0.673 | 0.547 | 1 | 8 | Sqstm1 | 0.126 | Young |
| 0.065208 | 0.085219 | 0.2 | 0.134 | 1 | 8 | Ticam2 | 0.066 | Young |
| 0.065316 | 0.241361 | 0.371 | 0.291 | 1 | 8 | Ep400 | 0.08 | Young |
| 0.065405 | 0.777285 | 0.138 | 0.084 | 1 | 8 | Ust | 0.054 | Young |
| 0.065652 | 0.507112 | 0.242 | 0.173 | 1 | 8 | Pfdn6 | 0.069 | Young |
| 0.065729 | 0.280559 | 0.268 | 0.19 | 1 | 8 | Polr2i | 0.078 | Young |
| 0.065754 | 0.738686 | 0.216 | 0.162 | 1 | 8 | Nhlrc3 | 0.054 | Young |
| 0.065808 | 1.060336 | 0.081 | 0.039 | 1 | 8 | Lsm11 | 0.042 | Young |
| 0.065815 | 0.227933 | 0.325 | 0.246 | 1 | 8 | Yme1l1 | 0.079 | Young |
| 0.065826 | 0.298838 | 0.662 | 0.609 | 1 | 8 | S100a11 | 0.053 | Young |
| 0.065874 | 0.893387 | 0.13 | 0.078 | 1 | 8 | Casp9 | 0.052 | Young |
| 0.065899 | 0.845619 | 0.109 | 0.061 | 1 | 8 | Tcf7l2 | 0.048 | Young |
| 0.06593 | 0.331383 | 0.161 | 0.101 | 1 | 8 | Mamdc2 | 0.06 | Young |
| 0.06611 | 0.107877 | 0.491 | 0.419 | 1 | 8 | Smim14 | 0.072 | Young |
| 0.0662 | 0.613103 | 0.049 | 0.017 | 1 | 8 | Gm10521 | 0.032 | Young |
| 0.066313 | 1.325116 | 0.073 | 0.034 | 1 | 8 | Dnajc27 | 0.039 | Young |
| 0.066369 | 0.522806 | 0.27 | 0.201 | 1 | 8 | Elp5 | 0.069 | Young |
| 0.066442 | 0.460529 | 0.234 | 0.168 | 1 | 8 | Fads1 | 0.066 | Young |
| 0.066462 | 0.321539 | 0.338 | 0.263 | 1 | 8 | Bcl2a1a | 0.075 | Young |
| 0.066615 | 0.384166 | 0.234 | 0.168 | 1 | 8 | Atp7a | 0.066 | Young |
| 0.066691 | 0.349848 | 0.257 | 0.19 | 1 | 8 | Arfgap2 | 0.067 | Young |
| 0.06671 | 0.222269 | 0.639 | 0.564 | 1 | 8 | Tcf25 | 0.075 | Young |
| 0.066788 | 0.032245 | 0.236 | 0.162 | 1 | 8 | Nipsnap3b | 0.074 | Young |
| 0.066845 | 0.582823 | 0.234 | 0.168 | 1 | 8 | Celf4 | 0.066 | Young |
| 0.066936 | 0.806914 | 0.049 | 0.017 | 1 | 8 | Ptpn21 | 0.032 | Young |
| 0.067085 | 0.244649 | 0.239 | 0.168 | 1 | 8 | Map1lc3a | 0.071 | Young |
| 0.067113 | 1.124329 | 0.073 | 0.034 | 1 | 8 | Zfp943 | 0.039 | Young |
| 0.067126 | 0.686913 | 0.21 | 0.151 | 1 | 8 | Ddx23 | 0.059 | Young |
| 0.067147 | 0.44364 | 0.151 | 0.095 | 1 | 8 | Bach2 | 0.056 | Young |
| 0.067183 | 0.152278 | 0.868 | 0.872 | 1 | 8 | Mgl2 | -0.004 | Young |
| 0.067184 | 0.135445 | 0.803 | 0.687 | 1 | 8 | Ccdc50 | 0.116 | Young |
| 0.067192 | 0.465175 | 0.234 | 0.168 | 1 | 8 | Zfp871 | 0.066 | Young |
| 0.067219 | 0.698166 | 0.169 | 0.112 | 1 | 8 | Tceanc2 | 0.057 | Young |
| 0.067459 | 0.275351 | 0.353 | 0.285 | 1 | 8 | Arih2 | 0.068 | Young |
| 0.067534 | 1.250606 | 0.057 | 0.022 | 1 | 8 | Zcchc10 | 0.035 | Young |
| 0.067551 | 0.585904 | 0.356 | 0.302 | 1 | 8 | Anapc13 | 0.054 | Young |
| 0.067564 | 0.148291 | 0.577 | 0.475 | 1 | 8 | Mxd4 | 0.102 | Young |
| 0.067699 | 0.477198 | 0.179 | 0.117 | 1 | 8 | Zeb2os | 0.062 | Young |
| 0.067716 | 0.205271 | 0.452 | 0.369 | 1 | 8 | Snw1 | 0.083 | Young |
| 0.06772 | 0.610131 | 0.14 | 0.084 | 1 | 8 | Fbxl6 | 0.056 | Young |
| 0.067814 | 0.505307 | 0.158 | 0.101 | 1 | 8 | Fmc1 | 0.057 | Young |
| 0.0679 | 0.282623 | 0.444 | 0.346 | 1 | 8 | H2-DMb2 | 0.098 | Young |
| 0.068064 | 0.200228 | 0.416 | 0.307 | 1 | 8 | Chp1 | 0.109 | Young |
| 0.068123 | 1.09699 | 0.073 | 0.034 | 1 | 8 | Fzd5 | 0.039 | Young |
| 0.068531 | 1.018317 | 0.073 | 0.034 | 1 | 8 | Hmgn5 | 0.039 | Young |

| | | | | | | | | |
|---|---|---|---|---|---|---|---|---|
| 0.068622 | 0.309907 | 0.351 | 0.279 | 1 | 8 | Imp3 | 0.072 | Young |
| 0.068707 | 0.327336 | 0.262 | 0.201 | 1 | 8 | Raf1 | 0.061 | Young |
| 0.068722 | 0.14033 | 0.53 | 0.419 | 1 | 8 | Hnrnpul1 | 0.111 | Young |
| 0.068747 | 1.155657 | 0.065 | 0.028 | 1 | 8 | Hist2h2be | 0.037 | Young |
| 0.068818 | 0.320997 | 0.242 | 0.168 | 1 | 8 | Stx6 | 0.074 | Young |
| 0.069052 | 2.169626 | 0.039 | 0.011 | 1 | 8 | Cd28 | 0.028 | Young |
| 0.069183 | 0.585412 | 0.065 | 0.028 | 1 | 8 | Ppip5k1 | 0.037 | Young |
| 0.069455 | 0.17945 | 0.125 | 0.073 | 1 | 8 | Kdelc2 | 0.052 | Young |
| 0.069555 | 1.195916 | 0.094 | 0.05 | 1 | 8 | Papolg | 0.044 | Young |
| 0.069621 | 1.012979 | 0.065 | 0.028 | 1 | 8 | Zfp143 | 0.037 | Young |
| 0.069651 | 0.625279 | 0.081 | 0.039 | 1 | 8 | Btbd2 | 0.042 | Young |
| 0.069864 | 0.706108 | 0.109 | 0.061 | 1 | 8 | Exoc6 | 0.048 | Young |
| 0.06995 | 2.000355 | 0.018 | 0 | 1 | 8 | Rims3 | 0.018 | Young |
| 0.06995 | 1.998043 | 0.018 | 0 | 1 | 8 | Gm12159 | 0.018 | Young |
| 0.06995 | 1.892935 | 0.018 | 0 | 1 | 8 | Cdk5rap1 | 0.018 | Young |
| 0.06995 | 1.885058 | 0.018 | 0 | 1 | 8 | Pxmp2 | 0.018 | Young |
| 0.06995 | 1.881557 | 0.018 | 0 | 1 | 8 | Acot1 | 0.018 | Young |
| 0.06995 | 1.782744 | 0.018 | 0 | 1 | 8 | Plcb4 | 0.018 | Young |
| 0.06995 | 1.758259 | 0.018 | 0 | 1 | 8 | Acad10 | 0.018 | Young |
| 0.06995 | 1.695002 | 0.018 | 0 | 1 | 8 | Ccl5 | 0.018 | Young |
| 0.06995 | 1.68567 | 0.018 | 0 | 1 | 8 | Atp9a | 0.018 | Young |
| 0.06995 | 1.682391 | 0.018 | 0 | 1 | 8 | Setbp1 | 0.018 | Young |
| 0.06995 | 1.656787 | 0.018 | 0 | 1 | 8 | Plpp1 | 0.018 | Young |
| 0.06995 | 1.632996 | 0.018 | 0 | 1 | 8 | Fut4 | 0.018 | Young |
| 0.06995 | 1.611602 | 0.018 | 0 | 1 | 8 | Kif5c | 0.018 | Young |
| 0.06995 | 1.586581 | 0.018 | 0 | 1 | 8 | Gm20324 | 0.018 | Young |
| 0.06995 | 1.537258 | 0.018 | 0 | 1 | 8 | Rdh12 | 0.018 | Young |
| 0.06999 | 0.356885 | 0.327 | 0.251 | 1 | 8 | Tpcn1 | 0.076 | Young |
| 0.07012 | 0.109845 | 0.426 | 0.335 | 1 | 8 | Arhgap26 | 0.091 | Young |
| 0.070205 | 1.98832 | 0.039 | 0.011 | 1 | 8 | Gm3448 | 0.028 | Young |
| 0.070447 | 0.731498 | 0.135 | 0.084 | 1 | 8 | Cep295 | 0.051 | Young |
| 0.070573 | 0.402434 | 0.205 | 0.14 | 1 | 8 | Lcor | 0.065 | Young |
| 0.07063 | 0.517283 | 0.179 | 0.117 | 1 | 8 | Rwdd4a | 0.062 | Young |
| 0.070824 | 0.303327 | 0.335 | 0.257 | 1 | 8 | Sde2 | 0.078 | Young |
| 0.07099 | 0.064395 | 0.592 | 0.48 | 1 | 8 | 1700020I1 | 0.112 | Young |
| 0.071081 | 1.593823 | 0.039 | 0.011 | 1 | 8 | Ptchd1 | 0.028 | Young |
| 0.071097 | 0.074871 | 0.587 | 0.453 | 1 | 8 | Abhd12 | 0.134 | Young |
| 0.071108 | 0.290527 | 0.242 | 0.173 | 1 | 8 | Wdr44 | 0.069 | Young |
| 0.071172 | 1.054132 | 0.065 | 0.028 | 1 | 8 | Smo | 0.037 | Young |
| 0.071174 | 0.1231 | 0.444 | 0.335 | 1 | 8 | Safb2 | 0.109 | Young |
| 0.071203 | 0.158378 | 0.657 | 0.525 | 1 | 8 | Atp5d | 0.132 | Young |
| 0.071285 | 0.133596 | 0.943 | 0.916 | 1 | 8 | Txnip | 0.027 | Young |
| 0.071433 | 0.305848 | 0.314 | 0.235 | 1 | 8 | Smarce1 | 0.079 | Young |
| 0.071451 | 0.147716 | 0.335 | 0.246 | 1 | 8 | Dscr3 | 0.089 | Young |
| 0.071663 | 0.316132 | 0.203 | 0.14 | 1 | 8 | Scamp5 | 0.063 | Young |

| | | | | | | | | |
|---|---|---|---|---|---|---|---|---|
| 0.071693 | 0.21255 | 0.369 | 0.285 | 1 | 8 | Osbpl8 | 0.084 | Young |
| 0.071722 | 0.103087 | 0.592 | 0.492 | 1 | 8 | Ppp1cb | 0.1 | Young |
| 0.071841 | 0.79798 | 0.081 | 0.039 | 1 | 8 | Ccdc170 | 0.042 | Young |
| 0.071965 | 1.393803 | 0.039 | 0.011 | 1 | 8 | Lss | 0.028 | Young |
| 0.071972 | 0.585411 | 0.19 | 0.128 | 1 | 8 | Arl13b | 0.062 | Young |
| 0.071977 | 0.181006 | 0.86 | 0.743 | 1 | 8 | Fcrls | 0.117 | Young |
| 0.071992 | 0.182269 | 0.634 | 0.514 | 1 | 8 | Cox5b | 0.12 | Young |
| 0.072002 | 0.296862 | 0.356 | 0.279 | 1 | 8 | Kxd1 | 0.077 | Young |
| 0.072262 | 1.510885 | 0.039 | 0.011 | 1 | 8 | Rbks | 0.028 | Young |
| 0.072302 | 0.642619 | 0.192 | 0.134 | 1 | 8 | Pdgfc | 0.058 | Young |
| 0.07244 | 0.668931 | 0.109 | 0.061 | 1 | 8 | Gbp5 | 0.048 | Young |
| 0.072449 | 0.538286 | 0.081 | 0.039 | 1 | 8 | Rpap3 | 0.042 | Young |
| 0.072478 | 0.175127 | 0.332 | 0.263 | 1 | 8 | 1500011K1 | 0.069 | Young |
| 0.0725 | 0.178296 | 0.153 | 0.095 | 1 | 8 | Brwd1 | 0.058 | Young |
| 0.072546 | 0.020208 | 0.652 | 0.553 | 1 | 8 | Fli1 | 0.099 | Young |
| 0.072623 | 0.841351 | 0.135 | 0.084 | 1 | 8 | Grap2 | 0.051 | Young |
| 0.072684 | 0.397672 | 0.195 | 0.134 | 1 | 8 | Ddx52 | 0.061 | Young |
| 0.072752 | 0.512335 | 0.065 | 0.028 | 1 | 8 | Tmco4 | 0.037 | Young |
| 0.07278 | 0.099654 | 0.19 | 0.128 | 1 | 8 | Sdccag8 | 0.062 | Young |
| 0.072794 | 0.511148 | 0.156 | 0.101 | 1 | 8 | Emc4 | 0.055 | Young |
| 0.072852 | 0.633873 | 0.244 | 0.184 | 1 | 8 | Tpd52l2 | 0.06 | Young |
| 0.073004 | 0.15177 | 0.418 | 0.33 | 1 | 8 | Kdm5a | 0.088 | Young |
| 0.073206 | 0.111443 | 0.366 | 0.291 | 1 | 8 | Poldip3 | 0.075 | Young |
| 0.073336 | 0.688512 | 0.125 | 0.073 | 1 | 8 | Itpkc | 0.052 | Young |
| 0.073459 | 1.227487 | 0.039 | 0.011 | 1 | 8 | Srr | 0.028 | Young |
| 0.073543 | 0.191785 | 0.449 | 0.341 | 1 | 8 | Srp72 | 0.108 | Young |
| 0.073668 | 0.1697 | 0.756 | 0.665 | 1 | 8 | Clic1 | 0.091 | Young |
| 0.073808 | 0.084358 | 0.343 | 0.257 | 1 | 8 | Atp13a2 | 0.086 | Young |
| 0.074024 | 0.302015 | 0.205 | 0.14 | 1 | 8 | Pycr2 | 0.065 | Young |
| 0.074144 | 0.490097 | 0.19 | 0.128 | 1 | 8 | Copg2 | 0.062 | Young |
| 0.074226 | 0.513188 | 0.117 | 0.067 | 1 | 8 | Zbtb34 | 0.05 | Young |
| 0.074246 | 1.13995 | 0.119 | 0.073 | 1 | 8 | Hdac7 | 0.046 | Young |
| 0.074328 | 0.272256 | 0.151 | 0.095 | 1 | 8 | Tex10 | 0.056 | Young |
| 0.074333 | 0.215068 | 0.231 | 0.162 | 1 | 8 | F630028O1 | 0.069 | Young |
| 0.07436 | 0.739276 | 0.065 | 0.028 | 1 | 8 | Rtl8b | 0.037 | Young |
| 0.074368 | 0.163401 | 0.195 | 0.134 | 1 | 8 | Cdk16 | 0.061 | Young |
| 0.074603 | 0.871308 | 0.094 | 0.05 | 1 | 8 | Lpcat2 | 0.044 | Young |
| 0.074673 | 0.502611 | 0.039 | 0.011 | 1 | 8 | Cdc6 | 0.028 | Young |
| 0.074807 | 0.330578 | 0.317 | 0.235 | 1 | 8 | Shoc2 | 0.082 | Young |
| 0.074824 | 0.199929 | 0.335 | 0.257 | 1 | 8 | Isy1 | 0.078 | Young |
| 0.074825 | 0.132581 | 0.205 | 0.14 | 1 | 8 | Mosmo | 0.065 | Young |
| 0.074957 | 0.322302 | 0.283 | 0.218 | 1 | 8 | BC004004 | 0.065 | Young |
| 0.075115 | 0.271295 | 0.306 | 0.223 | 1 | 8 | Lrpap1 | 0.083 | Young |
| 0.075286 | 0.937748 | 0.039 | 0.011 | 1 | 8 | Adgra2 | 0.028 | Young |
| 0.07536 | 0.224006 | 0.436 | 0.335 | 1 | 8 | Slc35b2 | 0.101 | Young |

| | | | | | | | | |
|---|---|---|---|---|---|---|---|---|
| 0.075373 | 0.447218 | 0.171 | 0.112 | 1 | 8 | Pfkl | 0.059 | Young |
| 0.07549 | 0.381027 | 0.473 | 0.413 | 1 | 8 | Ndufs5 | 0.06 | Young |
| 0.075929 | 0.116749 | 1 | 1 | 1 | 8 | Rpl37a | 0 | Young |
| 0.076212 | 0.455275 | 0.039 | 0.011 | 1 | 8 | Znhit3 | 0.028 | Young |
| 0.076578 | 0.320453 | 0.387 | 0.313 | 1 | 8 | S100a1 | 0.074 | Young |
| 0.07658 | 0.397427 | 0.187 | 0.123 | 1 | 8 | Clpx | 0.064 | Young |
| 0.07665 | 1.045687 | 0.119 | 0.073 | 1 | 8 | Gm20696 | 0.046 | Young |
| 0.076656 | 0.565184 | 0.187 | 0.128 | 1 | 8 | Mis18a | 0.059 | Young |
| 0.076714 | 2.090026 | 0.029 | 0.006 | 1 | 8 | Klhl42 | 0.023 | Young |
| 0.077197 | 0.744446 | 0.19 | 0.134 | 1 | 8 | Unc119b | 0.056 | Young |
| 0.077322 | 0.45092 | 0.158 | 0.101 | 1 | 8 | Pigyl | 0.057 | Young |
| 0.077457 | 1.96521 | 0.029 | 0.006 | 1 | 8 | Caprin2 | 0.023 | Young |
| 0.077457 | 1.761183 | 0.029 | 0.006 | 1 | 8 | Sh2d3c | 0.023 | Young |
| 0.077462 | 0.585441 | 0.221 | 0.162 | 1 | 8 | Hsd17b10 | 0.059 | Young |
| 0.077463 | 0.566227 | 0.039 | 0.011 | 1 | 8 | Med26 | 0.028 | Young |
| 0.077507 | 1.566297 | 0.055 | 0.022 | 1 | 8 | Mov10 | 0.033 | Young |
| 0.077624 | 0.187127 | 0.636 | 0.531 | 1 | 8 | Ndufa1 | 0.105 | Young |
| 0.077666 | 0.37762 | 0.309 | 0.235 | 1 | 8 | Map2k1 | 0.074 | Young |
| 0.077714 | 0.202765 | 0.914 | 0.832 | 1 | 8 | Fam46a | 0.082 | Young |
| 0.07778 | 0.136954 | 0.649 | 0.57 | 1 | 8 | Ndufc2 | 0.079 | Young |
| 0.077831 | 1.786975 | 0.029 | 0.006 | 1 | 8 | Mgp | 0.023 | Young |
| 0.077831 | 1.691308 | 0.029 | 0.006 | 1 | 8 | Tmem119 | 0.023 | Young |
| 0.077977 | 0.870538 | 0.101 | 0.056 | 1 | 8 | Traf3ip3 | 0.045 | Young |
| 0.078018 | 0.49923 | 0.161 | 0.106 | 1 | 8 | Srpk1 | 0.055 | Young |
| 0.078061 | 0.192188 | 0.268 | 0.19 | 1 | 8 | Pcif1 | 0.078 | Young |
| 0.078128 | 0.571739 | 0.13 | 0.078 | 1 | 8 | Ptcd2 | 0.052 | Young |
| 0.078684 | 0.356955 | 0.408 | 0.33 | 1 | 8 | 0610012G( | 0.078 | Young |
| 0.078862 | 0.46856 | 0.301 | 0.235 | 1 | 8 | Pnrc2 | 0.066 | Young |
| 0.078922 | 0.266951 | 0.603 | 0.514 | 1 | 8 | Fcho2 | 0.089 | Young |
| 0.07896 | 1.52689 | 0.029 | 0.006 | 1 | 8 | Cwf19l1 | 0.023 | Young |
| 0.079134 | 0.264703 | 0.244 | 0.173 | 1 | 8 | Ralgps2 | 0.071 | Young |
| 0.079261 | 0.237102 | 0.426 | 0.346 | 1 | 8 | Asph | 0.08 | Young |
| 0.079315 | 0.128769 | 0.268 | 0.19 | 1 | 8 | Lpin2 | 0.078 | Young |
| 0.079325 | 0.112819 | 0.964 | 0.944 | 1 | 8 | Mt1 | 0.02 | Young |
| 0.079368 | 1.129536 | 0.132 | 0.084 | 1 | 8 | Ypel2 | 0.048 | Young |
| 0.079711 | 1.197014 | 0.07 | 0.034 | 1 | 8 | Zfp865 | 0.036 | Young |
| 0.079721 | 1.236701 | 0.029 | 0.006 | 1 | 8 | Fgf11 | 0.023 | Young |
| 0.07997 | 0.142056 | 0.561 | 0.486 | 1 | 8 | Kdm2a | 0.075 | Young |
| 0.08006 | 0.695142 | 0.143 | 0.089 | 1 | 8 | Spn | 0.054 | Young |
| 0.080103 | 1.439569 | 0.029 | 0.006 | 1 | 8 | Chaf1a | 0.023 | Young |
| 0.080103 | 1.276154 | 0.029 | 0.006 | 1 | 8 | Zfp235 | 0.023 | Young |
| 0.080103 | 1.172927 | 0.029 | 0.006 | 1 | 8 | Ssx2ip | 0.023 | Young |
| 0.080103 | 1.034195 | 0.029 | 0.006 | 1 | 8 | Bcr | 0.023 | Young |
| 0.080131 | 0.325978 | 0.322 | 0.251 | 1 | 8 | Rabl6 | 0.071 | Young |
| 0.080216 | 1.259598 | 0.062 | 0.028 | 1 | 8 | Rubcnl | 0.034 | Young |

| | | | | | | | | |
|---|---|---|---|---|---|---|---|---|
| 0.080286 | 0.130535 | 0.231 | 0.162 | 1 | 8 | Mylip | 0.069 | Young |
| 0.080437 | 0.024892 | 0.59 | 0.469 | 1 | 8 | Eif3j1 | 0.121 | Young |
| 0.080487 | 1.14634 | 0.029 | 0.006 | 1 | 8 | AA467197 | 0.023 | Young |
| 0.080487 | 1.071116 | 0.029 | 0.006 | 1 | 8 | Ighmbp2 | 0.023 | Young |
| 0.080487 | 0.872304 | 0.029 | 0.006 | 1 | 8 | 1700066M | 0.023 | Young |
| 0.080593 | 0.212925 | 0.722 | 0.603 | 1 | 8 | Lgals1 | 0.119 | Young |
| 0.080725 | 0.990355 | 0.099 | 0.056 | 1 | 8 | Lrwd1 | 0.043 | Young |
| 0.080747 | 0.892097 | 0.086 | 0.045 | 1 | 8 | Gprc5a | 0.041 | Young |
| 0.080872 | 1.082427 | 0.029 | 0.006 | 1 | 8 | Pomt1 | 0.023 | Young |
| 0.080872 | 0.334876 | 0.029 | 0.006 | 1 | 8 | Zc4h2 | 0.023 | Young |
| 0.081065 | 0.447525 | 0.177 | 0.117 | 1 | 8 | Pcnx | 0.06 | Young |
| 0.081076 | 0.374648 | 0.377 | 0.307 | 1 | 8 | Mydgf | 0.07 | Young |
| 0.081089 | 0.163461 | 0.182 | 0.123 | 1 | 8 | Adgre4 | 0.059 | Young |
| 0.081196 | 0.250678 | 0.369 | 0.291 | 1 | 8 | Rbm33 | 0.078 | Young |
| 0.081239 | 1.341395 | 0.055 | 0.022 | 1 | 8 | Acad11 | 0.033 | Young |
| 0.081321 | 0.073465 | 0.878 | 0.804 | 1 | 8 | Atp5g2 | 0.074 | Young |
| 0.081498 | 0.079192 | 0.987 | 0.989 | 1 | 8 | Eif1 | -0.002 | Young |
| 0.081607 | 0.03465 | 0.634 | 0.514 | 1 | 8 | Usp9x | 0.12 | Young |
| 0.081795 | 0.897278 | 0.094 | 0.05 | 1 | 8 | Zc3h12c | 0.044 | Young |
| 0.081844 | 1.161224 | 0.07 | 0.034 | 1 | 8 | Gcdh | 0.036 | Young |
| 0.082057 | 1.381435 | 0.055 | 0.022 | 1 | 8 | Enho | 0.033 | Young |
| 0.082057 | 1.30454 | 0.055 | 0.022 | 1 | 8 | Tgif2 | 0.033 | Young |
| 0.082187 | 0.820472 | 0.047 | 0.017 | 1 | 8 | Slc52a3 | 0.03 | Young |
| 0.082397 | 1.100377 | 0.091 | 0.05 | 1 | 8 | Vps8 | 0.041 | Young |
| 0.082475 | 0.818925 | 0.086 | 0.045 | 1 | 8 | Sirt3 | 0.041 | Young |
| 0.082494 | 0.287088 | 0.145 | 0.089 | 1 | 8 | Rapgef1 | 0.056 | Young |
| 0.082646 | 0.132016 | 0.322 | 0.251 | 1 | 8 | Zdhhc9 | 0.071 | Young |
| 0.0829 | 0.472908 | 0.413 | 0.352 | 1 | 8 | Ankrd12 | 0.061 | Young |
| 0.082901 | 0.65751 | 0.125 | 0.073 | 1 | 8 | Ccdc122 | 0.052 | Young |
| 0.083025 | 0.562344 | 0.122 | 0.073 | 1 | 8 | Sigirr | 0.049 | Young |
| 0.083085 | 0.705715 | 0.047 | 0.017 | 1 | 8 | Rnf180 | 0.03 | Young |
| 0.083222 | 0.484879 | 0.257 | 0.19 | 1 | 8 | Tm9sf4 | 0.067 | Young |
| 0.083398 | 0.269148 | 0.358 | 0.285 | 1 | 8 | Exoc4 | 0.073 | Young |
| 0.083688 | 0.404269 | 0.047 | 0.017 | 1 | 8 | Rmdn1 | 0.03 | Young |
| 0.084058 | 0.316268 | 0.119 | 0.073 | 1 | 8 | Sertad3 | 0.046 | Young |
| 0.084084 | 0.167682 | 0.418 | 0.324 | 1 | 8 | Slc7a8 | 0.094 | Young |
| 0.084261 | 0.253716 | 0.403 | 0.33 | 1 | 8 | Ext1 | 0.073 | Young |
| 0.084552 | 0.75589 | 0.055 | 0.022 | 1 | 8 | Ntmt1 | 0.033 | Young |
| 0.08483 | 1.050369 | 0.062 | 0.028 | 1 | 8 | Zfp426 | 0.034 | Young |
| 0.084908 | 0.308488 | 0.21 | 0.151 | 1 | 8 | Exosc3 | 0.059 | Young |
| 0.084948 | 0.685034 | 0.153 | 0.101 | 1 | 8 | Camsap1 | 0.052 | Young |
| 0.084958 | 0.945264 | 0.091 | 0.05 | 1 | 8 | Sorl1 | 0.041 | Young |
| 0.085007 | 1.207228 | 0.07 | 0.034 | 1 | 8 | Tmem115 | 0.036 | Young |
| 0.08502 | 0.345047 | 0.317 | 0.246 | 1 | 8 | Msr1 | 0.071 | Young |
| 0.08511 | 0.253013 | 0.377 | 0.302 | 1 | 8 | Idh3g | 0.075 | Young |

| | | | | | | | | |
|---|---|---|---|---|---|---|---|---|
| 0.085255 | 0.719956 | 0.07 | 0.034 | 1 | 8 | Irgq | 0.036 | Young |
| 0.085472 | 0.314537 | 0.138 | 0.084 | 1 | 8 | Lias | 0.054 | Young |
| 0.085493 | 0.22974 | 0.966 | 0.922 | 1 | 8 | Zfp36l1 | 0.044 | Young |
| 0.08559 | 0.393732 | 0.309 | 0.246 | 1 | 8 | Rnf216 | 0.063 | Young |
| 0.085789 | 0.266194 | 0.335 | 0.263 | 1 | 8 | Samd9l | 0.072 | Young |
| 0.085901 | 0.635052 | 0.184 | 0.128 | 1 | 8 | Selenoh | 0.056 | Young |
| 0.086101 | 0.233208 | 0.67 | 0.598 | 1 | 8 | Wasf2 | 0.072 | Young |
| 0.086314 | 0.398261 | 0.335 | 0.257 | 1 | 8 | Sla | 0.078 | Young |
| 0.086732 | 0.275358 | 0.257 | 0.19 | 1 | 8 | Ophn1 | 0.067 | Young |
| 0.086895 | 0.419798 | 0.236 | 0.173 | 1 | 8 | Ipo5 | 0.063 | Young |
| 0.086968 | 0.372233 | 0.122 | 0.073 | 1 | 8 | Mitd1 | 0.049 | Young |
| 0.087042 | 0.374518 | 0.135 | 0.084 | 1 | 8 | Kctd5 | 0.051 | Young |
| 0.087084 | 0.059916 | 0.686 | 0.525 | 1 | 8 | Ssr1 | 0.161 | Young |
| 0.087284 | 0.269919 | 0.39 | 0.318 | 1 | 8 | Slc25a24 | 0.072 | Young |
| 0.087485 | 0.959945 | 0.062 | 0.028 | 1 | 8 | Cox18 | 0.034 | Young |
| 0.087485 | 0.748852 | 0.062 | 0.028 | 1 | 8 | Haus7 | 0.034 | Young |
| 0.087494 | 0.230687 | 0.268 | 0.196 | 1 | 8 | Pcmtd2 | 0.072 | Young |
| 0.087685 | 0.928152 | 0.055 | 0.022 | 1 | 8 | Coa4 | 0.033 | Young |
| 0.087741 | 0.327858 | 0.223 | 0.156 | 1 | 8 | Camk2g | 0.067 | Young |
| 0.087846 | 0.654553 | 0.171 | 0.117 | 1 | 8 | Cish | 0.054 | Young |
| 0.088179 | 0.158926 | 0.548 | 0.447 | 1 | 8 | Cwc15 | 0.101 | Young |
| 0.088269 | 1.013349 | 0.07 | 0.034 | 1 | 8 | Zfp68 | 0.036 | Young |
| 0.088321 | 0.257179 | 0.447 | 0.352 | 1 | 8 | Fes | 0.095 | Young |
| 0.088745 | 0.296202 | 0.418 | 0.346 | 1 | 8 | Maz | 0.072 | Young |
| 0.088775 | 0.257815 | 0.509 | 0.425 | 1 | 8 | Snx6 | 0.084 | Young |
| 0.088894 | 0.68343 | 0.106 | 0.061 | 1 | 8 | D430042O( | 0.045 | Young |
| 0.088976 | 0.226054 | 0.275 | 0.201 | 1 | 8 | Ppp1r11 | 0.074 | Young |
| 0.089053 | 0.333623 | 0.395 | 0.313 | 1 | 8 | Kmt5b | 0.082 | Young |
| 0.08911 | 0.0763 | 0.062 | 0.028 | 1 | 8 | Adck5 | 0.034 | Young |
| 0.089196 | 0.865429 | 0.125 | 0.078 | 1 | 8 | Mpp5 | 0.047 | Young |
| 0.089327 | 0.476252 | 0.592 | 0.525 | 1 | 8 | Lyz1 | 0.067 | Young |
| 0.089388 | 0.906257 | 0.125 | 0.078 | 1 | 8 | Rnf31 | 0.047 | Young |
| 0.089496 | 0.250468 | 0.41 | 0.33 | 1 | 8 | Acat1 | 0.08 | Young |
| 0.08957 | 0.33951 | 0.161 | 0.106 | 1 | 8 | Bid | 0.055 | Young |
| 0.089593 | 0.389737 | 0.273 | 0.207 | 1 | 8 | Cltb | 0.066 | Young |
| 0.089637 | 0.685257 | 0.104 | 0.061 | 1 | 8 | Vti1a | 0.043 | Young |
| 0.089727 | 0.821064 | 0.055 | 0.022 | 1 | 8 | Qrsl1 | 0.033 | Young |
| 0.089728 | 0.421524 | 0.106 | 0.061 | 1 | 8 | Gpatch11 | 0.045 | Young |
| 0.089809 | 0.095864 | 0.392 | 0.313 | 1 | 8 | Uqcc3 | 0.079 | Young |
| 0.089937 | 0.524466 | 0.106 | 0.061 | 1 | 8 | Adora2a | 0.045 | Young |
| 0.089975 | 0.673342 | 0.164 | 0.112 | 1 | 8 | Lyrm2 | 0.052 | Young |
| 0.090021 | 0.170641 | 0.93 | 0.821 | 1 | 8 | Rrbp1 | 0.109 | Young |
| 0.090059 | 0.508524 | 0.34 | 0.274 | 1 | 8 | Snx27 | 0.066 | Young |
| 0.090062 | 0.685801 | 0.205 | 0.151 | 1 | 8 | Maoa | 0.054 | Young |
| 0.090133 | 0.295923 | 0.268 | 0.201 | 1 | 8 | Sptan1 | 0.067 | Young |

| | | | | | | | | |
|---|---|---|---|---|---|---|---|---|
| 0.090288 | 0.107565 | 0.382 | 0.302 | 1 | 8 | Nacc2 | 0.08 | Young |
| 0.0903 | 0.523337 | 0.306 | 0.251 | 1 | 8 | Cklf | 0.055 | Young |
| 0.0904 | 0.198834 | 0.156 | 0.101 | 1 | 8 | Mfap1b | 0.055 | Young |
| 0.09062 | 0.506237 | 0.174 | 0.117 | 1 | 8 | Mapkapk3 | 0.057 | Young |
| 0.090669 | 1.525213 | 0.036 | 0.011 | 1 | 8 | Fastkd5 | 0.025 | Young |
| 0.090701 | 0.381842 | 0.205 | 0.145 | 1 | 8 | Nup210 | 0.06 | Young |
| 0.090724 | 0.274547 | 0.387 | 0.324 | 1 | 8 | Smc1a | 0.063 | Young |
| 0.090776 | 0.413403 | 0.309 | 0.251 | 1 | 8 | Inpp5k | 0.058 | Young |
| 0.090785 | 0.330223 | 0.192 | 0.134 | 1 | 8 | Trim24 | 0.058 | Young |
| 0.090892 | 0.236174 | 0.343 | 0.274 | 1 | 8 | Tcf12 | 0.069 | Young |
| 0.091038 | 1.423461 | 0.036 | 0.011 | 1 | 8 | Hist1h1e | 0.025 | Young |
| 0.091078 | 0.692674 | 0.083 | 0.045 | 1 | 8 | Dhx35 | 0.038 | Young |
| 0.091107 | 0.938268 | 0.07 | 0.034 | 1 | 8 | Entpd5 | 0.036 | Young |
| 0.091282 | 0.441928 | 0.195 | 0.14 | 1 | 8 | Wdr36 | 0.055 | Young |
| 0.091378 | 0.473425 | 0.127 | 0.078 | 1 | 8 | Ankrd46 | 0.049 | Young |
| 0.091408 | 1.664318 | 0.036 | 0.011 | 1 | 8 | Plpp5 | 0.025 | Young |
| 0.091408 | 1.572409 | 0.036 | 0.011 | 1 | 8 | Thg1l | 0.025 | Young |
| 0.091425 | 0.435868 | 0.151 | 0.101 | 1 | 8 | Fgfr1op | 0.05 | Young |
| 0.091437 | 0.238251 | 0.338 | 0.263 | 1 | 8 | Klhl5 | 0.075 | Young |
| 0.091779 | 1.513511 | 0.036 | 0.011 | 1 | 8 | Gm15445 | 0.025 | Young |
| 0.091809 | 0.130789 | 0.2 | 0.14 | 1 | 8 | Rbm12 | 0.06 | Young |
| 0.091822 | 0.203986 | 0.325 | 0.246 | 1 | 8 | Cd300lg | 0.079 | Young |
| 0.092132 | 0.243315 | 0.756 | 0.715 | 1 | 8 | Bach1 | 0.041 | Young |
| 0.092151 | 1.785934 | 0.036 | 0.011 | 1 | 8 | Osgin1 | 0.025 | Young |
| 0.092151 | 1.398843 | 0.036 | 0.011 | 1 | 8 | Stbd1 | 0.025 | Young |
| 0.092151 | 1.310476 | 0.036 | 0.011 | 1 | 8 | Sac3d1 | 0.025 | Young |
| 0.092604 | 0.080945 | 0.257 | 0.184 | 1 | 8 | Trim47 | 0.073 | Young |
| 0.092788 | 0.197487 | 0.501 | 0.413 | 1 | 8 | Ndufb4 | 0.088 | Young |
| 0.093056 | 0.130817 | 0.281 | 0.212 | 1 | 8 | Crybg3 | 0.069 | Young |
| 0.093156 | 0.250134 | 0.683 | 0.62 | 1 | 8 | Usf2 | 0.063 | Young |
| 0.093184 | 0.496753 | 0.226 | 0.168 | 1 | 8 | Nudt4 | 0.058 | Young |
| 0.093275 | 1.408888 | 0.036 | 0.011 | 1 | 8 | Adra2a | 0.025 | Young |
| 0.093275 | 1.17654 | 0.036 | 0.011 | 1 | 8 | Mtif3 | 0.025 | Young |
| 0.093313 | 0.294703 | 0.226 | 0.162 | 1 | 8 | Gcc2 | 0.064 | Young |
| 0.093424 | 0.233794 | 0.275 | 0.207 | 1 | 8 | Fbxl20 | 0.068 | Young |
| 0.093557 | 0.909302 | 0.106 | 0.061 | 1 | 8 | Tppp | 0.045 | Young |
| 0.093681 | 0.485541 | 0.361 | 0.307 | 1 | 8 | Nrd1 | 0.054 | Young |
| 0.09371 | 0.644709 | 0.125 | 0.078 | 1 | 8 | Lmnb1 | 0.047 | Young |
| 0.093721 | 1.805459 | 0.016 | 0 | 1 | 8 | Kif18a | 0.016 | Young |
| 0.093721 | 1.805195 | 0.016 | 0 | 1 | 8 | Glcci1 | 0.016 | Young |
| 0.093721 | 1.800935 | 0.016 | 0 | 1 | 8 | Cenpp | 0.016 | Young |
| 0.093721 | 1.747858 | 0.016 | 0 | 1 | 8 | Galnt12 | 0.016 | Young |
| 0.093721 | 1.670358 | 0.016 | 0 | 1 | 8 | Ttc30b | 0.016 | Young |
| 0.093721 | 1.645408 | 0.016 | 0 | 1 | 8 | Tox2 | 0.016 | Young |
| 0.093721 | 1.566375 | 0.016 | 0 | 1 | 8 | Aars2 | 0.016 | Young |

| | | | | | | | | |
|---|---|---|---|---|---|---|---|---|
| 0.093721 | 1.56379 | 0.016 | 0 | 1 | 8 | Agap2 | 0.016 | Young |
| 0.093721 | 1.541758 | 0.016 | 0 | 1 | 8 | Tshz2 | 0.016 | Young |
| 0.093721 | 1.538613 | 0.016 | 0 | 1 | 8 | Wdr74 | 0.016 | Young |
| 0.093721 | 1.523625 | 0.016 | 0 | 1 | 8 | Spats2l | 0.016 | Young |
| 0.093721 | 1.499924 | 0.016 | 0 | 1 | 8 | Cyp2j9 | 0.016 | Young |
| 0.093721 | 1.488485 | 0.016 | 0 | 1 | 8 | Trim46 | 0.016 | Young |
| 0.093721 | 1.48217 | 0.016 | 0 | 1 | 8 | Nlrc3 | 0.016 | Young |
| 0.093721 | 1.46993 | 0.016 | 0 | 1 | 8 | 1-Mar | 0.016 | Young |
| 0.093721 | 1.467675 | 0.016 | 0 | 1 | 8 | Gm29336 | 0.016 | Young |
| 0.093721 | 1.466317 | 0.016 | 0 | 1 | 8 | Tnfrsf23 | 0.016 | Young |
| 0.093721 | 1.466121 | 0.016 | 0 | 1 | 8 | Slc22a5 | 0.016 | Young |
| 0.093721 | 1.445021 | 0.016 | 0 | 1 | 8 | Cmtm8 | 0.016 | Young |
| 0.093721 | 1.439647 | 0.016 | 0 | 1 | 8 | Acy3 | 0.016 | Young |
| 0.093721 | 1.380843 | 0.016 | 0 | 1 | 8 | Lefty1 | 0.016 | Young |
| 0.093721 | 1.378331 | 0.016 | 0 | 1 | 8 | Gm40645 | 0.016 | Young |
| 0.093721 | 1.366869 | 0.016 | 0 | 1 | 8 | Gm28557 | 0.016 | Young |
| 0.093721 | 1.339379 | 0.016 | 0 | 1 | 8 | Gm26569 | 0.016 | Young |
| 0.093721 | 1.290503 | 0.016 | 0 | 1 | 8 | Gm26542 | 0.016 | Young |
| 0.093721 | 1.259674 | 0.016 | 0 | 1 | 8 | Ccdc125 | 0.016 | Young |
| 0.093967 | 0.483595 | 0.203 | 0.145 | 1 | 8 | Mb21d1 | 0.058 | Young |
| 0.094055 | 0.311742 | 0.395 | 0.335 | 1 | 8 | Wbp2 | 0.06 | Young |
| 0.094321 | 0.127475 | 0.844 | 0.782 | 1 | 8 | Ncl | 0.062 | Young |
| 0.094408 | 0.192623 | 0.236 | 0.168 | 1 | 8 | Wdr33 | 0.068 | Young |
| 0.09461 | 0.402891 | 0.177 | 0.123 | 1 | 8 | Ap5m1 | 0.054 | Young |
| 0.094717 | 0.949188 | 0.083 | 0.045 | 1 | 8 | Zmat3 | 0.038 | Young |
| 0.094763 | 0.233793 | 0.171 | 0.112 | 1 | 8 | Cyb5r1 | 0.059 | Young |
| 0.094792 | 1.180645 | 0.036 | 0.011 | 1 | 8 | Tnfrsf13c | 0.025 | Young |
| 0.095007 | 0.160088 | 0.803 | 0.693 | 1 | 8 | Tomm7 | 0.11 | Young |
| 0.095221 | 0.27402 | 0.366 | 0.291 | 1 | 8 | Stom | 0.075 | Young |
| 0.095305 | 0.314599 | 0.26 | 0.201 | 1 | 8 | Ezh1 | 0.059 | Young |
| 0.095446 | 0.291071 | 0.099 | 0.056 | 1 | 8 | Tdrd3 | 0.043 | Young |
| 0.095452 | 0.108117 | 0.364 | 0.285 | 1 | 8 | Trpm7 | 0.079 | Young |
| 0.095517 | 0.740944 | 0.242 | 0.19 | 1 | 8 | Cbx4 | 0.052 | Young |
| 0.095546 | 1.794921 | 0.044 | 0.017 | 1 | 8 | Map3k6 | 0.027 | Young |
| 0.095557 | 1.160955 | 0.036 | 0.011 | 1 | 8 | Lix1 | 0.025 | Young |
| 0.095557 | 1.119731 | 0.036 | 0.011 | 1 | 8 | Pcnx4 | 0.025 | Young |
| 0.095557 | 0.403135 | 0.036 | 0.011 | 1 | 8 | Zfp825 | 0.025 | Young |
| 0.095612 | 0.89637 | 0.091 | 0.05 | 1 | 8 | Dgkh | 0.041 | Young |
| 0.095872 | 0.39852 | 0.169 | 0.112 | 1 | 8 | Adcy9 | 0.057 | Young |
| 0.095935 | 0.172383 | 0.174 | 0.117 | 1 | 8 | Zcchc9 | 0.057 | Young |
| 0.096187 | 0.207401 | 0.19 | 0.128 | 1 | 8 | Sacm1l | 0.062 | Young |
| 0.09632 | 0.168017 | 0.535 | 0.453 | 1 | 8 | Nsd1 | 0.082 | Young |
| 0.096327 | 1.040955 | 0.036 | 0.011 | 1 | 8 | Fahd2a | 0.025 | Young |
| 0.096406 | 0.260107 | 0.338 | 0.268 | 1 | 8 | Med15 | 0.07 | Young |
| 0.096459 | 0.39305 | 0.171 | 0.117 | 1 | 8 | Plpp2 | 0.054 | Young |

| | | | | | | | | |
|---|---|---|---|---|---|---|---|---|
| 0.096715 | 0.882147 | 0.036 | 0.011 | 1 | 8 | Mtrf1 | 0.025 | Young |
| 0.096715 | 0.158295 | 0.319 | 0.246 | 1 | 8 | Cd80 | 0.073 | Young |
| 0.096933 | 1.534672 | 0.044 | 0.017 | 1 | 8 | Coq4 | 0.027 | Young |
| 0.097096 | 0.08799 | 0.61 | 0.52 | 1 | 8 | Mbtd1 | 0.09 | Young |
| 0.097284 | 0.427446 | 0.099 | 0.056 | 1 | 8 | Fbxo42 | 0.043 | Young |
| 0.097286 | 1.198395 | 0.068 | 0.034 | 1 | 8 | Dtx3 | 0.034 | Young |
| 0.097318 | 0.665377 | 0.249 | 0.19 | 1 | 8 | Lsr | 0.059 | Young |
| 0.097473 | 0.205752 | 0.423 | 0.352 | 1 | 8 | Slc4a7 | 0.071 | Young |
| 0.097646 | 1.339283 | 0.06 | 0.028 | 1 | 8 | Mthfsd | 0.032 | Young |
| 0.097658 | 0.333329 | 0.187 | 0.128 | 1 | 8 | Akap10 | 0.059 | Young |
| 0.097671 | 0.373761 | 0.283 | 0.218 | 1 | 8 | Ktn1 | 0.065 | Young |
| 0.097951 | 0.217314 | 0.543 | 0.486 | 1 | 8 | Zc3hav1 | 0.057 | Young |
| 0.097951 | 1.629938 | 0.052 | 0.022 | 1 | 8 | D3Ertd254 | 0.03 | Young |
| 0.097959 | 0.244185 | 0.291 | 0.223 | 1 | 8 | Slc38a9 | 0.068 | Young |
| 0.097983 | 1.493623 | 0.044 | 0.017 | 1 | 8 | Usp49 | 0.027 | Young |
| 0.097983 | 1.431716 | 0.044 | 0.017 | 1 | 8 | Zfp667 | 0.027 | Young |
| 0.098072 | 0.139215 | 0.192 | 0.134 | 1 | 8 | Phf12 | 0.058 | Young |
| 0.098116 | 0.195131 | 0.579 | 0.497 | 1 | 8 | Mapre2 | 0.082 | Young |
| 0.098152 | 0.179478 | 0.616 | 0.536 | 1 | 8 | Caprin1 | 0.08 | Young |
| 0.098266 | 0.422 | 0.151 | 0.101 | 1 | 8 | Nfyc | 0.05 | Young |
| 0.098274 | 0.609487 | 0.177 | 0.123 | 1 | 8 | Bckdha | 0.054 | Young |
| 0.098396 | 0.54792 | 0.104 | 0.061 | 1 | 8 | Fyco1 | 0.043 | Young |
| 0.098482 | 0.27493 | 0.231 | 0.168 | 1 | 8 | Zbtb1 | 0.063 | Young |
| 0.098594 | 0.571354 | 0.234 | 0.173 | 1 | 8 | Htatsf1 | 0.061 | Young |
| 0.099033 | 0.741473 | 0.125 | 0.078 | 1 | 8 | Prkab1 | 0.047 | Young |
| 0.099097 | 0.375599 | 0.301 | 0.24 | 1 | 8 | Cetn3 | 0.061 | Young |
| 0.099234 | 0.387619 | 0.127 | 0.078 | 1 | 8 | Anapc10 | 0.049 | Young |
| 0.099417 | 0.211598 | 0.223 | 0.156 | 1 | 8 | Fam98c | 0.067 | Young |
| 0.099431 | 0.148596 | 0.288 | 0.212 | 1 | 8 | Errfi1 | 0.076 | Young |
| 0.099472 | 0.216857 | 0.379 | 0.302 | 1 | 8 | Cacybp | 0.077 | Young |
| 0.099534 | 0.556824 | 0.257 | 0.196 | 1 | 8 | Fam76b | 0.061 | Young |
| 0.099881 | 0.177173 | 0.491 | 0.397 | 1 | 8 | Arpp19 | 0.094 | Young |
| 0.100243 | 0.188421 | 0.34 | 0.263 | 1 | 8 | Nap1l4 | 0.077 | Young |
| 0.100276 | 0.404251 | 0.27 | 0.207 | 1 | 8 | Txndc11 | 0.063 | Young |
| 0.100379 | 0.496866 | 0.343 | 0.285 | 1 | 8 | Supt4a | 0.058 | Young |
| 0.100459 | 0.139664 | 0.429 | 0.335 | 1 | 8 | Rnf111 | 0.094 | Young |
| 0.100533 | 0.22388 | 0.597 | 0.525 | 1 | 8 | Cggbp1 | 0.072 | Young |
| 0.100723 | 0.138983 | 0.47 | 0.408 | 1 | 8 | Hp1bp3 | 0.062 | Young |
| 0.100837 | 1.089857 | 0.088 | 0.05 | 1 | 8 | Psmg1 | 0.038 | Young |
| 0.101058 | 0.168438 | 0.304 | 0.24 | 1 | 8 | Atraid | 0.064 | Young |
| 0.101307 | 1.775283 | 0.026 | 0.006 | 1 | 8 | Bbs4 | 0.02 | Young |
| 0.101307 | 1.653261 | 0.026 | 0.006 | 1 | 8 | Adipoq | 0.02 | Young |
| 0.101307 | 1.543232 | 0.026 | 0.006 | 1 | 8 | Zfp472 | 0.02 | Young |
| 0.101529 | 0.073313 | 0.353 | 0.268 | 1 | 8 | Elovl5 | 0.085 | Young |
| 0.101589 | 0.677145 | 0.06 | 0.028 | 1 | 8 | Sdhaf3 | 0.032 | Young |

| | | | | | | | | |
|---|---|---|---|---|---|---|---|---|
| 0.101742 | 0.546125 | 0.075 | 0.039 | 1 | 8 | Ttc38 | 0.036 | Young |
| 0.101791 | 1.911669 | 0.026 | 0.006 | 1 | 8 | Col1a2 | 0.02 | Young |
| 0.101791 | 1.831902 | 0.026 | 0.006 | 1 | 8 | Cd101 | 0.02 | Young |
| 0.101791 | 1.663364 | 0.026 | 0.006 | 1 | 8 | Gm28707 | 0.02 | Young |
| 0.101877 | 1.363645 | 0.052 | 0.022 | 1 | 8 | Inpp5e | 0.03 | Young |
| 0.102075 | 0.459769 | 0.156 | 0.106 | 1 | 8 | Engase | 0.05 | Young |
| 0.102185 | 0.285885 | 0.208 | 0.151 | 1 | 8 | Mgat4a | 0.057 | Young |
| 0.102209 | 1.304968 | 0.052 | 0.022 | 1 | 8 | Zhx3 | 0.03 | Young |
| 0.102277 | 1.688893 | 0.026 | 0.006 | 1 | 8 | Rab27a | 0.02 | Young |
| 0.102277 | 1.355056 | 0.026 | 0.006 | 1 | 8 | Smco4 | 0.02 | Young |
| 0.102303 | 0.171794 | 0.304 | 0.246 | 1 | 8 | Ubp1 | 0.058 | Young |
| 0.102361 | 0.025978 | 0.465 | 0.369 | 1 | 8 | Dctn1 | 0.096 | Young |
| 0.102652 | 0.18093 | 0.161 | 0.106 | 1 | 8 | Chchd5 | 0.055 | Young |
| 0.102704 | 0.380361 | 0.249 | 0.184 | 1 | 8 | Jkamp | 0.065 | Young |
| 0.102764 | 1.734987 | 0.026 | 0.006 | 1 | 8 | Palb2 | 0.02 | Young |
| 0.102764 | 1.417325 | 0.026 | 0.006 | 1 | 8 | Cep152 | 0.02 | Young |
| 0.102942 | 0.726067 | 0.096 | 0.056 | 1 | 8 | Pmepa1 | 0.04 | Young |
| 0.103103 | 0.323882 | 0.338 | 0.279 | 1 | 8 | Cd180 | 0.059 | Young |
| 0.103139 | 0.393258 | 0.273 | 0.207 | 1 | 8 | Mrps25 | 0.066 | Young |
| 0.103336 | 0.10573 | 0.26 | 0.19 | 1 | 8 | Brd8 | 0.07 | Young |
| 0.103372 | 0.962391 | 0.044 | 0.017 | 1 | 8 | B3gnt8 | 0.027 | Young |
| 0.103549 | 1.218333 | 0.052 | 0.022 | 1 | 8 | Pigh | 0.03 | Young |
| 0.103609 | 0.281563 | 0.426 | 0.358 | 1 | 8 | Psma1 | 0.068 | Young |
| 0.10374 | 1.068178 | 0.044 | 0.017 | 1 | 8 | Nudt15 | 0.027 | Young |
| 0.10374 | 0.94623 | 0.044 | 0.017 | 1 | 8 | Gm11772 | 0.027 | Young |
| 0.103884 | 1.08447 | 0.068 | 0.034 | 1 | 8 | Rps6kb2 | 0.034 | Young |
| 0.103887 | 0.633628 | 0.109 | 0.067 | 1 | 8 | Edc3 | 0.042 | Young |
| 0.10392 | 0.30811 | 0.571 | 0.514 | 1 | 8 | Traf1 | 0.057 | Young |
| 0.103989 | 0.037164 | 0.306 | 0.229 | 1 | 8 | Wwp2 | 0.077 | Young |
| 0.104026 | 0.375377 | 0.145 | 0.095 | 1 | 8 | Pml | 0.05 | Young |
| 0.104333 | 0.421913 | 0.184 | 0.128 | 1 | 8 | St3gal3 | 0.056 | Young |
| 0.104533 | 0.478703 | 0.145 | 0.095 | 1 | 8 | Mphosph1 | 0.05 | Young |
| 0.104563 | 1.32545 | 0.052 | 0.022 | 1 | 8 | Prr7 | 0.03 | Young |
| 0.104563 | 1.154372 | 0.052 | 0.022 | 1 | 8 | Gbp9 | 0.03 | Young |
| 0.104653 | 0.054958 | 1 | 1 | 1 | 8 | Tmsb4x | 0 | Young |
| 0.104733 | 1.291557 | 0.026 | 0.006 | 1 | 8 | Ogg1 | 0.02 | Young |
| 0.10477 | 0.061258 | 0.068 | 0.034 | 1 | 8 | Dgcr14 | 0.034 | Young |
| 0.105157 | 0.539198 | 0.229 | 0.173 | 1 | 8 | Ccdc34 | 0.056 | Young |
| 0.10523 | 1.330962 | 0.026 | 0.006 | 1 | 8 | Gm42722 | 0.02 | Young |
| 0.10523 | 1.153613 | 0.026 | 0.006 | 1 | 8 | 4930512J1 | 0.02 | Young |
| 0.105327 | 0.206798 | 0.538 | 0.464 | 1 | 8 | Ifi203 | 0.074 | Young |
| 0.10549 | 0.676085 | 0.218 | 0.168 | 1 | 8 | Slc30a7 | 0.05 | Young |
| 0.105585 | 0.838684 | 0.052 | 0.022 | 1 | 8 | Wdr53 | 0.03 | Young |
| 0.105593 | 0.632313 | 0.044 | 0.017 | 1 | 8 | Fam131a | 0.027 | Young |
| 0.105663 | 0.99615 | 0.068 | 0.034 | 1 | 8 | Fmnl3 | 0.034 | Young |

| | | | | | | | | |
|---|---|---|---|---|---|---|---|---|
| 0.105881 | 0.202273 | 0.514 | 0.425 | 1 | 8 | Nars | 0.089 | Young |
| 0.10591 | 0.64444 | 0.096 | 0.056 | 1 | 8 | Carm1 | 0.04 | Young |
| 0.10623 | 0.169945 | 0.026 | 0.006 | 1 | 8 | Kctd7 | 0.02 | Young |
| 0.106255 | 0.452265 | 0.148 | 0.101 | 1 | 8 | Sun1 | 0.047 | Young |
| 0.106341 | 0.096031 | 0.044 | 0.017 | 1 | 8 | Fxn | 0.027 | Young |
| 0.106473 | 0.301796 | 0.351 | 0.285 | 1 | 8 | Ergic3 | 0.066 | Young |
| 0.106717 | 0.889615 | 0.044 | 0.017 | 1 | 8 | Glmn | 0.027 | Young |
| 0.106935 | 0.329493 | 0.06 | 0.028 | 1 | 8 | Cbr4 | 0.032 | Young |
| 0.107047 | 0.14177 | 0.187 | 0.128 | 1 | 8 | Stoml2 | 0.059 | Young |
| 0.107314 | 1.484116 | 0.073 | 0.039 | 1 | 8 | 4921524J1 | 0.034 | Young |
| 0.107401 | 0.982797 | 0.109 | 0.067 | 1 | 8 | Amz1 | 0.042 | Young |
| 0.107851 | 0.638327 | 0.044 | 0.017 | 1 | 8 | Dcaf4 | 0.027 | Young |
| 0.107877 | 0.842341 | 0.161 | 0.112 | 1 | 8 | Kat6b | 0.049 | Young |
| 0.107986 | 0.074596 | 0.694 | 0.592 | 1 | 8 | Arhgef6 | 0.102 | Young |
| 0.108014 | 0.111947 | 0.569 | 0.486 | 1 | 8 | Hdlbp | 0.083 | Young |
| 0.108071 | 0.785876 | 0.068 | 0.034 | 1 | 8 | Oraov1 | 0.034 | Young |
| 0.108184 | 0.113373 | 0.681 | 0.615 | 1 | 8 | Nav1 | 0.066 | Young |
| 0.108265 | 0.652371 | 0.117 | 0.073 | 1 | 8 | Inpp5a | 0.044 | Young |
| 0.108363 | 0.026247 | 0.875 | 0.771 | 1 | 8 | Snrnp70 | 0.104 | Young |
| 0.108431 | 0.12932 | 0.353 | 0.279 | 1 | 8 | Chd9 | 0.074 | Young |
| 0.108538 | 0.177727 | 0.442 | 0.363 | 1 | 8 | Chic2 | 0.079 | Young |
| 0.108596 | 0.158809 | 0.834 | 0.715 | 1 | 8 | Uqcr11 | 0.119 | Young |
| 0.108612 | 0.612214 | 0.044 | 0.017 | 1 | 8 | 4933433G1 | 0.027 | Young |
| 0.108673 | 0.321861 | 0.229 | 0.168 | 1 | 8 | Nsmaf | 0.061 | Young |
| 0.108687 | 0.116868 | 0.888 | 0.816 | 1 | 8 | Ehd4 | 0.072 | Young |
| 0.108845 | 0.656123 | 0.19 | 0.14 | 1 | 8 | Jade1 | 0.05 | Young |
| 0.108915 | 0.226725 | 0.234 | 0.168 | 1 | 8 | Atp13a1 | 0.066 | Young |
| 0.108916 | 0.093075 | 0.548 | 0.447 | 1 | 8 | Mrps33 | 0.101 | Young |
| 0.108936 | 0.111492 | 0.987 | 0.983 | 1 | 8 | Btg2 | 0.004 | Young |
| 0.109048 | 0.683979 | 0.052 | 0.022 | 1 | 8 | Plcl1 | 0.03 | Young |
| 0.109053 | 0.164186 | 0.745 | 0.637 | 1 | 8 | Ndufb7 | 0.108 | Young |
| 0.109067 | 0.396256 | 0.075 | 0.039 | 1 | 8 | Gm13373 | 0.036 | Young |
| 0.109235 | 0.282042 | 0.397 | 0.33 | 1 | 8 | Krcc1 | 0.067 | Young |
| 0.109246 | 0.468865 | 0.179 | 0.128 | 1 | 8 | Commd10 | 0.051 | Young |
| 0.110415 | 0.205966 | 0.405 | 0.341 | 1 | 8 | Rala | 0.064 | Young |
| 0.110431 | 0.427788 | 0.151 | 0.101 | 1 | 8 | Camsap2 | 0.05 | Young |
| 0.110443 | 0.062278 | 0.322 | 0.246 | 1 | 8 | Ints6l | 0.076 | Young |
| 0.11047 | 0.005121 | 0.512 | 0.43 | 1 | 8 | Eif3m | 0.082 | Young |
| 0.110511 | 0.757933 | 0.06 | 0.028 | 1 | 8 | Zmat1 | 0.032 | Young |
| 0.110522 | 0.011938 | 0.075 | 0.039 | 1 | 8 | Ubxn2b | 0.036 | Young |
| 0.110531 | 0.536894 | 0.174 | 0.123 | 1 | 8 | Gt(ROSA)2( | 0.051 | Young |
| 0.110646 | 0.099981 | 0.148 | 0.095 | 1 | 8 | Tmem159 | 0.053 | Young |
| 0.110675 | 0.423532 | 0.143 | 0.095 | 1 | 8 | Pde12 | 0.048 | Young |
| 0.111177 | 0.290524 | 0.301 | 0.229 | 1 | 8 | Atg101 | 0.072 | Young |
| 0.111292 | 0.152631 | 0.366 | 0.296 | 1 | 8 | Pik3r6 | 0.07 | Young |

| | | | | | | | | |
|---|---|---|---|---|---|---|---|---|
| 0.111399 | 0.46394 | 0.195 | 0.14 | 1 | 8 | Rexo1 | 0.055 | Young |
| 0.111403 | 0.390058 | 0.075 | 0.039 | 1 | 8 | Tnfrsf18 | 0.036 | Young |
| 0.111445 | 0.200873 | 0.21 | 0.156 | 1 | 8 | Dennd1c | 0.054 | Young |
| 0.111471 | 0.08545 | 0.99 | 0.978 | 1 | 8 | Nfkbia | 0.012 | Young |
| 0.111678 | 0.107291 | 0.434 | 0.358 | 1 | 8 | Pnn | 0.076 | Young |
| 0.111743 | 0.908721 | 0.109 | 0.067 | 1 | 8 | Nxt1 | 0.042 | Young |
| 0.111786 | 0.363556 | 0.332 | 0.263 | 1 | 8 | Clasp2 | 0.069 | Young |
| 0.111788 | 0.844417 | 0.135 | 0.089 | 1 | 8 | Mrpl27 | 0.046 | Young |
| 0.111809 | 0.129104 | 0.299 | 0.235 | 1 | 8 | Akna | 0.064 | Young |
| 0.111875 | 0.058123 | 0.696 | 0.592 | 1 | 8 | Ube2b | 0.104 | Young |
| 0.112085 | 0.230961 | 0.317 | 0.246 | 1 | 8 | Dhx36 | 0.071 | Young |
| 0.112171 | 0.343593 | 0.304 | 0.246 | 1 | 8 | Ankrd40 | 0.058 | Young |
| 0.112224 | 0.517533 | 0.132 | 0.084 | 1 | 8 | Ddr1 | 0.048 | Young |
| 0.112538 | 0.496443 | 0.187 | 0.134 | 1 | 8 | Foxo1 | 0.053 | Young |
| 0.112567 | 1.227295 | 0.086 | 0.05 | 1 | 8 | Adck1 | 0.036 | Young |
| 0.112858 | 0.269298 | 0.519 | 0.458 | 1 | 8 | Vps35 | 0.061 | Young |
| 0.112958 | 0.467988 | 0.151 | 0.101 | 1 | 8 | Hivep3 | 0.05 | Young |
| 0.113084 | 0.077057 | 0.213 | 0.151 | 1 | 8 | C5ar2 | 0.062 | Young |
| 0.113122 | 0.195877 | 0.704 | 0.62 | 1 | 8 | Slc38a2 | 0.084 | Young |
| 0.113263 | 0.130184 | 0.223 | 0.162 | 1 | 8 | Cks2 | 0.061 | Young |
| 0.113441 | 0.191625 | 0.317 | 0.251 | 1 | 8 | Usp3 | 0.066 | Young |
| 0.113475 | 0.365204 | 0.195 | 0.14 | 1 | 8 | Smg5 | 0.055 | Young |
| 0.113681 | 0.279365 | 0.623 | 0.553 | 1 | 8 | Tmem160 | 0.07 | Young |
| 0.113996 | 0.338206 | 0.088 | 0.05 | 1 | 8 | Tfec | 0.038 | Young |
| 0.114243 | 0.387274 | 0.356 | 0.302 | 1 | 8 | Stk40 | 0.054 | Young |
| 0.114283 | 0.691078 | 0.218 | 0.168 | 1 | 8 | Got2 | 0.05 | Young |
| 0.114443 | 0.356026 | 0.306 | 0.246 | 1 | 8 | Gabarapl1 | 0.06 | Young |
| 0.114695 | 0.554618 | 0.174 | 0.123 | 1 | 8 | Mrpl35 | 0.051 | Young |
| 0.114923 | 0.525906 | 0.13 | 0.084 | 1 | 8 | Hsf1 | 0.046 | Young |
| 0.115028 | 0.503675 | 0.229 | 0.173 | 1 | 8 | Prdx4 | 0.056 | Young |
| 0.115089 | 0.194539 | 0.309 | 0.251 | 1 | 8 | Ebp | 0.058 | Young |
| 0.115099 | 0.578007 | 0.151 | 0.101 | 1 | 8 | Stard3 | 0.05 | Young |
| 0.115153 | 0.163717 | 0.423 | 0.352 | 1 | 8 | Dyrk2 | 0.071 | Young |
| 0.115285 | 0.342152 | 0.221 | 0.162 | 1 | 8 | Ece1 | 0.059 | Young |
| 0.115292 | 0.412423 | 0.444 | 0.38 | 1 | 8 | Ttc3 | 0.064 | Young |
| 0.115591 | 0.206399 | 0.405 | 0.33 | 1 | 8 | Copb2 | 0.075 | Young |
| 0.115898 | 0.247993 | 0.374 | 0.307 | 1 | 8 | Psmb2 | 0.067 | Young |
| 0.115946 | 0.111669 | 0.803 | 0.715 | 1 | 8 | Ccdc88a | 0.088 | Young |
| 0.116062 | 1.719107 | 0.034 | 0.011 | 1 | 8 | D130040H: | 0.023 | Young |
| 0.116247 | 0.694416 | 0.153 | 0.106 | 1 | 8 | Yif1a | 0.047 | Young |
| 0.116352 | 0.169163 | 0.143 | 0.095 | 1 | 8 | Irgm1 | 0.048 | Young |
| 0.116352 | 0.203011 | 0.338 | 0.279 | 1 | 8 | Mdm4 | 0.059 | Young |
| 0.116359 | 0.210873 | 0.621 | 0.553 | 1 | 8 | Vasp | 0.068 | Young |
| 0.116527 | 0.092221 | 0.901 | 0.821 | 1 | 8 | Clta | 0.08 | Young |
| 0.116798 | 0.220904 | 0.327 | 0.257 | 1 | 8 | Map3k2 | 0.07 | Young |

| | | | | | | | | |
|---|---|---|---|---|---|---|---|---|
| 0.116937 | 0.269843 | 0.171 | 0.117 | 1 | 8 | Sesn3 | 0.054 | Young |
| 0.116989 | 2.120733 | 0.034 | 0.011 | 1 | 8 | Plac8 | 0.023 | Young |
| 0.117003 | 0.215854 | 0.452 | 0.385 | 1 | 8 | Smim11 | 0.067 | Young |
| 0.117015 | 0.304864 | 0.356 | 0.285 | 1 | 8 | Abcf1 | 0.071 | Young |
| 0.11756 | 0.231159 | 0.249 | 0.184 | 1 | 8 | Zcchc2 | 0.065 | Young |
| 0.117699 | 0.566605 | 0.242 | 0.19 | 1 | 8 | Kbtbd2 | 0.052 | Young |
| 0.117777 | 0.68689 | 0.164 | 0.117 | 1 | 8 | Scaf1 | 0.047 | Young |
| 0.117881 | 0.089098 | 0.239 | 0.173 | 1 | 8 | Cpeb2 | 0.066 | Young |
| 0.118199 | 0.692401 | 0.164 | 0.117 | 1 | 8 | Car3 | 0.047 | Young |
| 0.118217 | 0.451181 | 0.205 | 0.151 | 1 | 8 | Pvr | 0.054 | Young |
| 0.118422 | 0.177197 | 0.335 | 0.257 | 1 | 8 | Spag7 | 0.078 | Young |
| 0.118581 | 0.407471 | 0.117 | 0.073 | 1 | 8 | Cdc42ep4 | 0.044 | Young |
| 0.118848 | 0.023947 | 0.634 | 0.514 | 1 | 8 | Prkar2a | 0.12 | Young |
| 0.119026 | 0.757585 | 0.081 | 0.045 | 1 | 8 | B4gat1 | 0.036 | Young |
| 0.119291 | 0.230292 | 0.234 | 0.173 | 1 | 8 | Bag6 | 0.061 | Young |
| 0.119425 | 0.138106 | 0.351 | 0.274 | 1 | 8 | Rbbp7 | 0.077 | Young |
| 0.119568 | 0.952805 | 0.112 | 0.073 | 1 | 8 | Gna15 | 0.039 | Young |
| 0.119599 | 0.375591 | 0.148 | 0.101 | 1 | 8 | Cib2 | 0.047 | Young |
| 0.119623 | 0.692303 | 0.088 | 0.05 | 1 | 8 | Gngt2 | 0.038 | Young |
| 0.12008 | 0.366509 | 0.117 | 0.073 | 1 | 8 | Nmi | 0.044 | Young |
| 0.120204 | 0.360353 | 0.13 | 0.084 | 1 | 8 | Jpt2 | 0.046 | Young |
| 0.120281 | 1.129167 | 0.034 | 0.011 | 1 | 8 | Ctps | 0.023 | Young |
| 0.120306 | 0.270295 | 0.294 | 0.229 | 1 | 8 | Pip5k1a | 0.065 | Young |
| 0.120352 | 0.397355 | 0.073 | 0.039 | 1 | 8 | Pdpr | 0.034 | Young |
| 0.120544 | 0.271717 | 0.26 | 0.207 | 1 | 8 | Vps18 | 0.053 | Young |
| 0.120749 | 0.152795 | 0.423 | 0.352 | 1 | 8 | Zranb2 | 0.071 | Young |
| 0.12076 | 0.180036 | 0.512 | 0.43 | 1 | 8 | Pkn1 | 0.082 | Young |
| 0.12088 | 0.344661 | 0.304 | 0.24 | 1 | 8 | Chka | 0.064 | Young |
| 0.121013 | 1.303538 | 0.042 | 0.017 | 1 | 8 | Bphl | 0.025 | Young |
| 0.121096 | 1.315096 | 0.057 | 0.028 | 1 | 8 | Csrp2 | 0.029 | Young |
| 0.121235 | 1.223066 | 0.034 | 0.011 | 1 | 8 | Sag | 0.023 | Young |
| 0.121235 | 1.11324 | 0.034 | 0.011 | 1 | 8 | Rpgr | 0.023 | Young |
| 0.121258 | 0.038257 | 0.257 | 0.19 | 1 | 8 | Xpo7 | 0.067 | Young |
| 0.121305 | 0.33 | 0.377 | 0.313 | 1 | 8 | Ddx50 | 0.064 | Young |
| 0.121305 | 0.37201 | 0.252 | 0.19 | 1 | 8 | Tmem88 | 0.062 | Young |
| 0.121623 | 0.490101 | 0.109 | 0.067 | 1 | 8 | Stk35 | 0.042 | Young |
| 0.121714 | 1.525258 | 0.034 | 0.011 | 1 | 8 | Mad2l1 | 0.023 | Young |
| 0.121714 | 1.333157 | 0.034 | 0.011 | 1 | 8 | Tmem86b | 0.023 | Young |
| 0.121895 | 0.431511 | 0.151 | 0.106 | 1 | 8 | Inafm1 | 0.045 | Young |
| 0.121973 | 0.631022 | 0.231 | 0.179 | 1 | 8 | Sar1b | 0.052 | Young |
| 0.122055 | 0.187293 | 0.488 | 0.397 | 1 | 8 | Sec61a1 | 0.091 | Young |
| 0.122195 | 1.060223 | 0.034 | 0.011 | 1 | 8 | Klhl13 | 0.023 | Young |
| 0.122225 | 0.078449 | 0.896 | 0.844 | 1 | 8 | Gas6 | 0.052 | Young |
| 0.122259 | 0.121826 | 0.304 | 0.235 | 1 | 8 | Aagab | 0.069 | Young |
| 0.122267 | 0.425524 | 0.161 | 0.112 | 1 | 8 | Scaf4 | 0.049 | Young |

| | | | | | | | | |
|---|---|---|---|---|---|---|---|---|
| 0.122506 | 0.210003 | 0.46 | 0.374 | 1 | 8 | Pet100 | 0.086 | Young |
| 0.122677 | 1.104601 | 0.034 | 0.011 | 1 | 8 | B3gnt3 | 0.023 | Young |
| 0.12282 | 0.717168 | 0.119 | 0.078 | 1 | 8 | Cdr2 | 0.041 | Young |
| 0.123405 | 0.617047 | 0.135 | 0.089 | 1 | 8 | Gm38399 | 0.046 | Young |
| 0.12352 | 0.149176 | 0.462 | 0.385 | 1 | 8 | Dnajc8 | 0.077 | Young |
| 0.123755 | 0.411207 | 0.151 | 0.101 | 1 | 8 | Stx17 | 0.05 | Young |
| 0.123912 | 0.233658 | 0.47 | 0.391 | 1 | 8 | Arid4a | 0.079 | Young |
| 0.123931 | 0.260456 | 0.483 | 0.397 | 1 | 8 | Ubxn4 | 0.086 | Young |
| 0.124004 | 1.238201 | 0.057 | 0.028 | 1 | 8 | AU022252 | 0.029 | Young |
| 0.124011 | 0.395499 | 0.208 | 0.156 | 1 | 8 | Zmpste24 | 0.052 | Young |
| 0.124164 | 0.561797 | 0.187 | 0.14 | 1 | 8 | Selenbp1 | 0.047 | Young |
| 0.124177 | 0.14965 | 0.784 | 0.704 | 1 | 8 | Tpr | 0.08 | Young |
| 0.124204 | 0.821766 | 0.073 | 0.039 | 1 | 8 | Plk4 | 0.034 | Young |
| 0.12444 | 0.184326 | 0.356 | 0.291 | 1 | 8 | Tyk2 | 0.065 | Young |
| 0.124444 | 0.542215 | 0.164 | 0.117 | 1 | 8 | Tgfbr1 | 0.047 | Young |
| 0.124468 | 0.932812 | 0.065 | 0.034 | 1 | 8 | Rnf135 | 0.031 | Young |
| 0.124512 | 0.389706 | 0.171 | 0.123 | 1 | 8 | Ing3 | 0.048 | Young |
| 0.124522 | 0.830297 | 0.161 | 0.117 | 1 | 8 | Rnf167 | 0.044 | Young |
| 0.124652 | 1.041748 | 0.086 | 0.05 | 1 | 8 | Tlcd2 | 0.036 | Young |
| 0.124862 | 0.115771 | 0.452 | 0.38 | 1 | 8 | Pacsin2 | 0.072 | Young |
| 0.124974 | 1.509668 | 0.049 | 0.022 | 1 | 8 | Zfp81 | 0.027 | Young |
| 0.1254 | 0.266031 | 0.387 | 0.318 | 1 | 8 | Nbeal1 | 0.069 | Young |
| 0.125494 | 0.25862 | 0.384 | 0.313 | 1 | 8 | Clcn4 | 0.071 | Young |
| 0.125719 | 0.280103 | 0.551 | 0.48 | 1 | 8 | Ecm1 | 0.071 | Young |
| 0.12577 | 1.717219 | 0.042 | 0.017 | 1 | 8 | 2700038G2 | 0.025 | Young |
| 0.125797 | 0.155618 | 0.543 | 0.441 | 1 | 8 | Mt2 | 0.102 | Young |
| 0.125959 | 0.528296 | 0.101 | 0.061 | 1 | 8 | Mib2 | 0.04 | Young |
| 0.126082 | 0.540078 | 0.14 | 0.095 | 1 | 8 | Ppard | 0.045 | Young |
| 0.126174 | 1.094664 | 0.049 | 0.022 | 1 | 8 | Gpatch4 | 0.027 | Young |
| 0.126253 | 0.308658 | 0.356 | 0.285 | 1 | 8 | Arid1b | 0.071 | Young |
| 0.126412 | 1.961984 | 0.013 | 0 | 1 | 8 | Gm16754 | 0.013 | Young |
| 0.126412 | 1.712406 | 0.013 | 0 | 1 | 8 | Fam221a | 0.013 | Young |
| 0.126412 | 1.639166 | 0.013 | 0 | 1 | 8 | Nccrp1 | 0.013 | Young |
| 0.126412 | 1.632337 | 0.013 | 0 | 1 | 8 | Arhgap29 | 0.013 | Young |
| 0.126412 | 1.629579 | 0.013 | 0 | 1 | 8 | Uggt2 | 0.013 | Young |
| 0.126412 | 1.619747 | 0.013 | 0 | 1 | 8 | Kdm8 | 0.013 | Young |
| 0.126412 | 1.585603 | 0.013 | 0 | 1 | 8 | Dus4l | 0.013 | Young |
| 0.126412 | 1.569596 | 0.013 | 0 | 1 | 8 | 4933406P0 | 0.013 | Young |
| 0.126412 | 1.553021 | 0.013 | 0 | 1 | 8 | Mycl | 0.013 | Young |
| 0.126412 | 1.498805 | 0.013 | 0 | 1 | 8 | Gm19325 | 0.013 | Young |
| 0.126412 | 1.464805 | 0.013 | 0 | 1 | 8 | Ahcy | 0.013 | Young |
| 0.126412 | 1.43076 | 0.013 | 0 | 1 | 8 | Nrbp2 | 0.013 | Young |
| 0.126412 | 1.42421 | 0.013 | 0 | 1 | 8 | Catsper2 | 0.013 | Young |
| 0.126412 | 1.404905 | 0.013 | 0 | 1 | 8 | Tnnt3 | 0.013 | Young |
| 0.126412 | 1.384465 | 0.013 | 0 | 1 | 8 | Prima1 | 0.013 | Young |

| | | | | | | | | |
|---|---|---|---|---|---|---|---|---|
| 0.126412 | 1.374295 | 0.013 | 0 | 1 | 8 | Gm47709 | 0.013 | Young |
| 0.126412 | 1.374191 | 0.013 | 0 | 1 | 8 | Plat | 0.013 | Young |
| 0.126412 | 1.365121 | 0.013 | 0 | 1 | 8 | Zfp940 | 0.013 | Young |
| 0.126412 | 1.359502 | 0.013 | 0 | 1 | 8 | AI427809 | 0.013 | Young |
| 0.126412 | 1.338259 | 0.013 | 0 | 1 | 8 | Cdca7 | 0.013 | Young |
| 0.126412 | 1.334491 | 0.013 | 0 | 1 | 8 | 1700010I14 | 0.013 | Young |
| 0.126412 | 1.328948 | 0.013 | 0 | 1 | 8 | Dclk2 | 0.013 | Young |
| 0.126412 | 1.319091 | 0.013 | 0 | 1 | 8 | Frmd5 | 0.013 | Young |
| 0.126412 | 1.316859 | 0.013 | 0 | 1 | 8 | Zfp869 | 0.013 | Young |
| 0.126412 | 1.311653 | 0.013 | 0 | 1 | 8 | Xkr8 | 0.013 | Young |
| 0.126412 | 1.295051 | 0.013 | 0 | 1 | 8 | Tjp1 | 0.013 | Young |
| 0.126412 | 1.284489 | 0.013 | 0 | 1 | 8 | Lrrc3 | 0.013 | Young |
| 0.126412 | 1.281219 | 0.013 | 0 | 1 | 8 | Slc29a2 | 0.013 | Young |
| 0.126412 | 1.262744 | 0.013 | 0 | 1 | 8 | Rflnb | 0.013 | Young |
| 0.126412 | 1.242311 | 0.013 | 0 | 1 | 8 | Proser2 | 0.013 | Young |
| 0.126412 | 1.239219 | 0.013 | 0 | 1 | 8 | Rilpl1 | 0.013 | Young |
| 0.126412 | 1.238514 | 0.013 | 0 | 1 | 8 | Oas1g | 0.013 | Young |
| 0.126412 | 1.215076 | 0.013 | 0 | 1 | 8 | Gm45238 | 0.013 | Young |
| 0.126412 | 1.177501 | 0.013 | 0 | 1 | 8 | Gm45250 | 0.013 | Young |
| 0.126412 | 1.167138 | 0.013 | 0 | 1 | 8 | Tbc1d2 | 0.013 | Young |
| 0.126412 | 1.163539 | 0.013 | 0 | 1 | 8 | C130074G1 | 0.013 | Young |
| 0.126412 | 1.157788 | 0.013 | 0 | 1 | 8 | 4732471J0 | 0.013 | Young |
| 0.126412 | 1.131971 | 0.013 | 0 | 1 | 8 | Rapsn | 0.013 | Young |
| 0.12654 | 0.211773 | 0.416 | 0.341 | 1 | 8 | Emp1 | 0.075 | Young |
| 0.126549 | 0.900688 | 0.065 | 0.034 | 1 | 8 | Retsat | 0.031 | Young |
| 0.126624 | 0.195136 | 0.319 | 0.251 | 1 | 8 | Dusp16 | 0.068 | Young |
| 0.126651 | 0.23641 | 0.306 | 0.246 | 1 | 8 | Tmem173 | 0.06 | Young |
| 0.126824 | 0.415317 | 0.073 | 0.039 | 1 | 8 | Rpain | 0.034 | Young |
| 0.12684 | 0.121909 | 0.455 | 0.374 | 1 | 8 | Psmb5 | 0.081 | Young |
| 0.12686 | 0.092129 | 0.753 | 0.648 | 1 | 8 | Atrx | 0.105 | Young |
| 0.127159 | 0.249767 | 0.231 | 0.168 | 1 | 8 | Fez2 | 0.063 | Young |
| 0.127425 | 0.560971 | 0.239 | 0.184 | 1 | 8 | Gtf2i | 0.055 | Young |
| 0.127511 | 0.148409 | 0.691 | 0.592 | 1 | 8 | Igf1 | 0.099 | Young |
| 0.127535 | 1.39083 | 0.042 | 0.017 | 1 | 8 | Megf9 | 0.025 | Young |
| 0.127535 | 1.027769 | 0.042 | 0.017 | 1 | 8 | Tnfsfm13 | 0.025 | Young |
| 0.1276 | 0.591133 | 0.065 | 0.034 | 1 | 8 | Strada | 0.031 | Young |
| 0.127951 | 0.750314 | 0.065 | 0.034 | 1 | 8 | Znhit6 | 0.031 | Young |
| 0.12798 | 1.039409 | 0.042 | 0.017 | 1 | 8 | Adora3 | 0.025 | Young |
| 0.128199 | 0.39319 | 0.169 | 0.117 | 1 | 8 | Rxrb | 0.052 | Young |
| 0.128521 | 0.154322 | 0.416 | 0.352 | 1 | 8 | Mcfd2 | 0.064 | Young |
| 0.128567 | 0.121581 | 0.595 | 0.503 | 1 | 8 | Tlr2 | 0.092 | Young |
| 0.128727 | 0.554422 | 0.14 | 0.095 | 1 | 8 | Nckipsd | 0.045 | Young |
| 0.12878 | 0.553988 | 0.166 | 0.117 | 1 | 8 | Urm1 | 0.049 | Young |
| 0.129008 | 0.780529 | 0.049 | 0.022 | 1 | 8 | Aurkb | 0.027 | Young |
| 0.129316 | 0.568298 | 0.216 | 0.168 | 1 | 8 | Epb41 | 0.048 | Young |

| | | | | | | | | |
|---|---|---|---|---|---|---|---|---|
| 0.129333 | 0.069106 | 0.229 | 0.168 | 1 | 8 | Ylpm1 | 0.061 | Young |
| 0.129456 | 0.137317 | 0.312 | 0.24 | 1 | 8 | Maea | 0.072 | Young |
| 0.129583 | 0.475193 | 0.216 | 0.162 | 1 | 8 | 9930021J0 | 0.054 | Young |
| 0.129827 | 1.135864 | 0.049 | 0.022 | 1 | 8 | Rad9a | 0.027 | Young |
| 0.129981 | 0.031577 | 0.756 | 0.654 | 1 | 8 | Anp32b | 0.102 | Young |
| 0.129991 | 0.113598 | 0.462 | 0.391 | 1 | 8 | Elmsan1 | 0.071 | Young |
| 0.13017 | 0.071429 | 0.081 | 0.045 | 1 | 8 | Akr1c18 | 0.036 | Young |
| 0.130378 | 0.395749 | 0.205 | 0.151 | 1 | 8 | Sipa1l3 | 0.054 | Young |
| 0.130527 | 0.989391 | 0.078 | 0.045 | 1 | 8 | Cradd | 0.033 | Young |
| 0.130819 | 0.121309 | 0.777 | 0.715 | 1 | 8 | Mrpl33 | 0.062 | Young |
| 0.130848 | 0.231157 | 0.247 | 0.19 | 1 | 8 | Trappc6a | 0.057 | Young |
| 0.131151 | 0.718596 | 0.065 | 0.034 | 1 | 8 | Acvr1b | 0.031 | Young |
| 0.131156 | 0.382614 | 0.2 | 0.145 | 1 | 8 | Mgst1 | 0.055 | Young |
| 0.131415 | 0.497915 | 0.14 | 0.095 | 1 | 8 | Kansl1l | 0.045 | Young |
| 0.131471 | 0.653122 | 0.213 | 0.162 | 1 | 8 | Prr14 | 0.051 | Young |
| 0.131506 | 0.81868 | 0.138 | 0.095 | 1 | 8 | Tef | 0.043 | Young |
| 0.131644 | 0.085608 | 0.717 | 0.598 | 1 | 8 | Rnf13 | 0.119 | Young |
| 0.13175 | 0.179577 | 0.164 | 0.112 | 1 | 8 | Wipi2 | 0.052 | Young |
| 0.131779 | 0.349193 | 0.081 | 0.045 | 1 | 8 | Ift27 | 0.036 | Young |
| 0.13187 | 0.487434 | 0.065 | 0.034 | 1 | 8 | Frat2 | 0.031 | Young |
| 0.132288 | 0.066699 | 0.481 | 0.402 | 1 | 8 | Sptbn1 | 0.079 | Young |
| 0.132307 | 0.797069 | 0.049 | 0.022 | 1 | 8 | Abhd14b | 0.027 | Young |
| 0.132354 | 0.092668 | 0.255 | 0.19 | 1 | 8 | Filip1l | 0.065 | Young |
| 0.132516 | 0.188658 | 0.795 | 0.715 | 1 | 8 | Brk1 | 0.08 | Young |
| 0.132536 | 0.336425 | 0.073 | 0.039 | 1 | 8 | Mapk7 | 0.034 | Young |
| 0.132868 | 0.457214 | 0.205 | 0.156 | 1 | 8 | Dctn5 | 0.049 | Young |
| 0.133045 | 0.301009 | 0.4 | 0.352 | 1 | 8 | R3hdm2 | 0.048 | Young |
| 0.133072 | 0.157762 | 0.187 | 0.134 | 1 | 8 | Ctr9 | 0.053 | Young |
| 0.133094 | 0.346127 | 0.309 | 0.251 | 1 | 8 | Ei24 | 0.058 | Young |
| 0.133141 | 0.99575 | 0.122 | 0.084 | 1 | 8 | Bap1 | 0.038 | Young |
| 0.133142 | 1.073939 | 0.049 | 0.022 | 1 | 8 | Dnajb5 | 0.027 | Young |
| 0.133142 | 0.841526 | 0.049 | 0.022 | 1 | 8 | Mturn | 0.027 | Young |
| 0.133478 | 1.027909 | 0.078 | 0.045 | 1 | 8 | Kcne3 | 0.033 | Young |
| 0.133682 | 0.672728 | 0.065 | 0.034 | 1 | 8 | Rbm43 | 0.031 | Young |
| 0.133798 | 0.691801 | 0.21 | 0.162 | 1 | 8 | Med21 | 0.048 | Young |
| 0.13383 | 0.604557 | 0.057 | 0.028 | 1 | 8 | Slc19a1 | 0.029 | Young |
| 0.133883 | 0.206617 | 0.595 | 0.531 | 1 | 8 | Stap1 | 0.064 | Young |
| 0.133894 | 0.751019 | 0.094 | 0.056 | 1 | 8 | Car5b | 0.038 | Young |
| 0.133965 | 0.480929 | 0.114 | 0.073 | 1 | 8 | Kif1bp | 0.041 | Young |
| 0.133981 | 0.7931 | 0.049 | 0.022 | 1 | 8 | Zswim7 | 0.027 | Young |
| 0.134022 | 0.428217 | 0.135 | 0.089 | 1 | 8 | 2310033P0 | 0.046 | Young |
| 0.134023 | 1.718482 | 0.023 | 0.006 | 1 | 8 | Fhit | 0.017 | Young |
| 0.134184 | 0.58794 | 0.132 | 0.089 | 1 | 8 | Kif21b | 0.043 | Young |
| 0.134204 | 0.766619 | 0.086 | 0.05 | 1 | 8 | L1cam | 0.036 | Young |
| 0.134321 | 0.148744 | 0.351 | 0.296 | 1 | 8 | Nhp2 | 0.055 | Young |

| | | | | | | | | |
|---|---|---|---|---|---|---|---|---|
| 0.134328 | 0.617097 | 0.042 | 0.017 | 1 | 8 | Ap5b1 | 0.025 | Young |
| 0.134364 | 0.432386 | 0.203 | 0.151 | 1 | 8 | Nelfa | 0.052 | Young |
| 0.134423 | 0.387326 | 0.21 | 0.162 | 1 | 8 | Mink1 | 0.048 | Young |
| 0.134521 | 0.749616 | 0.086 | 0.05 | 1 | 8 | Ikbkg | 0.036 | Young |
| 0.134642 | 0.501071 | 0.226 | 0.173 | 1 | 8 | Kansl3 | 0.053 | Young |
| 0.134655 | 1.784883 | 0.023 | 0.006 | 1 | 8 | Rmdn2 | 0.017 | Young |
| 0.134655 | 1.60743 | 0.023 | 0.006 | 1 | 8 | Sox12 | 0.017 | Young |
| 0.134655 | 1.466095 | 0.023 | 0.006 | 1 | 8 | Gm4070 | 0.017 | Young |
| 0.135011 | 0.495008 | 0.143 | 0.095 | 1 | 8 | Decr1 | 0.048 | Young |
| 0.135013 | 0.126519 | 0.164 | 0.112 | 1 | 8 | Ctns | 0.052 | Young |
| 0.135119 | 0.024008 | 0.291 | 0.223 | 1 | 8 | Epc2 | 0.068 | Young |
| 0.135134 | 0.208721 | 0.33 | 0.263 | 1 | 8 | Cdc73 | 0.067 | Young |
| 0.135139 | 0.773496 | 0.078 | 0.045 | 1 | 8 | Lrch4 | 0.033 | Young |
| 0.135289 | 1.40118 | 0.023 | 0.006 | 1 | 8 | Nxpe3 | 0.017 | Young |
| 0.135289 | 1.274981 | 0.023 | 0.006 | 1 | 8 | Bbs12 | 0.017 | Young |
| 0.135414 | 0.115134 | 0.509 | 0.43 | 1 | 8 | Sbds | 0.079 | Young |
| 0.135433 | 0.333392 | 0.153 | 0.106 | 1 | 8 | Nom1 | 0.047 | Young |
| 0.135477 | 0.729753 | 0.086 | 0.05 | 1 | 8 | Pag1 | 0.036 | Young |
| 0.135579 | 0.166337 | 0.345 | 0.274 | 1 | 8 | Plcg2 | 0.071 | Young |
| 0.135902 | 0.314658 | 0.257 | 0.201 | 1 | 8 | Cir1 | 0.056 | Young |
| 0.135938 | 0.588242 | 0.187 | 0.14 | 1 | 8 | Tmem192 | 0.047 | Young |
| 0.136181 | 0.262885 | 0.384 | 0.313 | 1 | 8 | Map3k11 | 0.071 | Young |
| 0.136564 | 1.519744 | 0.023 | 0.006 | 1 | 8 | Pecr | 0.017 | Young |
| 0.136564 | 1.516234 | 0.023 | 0.006 | 1 | 8 | Myo1b | 0.017 | Young |
| 0.136564 | 1.457779 | 0.023 | 0.006 | 1 | 8 | Cldn5 | 0.017 | Young |
| 0.136564 | 1.456195 | 0.023 | 0.006 | 1 | 8 | 2510017J1 | 0.017 | Young |
| 0.136564 | 1.256383 | 0.023 | 0.006 | 1 | 8 | Fam187b | 0.017 | Young |
| 0.136564 | 1.092481 | 0.023 | 0.006 | 1 | 8 | Hist1h2ap | 0.017 | Young |
| 0.136584 | 0.425461 | 0.257 | 0.207 | 1 | 8 | 1810022KC | 0.05 | Young |
| 0.136599 | 0.087853 | 0.748 | 0.687 | 1 | 8 | Abracl | 0.061 | Young |
| 0.136601 | 0.114497 | 0.101 | 0.061 | 1 | 8 | Slc12a2 | 0.04 | Young |
| 0.13664 | 1.292131 | 0.07 | 0.039 | 1 | 8 | Ppox | 0.031 | Young |
| 0.13688 | 0.021801 | 0.358 | 0.285 | 1 | 8 | Trip11 | 0.073 | Young |
| 0.13695 | 0.938528 | 0.049 | 0.022 | 1 | 8 | Spp1 | 0.027 | Young |
| 0.137125 | 0.566076 | 0.156 | 0.112 | 1 | 8 | Chst12 | 0.044 | Young |
| 0.137205 | 1.083794 | 0.023 | 0.006 | 1 | 8 | 1110035H1 | 0.017 | Young |
| 0.13723 | 0.359541 | 0.164 | 0.117 | 1 | 8 | C2cd5 | 0.047 | Young |
| 0.13735 | 0.320189 | 0.306 | 0.251 | 1 | 8 | Rfk | 0.055 | Young |
| 0.137423 | 0.115465 | 0.397 | 0.307 | 1 | 8 | St3gal1 | 0.09 | Young |
| 0.137444 | 0.158957 | 0.519 | 0.447 | 1 | 8 | Yy1 | 0.072 | Young |
| 0.137474 | 0.761116 | 0.099 | 0.061 | 1 | 8 | Myef2 | 0.038 | Young |
| 0.137635 | 0.226001 | 0.486 | 0.419 | 1 | 8 | Baz2b | 0.067 | Young |
| 0.137767 | 0.28193 | 0.057 | 0.028 | 1 | 8 | Ttbk2 | 0.029 | Young |
| 0.137808 | 0.858411 | 0.049 | 0.022 | 1 | 8 | Sh2b2 | 0.027 | Young |
| 0.137848 | 1.40504 | 0.023 | 0.006 | 1 | 8 | Tmem177 | 0.017 | Young |

| | | | | | | | | |
|---|---|---|---|---|---|---|---|---|
| 0.137848 | 1.116785 | 0.023 | 0.006 | 1 | 8 | Zfp213 | 0.017 | Young |
| 0.137848 | 1.036088 | 0.023 | 0.006 | 1 | 8 | Gpt | 0.017 | Young |
| 0.137877 | 0.32945 | 0.19 | 0.145 | 1 | 8 | Shprh | 0.045 | Young |
| 0.138079 | 0.540781 | 0.132 | 0.089 | 1 | 8 | Cd72 | 0.043 | Young |
| 0.138253 | 0.189399 | 0.836 | 0.777 | 1 | 8 | Cdkn1a | 0.059 | Young |
| 0.138494 | 1.31057 | 0.023 | 0.006 | 1 | 8 | Dntt | 0.017 | Young |
| 0.138494 | 1.119812 | 0.023 | 0.006 | 1 | 8 | Gm39121 | 0.017 | Young |
| 0.138494 | 1.102832 | 0.023 | 0.006 | 1 | 8 | Npr1 | 0.017 | Young |
| 0.138494 | 1.073429 | 0.023 | 0.006 | 1 | 8 | Msrb3 | 0.017 | Young |
| 0.138494 | 1.068034 | 0.023 | 0.006 | 1 | 8 | Nrxn2 | 0.017 | Young |
| 0.138855 | 0.013913 | 0.634 | 0.547 | 1 | 8 | Anp32e | 0.087 | Young |
| 0.138865 | 0.156052 | 0.987 | 0.966 | 1 | 8 | Cd83 | 0.021 | Young |
| 0.139142 | 1.375965 | 0.023 | 0.006 | 1 | 8 | Phldb3 | 0.017 | Young |
| 0.139142 | 1.076141 | 0.023 | 0.006 | 1 | 8 | Zfp623 | 0.017 | Young |
| 0.139142 | 0.833994 | 0.023 | 0.006 | 1 | 8 | Zswim3 | 0.017 | Young |
| 0.139222 | 0.225609 | 0.317 | 0.251 | 1 | 8 | Kmt5a | 0.066 | Young |
| 0.139225 | 0.526638 | 0.14 | 0.095 | 1 | 8 | Prep | 0.045 | Young |
| 0.139489 | 0.100085 | 0.34 | 0.279 | 1 | 8 | Baz2a | 0.061 | Young |
| 0.139494 | 0.129967 | 0.366 | 0.302 | 1 | 8 | Lsm5 | 0.064 | Young |
| 0.13956 | 0.304846 | 0.41 | 0.346 | 1 | 8 | Zcrb1 | 0.064 | Young |
| 0.139702 | 0.344979 | 0.275 | 0.223 | 1 | 8 | Ifi27 | 0.052 | Young |
| 0.139792 | 0.765554 | 0.023 | 0.006 | 1 | 8 | Disp1 | 0.017 | Young |
| 0.139792 | 0.56664 | 0.023 | 0.006 | 1 | 8 | Ccr7 | 0.017 | Young |
| 0.139792 | 0.474138 | 0.023 | 0.006 | 1 | 8 | Spryd4 | 0.017 | Young |
| 0.139881 | 0.189671 | 0.244 | 0.19 | 1 | 8 | Cldnd1 | 0.054 | Young |
| 0.139897 | 0.724792 | 0.099 | 0.061 | 1 | 8 | Pde8a | 0.038 | Young |
| 0.140137 | 0.592343 | 0.112 | 0.073 | 1 | 8 | Abtb1 | 0.039 | Young |
| 0.140162 | 0.089698 | 0.491 | 0.413 | 1 | 8 | Cyth1 | 0.078 | Young |
| 0.140376 | 0.151835 | 0.499 | 0.43 | 1 | 8 | Ndufs2 | 0.069 | Young |
| 0.140982 | 0.164557 | 0.683 | 0.564 | 1 | 8 | Sec11c | 0.119 | Young |
| 0.141248 | 0.048986 | 0.584 | 0.497 | 1 | 8 | Herc1 | 0.087 | Young |
| 0.141606 | 0.166647 | 0.384 | 0.313 | 1 | 8 | Naga | 0.071 | Young |
| 0.141627 | 0.330103 | 0.283 | 0.223 | 1 | 8 | Cdc37l1 | 0.06 | Young |
| 0.141751 | 0.410893 | 0.171 | 0.123 | 1 | 8 | Ceacam1 | 0.048 | Young |
| 0.142079 | 1.23195 | 0.122 | 0.084 | 1 | 8 | Gm20186 | 0.038 | Young |
| 0.142328 | 0.602095 | 0.132 | 0.089 | 1 | 8 | Rps19bp1 | 0.043 | Young |
| 0.142573 | 0.365522 | 0.327 | 0.274 | 1 | 8 | Ufsp2 | 0.053 | Young |
| 0.142629 | 0.007895 | 0.455 | 0.374 | 1 | 8 | Borcs6 | 0.081 | Young |
| 0.142773 | 0.825544 | 0.07 | 0.039 | 1 | 8 | Zeb1 | 0.031 | Young |
| 0.142994 | 0.03401 | 0.317 | 0.257 | 1 | 8 | Ip6k1 | 0.06 | Young |
| 0.143001 | 0.310928 | 0.291 | 0.235 | 1 | 8 | Micu1 | 0.056 | Young |
| 0.143021 | 0.178727 | 0.309 | 0.251 | 1 | 8 | Herpud2 | 0.058 | Young |
| 0.143186 | 0.165877 | 0.592 | 0.497 | 1 | 8 | Zcchc11 | 0.095 | Young |
| 0.143579 | 0.316354 | 0.2 | 0.145 | 1 | 8 | Mzt1 | 0.055 | Young |
| 0.144382 | 0.663278 | 0.106 | 0.067 | 1 | 8 | Msantd2 | 0.039 | Young |

| | | | | | | | | |
|---|---|---|---|---|---|---|---|---|
| 0.144547 | 0.140126 | 0.504 | 0.425 | 1 | 8 | Baz1b | 0.079 | Young |
| 0.144773 | 0.092208 | 0.842 | 0.726 | 1 | 8 | Pip4k2a | 0.116 | Young |
| 0.14487 | 0.291955 | 0.416 | 0.341 | 1 | 8 | Ubn1 | 0.075 | Young |
| 0.145211 | 0.094402 | 0.405 | 0.324 | 1 | 8 | Bcas2 | 0.081 | Young |
| 0.14531 | 0.547972 | 0.177 | 0.128 | 1 | 8 | Fut8 | 0.049 | Young |
| 0.145359 | 0.691461 | 0.07 | 0.039 | 1 | 8 | Ttll3 | 0.031 | Young |
| 0.14563 | 0.713229 | 0.104 | 0.067 | 1 | 8 | Nadk2 | 0.037 | Young |
| 0.145857 | 0.598354 | 0.143 | 0.101 | 1 | 8 | Trim65 | 0.042 | Young |
| 0.145897 | 0.107512 | 0.34 | 0.274 | 1 | 8 | Slc35c2 | 0.066 | Young |
| 0.14591 | 0.587806 | 0.255 | 0.207 | 1 | 8 | Tmem229b | 0.048 | Young |
| 0.145977 | 0.587679 | 0.195 | 0.151 | 1 | 8 | 2610037D0 | 0.044 | Young |
| 0.145992 | 0.248671 | 0.119 | 0.078 | 1 | 8 | Mbtps2 | 0.041 | Young |
| 0.146318 | 0.28713 | 0.125 | 0.084 | 1 | 8 | 9330160F1 | 0.041 | Young |
| 0.146548 | 0.134473 | 0.852 | 0.737 | 1 | 8 | Eps8 | 0.115 | Young |
| 0.146795 | 0.160561 | 0.522 | 0.464 | 1 | 8 | Tcp1 | 0.058 | Young |
| 0.147159 | 1.156036 | 0.062 | 0.034 | 1 | 8 | Lrrc20 | 0.028 | Young |
| 0.147187 | 0.611619 | 0.13 | 0.089 | 1 | 8 | Hint3 | 0.041 | Young |
| 0.147582 | 0.04227 | 0.306 | 0.235 | 1 | 8 | Vps41 | 0.071 | Young |
| 0.148192 | 0.383358 | 0.213 | 0.162 | 1 | 8 | Prkar2b | 0.051 | Young |
| 0.148736 | 0.6199 | 0.07 | 0.039 | 1 | 8 | Kctd3 | 0.031 | Young |
| 0.14883 | 0.175534 | 0.179 | 0.128 | 1 | 8 | Zfp397 | 0.051 | Young |
| 0.148948 | 0.352195 | 0.403 | 0.346 | 1 | 8 | Cyb5b | 0.057 | Young |
| 0.149041 | 1.334782 | 0.031 | 0.011 | 1 | 8 | Armcx2 | 0.02 | Young |
| 0.149211 | 0.051203 | 0.665 | 0.57 | 1 | 8 | Cep170 | 0.095 | Young |
| 0.149376 | 0.771288 | 0.083 | 0.05 | 1 | 8 | 5730455P1 | 0.033 | Young |
| 0.14946 | 0.485839 | 0.091 | 0.056 | 1 | 8 | Naa16 | 0.035 | Young |
| 0.149564 | 0.911235 | 0.062 | 0.034 | 1 | 8 | Ppm1f | 0.028 | Young |
| 0.149644 | 1.14949 | 0.055 | 0.028 | 1 | 8 | 1700019L1 | 0.027 | Young |
| 0.149856 | 0.305923 | 0.216 | 0.162 | 1 | 8 | Cab39l | 0.054 | Young |
| 0.149916 | 0.043409 | 0.179 | 0.123 | 1 | 8 | Tbcel | 0.056 | Young |
| 0.149967 | 1.028258 | 0.062 | 0.034 | 1 | 8 | Batf2 | 0.028 | Young |
| 0.150018 | 0.6269 | 0.122 | 0.084 | 1 | 8 | Cwc25 | 0.038 | Young |
| 0.150205 | 1.7562 | 0.031 | 0.011 | 1 | 8 | Cdyl2 | 0.02 | Young |
| 0.150759 | 0.21479 | 0.195 | 0.145 | 1 | 8 | Gpd1l | 0.05 | Young |
| 0.150779 | 0.169415 | 0.722 | 0.676 | 1 | 8 | Wipf1 | 0.046 | Young |
| 0.15079 | 1.509949 | 0.031 | 0.011 | 1 | 8 | Btnl2 | 0.02 | Young |
| 0.15079 | 1.431384 | 0.031 | 0.011 | 1 | 8 | Zfp768 | 0.02 | Young |
| 0.150909 | 0.273863 | 0.423 | 0.374 | 1 | 8 | Osgep | 0.049 | Young |
| 0.151392 | 0.330778 | 0.221 | 0.168 | 1 | 8 | Wdr13 | 0.053 | Young |
| 0.151492 | 0.384087 | 0.21 | 0.162 | 1 | 8 | Ppm1k | 0.048 | Young |
| 0.151536 | 0.72504 | 0.099 | 0.061 | 1 | 8 | Leo1 | 0.038 | Young |
| 0.151777 | 0.185215 | 0.338 | 0.274 | 1 | 8 | Mrpl36 | 0.064 | Young |
| 0.151909 | 0.084649 | 0.535 | 0.447 | 1 | 8 | Cpsf6 | 0.088 | Young |
| 0.151926 | 0.177757 | 0.525 | 0.453 | 1 | 8 | Tmem234 | 0.072 | Young |
| 0.151964 | 1.278011 | 0.031 | 0.011 | 1 | 8 | Rsl1 | 0.02 | Young |

| | | | | | | | | |
|---|---|---|---|---|---|---|---|---|
| 0.152391 | 0.324112 | 0.247 | 0.196 | 1 | 8 | Phf20 | 0.051 | Young |
| 0.152647 | 0.149232 | 0.218 | 0.162 | 1 | 8 | Atl2 | 0.056 | Young |
| 0.152907 | 1.879849 | 0.039 | 0.017 | 1 | 8 | Hexdc | 0.022 | Young |
| 0.152983 | 0.232197 | 0.26 | 0.201 | 1 | 8 | 4833439L1 | 0.059 | Young |
| 0.153287 | 0.28349 | 0.156 | 0.112 | 1 | 8 | Ttc1 | 0.044 | Young |
| 0.153739 | 1.193853 | 0.031 | 0.011 | 1 | 8 | Col4a2 | 0.02 | Young |
| 0.153741 | 0.417304 | 0.169 | 0.123 | 1 | 8 | Nomo1 | 0.046 | Young |
| 0.153884 | 0.551243 | 0.151 | 0.106 | 1 | 8 | Pja1 | 0.045 | Young |
| 0.15405 | 0.707156 | 0.062 | 0.034 | 1 | 8 | Aasdh | 0.028 | Young |
| 0.154177 | 0.185426 | 0.688 | 0.615 | 1 | 8 | Ndufb9 | 0.073 | Young |
| 0.15421 | 0.571059 | 0.143 | 0.101 | 1 | 8 | Ube2l6 | 0.042 | Young |
| 0.154334 | 1.502049 | 0.031 | 0.011 | 1 | 8 | Mllt3 | 0.02 | Young |
| 0.154349 | 0.490435 | 0.164 | 0.117 | 1 | 8 | Slc25a46 | 0.047 | Young |
| 0.154363 | 0.507895 | 0.216 | 0.168 | 1 | 8 | Snx15 | 0.048 | Young |
| 0.154379 | 0.027505 | 0.997 | 1 | 1 | 8 | Mrc1 | -0.003 | Young |
| 0.154779 | 0.225067 | 0.247 | 0.19 | 1 | 8 | Bag3 | 0.057 | Young |
| 0.154792 | 0.1091 | 0.616 | 0.542 | 1 | 8 | Cd68 | 0.074 | Young |
| 0.154941 | 0.158716 | 0.33 | 0.274 | 1 | 8 | Taz | 0.056 | Young |
| 0.15507 | 0.386753 | 0.364 | 0.318 | 1 | 8 | Adipor2 | 0.046 | Young |
| 0.155116 | 0.326244 | 0.13 | 0.089 | 1 | 8 | Lonp1 | 0.041 | Young |
| 0.155137 | 0.177887 | 0.494 | 0.441 | 1 | 8 | Stard3nl | 0.053 | Young |
| 0.155177 | 1.141006 | 0.109 | 0.073 | 1 | 8 | Cbx6 | 0.036 | Young |
| 0.155244 | 0.269895 | 0.312 | 0.263 | 1 | 8 | Mllt10 | 0.049 | Young |
| 0.155277 | 0.224135 | 0.319 | 0.251 | 1 | 8 | Zzz3 | 0.068 | Young |
| 0.155305 | 0.049084 | 0.813 | 0.743 | 1 | 8 | Fnbp1 | 0.07 | Young |
| 0.155529 | 1.31526 | 0.031 | 0.011 | 1 | 8 | Adrb1 | 0.02 | Young |
| 0.155529 | 1.04326 | 0.031 | 0.011 | 1 | 8 | Hist3h2ba | 0.02 | Young |
| 0.155771 | 0.133785 | 0.818 | 0.749 | 1 | 8 | Sod1 | 0.069 | Young |
| 0.155783 | 1.180805 | 0.055 | 0.028 | 1 | 8 | Phc1 | 0.027 | Young |
| 0.155795 | 0.090237 | 0.844 | 0.788 | 1 | 8 | Foxp1 | 0.056 | Young |
| 0.155851 | 0.781335 | 0.135 | 0.095 | 1 | 8 | Apeh | 0.04 | Young |
| 0.155852 | 0.364633 | 0.125 | 0.084 | 1 | 8 | Coa6 | 0.041 | Young |
| 0.155984 | 0.460084 | 0.13 | 0.089 | 1 | 8 | Rnf215 | 0.041 | Young |
| 0.156119 | 0.656506 | 0.109 | 0.073 | 1 | 8 | Arid3b | 0.036 | Young |
| 0.156122 | 0.663602 | 0.062 | 0.034 | 1 | 8 | Apba3 | 0.028 | Young |
| 0.15613 | 1.395151 | 0.031 | 0.011 | 1 | 8 | Rsph3b | 0.02 | Young |
| 0.15613 | 0.951048 | 0.031 | 0.011 | 1 | 8 | Dnajc17 | 0.02 | Young |
| 0.156328 | 0.742151 | 0.104 | 0.067 | 1 | 8 | Rnf214 | 0.037 | Young |
| 0.156369 | 0.258421 | 0.127 | 0.084 | 1 | 8 | Nde1 | 0.043 | Young |
| 0.156408 | 0.121887 | 0.494 | 0.413 | 1 | 8 | Lsm6 | 0.081 | Young |
| 0.156607 | 1.570201 | 0.039 | 0.017 | 1 | 8 | Gm49359 | 0.022 | Young |
| 0.156643 | 0.990315 | 0.075 | 0.045 | 1 | 8 | Gstp3 | 0.03 | Young |
| 0.156732 | 0.950426 | 0.031 | 0.011 | 1 | 8 | Gm4673 | 0.02 | Young |
| 0.156766 | 0.704614 | 0.184 | 0.14 | 1 | 8 | Chm | 0.044 | Young |
| 0.156838 | 0.208572 | 0.33 | 0.263 | 1 | 8 | Rnf6 | 0.067 | Young |

| | | | | | | | | |
|---|---|---|---|---|---|---|---|---|
| 0.157205 | 0.291308 | 0.405 | 0.352 | 1 | 8 | Acadm | 0.053 | Young |
| 0.157336 | 0.624136 | 0.031 | 0.011 | 1 | 8 | Pclaf | 0.02 | Young |
| 0.157615 | 0.286581 | 0.366 | 0.318 | 1 | 8 | Lemd2 | 0.048 | Young |
| 0.157762 | 0.672822 | 0.184 | 0.14 | 1 | 8 | Mrpl58 | 0.044 | Young |
| 0.157901 | 0.118172 | 0.73 | 0.682 | 1 | 8 | Cnn2 | 0.048 | Young |
| 0.158213 | 1.248024 | 0.039 | 0.017 | 1 | 8 | Nmrk1 | 0.022 | Young |
| 0.158216 | 0.161727 | 0.338 | 0.274 | 1 | 8 | C1qbp | 0.064 | Young |
| 0.158471 | 0.910844 | 0.055 | 0.028 | 1 | 8 | Usp30 | 0.027 | Young |
| 0.158474 | 0.191992 | 0.545 | 0.475 | 1 | 8 | Kpna4 | 0.07 | Young |
| 0.158549 | 0.92645 | 0.031 | 0.011 | 1 | 8 | Mks1 | 0.02 | Young |
| 0.158549 | 0.700691 | 0.031 | 0.011 | 1 | 8 | Zbtb8a | 0.02 | Young |
| 0.158751 | 1.688711 | 0.039 | 0.017 | 1 | 8 | Prr5 | 0.022 | Young |
| 0.158854 | 0.149219 | 0.242 | 0.184 | 1 | 8 | Sos2 | 0.058 | Young |
| 0.158901 | 0.348597 | 0.169 | 0.123 | 1 | 8 | Polr2b | 0.046 | Young |
| 0.158964 | 0.128151 | 0.642 | 0.575 | 1 | 8 | Eif4h | 0.067 | Young |
| 0.15923 | 0.798058 | 0.119 | 0.084 | 1 | 8 | Aatf | 0.035 | Young |
| 0.159291 | 1.250739 | 0.039 | 0.017 | 1 | 8 | Tnfsf14 | 0.022 | Young |
| 0.159375 | 0.987222 | 0.055 | 0.028 | 1 | 8 | Eif5a2 | 0.027 | Young |
| 0.159375 | 0.675484 | 0.055 | 0.028 | 1 | 8 | Mlh3 | 0.027 | Young |
| 0.15961 | 0.374327 | 0.158 | 0.112 | 1 | 8 | Sh3glb2 | 0.046 | Young |
| 0.159769 | 0.991638 | 0.031 | 0.011 | 1 | 8 | Them6 | 0.02 | Young |
| 0.159769 | 0.728404 | 0.031 | 0.011 | 1 | 8 | Col4a1 | 0.02 | Young |
| 0.159777 | 0.189628 | 0.506 | 0.436 | 1 | 8 | Cuta | 0.07 | Young |
| 0.159889 | 0.197489 | 0.79 | 0.749 | 1 | 8 | Slamf9 | 0.041 | Young |
| 0.159906 | 0.571569 | 0.062 | 0.034 | 1 | 8 | Dcbld2 | 0.028 | Young |
| 0.159949 | 0.217139 | 0.319 | 0.263 | 1 | 8 | Rap2c | 0.056 | Young |
| 0.160361 | 0.024294 | 0.226 | 0.168 | 1 | 8 | Sart1 | 0.058 | Young |
| 0.160378 | 0.412558 | 0.299 | 0.251 | 1 | 8 | Pink1 | 0.048 | Young |
| 0.160433 | 0.083977 | 0.543 | 0.453 | 1 | 8 | Strn3 | 0.09 | Young |
| 0.160537 | 0.209169 | 0.296 | 0.246 | 1 | 8 | Timm8b | 0.05 | Young |
| 0.160582 | 0.183704 | 0.429 | 0.341 | 1 | 8 | Crebrf | 0.088 | Young |
| 0.160648 | 0.139473 | 0.483 | 0.425 | 1 | 8 | Trim8 | 0.058 | Young |
| 0.160739 | 0.617344 | 0.055 | 0.028 | 1 | 8 | Dis3l | 0.027 | Young |
| 0.160918 | 1.216141 | 0.039 | 0.017 | 1 | 8 | F830016BC | 0.022 | Young |
| 0.160974 | 0.084346 | 0.299 | 0.235 | 1 | 8 | Ndfip2 | 0.064 | Young |
| 0.161195 | 0.243649 | 0.055 | 0.028 | 1 | 8 | Rtn4rl1 | 0.027 | Young |
| 0.161223 | 1.234912 | 0.047 | 0.022 | 1 | 8 | Nme7 | 0.025 | Young |
| 0.161443 | 0.138668 | 0.803 | 0.726 | 1 | 8 | Rab14 | 0.077 | Young |
| 0.161543 | 0.179357 | 0.151 | 0.106 | 1 | 8 | Amz2 | 0.045 | Young |
| 0.161608 | 0.193267 | 0.397 | 0.335 | 1 | 8 | Piezo1 | 0.062 | Young |
| 0.161718 | 1.319106 | 0.047 | 0.022 | 1 | 8 | Klhl33 | 0.025 | Young |
| 0.161718 | 0.877874 | 0.047 | 0.022 | 1 | 8 | Aldh4a1 | 0.025 | Young |
| 0.161845 | 1.264497 | 0.068 | 0.039 | 1 | 8 | Snx7 | 0.029 | Young |
| 0.162184 | 0.313529 | 0.099 | 0.061 | 1 | 8 | Maip1 | 0.038 | Young |
| 0.162417 | 0.292044 | 0.226 | 0.179 | 1 | 8 | Zdhhc5 | 0.047 | Young |

| | | | | | | | | |
|---|---|---|---|---|---|---|---|---|
| 0.162497 | 0.177979 | 0.278 | 0.212 | 1 | 8 | Zfr | 0.066 | Young |
| 0.162731 | 0.388598 | 0.174 | 0.128 | 1 | 8 | Mta3 | 0.046 | Young |
| 0.162738 | 0.174196 | 0.452 | 0.38 | 1 | 8 | Gm26740 | 0.072 | Young |
| 0.162798 | 0.168795 | 0.203 | 0.151 | 1 | 8 | Nudt9 | 0.052 | Young |
| 0.162828 | 0.071742 | 0.621 | 0.542 | 1 | 8 | Nedd8 | 0.079 | Young |
| 0.16285 | 0.415496 | 0.031 | 0.011 | 1 | 8 | Wdr24 | 0.02 | Young |
| 0.163107 | 1.01441 | 0.039 | 0.017 | 1 | 8 | Zfp606 | 0.022 | Young |
| 0.163274 | 0.899389 | 0.169 | 0.128 | 1 | 8 | Zfp260 | 0.041 | Young |
| 0.163319 | 0.067013 | 0.444 | 0.352 | 1 | 8 | Cnih4 | 0.092 | Young |
| 0.163399 | 0.999605 | 0.088 | 0.056 | 1 | 8 | Npat | 0.032 | Young |
| 0.163471 | 0.452639 | 0.031 | 0.011 | 1 | 8 | Srfbp1 | 0.02 | Young |
| 0.163658 | 0.77462 | 0.039 | 0.017 | 1 | 8 | Glb1l | 0.022 | Young |
| 0.16389 | 0.307056 | 0.166 | 0.123 | 1 | 8 | Gga3 | 0.043 | Young |
| 0.163927 | 0.654192 | 0.138 | 0.101 | 1 | 8 | Def6 | 0.037 | Young |
| 0.163954 | 0.972809 | 0.055 | 0.028 | 1 | 8 | Acss2 | 0.027 | Young |
| 0.16397 | 0.240707 | 0.197 | 0.145 | 1 | 8 | Cetn2 | 0.052 | Young |
| 0.164108 | 0.178858 | 0.504 | 0.43 | 1 | 8 | Sri | 0.074 | Young |
| 0.164258 | 0.105643 | 0.626 | 0.542 | 1 | 8 | Tet3 | 0.084 | Young |
| 0.164349 | 0.455127 | 0.184 | 0.14 | 1 | 8 | Eef1e1 | 0.044 | Young |
| 0.164881 | 0.072589 | 0.325 | 0.257 | 1 | 8 | Nfatc2 | 0.068 | Young |
| 0.164978 | 0.067901 | 0.566 | 0.48 | 1 | 8 | Uvrag | 0.086 | Young |
| 0.164981 | 0.093769 | 0.392 | 0.33 | 1 | 8 | Rasa1 | 0.062 | Young |
| 0.165072 | 0.737529 | 0.158 | 0.117 | 1 | 8 | Rubcn | 0.041 | Young |
| 0.165122 | 0.613787 | 0.109 | 0.073 | 1 | 8 | Lrrc42 | 0.036 | Young |
| 0.165185 | 0.220893 | 0.301 | 0.235 | 1 | 8 | Fcgr1 | 0.066 | Young |
| 0.165244 | 0.286285 | 0.226 | 0.179 | 1 | 8 | Spast | 0.047 | Young |
| 0.165249 | 0.095085 | 0.462 | 0.38 | 1 | 8 | Tnrc6b | 0.082 | Young |
| 0.165563 | 0.100715 | 0.462 | 0.413 | 1 | 8 | Kpnb1 | 0.049 | Young |
| 0.165658 | 0.389773 | 0.281 | 0.235 | 1 | 8 | Tlr7 | 0.046 | Young |
| 0.165681 | 0.079684 | 0.623 | 0.531 | 1 | 8 | Emb | 0.092 | Young |
| 0.165724 | 0.828881 | 0.047 | 0.022 | 1 | 8 | Slx4 | 0.025 | Young |
| 0.165774 | 0.276304 | 0.091 | 0.056 | 1 | 8 | Gm26782 | 0.035 | Young |
| 0.165815 | 0.210706 | 0.192 | 0.14 | 1 | 8 | Fzr1 | 0.052 | Young |
| 0.16595 | 0.175173 | 0.434 | 0.369 | 1 | 8 | Slc15a3 | 0.065 | Young |
| 0.1661 | 0.077618 | 0.618 | 0.52 | 1 | 8 | Hook3 | 0.098 | Young |
| 0.16623 | 0.905106 | 0.047 | 0.022 | 1 | 8 | Vwf | 0.025 | Young |
| 0.166252 | 0.043841 | 0.104 | 0.067 | 1 | 8 | Cep290 | 0.037 | Young |
| 0.166259 | 0.642544 | 0.151 | 0.112 | 1 | 8 | Dtnbp1 | 0.039 | Young |
| 0.166298 | 0.218002 | 0.221 | 0.173 | 1 | 8 | Ncor2 | 0.048 | Young |
| 0.166395 | 1.135322 | 0.068 | 0.039 | 1 | 8 | Zc3h3 | 0.029 | Young |
| 0.166432 | 0.864829 | 0.039 | 0.017 | 1 | 8 | Nt5dc2 | 0.022 | Young |
| 0.166737 | 0.681201 | 0.047 | 0.022 | 1 | 8 | Platr25 | 0.025 | Young |
| 0.166783 | 0.114045 | 0.125 | 0.084 | 1 | 8 | Faap100 | 0.041 | Young |
| 0.167008 | 0.15417 | 0.366 | 0.318 | 1 | 8 | Utrn | 0.048 | Young |
| 0.167286 | 0.285193 | 0.423 | 0.363 | 1 | 8 | Zfp638 | 0.06 | Young |

| | | | | | | | | |
|---|---|---|---|---|---|---|---|---|
| 0.167486 | 0.217936 | 0.377 | 0.318 | 1 | 8 | Hps3 | 0.059 | Young |
| 0.167724 | 0.064358 | 0.984 | 0.961 | 1 | 8 | Pabpc1 | 0.023 | Young |
| 0.167778 | 0.014318 | 0.717 | 0.62 | 1 | 8 | Psme1 | 0.097 | Young |
| 0.168131 | 0.175897 | 0.332 | 0.263 | 1 | 8 | Pgrmc1 | 0.069 | Young |
| 0.168314 | 0.5327 | 0.091 | 0.056 | 1 | 8 | Dtx4 | 0.035 | Young |
| 0.168361 | 0.179703 | 0.117 | 0.078 | 1 | 8 | Gphn | 0.039 | Young |
| 0.168527 | 0.383624 | 0.249 | 0.201 | 1 | 8 | Espn | 0.048 | Young |
| 0.168578 | 0.346724 | 0.174 | 0.128 | 1 | 8 | Naa60 | 0.046 | Young |
| 0.168635 | 0.419794 | 0.104 | 0.067 | 1 | 8 | Nsun6 | 0.037 | Young |
| 0.168708 | 0.232394 | 0.138 | 0.095 | 1 | 8 | Gm43848 | 0.043 | Young |
| 0.168709 | 0.30959 | 0.392 | 0.335 | 1 | 8 | Stxbp6 | 0.057 | Young |
| 0.168736 | 0.401999 | 0.143 | 0.101 | 1 | 8 | Smim26 | 0.042 | Young |
| 0.168793 | 0.075511 | 0.33 | 0.263 | 1 | 8 | Pigx | 0.067 | Young |
| 0.168799 | 0.061093 | 0.67 | 0.564 | 1 | 8 | Hnrnpc | 0.106 | Young |
| 0.168927 | 0.350909 | 0.197 | 0.151 | 1 | 8 | Washc5 | 0.046 | Young |
| 0.169087 | 0.574292 | 0.083 | 0.05 | 1 | 8 | Stxbp1 | 0.033 | Young |
| 0.16909 | 0.288023 | 0.442 | 0.385 | 1 | 8 | Ppp4r3b | 0.057 | Young |
| 0.169156 | 0.358564 | 0.475 | 0.43 | 1 | 8 | Samd4b | 0.045 | Young |
| 0.169185 | 0.413734 | 0.265 | 0.218 | 1 | 8 | Zfp217 | 0.047 | Young |
| 0.16925 | 0.22583 | 0.486 | 0.408 | 1 | 8 | Tsc22d1 | 0.078 | Young |
| 0.169363 | 0.267965 | 0.096 | 0.061 | 1 | 8 | Hes6 | 0.035 | Young |
| 0.169856 | 0.137972 | 0.857 | 0.743 | 1 | 8 | Eef1g | 0.114 | Young |
| 0.169898 | 0.259369 | 0.348 | 0.296 | 1 | 8 | Fkbp15 | 0.052 | Young |
| 0.17007 | 0.155449 | 0.304 | 0.257 | 1 | 8 | Hlcs | 0.047 | Young |
| 0.170139 | 0.272387 | 0.265 | 0.207 | 1 | 8 | Cpq | 0.058 | Young |
| 0.170189 | 0.961102 | 0.068 | 0.039 | 1 | 8 | B230118H( | 0.029 | Young |
| 0.170315 | 0.663127 | 0.187 | 0.14 | 1 | 8 | Apba1 | 0.047 | Young |
| 0.170456 | 0.078736 | 0.483 | 0.408 | 1 | 8 | Rex1bd | 0.075 | Young |
| 0.170511 | 0.447911 | 0.143 | 0.101 | 1 | 8 | Rpia | 0.042 | Young |
| 0.170577 | 0.263139 | 0.101 | 0.067 | 1 | 8 | Ticam1 | 0.034 | Young |
| 0.170781 | 0.285529 | 0.062 | 0.034 | 1 | 8 | Klhl36 | 0.028 | Young |
| 0.170824 | 0.125032 | 0.338 | 0.268 | 1 | 8 | Efcab14 | 0.07 | Young |
| 0.171006 | 0.529282 | 0.055 | 0.028 | 1 | 8 | Pinx1 | 0.027 | Young |
| 0.171113 | 0.123111 | 0.109 | 0.073 | 1 | 8 | Tns4 | 0.036 | Young |
| 0.171467 | 0.641313 | 0.114 | 0.078 | 1 | 8 | Srxn1 | 0.036 | Young |
| 0.171635 | 0.210125 | 0.164 | 0.117 | 1 | 8 | Wasl | 0.047 | Young |
| 0.171831 | 0.144417 | 0.218 | 0.168 | 1 | 8 | Slc36a4 | 0.05 | Young |
| 0.171869 | 0.609319 | 0.096 | 0.061 | 1 | 8 | Kank2 | 0.035 | Young |
| 0.171888 | 0.281507 | 0.273 | 0.218 | 1 | 8 | Ephx1 | 0.055 | Young |
| 0.172086 | 0.34156 | 0.192 | 0.145 | 1 | 8 | Mfsd10 | 0.047 | Young |
| 0.172095 | 0.425887 | 0.182 | 0.14 | 1 | 8 | Sec16a | 0.042 | Young |
| 0.172112 | 1.966494 | 0.01 | 0 | 1 | 8 | Gm4262 | 0.01 | Young |
| 0.172112 | 1.805498 | 0.01 | 0 | 1 | 8 | Spag5 | 0.01 | Young |
| 0.172112 | 1.763472 | 0.01 | 0 | 1 | 8 | H2-M2 | 0.01 | Young |
| 0.172112 | 1.742806 | 0.01 | 0 | 1 | 8 | Abcb9 | 0.01 | Young |

| | | | | | | | | |
|---|---|---|---|---|---|---|---|---|
| 0.172112 | 1.741686 | 0.01 | 0 | 1 | 8 | Xkrx | 0.01 | Young |
| 0.172112 | 1.684976 | 0.01 | 0 | 1 | 8 | B230317F2 | 0.01 | Young |
| 0.172112 | 1.678754 | 0.01 | 0 | 1 | 8 | Gm34006 | 0.01 | Young |
| 0.172112 | 1.668064 | 0.01 | 0 | 1 | 8 | Cd34 | 0.01 | Young |
| 0.172112 | 1.607338 | 0.01 | 0 | 1 | 8 | Klk13 | 0.01 | Young |
| 0.172112 | 1.580589 | 0.01 | 0 | 1 | 8 | Swsap1 | 0.01 | Young |
| 0.172112 | 1.550237 | 0.01 | 0 | 1 | 8 | Meox2 | 0.01 | Young |
| 0.172112 | 1.547861 | 0.01 | 0 | 1 | 8 | Hdgfl3 | 0.01 | Young |
| 0.172112 | 1.546265 | 0.01 | 0 | 1 | 8 | Myb | 0.01 | Young |
| 0.172112 | 1.524128 | 0.01 | 0 | 1 | 8 | Efhd1 | 0.01 | Young |
| 0.172112 | 1.508809 | 0.01 | 0 | 1 | 8 | Pnp2 | 0.01 | Young |
| 0.172112 | 1.502759 | 0.01 | 0 | 1 | 8 | Itgb3 | 0.01 | Young |
| 0.172112 | 1.475334 | 0.01 | 0 | 1 | 8 | Zfr2 | 0.01 | Young |
| 0.172112 | 1.471382 | 0.01 | 0 | 1 | 8 | Ntng2 | 0.01 | Young |
| 0.172112 | 1.462776 | 0.01 | 0 | 1 | 8 | Hdc | 0.01 | Young |
| 0.172112 | 1.415888 | 0.01 | 0 | 1 | 8 | Pla1a | 0.01 | Young |
| 0.172112 | 1.400937 | 0.01 | 0 | 1 | 8 | Gm49207 | 0.01 | Young |
| 0.172112 | 1.391061 | 0.01 | 0 | 1 | 8 | Abhd3 | 0.01 | Young |
| 0.172112 | 1.37436 | 0.01 | 0 | 1 | 8 | Pcdh17 | 0.01 | Young |
| 0.172112 | 1.364559 | 0.01 | 0 | 1 | 8 | Zbtb3 | 0.01 | Young |
| 0.172112 | 1.35683 | 0.01 | 0 | 1 | 8 | Caskin2 | 0.01 | Young |
| 0.172112 | 1.336999 | 0.01 | 0 | 1 | 8 | Rab4b | 0.01 | Young |
| 0.172112 | 1.334896 | 0.01 | 0 | 1 | 8 | Il1bos | 0.01 | Young |
| 0.172112 | 1.333189 | 0.01 | 0 | 1 | 8 | Fcrla | 0.01 | Young |
| 0.172112 | 1.3328 | 0.01 | 0 | 1 | 8 | Nupr1l | 0.01 | Young |
| 0.172112 | 1.331585 | 0.01 | 0 | 1 | 8 | Gm10785 | 0.01 | Young |
| 0.172112 | 1.323423 | 0.01 | 0 | 1 | 8 | 2010015M | 0.01 | Young |
| 0.172112 | 1.317261 | 0.01 | 0 | 1 | 8 | Tnfsf10 | 0.01 | Young |
| 0.172112 | 1.304342 | 0.01 | 0 | 1 | 8 | Cacna2d4 | 0.01 | Young |
| 0.172112 | 1.301918 | 0.01 | 0 | 1 | 8 | Nat8f1 | 0.01 | Young |
| 0.172112 | 1.297947 | 0.01 | 0 | 1 | 8 | Zfpm1 | 0.01 | Young |
| 0.172112 | 1.297911 | 0.01 | 0 | 1 | 8 | Med18 | 0.01 | Young |
| 0.172112 | 1.285105 | 0.01 | 0 | 1 | 8 | Zfp442 | 0.01 | Young |
| 0.172112 | 1.276033 | 0.01 | 0 | 1 | 8 | Spata24 | 0.01 | Young |
| 0.172112 | 1.275466 | 0.01 | 0 | 1 | 8 | Ccdc14 | 0.01 | Young |
| 0.172112 | 1.251637 | 0.01 | 0 | 1 | 8 | Gm29554 | 0.01 | Young |
| 0.172112 | 1.247801 | 0.01 | 0 | 1 | 8 | Ptprb | 0.01 | Young |
| 0.172112 | 1.242519 | 0.01 | 0 | 1 | 8 | Zwilch | 0.01 | Young |
| 0.172112 | 1.229446 | 0.01 | 0 | 1 | 8 | Gm15228 | 0.01 | Young |
| 0.172112 | 1.228062 | 0.01 | 0 | 1 | 8 | Crisp1 | 0.01 | Young |
| 0.172112 | 1.218688 | 0.01 | 0 | 1 | 8 | Sat2 | 0.01 | Young |
| 0.172112 | 1.217296 | 0.01 | 0 | 1 | 8 | Rad9b | 0.01 | Young |
| 0.172112 | 1.199054 | 0.01 | 0 | 1 | 8 | Dscc1 | 0.01 | Young |
| 0.172112 | 1.194587 | 0.01 | 0 | 1 | 8 | Inmt | 0.01 | Young |
| 0.172112 | 1.193537 | 0.01 | 0 | 1 | 8 | Tnfaip8l3 | 0.01 | Young |

| | | | | | | | | |
|---|---|---|---|---|---|---|---|---|
| 0.172112 | 1.184828 | 0.01 | 0 | 1 | 8 | C330011M | 0.01 | Young |
| 0.172112 | 1.179451 | 0.01 | 0 | 1 | 8 | Grhl1 | 0.01 | Young |
| 0.172112 | 1.177398 | 0.01 | 0 | 1 | 8 | Cpxm1 | 0.01 | Young |
| 0.172112 | 1.15802 | 0.01 | 0 | 1 | 8 | Efna1 | 0.01 | Young |
| 0.172112 | 1.157903 | 0.01 | 0 | 1 | 8 | Cenpk | 0.01 | Young |
| 0.172112 | 1.155928 | 0.01 | 0 | 1 | 8 | Tespa1 | 0.01 | Young |
| 0.172112 | 1.121392 | 0.01 | 0 | 1 | 8 | Gbgt1 | 0.01 | Young |
| 0.172112 | 1.121145 | 0.01 | 0 | 1 | 8 | Gm9949 | 0.01 | Young |
| 0.172112 | 1.11831 | 0.01 | 0 | 1 | 8 | Bicdl1 | 0.01 | Young |
| 0.172112 | 1.115244 | 0.01 | 0 | 1 | 8 | Abca5 | 0.01 | Young |
| 0.172112 | 1.090115 | 0.01 | 0 | 1 | 8 | Gpr18 | 0.01 | Young |
| 0.172112 | 1.075178 | 0.01 | 0 | 1 | 8 | Ttc25 | 0.01 | Young |
| 0.172112 | 1.057274 | 0.01 | 0 | 1 | 8 | Tesmin | 0.01 | Young |
| 0.172112 | 1.056832 | 0.01 | 0 | 1 | 8 | Dsn1 | 0.01 | Young |
| 0.172112 | 1.0558 | 0.01 | 0 | 1 | 8 | D130017N( | 0.01 | Young |
| 0.172112 | 1.054587 | 0.01 | 0 | 1 | 8 | Zfp455 | 0.01 | Young |
| 0.172112 | 1.050655 | 0.01 | 0 | 1 | 8 | Gm15420 | 0.01 | Young |
| 0.172112 | 1.03124 | 0.01 | 0 | 1 | 8 | Cenpi | 0.01 | Young |
| 0.172112 | 1.030085 | 0.01 | 0 | 1 | 8 | Cd22 | 0.01 | Young |
| 0.172112 | 1.028884 | 0.01 | 0 | 1 | 8 | Plin4 | 0.01 | Young |
| 0.172112 | 1.026971 | 0.01 | 0 | 1 | 8 | A530010L1 | 0.01 | Young |
| 0.172112 | 1.024572 | 0.01 | 0 | 1 | 8 | 1110006O: | 0.01 | Young |
| 0.172112 | 1.023033 | 0.01 | 0 | 1 | 8 | Mafa | 0.01 | Young |
| 0.172112 | 1.011918 | 0.01 | 0 | 1 | 8 | A130051J0 | 0.01 | Young |
| 0.172112 | 0.99797 | 0.01 | 0 | 1 | 8 | Zfp964 | 0.01 | Young |
| 0.172112 | 0.990857 | 0.01 | 0 | 1 | 8 | Btbd11 | 0.01 | Young |
| 0.172112 | 0.984745 | 0.01 | 0 | 1 | 8 | Gm17767 | 0.01 | Young |
| 0.172112 | 0.981204 | 0.01 | 0 | 1 | 8 | Irs1 | 0.01 | Young |
| 0.172112 | 0.975244 | 0.01 | 0 | 1 | 8 | Mlph | 0.01 | Young |
| 0.172112 | 0.952421 | 0.01 | 0 | 1 | 8 | Tmem79 | 0.01 | Young |
| 0.172112 | 0.952374 | 0.01 | 0 | 1 | 8 | Smarcd3 | 0.01 | Young |
| 0.172112 | 0.88254 | 0.01 | 0 | 1 | 8 | Gm29776 | 0.01 | Young |
| 0.172112 | 0.876674 | 0.01 | 0 | 1 | 8 | Adcy4 | 0.01 | Young |
| 0.172112 | 0.867325 | 0.01 | 0 | 1 | 8 | Gm34961 | 0.01 | Young |
| 0.172112 | 0.864569 | 0.01 | 0 | 1 | 8 | AC146911. | 0.01 | Young |
| 0.172112 | 0.82057 | 0.01 | 0 | 1 | 8 | Gm42670 | 0.01 | Young |
| 0.172467 | 0.638119 | 0.109 | 0.073 | 1 | 8 | Decr2 | 0.036 | Young |
| 0.172736 | 0.222804 | 0.244 | 0.19 | 1 | 8 | Abi2 | 0.054 | Young |
| 0.172881 | 0.398173 | 0.145 | 0.101 | 1 | 8 | Vhl | 0.044 | Young |
| 0.173771 | 0.43186 | 0.231 | 0.179 | 1 | 8 | Setdb1 | 0.052 | Young |
| 0.173923 | 0.165615 | 0.062 | 0.034 | 1 | 8 | Tab3 | 0.028 | Young |
| 0.173952 | 0.353904 | 0.195 | 0.145 | 1 | 8 | Erlin2 | 0.05 | Young |
| 0.174477 | 0.109404 | 0.216 | 0.162 | 1 | 8 | Naip2 | 0.054 | Young |
| 0.174546 | 0.235515 | 0.262 | 0.207 | 1 | 8 | CAAA0111: | 0.055 | Young |
| 0.174762 | 0.254302 | 0.439 | 0.363 | 1 | 8 | AB124611 | 0.076 | Young |

| | | | | | | | | |
|---|---|---|---|---|---|---|---|---|
| 0.174942 | 0.184765 | 0.655 | 0.575 | 1 | 8 | Akr1b3 | 0.08 | Young |
| 0.17498 | 0.428134 | 0.091 | 0.056 | 1 | 8 | Scyl3 | 0.035 | Young |
| 0.175152 | 0.177958 | 0.525 | 0.458 | 1 | 8 | Nufip2 | 0.067 | Young |
| 0.17537 | 0.138811 | 0.777 | 0.737 | 1 | 8 | Rock2 | 0.04 | Young |
| 0.175404 | 0.065651 | 0.138 | 0.095 | 1 | 8 | Odf2 | 0.043 | Young |
| 0.175811 | 0.208398 | 0.405 | 0.335 | 1 | 8 | Uqcrc2 | 0.07 | Young |
| 0.175867 | 0.513594 | 0.122 | 0.084 | 1 | 8 | Zfp467 | 0.038 | Young |
| 0.175908 | 0.564683 | 0.184 | 0.145 | 1 | 8 | 4931406P1 | 0.039 | Young |
| 0.17593 | 0.21291 | 0.351 | 0.302 | 1 | 8 | Npepl1 | 0.049 | Young |
| 0.175936 | 0.132609 | 0.104 | 0.067 | 1 | 8 | Slc25a10 | 0.037 | Young |
| 0.176344 | 0.066203 | 0.278 | 0.223 | 1 | 8 | Ncoa2 | 0.055 | Young |
| 0.17641 | 0.014774 | 0.252 | 0.196 | 1 | 8 | Uba5 | 0.056 | Young |
| 0.176723 | 0.101508 | 0.221 | 0.162 | 1 | 8 | Arid3a | 0.059 | Young |
| 0.176898 | 0.120554 | 0.27 | 0.218 | 1 | 8 | Epb41l4aos | 0.052 | Young |
| 0.176907 | 0.17389 | 0.421 | 0.346 | 1 | 8 | Rere | 0.075 | Young |
| 0.177093 | 0.822164 | 0.068 | 0.039 | 1 | 8 | Dynlt1f | 0.029 | Young |
| 0.177093 | 0.483552 | 0.068 | 0.039 | 1 | 8 | Susd1 | 0.029 | Young |
| 0.177697 | 1.656721 | 0.021 | 0.006 | 1 | 8 | Zranb3 | 0.015 | Young |
| 0.177697 | 1.623604 | 0.021 | 0.006 | 1 | 8 | Tulp3 | 0.015 | Young |
| 0.177697 | 1.474542 | 0.021 | 0.006 | 1 | 8 | Dzip1 | 0.015 | Young |
| 0.177697 | 1.44625 | 0.021 | 0.006 | 1 | 8 | Lekr1 | 0.015 | Young |
| 0.177697 | 1.287593 | 0.021 | 0.006 | 1 | 8 | Acbd4 | 0.015 | Young |
| 0.17797 | 0.645703 | 0.068 | 0.039 | 1 | 8 | Nmral1 | 0.029 | Young |
| 0.178299 | 0.572918 | 0.135 | 0.095 | 1 | 8 | Prr12 | 0.04 | Young |
| 0.178415 | 0.087701 | 0.345 | 0.279 | 1 | 8 | Sh2b3 | 0.066 | Young |
| 0.178521 | 1.625728 | 0.021 | 0.006 | 1 | 8 | Ift88 | 0.015 | Young |
| 0.178521 | 0.979278 | 0.021 | 0.006 | 1 | 8 | P3h1 | 0.015 | Young |
| 0.178666 | 0.594761 | 0.197 | 0.156 | 1 | 8 | Rtn1 | 0.041 | Young |
| 0.178702 | 0.349235 | 0.197 | 0.151 | 1 | 8 | Zfp592 | 0.046 | Young |
| 0.179004 | 0.276933 | 0.156 | 0.112 | 1 | 8 | Apmap | 0.044 | Young |
| 0.179123 | 1.22702 | 0.052 | 0.028 | 1 | 8 | Rpgrip1 | 0.024 | Young |
| 0.179179 | 0.156315 | 0.704 | 0.648 | 1 | 8 | Fkbp1a | 0.056 | Young |
| 0.179348 | 1.49676 | 0.021 | 0.006 | 1 | 8 | Lbx1 | 0.015 | Young |
| 0.179348 | 1.385838 | 0.021 | 0.006 | 1 | 8 | Pecam1 | 0.015 | Young |
| 0.179496 | 0.739555 | 0.081 | 0.05 | 1 | 8 | Rbmx2 | 0.031 | Young |
| 0.179553 | 0.331182 | 0.096 | 0.061 | 1 | 8 | Cactin | 0.035 | Young |
| 0.179597 | 0.239757 | 0.156 | 0.112 | 1 | 8 | Gab1 | 0.044 | Young |
| 0.180178 | 1.657513 | 0.021 | 0.006 | 1 | 8 | Lepr | 0.015 | Young |
| 0.180178 | 1.512189 | 0.021 | 0.006 | 1 | 8 | Thnsl1 | 0.015 | Young |
| 0.180178 | 1.165232 | 0.021 | 0.006 | 1 | 8 | Vsig10l | 0.015 | Young |
| 0.180274 | 0.11891 | 0.491 | 0.43 | 1 | 8 | Usp7 | 0.061 | Young |
| 0.180623 | 0.470314 | 0.075 | 0.045 | 1 | 8 | Socs2 | 0.03 | Young |
| 0.180712 | 0.146985 | 0.138 | 0.095 | 1 | 8 | Ddt | 0.043 | Young |
| 0.180847 | 0.32905 | 0.13 | 0.089 | 1 | 8 | Eng | 0.041 | Young |
| 0.18097 | 0.339644 | 0.117 | 0.078 | 1 | 8 | Dzip3 | 0.039 | Young |

| | | | | | | | | |
|---|---|---|---|---|---|---|---|---|
| 0.18101 | 1.256579 | 0.021 | 0.006 | 1 | 8 | Gm8113 | 0.015 | Young |
| 0.18101 | 1.157485 | 0.021 | 0.006 | 1 | 8 | Clec4g | 0.015 | Young |
| 0.18101 | 1.09688 | 0.021 | 0.006 | 1 | 8 | Gm6133 | 0.015 | Young |
| 0.18101 | 0.980094 | 0.021 | 0.006 | 1 | 8 | Sardh | 0.015 | Young |
| 0.181019 | 0.517963 | 0.088 | 0.056 | 1 | 8 | Slc39a13 | 0.032 | Young |
| 0.181067 | 0.823835 | 0.068 | 0.039 | 1 | 8 | Mki67 | 0.029 | Young |
| 0.181206 | 0.35983 | 0.158 | 0.117 | 1 | 8 | Rai1 | 0.041 | Young |
| 0.181532 | 0.107165 | 0.514 | 0.441 | 1 | 8 | Lpp | 0.073 | Young |
| 0.181556 | 0.047958 | 0.356 | 0.296 | 1 | 8 | Irs2 | 0.06 | Young |
| 0.181846 | 1.233661 | 0.021 | 0.006 | 1 | 8 | Armc2 | 0.015 | Young |
| 0.181846 | 1.110207 | 0.021 | 0.006 | 1 | 8 | Tmem246 | 0.015 | Young |
| 0.181846 | 1.055304 | 0.021 | 0.006 | 1 | 8 | Cdo1 | 0.015 | Young |
| 0.181846 | 1.039355 | 0.021 | 0.006 | 1 | 8 | 9130011E1 | 0.015 | Young |
| 0.181846 | 1.031338 | 0.021 | 0.006 | 1 | 8 | Dph7 | 0.015 | Young |
| 0.181846 | 0.973792 | 0.021 | 0.006 | 1 | 8 | Usp27x | 0.015 | Young |
| 0.182082 | 0.230922 | 0.377 | 0.318 | 1 | 8 | Fam105a | 0.059 | Young |
| 0.182164 | 0.252045 | 0.109 | 0.073 | 1 | 8 | Arl11 | 0.036 | Young |
| 0.182227 | 0.560203 | 0.187 | 0.145 | 1 | 8 | Rfx7 | 0.042 | Young |
| 0.182684 | 1.265869 | 0.021 | 0.006 | 1 | 8 | Gnb5 | 0.015 | Young |
| 0.182684 | 0.828929 | 0.021 | 0.006 | 1 | 8 | Loxl3 | 0.015 | Young |
| 0.182684 | 0.779236 | 0.021 | 0.006 | 1 | 8 | 9130023H2 | 0.015 | Young |
| 0.182923 | 0.526107 | 0.096 | 0.061 | 1 | 8 | Syne3 | 0.035 | Young |
| 0.18301 | 0.081245 | 0.6 | 0.542 | 1 | 8 | Psma4 | 0.058 | Young |
| 0.18305 | 0.037206 | 0.655 | 0.575 | 1 | 8 | Pacs2 | 0.08 | Young |
| 0.183092 | 0.434387 | 0.101 | 0.067 | 1 | 8 | Ccdc134 | 0.034 | Young |
| 0.18342 | 0.210354 | 0.273 | 0.218 | 1 | 8 | Ssbp3 | 0.055 | Young |
| 0.183456 | 0.055186 | 0.283 | 0.223 | 1 | 8 | Fam107b | 0.06 | Young |
| 0.183526 | 1.116375 | 0.021 | 0.006 | 1 | 8 | Kcnip3 | 0.015 | Young |
| 0.183526 | 0.759285 | 0.021 | 0.006 | 1 | 8 | BC030499 | 0.015 | Young |
| 0.183529 | 0.218985 | 0.184 | 0.14 | 1 | 8 | Letm1 | 0.044 | Young |
| 0.18359 | 0.150929 | 0.273 | 0.207 | 1 | 8 | Fam103a1 | 0.066 | Young |
| 0.183772 | 0.248213 | 0.46 | 0.408 | 1 | 8 | Kif1b | 0.052 | Young |
| 0.184135 | 0.188162 | 0.179 | 0.134 | 1 | 8 | Hspbp1 | 0.045 | Young |
| 0.184172 | 0.175164 | 0.506 | 0.43 | 1 | 8 | Bmp2k | 0.076 | Young |
| 0.18437 | 0.711364 | 0.021 | 0.006 | 1 | 8 | Plxdc1 | 0.015 | Young |
| 0.184735 | 0.482189 | 0.119 | 0.084 | 1 | 8 | Bmf | 0.035 | Young |
| 0.184844 | 0.403422 | 0.122 | 0.084 | 1 | 8 | Plekhg2 | 0.038 | Young |
| 0.184862 | 0.20571 | 0.384 | 0.324 | 1 | 8 | Aldh9a1 | 0.06 | Young |
| 0.184917 | 0.261749 | 0.125 | 0.084 | 1 | 8 | Rprd1a | 0.041 | Young |
| 0.184948 | 0.190784 | 0.436 | 0.363 | 1 | 8 | Mlec | 0.073 | Young |
| 0.185111 | 1.588156 | 0.044 | 0.022 | 1 | 8 | Atp6v0c | 0.022 | Young |
| 0.185124 | 0.242171 | 0.169 | 0.128 | 1 | 8 | Slc41a1 | 0.041 | Young |
| 0.185596 | 0.421997 | 0.14 | 0.101 | 1 | 8 | Exosc9 | 0.039 | Young |
| 0.185806 | 0.083954 | 0.571 | 0.497 | 1 | 8 | Amfr | 0.074 | Young |
| 0.185915 | 0.5588 | 0.14 | 0.101 | 1 | 8 | Mob2 | 0.039 | Young |

| | | | | | | | | |
|---|---|---|---|---|---|---|---|---|
| 0.186052 | 0.622428 | 0.081 | 0.05 | 1 | 8 | Nek6 | 0.031 | Young |
| 0.18616 | 0.035959 | 0.143 | 0.101 | 1 | 8 | Nbdy | 0.042 | Young |
| 0.186923 | 0.426673 | 0.068 | 0.039 | 1 | 8 | Tmem42 | 0.029 | Young |
| 0.187056 | 0.179113 | 0.265 | 0.218 | 1 | 8 | Rcn2 | 0.047 | Young |
| 0.187065 | 0.542553 | 0.153 | 0.112 | 1 | 8 | Exosc10 | 0.041 | Young |
| 0.187308 | 0.088372 | 0.774 | 0.67 | 1 | 8 | Gpi1 | 0.104 | Young |
| 0.187402 | 0.711534 | 0.132 | 0.095 | 1 | 8 | D10Wsu10 | 0.037 | Young |
| 0.187443 | 0.106222 | 0.265 | 0.207 | 1 | 8 | Prr5l | 0.058 | Young |
| 0.187598 | 0.143653 | 0.647 | 0.553 | 1 | 8 | Shtn1 | 0.094 | Young |
| 0.187649 | 0.195501 | 0.291 | 0.235 | 1 | 8 | Pqbp1 | 0.056 | Young |
| 0.187929 | 0.338988 | 0.416 | 0.358 | 1 | 8 | Ptbp1 | 0.058 | Young |
| 0.188188 | 0.201258 | 0.283 | 0.229 | 1 | 8 | Mtfr1l | 0.054 | Young |
| 0.188659 | 0.585953 | 0.101 | 0.067 | 1 | 8 | Fbxl12 | 0.034 | Young |
| 0.188917 | 0.441646 | 0.229 | 0.184 | 1 | 8 | Mboat7 | 0.045 | Young |
| 0.189096 | 0.103036 | 0.268 | 0.207 | 1 | 8 | Mau2 | 0.061 | Young |
| 0.189399 | 1.017392 | 0.091 | 0.061 | 1 | 8 | Gdpd3 | 0.03 | Young |
| 0.189507 | 0.242763 | 0.268 | 0.212 | 1 | 8 | Rarg | 0.056 | Young |
| 0.189702 | 0.085742 | 0.504 | 0.43 | 1 | 8 | Rb1cc1 | 0.074 | Young |
| 0.190091 | 0.395806 | 0.221 | 0.173 | 1 | 8 | Sema6d | 0.048 | Young |
| 0.190199 | 0.298347 | 0.2 | 0.151 | 1 | 8 | Gpr155 | 0.049 | Young |
| 0.190269 | 0.091878 | 0.291 | 0.235 | 1 | 8 | Psmd12 | 0.056 | Young |
| 0.190343 | 0.743481 | 0.073 | 0.045 | 1 | 8 | Ermard | 0.028 | Young |
| 0.190505 | 0.610971 | 0.075 | 0.045 | 1 | 8 | Tfip11 | 0.03 | Young |
| 0.190603 | 0.122132 | 0.351 | 0.285 | 1 | 8 | Atp2c1 | 0.066 | Young |
| 0.190667 | 0.145916 | 0.122 | 0.084 | 1 | 8 | Trp53inp1 | 0.038 | Young |
| 0.19093 | 0.023188 | 0.275 | 0.223 | 1 | 8 | Gab2 | 0.052 | Young |
| 0.190961 | 0.062948 | 0.41 | 0.358 | 1 | 8 | Anapc11 | 0.052 | Young |
| 0.191673 | 0.249206 | 0.452 | 0.397 | 1 | 8 | Commd8 | 0.055 | Young |
| 0.192358 | 0.339833 | 0.288 | 0.246 | 1 | 8 | Cdk12 | 0.042 | Young |
| 0.19252 | 0.18947 | 0.416 | 0.369 | 1 | 8 | Sp1 | 0.047 | Young |
| 0.192871 | 0.084347 | 0.218 | 0.173 | 1 | 8 | Lage3 | 0.045 | Young |
| 0.19304 | 0.226807 | 0.205 | 0.156 | 1 | 8 | Lats1 | 0.049 | Young |
| 0.19321 | 0.622614 | 0.081 | 0.05 | 1 | 8 | Zfp595 | 0.031 | Young |
| 0.193282 | 0.304436 | 0.234 | 0.184 | 1 | 8 | 2310011J0 | 0.05 | Young |
| 0.1933 | 0.005483 | 0.26 | 0.207 | 1 | 8 | Exosc5 | 0.053 | Young |
| 0.193337 | 1.773879 | 0.036 | 0.017 | 1 | 8 | Smyd4 | 0.019 | Young |
| 0.193671 | 1.23319 | 0.044 | 0.022 | 1 | 8 | 2810454H( | 0.022 | Young |
| 0.193689 | 0.710703 | 0.06 | 0.034 | 1 | 8 | Cd3eap | 0.026 | Young |
| 0.19387 | 0.901716 | 0.13 | 0.095 | 1 | 8 | Rnf144b | 0.035 | Young |
| 0.194118 | 0.099807 | 0.335 | 0.279 | 1 | 8 | Ctdsp1 | 0.056 | Young |
| 0.194179 | 0.923432 | 0.052 | 0.028 | 1 | 8 | Eral1 | 0.024 | Young |
| 0.194186 | 0.404693 | 0.06 | 0.034 | 1 | 8 | Tnfrsf17 | 0.026 | Young |
| 0.194216 | 0.194006 | 0.481 | 0.43 | 1 | 8 | Tuba1a | 0.051 | Young |
| 0.194601 | 0.568211 | 0.231 | 0.19 | 1 | 8 | Ndufs8 | 0.041 | Young |
| 0.194834 | 1.153934 | 0.044 | 0.022 | 1 | 8 | Pomt2 | 0.022 | Young |

| | | | | | | | | |
|---|---|---|---|---|---|---|---|---|
| 0.194851 | 0.190857 | 0.364 | 0.313 | 1 | 8 | Ndufaf8 | 0.051 | Young |
| 0.195155 | 0.061776 | 0.003 | 0.011 | 1 | 8 | Sfxn5 | -0.008 | Young |
| 0.195262 | 0.499235 | 0.068 | 0.039 | 1 | 8 | Dlg4 | 0.029 | Young |
| 0.195324 | 0.096905 | 0.714 | 0.592 | 1 | 8 | Dnajc3 | 0.122 | Young |
| 0.195487 | 0.175071 | 0.46 | 0.402 | 1 | 8 | Pilra | 0.058 | Young |
| 0.195548 | 0.491442 | 0.229 | 0.179 | 1 | 8 | Aip | 0.05 | Young |
| 0.195584 | 1.475349 | 0.029 | 0.011 | 1 | 8 | Slc25a26 | 0.018 | Young |
| 0.195584 | 1.283478 | 0.029 | 0.011 | 1 | 8 | Stau2 | 0.018 | Young |
| 0.195639 | 0.155774 | 0.364 | 0.313 | 1 | 8 | Usp24 | 0.051 | Young |
| 0.19584 | 0.306396 | 0.262 | 0.218 | 1 | 8 | Sap18 | 0.044 | Young |
| 0.195989 | 0.020854 | 0.249 | 0.19 | 1 | 8 | Arhgap39 | 0.059 | Young |
| 0.195999 | 0.113977 | 0.556 | 0.48 | 1 | 8 | Hivep2 | 0.076 | Young |
| 0.196588 | 0.932434 | 0.044 | 0.022 | 1 | 8 | Armc3 | 0.022 | Young |
| 0.197016 | 0.302478 | 0.377 | 0.33 | 1 | 8 | Glipr2 | 0.047 | Young |
| 0.197063 | 1.401026 | 0.029 | 0.011 | 1 | 8 | Rbfox2 | 0.018 | Young |
| 0.197271 | 0.232446 | 0.187 | 0.145 | 1 | 8 | BC035044 | 0.042 | Young |
| 0.19733 | 0.163871 | 0.213 | 0.162 | 1 | 8 | Trabd | 0.051 | Young |
| 0.197514 | 0.26892 | 0.081 | 0.05 | 1 | 8 | Tada2b | 0.031 | Young |
| 0.197806 | 1.758584 | 0.029 | 0.011 | 1 | 8 | Slc7a4 | 0.018 | Young |
| 0.19783 | 0.215662 | 0.143 | 0.101 | 1 | 8 | Thumpd3 | 0.042 | Young |
| 0.197867 | 1.30248 | 0.036 | 0.017 | 1 | 8 | Fbxw9 | 0.019 | Young |
| 0.198205 | 0.335411 | 0.278 | 0.24 | 1 | 8 | Dlgap4 | 0.038 | Young |
| 0.198354 | 1.144735 | 0.044 | 0.022 | 1 | 8 | Wdr90 | 0.022 | Young |
| 0.19855 | 1.126443 | 0.029 | 0.011 | 1 | 8 | 3300005D( | 0.018 | Young |
| 0.199035 | 1.060826 | 0.052 | 0.028 | 1 | 8 | Tacc3 | 0.024 | Young |
| 0.199054 | 0.349438 | 0.068 | 0.039 | 1 | 8 | Gm35154 | 0.029 | Young |
| 0.199537 | 1.045099 | 0.044 | 0.022 | 1 | 8 | Wee1 | 0.022 | Young |
| 0.199558 | 0.254277 | 0.278 | 0.229 | 1 | 8 | Acp1 | 0.049 | Young |
| 0.199597 | 0.103719 | 0.151 | 0.106 | 1 | 8 | Plau | 0.045 | Young |
| 0.199796 | 0.442018 | 0.094 | 0.061 | 1 | 8 | Mrvi1 | 0.033 | Young |
| 0.200131 | 0.996703 | 0.044 | 0.022 | 1 | 8 | Gpr157 | 0.022 | Young |
| 0.200166 | 0.314042 | 0.101 | 0.067 | 1 | 8 | 2510009E( | 0.034 | Young |
| 0.200184 | 0.408186 | 0.106 | 0.073 | 1 | 8 | Lsg1 | 0.033 | Young |
| 0.200258 | 0.440979 | 0.249 | 0.201 | 1 | 8 | Pde3b | 0.048 | Young |
| 0.200609 | 0.448654 | 0.094 | 0.061 | 1 | 8 | Cog4 | 0.033 | Young |
| 0.200796 | 0.311951 | 0.029 | 0.011 | 1 | 8 | Layn | 0.018 | Young |
| 0.200943 | 0.276471 | 0.151 | 0.112 | 1 | 8 | Prorsd1 | 0.039 | Young |
| 0.201151 | 1.029808 | 0.036 | 0.017 | 1 | 8 | Atp2b4 | 0.019 | Young |
| 0.201342 | 0.315974 | 0.158 | 0.117 | 1 | 8 | Pccb | 0.041 | Young |
| 0.201499 | 0.049788 | 0.205 | 0.156 | 1 | 8 | Pdxdc1 | 0.049 | Young |
| 0.201549 | 0.911381 | 0.029 | 0.011 | 1 | 8 | Pcdhga12 | 0.018 | Young |
| 0.201549 | 0.597615 | 0.029 | 0.011 | 1 | 8 | Gm28151 | 0.018 | Young |
| 0.201661 | 0.218391 | 0.418 | 0.358 | 1 | 8 | Psmc6 | 0.06 | Young |
| 0.20192 | 1.110481 | 0.044 | 0.022 | 1 | 8 | Fam84b | 0.022 | Young |
| 0.202162 | 0.239734 | 0.348 | 0.296 | 1 | 8 | Mrps24 | 0.052 | Young |

| | | | | | | | | |
|---|---|---|---|---|---|---|---|---|
| 0.20247 | 0.578496 | 0.218 | 0.179 | 1 | 8 | Hlx | 0.039 | Young |
| 0.202475 | 1.241529 | 0.036 | 0.017 | 1 | 8 | Gtf2h3 | 0.019 | Young |
| 0.202519 | 0.586908 | 0.044 | 0.022 | 1 | 8 | Il18rap | 0.022 | Young |
| 0.202815 | 0.250871 | 0.127 | 0.089 | 1 | 8 | Aldh7a1 | 0.038 | Young |
| 0.203061 | 1.048933 | 0.029 | 0.011 | 1 | 8 | Tmem69 | 0.018 | Young |
| 0.203098 | 0.188876 | 0.234 | 0.19 | 1 | 8 | Carhsp1 | 0.044 | Young |
| 0.203279 | 0.363221 | 0.242 | 0.196 | 1 | 8 | Spg7 | 0.046 | Young |
| 0.203587 | 0.191172 | 0.158 | 0.117 | 1 | 8 | Pdlim2 | 0.041 | Young |
| 0.203799 | 0.491582 | 0.203 | 0.162 | 1 | 8 | Clec4n | 0.041 | Young |
| 0.203799 | 0.15388 | 0.205 | 0.156 | 1 | 8 | Clcc1 | 0.049 | Young |
| 0.203806 | 1.014347 | 0.036 | 0.017 | 1 | 8 | Eml6 | 0.019 | Young |
| 0.20382 | 0.084526 | 0.029 | 0.011 | 1 | 8 | Nphp3 | 0.018 | Young |
| 0.20398 | 0.606572 | 0.052 | 0.028 | 1 | 8 | Ccdc66 | 0.024 | Young |
| 0.2043 | 0.115929 | 0.161 | 0.117 | 1 | 8 | Plcb2 | 0.044 | Young |
| 0.204459 | 0.708858 | 0.073 | 0.045 | 1 | 8 | Zfp719 | 0.028 | Young |
| 0.204474 | 1.078196 | 0.036 | 0.017 | 1 | 8 | Zbed3 | 0.019 | Young |
| 0.204927 | 0.799788 | 0.044 | 0.022 | 1 | 8 | Dis3 | 0.022 | Young |
| 0.204964 | 0.120616 | 0.517 | 0.453 | 1 | 8 | Uqcrfs1 | 0.064 | Young |
| 0.205288 | 0.203342 | 0.218 | 0.168 | 1 | 8 | Ctnnbip1 | 0.05 | Young |
| 0.205324 | 0.154102 | 0.642 | 0.559 | 1 | 8 | Smarca5 | 0.083 | Young |
| 0.205506 | 0.502603 | 0.143 | 0.106 | 1 | 8 | Tmem238 | 0.037 | Young |
| 0.205657 | 0.870946 | 0.065 | 0.039 | 1 | 8 | Rab13 | 0.026 | Young |
| 0.205702 | 0.084042 | 0.561 | 0.508 | 1 | 8 | Ep300 | 0.053 | Young |
| 0.205814 | 1.202815 | 0.036 | 0.017 | 1 | 8 | Blm | 0.019 | Young |
| 0.20584 | 0.04357 | 0.47 | 0.397 | 1 | 8 | Lmbrd1 | 0.073 | Young |
| 0.206096 | 0.25765 | 0.369 | 0.324 | 1 | 8 | Rtf2 | 0.045 | Young |
| 0.20611 | 0.57598 | 0.029 | 0.011 | 1 | 8 | Sirpb1a | 0.018 | Young |
| 0.206208 | 0.074958 | 0.288 | 0.223 | 1 | 8 | Dusp2 | 0.065 | Young |
| 0.206327 | 0.2003 | 0.294 | 0.246 | 1 | 8 | Coro7 | 0.048 | Young |
| 0.206332 | 0.37778 | 0.081 | 0.05 | 1 | 8 | Ghdc | 0.031 | Young |
| 0.206417 | 0.159606 | 0.577 | 0.492 | 1 | 8 | Git2 | 0.085 | Young |
| 0.206544 | 0.110644 | 0.226 | 0.179 | 1 | 8 | Dnajc10 | 0.047 | Young |
| 0.206728 | 0.118512 | 0.761 | 0.665 | 1 | 8 | Scand1 | 0.096 | Young |
| 0.20674 | 0.105384 | 0.766 | 0.687 | 1 | 8 | Hspe1 | 0.079 | Young |
| 0.207146 | 0.111317 | 0.377 | 0.307 | 1 | 8 | Rab8a | 0.07 | Young |
| 0.207161 | 0.834523 | 0.036 | 0.017 | 1 | 8 | 1110019D1 | 0.019 | Young |
| 0.207206 | 0.532465 | 0.094 | 0.061 | 1 | 8 | Pank1 | 0.033 | Young |
| 0.20756 | 0.253447 | 0.177 | 0.134 | 1 | 8 | Zfp992 | 0.043 | Young |
| 0.207626 | 0.155434 | 0.506 | 0.43 | 1 | 8 | Cd209d | 0.076 | Young |
| 0.207836 | 0.927423 | 0.036 | 0.017 | 1 | 8 | 1810011H1 | 0.019 | Young |
| 0.207836 | 0.215258 | 0.036 | 0.017 | 1 | 8 | Wdr59 | 0.019 | Young |
| 0.207986 | 0.069307 | 0.522 | 0.48 | 1 | 8 | Dnajc5 | 0.042 | Young |
| 0.207987 | 0.144619 | 0.369 | 0.307 | 1 | 8 | Acsl5 | 0.062 | Young |
| 0.208514 | 0.688834 | 0.036 | 0.017 | 1 | 8 | Nipa1 | 0.019 | Young |
| 0.208571 | 0.456132 | 0.143 | 0.106 | 1 | 8 | Nop9 | 0.037 | Young |

| | | | | | | | | |
|---|---|---|---|---|---|---|---|---|
| 0.20858 | 0.339011 | 0.081 | 0.05 | 1 | 8 | Lin9 | 0.031 | Young |
| 0.208631 | 0.238018 | 0.387 | 0.341 | 1 | 8 | Rrp1 | 0.046 | Young |
| 0.20869 | 0.659216 | 0.099 | 0.067 | 1 | 8 | Dhcr7 | 0.032 | Young |
| 0.208728 | 0.233837 | 0.416 | 0.363 | 1 | 8 | Rab35 | 0.053 | Young |
| 0.209257 | 0.294849 | 0.143 | 0.106 | 1 | 8 | Ints3 | 0.037 | Young |
| 0.209278 | 0.319478 | 0.239 | 0.196 | 1 | 8 | Sash3 | 0.043 | Young |
| 0.209429 | 0.329105 | 0.239 | 0.19 | 1 | 8 | D10Jhu81e | 0.049 | Young |
| 0.209445 | 0.014348 | 0.169 | 0.123 | 1 | 8 | Padi2 | 0.046 | Young |
| 0.209578 | 0.21899 | 0.052 | 0.028 | 1 | 8 | Ssh3 | 0.024 | Young |
| 0.209653 | 0.74339 | 0.065 | 0.039 | 1 | 8 | 1810043G( | 0.026 | Young |
| 0.209758 | 0.26311 | 0.138 | 0.101 | 1 | 8 | Prmt2 | 0.037 | Young |
| 0.210248 | 0.506797 | 0.166 | 0.128 | 1 | 8 | Thap3 | 0.038 | Young |
| 0.21038 | 0.063841 | 0.234 | 0.184 | 1 | 8 | Ltbr | 0.05 | Young |
| 0.210432 | 0.310197 | 0.122 | 0.084 | 1 | 8 | Senp3 | 0.038 | Young |
| 0.210753 | 1.161148 | 0.057 | 0.034 | 1 | 8 | Trim37 | 0.023 | Young |
| 0.210856 | 0.068063 | 0.27 | 0.218 | 1 | 8 | Cacul1 | 0.052 | Young |
| 0.210976 | 0.231615 | 0.078 | 0.05 | 1 | 8 | Fam149b | 0.028 | Young |
| 0.210976 | 0.055759 | 0.281 | 0.223 | 1 | 8 | Mknk1 | 0.058 | Young |
| 0.211063 | 0.707245 | 0.122 | 0.089 | 1 | 8 | Nosip | 0.033 | Young |
| 0.211232 | 0.064918 | 0.345 | 0.291 | 1 | 8 | Rnf166 | 0.054 | Young |
| 0.211355 | 0.197004 | 0.187 | 0.145 | 1 | 8 | Patl1 | 0.042 | Young |
| 0.211404 | 0.336117 | 0.306 | 0.257 | 1 | 8 | Rnf10 | 0.049 | Young |
| 0.211538 | 0.268541 | 0.455 | 0.408 | 1 | 8 | St3gal5 | 0.047 | Young |
| 0.21168 | 0.7814 | 0.117 | 0.084 | 1 | 8 | Uba7 | 0.033 | Young |
| 0.212277 | 0.18908 | 0.044 | 0.022 | 1 | 8 | Amer1 | 0.022 | Young |
| 0.21257 | 0.292334 | 0.242 | 0.196 | 1 | 8 | Sf3a3 | 0.046 | Young |
| 0.212795 | 0.18739 | 0.099 | 0.067 | 1 | 8 | Xrcc1 | 0.032 | Young |
| 0.213472 | 0.112405 | 0.218 | 0.168 | 1 | 8 | Rbl2 | 0.05 | Young |
| 0.213666 | 0.192898 | 0.33 | 0.279 | 1 | 8 | Zranb1 | 0.051 | Young |
| 0.213798 | 0.234495 | 0.356 | 0.307 | 1 | 8 | Chd3 | 0.049 | Young |
| 0.214143 | 0.170073 | 0.044 | 0.022 | 1 | 8 | Rnf219 | 0.022 | Young |
| 0.214215 | 0.8857 | 0.065 | 0.039 | 1 | 8 | Gem | 0.026 | Young |
| 0.214379 | 0.051674 | 0.514 | 0.464 | 1 | 8 | Ywhaq | 0.05 | Young |
| 0.214397 | 0.085233 | 0.288 | 0.229 | 1 | 8 | Osbp | 0.059 | Young |
| 0.214824 | 0.281005 | 0.325 | 0.279 | 1 | 8 | Nudcd3 | 0.046 | Young |
| 0.215326 | 0.586442 | 0.143 | 0.106 | 1 | 8 | Nudt19 | 0.037 | Young |
| 0.215386 | 0.257202 | 0.156 | 0.117 | 1 | 8 | Zfp219 | 0.039 | Young |
| 0.215618 | 0.826608 | 0.078 | 0.05 | 1 | 8 | Gfm2 | 0.028 | Young |
| 0.215702 | 0.246527 | 0.325 | 0.279 | 1 | 8 | Actr1a | 0.046 | Young |
| 0.215756 | 0.072306 | 0.158 | 0.117 | 1 | 8 | Nop16 | 0.041 | Young |
| 0.216101 | 0.398447 | 0.299 | 0.251 | 1 | 8 | Synrg | 0.048 | Young |
| 0.216143 | 0.334342 | 0.294 | 0.257 | 1 | 8 | Mysm1 | 0.037 | Young |
| 0.216446 | 0.188645 | 0.397 | 0.352 | 1 | 8 | Mrpl24 | 0.045 | Young |
| 0.216556 | 0.688375 | 0.078 | 0.05 | 1 | 8 | Thap4 | 0.028 | Young |
| 0.216717 | 0.020724 | 0.41 | 0.341 | 1 | 8 | Bcl3 | 0.069 | Young |

| | | | | | | | | |
|---|---|---|---|---|---|---|---|---|
| 0.21677 | 0.581716 | 0.091 | 0.061 | 1 | 8 | Zbtb42 | 0.03 | Young |
| 0.216771 | 0.076255 | 0.161 | 0.117 | 1 | 8 | Slc37a2 | 0.044 | Young |
| 0.216832 | 0.776471 | 0.13 | 0.095 | 1 | 8 | Thoc3 | 0.035 | Young |
| 0.216902 | 0.351293 | 0.114 | 0.078 | 1 | 8 | Ermp1 | 0.036 | Young |
| 0.217169 | 0.381763 | 0.213 | 0.173 | 1 | 8 | Phf11b | 0.04 | Young |
| 0.217362 | 0.275773 | 0.073 | 0.045 | 1 | 8 | Ndrg1 | 0.028 | Young |
| 0.217666 | 0.18793 | 0.561 | 0.531 | 1 | 8 | Eloc | 0.03 | Young |
| 0.217684 | 0.02939 | 0.501 | 0.425 | 1 | 8 | Psmb10 | 0.076 | Young |
| 0.217796 | 0.06888 | 0.4 | 0.33 | 1 | 8 | Wdfy3 | 0.07 | Young |
| 0.217836 | 0.032665 | 0.59 | 0.503 | 1 | 8 | Scpep1 | 0.087 | Young |
| 0.217961 | 0.094226 | 0.291 | 0.235 | 1 | 8 | Slc1a5 | 0.056 | Young |
| 0.218235 | 1.074226 | 0.083 | 0.056 | 1 | 8 | 5430427O1 | 0.027 | Young |
| 0.21833 | 0.659568 | 0.065 | 0.039 | 1 | 8 | Klhl15 | 0.026 | Young |
| 0.218562 | 0.053117 | 0.366 | 0.302 | 1 | 8 | Rreb1 | 0.064 | Young |
| 0.21864 | 0.270145 | 0.099 | 0.067 | 1 | 8 | Rtca | 0.032 | Young |
| 0.218688 | 0.06529 | 0.361 | 0.302 | 1 | 8 | Max | 0.059 | Young |
| 0.218907 | 0.217049 | 0.265 | 0.212 | 1 | 8 | Nr2c2 | 0.053 | Young |
| 0.219297 | 0.087749 | 0.306 | 0.251 | 1 | 8 | Rchy1 | 0.055 | Young |
| 0.219472 | 0.140173 | 0.353 | 0.291 | 1 | 8 | Bscl2 | 0.062 | Young |
| 0.219474 | 0.505747 | 0.229 | 0.19 | 1 | 8 | Cass4 | 0.039 | Young |
| 0.219758 | 0.790691 | 0.104 | 0.073 | 1 | 8 | Nlrc4 | 0.031 | Young |
| 0.22009 | 0.011903 | 0.847 | 0.709 | 1 | 8 | Cfh | 0.138 | Young |
| 0.220332 | 0.050866 | 0.135 | 0.101 | 1 | 8 | Zfp518a | 0.034 | Young |
| 0.220493 | 0.375047 | 0.151 | 0.112 | 1 | 8 | Amn1 | 0.039 | Young |
| 0.220765 | 0.255236 | 0.257 | 0.207 | 1 | 8 | Kbtbd11 | 0.05 | Young |
| 0.220868 | 0.077998 | 0.431 | 0.369 | 1 | 8 | Cebpz | 0.062 | Young |
| 0.220919 | 0.069665 | 0.234 | 0.184 | 1 | 8 | Slc39a10 | 0.05 | Young |
| 0.221029 | 0.038893 | 0.081 | 0.05 | 1 | 8 | Smyd5 | 0.031 | Young |
| 0.2215 | 0.01777 | 0.081 | 0.05 | 1 | 8 | Dbr1 | 0.031 | Young |
| 0.221708 | 1.045219 | 0.057 | 0.034 | 1 | 8 | Phtf1 | 0.023 | Young |
| 0.221812 | 0.263697 | 0.405 | 0.358 | 1 | 8 | Med13 | 0.047 | Young |
| 0.22238 | 0.040605 | 0.426 | 0.369 | 1 | 8 | Polr2e | 0.057 | Young |
| 0.222529 | 0.289677 | 0.26 | 0.218 | 1 | 8 | Tmem5 | 0.042 | Young |
| 0.222783 | 0.130971 | 0.073 | 0.045 | 1 | 8 | Gm27010 | 0.028 | Young |
| 0.222825 | 1.086766 | 0.057 | 0.034 | 1 | 8 | 9530036M | 0.023 | Young |
| 0.222891 | 0.648416 | 0.117 | 0.084 | 1 | 8 | Cep68 | 0.033 | Young |
| 0.222925 | 0.689716 | 0.091 | 0.061 | 1 | 8 | Cables2 | 0.03 | Young |
| 0.223086 | 0.381209 | 0.26 | 0.218 | 1 | 8 | Pigp | 0.042 | Young |
| 0.223447 | 1.068321 | 0.07 | 0.045 | 1 | 8 | Prickle3 | 0.025 | Young |
| 0.223678 | 0.474653 | 0.078 | 0.05 | 1 | 8 | D930016D( | 0.028 | Young |
| 0.223947 | 0.967712 | 0.057 | 0.034 | 1 | 8 | Lamc1 | 0.023 | Young |
| 0.224097 | 0.214163 | 0.213 | 0.168 | 1 | 8 | Trim27 | 0.045 | Young |
| 0.224136 | 0.250153 | 0.226 | 0.184 | 1 | 8 | Pdcd2l | 0.042 | Young |
| 0.224311 | 0.316424 | 0.369 | 0.33 | 1 | 8 | Polr2m | 0.039 | Young |
| 0.224796 | 0.453121 | 0.177 | 0.14 | 1 | 8 | Brip1os | 0.037 | Young |

| | | | | | | | | |
|---|---|---|---|---|---|---|---|---|
| 0.225137 | 0.710655 | 0.065 | 0.039 | 1 | 8 | Efnb2 | 0.026 | Young |
| 0.22514 | 0.264854 | 0.332 | 0.279 | 1 | 8 | Jak2 | 0.053 | Young |
| 0.225606 | 0.509309 | 0.078 | 0.05 | 1 | 8 | Trp53bp1 | 0.028 | Young |
| 0.225764 | 0.461438 | 0.151 | 0.112 | 1 | 8 | Tctex1d2 | 0.039 | Young |
| 0.225829 | 0.079348 | 0.291 | 0.24 | 1 | 8 | Mettl9 | 0.051 | Young |
| 0.225877 | 0.244666 | 0.166 | 0.123 | 1 | 8 | Zfp655 | 0.043 | Young |
| 0.226115 | 0.34047 | 0.169 | 0.128 | 1 | 8 | Kmt2b | 0.041 | Young |
| 0.226365 | 0.119776 | 0.442 | 0.38 | 1 | 8 | Ctnnd1 | 0.062 | Young |
| 0.226769 | 0.750433 | 0.057 | 0.034 | 1 | 8 | Zc3h12d | 0.023 | Young |
| 0.227001 | 0.609191 | 0.07 | 0.045 | 1 | 8 | Accs | 0.025 | Young |
| 0.227059 | 0.480305 | 0.078 | 0.05 | 1 | 8 | Itga6 | 0.028 | Young |
| 0.227262 | 0.079695 | 0.065 | 0.039 | 1 | 8 | Rbbp9 | 0.026 | Young |
| 0.227336 | 0.581934 | 0.057 | 0.034 | 1 | 8 | Mcm7 | 0.023 | Young |
| 0.227652 | 0.354876 | 0.257 | 0.212 | 1 | 8 | Spopl | 0.045 | Young |
| 0.227652 | 1.685297 | 0.049 | 0.028 | 1 | 8 | Gm5086 | 0.021 | Young |
| 0.227832 | 0.205215 | 0.192 | 0.151 | 1 | 8 | Slc28a2 | 0.041 | Young |
| 0.228401 | 0.092145 | 0.127 | 0.089 | 1 | 8 | Arhgap23 | 0.038 | Young |
| 0.228791 | 0.079773 | 0.566 | 0.503 | 1 | 8 | Tfe3 | 0.063 | Young |
| 0.228865 | 0.546234 | 0.065 | 0.039 | 1 | 8 | Fbf1 | 0.026 | Young |
| 0.2289 | 0.229452 | 0.548 | 0.531 | 1 | 8 | Lyve1 | 0.017 | Young |
| 0.229328 | 0.150393 | 0.151 | 0.112 | 1 | 8 | Gpr137 | 0.039 | Young |
| 0.230464 | 0.162486 | 0.382 | 0.33 | 1 | 8 | Mrpl54 | 0.052 | Young |
| 0.230656 | 0.103444 | 0.306 | 0.246 | 1 | 8 | Tbc1d10b | 0.06 | Young |
| 0.231112 | 0.973113 | 0.07 | 0.045 | 1 | 8 | Tlr8 | 0.025 | Young |
| 0.233205 | 0.483219 | 0.049 | 0.028 | 1 | 8 | Eid2b | 0.021 | Young |
| 0.233306 | 0.395455 | 0.091 | 0.061 | 1 | 8 | Ndufaf7 | 0.03 | Young |
| 0.233352 | 0.499413 | 0.122 | 0.089 | 1 | 8 | Mrpl37 | 0.033 | Young |
| 0.233429 | 0.319501 | 0.262 | 0.212 | 1 | 8 | Ppdpf | 0.05 | Young |
| 0.233626 | 0.265269 | 0.125 | 0.089 | 1 | 8 | Mtpap | 0.036 | Young |
| 0.233781 | 0.106627 | 0.371 | 0.33 | 1 | 8 | Lrrfip1 | 0.041 | Young |
| 0.234138 | 0.816701 | 0.083 | 0.056 | 1 | 8 | Mto1 | 0.027 | Young |
| 0.234223 | 0.32058 | 0.057 | 0.034 | 1 | 8 | Pmm1 | 0.023 | Young |
| 0.234313 | 0.090366 | 0.522 | 0.469 | 1 | 8 | Gpcpd1 | 0.053 | Young |
| 0.234333 | 0.206135 | 0.255 | 0.207 | 1 | 8 | Erap1 | 0.048 | Young |
| 0.234418 | 0.196296 | 0.125 | 0.089 | 1 | 8 | Mtf1 | 0.036 | Young |
| 0.234847 | 0.532197 | 0.166 | 0.128 | 1 | 8 | Tshz1 | 0.038 | Young |
| 0.234905 | 0.238978 | 0.366 | 0.318 | 1 | 8 | Srp19 | 0.048 | Young |
| 0.234973 | 0.410986 | 0.283 | 0.246 | 1 | 8 | Hint2 | 0.037 | Young |
| 0.235146 | 0.230593 | 0.091 | 0.061 | 1 | 8 | Slc2a8 | 0.03 | Young |
| 0.235673 | 0.269033 | 0.377 | 0.33 | 1 | 8 | Aurkaip1 | 0.047 | Young |
| 0.235773 | 0.561528 | 0.145 | 0.112 | 1 | 8 | Hsdl1 | 0.033 | Young |
| 0.23624 | 1.66572 | 0.018 | 0.006 | 1 | 8 | Tbc1d19 | 0.012 | Young |
| 0.23624 | 1.553064 | 0.018 | 0.006 | 1 | 8 | Lrrc56 | 0.012 | Young |
| 0.23624 | 1.480197 | 0.018 | 0.006 | 1 | 8 | Ddb2 | 0.012 | Young |
| 0.23624 | 1.422616 | 0.018 | 0.006 | 1 | 8 | Clec9a | 0.012 | Young |

| | | | | | | | | |
|---|---|---|---|---|---|---|---|---|
| 0.23624 | 1.210297 | 0.018 | 0.006 | 1 | 8 | Ccdc94 | 0.012 | Young |
| 0.23624 | 1.20544 | 0.018 | 0.006 | 1 | 8 | Zfp882 | 0.012 | Young |
| 0.236295 | 0.460829 | 0.231 | 0.196 | 1 | 8 | Ccm2 | 0.035 | Young |
| 0.236332 | 0.829864 | 0.049 | 0.028 | 1 | 8 | Cbr1 | 0.021 | Young |
| 0.236507 | 0.231162 | 0.413 | 0.358 | 1 | 8 | Denr | 0.055 | Young |
| 0.236551 | 0.57749 | 0.057 | 0.034 | 1 | 8 | Tmem9 | 0.023 | Young |
| 0.236659 | 0.268884 | 0.301 | 0.257 | 1 | 8 | Tmem87b | 0.044 | Young |
| 0.236689 | 0.050712 | 0.104 | 0.073 | 1 | 8 | Sclt1 | 0.031 | Young |
| 0.237028 | 0.834441 | 0.083 | 0.056 | 1 | 8 | Xlr4b | 0.027 | Young |
| 0.237124 | 0.321488 | 0.384 | 0.341 | 1 | 8 | Ythdf1 | 0.043 | Young |
| 0.237308 | 0.291729 | 0.151 | 0.112 | 1 | 8 | Bad | 0.039 | Young |
| 0.237311 | 0.557265 | 0.099 | 0.067 | 1 | 8 | Clp1 | 0.032 | Young |
| 0.237347 | 0.299638 | 0.13 | 0.095 | 1 | 8 | Bend4 | 0.035 | Young |
| 0.237556 | 0.032226 | 0.27 | 0.218 | 1 | 8 | Rnps1 | 0.052 | Young |
| 0.237731 | 0.996916 | 0.062 | 0.039 | 1 | 8 | Gm38843 | 0.023 | Young |
| 0.237819 | 0.061006 | 0.496 | 0.425 | 1 | 8 | Nfic | 0.071 | Young |
| 0.237921 | 0.358013 | 0.078 | 0.05 | 1 | 8 | Man2b2 | 0.028 | Young |
| 0.238362 | 1.241208 | 0.042 | 0.022 | 1 | 8 | Zgpat | 0.02 | Young |
| 0.238392 | 1.459796 | 0.018 | 0.006 | 1 | 8 | Gm10384 | 0.012 | Young |
| 0.238392 | 1.165493 | 0.018 | 0.006 | 1 | 8 | Mboat1 | 0.012 | Young |
| 0.238392 | 1.010285 | 0.018 | 0.006 | 1 | 8 | Fuk | 0.012 | Young |
| 0.23856 | 0.453598 | 0.179 | 0.14 | 1 | 8 | Milr1 | 0.039 | Young |
| 0.239053 | 1.490304 | 0.042 | 0.022 | 1 | 8 | Mcrip2 | 0.02 | Young |
| 0.239075 | 0.357297 | 0.096 | 0.067 | 1 | 8 | Zbtb33 | 0.029 | Young |
| 0.239472 | 0.494466 | 0.174 | 0.134 | 1 | 8 | Zfp236 | 0.04 | Young |
| 0.239473 | 1.302226 | 0.018 | 0.006 | 1 | 8 | Arhgef40 | 0.012 | Young |
| 0.239473 | 1.133171 | 0.018 | 0.006 | 1 | 8 | Pter | 0.012 | Young |
| 0.239473 | 1.119843 | 0.018 | 0.006 | 1 | 8 | Gm12089 | 0.012 | Young |
| 0.239473 | 1.028382 | 0.018 | 0.006 | 1 | 8 | Cenps | 0.012 | Young |
| 0.239473 | 0.995181 | 0.018 | 0.006 | 1 | 8 | Uhrf1 | 0.012 | Young |
| 0.239473 | 0.99334 | 0.018 | 0.006 | 1 | 8 | Prps1l3 | 0.012 | Young |
| 0.239916 | 0.209309 | 0.273 | 0.223 | 1 | 8 | Sfswap | 0.05 | Young |
| 0.240051 | 0.290712 | 0.358 | 0.324 | 1 | 8 | Ssbp4 | 0.034 | Young |
| 0.240122 | 0.723142 | 0.049 | 0.028 | 1 | 8 | Sumf2 | 0.021 | Young |
| 0.240557 | 1.253288 | 0.018 | 0.006 | 1 | 8 | Parvg | 0.012 | Young |
| 0.240557 | 1.013954 | 0.018 | 0.006 | 1 | 8 | Ube2v1 | 0.012 | Young |
| 0.240557 | 0.995307 | 0.018 | 0.006 | 1 | 8 | Padi4 | 0.012 | Young |
| 0.240557 | 0.877318 | 0.018 | 0.006 | 1 | 8 | Napb | 0.012 | Young |
| 0.240665 | 0.258147 | 0.057 | 0.034 | 1 | 8 | Zfyve19 | 0.023 | Young |
| 0.240734 | 0.058899 | 0.738 | 0.665 | 1 | 8 | Abca9 | 0.073 | Young |
| 0.240834 | 0.309203 | 0.161 | 0.123 | 1 | 8 | 2810004N2 | 0.038 | Young |
| 0.241123 | 0.201641 | 0.304 | 0.268 | 1 | 8 | Glrx2 | 0.036 | Young |
| 0.241174 | 0.254446 | 0.216 | 0.173 | 1 | 8 | Ctsf | 0.043 | Young |
| 0.241452 | 0.303916 | 0.078 | 0.05 | 1 | 8 | Zfp866 | 0.028 | Young |
| 0.241645 | 0.702948 | 0.018 | 0.006 | 1 | 8 | Gsg1 | 0.012 | Young |

| | | | | | | | | |
|---|---|---|---|---|---|---|---|---|
| 0.241798 | 0.142093 | 0.2 | 0.162 | 1 | 8 | Gabpa | 0.038 | Young |
| 0.241862 | 0.039784 | 0.525 | 0.453 | 1 | 8 | Gtf2h5 | 0.072 | Young |
| 0.242034 | 0.304804 | 0.049 | 0.028 | 1 | 8 | Gfod1 | 0.021 | Young |
| 0.242154 | 0.512784 | 0.07 | 0.045 | 1 | 8 | Dnajc4 | 0.025 | Young |
| 0.242263 | 0.435508 | 0.099 | 0.067 | 1 | 8 | Miip | 0.032 | Young |
| 0.242484 | 0.429945 | 0.13 | 0.095 | 1 | 8 | Rnf41 | 0.035 | Young |
| 0.242673 | 0.734816 | 0.049 | 0.028 | 1 | 8 | Rhoj | 0.021 | Young |
| 0.242725 | 0.28586 | 0.265 | 0.218 | 1 | 8 | Ric8a | 0.047 | Young |
| 0.242736 | 0.961362 | 0.018 | 0.006 | 1 | 8 | Xrcc5 | 0.012 | Young |
| 0.242736 | 0.852558 | 0.018 | 0.006 | 1 | 8 | Rad54l | 0.012 | Young |
| 0.242736 | 0.439851 | 0.018 | 0.006 | 1 | 8 | Gm29367 | 0.012 | Young |
| 0.242922 | 0.930554 | 0.075 | 0.05 | 1 | 8 | Asgr2 | 0.025 | Young |
| 0.242953 | 0.303244 | 0.252 | 0.207 | 1 | 8 | Atp8a1 | 0.045 | Young |
| 0.243247 | 0.11375 | 0.27 | 0.223 | 1 | 8 | Slf2 | 0.047 | Young |
| 0.243439 | 0.033081 | 0.213 | 0.162 | 1 | 8 | Esco1 | 0.051 | Young |
| 0.243523 | 0.183221 | 0.278 | 0.229 | 1 | 8 | Rab5b | 0.049 | Young |
| 0.243633 | 0.296019 | 0.195 | 0.156 | 1 | 8 | Iars2 | 0.039 | Young |
| 0.243831 | 0.718273 | 0.018 | 0.006 | 1 | 8 | Meis1 | 0.012 | Young |
| 0.243831 | 0.079561 | 0.018 | 0.006 | 1 | 8 | Gm30411 | 0.012 | Young |
| 0.243971 | 0.24803 | 0.197 | 0.156 | 1 | 8 | Mak16 | 0.041 | Young |
| 0.244238 | 0.030801 | 0.239 | 0.196 | 1 | 8 | Relt | 0.043 | Young |
| 0.244248 | 0.35361 | 0.192 | 0.151 | 1 | 8 | Polr3k | 0.041 | Young |
| 0.244362 | 0.420992 | 0.083 | 0.056 | 1 | 8 | Ints10 | 0.027 | Young |
| 0.244367 | 0.382285 | 0.19 | 0.151 | 1 | 8 | Cep83 | 0.039 | Young |
| 0.244466 | 0.287734 | 0.117 | 0.084 | 1 | 8 | Trmt2a | 0.033 | Young |
| 0.244628 | 0.799634 | 0.042 | 0.022 | 1 | 8 | Sv2b | 0.02 | Young |
| 0.244718 | 0.209633 | 0.33 | 0.296 | 1 | 8 | Stat5b | 0.034 | Young |
| 0.244791 | 0.354975 | 0.153 | 0.117 | 1 | 8 | Prpsap1 | 0.036 | Young |
| 0.245093 | 0.23135 | 0.278 | 0.235 | 1 | 8 | Rbfa | 0.043 | Young |
| 0.245101 | 0.293446 | 0.122 | 0.089 | 1 | 8 | Yod1 | 0.033 | Young |
| 0.245159 | 0.167512 | 0.273 | 0.223 | 1 | 8 | Tnip2 | 0.05 | Young |
| 0.245243 | 0.613496 | 0.049 | 0.028 | 1 | 8 | Neurl1b | 0.021 | Young |
| 0.245261 | 0.495139 | 0.143 | 0.106 | 1 | 8 | Helb | 0.037 | Young |
| 0.245332 | 0.815971 | 0.042 | 0.022 | 1 | 8 | Pigg | 0.02 | Young |
| 0.245645 | 0.106094 | 0.205 | 0.162 | 1 | 8 | Virma | 0.043 | Young |
| 0.24571 | 0.070096 | 0.473 | 0.419 | 1 | 8 | Zc3h7a | 0.054 | Young |
| 0.245796 | 0.055522 | 0.377 | 0.335 | 1 | 8 | Setx | 0.042 | Young |
| 0.245888 | 0.731528 | 0.049 | 0.028 | 1 | 8 | Zkscan14 | 0.021 | Young |
| 0.246048 | 0.735121 | 0.148 | 0.117 | 1 | 8 | Tmem181a | 0.031 | Young |
| 0.246283 | 0.158053 | 0.086 | 0.056 | 1 | 8 | Xlr | 0.03 | Young |
| 0.24639 | 0.077499 | 0.208 | 0.162 | 1 | 8 | Slc25a20 | 0.046 | Young |
| 0.246807 | 0.635851 | 0.101 | 0.073 | 1 | 8 | Phf6 | 0.028 | Young |
| 0.247177 | 0.431807 | 0.348 | 0.313 | 1 | 8 | Pik3ca | 0.035 | Young |
| 0.247396 | 0.895692 | 0.088 | 0.061 | 1 | 8 | Eif2b1 | 0.027 | Young |
| 0.2474 | 0.114344 | 0.153 | 0.117 | 1 | 8 | Ncoa5 | 0.036 | Young |

| | | | | | | | | |
|---|---|---|---|---|---|---|---|---|
| 0.247512 | 0.057226 | 0.977 | 0.922 | 1 | 8 | Npc2 | 0.055 | Young |
| 0.247832 | 0.358314 | 0.049 | 0.028 | 1 | 8 | Wdr92 | 0.021 | Young |
| 0.247897 | 0.392269 | 0.213 | 0.173 | 1 | 8 | Oxsr1 | 0.04 | Young |
| 0.248023 | 0.038475 | 0.182 | 0.14 | 1 | 8 | Hk3 | 0.042 | Young |
| 0.248159 | 0.950016 | 0.042 | 0.022 | 1 | 8 | Riox1 | 0.02 | Young |
| 0.248226 | 0.197634 | 0.418 | 0.374 | 1 | 8 | Leprotl1 | 0.044 | Young |
| 0.248496 | 0.059015 | 0.514 | 0.453 | 1 | 8 | Nfe2l1 | 0.061 | Young |
| 0.248717 | 0.026881 | 0.4 | 0.341 | 1 | 8 | Sbf2 | 0.059 | Young |
| 0.248831 | 0.254284 | 0.122 | 0.089 | 1 | 8 | Pin1 | 0.033 | Young |
| 0.249089 | 0.620228 | 0.062 | 0.039 | 1 | 8 | Zc2hc1a | 0.023 | Young |
| 0.249133 | 0.591479 | 0.049 | 0.028 | 1 | 8 | Tango6 | 0.021 | Young |
| 0.249147 | 0.25572 | 0.262 | 0.207 | 1 | 8 | Srek1 | 0.055 | Young |
| 0.249173 | 0.427954 | 0.07 | 0.045 | 1 | 8 | Rad51b | 0.025 | Young |
| 0.249506 | 1.279851 | 0.026 | 0.011 | 1 | 8 | Itpr3 | 0.015 | Young |
| 0.249595 | 0.047758 | 0.099 | 0.067 | 1 | 8 | Mfng | 0.032 | Young |
| 0.249786 | 0.153661 | 0.779 | 0.715 | 1 | 8 | Rpl22l1 | 0.064 | Young |
| 0.249786 | 0.170931 | 0.049 | 0.028 | 1 | 8 | Gemin5 | 0.021 | Young |
| 0.24988 | 0.142485 | 0.314 | 0.263 | 1 | 8 | Ptov1 | 0.051 | Young |
| 0.249974 | 0.262657 | 0.117 | 0.084 | 1 | 8 | Mrpl40 | 0.033 | Young |
| 0.25016 | 0.079019 | 0.213 | 0.168 | 1 | 8 | Gprc5c | 0.045 | Young |
| 0.250429 | 0.092027 | 0.457 | 0.397 | 1 | 8 | Lrrc25 | 0.06 | Young |
| 0.251116 | 0.054414 | 0.831 | 0.76 | 1 | 8 | Ncf2 | 0.071 | Young |
| 0.251725 | 0.884362 | 0.042 | 0.022 | 1 | 8 | Nipal3 | 0.02 | Young |
| 0.251728 | 0.199306 | 0.192 | 0.151 | 1 | 8 | Usp38 | 0.041 | Young |
| 0.251984 | 0.270838 | 0.348 | 0.302 | 1 | 8 | Cdk14 | 0.046 | Young |
| 0.252256 | 1.132635 | 0.026 | 0.011 | 1 | 8 | Fkbpl | 0.015 | Young |
| 0.252355 | 0.960762 | 0.034 | 0.017 | 1 | 8 | Gdpd5 | 0.017 | Young |
| 0.252544 | 0.547308 | 0.138 | 0.106 | 1 | 8 | Prpf4 | 0.032 | Young |
| 0.252945 | 0.746868 | 0.106 | 0.078 | 1 | 8 | Fam173b | 0.028 | Young |
| 0.252992 | 0.161446 | 0.13 | 0.095 | 1 | 8 | Rbm15b | 0.035 | Young |
| 0.253011 | 0.515422 | 0.07 | 0.045 | 1 | 8 | Hebp2 | 0.025 | Young |
| 0.253156 | 1.314512 | 0.034 | 0.017 | 1 | 8 | Ss18l1 | 0.017 | Young |
| 0.253156 | 1.241242 | 0.034 | 0.017 | 1 | 8 | Tyw5 | 0.017 | Young |
| 0.253177 | 1.238173 | 0.026 | 0.011 | 1 | 8 | Homer2 | 0.015 | Young |
| 0.253177 | 1.234115 | 0.026 | 0.011 | 1 | 8 | Pms1 | 0.015 | Young |
| 0.25344 | 0.24663 | 0.109 | 0.078 | 1 | 8 | Zcchc14 | 0.031 | Young |
| 0.253896 | 0.377487 | 0.096 | 0.067 | 1 | 8 | Mrps11 | 0.029 | Young |
| 0.254101 | 1.656165 | 0.026 | 0.011 | 1 | 8 | Als2 | 0.015 | Young |
| 0.254223 | 0.010033 | 0.203 | 0.162 | 1 | 8 | Cisd1 | 0.041 | Young |
| 0.25435 | 0.356522 | 0.184 | 0.145 | 1 | 8 | Pofut2 | 0.039 | Young |
| 0.254557 | 0.17364 | 0.268 | 0.223 | 1 | 8 | D16Ertd47 | 0.045 | Young |
| 0.254763 | 0.897602 | 0.034 | 0.017 | 1 | 8 | Hdac6 | 0.017 | Young |
| 0.254794 | 0.27976 | 0.332 | 0.291 | 1 | 8 | Atp6v1c1 | 0.041 | Young |
| 0.254866 | 0.093585 | 0.475 | 0.441 | 1 | 8 | Med13l | 0.034 | Young |
| 0.254895 | 0.269162 | 0.083 | 0.056 | 1 | 8 | Rab2b | 0.027 | Young |

| | | | | | | | | |
|---|---|---|---|---|---|---|---|---|
| 0.255222 | 0.194717 | 0.07 | 0.045 | 1 | 8 | Elk1 | 0.025 | Young |
| 0.255245 | 0.219348 | 0.249 | 0.207 | 1 | 8 | Aqr | 0.042 | Young |
| 0.255777 | 0.509497 | 0.07 | 0.045 | 1 | 8 | Phlpp1 | 0.025 | Young |
| 0.255955 | 2.240418 | 0.026 | 0.011 | 1 | 8 | Ch25h | 0.015 | Young |
| 0.255955 | 1.195744 | 0.026 | 0.011 | 1 | 8 | Tal1 | 0.015 | Young |
| 0.256052 | 0.451317 | 0.042 | 0.022 | 1 | 8 | Osbpl1a | 0.02 | Young |
| 0.256098 | 0.060688 | 0.112 | 0.078 | 1 | 8 | Wdr12 | 0.034 | Young |
| 0.256195 | 0.704978 | 0.088 | 0.061 | 1 | 8 | Cyb561d1 | 0.027 | Young |
| 0.256237 | 0.584569 | 0.132 | 0.101 | 1 | 8 | Dctpp1 | 0.031 | Young |
| 0.256377 | 0.888908 | 0.034 | 0.017 | 1 | 8 | Sars2 | 0.017 | Young |
| 0.256396 | 0.157419 | 0.281 | 0.235 | 1 | 8 | Cuedc2 | 0.046 | Young |
| 0.256707 | 0.093506 | 0.527 | 0.453 | 1 | 8 | Eno1 | 0.074 | Young |
| 0.256778 | 0.605831 | 0.042 | 0.022 | 1 | 8 | E130311K1 | 0.02 | Young |
| 0.256886 | 0.938615 | 0.026 | 0.011 | 1 | 8 | Zkscan5 | 0.015 | Young |
| 0.256995 | 0.065195 | 0.283 | 0.229 | 1 | 8 | Uba6 | 0.054 | Young |
| 0.257186 | 0.183791 | 0.034 | 0.017 | 1 | 8 | 9430015G1 | 0.017 | Young |
| 0.257278 | 0.267038 | 0.203 | 0.162 | 1 | 8 | Tbrg1 | 0.041 | Young |
| 0.257505 | 0.397182 | 0.042 | 0.022 | 1 | 8 | Arvcf | 0.02 | Young |
| 0.257524 | 0.223639 | 0.296 | 0.251 | 1 | 8 | Lpcat1 | 0.045 | Young |
| 0.257668 | 0.057068 | 0.569 | 0.475 | 1 | 8 | Phip | 0.094 | Young |
| 0.257849 | 0.656955 | 0.156 | 0.123 | 1 | 8 | Brap | 0.033 | Young |
| 0.257998 | 1.139315 | 0.034 | 0.017 | 1 | 8 | Tut1 | 0.017 | Young |
| 0.258123 | 0.646682 | 0.114 | 0.084 | 1 | 8 | Exosc4 | 0.03 | Young |
| 0.258144 | 0.262832 | 0.127 | 0.095 | 1 | 8 | Hdgfl2 | 0.032 | Young |
| 0.258234 | 0.03946 | 0.042 | 0.022 | 1 | 8 | Prepl | 0.02 | Young |
| 0.258754 | 0.928225 | 0.026 | 0.011 | 1 | 8 | Gatsl3 | 0.015 | Young |
| 0.258754 | 0.917458 | 0.026 | 0.011 | 1 | 8 | Myom1 | 0.015 | Young |
| 0.258754 | 0.842096 | 0.026 | 0.011 | 1 | 8 | AC162302. | 0.015 | Young |
| 0.258754 | 0.434984 | 0.026 | 0.011 | 1 | 8 | Lcmt2 | 0.015 | Young |
| 0.258811 | 1.006736 | 0.034 | 0.017 | 1 | 8 | Vars2 | 0.017 | Young |
| 0.25926 | 0.141109 | 0.356 | 0.307 | 1 | 8 | Plekhm3 | 0.049 | Young |
| 0.259379 | 0.374554 | 0.14 | 0.106 | 1 | 8 | Gatsl2 | 0.034 | Young |
| 0.259523 | 0.353169 | 0.34 | 0.307 | 1 | 8 | Tgs1 | 0.033 | Young |
| 0.259692 | 0.925961 | 0.026 | 0.011 | 1 | 8 | Bend6 | 0.015 | Young |
| 0.259692 | 0.90297 | 0.026 | 0.011 | 1 | 8 | Zfp658 | 0.015 | Young |
| 0.259692 | 0.500038 | 0.026 | 0.011 | 1 | 8 | Oplah | 0.015 | Young |
| 0.259742 | 1.007109 | 0.055 | 0.034 | 1 | 8 | Snap47 | 0.021 | Young |
| 0.259765 | 0.180748 | 0.216 | 0.173 | 1 | 8 | Otud4 | 0.043 | Young |
| 0.259814 | 0.580588 | 0.234 | 0.196 | 1 | 8 | Fbxo18 | 0.038 | Young |
| 0.260065 | 0.077989 | 0.151 | 0.112 | 1 | 8 | Nqo2 | 0.039 | Young |
| 0.260279 | 0.153486 | 0.231 | 0.19 | 1 | 8 | Smim19 | 0.041 | Young |
| 0.260443 | 0.420452 | 0.034 | 0.017 | 1 | 8 | Ms4a4a | 0.017 | Young |
| 0.260632 | 0.846597 | 0.026 | 0.011 | 1 | 8 | Sox4 | 0.015 | Young |
| 0.260632 | 0.724489 | 0.026 | 0.011 | 1 | 8 | Haus5 | 0.015 | Young |
| 0.260632 | 0.706697 | 0.026 | 0.011 | 1 | 8 | Gm16062 | 0.015 | Young |

| | | | | | | | | |
|---|---|---|---|---|---|---|---|---|
| 0.260927 | 0.057776 | 0.439 | 0.374 | 1 | 8 | Ttyh3 | 0.065 | Young |
| 0.260935 | 0.196962 | 0.221 | 0.179 | 1 | 8 | Bloc1s2 | 0.042 | Young |
| 0.261013 | 0.809004 | 0.055 | 0.034 | 1 | 8 | Alkbh2 | 0.021 | Young |
| 0.261223 | 0.168733 | 0.556 | 0.531 | 1 | 8 | Ctnna1 | 0.025 | Young |
| 0.261311 | 0.070482 | 0.543 | 0.48 | 1 | 8 | Hsbp1 | 0.063 | Young |
| 0.261435 | 0.279228 | 0.268 | 0.223 | 1 | 8 | Lsm3 | 0.045 | Young |
| 0.261643 | 0.238615 | 0.213 | 0.173 | 1 | 8 | Socs7 | 0.04 | Young |
| 0.261742 | 0.240678 | 0.195 | 0.156 | 1 | 8 | Klhl28 | 0.039 | Young |
| 0.261891 | 0.079086 | 0.135 | 0.101 | 1 | 8 | Isca2 | 0.034 | Young |
| 0.2619 | 0.462785 | 0.042 | 0.022 | 1 | 8 | Slc7a6 | 0.02 | Young |
| 0.262167 | 0.009885 | 0.629 | 0.547 | 1 | 8 | Ncor1 | 0.082 | Young |
| 0.262358 | 0.342144 | 0.257 | 0.212 | 1 | 8 | Ssr2 | 0.045 | Young |
| 0.262537 | 0.106631 | 0.192 | 0.151 | 1 | 8 | Med14 | 0.041 | Young |
| 0.262904 | 0.395222 | 0.034 | 0.017 | 1 | 8 | Ets1 | 0.017 | Young |
| 0.262995 | 0.320181 | 0.114 | 0.084 | 1 | 8 | Map4k2 | 0.03 | Young |
| 0.263096 | 0.155012 | 0.738 | 0.687 | 1 | 8 | Hcls1 | 0.051 | Young |
| 0.263244 | 0.052999 | 0.223 | 0.184 | 1 | 8 | Parg | 0.039 | Young |
| 0.263355 | 0.077773 | 0.109 | 0.078 | 1 | 8 | Zdhhc16 | 0.031 | Young |
| 0.263466 | 0.619698 | 0.026 | 0.011 | 1 | 8 | Txk | 0.015 | Young |
| 0.263466 | 0.612749 | 0.026 | 0.011 | 1 | 8 | Slc7a5 | 0.015 | Young |
| 0.263466 | 0.599078 | 0.026 | 0.011 | 1 | 8 | Nlgn2 | 0.015 | Young |
| 0.263466 | 0.236464 | 0.026 | 0.011 | 1 | 8 | Epop | 0.015 | Young |
| 0.263812 | 0.734197 | 0.062 | 0.039 | 1 | 8 | Gys1 | 0.023 | Young |
| 0.264197 | 0.572864 | 0.088 | 0.061 | 1 | 8 | Dalrd3 | 0.027 | Young |
| 0.264345 | 0.28145 | 0.117 | 0.084 | 1 | 8 | Atg2b | 0.033 | Young |
| 0.264368 | 0.214168 | 0.306 | 0.257 | 1 | 8 | B3gat3 | 0.049 | Young |
| 0.264408 | 0.243705 | 0.244 | 0.207 | 1 | 8 | Crtc2 | 0.037 | Young |
| 0.264421 | 0.027786 | 0.156 | 0.117 | 1 | 8 | Tmem63b | 0.039 | Young |
| 0.264451 | 0.449181 | 0.166 | 0.134 | 1 | 8 | Tbc1d4 | 0.032 | Young |
| 0.264553 | 0.614041 | 0.034 | 0.017 | 1 | 8 | Ttll1 | 0.017 | Young |
| 0.264553 | 0.354034 | 0.034 | 0.017 | 1 | 8 | Fntb | 0.017 | Young |
| 0.26477 | 0.368608 | 0.07 | 0.045 | 1 | 8 | Ankrd52 | 0.025 | Young |
| 0.264859 | 0.296658 | 0.042 | 0.022 | 1 | 8 | Ccdc130 | 0.02 | Young |
| 0.264903 | 0.488261 | 0.127 | 0.095 | 1 | 8 | Dhx29 | 0.032 | Young |
| 0.265054 | 0.178062 | 0.26 | 0.212 | 1 | 8 | Phf8 | 0.048 | Young |
| 0.265095 | 0.824816 | 0.094 | 0.067 | 1 | 8 | Ankmy2 | 0.027 | Young |
| 0.26532 | 0.080354 | 0.286 | 0.24 | 1 | 8 | Tbc1d20 | 0.046 | Young |
| 0.265585 | 0.016986 | 0.564 | 0.469 | 1 | 8 | Blnk | 0.095 | Young |
| 0.265743 | 0.284019 | 0.223 | 0.184 | 1 | 8 | Kpna3 | 0.039 | Young |
| 0.265982 | 0.507276 | 0.153 | 0.123 | 1 | 8 | Sos1 | 0.03 | Young |
| 0.266222 | 0.586389 | 0.062 | 0.039 | 1 | 8 | Fig4 | 0.023 | Young |
| 0.266222 | 0.113451 | 0.062 | 0.039 | 1 | 8 | Pi4kb | 0.023 | Young |
| 0.266291 | 0.232576 | 0.177 | 0.14 | 1 | 8 | Dhps | 0.037 | Young |
| 0.266347 | 0.276988 | 0.042 | 0.022 | 1 | 8 | Wdr4 | 0.02 | Young |
| 0.266732 | 0.025028 | 0.088 | 0.061 | 1 | 8 | Zc3h10 | 0.027 | Young |

| | | | | | | | | |
|---|---|---|---|---|---|---|---|---|
| 0.266978 | 0.165271 | 0.413 | 0.363 | 1 | 8 | Arid5a | 0.05 | Young |
| 0.267708 | 0.082133 | 0.992 | 0.989 | 1 | 8 | Selenop | 0.003 | Young |
| 0.26804 | 0.307035 | 0.062 | 0.039 | 1 | 8 | Clcf1 | 0.023 | Young |
| 0.268211 | 0.31717 | 0.262 | 0.218 | 1 | 8 | Snapin | 0.044 | Young |
| 0.268281 | 0.050396 | 0.247 | 0.201 | 1 | 8 | Trim12a | 0.046 | Young |
| 0.268487 | 0.12385 | 0.229 | 0.184 | 1 | 8 | Trim26 | 0.045 | Young |
| 0.268953 | 0.63166 | 0.106 | 0.078 | 1 | 8 | Mtmr4 | 0.028 | Young |
| 0.269232 | 0.058588 | 0.868 | 0.799 | 1 | 8 | Slc6a6 | 0.069 | Young |
| 0.26926 | 0.174102 | 0.117 | 0.084 | 1 | 8 | Kif16b | 0.033 | Young |
| 0.269731 | 0.280974 | 0.174 | 0.14 | 1 | 8 | Emc8 | 0.034 | Young |
| 0.269795 | 0.176129 | 0.281 | 0.235 | 1 | 8 | Gas2l3 | 0.046 | Young |
| 0.269958 | 0.368011 | 0.351 | 0.318 | 1 | 8 | Ufc1 | 0.033 | Young |
| 0.269972 | 0.391026 | 0.255 | 0.218 | 1 | 8 | Magoh | 0.037 | Young |
| 0.270193 | 0.431852 | 0.114 | 0.084 | 1 | 8 | Cep57 | 0.03 | Young |
| 0.270199 | 0.342284 | 0.119 | 0.089 | 1 | 8 | Smarcd1 | 0.03 | Young |
| 0.270224 | 0.091407 | 0.135 | 0.101 | 1 | 8 | Pkib | 0.034 | Young |
| 0.270302 | 0.370133 | 0.145 | 0.112 | 1 | 8 | Pitpnm1 | 0.033 | Young |
| 0.270382 | 0.321661 | 0.034 | 0.017 | 1 | 8 | Gm10036 | 0.017 | Young |
| 0.27056 | 0.159438 | 0.288 | 0.246 | 1 | 8 | Anapc16 | 0.042 | Young |
| 0.270715 | 0.191684 | 0.481 | 0.436 | 1 | 8 | Stk10 | 0.045 | Young |
| 0.270774 | 0.464204 | 0.075 | 0.05 | 1 | 8 | Zbtb6 | 0.025 | Young |
| 0.270821 | 0.068738 | 0.81 | 0.726 | 1 | 8 | Stat3 | 0.084 | Young |
| 0.270865 | 0.099992 | 0.452 | 0.397 | 1 | 8 | Tnrc6a | 0.055 | Young |
| 0.271009 | 0.158744 | 0.494 | 0.447 | 1 | 8 | Sidt2 | 0.047 | Young |
| 0.271102 | 0.376832 | 0.114 | 0.084 | 1 | 8 | Fam53a | 0.03 | Young |
| 0.271784 | 0.428219 | 0.127 | 0.095 | 1 | 8 | Sntb2 | 0.032 | Young |
| 0.271816 | 0.301473 | 0.156 | 0.123 | 1 | 8 | Flot1 | 0.033 | Young |
| 0.271839 | 0.503738 | 0.161 | 0.128 | 1 | 8 | Anapc4 | 0.033 | Young |
| 0.271847 | 1.059096 | 0.068 | 0.045 | 1 | 8 | Il1a | 0.023 | Young |
| 0.271888 | 0.562748 | 0.075 | 0.05 | 1 | 8 | Zbed4 | 0.025 | Young |
| 0.272135 | 0.102586 | 0.551 | 0.48 | 1 | 8 | Pds5a | 0.071 | Young |
| 0.272335 | 0.388142 | 0.135 | 0.101 | 1 | 8 | Syap1 | 0.034 | Young |
| 0.272933 | 0.163211 | 0.613 | 0.559 | 1 | 8 | Atp5c1 | 0.054 | Young |
| 0.273209 | 0.139075 | 0.143 | 0.106 | 1 | 8 | Trim36 | 0.037 | Young |
| 0.273265 | 0.104297 | 0.208 | 0.168 | 1 | 8 | Nsmce1 | 0.04 | Young |
| 0.273311 | 0.167297 | 0.286 | 0.24 | 1 | 8 | Slc35a4 | 0.046 | Young |
| 0.273367 | 0.279785 | 0.132 | 0.101 | 1 | 8 | Coq5 | 0.031 | Young |
| 0.273982 | 0.03406 | 0.868 | 0.827 | 1 | 8 | Adgre1 | 0.041 | Young |
| 0.274248 | 0.729373 | 0.169 | 0.14 | 1 | 8 | Acadvl | 0.029 | Young |
| 0.27447 | 0.078847 | 0.481 | 0.408 | 1 | 8 | Akt2 | 0.073 | Young |
| 0.274958 | 0.450281 | 0.088 | 0.061 | 1 | 8 | Gnl2 | 0.027 | Young |
| 0.275115 | 0.3132 | 0.119 | 0.089 | 1 | 8 | Tpra1 | 0.03 | Young |
| 0.275385 | 0.568229 | 0.068 | 0.045 | 1 | 8 | Hacd2 | 0.023 | Young |
| 0.2754 | 0.133649 | 0.41 | 0.346 | 1 | 8 | Tmem164 | 0.064 | Young |
| 0.275488 | 0.222671 | 0.384 | 0.335 | 1 | 8 | Ppp4r2 | 0.049 | Young |

| | | | | | | | | |
|---|---|---|---|---|---|---|---|---|
| 0.27585 | 0.074715 | 0.551 | 0.492 | 1 | 8 | Hnrnpa1 | 0.059 | Young |
| 0.27594 | 0.6676 | 0.055 | 0.034 | 1 | 8 | Piga | 0.021 | Young |
| 0.276022 | 0.501301 | 0.106 | 0.078 | 1 | 8 | Crcp | 0.028 | Young |
| 0.276148 | 0.701827 | 0.081 | 0.056 | 1 | 8 | Sephs1 | 0.025 | Young |
| 0.276367 | 0.219338 | 0.452 | 0.413 | 1 | 8 | Polr2a | 0.039 | Young |
| 0.276368 | 0.113701 | 0.743 | 0.676 | 1 | 8 | Taok3 | 0.067 | Young |
| 0.276425 | 0.287606 | 0.151 | 0.117 | 1 | 8 | Prps2 | 0.034 | Young |
| 0.276936 | 0.489918 | 0.075 | 0.05 | 1 | 8 | Sparc | 0.025 | Young |
| 0.277054 | 0.412434 | 0.283 | 0.246 | 1 | 8 | Rictor | 0.037 | Young |
| 0.277501 | 0.364766 | 0.075 | 0.05 | 1 | 8 | Ifitm5 | 0.025 | Young |
| 0.277917 | 0.163269 | 0.688 | 0.626 | 1 | 8 | Lilr4b | 0.062 | Young |
| 0.278766 | 0.028231 | 1 | 0.983 | 1 | 8 | mt-Nd2 | 0.017 | Young |
| 0.278935 | 0.08097 | 0.148 | 0.112 | 1 | 8 | Timm17b | 0.036 | Young |
| 0.279071 | 0.112139 | 0.322 | 0.263 | 1 | 8 | Rab3d | 0.059 | Young |
| 0.279154 | 1.001751 | 0.047 | 0.028 | 1 | 8 | Mus81 | 0.019 | Young |
| 0.279364 | 0.271196 | 0.106 | 0.078 | 1 | 8 | Sharpin | 0.028 | Young |
| 0.27951 | 1.026654 | 0.073 | 0.05 | 1 | 8 | Ppp1r16b | 0.023 | Young |
| 0.279553 | 0.570839 | 0.068 | 0.045 | 1 | 8 | Wbp1 | 0.023 | Young |
| 0.279673 | 0.175902 | 0.109 | 0.078 | 1 | 8 | Man2c1 | 0.031 | Young |
| 0.279928 | 0.718721 | 0.055 | 0.034 | 1 | 8 | Nbas | 0.021 | Young |
| 0.280178 | 0.012673 | 0.281 | 0.24 | 1 | 8 | Nudc | 0.041 | Young |
| 0.280202 | 0.297595 | 0.234 | 0.196 | 1 | 8 | Clec2i | 0.038 | Young |
| 0.280327 | 0.039417 | 0.403 | 0.341 | 1 | 8 | Cul1 | 0.062 | Young |
| 0.280555 | 0.29007 | 0.127 | 0.095 | 1 | 8 | 1600010M | 0.032 | Young |
| 0.280775 | 0.187447 | 0.114 | 0.084 | 1 | 8 | Slc33a1 | 0.03 | Young |
| 0.281335 | 0.124014 | 0.894 | 0.821 | 1 | 8 | Ubl3 | 0.073 | Young |
| 0.281851 | 0.156104 | 0.216 | 0.179 | 1 | 8 | Trim30d | 0.037 | Young |
| 0.282216 | 0.341187 | 0.13 | 0.101 | 1 | 8 | Surf1 | 0.029 | Young |
| 0.282349 | 0.765278 | 0.117 | 0.089 | 1 | 8 | Alg10b | 0.028 | Young |
| 0.282712 | 0.223489 | 0.231 | 0.19 | 1 | 8 | Cdk2ap1 | 0.041 | Young |
| 0.282769 | 1.142922 | 0.047 | 0.028 | 1 | 8 | Negr1 | 0.019 | Young |
| 0.282872 | 0.15111 | 0.096 | 0.067 | 1 | 8 | Calcoco1 | 0.029 | Young |
| 0.282935 | 0.294778 | 0.379 | 0.341 | 1 | 8 | Rtcb | 0.038 | Young |
| 0.282936 | 0.197407 | 0.239 | 0.196 | 1 | 8 | Map3k7 | 0.043 | Young |
| 0.282936 | 0.03214 | 0.964 | 0.905 | 1 | 8 | Cybb | 0.059 | Young |
| 0.283162 | 0.209827 | 0.257 | 0.212 | 1 | 8 | Fkbp3 | 0.045 | Young |
| 0.283166 | 0.090522 | 0.571 | 0.514 | 1 | 8 | Gatad2b | 0.057 | Young |
| 0.283186 | 0.333413 | 0.197 | 0.156 | 1 | 8 | Slc35f5 | 0.041 | Young |
| 0.283763 | 0.698802 | 0.068 | 0.045 | 1 | 8 | Bud13 | 0.023 | Young |
| 0.283955 | 0.750151 | 0.055 | 0.034 | 1 | 8 | St7 | 0.021 | Young |
| 0.283955 | 0.516702 | 0.055 | 0.034 | 1 | 8 | Slc52a2 | 0.021 | Young |
| 0.284042 | 0.50214 | 0.179 | 0.145 | 1 | 8 | Ulk1 | 0.034 | Young |
| 0.284808 | 0.058835 | 0.397 | 0.346 | 1 | 8 | Cd38 | 0.051 | Young |
| 0.284949 | 0.462494 | 0.088 | 0.061 | 1 | 8 | Ofd1 | 0.027 | Young |
| 0.284968 | 0.299936 | 0.283 | 0.24 | 1 | 8 | Tcerg1 | 0.043 | Young |

| | | | | | | | | |
|---|---|---|---|---|---|---|---|---|
| 0.285015 | 0.281432 | 0.169 | 0.134 | 1 | 8 | Sigmar1 | 0.035 | Young |
| 0.285111 | 0.165298 | 0.288 | 0.24 | 1 | 8 | Parp1 | 0.048 | Young |
| 0.285306 | 0.430689 | 0.055 | 0.034 | 1 | 8 | Ulk3 | 0.021 | Young |
| 0.285481 | 0.032453 | 0.504 | 0.419 | 1 | 8 | Phf3 | 0.085 | Young |
| 0.285482 | 0.140867 | 0.088 | 0.061 | 1 | 8 | Fam160a2 | 0.027 | Young |
| 0.285795 | 0.221408 | 0.223 | 0.184 | 1 | 8 | Ube3c | 0.039 | Young |
| 0.285945 | 0.080116 | 0.644 | 0.609 | 1 | 8 | Fyb | 0.035 | Young |
| 0.285983 | 0.408374 | 0.055 | 0.034 | 1 | 8 | Lancl2 | 0.021 | Young |
| 0.286666 | 0.735097 | 0.169 | 0.14 | 1 | 8 | Cdk5rap3 | 0.029 | Young |
| 0.286705 | 0.110248 | 0.512 | 0.458 | 1 | 8 | Tprgl | 0.054 | Young |
| 0.286733 | 0.666665 | 0.125 | 0.095 | 1 | 8 | Tbc1d31 | 0.03 | Young |
| 0.286756 | 0.005846 | 0.717 | 0.637 | 1 | 8 | 9-Sep | 0.08 | Young |
| 0.286796 | 0.732151 | 0.068 | 0.045 | 1 | 8 | Knstrn | 0.023 | Young |
| 0.287882 | 1.067863 | 0.047 | 0.028 | 1 | 8 | Zfp52 | 0.019 | Young |
| 0.288016 | 0.436575 | 0.068 | 0.045 | 1 | 8 | Tnfrsf25 | 0.023 | Young |
| 0.288016 | 0.349193 | 0.068 | 0.045 | 1 | 8 | Dolk | 0.023 | Young |
| 0.288201 | 0.084933 | 0.314 | 0.263 | 1 | 8 | Rheb | 0.051 | Young |
| 0.288978 | 0.130803 | 0.54 | 0.475 | 1 | 8 | Rin2 | 0.065 | Young |
| 0.289019 | 0.289128 | 0.145 | 0.112 | 1 | 8 | Sema4c | 0.033 | Young |
| 0.289055 | 0.198417 | 0.392 | 0.352 | 1 | 8 | Glrx3 | 0.04 | Young |
| 0.289057 | 0.100623 | 0.366 | 0.313 | 1 | 8 | Lta4h | 0.053 | Young |
| 0.28923 | 0.170816 | 0.203 | 0.162 | 1 | 8 | Smpd1 | 0.041 | Young |
| 0.290063 | 0.0668 | 0.501 | 0.453 | 1 | 8 | Higd2a | 0.048 | Young |
| 0.290229 | 0.315262 | 0.119 | 0.089 | 1 | 8 | Usp40 | 0.03 | Young |
| 0.290444 | 0.130733 | 0.6 | 0.508 | 1 | 8 | Capn2 | 0.092 | Young |
| 0.290831 | 0.726887 | 0.047 | 0.028 | 1 | 8 | Gm20045 | 0.019 | Young |
| 0.291333 | 0.039887 | 0.496 | 0.441 | 1 | 8 | Ppp1r12c | 0.055 | Young |
| 0.291333 | 0.079588 | 0.449 | 0.408 | 1 | 8 | Aff4 | 0.041 | Young |
| 0.29144 | 0.094955 | 0.055 | 0.034 | 1 | 8 | Kctd13 | 0.021 | Young |
| 0.291695 | 0.031843 | 0.068 | 0.045 | 1 | 8 | Mad1l1 | 0.023 | Young |
| 0.291715 | 0.005451 | 0.501 | 0.447 | 1 | 8 | Rsu1 | 0.054 | Young |
| 0.291864 | 0.356903 | 0.119 | 0.089 | 1 | 8 | Papd7 | 0.03 | Young |
| 0.291884 | 0.011392 | 0.265 | 0.229 | 1 | 8 | Ebna1bp2 | 0.036 | Young |
| 0.292016 | 0.272204 | 0.106 | 0.078 | 1 | 8 | Mcm4 | 0.028 | Young |
| 0.292276 | 0.498165 | 0.112 | 0.084 | 1 | 8 | Wdr7 | 0.028 | Young |
| 0.292615 | 0.267107 | 0.117 | 0.089 | 1 | 8 | Glce | 0.028 | Young |
| 0.292706 | 0.093869 | 0.647 | 0.592 | 1 | 8 | Prdx5 | 0.055 | Young |
| 0.29296 | 0.797848 | 0.091 | 0.067 | 1 | 8 | Rgp1 | 0.024 | Young |
| 0.293066 | 0.27448 | 0.405 | 0.369 | 1 | 8 | Xrn2 | 0.036 | Young |
| 0.293193 | 0.055383 | 0.642 | 0.575 | 1 | 8 | Irf5 | 0.067 | Young |
| 0.293551 | 0.121236 | 0.088 | 0.061 | 1 | 8 | Acvr1 | 0.027 | Young |
| 0.294508 | 0.490897 | 0.117 | 0.089 | 1 | 8 | Usb1 | 0.028 | Young |
| 0.294596 | 0.071152 | 0.408 | 0.352 | 1 | 8 | Sf3b3 | 0.056 | Young |
| 0.294685 | 1.422322 | 0.039 | 0.022 | 1 | 8 | Hps6 | 0.017 | Young |
| 0.294786 | 0.119195 | 0.068 | 0.045 | 1 | 8 | Trip6 | 0.023 | Young |

| | | | | | | | | |
|---|---|---|---|---|---|---|---|---|
| 0.294813 | 0.132552 | 0.075 | 0.05 | 1 | 8 | Gm45606 | 0.025 | Young |
| 0.294978 | 0.203606 | 0.226 | 0.184 | 1 | 8 | Mrps36 | 0.042 | Young |
| 0.295395 | 0.211188 | 0.21 | 0.173 | 1 | 8 | Wwc2 | 0.037 | Young |
| 0.295577 | 0.005669 | 0.055 | 0.034 | 1 | 8 | Ulbp1 | 0.021 | Young |
| 0.295733 | 0.101062 | 0.369 | 0.33 | 1 | 8 | Emd | 0.039 | Young |
| 0.29586 | 0.342092 | 0.119 | 0.089 | 1 | 8 | Slc35b3 | 0.03 | Young |
| 0.296001 | 0.613575 | 0.104 | 0.078 | 1 | 8 | Tyw1 | 0.026 | Young |
| 0.296641 | 0.097156 | 0.288 | 0.251 | 1 | 8 | Kidins220 | 0.037 | Young |
| 0.297255 | 0.682723 | 0.127 | 0.101 | 1 | 8 | Hat1 | 0.026 | Young |
| 0.297274 | 0.203268 | 0.068 | 0.045 | 1 | 8 | Fmn1 | 0.023 | Young |
| 0.297298 | 0.137499 | 0.239 | 0.196 | 1 | 8 | Exoc2 | 0.043 | Young |
| 0.297689 | 0.452866 | 0.143 | 0.112 | 1 | 8 | Mum1 | 0.031 | Young |
| 0.297823 | 0.002368 | 0.262 | 0.212 | 1 | 8 | Ell2 | 0.05 | Young |
| 0.297983 | 0.29694 | 0.195 | 0.162 | 1 | 8 | Get4 | 0.033 | Young |
| 0.298088 | 0.050917 | 0.223 | 0.184 | 1 | 8 | Mrpl18 | 0.039 | Young |
| 0.298121 | 0.069151 | 0.153 | 0.117 | 1 | 8 | Ctc1 | 0.036 | Young |
| 0.298294 | 0.751997 | 0.047 | 0.028 | 1 | 8 | Calml4 | 0.019 | Young |
| 0.298393 | 0.043241 | 0.823 | 0.732 | 1 | 8 | Diaph1 | 0.091 | Young |
| 0.298488 | 0.237508 | 0.081 | 0.056 | 1 | 8 | Tdrkh | 0.025 | Young |
| 0.298573 | 0.198274 | 0.395 | 0.363 | 1 | 8 | Golga4 | 0.032 | Young |
| 0.298769 | 0.63756 | 0.109 | 0.084 | 1 | 8 | Ap2a1 | 0.025 | Young |
| 0.298795 | 0.327695 | 0.19 | 0.156 | 1 | 8 | Usf3 | 0.034 | Young |
| 0.299047 | 0.256219 | 0.047 | 0.028 | 1 | 8 | Zfp821 | 0.019 | Young |
| 0.299047 | 0.129453 | 0.047 | 0.028 | 1 | 8 | Cd209e | 0.019 | Young |
| 0.29909 | 0.02846 | 0.187 | 0.145 | 1 | 8 | Smim12 | 0.042 | Young |
| 0.299179 | 0.51137 | 0.119 | 0.089 | 1 | 8 | Fgd3 | 0.03 | Young |
| 0.299349 | 0.058678 | 0.532 | 0.486 | 1 | 8 | Adcy7 | 0.046 | Young |
| 0.299469 | 0.017675 | 0.094 | 0.067 | 1 | 8 | AI987944 | 0.027 | Young |
| 0.299802 | 0.3019 | 0.047 | 0.028 | 1 | 8 | Abca2 | 0.019 | Young |
| 0.300134 | 0.875878 | 0.073 | 0.05 | 1 | 8 | Smpd4 | 0.023 | Young |
| 0.300267 | 0.232424 | 0.236 | 0.196 | 1 | 8 | Sirt7 | 0.04 | Young |
| 0.300465 | 0.760556 | 0.091 | 0.067 | 1 | 8 | Zbtb16 | 0.024 | Young |
| 0.300558 | 0.588588 | 0.047 | 0.028 | 1 | 8 | Cyb5d2 | 0.019 | Young |
| 0.300975 | 0.065501 | 0.403 | 0.352 | 1 | 8 | Zfp445 | 0.051 | Young |
| 0.301006 | 0.12478 | 0.125 | 0.095 | 1 | 8 | Cntln | 0.03 | Young |
| 0.30134 | 0.07836 | 0.683 | 0.598 | 1 | 8 | Synj1 | 0.085 | Young |
| 0.301397 | 0.061436 | 0.868 | 0.844 | 1 | 8 | Sub1 | 0.024 | Young |
| 0.301622 | 0.370943 | 0.179 | 0.145 | 1 | 8 | Taf3 | 0.034 | Young |
| 0.301729 | 0.508054 | 0.291 | 0.257 | 1 | 8 | Rps6ka4 | 0.034 | Young |
| 0.301901 | 0.163021 | 0.286 | 0.24 | 1 | 8 | Bcl7c | 0.046 | Young |
| 0.30193 | 0.022858 | 0.387 | 0.324 | 1 | 8 | Timm10b | 0.063 | Young |
| 0.302023 | 0.271821 | 0.257 | 0.218 | 1 | 8 | Rapgef2 | 0.039 | Young |
| 0.302043 | 0.149499 | 0.119 | 0.089 | 1 | 8 | Rbms2 | 0.03 | Young |
| 0.302045 | 0.056736 | 0.969 | 0.939 | 1 | 8 | Ubc | 0.03 | Young |
| 0.30221 | 0.581237 | 0.135 | 0.106 | 1 | 8 | Las1l | 0.029 | Young |

| | | | | | | | | |
|---|---|---|---|---|---|---|---|---|
| 0.302518 | 0.29984 | 0.081 | 0.056 | 1 | 8 | Tmem94 | 0.025 | Young |
| 0.302657 | 0.048705 | 0.571 | 0.52 | 1 | 8 | Tle3 | 0.051 | Young |
| 0.302662 | 0.210713 | 0.574 | 0.52 | 1 | 8 | Fem1c | 0.054 | Young |
| 0.303525 | 0.162443 | 0.106 | 0.078 | 1 | 8 | Sugp1 | 0.028 | Young |
| 0.303593 | 0.240107 | 0.047 | 0.028 | 1 | 8 | Olfr1167 | 0.019 | Young |
| 0.303679 | 0.225746 | 0.268 | 0.223 | 1 | 8 | Ptpn22 | 0.045 | Young |
| 0.303685 | 0.157047 | 0.878 | 0.838 | 1 | 8 | S100a4 | 0.04 | Young |
| 0.30378 | 0.851876 | 0.039 | 0.022 | 1 | 8 | Rabl3 | 0.017 | Young |
| 0.304289 | 0.078921 | 0.922 | 0.849 | 1 | 8 | Prcp | 0.073 | Young |
| 0.304615 | 0.891756 | 0.039 | 0.022 | 1 | 8 | Tctn1 | 0.017 | Young |
| 0.304883 | 0.492102 | 0.104 | 0.078 | 1 | 8 | Vegfb | 0.026 | Young |
| 0.305046 | 0.125819 | 0.852 | 0.782 | 1 | 8 | Il6ra | 0.07 | Young |
| 0.305193 | 0.033134 | 1 | 0.994 | 1 | 8 | H2-Eb1 | 0.006 | Young |
| 0.305629 | 0.089004 | 0.351 | 0.296 | 1 | 8 | Ndufv2 | 0.055 | Young |
| 0.305653 | 0.034614 | 0.582 | 0.531 | 1 | 8 | Cysltr1 | 0.051 | Young |
| 0.305659 | 0.06364 | 0.127 | 0.095 | 1 | 8 | Taf12 | 0.032 | Young |
| 0.305884 | 0.264855 | 0.047 | 0.028 | 1 | 8 | Palm | 0.019 | Young |
| 0.306071 | 0.132365 | 0.286 | 0.24 | 1 | 8 | Bcor | 0.046 | Young |
| 0.306289 | 0.090429 | 0.229 | 0.184 | 1 | 8 | Snrnp27 | 0.045 | Young |
| 0.306292 | 0.727772 | 0.039 | 0.022 | 1 | 8 | 4930555A( | 0.017 | Young |
| 0.306491 | 0.207801 | 0.306 | 0.268 | 1 | 8 | Rgl2 | 0.038 | Young |
| 0.306649 | 0.243995 | 0.047 | 0.028 | 1 | 8 | Nrg2 | 0.019 | Young |
| 0.306677 | 0.182656 | 0.281 | 0.24 | 1 | 8 | Atad2b | 0.041 | Young |
| 0.306728 | 0.185723 | 0.138 | 0.106 | 1 | 8 | Kcnq1 | 0.032 | Young |
| 0.30682 | 0.571755 | 0.073 | 0.05 | 1 | 8 | Atg10 | 0.023 | Young |
| 0.307091 | 0.112597 | 0.548 | 0.503 | 1 | 8 | Psmb8 | 0.045 | Young |
| 0.307417 | 0.338978 | 0.047 | 0.028 | 1 | 8 | Cad | 0.019 | Young |
| 0.307598 | 0.247428 | 0.106 | 0.078 | 1 | 8 | Vipas39 | 0.028 | Young |
| 0.307847 | 0.161224 | 0.132 | 0.101 | 1 | 8 | Adam33 | 0.031 | Young |
| 0.307879 | 0.130276 | 0.353 | 0.307 | 1 | 8 | Madd | 0.046 | Young |
| 0.308033 | 0.133725 | 0.361 | 0.313 | 1 | 8 | Me2 | 0.048 | Young |
| 0.30813 | 0.082507 | 0.592 | 0.547 | 1 | 8 | Cd48 | 0.045 | Young |
| 0.308521 | 0.953152 | 0.052 | 0.034 | 1 | 8 | Plekha8 | 0.018 | Young |
| 0.308552 | 0.017754 | 0.909 | 0.849 | 1 | 8 | Glud1 | 0.06 | Young |
| 0.308765 | 0.554481 | 0.06 | 0.039 | 1 | 8 | 3-Sep | 0.021 | Young |
| 0.30882 | 0.12875 | 0.255 | 0.212 | 1 | 8 | Exoc6b | 0.043 | Young |
| 0.309282 | 0.179463 | 0.218 | 0.179 | 1 | 8 | Samd8 | 0.039 | Young |
| 0.309399 | 0.05864 | 0.649 | 0.592 | 1 | 8 | Per1 | 0.057 | Young |
| 0.309663 | 0.692153 | 0.039 | 0.022 | 1 | 8 | Prdm15 | 0.017 | Young |
| 0.309726 | 0.152986 | 0.047 | 0.028 | 1 | 8 | Setd6 | 0.019 | Young |
| 0.309893 | 0.207568 | 0.101 | 0.073 | 1 | 8 | 3110062M | 0.028 | Young |
| 0.309924 | 0.104196 | 0.068 | 0.045 | 1 | 8 | Fktn | 0.023 | Young |
| 0.309983 | 0.057731 | 0.322 | 0.268 | 1 | 8 | Rps27rt | 0.054 | Young |
| 0.31057 | 0.90746 | 0.065 | 0.045 | 1 | 8 | AW554918 | 0.02 | Young |
| 0.310654 | 0.168896 | 0.431 | 0.385 | 1 | 8 | Pin4 | 0.046 | Young |

| | | | | | | | | |
|---|---|---|---|---|---|---|---|---|
| 0.310771 | 0.76303 | 0.109 | 0.084 | 1 | 8 | Prkca | 0.025 | Young |
| 0.310805 | 0.089446 | 0.06 | 0.039 | 1 | 8 | Gmppb | 0.021 | Young |
| 0.311015 | 0.559733 | 0.177 | 0.145 | 1 | 8 | Dcaf13 | 0.032 | Young |
| 0.311186 | 0.144057 | 0.114 | 0.084 | 1 | 8 | Ino80b | 0.03 | Young |
| 0.311266 | 0.056243 | 0.543 | 0.475 | 1 | 8 | Jup | 0.068 | Young |
| 0.31133 | 0.352505 | 0.132 | 0.101 | 1 | 8 | Scmh1 | 0.031 | Young |
| 0.311487 | 0.126505 | 0.356 | 0.307 | 1 | 8 | Lman1 | 0.049 | Young |
| 0.311779 | 0.238092 | 0.236 | 0.196 | 1 | 8 | Itga5 | 0.04 | Young |
| 0.311788 | 0.206844 | 0.135 | 0.106 | 1 | 8 | Mrps34 | 0.029 | Young |
| 0.311806 | 0.128681 | 0.226 | 0.19 | 1 | 8 | Pak1 | 0.036 | Young |
| 0.312047 | 0.328669 | 0.047 | 0.028 | 1 | 8 | Smug1 | 0.019 | Young |
| 0.312126 | 0.046325 | 0.27 | 0.229 | 1 | 8 | Fkbp4 | 0.041 | Young |
| 0.312581 | 0.504309 | 0.182 | 0.151 | 1 | 8 | Sphk2 | 0.031 | Young |
| 0.31272 | 0.101073 | 0.306 | 0.257 | 1 | 8 | Camta2 | 0.049 | Young |
| 0.312886 | 0.065526 | 0.312 | 0.263 | 1 | 8 | Rnf44 | 0.049 | Young |
| 0.313061 | 0.087343 | 0.403 | 0.341 | 1 | 8 | Ppp1r2 | 0.062 | Young |
| 0.31312 | 0.033456 | 0.299 | 0.251 | 1 | 8 | Rab24 | 0.048 | Young |
| 0.313806 | 0.069358 | 0.816 | 0.749 | 1 | 8 | Nap1l1 | 0.067 | Young |
| 0.31399 | 0.165842 | 0.208 | 0.173 | 1 | 8 | Pes1 | 0.035 | Young |
| 0.314181 | 0.031447 | 0.647 | 0.57 | 1 | 8 | Uqcr10 | 0.077 | Young |
| 0.314204 | 0.463187 | 0.145 | 0.117 | 1 | 8 | Pld1 | 0.028 | Young |
| 0.314225 | 0.712534 | 0.06 | 0.039 | 1 | 8 | D030056L2 | 0.021 | Young |
| 0.314353 | 0.280354 | 0.169 | 0.134 | 1 | 8 | Calhm2 | 0.035 | Young |
| 0.314359 | 0.225674 | 0.244 | 0.207 | 1 | 8 | 2210016F1 | 0.037 | Young |
| 0.314405 | 0.113874 | 0.145 | 0.112 | 1 | 8 | Zfp512 | 0.033 | Young |
| 0.314408 | 0.049635 | 0.538 | 0.475 | 1 | 8 | Snap23 | 0.063 | Young |
| 0.314486 | 0.969558 | 0.031 | 0.017 | 1 | 8 | Kif9 | 0.014 | Young |
| 0.3146 | 0.228129 | 0.156 | 0.123 | 1 | 8 | D730003I1 | 0.033 | Young |
| 0.314631 | 0.07202 | 0.374 | 0.324 | 1 | 8 | Elk3 | 0.05 | Young |
| 0.314802 | 0.172295 | 0.327 | 0.291 | 1 | 8 | Cpeb4 | 0.036 | Young |
| 0.314821 | 0.604965 | 0.086 | 0.061 | 1 | 8 | Ano8 | 0.025 | Young |
| 0.315116 | 1.337879 | 0.016 | 0.006 | 1 | 8 | 2010007H( | 0.01 | Young |
| 0.315116 | 1.272639 | 0.016 | 0.006 | 1 | 8 | Lysmd4 | 0.01 | Young |
| 0.315116 | 1.238964 | 0.016 | 0.006 | 1 | 8 | Gm10282 | 0.01 | Young |
| 0.315116 | 1.224533 | 0.016 | 0.006 | 1 | 8 | Yes1 | 0.01 | Young |
| 0.315116 | 1.212226 | 0.016 | 0.006 | 1 | 8 | Rapgef5 | 0.01 | Young |
| 0.315116 | 1.188138 | 0.016 | 0.006 | 1 | 8 | Zfp420 | 0.01 | Young |
| 0.315116 | 1.158045 | 0.016 | 0.006 | 1 | 8 | Jmjd4 | 0.01 | Young |
| 0.315116 | 1.044172 | 0.016 | 0.006 | 1 | 8 | Stard10 | 0.01 | Young |
| 0.315116 | 1.02448 | 0.016 | 0.006 | 1 | 8 | Rsph3a | 0.01 | Young |
| 0.315116 | 1.002909 | 0.016 | 0.006 | 1 | 8 | B3galt4 | 0.01 | Young |
| 0.315116 | 0.939144 | 0.016 | 0.006 | 1 | 8 | Sirt6 | 0.01 | Young |
| 0.315438 | 0.056188 | 0.652 | 0.609 | 1 | 8 | Ankrd11 | 0.043 | Young |
| 0.315715 | 0.2089 | 0.304 | 0.263 | 1 | 8 | Bbip1 | 0.041 | Young |
| 0.315858 | 0.662668 | 0.091 | 0.067 | 1 | 8 | Rab40c | 0.024 | Young |

| | | | | | | | | |
|---|---|---|---|---|---|---|---|---|
| 0.316014 | 0.21898 | 0.081 | 0.056 | 1 | 8 | Spic | 0.025 | Young |
| 0.316096 | 0.694398 | 0.073 | 0.05 | 1 | 8 | Rpp30 | 0.023 | Young |
| 0.316296 | 0.447942 | 0.099 | 0.073 | 1 | 8 | Blcap | 0.026 | Young |
| 0.316509 | 0.193416 | 0.166 | 0.134 | 1 | 8 | Erh | 0.032 | Young |
| 0.316513 | 1.286111 | 0.016 | 0.006 | 1 | 8 | Fam135a | 0.01 | Young |
| 0.316513 | 1.283149 | 0.016 | 0.006 | 1 | 8 | Btla | 0.01 | Young |
| 0.316513 | 0.833894 | 0.016 | 0.006 | 1 | 8 | Chst1 | 0.01 | Young |
| 0.317133 | 0.089118 | 0.881 | 0.816 | 1 | 8 | Kdm6b | 0.065 | Young |
| 0.317304 | 0.096203 | 0.431 | 0.391 | 1 | 8 | Swap70 | 0.04 | Young |
| 0.317337 | 0.4794 | 0.039 | 0.022 | 1 | 8 | Cenpq | 0.017 | Young |
| 0.317722 | 0.012268 | 0.114 | 0.084 | 1 | 8 | Ndufaf2 | 0.03 | Young |
| 0.317914 | 1.702325 | 0.016 | 0.006 | 1 | 8 | Apold1 | 0.01 | Young |
| 0.317914 | 1.154134 | 0.016 | 0.006 | 1 | 8 | Zbtb32 | 0.01 | Young |
| 0.317914 | 1.017184 | 0.016 | 0.006 | 1 | 8 | 2700046G( | 0.01 | Young |
| 0.317914 | 0.869946 | 0.016 | 0.006 | 1 | 8 | Magee1 | 0.01 | Young |
| 0.317914 | 0.836889 | 0.016 | 0.006 | 1 | 8 | Gdf9 | 0.01 | Young |
| 0.317914 | 0.820099 | 0.016 | 0.006 | 1 | 8 | Gm33103 | 0.01 | Young |
| 0.318155 | 0.107933 | 0.652 | 0.587 | 1 | 8 | Capza1 | 0.065 | Young |
| 0.318369 | 0.008698 | 0.34 | 0.296 | 1 | 8 | Ncoa7 | 0.044 | Young |
| 0.318379 | 0.58842 | 0.091 | 0.067 | 1 | 8 | Lancl1 | 0.024 | Young |
| 0.318456 | 0.529213 | 0.099 | 0.073 | 1 | 8 | Sap30bp | 0.026 | Young |
| 0.318939 | 0.161048 | 0.101 | 0.073 | 1 | 8 | Ints14 | 0.028 | Young |
| 0.318948 | 0.322495 | 0.109 | 0.084 | 1 | 8 | H2-Ob | 0.025 | Young |
| 0.318959 | 0.231415 | 0.361 | 0.324 | 1 | 8 | Hsd17b4 | 0.037 | Young |
| 0.319112 | 0.848578 | 0.065 | 0.045 | 1 | 8 | Gm37233 | 0.02 | Young |
| 0.319119 | 0.163718 | 0.148 | 0.117 | 1 | 8 | Thumpd1 | 0.031 | Young |
| 0.319318 | 1.150774 | 0.016 | 0.006 | 1 | 8 | C030006K1 | 0.01 | Young |
| 0.319318 | 1.001422 | 0.016 | 0.006 | 1 | 8 | Dynlt1c | 0.01 | Young |
| 0.319318 | 0.911026 | 0.016 | 0.006 | 1 | 8 | Zfp870 | 0.01 | Young |
| 0.319318 | 0.869836 | 0.016 | 0.006 | 1 | 8 | Gm10791 | 0.01 | Young |
| 0.319318 | 0.844756 | 0.016 | 0.006 | 1 | 8 | Gm2814 | 0.01 | Young |
| 0.319318 | 0.810892 | 0.016 | 0.006 | 1 | 8 | Trim16 | 0.01 | Young |
| 0.319355 | 0.306455 | 0.197 | 0.168 | 1 | 8 | Ubap1 | 0.029 | Young |
| 0.319463 | 0.442533 | 0.109 | 0.084 | 1 | 8 | Ccdc85b | 0.025 | Young |
| 0.319587 | 0.0707 | 0.478 | 0.419 | 1 | 8 | Kitl | 0.059 | Young |
| 0.319786 | 0.220833 | 0.091 | 0.067 | 1 | 8 | Casp2 | 0.024 | Young |
| 0.31979 | 0.365389 | 0.208 | 0.173 | 1 | 8 | Arhgap11a | 0.035 | Young |
| 0.320079 | 0.085912 | 0.249 | 0.212 | 1 | 8 | Bicd2 | 0.037 | Young |
| 0.320099 | 1.245996 | 0.023 | 0.011 | 1 | 8 | Kyat3 | 0.012 | Young |
| 0.320099 | 1.123803 | 0.023 | 0.011 | 1 | 8 | Gpr171 | 0.012 | Young |
| 0.320279 | 0.89191 | 0.052 | 0.034 | 1 | 8 | Rnf26 | 0.018 | Young |
| 0.320442 | 0.392484 | 0.06 | 0.039 | 1 | 8 | Slc2a9 | 0.021 | Young |
| 0.320651 | 0.248638 | 0.317 | 0.279 | 1 | 8 | Mrps15 | 0.038 | Young |
| 0.320727 | 0.938554 | 0.016 | 0.006 | 1 | 8 | Zfp759 | 0.01 | Young |
| 0.320727 | 0.857101 | 0.016 | 0.006 | 1 | 8 | Jmjd8 | 0.01 | Young |

| | | | | | | | | |
|---|---|---|---|---|---|---|---|---|
| 0.320727 | 0.737399 | 0.016 | 0.006 | 1 | 8 | Slc22a21 | 0.01 | Young |
| 0.320727 | 0.690335 | 0.016 | 0.006 | 1 | 8 | Zfp354c | 0.01 | Young |
| 0.320788 | 0.143477 | 0.039 | 0.022 | 1 | 8 | Inafm2 | 0.017 | Young |
| 0.321024 | 0.801145 | 0.052 | 0.034 | 1 | 8 | Pik3ap1 | 0.018 | Young |
| 0.321024 | 0.664002 | 0.052 | 0.034 | 1 | 8 | Sorbs3 | 0.018 | Young |
| 0.321105 | 0.754884 | 0.065 | 0.045 | 1 | 8 | Zfhx2 | 0.02 | Young |
| 0.321241 | 0.836398 | 0.031 | 0.017 | 1 | 8 | Slc16a13 | 0.014 | Young |
| 0.321378 | 0.226554 | 0.257 | 0.218 | 1 | 8 | Usp12 | 0.039 | Young |
| 0.321531 | 0.012746 | 0.413 | 0.363 | 1 | 8 | Higd1a | 0.05 | Young |
| 0.321711 | 0.459732 | 0.247 | 0.218 | 1 | 8 | Atf2 | 0.029 | Young |
| 0.32214 | 0.828604 | 0.016 | 0.006 | 1 | 8 | Cdkl2 | 0.01 | Young |
| 0.32214 | 0.652542 | 0.016 | 0.006 | 1 | 8 | St8sia1 | 0.01 | Young |
| 0.32214 | 0.609315 | 0.016 | 0.006 | 1 | 8 | Pdss1 | 0.01 | Young |
| 0.32214 | 0.5801 | 0.016 | 0.006 | 1 | 8 | Gm49087 | 0.01 | Young |
| 0.32214 | 0.579035 | 0.016 | 0.006 | 1 | 8 | Gm15706 | 0.01 | Young |
| 0.32214 | 0.535667 | 0.016 | 0.006 | 1 | 8 | Gpr21 | 0.01 | Young |
| 0.322284 | 0.141182 | 0.174 | 0.14 | 1 | 8 | Ppme1 | 0.034 | Young |
| 0.322329 | 0.372482 | 0.151 | 0.123 | 1 | 8 | Pak4 | 0.028 | Young |
| 0.32236 | 1.109121 | 0.023 | 0.011 | 1 | 8 | Zfp748 | 0.012 | Young |
| 0.32236 | 0.9176 | 0.023 | 0.011 | 1 | 8 | Isg20 | 0.012 | Young |
| 0.322406 | 0.115429 | 0.273 | 0.229 | 1 | 8 | Inpp5f | 0.044 | Young |
| 0.322516 | 0.727872 | 0.052 | 0.034 | 1 | 8 | Trp53rka | 0.018 | Young |
| 0.323062 | 0.01024 | 0.19 | 0.151 | 1 | 8 | Ech1 | 0.039 | Young |
| 0.323188 | 0.241745 | 0.031 | 0.017 | 1 | 8 | Fut7 | 0.014 | Young |
| 0.323291 | 0.296194 | 0.197 | 0.162 | 1 | 8 | Aph1b | 0.035 | Young |
| 0.323392 | 0.310059 | 0.039 | 0.022 | 1 | 8 | Gins2 | 0.017 | Young |
| 0.323494 | 1.471756 | 0.023 | 0.011 | 1 | 8 | Xlr4c | 0.012 | Young |
| 0.323494 | 1.294263 | 0.023 | 0.011 | 1 | 8 | Gm47230 | 0.012 | Young |
| 0.323505 | 0.141818 | 0.255 | 0.218 | 1 | 8 | Zbtb44 | 0.037 | Young |
| 0.323557 | 0.454162 | 0.016 | 0.006 | 1 | 8 | Nfib | 0.01 | Young |
| 0.323557 | 0.427138 | 0.016 | 0.006 | 1 | 8 | Gmpr | 0.01 | Young |
| 0.323557 | 0.080204 | 0.016 | 0.006 | 1 | 8 | Aldh1a1 | 0.01 | Young |
| 0.323707 | 0.523818 | 0.104 | 0.078 | 1 | 8 | Nxt2 | 0.026 | Young |
| 0.323751 | 0.211807 | 0.2 | 0.168 | 1 | 8 | Polg | 0.032 | Young |
| 0.323899 | 0.076564 | 0.26 | 0.212 | 1 | 8 | Zdhhc3 | 0.048 | Young |
| 0.324165 | 0.523283 | 0.031 | 0.017 | 1 | 8 | Kazald1 | 0.014 | Young |
| 0.324165 | 0.306334 | 0.031 | 0.017 | 1 | 8 | Crtc1 | 0.014 | Young |
| 0.324263 | 0.229627 | 0.039 | 0.022 | 1 | 8 | Ankrd13d | 0.017 | Young |
| 0.324813 | 0.352751 | 0.296 | 0.257 | 1 | 8 | Maff | 0.039 | Young |
| 0.325771 | 0.953977 | 0.023 | 0.011 | 1 | 8 | Tle2 | 0.012 | Young |
| 0.326124 | 0.602805 | 0.031 | 0.017 | 1 | 8 | Gm7160 | 0.014 | Young |
| 0.326267 | 0.105662 | 0.052 | 0.034 | 1 | 8 | Supt7l | 0.018 | Young |
| 0.326384 | 1.189778 | 0.044 | 0.028 | 1 | 8 | Pradc1 | 0.016 | Young |
| 0.326386 | 0.054031 | 0.138 | 0.106 | 1 | 8 | Med29 | 0.032 | Young |
| 0.326735 | 0.308692 | 0.125 | 0.095 | 1 | 8 | Lmf1 | 0.03 | Young |

| | | | | | | | | |
|---|---|---|---|---|---|---|---|---|
| 0.326914 | 0.832802 | 0.023 | 0.011 | 1 | 8 | Fam234b | 0.012 | Young |
| 0.326975 | 0.34774 | 0.221 | 0.19 | 1 | 8 | Dpy30 | 0.031 | Young |
| 0.327106 | 0.118859 | 0.031 | 0.017 | 1 | 8 | Ipp | 0.014 | Young |
| 0.327659 | 0.314193 | 0.158 | 0.128 | 1 | 8 | Tm2d3 | 0.03 | Young |
| 0.328059 | 0.998985 | 0.023 | 0.011 | 1 | 8 | Nipsnap1 | 0.012 | Young |
| 0.328059 | 0.684664 | 0.023 | 0.011 | 1 | 8 | Platr17 | 0.012 | Young |
| 0.32809 | 0.61961 | 0.031 | 0.017 | 1 | 8 | BC049352 | 0.014 | Young |
| 0.328106 | 0.491719 | 0.073 | 0.05 | 1 | 8 | Hist1h4i | 0.023 | Young |
| 0.328137 | 0.162659 | 0.551 | 0.52 | 1 | 8 | Lst1 | 0.031 | Young |
| 0.328145 | 0.112593 | 0.122 | 0.095 | 1 | 8 | Prmt3 | 0.027 | Young |
| 0.328334 | 0.095444 | 0.364 | 0.324 | 1 | 8 | Tmem219 | 0.04 | Young |
| 0.328614 | 0.092731 | 0.265 | 0.218 | 1 | 8 | Wipf2 | 0.047 | Young |
| 0.329076 | 0.219038 | 0.031 | 0.017 | 1 | 8 | Cyfip2 | 0.014 | Young |
| 0.329152 | 0.235399 | 0.2 | 0.168 | 1 | 8 | AU040320 | 0.032 | Young |
| 0.329157 | 0.496444 | 0.065 | 0.045 | 1 | 8 | Sec22c | 0.02 | Young |
| 0.329206 | 0.765699 | 0.023 | 0.011 | 1 | 8 | Ak1 | 0.012 | Young |
| 0.329206 | 0.722078 | 0.023 | 0.011 | 1 | 8 | Usp54 | 0.012 | Young |
| 0.329206 | 0.68429 | 0.023 | 0.011 | 1 | 8 | Kctd21 | 0.012 | Young |
| 0.329288 | 0.297741 | 0.081 | 0.056 | 1 | 8 | 2300009A0 | 0.025 | Young |
| 0.329456 | 0.082913 | 0.091 | 0.067 | 1 | 8 | Rnf157 | 0.024 | Young |
| 0.329484 | 0.538848 | 0.127 | 0.101 | 1 | 8 | Mtif2 | 0.026 | Young |
| 0.329525 | 0.009613 | 0.257 | 0.218 | 1 | 8 | Zfp385a | 0.039 | Young |
| 0.329555 | 0.058186 | 0.694 | 0.615 | 1 | 8 | Lifr | 0.079 | Young |
| 0.329599 | 0.5429 | 0.078 | 0.056 | 1 | 8 | Ppih | 0.022 | Young |
| 0.32988 | 0.113198 | 0.416 | 0.358 | 1 | 8 | Etnk1 | 0.058 | Young |
| 0.329935 | 0.011007 | 0.46 | 0.419 | 1 | 8 | Entpd1 | 0.041 | Young |
| 0.330356 | 0.854416 | 0.023 | 0.011 | 1 | 8 | Hells | 0.012 | Young |
| 0.330485 | 0.342997 | 0.288 | 0.263 | 1 | 8 | Phf10 | 0.025 | Young |
| 0.330607 | 0.49061 | 0.091 | 0.067 | 1 | 8 | Alkbh3 | 0.024 | Young |
| 0.330912 | 0.014614 | 0.39 | 0.335 | 1 | 8 | Parp14 | 0.055 | Young |
| 0.331441 | 0.894041 | 0.083 | 0.061 | 1 | 8 | Mettl6 | 0.022 | Young |
| 0.331479 | 0.071161 | 0.218 | 0.179 | 1 | 8 | Grap | 0.039 | Young |
| 0.331509 | 0.833129 | 0.023 | 0.011 | 1 | 8 | Eif3j2 | 0.012 | Young |
| 0.331509 | 0.446241 | 0.023 | 0.011 | 1 | 8 | Treml2 | 0.012 | Young |
| 0.331685 | 0.555693 | 0.132 | 0.106 | 1 | 8 | Pole3 | 0.026 | Young |
| 0.331738 | 0.477789 | 0.109 | 0.084 | 1 | 8 | Sp3os | 0.025 | Young |
| 0.332155 | 0.171767 | 0.369 | 0.33 | 1 | 8 | Ttc39b | 0.039 | Young |
| 0.332463 | 0.237617 | 0.322 | 0.285 | 1 | 8 | Cnot4 | 0.037 | Young |
| 0.332664 | 0.616387 | 0.023 | 0.011 | 1 | 8 | Plppr4 | 0.012 | Young |
| 0.332713 | 0.191313 | 0.288 | 0.251 | 1 | 8 | Psme2b | 0.037 | Young |
| 0.332745 | 0.073763 | 0.322 | 0.268 | 1 | 8 | Nfya | 0.054 | Young |
| 0.333652 | 0.000669 | 0.112 | 0.084 | 1 | 8 | Bmi1 | 0.028 | Young |
| 0.33366 | 0.194766 | 0.122 | 0.095 | 1 | 8 | Wdtc1 | 0.027 | Young |
| 0.333822 | 0.626833 | 0.023 | 0.011 | 1 | 8 | Cep97 | 0.012 | Young |
| 0.334132 | 0.281227 | 0.338 | 0.307 | 1 | 8 | Lyl1 | 0.031 | Young |

| | | | | | | | | |
|---|---|---|---|---|---|---|---|---|
| 0.334545 | 0.349381 | 0.073 | 0.05 | 1 | 8 | Dgka | 0.023 | Young |
| 0.334608 | 1.201947 | 0.044 | 0.028 | 1 | 8 | Suox | 0.016 | Young |
| 0.334656 | 0.155573 | 0.592 | 0.559 | 1 | 8 | Chmp2a | 0.033 | Young |
| 0.334983 | 0.226923 | 0.023 | 0.011 | 1 | 8 | Rabl2 | 0.012 | Young |
| 0.335193 | 0.412027 | 0.073 | 0.05 | 1 | 8 | Maml2 | 0.023 | Young |
| 0.335268 | 0.245333 | 0.184 | 0.151 | 1 | 8 | Vps16 | 0.033 | Young |
| 0.33572 | 0.093149 | 0.322 | 0.274 | 1 | 8 | Rsbn1l | 0.048 | Young |
| 0.335804 | 0.119825 | 0.416 | 0.363 | 1 | 8 | Tnfsf12 | 0.053 | Young |
| 0.336031 | 0.158643 | 0.465 | 0.413 | 1 | 8 | Psmb3 | 0.052 | Young |
| 0.336082 | 0.018535 | 0.816 | 0.743 | 1 | 8 | Pak2 | 0.073 | Young |
| 0.336249 | 0.493557 | 0.096 | 0.073 | 1 | 8 | Scaper | 0.023 | Young |
| 0.336269 | 1.043448 | 0.044 | 0.028 | 1 | 8 | Coil | 0.016 | Young |
| 0.336492 | 0.12354 | 0.073 | 0.05 | 1 | 8 | Pde6d | 0.023 | Young |
| 0.337311 | 0.152791 | 0.023 | 0.011 | 1 | 8 | Il18r1 | 0.012 | Young |
| 0.33732 | 0.110733 | 0.119 | 0.089 | 1 | 8 | Aimp2 | 0.03 | Young |
| 0.337893 | 0.069968 | 0.522 | 0.458 | 1 | 8 | Tnpo3 | 0.064 | Young |
| 0.3379 | 0.10276 | 0.164 | 0.134 | 1 | 8 | Tle1 | 0.03 | Young |
| 0.338011 | 0.234113 | 0.218 | 0.184 | 1 | 8 | Itpr2 | 0.034 | Young |
| 0.33877 | 0.811158 | 0.044 | 0.028 | 1 | 8 | Rnf122 | 0.016 | Young |
| 0.338925 | 0.41988 | 0.109 | 0.084 | 1 | 8 | Pdzd11 | 0.025 | Young |
| 0.339032 | 0.263579 | 0.187 | 0.156 | 1 | 8 | Cyp27a1 | 0.031 | Young |
| 0.339078 | 0.214601 | 0.184 | 0.151 | 1 | 8 | Ltv1 | 0.033 | Young |
| 0.33923 | 0.404061 | 0.052 | 0.034 | 1 | 8 | Ccdc126 | 0.018 | Young |
| 0.339379 | 0.399815 | 0.127 | 0.101 | 1 | 8 | Pex5 | 0.026 | Young |
| 0.339901 | 0.255265 | 0.091 | 0.067 | 1 | 8 | Fam60a | 0.024 | Young |
| 0.340062 | 0.033994 | 0.429 | 0.369 | 1 | 8 | BC017643 | 0.06 | Young |
| 0.340202 | 0.02127 | 0.179 | 0.145 | 1 | 8 | Frmd4a | 0.034 | Young |
| 0.340207 | 0.06381 | 0.491 | 0.441 | 1 | 8 | Arl5b | 0.05 | Young |
| 0.340226 | 0.221079 | 0.148 | 0.117 | 1 | 8 | Mapk8ip3 | 0.031 | Young |
| 0.340444 | 0.83749 | 0.044 | 0.028 | 1 | 8 | Fbxw17 | 0.016 | Young |
| 0.340464 | 0.322954 | 0.317 | 0.296 | 1 | 8 | Ccdc47 | 0.021 | Young |
| 0.340695 | 0.317815 | 0.148 | 0.117 | 1 | 8 | Klhdc2 | 0.031 | Young |
| 0.340767 | 0.03684 | 0.397 | 0.352 | 1 | 8 | Heca | 0.045 | Young |
| 0.341157 | 0.013789 | 0.099 | 0.073 | 1 | 8 | Rgcc | 0.026 | Young |
| 0.341479 | 0.221495 | 0.065 | 0.045 | 1 | 8 | Mblac2 | 0.02 | Young |
| 0.341897 | 0.029992 | 0.275 | 0.229 | 1 | 8 | Fam20b | 0.046 | Young |
| 0.34225 | 0.597841 | 0.008 | 0.017 | 1 | 8 | Krtcap3 | -0.009 | Young |
| 0.342858 | 0.025734 | 0.099 | 0.073 | 1 | 8 | Dglucy | 0.026 | Young |
| 0.34298 | 0.049546 | 0.286 | 0.246 | 1 | 8 | Mrpl12 | 0.04 | Young |
| 0.343083 | 0.512 | 0.096 | 0.073 | 1 | 8 | Txlng | 0.023 | Young |
| 0.343272 | 0.42204 | 0.104 | 0.078 | 1 | 8 | Tomm6 | 0.026 | Young |
| 0.343343 | 0.256816 | 0.184 | 0.151 | 1 | 8 | Rabggtb | 0.033 | Young |
| 0.343485 | 0.123344 | 0.213 | 0.179 | 1 | 8 | Selenoi | 0.034 | Young |
| 0.343513 | 0.855015 | 0.057 | 0.039 | 1 | 8 | Surf2 | 0.018 | Young |
| 0.343657 | 0.589905 | 0.096 | 0.073 | 1 | 8 | Rnf34 | 0.023 | Young |

| | | | | | | | | |
|---|---|---|---|---|---|---|---|---|
| 0.343951 | 0.163619 | 0.143 | 0.112 | 1 | 8 | Zfp110 | 0.031 | Young |
| 0.343976 | 0.231181 | 0.117 | 0.089 | 1 | 8 | Tubgcp2 | 0.028 | Young |
| 0.344761 | 0.280967 | 0.13 | 0.101 | 1 | 8 | Fam241a | 0.029 | Young |
| 0.344868 | 0.078282 | 0.122 | 0.095 | 1 | 8 | Rfx1 | 0.027 | Young |
| 0.345085 | 0.007193 | 0.314 | 0.268 | 1 | 8 | Nub1 | 0.046 | Young |
| 0.345213 | 0.148493 | 0.195 | 0.162 | 1 | 8 | Asl | 0.033 | Young |
| 0.345758 | 0.030848 | 0.429 | 0.391 | 1 | 8 | Ptpn11 | 0.038 | Young |
| 0.345824 | 0.135202 | 0.504 | 0.447 | 1 | 8 | Cdc37 | 0.057 | Young |
| 0.345944 | 0.312534 | 0.078 | 0.056 | 1 | 8 | Atm | 0.022 | Young |
| 0.346069 | 0.393993 | 0.169 | 0.14 | 1 | 8 | Ddx47 | 0.029 | Young |
| 0.346124 | 0.22836 | 0.262 | 0.235 | 1 | 8 | Relb | 0.027 | Young |
| 0.34635 | 0.460368 | 0.065 | 0.045 | 1 | 8 | Fancg | 0.02 | Young |
| 0.346387 | 0.530559 | 0.091 | 0.067 | 1 | 8 | Ccs | 0.024 | Young |
| 0.346779 | 0.148999 | 0.184 | 0.151 | 1 | 8 | Xdh | 0.033 | Young |
| 0.346871 | 0.024713 | 0.774 | 0.715 | 1 | 8 | 5031439G( | 0.059 | Young |
| 0.347081 | 0.321959 | 0.239 | 0.207 | 1 | 8 | Ifi209 | 0.032 | Young |
| 0.347567 | 0.072009 | 0.255 | 0.218 | 1 | 8 | Deptor | 0.037 | Young |
| 0.347766 | 0.362472 | 0.138 | 0.112 | 1 | 8 | Bag5 | 0.026 | Young |
| 0.348339 | 0.071014 | 0.353 | 0.313 | 1 | 8 | Diaph2 | 0.04 | Young |
| 0.348504 | 0.314913 | 0.078 | 0.056 | 1 | 8 | Unc13d | 0.022 | Young |
| 0.348575 | 0.143054 | 0.275 | 0.235 | 1 | 8 | Tpm1 | 0.04 | Young |
| 0.348577 | 0.061084 | 0.052 | 0.034 | 1 | 8 | Dedd2 | 0.018 | Young |
| 0.348829 | 0.457621 | 0.187 | 0.162 | 1 | 8 | Tlr4 | 0.025 | Young |
| 0.348834 | 0.078739 | 0.849 | 0.771 | 1 | 8 | Nfkbid | 0.078 | Young |
| 0.349209 | 0.411077 | 0.109 | 0.084 | 1 | 8 | Prkch | 0.025 | Young |
| 0.349221 | 0.217674 | 0.231 | 0.196 | 1 | 8 | Tatdn2 | 0.035 | Young |
| 0.349277 | 0.09679 | 0.67 | 0.592 | 1 | 8 | Acer3 | 0.078 | Young |
| 0.349808 | 0.147651 | 0.273 | 0.235 | 1 | 8 | Pcgf5 | 0.038 | Young |
| 0.349881 | 0.023429 | 0.369 | 0.313 | 1 | 8 | Spg21 | 0.056 | Young |
| 0.35085 | 0.008186 | 0.805 | 0.737 | 1 | 8 | Srp9 | 0.068 | Young |
| 0.350869 | 0.629749 | 0.145 | 0.123 | 1 | 8 | Rhbdf2 | 0.022 | Young |
| 0.351094 | 0.024856 | 0.787 | 0.67 | 1 | 8 | Reep3 | 0.117 | Young |
| 0.351263 | 0.085739 | 0.925 | 0.883 | 1 | 8 | Atp5l | 0.042 | Young |
| 0.351453 | 0.581574 | 0.164 | 0.14 | 1 | 8 | Rnaseh2a | 0.024 | Young |
| 0.351454 | 0.67992 | 0.044 | 0.028 | 1 | 8 | Gm340 | 0.016 | Young |
| 0.35172 | 0.246814 | 0.078 | 0.056 | 1 | 8 | Slc12a4 | 0.022 | Young |
| 0.351794 | 0.349544 | 0.192 | 0.162 | 1 | 8 | S100pbp | 0.03 | Young |
| 0.351846 | 0.528989 | 0.114 | 0.089 | 1 | 8 | Rmi1 | 0.025 | Young |
| 0.35197 | 0.289186 | 0.065 | 0.045 | 1 | 8 | Kctd11 | 0.02 | Young |
| 0.35197 | 0.036738 | 0.065 | 0.045 | 1 | 8 | Zxdb | 0.02 | Young |
| 0.352124 | 0.168237 | 0.122 | 0.095 | 1 | 8 | Mpg | 0.027 | Young |
| 0.352205 | 0.194692 | 0.104 | 0.078 | 1 | 8 | Pgam5 | 0.026 | Young |
| 0.35231 | 0.868039 | 0.044 | 0.028 | 1 | 8 | Havcr2 | 0.016 | Young |
| 0.352519 | 0.053662 | 0.052 | 0.034 | 1 | 8 | Zfp777 | 0.018 | Young |
| 0.353023 | 0.249251 | 0.148 | 0.117 | 1 | 8 | Pogz | 0.031 | Young |

| | | | | | | | | |
|---|---|---|---|---|---|---|---|---|
| 0.353696 | 0.070079 | 0.283 | 0.24 | 1 | 8 | Psmd7 | 0.043 | Young |
| 0.353768 | 0.067004 | 0.319 | 0.279 | 1 | 8 | Tmpo | 0.04 | Young |
| 0.354026 | 0.559841 | 0.044 | 0.028 | 1 | 8 | Nepro | 0.016 | Young |
| 0.354266 | 0.259499 | 0.114 | 0.089 | 1 | 8 | Gins4 | 0.025 | Young |
| 0.354669 | 0.319677 | 0.096 | 0.073 | 1 | 8 | Irgm2 | 0.023 | Young |
| 0.355075 | 0.293618 | 0.114 | 0.089 | 1 | 8 | Pygl | 0.025 | Young |
| 0.355122 | 1.342966 | 0.049 | 0.034 | 1 | 8 | Armcx5 | 0.015 | Young |
| 0.355297 | 0.05232 | 0.566 | 0.531 | 1 | 8 | Fis1 | 0.035 | Young |
| 0.355413 | 0.041916 | 0.66 | 0.575 | 1 | 8 | Zbtb7a | 0.085 | Young |
| 0.355498 | 0.001014 | 0.509 | 0.469 | 1 | 8 | Cmtm7 | 0.04 | Young |
| 0.35551 | 0.035658 | 0.278 | 0.24 | 1 | 8 | Ntan1 | 0.038 | Young |
| 0.355712 | 0.035728 | 0.322 | 0.274 | 1 | 8 | Txndc9 | 0.048 | Young |
| 0.356314 | 0.182865 | 0.122 | 0.095 | 1 | 8 | Mfn2 | 0.027 | Young |
| 0.35661 | 0.386167 | 0.044 | 0.028 | 1 | 8 | Ccdc102a | 0.016 | Young |
| 0.356966 | 0.022814 | 0.655 | 0.553 | 1 | 8 | Ogdh | 0.102 | Young |
| 0.357093 | 0.032168 | 0.878 | 0.816 | 1 | 8 | Picalm | 0.062 | Young |
| 0.357143 | 0.484915 | 0.161 | 0.134 | 1 | 8 | Slc13a3 | 0.027 | Young |
| 0.357441 | 0.256175 | 0.187 | 0.162 | 1 | 8 | Hspb11 | 0.025 | Young |
| 0.357465 | 0.05485 | 0.348 | 0.302 | 1 | 8 | Paics | 0.046 | Young |
| 0.357972 | 0.171435 | 0.14 | 0.112 | 1 | 8 | Akt3 | 0.028 | Young |
| 0.358013 | 0.008436 | 0.423 | 0.38 | 1 | 8 | Kmt2c | 0.043 | Young |
| 0.358329 | 0.084762 | 0.875 | 0.838 | 1 | 8 | Plek | 0.037 | Young |
| 0.35834 | 0.289405 | 0.044 | 0.028 | 1 | 8 | Gstz1 | 0.016 | Young |
| 0.358669 | 0.098446 | 0.309 | 0.274 | 1 | 8 | Eri1 | 0.035 | Young |
| 0.359059 | 0.164543 | 0.543 | 0.492 | 1 | 8 | Snrpb2 | 0.051 | Young |
| 0.359076 | 0.306519 | 0.065 | 0.045 | 1 | 8 | Selenoo | 0.02 | Young |
| 0.35909 | 0.145167 | 0.226 | 0.19 | 1 | 8 | Prkce | 0.036 | Young |
| 0.359206 | 0.415409 | 0.044 | 0.028 | 1 | 8 | Hdhd5 | 0.016 | Young |
| 0.359475 | 0.060729 | 0.2 | 0.162 | 1 | 8 | Dusp7 | 0.038 | Young |
| 0.359628 | 0.168781 | 0.255 | 0.218 | 1 | 8 | Coa3 | 0.037 | Young |
| 0.360989 | 0.068805 | 0.379 | 0.33 | 1 | 8 | Coro1b | 0.049 | Young |
| 0.36129 | 0.261859 | 0.127 | 0.101 | 1 | 8 | Ttc19 | 0.026 | Young |
| 0.361814 | 0.448 | 0.044 | 0.028 | 1 | 8 | Tsen2 | 0.016 | Young |
| 0.362041 | 0.232332 | 0.418 | 0.374 | 1 | 8 | Slc29a1 | 0.044 | Young |
| 0.362477 | 1.181848 | 0.036 | 0.022 | 1 | 8 | Odf2l | 0.014 | Young |
| 0.362686 | 0.755028 | 0.044 | 0.028 | 1 | 8 | Vamp1 | 0.016 | Young |
| 0.36269 | 0.254229 | 0.203 | 0.173 | 1 | 8 | Csad | 0.03 | Young |
| 0.363372 | 0.383541 | 0.127 | 0.101 | 1 | 8 | Srprb | 0.026 | Young |
| 0.363438 | 0.926391 | 0.036 | 0.022 | 1 | 8 | Tepsin | 0.014 | Young |
| 0.363757 | 0.005832 | 0.174 | 0.14 | 1 | 8 | Tubgcp4 | 0.034 | Young |
| 0.364499 | 0.058933 | 0.187 | 0.151 | 1 | 8 | Ncbp1 | 0.036 | Young |
| 0.364506 | 0.287754 | 0.182 | 0.151 | 1 | 8 | BC029722 | 0.031 | Young |
| 0.364773 | 0.150191 | 0.078 | 0.056 | 1 | 8 | Nat10 | 0.022 | Young |
| 0.365014 | 0.11377 | 0.171 | 0.14 | 1 | 8 | Eif2b5 | 0.031 | Young |
| 0.365024 | 0.139733 | 0.325 | 0.285 | 1 | 8 | Slain2 | 0.04 | Young |

| | | | | | | | | |
|---|---|---|---|---|---|---|---|---|
| 0.365069 | 0.008498 | 0.41 | 0.363 | 1 | 8 | Saraf | 0.047 | Young |
| 0.365075 | 0.080838 | 0.582 | 0.52 | 1 | 8 | Nono | 0.062 | Young |
| 0.36554 | 0.491321 | 0.083 | 0.061 | 1 | 8 | Mast2 | 0.022 | Young |
| 0.365845 | 0.42333 | 0.109 | 0.084 | 1 | 8 | Pparg | 0.025 | Young |
| 0.36635 | 0.127837 | 0.244 | 0.207 | 1 | 8 | Trim35 | 0.037 | Young |
| 0.366675 | 0.050633 | 0.413 | 0.374 | 1 | 8 | Ncstn | 0.039 | Young |
| 0.366718 | 0.032769 | 0.187 | 0.151 | 1 | 8 | Oard1 | 0.036 | Young |
| 0.366965 | 0.508399 | 0.14 | 0.117 | 1 | 8 | Sart3 | 0.023 | Young |
| 0.367066 | 0.266979 | 0.044 | 0.028 | 1 | 8 | Slc25a33 | 0.016 | Young |
| 0.367354 | 0.273674 | 0.119 | 0.095 | 1 | 8 | Thoc5 | 0.024 | Young |
| 0.367743 | 0.001193 | 0.068 | 0.089 | 1 | 8 | Plscr1 | -0.021 | Young |
| 0.36801 | 0.221795 | 0.187 | 0.156 | 1 | 8 | Wdr48 | 0.031 | Young |
| 0.368032 | 0.078387 | 0.353 | 0.307 | 1 | 8 | Tbl1x | 0.046 | Young |
| 0.368119 | 0.113242 | 0.268 | 0.223 | 1 | 8 | Bckdk | 0.045 | Young |
| 0.368271 | 0.969066 | 0.036 | 0.022 | 1 | 8 | Pank4 | 0.014 | Young |
| 0.368271 | 0.86739 | 0.036 | 0.022 | 1 | 8 | Dyrk1b | 0.014 | Young |
| 0.368422 | 0.312261 | 0.138 | 0.112 | 1 | 8 | Rnf170 | 0.026 | Young |
| 0.368672 | 0.152182 | 0.275 | 0.246 | 1 | 8 | Srp54a | 0.029 | Young |
| 0.369225 | 0.087708 | 0.057 | 0.039 | 1 | 8 | Hyal2 | 0.018 | Young |
| 0.36974 | 0.21924 | 0.231 | 0.201 | 1 | 8 | Chml | 0.03 | Young |
| 0.369948 | 0.254179 | 0.343 | 0.307 | 1 | 8 | Sys1 | 0.036 | Young |
| 0.370167 | 0.128705 | 0.273 | 0.229 | 1 | 8 | Cops2 | 0.044 | Young |
| 0.370677 | 0.078707 | 0.083 | 0.061 | 1 | 8 | Cyb561d2 | 0.022 | Young |
| 0.370774 | 0.072667 | 0.057 | 0.039 | 1 | 8 | Rfxank | 0.018 | Young |
| 0.370823 | 0.770754 | 0.075 | 0.056 | 1 | 8 | Rac3 | 0.019 | Young |
| 0.370842 | 0.0897 | 0.468 | 0.419 | 1 | 8 | Ndufb6 | 0.049 | Young |
| 0.371322 | 0.25992 | 0.083 | 0.061 | 1 | 8 | Elp3 | 0.022 | Young |
| 0.371361 | 0.126854 | 0.065 | 0.045 | 1 | 8 | Bloc1s1 | 0.02 | Young |
| 0.371499 | 0.26729 | 0.132 | 0.106 | 1 | 8 | Pcgf3 | 0.026 | Young |
| 0.372188 | 0.020081 | 0.514 | 0.464 | 1 | 8 | Ptpro | 0.05 | Young |
| 0.372388 | 0.872304 | 0.049 | 0.034 | 1 | 8 | 6330418K0 | 0.015 | Young |
| 0.372614 | 0.188167 | 0.083 | 0.061 | 1 | 8 | Vps37c | 0.022 | Young |
| 0.372635 | 0.1144 | 0.171 | 0.14 | 1 | 8 | Insig2 | 0.031 | Young |
| 0.372956 | 0.415434 | 0.07 | 0.05 | 1 | 8 | Men1 | 0.02 | Young |
| 0.373143 | 0.882287 | 0.036 | 0.022 | 1 | 8 | Strip2 | 0.014 | Young |
| 0.374039 | 0.048859 | 0.416 | 0.369 | 1 | 8 | Lactb | 0.047 | Young |
| 0.374122 | 0.787501 | 0.036 | 0.022 | 1 | 8 | Ptch1 | 0.014 | Young |
| 0.374142 | 0.062479 | 0.151 | 0.117 | 1 | 8 | Gsr | 0.034 | Young |
| 0.374663 | 0.502834 | 0.057 | 0.039 | 1 | 8 | Dcaf6 | 0.018 | Young |
| 0.375016 | 0.06923 | 0.429 | 0.385 | 1 | 8 | Osbpl9 | 0.044 | Young |
| 0.375143 | 0.56335 | 0.106 | 0.084 | 1 | 8 | Rnaseh2b | 0.022 | Young |
| 0.375493 | 0.034057 | 0.325 | 0.285 | 1 | 8 | Mta2 | 0.04 | Young |
| 0.375735 | 0.740509 | 0.049 | 0.034 | 1 | 8 | Nsun4 | 0.015 | Young |
| 0.375857 | 0.316906 | 0.083 | 0.061 | 1 | 8 | Zfp830 | 0.022 | Young |
| 0.375912 | 0.048353 | 0.592 | 0.531 | 1 | 8 | Brd2 | 0.061 | Young |

| | | | | | | | | |
|---|---|---|---|---|---|---|---|---|
| 0.376085 | 0.633402 | 0.036 | 0.022 | 1 | 8 | Mfsd4b4 | 0.014 | Young |
| 0.376238 | 0.467629 | 0.088 | 0.067 | 1 | 8 | Iqsec2 | 0.021 | Young |
| 0.376254 | 0.067079 | 0.792 | 0.732 | 1 | 8 | 1810037I1 | 0.06 | Young |
| 0.376596 | 0.040727 | 0.208 | 0.179 | 1 | 8 | Ubac2 | 0.029 | Young |
| 0.376932 | 0.109492 | 0.291 | 0.263 | 1 | 8 | Otud5 | 0.028 | Young |
| 0.377184 | 0.089771 | 0.122 | 0.095 | 1 | 8 | Khk | 0.027 | Young |
| 0.377532 | 0.00921 | 0.634 | 0.57 | 1 | 8 | Laptm4a | 0.064 | Young |
| 0.377626 | 0.137697 | 0.475 | 0.447 | 1 | 8 | Zc3h12a | 0.028 | Young |
| 0.378055 | 0.824046 | 0.036 | 0.022 | 1 | 8 | Zfp35 | 0.014 | Young |
| 0.378055 | 0.406756 | 0.036 | 0.022 | 1 | 8 | Fam212a | 0.014 | Young |
| 0.378319 | 0.138128 | 0.221 | 0.19 | 1 | 8 | Fam98b | 0.031 | Young |
| 0.378464 | 0.49547 | 0.083 | 0.061 | 1 | 8 | Tsr1 | 0.022 | Young |
| 0.378823 | 0.067616 | 0.294 | 0.257 | 1 | 8 | Rnf114 | 0.037 | Young |
| 0.378975 | 0.335503 | 0.075 | 0.056 | 1 | 8 | Qsox2 | 0.019 | Young |
| 0.379485 | 0.124577 | 0.213 | 0.179 | 1 | 8 | Tlk2 | 0.034 | Young |
| 0.379503 | 0.020536 | 0.738 | 0.665 | 1 | 8 | Prrc2c | 0.073 | Young |
| 0.379629 | 0.501458 | 0.062 | 0.045 | 1 | 8 | Amd1 | 0.017 | Young |
| 0.379657 | 0.189822 | 0.021 | 0.034 | 1 | 8 | Pex26 | -0.013 | Young |
| 0.379665 | 0.138271 | 0.345 | 0.307 | 1 | 8 | Marf1 | 0.038 | Young |
| 0.379696 | 0.08188 | 0.782 | 0.704 | 1 | 8 | Creg1 | 0.078 | Young |
| 0.379759 | 0.444236 | 0.143 | 0.117 | 1 | 8 | Kifc3 | 0.026 | Young |
| 0.380306 | 0.271389 | 0.161 | 0.134 | 1 | 8 | Hs2st1 | 0.027 | Young |
| 0.380386 | 0.432072 | 0.275 | 0.246 | 1 | 8 | Rxra | 0.029 | Young |
| 0.381021 | 0.464479 | 0.036 | 0.022 | 1 | 8 | Zkscan17 | 0.014 | Young |
| 0.381022 | 0.206362 | 0.148 | 0.117 | 1 | 8 | Adk | 0.031 | Young |
| 0.381033 | 0.115135 | 0.605 | 0.564 | 1 | 8 | Pde4b | 0.041 | Young |
| 0.38135 | 0.093378 | 0.481 | 0.408 | 1 | 8 | Cdipt | 0.073 | Young |
| 0.381512 | 0.404941 | 0.106 | 0.084 | 1 | 8 | Timm29 | 0.022 | Young |
| 0.382013 | 0.25229 | 0.036 | 0.022 | 1 | 8 | Pard6a | 0.014 | Young |
| 0.382263 | 0.277401 | 0.112 | 0.089 | 1 | 8 | Megf8 | 0.023 | Young |
| 0.38281 | 0.08375 | 0.27 | 0.235 | 1 | 8 | Eif4e3 | 0.035 | Young |
| 0.382855 | 0.055368 | 0.096 | 0.073 | 1 | 8 | Fastk | 0.023 | Young |
| 0.382873 | 0.063122 | 0.436 | 0.397 | 1 | 8 | Kat6a | 0.039 | Young |
| 0.3831 | 0.178957 | 0.192 | 0.162 | 1 | 8 | Cnot2 | 0.03 | Young |
| 0.383457 | 0.36694 | 0.174 | 0.151 | 1 | 8 | Kank3 | 0.023 | Young |
| 0.38355 | 0.080822 | 0.514 | 0.458 | 1 | 8 | Eif3b | 0.056 | Young |
| 0.383585 | 0.015141 | 0.255 | 0.223 | 1 | 8 | Eif2s1 | 0.032 | Young |
| 0.384002 | 0.569569 | 0.036 | 0.022 | 1 | 8 | Trim14 | 0.014 | Young |
| 0.384002 | 0.49357 | 0.036 | 0.022 | 1 | 8 | Inpp1 | 0.014 | Young |
| 0.384534 | 0.233137 | 0.294 | 0.257 | 1 | 8 | Ncaph2 | 0.037 | Young |
| 0.384673 | 0.037419 | 0.153 | 0.123 | 1 | 8 | Zfp422 | 0.03 | Young |
| 0.385135 | 0.149562 | 0.088 | 0.067 | 1 | 8 | Zfp358 | 0.021 | Young |
| 0.385317 | 0.029889 | 0.14 | 0.112 | 1 | 8 | Eya3 | 0.028 | Young |
| 0.38554 | 0.119822 | 0.517 | 0.458 | 1 | 8 | Tubb2a | 0.059 | Young |
| 0.385867 | 0.293418 | 0.109 | 0.084 | 1 | 8 | Bloc1s6 | 0.025 | Young |

| | | | | | | | | |
|---|---|---|---|---|---|---|---|---|
| 0.385974 | 0.047662 | 0.161 | 0.128 | 1 | 8 | Ralgapb | 0.033 | Young |
| 0.386352 | 0.103239 | 0.083 | 0.061 | 1 | 8 | Ube2o | 0.022 | Young |
| 0.386391 | 0.348097 | 0.143 | 0.117 | 1 | 8 | Mvd | 0.026 | Young |
| 0.386518 | 0.670643 | 0.081 | 0.061 | 1 | 8 | Nit2 | 0.02 | Young |
| 0.386535 | 0.083533 | 0.273 | 0.235 | 1 | 8 | Rprd2 | 0.038 | Young |
| 0.386703 | 0.174199 | 0.21 | 0.179 | 1 | 8 | Dpagt1 | 0.031 | Young |
| 0.386996 | 0.45184 | 0.036 | 0.022 | 1 | 8 | Mapk1ip1 | 0.014 | Young |
| 0.387185 | 0.589571 | 0.062 | 0.045 | 1 | 8 | Bnip1 | 0.017 | Young |
| 0.387279 | 0.104133 | 0.057 | 0.039 | 1 | 8 | Tcta | 0.018 | Young |
| 0.387595 | 0.347859 | 0.049 | 0.034 | 1 | 8 | Wdr6 | 0.015 | Young |
| 0.387811 | 0.02479 | 0.514 | 0.441 | 1 | 8 | Aph1a | 0.073 | Young |
| 0.388451 | 0.2435 | 0.049 | 0.034 | 1 | 8 | Zbtb12 | 0.015 | Young |
| 0.388475 | 0.178181 | 0.057 | 0.039 | 1 | 8 | Clstn1 | 0.018 | Young |
| 0.388703 | 0.106317 | 0.899 | 0.81 | 1 | 8 | Erp29 | 0.089 | Young |
| 0.388734 | 0.048645 | 0.361 | 0.324 | 1 | 8 | Bank1 | 0.037 | Young |
| 0.38932 | 0.908785 | 0.075 | 0.056 | 1 | 8 | Mir155hg | 0.019 | Young |
| 0.390241 | 0.11964 | 0.429 | 0.385 | 1 | 8 | Pnpla8 | 0.044 | Young |
| 0.390366 | 0.072607 | 0.883 | 0.838 | 1 | 8 | Ifitm2 | 0.045 | Young |
| 0.390659 | 1.047858 | 0.055 | 0.039 | 1 | 8 | 4632415L0 | 0.016 | Young |
| 0.391409 | 0.586112 | 0.075 | 0.056 | 1 | 8 | Lin37 | 0.019 | Young |
| 0.391674 | 0.259805 | 0.184 | 0.156 | 1 | 8 | Eif2ak1 | 0.028 | Young |
| 0.391709 | 0.143726 | 0.283 | 0.246 | 1 | 8 | Kdelr2 | 0.037 | Young |
| 0.391955 | 0.495231 | 0.14 | 0.117 | 1 | 8 | Oas2 | 0.023 | Young |
| 0.392269 | 0.176424 | 0.135 | 0.112 | 1 | 8 | Cog8 | 0.023 | Young |
| 0.392489 | 0.087734 | 0.312 | 0.279 | 1 | 8 | Birc2 | 0.033 | Young |
| 0.39264 | 1.070693 | 0.029 | 0.017 | 1 | 8 | 4930522L1 | 0.012 | Young |
| 0.392805 | 0.537608 | 0.075 | 0.056 | 1 | 8 | Cdc25a | 0.019 | Young |
| 0.392968 | 0.381968 | 0.151 | 0.123 | 1 | 8 | Trip4 | 0.028 | Young |
| 0.393134 | 0.075721 | 0.14 | 0.112 | 1 | 8 | Pbx1 | 0.028 | Young |
| 0.393782 | 0.863288 | 0.029 | 0.017 | 1 | 8 | Thumpd2 | 0.012 | Young |
| 0.39417 | 0.198795 | 0.275 | 0.24 | 1 | 8 | Lrrc8d | 0.035 | Young |
| 0.394595 | 0.27368 | 0.081 | 0.061 | 1 | 8 | Bcl9 | 0.02 | Young |
| 0.394671 | 0.195384 | 0.361 | 0.318 | 1 | 8 | Nfkbie | 0.043 | Young |
| 0.394722 | 0.487774 | 0.179 | 0.156 | 1 | 8 | 4833420G1 | 0.023 | Young |
| 0.39479 | 0.061619 | 0.309 | 0.268 | 1 | 8 | Slc25a4 | 0.041 | Young |
| 0.395104 | 0.141926 | 0.166 | 0.14 | 1 | 8 | Trappc9 | 0.026 | Young |
| 0.395352 | 0.10701 | 0.766 | 0.749 | 1 | 8 | Aif1 | 0.017 | Young |
| 0.395509 | 0.035694 | 0.119 | 0.095 | 1 | 8 | Trappc2 | 0.024 | Young |
| 0.395608 | 0.095663 | 0.075 | 0.056 | 1 | 8 | Lig1 | 0.019 | Young |
| 0.395716 | 0.188667 | 0.255 | 0.218 | 1 | 8 | Tmem106b | 0.037 | Young |
| 0.396187 | 0.114051 | 0.26 | 0.218 | 1 | 8 | Atg5 | 0.042 | Young |
| 0.396701 | 0.270825 | 0.208 | 0.179 | 1 | 8 | Vars | 0.029 | Young |
| 0.397086 | 0.08816 | 0.187 | 0.156 | 1 | 8 | Kdm3b | 0.031 | Young |
| 0.39712 | 0.213578 | 0.442 | 0.413 | 1 | 8 | Ddb1 | 0.029 | Young |
| 0.397144 | 0.490002 | 0.062 | 0.045 | 1 | 8 | Pcca | 0.017 | Young |

| | | | | | | | | |
|---|---|---|---|---|---|---|---|---|
| 0.397155 | 0.185668 | 0.275 | 0.24 | 1 | 8 | Dnttip2 | 0.035 | Young |
| 0.397218 | 0.891864 | 0.029 | 0.017 | 1 | 8 | Per2 | 0.012 | Young |
| 0.397367 | 0.361547 | 0.187 | 0.162 | 1 | 8 | Gm2000 | 0.025 | Young |
| 0.397468 | 0.147676 | 0.049 | 0.067 | 1 | 8 | Mcph1 | -0.018 | Young |
| 0.397706 | 0.202232 | 0.127 | 0.101 | 1 | 8 | Tecpr1 | 0.026 | Young |
| 0.397792 | 0.111006 | 0.86 | 0.771 | 1 | 8 | Ddx3x | 0.089 | Young |
| 0.397841 | 0.527805 | 0.117 | 0.095 | 1 | 8 | Kctd17 | 0.022 | Young |
| 0.398077 | 0.114917 | 0.273 | 0.235 | 1 | 8 | Ddx18 | 0.038 | Young |
| 0.398331 | 0.029714 | 0.538 | 0.492 | 1 | 8 | Rassf1 | 0.046 | Young |
| 0.398368 | 0.687649 | 0.029 | 0.017 | 1 | 8 | Farp2 | 0.012 | Young |
| 0.398694 | 0.025415 | 0.569 | 0.514 | 1 | 8 | Etf1 | 0.055 | Young |
| 0.399009 | 0.22184 | 0.203 | 0.173 | 1 | 8 | Tbc1d9b | 0.03 | Young |
| 0.399172 | 0.129742 | 0.338 | 0.307 | 1 | 8 | Pdcl3 | 0.031 | Young |
| 0.399176 | 0.059359 | 0.322 | 0.279 | 1 | 8 | Pitpnc1 | 0.043 | Young |
| 0.399565 | 0.204389 | 0.395 | 0.374 | 1 | 8 | Ssu72 | 0.021 | Young |
| 0.399756 | 0.202554 | 0.431 | 0.408 | 1 | 8 | Snx17 | 0.023 | Young |
| 0.400401 | 0.028348 | 0.447 | 0.397 | 1 | 8 | Coa5 | 0.05 | Young |
| 0.400529 | 0.02171 | 0.184 | 0.156 | 1 | 8 | Zufsp | 0.028 | Young |
| 0.400552 | 0.47737 | 0.049 | 0.034 | 1 | 8 | Arl15 | 0.015 | Young |
| 0.400552 | 0.093744 | 0.049 | 0.034 | 1 | 8 | Surf6 | 0.015 | Young |
| 0.400667 | 0.084236 | 0.935 | 0.888 | 1 | 8 | Maf | 0.047 | Young |
| 0.400673 | 0.833742 | 0.029 | 0.017 | 1 | 8 | Cidec | 0.012 | Young |
| 0.400683 | 0.338235 | 0.088 | 0.067 | 1 | 8 | Rtl8a | 0.021 | Young |
| 0.400685 | 0.076861 | 0.361 | 0.318 | 1 | 8 | Ankrd13c | 0.043 | Young |
| 0.401425 | 0.660876 | 0.049 | 0.034 | 1 | 8 | Aldoc | 0.015 | Young |
| 0.401477 | 0.236153 | 0.265 | 0.229 | 1 | 8 | Jdp2 | 0.036 | Young |
| 0.401566 | 0.65469 | 0.109 | 0.089 | 1 | 8 | Mrpl50 | 0.02 | Young |
| 0.401644 | 0.047835 | 0.564 | 0.503 | 1 | 8 | Bst2 | 0.061 | Young |
| 0.401749 | 0.244874 | 0.119 | 0.095 | 1 | 8 | Alg1 | 0.024 | Young |
| 0.401828 | 0.623051 | 0.029 | 0.017 | 1 | 8 | Mpp7 | 0.012 | Young |
| 0.401899 | 0.014064 | 0.345 | 0.302 | 1 | 8 | Antxr2 | 0.043 | Young |
| 0.401964 | 0.375751 | 0.143 | 0.117 | 1 | 8 | E2f4 | 0.026 | Young |
| 0.402299 | 0.410295 | 0.049 | 0.034 | 1 | 8 | Gtf3c5 | 0.015 | Young |
| 0.402506 | 0.458656 | 0.112 | 0.089 | 1 | 8 | Pik3cb | 0.023 | Young |
| 0.402955 | 0.263699 | 0.192 | 0.162 | 1 | 8 | Osgin2 | 0.03 | Young |
| 0.402959 | 0.190422 | 0.4 | 0.38 | 1 | 8 | Cyld | 0.02 | Young |
| 0.402986 | 0.718277 | 0.029 | 0.017 | 1 | 8 | Col3a1 | 0.012 | Young |
| 0.402986 | 0.13177 | 0.029 | 0.017 | 1 | 8 | Cav1 | 0.012 | Young |
| 0.404928 | 0.1078 | 0.049 | 0.034 | 1 | 8 | Kremen1 | 0.015 | Young |
| 0.405448 | 0.315097 | 0.125 | 0.101 | 1 | 8 | Rhot2 | 0.024 | Young |
| 0.405531 | 0.730824 | 0.086 | 0.067 | 1 | 8 | Slc37a4 | 0.019 | Young |
| 0.405806 | 0.115953 | 0.049 | 0.034 | 1 | 8 | Gsdme | 0.015 | Young |
| 0.405856 | 0.037793 | 0.491 | 0.464 | 1 | 8 | Slc38a1 | 0.027 | Young |
| 0.406154 | 0.133538 | 0.405 | 0.369 | 1 | 8 | Axl | 0.036 | Young |
| 0.406211 | 0.53854 | 0.081 | 0.061 | 1 | 8 | Stk38l | 0.02 | Young |

| | | | | | | | | |
|---|---|---|---|---|---|---|---|---|
| 0.406233 | 0.126993 | 0.288 | 0.257 | 1 | 8 | Sp110 | 0.031 | Young |
| 0.40661 | 0.291766 | 0.088 | 0.067 | 1 | 8 | Ccdc86 | 0.021 | Young |
| 0.406986 | 0.007576 | 0.405 | 0.363 | 1 | 8 | Zmat2 | 0.042 | Young |
| 0.407036 | 0.00486 | 0.403 | 0.358 | 1 | 8 | Fbxl3 | 0.045 | Young |
| 0.407085 | 0.314515 | 0.158 | 0.134 | 1 | 8 | Fam219b | 0.024 | Young |
| 0.407094 | 0.164915 | 0.335 | 0.302 | 1 | 8 | Fam234a | 0.033 | Young |
| 0.407579 | 0.100244 | 0.725 | 0.687 | 1 | 8 | Tpm4 | 0.038 | Young |
| 0.407697 | 0.147073 | 0.125 | 0.101 | 1 | 8 | Zswim4 | 0.024 | Young |
| 0.407713 | 0.407201 | 0.174 | 0.151 | 1 | 8 | Stx8 | 0.023 | Young |
| 0.407822 | 0.064504 | 0.177 | 0.151 | 1 | 8 | Cask | 0.026 | Young |
| 0.407835 | 0.096883 | 0.151 | 0.123 | 1 | 8 | Nhlrc2 | 0.028 | Young |
| 0.408367 | 0.097079 | 0.075 | 0.056 | 1 | 8 | Pnkp | 0.019 | Young |
| 0.408408 | 0.031982 | 0.21 | 0.179 | 1 | 8 | Atxn3 | 0.031 | Young |
| 0.408666 | 0.236074 | 0.114 | 0.089 | 1 | 8 | Ttc17 | 0.025 | Young |
| 0.408803 | 0.518979 | 0.029 | 0.017 | 1 | 8 | Efnb1 | 0.012 | Young |
| 0.408803 | 0.41999 | 0.029 | 0.017 | 1 | 8 | Nme2 | 0.012 | Young |
| 0.408803 | 0.369886 | 0.029 | 0.017 | 1 | 8 | Galnt6 | 0.012 | Young |
| 0.40886 | 0.20459 | 0.166 | 0.14 | 1 | 8 | Zdhhc17 | 0.026 | Young |
| 0.408951 | 0.267103 | 0.099 | 0.078 | 1 | 8 | Sesn2 | 0.021 | Young |
| 0.409093 | 0.021398 | 0.584 | 0.531 | 1 | 8 | Mrps21 | 0.053 | Young |
| 0.409131 | 0.191304 | 0.242 | 0.207 | 1 | 8 | Trrap | 0.035 | Young |
| 0.409359 | 0.184333 | 0.143 | 0.117 | 1 | 8 | Dpm1 | 0.026 | Young |
| 0.409543 | 0.017815 | 0.405 | 0.369 | 1 | 8 | Atg3 | 0.036 | Young |
| 0.409973 | 0.550398 | 0.029 | 0.017 | 1 | 8 | 1110051M | 0.012 | Young |
| 0.410217 | 0.228424 | 0.049 | 0.034 | 1 | 8 | AU041133 | 0.015 | Young |
| 0.410398 | 0.290226 | 0.062 | 0.045 | 1 | 8 | Rpap2 | 0.017 | Young |
| 0.410479 | 0.078548 | 0.294 | 0.257 | 1 | 8 | Mtmr6 | 0.037 | Young |
| 0.410836 | 0.333398 | 0.099 | 0.078 | 1 | 8 | Mon1b | 0.021 | Young |
| 0.411092 | 0.008186 | 0.158 | 0.128 | 1 | 8 | Gm5547 | 0.03 | Young |
| 0.411144 | 0.277124 | 0.029 | 0.017 | 1 | 8 | Zswim9 | 0.012 | Young |
| 0.412318 | 0.136757 | 0.029 | 0.017 | 1 | 8 | Tmem132a | 0.012 | Young |
| 0.412332 | 0.505151 | 0.068 | 0.05 | 1 | 8 | Nedd4l | 0.018 | Young |
| 0.412421 | 1.256009 | 0.021 | 0.011 | 1 | 8 | Gba2 | 0.01 | Young |
| 0.412532 | 0.066396 | 0.049 | 0.067 | 1 | 8 | Gm20703 | -0.018 | Young |
| 0.412997 | 0.046082 | 0.881 | 0.777 | 1 | 8 | mt-Nd4l | 0.104 | Young |
| 0.413138 | 0.206522 | 0.081 | 0.061 | 1 | 8 | 4930503L1 | 0.02 | Young |
| 0.41316 | 0.017006 | 0.501 | 0.441 | 1 | 8 | Ankrd17 | 0.06 | Young |
| 0.413243 | 0.180661 | 0.119 | 0.095 | 1 | 8 | Bex3 | 0.024 | Young |
| 0.413493 | 0.080784 | 0.029 | 0.017 | 1 | 8 | Xpo5 | 0.012 | Young |
| 0.413702 | 0.060249 | 0.457 | 0.408 | 1 | 8 | Prpf4b | 0.049 | Young |
| 0.413723 | 0.047659 | 0.13 | 0.106 | 1 | 8 | Elac1 | 0.024 | Young |
| 0.413807 | 1.130568 | 0.021 | 0.011 | 1 | 8 | Map3k9 | 0.01 | Young |
| 0.414785 | 0.130713 | 0.512 | 0.464 | 1 | 8 | 2810474O1 | 0.048 | Young |
| 0.414987 | 0.62899 | 0.068 | 0.05 | 1 | 8 | Chd1l | 0.018 | Young |
| 0.415059 | 0.171996 | 0.125 | 0.101 | 1 | 8 | Srcap | 0.024 | Young |

| | | | | | | | | |
|---|---|---|---|---|---|---|---|---|
| 0.415196 | 1.003742 | 0.021 | 0.011 | 1 | 8 | Zfp28 | 0.01 | Young |
| 0.415196 | 0.924581 | 0.021 | 0.011 | 1 | 8 | Adamts6 | 0.01 | Young |
| 0.415196 | 0.843433 | 0.021 | 0.011 | 1 | 8 | Vmac | 0.01 | Young |
| 0.415642 | 0.297553 | 0.122 | 0.101 | 1 | 8 | Gosr1 | 0.021 | Young |
| 0.415747 | 0.697046 | 0.068 | 0.05 | 1 | 8 | Nudt7 | 0.018 | Young |
| 0.416588 | 1.178383 | 0.021 | 0.011 | 1 | 8 | Slc39a8 | 0.01 | Young |
| 0.416588 | 1.056734 | 0.021 | 0.011 | 1 | 8 | Plekha5 | 0.01 | Young |
| 0.417328 | 0.231739 | 0.081 | 0.061 | 1 | 8 | Tmem140 | 0.02 | Young |
| 0.417798 | 0.501292 | 0.094 | 0.073 | 1 | 8 | Ctdspl | 0.021 | Young |
| 0.417983 | 1.0045 | 0.021 | 0.011 | 1 | 8 | Zfp976 | 0.01 | Young |
| 0.417983 | 0.978104 | 0.021 | 0.011 | 1 | 8 | Zhx2 | 0.01 | Young |
| 0.417983 | 0.613933 | 0.021 | 0.011 | 1 | 8 | Pde2a | 0.01 | Young |
| 0.418159 | 0.099738 | 0.148 | 0.123 | 1 | 8 | Fbxl17 | 0.025 | Young |
| 0.418558 | 0.054952 | 0.221 | 0.184 | 1 | 8 | Asnsd1 | 0.037 | Young |
| 0.419063 | 0.15205 | 0.119 | 0.095 | 1 | 8 | Ubl7 | 0.024 | Young |
| 0.419267 | 0.110721 | 0.405 | 0.369 | 1 | 8 | Pqlc1 | 0.036 | Young |
| 0.419497 | 0.575081 | 0.042 | 0.028 | 1 | 8 | Snapc4 | 0.014 | Young |
| 0.419784 | 0.998732 | 0.073 | 0.056 | 1 | 8 | Zmym3 | 0.017 | Young |
| 0.420125 | 0.191329 | 0.452 | 0.425 | 1 | 8 | Pan3 | 0.027 | Young |
| 0.420628 | 0.044352 | 0.249 | 0.212 | 1 | 8 | Tnf | 0.037 | Young |
| 0.420762 | 0.441306 | 0.117 | 0.095 | 1 | 8 | Golga1 | 0.022 | Young |
| 0.42078 | 0.806599 | 0.021 | 0.011 | 1 | 8 | Tmem53 | 0.01 | Young |
| 0.42078 | 0.794337 | 0.021 | 0.011 | 1 | 8 | Mterf1a | 0.01 | Young |
| 0.42078 | 0.625646 | 0.021 | 0.011 | 1 | 8 | BC043934 | 0.01 | Young |
| 0.42078 | 0.485451 | 0.021 | 0.011 | 1 | 8 | Tyw3 | 0.01 | Young |
| 0.421449 | 0.621127 | 0.042 | 0.028 | 1 | 8 | Elovl6 | 0.014 | Young |
| 0.421484 | 0.619254 | 0.055 | 0.039 | 1 | 8 | Acd | 0.016 | Young |
| 0.421763 | 0.002335 | 0.377 | 0.352 | 1 | 8 | Zmynd11 | 0.025 | Young |
| 0.421863 | 0.019158 | 0.229 | 0.201 | 1 | 8 | Esf1 | 0.028 | Young |
| 0.421866 | 0.208835 | 0.112 | 0.089 | 1 | 8 | Fbxw8 | 0.023 | Young |
| 0.422034 | 1.256313 | 0.013 | 0.006 | 1 | 8 | Prc1 | 0.007 | Young |
| 0.422034 | 1.06176 | 0.013 | 0.006 | 1 | 8 | Kcnj10 | 0.007 | Young |
| 0.422034 | 1.015513 | 0.013 | 0.006 | 1 | 8 | E2f8 | 0.007 | Young |
| 0.422034 | 0.963356 | 0.013 | 0.006 | 1 | 8 | Rita1 | 0.007 | Young |
| 0.422034 | 0.924636 | 0.013 | 0.006 | 1 | 8 | Penk | 0.007 | Young |
| 0.422034 | 0.864986 | 0.013 | 0.006 | 1 | 8 | Fcrl1 | 0.007 | Young |
| 0.422034 | 0.768635 | 0.013 | 0.006 | 1 | 8 | Gm20506 | 0.007 | Young |
| 0.422034 | 0.717302 | 0.013 | 0.006 | 1 | 8 | Gng7 | 0.007 | Young |
| 0.422183 | 0.885973 | 0.021 | 0.011 | 1 | 8 | Fzd4 | 0.01 | Young |
| 0.422183 | 0.680483 | 0.021 | 0.011 | 1 | 8 | Mfsd3 | 0.01 | Young |
| 0.422183 | 0.671127 | 0.021 | 0.011 | 1 | 8 | Senp8 | 0.01 | Young |
| 0.422183 | 0.499356 | 0.021 | 0.011 | 1 | 8 | C130013H( | 0.01 | Young |
| 0.422183 | 0.131636 | 0.021 | 0.011 | 1 | 8 | 9230111E0 | 0.01 | Young |
| 0.422866 | 0.004025 | 0.229 | 0.196 | 1 | 8 | Ube4b | 0.033 | Young |
| 0.423041 | 0.11785 | 0.094 | 0.073 | 1 | 8 | Cisd3 | 0.021 | Young |

| | | | | | | | | |
|---|---|---|---|---|---|---|---|---|
| 0.423212 | 0.41622 | 0.182 | 0.156 | 1 | 8 | Lemd3 | 0.026 | Young |
| 0.423588 | 0.721771 | 0.021 | 0.011 | 1 | 8 | Clec1a | 0.01 | Young |
| 0.423588 | 0.544118 | 0.021 | 0.011 | 1 | 8 | Nat8l | 0.01 | Young |
| 0.423588 | 0.518598 | 0.021 | 0.011 | 1 | 8 | Zfp677 | 0.01 | Young |
| 0.423588 | 0.309835 | 0.021 | 0.011 | 1 | 8 | AW209491 | 0.01 | Young |
| 0.423788 | 0.124827 | 0.423 | 0.374 | 1 | 8 | Ripk1 | 0.049 | Young |
| 0.423843 | 1.350421 | 0.013 | 0.006 | 1 | 8 | Tspoap1 | 0.007 | Young |
| 0.423843 | 1.192228 | 0.013 | 0.006 | 1 | 8 | Cdk1 | 0.007 | Young |
| 0.423843 | 1.170378 | 0.013 | 0.006 | 1 | 8 | Ppp1r13l | 0.007 | Young |
| 0.423843 | 1.044649 | 0.013 | 0.006 | 1 | 8 | Ager | 0.007 | Young |
| 0.423843 | 1.018835 | 0.013 | 0.006 | 1 | 8 | Prdm9 | 0.007 | Young |
| 0.423843 | 0.898119 | 0.013 | 0.006 | 1 | 8 | Gm43569 | 0.007 | Young |
| 0.423843 | 0.893043 | 0.013 | 0.006 | 1 | 8 | Cobll1 | 0.007 | Young |
| 0.423843 | 0.877362 | 0.013 | 0.006 | 1 | 8 | Bik | 0.007 | Young |
| 0.423843 | 0.772861 | 0.013 | 0.006 | 1 | 8 | Cdh24 | 0.007 | Young |
| 0.423843 | 0.757445 | 0.013 | 0.006 | 1 | 8 | Wdpcp | 0.007 | Young |
| 0.423843 | 0.744979 | 0.013 | 0.006 | 1 | 8 | Pdzd4 | 0.007 | Young |
| 0.423843 | 0.735335 | 0.013 | 0.006 | 1 | 8 | Gm15859 | 0.007 | Young |
| 0.423843 | 0.685491 | 0.013 | 0.006 | 1 | 8 | Hyal3 | 0.007 | Young |
| 0.423843 | 0.653991 | 0.013 | 0.006 | 1 | 8 | Tmem156 | 0.007 | Young |
| 0.424051 | 0.083845 | 0.665 | 0.615 | 1 | 8 | Evi2a | 0.05 | Young |
| 0.424175 | 0.060945 | 0.488 | 0.447 | 1 | 8 | Cd300a | 0.041 | Young |
| 0.424383 | 0.082795 | 0.865 | 0.816 | 1 | 8 | Ndufa6 | 0.049 | Young |
| 0.424387 | 0.580683 | 0.042 | 0.028 | 1 | 8 | Impa2 | 0.014 | Young |
| 0.424418 | 0.257542 | 0.166 | 0.14 | 1 | 8 | Ikzf5 | 0.026 | Young |
| 0.424718 | 0.408402 | 0.158 | 0.134 | 1 | 8 | Dcaf17 | 0.024 | Young |
| 0.42482 | 0.205394 | 0.099 | 0.078 | 1 | 8 | Ppp2r1b | 0.021 | Young |
| 0.42494 | 0.447839 | 0.187 | 0.168 | 1 | 8 | Smim13 | 0.019 | Young |
| 0.424996 | 0.720281 | 0.021 | 0.011 | 1 | 8 | Slc22a17 | 0.01 | Young |
| 0.424996 | 0.604099 | 0.021 | 0.011 | 1 | 8 | P2ry14 | 0.01 | Young |
| 0.424996 | 0.58339 | 0.021 | 0.011 | 1 | 8 | 1700008J0 | 0.01 | Young |
| 0.425196 | 0.2187 | 0.112 | 0.089 | 1 | 8 | Nudt2 | 0.023 | Young |
| 0.425306 | 0.620667 | 0.091 | 0.073 | 1 | 8 | Slc46a3 | 0.018 | Young |
| 0.425655 | 1.542561 | 0.013 | 0.006 | 1 | 8 | Kynu | 0.007 | Young |
| 0.425655 | 1.099987 | 0.013 | 0.006 | 1 | 8 | Eaf2 | 0.007 | Young |
| 0.425655 | 0.97192 | 0.013 | 0.006 | 1 | 8 | Icos | 0.007 | Young |
| 0.425655 | 0.909648 | 0.013 | 0.006 | 1 | 8 | Cyp2e1 | 0.007 | Young |
| 0.425655 | 0.851618 | 0.013 | 0.006 | 1 | 8 | Mustn1 | 0.007 | Young |
| 0.425655 | 0.740483 | 0.013 | 0.006 | 1 | 8 | Satb2 | 0.007 | Young |
| 0.425655 | 0.690875 | 0.013 | 0.006 | 1 | 8 | Gm16861 | 0.007 | Young |
| 0.425655 | 0.647571 | 0.013 | 0.006 | 1 | 8 | Tmem120b | 0.007 | Young |
| 0.425655 | 0.645777 | 0.013 | 0.006 | 1 | 8 | Zfp59 | 0.007 | Young |
| 0.425655 | 0.60488 | 0.013 | 0.006 | 1 | 8 | Ankrd9 | 0.007 | Young |
| 0.425655 | 0.573954 | 0.013 | 0.006 | 1 | 8 | Traip | 0.007 | Young |
| 0.426407 | 0.580622 | 0.021 | 0.011 | 1 | 8 | Pdia5 | 0.01 | Young |

| | | | | | | | | |
|---|---|---|---|---|---|---|---|---|
| 0.426407 | 0.477983 | 0.021 | 0.011 | 1 | 8 | Zfp974 | 0.01 | Young |
| 0.426407 | 0.128507 | 0.021 | 0.011 | 1 | 8 | Zfyve9 | 0.01 | Young |
| 0.426459 | 0.043905 | 0.2 | 0.168 | 1 | 8 | Ern1 | 0.032 | Young |
| 0.42675 | 0.096233 | 0.151 | 0.123 | 1 | 8 | Idh3a | 0.028 | Young |
| 0.426925 | 0.169091 | 0.348 | 0.313 | 1 | 8 | Wbp11 | 0.035 | Young |
| 0.427019 | 0.084549 | 0.112 | 0.089 | 1 | 8 | Srm | 0.023 | Young |
| 0.427472 | 0.904842 | 0.013 | 0.006 | 1 | 8 | Gm27168 | 0.007 | Young |
| 0.427472 | 0.76713 | 0.013 | 0.006 | 1 | 8 | Rbp4 | 0.007 | Young |
| 0.427472 | 0.754519 | 0.013 | 0.006 | 1 | 8 | Ly6c1 | 0.007 | Young |
| 0.427472 | 0.665997 | 0.013 | 0.006 | 1 | 8 | Gm9917 | 0.007 | Young |
| 0.427472 | 0.611725 | 0.013 | 0.006 | 1 | 8 | Cerkl | 0.007 | Young |
| 0.427472 | 0.601118 | 0.013 | 0.006 | 1 | 8 | Cyb5d1 | 0.007 | Young |
| 0.427472 | 0.568629 | 0.013 | 0.006 | 1 | 8 | 2810405F1 | 0.007 | Young |
| 0.427472 | 0.559205 | 0.013 | 0.006 | 1 | 8 | Cadm3 | 0.007 | Young |
| 0.427472 | 0.556764 | 0.013 | 0.006 | 1 | 8 | Ehhadh | 0.007 | Young |
| 0.427472 | 0.547327 | 0.013 | 0.006 | 1 | 8 | 1700052K1 | 0.007 | Young |
| 0.427472 | 0.544588 | 0.013 | 0.006 | 1 | 8 | Nbl1 | 0.007 | Young |
| 0.427472 | 0.539542 | 0.013 | 0.006 | 1 | 8 | Zfp607b | 0.007 | Young |
| 0.427472 | 0.530978 | 0.013 | 0.006 | 1 | 8 | Zpbp | 0.007 | Young |
| 0.427472 | 0.508679 | 0.013 | 0.006 | 1 | 8 | Hoxc8 | 0.007 | Young |
| 0.427501 | 0.044827 | 0.345 | 0.307 | 1 | 8 | Pabpn1 | 0.038 | Young |
| 0.427821 | 0.326997 | 0.021 | 0.011 | 1 | 8 | Mr1 | 0.01 | Young |
| 0.427821 | 0.191327 | 0.021 | 0.011 | 1 | 8 | Wdr35 | 0.01 | Young |
| 0.42812 | 0.012744 | 0.465 | 0.419 | 1 | 8 | Dusp22 | 0.046 | Young |
| 0.428183 | 0.816311 | 0.096 | 0.078 | 1 | 8 | Ppcdc | 0.018 | Young |
| 0.428324 | 0.437345 | 0.042 | 0.028 | 1 | 8 | Maged2 | 0.014 | Young |
| 0.428441 | 0.34205 | 0.169 | 0.145 | 1 | 8 | Tsc1 | 0.024 | Young |
| 0.428622 | 0.378555 | 0.081 | 0.061 | 1 | 8 | Smim8 | 0.02 | Young |
| 0.42891 | 0.225287 | 0.213 | 0.184 | 1 | 8 | Mbd6 | 0.029 | Young |
| 0.429104 | 0.316848 | 0.299 | 0.268 | 1 | 8 | Prkab2 | 0.031 | Young |
| 0.429294 | 0.796312 | 0.013 | 0.006 | 1 | 8 | Lhfpl2 | 0.007 | Young |
| 0.429294 | 0.683252 | 0.013 | 0.006 | 1 | 8 | Arpin | 0.007 | Young |
| 0.429294 | 0.595041 | 0.013 | 0.006 | 1 | 8 | Alkbh4 | 0.007 | Young |
| 0.429294 | 0.591724 | 0.013 | 0.006 | 1 | 8 | Ift81 | 0.007 | Young |
| 0.429294 | 0.547399 | 0.013 | 0.006 | 1 | 8 | Gm14295 | 0.007 | Young |
| 0.429294 | 0.534771 | 0.013 | 0.006 | 1 | 8 | Nova1 | 0.007 | Young |
| 0.429294 | 0.516488 | 0.013 | 0.006 | 1 | 8 | Ehd2 | 0.007 | Young |
| 0.429294 | 0.496702 | 0.013 | 0.006 | 1 | 8 | 4933423P2 | 0.007 | Young |
| 0.429294 | 0.488134 | 0.013 | 0.006 | 1 | 8 | Ly6c2 | 0.007 | Young |
| 0.429294 | 0.466098 | 0.013 | 0.006 | 1 | 8 | Gm26857 | 0.007 | Young |
| 0.429294 | 0.430593 | 0.013 | 0.006 | 1 | 8 | Gm11696 | 0.007 | Young |
| 0.429294 | 0.425095 | 0.013 | 0.006 | 1 | 8 | Ankrd28 | 0.007 | Young |
| 0.429294 | 0.414325 | 0.013 | 0.006 | 1 | 8 | Ccdc15 | 0.007 | Young |
| 0.429294 | 0.412998 | 0.013 | 0.006 | 1 | 8 | Abi3 | 0.007 | Young |
| 0.429294 | 0.349084 | 0.013 | 0.006 | 1 | 8 | Trim13 | 0.007 | Young |

| | | | | | | | | |
|---|---|---|---|---|---|---|---|---|
| 0.429294 | 0.163233 | 0.013 | 0.006 | 1 | 8 | Angel1 | 0.007 | Young |
| 0.429313 | 0.694897 | 0.091 | 0.073 | 1 | 8 | Fam117a | 0.018 | Young |
| 0.429362 | 0.218828 | 0.223 | 0.196 | 1 | 8 | G6pdx | 0.027 | Young |
| 0.429541 | 0.010655 | 0.317 | 0.285 | 1 | 8 | Rbm42 | 0.032 | Young |
| 0.429572 | 0.258412 | 0.068 | 0.05 | 1 | 8 | Lrrc61 | 0.018 | Young |
| 0.429617 | 0.421708 | 0.195 | 0.168 | 1 | 8 | Tmem222 | 0.027 | Young |
| 0.430047 | 0.27329 | 0.081 | 0.061 | 1 | 8 | Rora | 0.02 | Young |
| 0.430053 | 0.242587 | 0.055 | 0.039 | 1 | 8 | Stk11ip | 0.016 | Young |
| 0.430459 | 0.055001 | 0.545 | 0.497 | 1 | 8 | Rab31 | 0.048 | Young |
| 0.430656 | 0.324942 | 0.021 | 0.011 | 1 | 8 | Ugt1a6a | 0.01 | Young |
| 0.43076 | 0.234096 | 0.081 | 0.061 | 1 | 8 | Fam122a | 0.02 | Young |
| 0.430916 | 0.174896 | 0.055 | 0.039 | 1 | 8 | Cbx8 | 0.016 | Young |
| 0.43112 | 0.422706 | 0.013 | 0.006 | 1 | 8 | Gm20517 | 0.007 | Young |
| 0.43112 | 0.419421 | 0.013 | 0.006 | 1 | 8 | Gm12216 | 0.007 | Young |
| 0.43112 | 0.243704 | 0.013 | 0.006 | 1 | 8 | Gm31105 | 0.007 | Young |
| 0.43112 | 0.167915 | 0.013 | 0.006 | 1 | 8 | Slc25a42 | 0.007 | Young |
| 0.43112 | 0.135914 | 0.013 | 0.006 | 1 | 8 | Trim6 | 0.007 | Young |
| 0.43112 | 0.086447 | 0.013 | 0.006 | 1 | 8 | Aplp1 | 0.007 | Young |
| 0.43129 | 0.420843 | 0.042 | 0.028 | 1 | 8 | Nedd4 | 0.014 | Young |
| 0.431474 | 0.117214 | 0.143 | 0.117 | 1 | 8 | Ubl4a | 0.026 | Young |
| 0.431779 | 0.126261 | 0.055 | 0.039 | 1 | 8 | Dnajc18 | 0.016 | Young |
| 0.432188 | 0.581895 | 0.081 | 0.061 | 1 | 8 | Fzd7 | 0.02 | Young |
| 0.432333 | 0.109047 | 0.312 | 0.274 | 1 | 8 | Bdp1 | 0.038 | Young |
| 0.433212 | 0.308604 | 0.099 | 0.078 | 1 | 8 | Cox6b2 | 0.021 | Young |
| 0.433507 | 0.111454 | 0.182 | 0.156 | 1 | 8 | Bcl2l1 | 0.026 | Young |
| 0.433919 | 0.015661 | 0.514 | 0.475 | 1 | 8 | Klf7 | 0.039 | Young |
| 0.434016 | 0.671038 | 0.091 | 0.073 | 1 | 8 | Caml | 0.018 | Young |
| 0.434072 | 0.230371 | 0.171 | 0.145 | 1 | 8 | Lpcat3 | 0.026 | Young |
| 0.434185 | 0.095743 | 0.382 | 0.346 | 1 | 8 | Mrps14 | 0.036 | Young |
| 0.434269 | 0.404657 | 0.042 | 0.028 | 1 | 8 | Gtpbp3 | 0.014 | Young |
| 0.435067 | 0.512901 | 0.114 | 0.095 | 1 | 8 | Dnajc30 | 0.019 | Young |
| 0.435163 | 0.103508 | 0.403 | 0.363 | 1 | 8 | Rdx | 0.04 | Young |
| 0.435167 | 0.190022 | 0.127 | 0.106 | 1 | 8 | Pigc | 0.021 | Young |
| 0.43541 | 0.376248 | 0.073 | 0.056 | 1 | 8 | Pias3 | 0.017 | Young |
| 0.435529 | 0.374692 | 0.104 | 0.084 | 1 | 8 | Champ1 | 0.02 | Young |
| 0.435594 | 0.320285 | 0.151 | 0.128 | 1 | 8 | Naxe | 0.023 | Young |
| 0.4356 | 0.230189 | 0.218 | 0.19 | 1 | 8 | Wdr61 | 0.028 | Young |
| 0.435623 | 0.20836 | 0.244 | 0.218 | 1 | 8 | Pilrb2 | 0.026 | Young |
| 0.436005 | 0.209266 | 0.205 | 0.179 | 1 | 8 | Ubfd1 | 0.026 | Young |
| 0.436487 | 0.272049 | 0.203 | 0.179 | 1 | 8 | Dmtf1 | 0.024 | Young |
| 0.4367 | 0.093328 | 0.138 | 0.117 | 1 | 8 | Prpf31 | 0.021 | Young |
| 0.437008 | 0.575476 | 0.005 | 0.011 | 1 | 8 | Chtf18 | -0.006 | Young |
| 0.437008 | 0.523002 | 0.005 | 0.011 | 1 | 8 | Adgra3 | -0.006 | Young |
| 0.437008 | 0.337155 | 0.005 | 0.011 | 1 | 8 | Man2c1os | -0.006 | Young |
| 0.437008 | 0.320079 | 0.005 | 0.011 | 1 | 8 | Ckap2 | -0.006 | Young |

| | | | | | | | | |
|---|---|---|---|---|---|---|---|---|
| 0.437008 | 0.318274 | 0.005 | 0.011 | 1 | 8 | Adgrl4 | -0.006 | Young |
| 0.437008 | 0.267916 | 0.005 | 0.011 | 1 | 8 | Ifit2 | -0.006 | Young |
| 0.437008 | 0.212249 | 0.005 | 0.011 | 1 | 8 | Hs3st1 | -0.006 | Young |
| 0.437125 | 0.003056 | 0.325 | 0.285 | 1 | 8 | Man1a2 | 0.04 | Young |
| 0.437391 | 0.274023 | 0.174 | 0.151 | 1 | 8 | Stam2 | 0.023 | Young |
| 0.437889 | 0.154782 | 0.242 | 0.207 | 1 | 8 | Dpf2 | 0.035 | Young |
| 0.438118 | 0.187253 | 0.031 | 0.045 | 1 | 8 | Katnb1 | -0.014 | Young |
| 0.438996 | 0.083829 | 0.086 | 0.067 | 1 | 8 | Tmem184c | 0.019 | Young |
| 0.439038 | 0.033968 | 0.647 | 0.603 | 1 | 8 | Abi1 | 0.044 | Young |
| 0.439672 | 0.298319 | 0.094 | 0.073 | 1 | 8 | Asb3 | 0.021 | Young |
| 0.440387 | 0.083507 | 0.099 | 0.078 | 1 | 8 | Cryzl2 | 0.021 | Young |
| 0.440469 | 0.175914 | 0.055 | 0.039 | 1 | 8 | Mccc1 | 0.016 | Young |
| 0.440505 | 0.245513 | 0.106 | 0.084 | 1 | 8 | Hace1 | 0.022 | Young |
| 0.441019 | 0.317385 | 0.094 | 0.073 | 1 | 8 | Nol9 | 0.021 | Young |
| 0.441271 | 0.120696 | 0.322 | 0.285 | 1 | 8 | Npm3 | 0.037 | Young |
| 0.441294 | 0.48962 | 0.068 | 0.05 | 1 | 8 | Vps33b | 0.018 | Young |
| 0.441294 | 0.07972 | 0.068 | 0.05 | 1 | 8 | Tnfsf13b | 0.018 | Young |
| 0.441308 | 0.452598 | 0.096 | 0.078 | 1 | 8 | Letmd1 | 0.018 | Young |
| 0.441693 | 0.238186 | 0.234 | 0.212 | 1 | 8 | Commd4 | 0.022 | Young |
| 0.441779 | 0.115174 | 0.543 | 0.492 | 1 | 8 | Atp2a2 | 0.051 | Young |
| 0.441886 | 0.066636 | 0.364 | 0.335 | 1 | 8 | Akirin1 | 0.029 | Young |
| 0.442134 | 0.017716 | 0.384 | 0.341 | 1 | 8 | Ncf4 | 0.043 | Young |
| 0.442223 | 0.098786 | 0.031 | 0.045 | 1 | 8 | Ccdc191 | -0.014 | Young |
| 0.442969 | 0.057304 | 0.231 | 0.196 | 1 | 8 | Dpp3 | 0.035 | Young |
| 0.443433 | 0.047022 | 0.166 | 0.14 | 1 | 8 | Dok3 | 0.026 | Young |
| 0.443469 | 0.181196 | 0.304 | 0.279 | 1 | 8 | Usp48 | 0.025 | Young |
| 0.443973 | 0.325196 | 0.055 | 0.039 | 1 | 8 | Ralgps1 | 0.016 | Young |
| 0.444074 | 0.08628 | 0.54 | 0.514 | 1 | 8 | Tbcb | 0.026 | Young |
| 0.444083 | 0.055869 | 0.184 | 0.156 | 1 | 8 | Pknox1 | 0.028 | Young |
| 0.444583 | 0.063616 | 0.642 | 0.587 | 1 | 8 | Dock8 | 0.055 | Young |
| 0.44487 | 0.062691 | 0.112 | 0.089 | 1 | 8 | Atxn1l | 0.023 | Young |
| 0.445103 | 0.060814 | 0.709 | 0.659 | 1 | 8 | Txn1 | 0.05 | Young |
| 0.445553 | 0.026222 | 0.169 | 0.14 | 1 | 8 | Ppt2 | 0.029 | Young |
| 0.446016 | 0.087459 | 0.086 | 0.067 | 1 | 8 | Ruvbl2 | 0.019 | Young |
| 0.44764 | 0.16244 | 0.169 | 0.145 | 1 | 8 | Mrto4 | 0.024 | Young |
| 0.448292 | 0.090784 | 0.099 | 0.078 | 1 | 8 | Eef2kmt | 0.021 | Young |
| 0.448871 | 0.356461 | 0.14 | 0.117 | 1 | 8 | Cd82 | 0.023 | Young |
| 0.448917 | 1.038011 | 0.034 | 0.022 | 1 | 8 | Rab3a | 0.012 | Young |
| 0.449089 | 0.351596 | 0.122 | 0.101 | 1 | 8 | Nat9 | 0.021 | Young |
| 0.449537 | 0.053669 | 0.756 | 0.682 | 1 | 8 | Atp5j2 | 0.074 | Young |
| 0.449833 | 0.597549 | 0.073 | 0.056 | 1 | 8 | Plekhn1 | 0.017 | Young |
| 0.450072 | 0.570318 | 0.143 | 0.123 | 1 | 8 | Sec23a | 0.02 | Young |
| 0.450124 | 0.058842 | 0.878 | 0.821 | 1 | 8 | Fus | 0.057 | Young |
| 0.450313 | 0.072267 | 0.501 | 0.453 | 1 | 8 | Psmb6 | 0.048 | Young |
| 0.450715 | 0.048247 | 0.106 | 0.084 | 1 | 8 | Trmt2b | 0.022 | Young |

| | | | | | | | | |
|---|---|---|---|---|---|---|---|---|
| 0.450752 | 0.090358 | 0.351 | 0.324 | 1 | 8 | Sugt1 | 0.027 | Young |
| 0.450882 | 0.033292 | 0.317 | 0.285 | 1 | 8 | Homer1 | 0.032 | Young |
| 0.452121 | 0.128455 | 0.158 | 0.134 | 1 | 8 | Nop14 | 0.024 | Young |
| 0.452388 | 0.360707 | 0.086 | 0.067 | 1 | 8 | Cul4b | 0.019 | Young |
| 0.452388 | 0.317509 | 0.086 | 0.067 | 1 | 8 | Kat5 | 0.019 | Young |
| 0.452539 | 0.116956 | 0.262 | 0.229 | 1 | 8 | Ddhd2 | 0.033 | Young |
| 0.452662 | 0.037903 | 0.449 | 0.397 | 1 | 8 | Rgs18 | 0.052 | Young |
| 0.453123 | 0.388335 | 0.091 | 0.073 | 1 | 8 | Fbxo6 | 0.018 | Young |
| 0.45324 | 0.093036 | 0.532 | 0.508 | 1 | 8 | Arrb1 | 0.024 | Young |
| 0.453417 | 0.001888 | 0.548 | 0.475 | 1 | 8 | 2-Mar | 0.073 | Young |
| 0.453515 | 0.179636 | 0.104 | 0.084 | 1 | 8 | Spryd7 | 0.02 | Young |
| 0.45376 | 0.28417 | 0.151 | 0.128 | 1 | 8 | Maged1 | 0.023 | Young |
| 0.453879 | 0.047347 | 0.613 | 0.57 | 1 | 8 | Limd1 | 0.043 | Young |
| 0.454086 | 0.566986 | 0.078 | 0.061 | 1 | 8 | Pyroxd1 | 0.017 | Young |
| 0.454439 | 0.027458 | 0.465 | 0.419 | 1 | 8 | Washc2 | 0.046 | Young |
| 0.454551 | 0.40158 | 0.034 | 0.022 | 1 | 8 | Nim1k | 0.012 | Young |
| 0.4549 | 0.107468 | 0.229 | 0.201 | 1 | 8 | Spata2 | 0.028 | Young |
| 0.456341 | 0.473933 | 0.13 | 0.112 | 1 | 8 | Skiv2l2 | 0.018 | Young |
| 0.456502 | 0.220065 | 0.169 | 0.145 | 1 | 8 | Map3k4 | 0.024 | Young |
| 0.456631 | 0.021997 | 0.436 | 0.391 | 1 | 8 | Zfand3 | 0.045 | Young |
| 0.456758 | 0.293968 | 0.073 | 0.056 | 1 | 8 | Fdxr | 0.017 | Young |
| 0.456812 | 0.280741 | 0.255 | 0.235 | 1 | 8 | Eftud2 | 0.02 | Young |
| 0.457105 | 0.020917 | 0.036 | 0.05 | 1 | 8 | Fndc7 | -0.014 | Young |
| 0.457152 | 0.513694 | 0.06 | 0.045 | 1 | 8 | 1110012L1 | 0.015 | Young |
| 0.457562 | 0.090813 | 0.197 | 0.168 | 1 | 8 | Cspp1 | 0.029 | Young |
| 0.457757 | 0.000812 | 0.4 | 0.352 | 1 | 8 | Sec11a | 0.048 | Young |
| 0.457983 | 0.109343 | 0.299 | 0.268 | 1 | 8 | Zmym5 | 0.031 | Young |
| 0.458003 | 0.47265 | 0.06 | 0.045 | 1 | 8 | Mcc | 0.015 | Young |
| 0.458076 | 0.104806 | 0.104 | 0.084 | 1 | 8 | Tmcc2 | 0.02 | Young |
| 0.458945 | 0.08436 | 0.164 | 0.14 | 1 | 8 | Sap30l | 0.024 | Young |
| 0.459022 | 0.113991 | 0.34 | 0.313 | 1 | 8 | Ss18 | 0.027 | Young |
| 0.459308 | 0.654419 | 0.078 | 0.061 | 1 | 8 | Ppp1r3b | 0.017 | Young |
| 0.459308 | 0.575446 | 0.078 | 0.061 | 1 | 8 | Ints8 | 0.017 | Young |
| 0.459707 | 0.438748 | 0.06 | 0.045 | 1 | 8 | Utp4 | 0.015 | Young |
| 0.459934 | 0.048751 | 0.756 | 0.709 | 1 | 8 | Slfn2 | 0.047 | Young |
| 0.460041 | 0.54294 | 0.109 | 0.089 | 1 | 8 | Rpl27-ps3 | 0.02 | Young |
| 0.460369 | 0.111305 | 0.335 | 0.302 | 1 | 8 | Igbp1 | 0.033 | Young |
| 0.461363 | 0.467266 | 0.034 | 0.022 | 1 | 8 | Hacl1 | 0.012 | Young |
| 0.461363 | 0.030085 | 0.034 | 0.022 | 1 | 8 | Mx1 | 0.012 | Young |
| 0.461416 | 0.628489 | 0.06 | 0.045 | 1 | 8 | Mamld1 | 0.015 | Young |
| 0.461604 | 0.205052 | 0.127 | 0.106 | 1 | 8 | Creb3l2 | 0.021 | Young |
| 0.461875 | 0.720849 | 0.047 | 0.034 | 1 | 8 | Wfs1 | 0.013 | Young |
| 0.462504 | 0.419287 | 0.034 | 0.022 | 1 | 8 | Tti1 | 0.012 | Young |
| 0.462964 | 0.329736 | 0.073 | 0.056 | 1 | 8 | Smad7 | 0.017 | Young |
| 0.463128 | 0.25983 | 0.06 | 0.045 | 1 | 8 | Dnal4 | 0.015 | Young |

| | | | | | | | | |
|---|---|---|---|---|---|---|---|---|
| 0.46395 | 0.030769 | 0.574 | 0.508 | 1 | 8 | Tmem258 | 0.066 | Young |
| 0.463979 | 0.123366 | 0.104 | 0.084 | 1 | 8 | Rev1 | 0.02 | Young |
| 0.464291 | 0.089303 | 0.439 | 0.425 | 1 | 8 | Elmo2 | 0.014 | Young |
| 0.464379 | 0.018125 | 0.449 | 0.408 | 1 | 8 | Mgat1 | 0.041 | Young |
| 0.464525 | 0.002705 | 0.462 | 0.419 | 1 | 8 | Dnajc19 | 0.043 | Young |
| 0.464637 | 0.042113 | 0.104 | 0.084 | 1 | 8 | Frs2 | 0.02 | Young |
| 0.464791 | 0.554974 | 0.034 | 0.022 | 1 | 8 | Tnks1bp1 | 0.012 | Young |
| 0.46493 | 0.651482 | 0.065 | 0.05 | 1 | 8 | Ccdc152 | 0.015 | Young |
| 0.465363 | 0.250958 | 0.151 | 0.128 | 1 | 8 | Ints12 | 0.023 | Young |
| 0.465694 | 0.522359 | 0.078 | 0.061 | 1 | 8 | Tspan33 | 0.017 | Young |
| 0.466364 | 0.417453 | 0.13 | 0.112 | 1 | 8 | Plxna4os1 | 0.018 | Young |
| 0.466562 | 0.525971 | 0.06 | 0.045 | 1 | 8 | Alg6 | 0.015 | Young |
| 0.466667 | 0.069276 | 0.117 | 0.095 | 1 | 8 | Abhd13 | 0.022 | Young |
| 0.46672 | 0.153675 | 0.047 | 0.034 | 1 | 8 | Mapkapk5 | 0.013 | Young |
| 0.467453 | 0.191326 | 0.086 | 0.067 | 1 | 8 | Ppm1d | 0.019 | Young |
| 0.46758 | 0.452237 | 0.101 | 0.084 | 1 | 8 | Bag4 | 0.017 | Young |
| 0.468598 | 0.105012 | 0.104 | 0.084 | 1 | 8 | Tfam | 0.02 | Young |
| 0.469387 | 0.235725 | 0.265 | 0.24 | 1 | 8 | Lmo1 | 0.025 | Young |
| 0.469405 | 0.21232 | 0.132 | 0.112 | 1 | 8 | Wdr81 | 0.02 | Young |
| 0.469411 | 0.53113 | 0.153 | 0.134 | 1 | 8 | Kctd12b | 0.019 | Young |
| 0.469691 | 0.068161 | 0.109 | 0.089 | 1 | 8 | Atg14 | 0.02 | Young |
| 0.470324 | 0.428903 | 0.138 | 0.117 | 1 | 8 | Vkorc1l1 | 0.021 | Young |
| 0.471122 | 0.548257 | 0.13 | 0.112 | 1 | 8 | Dusp8 | 0.018 | Young |
| 0.47117 | 0.003469 | 0.587 | 0.531 | 1 | 8 | Ythdc1 | 0.056 | Young |
| 0.471216 | 0.185862 | 0.127 | 0.106 | 1 | 8 | Plekha3 | 0.021 | Young |
| 0.471688 | 0.165886 | 0.034 | 0.022 | 1 | 8 | Ncapd2 | 0.012 | Young |
| 0.471845 | 0.083754 | 0.169 | 0.145 | 1 | 8 | Cryl1 | 0.024 | Young |
| 0.472268 | 0.05351 | 0.499 | 0.469 | 1 | 8 | Tex261 | 0.03 | Young |
| 0.472369 | 0.240233 | 0.132 | 0.112 | 1 | 8 | Rcbtb1 | 0.02 | Young |
| 0.472642 | 0.306945 | 0.171 | 0.151 | 1 | 8 | Nelfe | 0.02 | Young |
| 0.472827 | 0.129615 | 0.122 | 0.101 | 1 | 8 | Tpst1 | 0.021 | Young |
| 0.473473 | 0.37185 | 0.06 | 0.045 | 1 | 8 | Ttc27 | 0.015 | Young |
| 0.474 | 0.242105 | 0.034 | 0.022 | 1 | 8 | Adgrl1 | 0.012 | Young |
| 0.474108 | 0.080363 | 0.042 | 0.056 | 1 | 8 | Nop2 | -0.014 | Young |
| 0.474531 | 0.417954 | 0.047 | 0.034 | 1 | 8 | Atg4a | 0.013 | Young |
| 0.474849 | 0.165039 | 0.127 | 0.106 | 1 | 8 | Tmem86a | 0.021 | Young |
| 0.475519 | 0.007922 | 0.203 | 0.179 | 1 | 8 | Mast3 | 0.024 | Young |
| 0.475657 | 0.090486 | 0.197 | 0.173 | 1 | 8 | Gsto1 | 0.024 | Young |
| 0.475794 | 0.139298 | 0.14 | 0.117 | 1 | 8 | Tigd2 | 0.023 | Young |
| 0.475985 | 0.049568 | 0.143 | 0.117 | 1 | 8 | Cxcl10 | 0.026 | Young |
| 0.476395 | 0.190175 | 0.296 | 0.268 | 1 | 8 | Eif2s3y | 0.028 | Young |
| 0.476512 | 0.244325 | 0.109 | 0.089 | 1 | 8 | G6pc3 | 0.02 | Young |
| 0.477475 | 0.07616 | 0.574 | 0.531 | 1 | 8 | Psenen | 0.043 | Young |
| 0.477821 | 0.376525 | 0.06 | 0.045 | 1 | 8 | 2310057M | 0.015 | Young |
| 0.478064 | 0.070409 | 0.371 | 0.33 | 1 | 8 | Ppp2r2a | 0.041 | Young |

| | | | | | | | | |
|---|---|---|---|---|---|---|---|---|
| 0.478229 | 0.231809 | 0.078 | 0.061 | 1 | 8 | Tipin | 0.017 | Young |
| 0.478683 | 0.120832 | 0.073 | 0.056 | 1 | 8 | Toe1 | 0.017 | Young |
| 0.478741 | 0.035143 | 0.865 | 0.799 | 1 | 8 | Hnrnpa3 | 0.066 | Young |
| 0.479935 | 0.265477 | 0.114 | 0.095 | 1 | 8 | Cdc14b | 0.019 | Young |
| 0.480142 | 0.204492 | 0.205 | 0.184 | 1 | 8 | Slc35e1 | 0.021 | Young |
| 0.480154 | 0.191398 | 0.26 | 0.24 | 1 | 8 | Lsm8 | 0.02 | Young |
| 0.48107 | 0.168064 | 0.195 | 0.173 | 1 | 8 | Hck | 0.022 | Young |
| 0.481424 | 0.129349 | 0.047 | 0.034 | 1 | 8 | Pms2 | 0.013 | Young |
| 0.481607 | 0.154063 | 0.158 | 0.134 | 1 | 8 | Arsa | 0.024 | Young |
| 0.48206 | 0.0463 | 0.501 | 0.464 | 1 | 8 | Ap1s2 | 0.037 | Young |
| 0.482658 | 0.018345 | 0.073 | 0.056 | 1 | 8 | 3110002H1 | 0.017 | Young |
| 0.482683 | 0.005705 | 0.231 | 0.207 | 1 | 8 | B4galt3 | 0.024 | Young |
| 0.483211 | 0.318665 | 0.21 | 0.19 | 1 | 8 | Plod1 | 0.02 | Young |
| 0.483568 | 0.110322 | 0.262 | 0.229 | 1 | 8 | 5031425E2 | 0.033 | Young |
| 0.483736 | 0.09643 | 0.304 | 0.279 | 1 | 8 | Srrt | 0.025 | Young |
| 0.483944 | 0.235718 | 0.06 | 0.045 | 1 | 8 | Ppil1 | 0.015 | Young |
| 0.484215 | 0.044673 | 0.548 | 0.503 | 1 | 8 | Wdr83os | 0.045 | Young |
| 0.484346 | 0.366166 | 0.13 | 0.112 | 1 | 8 | Hectd4 | 0.018 | Young |
| 0.484823 | 0.276028 | 0.06 | 0.045 | 1 | 8 | Mmgt2 | 0.015 | Young |
| 0.484893 | 0.35839 | 0.065 | 0.05 | 1 | 8 | Cdk5rap2 | 0.015 | Young |
| 0.484943 | 0.905478 | 0.026 | 0.017 | 1 | 8 | Nck2 | 0.009 | Young |
| 0.48559 | 0.397275 | 0.14 | 0.123 | 1 | 8 | Chpf2 | 0.017 | Young |
| 0.485744 | 0.492758 | 0.112 | 0.095 | 1 | 8 | Nbeal2 | 0.017 | Young |
| 0.486076 | 0.115597 | 0.294 | 0.268 | 1 | 8 | Chd1 | 0.026 | Young |
| 0.486279 | 1.166295 | 0.026 | 0.017 | 1 | 8 | AW146154 | 0.009 | Young |
| 0.486687 | 0.094505 | 0.078 | 0.061 | 1 | 8 | Trpm4 | 0.017 | Young |
| 0.487257 | 0.020782 | 0.288 | 0.268 | 1 | 8 | Abhd17b | 0.02 | Young |
| 0.487356 | 0.030911 | 0.403 | 0.363 | 1 | 8 | Eprs | 0.04 | Young |
| 0.48747 | 0.114413 | 0.278 | 0.251 | 1 | 8 | Kif2a | 0.027 | Young |
| 0.488461 | 0.231614 | 0.2 | 0.179 | 1 | 8 | Dnmt1 | 0.021 | Young |
| 0.488682 | 0.034038 | 0.104 | 0.084 | 1 | 8 | 2310039H0 | 0.02 | Young |
| 0.488958 | 0.733752 | 0.026 | 0.017 | 1 | 8 | Zfp1 | 0.009 | Young |
| 0.488958 | 0.634123 | 0.026 | 0.017 | 1 | 8 | Zdhhc1 | 0.009 | Young |
| 0.489299 | 0.02742 | 0.844 | 0.777 | 1 | 8 | Clk1 | 0.067 | Young |
| 0.48932 | 0.326353 | 0.249 | 0.229 | 1 | 8 | Mospd3 | 0.02 | Young |
| 0.489715 | 0.124819 | 0.244 | 0.223 | 1 | 8 | Fam114a2 | 0.021 | Young |
| 0.489799 | 0.38171 | 0.174 | 0.156 | 1 | 8 | Gm37494 | 0.018 | Young |
| 0.490127 | 0.143787 | 0.208 | 0.184 | 1 | 8 | Swt1 | 0.024 | Young |
| 0.490162 | 0.012156 | 0.99 | 0.972 | 1 | 8 | Serinc3 | 0.018 | Young |
| 0.490283 | 0.003538 | 0.392 | 0.358 | 1 | 8 | Il4ra | 0.034 | Young |
| 0.490356 | 0.220723 | 0.317 | 0.291 | 1 | 8 | Nans | 0.026 | Young |
| 0.490363 | 0.076643 | 0.132 | 0.112 | 1 | 8 | Dera | 0.02 | Young |
| 0.490417 | 0.155333 | 0.184 | 0.162 | 1 | 8 | Usp1 | 0.022 | Young |
| 0.490527 | 0.829343 | 0.07 | 0.056 | 1 | 8 | Gna11 | 0.014 | Young |
| 0.490663 | 0.278048 | 0.218 | 0.201 | 1 | 8 | Maml1 | 0.017 | Young |

| | | | | | | | | |
|---|---|---|---|---|---|---|---|---|
| 0.491379 | 0.056452 | 0.343 | 0.296 | 1 | 8 | Flii | 0.047 | Young |
| 0.491474 | 0.278842 | 0.156 | 0.134 | 1 | 8 | Arhgap19 | 0.022 | Young |
| 0.491592 | 0.130181 | 0.257 | 0.235 | 1 | 8 | Zrsr2 | 0.022 | Young |
| 0.491644 | 0.618104 | 0.026 | 0.017 | 1 | 8 | Mdga1 | 0.009 | Young |
| 0.491644 | 0.602524 | 0.026 | 0.017 | 1 | 8 | Ctla2b | 0.009 | Young |
| 0.491973 | 0.186348 | 0.109 | 0.089 | 1 | 8 | Oxsm | 0.02 | Young |
| 0.492645 | 0.07043 | 0.735 | 0.715 | 1 | 8 | Pmaip1 | 0.02 | Young |
| 0.492851 | 0.007077 | 0.345 | 0.313 | 1 | 8 | BC005624 | 0.032 | Young |
| 0.492938 | 0.369309 | 0.106 | 0.089 | 1 | 8 | Metrn | 0.017 | Young |
| 0.49299 | 0.820038 | 0.026 | 0.017 | 1 | 8 | Immp2l | 0.009 | Young |
| 0.493346 | 0.325995 | 0.065 | 0.05 | 1 | 8 | Abcb10 | 0.015 | Young |
| 0.493434 | 0.115406 | 0.104 | 0.084 | 1 | 8 | Paqr7 | 0.02 | Young |
| 0.493578 | 0.516235 | 0.083 | 0.067 | 1 | 8 | Cks1b | 0.016 | Young |
| 0.493783 | 0.673257 | 0.052 | 0.039 | 1 | 8 | 0610009O2 | 0.013 | Young |
| 0.494131 | 0.110653 | 0.114 | 0.095 | 1 | 8 | Ppp5c | 0.019 | Young |
| 0.494155 | 0.167305 | 0.353 | 0.33 | 1 | 8 | Naa15 | 0.023 | Young |
| 0.494338 | 0.118056 | 0.026 | 0.017 | 1 | 8 | Mrps22 | 0.009 | Young |
| 0.495046 | 0.195109 | 0.065 | 0.05 | 1 | 8 | Dpysl3 | 0.015 | Young |
| 0.4951 | 0.293239 | 0.127 | 0.106 | 1 | 8 | Stoml1 | 0.021 | Young |
| 0.495342 | 0.202289 | 0.101 | 0.084 | 1 | 8 | Zc3h7b | 0.017 | Young |
| 0.49568 | 0.45359 | 0.052 | 0.039 | 1 | 8 | 1110004E0 | 0.013 | Young |
| 0.495688 | 0.066626 | 0.652 | 0.592 | 1 | 8 | Snrpf | 0.06 | Young |
| 0.495689 | 0.951168 | 0.026 | 0.017 | 1 | 8 | Trmt5 | 0.009 | Young |
| 0.495907 | 0.683791 | 0.039 | 0.028 | 1 | 8 | Nnt | 0.011 | Young |
| 0.495983 | 0.044597 | 0.377 | 0.341 | 1 | 8 | Ppp6r1 | 0.036 | Young |
| 0.496471 | 0.151 | 0.449 | 0.441 | 1 | 8 | Mtch1 | 0.008 | Young |
| 0.49663 | 0.555623 | 0.052 | 0.039 | 1 | 8 | Gstk1 | 0.013 | Young |
| 0.497581 | 0.400755 | 0.052 | 0.039 | 1 | 8 | Nmnat3 | 0.013 | Young |
| 0.497704 | 0.249407 | 0.156 | 0.134 | 1 | 8 | Hikeshi | 0.022 | Young |
| 0.498097 | 0.138383 | 0.101 | 0.084 | 1 | 8 | Ipo13 | 0.017 | Young |
| 0.49811 | 0.117057 | 0.083 | 0.067 | 1 | 8 | Ccdc167 | 0.016 | Young |
| 0.498395 | 0.622362 | 0.026 | 0.017 | 1 | 8 | Hdac8 | 0.009 | Young |
| 0.498532 | 0.651362 | 0.052 | 0.039 | 1 | 8 | Fbxl8 | 0.013 | Young |
| 0.498902 | 0.105831 | 0.177 | 0.156 | 1 | 8 | Tmem11 | 0.021 | Young |
| 0.499202 | 0.529741 | 0.039 | 0.028 | 1 | 8 | Rbm48 | 0.011 | Young |
| 0.499525 | 0.415068 | 0.078 | 0.061 | 1 | 8 | Tmem185a | 0.017 | Young |
| 0.500439 | 0.273851 | 0.052 | 0.039 | 1 | 8 | Mios | 0.013 | Young |
| 0.501109 | 0.443624 | 0.026 | 0.017 | 1 | 8 | Tasp1 | 0.009 | Young |
| 0.501109 | 0.393529 | 0.026 | 0.017 | 1 | 8 | Gm15448 | 0.009 | Young |
| 0.501109 | 0.293797 | 0.026 | 0.017 | 1 | 8 | Ifit1 | 0.009 | Young |
| 0.501424 | 0.029853 | 0.803 | 0.698 | 1 | 8 | Ostf1 | 0.105 | Young |
| 0.501546 | 0.029243 | 0.927 | 0.844 | 1 | 8 | Celf2 | 0.083 | Young |
| 0.501665 | 0.04106 | 0.405 | 0.38 | 1 | 8 | Map3k8 | 0.025 | Young |
| 0.50172 | 0.288148 | 0.096 | 0.078 | 1 | 8 | 1190002N1 | 0.018 | Young |
| 0.501733 | 0.015528 | 0.673 | 0.603 | 1 | 8 | Gnaq | 0.07 | Young |

| | | | | | | | | |
|---|---|---|---|---|---|---|---|---|
| 0.502562 | 0.006297 | 0.132 | 0.112 | 1 | 8 | Gtf3c1 | 0.02 | Young |
| 0.5026 | 0.140061 | 0.314 | 0.296 | 1 | 8 | Rraga | 0.018 | Young |
| 0.502901 | 0.027638 | 0.805 | 0.771 | 1 | 8 | Cbr2 | 0.034 | Young |
| 0.503613 | 0.548844 | 0.039 | 0.028 | 1 | 8 | Map4k3 | 0.011 | Young |
| 0.50375 | 0.116067 | 0.234 | 0.218 | 1 | 8 | Paxbp1 | 0.016 | Young |
| 0.504588 | 0.116098 | 0.358 | 0.324 | 1 | 8 | Rap2a | 0.034 | Young |
| 0.504719 | 0.405954 | 0.039 | 0.028 | 1 | 8 | Ankrd13b | 0.011 | Young |
| 0.504837 | 0.319152 | 0.125 | 0.106 | 1 | 8 | Alpk1 | 0.019 | Young |
| 0.505194 | 0.263045 | 0.026 | 0.017 | 1 | 8 | Tspyl4 | 0.009 | Young |
| 0.505312 | 0.631978 | 0.065 | 0.05 | 1 | 8 | Sort1 | 0.015 | Young |
| 0.50548 | 0.09879 | 0.195 | 0.173 | 1 | 8 | Cpd | 0.022 | Young |
| 0.50569 | 0.170989 | 0.192 | 0.173 | 1 | 8 | Uri1 | 0.019 | Young |
| 0.506073 | 0.073495 | 0.19 | 0.168 | 1 | 8 | Timm10 | 0.022 | Young |
| 0.506134 | 0.025808 | 0.169 | 0.145 | 1 | 8 | Sirt1 | 0.024 | Young |
| 0.506914 | 0.228823 | 0.229 | 0.207 | 1 | 8 | C1galt1 | 0.022 | Young |
| 0.507142 | 0.334339 | 0.052 | 0.039 | 1 | 8 | Acp5 | 0.013 | Young |
| 0.508627 | 0.011832 | 0.605 | 0.559 | 1 | 8 | Ywhag | 0.046 | Young |
| 0.509065 | 0.528934 | 0.052 | 0.039 | 1 | 8 | Serf1 | 0.013 | Young |
| 0.509156 | 0.615257 | 0.039 | 0.028 | 1 | 8 | Dynlt1a | 0.011 | Young |
| 0.509435 | 0.411497 | 0.07 | 0.056 | 1 | 8 | Extl2 | 0.014 | Young |
| 0.509654 | 1.218742 | 0.044 | 0.034 | 1 | 8 | 1700006J1 | 0.01 | Young |
| 0.509715 | 0.06658 | 0.023 | 0.034 | 1 | 8 | Igfbp6 | -0.011 | Young |
| 0.510029 | 0.215206 | 0.052 | 0.039 | 1 | 8 | Ick | 0.013 | Young |
| 0.510269 | 0.258405 | 0.039 | 0.028 | 1 | 8 | Mrpl47 | 0.011 | Young |
| 0.51035 | 0.001361 | 0.413 | 0.385 | 1 | 8 | Cdk11b | 0.028 | Young |
| 0.510406 | 0.09546 | 0.195 | 0.168 | 1 | 8 | Prelid2 | 0.027 | Young |
| 0.510459 | 0.132684 | 0.197 | 0.173 | 1 | 8 | Vkorc1 | 0.024 | Young |
| 0.511168 | 0.422074 | 0.094 | 0.078 | 1 | 8 | Ppp1r9a | 0.016 | Young |
| 0.51124 | 0.00811 | 0.273 | 0.251 | 1 | 8 | Commd3 | 0.022 | Young |
| 0.511908 | 0.343538 | 0.153 | 0.134 | 1 | 8 | Prps1 | 0.019 | Young |
| 0.512498 | 0.51655 | 0.039 | 0.028 | 1 | 8 | E130102H2 | 0.011 | Young |
| 0.512572 | 0.30427 | 0.143 | 0.123 | 1 | 8 | Snhg6 | 0.02 | Young |
| 0.512626 | 0.243078 | 0.309 | 0.285 | 1 | 8 | Erlin1 | 0.024 | Young |
| 0.512693 | 0.017517 | 0.101 | 0.084 | 1 | 8 | Bcas3 | 0.017 | Young |
| 0.512761 | 0.632702 | 0.07 | 0.056 | 1 | 8 | 1300002E1 | 0.014 | Young |
| 0.513139 | 0.08084 | 0.221 | 0.196 | 1 | 8 | Mrpl32 | 0.025 | Young |
| 0.513408 | 0.08078 | 0.052 | 0.039 | 1 | 8 | Tmem35b | 0.013 | Young |
| 0.513614 | 0.617725 | 0.039 | 0.028 | 1 | 8 | Abhd8 | 0.011 | Young |
| 0.513876 | 0.095068 | 0.096 | 0.078 | 1 | 8 | Ripk2 | 0.018 | Young |
| 0.51407 | 0.072889 | 0.416 | 0.385 | 1 | 8 | Svil | 0.031 | Young |
| 0.514086 | 0.32924 | 0.112 | 0.095 | 1 | 8 | Ccdc117 | 0.017 | Young |
| 0.514276 | 0.081872 | 0.447 | 0.397 | 1 | 8 | Snrpd1 | 0.05 | Young |
| 0.514794 | 0.069896 | 0.247 | 0.223 | 1 | 8 | H2-T22 | 0.024 | Young |
| 0.514819 | 0.472288 | 0.065 | 0.05 | 1 | 8 | Homer3 | 0.015 | Young |
| 0.514896 | 0.111343 | 0.226 | 0.201 | 1 | 8 | Oser1 | 0.025 | Young |

| | | | | | | | | |
|---|---|---|---|---|---|---|---|---|
| 0.515683 | 0.025892 | 0.927 | 0.883 | 1 | 8 | Morf4l1 | 0.044 | Young |
| 0.51585 | 0.376857 | 0.039 | 0.028 | 1 | 8 | BC017158 | 0.011 | Young |
| 0.51697 | 0.221649 | 0.039 | 0.028 | 1 | 8 | Kdm4b | 0.011 | Young |
| 0.517229 | 0.106529 | 0.083 | 0.067 | 1 | 8 | Thap7 | 0.016 | Young |
| 0.517293 | 0.073616 | 0.275 | 0.251 | 1 | 8 | Dync1li2 | 0.024 | Young |
| 0.517743 | 0.284413 | 0.14 | 0.123 | 1 | 8 | S1pr2 | 0.017 | Young |
| 0.518002 | 0.255172 | 0.083 | 0.067 | 1 | 8 | Il1rl2 | 0.016 | Young |
| 0.518015 | 0.018354 | 0.33 | 0.302 | 1 | 8 | 8-Sep | 0.028 | Young |
| 0.518726 | 0.191901 | 0.106 | 0.089 | 1 | 8 | Sfmbt1 | 0.017 | Young |
| 0.518822 | 0.239714 | 0.125 | 0.106 | 1 | 8 | Ripor2 | 0.019 | Young |
| 0.518924 | 0.077099 | 0.226 | 0.207 | 1 | 8 | Zcchc17 | 0.019 | Young |
| 0.519214 | 0.376046 | 0.039 | 0.028 | 1 | 8 | Lmnb2 | 0.011 | Young |
| 0.519214 | 0.354502 | 0.039 | 0.028 | 1 | 8 | Cpeb3 | 0.011 | Young |
| 0.519237 | 0.552449 | 0.057 | 0.045 | 1 | 8 | Ift172 | 0.012 | Young |
| 0.519407 | 0.53132 | 0.081 | 0.067 | 1 | 8 | Terf1 | 0.014 | Young |
| 0.519523 | 0.601468 | 0.13 | 0.112 | 1 | 8 | Zbtb21 | 0.018 | Young |
| 0.519716 | 0.005091 | 0.54 | 0.525 | 1 | 8 | Crebbp | 0.015 | Young |
| 0.519721 | 0.039843 | 0.101 | 0.084 | 1 | 8 | Anapc7 | 0.017 | Young |
| 0.520058 | 0.103037 | 0.223 | 0.201 | 1 | 8 | Agpat5 | 0.022 | Young |
| 0.520826 | 0.409854 | 0.117 | 0.101 | 1 | 8 | Cblb | 0.016 | Young |
| 0.521725 | 0.044529 | 0.273 | 0.24 | 1 | 8 | Cstf3 | 0.033 | Young |
| 0.521968 | 0.458771 | 0.07 | 0.056 | 1 | 8 | Haus2 | 0.014 | Young |
| 0.522017 | 0.551165 | 0.057 | 0.045 | 1 | 8 | Unc5a | 0.012 | Young |
| 0.522224 | 0.033618 | 0.029 | 0.039 | 1 | 8 | Helz2 | -0.01 | Young |
| 0.522589 | 0.282918 | 0.039 | 0.028 | 1 | 8 | 2810001G2 | 0.011 | Young |
| 0.523331 | 0.215694 | 0.361 | 0.352 | 1 | 8 | Grk3 | 0.009 | Young |
| 0.523584 | 0.038484 | 0.177 | 0.151 | 1 | 8 | Xpnpep1 | 0.026 | Young |
| 0.523874 | 0.659601 | 0.057 | 0.045 | 1 | 8 | Prdm10 | 0.012 | Young |
| 0.525095 | 0.004758 | 0.642 | 0.581 | 1 | 8 | Stard8 | 0.061 | Young |
| 0.525887 | 1.107405 | 0.018 | 0.011 | 1 | 8 | Gm9938 | 0.007 | Young |
| 0.526086 | 0.043323 | 0.01 | 0.017 | 1 | 8 | Gm43614 | -0.007 | Young |
| 0.526179 | 0.028367 | 0.545 | 0.486 | 1 | 8 | Srebf2 | 0.059 | Young |
| 0.526257 | 0.260885 | 0.075 | 0.061 | 1 | 8 | Sephs2 | 0.014 | Young |
| 0.526996 | 0.021476 | 0.216 | 0.19 | 1 | 8 | Myo18a | 0.026 | Young |
| 0.527051 | 0.072158 | 0.143 | 0.123 | 1 | 8 | N4bp2 | 0.02 | Young |
| 0.52753 | 0.222643 | 0.052 | 0.039 | 1 | 8 | Lyrm4 | 0.013 | Young |
| 0.527557 | 1.054828 | 0.018 | 0.011 | 1 | 8 | Dtwd2 | 0.007 | Young |
| 0.527557 | 0.978151 | 0.018 | 0.011 | 1 | 8 | Cmtr2 | 0.007 | Young |
| 0.527557 | 0.811677 | 0.018 | 0.011 | 1 | 8 | Fam92a | 0.007 | Young |
| 0.527645 | 0.067625 | 0.343 | 0.318 | 1 | 8 | Bmt2 | 0.025 | Young |
| 0.527895 | 0.296213 | 0.088 | 0.073 | 1 | 8 | Tpk1 | 0.015 | Young |
| 0.528168 | 0.112665 | 0.029 | 0.039 | 1 | 8 | Zfp688 | -0.01 | Young |
| 0.528254 | 0.214416 | 0.379 | 0.363 | 1 | 8 | Rnf145 | 0.016 | Young |
| 0.52836 | 0.158983 | 0.13 | 0.112 | 1 | 8 | Apex1 | 0.018 | Young |
| 0.52923 | 1.21256 | 0.018 | 0.011 | 1 | 8 | Gm48604 | 0.007 | Young |

| | | | | | | | | |
|---|---|---|---|---|---|---|---|---|
| 0.52923 | 0.984243 | 0.018 | 0.011 | 1 | 8 | C2 | 0.007 | Young |
| 0.529563 | 0.403949 | 0.07 | 0.056 | 1 | 8 | Cela1 | 0.014 | Young |
| 0.530211 | 0.049085 | 0.496 | 0.464 | 1 | 8 | Psmc3 | 0.032 | Young |
| 0.530355 | 0.103452 | 0.101 | 0.084 | 1 | 8 | Zfp780b | 0.017 | Young |
| 0.530411 | 0.395313 | 0.07 | 0.056 | 1 | 8 | Mrps31 | 0.014 | Young |
| 0.530867 | 0.123557 | 0.223 | 0.201 | 1 | 8 | Dph3 | 0.022 | Young |
| 0.530906 | 1.027795 | 0.018 | 0.011 | 1 | 8 | Cdc42bpa | 0.007 | Young |
| 0.530906 | 0.778822 | 0.018 | 0.011 | 1 | 8 | Gm26826 | 0.007 | Young |
| 0.530906 | 0.557452 | 0.018 | 0.011 | 1 | 8 | Gm12689 | 0.007 | Young |
| 0.531007 | 0.049994 | 0.319 | 0.291 | 1 | 8 | Rps6kb1 | 0.028 | Young |
| 0.531769 | 0.148936 | 0.01 | 0.017 | 1 | 8 | Lamb2 | -0.007 | Young |
| 0.531769 | 0.115766 | 0.01 | 0.017 | 1 | 8 | Phlda3 | -0.007 | Young |
| 0.531769 | 0.011884 | 0.01 | 0.017 | 1 | 8 | Cdkn3 | -0.007 | Young |
| 0.532275 | 0.584894 | 0.057 | 0.045 | 1 | 8 | Shmt2 | 0.012 | Young |
| 0.532275 | 0.132904 | 0.057 | 0.045 | 1 | 8 | Cry2 | 0.012 | Young |
| 0.532584 | 0.803883 | 0.018 | 0.011 | 1 | 8 | Uprt | 0.007 | Young |
| 0.532584 | 0.71212 | 0.018 | 0.011 | 1 | 8 | Ttll5 | 0.007 | Young |
| 0.532584 | 0.610266 | 0.018 | 0.011 | 1 | 8 | 2610301B2 | 0.007 | Young |
| 0.532584 | 0.49465 | 0.018 | 0.011 | 1 | 8 | Gm17435 | 0.007 | Young |
| 0.532829 | 0.27563 | 0.088 | 0.073 | 1 | 8 | Zfp64 | 0.015 | Young |
| 0.532948 | 0.048312 | 0.029 | 0.039 | 1 | 8 | Il15ra | -0.01 | Young |
| 0.53322 | 0.069142 | 0.904 | 0.838 | 1 | 8 | Rps27l | 0.066 | Young |
| 0.533572 | 0.204739 | 0.356 | 0.341 | 1 | 8 | Rbm27 | 0.015 | Young |
| 0.533585 | 0.635123 | 0.081 | 0.067 | 1 | 8 | Mcu | 0.014 | Young |
| 0.533585 | 0.536004 | 0.081 | 0.067 | 1 | 8 | Gde1 | 0.014 | Young |
| 0.533671 | 0.446392 | 0.01 | 0.017 | 1 | 8 | Dnah2 | -0.007 | Young |
| 0.533866 | 0.33257 | 0.044 | 0.034 | 1 | 8 | Oxnad1 | 0.01 | Young |
| 0.534265 | 0.567655 | 0.018 | 0.011 | 1 | 8 | Cep72 | 0.007 | Young |
| 0.534624 | 0.331605 | 0.21 | 0.196 | 1 | 8 | Hspa4l | 0.014 | Young |
| 0.535067 | 0.000183 | 0.156 | 0.134 | 1 | 8 | Gmeb1 | 0.022 | Young |
| 0.53551 | 0.371933 | 0.099 | 0.084 | 1 | 8 | Lix1l | 0.015 | Young |
| 0.535532 | 0.026517 | 0.909 | 0.804 | 1 | 8 | Emp3 | 0.105 | Young |
| 0.535948 | 0.625895 | 0.018 | 0.011 | 1 | 8 | Zfp438 | 0.007 | Young |
| 0.535948 | 0.614713 | 0.018 | 0.011 | 1 | 8 | 0610040B1 | 0.007 | Young |
| 0.535948 | 0.543546 | 0.018 | 0.011 | 1 | 8 | Lrfn1 | 0.007 | Young |
| 0.535948 | 0.518963 | 0.018 | 0.011 | 1 | 8 | Tarbp1 | 0.007 | Young |
| 0.535948 | 0.370163 | 0.018 | 0.011 | 1 | 8 | Sntg1 | 0.007 | Young |
| 0.536023 | 0.028119 | 0.164 | 0.14 | 1 | 8 | Zfp593 | 0.024 | Young |
| 0.536127 | 0.075393 | 0.603 | 0.581 | 1 | 8 | Tax1bp1 | 0.022 | Young |
| 0.536147 | 0.810164 | 0.031 | 0.022 | 1 | 8 | Zfp316 | 0.009 | Young |
| 0.53665 | 0.070777 | 0.569 | 0.564 | 1 | 8 | Mbnl2 | 0.005 | Young |
| 0.53674 | 0.003839 | 0.242 | 0.207 | 1 | 8 | Il17ra | 0.035 | Young |
| 0.537323 | 0.026319 | 0.535 | 0.48 | 1 | 8 | Zdhhc20 | 0.055 | Young |
| 0.537484 | 0.611274 | 0.01 | 0.017 | 1 | 8 | Fancf | -0.007 | Young |
| 0.537484 | 0.066068 | 0.01 | 0.017 | 1 | 8 | AC129328. | -0.007 | Young |

| | | | | | | | | |
|---|---|---|---|---|---|---|---|---|
| 0.537635 | 0.707449 | 0.018 | 0.011 | 1 | 8 | Wdr25 | 0.007 | Young |
| 0.537635 | 0.622278 | 0.018 | 0.011 | 1 | 8 | Asrgl1 | 0.007 | Young |
| 0.537635 | 0.51293 | 0.018 | 0.011 | 1 | 8 | Rpp40 | 0.007 | Young |
| 0.537635 | 0.479766 | 0.018 | 0.011 | 1 | 8 | Brca2 | 0.007 | Young |
| 0.537993 | 0.09425 | 0.273 | 0.251 | 1 | 8 | Zhx1 | 0.022 | Young |
| 0.538693 | 0.635661 | 0.031 | 0.022 | 1 | 8 | Fam206a | 0.009 | Young |
| 0.538856 | 0.387743 | 0.057 | 0.045 | 1 | 8 | Cc2d1a | 0.012 | Young |
| 0.538974 | 0.281836 | 0.19 | 0.173 | 1 | 8 | Zdhhc7 | 0.017 | Young |
| 0.539324 | 0.633957 | 0.018 | 0.011 | 1 | 8 | Card10 | 0.007 | Young |
| 0.539324 | 0.538034 | 0.018 | 0.011 | 1 | 8 | Rbp1 | 0.007 | Young |
| 0.539774 | 0.27984 | 0.14 | 0.123 | 1 | 8 | Trap1 | 0.017 | Young |
| 0.539969 | 0.714133 | 0.031 | 0.022 | 1 | 8 | Cenpw | 0.009 | Young |
| 0.540024 | 0.00171 | 0.901 | 0.855 | 1 | 8 | Dbi | 0.046 | Young |
| 0.54009 | 0.288033 | 0.169 | 0.151 | 1 | 8 | Arhgap6 | 0.018 | Young |
| 0.541009 | 0.067643 | 0.148 | 0.128 | 1 | 8 | Rabgap1l | 0.02 | Young |
| 0.541015 | 0.438729 | 0.018 | 0.011 | 1 | 8 | Il27 | 0.007 | Young |
| 0.541015 | 0.335557 | 0.018 | 0.011 | 1 | 8 | Slc19a2 | 0.007 | Young |
| 0.541015 | 0.277136 | 0.018 | 0.011 | 1 | 8 | Mfsd13a | 0.007 | Young |
| 0.541015 | 0.261376 | 0.018 | 0.011 | 1 | 8 | C920021L1 | 0.007 | Young |
| 0.541117 | 0.438502 | 0.166 | 0.151 | 1 | 8 | Nt5c2 | 0.015 | Young |
| 0.542346 | 0.170753 | 0.07 | 0.056 | 1 | 8 | Tars2 | 0.014 | Young |
| 0.542412 | 0.039522 | 0.499 | 0.458 | 1 | 8 | Atxn2l | 0.041 | Young |
| 0.542447 | 0.011118 | 0.449 | 0.402 | 1 | 8 | Dcaf12 | 0.047 | Young |
| 0.54271 | 0.34174 | 0.018 | 0.011 | 1 | 8 | Scml4 | 0.007 | Young |
| 0.54271 | 0.339966 | 0.018 | 0.011 | 1 | 8 | 1810009A1 | 0.007 | Young |
| 0.54271 | 0.281221 | 0.018 | 0.011 | 1 | 8 | Gm30025 | 0.007 | Young |
| 0.54271 | 0.063377 | 0.018 | 0.011 | 1 | 8 | Ccsap | 0.007 | Young |
| 0.543039 | 0.50227 | 0.086 | 0.073 | 1 | 8 | Ccdc69 | 0.013 | Young |
| 0.543152 | 0.054923 | 0.608 | 0.57 | 1 | 8 | Map1lc3b | 0.038 | Young |
| 0.543345 | 0.092362 | 0.603 | 0.57 | 1 | 8 | Srsf9 | 0.033 | Young |
| 0.543805 | 0.688073 | 0.031 | 0.022 | 1 | 8 | Haus1 | 0.009 | Young |
| 0.544282 | 0.352518 | 0.094 | 0.078 | 1 | 8 | Hpse | 0.016 | Young |
| 0.544407 | 0.268284 | 0.018 | 0.011 | 1 | 8 | Gm16867 | 0.007 | Young |
| 0.544407 | 0.223492 | 0.018 | 0.011 | 1 | 8 | Casp12 | 0.007 | Young |
| 0.544703 | 0.106831 | 0.327 | 0.296 | 1 | 8 | Gpr160 | 0.031 | Young |
| 0.54494 | 0.443769 | 0.099 | 0.084 | 1 | 8 | Slc39a3 | 0.015 | Young |
| 0.545547 | 0.70074 | 0.081 | 0.067 | 1 | 8 | Pstk | 0.014 | Young |
| 0.545865 | 0.175165 | 0.148 | 0.128 | 1 | 8 | Eif2ak3 | 0.02 | Young |
| 0.545893 | 0.090421 | 0.868 | 0.771 | 1 | 8 | Gnas | 0.097 | Young |
| 0.546053 | 0.174376 | 0.122 | 0.106 | 1 | 8 | Memo1 | 0.016 | Young |
| 0.546106 | 0.21299 | 0.018 | 0.011 | 1 | 8 | B130006D( | 0.007 | Young |
| 0.546106 | 0.19128 | 0.018 | 0.011 | 1 | 8 | Zfat | 0.007 | Young |
| 0.546106 | 0.189287 | 0.018 | 0.011 | 1 | 8 | Gm15564 | 0.007 | Young |
| 0.546106 | 0.161146 | 0.018 | 0.011 | 1 | 8 | Fchsd1 | 0.007 | Young |
| 0.546106 | 0.159796 | 0.018 | 0.011 | 1 | 8 | Gm3336 | 0.007 | Young |

| | | | | | | | | |
|---|---|---|---|---|---|---|---|---|
| 0.546146 | 0.093775 | 0.218 | 0.196 | 1 | 8 | Nr2f6 | 0.022 | Young |
| 0.546376 | 0.673302 | 0.125 | 0.112 | 1 | 8 | Ampd3 | 0.013 | Young |
| 0.546813 | 0.388641 | 0.075 | 0.061 | 1 | 8 | Cep162 | 0.014 | Young |
| 0.546906 | 0.174298 | 0.13 | 0.112 | 1 | 8 | Ankrd16 | 0.018 | Young |
| 0.547338 | 0.073457 | 0.919 | 0.877 | 1 | 8 | Cd36 | 0.042 | Young |
| 0.54737 | 0.052265 | 0.213 | 0.19 | 1 | 8 | Ube2j1 | 0.023 | Young |
| 0.547512 | 0.065997 | 0.231 | 0.207 | 1 | 8 | Pafah1b2 | 0.024 | Young |
| 0.548085 | 0.141278 | 0.249 | 0.229 | 1 | 8 | Znrd1 | 0.02 | Young |
| 0.548334 | 0.434472 | 0.156 | 0.14 | 1 | 8 | Actr1b | 0.016 | Young |
| 0.548884 | 0.481556 | 0.044 | 0.034 | 1 | 8 | Ms4a14 | 0.01 | Young |
| 0.54906 | 0.005035 | 0.325 | 0.291 | 1 | 8 | Atf7 | 0.034 | Young |
| 0.549134 | 0.11183 | 0.369 | 0.335 | 1 | 8 | Tm2d2 | 0.034 | Young |
| 0.549197 | 0.056002 | 0.151 | 0.128 | 1 | 8 | Smn1 | 0.023 | Young |
| 0.54922 | 0.384627 | 0.117 | 0.101 | 1 | 8 | Ipo9 | 0.016 | Young |
| 0.549673 | 0.056636 | 0.945 | 0.916 | 1 | 8 | Cyba | 0.029 | Young |
| 0.549964 | 0.230427 | 0.044 | 0.034 | 1 | 8 | Rundc1 | 0.01 | Young |
| 0.550118 | 0.175838 | 0.156 | 0.14 | 1 | 8 | Trmt6 | 0.016 | Young |
| 0.550163 | 0.109796 | 0.161 | 0.14 | 1 | 8 | Zfp598 | 0.021 | Young |
| 0.550191 | 0.03767 | 0.657 | 0.57 | 1 | 8 | Uqcrb | 0.087 | Young |
| 0.550229 | 1.066128 | 0.031 | 0.022 | 1 | 8 | Pfkm | 0.009 | Young |
| 0.550229 | 0.225442 | 0.031 | 0.022 | 1 | 8 | Cyp2s1 | 0.009 | Young |
| 0.551046 | 0.344185 | 0.044 | 0.034 | 1 | 8 | Gm26518 | 0.01 | Young |
| 0.551059 | 0.456576 | 0.145 | 0.128 | 1 | 8 | Yif1b | 0.017 | Young |
| 0.55148 | 0.290007 | 0.161 | 0.145 | 1 | 8 | Dcaf5 | 0.016 | Young |
| 0.552808 | 0.455216 | 0.031 | 0.022 | 1 | 8 | Ttc21b | 0.009 | Young |
| 0.552808 | 0.42849 | 0.031 | 0.022 | 1 | 8 | Praf2 | 0.009 | Young |
| 0.553023 | 0.008241 | 0.234 | 0.207 | 1 | 8 | Rhobtb1 | 0.027 | Young |
| 0.553048 | 0.236779 | 0.353 | 0.341 | 1 | 8 | Smc3 | 0.012 | Young |
| 0.553208 | 0.009647 | 0.135 | 0.117 | 1 | 8 | Tmem60 | 0.018 | Young |
| 0.553212 | 0.319718 | 0.044 | 0.034 | 1 | 8 | Rabep2 | 0.01 | Young |
| 0.553212 | 0.464933 | 0.044 | 0.034 | 1 | 8 | Pcyt1b | 0.01 | Young |
| 0.553304 | 0.699698 | 0.049 | 0.039 | 1 | 8 | Mapkbp1 | 0.01 | Young |
| 0.554284 | 0.11199 | 0.317 | 0.291 | 1 | 8 | Golga7 | 0.026 | Young |
| 0.554327 | 0.861417 | 0.049 | 0.039 | 1 | 8 | Cmss1 | 0.01 | Young |
| 0.554684 | 0.1953 | 0.122 | 0.106 | 1 | 8 | Isyna1 | 0.016 | Young |
| 0.555186 | 0.33026 | 0.099 | 0.084 | 1 | 8 | Gtf3c4 | 0.015 | Young |
| 0.555186 | 0.270417 | 0.099 | 0.084 | 1 | 8 | Slc25a19 | 0.015 | Young |
| 0.555383 | 0.130662 | 0.044 | 0.034 | 1 | 8 | Gm6563 | 0.01 | Young |
| 0.555501 | 0.114049 | 0.218 | 0.196 | 1 | 8 | Atrnl1 | 0.022 | Young |
| 0.555736 | 0.149002 | 0.421 | 0.397 | 1 | 8 | Mknk2 | 0.024 | Young |
| 0.556596 | 0.00924 | 0.99 | 0.955 | 1 | 8 | Oaz1 | 0.035 | Young |
| 0.556689 | 0.124796 | 0.031 | 0.022 | 1 | 8 | Fastkd1 | 0.009 | Young |
| 0.557604 | 0.065202 | 0.226 | 0.207 | 1 | 8 | Gle1 | 0.019 | Young |
| 0.557841 | 0.280998 | 0.088 | 0.073 | 1 | 8 | Adi1 | 0.015 | Young |
| 0.558119 | 0.006254 | 0.192 | 0.168 | 1 | 8 | Tmem41b | 0.024 | Young |

| | | | | | | | | |
|---|---|---|---|---|---|---|---|---|
| 0.558528 | 0.049189 | 0.374 | 0.335 | 1 | 8 | Kdm2b | 0.039 | Young |
| 0.558647 | 0.172777 | 0.044 | 0.034 | 1 | 8 | Apex2 | 0.01 | Young |
| 0.5592 | 0.137808 | 0.252 | 0.223 | 1 | 8 | Vav2 | 0.029 | Young |
| 0.559456 | 0.632017 | 0.049 | 0.039 | 1 | 8 | Cdkn1c | 0.01 | Young |
| 0.559817 | 0.094516 | 0.158 | 0.14 | 1 | 8 | Tmem245 | 0.018 | Young |
| 0.560485 | 0.775771 | 0.049 | 0.039 | 1 | 8 | Ankrd26 | 0.01 | Young |
| 0.561145 | 0.262807 | 0.187 | 0.168 | 1 | 8 | Igsf8 | 0.019 | Young |
| 0.561185 | 0.724077 | 0.068 | 0.056 | 1 | 8 | Rdh10 | 0.012 | Young |
| 0.561856 | 0.229507 | 0.075 | 0.061 | 1 | 8 | Zfp687 | 0.014 | Young |
| 0.561884 | 0.407509 | 0.031 | 0.022 | 1 | 8 | Dync2h1 | 0.009 | Young |
| 0.561884 | 0.274276 | 0.031 | 0.022 | 1 | 8 | Zfp747 | 0.009 | Young |
| 0.562039 | 0.269716 | 0.122 | 0.106 | 1 | 8 | Tjp2 | 0.016 | Young |
| 0.56277 | 0.022973 | 0.821 | 0.765 | 1 | 8 | Pomp | 0.056 | Young |
| 0.563926 | 0.061147 | 0.203 | 0.184 | 1 | 8 | Btaf1 | 0.019 | Young |
| 0.563938 | 0.220609 | 0.273 | 0.257 | 1 | 8 | Gsdmd | 0.016 | Young |
| 0.564283 | 0.021242 | 0.79 | 0.743 | 1 | 8 | C5ar1 | 0.047 | Young |
| 0.565202 | 0.276554 | 0.044 | 0.034 | 1 | 8 | Adat2 | 0.01 | Young |
| 0.56575 | 0.759524 | 0.073 | 0.061 | 1 | 8 | Supt3 | 0.012 | Young |
| 0.565776 | 0.297728 | 0.081 | 0.067 | 1 | 8 | Crip2 | 0.014 | Young |
| 0.566119 | 0.003204 | 0.706 | 0.654 | 1 | 8 | Nktr | 0.052 | Young |
| 0.566907 | 0.218179 | 0.112 | 0.095 | 1 | 8 | Cbll1 | 0.017 | Young |
| 0.567051 | 0.002478 | 0.296 | 0.268 | 1 | 8 | Snd1 | 0.028 | Young |
| 0.567093 | 0.026385 | 0.216 | 0.196 | 1 | 8 | Hps5 | 0.02 | Young |
| 0.567382 | 0.659125 | 0.068 | 0.056 | 1 | 8 | Ptcd3 | 0.012 | Young |
| 0.567386 | 0.072301 | 0.345 | 0.324 | 1 | 8 | Adrb2 | 0.021 | Young |
| 0.567431 | 0.174086 | 0.062 | 0.05 | 1 | 8 | Usp46 | 0.012 | Young |
| 0.567467 | 0.045249 | 0.426 | 0.385 | 1 | 8 | Mbp | 0.041 | Young |
| 0.567657 | 0.218062 | 0.322 | 0.296 | 1 | 8 | Maf1 | 0.026 | Young |
| 0.56799 | 1.405159 | 0.01 | 0.006 | 1 | 8 | Zfp30 | 0.004 | Young |
| 0.56799 | 1.109117 | 0.01 | 0.006 | 1 | 8 | AF529169 | 0.004 | Young |
| 0.56799 | 1.042886 | 0.01 | 0.006 | 1 | 8 | Cd300c | 0.004 | Young |
| 0.56799 | 1.003509 | 0.01 | 0.006 | 1 | 8 | Cystm1 | 0.004 | Young |
| 0.56799 | 0.994964 | 0.01 | 0.006 | 1 | 8 | Slc24a5 | 0.004 | Young |
| 0.56799 | 0.879803 | 0.01 | 0.006 | 1 | 8 | Gm26606 | 0.004 | Young |
| 0.56799 | 0.812727 | 0.01 | 0.006 | 1 | 8 | Mpzl1 | 0.004 | Young |
| 0.56799 | 0.801305 | 0.01 | 0.006 | 1 | 8 | Bgn | 0.004 | Young |
| 0.56799 | 0.727963 | 0.01 | 0.006 | 1 | 8 | 4833418N( | 0.004 | Young |
| 0.56799 | 0.711602 | 0.01 | 0.006 | 1 | 8 | Pidd1 | 0.004 | Young |
| 0.56799 | 0.659042 | 0.01 | 0.006 | 1 | 8 | Gm21988 | 0.004 | Young |
| 0.56799 | 0.635353 | 0.01 | 0.006 | 1 | 8 | Hist1h2ae | 0.004 | Young |
| 0.568238 | 0.235986 | 0.252 | 0.235 | 1 | 8 | Nedd9 | 0.017 | Young |
| 0.568411 | 0.054097 | 0.031 | 0.022 | 1 | 8 | 1700113A1 | 0.009 | Young |
| 0.568529 | 0.157523 | 0.182 | 0.162 | 1 | 8 | Fam53b | 0.02 | Young |
| 0.568797 | 0.429449 | 0.125 | 0.112 | 1 | 8 | Dohh | 0.013 | Young |
| 0.569069 | 0.199986 | 0.106 | 0.089 | 1 | 8 | Trappc11 | 0.017 | Young |

| | | | | | | | | |
|---|---|---|---|---|---|---|---|---|
| 0.569457 | 0.12715 | 0.242 | 0.223 | 1 | 8 | Trmt1 | 0.019 | Young |
| 0.56971 | 0.223846 | 0.13 | 0.112 | 1 | 8 | Afg3l2 | 0.018 | Young |
| 0.56972 | 0.223253 | 0.031 | 0.022 | 1 | 8 | 5430416N( | 0.009 | Young |
| 0.569974 | 0.031064 | 0.239 | 0.212 | 1 | 8 | Arpc5l | 0.027 | Young |
| 0.570151 | 0.13738 | 0.353 | 0.324 | 1 | 8 | Tifab | 0.029 | Young |
| 0.570208 | 0.215309 | 0.062 | 0.05 | 1 | 8 | Ncapd3 | 0.012 | Young |
| 0.57031 | 1.001133 | 0.01 | 0.006 | 1 | 8 | Krt7 | 0.004 | Young |
| 0.57031 | 0.94521 | 0.01 | 0.006 | 1 | 8 | Gm47389 | 0.004 | Young |
| 0.57031 | 0.869245 | 0.01 | 0.006 | 1 | 8 | Hykk | 0.004 | Young |
| 0.57031 | 0.790566 | 0.01 | 0.006 | 1 | 8 | 2810021J2 | 0.004 | Young |
| 0.57031 | 0.748675 | 0.01 | 0.006 | 1 | 8 | Npr3 | 0.004 | Young |
| 0.57031 | 0.728683 | 0.01 | 0.006 | 1 | 8 | Etnk2 | 0.004 | Young |
| 0.57031 | 0.684864 | 0.01 | 0.006 | 1 | 8 | Tmem178b | 0.004 | Young |
| 0.57031 | 0.639475 | 0.01 | 0.006 | 1 | 8 | Sncg | 0.004 | Young |
| 0.57031 | 0.577553 | 0.01 | 0.006 | 1 | 8 | Klhdc8b | 0.004 | Young |
| 0.57031 | 0.56338 | 0.01 | 0.006 | 1 | 8 | Notch4 | 0.004 | Young |
| 0.57031 | 0.527201 | 0.01 | 0.006 | 1 | 8 | Rftn2 | 0.004 | Young |
| 0.57031 | 0.52701 | 0.01 | 0.006 | 1 | 8 | Il17d | 0.004 | Young |
| 0.57031 | 0.517423 | 0.01 | 0.006 | 1 | 8 | Nap1l2 | 0.004 | Young |
| 0.57031 | 0.489457 | 0.01 | 0.006 | 1 | 8 | Ccnj | 0.004 | Young |
| 0.57031 | 0.480031 | 0.01 | 0.006 | 1 | 8 | Mpzl3 | 0.004 | Young |
| 0.57031 | 0.46422 | 0.01 | 0.006 | 1 | 8 | Ace | 0.004 | Young |
| 0.57031 | 0.457507 | 0.01 | 0.006 | 1 | 8 | Slc16a9 | 0.004 | Young |
| 0.57031 | 0.406958 | 0.01 | 0.006 | 1 | 8 | Gpr82 | 0.004 | Young |
| 0.57031 | 0.405606 | 0.01 | 0.006 | 1 | 8 | Xrcc3 | 0.004 | Young |
| 0.57031 | 0.405443 | 0.01 | 0.006 | 1 | 8 | Cdca7l | 0.004 | Young |
| 0.570573 | 0.016312 | 0.683 | 0.62 | 1 | 8 | St13 | 0.063 | Young |
| 0.571386 | 0.119854 | 0.268 | 0.246 | 1 | 8 | Smg7 | 0.022 | Young |
| 0.571661 | 0.003987 | 0.166 | 0.145 | 1 | 8 | Sdhaf4 | 0.021 | Young |
| 0.571811 | 0.334379 | 0.138 | 0.123 | 1 | 8 | Zfp945 | 0.015 | Young |
| 0.571953 | 0.054798 | 0.106 | 0.089 | 1 | 8 | Golim4 | 0.017 | Young |
| 0.572634 | 1.070363 | 0.01 | 0.006 | 1 | 8 | Cit | 0.004 | Young |
| 0.572634 | 1.055712 | 0.01 | 0.006 | 1 | 8 | Cep41 | 0.004 | Young |
| 0.572634 | 1.024511 | 0.01 | 0.006 | 1 | 8 | Peg10 | 0.004 | Young |
| 0.572634 | 0.863615 | 0.01 | 0.006 | 1 | 8 | Dock9 | 0.004 | Young |
| 0.572634 | 0.704684 | 0.01 | 0.006 | 1 | 8 | Mtus2 | 0.004 | Young |
| 0.572634 | 0.668687 | 0.01 | 0.006 | 1 | 8 | Cldn10 | 0.004 | Young |
| 0.572634 | 0.574994 | 0.01 | 0.006 | 1 | 8 | Gm3435 | 0.004 | Young |
| 0.572634 | 0.497128 | 0.01 | 0.006 | 1 | 8 | Hist2h3c1 | 0.004 | Young |
| 0.572634 | 0.49511 | 0.01 | 0.006 | 1 | 8 | Bbs10 | 0.004 | Young |
| 0.572634 | 0.476655 | 0.01 | 0.006 | 1 | 8 | Zfp850 | 0.004 | Young |
| 0.572634 | 0.432062 | 0.01 | 0.006 | 1 | 8 | Acot6 | 0.004 | Young |
| 0.572634 | 0.385423 | 0.01 | 0.006 | 1 | 8 | Col1a1 | 0.004 | Young |
| 0.572634 | 0.3452 | 0.01 | 0.006 | 1 | 8 | 9930111J2 | 0.004 | Young |
| 0.573145 | 0.396801 | 0.081 | 0.067 | 1 | 8 | Adprhl2 | 0.014 | Young |

| | | | | | | | | |
|---|---|---|---|---|---|---|---|---|
| 0.573921 | 0.295572 | 0.208 | 0.19 | 1 | 8 | Polr2h | 0.018 | Young |
| 0.574038 | 0.194787 | 0.187 | 0.168 | 1 | 8 | Mrpl11 | 0.019 | Young |
| 0.574505 | 0.68813 | 0.068 | 0.056 | 1 | 8 | Abhd6 | 0.012 | Young |
| 0.574515 | 0.295671 | 0.145 | 0.134 | 1 | 8 | Ap1m1 | 0.011 | Young |
| 0.574788 | 0.103497 | 0.156 | 0.14 | 1 | 8 | Cmc2 | 0.016 | Young |
| 0.574963 | 0.5529 | 0.01 | 0.006 | 1 | 8 | Fam171a1 | 0.004 | Young |
| 0.574963 | 0.459362 | 0.01 | 0.006 | 1 | 8 | Car7 | 0.004 | Young |
| 0.574963 | 0.448383 | 0.01 | 0.006 | 1 | 8 | Abcb4 | 0.004 | Young |
| 0.574963 | 0.396212 | 0.01 | 0.006 | 1 | 8 | Trpt1 | 0.004 | Young |
| 0.574963 | 0.394547 | 0.01 | 0.006 | 1 | 8 | Adamts4 | 0.004 | Young |
| 0.574963 | 0.38102 | 0.01 | 0.006 | 1 | 8 | Dok4 | 0.004 | Young |
| 0.574963 | 0.370969 | 0.01 | 0.006 | 1 | 8 | Ccdc157 | 0.004 | Young |
| 0.574963 | 0.349351 | 0.01 | 0.006 | 1 | 8 | Zfp428 | 0.004 | Young |
| 0.574963 | 0.343581 | 0.01 | 0.006 | 1 | 8 | Spsb4 | 0.004 | Young |
| 0.574963 | 0.313321 | 0.01 | 0.006 | 1 | 8 | Tmem216 | 0.004 | Young |
| 0.574963 | 0.305152 | 0.01 | 0.006 | 1 | 8 | Fbxo5 | 0.004 | Young |
| 0.574963 | 0.292403 | 0.01 | 0.006 | 1 | 8 | Fam72a | 0.004 | Young |
| 0.574963 | 0.29122 | 0.01 | 0.006 | 1 | 8 | Gm7324 | 0.004 | Young |
| 0.574963 | 0.249555 | 0.01 | 0.006 | 1 | 8 | Csf1 | 0.004 | Young |
| 0.574963 | 0.227374 | 0.01 | 0.006 | 1 | 8 | Camkk1 | 0.004 | Young |
| 0.574963 | 0.169067 | 0.01 | 0.006 | 1 | 8 | AC113595. | 0.004 | Young |
| 0.575154 | 0.267087 | 0.174 | 0.162 | 1 | 8 | Cdkn2aipnl | 0.012 | Young |
| 0.575628 | 0.195282 | 0.19 | 0.173 | 1 | 8 | Mtmr14 | 0.017 | Young |
| 0.575941 | 0.26002 | 0.257 | 0.246 | 1 | 8 | Ccdc59 | 0.011 | Young |
| 0.576023 | 0.460784 | 0.049 | 0.039 | 1 | 8 | Slc25a38 | 0.01 | Young |
| 0.576705 | 0.160901 | 0.164 | 0.151 | 1 | 8 | Fam219a | 0.013 | Young |
| 0.577297 | 0.242421 | 0.01 | 0.006 | 1 | 8 | Plcd1 | 0.004 | Young |
| 0.577297 | 0.155503 | 0.01 | 0.006 | 1 | 8 | Gm3604 | 0.004 | Young |
| 0.577297 | 0.09151 | 0.01 | 0.006 | 1 | 8 | Trim62 | 0.004 | Young |
| 0.577297 | 0.029622 | 0.01 | 0.006 | 1 | 8 | Cd177 | 0.004 | Young |
| 0.577297 | 0.001998 | 0.01 | 0.006 | 1 | 8 | Agbl3 | 0.004 | Young |
| 0.577322 | 0.06477 | 0.634 | 0.62 | 1 | 8 | Oxct1 | 0.014 | Young |
| 0.577608 | 0.017572 | 0.262 | 0.246 | 1 | 8 | Pip4k2b | 0.016 | Young |
| 0.577712 | 0.043406 | 0.049 | 0.061 | 1 | 8 | Zfp113 | -0.012 | Young |
| 0.578694 | 0.054925 | 0.218 | 0.196 | 1 | 8 | Faim | 0.022 | Young |
| 0.57891 | 0.002338 | 0.112 | 0.095 | 1 | 8 | Ppp2r5d | 0.017 | Young |
| 0.579371 | 0.220794 | 0.242 | 0.223 | 1 | 8 | Dopey1 | 0.019 | Young |
| 0.579512 | 0.203288 | 0.062 | 0.05 | 1 | 8 | Rabepk | 0.012 | Young |
| 0.580396 | 0.036431 | 0.19 | 0.173 | 1 | 8 | Pggt1b | 0.017 | Young |
| 0.580575 | 0.091754 | 0.821 | 0.793 | 1 | 8 | Tcf4 | 0.028 | Young |
| 0.580607 | 0.051522 | 0.016 | 0.022 | 1 | 8 | Tmc8 | -0.006 | Young |
| 0.581247 | 0.609508 | 0.049 | 0.039 | 1 | 8 | 2700097O( | 0.01 | Young |
| 0.581359 | 0.346258 | 0.216 | 0.201 | 1 | 8 | Apopt1 | 0.015 | Young |
| 0.581514 | 0.23306 | 0.148 | 0.134 | 1 | 8 | Cdc14a | 0.014 | Young |
| 0.581871 | 0.022099 | 0.418 | 0.402 | 1 | 8 | Rc3h1 | 0.016 | Young |

| | | | | | | | | |
|---|---|---|---|---|---|---|---|---|
| 0.583114 | 0.082934 | 0.07 | 0.084 | 1 | 8 | Gtdc1 | -0.014 | Young |
| 0.583159 | 0.125117 | 0.481 | 0.458 | 1 | 8 | Ciita | 0.023 | Young |
| 0.583819 | 0.454251 | 0.073 | 0.061 | 1 | 8 | Clcn6 | 0.012 | Young |
| 0.584883 | 0.555676 | 0.125 | 0.112 | 1 | 8 | Fkbp1b | 0.013 | Young |
| 0.585084 | 0.064412 | 0.395 | 0.369 | 1 | 8 | Twf1 | 0.026 | Young |
| 0.585278 | 0.016555 | 0.244 | 0.223 | 1 | 8 | 0610030E2 | 0.021 | Young |
| 0.585442 | 0.278503 | 0.049 | 0.039 | 1 | 8 | Nudt22 | 0.01 | Young |
| 0.585607 | 0.01107 | 0.016 | 0.022 | 1 | 8 | Gm7967 | -0.006 | Young |
| 0.586112 | 0.05899 | 0.356 | 0.33 | 1 | 8 | Tceal9 | 0.026 | Young |
| 0.586875 | 0.004058 | 0.392 | 0.358 | 1 | 8 | Sh3bp1 | 0.034 | Young |
| 0.587006 | 0.008476 | 0.055 | 0.067 | 1 | 8 | Zc3h8 | -0.012 | Young |
| 0.587233 | 0.387981 | 0.099 | 0.084 | 1 | 8 | Tfdp2 | 0.015 | Young |
| 0.587946 | 0.336397 | 0.062 | 0.05 | 1 | 8 | Mospd1 | 0.012 | Young |
| 0.588279 | 0.140061 | 0.213 | 0.196 | 1 | 8 | Ppm1h | 0.017 | Young |
| 0.588494 | 0.037206 | 0.543 | 0.525 | 1 | 8 | Ndufa3 | 0.018 | Young |
| 0.588618 | 0.093477 | 0.27 | 0.24 | 1 | 8 | Ubald1 | 0.03 | Young |
| 0.588654 | 0.045279 | 0.166 | 0.145 | 1 | 8 | Metap1 | 0.021 | Young |
| 0.588915 | 0.131123 | 0.374 | 0.352 | 1 | 8 | Tpp2 | 0.022 | Young |
| 0.589264 | 0.071658 | 0.61 | 0.592 | 1 | 8 | Sec62 | 0.018 | Young |
| 0.589683 | 0.247449 | 0.081 | 0.067 | 1 | 8 | Mthfd1l | 0.014 | Young |
| 0.589779 | 0.220651 | 0.068 | 0.056 | 1 | 8 | Gnpnat1 | 0.012 | Young |
| 0.58979 | 0.143306 | 0.291 | 0.274 | 1 | 8 | Osbpl11 | 0.017 | Young |
| 0.589987 | 0.170844 | 0.177 | 0.156 | 1 | 8 | Edem2 | 0.021 | Young |
| 0.590113 | 0.071969 | 0.506 | 0.531 | 1 | 8 | Psma2 | -0.025 | Young |
| 0.59075 | 0.119898 | 0.288 | 0.263 | 1 | 8 | Impdh2 | 0.025 | Young |
| 0.590794 | 0.040948 | 0.592 | 0.525 | 1 | 8 | Parvb | 0.067 | Young |
| 0.59108 | 0.348324 | 0.179 | 0.162 | 1 | 8 | Kdm3a | 0.017 | Young |
| 0.591681 | 0.074757 | 0.886 | 0.816 | 1 | 8 | Srrm2 | 0.07 | Young |
| 0.591694 | 0.660473 | 0.036 | 0.028 | 1 | 8 | Haao | 0.008 | Young |
| 0.591694 | 0.367033 | 0.036 | 0.028 | 1 | 8 | Slc36a3os | 0.008 | Young |
| 0.592481 | 0.011864 | 0.148 | 0.134 | 1 | 8 | As3mt | 0.014 | Young |
| 0.592688 | 0.049146 | 0.288 | 0.263 | 1 | 8 | Smarca2 | 0.025 | Young |
| 0.592922 | 0.757273 | 0.036 | 0.028 | 1 | 8 | Taf9b | 0.008 | Young |
| 0.593814 | 0.196451 | 0.132 | 0.117 | 1 | 8 | Tpgs1 | 0.015 | Young |
| 0.594468 | 0.730279 | 0.055 | 0.045 | 1 | 8 | Ap4e1 | 0.01 | Young |
| 0.594645 | 0.002204 | 0.694 | 0.637 | 1 | 8 | Tbxas1 | 0.057 | Young |
| 0.595247 | 0.147859 | 0.229 | 0.201 | 1 | 8 | Iws1 | 0.028 | Young |
| 0.595557 | 1.068885 | 0.023 | 0.017 | 1 | 8 | D11Wsu47 | 0.006 | Young |
| 0.59564 | 0.153611 | 0.091 | 0.106 | 1 | 8 | Ttc37 | -0.015 | Young |
| 0.596123 | 0.423013 | 0.068 | 0.056 | 1 | 8 | Dut | 0.012 | Young |
| 0.596765 | 0.054107 | 0.252 | 0.223 | 1 | 8 | Fundc1 | 0.029 | Young |
| 0.596861 | 0.072825 | 0.309 | 0.291 | 1 | 8 | Taf6l | 0.018 | Young |
| 0.596928 | 0.103477 | 0.081 | 0.095 | 1 | 8 | Psmf1 | -0.014 | Young |
| 0.597596 | 0.104144 | 0.21 | 0.184 | 1 | 8 | Snrnp48 | 0.026 | Young |
| 0.597741 | 0.192269 | 0.119 | 0.106 | 1 | 8 | Mfsd7a | 0.013 | Young |

| | | | | | | | | |
|---|---|---|---|---|---|---|---|---|
| 0.598179 | 0.291651 | 0.114 | 0.101 | 1 | 8 | Dnlz | 0.013 | Young |
| 0.598259 | 0.12399 | 0.252 | 0.229 | 1 | 8 | Sms | 0.023 | Young |
| 0.59851 | 0.002522 | 0.353 | 0.324 | 1 | 8 | Csnk1g3 | 0.029 | Young |
| 0.598639 | 0.811697 | 0.023 | 0.017 | 1 | 8 | Rwdd2b | 0.006 | Young |
| 0.5987 | 0.132659 | 0.216 | 0.196 | 1 | 8 | Psmc2 | 0.02 | Young |
| 0.599069 | 0.277963 | 0.184 | 0.168 | 1 | 8 | Smurf2 | 0.016 | Young |
| 0.599279 | 0.164054 | 0.055 | 0.067 | 1 | 8 | Nt5c3b | -0.012 | Young |
| 0.600182 | 0.749437 | 0.023 | 0.017 | 1 | 8 | Tbc1d7 | 0.006 | Young |
| 0.600273 | 0.157394 | 0.086 | 0.073 | 1 | 8 | Metap1d | 0.013 | Young |
| 0.600373 | 0.015613 | 0.317 | 0.296 | 1 | 8 | Gatad1 | 0.021 | Young |
| 0.601087 | 0.260525 | 0.086 | 0.073 | 1 | 8 | Incenp | 0.013 | Young |
| 0.601958 | 0.120305 | 0.184 | 0.168 | 1 | 8 | Zfp664 | 0.016 | Young |
| 0.602152 | 0.36566 | 0.073 | 0.061 | 1 | 8 | Pus10 | 0.012 | Young |
| 0.602193 | 0.035968 | 0.192 | 0.173 | 1 | 8 | Bola3 | 0.019 | Young |
| 0.602787 | 0.247991 | 0.036 | 0.028 | 1 | 8 | Pald1 | 0.008 | Young |
| 0.60341 | 0.20654 | 0.068 | 0.056 | 1 | 8 | Bod1 | 0.012 | Young |
| 0.603427 | 0.171883 | 0.049 | 0.039 | 1 | 8 | Eefsec | 0.01 | Young |
| 0.603539 | 0.317683 | 0.055 | 0.045 | 1 | 8 | Lmtk2 | 0.01 | Young |
| 0.603778 | 0.2025 | 0.021 | 0.028 | 1 | 8 | Afg1l | -0.007 | Young |
| 0.604025 | 0.2606 | 0.036 | 0.028 | 1 | 8 | Nkrf | 0.008 | Young |
| 0.604442 | 0.289509 | 0.143 | 0.128 | 1 | 8 | H2-Ke6 | 0.015 | Young |
| 0.604495 | 0.046412 | 0.148 | 0.128 | 1 | 8 | Foxk2 | 0.02 | Young |
| 0.604551 | 0.454873 | 0.055 | 0.045 | 1 | 8 | Zfp609 | 0.01 | Young |
| 0.604824 | 0.595384 | 0.023 | 0.017 | 1 | 8 | Cenpo | 0.006 | Young |
| 0.604982 | 0.011567 | 0.062 | 0.05 | 1 | 8 | Gmnn | 0.012 | Young |
| 0.605236 | 0.136845 | 0.096 | 0.112 | 1 | 8 | Rrn3 | -0.016 | Young |
| 0.605559 | 0.102016 | 0.049 | 0.039 | 1 | 8 | Card11 | 0.01 | Young |
| 0.605672 | 0.180077 | 0.338 | 0.341 | 1 | 8 | Ssfa2 | -0.003 | Young |
| 0.605879 | 0.053531 | 0.831 | 0.782 | 1 | 8 | Serp1 | 0.049 | Young |
| 0.605943 | 0.015553 | 0.13 | 0.145 | 1 | 8 | Phkb | -0.015 | Young |
| 0.606375 | 0.716201 | 0.023 | 0.017 | 1 | 8 | Map7d3 | 0.006 | Young |
| 0.606375 | 0.589358 | 0.023 | 0.017 | 1 | 8 | Pyroxd2 | 0.006 | Young |
| 0.606456 | 0.447329 | 0.083 | 0.073 | 1 | 8 | Spryd3 | 0.01 | Young |
| 0.606505 | 0.68613 | 0.036 | 0.028 | 1 | 8 | Disc1 | 0.008 | Young |
| 0.606505 | 0.367361 | 0.036 | 0.028 | 1 | 8 | Ppp1r18os | 0.008 | Young |
| 0.607148 | 0.14796 | 0.371 | 0.335 | 1 | 8 | Eif2a | 0.036 | Young |
| 0.608052 | 0.066937 | 0.184 | 0.162 | 1 | 8 | Mrpl9 | 0.022 | Young |
| 0.608247 | 0.102163 | 0.169 | 0.151 | 1 | 8 | Pabpc4 | 0.018 | Young |
| 0.608836 | 0.136663 | 0.436 | 0.419 | 1 | 8 | Mat2b | 0.017 | Young |
| 0.609322 | 0.282563 | 0.096 | 0.084 | 1 | 8 | Pld2 | 0.012 | Young |
| 0.609482 | 0.51028 | 0.023 | 0.017 | 1 | 8 | Gm30329 | 0.006 | Young |
| 0.609482 | 0.382281 | 0.023 | 0.017 | 1 | 8 | Tbc1d25 | 0.006 | Young |
| 0.609482 | 0.354098 | 0.023 | 0.017 | 1 | 8 | Atp6v1g2 | 0.006 | Young |
| 0.60977 | 0.126809 | 0.335 | 0.307 | 1 | 8 | Atxn7l1 | 0.028 | Young |
| 0.609803 | 0.05293 | 0.182 | 0.168 | 1 | 8 | Sdad1 | 0.014 | Young |

| | | | | | | | | |
|---|---|---|---|---|---|---|---|---|
| 0.61015 | 0.110451 | 0.132 | 0.117 | 1 | 8 | Tmx2 | 0.015 | Young |
| 0.610554 | 0.131019 | 0.156 | 0.14 | 1 | 8 | Acsl1 | 0.016 | Young |
| 0.611038 | 0.514218 | 0.023 | 0.017 | 1 | 8 | Acat2 | 0.006 | Young |
| 0.611363 | 0.015419 | 0.519 | 0.492 | 1 | 8 | Daglb | 0.027 | Young |
| 0.612364 | 0.33982 | 0.13 | 0.117 | 1 | 8 | Fam98a | 0.013 | Young |
| 0.612463 | 0.303218 | 0.106 | 0.095 | 1 | 8 | Nfs1 | 0.011 | Young |
| 0.612596 | 0.151241 | 0.023 | 0.017 | 1 | 8 | Magi2 | 0.006 | Young |
| 0.613299 | 0.193447 | 0.166 | 0.151 | 1 | 8 | Eef1akmt1 | 0.015 | Young |
| 0.613869 | 0.001674 | 0.091 | 0.078 | 1 | 8 | Qser1 | 0.013 | Young |
| 0.614156 | 0.574532 | 0.023 | 0.017 | 1 | 8 | Alg12 | 0.006 | Young |
| 0.614156 | 0.55143 | 0.023 | 0.017 | 1 | 8 | Faap24 | 0.006 | Young |
| 0.614243 | 0.035666 | 0.021 | 0.028 | 1 | 8 | Ehbp1 | -0.007 | Young |
| 0.614244 | 0.729211 | 0.06 | 0.05 | 1 | 8 | Gatb | 0.01 | Young |
| 0.614522 | 0.059447 | 0.278 | 0.246 | 1 | 8 | Slc50a1 | 0.032 | Young |
| 0.61522 | 0.392493 | 0.036 | 0.028 | 1 | 8 | Mtg2 | 0.008 | Young |
| 0.615643 | 0.119164 | 0.192 | 0.173 | 1 | 8 | Lyst | 0.019 | Young |
| 0.615731 | 0.07944 | 0.055 | 0.045 | 1 | 8 | Poc5 | 0.01 | Young |
| 0.615745 | 0.044176 | 0.021 | 0.028 | 1 | 8 | Gm32856 | -0.007 | Young |
| 0.615903 | 0.280536 | 0.109 | 0.095 | 1 | 8 | Rad17 | 0.014 | Young |
| 0.615981 | 0.088054 | 0.582 | 0.553 | 1 | 8 | Cox17 | 0.029 | Young |
| 0.616399 | 0.06373 | 0.075 | 0.089 | 1 | 8 | Nr2c1 | -0.014 | Young |
| 0.616563 | 0.399304 | 0.171 | 0.162 | 1 | 8 | Lin54 | 0.009 | Young |
| 0.617281 | 0.328108 | 0.023 | 0.017 | 1 | 8 | Aplf | 0.006 | Young |
| 0.617281 | 0.245613 | 0.023 | 0.017 | 1 | 8 | Gm16287 | 0.006 | Young |
| 0.617788 | 0.763347 | 0.042 | 0.034 | 1 | 8 | Zfp429 | 0.008 | Young |
| 0.617859 | 0.070723 | 0.548 | 0.52 | 1 | 8 | Pdlim5 | 0.028 | Young |
| 0.617973 | 0.318953 | 0.221 | 0.207 | 1 | 8 | Hmgcl | 0.014 | Young |
| 0.618531 | 0.053988 | 0.449 | 0.441 | 1 | 8 | Ube2k | 0.008 | Young |
| 0.618643 | 0.133247 | 0.26 | 0.246 | 1 | 8 | Fam69a | 0.014 | Young |
| 0.619051 | 0.167299 | 0.101 | 0.089 | 1 | 8 | Hps4 | 0.012 | Young |
| 0.619158 | 0.019799 | 0.026 | 0.034 | 1 | 8 | Siglecf | -0.008 | Young |
| 0.619403 | 0.161223 | 0.151 | 0.134 | 1 | 8 | Gm12840 | 0.017 | Young |
| 0.619624 | 0.056665 | 0.416 | 0.419 | 1 | 8 | 1110004F1 | -0.003 | Young |
| 0.619818 | 0.181075 | 0.055 | 0.045 | 1 | 8 | Taf5 | 0.01 | Young |
| 0.620259 | 0.247373 | 0.021 | 0.028 | 1 | 8 | Cdc25b | -0.007 | Young |
| 0.620412 | 0.468577 | 0.023 | 0.017 | 1 | 8 | Ada | 0.006 | Young |
| 0.620412 | 0.414177 | 0.023 | 0.017 | 1 | 8 | Ggnbp1 | 0.006 | Young |
| 0.620464 | 0.230014 | 0.148 | 0.134 | 1 | 8 | Mrpl55 | 0.014 | Young |
| 0.620729 | 0.061907 | 0.442 | 0.425 | 1 | 8 | Creb1 | 0.017 | Young |
| 0.621291 | 0.625824 | 0.042 | 0.034 | 1 | 8 | C130050O1 | 0.008 | Young |
| 0.621376 | 0.238885 | 0.153 | 0.14 | 1 | 8 | BC003965 | 0.013 | Young |
| 0.621981 | 0.057101 | 0.023 | 0.017 | 1 | 8 | Tubg1 | 0.006 | Young |
| 0.621981 | 0.024922 | 0.023 | 0.017 | 1 | 8 | Gm29291 | 0.006 | Young |
| 0.622232 | 0.615816 | 0.083 | 0.073 | 1 | 8 | Hspbap1 | 0.01 | Young |
| 0.622335 | 0.022102 | 0.462 | 0.43 | 1 | 8 | Arhgef1 | 0.032 | Young |

| | | | | | | | | |
|---|---|---|---|---|---|---|---|---|
| 0.622788 | 0.176648 | 0.239 | 0.223 | 1 | 8 | Atf1 | 0.016 | Young |
| 0.623277 | 0.162974 | 0.021 | 0.028 | 1 | 8 | Mfsd6l | -0.007 | Young |
| 0.623551 | 0.211227 | 0.023 | 0.017 | 1 | 8 | 9130221H1 | 0.006 | Young |
| 0.623708 | 0.006343 | 0.332 | 0.307 | 1 | 8 | Nmt1 | 0.025 | Young |
| 0.62397 | 0.064566 | 0.132 | 0.117 | 1 | 8 | Brms1 | 0.015 | Young |
| 0.62449 | 0.011852 | 0.556 | 0.525 | 1 | 8 | Snx9 | 0.031 | Young |
| 0.624989 | 0.386504 | 0.06 | 0.05 | 1 | 8 | Mrgbp | 0.01 | Young |
| 0.625446 | 0.14096 | 0.158 | 0.14 | 1 | 8 | Smc2 | 0.018 | Young |
| 0.625494 | 0.04642 | 0.109 | 0.095 | 1 | 8 | Slc9a1 | 0.014 | Young |
| 0.625658 | 0.020816 | 0.203 | 0.184 | 1 | 8 | Lcmt1 | 0.019 | Young |
| 0.625912 | 0.120417 | 0.101 | 0.089 | 1 | 8 | Tbc1d10a | 0.012 | Young |
| 0.626154 | 0.078736 | 0.444 | 0.402 | 1 | 8 | Cisd2 | 0.042 | Young |
| 0.626235 | 0.065234 | 0.109 | 0.095 | 1 | 8 | Racgap1 | 0.014 | Young |
| 0.626478 | 0.096689 | 0.169 | 0.151 | 1 | 8 | Proser1 | 0.018 | Young |
| 0.627 | 0.252899 | 0.055 | 0.045 | 1 | 8 | Polr3b | 0.01 | Young |
| 0.627717 | 0.077744 | 0.109 | 0.095 | 1 | 8 | Tbc1d8 | 0.014 | Young |
| 0.627765 | 0.176114 | 0.036 | 0.028 | 1 | 8 | Mcat | 0.008 | Young |
| 0.628035 | 0.237468 | 0.096 | 0.084 | 1 | 8 | Tada3 | 0.012 | Young |
| 0.628035 | 0.144909 | 0.096 | 0.084 | 1 | 8 | Tacc2 | 0.012 | Young |
| 0.628373 | 0.168014 | 0.151 | 0.134 | 1 | 8 | Klhl2 | 0.017 | Young |
| 0.628592 | 0.008813 | 0.158 | 0.14 | 1 | 8 | Pno1 | 0.018 | Young |
| 0.62936 | 0.155646 | 0.19 | 0.173 | 1 | 8 | Drg1 | 0.017 | Young |
| 0.630883 | 0.528223 | 0.06 | 0.05 | 1 | 8 | Zscan21 | 0.01 | Young |
| 0.631442 | 0.18902 | 0.132 | 0.117 | 1 | 8 | Il18 | 0.015 | Young |
| 0.631542 | 0.043144 | 0.091 | 0.078 | 1 | 8 | Pyurf | 0.013 | Young |
| 0.632564 | 0.115707 | 0.473 | 0.464 | 1 | 8 | Sumo1 | 0.009 | Young |
| 0.632757 | 0.1031 | 0.203 | 0.184 | 1 | 8 | Stx5a | 0.019 | Young |
| 0.632835 | 0.060092 | 0.026 | 0.034 | 1 | 8 | Map2k3os | -0.008 | Young |
| 0.632968 | 0.348842 | 0.125 | 0.112 | 1 | 8 | Zfp322a | 0.013 | Young |
| 0.633249 | 0.0296 | 0.197 | 0.173 | 1 | 8 | Usp6nl | 0.024 | Young |
| 0.633667 | 0.068879 | 0.125 | 0.112 | 1 | 8 | 2410089E0 | 0.013 | Young |
| 0.634205 | 0.521002 | 0.042 | 0.034 | 1 | 8 | Macrod1 | 0.008 | Young |
| 0.634209 | 0.243032 | 0.026 | 0.034 | 1 | 8 | Alms1 | -0.008 | Young |
| 0.634821 | 0.035922 | 0.914 | 0.844 | 1 | 8 | Fosb | 0.07 | Young |
| 0.634826 | 0.323902 | 0.06 | 0.05 | 1 | 8 | Sft2d3 | 0.01 | Young |
| 0.635529 | 0.080237 | 0.213 | 0.196 | 1 | 8 | Rbm14 | 0.017 | Young |
| 0.635781 | 0.140205 | 0.156 | 0.14 | 1 | 8 | Rpf2 | 0.016 | Young |
| 0.63633 | 0.205488 | 0.114 | 0.101 | 1 | 8 | Borcs8 | 0.013 | Young |
| 0.636805 | 0.078976 | 0.091 | 0.078 | 1 | 8 | Zfp654 | 0.013 | Young |
| 0.637174 | 0.171954 | 0.158 | 0.145 | 1 | 8 | Phactr4 | 0.013 | Young |
| 0.637321 | 0.036857 | 0.055 | 0.045 | 1 | 8 | Slc35d2 | 0.01 | Young |
| 0.637689 | 0.277203 | 0.244 | 0.235 | 1 | 8 | Tmem50b | 0.009 | Young |
| 0.637747 | 0.173769 | 0.042 | 0.034 | 1 | 8 | Gm42031 | 0.008 | Young |
| 0.637859 | 0.026769 | 0.164 | 0.145 | 1 | 8 | Ctps2 | 0.019 | Young |
| 0.638475 | 0.036151 | 0.294 | 0.268 | 1 | 8 | Gm8186 | 0.026 | Young |

| | | | | | | | | |
|---|---|---|---|---|---|---|---|---|
| 0.638574 | 0.101708 | 0.208 | 0.19 | 1 | 8 | Mllt6 | 0.018 | Young |
| 0.639434 | 0.030855 | 0.138 | 0.123 | 1 | 8 | Fbxo9 | 0.015 | Young |
| 0.640259 | 0.106238 | 0.177 | 0.162 | 1 | 8 | Nifk | 0.015 | Young |
| 0.640353 | 0.001314 | 0.161 | 0.145 | 1 | 8 | Rexo4 | 0.016 | Young |
| 0.640875 | 0.380812 | 0.112 | 0.101 | 1 | 8 | Apip | 0.011 | Young |
| 0.640956 | 0.062659 | 0.166 | 0.151 | 1 | 8 | Pgp | 0.015 | Young |
| 0.641126 | 0.168617 | 0.213 | 0.196 | 1 | 8 | Rnf146 | 0.017 | Young |
| 0.641297 | 0.442329 | 0.042 | 0.034 | 1 | 8 | Tlr11 | 0.008 | Young |
| 0.64187 | 0.173579 | 0.109 | 0.095 | 1 | 8 | Zfp398 | 0.014 | Young |
| 0.642238 | 0.031226 | 0.205 | 0.19 | 1 | 8 | 3110043O2 | 0.015 | Young |
| 0.642345 | 0.115906 | 0.192 | 0.173 | 1 | 8 | Mesd | 0.019 | Young |
| 0.642482 | 0.002744 | 0.036 | 0.045 | 1 | 8 | Phykpl | -0.009 | Young |
| 0.642819 | 0.127513 | 0.101 | 0.089 | 1 | 8 | Flrt3 | 0.012 | Young |
| 0.643039 | 0.102788 | 0.361 | 0.352 | 1 | 8 | Polb | 0.009 | Young |
| 0.643676 | 0.318115 | 0.132 | 0.123 | 1 | 8 | Xpo4 | 0.009 | Young |
| 0.644456 | 0.044282 | 0.551 | 0.514 | 1 | 8 | Nop10 | 0.037 | Young |
| 0.644752 | 0.108336 | 0.101 | 0.089 | 1 | 8 | Rnaset2a | 0.012 | Young |
| 0.644855 | 0.434352 | 0.042 | 0.034 | 1 | 8 | Tmem191c | 0.008 | Young |
| 0.644855 | 0.38592 | 0.042 | 0.034 | 1 | 8 | Rfx2 | 0.008 | Young |
| 0.645306 | 0.166681 | 0.236 | 0.223 | 1 | 8 | Pdzd8 | 0.013 | Young |
| 0.645876 | 0.378879 | 0.13 | 0.117 | 1 | 8 | Gm48065 | 0.013 | Young |
| 0.646043 | 0.438728 | 0.042 | 0.034 | 1 | 8 | Suv39h1 | 0.008 | Young |
| 0.646043 | 0.232279 | 0.042 | 0.034 | 1 | 8 | Limk2 | 0.008 | Young |
| 0.648921 | 0.174113 | 0.374 | 0.358 | 1 | 8 | Impact | 0.016 | Young |
| 0.649201 | 0.097509 | 0.179 | 0.162 | 1 | 8 | Kcnk13 | 0.017 | Young |
| 0.650443 | 0.038054 | 0.073 | 0.061 | 1 | 8 | AC127341. | 0.012 | Young |
| 0.650917 | 0.038669 | 0.213 | 0.24 | 1 | 8 | Dolpp1 | -0.027 | Young |
| 0.651171 | 0.586108 | 0.029 | 0.022 | 1 | 8 | Focad | 0.007 | Young |
| 0.651326 | 0.076756 | 0.208 | 0.196 | 1 | 8 | Mbd1 | 0.012 | Young |
| 0.651792 | 0.097256 | 0.494 | 0.464 | 1 | 8 | Tgif1 | 0.03 | Young |
| 0.651996 | 0.200723 | 0.036 | 0.045 | 1 | 8 | Atpaf1 | -0.009 | Young |
| 0.652353 | 0.230733 | 0.143 | 0.128 | 1 | 8 | Kpna6 | 0.015 | Young |
| 0.652695 | 0.236958 | 0.06 | 0.05 | 1 | 8 | Nlrp1b | 0.01 | Young |
| 0.653152 | 0.010466 | 0.231 | 0.212 | 1 | 8 | Seh1l | 0.019 | Young |
| 0.653218 | 0.327906 | 0.169 | 0.156 | 1 | 8 | Lrif1 | 0.013 | Young |
| 0.653453 | 0.07348 | 0.135 | 0.151 | 1 | 8 | Phrf1 | -0.016 | Young |
| 0.653639 | 0.026759 | 0.197 | 0.179 | 1 | 8 | Itfg1 | 0.018 | Young |
| 0.654339 | 0.051629 | 0.655 | 0.609 | 1 | 8 | Cast | 0.046 | Young |
| 0.655481 | 0.571699 | 0.029 | 0.022 | 1 | 8 | Vps51 | 0.007 | Young |
| 0.655543 | 0.339252 | 0.065 | 0.056 | 1 | 8 | Spata5 | 0.009 | Young |
| 0.655578 | 0.022155 | 0.036 | 0.045 | 1 | 8 | Fads6 | -0.009 | Young |
| 0.655847 | 0.2855 | 0.145 | 0.134 | 1 | 8 | Nubp1 | 0.011 | Young |
| 0.655922 | 0.298017 | 0.14 | 0.128 | 1 | 8 | Gla | 0.012 | Young |
| 0.655974 | 0.09774 | 0.106 | 0.095 | 1 | 8 | Git1 | 0.011 | Young |
| 0.656079 | 0.008688 | 0.626 | 0.559 | 1 | 8 | Twf2 | 0.067 | Young |

| | | | | | | | | |
|---|---|---|---|---|---|---|---|---|
| 0.656799 | 0.168715 | 0.112 | 0.101 | 1 | 8 | Pus1 | 0.011 | Young |
| 0.656956 | 0.013552 | 0.088 | 0.101 | 1 | 8 | Zer1 | -0.013 | Young |
| 0.657623 | 0.152976 | 0.125 | 0.112 | 1 | 8 | Ormdl3 | 0.013 | Young |
| 0.657768 | 0.016259 | 0.644 | 0.581 | 1 | 8 | Irf2 | 0.063 | Young |
| 0.657971 | 0.251712 | 0.042 | 0.034 | 1 | 8 | Cbr3 | 0.008 | Young |
| 0.658094 | 0.093785 | 0.301 | 0.279 | 1 | 8 | Por | 0.022 | Young |
| 0.658162 | 0.197264 | 0.145 | 0.134 | 1 | 8 | Ddit3 | 0.011 | Young |
| 0.658168 | 0.514595 | 0.138 | 0.128 | 1 | 8 | Traf7 | 0.01 | Young |
| 0.658697 | 0.034506 | 0.481 | 0.447 | 1 | 8 | Arl8b | 0.034 | Young |
| 0.658957 | 0.208948 | 0.096 | 0.084 | 1 | 8 | Slc35a1 | 0.012 | Young |
| 0.659112 | 0.148472 | 0.182 | 0.168 | 1 | 8 | Sec23ip | 0.014 | Young |
| 0.659324 | 0.048111 | 0.784 | 0.726 | 1 | 8 | Klf3 | 0.058 | Young |
| 0.659401 | 0.34159 | 0.099 | 0.112 | 1 | 8 | 1700037H( | -0.013 | Young |
| 0.659969 | 0.028247 | 0.161 | 0.145 | 1 | 8 | Suclg2 | 0.016 | Young |
| 0.66006 | 0.291739 | 0.192 | 0.184 | 1 | 8 | Stau1 | 0.008 | Young |
| 0.660212 | 0.144009 | 0.171 | 0.156 | 1 | 8 | Acot9 | 0.015 | Young |
| 0.660243 | 0.139592 | 0.244 | 0.229 | 1 | 8 | Trappc10 | 0.015 | Young |
| 0.660345 | 0.356776 | 0.065 | 0.056 | 1 | 8 | Gstcd | 0.009 | Young |
| 0.660367 | 0.141398 | 0.042 | 0.034 | 1 | 8 | Abcb8 | 0.008 | Young |
| 0.661159 | 0.071591 | 0.439 | 0.391 | 1 | 8 | Nlrp3 | 0.048 | Young |
| 0.661706 | 0.437913 | 0.06 | 0.05 | 1 | 8 | Pan2 | 0.01 | Young |
| 0.661779 | 0.235228 | 0.229 | 0.223 | 1 | 8 | Hypk | 0.006 | Young |
| 0.662072 | 0.122777 | 0.114 | 0.101 | 1 | 8 | Oasl2 | 0.013 | Young |
| 0.66269 | 0.528528 | 0.029 | 0.022 | 1 | 8 | Plekhh3 | 0.007 | Young |
| 0.66269 | 0.488162 | 0.029 | 0.022 | 1 | 8 | Amotl2 | 0.007 | Young |
| 0.662853 | 0.303955 | 0.083 | 0.073 | 1 | 8 | Wdr46 | 0.01 | Young |
| 0.662946 | 0.173056 | 0.047 | 0.056 | 1 | 8 | Zfp758 | -0.009 | Young |
| 0.663576 | 0.039777 | 0.164 | 0.151 | 1 | 8 | Tmem19 | 0.013 | Young |
| 0.66398 | 0.087858 | 0.255 | 0.24 | 1 | 8 | Cdk13 | 0.015 | Young |
| 0.664136 | 0.739253 | 0.029 | 0.022 | 1 | 8 | Prnp | 0.007 | Young |
| 0.664136 | 0.288629 | 0.029 | 0.022 | 1 | 8 | Slc30a6 | 0.007 | Young |
| 0.664222 | 0.030475 | 0.86 | 0.816 | 1 | 8 | Msn | 0.044 | Young |
| 0.66423 | 0.006354 | 0.301 | 0.285 | 1 | 8 | Rab43 | 0.016 | Young |
| 0.664519 | 0.021815 | 0.138 | 0.123 | 1 | 8 | Serpinb1a | 0.015 | Young |
| 0.66472 | 0.076333 | 0.06 | 0.05 | 1 | 8 | Abca6 | 0.01 | Young |
| 0.66518 | 0.422437 | 0.047 | 0.039 | 1 | 8 | Mrpl49 | 0.008 | Young |
| 0.666076 | 0.184113 | 0.151 | 0.14 | 1 | 8 | Lztfl1 | 0.011 | Young |
| 0.66671 | 0.178519 | 0.14 | 0.128 | 1 | 8 | Minpp1 | 0.012 | Young |
| 0.666798 | 0.384075 | 0.07 | 0.061 | 1 | 8 | Trim23 | 0.009 | Young |
| 0.666936 | 0.178199 | 0.127 | 0.117 | 1 | 8 | Fnip2 | 0.01 | Young |
| 0.667031 | 0.396426 | 0.029 | 0.022 | 1 | 8 | Gtf2ird2 | 0.007 | Young |
| 0.667031 | 0.258299 | 0.029 | 0.022 | 1 | 8 | Gm26601 | 0.007 | Young |
| 0.667422 | 0.011094 | 0.992 | 0.933 | 1 | 8 | Sem1 | 0.059 | Young |
| 0.66769 | 0.105858 | 0.164 | 0.151 | 1 | 8 | Ago1 | 0.013 | Young |
| 0.667901 | 0.051942 | 0.119 | 0.106 | 1 | 8 | Gnl3l | 0.013 | Young |

| | | | | | | | | |
|---|---|---|---|---|---|---|---|---|
| 0.668084 | 0.10745 | 0.151 | 0.134 | 1 | 8 | Mmd | 0.017 | Young |
| 0.668507 | 0.341707 | 0.197 | 0.19 | 1 | 8 | Tmem135 | 0.007 | Young |
| 0.668577 | 0.48406 | 0.047 | 0.039 | 1 | 8 | Mkks | 0.008 | Young |
| 0.668577 | 0.057201 | 0.042 | 0.05 | 1 | 8 | Pick1 | -0.008 | Young |
| 0.668697 | 0.37519 | 0.236 | 0.229 | 1 | 8 | Clip1 | 0.007 | Young |
| 0.668781 | 0.100939 | 0.042 | 0.034 | 1 | 8 | Kat8 | 0.008 | Young |
| 0.668844 | 0.192382 | 0.182 | 0.173 | 1 | 8 | Zbtb7b | 0.009 | Young |
| 0.669249 | 0.078739 | 0.338 | 0.324 | 1 | 8 | Gak | 0.014 | Young |
| 0.669711 | 0.030717 | 0.047 | 0.039 | 1 | 8 | Nr1d1 | 0.008 | Young |
| 0.669931 | 0.332376 | 0.029 | 0.022 | 1 | 8 | Gm26669 | 0.007 | Young |
| 0.669994 | 0.055767 | 0.236 | 0.223 | 1 | 8 | Xrn1 | 0.013 | Young |
| 0.670145 | 0.029702 | 0.291 | 0.274 | 1 | 8 | Dazap1 | 0.017 | Young |
| 0.670628 | 0.472179 | 0.112 | 0.101 | 1 | 8 | Rcan1 | 0.011 | Young |
| 0.67083 | 0.316895 | 0.088 | 0.078 | 1 | 8 | Galk1 | 0.01 | Young |
| 0.671132 | 0.120143 | 0.143 | 0.128 | 1 | 8 | Bcl2 | 0.015 | Young |
| 0.671215 | 0.051051 | 0.052 | 0.061 | 1 | 8 | Grwd1 | -0.009 | Young |
| 0.671265 | 0.006746 | 0.649 | 0.631 | 1 | 8 | Selenof | 0.018 | Young |
| 0.671383 | 0.234975 | 0.029 | 0.022 | 1 | 8 | Ddit4 | 0.007 | Young |
| 0.671383 | 0.054297 | 0.029 | 0.022 | 1 | 8 | Pex12 | 0.007 | Young |
| 0.671438 | 1.002945 | 0.016 | 0.011 | 1 | 8 | Cdkn2b | 0.005 | Young |
| 0.671438 | 0.8663 | 0.016 | 0.011 | 1 | 8 | Camk2n1 | 0.005 | Young |
| 0.67145 | 0.509423 | 0.07 | 0.061 | 1 | 8 | Prmt7 | 0.009 | Young |
| 0.671901 | 0.037282 | 0.192 | 0.173 | 1 | 8 | Abhd16a | 0.019 | Young |
| 0.671926 | 0.16634 | 0.065 | 0.056 | 1 | 8 | Tbkbp1 | 0.009 | Young |
| 0.67198 | 0.11978 | 0.047 | 0.039 | 1 | 8 | Acot11 | 0.008 | Young |
| 0.672343 | 0.366751 | 0.094 | 0.084 | 1 | 8 | Wdsub1 | 0.01 | Young |
| 0.672786 | 0.160362 | 0.06 | 0.05 | 1 | 8 | Med6 | 0.01 | Young |
| 0.673416 | 0.736901 | 0.016 | 0.011 | 1 | 8 | Trmu | 0.005 | Young |
| 0.673637 | 0.147477 | 0.208 | 0.196 | 1 | 8 | Mindy3 | 0.012 | Young |
| 0.673646 | 0.120698 | 0.127 | 0.117 | 1 | 8 | Bicra | 0.01 | Young |
| 0.67429 | 0.126473 | 0.029 | 0.022 | 1 | 8 | Acsf3 | 0.007 | Young |
| 0.675396 | 0.901956 | 0.016 | 0.011 | 1 | 8 | Zfp169 | 0.005 | Young |
| 0.675396 | 0.875301 | 0.016 | 0.011 | 1 | 8 | A930007I1 | 0.005 | Young |
| 0.675396 | 0.651711 | 0.016 | 0.011 | 1 | 8 | Spata1 | 0.005 | Young |
| 0.675396 | 0.624 | 0.016 | 0.011 | 1 | 8 | Tlcd1 | 0.005 | Young |
| 0.675396 | 0.607518 | 0.016 | 0.011 | 1 | 8 | Il1r1 | 0.005 | Young |
| 0.675396 | 0.50398 | 0.016 | 0.011 | 1 | 8 | Gtf2ird1 | 0.005 | Young |
| 0.676529 | 0.446304 | 0.042 | 0.05 | 1 | 8 | Nxn | -0.008 | Young |
| 0.676529 | 0.310285 | 0.047 | 0.039 | 1 | 8 | Atrip | 0.008 | Young |
| 0.677202 | 0.03671 | 0.029 | 0.022 | 1 | 8 | Zcwpw1 | 0.007 | Young |
| 0.677378 | 0.512794 | 0.016 | 0.011 | 1 | 8 | Ftx | 0.005 | Young |
| 0.677378 | 0.504263 | 0.016 | 0.011 | 1 | 8 | Eno3 | 0.005 | Young |
| 0.677378 | 0.35145 | 0.016 | 0.011 | 1 | 8 | Fam109a | 0.005 | Young |
| 0.677668 | 0.064255 | 0.047 | 0.039 | 1 | 8 | Kcnd1 | 0.008 | Young |
| 0.677975 | 0.004429 | 0.101 | 0.089 | 1 | 8 | Brcc3 | 0.012 | Young |

| | | | | | | | | |
|---|---|---|---|---|---|---|---|---|
| 0.678381 | 0.076868 | 0.184 | 0.201 | 1 | 8 | Areg | -0.017 | Young |
| 0.678484 | 0.120281 | 0.358 | 0.346 | 1 | 8 | Avpi1 | 0.012 | Young |
| 0.678919 | 0.355329 | 0.07 | 0.061 | 1 | 8 | Zfp942 | 0.009 | Young |
| 0.679003 | 0.066862 | 0.057 | 0.067 | 1 | 8 | Tsr2 | -0.01 | Young |
| 0.679356 | 0.427909 | 0.166 | 0.156 | 1 | 8 | Tmx4 | 0.01 | Young |
| 0.679363 | 0.642557 | 0.016 | 0.011 | 1 | 8 | Capsl | 0.005 | Young |
| 0.679363 | 0.463201 | 0.016 | 0.011 | 1 | 8 | Aplnr | 0.005 | Young |
| 0.679363 | 0.429176 | 0.016 | 0.011 | 1 | 8 | Pglyrp1 | 0.005 | Young |
| 0.679363 | 0.379637 | 0.016 | 0.011 | 1 | 8 | Akap17b | 0.005 | Young |
| 0.679452 | 0.125727 | 0.164 | 0.151 | 1 | 8 | Shkbp1 | 0.013 | Young |
| 0.679486 | 0.009048 | 0.571 | 0.531 | 1 | 8 | Lamtor1 | 0.04 | Young |
| 0.679553 | 0.352555 | 0.132 | 0.123 | 1 | 8 | Aqp9 | 0.009 | Young |
| 0.679673 | 0.251684 | 0.127 | 0.117 | 1 | 8 | Heatr6 | 0.01 | Young |
| 0.679948 | 0.020844 | 0.047 | 0.039 | 1 | 8 | Tab1 | 0.008 | Young |
| 0.680119 | 0.347348 | 0.029 | 0.022 | 1 | 8 | Pcgf1 | 0.007 | Young |
| 0.680156 | 0.414553 | 0.047 | 0.056 | 1 | 8 | Tecpr2 | -0.009 | Young |
| 0.680705 | 0.160863 | 0.122 | 0.112 | 1 | 8 | Dgcr6 | 0.01 | Young |
| 0.680828 | 0.060978 | 0.13 | 0.117 | 1 | 8 | Selenom | 0.013 | Young |
| 0.681349 | 0.627944 | 0.016 | 0.011 | 1 | 8 | Zfp763 | 0.005 | Young |
| 0.681349 | 0.626644 | 0.016 | 0.011 | 1 | 8 | Dact3 | 0.005 | Young |
| 0.681349 | 0.434657 | 0.016 | 0.011 | 1 | 8 | Ercc6l | 0.005 | Young |
| 0.681349 | 0.394141 | 0.016 | 0.011 | 1 | 8 | Klhl3 | 0.005 | Young |
| 0.681349 | 0.345288 | 0.016 | 0.011 | 1 | 8 | Napepld | 0.005 | Young |
| 0.681349 | 0.34087 | 0.016 | 0.011 | 1 | 8 | Hspa12a | 0.005 | Young |
| 0.681349 | 0.259289 | 0.016 | 0.011 | 1 | 8 | Eid2 | 0.005 | Young |
| 0.681349 | 0.130847 | 0.016 | 0.011 | 1 | 8 | Gm28187 | 0.005 | Young |
| 0.681494 | 0.175205 | 0.052 | 0.061 | 1 | 8 | Asf1a | -0.009 | Young |
| 0.682231 | 0.438347 | 0.042 | 0.05 | 1 | 8 | Zfp944 | -0.008 | Young |
| 0.682345 | 0.070771 | 0.125 | 0.14 | 1 | 8 | Armc10 | -0.015 | Young |
| 0.683041 | 0.001936 | 0.029 | 0.022 | 1 | 8 | Gale | 0.007 | Young |
| 0.683127 | 0.092699 | 0.182 | 0.168 | 1 | 8 | Sympk | 0.014 | Young |
| 0.683338 | 0.386603 | 0.016 | 0.011 | 1 | 8 | Pnpla1 | 0.005 | Young |
| 0.683338 | 0.257487 | 0.016 | 0.011 | 1 | 8 | Hist1h4c | 0.005 | Young |
| 0.683338 | 0.159422 | 0.016 | 0.011 | 1 | 8 | Enkur | 0.005 | Young |
| 0.683373 | 0.224433 | 0.047 | 0.039 | 1 | 8 | Rida | 0.008 | Young |
| 0.683429 | 0.032848 | 0.169 | 0.184 | 1 | 8 | Agap3 | -0.015 | Young |
| 0.683472 | 0.077506 | 0.2 | 0.184 | 1 | 8 | Fbxo22 | 0.016 | Young |
| 0.684044 | 0.035568 | 0.574 | 0.564 | 1 | 8 | Snrpg | 0.01 | Young |
| 0.685179 | 0.151309 | 0.187 | 0.173 | 1 | 8 | Knop1 | 0.014 | Young |
| 0.685329 | 0.406862 | 0.016 | 0.011 | 1 | 8 | Dhfr | 0.005 | Young |
| 0.685329 | 0.225989 | 0.016 | 0.011 | 1 | 8 | 2010013B2 | 0.005 | Young |
| 0.685329 | 0.159885 | 0.016 | 0.011 | 1 | 8 | Wdr62 | 0.005 | Young |
| 0.685329 | 0.094702 | 0.016 | 0.011 | 1 | 8 | Zfp772 | 0.005 | Young |
| 0.685329 | 0.071732 | 0.016 | 0.011 | 1 | 8 | Gm17089 | 0.005 | Young |
| 0.685329 | 0.008131 | 0.016 | 0.011 | 1 | 8 | Vcan | 0.005 | Young |

| | | | | | | | | |
|---|---|---|---|---|---|---|---|---|
| 0.68542 | 0.093769 | 0.099 | 0.089 | 1 | 8 | Pgghg | 0.01 | Young |
| 0.687218 | 0.007276 | 0.829 | 0.749 | 1 | 8 | Sf3b1 | 0.08 | Young |
| 0.687324 | 0.09814 | 0.387 | 0.38 | 1 | 8 | 2410015M | 0.007 | Young |
| 0.687904 | 0.183181 | 0.13 | 0.117 | 1 | 8 | Usp36 | 0.013 | Young |
| 0.687951 | 0.212151 | 0.047 | 0.039 | 1 | 8 | Leng9 | 0.008 | Young |
| 0.687951 | 0.046539 | 0.047 | 0.039 | 1 | 8 | Inpp5b | 0.008 | Young |
| 0.688238 | 0.194867 | 0.106 | 0.095 | 1 | 8 | R3hcc1l | 0.011 | Young |
| 0.688765 | 0.163988 | 0.083 | 0.073 | 1 | 8 | Prr3 | 0.01 | Young |
| 0.689106 | 0.034906 | 0.148 | 0.134 | 1 | 8 | Rnf181 | 0.014 | Young |
| 0.689124 | 0.080695 | 0.275 | 0.268 | 1 | 8 | Ing1 | 0.007 | Young |
| 0.689241 | 0.046175 | 0.07 | 0.061 | 1 | 8 | Tbcc | 0.009 | Young |
| 0.689964 | 0.013093 | 0.221 | 0.207 | 1 | 8 | Papd5 | 0.014 | Young |
| 0.691124 | 0.291229 | 0.07 | 0.061 | 1 | 8 | Nek1 | 0.009 | Young |
| 0.691392 | 0.207647 | 0.047 | 0.039 | 1 | 8 | Mfsd4a | 0.008 | Young |
| 0.691402 | 0.206526 | 0.065 | 0.056 | 1 | 8 | Mbd4 | 0.009 | Young |
| 0.692054 | 0.239796 | 0.143 | 0.134 | 1 | 8 | Gypc | 0.009 | Young |
| 0.692225 | 0.019587 | 0.309 | 0.291 | 1 | 8 | Ago3 | 0.018 | Young |
| 0.693716 | 0.110976 | 0.327 | 0.318 | 1 | 8 | Ddx54 | 0.009 | Young |
| 0.694162 | 0.168396 | 0.008 | 0.011 | 1 | 8 | Ccdc18 | -0.003 | Young |
| 0.694162 | 0.117847 | 0.008 | 0.011 | 1 | 8 | Ksr2 | -0.003 | Young |
| 0.694162 | 0.115338 | 0.008 | 0.011 | 1 | 8 | Zfp39 | -0.003 | Young |
| 0.694162 | 0.079862 | 0.008 | 0.011 | 1 | 8 | Kif22 | -0.003 | Young |
| 0.694162 | 0.061132 | 0.008 | 0.011 | 1 | 8 | Spata2l | -0.003 | Young |
| 0.695279 | 0.067529 | 0.184 | 0.173 | 1 | 8 | Sucla2 | 0.011 | Young |
| 0.695447 | 0.463321 | 0.086 | 0.078 | 1 | 8 | Acad9 | 0.008 | Young |
| 0.695648 | 0.07461 | 0.278 | 0.263 | 1 | 8 | Tor1a | 0.015 | Young |
| 0.695886 | 0.08802 | 0.169 | 0.156 | 1 | 8 | Smim20 | 0.013 | Young |
| 0.696688 | 0.285042 | 0.008 | 0.011 | 1 | 8 | Gm826 | -0.003 | Young |
| 0.696688 | 0.189616 | 0.008 | 0.011 | 1 | 8 | Cenpf | -0.003 | Young |
| 0.696688 | 0.02407 | 0.008 | 0.011 | 1 | 8 | Mcts2 | -0.003 | Young |
| 0.696729 | 0.121618 | 0.14 | 0.128 | 1 | 8 | Ndst2 | 0.012 | Young |
| 0.696779 | 0.080553 | 0.2 | 0.184 | 1 | 8 | Ccdc186 | 0.016 | Young |
| 0.696785 | 0.384094 | 0.07 | 0.061 | 1 | 8 | Zfp729a | 0.009 | Young |
| 0.697023 | 0.216084 | 0.052 | 0.061 | 1 | 8 | Actr5 | -0.009 | Young |
| 0.697332 | 0.020685 | 0.249 | 0.235 | 1 | 8 | Actr10 | 0.014 | Young |
| 0.697773 | 0.481292 | 0.034 | 0.028 | 1 | 8 | Zfp40 | 0.006 | Young |
| 0.698675 | 0.195655 | 0.07 | 0.061 | 1 | 8 | Tnfaip8l1 | 0.009 | Young |
| 0.699219 | 0.482493 | 0.008 | 0.011 | 1 | 8 | Spa17 | -0.003 | Young |
| 0.699219 | 0.38344 | 0.008 | 0.011 | 1 | 8 | Serpinh1 | -0.003 | Young |
| 0.699219 | 0.369033 | 0.008 | 0.011 | 1 | 8 | 6720489N1 | -0.003 | Young |
| 0.699219 | 0.159211 | 0.008 | 0.011 | 1 | 8 | Acta2 | -0.003 | Young |
| 0.699219 | 0.053445 | 0.008 | 0.011 | 1 | 8 | Ddias | -0.003 | Young |
| 0.699421 | 0.222881 | 0.117 | 0.106 | 1 | 8 | Fdps | 0.011 | Young |
| 0.699477 | 0.269175 | 0.075 | 0.067 | 1 | 8 | Dgkg | 0.008 | Young |
| 0.699503 | 0.009385 | 0.268 | 0.251 | 1 | 8 | Fto | 0.017 | Young |

| | | | | | | | | |
|---|---|---|---|---|---|---|---|---|
| 0.699687 | 0.037382 | 0.143 | 0.128 | 1 | 8 | Gart | 0.015 | Young |
| 0.699952 | 0.338719 | 0.138 | 0.128 | 1 | 8 | Mtx1 | 0.01 | Young |
| 0.700414 | 0.048956 | 0.2 | 0.218 | 1 | 8 | Mpdu1 | -0.018 | Young |
| 0.70059 | 0.414897 | 0.086 | 0.078 | 1 | 8 | Hs1bp3 | 0.008 | Young |
| 0.70092 | 0.115901 | 0.161 | 0.151 | 1 | 8 | Plcb3 | 0.01 | Young |
| 0.702456 | 0.456839 | 0.081 | 0.073 | 1 | 8 | Prkdc | 0.008 | Young |
| 0.702529 | 0.22839 | 0.21 | 0.201 | 1 | 8 | Acadsb | 0.009 | Young |
| 0.703177 | 0.595396 | 0.034 | 0.028 | 1 | 8 | Mettl25 | 0.006 | Young |
| 0.703315 | 0.092152 | 0.19 | 0.173 | 1 | 8 | Uchl5 | 0.017 | Young |
| 0.704172 | 0.059605 | 0.065 | 0.056 | 1 | 8 | Gtf2h4 | 0.009 | Young |
| 0.704244 | 0.087096 | 0.221 | 0.212 | 1 | 8 | Map3k5 | 0.009 | Young |
| 0.70453 | 0.286764 | 0.034 | 0.028 | 1 | 8 | Tfb1m | 0.006 | Young |
| 0.704965 | 0.370819 | 0.075 | 0.067 | 1 | 8 | Lrrc40 | 0.008 | Young |
| 0.705284 | 0.046524 | 0.197 | 0.184 | 1 | 8 | Ak3 | 0.013 | Young |
| 0.705936 | 0.045537 | 0.221 | 0.207 | 1 | 8 | Trnt1 | 0.014 | Young |
| 0.706144 | 0.225046 | 0.065 | 0.056 | 1 | 8 | Stk39 | 0.009 | Young |
| 0.706544 | 0.025897 | 0.226 | 0.212 | 1 | 8 | Elovl1 | 0.014 | Young |
| 0.707153 | 0.120838 | 0.088 | 0.078 | 1 | 8 | Gabbr1 | 0.01 | Young |
| 0.70724 | 0.236501 | 0.034 | 0.028 | 1 | 8 | Endov | 0.006 | Young |
| 0.707447 | 0.158404 | 0.057 | 0.05 | 1 | 8 | Spred3 | 0.007 | Young |
| 0.708071 | 0.034587 | 0.33 | 0.302 | 1 | 8 | Sun2 | 0.028 | Young |
| 0.708248 | 0.016617 | 0.223 | 0.212 | 1 | 8 | Sft2d1 | 0.011 | Young |
| 0.709288 | 0.04983 | 0.106 | 0.095 | 1 | 8 | Saysd1 | 0.011 | Young |
| 0.709576 | 0.04439 | 0.052 | 0.045 | 1 | 8 | Fam110b | 0.007 | Young |
| 0.709662 | 0.244115 | 0.195 | 0.184 | 1 | 8 | Ilvbl | 0.011 | Young |
| 0.710058 | 0.006949 | 0.07 | 0.061 | 1 | 8 | Qpctl | 0.009 | Young |
| 0.710441 | 0.074652 | 0.081 | 0.073 | 1 | 8 | Tapt1 | 0.008 | Young |
| 0.710941 | 0.006858 | 0.465 | 0.447 | 1 | 8 | Ndufs7 | 0.018 | Young |
| 0.711311 | 0.125114 | 0.034 | 0.028 | 1 | 8 | Egfl7 | 0.006 | Young |
| 0.711768 | 0.117379 | 0.138 | 0.128 | 1 | 8 | Gopc | 0.01 | Young |
| 0.711936 | 0.098899 | 0.216 | 0.207 | 1 | 8 | Phf2 | 0.009 | Young |
| 0.711962 | 0.133579 | 0.07 | 0.061 | 1 | 8 | Cadm1 | 0.009 | Young |
| 0.713489 | 0.125674 | 0.166 | 0.156 | 1 | 8 | Med11 | 0.01 | Young |
| 0.713634 | 0.18466 | 0.244 | 0.235 | 1 | 8 | Slc35f6 | 0.009 | Young |
| 0.714029 | 0.329328 | 0.034 | 0.028 | 1 | 8 | Fbxo10 | 0.006 | Young |
| 0.714029 | 0.12649 | 0.034 | 0.028 | 1 | 8 | Ube2e2 | 0.006 | Young |
| 0.71539 | 0.635714 | 0.034 | 0.028 | 1 | 8 | Tmtc4 | 0.006 | Young |
| 0.716041 | 0.041234 | 0.099 | 0.089 | 1 | 8 | Lztr1 | 0.01 | Young |
| 0.716614 | 0.013155 | 0.104 | 0.095 | 1 | 8 | Kctd6 | 0.009 | Young |
| 0.716751 | 0.118661 | 0.034 | 0.028 | 1 | 8 | Gm26853 | 0.006 | Young |
| 0.716853 | 0.03821 | 0.099 | 0.089 | 1 | 8 | Stambp | 0.01 | Young |
| 0.716907 | 0.01527 | 0.122 | 0.112 | 1 | 8 | Dpp7 | 0.01 | Young |
| 0.71741 | 0.239753 | 0.135 | 0.123 | 1 | 8 | Gadd45gip | 0.012 | Young |
| 0.718114 | 0.421367 | 0.034 | 0.028 | 1 | 8 | Abcb6 | 0.006 | Young |
| 0.718393 | 0.407266 | 0.164 | 0.156 | 1 | 8 | Gps2 | 0.008 | Young |

| | | | | | | | | |
|---|---|---|---|---|---|---|---|---|
| 0.718421 | 0.150562 | 0.127 | 0.117 | 1 | 8 | Ciapin1 | 0.01 | Young |
| 0.718752 | 0.087551 | 0.075 | 0.067 | 1 | 8 | Pdhx | 0.008 | Young |
| 0.71945 | 0.091836 | 0.086 | 0.095 | 1 | 8 | Mcm5 | -0.009 | Young |
| 0.722207 | 0.508236 | 0.034 | 0.028 | 1 | 8 | Uevld | 0.006 | Young |
| 0.722392 | 0.011366 | 0.112 | 0.101 | 1 | 8 | Timm9 | 0.011 | Young |
| 0.722548 | 0.084312 | 0.099 | 0.089 | 1 | 8 | Cep135 | 0.01 | Young |
| 0.72309 | 0.020486 | 0.161 | 0.151 | 1 | 8 | Zfp646 | 0.01 | Young |
| 0.723369 | 0.141711 | 0.075 | 0.067 | 1 | 8 | Exd2 | 0.008 | Young |
| 0.724899 | 0.453175 | 0.062 | 0.056 | 1 | 8 | Mipep | 0.006 | Young |
| 0.725166 | 0.149167 | 0.249 | 0.24 | 1 | 8 | Wbp4 | 0.009 | Young |
| 0.726002 | 0.291965 | 0.148 | 0.14 | 1 | 8 | Ergic1 | 0.008 | Young |
| 0.726349 | 0.38204 | 0.057 | 0.05 | 1 | 8 | Nt5m | 0.007 | Young |
| 0.72661 | 0.244515 | 0.013 | 0.017 | 1 | 8 | A530040E1 | -0.004 | Young |
| 0.728024 | 0.034387 | 0.13 | 0.117 | 1 | 8 | Cgnl1 | 0.013 | Young |
| 0.728327 | 0.1011 | 0.052 | 0.045 | 1 | 8 | Ppp2r3a | 0.007 | Young |
| 0.728354 | 0.056919 | 0.345 | 0.341 | 1 | 8 | Eapp | 0.004 | Young |
| 0.728974 | 0.078009 | 0.182 | 0.168 | 1 | 8 | Abcg3 | 0.014 | Young |
| 0.729398 | 0.043337 | 0.413 | 0.391 | 1 | 8 | Pdcd6 | 0.022 | Young |
| 0.730094 | 0.140696 | 0.081 | 0.073 | 1 | 8 | Aacs | 0.008 | Young |
| 0.730094 | 0.078649 | 0.081 | 0.073 | 1 | 8 | Maml3 | 0.008 | Young |
| 0.730682 | 0.095173 | 0.013 | 0.017 | 1 | 8 | Bbs5 | -0.004 | Young |
| 0.730682 | 0.089211 | 0.013 | 0.017 | 1 | 8 | A130010J1 | -0.004 | Young |
| 0.730682 | 0.019767 | 0.013 | 0.017 | 1 | 8 | Shf | -0.004 | Young |
| 0.730883 | 0.039326 | 0.371 | 0.352 | 1 | 8 | Rtf1 | 0.019 | Young |
| 0.731001 | 0.042385 | 0.049 | 0.056 | 1 | 8 | Pex1 | -0.007 | Young |
| 0.731173 | 0.633821 | 0.039 | 0.034 | 1 | 8 | Yars2 | 0.005 | Young |
| 0.731275 | 0.280489 | 0.091 | 0.084 | 1 | 8 | Kat2a | 0.007 | Young |
| 0.732721 | 0.187701 | 0.013 | 0.017 | 1 | 8 | Gm26802 | -0.004 | Young |
| 0.733159 | 0.070429 | 0.034 | 0.028 | 1 | 8 | B930036N1 | 0.006 | Young |
| 0.734071 | 0.190633 | 0.265 | 0.257 | 1 | 8 | Babam2 | 0.008 | Young |
| 0.734343 | 0.042089 | 0.086 | 0.078 | 1 | 8 | 2610524H0 | 0.008 | Young |
| 0.734786 | 0.08481 | 0.93 | 0.866 | 1 | 8 | mt-Nd5 | 0.064 | Young |
| 0.734997 | 0.014789 | 0.296 | 0.274 | 1 | 8 | Crem | 0.022 | Young |
| 0.735083 | 0.608105 | 0.021 | 0.017 | 1 | 8 | C330027C0 | 0.004 | Young |
| 0.735083 | 0.58318 | 0.021 | 0.017 | 1 | 8 | Slc12a5 | 0.004 | Young |
| 0.735649 | 0.435765 | 0.068 | 0.061 | 1 | 8 | Sergef | 0.007 | Young |
| 0.735649 | 0.427721 | 0.068 | 0.061 | 1 | 8 | Ddx31 | 0.007 | Young |
| 0.736443 | 0.025039 | 0.616 | 0.598 | 1 | 8 | Stx7 | 0.018 | Young |
| 0.736763 | 0.001186 | 0.997 | 0.994 | 1 | 8 | Rps26 | 0.003 | Young |
| 0.736805 | 0.744435 | 0.013 | 0.017 | 1 | 8 | Clspn | -0.004 | Young |
| 0.736805 | 0.226087 | 0.013 | 0.017 | 1 | 8 | Gm14305 | -0.004 | Young |
| 0.736805 | 0.181141 | 0.013 | 0.017 | 1 | 8 | 4632404H1 | -0.004 | Young |
| 0.73683 | 0.533174 | 0.021 | 0.017 | 1 | 8 | Gm10642 | 0.004 | Young |
| 0.736867 | 0.190278 | 0.07 | 0.061 | 1 | 8 | Plekhg1 | 0.009 | Young |
| 0.737717 | 0.01674 | 0.927 | 0.872 | 1 | 8 | Ube2d3 | 0.055 | Young |

| | | | | | | | | |
|---|---|---|---|---|---|---|---|---|
| 0.738063 | 0.178005 | 0.169 | 0.162 | 1 | 8 | Usp33 | 0.007 | Young |
| 0.738135 | 0.112553 | 0.286 | 0.296 | 1 | 8 | Ptgs2 | -0.01 | Young |
| 0.738579 | 0.774218 | 0.021 | 0.017 | 1 | 8 | Stac2 | 0.004 | Young |
| 0.738579 | 0.761242 | 0.021 | 0.017 | 1 | 8 | Simc1 | 0.004 | Young |
| 0.73885 | 0.248378 | 0.013 | 0.017 | 1 | 8 | Gca | -0.004 | Young |
| 0.739075 | 0.009324 | 0.719 | 0.665 | 1 | 8 | Taldo1 | 0.054 | Young |
| 0.739521 | 0.038414 | 0.901 | 0.86 | 1 | 8 | Hexa | 0.041 | Young |
| 0.739761 | 0.189071 | 0.091 | 0.084 | 1 | 8 | Hook2 | 0.007 | Young |
| 0.740724 | 0.047607 | 0.07 | 0.061 | 1 | 8 | 1190007I0 | 0.009 | Young |
| 0.741167 | 0.229922 | 0.057 | 0.05 | 1 | 8 | Zfp971 | 0.007 | Young |
| 0.741373 | 0.115957 | 0.221 | 0.212 | 1 | 8 | Wdr3 | 0.009 | Young |
| 0.74208 | 0.555945 | 0.021 | 0.017 | 1 | 8 | Tmem17 | 0.004 | Young |
| 0.743621 | 0.038075 | 0.065 | 0.073 | 1 | 8 | Gm44751 | -0.008 | Young |
| 0.743832 | 0.959189 | 0.021 | 0.017 | 1 | 8 | Gm29562 | 0.004 | Young |
| 0.743832 | 0.421644 | 0.021 | 0.017 | 1 | 8 | Zfp553 | 0.004 | Young |
| 0.743832 | 0.100219 | 0.018 | 0.022 | 1 | 8 | Hist1h4d | -0.004 | Young |
| 0.743832 | 0.051106 | 0.018 | 0.022 | 1 | 8 | Ajuba | -0.004 | Young |
| 0.743939 | 0.396732 | 0.039 | 0.034 | 1 | 8 | Cnr2 | 0.005 | Young |
| 0.743939 | 0.384886 | 0.039 | 0.034 | 1 | 8 | E330009J0 | 0.005 | Young |
| 0.744076 | 0.172498 | 0.132 | 0.123 | 1 | 8 | Tmem250- | 0.009 | Young |
| 0.744201 | 0.195632 | 0.096 | 0.106 | 1 | 8 | 2310061I0 | -0.01 | Young |
| 0.744308 | 0.245085 | 0.171 | 0.162 | 1 | 8 | Rrm1 | 0.009 | Young |
| 0.744731 | 0.156035 | 0.075 | 0.067 | 1 | 8 | Zdhhc12 | 0.008 | Young |
| 0.745153 | 0.284702 | 0.106 | 0.101 | 1 | 8 | Cdc16 | 0.005 | Young |
| 0.745186 | 0.055717 | 0.296 | 0.285 | 1 | 8 | Vti1b | 0.011 | Young |
| 0.745586 | 0.359466 | 0.021 | 0.017 | 1 | 8 | Arsi | 0.004 | Young |
| 0.745586 | 0.294055 | 0.021 | 0.017 | 1 | 8 | Taf1a | 0.004 | Young |
| 0.746092 | 0.032561 | 0.247 | 0.24 | 1 | 8 | Prdx3 | 0.007 | Young |
| 0.746429 | 0.008529 | 0.748 | 0.682 | 1 | 8 | Ncf1 | 0.066 | Young |
| 0.746501 | 0.467643 | 0.039 | 0.034 | 1 | 8 | Fbxo46 | 0.005 | Young |
| 0.747164 | 0.115499 | 0.029 | 0.034 | 1 | 8 | Lipt2 | -0.005 | Young |
| 0.747341 | 0.652872 | 0.021 | 0.017 | 1 | 8 | Gm26982 | 0.004 | Young |
| 0.747341 | 0.413833 | 0.021 | 0.017 | 1 | 8 | Rhof | 0.004 | Young |
| 0.747341 | 0.251274 | 0.021 | 0.017 | 1 | 8 | Nupl2 | 0.004 | Young |
| 0.747341 | 0.212043 | 0.018 | 0.022 | 1 | 8 | Gbp4 | -0.004 | Young |
| 0.747522 | 0.04812 | 0.857 | 0.855 | 1 | 8 | Atf3 | 0.002 | Young |
| 0.747783 | 0.036379 | 0.039 | 0.034 | 1 | 8 | Zbtb25 | 0.005 | Young |
| 0.747955 | 0.021227 | 0.023 | 0.028 | 1 | 8 | Laptm4b | -0.005 | Young |
| 0.748236 | 0.354181 | 0.062 | 0.056 | 1 | 8 | Rfc4 | 0.006 | Young |
| 0.748345 | 0.296572 | 0.086 | 0.078 | 1 | 8 | Slc20a2 | 0.008 | Young |
| 0.749097 | 0.14277 | 0.018 | 0.022 | 1 | 8 | Nars2 | -0.004 | Young |
| 0.749452 | 0.017356 | 0.21 | 0.223 | 1 | 8 | Chrac1 | -0.013 | Young |
| 0.749868 | 0.032931 | 0.034 | 0.039 | 1 | 8 | Poll | -0.005 | Young |
| 0.750031 | 0.359716 | 0.145 | 0.14 | 1 | 8 | Mdc1 | 0.005 | Young |
| 0.750147 | 0.100927 | 0.26 | 0.246 | 1 | 8 | Ppp2r2d | 0.014 | Young |

| | | | | | | | | |
|---|---|---|---|---|---|---|---|---|
| 0.750277 | 0.101753 | 0.062 | 0.056 | 1 | 8 | Snrnp35 | 0.006 | Young |
| 0.750535 | 0.011214 | 0.527 | 0.503 | 1 | 8 | Lman2 | 0.024 | Young |
| 0.750855 | 0.428158 | 0.018 | 0.022 | 1 | 8 | Ccnb2 | -0.004 | Young |
| 0.750855 | 0.261706 | 0.018 | 0.022 | 1 | 8 | Gm5113 | -0.004 | Young |
| 0.750855 | 0.129624 | 0.021 | 0.017 | 1 | 8 | Bbs2 | 0.004 | Young |
| 0.750855 | 0.103101 | 0.018 | 0.022 | 1 | 8 | Shpk | -0.004 | Young |
| 0.750878 | 0.545883 | 0.044 | 0.039 | 1 | 8 | Dffa | 0.005 | Young |
| 0.751364 | 0.219838 | 0.065 | 0.073 | 1 | 8 | Srbd1 | -0.008 | Young |
| 0.751368 | 0.087672 | 0.184 | 0.173 | 1 | 8 | Nrf1 | 0.011 | Young |
| 0.751633 | 0.136215 | 0.039 | 0.034 | 1 | 8 | Zfp346 | 0.005 | Young |
| 0.751917 | 0.059881 | 0.164 | 0.151 | 1 | 8 | Kri1 | 0.013 | Young |
| 0.752225 | 0.073938 | 0.055 | 0.061 | 1 | 8 | Clasrp | -0.006 | Young |
| 0.752614 | 0.086155 | 0.018 | 0.022 | 1 | 8 | Osgepl1 | -0.004 | Young |
| 0.752614 | 0.073544 | 0.021 | 0.017 | 1 | 8 | Polr3g | 0.004 | Young |
| 0.752918 | 0.073865 | 0.039 | 0.034 | 1 | 8 | Ndufaf1 | 0.005 | Young |
| 0.753341 | 0.090562 | 0.062 | 0.056 | 1 | 8 | Hmgxb4 | 0.006 | Young |
| 0.753813 | 0.078318 | 0.034 | 0.039 | 1 | 8 | Zfp74 | -0.005 | Young |
| 0.753945 | 0.056365 | 0.057 | 0.05 | 1 | 8 | Spidr | 0.007 | Young |
| 0.754203 | 0.187944 | 0.039 | 0.034 | 1 | 8 | 4930430F0 | 0.005 | Young |
| 0.754374 | 0.391456 | 0.021 | 0.017 | 1 | 8 | Klri2 | 0.004 | Young |
| 0.754374 | 0.334377 | 0.021 | 0.017 | 1 | 8 | Gpx7 | 0.004 | Young |
| 0.754374 | 0.150389 | 0.018 | 0.022 | 1 | 8 | Gm27003 | -0.004 | Young |
| 0.754494 | 0.125931 | 0.044 | 0.039 | 1 | 8 | Klf8 | 0.005 | Young |
| 0.754526 | 0.311915 | 0.039 | 0.045 | 1 | 8 | Tcim | -0.006 | Young |
| 0.755463 | 0.072644 | 0.231 | 0.218 | 1 | 8 | Ergic2 | 0.013 | Young |
| 0.755957 | 0.085497 | 0.075 | 0.067 | 1 | 8 | Dhx33 | 0.008 | Young |
| 0.75603 | 0.126857 | 0.319 | 0.318 | 1 | 8 | Rsf1 | 0.001 | Young |
| 0.756135 | 0.240614 | 0.018 | 0.022 | 1 | 8 | Stard6 | -0.004 | Young |
| 0.756135 | 0.20768 | 0.018 | 0.022 | 1 | 8 | Cryab | -0.004 | Young |
| 0.756135 | 0.026737 | 0.021 | 0.017 | 1 | 8 | Tesk2 | 0.004 | Young |
| 0.757624 | 0.041019 | 0.153 | 0.145 | 1 | 8 | Atg4d | 0.008 | Young |
| 0.757898 | 0.300416 | 0.021 | 0.017 | 1 | 8 | Arhgap27o | 0.004 | Young |
| 0.757898 | 0.086179 | 0.021 | 0.017 | 1 | 8 | Arhgef4 | 0.004 | Young |
| 0.758064 | 0.533449 | 0.039 | 0.034 | 1 | 8 | Zfp961 | 0.005 | Young |
| 0.758457 | 0.187869 | 0.062 | 0.056 | 1 | 8 | Dtx2 | 0.006 | Young |
| 0.758457 | 0.133863 | 0.062 | 0.056 | 1 | 8 | Cdkn2c | 0.006 | Young |
| 0.758765 | 0.087587 | 0.044 | 0.05 | 1 | 8 | Pqlc2 | -0.006 | Young |
| 0.758918 | 0.105731 | 0.229 | 0.223 | 1 | 8 | Akap8 | 0.006 | Young |
| 0.759167 | 0.318685 | 0.068 | 0.061 | 1 | 8 | Zmynd19 | 0.007 | Young |
| 0.759976 | 0.039064 | 0.029 | 0.034 | 1 | 8 | Hibch | -0.005 | Young |
| 0.760026 | 0.381043 | 0.143 | 0.134 | 1 | 8 | Slc11a2 | 0.009 | Young |
| 0.760255 | 0.03985 | 0.086 | 0.095 | 1 | 8 | Pdp2 | -0.009 | Young |
| 0.760641 | 0.101691 | 0.039 | 0.034 | 1 | 8 | A430046D1 | 0.005 | Young |
| 0.761742 | 0.477697 | 0.044 | 0.039 | 1 | 8 | Deaf1 | 0.005 | Young |
| 0.762043 | 0.469709 | 0.023 | 0.028 | 1 | 8 | Tmem221 | -0.005 | Young |

| | | | | | | | | |
|---|---|---|---|---|---|---|---|---|
| 0.762575 | 0.224033 | 0.039 | 0.034 | 1 | 8 | Plcxd2 | 0.005 | Young |
| 0.763043 | 0.11098 | 0.034 | 0.039 | 1 | 8 | Rarb | -0.005 | Young |
| 0.763109 | 0.273042 | 0.068 | 0.061 | 1 | 8 | Zfp369 | 0.007 | Young |
| 0.763291 | 0.145512 | 0.294 | 0.291 | 1 | 8 | Arl5a | 0.003 | Young |
| 0.763614 | 0.097864 | 0.023 | 0.028 | 1 | 8 | Amigo1 | -0.005 | Young |
| 0.763721 | 0.163314 | 0.234 | 0.223 | 1 | 8 | Suclg1 | 0.011 | Young |
| 0.763806 | 0.302864 | 0.096 | 0.089 | 1 | 8 | Syt11 | 0.007 | Young |
| 0.764365 | 0.106948 | 0.034 | 0.039 | 1 | 8 | Fuz | -0.005 | Young |
| 0.764586 | 0.166122 | 0.044 | 0.05 | 1 | 8 | 5730409E0 | -0.006 | Young |
| 0.764962 | 0.81375 | 0.018 | 0.022 | 1 | 8 | Zfp46 | -0.004 | Young |
| 0.764962 | 0.601732 | 0.018 | 0.022 | 1 | 8 | Bora | -0.004 | Young |
| 0.765048 | 0.271133 | 0.119 | 0.112 | 1 | 8 | Serhl | 0.007 | Young |
| 0.765373 | 0.656612 | 0.044 | 0.039 | 1 | 8 | Ube2c | 0.005 | Young |
| 0.765687 | 0.063742 | 0.034 | 0.039 | 1 | 8 | Nod2 | -0.005 | Young |
| 0.765803 | 0.10019 | 0.039 | 0.034 | 1 | 8 | Acyp2 | 0.005 | Young |
| 0.766118 | 0.266727 | 0.138 | 0.128 | 1 | 8 | Dctn6 | 0.01 | Young |
| 0.766757 | 0.475571 | 0.023 | 0.028 | 1 | 8 | Ubiad1 | -0.005 | Young |
| 0.767187 | 0.200975 | 0.06 | 0.067 | 1 | 8 | Fam46c | -0.007 | Young |
| 0.767542 | 0.113345 | 0.262 | 0.257 | 1 | 8 | Trim56 | 0.005 | Young |
| 0.767691 | 0.238151 | 0.062 | 0.056 | 1 | 8 | Alkbh7 | 0.006 | Young |
| 0.767715 | 0.072222 | 0.205 | 0.201 | 1 | 8 | Ostm1 | 0.004 | Young |
| 0.767797 | 0.289376 | 0.044 | 0.039 | 1 | 8 | Nmnat1 | 0.005 | Young |
| 0.768045 | 0.234786 | 0.068 | 0.061 | 1 | 8 | Mrpl46 | 0.007 | Young |
| 0.768658 | 0.446733 | 0.049 | 0.045 | 1 | 8 | Zfp65 | 0.004 | Young |
| 0.768845 | 0.142033 | 0.091 | 0.084 | 1 | 8 | Fra10ac1 | 0.007 | Young |
| 0.769358 | 0.073739 | 0.878 | 0.832 | 1 | 8 | Tnip3 | 0.046 | Young |
| 0.769805 | 0.494241 | 0.049 | 0.045 | 1 | 8 | Tatdn1 | 0.004 | Young |
| 0.769987 | 0.29389 | 0.029 | 0.034 | 1 | 8 | Hacd1 | -0.005 | Young |
| 0.770223 | 0.168499 | 0.044 | 0.039 | 1 | 8 | Iqcb1 | 0.005 | Young |
| 0.771311 | 0.346288 | 0.083 | 0.078 | 1 | 8 | Drosha | 0.005 | Young |
| 0.771344 | 0.10471 | 0.096 | 0.089 | 1 | 8 | Alg9 | 0.007 | Young |
| 0.772157 | 0.330491 | 0.078 | 0.073 | 1 | 8 | Dock5 | 0.005 | Young |
| 0.772189 | 0.033143 | 0.987 | 0.95 | 1 | 8 | Ppia | 0.037 | Young |
| 0.772764 | 0.178392 | 0.174 | 0.168 | 1 | 8 | Diablo | 0.006 | Young |
| 0.773056 | 0.366279 | 0.023 | 0.028 | 1 | 8 | Nthl1 | -0.005 | Young |
| 0.77337 | 0.354862 | 0.236 | 0.235 | 1 | 8 | Cdc123 | 0.001 | Young |
| 0.774673 | 0.0574 | 0.114 | 0.106 | 1 | 8 | Setd1a | 0.008 | Young |
| 0.774685 | 0.026321 | 0.216 | 0.207 | 1 | 8 | Dda1 | 0.009 | Young |
| 0.775074 | 0.19267 | 0.306 | 0.302 | 1 | 8 | Smc4 | 0.004 | Young |
| 0.775724 | 0.380295 | 0.029 | 0.034 | 1 | 8 | Phf19 | -0.005 | Young |
| 0.775963 | 0.090046 | 0.068 | 0.061 | 1 | 8 | Zfp874a | 0.007 | Young |
| 0.776264 | 0.014662 | 0.242 | 0.229 | 1 | 8 | Cmc1 | 0.013 | Young |
| 0.776594 | 0.231762 | 0.086 | 0.078 | 1 | 8 | Pithd1 | 0.008 | Young |
| 0.777379 | 0.015681 | 0.153 | 0.145 | 1 | 8 | Kif13a | 0.008 | Young |
| 0.777515 | 0.305384 | 0.073 | 0.067 | 1 | 8 | Zfp317 | 0.006 | Young |

| | | | | | | | | |
|---|---|---|---|---|---|---|---|---|
| 0.778905 | 0.086034 | 0.091 | 0.101 | 1 | 8 | Dvl3 | -0.01 | Young |
| 0.781169 | 0.558452 | 0.044 | 0.039 | 1 | 8 | Slc35g1 | 0.005 | Young |
| 0.781548 | 0.018697 | 0.351 | 0.341 | 1 | 8 | Ube2e3 | 0.01 | Young |
| 0.781796 | 0.050604 | 0.114 | 0.123 | 1 | 8 | B3galnt1 | -0.009 | Young |
| 0.782812 | 0.080484 | 0.158 | 0.168 | 1 | 8 | Gpatch2 | -0.01 | Young |
| 0.782997 | 0.456184 | 0.039 | 0.045 | 1 | 8 | Xrcc6 | -0.006 | Young |
| 0.782997 | 0.014417 | 0.039 | 0.045 | 1 | 8 | Tmem206 | -0.006 | Young |
| 0.783015 | 0.079126 | 0.216 | 0.207 | 1 | 8 | Cox16 | 0.009 | Young |
| 0.783106 | 0.138511 | 0.158 | 0.173 | 1 | 8 | Brwd3 | -0.015 | Young |
| 0.783607 | 0.305028 | 0.044 | 0.039 | 1 | 8 | Asb13 | 0.005 | Young |
| 0.784123 | 0.095834 | 0.213 | 0.207 | 1 | 8 | Ythdc2 | 0.006 | Young |
| 0.78476 | 0.628372 | 0.049 | 0.045 | 1 | 8 | 2010320M | 0.004 | Young |
| 0.784816 | 0.089774 | 0.239 | 0.229 | 1 | 8 | Oaz2 | 0.01 | Young |
| 0.786692 | 0.191257 | 0.166 | 0.162 | 1 | 8 | Nr1h2 | 0.004 | Young |
| 0.787751 | 0.154194 | 0.158 | 0.173 | 1 | 8 | Dpp9 | -0.015 | Young |
| 0.787889 | 0.015334 | 0.13 | 0.14 | 1 | 8 | Pias4 | -0.01 | Young |
| 0.788056 | 0.345016 | 0.073 | 0.067 | 1 | 8 | Numb | 0.006 | Young |
| 0.788075 | 0.391367 | 0.026 | 0.022 | 1 | 8 | Carmil1 | 0.004 | Young |
| 0.788162 | 0.155798 | 0.296 | 0.285 | 1 | 8 | Vps13a | 0.011 | Young |
| 0.788295 | 0.104849 | 0.101 | 0.095 | 1 | 8 | Pcyt2 | 0.006 | Young |
| 0.788295 | 0.104668 | 0.101 | 0.095 | 1 | 8 | Zfp266 | 0.006 | Young |
| 0.789119 | 0.032462 | 0.096 | 0.106 | 1 | 8 | Hic2 | -0.01 | Young |
| 0.789226 | 0.037021 | 0.288 | 0.302 | 1 | 8 | Map4k1 | -0.014 | Young |
| 0.789661 | 0.443683 | 0.026 | 0.022 | 1 | 8 | Hyal1 | 0.004 | Young |
| 0.790329 | 0.098413 | 0.132 | 0.128 | 1 | 8 | Pygb | 0.004 | Young |
| 0.790937 | 0.281365 | 0.073 | 0.067 | 1 | 8 | Whamm | 0.006 | Young |
| 0.790943 | 0.362328 | 0.044 | 0.05 | 1 | 8 | Cdc42bpg | -0.006 | Young |
| 0.791093 | 0.039979 | 0.514 | 0.497 | 1 | 8 | Gna13 | 0.017 | Young |
| 0.791551 | 0.111487 | 0.096 | 0.089 | 1 | 8 | Mfn1 | 0.007 | Young |
| 0.791898 | 0.149747 | 0.073 | 0.067 | 1 | 8 | Wiz | 0.006 | Young |
| 0.792835 | 0.364558 | 0.026 | 0.022 | 1 | 8 | Tex9 | 0.004 | Young |
| 0.792835 | 0.329113 | 0.026 | 0.022 | 1 | 8 | Sirt5 | 0.004 | Young |
| 0.792835 | 0.151087 | 0.026 | 0.022 | 1 | 8 | Sall2 | 0.004 | Young |
| 0.792835 | 0.059196 | 0.081 | 0.073 | 1 | 8 | Eif2d | 0.008 | Young |
| 0.792851 | 0.016337 | 0.143 | 0.151 | 1 | 8 | Grpel1 | -0.008 | Young |
| 0.79297 | 0.130437 | 0.195 | 0.196 | 1 | 8 | Braf | -0.001 | Young |
| 0.79345 | 0.048276 | 0.922 | 0.899 | 1 | 8 | Naca | 0.023 | Young |
| 0.794423 | 0.489076 | 0.026 | 0.022 | 1 | 8 | Crybb1 | 0.004 | Young |
| 0.794423 | 0.246697 | 0.026 | 0.022 | 1 | 8 | Gm43936 | 0.004 | Young |
| 0.794423 | 0.059075 | 0.026 | 0.022 | 1 | 8 | Cenpv | 0.004 | Young |
| 0.794687 | 0.258588 | 0.07 | 0.078 | 1 | 8 | Snx16 | -0.008 | Young |
| 0.795953 | 0.064656 | 0.842 | 0.782 | 1 | 8 | Ptms | 0.06 | Young |
| 0.796012 | 0.21904 | 0.026 | 0.022 | 1 | 8 | Fam149a | 0.004 | Young |
| 0.796318 | 0.347014 | 0.049 | 0.045 | 1 | 8 | Rnf152 | 0.004 | Young |
| 0.796318 | 0.339968 | 0.049 | 0.045 | 1 | 8 | Nsun3 | 0.004 | Young |

| | | | | | | | | |
|---|---|---|---|---|---|---|---|---|
| 0.797602 | 0.509585 | 0.026 | 0.022 | 1 | 8 | 8030462N1 | 0.004 | Young |
| 0.799193 | 0.180903 | 0.026 | 0.022 | 1 | 8 | Sap18b | 0.004 | Young |
| 0.802377 | 0.074383 | 0.026 | 0.022 | 1 | 8 | Trp53i13 | 0.004 | Young |
| 0.802398 | 0.026784 | 0.909 | 0.86 | 1 | 8 | Calm2 | 0.049 | Young |
| 0.803142 | 0.705557 | 0.031 | 0.028 | 1 | 8 | 6330403L0 | 0.003 | Young |
| 0.803196 | 0.034229 | 0.197 | 0.19 | 1 | 8 | Aldh16a1 | 0.007 | Young |
| 0.803859 | 0.15562 | 0.135 | 0.128 | 1 | 8 | Josd2 | 0.007 | Young |
| 0.80397 | 0.104971 | 0.026 | 0.022 | 1 | 8 | Bcs1l | 0.004 | Young |
| 0.804197 | 0.141848 | 0.112 | 0.106 | 1 | 8 | Ikbke | 0.006 | Young |
| 0.804592 | 0.675713 | 0.031 | 0.028 | 1 | 8 | 3222401L1 | 0.003 | Young |
| 0.804904 | 0.150108 | 0.073 | 0.067 | 1 | 8 | Mtg1 | 0.006 | Young |
| 0.805408 | 0.022655 | 0.223 | 0.235 | 1 | 8 | Cxcl1 | -0.012 | Young |
| 0.805564 | 0.154164 | 0.026 | 0.022 | 1 | 8 | Otud3 | 0.004 | Young |
| 0.805996 | 0.025072 | 0.13 | 0.123 | 1 | 8 | Fbxo8 | 0.007 | Young |
| 0.806771 | 0.182642 | 0.135 | 0.128 | 1 | 8 | Slc16a6 | 0.007 | Young |
| 0.807159 | 0.136649 | 0.026 | 0.022 | 1 | 8 | Pex11g | 0.004 | Young |
| 0.807374 | 0.211829 | 0.255 | 0.257 | 1 | 8 | Kctd20 | -0.002 | Young |
| 0.807624 | 0.118807 | 0.153 | 0.145 | 1 | 8 | Tbc1d17 | 0.008 | Young |
| 0.808285 | 0.087807 | 0.073 | 0.067 | 1 | 8 | Fkbp9 | 0.006 | Young |
| 0.808303 | 0.003798 | 0.257 | 0.246 | 1 | 8 | Setd5 | 0.011 | Young |
| 0.80866 | 0.010914 | 0.088 | 0.095 | 1 | 8 | Zdhhc4 | -0.007 | Young |
| 0.80965 | 0.011711 | 0.891 | 0.849 | 1 | 8 | Gdi2 | 0.042 | Young |
| 0.810261 | 0.183476 | 0.078 | 0.073 | 1 | 8 | Tbc1d12 | 0.005 | Young |
| 0.810653 | 0.002134 | 0.956 | 0.883 | 1 | 8 | Pid1 | 0.073 | Young |
| 0.811999 | 0.137961 | 0.19 | 0.184 | 1 | 8 | Ccdc9 | 0.006 | Young |
| 0.812992 | 0.071328 | 0.348 | 0.335 | 1 | 8 | Arhgef12 | 0.013 | Young |
| 0.813112 | 0.147438 | 0.174 | 0.168 | 1 | 8 | Ndufv1 | 0.006 | Young |
| 0.813519 | 0.156852 | 0.2 | 0.196 | 1 | 8 | Rsbn1 | 0.004 | Young |
| 0.81379 | 0.368675 | 0.055 | 0.05 | 1 | 8 | Gpank1 | 0.005 | Young |
| 0.814286 | 0.090892 | 0.499 | 0.497 | 1 | 8 | Ptger4 | 0.002 | Young |
| 0.814469 | 0.154155 | 0.213 | 0.212 | 1 | 8 | Gtf2b | 0.001 | Young |
| 0.816986 | 0.09065 | 0.13 | 0.14 | 1 | 8 | Coasy | -0.01 | Young |
| 0.817117 | 0.092527 | 0.062 | 0.067 | 1 | 8 | Btg3 | -0.005 | Young |
| 0.81732 | 0.019097 | 0.179 | 0.173 | 1 | 8 | Parp2 | 0.006 | Young |
| 0.81992 | 0.00442 | 0.088 | 0.095 | 1 | 8 | Orc2 | -0.007 | Young |
| 0.820734 | 0.082538 | 0.049 | 0.045 | 1 | 8 | Nectin4 | 0.004 | Young |
| 0.821754 | 0.098597 | 0.338 | 0.33 | 1 | 8 | Sptlc2 | 0.008 | Young |
| 0.821975 | 0.327402 | 0.088 | 0.084 | 1 | 8 | Ring1 | 0.004 | Young |
| 0.822761 | 0.029689 | 0.094 | 0.089 | 1 | 8 | Nostrin | 0.005 | Young |
| 0.82428 | 0.245813 | 0.127 | 0.123 | 1 | 8 | Med25 | 0.004 | Young |
| 0.824961 | 0.253306 | 0.031 | 0.028 | 1 | 8 | 5430403G1 | 0.003 | Young |
| 0.824961 | 0.197818 | 0.031 | 0.028 | 1 | 8 | Asb10 | 0.003 | Young |
| 0.825613 | 0.077391 | 0.197 | 0.196 | 1 | 8 | Abcb7 | 0.001 | Young |
| 0.82629 | 0.042022 | 0.062 | 0.067 | 1 | 8 | Pnpt1 | -0.005 | Young |
| 0.826421 | 0.325226 | 0.031 | 0.028 | 1 | 8 | Eef1akmt4 | 0.003 | Young |

| | | | | | | | | |
|---|---|---|---|---|---|---|---|---|
| 0.826574 | 0.019103 | 0.047 | 0.05 | 1 | 8 | Prim2 | -0.003 | Young |
| 0.8267 | 0.026151 | 0.073 | 0.067 | 1 | 8 | Tbcd | 0.006 | Young |
| 0.826823 | 0.020687 | 0.156 | 0.145 | 1 | 8 | Hmgn2 | 0.011 | Young |
| 0.826904 | 0.008765 | 0.805 | 0.749 | 1 | 8 | Cyfip1 | 0.056 | Young |
| 0.827595 | 0.001438 | 0.829 | 0.777 | 1 | 8 | Cox6b1 | 0.052 | Young |
| 0.827823 | 0.271927 | 0.187 | 0.184 | 1 | 8 | Cbx7 | 0.003 | Young |
| 0.827881 | 0.445701 | 0.031 | 0.028 | 1 | 8 | Gm33973 | 0.003 | Young |
| 0.827881 | 0.191039 | 0.031 | 0.028 | 1 | 8 | C2cd2 | 0.003 | Young |
| 0.827931 | 0.060835 | 0.478 | 0.458 | 1 | 8 | Il6 | 0.02 | Young |
| 0.829342 | 0.242421 | 0.031 | 0.028 | 1 | 8 | Zfp668 | 0.003 | Young |
| 0.829789 | 0.2183 | 0.122 | 0.117 | 1 | 8 | Kdm4c | 0.005 | Young |
| 0.83006 | 0.189826 | 0.099 | 0.095 | 1 | 8 | Slc7a6os | 0.004 | Young |
| 0.830201 | 0.076497 | 0.078 | 0.084 | 1 | 8 | E130309D( | -0.006 | Young |
| 0.830448 | 0.268241 | 0.055 | 0.05 | 1 | 8 | Zfp760 | 0.005 | Young |
| 0.830558 | 0.644142 | 0.036 | 0.034 | 1 | 8 | Kptn | 0.002 | Young |
| 0.830803 | 0.211591 | 0.031 | 0.028 | 1 | 8 | Cdk20 | 0.003 | Young |
| 0.832156 | 0.336319 | 0.088 | 0.084 | 1 | 8 | Lsm10 | 0.004 | Young |
| 0.83225 | 0.002675 | 0.951 | 0.905 | 1 | 8 | Arhgdib | 0.046 | Young |
| 0.833269 | 0.007022 | 0.2 | 0.207 | 1 | 8 | Commd2 | -0.007 | Young |
| 0.833835 | 0.030245 | 0.135 | 0.128 | 1 | 8 | G2e3 | 0.007 | Young |
| 0.834505 | 0.049844 | 0.499 | 0.492 | 1 | 8 | Klf10 | 0.007 | Young |
| 0.835191 | 0.157548 | 0.031 | 0.028 | 1 | 8 | Ftsj1 | 0.003 | Young |
| 0.835608 | 0.025294 | 0.078 | 0.073 | 1 | 8 | Rnf185 | 0.005 | Young |
| 0.835826 | 0.116038 | 0.184 | 0.179 | 1 | 8 | Sema4b | 0.005 | Young |
| 0.836266 | 0.00395 | 0.301 | 0.318 | 1 | 8 | Mybbp1a | -0.017 | Young |
| 0.836527 | 0.177087 | 0.13 | 0.123 | 1 | 8 | Lpar6 | 0.007 | Young |
| 0.836728 | 0.138058 | 0.06 | 0.056 | 1 | 8 | Bcorl1 | 0.004 | Young |
| 0.837007 | 0.018178 | 0.145 | 0.14 | 1 | 8 | Crot | 0.005 | Young |
| 0.837951 | 0.032251 | 0.117 | 0.123 | 1 | 8 | Gbe1 | -0.006 | Young |
| 0.838119 | 0.239469 | 0.031 | 0.028 | 1 | 8 | Gm48066 | 0.003 | Young |
| 0.838248 | 0.283963 | 0.055 | 0.05 | 1 | 8 | Orc5 | 0.005 | Young |
| 0.839868 | 0.037171 | 0.187 | 0.196 | 1 | 8 | Ppfia4 | -0.009 | Young |
| 0.840588 | 0.184065 | 0.088 | 0.084 | 1 | 8 | Pck2 | 0.004 | Young |
| 0.840694 | 0.121121 | 0.361 | 0.358 | 1 | 8 | Mtm1 | 0.003 | Young |
| 0.842185 | 0.008273 | 0.112 | 0.117 | 1 | 8 | Rab10os | -0.005 | Young |
| 0.842738 | 0.425563 | 0.036 | 0.034 | 1 | 8 | Sgce | 0.002 | Young |
| 0.843799 | 0.088446 | 0.291 | 0.313 | 1 | 8 | Tia1 | -0.022 | Young |
| 0.843905 | 0.582004 | 0.135 | 0.134 | 1 | 8 | Rfwd3 | 0.001 | Young |
| 0.845449 | 0.116688 | 0.031 | 0.028 | 1 | 8 | Runx2 | 0.003 | Young |
| 0.84559 | 0.2297 | 0.083 | 0.078 | 1 | 8 | Ddah2 | 0.005 | Young |
| 0.845758 | 0.026548 | 0.073 | 0.078 | 1 | 8 | Sgsm2 | -0.005 | Young |
| 0.845923 | 0.026447 | 0.088 | 0.084 | 1 | 8 | Gpr89 | 0.004 | Young |
| 0.846187 | 0.182549 | 0.073 | 0.067 | 1 | 8 | Mier2 | 0.006 | Young |
| 0.846206 | 0.671271 | 0.013 | 0.011 | 1 | 8 | Slc9a7 | 0.002 | Young |
| 0.846621 | 0.1149 | 0.197 | 0.19 | 1 | 8 | Frmd8 | 0.007 | Young |

| | | | | | | | | |
|---|---|---|---|---|---|---|---|---|
| 0.847283 | 0.145223 | 0.078 | 0.084 | 1 | 8 | Sp4 | -0.006 | Young |
| 0.848136 | 0.106163 | 0.101 | 0.095 | 1 | 8 | Xxylt1 | 0.006 | Young |
| 0.848474 | 0.669712 | 0.013 | 0.011 | 1 | 8 | Pex10 | 0.002 | Young |
| 0.848474 | 0.651035 | 0.013 | 0.011 | 1 | 8 | Epm2a | 0.002 | Young |
| 0.848474 | 0.618272 | 0.013 | 0.011 | 1 | 8 | Gnb4 | 0.002 | Young |
| 0.849738 | 0.140631 | 0.816 | 0.877 | 1 | 8 | Cxcl2 | -0.061 | Young |
| 0.850743 | 0.911472 | 0.013 | 0.011 | 1 | 8 | Plpp3 | 0.002 | Young |
| 0.850743 | 0.524412 | 0.013 | 0.011 | 1 | 8 | Ahrr | 0.002 | Young |
| 0.850743 | 0.509052 | 0.013 | 0.011 | 1 | 8 | Rgs5 | 0.002 | Young |
| 0.850743 | 0.36639 | 0.013 | 0.011 | 1 | 8 | D830025C( | 0.002 | Young |
| 0.850743 | 0.336485 | 0.013 | 0.011 | 1 | 8 | Gm44067 | 0.002 | Young |
| 0.850868 | 0.175002 | 0.114 | 0.112 | 1 | 8 | Bysl | 0.002 | Young |
| 0.850906 | 0.014632 | 0.571 | 0.525 | 1 | 8 | Lrp6 | 0.046 | Young |
| 0.851183 | 0.258556 | 0.278 | 0.279 | 1 | 8 | Slirp | -0.001 | Young |
| 0.853012 | 0.600927 | 0.013 | 0.011 | 1 | 8 | Retn | 0.002 | Young |
| 0.853012 | 0.52072 | 0.013 | 0.011 | 1 | 8 | H2-Q10 | 0.002 | Young |
| 0.853012 | 0.433846 | 0.013 | 0.011 | 1 | 8 | Cdkl5 | 0.002 | Young |
| 0.853012 | 0.29048 | 0.013 | 0.011 | 1 | 8 | Fndc10 | 0.002 | Young |
| 0.853012 | 0.279756 | 0.013 | 0.011 | 1 | 8 | B3gntl1 | 0.002 | Young |
| 0.853012 | 0.242661 | 0.013 | 0.011 | 1 | 8 | Mybpc3 | 0.002 | Young |
| 0.853306 | 0.235148 | 0.19 | 0.184 | 1 | 8 | Upf3a | 0.006 | Young |
| 0.853914 | 0.063161 | 0.088 | 0.095 | 1 | 8 | Hddc2 | -0.007 | Young |
| 0.854527 | 0.007779 | 0.042 | 0.045 | 1 | 8 | Clec16a | -0.003 | Young |
| 0.854534 | 0.058963 | 0.148 | 0.145 | 1 | 8 | Dph5 | 0.003 | Young |
| 0.854734 | 0.064004 | 0.047 | 0.05 | 1 | 8 | Smim5 | -0.003 | Young |
| 0.854957 | 0.271912 | 0.036 | 0.034 | 1 | 8 | Erc1 | 0.002 | Young |
| 0.855284 | 0.404557 | 0.013 | 0.011 | 1 | 8 | Gm10645 | 0.002 | Young |
| 0.855284 | 0.395357 | 0.013 | 0.011 | 1 | 8 | Tssk6 | 0.002 | Young |
| 0.855284 | 0.312362 | 0.013 | 0.011 | 1 | 8 | Llgl2 | 0.002 | Young |
| 0.855284 | 0.291825 | 0.013 | 0.011 | 1 | 8 | P3h3 | 0.002 | Young |
| 0.855284 | 0.243825 | 0.013 | 0.011 | 1 | 8 | Spata7 | 0.002 | Young |
| 0.855284 | 0.17006 | 0.013 | 0.011 | 1 | 8 | Atxn7l2 | 0.002 | Young |
| 0.855355 | 0.005207 | 0.265 | 0.279 | 1 | 8 | Clk3 | -0.014 | Young |
| 0.856067 | 0.127511 | 0.138 | 0.134 | 1 | 8 | Grcc10 | 0.004 | Young |
| 0.856176 | 0.121446 | 0.138 | 0.145 | 1 | 8 | Dcps | -0.007 | Young |
| 0.856631 | 0.048143 | 0.244 | 0.246 | 1 | 8 | Siah2 | -0.002 | Young |
| 0.857556 | 0.166334 | 0.013 | 0.011 | 1 | 8 | Cage1 | 0.002 | Young |
| 0.857556 | 0.125665 | 0.013 | 0.011 | 1 | 8 | 9330020H( | 0.002 | Young |
| 0.857556 | 0.049118 | 0.013 | 0.011 | 1 | 8 | Csrnp2 | 0.002 | Young |
| 0.857556 | 0.022594 | 0.013 | 0.011 | 1 | 8 | Ly6g6d | 0.002 | Young |
| 0.859511 | 0.004849 | 0.26 | 0.257 | 1 | 8 | Ireb2 | 0.003 | Young |
| 0.859829 | 0.351109 | 0.013 | 0.011 | 1 | 8 | Gm4631 | 0.002 | Young |
| 0.859829 | 0.223671 | 0.013 | 0.011 | 1 | 8 | Acsbg1 | 0.002 | Young |
| 0.859829 | 0.215966 | 0.013 | 0.011 | 1 | 8 | Arl16 | 0.002 | Young |
| 0.859829 | 0.170864 | 0.013 | 0.011 | 1 | 8 | Cacnb1 | 0.002 | Young |

| | | | | | | | | |
|---|---|---|---|---|---|---|---|---|
| 0.859829 | 0.105876 | 0.013 | 0.011 | 1 | 8 | Zfp993 | 0.002 | Young |
| 0.859829 | 0.096181 | 0.013 | 0.011 | 1 | 8 | Ffar4 | 0.002 | Young |
| 0.859829 | 0.091592 | 0.013 | 0.011 | 1 | 8 | Sell | 0.002 | Young |
| 0.859829 | 0.037512 | 0.013 | 0.011 | 1 | 8 | Mical3 | 0.002 | Young |
| 0.859829 | 0.017856 | 0.013 | 0.011 | 1 | 8 | Gm45894 | 0.002 | Young |
| 0.859829 | 0.003029 | 0.013 | 0.011 | 1 | 8 | Ass1 | 0.002 | Young |
| 0.860259 | 0.113908 | 0.081 | 0.078 | 1 | 8 | Tstd2 | 0.003 | Young |
| 0.861103 | 0.417438 | 0.042 | 0.039 | 1 | 8 | Fastkd2 | 0.003 | Young |
| 0.861147 | 0.169118 | 0.062 | 0.067 | 1 | 8 | Alg8 | -0.005 | Young |
| 0.86176 | 0.318009 | 0.036 | 0.034 | 1 | 8 | Xndc1 | 0.002 | Young |
| 0.862677 | 0.257733 | 0.088 | 0.095 | 1 | 8 | Trub2 | -0.007 | Young |
| 0.862708 | 0.203021 | 0.07 | 0.067 | 1 | 8 | Atg16l2 | 0.003 | Young |
| 0.862854 | 0.014626 | 0.223 | 0.212 | 1 | 8 | Dhx30 | 0.011 | Young |
| 0.863025 | 0.186956 | 0.083 | 0.078 | 1 | 8 | 1110034G2 | 0.005 | Young |
| 0.864302 | 0.279473 | 0.094 | 0.089 | 1 | 8 | Abr | 0.005 | Young |
| 0.86438 | 0.0739 | 0.013 | 0.011 | 1 | 8 | Gm40773 | 0.002 | Young |
| 0.864484 | 0.042319 | 0.036 | 0.034 | 1 | 8 | Gimap6 | 0.002 | Young |
| 0.864945 | 0.007459 | 0.104 | 0.101 | 1 | 8 | Tbc1d24 | 0.003 | Young |
| 0.865115 | 0.134679 | 0.099 | 0.095 | 1 | 8 | Reep4 | 0.004 | Young |
| 0.865847 | 0.105942 | 0.036 | 0.034 | 1 | 8 | Gm16973 | 0.002 | Young |
| 0.86642 | 0.00847 | 0.117 | 0.112 | 1 | 8 | Mthfs | 0.005 | Young |
| 0.867064 | 0.101276 | 0.088 | 0.095 | 1 | 8 | Tcp11l1 | -0.007 | Young |
| 0.867594 | 0.030597 | 0.122 | 0.128 | 1 | 8 | Ubqln4 | -0.006 | Young |
| 0.867762 | 0.053537 | 0.06 | 0.056 | 1 | 8 | 1110032A0 | 0.004 | Young |
| 0.867778 | 0.317393 | 0.073 | 0.078 | 1 | 8 | Zfp335 | -0.005 | Young |
| 0.868168 | 0.348784 | 0.07 | 0.067 | 1 | 8 | Rmnd5b | 0.003 | Young |
| 0.86912 | 0.012144 | 0.086 | 0.089 | 1 | 8 | Pym1 | -0.003 | Young |
| 0.870909 | 0.012833 | 0.086 | 0.089 | 1 | 8 | Zfp410 | -0.003 | Young |
| 0.872059 | 0.195598 | 0.057 | 0.061 | 1 | 8 | 5730480H0 | -0.004 | Young |
| 0.872265 | 0.254184 | 0.036 | 0.039 | 1 | 8 | Slc22a18 | -0.003 | Young |
| 0.873146 | 0.594702 | 0.047 | 0.045 | 1 | 8 | Large1 | 0.002 | Young |
| 0.874203 | 0.058608 | 0.081 | 0.078 | 1 | 8 | Otof | 0.003 | Young |
| 0.874796 | 0.093827 | 0.047 | 0.05 | 1 | 8 | Fkbp7 | -0.003 | Young |
| 0.875592 | 0.254344 | 0.182 | 0.184 | 1 | 8 | P3h2 | -0.002 | Young |
| 0.876266 | 0.051545 | 0.036 | 0.039 | 1 | 8 | Zfp111 | -0.003 | Young |
| 0.878314 | 0.007544 | 0.156 | 0.162 | 1 | 8 | Steap3 | -0.006 | Young |
| 0.878959 | 0.030559 | 0.042 | 0.039 | 1 | 8 | Tfpt | 0.003 | Young |
| 0.879098 | 0.110496 | 0.275 | 0.279 | 1 | 8 | Tcof1 | -0.004 | Young |
| 0.879168 | 0.693221 | 0.047 | 0.045 | 1 | 8 | 3110001I2 | 0.002 | Young |
| 0.880554 | 0.014502 | 0.418 | 0.397 | 1 | 8 | Pla2g15 | 0.021 | Young |
| 0.88063 | 0.231631 | 0.223 | 0.24 | 1 | 8 | Pank2 | -0.017 | Young |
| 0.881597 | 0.149598 | 0.07 | 0.067 | 1 | 8 | Btbd3 | 0.003 | Young |
| 0.881881 | 0.061683 | 0.031 | 0.034 | 1 | 8 | Coq6 | -0.003 | Young |
| 0.882039 | 0.073512 | 0.156 | 0.151 | 1 | 8 | Ssna1 | 0.005 | Young |
| 0.882067 | 0.051132 | 0.151 | 0.145 | 1 | 8 | Kdm5b | 0.006 | Young |

| | | | | | | | | |
|---|---|---|---|---|---|---|---|---|
| 0.88254 | 0.124046 | 0.148 | 0.145 | 1 | 8 | Pmm2 | 0.003 | Young |
| 0.882794 | 0.345076 | 0.042 | 0.039 | 1 | 8 | Rad1 | 0.003 | Young |
| 0.88282 | 0.205668 | 0.057 | 0.061 | 1 | 8 | Osbpl7 | -0.004 | Young |
| 0.884863 | 0.133592 | 0.104 | 0.101 | 1 | 8 | Orc4 | 0.003 | Young |
| 0.885995 | 0.256935 | 0.216 | 0.218 | 1 | 8 | Cdkn2d | -0.002 | Young |
| 0.886065 | 0.831443 | 0.018 | 0.017 | 1 | 8 | Chaf1b | 0.001 | Young |
| 0.886065 | 0.4538 | 0.018 | 0.017 | 1 | 8 | Cracr2a | 0.001 | Young |
| 0.886403 | 0.287272 | 0.047 | 0.045 | 1 | 8 | Galnt9 | 0.002 | Young |
| 0.886527 | 0.09859 | 0.104 | 0.101 | 1 | 8 | Pde4a | 0.003 | Young |
| 0.887639 | 0.098136 | 0.031 | 0.034 | 1 | 8 | Wrap53 | -0.003 | Young |
| 0.887983 | 0.644376 | 0.018 | 0.017 | 1 | 8 | Zcchc3 | 0.001 | Young |
| 0.889079 | 0.088536 | 0.031 | 0.034 | 1 | 8 | 2700049A( | -0.003 | Young |
| 0.889314 | 0.278361 | 0.117 | 0.112 | 1 | 8 | Unc45a | 0.005 | Young |
| 0.889624 | 0.517843 | 0.036 | 0.039 | 1 | 8 | Gtf2h2 | -0.003 | Young |
| 0.889655 | 0.076062 | 0.314 | 0.318 | 1 | 8 | Mark3 | -0.004 | Young |
| 0.889902 | 1.326854 | 0.018 | 0.017 | 1 | 8 | Zrsr1 | 0.001 | Young |
| 0.889902 | 0.512313 | 0.018 | 0.017 | 1 | 8 | Fam120c | 0.001 | Young |
| 0.89005 | 0.275891 | 0.117 | 0.117 | 1 | 8 | 1600002K( | 0 | Young |
| 0.89006 | 0.259031 | 0.062 | 0.067 | 1 | 8 | Mettl21a | -0.005 | Young |
| 0.890465 | 0.015992 | 0.094 | 0.089 | 1 | 8 | Nup155 | 0.005 | Young |
| 0.890569 | 0.183989 | 0.07 | 0.067 | 1 | 8 | Rfx3 | 0.003 | Young |
| 0.890581 | 0.119772 | 0.153 | 0.151 | 1 | 8 | Crkl | 0.002 | Young |
| 0.891531 | 0.005766 | 0.086 | 0.089 | 1 | 8 | Tmem209 | -0.003 | Young |
| 0.893295 | 0.494488 | 0.042 | 0.045 | 1 | 8 | Mpst | -0.003 | Young |
| 0.893383 | 0.001723 | 0.506 | 0.492 | 1 | 8 | Ggnbp2 | 0.014 | Young |
| 0.893742 | 0.816335 | 0.018 | 0.017 | 1 | 8 | Bub1b | 0.001 | Young |
| 0.893742 | 0.558111 | 0.018 | 0.017 | 1 | 8 | Ift122 | 0.001 | Young |
| 0.893742 | 0.347627 | 0.018 | 0.017 | 1 | 8 | Dph2 | 0.001 | Young |
| 0.895398 | 0.05941 | 0.265 | 0.279 | 1 | 8 | Mertk | -0.014 | Young |
| 0.895445 | 0.022728 | 0.46 | 0.453 | 1 | 8 | Prkd3 | 0.007 | Young |
| 0.895663 | 0.653242 | 0.018 | 0.017 | 1 | 8 | Gm36266 | 0.001 | Young |
| 0.895663 | 0.318577 | 0.018 | 0.017 | 1 | 8 | Psrc1 | 0.001 | Young |
| 0.897275 | 0.206552 | 0.052 | 0.05 | 1 | 8 | Qtrt1 | 0.002 | Young |
| 0.897584 | 0.3769 | 0.018 | 0.017 | 1 | 8 | Dlc1 | 0.001 | Young |
| 0.898054 | 0.114005 | 0.127 | 0.123 | 1 | 8 | Bola1 | 0.004 | Young |
| 0.898463 | 0.391878 | 0.081 | 0.078 | 1 | 8 | Phf11d | 0.003 | Young |
| 0.898821 | 0.054896 | 0.226 | 0.229 | 1 | 8 | Fam213b | -0.003 | Young |
| 0.899262 | 0.014452 | 0.192 | 0.184 | 1 | 8 | Arl6ip4 | 0.008 | Young |
| 0.899506 | 0.365433 | 0.018 | 0.017 | 1 | 8 | Bfsp1 | 0.001 | Young |
| 0.899692 | 0.074653 | 0.047 | 0.045 | 1 | 8 | Qtrt2 | 0.002 | Young |
| 0.900123 | 0.065622 | 0.244 | 0.251 | 1 | 8 | Mbtps1 | -0.007 | Young |
| 0.900367 | 0.049845 | 0.291 | 0.285 | 1 | 8 | Dnajc1 | 0.006 | Young |
| 0.900478 | 0.116423 | 0.096 | 0.095 | 1 | 8 | Fcgr4 | 0.001 | Young |
| 0.900615 | 0.209184 | 0.031 | 0.034 | 1 | 8 | Nubpl | -0.003 | Young |
| 0.901135 | 0.011166 | 0.883 | 0.827 | 1 | 8 | Akr1a1 | 0.056 | Young |

| | | | | | | | | |
|---|---|---|---|---|---|---|---|---|
| 0.901429 | 0.372911 | 0.018 | 0.017 | 1 | 8 | Smim10l2a | 0.001 | Young |
| 0.901429 | 0.335555 | 0.018 | 0.017 | 1 | 8 | Nckap5l | 0.001 | Young |
| 0.901429 | 0.176259 | 0.018 | 0.017 | 1 | 8 | Sult1e1 | 0.001 | Young |
| 0.901429 | 0.119064 | 0.018 | 0.017 | 1 | 8 | R3hcc1 | 0.001 | Young |
| 0.901478 | 0.398481 | 0.075 | 0.073 | 1 | 8 | Pgap1 | 0.002 | Young |
| 0.901569 | 0.012685 | 0.608 | 0.581 | 1 | 8 | Tmem256 | 0.027 | Young |
| 0.902414 | 0.123032 | 0.125 | 0.123 | 1 | 8 | Xab2 | 0.002 | Young |
| 0.90329 | 0.091472 | 0.042 | 0.039 | 1 | 8 | Trim7 | 0.003 | Young |
| 0.90329 | 0.027883 | 0.042 | 0.039 | 1 | 8 | Rnf25 | 0.003 | Young |
| 0.903352 | 0.014396 | 0.018 | 0.017 | 1 | 8 | Ska2 | 0.001 | Young |
| 0.903552 | 0.04568 | 0.07 | 0.067 | 1 | 8 | Stn1 | 0.003 | Young |
| 0.904008 | 0.109256 | 0.236 | 0.251 | 1 | 8 | Prdm2 | -0.015 | Young |
| 0.904166 | 0.47778 | 0.052 | 0.05 | 1 | 8 | Gmds | 0.002 | Young |
| 0.904184 | 0.025886 | 0.161 | 0.168 | 1 | 8 | Hdac5 | -0.007 | Young |
| 0.905276 | 0.34614 | 0.018 | 0.017 | 1 | 8 | Trp53rkb | 0.001 | Young |
| 0.905276 | 0.315767 | 0.018 | 0.017 | 1 | 8 | Uros | 0.001 | Young |
| 0.905276 | 0.250259 | 0.018 | 0.017 | 1 | 8 | Cdk5r1 | 0.001 | Young |
| 0.905276 | 0.185355 | 0.018 | 0.017 | 1 | 8 | Tubd1 | 0.001 | Young |
| 0.905276 | 0.16701 | 0.018 | 0.017 | 1 | 8 | 4933406I1[illegible] | 0.001 | Young |
| 0.906552 | 0.108355 | 0.07 | 0.067 | 1 | 8 | Ccdc71 | 0.003 | Young |
| 0.906953 | 0.211408 | 0.047 | 0.045 | 1 | 8 | Dmap1 | 0.002 | Young |
| 0.907178 | 0.573891 | 0.021 | 0.022 | 1 | 8 | Kbtbd7 | -0.001 | Young |
| 0.907178 | 0.032361 | 0.021 | 0.022 | 1 | 8 | Ncaph | -0.001 | Young |
| 0.9072 | 0.099073 | 0.018 | 0.017 | 1 | 8 | Zc3h11a | 0.001 | Young |
| 0.9072 | 0.047219 | 0.018 | 0.017 | 1 | 8 | Gm31243 | 0.001 | Young |
| 0.90748 | 0.220964 | 0.143 | 0.14 | 1 | 8 | Golga5 | 0.003 | Young |
| 0.907614 | 0.309543 | 0.052 | 0.05 | 1 | 8 | Pus7 | 0.002 | Young |
| 0.908217 | 0.169797 | 0.187 | 0.184 | 1 | 8 | Qrich1 | 0.003 | Young |
| 0.908938 | 0.245605 | 0.021 | 0.022 | 1 | 8 | Rab23 | -0.001 | Young |
| 0.908938 | 0.055483 | 0.021 | 0.022 | 1 | 8 | Gm14296 | -0.001 | Young |
| 0.909125 | 0.120249 | 0.018 | 0.017 | 1 | 8 | Adap2os | 0.001 | Young |
| 0.909125 | 0.047162 | 0.018 | 0.017 | 1 | 8 | Mppe1 | 0.001 | Young |
| 0.90925 | 0.251908 | 0.171 | 0.173 | 1 | 8 | Atg7 | -0.002 | Young |
| 0.909291 | 0.125747 | 0.091 | 0.095 | 1 | 8 | Snapc3 | -0.004 | Young |
| 0.909609 | 0.866225 | 0.026 | 0.028 | 1 | 8 | Ppp1r3d | -0.002 | Young |
| 0.909609 | 0.212829 | 0.026 | 0.028 | 1 | 8 | Tmem202 | -0.002 | Young |
| 0.909692 | 0.087156 | 0.075 | 0.073 | 1 | 8 | Med24 | 0.002 | Young |
| 0.909704 | 0.056634 | 0.114 | 0.112 | 1 | 8 | Pmf1 | 0.002 | Young |
| 0.910387 | 0.083851 | 0.138 | 0.134 | 1 | 8 | Fam8a1 | 0.004 | Young |
| 0.910692 | 0.134738 | 0.096 | 0.101 | 1 | 8 | Fam13b | -0.005 | Young |
| 0.910726 | 0.464967 | 0.031 | 0.034 | 1 | 8 | Ccl27a | -0.003 | Young |
| 0.911074 | 0.031761 | 0.351 | 0.352 | 1 | 8 | Cpsf2 | -0.001 | Young |
| 0.911631 | 0.107839 | 0.125 | 0.123 | 1 | 8 | Inf2 | 0.002 | Young |
| 0.912771 | 0.017804 | 0.221 | 0.218 | 1 | 8 | Ppie | 0.003 | Young |
| 0.912976 | 0.036336 | 0.018 | 0.017 | 1 | 8 | Gm14286 | 0.001 | Young |

| | | | | | | | | |
|---|---|---|---|---|---|---|---|---|
| 0.913411 | 0.585091 | 0.042 | 0.045 | 1 | 8 | Pcolce | -0.003 | Young |
| 0.914108 | 0.030959 | 0.109 | 0.112 | 1 | 8 | Slc9a8 | -0.003 | Young |
| 0.914131 | 0.021689 | 0.21 | 0.218 | 1 | 8 | Pold4 | -0.008 | Young |
| 0.914221 | 0.250897 | 0.047 | 0.045 | 1 | 8 | Cox15 | 0.002 | Young |
| 0.914222 | 0.643833 | 0.021 | 0.022 | 1 | 8 | Macrod2 | -0.001 | Young |
| 0.914222 | 0.345474 | 0.021 | 0.022 | 1 | 8 | Timp3 | -0.001 | Young |
| 0.914373 | 0.130617 | 0.14 | 0.145 | 1 | 8 | Phb | -0.005 | Young |
| 0.914455 | 0.033807 | 0.016 | 0.017 | 1 | 8 | Zscan22 | -0.001 | Young |
| 0.914707 | 0.184086 | 0.135 | 0.134 | 1 | 8 | Ccser2 | 0.001 | Young |
| 0.914903 | 0.006953 | 0.018 | 0.017 | 1 | 8 | Armc9 | 0.001 | Young |
| 0.914959 | 0.751966 | 0.023 | 0.022 | 1 | 8 | N4bp3 | 0.001 | Young |
| 0.914969 | 0.15924 | 0.096 | 0.101 | 1 | 8 | Pigu | -0.005 | Young |
| 0.915449 | 0.047593 | 0.135 | 0.134 | 1 | 8 | Ercc3 | 0.001 | Young |
| 0.915825 | 0.064305 | 0.096 | 0.101 | 1 | 8 | B230307C2 | -0.005 | Young |
| 0.916544 | 0.131387 | 0.104 | 0.101 | 1 | 8 | Synpo | 0.003 | Young |
| 0.916654 | 0.700403 | 0.023 | 0.022 | 1 | 8 | Dtwd1 | 0.001 | Young |
| 0.916883 | 0.102422 | 0.075 | 0.078 | 1 | 8 | Dtd2 | -0.003 | Young |
| 0.917191 | 0.022489 | 0.081 | 0.078 | 1 | 8 | Endog | 0.003 | Young |
| 0.919979 | 0.037277 | 0.213 | 0.212 | 1 | 8 | Dhcr24 | 0.001 | Young |
| 0.920235 | 0.160287 | 0.075 | 0.078 | 1 | 8 | Kctd18 | -0.003 | Young |
| 0.920544 | 0.149101 | 0.016 | 0.017 | 1 | 8 | 3110009E1 | -0.001 | Young |
| 0.921272 | 0.371165 | 0.021 | 0.022 | 1 | 8 | Optn | -0.001 | Young |
| 0.921423 | 0.173112 | 0.052 | 0.05 | 1 | 8 | Chst15 | 0.002 | Young |
| 0.922284 | 0.260428 | 0.13 | 0.128 | 1 | 8 | Gid4 | 0.002 | Young |
| 0.922575 | 0.077022 | 0.016 | 0.017 | 1 | 8 | Ccdc114 | -0.001 | Young |
| 0.923834 | 0.607675 | 0.026 | 0.028 | 1 | 8 | Cnbd2 | -0.002 | Young |
| 0.924519 | 0.066942 | 0.081 | 0.084 | 1 | 8 | Xrcc4 | -0.003 | Young |
| 0.924606 | 0.988884 | 0.016 | 0.017 | 1 | 8 | Chil3 | -0.001 | Young |
| 0.924606 | 0.502327 | 0.016 | 0.017 | 1 | 8 | Diaph3 | -0.001 | Young |
| 0.924606 | 0.20322 | 0.016 | 0.017 | 1 | 8 | Zfp202 | -0.001 | Young |
| 0.924606 | 0.144208 | 0.016 | 0.017 | 1 | 8 | Rhox8 | -0.001 | Young |
| 0.9248 | 0.230251 | 0.021 | 0.022 | 1 | 8 | Cenpe | -0.001 | Young |
| 0.926603 | 0.087265 | 0.2 | 0.201 | 1 | 8 | Ino80e | -0.001 | Young |
| 0.926638 | 0.242778 | 0.016 | 0.017 | 1 | 8 | Hhat | -0.001 | Young |
| 0.926829 | 0.609732 | 0.023 | 0.022 | 1 | 8 | Polr3a | 0.001 | Young |
| 0.926871 | 0.002973 | 0.509 | 0.497 | 1 | 8 | Leng8 | 0.012 | Young |
| 0.928329 | 0.529896 | 0.021 | 0.022 | 1 | 8 | Cmbl | -0.001 | Young |
| 0.928526 | 0.464839 | 0.023 | 0.022 | 1 | 8 | Mettl22 | 0.001 | Young |
| 0.928526 | 0.325826 | 0.023 | 0.022 | 1 | 8 | Capn10 | 0.001 | Young |
| 0.92867 | 0.585856 | 0.016 | 0.017 | 1 | 8 | Tefm | -0.001 | Young |
| 0.92867 | 0.368774 | 0.016 | 0.017 | 1 | 8 | Ankrd66 | -0.001 | Young |
| 0.92867 | 0.345784 | 0.016 | 0.017 | 1 | 8 | Nr3c2 | -0.001 | Young |
| 0.930703 | 0.606285 | 0.016 | 0.017 | 1 | 8 | Slc22a23 | -0.001 | Young |
| 0.931002 | 0.176601 | 0.156 | 0.156 | 1 | 8 | Ranbp3 | 0 | Young |
| 0.931536 | 0.024465 | 0.01 | 0.011 | 1 | 8 | Nadsyn1 | -0.001 | Young |

| | | | | | | | | |
|---|---|---|---|---|---|---|---|---|
| 0.931859 | 0.429227 | 0.021 | 0.022 | 1 | 8 | Nicn1 | -0.001 | Young |
| 0.931921 | 0.276702 | 0.023 | 0.022 | 1 | 8 | Spc25 | 0.001 | Young |
| 0.931921 | 0.198078 | 0.023 | 0.022 | 1 | 8 | Tmem220 | 0.001 | Young |
| 0.932736 | 0.066081 | 0.016 | 0.017 | 1 | 8 | Gm17259 | -0.001 | Young |
| 0.933541 | 0.48048 | 0.083 | 0.084 | 1 | 8 | Scn2b | -0.001 | Young |
| 0.933619 | 0.364736 | 0.023 | 0.022 | 1 | 8 | B630019K0 | 0.001 | Young |
| 0.934019 | 0.497985 | 0.01 | 0.011 | 1 | 8 | Dgat2 | -0.001 | Young |
| 0.934019 | 0.139304 | 0.01 | 0.011 | 1 | 8 | Zfp27 | -0.001 | Young |
| 0.934019 | 0.083905 | 0.01 | 0.011 | 1 | 8 | 9130230L2 | -0.001 | Young |
| 0.934019 | 0.007674 | 0.01 | 0.011 | 1 | 8 | Tgtp2 | -0.001 | Young |
| 0.934467 | 0.245022 | 0.112 | 0.112 | 1 | 8 | Thyn1 | 0 | Young |
| 0.935642 | 0.12227 | 0.091 | 0.095 | 1 | 8 | Clec5a | -0.004 | Young |
| 0.936504 | 0.418222 | 0.01 | 0.011 | 1 | 8 | Lgalsl | -0.001 | Young |
| 0.936504 | 0.144917 | 0.01 | 0.011 | 1 | 8 | Cd200r3 | -0.001 | Young |
| 0.936504 | 0.128801 | 0.01 | 0.011 | 1 | 8 | Gm30074 | -0.001 | Young |
| 0.936504 | 0.094161 | 0.01 | 0.011 | 1 | 8 | Sema7a | -0.001 | Young |
| 0.936504 | 0.084341 | 0.01 | 0.011 | 1 | 8 | Elavl4 | -0.001 | Young |
| 0.936504 | 0.048088 | 0.01 | 0.011 | 1 | 8 | Naprt | -0.001 | Young |
| 0.937016 | 0.110858 | 0.023 | 0.022 | 1 | 8 | 2900060B1 | 0.001 | Young |
| 0.937526 | 0.071018 | 0.148 | 0.145 | 1 | 8 | Nsmce2 | 0.003 | Young |
| 0.937642 | 0.025282 | 0.057 | 0.056 | 1 | 8 | Pwwp2b | 0.001 | Young |
| 0.938989 | 0.460184 | 0.01 | 0.011 | 1 | 8 | Ispd | -0.001 | Young |
| 0.938989 | 0.422704 | 0.01 | 0.011 | 1 | 8 | Itgb8 | -0.001 | Young |
| 0.938989 | 0.391725 | 0.01 | 0.011 | 1 | 8 | Celsr3 | -0.001 | Young |
| 0.938989 | 0.330224 | 0.01 | 0.011 | 1 | 8 | Mettl18 | -0.001 | Young |
| 0.938989 | 0.243228 | 0.01 | 0.011 | 1 | 8 | Fgfbp3 | -0.001 | Young |
| 0.938989 | 0.206998 | 0.01 | 0.011 | 1 | 8 | Cbx2 | -0.001 | Young |
| 0.938989 | 0.099397 | 0.01 | 0.011 | 1 | 8 | Pik3ip1 | -0.001 | Young |
| 0.938989 | 0.084996 | 0.01 | 0.011 | 1 | 8 | Cacna1s | -0.001 | Young |
| 0.939161 | 0.05427 | 0.091 | 0.095 | 1 | 8 | Rnf113a2 | -0.004 | Young |
| 0.940414 | 0.194795 | 0.023 | 0.022 | 1 | 8 | Trmo | 0.001 | Young |
| 0.940414 | 0.057585 | 0.023 | 0.022 | 1 | 8 | Elmod3 | 0.001 | Young |
| 0.941475 | 0.440473 | 0.01 | 0.011 | 1 | 8 | Jcad | -0.001 | Young |
| 0.941475 | 0.159684 | 0.01 | 0.011 | 1 | 8 | Smpd5 | -0.001 | Young |
| 0.941697 | 0.021508 | 0.151 | 0.151 | 1 | 8 | Zbtb22 | 0 | Young |
| 0.942113 | 0.236421 | 0.023 | 0.022 | 1 | 8 | Ttc8 | 0.001 | Young |
| 0.943676 | 0.561722 | 0.083 | 0.084 | 1 | 8 | 2010315B0 | -0.001 | Young |
| 0.943813 | 0.123599 | 0.023 | 0.022 | 1 | 8 | Gm26890 | 0.001 | Young |
| 0.943961 | 0.466353 | 0.01 | 0.011 | 1 | 8 | Anks6 | -0.001 | Young |
| 0.943961 | 0.459475 | 0.01 | 0.011 | 1 | 8 | Fmo2 | -0.001 | Young |
| 0.943961 | 0.426937 | 0.01 | 0.011 | 1 | 8 | Fcer2a | -0.001 | Young |
| 0.943961 | 0.344083 | 0.01 | 0.011 | 1 | 8 | Hist1h1d | -0.001 | Young |
| 0.943961 | 0.300973 | 0.01 | 0.011 | 1 | 8 | Fam221b | -0.001 | Young |
| 0.943961 | 0.29088 | 0.01 | 0.011 | 1 | 8 | Cldn34c1 | -0.001 | Young |
| 0.943961 | 0.273522 | 0.01 | 0.011 | 1 | 8 | Zfp651 | -0.001 | Young |

| | | | | | | | | |
|---|---|---|---|---|---|---|---|---|
| 0.943961 | 0.262953 | 0.01 | 0.011 | 1 | 8 | Cysltr2 | -0.001 | Young |
| 0.9445 | 0.092583 | 0.132 | 0.134 | 1 | 8 | Znrf1 | -0.002 | Young |
| 0.944541 | 0.039243 | 0.273 | 0.291 | 1 | 8 | Chtf8 | -0.018 | Young |
| 0.9446 | 0.373632 | 0.029 | 0.028 | 1 | 8 | Gm17745 | 0.001 | Young |
| 0.944966 | 0.197081 | 0.112 | 0.112 | 1 | 8 | Pprc1 | 0 | Young |
| 0.944986 | 0.11683 | 0.304 | 0.313 | 1 | 8 | Ldb1 | -0.009 | Young |
| 0.945774 | 0.012465 | 0.112 | 0.112 | 1 | 8 | Vamp5 | 0 | Young |
| 0.947101 | 0.303122 | 0.101 | 0.106 | 1 | 8 | Hagh | -0.005 | Young |
| 0.947673 | 0.274145 | 0.029 | 0.028 | 1 | 8 | Arhgap22 | 0.001 | Young |
| 0.947793 | 0.014151 | 0.226 | 0.218 | 1 | 8 | Dyrk1a | 0.008 | Young |
| 0.947918 | 0.05508 | 0.14 | 0.14 | 1 | 8 | Btbd9 | 0 | Young |
| 0.949076 | 0.110799 | 0.078 | 0.078 | 1 | 8 | Nemp1 | 0 | Young |
| 0.950018 | 0.104377 | 0.143 | 0.14 | 1 | 8 | Rinl | 0.003 | Young |
| 0.95015 | 0.01804 | 0.125 | 0.123 | 1 | 8 | Rrad | 0.002 | Young |
| 0.951019 | 0.113786 | 0.371 | 0.374 | 1 | 8 | Aldh3b1 | -0.003 | Young |
| 0.95161 | 0.082041 | 0.068 | 0.067 | 1 | 8 | Gstp1 | 0.001 | Young |
| 0.951951 | 0.163053 | 0.081 | 0.078 | 1 | 8 | Exosc1 | 0.003 | Young |
| 0.952006 | 0.012868 | 0.626 | 0.62 | 1 | 8 | Mia2 | 0.006 | Young |
| 0.952284 | 0.412548 | 0.029 | 0.028 | 1 | 8 | Tamm41 | 0.001 | Young |
| 0.95338 | 0.093752 | 0.135 | 0.134 | 1 | 8 | Gm5617 | 0.001 | Young |
| 0.954071 | 0.040922 | 0.171 | 0.179 | 1 | 8 | Fam208b | -0.008 | Young |
| 0.954752 | 0.013342 | 0.436 | 0.43 | 1 | 8 | Papd4 | 0.006 | Young |
| 0.954819 | 0.059222 | 0.351 | 0.352 | 1 | 8 | Txnl1 | -0.001 | Young |
| 0.955232 | 0.298699 | 0.07 | 0.073 | 1 | 8 | Paxx | -0.003 | Young |
| 0.955278 | 0.037278 | 0.057 | 0.056 | 1 | 8 | Pola1 | 0.001 | Young |
| 0.95536 | 0.289593 | 0.029 | 0.028 | 1 | 8 | Gpatch3 | 0.001 | Young |
| 0.956108 | 0.09438 | 0.174 | 0.184 | 1 | 8 | Ccpg1 | -0.01 | Young |
| 0.957767 | 0.136793 | 0.099 | 0.101 | 1 | 8 | Limk1 | -0.002 | Young |
| 0.958178 | 0.310891 | 0.062 | 0.061 | 1 | 8 | Tpgs2 | 0.001 | Young |
| 0.959082 | 0.802164 | 0.039 | 0.039 | 1 | 8 | Plxna3 | 0 | Young |
| 0.959973 | 0.233189 | 0.029 | 0.028 | 1 | 8 | Mtcp1 | 0.001 | Young |
| 0.959973 | 0.190923 | 0.029 | 0.028 | 1 | 8 | Ptar1 | 0.001 | Young |
| 0.961196 | 0.445368 | 0.07 | 0.073 | 1 | 8 | Fen1 | -0.003 | Young |
| 0.961512 | 0.112335 | 0.029 | 0.028 | 1 | 8 | Gm16845 | 0.001 | Young |
| 0.961758 | 0.104871 | 0.205 | 0.207 | 1 | 8 | Tk2 | -0.002 | Young |
| 0.962355 | 0.0465 | 0.049 | 0.05 | 1 | 8 | Isoc2b | -0.001 | Young |
| 0.962736 | 0.02345 | 0.192 | 0.196 | 1 | 8 | Apobr | -0.004 | Young |
| 0.963972 | 0.133764 | 0.083 | 0.084 | 1 | 8 | Ccdc112 | -0.001 | Young |
| 0.964179 | 0.112571 | 0.07 | 0.073 | 1 | 8 | Hrh1 | -0.003 | Young |
| 0.964584 | 0.921942 | 0.034 | 0.034 | 1 | 8 | Tbc1d8b | 0 | Young |
| 0.965999 | 0.010075 | 0.642 | 0.642 | 1 | 8 | Itpripl2 | 0 | Young |
| 0.966197 | 0.133215 | 0.078 | 0.078 | 1 | 8 | Paip2b | 0 | Young |
| 0.968316 | 0.240476 | 0.039 | 0.039 | 1 | 8 | Natd1 | 0 | Young |
| 0.969576 | 0.035263 | 0.19 | 0.19 | 1 | 8 | Pi4k2b | 0 | Young |
| 0.970347 | 0.033765 | 0.062 | 0.061 | 1 | 8 | Map4k5 | 0.001 | Young |

| | | | | | | | | |
|---|---|---|---|---|---|---|---|---|
| 0.971411 | 0.134291 | 0.099 | 0.101 | 1 | 8 | Hectd3 | -0.002 | Young |
| 0.971657 | 0.009841 | 0.112 | 0.112 | 1 | 8 | Mrpl19 | 0 | Young |
| 0.972464 | 0.088431 | 0.062 | 0.061 | 1 | 8 | Ccdc28a | 0.001 | Young |
| 0.973805 | 0.107007 | 0.348 | 0.363 | 1 | 8 | Banf1 | -0.015 | Young |
| 0.974496 | 0.417304 | 0.034 | 0.034 | 1 | 8 | L3mbtl2 | 0 | Young |
| 0.974604 | 0.1962 | 0.203 | 0.212 | 1 | 8 | Ncbp2 | -0.009 | Young |
| 0.974763 | 0.002347 | 0.078 | 0.078 | 1 | 8 | Gss | 0 | Young |
| 0.97484 | 0.149232 | 0.114 | 0.117 | 1 | 8 | Dhrs4 | -0.003 | Young |
| 0.97737 | 0.06356 | 0.073 | 0.073 | 1 | 8 | Capn15 | 0 | Young |
| 0.979736 | 0.166426 | 0.132 | 0.134 | 1 | 8 | Jade2 | -0.002 | Young |
| 0.979815 | 0.16844 | 0.055 | 0.056 | 1 | 8 | Donson | -0.001 | Young |
| 0.980796 | 0.246108 | 0.099 | 0.101 | 1 | 8 | Gipc1 | -0.002 | Young |
| 0.981172 | 0.104886 | 0.049 | 0.05 | 1 | 8 | Rfx5 | -0.001 | Young |
| 0.981259 | 0.011091 | 0.462 | 0.48 | 1 | 8 | Mrpl23 | -0.018 | Young |
| 0.982996 | 0.263263 | 0.034 | 0.034 | 1 | 8 | Zfp995 | 0 | Young |
| 0.983557 | 0.009749 | 0.268 | 0.274 | 1 | 8 | Ilkap | -0.006 | Young |
| 0.98386 | 0.067305 | 0.125 | 0.128 | 1 | 8 | Caap1 | -0.003 | Young |
| 0.985717 | 0.001439 | 0.117 | 0.117 | 1 | 8 | Mpv17l2 | 0 | Young |
| 0.98583 | 0.133888 | 0.034 | 0.034 | 1 | 8 | Szt2 | 0 | Young |
| 0.986231 | 0.051214 | 0.112 | 0.112 | 1 | 8 | Agfg2 | 0 | Young |
| 0.987246 | 0.164994 | 0.034 | 0.034 | 1 | 8 | Zfp934 | 0 | Young |
| 0.98758 | 0.318845 | 0.044 | 0.045 | 1 | 8 | Ank2 | -0.001 | Young |
| 0.98758 | 0.175922 | 0.044 | 0.045 | 1 | 8 | Gcfc2 | -0.001 | Young |
| 0.987727 | 0.013483 | 0.156 | 0.156 | 1 | 8 | Sin3a | 0 | Young |
| 0.988663 | 0.138631 | 0.034 | 0.034 | 1 | 8 | Wdhd1 | 0 | Young |
| 0.989431 | 0.303044 | 0.13 | 0.134 | 1 | 8 | Tprkb | -0.004 | Young |
| 0.990833 | 0.17439 | 0.127 | 0.128 | 1 | 8 | Mrs2 | -0.001 | Young |
| 0.992288 | 0.020902 | 0.213 | 0.218 | 1 | 8 | Bcap29 | -0.005 | Young |
| 0.992928 | 0.061838 | 0.27 | 0.274 | 1 | 8 | Slc30a5 | -0.004 | Young |
| 0.993262 | 0.09846 | 0.088 | 0.089 | 1 | 8 | Naglu | -0.001 | Young |
| 0.994394 | 0.027697 | 0.068 | 0.067 | 1 | 8 | L3mbtl3 | 0.001 | Young |
| 0.996658 | 0.038504 | 0.8 | 0.715 | 1 | 8 | Rap1a | 0.085 | Young |
| 0.996856 | 0.098272 | 0.088 | 0.089 | 1 | 8 | Rbl1 | -0.001 | Young |
| 0.997048 | 0.312154 | 0.073 | 0.073 | 1 | 8 | Tom1l2 | 0 | Young |
| 0.997378 | 0.015768 | 0.249 | 0.251 | 1 | 8 | Snx1 | -0.002 | Young |
| 1.03E-05 | 1.465439 | 0.379 | 0.191 | 0.31912 | 9 | Pcdh1 | 0.188 | Old |
| 4.21E-05 | 0.549171 | 0.768 | 0.574 | 1 | 9 | Clic4 | 0.194 | Old |
| 6.63E-05 | 0.855906 | 0.603 | 0.426 | 1 | 9 | Plcb4 | 0.177 | Old |
| 7.37E-05 | 0.610926 | 0.76 | 0.64 | 1 | 9 | Ly6a | 0.12 | Old |
| 0.000275 | 2.432023 | 0.134 | 0.022 | 1 | 9 | Slc7a1 | 0.112 | Old |
| 0.000389 | 0.234377 | 0.997 | 1 | 1 | 9 | Tmsb4x | -0.003 | Old |
| 0.000442 | 2.348732 | 0.149 | 0.037 | 1 | 9 | Scaf8 | 0.112 | Old |
| 0.000512 | 1.0603 | 0.273 | 0.125 | 1 | 9 | Fhod1 | 0.148 | Old |
| 0.000531 | 0.772059 | 0.515 | 0.338 | 1 | 9 | Ctla2a | 0.177 | Old |
| 0.000645 | 0.758632 | 0.173 | 0.051 | 1 | 9 | Ubd | 0.122 | Old |

| | | | | | | | | |
|---|---|---|---|---|---|---|---|---|
| 0.000689 | 1.004482 | 0.353 | 0.199 | 1 | 9 | Bcl6b | 0.154 | Old |
| 0.000699 | 0.366785 | 0.946 | 0.875 | 1 | 9 | Serinc3 | 0.071 | Old |
| 0.000758 | 0.792529 | 0.43 | 0.257 | 1 | 9 | H2-Q7 | 0.173 | Old |
| 0.001173 | 0.99787 | 0.384 | 0.243 | 1 | 9 | Ankrd12 | 0.141 | Old |
| 0.001258 | 0.882988 | 0.335 | 0.199 | 1 | 9 | Sbds | 0.136 | Old |
| 0.001382 | 0.502403 | 0.508 | 0.368 | 1 | 9 | Ywhaq | 0.14 | Old |
| 0.001522 | 0.777956 | 0.343 | 0.191 | 1 | 9 | Ptpn12 | 0.152 | Old |
| 0.002028 | 0.657905 | 0.567 | 0.434 | 1 | 9 | S100a13 | 0.133 | Old |
| 0.002186 | 1.295475 | 0.186 | 0.074 | 1 | 9 | Scd2 | 0.112 | Old |
| 0.002227 | 0.439267 | 0.639 | 0.5 | 1 | 9 | Tjp1 | 0.139 | Old |
| 0.002338 | 1.806367 | 0.149 | 0.051 | 1 | 9 | Plat | 0.098 | Old |
| 0.002375 | 1.670296 | 0.191 | 0.081 | 1 | 9 | Rab8a | 0.11 | Old |
| 0.002494 | 1.5664 | 0.18 | 0.074 | 1 | 9 | Kat6b | 0.106 | Old |
| 0.002598 | 0.453368 | 0.696 | 0.588 | 1 | 9 | Cd81 | 0.108 | Old |
| 0.003317 | 0.76567 | 0.204 | 0.088 | 1 | 9 | Vamp5 | 0.116 | Old |
| 0.003431 | 1.09137 | 0.276 | 0.154 | 1 | 9 | Rab21 | 0.122 | Old |
| 0.003436 | 0.926638 | 0.325 | 0.191 | 1 | 9 | Afap1l1 | 0.134 | Old |
| 0.003452 | 3.763926 | 0.075 | 0.007 | 1 | 9 | Sft2d3 | 0.068 | Old |
| 0.003523 | 2.82837 | 0.111 | 0.029 | 1 | 9 | Nsmaf | 0.082 | Old |
| 0.003542 | 0.98014 | 0.263 | 0.14 | 1 | 9 | Fam53b | 0.123 | Old |
| 0.003595 | 0.51049 | 0.472 | 0.324 | 1 | 9 | Rabac1 | 0.148 | Old |
| 0.003611 | 0.284891 | 0.866 | 0.794 | 1 | 9 | Epas1 | 0.072 | Old |
| 0.003742 | 4.586813 | 0.059 | 0 | 1 | 9 | Tdrd3 | 0.059 | Old |
| 0.00376 | 0.630392 | 0.263 | 0.14 | 1 | 9 | Psmb6 | 0.123 | Old |
| 0.004044 | 2.287472 | 0.121 | 0.037 | 1 | 9 | Fam192a | 0.084 | Old |
| 0.004502 | 1.447788 | 0.206 | 0.103 | 1 | 9 | Dync1li2 | 0.103 | Old |
| 0.004531 | 1.581226 | 0.165 | 0.066 | 1 | 9 | Pbld1 | 0.099 | Old |
| 0.004751 | 1.535119 | 0.206 | 0.103 | 1 | 9 | Psenen | 0.103 | Old |
| 0.004757 | 0.403878 | 0.688 | 0.566 | 1 | 9 | Morf4l1 | 0.122 | Old |
| 0.004858 | 0.764347 | 0.332 | 0.206 | 1 | 9 | Arhgef1 | 0.126 | Old |
| 0.004877 | 0.907175 | 0.387 | 0.25 | 1 | 9 | Ptp4a3 | 0.137 | Old |
| 0.004887 | 1.163853 | 0.111 | 0.029 | 1 | 9 | Hoxb3os | 0.082 | Old |
| 0.004957 | 1.784383 | 0.108 | 0.029 | 1 | 9 | Ddt | 0.079 | Old |
| 0.005172 | 0.581574 | 0.613 | 0.515 | 1 | 9 | Hnrnpa0 | 0.098 | Old |
| 0.005245 | 1.372121 | 0.131 | 0.044 | 1 | 9 | Vwa1 | 0.087 | Old |
| 0.005254 | 0.424375 | 0.732 | 0.625 | 1 | 9 | Egfl7 | 0.107 | Old |
| 0.005379 | 1.085832 | 0.162 | 0.066 | 1 | 9 | Hcfc1 | 0.096 | Old |
| 0.005696 | 4.459782 | 0.054 | 0 | 1 | 9 | Kcnj15 | 0.054 | Old |
| 0.005696 | 4.357141 | 0.054 | 0 | 1 | 9 | Dcps | 0.054 | Old |
| 0.005894 | 0.798161 | 0.356 | 0.228 | 1 | 9 | Adarb1 | 0.128 | Old |
| 0.005924 | 0.376189 | 0.552 | 0.419 | 1 | 9 | Palmd | 0.133 | Old |
| 0.005935 | 1.314542 | 0.152 | 0.059 | 1 | 9 | Pcif1 | 0.093 | Old |
| 0.005979 | 0.462094 | 0.34 | 0.206 | 1 | 9 | Tecpr1 | 0.134 | Old |
| 0.006231 | 1.292875 | 0.178 | 0.081 | 1 | 9 | Hoxa7 | 0.097 | Old |
| 0.006338 | 1.815454 | 0.147 | 0.059 | 1 | 9 | Pde8a | 0.088 | Old |

| | | | | | | | | |
|---|---|---|---|---|---|---|---|---|
| 0.006546 | 0.759212 | 0.472 | 0.368 | 1 | 9 | Rgl1 | 0.104 | Old |
| 0.00658 | 0.619388 | 0.201 | 0.096 | 1 | 9 | Raf1 | 0.105 | Old |
| 0.006586 | 3.698833 | 0.067 | 0.007 | 1 | 9 | Fam210b | 0.06 | Old |
| 0.006637 | 1.120215 | 0.214 | 0.11 | 1 | 9 | Ryk | 0.104 | Old |
| 0.006851 | 1.545925 | 0.106 | 0.029 | 1 | 9 | Vps53 | 0.077 | Old |
| 0.007009 | 0.567163 | 0.456 | 0.331 | 1 | 9 | H2-T23 | 0.125 | Old |
| 0.007026 | 2.571504 | 0.08 | 0.015 | 1 | 9 | Pth1r | 0.065 | Old |
| 0.007068 | 0.477323 | 0.668 | 0.551 | 1 | 9 | Sult1a1 | 0.117 | Old |
| 0.007127 | 1.984836 | 0.093 | 0.022 | 1 | 9 | Atoh8 | 0.071 | Old |
| 0.007322 | 1.023315 | 0.196 | 0.096 | 1 | 9 | Rnf6 | 0.1 | Old |
| 0.007432 | 1.032067 | 0.271 | 0.162 | 1 | 9 | Stx12 | 0.109 | Old |
| 0.007555 | 0.21411 | 0.807 | 0.779 | 1 | 9 | Zbtb20 | 0.028 | Old |
| 0.0076 | 0.815537 | 0.276 | 0.162 | 1 | 9 | Zbtb38 | 0.114 | Old |
| 0.00767 | 1.792261 | 0.124 | 0.044 | 1 | 9 | Zfp24 | 0.08 | Old |
| 0.007766 | 0.684338 | 0.361 | 0.235 | 1 | 9 | Rhoc | 0.126 | Old |
| 0.008072 | 0.685799 | 0.273 | 0.162 | 1 | 9 | Rasa1 | 0.111 | Old |
| 0.008157 | 0.703989 | 0.412 | 0.294 | 1 | 9 | Cast | 0.118 | Old |
| 0.008173 | 1.04763 | 0.25 | 0.147 | 1 | 9 | Grap | 0.103 | Old |
| 0.008484 | 2.611679 | 0.101 | 0.029 | 1 | 9 | Hoxd9 | 0.072 | Old |
| 0.008639 | 2.042949 | 0.09 | 0.022 | 1 | 9 | Bloc1s6 | 0.068 | Old |
| 0.008667 | 0.769492 | 0.304 | 0.199 | 1 | 9 | Chmp5 | 0.105 | Old |
| 0.008674 | 3.81316 | 0.049 | 0 | 1 | 9 | Csprs | 0.049 | Old |
| 0.0087 | 2.638262 | 0.064 | 0.007 | 1 | 9 | Medag | 0.057 | Old |
| 0.008751 | 1.561662 | 0.09 | 0.022 | 1 | 9 | Fktn | 0.068 | Old |
| 0.008869 | 0.453828 | 0.74 | 0.632 | 1 | 9 | Tm4sf1 | 0.108 | Old |
| 0.008989 | 0.218338 | 0.928 | 0.912 | 1 | 9 | Cav1 | 0.016 | Old |
| 0.009015 | 0.546094 | 0.278 | 0.169 | 1 | 9 | Cdc42se2 | 0.109 | Old |
| 0.009298 | 0.437772 | 0.49 | 0.36 | 1 | 9 | Rpl35 | 0.13 | Old |
| 0.009349 | 0.571332 | 0.49 | 0.368 | 1 | 9 | Tmod3 | 0.122 | Old |
| 0.009359 | 1.563867 | 0.204 | 0.11 | 1 | 9 | Ilk | 0.094 | Old |
| 0.009688 | 0.576678 | 0.448 | 0.316 | 1 | 9 | Nav1 | 0.132 | Old |
| 0.009747 | 1.540881 | 0.131 | 0.051 | 1 | 9 | Epha2 | 0.08 | Old |
| 0.009762 | 0.484632 | 0.247 | 0.14 | 1 | 9 | Dido1 | 0.107 | Old |
| 0.010032 | 0.24662 | 0.941 | 0.919 | 1 | 9 | Timp3 | 0.022 | Old |
| 0.010246 | 1.02949 | 0.271 | 0.162 | 1 | 9 | Zc3h13 | 0.109 | Old |
| 0.010592 | 2.19128 | 0.098 | 0.029 | 1 | 9 | Snf8 | 0.069 | Old |
| 0.010664 | 0.866262 | 0.312 | 0.206 | 1 | 9 | Gimap6 | 0.106 | Old |
| 0.010809 | 1.085409 | 0.075 | 0.015 | 1 | 9 | Immp1l | 0.06 | Old |
| 0.01099 | 1.014618 | 0.247 | 0.147 | 1 | 9 | 5031425E2 | 0.1 | Old |
| 0.011105 | 1.187767 | 0.088 | 0.022 | 1 | 9 | Riok2 | 0.066 | Old |
| 0.011301 | 0.634869 | 0.268 | 0.154 | 1 | 9 | Setd2 | 0.114 | Old |
| 0.011356 | 1.332615 | 0.119 | 0.044 | 1 | 9 | Pnpla6 | 0.075 | Old |
| 0.011901 | 0.909081 | 0.374 | 0.265 | 1 | 9 | Sema3g | 0.109 | Old |
| 0.012008 | 0.905459 | 0.291 | 0.184 | 1 | 9 | Emc7 | 0.107 | Old |
| 0.012095 | 0.484032 | 0.389 | 0.265 | 1 | 9 | Yes1 | 0.124 | Old |

| | | | | | | | | |
|---|---|---|---|---|---|---|---|---|
| 0.012243 | 3.397958 | 0.072 | 0.015 | 1 | 9 | Rpap3 | 0.057 | Old |
| 0.012394 | 0.271395 | 0.588 | 0.456 | 1 | 9 | Efnb1 | 0.132 | Old |
| 0.012557 | 1.796924 | 0.085 | 0.022 | 1 | 9 | Fam114a1 | 0.063 | Old |
| 0.012601 | 0.692404 | 0.338 | 0.228 | 1 | 9 | Abca1 | 0.11 | Old |
| 0.012703 | 2.698767 | 0.072 | 0.015 | 1 | 9 | 9530026P0 | 0.057 | Old |
| 0.012761 | 2.509048 | 0.072 | 0.015 | 1 | 9 | Tnfrsf25 | 0.057 | Old |
| 0.012827 | 0.695152 | 0.345 | 0.235 | 1 | 9 | Sp1 | 0.11 | Old |
| 0.012876 | 0.44248 | 0.451 | 0.324 | 1 | 9 | S100a10 | 0.127 | Old |
| 0.013178 | 1.676915 | 0.072 | 0.015 | 1 | 9 | Brip1 | 0.057 | Old |
| 0.013221 | 4.214196 | 0.044 | 0 | 1 | 9 | Dhx33 | 0.044 | Old |
| 0.013231 | 1.775121 | 0.144 | 0.066 | 1 | 9 | Tnks1bp1 | 0.078 | Old |
| 0.013285 | 1.203513 | 0.201 | 0.11 | 1 | 9 | Taf7 | 0.091 | Old |
| 0.013295 | 1.231182 | 0.108 | 0.037 | 1 | 9 | Prelp | 0.071 | Old |
| 0.013348 | 0.146548 | 0.961 | 0.993 | 1 | 9 | mt-Nd2 | -0.032 | Old |
| 0.013446 | 0.977255 | 0.224 | 0.132 | 1 | 9 | Grina | 0.092 | Old |
| 0.013471 | 1.913117 | 0.059 | 0.007 | 1 | 9 | Rbm48 | 0.052 | Old |
| 0.013639 | 0.570617 | 0.41 | 0.301 | 1 | 9 | Pxdn | 0.109 | Old |
| 0.013883 | 0.796143 | 0.351 | 0.243 | 1 | 9 | Zbtb4 | 0.108 | Old |
| 0.014137 | 2.670574 | 0.082 | 0.022 | 1 | 9 | Lias | 0.06 | Old |
| 0.014384 | 0.594836 | 0.425 | 0.294 | 1 | 9 | Rasgrf2 | 0.131 | Old |
| 0.014491 | 1.383471 | 0.173 | 0.088 | 1 | 9 | Npdc1 | 0.085 | Old |
| 0.014539 | 1.312182 | 0.116 | 0.044 | 1 | 9 | Ebna1bp2 | 0.072 | Old |
| 0.01458 | 0.112831 | 0.224 | 0.118 | 1 | 9 | Shroom4 | 0.106 | Old |
| 0.014588 | 3.56686 | 0.07 | 0.015 | 1 | 9 | Trp53bp1 | 0.055 | Old |
| 0.014685 | 2.322814 | 0.082 | 0.022 | 1 | 9 | Oas2 | 0.06 | Old |
| 0.015119 | 0.411104 | 0.399 | 0.279 | 1 | 9 | Thra | 0.12 | Old |
| 0.015177 | 3.531522 | 0.057 | 0.007 | 1 | 9 | Tshz3 | 0.05 | Old |
| 0.015318 | 0.585409 | 0.448 | 0.338 | 1 | 9 | Cbfa2t3 | 0.11 | Old |
| 0.015658 | 1.129602 | 0.116 | 0.044 | 1 | 9 | Ppm1g | 0.072 | Old |
| 0.015739 | 2.164359 | 0.121 | 0.051 | 1 | 9 | Irak2 | 0.07 | Old |
| 0.016022 | 0.939313 | 0.165 | 0.081 | 1 | 9 | Myo1d | 0.084 | Old |
| 0.016592 | 0.679115 | 0.32 | 0.213 | 1 | 9 | Arpp19 | 0.107 | Old |
| 0.016831 | 0.615667 | 0.247 | 0.147 | 1 | 9 | Clec1a | 0.1 | Old |
| 0.016837 | 1.50237 | 0.168 | 0.088 | 1 | 9 | Plxna4 | 0.08 | Old |
| 0.016838 | 0.265564 | 0.907 | 0.853 | 1 | 9 | Cldn5 | 0.054 | Old |
| 0.017014 | 0.38817 | 0.771 | 0.691 | 1 | 9 | H3f3b | 0.08 | Old |
| 0.017192 | 0.60967 | 0.33 | 0.228 | 1 | 9 | Syncrip | 0.102 | Old |
| 0.017384 | 2.846526 | 0.08 | 0.022 | 1 | 9 | AA467197 | 0.058 | Old |
| 0.017682 | 0.373437 | 0.686 | 0.588 | 1 | 9 | Ppp3ca | 0.098 | Old |
| 0.01791 | 1.456512 | 0.149 | 0.074 | 1 | 9 | Kansl3 | 0.075 | Old |
| 0.017962 | 0.478034 | 0.332 | 0.221 | 1 | 9 | Gapdh | 0.111 | Old |
| 0.01799 | 0.385265 | 0.559 | 0.456 | 1 | 9 | Atp5j | 0.103 | Old |
| 0.018016 | 0.887399 | 0.209 | 0.118 | 1 | 9 | Wsb2 | 0.091 | Old |
| 0.01812 | 0.381506 | 0.332 | 0.221 | 1 | 9 | Pcnp | 0.111 | Old |
| 0.018219 | 0.474777 | 0.593 | 0.471 | 1 | 9 | Ubb | 0.122 | Old |

| | | | | | | | | |
|---|---|---|---|---|---|---|---|---|
| 0.018461 | 3.626081 | 0.054 | 0.007 | 1 | 9 | Qpctl | 0.047 | Old |
| 0.018642 | 2.109548 | 0.111 | 0.044 | 1 | 9 | Ccdc117 | 0.067 | Old |
| 0.018859 | 1.37649 | 0.209 | 0.125 | 1 | 9 | Tes | 0.084 | Old |
| 0.01954 | 1.860739 | 0.09 | 0.029 | 1 | 9 | Hgsnat | 0.061 | Old |
| 0.019643 | 1.390933 | 0.149 | 0.074 | 1 | 9 | Zhx3 | 0.075 | Old |
| 0.019797 | 0.461659 | 0.521 | 0.404 | 1 | 9 | Vcp | 0.117 | Old |
| 0.019921 | 2.007756 | 0.09 | 0.029 | 1 | 9 | Rnf168 | 0.061 | Old |
| 0.020012 | 0.954106 | 0.278 | 0.191 | 1 | 9 | Nectin2 | 0.087 | Old |
| 0.020097 | 1.226168 | 0.067 | 0.015 | 1 | 9 | Vars | 0.052 | Old |
| 0.020188 | 3.611452 | 0.039 | 0 | 1 | 9 | Trap1 | 0.039 | Old |
| 0.02043 | 0.980715 | 0.054 | 0.007 | 1 | 9 | Ntn1 | 0.047 | Old |
| 0.020629 | 0.513329 | 0.322 | 0.213 | 1 | 9 | Srsf11 | 0.109 | Old |
| 0.020761 | 0.543562 | 0.343 | 0.243 | 1 | 9 | Lgals9 | 0.1 | Old |
| 0.020954 | 0.78166 | 0.24 | 0.147 | 1 | 9 | Prdm2 | 0.093 | Old |
| 0.021103 | 1.646551 | 0.09 | 0.029 | 1 | 9 | Spns1 | 0.061 | Old |
| 0.021504 | 1.231063 | 0.201 | 0.118 | 1 | 9 | Slc48a1 | 0.083 | Old |
| 0.021748 | 0.862458 | 0.144 | 0.066 | 1 | 9 | Csgalnact1 | 0.078 | Old |
| 0.022114 | 0.11894 | 0.173 | 0.088 | 1 | 9 | Hoxd8 | 0.085 | Old |
| 0.022279 | 0.493038 | 0.557 | 0.441 | 1 | 9 | Mef2c | 0.116 | Old |
| 0.022567 | 0.954911 | 0.196 | 0.11 | 1 | 9 | Trrap | 0.086 | Old |
| 0.022689 | 0.817113 | 0.175 | 0.096 | 1 | 9 | Phf1 | 0.079 | Old |
| 0.022774 | 1.03813 | 0.09 | 0.029 | 1 | 9 | Gimap9 | 0.061 | Old |
| 0.022781 | 0.362849 | 0.572 | 0.441 | 1 | 9 | Pdgfb | 0.131 | Old |
| 0.022784 | 0.698977 | 0.111 | 0.044 | 1 | 9 | Ikbip | 0.067 | Old |
| 0.022787 | 0.239503 | 0.655 | 0.574 | 1 | 9 | Fosb | 0.081 | Old |
| 0.022881 | 0.21884 | 0.918 | 0.846 | 1 | 9 | Cdh5 | 0.072 | Old |
| 0.022899 | 0.90222 | 0.077 | 0.022 | 1 | 9 | Lin54 | 0.055 | Old |
| 0.023121 | 0.816416 | 0.09 | 0.029 | 1 | 9 | Gramd1b | 0.061 | Old |
| 0.023145 | 1.418958 | 0.119 | 0.051 | 1 | 9 | Man1c1 | 0.068 | Old |
| 0.023364 | 1.574227 | 0.155 | 0.081 | 1 | 9 | Samd12 | 0.074 | Old |
| 0.023659 | 1.531619 | 0.077 | 0.022 | 1 | 9 | Aph1b | 0.055 | Old |
| 0.023747 | 2.682559 | 0.052 | 0.007 | 1 | 9 | Bola1 | 0.045 | Old |
| 0.024092 | 1.218888 | 0.147 | 0.074 | 1 | 9 | Zfp131 | 0.073 | Old |
| 0.024116 | 0.809887 | 0.232 | 0.147 | 1 | 9 | Ubxn2a | 0.085 | Old |
| 0.024157 | 0.711627 | 0.34 | 0.235 | 1 | 9 | Gm26532 | 0.105 | Old |
| 0.024308 | 1.325444 | 0.191 | 0.11 | 1 | 9 | Chp1 | 0.081 | Old |
| 0.024903 | 0.808641 | 0.064 | 0.015 | 1 | 9 | Dym | 0.049 | Old |
| 0.024972 | 4.303722 | 0.036 | 0 | 1 | 9 | Nxpe2 | 0.036 | Old |
| 0.024972 | 3.997338 | 0.036 | 0 | 1 | 9 | Aplp1 | 0.036 | Old |
| 0.024972 | 3.721637 | 0.036 | 0 | 1 | 9 | Tnfsfm13 | 0.036 | Old |
| 0.024972 | 3.658282 | 0.036 | 0 | 1 | 9 | Nxt2 | 0.036 | Old |
| 0.024972 | 3.5718 | 0.036 | 0 | 1 | 9 | Zfp959 | 0.036 | Old |
| 0.025039 | 0.374217 | 0.399 | 0.287 | 1 | 9 | Fmo1 | 0.112 | Old |
| 0.025164 | 0.897358 | 0.139 | 0.066 | 1 | 9 | Carhsp1 | 0.073 | Old |
| 0.025217 | 1.786798 | 0.098 | 0.037 | 1 | 9 | Rbl2 | 0.061 | Old |

| | | | | | | | | |
|---|---|---|---|---|---|---|---|---|
| 0.025313 | 1.882562 | 0.116 | 0.051 | 1 | 9 | Ilkap | 0.065 | Old |
| 0.025494 | 0.26141 | 0.822 | 0.684 | 1 | 9 | Ddx3x | 0.138 | Old |
| 0.025623 | 0.275535 | 0.647 | 0.493 | 1 | 9 | Hnrnpk | 0.154 | Old |
| 0.025641 | 0.418979 | 0.265 | 0.162 | 1 | 9 | Dusp6 | 0.103 | Old |
| 0.025943 | 0.496983 | 0.353 | 0.25 | 1 | 9 | Hip1 | 0.103 | Old |
| 0.02605 | 0.761909 | 0.302 | 0.206 | 1 | 9 | Elob | 0.096 | Old |
| 0.026174 | 0.607899 | 0.188 | 0.103 | 1 | 9 | Eif1a | 0.085 | Old |
| 0.026227 | 0.713897 | 0.361 | 0.265 | 1 | 9 | Rpl22l1 | 0.096 | Old |
| 0.026288 | 0.387377 | 0.322 | 0.221 | 1 | 9 | Auts2 | 0.101 | Old |
| 0.027109 | 0.260021 | 0.698 | 0.618 | 1 | 9 | Rps16 | 0.08 | Old |
| 0.027461 | 1.650015 | 0.106 | 0.044 | 1 | 9 | Plekhm2 | 0.062 | Old |
| 0.027675 | 0.861354 | 0.17 | 0.088 | 1 | 9 | Gpr182 | 0.082 | Old |
| 0.027702 | 1.981032 | 0.085 | 0.029 | 1 | 9 | Mmp11 | 0.056 | Old |
| 0.027898 | 0.753556 | 0.255 | 0.162 | 1 | 9 | Notch4 | 0.093 | Old |
| 0.028276 | 0.396406 | 0.485 | 0.404 | 1 | 9 | Cnbp | 0.081 | Old |
| 0.028575 | 0.559596 | 0.245 | 0.154 | 1 | 9 | Brd8 | 0.091 | Old |
| 0.028628 | 2.975378 | 0.093 | 0.037 | 1 | 9 | Chrac1 | 0.056 | Old |
| 0.02887 | 1.626872 | 0.175 | 0.103 | 1 | 9 | Zcrb1 | 0.072 | Old |
| 0.028878 | 0.361155 | 0.598 | 0.493 | 1 | 9 | Myl12a | 0.105 | Old |
| 0.029015 | 0.280559 | 0.693 | 0.596 | 1 | 9 | Fli1 | 0.097 | Old |
| 0.029025 | 2.727778 | 0.082 | 0.029 | 1 | 9 | Cul9 | 0.053 | Old |
| 0.029104 | 0.680121 | 0.075 | 0.022 | 1 | 9 | Nr1d2 | 0.053 | Old |
| 0.029146 | 1.077348 | 0.219 | 0.14 | 1 | 9 | Unc5b | 0.079 | Old |
| 0.029336 | 1.363181 | 0.186 | 0.11 | 1 | 9 | Jag2 | 0.076 | Old |
| 0.029772 | 0.721261 | 0.446 | 0.368 | 1 | 9 | Nus1 | 0.078 | Old |
| 0.029951 | 0.993415 | 0.191 | 0.11 | 1 | 9 | Fuca1 | 0.081 | Old |
| 0.030039 | 0.640078 | 0.162 | 0.088 | 1 | 9 | Mfsd14a | 0.074 | Old |
| 0.030177 | 0.612483 | 0.26 | 0.176 | 1 | 9 | Adcy4 | 0.084 | Old |
| 0.030228 | 2.437379 | 0.049 | 0.007 | 1 | 9 | Zfp768 | 0.042 | Old |
| 0.030239 | 0.384299 | 0.487 | 0.368 | 1 | 9 | Tomm7 | 0.119 | Old |
| 0.030431 | 0.509492 | 0.381 | 0.287 | 1 | 9 | Fmo2 | 0.094 | Old |
| 0.030532 | 1.69121 | 0.049 | 0.007 | 1 | 9 | Strada | 0.042 | Old |
| 0.030685 | 1.653603 | 0.049 | 0.007 | 1 | 9 | Zfp729b | 0.042 | Old |
| 0.0307 | 0.953131 | 0.147 | 0.074 | 1 | 9 | Arhgap1 | 0.073 | Old |
| 0.030762 | 0.762081 | 0.24 | 0.154 | 1 | 9 | Ifi27 | 0.086 | Old |
| 0.030838 | 1.398271 | 0.049 | 0.007 | 1 | 9 | Phospho2 | 0.042 | Old |
| 0.030838 | 1.132908 | 0.049 | 0.007 | 1 | 9 | Slc39a13 | 0.042 | Old |
| 0.030919 | 4.328889 | 0.034 | 0 | 1 | 9 | Osgin1 | 0.034 | Old |
| 0.030919 | 4.286452 | 0.034 | 0 | 1 | 9 | Mapt | 0.034 | Old |
| 0.030919 | 3.940344 | 0.034 | 0 | 1 | 9 | Clec16a | 0.034 | Old |
| 0.030919 | 3.73388 | 0.034 | 0 | 1 | 9 | Zbtb16 | 0.034 | Old |
| 0.030919 | 3.084353 | 0.034 | 0 | 1 | 9 | Iqsec2 | 0.034 | Old |
| 0.031059 | 0.201146 | 0.544 | 0.412 | 1 | 9 | Mapre1 | 0.132 | Old |
| 0.031074 | 0.225763 | 0.567 | 0.456 | 1 | 9 | Bst2 | 0.111 | Old |
| 0.031144 | 0.557695 | 0.477 | 0.375 | 1 | 9 | Adam15 | 0.102 | Old |

| | | | | | | | | |
|---|---|---|---|---|---|---|---|---|
| 0.03129 | 1.093883 | 0.162 | 0.088 | 1 | 9 | Cyth3 | 0.074 | Old |
| 0.031411 | 2.047836 | 0.093 | 0.037 | 1 | 9 | Thumpd1 | 0.056 | Old |
| 0.031543 | 0.673201 | 0.253 | 0.162 | 1 | 9 | Csnk2a2 | 0.091 | Old |
| 0.031571 | 0.921989 | 0.095 | 0.037 | 1 | 9 | Tnip1 | 0.058 | Old |
| 0.031576 | 1.282045 | 0.232 | 0.154 | 1 | 9 | Gtpbp2 | 0.078 | Old |
| 0.031744 | 0.149702 | 0.928 | 0.868 | 1 | 9 | Aplp2 | 0.06 | Old |
| 0.031819 | 0.512028 | 0.534 | 0.412 | 1 | 9 | Prdx1 | 0.122 | Old |
| 0.031964 | 0.17023 | 0.925 | 0.846 | 1 | 9 | Adgrf5 | 0.079 | Old |
| 0.032021 | 0.264582 | 0.722 | 0.588 | 1 | 9 | Wnk1 | 0.134 | Old |
| 0.032604 | 0.799495 | 0.335 | 0.25 | 1 | 9 | Man2a1 | 0.085 | Old |
| 0.032631 | 0.449804 | 0.572 | 0.463 | 1 | 9 | Sgms1 | 0.109 | Old |
| 0.032796 | 1.022064 | 0.18 | 0.103 | 1 | 9 | Cpsf7 | 0.077 | Old |
| 0.03322 | 0.707175 | 0.466 | 0.39 | 1 | 9 | Fabp5 | 0.076 | Old |
| 0.033399 | 0.177767 | 0.853 | 0.801 | 1 | 9 | Ly6c1 | 0.052 | Old |
| 0.033495 | 0.327272 | 0.536 | 0.426 | 1 | 9 | Lamp1 | 0.11 | Old |
| 0.033513 | 0.427319 | 0.34 | 0.243 | 1 | 9 | Cald1 | 0.097 | Old |
| 0.033518 | 0.77565 | 0.286 | 0.199 | 1 | 9 | Tnrc6a | 0.087 | Old |
| 0.033713 | 0.618419 | 0.224 | 0.14 | 1 | 9 | Lmcd1 | 0.084 | Old |
| 0.03385 | 1.057041 | 0.095 | 0.037 | 1 | 9 | Btbd10 | 0.058 | Old |
| 0.033869 | 2.496957 | 0.101 | 0.044 | 1 | 9 | Chmp1b | 0.057 | Old |
| 0.033918 | 0.254971 | 0.858 | 0.816 | 1 | 9 | Ddx5 | 0.042 | Old |
| 0.034007 | 0.324282 | 0.544 | 0.419 | 1 | 9 | Fzd4 | 0.125 | Old |
| 0.034034 | 2.777414 | 0.059 | 0.015 | 1 | 9 | Rufy1 | 0.044 | Old |
| 0.034174 | 0.455078 | 0.387 | 0.279 | 1 | 9 | Pgrmc1 | 0.108 | Old |
| 0.034232 | 0.715601 | 0.361 | 0.272 | 1 | 9 | Rad21 | 0.089 | Old |
| 0.034354 | 0.721418 | 0.186 | 0.103 | 1 | 9 | Apol10b | 0.083 | Old |
| 0.034635 | 1.809053 | 0.082 | 0.029 | 1 | 9 | Pitrm1 | 0.053 | Old |
| 0.03464 | 2.907415 | 0.059 | 0.015 | 1 | 9 | Ephb1 | 0.044 | Old |
| 0.034817 | 2.97445 | 0.046 | 0.007 | 1 | 9 | Itga9 | 0.039 | Old |
| 0.035523 | 2.901345 | 0.046 | 0.007 | 1 | 9 | Igkc | 0.039 | Old |
| 0.035545 | 0.927205 | 0.082 | 0.029 | 1 | 9 | Tef | 0.053 | Old |
| 0.035716 | 1.062552 | 0.121 | 0.059 | 1 | 9 | Creb5 | 0.062 | Old |
| 0.035846 | 0.474706 | 0.106 | 0.044 | 1 | 9 | Syde1 | 0.062 | Old |
| 0.036038 | 2.079717 | 0.059 | 0.015 | 1 | 9 | Elac1 | 0.044 | Old |
| 0.036038 | 1.26858 | 0.059 | 0.015 | 1 | 9 | Bop1 | 0.044 | Old |
| 0.036073 | 0.590955 | 0.222 | 0.14 | 1 | 9 | Ppm1a | 0.082 | Old |
| 0.03642 | 0.459447 | 0.214 | 0.132 | 1 | 9 | Zfp652 | 0.082 | Old |
| 0.036631 | 0.345254 | 0.448 | 0.338 | 1 | 9 | Rab14 | 0.11 | Old |
| 0.037187 | 1.622383 | 0.129 | 0.066 | 1 | 9 | Pigk | 0.063 | Old |
| 0.037841 | 0.621802 | 0.343 | 0.25 | 1 | 9 | Trim56 | 0.093 | Old |
| 0.0379 | 0.919949 | 0.046 | 0.007 | 1 | 9 | Zfp369 | 0.039 | Old |
| 0.037976 | 1.059338 | 0.059 | 0.015 | 1 | 9 | Rnf185 | 0.044 | Old |
| 0.038089 | 0.815106 | 0.046 | 0.007 | 1 | 9 | Kbtbd7 | 0.039 | Old |
| 0.038244 | 0.807412 | 0.204 | 0.125 | 1 | 9 | Hotairm1 | 0.079 | Old |
| 0.038325 | 4.045541 | 0.031 | 0 | 1 | 9 | Prkcz | 0.031 | Old |

| | | | | | | | | |
|---|---|---|---|---|---|---|---|---|
| 0.038325 | 3.933423 | 0.031 | 0 | 1 | 9 | Ppp1r18os | 0.031 | Old |
| 0.038325 | 3.433315 | 0.031 | 0 | 1 | 9 | Coq6 | 0.031 | Old |
| 0.038325 | 3.18086 | 0.031 | 0 | 1 | 9 | Gm26510 | 0.031 | Old |
| 0.038732 | 1.08216 | 0.093 | 0.037 | 1 | 9 | Zfp407 | 0.056 | Old |
| 0.038778 | 0.853679 | 0.178 | 0.103 | 1 | 9 | Nudcd3 | 0.075 | Old |
| 0.039004 | 1.232763 | 0.131 | 0.066 | 1 | 9 | Taz | 0.065 | Old |
| 0.039073 | 0.919272 | 0.276 | 0.199 | 1 | 9 | Erbin | 0.077 | Old |
| 0.039101 | 0.380609 | 0.603 | 0.493 | 1 | 9 | Dynll1 | 0.11 | Old |
| 0.039233 | 0.402567 | 0.271 | 0.176 | 1 | 9 | Mpzl1 | 0.095 | Old |
| 0.040302 | 0.489665 | 0.325 | 0.221 | 1 | 9 | Atp8a1 | 0.104 | Old |
| 0.040345 | 1.102323 | 0.147 | 0.081 | 1 | 9 | Ifih1 | 0.066 | Old |
| 0.04066 | 0.323049 | 0.657 | 0.559 | 1 | 9 | Id3 | 0.098 | Old |
| 0.040676 | 0.679425 | 0.407 | 0.316 | 1 | 9 | Scaf11 | 0.091 | Old |
| 0.041006 | 0.499469 | 0.456 | 0.368 | 1 | 9 | Tprgl | 0.088 | Old |
| 0.041035 | 2.804668 | 0.057 | 0.015 | 1 | 9 | Rabep2 | 0.042 | Old |
| 0.041217 | 1.531554 | 0.09 | 0.037 | 1 | 9 | Tbc1d12 | 0.053 | Old |
| 0.041304 | 0.295435 | 0.325 | 0.221 | 1 | 9 | Acvrl1 | 0.104 | Old |
| 0.041427 | 0.61893 | 0.198 | 0.118 | 1 | 9 | Ss18 | 0.08 | Old |
| 0.041429 | 0.45589 | 0.381 | 0.287 | 1 | 9 | Sdf4 | 0.094 | Old |
| 0.041587 | 0.183265 | 0.57 | 0.463 | 1 | 9 | Tmem245 | 0.107 | Old |
| 0.041858 | 0.027877 | 0.776 | 0.662 | 1 | 9 | Rpl23 | 0.114 | Old |
| 0.041868 | 0.264494 | 0.381 | 0.279 | 1 | 9 | Selenot | 0.102 | Old |
| 0.041968 | 0.421953 | 0.402 | 0.316 | 1 | 9 | Dennd5a | 0.086 | Old |
| 0.042222 | 0.195468 | 0.508 | 0.412 | 1 | 9 | Zcchc6 | 0.096 | Old |
| 0.042267 | 1.56442 | 0.101 | 0.044 | 1 | 9 | Ehd3 | 0.057 | Old |
| 0.042401 | 0.4355 | 0.397 | 0.294 | 1 | 9 | Med13l | 0.103 | Old |
| 0.04288 | 0.336119 | 0.425 | 0.324 | 1 | 9 | Tmem88 | 0.101 | Old |
| 0.043004 | 1.207907 | 0.07 | 0.022 | 1 | 9 | Pskh1 | 0.048 | Old |
| 0.043039 | 2.903966 | 0.044 | 0.007 | 1 | 9 | Acad12 | 0.037 | Old |
| 0.044065 | 0.418383 | 0.407 | 0.301 | 1 | 9 | Tnfaip1 | 0.106 | Old |
| 0.044125 | 3.344836 | 0.044 | 0.007 | 1 | 9 | Eid2 | 0.037 | Old |
| 0.04414 | 0.394932 | 0.423 | 0.316 | 1 | 9 | Capns1 | 0.107 | Old |
| 0.044303 | 0.963712 | 0.08 | 0.029 | 1 | 9 | Abca5 | 0.051 | Old |
| 0.044353 | 0.26504 | 0.405 | 0.301 | 1 | 9 | Synm | 0.104 | Old |
| 0.044454 | 0.419709 | 0.649 | 0.603 | 1 | 9 | Timp4 | 0.046 | Old |
| 0.044868 | 0.545408 | 0.204 | 0.125 | 1 | 9 | Wdr37 | 0.079 | Old |
| 0.04495 | 0.95738 | 0.08 | 0.029 | 1 | 9 | Snx19 | 0.051 | Old |
| 0.044961 | 1.814221 | 0.057 | 0.015 | 1 | 9 | Zkscan17 | 0.042 | Old |
| 0.045435 | 1.206832 | 0.147 | 0.081 | 1 | 9 | Rif1 | 0.066 | Old |
| 0.046138 | 1.88256 | 0.044 | 0.007 | 1 | 9 | Cklf | 0.037 | Old |
| 0.046403 | 0.159785 | 0.343 | 0.243 | 1 | 9 | Rpn2 | 0.1 | Old |
| 0.046404 | 0.38967 | 0.307 | 0.213 | 1 | 9 | Nop53 | 0.094 | Old |
| 0.046521 | 0.419414 | 0.216 | 0.14 | 1 | 9 | Cul1 | 0.076 | Old |
| 0.0466 | 0.784879 | 0.08 | 0.029 | 1 | 9 | Emc4 | 0.051 | Old |
| 0.046644 | 0.345104 | 0.58 | 0.5 | 1 | 9 | Meox2 | 0.08 | Old |

| | | | | | | | | |
|---|---|---|---|---|---|---|---|---|
| 0.046885 | 1.908421 | 0.067 | 0.022 | 1 | 9 | P4ha2 | 0.045 | Old |
| 0.046922 | 0.315281 | 0.642 | 0.566 | 1 | 9 | Hmgb1 | 0.076 | Old |
| 0.04703 | 0.514497 | 0.271 | 0.184 | 1 | 9 | Ifnar2 | 0.087 | Old |
| 0.047573 | 3.592674 | 0.028 | 0 | 1 | 9 | Rps6kl1 | 0.028 | Old |
| 0.047573 | 3.478251 | 0.028 | 0 | 1 | 9 | Bbs4 | 0.028 | Old |
| 0.047573 | 3.45588 | 0.028 | 0 | 1 | 9 | Zc3h10 | 0.028 | Old |
| 0.047573 | 3.417439 | 0.028 | 0 | 1 | 9 | Ccdc84 | 0.028 | Old |
| 0.047573 | 3.053681 | 0.028 | 0 | 1 | 9 | Dnajc27 | 0.028 | Old |
| 0.047634 | 0.986187 | 0.162 | 0.096 | 1 | 9 | Akap8 | 0.066 | Old |
| 0.047798 | 0.660079 | 0.332 | 0.243 | 1 | 9 | Cyb5r3 | 0.089 | Old |
| 0.047865 | 0.289997 | 0.881 | 0.816 | 1 | 9 | Pecam1 | 0.065 | Old |
| 0.047945 | 0.719708 | 0.219 | 0.14 | 1 | 9 | Pola2 | 0.079 | Old |
| 0.047999 | 1.626571 | 0.067 | 0.022 | 1 | 9 | Enpp5 | 0.045 | Old |
| 0.048137 | 0.240342 | 0.626 | 0.5 | 1 | 9 | Rpl28 | 0.126 | Old |
| 0.048203 | 0.348055 | 0.263 | 0.184 | 1 | 9 | Dynlt3 | 0.079 | Old |
| 0.048246 | 1.639817 | 0.098 | 0.044 | 1 | 9 | Cadm1 | 0.054 | Old |
| 0.04856 | 0.528282 | 0.353 | 0.272 | 1 | 9 | Lims2 | 0.081 | Old |
| 0.048695 | 0.657851 | 0.139 | 0.074 | 1 | 9 | Nbea | 0.065 | Old |
| 0.048888 | 0.226219 | 0.796 | 0.706 | 1 | 9 | Gng11 | 0.09 | Old |
| 0.048974 | 0.600613 | 0.227 | 0.147 | 1 | 9 | Mylk | 0.08 | Old |
| 0.049103 | 1.997996 | 0.088 | 0.037 | 1 | 9 | Ifit1 | 0.051 | Old |
| 0.049461 | 1.028416 | 0.077 | 0.029 | 1 | 9 | Epb41l4aos | 0.048 | Old |
| 0.049464 | 3.278697 | 0.054 | 0.015 | 1 | 9 | Sh2b1 | 0.039 | Old |
| 0.049532 | 0.449666 | 0.17 | 0.103 | 1 | 9 | Vps13a | 0.067 | Old |
| 0.049941 | 1.659739 | 0.088 | 0.037 | 1 | 9 | Scmh1 | 0.051 | Old |
| 0.049948 | 1.330115 | 0.108 | 0.051 | 1 | 9 | Lsm2 | 0.057 | Old |
| 0.050239 | 0.266984 | 0.42 | 0.331 | 1 | 9 | Eef1g | 0.089 | Old |
| 0.05028 | 1.43061 | 0.088 | 0.037 | 1 | 9 | Ptpn21 | 0.051 | Old |
| 0.050709 | 0.653054 | 0.32 | 0.25 | 1 | 9 | Sumo3 | 0.07 | Old |
| 0.05073 | 0.63703 | 0.26 | 0.176 | 1 | 9 | Rgs3 | 0.084 | Old |
| 0.050892 | 0.61134 | 0.309 | 0.221 | 1 | 9 | Fbln2 | 0.088 | Old |
| 0.050987 | 2.787317 | 0.054 | 0.015 | 1 | 9 | Crispld1 | 0.039 | Old |
| 0.051348 | 0.585332 | 0.389 | 0.309 | 1 | 9 | Uqcrb | 0.08 | Old |
| 0.051532 | 0.856198 | 0.307 | 0.228 | 1 | 9 | Kif26a | 0.079 | Old |
| 0.051761 | 0.779005 | 0.229 | 0.154 | 1 | 9 | Mecp2 | 0.075 | Old |
| 0.051877 | 0.138656 | 0.332 | 0.235 | 1 | 9 | Ptk2 | 0.097 | Old |
| 0.051939 | 1.101652 | 0.18 | 0.11 | 1 | 9 | Psme4 | 0.07 | Old |
| 0.052201 | 0.910898 | 0.077 | 0.029 | 1 | 9 | Gtdc1 | 0.048 | Old |
| 0.052295 | 0.29249 | 0.776 | 0.75 | 1 | 9 | Gnai2 | 0.026 | Old |
| 0.052388 | 0.656303 | 0.077 | 0.029 | 1 | 9 | Relb | 0.048 | Old |
| 0.052576 | 1.231006 | 0.077 | 0.029 | 1 | 9 | Med20 | 0.048 | Old |
| 0.052905 | 0.335076 | 0.557 | 0.485 | 1 | 9 | Eng | 0.072 | Old |
| 0.053142 | 1.081353 | 0.077 | 0.029 | 1 | 9 | Zufsp | 0.048 | Old |
| 0.053224 | 4.139754 | 0.041 | 0.007 | 1 | 9 | Mill2 | 0.034 | Old |
| 0.053224 | 3.246191 | 0.041 | 0.007 | 1 | 9 | Homer1 | 0.034 | Old |

| | | | | | | | | |
|---|---|---|---|---|---|---|---|---|
| 0.053688 | 2.043547 | 0.054 | 0.015 | 1 | 9 | Timm17b | 0.039 | Old |
| 0.053714 | 0.511731 | 0.077 | 0.029 | 1 | 9 | Aamdc | 0.048 | Old |
| 0.053991 | 0.178474 | 0.838 | 0.838 | 1 | 9 | Gpihbp1 | 0 | Old |
| 0.054021 | 2.904161 | 0.041 | 0.007 | 1 | 9 | Fgfbp3 | 0.034 | Old |
| 0.054097 | 1.14383 | 0.077 | 0.029 | 1 | 9 | Hoxd3os1 | 0.048 | Old |
| 0.054239 | 0.276301 | 0.34 | 0.243 | 1 | 9 | Ctsd | 0.097 | Old |
| 0.054333 | 0.73071 | 0.222 | 0.147 | 1 | 9 | Arpc1a | 0.075 | Old |
| 0.054466 | 0.42634 | 0.289 | 0.206 | 1 | 9 | Csrp1 | 0.083 | Old |
| 0.054552 | 0.639438 | 0.299 | 0.221 | 1 | 9 | Swi5 | 0.078 | Old |
| 0.054569 | 0.349184 | 0.265 | 0.184 | 1 | 9 | Inf2 | 0.081 | Old |
| 0.054729 | 0.408265 | 0.402 | 0.301 | 1 | 9 | Trp53i11 | 0.101 | Old |
| 0.054828 | 3.482057 | 0.041 | 0.007 | 1 | 9 | Zfp938 | 0.034 | Old |
| 0.054846 | 1.341127 | 0.116 | 0.059 | 1 | 9 | Arhgap21 | 0.057 | Old |
| 0.055206 | 1.578911 | 0.095 | 0.044 | 1 | 9 | Pmpca | 0.051 | Old |
| 0.055319 | 1.366344 | 0.054 | 0.015 | 1 | 9 | Fam149b | 0.039 | Old |
| 0.055555 | 1.624198 | 0.054 | 0.015 | 1 | 9 | Dtx3 | 0.039 | Old |
| 0.055679 | 1.070513 | 0.178 | 0.11 | 1 | 9 | Fbxw11 | 0.068 | Old |
| 0.055707 | 2.349253 | 0.064 | 0.022 | 1 | 9 | Map1s | 0.042 | Old |
| 0.055871 | 0.080586 | 0.415 | 0.316 | 1 | 9 | Tob2 | 0.099 | Old |
| 0.055923 | 2.353225 | 0.064 | 0.022 | 1 | 9 | Esm1 | 0.042 | Old |
| 0.056196 | 0.499267 | 0.211 | 0.132 | 1 | 9 | Exoc6 | 0.079 | Old |
| 0.056419 | 2.038541 | 0.075 | 0.029 | 1 | 9 | Myo7a | 0.046 | Old |
| 0.056679 | 1.678246 | 0.106 | 0.051 | 1 | 9 | Fam204a | 0.055 | Old |
| 0.056682 | 0.643494 | 0.332 | 0.25 | 1 | 9 | Adipor2 | 0.082 | Old |
| 0.056694 | 0.664613 | 0.353 | 0.272 | 1 | 9 | Tbl1x | 0.081 | Old |
| 0.056768 | 0.110885 | 0.17 | 0.103 | 1 | 9 | Cdc40 | 0.067 | Old |
| 0.056961 | 0.373828 | 0.539 | 0.434 | 1 | 9 | Sec62 | 0.105 | Old |
| 0.05703 | 1.461053 | 0.041 | 0.007 | 1 | 9 | Nprl2 | 0.034 | Old |
| 0.05731 | 1.272804 | 0.041 | 0.007 | 1 | 9 | Zfp595 | 0.034 | Old |
| 0.057361 | 0.411089 | 0.284 | 0.199 | 1 | 9 | Cd151 | 0.085 | Old |
| 0.057372 | 0.953446 | 0.16 | 0.096 | 1 | 9 | Fxr2 | 0.064 | Old |
| 0.057377 | 0.465308 | 0.173 | 0.103 | 1 | 9 | Rad23a | 0.07 | Old |
| 0.057592 | 1.141967 | 0.041 | 0.007 | 1 | 9 | Rfx3 | 0.034 | Old |
| 0.057824 | 0.230918 | 0.784 | 0.735 | 1 | 9 | Esam | 0.049 | Old |
| 0.057941 | 1.144497 | 0.16 | 0.096 | 1 | 9 | Sdhd | 0.064 | Old |
| 0.058034 | 0.624648 | 0.284 | 0.199 | 1 | 9 | Cltb | 0.085 | Old |
| 0.058628 | 0.743254 | 0.095 | 0.044 | 1 | 9 | Sympk | 0.051 | Old |
| 0.058752 | 0.57342 | 0.108 | 0.051 | 1 | 9 | Irf7 | 0.057 | Old |
| 0.059157 | 3.732772 | 0.026 | 0 | 1 | 9 | Aox1 | 0.026 | Old |
| 0.059157 | 3.713198 | 0.026 | 0 | 1 | 9 | Spats2 | 0.026 | Old |
| 0.059157 | 3.645647 | 0.026 | 0 | 1 | 9 | Slc25a27 | 0.026 | Old |
| 0.059157 | 3.605551 | 0.026 | 0 | 1 | 9 | Polr2d | 0.026 | Old |
| 0.059157 | 3.549943 | 0.026 | 0 | 1 | 9 | E2f6 | 0.026 | Old |
| 0.059157 | 3.459591 | 0.026 | 0 | 1 | 9 | Gpat3 | 0.026 | Old |
| 0.059157 | 3.435757 | 0.026 | 0 | 1 | 9 | Hhipl1 | 0.026 | Old |

| | | | | | | | | |
|---|---|---|---|---|---|---|---|---|
| 0.059157 | 3.428457 | 0.026 | 0 | 1 | 9 | Pdp2 | 0.026 | Old |
| 0.059157 | 3.349746 | 0.026 | 0 | 1 | 9 | Csrnp2 | 0.026 | Old |
| 0.059247 | 0.601498 | 0.312 | 0.235 | 1 | 9 | Ptprm | 0.077 | Old |
| 0.059637 | 0.966142 | 0.222 | 0.154 | 1 | 9 | Filip1l | 0.068 | Old |
| 0.059779 | 1.460251 | 0.085 | 0.037 | 1 | 9 | Tceal8 | 0.048 | Old |
| 0.060377 | 1.339747 | 0.085 | 0.037 | 1 | 9 | Nr1d1 | 0.048 | Old |
| 0.060415 | 0.49071 | 0.214 | 0.147 | 1 | 9 | Brd3 | 0.067 | Old |
| 0.060459 | 0.048001 | 0.129 | 0.066 | 1 | 9 | BC005561 | 0.063 | Old |
| 0.061206 | 1.144989 | 0.18 | 0.118 | 1 | 9 | Magi3 | 0.062 | Old |
| 0.061257 | 0.466281 | 0.008 | 0.029 | 1 | 9 | Lrrc1 | -0.021 | Old |
| 0.061447 | 0.513341 | 0.106 | 0.051 | 1 | 9 | Scaf4 | 0.055 | Old |
| 0.06161 | 0.993511 | 0.142 | 0.081 | 1 | 9 | AY036118 | 0.061 | Old |
| 0.06169 | 0.317738 | 0.49 | 0.397 | 1 | 9 | Pkn2 | 0.093 | Old |
| 0.06169 | 2.665823 | 0.052 | 0.015 | 1 | 9 | Mre11a | 0.037 | Old |
| 0.061997 | 1.595741 | 0.085 | 0.037 | 1 | 9 | Fnip2 | 0.048 | Old |
| 0.0621 | 1.75269 | 0.075 | 0.029 | 1 | 9 | Zmat3 | 0.046 | Old |
| 0.062165 | 0.122659 | 0.657 | 0.581 | 1 | 9 | Cav2 | 0.076 | Old |
| 0.062455 | 0.503581 | 0.216 | 0.14 | 1 | 9 | Nemf | 0.076 | Old |
| 0.062644 | 1.161507 | 0.113 | 0.059 | 1 | 9 | Dlgap4 | 0.054 | Old |
| 0.062758 | 0.256544 | 0.363 | 0.272 | 1 | 9 | 2010107E0 | 0.091 | Old |
| 0.062759 | 0.573519 | 0.075 | 0.029 | 1 | 9 | Unc13b | 0.046 | Old |
| 0.062876 | 0.71698 | 0.343 | 0.265 | 1 | 9 | Smc1a | 0.078 | Old |
| 0.062966 | 1.076132 | 0.149 | 0.088 | 1 | 9 | Dgcr2 | 0.061 | Old |
| 0.063122 | 1.025475 | 0.131 | 0.074 | 1 | 9 | Senp2 | 0.057 | Old |
| 0.063277 | 0.649072 | 0.126 | 0.066 | 1 | 9 | Ddit3 | 0.06 | Old |
| 0.063652 | 0.704053 | 0.085 | 0.037 | 1 | 9 | Fam53a | 0.048 | Old |
| 0.063894 | 0.785902 | 0.16 | 0.096 | 1 | 9 | Arap2 | 0.064 | Old |
| 0.063894 | 0.564044 | 0.16 | 0.096 | 1 | 9 | Clec14a | 0.064 | Old |
| 0.063999 | 0.36021 | 0.399 | 0.309 | 1 | 9 | Spred2 | 0.09 | Old |
| 0.064096 | 1.250379 | 0.075 | 0.029 | 1 | 9 | Pcsk7 | 0.046 | Old |
| 0.064123 | 0.809988 | 0.204 | 0.132 | 1 | 9 | Timm13 | 0.072 | Old |
| 0.064424 | 1.049868 | 0.155 | 0.096 | 1 | 9 | Hsd17b12 | 0.059 | Old |
| 0.06469 | 0.705399 | 0.142 | 0.081 | 1 | 9 | Chst1 | 0.061 | Old |
| 0.065034 | 0.613534 | 0.144 | 0.081 | 1 | 9 | Fzd6 | 0.063 | Old |
| 0.065429 | 0.29254 | 0.647 | 0.566 | 1 | 9 | Ywhab | 0.081 | Old |
| 0.065475 | 1.892624 | 0.052 | 0.015 | 1 | 9 | Noc4l | 0.037 | Old |
| 0.065486 | 0.570839 | 0.24 | 0.169 | 1 | 9 | Itch | 0.071 | Old |
| 0.065854 | 3.52302 | 0.039 | 0.007 | 1 | 9 | Macrod2 | 0.032 | Old |
| 0.065854 | 3.301755 | 0.039 | 0.007 | 1 | 9 | Ankmy2 | 0.032 | Old |
| 0.066018 | 1.39748 | 0.155 | 0.096 | 1 | 9 | Hoxa5 | 0.059 | Old |
| 0.066032 | 1.627604 | 0.052 | 0.015 | 1 | 9 | Fkbp7 | 0.037 | Old |
| 0.06618 | 3.095653 | 0.039 | 0.007 | 1 | 9 | Riox2 | 0.032 | Old |
| 0.066238 | 0.400233 | 0.281 | 0.199 | 1 | 9 | Dock6 | 0.082 | Old |
| 0.066304 | 0.505263 | 0.332 | 0.25 | 1 | 9 | Chmp2a | 0.082 | Old |
| 0.06633 | 1.436279 | 0.129 | 0.074 | 1 | 9 | Il17rd | 0.055 | Old |

| | | | | | | | | |
|---|---|---|---|---|---|---|---|---|
| 0.066512 | 0.296047 | 0.657 | 0.559 | 1 | 9 | Hnrnph1 | 0.098 | Old |
| 0.066616 | 0.307428 | 0.42 | 0.346 | 1 | 9 | Elf1 | 0.074 | Old |
| 0.066772 | 1.402164 | 0.093 | 0.044 | 1 | 9 | Chml | 0.049 | Old |
| 0.067371 | 0.667649 | 0.361 | 0.294 | 1 | 9 | Rassf1 | 0.067 | Old |
| 0.06747 | 0.314789 | 0.552 | 0.449 | 1 | 9 | Tmed5 | 0.103 | Old |
| 0.067494 | 2.852058 | 0.039 | 0.007 | 1 | 9 | Bnip1 | 0.032 | Old |
| 0.067494 | 2.073315 | 0.039 | 0.007 | 1 | 9 | Neo1 | 0.032 | Old |
| 0.067712 | 0.100479 | 0.778 | 0.699 | 1 | 9 | Rpl9 | 0.079 | Old |
| 0.067746 | 0.865378 | 0.271 | 0.206 | 1 | 9 | Setd5 | 0.065 | Old |
| 0.067822 | 0.706165 | 0.216 | 0.147 | 1 | 9 | Ppp1r13b | 0.069 | Old |
| 0.067938 | 1.861587 | 0.062 | 0.022 | 1 | 9 | Ctif | 0.04 | Old |
| 0.067968 | 0.408862 | 0.5 | 0.404 | 1 | 9 | Btf3 | 0.096 | Old |
| 0.06803 | 0.338093 | 0.696 | 0.654 | 1 | 9 | Itpkb | 0.042 | Old |
| 0.068088 | 0.190408 | 0.66 | 0.559 | 1 | 9 | Rpl23a | 0.101 | Old |
| 0.06824 | 1.084474 | 0.093 | 0.044 | 1 | 9 | Nudt9 | 0.049 | Old |
| 0.068664 | 1.544158 | 0.093 | 0.044 | 1 | 9 | Zfp639 | 0.049 | Old |
| 0.068666 | 1.879674 | 0.082 | 0.037 | 1 | 9 | Haus2 | 0.045 | Old |
| 0.068707 | 0.529075 | 0.5 | 0.412 | 1 | 9 | Il6st | 0.088 | Old |
| 0.069057 | 0.153912 | 0.392 | 0.301 | 1 | 9 | Tgm2 | 0.091 | Old |
| 0.069365 | 0.932736 | 0.113 | 0.059 | 1 | 9 | Usp33 | 0.054 | Old |
| 0.06939 | 0.296415 | 0.474 | 0.375 | 1 | 9 | Nisch | 0.099 | Old |
| 0.06945 | 1.137206 | 0.052 | 0.015 | 1 | 9 | Amot | 0.037 | Old |
| 0.069722 | 0.395699 | 0.106 | 0.051 | 1 | 9 | Fbxo8 | 0.055 | Old |
| 0.069765 | 0.232944 | 0.134 | 0.074 | 1 | 9 | Pdk1 | 0.06 | Old |
| 0.070057 | 0.926585 | 0.101 | 0.051 | 1 | 9 | Mettl16 | 0.05 | Old |
| 0.070169 | 0.932363 | 0.157 | 0.096 | 1 | 9 | Bcor | 0.061 | Old |
| 0.07057 | 1.477326 | 0.062 | 0.022 | 1 | 9 | Nsun4 | 0.04 | Old |
| 0.070666 | 0.97315 | 0.129 | 0.074 | 1 | 9 | Tmem205 | 0.055 | Old |
| 0.070834 | 0.586291 | 0.103 | 0.051 | 1 | 9 | Cpped1 | 0.052 | Old |
| 0.070838 | 0.387023 | 0.193 | 0.125 | 1 | 9 | Ing1 | 0.068 | Old |
| 0.070991 | 0.183872 | 0.601 | 0.5 | 1 | 9 | Mbnl2 | 0.101 | Old |
| 0.071154 | 0.717568 | 0.353 | 0.287 | 1 | 9 | Ewsr1 | 0.066 | Old |
| 0.071414 | 0.317317 | 0.647 | 0.537 | 1 | 9 | Msn | 0.11 | Old |
| 0.071504 | 0.705218 | 0.198 | 0.132 | 1 | 9 | Flot2 | 0.066 | Old |
| 0.071543 | 0.355713 | 0.381 | 0.294 | 1 | 9 | Slc43a3 | 0.087 | Old |
| 0.071637 | 0.480551 | 0.175 | 0.11 | 1 | 9 | Ercc1 | 0.065 | Old |
| 0.071747 | 0.301226 | 0.637 | 0.551 | 1 | 9 | Dst | 0.086 | Old |
| 0.071785 | 1.037385 | 0.129 | 0.074 | 1 | 9 | Got2 | 0.055 | Old |
| 0.071965 | 0.100821 | 0.116 | 0.059 | 1 | 9 | Fbrs | 0.057 | Old |
| 0.072589 | 0.498662 | 0.198 | 0.132 | 1 | 9 | Stip1 | 0.066 | Old |
| 0.072615 | 1.233896 | 0.16 | 0.103 | 1 | 9 | Smarcb1 | 0.057 | Old |
| 0.072646 | 0.733478 | 0.183 | 0.118 | 1 | 9 | Cd59a | 0.065 | Old |
| 0.072702 | 0.21281 | 0.5 | 0.426 | 1 | 9 | Hipk1 | 0.074 | Old |
| 0.073028 | 0.494291 | 0.26 | 0.199 | 1 | 9 | Ndufa5 | 0.061 | Old |
| 0.073616 | 0.098526 | 0.853 | 0.816 | 1 | 9 | Ly6e | 0.037 | Old |

| | | | | | | | | |
|---|---|---|---|---|---|---|---|---|
| 0.073724 | 3.957745 | 0.023 | 0 | 1 | 9 | Top3a | 0.023 | Old |
| 0.073724 | 3.910757 | 0.023 | 0 | 1 | 9 | Stk11ip | 0.023 | Old |
| 0.073724 | 3.521401 | 0.023 | 0 | 1 | 9 | Twist1 | 0.023 | Old |
| 0.073724 | 3.336782 | 0.023 | 0 | 1 | 9 | Cox10 | 0.023 | Old |
| 0.073724 | 3.302176 | 0.023 | 0 | 1 | 9 | E330009J0 | 0.023 | Old |
| 0.073724 | 3.264688 | 0.023 | 0 | 1 | 9 | Fut10 | 0.023 | Old |
| 0.073724 | 3.210974 | 0.023 | 0 | 1 | 9 | Ccdc125 | 0.023 | Old |
| 0.073724 | 3.175239 | 0.023 | 0 | 1 | 9 | BC024978 | 0.023 | Old |
| 0.073724 | 3.11853 | 0.023 | 0 | 1 | 9 | Ccdc167 | 0.023 | Old |
| 0.073724 | 3.014811 | 0.023 | 0 | 1 | 9 | Zfp446 | 0.023 | Old |
| 0.073724 | 2.978723 | 0.023 | 0 | 1 | 9 | Rnf135 | 0.023 | Old |
| 0.073724 | 2.968335 | 0.023 | 0 | 1 | 9 | Rtn4rl1 | 0.023 | Old |
| 0.073724 | 2.919389 | 0.023 | 0 | 1 | 9 | Adamts18 | 0.023 | Old |
| 0.073724 | 2.669753 | 0.023 | 0 | 1 | 9 | Dhh | 0.023 | Old |
| 0.073724 | 2.640448 | 0.023 | 0 | 1 | 9 | Qrsl1 | 0.023 | Old |
| 0.073868 | 0.697773 | 0.131 | 0.074 | 1 | 9 | Washc2 | 0.057 | Old |
| 0.074115 | 0.326679 | 0.062 | 0.022 | 1 | 9 | Prdm15 | 0.04 | Old |
| 0.074547 | 1.32686 | 0.072 | 0.029 | 1 | 9 | Lonp1 | 0.043 | Old |
| 0.074907 | 0.479342 | 0.335 | 0.25 | 1 | 9 | Degs1 | 0.085 | Old |
| 0.074943 | 0.177784 | 0.621 | 0.507 | 1 | 9 | Rpl34 | 0.114 | Old |
| 0.075025 | 0.482995 | 0.232 | 0.154 | 1 | 9 | Mafg | 0.078 | Old |
| 0.075237 | 0.534961 | 0.286 | 0.213 | 1 | 9 | Tmed9 | 0.073 | Old |
| 0.075294 | 0.930265 | 0.139 | 0.081 | 1 | 9 | Rsbn1 | 0.058 | Old |
| 0.075371 | 0.803047 | 0.111 | 0.059 | 1 | 9 | Tyk2 | 0.052 | Old |
| 0.075792 | 0.93 | 0.111 | 0.059 | 1 | 9 | Trim33 | 0.052 | Old |
| 0.075843 | 1.148178 | 0.101 | 0.051 | 1 | 9 | Acp1 | 0.05 | Old |
| 0.075982 | 0.772803 | 0.175 | 0.11 | 1 | 9 | Eif1ax | 0.065 | Old |
| 0.076151 | 0.973208 | 0.121 | 0.066 | 1 | 9 | Mitd1 | 0.055 | Old |
| 0.076208 | 2.885623 | 0.049 | 0.015 | 1 | 9 | Agk | 0.034 | Old |
| 0.076702 | 0.567963 | 0.245 | 0.169 | 1 | 9 | D8Ertd738 | 0.076 | Old |
| 0.076734 | 0.623234 | 0.18 | 0.118 | 1 | 9 | Skil | 0.062 | Old |
| 0.076852 | 2.150563 | 0.049 | 0.015 | 1 | 9 | Dalrd3 | 0.034 | Old |
| 0.076956 | 1.383807 | 0.101 | 0.051 | 1 | 9 | Exoc2 | 0.05 | Old |
| 0.077236 | 0.776525 | 0.229 | 0.162 | 1 | 9 | 2-Mar | 0.067 | Old |
| 0.077371 | 1.00084 | 0.155 | 0.096 | 1 | 9 | Fam114a2 | 0.059 | Old |
| 0.077486 | 0.765955 | 0.17 | 0.11 | 1 | 9 | Mknk2 | 0.06 | Old |
| 0.077573 | 1.797837 | 0.07 | 0.029 | 1 | 9 | Ints12 | 0.041 | Old |
| 0.077826 | 2.609739 | 0.049 | 0.015 | 1 | 9 | Sdhaf2 | 0.034 | Old |
| 0.0779 | 0.343001 | 0.412 | 0.324 | 1 | 9 | Irf1 | 0.088 | Old |
| 0.078292 | 0.783536 | 0.155 | 0.096 | 1 | 9 | Ipo5 | 0.059 | Old |
| 0.078304 | 0.257647 | 0.374 | 0.272 | 1 | 9 | Nxf1 | 0.102 | Old |
| 0.078315 | 0.792634 | 0.263 | 0.199 | 1 | 9 | Cct5 | 0.064 | Old |
| 0.078481 | 2.092796 | 0.049 | 0.015 | 1 | 9 | Fkrp | 0.034 | Old |
| 0.078734 | 1.195318 | 0.198 | 0.14 | 1 | 9 | Pkig | 0.058 | Old |
| 0.078935 | 1.976174 | 0.07 | 0.029 | 1 | 9 | Naa60 | 0.041 | Old |

| | | | | | | | | |
|---|---|---|---|---|---|---|---|---|
| 0.07897 | 0.309473 | 0.062 | 0.022 | 1 | 9 | Tmem237 | 0.04 | Old |
| 0.079012 | 0.683199 | 0.304 | 0.235 | 1 | 9 | Ghitm | 0.069 | Old |
| 0.079141 | 2.002913 | 0.049 | 0.015 | 1 | 9 | Aasdhppt | 0.034 | Old |
| 0.079317 | 0.972946 | 0.082 | 0.037 | 1 | 9 | Jade2 | 0.045 | Old |
| 0.079376 | 2.514926 | 0.059 | 0.022 | 1 | 9 | Fpgt | 0.037 | Old |
| 0.079704 | 0.576624 | 0.16 | 0.096 | 1 | 9 | Gcc2 | 0.064 | Old |
| 0.079977 | 2.359583 | 0.059 | 0.022 | 1 | 9 | Zscan21 | 0.037 | Old |
| 0.080138 | 0.370495 | 0.049 | 0.015 | 1 | 9 | Procr | 0.034 | Old |
| 0.080279 | 1.932818 | 0.059 | 0.022 | 1 | 9 | Sh3bgrl2 | 0.037 | Old |
| 0.080578 | 0.944048 | 0.137 | 0.081 | 1 | 9 | Bccip | 0.056 | Old |
| 0.080677 | 0.154729 | 0.072 | 0.029 | 1 | 9 | Csf2rb2 | 0.043 | Old |
| 0.08111 | 0.60857 | 0.155 | 0.096 | 1 | 9 | Bcar1 | 0.059 | Old |
| 0.081483 | 1.015304 | 0.129 | 0.074 | 1 | 9 | Mob2 | 0.055 | Old |
| 0.081824 | 1.024054 | 0.049 | 0.015 | 1 | 9 | Vangl2 | 0.034 | Old |
| 0.081874 | 0.266462 | 0.696 | 0.64 | 1 | 9 | Ece1 | 0.056 | Old |
| 0.082027 | 0.194237 | 0.75 | 0.684 | 1 | 9 | Ppp1r2 | 0.066 | Old |
| 0.082338 | 2.65282 | 0.036 | 0.007 | 1 | 9 | Nid2 | 0.029 | Old |
| 0.082474 | 0.315814 | 0.469 | 0.397 | 1 | 9 | Tpr | 0.072 | Old |
| 0.083014 | 0.92491 | 0.005 | 0.022 | 1 | 9 | Ercc6l | -0.017 | Old |
| 0.083037 | 1.787601 | 0.059 | 0.022 | 1 | 9 | Vps72 | 0.037 | Old |
| 0.083193 | 1.374737 | 0.049 | 0.015 | 1 | 9 | Klhl3 | 0.034 | Old |
| 0.083551 | 2.412151 | 0.036 | 0.007 | 1 | 9 | Tmem251 | 0.029 | Old |
| 0.083676 | 0.347404 | 0.101 | 0.051 | 1 | 9 | Jmy | 0.05 | Old |
| 0.083741 | 0.368498 | 0.242 | 0.169 | 1 | 9 | Marf1 | 0.073 | Old |
| 0.084106 | 0.539457 | 0.191 | 0.125 | 1 | 9 | Golga7 | 0.066 | Old |
| 0.08427 | 0.15457 | 0.152 | 0.088 | 1 | 9 | Ppm1l | 0.064 | Old |
| 0.084603 | 1.366376 | 0.059 | 0.022 | 1 | 9 | Coro7 | 0.037 | Old |
| 0.084643 | 0.381042 | 0.477 | 0.397 | 1 | 9 | Eif4a1 | 0.08 | Old |
| 0.084871 | 1.605473 | 0.07 | 0.029 | 1 | 9 | Fbxo6 | 0.041 | Old |
| 0.084921 | 0.297181 | 0.219 | 0.147 | 1 | 9 | Ei24 | 0.072 | Old |
| 0.085191 | 2.060636 | 0.036 | 0.007 | 1 | 9 | H2afx | 0.029 | Old |
| 0.085197 | 1.568528 | 0.124 | 0.074 | 1 | 9 | Usp8 | 0.05 | Old |
| 0.085236 | 1.781218 | 0.059 | 0.022 | 1 | 9 | Tspan3 | 0.037 | Old |
| 0.085296 | 0.450249 | 0.214 | 0.14 | 1 | 9 | Ptch1 | 0.074 | Old |
| 0.085417 | 1.377862 | 0.139 | 0.088 | 1 | 9 | Myd88 | 0.051 | Old |
| 0.085553 | 1.662337 | 0.059 | 0.022 | 1 | 9 | Alas1 | 0.037 | Old |
| 0.085571 | 0.94245 | 0.108 | 0.059 | 1 | 9 | Cpne8 | 0.049 | Old |
| 0.085579 | 0.44715 | 0.24 | 0.169 | 1 | 9 | Mgat1 | 0.071 | Old |
| 0.085611 | 0.9437 | 0.101 | 0.051 | 1 | 9 | Hps3 | 0.05 | Old |
| 0.085688 | 1.351927 | 0.08 | 0.037 | 1 | 9 | Gpatch2 | 0.043 | Old |
| 0.085962 | 1.385591 | 0.08 | 0.037 | 1 | 9 | Adpgk | 0.043 | Old |
| 0.086356 | 0.688993 | 0.137 | 0.081 | 1 | 9 | Ndst1 | 0.056 | Old |
| 0.086629 | 0.70073 | 0.013 | 0.037 | 1 | 9 | Ndnf | -0.024 | Old |
| 0.086632 | 1.561042 | 0.07 | 0.029 | 1 | 9 | Tesk1 | 0.041 | Old |
| 0.086761 | 0.31247 | 0.461 | 0.39 | 1 | 9 | Smad1 | 0.071 | Old |

| | | | | | | | | |
|---|---|---|---|---|---|---|---|---|
| 0.086856 | 0.750049 | 0.036 | 0.007 | 1 | 9 | Lrwd1 | 0.029 | Old |
| 0.086856 | 0.691894 | 0.036 | 0.007 | 1 | 9 | Ogfod2 | 0.029 | Old |
| 0.086935 | 0.69584 | 0.131 | 0.074 | 1 | 9 | Slc1a1 | 0.057 | Old |
| 0.086988 | 0.835388 | 0.144 | 0.088 | 1 | 9 | Tspyl1 | 0.056 | Old |
| 0.087191 | 0.097321 | 0.856 | 0.816 | 1 | 9 | Jun | 0.04 | Old |
| 0.087256 | 0.461244 | 0.307 | 0.235 | 1 | 9 | Sar1a | 0.072 | Old |
| 0.087618 | 0.41107 | 0.142 | 0.081 | 1 | 9 | Rxra | 0.061 | Old |
| 0.087694 | 0.230488 | 0.351 | 0.265 | 1 | 9 | Nudt4 | 0.086 | Old |
| 0.088147 | 0.696676 | 0.111 | 0.059 | 1 | 9 | Nop14 | 0.052 | Old |
| 0.088303 | 0.393579 | 0.399 | 0.331 | 1 | 9 | Lamc1 | 0.068 | Old |
| 0.088336 | 0.736351 | 0.119 | 0.066 | 1 | 9 | Cabin1 | 0.053 | Old |
| 0.088419 | 0.196436 | 0.286 | 0.213 | 1 | 9 | Fez2 | 0.073 | Old |
| 0.088459 | 1.432663 | 0.08 | 0.037 | 1 | 9 | Ranbp3 | 0.043 | Old |
| 0.088582 | 0.348753 | 0.101 | 0.051 | 1 | 9 | Faap20 | 0.05 | Old |
| 0.088663 | 2.441311 | 0.077 | 0.037 | 1 | 9 | Mrps5 | 0.04 | Old |
| 0.088732 | 0.040783 | 0.157 | 0.096 | 1 | 9 | Ncbp3 | 0.061 | Old |
| 0.088793 | 0.440304 | 0.356 | 0.294 | 1 | 9 | Rasal2 | 0.062 | Old |
| 0.088799 | 0.935497 | 0.119 | 0.066 | 1 | 9 | Kit | 0.053 | Old |
| 0.089026 | 1.006231 | 0.07 | 0.029 | 1 | 9 | Utp6 | 0.041 | Old |
| 0.089389 | 0.356815 | 0.43 | 0.36 | 1 | 9 | Tmed10 | 0.07 | Old |
| 0.089819 | 1.444228 | 0.098 | 0.051 | 1 | 9 | Ttc5 | 0.047 | Old |
| 0.090407 | 1.497554 | 0.088 | 0.044 | 1 | 9 | Mrpl27 | 0.044 | Old |
| 0.090623 | 0.7557 | 0.108 | 0.059 | 1 | 9 | Acsl1 | 0.049 | Old |
| 0.090856 | 0.88381 | 0.07 | 0.029 | 1 | 9 | Prkd1 | 0.041 | Old |
| 0.090961 | 0.159992 | 0.126 | 0.074 | 1 | 9 | Rrm2b | 0.052 | Old |
| 0.09136 | 2.658237 | 0.046 | 0.015 | 1 | 9 | Msmp | 0.031 | Old |
| 0.091905 | 0.196695 | 0.206 | 0.14 | 1 | 9 | Trp53inp2 | 0.066 | Old |
| 0.09209 | 0.362627 | 0.351 | 0.272 | 1 | 9 | Rapgef1 | 0.079 | Old |
| 0.092094 | 0.539837 | 0.255 | 0.184 | 1 | 9 | Pebp1 | 0.071 | Old |
| 0.092132 | 3.411845 | 0.021 | 0 | 1 | 9 | Capn10 | 0.021 | Old |
| 0.092132 | 3.34765 | 0.021 | 0 | 1 | 9 | Meis2 | 0.021 | Old |
| 0.092132 | 3.337154 | 0.021 | 0 | 1 | 9 | Rtel1 | 0.021 | Old |
| 0.092132 | 3.280739 | 0.021 | 0 | 1 | 9 | Trim16 | 0.021 | Old |
| 0.092132 | 3.265292 | 0.021 | 0 | 1 | 9 | Cep89 | 0.021 | Old |
| 0.092132 | 3.256354 | 0.021 | 0 | 1 | 9 | 5033430I1 | 0.021 | Old |
| 0.092132 | 3.192539 | 0.021 | 0 | 1 | 9 | Gm16638 | 0.021 | Old |
| 0.092132 | 3.165669 | 0.021 | 0 | 1 | 9 | 1500015A0 | 0.021 | Old |
| 0.092132 | 3.105336 | 0.021 | 0 | 1 | 9 | Zfp87 | 0.021 | Old |
| 0.092132 | 3.033024 | 0.021 | 0 | 1 | 9 | Ift57 | 0.021 | Old |
| 0.092132 | 3.022231 | 0.021 | 0 | 1 | 9 | Mettl1 | 0.021 | Old |
| 0.092132 | 3.015721 | 0.021 | 0 | 1 | 9 | Ppcs | 0.021 | Old |
| 0.092132 | 2.974467 | 0.021 | 0 | 1 | 9 | 4930403P2 | 0.021 | Old |
| 0.092132 | 2.957314 | 0.021 | 0 | 1 | 9 | Rcc1 | 0.021 | Old |
| 0.092132 | 2.936319 | 0.021 | 0 | 1 | 9 | Mettl25 | 0.021 | Old |
| 0.092132 | 2.808982 | 0.021 | 0 | 1 | 9 | Cyb5r2 | 0.021 | Old |

| | | | | | | | | |
|---|---|---|---|---|---|---|---|---|
| 0.092132 | 2.794895 | 0.021 | 0 | 1 | 9 | Wdr19 | 0.021 | Old |
| 0.092132 | 2.676259 | 0.021 | 0 | 1 | 9 | Dynlt1f | 0.021 | Old |
| 0.092132 | 2.656889 | 0.021 | 0 | 1 | 9 | Mmgt2 | 0.021 | Old |
| 0.092132 | 2.609071 | 0.021 | 0 | 1 | 9 | Tmed1 | 0.021 | Old |
| 0.092132 | 2.60454 | 0.021 | 0 | 1 | 9 | Gm36266 | 0.021 | Old |
| 0.092132 | 2.458247 | 0.021 | 0 | 1 | 9 | Hist2h3c1 | 0.021 | Old |
| 0.092132 | 2.446204 | 0.021 | 0 | 1 | 9 | Tgds | 0.021 | Old |
| 0.092132 | 2.335034 | 0.021 | 0 | 1 | 9 | H6pd | 0.021 | Old |
| 0.092453 | 0.416172 | 0.451 | 0.375 | 1 | 9 | Nova2 | 0.076 | Old |
| 0.092699 | 2.467687 | 0.085 | 0.044 | 1 | 9 | Rpp21 | 0.041 | Old |
| 0.092733 | 0.505337 | 0.222 | 0.154 | 1 | 9 | Kctd12b | 0.068 | Old |
| 0.09281 | 1.090395 | 0.134 | 0.081 | 1 | 9 | Tpi1 | 0.053 | Old |
| 0.092829 | 0.727918 | 0.175 | 0.118 | 1 | 9 | Dlst | 0.057 | Old |
| 0.092885 | 1.031505 | 0.152 | 0.096 | 1 | 9 | Fbrsl1 | 0.056 | Old |
| 0.093099 | 0.748444 | 0.152 | 0.096 | 1 | 9 | Abcg1 | 0.056 | Old |
| 0.09316 | 1.167672 | 0.088 | 0.044 | 1 | 9 | Zcchc8 | 0.044 | Old |
| 0.093242 | 0.3848 | 0.343 | 0.257 | 1 | 9 | Nfic | 0.086 | Old |
| 0.093315 | 1.776915 | 0.077 | 0.037 | 1 | 9 | Rassf8 | 0.04 | Old |
| 0.093315 | 1.520618 | 0.077 | 0.037 | 1 | 9 | Mphosph1 | 0.04 | Old |
| 0.093396 | 0.36502 | 0.601 | 0.544 | 1 | 9 | Jam2 | 0.057 | Old |
| 0.093613 | 1.654437 | 0.077 | 0.037 | 1 | 9 | Slc4a1ap | 0.04 | Old |
| 0.093637 | 0.453268 | 0.345 | 0.257 | 1 | 9 | Rpl27 | 0.088 | Old |
| 0.093666 | 2.432652 | 0.046 | 0.015 | 1 | 9 | Mest | 0.031 | Old |
| 0.093719 | 1.182627 | 0.134 | 0.081 | 1 | 9 | Scaf1 | 0.053 | Old |
| 0.093948 | 0.875222 | 0.134 | 0.081 | 1 | 9 | Trak2 | 0.053 | Old |
| 0.09413 | 0.568301 | 0.191 | 0.132 | 1 | 9 | Psma1 | 0.059 | Old |
| 0.094305 | 0.549847 | 0.162 | 0.103 | 1 | 9 | Stk25 | 0.059 | Old |
| 0.094495 | 0.417446 | 0.345 | 0.265 | 1 | 9 | Kctd10 | 0.08 | Old |
| 0.094503 | 0.529818 | 0.286 | 0.213 | 1 | 9 | Papola | 0.073 | Old |
| 0.094607 | 0.997215 | 0.201 | 0.14 | 1 | 9 | Nxn | 0.061 | Old |
| 0.094628 | 0.785523 | 0.116 | 0.066 | 1 | 9 | Sirt7 | 0.05 | Old |
| 0.094773 | 0.410784 | 0.216 | 0.147 | 1 | 9 | Rab5a | 0.069 | Old |
| 0.094894 | 0.836807 | 0.18 | 0.125 | 1 | 9 | Arsa | 0.055 | Old |
| 0.094938 | 0.438912 | 0.162 | 0.103 | 1 | 9 | Seh1l | 0.059 | Old |
| 0.095411 | 1.5096 | 0.088 | 0.044 | 1 | 9 | Cog7 | 0.044 | Old |
| 0.095553 | 0.427218 | 0.216 | 0.154 | 1 | 9 | Gng12 | 0.062 | Old |
| 0.095866 | 0.894602 | 0.116 | 0.066 | 1 | 9 | Spata2 | 0.05 | Old |
| 0.096118 | 2.608942 | 0.085 | 0.044 | 1 | 9 | Dapk3 | 0.041 | Old |
| 0.096434 | 0.745883 | 0.142 | 0.088 | 1 | 9 | 1810013L2 | 0.054 | Old |
| 0.096504 | 0.701466 | 0.384 | 0.324 | 1 | 9 | Pfdn5 | 0.06 | Old |
| 0.096569 | 0.280282 | 0.541 | 0.449 | 1 | 9 | Klf3 | 0.092 | Old |
| 0.096682 | 0.593637 | 0.304 | 0.235 | 1 | 9 | Litaf | 0.069 | Old |
| 0.096792 | 0.369659 | 0.255 | 0.184 | 1 | 9 | Psmd4 | 0.071 | Old |
| 0.096881 | 0.221519 | 0.585 | 0.485 | 1 | 9 | Atf4 | 0.1 | Old |
| 0.097212 | 1.882522 | 0.046 | 0.015 | 1 | 9 | Dnajc24 | 0.031 | Old |

| | | | | | | | | |
|---|---|---|---|---|---|---|---|---|
| 0.097347 | 1.424145 | 0.142 | 0.088 | 1 | 9 | Ccdc12 | 0.054 | Old |
| 0.097663 | 0.676971 | 0.134 | 0.081 | 1 | 9 | Impa1 | 0.053 | Old |
| 0.097957 | 0.67063 | 0.124 | 0.074 | 1 | 9 | Hist1h2bc | 0.05 | Old |
| 0.097979 | 0.444897 | 0.25 | 0.184 | 1 | 9 | Sh2b3 | 0.066 | Old |
| 0.098014 | 1.859199 | 0.046 | 0.015 | 1 | 9 | Camk2n2 | 0.031 | Old |
| 0.098073 | 0.227155 | 0.469 | 0.39 | 1 | 9 | Uqcr11 | 0.079 | Old |
| 0.098243 | 1.503381 | 0.113 | 0.066 | 1 | 9 | Vcpip1 | 0.047 | Old |
| 0.098576 | 1.07298 | 0.088 | 0.044 | 1 | 9 | Zfpl1 | 0.044 | Old |
| 0.098602 | 0.976931 | 0.183 | 0.125 | 1 | 9 | Mrpl24 | 0.058 | Old |
| 0.09908 | 0.490434 | 0.325 | 0.257 | 1 | 9 | Ppp6r3 | 0.068 | Old |
| 0.099429 | 0.946738 | 0.183 | 0.125 | 1 | 9 | Clcn4 | 0.058 | Old |
| 0.099479 | 0.337926 | 0.356 | 0.272 | 1 | 9 | Atxn1 | 0.084 | Old |
| 0.09958 | 1.346964 | 0.067 | 0.029 | 1 | 9 | Mthfd2l | 0.038 | Old |
| 0.100154 | 0.633358 | 0.17 | 0.11 | 1 | 9 | Cmpk2 | 0.06 | Old |
| 0.100273 | 0.890727 | 0.098 | 0.051 | 1 | 9 | Dmxl2 | 0.047 | Old |
| 0.100763 | 0.540266 | 0.268 | 0.199 | 1 | 9 | Hp1bp3 | 0.069 | Old |
| 0.100935 | 0.805707 | 0.124 | 0.074 | 1 | 9 | Exosc10 | 0.05 | Old |
| 0.101037 | 2.963527 | 0.034 | 0.007 | 1 | 9 | Trib3 | 0.027 | Old |
| 0.101207 | 0.794023 | 0.149 | 0.096 | 1 | 9 | Gm1673 | 0.053 | Old |
| 0.101305 | 0.99891 | 0.106 | 0.059 | 1 | 9 | Ctr9 | 0.047 | Old |
| 0.101917 | 0.216478 | 0.338 | 0.257 | 1 | 9 | H2-Q6 | 0.081 | Old |
| 0.102021 | 2.32743 | 0.034 | 0.007 | 1 | 9 | Nmb | 0.027 | Old |
| 0.102021 | 2.121771 | 0.034 | 0.007 | 1 | 9 | Cep135 | 0.027 | Old |
| 0.102056 | 1.774935 | 0.057 | 0.022 | 1 | 9 | Prmt2 | 0.035 | Old |
| 0.102234 | 0.804721 | 0.327 | 0.272 | 1 | 9 | Elf2 | 0.055 | Old |
| 0.102516 | 4.397808 | 0.034 | 0.007 | 1 | 9 | Cxcl13 | 0.027 | Old |
| 0.102516 | 2.725314 | 0.034 | 0.007 | 1 | 9 | Adgrg6 | 0.027 | Old |
| 0.102806 | 1.700844 | 0.057 | 0.022 | 1 | 9 | Pdp1 | 0.035 | Old |
| 0.103013 | 2.502672 | 0.034 | 0.007 | 1 | 9 | Lpin3 | 0.027 | Old |
| 0.103013 | 1.817308 | 0.034 | 0.007 | 1 | 9 | Ttc21b | 0.027 | Old |
| 0.103332 | 1.352672 | 0.067 | 0.029 | 1 | 9 | Mfsd4a | 0.038 | Old |
| 0.103332 | 1.254319 | 0.067 | 0.029 | 1 | 9 | Tada1 | 0.038 | Old |
| 0.103457 | 1.135433 | 0.095 | 0.051 | 1 | 9 | Mta1 | 0.044 | Old |
| 0.103511 | 2.510566 | 0.034 | 0.007 | 1 | 9 | Olfm2 | 0.027 | Old |
| 0.103511 | 2.31871 | 0.034 | 0.007 | 1 | 9 | Helq | 0.027 | Old |
| 0.103511 | 2.127153 | 0.034 | 0.007 | 1 | 9 | Cdpf1 | 0.027 | Old |
| 0.103559 | 1.346549 | 0.057 | 0.022 | 1 | 9 | Cpsf1 | 0.035 | Old |
| 0.103677 | 0.176766 | 0.91 | 0.846 | 1 | 9 | Calm1 | 0.064 | Old |
| 0.10371 | 0.156241 | 0.848 | 0.772 | 1 | 9 | Egr1 | 0.076 | Old |
| 0.104012 | 1.979848 | 0.034 | 0.007 | 1 | 9 | Pomt2 | 0.027 | Old |
| 0.104026 | 1.534619 | 0.067 | 0.029 | 1 | 9 | Pam16 | 0.038 | Old |
| 0.104076 | 0.400086 | 0.25 | 0.184 | 1 | 9 | Nudt3 | 0.066 | Old |
| 0.104285 | 0.165527 | 0.389 | 0.294 | 1 | 9 | Nfe2l2 | 0.095 | Old |
| 0.104295 | 0.398631 | 0.397 | 0.324 | 1 | 9 | Sumo2 | 0.073 | Old |
| 0.104338 | 0.993421 | 0.139 | 0.088 | 1 | 9 | Rprd1b | 0.051 | Old |

| | | | | | | | | |
|---|---|---|---|---|---|---|---|---|
| 0.104514 | 2.251194 | 0.034 | 0.007 | 1 | 9 | Irs2 | 0.027 | Old |
| 0.104514 | 1.652566 | 0.034 | 0.007 | 1 | 9 | Ncapd2 | 0.027 | Old |
| 0.104851 | 0.174113 | 0.894 | 0.838 | 1 | 9 | Itga1 | 0.056 | Old |
| 0.104902 | 0.576096 | 0.325 | 0.257 | 1 | 9 | Bri3 | 0.068 | Old |
| 0.105272 | 0.011042 | 0.108 | 0.059 | 1 | 9 | Mybbp1a | 0.049 | Old |
| 0.105335 | 0.080998 | 0.327 | 0.25 | 1 | 9 | Emc10 | 0.077 | Old |
| 0.105525 | 1.912315 | 0.034 | 0.007 | 1 | 9 | Dcun1d4 | 0.027 | Old |
| 0.105525 | 1.671871 | 0.034 | 0.007 | 1 | 9 | Hdhd5 | 0.027 | Old |
| 0.10558 | 0.726803 | 0.211 | 0.147 | 1 | 9 | Eps8 | 0.064 | Old |
| 0.105967 | 0.717417 | 0.116 | 0.066 | 1 | 9 | Nfx1 | 0.05 | Old |
| 0.106033 | 1.5012 | 0.034 | 0.007 | 1 | 9 | Adamtsl3 | 0.027 | Old |
| 0.106543 | 1.535899 | 0.034 | 0.007 | 1 | 9 | Kctd6 | 0.027 | Old |
| 0.107159 | 0.226904 | 0.601 | 0.507 | 1 | 9 | Slc9a3r2 | 0.094 | Old |
| 0.107308 | 1.386236 | 0.121 | 0.074 | 1 | 9 | Traf7 | 0.047 | Old |
| 0.107457 | 0.450117 | 0.077 | 0.037 | 1 | 9 | Zfp236 | 0.04 | Old |
| 0.107569 | 1.057912 | 0.034 | 0.007 | 1 | 9 | Zfp251 | 0.027 | Old |
| 0.107569 | 0.815232 | 0.034 | 0.007 | 1 | 9 | Mars | 0.027 | Old |
| 0.107569 | 0.629484 | 0.034 | 0.007 | 1 | 9 | Tmem129 | 0.027 | Old |
| 0.107569 | 0.003911 | 0.034 | 0.007 | 1 | 9 | Mxra8 | 0.027 | Old |
| 0.107613 | 1.528496 | 0.085 | 0.044 | 1 | 9 | Pi4ka | 0.041 | Old |
| 0.107783 | 0.877826 | 0.057 | 0.022 | 1 | 9 | Kif5c | 0.035 | Old |
| 0.107784 | 0.244834 | 0.237 | 0.176 | 1 | 9 | Capza1 | 0.061 | Old |
| 0.10787 | 0.405852 | 0.338 | 0.272 | 1 | 9 | Mtch1 | 0.066 | Old |
| 0.107909 | 1.309711 | 0.067 | 0.029 | 1 | 9 | Mrpl55 | 0.038 | Old |
| 0.108423 | 0.49199 | 0.477 | 0.419 | 1 | 9 | Tcf15 | 0.058 | Old |
| 0.108458 | 1.037305 | 0.131 | 0.081 | 1 | 9 | Gga1 | 0.05 | Old |
| 0.108474 | 0.37211 | 0.281 | 0.213 | 1 | 9 | Aida | 0.068 | Old |
| 0.108601 | 0.212947 | 0.611 | 0.507 | 1 | 9 | Eif3a | 0.104 | Old |
| 0.108637 | 0.537507 | 0.284 | 0.213 | 1 | 9 | Kdelr1 | 0.071 | Old |
| 0.108699 | 0.102486 | 0.003 | 0.015 | 1 | 9 | Rasgrp1 | -0.012 | Old |
| 0.108884 | 1.37214 | 0.085 | 0.044 | 1 | 9 | Zfp180 | 0.041 | Old |
| 0.109091 | 0.420815 | 0.152 | 0.096 | 1 | 9 | Cnppd1 | 0.056 | Old |
| 0.109156 | 0.122424 | 0.76 | 0.721 | 1 | 9 | Fus | 0.039 | Old |
| 0.109335 | 1.125332 | 0.165 | 0.11 | 1 | 9 | Asxl2 | 0.055 | Old |
| 0.109388 | 0.449375 | 0.095 | 0.051 | 1 | 9 | Rbm34 | 0.044 | Old |
| 0.110262 | 1.021297 | 0.121 | 0.074 | 1 | 9 | Tbc1d15 | 0.047 | Old |
| 0.110304 | 1.264009 | 0.113 | 0.066 | 1 | 9 | L1cam | 0.047 | Old |
| 0.110355 | 0.647567 | 0.142 | 0.088 | 1 | 9 | Fam91a1 | 0.054 | Old |
| 0.110392 | 0.561005 | 0.503 | 0.456 | 1 | 9 | S100a16 | 0.047 | Old |
| 0.110862 | 0.633167 | 0.142 | 0.088 | 1 | 9 | Tshz1 | 0.054 | Old |
| 0.110911 | 1.195172 | 0.095 | 0.051 | 1 | 9 | Slc38a9 | 0.044 | Old |
| 0.11094 | 0.876995 | 0.057 | 0.022 | 1 | 9 | Pcyox1l | 0.035 | Old |
| 0.111348 | 0.484155 | 0.247 | 0.176 | 1 | 9 | Atxn2 | 0.071 | Old |
| 0.111502 | 0.499457 | 0.077 | 0.037 | 1 | 9 | Peli2 | 0.04 | Old |
| 0.111657 | 1.467831 | 0.075 | 0.037 | 1 | 9 | Tmem71 | 0.038 | Old |

| | | | | | | | | |
|---|---|---|---|---|---|---|---|---|
| 0.111784 | 2.94475 | 0.044 | 0.015 | 1 | 9 | Ap4e1 | 0.029 | Old |
| 0.111834 | 1.045921 | 0.095 | 0.051 | 1 | 9 | Qsox2 | 0.044 | Old |
| 0.112143 | 0.45075 | 0.057 | 0.022 | 1 | 9 | Kcnc2 | 0.035 | Old |
| 0.112288 | 0.094289 | 0.68 | 0.559 | 1 | 9 | Rps11 | 0.121 | Old |
| 0.112406 | 0.188235 | 0.088 | 0.044 | 1 | 9 | Katna1 | 0.044 | Old |
| 0.112651 | 0.835715 | 0.142 | 0.088 | 1 | 9 | Sik3 | 0.054 | Old |
| 0.112702 | 1.625427 | 0.075 | 0.037 | 1 | 9 | Bcl2l13 | 0.038 | Old |
| 0.113102 | 0.638005 | 0.268 | 0.206 | 1 | 9 | Galnt1 | 0.062 | Old |
| 0.113226 | 0.326521 | 0.235 | 0.169 | 1 | 9 | Tcn2 | 0.066 | Old |
| 0.113732 | 0.037882 | 0.131 | 0.199 | 1 | 9 | Shoc2 | -0.068 | Old |
| 0.113826 | 0.755929 | 0.201 | 0.14 | 1 | 9 | Ttc39b | 0.061 | Old |
| 0.113988 | 0.884426 | 0.131 | 0.081 | 1 | 9 | Fbln5 | 0.05 | Old |
| 0.114049 | 0.527191 | 0.32 | 0.25 | 1 | 9 | Ggta1 | 0.07 | Old |
| 0.114107 | 1.590971 | 0.075 | 0.037 | 1 | 9 | Btrc | 0.038 | Old |
| 0.114558 | 2.61709 | 0.044 | 0.015 | 1 | 9 | Atat1 | 0.029 | Old |
| 0.114706 | 1.013543 | 0.162 | 0.11 | 1 | 9 | Tmx1 | 0.052 | Old |
| 0.115186 | 2.324733 | 0.054 | 0.022 | 1 | 9 | Nipal3 | 0.032 | Old |
| 0.115186 | 2.258319 | 0.054 | 0.022 | 1 | 9 | Hars2 | 0.032 | Old |
| 0.115199 | 0.407027 | 0.193 | 0.132 | 1 | 9 | Jak2 | 0.061 | Old |
| 0.115265 | 0.531017 | 0.222 | 0.162 | 1 | 9 | Txndc17 | 0.06 | Old |
| 0.115543 | 4.127509 | 0.018 | 0 | 1 | 9 | Pgf | 0.018 | Old |
| 0.115543 | 3.635646 | 0.018 | 0 | 1 | 9 | Stau2 | 0.018 | Old |
| 0.115543 | 3.600434 | 0.018 | 0 | 1 | 9 | Mfsd9 | 0.018 | Old |
| 0.115543 | 3.582157 | 0.018 | 0 | 1 | 9 | Abcb6 | 0.018 | Old |
| 0.115543 | 3.47535 | 0.018 | 0 | 1 | 9 | Slco2a1 | 0.018 | Old |
| 0.115543 | 3.347892 | 0.018 | 0 | 1 | 9 | Des | 0.018 | Old |
| 0.115543 | 3.308082 | 0.018 | 0 | 1 | 9 | Ly75 | 0.018 | Old |
| 0.115543 | 3.20353 | 0.018 | 0 | 1 | 9 | Atad5 | 0.018 | Old |
| 0.115543 | 3.150489 | 0.018 | 0 | 1 | 9 | Zfp334 | 0.018 | Old |
| 0.115543 | 3.138197 | 0.018 | 0 | 1 | 9 | Cadps2 | 0.018 | Old |
| 0.115543 | 3.069795 | 0.018 | 0 | 1 | 9 | Plppr2 | 0.018 | Old |
| 0.115543 | 3.035261 | 0.018 | 0 | 1 | 9 | Rdh5 | 0.018 | Old |
| 0.115543 | 3.030262 | 0.018 | 0 | 1 | 9 | Xlr | 0.018 | Old |
| 0.115543 | 3.013237 | 0.018 | 0 | 1 | 9 | Ovca2 | 0.018 | Old |
| 0.115543 | 2.998414 | 0.018 | 0 | 1 | 9 | Bphl | 0.018 | Old |
| 0.115543 | 2.987792 | 0.018 | 0 | 1 | 9 | Exoc3l4 | 0.018 | Old |
| 0.115543 | 2.959908 | 0.018 | 0 | 1 | 9 | Lmln | 0.018 | Old |
| 0.115543 | 2.93754 | 0.018 | 0 | 1 | 9 | Cadm3 | 0.018 | Old |
| 0.115543 | 2.917943 | 0.018 | 0 | 1 | 9 | Dffa | 0.018 | Old |
| 0.115543 | 2.910481 | 0.018 | 0 | 1 | 9 | Dync2li1 | 0.018 | Old |
| 0.115543 | 2.909513 | 0.018 | 0 | 1 | 9 | Map7d3 | 0.018 | Old |
| 0.115543 | 2.901792 | 0.018 | 0 | 1 | 9 | Tmem191c | 0.018 | Old |
| 0.115543 | 2.76392 | 0.018 | 0 | 1 | 9 | 1110035H1 | 0.018 | Old |
| 0.115543 | 2.743274 | 0.018 | 0 | 1 | 9 | Gm10419 | 0.018 | Old |
| 0.115543 | 2.73647 | 0.018 | 0 | 1 | 9 | Zfp57 | 0.018 | Old |

| | | | | | | | | |
|---|---|---|---|---|---|---|---|---|
| 0.115543 | 2.683747 | 0.018 | 0 | 1 | 9 | Rbms3 | 0.018 | Old |
| 0.115543 | 2.675293 | 0.018 | 0 | 1 | 9 | Stard10 | 0.018 | Old |
| 0.115543 | 2.667868 | 0.018 | 0 | 1 | 9 | Cdkl5 | 0.018 | Old |
| 0.115543 | 2.667098 | 0.018 | 0 | 1 | 9 | Naip2 | 0.018 | Old |
| 0.115543 | 2.663545 | 0.018 | 0 | 1 | 9 | Meis3 | 0.018 | Old |
| 0.115543 | 2.644532 | 0.018 | 0 | 1 | 9 | Nnt | 0.018 | Old |
| 0.115543 | 2.56505 | 0.018 | 0 | 1 | 9 | Gm10399 | 0.018 | Old |
| 0.115543 | 2.541549 | 0.018 | 0 | 1 | 9 | AC164424. | 0.018 | Old |
| 0.115543 | 2.526545 | 0.018 | 0 | 1 | 9 | Nup37 | 0.018 | Old |
| 0.115543 | 2.516223 | 0.018 | 0 | 1 | 9 | Lrrc61 | 0.018 | Old |
| 0.115543 | 2.514375 | 0.018 | 0 | 1 | 9 | Pcdhga9 | 0.018 | Old |
| 0.115543 | 2.490859 | 0.018 | 0 | 1 | 9 | AC130815. | 0.018 | Old |
| 0.115543 | 2.471191 | 0.018 | 0 | 1 | 9 | Fam120aos | 0.018 | Old |
| 0.115543 | 2.468076 | 0.018 | 0 | 1 | 9 | Prr5l | 0.018 | Old |
| 0.115543 | 2.421966 | 0.018 | 0 | 1 | 9 | Il27ra | 0.018 | Old |
| 0.115543 | 2.372692 | 0.018 | 0 | 1 | 9 | Rfc4 | 0.018 | Old |
| 0.115543 | 2.262415 | 0.018 | 0 | 1 | 9 | C1qtnf6 | 0.018 | Old |
| 0.115543 | 2.254794 | 0.018 | 0 | 1 | 9 | Ficd | 0.018 | Old |
| 0.115811 | 0.24419 | 0.693 | 0.588 | 1 | 9 | Rbm39 | 0.105 | Old |
| 0.115965 | 2.01255 | 0.044 | 0.015 | 1 | 9 | Pfkm | 0.029 | Old |
| 0.116026 | 2.439102 | 0.054 | 0.022 | 1 | 9 | Atg4d | 0.032 | Old |
| 0.116039 | 0.342615 | 0.327 | 0.265 | 1 | 9 | Ubl5 | 0.062 | Old |
| 0.116437 | 1.803314 | 0.044 | 0.015 | 1 | 9 | Col15a1 | 0.029 | Old |
| 0.116459 | 0.628793 | 0.188 | 0.132 | 1 | 9 | Rbm27 | 0.056 | Old |
| 0.116828 | 0.351029 | 0.149 | 0.096 | 1 | 9 | Dlg1 | 0.053 | Old |
| 0.117386 | 1.663645 | 0.044 | 0.015 | 1 | 9 | Klhl42 | 0.029 | Old |
| 0.117722 | 2.0841 | 0.054 | 0.022 | 1 | 9 | Ano10 | 0.032 | Old |
| 0.117722 | 2.038654 | 0.054 | 0.022 | 1 | 9 | Capn15 | 0.032 | Old |
| 0.117722 | 1.924073 | 0.054 | 0.022 | 1 | 9 | Manba | 0.032 | Old |
| 0.118648 | 1.502292 | 0.064 | 0.029 | 1 | 9 | 2310022A1 | 0.035 | Old |
| 0.118648 | 1.471172 | 0.064 | 0.029 | 1 | 9 | Acsf2 | 0.035 | Old |
| 0.118661 | 0.265542 | 0.729 | 0.676 | 1 | 9 | Tspan7 | 0.053 | Old |
| 0.118797 | 0.835228 | 0.188 | 0.132 | 1 | 9 | Gsk3a | 0.056 | Old |
| 0.118974 | 1.017091 | 0.139 | 0.088 | 1 | 9 | Insig2 | 0.051 | Old |
| 0.118974 | 0.911317 | 0.139 | 0.088 | 1 | 9 | Gpalpp1 | 0.051 | Old |
| 0.119114 | 0.975041 | 0.144 | 0.096 | 1 | 9 | Chm | 0.048 | Old |
| 0.119273 | 0.503955 | 0.113 | 0.066 | 1 | 9 | Rtp3 | 0.047 | Old |
| 0.119406 | 0.618526 | 0.206 | 0.147 | 1 | 9 | Ptp4a1 | 0.059 | Old |
| 0.119499 | 0.52802 | 0.191 | 0.132 | 1 | 9 | Btg3 | 0.059 | Old |
| 0.119601 | 0.388731 | 0.201 | 0.14 | 1 | 9 | Parva | 0.061 | Old |
| 0.119785 | 1.414061 | 0.044 | 0.015 | 1 | 9 | Nudt14 | 0.029 | Old |
| 0.119787 | 0.580144 | 0.178 | 0.125 | 1 | 9 | Tmem57 | 0.053 | Old |
| 0.119867 | 1.096966 | 0.111 | 0.066 | 1 | 9 | Glo1 | 0.045 | Old |
| 0.11987 | 1.951375 | 0.054 | 0.022 | 1 | 9 | Rbm12b1 | 0.032 | Old |
| 0.119957 | 1.524399 | 0.093 | 0.051 | 1 | 9 | Nup85 | 0.042 | Old |

| | | | | | | | | |
|---|---|---|---|---|---|---|---|---|
| 0.120341 | 0.138078 | 0.539 | 0.441 | 1 | 9 | Tie1 | 0.098 | Old |
| 0.120673 | 0.140393 | 0.485 | 0.397 | 1 | 9 | 1810037I1 | 0.088 | Old |
| 0.120755 | 0.959876 | 0.044 | 0.015 | 1 | 9 | Tyw1 | 0.029 | Old |
| 0.120819 | 0.458347 | 0.219 | 0.154 | 1 | 9 | Dstyk | 0.065 | Old |
| 0.120931 | 0.714452 | 0.186 | 0.132 | 1 | 9 | Mysm1 | 0.054 | Old |
| 0.12114 | 0.814245 | 0.103 | 0.059 | 1 | 9 | Il11ra1 | 0.044 | Old |
| 0.12121 | 0.733026 | 0.085 | 0.044 | 1 | 9 | Mrpl9 | 0.041 | Old |
| 0.121242 | 0.270673 | 0.044 | 0.015 | 1 | 9 | Mtg1 | 0.029 | Old |
| 0.121535 | 0.145866 | 0.384 | 0.294 | 1 | 9 | Pls3 | 0.09 | Old |
| 0.121638 | 0.353353 | 0.186 | 0.125 | 1 | 9 | Sun1 | 0.061 | Old |
| 0.121798 | 0.618014 | 0.165 | 0.11 | 1 | 9 | Hhex | 0.055 | Old |
| 0.121906 | 0.869313 | 0.085 | 0.044 | 1 | 9 | Hoxb3 | 0.041 | Old |
| 0.122228 | 0.246472 | 0.611 | 0.522 | 1 | 9 | Gda | 0.089 | Old |
| 0.122358 | 1.501455 | 0.101 | 0.059 | 1 | 9 | 1300002E1 | 0.042 | Old |
| 0.122371 | 0.082753 | 0.454 | 0.368 | 1 | 9 | Rock1 | 0.086 | Old |
| 0.122474 | 0.308304 | 0.229 | 0.162 | 1 | 9 | Plod1 | 0.067 | Old |
| 0.122696 | 1.300628 | 0.082 | 0.044 | 1 | 9 | Smg9 | 0.038 | Old |
| 0.123213 | 0.233253 | 0.461 | 0.382 | 1 | 9 | Amotl2 | 0.079 | Old |
| 0.12338 | 0.501732 | 0.276 | 0.206 | 1 | 9 | Exoc5 | 0.07 | Old |
| 0.123702 | 0.392142 | 0.044 | 0.015 | 1 | 9 | Tmem246 | 0.029 | Old |
| 0.123941 | 0.58296 | 0.214 | 0.154 | 1 | 9 | Pdcd5 | 0.06 | Old |
| 0.124198 | 1.017138 | 0.044 | 0.015 | 1 | 9 | Rtn1 | 0.029 | Old |
| 0.124637 | 1.286123 | 0.093 | 0.051 | 1 | 9 | Ap1ar | 0.042 | Old |
| 0.124702 | 1.471336 | 0.054 | 0.022 | 1 | 9 | Fam20a | 0.032 | Old |
| 0.125005 | 0.578138 | 0.157 | 0.103 | 1 | 9 | Ddx58 | 0.054 | Old |
| 0.125055 | 0.687614 | 0.064 | 0.029 | 1 | 9 | Pex14 | 0.035 | Old |
| 0.125328 | 3.290973 | 0.031 | 0.007 | 1 | 9 | Gtpbp8 | 0.024 | Old |
| 0.125328 | 3.06216 | 0.031 | 0.007 | 1 | 9 | Uxt | 0.024 | Old |
| 0.125328 | 2.511417 | 0.031 | 0.007 | 1 | 9 | Btd | 0.024 | Old |
| 0.125597 | 1.129056 | 0.054 | 0.022 | 1 | 9 | Mcm4 | 0.032 | Old |
| 0.125932 | 2.407738 | 0.031 | 0.007 | 1 | 9 | Prtg | 0.024 | Old |
| 0.126066 | 0.905665 | 0.129 | 0.081 | 1 | 9 | Tmem87b | 0.048 | Old |
| 0.126191 | 0.891679 | 0.165 | 0.11 | 1 | 9 | Zmat2 | 0.055 | Old |
| 0.12641 | 0.446647 | 0.42 | 0.353 | 1 | 9 | Ctbp1 | 0.067 | Old |
| 0.126507 | 0.871467 | 0.085 | 0.044 | 1 | 9 | Sdk2 | 0.041 | Old |
| 0.126698 | 1.007712 | 0.064 | 0.029 | 1 | 9 | Trim11 | 0.035 | Old |
| 0.126987 | 0.972248 | 0.126 | 0.081 | 1 | 9 | Mnt | 0.045 | Old |
| 0.127135 | 0.761806 | 0.155 | 0.103 | 1 | 9 | Uap1 | 0.052 | Old |
| 0.127145 | 2.386449 | 0.031 | 0.007 | 1 | 9 | Lrrc45 | 0.024 | Old |
| 0.127226 | 0.37382 | 0.32 | 0.25 | 1 | 9 | Cbx1 | 0.07 | Old |
| 0.127275 | 0.059571 | 0.995 | 0.993 | 1 | 9 | mt-Atp6 | 0.002 | Old |
| 0.127487 | 0.254165 | 0.531 | 0.478 | 1 | 9 | Tm9sf3 | 0.053 | Old |
| 0.127629 | 0.267987 | 0.175 | 0.118 | 1 | 9 | Psmd12 | 0.057 | Old |
| 0.127755 | 1.925806 | 0.031 | 0.007 | 1 | 9 | Idua | 0.024 | Old |
| 0.127885 | 0.301194 | 0.57 | 0.5 | 1 | 9 | Ctnnbip1 | 0.07 | Old |

| | | | | | | | | |
|---|---|---|---|---|---|---|---|---|
| 0.127912 | 0.489964 | 0.103 | 0.059 | 1 | 9 | Fam49a | 0.044 | Old |
| 0.127941 | 0.875611 | 0.064 | 0.029 | 1 | 9 | Rnf8 | 0.035 | Old |
| 0.128019 | 0.350057 | 0.201 | 0.14 | 1 | 9 | Cttn | 0.061 | Old |
| 0.128043 | 0.269224 | 0.562 | 0.463 | 1 | 9 | Nr3c1 | 0.099 | Old |
| 0.128191 | 0.718847 | 0.183 | 0.125 | 1 | 9 | Ube2z | 0.058 | Old |
| 0.128474 | 0.143962 | 0.379 | 0.301 | 1 | 9 | Ssr3 | 0.078 | Old |
| 0.128762 | 1.088426 | 0.093 | 0.051 | 1 | 9 | Exoc1 | 0.042 | Old |
| 0.128775 | 1.034416 | 0.064 | 0.029 | 1 | 9 | Ccm2 | 0.035 | Old |
| 0.128775 | 0.423482 | 0.064 | 0.029 | 1 | 9 | Csf1 | 0.035 | Old |
| 0.128803 | 0.212854 | 0.113 | 0.066 | 1 | 9 | Rbm15b | 0.047 | Old |
| 0.128981 | 2.16789 | 0.031 | 0.007 | 1 | 9 | Fzd1 | 0.024 | Old |
| 0.128981 | 2.152821 | 0.031 | 0.007 | 1 | 9 | Fam83d | 0.024 | Old |
| 0.128981 | 1.60531 | 0.031 | 0.007 | 1 | 9 | Rmnd1 | 0.024 | Old |
| 0.129614 | 0.670528 | 0.064 | 0.029 | 1 | 9 | Surf1 | 0.035 | Old |
| 0.130217 | 1.862811 | 0.031 | 0.007 | 1 | 9 | Usp49 | 0.024 | Old |
| 0.130696 | 0.66883 | 0.193 | 0.14 | 1 | 9 | Pigt | 0.053 | Old |
| 0.130839 | 1.53248 | 0.031 | 0.007 | 1 | 9 | Agbl3 | 0.024 | Old |
| 0.130839 | 1.417443 | 0.031 | 0.007 | 1 | 9 | Shb | 0.024 | Old |
| 0.130842 | 0.532385 | 0.17 | 0.118 | 1 | 9 | Luc7l | 0.052 | Old |
| 0.130968 | 0.110592 | 0.915 | 0.846 | 1 | 9 | Ptprb | 0.069 | Old |
| 0.131355 | 0.412461 | 0.113 | 0.066 | 1 | 9 | 2510009E0 | 0.047 | Old |
| 0.131462 | 1.108252 | 0.031 | 0.007 | 1 | 9 | Slc30a6 | 0.024 | Old |
| 0.131764 | 0.299481 | 0.479 | 0.412 | 1 | 9 | Limch1 | 0.067 | Old |
| 0.131857 | 1.496963 | 0.09 | 0.051 | 1 | 9 | Bcl3 | 0.039 | Old |
| 0.132022 | 0.54028 | 0.178 | 0.125 | 1 | 9 | Nhsl2 | 0.053 | Old |
| 0.132024 | 0.76044 | 0.129 | 0.081 | 1 | 9 | Ahsa1 | 0.048 | Old |
| 0.132088 | 0.914588 | 0.031 | 0.007 | 1 | 9 | Tmem101 | 0.024 | Old |
| 0.132246 | 0.068658 | 0.224 | 0.162 | 1 | 9 | Rrp1 | 0.062 | Old |
| 0.132546 | 0.31316 | 0.26 | 0.199 | 1 | 9 | Trim12c | 0.061 | Old |
| 0.133412 | 0.792995 | 0.111 | 0.066 | 1 | 9 | Cdip1 | 0.045 | Old |
| 0.134877 | 0.369901 | 0.32 | 0.25 | 1 | 9 | Prpf4b | 0.07 | Old |
| 0.135394 | 0.167842 | 0.616 | 0.551 | 1 | 9 | Myo10 | 0.065 | Old |
| 0.13582 | 1.486612 | 0.108 | 0.066 | 1 | 9 | E2f4 | 0.042 | Old |
| 0.13604 | 0.177007 | 0.265 | 0.191 | 1 | 9 | Tma7 | 0.074 | Old |
| 0.136076 | 0.065829 | 0.32 | 0.235 | 1 | 9 | Vezf1 | 0.085 | Old |
| 0.136341 | 0.977909 | 0.152 | 0.103 | 1 | 9 | Srpk2 | 0.049 | Old |
| 0.136376 | 0.794713 | 0.296 | 0.243 | 1 | 9 | Dctn1 | 0.053 | Old |
| 0.136383 | 1.309984 | 0.082 | 0.044 | 1 | 9 | Tatdn2 | 0.038 | Old |
| 0.136598 | 0.845825 | 0.093 | 0.051 | 1 | 9 | Mdc1 | 0.042 | Old |
| 0.136742 | 2.899488 | 0.052 | 0.022 | 1 | 9 | Lcmt1 | 0.03 | Old |
| 0.136783 | 0.660021 | 0.137 | 0.088 | 1 | 9 | Maoa | 0.049 | Old |
| 0.136842 | 0.095442 | 0.119 | 0.176 | 1 | 9 | Cir1 | -0.057 | Old |
| 0.137087 | 0.827059 | 0.137 | 0.088 | 1 | 9 | Herpud2 | 0.049 | Old |
| 0.137223 | 1.882474 | 0.041 | 0.015 | 1 | 9 | 2810006K2 | 0.026 | Old |
| 0.137259 | 0.861214 | 0.129 | 0.081 | 1 | 9 | Eaf1 | 0.048 | Old |

| | | | | | | | | |
|---|---|---|---|---|---|---|---|---|
| 0.137481 | 1.089125 | 0.134 | 0.088 | 1 | 9 | Cask | 0.046 | Old |
| 0.137836 | 0.591137 | 0.168 | 0.11 | 1 | 9 | Kif13b | 0.058 | Old |
| 0.137931 | 1.299003 | 0.082 | 0.044 | 1 | 9 | Adprm | 0.038 | Old |
| 0.13802 | 0.459812 | 0.379 | 0.324 | 1 | 9 | Srsf1 | 0.055 | Old |
| 0.138306 | 0.903405 | 0.198 | 0.147 | 1 | 9 | Banf1 | 0.051 | Old |
| 0.13838 | 1.985915 | 0.062 | 0.029 | 1 | 9 | Nectin3 | 0.033 | Old |
| 0.138676 | 0.710568 | 0.16 | 0.11 | 1 | 9 | Myef2 | 0.05 | Old |
| 0.138803 | 0.396264 | 0.152 | 0.103 | 1 | 9 | Mrpl20 | 0.049 | Old |
| 0.13883 | 1.514431 | 0.062 | 0.029 | 1 | 9 | Zfp771 | 0.033 | Old |
| 0.139009 | 0.945416 | 0.072 | 0.037 | 1 | 9 | Esrra | 0.035 | Old |
| 0.139017 | 0.424398 | 0.216 | 0.154 | 1 | 9 | Ide | 0.062 | Old |
| 0.1391 | 0.54051 | 0.082 | 0.044 | 1 | 9 | Armc5 | 0.038 | Old |
| 0.139427 | 1.024858 | 0.072 | 0.037 | 1 | 9 | Erg | 0.035 | Old |
| 0.139443 | 2.937822 | 0.041 | 0.015 | 1 | 9 | Ift46 | 0.026 | Old |
| 0.139701 | 1.963643 | 0.052 | 0.022 | 1 | 9 | Tbp | 0.03 | Old |
| 0.139834 | 0.124705 | 0.284 | 0.213 | 1 | 9 | Med13 | 0.071 | Old |
| 0.140002 | 2.293912 | 0.041 | 0.015 | 1 | 9 | Fam234a | 0.026 | Old |
| 0.140002 | 2.197921 | 0.041 | 0.015 | 1 | 9 | Gmpr2 | 0.026 | Old |
| 0.140002 | 2.163705 | 0.041 | 0.015 | 1 | 9 | Fam207a | 0.026 | Old |
| 0.140099 | 0.578133 | 0.235 | 0.176 | 1 | 9 | Sltm | 0.059 | Old |
| 0.140315 | 0.379884 | 0.423 | 0.346 | 1 | 9 | Strn3 | 0.077 | Old |
| 0.140651 | 0.629248 | 0.093 | 0.051 | 1 | 9 | Prg4 | 0.042 | Old |
| 0.141062 | 0.610713 | 0.206 | 0.147 | 1 | 9 | Zdhhc18 | 0.059 | Old |
| 0.141067 | 0.211214 | 0.082 | 0.044 | 1 | 9 | Thoc3 | 0.038 | Old |
| 0.141178 | 0.600357 | 0.157 | 0.103 | 1 | 9 | Pitpnm2 | 0.054 | Old |
| 0.141531 | 0.933632 | 0.072 | 0.037 | 1 | 9 | Oma1 | 0.035 | Old |
| 0.141595 | 0.466544 | 0.152 | 0.103 | 1 | 9 | Cdk4 | 0.049 | Old |
| 0.141621 | 1.114029 | 0.108 | 0.066 | 1 | 9 | Retreg3 | 0.042 | Old |
| 0.14166 | 0.207717 | 0.015 | 0.037 | 1 | 9 | AI467606 | -0.022 | Old |
| 0.141974 | 0.719113 | 0.119 | 0.074 | 1 | 9 | Crkl | 0.045 | Old |
| 0.142011 | 1.620613 | 0.062 | 0.029 | 1 | 9 | BC003965 | 0.033 | Old |
| 0.142257 | 1.683506 | 0.041 | 0.015 | 1 | 9 | Trip6 | 0.026 | Old |
| 0.142364 | 0.212365 | 0.415 | 0.346 | 1 | 9 | Lmo7 | 0.069 | Old |
| 0.14265 | 0.240507 | 0.482 | 0.404 | 1 | 9 | Arhgef15 | 0.078 | Old |
| 0.142889 | 0.250105 | 0.193 | 0.14 | 1 | 9 | Tal1 | 0.053 | Old |
| 0.142931 | 1.624715 | 0.062 | 0.029 | 1 | 9 | Prr12 | 0.033 | Old |
| 0.142952 | 0.961732 | 0.09 | 0.051 | 1 | 9 | Lysmd2 | 0.039 | Old |
| 0.142981 | 0.733713 | 0.175 | 0.125 | 1 | 9 | Lama5 | 0.05 | Old |
| 0.143102 | 0.69992 | 0.015 | 0.037 | 1 | 9 | Coro2a | -0.022 | Old |
| 0.143208 | 0.489647 | 0.17 | 0.118 | 1 | 9 | Txndc9 | 0.052 | Old |
| 0.143278 | 0.294585 | 0.418 | 0.338 | 1 | 9 | 7-Sep | 0.08 | Old |
| 0.143392 | 1.506527 | 0.062 | 0.029 | 1 | 9 | Sspn | 0.033 | Old |
| 0.143437 | 0.097995 | 0.309 | 0.243 | 1 | 9 | Cops9 | 0.066 | Old |
| 0.143468 | 0.97349 | 0.126 | 0.081 | 1 | 9 | Zfp950 | 0.045 | Old |
| 0.143854 | 1.22811 | 0.062 | 0.029 | 1 | 9 | Wdfy1 | 0.033 | Old |

| | | | | | | | | |
|---|---|---|---|---|---|---|---|---|
| 0.144067 | 0.228008 | 0.206 | 0.147 | 1 | 9 | Rraga | 0.059 | Old |
| 0.144222 | 0.420355 | 0.294 | 0.235 | 1 | 9 | Akap13 | 0.059 | Old |
| 0.144318 | 1.422681 | 0.062 | 0.029 | 1 | 9 | Vps26b | 0.033 | Old |
| 0.14474 | 0.202937 | 0.036 | 0.066 | 1 | 9 | Cysltr1 | -0.03 | Old |
| 0.144922 | 0.790534 | 0.144 | 0.096 | 1 | 9 | Prpf38a | 0.048 | Old |
| 0.145114 | 1.493545 | 0.041 | 0.015 | 1 | 9 | Unk | 0.026 | Old |
| 0.145339 | 0.367415 | 0.5 | 0.426 | 1 | 9 | Selenow | 0.074 | Old |
| 0.145422 | 0.628202 | 0.198 | 0.147 | 1 | 9 | Arf3 | 0.051 | Old |
| 0.145567 | 3.592467 | 0.015 | 0 | 1 | 9 | Lbp | 0.015 | Old |
| 0.145567 | 3.59088 | 0.015 | 0 | 1 | 9 | Klhdc1 | 0.015 | Old |
| 0.145567 | 3.342885 | 0.015 | 0 | 1 | 9 | Hyls1 | 0.015 | Old |
| 0.145567 | 3.191938 | 0.015 | 0 | 1 | 9 | Nov | 0.015 | Old |
| 0.145567 | 3.188424 | 0.015 | 0 | 1 | 9 | Ctu1 | 0.015 | Old |
| 0.145567 | 3.187532 | 0.015 | 0 | 1 | 9 | Gm3636 | 0.015 | Old |
| 0.145567 | 3.117647 | 0.015 | 0 | 1 | 9 | Ubiad1 | 0.015 | Old |
| 0.145567 | 3.05393 | 0.015 | 0 | 1 | 9 | Nap1l3 | 0.015 | Old |
| 0.145567 | 3.029357 | 0.015 | 0 | 1 | 9 | A430110L2 | 0.015 | Old |
| 0.145567 | 3.022209 | 0.015 | 0 | 1 | 9 | Trim59 | 0.015 | Old |
| 0.145567 | 3.018333 | 0.015 | 0 | 1 | 9 | Pcdhb20 | 0.015 | Old |
| 0.145567 | 3.012072 | 0.015 | 0 | 1 | 9 | Morn2 | 0.015 | Old |
| 0.145567 | 3.005923 | 0.015 | 0 | 1 | 9 | Tti1 | 0.015 | Old |
| 0.145567 | 3.004301 | 0.015 | 0 | 1 | 9 | Zfp105 | 0.015 | Old |
| 0.145567 | 2.989146 | 0.015 | 0 | 1 | 9 | Cacnb2 | 0.015 | Old |
| 0.145567 | 2.964578 | 0.015 | 0 | 1 | 9 | Brdt | 0.015 | Old |
| 0.145567 | 2.915911 | 0.015 | 0 | 1 | 9 | Zfp442 | 0.015 | Old |
| 0.145567 | 2.895996 | 0.015 | 0 | 1 | 9 | Pglyrp1 | 0.015 | Old |
| 0.145567 | 2.874196 | 0.015 | 0 | 1 | 9 | 4933408B1 | 0.015 | Old |
| 0.145567 | 2.870301 | 0.015 | 0 | 1 | 9 | Lama3 | 0.015 | Old |
| 0.145567 | 2.842531 | 0.015 | 0 | 1 | 9 | Ppp1r14a | 0.015 | Old |
| 0.145567 | 2.831471 | 0.015 | 0 | 1 | 9 | Zbtb8a | 0.015 | Old |
| 0.145567 | 2.758218 | 0.015 | 0 | 1 | 9 | Swsap1 | 0.015 | Old |
| 0.145567 | 2.729219 | 0.015 | 0 | 1 | 9 | Gm13715 | 0.015 | Old |
| 0.145567 | 2.718813 | 0.015 | 0 | 1 | 9 | D5Ertd615 | 0.015 | Old |
| 0.145567 | 2.686514 | 0.015 | 0 | 1 | 9 | Gnao1 | 0.015 | Old |
| 0.145567 | 2.645921 | 0.015 | 0 | 1 | 9 | Ttyh1 | 0.015 | Old |
| 0.145567 | 2.645526 | 0.015 | 0 | 1 | 9 | Zfp688 | 0.015 | Old |
| 0.145567 | 2.623952 | 0.015 | 0 | 1 | 9 | Parp3 | 0.015 | Old |
| 0.145567 | 2.617519 | 0.015 | 0 | 1 | 9 | Chaf1b | 0.015 | Old |
| 0.145567 | 2.611981 | 0.015 | 0 | 1 | 9 | Dnaaf5 | 0.015 | Old |
| 0.145567 | 2.611973 | 0.015 | 0 | 1 | 9 | Uhrf1bp1 | 0.015 | Old |
| 0.145567 | 2.61184 | 0.015 | 0 | 1 | 9 | 9130019O2 | 0.015 | Old |
| 0.145567 | 2.588701 | 0.015 | 0 | 1 | 9 | Gm15050 | 0.015 | Old |
| 0.145567 | 2.57597 | 0.015 | 0 | 1 | 9 | Gm16287 | 0.015 | Old |
| 0.145567 | 2.568031 | 0.015 | 0 | 1 | 9 | Gm10638 | 0.015 | Old |
| 0.145567 | 2.563792 | 0.015 | 0 | 1 | 9 | Angptl2 | 0.015 | Old |

| | | | | | | | | |
|---|---|---|---|---|---|---|---|---|
| 0.145567 | 2.563566 | 0.015 | 0 | 1 | 9 | Rspo1 | 0.015 | Old |
| 0.145567 | 2.562135 | 0.015 | 0 | 1 | 9 | Rad18 | 0.015 | Old |
| 0.145567 | 2.552621 | 0.015 | 0 | 1 | 9 | Gm5547 | 0.015 | Old |
| 0.145567 | 2.546681 | 0.015 | 0 | 1 | 9 | Ccdc3 | 0.015 | Old |
| 0.145567 | 2.521142 | 0.015 | 0 | 1 | 9 | Hoxd10 | 0.015 | Old |
| 0.145567 | 2.498741 | 0.015 | 0 | 1 | 9 | Zfp28 | 0.015 | Old |
| 0.145567 | 2.477 | 0.015 | 0 | 1 | 9 | Supv3l1 | 0.015 | Old |
| 0.145567 | 2.471839 | 0.015 | 0 | 1 | 9 | Paqr3 | 0.015 | Old |
| 0.145567 | 2.45456 | 0.015 | 0 | 1 | 9 | Fam174b | 0.015 | Old |
| 0.145567 | 2.454422 | 0.015 | 0 | 1 | 9 | Dusp28 | 0.015 | Old |
| 0.145567 | 2.431833 | 0.015 | 0 | 1 | 9 | Rangrf | 0.015 | Old |
| 0.145567 | 2.407512 | 0.015 | 0 | 1 | 9 | Lbx2 | 0.015 | Old |
| 0.145567 | 2.362527 | 0.015 | 0 | 1 | 9 | Cdh2 | 0.015 | Old |
| 0.145567 | 2.320055 | 0.015 | 0 | 1 | 9 | Srfbp1 | 0.015 | Old |
| 0.145567 | 2.296596 | 0.015 | 0 | 1 | 9 | Dph2 | 0.015 | Old |
| 0.145567 | 2.280976 | 0.015 | 0 | 1 | 9 | Dync2h1 | 0.015 | Old |
| 0.145567 | 2.233612 | 0.015 | 0 | 1 | 9 | Lrrc20 | 0.015 | Old |
| 0.145567 | 2.223799 | 0.015 | 0 | 1 | 9 | Nfatc4 | 0.015 | Old |
| 0.145567 | 2.210706 | 0.015 | 0 | 1 | 9 | Wdr74 | 0.015 | Old |
| 0.145567 | 2.101584 | 0.015 | 0 | 1 | 9 | Cwf19l1 | 0.015 | Old |
| 0.145567 | 2.050139 | 0.015 | 0 | 1 | 9 | Tut1 | 0.015 | Old |
| 0.145641 | 1.222025 | 0.09 | 0.051 | 1 | 9 | 2310022B0 | 0.039 | Old |
| 0.145691 | 1.667964 | 0.041 | 0.015 | 1 | 9 | Rbm45 | 0.026 | Old |
| 0.145716 | 1.496715 | 0.062 | 0.029 | 1 | 9 | Slitrk4 | 0.033 | Old |
| 0.145765 | 1.392554 | 0.052 | 0.022 | 1 | 9 | Cyp2j9 | 0.03 | Old |
| 0.14602 | 0.061983 | 0.015 | 0.037 | 1 | 9 | Ramp1 | -0.022 | Old |
| 0.146199 | 0.966117 | 0.116 | 0.074 | 1 | 9 | Prkaa1 | 0.042 | Old |
| 0.146342 | 0.62106 | 0.111 | 0.066 | 1 | 9 | Crlf2 | 0.045 | Old |
| 0.147094 | 1.068497 | 0.082 | 0.044 | 1 | 9 | Map2k5 | 0.038 | Old |
| 0.147312 | 1.598005 | 0.052 | 0.022 | 1 | 9 | Mtmr14 | 0.03 | Old |
| 0.147337 | 0.103705 | 0.82 | 0.75 | 1 | 9 | S1pr1 | 0.07 | Old |
| 0.147688 | 0.20136 | 0.477 | 0.397 | 1 | 9 | Ednrb | 0.08 | Old |
| 0.147781 | 1.022961 | 0.09 | 0.051 | 1 | 9 | Il16 | 0.039 | Old |
| 0.148015 | 1.121996 | 0.041 | 0.015 | 1 | 9 | Taf1b | 0.026 | Old |
| 0.148068 | 1.579837 | 0.062 | 0.029 | 1 | 9 | Fam135a | 0.033 | Old |
| 0.148499 | 0.376854 | 0.276 | 0.213 | 1 | 9 | Ankrd44 | 0.063 | Old |
| 0.148868 | 0.454157 | 0.255 | 0.199 | 1 | 9 | Prkd2 | 0.056 | Old |
| 0.148893 | 0.834698 | 0.041 | 0.015 | 1 | 9 | Ankrd26 | 0.026 | Old |
| 0.149394 | 0.22153 | 0.052 | 0.022 | 1 | 9 | Alkbh3 | 0.03 | Old |
| 0.149766 | 0.617376 | 0.152 | 0.103 | 1 | 9 | Chd2 | 0.049 | Old |
| 0.149918 | 0.998681 | 0.052 | 0.022 | 1 | 9 | Ubxn2b | 0.03 | Old |
| 0.149945 | 0.673174 | 0.09 | 0.051 | 1 | 9 | Rmdn3 | 0.039 | Old |
| 0.150134 | 0.614366 | 0.16 | 0.11 | 1 | 9 | Adgre5 | 0.05 | Old |
| 0.150206 | 0.315818 | 0.129 | 0.081 | 1 | 9 | Actr10 | 0.048 | Old |
| 0.150219 | 0.41833 | 0.327 | 0.272 | 1 | 9 | Btbd7 | 0.055 | Old |

| | | | | | | | | |
|---|---|---|---|---|---|---|---|---|
| 0.150367 | 0.015056 | 0.041 | 0.015 | 1 | 9 | Rab27a | 0.026 | Old |
| 0.150636 | 0.950037 | 0.072 | 0.037 | 1 | 9 | D10Wsu10 | 0.035 | Old |
| 0.150738 | 1.115759 | 0.09 | 0.051 | 1 | 9 | Gabpb2 | 0.039 | Old |
| 0.150871 | 0.242697 | 0.031 | 0.059 | 1 | 9 | Dock7 | -0.028 | Old |
| 0.15097 | 1.043003 | 0.052 | 0.022 | 1 | 9 | Map6 | 0.03 | Old |
| 0.150982 | 0.61892 | 0.175 | 0.125 | 1 | 9 | Cpt1a | 0.05 | Old |
| 0.15099 | 0.119612 | 0.021 | 0.044 | 1 | 9 | Nlrc5 | -0.023 | Old |
| 0.151343 | 0.103323 | 0.964 | 0.963 | 1 | 9 | mt-Nd4 | 0.001 | Old |
| 0.15141 | 1.144883 | 0.062 | 0.029 | 1 | 9 | Endod1 | 0.033 | Old |
| 0.151892 | 0.848277 | 0.062 | 0.029 | 1 | 9 | Casp9 | 0.033 | Old |
| 0.151973 | 0.940899 | 0.072 | 0.037 | 1 | 9 | Rbsn | 0.035 | Old |
| 0.152028 | 1.066579 | 0.052 | 0.022 | 1 | 9 | Sdhaf4 | 0.03 | Old |
| 0.15215 | 0.743388 | 0.041 | 0.015 | 1 | 9 | Diexf | 0.026 | Old |
| 0.152471 | 0.665501 | 0.126 | 0.081 | 1 | 9 | Nup153 | 0.045 | Old |
| 0.152559 | 0.993711 | 0.052 | 0.022 | 1 | 9 | Sigmar1 | 0.03 | Old |
| 0.152701 | 1.284375 | 0.098 | 0.059 | 1 | 9 | Atp9b | 0.039 | Old |
| 0.152893 | 0.106681 | 0.428 | 0.346 | 1 | 9 | Gatad2b | 0.082 | Old |
| 0.153591 | 0.496305 | 0.137 | 0.088 | 1 | 9 | Polr2e | 0.049 | Old |
| 0.153613 | 0.797342 | 0.149 | 0.103 | 1 | 9 | Tlk1 | 0.046 | Old |
| 0.153652 | 0.493034 | 0.119 | 0.074 | 1 | 9 | Zfp422 | 0.045 | Old |
| 0.153738 | 0.265946 | 0.162 | 0.11 | 1 | 9 | Hadh | 0.052 | Old |
| 0.153751 | 1.467343 | 0.08 | 0.044 | 1 | 9 | Nudt16l1 | 0.036 | Old |
| 0.153804 | 0.864202 | 0.134 | 0.088 | 1 | 9 | Nudt21 | 0.046 | Old |
| 0.154308 | 0.940116 | 0.165 | 0.118 | 1 | 9 | Smagp | 0.047 | Old |
| 0.154417 | 1.888373 | 0.119 | 0.081 | 1 | 9 | Asph | 0.038 | Old |
| 0.154621 | 0.271719 | 0.111 | 0.066 | 1 | 9 | Nsd2 | 0.045 | Old |
| 0.15464 | 0.185021 | 0.505 | 0.449 | 1 | 9 | 4-Sep | 0.056 | Old |
| 0.154775 | 0.743678 | 0.157 | 0.11 | 1 | 9 | Akirin1 | 0.047 | Old |
| 0.155092 | 0.390808 | 0.448 | 0.382 | 1 | 9 | Lbr | 0.066 | Old |
| 0.155338 | 0.23819 | 0.673 | 0.618 | 1 | 9 | Mylip | 0.055 | Old |
| 0.155562 | 0.555112 | 0.09 | 0.051 | 1 | 9 | Dpagt1 | 0.039 | Old |
| 0.155651 | 3.23391 | 0.028 | 0.007 | 1 | 9 | Mpv17l2 | 0.021 | Old |
| 0.155651 | 3.222679 | 0.028 | 0.007 | 1 | 9 | Rbm41 | 0.021 | Old |
| 0.155651 | 3.107801 | 0.028 | 0.007 | 1 | 9 | Hebp1 | 0.021 | Old |
| 0.155651 | 3.001158 | 0.028 | 0.007 | 1 | 9 | Dph7 | 0.021 | Old |
| 0.155651 | 2.877811 | 0.028 | 0.007 | 1 | 9 | Fkbp14 | 0.021 | Old |
| 0.155803 | 0.731961 | 0.098 | 0.059 | 1 | 9 | Wipi1 | 0.039 | Old |
| 0.155897 | 0.321528 | 0.376 | 0.324 | 1 | 9 | Ash1l | 0.052 | Old |
| 0.156004 | 0.562327 | 0.175 | 0.125 | 1 | 9 | Grk5 | 0.05 | Old |
| 0.156392 | 2.375826 | 0.028 | 0.007 | 1 | 9 | Dhx37 | 0.021 | Old |
| 0.156422 | 0.537627 | 0.289 | 0.228 | 1 | 9 | Ddx50 | 0.061 | Old |
| 0.156813 | 0.06999 | 0.041 | 0.074 | 1 | 9 | Pdxk | -0.033 | Old |
| 0.156956 | 0.223039 | 0.126 | 0.081 | 1 | 9 | Hmgcs1 | 0.045 | Old |
| 0.157273 | 0.713511 | 0.062 | 0.029 | 1 | 9 | Pcdhb14 | 0.033 | Old |
| 0.157381 | 0.38202 | 0.577 | 0.515 | 1 | 9 | Arhgap29 | 0.062 | Old |

| | | | | | | | | |
|---|---|---|---|---|---|---|---|---|
| 0.157474 | 0.266403 | 0.495 | 0.441 | 1 | 9 | Jcad | 0.054 | Old |
| 0.157684 | 0.414297 | 0.144 | 0.096 | 1 | 9 | Prkcg | 0.048 | Old |
| 0.157882 | 2.382257 | 0.028 | 0.007 | 1 | 9 | Tceal1 | 0.021 | Old |
| 0.157893 | 0.101582 | 0.108 | 0.066 | 1 | 9 | Akt3 | 0.042 | Old |
| 0.158072 | 0.940737 | 0.08 | 0.044 | 1 | 9 | Dpysl3 | 0.036 | Old |
| 0.158254 | 0.131185 | 0.515 | 0.419 | 1 | 9 | Anxa2 | 0.096 | Old |
| 0.15827 | 0.70134 | 0.108 | 0.066 | 1 | 9 | Lonp2 | 0.042 | Old |
| 0.158494 | 0.338451 | 0.052 | 0.022 | 1 | 9 | A230050P2 | 0.03 | Old |
| 0.158631 | 2.142119 | 0.028 | 0.007 | 1 | 9 | Higd1b | 0.021 | Old |
| 0.158631 | 2.116789 | 0.028 | 0.007 | 1 | 9 | Spata5 | 0.021 | Old |
| 0.158631 | 1.756112 | 0.028 | 0.007 | 1 | 9 | Itm2a | 0.021 | Old |
| 0.158648 | 0.464844 | 0.108 | 0.066 | 1 | 9 | AU019823 | 0.042 | Old |
| 0.158738 | 0.323627 | 0.423 | 0.346 | 1 | 9 | Kdm6b | 0.077 | Old |
| 0.159161 | 0.111675 | 0.271 | 0.206 | 1 | 9 | Nrip1 | 0.065 | Old |
| 0.159515 | 0.343049 | 0.631 | 0.588 | 1 | 9 | mt-Nd5 | 0.043 | Old |
| 0.15954 | 0.181732 | 0.655 | 0.574 | 1 | 9 | Mcl1 | 0.081 | Old |
| 0.159601 | 0.429538 | 0.397 | 0.346 | 1 | 9 | Arglu1 | 0.051 | Old |
| 0.159979 | 0.190651 | 0.206 | 0.147 | 1 | 9 | Pds5b | 0.059 | Old |
| 0.160012 | 0.869952 | 0.18 | 0.132 | 1 | 9 | Rere | 0.048 | Old |
| 0.1601 | 0.354585 | 0.286 | 0.228 | 1 | 9 | Cwc15 | 0.058 | Old |
| 0.160215 | 0.292918 | 0.33 | 0.257 | 1 | 9 | Cdc42bpb | 0.073 | Old |
| 0.160539 | 0.222202 | 0.229 | 0.169 | 1 | 9 | Pnpla8 | 0.06 | Old |
| 0.160816 | 0.314776 | 0.371 | 0.309 | 1 | 9 | Xist | 0.062 | Old |
| 0.160894 | 1.870403 | 0.028 | 0.007 | 1 | 9 | Hist1h4d | 0.021 | Old |
| 0.160894 | 1.750509 | 0.028 | 0.007 | 1 | 9 | Cep128 | 0.021 | Old |
| 0.160967 | 0.476147 | 0.309 | 0.25 | 1 | 9 | Aldoa | 0.059 | Old |
| 0.161176 | 0.809151 | 0.116 | 0.074 | 1 | 9 | Dll1 | 0.042 | Old |
| 0.16159 | 1.375183 | 0.095 | 0.059 | 1 | 9 | Tmc6 | 0.036 | Old |
| 0.161653 | 1.356421 | 0.028 | 0.007 | 1 | 9 | Def8 | 0.021 | Old |
| 0.161653 | 1.057705 | 0.028 | 0.007 | 1 | 9 | Kptn | 0.021 | Old |
| 0.162078 | 0.86527 | 0.108 | 0.066 | 1 | 9 | Cops3 | 0.042 | Old |
| 0.162151 | 0.454626 | 0.098 | 0.059 | 1 | 9 | Dld | 0.039 | Old |
| 0.162304 | 0.281824 | 0.479 | 0.426 | 1 | 9 | Arap3 | 0.053 | Old |
| 0.162384 | 1.276143 | 0.07 | 0.037 | 1 | 9 | Thsd7a | 0.033 | Old |
| 0.162384 | 0.855951 | 0.07 | 0.037 | 1 | 9 | Trp53bp2 | 0.033 | Old |
| 0.162416 | 1.193767 | 0.028 | 0.007 | 1 | 9 | Kctd13 | 0.021 | Old |
| 0.162484 | 0.351372 | 0.08 | 0.044 | 1 | 9 | Mrps28 | 0.036 | Old |
| 0.162742 | 1.885 | 0.059 | 0.029 | 1 | 9 | Sppl2b | 0.03 | Old |
| 0.163334 | 0.805433 | 0.312 | 0.265 | 1 | 9 | Psma3 | 0.047 | Old |
| 0.163427 | 0.191406 | 0.186 | 0.132 | 1 | 9 | Uba7 | 0.054 | Old |
| 0.163568 | 0.648389 | 0.17 | 0.118 | 1 | 9 | Abcc4 | 0.052 | Old |
| 0.163881 | 1.713782 | 0.077 | 0.044 | 1 | 9 | Neurl4 | 0.033 | Old |
| 0.163949 | 0.392828 | 0.028 | 0.007 | 1 | 9 | Hist2h3c2 | 0.021 | Old |
| 0.163949 | 0.546323 | 0.028 | 0.007 | 1 | 9 | Nr6a1 | 0.021 | Old |
| 0.164091 | 1.031419 | 0.088 | 0.051 | 1 | 9 | Ap1m1 | 0.037 | Old |

| | | | | | | | | |
|---|---|---|---|---|---|---|---|---|
| 0.165029 | 0.868677 | 0.18 | 0.132 | 1 | 9 | Ndufab1 | 0.048 | Old |
| 0.165264 | 0.955612 | 0.07 | 0.037 | 1 | 9 | Fads3 | 0.033 | Old |
| 0.165479 | 0.49982 | 0.134 | 0.088 | 1 | 9 | Stx5a | 0.046 | Old |
| 0.165667 | 0.491906 | 0.214 | 0.162 | 1 | 9 | Rsbn1l | 0.052 | Old |
| 0.16621 | 0.30637 | 0.198 | 0.147 | 1 | 9 | Vps37a | 0.051 | Old |
| 0.166239 | 0.101127 | 0.495 | 0.419 | 1 | 9 | Tmem182 | 0.076 | Old |
| 0.166421 | 0.773183 | 0.098 | 0.059 | 1 | 9 | Shc1 | 0.039 | Old |
| 0.166609 | 0.198185 | 0.24 | 0.184 | 1 | 9 | Inpp1 | 0.056 | Old |
| 0.167063 | 0.714791 | 0.178 | 0.132 | 1 | 9 | Snrnp200 | 0.046 | Old |
| 0.167344 | 2.362038 | 0.049 | 0.022 | 1 | 9 | Timm9 | 0.027 | Old |
| 0.167647 | 0.088222 | 0.704 | 0.632 | 1 | 9 | Rpl18a | 0.072 | Old |
| 0.167939 | 0.093727 | 0.116 | 0.074 | 1 | 9 | Tmem252 | 0.042 | Old |
| 0.168011 | 0.188989 | 0.608 | 0.529 | 1 | 9 | Sypl | 0.079 | Old |
| 0.168289 | 2.491415 | 0.039 | 0.015 | 1 | 9 | Endog | 0.024 | Old |
| 0.168512 | 1.803228 | 0.049 | 0.022 | 1 | 9 | Dnajc18 | 0.027 | Old |
| 0.168568 | 0.829467 | 0.232 | 0.184 | 1 | 9 | Rbck1 | 0.048 | Old |
| 0.168621 | 1.737697 | 0.103 | 0.066 | 1 | 9 | Gm26802 | 0.037 | Old |
| 0.168779 | 0.382172 | 0.387 | 0.338 | 1 | 9 | Baz2b | 0.049 | Old |
| 0.16892 | 0.508836 | 0.358 | 0.301 | 1 | 9 | Ppp1r12a | 0.057 | Old |
| 0.168952 | 2.264357 | 0.039 | 0.015 | 1 | 9 | Rnf126 | 0.024 | Old |
| 0.169051 | 0.245352 | 0.317 | 0.25 | 1 | 9 | Foxo1 | 0.067 | Old |
| 0.169152 | 0.523199 | 0.16 | 0.11 | 1 | 9 | Cfdp1 | 0.05 | Old |
| 0.169438 | 1.403736 | 0.095 | 0.059 | 1 | 9 | Csad | 0.036 | Old |
| 0.169449 | 0.998876 | 0.113 | 0.074 | 1 | 9 | Atg5 | 0.039 | Old |
| 0.169574 | 1.457235 | 0.059 | 0.029 | 1 | 9 | Cep164 | 0.03 | Old |
| 0.169768 | 0.474254 | 0.165 | 0.118 | 1 | 9 | Rap2c | 0.047 | Old |
| 0.170197 | 0.591174 | 0.168 | 0.118 | 1 | 9 | Iqsec1 | 0.05 | Old |
| 0.170276 | 1.947846 | 0.049 | 0.022 | 1 | 9 | Wdr70 | 0.027 | Old |
| 0.170372 | 0.193399 | 0.224 | 0.169 | 1 | 9 | S100a1 | 0.055 | Old |
| 0.170565 | 0.757636 | 0.278 | 0.235 | 1 | 9 | Gpi1 | 0.043 | Old |
| 0.170658 | 0.688581 | 0.08 | 0.044 | 1 | 9 | Adcy6 | 0.036 | Old |
| 0.170887 | 0.404734 | 0.278 | 0.221 | 1 | 9 | Hspe1 | 0.057 | Old |
| 0.171207 | 0.414028 | 0.399 | 0.353 | 1 | 9 | Zmiz1 | 0.046 | Old |
| 0.171274 | 1.478935 | 0.077 | 0.044 | 1 | 9 | Snhg12 | 0.033 | Old |
| 0.171495 | 0.942058 | 0.088 | 0.051 | 1 | 9 | Pyurf | 0.037 | Old |
| 0.171719 | 1.021023 | 0.059 | 0.029 | 1 | 9 | Zfp74 | 0.03 | Old |
| 0.172228 | 0.594491 | 0.206 | 0.162 | 1 | 9 | Cmtm8 | 0.044 | Old |
| 0.172298 | 1.641701 | 0.039 | 0.015 | 1 | 9 | Spg11 | 0.024 | Old |
| 0.172382 | 0.738072 | 0.088 | 0.051 | 1 | 9 | Ubac2 | 0.037 | Old |
| 0.172405 | 0.099665 | 0.564 | 0.463 | 1 | 9 | Gm10076 | 0.101 | Old |
| 0.172434 | 1.152744 | 0.137 | 0.096 | 1 | 9 | Cul2 | 0.041 | Old |
| 0.172688 | 1.289178 | 0.077 | 0.044 | 1 | 9 | Fam214b | 0.033 | Old |
| 0.173246 | 1.864703 | 0.049 | 0.022 | 1 | 9 | Trub2 | 0.027 | Old |
| 0.173373 | 0.263825 | 0.137 | 0.088 | 1 | 9 | Lgals8 | 0.049 | Old |
| 0.173444 | 0.209222 | 0.701 | 0.625 | 1 | 9 | Srsf5 | 0.076 | Old |

| | | | | | | | | |
|---|---|---|---|---|---|---|---|---|
| 0.173544 | 0.500257 | 0.149 | 0.103 | 1 | 9 | Mpdz | 0.046 | Old |
| 0.173655 | 0.045546 | 0.704 | 0.618 | 1 | 9 | Rasip1 | 0.086 | Old |
| 0.173884 | 0.807718 | 0.059 | 0.029 | 1 | 9 | Pigyl | 0.03 | Old |
| 0.17411 | 1.051608 | 0.095 | 0.059 | 1 | 9 | Gtf2b | 0.036 | Old |
| 0.174317 | 0.26291 | 0.668 | 0.632 | 1 | 9 | Itgb1 | 0.036 | Old |
| 0.174329 | 1.354568 | 0.039 | 0.015 | 1 | 9 | Pigg | 0.024 | Old |
| 0.174558 | 0.897465 | 0.175 | 0.132 | 1 | 9 | Rbm7 | 0.043 | Old |
| 0.174637 | 0.758599 | 0.07 | 0.037 | 1 | 9 | Tbck | 0.033 | Old |
| 0.175047 | 1.795994 | 0.049 | 0.022 | 1 | 9 | Pde6d | 0.027 | Old |
| 0.175123 | 0.251204 | 0.631 | 0.551 | 1 | 9 | Ddx6 | 0.08 | Old |
| 0.175541 | 1.535975 | 0.077 | 0.044 | 1 | 9 | Pgm1 | 0.033 | Old |
| 0.175647 | 0.915835 | 0.07 | 0.037 | 1 | 9 | Poc1b | 0.033 | Old |
| 0.175998 | 0.325336 | 0.106 | 0.066 | 1 | 9 | Mrpl12 | 0.04 | Old |
| 0.17607 | 0.829172 | 0.059 | 0.029 | 1 | 9 | Pold3 | 0.03 | Old |
| 0.176215 | 0.073724 | 0.992 | 1 | 1 | 9 | mt-Co2 | -0.008 | Old |
| 0.176255 | 1.599573 | 0.049 | 0.022 | 1 | 9 | Drosha | 0.027 | Old |
| 0.176356 | 0.471085 | 0.32 | 0.265 | 1 | 9 | Psmb8 | 0.055 | Old |
| 0.176378 | 1.112013 | 0.039 | 0.015 | 1 | 9 | Rnf113a2 | 0.024 | Old |
| 0.176604 | 0.571686 | 0.271 | 0.213 | 1 | 9 | Pcdh19 | 0.058 | Old |
| 0.176619 | 1.049273 | 0.059 | 0.029 | 1 | 9 | Vhl | 0.03 | Old |
| 0.176765 | 0.730104 | 0.276 | 0.221 | 1 | 9 | Casp8 | 0.055 | Old |
| 0.17681 | 0.448033 | 0.263 | 0.206 | 1 | 9 | Gdi1 | 0.057 | Old |
| 0.176815 | 0.549423 | 0.216 | 0.162 | 1 | 9 | Nfatc3 | 0.054 | Old |
| 0.177065 | 0.457156 | 0.039 | 0.015 | 1 | 9 | Cdc6 | 0.024 | Old |
| 0.177091 | 0.227282 | 0.629 | 0.551 | 1 | 9 | Anxa5 | 0.078 | Old |
| 0.177407 | 0.833003 | 0.129 | 0.088 | 1 | 9 | Mkrn1 | 0.041 | Old |
| 0.177417 | 0.581147 | 0.216 | 0.162 | 1 | 9 | Creb3l2 | 0.054 | Old |
| 0.177469 | 1.242892 | 0.049 | 0.022 | 1 | 9 | Ttll4 | 0.027 | Old |
| 0.177828 | 0.335853 | 0.07 | 0.11 | 1 | 9 | Plaur | -0.04 | Old |
| 0.177945 | 1.042026 | 0.077 | 0.044 | 1 | 9 | Tmx4 | 0.033 | Old |
| 0.178004 | 1.154246 | 0.095 | 0.059 | 1 | 9 | Faf2 | 0.036 | Old |
| 0.178807 | 0.41297 | 0.175 | 0.132 | 1 | 9 | Pitpnb | 0.043 | Old |
| 0.17883 | 0.249826 | 0.059 | 0.029 | 1 | 9 | Dolk | 0.03 | Old |
| 0.178925 | 1.050659 | 0.103 | 0.066 | 1 | 9 | Oser1 | 0.037 | Old |
| 0.179157 | 1.340264 | 0.077 | 0.044 | 1 | 9 | Arrdc3 | 0.033 | Old |
| 0.179279 | 0.448808 | 0.116 | 0.074 | 1 | 9 | 2010111I0 | 0.042 | Old |
| 0.179603 | 0.459475 | 0.088 | 0.051 | 1 | 9 | Tmem55a | 0.037 | Old |
| 0.179696 | 0.808169 | 0.131 | 0.088 | 1 | 9 | As3mt | 0.043 | Old |
| 0.179885 | 0.813599 | 0.162 | 0.118 | 1 | 9 | Psip1 | 0.044 | Old |
| 0.180196 | 0.703236 | 0.095 | 0.059 | 1 | 9 | Dip2a | 0.036 | Old |
| 0.180537 | 0.760645 | 0.232 | 0.184 | 1 | 9 | Stk10 | 0.048 | Old |
| 0.180601 | 0.307968 | 0.149 | 0.103 | 1 | 9 | Cnot2 | 0.046 | Old |
| 0.18104 | 1.066773 | 0.103 | 0.066 | 1 | 9 | Mpv17 | 0.037 | Old |
| 0.181048 | 0.21252 | 0.098 | 0.059 | 1 | 9 | Wdr61 | 0.039 | Old |
| 0.181078 | 0.572419 | 0.095 | 0.059 | 1 | 9 | Fbxo28 | 0.036 | Old |

| | | | | | | | | |
|---|---|---|---|---|---|---|---|---|
| 0.18115 | 0.944995 | 0.049 | 0.022 | 1 | 9 | 1700037H( | 0.027 | Old |
| 0.18115 | 0.907935 | 0.049 | 0.022 | 1 | 9 | Pex2 | 0.027 | Old |
| 0.181229 | 0.700601 | 0.039 | 0.015 | 1 | 9 | Gm14005 | 0.024 | Old |
| 0.181353 | 1.110427 | 0.077 | 0.044 | 1 | 9 | Zfp715 | 0.033 | Old |
| 0.181486 | 0.013223 | 0.098 | 0.059 | 1 | 9 | Snapin | 0.039 | Old |
| 0.181491 | 1.611669 | 0.093 | 0.059 | 1 | 9 | Ndfip1 | 0.034 | Old |
| 0.181623 | 0.748611 | 0.059 | 0.029 | 1 | 9 | Zfp160 | 0.03 | Old |
| 0.181639 | 0.76806 | 0.008 | 0.022 | 1 | 9 | Phpt1 | -0.014 | Old |
| 0.182185 | 0.696959 | 0.059 | 0.029 | 1 | 9 | Lgals4 | 0.03 | Old |
| 0.182335 | 0.40465 | 0.077 | 0.044 | 1 | 9 | Dchs1 | 0.033 | Old |
| 0.182738 | 0.723932 | 0.131 | 0.088 | 1 | 9 | Churc1 | 0.043 | Old |
| 0.182763 | 0.530482 | 0.139 | 0.096 | 1 | 9 | Mfap1b | 0.043 | Old |
| 0.182833 | 0.43115 | 0.088 | 0.051 | 1 | 9 | Pitpnm3 | 0.037 | Old |
| 0.183003 | 0.973904 | 0.113 | 0.074 | 1 | 9 | Yrdc | 0.039 | Old |
| 0.183321 | 1.224077 | 0.077 | 0.044 | 1 | 9 | St7l | 0.033 | Old |
| 0.183539 | 0.849468 | 0.106 | 0.066 | 1 | 9 | Trappc5 | 0.04 | Old |
| 0.183756 | 0.532969 | 0.173 | 0.125 | 1 | 9 | Psmd6 | 0.048 | Old |
| 0.184149 | 0.61004 | 0.188 | 0.14 | 1 | 9 | Anapc16 | 0.048 | Old |
| 0.18419 | 0.533759 | 0.095 | 0.059 | 1 | 9 | Pigq | 0.036 | Old |
| 0.184402 | 0.324638 | 0.049 | 0.081 | 1 | 9 | Dvl3 | -0.032 | Old |
| 0.1845 | 0.741103 | 0.242 | 0.191 | 1 | 9 | Nek7 | 0.051 | Old |
| 0.184519 | 3.326937 | 0.013 | 0 | 1 | 9 | Gm42595 | 0.013 | Old |
| 0.184519 | 3.274498 | 0.013 | 0 | 1 | 9 | Gcdh | 0.013 | Old |
| 0.184519 | 3.25422 | 0.013 | 0 | 1 | 9 | Phgdh | 0.013 | Old |
| 0.184519 | 3.058792 | 0.013 | 0 | 1 | 9 | Fndc10 | 0.013 | Old |
| 0.184519 | 3.04276 | 0.013 | 0 | 1 | 9 | Reln | 0.013 | Old |
| 0.184519 | 3.025147 | 0.013 | 0 | 1 | 9 | Cenpq | 0.013 | Old |
| 0.184519 | 3.005667 | 0.013 | 0 | 1 | 9 | Fstl3 | 0.013 | Old |
| 0.184519 | 2.96937 | 0.013 | 0 | 1 | 9 | Ogn | 0.013 | Old |
| 0.184519 | 2.879677 | 0.013 | 0 | 1 | 9 | Gm14418 | 0.013 | Old |
| 0.184519 | 2.843827 | 0.013 | 0 | 1 | 9 | Slc37a1 | 0.013 | Old |
| 0.184519 | 2.824384 | 0.013 | 0 | 1 | 9 | Irx5 | 0.013 | Old |
| 0.184519 | 2.820971 | 0.013 | 0 | 1 | 9 | Haus1 | 0.013 | Old |
| 0.184519 | 2.800671 | 0.013 | 0 | 1 | 9 | 1110051M | 0.013 | Old |
| 0.184519 | 2.791843 | 0.013 | 0 | 1 | 9 | C1qtnf4 | 0.013 | Old |
| 0.184519 | 2.78214 | 0.013 | 0 | 1 | 9 | Adamts6 | 0.013 | Old |
| 0.184519 | 2.705866 | 0.013 | 0 | 1 | 9 | Dand5 | 0.013 | Old |
| 0.184519 | 2.70138 | 0.013 | 0 | 1 | 9 | Gpr19 | 0.013 | Old |
| 0.184519 | 2.663854 | 0.013 | 0 | 1 | 9 | Prim1 | 0.013 | Old |
| 0.184519 | 2.632853 | 0.013 | 0 | 1 | 9 | Tpm2 | 0.013 | Old |
| 0.184519 | 2.631065 | 0.013 | 0 | 1 | 9 | Usp27x | 0.013 | Old |
| 0.184519 | 2.629699 | 0.013 | 0 | 1 | 9 | Slc25a35 | 0.013 | Old |
| 0.184519 | 2.58775 | 0.013 | 0 | 1 | 9 | Nudt6 | 0.013 | Old |
| 0.184519 | 2.556752 | 0.013 | 0 | 1 | 9 | Wdr55 | 0.013 | Old |
| 0.184519 | 2.496818 | 0.013 | 0 | 1 | 9 | Thg1l | 0.013 | Old |

| | | | | | | | | |
|---|---|---|---|---|---|---|---|---|
| 0.184519 | 2.469828 | 0.013 | 0 | 1 | 9 | Cd209b | 0.013 | Old |
| 0.184519 | 2.442382 | 0.013 | 0 | 1 | 9 | Kremen2 | 0.013 | Old |
| 0.184519 | 2.421081 | 0.013 | 0 | 1 | 9 | Atp6v0e2 | 0.013 | Old |
| 0.184519 | 2.384296 | 0.013 | 0 | 1 | 9 | Cacng1 | 0.013 | Old |
| 0.184519 | 2.356824 | 0.013 | 0 | 1 | 9 | Trim69 | 0.013 | Old |
| 0.184519 | 2.345787 | 0.013 | 0 | 1 | 9 | Gm9938 | 0.013 | Old |
| 0.184519 | 2.338365 | 0.013 | 0 | 1 | 9 | Gm26847 | 0.013 | Old |
| 0.184519 | 2.33079 | 0.013 | 0 | 1 | 9 | Ttc30b | 0.013 | Old |
| 0.184519 | 2.314876 | 0.013 | 0 | 1 | 9 | Zfp72 | 0.013 | Old |
| 0.184519 | 2.280397 | 0.013 | 0 | 1 | 9 | Dcaf4 | 0.013 | Old |
| 0.184519 | 2.278759 | 0.013 | 0 | 1 | 9 | Polm | 0.013 | Old |
| 0.184519 | 2.274992 | 0.013 | 0 | 1 | 9 | Mterf1a | 0.013 | Old |
| 0.184519 | 2.253674 | 0.013 | 0 | 1 | 9 | Mcoln2 | 0.013 | Old |
| 0.184519 | 2.188078 | 0.013 | 0 | 1 | 9 | Hdac8 | 0.013 | Old |
| 0.184519 | 2.186983 | 0.013 | 0 | 1 | 9 | Rpl9-ps6 | 0.013 | Old |
| 0.184519 | 2.182729 | 0.013 | 0 | 1 | 9 | P4htm | 0.013 | Old |
| 0.184519 | 2.177288 | 0.013 | 0 | 1 | 9 | Zscan20 | 0.013 | Old |
| 0.184519 | 2.176253 | 0.013 | 0 | 1 | 9 | Ttll5 | 0.013 | Old |
| 0.184519 | 2.169373 | 0.013 | 0 | 1 | 9 | 4732496C0 | 0.013 | Old |
| 0.184519 | 2.152419 | 0.013 | 0 | 1 | 9 | Slc19a2 | 0.013 | Old |
| 0.184519 | 2.148168 | 0.013 | 0 | 1 | 9 | Kank4 | 0.013 | Old |
| 0.184519 | 2.148068 | 0.013 | 0 | 1 | 9 | Cttnbp2 | 0.013 | Old |
| 0.184519 | 2.146183 | 0.013 | 0 | 1 | 9 | Sox12 | 0.013 | Old |
| 0.184519 | 2.145795 | 0.013 | 0 | 1 | 9 | Scn7a | 0.013 | Old |
| 0.184519 | 2.130798 | 0.013 | 0 | 1 | 9 | Sema3c | 0.013 | Old |
| 0.184519 | 2.123178 | 0.013 | 0 | 1 | 9 | Gm15446 | 0.013 | Old |
| 0.184519 | 2.114066 | 0.013 | 0 | 1 | 9 | Gm39822 | 0.013 | Old |
| 0.184519 | 2.088595 | 0.013 | 0 | 1 | 9 | Hcar1 | 0.013 | Old |
| 0.184519 | 2.069727 | 0.013 | 0 | 1 | 9 | 4632404H1 | 0.013 | Old |
| 0.184519 | 2.068457 | 0.013 | 0 | 1 | 9 | Mettl2 | 0.013 | Old |
| 0.184519 | 2.029038 | 0.013 | 0 | 1 | 9 | Tmem120b | 0.013 | Old |
| 0.184519 | 2.008797 | 0.013 | 0 | 1 | 9 | Aldh18a1 | 0.013 | Old |
| 0.184519 | 1.995573 | 0.013 | 0 | 1 | 9 | Slc4a11 | 0.013 | Old |
| 0.184519 | 1.951791 | 0.013 | 0 | 1 | 9 | Dglucy | 0.013 | Old |
| 0.184519 | 1.89056 | 0.013 | 0 | 1 | 9 | Dsn1 | 0.013 | Old |
| 0.184519 | 1.889472 | 0.013 | 0 | 1 | 9 | Ccdc134 | 0.013 | Old |
| 0.184519 | 1.88839 | 0.013 | 0 | 1 | 9 | Setbp1 | 0.013 | Old |
| 0.184519 | 1.869566 | 0.013 | 0 | 1 | 9 | Gtf2ird2 | 0.013 | Old |
| 0.184519 | 1.86617 | 0.013 | 0 | 1 | 9 | Golm1 | 0.013 | Old |
| 0.184519 | 1.786791 | 0.013 | 0 | 1 | 9 | Lhfp | 0.013 | Old |
| 0.184519 | 1.781634 | 0.013 | 0 | 1 | 9 | Ybey | 0.013 | Old |
| 0.184519 | 1.598119 | 0.013 | 0 | 1 | 9 | Hvcn1 | 0.013 | Old |
| 0.184519 | 1.425051 | 0.013 | 0 | 1 | 9 | Susd1 | 0.013 | Old |
| 0.184528 | 0.206069 | 0.572 | 0.507 | 1 | 9 | Nfe2l1 | 0.065 | Old |
| 0.184554 | 0.312095 | 0.338 | 0.279 | 1 | 9 | 7-Mar | 0.059 | Old |

| | | | | | | | | |
|---|---|---|---|---|---|---|---|---|
| 0.184808 | 0.975839 | 0.077 | 0.044 | 1 | 9 | Eci1 | 0.033 | Old |
| 0.18484 | 0.022918 | 0.564 | 0.478 | 1 | 9 | Tgfbr3 | 0.086 | Old |
| 0.184886 | 0.355746 | 0.049 | 0.022 | 1 | 9 | P2ry2 | 0.027 | Old |
| 0.185015 | 0.486911 | 0.232 | 0.184 | 1 | 9 | Tnfrsf1a | 0.048 | Old |
| 0.185296 | 0.302964 | 0.389 | 0.331 | 1 | 9 | Luzp1 | 0.058 | Old |
| 0.185313 | 0.178742 | 0.165 | 0.118 | 1 | 9 | Paf1 | 0.047 | Old |
| 0.185476 | 1.016205 | 0.113 | 0.074 | 1 | 9 | Unkl | 0.039 | Old |
| 0.185515 | 0.950135 | 0.049 | 0.022 | 1 | 9 | Cd46 | 0.027 | Old |
| 0.185515 | 0.830597 | 0.049 | 0.022 | 1 | 9 | Fdxr | 0.027 | Old |
| 0.18576 | 1.125116 | 0.103 | 0.066 | 1 | 9 | Ppp1r37 | 0.037 | Old |
| 0.185906 | 0.720009 | 0.152 | 0.11 | 1 | 9 | Stk40 | 0.042 | Old |
| 0.186144 | 0.673994 | 0.049 | 0.022 | 1 | 9 | Atg14 | 0.027 | Old |
| 0.186605 | 1.486574 | 0.067 | 0.037 | 1 | 9 | Trpm4 | 0.03 | Old |
| 0.186833 | 0.010241 | 0.356 | 0.287 | 1 | 9 | Dll4 | 0.069 | Old |
| 0.1869 | 0.609101 | 0.284 | 0.235 | 1 | 9 | Tmx3 | 0.049 | Old |
| 0.186995 | 0.292416 | 0.121 | 0.081 | 1 | 9 | Utp3 | 0.04 | Old |
| 0.187187 | 0.479362 | 0.255 | 0.206 | 1 | 9 | Helz | 0.049 | Old |
| 0.188183 | 0.17805 | 0.309 | 0.25 | 1 | 9 | Ndufa3 | 0.059 | Old |
| 0.188221 | 1.609436 | 0.067 | 0.037 | 1 | 9 | Smim4 | 0.03 | Old |
| 0.188618 | 0.790364 | 0.162 | 0.118 | 1 | 9 | Ift20 | 0.044 | Old |
| 0.188764 | 0.104824 | 0.049 | 0.081 | 1 | 9 | Idh2 | -0.032 | Old |
| 0.189232 | 0.653016 | 0.113 | 0.074 | 1 | 9 | Arl4d | 0.039 | Old |
| 0.189304 | 1.406677 | 0.067 | 0.037 | 1 | 9 | Mak16 | 0.03 | Old |
| 0.189329 | 0.408544 | 0.284 | 0.228 | 1 | 9 | Amotl1 | 0.056 | Old |
| 0.189847 | 1.332625 | 0.067 | 0.037 | 1 | 9 | Mrps26 | 0.03 | Old |
| 0.19013 | 0.843595 | 0.103 | 0.066 | 1 | 9 | Togaram1 | 0.037 | Old |
| 0.190457 | 0.426786 | 0.369 | 0.316 | 1 | 9 | Piezo1 | 0.053 | Old |
| 0.190654 | 0.296363 | 0.307 | 0.25 | 1 | 9 | Atp9a | 0.057 | Old |
| 0.190776 | 0.341333 | 0.059 | 0.029 | 1 | 9 | Kif1bp | 0.03 | Old |
| 0.190796 | 0.914742 | 0.155 | 0.11 | 1 | 9 | Camta2 | 0.045 | Old |
| 0.190937 | 0.934532 | 0.067 | 0.037 | 1 | 9 | Tmem243 | 0.03 | Old |
| 0.190986 | 0.117538 | 0.724 | 0.647 | 1 | 9 | Cd47 | 0.077 | Old |
| 0.1911 | 0.646272 | 0.144 | 0.103 | 1 | 9 | Creg1 | 0.041 | Old |
| 0.191181 | 0.349678 | 0.353 | 0.287 | 1 | 9 | Cep85l | 0.066 | Old |
| 0.192016 | 0.897957 | 0.175 | 0.132 | 1 | 9 | Gimap1 | 0.043 | Old |
| 0.192031 | 0.951719 | 0.067 | 0.037 | 1 | 9 | Adamts10 | 0.03 | Old |
| 0.192094 | 0.389242 | 0.278 | 0.228 | 1 | 9 | Atf7 | 0.05 | Old |
| 0.19215 | 0.153297 | 0.317 | 0.257 | 1 | 9 | Pxn | 0.06 | Old |
| 0.192788 | 0.868366 | 0.103 | 0.066 | 1 | 9 | Car5b | 0.037 | Old |
| 0.192939 | 0.486088 | 0.405 | 0.368 | 1 | 9 | Atp5k | 0.037 | Old |
| 0.193254 | 1.51024 | 0.075 | 0.044 | 1 | 9 | Tuba4a | 0.031 | Old |
| 0.193356 | 0.64424 | 0.235 | 0.184 | 1 | 9 | Ppm1f | 0.051 | Old |
| 0.193374 | 0.54267 | 0.155 | 0.11 | 1 | 9 | Qsox1 | 0.045 | Old |
| 0.19348 | 0.500011 | 0.157 | 0.11 | 1 | 9 | Camk2n1 | 0.047 | Old |
| 0.193597 | 3.070541 | 0.026 | 0.007 | 1 | 9 | Mthfd1 | 0.019 | Old |

| | | | | | | | | |
|---|---|---|---|---|---|---|---|---|
| 0.193597 | 2.668425 | 0.026 | 0.007 | 1 | 9 | Thap7 | 0.019 | Old |
| 0.193597 | 2.622234 | 0.026 | 0.007 | 1 | 9 | Nif3l1 | 0.019 | Old |
| 0.193597 | 2.570706 | 0.026 | 0.007 | 1 | 9 | Tmco6 | 0.019 | Old |
| 0.193597 | 2.426081 | 0.026 | 0.007 | 1 | 9 | Gm10138 | 0.019 | Old |
| 0.193597 | 2.394967 | 0.026 | 0.007 | 1 | 9 | Olfr920 | 0.019 | Old |
| 0.193597 | 2.344485 | 0.026 | 0.007 | 1 | 9 | Trmt44 | 0.019 | Old |
| 0.193597 | 2.242268 | 0.026 | 0.007 | 1 | 9 | Enho | 0.019 | Old |
| 0.193597 | 2.214386 | 0.026 | 0.007 | 1 | 9 | Rpap2 | 0.019 | Old |
| 0.193765 | 0.265456 | 0.095 | 0.059 | 1 | 9 | Tomm40 | 0.036 | Old |
| 0.193857 | 0.470053 | 0.273 | 0.228 | 1 | 9 | Prpf38b | 0.045 | Old |
| 0.193917 | 0.89242 | 0.077 | 0.044 | 1 | 9 | Dnajb2 | 0.033 | Old |
| 0.194031 | 0.064297 | 0.771 | 0.654 | 1 | 9 | Tmcc3 | 0.117 | Old |
| 0.194057 | 0.118951 | 0.918 | 0.89 | 1 | 9 | Rps21 | 0.028 | Old |
| 0.194185 | 0.099947 | 0.108 | 0.066 | 1 | 9 | Clpx | 0.042 | Old |
| 0.194234 | 1.210206 | 0.067 | 0.037 | 1 | 9 | Znhit6 | 0.03 | Old |
| 0.194278 | 0.146345 | 0.577 | 0.515 | 1 | 9 | Kank3 | 0.062 | Old |
| 0.194402 | 0.223183 | 0.304 | 0.243 | 1 | 9 | Epb41l2 | 0.061 | Old |
| 0.194509 | 0.279773 | 0.32 | 0.25 | 1 | 9 | Arf4 | 0.07 | Old |
| 0.194776 | 0.125407 | 0.696 | 0.61 | 1 | 9 | Macf1 | 0.086 | Old |
| 0.194781 | 0.650992 | 0.085 | 0.051 | 1 | 9 | Fbxo7 | 0.034 | Old |
| 0.195136 | 0.068494 | 0.309 | 0.25 | 1 | 9 | Impdh1 | 0.059 | Old |
| 0.195204 | 0.524404 | 0.088 | 0.051 | 1 | 9 | Zfp467 | 0.037 | Old |
| 0.195317 | 0.629975 | 0.268 | 0.228 | 1 | 9 | Rhoj | 0.04 | Old |
| 0.19542 | 2.461131 | 0.026 | 0.007 | 1 | 9 | Nek4 | 0.019 | Old |
| 0.19542 | 1.895313 | 0.026 | 0.007 | 1 | 9 | Atp1b2 | 0.019 | Old |
| 0.19542 | 1.870337 | 0.026 | 0.007 | 1 | 9 | P3h1 | 0.019 | Old |
| 0.195898 | 0.907494 | 0.067 | 0.037 | 1 | 9 | Stc1 | 0.03 | Old |
| 0.196336 | 2.220952 | 0.026 | 0.007 | 1 | 9 | Arl6 | 0.019 | Old |
| 0.196374 | 0.706379 | 0.103 | 0.066 | 1 | 9 | F2r | 0.037 | Old |
| 0.196465 | 1.640591 | 0.057 | 0.029 | 1 | 9 | Tmem238 | 0.028 | Old |
| 0.196848 | 1.217029 | 0.149 | 0.11 | 1 | 9 | Rragc | 0.039 | Old |
| 0.196864 | 0.082996 | 0.147 | 0.103 | 1 | 9 | Phf2 | 0.044 | Old |
| 0.196965 | 0.337743 | 0.229 | 0.184 | 1 | 9 | Mrps21 | 0.045 | Old |
| 0.19708 | 0.456972 | 0.111 | 0.074 | 1 | 9 | Asap2 | 0.037 | Old |
| 0.197256 | 2.041807 | 0.026 | 0.007 | 1 | 9 | Btf3l4 | 0.019 | Old |
| 0.197256 | 1.832207 | 0.026 | 0.007 | 1 | 9 | Rarres2 | 0.019 | Old |
| 0.197256 | 1.659616 | 0.026 | 0.007 | 1 | 9 | Syp | 0.019 | Old |
| 0.197419 | 0.442014 | 0.129 | 0.088 | 1 | 9 | Svbp | 0.041 | Old |
| 0.19768 | 1.757526 | 0.057 | 0.029 | 1 | 9 | Abi3 | 0.028 | Old |
| 0.197781 | 0.357583 | 0.402 | 0.36 | 1 | 9 | Mndal | 0.042 | Old |
| 0.198133 | 0.208414 | 0.067 | 0.037 | 1 | 9 | Cnnm3 | 0.03 | Old |
| 0.198174 | 0.163438 | 0.626 | 0.537 | 1 | 9 | Arhgap18 | 0.089 | Old |
| 0.198178 | 1.946047 | 0.026 | 0.007 | 1 | 9 | Fam136a | 0.019 | Old |
| 0.198178 | 1.798231 | 0.026 | 0.007 | 1 | 9 | Usp20 | 0.019 | Old |
| 0.198178 | 1.70185 | 0.026 | 0.007 | 1 | 9 | Coa7 | 0.019 | Old |

| | | | | | | | | |
|---|---|---|---|---|---|---|---|---|
| 0.198178 | 1.353579 | 0.026 | 0.007 | 1 | 9 | Ppm1n | 0.019 | Old |
| 0.198222 | 0.187821 | 0.534 | 0.478 | 1 | 9 | Wasf2 | 0.056 | Old |
| 0.198289 | 1.454867 | 0.057 | 0.029 | 1 | 9 | Zfp747 | 0.028 | Old |
| 0.198654 | 0.68244 | 0.216 | 0.169 | 1 | 9 | Arl2bp | 0.047 | Old |
| 0.198695 | 0.806255 | 0.067 | 0.037 | 1 | 9 | Farsb | 0.03 | Old |
| 0.198907 | 0.171472 | 0.613 | 0.566 | 1 | 9 | Gnb2 | 0.047 | Old |
| 0.19891 | 0.090143 | 0.428 | 0.368 | 1 | 9 | Ccnd2 | 0.06 | Old |
| 0.199223 | 1.614095 | 0.116 | 0.081 | 1 | 9 | Chmp1a | 0.035 | Old |
| 0.199266 | 0.076089 | 0.227 | 0.162 | 1 | 9 | Msrb3 | 0.065 | Old |
| 0.199822 | 0.577604 | 0.067 | 0.037 | 1 | 9 | Ftsj3 | 0.03 | Old |
| 0.199953 | 0.614954 | 0.147 | 0.103 | 1 | 9 | Cdc42ep4 | 0.044 | Old |
| 0.200033 | 1.448502 | 0.026 | 0.007 | 1 | 9 | Mex3a | 0.019 | Old |
| 0.200064 | 0.13685 | 0.253 | 0.199 | 1 | 9 | Snn | 0.054 | Old |
| 0.200387 | 1.001876 | 0.067 | 0.037 | 1 | 9 | Prkrip1 | 0.03 | Old |
| 0.200387 | 0.956971 | 0.067 | 0.037 | 1 | 9 | Zdhhc6 | 0.03 | Old |
| 0.20079 | 0.71226 | 0.229 | 0.184 | 1 | 9 | Rbfox2 | 0.045 | Old |
| 0.200811 | 1.880738 | 0.046 | 0.022 | 1 | 9 | AW554918 | 0.024 | Old |
| 0.200965 | 1.143109 | 0.026 | 0.007 | 1 | 9 | Klhl26 | 0.019 | Old |
| 0.200965 | 0.849226 | 0.026 | 0.007 | 1 | 9 | Amigo2 | 0.019 | Old |
| 0.200986 | 0.42992 | 0.242 | 0.184 | 1 | 9 | Cramp1l | 0.058 | Old |
| 0.201012 | 0.285098 | 0.41 | 0.346 | 1 | 9 | Prnp | 0.064 | Old |
| 0.201147 | 0.516491 | 0.106 | 0.066 | 1 | 9 | Zdhhc21 | 0.04 | Old |
| 0.201314 | 0.055316 | 0.621 | 0.574 | 1 | 9 | Prkar1a | 0.047 | Old |
| 0.201343 | 0.275322 | 0.554 | 0.515 | 1 | 9 | Ushbp1 | 0.039 | Old |
| 0.201468 | 0.932916 | 0.142 | 0.103 | 1 | 9 | Stat6 | 0.039 | Old |
| 0.201488 | 0.221397 | 0.039 | 0.066 | 1 | 9 | Med29 | -0.027 | Old |
| 0.201521 | 0.595806 | 0.121 | 0.081 | 1 | 9 | Klhl28 | 0.04 | Old |
| 0.2019 | 0.49985 | 0.026 | 0.007 | 1 | 9 | Timm8a1 | 0.019 | Old |
| 0.2019 | 0.464159 | 0.026 | 0.007 | 1 | 9 | AC126280. | 0.019 | Old |
| 0.202029 | 0.121446 | 0.438 | 0.375 | 1 | 9 | Tax1bp1 | 0.063 | Old |
| 0.20223 | 2.544586 | 0.036 | 0.015 | 1 | 9 | G6pc3 | 0.021 | Old |
| 0.202659 | 0.098879 | 0.067 | 0.037 | 1 | 9 | Slc6a8 | 0.03 | Old |
| 0.202685 | 1.041578 | 0.144 | 0.103 | 1 | 9 | Abhd2 | 0.041 | Old |
| 0.202738 | 0.527947 | 0.095 | 0.059 | 1 | 9 | Vps52 | 0.036 | Old |
| 0.202819 | 0.762697 | 0.111 | 0.074 | 1 | 9 | Patl1 | 0.037 | Old |
| 0.202819 | 0.963021 | 0.077 | 0.044 | 1 | 9 | Spon2 | 0.033 | Old |
| 0.202838 | 0.802184 | 0.026 | 0.007 | 1 | 9 | Slc26a10 | 0.019 | Old |
| 0.202838 | 0.453779 | 0.026 | 0.007 | 1 | 9 | Ccdc181 | 0.019 | Old |
| 0.202838 | 0.329202 | 0.026 | 0.007 | 1 | 9 | Tmem186 | 0.019 | Old |
| 0.203013 | 2.128879 | 0.036 | 0.015 | 1 | 9 | Repin1 | 0.021 | Old |
| 0.20328 | 0.284951 | 0.5 | 0.441 | 1 | 9 | Fermt2 | 0.059 | Old |
| 0.203697 | 0.140303 | 0.521 | 0.456 | 1 | 9 | Taok1 | 0.065 | Old |
| 0.20395 | 0.116195 | 0.191 | 0.14 | 1 | 9 | Gpr146 | 0.051 | Old |
| 0.204151 | 1.175281 | 0.093 | 0.059 | 1 | 9 | Gps2 | 0.034 | Old |
| 0.204241 | 1.893365 | 0.046 | 0.022 | 1 | 9 | Mrps11 | 0.024 | Old |

| | | | | | | | | |
|---|---|---|---|---|---|---|---|---|
| 0.204241 | 1.560925 | 0.046 | 0.022 | 1 | 9 | Ints14 | 0.024 | Old |
| 0.204297 | 0.180847 | 0.601 | 0.522 | 1 | 9 | Fryl | 0.079 | Old |
| 0.204376 | 0.690778 | 0.067 | 0.037 | 1 | 9 | Uchl1 | 0.03 | Old |
| 0.20491 | 1.658915 | 0.082 | 0.051 | 1 | 9 | Nceh1 | 0.031 | Old |
| 0.204932 | 1.537298 | 0.046 | 0.022 | 1 | 9 | Rnf40 | 0.024 | Old |
| 0.205022 | 0.494319 | 0.137 | 0.096 | 1 | 9 | Slc39a14 | 0.041 | Old |
| 0.205098 | 0.297661 | 0.155 | 0.11 | 1 | 9 | Gse1 | 0.045 | Old |
| 0.205373 | 2.614794 | 0.036 | 0.015 | 1 | 9 | Supt3 | 0.021 | Old |
| 0.205449 | 0.987899 | 0.142 | 0.103 | 1 | 9 | Mtmr10 | 0.039 | Old |
| 0.205625 | 1.430057 | 0.046 | 0.022 | 1 | 9 | Cpsf4 | 0.024 | Old |
| 0.205738 | 0.936437 | 0.216 | 0.176 | 1 | 9 | Txn2 | 0.04 | Old |
| 0.206073 | 0.303183 | 0.41 | 0.36 | 1 | 9 | Hsd17b11 | 0.05 | Old |
| 0.206164 | 2.247976 | 0.036 | 0.015 | 1 | 9 | Zbtb45 | 0.021 | Old |
| 0.206164 | 1.961069 | 0.036 | 0.015 | 1 | 9 | Zfp606 | 0.021 | Old |
| 0.206319 | 1.777408 | 0.046 | 0.022 | 1 | 9 | Nemp1 | 0.024 | Old |
| 0.206353 | 0.019223 | 0.634 | 0.588 | 1 | 9 | Scarb1 | 0.046 | Old |
| 0.206674 | 0.530155 | 0.232 | 0.184 | 1 | 9 | Tex261 | 0.048 | Old |
| 0.206958 | 2.020551 | 0.036 | 0.015 | 1 | 9 | C1s1 | 0.021 | Old |
| 0.207093 | 0.150713 | 0.095 | 0.059 | 1 | 9 | Clca2 | 0.036 | Old |
| 0.207176 | 0.121976 | 0.281 | 0.228 | 1 | 9 | Ext1 | 0.053 | Old |
| 0.207459 | 0.421004 | 0.144 | 0.103 | 1 | 9 | Casp4 | 0.041 | Old |
| 0.207754 | 1.980337 | 0.036 | 0.015 | 1 | 9 | Ptges2 | 0.021 | Old |
| 0.207894 | 0.837042 | 0.103 | 0.066 | 1 | 9 | Casp12 | 0.037 | Old |
| 0.207993 | 0.484676 | 0.121 | 0.081 | 1 | 9 | Acp2 | 0.04 | Old |
| 0.208227 | 0.893677 | 0.057 | 0.029 | 1 | 9 | Mcm6 | 0.028 | Old |
| 0.208227 | 0.449935 | 0.057 | 0.029 | 1 | 9 | Pcdhgc4 | 0.028 | Old |
| 0.208244 | 0.35269 | 0.497 | 0.456 | 1 | 9 | Pcmtd1 | 0.041 | Old |
| 0.208552 | 1.913406 | 0.036 | 0.015 | 1 | 9 | Saal1 | 0.021 | Old |
| 0.208582 | 0.319849 | 0.436 | 0.382 | 1 | 9 | Ndel1 | 0.054 | Old |
| 0.208988 | 0.509068 | 0.155 | 0.11 | 1 | 9 | Rex1bd | 0.045 | Old |
| 0.209092 | 0.555203 | 0.013 | 0.029 | 1 | 9 | Map4k1 | -0.016 | Old |
| 0.209593 | 0.507891 | 0.229 | 0.184 | 1 | 9 | Zfp664 | 0.045 | Old |
| 0.209813 | 0.538717 | 0.093 | 0.059 | 1 | 9 | Qrich1 | 0.034 | Old |
| 0.210005 | 0.268711 | 0.216 | 0.162 | 1 | 9 | Gm12002 | 0.054 | Old |
| 0.210141 | 0.583081 | 0.168 | 0.125 | 1 | 9 | Glmp | 0.043 | Old |
| 0.210174 | 0.638831 | 0.067 | 0.037 | 1 | 9 | Grpel1 | 0.03 | Old |
| 0.210282 | 0.663873 | 0.144 | 0.103 | 1 | 9 | 2310035C2 | 0.041 | Old |
| 0.210479 | 0.570439 | 0.085 | 0.051 | 1 | 9 | Nipsnap3b | 0.034 | Old |
| 0.210522 | 1.354138 | 0.046 | 0.022 | 1 | 9 | Ctgf | 0.024 | Old |
| 0.210522 | 0.477773 | 0.046 | 0.022 | 1 | 9 | Tpm3-rs7 | 0.024 | Old |
| 0.210629 | 0.3691 | 0.24 | 0.191 | 1 | 9 | Icam2 | 0.049 | Old |
| 0.210678 | 0.858955 | 0.124 | 0.088 | 1 | 9 | Lemd3 | 0.036 | Old |
| 0.210959 | 1.438317 | 0.036 | 0.015 | 1 | 9 | Wdr92 | 0.021 | Old |
| 0.210984 | 0.045001 | 0.871 | 0.831 | 1 | 9 | Hsp90ab1 | 0.04 | Old |
| 0.211358 | 0.102929 | 0.817 | 0.772 | 1 | 9 | Nrp1 | 0.045 | Old |

| | | | | | | | | |
|---|---|---|---|---|---|---|---|---|
| 0.211405 | 0.927198 | 0.057 | 0.029 | 1 | 9 | Rdm1 | 0.028 | Old |
| 0.211766 | 1.778915 | 0.036 | 0.015 | 1 | 9 | Polr3e | 0.021 | Old |
| 0.211766 | 1.767156 | 0.036 | 0.015 | 1 | 9 | Hdhd2 | 0.021 | Old |
| 0.211937 | 0.586018 | 0.046 | 0.022 | 1 | 9 | Dph6 | 0.024 | Old |
| 0.211938 | 0.012405 | 0.039 | 0.066 | 1 | 9 | Zfyve26 | -0.027 | Old |
| 0.212064 | 0.208991 | 0.314 | 0.265 | 1 | 9 | Aldh2 | 0.049 | Old |
| 0.212575 | 1.375041 | 0.036 | 0.015 | 1 | 9 | Xndc1 | 0.021 | Old |
| 0.212647 | 0.216666 | 0.046 | 0.022 | 1 | 9 | Upp1 | 0.024 | Old |
| 0.212687 | 0.159311 | 0.057 | 0.029 | 1 | 9 | Pde4dip | 0.028 | Old |
| 0.213095 | 0.260688 | 0.119 | 0.081 | 1 | 9 | Brwd1 | 0.038 | Old |
| 0.213219 | 0.292517 | 0.678 | 0.618 | 1 | 9 | Ptms | 0.06 | Old |
| 0.213386 | 1.057509 | 0.036 | 0.015 | 1 | 9 | Gtf3c5 | 0.021 | Old |
| 0.213386 | 0.835182 | 0.036 | 0.015 | 1 | 9 | Ubtd1 | 0.021 | Old |
| 0.213716 | 0.349587 | 0.175 | 0.132 | 1 | 9 | Tmbim4 | 0.043 | Old |
| 0.214072 | 1.034978 | 0.046 | 0.022 | 1 | 9 | Adgrl1 | 0.024 | Old |
| 0.214199 | 1.608682 | 0.036 | 0.015 | 1 | 9 | Sgip1 | 0.021 | Old |
| 0.214217 | 0.331499 | 0.018 | 0.037 | 1 | 9 | Lpxn | -0.019 | Old |
| 0.214324 | 0.888016 | 0.093 | 0.059 | 1 | 9 | Med17 | 0.034 | Old |
| 0.214424 | 0.412205 | 0.325 | 0.272 | 1 | 9 | Stat1 | 0.053 | Old |
| 0.214619 | 0.692087 | 0.057 | 0.029 | 1 | 9 | Tnfaip2 | 0.028 | Old |
| 0.214876 | 0.669314 | 0.101 | 0.066 | 1 | 9 | Usp1 | 0.035 | Old |
| 0.214987 | 0.814318 | 0.137 | 0.096 | 1 | 9 | Bet1 | 0.041 | Old |
| 0.215155 | 0.089137 | 0.018 | 0.037 | 1 | 9 | Tctex1d2 | -0.019 | Old |
| 0.215203 | 0.410812 | 0.085 | 0.051 | 1 | 9 | Ghr | 0.034 | Old |
| 0.215675 | 0.311171 | 0.451 | 0.404 | 1 | 9 | Dgkh | 0.047 | Old |
| 0.215677 | 0.314426 | 0.389 | 0.353 | 1 | 9 | Slc25a5 | 0.036 | Old |
| 0.215932 | 1.655037 | 0.064 | 0.037 | 1 | 9 | Hlx | 0.027 | Old |
| 0.215971 | 0.756085 | 0.082 | 0.051 | 1 | 9 | Arl6ip6 | 0.031 | Old |
| 0.21612 | 1.307637 | 0.09 | 0.059 | 1 | 9 | Kif3a | 0.031 | Old |
| 0.216129 | 0.087636 | 0.647 | 0.566 | 1 | 9 | Myadm | 0.081 | Old |
| 0.216244 | 0.518353 | 0.119 | 0.081 | 1 | 9 | 2410015M | 0.038 | Old |
| 0.216844 | 0.752862 | 0.168 | 0.125 | 1 | 9 | Mon2 | 0.043 | Old |
| 0.217218 | 0.428387 | 0.201 | 0.154 | 1 | 9 | Pla2g16 | 0.047 | Old |
| 0.217808 | 1.215991 | 0.101 | 0.066 | 1 | 9 | A4galt | 0.035 | Old |
| 0.217889 | 0.136273 | 0.067 | 0.037 | 1 | 9 | Dnajc30 | 0.03 | Old |
| 0.218368 | 1.657769 | 0.064 | 0.037 | 1 | 9 | Ctsf | 0.027 | Old |
| 0.218368 | 1.20149 | 0.064 | 0.037 | 1 | 9 | Zfp68 | 0.027 | Old |
| 0.21887 | 0.081516 | 0.111 | 0.074 | 1 | 9 | Msl3 | 0.037 | Old |
| 0.21898 | 1.404371 | 0.064 | 0.037 | 1 | 9 | Dusp26 | 0.027 | Old |
| 0.219138 | 0.015504 | 0.696 | 0.625 | 1 | 9 | Adgrl4 | 0.071 | Old |
| 0.219384 | 0.045626 | 0.418 | 0.346 | 1 | 9 | Gng5 | 0.072 | Old |
| 0.21985 | 0.230346 | 0.162 | 0.118 | 1 | 9 | Abhd17a | 0.044 | Old |
| 0.219957 | 0.698073 | 0.036 | 0.015 | 1 | 9 | Parp16 | 0.021 | Old |
| 0.220004 | 0.348669 | 0.384 | 0.346 | 1 | 9 | Txn1 | 0.038 | Old |
| 0.220178 | 0.536336 | 0.216 | 0.169 | 1 | 9 | Akap9 | 0.047 | Old |

| | | | | | | | | |
|---|---|---|---|---|---|---|---|---|
| 0.220273 | 0.565047 | 0.101 | 0.066 | 1 | 9 | Mrps12 | 0.035 | Old |
| 0.220295 | 1.507332 | 0.082 | 0.051 | 1 | 9 | Pdha1 | 0.031 | Old |
| 0.220491 | 0.680852 | 0.057 | 0.029 | 1 | 9 | Fra10ac1 | 0.028 | Old |
| 0.220849 | 0.436009 | 0.018 | 0.037 | 1 | 9 | A430033K( | -0.019 | Old |
| 0.221406 | 0.207683 | 0.222 | 0.169 | 1 | 9 | Chchd10 | 0.053 | Old |
| 0.221571 | 0.165639 | 0.023 | 0.044 | 1 | 9 | Mzt2 | -0.021 | Old |
| 0.221611 | 1.065506 | 0.175 | 0.14 | 1 | 9 | Stt3a | 0.035 | Old |
| 0.221832 | 0.180578 | 0.366 | 0.309 | 1 | 9 | Ube2h | 0.057 | Old |
| 0.221933 | 0.884302 | 0.082 | 0.051 | 1 | 9 | Fbxo21 | 0.031 | Old |
| 0.222187 | 0.418287 | 0.111 | 0.074 | 1 | 9 | Ppil2 | 0.037 | Old |
| 0.222232 | 0.0014 | 0.152 | 0.11 | 1 | 9 | Rchy1 | 0.042 | Old |
| 0.222323 | 0.751778 | 0.209 | 0.162 | 1 | 9 | Fkbp9 | 0.047 | Old |
| 0.222679 | 1.053865 | 0.064 | 0.037 | 1 | 9 | Casp3 | 0.027 | Old |
| 0.222838 | 0.125976 | 0.428 | 0.375 | 1 | 9 | Tmem59 | 0.053 | Old |
| 0.222874 | 0.611334 | 0.175 | 0.132 | 1 | 9 | Rab35 | 0.043 | Old |
| 0.222884 | 0.108288 | 0.995 | 1 | 1 | 9 | Cd36 | -0.005 | Old |
| 0.22295 | 1.271742 | 0.116 | 0.081 | 1 | 9 | Topbp1 | 0.035 | Old |
| 0.223141 | 0.540647 | 0.111 | 0.074 | 1 | 9 | Fbxo30 | 0.037 | Old |
| 0.223569 | 0.102295 | 0.273 | 0.213 | 1 | 9 | Cers4 | 0.06 | Old |
| 0.223599 | 0.319516 | 0.188 | 0.147 | 1 | 9 | Sike1 | 0.041 | Old |
| 0.223648 | 0.332021 | 0.284 | 0.228 | 1 | 9 | Fbxl3 | 0.056 | Old |
| 0.223744 | 0.766686 | 0.126 | 0.088 | 1 | 9 | Fam172a | 0.038 | Old |
| 0.223908 | 2.198792 | 0.054 | 0.029 | 1 | 9 | Coasy | 0.025 | Old |
| 0.224194 | 0.492068 | 0.126 | 0.088 | 1 | 9 | Lrpap1 | 0.038 | Old |
| 0.224819 | 1.168817 | 0.116 | 0.081 | 1 | 9 | Ppp1r7 | 0.035 | Old |
| 0.22488 | 0.24465 | 0.399 | 0.36 | 1 | 9 | Dstn | 0.039 | Old |
| 0.224942 | 0.325986 | 0.366 | 0.316 | 1 | 9 | Syne2 | 0.05 | Old |
| 0.224975 | 1.226004 | 0.09 | 0.059 | 1 | 9 | Nelfb | 0.031 | Old |
| 0.225024 | 0.348947 | 0.144 | 0.103 | 1 | 9 | Maged1 | 0.041 | Old |
| 0.225058 | 0.305292 | 0.139 | 0.096 | 1 | 9 | Micall1 | 0.043 | Old |
| 0.225159 | 0.189 | 0.466 | 0.404 | 1 | 9 | Map4 | 0.062 | Old |
| 0.225203 | 0.38779 | 0.075 | 0.044 | 1 | 9 | Ebpl | 0.031 | Old |
| 0.225411 | 0.2061 | 0.575 | 0.493 | 1 | 9 | Laptm4a | 0.082 | Old |
| 0.225549 | 0.547892 | 0.126 | 0.088 | 1 | 9 | Fas | 0.038 | Old |
| 0.226343 | 0.819123 | 0.082 | 0.051 | 1 | 9 | Hic1 | 0.031 | Old |
| 0.226444 | 0.121742 | 0.289 | 0.235 | 1 | 9 | Supt16 | 0.054 | Old |
| 0.226635 | 1.874603 | 0.054 | 0.029 | 1 | 9 | Pole3 | 0.025 | Old |
| 0.22666 | 1.248722 | 0.098 | 0.066 | 1 | 9 | Fkbp2 | 0.032 | Old |
| 0.226847 | 0.298287 | 0.441 | 0.39 | 1 | 9 | Git2 | 0.051 | Old |
| 0.226953 | 0.60034 | 0.075 | 0.044 | 1 | 9 | Eef2k | 0.031 | Old |
| 0.227172 | 1.030338 | 0.116 | 0.081 | 1 | 9 | Sgta | 0.035 | Old |
| 0.227957 | 0.525226 | 0.111 | 0.074 | 1 | 9 | Tnfrsf21 | 0.037 | Old |
| 0.227957 | 0.329649 | 0.539 | 0.507 | 1 | 9 | Spag9 | 0.032 | Old |
| 0.228263 | 0.866852 | 0.18 | 0.14 | 1 | 9 | Park7 | 0.04 | Old |
| 0.228742 | 0.265463 | 0.384 | 0.331 | 1 | 9 | Cyp4b1 | 0.053 | Old |

| | | | | | | | | |
|---|---|---|---|---|---|---|---|---|
| 0.228934 | 1.476936 | 0.072 | 0.044 | 1 | 9 | Pibf1 | 0.028 | Old |
| 0.228934 | 1.40231 | 0.072 | 0.044 | 1 | 9 | Fam173a | 0.028 | Old |
| 0.228942 | 1.177125 | 0.064 | 0.037 | 1 | 9 | Ormdl2 | 0.027 | Old |
| 0.229018 | 0.484109 | 0.191 | 0.147 | 1 | 9 | Ctdnep1 | 0.044 | Old |
| 0.229332 | 0.17936 | 0.093 | 0.059 | 1 | 9 | Rab11fip5 | 0.034 | Old |
| 0.229691 | 0.93983 | 0.082 | 0.051 | 1 | 9 | Copz2 | 0.031 | Old |
| 0.230339 | 0.44137 | 0.108 | 0.074 | 1 | 9 | Lcor | 0.034 | Old |
| 0.230591 | 0.287068 | 0.399 | 0.338 | 1 | 9 | Set | 0.061 | Old |
| 0.230841 | 0.985684 | 0.09 | 0.059 | 1 | 9 | Scyl2 | 0.031 | Old |
| 0.230841 | 1.430163 | 0.09 | 0.059 | 1 | 9 | Acot7 | 0.031 | Old |
| 0.231094 | 1.16554 | 0.139 | 0.103 | 1 | 9 | Aip | 0.036 | Old |
| 0.231274 | 0.273348 | 0.134 | 0.096 | 1 | 9 | Mlxip | 0.038 | Old |
| 0.231482 | 0.938187 | 0.064 | 0.037 | 1 | 9 | Slc39a3 | 0.027 | Old |
| 0.231991 | 0.628835 | 0.33 | 0.301 | 1 | 9 | Pomp | 0.029 | Old |
| 0.23212 | 0.878168 | 0.064 | 0.037 | 1 | 9 | Prdm1 | 0.027 | Old |
| 0.232248 | 0.178662 | 0.387 | 0.331 | 1 | 9 | Anxa7 | 0.056 | Old |
| 0.232302 | 0.349026 | 0.201 | 0.154 | 1 | 9 | Lman2 | 0.047 | Old |
| 0.233149 | 0.256742 | 0.307 | 0.25 | 1 | 9 | Tgfbr2 | 0.057 | Old |
| 0.233179 | 0.240618 | 0.085 | 0.051 | 1 | 9 | Epm2aip1 | 0.034 | Old |
| 0.2334 | 1.191862 | 0.064 | 0.037 | 1 | 9 | Pef1 | 0.027 | Old |
| 0.233544 | 0.9757 | 0.09 | 0.059 | 1 | 9 | Stk19 | 0.031 | Old |
| 0.233739 | 0.541606 | 0.263 | 0.221 | 1 | 9 | Srsf10 | 0.042 | Old |
| 0.233783 | 0.175029 | 0.567 | 0.5 | 1 | 9 | Sorbs1 | 0.067 | Old |
| 0.233927 | 0.273409 | 0.119 | 0.081 | 1 | 9 | Cpeb2 | 0.038 | Old |
| 0.234041 | 0.985333 | 0.064 | 0.037 | 1 | 9 | Ganc | 0.027 | Old |
| 0.234254 | 1.30293 | 0.054 | 0.029 | 1 | 9 | Exosc4 | 0.025 | Old |
| 0.234311 | 0.469933 | 0.175 | 0.132 | 1 | 9 | Zrsr2 | 0.043 | Old |
| 0.234581 | 0.178341 | 0.513 | 0.456 | 1 | 9 | Ltbp4 | 0.057 | Old |
| 0.234895 | 0.953704 | 0.131 | 0.096 | 1 | 9 | Fam222b | 0.035 | Old |
| 0.235216 | 0.267153 | 0.093 | 0.059 | 1 | 9 | Pot1a | 0.034 | Old |
| 0.235349 | 0.70892 | 0.082 | 0.051 | 1 | 9 | Pabpc4 | 0.031 | Old |
| 0.235749 | 1.060156 | 0.106 | 0.074 | 1 | 9 | Trappc2l | 0.032 | Old |
| 0.235787 | 0.712093 | 0.116 | 0.081 | 1 | 9 | Zfp397 | 0.035 | Old |
| 0.235897 | 3.3727 | 0.01 | 0 | 1 | 9 | Cela1 | 0.01 | Old |
| 0.235897 | 3.235803 | 0.01 | 0 | 1 | 9 | Cenpf | 0.01 | Old |
| 0.235897 | 3.074775 | 0.01 | 0 | 1 | 9 | Zfp418 | 0.01 | Old |
| 0.235897 | 3.007015 | 0.01 | 0 | 1 | 9 | Spag1 | 0.01 | Old |
| 0.235897 | 2.872869 | 0.01 | 0 | 1 | 9 | Sycp2 | 0.01 | Old |
| 0.235897 | 2.85203 | 0.01 | 0 | 1 | 9 | Il33 | 0.01 | Old |
| 0.235897 | 2.844886 | 0.01 | 0 | 1 | 9 | Map3k13 | 0.01 | Old |
| 0.235897 | 2.764377 | 0.01 | 0 | 1 | 9 | Rpusd1 | 0.01 | Old |
| 0.235897 | 2.76196 | 0.01 | 0 | 1 | 9 | Fam184a | 0.01 | Old |
| 0.235897 | 2.722498 | 0.01 | 0 | 1 | 9 | Peg3 | 0.01 | Old |
| 0.235897 | 2.717618 | 0.01 | 0 | 1 | 9 | Gm48242 | 0.01 | Old |
| 0.235897 | 2.697423 | 0.01 | 0 | 1 | 9 | Cox4i2 | 0.01 | Old |

| | | | | | | | | |
|---|---|---|---|---|---|---|---|---|
| 0.235897 | 2.69409 | 0.01 | 0 | 1 | 9 | Gnrh1 | 0.01 | Old |
| 0.235897 | 2.688429 | 0.01 | 0 | 1 | 9 | Chil3 | 0.01 | Old |
| 0.235897 | 2.67369 | 0.01 | 0 | 1 | 9 | Hsf5 | 0.01 | Old |
| 0.235897 | 2.628806 | 0.01 | 0 | 1 | 9 | Pard6b | 0.01 | Old |
| 0.235897 | 2.625552 | 0.01 | 0 | 1 | 9 | Aspa | 0.01 | Old |
| 0.235897 | 2.609042 | 0.01 | 0 | 1 | 9 | Pcdhga10 | 0.01 | Old |
| 0.235897 | 2.605814 | 0.01 | 0 | 1 | 9 | Patj | 0.01 | Old |
| 0.235897 | 2.601493 | 0.01 | 0 | 1 | 9 | Robo2 | 0.01 | Old |
| 0.235897 | 2.600344 | 0.01 | 0 | 1 | 9 | Cdkn2a | 0.01 | Old |
| 0.235897 | 2.58757 | 0.01 | 0 | 1 | 9 | Aars2 | 0.01 | Old |
| 0.235897 | 2.493701 | 0.01 | 0 | 1 | 9 | Kcns3 | 0.01 | Old |
| 0.235897 | 2.484633 | 0.01 | 0 | 1 | 9 | Myh11 | 0.01 | Old |
| 0.235897 | 2.46353 | 0.01 | 0 | 1 | 9 | Ccna2 | 0.01 | Old |
| 0.235897 | 2.460148 | 0.01 | 0 | 1 | 9 | Cenpt | 0.01 | Old |
| 0.235897 | 2.457104 | 0.01 | 0 | 1 | 9 | Rundc3b | 0.01 | Old |
| 0.235897 | 2.45353 | 0.01 | 0 | 1 | 9 | Polh | 0.01 | Old |
| 0.235897 | 2.443458 | 0.01 | 0 | 1 | 9 | AC163040. | 0.01 | Old |
| 0.235897 | 2.43466 | 0.01 | 0 | 1 | 9 | Gpt2 | 0.01 | Old |
| 0.235897 | 2.42953 | 0.01 | 0 | 1 | 9 | Trmt10b | 0.01 | Old |
| 0.235897 | 2.412525 | 0.01 | 0 | 1 | 9 | Aldh5a1 | 0.01 | Old |
| 0.235897 | 2.40713 | 0.01 | 0 | 1 | 9 | Lck | 0.01 | Old |
| 0.235897 | 2.386252 | 0.01 | 0 | 1 | 9 | Ch25h | 0.01 | Old |
| 0.235897 | 2.341048 | 0.01 | 0 | 1 | 9 | 4930579G2 | 0.01 | Old |
| 0.235897 | 2.339117 | 0.01 | 0 | 1 | 9 | Abhd3 | 0.01 | Old |
| 0.235897 | 2.328004 | 0.01 | 0 | 1 | 9 | Prkaa2 | 0.01 | Old |
| 0.235897 | 2.321142 | 0.01 | 0 | 1 | 9 | Tnni3 | 0.01 | Old |
| 0.235897 | 2.310976 | 0.01 | 0 | 1 | 9 | Zfp960 | 0.01 | Old |
| 0.235897 | 2.294442 | 0.01 | 0 | 1 | 9 | Igha | 0.01 | Old |
| 0.235897 | 2.284026 | 0.01 | 0 | 1 | 9 | Gm26641 | 0.01 | Old |
| 0.235897 | 2.27198 | 0.01 | 0 | 1 | 9 | Ccne2 | 0.01 | Old |
| 0.235897 | 2.269384 | 0.01 | 0 | 1 | 9 | Sgtb | 0.01 | Old |
| 0.235897 | 2.24759 | 0.01 | 0 | 1 | 9 | Gm15706 | 0.01 | Old |
| 0.235897 | 2.245744 | 0.01 | 0 | 1 | 9 | Cdk1 | 0.01 | Old |
| 0.235897 | 2.223451 | 0.01 | 0 | 1 | 9 | Fam160a1 | 0.01 | Old |
| 0.235897 | 2.213087 | 0.01 | 0 | 1 | 9 | Pcdhgc3 | 0.01 | Old |
| 0.235897 | 2.207718 | 0.01 | 0 | 1 | 9 | 2810405F1 | 0.01 | Old |
| 0.235897 | 2.193336 | 0.01 | 0 | 1 | 9 | Rcan2 | 0.01 | Old |
| 0.235897 | 2.188342 | 0.01 | 0 | 1 | 9 | Mcat | 0.01 | Old |
| 0.235897 | 2.181757 | 0.01 | 0 | 1 | 9 | Zfp184 | 0.01 | Old |
| 0.235897 | 2.166487 | 0.01 | 0 | 1 | 9 | Fam110a | 0.01 | Old |
| 0.235897 | 2.162039 | 0.01 | 0 | 1 | 9 | Abi3bp | 0.01 | Old |
| 0.235897 | 2.154125 | 0.01 | 0 | 1 | 9 | Pde4d | 0.01 | Old |
| 0.235897 | 2.148102 | 0.01 | 0 | 1 | 9 | Nek8 | 0.01 | Old |
| 0.235897 | 2.141742 | 0.01 | 0 | 1 | 9 | Syce2 | 0.01 | Old |
| 0.235897 | 2.128397 | 0.01 | 0 | 1 | 9 | Cacna1c | 0.01 | Old |

| | | | | | | | | |
|---|---|---|---|---|---|---|---|---|
| 0.235897 | 2.121169 | 0.01 | 0 | 1 | 9 | Clrn1 | 0.01 | Old |
| 0.235897 | 2.103845 | 0.01 | 0 | 1 | 9 | BC051077 | 0.01 | Old |
| 0.235897 | 2.088392 | 0.01 | 0 | 1 | 9 | Gm32401 | 0.01 | Old |
| 0.235897 | 2.085204 | 0.01 | 0 | 1 | 9 | Xrcc2 | 0.01 | Old |
| 0.235897 | 2.061926 | 0.01 | 0 | 1 | 9 | Iba57 | 0.01 | Old |
| 0.235897 | 2.053343 | 0.01 | 0 | 1 | 9 | Adck5 | 0.01 | Old |
| 0.235897 | 2.049555 | 0.01 | 0 | 1 | 9 | Sele | 0.01 | Old |
| 0.235897 | 2.030242 | 0.01 | 0 | 1 | 9 | Znrf3 | 0.01 | Old |
| 0.235897 | 2.029087 | 0.01 | 0 | 1 | 9 | Mybl1 | 0.01 | Old |
| 0.235897 | 2.019975 | 0.01 | 0 | 1 | 9 | Pus3 | 0.01 | Old |
| 0.235897 | 2.017317 | 0.01 | 0 | 1 | 9 | Gm28068 | 0.01 | Old |
| 0.235897 | 2.010723 | 0.01 | 0 | 1 | 9 | Sema5a | 0.01 | Old |
| 0.235897 | 1.980892 | 0.01 | 0 | 1 | 9 | Napb | 0.01 | Old |
| 0.235897 | 1.979018 | 0.01 | 0 | 1 | 9 | Hdx | 0.01 | Old |
| 0.235897 | 1.978526 | 0.01 | 0 | 1 | 9 | Sv2a | 0.01 | Old |
| 0.235897 | 1.972491 | 0.01 | 0 | 1 | 9 | Agtr1a | 0.01 | Old |
| 0.235897 | 1.952883 | 0.01 | 0 | 1 | 9 | Gm44777 | 0.01 | Old |
| 0.235897 | 1.936199 | 0.01 | 0 | 1 | 9 | Gm27224 | 0.01 | Old |
| 0.235897 | 1.935417 | 0.01 | 0 | 1 | 9 | Tmem14a | 0.01 | Old |
| 0.235897 | 1.927316 | 0.01 | 0 | 1 | 9 | Pelo | 0.01 | Old |
| 0.235897 | 1.926761 | 0.01 | 0 | 1 | 9 | A030005L1 | 0.01 | Old |
| 0.235897 | 1.909211 | 0.01 | 0 | 1 | 9 | Map9 | 0.01 | Old |
| 0.235897 | 1.872521 | 0.01 | 0 | 1 | 9 | Hist1h4h | 0.01 | Old |
| 0.235897 | 1.86772 | 0.01 | 0 | 1 | 9 | Ptchd1 | 0.01 | Old |
| 0.235897 | 1.862412 | 0.01 | 0 | 1 | 9 | Oxnad1 | 0.01 | Old |
| 0.235897 | 1.851098 | 0.01 | 0 | 1 | 9 | BC052040 | 0.01 | Old |
| 0.235897 | 1.829584 | 0.01 | 0 | 1 | 9 | Hgh1 | 0.01 | Old |
| 0.235897 | 1.8236 | 0.01 | 0 | 1 | 9 | Etfbkmt | 0.01 | Old |
| 0.235897 | 1.814459 | 0.01 | 0 | 1 | 9 | Prr15 | 0.01 | Old |
| 0.235897 | 1.813304 | 0.01 | 0 | 1 | 9 | Blm | 0.01 | Old |
| 0.235897 | 1.782663 | 0.01 | 0 | 1 | 9 | 2010310C0 | 0.01 | Old |
| 0.235897 | 1.780394 | 0.01 | 0 | 1 | 9 | Gm5141 | 0.01 | Old |
| 0.235897 | 1.777964 | 0.01 | 0 | 1 | 9 | Slc22a4 | 0.01 | Old |
| 0.235897 | 1.775875 | 0.01 | 0 | 1 | 9 | Lrig1 | 0.01 | Old |
| 0.235897 | 1.761323 | 0.01 | 0 | 1 | 9 | Mindy4 | 0.01 | Old |
| 0.235897 | 1.752539 | 0.01 | 0 | 1 | 9 | Nr2f1 | 0.01 | Old |
| 0.235897 | 1.734229 | 0.01 | 0 | 1 | 9 | Pcdhga5 | 0.01 | Old |
| 0.235897 | 1.723569 | 0.01 | 0 | 1 | 9 | Tmem255a | 0.01 | Old |
| 0.235897 | 1.715514 | 0.01 | 0 | 1 | 9 | 9130023H2 | 0.01 | Old |
| 0.235897 | 1.713623 | 0.01 | 0 | 1 | 9 | 4930404N1 | 0.01 | Old |
| 0.235897 | 1.705885 | 0.01 | 0 | 1 | 9 | Gm3558 | 0.01 | Old |
| 0.235897 | 1.680391 | 0.01 | 0 | 1 | 9 | Hck | 0.01 | Old |
| 0.235897 | 1.655359 | 0.01 | 0 | 1 | 9 | Mipol1 | 0.01 | Old |
| 0.235897 | 1.641158 | 0.01 | 0 | 1 | 9 | Slc24a2 | 0.01 | Old |
| 0.235897 | 1.639045 | 0.01 | 0 | 1 | 9 | AC158554. | 0.01 | Old |

| | | | | | | | | |
|---|---|---|---|---|---|---|---|---|
| 0.235897 | 1.624942 | 0.01 | 0 | 1 | 9 | Ulk4 | 0.01 | Old |
| 0.235897 | 1.623268 | 0.01 | 0 | 1 | 9 | Chmp4c | 0.01 | Old |
| 0.235897 | 1.606086 | 0.01 | 0 | 1 | 9 | Il18 | 0.01 | Old |
| 0.235897 | 1.582906 | 0.01 | 0 | 1 | 9 | 2210011C2 | 0.01 | Old |
| 0.235897 | 1.543001 | 0.01 | 0 | 1 | 9 | Hpse | 0.01 | Old |
| 0.235897 | 1.526321 | 0.01 | 0 | 1 | 9 | Fam20c | 0.01 | Old |
| 0.235897 | 1.462116 | 0.01 | 0 | 1 | 9 | Ggt5 | 0.01 | Old |
| 0.235897 | 1.458812 | 0.01 | 0 | 1 | 9 | Itgb4 | 0.01 | Old |
| 0.235897 | 1.448613 | 0.01 | 0 | 1 | 9 | A630089N( | 0.01 | Old |
| 0.235897 | 1.424463 | 0.01 | 0 | 1 | 9 | Fam221a | 0.01 | Old |
| 0.235897 | 1.396488 | 0.01 | 0 | 1 | 9 | Myom1 | 0.01 | Old |
| 0.235897 | 1.350609 | 0.01 | 0 | 1 | 9 | Pcdhb5 | 0.01 | Old |
| 0.235897 | 1.344616 | 0.01 | 0 | 1 | 9 | Padi2 | 0.01 | Old |
| 0.235897 | 1.335489 | 0.01 | 0 | 1 | 9 | Ppp1r9a | 0.01 | Old |
| 0.235897 | 1.305581 | 0.01 | 0 | 1 | 9 | Dnd1 | 0.01 | Old |
| 0.235897 | 1.248497 | 0.01 | 0 | 1 | 9 | Akr1b10 | 0.01 | Old |
| 0.235897 | 1.216551 | 0.01 | 0 | 1 | 9 | Slc2a4 | 0.01 | Old |
| 0.235897 | 1.143973 | 0.01 | 0 | 1 | 9 | Vtn | 0.01 | Old |
| 0.235897 | 1.085215 | 0.01 | 0 | 1 | 9 | Col27a1 | 0.01 | Old |
| 0.23592 | 0.576409 | 0.175 | 0.132 | 1 | 9 | Gle1 | 0.043 | Old |
| 0.236277 | 0.071 | 0.389 | 0.324 | 1 | 9 | Scp2 | 0.065 | Old |
| 0.23647 | 0.159831 | 0.258 | 0.213 | 1 | 9 | Cct2 | 0.045 | Old |
| 0.237048 | 0.120522 | 0.101 | 0.066 | 1 | 9 | Poglut1 | 0.035 | Old |
| 0.237266 | 0.505111 | 0.314 | 0.265 | 1 | 9 | Paip2 | 0.049 | Old |
| 0.237561 | 0.210423 | 0.42 | 0.368 | 1 | 9 | Nid1 | 0.052 | Old |
| 0.237777 | 1.27478 | 0.054 | 0.029 | 1 | 9 | Trmt13 | 0.025 | Old |
| 0.237919 | 0.121711 | 0.064 | 0.037 | 1 | 9 | Zfp524 | 0.027 | Old |
| 0.237935 | 0.264432 | 0.49 | 0.434 | 1 | 9 | Foxn3 | 0.056 | Old |
| 0.238067 | 0.197891 | 0.49 | 0.426 | 1 | 9 | Ppp1cb | 0.064 | Old |
| 0.238093 | 0.399762 | 0.101 | 0.066 | 1 | 9 | Nom1 | 0.035 | Old |
| 0.238645 | 0.160023 | 0.495 | 0.449 | 1 | 9 | Ywhah | 0.046 | Old |
| 0.238908 | 2.411512 | 0.044 | 0.022 | 1 | 9 | Dbndd2 | 0.022 | Old |
| 0.239287 | 0.306914 | 0.258 | 0.206 | 1 | 9 | Ids | 0.052 | Old |
| 0.239671 | 0.125916 | 0.469 | 0.397 | 1 | 9 | Sec14l1 | 0.072 | Old |
| 0.239733 | 0.194953 | 0.526 | 0.485 | 1 | 9 | Ctnna1 | 0.041 | Old |
| 0.239843 | 0.626456 | 0.211 | 0.169 | 1 | 9 | Emc3 | 0.042 | Old |
| 0.239852 | 0.43303 | 0.229 | 0.184 | 1 | 9 | Stard9 | 0.045 | Old |
| 0.240205 | 0.345756 | 0.119 | 0.081 | 1 | 9 | Trak1 | 0.038 | Old |
| 0.240458 | 0.391357 | 0.479 | 0.441 | 1 | 9 | Kif1b | 0.038 | Old |
| 0.240622 | 1.290859 | 0.054 | 0.029 | 1 | 9 | Ppp2r3a | 0.025 | Old |
| 0.240819 | 0.807939 | 0.206 | 0.169 | 1 | 9 | Dnttip2 | 0.037 | Old |
| 0.240833 | 0.436404 | 0.126 | 0.088 | 1 | 9 | Cops4 | 0.038 | Old |
| 0.240871 | 0.37731 | 0.17 | 0.125 | 1 | 9 | Prkce | 0.045 | Old |
| 0.241004 | 0.611775 | 0.307 | 0.257 | 1 | 9 | Arcn1 | 0.05 | Old |
| 0.241105 | 0.495929 | 0.082 | 0.051 | 1 | 9 | Parg | 0.031 | Old |

| | | | | | | | | |
|---|---|---|---|---|---|---|---|---|
| 0.241177 | 0.682322 | 0.188 | 0.147 | 1 | 9 | Sh3glb2 | 0.041 | Old |
| 0.241227 | 2.668046 | 0.023 | 0.007 | 1 | 9 | Bbs7 | 0.016 | Old |
| 0.241227 | 2.805105 | 0.023 | 0.007 | 1 | 9 | Zfp809 | 0.016 | Old |
| 0.241227 | 2.283646 | 0.023 | 0.007 | 1 | 9 | Zfp605 | 0.016 | Old |
| 0.241227 | 2.220948 | 0.023 | 0.007 | 1 | 9 | Accs | 0.016 | Old |
| 0.241227 | 1.818899 | 0.023 | 0.007 | 1 | 9 | Lair1 | 0.016 | Old |
| 0.241479 | 0.196152 | 0.477 | 0.412 | 1 | 9 | Uaca | 0.065 | Old |
| 0.241509 | 0.913704 | 0.165 | 0.125 | 1 | 9 | Ndufv3 | 0.04 | Old |
| 0.241701 | 0.181674 | 0.423 | 0.36 | 1 | 9 | Rab11a | 0.063 | Old |
| 0.241811 | 0.379505 | 0.201 | 0.162 | 1 | 9 | Kpna1 | 0.039 | Old |
| 0.242054 | 1.296761 | 0.054 | 0.029 | 1 | 9 | Crls1 | 0.025 | Old |
| 0.242058 | 0.450633 | 0.142 | 0.103 | 1 | 9 | Wipf2 | 0.039 | Old |
| 0.242085 | 1.962675 | 0.044 | 0.022 | 1 | 9 | Timm21 | 0.022 | Old |
| 0.242344 | 2.337951 | 0.023 | 0.007 | 1 | 9 | Sirt4 | 0.016 | Old |
| 0.242344 | 2.142905 | 0.023 | 0.007 | 1 | 9 | Scrn2 | 0.016 | Old |
| 0.242344 | 1.972493 | 0.023 | 0.007 | 1 | 9 | Tmem29 | 0.016 | Old |
| 0.242499 | 0.161605 | 0.064 | 0.037 | 1 | 9 | Atxn1l | 0.027 | Old |
| 0.242772 | 1.224902 | 0.054 | 0.029 | 1 | 9 | Slc25a45 | 0.025 | Old |
| 0.242817 | 0.245518 | 0.531 | 0.5 | 1 | 9 | Bmpr2 | 0.031 | Old |
| 0.242898 | 1.186767 | 0.09 | 0.059 | 1 | 9 | Eif2s1 | 0.031 | Old |
| 0.242955 | 0.972821 | 0.155 | 0.118 | 1 | 9 | Paics | 0.037 | Old |
| 0.242961 | 0.38303 | 0.142 | 0.103 | 1 | 9 | Mpp5 | 0.039 | Old |
| 0.243115 | 0.34412 | 0.286 | 0.235 | 1 | 9 | Nos3 | 0.051 | Old |
| 0.243465 | 2.637343 | 0.023 | 0.007 | 1 | 9 | Fen1 | 0.016 | Old |
| 0.243465 | 2.2784 | 0.023 | 0.007 | 1 | 9 | Ap4b1 | 0.016 | Old |
| 0.243465 | 1.813974 | 0.023 | 0.007 | 1 | 9 | Rmdn2 | 0.016 | Old |
| 0.243552 | 0.49512 | 0.309 | 0.265 | 1 | 9 | Nckap1 | 0.044 | Old |
| 0.243664 | 0.599382 | 0.278 | 0.228 | 1 | 9 | Ehd1 | 0.05 | Old |
| 0.243685 | 1.688338 | 0.044 | 0.022 | 1 | 9 | Mllt1 | 0.022 | Old |
| 0.244488 | 1.821164 | 0.044 | 0.022 | 1 | 9 | Telo2 | 0.022 | Old |
| 0.244589 | 2.309002 | 0.023 | 0.007 | 1 | 9 | C730034F0 | 0.016 | Old |
| 0.244589 | 2.253182 | 0.023 | 0.007 | 1 | 9 | Det1 | 0.016 | Old |
| 0.244589 | 2.018232 | 0.023 | 0.007 | 1 | 9 | D6Wsu163 | 0.016 | Old |
| 0.245117 | 0.242849 | 0.42 | 0.346 | 1 | 9 | Actr3 | 0.074 | Old |
| 0.245293 | 1.55723 | 0.044 | 0.022 | 1 | 9 | Gmpr | 0.022 | Old |
| 0.245658 | 0.853947 | 0.054 | 0.029 | 1 | 9 | Med24 | 0.025 | Old |
| 0.245683 | 0.593692 | 0.142 | 0.103 | 1 | 9 | Rnf215 | 0.039 | Old |
| 0.245716 | 1.612084 | 0.023 | 0.007 | 1 | 9 | Gcfc2 | 0.016 | Old |
| 0.24578 | 0.241203 | 0.082 | 0.051 | 1 | 9 | Xab2 | 0.031 | Old |
| 0.245812 | 0.425745 | 0.191 | 0.147 | 1 | 9 | Spty2d1 | 0.044 | Old |
| 0.246101 | 0.64326 | 0.113 | 0.081 | 1 | 9 | Tmem260 | 0.032 | Old |
| 0.246296 | 0.769971 | 0.149 | 0.11 | 1 | 9 | Gorasp2 | 0.039 | Old |
| 0.246384 | 0.893316 | 0.054 | 0.029 | 1 | 9 | Fbxw9 | 0.025 | Old |
| 0.246848 | 1.513394 | 0.023 | 0.007 | 1 | 9 | Cks1b | 0.016 | Old |
| 0.246858 | 0.243261 | 0.273 | 0.213 | 1 | 9 | Bcl2l1 | 0.06 | Old |

| | | | | | | | | |
|---|---|---|---|---|---|---|---|---|
| 0.246907 | 1.889783 | 0.044 | 0.022 | 1 | 9 | Bex3 | 0.022 | Old |
| 0.246907 | 1.507867 | 0.044 | 0.022 | 1 | 9 | Cldn12 | 0.022 | Old |
| 0.247183 | 0.366945 | 0.428 | 0.382 | 1 | 9 | Csf2rb | 0.046 | Old |
| 0.247186 | 0.068527 | 0.366 | 0.309 | 1 | 9 | Tnrc18 | 0.057 | Old |
| 0.247563 | 2.117532 | 0.034 | 0.015 | 1 | 9 | Svip | 0.019 | Old |
| 0.247625 | 0.126718 | 0.722 | 0.684 | 1 | 9 | Jup | 0.038 | Old |
| 0.247717 | 1.46707 | 0.044 | 0.022 | 1 | 9 | Cpeb3 | 0.022 | Old |
| 0.247983 | 1.428986 | 0.023 | 0.007 | 1 | 9 | Sesn2 | 0.016 | Old |
| 0.247983 | 1.402991 | 0.023 | 0.007 | 1 | 9 | Ntf3 | 0.016 | Old |
| 0.247983 | 1.36348 | 0.023 | 0.007 | 1 | 9 | Acat2 | 0.016 | Old |
| 0.247983 | 1.237676 | 0.023 | 0.007 | 1 | 9 | Nrbp2 | 0.016 | Old |
| 0.248009 | 1.678452 | 0.062 | 0.037 | 1 | 9 | Gstk1 | 0.025 | Old |
| 0.248056 | 0.157742 | 0.108 | 0.074 | 1 | 9 | Tbk1 | 0.034 | Old |
| 0.248223 | 0.508521 | 0.191 | 0.147 | 1 | 9 | Cers5 | 0.044 | Old |
| 0.248986 | 0.393911 | 0.126 | 0.088 | 1 | 9 | Ldlrap1 | 0.038 | Old |
| 0.249301 | 1.049658 | 0.054 | 0.029 | 1 | 9 | Rmnd5b | 0.025 | Old |
| 0.249431 | 2.187379 | 0.034 | 0.015 | 1 | 9 | Vps8 | 0.019 | Old |
| 0.249923 | 0.82127 | 0.082 | 0.051 | 1 | 9 | Katnbl1 | 0.031 | Old |
| 0.250034 | 0.441611 | 0.054 | 0.029 | 1 | 9 | Zfp574 | 0.025 | Old |
| 0.250159 | 1.485013 | 0.044 | 0.022 | 1 | 9 | Ehbp1 | 0.022 | Old |
| 0.250159 | 1.007787 | 0.044 | 0.022 | 1 | 9 | Slc9a1 | 0.022 | Old |
| 0.250264 | 0.235904 | 0.023 | 0.007 | 1 | 9 | 1700096K1 | 0.016 | Old |
| 0.250343 | 0.509555 | 0.101 | 0.066 | 1 | 9 | Prps1 | 0.035 | Old |
| 0.250369 | 2.236431 | 0.034 | 0.015 | 1 | 9 | Ruvbl2 | 0.019 | Old |
| 0.250413 | 0.150435 | 0.348 | 0.279 | 1 | 9 | Nfat5 | 0.069 | Old |
| 0.250663 | 0.17169 | 0.108 | 0.074 | 1 | 9 | Ugp2 | 0.034 | Old |
| 0.250977 | 1.02722 | 0.044 | 0.022 | 1 | 9 | 5730480H( | 0.022 | Old |
| 0.250977 | 0.789355 | 0.044 | 0.022 | 1 | 9 | Ccdc58 | 0.022 | Old |
| 0.251309 | 2.163419 | 0.034 | 0.015 | 1 | 9 | Vegfb | 0.019 | Old |
| 0.25141 | 0.29582 | 0.023 | 0.007 | 1 | 9 | Itfg2 | 0.016 | Old |
| 0.25141 | 0.043376 | 0.023 | 0.007 | 1 | 9 | Prr5 | 0.016 | Old |
| 0.251504 | 0.74874 | 0.054 | 0.029 | 1 | 9 | Orc3 | 0.025 | Old |
| 0.251796 | 1.141893 | 0.044 | 0.022 | 1 | 9 | Has3 | 0.022 | Old |
| 0.252183 | 0.664312 | 0.155 | 0.118 | 1 | 9 | Tug1 | 0.037 | Old |
| 0.252981 | 0.314841 | 0.054 | 0.029 | 1 | 9 | Tnfaip8l1 | 0.025 | Old |
| 0.253197 | 1.687565 | 0.034 | 0.015 | 1 | 9 | Enpp2 | 0.019 | Old |
| 0.253197 | 1.467416 | 0.034 | 0.015 | 1 | 9 | Exoc3l | 0.019 | Old |
| 0.253199 | 0.375182 | 0.098 | 0.066 | 1 | 9 | Rab24 | 0.032 | Old |
| 0.253201 | 0.004211 | 0.064 | 0.037 | 1 | 9 | Psen1 | 0.027 | Old |
| 0.253261 | 0.567461 | 0.18 | 0.14 | 1 | 9 | Flii | 0.04 | Old |
| 0.253416 | 0.233885 | 0.186 | 0.14 | 1 | 9 | Mertk | 0.046 | Old |
| 0.253571 | 0.291725 | 0.312 | 0.265 | 1 | 9 | Magi1 | 0.047 | Old |
| 0.253682 | 1.005289 | 0.072 | 0.044 | 1 | 9 | Pde4a | 0.028 | Old |
| 0.253817 | 0.204759 | 0.108 | 0.074 | 1 | 9 | Gnl3 | 0.034 | Old |
| 0.254145 | 1.677553 | 0.034 | 0.015 | 1 | 9 | Xrcc6 | 0.019 | Old |

| | | | | | | | | |
|---|---|---|---|---|---|---|---|---|
| 0.254145 | 1.319442 | 0.034 | 0.015 | 1 | 9 | Acot13 | 0.019 | Old |
| 0.25426 | 0.535405 | 0.139 | 0.103 | 1 | 9 | Agtpbp1 | 0.036 | Old |
| 0.254266 | 1.442313 | 0.044 | 0.022 | 1 | 9 | Tmem38b | 0.022 | Old |
| 0.254272 | 0.313492 | 0.271 | 0.221 | 1 | 9 | Tpp1 | 0.05 | Old |
| 0.254328 | 0.783375 | 0.088 | 0.059 | 1 | 9 | Abcd3 | 0.029 | Old |
| 0.254382 | 0.222911 | 0.119 | 0.081 | 1 | 9 | Mapk8 | 0.038 | Old |
| 0.254966 | 0.384024 | 0.072 | 0.044 | 1 | 9 | Gpr137b | 0.028 | Old |
| 0.255095 | 1.553419 | 0.034 | 0.015 | 1 | 9 | Dhrs4 | 0.019 | Old |
| 0.255207 | 0.751709 | 0.054 | 0.029 | 1 | 9 | Fitm2 | 0.025 | Old |
| 0.255254 | 0.292914 | 0.101 | 0.066 | 1 | 9 | Slc4a2 | 0.035 | Old |
| 0.255389 | 0.111853 | 0.08 | 0.051 | 1 | 9 | Stxbp1 | 0.029 | Old |
| 0.25561 | 0.522329 | 0.072 | 0.044 | 1 | 9 | Smad6 | 0.028 | Old |
| 0.25581 | 0.576611 | 0.131 | 0.096 | 1 | 9 | Snx6 | 0.035 | Old |
| 0.255968 | 0.348478 | 0.332 | 0.287 | 1 | 9 | Clic1 | 0.045 | Old |
| 0.25597 | 0.138278 | 0.09 | 0.059 | 1 | 9 | Bod1 | 0.031 | Old |
| 0.256047 | 1.490885 | 0.034 | 0.015 | 1 | 9 | Eml2 | 0.019 | Old |
| 0.256047 | 1.325009 | 0.034 | 0.015 | 1 | 9 | Tmem81 | 0.019 | Old |
| 0.256263 | 1.181894 | 0.062 | 0.037 | 1 | 9 | Zfp672 | 0.025 | Old |
| 0.256265 | 0.533705 | 0.116 | 0.081 | 1 | 9 | Gas2l1 | 0.035 | Old |
| 0.256672 | 0.729488 | 0.129 | 0.096 | 1 | 9 | Sf3b4 | 0.033 | Old |
| 0.256747 | 0.396545 | 0.325 | 0.279 | 1 | 9 | Atp6v1g1 | 0.046 | Old |
| 0.256752 | 1.072818 | 0.044 | 0.022 | 1 | 9 | Pard3b | 0.022 | Old |
| 0.256752 | 1.012839 | 0.044 | 0.022 | 1 | 9 | Dolpp1 | 0.022 | Old |
| 0.25703 | 0.416966 | 0.144 | 0.11 | 1 | 9 | Slc29a1 | 0.034 | Old |
| 0.257356 | 0.651332 | 0.124 | 0.088 | 1 | 9 | Rnps1 | 0.036 | Old |
| 0.257393 | 0.180237 | 0.131 | 0.169 | 1 | 9 | Igfbp4 | -0.038 | Old |
| 0.257551 | 0.039892 | 0.389 | 0.331 | 1 | 9 | Ndufa6 | 0.058 | Old |
| 0.257562 | 0.150454 | 0.448 | 0.382 | 1 | 9 | Sf3b2 | 0.066 | Old |
| 0.257564 | 0.790614 | 0.155 | 0.118 | 1 | 9 | Nf1 | 0.037 | Old |
| 0.257584 | 0.900025 | 0.044 | 0.022 | 1 | 9 | Urm1 | 0.022 | Old |
| 0.257815 | 0.11536 | 0.575 | 0.5 | 1 | 9 | Hnrnpu | 0.075 | Old |
| 0.25796 | 1.34958 | 0.034 | 0.015 | 1 | 9 | AW209491 | 0.019 | Old |
| 0.258025 | 0.179747 | 0.474 | 0.434 | 1 | 9 | Gfod1 | 0.04 | Old |
| 0.258506 | 0.075415 | 0.312 | 0.257 | 1 | 9 | Tram1 | 0.055 | Old |
| 0.258858 | 0.328121 | 0.191 | 0.147 | 1 | 9 | Mgat4a | 0.044 | Old |
| 0.25892 | 1.637275 | 0.034 | 0.015 | 1 | 9 | Hirip3 | 0.019 | Old |
| 0.258936 | 0.325634 | 0.046 | 0.074 | 1 | 9 | Cyp4f13 | -0.028 | Old |
| 0.258953 | 1.241913 | 0.111 | 0.081 | 1 | 9 | 2610037D0 | 0.03 | Old |
| 0.259017 | 0.972927 | 0.088 | 0.059 | 1 | 9 | Fundc1 | 0.029 | Old |
| 0.259088 | 0.170556 | 0.41 | 0.346 | 1 | 9 | 2-Mar | 0.064 | Old |
| 0.25921 | 0.407409 | 0.281 | 0.228 | 1 | 9 | Aaed1 | 0.053 | Old |
| 0.259255 | 1.024785 | 0.044 | 0.022 | 1 | 9 | Gmppa | 0.022 | Old |
| 0.259316 | 0.676526 | 0.098 | 0.066 | 1 | 9 | Nr2f6 | 0.032 | Old |
| 0.259641 | 0.11218 | 0.781 | 0.691 | 1 | 9 | Kitl | 0.09 | Old |
| 0.259758 | 1.393765 | 0.062 | 0.037 | 1 | 9 | Hsf2 | 0.025 | Old |

| | | | | | | | | |
|---|---|---|---|---|---|---|---|---|
| 0.259822 | 0.628111 | 0.106 | 0.074 | 1 | 9 | C2cd2l | 0.032 | Old |
| 0.259882 | 0.590644 | 0.034 | 0.015 | 1 | 9 | Rgl3 | 0.019 | Old |
| 0.259996 | 0.488038 | 0.173 | 0.132 | 1 | 9 | Efr3b | 0.041 | Old |
| 0.260044 | 0.61185 | 0.235 | 0.191 | 1 | 9 | Slco2b1 | 0.044 | Old |
| 0.260415 | 0.461248 | 0.139 | 0.103 | 1 | 9 | Ap3m1 | 0.036 | Old |
| 0.260718 | 0.050628 | 0.461 | 0.39 | 1 | 9 | C130074G1 | 0.071 | Old |
| 0.260791 | 1.144139 | 0.088 | 0.059 | 1 | 9 | Suclg2 | 0.029 | Old |
| 0.260966 | 0.166466 | 0.613 | 0.581 | 1 | 9 | AW112010 | 0.032 | Old |
| 0.261215 | 0.033085 | 0.09 | 0.059 | 1 | 9 | Bag4 | 0.031 | Old |
| 0.261246 | 0.240594 | 0.441 | 0.39 | 1 | 9 | Angptl4 | 0.051 | Old |
| 0.261775 | 0.329462 | 0.044 | 0.022 | 1 | 9 | Sephs2 | 0.022 | Old |
| 0.262186 | 0.305667 | 0.265 | 0.221 | 1 | 9 | Kat2b | 0.044 | Old |
| 0.262245 | 0.327291 | 0.286 | 0.235 | 1 | 9 | Mob4 | 0.051 | Old |
| 0.262566 | 0.387033 | 0.155 | 0.118 | 1 | 9 | Rprd2 | 0.037 | Old |
| 0.262784 | 0.828549 | 0.034 | 0.015 | 1 | 9 | L2hgdh | 0.019 | Old |
| 0.263464 | 0.142039 | 0.044 | 0.022 | 1 | 9 | Nqo2 | 0.022 | Old |
| 0.263485 | 0.205713 | 0.054 | 0.029 | 1 | 9 | Mafa | 0.025 | Old |
| 0.263657 | 0.342855 | 0.196 | 0.154 | 1 | 9 | Fam126a | 0.042 | Old |
| 0.263745 | 0.845196 | 0.224 | 0.191 | 1 | 9 | Sumo1 | 0.033 | Old |
| 0.263757 | 1.541267 | 0.034 | 0.015 | 1 | 9 | 4933427D1 | 0.019 | Old |
| 0.263757 | 0.805442 | 0.034 | 0.015 | 1 | 9 | Gm29007 | 0.019 | Old |
| 0.264294 | 0.273742 | 0.157 | 0.118 | 1 | 9 | Wtap | 0.039 | Old |
| 0.26433 | 0.131674 | 0.188 | 0.147 | 1 | 9 | Atf2 | 0.041 | Old |
| 0.264501 | 0.745693 | 0.162 | 0.125 | 1 | 9 | Ormdl3 | 0.037 | Old |
| 0.264732 | 1.07371 | 0.034 | 0.015 | 1 | 9 | Sec61a2 | 0.019 | Old |
| 0.264877 | 0.195609 | 0.613 | 0.581 | 1 | 9 | Igfbp7 | 0.032 | Old |
| 0.264933 | 0.491627 | 0.124 | 0.088 | 1 | 9 | Ppp6r1 | 0.036 | Old |
| 0.265161 | 0.459443 | 0.044 | 0.022 | 1 | 9 | Gm9889 | 0.022 | Old |
| 0.265272 | 0.223762 | 0.387 | 0.338 | 1 | 9 | Crebbp | 0.049 | Old |
| 0.265583 | 0.15016 | 0.616 | 0.559 | 1 | 9 | Prex2 | 0.057 | Old |
| 0.265779 | 0.094725 | 0.343 | 0.287 | 1 | 9 | Bmp6 | 0.056 | Old |
| 0.265784 | 0.743044 | 0.155 | 0.118 | 1 | 9 | Vps29 | 0.037 | Old |
| 0.265797 | 0.130782 | 0.68 | 0.625 | 1 | 9 | Btnl9 | 0.055 | Old |
| 0.265935 | 0.205441 | 0.064 | 0.096 | 1 | 9 | Ncbp2 | -0.032 | Old |
| 0.266013 | 0.326235 | 0.044 | 0.022 | 1 | 9 | Polg2 | 0.022 | Old |
| 0.266128 | 0.369076 | 0.157 | 0.118 | 1 | 9 | Dcaf7 | 0.039 | Old |
| 0.266132 | 1.018031 | 0.062 | 0.037 | 1 | 9 | Prkci | 0.025 | Old |
| 0.266534 | 1.087246 | 0.103 | 0.074 | 1 | 9 | Opa1 | 0.029 | Old |
| 0.266689 | 0.753167 | 0.034 | 0.015 | 1 | 9 | Oaf | 0.019 | Old |
| 0.266689 | 0.509545 | 0.034 | 0.015 | 1 | 9 | Phlda3 | 0.019 | Old |
| 0.266689 | 0.487284 | 0.034 | 0.015 | 1 | 9 | Cygb | 0.019 | Old |
| 0.266689 | 0.201173 | 0.034 | 0.015 | 1 | 9 | Arfip2 | 0.019 | Old |
| 0.267195 | 0.942553 | 0.08 | 0.051 | 1 | 9 | Abhd5 | 0.029 | Old |
| 0.267611 | 0.247159 | 0.134 | 0.096 | 1 | 9 | Slc38a10 | 0.038 | Old |
| 0.267671 | 0.600928 | 0.034 | 0.015 | 1 | 9 | 2900093K2 | 0.019 | Old |

| | | | | | | | | |
|---|---|---|---|---|---|---|---|---|
| 0.268104 | 1.605798 | 0.052 | 0.029 | 1 | 9 | Srxn1 | 0.023 | Old |
| 0.268112 | 0.220942 | 0.131 | 0.096 | 1 | 9 | Agmo | 0.035 | Old |
| 0.26833 | 0.345671 | 0.149 | 0.11 | 1 | 9 | Mut | 0.039 | Old |
| 0.268448 | 0.208315 | 0.397 | 0.353 | 1 | 9 | Snrpb | 0.044 | Old |
| 0.268572 | 0.214757 | 0.165 | 0.125 | 1 | 9 | Mrps14 | 0.04 | Old |
| 0.268586 | 0.310179 | 0.332 | 0.301 | 1 | 9 | Mdfic | 0.031 | Old |
| 0.268763 | 1.118908 | 0.103 | 0.074 | 1 | 9 | Tspan17 | 0.029 | Old |
| 0.268767 | 1.337454 | 0.07 | 0.044 | 1 | 9 | Fbxo34 | 0.026 | Old |
| 0.268991 | 0.176559 | 0.572 | 0.515 | 1 | 9 | Cox7c | 0.057 | Old |
| 0.26901 | 0.973625 | 0.16 | 0.125 | 1 | 9 | Pgm2l1 | 0.035 | Old |
| 0.269207 | 1.24089 | 0.095 | 0.066 | 1 | 9 | Acyp2 | 0.029 | Old |
| 0.269547 | 0.637278 | 0.098 | 0.066 | 1 | 9 | Vta1 | 0.032 | Old |
| 0.269552 | 0.173846 | 0.124 | 0.088 | 1 | 9 | Fam162a | 0.036 | Old |
| 0.269713 | 0.051335 | 0.054 | 0.081 | 1 | 9 | Gmfg | -0.027 | Old |
| 0.269785 | 1.111848 | 0.088 | 0.059 | 1 | 9 | Uba3 | 0.029 | Old |
| 0.270246 | 0.736984 | 0.106 | 0.074 | 1 | 9 | Pmepa1 | 0.032 | Old |
| 0.270288 | 0.475585 | 0.361 | 0.324 | 1 | 9 | Ppp2r5c | 0.037 | Old |
| 0.270334 | 0.155481 | 0.327 | 0.279 | 1 | 9 | Luc7l3 | 0.048 | Old |
| 0.2707 | 0.746278 | 0.098 | 0.066 | 1 | 9 | Irf3 | 0.032 | Old |
| 0.270962 | 0.14758 | 0.209 | 0.162 | 1 | 9 | Gcn1l1 | 0.047 | Old |
| 0.271537 | 0.744059 | 0.16 | 0.125 | 1 | 9 | Rnf145 | 0.035 | Old |
| 0.271568 | 1.143675 | 0.103 | 0.074 | 1 | 9 | Ccdc124 | 0.029 | Old |
| 0.271888 | 0.765844 | 0.062 | 0.037 | 1 | 9 | Pou6f1 | 0.025 | Old |
| 0.271918 | 0.879323 | 0.106 | 0.074 | 1 | 9 | Ssbp2 | 0.032 | Old |
| 0.271934 | 0.800016 | 0.137 | 0.103 | 1 | 9 | Snx11 | 0.034 | Old |
| 0.27219 | 0.082085 | 0.425 | 0.368 | 1 | 9 | Tob1 | 0.057 | Old |
| 0.272613 | 0.749009 | 0.062 | 0.037 | 1 | 9 | Dhps | 0.025 | Old |
| 0.272844 | 1.668173 | 0.052 | 0.029 | 1 | 9 | Nat9 | 0.023 | Old |
| 0.272848 | 1.063273 | 0.07 | 0.044 | 1 | 9 | Ogfrl1 | 0.026 | Old |
| 0.273018 | 0.363457 | 0.098 | 0.066 | 1 | 9 | Sorbs3 | 0.032 | Old |
| 0.273207 | 0.090995 | 0.381 | 0.338 | 1 | 9 | Eif2s2 | 0.043 | Old |
| 0.273429 | 0.071734 | 0.299 | 0.25 | 1 | 9 | Metap2 | 0.049 | Old |
| 0.273442 | 0.904915 | 0.088 | 0.059 | 1 | 9 | Acox3 | 0.029 | Old |
| 0.273442 | 0.620937 | 0.088 | 0.059 | 1 | 9 | Ercc5 | 0.029 | Old |
| 0.273639 | 1.703501 | 0.052 | 0.029 | 1 | 9 | Slc36a1 | 0.023 | Old |
| 0.273863 | 0.217733 | 0.505 | 0.456 | 1 | 9 | Nrp2 | 0.049 | Old |
| 0.27412 | 0.469262 | 0.005 | 0.015 | 1 | 9 | Bicdl1 | -0.01 | Old |
| 0.27412 | 0.09256 | 0.005 | 0.015 | 1 | 9 | 2900060B1 | -0.01 | Old |
| 0.275241 | 0.359503 | 0.642 | 0.669 | 1 | 9 | Aqp1 | -0.027 | Old |
| 0.275528 | 0.636945 | 0.062 | 0.037 | 1 | 9 | Rae1 | 0.025 | Old |
| 0.275625 | 1.088705 | 0.111 | 0.081 | 1 | 9 | Babam2 | 0.03 | Old |
| 0.275828 | 0.489911 | 0.271 | 0.235 | 1 | 9 | Stk38 | 0.036 | Old |
| 0.275967 | 0.172217 | 0.149 | 0.11 | 1 | 9 | Lnx2 | 0.039 | Old |
| 0.276041 | 0.861934 | 0.005 | 0.015 | 1 | 9 | Mir142hg | -0.01 | Old |
| 0.276041 | 0.641065 | 0.005 | 0.015 | 1 | 9 | Gpr21 | -0.01 | Old |

| | | | | | | | | |
|---|---|---|---|---|---|---|---|---|
| 0.276041 | 0.60649 | 0.005 | 0.015 | 1 | 9 | Rexo5 | -0.01 | Old |
| 0.276041 | 0.59046 | 0.005 | 0.015 | 1 | 9 | Mfap3l | -0.01 | Old |
| 0.276041 | 0.364877 | 0.005 | 0.015 | 1 | 9 | Foxp4 | -0.01 | Old |
| 0.276041 | 0.288382 | 0.005 | 0.015 | 1 | 9 | Gm14322 | -0.01 | Old |
| 0.276041 | 0.247022 | 0.005 | 0.015 | 1 | 9 | Gm45767 | -0.01 | Old |
| 0.276041 | 0.177769 | 0.005 | 0.015 | 1 | 9 | 4931414P1 | -0.01 | Old |
| 0.276041 | 0.140856 | 0.005 | 0.015 | 1 | 9 | Fam26f | -0.01 | Old |
| 0.276041 | 0.091984 | 0.005 | 0.015 | 1 | 9 | Gm1976 | -0.01 | Old |
| 0.276956 | 0.473663 | 0.271 | 0.235 | 1 | 9 | Ephx1 | 0.036 | Old |
| 0.276993 | 0.771064 | 0.062 | 0.037 | 1 | 9 | Arid3a | 0.025 | Old |
| 0.276998 | 0.038099 | 0.139 | 0.103 | 1 | 9 | Mapk14 | 0.036 | Old |
| 0.27764 | 1.503698 | 0.052 | 0.029 | 1 | 9 | Adk | 0.023 | Old |
| 0.277661 | 0.055538 | 0.683 | 0.632 | 1 | 9 | Dusp3 | 0.051 | Old |
| 0.277751 | 0.687955 | 0.088 | 0.059 | 1 | 9 | Gaa | 0.029 | Old |
| 0.277785 | 0.909254 | 0.188 | 0.154 | 1 | 9 | Oxr1 | 0.034 | Old |
| 0.277991 | 0.165684 | 0.716 | 0.654 | 1 | 9 | Tmbim6 | 0.062 | Old |
| 0.278445 | 1.359585 | 0.052 | 0.029 | 1 | 9 | Dnlz | 0.023 | Old |
| 0.278616 | 0.388907 | 0.309 | 0.257 | 1 | 9 | Rnf130 | 0.052 | Old |
| 0.278797 | 0.452351 | 0.307 | 0.265 | 1 | 9 | Ccnt1 | 0.042 | Old |
| 0.279243 | 0.507407 | 0.106 | 0.074 | 1 | 9 | Daam1 | 0.032 | Old |
| 0.279576 | 0.923302 | 0.152 | 0.118 | 1 | 9 | Fam8a1 | 0.034 | Old |
| 0.279611 | 0.978503 | 0.088 | 0.059 | 1 | 9 | Lrrc36 | 0.029 | Old |
| 0.27965 | 2.208749 | 0.077 | 0.051 | 1 | 9 | Ccl8 | 0.026 | Old |
| 0.279743 | 0.957379 | 0.07 | 0.044 | 1 | 9 | Rab20 | 0.026 | Old |
| 0.279761 | 0.432842 | 0.268 | 0.221 | 1 | 9 | Sema7a | 0.047 | Old |
| 0.27984 | 0.027427 | 0.149 | 0.11 | 1 | 9 | Dusp8 | 0.039 | Old |
| 0.27992 | 0.062741 | 0.24 | 0.191 | 1 | 9 | Cdc73 | 0.049 | Old |
| 0.28006 | 1.565234 | 0.052 | 0.029 | 1 | 9 | Gm11808 | 0.023 | Old |
| 0.280573 | 0.445065 | 0.322 | 0.279 | 1 | 9 | St8sia4 | 0.043 | Old |
| 0.280803 | 0.033955 | 0.273 | 0.221 | 1 | 9 | Srrm1 | 0.052 | Old |
| 0.280958 | 0.393014 | 0.405 | 0.368 | 1 | 9 | Rap1a | 0.037 | Old |
| 0.281081 | 0.072998 | 0.613 | 0.529 | 1 | 9 | Ppfibp1 | 0.084 | Old |
| 0.281609 | 0.18629 | 0.072 | 0.044 | 1 | 9 | Nsun6 | 0.028 | Old |
| 0.281681 | 1.185887 | 0.052 | 0.029 | 1 | 9 | Atl1 | 0.023 | Old |
| 0.281719 | 0.298162 | 0.186 | 0.147 | 1 | 9 | BC031181 | 0.039 | Old |
| 0.2818 | 0.596129 | 0.155 | 0.118 | 1 | 9 | Zdhhc17 | 0.037 | Old |
| 0.281828 | 0.272396 | 0.098 | 0.066 | 1 | 9 | Mid1ip1 | 0.032 | Old |
| 0.281849 | 0.552961 | 0.131 | 0.096 | 1 | 9 | Ganab | 0.035 | Old |
| 0.281881 | 0.403985 | 0.286 | 0.243 | 1 | 9 | S100a11 | 0.043 | Old |
| 0.282387 | 0.531589 | 0.165 | 0.125 | 1 | 9 | Map2k3 | 0.04 | Old |
| 0.282637 | 0.506731 | 0.08 | 0.051 | 1 | 9 | Avl9 | 0.029 | Old |
| 0.283243 | 0.038431 | 0.106 | 0.074 | 1 | 9 | Ube2w | 0.032 | Old |
| 0.283308 | 0.879434 | 0.052 | 0.029 | 1 | 9 | Fam78b | 0.023 | Old |
| 0.283777 | 0.674428 | 0.222 | 0.184 | 1 | 9 | Adh1 | 0.038 | Old |
| 0.28426 | 0.971299 | 0.142 | 0.11 | 1 | 9 | Tgfb1i1 | 0.032 | Old |

| | | | | | | | | |
|---|---|---|---|---|---|---|---|---|
| 0.284581 | 0.063565 | 0.108 | 0.074 | 1 | 9 | Uba6 | 0.034 | Old |
| 0.284933 | 0.227919 | 0.363 | 0.301 | 1 | 9 | Wdr1 | 0.062 | Old |
| 0.285344 | 0.864683 | 0.07 | 0.044 | 1 | 9 | Ago4 | 0.026 | Old |
| 0.285596 | 0.210811 | 0.278 | 0.235 | 1 | 9 | Aamp | 0.043 | Old |
| 0.285807 | 0.201292 | 0.438 | 0.39 | 1 | 9 | Nipbl | 0.048 | Old |
| 0.285891 | 0.614214 | 0.103 | 0.074 | 1 | 9 | Cog3 | 0.029 | Old |
| 0.286049 | 0.926574 | 0.07 | 0.044 | 1 | 9 | Ubap1 | 0.026 | Old |
| 0.286234 | 0.000328 | 0.116 | 0.154 | 1 | 9 | Rora | -0.038 | Old |
| 0.286359 | 0.751738 | 0.152 | 0.118 | 1 | 9 | Atp6ap2 | 0.034 | Old |
| 0.286652 | 0.324684 | 0.214 | 0.169 | 1 | 9 | Chd8 | 0.045 | Old |
| 0.287337 | 0.491669 | 0.152 | 0.118 | 1 | 9 | Mapk6 | 0.034 | Old |
| 0.287464 | 0.484381 | 0.07 | 0.044 | 1 | 9 | Cry1 | 0.026 | Old |
| 0.287584 | 0.098585 | 0.843 | 0.706 | 1 | 9 | Tns1 | 0.137 | Old |
| 0.287862 | 0.46509 | 0.106 | 0.074 | 1 | 9 | Naxd | 0.032 | Old |
| 0.288026 | 0.224456 | 0.253 | 0.206 | 1 | 9 | Acap2 | 0.047 | Old |
| 0.288599 | 0.191487 | 0.338 | 0.294 | 1 | 9 | Ptpra | 0.044 | Old |
| 0.28876 | 0.562029 | 0.17 | 0.132 | 1 | 9 | Irx3 | 0.038 | Old |
| 0.288789 | 0.952025 | 0.142 | 0.11 | 1 | 9 | Rabl6 | 0.032 | Old |
| 0.28887 | 0.087064 | 0.142 | 0.184 | 1 | 9 | Bcas2 | -0.042 | Old |
| 0.289014 | 0.084022 | 0.098 | 0.066 | 1 | 9 | Hfe | 0.032 | Old |
| 0.289042 | 0.402891 | 0.088 | 0.059 | 1 | 9 | Synpo | 0.029 | Old |
| 0.28907 | 0.514904 | 0.155 | 0.118 | 1 | 9 | Ubqln1 | 0.037 | Old |
| 0.28923 | 1.97558 | 0.041 | 0.022 | 1 | 9 | Gcsh | 0.019 | Old |
| 0.28923 | 1.886168 | 0.041 | 0.022 | 1 | 9 | Bcl6 | 0.019 | Old |
| 0.2893 | 0.419318 | 0.113 | 0.081 | 1 | 9 | Slc30a5 | 0.032 | Old |
| 0.290297 | 0.220363 | 0.724 | 0.669 | 1 | 9 | Dnaja1 | 0.055 | Old |
| 0.290711 | 0.87634 | 0.052 | 0.029 | 1 | 9 | Prss23 | 0.023 | Old |
| 0.291084 | 1.618678 | 0.041 | 0.022 | 1 | 9 | Map3k6 | 0.019 | Old |
| 0.291152 | 0.114386 | 0.74 | 0.691 | 1 | 9 | Ets1 | 0.049 | Old |
| 0.2914 | 0.291684 | 0.441 | 0.397 | 1 | 9 | Cnot6l | 0.044 | Old |
| 0.291735 | 0.936074 | 0.07 | 0.044 | 1 | 9 | Cdc16 | 0.026 | Old |
| 0.291811 | 0.392652 | 0.25 | 0.206 | 1 | 9 | Insr | 0.044 | Old |
| 0.292013 | 1.403576 | 0.041 | 0.022 | 1 | 9 | 2300009A( | 0.019 | Old |
| 0.292051 | 0.386705 | 0.17 | 0.132 | 1 | 9 | Pde12 | 0.038 | Old |
| 0.29247 | 0.11627 | 0.552 | 0.471 | 1 | 9 | Adgrg1 | 0.081 | Old |
| 0.292613 | 0.205984 | 0.492 | 0.449 | 1 | 9 | Ifngr1 | 0.043 | Old |
| 0.292766 | 1.866339 | 0.059 | 0.037 | 1 | 9 | Rnf121 | 0.022 | Old |
| 0.293015 | 0.755414 | 0.111 | 0.081 | 1 | 9 | Acer3 | 0.03 | Old |
| 0.293045 | 1.168201 | 0.077 | 0.051 | 1 | 9 | Mterf3 | 0.026 | Old |
| 0.293146 | 0.07963 | 0.41 | 0.346 | 1 | 9 | Hectd1 | 0.064 | Old |
| 0.293207 | 0.13733 | 0.134 | 0.096 | 1 | 9 | Rangap1 | 0.038 | Old |
| 0.293537 | 0.825044 | 0.103 | 0.074 | 1 | 9 | Smurf1 | 0.029 | Old |
| 0.293542 | 1.314668 | 0.059 | 0.037 | 1 | 9 | Tmco3 | 0.022 | Old |
| 0.293879 | 1.473229 | 0.041 | 0.022 | 1 | 9 | Msh3 | 0.019 | Old |
| 0.294042 | 0.524084 | 0.052 | 0.029 | 1 | 9 | Crbn | 0.023 | Old |

| | | | | | | | | |
|---|---|---|---|---|---|---|---|---|
| 0.294651 | 0.376029 | 0.25 | 0.213 | 1 | 9 | Clip1 | 0.037 | Old |
| 0.29512 | 0.292832 | 0.124 | 0.088 | 1 | 9 | Babam1 | 0.036 | Old |
| 0.295323 | 0.561777 | 0.219 | 0.184 | 1 | 9 | Tia1 | 0.035 | Old |
| 0.295717 | 0.571158 | 0.052 | 0.029 | 1 | 9 | Ppat | 0.023 | Old |
| 0.295734 | 0.501062 | 0.376 | 0.346 | 1 | 9 | Tmem140 | 0.03 | Old |
| 0.295928 | 0.056218 | 0.263 | 0.213 | 1 | 9 | Zcchc14 | 0.05 | Old |
| 0.29623 | 0.931146 | 0.095 | 0.066 | 1 | 9 | Atp11a | 0.029 | Old |
| 0.296383 | 0.173907 | 0.582 | 0.537 | 1 | 9 | Sgk1 | 0.045 | Old |
| 0.29646 | 0.96416 | 0.077 | 0.051 | 1 | 9 | Fbl | 0.026 | Old |
| 0.296692 | 1.721918 | 0.041 | 0.022 | 1 | 9 | Lrrc8b | 0.019 | Old |
| 0.296897 | 0.330369 | 0.24 | 0.199 | 1 | 9 | Plscr4 | 0.041 | Old |
| 0.29709 | 0.167696 | 0.433 | 0.375 | 1 | 9 | Rpl7a | 0.058 | Old |
| 0.297633 | 1.478389 | 0.041 | 0.022 | 1 | 9 | Sgsh | 0.019 | Old |
| 0.297633 | 1.369551 | 0.041 | 0.022 | 1 | 9 | Pold4 | 0.019 | Old |
| 0.297633 | 1.161729 | 0.041 | 0.022 | 1 | 9 | Rpp14 | 0.019 | Old |
| 0.29777 | 0.565495 | 0.186 | 0.147 | 1 | 9 | Lin7c | 0.039 | Old |
| 0.297791 | 0.273869 | 0.265 | 0.221 | 1 | 9 | Sec61g | 0.044 | Old |
| 0.298035 | 0.183948 | 0.088 | 0.059 | 1 | 9 | 4930503L1 | 0.029 | Old |
| 0.298141 | 0.462966 | 0.24 | 0.199 | 1 | 9 | Pja2 | 0.041 | Old |
| 0.298327 | 0.274459 | 0.219 | 0.176 | 1 | 9 | Arf6 | 0.043 | Old |
| 0.298518 | 0.337558 | 0.134 | 0.096 | 1 | 9 | Tmem43 | 0.038 | Old |
| 0.298681 | 0.123289 | 0.459 | 0.404 | 1 | 9 | Fam43a | 0.055 | Old |
| 0.298778 | 1.052591 | 0.111 | 0.081 | 1 | 9 | Dirc2 | 0.03 | Old |
| 0.299086 | 0.637423 | 0.052 | 0.029 | 1 | 9 | Tmtc4 | 0.023 | Old |
| 0.299086 | 0.299491 | 0.052 | 0.029 | 1 | 9 | Ptgr2 | 0.023 | Old |
| 0.299113 | 0.260362 | 0.201 | 0.154 | 1 | 9 | Tmem64 | 0.047 | Old |
| 0.299463 | 1.199192 | 0.144 | 0.118 | 1 | 9 | Lpcat1 | 0.026 | Old |
| 0.299464 | 0.318982 | 0.129 | 0.096 | 1 | 9 | Ythdf2 | 0.033 | Old |
| 0.29953 | 0.242322 | 0.24 | 0.191 | 1 | 9 | Appbp2 | 0.049 | Old |
| 0.299594 | 1.069421 | 0.134 | 0.103 | 1 | 9 | Gtf3c1 | 0.031 | Old |
| 0.300078 | 0.999887 | 0.183 | 0.154 | 1 | 9 | Psmd7 | 0.029 | Old |
| 0.300433 | 2.91789 | 0.031 | 0.015 | 1 | 9 | Sdhaf3 | 0.016 | Old |
| 0.300433 | 2.41411 | 0.031 | 0.015 | 1 | 9 | Tradd | 0.016 | Old |
| 0.300433 | 2.398474 | 0.031 | 0.015 | 1 | 9 | B230118H( | 0.016 | Old |
| 0.300761 | 0.680407 | 0.152 | 0.118 | 1 | 9 | Agfg1 | 0.034 | Old |
| 0.30103 | 0.090567 | 0.459 | 0.397 | 1 | 9 | Ndufa4 | 0.062 | Old |
| 0.301227 | 2.665715 | 0.021 | 0.007 | 1 | 9 | 1190007I0 | 0.014 | Old |
| 0.301227 | 2.481942 | 0.021 | 0.007 | 1 | 9 | Nupr1 | 0.014 | Old |
| 0.301227 | 2.31976 | 0.021 | 0.007 | 1 | 9 | Flad1 | 0.014 | Old |
| 0.301227 | 2.175505 | 0.021 | 0.007 | 1 | 9 | Alyref2 | 0.014 | Old |
| 0.301227 | 2.096725 | 0.021 | 0.007 | 1 | 9 | Dlg5 | 0.014 | Old |
| 0.301227 | 2.00178 | 0.021 | 0.007 | 1 | 9 | Zfp764 | 0.014 | Old |
| 0.301227 | 1.856457 | 0.021 | 0.007 | 1 | 9 | Naif1 | 0.014 | Old |
| 0.301465 | 0.057374 | 1 | 1 | 1 | 9 | Fabp4 | 0 | Old |
| 0.301534 | 1.876316 | 0.031 | 0.015 | 1 | 9 | Chst12 | 0.016 | Old |

| | | | | | | | | |
|---|---|---|---|---|---|---|---|---|
| 0.302272 | 0.874968 | 0.111 | 0.081 | 1 | 9 | Myzap | 0.03 | Old |
| 0.302372 | 0.927714 | 0.041 | 0.022 | 1 | 9 | Thoc5 | 0.019 | Old |
| 0.302597 | 2.435122 | 0.021 | 0.007 | 1 | 9 | Akap17b | 0.014 | Old |
| 0.302597 | 2.250745 | 0.021 | 0.007 | 1 | 9 | Txlnb | 0.014 | Old |
| 0.302597 | 2.179015 | 0.021 | 0.007 | 1 | 9 | Aacs | 0.014 | Old |
| 0.302597 | 1.939729 | 0.021 | 0.007 | 1 | 9 | Adck1 | 0.014 | Old |
| 0.302597 | 1.8681 | 0.021 | 0.007 | 1 | 9 | Ankrd33b | 0.014 | Old |
| 0.302917 | 0.272841 | 0.492 | 0.456 | 1 | 9 | Myl12b | 0.036 | Old |
| 0.302956 | 0.335239 | 0.026 | 0.044 | 1 | 9 | Lncpint | -0.018 | Old |
| 0.303135 | 0.643174 | 0.103 | 0.074 | 1 | 9 | Miga1 | 0.029 | Old |
| 0.303332 | 0.256763 | 0.07 | 0.044 | 1 | 9 | Trappc12 | 0.026 | Old |
| 0.303786 | 1.298559 | 0.116 | 0.088 | 1 | 9 | 1810030O( | 0.028 | Old |
| 0.303971 | 2.339775 | 0.021 | 0.007 | 1 | 9 | Gm17501 | 0.014 | Old |
| 0.303971 | 1.764991 | 0.021 | 0.007 | 1 | 9 | Abraxas1 | 0.014 | Old |
| 0.303971 | 1.762186 | 0.021 | 0.007 | 1 | 9 | Pold2 | 0.014 | Old |
| 0.303971 | 1.614849 | 0.021 | 0.007 | 1 | 9 | Wdr83 | 0.014 | Old |
| 0.303971 | 1.538546 | 0.021 | 0.007 | 1 | 9 | 9330160F1 | 0.014 | Old |
| 0.303971 | 1.514647 | 0.021 | 0.007 | 1 | 9 | Cdk5rap1 | 0.014 | Old |
| 0.303971 | 1.505669 | 0.021 | 0.007 | 1 | 9 | Mex3b | 0.014 | Old |
| 0.304144 | 0.401332 | 0.173 | 0.14 | 1 | 9 | Vgll4 | 0.033 | Old |
| 0.304277 | 0.138804 | 0.049 | 0.074 | 1 | 9 | Dda1 | -0.025 | Old |
| 0.304312 | 0.467908 | 0.144 | 0.11 | 1 | 9 | Hexb | 0.034 | Old |
| 0.304612 | 0.144142 | 0.366 | 0.316 | 1 | 9 | Ugcg | 0.05 | Old |
| 0.304853 | 2.223788 | 0.031 | 0.015 | 1 | 9 | Rnf157 | 0.016 | Old |
| 0.304853 | 2.216058 | 0.031 | 0.015 | 1 | 9 | Kat2a | 0.016 | Old |
| 0.305238 | 0.853115 | 0.041 | 0.022 | 1 | 9 | Ghdc | 0.019 | Old |
| 0.305283 | 0.635937 | 0.232 | 0.199 | 1 | 9 | Nin | 0.033 | Old |
| 0.305329 | 0.822547 | 0.245 | 0.213 | 1 | 9 | Pdcl3 | 0.032 | Old |
| 0.305338 | 0.83199 | 0.059 | 0.037 | 1 | 9 | Ttc13 | 0.022 | Old |
| 0.305348 | 1.869309 | 0.021 | 0.007 | 1 | 9 | Ldhb | 0.014 | Old |
| 0.305348 | 1.564786 | 0.021 | 0.007 | 1 | 9 | Mitf | 0.014 | Old |
| 0.305348 | 1.411106 | 0.021 | 0.007 | 1 | 9 | Zkscan14 | 0.014 | Old |
| 0.305348 | 1.399986 | 0.021 | 0.007 | 1 | 9 | Qtrt1 | 0.014 | Old |
| 0.305348 | 1.152359 | 0.021 | 0.007 | 1 | 9 | Ndufaf6 | 0.014 | Old |
| 0.305348 | 1.143283 | 0.021 | 0.007 | 1 | 9 | Prrx1 | 0.014 | Old |
| 0.305434 | 0.075866 | 0.186 | 0.147 | 1 | 9 | Anapc11 | 0.039 | Old |
| 0.305669 | 0.582533 | 0.01 | 0.022 | 1 | 9 | Ms4a6d | -0.012 | Old |
| 0.305865 | 0.52707 | 0.16 | 0.125 | 1 | 9 | Tmem110 | 0.035 | Old |
| 0.305965 | 2.290983 | 0.031 | 0.015 | 1 | 9 | Zfp35 | 0.016 | Old |
| 0.305965 | 2.212047 | 0.031 | 0.015 | 1 | 9 | Mbip | 0.016 | Old |
| 0.305965 | 1.741948 | 0.031 | 0.015 | 1 | 9 | Alox12 | 0.016 | Old |
| 0.306005 | 1.556904 | 0.067 | 0.044 | 1 | 9 | Dctn5 | 0.023 | Old |
| 0.306136 | 1.007794 | 0.059 | 0.037 | 1 | 9 | Rabif | 0.022 | Old |
| 0.306141 | 0.05099 | 0.015 | 0.029 | 1 | 9 | Rgs18 | -0.014 | Old |
| 0.306198 | 0.803798 | 0.041 | 0.022 | 1 | 9 | Tbcd | 0.019 | Old |

| | | | | | | | | |
|---|---|---|---|---|---|---|---|---|
| 0.306198 | 0.724121 | 0.041 | 0.022 | 1 | 9 | Mlph | 0.019 | Old |
| 0.306279 | 0.483493 | 0.07 | 0.044 | 1 | 9 | Trappc10 | 0.026 | Old |
| 0.30673 | 1.358336 | 0.021 | 0.007 | 1 | 9 | Col3a1 | 0.014 | Old |
| 0.30673 | 1.327571 | 0.021 | 0.007 | 1 | 9 | Slc52a3 | 0.014 | Old |
| 0.30673 | 1.200379 | 0.021 | 0.007 | 1 | 9 | Snapc2 | 0.014 | Old |
| 0.30673 | 1.123364 | 0.021 | 0.007 | 1 | 9 | Aven | 0.014 | Old |
| 0.30673 | 1.076924 | 0.021 | 0.007 | 1 | 9 | Sh3tc2 | 0.014 | Old |
| 0.306968 | 0.145191 | 0.456 | 0.39 | 1 | 9 | Cnn3 | 0.066 | Old |
| 0.306997 | 0.333287 | 0.291 | 0.257 | 1 | 9 | Tm6sf1 | 0.034 | Old |
| 0.307073 | 0.349912 | 0.263 | 0.228 | 1 | 9 | Papss2 | 0.035 | Old |
| 0.30708 | 2.213514 | 0.031 | 0.015 | 1 | 9 | Slc52a2 | 0.016 | Old |
| 0.30712 | 1.04668 | 0.026 | 0.044 | 1 | 9 | Carm1 | -0.018 | Old |
| 0.30716 | 1.026423 | 0.041 | 0.022 | 1 | 9 | Rpain | 0.019 | Old |
| 0.30716 | 0.85286 | 0.041 | 0.022 | 1 | 9 | Tbc1d16 | 0.019 | Old |
| 0.307176 | 0.218272 | 0.121 | 0.088 | 1 | 9 | Ppm1h | 0.033 | Old |
| 0.307876 | 0.417938 | 0.088 | 0.059 | 1 | 9 | Ugdh | 0.029 | Old |
| 0.308116 | 1.659302 | 0.021 | 0.007 | 1 | 9 | Mgst2 | 0.014 | Old |
| 0.308116 | 1.564647 | 0.021 | 0.007 | 1 | 9 | Cyp2u1 | 0.014 | Old |
| 0.308116 | 1.491805 | 0.021 | 0.007 | 1 | 9 | Ano8 | 0.014 | Old |
| 0.308116 | 1.103723 | 0.021 | 0.007 | 1 | 9 | Lepr | 0.014 | Old |
| 0.308116 | 1.006337 | 0.021 | 0.007 | 1 | 9 | Llgl2 | 0.014 | Old |
| 0.308124 | 0.871369 | 0.041 | 0.022 | 1 | 9 | Snx7 | 0.019 | Old |
| 0.308124 | 0.736478 | 0.041 | 0.022 | 1 | 9 | Deaf1 | 0.019 | Old |
| 0.308124 | 0.099471 | 0.041 | 0.022 | 1 | 9 | 5730409E0 | 0.019 | Old |
| 0.308158 | 0.295593 | 0.216 | 0.176 | 1 | 9 | Itpripl1 | 0.04 | Old |
| 0.308269 | 0.606317 | 0.077 | 0.051 | 1 | 9 | Hsf1 | 0.026 | Old |
| 0.308496 | 0.317561 | 0.402 | 0.368 | 1 | 9 | Pum2 | 0.034 | Old |
| 0.308526 | 0.775252 | 0.085 | 0.059 | 1 | 9 | Bap1 | 0.026 | Old |
| 0.308536 | 0.605328 | 0.059 | 0.037 | 1 | 9 | Bahcc1 | 0.022 | Old |
| 0.308839 | 0.193929 | 0.214 | 0.169 | 1 | 9 | Lzts2 | 0.045 | Old |
| 0.308976 | 0.044424 | 0.206 | 0.169 | 1 | 9 | Cnih1 | 0.037 | Old |
| 0.308992 | 0.026105 | 0.049 | 0.074 | 1 | 9 | Prpsap2 | -0.025 | Old |
| 0.309214 | 0.63696 | 0.173 | 0.14 | 1 | 9 | Pip5k1a | 0.033 | Old |
| 0.309334 | 0.143606 | 0.312 | 0.265 | 1 | 9 | Pdlim1 | 0.047 | Old |
| 0.309507 | 0.995189 | 0.021 | 0.007 | 1 | 9 | Aspn | 0.014 | Old |
| 0.309507 | 0.923578 | 0.021 | 0.007 | 1 | 9 | Myg1 | 0.014 | Old |
| 0.309507 | 0.892559 | 0.021 | 0.007 | 1 | 9 | Ogg1 | 0.014 | Old |
| 0.309783 | 0.228708 | 0.049 | 0.074 | 1 | 9 | Trmt2b | -0.025 | Old |
| 0.309843 | 0.319568 | 0.309 | 0.265 | 1 | 9 | Mindy2 | 0.044 | Old |
| 0.310081 | 0.575282 | 0.095 | 0.066 | 1 | 9 | Sms | 0.029 | Old |
| 0.310136 | 0.200802 | 0.021 | 0.037 | 1 | 9 | Efemp2 | -0.016 | Old |
| 0.310144 | 1.063888 | 0.059 | 0.037 | 1 | 9 | 6030458C1 | 0.022 | Old |
| 0.310386 | 0.541991 | 0.119 | 0.088 | 1 | 9 | Pdlim7 | 0.031 | Old |
| 0.310429 | 0.782839 | 0.173 | 0.14 | 1 | 9 | Mcam | 0.033 | Old |
| 0.310439 | 1.604806 | 0.031 | 0.015 | 1 | 9 | Zbtb24 | 0.016 | Old |

| | | | | | | | | |
|---|---|---|---|---|---|---|---|---|
| 0.310523 | 1.444021 | 0.067 | 0.044 | 1 | 9 | Hsd17b10 | 0.023 | Old |
| 0.310721 | 0.238883 | 0.095 | 0.066 | 1 | 9 | Tmem181a | 0.029 | Old |
| 0.310721 | 0.063865 | 0.08 | 0.11 | 1 | 9 | Mlx | -0.03 | Old |
| 0.310736 | 0.298535 | 0.18 | 0.14 | 1 | 9 | Sos2 | 0.04 | Old |
| 0.310901 | 0.940296 | 0.021 | 0.007 | 1 | 9 | Pctp | 0.014 | Old |
| 0.310901 | 0.782964 | 0.021 | 0.007 | 1 | 9 | Col5a3 | 0.014 | Old |
| 0.310901 | 0.708545 | 0.021 | 0.007 | 1 | 9 | Lrp4 | 0.014 | Old |
| 0.310901 | 0.561062 | 0.021 | 0.007 | 1 | 9 | Pcdhgb5 | 0.014 | Old |
| 0.310901 | 0.555845 | 0.021 | 0.007 | 1 | 9 | Hoxa4 | 0.014 | Old |
| 0.310901 | 0.523627 | 0.021 | 0.007 | 1 | 9 | Hlf | 0.014 | Old |
| 0.310901 | 0.442514 | 0.021 | 0.007 | 1 | 9 | Tgfb2 | 0.014 | Old |
| 0.310901 | 0.413569 | 0.021 | 0.007 | 1 | 9 | Pigo | 0.014 | Old |
| 0.310925 | 0.570936 | 0.198 | 0.162 | 1 | 9 | Hoxb4 | 0.036 | Old |
| 0.311756 | 1.141586 | 0.059 | 0.037 | 1 | 9 | Sumf1 | 0.022 | Old |
| 0.311756 | 0.223515 | 0.059 | 0.037 | 1 | 9 | Ranbp6 | 0.022 | Old |
| 0.311756 | 0.062581 | 0.059 | 0.037 | 1 | 9 | Rnaseh2a | 0.022 | Old |
| 0.311962 | 0.141196 | 0.01 | 0.022 | 1 | 9 | Jchain | -0.012 | Old |
| 0.311986 | 0.224005 | 0.258 | 0.213 | 1 | 9 | Sri | 0.045 | Old |
| 0.31206 | 0.004812 | 0.51 | 0.441 | 1 | 9 | Myct1 | 0.069 | Old |
| 0.312066 | 0.178794 | 0.423 | 0.36 | 1 | 9 | Tmem123 | 0.063 | Old |
| 0.312299 | 0.655827 | 0.021 | 0.007 | 1 | 9 | Zfp420 | 0.014 | Old |
| 0.312299 | 0.569561 | 0.021 | 0.007 | 1 | 9 | Rab4a | 0.014 | Old |
| 0.312299 | 0.410363 | 0.021 | 0.007 | 1 | 9 | 2610044O1 | 0.014 | Old |
| 0.312299 | 0.229903 | 0.021 | 0.007 | 1 | 9 | Gipc1 | 0.014 | Old |
| 0.312299 | 0.079267 | 0.021 | 0.007 | 1 | 9 | Dcxr | 0.014 | Old |
| 0.312326 | 0.26895 | 0.17 | 0.132 | 1 | 9 | Tpst2 | 0.038 | Old |
| 0.312471 | 0.701275 | 0.021 | 0.037 | 1 | 9 | Clec4e | -0.016 | Old |
| 0.312572 | 0.791095 | 0.085 | 0.059 | 1 | 9 | Sgce | 0.026 | Old |
| 0.312646 | 0.387888 | 0.093 | 0.066 | 1 | 9 | Aldh9a1 | 0.027 | Old |
| 0.312881 | 1.120557 | 0.101 | 0.074 | 1 | 9 | Tmem128 | 0.027 | Old |
| 0.312978 | 0.435737 | 0.07 | 0.044 | 1 | 9 | Mansc1 | 0.026 | Old |
| 0.313375 | 0.892744 | 0.059 | 0.037 | 1 | 9 | Minpp1 | 0.022 | Old |
| 0.313467 | 0.343992 | 0.312 | 0.272 | 1 | 9 | Taok2 | 0.04 | Old |
| 0.313493 | 0.324619 | 0.126 | 0.096 | 1 | 9 | Mapkap1 | 0.03 | Old |
| 0.313549 | 0.78051 | 0.01 | 0.022 | 1 | 9 | Phf7 | -0.012 | Old |
| 0.313549 | 0.482629 | 0.01 | 0.022 | 1 | 9 | Urb2 | -0.012 | Old |
| 0.313549 | 0.300896 | 0.01 | 0.022 | 1 | 9 | Slc16a10 | -0.012 | Old |
| 0.313623 | 0.180946 | 0.526 | 0.471 | 1 | 9 | Arhgdia | 0.055 | Old |
| 0.313642 | 0.365241 | 0.021 | 0.037 | 1 | 9 | Atg2b | -0.016 | Old |
| 0.313823 | 1.507289 | 0.031 | 0.015 | 1 | 9 | Ccdc61 | 0.016 | Old |
| 0.313823 | 1.115579 | 0.031 | 0.015 | 1 | 9 | Zfp874a | 0.016 | Old |
| 0.313823 | 0.413798 | 0.031 | 0.015 | 1 | 9 | Coq7 | 0.016 | Old |
| 0.313925 | 0.570233 | 0.16 | 0.125 | 1 | 9 | Acadm | 0.035 | Old |
| 0.313947 | 0.030973 | 0.041 | 0.022 | 1 | 9 | Vps33b | 0.019 | Old |
| 0.314045 | 0.281229 | 0.178 | 0.14 | 1 | 9 | Ceacam1 | 0.038 | Old |

| | | | | | | | | |
|---|---|---|---|---|---|---|---|---|
| 0.314057 | 0.53293 | 0.126 | 0.096 | 1 | 9 | Uimc1 | 0.03 | Old |
| 0.314321 | 1.430305 | 0.067 | 0.044 | 1 | 9 | Dis3l2 | 0.023 | Old |
| 0.314672 | 0.413609 | 0.211 | 0.176 | 1 | 9 | Ube2q1 | 0.035 | Old |
| 0.314773 | 0.276367 | 0.363 | 0.309 | 1 | 9 | Ube2b | 0.054 | Old |
| 0.3148 | 0.322313 | 0.17 | 0.132 | 1 | 9 | Tnks | 0.038 | Old |
| 0.314956 | 0.602596 | 0.031 | 0.015 | 1 | 9 | Kank1 | 0.016 | Old |
| 0.315019 | 0.63592 | 0.119 | 0.088 | 1 | 9 | Eml4 | 0.031 | Old |
| 0.315141 | 0.522121 | 0.01 | 0.022 | 1 | 9 | Bin2 | -0.012 | Old |
| 0.315141 | 0.367748 | 0.01 | 0.022 | 1 | 9 | Nme7 | -0.012 | Old |
| 0.315141 | 0.283263 | 0.01 | 0.022 | 1 | 9 | Ralgps1 | -0.012 | Old |
| 0.315141 | 0.239892 | 0.01 | 0.022 | 1 | 9 | Ftx | -0.012 | Old |
| 0.315141 | 0.194521 | 0.01 | 0.022 | 1 | 9 | Gm4631 | -0.012 | Old |
| 0.315158 | 0.106557 | 0.067 | 0.096 | 1 | 9 | Baiap2 | -0.029 | Old |
| 0.315275 | 0.492691 | 0.229 | 0.191 | 1 | 9 | Gna13 | 0.038 | Old |
| 0.315286 | 0.740095 | 0.134 | 0.103 | 1 | 9 | Lrrfip2 | 0.031 | Old |
| 0.315295 | 0.102143 | 0.938 | 0.912 | 1 | 9 | Actg1 | 0.026 | Old |
| 0.315385 | 0.144601 | 0.59 | 0.529 | 1 | 9 | Ifitm2 | 0.061 | Old |
| 0.315386 | 0.191519 | 0.113 | 0.081 | 1 | 9 | Ripply3 | 0.032 | Old |
| 0.316092 | 1.134284 | 0.031 | 0.015 | 1 | 9 | Cbfa2t2 | 0.016 | Old |
| 0.316092 | 0.925692 | 0.031 | 0.015 | 1 | 9 | BC002059 | 0.016 | Old |
| 0.316094 | 0.254193 | 0.271 | 0.228 | 1 | 9 | Fbxl7 | 0.043 | Old |
| 0.316155 | 0.284974 | 0.049 | 0.074 | 1 | 9 | Otulin | -0.025 | Old |
| 0.316304 | 0.248006 | 0.317 | 0.279 | 1 | 9 | Rpn1 | 0.038 | Old |
| 0.316611 | 0.126259 | 0.443 | 0.39 | 1 | 9 | Rdx | 0.053 | Old |
| 0.316795 | 0.332239 | 0.137 | 0.103 | 1 | 9 | Rab6b | 0.034 | Old |
| 0.316982 | 0.376673 | 0.144 | 0.11 | 1 | 9 | Impact | 0.034 | Old |
| 0.316999 | 0.141456 | 0.662 | 0.596 | 1 | 9 | Rpl26 | 0.066 | Old |
| 0.31712 | 0.268916 | 0.258 | 0.221 | 1 | 9 | Snw1 | 0.037 | Old |
| 0.317175 | 0.824976 | 0.021 | 0.037 | 1 | 9 | Mrpl44 | -0.016 | Old |
| 0.317231 | 0.652404 | 0.031 | 0.015 | 1 | 9 | B230217C1 | 0.016 | Old |
| 0.317231 | 0.624821 | 0.031 | 0.015 | 1 | 9 | Cplx2 | 0.016 | Old |
| 0.317371 | 0.143906 | 0.448 | 0.397 | 1 | 9 | Mtdh | 0.051 | Old |
| 0.317445 | 0.670142 | 0.059 | 0.037 | 1 | 9 | Ankrd16 | 0.022 | Old |
| 0.317726 | 0.175828 | 0.111 | 0.081 | 1 | 9 | 1810022K0 | 0.03 | Old |
| 0.317801 | 0.821815 | 0.031 | 0.015 | 1 | 9 | Ifitm5 | 0.016 | Old |
| 0.317878 | 0.082317 | 0.5 | 0.426 | 1 | 9 | Pcbp1 | 0.074 | Old |
| 0.317931 | 0.271694 | 0.044 | 0.066 | 1 | 9 | Slc12a9 | -0.022 | Old |
| 0.318032 | 1.539257 | 0.049 | 0.029 | 1 | 9 | Orc5 | 0.02 | Old |
| 0.318341 | 0.1689 | 0.423 | 0.382 | 1 | 9 | Pdcd6ip | 0.041 | Old |
| 0.318372 | 0.550326 | 0.031 | 0.015 | 1 | 9 | Ccser1 | 0.016 | Old |
| 0.318707 | 0.899438 | 0.085 | 0.059 | 1 | 9 | Carnmt1 | 0.026 | Old |
| 0.318789 | 0.163703 | 0.693 | 0.654 | 1 | 9 | F11r | 0.039 | Old |
| 0.318867 | 0.283258 | 0.5 | 0.471 | 1 | 9 | Ccnl1 | 0.029 | Old |
| 0.319393 | 1.14179 | 0.085 | 0.059 | 1 | 9 | Asb7 | 0.026 | Old |
| 0.319393 | 0.820329 | 0.085 | 0.059 | 1 | 9 | Zfyve16 | 0.026 | Old |

| | | | | | | | | |
|---|---|---|---|---|---|---|---|---|
| 0.319403 | 0.238589 | 0.273 | 0.221 | 1 | 9 | Plpp1 | 0.052 | Old |
| 0.319516 | 0.455237 | 0.031 | 0.015 | 1 | 9 | Def6 | 0.016 | Old |
| 0.319663 | 0.204393 | 0.077 | 0.051 | 1 | 9 | Actr8 | 0.026 | Old |
| 0.319698 | 0.204747 | 0.119 | 0.088 | 1 | 9 | Ipo7 | 0.031 | Old |
| 0.320277 | 0.57463 | 0.271 | 0.243 | 1 | 9 | Nol7 | 0.028 | Old |
| 0.320663 | 0.493158 | 0.031 | 0.015 | 1 | 9 | Nicn1 | 0.016 | Old |
| 0.320663 | 0.385087 | 0.031 | 0.015 | 1 | 9 | Notch3 | 0.016 | Old |
| 0.320726 | 0.449903 | 0.059 | 0.037 | 1 | 9 | Polk | 0.022 | Old |
| 0.320735 | 1.158576 | 0.049 | 0.029 | 1 | 9 | Rpl27-ps3 | 0.02 | Old |
| 0.32081 | 0.836132 | 0.082 | 0.059 | 1 | 9 | Cd247 | 0.023 | Old |
| 0.321092 | 0.355438 | 0.25 | 0.213 | 1 | 9 | Puf60 | 0.037 | Old |
| 0.321364 | 0.389076 | 0.111 | 0.081 | 1 | 9 | Eif6 | 0.03 | Old |
| 0.321528 | 0.540523 | 0.224 | 0.191 | 1 | 9 | Csnk2b | 0.033 | Old |
| 0.321549 | 0.557889 | 0.059 | 0.037 | 1 | 9 | Arhgef10 | 0.022 | Old |
| 0.32164 | 1.579693 | 0.049 | 0.029 | 1 | 9 | Sh3bp5l | 0.02 | Old |
| 0.321729 | 0.507659 | 0.273 | 0.243 | 1 | 9 | Esyt2 | 0.03 | Old |
| 0.32176 | 0.343495 | 0.147 | 0.11 | 1 | 9 | Tcf7l1 | 0.037 | Old |
| 0.321813 | 1.126157 | 0.031 | 0.015 | 1 | 9 | Kdm4b | 0.016 | Old |
| 0.321813 | 0.201096 | 0.031 | 0.015 | 1 | 9 | Pcdhb16 | 0.016 | Old |
| 0.321824 | 0.153533 | 0.041 | 0.022 | 1 | 9 | Tapt1 | 0.019 | Old |
| 0.321906 | 0.840071 | 0.108 | 0.081 | 1 | 9 | Vamp7 | 0.027 | Old |
| 0.322106 | 0.129013 | 0.49 | 0.449 | 1 | 9 | Plin2 | 0.041 | Old |
| 0.322148 | 0.575835 | 0.085 | 0.059 | 1 | 9 | Kremen1 | 0.026 | Old |
| 0.322375 | 0.716031 | 0.059 | 0.037 | 1 | 9 | Rnf152 | 0.022 | Old |
| 0.322454 | 0.230185 | 0.25 | 0.206 | 1 | 9 | Fis1 | 0.044 | Old |
| 0.322546 | 1.736724 | 0.049 | 0.029 | 1 | 9 | Dpm2 | 0.02 | Old |
| 0.322555 | 0.561643 | 0.077 | 0.051 | 1 | 9 | Txlng | 0.026 | Old |
| 0.322559 | 0.193343 | 0.224 | 0.184 | 1 | 9 | Pdcd10 | 0.04 | Old |
| 0.322631 | 0.211916 | 0.191 | 0.154 | 1 | 9 | Dnajc1 | 0.037 | Old |
| 0.322803 | 0.537299 | 0.17 | 0.132 | 1 | 9 | Snrpd1 | 0.038 | Old |
| 0.322883 | 0.693197 | 0.124 | 0.096 | 1 | 9 | Rabgap1 | 0.028 | Old |
| 0.322983 | 0.704863 | 0.101 | 0.074 | 1 | 9 | Opa3 | 0.027 | Old |
| 0.323177 | 0.057583 | 0.299 | 0.25 | 1 | 9 | Pde4b | 0.049 | Old |
| 0.323307 | 0.063173 | 0.17 | 0.132 | 1 | 9 | Pcgf3 | 0.038 | Old |
| 0.323826 | 0.036591 | 0.075 | 0.103 | 1 | 9 | Ulk1 | -0.028 | Old |
| 0.323907 | 0.141596 | 0.802 | 0.743 | 1 | 9 | Ablim1 | 0.059 | Old |
| 0.323937 | 0.710861 | 0.157 | 0.125 | 1 | 9 | Ube2g2 | 0.032 | Old |
| 0.324334 | 0.445421 | 0.314 | 0.279 | 1 | 9 | Eif3c | 0.035 | Old |
| 0.324391 | 0.931001 | 0.152 | 0.125 | 1 | 9 | Sec24c | 0.027 | Old |
| 0.324662 | 0.060093 | 0.508 | 0.441 | 1 | 9 | Heg1 | 0.067 | Old |
| 0.324759 | 0.058059 | 0.044 | 0.066 | 1 | 9 | Zfp622 | -0.022 | Old |
| 0.325277 | 0.16227 | 0.031 | 0.015 | 1 | 9 | Rrnad1 | 0.016 | Old |
| 0.325678 | 0.178675 | 0.232 | 0.184 | 1 | 9 | Mall | 0.048 | Old |
| 0.325898 | 0.714484 | 0.067 | 0.044 | 1 | 9 | Ap4m1 | 0.023 | Old |
| 0.326104 | 0.355906 | 0.059 | 0.037 | 1 | 9 | Commd1 | 0.022 | Old |

| | | | | | | | | |
|---|---|---|---|---|---|---|---|---|
| 0.326169 | 1.806803 | 0.057 | 0.037 | 1 | 9 | Micall2 | 0.02 | Old |
| 0.326187 | 1.498187 | 0.049 | 0.029 | 1 | 9 | Cd24a | 0.02 | Old |
| 0.326187 | 1.209873 | 0.049 | 0.029 | 1 | 9 | Zfp358 | 0.02 | Old |
| 0.326187 | 0.676022 | 0.049 | 0.029 | 1 | 9 | Rhobtb2 | 0.02 | Old |
| 0.326192 | 0.257363 | 0.077 | 0.051 | 1 | 9 | Ublcp1 | 0.026 | Old |
| 0.326192 | 0.08734 | 0.077 | 0.051 | 1 | 9 | Setd1a | 0.026 | Old |
| 0.32668 | 1.003647 | 0.067 | 0.044 | 1 | 9 | Dedd | 0.023 | Old |
| 0.326767 | 0.692083 | 0.173 | 0.14 | 1 | 9 | Naa50 | 0.033 | Old |
| 0.326846 | 0.94002 | 0.108 | 0.081 | 1 | 9 | Gab1 | 0.027 | Old |
| 0.327214 | 0.802245 | 0.245 | 0.221 | 1 | 9 | Leprot | 0.024 | Old |
| 0.327343 | 0.288326 | 0.044 | 0.066 | 1 | 9 | Cog1 | -0.022 | Old |
| 0.328083 | 0.166258 | 0.848 | 0.824 | 1 | 9 | Sh3bp5 | 0.024 | Old |
| 0.328189 | 0.252538 | 0.059 | 0.037 | 1 | 9 | Aldh6a1 | 0.022 | Old |
| 0.32859 | 0.066807 | 0.539 | 0.485 | 1 | 9 | Kif5b | 0.054 | Old |
| 0.329402 | 0.814466 | 0.168 | 0.14 | 1 | 9 | BC005624 | 0.028 | Old |
| 0.329855 | 1.14182 | 0.049 | 0.029 | 1 | 9 | Slc12a5 | 0.02 | Old |
| 0.330233 | 0.158403 | 0.392 | 0.338 | 1 | 9 | Dnajb6 | 0.054 | Old |
| 0.330372 | 0.533764 | 0.134 | 0.103 | 1 | 9 | Xpo1 | 0.031 | Old |
| 0.330705 | 0.094087 | 0.25 | 0.206 | 1 | 9 | Lamb2 | 0.044 | Old |
| 0.330776 | 0.796751 | 0.049 | 0.029 | 1 | 9 | Ddx49 | 0.02 | Old |
| 0.33094 | 0.081982 | 0.219 | 0.272 | 1 | 9 | Eps15 | -0.053 | Old |
| 0.331168 | 0.788276 | 0.093 | 0.066 | 1 | 9 | Cfap36 | 0.027 | Old |
| 0.331373 | 1.179311 | 0.082 | 0.059 | 1 | 9 | Prpf40b | 0.023 | Old |
| 0.331414 | 0.247104 | 0.436 | 0.39 | 1 | 9 | Mecom | 0.046 | Old |
| 0.332549 | 0.749031 | 0.149 | 0.118 | 1 | 9 | Uqcc3 | 0.031 | Old |
| 0.332623 | 1.06511 | 0.049 | 0.029 | 1 | 9 | Zfp511 | 0.02 | Old |
| 0.332856 | 0.485149 | 0.397 | 0.368 | 1 | 9 | Rapgef3 | 0.029 | Old |
| 0.333014 | 0.12382 | 0.469 | 0.412 | 1 | 9 | Tubb5 | 0.057 | Old |
| 0.333295 | 0.480602 | 0.101 | 0.074 | 1 | 9 | Ttbk2 | 0.027 | Old |
| 0.333338 | 0.492579 | 0.09 | 0.066 | 1 | 9 | Pfdn1 | 0.024 | Old |
| 0.333996 | 0.63799 | 0.124 | 0.096 | 1 | 9 | Fibp | 0.028 | Old |
| 0.334376 | 0.151385 | 0.76 | 0.728 | 1 | 9 | Cd300lg | 0.032 | Old |
| 0.334723 | 0.367971 | 0.149 | 0.118 | 1 | 9 | Eif3g | 0.031 | Old |
| 0.335357 | 0.744435 | 0.067 | 0.044 | 1 | 9 | Tipin | 0.023 | Old |
| 0.335357 | 0.694521 | 0.067 | 0.044 | 1 | 9 | Hectd4 | 0.023 | Old |
| 0.335407 | 1.167366 | 0.049 | 0.029 | 1 | 9 | Ankrd49 | 0.02 | Old |
| 0.335407 | 0.996687 | 0.049 | 0.029 | 1 | 9 | Nkiras1 | 0.02 | Old |
| 0.335407 | 0.978567 | 0.049 | 0.029 | 1 | 9 | Enoph1 | 0.02 | Old |
| 0.33554 | 0.150163 | 0.157 | 0.125 | 1 | 9 | Pptc7 | 0.032 | Old |
| 0.335954 | 0.156007 | 0.278 | 0.235 | 1 | 9 | Rbm8a | 0.043 | Old |
| 0.336338 | 0.794633 | 0.049 | 0.029 | 1 | 9 | Tmem70 | 0.02 | Old |
| 0.336396 | 0.266148 | 0.17 | 0.14 | 1 | 9 | Ap1g1 | 0.03 | Old |
| 0.336654 | 0.605789 | 0.155 | 0.125 | 1 | 9 | Kctd2 | 0.03 | Old |
| 0.336802 | 0.357575 | 0.16 | 0.125 | 1 | 9 | Snx27 | 0.035 | Old |
| 0.336831 | 0.339012 | 0.211 | 0.176 | 1 | 9 | Fto | 0.035 | Old |

| | | | | | | | | |
|---|---|---|---|---|---|---|---|---|
| 0.336874 | 0.140515 | 0.108 | 0.081 | 1 | 9 | Tars | 0.027 | Old |
| 0.336976 | 0.800881 | 0.116 | 0.088 | 1 | 9 | Zfp462 | 0.028 | Old |
| 0.337525 | 0.233654 | 0.394 | 0.36 | 1 | 9 | Ttc28 | 0.034 | Old |
| 0.337699 | 0.119526 | 0.763 | 0.706 | 1 | 9 | Ctnnb1 | 0.057 | Old |
| 0.338205 | 0.388042 | 0.049 | 0.029 | 1 | 9 | Pign | 0.02 | Old |
| 0.338313 | 0.150959 | 0.059 | 0.037 | 1 | 9 | Crcp | 0.022 | Old |
| 0.338485 | 0.467031 | 0.211 | 0.176 | 1 | 9 | Dhx36 | 0.035 | Old |
| 0.33853 | 0.290473 | 0.085 | 0.118 | 1 | 9 | Mocs2 | -0.033 | Old |
| 0.338714 | 0.504992 | 0.216 | 0.176 | 1 | 9 | Wdfy3 | 0.04 | Old |
| 0.33886 | 0.254906 | 0.366 | 0.331 | 1 | 9 | Slc3a2 | 0.035 | Old |
| 0.339004 | 0.083445 | 0.088 | 0.118 | 1 | 9 | Etv6 | -0.03 | Old |
| 0.339017 | 0.824594 | 0.075 | 0.051 | 1 | 9 | Phax | 0.024 | Old |
| 0.339059 | 0.682319 | 0.209 | 0.176 | 1 | 9 | Ubr4 | 0.033 | Old |
| 0.339141 | 0.359905 | 0.049 | 0.029 | 1 | 9 | Sdhaf1 | 0.02 | Old |
| 0.339384 | 0.503052 | 0.209 | 0.169 | 1 | 9 | Bag3 | 0.04 | Old |
| 0.339776 | 1.129808 | 0.075 | 0.051 | 1 | 9 | Zbed3 | 0.024 | Old |
| 0.339878 | 0.14509 | 0.397 | 0.346 | 1 | 9 | St13 | 0.051 | Old |
| 0.339968 | 0.38856 | 0.263 | 0.235 | 1 | 9 | Ndufs5 | 0.028 | Old |
| 0.340079 | 0.766486 | 0.049 | 0.029 | 1 | 9 | Rtca | 0.02 | Old |
| 0.340458 | 2.397329 | 0.039 | 0.022 | 1 | 9 | Cebpzos | 0.017 | Old |
| 0.340665 | 0.127433 | 0.101 | 0.132 | 1 | 9 | Mcrs1 | -0.031 | Old |
| 0.341262 | 0.786727 | 0.147 | 0.118 | 1 | 9 | Tlnrd1 | 0.029 | Old |
| 0.341262 | 0.765035 | 0.147 | 0.118 | 1 | 9 | Hdac1 | 0.029 | Old |
| 0.341493 | 0.378936 | 0.235 | 0.199 | 1 | 9 | Slc39a10 | 0.036 | Old |
| 0.341646 | 0.033784 | 0.235 | 0.287 | 1 | 9 | Ttc3 | -0.052 | Old |
| 0.341843 | 0.315402 | 0.134 | 0.103 | 1 | 9 | Mfsd5 | 0.031 | Old |
| 0.341959 | 0.872308 | 0.049 | 0.029 | 1 | 9 | Mb21d2 | 0.02 | Old |
| 0.343153 | 0.224404 | 0.18 | 0.147 | 1 | 9 | Arhgef7 | 0.033 | Old |
| 0.34324 | 0.525814 | 0.108 | 0.081 | 1 | 9 | Sgk3 | 0.027 | Old |
| 0.343307 | 0.069539 | 0.085 | 0.059 | 1 | 9 | Med14 | 0.026 | Old |
| 0.343613 | 0.838986 | 0.082 | 0.059 | 1 | 9 | Micu1 | 0.023 | Old |
| 0.343847 | 0.592776 | 0.049 | 0.029 | 1 | 9 | Socs1 | 0.02 | Old |
| 0.344385 | 0.201417 | 0.119 | 0.088 | 1 | 9 | Bbip1 | 0.031 | Old |
| 0.344793 | 0.570008 | 0.049 | 0.029 | 1 | 9 | Hivep3 | 0.02 | Old |
| 0.344793 | 0.082732 | 0.039 | 0.059 | 1 | 9 | Clec12a | -0.02 | Old |
| 0.344892 | 0.078726 | 0.296 | 0.265 | 1 | 9 | Brd4 | 0.031 | Old |
| 0.345002 | 0.18505 | 0.119 | 0.088 | 1 | 9 | Msx1 | 0.031 | Old |
| 0.345138 | 0.855981 | 0.057 | 0.037 | 1 | 9 | Acot2 | 0.02 | Old |
| 0.345167 | 0.268763 | 0.245 | 0.206 | 1 | 9 | Sav1 | 0.039 | Old |
| 0.345282 | 0.609964 | 0.227 | 0.199 | 1 | 9 | Csnk1d | 0.028 | Old |
| 0.345604 | 0.723054 | 0.162 | 0.132 | 1 | 9 | Mxra7 | 0.03 | Old |
| 0.34574 | 0.687007 | 0.049 | 0.029 | 1 | 9 | Galk2 | 0.02 | Old |
| 0.346395 | 0.154539 | 0.106 | 0.14 | 1 | 9 | Ube2a | -0.034 | Old |
| 0.34669 | 0.369102 | 0.049 | 0.029 | 1 | 9 | Asf1a | 0.02 | Old |
| 0.346736 | 0.093743 | 0.175 | 0.14 | 1 | 9 | Cand1 | 0.035 | Old |

| | | | | | | | | |
|---|---|---|---|---|---|---|---|---|
| 0.346896 | 0.994262 | 0.057 | 0.037 | 1 | 9 | Sugp1 | 0.02 | Old |
| 0.347062 | 0.355527 | 0.356 | 0.331 | 1 | 9 | Vat1 | 0.025 | Old |
| 0.34724 | 0.665154 | 0.155 | 0.125 | 1 | 9 | Rbm33 | 0.03 | Old |
| 0.347423 | 0.396975 | 0.075 | 0.051 | 1 | 9 | Ppp1r8 | 0.024 | Old |
| 0.347734 | 0.040609 | 0.673 | 0.61 | 1 | 9 | Neat1 | 0.063 | Old |
| 0.347742 | 0.787761 | 0.108 | 0.081 | 1 | 9 | Cish | 0.027 | Old |
| 0.347896 | 0.565335 | 0.09 | 0.066 | 1 | 9 | Ube2l6 | 0.024 | Old |
| 0.347905 | 1.8038 | 0.039 | 0.022 | 1 | 9 | Zfp426 | 0.017 | Old |
| 0.347928 | 0.165577 | 0.474 | 0.419 | 1 | 9 | Rpl10a | 0.055 | Old |
| 0.348001 | 1.020367 | 0.082 | 0.059 | 1 | 9 | Sart1 | 0.023 | Old |
| 0.348215 | 0.101081 | 0.43 | 0.375 | 1 | 9 | Rpl4 | 0.055 | Old |
| 0.348294 | 0.163721 | 0.093 | 0.066 | 1 | 9 | Ipo9 | 0.027 | Old |
| 0.348389 | 0.82357 | 0.108 | 0.081 | 1 | 9 | Ankrd28 | 0.027 | Old |
| 0.348593 | 0.716231 | 0.131 | 0.103 | 1 | 9 | Zcchc7 | 0.028 | Old |
| 0.348593 | 0.440936 | 0.049 | 0.029 | 1 | 9 | Cfap45 | 0.02 | Old |
| 0.34866 | 0.603291 | 0.057 | 0.037 | 1 | 9 | Zfand2a | 0.02 | Old |
| 0.34866 | 0.427673 | 0.057 | 0.037 | 1 | 9 | Rev3l | 0.02 | Old |
| 0.348718 | 0.029287 | 0.103 | 0.132 | 1 | 9 | Ndufs4 | -0.029 | Old |
| 0.348822 | 0.254644 | 0.436 | 0.382 | 1 | 9 | Cd2ap | 0.054 | Old |
| 0.349545 | 1.098782 | 0.057 | 0.037 | 1 | 9 | Axin1 | 0.02 | Old |
| 0.349548 | 0.185054 | 0.049 | 0.029 | 1 | 9 | Rbbp8 | 0.02 | Old |
| 0.350052 | 1.839065 | 0.039 | 0.022 | 1 | 9 | 2310009B1 | 0.017 | Old |
| 0.35032 | 0.049598 | 0.804 | 0.75 | 1 | 9 | Snrk | 0.054 | Old |
| 0.350357 | 0.371212 | 0.206 | 0.169 | 1 | 9 | Caskin2 | 0.037 | Old |
| 0.350521 | 0.209049 | 0.438 | 0.375 | 1 | 9 | Herc1 | 0.063 | Old |
| 0.350689 | 0.278043 | 0.157 | 0.125 | 1 | 9 | Psmc5 | 0.032 | Old |
| 0.351292 | 0.146096 | 0.144 | 0.11 | 1 | 9 | Ube2v2 | 0.034 | Old |
| 0.351318 | 0.8991 | 0.057 | 0.037 | 1 | 9 | Fam216a | 0.02 | Old |
| 0.351372 | 0.048438 | 0.668 | 0.603 | 1 | 9 | Arhgef12 | 0.065 | Old |
| 0.351785 | 0.484029 | 0.093 | 0.066 | 1 | 9 | Kdm3a | 0.027 | Old |
| 0.351835 | 0.400821 | 0.165 | 0.132 | 1 | 9 | Ivns1abp | 0.033 | Old |
| 0.351938 | 0.067066 | 0.057 | 0.081 | 1 | 9 | Msantd2 | -0.024 | Old |
| 0.352207 | 1.314398 | 0.039 | 0.022 | 1 | 9 | Zfp521 | 0.017 | Old |
| 0.352604 | 0.456736 | 0.162 | 0.132 | 1 | 9 | Coro1c | 0.03 | Old |
| 0.352974 | 0.197114 | 0.134 | 0.103 | 1 | 9 | Hdac4 | 0.031 | Old |
| 0.35324 | 0.525668 | 0.142 | 0.11 | 1 | 9 | Rnf10 | 0.032 | Old |
| 0.353288 | 1.173176 | 0.039 | 0.022 | 1 | 9 | Anapc15 | 0.017 | Old |
| 0.353288 | 1.074977 | 0.039 | 0.022 | 1 | 9 | Asb13 | 0.017 | Old |
| 0.35391 | 0.235587 | 0.209 | 0.169 | 1 | 9 | Lypla2 | 0.04 | Old |
| 0.354371 | 1.337161 | 0.039 | 0.022 | 1 | 9 | Tk2 | 0.017 | Old |
| 0.354371 | 1.28925 | 0.039 | 0.022 | 1 | 9 | Taf8 | 0.017 | Old |
| 0.354371 | 1.02845 | 0.039 | 0.022 | 1 | 9 | Mrpl49 | 0.017 | Old |
| 0.354436 | 0.003942 | 0.052 | 0.074 | 1 | 9 | Klf16 | -0.022 | Old |
| 0.354698 | 0.335188 | 0.149 | 0.118 | 1 | 9 | Cnot8 | 0.031 | Old |
| 0.35488 | 1.097931 | 0.057 | 0.037 | 1 | 9 | Tead4 | 0.02 | Old |

| | | | | | | | | |
|---|---|---|---|---|---|---|---|---|
| 0.355009 | 0.560637 | 0.253 | 0.221 | 1 | 9 | Sertad2 | 0.032 | Old |
| 0.356082 | 0.572116 | 0.155 | 0.125 | 1 | 9 | Rnf167 | 0.03 | Old |
| 0.356114 | 0.137712 | 0.119 | 0.154 | 1 | 9 | Dcaf12 | -0.035 | Old |
| 0.356543 | 1.389636 | 0.039 | 0.022 | 1 | 9 | Tubgcp6 | 0.017 | Old |
| 0.356543 | 1.297272 | 0.039 | 0.022 | 1 | 9 | Ppan | 0.017 | Old |
| 0.356543 | 0.788748 | 0.039 | 0.022 | 1 | 9 | Id4 | 0.017 | Old |
| 0.356543 | 0.108169 | 0.039 | 0.022 | 1 | 9 | Aga | 0.017 | Old |
| 0.356592 | 0.170498 | 0.214 | 0.176 | 1 | 9 | Zcchc11 | 0.038 | Old |
| 0.356918 | 0.163032 | 0.294 | 0.25 | 1 | 9 | Dhx15 | 0.044 | Old |
| 0.357133 | 0.913392 | 0.137 | 0.11 | 1 | 9 | Unc50 | 0.027 | Old |
| 0.357242 | 0.273066 | 0.049 | 0.029 | 1 | 9 | Mcu | 0.02 | Old |
| 0.357516 | 0.266524 | 0.108 | 0.081 | 1 | 9 | Rasa2 | 0.027 | Old |
| 0.357567 | 1.187864 | 0.057 | 0.037 | 1 | 9 | Pde7a | 0.02 | Old |
| 0.357632 | 1.09215 | 0.039 | 0.022 | 1 | 9 | Commd10 | 0.017 | Old |
| 0.357632 | 0.888816 | 0.039 | 0.022 | 1 | 9 | Haghl | 0.017 | Old |
| 0.35795 | 0.387648 | 0.25 | 0.221 | 1 | 9 | Ubald2 | 0.029 | Old |
| 0.358312 | 0.847533 | 0.137 | 0.11 | 1 | 9 | Napa | 0.027 | Old |
| 0.358444 | 0.1291 | 0.423 | 0.368 | 1 | 9 | Rbpms | 0.055 | Old |
| 0.359312 | 1.532108 | 0.064 | 0.044 | 1 | 9 | Pdrg1 | 0.02 | Old |
| 0.359366 | 1.087518 | 0.057 | 0.037 | 1 | 9 | Clcn6 | 0.02 | Old |
| 0.359694 | 0.003984 | 0.144 | 0.184 | 1 | 9 | Usp47 | -0.04 | Old |
| 0.360267 | 1.164089 | 0.057 | 0.037 | 1 | 9 | 2610008E1 | 0.02 | Old |
| 0.360792 | 0.60362 | 0.162 | 0.132 | 1 | 9 | Maf1 | 0.03 | Old |
| 0.361417 | 0.528208 | 0.116 | 0.088 | 1 | 9 | Map3k2 | 0.028 | Old |
| 0.361463 | 0.810418 | 0.075 | 0.051 | 1 | 9 | Ubald1 | 0.024 | Old |
| 0.362009 | 0.327821 | 0.039 | 0.022 | 1 | 9 | Wdr6 | 0.017 | Old |
| 0.362697 | 1.438719 | 0.064 | 0.044 | 1 | 9 | Gnpat | 0.02 | Old |
| 0.363532 | 1.594928 | 0.072 | 0.051 | 1 | 9 | Naxe | 0.021 | Old |
| 0.363734 | 0.337579 | 0.121 | 0.096 | 1 | 9 | Ssbp4 | 0.025 | Old |
| 0.363844 | 0.383544 | 0.224 | 0.191 | 1 | 9 | Dip2b | 0.033 | Old |
| 0.36386 | 0.188344 | 0.245 | 0.213 | 1 | 9 | Pros1 | 0.032 | Old |
| 0.363907 | 0.432571 | 0.147 | 0.118 | 1 | 9 | Smarca4 | 0.029 | Old |
| 0.363915 | 0.612281 | 0.193 | 0.162 | 1 | 9 | Chd7 | 0.031 | Old |
| 0.364517 | 0.038416 | 0.263 | 0.221 | 1 | 9 | Tsn | 0.042 | Old |
| 0.364584 | 0.267215 | 0.085 | 0.059 | 1 | 9 | Tirap | 0.026 | Old |
| 0.365205 | 0.740027 | 0.129 | 0.103 | 1 | 9 | Sars | 0.026 | Old |
| 0.365314 | 1.170981 | 0.039 | 0.022 | 1 | 9 | Mrpl16 | 0.017 | Old |
| 0.36549 | 0.420145 | 0.101 | 0.074 | 1 | 9 | Prpsap1 | 0.027 | Old |
| 0.36642 | 0.090348 | 0.039 | 0.022 | 1 | 9 | Rfxank | 0.017 | Old |
| 0.366667 | 0.380902 | 0.139 | 0.11 | 1 | 9 | Ndufa10 | 0.029 | Old |
| 0.366752 | 1.914624 | 0.028 | 0.015 | 1 | 9 | Znhit2 | 0.013 | Old |
| 0.366883 | 0.202997 | 0.322 | 0.287 | 1 | 9 | Ndufa7 | 0.035 | Old |
| 0.366956 | 1.105579 | 0.064 | 0.044 | 1 | 9 | 4921524J1 | 0.02 | Old |
| 0.367017 | 0.442507 | 0.075 | 0.051 | 1 | 9 | Fam57b | 0.024 | Old |
| 0.367262 | 0.644683 | 0.139 | 0.11 | 1 | 9 | Sncg | 0.029 | Old |

| | | | | | | | | |
|---|---|---|---|---|---|---|---|---|
| 0.367443 | 0.826642 | 0.183 | 0.154 | 1 | 9 | Deptor | 0.029 | Old |
| 0.367965 | 0.437264 | 0.134 | 0.103 | 1 | 9 | Ackr3 | 0.031 | Old |
| 0.368051 | 2.321343 | 0.028 | 0.015 | 1 | 9 | Hook2 | 0.013 | Old |
| 0.368051 | 1.59594 | 0.028 | 0.015 | 1 | 9 | Zbtb40 | 0.013 | Old |
| 0.368277 | 1.010602 | 0.129 | 0.103 | 1 | 9 | Tmem208 | 0.026 | Old |
| 0.368948 | 0.989566 | 0.082 | 0.059 | 1 | 9 | Tmem219 | 0.023 | Old |
| 0.368983 | 0.072513 | 0.982 | 0.978 | 1 | 9 | Actb | 0.004 | Old |
| 0.3691 | 0.340883 | 0.034 | 0.051 | 1 | 9 | Zfp637 | -0.017 | Old |
| 0.369349 | 0.0304 | 0.139 | 0.11 | 1 | 9 | Mbd3 | 0.029 | Old |
| 0.369352 | 1.839743 | 0.028 | 0.015 | 1 | 9 | Mterf4 | 0.013 | Old |
| 0.36971 | 0.237477 | 0.082 | 0.059 | 1 | 9 | Abcf3 | 0.023 | Old |
| 0.369946 | 0.021979 | 0.139 | 0.11 | 1 | 9 | Ddx21 | 0.029 | Old |
| 0.37044 | 0.868346 | 0.106 | 0.081 | 1 | 9 | Fbxl17 | 0.025 | Old |
| 0.370656 | 1.724012 | 0.028 | 0.015 | 1 | 9 | Praf2 | 0.013 | Old |
| 0.370774 | 0.542571 | 0.162 | 0.132 | 1 | 9 | Wbp11 | 0.03 | Old |
| 0.370811 | 1.972806 | 0.046 | 0.029 | 1 | 9 | Synj2 | 0.017 | Old |
| 0.371617 | 1.069201 | 0.072 | 0.051 | 1 | 9 | Ttc32 | 0.021 | Old |
| 0.371824 | 2.038938 | 0.046 | 0.029 | 1 | 9 | Tmem126b | 0.017 | Old |
| 0.371963 | 2.039949 | 0.028 | 0.015 | 1 | 9 | Osbpl5 | 0.013 | Old |
| 0.371963 | 1.860727 | 0.028 | 0.015 | 1 | 9 | Wfs1 | 0.013 | Old |
| 0.371963 | 1.441844 | 0.028 | 0.015 | 1 | 9 | Dbp | 0.013 | Old |
| 0.371963 | 1.392424 | 0.028 | 0.015 | 1 | 9 | Kbtbd11 | 0.013 | Old |
| 0.371963 | 1.3764 | 0.028 | 0.015 | 1 | 9 | Hs3st1 | 0.013 | Old |
| 0.372584 | 0.804956 | 0.113 | 0.088 | 1 | 9 | Kalrn | 0.025 | Old |
| 0.37284 | 1.974851 | 0.046 | 0.029 | 1 | 9 | Snx25 | 0.017 | Old |
| 0.373044 | 0.062392 | 0.747 | 0.691 | 1 | 9 | Hspb1 | 0.056 | Old |
| 0.373099 | 0.240666 | 0.039 | 0.022 | 1 | 9 | Slc35e2 | 0.017 | Old |
| 0.373105 | 0.360541 | 0.299 | 0.265 | 1 | 9 | Plec | 0.034 | Old |
| 0.37324 | 0.20126 | 0.284 | 0.243 | 1 | 9 | Tmed7 | 0.041 | Old |
| 0.373273 | 1.462881 | 0.028 | 0.015 | 1 | 9 | Ppp1r3d | 0.013 | Old |
| 0.373273 | 1.340553 | 0.028 | 0.015 | 1 | 9 | Mlkl | 0.013 | Old |
| 0.373515 | 0.002905 | 0.186 | 0.154 | 1 | 9 | Eif4a3 | 0.032 | Old |
| 0.373835 | 1.093596 | 0.064 | 0.044 | 1 | 9 | Nfil3 | 0.02 | Old |
| 0.373835 | 0.554791 | 0.064 | 0.044 | 1 | 9 | Tbc1d1 | 0.02 | Old |
| 0.373856 | 1.746473 | 0.046 | 0.029 | 1 | 9 | Wdr47 | 0.017 | Old |
| 0.374313 | 0.324695 | 0.052 | 0.074 | 1 | 9 | Myof | -0.022 | Old |
| 0.374585 | 1.809759 | 0.028 | 0.015 | 1 | 9 | 4930430F0 | 0.013 | Old |
| 0.375045 | 0.10399 | 0.126 | 0.096 | 1 | 9 | Mrpl51 | 0.03 | Old |
| 0.375073 | 0.043181 | 0.082 | 0.059 | 1 | 9 | Brix1 | 0.023 | Old |
| 0.375214 | 0.097735 | 0.116 | 0.147 | 1 | 9 | Sowahc | -0.031 | Old |
| 0.375332 | 0.005757 | 0.57 | 0.493 | 1 | 9 | Adamts1 | 0.077 | Old |
| 0.375342 | 0.405706 | 0.039 | 0.022 | 1 | 9 | Zc3h3 | 0.017 | Old |
| 0.375475 | 0.487042 | 0.034 | 0.051 | 1 | 9 | Prkcb | -0.017 | Old |
| 0.375546 | 0.454551 | 0.162 | 0.132 | 1 | 9 | Bdp1 | 0.03 | Old |
| 0.375719 | 0.691472 | 0.098 | 0.074 | 1 | 9 | Kat7 | 0.024 | Old |

| | | | | | | | | |
|---|---|---|---|---|---|---|---|---|
| 0.375901 | 1.692671 | 0.028 | 0.015 | 1 | 9 | Cluap1 | 0.013 | Old |
| 0.375901 | 1.363862 | 0.028 | 0.015 | 1 | 9 | Zxdc | 0.013 | Old |
| 0.375901 | 1.350406 | 0.028 | 0.015 | 1 | 9 | Scfd2 | 0.013 | Old |
| 0.375901 | 1.192016 | 0.028 | 0.015 | 1 | 9 | Slc2a8 | 0.013 | Old |
| 0.376435 | 0.375358 | 0.064 | 0.044 | 1 | 9 | Hmcn1 | 0.02 | Old |
| 0.376899 | 0.522553 | 0.144 | 0.118 | 1 | 9 | Adrm1 | 0.026 | Old |
| 0.377134 | 2.88251 | 0.018 | 0.007 | 1 | 9 | Fbxw17 | 0.011 | Old |
| 0.377134 | 2.470972 | 0.018 | 0.007 | 1 | 9 | Zdhhc13 | 0.011 | Old |
| 0.377134 | 2.347588 | 0.018 | 0.007 | 1 | 9 | A3galt2 | 0.011 | Old |
| 0.377134 | 2.209794 | 0.018 | 0.007 | 1 | 9 | Zfp30 | 0.011 | Old |
| 0.377134 | 2.122894 | 0.018 | 0.007 | 1 | 9 | Mfsd12 | 0.011 | Old |
| 0.377134 | 2.059266 | 0.018 | 0.007 | 1 | 9 | Rpusd3 | 0.011 | Old |
| 0.377134 | 2.009434 | 0.018 | 0.007 | 1 | 9 | Gm4707 | 0.011 | Old |
| 0.377134 | 1.845193 | 0.018 | 0.007 | 1 | 9 | Serf1 | 0.011 | Old |
| 0.377134 | 1.777811 | 0.018 | 0.007 | 1 | 9 | Dhx34 | 0.011 | Old |
| 0.377134 | 1.494809 | 0.018 | 0.007 | 1 | 9 | Snapc4 | 0.011 | Old |
| 0.377219 | 1.387059 | 0.028 | 0.015 | 1 | 9 | Snta1 | 0.013 | Old |
| 0.377219 | 1.344206 | 0.028 | 0.015 | 1 | 9 | Zfp280b | 0.013 | Old |
| 0.377219 | 1.03298 | 0.028 | 0.015 | 1 | 9 | Wdr4 | 0.013 | Old |
| 0.377661 | 0.443509 | 0.057 | 0.037 | 1 | 9 | Psmg3 | 0.02 | Old |
| 0.377839 | 0.020634 | 0.052 | 0.074 | 1 | 9 | Kpna3 | -0.022 | Old |
| 0.377872 | 0.695312 | 0.113 | 0.088 | 1 | 9 | Fblim1 | 0.025 | Old |
| 0.377941 | 1.082624 | 0.046 | 0.029 | 1 | 9 | Zfp787 | 0.017 | Old |
| 0.37854 | 1.049689 | 0.028 | 0.015 | 1 | 9 | Nt5m | 0.013 | Old |
| 0.37854 | 0.926408 | 0.028 | 0.015 | 1 | 9 | Jmjd8 | 0.013 | Old |
| 0.378729 | 0.084529 | 0.028 | 0.044 | 1 | 9 | Cd300a | -0.016 | Old |
| 0.37881 | 2.47694 | 0.018 | 0.007 | 1 | 9 | Tefm | 0.011 | Old |
| 0.37881 | 2.131745 | 0.018 | 0.007 | 1 | 9 | Rpgrip1 | 0.011 | Old |
| 0.37881 | 2.001 | 0.018 | 0.007 | 1 | 9 | Gdf11 | 0.011 | Old |
| 0.37881 | 1.873068 | 0.018 | 0.007 | 1 | 9 | Gm16181 | 0.011 | Old |
| 0.37881 | 1.763197 | 0.018 | 0.007 | 1 | 9 | Nit2 | 0.011 | Old |
| 0.37881 | 1.597254 | 0.018 | 0.007 | 1 | 9 | Mvk | 0.011 | Old |
| 0.37881 | 1.550519 | 0.018 | 0.007 | 1 | 9 | B630019K0 | 0.011 | Old |
| 0.37881 | 1.498266 | 0.018 | 0.007 | 1 | 9 | Acr | 0.011 | Old |
| 0.37881 | 1.403775 | 0.018 | 0.007 | 1 | 9 | Tm9sf1 | 0.011 | Old |
| 0.379161 | 0.25237 | 0.253 | 0.213 | 1 | 9 | Slc12a7 | 0.04 | Old |
| 0.379864 | 0.736347 | 0.028 | 0.015 | 1 | 9 | Ppp2r5b | 0.013 | Old |
| 0.379864 | 0.731503 | 0.028 | 0.015 | 1 | 9 | Utp14b | 0.013 | Old |
| 0.379864 | 0.624796 | 0.028 | 0.015 | 1 | 9 | Cyp39a1 | 0.013 | Old |
| 0.379993 | 1.221656 | 0.046 | 0.029 | 1 | 9 | Zbtb22 | 0.017 | Old |
| 0.380203 | 0.078128 | 0.899 | 0.875 | 1 | 9 | Itm2b | 0.024 | Old |
| 0.380454 | 0.009223 | 0.186 | 0.154 | 1 | 9 | Phf21a | 0.032 | Old |
| 0.380491 | 1.796115 | 0.018 | 0.007 | 1 | 9 | Tprn | 0.011 | Old |
| 0.380491 | 1.784254 | 0.018 | 0.007 | 1 | 9 | Zfp799 | 0.011 | Old |
| 0.380491 | 1.631215 | 0.018 | 0.007 | 1 | 9 | Gstm5 | 0.011 | Old |

| | | | | | | | | |
|---|---|---|---|---|---|---|---|---|
| 0.380491 | 1.626349 | 0.018 | 0.007 | 1 | 9 | Maged2 | 0.011 | Old |
| 0.380491 | 1.623818 | 0.018 | 0.007 | 1 | 9 | Kirrel3 | 0.011 | Old |
| 0.380491 | 1.618206 | 0.018 | 0.007 | 1 | 9 | Nxpe3 | 0.011 | Old |
| 0.380491 | 1.425374 | 0.018 | 0.007 | 1 | 9 | Kif23 | 0.011 | Old |
| 0.380491 | 1.42233 | 0.018 | 0.007 | 1 | 9 | Sh3pxd2b | 0.011 | Old |
| 0.380491 | 1.289526 | 0.018 | 0.007 | 1 | 9 | Coq3 | 0.011 | Old |
| 0.380491 | 1.284991 | 0.018 | 0.007 | 1 | 9 | Isyna1 | 0.011 | Old |
| 0.380491 | 1.246365 | 0.018 | 0.007 | 1 | 9 | Slc9a5 | 0.011 | Old |
| 0.380491 | 1.19834 | 0.018 | 0.007 | 1 | 9 | Pola1 | 0.011 | Old |
| 0.380491 | 1.111272 | 0.018 | 0.007 | 1 | 9 | Col4a3 | 0.011 | Old |
| 0.381191 | 0.829798 | 0.028 | 0.015 | 1 | 9 | Gmppb | 0.013 | Old |
| 0.381191 | 0.616197 | 0.028 | 0.015 | 1 | 9 | Gm13056 | 0.013 | Old |
| 0.381191 | 0.550928 | 0.028 | 0.015 | 1 | 9 | Riox1 | 0.013 | Old |
| 0.381191 | 0.259269 | 0.028 | 0.015 | 1 | 9 | Zfp949 | 0.013 | Old |
| 0.381259 | 0.465747 | 0.186 | 0.154 | 1 | 9 | Cbx7 | 0.032 | Old |
| 0.381261 | 0.665596 | 0.082 | 0.059 | 1 | 9 | Psen2 | 0.023 | Old |
| 0.381284 | 0.280078 | 0.209 | 0.176 | 1 | 9 | Psmb3 | 0.033 | Old |
| 0.381416 | 0.31869 | 0.098 | 0.074 | 1 | 9 | Slc25a24 | 0.024 | Old |
| 0.381435 | 0.521476 | 0.131 | 0.103 | 1 | 9 | Trappc8 | 0.028 | Old |
| 0.381668 | 0.829812 | 0.064 | 0.044 | 1 | 9 | Nrbf2 | 0.02 | Old |
| 0.381799 | 0.696702 | 0.206 | 0.176 | 1 | 9 | Mat2b | 0.03 | Old |
| 0.382132 | 0.641366 | 0.098 | 0.074 | 1 | 9 | Hipk2 | 0.024 | Old |
| 0.382177 | 2.399024 | 0.018 | 0.007 | 1 | 9 | Kcnj8 | 0.011 | Old |
| 0.382177 | 2.032204 | 0.018 | 0.007 | 1 | 9 | Emc9 | 0.011 | Old |
| 0.382177 | 1.795284 | 0.018 | 0.007 | 1 | 9 | Pus7l | 0.011 | Old |
| 0.382177 | 1.433563 | 0.018 | 0.007 | 1 | 9 | Mkks | 0.011 | Old |
| 0.382177 | 1.428584 | 0.018 | 0.007 | 1 | 9 | Lin9 | 0.011 | Old |
| 0.382177 | 1.374863 | 0.018 | 0.007 | 1 | 9 | Trabd2b | 0.011 | Old |
| 0.382177 | 1.074148 | 0.018 | 0.007 | 1 | 9 | Apol9a | 0.011 | Old |
| 0.382521 | 0.743715 | 0.028 | 0.015 | 1 | 9 | Cep97 | 0.013 | Old |
| 0.382521 | 0.63117 | 0.028 | 0.015 | 1 | 9 | Abcc9 | 0.013 | Old |
| 0.382539 | 0.379415 | 0.302 | 0.272 | 1 | 9 | Mbd2 | 0.03 | Old |
| 0.382664 | 0.492281 | 0.09 | 0.066 | 1 | 9 | Tor1a | 0.024 | Old |
| 0.382832 | 0.077604 | 0.451 | 0.412 | 1 | 9 | Atxn7l3b | 0.039 | Old |
| 0.382844 | 0.134428 | 0.142 | 0.11 | 1 | 9 | Pkd1 | 0.032 | Old |
| 0.383411 | 0.26158 | 0.09 | 0.066 | 1 | 9 | Mnat1 | 0.024 | Old |
| 0.383771 | 0.034198 | 0.869 | 0.787 | 1 | 9 | Myl6 | 0.082 | Old |
| 0.383853 | 0.940366 | 0.028 | 0.015 | 1 | 9 | Gin1 | 0.013 | Old |
| 0.383853 | 0.677755 | 0.028 | 0.015 | 1 | 9 | Gpc4 | 0.013 | Old |
| 0.383867 | 1.651777 | 0.018 | 0.007 | 1 | 9 | 0610040B1 | 0.011 | Old |
| 0.383867 | 1.582827 | 0.018 | 0.007 | 1 | 9 | Dhodh | 0.011 | Old |
| 0.383867 | 1.177301 | 0.018 | 0.007 | 1 | 9 | Rsl1 | 0.011 | Old |
| 0.383867 | 1.160152 | 0.018 | 0.007 | 1 | 9 | Zfp248 | 0.011 | Old |
| 0.383867 | 1.091071 | 0.018 | 0.007 | 1 | 9 | AW146154 | 0.011 | Old |
| 0.383867 | 0.986628 | 0.018 | 0.007 | 1 | 9 | Apol8 | 0.011 | Old |

| | | | | | | | | |
|---|---|---|---|---|---|---|---|---|
| 0.384059 | 0.598572 | 0.052 | 0.074 | 1 | 9 | Bin3 | -0.022 | Old |
| 0.384119 | 1.570531 | 0.046 | 0.029 | 1 | 9 | Abhd18 | 0.017 | Old |
| 0.38435 | 0.121079 | 0.544 | 0.493 | 1 | 9 | Tacc1 | 0.051 | Old |
| 0.384594 | 0.374189 | 0.126 | 0.096 | 1 | 9 | Abca8b | 0.03 | Old |
| 0.384812 | 0.04845 | 0.796 | 0.757 | 1 | 9 | Rps27 | 0.039 | Old |
| 0.385029 | 0.611263 | 0.168 | 0.14 | 1 | 9 | Cop1 | 0.028 | Old |
| 0.385065 | 0.330028 | 0.108 | 0.14 | 1 | 9 | Psmd14 | -0.032 | Old |
| 0.385175 | 0.075878 | 0.196 | 0.162 | 1 | 9 | Chd6 | 0.034 | Old |
| 0.385236 | 0.010588 | 0.126 | 0.096 | 1 | 9 | Dnpep | 0.03 | Old |
| 0.385562 | 1.866913 | 0.018 | 0.007 | 1 | 9 | Cep131 | 0.011 | Old |
| 0.385562 | 1.306265 | 0.018 | 0.007 | 1 | 9 | Gm45871 | 0.011 | Old |
| 0.385562 | 1.139485 | 0.018 | 0.007 | 1 | 9 | Mpi | 0.011 | Old |
| 0.385562 | 0.957775 | 0.018 | 0.007 | 1 | 9 | Tatdn3 | 0.011 | Old |
| 0.385562 | 0.90967 | 0.018 | 0.007 | 1 | 9 | Clec3b | 0.011 | Old |
| 0.385562 | 0.766654 | 0.018 | 0.007 | 1 | 9 | Ctdspl | 0.011 | Old |
| 0.385562 | 0.742391 | 0.018 | 0.007 | 1 | 9 | Zfp668 | 0.011 | Old |
| 0.385616 | 0.766426 | 0.072 | 0.051 | 1 | 9 | Naa25 | 0.021 | Old |
| 0.385654 | 0.311296 | 0.235 | 0.199 | 1 | 9 | Chd1 | 0.036 | Old |
| 0.385669 | 0.458866 | 0.206 | 0.176 | 1 | 9 | Clptm1 | 0.03 | Old |
| 0.385742 | 0.389129 | 0.242 | 0.199 | 1 | 9 | Igfbp3 | 0.043 | Old |
| 0.385986 | 0.416402 | 0.24 | 0.206 | 1 | 9 | Ndufb4 | 0.034 | Old |
| 0.385994 | 0.483995 | 0.286 | 0.257 | 1 | 9 | Scarb2 | 0.029 | Old |
| 0.386192 | 1.043169 | 0.046 | 0.029 | 1 | 9 | Rbm12b2 | 0.017 | Old |
| 0.386388 | 0.45974 | 0.152 | 0.125 | 1 | 9 | Mbtps1 | 0.027 | Old |
| 0.386527 | 0.697556 | 0.028 | 0.015 | 1 | 9 | Was | 0.013 | Old |
| 0.386527 | 0.561361 | 0.028 | 0.015 | 1 | 9 | Gjc1 | 0.013 | Old |
| 0.38676 | 0.056732 | 0.358 | 0.309 | 1 | 9 | Eif4ebp2 | 0.049 | Old |
| 0.386773 | 0.356153 | 0.229 | 0.199 | 1 | 9 | Fip1l1 | 0.03 | Old |
| 0.387231 | 1.433169 | 0.046 | 0.029 | 1 | 9 | Setdb1 | 0.017 | Old |
| 0.387231 | 1.246468 | 0.046 | 0.029 | 1 | 9 | Aldh7a1 | 0.017 | Old |
| 0.387261 | 1.013117 | 0.018 | 0.007 | 1 | 9 | Vars2 | 0.011 | Old |
| 0.387261 | 1.00195 | 0.018 | 0.007 | 1 | 9 | AC129328. | 0.011 | Old |
| 0.387261 | 0.921669 | 0.018 | 0.007 | 1 | 9 | Ddb2 | 0.011 | Old |
| 0.387261 | 0.768587 | 0.018 | 0.007 | 1 | 9 | Fbxo46 | 0.011 | Old |
| 0.387261 | 0.620104 | 0.018 | 0.007 | 1 | 9 | Chchd6 | 0.011 | Old |
| 0.387261 | 0.547595 | 0.018 | 0.007 | 1 | 9 | Gm27010 | 0.011 | Old |
| 0.387261 | 0.448078 | 0.018 | 0.007 | 1 | 9 | Pir | 0.011 | Old |
| 0.387261 | 0.429465 | 0.018 | 0.007 | 1 | 9 | Deup1 | 0.011 | Old |
| 0.387261 | 0.336582 | 0.018 | 0.007 | 1 | 9 | Spin4 | 0.011 | Old |
| 0.387261 | 0.240783 | 0.018 | 0.007 | 1 | 9 | Lcmt2 | 0.011 | Old |
| 0.387332 | 0.506523 | 0.124 | 0.096 | 1 | 9 | Polg | 0.028 | Old |
| 0.387453 | 0.289663 | 0.242 | 0.206 | 1 | 9 | Cept1 | 0.036 | Old |
| 0.38783 | 0.641232 | 0.064 | 0.044 | 1 | 9 | Ifi207 | 0.02 | Old |
| 0.38791 | 0.017187 | 0.08 | 0.103 | 1 | 9 | Abcc1 | -0.023 | Old |
| 0.38809 | 0.305125 | 0.028 | 0.044 | 1 | 9 | Map3k12 | -0.016 | Old |

| | | | | | | | | |
|---|---|---|---|---|---|---|---|---|
| 0.388527 | 1.915124 | 0.054 | 0.037 | 1 | 9 | Cep295 | 0.017 | Old |
| 0.388745 | 0.031322 | 0.325 | 0.279 | 1 | 9 | Hspa12b | 0.046 | Old |
| 0.38892 | 0.007932 | 0.294 | 0.25 | 1 | 9 | Diaph1 | 0.044 | Old |
| 0.388956 | 0.826294 | 0.072 | 0.051 | 1 | 9 | Zfp410 | 0.021 | Old |
| 0.388965 | 0.493932 | 0.018 | 0.007 | 1 | 9 | Fastkd2 | 0.011 | Old |
| 0.388965 | 0.282829 | 0.018 | 0.007 | 1 | 9 | Mars2 | 0.011 | Old |
| 0.388965 | 0.160709 | 0.018 | 0.007 | 1 | 9 | Fbxl22 | 0.011 | Old |
| 0.389112 | 0.202239 | 0.227 | 0.191 | 1 | 9 | Zyx | 0.036 | Old |
| 0.389212 | 0.5216 | 0.028 | 0.015 | 1 | 9 | Dnal4 | 0.013 | Old |
| 0.389212 | 0.123557 | 0.028 | 0.015 | 1 | 9 | Sirt3 | 0.013 | Old |
| 0.389309 | 0.652375 | 0.155 | 0.125 | 1 | 9 | Tm2d1 | 0.03 | Old |
| 0.389556 | 0.26517 | 0.436 | 0.419 | 1 | 9 | Rab2a | 0.017 | Old |
| 0.390358 | 1.172639 | 0.046 | 0.029 | 1 | 9 | Cep63 | 0.017 | Old |
| 0.390452 | 0.240988 | 0.374 | 0.324 | 1 | 9 | Glg1 | 0.05 | Old |
| 0.391062 | 0.335355 | 0.302 | 0.272 | 1 | 9 | Stag2 | 0.03 | Old |
| 0.391122 | 0.191105 | 0.394 | 0.338 | 1 | 9 | Tnks2 | 0.056 | Old |
| 0.391404 | 0.835767 | 0.046 | 0.029 | 1 | 9 | Cdkal1 | 0.017 | Old |
| 0.391678 | 0.301142 | 0.09 | 0.066 | 1 | 9 | Proser1 | 0.024 | Old |
| 0.391695 | 0.469868 | 0.273 | 0.25 | 1 | 9 | Irf2 | 0.023 | Old |
| 0.391964 | 0.585777 | 0.155 | 0.125 | 1 | 9 | Sema3f | 0.03 | Old |
| 0.392065 | 0.322258 | 0.317 | 0.279 | 1 | 9 | Mapk1 | 0.038 | Old |
| 0.392145 | 0.619627 | 0.129 | 0.103 | 1 | 9 | Rnf38 | 0.026 | Old |
| 0.392452 | 0.901824 | 0.046 | 0.029 | 1 | 9 | Gba | 0.017 | Old |
| 0.392894 | 0.301661 | 0.456 | 0.434 | 1 | 9 | Igf1r | 0.022 | Old |
| 0.392945 | 0.43225 | 0.214 | 0.184 | 1 | 9 | Ythdf1 | 0.03 | Old |
| 0.393015 | 0.184022 | 0.649 | 0.618 | 1 | 9 | Sema6a | 0.031 | Old |
| 0.393307 | 0.106701 | 0.245 | 0.206 | 1 | 9 | Fam160b1 | 0.039 | Old |
| 0.393501 | 0.747347 | 0.046 | 0.029 | 1 | 9 | Cep162 | 0.017 | Old |
| 0.393501 | 0.28303 | 0.046 | 0.029 | 1 | 9 | Rab40c | 0.017 | Old |
| 0.393718 | 0.392488 | 0.139 | 0.11 | 1 | 9 | Als2cl | 0.029 | Old |
| 0.393752 | 0.306127 | 0.101 | 0.074 | 1 | 9 | Sucla2 | 0.027 | Old |
| 0.393939 | 1.127886 | 0.088 | 0.066 | 1 | 9 | Dmac1 | 0.022 | Old |
| 0.394363 | 0.473475 | 0.144 | 0.118 | 1 | 9 | Acer2 | 0.026 | Old |
| 0.394552 | 0.721503 | 0.046 | 0.029 | 1 | 9 | Ikzf4 | 0.017 | Old |
| 0.394791 | 0.474567 | 0.137 | 0.11 | 1 | 9 | Pafah1b2 | 0.027 | Old |
| 0.395167 | 0.007472 | 0.124 | 0.096 | 1 | 9 | Krcc1 | 0.028 | Old |
| 0.395299 | 1.563024 | 0.054 | 0.037 | 1 | 9 | Nbeal2 | 0.017 | Old |
| 0.396123 | 0.412567 | 0.273 | 0.243 | 1 | 9 | Srp14 | 0.03 | Old |
| 0.396401 | 0.041628 | 0.394 | 0.346 | 1 | 9 | Tmem184b | 0.048 | Old |
| 0.396535 | 0.361861 | 0.072 | 0.051 | 1 | 9 | Foxo4 | 0.021 | Old |
| 0.396737 | 0.474648 | 0.064 | 0.044 | 1 | 9 | Wdr7 | 0.02 | Old |
| 0.396739 | 0.548931 | 0.113 | 0.088 | 1 | 9 | Mrpl42 | 0.025 | Old |
| 0.397276 | 0.963492 | 0.126 | 0.103 | 1 | 9 | Nacc2 | 0.023 | Old |
| 0.397715 | 0.565782 | 0.046 | 0.029 | 1 | 9 | Lsm11 | 0.017 | Old |
| 0.397735 | 0.876105 | 0.149 | 0.125 | 1 | 9 | Eif4e | 0.024 | Old |

| | | | | | | | | |
|---|---|---|---|---|---|---|---|---|
| 0.39809 | 0.364471 | 0.374 | 0.338 | 1 | 9 | Trim24 | 0.036 | Old |
| 0.398095 | 0.0349 | 0.235 | 0.199 | 1 | 9 | Tceal9 | 0.036 | Old |
| 0.398773 | 0.509712 | 0.046 | 0.029 | 1 | 9 | Spsb3 | 0.017 | Old |
| 0.399081 | 0.59721 | 0.072 | 0.051 | 1 | 9 | Prkaca | 0.021 | Old |
| 0.399832 | 0.684186 | 0.046 | 0.029 | 1 | 9 | Ptrh2 | 0.017 | Old |
| 0.399873 | 0.409029 | 0.129 | 0.103 | 1 | 9 | Prkcd | 0.026 | Old |
| 0.399932 | 0.517512 | 0.072 | 0.051 | 1 | 9 | Gnl2 | 0.021 | Old |
| 0.399973 | 0.218936 | 0.162 | 0.132 | 1 | 9 | Fam3c | 0.03 | Old |
| 0.400053 | 0.306562 | 0.09 | 0.066 | 1 | 9 | Gmcl1 | 0.024 | Old |
| 0.400893 | 0.755441 | 0.046 | 0.029 | 1 | 9 | Foxred1 | 0.017 | Old |
| 0.400893 | 0.665832 | 0.046 | 0.029 | 1 | 9 | Mgp | 0.017 | Old |
| 0.400893 | 0.536206 | 0.046 | 0.029 | 1 | 9 | AI606181 | 0.017 | Old |
| 0.401113 | 0.590296 | 0.144 | 0.118 | 1 | 9 | Arl5a | 0.026 | Old |
| 0.401791 | 0.391428 | 0.155 | 0.125 | 1 | 9 | Nub1 | 0.03 | Old |
| 0.402141 | 1.825654 | 0.054 | 0.037 | 1 | 9 | Nfkbib | 0.017 | Old |
| 0.402357 | 0.188358 | 0.253 | 0.213 | 1 | 9 | Ddx24 | 0.04 | Old |
| 0.402509 | 0.797995 | 0.098 | 0.074 | 1 | 9 | Arfrp1 | 0.024 | Old |
| 0.402627 | 0.05012 | 0.397 | 0.368 | 1 | 9 | Brd2 | 0.029 | Old |
| 0.402768 | 0.168415 | 0.08 | 0.059 | 1 | 9 | Ccdc97 | 0.021 | Old |
| 0.403045 | 0.018256 | 0.064 | 0.044 | 1 | 9 | Gm26624 | 0.02 | Old |
| 0.403124 | 1.351269 | 0.054 | 0.037 | 1 | 9 | Ak6 | 0.017 | Old |
| 0.403599 | 1.473935 | 0.036 | 0.022 | 1 | 9 | Nit1 | 0.014 | Old |
| 0.403599 | 0.994266 | 0.028 | 0.044 | 1 | 9 | Wdr91 | -0.016 | Old |
| 0.403711 | 0.064522 | 0.405 | 0.368 | 1 | 9 | B230219D2 | 0.037 | Old |
| 0.403777 | 0.207757 | 0.041 | 0.059 | 1 | 9 | Dse | -0.018 | Old |
| 0.403856 | 0.257145 | 0.106 | 0.081 | 1 | 9 | Atp6v1c1 | 0.025 | Old |
| 0.404009 | 0.16671 | 0.152 | 0.125 | 1 | 9 | Nmi | 0.027 | Old |
| 0.404087 | 0.736394 | 0.046 | 0.029 | 1 | 9 | 3110082I1 | 0.017 | Old |
| 0.404087 | 0.54151 | 0.046 | 0.029 | 1 | 9 | Rrp7a | 0.017 | Old |
| 0.404199 | 0.08763 | 0.245 | 0.206 | 1 | 9 | Setd7 | 0.039 | Old |
| 0.404397 | 0.719968 | 0.08 | 0.059 | 1 | 9 | C1qbp | 0.021 | Old |
| 0.404448 | 0.008802 | 0.173 | 0.14 | 1 | 9 | Cdk16 | 0.033 | Old |
| 0.404507 | 0.280296 | 0.216 | 0.184 | 1 | 9 | Cyb5b | 0.032 | Old |
| 0.404644 | 0.3283 | 0.211 | 0.176 | 1 | 9 | Hdac3 | 0.035 | Old |
| 0.405155 | 0.40655 | 0.046 | 0.029 | 1 | 9 | Acss2 | 0.017 | Old |
| 0.405392 | 0.231237 | 0.407 | 0.36 | 1 | 9 | Tmed2 | 0.047 | Old |
| 0.405508 | 0.02507 | 0.874 | 0.86 | 1 | 9 | Flt1 | 0.014 | Old |
| 0.405732 | 0.584543 | 0.219 | 0.191 | 1 | 9 | Znfx1 | 0.028 | Old |
| 0.406082 | 0.890319 | 0.054 | 0.037 | 1 | 9 | Mapk9 | 0.017 | Old |
| 0.406908 | 0.295979 | 0.023 | 0.037 | 1 | 9 | Tmem242 | -0.014 | Old |
| 0.407039 | 0.620355 | 0.07 | 0.096 | 1 | 9 | Mpdu1 | -0.026 | Old |
| 0.40723 | 1.764848 | 0.036 | 0.022 | 1 | 9 | Pus7 | 0.014 | Old |
| 0.407296 | 0.061696 | 0.046 | 0.029 | 1 | 9 | Tmem203 | 0.017 | Old |
| 0.407306 | 1.020534 | 0.134 | 0.11 | 1 | 9 | Arhgef5 | 0.024 | Old |
| 0.407596 | 0.174897 | 0.131 | 0.103 | 1 | 9 | Timm8b | 0.028 | Old |

| | | | | | | | | |
|---|---|---|---|---|---|---|---|---|
| 0.407665 | 0.947125 | 0.08 | 0.059 | 1 | 9 | Dcn | 0.021 | Old |
| 0.407936 | 0.363028 | 0.235 | 0.206 | 1 | 9 | Sel1l | 0.029 | Old |
| 0.407936 | 0.445864 | 0.235 | 0.206 | 1 | 9 | Cnot1 | 0.029 | Old |
| 0.408242 | 0.010699 | 0.023 | 0.037 | 1 | 9 | Glipr2 | -0.014 | Old |
| 0.408256 | 0.048503 | 0.887 | 0.809 | 1 | 9 | Rpl35a | 0.078 | Old |
| 0.408348 | 0.036299 | 0.129 | 0.103 | 1 | 9 | Dnajc19 | 0.026 | Old |
| 0.40847 | 0.246177 | 0.327 | 0.287 | 1 | 9 | Slc50a1 | 0.04 | Old |
| 0.408518 | 0.645243 | 0.245 | 0.221 | 1 | 9 | Bcap31 | 0.024 | Old |
| 0.408677 | 0.10944 | 0.059 | 0.081 | 1 | 9 | Tigd2 | -0.022 | Old |
| 0.408828 | 0.26404 | 0.17 | 0.14 | 1 | 9 | Cdk13 | 0.03 | Old |
| 0.40885 | 0.211124 | 0.188 | 0.228 | 1 | 9 | Uso1 | -0.04 | Old |
| 0.409006 | 0.284019 | 0.232 | 0.199 | 1 | 9 | Ldb2 | 0.033 | Old |
| 0.409053 | 1.246032 | 0.054 | 0.037 | 1 | 9 | Snx16 | 0.017 | Old |
| 0.409174 | 0.74755 | 0.152 | 0.125 | 1 | 9 | Jade1 | 0.027 | Old |
| 0.409662 | 1.929149 | 0.036 | 0.022 | 1 | 9 | Slc25a19 | 0.014 | Old |
| 0.409906 | 0.554214 | 0.149 | 0.125 | 1 | 9 | Tdrd7 | 0.024 | Old |
| 0.410127 | 0.555503 | 0.08 | 0.059 | 1 | 9 | Serhl | 0.021 | Old |
| 0.410294 | 0.002321 | 0.067 | 0.088 | 1 | 9 | Mrps7 | -0.021 | Old |
| 0.410303 | 0.373646 | 0.418 | 0.382 | 1 | 9 | Gsk3b | 0.036 | Old |
| 0.410501 | 0.008364 | 0.552 | 0.493 | 1 | 9 | Eef2 | 0.059 | Old |
| 0.410689 | 0.267984 | 0.098 | 0.074 | 1 | 9 | Kdm4a | 0.024 | Old |
| 0.410918 | 0.245163 | 0.023 | 0.037 | 1 | 9 | Dapk1 | -0.014 | Old |
| 0.411041 | 1.087768 | 0.054 | 0.037 | 1 | 9 | Gdpd5 | 0.017 | Old |
| 0.411636 | 0.16678 | 0.129 | 0.103 | 1 | 9 | Kifc3 | 0.026 | Old |
| 0.412037 | 0.881373 | 0.054 | 0.037 | 1 | 9 | Ino80c | 0.017 | Old |
| 0.412098 | 0.504689 | 0.273 | 0.25 | 1 | 9 | Spcs2 | 0.023 | Old |
| 0.412102 | 1.734863 | 0.036 | 0.022 | 1 | 9 | Ercc6 | 0.014 | Old |
| 0.412246 | 0.861317 | 0.095 | 0.074 | 1 | 9 | Nf2 | 0.021 | Old |
| 0.412304 | 0.02084 | 0.237 | 0.279 | 1 | 9 | Elavl1 | -0.042 | Old |
| 0.412348 | 1.510334 | 0.062 | 0.044 | 1 | 9 | Rbfa | 0.018 | Old |
| 0.412392 | 0.228362 | 0.119 | 0.096 | 1 | 9 | Qser1 | 0.023 | Old |
| 0.412597 | 0.878927 | 0.08 | 0.059 | 1 | 9 | Sacm1l | 0.021 | Old |
| 0.412658 | 0.070966 | 0.134 | 0.103 | 1 | 9 | Spry4 | 0.031 | Old |
| 0.412983 | 0.198703 | 0.263 | 0.228 | 1 | 9 | Ubn1 | 0.035 | Old |
| 0.413035 | 0.353208 | 0.054 | 0.037 | 1 | 9 | Dsel | 0.017 | Old |
| 0.413152 | 0.082956 | 0.25 | 0.213 | 1 | 9 | Tomm20 | 0.037 | Old |
| 0.413325 | 1.887513 | 0.036 | 0.022 | 1 | 9 | Rnf214 | 0.014 | Old |
| 0.413498 | 0.624856 | 0.103 | 0.081 | 1 | 9 | Slc44a2 | 0.022 | Old |
| 0.413758 | 0.191489 | 0.046 | 0.029 | 1 | 9 | Pde2a | 0.017 | Old |
| 0.414447 | 0.849113 | 0.119 | 0.096 | 1 | 9 | Ppp1r11 | 0.023 | Old |
| 0.414831 | 0.055262 | 0.121 | 0.096 | 1 | 9 | Afap1 | 0.025 | Old |
| 0.414952 | 0.259507 | 0.023 | 0.037 | 1 | 9 | Naip5 | -0.014 | Old |
| 0.415651 | 0.465827 | 0.16 | 0.132 | 1 | 9 | Snrnp27 | 0.028 | Old |
| 0.415778 | 2.181731 | 0.036 | 0.022 | 1 | 9 | Atp1b1 | 0.014 | Old |
| 0.415778 | 1.336931 | 0.036 | 0.022 | 1 | 9 | Fam13b | 0.014 | Old |

| | | | | | | | | |
|---|---|---|---|---|---|---|---|---|
| 0.416265 | 0.100948 | 0.129 | 0.103 | 1 | 9 | Aimp1 | 0.026 | Old |
| 0.416367 | 0.500079 | 0.25 | 0.228 | 1 | 9 | Acox1 | 0.022 | Old |
| 0.417008 | 1.566067 | 0.036 | 0.022 | 1 | 9 | Leng1 | 0.014 | Old |
| 0.417008 | 1.049935 | 0.036 | 0.022 | 1 | 9 | St7 | 0.014 | Old |
| 0.417825 | 0.301552 | 0.103 | 0.132 | 1 | 9 | Stard3nl | -0.029 | Old |
| 0.418066 | 0.005426 | 0.582 | 0.529 | 1 | 9 | Serf2 | 0.053 | Old |
| 0.41824 | 1.741431 | 0.036 | 0.022 | 1 | 9 | Cmklr1 | 0.014 | Old |
| 0.418467 | 0.306847 | 0.18 | 0.154 | 1 | 9 | Zmpste24 | 0.026 | Old |
| 0.41875 | 0.037713 | 0.131 | 0.103 | 1 | 9 | Ilf3 | 0.028 | Old |
| 0.418929 | 0.237472 | 0.121 | 0.096 | 1 | 9 | Tbc1d22a | 0.025 | Old |
| 0.418966 | 0.275259 | 0.165 | 0.132 | 1 | 9 | Flt4 | 0.033 | Old |
| 0.419245 | 0.120061 | 0.206 | 0.169 | 1 | 9 | Map1lc3a | 0.037 | Old |
| 0.419474 | 1.125986 | 0.036 | 0.022 | 1 | 9 | Acd | 0.014 | Old |
| 0.419725 | 0.106808 | 0.098 | 0.074 | 1 | 9 | Yipf1 | 0.024 | Old |
| 0.419756 | 0.43058 | 0.106 | 0.081 | 1 | 9 | Gadd45a | 0.025 | Old |
| 0.419758 | 0.328281 | 0.168 | 0.14 | 1 | 9 | Ndufb2 | 0.028 | Old |
| 0.420058 | 0.611386 | 0.054 | 0.037 | 1 | 9 | Gpr180 | 0.017 | Old |
| 0.420061 | 0.929362 | 0.08 | 0.059 | 1 | 9 | Gm8186 | 0.021 | Old |
| 0.420366 | 1.008483 | 0.023 | 0.037 | 1 | 9 | Dera | -0.014 | Old |
| 0.42071 | 1.212203 | 0.036 | 0.022 | 1 | 9 | Nars2 | 0.014 | Old |
| 0.421097 | 0.195014 | 0.24 | 0.206 | 1 | 9 | Dcaf8 | 0.034 | Old |
| 0.421342 | 0.02191 | 0.546 | 0.478 | 1 | 9 | Rpl36a | 0.068 | Old |
| 0.421379 | 0.252032 | 0.157 | 0.132 | 1 | 9 | Slc25a36 | 0.025 | Old |
| 0.421389 | 0.75861 | 0.095 | 0.074 | 1 | 9 | Nr1h3 | 0.021 | Old |
| 0.421557 | 0.316188 | 0.072 | 0.051 | 1 | 9 | Arhgap35 | 0.021 | Old |
| 0.421593 | 0.246749 | 0.129 | 0.103 | 1 | 9 | Rai1 | 0.026 | Old |
| 0.421948 | 1.302264 | 0.036 | 0.022 | 1 | 9 | Fastk | 0.014 | Old |
| 0.421948 | 0.959079 | 0.036 | 0.022 | 1 | 9 | Tspyl2 | 0.014 | Old |
| 0.421948 | 0.916491 | 0.036 | 0.022 | 1 | 9 | Dtx2 | 0.014 | Old |
| 0.422077 | 0.314279 | 0.054 | 0.037 | 1 | 9 | Zfp287 | 0.017 | Old |
| 0.422418 | 0.311494 | 0.302 | 0.265 | 1 | 9 | Ino80d | 0.037 | Old |
| 0.422562 | 0.047219 | 0.335 | 0.287 | 1 | 9 | Rtl8a | 0.048 | Old |
| 0.422686 | 0.176761 | 0.389 | 0.353 | 1 | 9 | Atox1 | 0.036 | Old |
| 0.423047 | 0.432726 | 0.144 | 0.118 | 1 | 9 | Fam20b | 0.026 | Old |
| 0.423047 | 0.31747 | 0.144 | 0.118 | 1 | 9 | Vps26a | 0.026 | Old |
| 0.423092 | 0.388199 | 0.235 | 0.206 | 1 | 9 | Psma2 | 0.029 | Old |
| 0.423189 | 1.287004 | 0.036 | 0.022 | 1 | 9 | Abcb4 | 0.014 | Old |
| 0.423189 | 1.040822 | 0.036 | 0.022 | 1 | 9 | Gpr1 | 0.014 | Old |
| 0.423419 | 0.572206 | 0.119 | 0.096 | 1 | 9 | Cdc26 | 0.023 | Old |
| 0.423821 | 0.09344 | 0.425 | 0.368 | 1 | 9 | Rsrc2 | 0.057 | Old |
| 0.424046 | 0.220929 | 0.273 | 0.243 | 1 | 9 | Prag1 | 0.03 | Old |
| 0.424102 | 0.748104 | 0.054 | 0.037 | 1 | 9 | Mrpl47 | 0.017 | Old |
| 0.424197 | 0.407882 | 0.072 | 0.051 | 1 | 9 | Cars | 0.021 | Old |
| 0.424322 | 0.447372 | 0.144 | 0.118 | 1 | 9 | Cd274 | 0.026 | Old |
| 0.424431 | 1.032242 | 0.036 | 0.022 | 1 | 9 | Lzic | 0.014 | Old |

| | | | | | | | | |
|---|---|---|---|---|---|---|---|---|
| 0.424431 | 0.692227 | 0.036 | 0.022 | 1 | 9 | Ppox | 0.014 | Old |
| 0.424751 | 0.004825 | 0.358 | 0.301 | 1 | 9 | Vcl | 0.057 | Old |
| 0.424912 | 0.346963 | 0.235 | 0.206 | 1 | 9 | Phc2 | 0.029 | Old |
| 0.424918 | 0.760889 | 0.103 | 0.081 | 1 | 9 | Trappc13 | 0.022 | Old |
| 0.425079 | 0.166402 | 0.072 | 0.051 | 1 | 9 | Srprb | 0.021 | Old |
| 0.425123 | 0.176409 | 0.054 | 0.074 | 1 | 9 | Tank | -0.02 | Old |
| 0.425443 | 0.094381 | 0.686 | 0.625 | 1 | 9 | Rpl17 | 0.061 | Old |
| 0.425676 | 0.864355 | 0.036 | 0.022 | 1 | 9 | Gm4070 | 0.014 | Old |
| 0.425676 | 0.439888 | 0.036 | 0.022 | 1 | 9 | Zfp719 | 0.014 | Old |
| 0.426392 | 1.119284 | 0.101 | 0.081 | 1 | 9 | Cyth2 | 0.02 | Old |
| 0.426923 | 0.631379 | 0.036 | 0.022 | 1 | 9 | Trim6 | 0.014 | Old |
| 0.426923 | 0.064091 | 0.036 | 0.022 | 1 | 9 | Armcx5 | 0.014 | Old |
| 0.427315 | 0.439744 | 0.142 | 0.118 | 1 | 9 | B4galt1 | 0.024 | Old |
| 0.427599 | 0.671766 | 0.119 | 0.096 | 1 | 9 | Virma | 0.023 | Old |
| 0.427627 | 0.392629 | 0.263 | 0.235 | 1 | 9 | Vps13d | 0.028 | Old |
| 0.427712 | 1.619939 | 0.044 | 0.029 | 1 | 9 | Zfp865 | 0.015 | Old |
| 0.428172 | 0.644686 | 0.036 | 0.022 | 1 | 9 | Grhpr | 0.014 | Old |
| 0.428561 | 0.197304 | 0.5 | 0.456 | 1 | 9 | Tinagl1 | 0.044 | Old |
| 0.428619 | 0.250219 | 0.072 | 0.051 | 1 | 9 | Ddx27 | 0.021 | Old |
| 0.428974 | 0.071009 | 0.497 | 0.463 | 1 | 9 | Sox18 | 0.034 | Old |
| 0.42919 | 0.986793 | 0.054 | 0.037 | 1 | 9 | Cfap43 | 0.017 | Old |
| 0.429326 | 1.119375 | 0.062 | 0.044 | 1 | 9 | Nrgn | 0.018 | Old |
| 0.429423 | 0.79375 | 0.036 | 0.022 | 1 | 9 | Usp39 | 0.014 | Old |
| 0.429423 | 0.537938 | 0.036 | 0.022 | 1 | 9 | Osbpl1a | 0.014 | Old |
| 0.429531 | 0.54972 | 0.214 | 0.191 | 1 | 9 | Gmfb | 0.023 | Old |
| 0.43021 | 0.406697 | 0.289 | 0.257 | 1 | 9 | Camk2d | 0.032 | Old |
| 0.430489 | 0.769734 | 0.085 | 0.066 | 1 | 9 | Mtx2 | 0.019 | Old |
| 0.430676 | 0.688625 | 0.036 | 0.022 | 1 | 9 | Ube2e2 | 0.014 | Old |
| 0.430676 | 0.628844 | 0.036 | 0.022 | 1 | 9 | Mrpl37 | 0.014 | Old |
| 0.431238 | 0.791096 | 0.062 | 0.044 | 1 | 9 | Slc29a3 | 0.018 | Old |
| 0.43137 | 0.44995 | 0.144 | 0.118 | 1 | 9 | Rnaseh2c | 0.026 | Old |
| 0.431428 | 0.637386 | 0.095 | 0.074 | 1 | 9 | Themis2 | 0.021 | Old |
| 0.431647 | 0.133294 | 0.412 | 0.368 | 1 | 9 | Myo1b | 0.044 | Old |
| 0.431828 | 0.236048 | 0.08 | 0.059 | 1 | 9 | Slc39a8 | 0.021 | Old |
| 0.431921 | 0.386242 | 0.175 | 0.147 | 1 | 9 | Api5 | 0.028 | Old |
| 0.431931 | 0.931177 | 0.036 | 0.022 | 1 | 9 | Pmm1 | 0.014 | Old |
| 0.432176 | 0.027291 | 0.062 | 0.081 | 1 | 9 | 4833403J1 | -0.019 | Old |
| 0.432195 | 0.835101 | 0.062 | 0.044 | 1 | 9 | Armcx1 | 0.018 | Old |
| 0.432286 | 0.183812 | 0.124 | 0.096 | 1 | 9 | Cmtm4 | 0.028 | Old |
| 0.432376 | 0.419517 | 0.139 | 0.11 | 1 | 9 | Angpt2 | 0.029 | Old |
| 0.432605 | 0.627992 | 0.126 | 0.103 | 1 | 9 | Ampd3 | 0.023 | Old |
| 0.432607 | 0.657703 | 0.088 | 0.066 | 1 | 9 | Snai2 | 0.022 | Old |
| 0.432676 | 0.564643 | 0.08 | 0.059 | 1 | 9 | Eml1 | 0.021 | Old |
| 0.432877 | 0.007218 | 0.165 | 0.132 | 1 | 9 | Fkbp3 | 0.033 | Old |
| 0.433154 | 0.649206 | 0.062 | 0.044 | 1 | 9 | Auh | 0.018 | Old |

| | | | | | | | | |
|---|---|---|---|---|---|---|---|---|
| 0.433208 | 0.672573 | 0.119 | 0.096 | 1 | 9 | St6galnac4 | 0.023 | Old |
| 0.433517 | 0.417813 | 0.018 | 0.029 | 1 | 9 | Fam58b | -0.011 | Old |
| 0.434114 | 0.681426 | 0.062 | 0.044 | 1 | 9 | Tmem161b | 0.018 | Old |
| 0.434448 | 0.453211 | 0.036 | 0.022 | 1 | 9 | Gpank1 | 0.014 | Old |
| 0.434774 | 0.002596 | 0.394 | 0.353 | 1 | 9 | Zbtb7a | 0.041 | Old |
| 0.434841 | 0.02205 | 0.356 | 0.316 | 1 | 9 | Msl1 | 0.04 | Old |
| 0.435075 | 0.313289 | 0.018 | 0.029 | 1 | 9 | Zfp825 | -0.011 | Old |
| 0.435075 | 0.228398 | 0.018 | 0.029 | 1 | 9 | Mblac2 | -0.011 | Old |
| 0.435248 | 0.095973 | 0.845 | 0.779 | 1 | 9 | Rps20 | 0.066 | Old |
| 0.435323 | 0.842349 | 0.119 | 0.096 | 1 | 9 | Triobp | 0.023 | Old |
| 0.435341 | 0.307198 | 0.054 | 0.037 | 1 | 9 | Ngrn | 0.017 | Old |
| 0.435475 | 0.11668 | 0.116 | 0.147 | 1 | 9 | Sec11a | -0.031 | Old |
| 0.435561 | 0.494878 | 0.121 | 0.096 | 1 | 9 | Rtp4 | 0.025 | Old |
| 0.435627 | 0.052653 | 0.214 | 0.25 | 1 | 9 | Ndfip2 | -0.036 | Old |
| 0.435707 | 0.784895 | 0.077 | 0.059 | 1 | 9 | Mical3 | 0.018 | Old |
| 0.43606 | 0.229371 | 0.291 | 0.25 | 1 | 9 | Sh3bgrl3 | 0.041 | Old |
| 0.436143 | 0.544236 | 0.103 | 0.081 | 1 | 9 | Zfp608 | 0.022 | Old |
| 0.436143 | 0.528559 | 0.103 | 0.081 | 1 | 9 | Hltf | 0.022 | Old |
| 0.436287 | 0.358945 | 0.394 | 0.368 | 1 | 9 | Elovl5 | 0.026 | Old |
| 0.436311 | 0.936247 | 0.07 | 0.051 | 1 | 9 | Cd68 | 0.019 | Old |
| 0.436405 | 0.208873 | 0.325 | 0.287 | 1 | 9 | Phf20l1 | 0.038 | Old |
| 0.436425 | 0.168013 | 0.178 | 0.147 | 1 | 9 | Stk11 | 0.031 | Old |
| 0.436673 | 0.183461 | 0.088 | 0.066 | 1 | 9 | Mms19 | 0.022 | Old |
| 0.437001 | 0.580117 | 0.062 | 0.044 | 1 | 9 | Parl | 0.018 | Old |
| 0.437408 | 0.303481 | 0.278 | 0.25 | 1 | 9 | Rras | 0.028 | Old |
| 0.437937 | 0.720511 | 0.036 | 0.051 | 1 | 9 | Gja5 | -0.015 | Old |
| 0.437966 | 0.769345 | 0.062 | 0.044 | 1 | 9 | Sh3gl1 | 0.018 | Old |
| 0.438091 | 0.39838 | 0.126 | 0.103 | 1 | 9 | Itpr3 | 0.023 | Old |
| 0.438161 | 0.198544 | 0.144 | 0.118 | 1 | 9 | Zhx1 | 0.026 | Old |
| 0.4382 | 0.461462 | 0.018 | 0.029 | 1 | 9 | 9930104L0 | -0.011 | Old |
| 0.438322 | 0.238163 | 0.054 | 0.074 | 1 | 9 | Pigs | -0.02 | Old |
| 0.438378 | 1.00952 | 0.041 | 0.059 | 1 | 9 | Memo1 | -0.018 | Old |
| 0.43934 | 0.148399 | 0.072 | 0.051 | 1 | 9 | Ints1 | 0.021 | Old |
| 0.439453 | 0.849096 | 0.041 | 0.059 | 1 | 9 | 4930578C1 | -0.018 | Old |
| 0.439507 | 0.228909 | 0.036 | 0.022 | 1 | 9 | Arrdc2 | 0.014 | Old |
| 0.439507 | 0.104283 | 0.036 | 0.022 | 1 | 9 | Zfp111 | 0.014 | Old |
| 0.439768 | 0.636756 | 0.018 | 0.029 | 1 | 9 | H2-DMb2 | -0.011 | Old |
| 0.439768 | 0.47161 | 0.018 | 0.029 | 1 | 9 | Cox15 | -0.011 | Old |
| 0.439842 | 0.149048 | 0.454 | 0.397 | 1 | 9 | Atp5e | 0.057 | Old |
| 0.439899 | 0.543583 | 0.062 | 0.044 | 1 | 9 | Hspa13 | 0.018 | Old |
| 0.440304 | 1.039496 | 0.093 | 0.074 | 1 | 9 | 0610037L1 | 0.019 | Old |
| 0.440508 | 0.278908 | 0.173 | 0.147 | 1 | 9 | Lrrc8d | 0.026 | Old |
| 0.440867 | 0.56398 | 0.062 | 0.044 | 1 | 9 | Phf11d | 0.018 | Old |
| 0.441338 | 0.078883 | 0.018 | 0.029 | 1 | 9 | 2700038G2 | -0.011 | Old |
| 0.441522 | 0.278464 | 0.247 | 0.221 | 1 | 9 | Myo18a | 0.026 | Old |

| | | | | | | | | |
|---|---|---|---|---|---|---|---|---|
| 0.441662 | 0.175544 | 0.302 | 0.272 | 1 | 9 | Tsc22d2 | 0.03 | Old |
| 0.441833 | 1.06919 | 0.144 | 0.125 | 1 | 9 | Tmtc1 | 0.019 | Old |
| 0.441935 | 0.202126 | 0.165 | 0.14 | 1 | 9 | Gpbp1l1 | 0.025 | Old |
| 0.441947 | 0.557348 | 0.291 | 0.272 | 1 | 9 | Sdha | 0.019 | Old |
| 0.442401 | 0.351769 | 0.088 | 0.066 | 1 | 9 | Chka | 0.022 | Old |
| 0.442507 | 0.728861 | 0.124 | 0.103 | 1 | 9 | Rbm28 | 0.021 | Old |
| 0.442526 | 1.234178 | 0.044 | 0.029 | 1 | 9 | Ccar2 | 0.015 | Old |
| 0.44269 | 0.325847 | 0.093 | 0.074 | 1 | 9 | Vps11 | 0.019 | Old |
| 0.442925 | 0.122381 | 0.126 | 0.103 | 1 | 9 | Skiv2l | 0.023 | Old |
| 0.442937 | 0.540738 | 0.235 | 0.213 | 1 | 9 | Vps35 | 0.022 | Old |
| 0.443223 | 0.196315 | 0.088 | 0.066 | 1 | 9 | Desi2 | 0.022 | Old |
| 0.443324 | 0.01888 | 0.036 | 0.022 | 1 | 9 | Eif2d | 0.014 | Old |
| 0.443621 | 0.41114 | 0.054 | 0.037 | 1 | 9 | Dxo | 0.017 | Old |
| 0.443792 | 2.368555 | 0.026 | 0.015 | 1 | 9 | Itpkc | 0.011 | Old |
| 0.443792 | 2.199195 | 0.026 | 0.015 | 1 | 9 | Rom1 | 0.011 | Old |
| 0.443824 | 0.799301 | 0.152 | 0.132 | 1 | 9 | Ifnar1 | 0.02 | Old |
| 0.444754 | 0.092155 | 0.062 | 0.044 | 1 | 9 | Atg9a | 0.018 | Old |
| 0.444803 | 1.339183 | 0.113 | 0.096 | 1 | 9 | Cgnl1 | 0.017 | Old |
| 0.445064 | 0.117581 | 0.175 | 0.147 | 1 | 9 | Zfp979 | 0.028 | Old |
| 0.445244 | 0.085148 | 0.454 | 0.426 | 1 | 9 | Rpl31 | 0.028 | Old |
| 0.445248 | 0.125814 | 0.067 | 0.088 | 1 | 9 | Aco1 | -0.021 | Old |
| 0.44526 | 0.281682 | 0.464 | 0.441 | 1 | 9 | Map4k4 | 0.023 | Old |
| 0.445291 | 0.093337 | 0.32 | 0.279 | 1 | 9 | Ecscr | 0.041 | Old |
| 0.445305 | 1.871954 | 0.026 | 0.015 | 1 | 9 | Dis3l | 0.011 | Old |
| 0.445553 | 0.503097 | 0.095 | 0.074 | 1 | 9 | Fam96b | 0.021 | Old |
| 0.445771 | 0.180055 | 0.32 | 0.279 | 1 | 9 | Psmb10 | 0.041 | Old |
| 0.445885 | 0.544965 | 0.201 | 0.176 | 1 | 9 | Ddx42 | 0.025 | Old |
| 0.446568 | 0.329163 | 0.235 | 0.206 | 1 | 9 | Kpna4 | 0.029 | Old |
| 0.446574 | 0.016988 | 0.178 | 0.147 | 1 | 9 | Exoc4 | 0.031 | Old |
| 0.446767 | 0.052194 | 0.281 | 0.235 | 1 | 9 | Smad7 | 0.046 | Old |
| 0.446821 | 2.288052 | 0.026 | 0.015 | 1 | 9 | 1600020E0 | 0.011 | Old |
| 0.447016 | 0.712002 | 0.041 | 0.059 | 1 | 9 | Utp23 | -0.018 | Old |
| 0.447142 | 1.49168 | 0.044 | 0.029 | 1 | 9 | Dbr1 | 0.015 | Old |
| 0.447142 | 1.412584 | 0.044 | 0.029 | 1 | 9 | Epor | 0.015 | Old |
| 0.4473 | 0.036372 | 0.049 | 0.066 | 1 | 9 | Utp15 | -0.017 | Old |
| 0.447311 | 0.207066 | 0.106 | 0.081 | 1 | 9 | Mrpl32 | 0.025 | Old |
| 0.447346 | 0.052957 | 0.088 | 0.066 | 1 | 9 | Snapc5 | 0.022 | Old |
| 0.447398 | 0.43004 | 0.147 | 0.125 | 1 | 9 | Maea | 0.022 | Old |
| 0.448212 | 0.855566 | 0.07 | 0.051 | 1 | 9 | Cpsf3 | 0.019 | Old |
| 0.448304 | 0.233129 | 0.103 | 0.081 | 1 | 9 | Itpa | 0.022 | Old |
| 0.44834 | 1.599939 | 0.026 | 0.015 | 1 | 9 | 1110034G2 | 0.011 | Old |
| 0.449243 | 0.393866 | 0.304 | 0.279 | 1 | 9 | Pdlim5 | 0.025 | Old |
| 0.449331 | 0.042418 | 0.147 | 0.118 | 1 | 9 | Vps36 | 0.029 | Old |
| 0.449842 | 0.234 | 0.451 | 0.412 | 1 | 9 | Cflar | 0.039 | Old |
| 0.449862 | 1.944928 | 0.026 | 0.015 | 1 | 9 | Entpd6 | 0.011 | Old |

| | | | | | | | | |
|---|---|---|---|---|---|---|---|---|
| 0.449862 | 1.767538 | 0.026 | 0.015 | 1 | 9 | Fam171a2 | 0.011 | Old |
| 0.450281 | 0.094417 | 0.085 | 0.11 | 1 | 9 | Foxn2 | -0.025 | Old |
| 0.45127 | 0.072445 | 0.595 | 0.544 | 1 | 9 | Dlc1 | 0.051 | Old |
| 0.451357 | 0.794654 | 0.168 | 0.147 | 1 | 9 | Ndufs7 | 0.021 | Old |
| 0.451386 | 1.74706 | 0.026 | 0.015 | 1 | 9 | Arhgef9 | 0.011 | Old |
| 0.451386 | 1.542028 | 0.026 | 0.015 | 1 | 9 | Zfp931 | 0.011 | Old |
| 0.451621 | 0.378869 | 0.24 | 0.213 | 1 | 9 | Mga | 0.027 | Old |
| 0.451766 | 0.130244 | 0.441 | 0.412 | 1 | 9 | Ndufc2 | 0.029 | Old |
| 0.451786 | 0.662418 | 0.044 | 0.029 | 1 | 9 | Urb1 | 0.015 | Old |
| 0.451838 | 0.059043 | 0.08 | 0.103 | 1 | 9 | Flot1 | -0.023 | Old |
| 0.452143 | 0.327467 | 0.103 | 0.081 | 1 | 9 | Cisd1 | 0.022 | Old |
| 0.452321 | 0.239662 | 0.088 | 0.066 | 1 | 9 | Alg10b | 0.022 | Old |
| 0.452498 | 0.430206 | 0.18 | 0.154 | 1 | 9 | Cnot4 | 0.026 | Old |
| 0.45259 | 1.532068 | 0.052 | 0.037 | 1 | 9 | Adamts4 | 0.015 | Old |
| 0.452914 | 2.416903 | 0.026 | 0.015 | 1 | 9 | Gm26870 | 0.011 | Old |
| 0.452914 | 1.489251 | 0.026 | 0.015 | 1 | 9 | Dclre1a | 0.011 | Old |
| 0.452914 | 1.401396 | 0.026 | 0.015 | 1 | 9 | Adam22 | 0.011 | Old |
| 0.452914 | 1.397355 | 0.026 | 0.015 | 1 | 9 | Fez1 | 0.011 | Old |
| 0.452951 | 1.210133 | 0.044 | 0.029 | 1 | 9 | Gfm2 | 0.015 | Old |
| 0.453663 | 1.41874 | 0.052 | 0.037 | 1 | 9 | Alkbh1 | 0.015 | Old |
| 0.453842 | 0.134107 | 0.188 | 0.162 | 1 | 9 | Trim8 | 0.026 | Old |
| 0.454024 | 0.894366 | 0.077 | 0.059 | 1 | 9 | 1600012H( | 0.018 | Old |
| 0.454024 | 0.244342 | 0.067 | 0.088 | 1 | 9 | Crnkl1 | -0.021 | Old |
| 0.454118 | 0.446298 | 0.036 | 0.051 | 1 | 9 | Pold1 | -0.015 | Old |
| 0.454118 | 0.303298 | 0.036 | 0.051 | 1 | 9 | Gpr155 | -0.015 | Old |
| 0.454349 | 0.139834 | 0.564 | 0.522 | 1 | 9 | Fcho2 | 0.042 | Old |
| 0.454445 | 1.351972 | 0.026 | 0.015 | 1 | 9 | Tbl2 | 0.011 | Old |
| 0.454445 | 1.274068 | 0.026 | 0.015 | 1 | 9 | Slfn3 | 0.011 | Old |
| 0.454445 | 1.163779 | 0.026 | 0.015 | 1 | 9 | Fntb | 0.011 | Old |
| 0.454696 | 0.530393 | 0.07 | 0.051 | 1 | 9 | Polr2j | 0.019 | Old |
| 0.455056 | 0.010279 | 0.206 | 0.176 | 1 | 9 | Cables2 | 0.03 | Old |
| 0.455162 | 0.086458 | 0.013 | 0.022 | 1 | 9 | Slc2a10 | -0.009 | Old |
| 0.455287 | 0.809023 | 0.044 | 0.029 | 1 | 9 | Ccdc66 | 0.015 | Old |
| 0.455287 | 0.483006 | 0.044 | 0.029 | 1 | 9 | Kat8 | 0.015 | Old |
| 0.455978 | 1.040208 | 0.026 | 0.015 | 1 | 9 | Lpcat4 | 0.011 | Old |
| 0.456196 | 0.326269 | 0.126 | 0.103 | 1 | 9 | Ckap5 | 0.023 | Old |
| 0.456355 | 0.858932 | 0.093 | 0.074 | 1 | 9 | Crebl2 | 0.019 | Old |
| 0.456457 | 0.387359 | 0.044 | 0.029 | 1 | 9 | Irgq | 0.015 | Old |
| 0.456457 | 0.365728 | 0.036 | 0.051 | 1 | 9 | Nup133 | -0.015 | Old |
| 0.456548 | 0.569126 | 0.126 | 0.103 | 1 | 9 | Fnip1 | 0.023 | Old |
| 0.456558 | 0.664991 | 0.07 | 0.051 | 1 | 9 | Sertad3 | 0.019 | Old |
| 0.456902 | 0.298648 | 0.131 | 0.162 | 1 | 9 | Vav3 | -0.031 | Old |
| 0.457166 | 0.938944 | 0.093 | 0.074 | 1 | 9 | Rcbtb1 | 0.019 | Old |
| 0.457486 | 0.649915 | 0.139 | 0.118 | 1 | 9 | Ggps1 | 0.021 | Old |
| 0.457514 | 0.88265 | 0.026 | 0.015 | 1 | 9 | Ptgfrn | 0.011 | Old |

| | | | | | | | | |
|---|---|---|---|---|---|---|---|---|
| 0.457618 | 0.830395 | 0.152 | 0.132 | 1 | 9 | Khsrp | 0.02 | Old |
| 0.457629 | 0.757036 | 0.044 | 0.029 | 1 | 9 | Faap100 | 0.015 | Old |
| 0.457629 | 0.461121 | 0.044 | 0.029 | 1 | 9 | Saysd1 | 0.015 | Old |
| 0.457949 | 0.007652 | 0.075 | 0.096 | 1 | 9 | Zpr1 | -0.021 | Old |
| 0.45797 | 1.607059 | 0.052 | 0.037 | 1 | 9 | Cmc4 | 0.015 | Old |
| 0.458148 | 0.005343 | 0.265 | 0.235 | 1 | 9 | Ndufs6 | 0.03 | Old |
| 0.458395 | 0.019048 | 0.546 | 0.493 | 1 | 9 | Etl4 | 0.053 | Old |
| 0.458448 | 0.75479 | 0.077 | 0.059 | 1 | 9 | Btbd9 | 0.018 | Old |
| 0.458803 | 0.710879 | 0.044 | 0.029 | 1 | 9 | Srgap1 | 0.015 | Old |
| 0.458823 | 0.163329 | 0.268 | 0.235 | 1 | 9 | Nono | 0.033 | Old |
| 0.458912 | 0.260859 | 0.013 | 0.022 | 1 | 9 | Trpm3 | -0.009 | Old |
| 0.458912 | 0.213706 | 0.013 | 0.022 | 1 | 9 | Plxna4os1 | -0.009 | Old |
| 0.459054 | 0.843211 | 0.026 | 0.015 | 1 | 9 | Ntmt1 | 0.011 | Old |
| 0.459054 | 0.600212 | 0.026 | 0.015 | 1 | 9 | Rcl1 | 0.011 | Old |
| 0.459203 | 0.081562 | 0.588 | 0.544 | 1 | 9 | Crip2 | 0.044 | Old |
| 0.459364 | 0.058308 | 0.216 | 0.184 | 1 | 9 | Rsf1 | 0.032 | Old |
| 0.459843 | 0.236424 | 0.088 | 0.066 | 1 | 9 | Atl3 | 0.022 | Old |
| 0.459978 | 0.813152 | 0.044 | 0.029 | 1 | 9 | Nudt19 | 0.015 | Old |
| 0.460596 | 0.963012 | 0.026 | 0.015 | 1 | 9 | Stambpl1 | 0.011 | Old |
| 0.460596 | 0.565611 | 0.026 | 0.015 | 1 | 9 | Ikzf1 | 0.011 | Old |
| 0.460596 | 0.549206 | 0.026 | 0.015 | 1 | 9 | Mmp14 | 0.011 | Old |
| 0.460596 | 0.361265 | 0.026 | 0.015 | 1 | 9 | Nup107 | 0.011 | Old |
| 0.460596 | 0.190439 | 0.026 | 0.015 | 1 | 9 | Gm17354 | 0.011 | Old |
| 0.460625 | 0.638726 | 0.108 | 0.088 | 1 | 9 | Hspa2 | 0.02 | Old |
| 0.461007 | 0.154021 | 0.361 | 0.331 | 1 | 9 | Lrrc8a | 0.03 | Old |
| 0.461155 | 0.774236 | 0.044 | 0.029 | 1 | 9 | Ncapd3 | 0.015 | Old |
| 0.461155 | 0.678419 | 0.044 | 0.029 | 1 | 9 | Kbtbd3 | 0.015 | Old |
| 0.461343 | 1.013528 | 0.085 | 0.066 | 1 | 9 | Polr3gl | 0.019 | Old |
| 0.46185 | 0.005887 | 0.113 | 0.14 | 1 | 9 | Enc1 | -0.027 | Old |
| 0.462141 | 0.26058 | 0.026 | 0.015 | 1 | 9 | Rad1 | 0.011 | Old |
| 0.462183 | 0.735139 | 0.214 | 0.191 | 1 | 9 | Eif4g3 | 0.023 | Old |
| 0.462211 | 0.519066 | 0.103 | 0.081 | 1 | 9 | Eci2 | 0.022 | Old |
| 0.462343 | 0.164675 | 0.649 | 0.581 | 1 | 9 | Utrn | 0.068 | Old |
| 0.462481 | 0.189786 | 0.062 | 0.044 | 1 | 9 | Rhno1 | 0.018 | Old |
| 0.46268 | 0.345826 | 0.013 | 0.022 | 1 | 9 | Aasdh | -0.009 | Old |
| 0.46268 | 0.167642 | 0.013 | 0.022 | 1 | 9 | Atp8b4 | -0.009 | Old |
| 0.46268 | 0.139074 | 0.013 | 0.022 | 1 | 9 | Celf4 | -0.009 | Old |
| 0.46268 | 0.119973 | 0.013 | 0.022 | 1 | 9 | Asxl3 | -0.009 | Old |
| 0.463313 | 0.498788 | 0.101 | 0.081 | 1 | 9 | Slc1a5 | 0.02 | Old |
| 0.463348 | 0.648181 | 0.119 | 0.096 | 1 | 9 | Klhl4 | 0.023 | Old |
| 0.463405 | 0.087656 | 0.173 | 0.147 | 1 | 9 | Nasp | 0.026 | Old |
| 0.463443 | 0.609479 | 0.17 | 0.147 | 1 | 9 | Pom121 | 0.023 | Old |
| 0.463477 | 0.140054 | 0.062 | 0.044 | 1 | 9 | Zer1 | 0.018 | Old |
| 0.463599 | 0.167991 | 0.43 | 0.39 | 1 | 9 | Mcf2l | 0.04 | Old |
| 0.463689 | 0.62379 | 0.026 | 0.015 | 1 | 9 | Armt1 | 0.011 | Old |

| | | | | | | | | |
|---|---|---|---|---|---|---|---|---|
| 0.463694 | 0.048506 | 0.309 | 0.279 | 1 | 9 | Atp6ap1 | 0.03 | Old |
| 0.463771 | 0.456822 | 0.229 | 0.206 | 1 | 9 | Hnrnph3 | 0.023 | Old |
| 0.463774 | 0.175282 | 0.17 | 0.14 | 1 | 9 | Tor1aip2 | 0.03 | Old |
| 0.464252 | 0.119384 | 0.144 | 0.118 | 1 | 9 | Dmwd | 0.026 | Old |
| 0.464472 | 1.138141 | 0.052 | 0.037 | 1 | 9 | Hoxb9 | 0.015 | Old |
| 0.46457 | 0.198591 | 0.013 | 0.022 | 1 | 9 | Tmem97 | -0.009 | Old |
| 0.464683 | 0.466826 | 0.077 | 0.059 | 1 | 9 | Map2k1 | 0.018 | Old |
| 0.464683 | 0.39498 | 0.077 | 0.059 | 1 | 9 | Ick | 0.018 | Old |
| 0.464683 | 0.261451 | 0.077 | 0.059 | 1 | 9 | Ica1 | 0.018 | Old |
| 0.46524 | 0.730911 | 0.026 | 0.015 | 1 | 9 | Ccdc137 | 0.011 | Old |
| 0.46524 | 0.697051 | 0.026 | 0.015 | 1 | 9 | Pdgfrb | 0.011 | Old |
| 0.46524 | 0.696781 | 0.026 | 0.015 | 1 | 9 | Pcnx4 | 0.011 | Old |
| 0.46524 | 0.337412 | 0.026 | 0.015 | 1 | 9 | Phka2 | 0.011 | Old |
| 0.465364 | 0.442239 | 0.24 | 0.221 | 1 | 9 | Tcf3 | 0.019 | Old |
| 0.465577 | 0.485645 | 0.077 | 0.059 | 1 | 9 | Mis12 | 0.018 | Old |
| 0.465929 | 0.024717 | 0.108 | 0.132 | 1 | 9 | Ankrd46 | -0.024 | Old |
| 0.466464 | 0.627235 | 0.013 | 0.022 | 1 | 9 | Aldh1a2 | -0.009 | Old |
| 0.466464 | 0.558152 | 0.013 | 0.022 | 1 | 9 | Gm38251 | -0.009 | Old |
| 0.466794 | 0.531197 | 0.026 | 0.015 | 1 | 9 | Hemk1 | 0.011 | Old |
| 0.466794 | 0.256444 | 0.026 | 0.015 | 1 | 9 | Rnaset2a | 0.011 | Old |
| 0.466794 | 0.231616 | 0.026 | 0.015 | 1 | 9 | Tmem98 | 0.011 | Old |
| 0.466794 | 0.167197 | 0.026 | 0.015 | 1 | 9 | Lzts1 | 0.011 | Old |
| 0.466794 | 0.095655 | 0.026 | 0.015 | 1 | 9 | Cdc25b | 0.011 | Old |
| 0.467369 | 0.258764 | 0.077 | 0.059 | 1 | 9 | Med31 | 0.018 | Old |
| 0.467428 | 1.294421 | 0.059 | 0.044 | 1 | 9 | Tmem39a | 0.015 | Old |
| 0.467743 | 0.12897 | 0.052 | 0.037 | 1 | 9 | Ddx41 | 0.015 | Old |
| 0.467779 | 0.272361 | 0.456 | 0.434 | 1 | 9 | Rrbp1 | 0.022 | Old |
| 0.468252 | 0.098992 | 0.044 | 0.029 | 1 | 9 | Dhx35 | 0.015 | Old |
| 0.46835 | 0.562906 | 0.026 | 0.015 | 1 | 9 | Plcl2 | 0.011 | Old |
| 0.46835 | 0.233011 | 0.026 | 0.015 | 1 | 9 | Eda2r | 0.011 | Old |
| 0.468455 | 0.192304 | 0.631 | 0.581 | 1 | 9 | Rpl3 | 0.05 | Old |
| 0.468767 | 0.646272 | 0.07 | 0.051 | 1 | 9 | Chchd7 | 0.019 | Old |
| 0.468836 | 0.900321 | 0.052 | 0.037 | 1 | 9 | Ccdc22 | 0.015 | Old |
| 0.468842 | 0.816781 | 0.101 | 0.081 | 1 | 9 | Ppp1r13l | 0.02 | Old |
| 0.468892 | 0.084648 | 0.134 | 0.11 | 1 | 9 | Abcc5 | 0.024 | Old |
| 0.469626 | 0.213943 | 0.232 | 0.206 | 1 | 9 | Psma6 | 0.026 | Old |
| 0.469772 | 0.450237 | 0.108 | 0.088 | 1 | 9 | 10-Sep | 0.02 | Old |
| 0.470006 | 0.357132 | 0.149 | 0.125 | 1 | 9 | Cant1 | 0.024 | Old |
| 0.470632 | 0.255874 | 0.044 | 0.029 | 1 | 9 | Nmrk1 | 0.015 | Old |
| 0.470675 | 0.200128 | 0.121 | 0.096 | 1 | 9 | Tanc2 | 0.025 | Old |
| 0.470826 | 0.130316 | 0.008 | 0.015 | 1 | 9 | Paqr7 | -0.007 | Old |
| 0.471572 | 0.444031 | 0.17 | 0.147 | 1 | 9 | 8-Sep | 0.023 | Old |
| 0.471594 | 0.203015 | 0.253 | 0.228 | 1 | 9 | Akt1 | 0.025 | Old |
| 0.471609 | 0.697842 | 0.085 | 0.066 | 1 | 9 | BC029722 | 0.019 | Old |
| 0.471824 | 0.222735 | 0.044 | 0.029 | 1 | 9 | Rps19bp1 | 0.015 | Old |

| | | | | | | | | |
|---|---|---|---|---|---|---|---|---|
| 0.471999 | 0.076459 | 0.399 | 0.353 | 1 | 9 | Nktr | 0.046 | Old |
| 0.472644 | 0.051883 | 0.175 | 0.147 | 1 | 9 | Actr1a | 0.028 | Old |
| 0.472729 | 0.956359 | 0.093 | 0.074 | 1 | 9 | Col28a1 | 0.019 | Old |
| 0.472824 | 0.079895 | 0.363 | 0.338 | 1 | 9 | Aes | 0.025 | Old |
| 0.473331 | 0.105546 | 0.247 | 0.287 | 1 | 9 | Srpr | -0.04 | Old |
| 0.473508 | 0.145897 | 0.284 | 0.243 | 1 | 9 | Kmt2c | 0.041 | Old |
| 0.47351 | 0.434865 | 0.07 | 0.051 | 1 | 9 | Rbl1 | 0.019 | Old |
| 0.473652 | 2.596711 | 0.015 | 0.007 | 1 | 9 | Gm14326 | 0.008 | Old |
| 0.473652 | 2.423878 | 0.015 | 0.007 | 1 | 9 | Thsd7b | 0.008 | Old |
| 0.473652 | 2.134802 | 0.015 | 0.007 | 1 | 9 | Farp2 | 0.008 | Old |
| 0.473652 | 2.133493 | 0.015 | 0.007 | 1 | 9 | Sap18b | 0.008 | Old |
| 0.473652 | 1.981987 | 0.015 | 0.007 | 1 | 9 | Dhcr24 | 0.008 | Old |
| 0.473652 | 1.863205 | 0.015 | 0.007 | 1 | 9 | Gm4673 | 0.008 | Old |
| 0.473652 | 1.829343 | 0.015 | 0.007 | 1 | 9 | Bbs5 | 0.008 | Old |
| 0.473652 | 1.712574 | 0.015 | 0.007 | 1 | 9 | Ecsit | 0.008 | Old |
| 0.473652 | 1.706847 | 0.015 | 0.007 | 1 | 9 | Gm11696 | 0.008 | Old |
| 0.473652 | 1.573435 | 0.015 | 0.007 | 1 | 9 | Mapkapk3 | 0.008 | Old |
| 0.473652 | 1.492015 | 0.015 | 0.007 | 1 | 9 | Timmdc1 | 0.008 | Old |
| 0.473652 | 1.426684 | 0.015 | 0.007 | 1 | 9 | Enpep | 0.008 | Old |
| 0.473652 | 1.375797 | 0.015 | 0.007 | 1 | 9 | Nradd | 0.008 | Old |
| 0.473652 | 1.292827 | 0.015 | 0.007 | 1 | 9 | Fam107a | 0.008 | Old |
| 0.473659 | 0.776109 | 0.075 | 0.059 | 1 | 9 | Utp14a | 0.016 | Old |
| 0.47367 | 0.587887 | 0.077 | 0.059 | 1 | 9 | Ptprk | 0.018 | Old |
| 0.474213 | 0.089991 | 0.044 | 0.029 | 1 | 9 | Gtf3c4 | 0.015 | Old |
| 0.474462 | 0.181977 | 0.07 | 0.051 | 1 | 9 | Rinl | 0.019 | Old |
| 0.474612 | 0.243855 | 0.268 | 0.243 | 1 | 9 | Mapk1ip1l | 0.025 | Old |
| 0.474965 | 0.042498 | 0.031 | 0.044 | 1 | 9 | Cdc25a | -0.013 | Old |
| 0.475308 | 0.413899 | 0.18 | 0.154 | 1 | 9 | Hecw2 | 0.026 | Old |
| 0.475311 | 0.316176 | 0.054 | 0.074 | 1 | 9 | Phb | -0.02 | Old |
| 0.475479 | 0.521658 | 0.077 | 0.059 | 1 | 9 | Rnf41 | 0.018 | Old |
| 0.475649 | 0.40249 | 0.008 | 0.015 | 1 | 9 | 1700037C1 | -0.007 | Old |
| 0.475649 | 0.254644 | 0.008 | 0.015 | 1 | 9 | Impa2 | -0.007 | Old |
| 0.475697 | 2.277976 | 0.015 | 0.007 | 1 | 9 | Ncapg2 | 0.008 | Old |
| 0.475697 | 2.231495 | 0.015 | 0.007 | 1 | 9 | Sdcbp2 | 0.008 | Old |
| 0.475697 | 2.184598 | 0.015 | 0.007 | 1 | 9 | Ptpdc1 | 0.008 | Old |
| 0.475697 | 2.090049 | 0.015 | 0.007 | 1 | 9 | Ccdc148 | 0.008 | Old |
| 0.475697 | 1.870507 | 0.015 | 0.007 | 1 | 9 | Zfp647 | 0.008 | Old |
| 0.475697 | 1.869548 | 0.015 | 0.007 | 1 | 9 | Lrfn3 | 0.008 | Old |
| 0.475697 | 1.769781 | 0.015 | 0.007 | 1 | 9 | Ccr1 | 0.008 | Old |
| 0.475697 | 1.4856 | 0.015 | 0.007 | 1 | 9 | Ss18l1 | 0.008 | Old |
| 0.475697 | 1.478099 | 0.015 | 0.007 | 1 | 9 | Gm42949 | 0.008 | Old |
| 0.475697 | 1.407356 | 0.015 | 0.007 | 1 | 9 | B3gnt5 | 0.008 | Old |
| 0.475697 | 1.398436 | 0.015 | 0.007 | 1 | 9 | Katnb1 | 0.008 | Old |
| 0.475697 | 1.268773 | 0.015 | 0.007 | 1 | 9 | Dynlt1a | 0.008 | Old |
| 0.47646 | 0.122402 | 0.111 | 0.088 | 1 | 9 | Inafm1 | 0.023 | Old |

| | | | | | | | | |
|---|---|---|---|---|---|---|---|---|
| 0.476526 | 0.888466 | 0.052 | 0.037 | 1 | 9 | Hinfp | 0.015 | Old |
| 0.476702 | 0.176732 | 0.211 | 0.184 | 1 | 9 | Azi2 | 0.027 | Old |
| 0.476873 | 0.571736 | 0.093 | 0.074 | 1 | 9 | Zdhhc8 | 0.019 | Old |
| 0.477547 | 0.438723 | 0.031 | 0.044 | 1 | 9 | Sass6 | -0.013 | Old |
| 0.477694 | 0.154407 | 0.216 | 0.257 | 1 | 9 | Nadk | -0.041 | Old |
| 0.477747 | 1.868755 | 0.015 | 0.007 | 1 | 9 | Htra3 | 0.008 | Old |
| 0.477747 | 1.793085 | 0.015 | 0.007 | 1 | 9 | H2afy2 | 0.008 | Old |
| 0.477747 | 1.574243 | 0.015 | 0.007 | 1 | 9 | Piga | 0.008 | Old |
| 0.477747 | 1.473457 | 0.015 | 0.007 | 1 | 9 | Zfand1 | 0.008 | Old |
| 0.477747 | 1.447321 | 0.015 | 0.007 | 1 | 9 | Mtrr | 0.008 | Old |
| 0.477747 | 1.322178 | 0.015 | 0.007 | 1 | 9 | Gm16486 | 0.008 | Old |
| 0.477747 | 1.14682 | 0.015 | 0.007 | 1 | 9 | Synpo2 | 0.008 | Old |
| 0.477747 | 1.057589 | 0.015 | 0.007 | 1 | 9 | Ccdc149 | 0.008 | Old |
| 0.477747 | 0.963115 | 0.015 | 0.007 | 1 | 9 | 4930524J0 | 0.008 | Old |
| 0.477747 | 0.92696 | 0.015 | 0.007 | 1 | 9 | Gm29994 | 0.008 | Old |
| 0.47807 | 0.849037 | 0.008 | 0.015 | 1 | 9 | Siglecg | -0.007 | Old |
| 0.47807 | 0.514514 | 0.008 | 0.015 | 1 | 9 | Serpinf1 | -0.007 | Old |
| 0.47807 | 0.486839 | 0.008 | 0.015 | 1 | 9 | Setd6 | -0.007 | Old |
| 0.47807 | 0.216962 | 0.008 | 0.015 | 1 | 9 | Mthfd1l | -0.007 | Old |
| 0.47807 | 0.182091 | 0.008 | 0.015 | 1 | 9 | Guca1a | -0.007 | Old |
| 0.47807 | 0.123062 | 0.008 | 0.015 | 1 | 9 | Pstpip1 | -0.007 | Old |
| 0.47807 | 0.121929 | 0.008 | 0.015 | 1 | 9 | Cnr2 | -0.007 | Old |
| 0.47807 | 0.041607 | 0.008 | 0.015 | 1 | 9 | Eefsec | -0.007 | Old |
| 0.47807 | 0.012088 | 0.008 | 0.015 | 1 | 9 | Gm45353 | -0.007 | Old |
| 0.478199 | 0.771428 | 0.067 | 0.088 | 1 | 9 | Bmp2k | -0.021 | Old |
| 0.478453 | 0.166921 | 0.095 | 0.118 | 1 | 9 | Rrm1 | -0.023 | Old |
| 0.478736 | 0.234655 | 0.242 | 0.213 | 1 | 9 | Dnajc8 | 0.029 | Old |
| 0.478846 | 0.189429 | 0.101 | 0.125 | 1 | 9 | Coa5 | -0.024 | Old |
| 0.478865 | 0.396609 | 0.204 | 0.176 | 1 | 9 | Nap1l4 | 0.028 | Old |
| 0.478865 | 0.165857 | 0.204 | 0.176 | 1 | 9 | Rer1 | 0.028 | Old |
| 0.479108 | 0.757837 | 0.077 | 0.059 | 1 | 9 | B230354K1 | 0.018 | Old |
| 0.479802 | 2.084098 | 0.015 | 0.007 | 1 | 9 | Trmo | 0.008 | Old |
| 0.479802 | 1.69411 | 0.015 | 0.007 | 1 | 9 | Plin4 | 0.008 | Old |
| 0.479802 | 1.420267 | 0.015 | 0.007 | 1 | 9 | A930004D1 | 0.008 | Old |
| 0.479802 | 0.988623 | 0.015 | 0.007 | 1 | 9 | Lss | 0.008 | Old |
| 0.479802 | 0.872802 | 0.015 | 0.007 | 1 | 9 | Arhgef17 | 0.008 | Old |
| 0.479802 | 0.826489 | 0.015 | 0.007 | 1 | 9 | Zfp628 | 0.008 | Old |
| 0.479802 | 0.797342 | 0.015 | 0.007 | 1 | 9 | Rita1 | 0.008 | Old |
| 0.480248 | 0.537595 | 0.152 | 0.132 | 1 | 9 | Leprotl1 | 0.02 | Old |
| 0.480419 | 0.107921 | 0.528 | 0.485 | 1 | 9 | Tgoln1 | 0.043 | Old |
| 0.480498 | 2.044921 | 0.008 | 0.015 | 1 | 9 | Rfc5 | -0.007 | Old |
| 0.480498 | 1.955463 | 0.008 | 0.015 | 1 | 9 | Setmar | -0.007 | Old |
| 0.480498 | 1.607787 | 0.008 | 0.015 | 1 | 9 | Gm43672 | -0.007 | Old |
| 0.480498 | 1.278752 | 0.008 | 0.015 | 1 | 9 | Ticam2 | -0.007 | Old |
| 0.480498 | 0.69968 | 0.008 | 0.015 | 1 | 9 | Fgd3 | -0.007 | Old |

| | | | | | | | | |
|---|---|---|---|---|---|---|---|---|
| 0.480498 | 0.662256 | 0.008 | 0.015 | 1 | 9 | Zfp27 | -0.007 | Old |
| 0.480498 | 0.584404 | 0.008 | 0.015 | 1 | 9 | Shq1 | -0.007 | Old |
| 0.480498 | 0.554602 | 0.008 | 0.015 | 1 | 9 | Gapt | -0.007 | Old |
| 0.480498 | 0.538878 | 0.008 | 0.015 | 1 | 9 | Tstd1 | -0.007 | Old |
| 0.480498 | 0.479245 | 0.008 | 0.015 | 1 | 9 | Zbtb49 | -0.007 | Old |
| 0.480498 | 0.456729 | 0.008 | 0.015 | 1 | 9 | Ccdc94 | -0.007 | Old |
| 0.480498 | 0.432428 | 0.008 | 0.015 | 1 | 9 | 2410080I0 | -0.007 | Old |
| 0.480498 | 0.352656 | 0.008 | 0.015 | 1 | 9 | Rab40b | -0.007 | Old |
| 0.480498 | 0.326016 | 0.008 | 0.015 | 1 | 9 | Zfp850 | -0.007 | Old |
| 0.480498 | 0.191164 | 0.008 | 0.015 | 1 | 9 | Fcgr4 | -0.007 | Old |
| 0.480498 | 0.020649 | 0.008 | 0.015 | 1 | 9 | Pqlc3 | -0.007 | Old |
| 0.480564 | 0.161085 | 0.387 | 0.346 | 1 | 9 | Eif5 | 0.041 | Old |
| 0.480939 | 0.341203 | 0.242 | 0.213 | 1 | 9 | Sntb2 | 0.029 | Old |
| 0.480952 | 0.845457 | 0.052 | 0.037 | 1 | 9 | Zfp994 | 0.015 | Old |
| 0.480952 | 0.666208 | 0.052 | 0.037 | 1 | 9 | Ap3s2 | 0.015 | Old |
| 0.481129 | 0.338504 | 0.085 | 0.066 | 1 | 9 | Srd5a3 | 0.019 | Old |
| 0.481314 | 0.022072 | 0.479 | 0.419 | 1 | 9 | Rasgrp3 | 0.06 | Old |
| 0.481862 | 1.788957 | 0.015 | 0.007 | 1 | 9 | Nphp1 | 0.008 | Old |
| 0.481862 | 1.442514 | 0.015 | 0.007 | 1 | 9 | Selenbp1 | 0.008 | Old |
| 0.481862 | 0.997033 | 0.015 | 0.007 | 1 | 9 | Frat2 | 0.008 | Old |
| 0.481862 | 0.963285 | 0.015 | 0.007 | 1 | 9 | Ift43 | 0.008 | Old |
| 0.481862 | 0.837512 | 0.015 | 0.007 | 1 | 9 | Stac | 0.008 | Old |
| 0.481862 | 0.786978 | 0.015 | 0.007 | 1 | 9 | B230369F2 | 0.008 | Old |
| 0.481862 | 0.762776 | 0.015 | 0.007 | 1 | 9 | Ramp3 | 0.008 | Old |
| 0.481862 | 0.741758 | 0.015 | 0.007 | 1 | 9 | Porcn | 0.008 | Old |
| 0.481862 | 0.725202 | 0.015 | 0.007 | 1 | 9 | Tgfa | 0.008 | Old |
| 0.481862 | 0.721722 | 0.015 | 0.007 | 1 | 9 | Alkbh2 | 0.008 | Old |
| 0.481862 | 0.516198 | 0.015 | 0.007 | 1 | 9 | Pag1 | 0.008 | Old |
| 0.481862 | 0.457593 | 0.015 | 0.007 | 1 | 9 | Cyb5d1 | 0.008 | Old |
| 0.481862 | 0.457046 | 0.015 | 0.007 | 1 | 9 | Nthl1 | 0.008 | Old |
| 0.482266 | 0.142623 | 0.222 | 0.199 | 1 | 9 | Ccnt2 | 0.023 | Old |
| 0.482421 | 0.418625 | 0.101 | 0.081 | 1 | 9 | Pex19 | 0.02 | Old |
| 0.482554 | 1.528958 | 0.034 | 0.022 | 1 | 9 | Mrps35 | 0.012 | Old |
| 0.483099 | 0.280486 | 0.387 | 0.375 | 1 | 9 | Nes | 0.012 | Old |
| 0.483544 | 0.590263 | 0.093 | 0.074 | 1 | 9 | Spast | 0.019 | Old |
| 0.483747 | 0.278217 | 0.273 | 0.243 | 1 | 9 | Trove2 | 0.03 | Old |
| 0.483757 | 0.623948 | 0.18 | 0.162 | 1 | 9 | Card10 | 0.018 | Old |
| 0.483927 | 0.903419 | 0.015 | 0.007 | 1 | 9 | 1700030K0 | 0.008 | Old |
| 0.483927 | 0.787195 | 0.015 | 0.007 | 1 | 9 | Tmem138 | 0.008 | Old |
| 0.483927 | 0.758618 | 0.015 | 0.007 | 1 | 9 | Gstcd | 0.008 | Old |
| 0.483927 | 0.622415 | 0.015 | 0.007 | 1 | 9 | Gm29666 | 0.008 | Old |
| 0.483927 | 0.588203 | 0.015 | 0.007 | 1 | 9 | Tdp1 | 0.008 | Old |
| 0.483927 | 0.541684 | 0.015 | 0.007 | 1 | 9 | Tamm41 | 0.008 | Old |
| 0.483927 | 0.391987 | 0.015 | 0.007 | 1 | 9 | Gm3468 | 0.008 | Old |
| 0.483927 | 0.357419 | 0.015 | 0.007 | 1 | 9 | Dars2 | 0.008 | Old |

| | | | | | | | | |
|---|---|---|---|---|---|---|---|---|
| 0.483927 | 0.122889 | 0.015 | 0.007 | 1 | 9 | Ano1 | 0.008 | Old |
| 0.484295 | 0.980099 | 0.178 | 0.162 | 1 | 9 | Mcfd2 | 0.016 | Old |
| 0.48458 | 0.306548 | 0.077 | 0.059 | 1 | 9 | Zmat1 | 0.018 | Old |
| 0.484745 | 0.159764 | 0.353 | 0.324 | 1 | 9 | Shisa5 | 0.029 | Old |
| 0.484853 | 0.107446 | 0.113 | 0.14 | 1 | 9 | Mrpl28 | -0.027 | Old |
| 0.484906 | 1.38255 | 0.059 | 0.044 | 1 | 9 | Nfkb2 | 0.015 | Old |
| 0.484906 | 0.615436 | 0.059 | 0.044 | 1 | 9 | Il15 | 0.015 | Old |
| 0.485299 | 0.284482 | 0.147 | 0.125 | 1 | 9 | Atraid | 0.022 | Old |
| 0.485312 | 1.731681 | 0.034 | 0.022 | 1 | 9 | Dzip3 | 0.012 | Old |
| 0.485399 | 0.426602 | 0.052 | 0.037 | 1 | 9 | Ercc4 | 0.015 | Old |
| 0.485888 | 0.047247 | 0.351 | 0.309 | 1 | 9 | 4930523C0 | 0.042 | Old |
| 0.485945 | 0.852293 | 0.059 | 0.044 | 1 | 9 | Ubash3b | 0.015 | Old |
| 0.485996 | 0.120698 | 0.015 | 0.007 | 1 | 9 | Adgrg3 | 0.008 | Old |
| 0.486257 | 0.042128 | 0.044 | 0.029 | 1 | 9 | Zfp275 | 0.015 | Old |
| 0.486584 | 0.523694 | 0.049 | 0.066 | 1 | 9 | Stard5 | -0.017 | Old |
| 0.486638 | 0.185859 | 0.265 | 0.235 | 1 | 9 | Creb1 | 0.03 | Old |
| 0.486647 | 0.120699 | 0.031 | 0.044 | 1 | 9 | Gfod2 | -0.013 | Old |
| 0.486694 | 1.762489 | 0.034 | 0.022 | 1 | 9 | Zfp982 | 0.012 | Old |
| 0.486704 | 0.224628 | 0.356 | 0.331 | 1 | 9 | Numa1 | 0.025 | Old |
| 0.487183 | 0.123373 | 0.211 | 0.184 | 1 | 9 | Cdc42ep2 | 0.027 | Old |
| 0.487308 | 0.060111 | 0.108 | 0.132 | 1 | 9 | Tiprl | -0.024 | Old |
| 0.487654 | 0.449723 | 0.124 | 0.103 | 1 | 9 | Lsm8 | 0.021 | Old |
| 0.48772 | 0.123243 | 0.513 | 0.507 | 1 | 9 | Pbrm1 | 0.006 | Old |
| 0.488153 | 1.05836 | 0.082 | 0.066 | 1 | 9 | Xpo4 | 0.016 | Old |
| 0.488153 | 0.730177 | 0.082 | 0.066 | 1 | 9 | Zfp704 | 0.016 | Old |
| 0.488364 | 0.641068 | 0.108 | 0.088 | 1 | 9 | Arih2 | 0.02 | Old |
| 0.489404 | 0.938037 | 0.098 | 0.081 | 1 | 9 | Grk6 | 0.017 | Old |
| 0.489423 | 0.489006 | 0.093 | 0.074 | 1 | 9 | Psmc2 | 0.019 | Old |
| 0.489465 | 1.237699 | 0.034 | 0.022 | 1 | 9 | Ccdc80 | 0.012 | Old |
| 0.489869 | 0.412543 | 0.052 | 0.037 | 1 | 9 | E4f1 | 0.015 | Old |
| 0.489948 | 0.057516 | 0.479 | 0.441 | 1 | 9 | Hnrnpm | 0.038 | Old |
| 0.490576 | 0.315622 | 0.031 | 0.044 | 1 | 9 | Hlcs | -0.013 | Old |
| 0.490727 | 0.042481 | 0.719 | 0.691 | 1 | 9 | Ifitm3 | 0.028 | Old |
| 0.49099 | 0.612806 | 0.052 | 0.037 | 1 | 9 | Cbr1 | 0.015 | Old |
| 0.491104 | 0.786024 | 0.067 | 0.051 | 1 | 9 | Polr3k | 0.016 | Old |
| 0.491104 | 1.18217 | 0.067 | 0.051 | 1 | 9 | Slc25a20 | 0.016 | Old |
| 0.491252 | 0.123438 | 0.392 | 0.368 | 1 | 9 | Cd34 | 0.024 | Old |
| 0.49132 | 0.529424 | 0.101 | 0.081 | 1 | 9 | Pygo2 | 0.02 | Old |
| 0.491494 | 0.264231 | 0.317 | 0.294 | 1 | 9 | Tmem204 | 0.023 | Old |
| 0.491718 | 0.381026 | 0.273 | 0.243 | 1 | 9 | U2af2 | 0.03 | Old |
| 0.491745 | 0.303843 | 0.454 | 0.426 | 1 | 9 | Anxa3 | 0.028 | Old |
| 0.492043 | 0.157787 | 0.201 | 0.176 | 1 | 9 | Psd3 | 0.025 | Old |
| 0.492182 | 0.414982 | 0.188 | 0.169 | 1 | 9 | Cnn2 | 0.019 | Old |
| 0.492204 | 1.004769 | 0.059 | 0.044 | 1 | 9 | Sec22a | 0.015 | Old |
| 0.492244 | 1.465969 | 0.034 | 0.022 | 1 | 9 | Rbm43 | 0.012 | Old |

| | | | | | | | | |
|---|---|---|---|---|---|---|---|---|
| 0.492542 | 0.116894 | 0.07 | 0.088 | 1 | 9 | Krt222 | -0.018 | Old |
| 0.492947 | 0.365276 | 0.101 | 0.081 | 1 | 9 | Otud5 | 0.02 | Old |
| 0.493507 | 0.552179 | 0.098 | 0.081 | 1 | 9 | Ncoa1 | 0.017 | Old |
| 0.493637 | 1.073847 | 0.034 | 0.022 | 1 | 9 | Plau | 0.012 | Old |
| 0.493686 | 0.518195 | 0.129 | 0.11 | 1 | 9 | Bola2 | 0.019 | Old |
| 0.493838 | 0.003019 | 0.168 | 0.14 | 1 | 9 | Rab5b | 0.028 | Old |
| 0.4943 | 0.236894 | 0.059 | 0.044 | 1 | 9 | Vps37b | 0.015 | Old |
| 0.494361 | 0.374083 | 0.052 | 0.037 | 1 | 9 | Mrps34 | 0.015 | Old |
| 0.494578 | 0.091433 | 0.101 | 0.081 | 1 | 9 | Maml1 | 0.02 | Old |
| 0.495032 | 1.598907 | 0.034 | 0.022 | 1 | 9 | Cptp | 0.012 | Old |
| 0.495032 | 1.177004 | 0.034 | 0.022 | 1 | 9 | Syne3 | 0.012 | Old |
| 0.495048 | 0.8026 | 0.067 | 0.051 | 1 | 9 | Terf2ip | 0.016 | Old |
| 0.495353 | 0.261802 | 0.247 | 0.221 | 1 | 9 | Bsdc1 | 0.026 | Old |
| 0.495517 | 0.108491 | 0.564 | 0.529 | 1 | 9 | Tpm4 | 0.035 | Old |
| 0.495596 | 0.507747 | 0.335 | 0.324 | 1 | 9 | Phip | 0.011 | Old |
| 0.495779 | 0.444299 | 0.17 | 0.147 | 1 | 9 | Rnf220 | 0.023 | Old |
| 0.49584 | 0.141335 | 0.031 | 0.044 | 1 | 9 | Ear2 | -0.013 | Old |
| 0.496401 | 0.481304 | 0.059 | 0.044 | 1 | 9 | Gm12216 | 0.015 | Old |
| 0.496429 | 1.472465 | 0.034 | 0.022 | 1 | 9 | Depdc5 | 0.012 | Old |
| 0.496429 | 1.245986 | 0.034 | 0.022 | 1 | 9 | Hes6 | 0.012 | Old |
| 0.496985 | 0.002048 | 0.149 | 0.125 | 1 | 9 | Ranbp1 | 0.024 | Old |
| 0.497168 | 0.104831 | 0.25 | 0.221 | 1 | 9 | Stub1 | 0.029 | Old |
| 0.497195 | 0.056596 | 0.134 | 0.11 | 1 | 9 | Fam219a | 0.024 | Old |
| 0.497321 | 0.026724 | 0.126 | 0.103 | 1 | 9 | Khnyn | 0.023 | Old |
| 0.497603 | 0.017645 | 0.946 | 0.963 | 1 | 9 | Sptbn1 | -0.017 | Old |
| 0.497828 | 1.143727 | 0.034 | 0.022 | 1 | 9 | Usp28 | 0.012 | Old |
| 0.498018 | 0.799534 | 0.067 | 0.051 | 1 | 9 | Inip | 0.016 | Old |
| 0.49901 | 0.897934 | 0.067 | 0.051 | 1 | 9 | Pld2 | 0.016 | Old |
| 0.499229 | 1.0631 | 0.034 | 0.022 | 1 | 9 | Cenpv | 0.012 | Old |
| 0.500007 | 0.537988 | 0.052 | 0.037 | 1 | 9 | Zfp532 | 0.015 | Old |
| 0.501673 | 0.428311 | 0.059 | 0.044 | 1 | 9 | 2610002M | 0.015 | Old |
| 0.50179 | 0.01832 | 0.149 | 0.125 | 1 | 9 | C330007P0 | 0.024 | Old |
| 0.501873 | 0.118236 | 0.428 | 0.375 | 1 | 9 | Plxna2 | 0.053 | Old |
| 0.501988 | 0.228816 | 0.131 | 0.11 | 1 | 9 | Dnm3 | 0.021 | Old |
| 0.502038 | 1.149163 | 0.034 | 0.022 | 1 | 9 | Sytl4 | 0.012 | Old |
| 0.502038 | 0.770593 | 0.034 | 0.022 | 1 | 9 | Nmnat1 | 0.012 | Old |
| 0.50215 | 0.126782 | 0.224 | 0.199 | 1 | 9 | Gnai3 | 0.025 | Old |
| 0.503097 | 0.292186 | 0.237 | 0.221 | 1 | 9 | Surf4 | 0.016 | Old |
| 0.503446 | 1.390325 | 0.034 | 0.022 | 1 | 9 | Cdk10 | 0.012 | Old |
| 0.503446 | 1.104486 | 0.034 | 0.022 | 1 | 9 | Ap1g2 | 0.012 | Old |
| 0.503446 | 0.656233 | 0.034 | 0.022 | 1 | 9 | Apba3 | 0.012 | Old |
| 0.503446 | 0.61621 | 0.034 | 0.022 | 1 | 9 | 0610009B2 | 0.012 | Old |
| 0.503857 | 0.160562 | 0.149 | 0.125 | 1 | 9 | Ppp2r2d | 0.024 | Old |
| 0.504076 | 0.734549 | 0.137 | 0.118 | 1 | 9 | Gemin7 | 0.019 | Old |
| 0.50442 | 0.601385 | 0.101 | 0.081 | 1 | 9 | Prkab1 | 0.02 | Old |

| | | | | | | | | |
|---|---|---|---|---|---|---|---|---|
| 0.504779 | 0.071145 | 0.155 | 0.132 | 1 | 9 | Tox4 | 0.023 | Old |
| 0.504851 | 0.771343 | 0.059 | 0.044 | 1 | 9 | Adgra3 | 0.015 | Old |
| 0.504855 | 0.477648 | 0.034 | 0.022 | 1 | 9 | Lrch2 | 0.012 | Old |
| 0.504983 | 0.602799 | 0.067 | 0.051 | 1 | 9 | Otud1 | 0.016 | Old |
| 0.505286 | 0.256 | 0.235 | 0.213 | 1 | 9 | Ep400 | 0.022 | Old |
| 0.505641 | 0.077104 | 0.209 | 0.243 | 1 | 9 | Pkm | -0.034 | Old |
| 0.505686 | 0.011159 | 0.052 | 0.037 | 1 | 9 | Asl | 0.015 | Old |
| 0.50586 | 0.566292 | 0.124 | 0.103 | 1 | 9 | Kmt5a | 0.021 | Old |
| 0.506267 | 0.642425 | 0.034 | 0.022 | 1 | 9 | Alg9 | 0.012 | Old |
| 0.506267 | 0.568598 | 0.034 | 0.022 | 1 | 9 | Inafm2 | 0.012 | Old |
| 0.506896 | 0.234395 | 0.101 | 0.081 | 1 | 9 | Lyar | 0.02 | Old |
| 0.506967 | 0.773679 | 0.082 | 0.066 | 1 | 9 | Lipo3 | 0.016 | Old |
| 0.50768 | 0.420186 | 0.034 | 0.022 | 1 | 9 | Mcm9 | 0.012 | Old |
| 0.507892 | 0.23315 | 0.191 | 0.169 | 1 | 9 | Pam | 0.022 | Old |
| 0.508428 | 0.950587 | 0.098 | 0.081 | 1 | 9 | Gtf2f1 | 0.017 | Old |
| 0.508512 | 0.205758 | 0.085 | 0.066 | 1 | 9 | Abcd4 | 0.019 | Old |
| 0.509006 | 0.712342 | 0.149 | 0.132 | 1 | 9 | Scamp3 | 0.017 | Old |
| 0.509111 | 0.378224 | 0.044 | 0.059 | 1 | 9 | Tbc1d22b | -0.015 | Old |
| 0.509122 | 0.085166 | 0.407 | 0.375 | 1 | 9 | Dab2ip | 0.032 | Old |
| 0.509132 | 0.032363 | 0.624 | 0.581 | 1 | 9 | Hnrnpf | 0.043 | Old |
| 0.509786 | 0.416375 | 0.129 | 0.11 | 1 | 9 | Zc3h14 | 0.019 | Old |
| 0.510171 | 0.679913 | 0.059 | 0.044 | 1 | 9 | Cwc22 | 0.015 | Old |
| 0.510431 | 0.474435 | 0.229 | 0.206 | 1 | 9 | Ppp1r12c | 0.023 | Old |
| 0.510735 | 0.137608 | 0.196 | 0.169 | 1 | 9 | Ccdc88a | 0.027 | Old |
| 0.510815 | 0.065058 | 0.673 | 0.618 | 1 | 9 | Crim1 | 0.055 | Old |
| 0.510994 | 0.507312 | 0.067 | 0.051 | 1 | 9 | Cenpc1 | 0.016 | Old |
| 0.51152 | 0.552291 | 0.121 | 0.103 | 1 | 9 | Rcor3 | 0.018 | Old |
| 0.511573 | 0.342902 | 0.219 | 0.191 | 1 | 9 | Synj2bp | 0.028 | Old |
| 0.511934 | 0.323005 | 0.034 | 0.022 | 1 | 9 | Ints2 | 0.012 | Old |
| 0.511934 | 0.022607 | 0.034 | 0.022 | 1 | 9 | Eva1c | 0.012 | Old |
| 0.512085 | 0.259703 | 0.232 | 0.206 | 1 | 9 | Plscr3 | 0.026 | Old |
| 0.512148 | 0.029302 | 0.214 | 0.184 | 1 | 9 | Rbpj | 0.03 | Old |
| 0.512287 | 1.663903 | 0.041 | 0.029 | 1 | 9 | Ptprr | 0.012 | Old |
| 0.512287 | 1.51903 | 0.041 | 0.029 | 1 | 9 | Gys1 | 0.012 | Old |
| 0.512287 | 1.395906 | 0.041 | 0.029 | 1 | 9 | Prr14l | 0.012 | Old |
| 0.512323 | 0.071182 | 0.322 | 0.294 | 1 | 9 | Clta | 0.028 | Old |
| 0.512411 | 0.868842 | 0.082 | 0.066 | 1 | 9 | Zfp318 | 0.016 | Old |
| 0.51278 | 0.332913 | 0.113 | 0.096 | 1 | 9 | Tmed4 | 0.017 | Old |
| 0.512911 | 0.418637 | 0.144 | 0.125 | 1 | 9 | Casp8ap2 | 0.019 | Old |
| 0.513575 | 0.318279 | 0.245 | 0.228 | 1 | 9 | Lmna | 0.017 | Old |
| 0.513803 | 0.054885 | 0.369 | 0.346 | 1 | 9 | Tmem109 | 0.023 | Old |
| 0.513873 | 0.177294 | 0.121 | 0.147 | 1 | 9 | Nae1 | -0.026 | Old |
| 0.514383 | 0.097487 | 0.173 | 0.206 | 1 | 9 | Mtmr3 | -0.033 | Old |
| 0.51478 | 0.007307 | 0.034 | 0.022 | 1 | 9 | Ogfod3 | 0.012 | Old |
| 0.514879 | 0.415197 | 0.16 | 0.14 | 1 | 9 | C1galt1 | 0.02 | Old |

| | | | | | | | | |
|---|---|---|---|---|---|---|---|---|
| 0.515021 | 0.416897 | 0.067 | 0.051 | 1 | 9 | Sacs | 0.016 | Old |
| 0.515711 | 0.241889 | 0.142 | 0.118 | 1 | 9 | Tspan6 | 0.024 | Old |
| 0.516446 | 0.102943 | 0.412 | 0.375 | 1 | 9 | Dock4 | 0.037 | Old |
| 0.516593 | 0.12091 | 0.059 | 0.044 | 1 | 9 | Zfp952 | 0.015 | Old |
| 0.516758 | 0.162526 | 0.134 | 0.162 | 1 | 9 | Trmt1l | -0.028 | Old |
| 0.517202 | 0.175873 | 0.312 | 0.272 | 1 | 9 | Tspan15 | 0.04 | Old |
| 0.517405 | 1.58877 | 0.041 | 0.029 | 1 | 9 | Pla2g12a | 0.012 | Old |
| 0.517405 | 1.324555 | 0.041 | 0.029 | 1 | 9 | Vipas39 | 0.012 | Old |
| 0.517707 | 0.584128 | 0.075 | 0.059 | 1 | 9 | Ndufaf4 | 0.016 | Old |
| 0.517787 | 0.043602 | 0.108 | 0.088 | 1 | 9 | Cops2 | 0.02 | Old |
| 0.517787 | 0.226351 | 0.098 | 0.118 | 1 | 9 | Clcc1 | -0.02 | Old |
| 0.517912 | 0.126987 | 0.399 | 0.368 | 1 | 9 | Lamp2 | 0.031 | Old |
| 0.518053 | 0.488346 | 0.067 | 0.051 | 1 | 9 | Fam3a | 0.016 | Old |
| 0.518172 | 0.327484 | 0.255 | 0.235 | 1 | 9 | Anp32b | 0.02 | Old |
| 0.5183 | 0.523639 | 0.044 | 0.059 | 1 | 9 | Apeh | -0.015 | Old |
| 0.51856 | 1.015433 | 0.111 | 0.096 | 1 | 9 | Tmem55b | 0.015 | Old |
| 0.519202 | 0.001703 | 0.415 | 0.382 | 1 | 9 | Bnip2 | 0.033 | Old |
| 0.519404 | 0.04263 | 0.351 | 0.316 | 1 | 9 | Zmynd11 | 0.035 | Old |
| 0.519663 | 0.081133 | 0.265 | 0.235 | 1 | 9 | Polr2m | 0.03 | Old |
| 0.519717 | 0.23958 | 0.082 | 0.066 | 1 | 9 | Ifngr2 | 0.016 | Old |
| 0.519816 | 0.069047 | 0.485 | 0.449 | 1 | 9 | Epn2 | 0.036 | Old |
| 0.519974 | 0.607965 | 0.106 | 0.088 | 1 | 9 | Fundc2 | 0.018 | Old |
| 0.519975 | 0.541711 | 0.041 | 0.029 | 1 | 9 | Tmem19 | 0.012 | Old |
| 0.520008 | 0.239447 | 0.129 | 0.154 | 1 | 9 | Capn1 | -0.025 | Old |
| 0.520056 | 0.129347 | 0.201 | 0.176 | 1 | 9 | Edf1 | 0.025 | Old |
| 0.520091 | 1.619147 | 0.049 | 0.037 | 1 | 9 | Scly | 0.012 | Old |
| 0.520194 | 0.241849 | 0.057 | 0.074 | 1 | 9 | Plek | -0.017 | Old |
| 0.520341 | 0.308216 | 0.131 | 0.11 | 1 | 9 | Slc25a46 | 0.021 | Old |
| 0.520497 | 0.473236 | 0.034 | 0.022 | 1 | 9 | Sfn | 0.012 | Old |
| 0.520683 | 0.622921 | 0.121 | 0.103 | 1 | 9 | Dpm1 | 0.018 | Old |
| 0.521262 | 1.091942 | 0.041 | 0.029 | 1 | 9 | Dcp1a | 0.012 | Old |
| 0.521282 | 0.041672 | 0.026 | 0.037 | 1 | 9 | Slc15a3 | -0.011 | Old |
| 0.521365 | 0.259933 | 0.124 | 0.103 | 1 | 9 | Ibtk | 0.021 | Old |
| 0.521931 | 0.184715 | 0.034 | 0.022 | 1 | 9 | Rabl3 | 0.012 | Old |
| 0.52255 | 1.234238 | 0.041 | 0.029 | 1 | 9 | Zscan29 | 0.012 | Old |
| 0.522613 | 0.042063 | 0.631 | 0.581 | 1 | 9 | H3f3a | 0.05 | Old |
| 0.5229 | 0.107046 | 0.204 | 0.176 | 1 | 9 | Mark2 | 0.028 | Old |
| 0.523484 | 0.388848 | 0.322 | 0.301 | 1 | 9 | Ostf1 | 0.021 | Old |
| 0.52359 | 0.37141 | 0.206 | 0.184 | 1 | 9 | Abca3 | 0.022 | Old |
| 0.523631 | 0.22476 | 0.155 | 0.132 | 1 | 9 | Aqr | 0.023 | Old |
| 0.523757 | 0.640567 | 0.121 | 0.103 | 1 | 9 | Vti1b | 0.018 | Old |
| 0.524247 | 0.446819 | 0.026 | 0.037 | 1 | 9 | Cyp27a1 | -0.011 | Old |
| 0.525041 | 0.420386 | 0.116 | 0.096 | 1 | 9 | Uba52 | 0.02 | Old |
| 0.525133 | 0.872382 | 0.041 | 0.029 | 1 | 9 | Tada2b | 0.012 | Old |
| 0.525133 | 0.594278 | 0.041 | 0.029 | 1 | 9 | Heatr3 | 0.012 | Old |

| | | | | | | | | |
|---|---|---|---|---|---|---|---|---|
| 0.52548 | 0.065444 | 0.348 | 0.324 | 1 | 9 | Ptger4 | 0.024 | Old |
| 0.52561 | 0.036086 | 0.271 | 0.309 | 1 | 9 | Rbbp6 | -0.038 | Old |
| 0.525732 | 0.251577 | 0.026 | 0.037 | 1 | 9 | Plekhn1 | -0.011 | Old |
| 0.525787 | 0.011169 | 0.41 | 0.375 | 1 | 9 | N4bp1 | 0.035 | Old |
| 0.526002 | 0.832843 | 0.049 | 0.037 | 1 | 9 | Rnaseh1 | 0.012 | Old |
| 0.526154 | 0.222409 | 0.082 | 0.066 | 1 | 9 | Golga3 | 0.016 | Old |
| 0.52642 | 0.056085 | 0.977 | 0.949 | 1 | 9 | mt-Cytb | 0.028 | Old |
| 0.526782 | 0.080298 | 0.621 | 0.566 | 1 | 9 | Gls | 0.055 | Old |
| 0.527109 | 1.809594 | 0.057 | 0.044 | 1 | 9 | Rab31 | 0.013 | Old |
| 0.527553 | 0.602471 | 0.229 | 0.213 | 1 | 9 | Efr3a | 0.016 | Old |
| 0.527721 | 0.874097 | 0.041 | 0.029 | 1 | 9 | Zfp523 | 0.012 | Old |
| 0.528377 | 1.110585 | 0.049 | 0.037 | 1 | 9 | Lyrm2 | 0.012 | Old |
| 0.528384 | 0.279859 | 0.121 | 0.103 | 1 | 9 | Pcnt | 0.018 | Old |
| 0.528709 | 0.415919 | 0.026 | 0.037 | 1 | 9 | Stxbp2 | -0.011 | Old |
| 0.528709 | 0.106747 | 0.026 | 0.037 | 1 | 9 | Igsf6 | -0.011 | Old |
| 0.529107 | 0.457909 | 0.317 | 0.294 | 1 | 9 | Huwe1 | 0.023 | Old |
| 0.529247 | 0.390298 | 0.067 | 0.051 | 1 | 9 | Taf5l | 0.016 | Old |
| 0.52949 | 0.552019 | 0.193 | 0.176 | 1 | 9 | Ubxn1 | 0.017 | Old |
| 0.529913 | 0.804101 | 0.17 | 0.154 | 1 | 9 | Snap23 | 0.016 | Old |
| 0.530269 | 0.146277 | 0.34 | 0.301 | 1 | 9 | Cebpd | 0.039 | Old |
| 0.530271 | 0.52949 | 0.067 | 0.051 | 1 | 9 | Bbs9 | 0.016 | Old |
| 0.530776 | 0.569674 | 0.082 | 0.066 | 1 | 9 | Sft2d1 | 0.016 | Old |
| 0.530776 | 0.54034 | 0.082 | 0.066 | 1 | 9 | Naga | 0.016 | Old |
| 0.531054 | 0.378783 | 0.09 | 0.074 | 1 | 9 | Fbxo38 | 0.016 | Old |
| 0.531354 | 0.443251 | 0.242 | 0.213 | 1 | 9 | Kif1c | 0.029 | Old |
| 0.531439 | 0.194535 | 0.296 | 0.272 | 1 | 9 | 2900097C1 | 0.024 | Old |
| 0.531617 | 0.66472 | 0.041 | 0.029 | 1 | 9 | Smyd2 | 0.012 | Old |
| 0.531617 | 0.039811 | 0.041 | 0.029 | 1 | 9 | Ston1 | 0.012 | Old |
| 0.531919 | 0.319052 | 0.265 | 0.243 | 1 | 9 | Zc3h7a | 0.022 | Old |
| 0.532288 | 0.431949 | 0.106 | 0.088 | 1 | 9 | Nolc1 | 0.018 | Old |
| 0.532425 | 0.022701 | 0.108 | 0.088 | 1 | 9 | Kif2a | 0.02 | Old |
| 0.532918 | 0.705221 | 0.041 | 0.029 | 1 | 9 | Trim62 | 0.012 | Old |
| 0.532929 | 0.572526 | 0.142 | 0.125 | 1 | 9 | Siah1a | 0.017 | Old |
| 0.533171 | 0.435116 | 0.075 | 0.059 | 1 | 9 | Fbxl12 | 0.016 | Old |
| 0.533572 | 0.872723 | 0.126 | 0.11 | 1 | 9 | Klhdc3 | 0.016 | Old |
| 0.533599 | 0.196295 | 0.196 | 0.169 | 1 | 9 | Epha4 | 0.027 | Old |
| 0.533778 | 0.160939 | 0.472 | 0.441 | 1 | 9 | Fscn1 | 0.031 | Old |
| 0.533941 | 0.624149 | 0.106 | 0.088 | 1 | 9 | S100pbp | 0.018 | Old |
| 0.533976 | 0.03952 | 0.317 | 0.272 | 1 | 9 | Sertad1 | 0.045 | Old |
| 0.534002 | 1.235507 | 0.088 | 0.074 | 1 | 9 | Ppp6r2 | 0.014 | Old |
| 0.534376 | 0.340275 | 0.067 | 0.051 | 1 | 9 | Ammecr1 | 0.016 | Old |
| 0.534385 | 0.068432 | 0.688 | 0.632 | 1 | 9 | Lpl | 0.056 | Old |
| 0.534949 | 0.85099 | 0.134 | 0.118 | 1 | 9 | Gabpa | 0.016 | Old |
| 0.535813 | 0.364443 | 0.155 | 0.132 | 1 | 9 | Yipf3 | 0.023 | Old |
| 0.536217 | 0.191796 | 0.247 | 0.228 | 1 | 9 | Ankrd40 | 0.019 | Old |

| | | | | | | | | |
|---|---|---|---|---|---|---|---|---|
| 0.536729 | 1.05081 | 0.049 | 0.037 | 1 | 9 | Klhdc8b | 0.012 | Old |
| 0.536729 | 0.949296 | 0.049 | 0.037 | 1 | 9 | Abhd17c | 0.012 | Old |
| 0.536772 | 0.227722 | 0.131 | 0.11 | 1 | 9 | Zfp945 | 0.021 | Old |
| 0.536926 | 0.022751 | 0.116 | 0.096 | 1 | 9 | Zc3h12c | 0.02 | Old |
| 0.537204 | 0.029516 | 0.052 | 0.066 | 1 | 9 | Trappc2 | -0.014 | Old |
| 0.537434 | 0.529001 | 0.162 | 0.147 | 1 | 9 | Prelid1 | 0.015 | Old |
| 0.537525 | 0.217508 | 0.131 | 0.11 | 1 | 9 | Bcl2 | 0.021 | Old |
| 0.537754 | 2.78744 | 0.023 | 0.015 | 1 | 9 | Fam206a | 0.008 | Old |
| 0.538141 | 0.478731 | 0.041 | 0.029 | 1 | 9 | Metap1d | 0.012 | Old |
| 0.538215 | 0.393484 | 0.082 | 0.066 | 1 | 9 | Ankrd27 | 0.016 | Old |
| 0.539127 | 1.397593 | 0.049 | 0.037 | 1 | 9 | Bmp4 | 0.012 | Old |
| 0.539315 | 0.642789 | 0.057 | 0.044 | 1 | 9 | Glod4 | 0.013 | Old |
| 0.539401 | 0.538778 | 0.211 | 0.199 | 1 | 9 | Ypel3 | 0.012 | Old |
| 0.539424 | 0.046604 | 0.039 | 0.051 | 1 | 9 | Gm17018 | -0.012 | Old |
| 0.539451 | 0.593492 | 0.041 | 0.029 | 1 | 9 | Nim1k | 0.012 | Old |
| 0.540639 | 0.298704 | 0.201 | 0.184 | 1 | 9 | Sec61a1 | 0.017 | Old |
| 0.540905 | 0.00628 | 0.093 | 0.074 | 1 | 9 | Eps15l1 | 0.019 | Old |
| 0.541558 | 0.155939 | 0.16 | 0.14 | 1 | 9 | Srsf4 | 0.02 | Old |
| 0.542074 | 0.104171 | 0.18 | 0.154 | 1 | 9 | Rnf216 | 0.026 | Old |
| 0.542226 | 0.119094 | 0.183 | 0.154 | 1 | 9 | Stxbp5 | 0.029 | Old |
| 0.542244 | 0.439805 | 0.106 | 0.088 | 1 | 9 | Pou2f1 | 0.018 | Old |
| 0.542589 | 0.374416 | 0.253 | 0.228 | 1 | 9 | Fndc3b | 0.025 | Old |
| 0.543014 | 2.118374 | 0.023 | 0.015 | 1 | 9 | S1pr4 | 0.008 | Old |
| 0.543014 | 1.80913 | 0.023 | 0.015 | 1 | 9 | Mettl8 | 0.008 | Old |
| 0.543014 | 1.520944 | 0.023 | 0.015 | 1 | 9 | Plk4 | 0.008 | Old |
| 0.543014 | 1.122662 | 0.023 | 0.015 | 1 | 9 | Disp1 | 0.008 | Old |
| 0.543262 | 0.080852 | 0.211 | 0.184 | 1 | 9 | Phactr4 | 0.027 | Old |
| 0.543389 | 0.009379 | 0.041 | 0.029 | 1 | 9 | C1qtnf1 | 0.012 | Old |
| 0.543536 | 0.352853 | 0.42 | 0.404 | 1 | 9 | Ndufa2 | 0.016 | Old |
| 0.543671 | 0.031943 | 0.067 | 0.051 | 1 | 9 | Trim21 | 0.016 | Old |
| 0.543895 | 0.374063 | 0.142 | 0.169 | 1 | 9 | Lmo2 | -0.027 | Old |
| 0.544371 | 0.197517 | 0.508 | 0.493 | 1 | 9 | Arpc3 | 0.015 | Old |
| 0.544766 | 0.512803 | 0.082 | 0.066 | 1 | 9 | Noc2l | 0.016 | Old |
| 0.544773 | 1.921465 | 0.023 | 0.015 | 1 | 9 | Rgs4 | 0.008 | Old |
| 0.544773 | 1.742261 | 0.023 | 0.015 | 1 | 9 | Gm13889 | 0.008 | Old |
| 0.544773 | 1.459493 | 0.023 | 0.015 | 1 | 9 | Zbtb5 | 0.008 | Old |
| 0.544773 | 1.211737 | 0.023 | 0.015 | 1 | 9 | Zfp790 | 0.008 | Old |
| 0.544909 | 1.165561 | 0.057 | 0.044 | 1 | 9 | Agpat1 | 0.013 | Old |
| 0.54492 | 0.324822 | 0.235 | 0.213 | 1 | 9 | Dnajc5 | 0.022 | Old |
| 0.544954 | 0.321114 | 0.155 | 0.132 | 1 | 9 | Hbp1 | 0.023 | Old |
| 0.545147 | 0.140867 | 0.049 | 0.037 | 1 | 9 | Tor2a | 0.012 | Old |
| 0.545466 | 0.476213 | 0.144 | 0.125 | 1 | 9 | Chd3 | 0.019 | Old |
| 0.545726 | 0.260396 | 0.106 | 0.125 | 1 | 9 | Mpp1 | -0.019 | Old |
| 0.546025 | 0.023344 | 0.773 | 0.75 | 1 | 9 | Rpl13 | 0.023 | Old |
| 0.546355 | 0.552241 | 0.049 | 0.037 | 1 | 9 | Alkbh6 | 0.012 | Old |

| | | | | | | | | |
|---|---|---|---|---|---|---|---|---|
| 0.546391 | 0.214192 | 0.175 | 0.154 | 1 | 9 | Cacybp | 0.021 | Old |
| 0.546534 | 1.101759 | 0.023 | 0.015 | 1 | 9 | Trim32 | 0.008 | Old |
| 0.546534 | 0.924789 | 0.023 | 0.015 | 1 | 9 | Galt | 0.008 | Old |
| 0.54675 | 0.953453 | 0.026 | 0.037 | 1 | 9 | Mob3b | -0.011 | Old |
| 0.54677 | 0.682088 | 0.064 | 0.051 | 1 | 9 | St8sia6 | 0.013 | Old |
| 0.546962 | 0.088277 | 0.039 | 0.051 | 1 | 9 | Tomm6 | -0.012 | Old |
| 0.547061 | 0.273737 | 0.175 | 0.154 | 1 | 9 | Mgat4b | 0.021 | Old |
| 0.547155 | 0.920862 | 0.057 | 0.044 | 1 | 9 | Mettl7a1 | 0.013 | Old |
| 0.54771 | 0.015188 | 0.24 | 0.213 | 1 | 9 | Cmtm6 | 0.027 | Old |
| 0.547717 | 0.852698 | 0.057 | 0.044 | 1 | 9 | Slf1 | 0.013 | Old |
| 0.54828 | 1.082041 | 0.057 | 0.044 | 1 | 9 | Pmvk | 0.013 | Old |
| 0.548299 | 1.526896 | 0.023 | 0.015 | 1 | 9 | Nt5dc1 | 0.008 | Old |
| 0.548299 | 0.91735 | 0.023 | 0.015 | 1 | 9 | Hist1h1e | 0.008 | Old |
| 0.548299 | 0.900229 | 0.023 | 0.015 | 1 | 9 | Tom1 | 0.008 | Old |
| 0.548299 | 0.767323 | 0.023 | 0.015 | 1 | 9 | Gm17103 | 0.008 | Old |
| 0.548475 | 0.500754 | 0.082 | 0.103 | 1 | 9 | Mzt1 | -0.021 | Old |
| 0.548645 | 0.061653 | 0.428 | 0.404 | 1 | 9 | Serpinb6a | 0.024 | Old |
| 0.548751 | 0.608662 | 0.072 | 0.059 | 1 | 9 | Cwc25 | 0.013 | Old |
| 0.548832 | 0.111845 | 0.17 | 0.147 | 1 | 9 | 1700025G( | 0.023 | Old |
| 0.548929 | 0.143644 | 0.183 | 0.162 | 1 | 9 | Wdr45b | 0.021 | Old |
| 0.54947 | 0.027007 | 0.082 | 0.066 | 1 | 9 | Cx3cl1 | 0.016 | Old |
| 0.549889 | 0.074725 | 0.134 | 0.118 | 1 | 9 | Mtch2 | 0.016 | Old |
| 0.549986 | 0.728505 | 0.049 | 0.037 | 1 | 9 | Ring1 | 0.012 | Old |
| 0.550066 | 1.547859 | 0.023 | 0.015 | 1 | 9 | Ift22 | 0.008 | Old |
| 0.550066 | 1.53263 | 0.023 | 0.015 | 1 | 9 | Zfp458 | 0.008 | Old |
| 0.550066 | 1.3337 | 0.023 | 0.015 | 1 | 9 | Kif3c | 0.008 | Old |
| 0.550066 | 0.793064 | 0.023 | 0.015 | 1 | 9 | Ints5.1 | 0.008 | Old |
| 0.550409 | 0.792114 | 0.142 | 0.125 | 1 | 9 | Evi5 | 0.017 | Old |
| 0.550641 | 0.26673 | 0.134 | 0.118 | 1 | 9 | Bcl2l11 | 0.016 | Old |
| 0.550773 | 0.361899 | 0.09 | 0.074 | 1 | 9 | Tubgcp3 | 0.016 | Old |
| 0.550843 | 0.134993 | 0.075 | 0.059 | 1 | 9 | Orc4 | 0.016 | Old |
| 0.550902 | 0.088181 | 0.137 | 0.118 | 1 | 9 | Tpd52l2 | 0.019 | Old |
| 0.55097 | 0.280268 | 0.24 | 0.213 | 1 | 9 | Arfgef1 | 0.027 | Old |
| 0.551186 | 0.596659 | 0.113 | 0.14 | 1 | 9 | Ube2e1 | -0.027 | Old |
| 0.551199 | 0.809381 | 0.049 | 0.037 | 1 | 9 | Zdhhc1 | 0.012 | Old |
| 0.551419 | 0.035515 | 0.147 | 0.125 | 1 | 9 | Sh3tc1 | 0.022 | Old |
| 0.551837 | 0.868851 | 0.023 | 0.015 | 1 | 9 | Grwd1 | 0.008 | Old |
| 0.551837 | 0.772487 | 0.023 | 0.015 | 1 | 9 | Zfp932 | 0.008 | Old |
| 0.551837 | 0.719865 | 0.023 | 0.015 | 1 | 9 | Sall2 | 0.008 | Old |
| 0.551837 | 0.694815 | 0.023 | 0.015 | 1 | 9 | Shroom2 | 0.008 | Old |
| 0.551837 | 0.588819 | 0.023 | 0.015 | 1 | 9 | Abhd8 | 0.008 | Old |
| 0.551837 | 0.576948 | 0.023 | 0.015 | 1 | 9 | Pyroxd1 | 0.008 | Old |
| 0.551999 | 0.229389 | 0.059 | 0.074 | 1 | 9 | Ddx55 | -0.015 | Old |
| 0.552641 | 0.011178 | 0.307 | 0.279 | 1 | 9 | Hdlbp | 0.028 | Old |
| 0.553399 | 0.279944 | 0.196 | 0.176 | 1 | 9 | Ccm2l | 0.02 | Old |

| | | | | | | | | |
|---|---|---|---|---|---|---|---|---|
| 0.55349 | 0.184488 | 0.09 | 0.074 | 1 | 9 | Cyp1a1 | 0.016 | Old |
| 0.55361 | 0.315065 | 0.023 | 0.015 | 1 | 9 | Fgfrl1 | 0.008 | Old |
| 0.553699 | 0.08147 | 0.098 | 0.081 | 1 | 9 | Npc1 | 0.017 | Old |
| 0.553738 | 0.249384 | 0.168 | 0.147 | 1 | 9 | Fmnl1 | 0.021 | Old |
| 0.554144 | 0.118862 | 0.204 | 0.184 | 1 | 9 | Gstm2 | 0.02 | Old |
| 0.554565 | 0.194244 | 0.322 | 0.301 | 1 | 9 | Ndufb11 | 0.021 | Old |
| 0.554769 | 0.127951 | 0.108 | 0.088 | 1 | 9 | Kdm4c | 0.02 | Old |
| 0.55529 | 0.066256 | 0.041 | 0.029 | 1 | 9 | Zc3h8 | 0.012 | Old |
| 0.555305 | 0.080372 | 0.09 | 0.074 | 1 | 9 | Lsm7 | 0.016 | Old |
| 0.555385 | 0.937595 | 0.023 | 0.015 | 1 | 9 | Tmem158 | 0.008 | Old |
| 0.555385 | 0.68011 | 0.023 | 0.015 | 1 | 9 | Rida | 0.008 | Old |
| 0.555385 | 0.251261 | 0.023 | 0.015 | 1 | 9 | Ap5z1 | 0.008 | Old |
| 0.555389 | 0.020424 | 0.716 | 0.691 | 1 | 9 | Cfl1 | 0.025 | Old |
| 0.555408 | 0.10047 | 0.106 | 0.125 | 1 | 9 | Cse1l | -0.019 | Old |
| 0.55541 | 0.084348 | 0.072 | 0.088 | 1 | 9 | Cyp20a1 | -0.016 | Old |
| 0.555728 | 0.280785 | 0.24 | 0.213 | 1 | 9 | Chmp3 | 0.027 | Old |
| 0.55662 | 0.238763 | 0.358 | 0.331 | 1 | 9 | H2-Q4 | 0.027 | Old |
| 0.556902 | 0.309207 | 0.137 | 0.118 | 1 | 9 | Prpf18 | 0.019 | Old |
| 0.557164 | 0.556535 | 0.023 | 0.015 | 1 | 9 | Tigar | 0.008 | Old |
| 0.557164 | 0.314845 | 0.023 | 0.015 | 1 | 9 | Aifm2 | 0.008 | Old |
| 0.557237 | 0.181365 | 0.067 | 0.051 | 1 | 9 | Zfp626 | 0.016 | Old |
| 0.557318 | 0.92155 | 0.057 | 0.044 | 1 | 9 | Cmc1 | 0.013 | Old |
| 0.557435 | 0.654044 | 0.134 | 0.118 | 1 | 9 | Ptdss1 | 0.016 | Old |
| 0.557475 | 0.989983 | 0.095 | 0.081 | 1 | 9 | Syvn1 | 0.014 | Old |
| 0.55754 | 0.167899 | 0.227 | 0.206 | 1 | 9 | Gabarapl2 | 0.021 | Old |
| 0.557702 | 0.24086 | 0.371 | 0.346 | 1 | 9 | Hint1 | 0.025 | Old |
| 0.557793 | 0.092671 | 0.075 | 0.059 | 1 | 9 | Pcdhga12 | 0.016 | Old |
| 0.558137 | 0.395077 | 0.129 | 0.11 | 1 | 9 | Letm1 | 0.019 | Old |
| 0.558441 | 0.46154 | 0.149 | 0.132 | 1 | 9 | Synrg | 0.017 | Old |
| 0.558945 | 0.493843 | 0.023 | 0.015 | 1 | 9 | Mbp | 0.008 | Old |
| 0.558945 | 0.355231 | 0.023 | 0.015 | 1 | 9 | Capn12 | 0.008 | Old |
| 0.558945 | 0.154263 | 0.023 | 0.015 | 1 | 9 | Fbxl8 | 0.008 | Old |
| 0.559253 | 0.596269 | 0.142 | 0.125 | 1 | 9 | Nop56 | 0.017 | Old |
| 0.559466 | 0.185007 | 0.106 | 0.125 | 1 | 9 | Stt3b | -0.019 | Old |
| 0.559639 | 0.796674 | 0.039 | 0.051 | 1 | 9 | Nfkbie | -0.012 | Old |
| 0.55973 | 0.087051 | 0.451 | 0.412 | 1 | 9 | Cdc42ep3 | 0.039 | Old |
| 0.559943 | 0.211056 | 0.021 | 0.029 | 1 | 9 | Pithd1 | -0.008 | Old |
| 0.559943 | 0.17512 | 0.021 | 0.029 | 1 | 9 | Ms4a7 | -0.008 | Old |
| 0.559943 | 0.091325 | 0.021 | 0.029 | 1 | 9 | Halr1 | -0.008 | Old |
| 0.559943 | 0.04545 | 0.021 | 0.029 | 1 | 9 | Surf6 | -0.008 | Old |
| 0.559951 | 0.191947 | 0.201 | 0.184 | 1 | 9 | Eny2 | 0.017 | Old |
| 0.560584 | 0.161142 | 0.149 | 0.125 | 1 | 9 | Pip5k1c | 0.024 | Old |
| 0.560729 | 0.195365 | 0.023 | 0.015 | 1 | 9 | Rai2 | 0.008 | Old |
| 0.561241 | 0.066052 | 0.423 | 0.397 | 1 | 9 | Selenok | 0.026 | Old |
| 0.561573 | 0.456498 | 0.052 | 0.066 | 1 | 9 | Lyst | -0.014 | Old |

| | | | | | | | | |
|---|---|---|---|---|---|---|---|---|
| 0.561582 | 0.304474 | 0.106 | 0.088 | 1 | 9 | Ptbp2 | 0.018 | Old |
| 0.561641 | 0.760159 | 0.064 | 0.051 | 1 | 9 | Car7 | 0.013 | Old |
| 0.561654 | 0.64153 | 0.021 | 0.029 | 1 | 9 | Rdh13 | -0.008 | Old |
| 0.561679 | 0.172972 | 0.09 | 0.074 | 1 | 9 | Mrpl15 | 0.016 | Old |
| 0.561761 | 0.080651 | 0.124 | 0.103 | 1 | 9 | Polr2g | 0.021 | Old |
| 0.562177 | 0.576 | 0.049 | 0.037 | 1 | 9 | Fiz1 | 0.012 | Old |
| 0.562516 | 0.262829 | 0.023 | 0.015 | 1 | 9 | Pik3r2 | 0.008 | Old |
| 0.562516 | 0.145527 | 0.023 | 0.015 | 1 | 9 | Ndrg2 | 0.008 | Old |
| 0.562588 | 0.5774 | 0.142 | 0.125 | 1 | 9 | Gtf3c2 | 0.017 | Old |
| 0.562694 | 0.245123 | 0.052 | 0.066 | 1 | 9 | Adcy9 | -0.014 | Old |
| 0.562919 | 0.051286 | 0.193 | 0.221 | 1 | 9 | Aftph | -0.028 | Old |
| 0.563368 | 0.028294 | 0.021 | 0.029 | 1 | 9 | Ggcx | -0.008 | Old |
| 0.563403 | 0.045827 | 0.049 | 0.037 | 1 | 9 | Ikbkap | 0.012 | Old |
| 0.563421 | 0.413328 | 0.245 | 0.228 | 1 | 9 | Gpr160 | 0.017 | Old |
| 0.564126 | 0.000621 | 0.144 | 0.169 | 1 | 9 | Ptpa | -0.025 | Old |
| 0.564345 | 0.565479 | 0.139 | 0.125 | 1 | 9 | Tmem168 | 0.014 | Old |
| 0.564888 | 0.202587 | 0.144 | 0.125 | 1 | 9 | Gripap1 | 0.019 | Old |
| 0.565285 | 0.509998 | 0.057 | 0.044 | 1 | 9 | Prdm4 | 0.013 | Old |
| 0.56562 | 0.227066 | 0.155 | 0.132 | 1 | 9 | Yipf4 | 0.023 | Old |
| 0.56575 | 0.021573 | 0.242 | 0.272 | 1 | 9 | Eif2s3x | -0.03 | Old |
| 0.565871 | 0.153331 | 0.173 | 0.154 | 1 | 9 | Bbx | 0.019 | Old |
| 0.565969 | 0.354262 | 0.175 | 0.154 | 1 | 9 | Atrn | 0.021 | Old |
| 0.566051 | 0.191271 | 0.291 | 0.265 | 1 | 9 | Zeb1 | 0.026 | Old |
| 0.566061 | 0.574995 | 0.052 | 0.066 | 1 | 9 | Exosc7 | -0.014 | Old |
| 0.566137 | 0.244134 | 0.121 | 0.103 | 1 | 9 | Nbdy | 0.018 | Old |
| 0.56661 | 0.071879 | 0.263 | 0.235 | 1 | 9 | Drap1 | 0.028 | Old |
| 0.566676 | 0.335494 | 0.142 | 0.125 | 1 | 9 | Cyld | 0.017 | Old |
| 0.566803 | 0.13535 | 0.021 | 0.029 | 1 | 9 | Mthfs | -0.008 | Old |
| 0.566861 | 0.41747 | 0.098 | 0.081 | 1 | 9 | Hoxb7 | 0.017 | Old |
| 0.567171 | 0.020963 | 0.09 | 0.074 | 1 | 9 | Nup214 | 0.016 | Old |
| 0.56732 | 0.225454 | 0.09 | 0.11 | 1 | 9 | Plbd2 | -0.02 | Old |
| 0.567331 | 0.455758 | 0.119 | 0.103 | 1 | 9 | 1810058I2 | 0.016 | Old |
| 0.567346 | 0.067847 | 0.08 | 0.096 | 1 | 9 | Pip4k2c | -0.016 | Old |
| 0.567421 | 0.735604 | 0.142 | 0.125 | 1 | 9 | Maz | 0.017 | Old |
| 0.567518 | 0.151192 | 0.113 | 0.096 | 1 | 9 | Klhl5 | 0.017 | Old |
| 0.567643 | 0.111121 | 0.314 | 0.287 | 1 | 9 | Trio | 0.027 | Old |
| 0.56765 | 0.054809 | 0.253 | 0.272 | 1 | 9 | Susd6 | -0.019 | Old |
| 0.567819 | 0.477477 | 0.17 | 0.154 | 1 | 9 | Tril | 0.016 | Old |
| 0.567951 | 0.048976 | 0.046 | 0.059 | 1 | 9 | Ttc1 | -0.013 | Old |
| 0.567951 | 0.017072 | 0.046 | 0.059 | 1 | 9 | Pias2 | -0.013 | Old |
| 0.568154 | 0.483492 | 0.204 | 0.184 | 1 | 9 | Tpp2 | 0.02 | Old |
| 0.568203 | 0.648708 | 0.111 | 0.096 | 1 | 9 | Taf2 | 0.015 | Old |
| 0.568524 | 0.525784 | 0.021 | 0.029 | 1 | 9 | E2f3 | -0.008 | Old |
| 0.568524 | 0.397495 | 0.021 | 0.029 | 1 | 9 | Mlh1 | -0.008 | Old |
| 0.568524 | 0.043784 | 0.021 | 0.029 | 1 | 9 | Fdps | -0.008 | Old |

| | | | | | | | | |
|---|---|---|---|---|---|---|---|---|
| 0.568714 | 0.092302 | 0.454 | 0.412 | 1 | 9 | Pltp | 0.042 | Old |
| 0.569054 | 0.013384 | 0.412 | 0.375 | 1 | 9 | Cdc42bpa | 0.037 | Old |
| 0.569087 | 0.056014 | 0.361 | 0.338 | 1 | 9 | Psme2 | 0.023 | Old |
| 0.569366 | 0.19869 | 0.222 | 0.206 | 1 | 9 | Zfr | 0.016 | Old |
| 0.569553 | 0.585445 | 0.049 | 0.037 | 1 | 9 | Pitpnm1 | 0.012 | Old |
| 0.569722 | 2.344825 | 0.031 | 0.022 | 1 | 9 | Zfp839 | 0.009 | Old |
| 0.569926 | 0.252407 | 0.09 | 0.074 | 1 | 9 | Gabpb1 | 0.016 | Old |
| 0.570248 | 0.242089 | 0.021 | 0.029 | 1 | 9 | Ddx19b | -0.008 | Old |
| 0.570347 | 0.081876 | 0.224 | 0.206 | 1 | 9 | Rnf7 | 0.018 | Old |
| 0.570562 | 0.348522 | 0.119 | 0.103 | 1 | 9 | Pgls | 0.016 | Old |
| 0.570619 | 0.281294 | 0.314 | 0.294 | 1 | 9 | Uqcrq | 0.02 | Old |
| 0.570846 | 0.327187 | 0.09 | 0.074 | 1 | 9 | Dmtn | 0.016 | Old |
| 0.57085 | 0.122713 | 0.072 | 0.088 | 1 | 9 | Upf2 | -0.016 | Old |
| 0.571009 | 0.406232 | 0.057 | 0.044 | 1 | 9 | Ctbs | 0.013 | Old |
| 0.57165 | 0.231872 | 0.113 | 0.096 | 1 | 9 | Ctbp2 | 0.017 | Old |
| 0.572381 | 0.189248 | 0.064 | 0.051 | 1 | 9 | Ldah | 0.013 | Old |
| 0.572593 | 0.259513 | 0.095 | 0.081 | 1 | 9 | Usf3 | 0.014 | Old |
| 0.57302 | 0.811942 | 0.072 | 0.059 | 1 | 9 | Timm29 | 0.013 | Old |
| 0.573259 | 0.403902 | 0.049 | 0.037 | 1 | 9 | Mlycd | 0.012 | Old |
| 0.573307 | 0.560358 | 0.057 | 0.044 | 1 | 9 | Cactin | 0.013 | Old |
| 0.573703 | 0.63579 | 0.021 | 0.029 | 1 | 9 | Tmem104 | -0.008 | Old |
| 0.573703 | 0.401299 | 0.021 | 0.029 | 1 | 9 | Neurl2 | -0.008 | Old |
| 0.573894 | 0.232578 | 0.046 | 0.059 | 1 | 9 | Utp18 | -0.013 | Old |
| 0.574042 | 0.748961 | 0.072 | 0.059 | 1 | 9 | Zfp282 | 0.013 | Old |
| 0.574042 | 0.054237 | 0.072 | 0.059 | 1 | 9 | Vkorc1 | 0.013 | Old |
| 0.574497 | 0.090566 | 0.049 | 0.037 | 1 | 9 | Tango6 | 0.012 | Old |
| 0.574762 | 0.073885 | 0.714 | 0.699 | 1 | 9 | Itga6 | 0.015 | Old |
| 0.574862 | 0.182383 | 0.487 | 0.485 | 1 | 9 | Kmt2e | 0.002 | Old |
| 0.575258 | 0.36813 | 0.129 | 0.11 | 1 | 9 | Palld | 0.019 | Old |
| 0.575645 | 0.067442 | 0.142 | 0.125 | 1 | 9 | Glrx5 | 0.017 | Old |
| 0.576178 | 0.885037 | 0.095 | 0.081 | 1 | 9 | Gosr1 | 0.014 | Old |
| 0.576204 | 0.649182 | 0.08 | 0.066 | 1 | 9 | Tbc1d9 | 0.014 | Old |
| 0.576761 | 0.217686 | 0.183 | 0.213 | 1 | 9 | Rnf187 | -0.03 | Old |
| 0.576957 | 0.597779 | 0.178 | 0.162 | 1 | 9 | Atxn7 | 0.016 | Old |
| 0.576976 | 0.03953 | 0.049 | 0.037 | 1 | 9 | Cd3eap | 0.012 | Old |
| 0.577084 | 0.060994 | 0.24 | 0.213 | 1 | 9 | Serpinb9 | 0.027 | Old |
| 0.577167 | 0.535199 | 0.021 | 0.029 | 1 | 9 | Elac2 | -0.008 | Old |
| 0.577505 | 0.076563 | 0.085 | 0.103 | 1 | 9 | Spg20 | -0.018 | Old |
| 0.577788 | 0.732715 | 0.064 | 0.051 | 1 | 9 | Mxi1 | 0.013 | Old |
| 0.577915 | 0.508389 | 0.057 | 0.044 | 1 | 9 | Guf1 | 0.013 | Old |
| 0.57798 | 0.20015 | 0.268 | 0.235 | 1 | 9 | Il4ra | 0.033 | Old |
| 0.578157 | 0.146873 | 0.08 | 0.066 | 1 | 9 | Bcorl1 | 0.014 | Old |
| 0.578424 | 0.307234 | 0.059 | 0.074 | 1 | 9 | Chmp7 | -0.015 | Old |
| 0.578649 | 0.691801 | 0.111 | 0.096 | 1 | 9 | Mrpl34 | 0.015 | Old |
| 0.578677 | 0.254222 | 0.34 | 0.324 | 1 | 9 | Khdrbs1 | 0.016 | Old |

| | | | | | | | | |
|---|---|---|---|---|---|---|---|---|
| 0.579038 | 1.538812 | 0.031 | 0.022 | 1 | 9 | Oxsm | 0.009 | Old |
| 0.579166 | 0.586538 | 0.072 | 0.059 | 1 | 9 | Med30 | 0.013 | Old |
| 0.579279 | 0.038255 | 0.098 | 0.081 | 1 | 9 | Zfp266 | 0.017 | Old |
| 0.579726 | 0.263985 | 0.034 | 0.044 | 1 | 9 | Rgs10 | -0.01 | Old |
| 0.579927 | 0.808504 | 0.088 | 0.074 | 1 | 9 | Wdr11 | 0.014 | Old |
| 0.580121 | 0.26865 | 0.098 | 0.118 | 1 | 9 | Inpp5d | -0.02 | Old |
| 0.580155 | 0.048078 | 0.142 | 0.125 | 1 | 9 | Timeless | 0.017 | Old |
| 0.580171 | 0.041643 | 0.098 | 0.081 | 1 | 9 | Naa38 | 0.017 | Old |
| 0.580225 | 0.085492 | 0.057 | 0.044 | 1 | 9 | Zfp423 | 0.013 | Old |
| 0.581092 | 0.046081 | 0.08 | 0.066 | 1 | 9 | Mmadhc | 0.014 | Old |
| 0.581118 | 0.027682 | 0.034 | 0.044 | 1 | 9 | Itsn1 | -0.01 | Old |
| 0.581501 | 0.070732 | 0.255 | 0.228 | 1 | 9 | Tbx3 | 0.027 | Old |
| 0.581576 | 0.700445 | 0.095 | 0.081 | 1 | 9 | G2e3 | 0.014 | Old |
| 0.581802 | 0.410564 | 0.088 | 0.074 | 1 | 9 | Zbtb14 | 0.014 | Old |
| 0.582511 | 0.245912 | 0.034 | 0.044 | 1 | 9 | Ahi1 | -0.01 | Old |
| 0.582847 | 0.098472 | 0.247 | 0.221 | 1 | 9 | Ccnd1 | 0.026 | Old |
| 0.583044 | 0.23072 | 0.42 | 0.397 | 1 | 9 | Elk4 | 0.023 | Old |
| 0.583052 | 0.450768 | 0.08 | 0.066 | 1 | 9 | Brwd3 | 0.014 | Old |
| 0.583305 | 0.167323 | 0.242 | 0.279 | 1 | 9 | Smarca2 | -0.037 | Old |
| 0.583428 | 0.023048 | 0.206 | 0.184 | 1 | 9 | Trib2 | 0.022 | Old |
| 0.58391 | 0.000948 | 0.155 | 0.176 | 1 | 9 | Brd7 | -0.021 | Old |
| 0.583987 | 0.012741 | 0.232 | 0.199 | 1 | 9 | Ube2k | 0.033 | Old |
| 0.584034 | 0.625774 | 0.08 | 0.066 | 1 | 9 | Eri1 | 0.014 | Old |
| 0.58447 | 0.031285 | 0.168 | 0.191 | 1 | 9 | Jmjd6 | -0.023 | Old |
| 0.584496 | 0.022359 | 0.567 | 0.515 | 1 | 9 | Wwtr1 | 0.052 | Old |
| 0.585194 | 0.364852 | 0.157 | 0.14 | 1 | 9 | Copg1 | 0.017 | Old |
| 0.585245 | 0.19389 | 0.227 | 0.206 | 1 | 9 | Synj1 | 0.021 | Old |
| 0.585288 | 1.062983 | 0.031 | 0.022 | 1 | 9 | Extl2 | 0.009 | Old |
| 0.585302 | 0.002312 | 0.034 | 0.044 | 1 | 9 | Sco2 | -0.01 | Old |
| 0.585346 | 0.271446 | 0.152 | 0.132 | 1 | 9 | Rest | 0.02 | Old |
| 0.58556 | 0.702022 | 0.088 | 0.074 | 1 | 9 | Arhgap26 | 0.014 | Old |
| 0.58609 | 0.196713 | 0.232 | 0.213 | 1 | 9 | Klhl9 | 0.019 | Old |
| 0.5867 | 1.601368 | 0.039 | 0.029 | 1 | 9 | Tns3 | 0.01 | Old |
| 0.586719 | 0.343069 | 0.09 | 0.11 | 1 | 9 | Atxn7l3 | -0.02 | Old |
| 0.586855 | 1.168657 | 0.031 | 0.022 | 1 | 9 | Donson | 0.009 | Old |
| 0.586855 | 0.852984 | 0.031 | 0.022 | 1 | 9 | Noc3l | 0.009 | Old |
| 0.586855 | 0.833488 | 0.031 | 0.022 | 1 | 9 | Myl9 | 0.009 | Old |
| 0.586855 | 0.769534 | 0.031 | 0.022 | 1 | 9 | Mta3 | 0.009 | Old |
| 0.58696 | 0.346997 | 0.149 | 0.132 | 1 | 9 | Zc3h18 | 0.017 | Old |
| 0.587071 | 0.021494 | 0.046 | 0.059 | 1 | 9 | Schip1 | -0.013 | Old |
| 0.587162 | 0.18609 | 0.265 | 0.25 | 1 | 9 | Atp5o.1 | 0.015 | Old |
| 0.587182 | 0.105398 | 0.057 | 0.044 | 1 | 9 | Agrn | 0.013 | Old |
| 0.587967 | 0.845088 | 0.08 | 0.066 | 1 | 9 | Irf2bp1 | 0.014 | Old |
| 0.5881 | 0.063526 | 0.034 | 0.044 | 1 | 9 | Dnajc25 | -0.01 | Old |
| 0.588225 | 0.234299 | 0.168 | 0.147 | 1 | 9 | Atad2b | 0.021 | Old |

| | | | | | | | | |
|---|---|---|---|---|---|---|---|---|
| 0.588425 | 1.162361 | 0.031 | 0.022 | 1 | 9 | Amigo1 | 0.009 | Old |
| 0.588425 | 0.915196 | 0.031 | 0.022 | 1 | 9 | A430105I1 | 0.009 | Old |
| 0.588425 | 0.903226 | 0.031 | 0.022 | 1 | 9 | P2rx7 | 0.009 | Old |
| 0.588425 | 0.880352 | 0.031 | 0.022 | 1 | 9 | 3110002H1 | 0.009 | Old |
| 0.588425 | 0.278688 | 0.031 | 0.022 | 1 | 9 | Hsd17b7 | 0.009 | Old |
| 0.588655 | 0.188117 | 0.173 | 0.154 | 1 | 9 | Lrrfip1 | 0.019 | Old |
| 0.588873 | 0.062714 | 0.155 | 0.176 | 1 | 9 | Dcun1d5 | -0.021 | Old |
| 0.589166 | 0.13255 | 0.5 | 0.471 | 1 | 9 | Galnt15 | 0.029 | Old |
| 0.589172 | 0.815098 | 0.108 | 0.096 | 1 | 9 | Snd1 | 0.012 | Old |
| 0.589617 | 0.328842 | 0.216 | 0.199 | 1 | 9 | Fam76a | 0.017 | Old |
| 0.589671 | 0.207821 | 0.369 | 0.353 | 1 | 9 | Ankrd11 | 0.016 | Old |
| 0.589766 | 0.467436 | 0.064 | 0.051 | 1 | 9 | Leo1 | 0.013 | Old |
| 0.589766 | 0.152807 | 0.064 | 0.051 | 1 | 9 | Orc6 | 0.013 | Old |
| 0.589996 | 0.728048 | 0.031 | 0.022 | 1 | 9 | Izumo4 | 0.009 | Old |
| 0.589996 | 0.604994 | 0.031 | 0.022 | 1 | 9 | Gmds | 0.009 | Old |
| 0.590273 | 0.412462 | 0.088 | 0.074 | 1 | 9 | Sec24a | 0.014 | Old |
| 0.590405 | 0.217394 | 0.191 | 0.169 | 1 | 9 | Swt1 | 0.022 | Old |
| 0.5907 | 0.016628 | 0.168 | 0.147 | 1 | 9 | Parp9 | 0.021 | Old |
| 0.590904 | 0.272676 | 0.034 | 0.044 | 1 | 9 | Atm | -0.01 | Old |
| 0.590904 | 0.033759 | 0.034 | 0.044 | 1 | 9 | Foxk2 | -0.01 | Old |
| 0.591188 | 1.038551 | 0.093 | 0.081 | 1 | 9 | Nkap | 0.012 | Old |
| 0.5915 | 0.257562 | 0.142 | 0.125 | 1 | 9 | Fam193b | 0.017 | Old |
| 0.59155 | 0.023624 | 0.072 | 0.059 | 1 | 9 | Fbxw8 | 0.013 | Old |
| 0.591569 | 0.720931 | 0.031 | 0.022 | 1 | 9 | Polr1b | 0.009 | Old |
| 0.591569 | 0.49941 | 0.031 | 0.022 | 1 | 9 | Ttll3 | 0.009 | Old |
| 0.591956 | 0.550167 | 0.064 | 0.051 | 1 | 9 | Tada3 | 0.013 | Old |
| 0.592729 | 0.437333 | 0.162 | 0.147 | 1 | 9 | Vbp1 | 0.015 | Old |
| 0.592748 | 0.105212 | 0.214 | 0.191 | 1 | 9 | Chic2 | 0.023 | Old |
| 0.592824 | 0.057284 | 0.995 | 1 | 1 | 9 | mt-Co1 | -0.005 | Old |
| 0.593143 | 0.846107 | 0.137 | 0.125 | 1 | 9 | Dnm1l | 0.012 | Old |
| 0.593144 | 0.857252 | 0.031 | 0.022 | 1 | 9 | Adsl | 0.009 | Old |
| 0.593144 | 0.538554 | 0.031 | 0.022 | 1 | 9 | Zfp954 | 0.009 | Old |
| 0.593713 | 0.508497 | 0.034 | 0.044 | 1 | 9 | Hax1 | -0.01 | Old |
| 0.593713 | 0.449925 | 0.034 | 0.044 | 1 | 9 | Cops7b | -0.01 | Old |
| 0.593942 | 0.263604 | 0.085 | 0.103 | 1 | 9 | Tmem80 | -0.018 | Old |
| 0.594246 | 0.105956 | 0.188 | 0.169 | 1 | 9 | Acadl | 0.019 | Old |
| 0.594316 | 0.320247 | 0.046 | 0.059 | 1 | 9 | Abhd13 | -0.013 | Old |
| 0.594316 | 0.185271 | 0.046 | 0.059 | 1 | 9 | Bclaf3 | -0.013 | Old |
| 0.594721 | 0.087618 | 0.031 | 0.022 | 1 | 9 | Mturn | 0.009 | Old |
| 0.595121 | 1.758236 | 0.039 | 0.029 | 1 | 9 | 2700097O( | 0.01 | Old |
| 0.595211 | 0.203508 | 0.258 | 0.235 | 1 | 9 | Kcmf1 | 0.023 | Old |
| 0.595288 | 0.152353 | 0.224 | 0.206 | 1 | 9 | Prkd3 | 0.018 | Old |
| 0.595375 | 0.396385 | 0.072 | 0.088 | 1 | 9 | Dnase2a | -0.016 | Old |
| 0.595856 | 1.367439 | 0.062 | 0.051 | 1 | 9 | C2cd5 | 0.011 | Old |
| 0.5963 | 0.840754 | 0.031 | 0.022 | 1 | 9 | Dgkq | 0.009 | Old |

| | | | | | | | | |
|---|---|---|---|---|---|---|---|---|
| 0.5963 | 0.649152 | 0.031 | 0.022 | 1 | 9 | Alad | 0.009 | Old |
| 0.5963 | 0.509645 | 0.031 | 0.022 | 1 | 9 | Sap30 | 0.009 | Old |
| 0.5963 | 0.50016 | 0.031 | 0.022 | 1 | 9 | Ptcd2 | 0.009 | Old |
| 0.596529 | 1.532884 | 0.039 | 0.029 | 1 | 9 | Slc16a11 | 0.01 | Old |
| 0.596529 | 1.307622 | 0.039 | 0.029 | 1 | 9 | Rab32 | 0.01 | Old |
| 0.596537 | 1.477953 | 0.046 | 0.037 | 1 | 9 | Rilpl1 | 0.009 | Old |
| 0.596661 | 0.110901 | 0.126 | 0.147 | 1 | 9 | Zfx | -0.021 | Old |
| 0.5971 | 2.738335 | 0.013 | 0.007 | 1 | 9 | Zfp438 | 0.006 | Old |
| 0.5971 | 2.681571 | 0.013 | 0.007 | 1 | 9 | Zfp213 | 0.006 | Old |
| 0.5971 | 2.399122 | 0.013 | 0.007 | 1 | 9 | Zfyve28 | 0.006 | Old |
| 0.5971 | 2.365884 | 0.013 | 0.007 | 1 | 9 | Clcf1 | 0.006 | Old |
| 0.5971 | 2.202444 | 0.013 | 0.007 | 1 | 9 | E030030I0( | 0.006 | Old |
| 0.5971 | 2.105205 | 0.013 | 0.007 | 1 | 9 | Zfp773 | 0.006 | Old |
| 0.5971 | 2.055901 | 0.013 | 0.007 | 1 | 9 | Tmem240 | 0.006 | Old |
| 0.5971 | 1.934972 | 0.013 | 0.007 | 1 | 9 | Zbp1 | 0.006 | Old |
| 0.5971 | 1.87997 | 0.013 | 0.007 | 1 | 9 | Umps | 0.006 | Old |
| 0.5971 | 1.754663 | 0.013 | 0.007 | 1 | 9 | Plcd3 | 0.006 | Old |
| 0.5971 | 1.753996 | 0.013 | 0.007 | 1 | 9 | Gdpd3 | 0.006 | Old |
| 0.5971 | 1.643824 | 0.013 | 0.007 | 1 | 9 | Mrm2 | 0.006 | Old |
| 0.5971 | 1.62894 | 0.013 | 0.007 | 1 | 9 | Mfsd3 | 0.006 | Old |
| 0.5971 | 1.494557 | 0.013 | 0.007 | 1 | 9 | Neto2 | 0.006 | Old |
| 0.5971 | 1.383702 | 0.013 | 0.007 | 1 | 9 | Tmlhe | 0.006 | Old |
| 0.5971 | 1.366019 | 0.013 | 0.007 | 1 | 9 | Poli | 0.006 | Old |
| 0.5971 | 1.264794 | 0.013 | 0.007 | 1 | 9 | Tedc2 | 0.006 | Old |
| 0.5971 | 0.997498 | 0.013 | 0.007 | 1 | 9 | A330069E1 | 0.006 | Old |
| 0.597776 | 0.195969 | 0.193 | 0.176 | 1 | 9 | Fkbp8 | 0.017 | Old |
| 0.597787 | 0.54876 | 0.072 | 0.059 | 1 | 9 | Fmo5 | 0.013 | Old |
| 0.597787 | 0.263014 | 0.072 | 0.059 | 1 | 9 | Tubgcp4 | 0.013 | Old |
| 0.597881 | 0.048541 | 0.031 | 0.022 | 1 | 9 | Tmem184c | 0.009 | Old |
| 0.59794 | 1.198225 | 0.039 | 0.029 | 1 | 9 | Per3 | 0.01 | Old |
| 0.59794 | 0.134192 | 0.034 | 0.044 | 1 | 9 | Zfp943 | -0.01 | Old |
| 0.597954 | 0.232951 | 0.046 | 0.059 | 1 | 9 | Stk38l | -0.013 | Old |
| 0.598546 | 1.049121 | 0.093 | 0.081 | 1 | 9 | Crtc3 | 0.012 | Old |
| 0.598868 | 0.115965 | 0.168 | 0.147 | 1 | 9 | Dbnl | 0.021 | Old |
| 0.599279 | 0.186395 | 0.095 | 0.081 | 1 | 9 | Igbp1 | 0.014 | Old |
| 0.599351 | 0.968276 | 0.039 | 0.029 | 1 | 9 | Smcr8 | 0.01 | Old |
| 0.599464 | 0.113352 | 0.031 | 0.022 | 1 | 9 | Cd209g | 0.009 | Old |
| 0.599517 | 0.026676 | 0.237 | 0.265 | 1 | 9 | Cpeb4 | -0.028 | Old |
| 0.59958 | 1.630798 | 0.013 | 0.007 | 1 | 9 | 9330020H0 | 0.006 | Old |
| 0.59958 | 1.447986 | 0.013 | 0.007 | 1 | 9 | Dnah2os | 0.006 | Old |
| 0.59958 | 1.361076 | 0.013 | 0.007 | 1 | 9 | Tmem62 | 0.006 | Old |
| 0.59958 | 1.225804 | 0.013 | 0.007 | 1 | 9 | Mrm1 | 0.006 | Old |
| 0.59958 | 1.114731 | 0.013 | 0.007 | 1 | 9 | Pex11g | 0.006 | Old |
| 0.59958 | 0.993273 | 0.013 | 0.007 | 1 | 9 | Taf1c | 0.006 | Old |
| 0.59958 | 0.92851 | 0.013 | 0.007 | 1 | 9 | Dhrs11 | 0.006 | Old |

| | | | | | | | | |
|---|---|---|---|---|---|---|---|---|
| 0.59958 | 0.8998 | 0.013 | 0.007 | 1 | 9 | BC017158 | 0.006 | Old |
| 0.59958 | 0.846342 | 0.013 | 0.007 | 1 | 9 | Nudt8.1 | 0.006 | Old |
| 0.59958 | 0.835368 | 0.013 | 0.007 | 1 | 9 | S100a14 | 0.006 | Old |
| 0.59958 | 0.817347 | 0.013 | 0.007 | 1 | 9 | Gm10762 | 0.006 | Old |
| 0.59958 | 0.730353 | 0.013 | 0.007 | 1 | 9 | 2900089D1 | 0.006 | Old |
| 0.59958 | 0.593355 | 0.013 | 0.007 | 1 | 9 | Afap1l2 | 0.006 | Old |
| 0.599644 | 0.432855 | 0.32 | 0.316 | 1 | 9 | Spaar | 0.004 | Old |
| 0.599787 | 0.22532 | 0.291 | 0.272 | 1 | 9 | Usp7 | 0.019 | Old |
| 0.600338 | 0.457252 | 0.072 | 0.088 | 1 | 9 | Wrn | -0.016 | Old |
| 0.600765 | 1.607653 | 0.039 | 0.029 | 1 | 9 | Tchp | 0.01 | Old |
| 0.600943 | 0.538478 | 0.126 | 0.11 | 1 | 9 | Sf3a3 | 0.016 | Old |
| 0.60118 | 0.15489 | 0.08 | 0.096 | 1 | 9 | Mrpl48 | -0.016 | Old |
| 0.601653 | 0.004103 | 0.206 | 0.184 | 1 | 9 | Yme1l1 | 0.022 | Old |
| 0.601686 | 0.26661 | 0.198 | 0.176 | 1 | 9 | Map4k2 | 0.022 | Old |
| 0.60183 | 0.304359 | 0.08 | 0.066 | 1 | 9 | Pbdc1 | 0.014 | Old |
| 0.601961 | 0.501371 | 0.072 | 0.059 | 1 | 9 | Vsig10 | 0.013 | Old |
| 0.602065 | 2.230415 | 0.013 | 0.007 | 1 | 9 | Zfp985 | 0.006 | Old |
| 0.602065 | 1.436164 | 0.013 | 0.007 | 1 | 9 | Cacna2d1 | 0.006 | Old |
| 0.602065 | 1.345086 | 0.013 | 0.007 | 1 | 9 | Gm42047 | 0.006 | Old |
| 0.602065 | 1.33542 | 0.013 | 0.007 | 1 | 9 | Ppp1r3b | 0.006 | Old |
| 0.602065 | 1.293964 | 0.013 | 0.007 | 1 | 9 | Alpk1 | 0.006 | Old |
| 0.602065 | 1.091255 | 0.013 | 0.007 | 1 | 9 | Usp2 | 0.006 | Old |
| 0.602065 | 1.077232 | 0.013 | 0.007 | 1 | 9 | Nptxr | 0.006 | Old |
| 0.602065 | 0.989946 | 0.013 | 0.007 | 1 | 9 | Kif22 | 0.006 | Old |
| 0.602065 | 0.899659 | 0.013 | 0.007 | 1 | 9 | Hoxa6 | 0.006 | Old |
| 0.602065 | 0.833549 | 0.013 | 0.007 | 1 | 9 | Stxbp4 | 0.006 | Old |
| 0.602065 | 0.556541 | 0.013 | 0.007 | 1 | 9 | Pcdhb19 | 0.006 | Old |
| 0.602065 | 0.46094 | 0.013 | 0.007 | 1 | 9 | Pgghg | 0.006 | Old |
| 0.602158 | 0.330785 | 0.059 | 0.074 | 1 | 9 | Sesn1 | -0.015 | Old |
| 0.602179 | 0.236004 | 0.039 | 0.029 | 1 | 9 | Oraov1 | 0.01 | Old |
| 0.602507 | 0.3483 | 0.119 | 0.103 | 1 | 9 | Sec23b | 0.016 | Old |
| 0.602804 | 0.316425 | 0.147 | 0.132 | 1 | 9 | Pikfyve | 0.015 | Old |
| 0.602962 | 0.094259 | 0.064 | 0.051 | 1 | 9 | Chmp6 | 0.013 | Old |
| 0.603095 | 0.207279 | 0.34 | 0.331 | 1 | 9 | Smarca5 | 0.009 | Old |
| 0.603361 | 0.089154 | 0.126 | 0.11 | 1 | 9 | Trnau1ap | 0.016 | Old |
| 0.603596 | 1.157277 | 0.039 | 0.029 | 1 | 9 | Eif2ak3 | 0.01 | Old |
| 0.603786 | 0.058844 | 0.17 | 0.154 | 1 | 9 | Ergic2 | 0.016 | Old |
| 0.604228 | 0.11281 | 0.183 | 0.162 | 1 | 9 | Socs4 | 0.021 | Old |
| 0.604481 | 0.175069 | 0.142 | 0.125 | 1 | 9 | Psmd3 | 0.017 | Old |
| 0.604555 | 1.067828 | 0.013 | 0.007 | 1 | 9 | Klra2 | 0.006 | Old |
| 0.604555 | 1.01219 | 0.013 | 0.007 | 1 | 9 | Fgf2 | 0.006 | Old |
| 0.604555 | 0.91637 | 0.013 | 0.007 | 1 | 9 | Fam83g | 0.006 | Old |
| 0.604555 | 0.90826 | 0.013 | 0.007 | 1 | 9 | Sdr39u1 | 0.006 | Old |
| 0.604555 | 0.868839 | 0.013 | 0.007 | 1 | 9 | Arhgap4 | 0.006 | Old |
| 0.604555 | 0.751111 | 0.013 | 0.007 | 1 | 9 | Trim14 | 0.006 | Old |

| | | | | | | | | |
|---|---|---|---|---|---|---|---|---|
| 0.604555 | 0.625895 | 0.013 | 0.007 | 1 | 9 | Mettl21a | 0.006 | Old |
| 0.604555 | 0.61309 | 0.013 | 0.007 | 1 | 9 | Gm17435 | 0.006 | Old |
| 0.604555 | 0.486349 | 0.013 | 0.007 | 1 | 9 | Bmpr1a | 0.006 | Old |
| 0.604938 | 0.228245 | 0.186 | 0.169 | 1 | 9 | Myh10 | 0.017 | Old |
| 0.605013 | 0.401476 | 0.039 | 0.029 | 1 | 9 | Cpt2 | 0.01 | Old |
| 0.605646 | 1.213353 | 0.07 | 0.059 | 1 | 9 | Sp2 | 0.011 | Old |
| 0.606098 | 0.192257 | 0.18 | 0.162 | 1 | 9 | Snx17 | 0.018 | Old |
| 0.606433 | 1.199423 | 0.039 | 0.029 | 1 | 9 | Dmac2 | 0.01 | Old |
| 0.607049 | 0.674558 | 0.013 | 0.007 | 1 | 9 | Pou3f1 | 0.006 | Old |
| 0.607049 | 0.663088 | 0.013 | 0.007 | 1 | 9 | Mcts2 | 0.006 | Old |
| 0.607049 | 0.621464 | 0.013 | 0.007 | 1 | 9 | Bst1 | 0.006 | Old |
| 0.607049 | 0.389259 | 0.013 | 0.007 | 1 | 9 | Ddx59 | 0.006 | Old |
| 0.607049 | 0.355358 | 0.013 | 0.007 | 1 | 9 | Lrsam1 | 0.006 | Old |
| 0.607049 | 0.329977 | 0.013 | 0.007 | 1 | 9 | 6430573F1 | 0.006 | Old |
| 0.607049 | 0.301015 | 0.013 | 0.007 | 1 | 9 | Hyal3 | 0.006 | Old |
| 0.607049 | 0.298406 | 0.013 | 0.007 | 1 | 9 | Cdca8 | 0.006 | Old |
| 0.607049 | 0.246433 | 0.013 | 0.007 | 1 | 9 | Cox6b2 | 0.006 | Old |
| 0.607049 | 0.218465 | 0.013 | 0.007 | 1 | 9 | Spred3 | 0.006 | Old |
| 0.607049 | 0.052436 | 0.013 | 0.007 | 1 | 9 | 6030408B1 | 0.006 | Old |
| 0.607759 | 0.261406 | 0.054 | 0.066 | 1 | 9 | Slc9a6 | -0.012 | Old |
| 0.607943 | 0.142123 | 0.186 | 0.206 | 1 | 9 | Cs | -0.02 | Old |
| 0.608078 | 0.166283 | 0.101 | 0.118 | 1 | 9 | Papss1 | -0.017 | Old |
| 0.608162 | 1.11795 | 0.046 | 0.037 | 1 | 9 | Itpk1 | 0.009 | Old |
| 0.608236 | 0.232432 | 0.322 | 0.294 | 1 | 9 | Rbx1 | 0.028 | Old |
| 0.608247 | 0.147317 | 0.072 | 0.059 | 1 | 9 | Exosc8 | 0.013 | Old |
| 0.608325 | 0.177168 | 0.119 | 0.103 | 1 | 9 | Tmtc3 | 0.016 | Old |
| 0.608486 | 0.146562 | 0.268 | 0.25 | 1 | 9 | 2610507B1 | 0.018 | Old |
| 0.608616 | 0.08551 | 0.119 | 0.14 | 1 | 9 | Ipo8 | -0.021 | Old |
| 0.608919 | 0.657176 | 0.165 | 0.147 | 1 | 9 | Odc1 | 0.018 | Old |
| 0.609 | 0.105736 | 0.031 | 0.022 | 1 | 9 | Dnmbp | 0.009 | Old |
| 0.609276 | 1.160452 | 0.039 | 0.029 | 1 | 9 | Zc3h6 | 0.01 | Old |
| 0.609276 | 0.019503 | 0.039 | 0.029 | 1 | 9 | Zfp777 | 0.01 | Old |
| 0.6093 | 0.059708 | 0.088 | 0.074 | 1 | 9 | Srp68 | 0.014 | Old |
| 0.609548 | 0.125235 | 0.013 | 0.007 | 1 | 9 | Slc9a9 | 0.006 | Old |
| 0.609548 | 0.109011 | 0.013 | 0.007 | 1 | 9 | Primpol | 0.006 | Old |
| 0.609548 | 0.072512 | 0.013 | 0.007 | 1 | 9 | Cdh4 | 0.006 | Old |
| 0.609826 | 0.078991 | 0.015 | 0.022 | 1 | 9 | Mthfsd | -0.007 | Old |
| 0.610505 | 0.142552 | 0.103 | 0.088 | 1 | 9 | Rit1 | 0.015 | Old |
| 0.610596 | 0.03723 | 0.031 | 0.022 | 1 | 9 | Gm42715 | 0.009 | Old |
| 0.610699 | 1.238912 | 0.039 | 0.029 | 1 | 9 | 4930555A( | 0.01 | Old |
| 0.610718 | 0.12473 | 0.126 | 0.147 | 1 | 9 | Ngdn | -0.021 | Old |
| 0.610826 | 0.009827 | 0.119 | 0.103 | 1 | 9 | Dennd1b | 0.016 | Old |
| 0.611097 | 0.049309 | 0.247 | 0.235 | 1 | 9 | Dynlrb1 | 0.012 | Old |
| 0.611394 | 0.094588 | 0.103 | 0.088 | 1 | 9 | Map3k20 | 0.015 | Old |
| 0.611404 | 0.52147 | 0.142 | 0.125 | 1 | 9 | Irgm2 | 0.017 | Old |

| | | | | | | | | |
|---|---|---|---|---|---|---|---|---|
| 0.611545 | 0.142851 | 0.101 | 0.118 | 1 | 9 | Slc35b1 | -0.017 | Old |
| 0.611753 | 0.112786 | 0.304 | 0.279 | 1 | 9 | Rapgef2 | 0.025 | Old |
| 0.611876 | 0.341301 | 0.015 | 0.022 | 1 | 9 | Sgsm3 | -0.007 | Old |
| 0.611876 | 0.175116 | 0.015 | 0.022 | 1 | 9 | Nudt7 | -0.007 | Old |
| 0.612059 | 0.990381 | 0.046 | 0.037 | 1 | 9 | Mcee | 0.009 | Old |
| 0.612124 | 0.883766 | 0.039 | 0.029 | 1 | 9 | Tmem255b | 0.01 | Old |
| 0.612124 | 0.311237 | 0.039 | 0.029 | 1 | 9 | Rfc3 | 0.01 | Old |
| 0.612177 | 0.14943 | 0.088 | 0.074 | 1 | 9 | Fubp3 | 0.014 | Old |
| 0.612453 | 0.58571 | 0.085 | 0.103 | 1 | 9 | Usp48 | -0.018 | Old |
| 0.612883 | 1.260698 | 0.054 | 0.044 | 1 | 9 | Sclt1 | 0.01 | Old |
| 0.613678 | 0.197829 | 0.289 | 0.272 | 1 | 9 | Larp4b | 0.017 | Old |
| 0.613929 | 0.416423 | 0.015 | 0.022 | 1 | 9 | Gata6 | -0.007 | Old |
| 0.61433 | 0.46364 | 0.072 | 0.088 | 1 | 9 | Tmem164 | -0.016 | Old |
| 0.614476 | 0.140774 | 0.232 | 0.213 | 1 | 9 | Anp32e | 0.019 | Old |
| 0.614563 | 0.208581 | 0.072 | 0.059 | 1 | 9 | Klhl7 | 0.013 | Old |
| 0.614871 | 0.075746 | 0.067 | 0.081 | 1 | 9 | Iah1 | -0.014 | Old |
| 0.614979 | 0.807227 | 0.039 | 0.029 | 1 | 9 | Cdc14b | 0.01 | Old |
| 0.614979 | 0.740428 | 0.039 | 0.029 | 1 | 9 | D930048N1 | 0.01 | Old |
| 0.614979 | 0.559155 | 0.039 | 0.029 | 1 | 9 | Prdm5 | 0.01 | Old |
| 0.615027 | 0.062981 | 0.43 | 0.426 | 1 | 9 | Nucks1 | 0.004 | Old |
| 0.615097 | 0.230446 | 0.322 | 0.301 | 1 | 9 | Sbf2 | 0.021 | Old |
| 0.615464 | 0.018544 | 0.198 | 0.221 | 1 | 9 | Ik | -0.023 | Old |
| 0.615589 | 0.072441 | 0.265 | 0.243 | 1 | 9 | Hnrnph2 | 0.022 | Old |
| 0.615906 | 0.245303 | 0.155 | 0.14 | 1 | 9 | Acaa2 | 0.015 | Old |
| 0.615985 | 0.381714 | 0.015 | 0.022 | 1 | 9 | Ltb4r1 | -0.007 | Old |
| 0.615985 | 0.170699 | 0.015 | 0.022 | 1 | 9 | Fam109a | -0.007 | Old |
| 0.616185 | 0.831216 | 0.085 | 0.074 | 1 | 9 | Ppp4r1 | 0.011 | Old |
| 0.616185 | 0.470093 | 0.085 | 0.074 | 1 | 9 | Unc45a | 0.011 | Old |
| 0.616186 | 0.929372 | 0.093 | 0.081 | 1 | 9 | Zfp110 | 0.012 | Old |
| 0.617158 | 0.592102 | 0.165 | 0.154 | 1 | 9 | Nbr1 | 0.011 | Old |
| 0.617167 | 0.162362 | 0.134 | 0.118 | 1 | 9 | Spns2 | 0.016 | Old |
| 0.617273 | 0.814067 | 0.046 | 0.037 | 1 | 9 | D230025D1 | 0.009 | Old |
| 0.617373 | 0.139861 | 0.155 | 0.132 | 1 | 9 | Malt1 | 0.023 | Old |
| 0.61784 | 0.59659 | 0.039 | 0.029 | 1 | 9 | Blzf1 | 0.01 | Old |
| 0.61784 | 0.316203 | 0.039 | 0.029 | 1 | 9 | Atf5 | 0.01 | Old |
| 0.618044 | 0.813332 | 0.015 | 0.022 | 1 | 9 | Ccdc130 | -0.007 | Old |
| 0.618044 | 0.804646 | 0.015 | 0.022 | 1 | 9 | Glmn | -0.007 | Old |
| 0.618044 | 0.468593 | 0.015 | 0.022 | 1 | 9 | Dna2 | -0.007 | Old |
| 0.618044 | 0.162051 | 0.015 | 0.022 | 1 | 9 | Afg1l | -0.007 | Old |
| 0.618122 | 0.010745 | 0.296 | 0.272 | 1 | 9 | Igf2r | 0.024 | Old |
| 0.618161 | 0.296509 | 0.101 | 0.088 | 1 | 9 | Csk | 0.013 | Old |
| 0.618992 | 0.522984 | 0.085 | 0.103 | 1 | 9 | Eif1ad | -0.018 | Old |
| 0.61924 | 0.320745 | 0.054 | 0.066 | 1 | 9 | Mtmr1 | -0.012 | Old |
| 0.619504 | 0.370766 | 0.216 | 0.199 | 1 | 9 | Snrpf | 0.017 | Old |
| 0.619536 | 0.195905 | 0.255 | 0.228 | 1 | 9 | Klc1 | 0.027 | Old |

| | | | | | | | | |
|---|---|---|---|---|---|---|---|---|
| 0.620009 | 0.023606 | 0.557 | 0.551 | 1 | 9 | Slc25a3 | 0.006 | Old |
| 0.620106 | 0.592627 | 0.015 | 0.022 | 1 | 9 | Gas2l3 | -0.007 | Old |
| 0.620106 | 0.542777 | 0.015 | 0.022 | 1 | 9 | Prom1 | -0.007 | Old |
| 0.620106 | 0.172276 | 0.015 | 0.022 | 1 | 9 | Cd82 | -0.007 | Old |
| 0.620138 | 0.80775 | 0.054 | 0.044 | 1 | 9 | Psmg2 | 0.01 | Old |
| 0.620236 | 0.293229 | 0.147 | 0.132 | 1 | 9 | Ttc7b | 0.015 | Old |
| 0.62031 | 0.058431 | 0.26 | 0.235 | 1 | 9 | H2-T22 | 0.025 | Old |
| 0.620408 | 0.048009 | 0.126 | 0.11 | 1 | 9 | Hsd17b4 | 0.016 | Old |
| 0.620706 | 0.328688 | 0.039 | 0.029 | 1 | 9 | Mettl5 | 0.01 | Old |
| 0.621145 | 0.108491 | 0.201 | 0.176 | 1 | 9 | Sorbs2 | 0.025 | Old |
| 0.621196 | 0.866033 | 0.046 | 0.037 | 1 | 9 | Abraxas2 | 0.009 | Old |
| 0.621215 | 0.158357 | 0.103 | 0.088 | 1 | 9 | Cope | 0.015 | Old |
| 0.62133 | 0.145831 | 0.075 | 0.088 | 1 | 9 | Gtpbp6 | -0.013 | Old |
| 0.621762 | 0.478097 | 0.147 | 0.132 | 1 | 9 | Acsl4 | 0.015 | Old |
| 0.622171 | 0.931816 | 0.015 | 0.022 | 1 | 9 | Zfp12 | -0.007 | Old |
| 0.622171 | 0.224627 | 0.015 | 0.022 | 1 | 9 | Rem1 | -0.007 | Old |
| 0.622506 | 0.867457 | 0.046 | 0.037 | 1 | 9 | Cox19 | 0.009 | Old |
| 0.622612 | 0.23986 | 0.111 | 0.096 | 1 | 9 | Gars | 0.015 | Old |
| 0.623578 | 0.280723 | 0.039 | 0.029 | 1 | 9 | Mettl6 | 0.01 | Old |
| 0.623746 | 0.097839 | 0.088 | 0.074 | 1 | 9 | Elp5 | 0.014 | Old |
| 0.62378 | 0.992033 | 0.054 | 0.044 | 1 | 9 | Tmem170b | 0.01 | Old |
| 0.623981 | 0.219073 | 0.028 | 0.037 | 1 | 9 | 2510046G1 | -0.009 | Old |
| 0.624121 | 0.187589 | 0.193 | 0.176 | 1 | 9 | Syngr2 | 0.017 | Old |
| 0.624239 | 1.422007 | 0.015 | 0.022 | 1 | 9 | Irf4 | -0.007 | Old |
| 0.624239 | 0.785084 | 0.015 | 0.022 | 1 | 9 | Zfp870 | -0.007 | Old |
| 0.624239 | 0.763013 | 0.015 | 0.022 | 1 | 9 | Gm34084 | -0.007 | Old |
| 0.624239 | 0.278766 | 0.015 | 0.022 | 1 | 9 | Pcgf6 | -0.007 | Old |
| 0.625129 | 0.351482 | 0.046 | 0.037 | 1 | 9 | Pelp1 | 0.009 | Old |
| 0.625714 | 0.160748 | 0.227 | 0.206 | 1 | 9 | Usp4 | 0.021 | Old |
| 0.625927 | 0.090866 | 0.119 | 0.103 | 1 | 9 | Sptlc1 | 0.016 | Old |
| 0.625949 | 0.895162 | 0.085 | 0.074 | 1 | 9 | Ecd | 0.011 | Old |
| 0.62631 | 0.697865 | 0.015 | 0.022 | 1 | 9 | Lipt2 | -0.007 | Old |
| 0.62631 | 0.451593 | 0.015 | 0.022 | 1 | 9 | S1pr2 | -0.007 | Old |
| 0.626456 | 0.260222 | 0.039 | 0.029 | 1 | 9 | Mapk11 | 0.01 | Old |
| 0.626493 | 0.018515 | 0.193 | 0.213 | 1 | 9 | Sft2d2 | -0.02 | Old |
| 0.626913 | 0.328885 | 0.142 | 0.125 | 1 | 9 | Cnot10 | 0.017 | Old |
| 0.627109 | 0.194183 | 0.028 | 0.037 | 1 | 9 | Kdm1b | -0.009 | Old |
| 0.627251 | 0.234382 | 0.108 | 0.096 | 1 | 9 | Crem | 0.012 | Old |
| 0.627464 | 0.157072 | 0.062 | 0.051 | 1 | 9 | Atxn10 | 0.011 | Old |
| 0.627826 | 0.126973 | 0.111 | 0.096 | 1 | 9 | Hbs1l | 0.015 | Old |
| 0.627897 | 0.552883 | 0.039 | 0.029 | 1 | 9 | B3glct | 0.01 | Old |
| 0.628254 | 0.475959 | 0.263 | 0.257 | 1 | 9 | Rc3h2 | 0.006 | Old |
| 0.628334 | 0.069881 | 0.075 | 0.088 | 1 | 9 | Bicra | -0.013 | Old |
| 0.628384 | 1.155897 | 0.015 | 0.022 | 1 | 9 | Fastkd1 | -0.007 | Old |
| 0.628435 | 0.300757 | 0.224 | 0.206 | 1 | 9 | Ano6 | 0.018 | Old |

| | | | | | | | | |
|---|---|---|---|---|---|---|---|---|
| 0.628731 | 0.10388 | 0.451 | 0.434 | 1 | 9 | Mef2a | 0.017 | Old |
| 0.629074 | 0.871046 | 0.046 | 0.037 | 1 | 9 | Igip | 0.009 | Old |
| 0.62909 | 0.024927 | 0.412 | 0.375 | 1 | 9 | Fcgrt | 0.037 | Old |
| 0.629208 | 0.482387 | 0.07 | 0.059 | 1 | 9 | Mrps36 | 0.011 | Old |
| 0.62921 | 0.176158 | 0.113 | 0.132 | 1 | 9 | Ifi47 | -0.019 | Old |
| 0.62987 | 0.653701 | 0.054 | 0.044 | 1 | 9 | Ptpn23 | 0.01 | Old |
| 0.629913 | 0.309758 | 0.198 | 0.184 | 1 | 9 | Stard4 | 0.014 | Old |
| 0.630109 | 0.149719 | 0.155 | 0.14 | 1 | 9 | Jag1 | 0.015 | Old |
| 0.63016 | 0.62737 | 0.116 | 0.103 | 1 | 9 | Crebzf | 0.013 | Old |
| 0.630332 | 0.699097 | 0.129 | 0.118 | 1 | 9 | Ankrd13c | 0.011 | Old |
| 0.630342 | 0.379505 | 0.075 | 0.088 | 1 | 9 | Rpf1 | -0.013 | Old |
| 0.630391 | 0.488913 | 0.046 | 0.037 | 1 | 9 | Plagl2 | 0.009 | Old |
| 0.630391 | 0.114808 | 0.041 | 0.051 | 1 | 9 | Tex2 | -0.01 | Old |
| 0.630411 | 0.109888 | 0.162 | 0.184 | 1 | 9 | Fam32a | -0.022 | Old |
| 0.630504 | 0.004696 | 0.129 | 0.11 | 1 | 9 | Ldlrad3 | 0.019 | Old |
| 0.630677 | 0.062471 | 0.564 | 0.544 | 1 | 9 | Sppl2a | 0.02 | Old |
| 0.630821 | 0.042445 | 0.343 | 0.316 | 1 | 9 | Hnrnpdl | 0.027 | Old |
| 0.631026 | 0.062077 | 0.173 | 0.191 | 1 | 9 | Rbbp4 | -0.018 | Old |
| 0.631091 | 0.715459 | 0.054 | 0.044 | 1 | 9 | Zfp513 | 0.01 | Old |
| 0.631184 | 0.060611 | 0.095 | 0.081 | 1 | 9 | Sigirr | 0.014 | Old |
| 0.631607 | 0.257251 | 0.304 | 0.294 | 1 | 9 | Xdh | 0.01 | Old |
| 0.631709 | 0.364872 | 0.046 | 0.037 | 1 | 9 | Eif2ak4 | 0.009 | Old |
| 0.631842 | 0.028096 | 0.119 | 0.103 | 1 | 9 | Cyhr1 | 0.016 | Old |
| 0.632229 | 0.180494 | 0.039 | 0.029 | 1 | 9 | Xpnpep3 | 0.01 | Old |
| 0.632302 | 0.189536 | 0.139 | 0.125 | 1 | 9 | Ube2e3 | 0.014 | Old |
| 0.632313 | 0.439064 | 0.054 | 0.044 | 1 | 9 | Dcp1b | 0.01 | Old |
| 0.632435 | 0.609566 | 0.124 | 0.11 | 1 | 9 | Nr5a2 | 0.014 | Old |
| 0.632597 | 0.062499 | 0.129 | 0.147 | 1 | 9 | Dnmt3a | -0.018 | Old |
| 0.633029 | 0.527676 | 0.046 | 0.037 | 1 | 9 | Igsf8 | 0.009 | Old |
| 0.633029 | 0.451727 | 0.046 | 0.037 | 1 | 9 | L3mbtl3 | 0.009 | Old |
| 0.633029 | 0.307925 | 0.046 | 0.037 | 1 | 9 | Tmem67 | 0.009 | Old |
| 0.633386 | 0.008375 | 0.028 | 0.037 | 1 | 9 | Hmbs | -0.009 | Old |
| 0.63428 | 0.039676 | 0.66 | 0.647 | 1 | 9 | Srgn | 0.013 | Old |
| 0.63433 | 0.899175 | 0.062 | 0.051 | 1 | 9 | Smarcal1 | 0.011 | Old |
| 0.63476 | 0.804877 | 0.054 | 0.044 | 1 | 9 | Zhx2 | 0.01 | Old |
| 0.634959 | 0.53275 | 0.028 | 0.037 | 1 | 9 | Srm | -0.009 | Old |
| 0.63517 | 0.18508 | 0.247 | 0.235 | 1 | 9 | Ctcf | 0.012 | Old |
| 0.635449 | 0.30763 | 0.173 | 0.154 | 1 | 9 | Szrd1 | 0.019 | Old |
| 0.635554 | 0.801089 | 0.142 | 0.132 | 1 | 9 | Pank2 | 0.01 | Old |
| 0.636045 | 0.136393 | 0.33 | 0.309 | 1 | 9 | Acin1 | 0.021 | Old |
| 0.636052 | 0.303581 | 0.175 | 0.162 | 1 | 9 | Ccdc186 | 0.013 | Old |
| 0.636112 | 0.018347 | 0.781 | 0.706 | 1 | 9 | App | 0.075 | Old |
| 0.63638 | 0.110951 | 0.075 | 0.088 | 1 | 9 | Imp4 | -0.013 | Old |
| 0.636534 | 0.330728 | 0.028 | 0.037 | 1 | 9 | Rab3ip | -0.009 | Old |
| 0.636534 | 0.054633 | 0.028 | 0.037 | 1 | 9 | Sharpin | -0.009 | Old |

| | | | | | | | | |
|---|---|---|---|---|---|---|---|---|
| 0.636708 | 0.032885 | 0.76 | 0.75 | 1 | 9 | Afdn | 0.01 | Old |
| 0.636891 | 0.071462 | 0.173 | 0.154 | 1 | 9 | Pgrmc2 | 0.019 | Old |
| 0.637211 | 0.368411 | 0.054 | 0.044 | 1 | 9 | Josd2 | 0.01 | Old |
| 0.637687 | 0.048846 | 0.16 | 0.176 | 1 | 9 | Pwwp2a | -0.016 | Old |
| 0.637884 | 0.018878 | 0.095 | 0.11 | 1 | 9 | Edem3 | -0.015 | Old |
| 0.638111 | 0.224065 | 0.028 | 0.037 | 1 | 9 | Acy3 | -0.009 | Old |
| 0.638192 | 0.103786 | 0.497 | 0.544 | 1 | 9 | Cdh13 | -0.047 | Old |
| 0.638959 | 0.822667 | 0.07 | 0.059 | 1 | 9 | Pofut2 | 0.011 | Old |
| 0.639264 | 0.105022 | 0.137 | 0.118 | 1 | 9 | Gsr | 0.019 | Old |
| 0.639409 | 0.253798 | 0.075 | 0.088 | 1 | 9 | Galnt11 | -0.013 | Old |
| 0.639689 | 0.304362 | 0.028 | 0.037 | 1 | 9 | Pno1 | -0.009 | Old |
| 0.639689 | 0.043629 | 0.028 | 0.037 | 1 | 9 | Bri3bp | -0.009 | Old |
| 0.639729 | 0.714121 | 0.085 | 0.074 | 1 | 9 | Cdk8 | 0.011 | Old |
| 0.639871 | 0.56635 | 0.077 | 0.066 | 1 | 9 | Enox2 | 0.011 | Old |
| 0.640075 | 0.723289 | 0.062 | 0.051 | 1 | 9 | Irak3 | 0.011 | Old |
| 0.640221 | 0.022316 | 0.121 | 0.103 | 1 | 9 | Dcaf1 | 0.018 | Old |
| 0.640888 | 0.199715 | 0.299 | 0.279 | 1 | 9 | Tmco1 | 0.02 | Old |
| 0.640895 | 0.633745 | 0.054 | 0.044 | 1 | 9 | Eps8l2 | 0.01 | Old |
| 0.64093 | 0.227749 | 0.039 | 0.029 | 1 | 9 | Tmem11 | 0.01 | Old |
| 0.640944 | 0.368814 | 0.111 | 0.096 | 1 | 9 | Hypk | 0.015 | Old |
| 0.64097 | 0.196501 | 0.041 | 0.051 | 1 | 9 | Polr1a | -0.01 | Old |
| 0.64097 | 0.14202 | 0.165 | 0.184 | 1 | 9 | Fbxo3 | -0.019 | Old |
| 0.641068 | 0.100805 | 0.247 | 0.228 | 1 | 9 | Rpl13a | 0.019 | Old |
| 0.641227 | 0.654614 | 0.062 | 0.051 | 1 | 9 | Mfsd10 | 0.011 | Old |
| 0.641269 | 0.015136 | 0.028 | 0.037 | 1 | 9 | Rrs1 | -0.009 | Old |
| 0.641708 | 0.607903 | 0.085 | 0.074 | 1 | 9 | Ptgs1 | 0.011 | Old |
| 0.641779 | 0.524712 | 0.139 | 0.125 | 1 | 9 | Stim1 | 0.014 | Old |
| 0.642138 | 0.417202 | 0.116 | 0.103 | 1 | 9 | Mcur1 | 0.013 | Old |
| 0.642421 | 0.095159 | 0.095 | 0.11 | 1 | 9 | Ptpn2 | -0.015 | Old |
| 0.642523 | 0.012005 | 0.147 | 0.132 | 1 | 9 | Strn4 | 0.015 | Old |
| 0.642977 | 0.534337 | 0.077 | 0.066 | 1 | 9 | Cep192 | 0.011 | Old |
| 0.64354 | 0.144555 | 0.093 | 0.081 | 1 | 9 | Dhrs1 | 0.012 | Old |
| 0.643627 | 0.912059 | 0.041 | 0.051 | 1 | 9 | BC037034 | -0.01 | Old |
| 0.643627 | 0.373243 | 0.041 | 0.051 | 1 | 9 | Tmcc2 | -0.01 | Old |
| 0.643979 | 0.118672 | 0.196 | 0.176 | 1 | 9 | U2af1 | 0.02 | Old |
| 0.644588 | 0.603445 | 0.054 | 0.044 | 1 | 9 | Taf12 | 0.01 | Old |
| 0.644805 | 0.031404 | 0.088 | 0.103 | 1 | 9 | Mfn1 | -0.015 | Old |
| 0.644872 | 0.006941 | 0.062 | 0.074 | 1 | 9 | D10Jhu81e | -0.012 | Old |
| 0.645214 | 0.178011 | 0.134 | 0.118 | 1 | 9 | Cwf19l2 | 0.016 | Old |
| 0.646287 | 0.028698 | 0.046 | 0.037 | 1 | 9 | Sephs1 | 0.009 | Old |
| 0.646469 | 0.49007 | 0.067 | 0.081 | 1 | 9 | Npm3 | -0.014 | Old |
| 0.646811 | 0.122877 | 0.222 | 0.206 | 1 | 9 | Ppp1r10 | 0.016 | Old |
| 0.646999 | 0.009168 | 0.062 | 0.051 | 1 | 9 | CAAA0111[illegible] | 0.011 | Old |
| 0.647084 | 0.13243 | 0.348 | 0.316 | 1 | 9 | Kdm2a | 0.032 | Old |
| 0.647508 | 0.006142 | 0.397 | 0.368 | 1 | 9 | Cox7a2 | 0.029 | Old |

| | | | | | | | | |
|---|---|---|---|---|---|---|---|---|
| 0.647751 | 0.297965 | 0.173 | 0.154 | 1 | 9 | Atp11b | 0.019 | Old |
| 0.647838 | 0.066119 | 0.32 | 0.294 | 1 | 9 | Adora2a | 0.026 | Old |
| 0.648146 | 0.129281 | 0.088 | 0.074 | 1 | 9 | Pml | 0.014 | Old |
| 0.648226 | 0.518792 | 0.101 | 0.088 | 1 | 9 | Hpf1 | 0.013 | Old |
| 0.64829 | 0.389578 | 0.054 | 0.044 | 1 | 9 | Mphosph6 | 0.01 | Old |
| 0.648655 | 0.601453 | 0.085 | 0.074 | 1 | 9 | Pogz | 0.011 | Old |
| 0.648655 | 0.25207 | 0.085 | 0.074 | 1 | 9 | Pik3c3 | 0.011 | Old |
| 0.649024 | 0.417339 | 0.168 | 0.154 | 1 | 9 | Ncstn | 0.014 | Old |
| 0.649313 | 0.543397 | 0.062 | 0.051 | 1 | 9 | 2810004N2 | 0.011 | Old |
| 0.64965 | 0.524228 | 0.085 | 0.074 | 1 | 9 | Supt4a | 0.011 | Old |
| 0.649744 | 0.346006 | 0.175 | 0.199 | 1 | 9 | Rp9 | -0.024 | Old |
| 0.649792 | 2.177868 | 0.021 | 0.015 | 1 | 9 | Dnajc12 | 0.006 | Old |
| 0.650287 | 0.275291 | 0.041 | 0.051 | 1 | 9 | Pigu | -0.01 | Old |
| 0.650287 | 0.093641 | 0.046 | 0.037 | 1 | 9 | Zfp617 | 0.009 | Old |
| 0.650472 | 0.289872 | 0.062 | 0.051 | 1 | 9 | Avpi1 | 0.011 | Old |
| 0.650989 | 0.636966 | 0.093 | 0.11 | 1 | 9 | Elmo2 | -0.017 | Old |
| 0.651124 | 0.015798 | 0.25 | 0.228 | 1 | 9 | Syne1 | 0.022 | Old |
| 0.651631 | 0.522769 | 0.062 | 0.051 | 1 | 9 | Ttc4 | 0.011 | Old |
| 0.651792 | 2.027722 | 0.021 | 0.015 | 1 | 9 | Zdhhc12 | 0.006 | Old |
| 0.651792 | 1.950883 | 0.021 | 0.015 | 1 | 9 | Fam72a | 0.006 | Old |
| 0.651792 | 1.77592 | 0.021 | 0.015 | 1 | 9 | Adnp | 0.006 | Old |
| 0.651809 | 0.498758 | 0.173 | 0.162 | 1 | 9 | Ahcyl1 | 0.011 | Old |
| 0.651875 | 0.837765 | 0.054 | 0.066 | 1 | 9 | Lrp1 | -0.012 | Old |
| 0.651894 | 0.49016 | 0.178 | 0.169 | 1 | 9 | Rcsd1 | 0.009 | Old |
| 0.652467 | 0.31116 | 0.124 | 0.11 | 1 | 9 | Cdyl | 0.014 | Old |
| 0.652479 | 0.134507 | 0.222 | 0.243 | 1 | 9 | Setx | -0.021 | Old |
| 0.652668 | 0.284987 | 0.286 | 0.272 | 1 | 9 | She | 0.014 | Old |
| 0.652952 | 0.059578 | 0.482 | 0.463 | 1 | 9 | 2900026A0 | 0.019 | Old |
| 0.65296 | 0.074341 | 0.046 | 0.037 | 1 | 9 | Cacfd1 | 0.009 | Old |
| 0.65296 | 0.028411 | 0.046 | 0.037 | 1 | 9 | Tfrc | 0.009 | Old |
| 0.653376 | 0.385706 | 0.077 | 0.066 | 1 | 9 | Cherp | 0.011 | Old |
| 0.653694 | 0.209828 | 0.131 | 0.118 | 1 | 9 | Mafk | 0.013 | Old |
| 0.653794 | 2.186057 | 0.021 | 0.015 | 1 | 9 | Acp6 | 0.006 | Old |
| 0.653794 | 1.562025 | 0.021 | 0.015 | 1 | 9 | Nod2 | 0.006 | Old |
| 0.653794 | 1.355802 | 0.021 | 0.015 | 1 | 9 | Commd9 | 0.006 | Old |
| 0.653794 | 1.35186 | 0.021 | 0.015 | 1 | 9 | Bhlhb9 | 0.006 | Old |
| 0.653794 | 1.056862 | 0.021 | 0.015 | 1 | 9 | Lins1 | 0.006 | Old |
| 0.653794 | 0.936622 | 0.021 | 0.015 | 1 | 9 | Pdcd2 | 0.006 | Old |
| 0.653794 | 0.886286 | 0.021 | 0.015 | 1 | 9 | Ercc2 | 0.006 | Old |
| 0.654148 | 0.194417 | 0.124 | 0.11 | 1 | 9 | Dcp2 | 0.014 | Old |
| 0.654251 | 0.609814 | 0.07 | 0.059 | 1 | 9 | Aktip | 0.011 | Old |
| 0.654478 | 0.464131 | 0.054 | 0.044 | 1 | 9 | Snip1 | 0.01 | Old |
| 0.654552 | 0.044659 | 0.709 | 0.691 | 1 | 9 | Klf7 | 0.018 | Old |
| 0.654606 | 0.032335 | 0.778 | 0.706 | 1 | 9 | Rpl27a | 0.072 | Old |
| 0.654681 | 0.460969 | 0.101 | 0.088 | 1 | 9 | Trim3 | 0.013 | Old |

| | | | | | | | | |
|---|---|---|---|---|---|---|---|---|
| 0.655343 | 0.503089 | 0.137 | 0.125 | 1 | 9 | Map3k7 | 0.012 | Old |
| 0.655605 | 0.051134 | 0.101 | 0.088 | 1 | 9 | Slc26a2 | 0.013 | Old |
| 0.655632 | 0.587445 | 0.085 | 0.074 | 1 | 9 | Emilin1 | 0.011 | Old |
| 0.655719 | 0.665551 | 0.054 | 0.044 | 1 | 9 | Cnot11 | 0.01 | Old |
| 0.655719 | 0.321809 | 0.054 | 0.044 | 1 | 9 | Parp11 | 0.01 | Old |
| 0.656277 | 0.497815 | 0.062 | 0.051 | 1 | 9 | Zbtb2 | 0.011 | Old |
| 0.656484 | 0.024352 | 0.101 | 0.118 | 1 | 9 | Tmem258 | -0.017 | Old |
| 0.657043 | 0.165681 | 0.276 | 0.257 | 1 | 9 | Brk1 | 0.019 | Old |
| 0.657362 | 0.120217 | 0.268 | 0.257 | 1 | 9 | Noct | 0.011 | Old |
| 0.657441 | 0.569131 | 0.062 | 0.051 | 1 | 9 | Bgn | 0.011 | Old |
| 0.657807 | 1.563987 | 0.021 | 0.015 | 1 | 9 | Rbm47 | 0.006 | Old |
| 0.657807 | 1.410099 | 0.021 | 0.015 | 1 | 9 | Tubd1 | 0.006 | Old |
| 0.657807 | 1.102687 | 0.021 | 0.015 | 1 | 9 | Usp54 | 0.006 | Old |
| 0.658078 | 0.234191 | 0.446 | 0.434 | 1 | 9 | Jak1 | 0.012 | Old |
| 0.658647 | 0.417871 | 0.07 | 0.059 | 1 | 9 | Prkra | 0.011 | Old |
| 0.65872 | 0.309668 | 0.168 | 0.191 | 1 | 9 | Ccdc59 | -0.023 | Old |
| 0.65927 | 0.114453 | 0.168 | 0.147 | 1 | 9 | Tspan9 | 0.021 | Old |
| 0.659514 | 0.321363 | 0.227 | 0.213 | 1 | 9 | Pnpla2 | 0.014 | Old |
| 0.659632 | 0.457129 | 0.085 | 0.074 | 1 | 9 | Wiz | 0.011 | Old |
| 0.659817 | 1.458317 | 0.021 | 0.015 | 1 | 9 | AI597479 | 0.006 | Old |
| 0.659817 | 1.13416 | 0.021 | 0.015 | 1 | 9 | Siah1b | 0.006 | Old |
| 0.659817 | 0.808458 | 0.021 | 0.015 | 1 | 9 | Ccdc69 | 0.006 | Old |
| 0.659916 | 0.155798 | 0.268 | 0.25 | 1 | 9 | Sec31a | 0.018 | Old |
| 0.66028 | 0.370141 | 0.116 | 0.103 | 1 | 9 | Hmg20a | 0.013 | Old |
| 0.660533 | 0.029372 | 0.175 | 0.154 | 1 | 9 | Tmem176a | 0.021 | Old |
| 0.660536 | 0.19037 | 0.175 | 0.162 | 1 | 9 | Ankhd1 | 0.013 | Old |
| 0.660637 | 0.106238 | 0.418 | 0.397 | 1 | 9 | Sema4c | 0.021 | Old |
| 0.660846 | 0.155202 | 0.379 | 0.368 | 1 | 9 | Map1lc3b | 0.011 | Old |
| 0.661183 | 0.279829 | 0.168 | 0.154 | 1 | 9 | Myo1e | 0.014 | Old |
| 0.661829 | 1.232394 | 0.021 | 0.015 | 1 | 9 | Txnrd3 | 0.006 | Old |
| 0.661829 | 1.156889 | 0.021 | 0.015 | 1 | 9 | Nav2 | 0.006 | Old |
| 0.661829 | 1.017582 | 0.021 | 0.015 | 1 | 9 | Zbtb12 | 0.006 | Old |
| 0.661829 | 0.712078 | 0.021 | 0.015 | 1 | 9 | Nt5c2 | 0.006 | Old |
| 0.661829 | 0.665023 | 0.021 | 0.015 | 1 | 9 | Trim68 | 0.006 | Old |
| 0.662059 | 0.664158 | 0.098 | 0.088 | 1 | 9 | Phrf1 | 0.01 | Old |
| 0.662885 | 0.077203 | 0.235 | 0.213 | 1 | 9 | Lmbrd1 | 0.022 | Old |
| 0.662913 | 0.371717 | 0.183 | 0.169 | 1 | 9 | Cox7b | 0.014 | Old |
| 0.662996 | 0.895223 | 0.098 | 0.088 | 1 | 9 | Kars | 0.01 | Old |
| 0.663055 | 0.373613 | 0.07 | 0.059 | 1 | 9 | Arfgap1 | 0.011 | Old |
| 0.663183 | 0.162633 | 0.049 | 0.059 | 1 | 9 | Fam57a | -0.01 | Old |
| 0.663226 | 0.391755 | 0.062 | 0.074 | 1 | 9 | Scpep1 | -0.012 | Old |
| 0.663842 | 0.115187 | 0.077 | 0.066 | 1 | 9 | Zfp217 | 0.011 | Old |
| 0.663844 | 0.475555 | 0.021 | 0.015 | 1 | 9 | Foxd2 | 0.006 | Old |
| 0.664369 | 0.205193 | 0.131 | 0.118 | 1 | 9 | Pdap1 | 0.013 | Old |
| 0.665133 | 0.731638 | 0.106 | 0.096 | 1 | 9 | 9030624J0 | 0.01 | Old |

| | | | | | | | | |
|---|---|---|---|---|---|---|---|---|
| 0.66545 | 0.085436 | 0.108 | 0.096 | 1 | 9 | Smpd1 | 0.012 | Old |
| 0.665678 | 0.334716 | 0.054 | 0.044 | 1 | 9 | Zfp386 | 0.01 | Old |
| 0.665861 | 0.588795 | 0.021 | 0.015 | 1 | 9 | Pcbp3 | 0.006 | Old |
| 0.665861 | 0.561443 | 0.021 | 0.015 | 1 | 9 | Trmt61a | 0.006 | Old |
| 0.665861 | 0.546138 | 0.021 | 0.015 | 1 | 9 | F830016B0 | 0.006 | Old |
| 0.665861 | 0.437945 | 0.021 | 0.015 | 1 | 9 | Lig4 | 0.006 | Old |
| 0.665861 | 0.244008 | 0.021 | 0.015 | 1 | 9 | Nlgn2 | 0.006 | Old |
| 0.665861 | 0.222085 | 0.021 | 0.015 | 1 | 9 | Mical1 | 0.006 | Old |
| 0.665861 | 0.155814 | 0.021 | 0.015 | 1 | 9 | Pnpla7 | 0.006 | Old |
| 0.666032 | 0.326162 | 0.255 | 0.243 | 1 | 9 | Glyr1 | 0.012 | Old |
| 0.666074 | 0.4223 | 0.054 | 0.066 | 1 | 9 | Ciao1 | -0.012 | Old |
| 0.666297 | 0.414041 | 0.036 | 0.044 | 1 | 9 | Pcolce | -0.008 | Old |
| 0.666395 | 0.246597 | 0.209 | 0.199 | 1 | 9 | Ssu72 | 0.01 | Old |
| 0.666757 | 0.547093 | 0.183 | 0.176 | 1 | 9 | Smim11 | 0.007 | Old |
| 0.667251 | 0.214801 | 0.307 | 0.294 | 1 | 9 | Copa | 0.013 | Old |
| 0.667262 | 1.02919 | 0.054 | 0.066 | 1 | 9 | Sh3kbp1 | -0.012 | Old |
| 0.667469 | 0.287215 | 0.186 | 0.169 | 1 | 9 | Pdpk1 | 0.017 | Old |
| 0.667493 | 0.034978 | 0.147 | 0.132 | 1 | 9 | Mob1b | 0.015 | Old |
| 0.667759 | 0.060056 | 0.237 | 0.213 | 1 | 9 | Abcb1b | 0.024 | Old |
| 0.667881 | 0.612693 | 0.021 | 0.015 | 1 | 9 | Farp1 | 0.006 | Old |
| 0.667881 | 0.259192 | 0.021 | 0.015 | 1 | 9 | Ebf2 | 0.006 | Old |
| 0.667881 | 0.171462 | 0.021 | 0.015 | 1 | 9 | Pex26 | 0.006 | Old |
| 0.667881 | 0.045453 | 0.021 | 0.015 | 1 | 9 | Nhsl1 | 0.006 | Old |
| 0.668134 | 0.035569 | 0.155 | 0.14 | 1 | 9 | Tbc1d20 | 0.015 | Old |
| 0.668249 | 0.338966 | 0.201 | 0.191 | 1 | 9 | Ddost | 0.01 | Old |
| 0.66849 | 0.027574 | 0.178 | 0.162 | 1 | 9 | Mosmo | 0.016 | Old |
| 0.668506 | 0.045042 | 0.124 | 0.11 | 1 | 9 | Por | 0.014 | Old |
| 0.668582 | 0.208123 | 0.07 | 0.059 | 1 | 9 | Cbx4 | 0.011 | Old |
| 0.669122 | 0.895003 | 0.062 | 0.051 | 1 | 9 | Vcam1 | 0.011 | Old |
| 0.669354 | 0.178848 | 0.124 | 0.11 | 1 | 9 | Mtmr2 | 0.014 | Old |
| 0.66969 | 0.3358 | 0.07 | 0.059 | 1 | 9 | Cryzl1 | 0.011 | Old |
| 0.669882 | 0.036706 | 0.204 | 0.221 | 1 | 9 | Vapb | -0.017 | Old |
| 0.669903 | 0.904692 | 0.021 | 0.015 | 1 | 9 | D3Ertd751 | 0.006 | Old |
| 0.669903 | 0.800761 | 0.021 | 0.015 | 1 | 9 | Ermard | 0.006 | Old |
| 0.669903 | 0.383694 | 0.021 | 0.015 | 1 | 9 | Fahd1 | 0.006 | Old |
| 0.669903 | 0.10518 | 0.021 | 0.015 | 1 | 9 | Cep57l1 | 0.006 | Old |
| 0.669942 | 0.035918 | 0.168 | 0.191 | 1 | 9 | Ap1s2 | -0.023 | Old |
| 0.671494 | 1.159947 | 0.028 | 0.022 | 1 | 9 | Tbc1d32 | 0.006 | Old |
| 0.671928 | 0.196858 | 0.021 | 0.015 | 1 | 9 | Fbxl19 | 0.006 | Old |
| 0.671928 | 0.000341 | 0.021 | 0.015 | 1 | 9 | Col6a3 | 0.006 | Old |
| 0.671956 | 0.058718 | 0.268 | 0.25 | 1 | 9 | Mdh2 | 0.018 | Old |
| 0.67196 | 0.194754 | 0.18 | 0.169 | 1 | 9 | Ypel5 | 0.011 | Old |
| 0.6722 | 0.30323 | 0.062 | 0.074 | 1 | 9 | Acads | -0.012 | Old |
| 0.672642 | 0.147874 | 0.062 | 0.051 | 1 | 9 | Tufm | 0.011 | Old |
| 0.673519 | 0.105576 | 0.036 | 0.044 | 1 | 9 | Etaa1 | -0.008 | Old |

| | | | | | | | | |
|---|---|---|---|---|---|---|---|---|
| 0.67394 | 0.111056 | 0.477 | 0.434 | 1 | 9 | Celf1 | 0.043 | Old |
| 0.673955 | 0.548933 | 0.021 | 0.015 | 1 | 9 | Ccdc191 | 0.006 | Old |
| 0.673955 | 0.075368 | 0.021 | 0.015 | 1 | 9 | Gm20696 | 0.006 | Old |
| 0.674127 | 0.369799 | 0.07 | 0.059 | 1 | 9 | Egflam | 0.011 | Old |
| 0.674127 | 0.037043 | 0.07 | 0.059 | 1 | 9 | Paip2b | 0.011 | Old |
| 0.675065 | 0.212076 | 0.196 | 0.184 | 1 | 9 | Dnajb14 | 0.012 | Old |
| 0.675472 | 0.059126 | 0.456 | 0.441 | 1 | 9 | Atp5a1 | 0.015 | Old |
| 0.675577 | 1.144124 | 0.067 | 0.059 | 1 | 9 | Timm44 | 0.008 | Old |
| 0.675697 | 0.340092 | 0.049 | 0.059 | 1 | 9 | Gmip | -0.01 | Old |
| 0.676401 | 0.089742 | 0.271 | 0.294 | 1 | 9 | Trim30a | -0.023 | Old |
| 0.676489 | 0.207284 | 0.077 | 0.066 | 1 | 9 | Kti12 | 0.011 | Old |
| 0.676657 | 1.42702 | 0.028 | 0.022 | 1 | 9 | Zmym1 | 0.006 | Old |
| 0.676698 | 0.905248 | 0.147 | 0.14 | 1 | 9 | Plekhg2 | 0.007 | Old |
| 0.676951 | 0.039986 | 0.727 | 0.699 | 1 | 9 | Marcks | 0.028 | Old |
| 0.67787 | 0.259377 | 0.121 | 0.11 | 1 | 9 | Cdkn2aip | 0.011 | Old |
| 0.677942 | 0.036334 | 0.209 | 0.199 | 1 | 9 | Gbp5 | 0.01 | Old |
| 0.678285 | 1.316926 | 0.044 | 0.037 | 1 | 9 | Lactb2 | 0.007 | Old |
| 0.678381 | 1.786052 | 0.028 | 0.022 | 1 | 9 | Rnd2 | 0.006 | Old |
| 0.678381 | 1.287918 | 0.028 | 0.022 | 1 | 9 | Tnfrsf11a | 0.006 | Old |
| 0.678381 | 1.070233 | 0.028 | 0.022 | 1 | 9 | Slc35c1 | 0.006 | Old |
| 0.678381 | 1.023965 | 0.028 | 0.022 | 1 | 9 | 4430402I1 | 0.006 | Old |
| 0.678381 | 0.986555 | 0.028 | 0.022 | 1 | 9 | Cmtr1 | 0.006 | Old |
| 0.679142 | 0.11104 | 0.168 | 0.184 | 1 | 9 | Mta2 | -0.016 | Old |
| 0.679261 | 0.268865 | 0.131 | 0.118 | 1 | 9 | Lsm5 | 0.013 | Old |
| 0.679372 | 0.179681 | 0.211 | 0.199 | 1 | 9 | Cdk9 | 0.012 | Old |
| 0.679665 | 0.197753 | 0.077 | 0.066 | 1 | 9 | Lsm14b | 0.011 | Old |
| 0.679763 | 0.532996 | 0.075 | 0.066 | 1 | 9 | Armc10 | 0.009 | Old |
| 0.67997 | 0.10603 | 0.01 | 0.015 | 1 | 9 | Zfp11 | -0.005 | Old |
| 0.680813 | 0.480844 | 0.152 | 0.14 | 1 | 9 | Scamp2 | 0.012 | Old |
| 0.680924 | 0.126298 | 0.124 | 0.14 | 1 | 9 | Npepl1 | -0.016 | Old |
| 0.681095 | 0.457385 | 0.093 | 0.081 | 1 | 9 | Rnf103 | 0.012 | Old |
| 0.68122 | 0.595745 | 0.062 | 0.074 | 1 | 9 | Adam9 | -0.012 | Old |
| 0.681341 | 0.043483 | 0.165 | 0.147 | 1 | 9 | Mau2 | 0.018 | Old |
| 0.681691 | 0.813376 | 0.052 | 0.044 | 1 | 9 | Ccdc9 | 0.008 | Old |
| 0.681834 | 1.450842 | 0.028 | 0.022 | 1 | 9 | Nop2 | 0.006 | Old |
| 0.681834 | 1.192958 | 0.028 | 0.022 | 1 | 9 | Gfer | 0.006 | Old |
| 0.681834 | 1.043632 | 0.028 | 0.022 | 1 | 9 | Asb6 | 0.006 | Old |
| 0.681834 | 1.014688 | 0.028 | 0.022 | 1 | 9 | Snx33 | 0.006 | Old |
| 0.681882 | 0.109847 | 0.139 | 0.125 | 1 | 9 | Ddx18 | 0.014 | Old |
| 0.682131 | 0.022992 | 0.124 | 0.11 | 1 | 9 | Smg6 | 0.014 | Old |
| 0.68259 | 0.381022 | 0.01 | 0.015 | 1 | 9 | Knstrn | -0.005 | Old |
| 0.68259 | 0.078315 | 0.01 | 0.015 | 1 | 9 | Smpd3 | -0.005 | Old |
| 0.68259 | 0.030063 | 0.01 | 0.015 | 1 | 9 | Atrip | -0.005 | Old |
| 0.68259 | 0.024536 | 0.01 | 0.015 | 1 | 9 | Zfp711 | -0.005 | Old |
| 0.68259 | 0.018829 | 0.01 | 0.015 | 1 | 9 | B230307C | -0.005 | Old |

| | | | | | | | | |
|---|---|---|---|---|---|---|---|---|
| 0.683564 | 1.331252 | 0.028 | 0.022 | 1 | 9 | Tstd3 | 0.006 | Old |
| 0.683564 | 0.915192 | 0.028 | 0.022 | 1 | 9 | Gnpnat1 | 0.006 | Old |
| 0.683592 | 0.23508 | 0.155 | 0.14 | 1 | 9 | Mbd6 | 0.015 | Old |
| 0.683733 | 0.055293 | 0.289 | 0.265 | 1 | 9 | Atp2b4 | 0.024 | Old |
| 0.684099 | 0.095025 | 0.714 | 0.669 | 1 | 9 | Actn4 | 0.045 | Old |
| 0.684369 | 0.154818 | 0.147 | 0.162 | 1 | 9 | Sdccag3 | -0.015 | Old |
| 0.685087 | 0.178946 | 0.131 | 0.118 | 1 | 9 | Ttll7 | 0.013 | Old |
| 0.68515 | 0.020527 | 0.964 | 0.912 | 1 | 9 | Cavin2 | 0.052 | Old |
| 0.685212 | 0.694539 | 0.01 | 0.015 | 1 | 9 | Nepro | -0.005 | Old |
| 0.685212 | 0.277348 | 0.01 | 0.015 | 1 | 9 | Ccdc120 | -0.005 | Old |
| 0.685212 | 0.271777 | 0.01 | 0.015 | 1 | 9 | Hps1 | -0.005 | Old |
| 0.685212 | 0.210433 | 0.01 | 0.015 | 1 | 9 | Sirt6 | -0.005 | Old |
| 0.685212 | 0.160203 | 0.01 | 0.015 | 1 | 9 | Gm46224 | -0.005 | Old |
| 0.685212 | 0.130456 | 0.01 | 0.015 | 1 | 9 | Ddx11 | -0.005 | Old |
| 0.685212 | 0.09735 | 0.01 | 0.015 | 1 | 9 | Rhbdd3 | -0.005 | Old |
| 0.685212 | 0.021789 | 0.01 | 0.015 | 1 | 9 | Noa1 | -0.005 | Old |
| 0.685252 | 0.018243 | 0.528 | 0.5 | 1 | 9 | Apbb2 | 0.028 | Old |
| 0.685294 | 0.86993 | 0.028 | 0.022 | 1 | 9 | Hs3st3b1 | 0.006 | Old |
| 0.685779 | 0.032738 | 0.09 | 0.103 | 1 | 9 | Nsmce3 | -0.013 | Old |
| 0.68597 | 0.125494 | 0.093 | 0.081 | 1 | 9 | Rnft1 | 0.012 | Old |
| 0.686006 | 0.140725 | 0.082 | 0.096 | 1 | 9 | Dnajc11 | -0.014 | Old |
| 0.686596 | 0.494846 | 0.036 | 0.044 | 1 | 9 | Notch2 | -0.008 | Old |
| 0.686596 | 0.4351 | 0.036 | 0.044 | 1 | 9 | Mtx3 | -0.008 | Old |
| 0.686667 | 0.019876 | 0.144 | 0.162 | 1 | 9 | Birc2 | -0.018 | Old |
| 0.687027 | 0.938687 | 0.028 | 0.022 | 1 | 9 | Lrrc25 | 0.006 | Old |
| 0.687098 | 0.21703 | 0.077 | 0.066 | 1 | 9 | Slc4a7 | 0.011 | Old |
| 0.687163 | 0.077665 | 0.108 | 0.096 | 1 | 9 | B3gat3 | 0.012 | Old |
| 0.687164 | 0.070393 | 0.183 | 0.162 | 1 | 9 | 2700081O1 | 0.021 | Old |
| 0.687244 | 0.07182 | 0.25 | 0.235 | 1 | 9 | Krit1 | 0.015 | Old |
| 0.687287 | 0.169236 | 0.07 | 0.081 | 1 | 9 | Lztr1 | -0.011 | Old |
| 0.687596 | 1.207655 | 0.059 | 0.051 | 1 | 9 | Frmd8 | 0.008 | Old |
| 0.687839 | 1.251073 | 0.01 | 0.015 | 1 | 9 | Pck1 | -0.005 | Old |
| 0.687839 | 1.201824 | 0.01 | 0.015 | 1 | 9 | Purg | -0.005 | Old |
| 0.687839 | 0.631902 | 0.01 | 0.015 | 1 | 9 | Ttc27 | -0.005 | Old |
| 0.687839 | 0.509735 | 0.01 | 0.015 | 1 | 9 | D130040H1 | -0.005 | Old |
| 0.687839 | 0.490739 | 0.01 | 0.015 | 1 | 9 | Krt10 | -0.005 | Old |
| 0.687839 | 0.423856 | 0.01 | 0.015 | 1 | 9 | Hykk | -0.005 | Old |
| 0.687839 | 0.417167 | 0.01 | 0.015 | 1 | 9 | Hpgds | -0.005 | Old |
| 0.688015 | 0.486514 | 0.062 | 0.074 | 1 | 9 | Strbp | -0.012 | Old |
| 0.68805 | 0.107673 | 0.598 | 0.588 | 1 | 9 | Srrm2 | 0.01 | Old |
| 0.688277 | 0.237168 | 0.258 | 0.287 | 1 | 9 | Hivep2 | -0.029 | Old |
| 0.688737 | 0.304813 | 0.113 | 0.103 | 1 | 9 | Eva1b | 0.01 | Old |
| 0.688761 | 0.917617 | 0.028 | 0.022 | 1 | 9 | AW011738 | 0.006 | Old |
| 0.688761 | 0.901453 | 0.028 | 0.022 | 1 | 9 | Cd209d | 0.006 | Old |
| 0.688799 | 1.032445 | 0.059 | 0.051 | 1 | 9 | Frs2 | 0.008 | Old |

| | | | | | | | | |
|---|---|---|---|---|---|---|---|---|
| 0.688985 | 0.082676 | 0.108 | 0.096 | 1 | 9 | Agpat3 | 0.012 | Old |
| 0.68904 | 0.308073 | 0.129 | 0.147 | 1 | 9 | Spopl | -0.018 | Old |
| 0.689429 | 0.090936 | 0.291 | 0.279 | 1 | 9 | Cyth1 | 0.012 | Old |
| 0.689515 | 0.091281 | 0.036 | 0.044 | 1 | 9 | Slc35e3 | -0.008 | Old |
| 0.689614 | 0.499805 | 0.023 | 0.029 | 1 | 9 | Mettl13 | -0.006 | Old |
| 0.690003 | 1.050618 | 0.059 | 0.051 | 1 | 9 | R3hcc1l | 0.008 | Old |
| 0.690105 | 0.074771 | 0.34 | 0.316 | 1 | 9 | Ralb | 0.024 | Old |
| 0.690446 | 0.082703 | 0.147 | 0.132 | 1 | 9 | Ssfa2 | 0.015 | Old |
| 0.69047 | 2.331578 | 0.01 | 0.015 | 1 | 9 | Mki67 | -0.005 | Old |
| 0.69047 | 1.619379 | 0.01 | 0.015 | 1 | 9 | Fam120c | -0.005 | Old |
| 0.69047 | 1.203257 | 0.01 | 0.015 | 1 | 9 | Adhfe1 | -0.005 | Old |
| 0.69047 | 1.047615 | 0.01 | 0.015 | 1 | 9 | Gsap | -0.005 | Old |
| 0.69047 | 0.970434 | 0.01 | 0.015 | 1 | 9 | Tbc1d19 | -0.005 | Old |
| 0.69047 | 0.842062 | 0.01 | 0.015 | 1 | 9 | Dennd1c | -0.005 | Old |
| 0.69047 | 0.744502 | 0.01 | 0.015 | 1 | 9 | 1110019D1 | -0.005 | Old |
| 0.69047 | 0.656593 | 0.01 | 0.015 | 1 | 9 | Tmem42 | -0.005 | Old |
| 0.69047 | 0.124291 | 0.01 | 0.015 | 1 | 9 | Ppfibp2 | -0.005 | Old |
| 0.690497 | 0.890093 | 0.028 | 0.022 | 1 | 9 | Ppcdc | 0.006 | Old |
| 0.690497 | 0.731565 | 0.028 | 0.022 | 1 | 9 | Mccc1 | 0.006 | Old |
| 0.690497 | 0.713928 | 0.028 | 0.022 | 1 | 9 | Bicd1 | 0.006 | Old |
| 0.690497 | 0.614112 | 0.028 | 0.022 | 1 | 9 | Gnptab | 0.006 | Old |
| 0.690497 | 0.369053 | 0.028 | 0.022 | 1 | 9 | Cstf1 | 0.006 | Old |
| 0.690694 | 0.947774 | 0.052 | 0.044 | 1 | 9 | Tmem198b | 0.008 | Old |
| 0.691097 | 1.030193 | 0.036 | 0.029 | 1 | 9 | Arid3b | 0.007 | Old |
| 0.691414 | 0.338185 | 0.023 | 0.029 | 1 | 9 | Haus4 | -0.006 | Old |
| 0.691539 | 0.32107 | 0.129 | 0.118 | 1 | 9 | Mrpl23 | 0.011 | Old |
| 0.691991 | 1.050215 | 0.088 | 0.081 | 1 | 9 | Tm2d3 | 0.007 | Old |
| 0.692099 | 0.428434 | 0.049 | 0.059 | 1 | 9 | Commd4 | -0.01 | Old |
| 0.692099 | 0.336253 | 0.049 | 0.059 | 1 | 9 | Mrpl54 | -0.01 | Old |
| 0.692407 | 0.751249 | 0.103 | 0.096 | 1 | 9 | Mrpl4 | 0.007 | Old |
| 0.692439 | 0.235428 | 0.036 | 0.044 | 1 | 9 | Cox18 | -0.008 | Old |
| 0.693152 | 0.313111 | 0.103 | 0.118 | 1 | 9 | Tbc1d23 | -0.015 | Old |
| 0.693216 | 0.164546 | 0.023 | 0.029 | 1 | 9 | Efcab2 | -0.006 | Old |
| 0.693494 | 0.052967 | 0.077 | 0.066 | 1 | 9 | Brip1os | 0.011 | Old |
| 0.693508 | 0.053891 | 0.332 | 0.301 | 1 | 9 | Tcf12 | 0.031 | Old |
| 0.693548 | 0.232277 | 0.162 | 0.147 | 1 | 9 | Gtpbp4 | 0.015 | Old |
| 0.693697 | 0.329558 | 0.067 | 0.059 | 1 | 9 | Gdap10 | 0.008 | Old |
| 0.694176 | 1.223749 | 0.036 | 0.029 | 1 | 9 | Ofd1 | 0.007 | Old |
| 0.694415 | 0.025374 | 0.168 | 0.154 | 1 | 9 | Rnf14 | 0.014 | Old |
| 0.694636 | 0.73262 | 0.049 | 0.059 | 1 | 9 | Apln | -0.01 | Old |
| 0.69502 | 0.305937 | 0.023 | 0.029 | 1 | 9 | Ero1lb | -0.006 | Old |
| 0.69502 | 0.048336 | 0.023 | 0.029 | 1 | 9 | Mto1 | -0.006 | Old |
| 0.69513 | 0.343858 | 0.121 | 0.11 | 1 | 9 | Taf3 | 0.011 | Old |
| 0.695386 | 0.28963 | 0.116 | 0.103 | 1 | 9 | Usp46 | 0.013 | Old |
| 0.695714 | 0.842464 | 0.028 | 0.022 | 1 | 9 | Stat5a | 0.006 | Old |

| | | | | | | | | |
|---|---|---|---|---|---|---|---|---|
| 0.695714 | 0.53547 | 0.028 | 0.022 | 1 | 9 | Selp | 0.006 | Old |
| 0.695714 | 0.306023 | 0.028 | 0.022 | 1 | 9 | Camk2b | 0.006 | Old |
| 0.695859 | 1.007728 | 0.052 | 0.044 | 1 | 9 | Slc35c2 | 0.008 | Old |
| 0.695957 | 0.139265 | 0.124 | 0.14 | 1 | 9 | Dok4 | -0.016 | Old |
| 0.696101 | 0.575996 | 0.098 | 0.088 | 1 | 9 | Desi1 | 0.01 | Old |
| 0.696426 | 0.623151 | 0.214 | 0.206 | 1 | 9 | Slc25a4 | 0.008 | Old |
| 0.696826 | 0.581184 | 0.023 | 0.029 | 1 | 9 | Tspan32 | -0.006 | Old |
| 0.696826 | 0.053404 | 0.023 | 0.029 | 1 | 9 | Cd200r1 | -0.006 | Old |
| 0.696886 | 0.331499 | 0.09 | 0.081 | 1 | 9 | Stx2 | 0.009 | Old |
| 0.697111 | 0.07928 | 0.335 | 0.324 | 1 | 9 | Arid1a | 0.011 | Old |
| 0.697456 | 0.511451 | 0.028 | 0.022 | 1 | 9 | Zc3h11a | 0.006 | Old |
| 0.697456 | 0.337425 | 0.028 | 0.022 | 1 | 9 | Tango2 | 0.006 | Old |
| 0.697456 | 0.193985 | 0.028 | 0.022 | 1 | 9 | Cbwd1 | 0.006 | Old |
| 0.697456 | 0.171257 | 0.028 | 0.022 | 1 | 9 | Clcn5 | 0.006 | Old |
| 0.69777 | 0.065815 | 0.077 | 0.066 | 1 | 9 | Abcf2 | 0.011 | Old |
| 0.698089 | 0.205069 | 0.075 | 0.066 | 1 | 9 | Rnf166 | 0.009 | Old |
| 0.698099 | 0.53376 | 0.082 | 0.074 | 1 | 9 | Plekha3 | 0.008 | Old |
| 0.698255 | 0.829844 | 0.067 | 0.059 | 1 | 9 | Haus3 | 0.008 | Old |
| 0.6983 | 0.118026 | 0.036 | 0.044 | 1 | 9 | Brpf3 | -0.008 | Old |
| 0.698446 | 0.908115 | 0.052 | 0.044 | 1 | 9 | D430042O( | 0.008 | Old |
| 0.698491 | 0.223114 | 0.129 | 0.147 | 1 | 9 | Tmem47 | -0.018 | Old |
| 0.698617 | 0.62446 | 0.082 | 0.074 | 1 | 9 | Nab2 | 0.008 | Old |
| 0.698633 | 0.467045 | 0.023 | 0.029 | 1 | 9 | Sccpdh | -0.006 | Old |
| 0.698633 | 0.228977 | 0.023 | 0.029 | 1 | 9 | Arhgap27o | -0.006 | Old |
| 0.698633 | 0.217301 | 0.023 | 0.029 | 1 | 9 | Mcm7 | -0.006 | Old |
| 0.69893 | 0.07905 | 0.711 | 0.684 | 1 | 9 | Sh3glb1 | 0.027 | Old |
| 0.6992 | 0.158154 | 0.028 | 0.022 | 1 | 9 | Adi1 | 0.006 | Old |
| 0.699205 | 1.304594 | 0.044 | 0.037 | 1 | 9 | Rab23 | 0.007 | Old |
| 0.699205 | 1.162854 | 0.044 | 0.037 | 1 | 9 | Osgin2 | 0.007 | Old |
| 0.699287 | 0.111441 | 0.662 | 0.662 | 1 | 9 | Gnb1 | 0 | Old |
| 0.699288 | 0.002461 | 0.247 | 0.265 | 1 | 9 | Smg1 | -0.018 | Old |
| 0.699768 | 0.195754 | 0.036 | 0.044 | 1 | 9 | Ints10 | -0.008 | Old |
| 0.699903 | 0.379109 | 0.139 | 0.132 | 1 | 9 | Rfc1 | 0.007 | Old |
| 0.699939 | 0.136285 | 0.106 | 0.118 | 1 | 9 | Apmap | -0.012 | Old |
| 0.700154 | 0.165945 | 0.131 | 0.118 | 1 | 9 | Cdk14 | 0.013 | Old |
| 0.700154 | 0.142821 | 0.131 | 0.118 | 1 | 9 | Rubcn | 0.013 | Old |
| 0.700348 | 1.03603 | 0.036 | 0.029 | 1 | 9 | Cd302 | 0.007 | Old |
| 0.700348 | 0.849046 | 0.036 | 0.029 | 1 | 9 | Akr7a5 | 0.007 | Old |
| 0.70036 | 0.537051 | 0.09 | 0.081 | 1 | 9 | Ahdc1 | 0.009 | Old |
| 0.700442 | 0.503912 | 0.023 | 0.029 | 1 | 9 | Casp1 | -0.006 | Old |
| 0.700442 | 0.285663 | 0.023 | 0.029 | 1 | 9 | Wdr76 | -0.006 | Old |
| 0.700442 | 0.100126 | 0.023 | 0.029 | 1 | 9 | Smyd4 | -0.006 | Old |
| 0.700946 | 0.110152 | 0.028 | 0.022 | 1 | 9 | Ccdc138 | 0.006 | Old |
| 0.700946 | 0.016553 | 0.028 | 0.022 | 1 | 9 | Gpsm1 | 0.006 | Old |
| 0.700992 | 0.35494 | 0.049 | 0.059 | 1 | 9 | Prmt5 | -0.01 | Old |

| | | | | | | | | |
|---|---|---|---|---|---|---|---|---|
| 0.701037 | 0.840168 | 0.052 | 0.044 | 1 | 9 | Vegfc | 0.008 | Old |
| 0.70151 | 0.022004 | 0.101 | 0.088 | 1 | 9 | Atg4b | 0.013 | Old |
| 0.701894 | 0.529201 | 0.036 | 0.029 | 1 | 9 | Acrbp | 0.007 | Old |
| 0.701897 | 0.136609 | 0.183 | 0.169 | 1 | 9 | Snapc3 | 0.014 | Old |
| 0.702007 | 0.075528 | 0.302 | 0.287 | 1 | 9 | Atp13a3 | 0.015 | Old |
| 0.702013 | 0.661772 | 0.044 | 0.037 | 1 | 9 | Zcchc4 | 0.007 | Old |
| 0.702045 | 0.097705 | 0.265 | 0.243 | 1 | 9 | Tcea1 | 0.022 | Old |
| 0.702088 | 0.624782 | 0.059 | 0.051 | 1 | 9 | Ifi204 | 0.008 | Old |
| 0.702253 | 0.468658 | 0.023 | 0.029 | 1 | 9 | Dis3 | -0.006 | Old |
| 0.702253 | 0.32161 | 0.023 | 0.029 | 1 | 9 | Klhl8 | -0.006 | Old |
| 0.702693 | 0.041531 | 0.028 | 0.022 | 1 | 9 | Fam13a | 0.006 | Old |
| 0.702707 | 0.845038 | 0.036 | 0.044 | 1 | 9 | 6430548M | -0.008 | Old |
| 0.703031 | 0.463323 | 0.423 | 0.404 | 1 | 9 | Tmsb10 | 0.019 | Old |
| 0.703418 | 0.818724 | 0.044 | 0.037 | 1 | 9 | Tagln | 0.007 | Old |
| 0.703442 | 0.537127 | 0.036 | 0.029 | 1 | 9 | Ctu2 | 0.007 | Old |
| 0.703514 | 0.058134 | 0.162 | 0.147 | 1 | 9 | Npr2 | 0.015 | Old |
| 0.703625 | 0.083781 | 0.198 | 0.184 | 1 | 9 | Lgals3bp | 0.014 | Old |
| 0.703714 | 0.001695 | 0.397 | 0.375 | 1 | 9 | Rab11b | 0.022 | Old |
| 0.704066 | 0.646688 | 0.023 | 0.029 | 1 | 9 | Uty | -0.006 | Old |
| 0.704109 | 0.45188 | 0.219 | 0.213 | 1 | 9 | Sec63 | 0.006 | Old |
| 0.704441 | 0.609424 | 0.028 | 0.022 | 1 | 9 | Ackr1 | 0.006 | Old |
| 0.704441 | 0.122084 | 0.028 | 0.022 | 1 | 9 | Aldh1l1 | 0.006 | Old |
| 0.704441 | 0.036457 | 0.028 | 0.022 | 1 | 9 | Tti2 | 0.006 | Old |
| 0.704512 | 0.208935 | 0.219 | 0.206 | 1 | 9 | Uhmk1 | 0.013 | Old |
| 0.704514 | 0.684624 | 0.059 | 0.051 | 1 | 9 | Cnpy4 | 0.008 | Old |
| 0.704824 | 1.014715 | 0.044 | 0.037 | 1 | 9 | Cln6 | 0.007 | Old |
| 0.704824 | 0.901452 | 0.044 | 0.037 | 1 | 9 | Ndst2 | 0.007 | Old |
| 0.70499 | 0.903322 | 0.036 | 0.029 | 1 | 9 | Gvin1 | 0.007 | Old |
| 0.70499 | 0.693715 | 0.036 | 0.029 | 1 | 9 | Khk | 0.007 | Old |
| 0.705354 | 0.180623 | 0.082 | 0.074 | 1 | 9 | Ninj1 | 0.008 | Old |
| 0.706231 | 0.772356 | 0.044 | 0.037 | 1 | 9 | Ino80b | 0.007 | Old |
| 0.706313 | 0.341739 | 0.085 | 0.074 | 1 | 9 | Rnpc3 | 0.011 | Old |
| 0.70654 | 0.967516 | 0.036 | 0.029 | 1 | 9 | Prkdc | 0.007 | Old |
| 0.706943 | 0.430737 | 0.059 | 0.051 | 1 | 9 | Mtmr4 | 0.008 | Old |
| 0.706988 | 0.724717 | 0.188 | 0.184 | 1 | 9 | Ehd4 | 0.004 | Old |
| 0.706988 | 0.015771 | 0.191 | 0.176 | 1 | 9 | Cux1 | 0.015 | Old |
| 0.707348 | 0.688171 | 0.188 | 0.184 | 1 | 9 | Ppp6c | 0.004 | Old |
| 0.707403 | 0.181729 | 0.067 | 0.059 | 1 | 9 | Syap1 | 0.008 | Old |
| 0.707696 | 0.756784 | 0.023 | 0.029 | 1 | 9 | Zbtb42 | -0.006 | Old |
| 0.707866 | 0.206625 | 0.232 | 0.257 | 1 | 9 | Serp1 | -0.025 | Old |
| 0.708091 | 0.585342 | 0.036 | 0.029 | 1 | 9 | Mrgbp | 0.007 | Old |
| 0.708267 | 0.062779 | 0.16 | 0.147 | 1 | 9 | Anapc1 | 0.013 | Old |
| 0.708592 | 0.040661 | 0.211 | 0.228 | 1 | 9 | Sp3 | -0.017 | Old |
| 0.708782 | 0.238709 | 0.441 | 0.441 | 1 | 9 | Sod1 | 0 | Old |
| 0.709322 | 0.242725 | 0.085 | 0.096 | 1 | 9 | Naaa | -0.011 | Old |

| | | | | | | | | |
|---|---|---|---|---|---|---|---|---|
| 0.709347 | 0.215833 | 0.209 | 0.228 | 1 | 9 | Hsbp1 | -0.019 | Old |
| 0.709643 | 0.708188 | 0.036 | 0.029 | 1 | 9 | Lin52 | 0.007 | Old |
| 0.709643 | 0.510131 | 0.036 | 0.029 | 1 | 9 | Syt11 | 0.007 | Old |
| 0.709643 | 0.393093 | 0.036 | 0.029 | 1 | 9 | Rpap1 | 0.007 | Old |
| 0.709643 | 0.042357 | 0.036 | 0.029 | 1 | 9 | Spire1 | 0.007 | Old |
| 0.709677 | 0.030478 | 0.34 | 0.331 | 1 | 9 | Cdk17 | 0.009 | Old |
| 0.709697 | 0.599223 | 0.067 | 0.059 | 1 | 9 | Pacs1 | 0.008 | Old |
| 0.709874 | 0.113822 | 0.152 | 0.14 | 1 | 9 | Rcor1 | 0.012 | Old |
| 0.709925 | 0.330146 | 0.121 | 0.11 | 1 | 9 | Slc10a7 | 0.011 | Old |
| 0.710133 | 0.48436 | 0.052 | 0.044 | 1 | 9 | Mical2 | 0.008 | Old |
| 0.710164 | 0.091354 | 0.621 | 0.603 | 1 | 9 | Arhgap31 | 0.018 | Old |
| 0.710625 | 0.273604 | 0.057 | 0.066 | 1 | 9 | Rprd1a | -0.009 | Old |
| 0.711128 | 0.043159 | 0.216 | 0.199 | 1 | 9 | Lima1 | 0.017 | Old |
| 0.711197 | 0.650085 | 0.036 | 0.029 | 1 | 9 | Man2c1 | 0.007 | Old |
| 0.711332 | 0.626241 | 0.023 | 0.029 | 1 | 9 | Zfp810 | -0.006 | Old |
| 0.711552 | 1.179679 | 0.036 | 0.044 | 1 | 9 | Dhdh | -0.008 | Old |
| 0.711709 | 0.0068 | 0.186 | 0.199 | 1 | 9 | Nup98 | -0.013 | Old |
| 0.711809 | 0.551143 | 0.059 | 0.051 | 1 | 9 | Dgcr8 | 0.008 | Old |
| 0.711823 | 0.236879 | 0.09 | 0.103 | 1 | 9 | AU040320 | -0.013 | Old |
| 0.711869 | 0.353727 | 0.044 | 0.037 | 1 | 9 | Agpat4 | 0.007 | Old |
| 0.711973 | 0.07507 | 0.268 | 0.243 | 1 | 9 | Slain2 | 0.025 | Old |
| 0.711973 | 0.211574 | 0.265 | 0.25 | 1 | 9 | Thrap3 | 0.015 | Old |
| 0.712238 | 0.104951 | 0.384 | 0.404 | 1 | 9 | Gdi2 | -0.02 | Old |
| 0.712288 | 0.030751 | 0.064 | 0.074 | 1 | 9 | Ralgapa2 | -0.01 | Old |
| 0.712418 | 0.019473 | 0.371 | 0.397 | 1 | 9 | Gnaq | -0.026 | Old |
| 0.712698 | 0.073018 | 0.503 | 0.478 | 1 | 9 | Ramp2 | 0.025 | Old |
| 0.71274 | 0.081457 | 0.052 | 0.044 | 1 | 9 | Cep250 | 0.008 | Old |
| 0.712751 | 0.552425 | 0.036 | 0.029 | 1 | 9 | Bloc1s5 | 0.007 | Old |
| 0.712751 | 0.422267 | 0.036 | 0.029 | 1 | 9 | Map3k10 | 0.007 | Old |
| 0.713153 | 0.863919 | 0.023 | 0.029 | 1 | 9 | Ankrd24 | -0.006 | Old |
| 0.713193 | 0.033975 | 0.116 | 0.103 | 1 | 9 | Pogk | 0.013 | Old |
| 0.713282 | 0.738823 | 0.044 | 0.037 | 1 | 9 | Anapc10 | 0.007 | Old |
| 0.713318 | 0.314349 | 0.09 | 0.081 | 1 | 9 | Mrpl43 | 0.009 | Old |
| 0.713766 | 0.404574 | 0.09 | 0.103 | 1 | 9 | Mrps24 | -0.013 | Old |
| 0.714509 | 0.668 | 0.036 | 0.044 | 1 | 9 | Aagab | -0.008 | Old |
| 0.7146 | 0.355701 | 0.198 | 0.191 | 1 | 9 | Sppl3 | 0.007 | Old |
| 0.714739 | 0.327685 | 0.09 | 0.103 | 1 | 9 | Fkbp15 | -0.013 | Old |
| 0.714976 | 1.258124 | 0.023 | 0.029 | 1 | 9 | Rars2 | -0.006 | Old |
| 0.714976 | 1.210892 | 0.023 | 0.029 | 1 | 9 | Zbed4 | -0.006 | Old |
| 0.715198 | 0.144564 | 0.361 | 0.36 | 1 | 9 | Itsn2 | 0.001 | Old |
| 0.715436 | 0.3057 | 0.134 | 0.125 | 1 | 9 | Dpp9 | 0.009 | Old |
| 0.715443 | 0.469928 | 0.067 | 0.059 | 1 | 9 | Patz1 | 0.008 | Old |
| 0.715864 | 0.289213 | 0.036 | 0.029 | 1 | 9 | Atpaf2 | 0.007 | Old |
| 0.715989 | 0.38462 | 0.036 | 0.044 | 1 | 9 | Pik3cd | -0.008 | Old |
| 0.71669 | 0.915441 | 0.103 | 0.096 | 1 | 9 | Zkscan1 | 0.007 | Old |

| | | | | | | | | |
|---|---|---|---|---|---|---|---|---|
| 0.716809 | 0.015624 | 0.077 | 0.088 | 1 | 9 | Thap11 | -0.011 | Old |
| 0.717513 | 0.231984 | 0.124 | 0.11 | 1 | 9 | 5031439G( | 0.014 | Old |
| 0.717831 | 0.108992 | 0.057 | 0.066 | 1 | 9 | Scaper | -0.009 | Old |
| 0.71791 | 0.71415 | 0.059 | 0.051 | 1 | 9 | Sbf1 | 0.008 | Old |
| 0.718632 | 0.45999 | 0.09 | 0.103 | 1 | 9 | Arpc5l | -0.013 | Old |
| 0.718898 | 0.315408 | 0.077 | 0.088 | 1 | 9 | Kri1 | -0.011 | Old |
| 0.718982 | 0.523061 | 0.036 | 0.029 | 1 | 9 | Dimt1 | 0.007 | Old |
| 0.718995 | 0.120414 | 0.152 | 0.14 | 1 | 9 | Trim44 | 0.012 | Old |
| 0.719132 | 0.755192 | 0.059 | 0.051 | 1 | 9 | Uchl5 | 0.008 | Old |
| 0.719142 | 0.043682 | 0.263 | 0.279 | 1 | 9 | Hnrnpr | -0.016 | Old |
| 0.71951 | 0.481258 | 0.103 | 0.096 | 1 | 9 | Rhbdf2 | 0.007 | Old |
| 0.719524 | 0.376663 | 0.201 | 0.191 | 1 | 9 | Snx9 | 0.01 | Old |
| 0.72034 | 0.307673 | 0.193 | 0.184 | 1 | 9 | Sdhc | 0.009 | Old |
| 0.720355 | 0.677942 | 0.059 | 0.051 | 1 | 9 | Taf11 | 0.008 | Old |
| 0.720355 | 0.19556 | 0.059 | 0.051 | 1 | 9 | Faf1 | 0.008 | Old |
| 0.720357 | 0.101221 | 0.044 | 0.037 | 1 | 9 | Gon7 | 0.007 | Old |
| 0.720579 | 0.730607 | 0.052 | 0.044 | 1 | 9 | Mien1 | 0.008 | Old |
| 0.721444 | 0.567092 | 0.057 | 0.066 | 1 | 9 | Arfip1 | -0.009 | Old |
| 0.721444 | 0.306824 | 0.057 | 0.066 | 1 | 9 | Caap1 | -0.009 | Old |
| 0.721775 | 0.38098 | 0.044 | 0.037 | 1 | 9 | Smim20 | 0.007 | Old |
| 0.721889 | 0.123792 | 0.052 | 0.044 | 1 | 9 | Mdn1 | 0.008 | Old |
| 0.722104 | 0.314316 | 0.036 | 0.029 | 1 | 9 | Zbtb17 | 0.007 | Old |
| 0.723072 | 0.292411 | 0.121 | 0.11 | 1 | 9 | Chd9 | 0.011 | Old |
| 0.72315 | 0.167864 | 0.106 | 0.096 | 1 | 9 | Vkorc1l1 | 0.01 | Old |
| 0.723216 | 0.012685 | 0.933 | 0.897 | 1 | 9 | Kdr | 0.036 | Old |
| 0.723517 | 0.031809 | 0.232 | 0.25 | 1 | 9 | Gatad2a | -0.018 | Old |
| 0.724351 | 0.261174 | 0.173 | 0.162 | 1 | 9 | Gbp9 | 0.011 | Old |
| 0.724593 | 0.003941 | 0.152 | 0.169 | 1 | 9 | Tsg101 | -0.017 | Old |
| 0.724613 | 0.074769 | 0.044 | 0.037 | 1 | 9 | D730003I1 | 0.007 | Old |
| 0.72467 | 0.528727 | 0.067 | 0.059 | 1 | 9 | Cog4 | 0.008 | Old |
| 0.724851 | 0.017576 | 0.108 | 0.096 | 1 | 9 | Bms1 | 0.012 | Old |
| 0.724949 | 0.145608 | 0.098 | 0.088 | 1 | 9 | Zbtb10 | 0.01 | Old |
| 0.725358 | 0.092462 | 0.075 | 0.066 | 1 | 9 | Usp3 | 0.009 | Old |
| 0.72559 | 0.005644 | 0.245 | 0.265 | 1 | 9 | Cox5b | -0.02 | Old |
| 0.725822 | 0.481555 | 0.052 | 0.044 | 1 | 9 | Ccdc32 | 0.008 | Old |
| 0.72633 | 0.006598 | 0.482 | 0.434 | 1 | 9 | Npm1 | 0.048 | Old |
| 0.727456 | 0.662973 | 0.044 | 0.037 | 1 | 9 | Mlst8 | 0.007 | Old |
| 0.727495 | 0.244619 | 0.34 | 0.331 | 1 | 9 | Calu | 0.009 | Old |
| 0.727507 | 0.105393 | 0.198 | 0.213 | 1 | 9 | Ppp1cc | -0.015 | Old |
| 0.727694 | 0.54272 | 0.147 | 0.14 | 1 | 9 | Ehmt1 | 0.007 | Old |
| 0.727734 | 0.103182 | 0.466 | 0.449 | 1 | 9 | Raph1 | 0.017 | Old |
| 0.727828 | 0.369413 | 0.106 | 0.096 | 1 | 9 | Naa20 | 0.01 | Old |
| 0.72822 | 0.031914 | 0.064 | 0.074 | 1 | 9 | Irak4 | -0.01 | Old |
| 0.728437 | 0.679904 | 0.134 | 0.125 | 1 | 9 | Timm10b | 0.009 | Old |
| 0.729197 | 0.076823 | 0.113 | 0.103 | 1 | 9 | Acbd6 | 0.01 | Old |

| | | | | | | | | |
|---|---|---|---|---|---|---|---|---|
| 0.729381 | 0.402148 | 0.09 | 0.081 | 1 | 9 | Clns1a | 0.009 | Old |
| 0.729927 | 0.029044 | 0.044 | 0.051 | 1 | 9 | Prickle2 | -0.007 | Old |
| 0.730166 | 0.001754 | 0.68 | 0.684 | 1 | 9 | Ywhaz | -0.004 | Old |
| 0.730342 | 0.360393 | 0.219 | 0.243 | 1 | 9 | Zkscan3 | -0.024 | Old |
| 0.730832 | 0.664425 | 0.103 | 0.096 | 1 | 9 | Eed | 0.007 | Old |
| 0.731078 | 0.469941 | 0.052 | 0.044 | 1 | 9 | Ubac1 | 0.008 | Old |
| 0.731393 | 0.354148 | 0.059 | 0.051 | 1 | 9 | Fh1 | 0.008 | Old |
| 0.731617 | 0.058777 | 0.067 | 0.059 | 1 | 9 | Scnm1 | 0.008 | Old |
| 0.732094 | 0.048006 | 0.198 | 0.184 | 1 | 9 | Polr2k | 0.014 | Old |
| 0.732162 | 0.008562 | 0.281 | 0.265 | 1 | 9 | Hmgb2 | 0.016 | Old |
| 0.73218 | 0.011989 | 0.49 | 0.522 | 1 | 9 | Zfand5 | -0.032 | Old |
| 0.732517 | 0.062908 | 0.106 | 0.096 | 1 | 9 | Bok | 0.01 | Old |
| 0.733088 | 0.084083 | 0.188 | 0.176 | 1 | 9 | Ythdf3 | 0.012 | Old |
| 0.733567 | 0.394517 | 0.126 | 0.14 | 1 | 9 | Ext2 | -0.014 | Old |
| 0.733682 | 0.331767 | 0.098 | 0.088 | 1 | 9 | Igtp | 0.01 | Old |
| 0.733938 | 0.325849 | 0.067 | 0.059 | 1 | 9 | Ppip5k2 | 0.008 | Old |
| 0.734168 | 0.053733 | 0.098 | 0.088 | 1 | 9 | Cops7a | 0.01 | Old |
| 0.734579 | 0.222961 | 0.044 | 0.037 | 1 | 9 | Dpt | 0.007 | Old |
| 0.734587 | 0.377473 | 0.186 | 0.176 | 1 | 9 | Stx4a | 0.01 | Old |
| 0.734726 | 0.519547 | 0.152 | 0.147 | 1 | 9 | Zscan26 | 0.005 | Old |
| 0.734866 | 0.035671 | 0.384 | 0.368 | 1 | 9 | Gbp7 | 0.016 | Old |
| 0.734928 | 0.042704 | 0.113 | 0.125 | 1 | 9 | BC028528 | -0.012 | Old |
| 0.73569 | 0.607674 | 0.077 | 0.088 | 1 | 9 | Sap30l | -0.011 | Old |
| 0.735741 | 0.448106 | 0.183 | 0.169 | 1 | 9 | Icam1 | 0.014 | Old |
| 0.735966 | 0.040438 | 0.286 | 0.309 | 1 | 9 | Tmcc1 | -0.023 | Old |
| 0.736261 | 0.23482 | 0.067 | 0.059 | 1 | 9 | Uxs1 | 0.008 | Old |
| 0.736346 | 0.388744 | 0.052 | 0.044 | 1 | 9 | Tpbgl | 0.008 | Old |
| 0.736447 | 0.093055 | 0.09 | 0.081 | 1 | 9 | Mrps15 | 0.009 | Old |
| 0.736447 | 0.043651 | 0.085 | 0.096 | 1 | 9 | Sox13 | -0.011 | Old |
| 0.736512 | 0.030982 | 0.363 | 0.338 | 1 | 9 | Baz1b | 0.025 | Old |
| 0.736767 | 0.549887 | 0.044 | 0.051 | 1 | 9 | Kbtbd4 | -0.007 | Old |
| 0.736767 | 0.456072 | 0.044 | 0.051 | 1 | 9 | Tmem185b | -0.007 | Old |
| 0.736806 | 0.030037 | 0.392 | 0.368 | 1 | 9 | Mrfap1 | 0.024 | Old |
| 0.737458 | 0.016681 | 0.085 | 0.096 | 1 | 9 | Asxl1 | -0.011 | Old |
| 0.737629 | 0.109738 | 0.121 | 0.11 | 1 | 9 | Oxct1 | 0.011 | Old |
| 0.738324 | 0.135586 | 0.17 | 0.162 | 1 | 9 | Tmem189 | 0.008 | Old |
| 0.738574 | 0.105958 | 0.075 | 0.066 | 1 | 9 | Ciz1 | 0.009 | Old |
| 0.738985 | 0.007854 | 0.052 | 0.044 | 1 | 9 | Gm26782 | 0.008 | Old |
| 0.740464 | 0.122103 | 0.299 | 0.316 | 1 | 9 | Grb2 | -0.017 | Old |
| 0.740881 | 0.791577 | 0.044 | 0.051 | 1 | 9 | Idi1 | -0.007 | Old |
| 0.741402 | 0.356483 | 0.106 | 0.118 | 1 | 9 | Unc119b | -0.012 | Old |
| 0.741595 | 0.125031 | 0.307 | 0.301 | 1 | 9 | Pum1 | 0.006 | Old |
| 0.741719 | 0.30762 | 0.18 | 0.169 | 1 | 9 | Zfp326 | 0.011 | Old |
| 0.74193 | 0.442454 | 0.126 | 0.118 | 1 | 9 | Trim34a | 0.008 | Old |
| 0.74197 | 0.22817 | 0.064 | 0.074 | 1 | 9 | Rras2 | -0.01 | Old |

| | | | | | | | | |
|---|---|---|---|---|---|---|---|---|
| 0.74202 | 0.179417 | 0.082 | 0.074 | 1 | 9 | Mlec | 0.008 | Old |
| 0.742254 | 0.237202 | 0.044 | 0.051 | 1 | 9 | Gm11730 | -0.007 | Old |
| 0.742575 | 0.219336 | 0.152 | 0.14 | 1 | 9 | Cdk19 | 0.012 | Old |
| 0.742582 | 0.689571 | 0.088 | 0.081 | 1 | 9 | Arhgap10 | 0.007 | Old |
| 0.74312 | 0.028581 | 0.064 | 0.074 | 1 | 9 | Wdr44 | -0.01 | Old |
| 0.743244 | 0.40464 | 0.067 | 0.059 | 1 | 9 | Wdr45 | 0.008 | Old |
| 0.743495 | 0.08243 | 0.327 | 0.316 | 1 | 9 | Rpl29 | 0.011 | Old |
| 0.744104 | 0.319738 | 0.075 | 0.066 | 1 | 9 | Arl6ip4 | 0.009 | Old |
| 0.74427 | 0.193308 | 0.064 | 0.074 | 1 | 9 | Tsc1 | -0.01 | Old |
| 0.74451 | 0.081087 | 0.209 | 0.199 | 1 | 9 | Usp19 | 0.01 | Old |
| 0.744803 | 0.070761 | 0.168 | 0.184 | 1 | 9 | Camk2g | -0.016 | Old |
| 0.744815 | 0.078584 | 0.173 | 0.162 | 1 | 9 | Cfl2 | 0.011 | Old |
| 0.745165 | 0.237322 | 0.121 | 0.11 | 1 | 9 | Rhog | 0.011 | Old |
| 0.745191 | 0.295692 | 0.082 | 0.074 | 1 | 9 | Cnp | 0.008 | Old |
| 0.745412 | 0.088845 | 0.072 | 0.081 | 1 | 9 | Slc25a28 | -0.009 | Old |
| 0.74553 | 0.077011 | 0.165 | 0.154 | 1 | 9 | Tmem14c | 0.011 | Old |
| 0.746195 | 0.160153 | 0.286 | 0.272 | 1 | 9 | Pparg | 0.014 | Old |
| 0.746518 | 0.161416 | 0.113 | 0.103 | 1 | 9 | Sae1 | 0.01 | Old |
| 0.746784 | 0.21023 | 0.08 | 0.088 | 1 | 9 | Tdg | -0.008 | Old |
| 0.746908 | 0.047576 | 0.031 | 0.037 | 1 | 9 | Tjap1 | -0.006 | Old |
| 0.747654 | 0.078953 | 0.405 | 0.382 | 1 | 9 | Hnrnpc | 0.023 | Old |
| 0.747792 | 0.404556 | 0.155 | 0.169 | 1 | 9 | Twistnb | -0.014 | Old |
| 0.748781 | 0.148193 | 0.255 | 0.243 | 1 | 9 | Peli1 | 0.012 | Old |
| 0.748877 | 0.098564 | 0.064 | 0.074 | 1 | 9 | Pts | -0.01 | Old |
| 0.748954 | 0.162033 | 0.134 | 0.147 | 1 | 9 | Vma21 | -0.013 | Old |
| 0.74901 | 0.066853 | 0.216 | 0.206 | 1 | 9 | Smc3 | 0.01 | Old |
| 0.749146 | 0.296522 | 0.134 | 0.125 | 1 | 9 | Plekha2 | 0.009 | Old |
| 0.749448 | 0.101252 | 0.198 | 0.213 | 1 | 9 | Rnf44 | -0.015 | Old |
| 0.749515 | 0.238402 | 0.348 | 0.338 | 1 | 9 | Ucp2 | 0.01 | Old |
| 0.750117 | 0.350528 | 0.211 | 0.206 | 1 | 9 | Fam168a | 0.005 | Old |
| 0.750158 | 0.043582 | 0.031 | 0.037 | 1 | 9 | Smpd4 | -0.006 | Old |
| 0.750272 | 0.12957 | 0.098 | 0.088 | 1 | 9 | Ice1 | 0.01 | Old |
| 0.750609 | 0.398771 | 0.111 | 0.125 | 1 | 9 | Prpf4 | -0.014 | Old |
| 0.751061 | 0.104919 | 0.289 | 0.309 | 1 | 9 | Cnot6 | -0.02 | Old |
| 0.751184 | 0.327584 | 0.064 | 0.074 | 1 | 9 | Ufsp2 | -0.01 | Old |
| 0.751577 | 0.214659 | 0.191 | 0.184 | 1 | 9 | Tle3 | 0.007 | Old |
| 0.75162 | 0.204586 | 0.26 | 0.287 | 1 | 9 | Psmd2 | -0.027 | Old |
| 0.751661 | 0.222264 | 0.085 | 0.096 | 1 | 9 | Fam212b | -0.011 | Old |
| 0.751739 | 0.271573 | 0.103 | 0.096 | 1 | 9 | Coa3 | 0.007 | Old |
| 0.752139 | 0.055562 | 0.276 | 0.265 | 1 | 9 | Tspan2 | 0.011 | Old |
| 0.752357 | 0.231082 | 0.119 | 0.11 | 1 | 9 | Atad1 | 0.009 | Old |
| 0.752397 | 0.030489 | 0.196 | 0.184 | 1 | 9 | Lrp5 | 0.012 | Old |
| 0.752694 | 0.363834 | 0.098 | 0.11 | 1 | 9 | Acacb | -0.012 | Old |
| 0.752844 | 0.66306 | 0.088 | 0.081 | 1 | 9 | Cdadc1 | 0.007 | Old |
| 0.753196 | 0.372554 | 0.317 | 0.309 | 1 | 9 | Oaz2 | 0.008 | Old |

| | | | | | | | | |
|---|---|---|---|---|---|---|---|---|
| 0.753413 | 0.474997 | 0.031 | 0.037 | 1 | 9 | Lclat1 | -0.006 | Old |
| 0.753545 | 0.221798 | 0.268 | 0.265 | 1 | 9 | Cul3 | 0.003 | Old |
| 0.753557 | 0.249151 | 0.139 | 0.154 | 1 | 9 | St6galnac2 | -0.015 | Old |
| 0.753888 | 0.05944 | 0.242 | 0.228 | 1 | 9 | Secisbp2l | 0.014 | Old |
| 0.754005 | 0.079989 | 0.624 | 0.632 | 1 | 9 | Tmem30a | -0.008 | Old |
| 0.754085 | 0.100677 | 0.052 | 0.059 | 1 | 9 | Kif3b | -0.007 | Old |
| 0.754085 | 0.063781 | 0.052 | 0.059 | 1 | 9 | Smim12 | -0.007 | Old |
| 0.75436 | 0.015083 | 0.807 | 0.779 | 1 | 9 | Rpl37 | 0.028 | Old |
| 0.754466 | 0.079161 | 0.263 | 0.287 | 1 | 9 | Taf15 | -0.024 | Old |
| 0.754983 | 0.044374 | 0.095 | 0.088 | 1 | 9 | Zadh2 | 0.007 | Old |
| 0.755042 | 0.090616 | 0.031 | 0.037 | 1 | 9 | Mast3 | -0.006 | Old |
| 0.755215 | 0.543982 | 0.139 | 0.132 | 1 | 9 | Slc30a9 | 0.007 | Old |
| 0.755931 | 0.994524 | 0.088 | 0.081 | 1 | 9 | Cdc123 | 0.007 | Old |
| 0.756003 | 2.657637 | 0.01 | 0.007 | 1 | 9 | Gzmm | 0.003 | Old |
| 0.756003 | 2.463155 | 0.01 | 0.007 | 1 | 9 | Tcf23 | 0.003 | Old |
| 0.756003 | 2.437618 | 0.01 | 0.007 | 1 | 9 | Efna5 | 0.003 | Old |
| 0.756003 | 2.329841 | 0.01 | 0.007 | 1 | 9 | Hist3h2ba | 0.003 | Old |
| 0.756003 | 2.164385 | 0.01 | 0.007 | 1 | 9 | BC048403 | 0.003 | Old |
| 0.756003 | 2.05642 | 0.01 | 0.007 | 1 | 9 | Arv1 | 0.003 | Old |
| 0.756003 | 1.773767 | 0.01 | 0.007 | 1 | 9 | Clmp | 0.003 | Old |
| 0.756003 | 1.768504 | 0.01 | 0.007 | 1 | 9 | Palb2 | 0.003 | Old |
| 0.756003 | 1.721457 | 0.01 | 0.007 | 1 | 9 | Pigb | 0.003 | Old |
| 0.756003 | 1.679679 | 0.01 | 0.007 | 1 | 9 | Slc41a2 | 0.003 | Old |
| 0.756003 | 1.546288 | 0.01 | 0.007 | 1 | 9 | Ephx2 | 0.003 | Old |
| 0.756003 | 1.502078 | 0.01 | 0.007 | 1 | 9 | Setd4 | 0.003 | Old |
| 0.756003 | 1.446075 | 0.01 | 0.007 | 1 | 9 | Tgtp1 | 0.003 | Old |
| 0.756003 | 1.413002 | 0.01 | 0.007 | 1 | 9 | Susd4 | 0.003 | Old |
| 0.756003 | 1.401466 | 0.01 | 0.007 | 1 | 9 | Arhgef25 | 0.003 | Old |
| 0.756003 | 1.28887 | 0.01 | 0.007 | 1 | 9 | Pfn2 | 0.003 | Old |
| 0.756003 | 1.261793 | 0.01 | 0.007 | 1 | 9 | Bdkrb2 | 0.003 | Old |
| 0.756003 | 1.226233 | 0.01 | 0.007 | 1 | 9 | Nsdhl | 0.003 | Old |
| 0.756003 | 1.195225 | 0.01 | 0.007 | 1 | 9 | Bend5 | 0.003 | Old |
| 0.756003 | 1.136621 | 0.01 | 0.007 | 1 | 9 | Meis1 | 0.003 | Old |
| 0.756003 | 1.116602 | 0.01 | 0.007 | 1 | 9 | Zfp428 | 0.003 | Old |
| 0.756003 | 1.06071 | 0.01 | 0.007 | 1 | 9 | Klf8 | 0.003 | Old |
| 0.756003 | 1.015765 | 0.01 | 0.007 | 1 | 9 | 1110002L0 | 0.003 | Old |
| 0.756003 | 1.012656 | 0.01 | 0.007 | 1 | 9 | Hes7 | 0.003 | Old |
| 0.756003 | 0.977665 | 0.01 | 0.007 | 1 | 9 | Gm36161 | 0.003 | Old |
| 0.756003 | 0.885195 | 0.01 | 0.007 | 1 | 9 | Slc39a11 | 0.003 | Old |
| 0.756003 | 0.45035 | 0.01 | 0.007 | 1 | 9 | Rhoh | 0.003 | Old |
| 0.756672 | 0.361576 | 0.031 | 0.037 | 1 | 9 | Wrb | -0.006 | Old |
| 0.756672 | 0.140624 | 0.031 | 0.037 | 1 | 9 | Cd86 | -0.006 | Old |
| 0.756672 | 0.017377 | 0.031 | 0.037 | 1 | 9 | Etv5 | -0.006 | Old |
| 0.757679 | 0.10138 | 0.807 | 0.794 | 1 | 9 | Rps15 | 0.013 | Old |
| 0.758098 | 0.041172 | 0.175 | 0.191 | 1 | 9 | Erap1 | -0.016 | Old |

| | | | | | | | | |
|---|---|---|---|---|---|---|---|---|
| 0.758304 | 0.80976 | 0.031 | 0.037 | 1 | 9 | Emilin2 | -0.006 | Old |
| 0.758454 | 0.819376 | 0.072 | 0.066 | 1 | 9 | Nol8 | 0.006 | Old |
| 0.758544 | 0.136904 | 0.075 | 0.066 | 1 | 9 | Lage3 | 0.009 | Old |
| 0.758977 | 1.131495 | 0.01 | 0.007 | 1 | 9 | Pkdcc | 0.003 | Old |
| 0.758977 | 1.096381 | 0.01 | 0.007 | 1 | 9 | Gins3 | 0.003 | Old |
| 0.758977 | 1.050931 | 0.01 | 0.007 | 1 | 9 | Actr3b | 0.003 | Old |
| 0.758977 | 0.812621 | 0.01 | 0.007 | 1 | 9 | 1810024B0 | 0.003 | Old |
| 0.758977 | 0.591402 | 0.01 | 0.007 | 1 | 9 | Pabpn1l | 0.003 | Old |
| 0.758977 | 0.520191 | 0.01 | 0.007 | 1 | 9 | A030001D2 | 0.003 | Old |
| 0.759635 | 0.334566 | 0.191 | 0.184 | 1 | 9 | Arid4b | 0.007 | Old |
| 0.759936 | 0.110671 | 0.031 | 0.037 | 1 | 9 | Ndc1 | -0.006 | Old |
| 0.761063 | 0.216143 | 0.134 | 0.125 | 1 | 9 | Rai14 | 0.009 | Old |
| 0.761113 | 0.153059 | 0.082 | 0.074 | 1 | 9 | Lemd2 | 0.008 | Old |
| 0.761555 | 0.36032 | 0.124 | 0.14 | 1 | 9 | Ddx54 | -0.016 | Old |
| 0.761569 | 0.503916 | 0.031 | 0.037 | 1 | 9 | Fgf11 | -0.006 | Old |
| 0.761794 | 0.111627 | 0.052 | 0.059 | 1 | 9 | Trp53rka | -0.007 | Old |
| 0.761859 | 0.155532 | 0.085 | 0.096 | 1 | 9 | Hoxb6 | -0.011 | Old |
| 0.761954 | 1.546273 | 0.01 | 0.007 | 1 | 9 | Chpf | 0.003 | Old |
| 0.761954 | 1.528455 | 0.01 | 0.007 | 1 | 9 | Adamtsl5 | 0.003 | Old |
| 0.761954 | 1.372165 | 0.01 | 0.007 | 1 | 9 | Ssx2ip | 0.003 | Old |
| 0.761954 | 1.323813 | 0.01 | 0.007 | 1 | 9 | Zranb3 | 0.003 | Old |
| 0.761954 | 1.135149 | 0.01 | 0.007 | 1 | 9 | Rpl36a-ps1 | 0.003 | Old |
| 0.761954 | 1.041833 | 0.01 | 0.007 | 1 | 9 | Ccdc152 | 0.003 | Old |
| 0.761954 | 1.033207 | 0.01 | 0.007 | 1 | 9 | Coa4 | 0.003 | Old |
| 0.761954 | 0.949615 | 0.01 | 0.007 | 1 | 9 | Serpine2 | 0.003 | Old |
| 0.761954 | 0.83024 | 0.01 | 0.007 | 1 | 9 | Camkk1 | 0.003 | Old |
| 0.761954 | 0.74972 | 0.01 | 0.007 | 1 | 9 | Gm28557 | 0.003 | Old |
| 0.761954 | 0.722116 | 0.01 | 0.007 | 1 | 9 | Adssl1 | 0.003 | Old |
| 0.761954 | 0.678916 | 0.01 | 0.007 | 1 | 9 | Gm11423 | 0.003 | Old |
| 0.761954 | 0.667481 | 0.01 | 0.007 | 1 | 9 | Tctn1 | 0.003 | Old |
| 0.761954 | 0.662404 | 0.01 | 0.007 | 1 | 9 | Gm15445 | 0.003 | Old |
| 0.761954 | 0.575401 | 0.01 | 0.007 | 1 | 9 | Gm43254 | 0.003 | Old |
| 0.761954 | 0.511129 | 0.01 | 0.007 | 1 | 9 | Ikzf2 | 0.003 | Old |
| 0.761954 | 0.483684 | 0.01 | 0.007 | 1 | 9 | Pex12 | 0.003 | Old |
| 0.761954 | 0.462523 | 0.01 | 0.007 | 1 | 9 | Tmc5 | 0.003 | Old |
| 0.761954 | 0.38595 | 0.01 | 0.007 | 1 | 9 | Gm5113 | 0.003 | Old |
| 0.761954 | 0.378577 | 0.01 | 0.007 | 1 | 9 | Pter | 0.003 | Old |
| 0.761954 | 0.298975 | 0.01 | 0.007 | 1 | 9 | Mpv17l | 0.003 | Old |
| 0.761954 | 0.289693 | 0.01 | 0.007 | 1 | 9 | Ndc80 | 0.003 | Old |
| 0.761954 | 0.223727 | 0.01 | 0.007 | 1 | 9 | A630072L1 | 0.003 | Old |
| 0.762696 | 0.128325 | 0.119 | 0.11 | 1 | 9 | Gtf2a2 | 0.009 | Old |
| 0.762713 | 0.110329 | 0.43 | 0.404 | 1 | 9 | Foxp1 | 0.026 | Old |
| 0.763204 | 0.606915 | 0.031 | 0.037 | 1 | 9 | Taf6 | -0.006 | Old |
| 0.763277 | 0.021363 | 0.201 | 0.184 | 1 | 9 | Stat2 | 0.017 | Old |
| 0.763671 | 0.703112 | 0.101 | 0.096 | 1 | 9 | Tuba1c | 0.005 | Old |

| | | | | | | | | |
|---|---|---|---|---|---|---|---|---|
| 0.764327 | 1.048454 | 0.057 | 0.051 | 1 | 9 | Aifm1 | 0.006 | Old |
| 0.764787 | 0.041247 | 0.322 | 0.346 | 1 | 9 | Eif4h | -0.024 | Old |
| 0.764839 | 0.513286 | 0.031 | 0.037 | 1 | 9 | Dok1 | -0.006 | Old |
| 0.764839 | 0.29953 | 0.031 | 0.037 | 1 | 9 | Ifit3b | -0.006 | Old |
| 0.764926 | 0.125529 | 0.085 | 0.096 | 1 | 9 | Mtfr1l | -0.011 | Old |
| 0.764934 | 0.719339 | 0.01 | 0.007 | 1 | 9 | Neil1 | 0.003 | Old |
| 0.764934 | 0.689448 | 0.01 | 0.007 | 1 | 9 | Cacna1a | 0.003 | Old |
| 0.764934 | 0.598079 | 0.01 | 0.007 | 1 | 9 | Ldlrad4 | 0.003 | Old |
| 0.764934 | 0.521134 | 0.01 | 0.007 | 1 | 9 | Pla2r1 | 0.003 | Old |
| 0.764934 | 0.442347 | 0.01 | 0.007 | 1 | 9 | Nav3 | 0.003 | Old |
| 0.764934 | 0.402756 | 0.01 | 0.007 | 1 | 9 | Tnfrsf9 | 0.003 | Old |
| 0.764934 | 0.401315 | 0.01 | 0.007 | 1 | 9 | Nat14 | 0.003 | Old |
| 0.764934 | 0.346829 | 0.01 | 0.007 | 1 | 9 | Eif3j2 | 0.003 | Old |
| 0.764934 | 0.322881 | 0.01 | 0.007 | 1 | 9 | Zfp51 | 0.003 | Old |
| 0.764934 | 0.305988 | 0.01 | 0.007 | 1 | 9 | Rgma | 0.003 | Old |
| 0.764934 | 0.136569 | 0.01 | 0.007 | 1 | 9 | Cspg4 | 0.003 | Old |
| 0.764934 | 0.101415 | 0.01 | 0.007 | 1 | 9 | Zfand4 | 0.003 | Old |
| 0.764934 | 0.052508 | 0.01 | 0.007 | 1 | 9 | Gcnt1 | 0.003 | Old |
| 0.764934 | 0.035862 | 0.01 | 0.007 | 1 | 9 | Gm7334 | 0.003 | Old |
| 0.765231 | 1.078738 | 0.072 | 0.066 | 1 | 9 | Ttpal | 0.006 | Old |
| 0.765336 | 1.330741 | 0.049 | 0.044 | 1 | 9 | Zfp654 | 0.005 | Old |
| 0.765501 | 0.29619 | 0.126 | 0.118 | 1 | 9 | Sun2 | 0.008 | Old |
| 0.765723 | 0.311492 | 0.119 | 0.132 | 1 | 9 | Socs5 | -0.013 | Old |
| 0.766257 | 0.163277 | 0.286 | 0.279 | 1 | 9 | Pten | 0.007 | Old |
| 0.766895 | 0.896715 | 0.095 | 0.088 | 1 | 9 | Inpp4a | 0.007 | Old |
| 0.766928 | 0.130682 | 0.399 | 0.419 | 1 | 9 | Lnpep | -0.02 | Old |
| 0.767256 | 0.25146 | 0.126 | 0.118 | 1 | 9 | Mfng | 0.008 | Old |
| 0.767964 | 0.082066 | 0.093 | 0.103 | 1 | 9 | Tm2d2 | -0.01 | Old |
| 0.768237 | 0.207209 | 0.052 | 0.059 | 1 | 9 | Fgfr1 | -0.007 | Old |
| 0.768486 | 0.041821 | 0.541 | 0.551 | 1 | 9 | Ctsl | -0.01 | Old |
| 0.769422 | 0.163883 | 0.101 | 0.11 | 1 | 9 | Otud7b | -0.009 | Old |
| 0.769444 | 0.021571 | 0.724 | 0.713 | 1 | 9 | Rbms1 | 0.011 | Old |
| 0.769457 | 0.926559 | 0.064 | 0.059 | 1 | 9 | Zdhhc7 | 0.005 | Old |
| 0.769528 | 0.583946 | 0.052 | 0.059 | 1 | 9 | Galnt4 | -0.007 | Old |
| 0.770152 | 0.11983 | 0.214 | 0.235 | 1 | 9 | Ywhag | -0.021 | Old |
| 0.77068 | 0.01315 | 0.039 | 0.044 | 1 | 9 | Vps39 | -0.005 | Old |
| 0.770781 | 1.046512 | 0.049 | 0.044 | 1 | 9 | Urod | 0.005 | Old |
| 0.770819 | 0.05393 | 0.052 | 0.059 | 1 | 9 | Erlin1 | -0.007 | Old |
| 0.770891 | 0.570813 | 0.072 | 0.066 | 1 | 9 | Parp6 | 0.006 | Old |
| 0.77173 | 0.010888 | 0.268 | 0.279 | 1 | 9 | Gata2 | -0.011 | Old |
| 0.772042 | 0.69498 | 0.064 | 0.074 | 1 | 9 | Zfp598 | -0.01 | Old |
| 0.772558 | 0.618301 | 0.08 | 0.074 | 1 | 9 | Rwdd4a | 0.006 | Old |
| 0.772722 | 0.757202 | 0.018 | 0.022 | 1 | 9 | Adgra2 | -0.004 | Old |
| 0.772722 | 0.050869 | 0.018 | 0.022 | 1 | 9 | Tslp | -0.004 | Old |
| 0.772851 | 0.044217 | 0.116 | 0.125 | 1 | 9 | Clk3 | -0.009 | Old |

| | | | | | | | | |
|---|---|---|---|---|---|---|---|---|
| 0.773159 | 0.762561 | 0.072 | 0.066 | 1 | 9 | Sart3 | 0.006 | Old |
| 0.773292 | 0.173193 | 0.211 | 0.199 | 1 | 9 | Cdk12 | 0.012 | Old |
| 0.77353 | 0.280553 | 0.119 | 0.11 | 1 | 9 | Tppp | 0.009 | Old |
| 0.774051 | 1.253497 | 0.044 | 0.051 | 1 | 9 | Batf3 | -0.007 | Old |
| 0.774849 | 0.344987 | 0.018 | 0.022 | 1 | 9 | Fdxacb1 | -0.004 | Old |
| 0.776048 | 0.215631 | 0.126 | 0.118 | 1 | 9 | Trim26 | 0.008 | Old |
| 0.776333 | 0.123021 | 0.245 | 0.235 | 1 | 9 | Cct4 | 0.01 | Old |
| 0.776564 | 0.516601 | 0.072 | 0.066 | 1 | 9 | Ccdc85c | 0.006 | Old |
| 0.776936 | 0.471901 | 0.116 | 0.11 | 1 | 9 | Dctn2 | 0.006 | Old |
| 0.776976 | 0.426392 | 0.018 | 0.022 | 1 | 9 | Cep95 | -0.004 | Old |
| 0.776976 | 0.285603 | 0.018 | 0.022 | 1 | 9 | Mpp4 | -0.004 | Old |
| 0.776976 | 0.127596 | 0.018 | 0.022 | 1 | 9 | Hoxa9 | -0.004 | Old |
| 0.776976 | 0.023615 | 0.018 | 0.022 | 1 | 9 | Tyw5 | -0.004 | Old |
| 0.777233 | 0.053808 | 0.09 | 0.081 | 1 | 9 | Adm | 0.009 | Old |
| 0.777305 | 0.326483 | 0.191 | 0.184 | 1 | 9 | Rtl8b | 0.007 | Old |
| 0.777834 | 1.001253 | 0.064 | 0.059 | 1 | 9 | Tlr2 | 0.005 | Old |
| 0.777834 | 0.591635 | 0.064 | 0.059 | 1 | 9 | Vps50 | 0.005 | Old |
| 0.779106 | 1.144767 | 0.018 | 0.022 | 1 | 9 | Ppp1r3e | -0.004 | Old |
| 0.779106 | 0.54855 | 0.018 | 0.022 | 1 | 9 | Cldn34c1 | -0.004 | Old |
| 0.779106 | 0.39324 | 0.018 | 0.022 | 1 | 9 | Sqle | -0.004 | Old |
| 0.779132 | 0.296845 | 0.139 | 0.154 | 1 | 9 | Arrdc4 | -0.015 | Old |
| 0.779874 | 0.46702 | 0.052 | 0.059 | 1 | 9 | Mvb12b | -0.007 | Old |
| 0.780141 | 0.044967 | 0.196 | 0.213 | 1 | 9 | Srek1 | -0.017 | Old |
| 0.780543 | 0.197202 | 0.103 | 0.096 | 1 | 9 | Hdac5 | 0.007 | Old |
| 0.781067 | 1.247191 | 0.041 | 0.037 | 1 | 9 | Kif16b | 0.004 | Old |
| 0.781067 | 0.215798 | 0.039 | 0.044 | 1 | 9 | Invs | -0.005 | Old |
| 0.782269 | 1.838498 | 0.018 | 0.015 | 1 | 9 | Kazald1 | 0.003 | Old |
| 0.782269 | 1.167822 | 0.018 | 0.015 | 1 | 9 | Slc25a38 | 0.003 | Old |
| 0.782554 | 0.183372 | 0.039 | 0.044 | 1 | 9 | Prep | -0.005 | Old |
| 0.782554 | 0.104018 | 0.039 | 0.044 | 1 | 9 | Ascc2 | -0.005 | Old |
| 0.78337 | 0.811979 | 0.018 | 0.022 | 1 | 9 | Zfp1 | -0.004 | Old |
| 0.78337 | 0.788644 | 0.018 | 0.022 | 1 | 9 | Atg10 | -0.004 | Old |
| 0.78337 | 0.778105 | 0.018 | 0.022 | 1 | 9 | Apba1 | -0.004 | Old |
| 0.78337 | 0.744841 | 0.018 | 0.022 | 1 | 9 | B4galnt1 | -0.004 | Old |
| 0.78337 | 0.56147 | 0.018 | 0.022 | 1 | 9 | Incenp | -0.004 | Old |
| 0.78337 | 0.495692 | 0.018 | 0.022 | 1 | 9 | Anpep | -0.004 | Old |
| 0.783704 | 0.006449 | 0.515 | 0.5 | 1 | 9 | Nfia | 0.015 | Old |
| 0.783884 | 0.316318 | 0.088 | 0.081 | 1 | 9 | Wwp2 | 0.007 | Old |
| 0.784042 | 1.471707 | 0.041 | 0.037 | 1 | 9 | Fbxo31 | 0.004 | Old |
| 0.784042 | 0.238734 | 0.039 | 0.044 | 1 | 9 | Cln3 | -0.005 | Old |
| 0.784443 | 0.868393 | 0.049 | 0.044 | 1 | 9 | Ppard | 0.005 | Old |
| 0.784516 | 1.9029 | 0.018 | 0.015 | 1 | 9 | Fign | 0.003 | Old |
| 0.784516 | 1.631529 | 0.018 | 0.015 | 1 | 9 | Cngb1 | 0.003 | Old |
| 0.784516 | 1.18967 | 0.018 | 0.015 | 1 | 9 | Tsen2 | 0.003 | Old |
| 0.784552 | 0.015753 | 0.603 | 0.596 | 1 | 9 | Sfpq | 0.007 | Old |

| | | | | | | | | |
|---|---|---|---|---|---|---|---|---|
| 0.784678 | 0.969963 | 0.057 | 0.051 | 1 | 9 | Pdk4 | 0.006 | Old |
| 0.785028 | 0.065387 | 0.358 | 0.346 | 1 | 9 | Sik1 | 0.012 | Old |
| 0.785033 | 0.070957 | 0.142 | 0.132 | 1 | 9 | Bvht | 0.01 | Old |
| 0.785286 | 0.150022 | 0.24 | 0.221 | 1 | 9 | Hspa4 | 0.019 | Old |
| 0.785411 | 0.050233 | 0.144 | 0.132 | 1 | 9 | Dmd | 0.012 | Old |
| 0.785468 | 0.140587 | 0.09 | 0.081 | 1 | 9 | Prps2 | 0.009 | Old |
| 0.785502 | 0.110052 | 0.106 | 0.118 | 1 | 9 | Sp110 | -0.012 | Old |
| 0.785504 | 1.169176 | 0.018 | 0.022 | 1 | 9 | Nat8l | -0.004 | Old |
| 0.785504 | 0.449437 | 0.018 | 0.022 | 1 | 9 | Brf2 | -0.004 | Old |
| 0.786533 | 0.247782 | 0.219 | 0.213 | 1 | 9 | Lama4 | 0.006 | Old |
| 0.786764 | 1.80977 | 0.018 | 0.015 | 1 | 9 | Chac1 | 0.003 | Old |
| 0.786764 | 1.717027 | 0.018 | 0.015 | 1 | 9 | Arc | 0.003 | Old |
| 0.786764 | 1.467789 | 0.018 | 0.015 | 1 | 9 | Fbxl15 | 0.003 | Old |
| 0.786764 | 1.413872 | 0.018 | 0.015 | 1 | 9 | Slc22a18 | 0.003 | Old |
| 0.786764 | 0.936916 | 0.018 | 0.015 | 1 | 9 | Oas1a | 0.003 | Old |
| 0.786805 | 0.58155 | 0.072 | 0.066 | 1 | 9 | St5 | 0.006 | Old |
| 0.786991 | 0.612892 | 0.116 | 0.11 | 1 | 9 | Mgrn1 | 0.006 | Old |
| 0.787224 | 1.263124 | 0.026 | 0.022 | 1 | 9 | Ampd2 | 0.004 | Old |
| 0.787231 | 0.160033 | 0.16 | 0.176 | 1 | 9 | Smim7 | -0.016 | Old |
| 0.787232 | 0.509682 | 0.057 | 0.051 | 1 | 9 | Eya3 | 0.006 | Old |
| 0.78764 | 0.919049 | 0.018 | 0.022 | 1 | 9 | Pigv | -0.004 | Old |
| 0.78764 | 0.309291 | 0.018 | 0.022 | 1 | 9 | Rcn3 | -0.004 | Old |
| 0.78838 | 0.712905 | 0.064 | 0.074 | 1 | 9 | Cd2bp2 | -0.01 | Old |
| 0.788511 | 0.89363 | 0.041 | 0.037 | 1 | 9 | Mrto4 | 0.004 | Old |
| 0.788757 | 0.100861 | 0.098 | 0.088 | 1 | 9 | Wdr5 | 0.01 | Old |
| 0.789015 | 1.871164 | 0.018 | 0.015 | 1 | 9 | Ass1 | 0.003 | Old |
| 0.789015 | 1.559065 | 0.018 | 0.015 | 1 | 9 | Ifrd2 | 0.003 | Old |
| 0.789494 | 1.848893 | 0.034 | 0.029 | 1 | 9 | Slc5a6 | 0.005 | Old |
| 0.789505 | 0.36064 | 0.152 | 0.147 | 1 | 9 | Rnf20 | 0.005 | Old |
| 0.789778 | 0.591078 | 0.018 | 0.022 | 1 | 9 | Focad | -0.004 | Old |
| 0.790002 | 1.101839 | 0.041 | 0.037 | 1 | 9 | Gm2000 | 0.004 | Old |
| 0.790002 | 0.832168 | 0.041 | 0.037 | 1 | 9 | Slc45a4 | 0.004 | Old |
| 0.790631 | 0.075822 | 0.348 | 0.324 | 1 | 9 | Mier1 | 0.024 | Old |
| 0.790985 | 1.147984 | 0.026 | 0.022 | 1 | 9 | Mettl26 | 0.004 | Old |
| 0.791007 | 0.452505 | 0.08 | 0.074 | 1 | 9 | Tmem86a | 0.006 | Old |
| 0.79114 | 0.002289 | 0.363 | 0.382 | 1 | 9 | Chd4 | -0.019 | Old |
| 0.79114 | 0.185293 | 0.059 | 0.066 | 1 | 9 | Iffo1 | -0.007 | Old |
| 0.791146 | 1.140852 | 0.034 | 0.029 | 1 | 9 | Fam118a | 0.005 | Old |
| 0.791267 | 1.808665 | 0.018 | 0.015 | 1 | 9 | Col5a2 | 0.003 | Old |
| 0.791267 | 1.527036 | 0.018 | 0.015 | 1 | 9 | Arl16 | 0.003 | Old |
| 0.791267 | 1.101692 | 0.018 | 0.015 | 1 | 9 | Cystm1 | 0.003 | Old |
| 0.791267 | 0.806622 | 0.018 | 0.015 | 1 | 9 | Card6 | 0.003 | Old |
| 0.791267 | 0.78514 | 0.018 | 0.015 | 1 | 9 | Zfp934 | 0.003 | Old |
| 0.791267 | 0.762605 | 0.018 | 0.015 | 1 | 9 | Snx32 | 0.003 | Old |
| 0.791267 | 0.608071 | 0.018 | 0.015 | 1 | 9 | Bloc1s3 | 0.003 | Old |

| | | | | | | | | |
|---|---|---|---|---|---|---|---|---|
| 0.791299 | 1.281147 | 0.049 | 0.044 | 1 | 9 | Mb21d1 | 0.005 | Old |
| 0.791546 | 0.007226 | 0.126 | 0.132 | 1 | 9 | Parp12 | -0.006 | Old |
| 0.791917 | 1.918885 | 0.018 | 0.022 | 1 | 9 | Atg4c | -0.004 | Old |
| 0.792348 | 0.24198 | 0.057 | 0.051 | 1 | 9 | Nat6 | 0.006 | Old |
| 0.792672 | 0.443341 | 0.049 | 0.044 | 1 | 9 | Kantr | 0.005 | Old |
| 0.792867 | 1.295892 | 0.026 | 0.022 | 1 | 9 | Btbd6 | 0.004 | Old |
| 0.79289 | 0.193175 | 0.095 | 0.088 | 1 | 9 | Bcl7c | 0.007 | Old |
| 0.79289 | 0.117908 | 0.095 | 0.088 | 1 | 9 | Mdp1 | 0.007 | Old |
| 0.792905 | 0.100668 | 0.178 | 0.169 | 1 | 9 | Npepps | 0.009 | Old |
| 0.792949 | 0.579097 | 0.116 | 0.11 | 1 | 9 | Arl13b | 0.006 | Old |
| 0.793185 | 0.459559 | 0.08 | 0.074 | 1 | 9 | Mrps9 | 0.006 | Old |
| 0.793458 | 0.44889 | 0.064 | 0.059 | 1 | 9 | Flvcr1 | 0.005 | Old |
| 0.793521 | 1.220949 | 0.018 | 0.015 | 1 | 9 | Lysmd1 | 0.003 | Old |
| 0.793521 | 0.919142 | 0.018 | 0.015 | 1 | 9 | Cp | 0.003 | Old |
| 0.793521 | 0.692501 | 0.018 | 0.015 | 1 | 9 | Cbr3 | 0.003 | Old |
| 0.793521 | 0.28372 | 0.018 | 0.015 | 1 | 9 | Fam173b | 0.003 | Old |
| 0.793894 | 0.179185 | 0.095 | 0.088 | 1 | 9 | Tmem50b | 0.007 | Old |
| 0.794244 | 0.27097 | 0.108 | 0.103 | 1 | 9 | Tmem147 | 0.005 | Old |
| 0.794452 | 1.248369 | 0.034 | 0.029 | 1 | 9 | 2310061I0 | 0.005 | Old |
| 0.794452 | 1.013983 | 0.034 | 0.029 | 1 | 9 | Eef1akmt1 | 0.005 | Old |
| 0.794534 | 0.287675 | 0.095 | 0.103 | 1 | 9 | Trip10 | -0.008 | Old |
| 0.794605 | 0.160874 | 0.126 | 0.118 | 1 | 9 | Zfyve9 | 0.008 | Old |
| 0.794618 | 0.133042 | 0.139 | 0.147 | 1 | 9 | Rnf19a | -0.008 | Old |
| 0.794663 | 0.061662 | 0.273 | 0.265 | 1 | 9 | mt-Atp8 | 0.008 | Old |
| 0.794942 | 0.37281 | 0.173 | 0.169 | 1 | 9 | Smpdl3a | 0.004 | Old |
| 0.795028 | 0.464378 | 0.124 | 0.118 | 1 | 9 | Adar | 0.006 | Old |
| 0.795777 | 0.807782 | 0.018 | 0.015 | 1 | 9 | Col6a2 | 0.003 | Old |
| 0.795777 | 0.632382 | 0.018 | 0.015 | 1 | 9 | Simc1 | 0.003 | Old |
| 0.795777 | 0.605761 | 0.018 | 0.015 | 1 | 9 | Cbx8 | 0.003 | Old |
| 0.795777 | 0.594519 | 0.018 | 0.015 | 1 | 9 | Frs3 | 0.003 | Old |
| 0.795777 | 0.544839 | 0.018 | 0.015 | 1 | 9 | Zfp58 | 0.003 | Old |
| 0.795777 | 0.292602 | 0.018 | 0.015 | 1 | 9 | Nckap5l | 0.003 | Old |
| 0.796279 | 0.453057 | 0.072 | 0.081 | 1 | 9 | Rab28 | -0.009 | Old |
| 0.796636 | 1.074876 | 0.026 | 0.022 | 1 | 9 | Fgfr3 | 0.004 | Old |
| 0.796636 | 1.007119 | 0.026 | 0.022 | 1 | 9 | Per2 | 0.004 | Old |
| 0.796796 | 0.483528 | 0.049 | 0.044 | 1 | 9 | Ell | 0.005 | Old |
| 0.797076 | 0.298616 | 0.064 | 0.059 | 1 | 9 | Dgka | 0.005 | Old |
| 0.797211 | 0.37541 | 0.101 | 0.11 | 1 | 9 | Commd8 | -0.009 | Old |
| 0.79743 | 0.207074 | 0.139 | 0.154 | 1 | 9 | Nr1h2 | -0.015 | Old |
| 0.797473 | 0.758732 | 0.057 | 0.051 | 1 | 9 | Ndufaf3 | 0.006 | Old |
| 0.797473 | 0.363767 | 0.057 | 0.051 | 1 | 9 | Endou | 0.006 | Old |
| 0.798034 | 0.731636 | 0.018 | 0.015 | 1 | 9 | Zfp429 | 0.003 | Old |
| 0.798034 | 0.700911 | 0.018 | 0.015 | 1 | 9 | Tmem150a | 0.003 | Old |
| 0.798034 | 0.472456 | 0.018 | 0.015 | 1 | 9 | Pmf1 | 0.003 | Old |
| 0.798034 | 0.337858 | 0.018 | 0.015 | 1 | 9 | Dpf3 | 0.003 | Old |

| | | | | | | | | |
|---|---|---|---|---|---|---|---|---|
| 0.798034 | 0.323857 | 0.018 | 0.015 | 1 | 9 | Nat2 | 0.003 | Old |
| 0.798034 | 0.189611 | 0.018 | 0.015 | 1 | 9 | Mpp3 | 0.003 | Old |
| 0.798034 | 0.134029 | 0.018 | 0.015 | 1 | 9 | Zswim7 | 0.003 | Old |
| 0.798965 | 0.655639 | 0.041 | 0.037 | 1 | 9 | Gpr89 | 0.004 | Old |
| 0.799417 | 1.148445 | 0.034 | 0.029 | 1 | 9 | Ccdc93 | 0.005 | Old |
| 0.799417 | 0.998745 | 0.034 | 0.029 | 1 | 9 | Pinx1 | 0.005 | Old |
| 0.799799 | 0.049556 | 0.173 | 0.184 | 1 | 9 | Fem1c | -0.011 | Old |
| 0.799806 | 0.013201 | 0.131 | 0.125 | 1 | 9 | Supt6 | 0.006 | Old |
| 0.800039 | 0.62092 | 0.057 | 0.051 | 1 | 9 | Clk2 | 0.006 | Old |
| 0.800159 | 0.639536 | 0.085 | 0.081 | 1 | 9 | Emc8 | 0.004 | Old |
| 0.800293 | 0.521586 | 0.018 | 0.015 | 1 | 9 | Lyrm4 | 0.003 | Old |
| 0.800293 | 0.339547 | 0.018 | 0.015 | 1 | 9 | Zscan22 | 0.003 | Old |
| 0.800293 | 0.331526 | 0.018 | 0.015 | 1 | 9 | Gm42722 | 0.003 | Old |
| 0.800293 | 0.306968 | 0.018 | 0.015 | 1 | 9 | Sema4g | 0.003 | Old |
| 0.800293 | 0.216286 | 0.018 | 0.015 | 1 | 9 | P2ry6 | 0.003 | Old |
| 0.800293 | 0.132032 | 0.018 | 0.015 | 1 | 9 | Gm20661 | 0.003 | Old |
| 0.800409 | 1.026446 | 0.026 | 0.022 | 1 | 9 | Clasrp | 0.004 | Old |
| 0.800409 | 0.982008 | 0.026 | 0.022 | 1 | 9 | Wdr81 | 0.004 | Old |
| 0.800409 | 0.545777 | 0.026 | 0.022 | 1 | 9 | C87436 | 0.004 | Old |
| 0.800461 | 0.5842 | 0.041 | 0.037 | 1 | 9 | Tnfrsf12a | 0.004 | Old |
| 0.800461 | 0.461307 | 0.039 | 0.044 | 1 | 9 | Zfand2b | -0.005 | Old |
| 0.800461 | 0.122615 | 0.039 | 0.044 | 1 | 9 | Zfp319 | -0.005 | Old |
| 0.800605 | 0.243041 | 0.134 | 0.125 | 1 | 9 | Trim35 | 0.009 | Old |
| 0.800688 | 0.317422 | 0.101 | 0.096 | 1 | 9 | Hist1h1c | 0.005 | Old |
| 0.800925 | 0.395028 | 0.049 | 0.044 | 1 | 9 | Zfp758 | 0.005 | Old |
| 0.801074 | 0.864165 | 0.034 | 0.029 | 1 | 9 | Tmem39b | 0.005 | Old |
| 0.801074 | 0.673186 | 0.034 | 0.029 | 1 | 9 | Ltv1 | 0.005 | Old |
| 0.801312 | 0.018113 | 0.276 | 0.265 | 1 | 9 | Nptn | 0.011 | Old |
| 0.801376 | 0.013384 | 0.575 | 0.566 | 1 | 9 | Csde1 | 0.009 | Old |
| 0.801958 | 0.598354 | 0.039 | 0.044 | 1 | 9 | Fam210a | -0.005 | Old |
| 0.802297 | 0.713919 | 0.026 | 0.022 | 1 | 9 | Gm49359 | 0.004 | Old |
| 0.802297 | 0.084938 | 0.026 | 0.022 | 1 | 9 | Rdh14 | 0.004 | Old |
| 0.802403 | 0.195785 | 0.111 | 0.103 | 1 | 9 | Mrps18c | 0.008 | Old |
| 0.802554 | 0.20654 | 0.018 | 0.015 | 1 | 9 | Rad9a | 0.003 | Old |
| 0.802585 | 0.021088 | 0.464 | 0.456 | 1 | 9 | Klf13 | 0.008 | Old |
| 0.802732 | 0.877106 | 0.034 | 0.029 | 1 | 9 | Klc4 | 0.005 | Old |
| 0.802959 | 0.340926 | 0.291 | 0.294 | 1 | 9 | Srp9 | -0.003 | Old |
| 0.803203 | 0.022398 | 0.129 | 0.14 | 1 | 9 | 4933434E2 | -0.011 | Old |
| 0.803382 | 0.200182 | 0.059 | 0.066 | 1 | 9 | Ppm1d | -0.007 | Old |
| 0.803456 | 0.940901 | 0.041 | 0.037 | 1 | 9 | Alg6 | 0.004 | Old |
| 0.803456 | 0.874386 | 0.041 | 0.037 | 1 | 9 | Zfp846 | 0.004 | Old |
| 0.803456 | 0.733976 | 0.039 | 0.044 | 1 | 9 | Nbas | -0.005 | Old |
| 0.803456 | 0.508986 | 0.039 | 0.044 | 1 | 9 | Kcnj2 | -0.005 | Old |
| 0.803456 | 0.335217 | 0.041 | 0.037 | 1 | 9 | Tmem175 | 0.004 | Old |
| 0.803544 | 0.02166 | 0.113 | 0.125 | 1 | 9 | Pcyt1a | -0.012 | Old |

| | | | | | | | | |
|---|---|---|---|---|---|---|---|---|
| 0.803807 | 0.007807 | 0.291 | 0.309 | 1 | 9 | Dusp11 | -0.018 | Old |
| 0.803861 | 0.011805 | 0.082 | 0.088 | 1 | 9 | Josd1 | -0.006 | Old |
| 0.804391 | 0.578803 | 0.034 | 0.029 | 1 | 9 | Nipsnap1 | 0.005 | Old |
| 0.804673 | 0.104879 | 0.108 | 0.103 | 1 | 9 | Camkk2 | 0.005 | Old |
| 0.804816 | 0.084855 | 0.018 | 0.015 | 1 | 9 | Zfp229 | 0.003 | Old |
| 0.804955 | 0.356993 | 0.039 | 0.044 | 1 | 9 | Isca2 | -0.005 | Old |
| 0.80504 | 0.382602 | 0.08 | 0.088 | 1 | 9 | Bmt2 | -0.008 | Old |
| 0.80506 | 0.573794 | 0.049 | 0.044 | 1 | 9 | Fbxl14 | 0.005 | Old |
| 0.805256 | 0.659675 | 0.134 | 0.132 | 1 | 9 | Usp6nl | 0.002 | Old |
| 0.805675 | 0.076416 | 0.072 | 0.066 | 1 | 9 | Nudt5 | 0.006 | Old |
| 0.805789 | 0.148888 | 0.454 | 0.449 | 1 | 9 | Eif4a2 | 0.005 | Old |
| 0.806051 | 0.538789 | 0.034 | 0.029 | 1 | 9 | Prmt9 | 0.005 | Old |
| 0.806231 | 1.00555 | 0.093 | 0.088 | 1 | 9 | Zwint | 0.005 | Old |
| 0.806437 | 0.232538 | 0.147 | 0.14 | 1 | 9 | Ahctf1 | 0.007 | Old |
| 0.806463 | 0.217637 | 0.057 | 0.051 | 1 | 9 | Nt5c3 | 0.006 | Old |
| 0.806871 | 0.280569 | 0.088 | 0.081 | 1 | 9 | Gstp1 | 0.007 | Old |
| 0.80754 | 0.124507 | 0.211 | 0.206 | 1 | 9 | Wbp2 | 0.005 | Old |
| 0.807631 | 0.152718 | 0.108 | 0.118 | 1 | 9 | Dr1 | -0.01 | Old |
| 0.807819 | 0.184927 | 0.046 | 0.051 | 1 | 9 | Zfp830 | -0.005 | Old |
| 0.807819 | 0.11937 | 0.049 | 0.044 | 1 | 9 | Ctc1 | 0.005 | Old |
| 0.807826 | 0.031389 | 0.474 | 0.471 | 1 | 9 | Fry | 0.003 | Old |
| 0.807954 | 0.399846 | 0.041 | 0.037 | 1 | 9 | AI480526 | 0.004 | Old |
| 0.807969 | 0.705892 | 0.026 | 0.022 | 1 | 9 | Lhfpl2 | 0.004 | Old |
| 0.807969 | 0.43328 | 0.026 | 0.022 | 1 | 9 | Snap47 | 0.004 | Old |
| 0.807969 | 0.27188 | 0.026 | 0.022 | 1 | 9 | Colec12 | 0.004 | Old |
| 0.807971 | 0.761117 | 0.072 | 0.066 | 1 | 9 | Capn7 | 0.006 | Old |
| 0.808548 | 0.16294 | 0.116 | 0.125 | 1 | 9 | Ube2j1 | -0.009 | Old |
| 0.80866 | 0.123126 | 0.137 | 0.147 | 1 | 9 | Cript | -0.01 | Old |
| 0.809182 | 0.234908 | 0.165 | 0.154 | 1 | 9 | Syt7 | 0.011 | Old |
| 0.8092 | 0.746405 | 0.049 | 0.044 | 1 | 9 | Aatk | 0.005 | Old |
| 0.809346 | 0.746381 | 0.018 | 0.015 | 1 | 9 | Zfp995 | 0.003 | Old |
| 0.809373 | 0.474506 | 0.034 | 0.029 | 1 | 9 | B4galt7 | 0.005 | Old |
| 0.809373 | 0.391399 | 0.034 | 0.029 | 1 | 9 | Syn2 | 0.005 | Old |
| 0.809455 | 0.531992 | 0.039 | 0.044 | 1 | 9 | Mkrn2 | -0.005 | Old |
| 0.809573 | 0.078362 | 0.08 | 0.074 | 1 | 9 | Trim30d | 0.006 | Old |
| 0.809628 | 0.069345 | 0.214 | 0.206 | 1 | 9 | Ccdc47 | 0.008 | Old |
| 0.809861 | 0.472946 | 0.026 | 0.022 | 1 | 9 | Pwp2 | 0.004 | Old |
| 0.809861 | 0.23607 | 0.026 | 0.022 | 1 | 9 | Mib2 | 0.004 | Old |
| 0.810581 | 0.066931 | 0.049 | 0.044 | 1 | 9 | Triap1 | 0.005 | Old |
| 0.810751 | 0.454317 | 0.059 | 0.066 | 1 | 9 | Reps1 | -0.007 | Old |
| 0.810793 | 0.087912 | 0.067 | 0.074 | 1 | 9 | Mrpl46 | -0.007 | Old |
| 0.811024 | 0.185692 | 0.095 | 0.088 | 1 | 9 | Dcaf17 | 0.007 | Old |
| 0.811035 | 0.464514 | 0.034 | 0.029 | 1 | 9 | Man2b2 | 0.005 | Old |
| 0.811035 | 0.113782 | 0.034 | 0.029 | 1 | 9 | Rpf2 | 0.005 | Old |
| 0.811755 | 0.539331 | 0.026 | 0.022 | 1 | 9 | Gm45716 | 0.004 | Old |

| | | | | | | | | |
|---|---|---|---|---|---|---|---|---|
| 0.811755 | 0.293252 | 0.026 | 0.022 | 1 | 9 | Tab1 | 0.004 | Old |
| 0.811819 | 0.518484 | 0.116 | 0.11 | 1 | 9 | Nup88 | 0.006 | Old |
| 0.811935 | 0.011632 | 0.296 | 0.287 | 1 | 9 | Cbx3 | 0.009 | Old |
| 0.811981 | 0.249572 | 0.059 | 0.066 | 1 | 9 | St3gal4 | -0.007 | Old |
| 0.812458 | 0.903453 | 0.041 | 0.037 | 1 | 9 | Kcnq1 | 0.004 | Old |
| 0.812458 | 0.388559 | 0.041 | 0.037 | 1 | 9 | Edrf1 | 0.004 | Old |
| 0.812698 | 0.069614 | 0.034 | 0.029 | 1 | 9 | Gtf2f2 | 0.005 | Old |
| 0.813121 | 0.103012 | 0.544 | 0.537 | 1 | 9 | Tanc1 | 0.007 | Old |
| 0.81317 | 0.1386 | 0.49 | 0.493 | 1 | 9 | Serpinh1 | -0.003 | Old |
| 0.813346 | 0.931407 | 0.049 | 0.044 | 1 | 9 | Polr1c | 0.005 | Old |
| 0.81365 | 0.549323 | 0.026 | 0.022 | 1 | 9 | Plekha7 | 0.004 | Old |
| 0.81365 | 0.174956 | 0.026 | 0.022 | 1 | 9 | Stap1 | 0.004 | Old |
| 0.813767 | 0.23589 | 0.209 | 0.199 | 1 | 9 | Tomm70a | 0.01 | Old |
| 0.813961 | 0.522357 | 0.041 | 0.037 | 1 | 9 | Cdk5rap2 | 0.004 | Old |
| 0.814362 | 0.639293 | 0.034 | 0.029 | 1 | 9 | Mpp2 | 0.005 | Old |
| 0.814362 | 0.421642 | 0.034 | 0.029 | 1 | 9 | Rab11fip3 | 0.005 | Old |
| 0.814362 | 0.299049 | 0.034 | 0.029 | 1 | 9 | Rabggta | 0.005 | Old |
| 0.814463 | 0.147424 | 0.116 | 0.125 | 1 | 9 | R3hdm4 | -0.009 | Old |
| 0.814729 | 0.703702 | 0.049 | 0.044 | 1 | 9 | Sprtn | 0.005 | Old |
| 0.814729 | 0.521357 | 0.049 | 0.044 | 1 | 9 | 2410004B1 | 0.005 | Old |
| 0.814729 | 0.490277 | 0.049 | 0.044 | 1 | 9 | Plekhh3 | 0.005 | Old |
| 0.814784 | 0.160297 | 0.191 | 0.206 | 1 | 9 | Emsy | -0.015 | Old |
| 0.815464 | 0.355538 | 0.041 | 0.037 | 1 | 9 | Rasl11a | 0.004 | Old |
| 0.815547 | 0.185267 | 0.026 | 0.022 | 1 | 9 | Amacr | 0.004 | Old |
| 0.815673 | 0.904434 | 0.059 | 0.066 | 1 | 9 | Mt2 | -0.007 | Old |
| 0.816112 | 0.220707 | 0.049 | 0.044 | 1 | 9 | Tfcp2 | 0.005 | Old |
| 0.816654 | 0.101769 | 0.157 | 0.169 | 1 | 9 | Pin4 | -0.012 | Old |
| 0.816969 | 0.19294 | 0.041 | 0.037 | 1 | 9 | Traf2 | 0.004 | Old |
| 0.817444 | 0.016441 | 0.026 | 0.022 | 1 | 9 | Pdss2 | 0.004 | Old |
| 0.817692 | 0.349031 | 0.034 | 0.029 | 1 | 9 | Lyrm9 | 0.005 | Old |
| 0.817692 | 0.342427 | 0.034 | 0.029 | 1 | 9 | Vav2 | 0.005 | Old |
| 0.817692 | 0.107873 | 0.034 | 0.029 | 1 | 9 | Ten1 | 0.005 | Old |
| 0.817795 | 0.181398 | 0.088 | 0.096 | 1 | 9 | Tubgcp5 | -0.008 | Old |
| 0.817943 | 0.000638 | 0.108 | 0.118 | 1 | 9 | Pmp22 | -0.01 | Old |
| 0.818012 | 0.059494 | 0.526 | 0.559 | 1 | 9 | Mxd4 | -0.033 | Old |
| 0.81831 | 0.111854 | 0.026 | 0.029 | 1 | 9 | Dtd2 | -0.003 | Old |
| 0.818446 | 0.446306 | 0.232 | 0.228 | 1 | 9 | Ptpn11 | 0.004 | Old |
| 0.818473 | 0.325528 | 0.041 | 0.037 | 1 | 9 | Acyp1 | 0.004 | Old |
| 0.818951 | 0.278963 | 0.108 | 0.103 | 1 | 9 | Ykt6 | 0.005 | Old |
| 0.819309 | 0.272231 | 0.095 | 0.103 | 1 | 9 | Lrrc47 | -0.008 | Old |
| 0.819442 | 0.063075 | 0.103 | 0.11 | 1 | 9 | Aprt | -0.007 | Old |
| 0.819955 | 0.041304 | 0.665 | 0.618 | 1 | 9 | Id1 | 0.047 | Old |
| 0.820518 | 0.675909 | 0.093 | 0.088 | 1 | 9 | Hip1r | 0.005 | Old |
| 0.820546 | 0.013184 | 0.08 | 0.074 | 1 | 9 | Ints13 | 0.006 | Old |
| 0.820613 | 0.140652 | 0.147 | 0.14 | 1 | 9 | Epc2 | 0.007 | Old |

| | | | | | | | | |
|---|---|---|---|---|---|---|---|---|
| 0.820709 | 0.210304 | 0.149 | 0.162 | 1 | 9 | Fndc3a | -0.013 | Old |
| 0.821485 | 0.423576 | 0.041 | 0.037 | 1 | 9 | Tab3 | 0.004 | Old |
| 0.821652 | 0.085755 | 0.046 | 0.051 | 1 | 9 | Caml | -0.005 | Old |
| 0.821816 | 0.040668 | 0.441 | 0.434 | 1 | 9 | Tln1 | 0.007 | Old |
| 0.821932 | 0.364863 | 0.057 | 0.051 | 1 | 9 | Chst2 | 0.006 | Old |
| 0.821973 | 0.548415 | 0.026 | 0.029 | 1 | 9 | Zbtb11os1 | -0.003 | Old |
| 0.821973 | 0.134098 | 0.026 | 0.029 | 1 | 9 | Selenoo | -0.003 | Old |
| 0.822515 | 0.039222 | 0.219 | 0.213 | 1 | 9 | Psma4 | 0.006 | Old |
| 0.822769 | 0.227923 | 0.108 | 0.103 | 1 | 9 | Camsap1 | 0.005 | Old |
| 0.822927 | 0.585686 | 0.157 | 0.154 | 1 | 9 | Chmp2b | 0.003 | Old |
| 0.822977 | 0.713642 | 0.077 | 0.074 | 1 | 9 | 0610010F0 | 0.003 | Old |
| 0.823039 | 0.043355 | 0.046 | 0.051 | 1 | 9 | Serpinb1a | -0.005 | Old |
| 0.823291 | 0.077854 | 0.095 | 0.103 | 1 | 9 | Adh5 | -0.008 | Old |
| 0.823632 | 0.20556 | 0.26 | 0.257 | 1 | 9 | Stag1 | 0.003 | Old |
| 0.824363 | 0.445371 | 0.034 | 0.029 | 1 | 9 | Rgs16 | 0.005 | Old |
| 0.824499 | 0.077165 | 0.041 | 0.037 | 1 | 9 | Jrkl | 0.004 | Old |
| 0.824749 | 0.681956 | 0.093 | 0.103 | 1 | 9 | Zfp655 | -0.01 | Old |
| 0.824788 | 0.015286 | 0.067 | 0.074 | 1 | 9 | Gpd1l | -0.007 | Old |
| 0.824945 | 0.062283 | 0.08 | 0.074 | 1 | 9 | Cbll1 | 0.006 | Old |
| 0.825591 | 0.014741 | 0.322 | 0.346 | 1 | 9 | Myo1c | -0.024 | Old |
| 0.825696 | 0.393536 | 0.149 | 0.169 | 1 | 9 | Cops8 | -0.02 | Old |
| 0.825813 | 0.383321 | 0.046 | 0.051 | 1 | 9 | Pomgnt1 | -0.005 | Old |
| 0.825813 | 0.193641 | 0.046 | 0.051 | 1 | 9 | Ctps2 | -0.005 | Old |
| 0.825969 | 0.008503 | 0.162 | 0.176 | 1 | 9 | Hras | -0.014 | Old |
| 0.826008 | 0.081482 | 0.041 | 0.037 | 1 | 9 | Polr3a | 0.004 | Old |
| 0.826033 | 0.965037 | 0.034 | 0.029 | 1 | 9 | Prmt3 | 0.005 | Old |
| 0.826156 | 0.011494 | 0.119 | 0.125 | 1 | 9 | Sncaip | -0.006 | Old |
| 0.826542 | 0.132092 | 0.201 | 0.213 | 1 | 9 | Becn1 | -0.012 | Old |
| 0.826865 | 0.306758 | 0.134 | 0.132 | 1 | 9 | Prdx6 | 0.002 | Old |
| 0.827087 | 0.014201 | 0.183 | 0.169 | 1 | 9 | Lrp10 | 0.014 | Old |
| 0.827476 | 0.628076 | 0.026 | 0.029 | 1 | 9 | Prmt7 | -0.003 | Old |
| 0.827476 | 0.233656 | 0.026 | 0.029 | 1 | 9 | Twnk | -0.003 | Old |
| 0.827703 | 0.17567 | 0.034 | 0.029 | 1 | 9 | E130307A1 | 0.005 | Old |
| 0.827986 | 0.204362 | 0.075 | 0.081 | 1 | 9 | Ankra2 | -0.006 | Old |
| 0.828248 | 0.088464 | 0.08 | 0.074 | 1 | 9 | Decr1 | 0.006 | Old |
| 0.828592 | 0.008685 | 0.858 | 0.86 | 1 | 9 | Eif1 | -0.002 | Old |
| 0.829049 | 0.538749 | 0.155 | 0.169 | 1 | 9 | Nek9 | -0.014 | Old |
| 0.829375 | 0.136952 | 0.034 | 0.029 | 1 | 9 | Gm26669 | 0.005 | Old |
| 0.829653 | 0.056501 | 0.562 | 0.529 | 1 | 9 | Rps7 | 0.033 | Old |
| 0.830433 | 0.25417 | 0.072 | 0.066 | 1 | 9 | Phlpp2 | 0.006 | Old |
| 0.830607 | 0.219901 | 0.129 | 0.14 | 1 | 9 | Esd | -0.011 | Old |
| 0.830674 | 0.197526 | 0.046 | 0.051 | 1 | 9 | Ppp1r12b | -0.005 | Old |
| 0.830989 | 0.097246 | 0.054 | 0.059 | 1 | 9 | Thoc6 | -0.005 | Old |
| 0.831718 | 0.107636 | 0.059 | 0.066 | 1 | 9 | Slc39a6 | -0.007 | Old |
| 0.832445 | 0.092709 | 0.075 | 0.081 | 1 | 9 | Smim13 | -0.006 | Old |

| | | | | | | | | |
|---|---|---|---|---|---|---|---|---|
| 0.832463 | 0.129929 | 0.137 | 0.147 | 1 | 9 | Egln2 | -0.01 | Old |
| 0.832759 | 0.032794 | 0.046 | 0.051 | 1 | 9 | Zfp984 | -0.005 | Old |
| 0.832883 | 0.538413 | 0.289 | 0.301 | 1 | 9 | Slc6a6 | -0.012 | Old |
| 0.832998 | 0.020153 | 0.108 | 0.118 | 1 | 9 | Akap1 | -0.01 | Old |
| 0.833696 | 0.301936 | 0.201 | 0.221 | 1 | 9 | Rad23b | -0.02 | Old |
| 0.833895 | 0.040012 | 0.16 | 0.154 | 1 | 9 | Atp6v0a2 | 0.006 | Old |
| 0.834118 | 0.293788 | 0.077 | 0.074 | 1 | 9 | Dennd3 | 0.003 | Old |
| 0.834353 | 0.484564 | 0.144 | 0.14 | 1 | 9 | Wdr82 | 0.004 | Old |
| 0.834804 | 0.193396 | 0.137 | 0.132 | 1 | 9 | Tapbpl | 0.005 | Old |
| 0.834827 | 0.620885 | 0.026 | 0.029 | 1 | 9 | Endov | -0.003 | Old |
| 0.835321 | 0.57067 | 0.067 | 0.074 | 1 | 9 | Tgif1 | -0.007 | Old |
| 0.835668 | 0.501615 | 0.137 | 0.132 | 1 | 9 | Lpcat3 | 0.005 | Old |
| 0.836667 | 0.149352 | 0.026 | 0.029 | 1 | 9 | Zfp185 | -0.003 | Old |
| 0.836667 | 0.192852 | 0.026 | 0.029 | 1 | 9 | Tmc8 | -0.003 | Old |
| 0.836791 | 0.052466 | 0.706 | 0.706 | 1 | 9 | Qk | 0 | Old |
| 0.836933 | 0.184371 | 0.049 | 0.044 | 1 | 9 | Ikbkg | 0.005 | Old |
| 0.837783 | 0.037712 | 0.343 | 0.331 | 1 | 9 | Snx3 | 0.012 | Old |
| 0.837911 | 0.109152 | 0.082 | 0.088 | 1 | 9 | Cacul1 | -0.006 | Old |
| 0.838341 | 0.177837 | 0.121 | 0.132 | 1 | 9 | Mum1 | -0.011 | Old |
| 0.838584 | 0.509258 | 0.077 | 0.074 | 1 | 9 | Gphn | 0.003 | Old |
| 0.838921 | 0.683564 | 0.113 | 0.11 | 1 | 9 | Suclg1 | 0.003 | Old |
| 0.839047 | 0.128775 | 0.147 | 0.14 | 1 | 9 | Plekhb2 | 0.007 | Old |
| 0.839143 | 0.387967 | 0.075 | 0.081 | 1 | 9 | Ebp | -0.006 | Old |
| 0.839463 | 0.45291 | 0.08 | 0.088 | 1 | 9 | Ncor2 | -0.008 | Old |
| 0.839609 | 0.068487 | 0.041 | 0.037 | 1 | 9 | D830026I1 | 0.004 | Old |
| 0.839701 | 0.275346 | 0.077 | 0.074 | 1 | 9 | Fam160a2 | 0.003 | Old |
| 0.840349 | 0.183452 | 0.026 | 0.029 | 1 | 9 | Zfp37 | -0.003 | Old |
| 0.84056 | 0.047046 | 0.116 | 0.11 | 1 | 9 | Cxcr4 | 0.006 | Old |
| 0.841032 | 0.444789 | 0.093 | 0.088 | 1 | 9 | Fam133b | 0.005 | Old |
| 0.841655 | 0.547859 | 0.085 | 0.081 | 1 | 9 | Pik3r4 | 0.004 | Old |
| 0.842192 | 0.963695 | 0.026 | 0.029 | 1 | 9 | Fbxo32 | -0.003 | Old |
| 0.842505 | 0.571132 | 0.046 | 0.051 | 1 | 9 | Dhdds | -0.005 | Old |
| 0.842505 | 0.467037 | 0.046 | 0.051 | 1 | 9 | Klhl12 | -0.005 | Old |
| 0.843055 | 0.658126 | 0.077 | 0.074 | 1 | 9 | Dnmt1 | 0.003 | Old |
| 0.843804 | 0.02527 | 0.196 | 0.206 | 1 | 9 | Sf3b3 | -0.01 | Old |
| 0.844035 | 0.67055 | 0.026 | 0.029 | 1 | 9 | Bmp2 | -0.003 | Old |
| 0.844035 | 0.352397 | 0.026 | 0.029 | 1 | 9 | Cysltr2 | -0.003 | Old |
| 0.844073 | 0.136199 | 0.157 | 0.147 | 1 | 9 | Rybp | 0.01 | Old |
| 0.844708 | 0.612874 | 0.067 | 0.074 | 1 | 9 | Slc25a1 | -0.007 | Old |
| 0.845294 | 0.412567 | 0.046 | 0.051 | 1 | 9 | Zfp317 | -0.005 | Old |
| 0.845322 | 0.256358 | 0.204 | 0.199 | 1 | 9 | Ndufb6 | 0.005 | Old |
| 0.845403 | 0.20671 | 0.082 | 0.088 | 1 | 9 | Odf2l | -0.006 | Old |
| 0.845403 | 0.130736 | 0.082 | 0.088 | 1 | 9 | Spry2 | -0.006 | Old |
| 0.845487 | 0.038529 | 0.072 | 0.066 | 1 | 9 | Ergic1 | 0.006 | Old |
| 0.845515 | 0.483373 | 0.211 | 0.213 | 1 | 9 | Vwa5a | -0.002 | Old |

| | | | | | | | | |
|---|---|---|---|---|---|---|---|---|
| 0.845883 | 0.421331 | 0.067 | 0.074 | 1 | 9 | Pdcd11 | -0.007 | Old |
| 0.845938 | 0.509571 | 0.085 | 0.081 | 1 | 9 | Atg2a | 0.004 | Old |
| 0.846361 | 0.069587 | 0.809 | 0.838 | 1 | 9 | Fos | -0.029 | Old |
| 0.846514 | 0.128034 | 0.25 | 0.243 | 1 | 9 | Rnf11 | 0.007 | Old |
| 0.846562 | 1.656202 | 0.046 | 0.044 | 1 | 9 | Polr3f | 0.002 | Old |
| 0.846958 | 0.030459 | 0.214 | 0.206 | 1 | 9 | Actn1 | 0.008 | Old |
| 0.847001 | 0.081771 | 0.296 | 0.294 | 1 | 9 | Ranbp2 | 0.002 | Old |
| 0.847364 | 0.027223 | 0.343 | 0.36 | 1 | 9 | Rab5c | -0.017 | Old |
| 0.847875 | 0.026689 | 0.211 | 0.206 | 1 | 9 | Rbm18 | 0.005 | Old |
| 0.84807 | 0.380481 | 0.201 | 0.199 | 1 | 9 | 4932438A1 | 0.002 | Old |
| 0.848085 | 0.673312 | 0.046 | 0.051 | 1 | 9 | Tmem230 | -0.005 | Old |
| 0.848662 | 0.521073 | 0.054 | 0.051 | 1 | 9 | Rgs19 | 0.003 | Old |
| 0.848911 | 0.162028 | 0.289 | 0.309 | 1 | 9 | Tnrc6c | -0.02 | Old |
| 0.849134 | 0.560258 | 0.121 | 0.132 | 1 | 9 | Slc35a2 | -0.011 | Old |
| 0.849269 | 0.435381 | 0.093 | 0.088 | 1 | 9 | Gulp1 | 0.005 | Old |
| 0.849409 | 1.516681 | 0.046 | 0.044 | 1 | 9 | Dnmt3b | 0.002 | Old |
| 0.849812 | 0.084335 | 0.191 | 0.184 | 1 | 9 | Ddit4 | 0.007 | Old |
| 0.850226 | 0.328169 | 0.085 | 0.081 | 1 | 9 | Trappc3 | 0.004 | Old |
| 0.85047 | 0.061933 | 0.054 | 0.059 | 1 | 9 | Chp2 | -0.005 | Old |
| 0.850487 | 0.048977 | 0.284 | 0.301 | 1 | 9 | Calm3 | -0.017 | Old |
| 0.850729 | 0.320227 | 0.152 | 0.147 | 1 | 9 | Hoxb8 | 0.005 | Old |
| 0.851016 | 0.109693 | 0.124 | 0.132 | 1 | 9 | Slf2 | -0.008 | Old |
| 0.851418 | 1.455867 | 0.026 | 0.029 | 1 | 9 | Slc2a1 | -0.003 | Old |
| 0.851699 | 0.305147 | 0.142 | 0.14 | 1 | 9 | Pacsin2 | 0.002 | Old |
| 0.851723 | 0.143453 | 0.242 | 0.243 | 1 | 9 | Cox17 | -0.001 | Old |
| 0.852131 | 0.202183 | 0.193 | 0.206 | 1 | 9 | Ost4 | -0.013 | Old |
| 0.853015 | 0.175795 | 0.216 | 0.213 | 1 | 9 | Iws1 | 0.003 | Old |
| 0.853057 | 0.255071 | 0.101 | 0.096 | 1 | 9 | Icmt | 0.005 | Old |
| 0.853393 | 0.238577 | 0.093 | 0.088 | 1 | 9 | Abr | 0.005 | Old |
| 0.854086 | 0.360023 | 0.09 | 0.088 | 1 | 9 | Morc3 | 0.002 | Old |
| 0.854461 | 0.156789 | 0.253 | 0.25 | 1 | 9 | Zc3hav1 | 0.003 | Old |
| 0.854518 | 0.199279 | 0.085 | 0.081 | 1 | 9 | Ngly1 | 0.004 | Old |
| 0.854613 | 0.054681 | 0.247 | 0.243 | 1 | 9 | Trip11 | 0.004 | Old |
| 0.854772 | 0.194933 | 0.16 | 0.169 | 1 | 9 | Nampt | -0.009 | Old |
| 0.854917 | 0.050865 | 0.374 | 0.368 | 1 | 9 | Tcf25 | 0.006 | Old |
| 0.855055 | 0.140713 | 0.082 | 0.088 | 1 | 9 | Ccnk | -0.006 | Old |
| 0.855114 | 1.82807 | 0.026 | 0.029 | 1 | 9 | Actr5 | -0.003 | Old |
| 0.855309 | 0.252092 | 0.121 | 0.118 | 1 | 9 | Hjurp | 0.003 | Old |
| 0.855399 | 0.08771 | 0.222 | 0.235 | 1 | 9 | Golga4 | -0.013 | Old |
| 0.856044 | 0.140709 | 0.155 | 0.147 | 1 | 9 | Sh3bp4 | 0.008 | Old |
| 0.85635 | 0.34281 | 0.144 | 0.14 | 1 | 9 | Mrpl17 | 0.004 | Old |
| 0.857039 | 0.271287 | 0.101 | 0.096 | 1 | 9 | Stx16 | 0.005 | Old |
| 0.857062 | 0.563871 | 0.07 | 0.066 | 1 | 9 | Tpcn1 | 0.004 | Old |
| 0.857161 | 0.735824 | 0.062 | 0.059 | 1 | 9 | Selenon | 0.003 | Old |
| 0.857198 | 0.319685 | 0.144 | 0.14 | 1 | 9 | P4ha1 | 0.004 | Old |

| | | | | | | | | |
|---|---|---|---|---|---|---|---|---|
| 0.857621 | 0.434964 | 0.077 | 0.074 | 1 | 9 | Rabgef1 | 0.003 | Old |
| 0.858096 | 0.252356 | 0.139 | 0.147 | 1 | 9 | Plpbp | -0.008 | Old |
| 0.858114 | 0.639832 | 0.106 | 0.103 | 1 | 9 | Snx4 | 0.003 | Old |
| 0.859628 | 0.558772 | 0.227 | 0.25 | 1 | 9 | Mkl2 | -0.023 | Old |
| 0.860599 | 0.17491 | 0.07 | 0.066 | 1 | 9 | Mplkip | 0.004 | Old |
| 0.86066 | 0.1354 | 0.16 | 0.154 | 1 | 9 | Mindy1 | 0.006 | Old |
| 0.861479 | 0.086453 | 0.111 | 0.118 | 1 | 9 | Pfkfb3 | -0.007 | Old |
| 0.861639 | 0.372975 | 0.062 | 0.066 | 1 | 9 | Rfc2 | -0.004 | Old |
| 0.861738 | 0.421653 | 0.129 | 0.14 | 1 | 9 | Osbpl11 | -0.011 | Old |
| 0.862333 | 0.715075 | 0.098 | 0.096 | 1 | 9 | Brpf1 | 0.002 | Old |
| 0.862333 | 0.608161 | 0.098 | 0.096 | 1 | 9 | Hdgfl2 | 0.002 | Old |
| 0.862549 | 0.240797 | 0.034 | 0.037 | 1 | 9 | Psmg1 | -0.003 | Old |
| 0.863139 | 0.32963 | 0.268 | 0.287 | 1 | 9 | Nrep | -0.019 | Old |
| 0.863167 | 0.271059 | 0.165 | 0.176 | 1 | 9 | Dhx40 | -0.011 | Old |
| 0.864569 | 0.804894 | 0.054 | 0.051 | 1 | 9 | Nfkbil1 | 0.003 | Old |
| 0.865201 | 0.074562 | 0.335 | 0.331 | 1 | 9 | Eif5b | 0.004 | Old |
| 0.865378 | 0.069797 | 0.119 | 0.125 | 1 | 9 | Trp53 | -0.006 | Old |
| 0.866107 | 0.281764 | 0.263 | 0.279 | 1 | 9 | Skap2 | -0.016 | Old |
| 0.866308 | 0.187111 | 0.093 | 0.088 | 1 | 9 | Fchsd2 | 0.005 | Old |
| 0.866556 | 0.159673 | 0.062 | 0.066 | 1 | 9 | Zbtb37 | -0.004 | Old |
| 0.867132 | 0.646083 | 0.062 | 0.059 | 1 | 9 | A630072M | 0.003 | Old |
| 0.86747 | 0.559129 | 0.034 | 0.037 | 1 | 9 | Zfp518b | -0.003 | Old |
| 0.86747 | 0.289515 | 0.034 | 0.037 | 1 | 9 | Lrrc57 | -0.003 | Old |
| 0.868135 | 0.10189 | 0.371 | 0.368 | 1 | 9 | Bzw1 | 0.003 | Old |
| 0.868431 | 0.125673 | 0.186 | 0.184 | 1 | 9 | Mff | 0.002 | Old |
| 0.868813 | 0.267205 | 0.103 | 0.11 | 1 | 9 | Pgap2 | -0.007 | Old |
| 0.869016 | 0.289697 | 0.062 | 0.066 | 1 | 9 | Dhx57 | -0.004 | Old |
| 0.869112 | 0.867703 | 0.034 | 0.037 | 1 | 9 | Champ1 | -0.003 | Old |
| 0.869112 | 0.427838 | 0.034 | 0.037 | 1 | 9 | Nol9 | -0.003 | Old |
| 0.869112 | 0.337271 | 0.034 | 0.037 | 1 | 9 | Coprs | -0.003 | Old |
| 0.869112 | 0.028213 | 0.034 | 0.037 | 1 | 9 | Zfp142 | -0.003 | Old |
| 0.870246 | 0.150228 | 0.062 | 0.066 | 1 | 9 | Gramd4 | -0.004 | Old |
| 0.870547 | 0.571201 | 0.075 | 0.081 | 1 | 9 | Mrpl11 | -0.006 | Old |
| 0.870754 | 0.253626 | 0.034 | 0.037 | 1 | 9 | Zfp335 | -0.003 | Old |
| 0.870754 | 0.070355 | 0.034 | 0.037 | 1 | 9 | Ice2 | -0.003 | Old |
| 0.87135 | 0.53671 | 0.054 | 0.059 | 1 | 9 | Slc25a17 | -0.005 | Old |
| 0.871477 | 0.103006 | 0.062 | 0.066 | 1 | 9 | Mir17hg | -0.004 | Old |
| 0.871486 | 0.053141 | 0.389 | 0.39 | 1 | 9 | Cox6b1 | -0.001 | Old |
| 0.871747 | 0.253857 | 0.302 | 0.316 | 1 | 9 | Atp6v1f | -0.014 | Old |
| 0.872078 | 0.007692 | 0.175 | 0.169 | 1 | 9 | Herc6 | 0.006 | Old |
| 0.872086 | 0.01991 | 0.312 | 0.324 | 1 | 9 | Ogdh | -0.012 | Old |
| 0.872397 | 0.736007 | 0.034 | 0.037 | 1 | 9 | Bcl9 | -0.003 | Old |
| 0.872471 | 0.912723 | 0.013 | 0.015 | 1 | 9 | Osgepl1 | -0.002 | Old |
| 0.872658 | 0.462055 | 0.054 | 0.059 | 1 | 9 | Pfdn6 | -0.005 | Old |
| 0.873146 | 0.781723 | 0.101 | 0.11 | 1 | 9 | Irf8 | -0.009 | Old |

| | | | | | | | | |
|---|---|---|---|---|---|---|---|---|
| 0.873315 | 0.01159 | 0.16 | 0.169 | 1 | 9 | Nlk | -0.009 | Old |
| 0.873374 | 0.330811 | 0.062 | 0.059 | 1 | 9 | Hspbp1 | 0.003 | Old |
| 0.873859 | 0.251672 | 0.137 | 0.132 | 1 | 9 | Rnf2 | 0.005 | Old |
| 0.874772 | 0.383759 | 0.07 | 0.066 | 1 | 9 | Wdr43 | 0.004 | Old |
| 0.874962 | 0.294703 | 0.085 | 0.081 | 1 | 9 | Slit3 | 0.004 | Old |
| 0.875051 | 0.562154 | 0.075 | 0.081 | 1 | 9 | Zfp212 | -0.006 | Old |
| 0.875079 | 0.530682 | 0.013 | 0.015 | 1 | 9 | 1700086O( | -0.002 | Old |
| 0.875113 | 0.721482 | 0.046 | 0.044 | 1 | 9 | Htatip2 | 0.002 | Old |
| 0.87553 | 0.290325 | 0.165 | 0.162 | 1 | 9 | Map2k2 | 0.003 | Old |
| 0.875684 | 0.966409 | 0.034 | 0.037 | 1 | 9 | Dnajc17 | -0.003 | Old |
| 0.875827 | 0.091743 | 0.041 | 0.044 | 1 | 9 | Cep44 | -0.003 | Old |
| 0.876212 | 0.338664 | 0.281 | 0.279 | 1 | 9 | Ppp4r3b | 0.002 | Old |
| 0.876239 | 0.102906 | 0.224 | 0.221 | 1 | 9 | Mfsd6 | 0.003 | Old |
| 0.876388 | 0.194617 | 0.121 | 0.118 | 1 | 9 | Sf3a2 | 0.003 | Old |
| 0.876411 | 0.16153 | 0.229 | 0.243 | 1 | 9 | Stom | -0.014 | Old |
| 0.876753 | 0.109461 | 0.157 | 0.154 | 1 | 9 | Kdm5c | 0.003 | Old |
| 0.877328 | 0.535755 | 0.034 | 0.037 | 1 | 9 | Gnat3 | -0.003 | Old |
| 0.877328 | 0.23451 | 0.034 | 0.037 | 1 | 9 | Nt5dc3 | -0.003 | Old |
| 0.877688 | 0.899028 | 0.013 | 0.015 | 1 | 9 | Bicc1 | -0.002 | Old |
| 0.877688 | 0.633513 | 0.013 | 0.015 | 1 | 9 | Bcl2a1a | -0.002 | Old |
| 0.877688 | 0.551364 | 0.013 | 0.015 | 1 | 9 | Gm42517 | -0.002 | Old |
| 0.877688 | 0.535967 | 0.013 | 0.015 | 1 | 9 | Tlr6 | -0.002 | Old |
| 0.877688 | 0.502344 | 0.013 | 0.015 | 1 | 9 | Bbs1 | -0.002 | Old |
| 0.877688 | 0.430629 | 0.013 | 0.015 | 1 | 9 | Nadsyn1 | -0.002 | Old |
| 0.877688 | 0.340138 | 0.013 | 0.015 | 1 | 9 | Lppos | -0.002 | Old |
| 0.877688 | 0.166421 | 0.013 | 0.015 | 1 | 9 | 9430015G1 | -0.002 | Old |
| 0.877688 | 0.129979 | 0.013 | 0.015 | 1 | 9 | 2210408I2 | -0.002 | Old |
| 0.877688 | 0.11509 | 0.013 | 0.015 | 1 | 9 | Adat2 | -0.002 | Old |
| 0.877688 | 0.107932 | 0.013 | 0.015 | 1 | 9 | Rrad | -0.002 | Old |
| 0.877688 | 0.04943 | 0.013 | 0.015 | 1 | 9 | Arhgef10l | -0.002 | Old |
| 0.877688 | 0.008133 | 0.013 | 0.015 | 1 | 9 | 1810043G( | -0.002 | Old |
| 0.877762 | 0.165435 | 0.229 | 0.228 | 1 | 9 | Cbx5 | 0.001 | Old |
| 0.877867 | 0.601037 | 0.054 | 0.051 | 1 | 9 | Gpatch11 | 0.003 | Old |
| 0.878649 | 0.11481 | 0.175 | 0.184 | 1 | 9 | Stat5b | -0.009 | Old |
| 0.878973 | 0.983031 | 0.034 | 0.037 | 1 | 9 | Slc27a4 | -0.003 | Old |
| 0.878973 | 0.532758 | 0.034 | 0.037 | 1 | 9 | Znhit3 | -0.003 | Old |
| 0.878973 | 0.436148 | 0.034 | 0.037 | 1 | 9 | Bace2 | -0.003 | Old |
| 0.878995 | 0.192905 | 0.077 | 0.074 | 1 | 9 | Mycbp | 0.003 | Old |
| 0.879002 | 0.038176 | 0.101 | 0.096 | 1 | 9 | Pspc1 | 0.005 | Old |
| 0.879277 | 0.028189 | 0.085 | 0.081 | 1 | 9 | Ubr7 | 0.004 | Old |
| 0.879411 | 0.782921 | 0.046 | 0.044 | 1 | 9 | Polr2h | 0.002 | Old |
| 0.879538 | 0.378674 | 0.103 | 0.103 | 1 | 9 | Srp54a | 0 | Old |
| 0.879817 | 0.385828 | 0.082 | 0.088 | 1 | 9 | Tbc1d10b | -0.006 | Old |
| 0.880046 | 0.103333 | 0.286 | 0.279 | 1 | 9 | Prkch | 0.007 | Old |
| 0.880298 | 1.655251 | 0.013 | 0.015 | 1 | 9 | Lsr | -0.002 | Old |

| | | | | | | | | |
|---|---|---|---|---|---|---|---|---|
| 0.880298 | 1.274606 | 0.013 | 0.015 | 1 | 9 | Alkbh4 | -0.002 | Old |
| 0.880298 | 0.315177 | 0.013 | 0.015 | 1 | 9 | Armc7 | -0.002 | Old |
| 0.880614 | 0.356258 | 0.116 | 0.125 | 1 | 9 | Mtmr6 | -0.009 | Old |
| 0.881007 | 0.834445 | 0.075 | 0.074 | 1 | 9 | Fam241a | 0.001 | Old |
| 0.881334 | 0.234208 | 0.062 | 0.066 | 1 | 9 | Agfg2 | -0.004 | Old |
| 0.881642 | 0.067484 | 0.227 | 0.235 | 1 | 9 | Epn1 | -0.008 | Old |
| 0.881863 | 0.525738 | 0.054 | 0.051 | 1 | 9 | Nfyc | 0.003 | Old |
| 0.881873 | 0.354223 | 0.07 | 0.066 | 1 | 9 | Acvr2b | 0.004 | Old |
| 0.882003 | 0.395967 | 0.119 | 0.118 | 1 | 9 | Cep120 | 0.001 | Old |
| 0.882264 | 0.891576 | 0.034 | 0.037 | 1 | 9 | Mlh3 | -0.003 | Old |
| 0.882278 | 0.707572 | 0.046 | 0.044 | 1 | 9 | Lman2l | 0.002 | Old |
| 0.88291 | 1.193838 | 0.013 | 0.015 | 1 | 9 | C5ar2 | -0.002 | Old |
| 0.88291 | 1.014288 | 0.013 | 0.015 | 1 | 9 | Ism1 | -0.002 | Old |
| 0.88291 | 0.859023 | 0.013 | 0.015 | 1 | 9 | Hist2h2be | -0.002 | Old |
| 0.88291 | 0.782563 | 0.013 | 0.015 | 1 | 9 | Gja4 | -0.002 | Old |
| 0.88291 | 0.756814 | 0.013 | 0.015 | 1 | 9 | Ighmbp2 | -0.002 | Old |
| 0.88291 | 0.745456 | 0.013 | 0.015 | 1 | 9 | Zfp119b | -0.002 | Old |
| 0.88291 | 0.71925 | 0.013 | 0.015 | 1 | 9 | Gm26982 | -0.002 | Old |
| 0.88291 | 0.424992 | 0.013 | 0.015 | 1 | 9 | Cd72 | -0.002 | Old |
| 0.88291 | 0.412141 | 0.013 | 0.015 | 1 | 9 | Dnajc4 | -0.002 | Old |
| 0.88291 | 0.306808 | 0.013 | 0.015 | 1 | 9 | Atp6v1g2 | -0.002 | Old |
| 0.88291 | 0.230211 | 0.013 | 0.015 | 1 | 9 | Atoh7 | -0.002 | Old |
| 0.88291 | 0.219999 | 0.013 | 0.015 | 1 | 9 | Nemp2 | -0.002 | Old |
| 0.88291 | 0.21001 | 0.013 | 0.015 | 1 | 9 | Gtf2h3 | -0.002 | Old |
| 0.882953 | 0.001126 | 0.106 | 0.103 | 1 | 9 | Sqor | 0.003 | Old |
| 0.883212 | 0.097319 | 0.129 | 0.125 | 1 | 9 | Dyrk2 | 0.004 | Old |
| 0.883353 | 0.792331 | 0.039 | 0.037 | 1 | 9 | Tsta3 | 0.002 | Old |
| 0.883666 | 0.165729 | 0.157 | 0.169 | 1 | 9 | Tet2 | -0.012 | Old |
| 0.883677 | 0.09852 | 0.113 | 0.11 | 1 | 9 | Chst11 | 0.003 | Old |
| 0.883802 | 0.285145 | 0.062 | 0.066 | 1 | 9 | Cblb | -0.004 | Old |
| 0.884447 | 0.556404 | 0.054 | 0.059 | 1 | 9 | Ciapin1 | -0.005 | Old |
| 0.884661 | 0.152242 | 0.119 | 0.125 | 1 | 9 | Agpat5 | -0.006 | Old |
| 0.884792 | 0.138476 | 0.041 | 0.044 | 1 | 9 | Pex5 | -0.003 | Old |
| 0.885146 | 0.504801 | 0.046 | 0.044 | 1 | 9 | Prpf31 | 0.002 | Old |
| 0.885523 | 1.132568 | 0.013 | 0.015 | 1 | 9 | Gm21781 | -0.002 | Old |
| 0.885523 | 0.693153 | 0.013 | 0.015 | 1 | 9 | Rilp | -0.002 | Old |
| 0.885523 | 0.341799 | 0.013 | 0.015 | 1 | 9 | Dennd6b | -0.002 | Old |
| 0.885523 | 0.24391 | 0.013 | 0.015 | 1 | 9 | Lekr1 | -0.002 | Old |
| 0.885787 | 0.148948 | 0.157 | 0.154 | 1 | 9 | Acat1 | 0.003 | Old |
| 0.886005 | 0.267004 | 0.09 | 0.096 | 1 | 9 | Zfp516 | -0.006 | Old |
| 0.886045 | 0.454282 | 0.111 | 0.118 | 1 | 9 | Uba5 | -0.007 | Old |
| 0.886128 | 0.036211 | 0.415 | 0.397 | 1 | 9 | Cd38 | 0.018 | Old |
| 0.886484 | 0.949403 | 0.039 | 0.037 | 1 | 9 | Zbtb7b | 0.002 | Old |
| 0.886581 | 0.851398 | 0.046 | 0.044 | 1 | 9 | Mcm3 | 0.002 | Old |
| 0.886893 | 0.130562 | 0.077 | 0.074 | 1 | 9 | 3110062M | 0.003 | Old |

| | | | | | | | | |
|---|---|---|---|---|---|---|---|---|
| 0.887349 | 0.131798 | 0.103 | 0.11 | 1 | 9 | Crlf3 | -0.007 | Old |
| 0.887378 | 0.367439 | 0.082 | 0.088 | 1 | 9 | Ermp1 | -0.006 | Old |
| 0.887799 | 0.207368 | 0.07 | 0.066 | 1 | 9 | Gypc | 0.004 | Old |
| 0.888009 | 0.162338 | 0.175 | 0.184 | 1 | 9 | Bag1 | -0.009 | Old |
| 0.888137 | 1.827717 | 0.013 | 0.015 | 1 | 9 | Ccdc122 | -0.002 | Old |
| 0.888137 | 1.27778 | 0.013 | 0.015 | 1 | 9 | Dpp7 | -0.002 | Old |
| 0.888137 | 1.227354 | 0.013 | 0.015 | 1 | 9 | Naprt | -0.002 | Old |
| 0.888137 | 1.02623 | 0.013 | 0.015 | 1 | 9 | Pcgf1 | -0.002 | Old |
| 0.888137 | 0.928439 | 0.013 | 0.015 | 1 | 9 | Cpm | -0.002 | Old |
| 0.888137 | 0.813111 | 0.013 | 0.015 | 1 | 9 | Siglece | -0.002 | Old |
| 0.888137 | 0.434755 | 0.013 | 0.015 | 1 | 9 | Armc9 | -0.002 | Old |
| 0.888294 | 0.11873 | 0.214 | 0.213 | 1 | 9 | Tubb2a | 0.001 | Old |
| 0.888388 | 0.20143 | 0.062 | 0.059 | 1 | 9 | Nhlrc2 | 0.003 | Old |
| 0.888531 | 0.695442 | 0.054 | 0.051 | 1 | 9 | Slc9a8 | 0.003 | Old |
| 0.888531 | 0.460148 | 0.054 | 0.051 | 1 | 9 | Mtm1 | 0.003 | Old |
| 0.888853 | 0.402863 | 0.034 | 0.037 | 1 | 9 | Kmt5c | -0.003 | Old |
| 0.888887 | 0.114327 | 0.111 | 0.118 | 1 | 9 | Rnf122 | -0.007 | Old |
| 0.889304 | 0.167251 | 0.106 | 0.103 | 1 | 9 | Stam | 0.003 | Old |
| 0.890397 | 0.00803 | 0.077 | 0.081 | 1 | 9 | Hikeshi | -0.004 | Old |
| 0.890888 | 0.50012 | 0.046 | 0.044 | 1 | 9 | Herc3 | 0.002 | Old |
| 0.890888 | 0.01523 | 0.046 | 0.044 | 1 | 9 | Med27 | 0.002 | Old |
| 0.890894 | 0.178838 | 0.062 | 0.059 | 1 | 9 | Mknk1 | 0.003 | Old |
| 0.890959 | 0.148254 | 0.175 | 0.169 | 1 | 9 | Nos2 | 0.006 | Old |
| 0.891008 | 0.6004 | 0.054 | 0.059 | 1 | 9 | Mfsd11 | -0.005 | Old |
| 0.891184 | 1.097294 | 0.039 | 0.037 | 1 | 9 | Usb1 | 0.002 | Old |
| 0.891357 | 0.610811 | 0.103 | 0.103 | 1 | 9 | Polb | 0 | Old |
| 0.891403 | 0.294174 | 0.155 | 0.154 | 1 | 9 | Golgb1 | 0.001 | Old |
| 0.891703 | 0.462777 | 0.082 | 0.088 | 1 | 9 | Gab2 | -0.006 | Old |
| 0.891846 | 0.130466 | 0.204 | 0.206 | 1 | 9 | Ptpn1 | -0.002 | Old |
| 0.892535 | 0.490647 | 0.054 | 0.051 | 1 | 9 | Ecm1 | 0.003 | Old |
| 0.892752 | 0.620151 | 0.039 | 0.037 | 1 | 9 | Tada2a | 0.002 | Old |
| 0.893326 | 0.081911 | 0.085 | 0.081 | 1 | 9 | Rbms2 | 0.004 | Old |
| 0.894319 | 0.93128 | 0.039 | 0.037 | 1 | 9 | Fgd6 | 0.002 | Old |
| 0.894593 | 0.026643 | 0.235 | 0.243 | 1 | 9 | Crybg3 | -0.008 | Old |
| 0.894713 | 0.026783 | 0.18 | 0.176 | 1 | 9 | Acbd5 | 0.004 | Old |
| 0.894821 | 0.017058 | 0.137 | 0.132 | 1 | 9 | Rab3d | 0.005 | Old |
| 0.894949 | 0.250757 | 0.082 | 0.088 | 1 | 9 | Ccz1 | -0.006 | Old |
| 0.89528 | 0.041014 | 0.152 | 0.147 | 1 | 9 | Cped1 | 0.005 | Old |
| 0.895469 | 1.319788 | 0.031 | 0.029 | 1 | 9 | Zswim4 | 0.002 | Old |
| 0.895888 | 0.967784 | 0.039 | 0.037 | 1 | 9 | Cep85 | 0.002 | Old |
| 0.896768 | 0.27158 | 0.041 | 0.044 | 1 | 9 | Zfhx2 | -0.003 | Old |
| 0.896874 | 0.162552 | 0.188 | 0.191 | 1 | 9 | Guk1 | -0.003 | Old |
| 0.897365 | 0.021704 | 0.198 | 0.199 | 1 | 9 | Ndufa1 | -0.001 | Old |
| 0.897421 | 0.308331 | 0.113 | 0.11 | 1 | 9 | Mindy3 | 0.003 | Old |
| 0.897456 | 0.888202 | 0.039 | 0.037 | 1 | 9 | Sesn3 | 0.002 | Old |

| | | | | | | | | |
|---|---|---|---|---|---|---|---|---|
| 0.897456 | 0.622791 | 0.039 | 0.037 | 1 | 9 | Coq8b | 0.002 | Old |
| 0.897556 | 0.159231 | 0.129 | 0.125 | 1 | 9 | Taf1d | 0.004 | Old |
| 0.898073 | 0.452899 | 0.046 | 0.044 | 1 | 9 | Lst1 | 0.002 | Old |
| 0.898222 | 0.012254 | 0.077 | 0.081 | 1 | 9 | Zfp512 | -0.004 | Old |
| 0.898861 | 0.132556 | 0.196 | 0.206 | 1 | 9 | Rwdd1 | -0.01 | Old |
| 0.898986 | 0.046538 | 0.505 | 0.515 | 1 | 9 | Arf1 | -0.01 | Old |
| 0.899866 | 0.431004 | 0.062 | 0.066 | 1 | 9 | Dscr3 | -0.004 | Old |
| 0.900344 | 0.224469 | 0.07 | 0.074 | 1 | 9 | Dhx8 | -0.004 | Old |
| 0.900347 | 0.175074 | 0.134 | 0.132 | 1 | 9 | Vps4a | 0.002 | Old |
| 0.900595 | 0.325415 | 0.039 | 0.037 | 1 | 9 | Nedd1 | 0.002 | Old |
| 0.90117 | 0.017738 | 0.049 | 0.051 | 1 | 9 | Gpatch2l | -0.002 | Old |
| 0.901266 | 0.706117 | 0.041 | 0.044 | 1 | 9 | E130309D( | -0.003 | Old |
| 0.901279 | 0.028272 | 0.366 | 0.36 | 1 | 9 | Trip12 | 0.006 | Old |
| 0.901578 | 0.243031 | 0.077 | 0.081 | 1 | 9 | Atp2a3 | -0.004 | Old |
| 0.90159 | 0.034691 | 0.077 | 0.074 | 1 | 9 | Rpe | 0.003 | Old |
| 0.902388 | 0.659829 | 0.046 | 0.044 | 1 | 9 | Ftsj1 | 0.002 | Old |
| 0.902565 | 0.760311 | 0.067 | 0.066 | 1 | 9 | 0610030E2 | 0.001 | Old |
| 0.903117 | 0.110428 | 0.113 | 0.11 | 1 | 9 | Mfap1a | 0.003 | Old |
| 0.903292 | 0.40732 | 0.106 | 0.11 | 1 | 9 | Fam98b | -0.004 | Old |
| 0.904423 | 0.144188 | 0.07 | 0.066 | 1 | 9 | Pdpr | 0.004 | Old |
| 0.904741 | 0.051447 | 0.126 | 0.132 | 1 | 9 | Tcof1 | -0.006 | Old |
| 0.904817 | 0.752028 | 0.062 | 0.066 | 1 | 9 | Casp2 | -0.004 | Old |
| 0.905825 | 1.337417 | 0.059 | 0.059 | 1 | 9 | Snx8 | 0 | Old |
| 0.906042 | 0.02962 | 0.121 | 0.125 | 1 | 9 | 4931406C( | -0.004 | Old |
| 0.906212 | 0.396115 | 0.07 | 0.074 | 1 | 9 | Ola1 | -0.004 | Old |
| 0.90647 | 0.036394 | 0.951 | 0.912 | 1 | 9 | H2-K1 | 0.039 | Old |
| 0.906585 | 0.100011 | 0.361 | 0.353 | 1 | 9 | Gm26917 | 0.008 | Old |
| 0.906706 | 0.395095 | 0.046 | 0.044 | 1 | 9 | Ccdc28b | 0.002 | Old |
| 0.906866 | 0.55644 | 0.082 | 0.088 | 1 | 9 | Kdm2b | -0.006 | Old |
| 0.907003 | 0.036319 | 0.296 | 0.301 | 1 | 9 | Gna11 | -0.005 | Old |
| 0.90711 | 0.034388 | 0.327 | 0.331 | 1 | 9 | Enpp4 | -0.004 | Old |
| 0.907384 | 0.678638 | 0.067 | 0.066 | 1 | 9 | Gnl3l | 0.001 | Old |
| 0.907518 | 0.047966 | 0.278 | 0.279 | 1 | 9 | Sbno1 | -0.001 | Old |
| 0.907708 | 1.168438 | 0.031 | 0.029 | 1 | 9 | 2310039H( | 0.002 | Old |
| 0.907708 | 0.845531 | 0.031 | 0.029 | 1 | 9 | Gipc3 | 0.002 | Old |
| 0.907711 | 0.009104 | 0.312 | 0.309 | 1 | 9 | Ep300 | 0.003 | Old |
| 0.908022 | 0.201254 | 0.157 | 0.154 | 1 | 9 | Smu1 | 0.003 | Old |
| 0.908333 | 0.091131 | 0.188 | 0.184 | 1 | 9 | Phf14 | 0.004 | Old |
| 0.909459 | 0.821936 | 0.031 | 0.029 | 1 | 9 | Nprl3 | 0.002 | Old |
| 0.909624 | 0.220696 | 0.155 | 0.154 | 1 | 9 | Sys1 | 0.001 | Old |
| 0.90995 | 0.164832 | 0.268 | 0.279 | 1 | 9 | Colgalt1 | -0.011 | Old |
| 0.91027 | 0.136559 | 0.041 | 0.044 | 1 | 9 | Mxd1 | -0.003 | Old |
| 0.910662 | 0.954931 | 0.088 | 0.096 | 1 | 9 | Shtn1 | -0.008 | Old |
| 0.910861 | 0.295261 | 0.142 | 0.14 | 1 | 9 | Mphosph8 | 0.002 | Old |
| 0.911255 | 0.303507 | 0.054 | 0.051 | 1 | 9 | Cnpy3 | 0.003 | Old |

| | | | | | | | | |
|---|---|---|---|---|---|---|---|---|
| 0.911366 | 0.369977 | 0.082 | 0.081 | 1 | 9 | Wdr89 | 0.001 | Old |
| 0.912086 | 0.202173 | 0.07 | 0.074 | 1 | 9 | Nepn | -0.004 | Old |
| 0.912113 | 0.391146 | 0.155 | 0.154 | 1 | 9 | Map2k7 | 0.001 | Old |
| 0.912113 | 0.073823 | 0.155 | 0.154 | 1 | 9 | Gclm | 0.001 | Old |
| 0.912251 | 0.870601 | 0.062 | 0.066 | 1 | 9 | Rbm14 | -0.004 | Old |
| 0.912269 | 0.205083 | 0.173 | 0.184 | 1 | 9 | Irak1 | -0.011 | Old |
| 0.912465 | 0.048575 | 0.253 | 0.243 | 1 | 9 | Clock | 0.01 | Old |
| 0.912778 | 0.262566 | 0.077 | 0.081 | 1 | 9 | Smarcd1 | -0.004 | Old |
| 0.912961 | 0.674692 | 0.031 | 0.029 | 1 | 9 | Vegfa | 0.002 | Old |
| 0.913047 | 0.47442 | 0.075 | 0.074 | 1 | 9 | Lurap1l | 0.001 | Old |
| 0.913274 | 0.594261 | 0.041 | 0.044 | 1 | 9 | Smox | -0.003 | Old |
| 0.913274 | 0.503282 | 0.041 | 0.044 | 1 | 9 | Arsk | -0.003 | Old |
| 0.913274 | 0.488945 | 0.041 | 0.044 | 1 | 9 | Sestd1 | -0.003 | Old |
| 0.913368 | 0.077338 | 0.222 | 0.228 | 1 | 9 | Scamp1 | -0.006 | Old |
| 0.913909 | 0.032269 | 0.046 | 0.044 | 1 | 9 | Erbb2 | 0.002 | Old |
| 0.913941 | 0.359971 | 0.07 | 0.066 | 1 | 9 | Kin | 0.004 | Old |
| 0.913993 | 0.370769 | 0.106 | 0.11 | 1 | 9 | Il3ra | -0.004 | Old |
| 0.914115 | 0.205346 | 0.09 | 0.096 | 1 | 9 | Tmem183a | -0.006 | Old |
| 0.91453 | 0.302514 | 0.126 | 0.125 | 1 | 9 | Erp44 | 0.001 | Old |
| 0.914712 | 0.656774 | 0.031 | 0.029 | 1 | 9 | Scamp5 | 0.002 | Old |
| 0.914712 | 0.517571 | 0.031 | 0.029 | 1 | 9 | Tmem115 | 0.002 | Old |
| 0.914712 | 0.397612 | 0.031 | 0.029 | 1 | 9 | Stambp | 0.002 | Old |
| 0.915043 | 1.667821 | 0.023 | 0.022 | 1 | 9 | Antxr2 | 0.001 | Old |
| 0.915826 | 0.771512 | 0.067 | 0.066 | 1 | 9 | Lsm1 | 0.001 | Old |
| 0.915971 | 0.977232 | 0.062 | 0.066 | 1 | 9 | Lgalsl | -0.004 | Old |
| 0.916224 | 0.047923 | 0.443 | 0.449 | 1 | 9 | Shank3 | -0.006 | Old |
| 0.916294 | 0.058594 | 0.16 | 0.169 | 1 | 9 | Arid2 | -0.009 | Old |
| 0.91642 | 0.391305 | 0.049 | 0.051 | 1 | 9 | Fcgr3 | -0.002 | Old |
| 0.916464 | 0.139519 | 0.031 | 0.029 | 1 | 9 | Uck1 | 0.002 | Old |
| 0.916613 | 0.152816 | 0.054 | 0.051 | 1 | 9 | Prdx3 | 0.003 | Old |
| 0.91696 | 0.057962 | 0.325 | 0.331 | 1 | 9 | Gapvd1 | -0.006 | Old |
| 0.917059 | 1.237494 | 0.023 | 0.022 | 1 | 9 | Tpk1 | 0.001 | Old |
| 0.917059 | 1.048828 | 0.023 | 0.022 | 1 | 9 | Rnf24 | 0.001 | Old |
| 0.917494 | 0.297524 | 0.093 | 0.096 | 1 | 9 | Iars | -0.003 | Old |
| 0.917808 | 0.519871 | 0.049 | 0.051 | 1 | 9 | Tcaf2 | -0.002 | Old |
| 0.918217 | 0.255338 | 0.031 | 0.029 | 1 | 9 | Cnnm4 | 0.002 | Old |
| 0.918554 | 0.226041 | 0.085 | 0.088 | 1 | 9 | Tbc1d5 | -0.003 | Old |
| 0.919075 | 1.007901 | 0.023 | 0.022 | 1 | 9 | Pdcd2l | 0.001 | Old |
| 0.919457 | 0.046088 | 0.039 | 0.037 | 1 | 9 | Gm46430 | 0.002 | Old |
| 0.919676 | 0.023824 | 0.046 | 0.044 | 1 | 9 | Tex10 | 0.002 | Old |
| 0.919969 | 0.006981 | 0.031 | 0.029 | 1 | 9 | Ifit3 | 0.002 | Old |
| 0.919969 | 0.938866 | 0.031 | 0.029 | 1 | 9 | Eid2b | 0.002 | Old |
| 0.919969 | 0.306718 | 0.031 | 0.029 | 1 | 9 | D17H6S53I | 0.002 | Old |
| 0.920144 | 0.287652 | 0.09 | 0.088 | 1 | 9 | Ndrg3 | 0.002 | Old |
| 0.920672 | 0.16285 | 0.098 | 0.103 | 1 | 9 | Etfa | -0.005 | Old |

| | | | | | | | | |
|---|---|---|---|---|---|---|---|---|
| 0.920829 | 0.180731 | 0.021 | 0.022 | 1 | 9 | Mr1 | -0.001 | Old |
| 0.921031 | 0.252942 | 0.039 | 0.037 | 1 | 9 | Mgmt | 0.002 | Old |
| 0.921091 | 1.366962 | 0.023 | 0.022 | 1 | 9 | Sema4d | 0.001 | Old |
| 0.921493 | 0.042826 | 0.291 | 0.287 | 1 | 9 | Nsd1 | 0.004 | Old |
| 0.921679 | 0.30763 | 0.098 | 0.103 | 1 | 9 | Man1b1 | -0.005 | Old |
| 0.922012 | 0.09836 | 0.098 | 0.096 | 1 | 9 | Pop5 | 0.002 | Old |
| 0.922293 | 1.175438 | 0.041 | 0.044 | 1 | 9 | Cyp51 | -0.003 | Old |
| 0.922332 | 0.417852 | 0.116 | 0.118 | 1 | 9 | Thap12 | -0.002 | Old |
| 0.922653 | 0.155433 | 0.168 | 0.176 | 1 | 9 | Spin1 | -0.008 | Old |
| 0.922855 | 0.000949 | 0.085 | 0.088 | 1 | 9 | Xxylt1 | -0.003 | Old |
| 0.922883 | 0.352415 | 0.119 | 0.118 | 1 | 9 | Atf6 | 0.001 | Old |
| 0.923108 | 1.077527 | 0.023 | 0.022 | 1 | 9 | Map3k14 | 0.001 | Old |
| 0.923108 | 0.774163 | 0.023 | 0.022 | 1 | 9 | Ctnnal1 | 0.001 | Old |
| 0.923363 | 0.516763 | 0.049 | 0.051 | 1 | 9 | Coq5 | -0.002 | Old |
| 0.923638 | 0.149153 | 0.129 | 0.125 | 1 | 9 | Ankrd50 | 0.004 | Old |
| 0.923761 | 0.319054 | 0.137 | 0.132 | 1 | 9 | Ddah2 | 0.005 | Old |
| 0.925039 | 0.605943 | 0.021 | 0.022 | 1 | 9 | Gk | -0.001 | Old |
| 0.925125 | 1.078355 | 0.023 | 0.022 | 1 | 9 | Cad | 0.001 | Old |
| 0.925125 | 0.79131 | 0.023 | 0.022 | 1 | 9 | Phf11b | 0.001 | Old |
| 0.92523 | 0.398458 | 0.031 | 0.029 | 1 | 9 | Platr25 | 0.002 | Old |
| 0.92523 | 0.391627 | 0.031 | 0.029 | 1 | 9 | P3h4 | 0.002 | Old |
| 0.925366 | 0.119624 | 0.157 | 0.154 | 1 | 9 | Abce1 | 0.003 | Old |
| 0.925518 | 0.263593 | 0.137 | 0.132 | 1 | 9 | Spaca6 | 0.005 | Old |
| 0.925755 | 0.086535 | 0.039 | 0.037 | 1 | 9 | Hmgxb3 | 0.002 | Old |
| 0.925755 | 0.017186 | 0.039 | 0.037 | 1 | 9 | Rdh11 | 0.002 | Old |
| 0.92586 | 0.455752 | 0.124 | 0.132 | 1 | 9 | Eif2a | -0.008 | Old |
| 0.926985 | 0.518592 | 0.031 | 0.029 | 1 | 9 | Pick1 | 0.002 | Old |
| 0.926985 | 0.441876 | 0.031 | 0.029 | 1 | 9 | Mrps2 | 0.002 | Old |
| 0.926985 | 0.044902 | 0.031 | 0.029 | 1 | 9 | Gem | 0.002 | Old |
| 0.927143 | 1.159471 | 0.023 | 0.022 | 1 | 9 | Zfp40 | 0.001 | Old |
| 0.927143 | 0.742227 | 0.023 | 0.022 | 1 | 9 | Ttc38 | 0.001 | Old |
| 0.927144 | 0.793303 | 0.021 | 0.022 | 1 | 9 | Nudt2 | -0.001 | Old |
| 0.927144 | 0.35203 | 0.021 | 0.022 | 1 | 9 | Adap2 | -0.001 | Old |
| 0.927144 | 0.229732 | 0.021 | 0.022 | 1 | 9 | Ptar1 | -0.001 | Old |
| 0.927144 | 0.205449 | 0.021 | 0.022 | 1 | 9 | Plcd1 | -0.001 | Old |
| 0.927961 | 0.175782 | 0.353 | 0.36 | 1 | 9 | Sf1 | -0.007 | Old |
| 0.927964 | 0.024496 | 0.307 | 0.301 | 1 | 9 | Mib1 | 0.006 | Old |
| 0.928739 | 0.895248 | 0.031 | 0.029 | 1 | 9 | Ctns | 0.002 | Old |
| 0.928739 | 0.495944 | 0.031 | 0.029 | 1 | 9 | Eef1akmt2 | 0.002 | Old |
| 0.928921 | 0.13751 | 0.049 | 0.051 | 1 | 9 | Slc25a40 | -0.002 | Old |
| 0.929123 | 0.69857 | 0.075 | 0.074 | 1 | 9 | Ythdc2 | 0.001 | Old |
| 0.929162 | 1.217772 | 0.023 | 0.022 | 1 | 9 | Bcs1l | 0.001 | Old |
| 0.929162 | 0.743841 | 0.023 | 0.022 | 1 | 9 | Mpg | 0.001 | Old |
| 0.929162 | 0.610239 | 0.023 | 0.022 | 1 | 9 | Ttf2 | 0.001 | Old |
| 0.929251 | 0.747883 | 0.021 | 0.022 | 1 | 9 | Snupn | -0.001 | Old |

| | | | | | | | | |
|---|---|---|---|---|---|---|---|---|
| 0.929251 | 0.554288 | 0.021 | 0.022 | 1 | 9 | Ntpcr | -0.001 | Old |
| 0.929251 | 0.48256 | 0.021 | 0.022 | 1 | 9 | Cyb561d2 | -0.001 | Old |
| 0.929617 | 0.014463 | 0.299 | 0.294 | 1 | 9 | Cpsf6 | 0.005 | Old |
| 0.92971 | 0.367538 | 0.111 | 0.11 | 1 | 9 | Rftn1 | 0.001 | Old |
| 0.929741 | 0.439601 | 0.144 | 0.154 | 1 | 9 | Ubqln2 | -0.01 | Old |
| 0.929754 | 0.076268 | 0.536 | 0.529 | 1 | 9 | Serbp1 | 0.007 | Old |
| 0.930311 | 0.420212 | 0.049 | 0.051 | 1 | 9 | Phtf1 | -0.002 | Old |
| 0.930482 | 0.169103 | 0.039 | 0.037 | 1 | 9 | Eif2b2 | 0.002 | Old |
| 0.931092 | 0.183765 | 0.178 | 0.176 | 1 | 9 | Arnt | 0.002 | Old |
| 0.931181 | 1.12889 | 0.023 | 0.022 | 1 | 9 | Lrrc40 | 0.001 | Old |
| 0.931181 | 0.474074 | 0.023 | 0.022 | 1 | 9 | Pla2g2d | 0.001 | Old |
| 0.931181 | 0.37841 | 0.023 | 0.022 | 1 | 9 | Elk1 | 0.001 | Old |
| 0.931323 | 1.326517 | 0.041 | 0.044 | 1 | 9 | Ivd | -0.003 | Old |
| 0.931357 | 0.738085 | 0.021 | 0.022 | 1 | 9 | Dtwd1 | -0.001 | Old |
| 0.931357 | 0.562289 | 0.021 | 0.022 | 1 | 9 | Gm15564 | -0.001 | Old |
| 0.931357 | 0.166598 | 0.021 | 0.022 | 1 | 9 | Zfp944 | -0.001 | Old |
| 0.931375 | 0.605872 | 0.059 | 0.059 | 1 | 9 | Kctd11 | 0 | Old |
| 0.931491 | 0.032716 | 0.433 | 0.426 | 1 | 9 | Akr1a1 | 0.007 | Old |
| 0.931528 | 0.657536 | 0.067 | 0.066 | 1 | 9 | Arfgef2 | 0.001 | Old |
| 0.931611 | 0.035564 | 0.057 | 0.059 | 1 | 9 | Eml3 | -0.002 | Old |
| 0.931828 | 0.136344 | 0.214 | 0.221 | 1 | 9 | Rab18 | -0.007 | Old |
| 0.932675 | 0.496848 | 0.124 | 0.125 | 1 | 9 | Denr | -0.001 | Old |
| 0.932738 | 0.677351 | 0.157 | 0.162 | 1 | 9 | Tgs1 | -0.005 | Old |
| 0.932867 | 0.097778 | 0.219 | 0.228 | 1 | 9 | Lrrc32 | -0.009 | Old |
| 0.93291 | 0.298116 | 0.057 | 0.059 | 1 | 9 | Fam45a | -0.002 | Old |
| 0.933026 | 0.045882 | 0.134 | 0.132 | 1 | 9 | Cmtm3 | 0.002 | Old |
| 0.933092 | 0.387151 | 0.049 | 0.051 | 1 | 9 | Smg8 | -0.002 | Old |
| 0.9332 | 1.024794 | 0.023 | 0.022 | 1 | 9 | Fam69a | 0.001 | Old |
| 0.9332 | 0.419152 | 0.023 | 0.022 | 1 | 9 | Gpatch4 | 0.001 | Old |
| 0.9332 | 0.325726 | 0.023 | 0.022 | 1 | 9 | Rrp9 | 0.001 | Old |
| 0.933464 | 0.553117 | 0.021 | 0.022 | 1 | 9 | Tatdn1 | -0.001 | Old |
| 0.933634 | 0.069067 | 0.039 | 0.037 | 1 | 9 | Serpinb6b | 0.002 | Old |
| 0.93398 | 0.105833 | 0.232 | 0.243 | 1 | 9 | Helz2 | -0.011 | Old |
| 0.934005 | 0.269826 | 0.031 | 0.029 | 1 | 9 | Cd37 | 0.002 | Old |
| 0.934005 | 0.071615 | 0.031 | 0.029 | 1 | 9 | Asnsd1 | 0.002 | Old |
| 0.934105 | 0.5513 | 0.077 | 0.081 | 1 | 9 | Stoml2 | -0.004 | Old |
| 0.934329 | 0.07738 | 0.296 | 0.294 | 1 | 9 | Top2b | 0.002 | Old |
| 0.93487 | 0.122589 | 0.381 | 0.382 | 1 | 9 | Wdr26 | -0.001 | Old |
| 0.935006 | 0.108527 | 0.119 | 0.118 | 1 | 9 | Sema6b | 0.001 | Old |
| 0.935211 | 0.089868 | 0.039 | 0.037 | 1 | 9 | Gpsm3 | 0.002 | Old |
| 0.93522 | 0.396608 | 0.023 | 0.022 | 1 | 9 | Cep41 | 0.001 | Old |
| 0.93522 | 0.153232 | 0.023 | 0.022 | 1 | 9 | Ankrd52 | 0.001 | Old |
| 0.93551 | 0.027295 | 0.057 | 0.059 | 1 | 9 | Bach2 | -0.002 | Old |
| 0.935957 | 0.214726 | 0.113 | 0.118 | 1 | 9 | Crybb3 | -0.005 | Old |
| 0.936016 | 0.044688 | 0.356 | 0.36 | 1 | 9 | Thoc2 | -0.004 | Old |

| | | | | | | | | |
|---|---|---|---|---|---|---|---|---|
| 0.93621 | 0.077523 | 0.155 | 0.154 | 1 | 9 | Golph3 | 0.001 | Old |
| 0.936376 | 0.0021 | 0.186 | 0.191 | 1 | 9 | Ncoa7 | -0.005 | Old |
| 0.936805 | 0.583249 | 0.07 | 0.074 | 1 | 9 | Foxo3 | -0.004 | Old |
| 0.93681 | 0.13051 | 0.057 | 0.059 | 1 | 9 | Lsg1 | -0.002 | Old |
| 0.936931 | 0.04832 | 0.593 | 0.581 | 1 | 9 | Col4a2 | 0.012 | Old |
| 0.937015 | 2.433477 | 0.015 | 0.015 | 1 | 9 | Fam151b | 0 | Old |
| 0.937015 | 2.278515 | 0.015 | 0.015 | 1 | 9 | Acta2 | 0 | Old |
| 0.937015 | 1.942814 | 0.015 | 0.015 | 1 | 9 | Intu | 0 | Old |
| 0.937015 | 1.25811 | 0.015 | 0.015 | 1 | 9 | Zfp324 | 0 | Old |
| 0.937015 | 1.013029 | 0.015 | 0.015 | 1 | 9 | Isg20 | 0 | Old |
| 0.937015 | 1.006122 | 0.015 | 0.015 | 1 | 9 | Gpr35 | 0 | Old |
| 0.937015 | 0.715808 | 0.015 | 0.015 | 1 | 9 | Fbxo10 | 0 | Old |
| 0.937173 | 0.097738 | 0.075 | 0.074 | 1 | 9 | Map3k5 | 0.001 | Old |
| 0.93724 | 0.473104 | 0.023 | 0.022 | 1 | 9 | Trub1 | 0.001 | Old |
| 0.93724 | 0.430788 | 0.023 | 0.022 | 1 | 9 | Mtrf1 | 0.001 | Old |
| 0.93724 | 0.401134 | 0.023 | 0.022 | 1 | 9 | Nat10 | 0.001 | Old |
| 0.93724 | 0.209048 | 0.023 | 0.022 | 1 | 9 | Sag | 0.001 | Old |
| 0.937265 | 0.447009 | 0.049 | 0.051 | 1 | 9 | Rnf170 | -0.002 | Old |
| 0.937361 | 0.645899 | 0.116 | 0.118 | 1 | 9 | Cstf2t | -0.002 | Old |
| 0.937477 | 0.088564 | 0.077 | 0.081 | 1 | 9 | Plvap | -0.004 | Old |
| 0.93768 | 1.201625 | 0.021 | 0.022 | 1 | 9 | Mthfd2 | -0.001 | Old |
| 0.93768 | 1.100159 | 0.021 | 0.022 | 1 | 9 | Itgb3bp | -0.001 | Old |
| 0.937951 | 0.193994 | 0.211 | 0.213 | 1 | 9 | Cdc37 | -0.002 | Old |
| 0.938357 | 0.276754 | 0.111 | 0.11 | 1 | 9 | Naa35 | 0.001 | Old |
| 0.93926 | 0.36121 | 0.023 | 0.022 | 1 | 9 | Gm26660 | 0.001 | Old |
| 0.93926 | 0.217646 | 0.023 | 0.022 | 1 | 9 | Alg3 | 0.001 | Old |
| 0.93948 | 1.467802 | 0.015 | 0.015 | 1 | 9 | Cox11 | 0 | Old |
| 0.93948 | 1.189707 | 0.015 | 0.015 | 1 | 9 | Ift88 | 0 | Old |
| 0.939844 | 0.341851 | 0.103 | 0.103 | 1 | 9 | Ube2d1 | 0 | Old |
| 0.940342 | 0.659672 | 0.07 | 0.074 | 1 | 9 | Fem1a | -0.004 | Old |
| 0.941031 | 0.010997 | 0.031 | 0.029 | 1 | 9 | Mypopos | 0.002 | Old |
| 0.941281 | 0.110823 | 0.023 | 0.022 | 1 | 9 | Fuom | 0.001 | Old |
| 0.941513 | 0.084937 | 0.317 | 0.316 | 1 | 9 | Cyfip1 | 0.001 | Old |
| 0.941868 | 1.424313 | 0.041 | 0.044 | 1 | 9 | Fer | -0.003 | Old |
| 0.941897 | 1.019935 | 0.021 | 0.022 | 1 | 9 | AC127341. | -0.001 | Old |
| 0.941946 | 1.242624 | 0.015 | 0.015 | 1 | 9 | Zfp408 | 0 | Old |
| 0.941946 | 1.054617 | 0.015 | 0.015 | 1 | 9 | Msrb2 | 0 | Old |
| 0.941946 | 0.903681 | 0.015 | 0.015 | 1 | 9 | Serpine1 | 0 | Old |
| 0.941946 | 0.851956 | 0.015 | 0.015 | 1 | 9 | Fhod3 | 0 | Old |
| 0.941946 | 0.8474 | 0.015 | 0.015 | 1 | 9 | Tcea2 | 0 | Old |
| 0.941946 | 0.797204 | 0.015 | 0.015 | 1 | 9 | Msr1 | 0 | Old |
| 0.941946 | 0.713951 | 0.015 | 0.015 | 1 | 9 | Tmem86b | 0 | Old |
| 0.941946 | 0.708848 | 0.015 | 0.015 | 1 | 9 | Gsto1 | 0 | Old |
| 0.941946 | 0.399335 | 0.015 | 0.015 | 1 | 9 | Sardh | 0 | Old |
| 0.942 | 0.029687 | 0.188 | 0.191 | 1 | 9 | Fbxo11 | -0.003 | Old |

| | | | | | | | | |
|---|---|---|---|---|---|---|---|---|
| 0.942811 | 0.060079 | 0.142 | 0.14 | 1 | 9 | Atg3 | 0.002 | Old |
| 0.942874 | 0.167127 | 0.119 | 0.125 | 1 | 9 | Itfg1 | -0.006 | Old |
| 0.943313 | 0.231057 | 0.057 | 0.059 | 1 | 9 | Msmo1 | -0.002 | Old |
| 0.943314 | 0.076861 | 0.085 | 0.088 | 1 | 9 | Zcchc17 | -0.003 | Old |
| 0.943353 | 0.145435 | 0.271 | 0.272 | 1 | 9 | Fyttd1 | -0.001 | Old |
| 0.943815 | 0.354213 | 0.103 | 0.103 | 1 | 9 | Rnf139 | 0 | Old |
| 0.944007 | 0.997613 | 0.021 | 0.022 | 1 | 9 | Chst14 | -0.001 | Old |
| 0.944078 | 0.255515 | 0.075 | 0.074 | 1 | 9 | Atp7a | 0.001 | Old |
| 0.944179 | 0.043369 | 0.108 | 0.11 | 1 | 9 | Tecr | -0.002 | Old |
| 0.944412 | 1.315697 | 0.015 | 0.015 | 1 | 9 | Pcp4l1 | 0 | Old |
| 0.944614 | 0.174597 | 0.057 | 0.059 | 1 | 9 | Zfyve27 | -0.002 | Old |
| 0.945324 | 0.542796 | 0.023 | 0.022 | 1 | 9 | Dlg4 | 0.001 | Old |
| 0.945324 | 0.351666 | 0.023 | 0.022 | 1 | 9 | Borcs7 | 0.001 | Old |
| 0.945384 | 0.001781 | 0.464 | 0.485 | 1 | 9 | Tra2b | -0.021 | Old |
| 0.945631 | 0.05291 | 0.229 | 0.235 | 1 | 9 | Zfp445 | -0.006 | Old |
| 0.945915 | 0.249635 | 0.057 | 0.059 | 1 | 9 | Fgd4 | -0.002 | Old |
| 0.945925 | 0.249896 | 0.064 | 0.066 | 1 | 9 | Pycr2 | -0.002 | Old |
| 0.946412 | 0.151918 | 0.028 | 0.029 | 1 | 9 | Ndufaf7 | -0.001 | Old |
| 0.946879 | 1.85178 | 0.015 | 0.015 | 1 | 9 | Galns | 0 | Old |
| 0.946879 | 1.640815 | 0.015 | 0.015 | 1 | 9 | Bpgm | 0 | Old |
| 0.946879 | 1.3906 | 0.015 | 0.015 | 1 | 9 | Cdkl2 | 0 | Old |
| 0.946879 | 1.241337 | 0.015 | 0.015 | 1 | 9 | Tbccd1 | 0 | Old |
| 0.946879 | 0.882849 | 0.015 | 0.015 | 1 | 9 | 3110040N1 | 0 | Old |
| 0.946879 | 0.568145 | 0.015 | 0.015 | 1 | 9 | Tspyl4 | 0 | Old |
| 0.946879 | 0.517237 | 0.015 | 0.015 | 1 | 9 | Zswim1 | 0 | Old |
| 0.946879 | 0.40439 | 0.015 | 0.015 | 1 | 9 | Acbd4 | 0 | Old |
| 0.946879 | 0.327332 | 0.015 | 0.015 | 1 | 9 | Aldh4a1 | 0 | Old |
| 0.946879 | 0.258816 | 0.015 | 0.015 | 1 | 9 | Arhgap19 | 0 | Old |
| 0.946879 | 0.071492 | 0.015 | 0.015 | 1 | 9 | Vav1 | 0 | Old |
| 0.947092 | 0.276592 | 0.126 | 0.132 | 1 | 9 | Fam193a | -0.006 | Old |
| 0.947217 | 0.326574 | 0.057 | 0.059 | 1 | 9 | Msh2 | -0.002 | Old |
| 0.947349 | 0.382339 | 0.101 | 0.103 | 1 | 9 | H2-Ke6 | -0.002 | Old |
| 0.947759 | 0.057774 | 0.131 | 0.132 | 1 | 9 | Pdia4 | -0.001 | Old |
| 0.94816 | 0.209805 | 0.144 | 0.147 | 1 | 9 | Eogt | -0.003 | Old |
| 0.948272 | 0.347884 | 0.113 | 0.118 | 1 | 9 | Emc6 | -0.005 | Old |
| 0.948467 | 0.66959 | 0.067 | 0.066 | 1 | 9 | Dcun1d3 | 0.001 | Old |
| 0.948588 | 0.191086 | 0.209 | 0.206 | 1 | 9 | Setd1b | 0.003 | Old |
| 0.948968 | 0.147348 | 0.134 | 0.132 | 1 | 9 | Trmt112 | 0.002 | Old |
| 0.949305 | 0.460186 | 0.059 | 0.059 | 1 | 9 | Pstk | 0 | Old |
| 0.949346 | 0.753089 | 0.015 | 0.015 | 1 | 9 | Nkapl | 0 | Old |
| 0.949346 | 0.559452 | 0.015 | 0.015 | 1 | 9 | Tsr1 | 0 | Old |
| 0.949346 | 0.112873 | 0.015 | 0.015 | 1 | 9 | Gpr157 | 0 | Old |
| 0.949959 | 0.136573 | 0.276 | 0.272 | 1 | 9 | Sipa1 | 0.004 | Old |
| 0.95004 | 0.344879 | 0.028 | 0.029 | 1 | 9 | Adprhl2 | -0.001 | Old |
| 0.95004 | 0.168356 | 0.028 | 0.029 | 1 | 9 | Dut | -0.001 | Old |

| | | | | | | | | |
|---|---|---|---|---|---|---|---|---|
| 0.950766 | 0.200767 | 0.103 | 0.103 | 1 | 9 | Jkamp | 0 | Old |
| 0.950861 | 0.254241 | 0.111 | 0.11 | 1 | 9 | Dhx38 | 0.001 | Old |
| 0.951186 | 0.888045 | 0.049 | 0.051 | 1 | 9 | Umad1 | -0.002 | Old |
| 0.951367 | 0.636514 | 0.052 | 0.051 | 1 | 9 | Elp3 | 0.001 | Old |
| 0.951814 | 0.319812 | 0.015 | 0.015 | 1 | 9 | Haus5 | 0 | Old |
| 0.951814 | 0.276632 | 0.015 | 0.015 | 1 | 9 | Tnxb | 0 | Old |
| 0.951814 | 0.124689 | 0.015 | 0.015 | 1 | 9 | Cmss1 | 0 | Old |
| 0.951855 | 0.387728 | 0.028 | 0.029 | 1 | 9 | Tsr2 | -0.001 | Old |
| 0.951855 | 0.187544 | 0.028 | 0.029 | 1 | 9 | Zdhhc14 | -0.001 | Old |
| 0.952064 | 0.217113 | 0.113 | 0.118 | 1 | 9 | Tmem165 | -0.005 | Old |
| 0.952973 | 0.00717 | 0.598 | 0.574 | 1 | 9 | Ncor1 | 0.024 | Old |
| 0.95315 | 0.57172 | 0.059 | 0.059 | 1 | 9 | Isca1 | 0 | Old |
| 0.953295 | 0.46786 | 0.144 | 0.147 | 1 | 9 | Uqcc2 | -0.003 | Old |
| 0.95367 | 0.1283 | 0.028 | 0.029 | 1 | 9 | Chd3os | -0.001 | Old |
| 0.954223 | 0.018426 | 0.838 | 0.801 | 1 | 9 | Cd93 | 0.037 | Old |
| 0.954282 | 0.327787 | 0.015 | 0.015 | 1 | 9 | Msantd3 | 0 | Old |
| 0.954518 | 0.057797 | 0.064 | 0.066 | 1 | 9 | Ddx1 | -0.002 | Old |
| 0.955135 | 0.123602 | 0.232 | 0.243 | 1 | 9 | Gnb4 | -0.011 | Old |
| 0.955479 | 0.683167 | 0.077 | 0.081 | 1 | 9 | Mrpl38 | -0.004 | Old |
| 0.955485 | 0.677378 | 0.028 | 0.029 | 1 | 9 | Pcdhgb4 | -0.001 | Old |
| 0.955485 | 0.600224 | 0.028 | 0.029 | 1 | 9 | Mrps27 | -0.001 | Old |
| 0.955485 | 0.488855 | 0.028 | 0.029 | 1 | 9 | Gab3 | -0.001 | Old |
| 0.955625 | 0.436465 | 0.16 | 0.162 | 1 | 9 | Uba2 | -0.002 | Old |
| 0.956659 | 0.120477 | 0.142 | 0.14 | 1 | 9 | Fam208b | 0.002 | Old |
| 0.956751 | 0.127846 | 0.015 | 0.015 | 1 | 9 | Armc6 | 0 | Old |
| 0.956751 | 0.074308 | 0.015 | 0.015 | 1 | 9 | C030006K1 | 0 | Old |
| 0.957104 | 0.070742 | 0.369 | 0.39 | 1 | 9 | Pura | -0.021 | Old |
| 0.957233 | 0.300534 | 0.178 | 0.184 | 1 | 9 | Gigyf2 | -0.006 | Old |
| 0.9573 | 1.01436 | 0.028 | 0.029 | 1 | 9 | Fam117a | -0.001 | Old |
| 0.958121 | 0.242356 | 0.211 | 0.213 | 1 | 9 | Pttg1 | -0.002 | Old |
| 0.958153 | 0.743347 | 0.049 | 0.051 | 1 | 9 | Lrrc42 | -0.002 | Old |
| 0.958644 | 0.059264 | 0.082 | 0.081 | 1 | 9 | Hyou1 | 0.001 | Old |
| 0.959115 | 0.901768 | 0.028 | 0.029 | 1 | 9 | C230037L1 | -0.001 | Old |
| 0.959115 | 0.82629 | 0.028 | 0.029 | 1 | 9 | Lancl1 | -0.001 | Old |
| 0.959562 | 0.104042 | 0.059 | 0.059 | 1 | 9 | Fmnl2 | 0 | Old |
| 0.960931 | 0.222548 | 0.028 | 0.029 | 1 | 9 | Suox | -0.001 | Old |
| 0.961028 | 0.242865 | 0.093 | 0.096 | 1 | 9 | Galnt10 | -0.003 | Old |
| 0.961028 | 0.057348 | 0.093 | 0.096 | 1 | 9 | Spg7 | -0.003 | Old |
| 0.961198 | 0.119847 | 0.139 | 0.14 | 1 | 9 | Ufl1 | -0.001 | Old |
| 0.961229 | 0.060869 | 0.474 | 0.5 | 1 | 9 | Nbeal1 | -0.026 | Old |
| 0.961353 | 0.035782 | 0.291 | 0.287 | 1 | 9 | Cdipt | 0.004 | Old |
| 0.96169 | 0.037447 | 0.015 | 0.015 | 1 | 9 | Scarf2 | 0 | Old |
| 0.962378 | 0.02987 | 0.358 | 0.36 | 1 | 9 | Prr13 | -0.002 | Old |
| 0.962747 | 0.868411 | 0.028 | 0.029 | 1 | 9 | Msh6 | -0.001 | Old |
| 0.962747 | 0.427979 | 0.028 | 0.029 | 1 | 9 | Taf4b | -0.001 | Old |

| | | | | | | | | |
|---|---|---|---|---|---|---|---|---|
| 0.962748 | 0.048203 | 0.518 | 0.537 | 1 | 9 | Mcc | -0.019 | Old |
| 0.963005 | 0.14875 | 0.067 | 0.066 | 1 | 9 | Bmp1 | 0.001 | Old |
| 0.963116 | 0.099692 | 0.064 | 0.066 | 1 | 9 | Fech | -0.002 | Old |
| 0.963686 | 0.514833 | 0.052 | 0.051 | 1 | 9 | Gzf1 | 0.001 | Old |
| 0.963686 | 0.31042 | 0.052 | 0.051 | 1 | 9 | Rabepk | 0.001 | Old |
| 0.964227 | 0.069563 | 0.142 | 0.147 | 1 | 9 | Zfp800 | -0.005 | Old |
| 0.964563 | 0.959801 | 0.028 | 0.029 | 1 | 9 | Nudt16 | -0.001 | Old |
| 0.965056 | 0.775982 | 0.052 | 0.051 | 1 | 9 | Slc38a6 | 0.001 | Old |
| 0.965448 | 0.596364 | 0.057 | 0.059 | 1 | 9 | Zdhhc9 | -0.002 | Old |
| 0.96597 | 0.502603 | 0.085 | 0.088 | 1 | 9 | Dusp16 | -0.003 | Old |
| 0.966286 | 0.171525 | 0.129 | 0.132 | 1 | 9 | Wwp1 | -0.003 | Old |
| 0.966379 | 0.815857 | 0.028 | 0.029 | 1 | 9 | Pop4 | -0.001 | Old |
| 0.966425 | 0.631737 | 0.052 | 0.051 | 1 | 9 | Ttc9c | 0.001 | Old |
| 0.967795 | 0.009559 | 0.052 | 0.051 | 1 | 9 | Copg2 | 0.001 | Old |
| 0.968031 | 0.280357 | 0.064 | 0.066 | 1 | 9 | Gadd45gip | -0.002 | Old |
| 0.968195 | 0.960891 | 0.028 | 0.029 | 1 | 9 | Lmf1 | -0.001 | Old |
| 0.968195 | 0.421884 | 0.028 | 0.029 | 1 | 9 | Optn | -0.001 | Old |
| 0.968389 | 0.238695 | 0.036 | 0.037 | 1 | 9 | Cdan1 | -0.001 | Old |
| 0.968543 | 0.427627 | 0.059 | 0.059 | 1 | 9 | Usp5 | 0 | Old |
| 0.969066 | 0.164715 | 0.067 | 0.066 | 1 | 9 | Ube2q2 | 0.001 | Old |
| 0.969354 | 0.158859 | 0.196 | 0.199 | 1 | 9 | Plpp6 | -0.003 | Old |
| 0.969826 | 0.100365 | 0.059 | 0.059 | 1 | 9 | Rogdi | 0 | Old |
| 0.970012 | 0.901777 | 0.028 | 0.029 | 1 | 9 | Zfp788 | -0.001 | Old |
| 0.970012 | 0.544564 | 0.028 | 0.029 | 1 | 9 | Inpp5e | -0.001 | Old |
| 0.970012 | 0.52887 | 0.028 | 0.029 | 1 | 9 | Rint1 | -0.001 | Old |
| 0.970051 | 0.482507 | 0.088 | 0.088 | 1 | 9 | Ilvbl | 0 | Old |
| 0.970535 | 0.399237 | 0.052 | 0.051 | 1 | 9 | Gbe1 | 0.001 | Old |
| 0.971099 | 0.049387 | 0.175 | 0.176 | 1 | 9 | Pja1 | -0.001 | Old |
| 0.971484 | 0.028786 | 0.534 | 0.537 | 1 | 9 | Sp100 | -0.003 | Old |
| 0.971718 | 0.348944 | 0.064 | 0.066 | 1 | 9 | Dopey2 | -0.002 | Old |
| 0.971828 | 1.170286 | 0.028 | 0.029 | 1 | 9 | Lyl1 | -0.001 | Old |
| 0.971828 | 1.155923 | 0.028 | 0.029 | 1 | 9 | Lacc1 | -0.001 | Old |
| 0.972124 | 0.134615 | 0.08 | 0.081 | 1 | 9 | Twsg1 | -0.001 | Old |
| 0.972218 | 0.074283 | 0.294 | 0.287 | 1 | 9 | Psmd11 | 0.007 | Old |
| 0.972325 | 0.041728 | 0.157 | 0.162 | 1 | 9 | Srrt | -0.005 | Old |
| 0.972449 | 0.063649 | 0.227 | 0.228 | 1 | 9 | Ensa | -0.001 | Old |
| 0.972454 | 0.476169 | 0.093 | 0.096 | 1 | 9 | Rps27rt | -0.003 | Old |
| 0.972622 | 0.181897 | 0.229 | 0.235 | 1 | 9 | 4930402H2 | -0.006 | Old |
| 0.97269 | 0.025034 | 0.201 | 0.206 | 1 | 9 | Dpm3 | -0.005 | Old |
| 0.973268 | 0.791611 | 0.057 | 0.059 | 1 | 9 | Dgcr6 | -0.002 | Old |
| 0.973502 | 0.067966 | 0.26 | 0.265 | 1 | 9 | Uqcrc2 | -0.005 | Old |
| 0.974645 | 0.297236 | 0.052 | 0.051 | 1 | 9 | Arfgap3 | 0.001 | Old |
| 0.974821 | 0.009386 | 0.312 | 0.324 | 1 | 9 | Smim10l1 | -0.012 | Old |
| 0.974871 | 0.303923 | 0.036 | 0.037 | 1 | 9 | Sphk2 | -0.001 | Old |
| 0.975027 | 0.049403 | 0.196 | 0.199 | 1 | 9 | Ikbkb | -0.003 | Old |

| | | | | | | | | |
|---|---|---|---|---|---|---|---|---|
| 0.975129 | 0.134272 | 0.067 | 0.066 | 1 | 9 | Gprin3 | 0.001 | Old |
| 0.975406 | 0.137137 | 0.196 | 0.199 | 1 | 9 | Cuta | -0.003 | Old |
| 0.975592 | 0.769525 | 0.044 | 0.044 | 1 | 9 | Selenoh | 0 | Old |
| 0.975875 | 0.556748 | 0.057 | 0.059 | 1 | 9 | Wdr36 | -0.002 | Old |
| 0.975904 | 0.072175 | 0.242 | 0.243 | 1 | 9 | AU020206 | -0.001 | Old |
| 0.976015 | 0.706168 | 0.052 | 0.051 | 1 | 9 | Kctd18 | 0.001 | Old |
| 0.976347 | 0.000654 | 0.075 | 0.074 | 1 | 9 | Tmem63b | 0.001 | Old |
| 0.976492 | 0.271142 | 0.036 | 0.037 | 1 | 9 | Thap4 | -0.001 | Old |
| 0.976553 | 0.054924 | 0.147 | 0.147 | 1 | 9 | Gpat4 | 0 | Old |
| 0.977111 | 0.13338 | 0.539 | 0.566 | 1 | 9 | Luc7l2 | -0.027 | Old |
| 0.977528 | 0.079806 | 0.059 | 0.059 | 1 | 9 | Orai3 | 0 | Old |
| 0.977622 | 0.030013 | 0.412 | 0.404 | 1 | 9 | Pdia3 | 0.008 | Old |
| 0.977863 | 0.07637 | 0.428 | 0.426 | 1 | 9 | Wwc2 | 0.002 | Old |
| 0.977864 | 0.444128 | 0.064 | 0.066 | 1 | 9 | Phkb | -0.002 | Old |
| 0.97796 | 0.082647 | 0.222 | 0.221 | 1 | 9 | Kcna5 | 0.001 | Old |
| 0.978432 | 0.085716 | 0.196 | 0.199 | 1 | 9 | Arid5a | -0.003 | Old |
| 0.978755 | 0.343494 | 0.052 | 0.051 | 1 | 9 | Zfp362 | 0.001 | Old |
| 0.979097 | 0.706721 | 0.028 | 0.029 | 1 | 9 | Pdzd11 | -0.001 | Old |
| 0.979589 | 0.009805 | 0.273 | 0.272 | 1 | 9 | Cracr2b | 0.001 | Old |
| 0.979734 | 0.01967 | 0.036 | 0.037 | 1 | 9 | Rab10os | -0.001 | Old |
| 0.980702 | 0.039036 | 0.196 | 0.199 | 1 | 9 | Twf1 | -0.003 | Old |
| 0.981355 | 0.098723 | 0.036 | 0.037 | 1 | 9 | Thsd1 | -0.001 | Old |
| 0.981508 | 0.654168 | 0.044 | 0.044 | 1 | 9 | Xrcc4 | 0 | Old |
| 0.981508 | 0.317667 | 0.044 | 0.044 | 1 | 9 | Nob1 | 0 | Old |
| 0.981781 | 0.039895 | 0.126 | 0.125 | 1 | 9 | Ccdc25 | 0.001 | Old |
| 0.982157 | 0.290749 | 0.08 | 0.081 | 1 | 9 | Cdkn2aipnl | -0.001 | Old |
| 0.982171 | 1.11453 | 0.085 | 0.088 | 1 | 9 | Ccdc34 | -0.003 | Old |
| 0.98258 | 0.05948 | 0.165 | 0.169 | 1 | 9 | Samd8 | -0.004 | Old |
| 0.982729 | 0.093954 | 0.178 | 0.184 | 1 | 9 | Tmf1 | -0.006 | Old |
| 0.982731 | 1.332559 | 0.028 | 0.029 | 1 | 9 | Slx4ip | -0.001 | Old |
| 0.982866 | 0.048923 | 0.052 | 0.051 | 1 | 9 | Sec24d | 0.001 | Old |
| 0.982976 | 2.216613 | 0.036 | 0.037 | 1 | 9 | Rgs1 | -0.001 | Old |
| 0.982976 | 0.589403 | 0.036 | 0.037 | 1 | 9 | Akna | -0.001 | Old |
| 0.982987 | 0.869759 | 0.044 | 0.044 | 1 | 9 | Gmeb2 | 0 | Old |
| 0.982987 | 0.723803 | 0.044 | 0.044 | 1 | 9 | Atmin | 0 | Old |
| 0.983886 | 0.350695 | 0.093 | 0.096 | 1 | 9 | Gskip | -0.003 | Old |
| 0.984234 | 0.041109 | 0.072 | 0.074 | 1 | 9 | Dubr | -0.002 | Old |
| 0.984293 | 0.007773 | 0.399 | 0.404 | 1 | 9 | Atp1b3 | -0.005 | Old |
| 0.984294 | 0.016616 | 0.353 | 0.346 | 1 | 9 | Kmt2a | 0.007 | Old |
| 0.984597 | 0.316547 | 0.036 | 0.037 | 1 | 9 | Mamstr | -0.001 | Old |
| 0.984597 | 0.189941 | 0.036 | 0.037 | 1 | 9 | Mroh1 | -0.001 | Old |
| 0.984625 | 0.215813 | 0.178 | 0.176 | 1 | 9 | Pak1ip1 | 0.002 | Old |
| 0.984659 | 0.183992 | 0.201 | 0.206 | 1 | 9 | Hprt | -0.005 | Old |
| 0.984869 | 0.074028 | 0.124 | 0.125 | 1 | 9 | Trip4 | -0.001 | Old |
| 0.985242 | 0.050832 | 0.196 | 0.199 | 1 | 9 | Kdm6a | -0.003 | Old |

| | | | | | | | | |
|---|---|---|---|---|---|---|---|---|
| 0.985502 | 0.258663 | 0.08 | 0.081 | 1 | 9 | Atp13a1 | -0.001 | Old |
| 0.985607 | 0.154092 | 0.052 | 0.051 | 1 | 9 | Ppp2r3c | 0.001 | Old |
| 0.985608 | 0.052729 | 0.139 | 0.14 | 1 | 9 | Glrx3 | -0.001 | Old |
| 0.985773 | 0.308665 | 0.142 | 0.147 | 1 | 9 | Pgk1 | -0.005 | Old |
| 0.985946 | 0.149523 | 0.044 | 0.044 | 1 | 9 | Tstd2 | 0 | Old |
| 0.986218 | 0.578818 | 0.036 | 0.037 | 1 | 9 | Stk39 | -0.001 | Old |
| 0.986608 | 0.100404 | 0.095 | 0.096 | 1 | 9 | Heatr5a | -0.001 | Old |
| 0.987567 | 0.316059 | 0.144 | 0.147 | 1 | 9 | Tmem173 | -0.003 | Old |
| 0.987592 | 0.136702 | 0.232 | 0.235 | 1 | 9 | Rnf213 | -0.003 | Old |
| 0.987779 | 4.47E-06 | 0.178 | 0.176 | 1 | 9 | Kbtbd2 | 0.002 | Old |
| 0.987856 | 0.378372 | 0.108 | 0.11 | 1 | 9 | Stx6 | -0.002 | Old |
| 0.987998 | 0.567353 | 0.121 | 0.125 | 1 | 9 | Polr2i | -0.004 | Old |
| 0.988654 | 0.825976 | 0.085 | 0.088 | 1 | 9 | Cstf3 | -0.003 | Old |
| 0.988848 | 0.380911 | 0.08 | 0.081 | 1 | 9 | Smc2 | -0.001 | Old |
| 0.989009 | 0.212774 | 0.178 | 0.184 | 1 | 9 | Clcn3 | -0.006 | Old |
| 0.989723 | 0.116349 | 0.131 | 0.132 | 1 | 9 | Armc1 | -0.001 | Old |
| 0.990073 | 0.813827 | 0.072 | 0.074 | 1 | 9 | Golga5 | -0.002 | Old |
| 0.990073 | 0.581281 | 0.072 | 0.074 | 1 | 9 | Cpox | -0.002 | Old |
| 0.990293 | 0.202754 | 0.464 | 0.463 | 1 | 9 | Cyr61 | 0.001 | Old |
| 0.990348 | 0.045195 | 0.137 | 0.14 | 1 | 9 | Cldnd1 | -0.003 | Old |
| 0.990835 | 0.129697 | 0.358 | 0.368 | 1 | 9 | Fam208a | -0.01 | Old |
| 0.99109 | 0.102042 | 0.052 | 0.051 | 1 | 9 | Lmnb1 | 0.001 | Old |
| 0.991163 | 0.814119 | 0.093 | 0.096 | 1 | 9 | Fbxo9 | -0.003 | Old |
| 0.991863 | 0.0031 | 0.044 | 0.044 | 1 | 9 | 2210016F1 | 0 | Old |
| 0.992193 | 0.041805 | 0.08 | 0.081 | 1 | 9 | Il2rg | -0.001 | Old |
| 0.992461 | 0.143916 | 0.052 | 0.051 | 1 | 9 | Nek6 | 0.001 | Old |
| 0.992788 | 0.027772 | 0.095 | 0.096 | 1 | 9 | Oasl2 | -0.001 | Old |
| 0.99298 | 0.214601 | 0.137 | 0.14 | 1 | 9 | Sar1b | -0.003 | Old |
| 0.993292 | 0.128738 | 0.423 | 0.434 | 1 | 9 | Matr3 | -0.011 | Old |
| 0.993542 | 0.067215 | 0.193 | 0.199 | 1 | 9 | Plekho1 | -0.006 | Old |
| 0.993576 | 0.68797 | 0.072 | 0.074 | 1 | 9 | Pld3 | -0.002 | Old |
| 0.993819 | 0.078462 | 0.095 | 0.096 | 1 | 9 | Erlec1 | -0.001 | Old |
| 0.993831 | 0.00776 | 0.052 | 0.051 | 1 | 9 | Wdr75 | 0.001 | Old |
| 0.994325 | 0.648098 | 0.036 | 0.037 | 1 | 9 | Atg4a | -0.001 | Old |
| 0.994325 | 0.562146 | 0.036 | 0.037 | 1 | 9 | Foxf1 | -0.001 | Old |
| 0.994325 | 0.529404 | 0.036 | 0.037 | 1 | 9 | Airn | -0.001 | Old |
| 0.994822 | 0.322038 | 0.044 | 0.044 | 1 | 9 | Fancb | 0 | Old |
| 0.99508 | 0.619992 | 0.064 | 0.066 | 1 | 9 | Ccdc115 | -0.002 | Old |
| 0.995205 | 0.270748 | 0.17 | 0.176 | 1 | 9 | Zfp106 | -0.006 | Old |
| 0.995539 | 0.128384 | 0.08 | 0.081 | 1 | 9 | Zfp329 | -0.001 | Old |
| 0.995884 | 0.090031 | 0.201 | 0.206 | 1 | 9 | Ndufa11 | -0.005 | Old |
| 0.995946 | 0.609965 | 0.036 | 0.037 | 1 | 9 | Glb1 | -0.001 | Old |
| 0.995946 | 0.148416 | 0.036 | 0.037 | 1 | 9 | Poc5 | -0.001 | Old |
| 0.996255 | 0.14029 | 0.088 | 0.088 | 1 | 9 | Pdcd7 | 0 | Old |
| 0.99631 | 0.61055 | 0.064 | 0.066 | 1 | 9 | Rnf169 | -0.002 | Old |

| | | | | | | | | |
|---|---|---|---|---|---|---|---|---|
| 0.996465 | 0.152469 | 0.232 | 0.243 | 1 | 9 | Rela | -0.011 | Old |
| 0.996654 | 0.543642 | 0.08 | 0.081 | 1 | 9 | Slc36a4 | -0.001 | Old |
| 0.99708 | 0.598193 | 0.072 | 0.074 | 1 | 9 | Tfdp2 | -0.002 | Old |
| 0.997712 | 0.109913 | 0.456 | 0.471 | 1 | 9 | Adipor1 | -0.015 | Old |
| 0.997781 | 0.662787 | 0.044 | 0.044 | 1 | 9 | Rnase4 | 0 | Old |
| 0.997944 | 0.313023 | 0.052 | 0.051 | 1 | 9 | Shkbp1 | 0.001 | Old |
| 0.998072 | 0.007089 | 0.307 | 0.309 | 1 | 9 | Nfix | -0.002 | Old |
| 0.998248 | 0.229774 | 0.072 | 0.074 | 1 | 9 | Arhgef11 | -0.002 | Old |
| 0.998387 | 0.016422 | 0.168 | 0.169 | 1 | 9 | Os9 | -0.001 | Old |
| 0.998495 | 0.333149 | 0.101 | 0.103 | 1 | 9 | Fbxl18 | -0.002 | Old |
| 0.998709 | 0.062639 | 0.302 | 0.309 | 1 | 9 | P4hb | -0.007 | Old |
| 0.999028 | 0.146001 | 0.108 | 0.11 | 1 | 9 | Acadsb | -0.002 | Old |
| 0.999189 | 0.787321 | 0.036 | 0.037 | 1 | 9 | Zfp120 | -0.001 | Old |
| 0.999189 | 0.681055 | 0.036 | 0.037 | 1 | 9 | Narf | -0.001 | Old |
| 0.99926 | 0.377107 | 0.044 | 0.044 | 1 | 9 | Zfp84 | 0 | Old |
| 0.99926 | 0.266791 | 0.044 | 0.044 | 1 | 9 | Dnajb5 | 0 | Old |
| 0.99926 | 0.150496 | 0.044 | 0.044 | 1 | 9 | Slc16a2 | 0 | Old |
| 0.99926 | 0.068567 | 0.044 | 0.044 | 1 | 9 | Rptor | 0 | Old |
| 0.999267 | 0.258119 | 0.211 | 0.221 | 1 | 9 | Prkar2a | -0.01 | Old |
| 0.999358 | 0.040241 | 0.059 | 0.059 | 1 | 9 | Gm28875 | 0 | Old |
| 0.999514 | 0.564311 | 0.108 | 0.11 | 1 | 9 | Usp12 | -0.002 | Old |
| 0.999553 | 0.345468 | 0.131 | 0.132 | 1 | 9 | Baz1a | -0.001 | Old |
| 7.39E-07 | 0.473698 | 1 | 1 | 0.022934 | 9 | Gm42418 | 0 | Young |
| 3.47E-05 | 2.324579 | 0.14 | 0.039 | 1 | 9 | Fcgr2b | 0.101 | Young |
| 9.07E-05 | 2.76634 | 0.066 | 0.008 | 1 | 9 | Cryzl2 | 0.058 | Young |
| 0.00019 | 2.079864 | 0.088 | 0.018 | 1 | 9 | Emb | 0.07 | Young |
| 0.000209 | 1.197915 | 0.316 | 0.17 | 1 | 9 | Tkt | 0.146 | Young |
| 0.00023 | 1.748985 | 0.103 | 0.026 | 1 | 9 | Cdo1 | 0.077 | Young |
| 0.000287 | 2.010671 | 0.125 | 0.039 | 1 | 9 | Cfd | 0.086 | Young |
| 0.000308 | 1.764201 | 0.096 | 0.023 | 1 | 9 | Dennd4b | 0.073 | Young |
| 0.000323 | 2.478049 | 0.221 | 0.103 | 1 | 9 | Scd1 | 0.118 | Young |
| 0.000354 | 3.066152 | 0.051 | 0.005 | 1 | 9 | Pira2 | 0.046 | Young |
| 0.000354 | 2.897616 | 0.051 | 0.005 | 1 | 9 | Pilra | 0.046 | Young |
| 0.000376 | 1.115516 | 0.434 | 0.304 | 1 | 9 | H2-Aa | 0.13 | Young |
| 0.000493 | 2.045076 | 0.478 | 0.34 | 1 | 9 | Retnla | 0.138 | Young |
| 0.000519 | 2.255838 | 0.081 | 0.018 | 1 | 9 | Rab7b | 0.063 | Young |
| 0.000679 | 3.301052 | 0.088 | 0.023 | 1 | 9 | Ccl24 | 0.065 | Young |
| 0.00071 | 3.331087 | 0.029 | 0 | 1 | 9 | Pafah1b3 | 0.029 | Young |
| 0.00071 | 2.343362 | 0.029 | 0 | 1 | 9 | Plpp2 | 0.029 | Young |
| 0.00071 | 2.198545 | 0.029 | 0 | 1 | 9 | Ptgir | 0.029 | Young |
| 0.00086 | 1.944098 | 0.125 | 0.044 | 1 | 9 | Rps12-ps3 | 0.081 | Young |
| 0.001031 | 1.84586 | 0.088 | 0.023 | 1 | 9 | Pik3r5 | 0.065 | Young |
| 0.001037 | 1.322013 | 0.125 | 0.044 | 1 | 9 | Trf | 0.081 | Young |
| 0.001169 | 1.267049 | 0.132 | 0.049 | 1 | 9 | Metap1 | 0.083 | Young |
| 0.001234 | 1.439947 | 0.176 | 0.08 | 1 | 9 | Nfkbid | 0.096 | Young |

| | | | | | | | | |
|---|---|---|---|---|---|---|---|---|
| 0.00124 | 1.005058 | 0.206 | 0.098 | 1 | 9 | Trps1 | 0.108 | Young |
| 0.001272 | 1.554233 | 0.081 | 0.021 | 1 | 9 | Spryd7 | 0.06 | Young |
| 0.001291 | 3.682199 | 0.037 | 0.003 | 1 | 9 | AI661453 | 0.034 | Young |
| 0.001291 | 2.382019 | 0.037 | 0.003 | 1 | 9 | Tldc1 | 0.034 | Young |
| 0.001308 | 3.10622 | 0.037 | 0.003 | 1 | 9 | Bend4 | 0.034 | Young |
| 0.001378 | 1.520655 | 0.051 | 0.008 | 1 | 9 | Arl5c | 0.043 | Young |
| 0.001421 | 3.24189 | 0.044 | 0.005 | 1 | 9 | Arhgap32 | 0.039 | Young |
| 0.001436 | 2.798146 | 0.044 | 0.005 | 1 | 9 | Cd2 | 0.039 | Young |
| 0.001452 | 2.770115 | 0.044 | 0.005 | 1 | 9 | Sh3bp2 | 0.039 | Young |
| 0.001452 | 2.252194 | 0.044 | 0.005 | 1 | 9 | Tubb2b | 0.039 | Young |
| 0.001499 | 1.847114 | 0.044 | 0.005 | 1 | 9 | Arhgap6 | 0.039 | Young |
| 0.001527 | 1.814063 | 0.11 | 0.039 | 1 | 9 | Syk | 0.071 | Young |
| 0.001613 | 1.644328 | 0.199 | 0.101 | 1 | 9 | Mmd | 0.098 | Young |
| 0.001661 | 0.925422 | 0.14 | 0.054 | 1 | 9 | Tvp23b | 0.086 | Young |
| 0.001852 | 1.603037 | 0.088 | 0.026 | 1 | 9 | Nagk | 0.062 | Young |
| 0.002122 | 1.533519 | 0.287 | 0.178 | 1 | 9 | Car3 | 0.109 | Young |
| 0.002144 | 2.133766 | 0.096 | 0.031 | 1 | 9 | Ccl4 | 0.065 | Young |
| 0.00229 | 1.629041 | 0.096 | 0.031 | 1 | 9 | Nup93 | 0.065 | Young |
| 0.002398 | 1.780793 | 0.066 | 0.015 | 1 | 9 | Rrp1b | 0.051 | Young |
| 0.002401 | 2.077454 | 0.081 | 0.023 | 1 | 9 | Poldip2 | 0.058 | Young |
| 0.002416 | 1.578315 | 0.066 | 0.015 | 1 | 9 | Slamf7 | 0.051 | Young |
| 0.0028 | 0.784205 | 0.353 | 0.232 | 1 | 9 | Reep5 | 0.121 | Young |
| 0.002864 | 1.227649 | 0.081 | 0.023 | 1 | 9 | Evi2a | 0.058 | Young |
| 0.003083 | 2.34575 | 0.059 | 0.013 | 1 | 9 | Gm6377 | 0.046 | Young |
| 0.003279 | 1.810661 | 0.213 | 0.121 | 1 | 9 | Mafb | 0.092 | Young |
| 0.003303 | 1.655054 | 0.088 | 0.028 | 1 | 9 | Bmf | 0.06 | Young |
| 0.003309 | 1.145585 | 0.235 | 0.131 | 1 | 9 | Rnpep | 0.104 | Young |
| 0.003405 | 3.98799 | 0.022 | 0 | 1 | 9 | Aqp9 | 0.022 | Young |
| 0.003405 | 3.769537 | 0.022 | 0 | 1 | 9 | Lgals12 | 0.022 | Young |
| 0.003405 | 3.466906 | 0.022 | 0 | 1 | 9 | Egfr | 0.022 | Young |
| 0.003405 | 3.041301 | 0.022 | 0 | 1 | 9 | Frat1 | 0.022 | Young |
| 0.003405 | 2.918706 | 0.022 | 0 | 1 | 9 | Gm9465 | 0.022 | Young |
| 0.003405 | 2.602363 | 0.022 | 0 | 1 | 9 | Gpatch3 | 0.022 | Young |
| 0.003405 | 2.571861 | 0.022 | 0 | 1 | 9 | Pcdhga11 | 0.022 | Young |
| 0.003405 | 2.468292 | 0.022 | 0 | 1 | 9 | Igsf21 | 0.022 | Young |
| 0.003405 | 1.93623 | 0.022 | 0 | 1 | 9 | Nmnat3 | 0.022 | Young |
| 0.003405 | 1.733335 | 0.022 | 0 | 1 | 9 | Kif9 | 0.022 | Young |
| 0.003538 | 1.259296 | 0.169 | 0.082 | 1 | 9 | Fcer1g | 0.087 | Young |
| 0.00358 | 1.178786 | 0.14 | 0.059 | 1 | 9 | Ptafr | 0.081 | Young |
| 0.003604 | 1.38212 | 0.074 | 0.021 | 1 | 9 | Mavs | 0.053 | Young |
| 0.003726 | 1.403169 | 0.132 | 0.057 | 1 | 9 | Ccr2 | 0.075 | Young |
| 0.003948 | 2.530521 | 0.051 | 0.01 | 1 | 9 | Ptpn7 | 0.041 | Young |
| 0.003968 | 1.833446 | 0.096 | 0.034 | 1 | 9 | Abcd1 | 0.062 | Young |
| 0.00417 | 1.057121 | 0.096 | 0.034 | 1 | 9 | 3110043O2 | 0.062 | Young |
| 0.004818 | 1.987017 | 0.081 | 0.026 | 1 | 9 | Ttc7 | 0.055 | Young |

| | | | | | | | | |
|---|---|---|---|---|---|---|---|---|
| 0.004881 | 2.37086 | 0.044 | 0.008 | 1 | 9 | Ccdc112 | 0.036 | Young |
| 0.00491 | 1.115416 | 0.206 | 0.111 | 1 | 9 | Mt1 | 0.095 | Young |
| 0.00497 | 2.337546 | 0.044 | 0.008 | 1 | 9 | Tmem37 | 0.036 | Young |
| 0.005127 | 1.508182 | 0.118 | 0.049 | 1 | 9 | Pid1 | 0.069 | Young |
| 0.005154 | 2.185367 | 0.044 | 0.008 | 1 | 9 | Dusp19 | 0.036 | Young |
| 0.005295 | 1.342887 | 0.044 | 0.008 | 1 | 9 | Ryr1 | 0.036 | Young |
| 0.005358 | 1.301729 | 0.11 | 0.044 | 1 | 9 | Itgam | 0.066 | Young |
| 0.005552 | 3.542301 | 0.029 | 0.003 | 1 | 9 | Atp1a2 | 0.026 | Young |
| 0.005614 | 1.883863 | 0.14 | 0.064 | 1 | 9 | Wfdc17 | 0.076 | Young |
| 0.005619 | 3.461613 | 0.029 | 0.003 | 1 | 9 | Gm12355 | 0.026 | Young |
| 0.005619 | 3.457028 | 0.029 | 0.003 | 1 | 9 | Mapk8ip1 | 0.026 | Young |
| 0.005619 | 2.972775 | 0.029 | 0.003 | 1 | 9 | Mmp28 | 0.026 | Young |
| 0.005619 | 2.839731 | 0.029 | 0.003 | 1 | 9 | Boc | 0.026 | Young |
| 0.005687 | 2.667105 | 0.029 | 0.003 | 1 | 9 | Pex11a | 0.026 | Young |
| 0.005687 | 2.4332 | 0.029 | 0.003 | 1 | 9 | Ptgr1 | 0.026 | Young |
| 0.005687 | 2.418055 | 0.029 | 0.003 | 1 | 9 | Slc25a42 | 0.026 | Young |
| 0.005687 | 2.082342 | 0.029 | 0.003 | 1 | 9 | P2ry10 | 0.026 | Young |
| 0.005746 | 2.633104 | 0.037 | 0.005 | 1 | 9 | Clec2i | 0.032 | Young |
| 0.005755 | 2.614102 | 0.029 | 0.003 | 1 | 9 | Plcl1 | 0.026 | Young |
| 0.005755 | 2.080032 | 0.029 | 0.003 | 1 | 9 | Gm10643 | 0.026 | Young |
| 0.005804 | 2.374818 | 0.037 | 0.005 | 1 | 9 | Zfp953 | 0.032 | Young |
| 0.005804 | 2.370487 | 0.037 | 0.005 | 1 | 9 | Rnase6 | 0.032 | Young |
| 0.005848 | 1.187187 | 0.228 | 0.131 | 1 | 9 | Cd83 | 0.097 | Young |
| 0.005868 | 0.801173 | 0.081 | 0.026 | 1 | 9 | Sema4a | 0.055 | Young |
| 0.006619 | 1.99665 | 0.059 | 0.015 | 1 | 9 | Ms4a8a | 0.044 | Young |
| 0.006689 | 1.658105 | 0.096 | 0.036 | 1 | 9 | Adgre1 | 0.06 | Young |
| 0.00681 | 2.142717 | 0.059 | 0.015 | 1 | 9 | Adcy7 | 0.044 | Young |
| 0.006957 | 2.281113 | 0.059 | 0.015 | 1 | 9 | Sh3bp1 | 0.044 | Young |
| 0.007093 | 1.025069 | 0.162 | 0.082 | 1 | 9 | Tm9sf4 | 0.08 | Young |
| 0.007196 | 1.8619 | 0.074 | 0.023 | 1 | 9 | Zfp280c | 0.051 | Young |
| 0.007196 | 1.503516 | 0.074 | 0.023 | 1 | 9 | Cd163 | 0.051 | Young |
| 0.007258 | 1.291198 | 0.059 | 0.015 | 1 | 9 | 1110012L1 | 0.044 | Young |
| 0.007327 | 0.866652 | 0.11 | 0.044 | 1 | 9 | Inpp5f | 0.066 | Young |
| 0.007378 | 1.125378 | 0.096 | 0.036 | 1 | 9 | Mrpl19 | 0.06 | Young |
| 0.007418 | 1.301971 | 0.132 | 0.062 | 1 | 9 | Nsun2 | 0.07 | Young |
| 0.007624 | 0.015687 | 0.059 | 0.015 | 1 | 9 | Tmub1 | 0.044 | Young |
| 0.00775 | 1.60535 | 0.11 | 0.046 | 1 | 9 | Necap1 | 0.064 | Young |
| 0.007927 | 1.070058 | 0.176 | 0.093 | 1 | 9 | Itgb5 | 0.083 | Young |
| 0.008168 | 1.23249 | 0.162 | 0.082 | 1 | 9 | Arhgdib | 0.08 | Young |
| 0.008652 | 0.925808 | 0.243 | 0.144 | 1 | 9 | Blmh | 0.099 | Young |
| 0.008891 | 0.995411 | 0.103 | 0.041 | 1 | 9 | Clec4a3 | 0.062 | Young |
| 0.009204 | 1.120454 | 0.081 | 0.028 | 1 | 9 | Akap10 | 0.053 | Young |
| 0.009381 | 2.2266 | 0.066 | 0.021 | 1 | 9 | Hectd3 | 0.045 | Young |
| 0.009435 | 2.487292 | 0.051 | 0.013 | 1 | 9 | Il1rl1 | 0.038 | Young |
| 0.009576 | 1.754431 | 0.051 | 0.013 | 1 | 9 | Il17ra | 0.038 | Young |

| | | | | | | | | |
|---|---|---|---|---|---|---|---|---|
| 0.009618 | 1.975342 | 0.066 | 0.021 | 1 | 9 | Tmem132a | 0.045 | Young |
| 0.009678 | 2.142377 | 0.066 | 0.021 | 1 | 9 | Arhgap45 | 0.045 | Young |
| 0.00979 | 1.94033 | 0.051 | 0.013 | 1 | 9 | Sp3os | 0.038 | Young |
| 0.010008 | 1.976732 | 0.051 | 0.013 | 1 | 9 | Rassf5 | 0.038 | Young |
| 0.010082 | 1.365541 | 0.051 | 0.013 | 1 | 9 | Cc2d1a | 0.038 | Young |
| 0.010298 | 1.623568 | 0.066 | 0.021 | 1 | 9 | C3ar1 | 0.045 | Young |
| 0.010382 | 0.307781 | 0.912 | 0.923 | 1 | 9 | Psap | -0.011 | Young |
| 0.010426 | 1.540392 | 0.066 | 0.021 | 1 | 9 | Nubp2 | 0.045 | Young |
| 0.010855 | 1.579454 | 0.088 | 0.034 | 1 | 9 | Tln2 | 0.054 | Young |
| 0.011071 | 0.570365 | 0.868 | 0.83 | 1 | 9 | Klf2 | 0.038 | Young |
| 0.011293 | 0.425109 | 0.066 | 0.021 | 1 | 9 | AI413582 | 0.045 | Young |
| 0.011485 | 1.602369 | 0.14 | 0.07 | 1 | 9 | Top3b | 0.07 | Young |
| 0.011584 | 2.292264 | 0.081 | 0.031 | 1 | 9 | Me1 | 0.05 | Young |
| 0.011702 | 1.931114 | 0.074 | 0.026 | 1 | 9 | Dock10 | 0.048 | Young |
| 0.011872 | 1.439981 | 0.125 | 0.059 | 1 | 9 | Erlin2 | 0.066 | Young |
| 0.012022 | 0.542114 | 0.412 | 0.312 | 1 | 9 | Fam198b | 0.1 | Young |
| 0.012333 | 0.88821 | 0.147 | 0.075 | 1 | 9 | Dusp22 | 0.072 | Young |
| 0.012365 | 1.07515 | 0.191 | 0.111 | 1 | 9 | Serinc5 | 0.08 | Young |
| 0.012442 | 0.926248 | 0.118 | 0.054 | 1 | 9 | Tep1 | 0.064 | Young |
| 0.012448 | 1.713005 | 0.074 | 0.026 | 1 | 9 | Nfam1 | 0.048 | Young |
| 0.012629 | 1.131376 | 0.14 | 0.07 | 1 | 9 | C1qc | 0.07 | Young |
| 0.012887 | 2.791933 | 0.044 | 0.01 | 1 | 9 | AB124611 | 0.034 | Young |
| 0.013117 | 1.138118 | 0.243 | 0.157 | 1 | 9 | Ppp3r1 | 0.086 | Young |
| 0.013258 | 0.61358 | 0.265 | 0.165 | 1 | 9 | Tppp3 | 0.1 | Young |
| 0.013295 | 1.800408 | 0.044 | 0.01 | 1 | 9 | Glrx | 0.034 | Young |
| 0.013306 | 2.043819 | 0.059 | 0.018 | 1 | 9 | Fdft1 | 0.041 | Young |
| 0.013391 | 2.118928 | 0.059 | 0.018 | 1 | 9 | Ptpro | 0.041 | Young |
| 0.013399 | 1.905329 | 0.044 | 0.01 | 1 | 9 | 1600002K0 | 0.034 | Young |
| 0.013503 | 1.829098 | 0.044 | 0.01 | 1 | 9 | Tmem141 | 0.034 | Young |
| 0.013691 | 0.644854 | 0.118 | 0.054 | 1 | 9 | Yif1b | 0.064 | Young |
| 0.013715 | 1.296043 | 0.044 | 0.01 | 1 | 9 | Kiz | 0.034 | Young |
| 0.013929 | 1.107233 | 0.044 | 0.01 | 1 | 9 | Dock5 | 0.034 | Young |
| 0.01406 | 1.814098 | 0.096 | 0.041 | 1 | 9 | Plbd1 | 0.055 | Young |
| 0.014442 | 2.101357 | 0.088 | 0.036 | 1 | 9 | Utp20 | 0.052 | Young |
| 0.014634 | 0.973419 | 0.059 | 0.018 | 1 | 9 | Lrmp | 0.041 | Young |
| 0.014654 | 1.992056 | 0.088 | 0.036 | 1 | 9 | Arl14ep | 0.052 | Young |
| 0.014831 | 1.191209 | 0.081 | 0.031 | 1 | 9 | Ccp110 | 0.05 | Young |
| 0.014839 | 1.04193 | 0.169 | 0.093 | 1 | 9 | Ppm1b | 0.076 | Young |
| 0.015111 | 0.524829 | 0.103 | 0.044 | 1 | 9 | Zcchc9 | 0.059 | Young |
| 0.015152 | 1.018591 | 0.125 | 0.059 | 1 | 9 | Tgfbrap1 | 0.066 | Young |
| 0.015177 | 1.123167 | 0.081 | 0.031 | 1 | 9 | Fcrls | 0.05 | Young |
| 0.015305 | 1.323823 | 0.088 | 0.036 | 1 | 9 | Cd300ld | 0.052 | Young |
| 0.015597 | 0.575688 | 0.272 | 0.173 | 1 | 9 | Pfkp | 0.099 | Young |
| 0.016398 | 1.285003 | 0.199 | 0.121 | 1 | 9 | C1qa | 0.078 | Young |
| 0.016736 | 1.249969 | 0.176 | 0.103 | 1 | 9 | Laptm5 | 0.073 | Young |

| | | | | | | | | |
|---|---|---|---|---|---|---|---|---|
| 0.016784 | 1.407208 | 0.11 | 0.052 | 1 | 9 | Mpeg1 | 0.058 | Young |
| 0.016945 | 1.786323 | 0.015 | 0 | 1 | 9 | Map3k9 | 0.015 | Young |
| 0.016945 | 3.938231 | 0.015 | 0 | 1 | 9 | Adam33 | 0.015 | Young |
| 0.016945 | 3.83207 | 0.015 | 0 | 1 | 9 | Rnf32 | 0.015 | Young |
| 0.016945 | 3.318182 | 0.015 | 0 | 1 | 9 | Aif1l | 0.015 | Young |
| 0.016945 | 3.307159 | 0.015 | 0 | 1 | 9 | Klrd1 | 0.015 | Young |
| 0.016945 | 3.276014 | 0.015 | 0 | 1 | 9 | Olfr1322 | 0.015 | Young |
| 0.016945 | 2.739233 | 0.015 | 0 | 1 | 9 | Dzank1 | 0.015 | Young |
| 0.016945 | 2.676701 | 0.015 | 0 | 1 | 9 | E2f8 | 0.015 | Young |
| 0.016945 | 2.675148 | 0.015 | 0 | 1 | 9 | Ndor1 | 0.015 | Young |
| 0.016945 | 2.649764 | 0.015 | 0 | 1 | 9 | Samd15 | 0.015 | Young |
| 0.016945 | 2.645286 | 0.015 | 0 | 1 | 9 | Aldh1a7 | 0.015 | Young |
| 0.016945 | 2.635846 | 0.015 | 0 | 1 | 9 | Plin1 | 0.015 | Young |
| 0.016945 | 2.626365 | 0.015 | 0 | 1 | 9 | Klri2 | 0.015 | Young |
| 0.016945 | 2.563341 | 0.015 | 0 | 1 | 9 | Gm9968 | 0.015 | Young |
| 0.016945 | 2.515762 | 0.015 | 0 | 1 | 9 | 0610043K1 | 0.015 | Young |
| 0.016945 | 2.462793 | 0.015 | 0 | 1 | 9 | Gm44148 | 0.015 | Young |
| 0.016945 | 2.384345 | 0.015 | 0 | 1 | 9 | Gm26737 | 0.015 | Young |
| 0.016945 | 2.321052 | 0.015 | 0 | 1 | 9 | 1190005I0 | 0.015 | Young |
| 0.016945 | 2.305194 | 0.015 | 0 | 1 | 9 | CT025521.1 | 0.015 | Young |
| 0.016945 | 2.289937 | 0.015 | 0 | 1 | 9 | Capn11 | 0.015 | Young |
| 0.016945 | 2.276827 | 0.015 | 0 | 1 | 9 | N4bp2l1 | 0.015 | Young |
| 0.016945 | 2.251914 | 0.015 | 0 | 1 | 9 | Gm14698 | 0.015 | Young |
| 0.016945 | 2.165748 | 0.015 | 0 | 1 | 9 | Tnfsf13b | 0.015 | Young |
| 0.016945 | 2.161357 | 0.015 | 0 | 1 | 9 | Gng10 | 0.015 | Young |
| 0.016945 | 2.049918 | 0.015 | 0 | 1 | 9 | 2810457G0 | 0.015 | Young |
| 0.016945 | 2.048077 | 0.015 | 0 | 1 | 9 | Sdr42e1 | 0.015 | Young |
| 0.016945 | 1.968146 | 0.015 | 0 | 1 | 9 | Scgb3a1 | 0.015 | Young |
| 0.016945 | 1.86838 | 0.015 | 0 | 1 | 9 | 3632451O0 | 0.015 | Young |
| 0.016945 | 1.861081 | 0.015 | 0 | 1 | 9 | Gm14964 | 0.015 | Young |
| 0.016945 | 1.796043 | 0.015 | 0 | 1 | 9 | Depdc7 | 0.015 | Young |
| 0.016945 | 1.642075 | 0.015 | 0 | 1 | 9 | Zfp941 | 0.015 | Young |
| 0.016945 | 1.551184 | 0.015 | 0 | 1 | 9 | Cstad | 0.015 | Young |
| 0.016945 | 1.464942 | 0.015 | 0 | 1 | 9 | Gm15890 | 0.015 | Young |
| 0.016945 | 1.434067 | 0.015 | 0 | 1 | 9 | Ipcef1 | 0.015 | Young |
| 0.016945 | 1.409851 | 0.015 | 0 | 1 | 9 | 2610016A1 | 0.015 | Young |
| 0.016945 | 1.385045 | 0.015 | 0 | 1 | 9 | Gm26882 | 0.015 | Young |
| 0.016945 | 1.325856 | 0.015 | 0 | 1 | 9 | Msantd1 | 0.015 | Young |
| 0.016945 | 1.066433 | 0.015 | 0 | 1 | 9 | St8sia2 | 0.015 | Young |
| 0.016945 | 1.002865 | 0.015 | 0 | 1 | 9 | Gm4285 | 0.015 | Young |
| 0.016945 | 1.000978 | 0.015 | 0 | 1 | 9 | Gm29340 | 0.015 | Young |
| 0.016945 | 0.93439 | 0.015 | 0 | 1 | 9 | P2ry12 | 0.015 | Young |
| 0.016945 | 0.930399 | 0.015 | 0 | 1 | 9 | Ccdc170 | 0.015 | Young |
| 0.017167 | 1.07995 | 0.088 | 0.036 | 1 | 9 | Tnip3 | 0.052 | Young |
| 0.017415 | 0.853761 | 0.25 | 0.165 | 1 | 9 | Clasp2 | 0.085 | Young |

| | | | | | | | | |
|---|---|---|---|---|---|---|---|---|
| 0.017461 | 2.348796 | 0.037 | 0.008 | 1 | 9 | Espl1 | 0.029 | Young |
| 0.017461 | 2.328221 | 0.037 | 0.008 | 1 | 9 | Ssh3 | 0.029 | Young |
| 0.017608 | 2.284646 | 0.037 | 0.008 | 1 | 9 | Gm26890 | 0.029 | Young |
| 0.017659 | 1.166913 | 0.096 | 0.041 | 1 | 9 | H2-DMa | 0.055 | Young |
| 0.017765 | 0.730963 | 0.221 | 0.137 | 1 | 9 | Abhd17b | 0.084 | Young |
| 0.017906 | 1.606896 | 0.037 | 0.008 | 1 | 9 | Cables1 | 0.029 | Young |
| 0.017906 | 0.22383 | 0.037 | 0.008 | 1 | 9 | Hk3 | 0.029 | Young |
| 0.018056 | 1.091604 | 0.037 | 0.008 | 1 | 9 | Akip1 | 0.029 | Young |
| 0.018258 | 0.793128 | 0.154 | 0.082 | 1 | 9 | Hcfc2 | 0.072 | Young |
| 0.018316 | 1.907554 | 0.066 | 0.023 | 1 | 9 | Trpv4 | 0.043 | Young |
| 0.01836 | 0.405652 | 0.037 | 0.008 | 1 | 9 | Slc13a3 | 0.029 | Young |
| 0.018475 | 1.067059 | 0.118 | 0.057 | 1 | 9 | Rhoq | 0.061 | Young |
| 0.018592 | 1.041573 | 0.14 | 0.072 | 1 | 9 | Med8 | 0.068 | Young |
| 0.018627 | 1.216308 | 0.066 | 0.023 | 1 | 9 | 9130401M | 0.043 | Young |
| 0.018647 | 1.347388 | 0.125 | 0.062 | 1 | 9 | Sgpp1 | 0.063 | Young |
| 0.018825 | 0.523827 | 0.037 | 0.008 | 1 | 9 | Slc16a7 | 0.029 | Young |
| 0.018966 | 1.518011 | 0.103 | 0.046 | 1 | 9 | Cmc2 | 0.057 | Young |
| 0.019011 | 0.543314 | 0.346 | 0.237 | 1 | 9 | Tsnax | 0.109 | Young |
| 0.019018 | 1.9704 | 0.118 | 0.059 | 1 | 9 | Edc4 | 0.059 | Young |
| 0.019266 | 1.292227 | 0.14 | 0.072 | 1 | 9 | Apaf1 | 0.068 | Young |
| 0.019291 | 0.686938 | 0.103 | 0.046 | 1 | 9 | Trappc9 | 0.057 | Young |
| 0.019342 | 1.787667 | 0.051 | 0.015 | 1 | 9 | 2610035D1 | 0.036 | Young |
| 0.019483 | 1.196284 | 0.147 | 0.08 | 1 | 9 | Anapc2 | 0.067 | Young |
| 0.019649 | 2.30417 | 0.081 | 0.034 | 1 | 9 | Slc15a2 | 0.047 | Young |
| 0.020066 | 1.98827 | 0.074 | 0.028 | 1 | 9 | Clec4b1 | 0.046 | Young |
| 0.020278 | 0.919073 | 0.243 | 0.157 | 1 | 9 | Ap3s1 | 0.086 | Young |
| 0.020779 | 0.502501 | 0.537 | 0.459 | 1 | 9 | Hspa1a | 0.078 | Young |
| 0.021268 | 0.853903 | 0.11 | 0.052 | 1 | 9 | Pdlim4 | 0.058 | Young |
| 0.021309 | 1.383454 | 0.11 | 0.054 | 1 | 9 | Fam46a | 0.056 | Young |
| 0.021599 | 0.282731 | 0.051 | 0.015 | 1 | 9 | Zfyve21 | 0.036 | Young |
| 0.021673 | 1.053598 | 0.074 | 0.028 | 1 | 9 | Arhgap9 | 0.046 | Young |
| 0.021864 | 1.845842 | 0.088 | 0.039 | 1 | 9 | Gga3 | 0.049 | Young |
| 0.021958 | 3.396332 | 0.029 | 0.005 | 1 | 9 | Ppp1r1a | 0.024 | Young |
| 0.021958 | 2.569093 | 0.029 | 0.005 | 1 | 9 | Runx3 | 0.024 | Young |
| 0.02202 | 0.980229 | 0.051 | 0.015 | 1 | 9 | Cks2 | 0.036 | Young |
| 0.022162 | 0.876548 | 0.051 | 0.015 | 1 | 9 | Il1rap | 0.036 | Young |
| 0.022371 | 0.786737 | 0.287 | 0.204 | 1 | 9 | Dab2 | 0.083 | Young |
| 0.022375 | 1.894634 | 0.029 | 0.005 | 1 | 9 | Bmx | 0.024 | Young |
| 0.022375 | 1.684218 | 0.029 | 0.005 | 1 | 9 | Ssu2 | 0.024 | Young |
| 0.022382 | 0.422535 | 0.243 | 0.155 | 1 | 9 | Arhgef3 | 0.088 | Young |
| 0.022585 | 0.312039 | 0.029 | 0.005 | 1 | 9 | Itgal | 0.024 | Young |
| 0.022585 | 2.075932 | 0.029 | 0.005 | 1 | 9 | Steap3 | 0.024 | Young |
| 0.022585 | 1.287097 | 0.029 | 0.005 | 1 | 9 | Pycrl | 0.024 | Young |
| 0.022585 | 1.204715 | 0.029 | 0.005 | 1 | 9 | Rrm2 | 0.024 | Young |
| 0.022797 | 0.543699 | 0.449 | 0.358 | 1 | 9 | Irf2bp2 | 0.091 | Young |

| | | | | | | | | |
|---|---|---|---|---|---|---|---|---|
| 0.022917 | 0.822499 | 0.206 | 0.126 | 1 | 9 | Slc25a51 | 0.08 | Young |
| 0.023059 | 1.521684 | 0.11 | 0.054 | 1 | 9 | Cd53 | 0.056 | Young |
| 0.023228 | 0.153868 | 0.029 | 0.005 | 1 | 9 | Plxdc2 | 0.024 | Young |
| 0.023274 | 2.673127 | 0.059 | 0.021 | 1 | 9 | Ppfia4 | 0.038 | Young |
| 0.023355 | 1.431421 | 0.081 | 0.034 | 1 | 9 | Ly86 | 0.047 | Young |
| 0.024045 | 0.296064 | 0.118 | 0.057 | 1 | 9 | Rps6ka1 | 0.061 | Young |
| 0.024093 | 0.585456 | 0.39 | 0.294 | 1 | 9 | Sbno2 | 0.096 | Young |
| 0.024375 | 1.287349 | 0.118 | 0.059 | 1 | 9 | Lilr4b | 0.059 | Young |
| 0.024553 | 2.012461 | 0.022 | 0.003 | 1 | 9 | Slc25a33 | 0.019 | Young |
| 0.024832 | 3.597833 | 0.022 | 0.003 | 1 | 9 | Acy1 | 0.019 | Young |
| 0.024832 | 3.525699 | 0.022 | 0.003 | 1 | 9 | Cst6 | 0.019 | Young |
| 0.024832 | 3.287233 | 0.022 | 0.003 | 1 | 9 | Runx2 | 0.019 | Young |
| 0.024832 | 3.212456 | 0.022 | 0.003 | 1 | 9 | Hist1h2ap | 0.019 | Young |
| 0.024832 | 2.650099 | 0.022 | 0.003 | 1 | 9 | Bcl11a | 0.019 | Young |
| 0.024832 | 2.370579 | 0.022 | 0.003 | 1 | 9 | Itgax | 0.019 | Young |
| 0.024832 | 2.32953 | 0.022 | 0.003 | 1 | 9 | Vasn | 0.019 | Young |
| 0.024832 | 2.312611 | 0.022 | 0.003 | 1 | 9 | Tmem206 | 0.019 | Young |
| 0.024832 | 2.115288 | 0.022 | 0.003 | 1 | 9 | Slc14a1 | 0.019 | Young |
| 0.025115 | 3.144738 | 0.022 | 0.003 | 1 | 9 | Fxyd1 | 0.019 | Young |
| 0.025115 | 2.291287 | 0.022 | 0.003 | 1 | 9 | D030040B2 | 0.019 | Young |
| 0.025115 | 2.149095 | 0.022 | 0.003 | 1 | 9 | D730045B0 | 0.019 | Young |
| 0.025115 | 1.85759 | 0.022 | 0.003 | 1 | 9 | Skint3 | 0.019 | Young |
| 0.025115 | 1.734579 | 0.022 | 0.003 | 1 | 9 | AC163018. | 0.019 | Young |
| 0.025115 | 1.673248 | 0.022 | 0.003 | 1 | 9 | Abcb8 | 0.019 | Young |
| 0.025238 | 1.109703 | 0.088 | 0.039 | 1 | 9 | Selplg | 0.049 | Young |
| 0.0254 | 1.689154 | 0.022 | 0.003 | 1 | 9 | Aldh1a1 | 0.019 | Young |
| 0.0254 | 1.496426 | 0.022 | 0.003 | 1 | 9 | Cdkl1 | 0.019 | Young |
| 0.0254 | 1.396889 | 0.022 | 0.003 | 1 | 9 | Fgd1 | 0.019 | Young |
| 0.0254 | 1.261494 | 0.022 | 0.003 | 1 | 9 | D830030K2 | 0.019 | Young |
| 0.0254 | 1.035033 | 0.022 | 0.003 | 1 | 9 | F2rl1 | 0.019 | Young |
| 0.0254 | 0.998618 | 0.022 | 0.003 | 1 | 9 | Cd300lb | 0.019 | Young |
| 0.0254 | 0.760367 | 0.022 | 0.003 | 1 | 9 | 6720489N1 | 0.019 | Young |
| 0.025511 | 1.680101 | 0.059 | 0.021 | 1 | 9 | Iqce | 0.038 | Young |
| 0.025581 | 0.489899 | 0.147 | 0.08 | 1 | 9 | H2-M3 | 0.067 | Young |
| 0.026367 | 0.740899 | 0.228 | 0.147 | 1 | 9 | Gltp | 0.081 | Young |
| 0.026462 | 0.614283 | 0.184 | 0.108 | 1 | 9 | Gm47283 | 0.076 | Young |
| 0.026614 | 0.691156 | 0.088 | 0.039 | 1 | 9 | Itga4 | 0.049 | Young |
| 0.026732 | 0.603858 | 0.088 | 0.039 | 1 | 9 | Trim5 | 0.049 | Young |
| 0.027316 | 0.493596 | 0.14 | 0.075 | 1 | 9 | Zfp592 | 0.065 | Young |
| 0.027689 | 0.807765 | 0.088 | 0.039 | 1 | 9 | Nans | 0.049 | Young |
| 0.028294 | 2.032089 | 0.044 | 0.013 | 1 | 9 | Sco1 | 0.031 | Young |
| 0.028294 | 1.812653 | 0.044 | 0.013 | 1 | 9 | Kdm6bos | 0.031 | Young |
| 0.028348 | 1.15371 | 0.184 | 0.113 | 1 | 9 | Dpf2 | 0.071 | Young |
| 0.028611 | 0.821283 | 0.176 | 0.103 | 1 | 9 | Fnbp1 | 0.073 | Young |
| 0.028616 | 0.794147 | 0.184 | 0.108 | 1 | 9 | Bak1 | 0.076 | Young |

| | | | | | | | | |
|---|---|---|---|---|---|---|---|---|
| 0.028625 | 2.356255 | 0.074 | 0.031 | 1 | 9 | Pcid2 | 0.043 | Young |
| 0.029313 | 1.965408 | 0.11 | 0.057 | 1 | 9 | Tcaf1 | 0.053 | Young |
| 0.029857 | 1.668102 | 0.044 | 0.013 | 1 | 9 | Cutc | 0.031 | Young |
| 0.029857 | 1.578971 | 0.044 | 0.013 | 1 | 9 | Dusp18 | 0.031 | Young |
| 0.030057 | 1.089588 | 0.044 | 0.013 | 1 | 9 | Plekha5 | 0.031 | Young |
| 0.030517 | 1.281144 | 0.066 | 0.026 | 1 | 9 | Mad2l1bp | 0.04 | Young |
| 0.030611 | 1.933047 | 0.074 | 0.031 | 1 | 9 | Camk1d | 0.043 | Young |
| 0.03117 | 0.795819 | 0.309 | 0.229 | 1 | 9 | Gbp4 | 0.08 | Young |
| 0.031306 | 1.134744 | 0.066 | 0.026 | 1 | 9 | Cdk5rap3 | 0.04 | Young |
| 0.031306 | 0.737653 | 0.066 | 0.026 | 1 | 9 | Cd48 | 0.04 | Young |
| 0.031918 | 0.917024 | 0.14 | 0.077 | 1 | 9 | Mynn | 0.063 | Young |
| 0.0321 | 1.429587 | 0.074 | 0.031 | 1 | 9 | 2410002F2 | 0.043 | Young |
| 0.032253 | 1.417333 | 0.074 | 0.031 | 1 | 9 | Sepsecs | 0.043 | Young |
| 0.032666 | 0.820228 | 0.162 | 0.098 | 1 | 9 | Fn1 | 0.064 | Young |
| 0.033007 | 0.60057 | 0.169 | 0.101 | 1 | 9 | Pkn3 | 0.068 | Young |
| 0.033024 | 1.298919 | 0.074 | 0.031 | 1 | 9 | E2f1 | 0.043 | Young |
| 0.033437 | 1.156605 | 0.081 | 0.036 | 1 | 9 | Whamm | 0.045 | Young |
| 0.033478 | 1.121978 | 0.096 | 0.046 | 1 | 9 | Hsdl2 | 0.05 | Young |
| 0.033729 | 0.869264 | 0.132 | 0.072 | 1 | 9 | Il6ra | 0.06 | Young |
| 0.033949 | 0.891799 | 0.118 | 0.062 | 1 | 9 | Tdp2 | 0.056 | Young |
| 0.034385 | 1.255251 | 0.11 | 0.057 | 1 | 9 | Phf6 | 0.053 | Young |
| 0.034753 | 0.769239 | 0.132 | 0.072 | 1 | 9 | Wdr20 | 0.06 | Young |
| 0.034776 | 1.478818 | 0.074 | 0.031 | 1 | 9 | Ldlr | 0.043 | Young |
| 0.035215 | 0.657585 | 0.618 | 0.593 | 1 | 9 | Arpc2 | 0.025 | Young |
| 0.035218 | 0.224052 | 0.11 | 0.054 | 1 | 9 | Klhdc4 | 0.056 | Young |
| 0.035379 | 2.24812 | 0.051 | 0.018 | 1 | 9 | Rundc1 | 0.033 | Young |
| 0.035467 | 0.612702 | 0.331 | 0.242 | 1 | 9 | Hnrnpd | 0.089 | Young |
| 0.03613 | 1.123493 | 0.243 | 0.17 | 1 | 9 | Tmem176b | 0.073 | Young |
| 0.036165 | 0.491507 | 0.449 | 0.366 | 1 | 9 | 8430408G2 | 0.083 | Young |
| 0.036941 | 0.969411 | 0.088 | 0.041 | 1 | 9 | Clec4a1 | 0.047 | Young |
| 0.037085 | 1.005184 | 0.103 | 0.052 | 1 | 9 | Flt3l | 0.051 | Young |
| 0.037373 | 0.68565 | 0.154 | 0.093 | 1 | 9 | Plekhj1 | 0.061 | Young |
| 0.037393 | 0.40775 | 0.11 | 0.057 | 1 | 9 | Slc35b2 | 0.053 | Young |
| 0.037731 | 0.511994 | 0.162 | 0.093 | 1 | 9 | Anxa6 | 0.069 | Young |
| 0.037976 | 0.965501 | 0.081 | 0.036 | 1 | 9 | Ptpn6 | 0.045 | Young |
| 0.038202 | 1.627774 | 0.118 | 0.064 | 1 | 9 | Rgs2 | 0.054 | Young |
| 0.03833 | 0.401978 | 0.221 | 0.142 | 1 | 9 | Clk4 | 0.079 | Young |
| 0.038578 | 1.364756 | 0.051 | 0.018 | 1 | 9 | G6pdx | 0.033 | Young |
| 0.038715 | 0.941156 | 0.309 | 0.222 | 1 | 9 | Atf7ip | 0.087 | Young |
| 0.038978 | 0.854873 | 0.081 | 0.036 | 1 | 9 | Wdr3 | 0.045 | Young |
| 0.039071 | 0.226958 | 0.691 | 0.644 | 1 | 9 | Cdc42 | 0.047 | Young |
| 0.039472 | 1.432621 | 0.051 | 0.018 | 1 | 9 | Il1rn | 0.033 | Young |
| 0.039611 | 0.766773 | 0.118 | 0.062 | 1 | 9 | Slc35e1 | 0.056 | Young |
| 0.03965 | 3.033482 | 0.037 | 0.01 | 1 | 9 | Spa17 | 0.027 | Young |
| 0.039932 | 1.874511 | 0.037 | 0.01 | 1 | 9 | Camk2a | 0.027 | Young |

| | | | | | | | | |
|---|---|---|---|---|---|---|---|---|
| 0.040125 | 0.804111 | 0.103 | 0.052 | 1 | 9 | Mfn2 | 0.051 | Young |
| 0.040175 | 0.292377 | 0.081 | 0.036 | 1 | 9 | Mrpl22 | 0.045 | Young |
| 0.040216 | 2.704407 | 0.037 | 0.01 | 1 | 9 | Icosl | 0.027 | Young |
| 0.040894 | 2.980354 | 0.059 | 0.023 | 1 | 9 | Gatm | 0.036 | Young |
| 0.041368 | 1.43925 | 0.037 | 0.01 | 1 | 9 | Smim24 | 0.027 | Young |
| 0.041368 | 1.220814 | 0.037 | 0.01 | 1 | 9 | Iqcc | 0.027 | Young |
| 0.041387 | 0.927941 | 0.169 | 0.103 | 1 | 9 | Ap1b1 | 0.066 | Young |
| 0.041547 | 0.45731 | 0.051 | 0.018 | 1 | 9 | Chchd5 | 0.033 | Young |
| 0.041905 | 0.908154 | 0.176 | 0.111 | 1 | 9 | Cnep1r1 | 0.065 | Young |
| 0.042444 | 1.018594 | 0.169 | 0.103 | 1 | 9 | Eif3i | 0.066 | Young |
| 0.042547 | 0.694272 | 0.037 | 0.01 | 1 | 9 | Napsa | 0.027 | Young |
| 0.04257 | 0.357275 | 0.596 | 0.51 | 1 | 9 | Nfkbia | 0.086 | Young |
| 0.042616 | 1.635164 | 0.059 | 0.023 | 1 | 9 | Arhgap15 | 0.036 | Young |
| 0.042702 | 0.792849 | 0.221 | 0.155 | 1 | 9 | Efhd2 | 0.066 | Young |
| 0.043057 | 1.522529 | 0.059 | 0.023 | 1 | 9 | Rab13 | 0.036 | Young |
| 0.043057 | 1.380748 | 0.059 | 0.023 | 1 | 9 | Blnk | 0.036 | Young |
| 0.043382 | 1.32424 | 0.11 | 0.059 | 1 | 9 | Zfp518a | 0.051 | Young |
| 0.044061 | 0.408235 | 0.037 | 0.01 | 1 | 9 | Rft1 | 0.027 | Young |
| 0.044174 | 1.305885 | 0.059 | 0.023 | 1 | 9 | AC163032. | 0.036 | Young |
| 0.044384 | 1.530386 | 0.147 | 0.09 | 1 | 9 | Cirbp | 0.057 | Young |
| 0.044627 | 1.544784 | 0.059 | 0.023 | 1 | 9 | Mus81 | 0.036 | Young |
| 0.044627 | 0.963219 | 0.059 | 0.023 | 1 | 9 | Ankrd39 | 0.036 | Young |
| 0.045924 | 1.607491 | 0.074 | 0.034 | 1 | 9 | Gm21188 | 0.04 | Young |
| 0.04696 | 1.522888 | 0.066 | 0.028 | 1 | 9 | Slc7a7 | 0.038 | Young |
| 0.047064 | 0.651353 | 0.199 | 0.129 | 1 | 9 | Rheb | 0.07 | Young |
| 0.04732 | 0.569933 | 0.11 | 0.059 | 1 | 9 | Atp8b1 | 0.051 | Young |
| 0.04777 | 1.707369 | 0.074 | 0.034 | 1 | 9 | Fasn | 0.04 | Young |
| 0.047788 | 0.623362 | 0.25 | 0.178 | 1 | 9 | Cep350 | 0.072 | Young |
| 0.04807 | 0.887062 | 0.066 | 0.028 | 1 | 9 | Alcam | 0.038 | Young |
| 0.048164 | 1.37969 | 0.096 | 0.049 | 1 | 9 | Arrb2 | 0.047 | Young |
| 0.048295 | 0.843757 | 0.066 | 0.028 | 1 | 9 | Hs6st1 | 0.038 | Young |
| 0.049136 | 1.070384 | 0.088 | 0.044 | 1 | 9 | Bet1l | 0.044 | Young |
| 0.049149 | 0.742579 | 0.243 | 0.17 | 1 | 9 | Stk17b | 0.073 | Young |
| 0.049384 | 0.237403 | 0.059 | 0.023 | 1 | 9 | Ciita | 0.036 | Young |
| 0.049968 | 0.83795 | 0.147 | 0.088 | 1 | 9 | Fam220a | 0.059 | Young |
| 0.049971 | 0.420711 | 0.331 | 0.242 | 1 | 9 | Dag1 | 0.089 | Young |
| 0.050323 | 0.894171 | 0.081 | 0.039 | 1 | 9 | Gart | 0.042 | Young |
| 0.05034 | 2.610365 | 0.044 | 0.015 | 1 | 9 | Pnpt1 | 0.029 | Young |
| 0.050361 | 1.136309 | 0.096 | 0.049 | 1 | 9 | S100a4 | 0.047 | Young |
| 0.050454 | 0.501912 | 0.206 | 0.134 | 1 | 9 | Dnajc10 | 0.072 | Young |
| 0.050758 | 0.896999 | 0.14 | 0.082 | 1 | 9 | BC004004 | 0.058 | Young |
| 0.050874 | 0.691851 | 0.118 | 0.064 | 1 | 9 | Gpd2 | 0.054 | Young |
| 0.050884 | 1.500533 | 0.088 | 0.044 | 1 | 9 | Gramd1a | 0.044 | Young |
| 0.051383 | 0.928198 | 0.125 | 0.072 | 1 | 9 | Cotl1 | 0.053 | Young |
| 0.051695 | 0.483167 | 0.338 | 0.245 | 1 | 9 | Smim14 | 0.093 | Young |

| | | | | | | | | |
|---|---|---|---|---|---|---|---|---|
| 0.051852 | 2.191265 | 0.044 | 0.015 | 1 | 9 | Zkscan6 | 0.029 | Young |
| 0.051877 | 0.937405 | 0.088 | 0.044 | 1 | 9 | Rbbp5 | 0.044 | Young |
| 0.051895 | 0.432774 | 0.478 | 0.405 | 1 | 9 | Ptprg | 0.073 | Young |
| 0.051965 | 1.219092 | 0.206 | 0.144 | 1 | 9 | Asap1 | 0.062 | Young |
| 0.052063 | 1.093371 | 0.096 | 0.049 | 1 | 9 | Akr1e1 | 0.047 | Young |
| 0.052211 | 0.553635 | 0.235 | 0.162 | 1 | 9 | Sfswap | 0.073 | Young |
| 0.052467 | 1.128888 | 0.044 | 0.015 | 1 | 9 | Gdpd1 | 0.029 | Young |
| 0.052597 | 1.344915 | 0.103 | 0.054 | 1 | 9 | Vwf | 0.049 | Young |
| 0.052731 | 0.669805 | 0.169 | 0.106 | 1 | 9 | Snx5 | 0.063 | Young |
| 0.052777 | 2.908403 | 0.044 | 0.015 | 1 | 9 | Creb3l1 | 0.029 | Young |
| 0.053089 | 2.203588 | 0.044 | 0.015 | 1 | 9 | Nlrp3 | 0.029 | Young |
| 0.053306 | 0.498002 | 0.169 | 0.103 | 1 | 9 | Ube2j2 | 0.066 | Young |
| 0.053409 | 0.517668 | 0.279 | 0.201 | 1 | 9 | Thrsp | 0.078 | Young |
| 0.05343 | 0.960451 | 0.132 | 0.077 | 1 | 9 | D16Ertd47 | 0.055 | Young |
| 0.053711 | 0.265384 | 0.066 | 0.028 | 1 | 9 | Me2 | 0.038 | Young |
| 0.053716 | 1.769878 | 0.044 | 0.015 | 1 | 9 | Fam213b | 0.029 | Young |
| 0.053818 | 0.588772 | 0.294 | 0.224 | 1 | 9 | Pabpn1 | 0.07 | Young |
| 0.05388 | 0.300359 | 0.11 | 0.059 | 1 | 9 | H2-DMb1 | 0.051 | Young |
| 0.054032 | 1.68079 | 0.044 | 0.015 | 1 | 9 | Rnf150 | 0.029 | Young |
| 0.054316 | 1.68381 | 0.11 | 0.062 | 1 | 9 | Csrp2 | 0.048 | Young |
| 0.054326 | 0.757483 | 0.088 | 0.044 | 1 | 9 | Gdap2 | 0.044 | Young |
| 0.054535 | 0.729144 | 0.088 | 0.044 | 1 | 9 | Mid1 | 0.044 | Young |
| 0.054669 | 2.194395 | 0.044 | 0.015 | 1 | 9 | Mfsd8 | 0.029 | Young |
| 0.054798 | 1.317713 | 0.103 | 0.057 | 1 | 9 | Zfp46 | 0.046 | Young |
| 0.054812 | 0.994958 | 0.206 | 0.139 | 1 | 9 | Dync1li1 | 0.067 | Young |
| 0.05498 | 0.79984 | 0.221 | 0.149 | 1 | 9 | Mfsd1 | 0.072 | Young |
| 0.055076 | 0.652878 | 0.132 | 0.077 | 1 | 9 | Tbpl1 | 0.055 | Young |
| 0.055555 | 0.616805 | 0.074 | 0.034 | 1 | 9 | Itgb2 | 0.04 | Young |
| 0.055636 | 2.172502 | 0.044 | 0.015 | 1 | 9 | Fgr | 0.029 | Young |
| 0.056025 | 0.774418 | 0.103 | 0.054 | 1 | 9 | Sfmbt1 | 0.049 | Young |
| 0.056026 | 0.347991 | 0.588 | 0.523 | 1 | 9 | Iqgap1 | 0.065 | Young |
| 0.056386 | 2.3152 | 0.029 | 0.008 | 1 | 9 | Uros | 0.021 | Young |
| 0.056617 | 0.782755 | 0.044 | 0.015 | 1 | 9 | Lat2 | 0.029 | Young |
| 0.056617 | 0.523632 | 0.044 | 0.015 | 1 | 9 | Smug1 | 0.029 | Young |
| 0.056815 | 2.61652 | 0.029 | 0.008 | 1 | 9 | Fastkd5 | 0.021 | Young |
| 0.056815 | 2.446711 | 0.029 | 0.008 | 1 | 9 | Sez6l2 | 0.021 | Young |
| 0.056815 | 2.285476 | 0.029 | 0.008 | 1 | 9 | 1600010M | 0.021 | Young |
| 0.056947 | 1.018098 | 0.044 | 0.015 | 1 | 9 | Ccl3 | 0.029 | Young |
| 0.057247 | 2.200135 | 0.029 | 0.008 | 1 | 9 | Nbl1 | 0.021 | Young |
| 0.057247 | 1.944718 | 0.029 | 0.008 | 1 | 9 | Spout1 | 0.021 | Young |
| 0.057272 | 1.00202 | 0.25 | 0.188 | 1 | 9 | H2-Eb1 | 0.062 | Young |
| 0.057495 | 0.558139 | 0.154 | 0.095 | 1 | 9 | Exoc3 | 0.059 | Young |
| 0.057612 | 1.330309 | 0.044 | 0.015 | 1 | 9 | Sp140 | 0.029 | Young |
| 0.057681 | 2.003565 | 0.029 | 0.008 | 1 | 9 | Fam78a | 0.021 | Young |
| 0.057681 | 1.859296 | 0.029 | 0.008 | 1 | 9 | Tmie | 0.021 | Young |

| | | | | | | | | |
|---|---|---|---|---|---|---|---|---|
| 0.057681 | 1.743262 | 0.029 | 0.008 | 1 | 9 | Rhebl1 | 0.021 | Young |
| 0.058119 | 2.028371 | 0.029 | 0.008 | 1 | 9 | Slc37a4 | 0.021 | Young |
| 0.058119 | 1.985808 | 0.029 | 0.008 | 1 | 9 | Elovl6 | 0.021 | Young |
| 0.058283 | 0.307684 | 0.044 | 0.015 | 1 | 9 | Ppih | 0.029 | Young |
| 0.058483 | 0.508077 | 0.125 | 0.072 | 1 | 9 | Insig1 | 0.053 | Young |
| 0.058508 | 0.527507 | 0.096 | 0.049 | 1 | 9 | Alg5 | 0.047 | Young |
| 0.058559 | 1.849947 | 0.029 | 0.008 | 1 | 9 | Adig | 0.021 | Young |
| 0.058559 | 1.740587 | 0.029 | 0.008 | 1 | 9 | N6amt1 | 0.021 | Young |
| 0.058559 | 1.480869 | 0.029 | 0.008 | 1 | 9 | Rsad1 | 0.021 | Young |
| 0.058559 | 0.635251 | 0.029 | 0.008 | 1 | 9 | Etv1 | 0.021 | Young |
| 0.058812 | 1.423535 | 0.11 | 0.062 | 1 | 9 | Terf1 | 0.048 | Young |
| 0.059001 | 2.97634 | 0.029 | 0.008 | 1 | 9 | 1500011B0 | 0.021 | Young |
| 0.059001 | 1.951418 | 0.029 | 0.008 | 1 | 9 | Fars2 | 0.021 | Young |
| 0.059001 | 1.778769 | 0.029 | 0.008 | 1 | 9 | Slc27a3 | 0.021 | Young |
| 0.059001 | 1.144359 | 0.029 | 0.008 | 1 | 9 | Plcb2 | 0.021 | Young |
| 0.059001 | 1.026349 | 0.029 | 0.008 | 1 | 9 | Qpct | 0.021 | Young |
| 0.059001 | 0.792288 | 0.029 | 0.008 | 1 | 9 | Slc37a2 | 0.021 | Young |
| 0.059006 | 1.24409 | 0.11 | 0.062 | 1 | 9 | Larp7 | 0.048 | Young |
| 0.059069 | 0.524508 | 0.088 | 0.044 | 1 | 9 | Nek1 | 0.044 | Young |
| 0.059572 | 1.607151 | 0.125 | 0.075 | 1 | 9 | F13a1 | 0.05 | Young |
| 0.059895 | 1.112193 | 0.029 | 0.008 | 1 | 9 | Hdac10 | 0.021 | Young |
| 0.059895 | 1.060778 | 0.029 | 0.008 | 1 | 9 | Grk4 | 0.021 | Young |
| 0.059895 | 1.055338 | 0.029 | 0.008 | 1 | 9 | Ifitm1 | 0.021 | Young |
| 0.059895 | 0.47022 | 0.029 | 0.008 | 1 | 9 | Igfbp6 | 0.021 | Young |
| 0.059898 | 0.712666 | 0.25 | 0.178 | 1 | 9 | R3hdm1 | 0.072 | Young |
| 0.060485 | 0.704779 | 0.162 | 0.101 | 1 | 9 | Atg16l1 | 0.061 | Young |
| 0.0608 | 0.562272 | 0.029 | 0.008 | 1 | 9 | Gm13212 | 0.021 | Young |
| 0.060849 | 0.286078 | 0.831 | 0.812 | 1 | 9 | Rps3a1 | 0.019 | Young |
| 0.060932 | 1.275044 | 0.051 | 0.021 | 1 | 9 | Ccdc163 | 0.03 | Young |
| 0.061397 | 1.00223 | 0.184 | 0.121 | 1 | 9 | Txndc15 | 0.063 | Young |
| 0.061969 | 1.365118 | 0.125 | 0.075 | 1 | 9 | Socs6 | 0.05 | Young |
| 0.062328 | 0.726369 | 0.118 | 0.067 | 1 | 9 | Gpcpd1 | 0.051 | Young |
| 0.062383 | 1.13409 | 0.206 | 0.144 | 1 | 9 | Adam17 | 0.062 | Young |
| 0.06252 | 1.393954 | 0.051 | 0.021 | 1 | 9 | Akap5 | 0.03 | Young |
| 0.06252 | 1.360195 | 0.051 | 0.021 | 1 | 9 | Suv39h1 | 0.03 | Young |
| 0.0632 | 0.991639 | 0.11 | 0.062 | 1 | 9 | Alox5ap | 0.048 | Young |
| 0.063488 | 1.355285 | 0.051 | 0.021 | 1 | 9 | Pwwp2b | 0.03 | Young |
| 0.06356 | 0.454044 | 0.235 | 0.16 | 1 | 9 | St3gal1 | 0.075 | Young |
| 0.063835 | 0.394218 | 0.493 | 0.423 | 1 | 9 | Lpar6 | 0.07 | Young |
| 0.064141 | 1.208461 | 0.051 | 0.021 | 1 | 9 | Ncdn | 0.03 | Young |
| 0.064141 | 1.037832 | 0.051 | 0.021 | 1 | 9 | Polr3d | 0.03 | Young |
| 0.064203 | 0.166628 | 0.243 | 0.162 | 1 | 9 | Cap1 | 0.081 | Young |
| 0.064646 | 0.664842 | 0.162 | 0.101 | 1 | 9 | Gpr107 | 0.061 | Young |
| 0.064799 | 0.965972 | 0.051 | 0.021 | 1 | 9 | Retn | 0.03 | Young |
| 0.065455 | 1.543196 | 0.096 | 0.052 | 1 | 9 | Gpr132 | 0.044 | Young |

| | | | | | | | | |
|---|---|---|---|---|---|---|---|---|
| 0.065456 | 0.612594 | 0.213 | 0.144 | 1 | 9 | Tmem106b | 0.069 | Young |
| 0.065883 | 0.424272 | 0.603 | 0.541 | 1 | 9 | Hspa1b | 0.062 | Young |
| 0.066084 | 0.820435 | 0.147 | 0.09 | 1 | 9 | Rexo1 | 0.057 | Young |
| 0.066132 | 1.057623 | 0.051 | 0.021 | 1 | 9 | Fancg | 0.03 | Young |
| 0.066242 | 1.301297 | 0.154 | 0.101 | 1 | 9 | Mex3c | 0.053 | Young |
| 0.066322 | 0.807581 | 0.235 | 0.173 | 1 | 9 | Foxj2 | 0.062 | Young |
| 0.06672 | 0.415484 | 0.199 | 0.131 | 1 | 9 | Fam126b | 0.068 | Young |
| 0.066881 | 1.175433 | 0.059 | 0.026 | 1 | 9 | Rngtt | 0.033 | Young |
| 0.067162 | 1.93084 | 0.074 | 0.036 | 1 | 9 | Tnip2 | 0.038 | Young |
| 0.06723 | 1.285741 | 0.118 | 0.07 | 1 | 9 | Spint2 | 0.048 | Young |
| 0.06801 | 0.643677 | 0.096 | 0.052 | 1 | 9 | Dgat1 | 0.044 | Young |
| 0.06824 | 0.39357 | 0.118 | 0.067 | 1 | 9 | Celf2 | 0.051 | Young |
| 0.068284 | 0.678536 | 0.368 | 0.291 | 1 | 9 | Ric1 | 0.077 | Young |
| 0.068513 | 0.903524 | 0.11 | 0.062 | 1 | 9 | Ppp1r14b | 0.048 | Young |
| 0.068518 | 0.602157 | 0.051 | 0.021 | 1 | 9 | Ifitm6 | 0.03 | Young |
| 0.068518 | 0.521911 | 0.051 | 0.021 | 1 | 9 | Phc1 | 0.03 | Young |
| 0.068524 | 0.953911 | 0.066 | 0.031 | 1 | 9 | Ncf1 | 0.035 | Young |
| 0.068596 | 1.238421 | 0.147 | 0.095 | 1 | 9 | Zdhhc5 | 0.052 | Young |
| 0.068865 | 0.527691 | 0.051 | 0.021 | 1 | 9 | Mrps18b | 0.03 | Young |
| 0.068901 | 1.384174 | 0.118 | 0.07 | 1 | 9 | Ccl7 | 0.048 | Young |
| 0.069053 | 0.211018 | 0.228 | 0.155 | 1 | 9 | Pcmtd2 | 0.073 | Young |
| 0.069084 | 1.361045 | 0.059 | 0.026 | 1 | 9 | Rab3il1 | 0.033 | Young |
| 0.069084 | 1.139548 | 0.059 | 0.026 | 1 | 9 | Mad2l2 | 0.033 | Young |
| 0.069248 | 0.749695 | 0.301 | 0.237 | 1 | 9 | Ap2a2 | 0.064 | Young |
| 0.069719 | 0.398135 | 0.301 | 0.229 | 1 | 9 | Pik3r1 | 0.072 | Young |
| 0.069724 | 1.081153 | 0.059 | 0.026 | 1 | 9 | Grk3 | 0.033 | Young |
| 0.070046 | 0.938457 | 0.059 | 0.026 | 1 | 9 | Cxcl1 | 0.033 | Young |
| 0.070485 | 1.007193 | 0.176 | 0.119 | 1 | 9 | Spag7 | 0.057 | Young |
| 0.070608 | 1.223577 | 0.066 | 0.031 | 1 | 9 | Entpd7 | 0.035 | Young |
| 0.070866 | 0.67829 | 0.243 | 0.173 | 1 | 9 | Nab1 | 0.07 | Young |
| 0.071019 | 0.214346 | 0.059 | 0.026 | 1 | 9 | Btk | 0.033 | Young |
| 0.071346 | 0.566191 | 0.059 | 0.026 | 1 | 9 | Rcn1 | 0.033 | Young |
| 0.071581 | 0.816959 | 0.235 | 0.168 | 1 | 9 | Cisd2 | 0.067 | Young |
| 0.071608 | 0.006539 | 0.272 | 0.366 | 1 | 9 | Rab6a | -0.094 | Young |
| 0.071674 | 0.807386 | 0.059 | 0.026 | 1 | 9 | Stard6 | 0.033 | Young |
| 0.071872 | 0.488139 | 0.11 | 0.062 | 1 | 9 | Rabggtb | 0.048 | Young |
| 0.072291 | 1.095196 | 0.088 | 0.046 | 1 | 9 | Bad | 0.042 | Young |
| 0.072484 | 1.014648 | 0.074 | 0.036 | 1 | 9 | Cep290 | 0.038 | Young |
| 0.072823 | 1.590711 | 0.103 | 0.059 | 1 | 9 | Msrb1 | 0.044 | Young |
| 0.072868 | 0.51796 | 0.287 | 0.216 | 1 | 9 | Csrnp1 | 0.071 | Young |
| 0.073106 | 0.640175 | 0.228 | 0.16 | 1 | 9 | Bhlhe40 | 0.068 | Young |
| 0.073127 | 0.759235 | 0.199 | 0.137 | 1 | 9 | Pigm | 0.062 | Young |
| 0.073592 | 0.433021 | 0.221 | 0.149 | 1 | 9 | Osbp | 0.072 | Young |
| 0.073594 | 0.075964 | 0.176 | 0.258 | 1 | 9 | Cavin3 | -0.082 | Young |
| 0.074005 | 0.345637 | 0.059 | 0.026 | 1 | 9 | Cytip | 0.033 | Young |

| | | | | | | | | |
|---|---|---|---|---|---|---|---|---|
| 0.074343 | 0.642456 | 0.059 | 0.026 | 1 | 9 | C030034I2 | 0.033 | Young |
| 0.074588 | 1.039795 | 0.14 | 0.088 | 1 | 9 | Slc23a2 | 0.052 | Young |
| 0.074617 | 0.580836 | 0.066 | 0.031 | 1 | 9 | Myo1f | 0.035 | Young |
| 0.074954 | 1.328457 | 0.081 | 0.041 | 1 | 9 | Tbc1d10a | 0.04 | Young |
| 0.075175 | 0.959524 | 0.257 | 0.196 | 1 | 9 | Manf | 0.061 | Young |
| 0.075235 | 1.20062 | 0.081 | 0.041 | 1 | 9 | Chsy1 | 0.04 | Young |
| 0.075337 | 0.684549 | 0.235 | 0.17 | 1 | 9 | Kmt5b | 0.065 | Young |
| 0.075366 | 0.176198 | 0.11 | 0.062 | 1 | 9 | Xpa | 0.048 | Young |
| 0.075379 | 0.544593 | 0.434 | 0.361 | 1 | 9 | Xbp1 | 0.073 | Young |
| 0.075459 | 0.944724 | 0.088 | 0.046 | 1 | 9 | Nfrkb | 0.042 | Young |
| 0.075517 | 0.586718 | 0.081 | 0.041 | 1 | 9 | Dok2 | 0.04 | Young |
| 0.075575 | 1.002575 | 0.169 | 0.113 | 1 | 9 | Rel | 0.056 | Young |
| 0.075907 | 0.556839 | 0.184 | 0.121 | 1 | 9 | Ogfr | 0.063 | Young |
| 0.076084 | 0.79498 | 0.081 | 0.041 | 1 | 9 | Sdad1 | 0.04 | Young |
| 0.076208 | 0.716297 | 0.066 | 0.031 | 1 | 9 | Fam89a | 0.035 | Young |
| 0.07635 | 0.525208 | 0.14 | 0.085 | 1 | 9 | Vwa3a | 0.055 | Young |
| 0.076529 | 0.527979 | 0.162 | 0.103 | 1 | 9 | Ndufs3 | 0.059 | Young |
| 0.07674 | 0.408173 | 0.265 | 0.191 | 1 | 9 | Arl6ip5 | 0.074 | Young |
| 0.076858 | 0.867788 | 0.235 | 0.17 | 1 | 9 | Stxbp3 | 0.065 | Young |
| 0.077183 | 0.140189 | 0.272 | 0.193 | 1 | 9 | Sin3b | 0.079 | Young |
| 0.077534 | 0.264878 | 0.074 | 0.036 | 1 | 9 | Slc18b1 | 0.038 | Young |
| 0.078146 | 0.758658 | 0.074 | 0.036 | 1 | 9 | Tmem131l | 0.038 | Young |
| 0.078298 | 0.418276 | 0.478 | 0.394 | 1 | 9 | Dbi | 0.084 | Young |
| 0.078372 | 0.490777 | 0.419 | 0.338 | 1 | 9 | Nras | 0.081 | Young |
| 0.078922 | 0.630131 | 0.338 | 0.271 | 1 | 9 | Ssr1 | 0.067 | Young |
| 0.07897 | 0.339433 | 0.081 | 0.041 | 1 | 9 | Cdk7 | 0.04 | Young |
| 0.079069 | 0.514382 | 0.654 | 0.616 | 1 | 9 | Cd200 | 0.038 | Young |
| 0.07924 | 2.119985 | 0.037 | 0.013 | 1 | 9 | Slc38a1 | 0.024 | Young |
| 0.07924 | 1.418659 | 0.037 | 0.013 | 1 | 9 | Zfp553 | 0.024 | Young |
| 0.079601 | 3.049149 | 0.022 | 0.005 | 1 | 9 | Aptx | 0.017 | Young |
| 0.079601 | 2.793395 | 0.022 | 0.005 | 1 | 9 | Arxes2 | 0.017 | Young |
| 0.079601 | 2.045072 | 0.022 | 0.005 | 1 | 9 | Mblac1 | 0.017 | Young |
| 0.079717 | 2.793938 | 0.037 | 0.013 | 1 | 9 | Sdc1 | 0.024 | Young |
| 0.079717 | 1.351546 | 0.037 | 0.013 | 1 | 9 | Oas1c | 0.024 | Young |
| 0.079757 | 1.281453 | 0.125 | 0.077 | 1 | 9 | Lmf2 | 0.048 | Young |
| 0.079763 | 0.521098 | 0.25 | 0.178 | 1 | 9 | Pan3 | 0.072 | Young |
| 0.079929 | 1.605678 | 0.096 | 0.054 | 1 | 9 | Fam98c | 0.042 | Young |
| 0.080195 | 1.728883 | 0.037 | 0.013 | 1 | 9 | Gm16143 | 0.024 | Young |
| 0.080275 | 2.677607 | 0.022 | 0.005 | 1 | 9 | Ust | 0.017 | Young |
| 0.080275 | 2.574455 | 0.022 | 0.005 | 1 | 9 | Mss51 | 0.017 | Young |
| 0.080275 | 2.574175 | 0.022 | 0.005 | 1 | 9 | Pigw | 0.017 | Young |
| 0.080275 | 2.552651 | 0.022 | 0.005 | 1 | 9 | Adgre4 | 0.017 | Young |
| 0.080275 | 2.523719 | 0.022 | 0.005 | 1 | 9 | Il20rb | 0.017 | Young |
| 0.080275 | 1.948828 | 0.022 | 0.005 | 1 | 9 | Bend3 | 0.017 | Young |
| 0.080321 | 0.370974 | 0.074 | 0.036 | 1 | 9 | Zdhhc4 | 0.038 | Young |

| | | | | | | | | |
|---|---|---|---|---|---|---|---|---|
| 0.080636 | 0.658932 | 0.074 | 0.036 | 1 | 9 | Coq2 | 0.038 | Young |
| 0.080952 | 2.893632 | 0.022 | 0.005 | 1 | 9 | H2-Oa | 0.017 | Young |
| 0.080952 | 2.374053 | 0.022 | 0.005 | 1 | 9 | Pcdhb21 | 0.017 | Young |
| 0.080952 | 2.34611 | 0.022 | 0.005 | 1 | 9 | Tec | 0.017 | Young |
| 0.080952 | 2.263357 | 0.022 | 0.005 | 1 | 9 | Lrp12 | 0.017 | Young |
| 0.080952 | 2.145186 | 0.022 | 0.005 | 1 | 9 | Zeb2os | 0.017 | Young |
| 0.080952 | 1.877426 | 0.022 | 0.005 | 1 | 9 | Kcnk13 | 0.017 | Young |
| 0.080952 | 1.60742 | 0.022 | 0.005 | 1 | 9 | Map2k3os | 0.017 | Young |
| 0.080952 | 1.251161 | 0.022 | 0.005 | 1 | 9 | Cebpe | 0.017 | Young |
| 0.081219 | 0.407672 | 0.096 | 0.052 | 1 | 9 | Creld2 | 0.044 | Young |
| 0.081574 | 0.786275 | 0.287 | 0.206 | 1 | 9 | Ndufc1 | 0.081 | Young |
| 0.081577 | 0.658467 | 0.228 | 0.162 | 1 | 9 | Mark3 | 0.066 | Young |
| 0.081635 | 2.304746 | 0.022 | 0.005 | 1 | 9 | Morc4 | 0.017 | Young |
| 0.081635 | 1.942175 | 0.022 | 0.005 | 1 | 9 | Gstm4 | 0.017 | Young |
| 0.081635 | 1.931377 | 0.022 | 0.005 | 1 | 9 | Tagap | 0.017 | Young |
| 0.081635 | 1.643208 | 0.022 | 0.005 | 1 | 9 | Bbs10 | 0.017 | Young |
| 0.081635 | 1.569982 | 0.022 | 0.005 | 1 | 9 | Adat1 | 0.017 | Young |
| 0.081635 | 1.556757 | 0.022 | 0.005 | 1 | 9 | Pak4 | 0.017 | Young |
| 0.081635 | 1.484841 | 0.022 | 0.005 | 1 | 9 | Fcgr1 | 0.017 | Young |
| 0.081635 | 1.465876 | 0.022 | 0.005 | 1 | 9 | F8a | 0.017 | Young |
| 0.081635 | 0.754262 | 0.022 | 0.005 | 1 | 9 | Hps6 | 0.017 | Young |
| 0.081644 | 1.244887 | 0.037 | 0.013 | 1 | 9 | Frrs1 | 0.024 | Young |
| 0.081644 | 1.133104 | 0.037 | 0.013 | 1 | 9 | Chid1 | 0.024 | Young |
| 0.081644 | 1.070784 | 0.037 | 0.013 | 1 | 9 | Cdyl2 | 0.024 | Young |
| 0.082132 | 1.65322 | 0.037 | 0.013 | 1 | 9 | Pak1 | 0.024 | Young |
| 0.082132 | 0.718173 | 0.037 | 0.013 | 1 | 9 | Dhrs13 | 0.024 | Young |
| 0.082322 | 1.981821 | 0.022 | 0.005 | 1 | 9 | Nsl1 | 0.017 | Young |
| 0.082322 | 1.495084 | 0.022 | 0.005 | 1 | 9 | Fam161b | 0.017 | Young |
| 0.082322 | 1.411607 | 0.022 | 0.005 | 1 | 9 | B430010I2 | 0.017 | Young |
| 0.082322 | 1.396689 | 0.022 | 0.005 | 1 | 9 | Col5a1 | 0.017 | Young |
| 0.082322 | 1.383355 | 0.022 | 0.005 | 1 | 9 | Gpr34 | 0.017 | Young |
| 0.082322 | 1.324326 | 0.022 | 0.005 | 1 | 9 | Rab11fip1 | 0.017 | Young |
| 0.082322 | 1.162414 | 0.022 | 0.005 | 1 | 9 | 1110032A0 | 0.017 | Young |
| 0.082322 | 1.109627 | 0.022 | 0.005 | 1 | 9 | Fam213a | 0.017 | Young |
| 0.082322 | 1.104047 | 0.022 | 0.005 | 1 | 9 | Ddr1 | 0.017 | Young |
| 0.082322 | 0.762708 | 0.022 | 0.005 | 1 | 9 | AI839979 | 0.017 | Young |
| 0.082322 | 0.1656 | 0.022 | 0.005 | 1 | 9 | Plxna3 | 0.017 | Young |
| 0.082369 | 1.496326 | 0.118 | 0.072 | 1 | 9 | Pmpcb | 0.046 | Young |
| 0.082457 | 0.620399 | 0.272 | 0.206 | 1 | 9 | Cd44 | 0.066 | Young |
| 0.082506 | 0.178358 | 0.066 | 0.031 | 1 | 9 | Sfi1 | 0.035 | Young |
| 0.082775 | 0.498059 | 0.147 | 0.093 | 1 | 9 | Polr2f | 0.054 | Young |
| 0.082849 | 0.186382 | 0.015 | 0.049 | 1 | 9 | Myc | -0.034 | Young |
| 0.082854 | 0.190576 | 0.081 | 0.041 | 1 | 9 | Ppa2 | 0.04 | Young |
| 0.082933 | 0.837208 | 0.103 | 0.059 | 1 | 9 | Rev1 | 0.044 | Young |
| 0.083014 | 0.842135 | 0.022 | 0.005 | 1 | 9 | AU041133 | 0.017 | Young |

| | | | | | | | | |
|---|---|---|---|---|---|---|---|---|
| 0.083014 | 0.669569 | 0.022 | 0.005 | 1 | 9 | Tox2 | 0.017 | Young |
| 0.083115 | 0.713993 | 0.037 | 0.013 | 1 | 9 | Wdhd1 | 0.024 | Young |
| 0.083115 | 0.667896 | 0.037 | 0.013 | 1 | 9 | Alms1 | 0.024 | Young |
| 0.083115 | 0.512861 | 0.037 | 0.013 | 1 | 9 | Disc1 | 0.024 | Young |
| 0.083115 | 0.008318 | 0.037 | 0.013 | 1 | 9 | Cenpl | 0.024 | Young |
| 0.083347 | 0.712778 | 0.169 | 0.113 | 1 | 9 | Dyrk1a | 0.056 | Young |
| 0.083528 | 0.893929 | 0.118 | 0.07 | 1 | 9 | Ncln | 0.048 | Young |
| 0.08371 | 0.00065 | 0.022 | 0.005 | 1 | 9 | Top1mt | 0.017 | Young |
| 0.083714 | 1.367027 | 0.088 | 0.049 | 1 | 9 | Lsp1 | 0.039 | Young |
| 0.083998 | 0.497047 | 0.162 | 0.106 | 1 | 9 | Hmgcl | 0.056 | Young |
| 0.084107 | 0.99914 | 0.037 | 0.013 | 1 | 9 | Cers6 | 0.024 | Young |
| 0.084606 | 0.285608 | 0.037 | 0.013 | 1 | 9 | Ift74 | 0.024 | Young |
| 0.085163 | 1.813657 | 0.088 | 0.049 | 1 | 9 | Rab29 | 0.039 | Young |
| 0.085165 | 0.054903 | 0.147 | 0.229 | 1 | 9 | Gpr4 | -0.082 | Young |
| 0.08539 | 0.399772 | 0.463 | 0.407 | 1 | 9 | Dhrs3 | 0.056 | Young |
| 0.086796 | 0.627884 | 0.14 | 0.088 | 1 | 9 | Tnfrsf1b | 0.052 | Young |
| 0.086887 | 0.730737 | 0.176 | 0.121 | 1 | 9 | Atf1 | 0.055 | Young |
| 0.087334 | 0.483775 | 0.125 | 0.075 | 1 | 9 | Polr2l | 0.05 | Young |
| 0.08753 | 0.882418 | 0.162 | 0.108 | 1 | 9 | Il1b | 0.054 | Young |
| 0.087706 | 0.583833 | 0.154 | 0.101 | 1 | 9 | Trim41 | 0.053 | Young |
| 0.08796 | 0.640061 | 0.14 | 0.088 | 1 | 9 | Apopt1 | 0.052 | Young |
| 0.088038 | 0.585454 | 0.154 | 0.101 | 1 | 9 | Zfp395 | 0.053 | Young |
| 0.088399 | 0.864212 | 0.14 | 0.09 | 1 | 9 | Knop1 | 0.05 | Young |
| 0.08919 | 0.366954 | 0.11 | 0.064 | 1 | 9 | Slc10a6 | 0.046 | Young |
| 0.08934 | 2.493541 | 0.044 | 0.018 | 1 | 9 | Fgd2 | 0.026 | Young |
| 0.089632 | 1.095847 | 0.088 | 0.049 | 1 | 9 | Pprc1 | 0.039 | Young |
| 0.08966 | 0.361593 | 0.132 | 0.08 | 1 | 9 | Tfam | 0.052 | Young |
| 0.090265 | 2.138064 | 0.044 | 0.018 | 1 | 9 | Tiam1 | 0.026 | Young |
| 0.090265 | 1.563191 | 0.044 | 0.018 | 1 | 9 | Gm32688 | 0.026 | Young |
| 0.09044 | 0.58816 | 0.294 | 0.219 | 1 | 9 | Senp6 | 0.075 | Young |
| 0.090503 | 0.280023 | 0.154 | 0.098 | 1 | 9 | Tmem263 | 0.056 | Young |
| 0.090879 | 0.677778 | 0.147 | 0.095 | 1 | 9 | Ostc | 0.052 | Young |
| 0.091163 | 0.993933 | 0.088 | 0.049 | 1 | 9 | Cyth4 | 0.039 | Young |
| 0.091381 | 1.238723 | 0.14 | 0.093 | 1 | 9 | Ctsc | 0.047 | Young |
| 0.091666 | 1.680935 | 0.044 | 0.018 | 1 | 9 | Ccdc71 | 0.026 | Young |
| 0.091666 | 1.323938 | 0.044 | 0.018 | 1 | 9 | Col1a2 | 0.026 | Young |
| 0.091782 | 0.971866 | 0.088 | 0.049 | 1 | 9 | Med22 | 0.039 | Young |
| 0.091884 | 1.696264 | 0.103 | 0.062 | 1 | 9 | Prickle1 | 0.041 | Young |
| 0.091988 | 0.298991 | 0.257 | 0.183 | 1 | 9 | Ifrd1 | 0.074 | Young |
| 0.092021 | 0.67181 | 0.294 | 0.232 | 1 | 9 | Map4k5 | 0.062 | Young |
| 0.092137 | 1.637523 | 0.044 | 0.018 | 1 | 9 | Hpgd | 0.026 | Young |
| 0.092576 | 1.36279 | 0.081 | 0.044 | 1 | 9 | Aar2 | 0.037 | Young |
| 0.092609 | 1.134757 | 0.044 | 0.018 | 1 | 9 | Dmpk | 0.026 | Young |
| 0.092834 | 0.308583 | 0.684 | 0.642 | 1 | 9 | Dock9 | 0.042 | Young |
| 0.093084 | 1.204197 | 0.044 | 0.018 | 1 | 9 | Lipg | 0.026 | Young |

| | | | | | | | | |
|---|---|---|---|---|---|---|---|---|
| 0.09323 | 0.667644 | 0.147 | 0.095 | 1 | 9 | Gcc1 | 0.052 | Young |
| 0.093737 | 1.441276 | 0.051 | 0.023 | 1 | 9 | Tyms | 0.028 | Young |
| 0.09404 | 1.168561 | 0.044 | 0.018 | 1 | 9 | Ska2 | 0.026 | Young |
| 0.09404 | 1.091582 | 0.044 | 0.018 | 1 | 9 | Wdr73 | 0.026 | Young |
| 0.09404 | 1.083371 | 0.044 | 0.018 | 1 | 9 | Mpp7 | 0.026 | Young |
| 0.094965 | 0.615781 | 0.125 | 0.077 | 1 | 9 | Cxxc5 | 0.048 | Young |
| 0.095003 | 0.906825 | 0.044 | 0.018 | 1 | 9 | Fxyd2 | 0.026 | Young |
| 0.095022 | 1.925192 | 0.059 | 0.028 | 1 | 9 | Prr3 | 0.031 | Young |
| 0.095041 | 1.915006 | 0.051 | 0.023 | 1 | 9 | Dus3l | 0.028 | Young |
| 0.095324 | 0.362877 | 0.838 | 0.832 | 1 | 9 | Rps9 | 0.006 | Young |
| 0.095367 | 1.431524 | 0.14 | 0.093 | 1 | 9 | Mtr | 0.047 | Young |
| 0.095488 | 1.13099 | 0.044 | 0.018 | 1 | 9 | Rasgef1b | 0.026 | Young |
| 0.095488 | 0.514162 | 0.044 | 0.018 | 1 | 9 | 1110065P2 | 0.026 | Young |
| 0.095807 | 0.173568 | 0.971 | 0.948 | 1 | 9 | Txnip | 0.023 | Young |
| 0.095933 | 1.341101 | 0.066 | 0.034 | 1 | 9 | Cln5 | 0.032 | Young |
| 0.095974 | 0.622148 | 0.044 | 0.018 | 1 | 9 | Pot1b | 0.026 | Young |
| 0.096463 | 0.512816 | 0.044 | 0.018 | 1 | 9 | 2810001G2 | 0.026 | Young |
| 0.096574 | 0.725756 | 0.096 | 0.054 | 1 | 9 | Rbbp9 | 0.042 | Young |
| 0.096602 | 0.532752 | 0.147 | 0.095 | 1 | 9 | Arrb1 | 0.052 | Young |
| 0.096804 | 1.340627 | 0.051 | 0.023 | 1 | 9 | Usp43 | 0.028 | Young |
| 0.096953 | 0.771884 | 0.044 | 0.018 | 1 | 9 | Nup188 | 0.026 | Young |
| 0.096983 | 0.219113 | 0.11 | 0.064 | 1 | 9 | Msantd4 | 0.046 | Young |
| 0.097051 | 1.25629 | 0.059 | 0.028 | 1 | 9 | Pla2g7 | 0.031 | Young |
| 0.097072 | 2.149187 | 0.066 | 0.034 | 1 | 9 | Cyb561 | 0.032 | Young |
| 0.097116 | 0.565573 | 0.14 | 0.09 | 1 | 9 | Ctso | 0.05 | Young |
| 0.097174 | 0.72463 | 0.088 | 0.049 | 1 | 9 | Ric8b | 0.039 | Young |
| 0.097622 | 0.745683 | 0.125 | 0.077 | 1 | 9 | Ube3b | 0.048 | Young |
| 0.097695 | 1.53278 | 0.051 | 0.023 | 1 | 9 | Cd209a | 0.028 | Young |
| 0.09778 | 0.548105 | 0.213 | 0.152 | 1 | 9 | Ip6k1 | 0.061 | Young |
| 0.097874 | 0.043513 | 0.074 | 0.129 | 1 | 9 | Rlf | -0.055 | Young |
| 0.09806 | 0.814339 | 0.287 | 0.222 | 1 | 9 | Cdv3 | 0.065 | Young |
| 0.09843 | 0.405932 | 0.265 | 0.196 | 1 | 9 | Ccnd3 | 0.069 | Young |
| 0.098808 | 0.492539 | 0.088 | 0.049 | 1 | 9 | Hacd3 | 0.039 | Young |
| 0.098917 | 0.436591 | 0.382 | 0.317 | 1 | 9 | Ldha | 0.065 | Young |
| 0.099043 | 1.478092 | 0.051 | 0.023 | 1 | 9 | Setdb2 | 0.028 | Young |
| 0.099102 | 0.932839 | 0.074 | 0.039 | 1 | 9 | Hilpda | 0.035 | Young |
| 0.099313 | 0.956723 | 0.081 | 0.044 | 1 | 9 | Dock2 | 0.037 | Young |
| 0.099436 | 0.126418 | 0.044 | 0.018 | 1 | 9 | Agl | 0.026 | Young |
| 0.099468 | 0.817345 | 0.074 | 0.039 | 1 | 9 | Atg7 | 0.035 | Young |
| 0.099918 | 1.004843 | 0.199 | 0.149 | 1 | 9 | Unc93b1 | 0.05 | Young |
| 0.100163 | 1.485587 | 0.066 | 0.034 | 1 | 9 | Zfp101 | 0.032 | Young |
| 0.100202 | 0.825671 | 0.074 | 0.039 | 1 | 9 | Cd84 | 0.035 | Young |
| 0.100323 | 1.000236 | 0.147 | 0.095 | 1 | 9 | Socs7 | 0.052 | Young |
| 0.100357 | 0.573258 | 0.081 | 0.044 | 1 | 9 | Csf2ra | 0.037 | Young |
| 0.100379 | 1.030488 | 0.11 | 0.067 | 1 | 9 | Rbmxl1 | 0.043 | Young |

| | | | | | | | | |
|---|---|---|---|---|---|---|---|---|
| 0.10075 | 0.393489 | 0.691 | 0.678 | 1 | 9 | Rps28 | 0.013 | Young |
| 0.101077 | 0.071237 | 0.14 | 0.216 | 1 | 9 | Pcdh12 | -0.076 | Young |
| 0.10168 | 0.843233 | 0.118 | 0.075 | 1 | 9 | Cc2d1b | 0.043 | Young |
| 0.101737 | 0.944652 | 0.066 | 0.034 | 1 | 9 | Bin1 | 0.032 | Young |
| 0.101843 | 0.551273 | 0.147 | 0.095 | 1 | 9 | Dap | 0.052 | Young |
| 0.102168 | 0.516729 | 0.309 | 0.25 | 1 | 9 | Tulp4 | 0.059 | Young |
| 0.102196 | 1.022975 | 0.147 | 0.098 | 1 | 9 | Gm2a | 0.049 | Young |
| 0.102875 | 0.592583 | 0.257 | 0.196 | 1 | 9 | Mfhas1 | 0.061 | Young |
| 0.103478 | 0.179743 | 0.912 | 0.889 | 1 | 9 | Ubc | 0.023 | Young |
| 0.103541 | 0.620741 | 0.081 | 0.044 | 1 | 9 | Nip7 | 0.037 | Young |
| 0.103646 | 1.321643 | 0.051 | 0.023 | 1 | 9 | St3gal3 | 0.028 | Young |
| 0.103646 | 0.738157 | 0.051 | 0.023 | 1 | 9 | Ptdss2 | 0.028 | Young |
| 0.103697 | 0.712318 | 0.265 | 0.204 | 1 | 9 | Zfand3 | 0.061 | Young |
| 0.103778 | 1.066368 | 0.059 | 0.028 | 1 | 9 | Malsu1 | 0.031 | Young |
| 0.103849 | 0.611536 | 0.103 | 0.059 | 1 | 9 | Gm10073 | 0.044 | Young |
| 0.104094 | 0.658795 | 0.14 | 0.09 | 1 | 9 | Nsmce1 | 0.05 | Young |
| 0.104103 | 1.023054 | 0.169 | 0.116 | 1 | 9 | Coro1b | 0.053 | Young |
| 0.104645 | 0.936953 | 0.059 | 0.028 | 1 | 9 | Mtif3 | 0.031 | Young |
| 0.10563 | 0.533378 | 0.169 | 0.113 | 1 | 9 | Ppfia1 | 0.056 | Young |
| 0.106489 | 0.184774 | 0.051 | 0.023 | 1 | 9 | Ormdl1 | 0.028 | Young |
| 0.106496 | 3.570447 | 0.015 | 0.003 | 1 | 9 | Isoc2b | 0.012 | Young |
| 0.106496 | 2.768018 | 0.015 | 0.003 | 1 | 9 | Tmem69 | 0.012 | Young |
| 0.106496 | 2.595806 | 0.015 | 0.003 | 1 | 9 | Ankef1 | 0.012 | Young |
| 0.106496 | 2.518583 | 0.015 | 0.003 | 1 | 9 | Lrrc71 | 0.012 | Young |
| 0.106496 | 2.454976 | 0.015 | 0.003 | 1 | 9 | Tlr1 | 0.012 | Young |
| 0.106496 | 2.408138 | 0.015 | 0.003 | 1 | 9 | Otub2 | 0.012 | Young |
| 0.106496 | 2.371298 | 0.015 | 0.003 | 1 | 9 | AC174678. | 0.012 | Young |
| 0.106496 | 2.150904 | 0.015 | 0.003 | 1 | 9 | Shcbp1 | 0.012 | Young |
| 0.106496 | 2.000395 | 0.015 | 0.003 | 1 | 9 | Il18rap | 0.012 | Young |
| 0.106496 | 1.955868 | 0.015 | 0.003 | 1 | 9 | Kcne4 | 0.012 | Young |
| 0.106496 | 1.752076 | 0.015 | 0.003 | 1 | 9 | C130013H( | 0.012 | Young |
| 0.106496 | 1.523427 | 0.015 | 0.003 | 1 | 9 | Ttc39c | 0.012 | Young |
| 0.106618 | 0.495615 | 0.074 | 0.039 | 1 | 9 | Zfp367 | 0.035 | Young |
| 0.107005 | 0.496358 | 0.074 | 0.039 | 1 | 9 | Eif5a2 | 0.035 | Young |
| 0.107306 | 0.321266 | 0.088 | 0.049 | 1 | 9 | Tmem209 | 0.039 | Young |
| 0.107523 | 0.697505 | 0.103 | 0.062 | 1 | 9 | Cd52 | 0.041 | Young |
| 0.107593 | 3.095068 | 0.015 | 0.003 | 1 | 9 | Ifi211 | 0.012 | Young |
| 0.107593 | 2.546433 | 0.015 | 0.003 | 1 | 9 | Ebi3 | 0.012 | Young |
| 0.107593 | 2.349745 | 0.015 | 0.003 | 1 | 9 | 6030443J0 | 0.012 | Young |
| 0.107593 | 2.322702 | 0.015 | 0.003 | 1 | 9 | Atp1a3 | 0.012 | Young |
| 0.107593 | 2.314064 | 0.015 | 0.003 | 1 | 9 | Cmbl | 0.012 | Young |
| 0.107593 | 2.220421 | 0.015 | 0.003 | 1 | 9 | G730003C: | 0.012 | Young |
| 0.107593 | 2.2048 | 0.015 | 0.003 | 1 | 9 | Kcng2 | 0.012 | Young |
| 0.107593 | 1.932769 | 0.015 | 0.003 | 1 | 9 | Gm12689 | 0.012 | Young |
| 0.107593 | 1.932113 | 0.015 | 0.003 | 1 | 9 | AV099323 | 0.012 | Young |

| | | | | | | | | |
|---|---|---|---|---|---|---|---|---|
| 0.107593 | 1.811286 | 0.015 | 0.003 | 1 | 9 | Gm3448 | 0.012 | Young |
| 0.107593 | 1.760035 | 0.015 | 0.003 | 1 | 9 | Mettl15 | 0.012 | Young |
| 0.107593 | 1.757247 | 0.015 | 0.003 | 1 | 9 | Gm8281 | 0.012 | Young |
| 0.107593 | 1.63225 | 0.015 | 0.003 | 1 | 9 | Gm26664 | 0.012 | Young |
| 0.107593 | 1.614194 | 0.015 | 0.003 | 1 | 9 | Dpp4 | 0.012 | Young |
| 0.107593 | 1.592256 | 0.015 | 0.003 | 1 | 9 | Gm26910 | 0.012 | Young |
| 0.107593 | 1.573471 | 0.015 | 0.003 | 1 | 9 | Sars2 | 0.012 | Young |
| 0.107593 | 1.52903 | 0.015 | 0.003 | 1 | 9 | Cnnm2 | 0.012 | Young |
| 0.107593 | 1.361964 | 0.015 | 0.003 | 1 | 9 | Rfx2 | 0.012 | Young |
| 0.107593 | 1.353608 | 0.015 | 0.003 | 1 | 9 | Pvt1 | 0.012 | Young |
| 0.107593 | 1.279792 | 0.015 | 0.003 | 1 | 9 | Klrb1b | 0.012 | Young |
| 0.107593 | 1.154733 | 0.015 | 0.003 | 1 | 9 | Kcne3 | 0.012 | Young |
| 0.107593 | 1.054072 | 0.015 | 0.003 | 1 | 9 | Tlr13 | 0.012 | Young |
| 0.107593 | 1.050916 | 0.015 | 0.003 | 1 | 9 | Gm35154 | 0.012 | Young |
| 0.107663 | 0.78817 | 0.184 | 0.131 | 1 | 9 | Stxbp6 | 0.053 | Young |
| 0.107718 | 0.832164 | 0.125 | 0.08 | 1 | 9 | Fbxo18 | 0.045 | Young |
| 0.107722 | 0.646015 | 0.059 | 0.028 | 1 | 9 | St3gal5 | 0.031 | Young |
| 0.107722 | 0.489798 | 0.059 | 0.028 | 1 | 9 | Tmem192 | 0.031 | Young |
| 0.107823 | 0.633165 | 0.176 | 0.121 | 1 | 9 | Lrch1 | 0.055 | Young |
| 0.10801 | 0.390007 | 0.088 | 0.049 | 1 | 9 | Slc8b1 | 0.039 | Young |
| 0.108257 | 0.236826 | 0.824 | 0.796 | 1 | 9 | Rpl38 | 0.028 | Young |
| 0.108414 | 0.504703 | 0.14 | 0.09 | 1 | 9 | Elf4 | 0.05 | Young |
| 0.108699 | 1.297828 | 0.015 | 0.003 | 1 | 9 | Slc2a13 | 0.012 | Young |
| 0.108699 | 1.20508 | 0.015 | 0.003 | 1 | 9 | Gm19325 | 0.012 | Young |
| 0.108699 | 1.133185 | 0.015 | 0.003 | 1 | 9 | Atp10a | 0.012 | Young |
| 0.108699 | 1.002339 | 0.015 | 0.003 | 1 | 9 | Nnat | 0.012 | Young |
| 0.108699 | 1.001747 | 0.015 | 0.003 | 1 | 9 | Mrm3 | 0.012 | Young |
| 0.108699 | 0.940674 | 0.015 | 0.003 | 1 | 9 | Fam109b | 0.012 | Young |
| 0.108699 | 0.883493 | 0.015 | 0.003 | 1 | 9 | Pdgfc | 0.012 | Young |
| 0.108699 | 0.844801 | 0.015 | 0.003 | 1 | 9 | Traf4 | 0.012 | Young |
| 0.108699 | 0.828312 | 0.015 | 0.003 | 1 | 9 | Asb4 | 0.012 | Young |
| 0.108699 | 0.811475 | 0.015 | 0.003 | 1 | 9 | Ccsap | 0.012 | Young |
| 0.108699 | 0.792872 | 0.015 | 0.003 | 1 | 9 | Gm38843 | 0.012 | Young |
| 0.108699 | 0.785038 | 0.015 | 0.003 | 1 | 9 | Gna15 | 0.012 | Young |
| 0.108699 | 0.758627 | 0.015 | 0.003 | 1 | 9 | Gm13373 | 0.012 | Young |
| 0.108699 | 0.747682 | 0.015 | 0.003 | 1 | 9 | Smpd5 | 0.012 | Young |
| 0.108699 | 0.712444 | 0.015 | 0.003 | 1 | 9 | Mag | 0.012 | Young |
| 0.108699 | 0.618443 | 0.015 | 0.003 | 1 | 9 | Tdrkh | 0.012 | Young |
| 0.108699 | 0.580313 | 0.015 | 0.003 | 1 | 9 | Fkbpl | 0.012 | Young |
| 0.108699 | 0.422928 | 0.015 | 0.003 | 1 | 9 | Dlg3 | 0.012 | Young |
| 0.108699 | 0.230473 | 0.015 | 0.003 | 1 | 9 | Mir155hg | 0.012 | Young |
| 0.108699 | 0.143942 | 0.015 | 0.003 | 1 | 9 | Lrrd1 | 0.012 | Young |
| 0.108699 | 0.130762 | 0.015 | 0.003 | 1 | 9 | Gm26778 | 0.012 | Young |
| 0.108699 | 0.03597 | 0.015 | 0.003 | 1 | 9 | Smyd5 | 0.012 | Young |
| 0.109228 | 1.123285 | 0.147 | 0.098 | 1 | 9 | Sin3a | 0.049 | Young |

| | | | | | | | | |
|---|---|---|---|---|---|---|---|---|
| 0.109813 | 0.600235 | 0.132 | 0.085 | 1 | 9 | Ccng2 | 0.047 | Young |
| 0.110866 | 0.216197 | 0.287 | 0.216 | 1 | 9 | Pnn | 0.071 | Young |
| 0.111024 | 0.609806 | 0.221 | 0.16 | 1 | 9 | Klf11 | 0.061 | Young |
| 0.111863 | 1.228135 | 0.154 | 0.108 | 1 | 9 | Twf2 | 0.046 | Young |
| 0.11199 | 0.905466 | 0.11 | 0.067 | 1 | 9 | Ctsh | 0.043 | Young |
| 0.112458 | 0.980977 | 0.066 | 0.034 | 1 | 9 | Rilpl2 | 0.032 | Young |
| 0.112577 | 0.365076 | 0.228 | 0.165 | 1 | 9 | Wdr83os | 0.063 | Young |
| 0.112703 | 0.373096 | 0.059 | 0.028 | 1 | 9 | Trim39 | 0.031 | Young |
| 0.112738 | 0.413915 | 0.103 | 0.062 | 1 | 9 | Mrps23 | 0.041 | Young |
| 0.113442 | 0.280817 | 0.14 | 0.09 | 1 | 9 | Banp | 0.05 | Young |
| 0.113584 | 0.374712 | 0.728 | 0.719 | 1 | 9 | Hspg2 | 0.009 | Young |
| 0.113651 | 2.336932 | 0.029 | 0.01 | 1 | 9 | Dok3 | 0.019 | Young |
| 0.113762 | 0.495223 | 0.074 | 0.039 | 1 | 9 | Sgcb | 0.035 | Young |
| 0.11409 | 0.240117 | 0.368 | 0.294 | 1 | 9 | Ppig | 0.074 | Young |
| 0.114166 | 0.323934 | 0.118 | 0.072 | 1 | 9 | Nt5c | 0.046 | Young |
| 0.114361 | 0.169442 | 0.213 | 0.152 | 1 | 9 | Araf | 0.061 | Young |
| 0.11439 | 0.412488 | 0.618 | 0.588 | 1 | 9 | Ftl1 | 0.03 | Young |
| 0.115789 | 2.652675 | 0.029 | 0.01 | 1 | 9 | Ankzf1 | 0.019 | Young |
| 0.116018 | 1.248073 | 0.096 | 0.057 | 1 | 9 | Ppm1k | 0.039 | Young |
| 0.116508 | 0.382335 | 0.132 | 0.085 | 1 | 9 | Pgp | 0.047 | Young |
| 0.116548 | 0.375627 | 0.265 | 0.196 | 1 | 9 | Bach1 | 0.069 | Young |
| 0.117232 | 2.281633 | 0.029 | 0.01 | 1 | 9 | Cpq | 0.019 | Young |
| 0.117232 | 2.257423 | 0.029 | 0.01 | 1 | 9 | Il1r1 | 0.019 | Young |
| 0.117232 | 1.833817 | 0.029 | 0.01 | 1 | 9 | Trem2 | 0.019 | Young |
| 0.117232 | 1.732822 | 0.029 | 0.01 | 1 | 9 | F630028O1 | 0.019 | Young |
| 0.117459 | 1.560768 | 0.11 | 0.07 | 1 | 9 | Tmem41b | 0.04 | Young |
| 0.11765 | 0.471283 | 0.294 | 0.222 | 1 | 9 | Amfr | 0.072 | Young |
| 0.117959 | 2.216024 | 0.029 | 0.01 | 1 | 9 | Tpmt | 0.019 | Young |
| 0.117959 | 2.193497 | 0.029 | 0.01 | 1 | 9 | Tbc1d8 | 0.019 | Young |
| 0.118005 | 0.298156 | 0.132 | 0.085 | 1 | 9 | Fbxo33 | 0.047 | Young |
| 0.118006 | 0.905431 | 0.14 | 0.093 | 1 | 9 | Rars | 0.047 | Young |
| 0.118443 | 0.692487 | 0.14 | 0.093 | 1 | 9 | Lsm12 | 0.047 | Young |
| 0.118689 | 2.484859 | 0.029 | 0.01 | 1 | 9 | Zfp385a | 0.019 | Young |
| 0.118689 | 1.28084 | 0.029 | 0.01 | 1 | 9 | Hk2 | 0.019 | Young |
| 0.118689 | 1.248416 | 0.029 | 0.01 | 1 | 9 | Gm16973 | 0.019 | Young |
| 0.118689 | 1.129338 | 0.029 | 0.01 | 1 | 9 | Rad52 | 0.019 | Young |
| 0.118841 | 0.505563 | 0.103 | 0.062 | 1 | 9 | Gm43848 | 0.041 | Young |
| 0.119213 | 0.354945 | 0.132 | 0.085 | 1 | 9 | Prrc1 | 0.047 | Young |
| 0.119341 | 1.897227 | 0.074 | 0.041 | 1 | 9 | Rbmx2 | 0.033 | Young |
| 0.119423 | 1.472059 | 0.029 | 0.01 | 1 | 9 | Lyve1 | 0.019 | Young |
| 0.119423 | 1.409845 | 0.029 | 0.01 | 1 | 9 | Surf2 | 0.019 | Young |
| 0.119786 | 0.73993 | 0.162 | 0.113 | 1 | 9 | Mrrf | 0.049 | Young |
| 0.119835 | 0.041558 | 0.14 | 0.09 | 1 | 9 | Erf | 0.05 | Young |
| 0.120007 | 0.181185 | 0.64 | 0.585 | 1 | 9 | Sdc3 | 0.055 | Young |
| 0.12016 | 1.215163 | 0.029 | 0.01 | 1 | 9 | Cda | 0.019 | Young |

| | | | | | | | | |
|---|---|---|---|---|---|---|---|---|
| 0.12016 | 0.987105 | 0.029 | 0.01 | 1 | 9 | Zfp113 | 0.019 | Young |
| 0.12016 | 0.935909 | 0.029 | 0.01 | 1 | 9 | Kif18a | 0.019 | Young |
| 0.12016 | 0.718901 | 0.029 | 0.01 | 1 | 9 | Slc35d2 | 0.019 | Young |
| 0.12016 | 0.703323 | 0.029 | 0.01 | 1 | 9 | Ddx51 | 0.019 | Young |
| 0.120632 | 0.302283 | 0.184 | 0.126 | 1 | 9 | Atp6v1h | 0.058 | Young |
| 0.120677 | 0.820508 | 0.096 | 0.057 | 1 | 9 | Dclre1b | 0.039 | Young |
| 0.120901 | 0.974718 | 0.029 | 0.01 | 1 | 9 | Pilrb1 | 0.019 | Young |
| 0.120901 | 0.397065 | 0.029 | 0.01 | 1 | 9 | Pfkfb2 | 0.019 | Young |
| 0.120901 | 0.097847 | 0.029 | 0.01 | 1 | 9 | Kif19a | 0.019 | Young |
| 0.12102 | 1.040393 | 0.081 | 0.046 | 1 | 9 | Tbc1d2b | 0.035 | Young |
| 0.121645 | 1.055759 | 0.029 | 0.01 | 1 | 9 | Cdh24 | 0.019 | Young |
| 0.121645 | 0.990444 | 0.029 | 0.01 | 1 | 9 | Ccl27a | 0.019 | Young |
| 0.121742 | 0.351322 | 0.206 | 0.149 | 1 | 9 | Arid1b | 0.057 | Young |
| 0.121813 | 0.138575 | 0.176 | 0.121 | 1 | 9 | Ifi27l2a | 0.055 | Young |
| 0.122393 | 0.460833 | 0.029 | 0.01 | 1 | 9 | Borcs8 | 0.019 | Young |
| 0.122713 | 0.124201 | 0.015 | 0.044 | 1 | 9 | Eri3 | -0.029 | Young |
| 0.123137 | 0.130132 | 0.368 | 0.281 | 1 | 9 | Smchd1 | 0.087 | Young |
| 0.123706 | 0.178846 | 0.221 | 0.157 | 1 | 9 | Mgea5 | 0.064 | Young |
| 0.123987 | 0.757615 | 0.096 | 0.057 | 1 | 9 | Csf1r | 0.039 | Young |
| 0.124574 | 1.004933 | 0.088 | 0.052 | 1 | 9 | Cluh | 0.036 | Young |
| 0.124618 | 0.45067 | 0.14 | 0.209 | 1 | 9 | 2810403A( | -0.069 | Young |
| 0.124963 | 0.711143 | 0.088 | 0.052 | 1 | 9 | Mpp6 | 0.036 | Young |
| 0.125491 | 0.36218 | 0.199 | 0.142 | 1 | 9 | Dynll2 | 0.057 | Young |
| 0.126015 | 0.750993 | 0.176 | 0.126 | 1 | 9 | Lamtor1 | 0.05 | Young |
| 0.126285 | 0.872474 | 0.081 | 0.046 | 1 | 9 | Gng2 | 0.035 | Young |
| 0.126994 | 1.588041 | 0.074 | 0.041 | 1 | 9 | Lcp2 | 0.033 | Young |
| 0.127661 | 0.806352 | 0.235 | 0.178 | 1 | 9 | Arl8b | 0.057 | Young |
| 0.127869 | 0.690955 | 0.162 | 0.113 | 1 | 9 | Pf4 | 0.049 | Young |
| 0.128108 | 0.819027 | 0.169 | 0.121 | 1 | 9 | Emd | 0.048 | Young |
| 0.128358 | 0.810757 | 0.081 | 0.046 | 1 | 9 | Rfx1 | 0.035 | Young |
| 0.128407 | 0.429257 | 0.125 | 0.08 | 1 | 9 | Fam181b | 0.045 | Young |
| 0.128496 | 0.375243 | 0.162 | 0.111 | 1 | 9 | Fnbp4 | 0.051 | Young |
| 0.128798 | 0.798442 | 0.147 | 0.101 | 1 | 9 | Stau1 | 0.046 | Young |
| 0.128903 | 0.232325 | 0.176 | 0.121 | 1 | 9 | Arntl | 0.055 | Young |
| 0.129331 | 0.691684 | 0.213 | 0.155 | 1 | 9 | Cerk | 0.058 | Young |
| 0.129614 | 1.075745 | 0.081 | 0.046 | 1 | 9 | Heph | 0.035 | Young |
| 0.130072 | 1.18054 | 0.074 | 0.041 | 1 | 9 | Gnpda1 | 0.033 | Young |
| 0.130456 | 0.473405 | 0.081 | 0.046 | 1 | 9 | Zw10 | 0.035 | Young |
| 0.131276 | 0.1408 | 0.037 | 0.075 | 1 | 9 | Uhrf1bp1l | -0.038 | Young |
| 0.131408 | 1.474607 | 0.074 | 0.041 | 1 | 9 | AI987944 | 0.033 | Young |
| 0.131475 | 1.063099 | 0.154 | 0.108 | 1 | 9 | Abl1 | 0.046 | Young |
| 0.131478 | 0.278387 | 0.662 | 0.611 | 1 | 9 | Rsrp1 | 0.051 | Young |
| 0.131613 | 0.756544 | 0.103 | 0.064 | 1 | 9 | Txndc11 | 0.039 | Young |
| 0.132486 | 2.184988 | 0.044 | 0.021 | 1 | 9 | Zfp593 | 0.023 | Young |
| 0.132545 | 0.407692 | 0.088 | 0.052 | 1 | 9 | Abhd12 | 0.036 | Young |

| | | | | | | | | |
|---|---|---|---|---|---|---|---|---|
| 0.132948 | 0.445782 | 0.14 | 0.093 | 1 | 9 | Tnfaip3 | 0.047 | Young |
| 0.133009 | 0.681549 | 0.081 | 0.046 | 1 | 9 | Exd2 | 0.035 | Young |
| 0.133242 | 1.406485 | 0.037 | 0.015 | 1 | 9 | Tle2 | 0.022 | Young |
| 0.133928 | 1.733189 | 0.037 | 0.015 | 1 | 9 | Uevld | 0.022 | Young |
| 0.134118 | 0.21441 | 0.64 | 0.603 | 1 | 9 | Mat2a | 0.037 | Young |
| 0.134461 | 0.01259 | 0.125 | 0.08 | 1 | 9 | Slc30a7 | 0.045 | Young |
| 0.134617 | 1.684141 | 0.037 | 0.015 | 1 | 9 | Fam105a | 0.022 | Young |
| 0.134617 | 1.508566 | 0.037 | 0.015 | 1 | 9 | Tnfrsf13b | 0.022 | Young |
| 0.134617 | 1.45905 | 0.037 | 0.015 | 1 | 9 | Rttn | 0.022 | Young |
| 0.135302 | 2.384618 | 0.051 | 0.026 | 1 | 9 | Ttll12 | 0.025 | Young |
| 0.135309 | 1.167345 | 0.037 | 0.015 | 1 | 9 | Ndufa4l2 | 0.022 | Young |
| 0.135309 | 0.726332 | 0.037 | 0.015 | 1 | 9 | Lilrb4a | 0.022 | Young |
| 0.135417 | 0.953527 | 0.118 | 0.077 | 1 | 9 | Zmiz2 | 0.041 | Young |
| 0.135521 | 0.747818 | 0.14 | 0.093 | 1 | 9 | Bambi | 0.047 | Young |
| 0.135719 | 0.494128 | 0.331 | 0.273 | 1 | 9 | N4bp2l2 | 0.058 | Young |
| 0.136004 | 1.269836 | 0.037 | 0.015 | 1 | 9 | Gfra2 | 0.022 | Young |
| 0.136004 | 1.224125 | 0.037 | 0.015 | 1 | 9 | Zfp182 | 0.022 | Young |
| 0.136701 | 1.273782 | 0.037 | 0.015 | 1 | 9 | Cyp2e1 | 0.022 | Young |
| 0.136747 | 0.464295 | 0.493 | 0.425 | 1 | 9 | Tcim | 0.068 | Young |
| 0.137287 | 0.076612 | 0.037 | 0.075 | 1 | 9 | Med19 | -0.038 | Young |
| 0.137401 | 1.446356 | 0.037 | 0.015 | 1 | 9 | Gata3 | 0.022 | Young |
| 0.13756 | 2.094956 | 0.051 | 0.026 | 1 | 9 | Slc11a1 | 0.025 | Young |
| 0.137723 | 1.408509 | 0.066 | 0.036 | 1 | 9 | Metrnl | 0.03 | Young |
| 0.138251 | 1.074028 | 0.074 | 0.041 | 1 | 9 | Bbc3 | 0.033 | Young |
| 0.138251 | 0.760666 | 0.074 | 0.041 | 1 | 9 | 0610009O2 | 0.033 | Young |
| 0.138674 | 0.782805 | 0.081 | 0.046 | 1 | 9 | Capg | 0.035 | Young |
| 0.138804 | 0.759937 | 0.147 | 0.103 | 1 | 9 | Ube2g1 | 0.044 | Young |
| 0.13881 | 1.196817 | 0.037 | 0.015 | 1 | 9 | Adipoq | 0.022 | Young |
| 0.13881 | 0.972173 | 0.037 | 0.015 | 1 | 9 | Gnai1 | 0.022 | Young |
| 0.139204 | 0.946959 | 0.066 | 0.036 | 1 | 9 | Nol12 | 0.03 | Young |
| 0.139255 | 0.328451 | 0.176 | 0.124 | 1 | 9 | C2cd2 | 0.052 | Young |
| 0.139287 | 2.048824 | 0.044 | 0.021 | 1 | 9 | Rbm38 | 0.023 | Young |
| 0.139289 | 1.120084 | 0.096 | 0.059 | 1 | 9 | Ttc33 | 0.037 | Young |
| 0.139381 | 1.264915 | 0.059 | 0.031 | 1 | 9 | Gm26601 | 0.028 | Young |
| 0.139518 | 1.272061 | 0.037 | 0.015 | 1 | 9 | Hdc | 0.022 | Young |
| 0.139518 | 0.773073 | 0.037 | 0.015 | 1 | 9 | Prmt6 | 0.022 | Young |
| 0.14023 | 0.639306 | 0.037 | 0.015 | 1 | 9 | Dffb | 0.022 | Young |
| 0.140552 | 1.371122 | 0.044 | 0.021 | 1 | 9 | Ccl12 | 0.023 | Young |
| 0.140724 | 0.685902 | 0.213 | 0.16 | 1 | 9 | Psmb7 | 0.053 | Young |
| 0.141143 | 1.029888 | 0.14 | 0.095 | 1 | 9 | Slc35a4 | 0.045 | Young |
| 0.141187 | 1.146565 | 0.044 | 0.021 | 1 | 9 | Atg16l2 | 0.023 | Young |
| 0.141537 | 0.964838 | 0.074 | 0.041 | 1 | 9 | Gtf3a | 0.033 | Young |
| 0.141537 | 0.649801 | 0.074 | 0.041 | 1 | 9 | Mcub | 0.033 | Young |
| 0.141581 | 0.90217 | 0.051 | 0.026 | 1 | 9 | Drg2 | 0.025 | Young |
| 0.14166 | 0.81753 | 0.037 | 0.015 | 1 | 9 | Ulk3 | 0.022 | Young |

| | | | | | | | | |
|---|---|---|---|---|---|---|---|---|
| 0.14166 | 0.315189 | 0.037 | 0.015 | 1 | 9 | Mtg2 | 0.022 | Young |
| 0.142163 | 1.315378 | 0.051 | 0.026 | 1 | 9 | Nufip1 | 0.025 | Young |
| 0.142233 | 0.838328 | 0.25 | 0.201 | 1 | 9 | Lyz2 | 0.049 | Young |
| 0.142593 | 1.038314 | 0.059 | 0.031 | 1 | 9 | Mrpl58 | 0.028 | Young |
| 0.142635 | 0.224129 | 0.801 | 0.814 | 1 | 9 | mt-Nd3 | -0.013 | Young |
| 0.142747 | 1.352804 | 0.051 | 0.026 | 1 | 9 | Cog2 | 0.025 | Young |
| 0.143102 | 0.579517 | 0.037 | 0.015 | 1 | 9 | Slc25a53 | 0.022 | Young |
| 0.143156 | 0.384883 | 0.081 | 0.046 | 1 | 9 | Got1 | 0.035 | Young |
| 0.143319 | 0.810073 | 0.176 | 0.129 | 1 | 9 | Zswim8 | 0.047 | Young |
| 0.143332 | 1.750452 | 0.051 | 0.026 | 1 | 9 | Ttc23 | 0.025 | Young |
| 0.143332 | 1.603225 | 0.051 | 0.026 | 1 | 9 | Tbl3 | 0.025 | Young |
| 0.143441 | 0.65227 | 0.074 | 0.041 | 1 | 9 | E2f7 | 0.033 | Young |
| 0.143444 | 0.051554 | 0.132 | 0.193 | 1 | 9 | Spata13 | -0.061 | Young |
| 0.143611 | 0.117725 | 0.081 | 0.046 | 1 | 9 | Phkg2 | 0.035 | Young |
| 0.143721 | 1.019969 | 0.066 | 0.036 | 1 | 9 | Pex16 | 0.03 | Young |
| 0.143752 | 0.868627 | 0.044 | 0.021 | 1 | 9 | Cds1 | 0.023 | Young |
| 0.14421 | 0.305921 | 0.162 | 0.111 | 1 | 9 | Pqlc1 | 0.051 | Young |
| 0.14428 | 0.541739 | 0.176 | 0.126 | 1 | 9 | Idnk | 0.05 | Young |
| 0.144399 | 1.160728 | 0.044 | 0.021 | 1 | 9 | Ddr2 | 0.023 | Young |
| 0.144508 | 1.158065 | 0.051 | 0.026 | 1 | 9 | Rbak | 0.025 | Young |
| 0.144925 | 0.535908 | 0.147 | 0.101 | 1 | 9 | Hmox1 | 0.046 | Young |
| 0.144995 | 0.567241 | 0.309 | 0.253 | 1 | 9 | Ppp1r9b | 0.056 | Young |
| 0.145219 | 0.271443 | 0.118 | 0.075 | 1 | 9 | Ncoa4 | 0.043 | Young |
| 0.145239 | 0.06196 | 0.11 | 0.168 | 1 | 9 | Ikzf5 | -0.058 | Young |
| 0.145324 | 0.864476 | 0.25 | 0.191 | 1 | 9 | Jarid2 | 0.059 | Young |
| 0.145369 | 0.308602 | 0.434 | 0.371 | 1 | 9 | Iigp1 | 0.063 | Young |
| 0.14556 | 1.05723 | 0.11 | 0.072 | 1 | 9 | Srebf2 | 0.038 | Young |
| 0.145641 | 0.759908 | 0.11 | 0.07 | 1 | 9 | Pip4k2b | 0.04 | Young |
| 0.145692 | 1.323908 | 0.051 | 0.026 | 1 | 9 | Jpt2 | 0.025 | Young |
| 0.146185 | 0.777366 | 0.14 | 0.095 | 1 | 9 | Cops6 | 0.045 | Young |
| 0.146287 | 1.112038 | 0.051 | 0.026 | 1 | 9 | Rnf34 | 0.025 | Young |
| 0.146334 | 0.830752 | 0.074 | 0.041 | 1 | 9 | Men1 | 0.033 | Young |
| 0.146353 | 0.753163 | 0.044 | 0.021 | 1 | 9 | Sfxn2 | 0.023 | Young |
| 0.146401 | 0.469591 | 0.257 | 0.193 | 1 | 9 | Rexo2 | 0.064 | Young |
| 0.146411 | 0.881827 | 0.059 | 0.031 | 1 | 9 | Cryl1 | 0.028 | Young |
| 0.146808 | 0.833982 | 0.184 | 0.137 | 1 | 9 | Tomm22 | 0.047 | Young |
| 0.146849 | 0.138576 | 0.015 | 0.041 | 1 | 9 | Rrp8 | -0.026 | Young |
| 0.14691 | 0.671577 | 0.221 | 0.168 | 1 | 9 | Ier5l | 0.053 | Young |
| 0.147009 | 0.561437 | 0.044 | 0.021 | 1 | 9 | Syt15 | 0.023 | Young |
| 0.147517 | 0.949468 | 0.059 | 0.031 | 1 | 9 | Slamf9 | 0.028 | Young |
| 0.147798 | 0.321834 | 0.074 | 0.041 | 1 | 9 | Scyl1 | 0.033 | Young |
| 0.14783 | 0.771146 | 0.066 | 0.036 | 1 | 9 | Ptpn4 | 0.03 | Young |
| 0.14783 | 0.492346 | 0.066 | 0.036 | 1 | 9 | Snhg6 | 0.03 | Young |
| 0.148082 | 0.946135 | 0.051 | 0.026 | 1 | 9 | Slc22a23 | 0.025 | Young |
| 0.148166 | 0.079811 | 0.169 | 0.232 | 1 | 9 | Plscr2 | -0.063 | Young |

| | | | | | | | | |
|---|---|---|---|---|---|---|---|---|
| 0.148508 | 0.506141 | 0.294 | 0.232 | 1 | 9 | Pik3ca | 0.062 | Young |
| 0.148989 | 0.658784 | 0.044 | 0.021 | 1 | 9 | Smyd3 | 0.023 | Young |
| 0.148989 | 0.489741 | 0.044 | 0.021 | 1 | 9 | Rph3al | 0.023 | Young |
| 0.149394 | 0.601049 | 0.066 | 0.036 | 1 | 9 | Eipr1 | 0.03 | Young |
| 0.149626 | 1.133079 | 0.132 | 0.09 | 1 | 9 | Prdx5 | 0.042 | Young |
| 0.149679 | 0.602086 | 0.272 | 0.211 | 1 | 9 | Lman1 | 0.061 | Young |
| 0.149746 | 0.837074 | 0.059 | 0.031 | 1 | 9 | Haus8 | 0.028 | Young |
| 0.149746 | 0.768387 | 0.059 | 0.031 | 1 | 9 | Zfp90 | 0.028 | Young |
| 0.150207 | 0.919147 | 0.11 | 0.072 | 1 | 9 | Hspa4l | 0.038 | Young |
| 0.150308 | 0.647134 | 0.059 | 0.031 | 1 | 9 | Adnp2 | 0.028 | Young |
| 0.150321 | 0.156097 | 0.044 | 0.021 | 1 | 9 | Cd300c2 | 0.023 | Young |
| 0.150871 | 0.200038 | 0.059 | 0.031 | 1 | 9 | Ift52 | 0.028 | Young |
| 0.150879 | 0.55277 | 0.257 | 0.201 | 1 | 9 | Zcchc24 | 0.056 | Young |
| 0.150902 | 0.156023 | 0.559 | 0.51 | 1 | 9 | Kctd12 | 0.049 | Young |
| 0.151006 | 0.504127 | 0.176 | 0.126 | 1 | 9 | Dazap1 | 0.05 | Young |
| 0.151111 | 0.412751 | 0.051 | 0.026 | 1 | 9 | Xylt1 | 0.025 | Young |
| 0.151202 | 0.450065 | 0.449 | 0.387 | 1 | 9 | Thbd | 0.062 | Young |
| 0.151227 | 0.164116 | 0.875 | 0.817 | 1 | 9 | Junb | 0.058 | Young |
| 0.151538 | 0.03897 | 0.243 | 0.178 | 1 | 9 | Kctd20 | 0.065 | Young |
| 0.15155 | 0.791732 | 0.118 | 0.077 | 1 | 9 | Ddx23 | 0.041 | Young |
| 0.152197 | 0.868467 | 0.213 | 0.162 | 1 | 9 | Scoc | 0.051 | Young |
| 0.152673 | 0.724677 | 0.25 | 0.193 | 1 | 9 | Pnp | 0.057 | Young |
| 0.152951 | 0.519983 | 0.051 | 0.026 | 1 | 9 | Reep4 | 0.025 | Young |
| 0.153031 | 0.701625 | 0.132 | 0.09 | 1 | 9 | Pik3cb | 0.042 | Young |
| 0.153419 | 0.627187 | 0.279 | 0.224 | 1 | 9 | Phc3 | 0.055 | Young |
| 0.15382 | 0.576646 | 0.154 | 0.108 | 1 | 9 | Trabd | 0.046 | Young |
| 0.155126 | 0.717975 | 0.088 | 0.054 | 1 | 9 | Cmtm7 | 0.034 | Young |
| 0.155203 | 0.564016 | 0.272 | 0.216 | 1 | 9 | Fxr1 | 0.056 | Young |
| 0.155449 | 0.398518 | 0.419 | 0.353 | 1 | 9 | Jmjd1c | 0.066 | Young |
| 0.155466 | 0.216457 | 0.875 | 0.858 | 1 | 9 | Zfp36l1 | 0.017 | Young |
| 0.155588 | 0.931698 | 0.176 | 0.129 | 1 | 9 | Evl | 0.047 | Young |
| 0.155664 | 0.171074 | 0.61 | 0.549 | 1 | 9 | Csnk1a1 | 0.061 | Young |
| 0.155939 | 1.336864 | 0.081 | 0.049 | 1 | 9 | Sdc4 | 0.032 | Young |
| 0.156686 | 0.234586 | 0.176 | 0.126 | 1 | 9 | Tspan4 | 0.05 | Young |
| 0.157724 | 0.519663 | 0.154 | 0.108 | 1 | 9 | Idh1 | 0.046 | Young |
| 0.158344 | 1.153185 | 0.162 | 0.121 | 1 | 9 | Samd5 | 0.041 | Young |
| 0.158509 | 0.447831 | 0.044 | 0.08 | 1 | 9 | Ppid | -0.036 | Young |
| 0.159207 | 0.128618 | 0.051 | 0.026 | 1 | 9 | Tfpt | 0.025 | Young |
| 0.159546 | 0.442813 | 0.191 | 0.139 | 1 | 9 | Poldip3 | 0.052 | Young |
| 0.159576 | 0.412857 | 0.14 | 0.095 | 1 | 9 | Senp5 | 0.045 | Young |
| 0.159985 | 0.450729 | 0.162 | 0.111 | 1 | 9 | Tmem5 | 0.051 | Young |
| 0.160328 | 0.19606 | 0.338 | 0.273 | 1 | 9 | Ran | 0.065 | Young |
| 0.16039 | 0.518712 | 0.176 | 0.129 | 1 | 9 | Zfp740 | 0.047 | Young |
| 0.161075 | 0.965486 | 0.11 | 0.072 | 1 | 9 | Ralgapb | 0.038 | Young |
| 0.161347 | 0.009628 | 0.059 | 0.098 | 1 | 9 | Elp2 | -0.039 | Young |

| | | | | | | | | |
|---|---|---|---|---|---|---|---|---|
| 0.162029 | 0.416575 | 0.25 | 0.196 | 1 | 9 | Prcp | 0.054 | Young |
| 0.16241 | 0.565435 | 0.309 | 0.253 | 1 | 9 | Arhgef2 | 0.056 | Young |
| 0.162487 | 0.696494 | 0.243 | 0.196 | 1 | 9 | Mdm4 | 0.047 | Young |
| 0.162636 | 0.63123 | 0.147 | 0.106 | 1 | 9 | Tcp11l2 | 0.041 | Young |
| 0.162748 | 0.631309 | 0.287 | 0.229 | 1 | 9 | Cuedc1 | 0.058 | Young |
| 0.163065 | 0.514107 | 0.213 | 0.16 | 1 | 9 | Setd3 | 0.053 | Young |
| 0.163234 | 0.410061 | 0.603 | 0.585 | 1 | 9 | Tuba1b | 0.018 | Young |
| 0.163603 | 1.696788 | 0.103 | 0.067 | 1 | 9 | Crot | 0.036 | Young |
| 0.163706 | 0.702056 | 0.147 | 0.106 | 1 | 9 | Nipa2 | 0.041 | Young |
| 0.163768 | 0.357687 | 0.118 | 0.077 | 1 | 9 | Marcksl1 | 0.041 | Young |
| 0.163949 | 1.574886 | 0.007 | 0.028 | 1 | 9 | Ncaph | -0.021 | Young |
| 0.164332 | 0.185918 | 0.574 | 0.497 | 1 | 9 | Tsc22d1 | 0.077 | Young |
| 0.1647 | 1.044682 | 0.081 | 0.049 | 1 | 9 | Zmat5 | 0.032 | Young |
| 0.164884 | 0.441975 | 0.088 | 0.054 | 1 | 9 | Bcap29 | 0.034 | Young |
| 0.165444 | 1.141883 | 0.213 | 0.165 | 1 | 9 | Limd1 | 0.048 | Young |
| 0.165707 | 0.491875 | 0.382 | 0.343 | 1 | 9 | Cox8a | 0.039 | Young |
| 0.16713 | 0.32043 | 0.353 | 0.296 | 1 | 9 | Glul | 0.057 | Young |
| 0.16745 | 0.884929 | 0.074 | 0.044 | 1 | 9 | Tbkbp1 | 0.03 | Young |
| 0.16766 | 0.209654 | 0.213 | 0.152 | 1 | 9 | Psmd8 | 0.061 | Young |
| 0.167955 | 0.357093 | 0.162 | 0.116 | 1 | 9 | Lcp1 | 0.046 | Young |
| 0.168151 | 0.346959 | 0.228 | 0.17 | 1 | 9 | Per1 | 0.058 | Young |
| 0.168199 | 0.458354 | 0.169 | 0.121 | 1 | 9 | Uvrag | 0.048 | Young |
| 0.16839 | 0.613788 | 0.176 | 0.131 | 1 | 9 | Chuk | 0.045 | Young |
| 0.168403 | 0.206155 | 0.081 | 0.129 | 1 | 9 | Gkap1 | -0.048 | Young |
| 0.168412 | 0.006913 | 0.118 | 0.168 | 1 | 9 | Iffo2 | -0.05 | Young |
| 0.168607 | 0.330078 | 0.147 | 0.103 | 1 | 9 | Cul4a | 0.044 | Young |
| 0.168631 | 0.695007 | 0.213 | 0.165 | 1 | 9 | Nuak1 | 0.048 | Young |
| 0.169216 | 0.622757 | 0.081 | 0.049 | 1 | 9 | Atr | 0.032 | Young |
| 0.170109 | 1.216221 | 0.074 | 0.044 | 1 | 9 | Zfyve19 | 0.03 | Young |
| 0.170629 | 0.76899 | 0.103 | 0.067 | 1 | 9 | Igf1 | 0.036 | Young |
| 0.170941 | 0.215678 | 0.257 | 0.201 | 1 | 9 | Gpatch8 | 0.056 | Young |
| 0.171253 | 1.486957 | 0.081 | 0.049 | 1 | 9 | Hsdl1 | 0.032 | Young |
| 0.171297 | 1.265717 | 0.088 | 0.057 | 1 | 9 | Taf9b | 0.031 | Young |
| 0.171585 | 0.88867 | 0.096 | 0.062 | 1 | 9 | Lgals3 | 0.034 | Young |
| 0.171765 | 0.551978 | 0.081 | 0.049 | 1 | 9 | Blcap | 0.032 | Young |
| 0.172092 | 0.392126 | 0.11 | 0.072 | 1 | 9 | Spg21 | 0.038 | Young |
| 0.172474 | 2.719329 | 0.022 | 0.008 | 1 | 9 | Mcph1 | 0.014 | Young |
| 0.172474 | 2.20698 | 0.022 | 0.008 | 1 | 9 | Zfp59 | 0.014 | Young |
| 0.172871 | 0.412492 | 0.103 | 0.067 | 1 | 9 | Scfd1 | 0.036 | Young |
| 0.172967 | 0.337899 | 0.265 | 0.204 | 1 | 9 | Hipk3 | 0.061 | Young |
| 0.17297 | 1.389369 | 0.066 | 0.039 | 1 | 9 | Tbc1d4 | 0.027 | Young |
| 0.173197 | 0.1587 | 0.176 | 0.124 | 1 | 9 | Htt | 0.052 | Young |
| 0.173309 | 0.571098 | 0.081 | 0.049 | 1 | 9 | Rhot2 | 0.032 | Young |
| 0.173583 | 1.008165 | 0.14 | 0.098 | 1 | 9 | Pigp | 0.042 | Young |
| 0.1736 | 2.542089 | 0.022 | 0.008 | 1 | 9 | Mrps22 | 0.014 | Young |

| | | | | | | | | |
|---|---|---|---|---|---|---|---|---|
| 0.173711 | 0.488596 | 0.206 | 0.152 | 1 | 9 | Gsn | 0.054 | Young |
| 0.173711 | 0.042141 | 0.125 | 0.18 | 1 | 9 | Vdac3 | -0.055 | Young |
| 0.174031 | 0.603773 | 0.235 | 0.186 | 1 | 9 | Inpp5a | 0.049 | Young |
| 0.174134 | 0.758013 | 0.449 | 0.428 | 1 | 9 | Zfp36l2 | 0.021 | Young |
| 0.174732 | 2.907494 | 0.022 | 0.008 | 1 | 9 | Tusc5 | 0.014 | Young |
| 0.174732 | 2.401456 | 0.022 | 0.008 | 1 | 9 | Gm37305 | 0.014 | Young |
| 0.174732 | 2.357185 | 0.022 | 0.008 | 1 | 9 | Ptpn22 | 0.014 | Young |
| 0.174732 | 1.864177 | 0.022 | 0.008 | 1 | 9 | Chn1 | 0.014 | Young |
| 0.174732 | 1.841063 | 0.022 | 0.008 | 1 | 9 | Taf1a | 0.014 | Young |
| 0.174732 | 1.447597 | 0.022 | 0.008 | 1 | 9 | Tmem8 | 0.014 | Young |
| 0.174732 | 1.4167 | 0.022 | 0.008 | 1 | 9 | Itga7 | 0.014 | Young |
| 0.174808 | 0.62653 | 0.169 | 0.124 | 1 | 9 | Pcmt1 | 0.045 | Young |
| 0.17525 | 0.3944 | 0.132 | 0.088 | 1 | 9 | Taok3 | 0.044 | Young |
| 0.175521 | 0.237394 | 0.074 | 0.044 | 1 | 9 | Unc119 | 0.03 | Young |
| 0.175847 | 0.843039 | 0.309 | 0.268 | 1 | 9 | Cebpb | 0.041 | Young |
| 0.175869 | 2.708244 | 0.022 | 0.008 | 1 | 9 | Zfp958 | 0.014 | Young |
| 0.175869 | 2.127467 | 0.022 | 0.008 | 1 | 9 | A430106G1 | 0.014 | Young |
| 0.175869 | 1.928869 | 0.022 | 0.008 | 1 | 9 | Gm45733 | 0.014 | Young |
| 0.175869 | 1.236205 | 0.022 | 0.008 | 1 | 9 | Tmem154 | 0.014 | Young |
| 0.175869 | 1.016504 | 0.022 | 0.008 | 1 | 9 | Kpna2 | 0.014 | Young |
| 0.175869 | 0.152958 | 0.022 | 0.008 | 1 | 9 | Cdca7 | 0.014 | Young |
| 0.175904 | 0.669486 | 0.081 | 0.049 | 1 | 9 | Sap30bp | 0.032 | Young |
| 0.176934 | 0.078423 | 0.404 | 0.505 | 1 | 9 | Mast4 | -0.101 | Young |
| 0.177012 | 2.442569 | 0.022 | 0.008 | 1 | 9 | Ccl22 | 0.014 | Young |
| 0.177012 | 2.05462 | 0.022 | 0.008 | 1 | 9 | Kdelc1 | 0.014 | Young |
| 0.177012 | 2.021505 | 0.022 | 0.008 | 1 | 9 | Zfp873 | 0.014 | Young |
| 0.177012 | 1.227732 | 0.022 | 0.008 | 1 | 9 | Pcx | 0.014 | Young |
| 0.177012 | 0.908198 | 0.022 | 0.008 | 1 | 9 | Tet1 | 0.014 | Young |
| 0.177139 | 0.046214 | 0.272 | 0.211 | 1 | 9 | Nsa2 | 0.061 | Young |
| 0.178161 | 2.06781 | 0.022 | 0.008 | 1 | 9 | Txnl4b | 0.014 | Young |
| 0.178161 | 1.345558 | 0.022 | 0.008 | 1 | 9 | Hspbap1 | 0.014 | Young |
| 0.178161 | 1.233431 | 0.022 | 0.008 | 1 | 9 | Col6a1 | 0.014 | Young |
| 0.178161 | 1.062857 | 0.022 | 0.008 | 1 | 9 | B3gnt2 | 0.014 | Young |
| 0.178161 | 1.054465 | 0.022 | 0.008 | 1 | 9 | Gm48551 | 0.014 | Young |
| 0.178161 | 0.613416 | 0.022 | 0.008 | 1 | 9 | Mvd | 0.014 | Young |
| 0.178183 | 0.917404 | 0.066 | 0.039 | 1 | 9 | Dock8 | 0.027 | Young |
| 0.178275 | 0.49755 | 0.074 | 0.044 | 1 | 9 | Erh | 0.03 | Young |
| 0.178375 | 0.528792 | 0.118 | 0.08 | 1 | 9 | Sirpa | 0.038 | Young |
| 0.17877 | 1.013242 | 0.066 | 0.039 | 1 | 9 | AI837181 | 0.027 | Young |
| 0.179115 | 0.434411 | 0.154 | 0.111 | 1 | 9 | Rbm17 | 0.043 | Young |
| 0.179233 | 0.683491 | 0.118 | 0.08 | 1 | 9 | Rps6ka2 | 0.038 | Young |
| 0.179264 | 0.690964 | 0.103 | 0.067 | 1 | 9 | Med12 | 0.036 | Young |
| 0.179315 | 2.386271 | 0.022 | 0.008 | 1 | 9 | Nup210 | 0.014 | Young |
| 0.179315 | 1.892771 | 0.022 | 0.008 | 1 | 9 | Gm15984 | 0.014 | Young |
| 0.179315 | 1.808397 | 0.022 | 0.008 | 1 | 9 | Capn5 | 0.014 | Young |

| | | | | | | | | |
|---|---|---|---|---|---|---|---|---|
| 0.179315 | 0.964043 | 0.022 | 0.008 | 1 | 9 | Thumpd2 | 0.014 | Young |
| 0.179315 | 0.921087 | 0.022 | 0.008 | 1 | 9 | Pgm3 | 0.014 | Young |
| 0.179315 | 0.23598 | 0.022 | 0.008 | 1 | 9 | C1ra | 0.014 | Young |
| 0.179358 | 1.039887 | 0.066 | 0.039 | 1 | 9 | Abtb1 | 0.027 | Young |
| 0.180474 | 0.45874 | 0.022 | 0.008 | 1 | 9 | Ift140 | 0.014 | Young |
| 0.180474 | 0.428856 | 0.022 | 0.008 | 1 | 9 | Ly9 | 0.014 | Young |
| 0.180502 | 1.022022 | 0.074 | 0.044 | 1 | 9 | Cdkn2d | 0.03 | Young |
| 0.180697 | 0.380739 | 0.279 | 0.227 | 1 | 9 | Aco2 | 0.052 | Young |
| 0.180744 | 0.922942 | 0.331 | 0.302 | 1 | 9 | H2-Ab1 | 0.029 | Young |
| 0.181229 | 0.260656 | 0.015 | 0.039 | 1 | 9 | Zfp942 | -0.024 | Young |
| 0.181903 | 0.519864 | 0.221 | 0.168 | 1 | 9 | Map1b | 0.053 | Young |
| 0.18225 | 0.330837 | 0.081 | 0.049 | 1 | 9 | Dnajc16 | 0.032 | Young |
| 0.182365 | 0.349559 | 0.257 | 0.196 | 1 | 9 | Rnpepl1 | 0.061 | Young |
| 0.182749 | 0.692635 | 0.074 | 0.044 | 1 | 9 | Cep104 | 0.03 | Young |
| 0.182788 | 0.610806 | 0.221 | 0.17 | 1 | 9 | Sf3b6 | 0.051 | Young |
| 0.183012 | 1.147525 | 0.059 | 0.034 | 1 | 9 | Pou5f2 | 0.025 | Young |
| 0.183423 | 0.018782 | 0.066 | 0.108 | 1 | 9 | Fam212a | -0.042 | Young |
| 0.183437 | 0.376035 | 0.272 | 0.227 | 1 | 9 | Flna | 0.045 | Young |
| 0.18388 | 0.825949 | 0.074 | 0.044 | 1 | 9 | Nostrin | 0.03 | Young |
| 0.18388 | 0.770614 | 0.074 | 0.044 | 1 | 9 | Daglb | 0.03 | Young |
| 0.184713 | 1.241274 | 0.066 | 0.039 | 1 | 9 | Zfp568 | 0.027 | Young |
| 0.185315 | 0.895361 | 0.066 | 0.039 | 1 | 9 | Mtfr1 | 0.027 | Young |
| 0.185327 | 0.811765 | 0.118 | 0.08 | 1 | 9 | Gfm1 | 0.038 | Young |
| 0.185465 | 0.248522 | 0.015 | 0.039 | 1 | 9 | Tmem94 | -0.024 | Young |
| 0.185465 | 0.083364 | 0.015 | 0.039 | 1 | 9 | Zxdb | -0.024 | Young |
| 0.185768 | 0.38194 | 0.272 | 0.222 | 1 | 9 | Far1 | 0.05 | Young |
| 0.185919 | 0.73833 | 0.066 | 0.039 | 1 | 9 | Bcas3 | 0.027 | Young |
| 0.186508 | 0.132246 | 0.037 | 0.07 | 1 | 9 | Camta1 | -0.033 | Young |
| 0.18673 | 0.166331 | 0.029 | 0.059 | 1 | 9 | Scn1b | -0.03 | Young |
| 0.18673 | 0.017701 | 0.029 | 0.059 | 1 | 9 | Frmd6 | -0.03 | Young |
| 0.186782 | 0.593762 | 0.184 | 0.137 | 1 | 9 | Cspp1 | 0.047 | Young |
| 0.18702 | 0.443494 | 0.147 | 0.106 | 1 | 9 | Tdrp | 0.041 | Young |
| 0.188391 | 0.372275 | 0.199 | 0.152 | 1 | 9 | Dnaja4 | 0.047 | Young |
| 0.188491 | 0.40976 | 0.301 | 0.245 | 1 | 9 | Cct8 | 0.056 | Young |
| 0.188792 | 0.698146 | 0.243 | 0.201 | 1 | 9 | Ssbp3 | 0.042 | Young |
| 0.189421 | 0.5985 | 0.096 | 0.062 | 1 | 9 | Tbce | 0.034 | Young |
| 0.190121 | 0.037101 | 0.213 | 0.162 | 1 | 9 | Tmem127 | 0.051 | Young |
| 0.190442 | 0.19951 | 0.64 | 0.606 | 1 | 9 | Btg1 | 0.034 | Young |
| 0.190479 | 0.353877 | 0.228 | 0.178 | 1 | 9 | Camsap2 | 0.05 | Young |
| 0.190772 | 0.866769 | 0.059 | 0.034 | 1 | 9 | Pgpep1 | 0.025 | Young |
| 0.190801 | 1.406443 | 0.066 | 0.039 | 1 | 9 | Traf1 | 0.027 | Young |
| 0.190854 | 1.117179 | 0.103 | 0.07 | 1 | 9 | Ddhd1 | 0.033 | Young |
| 0.190911 | 0.256972 | 0.169 | 0.121 | 1 | 9 | Fdx1 | 0.048 | Young |
| 0.192634 | 0.291312 | 0.191 | 0.25 | 1 | 9 | Dennd6a | -0.059 | Young |
| 0.19275 | 0.577883 | 0.059 | 0.034 | 1 | 9 | Atpaf1 | 0.025 | Young |

| | | | | | | | | |
|---|---|---|---|---|---|---|---|---|
| 0.193116 | 0.142999 | 0.074 | 0.044 | 1 | 9 | Lap3 | 0.03 | Young |
| 0.193374 | 0.528823 | 0.272 | 0.224 | 1 | 9 | Erp29 | 0.048 | Young |
| 0.193624 | 2.044404 | 0.029 | 0.013 | 1 | 9 | Tgtp2 | 0.016 | Young |
| 0.194037 | 0.65069 | 0.228 | 0.18 | 1 | 9 | Nme1 | 0.048 | Young |
| 0.194628 | 2.21481 | 0.029 | 0.013 | 1 | 9 | Scnn1b | 0.016 | Young |
| 0.194977 | 0.277693 | 0.162 | 0.116 | 1 | 9 | D5Ertd579 | 0.046 | Young |
| 0.195185 | 0.761049 | 0.132 | 0.093 | 1 | 9 | Tmx2 | 0.039 | Young |
| 0.195199 | 2.046852 | 0.051 | 0.028 | 1 | 9 | Cobll1 | 0.023 | Young |
| 0.195294 | 0.329528 | 0.125 | 0.085 | 1 | 9 | Taf13 | 0.04 | Young |
| 0.19541 | 0.446439 | 0.25 | 0.196 | 1 | 9 | Psmd13 | 0.054 | Young |
| 0.195446 | 0.017731 | 0.132 | 0.188 | 1 | 9 | Palm | -0.056 | Young |
| 0.195503 | 0.709658 | 0.235 | 0.188 | 1 | 9 | Ppib | 0.047 | Young |
| 0.196458 | 0.732585 | 0.11 | 0.075 | 1 | 9 | Atf6b | 0.035 | Young |
| 0.196509 | 0.253996 | 0.103 | 0.067 | 1 | 9 | Naa16 | 0.036 | Young |
| 0.196626 | 0.595211 | 0.324 | 0.271 | 1 | 9 | Sh2d3c | 0.053 | Young |
| 0.196648 | 2.357539 | 0.029 | 0.013 | 1 | 9 | Zfp512b | 0.016 | Young |
| 0.196689 | 0.486638 | 0.25 | 0.196 | 1 | 9 | Trim12a | 0.054 | Young |
| 0.197423 | 0.850045 | 0.059 | 0.034 | 1 | 9 | Clip2 | 0.025 | Young |
| 0.19751 | 0.285435 | 0.118 | 0.08 | 1 | 9 | Grcc10 | 0.038 | Young |
| 0.197664 | 2.013567 | 0.029 | 0.013 | 1 | 9 | Lrrc3b | 0.016 | Young |
| 0.197664 | 1.699936 | 0.029 | 0.013 | 1 | 9 | Gm15859 | 0.016 | Young |
| 0.197784 | 0.523584 | 0.088 | 0.057 | 1 | 9 | Tusc2 | 0.031 | Young |
| 0.197864 | 0.303178 | 0.699 | 0.737 | 1 | 9 | Hnrnpa2b1 | -0.038 | Young |
| 0.198097 | 1.534054 | 0.059 | 0.034 | 1 | 9 | Nckap1l | 0.025 | Young |
| 0.198388 | 2.957115 | 0.044 | 0.023 | 1 | 9 | Klhl6 | 0.021 | Young |
| 0.198684 | 0.86674 | 0.029 | 0.013 | 1 | 9 | Hacl1 | 0.016 | Young |
| 0.198773 | 0.496266 | 0.059 | 0.034 | 1 | 9 | Aatf | 0.025 | Young |
| 0.198773 | 0.472744 | 0.059 | 0.034 | 1 | 9 | Arel1 | 0.025 | Young |
| 0.198997 | 0.154603 | 0.103 | 0.067 | 1 | 9 | Wls | 0.036 | Young |
| 0.199184 | 1.663321 | 0.044 | 0.023 | 1 | 9 | Dvl2 | 0.021 | Young |
| 0.199294 | 0.366299 | 0.235 | 0.191 | 1 | 9 | Kmt2d | 0.044 | Young |
| 0.19945 | 0.527792 | 0.059 | 0.034 | 1 | 9 | Cenpj | 0.025 | Young |
| 0.19955 | 1.028688 | 0.051 | 0.028 | 1 | 9 | Cwc27 | 0.023 | Young |
| 0.199707 | 1.225762 | 0.029 | 0.013 | 1 | 9 | Pik3cg | 0.016 | Young |
| 0.199718 | 0.890864 | 0.132 | 0.095 | 1 | 9 | Ccl9 | 0.037 | Young |
| 0.199939 | 0.26252 | 0.088 | 0.057 | 1 | 9 | Mmgt1 | 0.031 | Young |
| 0.200085 | 0.439107 | 0.206 | 0.157 | 1 | 9 | Ssh1 | 0.049 | Young |
| 0.20021 | 1.039436 | 0.088 | 0.057 | 1 | 9 | Cdk6 | 0.031 | Young |
| 0.200321 | 0.34136 | 0.382 | 0.325 | 1 | 9 | Ebf3 | 0.057 | Young |
| 0.200481 | 0.535052 | 0.088 | 0.057 | 1 | 9 | Ccnc | 0.031 | Young |
| 0.200735 | 1.14127 | 0.029 | 0.013 | 1 | 9 | Tifab | 0.016 | Young |
| 0.200735 | 1.064759 | 0.029 | 0.013 | 1 | 9 | Aoah | 0.016 | Young |
| 0.200739 | 0.138192 | 0.213 | 0.162 | 1 | 9 | Stx7 | 0.051 | Young |
| 0.200784 | 1.340881 | 0.044 | 0.023 | 1 | 9 | Zfp874b | 0.021 | Young |
| 0.201016 | 1.08029 | 0.051 | 0.028 | 1 | 9 | Tmem267 | 0.023 | Young |

| | | | | | | | | |
|---|---|---|---|---|---|---|---|---|
| 0.20107 | 0.318407 | 0.574 | 0.554 | 1 | 9 | Car4 | 0.02 | Young |
| 0.201386 | 2.094052 | 0.037 | 0.018 | 1 | 9 | Tlcd2 | 0.019 | Young |
| 0.201587 | 1.445742 | 0.044 | 0.023 | 1 | 9 | Ern1 | 0.021 | Young |
| 0.201599 | 0.516842 | 0.228 | 0.178 | 1 | 9 | Eif3l | 0.05 | Young |
| 0.201766 | 1.43653 | 0.029 | 0.013 | 1 | 9 | Wrap53 | 0.016 | Young |
| 0.201766 | 0.70607 | 0.029 | 0.013 | 1 | 9 | Stradb | 0.016 | Young |
| 0.202489 | 1.213706 | 0.051 | 0.028 | 1 | 9 | Trim23 | 0.023 | Young |
| 0.202489 | 0.663139 | 0.051 | 0.028 | 1 | 9 | Klhl15 | 0.023 | Young |
| 0.20255 | 0.191863 | 0.147 | 0.103 | 1 | 9 | Edem1 | 0.044 | Young |
| 0.202657 | 1.012571 | 0.088 | 0.057 | 1 | 9 | Cln8 | 0.031 | Young |
| 0.202801 | 1.276291 | 0.029 | 0.013 | 1 | 9 | Dnal1 | 0.016 | Young |
| 0.202801 | 1.2106 | 0.029 | 0.013 | 1 | 9 | Agbl5 | 0.016 | Young |
| 0.202801 | 0.577888 | 0.029 | 0.013 | 1 | 9 | Ddx28 | 0.016 | Young |
| 0.202838 | 0.238811 | 0.007 | 0.026 | 1 | 9 | Slc46a3 | -0.019 | Young |
| 0.202845 | 1.461832 | 0.096 | 0.064 | 1 | 9 | Usp40 | 0.032 | Young |
| 0.203036 | 0.651845 | 0.176 | 0.134 | 1 | 9 | Rtcb | 0.042 | Young |
| 0.203551 | 0.049451 | 0.059 | 0.034 | 1 | 9 | Vrk2 | 0.025 | Young |
| 0.203553 | 0.607367 | 0.199 | 0.149 | 1 | 9 | Ccni | 0.05 | Young |
| 0.203798 | 0.476479 | 0.162 | 0.119 | 1 | 9 | 1110008F1 | 0.043 | Young |
| 0.20384 | 1.271096 | 0.029 | 0.013 | 1 | 9 | Gm20604 | 0.016 | Young |
| 0.20384 | 0.839722 | 0.029 | 0.013 | 1 | 9 | Map2k6 | 0.016 | Young |
| 0.20384 | 0.527728 | 0.029 | 0.013 | 1 | 9 | Sumf2 | 0.016 | Young |
| 0.204064 | 0.094757 | 0.066 | 0.039 | 1 | 9 | Ell2 | 0.027 | Young |
| 0.204087 | 1.619031 | 0.037 | 0.018 | 1 | 9 | Recql5 | 0.019 | Young |
| 0.204376 | 0.679995 | 0.081 | 0.052 | 1 | 9 | Idh3a | 0.029 | Young |
| 0.204714 | 1.208737 | 0.051 | 0.028 | 1 | 9 | Dtd1 | 0.023 | Young |
| 0.204714 | 0.764919 | 0.051 | 0.028 | 1 | 9 | Slfn9 | 0.023 | Young |
| 0.204823 | 1.728447 | 0.044 | 0.023 | 1 | 9 | Wdfy4 | 0.021 | Young |
| 0.204882 | 0.911073 | 0.029 | 0.013 | 1 | 9 | Batf2 | 0.016 | Young |
| 0.204882 | 0.650253 | 0.029 | 0.013 | 1 | 9 | Nudt22 | 0.016 | Young |
| 0.20495 | 0.751261 | 0.081 | 0.052 | 1 | 9 | Rarres1 | 0.029 | Young |
| 0.205017 | 0.488883 | 0.272 | 0.222 | 1 | 9 | Vamp8 | 0.05 | Young |
| 0.205095 | 0.381734 | 0.243 | 0.186 | 1 | 9 | Retreg2 | 0.057 | Young |
| 0.205826 | 0.64987 | 0.059 | 0.098 | 1 | 9 | Traf6 | -0.039 | Young |
| 0.205929 | 0.773685 | 0.029 | 0.013 | 1 | 9 | Fancc | 0.016 | Young |
| 0.206103 | 0.693743 | 0.081 | 0.052 | 1 | 9 | Rfwd3 | 0.029 | Young |
| 0.206252 | 0.381442 | 0.14 | 0.101 | 1 | 9 | Nfya | 0.039 | Young |
| 0.206455 | 1.313446 | 0.044 | 0.023 | 1 | 9 | Cyp4f18 | 0.021 | Young |
| 0.206669 | 0.027842 | 0.478 | 0.539 | 1 | 9 | C1qtnf9 | -0.061 | Young |
| 0.206815 | 1.614618 | 0.037 | 0.018 | 1 | 9 | Zfp770 | 0.019 | Young |
| 0.206847 | 0.538829 | 0.324 | 0.271 | 1 | 9 | Acly | 0.053 | Young |
| 0.206979 | 0.478929 | 0.029 | 0.013 | 1 | 9 | Fxn | 0.016 | Young |
| 0.207275 | 1.508644 | 0.044 | 0.023 | 1 | 9 | Gorab | 0.021 | Young |
| 0.207275 | 0.9288 | 0.044 | 0.023 | 1 | 9 | Cdr2l | 0.021 | Young |
| 0.207275 | 0.835456 | 0.044 | 0.023 | 1 | 9 | Dcbld2 | 0.021 | Young |

| | | | | | | | | |
|---|---|---|---|---|---|---|---|---|
| 0.207513 | 0.971402 | 0.206 | 0.16 | 1 | 9 | Slc39a7 | 0.046 | Young |
| 0.207828 | 0.684997 | 0.169 | 0.124 | 1 | 9 | Atp6v1d | 0.045 | Young |
| 0.207993 | 0.350497 | 0.191 | 0.142 | 1 | 9 | Mgat2 | 0.049 | Young |
| 0.208034 | 0.334536 | 0.029 | 0.013 | 1 | 9 | Samsn1 | 0.016 | Young |
| 0.208034 | 0.083753 | 0.029 | 0.013 | 1 | 9 | Serpinb8 | 0.016 | Young |
| 0.208434 | 0.639673 | 0.103 | 0.07 | 1 | 9 | 1700017B0 | 0.033 | Young |
| 0.208648 | 1.236118 | 0.037 | 0.018 | 1 | 9 | Borcs5 | 0.019 | Young |
| 0.208887 | 1.073169 | 0.191 | 0.149 | 1 | 9 | Ckb | 0.042 | Young |
| 0.209005 | 1.434794 | 0.081 | 0.052 | 1 | 9 | Snx15 | 0.029 | Young |
| 0.209114 | 0.250703 | 0.059 | 0.034 | 1 | 9 | Katnal1 | 0.025 | Young |
| 0.209202 | 0.728534 | 0.103 | 0.07 | 1 | 9 | Emp3 | 0.033 | Young |
| 0.209304 | 0.04396 | 0.118 | 0.08 | 1 | 9 | Uspl1 | 0.038 | Young |
| 0.209568 | 1.345401 | 0.037 | 0.018 | 1 | 9 | Gm12840 | 0.019 | Young |
| 0.209747 | 0.631351 | 0.044 | 0.023 | 1 | 9 | Haus6 | 0.021 | Young |
| 0.209748 | 1.189716 | 0.044 | 0.023 | 1 | 9 | Ulbp1 | 0.021 | Young |
| 0.210062 | 0.191993 | 0.11 | 0.075 | 1 | 9 | Casp7 | 0.035 | Young |
| 0.210174 | 0.745209 | 0.081 | 0.052 | 1 | 9 | Traf3 | 0.029 | Young |
| 0.210522 | 0.135306 | 0.059 | 0.034 | 1 | 9 | Fam118b | 0.025 | Young |
| 0.210645 | 0.760867 | 0.25 | 0.216 | 1 | 9 | Lcorl | 0.034 | Young |
| 0.210761 | 0.25069 | 0.037 | 0.067 | 1 | 9 | Mea1 | -0.03 | Young |
| 0.211302 | 0.007757 | 0.728 | 0.771 | 1 | 9 | Apoe | -0.043 | Young |
| 0.211419 | 1.158152 | 0.037 | 0.018 | 1 | 9 | Atad3a | 0.019 | Young |
| 0.211419 | 0.736217 | 0.037 | 0.018 | 1 | 9 | Zfp868 | 0.019 | Young |
| 0.21226 | 0.409179 | 0.051 | 0.028 | 1 | 9 | Ftl1-ps1 | 0.023 | Young |
| 0.212349 | 1.14866 | 0.037 | 0.018 | 1 | 9 | Dtnb | 0.019 | Young |
| 0.212349 | 1.116238 | 0.037 | 0.018 | 1 | 9 | Mecr | 0.019 | Young |
| 0.212592 | 0.326664 | 0.39 | 0.33 | 1 | 9 | Wsb1 | 0.06 | Young |
| 0.212637 | 0.207335 | 0.066 | 0.103 | 1 | 9 | Mid2 | -0.037 | Young |
| 0.212647 | 0.090424 | 0.059 | 0.034 | 1 | 9 | Clec4a2 | 0.025 | Young |
| 0.21277 | 0.436633 | 0.228 | 0.186 | 1 | 9 | Sh3pxd2a | 0.042 | Young |
| 0.213281 | 0.749743 | 0.037 | 0.018 | 1 | 9 | Sla | 0.019 | Young |
| 0.213281 | 0.099496 | 0.037 | 0.018 | 1 | 9 | Ndufaf5 | 0.019 | Young |
| 0.213358 | 0.800072 | 0.059 | 0.034 | 1 | 9 | Sdf2l1 | 0.025 | Young |
| 0.213697 | 0.366074 | 0.272 | 0.222 | 1 | 9 | Grk2 | 0.05 | Young |
| 0.213793 | 0.171391 | 0.051 | 0.028 | 1 | 9 | Pi4k2b | 0.023 | Young |
| 0.21392 | 0.144437 | 0.36 | 0.438 | 1 | 9 | Cxcl9 | -0.078 | Young |
| 0.214217 | 0.65573 | 0.037 | 0.018 | 1 | 9 | Dapp1 | 0.019 | Young |
| 0.214217 | 0.004229 | 0.037 | 0.018 | 1 | 9 | Dusp12 | 0.019 | Young |
| 0.21436 | 0.661935 | 0.221 | 0.173 | 1 | 9 | Tsc22d3 | 0.048 | Young |
| 0.214619 | 0.200601 | 0.029 | 0.057 | 1 | 9 | Hist3h2a | -0.028 | Young |
| 0.214936 | 0.068451 | 0.088 | 0.057 | 1 | 9 | Sfxn3 | 0.031 | Young |
| 0.215121 | 0.278165 | 0.243 | 0.191 | 1 | 9 | Adss | 0.052 | Young |
| 0.215155 | 0.471971 | 0.037 | 0.018 | 1 | 9 | Tsga10 | 0.019 | Young |
| 0.215436 | 0.07193 | 0.213 | 0.165 | 1 | 9 | AC160336. | 0.048 | Young |
| 0.216097 | 1.057268 | 0.037 | 0.018 | 1 | 9 | Nme6 | 0.019 | Young |

| | | | | | | | | |
|---|---|---|---|---|---|---|---|---|
| 0.216097 | 0.489498 | 0.037 | 0.018 | 1 | 9 | Zmym6 | 0.019 | Young |
| 0.216107 | 0.185864 | 0.051 | 0.028 | 1 | 9 | Myo5a | 0.023 | Young |
| 0.216446 | 0.112012 | 0.044 | 0.023 | 1 | 9 | Tbrg4 | 0.021 | Young |
| 0.216719 | 1.226371 | 0.074 | 0.046 | 1 | 9 | Tnfrsf22 | 0.028 | Young |
| 0.217294 | 0.358545 | 0.044 | 0.023 | 1 | 9 | Tmem51 | 0.021 | Young |
| 0.217331 | 0.370948 | 0.176 | 0.131 | 1 | 9 | Lztfl1 | 0.045 | Young |
| 0.21737 | 0.343747 | 0.154 | 0.111 | 1 | 9 | Vps54 | 0.043 | Young |
| 0.217989 | 0.221464 | 0.037 | 0.018 | 1 | 9 | Dusp10 | 0.019 | Young |
| 0.218118 | 0.6453 | 0.169 | 0.129 | 1 | 9 | Arhgef6 | 0.04 | Young |
| 0.218451 | 0.017121 | 0.125 | 0.178 | 1 | 9 | Osbpl8 | -0.053 | Young |
| 0.21861 | 1.132385 | 0.074 | 0.046 | 1 | 9 | Ccdc90b | 0.028 | Young |
| 0.218647 | 0.491717 | 0.11 | 0.075 | 1 | 9 | Borcs6 | 0.035 | Young |
| 0.218839 | 0.304002 | 0.419 | 0.361 | 1 | 9 | Dync1i2 | 0.058 | Young |
| 0.218939 | 0.779601 | 0.037 | 0.018 | 1 | 9 | Pclaf | 0.019 | Young |
| 0.219096 | 0.004994 | 0.103 | 0.149 | 1 | 9 | Hdgfl3 | -0.046 | Young |
| 0.21916 | 0.400295 | 0.11 | 0.075 | 1 | 9 | Arhgap25 | 0.035 | Young |
| 0.219208 | 0.877337 | 0.14 | 0.103 | 1 | 9 | Fnta | 0.037 | Young |
| 0.219405 | 0.513718 | 0.096 | 0.064 | 1 | 9 | Cib1 | 0.032 | Young |
| 0.219441 | 1.163956 | 0.088 | 0.059 | 1 | 9 | Pepd | 0.029 | Young |
| 0.219632 | 0.886697 | 0.147 | 0.111 | 1 | 9 | Sipa1l1 | 0.036 | Young |
| 0.219877 | 2.030175 | 0.074 | 0.046 | 1 | 9 | Tbc1d8b | 0.028 | Young |
| 0.219957 | 0.062115 | 0.015 | 0.036 | 1 | 9 | Stoml1 | -0.021 | Young |
| 0.220257 | 0.173665 | 0.316 | 0.263 | 1 | 9 | Tet3 | 0.053 | Young |
| 0.220405 | 0.324495 | 0.154 | 0.111 | 1 | 9 | Pole4 | 0.043 | Young |
| 0.22058 | 0.806455 | 0.169 | 0.129 | 1 | 9 | Trafd1 | 0.04 | Young |
| 0.221081 | 0.164841 | 0.051 | 0.085 | 1 | 9 | Ccdc107 | -0.034 | Young |
| 0.221149 | 0.654378 | 0.074 | 0.046 | 1 | 9 | Ms4a6c | 0.028 | Young |
| 0.221297 | 0.520326 | 0.199 | 0.157 | 1 | 9 | Ppp4r3a | 0.042 | Young |
| 0.22176 | 0.322798 | 0.544 | 0.534 | 1 | 9 | Pabpc1 | 0.01 | Young |
| 0.221842 | 0.201199 | 0.14 | 0.101 | 1 | 9 | Soga1 | 0.039 | Young |
| 0.222058 | 1.271913 | 0.081 | 0.054 | 1 | 9 | Rsrc1 | 0.027 | Young |
| 0.222387 | 0.332043 | 0.346 | 0.289 | 1 | 9 | Ubtf | 0.057 | Young |
| 0.222828 | 0.613543 | 0.199 | 0.157 | 1 | 9 | Ppil4 | 0.042 | Young |
| 0.222874 | 0.575958 | 0.154 | 0.113 | 1 | 9 | Snx10 | 0.041 | Young |
| 0.223216 | 0.225929 | 0.14 | 0.101 | 1 | 9 | Riok1 | 0.039 | Young |
| 0.223677 | 0.066993 | 0.132 | 0.093 | 1 | 9 | Tm7sf3 | 0.039 | Young |
| 0.224009 | 0.429056 | 0.118 | 0.082 | 1 | 9 | Ndufaf8 | 0.036 | Young |
| 0.22404 | 0.797054 | 0.147 | 0.111 | 1 | 9 | Zeb2 | 0.036 | Young |
| 0.22439 | 0.874639 | 0.096 | 0.064 | 1 | 9 | Dguok | 0.032 | Young |
| 0.224998 | 0.957817 | 0.074 | 0.046 | 1 | 9 | Tfb2m | 0.028 | Young |
| 0.225063 | 0.7062 | 0.154 | 0.113 | 1 | 9 | Inpp5k | 0.041 | Young |
| 0.225591 | 1.827568 | 0.118 | 0.088 | 1 | 9 | Ccl2 | 0.03 | Young |
| 0.225951 | 1.559042 | 0.066 | 0.041 | 1 | 9 | Il18bp | 0.025 | Young |
| 0.226291 | 0.213607 | 0.074 | 0.046 | 1 | 9 | Zfp729a | 0.028 | Young |
| 0.227149 | 0.086949 | 0.088 | 0.057 | 1 | 9 | Acaca | 0.031 | Young |

| | | | | | | | | |
|---|---|---|---|---|---|---|---|---|
| 0.227312 | 0.595915 | 0.184 | 0.142 | 1 | 9 | Fem1b | 0.042 | Young |
| 0.227502 | 0.647945 | 0.162 | 0.121 | 1 | 9 | Preb | 0.041 | Young |
| 0.22759 | 0.680407 | 0.074 | 0.046 | 1 | 9 | Tfeb | 0.028 | Young |
| 0.228241 | 0.81689 | 0.074 | 0.046 | 1 | 9 | Rdh10 | 0.028 | Young |
| 0.228249 | 0.156842 | 0.103 | 0.07 | 1 | 9 | Slirp | 0.033 | Young |
| 0.228715 | 0.146911 | 0.868 | 0.861 | 1 | 9 | Rplp2 | 0.007 | Young |
| 0.230076 | 1.040906 | 0.066 | 0.041 | 1 | 9 | Wdr77 | 0.025 | Young |
| 0.230203 | 0.546967 | 0.074 | 0.046 | 1 | 9 | Dpp3 | 0.028 | Young |
| 0.230353 | 0.578868 | 0.243 | 0.204 | 1 | 9 | Nrros | 0.039 | Young |
| 0.231209 | 0.212645 | 0.279 | 0.229 | 1 | 9 | Cct7 | 0.05 | Young |
| 0.231596 | 0.642892 | 0.14 | 0.103 | 1 | 9 | Cfap20 | 0.037 | Young |
| 0.231665 | 0.92032 | 0.088 | 0.059 | 1 | 9 | Ruvbl1 | 0.029 | Young |
| 0.231881 | 1.217562 | 0.118 | 0.085 | 1 | 9 | Ints3 | 0.033 | Young |
| 0.231896 | 0.470661 | 0.412 | 0.369 | 1 | 9 | Ralbp1 | 0.043 | Young |
| 0.231995 | 0.593407 | 0.11 | 0.077 | 1 | 9 | Vps41 | 0.033 | Young |
| 0.232259 | 0.967535 | 0.088 | 0.059 | 1 | 9 | Slc25a16 | 0.029 | Young |
| 0.232283 | 0.175285 | 0.176 | 0.134 | 1 | 9 | Nfatc1 | 0.042 | Young |
| 0.23298 | 0.298212 | 0.647 | 0.634 | 1 | 9 | Podxl | 0.013 | Young |
| 0.233007 | 0.287171 | 0.397 | 0.345 | 1 | 9 | Snu13 | 0.052 | Young |
| 0.233198 | 0.075032 | 0.103 | 0.07 | 1 | 9 | Clec10a | 0.033 | Young |
| 0.233411 | 0.657391 | 0.103 | 0.072 | 1 | 9 | Tbc1d17 | 0.031 | Young |
| 0.2335 | 0.455314 | 0.074 | 0.046 | 1 | 9 | Ncf2 | 0.028 | Young |
| 0.233615 | 0.573021 | 0.169 | 0.129 | 1 | 9 | Mtss1l | 0.04 | Young |
| 0.234142 | 0.502841 | 0.199 | 0.155 | 1 | 9 | 1110004F1 | 0.044 | Young |
| 0.234808 | 0.311863 | 0.537 | 0.497 | 1 | 9 | Ebf1 | 0.04 | Young |
| 0.234905 | 0.653394 | 0.169 | 0.129 | 1 | 9 | Thoc7 | 0.04 | Young |
| 0.237553 | 0.436921 | 0.199 | 0.155 | 1 | 9 | Zfp451 | 0.044 | Young |
| 0.237641 | 0.518857 | 0.132 | 0.095 | 1 | 9 | Dennd1a | 0.037 | Young |
| 0.237924 | 0.578522 | 0.213 | 0.162 | 1 | 9 | Utp11 | 0.051 | Young |
| 0.238016 | 0.527182 | 0.221 | 0.175 | 1 | 9 | Safb2 | 0.046 | Young |
| 0.238637 | 0.850639 | 0.228 | 0.188 | 1 | 9 | Eif3h | 0.04 | Young |
| 0.239647 | 0.352806 | 0.14 | 0.103 | 1 | 9 | Nck2 | 0.037 | Young |
| 0.239933 | 0.310421 | 0.309 | 0.255 | 1 | 9 | Cdc37l1 | 0.054 | Young |
| 0.240027 | 0.460766 | 0.11 | 0.077 | 1 | 9 | Jdp2 | 0.033 | Young |
| 0.240617 | 0.646738 | 0.103 | 0.072 | 1 | 9 | Usp53 | 0.031 | Young |
| 0.240719 | 0.437971 | 0.118 | 0.082 | 1 | 9 | Dvl1 | 0.036 | Young |
| 0.240863 | 0.014665 | 0.507 | 0.58 | 1 | 9 | Elk3 | -0.073 | Young |
| 0.240871 | 0.229792 | 0.191 | 0.147 | 1 | 9 | Ylpm1 | 0.044 | Young |
| 0.241298 | 0.523565 | 0.088 | 0.059 | 1 | 9 | Snx14 | 0.029 | Young |
| 0.241381 | 0.004196 | 0.706 | 0.786 | 1 | 9 | Fau | -0.08 | Young |
| 0.241401 | 1.065199 | 0.059 | 0.036 | 1 | 9 | Thap1 | 0.023 | Young |
| 0.241401 | 1.021594 | 0.059 | 0.036 | 1 | 9 | Cars2 | 0.023 | Young |
| 0.241548 | 0.37503 | 0.074 | 0.046 | 1 | 9 | Siae | 0.028 | Young |
| 0.241739 | 0.293955 | 0.103 | 0.072 | 1 | 9 | Wdfy2 | 0.031 | Young |
| 0.242271 | 0.937686 | 0.147 | 0.113 | 1 | 9 | Cxcl2 | 0.034 | Young |

| | | | | | | | | |
|---|---|---|---|---|---|---|---|---|
| 0.24231 | 0.763935 | 0.169 | 0.129 | 1 | 9 | Btbd1 | 0.04 | Young |
| 0.242434 | 0.340626 | 0.184 | 0.142 | 1 | 9 | Smndc1 | 0.042 | Young |
| 0.242595 | 0.432318 | 0.243 | 0.196 | 1 | 9 | Brd1 | 0.047 | Young |
| 0.243505 | 0.271655 | 0.169 | 0.131 | 1 | 9 | Xrn2 | 0.038 | Young |
| 0.24402 | 1.029751 | 0.096 | 0.067 | 1 | 9 | Cfp | 0.029 | Young |
| 0.24504 | 0.09871 | 0.103 | 0.07 | 1 | 9 | Sgpl1 | 0.033 | Young |
| 0.245658 | 1.250289 | 0.066 | 0.041 | 1 | 9 | Rpl10-ps3 | 0.025 | Young |
| 0.245658 | 0.587914 | 0.029 | 0.054 | 1 | 9 | Fpgs | -0.025 | Young |
| 0.245765 | 0.40145 | 0.118 | 0.082 | 1 | 9 | Srebf1 | 0.036 | Young |
| 0.245904 | 0.658557 | 0.125 | 0.093 | 1 | 9 | Ankle2 | 0.032 | Young |
| 0.246265 | 0.816748 | 0.103 | 0.072 | 1 | 9 | Slc25a12 | 0.031 | Young |
| 0.246475 | 0.395279 | 0.037 | 0.064 | 1 | 9 | Ppp2r5d | -0.027 | Young |
| 0.246475 | 0.164949 | 0.037 | 0.064 | 1 | 9 | Wrnip1 | -0.027 | Young |
| 0.247055 | 0.072913 | 0.184 | 0.137 | 1 | 9 | Selenos | 0.047 | Young |
| 0.247523 | 0.175879 | 0.096 | 0.064 | 1 | 9 | Commd6 | 0.032 | Young |
| 0.247543 | 0.999679 | 0.059 | 0.036 | 1 | 9 | Cfh | 0.023 | Young |
| 0.247615 | 0.335633 | 0.132 | 0.095 | 1 | 9 | Mrpl14 | 0.037 | Young |
| 0.247839 | 0.405857 | 0.066 | 0.041 | 1 | 9 | Papolg | 0.025 | Young |
| 0.248498 | 0.396435 | 0.243 | 0.204 | 1 | 9 | Cdc5l | 0.039 | Young |
| 0.24855 | 0.421602 | 0.103 | 0.072 | 1 | 9 | Psme3 | 0.031 | Young |
| 0.248569 | 0.271275 | 0.066 | 0.041 | 1 | 9 | Map3k8 | 0.025 | Young |
| 0.248596 | 0.088201 | 0.213 | 0.165 | 1 | 9 | Pbxip1 | 0.048 | Young |
| 0.248706 | 0.326516 | 0.088 | 0.059 | 1 | 9 | Slfn2 | 0.029 | Young |
| 0.248887 | 0.262414 | 0.206 | 0.162 | 1 | 9 | Fam174a | 0.044 | Young |
| 0.249031 | 0.47242 | 0.243 | 0.193 | 1 | 9 | Copb1 | 0.05 | Young |
| 0.249377 | 0.344486 | 0.184 | 0.139 | 1 | 9 | Pbx2 | 0.045 | Young |
| 0.249706 | 0.349824 | 0.125 | 0.17 | 1 | 9 | Cyp26b1 | -0.045 | Young |
| 0.249921 | 0.507551 | 0.11 | 0.077 | 1 | 9 | Sc5d | 0.033 | Young |
| 0.250263 | 0.505502 | 0.081 | 0.054 | 1 | 9 | Pld4 | 0.027 | Young |
| 0.250282 | 0.580024 | 0.154 | 0.119 | 1 | 9 | Agap3 | 0.035 | Young |
| 0.250764 | 0.917522 | 0.176 | 0.139 | 1 | 9 | Arf2 | 0.037 | Young |
| 0.251408 | 0.231756 | 0.221 | 0.173 | 1 | 9 | Cd9 | 0.048 | Young |
| 0.25141 | 0.347744 | 0.007 | 0.023 | 1 | 9 | Tasp1 | -0.016 | Young |
| 0.25141 | 0.007619 | 0.007 | 0.023 | 1 | 9 | Zfp579 | -0.016 | Young |
| 0.25179 | 0.438225 | 0.118 | 0.085 | 1 | 9 | Foxk1 | 0.033 | Young |
| 0.251846 | 0.067014 | 0.037 | 0.064 | 1 | 9 | Zc3hc1 | -0.027 | Young |
| 0.25222 | 1.14813 | 0.059 | 0.036 | 1 | 9 | Dcakd | 0.023 | Young |
| 0.25222 | 0.904159 | 0.059 | 0.036 | 1 | 9 | Zfp41 | 0.023 | Young |
| 0.25222 | 0.257106 | 0.25 | 0.196 | 1 | 9 | Ddi2 | 0.054 | Young |
| 0.252242 | 0.630442 | 0.066 | 0.041 | 1 | 9 | Manbal | 0.025 | Young |
| 0.252599 | 0.445631 | 0.154 | 0.116 | 1 | 9 | Zc3h4 | 0.038 | Young |
| 0.252755 | 0.209552 | 0.574 | 0.544 | 1 | 9 | Hspa5 | 0.03 | Young |
| 0.253317 | 0.333526 | 0.287 | 0.242 | 1 | 9 | Vdac2 | 0.045 | Young |
| 0.253334 | 1.01429 | 0.162 | 0.126 | 1 | 9 | Mrc1 | 0.036 | Young |
| 0.253721 | 0.230618 | 0.066 | 0.041 | 1 | 9 | Il13ra1 | 0.025 | Young |

| | | | | | | | | |
|---|---|---|---|---|---|---|---|---|
| 0.254581 | 0.741373 | 0.059 | 0.036 | 1 | 9 | Gpn1 | 0.023 | Young |
| 0.254948 | 0.75367 | 0.11 | 0.08 | 1 | 9 | Ahsa2 | 0.03 | Young |
| 0.255372 | 1.006201 | 0.059 | 0.036 | 1 | 9 | Coa6 | 0.023 | Young |
| 0.255372 | 0.664993 | 0.059 | 0.036 | 1 | 9 | B4galt5 | 0.023 | Young |
| 0.255389 | 0.063989 | 0.096 | 0.064 | 1 | 9 | Pxmp4 | 0.032 | Young |
| 0.255523 | 1.102027 | 0.051 | 0.031 | 1 | 9 | Fermt3 | 0.02 | Young |
| 0.256049 | 0.389256 | 0.272 | 0.229 | 1 | 9 | Magt1 | 0.043 | Young |
| 0.256674 | 0.361111 | 0.11 | 0.077 | 1 | 9 | Entpd1 | 0.033 | Young |
| 0.257035 | 0.449593 | 0.309 | 0.255 | 1 | 9 | Nxpe4 | 0.054 | Young |
| 0.257522 | 1.090676 | 0.081 | 0.054 | 1 | 9 | Pdik1l | 0.027 | Young |
| 0.257564 | 0.166823 | 0.176 | 0.134 | 1 | 9 | Ndufa12 | 0.042 | Young |
| 0.257754 | 0.887888 | 0.059 | 0.036 | 1 | 9 | Folr2 | 0.023 | Young |
| 0.258004 | 0.410445 | 0.221 | 0.18 | 1 | 9 | Chmp4b | 0.041 | Young |
| 0.258068 | 0.682938 | 0.132 | 0.098 | 1 | 9 | Pip4k2a | 0.034 | Young |
| 0.258342 | 0.01139 | 0.691 | 0.755 | 1 | 9 | Klf4 | -0.064 | Young |
| 0.25846 | 0.04283 | 0.096 | 0.064 | 1 | 9 | Abhd4 | 0.032 | Young |
| 0.258937 | 1.319964 | 0.051 | 0.031 | 1 | 9 | Pld1 | 0.02 | Young |
| 0.258965 | 1.207568 | 0.096 | 0.067 | 1 | 9 | Stx17 | 0.029 | Young |
| 0.259133 | 0.045333 | 0.074 | 0.108 | 1 | 9 | Sec23a | -0.034 | Young |
| 0.25935 | 1.21875 | 0.059 | 0.036 | 1 | 9 | Flcn | 0.023 | Young |
| 0.259524 | 0.642351 | 0.11 | 0.077 | 1 | 9 | Zfp710 | 0.033 | Young |
| 0.259524 | 0.540621 | 0.11 | 0.077 | 1 | 9 | Cnpy2 | 0.033 | Young |
| 0.259623 | 0.345354 | 0.184 | 0.144 | 1 | 9 | Cbfb | 0.04 | Young |
| 0.259795 | 1.539626 | 0.051 | 0.031 | 1 | 9 | Prune1 | 0.02 | Young |
| 0.260077 | 0.349124 | 0.272 | 0.224 | 1 | 9 | Skp1a | 0.048 | Young |
| 0.260151 | 1.102617 | 0.059 | 0.036 | 1 | 9 | Phka1 | 0.023 | Young |
| 0.260197 | 0.610755 | 0.081 | 0.054 | 1 | 9 | Dcbld1 | 0.027 | Young |
| 0.260587 | 0.972818 | 0.11 | 0.08 | 1 | 9 | Dcaf10 | 0.03 | Young |
| 0.260757 | 0.016922 | 0.088 | 0.059 | 1 | 9 | Prpf19 | 0.029 | Young |
| 0.260933 | 0.019439 | 0.022 | 0.044 | 1 | 9 | Abt1 | -0.022 | Young |
| 0.261017 | 1.334916 | 0.118 | 0.088 | 1 | 9 | Nol11 | 0.03 | Young |
| 0.261086 | 0.957199 | 0.051 | 0.031 | 1 | 9 | Lig1 | 0.02 | Young |
| 0.261375 | 0.572002 | 0.206 | 0.162 | 1 | 9 | Mettl9 | 0.044 | Young |
| 0.261758 | 0.713016 | 0.059 | 0.036 | 1 | 9 | Pank4 | 0.023 | Young |
| 0.261758 | 0.338801 | 0.059 | 0.036 | 1 | 9 | Tarbp2 | 0.023 | Young |
| 0.262382 | 1.744967 | 0.051 | 0.031 | 1 | 9 | Ctdp1 | 0.02 | Young |
| 0.262768 | 0.000476 | 0.044 | 0.072 | 1 | 9 | 2200002D( | -0.028 | Young |
| 0.263391 | 0.790643 | 0.147 | 0.113 | 1 | 9 | Ccl6 | 0.034 | Young |
| 0.263485 | 0.089413 | 0.066 | 0.041 | 1 | 9 | Vps18 | 0.025 | Young |
| 0.263537 | 0.677966 | 0.213 | 0.173 | 1 | 9 | Eif3d | 0.04 | Young |
| 0.264181 | 0.524214 | 0.059 | 0.036 | 1 | 9 | Xpc | 0.023 | Young |
| 0.264362 | 0.455578 | 0.294 | 0.25 | 1 | 9 | Rps6ka3 | 0.044 | Young |
| 0.264403 | 0.808487 | 0.118 | 0.085 | 1 | 9 | Ptpn18 | 0.033 | Young |
| 0.264458 | 0.116322 | 0.184 | 0.139 | 1 | 9 | Spcs3 | 0.045 | Young |
| 0.2646 | 0.014967 | 0.162 | 0.206 | 1 | 9 | Ptbp1 | -0.044 | Young |

| | | | | | | | | |
|---|---|---|---|---|---|---|---|---|
| 0.265161 | 0.228489 | 0.022 | 0.044 | 1 | 9 | Uchl3 | -0.022 | Young |
| 0.265898 | 0.510944 | 0.132 | 0.098 | 1 | 9 | Tpd52 | 0.034 | Young |
| 0.26662 | 0.333295 | 0.059 | 0.036 | 1 | 9 | Ccdc157 | 0.023 | Young |
| 0.26724 | 0.268921 | 0.228 | 0.18 | 1 | 9 | Rbm26 | 0.048 | Young |
| 0.267509 | 0.329601 | 0.176 | 0.137 | 1 | 9 | Smim15 | 0.039 | Young |
| 0.268254 | 0.354118 | 0.059 | 0.036 | 1 | 9 | Plekhm3 | 0.023 | Young |
| 0.268375 | 0.804355 | 0.154 | 0.121 | 1 | 9 | Fbxo22 | 0.033 | Young |
| 0.268415 | 3.070591 | 0.015 | 0.005 | 1 | 9 | Ikbke | 0.01 | Young |
| 0.268415 | 2.590182 | 0.015 | 0.005 | 1 | 9 | Gas8 | 0.01 | Young |
| 0.268415 | 2.49365 | 0.015 | 0.005 | 1 | 9 | Zfp955b | 0.01 | Young |
| 0.268415 | 2.443477 | 0.015 | 0.005 | 1 | 9 | Gm39469 | 0.01 | Young |
| 0.268415 | 2.271913 | 0.015 | 0.005 | 1 | 9 | Mc1r | 0.01 | Young |
| 0.268415 | 2.084564 | 0.015 | 0.005 | 1 | 9 | Gm43597 | 0.01 | Young |
| 0.268415 | 1.949945 | 0.015 | 0.005 | 1 | 9 | Gm26825 | 0.01 | Young |
| 0.268415 | 1.901809 | 0.015 | 0.005 | 1 | 9 | Fam177a | 0.01 | Young |
| 0.268415 | 1.766464 | 0.015 | 0.005 | 1 | 9 | Gm11716 | 0.01 | Young |
| 0.268415 | 1.662409 | 0.015 | 0.005 | 1 | 9 | Gm26652 | 0.01 | Young |
| 0.268415 | 1.309129 | 0.015 | 0.005 | 1 | 9 | Gm16104 | 0.01 | Young |
| 0.268415 | 1.219736 | 0.015 | 0.005 | 1 | 9 | Bub1b | 0.01 | Young |
| 0.268415 | 1.073575 | 0.015 | 0.005 | 1 | 9 | Naip6 | 0.01 | Young |
| 0.269026 | 0.719786 | 0.081 | 0.054 | 1 | 9 | Slc17a5 | 0.027 | Young |
| 0.269132 | 1.465104 | 0.132 | 0.101 | 1 | 9 | Stk24 | 0.031 | Young |
| 0.26955 | 0.24574 | 0.154 | 0.116 | 1 | 9 | Dctn3 | 0.038 | Young |
| 0.269615 | 0.636956 | 0.235 | 0.191 | 1 | 9 | Usp15 | 0.044 | Young |
| 0.269883 | 0.33211 | 0.118 | 0.088 | 1 | 9 | Mtpap | 0.03 | Young |
| 0.270205 | 0.343956 | 0.154 | 0.119 | 1 | 9 | Ddx3y | 0.035 | Young |
| 0.270249 | 0.674568 | 0.051 | 0.031 | 1 | 9 | Mmp9 | 0.02 | Young |
| 0.270249 | 0.603246 | 0.051 | 0.031 | 1 | 9 | Tbxas1 | 0.02 | Young |
| 0.270249 | 0.254908 | 0.051 | 0.031 | 1 | 9 | 2700049A( | 0.02 | Young |
| 0.270304 | 0.371604 | 0.25 | 0.209 | 1 | 9 | Dhx9 | 0.041 | Young |
| 0.270305 | 0.118272 | 0.199 | 0.25 | 1 | 9 | Ubxn4 | -0.051 | Young |
| 0.270308 | 2.433901 | 0.015 | 0.005 | 1 | 9 | Agt | 0.01 | Young |
| 0.270308 | 2.283278 | 0.015 | 0.005 | 1 | 9 | Exog | 0.01 | Young |
| 0.270308 | 1.788281 | 0.015 | 0.005 | 1 | 9 | Slc25a26 | 0.01 | Young |
| 0.270308 | 1.736656 | 0.015 | 0.005 | 1 | 9 | Exd1 | 0.01 | Young |
| 0.270308 | 1.736137 | 0.015 | 0.005 | 1 | 9 | Kif20b | 0.01 | Young |
| 0.270308 | 1.694466 | 0.015 | 0.005 | 1 | 9 | Cass4 | 0.01 | Young |
| 0.270308 | 1.599649 | 0.015 | 0.005 | 1 | 9 | Ppif | 0.01 | Young |
| 0.270308 | 1.584889 | 0.015 | 0.005 | 1 | 9 | Col11a2 | 0.01 | Young |
| 0.270308 | 1.565777 | 0.015 | 0.005 | 1 | 9 | 4930539J0 | 0.01 | Young |
| 0.270308 | 1.514505 | 0.015 | 0.005 | 1 | 9 | Pcdhga7 | 0.01 | Young |
| 0.270308 | 1.425434 | 0.015 | 0.005 | 1 | 9 | Rbp1 | 0.01 | Young |
| 0.270308 | 1.409132 | 0.015 | 0.005 | 1 | 9 | Vcpkmt | 0.01 | Young |
| 0.270308 | 1.357157 | 0.015 | 0.005 | 1 | 9 | G0s2 | 0.01 | Young |
| 0.270308 | 1.309238 | 0.015 | 0.005 | 1 | 9 | Gm35339 | 0.01 | Young |

| | | | | | | | | |
|---|---|---|---|---|---|---|---|---|
| 0.270308 | 1.292632 | 0.015 | 0.005 | 1 | 9 | Rad51c | 0.01 | Young |
| 0.270308 | 1.19911 | 0.015 | 0.005 | 1 | 9 | Lpar1 | 0.01 | Young |
| 0.270308 | 1.193923 | 0.015 | 0.005 | 1 | 9 | Gm12758 | 0.01 | Young |
| 0.270402 | 0.527103 | 0.081 | 0.054 | 1 | 9 | Vezt | 0.027 | Young |
| 0.270441 | 0.730252 | 0.074 | 0.049 | 1 | 9 | Ubl7 | 0.025 | Young |
| 0.270954 | 0.633511 | 0.11 | 0.08 | 1 | 9 | Wdr48 | 0.03 | Young |
| 0.271039 | 0.210021 | 0.154 | 0.116 | 1 | 9 | Mcts1 | 0.038 | Young |
| 0.271049 | 0.26013 | 0.279 | 0.237 | 1 | 9 | Epb41 | 0.042 | Young |
| 0.271928 | 0.166112 | 0.059 | 0.09 | 1 | 9 | Cd320 | -0.031 | Young |
| 0.272019 | 1.167155 | 0.051 | 0.031 | 1 | 9 | Sulf2 | 0.02 | Young |
| 0.272209 | 2.108117 | 0.015 | 0.005 | 1 | 9 | Sod3 | 0.01 | Young |
| 0.272209 | 1.882447 | 0.015 | 0.005 | 1 | 9 | Orai2 | 0.01 | Young |
| 0.272209 | 1.74777 | 0.015 | 0.005 | 1 | 9 | Mcemp1 | 0.01 | Young |
| 0.272209 | 1.460849 | 0.015 | 0.005 | 1 | 9 | AC163354. | 0.01 | Young |
| 0.272209 | 1.30483 | 0.015 | 0.005 | 1 | 9 | Guca1b | 0.01 | Young |
| 0.272209 | 1.298774 | 0.015 | 0.005 | 1 | 9 | Mamdc2 | 0.01 | Young |
| 0.272209 | 1.243784 | 0.015 | 0.005 | 1 | 9 | Fggy | 0.01 | Young |
| 0.272209 | 1.154177 | 0.015 | 0.005 | 1 | 9 | Gm49201 | 0.01 | Young |
| 0.272209 | 1.130737 | 0.015 | 0.005 | 1 | 9 | Cd27 | 0.01 | Young |
| 0.272209 | 1.121444 | 0.015 | 0.005 | 1 | 9 | Proser3 | 0.01 | Young |
| 0.272209 | 1.044024 | 0.015 | 0.005 | 1 | 9 | Gprc5c | 0.01 | Young |
| 0.272209 | 1.040291 | 0.015 | 0.005 | 1 | 9 | D11Wsu47 | 0.01 | Young |
| 0.272209 | 1.034448 | 0.015 | 0.005 | 1 | 9 | Acss3 | 0.01 | Young |
| 0.272209 | 1.03153 | 0.015 | 0.005 | 1 | 9 | D130017N( | 0.01 | Young |
| 0.272209 | 1.013778 | 0.015 | 0.005 | 1 | 9 | Pcsk5 | 0.01 | Young |
| 0.272209 | 0.861483 | 0.015 | 0.005 | 1 | 9 | Zfp354c | 0.01 | Young |
| 0.272209 | 0.858432 | 0.015 | 0.005 | 1 | 9 | Hapln3 | 0.01 | Young |
| 0.272209 | 0.858177 | 0.015 | 0.005 | 1 | 9 | Flt3 | 0.01 | Young |
| 0.272209 | 0.858161 | 0.015 | 0.005 | 1 | 9 | Pon3 | 0.01 | Young |
| 0.272209 | 0.838911 | 0.015 | 0.005 | 1 | 9 | Zfp61 | 0.01 | Young |
| 0.272209 | 0.818562 | 0.015 | 0.005 | 1 | 9 | 2010320M | 0.01 | Young |
| 0.272209 | 0.789234 | 0.015 | 0.005 | 1 | 9 | Kif21b | 0.01 | Young |
| 0.272209 | 0.760607 | 0.015 | 0.005 | 1 | 9 | Fam83e | 0.01 | Young |
| 0.272209 | 0.730782 | 0.015 | 0.005 | 1 | 9 | Alg8 | 0.01 | Young |
| 0.272209 | 0.495592 | 0.015 | 0.005 | 1 | 9 | Prps1l3 | 0.01 | Young |
| 0.272209 | 0.435963 | 0.015 | 0.005 | 1 | 9 | Gm14167 | 0.01 | Young |
| 0.272209 | 0.261776 | 0.015 | 0.005 | 1 | 9 | Batf | 0.01 | Young |
| 0.272229 | 0.023303 | 0.184 | 0.139 | 1 | 9 | Tle4 | 0.045 | Young |
| 0.273559 | 0.19302 | 0.75 | 0.737 | 1 | 9 | Rgcc | 0.013 | Young |
| 0.2739 | 0.023797 | 0.176 | 0.222 | 1 | 9 | Gigyf1 | -0.046 | Young |
| 0.27412 | 0.955092 | 0.015 | 0.005 | 1 | 9 | Pcdhb7 | 0.01 | Young |
| 0.27412 | 0.783462 | 0.015 | 0.005 | 1 | 9 | Fzd7 | 0.01 | Young |
| 0.27412 | 0.621008 | 0.015 | 0.005 | 1 | 9 | Med18 | 0.01 | Young |
| 0.27412 | 0.618863 | 0.015 | 0.005 | 1 | 9 | Siglec1 | 0.01 | Young |
| 0.27412 | 0.571803 | 0.015 | 0.005 | 1 | 9 | Eno3 | 0.01 | Young |

| | | | | | | | | |
|---|---|---|---|---|---|---|---|---|
| 0.27412 | 0.57082 | 0.015 | 0.005 | 1 | 9 | Traip | 0.01 | Young |
| 0.27412 | 0.564344 | 0.015 | 0.005 | 1 | 9 | Pdzd4 | 0.01 | Young |
| 0.27412 | 0.494138 | 0.015 | 0.005 | 1 | 9 | Map7 | 0.01 | Young |
| 0.27412 | 0.466216 | 0.015 | 0.005 | 1 | 9 | Rbmxl2 | 0.01 | Young |
| 0.27412 | 0.408206 | 0.015 | 0.005 | 1 | 9 | Ccl11 | 0.01 | Young |
| 0.27412 | 0.275221 | 0.015 | 0.005 | 1 | 9 | A930024N1 | 0.01 | Young |
| 0.274668 | 0.298513 | 0.103 | 0.072 | 1 | 9 | Siva1 | 0.031 | Young |
| 0.274859 | 0.035229 | 0.059 | 0.036 | 1 | 9 | Pycard | 0.023 | Young |
| 0.275829 | 0.456556 | 0.044 | 0.026 | 1 | 9 | Sez6 | 0.018 | Young |
| 0.275955 | 0.448106 | 0.081 | 0.054 | 1 | 9 | Snhg20 | 0.027 | Young |
| 0.276041 | 0.581713 | 0.015 | 0.005 | 1 | 9 | Trmu | 0.01 | Young |
| 0.276041 | 0.400324 | 0.015 | 0.005 | 1 | 9 | Slc36a3os | 0.01 | Young |
| 0.276041 | 0.378471 | 0.015 | 0.005 | 1 | 9 | Ccnj | 0.01 | Young |
| 0.276041 | 0.309199 | 0.015 | 0.005 | 1 | 9 | Tsku | 0.01 | Young |
| 0.276041 | 0.284314 | 0.015 | 0.005 | 1 | 9 | 1700084E1 | 0.01 | Young |
| 0.276041 | 0.265069 | 0.015 | 0.005 | 1 | 9 | 2610524H0 | 0.01 | Young |
| 0.276041 | 0.26411 | 0.015 | 0.005 | 1 | 9 | Fcor | 0.01 | Young |
| 0.276041 | 0.255617 | 0.015 | 0.005 | 1 | 9 | Slamf8 | 0.01 | Young |
| 0.276041 | 0.248143 | 0.015 | 0.005 | 1 | 9 | Gm26735 | 0.01 | Young |
| 0.276041 | 0.232676 | 0.015 | 0.005 | 1 | 9 | Foxred2 | 0.01 | Young |
| 0.276041 | 0.186819 | 0.015 | 0.005 | 1 | 9 | 5830454E0 | 0.01 | Young |
| 0.276041 | 0.099632 | 0.015 | 0.005 | 1 | 9 | Zfp2 | 0.01 | Young |
| 0.276777 | 0.101553 | 0.051 | 0.08 | 1 | 9 | Pmm2 | -0.029 | Young |
| 0.276803 | 0.953914 | 0.044 | 0.026 | 1 | 9 | Eid3 | 0.018 | Young |
| 0.276962 | 0.659126 | 0.147 | 0.113 | 1 | 9 | Zc3h7b | 0.034 | Young |
| 0.277249 | 0.29984 | 0.147 | 0.111 | 1 | 9 | Tiparp | 0.036 | Young |
| 0.277377 | 0.620196 | 0.051 | 0.031 | 1 | 9 | Vps33a | 0.02 | Young |
| 0.277722 | 0.453877 | 0.096 | 0.067 | 1 | 9 | Tmem135 | 0.029 | Young |
| 0.27778 | 1.001216 | 0.044 | 0.026 | 1 | 9 | Dennd2c | 0.018 | Young |
| 0.278097 | 0.628504 | 0.199 | 0.157 | 1 | 9 | Riok3 | 0.042 | Young |
| 0.278193 | 0.386959 | 0.184 | 0.144 | 1 | 9 | Lamtor2 | 0.04 | Young |
| 0.278277 | 1.146188 | 0.051 | 0.031 | 1 | 9 | Samd1 | 0.02 | Young |
| 0.278887 | 0.222038 | 0.11 | 0.077 | 1 | 9 | Alg14 | 0.033 | Young |
| 0.279178 | 0.375055 | 0.051 | 0.031 | 1 | 9 | Faim | 0.02 | Young |
| 0.279235 | 0.639806 | 0.103 | 0.075 | 1 | 9 | Tyrobp | 0.028 | Young |
| 0.27974 | 0.865203 | 0.044 | 0.026 | 1 | 9 | Qtrt2 | 0.018 | Young |
| 0.280053 | 0.284963 | 0.449 | 0.43 | 1 | 9 | Calr | 0.019 | Young |
| 0.28017 | 1.025255 | 0.088 | 0.062 | 1 | 9 | Maf | 0.026 | Young |
| 0.280626 | 0.037254 | 0.184 | 0.139 | 1 | 9 | Lta4h | 0.045 | Young |
| 0.280679 | 1.087414 | 0.074 | 0.049 | 1 | 9 | Apobec1 | 0.025 | Young |
| 0.28078 | 0.564873 | 0.272 | 0.245 | 1 | 9 | Ier3 | 0.027 | Young |
| 0.280871 | 0.843303 | 0.14 | 0.108 | 1 | 9 | Lrrk1 | 0.032 | Young |
| 0.280988 | 0.72805 | 0.051 | 0.031 | 1 | 9 | Irf5 | 0.02 | Young |
| 0.280988 | 0.133971 | 0.051 | 0.031 | 1 | 9 | Usp30 | 0.02 | Young |
| 0.281018 | 0.388424 | 0.162 | 0.124 | 1 | 9 | Use1 | 0.038 | Young |

| | | | | | | | | |
|---|---|---|---|---|---|---|---|---|
| 0.281018 | 0.34842 | 0.162 | 0.124 | 1 | 9 | Trappc4 | 0.038 | Young |
| 0.281111 | 0.158693 | 0.272 | 0.229 | 1 | 9 | Bod1l | 0.043 | Young |
| 0.281113 | 0.971434 | 0.36 | 0.343 | 1 | 9 | Cd74 | 0.017 | Young |
| 0.281421 | 0.673315 | 0.074 | 0.049 | 1 | 9 | Plcb3 | 0.025 | Young |
| 0.281709 | 1.131182 | 0.044 | 0.026 | 1 | 9 | Rps6kc1 | 0.018 | Young |
| 0.281709 | 0.730574 | 0.044 | 0.026 | 1 | 9 | Acad11 | 0.018 | Young |
| 0.281709 | 0.710627 | 0.044 | 0.026 | 1 | 9 | Wdr12 | 0.018 | Young |
| 0.281894 | 0.020295 | 0.169 | 0.126 | 1 | 9 | Man2b1 | 0.043 | Young |
| 0.281896 | 0.19079 | 0.051 | 0.031 | 1 | 9 | Zbtb43 | 0.02 | Young |
| 0.281978 | 0.035623 | 0.301 | 0.363 | 1 | 9 | Mfge8 | -0.062 | Young |
| 0.282163 | 0.705042 | 0.074 | 0.049 | 1 | 9 | Plekha8 | 0.025 | Young |
| 0.282387 | 0.845357 | 0.221 | 0.18 | 1 | 9 | 9530068E0 | 0.041 | Young |
| 0.282698 | 0.920347 | 0.044 | 0.026 | 1 | 9 | Tfdp1 | 0.018 | Young |
| 0.282938 | 0.916179 | 0.103 | 0.075 | 1 | 9 | B3galnt2 | 0.028 | Young |
| 0.282967 | 0.495385 | 0.213 | 0.173 | 1 | 9 | Ubr1 | 0.04 | Young |
| 0.283075 | 0.199928 | 0.279 | 0.227 | 1 | 9 | Plekhg1 | 0.052 | Young |
| 0.283401 | 0.527544 | 0.11 | 0.08 | 1 | 9 | Cybb | 0.03 | Young |
| 0.283688 | 0.929464 | 0.044 | 0.026 | 1 | 9 | Zfp472 | 0.018 | Young |
| 0.283704 | 0.527572 | 0.162 | 0.126 | 1 | 9 | Frmd4b | 0.036 | Young |
| 0.2842 | 0.858459 | 0.088 | 0.062 | 1 | 9 | Fut8 | 0.026 | Young |
| 0.284607 | 0.851387 | 0.11 | 0.08 | 1 | 9 | Akt1s1 | 0.03 | Young |
| 0.284681 | 1.415044 | 0.044 | 0.026 | 1 | 9 | Nop9 | 0.018 | Young |
| 0.284681 | 0.762272 | 0.044 | 0.026 | 1 | 9 | Zdhhc16 | 0.018 | Young |
| 0.284681 | 0.291443 | 0.044 | 0.026 | 1 | 9 | Cd40 | 0.018 | Young |
| 0.285444 | 0.216662 | 0.581 | 0.546 | 1 | 9 | Rassf3 | 0.035 | Young |
| 0.285676 | 1.55435 | 0.044 | 0.026 | 1 | 9 | Efl1 | 0.018 | Young |
| 0.285676 | 1.103594 | 0.044 | 0.026 | 1 | 9 | Mettl17 | 0.018 | Young |
| 0.285676 | 0.297917 | 0.044 | 0.026 | 1 | 9 | Slc25a22 | 0.018 | Young |
| 0.285896 | 0.939884 | 0.074 | 0.049 | 1 | 9 | Mllt3 | 0.025 | Young |
| 0.285896 | 0.77526 | 0.074 | 0.049 | 1 | 9 | Bcat2 | 0.025 | Young |
| 0.286054 | 0.834198 | 0.154 | 0.124 | 1 | 9 | Epc1 | 0.03 | Young |
| 0.286647 | 0.492449 | 0.037 | 0.062 | 1 | 9 | Cradd | -0.025 | Young |
| 0.286724 | 0.130081 | 0.184 | 0.142 | 1 | 9 | Hdac2 | 0.042 | Young |
| 0.287146 | 0.261483 | 0.566 | 0.526 | 1 | 9 | Slc38a2 | 0.04 | Young |
| 0.287158 | 0.232474 | 0.125 | 0.09 | 1 | 9 | 2310011J0 | 0.035 | Young |
| 0.287444 | 0.086655 | 0.125 | 0.168 | 1 | 9 | Fbxl20 | -0.043 | Young |
| 0.287474 | 0.184878 | 0.265 | 0.216 | 1 | 9 | Acvr2a | 0.049 | Young |
| 0.288152 | 0.316782 | 0.074 | 0.049 | 1 | 9 | Diablo | 0.025 | Young |
| 0.288317 | 0.510746 | 0.169 | 0.134 | 1 | 9 | Rsl1d1 | 0.035 | Young |
| 0.288676 | 0.38544 | 0.044 | 0.026 | 1 | 9 | Ipo13 | 0.018 | Young |
| 0.288906 | 0.246438 | 0.037 | 0.062 | 1 | 9 | Fgfr1op | -0.025 | Young |
| 0.288906 | 0.170724 | 0.037 | 0.062 | 1 | 9 | Psmf1 | -0.025 | Young |
| 0.289463 | 0.182265 | 0.066 | 0.095 | 1 | 9 | Brap | -0.029 | Young |
| 0.289559 | 0.017458 | 0.176 | 0.134 | 1 | 9 | Zzef1 | 0.042 | Young |
| 0.289643 | 1.18219 | 0.037 | 0.021 | 1 | 9 | Basp1 | 0.016 | Young |

| | | | | | | | | |
|---|---|---|---|---|---|---|---|---|
| 0.290524 | 0.291877 | 0.257 | 0.219 | 1 | 9 | Mapre2 | 0.038 | Young |
| 0.29058 | 0.917951 | 0.118 | 0.088 | 1 | 9 | Mfap3 | 0.03 | Young |
| 0.290622 | 0.756642 | 0.228 | 0.191 | 1 | 9 | Rtraf | 0.037 | Young |
| 0.290685 | 0.371344 | 0.11 | 0.08 | 1 | 9 | Lrrc41 | 0.03 | Young |
| 0.290757 | 1.148875 | 0.037 | 0.021 | 1 | 9 | Bcdin3d | 0.016 | Young |
| 0.290757 | 0.839797 | 0.037 | 0.021 | 1 | 9 | Gpn3 | 0.016 | Young |
| 0.290843 | 0.169957 | 0.75 | 0.771 | 1 | 9 | Col4a1 | -0.021 | Young |
| 0.291436 | 0.913778 | 0.11 | 0.082 | 1 | 9 | Fam199x | 0.028 | Young |
| 0.291697 | 0.698512 | 0.044 | 0.026 | 1 | 9 | Paxx | 0.018 | Young |
| 0.29184 | 0.425624 | 0.287 | 0.242 | 1 | 9 | Ppp1r18 | 0.045 | Young |
| 0.291874 | 1.430343 | 0.037 | 0.021 | 1 | 9 | Hdac6 | 0.016 | Young |
| 0.292013 | 0.043578 | 0.051 | 0.031 | 1 | 9 | Apbb1ip | 0.02 | Young |
| 0.292095 | 0.069554 | 0.272 | 0.219 | 1 | 9 | Pik3c2a | 0.053 | Young |
| 0.292236 | 0.393996 | 0.169 | 0.131 | 1 | 9 | Emc2 | 0.038 | Young |
| 0.292369 | 0.506497 | 0.066 | 0.044 | 1 | 9 | Phyh | 0.022 | Young |
| 0.292709 | 1.37996 | 0.044 | 0.026 | 1 | 9 | R3hcc1 | 0.018 | Young |
| 0.292709 | 0.669897 | 0.044 | 0.026 | 1 | 9 | Gpx3 | 0.018 | Young |
| 0.292994 | 0.681836 | 0.037 | 0.021 | 1 | 9 | Pced1a | 0.016 | Young |
| 0.293168 | 1.350905 | 0.081 | 0.057 | 1 | 9 | Adam19 | 0.024 | Young |
| 0.293171 | 1.181897 | 0.066 | 0.044 | 1 | 9 | Cdk5 | 0.022 | Young |
| 0.293202 | 2.587918 | 0.029 | 0.015 | 1 | 9 | Dram1 | 0.014 | Young |
| 0.293461 | 0.719419 | 0.037 | 0.062 | 1 | 9 | Fam98a | -0.025 | Young |
| 0.294116 | 0.829962 | 0.037 | 0.021 | 1 | 9 | 1-Mar | 0.016 | Young |
| 0.294131 | 0.613534 | 0.118 | 0.088 | 1 | 9 | Phlda1 | 0.03 | Young |
| 0.294372 | 0.300043 | 0.221 | 0.18 | 1 | 9 | Emc1 | 0.041 | Young |
| 0.29448 | 2.596241 | 0.029 | 0.015 | 1 | 9 | Rasgrp4 | 0.014 | Young |
| 0.294739 | 0.44261 | 0.044 | 0.026 | 1 | 9 | H2-Q10 | 0.018 | Young |
| 0.294739 | 0.063688 | 0.044 | 0.026 | 1 | 9 | Gm17745 | 0.018 | Young |
| 0.294805 | 0.562098 | 0.14 | 0.108 | 1 | 9 | Gadd45b | 0.032 | Young |
| 0.295056 | 0.278346 | 0.257 | 0.209 | 1 | 9 | Atp5c1 | 0.048 | Young |
| 0.295242 | 1.296885 | 0.037 | 0.021 | 1 | 9 | Creld1 | 0.016 | Young |
| 0.295708 | 1.168788 | 0.11 | 0.082 | 1 | 9 | Rnf141 | 0.028 | Young |
| 0.295758 | 0.951375 | 0.044 | 0.026 | 1 | 9 | Pcyt2 | 0.018 | Young |
| 0.296191 | 0.188133 | 0.125 | 0.093 | 1 | 9 | Odf2 | 0.032 | Young |
| 0.29637 | 0.662166 | 0.037 | 0.021 | 1 | 9 | Etfrf1 | 0.016 | Young |
| 0.296525 | 0.763282 | 0.074 | 0.049 | 1 | 9 | Pde9a | 0.025 | Young |
| 0.296669 | 0.078277 | 0.103 | 0.137 | 1 | 9 | Dhx30 | -0.034 | Young |
| 0.2968 | 0.206337 | 0.103 | 0.075 | 1 | 9 | Nmd3 | 0.028 | Young |
| 0.297045 | 2.913118 | 0.029 | 0.015 | 1 | 9 | Col1a1 | 0.014 | Young |
| 0.297045 | 1.405998 | 0.029 | 0.015 | 1 | 9 | Rnaset2b | 0.014 | Young |
| 0.29752 | 0.037633 | 0.603 | 0.678 | 1 | 9 | Ndrg1 | -0.075 | Young |
| 0.297922 | 1.4885 | 0.022 | 0.01 | 1 | 9 | Zfp455 | 0.012 | Young |
| 0.298017 | 1.253741 | 0.066 | 0.044 | 1 | 9 | 1110008P1 | 0.022 | Young |
| 0.298333 | 1.406952 | 0.029 | 0.015 | 1 | 9 | Arntl2 | 0.014 | Young |
| 0.298369 | 0.99167 | 0.154 | 0.124 | 1 | 9 | Ehbp1l1 | 0.03 | Young |

| | | | | | | | | |
|---|---|---|---|---|---|---|---|---|
| 0.298462 | 0.291393 | 0.397 | 0.371 | 1 | 9 | Nfkbiz | 0.026 | Young |
| 0.298636 | 0.78544 | 0.037 | 0.021 | 1 | 9 | Coq9 | 0.016 | Young |
| 0.298636 | 0.567159 | 0.037 | 0.021 | 1 | 9 | Mocs1 | 0.016 | Young |
| 0.298829 | 0.877314 | 0.066 | 0.044 | 1 | 9 | Cox16 | 0.022 | Young |
| 0.298837 | 0.345885 | 0.074 | 0.049 | 1 | 9 | Cd55 | 0.025 | Young |
| 0.299461 | 2.448307 | 0.022 | 0.01 | 1 | 9 | F8 | 0.012 | Young |
| 0.299461 | 1.961057 | 0.022 | 0.01 | 1 | 9 | Tubg1 | 0.012 | Young |
| 0.299461 | 1.436664 | 0.022 | 0.01 | 1 | 9 | Mbd4 | 0.012 | Young |
| 0.299625 | 1.87564 | 0.029 | 0.015 | 1 | 9 | Loxl2 | 0.014 | Young |
| 0.299625 | 1.03804 | 0.029 | 0.015 | 1 | 9 | Fn3krp | 0.014 | Young |
| 0.299773 | 0.376175 | 0.037 | 0.021 | 1 | 9 | Sergef | 0.016 | Young |
| 0.299933 | 0.075226 | 0.103 | 0.142 | 1 | 9 | Nfyb | -0.039 | Young |
| 0.300114 | 0.227631 | 0.118 | 0.088 | 1 | 9 | Pacs2 | 0.03 | Young |
| 0.300459 | 0.737674 | 0.066 | 0.044 | 1 | 9 | Alg2 | 0.022 | Young |
| 0.300748 | 0.481461 | 0.096 | 0.067 | 1 | 9 | Eif2ak1 | 0.029 | Young |
| 0.300913 | 0.484492 | 0.037 | 0.021 | 1 | 9 | Relt | 0.016 | Young |
| 0.300921 | 1.197298 | 0.029 | 0.015 | 1 | 9 | Slc25a10 | 0.014 | Young |
| 0.300921 | 1.084038 | 0.029 | 0.015 | 1 | 9 | Ifi213 | 0.014 | Young |
| 0.301005 | 1.866782 | 0.022 | 0.01 | 1 | 9 | Gm30918 | 0.012 | Young |
| 0.301005 | 1.482909 | 0.022 | 0.01 | 1 | 9 | Harbi1 | 0.012 | Young |
| 0.301005 | 1.163022 | 0.022 | 0.01 | 1 | 9 | Gm11772 | 0.012 | Young |
| 0.301005 | 0.787402 | 0.022 | 0.01 | 1 | 9 | Pomc | 0.012 | Young |
| 0.301106 | 0.341589 | 0.066 | 0.093 | 1 | 9 | Smim26 | -0.027 | Young |
| 0.301498 | 0.322954 | 0.316 | 0.273 | 1 | 9 | Pim1 | 0.043 | Young |
| 0.302055 | 0.636544 | 0.037 | 0.021 | 1 | 9 | Aldh3b1 | 0.016 | Young |
| 0.302125 | 1.371694 | 0.059 | 0.039 | 1 | 9 | Slc35d1 | 0.02 | Young |
| 0.302221 | 1.735112 | 0.029 | 0.015 | 1 | 9 | Bace1 | 0.014 | Young |
| 0.302221 | 1.671518 | 0.029 | 0.015 | 1 | 9 | Fancf | 0.014 | Young |
| 0.302378 | 0.397369 | 0.118 | 0.085 | 1 | 9 | Ralgds | 0.033 | Young |
| 0.302554 | 2.67895 | 0.022 | 0.01 | 1 | 9 | Gpr183 | 0.012 | Young |
| 0.302554 | 1.419435 | 0.022 | 0.01 | 1 | 9 | Krba1 | 0.012 | Young |
| 0.302554 | 1.201915 | 0.022 | 0.01 | 1 | 9 | Smim1 | 0.012 | Young |
| 0.302554 | 0.968166 | 0.022 | 0.01 | 1 | 9 | Dpep2 | 0.012 | Young |
| 0.302554 | 0.243065 | 0.022 | 0.01 | 1 | 9 | Marveld2 | 0.012 | Young |
| 0.302779 | 0.139632 | 0.096 | 0.067 | 1 | 9 | Nudt13 | 0.029 | Young |
| 0.302848 | 0.262668 | 0.787 | 0.776 | 1 | 9 | Rps26 | 0.011 | Young |
| 0.302857 | 0.700321 | 0.125 | 0.095 | 1 | 9 | Tnfaip8 | 0.03 | Young |
| 0.302915 | 0.636123 | 0.066 | 0.044 | 1 | 9 | Gm16536 | 0.022 | Young |
| 0.303475 | 0.11371 | 0.816 | 0.791 | 1 | 9 | Rflnb | 0.025 | Young |
| 0.303524 | 1.4512 | 0.029 | 0.015 | 1 | 9 | Lyz1 | 0.014 | Young |
| 0.303524 | 1.445069 | 0.029 | 0.015 | 1 | 9 | A130010J1 | 0.014 | Young |
| 0.303524 | 1.424657 | 0.029 | 0.015 | 1 | 9 | Smim8 | 0.014 | Young |
| 0.303524 | 1.182781 | 0.029 | 0.015 | 1 | 9 | Cd1d1 | 0.014 | Young |
| 0.303524 | 1.13863 | 0.029 | 0.015 | 1 | 9 | Gm7160 | 0.014 | Young |
| 0.303524 | 0.88162 | 0.029 | 0.015 | 1 | 9 | Ddx20 | 0.014 | Young |

| | | | | | | | | |
|---|---|---|---|---|---|---|---|---|
| 0.303733 | 0.154141 | 0.11 | 0.08 | 1 | 9 | Cuedc2 | 0.03 | Young |
| 0.303898 | 0.677634 | 0.103 | 0.075 | 1 | 9 | Ube4a | 0.028 | Young |
| 0.304067 | 0.614902 | 0.081 | 0.057 | 1 | 9 | Pop7 | 0.024 | Young |
| 0.304109 | 1.427564 | 0.022 | 0.01 | 1 | 9 | Rasl11b | 0.012 | Young |
| 0.304109 | 1.361046 | 0.022 | 0.01 | 1 | 9 | Npl | 0.012 | Young |
| 0.304109 | 0.930556 | 0.022 | 0.01 | 1 | 9 | Apoo | 0.012 | Young |
| 0.304109 | 0.872974 | 0.022 | 0.01 | 1 | 9 | Mccc2 | 0.012 | Young |
| 0.304109 | 0.776107 | 0.022 | 0.01 | 1 | 9 | Rap1gap2 | 0.012 | Young |
| 0.304314 | 0.043795 | 0.926 | 0.93 | 1 | 9 | Eef1a1 | -0.004 | Young |
| 0.30435 | 0.219541 | 0.037 | 0.021 | 1 | 9 | Pdk3 | 0.016 | Young |
| 0.304489 | 0.426021 | 0.279 | 0.253 | 1 | 9 | Itpripl2 | 0.026 | Young |
| 0.30456 | 0.544417 | 0.066 | 0.044 | 1 | 9 | Glt8d1 | 0.022 | Young |
| 0.304738 | 0.799662 | 0.059 | 0.039 | 1 | 9 | Ss18l2 | 0.02 | Young |
| 0.304831 | 0.792214 | 0.029 | 0.015 | 1 | 9 | Tmem106c | 0.014 | Young |
| 0.305059 | 0.162863 | 0.074 | 0.049 | 1 | 9 | Ptprj | 0.025 | Young |
| 0.305378 | 0.525111 | 0.169 | 0.131 | 1 | 9 | Fam217b | 0.038 | Young |
| 0.305501 | 0.298236 | 0.037 | 0.021 | 1 | 9 | Lsm10 | 0.016 | Young |
| 0.305669 | 1.0929 | 0.022 | 0.01 | 1 | 9 | Alg13 | 0.012 | Young |
| 0.305669 | 0.899513 | 0.022 | 0.01 | 1 | 9 | Tmem216 | 0.012 | Young |
| 0.305669 | 0.756835 | 0.022 | 0.01 | 1 | 9 | Pnpo | 0.012 | Young |
| 0.305669 | 0.481309 | 0.022 | 0.01 | 1 | 9 | Tmem17 | 0.012 | Young |
| 0.305843 | 0.221292 | 0.074 | 0.049 | 1 | 9 | Arsb | 0.025 | Young |
| 0.305952 | 0.373745 | 0.103 | 0.137 | 1 | 9 | Mydgf | -0.034 | Young |
| 0.306058 | 1.19001 | 0.096 | 0.07 | 1 | 9 | Ranbp10 | 0.026 | Young |
| 0.306142 | 1.290582 | 0.029 | 0.015 | 1 | 9 | Il10 | 0.014 | Young |
| 0.306142 | 0.618446 | 0.029 | 0.015 | 1 | 9 | Arl10 | 0.014 | Young |
| 0.306312 | 0.008513 | 0.235 | 0.281 | 1 | 9 | Hspa9 | -0.046 | Young |
| 0.306751 | 0.300276 | 0.324 | 0.271 | 1 | 9 | Hdgf | 0.053 | Young |
| 0.307235 | 1.607882 | 0.022 | 0.01 | 1 | 9 | Hcst | 0.012 | Young |
| 0.307235 | 1.211466 | 0.022 | 0.01 | 1 | 9 | Itgb7 | 0.012 | Young |
| 0.307235 | 0.847931 | 0.022 | 0.01 | 1 | 9 | Cep78 | 0.012 | Young |
| 0.307235 | 0.7882 | 0.022 | 0.01 | 1 | 9 | Gpt | 0.012 | Young |
| 0.307235 | 0.599449 | 0.022 | 0.01 | 1 | 9 | Cd180 | 0.012 | Young |
| 0.307235 | 0.549965 | 0.022 | 0.01 | 1 | 9 | Dhx58 | 0.012 | Young |
| 0.307235 | 0.027118 | 0.022 | 0.01 | 1 | 9 | Unc13d | 0.012 | Young |
| 0.307399 | 0.402386 | 0.118 | 0.088 | 1 | 9 | Tbc1d14 | 0.03 | Young |
| 0.307456 | 1.589956 | 0.029 | 0.015 | 1 | 9 | Cmah | 0.014 | Young |
| 0.307456 | 1.435841 | 0.029 | 0.015 | 1 | 9 | Nkrf | 0.014 | Young |
| 0.307456 | 0.868298 | 0.029 | 0.015 | 1 | 9 | Ccdc126 | 0.014 | Young |
| 0.307456 | 0.833811 | 0.029 | 0.015 | 1 | 9 | Hexdc | 0.014 | Young |
| 0.307456 | 0.805401 | 0.029 | 0.015 | 1 | 9 | Traf3ip1 | 0.014 | Young |
| 0.307564 | 0.635417 | 0.125 | 0.095 | 1 | 9 | Eif2b5 | 0.03 | Young |
| 0.308072 | 0.352811 | 0.14 | 0.108 | 1 | 9 | Commd3 | 0.032 | Young |
| 0.308203 | 0.100106 | 0.037 | 0.062 | 1 | 9 | Rnf217 | -0.025 | Young |
| 0.308484 | 0.761607 | 0.029 | 0.052 | 1 | 9 | Zrsr1 | -0.023 | Young |

| | | | | | | | | |
|---|---|---|---|---|---|---|---|---|
| 0.308624 | 0.430723 | 0.118 | 0.088 | 1 | 9 | Ksr1 | 0.03 | Young |
| 0.308774 | 2.315514 | 0.029 | 0.015 | 1 | 9 | Dock11 | 0.014 | Young |
| 0.308774 | 1.661345 | 0.029 | 0.015 | 1 | 9 | Hba-a2 | 0.014 | Young |
| 0.308774 | 1.045236 | 0.029 | 0.015 | 1 | 9 | Atp5s | 0.014 | Young |
| 0.308805 | 1.071774 | 0.022 | 0.01 | 1 | 9 | Mylpf | 0.012 | Young |
| 0.308805 | 0.464851 | 0.022 | 0.01 | 1 | 9 | Arhgap39 | 0.012 | Young |
| 0.308805 | 0.290816 | 0.022 | 0.01 | 1 | 9 | Lilra6 | 0.012 | Young |
| 0.308805 | 0.200618 | 0.022 | 0.01 | 1 | 9 | Eef1akmt4 | 0.012 | Young |
| 0.309271 | 0.986882 | 0.118 | 0.09 | 1 | 9 | Strip1 | 0.028 | Young |
| 0.309293 | 0.307089 | 0.529 | 0.521 | 1 | 9 | Pfn1 | 0.008 | Young |
| 0.309569 | 0.10369 | 0.324 | 0.286 | 1 | 9 | Mycbp2 | 0.038 | Young |
| 0.309904 | 0.222113 | 0.147 | 0.113 | 1 | 9 | Zfp260 | 0.034 | Young |
| 0.310096 | 0.752005 | 0.029 | 0.015 | 1 | 9 | Nup210l | 0.014 | Young |
| 0.310096 | 0.055595 | 0.029 | 0.015 | 1 | 9 | Zdhhc24 | 0.014 | Young |
| 0.310287 | 1.234532 | 0.103 | 0.077 | 1 | 9 | Cry2 | 0.026 | Young |
| 0.310381 | 0.460512 | 0.022 | 0.01 | 1 | 9 | Tarbp1 | 0.012 | Young |
| 0.310381 | 0.458707 | 0.022 | 0.01 | 1 | 9 | Gm9856 | 0.012 | Young |
| 0.310381 | 0.115894 | 0.022 | 0.01 | 1 | 9 | Tacc3 | 0.012 | Young |
| 0.310381 | 0.0499 | 0.022 | 0.01 | 1 | 9 | Pdss1 | 0.012 | Young |
| 0.31053 | 0.480924 | 0.125 | 0.095 | 1 | 9 | Ndufa9 | 0.03 | Young |
| 0.310662 | 0.409983 | 0.272 | 0.232 | 1 | 9 | Smarcc2 | 0.04 | Young |
| 0.310811 | 0.754907 | 0.096 | 0.07 | 1 | 9 | Rad50 | 0.026 | Young |
| 0.311026 | 0.122255 | 0.022 | 0.041 | 1 | 9 | Helb | -0.019 | Young |
| 0.311401 | 0.414974 | 0.316 | 0.276 | 1 | 9 | Phactr2 | 0.04 | Young |
| 0.311422 | 0.462579 | 0.029 | 0.015 | 1 | 9 | Snx30 | 0.014 | Young |
| 0.311422 | 0.438031 | 0.029 | 0.015 | 1 | 9 | Abcg3 | 0.014 | Young |
| 0.311782 | 0.766026 | 0.059 | 0.039 | 1 | 9 | Mrps18a | 0.02 | Young |
| 0.311962 | 0.285132 | 0.022 | 0.01 | 1 | 9 | Sac3d1 | 0.012 | Young |
| 0.311962 | 0.28176 | 0.022 | 0.01 | 1 | 9 | Zfp692 | 0.012 | Young |
| 0.312004 | 0.249747 | 0.169 | 0.134 | 1 | 9 | Pqbp1 | 0.035 | Young |
| 0.312067 | 0.585209 | 0.088 | 0.062 | 1 | 9 | Cdc34 | 0.026 | Young |
| 0.312086 | 0.498569 | 0.029 | 0.015 | 1 | 9 | Polrmt | 0.014 | Young |
| 0.312299 | 0.733136 | 0.007 | 0.021 | 1 | 9 | Wdr53 | -0.014 | Young |
| 0.312299 | 0.22541 | 0.007 | 0.021 | 1 | 9 | Fastkd3 | -0.014 | Young |
| 0.312299 | 0.104531 | 0.007 | 0.021 | 1 | 9 | Wdr90 | -0.014 | Young |
| 0.312751 | 0.961912 | 0.029 | 0.015 | 1 | 9 | Aph1c | 0.014 | Young |
| 0.312751 | 0.624301 | 0.029 | 0.015 | 1 | 9 | Slc22a5 | 0.014 | Young |
| 0.313368 | 0.161984 | 0.221 | 0.271 | 1 | 9 | Eloc | -0.05 | Young |
| 0.313462 | 0.024997 | 0.287 | 0.237 | 1 | 9 | Ccdc50 | 0.05 | Young |
| 0.313506 | 0.256544 | 0.088 | 0.062 | 1 | 9 | Rnf149 | 0.026 | Young |
| 0.313549 | 0.067957 | 0.022 | 0.01 | 1 | 9 | Poll | 0.012 | Young |
| 0.313559 | 0.693244 | 0.059 | 0.039 | 1 | 9 | Fam60a | 0.02 | Young |
| 0.313947 | 0.434921 | 0.022 | 0.041 | 1 | 9 | Dus1l | -0.019 | Young |
| 0.314084 | 0.321892 | 0.029 | 0.015 | 1 | 9 | Gm20186 | 0.014 | Young |
| 0.314179 | 0.198609 | 0.118 | 0.088 | 1 | 9 | Snrpa1 | 0.03 | Young |

| | | | | | | | | |
|---|---|---|---|---|---|---|---|---|
| 0.314186 | 0.249172 | 0.037 | 0.059 | 1 | 9 | Fzd5 | -0.022 | Young |
| 0.314607 | 0.128197 | 0.154 | 0.119 | 1 | 9 | Ctdspl2 | 0.035 | Young |
| 0.314645 | 1.133502 | 0.088 | 0.064 | 1 | 9 | Tgfbi | 0.024 | Young |
| 0.315088 | 0.845117 | 0.103 | 0.075 | 1 | 9 | Eftud2 | 0.028 | Young |
| 0.315141 | 0.135457 | 0.022 | 0.01 | 1 | 9 | 9130008F2 | 0.012 | Young |
| 0.315232 | 0.587353 | 0.081 | 0.057 | 1 | 9 | Gyg | 0.024 | Young |
| 0.315343 | 1.518769 | 0.059 | 0.039 | 1 | 9 | Fbn1 | 0.02 | Young |
| 0.315395 | 0.592039 | 0.066 | 0.044 | 1 | 9 | Usp50 | 0.022 | Young |
| 0.315421 | 0.346588 | 0.029 | 0.015 | 1 | 9 | Col18a1 | 0.014 | Young |
| 0.315421 | 0.325996 | 0.029 | 0.015 | 1 | 9 | Gm26740 | 0.014 | Young |
| 0.31545 | 0.392565 | 0.147 | 0.113 | 1 | 9 | Pik3r3 | 0.034 | Young |
| 0.315961 | 0.31996 | 0.125 | 0.16 | 1 | 9 | Slmap | -0.035 | Young |
| 0.316685 | 0.168903 | 0.618 | 0.544 | 1 | 9 | Son | 0.074 | Young |
| 0.316762 | 0.261778 | 0.029 | 0.015 | 1 | 9 | Frg2f1 | 0.014 | Young |
| 0.317122 | 0.815951 | 0.125 | 0.095 | 1 | 9 | Pck2 | 0.03 | Young |
| 0.31776 | 0.324721 | 0.103 | 0.075 | 1 | 9 | Vamp4 | 0.028 | Young |
| 0.318196 | 0.338593 | 0.184 | 0.147 | 1 | 9 | Mbtd1 | 0.037 | Young |
| 0.318255 | 0.648779 | 0.081 | 0.057 | 1 | 9 | Orc2 | 0.024 | Young |
| 0.318338 | 0.580479 | 0.11 | 0.082 | 1 | 9 | Nfu1 | 0.028 | Young |
| 0.31843 | 0.643446 | 0.103 | 0.075 | 1 | 9 | Ddx39 | 0.028 | Young |
| 0.318982 | 0.545409 | 0.11 | 0.082 | 1 | 9 | Zfp646 | 0.028 | Young |
| 0.319089 | 0.250201 | 0.096 | 0.131 | 1 | 9 | C1d | -0.035 | Young |
| 0.319516 | 0.925976 | 0.015 | 0.031 | 1 | 9 | Nuak2 | -0.016 | Young |
| 0.319629 | 0.332933 | 0.066 | 0.044 | 1 | 9 | Pla2g15 | 0.022 | Young |
| 0.319802 | 0.195363 | 0.191 | 0.237 | 1 | 9 | Tbcb | -0.046 | Young |
| 0.319843 | 0.880428 | 0.022 | 0.041 | 1 | 9 | Ddah1 | -0.019 | Young |
| 0.320419 | 0.029547 | 0.066 | 0.095 | 1 | 9 | Mrps16 | -0.029 | Young |
| 0.320534 | 1.333845 | 0.081 | 0.057 | 1 | 9 | Fads1 | 0.024 | Young |
| 0.321726 | 0.25621 | 0.5 | 0.454 | 1 | 9 | Bsg | 0.046 | Young |
| 0.322059 | 1.007589 | 0.081 | 0.057 | 1 | 9 | Isy1 | 0.024 | Young |
| 0.322151 | 0.342806 | 0.309 | 0.273 | 1 | 9 | Sqstm1 | 0.036 | Young |
| 0.322188 | 0.218902 | 0.515 | 0.474 | 1 | 9 | Arf5 | 0.041 | Young |
| 0.322203 | 0.48295 | 0.221 | 0.183 | 1 | 9 | Eea1 | 0.038 | Young |
| 0.322431 | 0.422857 | 0.324 | 0.281 | 1 | 9 | Srgap2 | 0.043 | Young |
| 0.322476 | 0.257834 | 0.132 | 0.101 | 1 | 9 | Svil | 0.031 | Young |
| 0.322867 | 0.895606 | 0.11 | 0.082 | 1 | 9 | Ttc37 | 0.028 | Young |
| 0.322924 | 0.002185 | 0.154 | 0.121 | 1 | 9 | Dgkd | 0.033 | Young |
| 0.323043 | 0.384822 | 0.066 | 0.044 | 1 | 9 | Npat | 0.022 | Young |
| 0.323086 | 0.093146 | 0.184 | 0.144 | 1 | 9 | Heca | 0.04 | Young |
| 0.323153 | 0.104059 | 0.412 | 0.474 | 1 | 9 | Rpl15 | -0.062 | Young |
| 0.323454 | 1.109645 | 0.059 | 0.039 | 1 | 9 | Itgav | 0.02 | Young |
| 0.32352 | 0.111061 | 0.029 | 0.015 | 1 | 9 | Pfkfb4 | 0.014 | Young |
| 0.3239 | 0.173076 | 0.066 | 0.044 | 1 | 9 | Fzr1 | 0.022 | Young |
| 0.324006 | 1.356413 | 0.088 | 0.064 | 1 | 9 | Kdm3b | 0.024 | Young |
| 0.32412 | 0.875958 | 0.015 | 0.031 | 1 | 9 | Gemin5 | -0.016 | Young |

| | | | | | | | | |
|---|---|---|---|---|---|---|---|---|
| 0.32412 | 0.730979 | 0.015 | 0.031 | 1 | 9 | Nr2c2ap | -0.016 | Young |
| 0.324215 | 0.506013 | 0.243 | 0.206 | 1 | 9 | Tuba1a | 0.037 | Young |
| 0.324428 | 1.391185 | 0.051 | 0.034 | 1 | 9 | Alg11 | 0.017 | Young |
| 0.324649 | 0.017468 | 0.206 | 0.26 | 1 | 9 | Arih1 | -0.054 | Young |
| 0.324849 | 0.312304 | 0.118 | 0.088 | 1 | 9 | Calcoco1 | 0.03 | Young |
| 0.324853 | 0.869749 | 0.191 | 0.16 | 1 | 9 | Wars | 0.031 | Young |
| 0.325277 | 0.280558 | 0.015 | 0.031 | 1 | 9 | B630019A1 | -0.016 | Young |
| 0.325277 | 0.12526 | 0.015 | 0.031 | 1 | 9 | Kyat3 | -0.016 | Young |
| 0.325402 | 1.267652 | 0.051 | 0.034 | 1 | 9 | Ddx56 | 0.017 | Young |
| 0.326062 | 0.380176 | 0.132 | 0.101 | 1 | 9 | Cdk2ap1 | 0.031 | Young |
| 0.326298 | 0.732283 | 0.074 | 0.052 | 1 | 9 | Papd7 | 0.022 | Young |
| 0.326618 | 0.069171 | 0.11 | 0.147 | 1 | 9 | Sirt1 | -0.037 | Young |
| 0.326756 | 0.139016 | 0.118 | 0.088 | 1 | 9 | Pdxdc1 | 0.03 | Young |
| 0.326812 | 0.819127 | 0.022 | 0.041 | 1 | 9 | Gm4951 | -0.019 | Young |
| 0.326983 | 0.528224 | 0.235 | 0.204 | 1 | 9 | Zbtb44 | 0.031 | Young |
| 0.327435 | 0.403456 | 0.081 | 0.057 | 1 | 9 | Med7 | 0.024 | Young |
| 0.327474 | 0.317278 | 0.096 | 0.07 | 1 | 9 | Sec16a | 0.026 | Young |
| 0.327912 | 0.783653 | 0.074 | 0.052 | 1 | 9 | Mrpl41 | 0.022 | Young |
| 0.328018 | 0.59374 | 0.059 | 0.039 | 1 | 9 | Chkb | 0.02 | Young |
| 0.328093 | 0.681491 | 0.11 | 0.082 | 1 | 9 | Bmi1 | 0.028 | Young |
| 0.328337 | 1.222752 | 0.051 | 0.034 | 1 | 9 | Mov10 | 0.017 | Young |
| 0.328592 | 0.728694 | 0.132 | 0.103 | 1 | 9 | Idh3g | 0.029 | Young |
| 0.328936 | 0.358278 | 0.059 | 0.039 | 1 | 9 | Iqgap2 | 0.02 | Young |
| 0.328938 | 0.24624 | 0.243 | 0.198 | 1 | 9 | Mllt10 | 0.045 | Young |
| 0.329406 | 0.575382 | 0.265 | 0.235 | 1 | 9 | Msl2 | 0.03 | Young |
| 0.329597 | 0.335311 | 0.096 | 0.07 | 1 | 9 | Hace1 | 0.026 | Young |
| 0.329611 | 0.281086 | 0.287 | 0.242 | 1 | 9 | Smdt1 | 0.045 | Young |
| 0.330343 | 0.226624 | 0.074 | 0.052 | 1 | 9 | Elof1 | 0.022 | Young |
| 0.33058 | 0.118854 | 0.081 | 0.111 | 1 | 9 | Prkag1 | -0.03 | Young |
| 0.330776 | 0.144905 | 0.059 | 0.039 | 1 | 9 | Abcb7 | 0.02 | Young |
| 0.330884 | 0.103721 | 0.132 | 0.101 | 1 | 9 | Lix1l | 0.031 | Young |
| 0.331155 | 0.832078 | 0.074 | 0.052 | 1 | 9 | Stk35 | 0.022 | Young |
| 0.331263 | 0.972657 | 0.147 | 0.121 | 1 | 9 | Rpl36al | 0.026 | Young |
| 0.33133 | 0.078683 | 0.051 | 0.077 | 1 | 9 | Rpp25l | -0.026 | Young |
| 0.331757 | 0.71548 | 0.125 | 0.098 | 1 | 9 | Ctss | 0.027 | Young |
| 0.332049 | 0.40046 | 0.11 | 0.082 | 1 | 9 | Scrib | 0.028 | Young |
| 0.332089 | 0.131905 | 0.044 | 0.07 | 1 | 9 | Cxxc1 | -0.026 | Young |
| 0.332522 | 0.500909 | 0.324 | 0.296 | 1 | 9 | Cggbp1 | 0.028 | Young |
| 0.333226 | 0.556361 | 0.037 | 0.059 | 1 | 9 | Manea | -0.022 | Young |
| 0.333396 | 0.060755 | 0.272 | 0.34 | 1 | 9 | Hrct1 | -0.068 | Young |
| 0.334007 | 0.045406 | 0.169 | 0.214 | 1 | 9 | Fbxw2 | -0.045 | Young |
| 0.334039 | 0.376991 | 0.11 | 0.082 | 1 | 9 | Thap2 | 0.028 | Young |
| 0.334477 | 0.294944 | 0.059 | 0.039 | 1 | 9 | Tsc2 | 0.02 | Young |
| 0.334482 | 0.088511 | 0.162 | 0.206 | 1 | 9 | Rab22a | -0.044 | Young |
| 0.334916 | 0.141871 | 0.037 | 0.059 | 1 | 9 | Mrps25 | -0.022 | Young |

| | | | | | | | | |
|---|---|---|---|---|---|---|---|---|
| 0.335179 | 0.601939 | 0.14 | 0.108 | 1 | 9 | Cpsf2 | 0.032 | Young |
| 0.335252 | 0.764042 | 0.051 | 0.034 | 1 | 9 | Epsti1 | 0.017 | Young |
| 0.335407 | 2.079046 | 0.059 | 0.039 | 1 | 9 | Fgl2 | 0.02 | Young |
| 0.335643 | 0.335171 | 0.287 | 0.25 | 1 | 9 | Ankrd17 | 0.037 | Young |
| 0.335927 | 0.23905 | 0.243 | 0.201 | 1 | 9 | Dnajb1 | 0.042 | Young |
| 0.336232 | 0.067631 | 0.309 | 0.361 | 1 | 9 | Eif3e | -0.052 | Young |
| 0.336338 | 0.051721 | 0.059 | 0.039 | 1 | 9 | Yif1a | 0.02 | Young |
| 0.336539 | 0.280229 | 0.39 | 0.351 | 1 | 9 | Pdia6 | 0.039 | Young |
| 0.336712 | 0.922293 | 0.14 | 0.113 | 1 | 9 | Tlr3 | 0.027 | Young |
| 0.337116 | 0.471867 | 0.147 | 0.113 | 1 | 9 | Yipf6 | 0.034 | Young |
| 0.337477 | 0.527271 | 0.154 | 0.121 | 1 | 9 | 0610012G( | 0.033 | Young |
| 0.337701 | 0.387233 | 0.118 | 0.088 | 1 | 9 | Kifap3 | 0.03 | Young |
| 0.338004 | 0.166067 | 0.36 | 0.32 | 1 | 9 | Trir | 0.04 | Young |
| 0.338647 | 0.515241 | 0.301 | 0.271 | 1 | 9 | Ppp1ca | 0.03 | Young |
| 0.338705 | 0.023569 | 0.419 | 0.495 | 1 | 9 | Uqcrh | -0.076 | Young |
| 0.338986 | 0.312069 | 0.294 | 0.255 | 1 | 9 | Fam102b | 0.039 | Young |
| 0.339245 | 1.333769 | 0.051 | 0.034 | 1 | 9 | Tecpr2 | 0.017 | Young |
| 0.339245 | 0.841352 | 0.051 | 0.034 | 1 | 9 | Wee1 | 0.017 | Young |
| 0.339354 | 1.404292 | 0.074 | 0.052 | 1 | 9 | Tmem250- | 0.022 | Young |
| 0.339354 | 0.478933 | 0.074 | 0.052 | 1 | 9 | B4gat1 | 0.022 | Young |
| 0.339509 | 0.006927 | 0.051 | 0.077 | 1 | 9 | Dap3 | -0.026 | Young |
| 0.339537 | 0.092744 | 0.272 | 0.222 | 1 | 9 | Ube2l3 | 0.05 | Young |
| 0.340181 | 0.908715 | 0.074 | 0.052 | 1 | 9 | Tubb6 | 0.022 | Young |
| 0.340248 | 0.747344 | 0.051 | 0.034 | 1 | 9 | Dclre1c | 0.017 | Young |
| 0.341009 | 0.445668 | 0.074 | 0.052 | 1 | 9 | Exoc7 | 0.022 | Young |
| 0.341009 | 0.427104 | 0.074 | 0.052 | 1 | 9 | Lrig2 | 0.022 | Young |
| 0.341041 | 0.470258 | 0.191 | 0.157 | 1 | 9 | H2afj | 0.034 | Young |
| 0.341055 | 0.067717 | 0.191 | 0.152 | 1 | 9 | Tnfrsf10b | 0.039 | Young |
| 0.341118 | 0.085499 | 0.588 | 0.513 | 1 | 9 | Atrx | 0.075 | Young |
| 0.341152 | 0.123606 | 0.059 | 0.085 | 1 | 9 | Bpnt1 | -0.026 | Young |
| 0.341195 | 0.368273 | 0.162 | 0.129 | 1 | 9 | Ndufs1 | 0.033 | Young |
| 0.341262 | 0.550144 | 0.162 | 0.131 | 1 | 9 | Dnajb11 | 0.031 | Young |
| 0.341709 | 0.222421 | 0.287 | 0.242 | 1 | 9 | Uhrf2 | 0.045 | Young |
| 0.341753 | 0.158144 | 0.162 | 0.129 | 1 | 9 | Stox2 | 0.033 | Young |
| 0.341861 | 0.143675 | 0.125 | 0.095 | 1 | 9 | Ate1 | 0.03 | Young |
| 0.342275 | 0.454843 | 0.074 | 0.103 | 1 | 9 | AW549877 | -0.029 | Young |
| 0.342475 | 0.604288 | 0.191 | 0.162 | 1 | 9 | Pcna | 0.029 | Young |
| 0.342587 | 0.614044 | 0.096 | 0.072 | 1 | 9 | Edc3 | 0.024 | Young |
| 0.342817 | 0.10503 | 0.515 | 0.585 | 1 | 9 | Apold1 | -0.07 | Young |
| 0.342902 | 0.116718 | 0.029 | 0.049 | 1 | 9 | Nol10 | -0.02 | Young |
| 0.343269 | 0.402981 | 0.088 | 0.064 | 1 | 9 | Chpf2 | 0.024 | Young |
| 0.343269 | 0.858178 | 0.088 | 0.064 | 1 | 9 | 1110059E2 | 0.024 | Young |
| 0.343847 | 0.234125 | 0.059 | 0.039 | 1 | 9 | Polr3c | 0.02 | Young |
| 0.344045 | 0.265634 | 0.301 | 0.263 | 1 | 9 | Kdm7a | 0.038 | Young |
| 0.344278 | 0.764722 | 0.051 | 0.034 | 1 | 9 | Heatr1 | 0.017 | Young |

| | | | | | | | | |
|---|---|---|---|---|---|---|---|---|
| 0.34478 | 0.528889 | 0.088 | 0.064 | 1 | 9 | Esco1 | 0.024 | Young |
| 0.344793 | 0.003378 | 0.059 | 0.039 | 1 | 9 | Pgs1 | 0.02 | Young |
| 0.344991 | 0.156833 | 0.044 | 0.067 | 1 | 9 | Timm22 | -0.023 | Young |
| 0.345267 | 0.62049 | 0.118 | 0.09 | 1 | 9 | Ppp5c | 0.028 | Young |
| 0.345291 | 1.476638 | 0.051 | 0.034 | 1 | 9 | Tnk2 | 0.017 | Young |
| 0.34574 | 0.024511 | 0.029 | 0.049 | 1 | 9 | Sirt5 | -0.02 | Young |
| 0.346005 | 0.156109 | 0.074 | 0.052 | 1 | 9 | Prcc | 0.022 | Young |
| 0.346005 | 0.075058 | 0.074 | 0.052 | 1 | 9 | Dnajc9 | 0.022 | Young |
| 0.3467 | 0.15436 | 0.221 | 0.178 | 1 | 9 | Tspan31 | 0.043 | Young |
| 0.347067 | 0.329176 | 0.235 | 0.198 | 1 | 9 | Fam117b | 0.037 | Young |
| 0.347157 | 0.301101 | 0.11 | 0.082 | 1 | 9 | Prrg2 | 0.028 | Young |
| 0.347366 | 0.61917 | 0.162 | 0.129 | 1 | 9 | Zmym4 | 0.033 | Young |
| 0.347477 | 0.359339 | 0.132 | 0.103 | 1 | 9 | Smurf2 | 0.029 | Young |
| 0.347545 | 0.103563 | 0.912 | 0.933 | 1 | 9 | Tpt1 | -0.021 | Young |
| 0.347648 | 0.04761 | 0.162 | 0.201 | 1 | 9 | Vamp2 | -0.039 | Young |
| 0.347681 | 0.477428 | 0.074 | 0.052 | 1 | 9 | Mphosph9 | 0.022 | Young |
| 0.347731 | 0.202434 | 0.412 | 0.384 | 1 | 9 | Cox7a2l | 0.028 | Young |
| 0.34868 | 0.057789 | 0.831 | 0.794 | 1 | 9 | Ybx1 | 0.037 | Young |
| 0.349047 | 0.427353 | 0.301 | 0.271 | 1 | 9 | Prpf40a | 0.03 | Young |
| 0.350649 | 0.56418 | 0.316 | 0.291 | 1 | 9 | Ddx39b | 0.025 | Young |
| 0.350911 | 0.818693 | 0.14 | 0.111 | 1 | 9 | Znhit1 | 0.029 | Young |
| 0.351027 | 0.602437 | 0.184 | 0.155 | 1 | 9 | Brox | 0.029 | Young |
| 0.351122 | 0.300077 | 0.103 | 0.142 | 1 | 9 | Rassf9 | -0.039 | Young |
| 0.351282 | 0.335682 | 0.059 | 0.085 | 1 | 9 | Foxc1 | -0.026 | Young |
| 0.351405 | 0.507994 | 0.051 | 0.034 | 1 | 9 | 2810454H( | 0.017 | Young |
| 0.351696 | 0.222226 | 0.301 | 0.26 | 1 | 9 | Wapl | 0.041 | Young |
| 0.351701 | 0.31705 | 0.324 | 0.281 | 1 | 9 | Hnrnpul1 | 0.043 | Young |
| 0.352431 | 1.313611 | 0.051 | 0.034 | 1 | 9 | Btbd19 | 0.017 | Young |
| 0.352483 | 0.45764 | 0.103 | 0.077 | 1 | 9 | Rara | 0.026 | Young |
| 0.353385 | 0.099453 | 0.853 | 0.809 | 1 | 9 | Zfp36 | 0.044 | Young |
| 0.353566 | 0.564836 | 0.147 | 0.119 | 1 | 9 | Mif4gd | 0.028 | Young |
| 0.353729 | 0.462086 | 0.228 | 0.198 | 1 | 9 | Btbd3 | 0.03 | Young |
| 0.353975 | 0.197344 | 0.463 | 0.554 | 1 | 9 | Tmem50a | -0.091 | Young |
| 0.354946 | 0.029682 | 0.294 | 0.242 | 1 | 9 | Abi1 | 0.052 | Young |
| 0.354949 | 0.454219 | 0.191 | 0.155 | 1 | 9 | Nucb1 | 0.036 | Young |
| 0.35578 | 0.036267 | 0.699 | 0.727 | 1 | 9 | Crip1 | -0.028 | Young |
| 0.357156 | 0.812353 | 0.096 | 0.072 | 1 | 9 | Sh3rf1 | 0.024 | Young |
| 0.357588 | 0.19961 | 0.051 | 0.034 | 1 | 9 | Cidec | 0.017 | Young |
| 0.357894 | 0.694705 | 0.096 | 0.072 | 1 | 9 | Pvr | 0.024 | Young |
| 0.358466 | 0.93908 | 0.066 | 0.046 | 1 | 9 | Spats2l | 0.02 | Young |
| 0.358625 | 0.06728 | 0.051 | 0.034 | 1 | 9 | Atg13 | 0.017 | Young |
| 0.358753 | 0.649784 | 0.132 | 0.103 | 1 | 9 | Rap1gds1 | 0.029 | Young |
| 0.359127 | 0.00943 | 0.059 | 0.082 | 1 | 9 | Stk4 | -0.023 | Young |
| 0.359303 | 1.557675 | 0.044 | 0.028 | 1 | 9 | Pfas | 0.016 | Young |
| 0.359303 | 0.624122 | 0.044 | 0.028 | 1 | 9 | Rac2 | 0.016 | Young |

| | | | | | | | | |
|---|---|---|---|---|---|---|---|---|
| 0.35935 | 0.432515 | 0.088 | 0.064 | 1 | 9 | Tmub2 | 0.024 | Young |
| 0.359556 | 0.349729 | 0.074 | 0.052 | 1 | 9 | Ppm1m | 0.022 | Young |
| 0.359851 | 0.069846 | 0.912 | 0.936 | 1 | 9 | Jund | -0.024 | Young |
| 0.360555 | 0.198597 | 0.485 | 0.466 | 1 | 9 | Atp2b1 | 0.019 | Young |
| 0.360806 | 0.038394 | 0.757 | 0.825 | 1 | 9 | Rps23 | -0.068 | Young |
| 0.360947 | 0.168172 | 0.632 | 0.603 | 1 | 9 | Ube2d3 | 0.029 | Young |
| 0.360972 | 0.225195 | 0.132 | 0.103 | 1 | 9 | Ireb2 | 0.029 | Young |
| 0.361273 | 0.212353 | 0.074 | 0.052 | 1 | 9 | Aspscr1 | 0.022 | Young |
| 0.361555 | 0.49255 | 0.044 | 0.028 | 1 | 9 | Bloc1s2 | 0.016 | Young |
| 0.362009 | 0.404087 | 0.022 | 0.039 | 1 | 9 | Nfatc2ip | -0.017 | Young |
| 0.362074 | 0.842127 | 0.066 | 0.046 | 1 | 9 | Smn1 | 0.02 | Young |
| 0.362133 | 0.679136 | 0.074 | 0.052 | 1 | 9 | Vps45 | 0.022 | Young |
| 0.362466 | 0.729568 | 0.088 | 0.064 | 1 | 9 | Mrps6 | 0.024 | Young |
| 0.362471 | 0.873829 | 0.125 | 0.101 | 1 | 9 | Rusc1 | 0.024 | Young |
| 0.362523 | 0.291713 | 0.147 | 0.119 | 1 | 9 | Ptov1 | 0.028 | Young |
| 0.362756 | 0.410442 | 0.529 | 0.526 | 1 | 9 | Pkp4 | 0.003 | Young |
| 0.36298 | 0.685014 | 0.066 | 0.046 | 1 | 9 | Slc7a6os | 0.02 | Young |
| 0.363254 | 0.970937 | 0.147 | 0.121 | 1 | 9 | 2610001J0 | 0.026 | Young |
| 0.363812 | 0.318754 | 0.162 | 0.134 | 1 | 9 | Myo9a | 0.028 | Young |
| 0.363816 | 1.227311 | 0.044 | 0.028 | 1 | 9 | Srr | 0.016 | Young |
| 0.363816 | 0.455908 | 0.044 | 0.028 | 1 | 9 | Zfp946 | 0.016 | Young |
| 0.363839 | 0.137964 | 0.051 | 0.034 | 1 | 9 | Letmd1 | 0.017 | Young |
| 0.364191 | 0.036986 | 0.522 | 0.598 | 1 | 9 | Ywhae | -0.076 | Young |
| 0.364315 | 0.572823 | 0.154 | 0.126 | 1 | 9 | Fam13c | 0.028 | Young |
| 0.364887 | 0.483408 | 0.051 | 0.034 | 1 | 9 | Tbc1d24 | 0.017 | Young |
| 0.365042 | 0.809565 | 0.029 | 0.049 | 1 | 9 | Pccb | -0.02 | Young |
| 0.365065 | 1.623019 | 0.059 | 0.041 | 1 | 9 | Exosc5 | 0.018 | Young |
| 0.365184 | 0.380145 | 0.235 | 0.201 | 1 | 9 | Furin | 0.034 | Young |
| 0.365681 | 0.106597 | 0.132 | 0.168 | 1 | 9 | Rbbp7 | -0.036 | Young |
| 0.365705 | 0.847653 | 0.066 | 0.046 | 1 | 9 | Trmt6 | 0.02 | Young |
| 0.365776 | 0.307207 | 0.154 | 0.124 | 1 | 9 | Zmynd8 | 0.03 | Young |
| 0.366616 | 0.539012 | 0.066 | 0.046 | 1 | 9 | Slc31a1 | 0.02 | Young |
| 0.366616 | 0.46638 | 0.066 | 0.046 | 1 | 9 | Nup155 | 0.02 | Young |
| 0.367009 | 0.233563 | 0.029 | 0.049 | 1 | 9 | Fam71a | -0.02 | Young |
| 0.367471 | 0.43904 | 0.162 | 0.129 | 1 | 9 | Plcb1 | 0.033 | Young |
| 0.367959 | 0.091274 | 0.088 | 0.064 | 1 | 9 | Dcaf6 | 0.024 | Young |
| 0.368073 | 1.177541 | 0.118 | 0.093 | 1 | 9 | Rcan1 | 0.025 | Young |
| 0.368365 | 1.379813 | 0.044 | 0.028 | 1 | 9 | Hmgn3 | 0.016 | Young |
| 0.368498 | 0.214497 | 0.184 | 0.147 | 1 | 9 | Ehmt2 | 0.037 | Young |
| 0.368747 | 0.231096 | 0.125 | 0.098 | 1 | 9 | Sec11c | 0.027 | Young |
| 0.368862 | 0.43602 | 0.213 | 0.18 | 1 | 9 | Coq10b | 0.033 | Young |
| 0.3691 | 0.264685 | 0.051 | 0.034 | 1 | 9 | Eef2kmt | 0.017 | Young |
| 0.369462 | 0.163782 | 0.191 | 0.155 | 1 | 9 | Senp1 | 0.036 | Young |
| 0.369508 | 0.673316 | 0.044 | 0.028 | 1 | 9 | Gdf15 | 0.016 | Young |
| 0.370231 | 0.128279 | 0.515 | 0.451 | 1 | 9 | Srsf2 | 0.064 | Young |

| | | | | | | | | |
|---|---|---|---|---|---|---|---|---|
| 0.370256 | 0.211496 | 0.088 | 0.119 | 1 | 9 | Csnk1g1 | -0.031 | Young |
| 0.370544 | 0.302605 | 0.154 | 0.124 | 1 | 9 | Srpk1 | 0.03 | Young |
| 0.370653 | 0.055354 | 0.044 | 0.028 | 1 | 9 | Snx20 | 0.016 | Young |
| 0.370666 | 0.002613 | 0.206 | 0.247 | 1 | 9 | Rictor | -0.041 | Young |
| 0.370808 | 0.004611 | 0.074 | 0.052 | 1 | 9 | Isg20l2 | 0.022 | Young |
| 0.371237 | 0.0056 | 0.059 | 0.082 | 1 | 9 | Jak3 | -0.023 | Young |
| 0.371802 | 0.475939 | 0.14 | 0.113 | 1 | 9 | Casd1 | 0.027 | Young |
| 0.37194 | 0.091165 | 0.132 | 0.106 | 1 | 9 | Cab39l | 0.026 | Young |
| 0.372108 | 1.556243 | 0.074 | 0.054 | 1 | 9 | Parp1 | 0.02 | Young |
| 0.372117 | 0.024827 | 0.551 | 0.621 | 1 | 9 | Ppp2r5a | -0.07 | Young |
| 0.372677 | 0.234885 | 0.103 | 0.077 | 1 | 9 | Ppp1r21 | 0.026 | Young |
| 0.372919 | 1.047931 | 0.103 | 0.082 | 1 | 9 | Gid4 | 0.021 | Young |
| 0.372948 | 0.435828 | 0.081 | 0.059 | 1 | 9 | Mbtps2 | 0.022 | Young |
| 0.372949 | 0.255542 | 0.044 | 0.028 | 1 | 9 | Fkbp1b | 0.016 | Young |
| 0.373072 | 0.185035 | 0.118 | 0.149 | 1 | 9 | Hoxb5 | -0.031 | Young |
| 0.373159 | 0.166133 | 0.228 | 0.196 | 1 | 9 | Srf | 0.032 | Young |
| 0.374101 | 0.412667 | 0.044 | 0.028 | 1 | 9 | Asns | 0.016 | Young |
| 0.374101 | 0.133354 | 0.044 | 0.028 | 1 | 9 | Narfl | 0.016 | Young |
| 0.374763 | 1.324621 | 0.059 | 0.041 | 1 | 9 | Kazn | 0.018 | Young |
| 0.375145 | 0.957434 | 0.096 | 0.072 | 1 | 9 | Ccnh | 0.024 | Young |
| 0.375396 | 0.471163 | 0.074 | 0.103 | 1 | 9 | Ptpn14 | -0.029 | Young |
| 0.375759 | 0.159471 | 0.096 | 0.124 | 1 | 9 | Hbegf | -0.028 | Young |
| 0.375906 | 0.698148 | 0.096 | 0.072 | 1 | 9 | Nabp2 | 0.024 | Young |
| 0.376214 | 0.06486 | 0.154 | 0.193 | 1 | 9 | Rgmb | -0.039 | Young |
| 0.376559 | 0.620428 | 0.154 | 0.124 | 1 | 9 | Rnf146 | 0.03 | Young |
| 0.37664 | 0.55801 | 0.375 | 0.348 | 1 | 9 | Selenof | 0.027 | Young |
| 0.376733 | 0.357059 | 0.066 | 0.046 | 1 | 9 | H2-T24 | 0.02 | Young |
| 0.376733 | 0.351756 | 0.066 | 0.046 | 1 | 9 | Pum3 | 0.02 | Young |
| 0.376766 | 0.659872 | 0.132 | 0.108 | 1 | 9 | Rbmx | 0.024 | Young |
| 0.377025 | 1.302401 | 0.088 | 0.067 | 1 | 9 | 6720427I0 | 0.021 | Young |
| 0.377097 | 0.212628 | 0.081 | 0.059 | 1 | 9 | Rufy2 | 0.022 | Young |
| 0.37754 | 0.876548 | 0.125 | 0.101 | 1 | 9 | Akr1c14 | 0.024 | Young |
| 0.377593 | 0.111191 | 0.022 | 0.039 | 1 | 9 | Rrp36 | -0.017 | Young |
| 0.377758 | 0.186334 | 0.14 | 0.175 | 1 | 9 | Armc8 | -0.035 | Young |
| 0.378313 | 0.241648 | 0.14 | 0.108 | 1 | 9 | 4833420G1 | 0.032 | Young |
| 0.378468 | 0.298264 | 0.228 | 0.193 | 1 | 9 | Aph1a | 0.035 | Young |
| 0.379142 | 0.142825 | 0.522 | 0.495 | 1 | 9 | Capzb | 0.027 | Young |
| 0.379496 | 2.724591 | 0.037 | 0.023 | 1 | 9 | Igfbp5 | 0.014 | Young |
| 0.379891 | 0.836411 | 0.044 | 0.028 | 1 | 9 | Ankrd54 | 0.016 | Young |
| 0.380009 | 0.090357 | 0.147 | 0.183 | 1 | 9 | Tmem65 | -0.036 | Young |
| 0.380152 | 0.008981 | 0.257 | 0.222 | 1 | 9 | Psmb9 | 0.035 | Young |
| 0.381388 | 0.372572 | 0.066 | 0.046 | 1 | 9 | Mboat2 | 0.02 | Young |
| 0.382106 | 0.219935 | 0.257 | 0.224 | 1 | 9 | Polr1d | 0.033 | Young |
| 0.382223 | 0.790681 | 0.044 | 0.028 | 1 | 9 | Tspan5 | 0.016 | Young |
| 0.382284 | 0.375617 | 0.169 | 0.139 | 1 | 9 | Dusp7 | 0.03 | Young |

| | | | | | | | | |
|---|---|---|---|---|---|---|---|---|
| 0.382954 | 0.228506 | 0.081 | 0.059 | 1 | 9 | Bcar3 | 0.022 | Young |
| 0.383244 | 0.233313 | 0.338 | 0.312 | 1 | 9 | Atp6v0b | 0.026 | Young |
| 0.383341 | 1.118204 | 0.037 | 0.023 | 1 | 9 | Tars2 | 0.014 | Young |
| 0.383392 | 0.095897 | 0.044 | 0.028 | 1 | 9 | Tmem218 | 0.016 | Young |
| 0.383423 | 0.996561 | 0.074 | 0.054 | 1 | 9 | Eif2b4 | 0.02 | Young |
| 0.383613 | 0.613768 | 0.132 | 0.106 | 1 | 9 | Bag6 | 0.026 | Young |
| 0.384038 | 0.442801 | 0.228 | 0.196 | 1 | 9 | Kidins220 | 0.032 | Young |
| 0.384079 | 0.082694 | 0.11 | 0.082 | 1 | 9 | Plekhm1 | 0.028 | Young |
| 0.384198 | 0.325038 | 0.037 | 0.057 | 1 | 9 | Zfp948 | -0.02 | Young |
| 0.384563 | 0.123308 | 0.044 | 0.028 | 1 | 9 | Decr2 | 0.016 | Young |
| 0.384637 | 0.501003 | 0.081 | 0.059 | 1 | 9 | Mrpl35 | 0.022 | Young |
| 0.384637 | 0.109736 | 0.081 | 0.059 | 1 | 9 | Ttc19 | 0.022 | Young |
| 0.384775 | 0.394796 | 0.125 | 0.098 | 1 | 9 | Ube4b | 0.027 | Young |
| 0.384982 | 0.832114 | 0.088 | 0.067 | 1 | 9 | Ezr | 0.021 | Young |
| 0.38515 | 0.126768 | 0.574 | 0.539 | 1 | 9 | Cpd | 0.035 | Young |
| 0.385773 | 0.127974 | 0.191 | 0.157 | 1 | 9 | Mia2 | 0.034 | Young |
| 0.385918 | 1.574877 | 0.037 | 0.023 | 1 | 9 | Gnb5 | 0.014 | Young |
| 0.385918 | 0.961016 | 0.037 | 0.023 | 1 | 9 | Ptcd1 | 0.014 | Young |
| 0.386451 | 0.169808 | 0.301 | 0.265 | 1 | 9 | Rlim | 0.036 | Young |
| 0.386667 | 0.094319 | 0.096 | 0.072 | 1 | 9 | Pih1d1 | 0.024 | Young |
| 0.386912 | 0.626705 | 0.044 | 0.028 | 1 | 9 | Tifa | 0.016 | Young |
| 0.38721 | 1.895303 | 0.037 | 0.023 | 1 | 9 | N4bp2 | 0.014 | Young |
| 0.387512 | 0.424999 | 0.059 | 0.082 | 1 | 9 | Pdgfd | -0.023 | Young |
| 0.388135 | 0.308482 | 0.228 | 0.196 | 1 | 9 | Ndufb3 | 0.032 | Young |
| 0.388279 | 0.339302 | 0.088 | 0.116 | 1 | 9 | Htatsf1 | -0.028 | Young |
| 0.388965 | 1.16024 | 0.007 | 0.018 | 1 | 9 | Top2a | -0.011 | Young |
| 0.388965 | 0.371834 | 0.007 | 0.018 | 1 | 9 | Bbs2 | -0.011 | Young |
| 0.388965 | 0.336781 | 0.007 | 0.018 | 1 | 9 | Cdc42bpg | -0.011 | Young |
| 0.388965 | 0.142852 | 0.007 | 0.018 | 1 | 9 | Pabpc1l | -0.011 | Young |
| 0.388965 | 0.133011 | 0.007 | 0.018 | 1 | 9 | Klhl36 | -0.011 | Young |
| 0.388965 | 0.104852 | 0.007 | 0.018 | 1 | 9 | Apol9b | -0.011 | Young |
| 0.389084 | 0.083995 | 0.059 | 0.082 | 1 | 9 | Gar1 | -0.023 | Young |
| 0.389369 | 0.390025 | 0.118 | 0.093 | 1 | 9 | Dnajc13 | 0.025 | Young |
| 0.389491 | 0.911706 | 0.147 | 0.124 | 1 | 9 | B4galt4 | 0.023 | Young |
| 0.389802 | 1.196748 | 0.037 | 0.023 | 1 | 9 | Gtf2h2 | 0.014 | Young |
| 0.390193 | 0.456301 | 0.132 | 0.106 | 1 | 9 | Upf3b | 0.026 | Young |
| 0.391102 | 0.880812 | 0.037 | 0.023 | 1 | 9 | Rpusd4 | 0.014 | Young |
| 0.391293 | 0.004788 | 0.213 | 0.178 | 1 | 9 | Jpt1 | 0.035 | Young |
| 0.3913 | 0.196838 | 0.103 | 0.077 | 1 | 9 | Mtmr11 | 0.026 | Young |
| 0.391421 | 1.375934 | 0.088 | 0.067 | 1 | 9 | Brms1l | 0.021 | Young |
| 0.391636 | 0.06724 | 0.044 | 0.028 | 1 | 9 | Tex30 | 0.016 | Young |
| 0.39231 | 1.328746 | 0.125 | 0.103 | 1 | 9 | Nploc4 | 0.022 | Young |
| 0.392405 | 1.327545 | 0.037 | 0.023 | 1 | 9 | Metrn | 0.014 | Young |
| 0.392405 | 1.094764 | 0.037 | 0.023 | 1 | 9 | Gm28151 | 0.014 | Young |
| 0.392405 | 0.924681 | 0.037 | 0.023 | 1 | 9 | Mthfsl | 0.014 | Young |

| | | | | | | | | |
|---|---|---|---|---|---|---|---|---|
| 0.392698 | 0.481754 | 0.147 | 0.121 | 1 | 9 | Napg | 0.026 | Young |
| 0.392873 | 0.211946 | 0.118 | 0.093 | 1 | 9 | Ipo11 | 0.025 | Young |
| 0.393498 | 0.68666 | 0.169 | 0.137 | 1 | 9 | Clptm1l | 0.032 | Young |
| 0.39371 | 0.895353 | 0.037 | 0.023 | 1 | 9 | Glcci1 | 0.014 | Young |
| 0.394004 | 0.114542 | 0.228 | 0.188 | 1 | 9 | Pias1 | 0.04 | Young |
| 0.394052 | 0.032805 | 0.044 | 0.064 | 1 | 9 | 0610010K1 | -0.02 | Young |
| 0.394363 | 0.014754 | 0.162 | 0.129 | 1 | 9 | Erc1 | 0.033 | Young |
| 0.394467 | 0.631967 | 0.096 | 0.072 | 1 | 9 | Aars | 0.024 | Young |
| 0.394832 | 0.320074 | 0.081 | 0.059 | 1 | 9 | Pisd | 0.022 | Young |
| 0.395018 | 1.220261 | 0.037 | 0.023 | 1 | 9 | Steap4 | 0.014 | Young |
| 0.395018 | 1.020309 | 0.037 | 0.023 | 1 | 9 | Supt7l | 0.014 | Young |
| 0.395018 | 0.633959 | 0.037 | 0.023 | 1 | 9 | Prim2 | 0.014 | Young |
| 0.395478 | 0.74296 | 0.088 | 0.067 | 1 | 9 | Nedd4l | 0.021 | Young |
| 0.395596 | 0.200037 | 0.257 | 0.219 | 1 | 9 | Ppp2r2a | 0.038 | Young |
| 0.395982 | 0.27412 | 0.147 | 0.116 | 1 | 9 | Orai1 | 0.031 | Young |
| 0.396329 | 0.494425 | 0.037 | 0.023 | 1 | 9 | Rhpn2 | 0.014 | Young |
| 0.397377 | 0.163832 | 0.103 | 0.077 | 1 | 9 | Pcbd2 | 0.026 | Young |
| 0.397635 | 0.588559 | 0.074 | 0.054 | 1 | 9 | Rce1 | 0.02 | Young |
| 0.398103 | 0.472857 | 0.11 | 0.088 | 1 | 9 | Ercc6l2 | 0.022 | Young |
| 0.398453 | 0.643803 | 0.125 | 0.098 | 1 | 9 | Psmg4 | 0.027 | Young |
| 0.398635 | 0.090068 | 0.206 | 0.173 | 1 | 9 | Qdpr | 0.033 | Young |
| 0.399191 | 0.220697 | 0.11 | 0.085 | 1 | 9 | Atl2 | 0.025 | Young |
| 0.399228 | 0.296657 | 0.118 | 0.093 | 1 | 9 | Psmd5 | 0.025 | Young |
| 0.39948 | 0.133546 | 0.088 | 0.113 | 1 | 9 | Hivep1 | -0.025 | Young |
| 0.399697 | 0.432949 | 0.059 | 0.041 | 1 | 9 | Atrnl1 | 0.018 | Young |
| 0.399823 | 0.605847 | 0.125 | 0.101 | 1 | 9 | Tnpo2 | 0.024 | Young |
| 0.399832 | 0.02182 | 0.029 | 0.046 | 1 | 9 | Pramef8 | -0.017 | Young |
| 0.400193 | 0.077125 | 0.779 | 0.845 | 1 | 9 | Rps24 | -0.066 | Young |
| 0.400277 | 0.695766 | 0.037 | 0.023 | 1 | 9 | Wscd1 | 0.014 | Young |
| 0.400527 | 0.199759 | 0.449 | 0.43 | 1 | 9 | Mtss1 | 0.019 | Young |
| 0.400632 | 0.27946 | 0.191 | 0.16 | 1 | 9 | Dicer1 | 0.031 | Young |
| 0.401038 | 0.017655 | 0.265 | 0.322 | 1 | 9 | Pxdc1 | -0.057 | Young |
| 0.401278 | 0.006226 | 0.228 | 0.273 | 1 | 9 | Cyp2d22 | -0.045 | Young |
| 0.401598 | 0.385437 | 0.037 | 0.023 | 1 | 9 | 2-Sep | 0.014 | Young |
| 0.401637 | 0.825254 | 0.081 | 0.062 | 1 | 9 | Bloc1s4 | 0.019 | Young |
| 0.401681 | 0.271139 | 0.375 | 0.332 | 1 | 9 | Eif4g1 | 0.043 | Young |
| 0.401888 | 0.464606 | 0.125 | 0.101 | 1 | 9 | Ubfd1 | 0.024 | Young |
| 0.402287 | 0.442241 | 0.037 | 0.057 | 1 | 9 | Tmem9 | -0.02 | Young |
| 0.40229 | 0.398792 | 0.375 | 0.353 | 1 | 9 | Gbp2 | 0.022 | Young |
| 0.402755 | 0.564887 | 0.059 | 0.041 | 1 | 9 | Pirb | 0.018 | Young |
| 0.403257 | 0.796572 | 0.088 | 0.067 | 1 | 9 | Med4 | 0.021 | Young |
| 0.4038 | 0.096424 | 0.39 | 0.348 | 1 | 9 | Efna1 | 0.042 | Young |
| 0.404226 | 0.380384 | 0.346 | 0.317 | 1 | 9 | Glud1 | 0.029 | Young |
| 0.404494 | 0.414735 | 0.088 | 0.067 | 1 | 9 | Ino80e | 0.021 | Young |
| 0.404494 | 0.764078 | 0.154 | 0.129 | 1 | 9 | Gngt2 | 0.025 | Young |

| | | | | | | | | |
|---|---|---|---|---|---|---|---|---|
| 0.40498 | 0.401305 | 0.221 | 0.186 | 1 | 9 | Ifi35 | 0.035 | Young |
| 0.405261 | 0.871299 | 0.103 | 0.08 | 1 | 9 | Strn | 0.023 | Young |
| 0.405577 | 0.569284 | 0.037 | 0.023 | 1 | 9 | Prkar2b | 0.014 | Young |
| 0.405577 | 0.533721 | 0.037 | 0.023 | 1 | 9 | Vps9d1 | 0.014 | Young |
| 0.405577 | 0.281526 | 0.037 | 0.023 | 1 | 9 | Tor4a | 0.014 | Young |
| 0.405577 | 0.222572 | 0.037 | 0.023 | 1 | 9 | Lrrc14 | 0.014 | Young |
| 0.405577 | 0.047283 | 0.037 | 0.023 | 1 | 9 | Sox6 | 0.014 | Young |
| 0.405675 | 0.086366 | 0.331 | 0.281 | 1 | 9 | Mlf2 | 0.05 | Young |
| 0.405777 | 0.449224 | 0.14 | 0.113 | 1 | 9 | Rcbtb2 | 0.027 | Young |
| 0.405809 | 0.116839 | 0.213 | 0.253 | 1 | 9 | Map3k11 | -0.04 | Young |
| 0.406791 | 0.61503 | 0.103 | 0.08 | 1 | 9 | Chn2 | 0.023 | Young |
| 0.40704 | 0.337542 | 0.176 | 0.147 | 1 | 9 | Lysmd3 | 0.029 | Young |
| 0.407937 | 0.013807 | 0.147 | 0.183 | 1 | 9 | Rgs5 | -0.036 | Young |
| 0.408156 | 0.05639 | 0.081 | 0.106 | 1 | 9 | Ddx60 | -0.025 | Young |
| 0.408246 | 0.603374 | 0.154 | 0.129 | 1 | 9 | Fam49b | 0.025 | Young |
| 0.408632 | 0.209683 | 0.088 | 0.067 | 1 | 9 | Ptprc | 0.021 | Young |
| 0.408815 | 0.354935 | 0.125 | 0.101 | 1 | 9 | Taf4 | 0.024 | Young |
| 0.409443 | 0.014177 | 0.029 | 0.046 | 1 | 9 | Pcgf2 | -0.017 | Young |
| 0.409579 | 0.206422 | 0.037 | 0.023 | 1 | 9 | Zscan12 | 0.014 | Young |
| 0.409622 | 0.010916 | 0.154 | 0.188 | 1 | 9 | Man1a | -0.034 | Young |
| 0.409737 | 0.388239 | 0.14 | 0.113 | 1 | 9 | Sidt2 | 0.027 | Young |
| 0.409837 | 0.232707 | 0.338 | 0.312 | 1 | 9 | Yy1 | 0.026 | Young |
| 0.410109 | 0.041437 | 0.14 | 0.111 | 1 | 9 | Nsf | 0.029 | Young |
| 0.41035 | 0.087473 | 0.096 | 0.072 | 1 | 9 | Rpa3 | 0.024 | Young |
| 0.41054 | 2.371004 | 0.029 | 0.018 | 1 | 9 | Pop1 | 0.011 | Young |
| 0.410918 | 0.13092 | 0.037 | 0.023 | 1 | 9 | Krt80 | 0.014 | Young |
| 0.410977 | 0.194502 | 0.059 | 0.041 | 1 | 9 | Tpgs2 | 0.018 | Young |
| 0.411242 | 1.157068 | 0.074 | 0.054 | 1 | 9 | Rapgef4 | 0.02 | Young |
| 0.411242 | 0.039734 | 0.044 | 0.064 | 1 | 9 | Zfp760 | -0.02 | Young |
| 0.411807 | 1.484181 | 0.066 | 0.049 | 1 | 9 | Zc2hc1a | 0.017 | Young |
| 0.412049 | 1.299327 | 0.029 | 0.018 | 1 | 9 | Paox | 0.011 | Young |
| 0.412159 | 0.089023 | 0.074 | 0.054 | 1 | 9 | Prkab2 | 0.02 | Young |
| 0.41226 | 0.663956 | 0.037 | 0.023 | 1 | 9 | 1-Sep | 0.014 | Young |
| 0.41226 | 0.179233 | 0.037 | 0.023 | 1 | 9 | Zfp7 | 0.014 | Young |
| 0.412795 | 0.284771 | 0.088 | 0.067 | 1 | 9 | Snrnp40 | 0.021 | Young |
| 0.412813 | 0.482054 | 0.11 | 0.088 | 1 | 9 | Usp37 | 0.022 | Young |
| 0.412897 | 0.131852 | 0.647 | 0.642 | 1 | 9 | Rhoa | 0.005 | Young |
| 0.413285 | 0.398387 | 0.154 | 0.126 | 1 | 9 | Ypel2 | 0.028 | Young |
| 0.413371 | 0.192461 | 0.176 | 0.216 | 1 | 9 | Lsm6 | -0.04 | Young |
| 0.413551 | 0.311576 | 0.184 | 0.152 | 1 | 9 | Alyref | 0.032 | Young |
| 0.413561 | 1.98524 | 0.029 | 0.018 | 1 | 9 | Them6 | 0.011 | Young |
| 0.413617 | 0.351138 | 0.191 | 0.157 | 1 | 9 | Krtcap2 | 0.034 | Young |
| 0.414101 | 0.332696 | 0.213 | 0.18 | 1 | 9 | Keap1 | 0.033 | Young |
| 0.414557 | 0.50577 | 0.154 | 0.126 | 1 | 9 | Aldh3a2 | 0.028 | Young |
| 0.414699 | 1.146942 | 0.066 | 0.049 | 1 | 9 | Dcaf15 | 0.017 | Young |

| | | | | | | | | |
|---|---|---|---|---|---|---|---|---|
| 0.415406 | 0.982257 | 0.051 | 0.036 | 1 | 9 | Zfp81 | 0.015 | Young |
| 0.415812 | 0.869839 | 0.125 | 0.101 | 1 | 9 | Rbm4b | 0.024 | Young |
| 0.415994 | 0.32484 | 0.316 | 0.289 | 1 | 9 | Raly | 0.027 | Young |
| 0.416431 | 0.254712 | 0.184 | 0.155 | 1 | 9 | Cat | 0.029 | Young |
| 0.416516 | 1.687707 | 0.051 | 0.036 | 1 | 9 | Rfng | 0.015 | Young |
| 0.416634 | 0.771834 | 0.066 | 0.049 | 1 | 9 | Nsmf | 0.017 | Young |
| 0.416829 | 0.542835 | 0.118 | 0.095 | 1 | 9 | Rfx7 | 0.023 | Young |
| 0.416835 | 0.120071 | 0.279 | 0.32 | 1 | 9 | Ndufa8 | -0.041 | Young |
| 0.416845 | 0.652273 | 0.103 | 0.082 | 1 | 9 | Zfp503 | 0.021 | Young |
| 0.417017 | 0.235695 | 0.228 | 0.196 | 1 | 9 | Vps13c | 0.032 | Young |
| 0.417436 | 0.167157 | 0.449 | 0.43 | 1 | 9 | Rap1b | 0.019 | Young |
| 0.417622 | 0.309561 | 0.294 | 0.26 | 1 | 9 | Clstn1 | 0.034 | Young |
| 0.417925 | 0.667262 | 0.125 | 0.101 | 1 | 9 | Phf8 | 0.024 | Young |
| 0.41833 | 0.343917 | 0.096 | 0.075 | 1 | 9 | Sipa1l3 | 0.021 | Young |
| 0.418729 | 0.286247 | 0.14 | 0.113 | 1 | 9 | Zcchc2 | 0.027 | Young |
| 0.418867 | 0.527828 | 0.162 | 0.137 | 1 | 9 | Magoh | 0.025 | Young |
| 0.418926 | 0.770463 | 0.081 | 0.062 | 1 | 9 | Hira | 0.019 | Young |
| 0.419134 | 1.099507 | 0.096 | 0.075 | 1 | 9 | Gnl1 | 0.021 | Young |
| 0.41919 | 0.337951 | 0.029 | 0.046 | 1 | 9 | Tcaim | -0.017 | Young |
| 0.419643 | 1.109978 | 0.029 | 0.018 | 1 | 9 | Lyrm1 | 0.011 | Young |
| 0.419643 | 0.995488 | 0.029 | 0.018 | 1 | 9 | Pou2f2 | 0.011 | Young |
| 0.41989 | 0.213204 | 0.235 | 0.201 | 1 | 9 | Cycs | 0.034 | Young |
| 0.420104 | 0.180842 | 0.11 | 0.085 | 1 | 9 | Abca7 | 0.025 | Young |
| 0.420186 | 0.210637 | 0.603 | 0.58 | 1 | 9 | Ptp4a2 | 0.023 | Young |
| 0.42052 | 1.413242 | 0.066 | 0.049 | 1 | 9 | Ly96 | 0.017 | Young |
| 0.42052 | 0.919233 | 0.066 | 0.049 | 1 | 9 | Hmces | 0.017 | Young |
| 0.42052 | 0.833945 | 0.066 | 0.049 | 1 | 9 | Mgl2 | 0.017 | Young |
| 0.42052 | 0.644539 | 0.066 | 0.049 | 1 | 9 | Plxnb2 | 0.017 | Young |
| 0.420973 | 1.512986 | 0.051 | 0.036 | 1 | 9 | Ap3m2 | 0.015 | Young |
| 0.421066 | 0.143894 | 0.184 | 0.147 | 1 | 9 | Arl4a | 0.037 | Young |
| 0.422436 | 1.154975 | 0.081 | 0.062 | 1 | 9 | Itih5 | 0.019 | Young |
| 0.422469 | 0.480414 | 0.029 | 0.046 | 1 | 9 | Zfyve1 | -0.017 | Young |
| 0.422703 | 0.935792 | 0.029 | 0.018 | 1 | 9 | Abhd11 | 0.011 | Young |
| 0.423429 | 0.111106 | 0.169 | 0.139 | 1 | 9 | Zyg11b | 0.03 | Young |
| 0.424197 | 0.569671 | 0.081 | 0.062 | 1 | 9 | Rhbdd1 | 0.019 | Young |
| 0.424238 | 0.830165 | 0.029 | 0.018 | 1 | 9 | Cd33 | 0.011 | Young |
| 0.424238 | 0.794753 | 0.029 | 0.018 | 1 | 9 | 4732440D( | 0.011 | Young |
| 0.424259 | 0.337063 | 0.272 | 0.33 | 1 | 9 | Dach1 | -0.058 | Young |
| 0.424335 | 0.6863 | 0.051 | 0.036 | 1 | 9 | Amdhd2 | 0.015 | Young |
| 0.424356 | 0.497155 | 0.228 | 0.198 | 1 | 9 | Atp5g1 | 0.03 | Young |
| 0.424793 | 0.232056 | 0.434 | 0.397 | 1 | 9 | Rab10 | 0.037 | Young |
| 0.424869 | 0.031328 | 0.206 | 0.168 | 1 | 9 | Elmsan1 | 0.038 | Young |
| 0.42495 | 0.199617 | 0.191 | 0.162 | 1 | 9 | Cox20 | 0.029 | Young |
| 0.425123 | 0.279156 | 0.074 | 0.054 | 1 | 9 | Slc35a5 | 0.02 | Young |
| 0.425123 | 0.247305 | 0.044 | 0.064 | 1 | 9 | Zfp398 | -0.02 | Young |

| | | | | | | | | |
|---|---|---|---|---|---|---|---|---|
| 0.425777 | 1.071548 | 0.029 | 0.018 | 1 | 9 | Fcna | 0.011 | Young |
| 0.425777 | 0.962398 | 0.029 | 0.018 | 1 | 9 | Reps2 | 0.011 | Young |
| 0.425943 | 0.057107 | 0.272 | 0.235 | 1 | 9 | Stard8 | 0.037 | Young |
| 0.426058 | 0.318737 | 0.074 | 0.054 | 1 | 9 | Kdelc2 | 0.02 | Young |
| 0.426469 | 0.237167 | 0.147 | 0.119 | 1 | 9 | Tlk2 | 0.028 | Young |
| 0.426585 | 0.720459 | 0.051 | 0.036 | 1 | 9 | Rfx5 | 0.015 | Young |
| 0.426738 | 0.685956 | 0.221 | 0.198 | 1 | 9 | Eif3m | 0.023 | Young |
| 0.427318 | 0.734999 | 0.029 | 0.018 | 1 | 9 | D2hgdh | 0.011 | Young |
| 0.427318 | 0.73379 | 0.029 | 0.018 | 1 | 9 | Zfp930 | 0.011 | Young |
| 0.427517 | 0.339952 | 0.118 | 0.144 | 1 | 9 | Ago3 | -0.026 | Young |
| 0.427712 | 0.970551 | 0.051 | 0.036 | 1 | 9 | Nadk2 | 0.015 | Young |
| 0.427712 | 0.555782 | 0.051 | 0.036 | 1 | 9 | Rnasel | 0.015 | Young |
| 0.42784 | 0.517606 | 0.11 | 0.088 | 1 | 9 | Usf1 | 0.022 | Young |
| 0.427932 | 0.075231 | 0.044 | 0.064 | 1 | 9 | 3222401L1 | -0.02 | Young |
| 0.427974 | 0.059128 | 0.14 | 0.175 | 1 | 9 | Sec24b | -0.035 | Young |
| 0.42816 | 0.419717 | 0.147 | 0.121 | 1 | 9 | Sap18 | 0.026 | Young |
| 0.428172 | 0.459208 | 0.022 | 0.036 | 1 | 9 | Zfp346 | -0.014 | Young |
| 0.428619 | 1.067266 | 0.081 | 0.062 | 1 | 9 | Rnf138 | 0.019 | Young |
| 0.428841 | 0.351264 | 0.051 | 0.036 | 1 | 9 | Ado | 0.015 | Young |
| 0.428863 | 0.493987 | 0.029 | 0.018 | 1 | 9 | Hacd1 | 0.011 | Young |
| 0.429018 | 0.225907 | 0.213 | 0.183 | 1 | 9 | Kdm5a | 0.03 | Young |
| 0.429321 | 0.192382 | 0.081 | 0.106 | 1 | 9 | Eif4e2 | -0.025 | Young |
| 0.429322 | 0.568739 | 0.14 | 0.116 | 1 | 9 | Tor1b | 0.024 | Young |
| 0.429405 | 0.175213 | 0.176 | 0.211 | 1 | 9 | Ist1 | -0.035 | Young |
| 0.430086 | 0.084855 | 0.14 | 0.17 | 1 | 9 | Smap2 | -0.03 | Young |
| 0.430251 | 0.340308 | 0.338 | 0.312 | 1 | 9 | Gas6 | 0.026 | Young |
| 0.430411 | 0.90294 | 0.029 | 0.018 | 1 | 9 | Smpd2 | 0.011 | Young |
| 0.430411 | 0.574254 | 0.029 | 0.018 | 1 | 9 | Tmem170 | 0.011 | Young |
| 0.430411 | 0.500942 | 0.029 | 0.018 | 1 | 9 | Zkscan5 | 0.011 | Young |
| 0.430627 | 0.05653 | 0.154 | 0.186 | 1 | 9 | Esyt1 | -0.032 | Young |
| 0.431727 | 0.308981 | 0.463 | 0.43 | 1 | 9 | Egln1 | 0.033 | Young |
| 0.431732 | 0.261846 | 0.184 | 0.152 | 1 | 9 | Crtc2 | 0.032 | Young |
| 0.431931 | 0.565327 | 0.022 | 0.036 | 1 | 9 | Pcbp4 | -0.014 | Young |
| 0.431963 | 0.740161 | 0.029 | 0.018 | 1 | 9 | Lca5 | 0.011 | Young |
| 0.432151 | 0.301261 | 0.059 | 0.082 | 1 | 9 | Gpaa1 | -0.023 | Young |
| 0.432176 | 0.221772 | 0.081 | 0.062 | 1 | 9 | Tmem63a | 0.019 | Young |
| 0.433188 | 0.331255 | 0.022 | 0.036 | 1 | 9 | Entpd5 | -0.014 | Young |
| 0.433188 | 0.128677 | 0.022 | 0.036 | 1 | 9 | Cep70 | -0.014 | Young |
| 0.433404 | 0.19202 | 0.169 | 0.142 | 1 | 9 | Sec13 | 0.027 | Young |
| 0.433517 | 0.3503 | 0.029 | 0.018 | 1 | 9 | Plcg2 | 0.011 | Young |
| 0.433766 | 0.499573 | 0.096 | 0.075 | 1 | 9 | Otub1 | 0.021 | Young |
| 0.434375 | 0.058606 | 0.059 | 0.08 | 1 | 9 | Usp42 | -0.021 | Young |
| 0.434448 | 0.197162 | 0.022 | 0.036 | 1 | 9 | Pde5a | -0.014 | Young |
| 0.434705 | 0.586385 | 0.11 | 0.088 | 1 | 9 | Akap7 | 0.022 | Young |
| 0.434959 | 0.41809 | 0.368 | 0.348 | 1 | 9 | Grn | 0.02 | Young |

| | | | | | | | | |
|---|---|---|---|---|---|---|---|---|
| 0.435075 | 0.509389 | 0.029 | 0.018 | 1 | 9 | Zmynd19 | 0.011 | Young |
| 0.435341 | 0.063384 | 0.037 | 0.054 | 1 | 9 | Iqcb1 | -0.017 | Young |
| 0.43541 | 0.528575 | 0.096 | 0.075 | 1 | 9 | Slc25a11 | 0.021 | Young |
| 0.43571 | 0.370812 | 0.022 | 0.036 | 1 | 9 | Kctd9 | -0.014 | Young |
| 0.435814 | 0.005277 | 0.537 | 0.564 | 1 | 9 | Rac1 | -0.027 | Young |
| 0.435843 | 0.589413 | 0.118 | 0.095 | 1 | 9 | Sec23ip | 0.023 | Young |
| 0.436371 | 0.094782 | 0.037 | 0.054 | 1 | 9 | Ppip5k1 | -0.017 | Young |
| 0.436422 | 0.130438 | 0.074 | 0.054 | 1 | 9 | Ndufaf2 | 0.02 | Young |
| 0.436636 | 0.023771 | 0.029 | 0.018 | 1 | 9 | Nrm | 0.011 | Young |
| 0.437158 | 1.265296 | 0.044 | 0.031 | 1 | 9 | Pde10a | 0.013 | Young |
| 0.437257 | 0.129785 | 0.243 | 0.278 | 1 | 9 | Tcf7l2 | -0.035 | Young |
| 0.437777 | 0.199933 | 0.11 | 0.088 | 1 | 9 | Usp36 | 0.022 | Young |
| 0.437937 | 0.428072 | 0.051 | 0.036 | 1 | 9 | Runx1 | 0.015 | Young |
| 0.438536 | 1.298958 | 0.074 | 0.057 | 1 | 9 | Slc35f5 | 0.017 | Young |
| 0.438708 | 0.579051 | 0.096 | 0.075 | 1 | 9 | Vopp1 | 0.021 | Young |
| 0.438763 | 0.112826 | 0.154 | 0.186 | 1 | 9 | Cdk2ap2 | -0.032 | Young |
| 0.438812 | 0.931 | 0.118 | 0.095 | 1 | 9 | Hacd4 | 0.023 | Young |
| 0.438812 | 0.525662 | 0.118 | 0.095 | 1 | 9 | Clcn7 | 0.023 | Young |
| 0.439317 | 0.320438 | 0.118 | 0.149 | 1 | 9 | Sarnp | -0.031 | Young |
| 0.439388 | 0.115466 | 0.118 | 0.093 | 1 | 9 | Prrg1 | 0.025 | Young |
| 0.43947 | 0.362768 | 0.037 | 0.054 | 1 | 9 | Rad51d | -0.017 | Young |
| 0.439768 | 0.059109 | 0.029 | 0.018 | 1 | 9 | Inpp5b | 0.011 | Young |
| 0.439824 | 0.030861 | 0.846 | 0.881 | 1 | 9 | Slfn5 | -0.035 | Young |
| 0.440226 | 0.010979 | 0.103 | 0.131 | 1 | 9 | Ankrd10 | -0.028 | Young |
| 0.440265 | 0.53229 | 0.066 | 0.049 | 1 | 9 | Cebpa | 0.017 | Young |
| 0.44044 | 0.008005 | 0.397 | 0.428 | 1 | 9 | G3bp1 | -0.031 | Young |
| 0.440584 | 0.456766 | 0.397 | 0.369 | 1 | 9 | Ptges3 | 0.028 | Young |
| 0.440777 | 0.31557 | 0.022 | 0.036 | 1 | 9 | Exoc3l2 | -0.014 | Young |
| 0.440857 | 0.343905 | 0.176 | 0.149 | 1 | 9 | Cep170 | 0.027 | Young |
| 0.441207 | 1.166242 | 0.088 | 0.07 | 1 | 9 | Rnh1 | 0.018 | Young |
| 0.442049 | 0.048289 | 0.022 | 0.036 | 1 | 9 | Cyb5d2 | -0.014 | Young |
| 0.442154 | 0.513551 | 0.397 | 0.369 | 1 | 9 | Sash1 | 0.028 | Young |
| 0.442176 | 0.61815 | 0.147 | 0.121 | 1 | 9 | Pex6 | 0.026 | Young |
| 0.442931 | 0.252133 | 0.213 | 0.242 | 1 | 9 | Zfand6 | -0.029 | Young |
| 0.443272 | 0.636176 | 0.066 | 0.049 | 1 | 9 | 1110004E0 | 0.017 | Young |
| 0.443336 | 1.762791 | 0.044 | 0.031 | 1 | 9 | Nfatc2 | 0.013 | Young |
| 0.443678 | 0.305729 | 0.051 | 0.036 | 1 | 9 | Zbtb6 | 0.015 | Young |
| 0.443852 | 0.572148 | 0.081 | 0.062 | 1 | 9 | Chst7 | 0.019 | Young |
| 0.443913 | 0.0766 | 0.404 | 0.472 | 1 | 9 | Rpl11 | -0.068 | Young |
| 0.444015 | 1.985328 | 0.022 | 0.013 | 1 | 9 | Mttp | 0.009 | Young |
| 0.444015 | 1.677672 | 0.022 | 0.013 | 1 | 9 | Unc5a | 0.009 | Young |
| 0.444015 | 0.899062 | 0.022 | 0.013 | 1 | 9 | Aaas | 0.009 | Young |
| 0.444025 | 0.412776 | 0.507 | 0.505 | 1 | 9 | Snrnp70 | 0.002 | Young |
| 0.444277 | 0.061341 | 0.147 | 0.119 | 1 | 9 | Inppl1 | 0.028 | Young |
| 0.444568 | 0.288386 | 0.176 | 0.149 | 1 | 9 | Snrpd2 | 0.027 | Young |

| | | | | | | | | |
|---|---|---|---|---|---|---|---|---|
| 0.444645 | 0.699526 | 0.088 | 0.07 | 1 | 9 | Wipi2 | 0.018 | Young |
| 0.444663 | 0.351678 | 0.037 | 0.054 | 1 | 9 | Gas1 | -0.017 | Young |
| 0.444663 | 0.008256 | 0.037 | 0.054 | 1 | 9 | Pawr | -0.017 | Young |
| 0.444799 | 0.559978 | 0.243 | 0.224 | 1 | 9 | Pcm1 | 0.019 | Young |
| 0.445085 | 0.51474 | 0.103 | 0.082 | 1 | 9 | Creb3 | 0.021 | Young |
| 0.445694 | 1.425703 | 0.096 | 0.077 | 1 | 9 | Hic2 | 0.019 | Young |
| 0.445862 | 2.221408 | 0.022 | 0.013 | 1 | 9 | Fam122b | 0.009 | Young |
| 0.445862 | 1.956783 | 0.022 | 0.013 | 1 | 9 | A630001G2 | 0.009 | Young |
| 0.445862 | 0.172288 | 0.022 | 0.013 | 1 | 9 | Gorasp1 | 0.009 | Young |
| 0.446303 | 0.313793 | 0.287 | 0.247 | 1 | 9 | Phf3 | 0.04 | Young |
| 0.446404 | 0.245971 | 0.103 | 0.129 | 1 | 9 | Gtf2h1 | -0.026 | Young |
| 0.446432 | 1.02469 | 0.154 | 0.139 | 1 | 9 | Cetn3 | 0.015 | Young |
| 0.44666 | 0.07925 | 0.147 | 0.119 | 1 | 9 | Xpo7 | 0.028 | Young |
| 0.447159 | 0.389293 | 0.022 | 0.036 | 1 | 9 | Cul4b | -0.014 | Young |
| 0.447159 | 0.225592 | 0.022 | 0.036 | 1 | 9 | Spindoc | -0.014 | Young |
| 0.447263 | 0.273776 | 0.368 | 0.338 | 1 | 9 | Psmb1 | 0.03 | Young |
| 0.447713 | 2.263962 | 0.022 | 0.013 | 1 | 9 | Rin1 | 0.009 | Young |
| 0.447713 | 1.51058 | 0.022 | 0.013 | 1 | 9 | Msto1 | 0.009 | Young |
| 0.447713 | 1.171326 | 0.022 | 0.013 | 1 | 9 | Gm48604 | 0.009 | Young |
| 0.447713 | 0.983048 | 0.022 | 0.013 | 1 | 9 | Atxn7l2 | 0.009 | Young |
| 0.447776 | 0.144063 | 0.191 | 0.162 | 1 | 9 | Xrn1 | 0.029 | Young |
| 0.4481 | 0.578995 | 0.088 | 0.07 | 1 | 9 | Kcnk6 | 0.018 | Young |
| 0.4481 | 0.242827 | 0.059 | 0.08 | 1 | 9 | Slc16a1 | -0.021 | Young |
| 0.44831 | 0.655963 | 0.066 | 0.049 | 1 | 9 | Mis18a | 0.017 | Young |
| 0.448314 | 1.998658 | 0.044 | 0.031 | 1 | 9 | Mpnd | 0.013 | Young |
| 0.448339 | 0.045725 | 0.213 | 0.253 | 1 | 9 | Pbx1 | -0.04 | Young |
| 0.448547 | 0.324847 | 0.375 | 0.351 | 1 | 9 | Nufip2 | 0.024 | Young |
| 0.448638 | 0.207595 | 0.096 | 0.121 | 1 | 9 | Gak | -0.025 | Young |
| 0.449001 | 1.116298 | 0.096 | 0.077 | 1 | 9 | Mvb12a | 0.019 | Young |
| 0.449321 | 0.131743 | 0.066 | 0.049 | 1 | 9 | Elmod2 | 0.017 | Young |
| 0.449569 | 1.292332 | 0.022 | 0.013 | 1 | 9 | Pqlc2 | 0.009 | Young |
| 0.449569 | 1.025415 | 0.022 | 0.013 | 1 | 9 | Zswim9 | 0.009 | Young |
| 0.449569 | 0.79994 | 0.022 | 0.013 | 1 | 9 | Arhgap11a | 0.009 | Young |
| 0.449569 | 0.696484 | 0.022 | 0.013 | 1 | 9 | Slc7a6 | 0.009 | Young |
| 0.449708 | 0.270305 | 0.096 | 0.121 | 1 | 9 | Lars | -0.025 | Young |
| 0.450815 | 1.703998 | 0.044 | 0.031 | 1 | 9 | Edn1 | 0.013 | Young |
| 0.450815 | 1.026798 | 0.044 | 0.031 | 1 | 9 | Hs1bp3 | 0.013 | Young |
| 0.45082 | 0.116838 | 0.206 | 0.242 | 1 | 9 | Ier3ip1 | -0.036 | Young |
| 0.451429 | 1.34547 | 0.022 | 0.013 | 1 | 9 | Gbp6 | 0.009 | Young |
| 0.451429 | 0.893129 | 0.022 | 0.013 | 1 | 9 | Pde7b | 0.009 | Young |
| 0.451429 | 0.855744 | 0.022 | 0.013 | 1 | 9 | Gm45605 | 0.009 | Young |
| 0.451429 | 0.850116 | 0.022 | 0.013 | 1 | 9 | Pole2 | 0.009 | Young |
| 0.451429 | 0.657317 | 0.022 | 0.013 | 1 | 9 | Epm2a | 0.009 | Young |
| 0.451429 | 0.580558 | 0.022 | 0.013 | 1 | 9 | AL590144.1 | 0.009 | Young |
| 0.451845 | 0.575593 | 0.125 | 0.101 | 1 | 9 | Plaa | 0.024 | Young |

| | | | | | | | | |
|---|---|---|---|---|---|---|---|---|
| 0.452068 | 1.444156 | 0.044 | 0.031 | 1 | 9 | Dtnbp1 | 0.013 | Young |
| 0.452302 | 0.500102 | 0.022 | 0.036 | 1 | 9 | Med6 | -0.014 | Young |
| 0.452311 | 0.552839 | 0.103 | 0.082 | 1 | 9 | Akt2 | 0.021 | Young |
| 0.452914 | 0.085439 | 0.154 | 0.186 | 1 | 9 | Gtf2i | -0.032 | Young |
| 0.452989 | 0.056615 | 0.471 | 0.436 | 1 | 9 | Eef1b2 | 0.035 | Young |
| 0.453253 | 0.031518 | 0.544 | 0.606 | 1 | 9 | Rps17 | -0.062 | Young |
| 0.453293 | 1.988008 | 0.022 | 0.013 | 1 | 9 | BC035044 | 0.009 | Young |
| 0.453293 | 1.803819 | 0.022 | 0.013 | 1 | 9 | Dcstamp | 0.009 | Young |
| 0.453293 | 1.040082 | 0.022 | 0.013 | 1 | 9 | Elp4 | 0.009 | Young |
| 0.453293 | 0.992629 | 0.022 | 0.013 | 1 | 9 | Rab30 | 0.009 | Young |
| 0.453293 | 0.274692 | 0.022 | 0.013 | 1 | 9 | Tyw3 | 0.009 | Young |
| 0.453313 | 0.028647 | 0.588 | 0.567 | 1 | 9 | Ddx17 | 0.021 | Young |
| 0.453505 | 0.603834 | 0.162 | 0.139 | 1 | 9 | Znrf2 | 0.023 | Young |
| 0.454118 | 0.34288 | 0.051 | 0.036 | 1 | 9 | Gpn2 | 0.015 | Young |
| 0.454198 | 0.361963 | 0.154 | 0.129 | 1 | 9 | Slc25a30 | 0.025 | Young |
| 0.455115 | 0.316524 | 0.074 | 0.095 | 1 | 9 | Grsf1 | -0.021 | Young |
| 0.455162 | 0.536418 | 0.022 | 0.013 | 1 | 9 | 2900076A( | 0.009 | Young |
| 0.455162 | 0.334326 | 0.022 | 0.013 | 1 | 9 | Tbc1d25 | 0.009 | Young |
| 0.455162 | 0.039776 | 0.022 | 0.013 | 1 | 9 | Adam8 | 0.009 | Young |
| 0.455593 | 0.772712 | 0.074 | 0.057 | 1 | 9 | Ank | 0.017 | Young |
| 0.455605 | 0.378385 | 0.221 | 0.186 | 1 | 9 | Rps6kb1 | 0.035 | Young |
| 0.455841 | 1.60191 | 0.044 | 0.031 | 1 | 9 | Pus1 | 0.013 | Young |
| 0.455927 | 0.511686 | 0.088 | 0.07 | 1 | 9 | Plekhf2 | 0.018 | Young |
| 0.456802 | 0.380576 | 0.088 | 0.07 | 1 | 9 | Coro1a | 0.018 | Young |
| 0.456833 | 0.018904 | 0.544 | 0.595 | 1 | 9 | Rplp0 | -0.051 | Young |
| 0.456985 | 0.190019 | 0.404 | 0.376 | 1 | 9 | Arhgap5 | 0.028 | Young |
| 0.457035 | 0.695537 | 0.022 | 0.013 | 1 | 9 | Dgat2 | 0.009 | Young |
| 0.457035 | 0.653453 | 0.022 | 0.013 | 1 | 9 | Racgap1 | 0.009 | Young |
| 0.457035 | 0.375787 | 0.022 | 0.013 | 1 | 9 | Alox5 | 0.009 | Young |
| 0.457102 | 0.793198 | 0.044 | 0.031 | 1 | 9 | Nxt1 | 0.013 | Young |
| 0.457102 | 0.733029 | 0.044 | 0.031 | 1 | 9 | Chd1l | 0.013 | Young |
| 0.457267 | 0.403882 | 0.037 | 0.054 | 1 | 9 | Gtpbp10 | -0.017 | Young |
| 0.457277 | 0.331499 | 0.206 | 0.258 | 1 | 9 | Eepd1 | -0.052 | Young |
| 0.457314 | 0.219783 | 0.581 | 0.588 | 1 | 9 | Rack1 | -0.007 | Young |
| 0.458042 | 0.386987 | 0.11 | 0.088 | 1 | 9 | Dkc1 | 0.022 | Young |
| 0.458584 | 0.525743 | 0.265 | 0.25 | 1 | 9 | Ctsz | 0.015 | Young |
| 0.458773 | 0.019562 | 0.14 | 0.175 | 1 | 9 | Msi2 | -0.035 | Young |
| 0.458789 | 0.252947 | 0.184 | 0.157 | 1 | 9 | Iscu | 0.027 | Young |
| 0.458912 | 1.375015 | 0.022 | 0.013 | 1 | 9 | Mtfmt | 0.009 | Young |
| 0.458912 | 0.631487 | 0.022 | 0.013 | 1 | 9 | Pgap1 | 0.009 | Young |
| 0.458912 | 0.512402 | 0.022 | 0.013 | 1 | 9 | Ercc8 | 0.009 | Young |
| 0.458912 | 0.284585 | 0.022 | 0.013 | 1 | 9 | Acvr1b | 0.009 | Young |
| 0.458912 | 0.175882 | 0.022 | 0.013 | 1 | 9 | Mrgpre | 0.009 | Young |
| 0.458912 | 0.13499 | 0.022 | 0.013 | 1 | 9 | Pradc1 | 0.009 | Young |
| 0.458988 | 0.055617 | 0.147 | 0.178 | 1 | 9 | Pcnx | -0.031 | Young |

| | | | | | | | | |
|---|---|---|---|---|---|---|---|---|
| 0.459054 | 0.722918 | 0.015 | 0.026 | 1 | 9 | Pdia5 | -0.011 | Young |
| 0.459054 | 0.356565 | 0.015 | 0.026 | 1 | 9 | 3110001I2: | -0.011 | Young |
| 0.459499 | 0.090493 | 0.044 | 0.062 | 1 | 9 | Timm50 | -0.018 | Young |
| 0.45963 | 1.025755 | 0.044 | 0.031 | 1 | 9 | Alg1 | 0.013 | Young |
| 0.45963 | 0.867711 | 0.044 | 0.031 | 1 | 9 | Mon1b | 0.013 | Young |
| 0.45977 | 0.995874 | 0.118 | 0.101 | 1 | 9 | Bend7 | 0.017 | Young |
| 0.459843 | 0.749492 | 0.096 | 0.077 | 1 | 9 | Nde1 | 0.019 | Young |
| 0.459848 | 0.087393 | 0.154 | 0.126 | 1 | 9 | Cic | 0.028 | Young |
| 0.460009 | 0.391394 | 0.228 | 0.198 | 1 | 9 | Safb | 0.03 | Young |
| 0.460683 | 0.053929 | 0.118 | 0.095 | 1 | 9 | 8-Mar | 0.023 | Young |
| 0.460683 | 0.029706 | 0.066 | 0.088 | 1 | 9 | Ttl | -0.022 | Young |
| 0.460704 | 0.469417 | 0.147 | 0.124 | 1 | 9 | Hcls1 | 0.023 | Young |
| 0.460794 | 0.208485 | 0.022 | 0.013 | 1 | 9 | Sash3 | 0.009 | Young |
| 0.460794 | 0.008743 | 0.022 | 0.013 | 1 | 9 | Srrd | 0.009 | Young |
| 0.460794 | 0.002903 | 0.022 | 0.013 | 1 | 9 | Zfp623 | 0.009 | Young |
| 0.460794 | 0.002458 | 0.022 | 0.013 | 1 | 9 | Gm20707 | 0.009 | Young |
| 0.460898 | 0.676532 | 0.044 | 0.031 | 1 | 9 | Vmac | 0.013 | Young |
| 0.460898 | 0.559981 | 0.044 | 0.031 | 1 | 9 | Zfp597 | 0.013 | Young |
| 0.461048 | 0.147113 | 0.14 | 0.116 | 1 | 9 | Ttyh3 | 0.024 | Young |
| 0.461155 | 0.182527 | 0.029 | 0.044 | 1 | 9 | Ippk | -0.015 | Young |
| 0.461275 | 0.047157 | 0.14 | 0.17 | 1 | 9 | Irgm1 | -0.03 | Young |
| 0.461485 | 0.221203 | 0.515 | 0.505 | 1 | 9 | Hdac7 | 0.01 | Young |
| 0.461583 | 0.747911 | 0.132 | 0.111 | 1 | 9 | Gid8 | 0.021 | Young |
| 0.462164 | 0.022771 | 0.081 | 0.062 | 1 | 9 | Mgst3 | 0.019 | Young |
| 0.462171 | 0.422998 | 0.449 | 0.43 | 1 | 9 | Sat1 | 0.019 | Young |
| 0.462257 | 0.073335 | 0.162 | 0.191 | 1 | 9 | Ulk2 | -0.029 | Young |
| 0.462348 | 0.098312 | 0.272 | 0.232 | 1 | 9 | Cmpk1 | 0.04 | Young |
| 0.462484 | 0.091096 | 0.912 | 0.91 | 1 | 9 | Rpl41 | 0.002 | Young |
| 0.46268 | 0.380549 | 0.022 | 0.013 | 1 | 9 | Bcl2a1d | 0.009 | Young |
| 0.46268 | 0.04306 | 0.022 | 0.013 | 1 | 9 | 1700052K1 | 0.009 | Young |
| 0.46268 | 0.03963 | 0.022 | 0.013 | 1 | 9 | Mctp1 | 0.009 | Young |
| 0.463683 | 0.416111 | 0.103 | 0.082 | 1 | 9 | Tle1 | 0.021 | Young |
| 0.463831 | 0.139924 | 0.059 | 0.08 | 1 | 9 | Tmem87a | -0.021 | Young |
| 0.46426 | 0.554953 | 0.074 | 0.057 | 1 | 9 | Tmem167b | 0.017 | Young |
| 0.464281 | 0.409888 | 0.191 | 0.168 | 1 | 9 | Pa2g4 | 0.023 | Young |
| 0.464345 | 0.053987 | 0.368 | 0.423 | 1 | 9 | Tek | -0.055 | Young |
| 0.464711 | 0.356995 | 0.044 | 0.031 | 1 | 9 | Fbxw5 | 0.013 | Young |
| 0.464942 | 0.112768 | 0.154 | 0.129 | 1 | 9 | Anks1 | 0.025 | Young |
| 0.465464 | 0.237923 | 0.235 | 0.201 | 1 | 9 | Etfb | 0.034 | Young |
| 0.466031 | 3.101922 | 0.015 | 0.008 | 1 | 9 | Jpx | 0.007 | Young |
| 0.466031 | 2.607941 | 0.015 | 0.008 | 1 | 9 | Fadd | 0.007 | Young |
| 0.466031 | 1.812357 | 0.015 | 0.008 | 1 | 9 | Glyctk | 0.007 | Young |
| 0.466031 | 1.030988 | 0.015 | 0.008 | 1 | 9 | Pilrb2 | 0.007 | Young |
| 0.466031 | 0.896393 | 0.015 | 0.008 | 1 | 9 | Hr | 0.007 | Young |
| 0.466198 | 0.837177 | 0.074 | 0.057 | 1 | 9 | Xylt2 | 0.017 | Young |

| | | | | | | | | |
|---|---|---|---|---|---|---|---|---|
| 0.466275 | 0.221452 | 0.794 | 0.812 | 1 | 9 | Dusp1 | -0.018 | Young |
| 0.466834 | 0.561234 | 0.213 | 0.193 | 1 | 9 | Dpp8 | 0.02 | Young |
| 0.466834 | 0.530074 | 0.213 | 0.193 | 1 | 9 | Stx3 | 0.02 | Young |
| 0.466873 | 0.15443 | 0.125 | 0.103 | 1 | 9 | Snrnp48 | 0.022 | Young |
| 0.467169 | 0.491644 | 0.074 | 0.057 | 1 | 9 | Cinp | 0.017 | Young |
| 0.467538 | 0.293604 | 0.199 | 0.175 | 1 | 9 | Srp19 | 0.024 | Young |
| 0.468425 | 1.968767 | 0.015 | 0.008 | 1 | 9 | Lum | 0.007 | Young |
| 0.468425 | 1.71318 | 0.015 | 0.008 | 1 | 9 | Gm36660 | 0.007 | Young |
| 0.468425 | 1.66018 | 0.015 | 0.008 | 1 | 9 | Cdc20 | 0.007 | Young |
| 0.468425 | 1.372312 | 0.015 | 0.008 | 1 | 9 | Nusap1 | 0.007 | Young |
| 0.468425 | 1.369158 | 0.015 | 0.008 | 1 | 9 | AC161165. | 0.007 | Young |
| 0.468425 | 1.036244 | 0.015 | 0.008 | 1 | 9 | Gemin8 | 0.007 | Young |
| 0.468425 | 0.721532 | 0.015 | 0.008 | 1 | 9 | Akr1c13 | 0.007 | Young |
| 0.468469 | 0.687334 | 0.11 | 0.093 | 1 | 9 | Nelfa | 0.017 | Young |
| 0.468477 | 0.561136 | 0.044 | 0.062 | 1 | 9 | Tmem60 | -0.018 | Young |
| 0.469411 | 0.201045 | 0.471 | 0.43 | 1 | 9 | Atp5g3 | 0.041 | Young |
| 0.469496 | 0.0614 | 0.846 | 0.866 | 1 | 9 | Rps12 | -0.02 | Young |
| 0.46964 | 0.439062 | 0.118 | 0.144 | 1 | 9 | Cul5 | -0.026 | Young |
| 0.469667 | 0.298645 | 0.221 | 0.193 | 1 | 9 | Kcnq1ot1 | 0.028 | Young |
| 0.469726 | 0.297045 | 0.382 | 0.361 | 1 | 9 | Atp5g2 | 0.021 | Young |
| 0.469822 | 0.322109 | 0.044 | 0.031 | 1 | 9 | Arhgap30 | 0.013 | Young |
| 0.46991 | 0.502904 | 0.015 | 0.026 | 1 | 9 | Cbr4 | -0.011 | Young |
| 0.46993 | 0.616746 | 0.059 | 0.044 | 1 | 9 | Nagpa | 0.015 | Young |
| 0.470093 | 0.557914 | 0.176 | 0.152 | 1 | 9 | Rnf13 | 0.024 | Young |
| 0.470367 | 0.350575 | 0.154 | 0.129 | 1 | 9 | Tap1 | 0.025 | Young |
| 0.470826 | 2.414646 | 0.015 | 0.008 | 1 | 9 | Gm9844 | 0.007 | Young |
| 0.470826 | 1.845646 | 0.015 | 0.008 | 1 | 9 | Prodh | 0.007 | Young |
| 0.470826 | 1.688189 | 0.015 | 0.008 | 1 | 9 | Otud3 | 0.007 | Young |
| 0.470826 | 1.020994 | 0.015 | 0.008 | 1 | 9 | 4833413G1 | 0.007 | Young |
| 0.470826 | 1.006743 | 0.015 | 0.008 | 1 | 9 | Clspn | 0.007 | Young |
| 0.470826 | 1.005056 | 0.015 | 0.008 | 1 | 9 | 1700066M | 0.007 | Young |
| 0.470826 | 0.970097 | 0.015 | 0.008 | 1 | 9 | Glis2 | 0.007 | Young |
| 0.470826 | 0.957069 | 0.015 | 0.008 | 1 | 9 | Thop1 | 0.007 | Young |
| 0.470826 | 0.939639 | 0.015 | 0.008 | 1 | 9 | 5430403G1 | 0.007 | Young |
| 0.470826 | 0.937108 | 0.015 | 0.008 | 1 | 9 | Hoxa2 | 0.007 | Young |
| 0.470826 | 0.760475 | 0.015 | 0.008 | 1 | 9 | Zfp991 | 0.007 | Young |
| 0.470826 | 0.755813 | 0.015 | 0.008 | 1 | 9 | AI504432 | 0.007 | Young |
| 0.470826 | 0.754788 | 0.015 | 0.008 | 1 | 9 | Klrb1a | 0.007 | Young |
| 0.470826 | 0.729057 | 0.015 | 0.008 | 1 | 9 | Cd226 | 0.007 | Young |
| 0.470826 | 0.463823 | 0.015 | 0.008 | 1 | 9 | Smo | 0.007 | Young |
| 0.471105 | 0.303496 | 0.044 | 0.031 | 1 | 9 | Cit | 0.013 | Young |
| 0.471296 | 0.203414 | 0.691 | 0.68 | 1 | 9 | Rpl8 | 0.011 | Young |
| 0.471472 | 1.152021 | 0.015 | 0.026 | 1 | 9 | Zfp933 | -0.011 | Young |
| 0.471472 | 0.636656 | 0.015 | 0.026 | 1 | 9 | 2510017J1 | -0.011 | Young |
| 0.471766 | 0.220846 | 0.176 | 0.206 | 1 | 9 | Rin3 | -0.03 | Young |

| | | | | | | | | |
|---|---|---|---|---|---|---|---|---|
| 0.472204 | 0.171 | 0.412 | 0.371 | 1 | 9 | Fubp1 | 0.041 | Young |
| 0.472556 | 0.54705 | 0.103 | 0.126 | 1 | 9 | Abl2 | -0.023 | Young |
| 0.472814 | 0.492406 | 0.11 | 0.09 | 1 | 9 | Megf8 | 0.02 | Young |
| 0.472941 | 0.320217 | 0.074 | 0.095 | 1 | 9 | Ing5 | -0.021 | Young |
| 0.473234 | 2.197707 | 0.015 | 0.008 | 1 | 9 | Noxo1 | 0.007 | Young |
| 0.473234 | 2.127234 | 0.015 | 0.008 | 1 | 9 | Foxm1 | 0.007 | Young |
| 0.473234 | 1.579892 | 0.015 | 0.008 | 1 | 9 | Ak1 | 0.007 | Young |
| 0.473234 | 1.408098 | 0.015 | 0.008 | 1 | 9 | Tmem177 | 0.007 | Young |
| 0.473234 | 1.3577 | 0.015 | 0.008 | 1 | 9 | Slc25a13 | 0.007 | Young |
| 0.473234 | 1.313376 | 0.015 | 0.008 | 1 | 9 | Thrb | 0.007 | Young |
| 0.473234 | 0.916758 | 0.015 | 0.008 | 1 | 9 | Pde3b | 0.007 | Young |
| 0.473234 | 0.912972 | 0.015 | 0.008 | 1 | 9 | Tnfrsf26 | 0.007 | Young |
| 0.473234 | 0.886806 | 0.015 | 0.008 | 1 | 9 | Oplah | 0.007 | Young |
| 0.473234 | 0.876345 | 0.015 | 0.008 | 1 | 9 | Sbk1 | 0.007 | Young |
| 0.473234 | 0.867835 | 0.015 | 0.008 | 1 | 9 | Mettl27 | 0.007 | Young |
| 0.473234 | 0.796666 | 0.015 | 0.008 | 1 | 9 | Adam12 | 0.007 | Young |
| 0.473234 | 0.729883 | 0.015 | 0.008 | 1 | 9 | Vsig10l | 0.007 | Young |
| 0.473234 | 0.696452 | 0.015 | 0.008 | 1 | 9 | Hoxd3 | 0.007 | Young |
| 0.473234 | 0.695216 | 0.015 | 0.008 | 1 | 9 | Xkr8 | 0.007 | Young |
| 0.473234 | 0.620559 | 0.015 | 0.008 | 1 | 9 | Clec4n | 0.007 | Young |
| 0.473234 | 0.617639 | 0.015 | 0.008 | 1 | 9 | Zfp316 | 0.007 | Young |
| 0.473234 | 0.600254 | 0.015 | 0.008 | 1 | 9 | Sirpb1b | 0.007 | Young |
| 0.473234 | 0.531401 | 0.015 | 0.008 | 1 | 9 | B930036N1 | 0.007 | Young |
| 0.473234 | 0.484412 | 0.015 | 0.008 | 1 | 9 | Pigl | 0.007 | Young |
| 0.473234 | 0.368257 | 0.015 | 0.008 | 1 | 9 | Apon | 0.007 | Young |
| 0.473594 | 0.575248 | 0.088 | 0.07 | 1 | 9 | Cd14 | 0.018 | Young |
| 0.473649 | 0.305171 | 0.301 | 0.273 | 1 | 9 | Mia3 | 0.028 | Young |
| 0.473663 | 0.295539 | 0.206 | 0.178 | 1 | 9 | Angel2 | 0.028 | Young |
| 0.473825 | 0.293172 | 0.162 | 0.139 | 1 | 9 | Pear1 | 0.023 | Young |
| 0.473981 | 0.068433 | 0.066 | 0.049 | 1 | 9 | Dusp23 | 0.017 | Young |
| 0.474605 | 0.683571 | 0.015 | 0.026 | 1 | 9 | Hscb | -0.011 | Young |
| 0.47541 | 0.093543 | 0.029 | 0.044 | 1 | 9 | Vwa8 | -0.015 | Young |
| 0.475587 | 0.303686 | 0.272 | 0.25 | 1 | 9 | M6pr | 0.022 | Young |
| 0.475649 | 0.714035 | 0.015 | 0.008 | 1 | 9 | Slc2a3 | 0.007 | Young |
| 0.475649 | 0.674857 | 0.015 | 0.008 | 1 | 9 | Psat1 | 0.007 | Young |
| 0.475649 | 0.637228 | 0.015 | 0.008 | 1 | 9 | Rgs14 | 0.007 | Young |
| 0.475649 | 0.632978 | 0.015 | 0.008 | 1 | 9 | Tbx3os1 | 0.007 | Young |
| 0.475649 | 0.568725 | 0.015 | 0.008 | 1 | 9 | Gm15952 | 0.007 | Young |
| 0.475649 | 0.49761 | 0.015 | 0.008 | 1 | 9 | Apobr | 0.007 | Young |
| 0.475649 | 0.473389 | 0.015 | 0.008 | 1 | 9 | Mcm8 | 0.007 | Young |
| 0.475649 | 0.417014 | 0.015 | 0.008 | 1 | 9 | Ddx31 | 0.007 | Young |
| 0.475649 | 0.388257 | 0.015 | 0.008 | 1 | 9 | Zfp973 | 0.007 | Young |
| 0.475649 | 0.37924 | 0.015 | 0.008 | 1 | 9 | Hhat | 0.007 | Young |
| 0.475649 | 0.296381 | 0.015 | 0.008 | 1 | 9 | Srms | 0.007 | Young |
| 0.475649 | 0.247375 | 0.015 | 0.008 | 1 | 9 | Rtn4ip1 | 0.007 | Young |

| | | | | | | | | |
|---|---|---|---|---|---|---|---|---|
| 0.475649 | 0.243261 | 0.015 | 0.008 | 1 | 9 | Evi5l | 0.007 | Young |
| 0.475649 | 0.233174 | 0.015 | 0.008 | 1 | 9 | Sfxn5 | 0.007 | Young |
| 0.475649 | 0.112037 | 0.015 | 0.008 | 1 | 9 | Fam189b | 0.007 | Young |
| 0.475649 | 0.07344 | 0.015 | 0.008 | 1 | 9 | Fuz | 0.007 | Young |
| 0.475649 | 0.024531 | 0.015 | 0.008 | 1 | 9 | Mrvi1 | 0.007 | Young |
| 0.47634 | 0.005399 | 0.934 | 0.941 | 1 | 9 | Tcf4 | -0.007 | Young |
| 0.476356 | 0.310061 | 0.081 | 0.103 | 1 | 9 | Tmem2 | -0.022 | Young |
| 0.476921 | 0.341588 | 0.132 | 0.108 | 1 | 9 | Herc4 | 0.024 | Young |
| 0.477149 | 0.136657 | 0.162 | 0.139 | 1 | 9 | Asah1 | 0.023 | Young |
| 0.477889 | 0.124782 | 0.199 | 0.168 | 1 | 9 | Appl1 | 0.031 | Young |
| 0.477921 | 0.740268 | 0.074 | 0.057 | 1 | 9 | Tram2 | 0.017 | Young |
| 0.47807 | 0.211436 | 0.015 | 0.008 | 1 | 9 | C1qtnf12 | 0.007 | Young |
| 0.47828 | 0.354858 | 0.051 | 0.07 | 1 | 9 | Hs2st1 | -0.019 | Young |
| 0.478494 | 0.277862 | 0.25 | 0.227 | 1 | 9 | Cyb5a | 0.023 | Young |
| 0.478807 | 0.387642 | 0.243 | 0.222 | 1 | 9 | Cers2 | 0.021 | Young |
| 0.479843 | 1.163044 | 0.059 | 0.044 | 1 | 9 | Eef1e1 | 0.015 | Young |
| 0.479865 | 0.303966 | 0.088 | 0.07 | 1 | 9 | Dcaf11 | 0.018 | Young |
| 0.480024 | 0.238821 | 0.831 | 0.817 | 1 | 9 | Rps14 | 0.014 | Young |
| 0.480215 | 0.012729 | 0.029 | 0.044 | 1 | 9 | Aldh16a1 | -0.015 | Young |
| 0.480299 | 0.312094 | 0.088 | 0.111 | 1 | 9 | Tbcel | -0.023 | Young |
| 0.480473 | 0.57836 | 0.243 | 0.224 | 1 | 9 | Higd2a | 0.019 | Young |
| 0.480498 | 0.277481 | 0.015 | 0.008 | 1 | 9 | Tfpi2 | 0.007 | Young |
| 0.480498 | 0.143857 | 0.015 | 0.008 | 1 | 9 | Clec12b | 0.007 | Young |
| 0.480498 | 0.028372 | 0.015 | 0.008 | 1 | 9 | Mmp19 | 0.007 | Young |
| 0.480952 | 0.470961 | 0.059 | 0.044 | 1 | 9 | Parn | 0.015 | Young |
| 0.481476 | 0.005807 | 0.559 | 0.631 | 1 | 9 | Sem1 | -0.072 | Young |
| 0.481684 | 0.27025 | 0.647 | 0.626 | 1 | 9 | Hsp90aa1 | 0.021 | Young |
| 0.482319 | 0.358937 | 0.081 | 0.103 | 1 | 9 | Samm50 | -0.022 | Young |
| 0.482465 | 0.082418 | 0.316 | 0.286 | 1 | 9 | D17Wsu92 | 0.03 | Young |
| 0.48278 | 0.383783 | 0.221 | 0.193 | 1 | 9 | Gxylt1 | 0.028 | Young |
| 0.483071 | 0.175668 | 0.566 | 0.557 | 1 | 9 | Hnrnpa3 | 0.009 | Young |
| 0.483336 | 0.329779 | 0.081 | 0.064 | 1 | 9 | Zfp26 | 0.017 | Young |
| 0.483835 | 0.011853 | 0.029 | 0.044 | 1 | 9 | Mgme1 | -0.015 | Young |
| 0.483927 | 0.4127 | 0.007 | 0.015 | 1 | 9 | Ccdc166 | -0.008 | Young |
| 0.483927 | 0.067889 | 0.007 | 0.015 | 1 | 9 | Tfap4 | -0.008 | Young |
| 0.483927 | 0.012225 | 0.007 | 0.015 | 1 | 9 | F2rl3 | -0.008 | Young |
| 0.484038 | 0.615115 | 0.044 | 0.031 | 1 | 9 | Gtf2h4 | 0.013 | Young |
| 0.484125 | 0.011432 | 0.103 | 0.126 | 1 | 9 | Fam76b | -0.023 | Young |
| 0.484163 | 0.470279 | 0.265 | 0.247 | 1 | 9 | Caprin1 | 0.018 | Young |
| 0.484646 | 0.105412 | 0.566 | 0.526 | 1 | 9 | mt-Nd4l | 0.04 | Young |
| 0.484795 | 0.124765 | 0.199 | 0.173 | 1 | 9 | Dgkz | 0.026 | Young |
| 0.484906 | 1.185173 | 0.066 | 0.052 | 1 | 9 | Zbtb33 | 0.014 | Young |
| 0.485116 | 0.020954 | 0.081 | 0.103 | 1 | 9 | Prickle3 | -0.022 | Young |
| 0.485996 | 2.068407 | 0.007 | 0.015 | 1 | 9 | Cdkl3 | -0.008 | Young |
| 0.485996 | 1.565988 | 0.007 | 0.015 | 1 | 9 | Alkbh7 | -0.008 | Young |

| | | | | | | | | |
|---|---|---|---|---|---|---|---|---|
| 0.485996 | 1.553462 | 0.007 | 0.015 | 1 | 9 | Bank1 | -0.008 | Young |
| 0.485996 | 1.347314 | 0.007 | 0.015 | 1 | 9 | Gpx8 | -0.008 | Young |
| 0.485996 | 1.246785 | 0.007 | 0.015 | 1 | 9 | Gpr137c | -0.008 | Young |
| 0.485996 | 1.168431 | 0.007 | 0.015 | 1 | 9 | Skp2 | -0.008 | Young |
| 0.485996 | 0.9516 | 0.007 | 0.015 | 1 | 9 | Wdyhv1 | -0.008 | Young |
| 0.485996 | 0.911023 | 0.007 | 0.015 | 1 | 9 | Tll1 | -0.008 | Young |
| 0.485996 | 0.865127 | 0.007 | 0.015 | 1 | 9 | A930033H1 | -0.008 | Young |
| 0.485996 | 0.413358 | 0.007 | 0.015 | 1 | 9 | Tsen54 | -0.008 | Young |
| 0.485996 | 0.245565 | 0.007 | 0.015 | 1 | 9 | Kcnk7 | -0.008 | Young |
| 0.485996 | 0.043156 | 0.007 | 0.015 | 1 | 9 | Kdm5d | -0.008 | Young |
| 0.485996 | 0.041175 | 0.007 | 0.015 | 1 | 9 | Ifi209 | -0.008 | Young |
| 0.486018 | 0.267065 | 0.25 | 0.224 | 1 | 9 | Rtf1 | 0.026 | Young |
| 0.486221 | 0.2931 | 0.206 | 0.186 | 1 | 9 | Kansl1 | 0.02 | Young |
| 0.486257 | 0.522897 | 0.029 | 0.044 | 1 | 9 | Pex11b | -0.015 | Young |
| 0.486647 | 0.635144 | 0.044 | 0.031 | 1 | 9 | Tprkb | 0.013 | Young |
| 0.487471 | 0.553471 | 0.029 | 0.044 | 1 | 9 | Clp1 | -0.015 | Young |
| 0.487851 | 0.199305 | 0.147 | 0.124 | 1 | 9 | Slc43a2 | 0.023 | Young |
| 0.488026 | 0.436579 | 0.066 | 0.052 | 1 | 9 | Mtx1 | 0.014 | Young |
| 0.488484 | 0.322261 | 0.184 | 0.165 | 1 | 9 | Mapk12 | 0.019 | Young |
| 0.489069 | 0.817505 | 0.066 | 0.052 | 1 | 9 | Abcb10 | 0.014 | Young |
| 0.489099 | 0.4926 | 0.338 | 0.299 | 1 | 9 | Ttc14 | 0.039 | Young |
| 0.489304 | 0.129129 | 0.338 | 0.309 | 1 | 9 | Efcab14 | 0.029 | Young |
| 0.489797 | 0.306854 | 0.044 | 0.062 | 1 | 9 | Golt1b | -0.018 | Young |
| 0.489804 | 0.618703 | 0.074 | 0.057 | 1 | 9 | Snap29 | 0.017 | Young |
| 0.489894 | 0.336808 | 0.162 | 0.139 | 1 | 9 | Slc5a3 | 0.023 | Young |
| 0.48993 | 0.080501 | 0.081 | 0.103 | 1 | 9 | Dpy30 | -0.022 | Young |
| 0.489966 | 0.480614 | 0.103 | 0.126 | 1 | 9 | St6galnac6 | -0.023 | Young |
| 0.490332 | 0.110584 | 0.081 | 0.103 | 1 | 9 | Uri1 | -0.022 | Young |
| 0.490341 | 0.468258 | 0.243 | 0.216 | 1 | 9 | Vapa | 0.027 | Young |
| 0.49038 | 0.225025 | 0.147 | 0.178 | 1 | 9 | Irf9 | -0.031 | Young |
| 0.490507 | 0.087982 | 0.11 | 0.09 | 1 | 9 | Ak2 | 0.02 | Young |
| 0.490507 | 0.309691 | 0.11 | 0.09 | 1 | 9 | Atxn3 | 0.02 | Young |
| 0.490619 | 1.123429 | 0.096 | 0.077 | 1 | 9 | Dlat | 0.019 | Young |
| 0.491326 | 0.380265 | 0.154 | 0.134 | 1 | 9 | Naa15 | 0.02 | Young |
| 0.49153 | 0.674177 | 0.132 | 0.162 | 1 | 9 | Pik3r6 | -0.03 | Young |
| 0.491668 | 0.540638 | 0.243 | 0.224 | 1 | 9 | Spen | 0.019 | Young |
| 0.491917 | 0.019357 | 0.088 | 0.111 | 1 | 9 | Ufc1 | -0.023 | Young |
| 0.492709 | 0.097771 | 0.926 | 0.869 | 1 | 9 | Rplp1 | 0.057 | Young |
| 0.493559 | 1.768199 | 0.037 | 0.026 | 1 | 9 | Psph | 0.011 | Young |
| 0.494041 | 0.017112 | 0.154 | 0.183 | 1 | 9 | Cbx6 | -0.029 | Young |
| 0.494124 | 1.414378 | 0.051 | 0.039 | 1 | 9 | Tinf2 | 0.012 | Young |
| 0.4943 | 0.671605 | 0.066 | 0.052 | 1 | 9 | AC165953. | 0.014 | Young |
| 0.494361 | 1.568846 | 0.059 | 0.044 | 1 | 9 | Wbp1 | 0.015 | Young |
| 0.494784 | 0.530098 | 0.125 | 0.106 | 1 | 9 | Elovl1 | 0.019 | Young |
| 0.49494 | 0.415138 | 0.132 | 0.111 | 1 | 9 | Eloa | 0.021 | Young |

| | | | | | | | | |
|---|---|---|---|---|---|---|---|---|
| 0.495043 | 0.057883 | 0.118 | 0.098 | 1 | 9 | Coro6 | 0.02 | Young |
| 0.495197 | 0.155833 | 0.206 | 0.242 | 1 | 9 | Ezh1 | -0.036 | Young |
| 0.495281 | 0.230052 | 0.088 | 0.07 | 1 | 9 | Gtf3c6 | 0.018 | Young |
| 0.496615 | 0.012062 | 0.037 | 0.052 | 1 | 9 | Yae1d1 | -0.015 | Young |
| 0.497453 | 0.124766 | 0.066 | 0.052 | 1 | 9 | Utp4 | 0.014 | Young |
| 0.497807 | 0.262673 | 0.316 | 0.371 | 1 | 9 | Sox7 | -0.055 | Young |
| 0.497979 | 0.735034 | 0.066 | 0.052 | 1 | 9 | Kat14 | 0.014 | Young |
| 0.498029 | 0.493638 | 0.088 | 0.07 | 1 | 9 | Mob3c | 0.018 | Young |
| 0.498077 | 0.523628 | 0.176 | 0.155 | 1 | 9 | Phb2 | 0.021 | Young |
| 0.498104 | 0.192518 | 0.125 | 0.101 | 1 | 9 | Trnt1 | 0.024 | Young |
| 0.498595 | 0.566061 | 0.051 | 0.07 | 1 | 9 | Fcf1 | -0.019 | Young |
| 0.498666 | 0.191545 | 0.11 | 0.09 | 1 | 9 | Glrx2 | 0.02 | Young |
| 0.498677 | 0.057337 | 0.257 | 0.284 | 1 | 9 | Sfr1 | -0.027 | Young |
| 0.498699 | 0.05173 | 0.191 | 0.165 | 1 | 9 | Zzz3 | 0.026 | Young |
| 0.498947 | 0.403855 | 0.088 | 0.07 | 1 | 9 | Marveld1 | 0.018 | Young |
| 0.499329 | 1.473034 | 0.037 | 0.026 | 1 | 9 | Ctla2b | 0.011 | Young |
| 0.499329 | 0.796504 | 0.037 | 0.026 | 1 | 9 | Zfp235 | 0.011 | Young |
| 0.499866 | 0.189287 | 0.088 | 0.07 | 1 | 9 | Snapc1 | 0.018 | Young |
| 0.500268 | 0.302917 | 0.301 | 0.268 | 1 | 9 | Atxn2l | 0.033 | Young |
| 0.500306 | 0.207316 | 0.11 | 0.09 | 1 | 9 | Rnf181 | 0.02 | Young |
| 0.500379 | 0.646675 | 0.228 | 0.206 | 1 | 9 | Dnaja2 | 0.022 | Young |
| 0.500449 | 0.912539 | 0.11 | 0.093 | 1 | 9 | Med10 | 0.017 | Young |
| 0.500633 | 0.288411 | 0.022 | 0.034 | 1 | 9 | Epg5 | -0.012 | Young |
| 0.501263 | 0.266368 | 0.081 | 0.103 | 1 | 9 | Zbtb41 | -0.022 | Young |
| 0.501315 | 0.252505 | 0.316 | 0.291 | 1 | 9 | Pnkd | 0.025 | Young |
| 0.50139 | 0.102887 | 0.132 | 0.155 | 1 | 9 | Slc25a39 | -0.023 | Young |
| 0.502099 | 0.019491 | 0.199 | 0.229 | 1 | 9 | Mrpl52 | -0.03 | Young |
| 0.502227 | 0.536849 | 0.037 | 0.026 | 1 | 9 | Abcb9 | 0.011 | Young |
| 0.502886 | 0.223651 | 0.36 | 0.332 | 1 | 9 | Atf3 | 0.028 | Young |
| 0.503223 | 0.512817 | 0.081 | 0.064 | 1 | 9 | Crat | 0.017 | Young |
| 0.503411 | 0.108423 | 0.059 | 0.044 | 1 | 9 | Soat1 | 0.015 | Young |
| 0.503596 | 0.760781 | 0.11 | 0.09 | 1 | 9 | Sgf29 | 0.02 | Young |
| 0.503647 | 0.043371 | 0.184 | 0.16 | 1 | 9 | Dmxl1 | 0.024 | Young |
| 0.503768 | 0.430082 | 0.103 | 0.085 | 1 | 9 | Pknox1 | 0.018 | Young |
| 0.503768 | 0.366458 | 0.103 | 0.085 | 1 | 9 | Washc1 | 0.018 | Young |
| 0.5049 | 0.25107 | 0.294 | 0.278 | 1 | 9 | Dad1 | 0.016 | Young |
| 0.505135 | 0.183704 | 0.037 | 0.026 | 1 | 9 | Mbnl3 | 0.011 | Young |
| 0.505135 | 0.087027 | 0.037 | 0.026 | 1 | 9 | Ccdc86 | 0.011 | Young |
| 0.505368 | 0.377861 | 0.14 | 0.119 | 1 | 9 | Fat4 | 0.021 | Young |
| 0.505537 | 0.370927 | 0.125 | 0.106 | 1 | 9 | Ddhd2 | 0.019 | Young |
| 0.50562 | 0.309502 | 0.228 | 0.201 | 1 | 9 | Prdx2 | 0.027 | Young |
| 0.505649 | 0.187612 | 0.228 | 0.204 | 1 | 9 | Pdcd6 | 0.024 | Young |
| 0.505771 | 0.437419 | 0.088 | 0.072 | 1 | 9 | Klhdc7a | 0.016 | Young |
| 0.505892 | 0.249582 | 0.074 | 0.057 | 1 | 9 | Washc3 | 0.017 | Young |
| 0.505913 | 0.404738 | 0.066 | 0.052 | 1 | 9 | 2410089E0 | 0.014 | Young |

| | | | | | | | | |
|---|---|---|---|---|---|---|---|---|
| 0.506099 | 0.599978 | 0.081 | 0.064 | 1 | 9 | Btaf1 | 0.017 | Young |
| 0.506163 | 0.664819 | 0.051 | 0.039 | 1 | 9 | Ptcd3 | 0.012 | Young |
| 0.506267 | 0.440389 | 0.022 | 0.034 | 1 | 9 | Fbxl6 | -0.012 | Young |
| 0.506549 | 0.119781 | 0.14 | 0.116 | 1 | 9 | Dot1l | 0.024 | Young |
| 0.506976 | 0.469562 | 0.066 | 0.052 | 1 | 9 | Ccr5 | 0.014 | Young |
| 0.506976 | 0.00569 | 0.044 | 0.059 | 1 | 9 | Slco3a1 | -0.015 | Young |
| 0.507163 | 0.804634 | 0.103 | 0.085 | 1 | 9 | Lrrc59 | 0.018 | Young |
| 0.507283 | 0.03766 | 0.132 | 0.16 | 1 | 9 | Trappc6b | -0.028 | Young |
| 0.507801 | 0.503903 | 0.11 | 0.093 | 1 | 9 | Acadvl | 0.017 | Young |
| 0.507923 | 0.11859 | 0.074 | 0.057 | 1 | 9 | Txndc16 | 0.017 | Young |
| 0.507923 | 0.013098 | 0.074 | 0.057 | 1 | 9 | Nabp1 | 0.017 | Young |
| 0.507968 | 0.0809 | 0.037 | 0.052 | 1 | 9 | Xpnpep1 | -0.015 | Young |
| 0.508014 | 0.314033 | 0.103 | 0.085 | 1 | 9 | Ccdc71l | 0.018 | Young |
| 0.50804 | 0.275538 | 0.066 | 0.052 | 1 | 9 | Mtmr9 | 0.014 | Young |
| 0.508051 | 2.197413 | 0.037 | 0.026 | 1 | 9 | Rcan3 | 0.011 | Young |
| 0.508051 | 0.946844 | 0.037 | 0.026 | 1 | 9 | Jade3 | 0.011 | Young |
| 0.508051 | 0.496194 | 0.037 | 0.026 | 1 | 9 | Mettl14 | 0.011 | Young |
| 0.508051 | 0.366419 | 0.037 | 0.026 | 1 | 9 | Nop16 | 0.011 | Young |
| 0.508051 | 0.011846 | 0.037 | 0.026 | 1 | 9 | Pwp1 | 0.011 | Young |
| 0.509221 | 0.156151 | 0.257 | 0.227 | 1 | 9 | Mcrip1 | 0.03 | Young |
| 0.509379 | 0.134406 | 0.11 | 0.09 | 1 | 9 | Ash2l | 0.02 | Young |
| 0.509485 | 0.259352 | 0.485 | 0.472 | 1 | 9 | Pafah1b1 | 0.013 | Young |
| 0.509512 | 1.136779 | 0.037 | 0.026 | 1 | 9 | Slc27a1 | 0.011 | Young |
| 0.509512 | 0.373776 | 0.037 | 0.026 | 1 | 9 | Wars2 | 0.011 | Young |
| 0.509512 | 0.10499 | 0.037 | 0.026 | 1 | 9 | Bcl2a1b | 0.011 | Young |
| 0.509718 | 0.475595 | 0.103 | 0.085 | 1 | 9 | Bckdk | 0.018 | Young |
| 0.509866 | 0.325618 | 0.103 | 0.082 | 1 | 9 | Csgalnact2 | 0.021 | Young |
| 0.509902 | 0.663619 | 0.125 | 0.108 | 1 | 9 | Tsfm | 0.017 | Young |
| 0.510019 | 0.393591 | 0.118 | 0.101 | 1 | 9 | Osgep | 0.017 | Young |
| 0.510162 | 0.284693 | 0.449 | 0.436 | 1 | 9 | Tnrc6b | 0.013 | Young |
| 0.510571 | 0.476073 | 0.074 | 0.095 | 1 | 9 | Axl | -0.021 | Young |
| 0.510846 | 0.2732 | 0.096 | 0.077 | 1 | 9 | Rpia | 0.019 | Young |
| 0.510976 | 0.378538 | 0.037 | 0.026 | 1 | 9 | Tmigd1 | 0.011 | Young |
| 0.511454 | 0.188577 | 0.184 | 0.16 | 1 | 9 | Mcmbp | 0.024 | Young |
| 0.511521 | 0.017193 | 0.397 | 0.438 | 1 | 9 | Rnf144a | -0.041 | Young |
| 0.511754 | 0.359513 | 0.206 | 0.175 | 1 | 9 | Hcfc1r1 | 0.031 | Young |
| 0.511889 | 0.028323 | 0.088 | 0.07 | 1 | 9 | 2310001H1 | 0.018 | Young |
| 0.511934 | 0.055703 | 0.022 | 0.034 | 1 | 9 | Cenpa | -0.012 | Young |
| 0.512118 | 0.265157 | 0.191 | 0.168 | 1 | 9 | Dopey1 | 0.023 | Young |
| 0.512442 | 0.408921 | 0.037 | 0.026 | 1 | 9 | Igsf9 | 0.011 | Young |
| 0.512459 | 0.270674 | 0.162 | 0.139 | 1 | 9 | Sec22b | 0.023 | Young |
| 0.512637 | 0.486627 | 0.132 | 0.113 | 1 | 9 | Cmip | 0.019 | Young |
| 0.513203 | 0.76816 | 0.118 | 0.101 | 1 | 9 | Chfr | 0.017 | Young |
| 0.513209 | 0.04736 | 0.257 | 0.227 | 1 | 9 | Arpc4 | 0.03 | Young |
| 0.51325 | 0.248159 | 0.162 | 0.134 | 1 | 9 | Acsl5 | 0.028 | Young |

| | | | | | | | | |
|---|---|---|---|---|---|---|---|---|
| 0.513356 | 1.103345 | 0.022 | 0.034 | 1 | 9 | Abhd14b | -0.012 | Young |
| 0.51346 | 0.391322 | 0.051 | 0.039 | 1 | 9 | Megf9 | 0.012 | Young |
| 0.513804 | 0.002635 | 0.338 | 0.304 | 1 | 9 | Ube2r2 | 0.034 | Young |
| 0.51381 | 0.652933 | 0.081 | 0.064 | 1 | 9 | Naa10 | 0.017 | Young |
| 0.51391 | 0.678639 | 0.037 | 0.026 | 1 | 9 | Stn1 | 0.011 | Young |
| 0.514682 | 0.626194 | 0.051 | 0.039 | 1 | 9 | Slc33a1 | 0.012 | Young |
| 0.514955 | 0.108357 | 0.125 | 0.149 | 1 | 9 | Rusc2 | -0.024 | Young |
| 0.515205 | 0.052422 | 0.162 | 0.183 | 1 | 9 | Git1 | -0.021 | Young |
| 0.51538 | 0.180086 | 0.037 | 0.026 | 1 | 9 | Fam46c | 0.011 | Young |
| 0.51552 | 0.104109 | 0.066 | 0.052 | 1 | 9 | Blvra | 0.014 | Young |
| 0.515925 | 0.283514 | 0.162 | 0.139 | 1 | 9 | Impdh2 | 0.023 | Young |
| 0.516306 | 0.642745 | 0.147 | 0.126 | 1 | 9 | Sde2 | 0.021 | Young |
| 0.516337 | 0.374181 | 0.279 | 0.258 | 1 | 9 | Plcg1 | 0.021 | Young |
| 0.516643 | 0.0011 | 0.11 | 0.131 | 1 | 9 | Tmed3 | -0.021 | Young |
| 0.516852 | 0.325378 | 0.037 | 0.026 | 1 | 9 | Mob3a | 0.011 | Young |
| 0.516852 | 0.167725 | 0.037 | 0.026 | 1 | 9 | Pms2 | 0.011 | Young |
| 0.517129 | 0.606365 | 0.051 | 0.039 | 1 | 9 | Gpr137 | 0.012 | Young |
| 0.517634 | 0.103737 | 0.022 | 0.034 | 1 | 9 | Trmt10c | -0.012 | Young |
| 0.517668 | 0.894048 | 0.066 | 0.052 | 1 | 9 | Pcnx3 | 0.014 | Young |
| 0.518355 | 0.663496 | 0.051 | 0.039 | 1 | 9 | Zfp507 | 0.012 | Young |
| 0.518878 | 0.453457 | 0.096 | 0.077 | 1 | 9 | Ncbp1 | 0.019 | Young |
| 0.519064 | 0.1349 | 0.022 | 0.034 | 1 | 9 | Klc2 | -0.012 | Young |
| 0.51927 | 0.460057 | 0.154 | 0.134 | 1 | 9 | Parp10 | 0.02 | Young |
| 0.519298 | 0.064478 | 0.191 | 0.162 | 1 | 9 | Copz1 | 0.029 | Young |
| 0.519583 | 0.306152 | 0.051 | 0.039 | 1 | 9 | Gins4 | 0.012 | Young |
| 0.519598 | 0.069096 | 0.294 | 0.325 | 1 | 9 | Rb1cc1 | -0.031 | Young |
| 0.51982 | 0.284511 | 0.066 | 0.052 | 1 | 9 | Zfp629 | 0.014 | Young |
| 0.51985 | 0.226791 | 0.162 | 0.137 | 1 | 9 | Gusb | 0.025 | Young |
| 0.520607 | 0.222854 | 0.081 | 0.064 | 1 | 9 | Rexo4 | 0.017 | Young |
| 0.520742 | 0.165664 | 0.14 | 0.162 | 1 | 9 | Washc4 | -0.022 | Young |
| 0.521056 | 0.351171 | 0.522 | 0.539 | 1 | 9 | Pim3 | -0.017 | Young |
| 0.521292 | 0.005921 | 0.588 | 0.629 | 1 | 9 | Rpl5 | -0.041 | Young |
| 0.521931 | 0.078872 | 0.022 | 0.034 | 1 | 9 | Nr2c1 | -0.012 | Young |
| 0.521977 | 0.265767 | 0.066 | 0.052 | 1 | 9 | Amn1 | 0.014 | Young |
| 0.522042 | 0.322421 | 0.051 | 0.039 | 1 | 9 | Cdc7 | 0.012 | Young |
| 0.52205 | 0.643584 | 0.066 | 0.085 | 1 | 9 | Rad17 | -0.019 | Young |
| 0.522566 | 0.837575 | 0.14 | 0.121 | 1 | 9 | Med15 | 0.019 | Young |
| 0.522618 | 0.510081 | 0.074 | 0.059 | 1 | 9 | Arl4c | 0.015 | Young |
| 0.522763 | 0.130499 | 0.037 | 0.026 | 1 | 9 | Hmgn2 | 0.011 | Young |
| 0.522867 | 0.185241 | 0.074 | 0.093 | 1 | 9 | Hspa14 | -0.019 | Young |
| 0.522927 | 0.728448 | 0.037 | 0.052 | 1 | 9 | Mrpl39 | -0.015 | Young |
| 0.524254 | 0.09301 | 0.096 | 0.116 | 1 | 9 | Atp8b2 | -0.02 | Young |
| 0.524318 | 0.44676 | 0.103 | 0.085 | 1 | 9 | Lrpprc | 0.018 | Young |
| 0.52442 | 0.156566 | 0.301 | 0.276 | 1 | 9 | Ap2m1 | 0.025 | Young |
| 0.524508 | 0.575845 | 0.051 | 0.039 | 1 | 9 | B4galt6 | 0.012 | Young |

| | | | | | | | | |
|---|---|---|---|---|---|---|---|---|
| 0.524508 | 0.279401 | 0.051 | 0.039 | 1 | 9 | Hrh2 | 0.012 | Young |
| 0.524508 | 0.058064 | 0.051 | 0.039 | 1 | 9 | Usp21 | 0.012 | Young |
| 0.524613 | 0.302906 | 0.074 | 0.093 | 1 | 9 | Brd9 | -0.019 | Young |
| 0.524805 | 0.083375 | 0.022 | 0.034 | 1 | 9 | Calhm2 | -0.012 | Young |
| 0.524972 | 0.020557 | 0.309 | 0.353 | 1 | 9 | Csnk1e | -0.044 | Young |
| 0.525743 | 0.38696 | 0.051 | 0.039 | 1 | 9 | Cep170b | 0.012 | Young |
| 0.52605 | 0.224835 | 0.265 | 0.245 | 1 | 9 | Vamp3 | 0.02 | Young |
| 0.526144 | 0.087045 | 0.316 | 0.356 | 1 | 9 | Tspo | -0.04 | Young |
| 0.526839 | 0.00593 | 0.132 | 0.111 | 1 | 9 | Prmt1 | 0.021 | Young |
| 0.527115 | 0.03995 | 0.118 | 0.098 | 1 | 9 | Rnf115 | 0.02 | Young |
| 0.527203 | 0.244268 | 0.074 | 0.059 | 1 | 9 | Galc | 0.015 | Young |
| 0.527219 | 0.02377 | 0.037 | 0.026 | 1 | 9 | Mthfr | 0.011 | Young |
| 0.529003 | 0.107164 | 0.294 | 0.335 | 1 | 9 | Npr1 | -0.041 | Young |
| 0.529015 | 0.428132 | 0.132 | 0.113 | 1 | 9 | mt-Nd6 | 0.019 | Young |
| 0.529031 | 0.182757 | 0.147 | 0.124 | 1 | 9 | Paip1 | 0.023 | Young |
| 0.529128 | 0.070357 | 0.206 | 0.224 | 1 | 9 | Ybx3 | -0.018 | Young |
| 0.529132 | 0.154412 | 0.022 | 0.034 | 1 | 9 | Prdm10 | -0.012 | Young |
| 0.529261 | 0.226149 | 0.221 | 0.247 | 1 | 9 | Atp2a2 | -0.026 | Young |
| 0.529414 | 0.110752 | 0.081 | 0.064 | 1 | 9 | Lpin1 | 0.017 | Young |
| 0.529456 | 0.501229 | 0.051 | 0.039 | 1 | 9 | Coq10a | 0.012 | Young |
| 0.529562 | 0.220145 | 0.066 | 0.052 | 1 | 9 | Brcc3 | 0.014 | Young |
| 0.529784 | 0.161651 | 0.132 | 0.113 | 1 | 9 | Pdcd4 | 0.019 | Young |
| 0.53026 | 0.362711 | 0.066 | 0.085 | 1 | 9 | Atg101 | -0.019 | Young |
| 0.530579 | 0.613095 | 0.022 | 0.034 | 1 | 9 | Echdc1 | -0.012 | Young |
| 0.530651 | 0.044525 | 0.066 | 0.052 | 1 | 9 | Cgrrf1 | 0.014 | Young |
| 0.530771 | 0.066129 | 0.265 | 0.286 | 1 | 9 | Prkx | -0.021 | Young |
| 0.530982 | 0.128428 | 0.125 | 0.103 | 1 | 9 | Psma5 | 0.022 | Young |
| 0.53127 | 0.51955 | 0.154 | 0.137 | 1 | 9 | Usp14 | 0.017 | Young |
| 0.531327 | 0.028989 | 0.75 | 0.755 | 1 | 9 | Btg2 | -0.005 | Young |
| 0.532569 | 1.280193 | 0.118 | 0.103 | 1 | 9 | Eln | 0.015 | Young |
| 0.532658 | 0.314033 | 0.287 | 0.265 | 1 | 9 | Nap1l1 | 0.022 | Young |
| 0.533183 | 0.195575 | 0.051 | 0.039 | 1 | 9 | Il6 | 0.012 | Young |
| 0.533348 | 0.584782 | 0.074 | 0.059 | 1 | 9 | Ccrl2 | 0.015 | Young |
| 0.533552 | 0.777554 | 0.11 | 0.093 | 1 | 9 | Nup54 | 0.017 | Young |
| 0.534077 | 1.557123 | 0.059 | 0.046 | 1 | 9 | Arhgap28 | 0.013 | Young |
| 0.534145 | 1.101951 | 0.081 | 0.067 | 1 | 9 | Ago1 | 0.014 | Young |
| 0.534182 | 0.235111 | 0.25 | 0.227 | 1 | 9 | Snrpe | 0.023 | Young |
| 0.534796 | 0.374994 | 0.206 | 0.18 | 1 | 9 | Slbp | 0.026 | Young |
| 0.534965 | 0.195214 | 0.221 | 0.196 | 1 | 9 | Ndufs2 | 0.025 | Young |
| 0.535874 | 0.271966 | 0.279 | 0.263 | 1 | 9 | Rasgrp2 | 0.016 | Young |
| 0.535934 | 0.163503 | 0.463 | 0.454 | 1 | 9 | Naca | 0.009 | Young |
| 0.536109 | 0.22293 | 0.066 | 0.052 | 1 | 9 | Renbp | 0.014 | Young |
| 0.536398 | 0.361426 | 0.184 | 0.165 | 1 | 9 | Hexa | 0.019 | Young |
| 0.536402 | 0.503881 | 0.088 | 0.072 | 1 | 9 | Gstt1 | 0.016 | Young |
| 0.536833 | 0.896797 | 0.029 | 0.041 | 1 | 9 | Cnksr3 | -0.012 | Young |

| | | | | | | | | |
|---|---|---|---|---|---|---|---|---|
| 0.537204 | 0.415302 | 0.066 | 0.052 | 1 | 9 | Hps5 | 0.014 | Young |
| 0.537204 | 0.107447 | 0.044 | 0.059 | 1 | 9 | Mrps10 | -0.015 | Young |
| 0.537305 | 0.085503 | 0.081 | 0.064 | 1 | 9 | Yars | 0.017 | Young |
| 0.53793 | 1.089817 | 0.029 | 0.021 | 1 | 9 | Zfp276 | 0.008 | Young |
| 0.537976 | 0.079933 | 0.551 | 0.549 | 1 | 9 | Gpx4 | 0.002 | Young |
| 0.538051 | 0.268925 | 0.059 | 0.075 | 1 | 9 | Gpkow | -0.016 | Young |
| 0.538564 | 0.307181 | 0.118 | 0.139 | 1 | 9 | Thbs1 | -0.021 | Young |
| 0.538843 | 0.145361 | 0.654 | 0.652 | 1 | 9 | Pcbp2 | 0.002 | Young |
| 0.539299 | 0.390139 | 0.147 | 0.129 | 1 | 9 | St3gal2 | 0.018 | Young |
| 0.539412 | 0.039148 | 0.228 | 0.25 | 1 | 9 | Zdhhc20 | -0.022 | Young |
| 0.539424 | 0.080947 | 0.051 | 0.039 | 1 | 9 | Bysl | 0.012 | Young |
| 0.539684 | 0.754508 | 0.257 | 0.237 | 1 | 9 | Gopc | 0.02 | Young |
| 0.539869 | 0.65942 | 0.243 | 0.235 | 1 | 9 | Ppic | 0.008 | Young |
| 0.540024 | 0.066345 | 0.618 | 0.637 | 1 | 9 | Tpm3 | -0.019 | Young |
| 0.540495 | 0.990948 | 0.066 | 0.052 | 1 | 9 | Ercc3 | 0.014 | Young |
| 0.540495 | 0.19465 | 0.066 | 0.052 | 1 | 9 | Amz2 | 0.014 | Young |
| 0.541031 | 0.318999 | 0.272 | 0.25 | 1 | 9 | Aplnr | 0.022 | Young |
| 0.541288 | 1.242754 | 0.029 | 0.021 | 1 | 9 | Tfpi | 0.008 | Young |
| 0.541541 | 0.369771 | 0.176 | 0.152 | 1 | 9 | Pgam1 | 0.024 | Young |
| 0.541724 | 0.919145 | 0.11 | 0.095 | 1 | 9 | Ndufs8 | 0.015 | Young |
| 0.542185 | 0.422827 | 0.169 | 0.196 | 1 | 9 | Chordc1 | -0.027 | Young |
| 0.543279 | 0.39111 | 0.169 | 0.147 | 1 | 9 | Vmp1 | 0.022 | Young |
| 0.543487 | 0.332294 | 0.176 | 0.157 | 1 | 9 | Abcf1 | 0.019 | Young |
| 0.543797 | 0.131443 | 0.066 | 0.052 | 1 | 9 | Pmaip1 | 0.014 | Young |
| 0.544237 | 0.236449 | 0.176 | 0.155 | 1 | 9 | Cdc42se1 | 0.021 | Young |
| 0.544543 | 0.305665 | 0.11 | 0.093 | 1 | 9 | Kif13a | 0.017 | Young |
| 0.544656 | 1.208097 | 0.029 | 0.021 | 1 | 9 | Tbc1d2 | 0.008 | Young |
| 0.544656 | 1.088771 | 0.029 | 0.021 | 1 | 9 | Tma16 | 0.008 | Young |
| 0.544656 | 0.731672 | 0.029 | 0.021 | 1 | 9 | Osbpl6 | 0.008 | Young |
| 0.544718 | 0.294705 | 0.426 | 0.402 | 1 | 9 | Ier5 | 0.024 | Young |
| 0.544747 | 0.003363 | 0.088 | 0.106 | 1 | 9 | Dcun1d1 | -0.018 | Young |
| 0.544775 | 0.683127 | 0.118 | 0.101 | 1 | 9 | Zfc3h1 | 0.017 | Young |
| 0.544855 | 0.547254 | 0.096 | 0.08 | 1 | 9 | Skiv2l2 | 0.016 | Young |
| 0.545008 | 0.02898 | 0.081 | 0.098 | 1 | 9 | Get4 | -0.017 | Young |
| 0.545451 | 0.890038 | 0.044 | 0.034 | 1 | 9 | Gatsl2 | 0.01 | Young |
| 0.545865 | 0.151885 | 0.088 | 0.072 | 1 | 9 | Cops5 | 0.016 | Young |
| 0.546345 | 0.821493 | 0.029 | 0.021 | 1 | 9 | Trim36 | 0.008 | Young |
| 0.546437 | 0.098771 | 0.426 | 0.402 | 1 | 9 | Slc28a2 | 0.024 | Young |
| 0.54667 | 0.261807 | 0.11 | 0.093 | 1 | 9 | Armcx3 | 0.017 | Young |
| 0.546802 | 0.883944 | 0.044 | 0.034 | 1 | 9 | Fyb | 0.01 | Young |
| 0.546846 | 0.678054 | 0.346 | 0.338 | 1 | 9 | Ubn2 | 0.008 | Young |
| 0.547276 | 0.093645 | 0.537 | 0.521 | 1 | 9 | Srsf3 | 0.016 | Young |
| 0.547284 | 0.21793 | 0.169 | 0.204 | 1 | 9 | Tspan18 | -0.035 | Young |
| 0.547342 | 0.375251 | 0.029 | 0.041 | 1 | 9 | Slc35b4 | -0.012 | Young |
| 0.547768 | 0.7556 | 0.088 | 0.072 | 1 | 9 | Anapc13 | 0.016 | Young |

| | | | | | | | | |
|---|---|---|---|---|---|---|---|---|
| 0.547812 | 0.386563 | 0.169 | 0.149 | 1 | 9 | Hadha | 0.02 | Young |
| 0.547827 | 0.169328 | 0.257 | 0.296 | 1 | 9 | Znrf1 | -0.039 | Young |
| 0.548035 | 1.600935 | 0.029 | 0.021 | 1 | 9 | Yipf2 | 0.008 | Young |
| 0.548035 | 1.123399 | 0.029 | 0.021 | 1 | 9 | Ppm1j | 0.008 | Young |
| 0.548035 | 0.971807 | 0.029 | 0.021 | 1 | 9 | Ppp1r16a | 0.008 | Young |
| 0.548234 | 1.062554 | 0.081 | 0.101 | 1 | 9 | Med21 | -0.02 | Young |
| 0.548525 | 0.147181 | 0.375 | 0.356 | 1 | 9 | Nfkb1 | 0.019 | Young |
| 0.548869 | 0.621153 | 0.074 | 0.059 | 1 | 9 | Rassf2 | 0.015 | Young |
| 0.549294 | 0.220973 | 0.235 | 0.209 | 1 | 9 | Dnm2 | 0.026 | Young |
| 0.549729 | 0.748333 | 0.029 | 0.021 | 1 | 9 | Zfp64 | 0.008 | Young |
| 0.549729 | 0.595795 | 0.029 | 0.021 | 1 | 9 | Ripk3 | 0.008 | Young |
| 0.549729 | 0.567941 | 0.029 | 0.021 | 1 | 9 | Gemin2 | 0.008 | Young |
| 0.549729 | 0.461037 | 0.029 | 0.021 | 1 | 9 | Zfpm2 | 0.008 | Young |
| 0.549985 | 0.306059 | 0.029 | 0.041 | 1 | 9 | Ifi208 | -0.012 | Young |
| 0.550627 | 0.499692 | 0.088 | 0.072 | 1 | 9 | Dip2c | 0.016 | Young |
| 0.550955 | 0.522227 | 0.074 | 0.059 | 1 | 9 | Exosc9 | 0.015 | Young |
| 0.551252 | 0.189003 | 0.235 | 0.211 | 1 | 9 | Med28 | 0.024 | Young |
| 0.551289 | 0.202187 | 0.191 | 0.216 | 1 | 9 | Tjp2 | -0.025 | Young |
| 0.551425 | 1.291483 | 0.029 | 0.021 | 1 | 9 | Mtrf1l | 0.008 | Young |
| 0.551425 | 0.929261 | 0.029 | 0.021 | 1 | 9 | Ptgs2 | 0.008 | Young |
| 0.551425 | 0.669842 | 0.029 | 0.021 | 1 | 9 | Acsf3 | 0.008 | Young |
| 0.551425 | 0.590175 | 0.029 | 0.021 | 1 | 9 | Clmn | 0.008 | Young |
| 0.551425 | 0.510915 | 0.029 | 0.021 | 1 | 9 | A330009N1 | 0.008 | Young |
| 0.551999 | 0.238089 | 0.074 | 0.059 | 1 | 9 | Zfp746 | 0.015 | Young |
| 0.552008 | 0.758465 | 0.154 | 0.144 | 1 | 9 | Strap | 0.01 | Young |
| 0.552133 | 0.165301 | 0.434 | 0.485 | 1 | 9 | 4931406P1 | -0.051 | Young |
| 0.553123 | 0.558029 | 0.029 | 0.021 | 1 | 9 | Hcn4 | 0.008 | Young |
| 0.553224 | 0.573925 | 0.221 | 0.198 | 1 | 9 | Grrp1 | 0.023 | Young |
| 0.553366 | 0.325354 | 0.147 | 0.129 | 1 | 9 | Mbd1 | 0.018 | Young |
| 0.553561 | 0.712197 | 0.125 | 0.108 | 1 | 9 | Mospd3 | 0.017 | Young |
| 0.55361 | 1.760067 | 0.015 | 0.023 | 1 | 9 | Nubp1 | -0.008 | Young |
| 0.553707 | 0.320365 | 0.287 | 0.263 | 1 | 9 | Ccdc85b | 0.024 | Young |
| 0.55381 | 0.778876 | 0.059 | 0.046 | 1 | 9 | Ift27 | 0.013 | Young |
| 0.553827 | 0.01672 | 0.176 | 0.206 | 1 | 9 | Grb10 | -0.03 | Young |
| 0.553898 | 0.301582 | 0.147 | 0.126 | 1 | 9 | Rnf114 | 0.021 | Young |
| 0.553961 | 0.138054 | 0.029 | 0.041 | 1 | 9 | Gm20045 | -0.012 | Young |
| 0.554588 | 0.771278 | 0.213 | 0.198 | 1 | 9 | Psmc1 | 0.015 | Young |
| 0.554814 | 0.066569 | 0.184 | 0.16 | 1 | 9 | Eprs | 0.024 | Young |
| 0.554946 | 0.855435 | 0.044 | 0.034 | 1 | 9 | Pcdhb17 | 0.01 | Young |
| 0.554946 | 0.775622 | 0.044 | 0.034 | 1 | 9 | Ift80 | 0.01 | Young |
| 0.555139 | 0.315385 | 0.051 | 0.067 | 1 | 9 | Rnf5 | -0.016 | Young |
| 0.555171 | 0.058746 | 0.213 | 0.191 | 1 | 9 | Rgp1 | 0.022 | Young |
| 0.555206 | 0.027055 | 0.125 | 0.103 | 1 | 9 | Yeats4 | 0.022 | Young |
| 0.556528 | 2.397194 | 0.029 | 0.021 | 1 | 9 | 4930522L1 | 0.008 | Young |
| 0.55693 | 0.103803 | 0.272 | 0.291 | 1 | 9 | Ski | -0.019 | Young |

| | | | | | | | | |
|---|---|---|---|---|---|---|---|---|
| 0.557029 | 0.023967 | 0.096 | 0.116 | 1 | 9 | Tcf20 | -0.02 | Young |
| 0.557164 | 0.17977 | 0.015 | 0.023 | 1 | 9 | Gm45902 | -0.008 | Young |
| 0.557197 | 0.193859 | 0.25 | 0.224 | 1 | 9 | Sugt1 | 0.026 | Young |
| 0.557373 | 0.154182 | 0.154 | 0.18 | 1 | 9 | Slc44a1 | -0.026 | Young |
| 0.557373 | 0.20805 | 0.191 | 0.168 | 1 | 9 | Mrps33 | 0.023 | Young |
| 0.557444 | 0.165273 | 0.235 | 0.206 | 1 | 9 | Zranb1 | 0.029 | Young |
| 0.558234 | 0.49642 | 0.029 | 0.021 | 1 | 9 | Ms4a6b | 0.008 | Young |
| 0.558521 | 0.183245 | 0.301 | 0.278 | 1 | 9 | Kras | 0.023 | Young |
| 0.559018 | 0.142231 | 0.213 | 0.242 | 1 | 9 | U2surp | -0.029 | Young |
| 0.55904 | 1.207882 | 0.044 | 0.034 | 1 | 9 | Tmem126a | 0.01 | Young |
| 0.55904 | 0.480016 | 0.044 | 0.034 | 1 | 9 | Pcca | 0.01 | Young |
| 0.559336 | 0.176913 | 0.066 | 0.052 | 1 | 9 | Fbf1 | 0.014 | Young |
| 0.559474 | 0.041342 | 0.081 | 0.101 | 1 | 9 | Ddx52 | -0.02 | Young |
| 0.560213 | 0.327453 | 0.088 | 0.072 | 1 | 9 | Adrb2 | 0.016 | Young |
| 0.560213 | 0.267147 | 0.088 | 0.072 | 1 | 9 | Ogfod1 | 0.016 | Young |
| 0.560253 | 0.281325 | 0.162 | 0.188 | 1 | 9 | Dapk2 | -0.026 | Young |
| 0.560408 | 0.759332 | 0.044 | 0.034 | 1 | 9 | Gamt | 0.01 | Young |
| 0.560408 | 0.604606 | 0.044 | 0.034 | 1 | 9 | Mtif2 | 0.01 | Young |
| 0.560784 | 0.091939 | 0.059 | 0.075 | 1 | 9 | Fopnl | -0.016 | Young |
| 0.560809 | 0.277573 | 0.176 | 0.206 | 1 | 9 | Abracl | -0.03 | Young |
| 0.56097 | 0.346864 | 0.132 | 0.113 | 1 | 9 | Smarcad1 | 0.019 | Young |
| 0.561654 | 0.005182 | 0.029 | 0.021 | 1 | 9 | Trpv2 | 0.008 | Young |
| 0.561777 | 1.108712 | 0.044 | 0.034 | 1 | 9 | Fkbp5 | 0.01 | Young |
| 0.561777 | 0.940552 | 0.044 | 0.034 | 1 | 9 | Ints7 | 0.01 | Young |
| 0.561959 | 0.392095 | 0.074 | 0.093 | 1 | 9 | Slc16a13 | -0.019 | Young |
| 0.562533 | 0.589646 | 0.11 | 0.093 | 1 | 9 | 2510002D2 | 0.017 | Young |
| 0.562746 | 0.765721 | 0.088 | 0.075 | 1 | 9 | C1qb | 0.013 | Young |
| 0.562746 | 0.221013 | 0.066 | 0.082 | 1 | 9 | Tbc1d13 | -0.016 | Young |
| 0.563204 | 0.241014 | 0.25 | 0.227 | 1 | 9 | Mllt6 | 0.023 | Young |
| 0.563368 | 0.047614 | 0.029 | 0.021 | 1 | 9 | Mocos | 0.008 | Young |
| 0.564053 | 0.116574 | 0.235 | 0.268 | 1 | 9 | Nr2f2 | -0.033 | Young |
| 0.564272 | 0.216941 | 0.338 | 0.32 | 1 | 9 | Minos1 | 0.018 | Young |
| 0.564306 | 0.768125 | 0.015 | 0.023 | 1 | 9 | Hibch | -0.008 | Young |
| 0.564306 | 0.612782 | 0.015 | 0.023 | 1 | 9 | Zfp65 | -0.008 | Young |
| 0.564395 | 0.268508 | 0.25 | 0.229 | 1 | 9 | Syf2 | 0.021 | Young |
| 0.5644 | 0.352207 | 0.059 | 0.046 | 1 | 9 | Wtip | 0.013 | Young |
| 0.5644 | 0.315733 | 0.059 | 0.046 | 1 | 9 | Timm10 | 0.013 | Young |
| 0.565069 | 0.228451 | 0.184 | 0.16 | 1 | 9 | Smg7 | 0.024 | Young |
| 0.565178 | 0.088246 | 0.235 | 0.206 | 1 | 9 | Nr4a2 | 0.029 | Young |
| 0.565284 | 0.05104 | 0.456 | 0.492 | 1 | 9 | Rpl14 | -0.036 | Young |
| 0.565572 | 0.098937 | 0.338 | 0.309 | 1 | 9 | Apc | 0.029 | Young |
| 0.565583 | 0.596875 | 0.059 | 0.046 | 1 | 9 | Dbt | 0.013 | Young |
| 0.565583 | 0.339549 | 0.059 | 0.046 | 1 | 9 | Fsd1l | 0.013 | Young |
| 0.565957 | 0.337374 | 0.096 | 0.08 | 1 | 9 | Eva1a | 0.016 | Young |
| 0.565979 | 0.010987 | 0.081 | 0.098 | 1 | 9 | Chchd3 | -0.017 | Young |

| | | | | | | | | |
|---|---|---|---|---|---|---|---|---|
| 0.565992 | 0.536381 | 0.11 | 0.093 | 1 | 9 | Prpf6 | 0.017 | Young |
| 0.566034 | 0.274235 | 0.243 | 0.211 | 1 | 9 | Herc2 | 0.032 | Young |
| 0.566098 | 0.544476 | 0.015 | 0.023 | 1 | 9 | Vamp1 | -0.008 | Young |
| 0.566098 | 0.345873 | 0.015 | 0.023 | 1 | 9 | Slc2a9 | -0.008 | Young |
| 0.566423 | 0.381902 | 0.272 | 0.24 | 1 | 9 | Crebrf | 0.032 | Young |
| 0.566533 | 0.280275 | 0.088 | 0.108 | 1 | 9 | Usp31 | -0.02 | Young |
| 0.566766 | 1.006913 | 0.059 | 0.046 | 1 | 9 | Abhd10 | 0.013 | Young |
| 0.56679 | 0.485037 | 0.081 | 0.067 | 1 | 9 | Dnaja3 | 0.014 | Young |
| 0.566803 | 0.482964 | 0.029 | 0.021 | 1 | 9 | Sssca1 | 0.008 | Young |
| 0.566803 | 0.000188 | 0.029 | 0.021 | 1 | 9 | Gm26853 | 0.008 | Young |
| 0.567132 | 0.035034 | 0.956 | 0.961 | 1 | 9 | mt-Nd1 | -0.005 | Young |
| 0.567271 | 0.84116 | 0.044 | 0.034 | 1 | 9 | Shf | 0.01 | Young |
| 0.567271 | 0.366096 | 0.044 | 0.034 | 1 | 9 | Usp18 | 0.01 | Young |
| 0.567317 | 0.225376 | 0.029 | 0.041 | 1 | 9 | Dpy19l3 | -0.012 | Young |
| 0.5679 | 0.40499 | 0.176 | 0.157 | 1 | 9 | Snx2 | 0.019 | Young |
| 0.568274 | 0.542229 | 0.096 | 0.08 | 1 | 9 | Rnmt | 0.016 | Young |
| 0.56843 | 0.059868 | 0.441 | 0.472 | 1 | 9 | Prrc2c | -0.031 | Young |
| 0.568649 | 0.718861 | 0.044 | 0.034 | 1 | 9 | Pgm5 | 0.01 | Young |
| 0.568909 | 0.016743 | 0.088 | 0.072 | 1 | 9 | Snrpa | 0.016 | Young |
| 0.56907 | 0.177625 | 0.257 | 0.245 | 1 | 9 | Fyn | 0.012 | Young |
| 0.569197 | 0.353834 | 0.184 | 0.16 | 1 | 9 | Fam111a | 0.024 | Young |
| 0.569691 | 1.318815 | 0.015 | 0.023 | 1 | 9 | Ticam1 | -0.008 | Young |
| 0.569879 | 0.318325 | 0.088 | 0.072 | 1 | 9 | Ambra1 | 0.016 | Young |
| 0.570006 | 0.121975 | 0.029 | 0.041 | 1 | 9 | Lancl3 | -0.012 | Young |
| 0.570028 | 0.569744 | 0.044 | 0.034 | 1 | 9 | Ints4 | 0.01 | Young |
| 0.570028 | 0.372493 | 0.044 | 0.034 | 1 | 9 | Apool | 0.01 | Young |
| 0.570325 | 0.573042 | 0.059 | 0.046 | 1 | 9 | Tmem229b | 0.013 | Young |
| 0.570335 | 0.3281 | 0.176 | 0.157 | 1 | 9 | Bcl9l | 0.019 | Young |
| 0.570595 | 0.578921 | 0.096 | 0.08 | 1 | 9 | Ophn1 | 0.016 | Young |
| 0.570787 | 1.055828 | 0.037 | 0.049 | 1 | 9 | Sil1 | -0.012 | Young |
| 0.570965 | 0.553512 | 0.051 | 0.067 | 1 | 9 | Fam122a | -0.016 | Young |
| 0.571409 | 0.95192 | 0.044 | 0.034 | 1 | 9 | Epb41l5 | 0.01 | Young |
| 0.571513 | 0.103167 | 0.059 | 0.046 | 1 | 9 | Ptprs | 0.013 | Young |
| 0.571822 | 0.138102 | 0.088 | 0.072 | 1 | 9 | Gabbr1 | 0.016 | Young |
| 0.572091 | 0.25613 | 0.132 | 0.113 | 1 | 9 | Cetn2 | 0.019 | Young |
| 0.572258 | 0.229963 | 0.632 | 0.629 | 1 | 9 | Rpl6 | 0.003 | Young |
| 0.572323 | 0.753383 | 0.088 | 0.075 | 1 | 9 | Smc5 | 0.013 | Young |
| 0.572576 | 0.24625 | 0.257 | 0.237 | 1 | 9 | 9-Sep | 0.02 | Young |
| 0.572743 | 0.135891 | 0.397 | 0.366 | 1 | 9 | Psma7 | 0.031 | Young |
| 0.572791 | 0.856231 | 0.044 | 0.034 | 1 | 9 | Ggct | 0.01 | Young |
| 0.572931 | 0.050233 | 0.868 | 0.899 | 1 | 9 | Tspan13 | -0.031 | Young |
| 0.573358 | 0.06138 | 0.103 | 0.121 | 1 | 9 | Ilf2 | -0.018 | Young |
| 0.573401 | 0.100563 | 0.125 | 0.108 | 1 | 9 | Psmc4 | 0.017 | Young |
| 0.573847 | 0.089232 | 0.449 | 0.492 | 1 | 9 | Gja1 | -0.043 | Young |
| 0.575425 | 0.078746 | 0.272 | 0.307 | 1 | 9 | Herpud1 | -0.035 | Young |

| | | | | | | | | |
|---|---|---|---|---|---|---|---|---|
| 0.575801 | 0.217727 | 0.235 | 0.265 | 1 | 9 | Cdkn1c | -0.03 | Young |
| 0.57662 | 0.256798 | 0.147 | 0.126 | 1 | 9 | Myo9b | 0.021 | Young |
| 0.576947 | 0.828972 | 0.044 | 0.034 | 1 | 9 | Htra2 | 0.01 | Young |
| 0.576947 | 0.183747 | 0.044 | 0.034 | 1 | 9 | Tsen15 | 0.01 | Young |
| 0.57714 | 0.663255 | 0.088 | 0.075 | 1 | 9 | Plscr1 | 0.013 | Young |
| 0.57739 | 1.21228 | 0.051 | 0.041 | 1 | 9 | Gne | 0.01 | Young |
| 0.57749 | 0.428163 | 0.125 | 0.108 | 1 | 9 | 1500011K1 | 0.017 | Young |
| 0.577525 | 0.23199 | 0.103 | 0.088 | 1 | 9 | Ecm2 | 0.015 | Young |
| 0.57798 | 0.099426 | 0.235 | 0.268 | 1 | 9 | Trpc4ap | -0.033 | Young |
| 0.57831 | 0.322187 | 0.125 | 0.108 | 1 | 9 | Rtf2 | 0.017 | Young |
| 0.578336 | 0.146388 | 0.044 | 0.034 | 1 | 9 | Tmem185a | 0.01 | Young |
| 0.578669 | 0.057308 | 0.059 | 0.046 | 1 | 9 | Ccdc91 | 0.013 | Young |
| 0.579866 | 0.670945 | 0.059 | 0.046 | 1 | 9 | Golga1 | 0.013 | Young |
| 0.579929 | 0.044514 | 0.11 | 0.093 | 1 | 9 | Mrpl36 | 0.017 | Young |
| 0.580064 | 0.792146 | 0.243 | 0.232 | 1 | 9 | Fgfr1op2 | 0.011 | Young |
| 0.580171 | 0.156085 | 0.081 | 0.098 | 1 | 9 | Ccdc174 | -0.017 | Young |
| 0.580704 | 0.347736 | 0.037 | 0.049 | 1 | 9 | Trp53i13 | -0.012 | Young |
| 0.580799 | 0.244518 | 0.096 | 0.116 | 1 | 9 | Bfar | -0.02 | Young |
| 0.580922 | 0.645372 | 0.081 | 0.067 | 1 | 9 | Thoc1 | 0.014 | Young |
| 0.581008 | 0.020397 | 0.066 | 0.082 | 1 | 9 | Ighm | -0.016 | Young |
| 0.581382 | 0.299505 | 0.044 | 0.057 | 1 | 9 | Pdhx | -0.013 | Young |
| 0.581585 | 0.295903 | 0.154 | 0.137 | 1 | 9 | Lamtor3 | 0.017 | Young |
| 0.581956 | 0.087404 | 0.081 | 0.098 | 1 | 9 | Cdr2 | -0.017 | Young |
| 0.582263 | 0.019753 | 0.059 | 0.046 | 1 | 9 | Atp13a2 | 0.013 | Young |
| 0.582511 | 0.126046 | 0.044 | 0.034 | 1 | 9 | Wrap73 | 0.01 | Young |
| 0.582706 | 0.907617 | 0.051 | 0.067 | 1 | 9 | Dhx29 | -0.016 | Young |
| 0.58279 | 0.122255 | 0.118 | 0.139 | 1 | 9 | Lrba | -0.021 | Young |
| 0.582946 | 0.066989 | 0.088 | 0.075 | 1 | 9 | Rps6ka4 | 0.013 | Young |
| 0.583906 | 0.079886 | 0.044 | 0.034 | 1 | 9 | Appl2 | 0.01 | Young |
| 0.583911 | 0.242809 | 0.11 | 0.129 | 1 | 9 | Fam129b | -0.019 | Young |
| 0.583978 | 0.61458 | 0.125 | 0.111 | 1 | 9 | Gfpt1 | 0.014 | Young |
| 0.584418 | 0.165247 | 0.368 | 0.332 | 1 | 9 | Tm9sf2 | 0.036 | Young |
| 0.584488 | 0.151082 | 0.118 | 0.101 | 1 | 9 | Net1 | 0.017 | Young |
| 0.584729 | 0.572701 | 0.074 | 0.09 | 1 | 9 | Plpp5 | -0.016 | Young |
| 0.58482 | 0.083149 | 0.375 | 0.402 | 1 | 9 | Ogt | -0.027 | Young |
| 0.58517 | 0.110885 | 0.287 | 0.263 | 1 | 9 | Rsu1 | 0.024 | Young |
| 0.585224 | 0.594817 | 0.154 | 0.139 | 1 | 9 | Plk3 | 0.015 | Young |
| 0.585224 | 0.189661 | 0.14 | 0.121 | 1 | 9 | Zbtb1 | 0.019 | Young |
| 0.585288 | 0.083951 | 0.022 | 0.031 | 1 | 9 | Ezh2 | -0.009 | Young |
| 0.585356 | 0.378996 | 0.331 | 0.327 | 1 | 9 | Ralgapa1 | 0.004 | Young |
| 0.585535 | 0.303239 | 0.081 | 0.098 | 1 | 9 | Chpt1 | -0.017 | Young |
| 0.585692 | 0.570487 | 0.037 | 0.049 | 1 | 9 | 9430038I0 | -0.012 | Young |
| 0.585692 | 0.351103 | 0.037 | 0.049 | 1 | 9 | Aen | -0.012 | Young |
| 0.586592 | 0.696373 | 0.096 | 0.082 | 1 | 9 | Lpin2 | 0.014 | Young |
| 0.587262 | 0.03885 | 0.154 | 0.173 | 1 | 9 | Hmgn1 | -0.019 | Young |

| | | | | | | | | |
|---|---|---|---|---|---|---|---|---|
| 0.587318 | 0.392828 | 0.066 | 0.082 | 1 | 9 | Acsl3 | -0.016 | Young |
| 0.587509 | 0.368142 | 0.375 | 0.358 | 1 | 9 | Rps27l | 0.017 | Young |
| 0.588095 | 1.127733 | 0.081 | 0.07 | 1 | 9 | Ctnnbl1 | 0.011 | Young |
| 0.588319 | 0.226611 | 0.118 | 0.101 | 1 | 9 | Enpp3 | 0.017 | Young |
| 0.588464 | 0.506418 | 0.066 | 0.054 | 1 | 9 | Gm20342 | 0.012 | Young |
| 0.588586 | 0.321372 | 0.169 | 0.149 | 1 | 9 | Rgs12 | 0.02 | Young |
| 0.58883 | 0.009795 | 0.184 | 0.204 | 1 | 9 | Taf1 | -0.02 | Young |
| 0.588867 | 0.037741 | 0.434 | 0.472 | 1 | 9 | Serinc1 | -0.038 | Young |
| 0.589328 | 0.205987 | 0.412 | 0.402 | 1 | 9 | Spop | 0.01 | Young |
| 0.589387 | 0.055794 | 0.294 | 0.268 | 1 | 9 | Usf2 | 0.026 | Young |
| 0.589602 | 0.692197 | 0.118 | 0.144 | 1 | 9 | Mospd1 | -0.026 | Young |
| 0.589641 | 0.020961 | 0.199 | 0.175 | 1 | 9 | Csnk1g2 | 0.024 | Young |
| 0.589773 | 0.494689 | 0.11 | 0.095 | 1 | 9 | Hyal2 | 0.015 | Young |
| 0.590032 | 0.133429 | 0.096 | 0.111 | 1 | 9 | Gt(ROSA)2( | -0.015 | Young |
| 0.590729 | 0.20288 | 0.088 | 0.075 | 1 | 9 | Extl3 | 0.013 | Young |
| 0.590925 | 0.399623 | 0.066 | 0.08 | 1 | 9 | Smim27 | -0.014 | Young |
| 0.591053 | 0.023086 | 0.132 | 0.155 | 1 | 9 | Ipmk | -0.023 | Young |
| 0.591117 | 0.167854 | 0.081 | 0.067 | 1 | 9 | Phldb1 | 0.014 | Young |
| 0.591525 | 0.12642 | 0.191 | 0.219 | 1 | 9 | Ssrp1 | -0.028 | Young |
| 0.591879 | 0.233454 | 0.353 | 0.325 | 1 | 9 | Vsir | 0.028 | Young |
| 0.591956 | 0.082979 | 0.051 | 0.064 | 1 | 9 | Gsdmd | -0.013 | Young |
| 0.59236 | 0.03652 | 0.382 | 0.415 | 1 | 9 | Tap2 | -0.033 | Young |
| 0.592658 | 0.062283 | 0.25 | 0.227 | 1 | 9 | Rbm5 | 0.023 | Young |
| 0.592896 | 0.90787 | 0.199 | 0.178 | 1 | 9 | Arhgef28 | 0.021 | Young |
| 0.593536 | 1.005383 | 0.022 | 0.015 | 1 | 9 | Axin2 | 0.007 | Young |
| 0.593967 | 0.752975 | 0.051 | 0.041 | 1 | 9 | Paqr8 | 0.01 | Young |
| 0.59411 | 0.961011 | 0.066 | 0.054 | 1 | 9 | Ap2a1 | 0.012 | Young |
| 0.59411 | 0.025932 | 0.456 | 0.518 | 1 | 9 | Rps2 | -0.062 | Young |
| 0.59415 | 0.089562 | 0.051 | 0.064 | 1 | 9 | Nfs1 | -0.013 | Young |
| 0.595252 | 1.178266 | 0.051 | 0.041 | 1 | 9 | Mast2 | 0.01 | Young |
| 0.595288 | 0.211984 | 0.235 | 0.214 | 1 | 9 | Scarf1 | 0.021 | Young |
| 0.595297 | 0.144366 | 0.434 | 0.415 | 1 | 9 | Clec2d | 0.019 | Young |
| 0.595348 | 0.117078 | 0.044 | 0.057 | 1 | 9 | Fbxo42 | -0.013 | Young |
| 0.595562 | 1.739765 | 0.022 | 0.015 | 1 | 9 | Gga2 | 0.007 | Young |
| 0.595563 | 0.023043 | 0.221 | 0.247 | 1 | 9 | Tmem234 | -0.026 | Young |
| 0.595887 | 0.00933 | 0.25 | 0.273 | 1 | 9 | Cpne3 | -0.023 | Young |
| 0.596464 | 0.929359 | 0.074 | 0.062 | 1 | 9 | Fbxl4 | 0.012 | Young |
| 0.596522 | 0.451037 | 0.132 | 0.119 | 1 | 9 | Adprh | 0.013 | Young |
| 0.596874 | 0.053122 | 0.118 | 0.101 | 1 | 9 | Ing3 | 0.017 | Young |
| 0.597189 | 0.145125 | 0.368 | 0.356 | 1 | 9 | Adgrl2 | 0.012 | Young |
| 0.597436 | 0.669834 | 0.162 | 0.147 | 1 | 9 | Golim4 | 0.015 | Young |
| 0.597824 | 0.80278 | 0.051 | 0.041 | 1 | 9 | Pex7 | 0.01 | Young |
| 0.597881 | 0.134912 | 0.022 | 0.031 | 1 | 9 | Gm5617 | -0.009 | Young |
| 0.59794 | 0.292773 | 0.044 | 0.034 | 1 | 9 | Dohh | 0.01 | Young |
| 0.59803 | 0.027773 | 0.176 | 0.198 | 1 | 9 | Lrch3 | -0.022 | Young |

| | | | | | | | | |
|---|---|---|---|---|---|---|---|---|
| 0.598303 | 0.106347 | 0.125 | 0.106 | 1 | 9 | Ap3d1 | 0.019 | Young |
| 0.598896 | 0.185579 | 0.074 | 0.059 | 1 | 9 | 1110059G1 | 0.015 | Young |
| 0.599622 | 2.229572 | 0.022 | 0.015 | 1 | 9 | Numbl | 0.007 | Young |
| 0.599622 | 1.315568 | 0.022 | 0.015 | 1 | 9 | Olfr1396 | 0.007 | Young |
| 0.599702 | 0.177549 | 0.096 | 0.113 | 1 | 9 | 9530082P2 | -0.017 | Young |
| 0.599708 | 0.06115 | 0.074 | 0.09 | 1 | 9 | Yaf2 | -0.016 | Young |
| 0.600343 | 0.348971 | 0.132 | 0.113 | 1 | 9 | Aup1 | 0.019 | Young |
| 0.600747 | 0.513007 | 0.074 | 0.062 | 1 | 9 | Cipc | 0.012 | Young |
| 0.600747 | 0.503882 | 0.074 | 0.062 | 1 | 9 | Colgalt2 | 0.012 | Young |
| 0.6009 | 0.14553 | 0.449 | 0.438 | 1 | 9 | Gabarap | 0.011 | Young |
| 0.60098 | 0.13642 | 0.25 | 0.227 | 1 | 9 | Eif3b | 0.023 | Young |
| 0.601049 | 0.853639 | 0.022 | 0.031 | 1 | 9 | Crtc1 | -0.009 | Young |
| 0.601049 | 0.618036 | 0.022 | 0.031 | 1 | 9 | Phf21b | -0.009 | Young |
| 0.601212 | 0.233255 | 0.044 | 0.057 | 1 | 9 | Gpatch1 | -0.013 | Young |
| 0.601656 | 1.883901 | 0.022 | 0.015 | 1 | 9 | Zfp961 | 0.007 | Young |
| 0.601656 | 1.869679 | 0.022 | 0.015 | 1 | 9 | Pkmyt1 | 0.007 | Young |
| 0.601656 | 1.372093 | 0.022 | 0.015 | 1 | 9 | 9130221H1 | 0.007 | Young |
| 0.601772 | 0.105555 | 0.228 | 0.253 | 1 | 9 | Calcrl | -0.025 | Young |
| 0.601813 | 0.35778 | 0.25 | 0.235 | 1 | 9 | Gimap8 | 0.015 | Young |
| 0.601893 | 0.130293 | 0.184 | 0.165 | 1 | 9 | Add3 | 0.019 | Young |
| 0.601931 | 0.1119 | 0.169 | 0.147 | 1 | 9 | Gprc5b | 0.022 | Young |
| 0.601961 | 0.608434 | 0.059 | 0.072 | 1 | 9 | Oxa1l | -0.013 | Young |
| 0.601997 | 0.182371 | 0.14 | 0.16 | 1 | 9 | Fmr1 | -0.02 | Young |
| 0.602425 | 0.583948 | 0.081 | 0.067 | 1 | 9 | Arl3 | 0.014 | Young |
| 0.602427 | 0.070889 | 0.272 | 0.304 | 1 | 9 | Rab7 | -0.032 | Young |
| 0.602536 | 0.405669 | 0.074 | 0.09 | 1 | 9 | Kxd1 | -0.016 | Young |
| 0.603194 | 0.26699 | 0.066 | 0.054 | 1 | 9 | Olfm1 | 0.012 | Young |
| 0.603194 | 0.154531 | 0.066 | 0.054 | 1 | 9 | Nbn | 0.012 | Young |
| 0.603694 | 1.906818 | 0.022 | 0.015 | 1 | 9 | Fam167b | 0.007 | Young |
| 0.603694 | 1.657353 | 0.022 | 0.015 | 1 | 9 | Adamts7 | 0.007 | Young |
| 0.603694 | 1.387551 | 0.022 | 0.015 | 1 | 9 | Spata7 | 0.007 | Young |
| 0.603694 | 1.289799 | 0.022 | 0.015 | 1 | 9 | Chac2 | 0.007 | Young |
| 0.603694 | 0.857832 | 0.022 | 0.015 | 1 | 9 | Stox1 | 0.007 | Young |
| 0.603853 | 0.275631 | 0.11 | 0.095 | 1 | 9 | Acot9 | 0.015 | Young |
| 0.604201 | 0.155683 | 0.243 | 0.227 | 1 | 9 | Lnx1 | 0.016 | Young |
| 0.604224 | 0.259846 | 0.022 | 0.031 | 1 | 9 | Als2 | -0.009 | Young |
| 0.605174 | 0.347699 | 0.051 | 0.064 | 1 | 9 | Mogs | -0.013 | Young |
| 0.605234 | 0.085787 | 0.118 | 0.137 | 1 | 9 | Kansl2 | -0.019 | Young |
| 0.605412 | 0.233492 | 0.434 | 0.425 | 1 | 9 | Psme1 | 0.009 | Young |
| 0.60546 | 0.026437 | 0.581 | 0.552 | 1 | 9 | Plxnd1 | 0.029 | Young |
| 0.605526 | 0.235787 | 0.081 | 0.067 | 1 | 9 | Ints11 | 0.014 | Young |
| 0.605589 | 0.089189 | 0.162 | 0.139 | 1 | 9 | Ubp1 | 0.023 | Young |
| 0.605646 | 0.220303 | 0.25 | 0.286 | 1 | 9 | Tnpo3 | -0.036 | Young |
| 0.605735 | 1.982877 | 0.022 | 0.015 | 1 | 9 | Cib2 | 0.007 | Young |
| 0.605735 | 1.653443 | 0.022 | 0.015 | 1 | 9 | Gatb | 0.007 | Young |

| | | | | | | | | |
|---|---|---|---|---|---|---|---|---|
| 0.605735 | 1.12657 | 0.022 | 0.015 | 1 | 9 | Tspyl3 | 0.007 | Young |
| 0.605735 | 0.628557 | 0.022 | 0.015 | 1 | 9 | Mmaa | 0.007 | Young |
| 0.605735 | 0.448843 | 0.022 | 0.015 | 1 | 9 | Mipep | 0.007 | Young |
| 0.60596 | 0.155879 | 0.206 | 0.227 | 1 | 9 | Psmb5 | -0.021 | Young |
| 0.606121 | 0.413704 | 0.074 | 0.062 | 1 | 9 | Cfap97 | 0.012 | Young |
| 0.606282 | 0.049478 | 0.051 | 0.064 | 1 | 9 | Meox1 | -0.013 | Young |
| 0.606378 | 0.874279 | 0.081 | 0.07 | 1 | 9 | Nmt2 | 0.011 | Young |
| 0.606615 | 0.01762 | 0.375 | 0.412 | 1 | 9 | H2afz | -0.037 | Young |
| 0.606616 | 0.3534 | 0.066 | 0.054 | 1 | 9 | Brf1 | 0.012 | Young |
| 0.606898 | 1.281528 | 0.037 | 0.028 | 1 | 9 | Zfp157 | 0.009 | Young |
| 0.607049 | 0.174627 | 0.007 | 0.013 | 1 | 9 | Map10 | -0.006 | Young |
| 0.607049 | 0.084617 | 0.007 | 0.013 | 1 | 9 | Sox11 | -0.006 | Young |
| 0.607049 | 0.082051 | 0.007 | 0.013 | 1 | 9 | Zfp189 | -0.006 | Young |
| 0.607396 | 0.362959 | 0.176 | 0.16 | 1 | 9 | Zfp281 | 0.016 | Young |
| 0.60821 | 0.535439 | 0.096 | 0.082 | 1 | 9 | Nipsnap2 | 0.014 | Young |
| 0.608443 | 0.757798 | 0.037 | 0.028 | 1 | 9 | Osbpl7 | 0.009 | Young |
| 0.608452 | 0.207865 | 0.103 | 0.088 | 1 | 9 | Zdhhc2 | 0.015 | Young |
| 0.609 | 0.775307 | 0.022 | 0.031 | 1 | 9 | Ttc8 | -0.009 | Young |
| 0.609055 | 0.282553 | 0.051 | 0.064 | 1 | 9 | Snx21 | -0.013 | Young |
| 0.60946 | 0.46894 | 0.051 | 0.041 | 1 | 9 | Dedd2 | 0.01 | Young |
| 0.609548 | 3.365991 | 0.007 | 0.013 | 1 | 9 | Hbb-bt | -0.006 | Young |
| 0.609548 | 1.336306 | 0.007 | 0.013 | 1 | 9 | Spock2 | -0.006 | Young |
| 0.609548 | 1.24313 | 0.007 | 0.013 | 1 | 9 | Ehhadh | -0.006 | Young |
| 0.609548 | 0.987896 | 0.007 | 0.013 | 1 | 9 | Pard6a | -0.006 | Young |
| 0.609548 | 0.811901 | 0.007 | 0.013 | 1 | 9 | Rarb | -0.006 | Young |
| 0.609548 | 0.761518 | 0.007 | 0.013 | 1 | 9 | Tcte2 | -0.006 | Young |
| 0.609548 | 0.496087 | 0.007 | 0.013 | 1 | 9 | Gm11837 | -0.006 | Young |
| 0.609548 | 0.492445 | 0.007 | 0.013 | 1 | 9 | Jmjd4 | -0.006 | Young |
| 0.609548 | 0.47309 | 0.007 | 0.013 | 1 | 9 | Naglu | -0.006 | Young |
| 0.609548 | 0.310574 | 0.007 | 0.013 | 1 | 9 | Tmem220 | -0.006 | Young |
| 0.609548 | 0.29866 | 0.007 | 0.013 | 1 | 9 | Zfp619 | -0.006 | Young |
| 0.609548 | 0.287349 | 0.007 | 0.013 | 1 | 9 | Matk | -0.006 | Young |
| 0.609548 | 0.272675 | 0.007 | 0.013 | 1 | 9 | Zbtb48 | -0.006 | Young |
| 0.609548 | 0.248146 | 0.007 | 0.013 | 1 | 9 | Park2 | -0.006 | Young |
| 0.609548 | 0.034106 | 0.007 | 0.013 | 1 | 9 | D830025C( | -0.006 | Young |
| 0.609548 | 0.032372 | 0.007 | 0.013 | 1 | 9 | Olfr1167 | -0.006 | Young |
| 0.60966 | 0.203386 | 0.199 | 0.18 | 1 | 9 | Gm16286 | 0.019 | Young |
| 0.609826 | 1.509755 | 0.022 | 0.015 | 1 | 9 | Ankrd29 | 0.007 | Young |
| 0.610048 | 0.449408 | 0.066 | 0.054 | 1 | 9 | Pias3 | 0.012 | Young |
| 0.610473 | 0.821228 | 0.059 | 0.049 | 1 | 9 | Aif1 | 0.01 | Young |
| 0.610589 | 0.116529 | 0.14 | 0.121 | 1 | 9 | Pdhb | 0.019 | Young |
| 0.610596 | 0.612531 | 0.022 | 0.031 | 1 | 9 | Ppp3cc | -0.009 | Young |
| 0.610648 | 0.507517 | 0.044 | 0.057 | 1 | 9 | Afg3l2 | -0.013 | Young |
| 0.610738 | 0.106181 | 0.096 | 0.08 | 1 | 9 | Ahcyl2 | 0.016 | Young |
| 0.610818 | 0.102006 | 0.206 | 0.232 | 1 | 9 | Ephb4 | -0.026 | Young |

| | | | | | | | | |
|---|---|---|---|---|---|---|---|---|
| 0.611139 | 0.25383 | 0.191 | 0.173 | 1 | 9 | Rbm22 | 0.018 | Young |
| 0.611217 | 0.053643 | 0.074 | 0.088 | 1 | 9 | Pcyox1 | -0.014 | Young |
| 0.611537 | 0.660481 | 0.037 | 0.028 | 1 | 9 | Vps37c | 0.009 | Young |
| 0.611777 | 0.192358 | 0.088 | 0.106 | 1 | 9 | Anxa4 | -0.018 | Young |
| 0.611844 | 0.169356 | 0.471 | 0.546 | 1 | 9 | Tra2a | -0.075 | Young |
| 0.611876 | 0.396848 | 0.022 | 0.015 | 1 | 9 | Trp53rkb | 0.007 | Young |
| 0.611876 | 0.080973 | 0.022 | 0.015 | 1 | 9 | Med9 | 0.007 | Young |
| 0.612124 | 0.274849 | 0.029 | 0.039 | 1 | 9 | Nol6 | -0.01 | Young |
| 0.612194 | 0.097967 | 0.022 | 0.031 | 1 | 9 | Adck2 | -0.009 | Young |
| 0.612273 | 0.272901 | 0.375 | 0.366 | 1 | 9 | Rab1a | 0.009 | Young |
| 0.612956 | 0.040061 | 0.074 | 0.09 | 1 | 9 | Cog5 | -0.016 | Young |
| 0.613086 | 0.991227 | 0.037 | 0.028 | 1 | 9 | Tepsin | 0.009 | Young |
| 0.613086 | 0.231683 | 0.037 | 0.028 | 1 | 9 | Gucd1 | 0.009 | Young |
| 0.613361 | 0.620069 | 0.051 | 0.041 | 1 | 9 | Dusp2 | 0.01 | Young |
| 0.613361 | 0.590386 | 0.051 | 0.041 | 1 | 9 | Brms1 | 0.01 | Young |
| 0.613543 | 0.087611 | 0.088 | 0.106 | 1 | 9 | Dph3 | -0.018 | Young |
| 0.613613 | 0.391378 | 0.103 | 0.09 | 1 | 9 | Aig1 | 0.013 | Young |
| 0.613659 | 0.359416 | 0.316 | 0.294 | 1 | 9 | Usp34 | 0.022 | Young |
| 0.613929 | 0.463053 | 0.022 | 0.015 | 1 | 9 | Rps6ka5 | 0.007 | Young |
| 0.613929 | 0.174665 | 0.022 | 0.015 | 1 | 9 | Abca2 | 0.007 | Young |
| 0.614247 | 0.12786 | 0.235 | 0.26 | 1 | 9 | Pds5a | -0.025 | Young |
| 0.61427 | 0.076403 | 0.375 | 0.407 | 1 | 9 | Bnip3l | -0.032 | Young |
| 0.614292 | 0.197065 | 0.206 | 0.232 | 1 | 9 | Ap2b1 | -0.026 | Young |
| 0.614667 | 0.114159 | 0.338 | 0.312 | 1 | 9 | Lpp | 0.026 | Young |
| 0.614859 | 0.098492 | 0.096 | 0.082 | 1 | 9 | Slc12a2 | 0.014 | Young |
| 0.614979 | 0.607189 | 0.029 | 0.039 | 1 | 9 | Tceanc2 | -0.01 | Young |
| 0.615811 | 0.560544 | 0.096 | 0.082 | 1 | 9 | Nup50 | 0.014 | Young |
| 0.615985 | 0.624615 | 0.022 | 0.015 | 1 | 9 | Fkbp10 | 0.007 | Young |
| 0.615985 | 0.446052 | 0.022 | 0.015 | 1 | 9 | Gprc5a | 0.007 | Young |
| 0.615985 | 0.118919 | 0.022 | 0.015 | 1 | 9 | Coil | 0.007 | Young |
| 0.61612 | 0.695531 | 0.118 | 0.103 | 1 | 9 | Pggt1b | 0.015 | Young |
| 0.61619 | 0.846708 | 0.037 | 0.028 | 1 | 9 | Mn1 | 0.009 | Young |
| 0.61619 | 0.583494 | 0.037 | 0.028 | 1 | 9 | Cdkn2c | 0.009 | Young |
| 0.616366 | 0.394654 | 0.265 | 0.25 | 1 | 9 | Srsf7 | 0.015 | Young |
| 0.616801 | 0.008422 | 0.316 | 0.286 | 1 | 9 | Vasp | 0.03 | Young |
| 0.616838 | 0.099335 | 0.147 | 0.129 | 1 | 9 | Yap1 | 0.018 | Young |
| 0.616931 | 0.001091 | 0.074 | 0.062 | 1 | 9 | Ostm1 | 0.012 | Young |
| 0.616998 | 0.283642 | 0.022 | 0.031 | 1 | 9 | Gm3336 | -0.009 | Young |
| 0.617189 | 0.000109 | 0.11 | 0.095 | 1 | 9 | Arhgap12 | 0.015 | Young |
| 0.617273 | 0.423592 | 0.051 | 0.041 | 1 | 9 | L3mbtl2 | 0.01 | Young |
| 0.617377 | 0.110801 | 0.279 | 0.299 | 1 | 9 | Pkd2 | -0.02 | Young |
| 0.617745 | 0.305022 | 0.037 | 0.028 | 1 | 9 | Slc10a3 | 0.009 | Young |
| 0.617768 | 0.214104 | 0.044 | 0.057 | 1 | 9 | Asb8 | -0.013 | Young |
| 0.618017 | 0.644625 | 0.074 | 0.062 | 1 | 9 | Mcoln1 | 0.012 | Young |
| 0.618039 | 0.026059 | 0.25 | 0.273 | 1 | 9 | Parvb | -0.023 | Young |

| | | | | | | | | |
|---|---|---|---|---|---|---|---|---|
| 0.618044 | 0.782663 | 0.022 | 0.015 | 1 | 9 | Upb1 | 0.007 | Young |
| 0.618044 | 0.066929 | 0.022 | 0.015 | 1 | 9 | Magee1 | 0.007 | Young |
| 0.618339 | 0.05132 | 0.088 | 0.075 | 1 | 9 | Bscl2 | 0.013 | Young |
| 0.618579 | 0.745299 | 0.051 | 0.041 | 1 | 9 | Hdac9 | 0.01 | Young |
| 0.618603 | 1.574234 | 0.022 | 0.031 | 1 | 9 | 4632415L0 | -0.009 | Young |
| 0.618755 | 0.051013 | 0.118 | 0.134 | 1 | 9 | Dnajc21 | -0.016 | Young |
| 0.618782 | 0.079468 | 0.191 | 0.173 | 1 | 9 | Uqcrfs1 | 0.018 | Young |
| 0.619216 | 0.227829 | 0.272 | 0.258 | 1 | 9 | H2afy | 0.014 | Young |
| 0.619302 | 0.949869 | 0.037 | 0.028 | 1 | 9 | Upp2 | 0.009 | Young |
| 0.619302 | 0.374214 | 0.037 | 0.028 | 1 | 9 | Mgat5 | 0.009 | Young |
| 0.619384 | 0.096664 | 0.257 | 0.24 | 1 | 9 | Ankrd13a | 0.017 | Young |
| 0.620106 | 0.097257 | 0.022 | 0.015 | 1 | 9 | Pomk | 0.007 | Young |
| 0.620138 | 0.809811 | 0.059 | 0.049 | 1 | 9 | Mex3d | 0.01 | Young |
| 0.62021 | 0.243426 | 0.022 | 0.031 | 1 | 9 | E130308A1 | -0.009 | Young |
| 0.620759 | 0.275315 | 0.206 | 0.186 | 1 | 9 | R3hdm2 | 0.02 | Young |
| 0.621196 | 0.640869 | 0.051 | 0.041 | 1 | 9 | Nudt12 | 0.01 | Young |
| 0.621196 | 0.400655 | 0.051 | 0.041 | 1 | 9 | Gnptg | 0.01 | Young |
| 0.621196 | 0.081858 | 0.051 | 0.041 | 1 | 9 | Srcap | 0.01 | Young |
| 0.621375 | 0.012833 | 0.39 | 0.433 | 1 | 9 | Ctnnd1 | -0.043 | Young |
| 0.622364 | 0.100271 | 0.184 | 0.165 | 1 | 9 | Usp22 | 0.019 | Young |
| 0.62242 | 0.117459 | 0.037 | 0.028 | 1 | 9 | Tmem201 | 0.009 | Young |
| 0.622712 | 0.438604 | 0.221 | 0.198 | 1 | 9 | Hmg20b | 0.023 | Young |
| 0.622795 | 0.30732 | 0.081 | 0.098 | 1 | 9 | Bicral | -0.017 | Young |
| 0.62292 | 0.009169 | 0.199 | 0.178 | 1 | 9 | Pxk | 0.021 | Young |
| 0.623008 | 0.061208 | 0.088 | 0.103 | 1 | 9 | Arfgap2 | -0.015 | Young |
| 0.623458 | 1.074168 | 0.074 | 0.062 | 1 | 9 | Rpa2 | 0.012 | Young |
| 0.623647 | 0.066748 | 0.206 | 0.235 | 1 | 9 | Zfp292 | -0.029 | Young |
| 0.623787 | 0.363827 | 0.184 | 0.165 | 1 | 9 | Atp6v0a1 | 0.019 | Young |
| 0.623981 | 0.491615 | 0.037 | 0.028 | 1 | 9 | Rmi1 | 0.009 | Young |
| 0.623981 | 0.476664 | 0.037 | 0.028 | 1 | 9 | Clu | 0.009 | Young |
| 0.624413 | 0.471258 | 0.096 | 0.082 | 1 | 9 | 5730455P1 | 0.014 | Young |
| 0.624907 | 0.682947 | 0.132 | 0.119 | 1 | 9 | Exoc6b | 0.013 | Young |
| 0.624907 | 0.119348 | 0.096 | 0.113 | 1 | 9 | Bag5 | -0.017 | Young |
| 0.625246 | 0.307992 | 0.051 | 0.064 | 1 | 9 | Morc2a | -0.013 | Young |
| 0.625507 | 0.274841 | 0.243 | 0.227 | 1 | 9 | Prkacb | 0.016 | Young |
| 0.625609 | 0.007643 | 0.147 | 0.165 | 1 | 9 | Neurl1b | -0.018 | Young |
| 0.625623 | 0.340514 | 0.404 | 0.394 | 1 | 9 | Hadhb | 0.01 | Young |
| 0.62565 | 0.197986 | 0.191 | 0.17 | 1 | 9 | Col4a3bp | 0.021 | Young |
| 0.625966 | 0.744366 | 0.11 | 0.098 | 1 | 9 | Errfi1 | 0.012 | Young |
| 0.626314 | 0.616606 | 0.162 | 0.149 | 1 | 9 | Cdh23 | 0.013 | Young |
| 0.626323 | 0.956117 | 0.066 | 0.057 | 1 | 9 | Lrrc28 | 0.009 | Young |
| 0.626349 | 0.552721 | 0.125 | 0.111 | 1 | 9 | Mdm2 | 0.014 | Young |
| 0.627047 | 0.078952 | 0.243 | 0.265 | 1 | 9 | Osbpl9 | -0.022 | Young |
| 0.627109 | 0.814775 | 0.037 | 0.028 | 1 | 9 | Mios | 0.009 | Young |
| 0.627109 | 0.5798 | 0.037 | 0.028 | 1 | 9 | Mad1l1 | 0.009 | Young |

| | | | | | | | | |
|---|---|---|---|---|---|---|---|---|
| 0.627109 | 0.013965 | 0.037 | 0.028 | 1 | 9 | Odf3b | 0.009 | Young |
| 0.62728 | 0.096013 | 0.059 | 0.072 | 1 | 9 | Ppp2r3d | -0.013 | Young |
| 0.62764 | 0.057566 | 0.191 | 0.168 | 1 | 9 | Csnk1g3 | 0.023 | Young |
| 0.627897 | 0.302303 | 0.029 | 0.039 | 1 | 9 | Prr7 | -0.01 | Young |
| 0.628122 | 0.222763 | 0.316 | 0.291 | 1 | 9 | Csnk2a1 | 0.025 | Young |
| 0.628188 | 0.709386 | 0.132 | 0.119 | 1 | 9 | Timp2 | 0.013 | Young |
| 0.628405 | 0.164445 | 0.676 | 0.724 | 1 | 9 | Hsp90b1 | -0.048 | Young |
| 0.628676 | 0.305507 | 0.037 | 0.028 | 1 | 9 | Mapkbp1 | 0.009 | Young |
| 0.628941 | 0.227883 | 0.36 | 0.353 | 1 | 9 | Eif3f | 0.007 | Young |
| 0.629176 | 0.19024 | 0.346 | 0.379 | 1 | 9 | Romo1 | -0.033 | Young |
| 0.629573 | 0.192844 | 0.287 | 0.268 | 1 | 9 | Smc6 | 0.019 | Young |
| 0.630014 | 0.357139 | 0.074 | 0.062 | 1 | 9 | 2210016L2 | 0.012 | Young |
| 0.630052 | 0.211789 | 0.5 | 0.497 | 1 | 9 | Mapk3 | 0.003 | Young |
| 0.630159 | 0.082045 | 0.64 | 0.688 | 1 | 9 | Selenop | -0.048 | Young |
| 0.630504 | 0.141219 | 0.11 | 0.129 | 1 | 9 | Pald1 | -0.019 | Young |
| 0.63111 | 0.477522 | 0.074 | 0.062 | 1 | 9 | Bola3 | 0.012 | Young |
| 0.631525 | 0.216367 | 0.118 | 0.134 | 1 | 9 | Znrd1 | -0.016 | Young |
| 0.631663 | 0.166431 | 0.235 | 0.214 | 1 | 9 | Eif1b | 0.021 | Young |
| 0.631814 | 0.324046 | 0.037 | 0.028 | 1 | 9 | Exosc1 | 0.009 | Young |
| 0.632597 | 0.496529 | 0.147 | 0.129 | 1 | 9 | Mdh1 | 0.018 | Young |
| 0.633065 | 0.291472 | 0.096 | 0.082 | 1 | 9 | Madd | 0.014 | Young |
| 0.633386 | 0.766757 | 0.037 | 0.028 | 1 | 9 | Ift172 | 0.009 | Young |
| 0.633676 | 0.867229 | 0.029 | 0.039 | 1 | 9 | Cxcl10 | -0.01 | Young |
| 0.634333 | 0.497065 | 0.14 | 0.126 | 1 | 9 | Tcirg1 | 0.014 | Young |
| 0.634749 | 0.211933 | 0.059 | 0.072 | 1 | 9 | Bik | -0.013 | Young |
| 0.634959 | 0.218983 | 0.037 | 0.028 | 1 | 9 | Med11 | 0.009 | Young |
| 0.635168 | 0.09183 | 0.118 | 0.101 | 1 | 9 | Ccndbp1 | 0.017 | Young |
| 0.635191 | 0.154397 | 0.279 | 0.258 | 1 | 9 | Rc3h1 | 0.021 | Young |
| 0.635398 | 0.072038 | 0.103 | 0.088 | 1 | 9 | Sra1 | 0.015 | Young |
| 0.635958 | 0.201307 | 0.257 | 0.232 | 1 | 9 | Tnpo1 | 0.025 | Young |
| 0.636053 | 0.019407 | 0.221 | 0.237 | 1 | 9 | Larp1 | -0.016 | Young |
| 0.636087 | 0.093796 | 0.331 | 0.314 | 1 | 9 | Dgke | 0.017 | Young |
| 0.636385 | 0.217772 | 0.074 | 0.088 | 1 | 9 | Kpna6 | -0.014 | Young |
| 0.63643 | 0.493328 | 0.11 | 0.131 | 1 | 9 | Chchd1 | -0.021 | Young |
| 0.636534 | 0.286329 | 0.037 | 0.028 | 1 | 9 | Lamb1 | 0.009 | Young |
| 0.636534 | 0.234945 | 0.037 | 0.028 | 1 | 9 | Nkiras2 | 0.009 | Young |
| 0.636573 | 0.101354 | 0.029 | 0.039 | 1 | 9 | Slc25a44 | -0.01 | Young |
| 0.636684 | 0.05581 | 0.515 | 0.466 | 1 | 9 | Clk1 | 0.049 | Young |
| 0.637148 | 0.535688 | 0.11 | 0.098 | 1 | 9 | Blvrb | 0.012 | Young |
| 0.637211 | 0.53262 | 0.059 | 0.049 | 1 | 9 | Amd2 | 0.01 | Young |
| 0.637651 | 0.012256 | 0.051 | 0.064 | 1 | 9 | Maip1 | -0.013 | Young |
| 0.637667 | 0.245478 | 0.434 | 0.508 | 1 | 9 | Ctsb | -0.074 | Young |
| 0.638093 | 0.10437 | 0.184 | 0.165 | 1 | 9 | Dnajc7 | 0.019 | Young |
| 0.63819 | 0.579912 | 0.243 | 0.227 | 1 | 9 | Smad4 | 0.016 | Young |
| 0.63819 | 0.100793 | 0.243 | 0.227 | 1 | 9 | Ube2s | 0.016 | Young |

| | | | | | | | | |
|---|---|---|---|---|---|---|---|---|
| 0.638318 | 0.123531 | 0.051 | 0.041 | 1 | 9 | Gatd1 | 0.01 | Young |
| 0.638318 | 0.450037 | 0.037 | 0.046 | 1 | 9 | Snai1 | -0.009 | Young |
| 0.638438 | 0.924678 | 0.059 | 0.049 | 1 | 9 | Spice1 | 0.01 | Young |
| 0.638438 | 0.596955 | 0.059 | 0.049 | 1 | 9 | Snx29 | 0.01 | Young |
| 0.639243 | 0.00064 | 0.088 | 0.103 | 1 | 9 | Mustn1 | -0.015 | Young |
| 0.639904 | 0.060986 | 0.074 | 0.062 | 1 | 9 | Arhgef26 | 0.012 | Young |
| 0.640094 | 0.289622 | 0.103 | 0.088 | 1 | 9 | Rab3gap2 | 0.015 | Young |
| 0.640895 | 0.936253 | 0.059 | 0.049 | 1 | 9 | Pik3ip1 | 0.01 | Young |
| 0.64097 | 0.237949 | 0.051 | 0.041 | 1 | 9 | Mrps30 | 0.01 | Young |
| 0.641734 | 0.160344 | 0.191 | 0.214 | 1 | 9 | Tbrg1 | -0.023 | Young |
| 0.641973 | 0.929575 | 0.044 | 0.036 | 1 | 9 | Limk2 | 0.008 | Young |
| 0.642421 | 0.296358 | 0.11 | 0.095 | 1 | 9 | Lfng | 0.015 | Young |
| 0.64289 | 0.266998 | 0.176 | 0.198 | 1 | 9 | Rbm42 | -0.022 | Young |
| 0.642891 | 0.423341 | 0.103 | 0.119 | 1 | 9 | Fbxl5 | -0.016 | Young |
| 0.643253 | 0.245851 | 0.132 | 0.116 | 1 | 9 | Tmem106a | 0.016 | Young |
| 0.643356 | 0.293246 | 0.059 | 0.049 | 1 | 9 | Large1 | 0.01 | Young |
| 0.643709 | 0.81501 | 0.096 | 0.082 | 1 | 9 | Plod2 | 0.014 | Young |
| 0.643775 | 0.10094 | 0.603 | 0.621 | 1 | 9 | Cox4i1 | -0.018 | Young |
| 0.644136 | 0.519011 | 0.103 | 0.09 | 1 | 9 | Ap1s1 | 0.013 | Young |
| 0.64432 | 0.139717 | 0.074 | 0.062 | 1 | 9 | Ube2o | 0.012 | Young |
| 0.644433 | 0.197995 | 0.037 | 0.028 | 1 | 9 | Tmtc2 | 0.009 | Young |
| 0.644478 | 0.28076 | 0.118 | 0.137 | 1 | 9 | Mier3 | -0.019 | Young |
| 0.644701 | 0.148177 | 0.132 | 0.119 | 1 | 9 | Slc20a1 | 0.013 | Young |
| 0.644968 | 0.03183 | 0.316 | 0.291 | 1 | 9 | Morf4l2 | 0.025 | Young |
| 0.6453 | 0.451258 | 0.029 | 0.039 | 1 | 9 | Wdr46 | -0.01 | Young |
| 0.645725 | 0.323015 | 0.213 | 0.206 | 1 | 9 | Atpif1 | 0.007 | Young |
| 0.645821 | 0.299698 | 0.044 | 0.054 | 1 | 9 | Uckl1 | -0.01 | Young |
| 0.646109 | 0.964838 | 0.088 | 0.077 | 1 | 9 | Wdr13 | 0.011 | Young |
| 0.646217 | 0.251891 | 0.162 | 0.142 | 1 | 9 | Sos1 | 0.02 | Young |
| 0.646932 | 0.27371 | 0.154 | 0.17 | 1 | 9 | Tmem33 | -0.016 | Young |
| 0.647108 | 0.879676 | 0.11 | 0.101 | 1 | 9 | Slc41a1 | 0.009 | Young |
| 0.64751 | 0.383194 | 0.088 | 0.106 | 1 | 9 | Lpar4 | -0.018 | Young |
| 0.648079 | 0.042354 | 0.154 | 0.134 | 1 | 9 | Dnajc14 | 0.02 | Young |
| 0.64829 | 0.559263 | 0.059 | 0.049 | 1 | 9 | Arap1 | 0.01 | Young |
| 0.648772 | 0.049549 | 0.059 | 0.07 | 1 | 9 | Fignl2 | -0.011 | Young |
| 0.648795 | 0.089617 | 0.11 | 0.095 | 1 | 9 | Atn1 | 0.015 | Young |
| 0.64896 | 0.401261 | 0.346 | 0.327 | 1 | 9 | Ccny | 0.019 | Young |
| 0.649271 | 0.368031 | 0.338 | 0.332 | 1 | 9 | Casz1 | 0.006 | Young |
| 0.649525 | 0.352858 | 0.059 | 0.049 | 1 | 9 | Zfp827 | 0.01 | Young |
| 0.649681 | 0.271797 | 0.029 | 0.039 | 1 | 9 | Nt5c3b | -0.01 | Young |
| 0.64973 | 0.19155 | 0.103 | 0.119 | 1 | 9 | Gm37494 | -0.016 | Young |
| 0.650443 | 0.583013 | 0.184 | 0.17 | 1 | 9 | Aff1 | 0.014 | Young |
| 0.650621 | 0.508335 | 0.11 | 0.095 | 1 | 9 | Krr1 | 0.015 | Young |
| 0.651623 | 0.82031 | 0.037 | 0.046 | 1 | 9 | Golph3l | -0.009 | Young |
| 0.651911 | 0.036843 | 0.088 | 0.101 | 1 | 9 | Nod1 | -0.013 | Young |

| | | | | | | | | |
|---|---|---|---|---|---|---|---|---|
| 0.651963 | 0.087397 | 0.118 | 0.103 | 1 | 9 | Nrf1 | 0.015 | Young |
| 0.652 | 0.825104 | 0.059 | 0.049 | 1 | 9 | Med16 | 0.01 | Young |
| 0.652142 | 0.294388 | 0.088 | 0.077 | 1 | 9 | Mrpl3 | 0.011 | Young |
| 0.653239 | 0.450554 | 0.044 | 0.054 | 1 | 9 | Ccs | -0.01 | Young |
| 0.653331 | 0.199421 | 0.25 | 0.235 | 1 | 9 | Psmc6 | 0.015 | Young |
| 0.653689 | 0.208512 | 0.301 | 0.289 | 1 | 9 | Phldb2 | 0.012 | Young |
| 0.65406 | 0.166519 | 0.074 | 0.088 | 1 | 9 | Ppp2cb | -0.014 | Young |
| 0.654427 | 0.212717 | 0.096 | 0.082 | 1 | 9 | Hint2 | 0.014 | Young |
| 0.654449 | 0.254229 | 0.125 | 0.111 | 1 | 9 | Atad2 | 0.014 | Young |
| 0.654804 | 0.700982 | 0.044 | 0.036 | 1 | 9 | Cd209f | 0.008 | Young |
| 0.655505 | 0.326734 | 0.118 | 0.103 | 1 | 9 | Senp7 | 0.015 | Young |
| 0.655713 | 0.094316 | 0.25 | 0.281 | 1 | 9 | Map4k3 | -0.031 | Young |
| 0.656177 | 0.615623 | 0.088 | 0.077 | 1 | 9 | Rxrb | 0.011 | Young |
| 0.656196 | 0.088598 | 0.287 | 0.312 | 1 | 9 | Enah | -0.025 | Young |
| 0.656328 | 0.519843 | 0.081 | 0.07 | 1 | 9 | Ppwd1 | 0.011 | Young |
| 0.656767 | 0.091393 | 0.235 | 0.25 | 1 | 9 | Ddb1 | -0.015 | Young |
| 0.656864 | 0.055505 | 0.368 | 0.343 | 1 | 9 | Osmr | 0.025 | Young |
| 0.656961 | 0.430504 | 0.059 | 0.049 | 1 | 9 | Cxcl16 | 0.01 | Young |
| 0.656961 | 0.356044 | 0.059 | 0.049 | 1 | 9 | Ric3 | 0.01 | Young |
| 0.656961 | 0.32141 | 0.059 | 0.049 | 1 | 9 | Bckdha | 0.01 | Young |
| 0.656961 | 0.287806 | 0.044 | 0.054 | 1 | 9 | Cyp2j6 | -0.01 | Young |
| 0.65767 | 0.700449 | 0.044 | 0.036 | 1 | 9 | Sugp2 | 0.008 | Young |
| 0.658317 | 0.320211 | 0.037 | 0.046 | 1 | 9 | Eif2b1 | -0.009 | Young |
| 0.658605 | 0.02182 | 0.051 | 0.062 | 1 | 9 | Ubap2 | -0.011 | Young |
| 0.658938 | 0.106222 | 0.301 | 0.281 | 1 | 9 | Usmg5 | 0.02 | Young |
| 0.659105 | 0.521559 | 0.044 | 0.036 | 1 | 9 | Ap5s1 | 0.008 | Young |
| 0.659304 | 0.113861 | 0.125 | 0.139 | 1 | 9 | Mtf2 | -0.014 | Young |
| 0.65977 | 0.974648 | 0.066 | 0.057 | 1 | 9 | St6galnac3 | 0.009 | Young |
| 0.65983 | 0.160438 | 0.279 | 0.273 | 1 | 9 | Ripor1 | 0.006 | Young |
| 0.660102 | 0.648171 | 0.103 | 0.09 | 1 | 9 | Anxa1 | 0.013 | Young |
| 0.660299 | 0.388326 | 0.287 | 0.273 | 1 | 9 | Dysf | 0.014 | Young |
| 0.660541 | 0.442935 | 0.044 | 0.036 | 1 | 9 | Atic | 0.008 | Young |
| 0.660564 | 0.678716 | 0.081 | 0.07 | 1 | 9 | Nucb2 | 0.011 | Young |
| 0.660691 | 0.016643 | 0.059 | 0.049 | 1 | 9 | Zkscan8 | 0.01 | Young |
| 0.660763 | 0.111388 | 0.066 | 0.08 | 1 | 9 | Bcl7a | -0.014 | Young |
| 0.66085 | 0.069933 | 0.059 | 0.07 | 1 | 9 | Ptpn9 | -0.011 | Young |
| 0.66097 | 0.272147 | 0.169 | 0.155 | 1 | 9 | Plekha1 | 0.014 | Young |
| 0.661128 | 0.330225 | 0.11 | 0.098 | 1 | 9 | Gch1 | 0.012 | Young |
| 0.661498 | 0.362666 | 0.125 | 0.113 | 1 | 9 | Asah2 | 0.012 | Young |
| 0.662614 | 0.075259 | 0.191 | 0.173 | 1 | 9 | Phf20 | 0.018 | Young |
| 0.662962 | 0.298122 | 0.265 | 0.245 | 1 | 9 | Sptssa | 0.02 | Young |
| 0.663183 | 0.677424 | 0.044 | 0.054 | 1 | 9 | Nelfe | -0.01 | Young |
| 0.663183 | 0.216104 | 0.059 | 0.049 | 1 | 9 | Jagn1 | 0.01 | Young |
| 0.663416 | 0.40689 | 0.044 | 0.036 | 1 | 9 | Zmym3 | 0.008 | Young |
| 0.663874 | 0.421031 | 0.066 | 0.08 | 1 | 9 | Pfkl | -0.014 | Young |

| | | | | | | | | |
|---|---|---|---|---|---|---|---|---|
| 0.663885 | 0.060045 | 0.868 | 0.835 | 1 | 9 | Cst3 | 0.033 | Young |
| 0.664856 | 0.566846 | 0.044 | 0.036 | 1 | 9 | Them4 | 0.008 | Young |
| 0.664875 | 0.003733 | 0.088 | 0.101 | 1 | 9 | Tbl1xr1 | -0.013 | Young |
| 0.665039 | 0.392775 | 0.037 | 0.046 | 1 | 9 | Ace | -0.009 | Young |
| 0.665295 | 0.124795 | 0.529 | 0.526 | 1 | 9 | Ubl3 | 0.003 | Young |
| 0.665305 | 0.290554 | 0.11 | 0.095 | 1 | 9 | Gps1 | 0.015 | Young |
| 0.665359 | 0.011001 | 0.228 | 0.206 | 1 | 9 | Il10rb | 0.022 | Young |
| 0.665861 | 0.148534 | 0.015 | 0.021 | 1 | 9 | Lmnb2 | -0.006 | Young |
| 0.666201 | 0.119501 | 0.096 | 0.082 | 1 | 9 | Fam96a | 0.014 | Young |
| 0.666297 | 0.191613 | 0.044 | 0.036 | 1 | 9 | Abca9 | 0.008 | Young |
| 0.666646 | 0.540756 | 0.125 | 0.113 | 1 | 9 | Cyc1 | 0.012 | Young |
| 0.666887 | 0.35373 | 0.096 | 0.085 | 1 | 9 | Stam2 | 0.011 | Young |
| 0.66694 | 0.270892 | 0.081 | 0.07 | 1 | 9 | Pfdn4 | 0.011 | Young |
| 0.667093 | 0.251914 | 0.243 | 0.232 | 1 | 9 | Etv3 | 0.011 | Young |
| 0.667505 | 0.306167 | 0.213 | 0.198 | 1 | 9 | Chtop | 0.015 | Young |
| 0.667699 | 0.062446 | 0.176 | 0.193 | 1 | 9 | Acbd3 | -0.017 | Young |
| 0.667881 | 0.969428 | 0.015 | 0.021 | 1 | 9 | 2610021A0 | -0.006 | Young |
| 0.667881 | 0.488636 | 0.015 | 0.021 | 1 | 9 | Bid | -0.006 | Young |
| 0.668075 | 0.108165 | 0.551 | 0.619 | 1 | 9 | Rpl18 | -0.068 | Young |
| 0.668177 | 0.004115 | 0.044 | 0.054 | 1 | 9 | D1Ertd622 | -0.01 | Young |
| 0.668185 | 0.057149 | 0.824 | 0.786 | 1 | 9 | Plpp3 | 0.038 | Young |
| 0.668427 | 0.055768 | 0.169 | 0.152 | 1 | 9 | Gtf2a1 | 0.017 | Young |
| 0.668573 | 0.126918 | 0.228 | 0.214 | 1 | 9 | Ddrgk1 | 0.014 | Young |
| 0.668582 | 0.261423 | 0.059 | 0.07 | 1 | 9 | Abhd6 | -0.011 | Young |
| 0.669085 | 1.164664 | 0.037 | 0.046 | 1 | 9 | Ints8 | -0.009 | Young |
| 0.669085 | 0.596474 | 0.037 | 0.046 | 1 | 9 | Hus1 | -0.009 | Young |
| 0.669182 | 0.935938 | 0.044 | 0.036 | 1 | 9 | Pym1 | 0.008 | Young |
| 0.669423 | 0.149634 | 0.338 | 0.332 | 1 | 9 | Ccdc85a | 0.006 | Young |
| 0.669533 | 2.07147 | 0.015 | 0.01 | 1 | 9 | Elp6 | 0.005 | Young |
| 0.669533 | 2.012885 | 0.015 | 0.01 | 1 | 9 | Fam124a | 0.005 | Young |
| 0.669533 | 1.853365 | 0.015 | 0.01 | 1 | 9 | Apex2 | 0.005 | Young |
| 0.669533 | 1.609001 | 0.015 | 0.01 | 1 | 9 | Cd276 | 0.005 | Young |
| 0.669533 | 1.546749 | 0.015 | 0.01 | 1 | 9 | Sec22c | 0.005 | Young |
| 0.669533 | 1.450664 | 0.015 | 0.01 | 1 | 9 | Zbtb26 | 0.005 | Young |
| 0.669533 | 1.44858 | 0.015 | 0.01 | 1 | 9 | Zfp14 | 0.005 | Young |
| 0.669533 | 1.309566 | 0.015 | 0.01 | 1 | 9 | 4930425P0 | 0.005 | Young |
| 0.669533 | 0.641259 | 0.015 | 0.01 | 1 | 9 | Tnfaip8l2 | 0.005 | Young |
| 0.670112 | 0.302093 | 0.066 | 0.08 | 1 | 9 | Stmn1 | -0.014 | Young |
| 0.670627 | 1.334378 | 0.044 | 0.036 | 1 | 9 | Lrrc49 | 0.008 | Young |
| 0.670627 | 0.928287 | 0.044 | 0.036 | 1 | 9 | Pex1 | 0.008 | Young |
| 0.671046 | 0.176188 | 0.184 | 0.198 | 1 | 9 | Ap2s1 | -0.014 | Young |
| 0.671186 | 0.191273 | 0.11 | 0.098 | 1 | 9 | Thap3 | 0.012 | Young |
| 0.671468 | 0.779345 | 0.066 | 0.057 | 1 | 9 | Med25 | 0.009 | Young |
| 0.671707 | 1.251927 | 0.029 | 0.023 | 1 | 9 | Ltbp1 | 0.006 | Young |
| 0.671931 | 0.198908 | 0.176 | 0.162 | 1 | 9 | Dnajb9 | 0.014 | Young |

| | | | | | | | | |
|---|---|---|---|---|---|---|---|---|
| 0.671985 | 0.101697 | 0.162 | 0.147 | 1 | 9 | Rab8b | 0.015 | Young |
| 0.672072 | 0.568358 | 0.257 | 0.237 | 1 | 9 | Cd164 | 0.02 | Young |
| 0.672072 | 0.789139 | 0.044 | 0.036 | 1 | 9 | Spi1 | 0.008 | Young |
| 0.672072 | 0.024929 | 0.044 | 0.036 | 1 | 9 | Rps6kb2 | 0.008 | Young |
| 0.672136 | 1.767665 | 0.015 | 0.01 | 1 | 9 | Mad2l1 | 0.005 | Young |
| 0.672136 | 1.44228 | 0.015 | 0.01 | 1 | 9 | Gdpgp1 | 0.005 | Young |
| 0.672136 | 1.289846 | 0.015 | 0.01 | 1 | 9 | Rnf123 | 0.005 | Young |
| 0.672136 | 1.149664 | 0.015 | 0.01 | 1 | 9 | 1700008J0 | 0.005 | Young |
| 0.672136 | 0.921232 | 0.015 | 0.01 | 1 | 9 | Fam217a | 0.005 | Young |
| 0.672136 | 0.797736 | 0.015 | 0.01 | 1 | 9 | Polr1e | 0.005 | Young |
| 0.67224 | 0.443549 | 0.206 | 0.193 | 1 | 9 | Rapgef6 | 0.013 | Young |
| 0.672257 | 0.083318 | 0.25 | 0.229 | 1 | 9 | Ppp4r2 | 0.021 | Young |
| 0.672487 | 0.269172 | 0.125 | 0.108 | 1 | 9 | Camk1 | 0.017 | Young |
| 0.672971 | 0.085327 | 0.243 | 0.224 | 1 | 9 | Snrpd3 | 0.019 | Young |
| 0.673128 | 0.698507 | 0.257 | 0.245 | 1 | 9 | Polr2a | 0.012 | Young |
| 0.67349 | 1.79704 | 0.029 | 0.023 | 1 | 9 | Fam35a | 0.006 | Young |
| 0.673519 | 0.547007 | 0.044 | 0.036 | 1 | 9 | Armcx4 | 0.008 | Young |
| 0.673519 | 0.092571 | 0.044 | 0.036 | 1 | 9 | Stmn2 | 0.008 | Young |
| 0.673864 | 0.058309 | 0.441 | 0.469 | 1 | 9 | Dpysl2 | -0.028 | Young |
| 0.673955 | 0.849422 | 0.015 | 0.021 | 1 | 9 | Zfp653 | -0.006 | Young |
| 0.674329 | 0.108291 | 0.346 | 0.32 | 1 | 9 | Srsf6 | 0.026 | Young |
| 0.674459 | 0.333621 | 0.088 | 0.077 | 1 | 9 | Rp2 | 0.011 | Young |
| 0.674691 | 0.422346 | 0.051 | 0.044 | 1 | 9 | Acad9 | 0.007 | Young |
| 0.674744 | 1.645353 | 0.015 | 0.01 | 1 | 9 | Cd109 | 0.005 | Young |
| 0.674744 | 1.615338 | 0.015 | 0.01 | 1 | 9 | Lag3 | 0.005 | Young |
| 0.674744 | 1.406812 | 0.015 | 0.01 | 1 | 9 | Rtl5 | 0.005 | Young |
| 0.674744 | 1.123433 | 0.015 | 0.01 | 1 | 9 | 4930481A1 | 0.005 | Young |
| 0.674744 | 0.976133 | 0.015 | 0.01 | 1 | 9 | Faap24 | 0.005 | Young |
| 0.674744 | 0.973598 | 0.015 | 0.01 | 1 | 9 | Mnd1 | 0.005 | Young |
| 0.674744 | 0.954014 | 0.015 | 0.01 | 1 | 9 | Efna2 | 0.005 | Young |
| 0.674744 | 0.796634 | 0.015 | 0.01 | 1 | 9 | Natd1 | 0.005 | Young |
| 0.674744 | 0.688211 | 0.015 | 0.01 | 1 | 9 | Nek3 | 0.005 | Young |
| 0.674744 | 0.591842 | 0.015 | 0.01 | 1 | 9 | Bora | 0.005 | Young |
| 0.674744 | 0.437517 | 0.015 | 0.01 | 1 | 9 | Zfp39 | 0.005 | Young |
| 0.674744 | 0.368793 | 0.015 | 0.01 | 1 | 9 | Gm45606 | 0.005 | Young |
| 0.674891 | 0.367863 | 0.228 | 0.214 | 1 | 9 | Ccdc82 | 0.014 | Young |
| 0.674968 | 0.079068 | 0.044 | 0.036 | 1 | 9 | Slc35b3 | 0.008 | Young |
| 0.675085 | 0.324486 | 0.096 | 0.082 | 1 | 9 | Smim19 | 0.014 | Young |
| 0.675454 | 0.408144 | 0.162 | 0.186 | 1 | 9 | Yipf5 | -0.024 | Young |
| 0.675623 | 0.306607 | 0.147 | 0.134 | 1 | 9 | Psmb2 | 0.013 | Young |
| 0.675772 | 0.30536 | 0.338 | 0.322 | 1 | 9 | Trim25 | 0.016 | Young |
| 0.675984 | 0.01332 | 0.015 | 0.021 | 1 | 9 | Rbm4 | -0.006 | Young |
| 0.676011 | 0.086204 | 0.154 | 0.17 | 1 | 9 | Srsf9 | -0.016 | Young |
| 0.676136 | 0.125519 | 0.11 | 0.126 | 1 | 9 | Usp16 | -0.016 | Young |
| 0.676225 | 0.124795 | 0.103 | 0.09 | 1 | 9 | Scai | 0.013 | Young |

| | | | | | | | | |
|---|---|---|---|---|---|---|---|---|
| 0.676489 | 0.359725 | 0.066 | 0.077 | 1 | 9 | Commd2 | -0.011 | Young |
| 0.676827 | 0.331432 | 0.279 | 0.278 | 1 | 9 | Eif3k | 0.001 | Young |
| 0.67706 | 0.571518 | 0.029 | 0.023 | 1 | 9 | Prpf3 | 0.006 | Young |
| 0.677078 | 0.197362 | 0.118 | 0.106 | 1 | 9 | Kdm5b | 0.012 | Young |
| 0.677238 | 0.126882 | 0.721 | 0.74 | 1 | 9 | Aqp7 | -0.019 | Young |
| 0.677347 | 0.055585 | 0.051 | 0.062 | 1 | 9 | Maml3 | -0.011 | Young |
| 0.677355 | 2.103416 | 0.015 | 0.01 | 1 | 9 | Slc16a3 | 0.005 | Young |
| 0.677355 | 0.918558 | 0.015 | 0.01 | 1 | 9 | Mms22l | 0.005 | Young |
| 0.677355 | 0.723785 | 0.015 | 0.01 | 1 | 9 | Alg12 | 0.005 | Young |
| 0.677355 | 0.194973 | 0.015 | 0.01 | 1 | 9 | Usp35 | 0.005 | Young |
| 0.678016 | 0.078005 | 0.015 | 0.021 | 1 | 9 | Gstm7 | -0.006 | Young |
| 0.678057 | 0.109005 | 0.096 | 0.082 | 1 | 9 | Klhdc10 | 0.014 | Young |
| 0.678525 | 0.538961 | 0.066 | 0.057 | 1 | 9 | Klhl21 | 0.009 | Young |
| 0.678525 | 0.284204 | 0.066 | 0.057 | 1 | 9 | Fes | 0.009 | Young |
| 0.678848 | 1.01756 | 0.029 | 0.023 | 1 | 9 | Lig3 | 0.006 | Young |
| 0.678944 | 0.128217 | 0.184 | 0.204 | 1 | 9 | Aebp2 | -0.02 | Young |
| 0.679076 | 0.312041 | 0.147 | 0.165 | 1 | 9 | Ube2i | -0.018 | Young |
| 0.679612 | 0.461553 | 0.096 | 0.085 | 1 | 9 | Mrs2 | 0.011 | Young |
| 0.679665 | 1.216399 | 0.081 | 0.072 | 1 | 9 | Ssna1 | 0.009 | Young |
| 0.67997 | 1.217952 | 0.015 | 0.01 | 1 | 9 | Cd244 | 0.005 | Young |
| 0.67997 | 1.210643 | 0.015 | 0.01 | 1 | 9 | Cdon | 0.005 | Young |
| 0.67997 | 1.080949 | 0.015 | 0.01 | 1 | 9 | Zfp169 | 0.005 | Young |
| 0.67997 | 0.828517 | 0.015 | 0.01 | 1 | 9 | F10 | 0.005 | Young |
| 0.67997 | 0.291331 | 0.015 | 0.01 | 1 | 9 | Cpne7 | 0.005 | Young |
| 0.67997 | 0.237057 | 0.015 | 0.01 | 1 | 9 | Layn | 0.005 | Young |
| 0.67997 | 0.171715 | 0.015 | 0.01 | 1 | 9 | Zfp691 | 0.005 | Young |
| 0.67997 | 0.101846 | 0.015 | 0.01 | 1 | 9 | Retsat | 0.005 | Young |
| 0.67997 | 0.067321 | 0.015 | 0.01 | 1 | 9 | Pacsin3 | 0.005 | Young |
| 0.67997 | 0.057317 | 0.015 | 0.01 | 1 | 9 | Mllt11 | 0.005 | Young |
| 0.68005 | 0.496751 | 0.015 | 0.021 | 1 | 9 | Reck | -0.006 | Young |
| 0.680356 | 0.109385 | 0.272 | 0.294 | 1 | 9 | Klf10 | -0.022 | Young |
| 0.680728 | 0.252607 | 0.059 | 0.049 | 1 | 9 | E2f5 | 0.01 | Young |
| 0.680976 | 0.345719 | 0.294 | 0.32 | 1 | 9 | Flnb | -0.026 | Young |
| 0.681189 | 0.110593 | 0.382 | 0.369 | 1 | 9 | Pnisr | 0.013 | Young |
| 0.681276 | 0.066093 | 0.11 | 0.124 | 1 | 9 | Polr2b | -0.014 | Young |
| 0.681279 | 0.375381 | 0.015 | 0.01 | 1 | 9 | Cep152 | 0.005 | Young |
| 0.681621 | 0.328757 | 0.088 | 0.077 | 1 | 9 | Slc25a37 | 0.011 | Young |
| 0.682065 | 0.090968 | 0.206 | 0.188 | 1 | 9 | Mepce | 0.018 | Young |
| 0.682086 | 0.882694 | 0.015 | 0.021 | 1 | 9 | Zfp976 | -0.006 | Young |
| 0.682086 | 0.808271 | 0.015 | 0.021 | 1 | 9 | Thada | -0.006 | Young |
| 0.682135 | 0.11386 | 0.309 | 0.291 | 1 | 9 | 1190002N1 | 0.018 | Young |
| 0.682429 | 0.961986 | 0.029 | 0.023 | 1 | 9 | Six5 | 0.006 | Young |
| 0.682429 | 0.496632 | 0.029 | 0.023 | 1 | 9 | Ltc4s | 0.006 | Young |
| 0.682429 | 0.227877 | 0.029 | 0.023 | 1 | 9 | Ccl25 | 0.006 | Young |
| 0.68259 | 0.760121 | 0.015 | 0.01 | 1 | 9 | Cx3cr1 | 0.005 | Young |

| | | | | | | | | |
|---|---|---|---|---|---|---|---|---|
| 0.68259 | 0.563885 | 0.015 | 0.01 | 1 | 9 | Nup43 | 0.005 | Young |
| 0.68259 | 0.181254 | 0.015 | 0.01 | 1 | 9 | Fam149a | 0.005 | Young |
| 0.682688 | 0.199063 | 0.11 | 0.098 | 1 | 9 | Zfp703 | 0.012 | Young |
| 0.682905 | 0.130814 | 0.206 | 0.224 | 1 | 9 | Rbm6 | -0.018 | Young |
| 0.683477 | 0.517219 | 0.081 | 0.095 | 1 | 9 | Fzd8 | -0.014 | Young |
| 0.683483 | 0.19632 | 0.441 | 0.518 | 1 | 9 | Robo4 | -0.077 | Young |
| 0.683526 | 0.050224 | 0.096 | 0.108 | 1 | 9 | Gcnt2 | -0.012 | Young |
| 0.683673 | 0.235805 | 0.088 | 0.077 | 1 | 9 | Echs1 | 0.011 | Young |
| 0.683682 | 0.25274 | 0.044 | 0.036 | 1 | 9 | Tulp3 | 0.008 | Young |
| 0.683864 | 0.058936 | 0.103 | 0.09 | 1 | 9 | Nvl | 0.013 | Young |
| 0.684152 | 0.169133 | 0.059 | 0.07 | 1 | 9 | Rffl | -0.011 | Young |
| 0.684223 | 0.551783 | 0.029 | 0.023 | 1 | 9 | Wdr41 | 0.006 | Young |
| 0.684428 | 0.702707 | 0.066 | 0.057 | 1 | 9 | Lonrf1 | 0.009 | Young |
| 0.684428 | 0.346487 | 0.066 | 0.057 | 1 | 9 | Trim65 | 0.009 | Young |
| 0.684428 | 0.271184 | 0.051 | 0.062 | 1 | 9 | Trappc1 | -0.011 | Young |
| 0.684509 | 0.364536 | 0.088 | 0.101 | 1 | 9 | Egr2 | -0.013 | Young |
| 0.684626 | 0.50045 | 0.272 | 0.271 | 1 | 9 | Tbca | 0.001 | Young |
| 0.684637 | 0.326092 | 0.206 | 0.191 | 1 | 9 | Rmnd5a | 0.015 | Young |
| 0.685212 | 0.130323 | 0.015 | 0.01 | 1 | 9 | Pkib | 0.005 | Young |
| 0.685212 | 0.169396 | 0.015 | 0.01 | 1 | 9 | Tex9 | 0.005 | Young |
| 0.685212 | 0.029844 | 0.015 | 0.01 | 1 | 9 | Pafah2 | 0.005 | Young |
| 0.685294 | 0.081805 | 0.022 | 0.028 | 1 | 9 | Slfn8 | -0.006 | Young |
| 0.68545 | 0.162028 | 0.088 | 0.101 | 1 | 9 | Ing4 | -0.013 | Young |
| 0.685773 | 0.504425 | 0.044 | 0.054 | 1 | 9 | Epb41l4a | -0.01 | Young |
| 0.686018 | 0.834542 | 0.029 | 0.023 | 1 | 9 | Rnf219 | 0.006 | Young |
| 0.686018 | 0.741916 | 0.029 | 0.023 | 1 | 9 | Pigf | 0.006 | Young |
| 0.686018 | 0.700208 | 0.029 | 0.023 | 1 | 9 | Rccd1 | 0.006 | Young |
| 0.686112 | 0.114327 | 0.5 | 0.485 | 1 | 9 | Dync1h1 | 0.015 | Young |
| 0.686237 | 0.379743 | 0.228 | 0.206 | 1 | 9 | Ctdsp1 | 0.022 | Young |
| 0.686403 | 0.281483 | 0.235 | 0.227 | 1 | 9 | Sec61b | 0.008 | Young |
| 0.686596 | 0.890865 | 0.044 | 0.036 | 1 | 9 | Bloc1s1 | 0.008 | Young |
| 0.687049 | 0.374629 | 0.147 | 0.137 | 1 | 9 | Shprh | 0.01 | Young |
| 0.687488 | 0.204283 | 0.096 | 0.085 | 1 | 9 | Zbtb21 | 0.011 | Young |
| 0.68782 | 0.206619 | 0.235 | 0.222 | 1 | 9 | Cenpb | 0.013 | Young |
| 0.687924 | 0.001773 | 0.081 | 0.093 | 1 | 9 | Mrpl21 | -0.012 | Young |
| 0.688016 | 0.81684 | 0.051 | 0.044 | 1 | 9 | Zfp821 | 0.007 | Young |
| 0.688055 | 0.008294 | 0.044 | 0.036 | 1 | 9 | Vac14 | 0.008 | Young |
| 0.688163 | 0.250847 | 0.066 | 0.077 | 1 | 9 | Fam120b | -0.011 | Young |
| 0.689567 | 0.240052 | 0.272 | 0.258 | 1 | 9 | Hnrnpul2 | 0.014 | Young |
| 0.68991 | 0.022512 | 0.404 | 0.42 | 1 | 9 | Lifr | -0.016 | Young |
| 0.690162 | 0.025031 | 0.507 | 0.539 | 1 | 9 | Sdcbp | -0.032 | Young |
| 0.69047 | 0.009165 | 0.015 | 0.01 | 1 | 9 | Cep112 | 0.005 | Young |
| 0.690635 | 0.208689 | 0.206 | 0.219 | 1 | 9 | Copb2 | -0.013 | Young |
| 0.690693 | 0.555046 | 0.051 | 0.044 | 1 | 9 | Snrnp25 | 0.007 | Young |
| 0.690746 | 0.243549 | 0.235 | 0.224 | 1 | 9 | Ndufb9 | 0.011 | Young |

| | | | | | | | | |
|---|---|---|---|---|---|---|---|---|
| 0.690872 | 0.034325 | 0.088 | 0.077 | 1 | 9 | Smg5 | 0.011 | Young |
| 0.691026 | 0.21594 | 0.206 | 0.193 | 1 | 9 | Dtx3l | 0.013 | Young |
| 0.691094 | 0.083137 | 0.176 | 0.188 | 1 | 9 | Bcl10 | -0.012 | Young |
| 0.691479 | 0.475362 | 0.11 | 0.098 | 1 | 9 | Arrdc1 | 0.012 | Young |
| 0.691521 | 0.128699 | 0.279 | 0.263 | 1 | 9 | Zfp644 | 0.016 | Young |
| 0.691753 | 0.396149 | 0.14 | 0.157 | 1 | 9 | Nr2c2 | -0.017 | Young |
| 0.691808 | 0.146852 | 0.125 | 0.111 | 1 | 9 | Actr1b | 0.014 | Young |
| 0.691837 | 0.117158 | 0.081 | 0.093 | 1 | 9 | Bahd1 | -0.012 | Young |
| 0.691838 | 0.402408 | 0.118 | 0.106 | 1 | 9 | Xpot | 0.012 | Young |
| 0.692033 | 0.411275 | 0.051 | 0.044 | 1 | 9 | Cep57 | 0.007 | Young |
| 0.692234 | 1.027418 | 0.022 | 0.028 | 1 | 9 | Nup160 | -0.006 | Young |
| 0.692426 | 0.214444 | 0.066 | 0.077 | 1 | 9 | Tusc3 | -0.011 | Young |
| 0.692464 | 0.112254 | 0.382 | 0.353 | 1 | 9 | Rnf4 | 0.029 | Young |
| 0.692968 | 0.326753 | 0.574 | 0.572 | 1 | 9 | Rps13 | 0.002 | Young |
| 0.693011 | 0.380157 | 0.169 | 0.193 | 1 | 9 | Arhgap23 | -0.024 | Young |
| 0.693216 | 0.226759 | 0.029 | 0.023 | 1 | 9 | Tmem120a | 0.006 | Young |
| 0.693321 | 0.302406 | 0.213 | 0.235 | 1 | 9 | Txnl1 | -0.022 | Young |
| 0.693567 | 0.66686 | 0.132 | 0.119 | 1 | 9 | Uqcrc1 | 0.013 | Young |
| 0.694022 | 0.350552 | 0.265 | 0.25 | 1 | 9 | Ago2 | 0.015 | Young |
| 0.694271 | 0.152195 | 0.184 | 0.173 | 1 | 9 | Atp6v1b2 | 0.011 | Young |
| 0.694842 | 0.057816 | 0.279 | 0.289 | 1 | 9 | Parp14 | -0.01 | Young |
| 0.69502 | 0.380778 | 0.029 | 0.023 | 1 | 9 | Ccdc77 | 0.006 | Young |
| 0.69502 | 0.21926 | 0.029 | 0.023 | 1 | 9 | Ncf4 | 0.006 | Young |
| 0.695714 | 0.750497 | 0.022 | 0.028 | 1 | 9 | Arpin | -0.006 | Young |
| 0.696058 | 0.454146 | 0.051 | 0.044 | 1 | 9 | Itga3 | 0.007 | Young |
| 0.696291 | 0.211579 | 0.066 | 0.057 | 1 | 9 | Mboat7 | 0.009 | Young |
| 0.696336 | 0.213673 | 0.25 | 0.229 | 1 | 9 | Scand1 | 0.021 | Young |
| 0.696481 | 0.060341 | 0.074 | 0.064 | 1 | 9 | Mtap | 0.01 | Young |
| 0.696579 | 0.11375 | 0.61 | 0.613 | 1 | 9 | Rhob | -0.003 | Young |
| 0.696826 | 0.315332 | 0.029 | 0.023 | 1 | 9 | Ppil1 | 0.006 | Young |
| 0.696826 | 0.088179 | 0.029 | 0.023 | 1 | 9 | Letm2 | 0.006 | Young |
| 0.697065 | 0.452954 | 0.088 | 0.077 | 1 | 9 | Rhot1 | 0.011 | Young |
| 0.69721 | 0.633328 | 0.103 | 0.113 | 1 | 9 | Emp1 | -0.01 | Young |
| 0.697256 | 0.094937 | 0.846 | 0.884 | 1 | 9 | Hspa8 | -0.038 | Young |
| 0.697456 | 0.557856 | 0.022 | 0.028 | 1 | 9 | Gm26518 | -0.006 | Young |
| 0.697482 | 0.505454 | 0.066 | 0.057 | 1 | 9 | Klhdc2 | 0.009 | Young |
| 0.697482 | 0.469382 | 0.066 | 0.057 | 1 | 9 | Vps16 | 0.009 | Young |
| 0.697521 | 0.098586 | 0.324 | 0.302 | 1 | 9 | H1f0 | 0.022 | Young |
| 0.697723 | 0.243146 | 0.081 | 0.093 | 1 | 9 | Slc16a6 | -0.012 | Young |
| 0.698253 | 0.036799 | 0.169 | 0.183 | 1 | 9 | Slc35a3 | -0.014 | Young |
| 0.698633 | 0.07751 | 0.029 | 0.023 | 1 | 9 | Slc7a8 | 0.006 | Young |
| 0.6987 | 0.172175 | 0.132 | 0.119 | 1 | 9 | Cstf2 | 0.013 | Young |
| 0.699134 | 0.122727 | 0.088 | 0.077 | 1 | 9 | Cndp2 | 0.011 | Young |
| 0.699612 | 0.737661 | 0.154 | 0.147 | 1 | 9 | Trim47 | 0.007 | Young |
| 0.699967 | 1.022789 | 0.081 | 0.095 | 1 | 9 | Ccnyl1 | -0.014 | Young |

| | | | | | | | | |
|---|---|---|---|---|---|---|---|---|
| 0.700261 | 0.045217 | 0.478 | 0.474 | 1 | 9 | Pitpnc1 | 0.004 | Young |
| 0.700269 | 0.020053 | 0.397 | 0.415 | 1 | 9 | Midn | -0.018 | Young |
| 0.700414 | 0.306413 | 0.132 | 0.119 | 1 | 9 | 2510039O1 | 0.013 | Young |
| 0.700609 | 0.389217 | 0.169 | 0.196 | 1 | 9 | Xaf1 | -0.027 | Young |
| 0.700673 | 0.067235 | 0.081 | 0.093 | 1 | 9 | Gm525 | -0.012 | Young |
| 0.700692 | 0.654268 | 0.235 | 0.224 | 1 | 9 | Vps4b | 0.011 | Young |
| 0.700946 | 0.05278 | 0.022 | 0.028 | 1 | 9 | Rep15 | -0.006 | Young |
| 0.701439 | 0.490629 | 0.051 | 0.044 | 1 | 9 | Kctd3 | 0.007 | Young |
| 0.701519 | 0.372254 | 0.272 | 0.265 | 1 | 9 | Eif3j1 | 0.007 | Young |
| 0.701837 | 0.45368 | 0.103 | 0.093 | 1 | 9 | Smad2 | 0.01 | Young |
| 0.702055 | 0.388518 | 0.081 | 0.072 | 1 | 9 | Las1l | 0.009 | Young |
| 0.702677 | 0.742788 | 0.191 | 0.186 | 1 | 9 | Tcp1 | 0.005 | Young |
| 0.703216 | 0.227645 | 0.191 | 0.18 | 1 | 9 | Bub3 | 0.011 | Young |
| 0.703239 | 0.267935 | 0.074 | 0.064 | 1 | 9 | Hacd2 | 0.01 | Young |
| 0.703418 | 0.321109 | 0.037 | 0.044 | 1 | 9 | Tpgs1 | -0.007 | Young |
| 0.703436 | 0.03255 | 0.551 | 0.598 | 1 | 9 | Ablim3 | -0.047 | Young |
| 0.703884 | 0.069742 | 0.993 | 0.997 | 1 | 9 | mt-Co3 | -0.004 | Young |
| 0.703925 | 0.05613 | 0.103 | 0.113 | 1 | 9 | Tmem248 | -0.01 | Young |
| 0.704178 | 0.818701 | 0.044 | 0.036 | 1 | 9 | Zcchc10 | 0.008 | Young |
| 0.704441 | 0.353032 | 0.022 | 0.028 | 1 | 9 | Sulf1 | -0.006 | Young |
| 0.704441 | 0.117042 | 0.022 | 0.028 | 1 | 9 | Tcta | -0.006 | Young |
| 0.705016 | 0.090537 | 0.485 | 0.469 | 1 | 9 | Sptan1 | 0.016 | Young |
| 0.705025 | 0.211425 | 0.125 | 0.111 | 1 | 9 | Uvssa | 0.014 | Young |
| 0.705484 | 0.830566 | 0.051 | 0.044 | 1 | 9 | Scyl3 | 0.007 | Young |
| 0.705484 | 0.377636 | 0.051 | 0.044 | 1 | 9 | Gm11131 | 0.007 | Young |
| 0.705529 | 0.001538 | 0.228 | 0.237 | 1 | 9 | Cct3 | -0.009 | Young |
| 0.705652 | 0.073447 | 0.537 | 0.526 | 1 | 9 | Picalm | 0.011 | Young |
| 0.705712 | 0.372543 | 0.199 | 0.191 | 1 | 9 | Yeats2 | 0.008 | Young |
| 0.706017 | 0.092407 | 0.243 | 0.222 | 1 | 9 | Fam84b | 0.021 | Young |
| 0.706192 | 0.355248 | 0.022 | 0.028 | 1 | 9 | Stx11 | -0.006 | Young |
| 0.706192 | 0.172949 | 0.022 | 0.028 | 1 | 9 | Dexi | -0.006 | Young |
| 0.706586 | 0.180784 | 0.081 | 0.093 | 1 | 9 | Drg1 | -0.012 | Young |
| 0.706721 | 0.187684 | 0.331 | 0.363 | 1 | 9 | Cr1l | -0.032 | Young |
| 0.707315 | 0.076359 | 0.11 | 0.098 | 1 | 9 | Mrpl57 | 0.012 | Young |
| 0.708163 | 0.180175 | 0.088 | 0.101 | 1 | 9 | Txndc5 | -0.013 | Young |
| 0.708229 | 0.022701 | 0.066 | 0.057 | 1 | 9 | Samd4 | 0.009 | Young |
| 0.70836 | 0.072171 | 0.154 | 0.144 | 1 | 9 | Supt20 | 0.01 | Young |
| 0.708369 | 0.180031 | 0.074 | 0.085 | 1 | 9 | Bcl2l2 | -0.011 | Young |
| 0.70934 | 0.035539 | 0.265 | 0.25 | 1 | 9 | Atp6v0e | 0.015 | Young |
| 0.709684 | 0.096677 | 0.125 | 0.139 | 1 | 9 | Nenf | -0.014 | Young |
| 0.709697 | 0.694659 | 0.022 | 0.028 | 1 | 9 | Sord | -0.006 | Young |
| 0.709975 | 0.47845 | 0.176 | 0.173 | 1 | 9 | Chst15 | 0.003 | Young |
| 0.710861 | 0.108986 | 0.478 | 0.461 | 1 | 9 | Capza2 | 0.017 | Young |
| 0.711488 | 0.149798 | 0.191 | 0.178 | 1 | 9 | Zfp280d | 0.013 | Young |
| 0.711527 | 0.558088 | 0.096 | 0.088 | 1 | 9 | Fam53c | 0.008 | Young |

| | | | | | | | | |
|---|---|---|---|---|---|---|---|---|
| 0.711916 | 0.102433 | 0.103 | 0.093 | 1 | 9 | Anapc4 | 0.01 | Young |
| 0.71219 | 0.152159 | 0.338 | 0.332 | 1 | 9 | Dennd5b | 0.006 | Young |
| 0.712249 | 0.270248 | 0.14 | 0.152 | 1 | 9 | Lgals1 | -0.012 | Young |
| 0.712298 | 0.340253 | 0.103 | 0.116 | 1 | 9 | Wdr33 | -0.013 | Young |
| 0.712479 | 0.015315 | 0.184 | 0.198 | 1 | 9 | Fkbp4 | -0.014 | Young |
| 0.712604 | 0.095442 | 0.5 | 0.479 | 1 | 9 | Ctdsp2 | 0.021 | Young |
| 0.712636 | 0.167509 | 0.088 | 0.077 | 1 | 9 | Mcm3ap | 0.011 | Young |
| 0.712844 | 0.01733 | 0.228 | 0.206 | 1 | 9 | Bicd2 | 0.022 | Young |
| 0.713208 | 0.647234 | 0.022 | 0.028 | 1 | 9 | Abhd14a | -0.006 | Young |
| 0.713599 | 0.889271 | 0.051 | 0.044 | 1 | 9 | Sik2 | 0.007 | Young |
| 0.713599 | 0.466301 | 0.051 | 0.044 | 1 | 9 | Psmd10 | 0.007 | Young |
| 0.713599 | 0.261801 | 0.051 | 0.044 | 1 | 9 | Kank2 | 0.007 | Young |
| 0.713678 | 0.072623 | 0.074 | 0.082 | 1 | 9 | Dnajc2 | -0.008 | Young |
| 0.713806 | 0.103313 | 0.147 | 0.131 | 1 | 9 | Rab1b | 0.016 | Young |
| 0.714051 | 0.235975 | 0.176 | 0.165 | 1 | 9 | Uggt1 | 0.011 | Young |
| 0.714062 | 0.102463 | 0.184 | 0.17 | 1 | 9 | Atp2c1 | 0.014 | Young |
| 0.714298 | 0.492745 | 0.125 | 0.116 | 1 | 9 | Rnf144b | 0.009 | Young |
| 0.714307 | 0.198199 | 0.029 | 0.036 | 1 | 9 | Lrig3 | -0.007 | Young |
| 0.714337 | 0.213062 | 0.257 | 0.242 | 1 | 9 | Dock1 | 0.015 | Young |
| 0.714659 | 1.30076 | 0.037 | 0.031 | 1 | 9 | Pnkp | 0.006 | Young |
| 0.715637 | 0.24711 | 0.257 | 0.242 | 1 | 9 | Psmc3 | 0.015 | Young |
| 0.715864 | 0.198238 | 0.029 | 0.036 | 1 | 9 | Pced1b | -0.007 | Young |
| 0.716048 | 0.465715 | 0.081 | 0.072 | 1 | 9 | Afg3l1 | 0.009 | Young |
| 0.716475 | 0.102574 | 0.551 | 0.549 | 1 | 9 | Purb | 0.002 | Young |
| 0.716517 | 0.054443 | 0.096 | 0.108 | 1 | 9 | Dennd4a | -0.012 | Young |
| 0.717423 | 0.189189 | 0.029 | 0.036 | 1 | 9 | Dgcr14 | -0.007 | Young |
| 0.717664 | 0.076885 | 0.228 | 0.247 | 1 | 9 | Kpnb1 | -0.019 | Young |
| 0.718863 | 0.859092 | 0.066 | 0.059 | 1 | 9 | Erich1 | 0.007 | Young |
| 0.718982 | 0.134374 | 0.029 | 0.036 | 1 | 9 | Tbcc | -0.007 | Young |
| 0.719035 | 0.055825 | 0.066 | 0.057 | 1 | 9 | Trappc6a | 0.009 | Young |
| 0.719101 | 0.384294 | 0.074 | 0.064 | 1 | 9 | Nhlrc3 | 0.01 | Young |
| 0.71923 | 0.163085 | 0.279 | 0.268 | 1 | 9 | Nedd8 | 0.011 | Young |
| 0.719464 | 0.353055 | 0.316 | 0.314 | 1 | 9 | Ubap2l | 0.002 | Young |
| 0.719466 | 1.025318 | 0.037 | 0.031 | 1 | 9 | Engase | 0.006 | Young |
| 0.719561 | 0.364221 | 0.206 | 0.193 | 1 | 9 | Nop10 | 0.013 | Young |
| 0.720543 | 0.876174 | 0.029 | 0.036 | 1 | 9 | Rhbdd2 | -0.007 | Young |
| 0.720728 | 0.14208 | 0.125 | 0.113 | 1 | 9 | Sema6d | 0.012 | Young |
| 0.721026 | 0.123012 | 0.507 | 0.523 | 1 | 9 | Atp5l | -0.016 | Young |
| 0.72107 | 0.979589 | 0.037 | 0.031 | 1 | 9 | Rnf25 | 0.006 | Young |
| 0.721296 | 0.243105 | 0.221 | 0.204 | 1 | 9 | Socs2 | 0.017 | Young |
| 0.721317 | 0.223143 | 0.382 | 0.361 | 1 | 9 | Hnrnpl | 0.021 | Young |
| 0.721747 | 0.030538 | 0.051 | 0.044 | 1 | 9 | Prkag2 | 0.007 | Young |
| 0.722076 | 0.164682 | 0.125 | 0.139 | 1 | 9 | Rab3gap1 | -0.014 | Young |
| 0.722494 | 0.427547 | 0.088 | 0.101 | 1 | 9 | Hibadh | -0.013 | Young |
| 0.722516 | 0.690265 | 0.074 | 0.064 | 1 | 9 | Ccdc43 | 0.01 | Young |

| | | | | | | | | |
|---|---|---|---|---|---|---|---|---|
| 0.722676 | 1.215434 | 0.037 | 0.031 | 1 | 9 | Toe1 | 0.006 | Young |
| 0.72315 | 0.390376 | 0.11 | 0.101 | 1 | 9 | Tfg | 0.009 | Young |
| 0.723239 | 0.042086 | 0.176 | 0.191 | 1 | 9 | Slc12a6 | -0.015 | Young |
| 0.723343 | 0.308746 | 0.096 | 0.085 | 1 | 9 | Ero1l | 0.011 | Young |
| 0.723499 | 0.941891 | 0.059 | 0.052 | 1 | 9 | Med23 | 0.007 | Young |
| 0.723656 | 0.053829 | 0.074 | 0.064 | 1 | 9 | Xrcc1 | 0.01 | Young |
| 0.723667 | 0.211519 | 0.029 | 0.036 | 1 | 9 | Gm16503 | -0.007 | Young |
| 0.72372 | 0.134923 | 0.566 | 0.564 | 1 | 9 | Cltc | 0.002 | Young |
| 0.723906 | 0.522259 | 0.199 | 0.191 | 1 | 9 | Gatad1 | 0.008 | Young |
| 0.724085 | 0.40519 | 0.11 | 0.101 | 1 | 9 | Trappc11 | 0.009 | Young |
| 0.724283 | 1.154737 | 0.037 | 0.031 | 1 | 9 | Bmyc | 0.006 | Young |
| 0.724283 | 0.549634 | 0.037 | 0.031 | 1 | 9 | Spidr | 0.006 | Young |
| 0.724433 | 0.01881 | 0.184 | 0.168 | 1 | 9 | Hmbox1 | 0.016 | Young |
| 0.724613 | 0.293392 | 0.037 | 0.044 | 1 | 9 | Wdr59 | -0.007 | Young |
| 0.724831 | 0.260207 | 0.125 | 0.116 | 1 | 9 | Ube2m | 0.009 | Young |
| 0.724897 | 0.792321 | 0.088 | 0.08 | 1 | 9 | Map3k4 | 0.008 | Young |
| 0.725012 | 0.034628 | 0.316 | 0.335 | 1 | 9 | Smtn | -0.019 | Young |
| 0.725177 | 0.017844 | 0.551 | 0.59 | 1 | 9 | Rps6 | -0.039 | Young |
| 0.725354 | 0.727549 | 0.096 | 0.085 | 1 | 9 | Ddx10 | 0.011 | Young |
| 0.725494 | 0.095303 | 0.14 | 0.129 | 1 | 9 | Trim2 | 0.011 | Young |
| 0.725606 | 0.049689 | 0.397 | 0.376 | 1 | 9 | Dnajc3 | 0.021 | Young |
| 0.726014 | 0.532337 | 0.029 | 0.036 | 1 | 9 | Rab3a | -0.007 | Young |
| 0.726122 | 0.102817 | 0.176 | 0.193 | 1 | 9 | Gpbp1 | -0.017 | Young |
| 0.726334 | 0.620561 | 0.14 | 0.129 | 1 | 9 | Grasp | 0.011 | Young |
| 0.727078 | 0.004302 | 0.074 | 0.064 | 1 | 9 | Nr3c2 | 0.01 | Young |
| 0.727135 | 0.277803 | 0.044 | 0.052 | 1 | 9 | Slc37a3 | -0.008 | Young |
| 0.727197 | 0.340094 | 0.051 | 0.044 | 1 | 9 | Zbtb39 | 0.007 | Young |
| 0.728122 | 0.077507 | 0.632 | 0.66 | 1 | 9 | Tagln2 | -0.028 | Young |
| 0.728374 | 0.595402 | 0.081 | 0.09 | 1 | 9 | Rbm15 | -0.009 | Young |
| 0.728787 | 0.189123 | 0.507 | 0.492 | 1 | 9 | Top1 | 0.015 | Young |
| 0.729192 | 0.071398 | 0.831 | 0.817 | 1 | 9 | Nfib | 0.014 | Young |
| 0.729563 | 1.509477 | 0.066 | 0.059 | 1 | 9 | Pigx | 0.007 | Young |
| 0.72993 | 0.165545 | 0.029 | 0.036 | 1 | 9 | Mrpl1 | -0.007 | Young |
| 0.730506 | 0.313009 | 0.074 | 0.064 | 1 | 9 | Zfp62 | 0.01 | Young |
| 0.730557 | 0.015845 | 0.191 | 0.178 | 1 | 9 | Eef1d | 0.013 | Young |
| 0.730724 | 1.189258 | 0.037 | 0.031 | 1 | 9 | Syn3 | 0.006 | Young |
| 0.730724 | 0.255826 | 0.037 | 0.031 | 1 | 9 | Msra | 0.006 | Young |
| 0.730987 | 0.169329 | 0.301 | 0.291 | 1 | 9 | Mtpn | 0.01 | Young |
| 0.731499 | 0.277417 | 0.029 | 0.036 | 1 | 9 | Recql | -0.007 | Young |
| 0.731499 | 0.04943 | 0.029 | 0.036 | 1 | 9 | Pkia | -0.007 | Young |
| 0.731778 | 0.346812 | 0.11 | 0.098 | 1 | 9 | Ppdpf | 0.012 | Young |
| 0.732775 | 0.39476 | 0.184 | 0.175 | 1 | 9 | Polr2c | 0.009 | Young |
| 0.733069 | 1.082028 | 0.029 | 0.036 | 1 | 9 | Eif2s3y | -0.007 | Young |
| 0.73371 | 0.268262 | 0.044 | 0.052 | 1 | 9 | Pgam5 | -0.008 | Young |
| 0.733714 | 0.007737 | 0.279 | 0.26 | 1 | 9 | Tspan12 | 0.019 | Young |

| | | | | | | | | |
|---|---|---|---|---|---|---|---|---|
| 0.733939 | 0.08955 | 0.074 | 0.064 | 1 | 9 | Heatr6 | 0.01 | Young |
| 0.734195 | 0.093467 | 0.309 | 0.296 | 1 | 9 | 9930111J2 | 0.013 | Young |
| 0.734316 | 0.231697 | 0.103 | 0.116 | 1 | 9 | Ppa1 | -0.013 | Young |
| 0.734562 | 0.097815 | 0.191 | 0.206 | 1 | 9 | Anxa11 | -0.015 | Young |
| 0.734768 | 0.031316 | 0.103 | 0.116 | 1 | 9 | Topors | -0.013 | Young |
| 0.734769 | 0.165042 | 0.787 | 0.796 | 1 | 9 | Rps27a | -0.009 | Young |
| 0.734968 | 0.091356 | 0.632 | 0.668 | 1 | 9 | Emcn | -0.036 | Young |
| 0.735085 | 0.870438 | 0.074 | 0.064 | 1 | 9 | Rnf31 | 0.01 | Young |
| 0.735271 | 0.278483 | 0.169 | 0.155 | 1 | 9 | Vps28 | 0.014 | Young |
| 0.735292 | 0.310524 | 0.088 | 0.08 | 1 | 9 | Wdsub1 | 0.008 | Young |
| 0.735504 | 0.064471 | 0.485 | 0.459 | 1 | 9 | Arid5b | 0.026 | Young |
| 0.73558 | 0.116348 | 0.081 | 0.072 | 1 | 9 | Nosip | 0.009 | Young |
| 0.735642 | 0.142377 | 0.434 | 0.451 | 1 | 9 | Lrrc58 | -0.017 | Young |
| 0.736006 | 0.707815 | 0.037 | 0.044 | 1 | 9 | Parp2 | -0.007 | Young |
| 0.736006 | 0.382457 | 0.037 | 0.044 | 1 | 9 | Magohb | -0.007 | Young |
| 0.736193 | 0.314741 | 0.059 | 0.052 | 1 | 9 | Bnip3 | 0.007 | Young |
| 0.736604 | 0.411365 | 0.162 | 0.152 | 1 | 9 | Arhgap27 | 0.01 | Young |
| 0.736729 | 0.192902 | 0.404 | 0.415 | 1 | 9 | Rpl10 | -0.011 | Young |
| 0.737385 | 0.630366 | 0.118 | 0.108 | 1 | 9 | Ggh | 0.01 | Young |
| 0.738421 | 0.413182 | 0.074 | 0.085 | 1 | 9 | Cnst | -0.011 | Young |
| 0.738549 | 0.053831 | 0.103 | 0.093 | 1 | 9 | Abi2 | 0.01 | Young |
| 0.738853 | 0.809484 | 0.081 | 0.072 | 1 | 9 | Nol4l | 0.009 | Young |
| 0.738853 | 0.280954 | 0.081 | 0.072 | 1 | 9 | Usp45 | 0.009 | Young |
| 0.739847 | 0.371876 | 0.147 | 0.139 | 1 | 9 | Crtap | 0.008 | Young |
| 0.740306 | 0.745243 | 0.044 | 0.052 | 1 | 9 | Ppt2 | -0.008 | Young |
| 0.740419 | 0.303969 | 0.081 | 0.093 | 1 | 9 | Mtmr12 | -0.012 | Young |
| 0.74042 | 0.443919 | 0.037 | 0.031 | 1 | 9 | Aimp2 | 0.006 | Young |
| 0.740501 | 0.091923 | 0.382 | 0.353 | 1 | 9 | Pcdh17 | 0.029 | Young |
| 0.740957 | 0.093102 | 0.184 | 0.201 | 1 | 9 | Cdc42ep1 | -0.017 | Young |
| 0.741155 | 0.066374 | 0.654 | 0.696 | 1 | 9 | Rpl19 | -0.042 | Young |
| 0.741291 | 0.314773 | 0.059 | 0.052 | 1 | 9 | Nup205 | 0.007 | Young |
| 0.741291 | 0.248474 | 0.059 | 0.052 | 1 | 9 | Farsa | 0.007 | Young |
| 0.741629 | 0.106738 | 0.162 | 0.149 | 1 | 9 | Sod2 | 0.013 | Young |
| 0.741654 | 0.174104 | 0.618 | 0.647 | 1 | 9 | Xiap | -0.029 | Young |
| 0.742041 | 0.396867 | 0.037 | 0.031 | 1 | 9 | Cchcr1 | 0.006 | Young |
| 0.742545 | 0.433486 | 0.074 | 0.064 | 1 | 9 | Ints9 | 0.01 | Young |
| 0.74271 | 0.318735 | 0.147 | 0.134 | 1 | 9 | Trib1 | 0.013 | Young |
| 0.742714 | 0.268436 | 0.132 | 0.124 | 1 | 9 | Eno1 | 0.008 | Young |
| 0.742875 | 0.066985 | 0.228 | 0.242 | 1 | 9 | Tead1 | -0.014 | Young |
| 0.743133 | 0.038145 | 0.132 | 0.144 | 1 | 9 | Larp1b | -0.012 | Young |
| 0.743318 | 0.828655 | 0.147 | 0.142 | 1 | 9 | Sf3b5 | 0.005 | Young |
| 0.743662 | 0.97779 | 0.037 | 0.031 | 1 | 9 | Cdt1 | 0.006 | Young |
| 0.743844 | 0.457504 | 0.059 | 0.052 | 1 | 9 | Dtymk | 0.007 | Young |
| 0.744041 | 0.164514 | 0.081 | 0.09 | 1 | 9 | Pias4 | -0.009 | Young |
| 0.744359 | 0.014356 | 0.419 | 0.397 | 1 | 9 | Hnrnpab | 0.022 | Young |

| | | | | | | | | |
|---|---|---|---|---|---|---|---|---|
| 0.744424 | 0.27876 | 0.176 | 0.17 | 1 | 9 | Atp11c | 0.006 | Young |
| 0.744589 | 0.831981 | 0.037 | 0.044 | 1 | 9 | Mul1 | -0.007 | Young |
| 0.744766 | 0.090967 | 0.191 | 0.201 | 1 | 9 | AI314180 | -0.01 | Young |
| 0.744898 | 0.051608 | 0.728 | 0.719 | 1 | 9 | Reep3 | 0.009 | Young |
| 0.745321 | 0.06944 | 0.294 | 0.322 | 1 | 9 | Rbm3 | -0.028 | Young |
| 0.745412 | 0.050802 | 0.081 | 0.072 | 1 | 9 | Inhbb | 0.009 | Young |
| 0.746005 | 0.356267 | 0.125 | 0.137 | 1 | 9 | Mpc2 | -0.012 | Young |
| 0.746198 | 0.498 | 0.051 | 0.059 | 1 | 9 | Vrk1 | -0.008 | Young |
| 0.74622 | 0.047377 | 0.824 | 0.869 | 1 | 9 | Ahnak | -0.045 | Young |
| 0.746593 | 0.353767 | 0.103 | 0.095 | 1 | 9 | Fstl1 | 0.008 | Young |
| 0.746643 | 0.429299 | 0.11 | 0.101 | 1 | 9 | Nudc | 0.009 | Young |
| 0.746643 | 0.019226 | 0.11 | 0.101 | 1 | 9 | Stap2 | 0.009 | Young |
| 0.747346 | 1.240817 | 0.022 | 0.018 | 1 | 9 | Vash1 | 0.004 | Young |
| 0.747401 | 0.059718 | 0.14 | 0.152 | 1 | 9 | Tacc2 | -0.012 | Young |
| 0.747436 | 0.358245 | 0.051 | 0.059 | 1 | 9 | Lancl2 | -0.008 | Young |
| 0.747588 | 0.054431 | 0.11 | 0.101 | 1 | 9 | Slc31a2 | 0.009 | Young |
| 0.747679 | 0.130692 | 0.059 | 0.052 | 1 | 9 | Stard3 | 0.007 | Young |
| 0.747791 | 0.268365 | 0.316 | 0.307 | 1 | 9 | Lats2 | 0.009 | Young |
| 0.748367 | 0.770506 | 0.074 | 0.082 | 1 | 9 | Dars | -0.008 | Young |
| 0.748927 | 0.321191 | 0.199 | 0.224 | 1 | 9 | Wasl | -0.025 | Young |
| 0.749126 | 0.083618 | 0.265 | 0.255 | 1 | 9 | Tnfsf10 | 0.01 | Young |
| 0.749269 | 0.063364 | 0.368 | 0.379 | 1 | 9 | Ssb | -0.011 | Young |
| 0.749292 | 0.134296 | 0.088 | 0.098 | 1 | 9 | Zfp384 | -0.01 | Young |
| 0.749451 | 1.247158 | 0.022 | 0.018 | 1 | 9 | Oasl1 | 0.004 | Young |
| 0.749599 | 0.136899 | 0.059 | 0.052 | 1 | 9 | Mark4 | 0.007 | Young |
| 0.749644 | 0.192271 | 0.603 | 0.634 | 1 | 9 | Rpl21 | -0.031 | Young |
| 0.749795 | 0.386833 | 0.081 | 0.072 | 1 | 9 | Trmt1 | 0.009 | Young |
| 0.750029 | 1.230088 | 0.044 | 0.039 | 1 | 9 | Sh3d19 | 0.005 | Young |
| 0.750153 | 0.211198 | 0.382 | 0.387 | 1 | 9 | Itpr2 | -0.005 | Young |
| 0.750158 | 0.090626 | 0.037 | 0.031 | 1 | 9 | Tlr4 | 0.006 | Young |
| 0.750239 | 0.700107 | 0.059 | 0.052 | 1 | 9 | 2310033PC | 0.007 | Young |
| 0.750633 | 0.039634 | 0.132 | 0.121 | 1 | 9 | Tfe3 | 0.011 | Young |
| 0.750645 | 0.208484 | 0.331 | 0.371 | 1 | 9 | Npc2 | -0.04 | Young |
| 0.750898 | 0.445568 | 0.044 | 0.052 | 1 | 9 | Nsmce2 | -0.008 | Young |
| 0.750932 | 0.089312 | 0.169 | 0.155 | 1 | 9 | Lamtor4 | 0.014 | Young |
| 0.751121 | 0.65264 | 0.066 | 0.059 | 1 | 9 | Osbpl2 | 0.007 | Young |
| 0.751557 | 1.995269 | 0.022 | 0.018 | 1 | 9 | Zfp69 | 0.004 | Young |
| 0.751557 | 1.166286 | 0.022 | 0.018 | 1 | 9 | Kyat1 | 0.004 | Young |
| 0.751785 | 1.095544 | 0.037 | 0.031 | 1 | 9 | Gpr65 | 0.006 | Young |
| 0.752465 | 0.134035 | 0.515 | 0.559 | 1 | 9 | Dazap2 | -0.044 | Young |
| 0.752802 | 0.672786 | 0.059 | 0.052 | 1 | 9 | Rap2b | 0.007 | Young |
| 0.75323 | 0.33736 | 0.162 | 0.152 | 1 | 9 | Slc25a32 | 0.01 | Young |
| 0.753265 | 0.243469 | 0.11 | 0.101 | 1 | 9 | Jtb | 0.009 | Young |
| 0.753403 | 0.263888 | 0.184 | 0.173 | 1 | 9 | B3gnt3 | 0.011 | Young |
| 0.753557 | 0.391219 | 0.132 | 0.147 | 1 | 9 | Fam104a | -0.015 | Young |

| | | | | | | | | |
|---|---|---|---|---|---|---|---|---|
| 0.753666 | 2.108118 | 0.022 | 0.018 | 1 | 9 | Zfp146 | 0.004 | Young |
| 0.754062 | 0.291496 | 0.125 | 0.116 | 1 | 9 | Cd63 | 0.009 | Young |
| 0.754084 | 0.341325 | 0.169 | 0.16 | 1 | 9 | Braf | 0.009 | Young |
| 0.754272 | 0.06775 | 0.176 | 0.191 | 1 | 9 | Max | -0.015 | Young |
| 0.754396 | 0.369869 | 0.11 | 0.124 | 1 | 9 | Dpy19l1 | -0.014 | Young |
| 0.75444 | 1.520216 | 0.044 | 0.039 | 1 | 9 | Rnd3 | 0.005 | Young |
| 0.75444 | 0.859613 | 0.044 | 0.039 | 1 | 9 | Urgcp | 0.005 | Young |
| 0.754504 | 0.3552 | 0.176 | 0.193 | 1 | 9 | Ptpre | -0.017 | Young |
| 0.755042 | 0.003938 | 0.037 | 0.031 | 1 | 9 | Sdccag8 | 0.006 | Young |
| 0.755745 | 0.026496 | 0.191 | 0.178 | 1 | 9 | Hpcal1 | 0.013 | Young |
| 0.755776 | 1.098169 | 0.022 | 0.018 | 1 | 9 | Gm14296 | 0.004 | Young |
| 0.755776 | 0.9998 | 0.022 | 0.018 | 1 | 9 | Zfp992 | 0.004 | Young |
| 0.756101 | 0.880136 | 0.059 | 0.067 | 1 | 9 | Arhgef18 | -0.008 | Young |
| 0.756101 | 0.030452 | 0.059 | 0.067 | 1 | 9 | Lnpk | -0.008 | Young |
| 0.756201 | 0.06832 | 0.213 | 0.229 | 1 | 9 | Pcf11 | -0.016 | Young |
| 0.756211 | 0.802487 | 0.044 | 0.052 | 1 | 9 | Alpl | -0.008 | Young |
| 0.756679 | 0.551113 | 0.154 | 0.147 | 1 | 9 | Eapp | 0.007 | Young |
| 0.75712 | 0.004602 | 0.301 | 0.278 | 1 | 9 | Cab39 | 0.023 | Young |
| 0.757144 | 0.590101 | 0.066 | 0.059 | 1 | 9 | Clpp | 0.007 | Young |
| 0.757144 | 0.514469 | 0.066 | 0.059 | 1 | 9 | Cbr2 | 0.007 | Young |
| 0.757409 | 0.058952 | 0.412 | 0.433 | 1 | 9 | Rtn3 | -0.021 | Young |
| 0.757907 | 0.234664 | 0.176 | 0.162 | 1 | 9 | Ckap4 | 0.014 | Young |
| 0.758778 | 0.039559 | 0.162 | 0.149 | 1 | 9 | Cstb | 0.013 | Young |
| 0.758885 | 0.18841 | 0.074 | 0.067 | 1 | 9 | Slc22a15 | 0.007 | Young |
| 0.759686 | 0.07683 | 0.081 | 0.072 | 1 | 9 | Vps13b | 0.009 | Young |
| 0.759851 | 0.257652 | 0.051 | 0.059 | 1 | 9 | Rrn3 | -0.008 | Young |
| 0.759905 | 0.005052 | 0.096 | 0.106 | 1 | 9 | Mink1 | -0.01 | Young |
| 0.760002 | 0.335168 | 0.022 | 0.018 | 1 | 9 | Rpp30 | 0.004 | Young |
| 0.760042 | 0.437103 | 0.132 | 0.124 | 1 | 9 | Arl15 | 0.008 | Young |
| 0.760085 | 0.023515 | 0.088 | 0.098 | 1 | 9 | Ap5m1 | -0.01 | Young |
| 0.760369 | 0.024061 | 0.199 | 0.186 | 1 | 9 | 4833439L1 | 0.013 | Young |
| 0.760758 | 0.758333 | 0.081 | 0.075 | 1 | 9 | Tmem68 | 0.006 | Young |
| 0.760766 | 0.283665 | 0.066 | 0.059 | 1 | 9 | Gas7 | 0.007 | Young |
| 0.760773 | 0.233071 | 0.147 | 0.162 | 1 | 9 | Papd5 | -0.015 | Young |
| 0.761108 | 0.007608 | 0.618 | 0.626 | 1 | 9 | Rpl24 | -0.008 | Young |
| 0.76178 | 0.052825 | 0.346 | 0.335 | 1 | 9 | Fosl2 | 0.011 | Young |
| 0.761888 | 0.586247 | 0.066 | 0.075 | 1 | 9 | Nhp2 | -0.009 | Young |
| 0.762118 | 2.05596 | 0.022 | 0.018 | 1 | 9 | Hmgb3 | 0.004 | Young |
| 0.762118 | 0.298308 | 0.022 | 0.018 | 1 | 9 | Nphp3 | 0.004 | Young |
| 0.762315 | 0.039612 | 0.074 | 0.067 | 1 | 9 | 4930453N2 | 0.007 | Young |
| 0.762465 | 0.066378 | 0.324 | 0.314 | 1 | 9 | Zfp366 | 0.01 | Young |
| 0.762499 | 0.191839 | 0.449 | 0.443 | 1 | 9 | Rps18 | 0.006 | Young |
| 0.763004 | 0.000649 | 0.066 | 0.075 | 1 | 9 | Tex264 | -0.009 | Young |
| 0.763143 | 0.344318 | 0.059 | 0.067 | 1 | 9 | Ccng1 | -0.008 | Young |
| 0.763281 | 0.121065 | 0.228 | 0.219 | 1 | 9 | Lmo1 | 0.009 | Young |

| | | | | | | | | |
|---|---|---|---|---|---|---|---|---|
| 0.763285 | 0.137767 | 0.044 | 0.039 | 1 | 9 | Klhl22 | 0.005 | Young |
| 0.763523 | 0.420951 | 0.118 | 0.129 | 1 | 9 | Ssr4 | -0.011 | Young |
| 0.763523 | 0.160297 | 0.132 | 0.124 | 1 | 9 | Ltbr | 0.008 | Young |
| 0.763575 | 0.018321 | 0.596 | 0.606 | 1 | 9 | Pdzd2 | -0.01 | Young |
| 0.764169 | 0.074004 | 0.154 | 0.168 | 1 | 9 | Trim28 | -0.014 | Young |
| 0.764235 | 0.526178 | 0.022 | 0.018 | 1 | 9 | Zfp566 | 0.004 | Young |
| 0.764235 | 0.427995 | 0.022 | 0.018 | 1 | 9 | Amz1 | 0.004 | Young |
| 0.764835 | 0.336671 | 0.051 | 0.059 | 1 | 9 | Nifk | -0.008 | Young |
| 0.764934 | 0.602542 | 0.007 | 0.01 | 1 | 9 | Sema4b | -0.003 | Young |
| 0.764934 | 0.213338 | 0.007 | 0.01 | 1 | 9 | Mier2 | -0.003 | Young |
| 0.764934 | 0.211439 | 0.007 | 0.01 | 1 | 9 | Pank1 | -0.003 | Young |
| 0.764934 | 0.157066 | 0.007 | 0.01 | 1 | 9 | Gm15441 | -0.003 | Young |
| 0.764934 | 0.029664 | 0.007 | 0.01 | 1 | 9 | Hspb6 | -0.003 | Young |
| 0.764934 | 0.014285 | 0.007 | 0.01 | 1 | 9 | Eda | -0.003 | Young |
| 0.764972 | 0.397772 | 0.176 | 0.17 | 1 | 9 | Man1a2 | 0.006 | Young |
| 0.765012 | 0.181469 | 0.206 | 0.198 | 1 | 9 | Cebpz | 0.008 | Young |
| 0.76507 | 0.642535 | 0.074 | 0.064 | 1 | 9 | Prex1 | 0.01 | Young |
| 0.765191 | 0.011474 | 0.279 | 0.26 | 1 | 9 | Aff4 | 0.019 | Young |
| 0.7662 | 0.058706 | 0.353 | 0.381 | 1 | 9 | Atp5b | -0.028 | Young |
| 0.76624 | 0.33353 | 0.044 | 0.039 | 1 | 9 | Rab2b | 0.005 | Young |
| 0.766255 | 0.009908 | 0.882 | 0.915 | 1 | 9 | Malat1 | -0.033 | Young |
| 0.766354 | 0.311592 | 0.022 | 0.018 | 1 | 9 | Mapkapk5 | 0.004 | Young |
| 0.766354 | 0.157204 | 0.022 | 0.018 | 1 | 9 | Pdlim2 | 0.004 | Young |
| 0.766354 | 0.069838 | 0.022 | 0.018 | 1 | 9 | Bckdhb | 0.004 | Young |
| 0.766873 | 0.807954 | 0.044 | 0.052 | 1 | 9 | Sufu | -0.008 | Young |
| 0.76782 | 0.170976 | 0.088 | 0.08 | 1 | 9 | Hey1 | 0.008 | Young |
| 0.767918 | 1.971164 | 0.007 | 0.01 | 1 | 9 | Hoxc8 | -0.003 | Young |
| 0.767918 | 1.813748 | 0.007 | 0.01 | 1 | 9 | Tcp11l1 | -0.003 | Young |
| 0.767918 | 1.086943 | 0.007 | 0.01 | 1 | 9 | Gm4632 | -0.003 | Young |
| 0.767918 | 1.074907 | 0.007 | 0.01 | 1 | 9 | Ubtd2 | -0.003 | Young |
| 0.767918 | 1.043943 | 0.007 | 0.01 | 1 | 9 | Gm10031 | -0.003 | Young |
| 0.767918 | 0.919447 | 0.007 | 0.01 | 1 | 9 | Daam2 | -0.003 | Young |
| 0.767918 | 0.902643 | 0.007 | 0.01 | 1 | 9 | Pard3 | -0.003 | Young |
| 0.767918 | 0.83171 | 0.007 | 0.01 | 1 | 9 | 2700062CC | -0.003 | Young |
| 0.767918 | 0.77139 | 0.007 | 0.01 | 1 | 9 | Wdr66 | -0.003 | Young |
| 0.767918 | 0.688718 | 0.007 | 0.01 | 1 | 9 | Zgrf1 | -0.003 | Young |
| 0.767918 | 0.621869 | 0.007 | 0.01 | 1 | 9 | Gm14703 | -0.003 | Young |
| 0.767918 | 0.605454 | 0.007 | 0.01 | 1 | 9 | Spsb2 | -0.003 | Young |
| 0.767918 | 0.408849 | 0.007 | 0.01 | 1 | 9 | Rac3 | -0.003 | Young |
| 0.767918 | 0.375177 | 0.007 | 0.01 | 1 | 9 | Npy1r | -0.003 | Young |
| 0.767918 | 0.367139 | 0.007 | 0.01 | 1 | 9 | Zswim3 | -0.003 | Young |
| 0.767918 | 0.333397 | 0.007 | 0.01 | 1 | 9 | Zfp955a | -0.003 | Young |
| 0.767918 | 0.268885 | 0.007 | 0.01 | 1 | 9 | Ttc30a1 | -0.003 | Young |
| 0.767918 | 0.119859 | 0.007 | 0.01 | 1 | 9 | Cnbd2 | -0.003 | Young |
| 0.768475 | 1.28564 | 0.022 | 0.018 | 1 | 9 | Dctpp1 | 0.004 | Young |

| | | | | | | | | |
|---|---|---|---|---|---|---|---|---|
| 0.768475 | 0.23236 | 0.022 | 0.018 | 1 | 9 | Htra1 | 0.004 | Young |
| 0.768987 | 0.700479 | 0.103 | 0.095 | 1 | 9 | Phf10 | 0.008 | Young |
| 0.769111 | 0.07464 | 0.691 | 0.755 | 1 | 9 | Chchd2 | -0.064 | Young |
| 0.769337 | 0.066502 | 0.382 | 0.389 | 1 | 9 | Lrrc8c | -0.007 | Young |
| 0.769385 | 0.310757 | 0.265 | 0.253 | 1 | 9 | Saraf | 0.012 | Young |
| 0.769616 | 0.083254 | 0.081 | 0.072 | 1 | 9 | Fam50a | 0.009 | Young |
| 0.770448 | 0.519807 | 0.066 | 0.059 | 1 | 9 | Nup62 | 0.007 | Young |
| 0.770568 | 0.430534 | 0.14 | 0.131 | 1 | 9 | Eif4enif1 | 0.009 | Young |
| 0.770598 | 1.029245 | 0.022 | 0.018 | 1 | 9 | Dcun1d2 | 0.004 | Young |
| 0.770598 | 0.525599 | 0.022 | 0.018 | 1 | 9 | Slc22a17 | 0.004 | Young |
| 0.770598 | 0.123169 | 0.022 | 0.018 | 1 | 9 | Epb41l1 | 0.004 | Young |
| 0.770598 | 0.013977 | 0.022 | 0.018 | 1 | 9 | Chic1 | 0.004 | Young |
| 0.770776 | 0.362486 | 0.11 | 0.103 | 1 | 9 | Plgrkt | 0.007 | Young |
| 0.770924 | 0.048877 | 0.103 | 0.093 | 1 | 9 | Ccpg1 | 0.01 | Young |
| 0.770973 | 0.202193 | 0.456 | 0.446 | 1 | 9 | Cox5a | 0.01 | Young |
| 0.77135 | 0.168254 | 0.176 | 0.165 | 1 | 9 | Rreb1 | 0.011 | Young |
| 0.77211 | 0.55542 | 0.059 | 0.052 | 1 | 9 | Pigc | 0.007 | Young |
| 0.772161 | 0.173656 | 0.044 | 0.039 | 1 | 9 | Btbd2 | 0.005 | Young |
| 0.77257 | 0.032282 | 0.426 | 0.41 | 1 | 9 | Pttg1ip | 0.016 | Young |
| 0.772632 | 0.013122 | 0.176 | 0.162 | 1 | 9 | Akirin2 | 0.014 | Young |
| 0.772697 | 0.084838 | 0.206 | 0.193 | 1 | 9 | Ints6 | 0.013 | Young |
| 0.772722 | 0.414482 | 0.022 | 0.018 | 1 | 9 | Milr1 | 0.004 | Young |
| 0.772722 | 0.087459 | 0.022 | 0.018 | 1 | 9 | E130102H2 | 0.004 | Young |
| 0.772874 | 0.34171 | 0.066 | 0.059 | 1 | 9 | Foxj3 | 0.007 | Young |
| 0.773107 | 0.027054 | 0.088 | 0.08 | 1 | 9 | Neurl3 | 0.008 | Young |
| 0.77315 | 0.004635 | 0.39 | 0.418 | 1 | 9 | Ppp2ca | -0.028 | Young |
| 0.77353 | 0.030865 | 0.11 | 0.119 | 1 | 9 | Mrpl30 | -0.009 | Young |
| 0.774193 | 0.076146 | 0.096 | 0.106 | 1 | 9 | Tor3a | -0.01 | Young |
| 0.774849 | 0.248339 | 0.022 | 0.018 | 1 | 9 | Armcx6 | 0.004 | Young |
| 0.774849 | 0.207583 | 0.022 | 0.018 | 1 | 9 | Smarcd3 | 0.004 | Young |
| 0.774849 | 0.198681 | 0.022 | 0.018 | 1 | 9 | Ccdc14 | 0.004 | Young |
| 0.774997 | 0.086406 | 0.596 | 0.619 | 1 | 9 | Rps15a | -0.023 | Young |
| 0.775219 | 0.108232 | 0.176 | 0.165 | 1 | 9 | Gosr2 | 0.011 | Young |
| 0.775222 | 0.140401 | 0.103 | 0.111 | 1 | 9 | Plin3 | -0.008 | Young |
| 0.775248 | 0.312793 | 0.199 | 0.219 | 1 | 9 | Gimap4 | -0.02 | Young |
| 0.775547 | 0.05476 | 0.11 | 0.121 | 1 | 9 | 9930021J0 | -0.011 | Young |
| 0.775547 | 0.210914 | 0.199 | 0.216 | 1 | 9 | Mtus1 | -0.017 | Young |
| 0.77572 | 0.081126 | 0.096 | 0.103 | 1 | 9 | Iars2 | -0.007 | Young |
| 0.775749 | 0.25844 | 0.309 | 0.317 | 1 | 9 | Tab2 | -0.008 | Young |
| 0.776679 | 0.049272 | 0.449 | 0.477 | 1 | 9 | Lasp1 | -0.028 | Young |
| 0.776855 | 0.001266 | 0.191 | 0.18 | 1 | 9 | Spred1 | 0.011 | Young |
| 0.776976 | 0.936613 | 0.022 | 0.018 | 1 | 9 | Kcnn4 | 0.004 | Young |
| 0.776976 | 0.509709 | 0.022 | 0.018 | 1 | 9 | Nectin1 | 0.004 | Young |
| 0.777553 | 0.822439 | 0.066 | 0.075 | 1 | 9 | Rab9 | -0.009 | Young |
| 0.777987 | 0.177327 | 0.324 | 0.345 | 1 | 9 | Eif5a | -0.021 | Young |

| | | | | | | | | |
|---|---|---|---|---|---|---|---|---|
| 0.77824 | 1.035132 | 0.081 | 0.075 | 1 | 9 | Ino80 | 0.006 | Young |
| 0.77897 | 0.338161 | 0.051 | 0.046 | 1 | 9 | Trmt11 | 0.005 | Young |
| 0.779088 | 0.185082 | 0.162 | 0.155 | 1 | 9 | Kctd17 | 0.007 | Young |
| 0.779243 | 0.656894 | 0.118 | 0.129 | 1 | 9 | Suds3 | -0.011 | Young |
| 0.779289 | 0.267475 | 0.081 | 0.09 | 1 | 9 | Rsl24d1 | -0.009 | Young |
| 0.779532 | 0.367012 | 0.074 | 0.067 | 1 | 9 | Pygm | 0.007 | Young |
| 0.779576 | 0.938847 | 0.059 | 0.054 | 1 | 9 | Ccdc102a | 0.005 | Young |
| 0.779581 | 0.723931 | 0.044 | 0.039 | 1 | 9 | Anapc7 | 0.005 | Young |
| 0.77978 | 0.272563 | 0.191 | 0.204 | 1 | 9 | Egr3 | -0.013 | Young |
| 0.780166 | 0.570421 | 0.066 | 0.059 | 1 | 9 | Mettl23 | 0.007 | Young |
| 0.780662 | 0.85848 | 0.088 | 0.082 | 1 | 9 | Tmem161a | 0.006 | Young |
| 0.780871 | 0.124601 | 0.096 | 0.088 | 1 | 9 | Ufd1 | 0.008 | Young |
| 0.780887 | 0.249333 | 0.096 | 0.106 | 1 | 9 | Agtrap | -0.01 | Young |
| 0.780945 | 0.205613 | 0.11 | 0.121 | 1 | 9 | Cox14 | -0.011 | Young |
| 0.781067 | 0.705606 | 0.044 | 0.039 | 1 | 9 | Lmbr1l | 0.005 | Young |
| 0.781099 | 0.524146 | 0.051 | 0.059 | 1 | 9 | Cdc14a | -0.008 | Young |
| 0.781099 | 0.428812 | 0.051 | 0.059 | 1 | 9 | Cdc23 | -0.008 | Young |
| 0.781837 | 0.521476 | 0.074 | 0.067 | 1 | 9 | Tmem159 | 0.007 | Young |
| 0.781837 | 0.340836 | 0.074 | 0.067 | 1 | 9 | Fam92a | 0.007 | Young |
| 0.782045 | 0.237044 | 0.235 | 0.258 | 1 | 9 | Tns2 | -0.023 | Young |
| 0.782176 | 0.28116 | 0.353 | 0.384 | 1 | 9 | Klhl24 | -0.031 | Young |
| 0.782185 | 0.227489 | 0.309 | 0.317 | 1 | 9 | Rell1 | -0.008 | Young |
| 0.782554 | 0.29114 | 0.044 | 0.039 | 1 | 9 | Nipa1 | 0.005 | Young |
| 0.782696 | 0.136929 | 0.184 | 0.196 | 1 | 9 | 2810474O1 | -0.012 | Young |
| 0.783104 | 0.509582 | 0.11 | 0.103 | 1 | 9 | Hmox2 | 0.007 | Young |
| 0.783156 | 0.399936 | 0.191 | 0.209 | 1 | 9 | Atp6v1a | -0.018 | Young |
| 0.78337 | 0.272322 | 0.397 | 0.438 | 1 | 9 | Rpl12 | -0.041 | Young |
| 0.783953 | 0.068261 | 0.36 | 0.387 | 1 | 9 | Apobec3 | -0.027 | Young |
| 0.784591 | 0.368456 | 0.191 | 0.186 | 1 | 9 | Casc3 | 0.005 | Young |
| 0.784702 | 0.107883 | 0.206 | 0.193 | 1 | 9 | Mvp | 0.013 | Young |
| 0.785096 | 0.373232 | 0.154 | 0.17 | 1 | 9 | Nmt1 | -0.016 | Young |
| 0.785502 | 0.171194 | 0.103 | 0.111 | 1 | 9 | Acvr1 | -0.008 | Young |
| 0.785753 | 0.144594 | 0.118 | 0.126 | 1 | 9 | Lypla1 | -0.008 | Young |
| 0.785813 | 0.673887 | 0.051 | 0.046 | 1 | 9 | Pex13 | 0.005 | Young |
| 0.786456 | 0.424872 | 0.176 | 0.17 | 1 | 9 | Ncoa3 | 0.006 | Young |
| 0.786714 | 0.041226 | 0.147 | 0.137 | 1 | 9 | Zfp871 | 0.01 | Young |
| 0.786943 | 0.194314 | 0.184 | 0.175 | 1 | 9 | Art3 | 0.009 | Young |
| 0.787183 | 1.523887 | 0.051 | 0.046 | 1 | 9 | Ttc17 | 0.005 | Young |
| 0.787465 | 0.092764 | 0.897 | 0.918 | 1 | 9 | Ptma | -0.021 | Young |
| 0.788124 | 0.725119 | 0.081 | 0.075 | 1 | 9 | Rab43 | 0.006 | Young |
| 0.788406 | 0.162482 | 0.118 | 0.126 | 1 | 9 | Sfxn1 | -0.008 | Young |
| 0.788511 | 0.632153 | 0.044 | 0.039 | 1 | 9 | Scrn3 | 0.005 | Young |
| 0.789789 | 0.234442 | 0.059 | 0.054 | 1 | 9 | Mapre3 | 0.005 | Young |
| 0.789916 | 0.502104 | 0.074 | 0.067 | 1 | 9 | Hars | 0.007 | Young |
| 0.790002 | 0.230691 | 0.044 | 0.039 | 1 | 9 | Zfp219 | 0.005 | Young |

| | | | | | | | | |
|---|---|---|---|---|---|---|---|---|
| 0.790175 | 0.032992 | 0.176 | 0.191 | 1 | 9 | Imp3 | -0.015 | Young |
| 0.790506 | 0.268726 | 0.287 | 0.286 | 1 | 9 | Nck1 | 0.001 | Young |
| 0.790628 | 0.372153 | 0.081 | 0.09 | 1 | 9 | Lmtk2 | -0.009 | Young |
| 0.790653 | 0.144056 | 0.118 | 0.129 | 1 | 9 | Rspry1 | -0.011 | Young |
| 0.790903 | 0.045785 | 0.36 | 0.381 | 1 | 9 | Tubb4b | -0.021 | Young |
| 0.790999 | 0.048177 | 0.684 | 0.701 | 1 | 9 | Cdkn1a | -0.017 | Young |
| 0.791022 | 0.653888 | 0.044 | 0.052 | 1 | 9 | Mrpl50 | -0.008 | Young |
| 0.791625 | 0.248258 | 0.279 | 0.276 | 1 | 9 | Eif4b | 0.003 | Young |
| 0.79164 | 0.159388 | 0.235 | 0.224 | 1 | 9 | Bud31 | 0.011 | Young |
| 0.791992 | 0.005047 | 0.64 | 0.662 | 1 | 9 | Tapbp | -0.022 | Young |
| 0.792458 | 0.379886 | 0.162 | 0.152 | 1 | 9 | Gnpda2 | 0.01 | Young |
| 0.792466 | 0.153005 | 0.588 | 0.601 | 1 | 9 | Vim | -0.013 | Young |
| 0.792679 | 0.341272 | 0.125 | 0.116 | 1 | 9 | Tmem9b | 0.009 | Young |
| 0.792768 | 0.201268 | 0.184 | 0.178 | 1 | 9 | Fhl1 | 0.006 | Young |
| 0.792987 | 0.940131 | 0.044 | 0.039 | 1 | 9 | Galk1 | 0.005 | Young |
| 0.793224 | 0.043876 | 0.206 | 0.219 | 1 | 9 | Stim2 | -0.013 | Young |
| 0.793578 | 0.097065 | 0.11 | 0.103 | 1 | 9 | Phf23 | 0.007 | Young |
| 0.794022 | 0.175145 | 0.162 | 0.173 | 1 | 9 | Cnih4 | -0.011 | Young |
| 0.794046 | 0.731443 | 0.051 | 0.046 | 1 | 9 | A430005L1 | 0.005 | Young |
| 0.794217 | 0.236292 | 0.118 | 0.111 | 1 | 9 | Nsrp1 | 0.007 | Young |
| 0.794367 | 0.243697 | 0.522 | 0.539 | 1 | 9 | Fnbp1l | -0.017 | Young |
| 0.794505 | 0.072292 | 0.14 | 0.131 | 1 | 9 | Ergic3 | 0.009 | Young |
| 0.794534 | 0.631421 | 0.103 | 0.095 | 1 | 9 | Actl6a | 0.008 | Young |
| 0.79491 | 0.225022 | 0.051 | 0.057 | 1 | 9 | St6gal1 | -0.006 | Young |
| 0.79491 | 0.007059 | 0.235 | 0.232 | 1 | 9 | Adamts5 | 0.003 | Young |
| 0.795341 | 0.244156 | 0.221 | 0.214 | 1 | 9 | Coro2b | 0.007 | Young |
| 0.795446 | 0.093054 | 0.243 | 0.253 | 1 | 9 | Psmd1 | -0.01 | Young |
| 0.795504 | 0.020994 | 0.14 | 0.149 | 1 | 9 | Nop58 | -0.009 | Young |
| 0.795777 | 0.045453 | 0.015 | 0.018 | 1 | 9 | Rpgrip1l | -0.003 | Young |
| 0.795835 | 0.168903 | 0.081 | 0.075 | 1 | 9 | Ppie | 0.006 | Young |
| 0.795974 | 0.224345 | 0.044 | 0.039 | 1 | 9 | Fam214a | 0.005 | Young |
| 0.79603 | 0.034703 | 0.25 | 0.265 | 1 | 9 | Zfp638 | -0.015 | Young |
| 0.796181 | 0.176726 | 0.154 | 0.147 | 1 | 9 | Cdk2 | 0.007 | Young |
| 0.796371 | 0.250794 | 0.309 | 0.309 | 1 | 9 | Efnb2 | 0 | Young |
| 0.796509 | 0.079485 | 0.103 | 0.095 | 1 | 9 | Hccs | 0.008 | Young |
| 0.797512 | 0.242609 | 0.147 | 0.16 | 1 | 9 | Med1 | -0.013 | Young |
| 0.797791 | 0.130233 | 0.301 | 0.317 | 1 | 9 | Ccser2 | -0.016 | Young |
| 0.798018 | 0.276814 | 0.074 | 0.067 | 1 | 9 | Tbc1d9b | 0.007 | Young |
| 0.798034 | 1.329079 | 0.015 | 0.018 | 1 | 9 | Cryz | -0.003 | Young |
| 0.798034 | 0.511527 | 0.015 | 0.018 | 1 | 9 | Asrgl1 | -0.003 | Young |
| 0.798034 | 0.476825 | 0.015 | 0.018 | 1 | 9 | 2610301B2 | -0.003 | Young |
| 0.798172 | 1.114585 | 0.051 | 0.046 | 1 | 9 | Mtor | 0.005 | Young |
| 0.798172 | 0.109075 | 0.044 | 0.049 | 1 | 9 | Fbxo4 | -0.005 | Young |
| 0.798231 | 1.684032 | 0.029 | 0.026 | 1 | 9 | Neu3 | 0.003 | Young |
| 0.798639 | 0.056951 | 0.103 | 0.111 | 1 | 9 | Card19 | -0.008 | Young |

| | | | | | | | | |
|---|---|---|---|---|---|---|---|---|
| 0.798916 | 0.099716 | 0.162 | 0.152 | 1 | 9 | Txlna | 0.01 | Young |
| 0.799392 | 0.155135 | 0.301 | 0.294 | 1 | 9 | Gstt2 | 0.007 | Young |
| 0.79958 | 0.36889 | 0.103 | 0.111 | 1 | 9 | Ccdc127 | -0.008 | Young |
| 0.799602 | 0.192663 | 0.184 | 0.178 | 1 | 9 | Gbp3 | 0.006 | Young |
| 0.800172 | 0.103625 | 0.368 | 0.353 | 1 | 9 | Atp5f1 | 0.015 | Young |
| 0.80028 | 0.138568 | 0.669 | 0.683 | 1 | 9 | Rpl39 | -0.014 | Young |
| 0.800293 | 0.570218 | 0.015 | 0.018 | 1 | 9 | Tmem268 | -0.003 | Young |
| 0.800293 | 0.404244 | 0.015 | 0.018 | 1 | 9 | Mfsd13a | -0.003 | Young |
| 0.800293 | 0.331528 | 0.015 | 0.018 | 1 | 9 | Szt2 | -0.003 | Young |
| 0.800293 | 0.086558 | 0.015 | 0.018 | 1 | 9 | Eral1 | -0.003 | Young |
| 0.800293 | 0.074807 | 0.015 | 0.018 | 1 | 9 | Ajuba | -0.003 | Young |
| 0.801074 | 0.358109 | 0.029 | 0.034 | 1 | 9 | Comtd1 | -0.005 | Young |
| 0.801323 | 1.077219 | 0.059 | 0.054 | 1 | 9 | Habp4 | 0.005 | Young |
| 0.801329 | 0.001255 | 0.096 | 0.088 | 1 | 9 | Llgl1 | 0.008 | Young |
| 0.801872 | 0.726821 | 0.029 | 0.026 | 1 | 9 | Elmod3 | 0.003 | Young |
| 0.801872 | 0.427317 | 0.029 | 0.026 | 1 | 9 | Dbf4 | 0.003 | Young |
| 0.802441 | 0.246253 | 0.103 | 0.095 | 1 | 9 | Micu2 | 0.008 | Young |
| 0.802656 | 0.183979 | 0.61 | 0.606 | 1 | 9 | Arl6ip1 | 0.004 | Young |
| 0.802758 | 0.145563 | 0.279 | 0.273 | 1 | 9 | Tpm1 | 0.006 | Young |
| 0.802831 | 0.025631 | 0.169 | 0.16 | 1 | 9 | Necap2 | 0.009 | Young |
| 0.803003 | 0.042456 | 0.441 | 0.456 | 1 | 9 | Abcg2 | -0.015 | Young |
| 0.803295 | 0.188449 | 0.434 | 0.428 | 1 | 9 | Tpst1 | 0.006 | Young |
| 0.803333 | 0.428138 | 0.176 | 0.178 | 1 | 9 | Ahr | -0.002 | Young |
| 0.803358 | 0.07494 | 0.191 | 0.201 | 1 | 9 | Pkn1 | -0.01 | Young |
| 0.803818 | 0.122576 | 0.074 | 0.067 | 1 | 9 | Prdx4 | 0.007 | Young |
| 0.803892 | 0.970234 | 0.059 | 0.054 | 1 | 9 | Cyp4f16 | 0.005 | Young |
| 0.803892 | 0.498584 | 0.059 | 0.054 | 1 | 9 | Pi4kb | 0.005 | Young |
| 0.803959 | 0.161685 | 0.088 | 0.095 | 1 | 9 | Lactb | -0.007 | Young |
| 0.80409 | 0.059504 | 0.11 | 0.103 | 1 | 9 | Zfp322a | 0.007 | Young |
| 0.804186 | 0.236034 | 0.022 | 0.026 | 1 | 9 | Zfp53 | -0.004 | Young |
| 0.804227 | 0.049518 | 0.176 | 0.17 | 1 | 9 | Arl1 | 0.006 | Young |
| 0.80446 | 0.202841 | 0.125 | 0.116 | 1 | 9 | Rb1 | 0.009 | Young |
| 0.804703 | 0.538347 | 0.096 | 0.09 | 1 | 9 | Ifi44 | 0.006 | Young |
| 0.804816 | 0.351458 | 0.015 | 0.018 | 1 | 9 | Gucy1a1 | -0.003 | Young |
| 0.804816 | 0.07292 | 0.015 | 0.018 | 1 | 9 | Ttll1 | -0.003 | Young |
| 0.804816 | 0.01602 | 0.015 | 0.018 | 1 | 9 | Nsg1 | -0.003 | Young |
| 0.805177 | 0.516095 | 0.059 | 0.054 | 1 | 9 | Cul7 | 0.005 | Young |
| 0.805341 | 0.236658 | 0.228 | 0.24 | 1 | 9 | Anapc5 | -0.012 | Young |
| 0.805534 | 1.171721 | 0.066 | 0.062 | 1 | 9 | Ubqln4 | 0.004 | Young |
| 0.805534 | 0.468872 | 0.066 | 0.062 | 1 | 9 | P2rx4 | 0.004 | Young |
| 0.805696 | 0.132262 | 0.147 | 0.137 | 1 | 9 | Phf12 | 0.01 | Young |
| 0.805979 | 0.133896 | 0.088 | 0.082 | 1 | 9 | Nr4a3 | 0.006 | Young |
| 0.80601 | 0.158179 | 0.184 | 0.175 | 1 | 9 | Nrbp1 | 0.009 | Young |
| 0.806439 | 0.489917 | 0.051 | 0.046 | 1 | 9 | Mrpl40 | 0.005 | Young |
| 0.80708 | 1.065998 | 0.015 | 0.018 | 1 | 9 | Hbb-bs | -0.003 | Young |

| | | | | | | | | |
|---|---|---|---|---|---|---|---|---|
| 0.80708 | 0.567399 | 0.015 | 0.018 | 1 | 9 | Peg12 | -0.003 | Young |
| 0.80708 | 0.492155 | 0.015 | 0.018 | 1 | 9 | Prune2 | -0.003 | Young |
| 0.80708 | 0.245838 | 0.015 | 0.018 | 1 | 9 | Gm38394 | -0.003 | Young |
| 0.80708 | 0.002063 | 0.015 | 0.018 | 1 | 9 | Thyn1 | -0.003 | Young |
| 0.807093 | 0.087104 | 0.169 | 0.178 | 1 | 9 | Cttnbp2nl | -0.009 | Young |
| 0.807168 | 0.159094 | 0.324 | 0.348 | 1 | 9 | Usp9x | -0.024 | Young |
| 0.807342 | 0.816041 | 0.029 | 0.026 | 1 | 9 | Hddc2 | 0.003 | Young |
| 0.807711 | 0.557511 | 0.029 | 0.034 | 1 | 9 | Naa40 | -0.005 | Young |
| 0.807819 | 0.29891 | 0.051 | 0.046 | 1 | 9 | Lsm3 | 0.005 | Young |
| 0.807819 | 0.662033 | 0.051 | 0.046 | 1 | 9 | Tomm40l | 0.005 | Young |
| 0.807826 | 0.235737 | 0.64 | 0.662 | 1 | 9 | Rps4x | -0.022 | Young |
| 0.80866 | 0.024362 | 0.154 | 0.162 | 1 | 9 | Gbf1 | -0.008 | Young |
| 0.809167 | 1.309553 | 0.029 | 0.026 | 1 | 9 | Zfp687 | 0.003 | Young |
| 0.809167 | 1.304185 | 0.029 | 0.026 | 1 | 9 | Gtf2ird1 | 0.003 | Young |
| 0.809167 | 0.832931 | 0.029 | 0.026 | 1 | 9 | Uap1l1 | 0.003 | Young |
| 0.809167 | 0.678526 | 0.029 | 0.026 | 1 | 9 | Ptchd4 | 0.003 | Young |
| 0.809167 | 0.090253 | 0.029 | 0.026 | 1 | 9 | Zc4h2 | 0.003 | Young |
| 0.8092 | 0.331191 | 0.051 | 0.046 | 1 | 9 | Tmem214 | 0.005 | Young |
| 0.809346 | 0.107095 | 0.015 | 0.018 | 1 | 9 | Agpat2 | -0.003 | Young |
| 0.809859 | 0.13898 | 0.125 | 0.119 | 1 | 9 | Rufy3 | 0.006 | Young |
| 0.810457 | 0.134212 | 0.279 | 0.265 | 1 | 9 | Lrp6 | 0.014 | Young |
| 0.810472 | 0.093737 | 0.434 | 0.443 | 1 | 9 | Pnrc1 | -0.009 | Young |
| 0.810581 | 0.056055 | 0.051 | 0.046 | 1 | 9 | Chchd4 | 0.005 | Young |
| 0.810669 | 0.029895 | 0.074 | 0.08 | 1 | 9 | Akap8l | -0.006 | Young |
| 0.810956 | 0.414902 | 0.037 | 0.041 | 1 | 9 | Fbxo45 | -0.004 | Young |
| 0.810994 | 0.596762 | 0.029 | 0.026 | 1 | 9 | Nup35 | 0.003 | Young |
| 0.811358 | 0.720099 | 0.118 | 0.113 | 1 | 9 | Gns | 0.005 | Young |
| 0.811697 | 0.232737 | 0.441 | 0.428 | 1 | 9 | Ncl | 0.013 | Young |
| 0.811705 | 0.037019 | 0.088 | 0.098 | 1 | 9 | Tom1l2 | -0.01 | Young |
| 0.811755 | 0.231196 | 0.022 | 0.026 | 1 | 9 | Dph5 | -0.004 | Young |
| 0.811991 | 0.723145 | 0.066 | 0.072 | 1 | 9 | Zfp263 | -0.006 | Young |
| 0.812342 | 0.408 | 0.088 | 0.082 | 1 | 9 | Tmem222 | 0.006 | Young |
| 0.812458 | 0.656121 | 0.037 | 0.041 | 1 | 9 | Fuca2 | -0.004 | Young |
| 0.812698 | 0.415682 | 0.029 | 0.034 | 1 | 9 | Zfp414 | -0.005 | Young |
| 0.812716 | 0.496532 | 0.11 | 0.103 | 1 | 9 | Rbm12 | 0.007 | Young |
| 0.812821 | 1.317675 | 0.029 | 0.026 | 1 | 9 | Rmdn1 | 0.003 | Young |
| 0.812821 | 1.014401 | 0.029 | 0.026 | 1 | 9 | Emp2 | 0.003 | Young |
| 0.812951 | 0.077916 | 0.221 | 0.216 | 1 | 9 | Stab1 | 0.005 | Young |
| 0.813211 | 0.245548 | 0.066 | 0.059 | 1 | 9 | Pi4k2a | 0.007 | Young |
| 0.81365 | 0.700846 | 0.022 | 0.026 | 1 | 9 | Tysnd1 | -0.004 | Young |
| 0.81365 | 0.351379 | 0.022 | 0.026 | 1 | 9 | B9d2 | -0.004 | Young |
| 0.813882 | 1.943311 | 0.015 | 0.018 | 1 | 9 | Kctd7 | -0.003 | Young |
| 0.813882 | 1.093157 | 0.015 | 0.018 | 1 | 9 | Lgr4 | -0.003 | Young |
| 0.813882 | 0.350162 | 0.015 | 0.018 | 1 | 9 | Fchsd1 | -0.003 | Young |
| 0.814206 | 0.068665 | 0.301 | 0.284 | 1 | 9 | Peak1 | 0.017 | Young |

| | | | | | | | | |
|---|---|---|---|---|---|---|---|---|
| 0.81465 | 1.0798 | 0.029 | 0.026 | 1 | 9 | Homez | 0.003 | Young |
| 0.81465 | 0.687301 | 0.029 | 0.026 | 1 | 9 | Srbd1 | 0.003 | Young |
| 0.81465 | 0.639809 | 0.029 | 0.026 | 1 | 9 | Cc2d2a | 0.003 | Young |
| 0.814701 | 0.149397 | 0.096 | 0.088 | 1 | 9 | Bcr | 0.008 | Young |
| 0.814729 | 0.577322 | 0.051 | 0.046 | 1 | 9 | Stx8 | 0.005 | Young |
| 0.814915 | 0.008257 | 0.213 | 0.201 | 1 | 9 | Trpm7 | 0.012 | Young |
| 0.815179 | 0.203597 | 0.125 | 0.137 | 1 | 9 | Eif2ak2 | -0.012 | Young |
| 0.815478 | 0.772691 | 0.059 | 0.054 | 1 | 9 | Wdtc1 | 0.005 | Young |
| 0.815512 | 0.206406 | 0.235 | 0.245 | 1 | 9 | Taldo1 | -0.01 | Young |
| 0.815528 | 0.213497 | 0.088 | 0.082 | 1 | 9 | Selenoi | 0.006 | Young |
| 0.815547 | 0.373799 | 0.022 | 0.026 | 1 | 9 | Dclk2 | -0.004 | Young |
| 0.815547 | 0.161637 | 0.022 | 0.026 | 1 | 9 | Zfp60 | -0.004 | Young |
| 0.815754 | 0.707938 | 0.081 | 0.075 | 1 | 9 | Ndufv1 | 0.006 | Young |
| 0.815822 | 0.247859 | 0.287 | 0.299 | 1 | 9 | Pitpna | -0.012 | Young |
| 0.816043 | 0.082054 | 0.14 | 0.131 | 1 | 9 | Hsph1 | 0.009 | Young |
| 0.816112 | 0.300079 | 0.051 | 0.046 | 1 | 9 | Txnl4a | 0.005 | Young |
| 0.816112 | 0.18293 | 0.051 | 0.046 | 1 | 9 | Rpa1 | 0.005 | Young |
| 0.816324 | 0.082145 | 0.081 | 0.088 | 1 | 9 | Cenpx | -0.007 | Young |
| 0.816479 | 1.433354 | 0.029 | 0.026 | 1 | 9 | Rrp15 | 0.003 | Young |
| 0.816514 | 0.07092 | 0.375 | 0.405 | 1 | 9 | Npr3 | -0.03 | Young |
| 0.816768 | 0.095514 | 0.051 | 0.057 | 1 | 9 | Hat1 | -0.006 | Young |
| 0.81689 | 0.014514 | 0.257 | 0.247 | 1 | 9 | Akap2 | 0.01 | Young |
| 0.817449 | 0.443346 | 0.353 | 0.343 | 1 | 9 | Lims1 | 0.01 | Young |
| 0.81746 | 0.434642 | 0.096 | 0.103 | 1 | 9 | Lsm4 | -0.007 | Young |
| 0.81746 | 0.14579 | 0.096 | 0.103 | 1 | 9 | Rpl7l1 | -0.007 | Young |
| 0.817657 | 1.291101 | 0.066 | 0.062 | 1 | 9 | Fam160b2 | 0.004 | Young |
| 0.817848 | 0.095156 | 0.154 | 0.144 | 1 | 9 | Itga5 | 0.01 | Young |
| 0.818055 | 0.021038 | 0.632 | 0.647 | 1 | 9 | Cxcl12 | -0.015 | Young |
| 0.818107 | 0.139199 | 0.14 | 0.147 | 1 | 9 | Akap11 | -0.007 | Young |
| 0.818112 | 0.250708 | 0.184 | 0.198 | 1 | 9 | Fmnl3 | -0.014 | Young |
| 0.81831 | 0.456892 | 0.029 | 0.026 | 1 | 9 | Traf5 | 0.003 | Young |
| 0.81831 | 0.06114 | 0.029 | 0.026 | 1 | 9 | Gss | 0.003 | Young |
| 0.818358 | 0.164062 | 0.118 | 0.111 | 1 | 9 | Ppme1 | 0.007 | Young |
| 0.819342 | 0.518234 | 0.022 | 0.026 | 1 | 9 | Fam124b | -0.004 | Young |
| 0.819342 | 0.144766 | 0.022 | 0.026 | 1 | 9 | Zfp820 | -0.004 | Young |
| 0.819349 | 0.020532 | 0.051 | 0.057 | 1 | 9 | Scap | -0.006 | Young |
| 0.819359 | 0.614403 | 0.029 | 0.034 | 1 | 9 | Cyb561d1 | -0.005 | Young |
| 0.819359 | 0.447825 | 0.029 | 0.034 | 1 | 9 | Grpel2 | -0.005 | Young |
| 0.820141 | 0.775637 | 0.029 | 0.026 | 1 | 9 | Zfp580 | 0.003 | Young |
| 0.820141 | 0.325658 | 0.029 | 0.026 | 1 | 9 | Eif2b3 | 0.003 | Young |
| 0.820141 | 0.307105 | 0.029 | 0.026 | 1 | 9 | 2010315B0 | 0.003 | Young |
| 0.820141 | 0.004558 | 0.029 | 0.026 | 1 | 9 | Phykpl | 0.003 | Young |
| 0.820266 | 0.688289 | 0.051 | 0.046 | 1 | 9 | Pin1 | 0.005 | Young |
| 0.820266 | 0.019174 | 0.051 | 0.046 | 1 | 9 | Exosc2 | 0.005 | Young |
| 0.820818 | 0.026434 | 0.368 | 0.394 | 1 | 9 | Rapgef5 | -0.026 | Young |

| | | | | | | | | |
|---|---|---|---|---|---|---|---|---|
| 0.820832 | 0.084334 | 0.375 | 0.371 | 1 | 9 | Atp5j2 | 0.004 | Young |
| 0.820844 | 0.272729 | 0.088 | 0.082 | 1 | 9 | Golga2 | 0.006 | Young |
| 0.821026 | 0.159379 | 0.029 | 0.034 | 1 | 9 | Leng9 | -0.005 | Young |
| 0.821171 | 0.060829 | 0.154 | 0.144 | 1 | 9 | Mapkapk2 | 0.01 | Young |
| 0.821241 | 0.081684 | 0.022 | 0.026 | 1 | 9 | Apex1 | -0.004 | Young |
| 0.821284 | 0.091314 | 0.074 | 0.067 | 1 | 9 | Secisbp2 | 0.007 | Young |
| 0.821485 | 0.584759 | 0.037 | 0.041 | 1 | 9 | Hagh | -0.004 | Young |
| 0.821621 | 0.038549 | 0.125 | 0.119 | 1 | 9 | 1700123O2 | 0.006 | Young |
| 0.821652 | 0.174869 | 0.051 | 0.046 | 1 | 9 | Zfpm1 | 0.005 | Young |
| 0.821932 | 0.508449 | 0.059 | 0.054 | 1 | 9 | Naf1 | 0.005 | Young |
| 0.821973 | 0.850402 | 0.029 | 0.026 | 1 | 9 | Gm6563 | 0.003 | Young |
| 0.821973 | 0.776434 | 0.029 | 0.026 | 1 | 9 | Amer1 | 0.003 | Young |
| 0.821973 | 0.127756 | 0.029 | 0.026 | 1 | 9 | Gpx7 | 0.003 | Young |
| 0.822053 | 0.216325 | 0.096 | 0.09 | 1 | 9 | Cyb5r4 | 0.006 | Young |
| 0.822053 | 0.209517 | 0.096 | 0.09 | 1 | 9 | Mrps17 | 0.006 | Young |
| 0.822438 | 0.15256 | 0.147 | 0.157 | 1 | 9 | Rap2a | -0.01 | Young |
| 0.822553 | 0.318372 | 0.265 | 0.263 | 1 | 9 | Impad1 | 0.002 | Young |
| 0.822694 | 0.07355 | 0.029 | 0.034 | 1 | 9 | Carf | -0.005 | Young |
| 0.822762 | 0.25121 | 0.191 | 0.186 | 1 | 9 | Xpr1 | 0.005 | Young |
| 0.822863 | 0.156105 | 0.176 | 0.168 | 1 | 9 | Ctsa | 0.008 | Young |
| 0.822985 | 0.111761 | 0.294 | 0.291 | 1 | 9 | Smco4 | 0.003 | Young |
| 0.823354 | 0.043581 | 0.544 | 0.513 | 1 | 9 | Mprip | 0.031 | Young |
| 0.823402 | 0.553017 | 0.162 | 0.157 | 1 | 9 | Bcl7b | 0.005 | Young |
| 0.823589 | 0.182839 | 0.191 | 0.18 | 1 | 9 | Ripk1 | 0.011 | Young |
| 0.823954 | 0.075735 | 0.118 | 0.111 | 1 | 9 | Cecr2 | 0.007 | Young |
| 0.824426 | 0.295236 | 0.051 | 0.046 | 1 | 9 | Fam102a | 0.005 | Young |
| 0.824499 | 1.288895 | 0.037 | 0.041 | 1 | 9 | AU022252 | -0.004 | Young |
| 0.8248 | 0.228479 | 0.103 | 0.098 | 1 | 9 | Rad54l2 | 0.005 | Young |
| 0.825009 | 0.977997 | 0.037 | 0.034 | 1 | 9 | Eps8l1 | 0.003 | Young |
| 0.825043 | 0.365258 | 0.022 | 0.026 | 1 | 9 | Zfp141 | -0.004 | Young |
| 0.825103 | 0.133732 | 0.088 | 0.082 | 1 | 9 | Phf5a | 0.006 | Young |
| 0.825641 | 0.454442 | 0.029 | 0.026 | 1 | 9 | Specc1 | 0.003 | Young |
| 0.825813 | 0.027512 | 0.066 | 0.072 | 1 | 9 | Srek1ip1 | -0.006 | Young |
| 0.825813 | 0.720012 | 0.074 | 0.07 | 1 | 9 | Tead2 | 0.004 | Young |
| 0.825928 | 0.497115 | 0.228 | 0.216 | 1 | 9 | Azin1 | 0.012 | Young |
| 0.826033 | 0.064286 | 0.029 | 0.034 | 1 | 9 | Lin37 | -0.005 | Young |
| 0.82614 | 0.155038 | 0.228 | 0.219 | 1 | 9 | Gadd45g | 0.009 | Young |
| 0.82671 | 0.194549 | 0.169 | 0.16 | 1 | 9 | Ammecr1l | 0.009 | Young |
| 0.826946 | 0.21909 | 0.022 | 0.026 | 1 | 9 | Taf5 | -0.004 | Young |
| 0.82708 | 0.091979 | 0.625 | 0.665 | 1 | 9 | Rpl22 | -0.04 | Young |
| 0.827516 | 0.966969 | 0.037 | 0.041 | 1 | 9 | Wwox | -0.004 | Young |
| 0.82769 | 0.325958 | 0.118 | 0.111 | 1 | 9 | Dmtf1 | 0.007 | Young |
| 0.827703 | 0.269951 | 0.029 | 0.034 | 1 | 9 | Efhd1 | -0.005 | Young |
| 0.827839 | 0.03891 | 0.434 | 0.43 | 1 | 9 | Unc45b | 0.004 | Young |
| 0.827986 | 0.266834 | 0.081 | 0.075 | 1 | 9 | Rasa3 | 0.006 | Young |

| | | | | | | | | |
|---|---|---|---|---|---|---|---|---|
| 0.828259 | 1.019764 | 0.037 | 0.034 | 1 | 9 | Snhg9 | 0.003 | Young |
| 0.828604 | 0.958637 | 0.066 | 0.062 | 1 | 9 | Kctd5 | 0.004 | Young |
| 0.828849 | 0.773612 | 0.022 | 0.026 | 1 | 9 | B3galt6 | -0.004 | Young |
| 0.828849 | 0.370857 | 0.022 | 0.026 | 1 | 9 | Serping1 | -0.004 | Young |
| 0.828914 | 0.094844 | 0.132 | 0.126 | 1 | 9 | Fxyd5 | 0.006 | Young |
| 0.828927 | 0.076093 | 0.309 | 0.296 | 1 | 9 | Rnf125 | 0.013 | Young |
| 0.829026 | 0.799557 | 0.037 | 0.041 | 1 | 9 | Ctps | -0.004 | Young |
| 0.829026 | 0.213127 | 0.037 | 0.041 | 1 | 9 | Uqcc1 | -0.004 | Young |
| 0.829148 | 0.429536 | 0.096 | 0.106 | 1 | 9 | Ints6l | -0.01 | Young |
| 0.829278 | 0.864429 | 0.074 | 0.07 | 1 | 9 | Dcaf13 | 0.004 | Young |
| 0.829313 | 0.181848 | 0.029 | 0.026 | 1 | 9 | Rassf10 | 0.003 | Young |
| 0.829375 | 0.302316 | 0.029 | 0.034 | 1 | 9 | Gtpbp3 | -0.005 | Young |
| 0.829452 | 0.301512 | 0.213 | 0.209 | 1 | 9 | Atp5d | 0.004 | Young |
| 0.829885 | 1.605403 | 0.037 | 0.034 | 1 | 9 | Armcx2 | 0.003 | Young |
| 0.830006 | 0.017824 | 0.287 | 0.294 | 1 | 9 | Hexim1 | -0.007 | Young |
| 0.830536 | 0.224821 | 0.037 | 0.041 | 1 | 9 | Ppp2r1b | -0.004 | Young |
| 0.830754 | 0.307836 | 0.022 | 0.026 | 1 | 9 | Cep19 | -0.004 | Young |
| 0.830989 | 0.056649 | 0.051 | 0.057 | 1 | 9 | Nomo1 | -0.006 | Young |
| 0.831047 | 0.680786 | 0.029 | 0.034 | 1 | 9 | Shroom3 | -0.005 | Young |
| 0.83115 | 0.706955 | 0.029 | 0.026 | 1 | 9 | Nsun3 | 0.003 | Young |
| 0.831494 | 0.021493 | 0.14 | 0.147 | 1 | 9 | Ube3c | -0.007 | Young |
| 0.831637 | 0.42075 | 0.051 | 0.057 | 1 | 9 | Gatsl3 | -0.006 | Young |
| 0.831768 | 0.182015 | 0.39 | 0.423 | 1 | 9 | Bptf | -0.033 | Young |
| 0.832139 | 0.146363 | 0.081 | 0.088 | 1 | 9 | Snx1 | -0.007 | Young |
| 0.83229 | 0.067351 | 0.176 | 0.165 | 1 | 9 | Pnrc2 | 0.011 | Young |
| 0.83229 | 0.184571 | 0.162 | 0.17 | 1 | 9 | Kdm1a | -0.008 | Young |
| 0.83229 | 1.226524 | 0.074 | 0.082 | 1 | 9 | Vti1a | -0.008 | Young |
| 0.832514 | 0.608259 | 0.125 | 0.119 | 1 | 9 | Abhd16a | 0.006 | Young |
| 0.832656 | 0.099125 | 0.074 | 0.08 | 1 | 9 | Hmgn5 | -0.006 | Young |
| 0.832659 | 0.308909 | 0.022 | 0.026 | 1 | 9 | Nubpl | -0.004 | Young |
| 0.832699 | 0.053557 | 0.154 | 0.162 | 1 | 9 | Dhrs7 | -0.008 | Young |
| 0.832715 | 0.008387 | 0.449 | 0.477 | 1 | 9 | Sf3b1 | -0.028 | Young |
| 0.832719 | 0.055002 | 0.029 | 0.034 | 1 | 9 | Ubxn8 | -0.005 | Young |
| 0.832871 | 0.754581 | 0.066 | 0.062 | 1 | 9 | Dhx16 | 0.004 | Young |
| 0.833139 | 1.383324 | 0.037 | 0.034 | 1 | 9 | Il15ra | 0.003 | Young |
| 0.833195 | 0.474171 | 0.088 | 0.085 | 1 | 9 | Alkbh8 | 0.003 | Young |
| 0.833195 | 0.321954 | 0.081 | 0.088 | 1 | 9 | Galnt18 | -0.007 | Young |
| 0.833375 | 0.153708 | 0.14 | 0.131 | 1 | 9 | Stk3 | 0.009 | Young |
| 0.833481 | 0.18176 | 0.059 | 0.064 | 1 | 9 | Slc35a1 | -0.005 | Young |
| 0.833482 | 0.089966 | 0.324 | 0.351 | 1 | 9 | Zfp91 | -0.027 | Young |
| 0.833715 | 0.404936 | 0.206 | 0.198 | 1 | 9 | Tollip | 0.008 | Young |
| 0.834767 | 0.863428 | 0.037 | 0.034 | 1 | 9 | Mcm5 | 0.003 | Young |
| 0.834868 | 0.063095 | 0.243 | 0.255 | 1 | 9 | Plekho2 | -0.012 | Young |
| 0.834877 | 0.520798 | 0.059 | 0.054 | 1 | 9 | Acss1 | 0.005 | Young |
| 0.834877 | 0.342279 | 0.059 | 0.054 | 1 | 9 | Runx1t1 | 0.005 | Young |

| | | | | | | | | |
|---|---|---|---|---|---|---|---|---|
| 0.834877 | 0.25192 | 0.051 | 0.057 | 1 | 9 | Mrpl45 | -0.006 | Young |
| 0.83507 | 0.838735 | 0.037 | 0.041 | 1 | 9 | B3galnt1 | -0.004 | Young |
| 0.835165 | 0.015029 | 0.507 | 0.495 | 1 | 9 | Stat3 | 0.012 | Young |
| 0.835227 | 0.059874 | 0.515 | 0.515 | 1 | 9 | Cox6c | 0 | Young |
| 0.835909 | 0.125341 | 0.11 | 0.116 | 1 | 9 | Ginm1 | -0.006 | Young |
| 0.836067 | 0.378928 | 0.029 | 0.034 | 1 | 9 | Fah | -0.005 | Young |
| 0.836174 | 0.361674 | 0.059 | 0.054 | 1 | 9 | Dhx32 | 0.005 | Young |
| 0.836406 | 0.515769 | 0.096 | 0.09 | 1 | 9 | Ubxn7 | 0.006 | Young |
| 0.836493 | 0.093491 | 0.074 | 0.067 | 1 | 9 | Zfp560 | 0.007 | Young |
| 0.836558 | 0.114471 | 0.213 | 0.227 | 1 | 9 | Rgl2 | -0.014 | Young |
| 0.836624 | 0.062696 | 0.272 | 0.265 | 1 | 9 | Sox4 | 0.007 | Young |
| 0.837373 | 0.362255 | 0.066 | 0.072 | 1 | 9 | Lxn | -0.006 | Young |
| 0.837434 | 0.188783 | 0.096 | 0.09 | 1 | 9 | Terf2 | 0.006 | Young |
| 0.837742 | 0.586678 | 0.029 | 0.034 | 1 | 9 | Clpb | -0.005 | Young |
| 0.837939 | 0.280636 | 0.154 | 0.16 | 1 | 9 | Selenom | -0.006 | Young |
| 0.838364 | 0.489755 | 0.066 | 0.062 | 1 | 9 | Styx | 0.004 | Young |
| 0.839489 | 0.580082 | 0.096 | 0.09 | 1 | 9 | Mrpl18 | 0.006 | Young |
| 0.840291 | 0.869573 | 0.022 | 0.026 | 1 | 9 | Dhcr7 | -0.004 | Young |
| 0.840848 | 0.02525 | 0.066 | 0.072 | 1 | 9 | Hmgcr | -0.006 | Young |
| 0.841117 | 0.358322 | 0.154 | 0.165 | 1 | 9 | Lrif1 | -0.011 | Young |
| 0.841123 | 0.768589 | 0.037 | 0.041 | 1 | 9 | Fance | -0.004 | Young |
| 0.841123 | 0.709752 | 0.037 | 0.041 | 1 | 9 | Acap3 | -0.004 | Young |
| 0.841613 | 0.142192 | 0.154 | 0.147 | 1 | 9 | Atp6v0d1 | 0.007 | Young |
| 0.842007 | 0.390079 | 0.066 | 0.072 | 1 | 9 | Qars | -0.006 | Young |
| 0.842654 | 0.024686 | 0.132 | 0.139 | 1 | 9 | Ncoa2 | -0.007 | Young |
| 0.842919 | 0.417302 | 0.037 | 0.034 | 1 | 9 | Trmt10a | 0.003 | Young |
| 0.843166 | 0.337794 | 0.154 | 0.162 | 1 | 9 | Mgst1 | -0.008 | Young |
| 0.843329 | 0.143596 | 0.368 | 0.361 | 1 | 9 | Mob1a | 0.007 | Young |
| 0.843567 | 0.405434 | 0.11 | 0.103 | 1 | 9 | Paxbp1 | 0.007 | Young |
| 0.843734 | 0.124884 | 0.118 | 0.126 | 1 | 9 | Chek2 | -0.008 | Young |
| 0.844551 | 0.351324 | 0.037 | 0.034 | 1 | 9 | Bud13 | 0.003 | Young |
| 0.844581 | 0.910723 | 0.044 | 0.041 | 1 | 9 | Sned1 | 0.003 | Young |
| 0.844932 | 0.268257 | 0.221 | 0.216 | 1 | 9 | Hif1a | 0.005 | Young |
| 0.845294 | 0.529332 | 0.051 | 0.046 | 1 | 9 | Mief1 | 0.005 | Young |
| 0.845363 | 0.197374 | 0.36 | 0.366 | 1 | 9 | Adam10 | -0.006 | Young |
| 0.84567 | 0.624902 | 0.037 | 0.041 | 1 | 9 | Cyp4v3 | -0.004 | Young |
| 0.8457 | 0.470543 | 0.066 | 0.062 | 1 | 9 | C2cd3 | 0.004 | Young |
| 0.845752 | 0.339246 | 0.132 | 0.126 | 1 | 9 | Wbp1l | 0.006 | Young |
| 0.84585 | 0.117707 | 0.5 | 0.539 | 1 | 9 | Clic5 | -0.039 | Young |
| 0.846184 | 0.456509 | 0.037 | 0.034 | 1 | 9 | Zfp738 | 0.003 | Young |
| 0.84659 | 0.02387 | 0.265 | 0.271 | 1 | 9 | Ccnl2 | -0.006 | Young |
| 0.846924 | 0.563429 | 0.066 | 0.062 | 1 | 9 | Ssbp1 | 0.004 | Young |
| 0.847208 | 0.046426 | 0.088 | 0.093 | 1 | 9 | Prelid3b | -0.005 | Young |
| 0.847327 | 0.102645 | 0.294 | 0.314 | 1 | 9 | Kat6a | -0.02 | Young |
| 0.847547 | 0.813415 | 0.044 | 0.041 | 1 | 9 | Mapk7 | 0.003 | Young |

| | | | | | | | | |
|---|---|---|---|---|---|---|---|---|
| 0.847588 | 0.219316 | 0.088 | 0.095 | 1 | 9 | Dctn6 | -0.007 | Young |
| 0.848079 | 0.092584 | 0.169 | 0.178 | 1 | 9 | Neu1 | -0.009 | Young |
| 0.848604 | 0.0498 | 0.301 | 0.296 | 1 | 9 | Cbl | 0.005 | Young |
| 0.848609 | 0.187155 | 0.088 | 0.095 | 1 | 9 | Tmem44 | -0.007 | Young |
| 0.849064 | 0.419563 | 0.162 | 0.165 | 1 | 9 | Tomm5 | -0.003 | Young |
| 0.849375 | 0.332408 | 0.132 | 0.129 | 1 | 9 | Phactr1 | 0.003 | Young |
| 0.849464 | 0.247165 | 0.338 | 0.335 | 1 | 9 | Socs3 | 0.003 | Young |
| 0.85009 | 0.041706 | 0.287 | 0.278 | 1 | 9 | Alkbh5 | 0.009 | Young |
| 0.850143 | 0.130585 | 0.169 | 0.175 | 1 | 9 | Nrd1 | -0.006 | Young |
| 0.851087 | 0.641515 | 0.037 | 0.034 | 1 | 9 | Tnf | 0.003 | Young |
| 0.851087 | 0.412356 | 0.037 | 0.034 | 1 | 9 | Postn | 0.003 | Young |
| 0.851382 | 0.127421 | 0.721 | 0.74 | 1 | 9 | Rpl7 | -0.019 | Young |
| 0.851583 | 0.283951 | 0.125 | 0.131 | 1 | 9 | Ubxn6 | -0.006 | Young |
| 0.851669 | 0.045041 | 0.779 | 0.809 | 1 | 9 | Sparcl1 | -0.03 | Young |
| 0.851792 | 0.229559 | 0.176 | 0.186 | 1 | 9 | Aak1 | -0.01 | Young |
| 0.852238 | 0.029653 | 0.934 | 0.966 | 1 | 9 | Cavin1 | -0.032 | Young |
| 0.852722 | 0.05049 | 0.037 | 0.034 | 1 | 9 | Samd10 | 0.003 | Young |
| 0.853619 | 0.275283 | 0.199 | 0.196 | 1 | 9 | Ndufb5 | 0.003 | Young |
| 0.853799 | 0.439749 | 0.103 | 0.101 | 1 | 9 | Kansl1l | 0.002 | Young |
| 0.853921 | 0.044222 | 0.301 | 0.314 | 1 | 9 | Cebpg | -0.013 | Young |
| 0.854185 | 0.235474 | 0.176 | 0.17 | 1 | 9 | Snx13 | 0.006 | Young |
| 0.854195 | 0.058447 | 0.14 | 0.147 | 1 | 9 | Psme2b | -0.007 | Young |
| 0.85424 | 0.031924 | 0.596 | 0.601 | 1 | 9 | Anp32a | -0.005 | Young |
| 0.854275 | 0.230358 | 0.066 | 0.062 | 1 | 9 | Gm38399 | 0.004 | Young |
| 0.855118 | 0.377222 | 0.25 | 0.271 | 1 | 9 | St3gal6 | -0.021 | Young |
| 0.85519 | 0.092217 | 0.375 | 0.389 | 1 | 9 | Wac | -0.014 | Young |
| 0.85548 | 0.046817 | 0.11 | 0.116 | 1 | 9 | Map2k4 | -0.006 | Young |
| 0.855954 | 0.660226 | 0.279 | 0.281 | 1 | 9 | Slk | -0.002 | Young |
| 0.855995 | 0.88017 | 0.037 | 0.034 | 1 | 9 | Zfp143 | 0.003 | Young |
| 0.856146 | 0.14178 | 0.272 | 0.265 | 1 | 9 | Ube2d2a | 0.007 | Young |
| 0.856249 | 0.15393 | 0.14 | 0.149 | 1 | 9 | Mmp15 | -0.009 | Young |
| 0.856341 | 0.163815 | 0.257 | 0.26 | 1 | 9 | Fam189a2 | -0.003 | Young |
| 0.856552 | 0.322382 | 0.169 | 0.186 | 1 | 9 | Isg15 | -0.017 | Young |
| 0.856714 | 0.060185 | 0.265 | 0.26 | 1 | 9 | Arl5b | 0.005 | Young |
| 0.856728 | 0.156655 | 0.066 | 0.062 | 1 | 9 | Cog6 | 0.004 | Young |
| 0.857086 | 0.329444 | 0.368 | 0.371 | 1 | 9 | Tardbp | -0.003 | Young |
| 0.857105 | 0.266431 | 0.074 | 0.07 | 1 | 9 | Cmas | 0.004 | Young |
| 0.857581 | 0.44113 | 0.088 | 0.085 | 1 | 9 | Kcnb1 | 0.003 | Young |
| 0.859183 | 0.675417 | 0.066 | 0.062 | 1 | 9 | Pex3 | 0.004 | Young |
| 0.859432 | 1.006194 | 0.044 | 0.041 | 1 | 9 | Cnot9 | 0.003 | Young |
| 0.859432 | 0.746966 | 0.066 | 0.072 | 1 | 9 | Edem2 | -0.006 | Young |
| 0.859593 | 0.79778 | 0.051 | 0.057 | 1 | 9 | Ap4s1 | -0.006 | Young |
| 0.859593 | 0.19276 | 0.051 | 0.057 | 1 | 9 | Gtf3c3 | -0.006 | Young |
| 0.860213 | 0.083995 | 0.647 | 0.657 | 1 | 9 | Rock2 | -0.01 | Young |
| 0.860329 | 0.015106 | 0.331 | 0.345 | 1 | 9 | Mospd2 | -0.014 | Young |

| | | | | | | | | |
|---|---|---|---|---|---|---|---|---|
| 0.860685 | 0.130648 | 0.147 | 0.152 | 1 | 9 | Rnf111 | -0.005 | Young |
| 0.860909 | 0.128074 | 0.037 | 0.034 | 1 | 9 | Cdca4 | 0.003 | Young |
| 0.861586 | 0.111514 | 0.301 | 0.307 | 1 | 9 | Tgfb1 | -0.006 | Young |
| 0.862055 | 1.883254 | 0.015 | 0.013 | 1 | 9 | Ret | 0.002 | Young |
| 0.862204 | 0.265459 | 0.235 | 0.237 | 1 | 9 | Filip1 | -0.002 | Young |
| 0.862409 | 1.145536 | 0.044 | 0.041 | 1 | 9 | Pus10 | 0.003 | Young |
| 0.862409 | 0.486575 | 0.044 | 0.041 | 1 | 9 | 2810402E2 | 0.003 | Young |
| 0.862549 | 0.514175 | 0.037 | 0.034 | 1 | 9 | Micu3 | 0.003 | Young |
| 0.862688 | 0.612402 | 0.103 | 0.101 | 1 | 9 | Pik3c2b | 0.002 | Young |
| 0.863076 | 0.107613 | 0.191 | 0.201 | 1 | 9 | Gabarapl1 | -0.01 | Young |
| 0.863173 | 0.001178 | 0.213 | 0.219 | 1 | 9 | Snrpc | -0.006 | Young |
| 0.863372 | 0.032556 | 0.301 | 0.314 | 1 | 9 | Abcb1a | -0.013 | Young |
| 0.863546 | 0.002809 | 0.118 | 0.121 | 1 | 9 | Gon4l | -0.003 | Young |
| 0.863576 | 0.233074 | 0.11 | 0.106 | 1 | 9 | Ppt1 | 0.004 | Young |
| 0.863796 | 0.008468 | 0.463 | 0.503 | 1 | 9 | S100a6 | -0.04 | Young |
| 0.864189 | 0.080685 | 0.037 | 0.034 | 1 | 9 | Mapk1ip1 | 0.003 | Young |
| 0.864309 | 0.427963 | 0.213 | 0.211 | 1 | 9 | Tspan14 | 0.002 | Young |
| 0.864564 | 0.150222 | 0.206 | 0.211 | 1 | 9 | Hnrnpa1 | -0.005 | Young |
| 0.864657 | 2.152935 | 0.015 | 0.013 | 1 | 9 | Klhl25 | 0.002 | Young |
| 0.864657 | 1.987694 | 0.015 | 0.013 | 1 | 9 | Spsb1 | 0.002 | Young |
| 0.864657 | 1.217805 | 0.015 | 0.013 | 1 | 9 | Rnf26 | 0.002 | Young |
| 0.864657 | 0.815392 | 0.015 | 0.013 | 1 | 9 | Ankrd13d | 0.002 | Young |
| 0.864893 | 0.276968 | 0.39 | 0.418 | 1 | 9 | Rtn4 | -0.028 | Young |
| 0.865318 | 0.487017 | 0.066 | 0.07 | 1 | 9 | Dhrs7b | -0.004 | Young |
| 0.865439 | 0.139119 | 0.235 | 0.227 | 1 | 9 | Elmo1 | 0.008 | Young |
| 0.865638 | 0.103066 | 0.581 | 0.624 | 1 | 9 | Mbnl1 | -0.043 | Young |
| 0.865652 | 0.611286 | 0.059 | 0.057 | 1 | 9 | Bzw2 | 0.002 | Young |
| 0.865691 | 0.511806 | 0.147 | 0.142 | 1 | 9 | Slu7 | 0.005 | Young |
| 0.866014 | 0.274246 | 0.147 | 0.144 | 1 | 9 | Ppp2r5e | 0.003 | Young |
| 0.866121 | 0.42433 | 0.051 | 0.057 | 1 | 9 | Tmed8 | -0.006 | Young |
| 0.866121 | 0.341556 | 0.059 | 0.054 | 1 | 9 | Erg28 | 0.005 | Young |
| 0.866499 | 0.007792 | 0.066 | 0.07 | 1 | 9 | Zfp609 | -0.004 | Young |
| 0.866825 | 0.229821 | 0.088 | 0.093 | 1 | 9 | D930016D( | -0.005 | Young |
| 0.866881 | 0.304923 | 0.088 | 0.082 | 1 | 9 | Otud6b | 0.006 | Young |
| 0.867203 | 0.190046 | 0.118 | 0.124 | 1 | 9 | Flywch1 | -0.006 | Young |
| 0.86726 | 0.827569 | 0.015 | 0.013 | 1 | 9 | Gm10053 | 0.002 | Young |
| 0.86726 | 0.787173 | 0.015 | 0.013 | 1 | 9 | Zfp975 | 0.002 | Young |
| 0.86726 | 0.657614 | 0.015 | 0.013 | 1 | 9 | Fhl3 | 0.002 | Young |
| 0.86726 | 0.609265 | 0.015 | 0.013 | 1 | 9 | Pla2g6 | 0.002 | Young |
| 0.86726 | 0.604746 | 0.015 | 0.013 | 1 | 9 | Tarsl2 | 0.002 | Young |
| 0.86726 | 0.382931 | 0.015 | 0.013 | 1 | 9 | Fuk | 0.002 | Young |
| 0.867785 | 0.332335 | 0.066 | 0.062 | 1 | 9 | Tgfbr1 | 0.004 | Young |
| 0.867785 | 0.288658 | 0.066 | 0.062 | 1 | 9 | Stk16 | 0.004 | Young |
| 0.86792 | 0.232693 | 0.103 | 0.108 | 1 | 9 | Usp32 | -0.005 | Young |
| 0.868119 | 0.013723 | 0.618 | 0.626 | 1 | 9 | BC005537 | -0.008 | Young |

| | | | | | | | | |
|---|---|---|---|---|---|---|---|---|
| 0.868368 | 0.323579 | 0.044 | 0.041 | 1 | 9 | Slc15a4 | 0.003 | Young |
| 0.868977 | 0.031528 | 0.316 | 0.338 | 1 | 9 | Zfp207 | -0.022 | Young |
| 0.868978 | 0.07766 | 0.816 | 0.832 | 1 | 9 | Gnas | -0.016 | Young |
| 0.869128 | 0.192985 | 0.375 | 0.394 | 1 | 9 | Mef2d | -0.019 | Young |
| 0.869215 | 0.4866 | 0.081 | 0.077 | 1 | 9 | Lmbrd2 | 0.004 | Young |
| 0.869321 | 0.220688 | 0.235 | 0.25 | 1 | 9 | Ubr5 | -0.015 | Young |
| 0.869377 | 0.034456 | 0.404 | 0.41 | 1 | 9 | G3bp2 | -0.006 | Young |
| 0.869533 | 0.130523 | 0.169 | 0.173 | 1 | 9 | Pfdn2 | -0.004 | Young |
| 0.869858 | 1.038082 | 0.044 | 0.041 | 1 | 9 | Cpne2 | 0.003 | Young |
| 0.869865 | 1.458687 | 0.015 | 0.013 | 1 | 9 | Zfp9 | 0.002 | Young |
| 0.869865 | 1.310131 | 0.015 | 0.013 | 1 | 9 | Cep76 | 0.002 | Young |
| 0.869865 | 1.017776 | 0.015 | 0.013 | 1 | 9 | C330018D2 | 0.002 | Young |
| 0.869865 | 0.962475 | 0.015 | 0.013 | 1 | 9 | Zfp974 | 0.002 | Young |
| 0.869865 | 0.902789 | 0.015 | 0.013 | 1 | 9 | Fig4 | 0.002 | Young |
| 0.869865 | 0.588873 | 0.015 | 0.013 | 1 | 9 | Crocc | 0.002 | Young |
| 0.869865 | 0.520465 | 0.015 | 0.013 | 1 | 9 | Pomt1 | 0.002 | Young |
| 0.869865 | 0.384849 | 0.015 | 0.013 | 1 | 9 | Cpe | 0.002 | Young |
| 0.869865 | 0.086799 | 0.015 | 0.013 | 1 | 9 | Asna1 | 0.002 | Young |
| 0.87088 | 0.240462 | 0.125 | 0.119 | 1 | 9 | Chtf8 | 0.006 | Young |
| 0.871188 | 0.10535 | 0.176 | 0.183 | 1 | 9 | Frmd4a | -0.007 | Young |
| 0.871272 | 0.320448 | 0.221 | 0.245 | 1 | 9 | Cyba | -0.024 | Young |
| 0.87135 | 1.295302 | 0.044 | 0.041 | 1 | 9 | Homer3 | 0.003 | Young |
| 0.87135 | 0.761331 | 0.044 | 0.041 | 1 | 9 | Mri1 | 0.003 | Young |
| 0.87135 | 0.480901 | 0.044 | 0.041 | 1 | 9 | Ascc1 | 0.003 | Young |
| 0.87135 | 0.507088 | 0.059 | 0.054 | 1 | 9 | Fmc1 | 0.005 | Young |
| 0.871511 | 0.272097 | 0.132 | 0.129 | 1 | 9 | Kmt2b | 0.003 | Young |
| 0.87211 | 0.285009 | 0.184 | 0.183 | 1 | 9 | Plekhg5 | 0.001 | Young |
| 0.872248 | 0.054242 | 0.235 | 0.235 | 1 | 9 | Wipf1 | 0 | Young |
| 0.872267 | 0.045258 | 0.088 | 0.082 | 1 | 9 | Dnajc15 | 0.006 | Young |
| 0.872327 | 0.041132 | 0.118 | 0.113 | 1 | 9 | Plp2 | 0.005 | Young |
| 0.872432 | 0.19726 | 0.206 | 0.219 | 1 | 9 | Ccar1 | -0.013 | Young |
| 0.872471 | 1.026735 | 0.015 | 0.013 | 1 | 9 | Itga2b | 0.002 | Young |
| 0.872471 | 0.544132 | 0.015 | 0.013 | 1 | 9 | Adora2b | 0.002 | Young |
| 0.872471 | 0.466179 | 0.015 | 0.013 | 1 | 9 | Trit1 | 0.002 | Young |
| 0.872471 | 0.445475 | 0.015 | 0.013 | 1 | 9 | Zfp712 | 0.002 | Young |
| 0.872471 | 0.424971 | 0.015 | 0.013 | 1 | 9 | Hyal1 | 0.002 | Young |
| 0.872471 | 0.334439 | 0.015 | 0.013 | 1 | 9 | Rpgr | 0.002 | Young |
| 0.872471 | 0.321032 | 0.015 | 0.013 | 1 | 9 | Lysmd4 | 0.002 | Young |
| 0.872471 | 0.305217 | 0.015 | 0.013 | 1 | 9 | Trdmt1 | 0.002 | Young |
| 0.872471 | 0.091342 | 0.015 | 0.013 | 1 | 9 | 1700109H0 | 0.002 | Young |
| 0.872658 | 0.356915 | 0.059 | 0.054 | 1 | 9 | Slc12a4 | 0.005 | Young |
| 0.87278 | 0.50646 | 0.11 | 0.108 | 1 | 9 | Cntln | 0.002 | Young |
| 0.873323 | 0.25994 | 0.265 | 0.281 | 1 | 9 | Mif | -0.016 | Young |
| 0.873693 | 0.047515 | 0.309 | 0.327 | 1 | 9 | Ndufb7 | -0.018 | Young |
| 0.87395 | 0.456832 | 0.346 | 0.353 | 1 | 9 | Lyn | -0.007 | Young |

| | | | | | | | | |
|---|---|---|---|---|---|---|---|---|
| 0.87415 | 0.374426 | 0.316 | 0.32 | 1 | 9 | Atp1a1 | -0.004 | Young |
| 0.874356 | 0.013552 | 0.324 | 0.309 | 1 | 9 | Rab12 | 0.015 | Young |
| 0.874459 | 0.064228 | 0.346 | 0.356 | 1 | 9 | Itpr1 | -0.01 | Young |
| 0.874577 | 1.181502 | 0.037 | 0.041 | 1 | 9 | Nynrin | -0.004 | Young |
| 0.87463 | 0.189661 | 0.316 | 0.325 | 1 | 9 | Fam120a | -0.009 | Young |
| 0.87505 | 0.300631 | 0.147 | 0.142 | 1 | 9 | Ascc3 | 0.005 | Young |
| 0.875079 | 1.461191 | 0.015 | 0.013 | 1 | 9 | Gm45051 | 0.002 | Young |
| 0.875079 | 1.142108 | 0.015 | 0.013 | 1 | 9 | Gm10605 | 0.002 | Young |
| 0.875079 | 1.069131 | 0.015 | 0.013 | 1 | 9 | Ccdc28a | 0.002 | Young |
| 0.875079 | 0.893321 | 0.015 | 0.013 | 1 | 9 | BC051226 | 0.002 | Young |
| 0.875079 | 0.834742 | 0.015 | 0.013 | 1 | 9 | Zgpat | 0.002 | Young |
| 0.875079 | 0.699417 | 0.015 | 0.013 | 1 | 9 | Tmem35b | 0.002 | Young |
| 0.875079 | 0.374858 | 0.015 | 0.013 | 1 | 9 | Bivm | 0.002 | Young |
| 0.875079 | 0.314842 | 0.015 | 0.013 | 1 | 9 | Nnmt | 0.002 | Young |
| 0.875079 | 0.309759 | 0.015 | 0.013 | 1 | 9 | Zfp867 | 0.002 | Young |
| 0.875079 | 0.139434 | 0.015 | 0.013 | 1 | 9 | A830010M | 0.002 | Young |
| 0.875079 | 0.075599 | 0.015 | 0.013 | 1 | 9 | 2310040G2 | 0.002 | Young |
| 0.875171 | 0.391725 | 0.066 | 0.062 | 1 | 9 | Ttf1 | 0.004 | Young |
| 0.875569 | 0.074373 | 0.441 | 0.456 | 1 | 9 | Calm2 | -0.015 | Young |
| 0.87561 | 0.320132 | 0.118 | 0.113 | 1 | 9 | Man2a2 | 0.005 | Young |
| 0.875915 | 0.114701 | 0.382 | 0.366 | 1 | 9 | Crk | 0.016 | Young |
| 0.876295 | 0.478016 | 0.051 | 0.049 | 1 | 9 | Gprasp1 | 0.002 | Young |
| 0.876741 | 0.031087 | 0.382 | 0.405 | 1 | 9 | Tor1aip1 | -0.023 | Young |
| 0.877084 | 0.130675 | 0.331 | 0.348 | 1 | 9 | Sox17 | -0.017 | Young |
| 0.877119 | 0.356459 | 0.081 | 0.085 | 1 | 9 | Hoxa3 | -0.004 | Young |
| 0.877306 | 0.164658 | 0.125 | 0.119 | 1 | 9 | Pcgf5 | 0.006 | Young |
| 0.877578 | 0.056391 | 0.169 | 0.175 | 1 | 9 | Fut11 | -0.006 | Young |
| 0.877641 | 0.296824 | 0.14 | 0.137 | 1 | 9 | Rnd1 | 0.003 | Young |
| 0.877688 | 0.046349 | 0.015 | 0.013 | 1 | 9 | Pcdhgc5 | 0.002 | Young |
| 0.877688 | 0.03065 | 0.015 | 0.013 | 1 | 9 | Gm15987 | 0.002 | Young |
| 0.878123 | 0.204413 | 0.081 | 0.077 | 1 | 9 | Dpy19l4 | 0.004 | Young |
| 0.878458 | 0.635254 | 0.132 | 0.147 | 1 | 9 | Cd99l2 | -0.015 | Young |
| 0.878511 | 0.050749 | 0.221 | 0.211 | 1 | 9 | Samd4b | 0.01 | Young |
| 0.878564 | 0.40073 | 0.206 | 0.204 | 1 | 9 | Lmo4 | 0.002 | Young |
| 0.878578 | 0.024234 | 0.485 | 0.518 | 1 | 9 | Rps3 | -0.033 | Young |
| 0.878813 | 0.984895 | 0.044 | 0.041 | 1 | 9 | Ip6k2 | 0.003 | Young |
| 0.879311 | 0.052272 | 0.14 | 0.144 | 1 | 9 | Wbp4 | -0.004 | Young |
| 0.879485 | 0.682207 | 0.162 | 0.17 | 1 | 9 | Pink1 | -0.008 | Young |
| 0.880003 | 0.072505 | 0.831 | 0.858 | 1 | 9 | Fth1 | -0.027 | Young |
| 0.880008 | 0.393258 | 0.11 | 0.106 | 1 | 9 | Tcerg1 | 0.004 | Young |
| 0.880038 | 0.496117 | 0.206 | 0.204 | 1 | 9 | Sptlc2 | 0.002 | Young |
| 0.880222 | 0.111151 | 0.184 | 0.193 | 1 | 9 | Etf1 | -0.009 | Young |
| 0.880298 | 0.05753 | 0.015 | 0.013 | 1 | 9 | Pcdhga6 | 0.002 | Young |
| 0.880353 | 0.568929 | 0.074 | 0.08 | 1 | 9 | Vrk3 | -0.006 | Young |
| 0.880655 | 0.085355 | 0.471 | 0.464 | 1 | 9 | Pak2 | 0.007 | Young |

| | | | | | | | | |
|---|---|---|---|---|---|---|---|---|
| 0.881633 | 0.228406 | 0.162 | 0.16 | 1 | 9 | Tmem131 | 0.002 | Young |
| 0.881863 | 0.067525 | 0.154 | 0.16 | 1 | 9 | Tnfsf12 | -0.006 | Young |
| 0.882213 | 0.110082 | 0.191 | 0.183 | 1 | 9 | Gclc | 0.008 | Young |
| 0.882504 | 0.480807 | 0.103 | 0.101 | 1 | 9 | Kdsr | 0.002 | Young |
| 0.882629 | 0.218579 | 0.397 | 0.42 | 1 | 9 | Gpx1 | -0.023 | Young |
| 0.882893 | 0.140439 | 0.265 | 0.258 | 1 | 9 | Lipa | 0.007 | Young |
| 0.883297 | 0.329214 | 0.044 | 0.041 | 1 | 9 | Rhbdf1 | 0.003 | Young |
| 0.883362 | 0.336411 | 0.301 | 0.312 | 1 | 9 | Tmpo | -0.011 | Young |
| 0.88348 | 0.395388 | 0.199 | 0.196 | 1 | 9 | Tial1 | 0.003 | Young |
| 0.884237 | 0.199043 | 0.301 | 0.304 | 1 | 9 | Capn2 | -0.003 | Young |
| 0.884687 | 0.163438 | 0.191 | 0.188 | 1 | 9 | Emg1 | 0.003 | Young |
| 0.885228 | 0.205716 | 0.279 | 0.273 | 1 | 9 | Eid1 | 0.006 | Young |
| 0.885373 | 0.232394 | 0.485 | 0.531 | 1 | 9 | Rpl32 | -0.046 | Young |
| 0.885757 | 0.034597 | 0.081 | 0.085 | 1 | 9 | Ing2 | -0.004 | Young |
| 0.886209 | 0.006516 | 0.184 | 0.191 | 1 | 9 | Id2 | -0.007 | Young |
| 0.886881 | 0.458147 | 0.118 | 0.113 | 1 | 9 | Zbtb11 | 0.005 | Young |
| 0.88729 | 0.226902 | 0.235 | 0.227 | 1 | 9 | Slc30a4 | 0.008 | Young |
| 0.887378 | 0.118013 | 0.088 | 0.082 | 1 | 9 | Fam219b | 0.006 | Young |
| 0.887421 | 0.232489 | 0.118 | 0.121 | 1 | 9 | Aurkaip1 | -0.003 | Young |
| 0.887513 | 0.314005 | 0.103 | 0.098 | 1 | 9 | Ap3b1 | 0.005 | Young |
| 0.888163 | 0.377274 | 0.074 | 0.08 | 1 | 9 | Col8a1 | -0.006 | Young |
| 0.888327 | 0.100246 | 0.103 | 0.106 | 1 | 9 | Map3k3 | -0.003 | Young |
| 0.888531 | 0.57845 | 0.051 | 0.054 | 1 | 9 | Dnajb12 | -0.003 | Young |
| 0.88859 | 0.074992 | 0.088 | 0.09 | 1 | 9 | Sp4 | -0.002 | Young |
| 0.889099 | 0.285688 | 0.199 | 0.196 | 1 | 9 | Arl8a | 0.003 | Young |
| 0.88966 | 0.022157 | 0.213 | 0.204 | 1 | 9 | Dctn4 | 0.009 | Young |
| 0.889778 | 0.045909 | 0.228 | 0.222 | 1 | 9 | 11-Sep | 0.006 | Young |
| 0.89066 | 0.454334 | 0.096 | 0.093 | 1 | 9 | Fbxw7 | 0.003 | Young |
| 0.890681 | 0.353558 | 0.088 | 0.093 | 1 | 9 | Oard1 | -0.005 | Young |
| 0.891036 | 0.240773 | 0.404 | 0.423 | 1 | 9 | Arpc5 | -0.019 | Young |
| 0.891184 | 0.473774 | 0.037 | 0.039 | 1 | 9 | Trmt2a | -0.002 | Young |
| 0.891483 | 0.386095 | 0.051 | 0.049 | 1 | 9 | Stx18 | 0.002 | Young |
| 0.891936 | 0.021272 | 0.147 | 0.152 | 1 | 9 | Rarg | -0.005 | Young |
| 0.892191 | 0.221055 | 0.206 | 0.201 | 1 | 9 | Slc30a1 | 0.005 | Young |
| 0.892502 | 0.461493 | 0.154 | 0.152 | 1 | 9 | Higd1a | 0.002 | Young |
| 0.892968 | 0.066253 | 0.338 | 0.353 | 1 | 9 | Rbp7 | -0.015 | Young |
| 0.893402 | 0.060231 | 0.059 | 0.062 | 1 | 9 | Snx12 | -0.003 | Young |
| 0.894017 | 0.389828 | 0.059 | 0.057 | 1 | 9 | Pygb | 0.002 | Young |
| 0.894803 | 0.202991 | 0.074 | 0.077 | 1 | 9 | Tnfsf9 | -0.003 | Young |
| 0.894856 | 0.303385 | 0.132 | 0.131 | 1 | 9 | Slc39a9 | 0.001 | Young |
| 0.895696 | 0.114081 | 0.132 | 0.137 | 1 | 9 | Ankib1 | -0.005 | Young |
| 0.895829 | 0.290937 | 0.147 | 0.155 | 1 | 9 | Derl1 | -0.008 | Young |
| 0.896603 | 0.523324 | 0.059 | 0.057 | 1 | 9 | Phf13 | 0.002 | Young |
| 0.896603 | 0.248493 | 0.059 | 0.057 | 1 | 9 | Fbxo25 | 0.002 | Young |
| 0.897306 | 0.145959 | 0.154 | 0.157 | 1 | 9 | Hk1 | -0.003 | Young |

| | | | | | | | | |
|---|---|---|---|---|---|---|---|---|
| 0.897795 | 0.308867 | 0.125 | 0.121 | 1 | 9 | Lats1 | 0.004 | Young |
| 0.89804 | 0.487428 | 0.088 | 0.09 | 1 | 9 | Dnajb4 | -0.002 | Young |
| 0.89841 | 0.289863 | 0.103 | 0.101 | 1 | 9 | Galnt7 | 0.002 | Young |
| 0.89919 | 0.241451 | 0.059 | 0.057 | 1 | 9 | Slx1b | 0.002 | Young |
| 0.899352 | 0.352067 | 0.125 | 0.129 | 1 | 9 | Ech1 | -0.004 | Young |
| 0.899931 | 0.110934 | 0.235 | 0.232 | 1 | 9 | Ppp2r1a | 0.003 | Young |
| 0.899958 | 0.22524 | 0.154 | 0.162 | 1 | 9 | Tmem167 | -0.008 | Young |
| 0.900073 | 0.334377 | 0.14 | 0.134 | 1 | 9 | Ufm1 | 0.006 | Young |
| 0.900227 | 0.170414 | 0.294 | 0.291 | 1 | 9 | Dek | 0.003 | Young |
| 0.900776 | 0.294664 | 0.265 | 0.278 | 1 | 9 | Tmbim1 | -0.013 | Young |
| 0.90093 | 0.170238 | 0.059 | 0.062 | 1 | 9 | Gba2 | -0.003 | Young |
| 0.901047 | 0.09262 | 0.331 | 0.33 | 1 | 9 | Smad5 | 0.001 | Young |
| 0.901348 | 0.128165 | 0.11 | 0.106 | 1 | 9 | Hid1 | 0.004 | Young |
| 0.901825 | 0.272574 | 0.14 | 0.134 | 1 | 9 | Commd7 | 0.006 | Young |
| 0.901834 | 0.542319 | 0.103 | 0.108 | 1 | 9 | Ndufv2 | -0.005 | Young |
| 0.901888 | 0.408655 | 0.051 | 0.054 | 1 | 9 | Rbm10 | -0.003 | Young |
| 0.901915 | 0.743804 | 0.022 | 0.021 | 1 | 9 | Hap1 | 0.001 | Young |
| 0.901915 | 0.633158 | 0.022 | 0.021 | 1 | 9 | Actr6 | 0.001 | Young |
| 0.90203 | 0.308049 | 0.213 | 0.219 | 1 | 9 | Nsd3 | -0.006 | Young |
| 0.902184 | 0.139983 | 0.353 | 0.374 | 1 | 9 | Adamts9 | -0.021 | Young |
| 0.902447 | 0.087945 | 0.088 | 0.085 | 1 | 9 | Cxxc4 | 0.003 | Young |
| 0.902766 | 0.290027 | 0.044 | 0.041 | 1 | 9 | Slc11a2 | 0.003 | Young |
| 0.904154 | 0.294392 | 0.147 | 0.155 | 1 | 9 | Taf10 | -0.008 | Young |
| 0.904423 | 0.274497 | 0.066 | 0.07 | 1 | 9 | Rab33b | -0.004 | Young |
| 0.904794 | 0.328101 | 0.176 | 0.175 | 1 | 9 | Gspt1 | 0.001 | Young |
| 0.904912 | 0.385981 | 0.162 | 0.162 | 1 | 9 | Spata6 | 0 | Young |
| 0.905081 | 0.499231 | 0.228 | 0.24 | 1 | 9 | Spcs1 | -0.012 | Young |
| 0.905267 | 0.191766 | 0.044 | 0.046 | 1 | 9 | Fgf9 | -0.002 | Young |
| 0.905612 | 0.126796 | 0.066 | 0.07 | 1 | 9 | Cep83 | -0.004 | Young |
| 0.905665 | 0.389211 | 0.088 | 0.085 | 1 | 9 | Pgd | 0.003 | Young |
| 0.905773 | 0.402437 | 0.169 | 0.162 | 1 | 9 | Srp72 | 0.007 | Young |
| 0.906114 | 1.101159 | 0.022 | 0.021 | 1 | 9 | Nsun5 | 0.001 | Young |
| 0.906128 | 0.085014 | 0.096 | 0.093 | 1 | 9 | Cntrl | 0.003 | Young |
| 0.906187 | 0.069057 | 0.11 | 0.116 | 1 | 9 | Klhl20 | -0.006 | Young |
| 0.906739 | 0.55395 | 0.088 | 0.085 | 1 | 9 | Rassf4 | 0.003 | Young |
| 0.906918 | 0.273415 | 0.11 | 0.113 | 1 | 9 | Ric8a | -0.003 | Young |
| 0.907239 | 0.020486 | 0.051 | 0.054 | 1 | 9 | Senp3 | -0.003 | Young |
| 0.907546 | 0.187311 | 0.206 | 0.216 | 1 | 9 | Ube2n | -0.01 | Young |
| 0.907812 | 0.662474 | 0.088 | 0.085 | 1 | 9 | Exoc8 | 0.003 | Young |
| 0.907812 | 0.450348 | 0.088 | 0.085 | 1 | 9 | Ccdc6 | 0.003 | Young |
| 0.908072 | 0.025449 | 0.14 | 0.137 | 1 | 9 | Comt | 0.003 | Young |
| 0.908214 | 1.206037 | 0.022 | 0.021 | 1 | 9 | Rbm19 | 0.001 | Young |
| 0.908214 | 0.195121 | 0.022 | 0.021 | 1 | 9 | 1700021F0 | 0.001 | Young |
| 0.908261 | 0.55294 | 0.169 | 0.168 | 1 | 9 | Gna12 | 0.001 | Young |
| 0.908326 | 0.250608 | 0.647 | 0.688 | 1 | 9 | Rps10 | -0.041 | Young |

| | | | | | | | | |
|---|---|---|---|---|---|---|---|---|
| 0.908414 | 0.266497 | 0.338 | 0.34 | 1 | 9 | Ppp1r15a | -0.002 | Young |
| 0.908491 | 0.018474 | 0.868 | 0.887 | 1 | 9 | Mgll | -0.019 | Young |
| 0.908727 | 0.097849 | 0.176 | 0.175 | 1 | 9 | Sirt2 | 0.001 | Young |
| 0.908847 | 0.090717 | 0.154 | 0.157 | 1 | 9 | Gimap5 | -0.003 | Young |
| 0.909098 | 0.26998 | 0.096 | 0.101 | 1 | 9 | Sf3a1 | -0.005 | Young |
| 0.909517 | 0.009764 | 0.184 | 0.175 | 1 | 9 | Klhl2 | 0.009 | Young |
| 0.909855 | 0.636214 | 0.081 | 0.08 | 1 | 9 | Dcaf5 | 0.001 | Young |
| 0.90989 | 0.12445 | 0.301 | 0.309 | 1 | 9 | Ggnbp2 | -0.008 | Young |
| 0.910315 | 1.137814 | 0.022 | 0.021 | 1 | 9 | Xpo5 | 0.001 | Young |
| 0.910315 | 0.511079 | 0.022 | 0.021 | 1 | 9 | 8030462N1 | 0.001 | Young |
| 0.910315 | 0.195034 | 0.022 | 0.021 | 1 | 9 | Rnaseh2b | 0.001 | Young |
| 0.910456 | 0.226817 | 0.206 | 0.216 | 1 | 9 | Baz2a | -0.01 | Young |
| 0.910464 | 0.053659 | 0.088 | 0.093 | 1 | 9 | Pofut1 | -0.005 | Young |
| 0.910982 | 0.007434 | 0.059 | 0.062 | 1 | 9 | Gan | -0.003 | Young |
| 0.911483 | 0.375819 | 0.169 | 0.175 | 1 | 9 | Snrpb2 | -0.006 | Young |
| 0.912001 | 0.101064 | 0.074 | 0.072 | 1 | 9 | Fyco1 | 0.002 | Young |
| 0.912142 | 0.23044 | 0.059 | 0.057 | 1 | 9 | Gtf2e2 | 0.002 | Young |
| 0.912417 | 0.923786 | 0.022 | 0.021 | 1 | 9 | Mcm2 | 0.001 | Young |
| 0.912417 | 0.662769 | 0.022 | 0.021 | 1 | 9 | Phkg1 | 0.001 | Young |
| 0.912417 | 0.553711 | 0.022 | 0.021 | 1 | 9 | Brat1 | 0.001 | Young |
| 0.91255 | 0.239975 | 0.088 | 0.093 | 1 | 9 | Fam103a1 | -0.005 | Young |
| 0.912832 | 0.648044 | 0.081 | 0.085 | 1 | 9 | Mtf1 | -0.004 | Young |
| 0.913095 | 0.132632 | 0.456 | 0.466 | 1 | 9 | Hook3 | -0.01 | Young |
| 0.913223 | 0.264727 | 0.14 | 0.134 | 1 | 9 | Ncaph2 | 0.006 | Young |
| 0.914237 | 0.087741 | 0.176 | 0.175 | 1 | 9 | Mfsd14b | 0.001 | Young |
| 0.914519 | 0.99252 | 0.022 | 0.021 | 1 | 9 | Dyrk1b | 0.001 | Young |
| 0.914519 | 0.401263 | 0.022 | 0.021 | 1 | 9 | Slc26a11 | 0.001 | Young |
| 0.91469 | 0.197741 | 0.243 | 0.24 | 1 | 9 | Mrpl33 | 0.003 | Young |
| 0.914737 | 0.670176 | 0.037 | 0.039 | 1 | 9 | Acot8 | -0.002 | Young |
| 0.914901 | 0.237806 | 0.096 | 0.098 | 1 | 9 | Usp38 | -0.002 | Young |
| 0.91642 | 0.44041 | 0.051 | 0.049 | 1 | 9 | Nrde2 | 0.002 | Young |
| 0.916507 | 1.050434 | 0.081 | 0.08 | 1 | 9 | Usp24 | 0.001 | Young |
| 0.916575 | 0.514787 | 0.206 | 0.206 | 1 | 9 | Txnrd1 | 0 | Young |
| 0.916613 | 0.629922 | 0.051 | 0.054 | 1 | 9 | Gde1 | -0.003 | Young |
| 0.916613 | 0.279987 | 0.051 | 0.054 | 1 | 9 | Klhl18 | -0.003 | Young |
| 0.916622 | 1.284326 | 0.022 | 0.021 | 1 | 9 | Gla | 0.001 | Young |
| 0.916622 | 1.136413 | 0.022 | 0.021 | 1 | 9 | Eml6 | 0.001 | Young |
| 0.916622 | 0.706897 | 0.022 | 0.021 | 1 | 9 | Fam107b | 0.001 | Young |
| 0.916622 | 0.428045 | 0.022 | 0.021 | 1 | 9 | Dus2 | 0.001 | Young |
| 0.916622 | 0.161329 | 0.022 | 0.021 | 1 | 9 | Hspb11 | 0.001 | Young |
| 0.916622 | 0.084687 | 0.022 | 0.021 | 1 | 9 | Gatc | 0.001 | Young |
| 0.916622 | 0.064086 | 0.022 | 0.021 | 1 | 9 | Ipo4 | 0.001 | Young |
| 0.917166 | 0.044432 | 0.213 | 0.209 | 1 | 9 | Zfp706 | 0.004 | Young |
| 0.917329 | 0.135308 | 0.059 | 0.057 | 1 | 9 | Mrpl10 | 0.002 | Young |
| 0.91787 | 0.103224 | 0.118 | 0.121 | 1 | 9 | Zfhx3 | -0.003 | Young |

| | | | | | | | | |
|---|---|---|---|---|---|---|---|---|
| 0.917883 | 0.435833 | 0.037 | 0.039 | 1 | 9 | A530017D2 | -0.002 | Young |
| 0.917953 | 0.429858 | 0.051 | 0.054 | 1 | 9 | Pes1 | -0.003 | Young |
| 0.918011 | 0.210315 | 0.353 | 0.358 | 1 | 9 | Irf2bpl | -0.005 | Young |
| 0.918217 | 0.307901 | 0.029 | 0.031 | 1 | 9 | Tbc1d31 | -0.002 | Young |
| 0.918234 | 0.370718 | 0.044 | 0.046 | 1 | 9 | Ethe1 | -0.002 | Young |
| 0.918725 | 0.189638 | 0.022 | 0.021 | 1 | 9 | Sh3d21 | 0.001 | Young |
| 0.918725 | 0.054886 | 0.022 | 0.021 | 1 | 9 | Med26 | 0.001 | Young |
| 0.918812 | 0.283164 | 0.088 | 0.093 | 1 | 9 | Derl2 | -0.005 | Young |
| 0.919012 | 0.572426 | 0.11 | 0.108 | 1 | 9 | Zc3h12a | 0.002 | Young |
| 0.919242 | 0.563109 | 0.029 | 0.028 | 1 | 9 | Rcc1l | 0.001 | Young |
| 0.919457 | 0.61173 | 0.037 | 0.039 | 1 | 9 | Hps4 | -0.002 | Young |
| 0.919457 | 0.03644 | 0.037 | 0.039 | 1 | 9 | Zfp866 | -0.002 | Young |
| 0.919676 | 0.349001 | 0.044 | 0.046 | 1 | 9 | Gtpbp1 | -0.002 | Young |
| 0.919896 | 0.072485 | 0.066 | 0.07 | 1 | 9 | Vangl1 | -0.004 | Young |
| 0.919961 | 0.031337 | 0.324 | 0.314 | 1 | 9 | Dram2 | 0.01 | Young |
| 0.920153 | 0.134055 | 0.132 | 0.131 | 1 | 9 | Hnrnpll | 0.001 | Young |
| 0.920544 | 0.153683 | 0.132 | 0.129 | 1 | 9 | Prdm16 | 0.003 | Young |
| 0.920829 | 0.09248 | 0.022 | 0.021 | 1 | 9 | Peg10 | 0.001 | Young |
| 0.920956 | 0.200417 | 0.228 | 0.237 | 1 | 9 | H13 | -0.009 | Young |
| 0.921304 | 0.193434 | 0.434 | 0.446 | 1 | 9 | Rsad2 | -0.012 | Young |
| 0.921723 | 0.241037 | 0.029 | 0.031 | 1 | 9 | Prorsd1 | -0.002 | Young |
| 0.922055 | 0.722614 | 0.081 | 0.08 | 1 | 9 | Trim27 | 0.001 | Young |
| 0.922332 | 0.173972 | 0.118 | 0.116 | 1 | 9 | Upf3a | 0.002 | Young |
| 0.922341 | 0.142646 | 0.184 | 0.18 | 1 | 9 | Tssc4 | 0.004 | Young |
| 0.922393 | 0.375627 | 0.191 | 0.191 | 1 | 9 | Gtf2h5 | 0 | Young |
| 0.922412 | 0.141229 | 0.132 | 0.134 | 1 | 9 | Akr1b3 | -0.002 | Young |
| 0.922431 | 0.288585 | 0.213 | 0.224 | 1 | 9 | Cited2 | -0.011 | Young |
| 0.922519 | 0.239324 | 0.059 | 0.057 | 1 | 9 | Zfp780b | 0.002 | Young |
| 0.922934 | 0.250658 | 0.022 | 0.021 | 1 | 9 | Cdc42ep5 | 0.001 | Young |
| 0.922934 | 0.211908 | 0.022 | 0.021 | 1 | 9 | Rrp12 | 0.001 | Young |
| 0.922947 | 0.050969 | 0.397 | 0.392 | 1 | 9 | Gstm1 | 0.005 | Young |
| 0.923127 | 0.217103 | 0.316 | 0.317 | 1 | 9 | Nedd9 | -0.001 | Young |
| 0.923307 | 0.237441 | 0.088 | 0.09 | 1 | 9 | Mrpl13 | -0.002 | Young |
| 0.923363 | 0.25676 | 0.051 | 0.049 | 1 | 9 | Sybu | 0.002 | Young |
| 0.923375 | 0.056873 | 0.147 | 0.144 | 1 | 9 | Tomm34 | 0.003 | Young |
| 0.923476 | 0.312393 | 0.029 | 0.031 | 1 | 9 | Zfp667 | -0.002 | Young |
| 0.923778 | 0.1903 | 0.14 | 0.137 | 1 | 9 | Gmps | 0.003 | Young |
| 0.923862 | 0.268809 | 0.066 | 0.064 | 1 | 9 | Wdcp | 0.002 | Young |
| 0.924433 | 0.233827 | 0.206 | 0.204 | 1 | 9 | 6-Mar | 0.002 | Young |
| 0.924907 | 0.065097 | 0.875 | 0.851 | 1 | 9 | H2-D1 | 0.024 | Young |
| 0.925039 | 0.233011 | 0.022 | 0.021 | 1 | 9 | Gmnn | 0.001 | Young |
| 0.925082 | 0.378683 | 0.147 | 0.144 | 1 | 9 | Clint1 | 0.003 | Young |
| 0.92523 | 0.118098 | 0.029 | 0.031 | 1 | 9 | P2ry10b | -0.002 | Young |
| 0.925501 | 0.142681 | 0.221 | 0.209 | 1 | 9 | Itm2c | 0.012 | Young |
| 0.925755 | 0.049074 | 0.037 | 0.039 | 1 | 9 | Hoxb2 | -0.002 | Young |

| | | | | | | | | |
|---|---|---|---|---|---|---|---|---|
| 0.925943 | 0.27351 | 0.074 | 0.072 | 1 | 9 | Arhgef40 | 0.002 | Young |
| 0.926234 | 0.024125 | 0.235 | 0.235 | 1 | 9 | Larp4 | 0 | Young |
| 0.92634 | 0.854005 | 0.103 | 0.101 | 1 | 9 | Ak3 | 0.002 | Young |
| 0.926657 | 0.122331 | 0.735 | 0.781 | 1 | 9 | Klf6 | -0.046 | Young |
| 0.926891 | 0.367623 | 0.044 | 0.046 | 1 | 9 | Dck | -0.002 | Young |
| 0.926985 | 0.055061 | 0.029 | 0.031 | 1 | 9 | Cby1 | -0.002 | Young |
| 0.927341 | 0.650809 | 0.051 | 0.054 | 1 | 9 | Pan2 | -0.003 | Young |
| 0.927544 | 0.548114 | 0.118 | 0.119 | 1 | 9 | Sipa1l2 | -0.001 | Young |
| 0.927842 | 0.558805 | 0.096 | 0.093 | 1 | 9 | Rfk | 0.003 | Young |
| 0.927882 | 0.858919 | 0.088 | 0.088 | 1 | 9 | Meaf6 | 0 | Young |
| 0.928107 | 0.360622 | 0.125 | 0.124 | 1 | 9 | Aggf1 | 0.001 | Young |
| 0.928155 | 0.055698 | 0.14 | 0.134 | 1 | 9 | Vdac1 | 0.006 | Young |
| 0.92829 | 1.17694 | 0.029 | 0.028 | 1 | 9 | D030056L2 | 0.001 | Young |
| 0.928335 | 0.872379 | 0.044 | 0.046 | 1 | 9 | Zbtb8os | -0.002 | Young |
| 0.928683 | 0.341953 | 0.051 | 0.054 | 1 | 9 | Nckipsd | -0.003 | Young |
| 0.928906 | 0.014015 | 0.037 | 0.039 | 1 | 9 | Thumpd3 | -0.002 | Young |
| 0.929049 | 0.345434 | 0.132 | 0.131 | 1 | 9 | Idh3b | 0.001 | Young |
| 0.929461 | 0.115547 | 0.353 | 0.376 | 1 | 9 | Prrc2b | -0.023 | Young |
| 0.929651 | 0.369758 | 0.11 | 0.108 | 1 | 9 | Oxsr1 | 0.002 | Young |
| 0.929781 | 0.096361 | 0.61 | 0.616 | 1 | 9 | Ier2 | -0.006 | Young |
| 0.92995 | 0.054283 | 0.324 | 0.33 | 1 | 9 | Cds2 | -0.006 | Young |
| 0.929985 | 0.548286 | 0.066 | 0.064 | 1 | 9 | Mageh1 | 0.002 | Young |
| 0.930482 | 0.049037 | 0.037 | 0.039 | 1 | 9 | Zfp618 | -0.002 | Young |
| 0.93049 | 0.132085 | 0.199 | 0.204 | 1 | 9 | AC149090. | -0.005 | Young |
| 0.931211 | 0.741895 | 0.066 | 0.064 | 1 | 9 | Hopx | 0.002 | Young |
| 0.931223 | 0.655176 | 0.044 | 0.046 | 1 | 9 | Asb3 | -0.002 | Young |
| 0.931234 | 0.096273 | 0.846 | 0.892 | 1 | 9 | Rps8 | -0.046 | Young |
| 0.931612 | 0.4323 | 0.169 | 0.165 | 1 | 9 | Dennd4c | 0.004 | Young |
| 0.932058 | 0.413349 | 0.037 | 0.039 | 1 | 9 | Rasd1 | -0.002 | Young |
| 0.932058 | 0.217028 | 0.037 | 0.039 | 1 | 9 | Bcl2l12 | -0.002 | Young |
| 0.932058 | 0.164664 | 0.037 | 0.039 | 1 | 9 | Casp6 | -0.002 | Young |
| 0.93225 | 0.270015 | 0.029 | 0.031 | 1 | 9 | Pgm2 | -0.002 | Young |
| 0.93225 | 0.079471 | 0.029 | 0.031 | 1 | 9 | Il17rc | -0.002 | Young |
| 0.932882 | 0.091294 | 0.154 | 0.155 | 1 | 9 | Kdelr2 | -0.001 | Young |
| 0.933327 | 0.195117 | 0.081 | 0.082 | 1 | 9 | BC017643 | -0.001 | Young |
| 0.933511 | 0.038352 | 0.647 | 0.68 | 1 | 9 | Eif4g2 | -0.033 | Young |
| 0.933723 | 0.572057 | 0.029 | 0.028 | 1 | 9 | Fam131a | 0.001 | Young |
| 0.933774 | 0.312357 | 0.39 | 0.407 | 1 | 9 | Psmb4 | -0.017 | Young |
| 0.934005 | 0.308909 | 0.029 | 0.031 | 1 | 9 | Tubgcp2 | -0.002 | Young |
| 0.934207 | 0.228426 | 0.066 | 0.07 | 1 | 9 | Atxn7l1 | -0.004 | Young |
| 0.934291 | 0.08647 | 0.618 | 0.631 | 1 | 9 | Mmrn2 | -0.013 | Young |
| 0.934444 | 0.105671 | 0.176 | 0.173 | 1 | 9 | Llph | 0.003 | Young |
| 0.934698 | 0.083736 | 0.147 | 0.152 | 1 | 9 | Rcn2 | -0.005 | Young |
| 0.935021 | 0.011461 | 0.176 | 0.178 | 1 | 9 | Crybg1 | -0.002 | Young |
| 0.935029 | 0.279958 | 0.154 | 0.165 | 1 | 9 | Ncoa6 | -0.011 | Young |

| | | | | | | | | |
|---|---|---|---|---|---|---|---|---|
| 0.935191 | 0.463901 | 0.132 | 0.137 | 1 | 9 | Cnot3 | -0.005 | Young |
| 0.935368 | 0.030028 | 0.5 | 0.536 | 1 | 9 | Actr2 | -0.036 | Young |
| 0.935389 | 0.565538 | 0.125 | 0.126 | 1 | 9 | Laptm4b | -0.001 | Young |
| 0.935535 | 1.648394 | 0.029 | 0.028 | 1 | 9 | Gm15417 | 0.001 | Young |
| 0.935535 | 1.241252 | 0.029 | 0.028 | 1 | 9 | Miip | 0.001 | Young |
| 0.935557 | 0.755254 | 0.044 | 0.046 | 1 | 9 | Aste1 | -0.002 | Young |
| 0.935557 | 0.547748 | 0.044 | 0.046 | 1 | 9 | Tpra1 | -0.002 | Young |
| 0.935761 | 0.054441 | 0.029 | 0.031 | 1 | 9 | Tusc1 | -0.002 | Young |
| 0.936738 | 0.64519 | 0.051 | 0.054 | 1 | 9 | Lrg1 | -0.003 | Young |
| 0.936788 | 0.386392 | 0.037 | 0.039 | 1 | 9 | Parp8 | -0.002 | Young |
| 0.936788 | 0.330701 | 0.037 | 0.039 | 1 | 9 | C4b | -0.002 | Young |
| 0.937002 | 0.787974 | 0.044 | 0.046 | 1 | 9 | Isoc1 | -0.002 | Young |
| 0.93724 | 0.415842 | 0.022 | 0.023 | 1 | 9 | Gm38910 | -0.001 | Young |
| 0.937477 | 0.464645 | 0.088 | 0.088 | 1 | 9 | Cep68 | 0 | Young |
| 0.937518 | 0.445755 | 0.029 | 0.031 | 1 | 9 | Mon1a | -0.002 | Young |
| 0.937518 | 0.350974 | 0.029 | 0.031 | 1 | 9 | Mamld1 | -0.002 | Young |
| 0.937578 | 0.342928 | 0.162 | 0.16 | 1 | 9 | Plod3 | 0.002 | Young |
| 0.937638 | 0.332004 | 0.272 | 0.276 | 1 | 9 | Cdkn1b | -0.004 | Young |
| 0.937874 | 0.283916 | 0.154 | 0.155 | 1 | 9 | Zbtb18 | -0.001 | Young |
| 0.938159 | 0.026101 | 0.125 | 0.124 | 1 | 9 | Upf1 | 0.001 | Young |
| 0.938619 | 0.052049 | 0.956 | 0.961 | 1 | 9 | Rpl37a | -0.005 | Young |
| 0.938656 | 0.149995 | 0.051 | 0.049 | 1 | 9 | Paxip1 | 0.002 | Young |
| 0.938744 | 0.525878 | 0.074 | 0.072 | 1 | 9 | Paqr4 | 0.002 | Young |
| 0.938946 | 0.028946 | 0.397 | 0.41 | 1 | 9 | Prrc2a | -0.013 | Young |
| 0.939166 | 0.071248 | 0.118 | 0.121 | 1 | 9 | Ntan1 | -0.003 | Young |
| 0.939238 | 0.462496 | 0.096 | 0.093 | 1 | 9 | Zfp330 | 0.003 | Young |
| 0.939274 | 0.866388 | 0.029 | 0.031 | 1 | 9 | Tmem41a | -0.002 | Young |
| 0.939334 | 0.221451 | 0.11 | 0.108 | 1 | 9 | Ocrl | 0.002 | Young |
| 0.939593 | 0.377346 | 0.272 | 0.281 | 1 | 9 | Uba1 | -0.009 | Young |
| 0.939601 | 0.030803 | 0.228 | 0.232 | 1 | 9 | Ndufa13 | -0.004 | Young |
| 0.93994 | 0.454189 | 0.206 | 0.211 | 1 | 9 | Nsmce4a | -0.005 | Young |
| 0.940972 | 0.317433 | 0.029 | 0.028 | 1 | 9 | Bud23 | 0.001 | Young |
| 0.941258 | 0.064246 | 0.684 | 0.67 | 1 | 9 | Ets2 | 0.014 | Young |
| 0.941281 | 0.127389 | 0.022 | 0.023 | 1 | 9 | Wdr18 | -0.001 | Young |
| 0.941439 | 0.105883 | 0.051 | 0.049 | 1 | 9 | Tmem241 | 0.002 | Young |
| 0.941539 | 0.604106 | 0.118 | 0.119 | 1 | 9 | Rasa4 | -0.001 | Young |
| 0.941879 | 0.003499 | 0.132 | 0.134 | 1 | 9 | Timm17a | -0.002 | Young |
| 0.942113 | 0.537212 | 0.051 | 0.054 | 1 | 9 | Slc25a23 | -0.003 | Young |
| 0.942197 | 0.127631 | 0.25 | 0.253 | 1 | 9 | Uqcr10 | -0.003 | Young |
| 0.942413 | 0.132535 | 0.493 | 0.497 | 1 | 9 | Hes1 | -0.004 | Young |
| 0.942785 | 0.627666 | 0.029 | 0.028 | 1 | 9 | P3h2 | 0.001 | Young |
| 0.942785 | 0.09857 | 0.029 | 0.028 | 1 | 9 | Tmem199 | 0.001 | Young |
| 0.942927 | 0.060311 | 0.074 | 0.075 | 1 | 9 | Daxx | -0.001 | Young |
| 0.943089 | 0.995238 | 0.044 | 0.044 | 1 | 9 | Gtf2e1 | 0 | Young |
| 0.943089 | 0.851392 | 0.044 | 0.044 | 1 | 9 | Etfdh | 0 | Young |

| | | | | | | | | |
|---|---|---|---|---|---|---|---|---|
| 0.943303 | 0.356629 | 0.022 | 0.023 | 1 | 9 | Aim2 | -0.001 | Young |
| 0.943588 | 0.354264 | 0.074 | 0.077 | 1 | 9 | Lhx6 | -0.003 | Young |
| 0.943734 | 0.006928 | 0.478 | 0.469 | 1 | 9 | Tshz2 | 0.009 | Young |
| 0.943808 | 0.222693 | 0.301 | 0.309 | 1 | 9 | Lipe | -0.008 | Young |
| 0.943826 | 0.035797 | 0.154 | 0.152 | 1 | 9 | Eif4e3 | 0.002 | Young |
| 0.943878 | 0.205763 | 0.088 | 0.088 | 1 | 9 | Ddx47 | 0 | Young |
| 0.944008 | 0.343211 | 0.11 | 0.116 | 1 | 9 | Agps | -0.006 | Young |
| 0.944049 | 0.092867 | 0.559 | 0.582 | 1 | 9 | Fkbp1a | -0.023 | Young |
| 0.944104 | 1.427624 | 0.037 | 0.036 | 1 | 9 | Slc38a7 | 0.001 | Young |
| 0.944282 | 0.329975 | 0.081 | 0.08 | 1 | 9 | Rab11fip2 | 0.001 | Young |
| 0.944396 | 0.199846 | 0.096 | 0.098 | 1 | 9 | Trp53inp1 | -0.002 | Young |
| 0.944599 | 0.47947 | 0.029 | 0.028 | 1 | 9 | Ptrhd1 | 0.001 | Young |
| 0.944663 | 0.130112 | 0.154 | 0.152 | 1 | 9 | Prkcsh | 0.002 | Young |
| 0.945003 | 0.277291 | 0.059 | 0.062 | 1 | 9 | 1600014C1 | -0.003 | Young |
| 0.945324 | 0.952171 | 0.022 | 0.023 | 1 | 9 | Adamts15 | -0.001 | Young |
| 0.94583 | 0.493206 | 0.221 | 0.227 | 1 | 9 | Nrarp | -0.006 | Young |
| 0.946145 | 0.133089 | 0.051 | 0.054 | 1 | 9 | Vsig2 | -0.003 | Young |
| 0.946254 | 0.45082 | 0.037 | 0.039 | 1 | 9 | Ripk2 | -0.002 | Young |
| 0.946254 | 0.263563 | 0.037 | 0.039 | 1 | 9 | Mrps31 | -0.002 | Young |
| 0.946377 | 0.076303 | 0.118 | 0.113 | 1 | 9 | Taf6l | 0.005 | Young |
| 0.946412 | 0.481362 | 0.029 | 0.028 | 1 | 9 | Tnfrsf23 | 0.001 | Young |
| 0.946507 | 0.176283 | 0.081 | 0.08 | 1 | 9 | Gpr108 | 0.001 | Young |
| 0.946635 | 0.397639 | 0.14 | 0.134 | 1 | 9 | Lpgat1 | 0.006 | Young |
| 0.946879 | 0.00979 | 0.015 | 0.015 | 1 | 9 | B130006D( | 0 | Young |
| 0.94734 | 0.649575 | 0.037 | 0.036 | 1 | 9 | 2310057M | 0.001 | Young |
| 0.94734 | 0.360053 | 0.037 | 0.036 | 1 | 9 | Tfip11 | 0.001 | Young |
| 0.947346 | 0.394603 | 0.022 | 0.023 | 1 | 9 | Ncald | -0.001 | Young |
| 0.947346 | 0.230762 | 0.022 | 0.023 | 1 | 9 | Nudt18 | -0.001 | Young |
| 0.947829 | 0.0302 | 0.162 | 0.16 | 1 | 9 | Ankfy1 | 0.002 | Young |
| 0.948062 | 0.905982 | 0.029 | 0.031 | 1 | 9 | Aqp11 | -0.002 | Young |
| 0.948062 | 0.511793 | 0.029 | 0.031 | 1 | 9 | Ar | -0.002 | Young |
| 0.948212 | 0.44864 | 0.103 | 0.106 | 1 | 9 | BC003331 | -0.003 | Young |
| 0.948226 | 0.955576 | 0.029 | 0.028 | 1 | 9 | Haus7 | 0.001 | Young |
| 0.948226 | 0.055643 | 0.029 | 0.028 | 1 | 9 | D3Ertd254 | 0.001 | Young |
| 0.948373 | 0.197419 | 0.272 | 0.271 | 1 | 9 | Arhgap17 | 0.001 | Young |
| 0.949097 | 0.140833 | 0.051 | 0.049 | 1 | 9 | Siah2 | 0.002 | Young |
| 0.949109 | 0.111828 | 0.316 | 0.325 | 1 | 9 | Plk2 | -0.009 | Young |
| 0.949137 | 0.358177 | 0.125 | 0.124 | 1 | 9 | Cnot7 | 0.001 | Young |
| 0.949137 | 0.085402 | 0.125 | 0.124 | 1 | 9 | Jam3 | 0.001 | Young |
| 0.949369 | 0.858119 | 0.022 | 0.023 | 1 | 9 | Slc41a3 | -0.001 | Young |
| 0.949369 | 0.620364 | 0.022 | 0.023 | 1 | 9 | 5430416N( | -0.001 | Young |
| 0.94982 | 0.574645 | 0.029 | 0.031 | 1 | 9 | Uck2 | -0.002 | Young |
| 0.949893 | 0.143148 | 0.199 | 0.204 | 1 | 9 | Cyb5r1 | -0.005 | Young |
| 0.95004 | 0.824069 | 0.029 | 0.028 | 1 | 9 | Nudt1 | 0.001 | Young |
| 0.950576 | 0.290375 | 0.037 | 0.036 | 1 | 9 | Gm29107 | 0.001 | Young |

| | | | | | | | | |
|---|---|---|---|---|---|---|---|---|
| 0.950959 | 0.533887 | 0.081 | 0.08 | 1 | 9 | Maff | 0.001 | Young |
| 0.950963 | 0.174949 | 0.176 | 0.18 | 1 | 9 | Nsfl1c | -0.004 | Young |
| 0.950987 | 0.105917 | 0.074 | 0.075 | 1 | 9 | Dusp5 | -0.001 | Young |
| 0.951391 | 0.801358 | 0.022 | 0.023 | 1 | 9 | Cnrip1 | -0.001 | Young |
| 0.951391 | 0.751199 | 0.022 | 0.023 | 1 | 9 | Dmap1 | -0.001 | Young |
| 0.951391 | 0.482489 | 0.022 | 0.023 | 1 | 9 | Apbb1 | -0.001 | Young |
| 0.951391 | 0.133645 | 0.022 | 0.023 | 1 | 9 | Panx1 | -0.001 | Young |
| 0.951562 | 0.144864 | 0.221 | 0.227 | 1 | 9 | Supt5 | -0.006 | Young |
| 0.951578 | 0.402511 | 0.029 | 0.031 | 1 | 9 | Ppil3 | -0.002 | Young |
| 0.951582 | 0.184303 | 0.551 | 0.552 | 1 | 9 | Cox6a1 | -0.001 | Young |
| 0.951618 | 0.226865 | 0.14 | 0.139 | 1 | 9 | Smap1 | 0.001 | Young |
| 0.951814 | 0.995689 | 0.015 | 0.015 | 1 | 9 | Mpst | 0 | Young |
| 0.951814 | 0.477928 | 0.015 | 0.015 | 1 | 9 | Sgsm2 | 0 | Young |
| 0.951814 | 0.301111 | 0.015 | 0.015 | 1 | 9 | E430018J2 | 0 | Young |
| 0.951814 | 0.288941 | 0.015 | 0.015 | 1 | 9 | Ptk2b | 0 | Young |
| 0.951814 | 0.153283 | 0.015 | 0.015 | 1 | 9 | Zfp383 | 0 | Young |
| 0.951814 | 0.018794 | 0.015 | 0.015 | 1 | 9 | Mettl3 | 0 | Young |
| 0.951855 | 0.800813 | 0.029 | 0.028 | 1 | 9 | Nudcd1 | 0.001 | Young |
| 0.951855 | 0.176341 | 0.029 | 0.028 | 1 | 9 | Prelid2 | 0.001 | Young |
| 0.951855 | 0.017442 | 0.029 | 0.028 | 1 | 9 | Ebag9 | 0.001 | Young |
| 0.952031 | 0.591777 | 0.154 | 0.155 | 1 | 9 | Tmem259 | -0.001 | Young |
| 0.952258 | 0.025959 | 0.691 | 0.735 | 1 | 9 | Myh9 | -0.044 | Young |
| 0.952306 | 0.34267 | 0.301 | 0.322 | 1 | 9 | Slc39a1 | -0.021 | Young |
| 0.952868 | 0.383142 | 0.051 | 0.054 | 1 | 9 | Zfp444 | -0.003 | Young |
| 0.952942 | 0.383315 | 0.213 | 0.216 | 1 | 9 | Zbtb46 | -0.003 | Young |
| 0.95329 | 0.360345 | 0.066 | 0.064 | 1 | 9 | Itgb1bp1 | 0.002 | Young |
| 0.953414 | 1.811329 | 0.022 | 0.023 | 1 | 9 | Hba-a1 | -0.001 | Young |
| 0.953414 | 0.289824 | 0.022 | 0.023 | 1 | 9 | Yars2 | -0.001 | Young |
| 0.953416 | 0.238841 | 0.382 | 0.392 | 1 | 9 | Mkln1 | -0.01 | Young |
| 0.953479 | 0.163132 | 0.199 | 0.196 | 1 | 9 | Ndufb8 | 0.003 | Young |
| 0.95367 | 0.832859 | 0.029 | 0.028 | 1 | 9 | Pigh | 0.001 | Young |
| 0.953981 | 0.340765 | 0.221 | 0.219 | 1 | 9 | Pank3 | 0.002 | Young |
| 0.954171 | 0.520839 | 0.081 | 0.085 | 1 | 9 | Yod1 | -0.004 | Young |
| 0.954282 | 0.58648 | 0.015 | 0.015 | 1 | 9 | Gm16740 | 0 | Young |
| 0.954282 | 0.227874 | 0.015 | 0.015 | 1 | 9 | 2410131K1 | 0 | Young |
| 0.954282 | 0.227428 | 0.015 | 0.015 | 1 | 9 | Zbtb34 | 0 | Young |
| 0.954667 | 0.015614 | 0.331 | 0.335 | 1 | 9 | Ccdc88c | -0.004 | Young |
| 0.95469 | 0.0522 | 0.449 | 0.456 | 1 | 9 | Arpc1b | -0.007 | Young |
| 0.954859 | 0.123584 | 0.228 | 0.232 | 1 | 9 | Tsen34 | -0.004 | Young |
| 0.955336 | 0.12504 | 0.279 | 0.299 | 1 | 9 | Cldn15 | -0.02 | Young |
| 0.955384 | 0.40104 | 0.125 | 0.126 | 1 | 9 | Suco | -0.001 | Young |
| 0.955485 | 1.174338 | 0.029 | 0.028 | 1 | 9 | Mkl1 | 0.001 | Young |
| 0.955693 | 0.097915 | 0.706 | 0.727 | 1 | 9 | Ppia | -0.021 | Young |
| 0.955746 | 0.119839 | 0.066 | 0.064 | 1 | 9 | Plxna1 | 0.002 | Young |
| 0.956036 | 0.119517 | 0.162 | 0.16 | 1 | 9 | Nars | 0.002 | Young |

| | | | | | | | | |
|---|---|---|---|---|---|---|---|---|
| 0.956138 | 0.096181 | 0.691 | 0.711 | 1 | 9 | Cyyr1 | -0.02 | Young |
| 0.956176 | 0.129603 | 0.118 | 0.116 | 1 | 9 | Sdhb | 0.002 | Young |
| 0.956369 | 0.201348 | 0.213 | 0.219 | 1 | 9 | Ppp1r15b | -0.006 | Young |
| 0.956747 | 0.004164 | 0.074 | 0.075 | 1 | 9 | Ssr2 | -0.001 | Young |
| 0.956748 | 0.209496 | 0.125 | 0.126 | 1 | 9 | Lbh | -0.001 | Young |
| 0.956751 | 0.68342 | 0.015 | 0.015 | 1 | 9 | Smpdl3b | 0 | Young |
| 0.956751 | 0.187629 | 0.015 | 0.015 | 1 | 9 | Ptger2 | 0 | Young |
| 0.957051 | 0.979347 | 0.037 | 0.036 | 1 | 9 | Kat5 | 0.001 | Young |
| 0.957072 | 0.079981 | 0.169 | 0.17 | 1 | 9 | Prpf39 | -0.001 | Young |
| 0.957112 | 0.111771 | 0.088 | 0.09 | 1 | 9 | Plekhg3 | -0.002 | Young |
| 0.957213 | 0.102491 | 0.485 | 0.495 | 1 | 9 | Canx | -0.01 | Young |
| 0.957461 | 1.231053 | 0.022 | 0.023 | 1 | 9 | Phtf2 | -0.001 | Young |
| 0.95758 | 0.098083 | 0.412 | 0.423 | 1 | 9 | Ktn1 | -0.011 | Young |
| 0.957874 | 0.325231 | 0.169 | 0.17 | 1 | 9 | Rabep1 | -0.001 | Young |
| 0.95817 | 0.089539 | 0.088 | 0.09 | 1 | 9 | Magix | -0.002 | Young |
| 0.958388 | 0.908538 | 0.074 | 0.077 | 1 | 9 | Ncoa5 | -0.003 | Young |
| 0.95867 | 1.033089 | 0.037 | 0.036 | 1 | 9 | Apip | 0.001 | Young |
| 0.958693 | 0.162912 | 0.243 | 0.247 | 1 | 9 | Atp6v1e1 | -0.004 | Young |
| 0.959097 | 0.163656 | 0.294 | 0.299 | 1 | 9 | Ppp1r16b | -0.005 | Young |
| 0.959115 | 0.389005 | 0.029 | 0.028 | 1 | 9 | Pdgfa | 0.001 | Young |
| 0.95922 | 0.646096 | 0.015 | 0.015 | 1 | 9 | Shmt2 | 0 | Young |
| 0.95922 | 0.352186 | 0.015 | 0.015 | 1 | 9 | Ndufaf1 | 0 | Young |
| 0.95922 | 0.325705 | 0.015 | 0.015 | 1 | 9 | Ccdc15 | 0 | Young |
| 0.959332 | 0.785996 | 0.044 | 0.044 | 1 | 9 | Spef1 | 0 | Young |
| 0.960051 | 0.032603 | 0.529 | 0.554 | 1 | 9 | Klf9 | -0.025 | Young |
| 0.960577 | 0.017842 | 0.39 | 0.389 | 1 | 9 | Map7d1 | 0.001 | Young |
| 0.960638 | 0.1556 | 0.176 | 0.173 | 1 | 9 | N4bp3 | 0.003 | Young |
| 0.960663 | 0.558396 | 0.088 | 0.093 | 1 | 9 | Rabgap1l | -0.005 | Young |
| 0.960867 | 0.217183 | 0.096 | 0.095 | 1 | 9 | Stard13 | 0.001 | Young |
| 0.961014 | 0.093983 | 0.338 | 0.348 | 1 | 9 | Ube3a | -0.01 | Young |
| 0.961142 | 0.365712 | 0.132 | 0.131 | 1 | 9 | Papd4 | 0.001 | Young |
| 0.961379 | 0.025661 | 0.603 | 0.624 | 1 | 9 | Rpl36 | -0.021 | Young |
| 0.96169 | 1.659867 | 0.015 | 0.015 | 1 | 9 | Adal | 0 | Young |
| 0.96169 | 0.329022 | 0.015 | 0.015 | 1 | 9 | Mettl18 | 0 | Young |
| 0.96169 | 0.233762 | 0.015 | 0.015 | 1 | 9 | Fbxo5 | 0 | Young |
| 0.96169 | 0.070968 | 0.015 | 0.015 | 1 | 9 | Tmem143 | 0 | Young |
| 0.961887 | 0.475965 | 0.066 | 0.064 | 1 | 9 | Hif1an | 0.002 | Young |
| 0.961896 | 0.355696 | 0.096 | 0.095 | 1 | 9 | Mrpl2 | 0.001 | Young |
| 0.961909 | 0.212711 | 0.037 | 0.036 | 1 | 9 | Slc35f6 | 0.001 | Young |
| 0.962159 | 0.300056 | 0.125 | 0.121 | 1 | 9 | Fam129a | 0.004 | Young |
| 0.962185 | 0.26328 | 0.184 | 0.188 | 1 | 9 | Lsm14a | -0.004 | Young |
| 0.9622 | 0.218828 | 0.25 | 0.263 | 1 | 9 | Fam171a1 | -0.013 | Young |
| 0.962362 | 0.673231 | 0.118 | 0.121 | 1 | 9 | Zmym2 | -0.003 | Young |
| 0.962687 | 0.608454 | 0.059 | 0.062 | 1 | 9 | Hint3 | -0.003 | Young |
| 0.962714 | 0.181598 | 0.147 | 0.144 | 1 | 9 | Specc1l | 0.003 | Young |

| | | | | | | | | |
|---|---|---|---|---|---|---|---|---|
| 0.962812 | 0.215884 | 0.294 | 0.314 | 1 | 9 | Spry1 | -0.02 | Young |
| 0.962893 | 0.59406 | 0.081 | 0.085 | 1 | 9 | Ocel1 | -0.004 | Young |
| 0.963116 | 0.512153 | 0.066 | 0.064 | 1 | 9 | Cdc27 | 0.002 | Young |
| 0.963291 | 0.31762 | 0.118 | 0.121 | 1 | 9 | Rnf19b | -0.003 | Young |
| 0.963381 | 0.294023 | 0.103 | 0.101 | 1 | 9 | Xpo6 | 0.002 | Young |
| 0.963529 | 0.019681 | 0.037 | 0.036 | 1 | 9 | Nqo1 | 0.001 | Young |
| 0.963534 | 0.687075 | 0.022 | 0.023 | 1 | 9 | Heatr5b | -0.001 | Young |
| 0.963954 | 0.320146 | 0.096 | 0.095 | 1 | 9 | Mapk8ip3 | 0.001 | Young |
| 0.963954 | 0.163478 | 0.096 | 0.095 | 1 | 9 | Ranbp9 | 0.001 | Young |
| 0.964563 | 0.189076 | 0.029 | 0.028 | 1 | 9 | Slc35e4 | 0.001 | Young |
| 0.964769 | 0.065479 | 0.176 | 0.175 | 1 | 9 | Slc25a25 | 0.001 | Young |
| 0.964903 | 0.430803 | 0.118 | 0.119 | 1 | 9 | Snx18 | -0.001 | Young |
| 0.965149 | 0.028577 | 0.037 | 0.036 | 1 | 9 | Tmem18 | 0.001 | Young |
| 0.965243 | 0.37756 | 0.044 | 0.044 | 1 | 9 | Asb1 | 0 | Young |
| 0.965284 | 0.261362 | 0.147 | 0.144 | 1 | 9 | Tmem160 | 0.003 | Young |
| 0.965704 | 0.021368 | 0.36 | 0.371 | 1 | 9 | Usp25 | -0.011 | Young |
| 0.96579 | 0.075472 | 0.14 | 0.137 | 1 | 9 | Spr | 0.003 | Young |
| 0.965977 | 0.372412 | 0.169 | 0.165 | 1 | 9 | Ociad1 | 0.004 | Young |
| 0.966013 | 0.312901 | 0.096 | 0.095 | 1 | 9 | Usp10 | 0.001 | Young |
| 0.966188 | 0.43657 | 0.14 | 0.142 | 1 | 9 | Bax | -0.002 | Young |
| 0.966359 | 0.528029 | 0.081 | 0.082 | 1 | 9 | Zfp277 | -0.001 | Young |
| 0.966379 | 0.098568 | 0.029 | 0.028 | 1 | 9 | Cisd3 | 0.001 | Young |
| 0.966447 | 0.34286 | 0.169 | 0.173 | 1 | 9 | Ldb1 | -0.004 | Young |
| 0.96663 | 1.738489 | 0.015 | 0.015 | 1 | 9 | Myrip | 0 | Young |
| 0.96663 | 1.625481 | 0.015 | 0.015 | 1 | 9 | Gk5 | 0 | Young |
| 0.96663 | 1.263181 | 0.015 | 0.015 | 1 | 9 | Wdr24 | 0 | Young |
| 0.96663 | 1.17441 | 0.015 | 0.015 | 1 | 9 | Zfp748 | 0 | Young |
| 0.96663 | 1.169932 | 0.015 | 0.015 | 1 | 9 | C77080 | 0 | Young |
| 0.966641 | 0.489129 | 0.066 | 0.067 | 1 | 9 | Lrrk2 | -0.001 | Young |
| 0.967391 | 0.116083 | 0.132 | 0.131 | 1 | 9 | Pdcl | 0.001 | Young |
| 0.967584 | 0.235984 | 0.184 | 0.188 | 1 | 9 | Rfxap | -0.004 | Young |
| 0.967666 | 0.220264 | 0.081 | 0.08 | 1 | 9 | Ddx19a | 0.001 | Young |
| 0.967848 | 0.463142 | 0.162 | 0.165 | 1 | 9 | Pon2 | -0.003 | Young |
| 0.967854 | 0.050394 | 0.066 | 0.067 | 1 | 9 | Nelfcd | -0.001 | Young |
| 0.967949 | 0.063515 | 0.853 | 0.892 | 1 | 9 | Sparc | -0.039 | Young |
| 0.968192 | 0.047216 | 0.11 | 0.111 | 1 | 9 | Stard7 | -0.001 | Young |
| 0.968235 | 0.030895 | 0.941 | 0.946 | 1 | 9 | Rps29 | -0.005 | Young |
| 0.968284 | 0.107076 | 0.132 | 0.131 | 1 | 9 | Rhobtb1 | 0.001 | Young |
| 0.96848 | 0.035479 | 0.132 | 0.134 | 1 | 9 | Smarcc1 | -0.002 | Young |
| 0.968977 | 0.095303 | 0.596 | 0.634 | 1 | 9 | Rps5 | -0.038 | Young |
| 0.969365 | 0.061534 | 0.699 | 0.691 | 1 | 9 | Nedd4 | 0.008 | Young |
| 0.969678 | 0.668401 | 0.044 | 0.044 | 1 | 9 | Nln | 0 | Young |
| 0.969866 | 0.500237 | 0.11 | 0.113 | 1 | 9 | 5-Mar | -0.003 | Young |
| 0.970145 | 0.167512 | 0.162 | 0.162 | 1 | 9 | Prr14 | 0 | Young |
| 0.970241 | 0.005849 | 0.338 | 0.345 | 1 | 9 | Diaph2 | -0.007 | Young |

| | | | | | | | | |
|---|---|---|---|---|---|---|---|---|
| 0.970535 | 0.344257 | 0.051 | 0.052 | 1 | 9 | Mras | -0.001 | Young |
| 0.97067 | 0.272189 | 0.184 | 0.188 | 1 | 9 | Pet100 | -0.004 | Young |
| 0.971099 | 0.063271 | 0.426 | 0.423 | 1 | 9 | Sub1 | 0.003 | Young |
| 0.97163 | 0.156719 | 0.037 | 0.036 | 1 | 9 | Limd2 | 0.001 | Young |
| 0.972127 | 0.09484 | 0.691 | 0.729 | 1 | 9 | Rpl30 | -0.038 | Young |
| 0.972456 | 0.012353 | 0.25 | 0.253 | 1 | 9 | Tmem256 | -0.003 | Young |
| 0.972622 | 0.190724 | 0.404 | 0.436 | 1 | 9 | Sh3bgrl | -0.032 | Young |
| 0.973202 | 0.670957 | 0.184 | 0.193 | 1 | 9 | Lgmn | -0.009 | Young |
| 0.97325 | 0.3172 | 0.037 | 0.036 | 1 | 9 | Myo1g | 0.001 | Young |
| 0.974433 | 0.271048 | 0.169 | 0.173 | 1 | 9 | Sdf2 | -0.004 | Young |
| 0.974514 | 0.268895 | 0.221 | 0.227 | 1 | 9 | Myo6 | -0.006 | Young |
| 0.974871 | 0.400052 | 0.037 | 0.036 | 1 | 9 | Il10ra | 0.001 | Young |
| 0.974968 | 0.039842 | 0.324 | 0.32 | 1 | 9 | Birc6 | 0.004 | Young |
| 0.975171 | 0.075816 | 0.162 | 0.165 | 1 | 9 | Glce | -0.003 | Young |
| 0.975592 | 0.531562 | 0.044 | 0.044 | 1 | 9 | C3 | 0 | Young |
| 0.975706 | 0.096161 | 0.581 | 0.577 | 1 | 9 | Ehd2 | 0.004 | Young |
| 0.975872 | 0.179064 | 0.184 | 0.183 | 1 | 9 | 1110038F1 | 0.001 | Young |
| 0.976015 | 0.53363 | 0.051 | 0.052 | 1 | 9 | Anks3 | -0.001 | Young |
| 0.976223 | 0.037406 | 0.301 | 0.309 | 1 | 9 | Hspd1 | -0.008 | Young |
| 0.976355 | 0.049786 | 0.397 | 0.407 | 1 | 9 | Fgd5 | -0.01 | Young |
| 0.976438 | 0.067445 | 0.346 | 0.343 | 1 | 9 | Ythdc1 | 0.003 | Young |
| 0.976536 | 0.172466 | 0.176 | 0.173 | 1 | 9 | Tax1bp3 | 0.003 | Young |
| 0.976805 | 0.253581 | 0.103 | 0.106 | 1 | 9 | Naa30 | -0.003 | Young |
| 0.977091 | 0.218869 | 0.301 | 0.304 | 1 | 9 | Lars2 | -0.003 | Young |
| 0.977139 | 0.170646 | 0.301 | 0.307 | 1 | 9 | Map3k1 | -0.006 | Young |
| 0.977385 | 0.420338 | 0.051 | 0.052 | 1 | 9 | Pphln1 | -0.001 | Young |
| 0.977385 | 0.062626 | 0.081 | 0.082 | 1 | 9 | Slc20a2 | -0.001 | Young |
| 0.97791 | 0.154804 | 0.515 | 0.552 | 1 | 9 | Rps25 | -0.037 | Young |
| 0.978126 | 0.007365 | 0.176 | 0.173 | 1 | 9 | Snrpg | 0.003 | Young |
| 0.978127 | 0.125718 | 0.485 | 0.508 | 1 | 9 | 1700020I1 | -0.023 | Young |
| 0.978488 | 0.124824 | 0.081 | 0.082 | 1 | 9 | Psmd9 | -0.001 | Young |
| 0.978492 | 0.081225 | 0.147 | 0.152 | 1 | 9 | Phlpp1 | -0.005 | Young |
| 0.97855 | 0.423425 | 0.044 | 0.044 | 1 | 9 | Fbxw4 | 0 | Young |
| 0.979049 | 0.261446 | 0.125 | 0.126 | 1 | 9 | Zdhhc3 | -0.001 | Young |
| 0.979137 | 0.013518 | 0.647 | 0.675 | 1 | 9 | Oaz1 | -0.028 | Young |
| 0.979326 | 0.006315 | 0.191 | 0.191 | 1 | 9 | Parp4 | 0 | Young |
| 0.979368 | 0.080333 | 0.125 | 0.124 | 1 | 9 | Ltn1 | 0.001 | Young |
| 0.979717 | 0.048068 | 0.176 | 0.173 | 1 | 9 | Tsc22d4 | 0.003 | Young |
| 0.979807 | 0.095527 | 0.176 | 0.175 | 1 | 9 | Samd9l | 0.001 | Young |
| 0.979995 | 0.069447 | 0.368 | 0.371 | 1 | 9 | Leng8 | -0.003 | Young |
| 0.980256 | 0.079758 | 0.221 | 0.227 | 1 | 9 | Immt | -0.006 | Young |
| 0.980753 | 0.236621 | 0.103 | 0.106 | 1 | 9 | Gm49336 | -0.003 | Young |
| 0.981508 | 0.181618 | 0.044 | 0.044 | 1 | 9 | Exosc3 | 0 | Young |
| 0.981631 | 0.381621 | 0.176 | 0.183 | 1 | 9 | Oat | -0.007 | Young |
| 0.982136 | 0.526779 | 0.235 | 0.242 | 1 | 9 | H2afv | -0.007 | Young |

| | | | | | | | | |
|---|---|---|---|---|---|---|---|---|
| 0.982215 | 0.126827 | 0.199 | 0.196 | 1 | 9 | Rala | 0.003 | Young |
| 0.982358 | 0.046521 | 0.346 | 0.345 | 1 | 9 | Etnk1 | 0.001 | Young |
| 0.983087 | 0.053659 | 0.235 | 0.237 | 1 | 9 | Zmym5 | -0.002 | Young |
| 0.98322 | 0.302508 | 0.169 | 0.173 | 1 | 9 | Clasp1 | -0.004 | Young |
| 0.983469 | 0.199345 | 0.132 | 0.131 | 1 | 9 | Birc3 | 0.001 | Young |
| 0.983593 | 0.033678 | 0.713 | 0.763 | 1 | 9 | Ptbp3 | -0.05 | Young |
| 0.983897 | 0.214192 | 0.228 | 0.227 | 1 | 9 | Smarce1 | 0.001 | Young |
| 0.984467 | 0.057905 | 0.044 | 0.044 | 1 | 9 | Polr3b | 0 | Young |
| 0.985015 | 0.110175 | 0.272 | 0.278 | 1 | 9 | Zfp148 | -0.006 | Young |
| 0.985052 | 0.239207 | 0.154 | 0.157 | 1 | 9 | Ppp4c | -0.003 | Young |
| 0.985576 | 0.725542 | 0.074 | 0.075 | 1 | 9 | Tnc | -0.001 | Young |
| 0.985576 | 0.226138 | 0.074 | 0.075 | 1 | 9 | Lrrcc1 | -0.001 | Young |
| 0.986318 | 0.081022 | 0.368 | 0.366 | 1 | 9 | Swap70 | 0.002 | Young |
| 0.986586 | 0.124273 | 0.221 | 0.219 | 1 | 9 | Smarcd2 | 0.002 | Young |
| 0.986608 | 0.221578 | 0.096 | 0.095 | 1 | 9 | Txndc12 | 0.001 | Young |
| 0.98673 | 0.221092 | 0.074 | 0.075 | 1 | 9 | Slc9a3r1 | -0.001 | Young |
| 0.986905 | 0.152742 | 0.213 | 0.219 | 1 | 9 | Fam168b | -0.006 | Young |
| 0.987256 | 0.256372 | 0.397 | 0.402 | 1 | 9 | Rbm25 | -0.005 | Young |
| 0.987312 | 0.157128 | 0.081 | 0.082 | 1 | 9 | C1galt1c1 | -0.001 | Young |
| 0.987371 | 0.468301 | 0.199 | 0.209 | 1 | 9 | Ubr3 | -0.01 | Young |
| 0.987395 | 0.04906 | 0.941 | 0.928 | 1 | 9 | B2m | 0.013 | Young |
| 0.988115 | 0.072457 | 0.221 | 0.224 | 1 | 9 | Arid4a | -0.003 | Young |
| 0.988258 | 0.231234 | 0.096 | 0.098 | 1 | 9 | Chrnb1 | -0.002 | Young |
| 0.988349 | 0.451898 | 0.051 | 0.052 | 1 | 9 | Plrg1 | -0.001 | Young |
| 0.988924 | 0.060782 | 0.522 | 0.539 | 1 | 9 | Samhd1 | -0.017 | Young |
| 0.988983 | 0.059203 | 0.199 | 0.193 | 1 | 9 | Ddx46 | 0.006 | Young |
| 0.989011 | 0.066907 | 0.25 | 0.255 | 1 | 9 | Ubr2 | -0.005 | Young |
| 0.989037 | 0.098724 | 0.074 | 0.075 | 1 | 9 | Gstt3 | -0.001 | Young |
| 0.989068 | 0.188878 | 0.125 | 0.126 | 1 | 9 | Zc3h15 | -0.001 | Young |
| 0.989537 | 0.009308 | 0.426 | 0.433 | 1 | 9 | Bcam | -0.007 | Young |
| 0.989672 | 0.643473 | 0.154 | 0.16 | 1 | 9 | Rin2 | -0.006 | Young |
| 0.989686 | 0.319704 | 0.066 | 0.067 | 1 | 9 | Nudcd2 | -0.001 | Young |
| 0.989686 | 0.308796 | 0.066 | 0.067 | 1 | 9 | Gm20300 | -0.001 | Young |
| 0.989719 | 0.6494 | 0.051 | 0.052 | 1 | 9 | Gm9917 | -0.001 | Young |
| 0.990042 | 0.168916 | 0.206 | 0.211 | 1 | 9 | Zranb2 | -0.005 | Young |
| 0.990191 | 0.447983 | 0.074 | 0.075 | 1 | 9 | Arl2 | -0.001 | Young |
| 0.990244 | 0.30288 | 0.221 | 0.222 | 1 | 9 | Otud4 | -0.001 | Young |
| 0.990348 | 0.032547 | 0.463 | 0.479 | 1 | 9 | Nr4a1 | -0.016 | Young |
| 0.990545 | 0.068737 | 0.162 | 0.16 | 1 | 9 | Lamtor5 | 0.002 | Young |
| 0.990622 | 0.173867 | 0.081 | 0.082 | 1 | 9 | Numb | -0.001 | Young |
| 0.992136 | 0.075213 | 0.603 | 0.603 | 1 | 9 | Notch1 | 0 | Young |
| 0.992228 | 0.0967 | 0.162 | 0.162 | 1 | 9 | Esf1 | 0 | Young |
| 0.992461 | 0.481876 | 0.051 | 0.052 | 1 | 9 | Ppp1r35 | -0.001 | Young |
| 0.992461 | 0.137856 | 0.051 | 0.052 | 1 | 9 | Acad8 | -0.001 | Young |
| 0.992937 | 0.280531 | 0.059 | 0.059 | 1 | 9 | Washc5 | 0 | Young |

| | | | | | | | | |
|---|---|---|---|---|---|---|---|---|
| 0.992937 | 0.113815 | 0.059 | 0.059 | 1 | 9 | Prrg3 | 0 | Young |
| 0.992979 | 0.035989 | 0.118 | 0.119 | 1 | 9 | Trim37 | -0.001 | Young |
| 0.993819 | 0.703051 | 0.096 | 0.095 | 1 | 9 | Smc4 | 0.001 | Young |
| 0.993831 | 0.837537 | 0.051 | 0.052 | 1 | 9 | Ubl4a | -0.001 | Young |
| 0.993831 | 0.154802 | 0.051 | 0.052 | 1 | 9 | Spryd3 | -0.001 | Young |
| 0.993831 | 0.019726 | 0.051 | 0.052 | 1 | 9 | Amd1 | -0.001 | Young |
| 0.993894 | 0.51185 | 0.162 | 0.165 | 1 | 9 | Prpf8 | -0.003 | Young |
| 0.994033 | 0.235577 | 0.375 | 0.397 | 1 | 9 | Akap12 | -0.022 | Young |
| 0.994116 | 0.068114 | 0.088 | 0.088 | 1 | 9 | Mpc1 | 0 | Young |
| 0.994221 | 0.184106 | 0.059 | 0.059 | 1 | 9 | Cog8 | 0 | Young |
| 0.995223 | 0.080383 | 0.706 | 0.729 | 1 | 9 | Rpsa | -0.023 | Young |
| 0.995692 | 0.045796 | 0.228 | 0.224 | 1 | 9 | Suz12 | 0.004 | Young |
| 0.995961 | 0.590246 | 0.074 | 0.075 | 1 | 9 | Hmgxb4 | -0.001 | Young |
| 0.996222 | 0.233754 | 0.412 | 0.438 | 1 | 9 | Pea15a | -0.026 | Young |
| 0.996573 | 0.09841 | 0.051 | 0.052 | 1 | 9 | Dnttip1 | -0.001 | Young |
| 0.996967 | 0.487913 | 0.066 | 0.067 | 1 | 9 | Dnase1l1 | -0.001 | Young |
| 0.996995 | 0.213768 | 0.382 | 0.381 | 1 | 9 | Ifi203 | 0.001 | Young |
| 0.997108 | 0.358664 | 0.11 | 0.111 | 1 | 9 | Mesd | -0.001 | Young |
| 0.997108 | 0.023847 | 0.11 | 0.111 | 1 | 9 | B4galt3 | -0.001 | Young |
| 0.997409 | 0.009648 | 0.206 | 0.209 | 1 | 9 | Ppp3cb | -0.003 | Young |
| 0.997676 | 0.188275 | 0.118 | 0.121 | 1 | 9 | Ifit2 | -0.003 | Young |
| 0.99794 | 0.202377 | 0.096 | 0.095 | 1 | 9 | Pbx3 | 0.001 | Young |
| 0.998074 | 0.162432 | 0.059 | 0.059 | 1 | 9 | Eif4ebp1 | 0 | Young |
| 0.998269 | 0.31592 | 0.074 | 0.075 | 1 | 9 | 3830406C1 | -0.001 | Young |
| 0.998345 | 0.525015 | 0.081 | 0.082 | 1 | 9 | Rcc2 | -0.001 | Young |
| 0.998345 | 0.256956 | 0.081 | 0.082 | 1 | 9 | Lmbr1 | -0.001 | Young |
| 0.998507 | 0.272095 | 0.103 | 0.103 | 1 | 9 | Acaa1a | 0 | Young |
| 0.998658 | 0.029534 | 0.272 | 0.271 | 1 | 9 | Ndufb10 | 0.001 | Young |
| 0.999193 | 0.05569 | 0.169 | 0.168 | 1 | 9 | Frg1 | 0.001 | Young |
| 0.999205 | 0.018064 | 0.713 | 0.742 | 1 | 9 | Rps19 | -0.029 | Young |
| 0.999315 | 0.265574 | 0.051 | 0.052 | 1 | 9 | C5ar1 | -0.001 | Young |
| 0.999315 | 0.002475 | 0.051 | 0.052 | 1 | 9 | Gmeb1 | -0.001 | Young |
| 0.999358 | 0.437918 | 0.059 | 0.059 | 1 | 9 | Nfxl1 | 0 | Young |
| 0.999395 | 0.044549 | 0.368 | 0.374 | 1 | 9 | Bclaf1 | -0.006 | Young |
| 0.999432 | 0.000978 | 0.493 | 0.469 | 1 | 9 | Ssh2 | 0.024 | Young |
| 0.999532 | 0.208841 | 0.118 | 0.119 | 1 | 9 | Maml2 | -0.001 | Young |
| 0.999553 | 0.380273 | 0.132 | 0.131 | 1 | 9 | Pdzd8 | 0.001 | Young |
| 0.999642 | 0.135533 | 0.228 | 0.227 | 1 | 9 | Agap1 | 0.001 | Young |
